# Immunomodulatory Functions of Intercalated Cells in Kidney Autoimmunity

**DOI:** 10.1101/2025.11.14.688275

**Authors:** Maria C. Avenatti, Maia L. Elizagaray, Micah C. Purba, Ferran Barrachina, Angela Chen, Isinsu Bastepe, Katherine Radovanovic, Maria A. Battistone

**Author notes:** Correspondence: Maria Agustina Battistone, PhD, MGH CNY, 149 13th St. Suite 4.325H, Boston, MA 02129. M.C.A. and M.A.B. were involved in the study design and conceptualization. M.C.A., M.C.P., A.C., M.L.E., F.B., I.B., and K.R. performed the experiments and analyzed the data. M.C.A. and M.A.B. were involved in data interpretation. M.C.A. and M.A.B. wrote the original manuscript. All authors contributed to the writing of the manuscript, made critical comments, and approved the final version. **Competing Interest Statement:** The authors declare no financial or non-financial competing interests.

## Abstract

Various autoimmune diseases frequently cause both acute and chronic kidney injuries through complex mechanisms involving autoantibodies and cellular immune responses that result in tissue damage. Intercalated cells (ICs), specialized renal tubular epithelial cells responsible for proton secretion, are strategically positioned at the epithelial-immune interface, making them ideal sensors of stress signals and potential triggers of immune responses. This study investigates the molecular mechanisms by which ICs interact with immune cells to maintain renal immune homeostasis and contribute to the development of autoimmune kidney disease. We depleted Foxp3⁺ regulatory T cells (Tregs) by injecting diphtheria toxin (DT) into male and female Foxp3-DTR mice. Two weeks after depletion, we observed autoimmune inflammation marked by increased renal immune infiltration, including neutrophils, macrophages, and subsets of T and B cells, along with the formation of ectopic lymphoid-like structures and enhanced antigen presentation. We found higher levels of renal autoantibodies in urine and serum, with antibodies depositing in glomeruli and tubules. Our analysis identified several renal antigens targeted by autoantibodies, suggesting their potential role in antibody-mediated renal injury. Kidney damage included smaller glomeruli, proximal tubular injury, an increased urine albumin/creatinine ratio, and decreased urine output. Disruption of immune tolerance led to the upregulation of inflammasome-related genes and IL-33 in ICs, which acts as a key alarmin signaling damage and promoting activation and expansion of Tregs. Our findings uncover a novel IC-Treg interaction mediated by the IL-33 pathway, revealing immune-regulating mechanisms that support renal immune tolerance. Although Tregs were initially depleted, a significant rebound in their numbers and function occurred. Understanding the cellular and molecular mechanisms behind autoimmune renal injury is crucial for developing targeted therapies and identifying appropriate biomarkers.

## Introduction

Chronic kidney disease (CKD), often arising from repeated or unresolved acute kidney injury (AKI), is a primary global health concern, impacting approximately 10–15% of the adult population worldwide^1–3^. Patients suffering from autoimmune conditions, such as systemic lupus erythematosus or vasculitis, are more vulnerable due to immune-mediated damage to the renal glomeruli and tubules, which leads to faster progression of kidney dysfunction^4–9^. Early treatment of autoimmune-related kidney injury is therefore crucial to prevent progression to end-stage renal disease and to improve patient outcomes. Classifying kidney disorder subtypes with targetable immune pathways could enable earlier intervention and more precise therapies. Importantly, there is an urgent need for early biomarkers and treatments that reduce inflammation and support kidney recovery.

Regulatory T cells (Tregs) play a crucial role in suppressing harmful immune responses caused by autoreactive cells and excess inflammation, thereby maintaining tissue homeostasis^10–15^. Dysregulation of this process often contributes to kidney diseases^16–22^. These include post-transplant kidney injury and primary immune-mediated disorders, such as membranous nephropathy and anti-glomerular basement membrane glomerulonephritis, as well as secondary immune-driven conditions, including anti-neutrophil cytoplasmic antibody-associated vasculitis, IgA nephropathy, and lupus nephritis. However, the complex mechanisms by which they prevent and reduce the severe damage caused by excessive inflammation in the kidney are still being studied.

The role of mucosal immunity, especially in epithelial cells such as renal intercalated cells (ICs), in coordinating immune responses in the kidney is gaining increasing interest^23–27^. ICs, which participate in proton secretion in renal tubules^28, 29^, are also crucial in initiating and regulating inflammation during ischemia-reperfusion injury^23, 24, 30^. These cells are situated at the boundary between the renal epithelial barrier and the immune system, making them well-positioned to detect and respond to stress signals^25, 31, 32^. However, the signaling pathways that enable these tissue-specific epithelial cells to communicate with the immune system, mainly with mononuclear phagocytes (MPs), are still largely unknown. This knowledge gap is significant, as understanding how ICs contribute to renal inflammation could lead to the development of new therapies for preventing kidney damage in conditions such as AKI. We have demonstrated that ICs play a crucial role in initiating renal inflammation through cell-to-cell communication involving a danger-associated molecular pattern (DAMP), specifically uridine diphosphate-glucose (UDP-Glc), which binds to the P2Y14 receptor. Consequently, ICs release pro-inflammatory mediators, attract neutrophils and monocytes, and worsen kidney injury in a pre-clinical model of AKI. Notably, our findings also reveal that urinary UDP-Glc levels are linked to AKI in a diverse group of ICU patients, indicating that UDP-Glc may be an early marker of AKI ^23, 24^. Although the exact ways by which ICs communicate with immunocytes to trigger immune activation remain unclear, this study advances our understanding by exploring a previously overlooked role of ICs in the development of autoimmune AKI.

Here, we explored the complex cellular and molecular mechanisms behind cell-cell communication between ICs, immune cells, and other immune components in autoimmune-related kidney injury. Specifically, it highlights how their interactions monitor the renal epithelial barrier and regulate the balance between inflammation and tolerance. Treg depletion resulted in autoimmune renal inflammation characterized by increased infiltration of neutrophils, macrophages, T and B cells, formation of B-cell clusters, and enhanced antigen presentation. We detected higher levels of renal autoantibodies in both urine and serum, with antibody deposits present in the glomeruli and tubules, ultimately leading to kidney damage. We identified the renal antigens targeted by these autoantibodies, emphasizing the potential use of specific antibody detection as a biomarker for autoimmune renal diseases. Disruption of tolerance caused ICs to upregulate inflammasome-related genes and IL-33, a key alarmin that signals damage and activates Tregs. Our findings reveal a novel IC-Treg crosstalk via the IL-33 pathway, uncovering immunoregulatory mechanisms that promote renal immune tolerance. The molecular foundation of this pathway offers potential therapeutic targets for autoimmune kidney injury.

## Methods

### Experimental Animals

Female C57BL/6-Tg (Foxp3-DTR/EGFP)23.2Spar/Mmjax mice and male C57BL/6 wild-type (WT) mice were obtained from Jackson Laboratory (Bar Harbor, ME). The breeding program was at Massachusetts General Hospital (MGH) and generated Foxp3-DTR or WT (control) male and female offspring. These transgenic mice express the diphtheria toxin receptor (DTR) fusion protein controlled by the endogenous forkhead box P3 (Foxp3) promoter/enhancer regions. B1-EGFP mice, expressing EGFP under the control of intercalated cell (IC)-specific V-ATPase B1 subunit (ATP6V1B1), were crossbred with Foxp3-DTR mice to generate B1-EGFP Foxp3-DTR offspring^33, 34^. Male and female mice, aged between 6-15 weeks, were utilized throughout this study. All experimental procedures were strictly followed in accordance with the National Institutes of Health (NIH) Guide for the Care and Use of Laboratory Animals and received approval from MGH’s Research Animal Care Subcommittee.

### Regulatory T Cell In Vivo Depletion

To deplete Tregs, six-week-old Foxp3-DTR and WT male and female littermates received intraperitoneal injections of diphtheria toxin (DT) from *Corynebacterium diphtheriae* (D0564, Sigma-Aldrich, St. Louis, MO) dissolved in phosphate-buffered saline (PBS) at 40 µg/kg of body weight. DT injections were administered on days 0, 2, and 4, and mice were analyzed at 2 weeks and 8 weeks post-injection^34^. WT mice given DT served as the control group. To confirm Treg depletion kinetics, some Foxp3-DTR and WT mice received a single DT injection and were analyzed 24 h later^34^.

### In vivo staining of CD45^+^ cells

To stain circulating CD45⁺ cells *in vivo*, mice were intravenously (i.v.) injected with 3 µg of anti-mouse CD45-PE or CD45-PE-Fire-700 (diluted in 100 µL saline) 10 min before euthanasia, following a published protocol^34^. After tissue collection, samples were processed for flow cytometric staining and fixation. The method for differentiating circulating from tissue-resident CD45⁺ cells is described in Figure 1.

**Figure 1.**
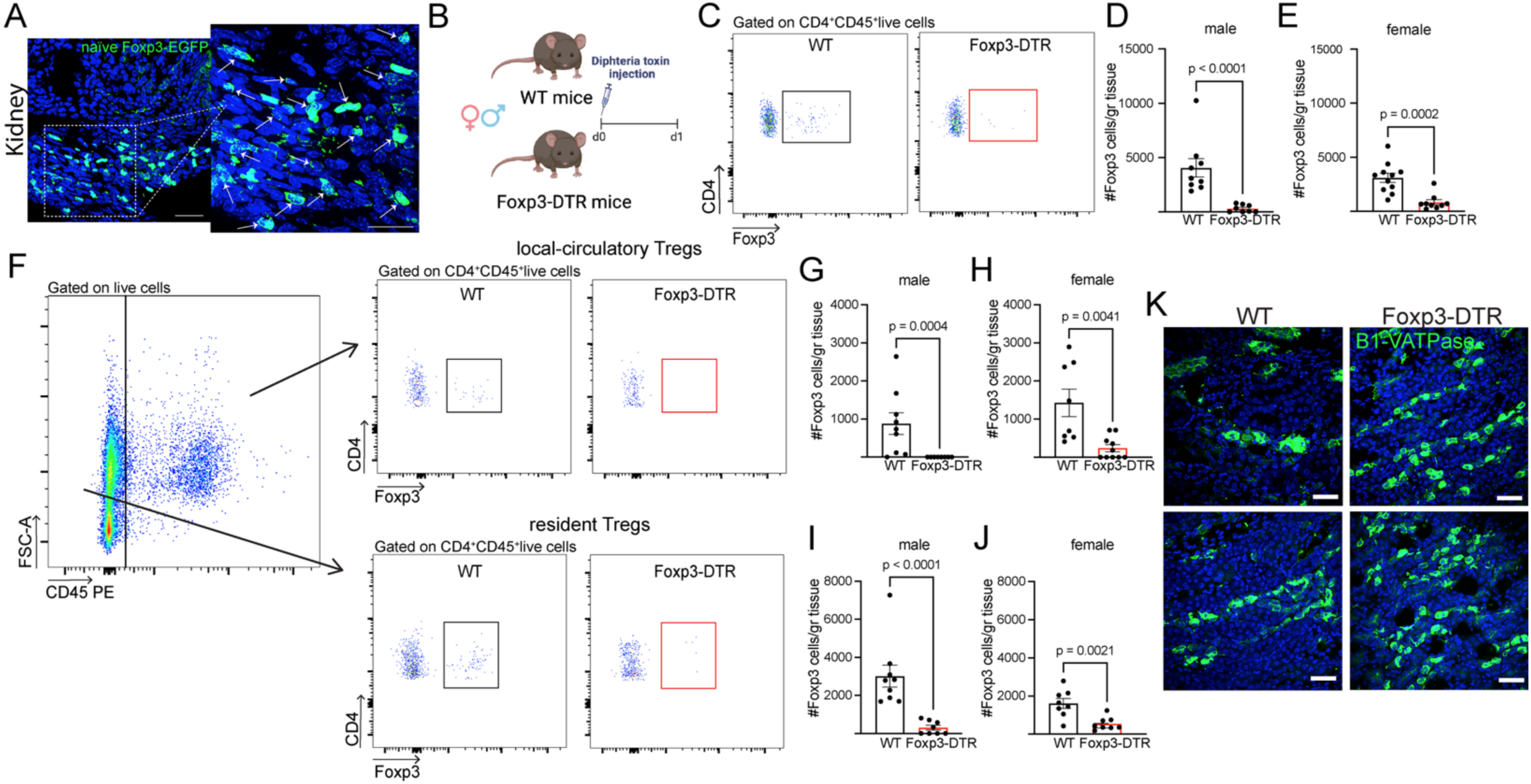
Treg population in the kidneys of Foxp3-DTR mice 1 day after Treg depletion. **A)** Confocal microscopy images showing Foxp3-EGFP^+^ Tregs (arrows) in the kidneys of naïve Foxp3-EGFP transgenic mice. *n* =3 kidneys. **B)** Diagram of regulatory T cell (Treg) depletion protocol and subsequent analysis 1 day after diphtheria toxin (DT) injection. The figure was created with BioRender.com. **C)** Flow cytometry gating strategy for identifying Tregs (Foxp3^+^CD4^+^CD45^+^) in the kidneys. Absolute count (#) of total Tregs per gram of tissue in male **(D)** and female **(E)** kidneys. Male: *n*=9 (WT), *n*=8 (Foxp3-DTR). Female: *n*=11 (WT), *n*=9 (Foxp3-DTR). **F)** Flow cytometry gating strategy for identifying tissue-resident Tregs (CD45-PE^-^Foxp3^+^CD4^+^BV711CD45^+^) and circulatory Tregs (CD45-PE^+^Foxp3^+^CD4^+^BV711CD45^-^). Absolute count (#) of circulatory Tregs per gram of tissue in male **(G)** and female **(H)** kidneys. Absolute count (#) of tissue-resident Tregs per gram of tissue in male **(I)** and female **(J)** kidneys. Male: *n*=9 (WT), *n*=8 (Foxp3-DTR). Female: *n*=8 (WT), *n*=10 (Foxp3-DTR). **K)** Confocal microscopy images showing B1V-ATPase^+^ cells (intercalated cells (IC); green) in kidney sections of 1-day DT-injected WT and Foxp3-DTR mice. *n*=3 kidneys per group. DAPI (blue, cell nuclei). Bars: 20 μm. Data are means ± SEM, and two-sided Mann–Whitney test **(D, E, G-I)** or Student’s t-test **(J)** were performed. *n*= number of samples from different mice.

### Mouse Kidney Collection

Under isoflurane anesthesia (2%, mixed with oxygen, Baxter, Deerfield, IL), mice underwent cardiac perfusion with PBS through the left ventricle of the heart until the organs were free of blood, ensuring that they were adequately cleared for subsequent ELISA analyses. Following perfusion, the kidneys were carefully dissected for further examination. Tissues designated for fixation continued perfusion with 4% paraformaldehyde solution (PFA; 15714-S, Paraformaldehyde 32% Solution EM grade, Electron Microscopy Sciences) for 5 min, followed by 4-hour post-collection immersion in 4% PFA^35^. For *in vivo* CD45 analysis, PBS perfusion was omitted, and kidneys with spleen were directly collected following euthanasia.

### Mouse Serum Collection and Biochemical Analysis of Renal Biomarkers

Whole blood was obtained via left ventricular puncture using a 1 mL syringe with a 25G × 5/8” BD PrecisionGlide™ needle (Becton, Dickinson and Company, Franklin Lakes, NJ) and collected in BD Microtainer® collection tubes (365967, Becton, Dickinson and Company, Franklin Lakes, NJ). Following a 30-min incubation period at room temperature to promote clotting, samples underwent centrifugation at 4,000 *g* for 10 minutes to separate and recover the serum fraction. Serum BUN and creatinine levels were determined with a Heska DriChem 7000 chemistry analyzer (http://www.heska.com/product/element-dc5x). Urinalysis for creatinine and albumin was performed using the DCA Vantage system (Siemens Healthineers) at the Center for Comparative Medicine of Massachusetts General Hospital.

### Tissue Homogenization and Protein Extraction

Previously cleaned, perfused, and frozen tissues were thawed in PBS containing protease inhibitor cocktail (#04693159001, cOmplete™ Mini EDTA-free Protease Inhibitor Cocktail, Roche, Basel, Switzerland). Tissue samples underwent mechanical disruption using scissors, followed by pestle homogenization, and were then maintained on ice for 30 min. Following centrifugation at 10,000g for 20 minutes, the resulting supernatant was collected for immediate ELISA analysis or stored at -80°C for subsequent use^34^.

### Immunofluorescence Microscopy

Following established protocols^35^, fixed tissue specimens were cryoprotected in 30% sucrose-PBS solution (containing 0.02% sodium azide) for 48 h at 4°C, then embedded in Tissue-Tek OCT compound (Sakura Finetek, Torrance, CA). Cryosections of 5, 10, or 16 μm thickness were generated using a Reichert Frigocut microtome and mounted on Fisherbrand Superfrost Plus slides (Fisher Scientific, Pittsburgh, PA).

The primary antibody panel included rat anti-mouse-F4/80^33^, rabbit anti-V-ATPase B1 subunit^35^, rat anti MHC Class II (I-A/I-E)^33^ (0.2 μg/mL, M5/114.15.2, 12-5321-81, Thermo Fisher Scientific), rat anti AQP1^23^ (200 µg/mL, sc-32737, Santa Cruz Biotechnology), goat anti-mouse-IgG1^34^ (1mg/mL, A10551, Invitrogen), goat anti-mouse-IgG2c^34^ (0.8mg/mL, 115-035-208, Jackson ImmunoResearch), donkey anti-mouse-IgM^34^ (0.8mg/mL, 715-035-020, Jackson ImmunoResearch), Armenian Hamster anti-mouse-CD3^34^ (0.5mg/mL, 100303, Bioegend), rat anti-mouse-IgD^34^ (0.5mg/mL, 405702, Biolegend), goat anti-mouse-CD20^34^ (0.5mg/mL, sc-7735, Santa Cruz), goat anti-mouse IL-33 (0.2 mg/mL; Polyclonal; AF3626, R&D Systems), rat anti-mouse CD138 (0.2 mg/ml; Clone 281-2, 563192, BD Biosciences) and Alexa Fluor 488 rat anti-mouse-CD45 (0.5mg/ml; Clone 30-F11, 103121, BD Biosciences).

Secondary antibodies comprised Alexa Fluor 488 goat anti-rabbit IgG^34^ (1:800 dilution; 111-545-144, Jackson ImmunoResearch), DyLight 649 goat anti-hamster^34^ (STAR104D649, AbD Serotec), Cy3 donkey anti-rabbit IgG^34^ (7.5 μg/ml; 711-165-152, Jackson ImmunoResearch), Cy3 donkey anti-rat IgG^34^ (1:800 dilution; 712-166-153, Jackson ImmunoResearch), Alexa Fluor 647 donkey anti-mouse^34^ (1:800 dilution; 715-606-151, Jackson ImmunoResearch), Cy3 donkey anti-goat IgG^34^ (1:800 dilution; 705-166-147, Jackson ImmunoResearch), Cy3 donkey anti-goat IgG(H+L)^34^ (1:800 dilution; 705-165-147, Jackson ImmunoResearch), Alexa Fluor 488 Donkey Anti-Goat IgG (H+L)^34^ (1.5mg/ml; 705-545-147, Jackson ImmunoResearch).

Antigen retrieval protocols were tailored to specific antibodies: 4-min treatment with 1% SDS in PBS (for V-ATPase B1, MHCII, CD45, CD20, CD3, CD138, IL-33, IgM, IgG1, IgG2c, and AQP1), or 1% SDS plus 0.1% Triton X-100 in PBS (for F4/80 and IgG antibodies). Slides underwent 30-minute blocking in 1% BSA-PBS at room temperature, followed by 18-hour primary antibody incubation at 4°C. All antibodies were diluted in DAKO antibody diluent (Dako North America, Cat: S0809). Slides were mounted using SlowFade Diamond Antifade Mounting medium (Thermo Fisher Scientific, S36963) with DAPI nuclear counterstain. Negative control slides received secondary antibody treatment only.

Images were taken using a Nikon CSA-W1 SoRa spinning disk confocal microscope (Nikon, 173 Yokogawa Electric Corporation, Tokyo, Japan) at the Molecular Imaging Core (MGH, Charlestown, MA). Quantification of tubular damage was performed using Fiji software as previously reported^23^

Renal tubular damage was evaluated using a three-tier scoring system applied to AQP1-immunostained kidney sections as previously described^23^. Two blinded investigators independently assessed tubular integrity by categorizing structures as intact, moderate injury, or severe injury based on AQP1 subcellular localization patterns in immunolabeled tissue. For AQP1 analysis, intact proximal tubules displayed characteristic apical brush border and basolateral membrane immunoreactivity. Moderated damage was defined by preserved basolateral but absent apical AQP1 expression, while severe damage showed complete loss of polarized AQP1 distribution.

### H&E evaluation

Tissue sections, cut at 5 μm thickness, were stained using Harris Modified Hematoxylin (Sigma-Aldrich, HHS32), followed by counterstaining with Eosin Y (Sigma-Aldrich, HT110132). The prepared slides were then digitized with a NanoZoomer 2.0RS scanner (Hamamatsu, Japan). Measurements of glomerular length were taken in two directions using Fiji software.

### Flow Cytometry analysis

The kidneys underwent mechanical and enzymatic processing using collagenase types I (0.5 mg/mL) and collagenase type II (0.5 mg/mL) for 30 min at 37°C in RPMI 1640 medium, as previously reported^23^. Cell suspensions were filtered through a 70 μm nylon mesh strainer for kidney samples. The filtrates were washed with PBS supplemented with 2% fetal bovine serum (FBS) and 2 mM EDTA, then centrifuged at 400 × g for 5 min. Erythrocytes were lysed using ACK buffer (#A10492-01, Gibco, Grand Island, NY) for 1 min, followed by another centrifugation step. The cell pellets were then incubated for 30 min at 4°C with a comprehensive panel of fluorochrome-conjugated anti-mouse antibodies (1:400 dilution, in 2% FBS in PBS with BD Horizon Brilliant Stain buffer) as described in detail in the Suppl. Table 1. Cell viability was assessed using DAPI (62248, ThermoScientific) or LIVE/DEAD™ Fixable Blue Dead Cell Stain Kit, for UV excitation (L23105, Invitrogen) or LIVE/DEAD™ Fixable Yellow Dead Cell Stain Kit, for 405 nm excitation (L34967, Invitrogen).

For the assessment of Foxp3 nuclear expression, CD152, IL-33 production, and intracellular IgM detection, cell preparations underwent fixation for 20 min using the 1x Fixation/Permeabilization working solution, followed by washing with 1x Permeabilization Buffer according to the manufacturer’s protocol (#00-5523-00, Thermo Fisher Scientific). The permeabilized cells were then stained with 1:100 dilution overnight with anti-Foxp3-APC, anti-CD152 BV421, anti-IL-33 and anti-IgM BV786, all resuspended in 1x Permeabilization Buffer (00-5523-00, Thermo Fisher Scientific) enriched with BD Horizon Brilliant Stain buffer (563794, BD Biosciences). For IL-33 detection, cells were subsequently incubated for 1 h with a donkey anti-goat Cy3 secondary Ab (#705-166-147, Jackson Immunoresearch Laboratories)^36^ at 1:100 dilution in 2% FBS in PBS. Following antibody incubation, cells were rinsed with PBS containing 2% FBS and passed through a 40 μm mesh filter.

Data acquisition was performed using a BD FACSAria II (BD Biosciences), BD LSRFortessa X-20 (BD Biosciences), or Cytek Aurora System flow cytometer, and subsequently analyzed with FlowJo software v10.8.1 (BD Biosciences). Gating parameters were defined through comprehensive control experiments, which incorporated unstained cellular preparations from each tissue and a fluorescence minus one (FMO) control, as per standard protocols. Complete gating strategies are provided in the Suppl. Fig. 1 and in previous publications^33, 34^. Tissue-infiltrating immune cells were analyzed following whole-body perfusion or *in vivo* CD45^PE^ staining, with immune populations identified by gating on CD45^PE^-negative cells.

Advanced computational analysis employed t-distributed Stochastic Neighbor Embedding (t-SNE) alongside FlowSOM algorithms to facilitate comprehensive cellular population characterization. The t-SNE algorithm enabled visualization of complex multi-parameter datasets through dimensionality reduction techniques, while FlowSOM provided automated clustering based on cellular phenotypic characteristics to identify discrete population subsets.

### Antibody level via ELISA

Kidney tissues from WT mice were homogenized using RIPA buffer (R26200-125.0, Research Products International, Mt. Prospect, IL) containing both a protease inhibitor cocktail (cOmplete Mini EDTA-free, #04693159001, Roche, Basel, Switzerland) and a phosphatase inhibitor cocktail (PhosSTOP, #4906837001, Roche). After incubation for 20 min at 4°C, samples were centrifuged at 10,000 × g for 10 minutes, and the resulting supernatants, containing soluble proteins, were collected for subsequent analysis. Microplate preparation entailed overnight coating at 4°C with kidney-derived proteins at a 6 µg/mL concentration in PBS buffer (pH 7.4). Following blocking with 3% BSA in PBS (1 h, 37°C), wells were incubated for 2 h at 37°C with biological samples diluted in 1% BSA-PBS: mouse serum (1:25), mouse urine (1:5) and tissue homogenates (1:3). Following washes, antigen-bound isotypes were detected using HRP-conjugated secondary antibodies: IgG (1:5000; 115-035-205, Jackson ImmunoResearch), IgA (1:3000; 62-6720, Invitrogen), IgG1 (1:2000; A10551, Invitrogen), IgG2b (1:5000; M32407, Invitrogen), IgG2c (1:5000; 115-035-208, Jackson ImmunoResearch), IgG3 (1:2000; M32707, Invitrogen), IgM (1:5000; 715-035-020, Jackson ImmunoResearch) and IgA (1:10000; 109-035-011, Jackson ImmunoResearch) —all incubated for 1 hour at room temperature. Signal development employed 3,3’,5,5’-tetramethylbenzidine (TMB) substrate (TMB substrate kit, 34021, Thermo Fisher Scientific) with acid quenching (H₂SO₄2N). Spectrophotometric analysis was conducted at 450 nm wavelength using a Promega GloMax Discover plate reader (Madison, WI).

### Immunoprecipitation of renal protein antigens

To investigate the renal antigens that autoantibodies recognized, serum from both WT and Foxp3-DTR mice were collected 2 weeks after DT treatment. These samples were incubated with Dynabeads® Protein G (Immunoprecipitation Kit, 10006D, Invitrogen) to capture specific antibodies, following the manufacturer’s protocol. After thorough washing, the beads were then exposed to renal protein extracts from WT mice, prepared as previously described^34^. The resulting complexes were eluted and separated by electrophoresis on a 10% NuPAGE™ Bis-Tris gel (NP0302BOX, Invitrogen), followed by staining with Coomassie R-250 (ImperialTM Protein Stain, 24615, Thermo Scientific) to visualize protein bands. Bands of interest were excised and sent for Mass Spectrometry analysis at the Taplin Biological Mass Spectrometry Facility, Harvard Medical School, Boston, MA.

### Renal Proteomic Data Interpretation

Datasets from each experimental group were compared using the VENNY 2.1 online Venn diagram tool. Protein abundance was determined by summing the intensities of all spectral peaks corresponding to each protein, and differences in protein expression across groups were visualized using heatmaps generated with the Morpheus web-based software (https://software.broadinstitute.org/morpheus)

### Isolation of EGFP^+^ Intercalated Cells (ICs) from Kidney, RNA Extraction, and RNA-seq Analysis

EGFP-positive (EGFP^+^) live ICs were isolated from the renal medulla of B1-VATPase^EGFP^ Foxp3-DTR and B1-VATPase^EGFP^ mice using double fluorescence-activated cell sorting (FACS) at the HSCI-CRM Flow Cytometry Core (Boston, MA). DAPI was employed to distinguish viable cells.

RNA was extracted from the sorted ICs using the PicoRNA kit (Thermo Fisher Scientific). To prevent DNA contamination, samples were treated with an RNase-free DNase set (Qiagen, Hilden, Germany). RNA quality and quantity were assessed using a Bioanalyzer (Agilent RNA 6000 Pico Kit, Agilent Technologies, Santa Clara, CA).

RNA-seq libraries were prepared using the Clontech SMARTER Kit v4, followed by sequencing on an Illumina HiSeq2500 platform. Transcriptome mapping was performed using STAR, with the Ensembl mm9 reference genome utilized for alignment. Gene-level read counts were generated using HTSeq. Differential expression analysis was conducted with the EdgeR package, considering only genes with counts per million (CPM) greater than 1. Differentially expressed genes were defined as those exhibiting a more than 2-fold change in expression and a P-value of less than 0.05.

Volcano plots were generated using Multiplot Studio software, and heatmaps were constructed using Morpheus. The complete RNA-seq dataset for kidney ICs will be deposited in the Gene Expression Omnibus.

### Statistical Methods

Statistical evaluations were conducted using GraphPad Prism version 10.4.1 (GraphPad Software, La Jolla, CA; https://www.graphpad.com). We assessed normal distribution using the Shapiro-Wilk normality test, and variance homogeneity was examined via an F-test (for two-group comparisons). For parametric data, we employed a two-tailed Student’s t-test or one-way ANOVA with subsequent Tukey’s multiple comparison tests. Non-parametric analyses employed a two-tailed Mann-Whitney test or a Kruskal-Wallis test, followed by Dunn’s post hoc analysis. Statistical significance was established at p<0.05, with results presented as means ± SEM.

## Results

### Tregs sustain immunity in the renal mucosa

Flow cytometry analysis and confocal microscopy revealed that the resting kidney contains a significant population of regulatory T cells (Tregs; Foxp3^+^CD4^+^CD45^+^ cells) in naïve and WT kidneys (Fig. 1A-E). To evaluate the functional importance of these cells, we used transgenic Foxp3-DTR/EGFP mice, in which Tregs can be depleted by administering diphtheria toxin (DT) (Fig. 1B). The total number of Tregs per gram of kidney tissue decreased 24 h after DT treatment in both female and male mice (Fig. 1C–E). To distinguish local Tregs from circulating ones, we performed *in vivo* intravascular labeling of CD45^+^ immune cells. Mice received an intravenous injection of anti-CD45-PE antibody 10 min before euthanasia, 24 h after DT^34^. Flow cytometry analysis then included staining with a second anti-CD45 antibody conjugated to BV711, which allowed us to differentiate circulating (CD45-PE⁺) from tissue-resident (CD45-BV711⁺ PE⁻) immune cells (Fig. 1F). Circulating Tregs decreased after DT injection in both male and female tissues (Fig. 1G-H). Quantification revealed a reduction in renal Tregs in male and female Foxp3-DTR mice compared to controls (Fig. 1I-J), confirming the effective depletion of Tregs. Kidney epithelial cells, including intercalated cells (ICs) remained unaffected after early Treg depletion (Fig. 1K).

### Treg ablation triggers kidney immune infiltration

Two weeks following Treg depletion (Fig. 2A), there was an increased infiltration of CD45^+^ immune cells (Fig. 2B-E), monocytes (CD11b^+^Ly6G^-^Ly6C^+^; Fig. 2F-H), macrophage-like mononuclear phagocytes (F4/80^+^CD64^+^; MPs; Fig. 2I-L), and MHCII^+^ cells (Fig. 2M-P) in the kidneys of both males and females. Notably, the proportion of neutrophils (CD11b^+^Ly6G^+^) increased in male kidneys but not in female kidneys (Fig. 2Q-S). Confocal microscopy showed many F4/80^+^ MPs (Fig. 2L, red) in the cortex and medulla of kidneys in DT-injected Foxp3-DTR mice. Infiltrating MHCII^+^ cells are located in the cortex and medulla (Fig. 2P). Interestingly, intercalated cells (ICs, B1 subunit V-ATPase+ cells) closely interact with MHCII^+^ cells in the kidney (Fig. 2T-U). This cell-cell interaction persists even after tolerance ablation.

**Figure 2.**
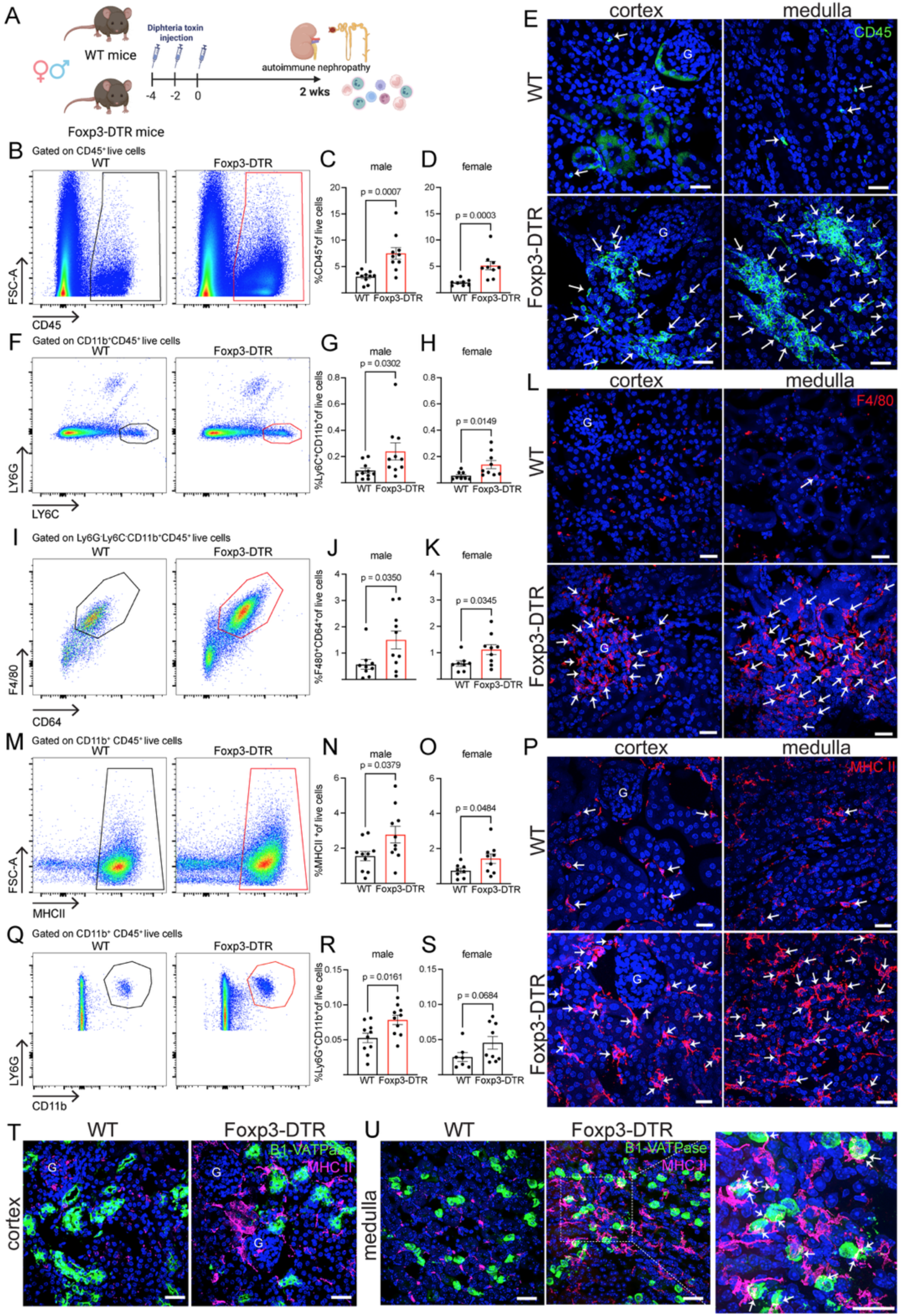
Renal recruitment of proinflammatory immune cells following Treg depletion in Foxp3-DTR mice after 2 weeks. **A)** Diagram of the Treg depletion protocol and subsequent analysis 2 weeks after DT injection. The panel was created with BioRender.com. Flow cytometry gating strategy and relative abundance of CD45^+^ immune cells **(B-D)**, monocytes (Ly6C^+^Ly6G^−^CD11b^+^CD45^+^) **(F-H)**, F4/80^+^ mononuclear phagocytes (MPs; F4/80^+^Ly6G^-^Ly6C^-^CD11b^+^CD45^+^) **(I-K)**, MHCII^+^MPs (MHCII^+^CD11b^+^CD45^+^) **(M-O),** and neutrophils (Ly6G^+^CD11b^+^CD45^+^) **(Q-S)** in male and female kidneys. Male: *n*=10 (WT), *n*=10 (Foxp3-DTR). Female: *n*=8 (WT), *n*=9 (Foxp3-DTR). Confocal microscopy images showing infiltration of CD45^+^ immune cells (arrows, green) **(E)**, F4/80^+^MPs (arrows, red) **(L)**, and MHCII^+^ MPs (arrows, red) **(P)** in the cortex and renal medulla of WT and Foxp3-DTR mice after DT. **Q)** Confocal microscopy images showing B1V-ATPase^+^ cells (intercalated cells (ICs); green), and MCHII^+^ MP (MP, pink), DT-injected WT and Foxp3-DTR mice. Arrows indicate MP-IC interaction. DAPI (blue, cell nuclei). Bars: 20 μm. G: Glomeruli. Data were analyzed using a two-sided Mann–Whitney test (**C, D, G, H, J, S)** or a Student’s t-test **(K, N, O, R).** Data are presented as means ± SEM. *n=* number of samples from different mice.

t-SNE (t-distributed Stochastic Neighbor Embedding) and FlowSOM (a clustering algorithm) analysis of the immune cell node (CD45^+^) revealed multiple lymphocyte subtypes. They revealed a shift in the renal immune cell landscape after Treg depletion (Fig. 3A, Suppl. Fig. 2A). Interestingly, 2 weeks after DT injections, an increase in renal Tregs was observed, suggesting that these cells repopulate the kidney to help counteract the heightened proinflammatory environment (Fig. 3B-C). Additionally, the number of CD4⁺Foxp3⁻ cells increased in both male and female kidneys (Fig. 3D). We also detected higher levels of follicular T cells and follicular regulatory T cells (Fig. 3F-H). Notably, the expression of CD69, PD-1, CD25, CD152, and LAG3 on Tregs was also elevated, indicating these repopulating cells are functionally active and have enhanced suppressive capacity within the tissue (Fig. 3I-R). Treg ablation also led to infiltration of CD8⁺ T cells and activated CD8⁺ T cells into the kidneys (Fig. 3S-V). An increase in B cell abundance was observed in the kidney 2 weeks after tolerance breakdown (Fig. 4A–H). Remarkably, naïve mature (Fig. 4H-J) and plasma B cells (Fig. 4K–M) were also elevated, suggesting B cell maturation and autoantibody production may be occurring locally within the renal tissue. After Treg depletion, both germinal center (GC)-like activity and follicular dendritic cell populations increased, indicating activation of local B cell responses (Fig. 4N–O, R-S). Furthermore, memory B cells and regulatory B cells (Bregs) were elevated 2 weeks after DT treatment (Fig. 4P-Q, Suppl. Fig. 2B). Confocal microscopy revealed B-cell clusters resembling ectopic lymphoid-like structures in the kidney following depletion (Fig. 4G, H, M). These structures contained both IgD⁺ B cells and CD138⁺ plasma cells. Notably, the ectopic clusters were located within a highly vascularized region of the medulla, in agreement with prior studies that have reported similar formations in this anatomical site^37, 38^.

**Figure 3.**
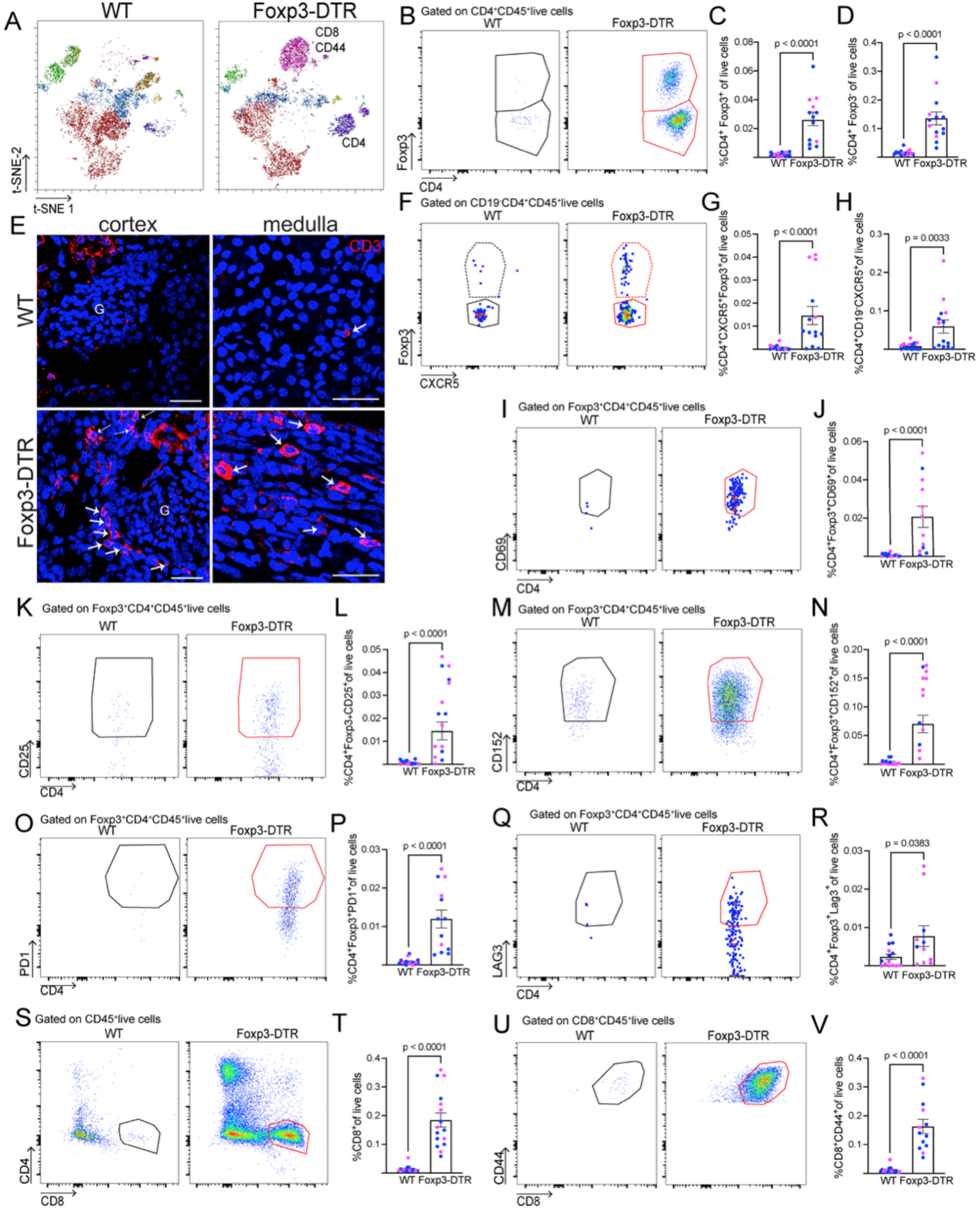
Renal T cell profiling in Foxp3-DTR mice 2 weeks after Treg depletion. **A)** t-SNE and FlowSOM analysis of kidneys from WT and Foxp3-DTR mice: Magenta: CD8/CD44 and purple: CD4. *n*=3. The most relevant populations that relatively increased in the Foxp3-DTR samples are highlighted. Flow cytometry gating strategy and relative abundance of Tregs (Foxp3^+^CD4^+^CD45+) (**B-C)**, and conventional T cells (Foxp3^-^CD4^+^CD45^+^) **(D)** in male and female kidneys of WT and Foxp3-DTR mice. Male (blue dots): *n* = 11 (WT), *n =* 10 (Foxp3-DTR). Female (pink dots): *n* = 5 (WT), *n=* 4 (Foxp3-DTR). **E)** Confocal microscopy images showing infiltration of CD3 (red) in the cortex and renal medulla of DT-treated Foxp3-DTR and WT mice. Arrows: T cells. *n*=3 per group. **F)** Flow cytometry gating strategy and relative abundance of T follicular regulatory cells (Tfr, CD4^+^CD19^−^CD8^−^CXCR5^+^Foxp3^+^) **(G)** and T follicular helper cells (Tfh, CD4^+^CD19^−^CD8^−^CXCR5^+^) **(H)** in male and female kidneys of WT and Foxp3-DTR mice. Male: *n* = 11 (WT), *n =* 10 (Foxp3-DTR). Female: *n* = 5 (WT), *n* = 4 (Foxp3-DTR). Flow cytometry gating strategies **(I, K, M, O, Q, S, U)** and relative abundance of CD69^+^**(J)**, CD25^+^**(L)**, CD152^+^**(N)**, PD1^+^**(P)**, Lag3^+^**(R)**, CD8^+^ T cells **(T)** and activated CD8^+^ T cells (CD4^−^CD8^+^CD44^+^) **(V)** in male and female kidneys of DT-treated WT and Foxp3-DTR mice. Male(blue dots): *n* = 6 (WT), *n =* 4 (Foxp3-DTR). Female(pink dots): *n* = 8 (WT), *n* = 8 (Foxp3-DTR). DAPI (blue, cell nuclei). *n*= number of samples from different mice. Bars: 20 μm. G: Glomeruli. Data were analyzed using the two-sided Mann–Whitney test. Data are shown as means ± SEM.

**Figure 4.**
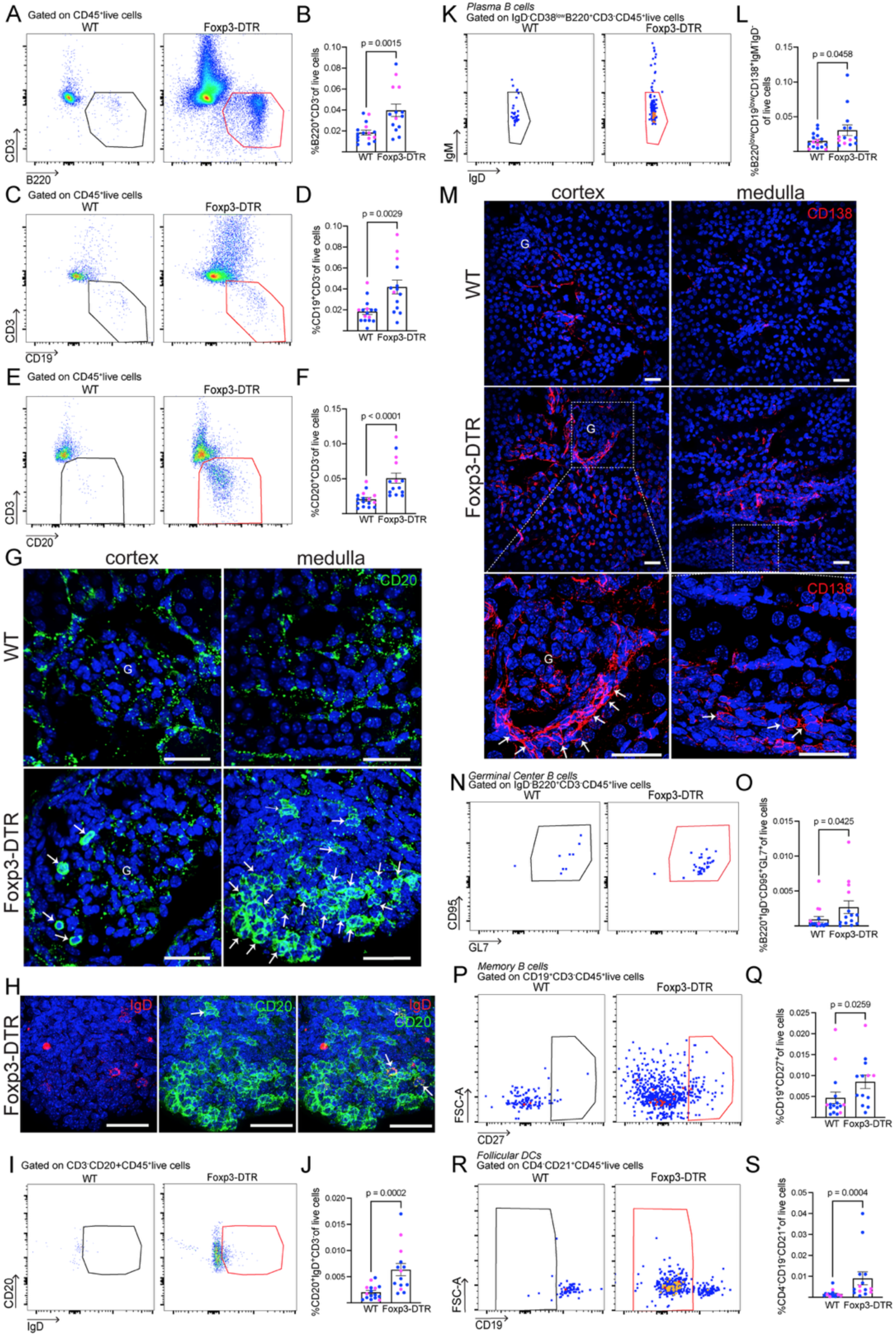
B cell analysis in the mouse kidneys 2 weeks after Treg depletion. Flow cytometry gating strategy and relative abundance of total B cells: CD3^−^B220**^+^ (A-B)**, CD3^−^CD19^+^ **(C-D),** and CD3^−^CD20^+^ **(E-F)** of male and female kidneys. Male (blue dots): *n =* 11 (WT), *n*= 10 (Foxp3-DTR). Female (pink dots): *n=* 5 (WT), *n=* 4 (Foxp3-DTR). **G)** Immunolabeling of CD20 (arrows, green) in the kidneys of DT-injected WT and Foxp3-DTR mice. Non-specific dots are visible in both genotypes. *n=* 3. **H)** Immunolabeling of IgD (red), and CD20 (green), in the DT-injected Foxp3-DTR distal epididymis. Non-specific background visible in both genotypes. Arrows: double-positive cells. *n =* 3. Flow cytometry gating strategy **(I)** and relative abundance of IgD^+^CD20^+^B cells **(J)** in male and female kidneys. Flow cytometry gating strategy **(K)** and relative abundance of Plasma B cells (B220^low^CD19^low^IgD⁻IgM^-^CD138⁺) in male and female kidneys. Male: *n =* 11 (WT), *n*= 10 (Foxp3-DTR). Female: *n=* 5 (WT), *n=* 4 (Foxp3-DTR). **M)** Immunolabeling of CD138 (arrows, red) in the kidneys of DT-injected WT and Foxp3-DTR mice. Non-specific background visible in both genotypes*. n=* 3. Flow cytometry gating strategy **(N, P, R)** and relative abundance of Germinal center (GC) B cells (CD3⁻B220⁺IgD^-^CD95^+^GL7⁺) **(O)**, Memory B cells (CD19^+^CD3^-^CD27^+^) (**Q**), follicular dendritic cells (DC, CD4^−^CD19^−^CD21^+^) **(S)** in male and female kidneys. Male: *n=* 11 (WT), *n*= 10 (Foxp3-DTR). Female: *n=* 5 (WT), *n=* 4 (Foxp3-DTR). DAPI (blue). Bars: 20 μm. G=Glomeruli. Data are means ± SEM, and a two-sided Mann–Whitney test was performed. *n=* number of samples from different mice.

### Kidney-specific autoantibody production after Treg depletion

Ablation of Tregs caused significant deposition of immunoglobulins (Ig), including IgG and IgM, not only in the glomeruli but also throughout the peritubular interstitial areas of the cortex and medulla (Fig. 5A, D). ELISA confirmed increased levels of total IgG and IgM autoantibodies in the kidneys of Treg-depleted male and female mice compared to DT-injected WT (Fig. 5B, C, E, F). Further isotyping of the autoantibodies revealed higher levels of IgG1 (Fig. 5G-I) and IgG2c (Fig. 5J-L) after Treg depletion, while IgG2b levels increased only in males and IgG3 decreased only in females, indicating sex-specific autoantibody responses (Suppl. Fig. 3A-D). In contrast, IgA levels did not increase in either sex after DT treatment (Suppl. Fig. 3E, F). Within the glomerulus, IgM and IgG antibodies display distinct localization patterns; IgM appears more punctate and concentrated in specific areas, while IgG is more diffusely distributed across the glomerular structure (Fig. 5M). This spatial difference suggests that these immunoglobulins may target different antigens or operate through different mechanisms of immune complex deposition and recognition. We also examined the humoral response in serum and urine following depletion of Tregs. In Treg-ablated mice, serum levels of IgG1, IgG2b, and IgG2c against renal antigens were increased in both sexes, while IgM was observed only in males (Suppl. Fig. 3G-R). Additionally, urinary levels of IgG and IgM were elevated in both male and female Treg-depleted mice, whereas IgA elevation was male-specific (Fig. 5N-S).

**Figure 5.**
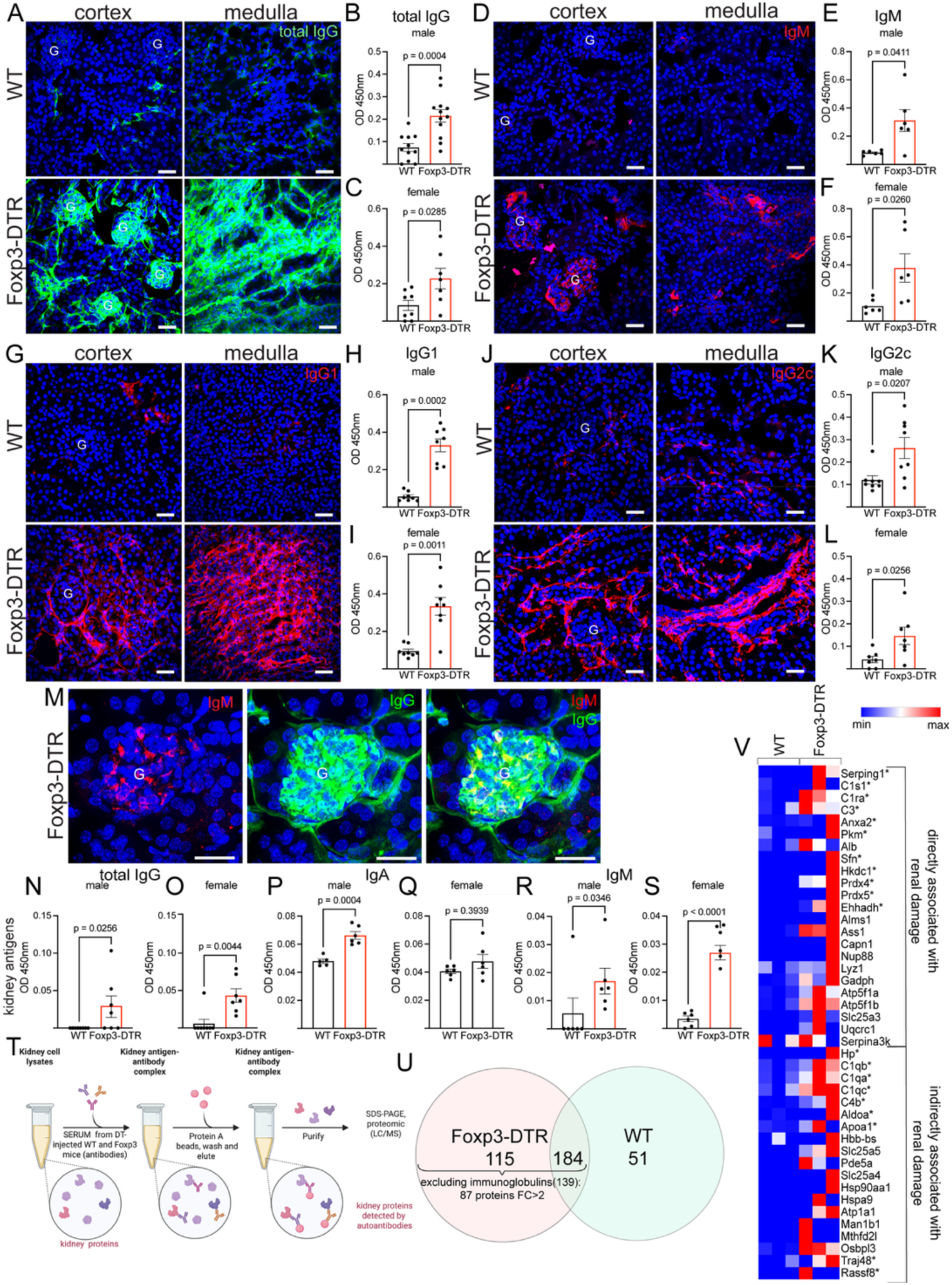
Isotypification of autoantibody levels in male and female kidneys of WT and Foxp3-DTR mice 2 weeks after Treg depletion. Immunolabeling of total IgG (green) (**A**), IgM (red) (**D**), IgG1(red) (**G**), and IgG2c (red) **(J)** in cortex and renal medulla of DT-injected WT and Foxp3-DTR mice. *n=* 5 kidneys from different mice. Isotypification of autoantibody levels of total IgG **(B, C),** IgM **(E, F),** IgG1 **(H, I)**, and IgG2c **(K, L)** in male and female kidney homogenates against kidney antigens. *n =* 8 (male WT and Foxp3-DTR: total IgG, IgG1, IgG2c), *n=* 6 (male WT and Foxp3-DTR: IgM). *n=* 8 (female WT: totIgG, IgG1; female Foxp3-DTR: IgG1), *n=* 6 (female WT: IgM; Foxp3-DTR: IgM), *n =* 7 (female WT: IgG2c; female Foxp3-DTR: totIgG, IgG2c). **M)** Immunolabeling of IgM (red) and total IgG (green), in Foxp3-DTR. *n*= 3 kidneys from different mice. Localized IgM deposition (red) and diffuse IgG deposition can be observed in the glomeruli. Urine total IgG, IgA, and IgM levels of DT-injected Foxp3-DTR and WT mice against kidney antigens **(N-S)**. N-O: *n*=8 (male and female WT), *n=*7 (male and female Foxp3-DTR). P-Q: *n*=5 (male WT), *n*=6 (female WT; male and female Foxp3-DTR). R-S: *n*=6 (male and female WT, Foxp3-DTR). Nuclei are labeled with DAPI (blue). *n* = number of samples from different mice. Bars: 20 μm. G: Glomeruli. Data were analyzed using two-sided Student’s t-test **(B, C, N, P, S)** or Mann–Whitney test **(E, F, H, I, K, L, O, Q, R)**. Data are shown as means ± SEM. **T)** Kidney protein immunoprecipitation strategy with serum protocol diagram. The figure was created with BioRender.com. **U)** Venn diagram showing the serum immunoprecipitated kidney proteins detected in the DT-treated WT and Foxp3-DTR samples by proteomics. See also Suppl. Data 1. *n=*3 independent experiments. **V)** Heat-map of the most representative renal injury response-associated proteins (2-fold-changed detected with Treg-depleted serum). * Indicates associated with inflammation.

Immunoprecipitation of renal proteins using serum from DT-treated WT and Foxp3-DTR mice (Fig. 5T) confirmed the presence of autoantibodies following depletion (Suppl. Table 2 in red immunoglobulins). We detected 87 proteins (excluding immunoglobulins) with a ≥2-fold increase or uniquely detected in Foxp3-DTR samples versus WT after immunoprecipitation (Fig. 5U). In Treg-ablated serum, we detected 138 immunoglobulin (Ig)-related proteins. These confirmed the presence of autoantibodies against renal antigens, including IgM, IgG2b, IgG1, and IgG2c, in the serum following Treg ablation. We identified 8 complement proteins in the immunoprecipitate, several of which are involved in mediating. renal injury, consistent with the presence of anti-complement antibodies in autoimmune nephropathies^39, 40^. Moreover, we detected autoantibodies against Serping 1, a key regulator of complement activation, suggesting dysregulation of complement pathways that could drive excessive glomerular complement deposition and inflammation —a hallmark mechanism in autoimmune kidney diseases. EnrichR pathway analysis of proteins showing at least a two-fold increase in the immunoprecipitated derived from Treg-depleted serum revealed significant enrichment in pathways associated with nephritis, systemic lupus erythematosus, and complement deficiency–related disorders, consistent with the autoimmune phenotype observed in our mouse model. We identified specific renal antigens, several of which are proteins whose dysregulation is associated with kidney injury (Fig. 5V), including Ehhadh^41^ (enoyl-Coenzyme A hydratase/3-hydroxyacyl CoA dehydrogenase), Serping1^42^, Slc25a3^43^ (mitochondrial phosphate carrier protein), Uqcrc1^44^ (ubiquinol-cytochrome c reductase core protein 1), Prdx4^45^ (Peroxiredoxin 4), Ass1^46^ (enzyme argininosuccinate synthase 1), Gapdh^47^ (glyceraldehyde-3-phosphate dehydrogenase), Atp5f1a^48^ (ATP synthase F1 subunit alpha), Atp5f1b^49^ (ATP synthase F1 subunit beta) and Serpina3k^50^, Pkm^51^ (pyruvate kinase, muscle type), Lyz1^52^ (C-type lysozyme), and Capn1^53^ (calpain-1). Additional proteins are indirectly linked to renal damage due to their roles in oxidative stress, mitochondrial dysfunction, and cytoskeletal regulation, which are also observed in other organs (Fig. 5V). For example: Slc25a5^54, 55^ (mitochondrial ADP/ATP translocase 2), Man1b1^56^ (mannosidase alpha class 1B member 1), Mthfd2l^57^ (methylenetetrahydrofolate dehydrogenase 2-like), and Osbpl3^58^ (oxysterol-binding protein 3). The serum autoantibodies recognize renal proteins that function as immune modulators, such as Serping1^42^, Aldoa^59^ (fructose-bisphosphate aldolase protein family), Slc25a3^43, 55^ (mitochondrial phosphate carrier protein), Apoa1 (apolipoprotein A-I), Traj48^60^ (T cell receptor alpha), Rassf8^61^ (Ras association domain family member 8), and Anxa2^62^ (annexin A2). Moreover, some of the proteins participate in the IF-1 signaling pathway (Hkdc1, Aldoa, Gapdh), Neutrophil extracellular trap formation (C3, H2ac4, Slc25a5, Slc25a4) and phagocytosis and bacterial infection (Tubb1, Tubb4b, Gapdh, Tuba4a, Hsp90aa1, Anxa2).

### Treg depletion induces kidney injury

Furthermore, 2 weeks after Treg depletion in both female and male mice, we observed a reduction in glomerular length (Fig. 6A-C), accompanied by a decrease in kidney weight (Fig. 6D-E). Notably, serum creatinine (sCr) levels were elevated in Treg-depleted mice (Fig. 6F-G), indicating impaired renal function. These parameters collectively suggest an early onset of kidney injury following Treg ablation. No effect was seen in BUN (Suppl. Fig. 4A-B). Interestingly, microalbuminuria increased in female Treg-depleted mice but not in males at this time point (Fig. 6H-I). Treg depletion caused decreased urinary output (Fig. 6J-K) and a reduction in urine osmolarity in male Treg-ablated mice (Fig. 6L-M). However, by 8 weeks post-depletion, both sexes showed a significant increase in this urinary marker of tubular damage, indicating progressive renal injury over time (Suppl. Fig.4G-H). No change in sCr or BUN at this time point (Suppl. Fig 4C-F). To assess proximal tubule (PT) damage, we labeled kidney sections for aquaporin 1 (AQP1), which is located in the brush border and basolateral membrane of intact PT cells. At week 2 after Treg depletion, we observed relocalization of AQP1 in some PTs from the apical membrane to the intracellular compartment (Fig. 6N-O, arrowheads). Note that the basolateral staining was still visible. After Treg ablation, while some PT cells showed a complete loss of AQP1 polarity with absent apical and basolateral staining (Fig. 6O; very damaged PTs), some cells showed partial redistribution with basolateral staining still visible (Fig. 6O; moderately damaged PTs). We quantified the number of intact, moderately damaged, and significantly damaged PTs based on these AQP1 staining patterns and found a reduction in the number of intact PTs in the Treg-depleted kidney compared to the control group (Fig. 6P). Neutrophil Gelatinase-Associated Lipocalin (NGAL) transcript (*Lcn2*) was increasingly detected post-depletion, suggesting that it may serve as a biomarker of autoimmune renal injury (Fig. 6Q).

**Figure 6.**
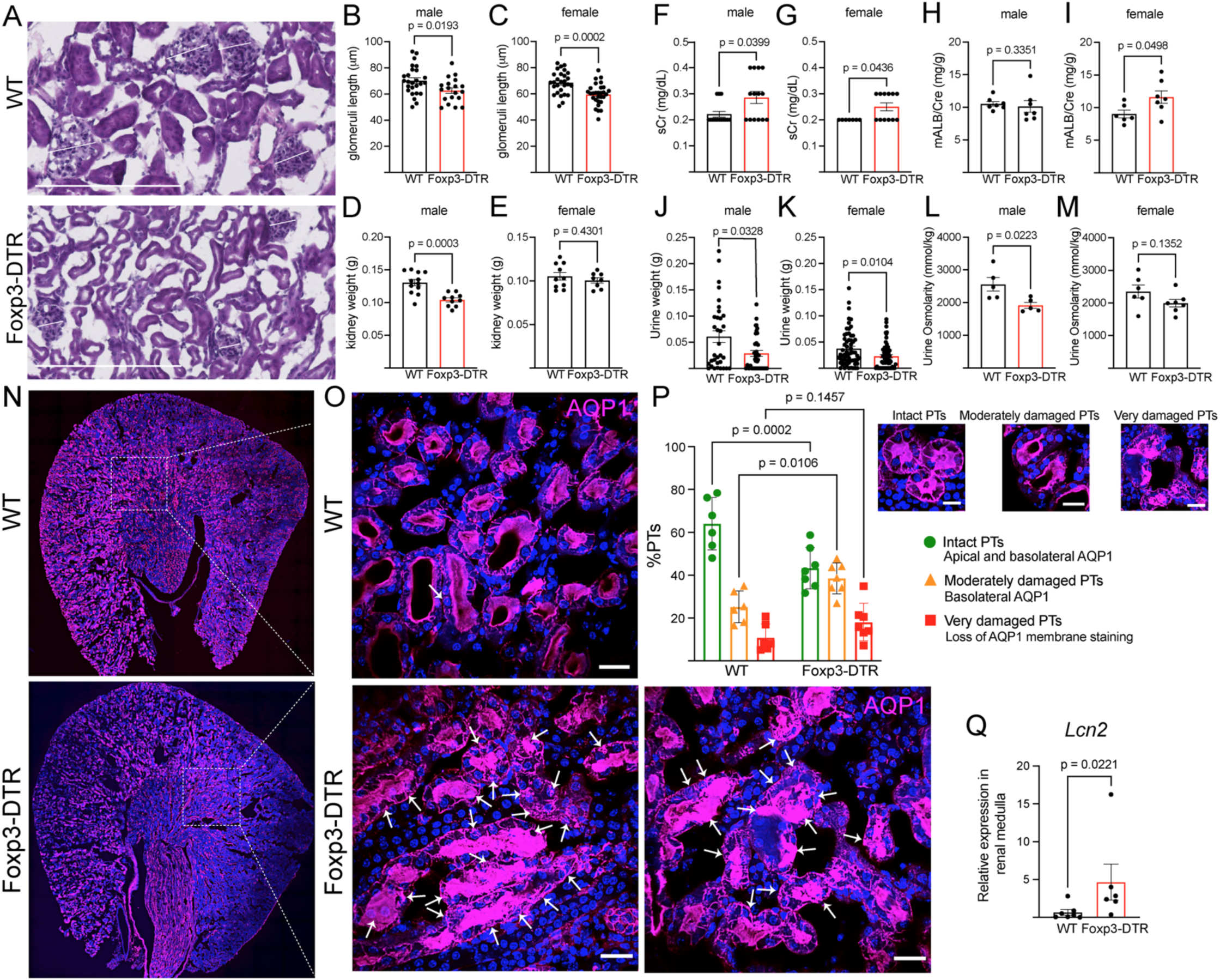
Treg depletion exacerbates renal inflammation and promotes kidney injury after 2 weeks of DT injection. Kidney sections stained using H&E **(A)** and quantification of the glomeruli length (μm) of male **(B)** and female **(C)** kidneys in WT and Foxp3-DTR mice. *n*= 3 (male and female WT), *n*=3 (male and female Foxp3-DTR). Each area quantified is represented as a dot. Kidney weight (g) of male **(D)** and female **(E)** in WT and Foxp3-DTR mice. Male: *n*=12 (WT), *n*=10 (Foxp3-DTR). Female: *n*=10 (WT), *n*=8 (Foxp3-DTR). Serum creatinine (sCr) of male **(F)** and female **(G)** mice after Treg depletion. Male: *n=*14 (WT, Foxp3-DTR). Female: *n=*7(WT), *n=*12 (Foxp3-DTR). Urine microalbumin/creatinine ratio (mALB/Cre) of male **(H)** and female **(I)** mice after Treg depletion. Each dot represents a pool of 2-3 urine samples. Urine weight (g) was measured in male **(J)** and female **(K)** mice. Urine osmolarity of WT and Foxp3-DTR male **(L)** and female **(M)** mice. Male: *n=*5 (WT, Foxp3-DTR). Female: *n=*6(WT), *n=*7 (Foxp3-DTR). **N-P)** Quantification of the percentage of intact tubules (green bars), moderately damaged tubules with detectable cellular structures (orange bars), and significantly damaged tubules with a complete loss of cell architecture (red bars). WT (*n*=8), Foxp3-DTR (*n*=8), by 2-way ANOVA. Between 1100 and 1500, tubules were analyzed in each group. Arrows indicate AQP1^+^ cell damage. **Q)** Relative quantification of *Lcn2* transcript in the kidney by qPCR. Samples were normalized to their internal control (*Gapdh*) and expressed as relative values compared to the control group*. n=*7 (WT); *n=*6 (Foxp3-DTR). Results are expressed as mean ± SEM. DAPI (blue). Bars: 250 μm(A) 20 μm(N-O) Data were analyzed using two-sided Student’s t-test **(B-E, I, L, M)** or Mann–Whitney test **(F-H, J, K, Q)**. *n* = number of samples from different mice.

### Treg depletion reprograms IC transcriptomes

The specific role of proton-secreting renal ICs in tissue homeostasis and autoimmunity remains poorly understood. To investigate this, we crossed B1-V-ATPase-EGFP mice with Foxp3-DTR mice, enabling the depletion of Tregs and the isolation of ICs. Two weeks after Treg ablation, we isolated EGFP-positive CD45⁻ ICs using fluorescence-activated cell sorting (FACS) for transcriptomic analysis via RNA sequencing (Fig. 7A, Suppl. Table 3). The complete transcriptome dataset is presented in the Suppl. Table 3. A volcano plot (fold change (FC) versus P value) comparing gene expression profiles of ICs, Treg-depleted versus WT, shows increased expression of several pro-inflammatory genes (Fig. 7B). The full list of upregulated genes is provided in the Suppl. Table 4, while downregulated genes are listed in the Suppl. Table 5. These include genes involved in metabolic and peroxidative pathways.

**Figure 7.**
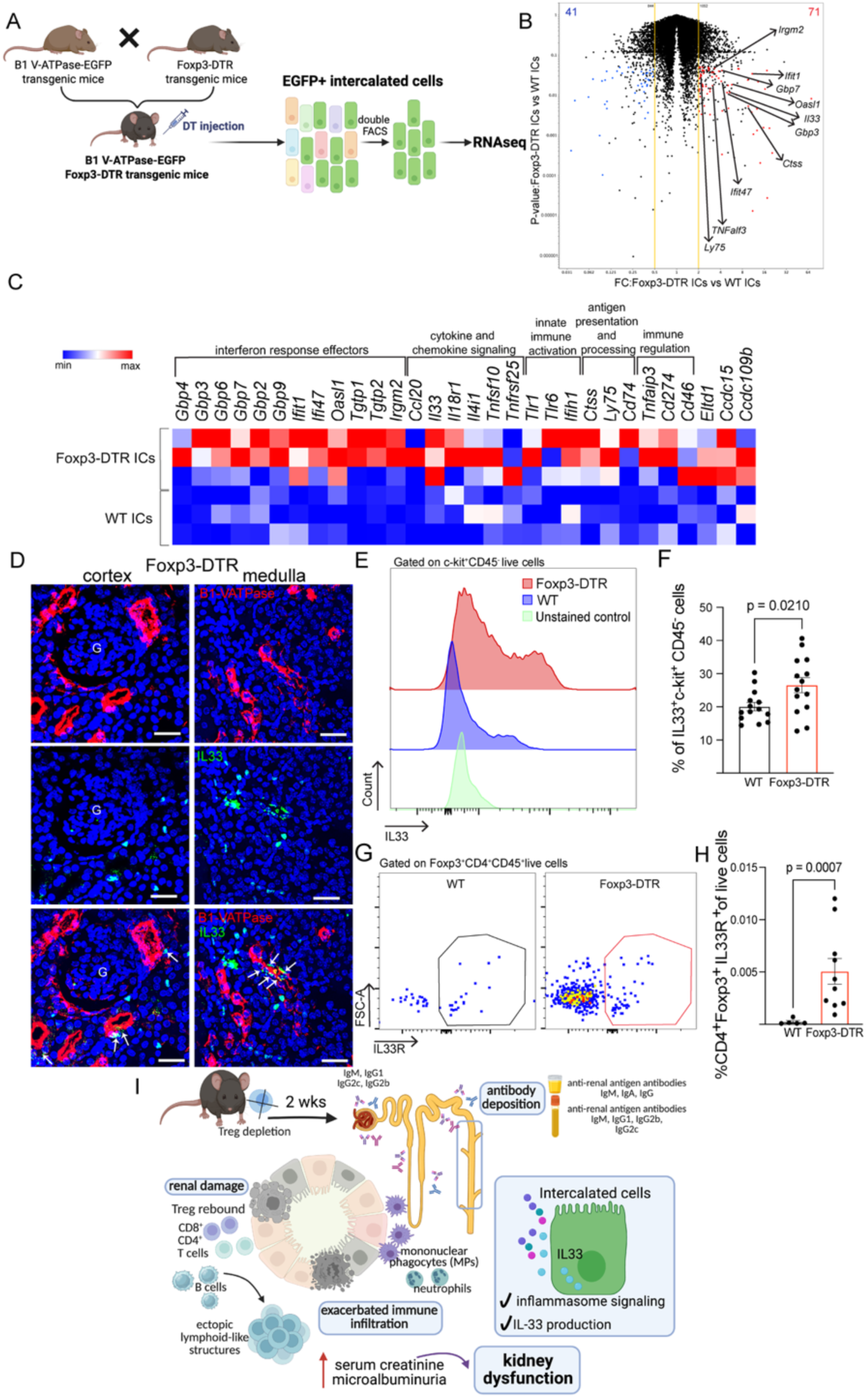
Treg depletion reshapes the functional landscape of intercalated cells (ICs). **A)** B1 V-ATPase-EGFP and Foxp3-DTR mice were crossed to generate the double transgenic mice. After 2 weeks of DT-induced Treg depletion, EGFP^+^ ICs were FACS-sorted and analyzed by RNA sequencing. **B)** Volcano plots (fold change (FC) versus P value) of gene expression profiles of ICs, isolated by FACS 2 weeks after Treg depletion (EGFP^+^ Foxp3-DTR ICs) versus EGFP^+^ WT ICs. Each sample of RNA (*n*=3) was obtained from a pool of two kidneys from two mice in each group. Several pro-inflammatory-associated genes were up-regulated after Treg depletion compared to CTR. The yellow line represents ± 2-fold change, and the black dots denote transcripts that were not significantly differentially expressed. Data were analyzed using Student’s t-test, two-tailed, and a value of P<0.05 was considered significant. Genes up-regulated after Treg ablation are shown in red, and genes down-regulated are shown in blue. **C)** Heatmap of immune-related genes up-regulated in the ICs after Treg depletion. **D)** Immunolabeling of B1VATP-ase (red) and IL33 (green) in the renal cortex and medulla of DT-injected Foxp3-DTR mice. Arrows: IL-33 expression in intercalated cells. **E)** Histogram overlay showing intracellular cytokine staining for IL-33 ICs cells. Foxp3-DTR cells (red) display elevated IL-33 expression compared to WT (blue), with both showing positive staining above the unstained control (green). Data are representative of independent experiments. **F)** Percentage of IL33^+^c-kit^+^CD45^-^cells in the kidneys from DT-treated WT and Foxp3-DTR mice. *n=*14 Flow cytometry gating strategy **(G)** and relative abundance of CD4^+^Foxp3^+^IL33R^+^**(H)** in the kidneys. *n*=5(WT); *n=*10 (Foxp3-DTR) **I)** Graphical representation of the key findings at 2 after Treg depletion in the kidneys. The figure was created with BioRender.com. Data were analyzed using two-sided Student’s t-test **(F)** or Mann–Whitney test **(H).** *n* = number of samples from different mice.

The transcriptomic profile of ICs from Treg-depleted mice (Fig. 7B-C) revealed upregulation of inflammasome-related genes, including *Gbp3*, *Gbp7*, *Ifit1*, and *Irgm2.* Guanylate-binding proteins (GBPs), such as GBP3 and GBP7, are interferon (IFN)-inducible GTPases that play crucial roles in defense mechanisms and innate immune response, specifically regulating inflammasome activation^63^. *Ifit1* is a critical regulatory node in IFN signaling, modulating downstream pathways^64^. *Irgm2* is a key regulator of the non-canonical inflammasome pathway, further implicating ICs in immune surveillance and inflammatory signaling. Moreover, ICs exhibited upregulation of *Ctss* (cathepsin S) following Treg ablation. CTSS is a lysosomal cysteine protease that processes pro-inflammatory cytokines into their biologically active forms^65^. CTSS is overexpressed in the kidneys of patients with CKD, such as IgA nephropathy and lupus nephritis. High CTSS levels in renal tissue and serum correlate with disease severity, glomerular injury, proteinuria, and impaired renal function^66^.

Additionally, ICs upregulated *Tnfaip3*, which is known to limit inflammation and injury in the kidney^67^, particularly during stress conditions such as IRI^68^. In the mouse renal ischemia-reperfusion injury model, renal TNFAIP3 variants exhibit increased NF-κB activation and inflammation in the kidney, as well as an increase in the numbers of Tregs, which may provide protective effects by promoting immune regulation and limiting tissue damage^68^. ICs upregulate *Ly75*, suggesting a role in promoting immune tolerance through antigen presentation that supports Treg function and helps maintain renal immune quiescence^69, 70^. These findings suggest that ICs play an underappreciated role in shaping the inflammatory microenvironment that emerges upon Treg depletion. Notably, ICs upregulated other genes associated with T-cell responses, including *Cd274* (PD-1L), *Ccl20*, *Tlr1,* and *Tlr6.* Interestingly, PD-1^+^ Tregs increased after Treg depletion (Fig. 3O-P). It was previously reported that ICs expressed TLR receptors^31,32^.

Interestingly, IL-33 expression and production were elevated in ICs 2 weeks after Treg ablation, suggesting that ICs may trigger tissue repair responses and modulate local lymphocyte activity (Fig.7C, D-F). IL-33, a member of the IL-1 cytokine family, is known to act as an alarmin released upon cellular stress or damage^36^. In the kidney, IL-33 regulates inflammation and promotes tissue remodeling and fibrosis, particularly by influencing the recruitment and activation of Tregs^71–78^. Intracellular IL-33 levels in ICs increased 2 weeks after Treg depletion (Fig. 7D–F). IF analyses further confirmed the localization of IL-33 within cortical and medullary ICs (Fig. 7F). This suggests that IC-mediated IL-33 signaling may affect the renal Treg function, potentially promoting tissue repair. Moreover, renal Tregs upregulated ST2, the receptor for IL-33 (Fig. 7G-H). These findings highlight a potential feedback loop in which ICs, via IL-33/ST2 axis signaling, orchestrate immune responses and tissue homeostasis following autoimmune perturbation.

### Tolerance disruption triggers kidney autoimmunity, impairing renal function

The loss of immune tolerance led to chronic inflammation and localized autoimmune activity in the kidney, characterized by a distinct immune landscape of myeloid and lymphocyte cells, along with the accumulation of autoantibodies in the serum, urine, and renal tissue. This immune imbalance damages kidney function and contributes to ongoing nephropathy. The presence of ectopic lymphoid-like structures may further worsen inflammation by promoting local autoantibody production. Our research highlights key molecular factors involved in interactions between ICs and immune cells, showing that kidney disorders stem from immune dysregulation marked by pro-inflammatory immune cell infiltration and autoantibody development (Fig. 7I). ICs contribute not only to pro-inflammatory responses but also to tissue repair through the secretion of IL-33, a crucial cytokine involved in orchestrating reparative processes. Renal Tregs showed a significant rebound, indicating that activated Tregs repopulate the kidney 2 weeks after DT to help counteract increased inflammation. This study offers a detailed characterization of epithelial ICs and immune cell diversity in the kidney, providing a foundation for developing targeted therapies to treat renal damage and chronic inflammation.

## Discussion

Autoimmune diseases induce both glomerulonephritis and tubulointerstitial nephritis, ultimately leading to AKI and CKD^1–3^. CKD dramatically increases the risk of end-stage renal disease, cardiovascular events, infections, and premature mortality, leading to significant social and economic burdens. Improving mechanistic understanding and developing therapies are crucial for mitigating its impact and enhancing public health. To study autoimmune-mediated kidney injury, we used Treg-depleted mice^15, 33, 34^, which allowed for a detailed examination of cellular and molecular mechanisms. This model offered essential insights into how ICs, along with innate and adaptive immune cells—including MPs, B cells, and T cells—contribute to kidney damage and dysfunction. Notably, we found that ICs function as active immunoregulatory cells rather than passive targets in the context of autoimmune disease. These cells produce inflammatory cytokines and chemokines, influencing local immune responses by activating inflammasome signaling pathways. Additionally, we discovered a new positive feedback loop between ICs and Tregs that controls inflammation, involving the cytokine IL-33.

Specialized epithelial ICs play a central role in organizing the inflammatory microenvironment and both initiating and sustaining autoimmune-mediated kidney injury. Kidneys are a frequent target of systemic autoimmunity^2, 20^; in our mouse model, autoimmune responses directly impact kidney structure and function through failures in tolerance mechanisms, leading to persistent immune activation, infiltration of autoreactive lymphocytes, and sustained local inflammation. The combination of autoantibody deposition and cytokine production leads to tissue damage^2, 20, 79, 80^. Among the most well-known autoimmune kidney diseases are lupus nephritis, anti-GBM disease, and ANCA-associated vasculitis, each affecting different segments of the nephrons. For example, in lupus nephritis, immune complex deposition and complement activation led to glomerular inflammation and fibrosis^4, 5, 20, 81^. In anti-GBM disease, autoantibodies attack the glomerular basement membrane, triggering rapid and severe glomerulonephritis^82^. These examples highlight the significance of the epithelial microenvironment and intercellular communication between tissue-resident immune cells and epithelial cells in driving local inflammation and hindering tissue repair. Understanding these cellular and molecular interactions is essential for developing new targeted therapies. Current strategies modulating immune responses—such as B cell depletion, checkpoint inhibitors, or cytokine blockade—are being studied to prevent or reverse renal autoimmunity without causing widespread immunosuppression^83–85^. The current study suggests that ICs may serve as promising therapeutic targets for mitigating kidney injury. Furthermore, we identified multiple renal antigens that are targeted by autoantibodies, implicating these proteins in the development of antibody-mediated kidney damage. These results support the concept that the loss of tolerance to kidney-specific antigens is a key driver of tissue injury in autoimmune nephritis.

Treg ablation triggered an intense and persistent renal inflammatory response, leading to severe autoimmune signs characterized by infiltration of both myeloid and lymphoid cells and deposition of autoantibodies. Two weeks after DT, renal Tregs showed a notable^23, 25, 31, 86^ rebound, indicating that Tregs repopulate the kidney —a pattern consistent with previous findings in secondary lymphoid organs, such as the spleen^87^. Despite this recovery, inflammation worsened substantially, evidenced by a large influx of F4/80⁺ and MHCII⁺ MPs, neutrophils, and monocytes. These innate immune cells are recognized for their robust phagocytic ability, antigen-presenting roles, and the release of pro-inflammatory and cytotoxic mediators, which further amplify immune responses^88, 89^. MPs are essential effectors in disrupting immunological tolerance and are known to promote pro-inflammatory responses in various organ systems following Treg depletion^33, 34, 90^. Specifically in the kidney, MPs have been linked to mediating tubular-interstitial injury in murine models of AKI^91–93^. MPs may not only cause direct kidney injury but also act as mediators between the innate and adaptive immune systems, thereby maintaining ongoing inflammation and kidney dysfunction. After Treg ablation, activated T and B lymphocytes also infiltrated the renal interstitium. Most likely, autoreactive CD4^+^ and CD8^+^ subsets recognize renal self-antigens, resulting in cellular damage and increased inflammation. Remarkably, in our mouse model, the complex immune responses were triggered independently, without the need for a secondary stimulus. This finding differs from traditional nephropathy models, which typically rely on exogenous antigens, such as sheep IgG, to induce renal injury^94^. Our study suggests that the intrinsic breakdown of immune mechanisms may be sufficient to initiate and sustain renal inflammation.

Intrarenal B cells contribute to autoimmune kidney disease by producing autoantibodies that target renal components—including glomerular, tubular, and nuclear antigens—leading to immune complex deposition and complement activation^2, 80, 84^. This triggers chronic inflammation that forms tertiary lymphoid-like structures, similar to those observed in our mouse model, which drive diseases such as lupus nephritis, transplant rejection, and ANCA-associated vasculitis^38, 95^. Both memory and plasma B cells participate in this process, driving continuous tissue damage^96^. Current B-cell therapies, such as anti-CD20 antibodies, can reduce relapse rates but often do not lead to lasting remission unless plasma cells are also depleted^97, 98^. Anti-CD38 therapies, which target plasma cells, offer additional therapeutic potential. Broad B-cell depletion with anti-CD19 CAR-T therapy may help reverse systemic lupus erythematosus and related autoimmune diseases, while BAFF/APRIL blockade shows promise for IgA nephropathy. These emerging strategies offer hope for more effective and targeted treatment of B–cell–mediated kidney disorders^98^. Conversely, Bregs are often reduced or dysfunctional in autoimmune nephropathy, and this is linked to higher disease activity. Restoring Bregs can help promote immune tolerance and remission^38^. Effective treatment may therefore need to eliminate harmful B-cell subsets while maintaining Bregs selectively. Our research suggests that ICs serve as kidney-specific immune regulators, capable of modulating local T- and B-cell responses. Moreover, the increase in follicular regulatory T cells suggests a compensatory response that reduces inflammation and autoantibody production, consistent with their protective role in kidney allograft rejection^99^.

Tertiary lymphoid structures (TLSs), composed of T and B lymphocytes and stromal fibroblasts, resemble secondary lymphoid organs and can initiate adaptive immune responses and antibody production^37, 95, 100–102^. TLSs have been found in aged kidneys and in renal autoimmune and inflammatory diseases, including allograft rejection and IgA nephropathy^37, 38, 103–107^. Some develop GCs that promote B-cell proliferation, class switching, and the differentiation of plasma cells. After Treg depletion, we observed the formation of ectopic lymphoid-like structures that lacked full maturation in the medullary region of the kidney. We found increased numbers of follicular T cells, GC B cells, memory B cells, and follicular DCs in both tissues, suggesting the presence of more mature ectopic structures. Notably, GC B cells appeared in the Treg-depleted kidney, suggesting the start of *in situ* B cell maturation. Bregs were also elevated, possibly reflecting a compensatory anti-inflammatory response. Interestingly, in the epididymis, Treg loss led to the formation of mature TLSs 8 weeks after DT treatment^34^.

Furthermore, our research highlights the crucial connection between inflammasome activation in ICs and the progression of autoimmune nephropathy. Previous studies have shown that P2Y14 in ICs plays a key role in AKI, using both mouse models and human samples^23, 24, 30^. P2Y14, a G protein–coupled receptor activated by UDP-sugars, is highly expressed in ICs and encourages inflammatory signaling. Its activation can activate the NLRP3 inflammasome, a multiprotein complex responsible for releasing pro-inflammatory cytokines and involved in the development of AKI^27, 108^. In cases of AKI and autoimmune nephropathies, this signaling pathway may worsen kidney damage by promoting immune activation within ICs. Exploring this pathway reveals potential therapeutic targets to reduce kidney inflammation and preserve renal function.

Our study shows that renal ICs actively influence Treg dynamics, suggesting a local mechanism for immune regulation in nephritis. Tregs inhibit harmful immune activation by restraining effector T cells, thereby decreasing inflammation and supporting tissue repair^10, 12, 13, 15, 20, 21, 109^. Impaired Treg function or depletion leads to excessive immune responses, contributing to autoimmune kidney diseases, such as lupus nephritis and glomerulonephritis. Therefore, enhancing Treg activity offers a promising therapeutic approach to protect kidney tissue. In experimental glomerulonephritis, Tregs localize near glomeruli and tubulointerstitial regions, and their loss worsens disease severity^20–22, 110, 111^.

IL-33 has emerged as a key alarmin cytokine in renal injury, showing a rapid increase after tubular epithelial cell damage, glomerulonephritis, and renal allograft fibrosis^71–73, 76, 78, 112–118^. In AKI and CKD models, IL-33 is released into the local tissue environment, acting as a potent immunomodulator^112, 113, 78^. It promotes the recruitment and activation Tregs and macrophages, which aid in tissue repair and limit fibrosis in the injured kidney^74, 77, 78, 119^. Interestingly, IL-33 also activates B cells and stimulates the production of proinflammatory cytokines, including IL-8, TNF-α, and CCL2. Systemic inhibition of IL-33 signaling in diabetic mice decreased endothelial inflammation, enhanced glomerular integrity, and slowed the progression of kidney disease^117^. An ongoing trial is evaluating IL-33–targeted therapy as a treatment for diabetic kidney disease^117^. Our study extends these findings by identifying renal ICs as previously unrecognized sources of IL-33 during autoimmune-mediated renal inflammation. This discovery reveals a new aspect of local immune regulation, suggesting that IL-33 from ICs may influence disease progression by directing the recruitment and activation of immune cells within the kidney microenvironment.

IL-33 is expressed by endothelial, epithelial, and fibroblastic stromal cells in various organs ^36, 75, 120^. The role of IL-33 varies depending on the severity and type of damage. While IL-33 can exacerbate inflammation in certain situations, in others, it supports tissue repair by promoting anti-inflammatory immune responses and resolving fibrosis^10, 36, 75^. Overall, IL-33 acts as a context-dependent cytokine, integrating signals from tissue injury and environmental stress to coordinate immune responses. The presence of IL-33-producing special epithelial cells in kidney tissue, therefore, makes them promising targets for therapies aimed at modulating immune responses and promoting tissue repair. ICs producing proinflammatory mediators and IL-33 play central roles in both inflammation and tissue repair. Modulating the IL-33/ST2 axis represents a promising therapeutic approach for autoimmune diseases, although interventions must be precisely timed and tissue-specific to balance immune suppression with the preservation of protective immunity.

By understanding how epithelial cells function as immune modulators, we can identify new biomarkers for active inflammation and develop targeted, organ-specific strategies to mitigate renal injury. ICs are not just proton-secreting epithelial cells but may also play an active role in autoimmune responses. Urinary biomarkers based on these renal epithelial-related molecules could allow for earlier clinical assessment and targeted treatment for high-risk patients, thereby improving their prognosis. Classifying patients who are at high risk of developing autoimmune-related renal injury could enhance treatment outcomes. Additionally, developing new therapies that selectively target IC-related molecules offers promise for achieving more precise, tissue-specific control of immune responses. This approach could lessen dependence on broad immunosuppressive agents and decrease their side effects. Further studies are necessary to validate these biomarkers and support the translation of IC-targeting therapies into clinical practice.

## Acknowledgments

The authors thank the Microscopy Core of the Program in Membrane Biology (PMB) (MGH, Boston, MA), the MGB Nikon for Excellence Molecular Imaging Core (MGH, Charlestown, MA), and the MGH Pathology Flow Cytometry Facility (MGH, Boston, MA). The National Institutes of Health supported this work (grant HD104672 to M.A.B.), the MGH Physician/Scientist Development and Claflin Distinguished Scholar Awards (to M.A.B.).

## Ethics approval

All animal procedures were approved by the Massachusetts General Hospital (MGH) Subcommittee on Research Animal Care and followed the NIH Guide for the Care and Use of Laboratory Animals (National Academies Press, 2011; protocol 2015N000016).

