## Supplementary material for "Immunomodulatory Functions of Intercalated Cells in Kidney Autoimmunity": Supplementary Material.pdf

**Supplementary Table 1.** List of antibodies used in this study.

**Supplementary Table 2.** Proteomics raw data and analysis of kidney proteins recognized by serum of WT and Foxp3-DTR mice after 2 weeks of DT treatment. In red are indicated the detected immunoglobulins.

**Supplementary Table 3.** The complete transcriptome dataset of renal ICs from WT and Foxp3-DTR mice after 2 weeks of DT treatment.

**Supplementary Table 4.** List of transcripts upregulated (<2-fold increase and p<0.05) in ICs after Treg depletion compared to control.

**Supplementary Table 5.** List of transcripts downregulated in ICs (<2-fold increase and p<0.05) after Treg depletion compared to control.

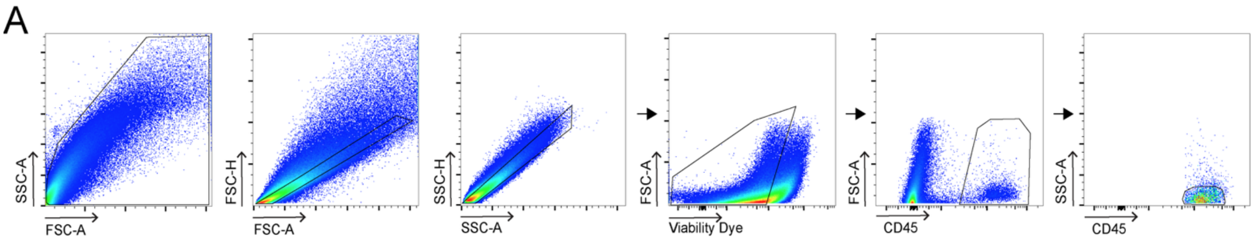


**Supplementary Figure 1. Flow cytometry gating strategy used to analyze lymphoid immune cells in the kidney.** SSC-H: side scatter height, FSC-H: forward scatter height, SSC-A: side scatter area, FSC-A: forward scatter area, Viability dye: LIVE/DEAD™ Fixable Yellow Dead Cell, CD45: pan immune cell marker.

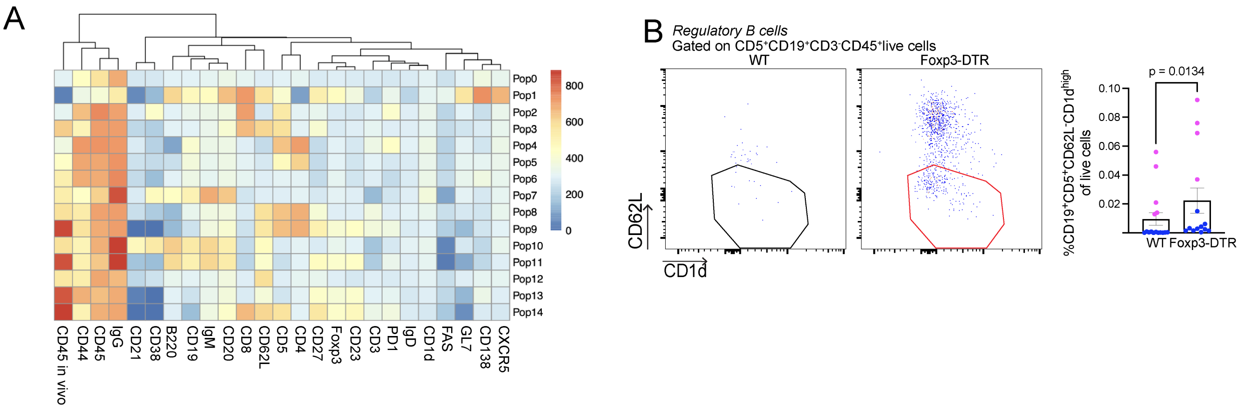


**Supplementary Figure 2. Renal B cells infiltration after Treg depletion. A)** The relative marker expression heatmap of the t-SNE and FlowSOM analysis of the $CD45^+$ cell population of kidneys from DT-treated Foxp3-DTR and WT mice. **B)** Flow cytometry gating strategy and relative abundance of Regulatory B cells ($CD3^-CD19^+CD5^+CD62L^-CD1d^{hi}$) in the kidneys of DT-treated WT and Foxp3-DTR mice. Male: *n*=9 (WT), *n*=8 (Foxp3-DTR). Female: *n*=8 (WT), *n*=10 (Foxp3-DTR). *n* = number of samples from different mice. Data are means ± SEM, and a two-sided Mann-Whitney test was performed.

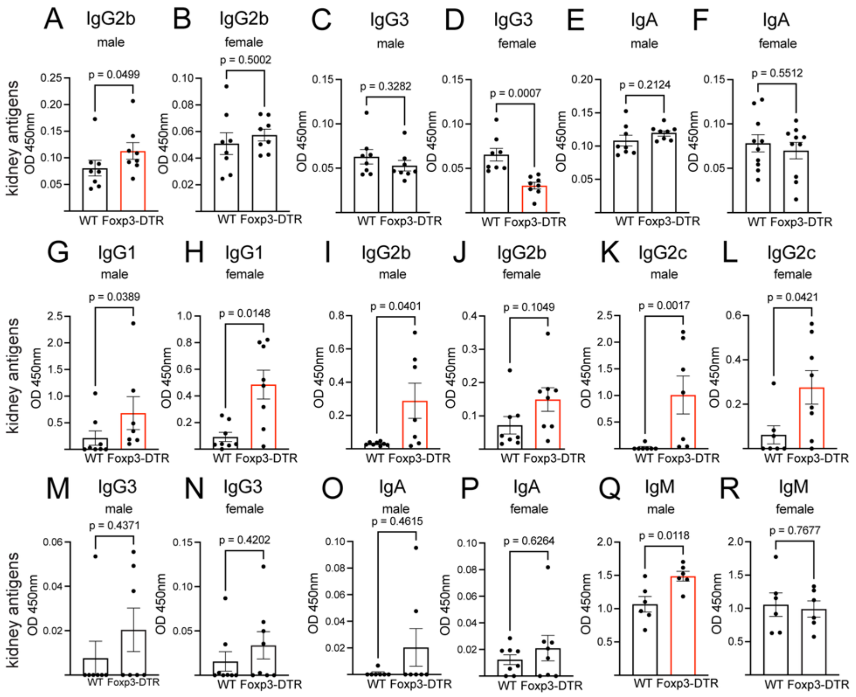


**Supplementary Figure 3. Autoantibody isotype profiling of Foxp3-DTR mice 2 weeks after DT treatment by ELISA. A)** Isotype levels (IgG2b, IgG3, and IgA) in kidney homogenates against kidney antigens (**A-F**). *n*=7 (WT male: IgG2b, IgA; WT female: IgG2b), *n*=8 (WT: IgG3; Foxp3-DTR male: IgG2b, IgG3, IgA; Foxp3-DTR female: IgG2b, IgG3), *n*=10 (WT and Foxp3-DTR female: IgA). **G-R)** Serum levels of IgG1, IgG2b, IgG2c, IgG3, IgA, and IgM in DT-injected Foxp3-DTR and WT male and female mice against kidney antigens, with *n*=5-10. *n* = number of samples from different mice. Data were analyzed using Student's t-test **(B-D, F, Q, R)** or Mann-Whitney test **(A, E, G-P)**. Data are shown as means ± SEM.

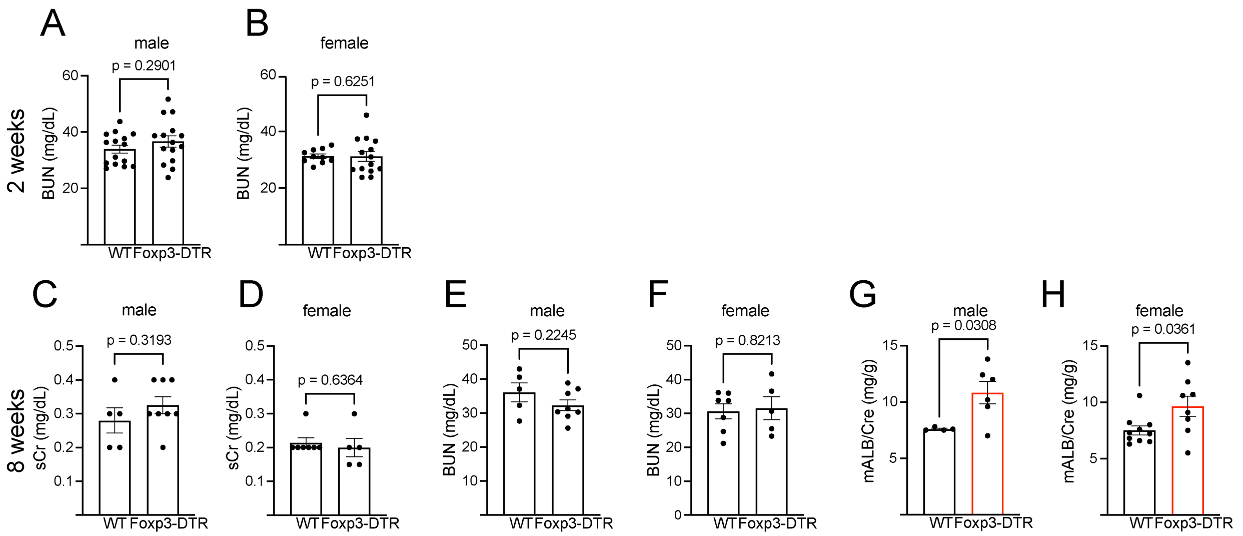


**Supplementary Figure 4. Kidney function parameters post-Treg depletion. A)** Serum Blood Urea Nitrogen (BUN) 2 weeks post-Treg depletion *n*=14(WT male; Foxp3-DTR female); *n*=15(Foxp3-DTR male); *n*=10 (WT female). Serum Creatinine (SCr) (**A**), BUN (**B**) 8 weeks post-Treg depletion. *n*=5(WT male; Foxp3-DTR female); *n*=8(Foxp3-DTR male); *n*=7(WT female). **C)** Urine albumin-creatinine ratio (mALB/cre) 8 weeks post-Treg depletion. *n*=4(WT male); *n*=6(Foxp3-DTR male); *n*=10 (WT female); *n*=8 (Foxp3-DTR female). *n* = number of samples from different mice. Data were analyzed using Student's t-test **(A, C, E, F, G)** or Mann-Whitney test **(B, D, H)**. Data are shown as means ± SEM.

**Supplementary Table 1**

**Format**: Fluorochrome – Species – Anti- – Antibody

| | |
|---|---|
| BV711 Rat anti-mouse CD45 | 2 mg/ml; Clone 30-F11, 563709, BD Biosciences |
| PE/Fire™ 700 Rat anti-mouse CD45 | 0.2 mg/ml; Clone 30-F11, 103177, BioLegend |
| BUV395 Rat anti-mouse CD45 | 0.2 mg/ml; Clone 30-F11; 565967, BD Biosciences |
| PE Rat anti-mouse Ly-6G | 0.2 mg/ml; Clone 1A8, 551461, BD Biosciences |
| FITC Rat anti-mouse Ly-6C | 0.5 mg/ml; Clone AL-21; 553104, BD Biosciences |
| PE/Cyanine7 Rat anti-mouse F4/80 | 0.2 mg/ml; Clone BM8, 123113, BioLegend |
| APC/Cyanine7 Rat anti-mouse/human CD11b | 0.2 mg/ml; Clone M1/70, 101225, BioLegend |
| Alexa Fluor™ 700 Rat anti-mouse MHC Class II (I-A/I-E) | 0.2 mg/ml; Clone M5/114.15.2, 56-5321-82, Thermo Fisher Scientific |
| Alexa Fluor® 647 Mouse anti-mouse CD64 a and b Alloantigens | 0.2mg/ml; Clone 184 X54-5/7.1; 558539, BD Biosciences |
| BV421 Armenian Hamster anti-mouse CD11c | 0.05mg/mL; Clone N418, 117329, BioLegend |
| BV605 Rat anti-mouse CD206 (MMR) | 0.1mg/ml; Clone: C068C2, 141721, Biolegend |
| PE Rat anti-mouse CD4 | 0.2 mg/ml; Clone RM4-5, 100511, BioLegend |
| APC/Cyanine7 Rat anti-mouse CD279 (PD-1) | 0.2 mg/ml; Clone 29F.1A12, 135223, BioLegend |
| BV786 Rat Anti-Mouse CD25 | 0.2mg/ml; Clone 3C7, 564368, BD Biosciences |
| BV650 Rat anti-mouse CD223 (LAG-3) | 0.2mg/ml; Clone C9B7w, 125227, BioLegend |
| PE/Cyanine7 Armenian Hamster anti-mouse CD69 | 0.2mg/ml; Clone H1.2F3, 104511, BioLegend |
| BV421 Armenian Hamster anti-mouse CD152 | 0.2mg/ml; Clone UC10-4B9, 106311 BioLegend |
| BUV737 Rat Anti-Mouse CD44 | 0.2 mg/ml; Clone IM7; 612799, BD Biosciences |
| BUV805 Rat Anti-Mouse CD1d | 0.2 mg/ml; Clone IB1; 741965, BD Biosciences |
| BV480 Rat Anti-Mouse CD4 | 0.2 mg/ml; Clone RM4-5; 565634, BD Biosciences |
| BV750 Rat Anti-Mouse CD19 | 0.2 mg/ml; Clone 1D3; 747332, BD Biosciences |
| BUV615 Mouse Anti-Mouse IgD[b] | 0.2 mg/ml; Clone 217-170, 751474, BD Biosciences |
| BV510 Rat Anti-Mouse CD138 | 0.2 mg/ml; Clone 281-2, 563192, BD Biosciences |
| BV421 Rat anti-mouse CD3 | 0.2 mg/ml; Clone 17A2, 100227, BioLegend |
| PE Rat anti-mouse CD20 | 0.2 mg/ml; Clone SA275A11, 150409, BioLegend |
| Pacific Blue™ Rat anti-mouse/human CD45R/B220 | 0.5 mg/ml; Clone RA3-6B2, 103230, BioLegend |
| APC/Fire™ 810 Mouse anti-mouse CD95 (Fas) | 0.2 mg/ml; Clone SA367H8, 152623, BioLegend |
| Alexa Fluor® 594 Rat anti-mouse CD5 | 0.5 mg/ml; Clone 53-7.3, 100632, BioLegend |
| PerCP/Cyanine5.5 Rat anti-mouse/human GL7 | 0.2 mg/ml; Clone GL7, 144609, BioLegend |
| FITC Rat anti-mouse CD8a | 0.5 mg/ml; Clone 53-6.7, 100706, BioLegend |
| BV605 Rat anti-mouse CD185 (CXCR5) | 0.1 mg/ml; Clone L138D7, 145513, BioLegend |
| PE/Cyanine5 Rat anti-mouse CD62L | 0.2 mg/ml; Clone MEL-14, 104410, BioLegend |
| Alexa Fluor® 647 Rat anti-mouse CD21/CD35 (CR2/CR1) | 0.5 mg/ml; Clone 7E9, 123423, BioLegend |

| | |
|---|---|
| Alexa Fluor® 700 Rat anti-mouse CD38 | 0.5 mg/ml; Clone 90, 102741, BioLegend |
| Goat anti-mouse IL-33 (unconjugated) | 0.2 mg/mL; Polyclonal; Cat AF3626, R&D Systems |
| eFluor™ 450 Rat anti-mouse IL-33R(ST2) | 0.2 mg/mL; Clone: RMST2-2; 48-9335-82, Invitrogen |
| PerCP Rabbit anti-mouse CD23 (FCER2) | 0.1 mg/ml; Clone #011, 50695-R011, SinoBiological Inc |
| PerCP-eFluor™ 710 Armenian Hamster anti-human/mouse/rat CD27 | 0.2 mg/ml; Clone LG.7F9, 46-0271-80, Invitrogen |
| APC Rat anti-bovine/dog/cat/mouse/pig/rat FOXP3 | 0.2mg/ml; Clone FJK-16s, 17-5773-82 ThermoFisher Scientific |
| BV786 Rat Anti-Mouse IgM | 0.2 mg/ml; Clone R6-60.2, 564028, BD Biosciences |
| PE/Cyanine7 Goat anti-mouse IgG | 0.2 mg/ml; Clone Poly4053, 405315, BioLegend |

Suppl. Table 2. List of proteins detected in the immunoprecipitated using serum from WT and Treg depleted mice.

[illegible]

Suppl. Table 2 Complete list of genes expressed by IC after Treg depletion and control

| ID | Name | Length | B1.eGFP.r1 | B1.eGFP.r2 | B1.eGFP.r3 | B1.Foxp3.r1 | B1.Foxp3.r2 | B1.Foxp3.r3 |
|---|---|---|---|---|---|---|---|---|
| ENSMUSG00 | Gnai3 | 3253 | 70.2350896 | 76.6568823 | 89.4499639 | 90.8207134 | 78.7434389 | 86.6365962 |
| ENSMUSG00 | Pbsn | 1107 | 0 | 0 | 0 | 0 | 0 | 0 |
| ENSMUSG00 | Cdc45 | 2338 | 2.94170151 | 0.70074381 | 1.19106763 | 0.61731471 | 0.45832074 | 0.20998439 |
| ENSMUSG00 | H19 | 2893 | 0 | 0 | 0 | 0 | 0 | 0 |
| ENSMUSG00 | Scml2 | 6101 | 1.92228598 | 1.30319013 | 5.10773633 | 1.31293458 | 2.3418104 | 4.24252297 |
| ENSMUSG00 | Apoh | 1584 | 0.40157051 | 0 | 0 | 0 | 0.03221362 | 0 |
| ENSMUSG00 | Narf | 4795 | 15.0979625 | 20.7920227 | 24.8525899 | 21.4234788 | 17.505399 | 16.0121188 |
| ENSMUSG00 | Cav2 | 3515 | 244.12014 | 435.706882 | 337.505907 | 464.457521 | 291.438271 | 391.575361 |
| ENSMUSG00 | Klf6 | 4213 | 23.1191377 | 32.9744868 | 39.0189068 | 56.8679743 | 32.301767 | 36.253971 |
| ENSMUSG00 | Scmh1 | 5784 | 8.93536042 | 15.4123302 | 5.17369428 | 12.5388848 | 15.3502583 | 6.93654812 |
| ENSMUSG00 | Cox5a | 685 | 143.003655 | 140.338616 | 186.615664 | 140.27224 | 78.8860321 | 276.767909 |
| ENSMUSG00 | Tbx2 | 3619 | 65.7355058 | 73.2583047 | 54.9622254 | 68.1261703 | 74.8405645 | 59.319891 |
| ENSMUSG00 | Tbx4 | 3391 | 0 | 0 | 0 | 0 | 0 | 0 |
| ENSMUSG00 | Zfy2 | 2807 | 0 | 0 | 0 | 0 | 0 | 0 |
| ENSMUSG00 | Ngfr | 3440 | 0 | 0 | 0 | 0 | 0 | 0 |
| ENSMUSG00 | Wnt3 | 3071 | 0 | 0 | 0 | 0 | 0.14954001 | 0 |
| ENSMUSG00 | Wnt9a | 3462 | 1.69954105 | 0.58458422 | 0.33196069 | 0.11464543 | 0.81064433 | 0.49633226 |
| ENSMUSG00 | Fer | 4416 | 21.795294 | 14.1307615 | 17.2863639 | 20.0837105 | 18.1411708 | 20.2336318 |
| ENSMUSG00 | Xpo6 | 9659 | 9.48715008 | 9.7380614 | 9.15704399 | 10.2504535 | 10.6342369 | 6.81936575 |
| ENSMUSG00 | Tfe3 | 8492 | 4.16655413 | 4.590536 | 4.01834937 | 4.97976404 | 4.87911177 | 3.06727278 |
| ENSMUSG00 | Axin2 | 5492 | 0.03619399 | 0.00877393 | 0.1448713 | 0.12482863 | 0 | 0.03972999 |
| ENSMUSG00 | Baat1 | 6223 | 7.49368129 | 9.12932915 | 5.61135401 | 2.41783908 | 6.74830585 | 2.98912164 |
| ENSMUSG00 | Gna12 | 4017 | 12.1730747 | 9.2726213 | 11.905998 | 16.7340923 | 20.6163326 | 7.08176468 |
| ENSMUSG00 | Slc22a18 | 1807 | 6.77625235 | 7.83996347 | 7.85212596 | 6.36976879 | 3.72743863 | 4.86022902 |
| ENSMUSG00 | Itgb2l | 3359 | 0.41424288 | 0.80334646 | 0.82903129 | 0.96677107 | 1.12412974 | 0.72266832 |
| ENSMUSG00 | Igsf5 | 2802 | 43.4160494 | 39.5018764 | 36.96107 | 42.4434043 | 37.987516 | 29.406383 |
| ENSMUSG00 | Pih1d2 | 2882 | 2.0139834 | 1.8057376 | 1.088942 | 1.21441996 | 1.50494169 | 1.58045273 |
| ENSMUSG00 | Dlat | 4667 | 77.5346869 | 53.9684022 | 79.7944134 | 66.5356921 | 51.0591762 | 79.4219729 |
| ENSMUSG00 | Sdhd | 1283 | 630.014365 | 533.506166 | 737.202679 | 494.152459 | 408.608718 | 648.171279 |
| ENSMUSG00 | Fgf23 | 1811 | 0 | 0 | 0 | 0 | 0 | 0 |
| ENSMUSG00 | Fgf6 | 1229 | 0 | 0 | 0 | 0 | 0 | 0 |
| ENSMUSG00 | Ccnd2 | 5696 | 0.46762942 | 0.37222673 | 0.45784915 | 0.74115155 | 0.1522908 | 0.38307078 |
| ENSMUSG00 | Gpr107 | 10361 | 8.40309838 | 7.70164542 | 6.55284537 | 7.5047588 | 7.6876916 | 5.75713117 |
| ENSMUSG00 | Nalcn | 7257 | 0 | 0 | 0 | 0 | 0.02812533 | 0 |
| ENSMUSG00 | Btbd17 | 3162 | 0.38975961 | 0 | 0 | 0.04564459 | 0.09682424 | 0 |
| ENSMUSG00 | Slfn4 | 3572 | 0 | 0 | 0 | 0 | 0 | 0 |
| ENSMUSG00 | Th | 2574 | 0 | 0 | 0 | 0 | 0 | 0 |
| ENSMUSG00 | Ins2 | 822 | 0 | 0 | 0 | 0 | 0 | 0 |
| ENSMUSG00 | Scnn1g | 2978 | 288.340706 | 428.305952 | 94.6821882 | 318.510674 | 419.51974 | 150.065468 |
| ENSMUSG00 | Drp2 | 8620 | 0 | 0 | 0 | 0 | 0 | 0 |
| ENSMUSG00 | Tspan32 | 2962 | 0 | 0 | 0.16415269 | 0 | 0.20672401 | 0.25782914 |
| ENSMUSG00 | Lhx2 | 4702 | 0 | 0 | 0 | 0 | 0 | 0 |
| ENSMUSG00 | Clec2g | 3079 | 0 | 0 | 0 | 0 | 0 | 0 |
| ENSMUSG00 | Gmpr | 1861 | 34.350786 | 43.6034221 | 37.4326193 | 29.7226084 | 45.7345486 | 36.9915043 |
| ENSMUSG00 | Glra1 | 3270 | 0 | 0 | 0 | 0 | 0 | 0 |
| ENSMUSG00 | Mid2 | 8103 | 2.98301026 | 2.92579656 | 2.36746882 | 5.85114236 | 1.87657164 | 1.2083914 |
| ENSMUSG00 | Trim25 | 6232 | 3.92961746 | 4.23719036 | 3.79460623 | 3.90232638 | 3.98745909 | 3.86011422 |
| ENSMUSG00 | Dgke | 11704 | 1.61684969 | 1.90621349 | 1.35581525 | 1.34105343 | 1.70465763 | 2.49582524 |
| ENSMUSG00 | Scpep1 | 2786 | 12.1150047 | 14.7188307 | 20.4191572 | 17.574779 | 12.9672197 | 16.8288244 |
| ENSMUSG00 | Mnt | 2671 | 0.99723594 | 1.20872019 | 1.42319674 | 0.97263467 | 0.80236158 | 1.84826271 |
| ENSMUSG00 | Itgb2 | 3148 | 0 | 0 | 0 | 0 | 0 | 0 |
| ENSMUSG00 | Hddc2 | 2549 | 7.36154837 | 6.99450121 | 7.17911456 | 9.63980868 | 6.22565825 | 12.4014543 |
| ENSMUSG00 | Tpd52l1 | 1274 | 103.72623 | 130.64048 | 81.7074885 | 99.2681052 | 91.2787367 | 117.169624 |
| ENSMUSG00 | Pemt | 2147 | 0.29626814 | 1.27928608 | 1.42055294 | 0 | 0.66545809 | 1.60065419 |
| ENSMUSG00 | Cdh1 | 3837 | 267.823668 | 228.147796 | 217.737564 | 263.172296 | 302.594255 | 194.960003 |
| ENSMUSG00 | Cdh4 | 7257 | 1.85711835 | 2.43023809 | 0.1461822 | 1.45183368 | 1.0125118 | 0 |
| ENSMUSG00 | Clmt1 | 2658 | 364.021401 | 242.237487 | 460.510699 | 243.682762 | 244.323809 | 312.550127 |
| ENSMUSG00 | Bcl6b | 3710 | 0.51435662 | 0.31171822 | 0.40508428 | 0.40847598 | 0.97651555 | 0.26465957 |
| ENSMUSG00 | Clec10a | 1990 | 0 | 0 | 0 | 0 | 0 | 0 |
| ENSMUSG00 | Alox12 | 3971 | 24.2777739 | 13.6271404 | 36.8107517 | 19.7447199 | 14.5844716 | 26.3886084 |
| ENSMUSG00 | Arvcf | 5943 | 8.60264983 | 6.92431795 | 5.8385406 | 6.82419957 | 5.90714228 | 6.70507285 |
| ENSMUSG00 | Comt | 3219 | 25.3674299 | 25.0587462 | 31.1019492 | 21.902552 | 22.6361183 | 30.773995 |
| ENSMUSG00 | Rtcd1 | 1696 | 16.760123 | 18.3824458 | 15.7677565 | 11.4883868 | 11.583228 | 13.3960935 |
| ENSMUSG00 | Dbt | 3285 | 44.6084326 | 35.3660615 | 44.1076534 | 39.3442568 | 43.5549339 | 36.3744885 |
| ENSMUSG00 | Dazap2 | 1834 | 416.435407 | 432.285459 | 444.283955 | 383.937349 | 390.237703 | 403.736045 |
| ENSMUSG00 | Mcts1 | 1781 | 27.054263 | 22.8892364 | 24.9922829 | 28.2213858 | 17.1042933 | 51.7927175 |
| ENSMUSG00 | Rem1 | 2112 | 0 | 0 | 0 | 0 | 0.04832043 | 0 |
| ENSMUSG00 | Rnf17 | 5120 | 0.61341464 | 0.23528536 | 0.31942738 | 0.17618186 | 0.62786361 | 0.47410994 |
| ENSMUSG00 | Trappc10 | 5025 | 5.90200765 | 9.22494674 | 7.9871191 | 6.36910915 | 5.70683049 | 7.1918203 |
| ENSMUSG00 | Ccm2 | 3312 | 8.01831551 | 7.85648513 | 7.47374157 | 6.78717202 | 8.18085917 | 6.37396461 |
| ENSMUSG00 | Wap | 2761 | 0 | 0 | 0.09605616 | 0.01306847 | 0 | 0 |
| ENSMUSG00 | Tbrg4 | 3249 | 2.06792127 | 1.97254442 | 1.45570859 | 1.22161431 | 1.58623083 | 2.3169592 |
| ENSMUSG00 | Tmprss2 | 3145 | 342.996887 | 396.844845 | 330.241878 | 361.176103 | 351.067956 | 257.80388 |
| ENSMUSG00 | Mx1 | 4963 | 0 | 0 | 0 | 0 | 0.13365764 | 0.25279738 |
| ENSMUSG00 | Fap | 5810 | 0.28738902 | 0.29857348 | 0.22823671 | 0 | 0 | 0 |
| ENSMUSG00 | Gcg | 1487 | 0 | 0 | 0 | 0 | 0 | 0 |
| ENSMUSG00 | Ndufa9 | 1311 | 226.797284 | 240.233857 | 263.862413 | 195.189824 | 193.830169 | 218.883661 |
| ENSMUSG00 | Egfl6 | 2692 | 3.39664209 | 2.04058485 | 0.95234283 | 0.0804206 | 3.60141583 | 2.59372498 |
| ENSMUSG00 | Lck | 3548 | 0.22410079 | 0.58399578 | 0.03737473 | 0 | 0.86290376 | 0 |
| ENSMUSG00 | Tssk3 | 1278 | 0 | 0.263932 | 0 | 0.08469964 | 0.27948719 | 0.19207492 |
| ENSMUSG00 | Cttnbp2 | 9995 | 4.26789701 | 4.58964411 | 2.17139619 | 6.27058643 | 6.41722405 | 5.61047107 |
| ENSMUSG00 | Galnt1 | 5503 | 18.6243192 | 18.8700314 | 23.3097853 | 19.4081141 | 16.5791677 | 21.7086898 |
| ENSMUSG00 | Myf5 | 2080 | 0 | 0 | 0 | 0 | 0 | 0 |
| ENSMUSG00 | Mkrn2 | 2314 | 17.0773326 | 14.764126 | 22.9223041 | 15.9671623 | 18.2583575 | 11.7867931 |
| ENSMUSG00 | Pparg | 1975 | 0.06038807 | 0.39037118 | 0.04476136 | 0.09134695 | 0 | 0 |
| ENSMUSG00 | Raf1 | 6697 | 27.734449 | 23.4996148 | 18.6918948 | 21.4056985 | 23.2921653 | 15.561654 |
| ENSMUSG00 | 1-Sep | 2354 | 0.21955023 | 0.18423024 | 0.35676934 | 0.16860769 | 0.13005873 | 0.70677697 |
| ENSMUSG00 | Pdgfb | 2911 | 0.54627936 | 1.04285327 | 0 | 0.12395069 | 0.42069152 | 0 |
| ENSMUSG00 | Acvrl1 | 4297 | 2.74781888 | 2.10822693 | 2.24249518 | 1.2511577 | 2.50560049 | 0.95845253 |
| ENSMUSG00 | Grasp | 2095 | 0.31725072 | 0.26436452 | 0.6393284 | 0.61186509 | 0.89073474 | 0.13603311 |
| ENSMUSG00 | Acvr1b | 3268 | 1.94641275 | 1.79888187 | 2.70513115 | 1.61198853 | 1.32718542 | 2.54552172 |
| ENSMUSG00 | Tom1l2 | 12272 | 22.8030335 | 15.1760592 | 18.7620111 | 15.9064424 | 23.3302636 | 14.3863054 |
| ENSMUSG00 | Gpa33 | 2997 | 0.02653018 | 0 | 0 | 0 | 0 | 0 |
| ENSMUSG00 | Zfp385a | 3711 | 0.05356438 | 0.01298476 | 0 | 0.09722998 | 0.11000027 | 0 |
| ENSMUSG00 | Itga5 | 4349 | 0.21939102 | 0.18835813 | 0.23376463 | 0.17422923 | 0.61011073 | 1.04106464 |
| ENSMUSG00 | Gabra2 | 2382 | 0 | 0 | 0 | 0 | 0 | 0 |
| ENSMUSG00 | Wdr77 | 6811 | 10.955959 | 9.02036613 | 9.55294566 | 7.82457492 | 8.06113352 | 9.46663444 |
| ENSMUSG00 | Adora3 | 2521 | 0 | 0 | 0 | 0 | 0 | 0.04327591 |
| ENSMUSG00 | Atp5f1 | 2743 | 271.664866 | 230.479811 | 285.402421 | 231.317081 | 180.945519 | 308.691643 |
| ENSMUSG00 | Sox9 | 1698 | 0.51508868 | 1.58918773 | 1.04126839 | 0.84998927 | 1.35228911 | 1.07620779 |
| ENSMUSG00 | Hnrnpd | 5344 | 28.6188873 | 21.2979746 | 19.3962054 | 25.5086011 | 20.3666279 | 40.9170627 |
| ENSMUSG00 | Dynll1c | 6502 | 0 | 0 | 0 | 0 | 0 | 0 |
| ENSMUSG00 | C1d | 4056 | 14.99652 | 19.1510219 | 14.2108489 | 16.3152046 | 11.6243519 | 22.0160673 |
| ENSMUSG00 | Gm2a | 4123 | 29.6213524 | 41.162418 | 32.3553631 | 29.1159258 | 30.6306767 | 24.0331711 |
| ENSMUSG00 | Krit1 | 4763 | 27.4273586 | 29.5815992 | 32.5736865 | 42.483331 | 29.2359814 | 49.057702 |
| ENSMUSG00 | Clcn4-2 | 4759 | 52.5951892 | 47.1941599 | 73.0319587 | 54.528696 | 56.3874155 | 53.9073874 |
| ENSMUSG00 | Vmn2r88 | 3174 | 0 | 0 | 0 | 0 | 0 | 0 |
| ENSMUSG00 | Grm6 | 5410 | 0 | 0 | 0 | 0 | 0 | 0 |
| ENSMUSG00 | Sema4f | 5143 | 0 | 0 | 0 | 0 | 0 | 0 |
| ENSMUSG00 | Hk2 | 6685 | 0 | 0.14416288 | 0.01322419 | 0.22129601 | 0 | 0 |
| ENSMUSG00 | Myo18a | 10638 | 15.0531186 | 14.9161453 | 11.5386838 | 19.9065162 | 22.3234398 | 11.2195734 |
| ENSMUSG00 | Sez6 | 5750 | 0.05531197 | 0 | 0 | 0 | 0 | 0 |
| ENSMUSG00 | Haao | 1212 | 0 | 0.51685128 | 0.1094105 | 0.11908266 | 0.63151455 | 1.6202756 |
| ENSMUSG00 | Cd52 | 1183 | 0 | 0 | 0.52309873 | 0 | 0 | 0.02305549 |
| ENSMUSG00 | Abhd15 | 1892 | 0 | 0.07640556 | 0 | 0.09535424 | 0 | 0 |
| ENSMUSG00 | Hoxb6 | 2073 | 13.2326491 | 13.7376687 | 15.8000799 | 16.3787766 | 13.8088745 | 18.4593916 |
| ENSMUSG00 | Loxl3 | 4183 | 0.00950406 | 0.02303918 | 0.14793827 | 0.47442325 | 0.07319107 | 0.3651398 |
| ENSMUSG00 | Tcl1b5 | 1035 | 0 | 0 | 0 | 0 | 0 | 0 |
| ENSMUSG00 | Btn1a1 | 3925 | 0 | 0 | 0 | 0 | 0 | 0 |
| ENSMUSG00 | Kat2b | 4198 | 15.0385191 | 10.2387581 | 20.153008 | 15.8149028 | 13.4676571 | 16.5674917 |
| ENSMUSG00 | Rab5b | 3160 | 10.4421041 | 8.95108909 | 10.4210041 | 7.83300081 | 10.17298 | 10.3833516 |
| ENSMUSG00 | Cryba1 | 1908 | 0 | 0 | 0 | 0 | 0 | 0 |
| ENSMUSG00 | Dnmt3l | 2970 | 0 | 0 | 0 | 0 | 0.05154179 | 0 |
| ENSMUSG00 | Aire | 4111 | 0 | 0.15237746 | 0 | 0 | 0 | 0 |
| ENSMUSG00 | Icosl | 2834 | 2.14629094 | 4.40377153 | 1.55969806 | 3.9214078 | 5.50955222 | 2.09804917 |
| ENSMUSG00 | Spg7 | 3939 | 31.136242 | 38.2898106 | 35.9427528 | 25.6485719 | 28.7322929 | 20.987416 |
| ENSMUSG00 | Sult5a1 | 2053 | 2.14946826 | 1.5491014 | 8.78438965 | 1.77510302 | 0.49709085 | 3.41431188 |
| ENSMUSG00 | Rpl13 | 2348 | 2.62440353 | 3.77610961 | 3.42620759 | 3.62664502 | 2.80340821 | 3.8913987 |
| ENSMUSG00 | Chmp1a | 3395 | 29.6731628 | 28.8976719 | 26.1956136 | 22.7864222 | 31.9384532 | 16.0675342 |
| ENSMUSG00 | Rpa1 | 6612 | 8.35153697 | 6.34032134 | 9.0248772 | 9.29336386 | 10.0787125 | 9.38854793 |
| ENSMUSG00 | Serpinf1 | 2712 | 0 | 0.05330359 | 0.03259723 | 0 | 0 | 0 |
| ENSMUSG00 | Tubgcp3 | 9566 | 5.23645259 | 6.94638341 | 6.0808724 | 8.23406891 | 9.66546027 | 6.9027704 |
| ENSMUSG00 | Oprm1 | 8203 | 0.48949208 | 0.08223945 | 0.29636735 | 0.71697833 | 0.47897488 | 0.40564501 |
| ENSMUSG00 | Polr3d | 2040 | 18.4161417 | 14.8811071 | 11.7221554 | 15.8831745 | 12.2813483 | 14.9074621 |
| ENSMUSG00 | Tcf7 | 4136 | 0 | 0.5592237 | 0 | 0.18320187 | 0.16038271 | 0 |
| ENSMUSG00 | Ddx3x | 3287 | 111.102478 | 96.6514186 | 99.1213218 | 115.743559 | 96.6349831 | 121.62813 |
| ENSMUSG00 | Il12rb1 | 2991 | 0 | 0 | 0 | 0 | 0 | 0 |
| ENSMUSG00 | Slc5a5 | 2913 | 0.01364761 | 0.03308372 | 0.80422166 | 0.32205052 | 0.22771812 | 0.31834457 |
| ENSMUSG00 | Kcnn3 | 3072 | 0.19411856 | 0.36077089 | 0.05755448 | 0.42283646 | 0 | 0.07102771 |
| ENSMUSG00 | Usp32 | 8579 | 8.22079758 | 6.37505675 | 4.97715122 | 9.82908708 | 6.89352717 | 8.22785481 |
| ENSMUSG00 | Car4 | 2211 | 0.26971154 | 0.80637646 | 0.25989324 | 0.97915996 | 0.43848989 | 0.90051952 |
| ENSMUSG00 | Txnrd3 | 2820 | 11.3345413 | 12.5079701 | 12.1006464 | 9.51951813 | 9.13770205 | 6.22867648 |
| ENSMUSG00 | Fasl | 2382 | 0 | 0 | 0 | 0 | 0 | 0 |
| ENSMUSG00 | Znf512b | 9357 | 6.00347269 | 5.29396842 | 6.00411911 | 6.12357451 | 7.6073308 | 7.75944103 |
| ENSMUSG00 | Dnajc5 | 2094 | 113.931537 | 104.795156 | 103.876442 | 108.952611 | 127.370996 | 99.7986084 |
| ENSMUSG00 | Tpd52l2 | 5140 | 22.1903645 | 25.2556956 | 26.5555041 | 23.0040584 | 22.7037591 | 22.8703687 |
| ENSMUSG00 | Fmr1 | 8279 | 17.7816819 | 20.7494463 | 16.700491 | 18.9976606 | 16.616392 | 21.8157581 |
| ENSMUSG00 | Bcl11a | 13496 | 0.08542597 | 0.14995781 | 0 | 0 | 0.04158937 | 0 |
| ENSMUSG00 | Il4 | 1220 | 0.1629323 | 0 | 0 | 0 | 0.1254747 | 0 |
| ENSMUSG00 | Pxmp4 | 3109 | 12.4036076 | 10.6013273 | 20.0607271 | 12.8939053 | 14.4593879 | 10.9221376 |
| ENSMUSG00 | Dlg3 | 6526 | 6.75587311 | 8.07783753 | 10.6813219 | 8.60860302 | 8.48354547 | 5.47498888 |
| ENSMUSG00 | Gnb1l | 4178 | 1.0276668 | 2.27207494 | 1.37535653 | 1.03634403 | 1.86860591 | 1.4818916 |
| ENSMUSG00 | Dbh | 2398 | 0 | 0 | 0 | 0 | 0 | 0 |
| ENSMUSG00 | Mmp11 | 2264 | 2.87981396 | 2.72432182 | 4.39285101 | 4.12776039 | 4.59778298 | 3.74664877 |
| ENSMUSG00 | Smarcb1 | 7032 | 9.97279119 | 10.1895961 | 10.3087347 | 9.74399923 | 8.2504251 | 9.59577041 |
| ENSMUSG00 | Vpreb3 | 856 | 0.13932996 | 0 | 0.05163767 | 0 | 0.35766151 | 0 |
| ENSMUSG00 | Hip1r | 4878 | 38.8426856 | 31.1858544 | 42.5707787 | 32.2208663 | 38.9340244 | 34.1016861 |
| ENSMUSG00 | Nsun5 | 3239 | 5.51102524 | 8.03355318 | 6.14104025 | 8.8673379 | 5.46654906 | 4.68190782 |
| ENSMUSG00 | Top1mt | 1889 | 3.87242372 | 9.38730053 | 8.47065493 | 5.34831788 | 6.15882143 | 4.38935433 |
| ENSMUSG00 | Hoxa10 | 4974 | 8.88783561 | 8.37014194 | 7.215902 | 7.69663234 | 7.05793055 | 8.57061945 |
| ENSMUSG00 | Hoxa4 | 1982 | 0.10029132 | 0.36468044 | 0 | 0.18204866 | 0.3346836 | 0.20641755 |
| ENSMUSG00 | Snrpn | 2185 | 17.4851335 | 18.8555643 | 16.8310908 | 22.4914163 | 14.0818785 | 23.8793524 |
| ENSMUSG00 | Mmp14 | 2587 | 3.39619683 | 4.56346282 | 2.13576744 | 2.98475359 | 5.77917592 | 1.96099068 |
| ENSMUSG00 | Slc7a7 | 2259 | 0.77434314 | 0.98122016 | 0.07826798 | 0.55904009 | 1.60375063 | 0.77272108 |
| ENSMUSG00 | Oxa1l | 2524 | 23.6737271 | 19.8358611 | 27.4247568 | 20.5570444 | 21.1868629 | 24.0111919 |
| ENSMUSG00 | Heatr6 | 6626 | 5.35792999 | 4.47975375 | 4.3761559 | 3.66483791 | 7.94736188 | 4.0010488 |
| ENSMUSG00 | Ccl3 | 763 | 0.31262501 | 0.44207745 | 0 | 0 | 0.33437992 | 0.32171921 |
| ENSMUSG00 | Expi | 823 | 0 | 0 | 0 | 0 | 0 | 0 |
| ENSMUSG00 | Icam4 | 904 | 0 | 0 | 0 | 0 | 0 | 0 |
| ENSMUSG00 | Ilf2 | 1959 | 58.2836856 | 56.2298146 | 55.4384524 | 48.9013927 | 42.639192 | 47.2259195 |
| ENSMUSG00 | 2500003M10 | 4779 | 21.1962679 | 17.3225168 | 20.7551655 | 18.512905 | 17.937709 | 19.8153479 |
| ENSMUSG00 | Snapin | 3836 | 37.0194412 | 40.1595557 | 33.9118936 | 36.5617589 | 42.5663198 | 49.1668745 |
| ENSMUSG00 | S100a4 | 619 | 0.32112666 | 0.15569125 | 0 | 0.23316345 | 0.08243356 | 0 |
| ENSMUSG00 | S100a3 | 709 | 0 | 0 | 0 | 0 | 0 | 0 |
| ENSMUSG00 | S100a5 | 429 | 0 | 0 | 0 | 0 | 0 | 0 |
| ENSMUSG00 | S100a6 | 716 | 1.49915918 | 1.480589 | 1.17295394 | 0.50393917 | 1.2115201 | 0.45711686 |
| ENSMUSG00 | Scn4a | 7015 | 0.18135073 | 0.19920268 | 0.01890314 | 0.03600489 | 0.21821686 | 0 |
| ENSMUSG00 | Icam2 | 1897 | 1.42507785 | 1.7780975 | 1.46795789 | 1.06515261 | 1.21043063 | 2.08477738 |
| ENSMUSG00 | Mapk7 | 4299 | 6.50107067 | 5.41382916 | 5.63447545 | 6.01787065 | 5.04447772 | 3.69879383 |
| ENSMUSG00 | Epn2 | 5209 | 7.28863196 | 8.07578499 | 12.0686195 | 11.7964526 | 10.7753913 | 6.43511838 |
| ENSMUSG00 | B9d1 | 1654 | 5.48019925 | 9.55571525 | 5.10432407 | 7.9406676 | 3.64033395 | 8.95412885 |
| ENSMUSG00 | Sec24b | 6548 | 12.9259954 | 9.58136037 | 7.77650018 | 9.97380889 | 12.2111074 | 7.56842082 |
| ENSMUSG00 | N4bp3 | 2721 | 4.66078584 | 4.39185507 | 2.66413166 | 4.25664693 | 2.77541478 | 3.97943101 |
| ENSMUSG00 | Rmnd5b | 2624 | 19.7110824 | 22.3119387 | 19.7762819 | 20.4061563 | 18.0070213 | 14.6351723 |
| ENSMUSG00 | Nhp2 | 1349 | 8.90004086 | 10.6803159 | 10.9112037 | 7.97068144 | 5.67380013 | 12.4141056 |
| ENSMUSG00 | 1300018I17R | 5563 | 13.2351519 | 13.4953221 | 9.16931994 | 10.6631101 | 11.6948821 | 11.1393098 |
| ENSMUSG00 | Zfp276 | 5861 | 8.47881772 | 8.39419168 | 6.71964547 | 11.1798316 | 10.3341253 | 6.98037176 |
| ENSMUSG00 | C1ql4 | 872 | 0 | 0 | 0.10138037 | 0 | 0.46813189 | 0 |
| ENSMUSG00 | Mfsd10 | 2961 | 8.04239542 | 6.50948222 | 8.86723902 | 7.54298736 | 10.4775537 | 8.07830416 |
| ENSMUSG00 | Luzp1 | 4366 | 23.1466861 | 27.338025 | 20.0052317 | 33.3548172 | 29.0076649 | 24.2010885 |
| ENSMUSG00 | Slc13a2 | 2400 | 0 | 0 | 0.20259178 | 0 | 0.06378297 | 0.11364433 |
| ENSMUSG00 | Kctd10 | 4695 | 4.62332106 | 4.31060824 | 5.65821249 | 4.2499192 | 3.35828544 | 3.50300479 |
| ENSMUSG00 | Poldip2 | 3524 | 42.0117507 | 38.1225314 | 42.9851635 | 32.7236704 | 38.1973835 | 33.6056991 |
| ENSMUSG00 | Sebox | 1432 | 0 | 0 | 0 | 0 | 0 | 0 |
| ENSMUSG00 | Ift20 | 1668 | 31.1513268 | 23.1687809 | 36.887869 | 27.8619145 | 19.5478742 | 43.3810659 |
| ENSMUSG00 | Col6a1 | 4358 | 0 | 0 | 0.01014269 | 0 | 0.05854334 | 0 |
| ENSMUSG00 | Pcbp3 | 3578 | 6.26665484 | 9.30598981 | 3.66907417 | 5.5867671 | 7.24466152 | 6.52517921 |
| ENSMUSG00 | Lgals9 | 2688 | 1.4050486 | 1.36241429 | 0.44399173 | 3.47665533 | 1.04406647 | 0.2638172 |
| ENSMUSG00 | Araf | 4311 | 60.8366748 | 66.2046618 | 55.3984129 | 79.2700171 | 63.8924557 | 65.5261978 |
| ENSMUSG00 | Cfp | 1835 | 1.06159049 | 0.94534709 | 0.93943972 | 0.45225451 | 1.58501549 | 1.82821833 |
| ENSMUSG00 | Timp1 | 985 | 0 | 0.83164418 | 0 | 0.14652607 | 0.62164113 | 0 |
| ENSMUSG00 | Uxt | 1631 | 3.58311198 | 2.80669038 | 3.65864427 | 2.67684083 | 2.31511454 | 5.6188098 |
| ENSMUSG00 | Cnnm3 | 5605 | 9.17816089 | 9.14725681 | 6.49029738 | 10.4158337 | 10.7423949 | 5.72743094 |
| ENSMUSG00 | Lman2l | 7302 | 11.6511542 | 13.2179463 | 15.0729374 | 14.6512273 | 14.339376 | 9.38289425 |
| ENSMUSG00 | Mcm3ap | 6640 | 5.67593306 | 4.43402352 | 4.10730981 | 4.08096618 | 8.04587583 | 3.1587647 |
| ENSMUSG00 | Pcnt | 10004 | 2.25323445 | 2.59621075 | 2.83662367 | 2.8854094 | 5.26891706 | 3.27982717 |
| ENSMUSG00 | Ftcd | 3285 | 0 | 0 | 0 | 0 | 0 | 0 |
| ENSMUSG00 | Mxd1 | 4656 | 5.58421054 | 5.52653782 | 5.25941173 | 6.09116989 | 7.38655011 | 4.20015385 |
| ENSMUSG00 | Gmcl1 | 3808 | 40.4131999 | 39.0882142 | 40.7775931 | 42.6673967 | 40.3467483 | 39.4651424 |
| ENSMUSG00 | Snrnp27 | 1936 | 18.4403004 | 20.6584437 | 24.9776944 | 18.4137707 | 14.4170597 | 34.3750619 |
| ENSMUSG00 | Oas1c | 3433 | 2.62874864 | 1.60013236 | 0.66952981 | 0.87235936 | 1.09989858 | 0.78653926 |
| ENSMUSG00 | Oas1h | 1981 | 0 | 0 | 0 | 0 | 0 | 0 |
| ENSMUSG00 | Ocrl | 9714 | 7.47717345 | 8.15508656 | 4.21814994 | 4.4536104 | 6.99682238 | 4.90235949 |
| ENSMUSG00 | Calm1 | 4878 | 101.206141 | 104.917631 | 99.2412023 | 86.9430815 | 74.1336461 | 114.091639 |
| ENSMUSG00 | Agpat3 | 9772 | 12.6158151 | 15.1729106 | 10.1277043 | 13.5547672 | 15.1690157 | 10.7624856 |
| ENSMUSG00 | Slc26a3 | 5215 | 0.03811647 | 0 | 0.01695181 | 0 | 0 | 0 |
| ENSMUSG00 | Sema6b | 3959 | 0 | 0 | 0 | 0 | 0.27066277 | 0.07578202 |
| ENSMUSG00 | Uhrf1 | 4188 | 1.58528301 | 2.22062639 | 0.58049221 | 0.56001263 | 1.23057879 | 0.48193011 |
| ENSMUSG00 | Dpp9 | 3611 | 21.6667919 | 18.2817573 | 12.9508585 | 20.0944313 | 17.4091651 | 11.7074746 |
| ENSMUSG00 | Ramp2 | 3294 | 0.72414354 | 0.70217038 | 0.28179681 | 0.16430804 | 0.5111932 | 0.17388204 |
| ENSMUSG00 | Lsr | 2351 | 15.2866671 | 18.7334786 | 17.0339727 | 15.8539992 | 19.2515506 | 10.4179609 |
| ENSMUSG00 | Gramd1a | 2663 | 30.096526 | 20.9356791 | 19.6360422 | 24.6599028 | 37.3069297 | 21.9897298 |
| ENSMUSG00 | Hpn | 3272 | 60.1192291 | 48.2895305 | 69.9637362 | 45.6429041 | 40.4998466 | 47.4055237 |
| ENSMUSG00 | Gabrg1 | 5023 | 0.28492879 | 3.32882648 | 0.76559035 | 0.53875241 | 1.32061096 | 0.44525591 |
| ENSMUSG00 | Ckb | 1470 | 17.1732858 | 15.3409897 | 27.1224914 | 18.3601152 | 16.6269619 | 15.9009291 |
| ENSMUSG00 | Sp1 | 8316 | 5.12958521 | 6.65200043 | 4.16717712 | 3.01551478 | 5.24008237 | 6.10694785 |
| ENSMUSG00 | Itgb7 | 2727 | 0.11662774 | 0 | 0.34038823 | 0 | 0 | 0 |
| ENSMUSG00 | Myg1 | 3050 | 11.3922262 | 11.6279415 | 10.2606245 | 13.1078378 | 9.05090797 | 8.9872172 |
| ENSMUSG00 | Rarg | 5378 | 0.99795275 | 0.80639267 | 0.37807452 | 1.12043537 | 1.13855803 | 1.29830934 |
| ENSMUSG00 | Pfdn5 | 1501 | 12.6073343 | 14.7352278 | 20.702129 | 15.6491745 | 12.3741511 | 20.660403 |
| ENSMUSG00 | Efnb2 | 7055 | 4.13051272 | 6.1197806 | 5.23780875 | 6.78168548 | 5.6920989 | 8.20365478 |
| ENSMUSG00 | Rrp15 | 3689 | 3.89041162 | 2.71693683 | 3.01948074 | 2.74845609 | 3.61016104 | 2.58033318 |
| ENSMUSG00 | Rnd2 | 1983 | 1.08260007 | 2.30847807 | 1.24826183 | 1.47385053 | 2.18721228 | 1.5267196 |
| ENSMUSG00 | Srr | 6885 | 20.8506957 | 9.58829713 | 10.7727949 | 8.5580216 | 10.6795944 | 5.79165179 |
| ENSMUSG00 | Sync | 2356 | 1.13056757 | 1.55440136 | 2.42021976 | 2.69543287 | 2.57730846 | 1.92172755 |
| ENSMUSG00 | Fndc5 | 2757 | 0.519114 | 0.12234498 | 0.68940125 | 0.1308743 | 0.55523804 | 0.13850016 |
| ENSMUSG00 | Acp5 | 1605 | 1.01556056 | 0.72054493 | 2.36844766 | 0.38217742 | 0.69942696 | 1.03660623 |
| ENSMUSG00 | Cnn1 | 2361 | 0.03367682 | 0 | 0 | 0 | 0.08644875 | 0.01155216 |
| ENSMUSG00 | Fbxo9 | 3675 | 43.2063644 | 40.5159473 | 53.5594033 | 43.0726338 | 39.5714751 | 36.6333658 |
| ENSMUSG00 | Ccdc132 | 10045 | 9.73603604 | 10.937291 | 9.36401413 | 12.2596335 | 11.3888636 | 10.459125 |
| ENSMUSG00 | Apbb3 | 2048 | 13.0835908 | 8.7055584 | 5.6115621 | 11.3989662 | 11.8845612 | 8.62986633 |
| ENSMUSG00 | Hars | 1974 | 41.5277613 | 34.8094562 | 42.6567938 | 40.688263 | 36.0338237 | 34.003489 |
| ENSMUSG00 | Zmat2 | 2251 | 22.8536614 | 23.2048438 | 24.9776741 | 18.5780052 | 19.9934533 | 26.6445721 |
| ENSMUSG00 | Ube2c | 2770 | 0.11481727 | 0.34791655 | 0 | 0.48196233 | 0.03684215 | 0.12800372 |
| ENSMUSG00 | Smg5 | 4425 | 6.70227988 | 5.9892753 | 7.35199016 | 7.22456304 | 9.13285642 | 7.3718573 |
| ENSMUSG00 | Cct3 | 4575 | 55.4751885 | 48.3760282 | 52.9166108 | 47.5178838 | 39.2707911 | 38.8283553 |
| ENSMUSG00 | Rhbg | 2327 | 368.06707 | 337.201557 | 303.904292 | 368.123687 | 337.997661 | 275.969652 |
| ENSMUSG00 | 0610031J06R | 2546 | 39.1308854 | 43.96587 | 40.6254174 | 31.1076543 | 28.8401237 | 33.702284 |
| ENSMUSG00 | Mef2d | 3903 | 10.827588 | 12.1608111 | 10.090659 | 13.9317553 | 12.8121569 | 13.6967187 |
| ENSMUSG00 | Tmem79 | 3547 | 1.53552321 | 1.8747474 | 1.5327958 | 1.38346717 | 2.0715529 | 1.46100408 |
| ENSMUSG00 | Snd1 | 7518 | 16.8053893 | 15.3827429 | 15.1455105 | 14.8302098 | 17.2327704 | 10.8801068 |
| ENSMUSG00 | Col18a1 | 7962 | 104.047313 | 94.9627561 | 95.4653218 | 94.1568248 | 118.279678 | 80.2515667 |
| ENSMUSG00 | Slc19a1 | 4637 | 5.20413558 | 3.82415586 | 2.65954587 | 4.98005359 | 3.07016581 | 1.95281006 |
| ENSMUSG00 | Kpnb1 | 6740 | 6.37621282 | 9.22977697 | 8.4665548 | 10.3802796 | 9.12266808 | 10.5780277 |
| ENSMUSG00 | Npepps | 6811 | 9.4733732 | 8.68077588 | 11.4739184 | 7.27362313 | 8.36829567 | 7.7126641 |
| ENSMUSG00 | Tbx21 | 2482 | 0 | 0 | 0 | 0 | 0 | 0 |
| ENSMUSG00 | Mrpl10 | 1065 | 50.9542074 | 47.8697237 | 62.1731088 | 48.1771524 | 42.1626392 | 45.9187119 |
| ENSMUSG00 | Cyp51 | 3896 | 21.0920375 | 17.9214976 | 25.1641909 | 21.4213986 | 24.8845262 | 19.3428717 |
| ENSMUSG00 | Tcf25 | 3984 | 88.2324193 | 84.5197936 | 88.1374099 | 93.2119484 | 99.2222223 | 65.1401086 |
| ENSMUSG00 | Tubb6 | 1753 | 1.33803386 | 0.93459158 | 1.1094587 | 0.90565314 | 1.45540148 | 0.65347111 |
| ENSMUSG00 | Def8 | 4028 | 15.1007659 | 15.049291 | 9.39344032 | 10.0148277 | 14.2640763 | 4.80082404 |
| ENSMUSG00 | Meox1 | 2225 | 0.23227921 | 0.19491145 | 0.11919598 | 0.37298293 | 0.16053242 | 0 |
| ENSMUSG00 | Sost | 2064 | 0 | 0.11673072 | 0 | 0 | 0 | 0 |
| ENSMUSG00 | Nkx2-1 | 2829 | 0 | 0 | 0 | 0 | 0 | 0 |
| ENSMUSG00 | Pax9 | 5052 | 0 | 0.07630474 | 0.01749875 | 0.34282228 | 0 | 0 |
| ENSMUSG00 | Irx2 | 3777 | 0 | 0 | 0 | 0 | 0 | 0 |
| ENSMUSG00 | Col1a1 | 6702 | 0 | 0.0359493 | 0 | 0 | 0 | 0 |
| ENSMUSG00 | Itga3 | 6963 | 14.1996094 | 14.3251379 | 13.3627574 | 11.6024268 | 13.0515547 | 11.5828103 |
| ENSMUSG00 | Sgca | 2007 | 0 | 0.24009189 | 0 | 0 | 0 | 0 |
| ENSMUSG00 | Dlx3 | 2634 | 0.24149115 | 0 | 0.13425009 | 0 | 0.34869961 | 0 |
| ENSMUSG00 | Foxm1 | 5205 | 0.25205204 | 0.20366988 | 0.39913288 | 0.18716911 | 0.1274434 | 0.10480169 |
| ENSMUSG00 | Itfg2 | 3622 | 7.0356883 | 6.47896116 | 8.29852381 | 6.63463326 | 7.25526879 | 7.77876928 |
| ENSMUSG00 | Nrip2 | 3639 | 1.33283007 | 0.47670017 | 1.46975076 | 1.63603664 | 1.14981116 | 0.96686685 |
| ENSMUSG00 | Tulp3 | 6894 | 0.63433462 | 1.28609013 | 0.64116395 | 1.3974334 | 0.71795162 | 0.53805497 |
| ENSMUSG00 | Gtf2h4 | 2952 | 4.41727561 | 3.72171708 | 3.71343397 | 4.2046827 | 4.47690763 | 3.51096305 |
| ENSMUSG00 | Tubb5 | 3753 | 6.13339952 | 11.2345156 | 7.69086157 | 3.50917832 | 3.86130848 | 7.52905043 |
| ENSMUSG00 | Ell2 | 3635 | 6.89022083 | 7.56931457 | 5.54499049 | 8.20903737 | 9.08227377 | 6.2427785 |
| ENSMUSG00 | Jup | 6380 | 48.6412666 | 46.4115497 | 32.2922869 | 43.7508929 | 51.0983566 | 24.671151 |
| ENSMUSG00 | Fkbp10 | 3185 | 0.59914068 | 0.2420669 | 0.22205008 | 0.03398623 | 0 | 0 |
| ENSMUSG00 | Klhl10 | 2453 | 0.11344817 | 0 | 0.36039008 | 0.82372381 | 0.06240486 | 0 |
| ENSMUSG00 | Mnx1 | 3262 | 0 | 0 | 0 | 0 | 0 | 0 |
| ENSMUSG00 | Nom1 | 4884 | 11.2494009 | 8.02120751 | 11.7382863 | 9.81839418 | 12.2655539 | 9.80077639 |
| ENSMUSG00 | Erglc1 | 6555 | 18.922517 | 25.9567242 | 14.7878935 | 16.8217739 | 19.5153508 | 12.3203977 |
| ENSMUSG00 | Tnk1 | 4467 | 5.90947819 | 4.83266758 | 4.66063758 | 5.67038175 | 6.69385633 | 7.09494757 |
| ENSMUSG00 | Acap1 | 4708 | 0.01688848 | 0.17399522 | 0 | 0 | 0.01083823 | 0.01737976 |
| ENSMUSG00 | Tcea3 | 1388 | 2.14817078 | 2.39543552 | 3.34379936 | 1.19580263 | 2.27927615 | 1.13971836 |
| ENSMUSG00 | Csn3 | 4218 | 0 | 0 | 0 | 0.0085543 | 0 | 0 |
| ENSMUSG00 | Ifrd1 | 5303 | 13.5092167 | 10.0225619 | 9.27713582 | 9.64138357 | 9.86272586 | 13.2027153 |
| ENSMUSG00 | Stk38l | 4568 | 5.77881766 | 8.72377138 | 7.00572118 | 5.76617612 | 6.66872731 | 7.71428063 |
| ENSMUSG00 | Brpf1 | 6772 | 8.97609712 | 7.95517459 | 5.86137847 | 9.13772982 | 14.9115768 | 5.28013173 |
| ENSMUSG00 | Akr1b3 | 3257 | 4.30877637 | 3.63950407 | 2.18498518 | 3.06869092 | 2.92966911 | 1.83394105 |
| ENSMUSG00 | Hoxc13 | 2459 | 0 | 0 | 0 | 0 | 0 | 0 |
| ENSMUSG00 | Hoxc11 | 2019 | 0 | 0 | 0 | 0 | 0 | 0 |
| ENSMUSG00 | Hoxc8 | 2297 | 1.10768426 | 0.8181416 | 0.5195689 | 2.46620853 | 0.84414553 | 3.0278768 |
| ENSMUSG00 | Hoxc6 | 2498 | 0.5251925 | 0.67515031 | 0.14155915 | 0.82332928 | 0.67408743 | 0.44766221 |
| ENSMUSG00 | Gstt1 | 1771 | 167.237905 | 215.737041 | 145.858594 | 162.297892 | 136.800244 | 148.200934 |
| ENSMUSG00 | Gstt3 | 2457 | 68.9127357 | 84.8999219 | 76.8899784 | 71.547302 | 74.3485655 | 90.9376657 |
| ENSMUSG00 | Ddt | 1043 | 14.2555606 | 15.4769493 | 14.7480742 | 14.6680603 | 8.36578165 | 17.154519 |
| ENSMUSG00 | Tat | 2407 | 0 | 0 | 0 | 0 | 0 | 0 |
| ENSMUSG00 | Marveld3 | 3280 | 14.6780143 | 11.1945332 | 14.1230279 | 9.33952921 | 11.2009118 | 12.4981045 |
| ENSMUSG00 | Ddx18 | 2469 | 3.94495453 | 3.18120294 | 3.90279537 | 4.96877081 | 3.88536195 | 3.47975349 |
| ENSMUSG00 | Ubl3 | 3657 | 79.9131904 | 77.0693191 | 94.2658391 | 69.02652 | 69.1374603 | 75.8573573 |
| ENSMUSG00 | Gramd3 | 3197 | 24.7834447 | 26.9494397 | 23.6010466 | 23.780065 | 25.4573254 | 23.938892 |
| ENSMUSG00 | Eef1e1 | 1028 | 21.6953546 | 24.6088348 | 30.184527 | 30.5012614 | 12.4587746 | 20.2967889 |
| ENSMUSG00 | Akt1 | 4183 | 43.7471855 | 36.7129312 | 35.547454 | 27.3181532 | 41.5115364 | 23.4145899 |
| ENSMUSG00 | Cldn15 | 2117 | 1.12674957 | 0.22761664 | 0.9813352 | 0.698802 | 1.68722074 | 2.92458342 |
| ENSMUSG00 | Il16 | 8831 | 0.04051629 | 0.16915182 | 0.0050053 | 0.05311591 | 0.47380397 | 0.15442554 |
| ENSMUSG00 | Tcirg1 | 4391 | 4.30058147 | 3.55554709 | 1.8119635 | 2.50626818 | 3.49782262 | 3.29208809 |
| ENSMUSG00 | Naglu | 2651 | 14.8614414 | 17.3223989 | 18.0242144 | 12.5627035 | 17.9968545 | 8.88920721 |
| ENSMUSG00 | Coasy | 2930 | 25.7528676 | 22.3170655 | 22.0254917 | 22.3019053 | 24.6946761 | 19.557685 |
| ENSMUSG00 | Smo | 3767 | 5.11850492 | 5.58998546 | 2.38199473 | 3.02679216 | 5.82461949 | 3.72157276 |
| ENSMUSG00 | Tspan33 | 5497 | 4.58522369 | 4.58459328 | 1.25440923 | 2.46804598 | 4.33496771 | 1.06181059 |
| ENSMUSG00 | Crnkl1 | 3830 | 14.6462097 | 16.9470333 | 13.8376005 | 15.1017017 | 16.6535169 | 16.6710001 |
| ENSMUSG00 | Rin2 | 5384 | 14.9304572 | 17.5239699 | 13.3245898 | 18.1750566 | 19.6182389 | 17.416458 |
| ENSMUSG00 | Folh1 | 2652 | 4.9919212 | 1.5626071 | 0.783366 | 4.68032553 | 2.52053365 | 2.75626068 |
| ENSMUSG00 | Chordc1 | 2194 | 86.7952378 | 81.6797531 | 100.229794 | 136.48354 | 60.4920709 | 110.56547 |
| ENSMUSG00 | D10Wsu52e | 2603 | 56.7695432 | 48.2975135 | 58.6867343 | 49.1259185 | 54.260856 | 40.3932902 |
| ENSMUSG00 | Pwp1 | 2845 | 29.8480515 | 25.2703591 | 34.5846406 | 34.1923346 | 24.9302856 | 23.6987375 |
| ENSMUSG00 | Fbxo7 | 6326 | 7.01345499 | 9.30822515 | 7.66509987 | 9.2686251 | 6.63842986 | 5.47993463 |
| ENSMUSG00 | Capns1 | 2928 | 14.0121776 | 17.8888875 | 15.9718408 | 17.6220368 | 11.2578684 | 17.9781604 |
| ENSMUSG00 | Lrp3 | 4942 | 0.10457735 | 0.01950079 | 0.0447206 | 0.48187271 | 0.15487569 | 0 |
| ENSMUSG00 | Dsg4 | 3717 | 0 | 0 | 0 | 0 | 0 | 0 |
| ENSMUSG00 | Evx2 | 7406 | 0 | 0 | 0 | 0 | 0 | 0 |
| ENSMUSG00 | Hoxd13 | 2481 | 0 | 0 | 0 | 0 | 0 | 0 |
| ENSMUSG00 | Hoxd12 | 2617 | 0 | 0 | 0 | 0 | 0 | 0 |
| ENSMUSG00 | Folr1 | 2131 | 2.74240058 | 1.58284887 | 2.4890761 | 2.37047688 | 3.71144451 | 2.08623472 |
| ENSMUSG00 | Clpb | 10260 | 16.4020418 | 13.0047035 | 15.9359276 | 11.4470814 | 13.2439804 | 10.713138 |
| ENSMUSG00 | 7-Sep | 3625 | 77.7233353 | 92.3451624 | 88.3792988 | 104.065593 | 63.8498319 | 135.711081 |
| ENSMUSG00 | Zdhhc4 | 3175 | 17.905618 | 20.7163759 | 22.9431929 | 16.7852535 | 22.0819655 | 18.6154782 |
| ENSMUSG00 | Rac1 | 4266 | 63.2210923 | 55.4042426 | 59.4124163 | 54.7912528 | 51.4330558 | 63.122952 |
| ENSMUSG00 | Nup214 | 9661 | 4.66234959 | 4.85804727 | 4.23671531 | 5.40054201 | 7.66900918 | 3.94114444 |
| ENSMUSG00 | Aif1l | 4009 | 448.922575 | 431.43032 | 354.000642 | 430.392459 | 404.863554 | 340.371714 |
| ENSMUSG00 | Cpa3 | 1441 | 0 | 0 | 0 | 0 | 0 | 0 |
| ENSMUSG00 | Ltbp1 | 11195 | 0.14914964 | 0.37016829 | 0.19741779 | 0.00966915 | 0.2233401 | 0 |
| ENSMUSG00 | Ugp2 | 3514 | 44.1337876 | 41.7551839 | 44.4031035 | 51.556046 | 34.0224242 | 50.8857527 |
| ENSMUSG00 | Kcnh6 | 3759 | 0 | 0.03845686 | 0.02351787 | 0 | 0 | 0 |
| ENSMUSG00 | Trmt1 | 6253 | 5.10533358 | 6.17261157 | 6.36908053 | 5.87422379 | 7.1647462 | 4.07832843 |
| ENSMUSG00 | Nacc1 | 4294 | 3.31449977 | 2.98500084 | 2.98521995 | 3.48720272 | 4.01651491 | 3.34739983 |
| ENSMUSG00 | Nfix | 6243 | 3.31773272 | 4.98613513 | 2.64092383 | 3.36372736 | 3.8169658 | 3.03197175 |
| ENSMUSG00 | Slc1a5 | 5242 | 41.5680635 | 36.4293915 | 30.7777045 | 43.7775283 | 41.0196772 | 44.1222703 |
| ENSMUSG00 | Uba1 | 4895 | 53.9846127 | 51.5235625 | 52.3198117 | 56.7803859 | 55.2481699 | 40.0510944 |
| ENSMUSG00 | Vwf | 16008 | 0.06705385 | 0 | 0.17671901 | 0.06085802 | 0 | 0.0136305 |
| ENSMUSG00 | Siae | 3423 | 12.5085167 | 13.2326192 | 18.1946821 | 16.2016075 | 14.2211985 | 13.0915081 |
| ENSMUSG00 | Vsig2 | 1310 | 0.39452004 | 0.47818607 | 0 | 0 | 0.07790286 | 0.39558637 |
| ENSMUSG00 | Esam | 2857 | 5.2877433 | 1.83840469 | 1.23771349 | 3.59936391 | 4.12568877 | 1.39380376 |
| ENSMUSG00 | Spa17 | 572 | 5.56020706 | 10.3617699 | 4.09562531 | 4.41563482 | 6.69052131 | 13.7326855 |
| ENSMUSG00 | Fam60a | 2532 | 25.9698123 | 18.8597015 | 21.7692331 | 20.9908577 | 19.0845094 | 23.7629756 |
| ENSMUSG00 | Emd | 2791 | 3.94563529 | 4.60973846 | 4.75118341 | 4.79628754 | 2.02935425 | 6.49861523 |
| ENSMUSG00 | Taco1 | 1378 | 8.02033642 | 4.33608043 | 4.07375476 | 3.90146926 | 5.96171735 | 6.21497585 |
| ENSMUSG00 | Grik3 | 8962 | 0.01330802 | 0.2957213 | 0.00986428 | 0.03220892 | 0.17650275 | 0.01217346 |
| ENSMUSG00 | Gria3 | 6707 | 0.01778238 | 0.043107 | 0.36906265 | 0.00537976 | 0.16737443 | 0.02033297 |
| ENSMUSG00 | Npas1 | 2079 | 0.07648962 | 0.13906621 | 0.14882775 | 0 | 0.09817485 | 0.27550141 |
| ENSMUSG00 | Sipa1l2 | 6344 | 4.3929054 | 3.82817888 | 6.06869566 | 2.59353914 | 4.00553833 | 5.43859058 |
| ENSMUSG00 | Ap4e1 | 7188 | 2.01321576 | 4.30379742 | 2.84716935 | 2.13841833 | 3.08089136 | 2.34877597 |
| ENSMUSG00 | Blvra | 2047 | 18.5667999 | 17.7727229 | 16.1519192 | 13.7488983 | 12.5135519 | 17.4014064 |
| ENSMUSG00 | Pdzd4 | 3962 | 0.03010259 | 0.10945936 | 0.16734671 | 0.17303353 | 0.06439472 | 0.12391305 |
| ENSMUSG00 | Srpk3 | 2192 | 0 | 2.39613239 | 0.38313641 | 1.8600689 | 0 | 1.04519603 |
| ENSMUSG00 | Idh3g | 2398 | 208.094575 | 188.606316 | 233.285457 | 175.595271 | 166.506001 | 193.299622 |
| ENSMUSG00 | Pnck | 2254 | 0.33511718 | 0.21378191 | 0.37259761 | 0.24012008 | 0.97344018 | 0.30251375 |
| ENSMUSG00 | Ssr4 | 1639 | 43.1270557 | 40.5718671 | 46.5211588 | 41.05736 | 31.2571576 | 53.9335606 |
| ENSMUSG00 | Bcap31 | 1211 | 173.696323 | 204.802327 | 202.905075 | 148.946441 | 174.821167 | 202.026023 |
| ENSMUSG00 | Fam98a | 3471 | 13.217466 | 11.6058386 | 8.69774093 | 14.7613089 | 9.21738336 | 12.2346912 |
| ENSMUSG00 | Ltbp2 | 8635 | 0 | 0.00558036 | 0 | 0 | 0 | 0.11371014 |
| ENSMUSG00 | Mll1 | 16633 | 3.58523542 | 3.58073964 | 4.3768674 | 6.16299456 | 4.14763784 | 4.23721925 |
| ENSMUSG00 | Ift46 | 5301 | 13.7318025 | 10.399036 | 30.2517047 | 10.3324927 | 13.6108839 | 24.1052037 |
| ENSMUSG00 | Tmem25 | 4167 | 28.9364945 | 26.099544 | 23.0503011 | 25.500749 | 31.9113836 | 21.9925097 |
| ENSMUSG00 | Cd3g | 1226 | 0 | 0 | 0 | 0 | 0 | 0 |
| ENSMUSG00 | Trappc6a | 1430 | 9.89716857 | 11.692794 | 11.9314066 | 9.28544922 | 7.88589445 | 16.9179056 |
| ENSMUSG00 | Supt6h | 7798 | 8.40177377 | 9.26900016 | 7.42554685 | 10.1888513 | 11.595119 | 6.0404337 |
| ENSMUSG00 | Spag5 | 5247 | 2.76553276 | 0.7255058 | 0.63181594 | 0.90772439 | 1.62405275 | 1.53345123 |
| ENSMUSG00 | Foxn1 | 3205 | 0 | 0 | 0 | 0 | 0 | 0 |
| ENSMUSG00 | Unc119 | 2732 | 2.47380369 | 3.98614053 | 1.45613685 | 2.13956487 | 2.57746701 | 2.97505216 |
| ENSMUSG00 | Rab34 | 2266 | 19.0531562 | 24.5397856 | 17.3413232 | 17.1174748 | 19.4107396 | 17.2001145 |
| ENSMUSG00 | Sdf2 | 1511 | 36.5718848 | 33.4849532 | 39.3457835 | 33.0493379 | 26.9484103 | 34.4588262 |
| ENSMUSG00 | Ccne1 | 2651 | 0 | 0 | 0 | 0 | 0 | 0 |
| ENSMUSG00 | Hsf2bp | 3447 | 0 | 0 | 0.12823279 | 0.062806 | 0 | 0 |
| ENSMUSG00 | Bbc3 | 2003 | 0.83361467 | 0.72171406 | 0.50755986 | 0.37829403 | 0.56044946 | 1.29360496 |
| ENSMUSG00 | Mybpc3 | 4131 | 0.13473172 | 0 | 0 | 0 | 0 | 0 |
| ENSMUSG00 | Psmc3 | 2869 | 85.8435698 | 96.8767509 | 92.0396689 | 85.5455095 | 76.4773148 | 80.1317303 |
| ENSMUSG00 | Acp2 | 4354 | 18.6542138 | 17.2869138 | 21.3801289 | 23.9911619 | 18.6456049 | 16.1680854 |
| ENSMUSG00 | Rapsn | 2155 | 3.4313315 | 4.4496993 | 1.88703924 | 2.51151122 | 2.69930712 | 2.75910504 |
| ENSMUSG00 | Slc39a13 | 4475 | 14.125411 | 14.5689958 | 11.3788879 | 13.9570992 | 15.9065462 | 12.4457684 |
| ENSMUSG00 | Celf2 | 18155 | 0.89343079 | 1.60311821 | 0.58919559 | 0.84863856 | 1.36313921 | 1.48279891 |
| ENSMUSG00 | Nr1h3 | 2896 | 1.30413351 | 1.43095098 | 1.86209422 | 2.18037217 | 1.32147037 | 1.03598423 |
| ENSMUSG00 | Ddb2 | 2775 | 6.93392885 | 6.78951311 | 6.833366 | 4.08279711 | 10.6649722 | 8.10867653 |
| ENSMUSG00 | Slpi1 | 2803 | 0 | 0.08595512 | 0 | 0 | 0 | 0 |
| ENSMUSG00 | Sf3a1 | 6327 | 8.85968508 | 8.0653457 | 7.46827409 | 10.6244427 | 8.7987634 | 9.16051974 |
| ENSMUSG00 | Stat6 | 4430 | 34.3799652 | 28.5855463 | 36.5089263 | 49.7166591 | 31.1226337 | 32.8219192 |
| ENSMUSG00 | Clgn | 4109 | 0 | 0 | 0 | 0 | 0 | 0 |
| ENSMUSG00 | Napsa | 2149 | 4.29188994 | 3.9464001 | 30.6882968 | 2.88790677 | 2.0420048 | 20.2814674 |
| ENSMUSG00 | Vrk3 | 3169 | 20.323092 | 17.56243 | 22.833202 | 16.0655616 | 19.2737683 | 17.3941451 |
| ENSMUSG00 | 1500002O20 | 3855 | 5.49667214 | 5.44988036 | 5.136816 | 5.92475595 | 6.79028036 | 4.87476691 |
| ENSMUSG00 | Paxip1 | 5031 | 12.8093091 | 9.76946352 | 13.2052018 | 10.8152898 | 16.0756919 | 7.60610592 |
| ENSMUSG00 | Rmnd5a | 9948 | 11.7691888 | 14.7397812 | 14.7739373 | 14.4828713 | 12.8386629 | 14.2048559 |
| ENSMUSG00 | Mov10 | 4201 | 3.18914471 | 2.79873646 | 2.43052267 | 3.2895556 | 5.01639924 | 2.14249725 |
| ENSMUSG00 | Ppm1j | 1711 | 0.76676263 | 1.07018399 | 0.31000708 | 0.46394213 | 1.28236947 | 0.78109721 |
| ENSMUSG00 | Rhoc | 910 | 1.04849619 | 1.58856403 | 0.92289562 | 1.38777094 | 2.41113644 | 0.92913606 |
| ENSMUSG00 | 4930432K09 | 1525 | 0 | 0 | 0 | 0 | 0 | 0 |
| ENSMUSG00 | Tead3 | 5375 | 0.75442958 | 0.72615848 | 0.55920471 | 1.03379253 | 0.67402283 | 0.76115272 |
| ENSMUSG00 | Ppard | 4719 | 0.46335059 | 0.76583665 | 0.33720414 | 0.37465992 | 0.45414448 | 0.39302299 |
| ENSMUSG00 | Def6 | 2612 | 9.54314176 | 7.08407113 | 4.82294229 | 7.5700461 | 6.21224637 | 5.80578079 |
| ENSMUSG00 | Peg3 | 13121 | 8.25045145 | 3.24646101 | 10.9620301 | 4.99665231 | 5.37447233 | 11.9857915 |
| ENSMUSG00 | Zim1 | 5888 | 0 | 0 | 0 | 0 | 0 | 0 |
| ENSMUSG00 | Metrn | 1771 | 0.02244804 | 0.13604303 | 0.12479346 | 0.14261678 | 0.23049746 | 0.0308014 |
| ENSMUSG00 | Lmf1 | 14147 | 2.76520766 | 3.15407121 | 2.77765169 | 3.70588893 | 3.09108674 | 2.14773084 |
| ENSMUSG00 | Narfl | 3347 | 25.3237778 | 16.8731731 | 27.6674219 | 18.9950889 | 18.035316 | 18.5144847 |
| ENSMUSG00 | Angptl4 | 3850 | 0.19619588 | 0.41302665 | 0.30998695 | 0 | 0 | 0.26920423 |
| ENSMUSG00 | Rab11b-ps2 | 1544 | 1.10717983 | 0.81142973 | 0.8588441 | 0.3972764 | 0.59486708 | 0.38862828 |
| ENSMUSG00 | Dbl4 | 7708 | 0.63955366 | 0.75642962 | 0.37847971 | 0.76770308 | 1.26440552 | 0.62985026 |
| ENSMUSG00 | Daxx | 3002 | 6.70095707 | 9.05301578 | 8.51054806 | 6.62265374 | 7.25791555 | 7.37741741 |
| ENSMUSG00 | Cd320 | 2774 | 15.234346 | 13.1149113 | 17.1772122 | 14.7632014 | 11.2942303 | 13.7258098 |
| ENSMUSG00 | Ipo4 | 5645 | 13.2048762 | 12.2408075 | 10.688311 | 14.0301307 | 13.0797459 | 6.53721646 |
| ENSMUSG00 | Tm9sf1 | 4075 | 26.7800722 | 23.0230682 | 28.5495094 | 23.5972144 | 31.3170469 | 16.793146 |
| ENSMUSG00 | Rec8 | 2812 | 11.9464371 | 17.2045476 | 4.07122238 | 17.6304158 | 17.2749324 | 8.06017966 |
| ENSMUSG00 | Irf9 | 3689 | 10.1409344 | 14.1855452 | 10.1967385 | 11.3361587 | 24.5656905 | 23.2525726 |
| ENSMUSG00 | Gmpr2 | 1533 | 24.1437393 | 20.5884668 | 20.4718255 | 18.8765817 | 15.4110972 | 16.7775504 |
| ENSMUSG00 | Mdp1 | 1727 | 34.6910881 | 38.7277254 | 39.9275478 | 38.0668704 | 37.6714703 | 30.7649087 |
| ENSMUSG00 | Dhrs1 | 3150 | 26.5919992 | 27.7033799 | 24.4443208 | 23.6423301 | 25.3026029 | 22.6682557 |
| ENSMUSG00 | Ncan | 7180 | 0 | 0 | 0 | 0 | 0 | 0 |
| ENSMUSG00 | Tmem161a | 3697 | 16.6893443 | 14.7283418 | 10.4735771 | 9.06687026 | 12.960175 | 8.35133666 |
| ENSMUSG00 | Armc6 | 3933 | 2.24401644 | 4.03085163 | 1.16882575 | 2.31189301 | 1.19359943 | 0.66574253 |
| ENSMUSG00 | 2310045N01 | 2858 | 7.77582701 | 6.89582045 | 8.98574905 | 5.61809301 | 6.83803427 | 7.24333492 |
| ENSMUSG00 | Slc25a42 | 6877 | 2.67079133 | 2.92187675 | 2.46815584 | 3.23201097 | 4.33319812 | 3.02610873 |
| ENSMUSG00 | Snx9 | 3288 | 23.2753345 | 21.3820009 | 31.0676579 | 25.7556443 | 23.4802027 | 27.2165728 |
| ENSMUSG00 | Ranbp3 | 2463 | 23.6951057 | 25.3942355 | 23.4558704 | 17.5502596 | 24.1147793 | 16.466662 |
| ENSMUSG00 | Ndufa11 | 2657 | 23.3116442 | 26.7682306 | 33.4882612 | 20.953931 | 16.8423529 | 46.2139352 |
| ENSMUSG00 | Bmp8b | 3870 | 0 | 0.13696405 | 0 | 0 | 0 | 0 |
| ENSMUSG00 | Nr2f6 | 3217 | 14.4711432 | 14.3795413 | 12.6271351 | 11.7880724 | 12.070585 | 7.80002117 |
| ENSMUSG00 | Use1 | 2334 | 13.6606236 | 10.3433623 | 12.7454329 | 11.9655109 | 9.55378159 | 17.8675764 |

| | | | | | | | | |
|---|---|---|---|---|---|---|---|---|
| ENSMUSG00 | Ocel1 | 7775 | 3.74800865 | 2.96245908 | 3.14387385 | 4.34376768 | 4.20024187 | 3.42380037 |
| ENSMUSG00 | Dyrk1b | 2829 | 6.57672848 | 4.99067782 | 6.73417969 | 7.55057277 | 6.25880255 | 5.12905906 |
| ENSMUSG00 | Braf | 10697 | 5.1659454 | 5.98220017 | 6.36354476 | 8.36865967 | 7.72765531 | 7.37641688 |
| ENSMUSG00 | Ndufb2 | 4711 | 14.6329873 | 8.20325336 | 13.3609477 | 10.2555418 | 12.867615 | 18.1734436 |
| ENSMUSG00 | Hltf | 6079 | 13.9624858 | 9.40107261 | 14.3243347 | 14.9159694 | 15.3104309 | 11.6519556 |
| ENSMUSG00 | Prpf6 | 3837 | 36.9476267 | 26.9753656 | 36.7599899 | 35.6118589 | 39.5098678 | 22.8888033 |
| ENSMUSG00 | Rgs19 | 2429 | 4.73006746 | 6.26880021 | 8.46185961 | 5.94187641 | 4.93668107 | 5.40102984 |
| ENSMUSG00 | Rgs20 | 4922 | 0 | 0 | 0 | 0 | 0 | 0 |
| ENSMUSG00 | Abhd3 | 2775 | 11.5469972 | 21.6882401 | 7.62979561 | 14.2767874 | 11.7498574 | 5.55321484 |
| ENSMUSG00 | Snrpd1 | 833 | 23.5287538 | 19.2630075 | 17.5109342 | 30.27773 | 12.4349992 | 39.487653 |
| ENSMUSG00 | Tchp | 3978 | 6.26613531 | 2.70125103 | 3.20013343 | 4.8708039 | 6.34943591 | 4.13439102 |
| ENSMUSG00 | Tiam1 | 10002 | 1.07318334 | 0.57812168 | 0.49938095 | 0.67099183 | 0.67341348 | 0.62446435 |
| ENSMUSG00 | Tsc2 | 6299 | 17.280601 | 20.3333169 | 22.5605533 | 24.1501667 | 22.276954 | 10.9808676 |
| ENSMUSG00 | Rpl3l | 1584 | 0.72784655 | 0.82135981 | 0.13952602 | 1.07061622 | 0.74091329 | 0.41325211 |
| ENSMUSG00 | Slc9a3r2 | 3350 | 6.65755957 | 4.97686835 | 2.13752196 | 3.942098 | 9.06289382 | 4.34765891 |
| ENSMUSG00 | Puf60 | 1900 | 96.3756543 | 97.6409486 | 121.066522 | 91.7433458 | 93.1902758 | 96.8249693 |
| ENSMUSG00 | Golga2 | 5394 | 22.2362411 | 17.4378819 | 20.4701898 | 17.8002077 | 24.6712622 | 20.4079429 |
| ENSMUSG00 | Uck1 | 3952 | 7.01153085 | 7.43768464 | 8.44443104 | 7.49578911 | 7.95350392 | 6.42527559 |
| ENSMUSG00 | Scin | 3848 | 33.7219045 | 53.8966858 | 14.7952115 | 37.7604972 | 48.2151514 | 24.4961417 |
| ENSMUSG00 | Ikzf4 | 8382 | 0.05217255 | 0.3794208 | 0.30585861 | 0.21523529 | 1.12620849 | 0.16269768 |
| ENSMUSG00 | 1810046J19R | 743 | 35.6354644 | 48.3810038 | 54.553284 | 42.8322521 | 30.0801516 | 57.412565 |
| ENSMUSG00 | Pon1 | 2890 | 0 | 0.20008211 | 0.07647378 | 0 | 0 | 0 |
| ENSMUSG00 | Axl | 6694 | 0.00593897 | 0 | 0.09244485 | 0 | 0.15245407 | 0 |
| ENSMUSG00 | Tgfb1 | 2551 | 0.79479793 | 1.11446495 | 1.12627197 | 0.49504961 | 0.42005249 | 0.85533953 |
| ENSMUSG00 | Ccdc97 | 3902 | 9.73000612 | 10.6449803 | 8.4959975 | 8.55353439 | 7.3754167 | 9.64607949 |
| ENSMUSG00 | Zfp40 | 7290 | 14.6424506 | 13.0810657 | 8.7251649 | 21.4858924 | 8.3294084 | 13.8543195 |
| ENSMUSG00 | Akap8l | 2075 | 52.2281638 | 47.0949902 | 38.8976219 | 39.9249996 | 64.305529 | 59.438997 |
| ENSMUSG00 | Shh | 2689 | 0.1034914 | 0.4838356 | 0 | 0.52331712 | 0.1138558 | 0.04057217 |
| ENSMUSG00 | Pdcd2l | 1466 | 9.49141758 | 10.7482719 | 13.1761292 | 9.5989068 | 9.85024854 | 12.5396363 |
| ENSMUSG00 | Gtf2f1 | 1718 | 40.1025889 | 43.5024283 | 30.4112797 | 40.7025625 | 41.0171275 | 42.1185203 |
| ENSMUSG00 | Clpp | 1201 | 55.6444319 | 58.4576571 | 70.4064327 | 59.756192 | 41.0845174 | 63.8603544 |
| ENSMUSG00 | Alkbh7 | 1250 | 11.7676222 | 12.9139665 | 15.948025 | 9.81431611 | 6.1639862 | 17.9358028 |
| ENSMUSG00 | Pspn | 472 | 0.33691085 | 0.40835968 | 0.74918378 | 0.7644501 | 0.10810673 | 1.0979196 |
| ENSMUSG00 | Dennd1c | 2496 | 4.96943506 | 4.78775547 | 3.80745042 | 4.14885688 | 4.25219799 | 3.04872771 |
| ENSMUSG00 | Med6 | 1838 | 16.5034993 | 17.6438931 | 17.4113897 | 13.1724983 | 16.3795222 | 19.6620767 |
| ENSMUSG00 | Prkd1 | 3759 | 0.62398865 | 3.87132363 | 1.21117048 | 1.65100071 | 1.46604113 | 1.16818753 |
| ENSMUSG00 | Lcp2 | 4725 | 13.2181714 | 8.1177583 | 20.5059132 | 13.8524505 | 8.30461016 | 22.0102009 |
| ENSMUSG00 | Cse1l | 5087 | 32.2764172 | 27.6311877 | 27.9530822 | 31.9681098 | 26.4811544 | 24.0254882 |
| ENSMUSG00 | Naa20 | 3592 | 18.0294203 | 16.2320699 | 20.2427705 | 16.2429471 | 13.6373388 | 25.2017059 |
| ENSMUSG00 | Prkra | 2020 | 11.41494 | 12.6429774 | 15.1642957 | 8.34174 | 11.3167408 | 11.6254774 |
| ENSMUSG00 | Fkbp7 | 1768 | 0.4272365 | 1.68979605 | 1.55006463 | 0.81633585 | 1.12558182 | 1.65067104 |
| ENSMUSG00 | Plekha3 | 3118 | 40.2909937 | 36.5648243 | 42.6566214 | 39.1602433 | 41.8946511 | 48.0936391 |
| ENSMUSG00 | Ykt6 | 2977 | 17.1468045 | 18.8731581 | 16.718601 | 13.635304 | 12.6151873 | 12.7532072 |
| ENSMUSG00 | Baz1b | 10776 | 4.784972 | 5.6655737 | 4.73356782 | 5.93331319 | 6.31673955 | 5.24181309 |
| ENSMUSG00 | Pex6 | 4686 | 21.7696464 | 15.3114836 | 16.016801 | 15.2228344 | 19.0450558 | 10.8202207 |
| ENSMUSG00 | Mrpl2 | 1209 | 51.8892872 | 53.0887849 | 54.9141176 | 42.5284644 | 36.2966776 | 50.2629415 |
| ENSMUSG00 | Mea1 | 836 | 41.7527654 | 54.3538456 | 58.1603198 | 47.6921761 | 34.7907108 | 63.6517 |
| ENSMUSG00 | Gnmt | 1201 | 14.4324642 | 10.1107272 | 19.0277709 | 9.46365031 | 8.70974775 | 15.3519202 |
| ENSMUSG00 | Grin2d | 6606 | 0.13841599 | 0.05106041 | 0.07360283 | 0.10924022 | 0.18538193 | 0.16927948 |
| ENSMUSG00 | Kdelr1 | 2143 | 21.8349046 | 20.7541232 | 20.4198902 | 16.3488872 | 12.5482501 | 18.4800636 |
| ENSMUSG00 | Tmem143 | 5232 | 3.10779655 | 4.13526616 | 2.92313411 | 0.88963757 | 0.99478026 | 2.28331269 |
| ENSMUSG00 | Pdxk-ps | 2639 | 0 | 0.29214971 | 0 | 0 | 0.05800649 | 0 |
| ENSMUSG00 | Ggct | 4786 | 1.22937967 | 1.72166352 | 0.75732368 | 1.46258183 | 1.27939095 | 1.95470148 |
| ENSMUSG00 | Jag2 | 4028 | 0.55270777 | 1.01684399 | 0.51576133 | 0.16124052 | 0.35470172 | 0.11511144 |
| ENSMUSG00 | Btbd6 | 2083 | 15.6120898 | 14.226914 | 20.4987904 | 15.0875952 | 16.094253 | 16.8387835 |
| ENSMUSG00 | Nudt14 | 788 | 21.9462361 | 21.2191566 | 30.4588842 | 16.4383934 | 14.8934853 | 27.8630515 |
| ENSMUSG00 | Epdr1 | 2383 | 6.55640111 | 5.90450739 | 2.96781153 | 6.54110081 | 1.75583836 | 0.89274942 |
| ENSMUSG00 | Flii | 5313 | 30.5592664 | 31.0722286 | 32.6293296 | 30.5471553 | 40.020122 | 22.3874839 |
| ENSMUSG00 | Top3a | 4995 | 3.21545828 | 2.72043577 | 3.94675115 | 3.00503114 | 6.4459746 | 3.40728226 |
| ENSMUSG00 | Atg4d | 2531 | 38.7973279 | 38.0389219 | 39.7135484 | 30.7787966 | 43.6476111 | 25.6150998 |
| ENSMUSG00 | Qtrt1 | 1309 | 6.71196424 | 10.3072604 | 8.71205156 | 9.23413668 | 7.36744464 | 10.3347755 |
| ENSMUSG00 | Plin4 | 5746 | 0 | 0 | 0.00769263 | 0 | 0 | 0.06645405 |
| ENSMUSG00 | Hdgfrp2 | 2454 | 22.9719932 | 22.3652639 | 16.5892003 | 19.011444 | 25.1389113 | 15.1822157 |
| ENSMUSG00 | Chaf1a | 2830 | 0.96930324 | 0.93648563 | 1.07771278 | 1.01998712 | 1.49653328 | 0.06746377 |
| ENSMUSG00 | Adprh | 1579 | 20.0665725 | 20.1107444 | 17.3560118 | 15.7444767 | 16.5456013 | 15.5460262 |
| ENSMUSG00 | Tmem39a | 3017 | 6.16690914 | 8.00179235 | 7.25220825 | 11.3018005 | 7.64465425 | 5.25242473 |
| ENSMUSG00 | 4930455C21 | 1776 | 2.95479923 | 4.47678094 | 2.63817306 | 3.51474871 | 3.73503877 | 3.67040472 |
| ENSMUSG00 | Pla1a | 1961 | 0.36491517 | 0 | 0.09016184 | 0 | 0 | 0 |
| ENSMUSG00 | Mcm2 | 3354 | 16.025465 | 14.8553313 | 17.2247492 | 13.5226983 | 15.0005983 | 11.994661 |
| ENSMUSG00 | Tpra1 | 2339 | 36.3221299 | 37.8033866 | 33.9214657 | 38.0874596 | 38.7660837 | 25.9802891 |
| ENSMUSG00 | Nab1 | 4162 | 19.6007319 | 18.0380771 | 21.9522368 | 18.4484841 | 21.0627856 | 19.6138864 |
| ENSMUSG00 | Cd97 | 4133 | 4.90570894 | 4.72187493 | 5.53993581 | 4.67940379 | 5.39523983 | 3.54378932 |
| ENSMUSG00 | Il17ra | 3957 | 5.24446824 | 4.2621316 | 4.34534166 | 2.89969425 | 4.95176354 | 3.86683665 |
| ENSMUSG00 | Lamb1 | 9879 | 12.0888211 | 14.0574275 | 4.21481285 | 10.4020309 | 16.6265719 | 3.46213155 |
| ENSMUSG00 | Kcnn1 | 4039 | 0.61025991 | 2.08780079 | 0.78795066 | 0.03573364 | 1.09910738 | 0.29037125 |
| ENSMUSG00 | Arrdc2 | 2135 | 14.5056297 | 7.74142841 | 14.0369319 | 5.40808162 | 11.8304712 | 11.446406 |
| ENSMUSG00 | Ppp1r17 | 1733 | 0 | 0 | 0 | 0 | 0 | 0 |
| ENSMUSG00 | Cd36 | 3910 | 3.05029262 | 2.34154067 | 0 | 0.71979526 | 2.6622457 | 2.09268332 |
| ENSMUSG00 | Map2k7 | 5614 | 6.71325179 | 6.52328038 | 6.82632666 | 7.9825257 | 8.92552566 | 10.1296086 |
| ENSMUSG00 | Timm44 | 4159 | 16.6707281 | 16.4406255 | 19.502376 | 15.8504197 | 20.4399957 | 15.3719054 |
| ENSMUSG00 | Ap2a2 | 4617 | 34.0035505 | 26.5197638 | 31.7751607 | 24.7893102 | 32.6139994 | 21.3376645 |
| ENSMUSG00 | Pnkp | 3116 | 10.0664551 | 12.7270353 | 9.68865814 | 7.28357061 | 10.3493805 | 10.7312926 |
| ENSMUSG00 | Med25 | 2521 | 24.7742403 | 19.1331331 | 23.3896305 | 21.3543873 | 22.0216965 | 21.9625219 |
| ENSMUSG00 | Bcam | 3244 | 77.8932904 | 103.146318 | 50.0336521 | 67.7817446 | 101.675245 | 43.0979657 |
| ENSMUSG00 | Clptm1 | 4116 | 75.4639381 | 70.6757899 | 80.8111926 | 66.0189206 | 72.4733219 | 63.8860051 |
| ENSMUSG00 | Relb | 3316 | 1.79834803 | 1.17705121 | 1.95949063 | 1.99125879 | 6.89379263 | 1.39005248 |
| ENSMUSG00 | Tomm40 | 4759 | 12.2465927 | 12.0693925 | 12.7617845 | 10.5842686 | 8.18094658 | 7.74281102 |
| ENSMUSG00 | Apoe | 2810 | 2.27780511 | 2.93234221 | 1.46290797 | 2.04165305 | 3.79520019 | 2.05773079 |
| ENSMUSG00 | Apoc2 | 1189 | 0 | 0 | 0 | 0 | 0 | 0 |
| ENSMUSG00 | Hbp1 | 6805 | 45.8663156 | 33.7057259 | 54.4322475 | 36.5168318 | 49.2567616 | 52.565304 |
| ENSMUSG00 | Prkar2b | 3885 | 0.23536063 | 0.4713217 | 0 | 0.78943984 | 0.65671011 | 0 |
| ENSMUSG00 | Cdkn1b | 4550 | 2.54260326 | 2.22398964 | 2.2149495 | 2.73589129 | 2.19805927 | 3.63861671 |
| ENSMUSG00 | Klf4 | 4212 | 0.31147456 | 0.49193186 | 0.54570177 | 0.17132975 | 0.55726811 | 0.6540215 |
| ENSMUSG00 | Ap1m1 | 3045 | 21.9601702 | 17.7553984 | 17.4484779 | 19.4215011 | 18.6510201 | 13.1222813 |
| ENSMUSG00 | Rab8a | 2209 | 51.4715592 | 51.6766326 | 48.0037218 | 46.6175442 | 37.4670808 | 42.6714139 |
| ENSMUSG00 | Hmgn2 | 3092 | 3.25295491 | 3.69346274 | 4.67464511 | 4.3760565 | 3.92764471 | 7.61255293 |
| ENSMUSG00 | Fam32a | 1906 | 73.7749919 | 80.4708528 | 92.6708104 | 65.1786355 | 64.5191845 | 73.7960727 |
| ENSMUSG00 | Elf3 | 1991 | 64.0360753 | 66.1928274 | 87.2492431 | 61.1636867 | 65.9421723 | 56.1931543 |
| ENSMUSG00 | Cyp2c29 | 3087 | 0 | 0 | 0.05727482 | 0 | 0 | 0 |
| ENSMUSG00 | Stard3nl | 1274 | 21.1571553 | 29.9936017 | 25.3969773 | 19.7403336 | 22.749593 | 20.2525971 |
| ENSMUSG00 | Stk11 | 5550 | 6.06718775 | 6.51168137 | 6.46700838 | 7.29442414 | 5.94848021 | 7.19460754 |
| ENSMUSG00 | Efna2 | 2127 | 0.54203523 | 0.38512906 | 0.70656449 | 0.52587841 | 0.04797967 | 0.76938334 |
| ENSMUSG00 | Atp5d | 1920 | 115.394423 | 124.983584 | 145.58982 | 98.2484004 | 78.027833 | 146.416514 |
| ENSMUSG00 | Ppp6c | 3267 | 34.6810895 | 32.0357981 | 26.3019231 | 25.8107555 | 24.4901614 | 19.310144 |
| ENSMUSG00 | Cdk12 | 11101 | 2.75398293 | 2.27454245 | 2.91466977 | 3.62738147 | 3.32790704 | 3.09576123 |
| ENSMUSG00 | Lipe | 5029 | 1.62848061 | 1.68638175 | 1.8721401 | 1.13361763 | 3.19612416 | 2.27785812 |
| ENSMUSG00 | Pafah1b2 | 6876 | 35.0780251 | 24.0091261 | 33.8006531 | 29.5593596 | 30.9527361 | 32.6058078 |
| ENSMUSG00 | Tbc1d8 | 4416 | 2.61975653 | 2.65156373 | 1.60151605 | 2.27964004 | 2.04521479 | 1.46378838 |
| ENSMUSG00 | D1Bwg0212e | 3492 | 6.2502173 | 9.21779592 | 7.72139869 | 8.59686742 | 8.25598578 | 11.1847891 |
| ENSMUSG00 | Slc2a3 | 7010 | 0.11342505 | 0.21309268 | 0.02522216 | 0.00514723 | 0 | 0.19065012 |
| ENSMUSG00 | Foxj2 | 5272 | 7.51825001 | 9.27716972 | 9.07177431 | 13.934568 | 10.5498387 | 10.3055921 |
| ENSMUSG00 | Sri | 3725 | 115.210849 | 131.118867 | 115.18585 | 141.005724 | 101.696594 | 165.002415 |
| ENSMUSG00 | Dgcr2 | 8258 | 11.9150901 | 10.4098587 | 12.3217053 | 7.40603844 | 9.70724587 | 6.73443805 |
| ENSMUSG00 | Mical3 | 3295 | 5.69486701 | 3.15880774 | 5.68788511 | 4.63208037 | 5.79176466 | 2.65710446 |
| ENSMUSG00 | Irf3 | 3282 | 16.2680105 | 16.6787929 | 15.4073456 | 13.6544483 | 16.4957297 | 15.3326354 |
| ENSMUSG00 | Bcl2l12 | 1877 | 0.82603289 | 1.25793056 | 1.88393577 | 1.2495114 | 1.41362362 | 2.17964618 |
| ENSMUSG00 | Zfp959 | 1616 | 23.9861346 | 31.6670802 | 29.2126042 | 43.5619238 | 24.0607044 | 62.194954 |
| ENSMUSG00 | Mpnd | 2282 | 61.7586233 | 64.4879029 | 60.0463249 | 53.8226466 | 55.6773338 | 53.5931124 |
| ENSMUSG00 | Sh3gl1 | 2060 | 21.7883192 | 27.7422915 | 27.3150224 | 22.8578001 | 32.4488118 | 21.2371463 |
| ENSMUSG00 | Ebi3 | 1165 | 0 | 0.12408526 | 0.30353197 | 0.09291514 | 0.26279679 | 0.3043522 |
| ENSMUSG00 | Ccdc94 | 1303 | 4.39354505 | 5.21434255 | 4.24039169 | 5.45522853 | 6.30487069 | 5.08652136 |
| ENSMUSG00 | Ranbp2 | 9448 | 12.7328624 | 11.5671942 | 15.2002316 | 18.3465434 | 15.4623745 | 18.34283 |
| ENSMUSG00 | Edar | 3625 | 0 | 0.02658562 | 0 | 0 | 0 | 0 |
| ENSMUSG00 | Grk5 | 3149 | 0.60599018 | 0.8569199 | 0.82817045 | 0.80207784 | 1.39354345 | 0.38110007 |
| ENSMUSG00 | Dvl3 | 4272 | 11.4278394 | 11.1780815 | 9.91230468 | 10.9462382 | 11.7532663 | 9.49377073 |
| ENSMUSG00 | Abcf3 | 3538 | 35.395637 | 24.0796014 | 31.3835584 | 23.7623414 | 27.3592524 | 22.1558262 |
| ENSMUSG00 | Eif2b5 | 3193 | 18.0785962 | 21.7163452 | 23.6582993 | 19.6287226 | 22.1172891 | 12.5482133 |
| ENSMUSG00 | Cyth2 | 3989 | 19.6335665 | 16.4044092 | 22.1507857 | 13.7489866 | 18.9701969 | 16.7654589 |
| ENSMUSG00 | Sult2b1 | 2077 | 1.51212468 | 3.24800284 | 1.23433168 | 0.86860964 | 1.79341619 | 1.35256998 |
| ENSMUSG00 | Car11 | 1918 | 0.72546497 | 1.33153359 | 0.46091599 | 1.5614232 | 0.98434615 | 0.8390009 |
| ENSMUSG00 | Dlgap1 | 15584 | 0.06377612 | 0.02473637 | 0.00850908 | 0.02083794 | 0.02619424 | 0 |
| ENSMUSG00 | Plag1 | 5429 | 1.08377438 | 0.32840272 | 0.3989483 | 0.01329234 | 0.32895987 | 0.04019103 |
| ENSMUSG00 | Hck | 2079 | 0 | 0 | 0 | 0 | 0 | 0 |
| ENSMUSG00 | Mrpl4 | 1276 | 46.267154 | 57.6651232 | 64.6399993 | 52.5395288 | 40.5491733 | 60.2564326 |
| ENSMUSG00 | Keap1 | 4465 | 12.3940938 | 15.1736031 | 9.83033149 | 11.8468706 | 14.0451099 | 8.66802085 |
| ENSMUSG00 | Ap1m2 | 1727 | 83.0560357 | 57.4219444 | 88.4804698 | 64.5172863 | 62.7266913 | 61.7825065 |
| ENSMUSG00 | Glg1 | 7600 | 36.7476646 | 33.4134934 | 25.4742201 | 30.2044694 | 37.7595181 | 20.5851751 |
| ENSMUSG00 | Atp8b3 | 5471 | 0 | 0 | 0 | 0 | 0 | 0 |
| ENSMUSG00 | Btbd2 | 4393 | 8.79634124 | 8.40218864 | 5.34285878 | 7.90141744 | 8.3514601 | 4.97313591 |
| ENSMUSG00 | Csnk1g2 | 2499 | 48.5847289 | 37.1376901 | 31.4312425 | 31.8370981 | 44.9211792 | 20.4205082 |
| ENSMUSG00 | Fam108a | 1879 | 29.5397431 | 34.1331317 | 38.2267136 | 28.8041867 | 20.3399444 | 27.695589 |
| ENSMUSG00 | Mob3a | 2640 | 1.7920084 | 1.35068058 | 1.18876169 | 1.72209758 | 1.43028478 | 0.87816073 |
| ENSMUSG00 | Cacnb3 | 3106 | 8.07653194 | 10.332315 | 6.13361054 | 8.55002729 | 8.13202062 | 4.86482619 |
| ENSMUSG00 | Ccdc65 | 1904 | 0.93959907 | 1.69563636 | 0.25536779 | 0.11370392 | 1.17918095 | 0.15757407 |
| ENSMUSG00 | Fkbp11 | 725 | 3.45461417 | 3.45613102 | 3.04840297 | 2.83679523 | 3.58944161 | 6.73401438 |
| ENSMUSG00 | Ddx23 | 5089 | 0.42185026 | 1.05103067 | 1.03360568 | 1.09189131 | 0.94251903 | 0.87896263 |
| ENSMUSG00 | Pld3 | 3199 | 70.2524949 | 69.1239709 | 62.371716 | 53.0120692 | 57.7894841 | 42.391218 |
| ENSMUSG00 | Grik5 | 3816 | 3.10459465 | 4.15443906 | 1.85332675 | 2.8933715 | 3.04874573 | 2.1871173 |
| ENSMUSG00 | Cd79a | 1452 | 0 | 0 | 0.03044204 | 0.27334882 | 0 | 0.31933118 |
| ENSMUSG00 | Rabac1 | 1112 | 104.751401 | 129.349397 | 135.03027 | 101.399631 | 72.6850533 | 134.926071 |
| ENSMUSG00 | Etv3 | 6118 | 4.24327048 | 4.16649688 | 3.75693991 | 5.37279218 | 4.22856694 | 3.83396367 |
| ENSMUSG00 | Prkcsh | 1995 | 38.5996821 | 40.9645142 | 26.6983563 | 23.2227294 | 35.9102916 | 19.3725132 |
| ENSMUSG00 | Elavl3 | 5020 | 0.04751651 | 0.16318118 | 0.03522059 | 0.05031361 | 0.05082308 | 0.13582988 |
| ENSMUSG00 | Rab3b | 3536 | 0.16864599 | 0.13627388 | 0 | 0 | 0 | 0.08484758 |
| ENSMUSG00 | St8sia6 | 9750 | 0.17533186 | 0 | 0 | 0 | 0 | 0 |
| ENSMUSG00 | Fcgrt | 1731 | 50.5957964 | 48.7988637 | 64.374839 | 47.5465878 | 48.9334396 | 35.1056593 |
| ENSMUSG00 | Nosip | 2987 | 15.026427 | 11.8086628 | 15.2124187 | 14.2540384 | 11.8725582 | 17.8330668 |
| ENSMUSG00 | Pih1d1 | 1997 | 5.71348168 | 7.72141286 | 9.73901398 | 6.81168292 | 7.66545456 | 6.30990653 |
| ENSMUSG00 | Rps11 | 748 | 25.1926441 | 30.9217811 | 40.1244003 | 23.2507292 | 18.2139604 | 53.382449 |
| ENSMUSG00 | Supt5h | 3479 | 31.8820899 | 34.4742899 | 32.7035194 | 33.8314542 | 37.1807596 | 26.6082569 |
| ENSMUSG00 | Dll3 | 2750 | 0 | 0 | 0 | 0 | 0 | 0 |
| ENSMUSG00 | Paf1 | 3665 | 23.1156696 | 22.9296467 | 23.7471839 | 20.9502294 | 23.3064538 | 18.8503851 |
| ENSMUSG00 | Timm50 | 3021 | 20.3975488 | 23.3515231 | 19.8403506 | 21.2837482 | 20.0659829 | 20.2415562 |
| ENSMUSG00 | Med29 | 1277 | 3.05092959 | 4.15075069 | 3.91136121 | 3.36238316 | 2.83701855 | 5.65996821 |
| ENSMUSG00 | Bicd1 | 14252 | 5.19119767 | 6.00809063 | 4.52191181 | 8.66607062 | 7.26083989 | 5.10203398 |
| ENSMUSG00 | Ncstn | 4055 | 77.9325067 | 62.4225353 | 80.3699602 | 59.4575146 | 67.3850167 | 63.9255663 |
| ENSMUSG00 | Pex19 | 3899 | 43.1202686 | 37.9411073 | 35.3025235 | 33.5186369 | 39.5097791 | 41.4820751 |
| ENSMUSG00 | Phyhip | 2904 | 0.13689904 | 0.34845568 | 0.19787326 | 0.48457291 | 0.07028426 | 0.22541024 |
| ENSMUSG00 | Crhr2 | 4057 | 0 | 0 | 0 | 0 | 0 | 0 |
| ENSMUSG00 | Inmt | 1597 | 20.1142318 | 23.6557111 | 1.99281947 | 15.6347995 | 13.0361687 | 7.6341664 |
| ENSMUSG00 | Cyp4f18 | 2096 | 0.15173848 | 0.02298972 | 0 | 0.03442943 | 0.24344645 | 0.20820335 |
| ENSMUSG00 | Impdh1 | 4059 | 2.9089376 | 2.56424526 | 2.10173829 | 2.400136 | 1.87310421 | 2.3988781 |
| ENSMUSG00 | Psg18 | 2531 | 0 | 0 | 0 | 0 | 0 | 0 |
| ENSMUSG00 | Dusp3 | 5071 | 2.92423471 | 2.27106284 | 1.79561815 | 2.23422638 | 1.499296 | 1.85022203 |
| ENSMUSG00 | Prodh | 3444 | 21.4130129 | 19.2102163 | 14.8237888 | 14.5417764 | 18.3866819 | 14.3896689 |
| ENSMUSG00 | Dgcr14 | 2809 | 13.8698366 | 11.7678531 | 11.565808 | 13.9755302 | 14.1144514 | 11.8167447 |
| ENSMUSG00 | Slc25a1 | 2363 | 50.8257751 | 36.8280637 | 53.348987 | 31.058349 | 31.4407123 | 37.5473557 |
| ENSMUSG00 | Dgcr6 | 3046 | 14.9311457 | 12.8296798 | 16.9783836 | 11.6443368 | 11.8938696 | 15.6699339 |
| ENSMUSG00 | Ddr1 | 5540 | 37.8609955 | 42.9937876 | 32.0024535 | 36.3621037 | 37.192149 | 28.7860678 |
| ENSMUSG00 | Ier3 | 1129 | 53.2068477 | 133.547721 | 63.4642936 | 82.5508601 | 87.1832409 | 61.7967469 |
| ENSMUSG00 | Fosb | 3747 | 0.36073828 | 0.46296021 | 0.71959232 | 1.24221611 | 0.65366054 | 1.25927745 |
| ENSMUSG00 | Klc4 | 2438 | 34.7819688 | 36.1892435 | 31.129846 | 33.5808692 | 41.3568985 | 29.6351597 |
| ENSMUSG00 | Ercc1 | 2664 | 8.65547248 | 8.82694586 | 9.15894044 | 9.75190393 | 8.31285552 | 9.78774592 |
| ENSMUSG00 | Cyp17a1 | 2158 | 0 | 0 | 0 | 0 | 0 | 0 |
| ENSMUSG00 | As3mt | 1740 | 34.4547498 | 30.6565468 | 37.5207598 | 23.3289081 | 34.4281409 | 29.5945511 |
| ENSMUSG00 | Homer3 | 3396 | 1.53355947 | 1.16351244 | 0.74190373 | 1.14748552 | 1.53259433 | 1.90344214 |
| ENSMUSG00 | Crtc1 | 11168 | 1.95075245 | 2.06242115 | 1.80480305 | 2.05481557 | 2.26164542 | 2.0416904 |
| ENSMUSG00 | Rnf215 | 2572 | 11.1908895 | 15.4189121 | 10.6895592 | 14.4636811 | 14.5817987 | 10.3923587 |
| ENSMUSG00 | Sec14l2 | 4153 | 0.40205398 | 0.09282243 | 0.28737052 | 0.5039149 | 0.39317217 | 0.03283727 |
| ENSMUSG00 | Aven | 5112 | 3.04854232 | 3.09177485 | 3.50190658 | 3.48680164 | 2.19597079 | 4.18296502 |
| ENSMUSG00 | Cp | 9266 | 9.09579308 | 4.92473135 | 13.1231675 | 5.33481556 | 7.63798654 | 15.4122611 |
| ENSMUSG00 | Crot | 5437 | 73.3249878 | 70.9103777 | 80.90799 | 73.119729 | 62.7764259 | 81.7737529 |
| ENSMUSG00 | Rps6ka1 | 4802 | 52.2235675 | 38.9946927 | 61.1020062 | 43.6185482 | 57.837685 | 36.8849453 |
| ENSMUSG00 | Calb2 | 1730 | 0 | 0 | 0 | 0 | 0 | 0 |
| ENSMUSG00 | Snrnp200 | 7214 | 19.4864679 | 16.7790883 | 14.8585347 | 17.7959405 | 21.8916874 | 9.9775122 |
| ENSMUSG00 | Ciao1 | 3485 | 21.3892471 | 23.9756932 | 23.7561125 | 22.7259936 | 22.0943475 | 23.7762852 |
| ENSMUSG00 | Has1 | 2085 | 0 | 0.13866602 | 0 | 0 | 0.31815007 | 0.11773226 |
| ENSMUSG00 | Taf6l | 4101 | 3.32507433 | 4.97021825 | 3.8155212 | 3.29938227 | 3.54609049 | 1.59617494 |
| ENSMUSG00 | Insig2 | 3532 | 54.2304375 | 47.1086593 | 55.4275093 | 54.4193237 | 49.4227723 | 59.7926194 |
| ENSMUSG00 | Kpna6 | 7408 | 6.43987265 | 4.75489865 | 5.45362912 | 4.86094498 | 3.91238951 | 4.7016353 |
| ENSMUSG00 | Man1a | 9592 | 1.89410494 | 1.72309734 | 2.63128152 | 2.76483556 | 3.18648865 | 2.72120827 |
| ENSMUSG00 | Itpkc | 4111 | 5.66692084 | 4.98157089 | 5.64484738 | 3.88818918 | 5.69717988 | 2.12305633 |
| ENSMUSG00 | Adck4 | 3157 | 9.94831473 | 11.7375274 | 8.55474377 | 6.33178735 | 8.35623576 | 5.66745751 |
| ENSMUSG00 | Brd8 | 8817 | 17.5308277 | 16.0293563 | 18.759589 | 17.8711907 | 19.3410625 | 20.5279013 |
| ENSMUSG00 | Kif20a | 3873 | 0.41059107 | 1.14463018 | 0.44509989 | 1.12727276 | 0.68509464 | 0.96478843 |
| ENSMUSG00 | Farsa | 4608 | 14.1404585 | 9.27547183 | 8.75787384 | 7.04727432 | 9.17987534 | 5.60526983 |
| ENSMUSG00 | Gcdh | 2496 | 38.5768325 | 42.8388282 | 39.2787211 | 33.7112692 | 39.067069 | 54.9645174 |
| ENSMUSG00 | Mast2 | 8848 | 3.91354244 | 5.10837282 | 3.79172681 | 3.70689178 | 5.91117035 | 2.74965848 |
| ENSMUSG00 | Dnase2a | 3756 | 8.65814245 | 8.5955581 | 8.46142842 | 10.7400761 | 8.05608117 | 11.8073918 |
| ENSMUSG00 | Rad23a | 3937 | 13.8644335 | 10.6972188 | 14.6965234 | 10.5854106 | 12.9607254 | 9.51183127 |
| ENSMUSG00 | Calr | 2936 | 114.337629 | 109.551942 | 146.245471 | 128.069137 | 105.771977 | 148.291916 |
| ENSMUSG00 | Syce2 | 2023 | 2.14204023 | 1.73880884 | 2.84045457 | 1.9084423 | 0.73147054 | 2.29198649 |
| ENSMUSG00 | Nfat5 | 18323 | 6.14677052 | 6.76656201 | 5.76797504 | 10.5491193 | 7.0845986 | 7.62135856 |
| ENSMUSG00 | Nob1 | 2988 | 9.30022787 | 11.2725311 | 10.6066672 | 8.47710685 | 7.78715776 | 7.74059374 |
| ENSMUSG00 | Nqo1 | 1547 | 29.9900748 | 172.779699 | 10.9718861 | 108.152838 | 28.3003429 | 9.99658724 |
| ENSMUSG00 | Ppfia3 | 6091 | 0 | 0.07911089 | 0.14513821 | 0.03554298 | 0.22618817 | 0.08507932 |
| ENSMUSG00 | Gys1 | 5066 | 17.5313351 | 15.9321537 | 19.9981493 | 20.191985 | 21.3734642 | 16.2108051 |
| ENSMUSG00 | Ruvbl2 | 2662 | 8.24381114 | 10.1368924 | 9.87982592 | 9.94899349 | 8.26159578 | 7.1926359 |
| ENSMUSG00 | Lin7b | 1563 | 3.30659787 | 2.71299226 | 1.52712701 | 3.02415089 | 2.02407889 | 2.8094798 |
| ENSMUSG00 | Bax | 1438 | 23.8311712 | 30.0913943 | 27.9719591 | 26.1958654 | 18.6292401 | 32.6423186 |
| ENSMUSG00 | Il7r | 3200 | 0 | 0 | 0 | 0 | 0 | 0 |
| ENSMUSG00 | Tfam | 4682 | 14.655694 | 13.3382377 | 15.5962077 | 13.3708559 | 16.7181676 | 13.2120636 |
| ENSMUSG00 | Zfp81 | 7454 | 5.19477301 | 3.93688533 | 6.20865705 | 7.77888993 | 7.13985913 | 13.5238887 |
| ENSMUSG00 | Efnb3 | 3178 | 0 | 0.07581253 | 0 | 0 | 0.09633677 | 0 |
| ENSMUSG00 | Mmd | 5907 | 2.07964169 | 6.86047026 | 1.09251212 | 2.22954905 | 3.28254999 | 2.12397732 |
| ENSMUSG00 | Hlf | 11048 | 0.26268556 | 0.21807767 | 0.07601693 | 0.01959561 | 0.06927911 | 0.35549856 |
| ENSMUSG00 | Fam162a | 630 | 142.488691 | 129.33853 | 171.755733 | 102.461552 | 77.5114947 | 172.99914 |
| ENSMUSG00 | Rpl8 | 856 | 67.8536881 | 66.2000654 | 73.0672989 | 55.8934475 | 43.2174328 | 65.9561954 |
| ENSMUSG00 | Grm3 | 3571 | 0.02226574 | 0 | 0 | 0 | 0 | 0 |
| ENSMUSG00 | Ssbp2 | 5785 | 6.16433293 | 10.9366981 | 5.76877986 | 8.41394608 | 8.89984671 | 8.27904348 |
| ENSMUSG00 | Fancl | 3142 | 5.4407564 | 5.56705236 | 4.83941248 | 6.99362353 | 3.96258297 | 5.25180037 |
| ENSMUSG00 | Fam5b | 4091 | 0 | 0 | 0.05402327 | 0 | 0 | 0 |
| ENSMUSG00 | Gstm5 | 1432 | 7.44027149 | 11.3736155 | 9.63056915 | 7.05514837 | 4.95297922 | 9.92324515 |
| ENSMUSG00 | Gstm7 | 1862 | 3.58696065 | 4.70995299 | 8.56974507 | 2.96485113 | 2.98704348 | 7.04570434 |
| ENSMUSG00 | Gstm3 | 1384 | 0.25852552 | 0 | 0 | 0 | 0 | 0 |
| ENSMUSG00 | Stat3 | 9647 | 30.7263256 | 27.1876028 | 26.7630315 | 24.4536547 | 33.2012606 | 20.7266902 |
| ENSMUSG00 | Stat5a | 5012 | 3.75186398 | 2.06705608 | 1.37579559 | 1.38223315 | 2.16851916 | 1.85567677 |
| ENSMUSG00 | Ptrf | 4805 | 4.27754078 | 5.11448189 | 4.12121242 | 2.93612475 | 3.34512141 | 3.28657984 |
| ENSMUSG00 | Map3k11 | 3789 | 10.3139713 | 9.98320324 | 8.8543676 | 10.0370744 | 10.4099732 | 7.60148299 |
| ENSMUSG00 | Akt2 | 5865 | 7.68673741 | 8.41311454 | 9.55634048 | 7.81931434 | 7.27332484 | 6.31060281 |
| ENSMUSG00 | Dnaja3 | 3897 | 48.8246676 | 49.6455133 | 55.1587279 | 46.2946427 | 46.9149357 | 42.4482133 |
| ENSMUSG00 | Hmox2 | 2274 | 29.073599 | 36.7861317 | 38.6814721 | 28.70379 | 22.169771 | 28.5819988 |
| ENSMUSG00 | 5730403B10 | 5739 | 25.2913851 | 24.0302487 | 26.9108276 | 21.6592862 | 26.3534027 | 20.5165739 |
| ENSMUSG00 | B230120H23 | 10280 | 6.77931492 | 4.26553136 | 8.68557616 | 4.76297027 | 8.04112148 | 5.63799693 |
| ENSMUSG00 | Cwc15 | 1104 | 75.9099211 | 69.093422 | 68.7851144 | 59.5484467 | 60.824919 | 82.1450865 |
| ENSMUSG00 | Col5a3 | 6742 | 0 | 0 | 0 | 0 | 0 | 0 |
| ENSMUSG00 | Dnmt1 | 5225 | 4.30652669 | 5.42270392 | 3.67150237 | 5.53142922 | 6.87513274 | 4.80242452 |
| ENSMUSG00 | Ppan | 2474 | 10.5575387 | 14.1793573 | 11.3809919 | 11.0988019 | 13.9012843 | 8.01481921 |
| ENSMUSG00 | Angptl2 | 6092 | 4.59419867 | 2.50740351 | 3.06916934 | 3.5359456 | 2.00185552 | 1.46401954 |
| ENSMUSG00 | Cacna1e | 12649 | 1.40176649 | 0.89142442 | 2.43915617 | 1.74576735 | 0.55669538 | 1.88673487 |
| ENSMUSG00 | Cacna1b | 13098 | 0 | 0 | 0 | 0 | 0 | 0 |
| ENSMUSG00 | Rgmtd2 | 7073 | 2.1696049 | 1.15135144 | 2.36226448 | 2.39254615 | 1.45727809 | 2.05533476 |
| ENSMUSG00 | Etv1 | 5276 | 0.8137968 | 0.94984647 | 2.22014564 | 0.37613959 | 0.7350274 | 1.34408761 |
| ENSMUSG00 | Kifc2 | 3199 | 14.6644161 | 12.5775802 | 10.0866975 | 14.7531461 | 13.0157933 | 10.0265632 |
| ENSMUSG00 | Psap | 2867 | 314.923864 | 318.86752 | 289.817316 | 237.975368 | 264.600324 | 187.088614 |
| ENSMUSG00 | Ikbkg | 8205 | 15.4418911 | 15.727397 | 14.6477527 | 15.4838365 | 16.1878923 | 14.5963365 |
| ENSMUSG00 | Pax2 | 3611 | 1.66244186 | 1.20099136 | 1.30977491 | 0.92928002 | 1.38481995 | 1.57862077 |
| ENSMUSG00 | Wars2 | 5972 | 2.85584413 | 2.54164924 | 3.16784809 | 2.58592014 | 1.48670285 | 3.67193737 |
| ENSMUSG00 | Atn1 | 5070 | 2.15636236 | 2.19547695 | 2.31907105 | 2.43393673 | 1.90216867 | 2.81891735 |
| ENSMUSG00 | Phb2 | 2313 | 51.6666556 | 48.7905956 | 56.0310435 | 50.1841493 | 40.7019729 | 47.5802721 |
| ENSMUSG00 | Ptpn6 | 2742 | 11.7149629 | 12.2135585 | 10.7844759 | 11.0009443 | 14.6454259 | 10.6134355 |
| ENSMUSG00 | Eno2 | 5770 | 0.81302369 | 1.00214438 | 0.6588143 | 1.41951891 | 0.7251582 | 0.46324344 |
| ENSMUSG00 | Emg1 | 1377 | 27.1099464 | 24.7406061 | 22.566373 | 26.3867965 | 19.3062758 | 20.6590042 |
| ENSMUSG00 | Lpcat3 | 3370 | 48.9098582 | 43.038929 | 50.3533755 | 43.6410723 | 41.8658543 | 37.9902542 |
| ENSMUSG00 | Atp6v1f | 1059 | 213.455772 | 218.226796 | 298.393745 | 200.853307 | 140.455038 | 352.715944 |
| ENSMUSG00 | Il12b | 2014 | 0 | 0 | 0 | 0 | 0 | 0 |
| ENSMUSG00 | Clcn5 | 11787 | 15.8556472 | 12.0558088 | 22.7065546 | 15.6946333 | 17.0953727 | 22.1723588 |
| ENSMUSG00 | Clcn3 | 18308 | 22.6832942 | 17.4290267 | 31.9659385 | 22.0319628 | 20.5883678 | 26.9379521 |
| ENSMUSG00 | Hif3a | 7124 | 0.011161 | 0 | 0.00620464 | 0 | 0.02148781 | 0 |
| ENSMUSG00 | Gpx6 | 1270 | 8.51455961 | 9.52346219 | 4.97705791 | 4.91511315 | 10.3258099 | 6.50725645 |
| ENSMUSG00 | Gpx5 | 1669 | 0 | 0 | 0 | 0 | 0 | 0 |
| ENSMUSG00 | Pde1c | 13929 | 2.97117203 | 1.7089599 | 4.10951157 | 2.54638881 | 4.00034478 | 3.45803812 |
| ENSMUSG00 | Utp20 | 8769 | 0.87499233 | 0.99461125 | 0.96781319 | 1.53479332 | 1.06486792 | 1.36233231 |
| ENSMUSG00 | Spic | 3907 | 0 | 0 | 0 | 0 | 0 | 0 |
| ENSMUSG00 | 9330159F19 | 5638 | 0 | 0 | 0 | 0 | 0 | 0 |
| ENSMUSG00 | Cul3 | 5320 | 69.3553787 | 65.4411738 | 60.7692255 | 73.9885383 | 55.0451825 | 95.9585381 |
| ENSMUSG00 | Sst | 597 | 0 | 0 | 0 | 0 | 0 | 0 |
| ENSMUSG00 | Il11 | 1828 | 0.23922882 | 0.10544079 | 0 | 0.33555512 | 0.02791377 | 0 |
| ENSMUSG00 | Large | 7868 | 0.44464696 | 0.47769986 | 0.20224534 | 0.15133547 | 0.59016271 | 0.29118832 |
| ENSMUSG00 | Ddx56 | 2519 | 12.3732818 | 14.8059969 | 15.3188602 | 13.2210112 | 16.6914385 | 12.1160466 |
| ENSMUSG00 | Tmed4 | 2308 | 74.653489 | 87.7711451 | 98.8029931 | 75.7285198 | 65.6179738 | 86.0072896 |
| ENSMUSG00 | Emid2 | 6701 | 1.99341016 | 4.80354326 | 1.00923474 | 3.21459194 | 3.36571528 | 2.37294653 |
| ENSMUSG00 | Bid | 3541 | 3.74988153 | 3.44286074 | 3.45775502 | 4.259332 | 3.28551643 | 4.33652129 |
| ENSMUSG00 | Ralb | 4728 | 9.72865714 | 8.93813658 | 10.7980391 | 6.90656732 | 13.3070054 | 8.45698417 |
| ENSMUSG00 | Ppp1cc | 3676 | 3.62298313 | 2.80519549 | 3.88389426 | 4.01456915 | 3.4841187 | 3.84337952 |
| ENSMUSG00 | Dnajb11 | 3246 | 42.7806203 | 42.1891154 | 51.8275461 | 43.5518948 | 41.9089088 | 44.7687239 |
| ENSMUSG00 | Tbccd1 | 2694 | 14.1962778 | 8.51401131 | 15.6035459 | 13.0854334 | 18.959689 | 9.95210482 |
| ENSMUSG00 | Clec11a | 1948 | 1.61226025 | 0.8163001 | 1.81527076 | 1.44476359 | 0.62866171 | 3.43033194 |
| ENSMUSG00 | Zfp324 | 5226 | 4.92949701 | 6.50046722 | 4.15290948 | 3.50740119 | 5.42875334 | 2.82349404 |
| ENSMUSG00 | Gab2 | 5987 | 4.91382255 | 4.76886472 | 3.69886811 | 4.05599064 | 4.49154836 | 3.59895858 |
| ENSMUSG00 | Coro1c | 9649 | 4.48686064 | 2.95640727 | 4.5763956 | 3.73945948 | 4.17243347 | 3.97431265 |
| ENSMUSG00 | Tax1bp1 | 6482 | 84.0250051 | 69.4919564 | 81.2571986 | 80.714231 | 95.7551428 | 86.2757179 |
| ENSMUSG00 | Psg17 | 2095 | 0 | 0 | 0 | 0 | 0 | 0 |
| ENSMUSG00 | Psg19 | 3082 | 0 | 0 | 0 | 0 | 0 | 0 |
| ENSMUSG00 | Ctse | 3221 | 6.78842418 | 4.39826575 | 11.1019221 | 1.62430812 | 5.16442053 | 5.92742859 |
| ENSMUSG00 | Ndrg2 | 2141 | 150.536049 | 132.630875 | 136.899776 | 110.167363 | 153.293624 | 125.914308 |
| ENSMUSG00 | Mettl17 | 3719 | 11.4594986 | 8.86247013 | 9.68660986 | 8.58634294 | 15.4080721 | 10.7734458 |
| ENSMUSG00 | Arhgef40 | 5620 | 0.32540073 | 0.9260028 | 0.31460387 | 0.8667395 | 2.278936 | 0.99974656 |
| ENSMUSG00 | Pnpla6 | 4461 | 14.1430055 | 12.0655135 | 13.7133716 | 12.5450014 | 14.8126332 | 7.74034819 |
| ENSMUSG00 | Mcoln1 | 3350 | 25.1468248 | 26.5385032 | 32.3003319 | 19.9366759 | 27.8131828 | 22.4873294 |
| ENSMUSG00 | Arhgef18 | 8150 | 22.5118457 | 22.6091969 | 24.5686318 | 23.1721989 | 25.0811855 | 19.3332013 |
| ENSMUSG00 | Pkn2 | 7617 | 30.4129165 | 23.6029428 | 39.3156737 | 37.9152796 | 41.1721289 | 39.1555967 |
| ENSMUSG00 | Cd33 | 2564 | 0 | 0 | 0 | 0 | 0 | 0 |
| ENSMUSG00 | Etfb | 1115 | 162.088264 | 167.075144 | 184.537739 | 152.25652 | 111.617337 | 187.033087 |
| ENSMUSG00 | Nkg7 | 1204 | 0 | 0 | 0 | 0 | 0 | 0 |
| ENSMUSG00 | Lim2 | 1850 | 0 | 0 | 0 | 0 | 0 | 0 |
| ENSMUSG00 | Stxbp2 | 4119 | 59.6283949 | 60.937892 | 58.538736 | 51.2892379 | 65.6566623 | 49.3645146 |
| ENSMUSG00 | Pcp2 | 966 | 0.24692845 | 0 | 0 | 0 | 0.26411168 | 0 |
| ENSMUSG00 | Sgce | 3352 | 3.77155215 | 5.26140747 | 1.37141756 | 5.8880902 | 6.53052384 | 2.47359497 |
| ENSMUSG00 | Chn2 | 6308 | 2.49574672 | 2.02431946 | 0.9459811 | 1.14972907 | 2.35394346 | 0.53615334 |
| ENSMUSG00 | Wwox | 10339 | 1.79570649 | 4.38566999 | 3.1893389 | 2.59299343 | 2.7243021 | 2.25024347 |
| ENSMUSG00 | Slbp | 3912 | 19.9590398 | 21.1123624 | 22.1009215 | 20.2915383 | 18.7044537 | 27.8114867 |
| ENSMUSG00 | Tyr | 5394 | 0 | 0 | 0 | 0 | 0 | 0 |
| ENSMUSG00 | Ghrhr | 1585 | 0.17557626 | 0 | 0 | 0 | 0 | 0 |
| ENSMUSG00 | Aqp1 | 2623 | 56.8520043 | 19.1239368 | 7.17879723 | 1.66447861 | 22.9161535 | 9.78476383 |
| ENSMUSG00 | Arid3b | 6532 | 3.06139111 | 2.55981253 | 1.81354775 | 3.3364291 | 2.35133789 | 2.54290498 |
| ENSMUSG00 | Cnn2 | 2570 | 2.32035879 | 2.84993743 | 1.68551775 | 1.74092355 | 2.22371755 | 1.33720021 |
| ENSMUSG00 | Polr2e | 2954 | 11.9104943 | 17.11157 | 15.7714091 | 13.411674 | 11.2106019 | 12.2246521 |
| ENSMUSG00 | Abca13 | 19682 | 5.14062077 | 6.75716799 | 1.48671985 | 7.64832282 | 5.63099727 | 4.23767132 |
| ENSMUSG00 | Myo9b | 7346 | 5.90974472 | 5.10988816 | 5.55381175 | 4.64655787 | 4.89703171 | 4.91582118 |
| ENSMUSG00 | Hdac9 | 9564 | 0 | 0.03022989 | 0.34200506 | 0 | 0.25609223 | 0.07699867 |
| ENSMUSG00 | Ly9 | 3789 | 0 | 0 | 0 | 0 | 0 | 0 |
| ENSMUSG00 | Cd244 | 2243 | 0 | 0 | 0 | 0 | 0 | 0 |
| ENSMUSG00 | Emr1 | 3274 | 0 | 0 | 0 | 0 | 0 | 0 |
| ENSMUSG00 | Mtfp1 | 1733 | 16.4252303 | 17.3226506 | 24.7662952 | 14.65768 | 11.6598066 | 15.5023195 |
| ENSMUSG00 | Rab23 | 8551 | 2.55707102 | 1.89342119 | 2.16072628 | 2.33767427 | 2.94784583 | 2.02861309 |
| ENSMUSG00 | Rab11a | 3332 | 139.454399 | 125.55663 | 127.0604 | 119.226684 | 98.7602334 | 168.968518 |
| ENSMUSG00 | Eif2b2 | 2001 | 12.2981721 | 16.3511216 | 14.0270716 | 10.5667557 | 14.739253 | 13.7122874 |
| ENSMUSG00 | Dlst | 2765 | 143.335402 | 110.384421 | 128.736869 | 128.459908 | 107.109253 | 87.2689814 |
| ENSMUSG00 | Pgf | 1575 | 0 | 0 | 0 | 0 | 0 | 0 |
| ENSMUSG00 | Ulk2 | 6287 | 14.1202462 | 15.9573998 | 7.03770397 | 16.3106681 | 14.4305545 | 11.9952883 |
| ENSMUSG00 | Ccl24 | 1286 | 0 | 0 | 0 | 0 | 0 | 0 |
| ENSMUSG00 | Dgkq | 8837 | 3.22110717 | 4.55854539 | 3.25123888 | 6.99429017 | 5.20253079 | 3.12345082 |
| ENSMUSG00 | Tmed11 | 758 | 0 | 0 | 0 | 0 | 0 | 0 |
| ENSMUSG00 | Grap | 1787 | 0.0444941 | 0.83591478 | 0.34629312 | 0.34325392 | 0.31409633 | 0.06105124 |
| ENSMUSG00 | Pou1f1 | 3896 | 0 | 0 | 0 | 0.01851311 | 0 | 0.02798834 |
| ENSMUSG00 | Chmp2b | 1832 | 80.8129964 | 84.9575372 | 103.386953 | 94.498717 | 70.6347648 | 99.6000746 |
| ENSMUSG00 | Plod3 | 4812 | 4.06477482 | 5.91816029 | 6.1636402 | 4.6039849 | 6.26695514 | 3.65022188 |
| ENSMUSG00 | Ap1s1 | 2640 | 21.0071573 | 21.2458404 | 15.5376175 | 13.4624295 | 13.1431574 | 17.1086374 |
| ENSMUSG00 | Mapk13 | 1693 | 4.01546791 | 9.8763706 | 3.18524799 | 5.37074732 | 6.0279239 | 8.32899498 |
| ENSMUSG00 | Srpk1 | 9869 | 4.81384124 | 5.55640604 | 6.09572483 | 6.70894231 | 6.05967296 | 7.17448206 |
| ENSMUSG00 | Pax3 | 7743 | 0 | 0 | 0 | 0 | 0 | 0.00352249 |
| ENSMUSG00 | Lbr | 3537 | 9.85738095 | 10.2448981 | 10.3100143 | 10.0482934 | 13.8782509 | 11.0687564 |
| ENSMUSG00 | Crabp2 | 1318 | 0.60326981 | 0.84088632 | 0.16768529 | 0.24638725 | 0.038715 | 0 |
| ENSMUSG00 | Nes | 7307 | 0 | 0.13848574 | 0.0907387 | 0.26171457 | 0.18854689 | 0.00746535 |
| ENSMUSG00 | Bcan | 3253 | 0 | 0.13331632 | 0 | 0.32166594 | 0 | 0 |
| ENSMUSG00 | Hapln2 | 3096 | 0 | 0 | 0 | 0 | 0 | 0 |
| ENSMUSG00 | Prcc | 2095 | 3.6434617 | 3.42710257 | 2.29976176 | 4.06461218 | 3.45858967 | 2.08302734 |
| ENSMUSG00 | Rrnad1 | 5313 | 3.99575129 | 4.73429753 | 5.19140788 | 7.02896971 | 5.86808124 | 5.15410085 |
| ENSMUSG00 | Hdgf | 2863 | 45.0599142 | 53.2188369 | 42.4263586 | 39.5478364 | 35.4671631 | 34.7339625 |
| ENSMUSG00 | Slc25a18 | 1372 | 0 | 0.07024263 | 0 | 0.0788966 | 0.29752969 | 0.05963842 |
| ENSMUSG00 | Thop1 | 3685 | 3.87305767 | 4.49827303 | 3.38261051 | 3.82851544 | 6.02346635 | 2.27226981 |
| ENSMUSG00 | Apba3 | 2227 | 28.1340985 | 32.1315072 | 21.6146417 | 19.3452902 | 26.4869737 | 13.5944542 |
| ENSMUSG00 | Matk | 3814 | 1.17786295 | 0.6190707 | 0.22019796 | 0.71899197 | 0.41473981 | 0.51488569 |
| ENSMUSG00 | Pias4 | 4551 | 5.36362668 | 6.24697888 | 5.88580902 | 4.82045332 | 6.76091071 | 5.26794285 |
| ENSMUSG00 | Map2k1 | 3649 | 47.0877464 | 39.4444787 | 32.1853376 | 35.5381935 | 34.1341144 | 37.7763844 |
| ENSMUSG00 | Sgta | 1957 | 52.5536168 | 52.4461532 | 56.1727049 | 36.8011042 | 50.896007 | 38.4799586 |
| ENSMUSG00 | Itgb1bp3 | 1833 | 0 | 0 | 0 | 0 | 0 | 0 |
| ENSMUSG00 | 5730437N04 | 999 | 21.5292442 | 23.5385223 | 34.8216721 | 25.2465957 | 15.5275458 | 33.2537644 |
| ENSMUSG00 | Dtx2 | 7175 | 0.90315586 | 0.98051854 | 1.08425427 | 1.39802207 | 0.9245197 | 0.85530228 |
| ENSMUSG00 | Zp3 | 1589 | 0 | 0 | 0 | 0 | 0 | 0 |
| ENSMUSG00 | Hspb1 | 1408 | 3.10589692 | 2.19029283 | 2.32310823 | 3.45957103 | 2.50058234 | 2.15020238 |
| ENSMUSG00 | Rasa4 | 6211 | 0.55687117 | 0.27929672 | 0.12810066 | 0.34275328 | 0.41898972 | 0.43913443 |
| ENSMUSG00 | Syt5 | 1739 | 0 | 0.72044134 | 0 | 0 | 0 | 0 |
| ENSMUSG00 | Hnrnpa2b1 | 4614 | 327.504511 | 342.871496 | 343.02891 | 380.636635 | 301.18169 | 524.20036 |
| ENSMUSG00 | Fxyd4 | 1191 | 156.451669 | 277.021469 | 91.7809889 | 159.021538 | 131.871692 | 171.18214 |
| ENSMUSG00 | Ccdc130 | 3310 | 6.73801346 | 7.54095983 | 8.03912674 | 7.94677053 | 10.7139973 | 7.73742787 |
| ENSMUSG00 | Mri1 | 3388 | 7.42775064 | 6.51398775 | 6.22322288 | 5.7083754 | 7.19911678 | 4.92682386 |
| ENSMUSG00 | Prkacb | 8580 | 20.6376352 | 19.8642711 | 24.0688824 | 24.647653 | 18.1446938 | 19.3688085 |
| ENSMUSG00 | Sgsh | 7753 | 1.49730431 | 1.67188868 | 1.15735511 | 1.5730338 | 0.90824711 | 1.0061327 |
| ENSMUSG00 | Chd5 | 9686 | 0 | 0 | 0 | 0 | 0 | 0 |
| ENSMUSG00 | Cstb | 596 | 54.8972491 | 67.1052802 | 88.9228352 | 71.3165242 | 39.3827733 | 113.949416 |
| ENSMUSG00 | Sh2b2 | 2839 | 0.05601336 | 0.32248764 | 0 | 0 | 0 | 0.02882139 |
| ENSMUSG00 | Pex5 | 5279 | 18.1494048 | 21.4871917 | 13.3216769 | 13.5401648 | 19.0805202 | 13.3144053 |
| ENSMUSG00 | Jkamp | 2683 | 20.640844 | 19.1811853 | 24.6627503 | 17.4291203 | 14.7012257 | 19.9959804 |
| ENSMUSG00 | Cd44 | 8637 | 0 | 0 | 0.16376739 | 0 | 0 | 0 |
| ENSMUSG00 | Slc1a2 | 13209 | 0.00300973 | 0 | 0.00669269 | 0.00819488 | 0.08112301 | 0 |
| ENSMUSG00 | Eif2ak4 | 7651 | 5.70533494 | 3.16162514 | 6.2509991 | 3.24460157 | 5.78890266 | 4.662819 |
| ENSMUSG00 | Wdr1 | 2933 | 129.811878 | 138.973445 | 112.682298 | 123.500731 | 107.463322 | 79.4898112 |
| ENSMUSG00 | Slc2a9 | 7127 | 5.845902 | 10.2092785 | 2.0342647 | 5.06778821 | 8.44115295 | 3.37919271 |
| ENSMUSG00 | Wisp1 | 5759 | 0 | 6.13312417 | 1.32014534 | 3.20158443 | 5.06808247 | 1.31187273 |
| ENSMUSG00 | Ndrg1 | 5894 | 71.039145 | 127.349883 | 44.0818516 | 92.9232022 | 104.225745 | 55.7478078 |
| ENSMUSG00 | 4930550C14 | 1351 | 26.5428893 | 41.4096864 | 14.7884775 | 40.5422232 | 37.7315688 | 29.8789534 |
| ENSMUSG00 | Man2b1 | 4301 | 24.9569396 | 23.1913358 | 22.8973974 | 23.5485466 | 19.3617869 | 14.6234174 |
| ENSMUSG00 | Klf5 | 1717 | 7.03882707 | 7.18446862 | 5.32893506 | 5.98915707 | 8.97493622 | 6.57641273 |
| ENSMUSG00 | Wdr83 | 2322 | 14.0394031 | 17.5147964 | 18.7696026 | 16.6735718 | 12.3940034 | 19.5691426 |
| ENSMUSG00 | Prdx2 | 3195 | 87.8477911 | 97.4436941 | 113.555157 | 86.5517337 | 67.3005157 | 92.3686969 |
| ENSMUSG00 | Polr2a | 8097 | 9.35336426 | 9.40281321 | 8.57615109 | 6.67542333 | 10.3539997 | 9.75176987 |
| ENSMUSG00 | Shbg | 1498 | 0 | 0 | 0 | 0 | 0 | 0 |
| ENSMUSG00 | Senp3 | 4111 | 5.87967214 | 5.27460447 | 4.37610073 | 4.03739734 | 4.79109245 | 3.39689012 |
| ENSMUSG00 | Corin | 7030 | 1.28936694 | 1.32289948 | 1.01230394 | 0.09238646 | 0.54437812 | 0 |
| ENSMUSG00 | Plekha8 | 6990 | 3.24186322 | 4.91515497 | 3.13649702 | 3.28816343 | 3.98575124 | 2.52066051 |
| ENSMUSG00 | G6pc2 | 1968 | 0 | 0.58763954 | 0.53904687 | 0 | 0 | 0.88698014 |
| ENSMUSG00 | Spc25 | 3830 | 22.8256662 | 24.9110066 | 19.9427636 | 18.8041151 | 16.6401941 | 22.9733646 |
| ENSMUSG00 | Dnahc2 | 14507 | 0 | 0 | 0 | 0.15918183 | 0.17235076 | 0.06580357 |
| ENSMUSG00 | Ripk4 | 3944 | 22.9118172 | 15.8707374 | 18.3464546 | 15.7355772 | 22.3305083 | 15.9954971 |
| ENSMUSG00 | Ufd1l | 4187 | 22.0093632 | 18.7359751 | 20.9660521 | 21.6819737 | 25.8361397 | 23.7244414 |
| ENSMUSG00 | Zfp287 | 6994 | 1.38126705 | 1.16435641 | 1.36511268 | 1.07307196 | 2.17412925 | 1.97325814 |
| ENSMUSG00 | Prlr | 15933 | 2.51762253 | 2.75817707 | 6.72751329 | 3.39918087 | 3.21536944 | 6.25846238 |
| ENSMUSG00 | Letm1 | 9714 | 24.4287074 | 19.1674297 | 18.0511335 | 19.9725297 | 21.0535016 | 12.8932616 |
| ENSMUSG00 | Ubqln1 | 3651 | 110.903744 | 96.1485158 | 121.091984 | 112.881159 | 90.3967677 | 96.1447842 |
| ENSMUSG00 | Fgfr4 | 3534 | 0.05624714 | 0.49086359 | 0 | 0.24503935 | 0.02887741 | 0 |
| ENSMUSG00 | Cadm3 | 7359 | 0 | 0 | 0 | 0 | 0 | 0 |
| ENSMUSG00 | Fcer1a | 1021 | 0 | 0 | 0 | 0 | 0 | 0 |
| ENSMUSG00 | Txn2 | 4387 | 56.8103732 | 53.1731403 | 58.7712219 | 48.2464721 | 38.7321248 | 54.6487529 |
| ENSMUSG00 | Casp14 | 2322 | 0 | 0 | 0.11421665 | 0 | 0 | 0 |
| ENSMUSG00 | Slc1a6 | 2079 | 0 | 0 | 0 | 0 | 0 | 0 |
| ENSMUSG00 | Slc1a3 | 9684 | 0.68147561 | 1.77141402 | 0.7987735 | 0.89422663 | 1.3647079 | 0.69003373 |
| ENSMUSG00 | Crbn | 5166 | 22.3172461 | 24.1864972 | 23.7095058 | 26.2478365 | 18.5397808 | 19.0331155 |
| ENSMUSG00 | Il5ra | 3774 | 0 | 0 | 0.2459562 | 0.90826556 | 1.1357222 | 0 |
| ENSMUSG00 | Msh6 | 4244 | 6.02327379 | 6.40366479 | 9.0299241 | 10.2617879 | 10.7968156 | 7.97545413 |
| ENSMUSG00 | Fbxo11 | 4878 | 14.0016227 | 15.9633642 | 13.1481907 | 16.7613598 | 12.4375484 | 22.0019896 |
| ENSMUSG00 | Mlxipl | 4673 | 0.46791171 | 0.60838863 | 0.18917973 | 0.94200929 | 0.98274638 | 0.51362471 |
| ENSMUSG00 | Tbl2 | 6138 | 4.91600028 | 5.5346109 | 5.14175887 | 5.73738603 | 6.06031736 | 4.7012982 |
| ENSMUSG00 | Wbscr22 | 3098 | 12.3578204 | 13.6875627 | 10.4155408 | 10.4123137 | 10.7389274 | 7.97637933 |
| ENSMUSG00 | Nid1 | 6026 | 0.6597325 | 0.53576031 | 0.71151324 | 0.62871136 | 1.80362065 | 0.00905232 |
| ENSMUSG00 | Mcm5 | 3479 | 1.62267267 | 1.93909224 | 1.35947031 | 0.39411259 | 1.76003596 | 0.79181908 |
| ENSMUSG00 | Hmox1 | 2153 | 23.1922357 | 23.7463145 | 25.5397551 | 28.7248603 | 15.4998838 | 13.2509395 |
| ENSMUSG00 | Mprip | 13343 | 29.8039475 | 26.0957229 | 26.5582085 | 31.6688019 | 42.0777346 | 28.3191825 |
| ENSMUSG00 | Cic | 10261 | 1.65437971 | 1.09418585 | 1.14586203 | 1.04086202 | 1.34764135 | 1.25727555 |
| ENSMUSG00 | Pafah1b3 | 2083 | 8.14958722 | 10.0166728 | 14.3236889 | 7.18869346 | 7.20199448 | 12.3737562 |
| ENSMUSG00 | Il27ra | 2769 | 0.1005014 | 0 | 0 | 0 | 0.81081998 | 0 |
| ENSMUSG00 | Prkaca | 2304 | 28.0393472 | 27.1048737 | 28.4510995 | 31.6814132 | 26.6869717 | 31.0982974 |
| ENSMUSG00 | Asf1b | 2557 | 0.74628982 | 1.92218111 | 1.10634257 | 0.52211015 | 0.71840029 | 0.55466611 |
| ENSMUSG00 | Myl10 | 816 | 0 | 0 | 0 | 0 | 0 | 0 |
| ENSMUSG00 | Ddx39 | 5211 | 13.1450188 | 12.742452 | 13.1562193 | 15.2540288 | 11.5350356 | 15.5189609 |
| ENSMUSG00 | Dnajb1 | 2467 | 22.141885 | 31.1151205 | 34.9207102 | 34.2830614 | 28.0055585 | 27.2746393 |
| ENSMUSG00 | Msh4 | 4441 | 0 | 0 | 0 | 0 | 0 | 0 |
| ENSMUSG00 | Usp40 | 5548 | 11.6371486 | 12.5764182 | 12.9386794 | 14.4120063 | 13.9982235 | 11.7544453 |
| ENSMUSG00 | Evx1 | 3538 | 0 | 0 | 0 | 0 | 0 | 0 |
| ENSMUSG00 | Kbtbd4 | 2016 | 35.0818452 | 43.1909231 | 46.109363 | 31.8223587 | 38.1938498 | 41.3178315 |

| | | | | | | | | |
|---|---|---|---|---|---|---|---|---|
| ENSMUSG00 | Celf1 | 8674 | 22.3848013 | 17.7602093 | 17.4840354 | 23.4404336 | 23.795445 | 22.0046029 |
| ENSMUSG00 | Ndufs3 | 2411 | 43.6139552 | 37.0741809 | 50.3068674 | 40.9608278 | 28.1057765 | 52.9768294 |
| ENSMUSG00 | Por | 4002 | 18.3578531 | 18.3739407 | 13.1213867 | 17.8246382 | 21.7646236 | 9.92300718 |
| ENSMUSG00 | Igf1r | 4454 | 2.44566719 | 2.6289415 | 0.75422992 | 3.3700338 | 1.64970771 | 1.48191798 |
| ENSMUSG00 | Insr | 9647 | 11.1638433 | 11.8930153 | 8.09626377 | 11.2842882 | 11.1499534 | 7.37916538 |
| ENSMUSG00 | Fcer2a | 2970 | 0 | 0 | 0 | 0 | 0 | 0 |
| ENSMUSG00 | Cyp2a5 | 2148 | 0 | 0.31406433 | 0 | 0 | 0 | 0 |
| ENSMUSG00 | Atp4a | 3771 | 79.6163853 | 83.2752701 | 49.3710323 | 72.0684061 | 105.855035 | 38.8759443 |
| ENSMUSG00 | Trim28 | 3829 | 16.1451481 | 21.2931471 | 15.9306721 | 21.4098734 | 15.7250257 | 15.1937335 |
| ENSMUSG00 | Ube2m | 2676 | 16.6539214 | 18.7991949 | 20.9777058 | 19.9422062 | 12.0892086 | 27.3154085 |
| ENSMUSG00 | Adcy9 | 6730 | 1.25823438 | 1.00239256 | 0.74217062 | 0.44499401 | 0.29569519 | 1.09017503 |
| ENSMUSG00 | Mef2c | 7382 | 0.7647356 | 0.28721261 | 0.38321836 | 0.37636378 | 0.26957852 | 0.18104272 |
| ENSMUSG00 | Ctr9 | 5287 | 24.2503168 | 24.9909636 | 17.6406069 | 25.7699641 | 36.9837473 | 17.4264671 |
| ENSMUSG00 | Eif4g2 | 12073 | 94.8000106 | 81.2140913 | 103.286523 | 108.407521 | 96.0173284 | 122.556275 |
| ENSMUSG00 | Mrvi1 | 9724 | 0 | 0 | 0 | 0 | 0 | 0 |
| ENSMUSG00 | Pcyt1a | 5849 | 16.1767898 | 12.8271996 | 13.3232774 | 17.2298086 | 13.1120949 | 9.76925444 |
| ENSMUSG00 | Zfp592 | 5829 | 14.424917 | 12.4826776 | 11.5490491 | 16.1066186 | 13.2446229 | 11.0240264 |
| ENSMUSG00 | Psmd4 | 2517 | 39.2974317 | 43.7257981 | 45.0096637 | 33.6593725 | 32.9836764 | 40.3972408 |
| ENSMUSG00 | Tmod4 | 1770 | 0.31445013 | 0.08167194 | 0.17480955 | 0.36693605 | 0 | 0.0462282 |
| ENSMUSG00 | Insrr | 6185 | 315.614164 | 179.883066 | 394.629195 | 243.81778 | 283.437699 | 285.433399 |
| ENSMUSG00 | Cabp5 | 2595 | 0 | 0 | 0 | 0 | 0 | 0 |
| ENSMUSG00 | Snx6 | 1914 | 79.884837 | 70.6432375 | 70.5520535 | 64.1898441 | 60.0906223 | 68.4288494 |
| ENSMUSG00 | Mthfd2 | 2965 | 4.70629803 | 7.18327401 | 6.2613066 | 6.04815384 | 5.71357733 | 5.14216639 |
| ENSMUSG00 | Kit | 5973 | 93.2087308 | 50.2920275 | 171.049958 | 88.5650184 | 143.442805 | 119.984668 |
| ENSMUSG00 | Tomm40l | 4632 | 2.90098282 | 4.66051945 | 2.99641164 | 3.20158038 | 2.69893395 | 2.37298783 |
| ENSMUSG00 | Nr1i3 | 3574 | 0.11123526 | 0.08089498 | 0.03710283 | 0.14133985 | 0.01427711 | 0.20604792 |
| ENSMUSG00 | Apoa2 | 572 | 0.49651812 | 0.08424203 | 0 | 0.37848299 | 0.71365561 | 0.28609762 |
| ENSMUSG00 | Pan2 | 4418 | 13.11085 | 10.1542729 | 11.9058835 | 12.1525763 | 18.7335404 | 10.0998891 |
| ENSMUSG00 | Cs | 2817 | 219.113806 | 204.360463 | 242.52172 | 203.222477 | 193.182924 | 168.043535 |
| ENSMUSG00 | Ampd3 | 5623 | 26.3009759 | 30.2589947 | 20.682029 | 31.6094881 | 25.3634573 | 29.8163271 |
| ENSMUSG00 | Bcas2 | 4548 | 18.7501112 | 18.287115 | 19.2143895 | 20.8415855 | 18.9273298 | 25.5115029 |
| ENSMUSG00 | Sh2d1a | 2724 | 0 | 0 | 0 | 0 | 0 | 0 |
| ENSMUSG00 | Ctcf | 4628 | 22.7812304 | 24.0203375 | 22.225084 | 26.819843 | 19.7027082 | 24.3397278 |
| ENSMUSG00 | Pard6a | 1424 | 3.62936269 | 5.95562768 | 4.00423999 | 4.53559408 | 4.29997549 | 5.51621917 |
| ENSMUSG00 | Agrp | 654 | 0.60788197 | 0.36839788 | 1.21656449 | 0.05517132 | 0.46813189 | 0.25022605 |
| ENSMUSG00 | Pvalb | 1613 | 0.59152606 | 2.83801116 | 0.93171895 | 1.61060583 | 1.83479839 | 2.0798392 |
| ENSMUSG00 | Tfap4 | 2106 | 0.07550899 | 0.09152221 | 0 | 0.01713298 | 0.02422905 | 0.18131289 |
| ENSMUSG00 | Ranbp1 | 2049 | 30.0736919 | 26.927026 | 29.6189021 | 29.8834698 | 20.3458024 | 34.5425519 |
| ENSMUSG00 | Cd247 | 5557 | 0.11446602 | 0 | 0.02386279 | 0.03895848 | 0.00918236 | 0.02944895 |
| ENSMUSG00 | Rfx5 | 5528 | 2.91262113 | 2.2140659 | 2.81458914 | 4.96715555 | 4.3660412 | 4.24809414 |
| ENSMUSG00 | Psmb4 | 1495 | 33.4531067 | 35.487139 | 40.5651696 | 39.4851002 | 25.4961222 | 38.8047878 |
| ENSMUSG00 | Mmp8 | 2443 | 0 | 0 | 0 | 0 | 0 | 0 |
| ENSMUSG00 | Slc30a4 | 7410 | 3.66437155 | 6.46387631 | 6.01884745 | 4.50416885 | 4.1661211 | 7.68548539 |
| ENSMUSG00 | Sqrdl | 3886 | 21.7806017 | 29.8220261 | 22.7037773 | 27.5954288 | 21.8496885 | 16.6483387 |
| ENSMUSG00 | Pldn | 4082 | 19.4005187 | 15.3814145 | 17.6504175 | 11.261275 | 15.8129264 | 17.6797392 |
| ENSMUSG00 | Metap1 | 2538 | 22.1646986 | 32.7888044 | 30.8263444 | 30.1678875 | 21.5726167 | 30.7672546 |
| ENSMUSG00 | Gpr108 | 2092 | 47.0718572 | 54.4285673 | 37.4405668 | 43.1363257 | 49.1726452 | 33.1024422 |
| ENSMUSG00 | Tnfsf14 | 1855 | 0 | 0 | 0 | 0 | 0 | 0 |
| ENSMUSG00 | Gata6 | 3187 | 0 | 0 | 0 | 0 | 0 | 0.17971993 |
| ENSMUSG00 | Rsl1d1 | 3011 | 40.534482 | 40.9528746 | 39.210602 | 41.6183795 | 36.757293 | 43.7879795 |
| ENSMUSG00 | Cnga2 | 3423 | 0 | 0 | 0 | 0 | 0 | 0 |
| ENSMUSG00 | Apc | 15806 | 7.62613039 | 8.17639111 | 8.48465087 | 10.3251352 | 10.3628158 | 8.14995073 |
| ENSMUSG00 | Reep5 | 2889 | 21.9350073 | 18.6807945 | 35.9704164 | 22.7807716 | 22.3604679 | 30.5883805 |
| ENSMUSG00 | Ergic3 | 2230 | 54.445398 | 53.5452931 | 67.0361583 | 52.4403167 | 54.5730522 | 49.4857356 |
| ENSMUSG00 | Uqcc | 6351 | 22.541251 | 15.4096464 | 19.8494184 | 17.2257533 | 16.7115197 | 13.1026491 |
| ENSMUSG00 | Spo11 | 3020 | 0 | 0 | 0 | 0 | 0 | 0.55994293 |
| ENSMUSG00 | Ncoa2 | 17932 | 4.20567402 | 3.88565667 | 5.18629699 | 6.77494111 | 5.86752102 | 4.40482686 |
| ENSMUSG00 | Prl4a1 | 860 | 0 | 0 | 0 | 0 | 0 | 0 |
| ENSMUSG00 | Trh | 1381 | 0 | 0.24424699 | 0 | 0 | 0 | 0 |
| ENSMUSG00 | Nr2c2 | 9134 | 3.92592987 | 4.31536125 | 4.4182486 | 5.44746435 | 6.36852074 | 4.64928983 |
| ENSMUSG00 | Nr2c1 | 4591 | 6.43396254 | 6.17155913 | 6.98025184 | 6.138113 | 6.05736764 | 5.55473485 |
| ENSMUSG00 | Smpd4 | 5463 | 8.19415542 | 6.66830501 | 6.77227578 | 6.94825386 | 9.06014727 | 6.76498923 |
| ENSMUSG00 | Pex1 | 7883 | 4.22115149 | 4.68844363 | 3.31948383 | 2.95229211 | 3.76726509 | 3.90972249 |
| ENSMUSG00 | Otx1 | 6170 | 0 | 0 | 0.03581997 | 0 | 0 | 0 |
| ENSMUSG00 | Kctd20 | 4368 | 14.7535792 | 10.6455853 | 13.792837 | 11.093907 | 15.2331488 | 8.39219669 |
| ENSMUSG00 | Itgae | 4550 | 0 | 0.07413299 | 0 | 0 | 0.30279388 | 0.04795541 |
| ENSMUSG00 | Ctns | 4819 | 8.39823182 | 6.1095489 | 8.42944464 | 5.6455409 | 5.55900546 | 5.99940187 |
| ENSMUSG00 | P2rx5 | 2530 | 0.09428177 | 0.03809205 | 0.26206626 | 0.89848569 | 0.54455026 | 0.03234147 |
| ENSMUSG00 | Shpk | 2977 | 9.84205211 | 8.54633572 | 8.09204046 | 8.15694188 | 6.05049065 | 7.08206118 |
| ENSMUSG00 | Trpv1 | 3636 | 0 | 0.17228376 | 0 | 0 | 0.16840388 | 0 |
| ENSMUSG00 | Ephb3 | 6757 | 5.00694301 | 4.79225827 | 4.06235674 | 2.75541438 | 6.06396031 | 3.67321618 |
| ENSMUSG00 | Tuft1 | 4536 | 8.65049807 | 9.05089257 | 6.09042149 | 6.11708383 | 10.3942617 | 5.0688979 |
| ENSMUSG00 | Rcn1 | 4287 | 21.4681449 | 23.233351 | 26.3334516 | 19.9053033 | 20.2224895 | 30.341207 |
| ENSMUSG00 | Dnase1 | 2259 | 1.83026559 | 2.34639603 | 1.3501227 | 3.57785656 | 3.13973716 | 4.58803139 |
| ENSMUSG00 | Trap1 | 2787 | 111.263993 | 91.7911092 | 100.837933 | 91.2990951 | 85.5566036 | 83.507175 |
| ENSMUSG00 | Nat15 | 3756 | 21.200806 | 18.8075943 | 17.6642615 | 16.4079159 | 17.103889 | 13.7317207 |
| ENSMUSG00 | 1700037C18l | 3362 | 3.14543659 | 3.2391838 | 3.15539629 | 2.60795265 | 3.29349303 | 3.91028439 |
| ENSMUSG00 | Ankrd13d | 2653 | 3.16185691 | 5.53971536 | 2.93234993 | 6.20181391 | 4.98146677 | 1.70659258 |
| ENSMUSG00 | Tyrp1 | 2251 | 0 | 0 | 0 | 0 | 0 | 0 |
| ENSMUSG00 | Tpr | 9887 | 16.6106898 | 18.9295177 | 17.1853833 | 19.5573659 | 17.3098477 | 23.5449627 |
| ENSMUSG00 | Pdc | 1313 | 0 | 0 | 0 | 0 | 0 | 0 |
| ENSMUSG00 | BC003331 | 5645 | 20.8883534 | 25.9925095 | 20.5544443 | 26.0084746 | 22.6793936 | 31.2124304 |
| ENSMUSG00 | Prg4 | 4527 | 0.1317279 | 0.29803852 | 0.17575286 | 0.10361533 | 0.05635783 | 0.2289455 |
| ENSMUSG00 | Dhx34 | 4778 | 3.62775 | 3.29781636 | 2.82159107 | 3.82871423 | 4.23973864 | 1.55267934 |
| ENSMUSG00 | Kptn | 2933 | 8.03784519 | 6.17732774 | 9.87119235 | 8.42693505 | 8.05496488 | 6.36067277 |
| ENSMUSG00 | Napa | 3943 | 63.3587217 | 71.613632 | 79.6821456 | 71.7431471 | 62.4403407 | 54.8467194 |
| ENSMUSG00 | Sra1 | 2217 | 38.0698625 | 36.2539402 | 35.6087017 | 38.1327155 | 27.1588288 | 42.3575025 |
| ENSMUSG00 | Calcoco2 | 2535 | 0 | 0 | 0 | 0 | 0 | 0 |
| ENSMUSG00 | Atp5g1 | 5133 | 7.86125321 | 7.94189169 | 9.62744214 | 6.10857134 | 5.58675691 | 9.29877629 |
| ENSMUSG00 | Snf8 | 2785 | 7.56567493 | 7.35340679 | 10.33228 | 8.88771366 | 7.1455248 | 12.2907261 |
| ENSMUSG00 | Tbcb | 1589 | 34.326318 | 36.0261128 | 50.0991311 | 34.0610867 | 25.1760092 | 40.9720352 |
| ENSMUSG00 | Inpp5k | 5181 | 32.0591387 | 33.370576 | 31.2509846 | 35.838294 | 43.0193417 | 28.6644298 |
| ENSMUSG00 | Crkl | 3860 | 18.5903218 | 22.8948018 | 20.772576 | 17.6016813 | 20.1065072 | 15.8489678 |
| ENSMUSG00 | 2310043J07R | 1288 | 0 | 0 | 0 | 0 | 0 | 0 |
| ENSMUSG00 | Eps8l1 | 4944 | 5.17850515 | 4.30793031 | 3.49573637 | 4.43727408 | 3.62262498 | 3.53621435 |
| ENSMUSG00 | Clint1 | 4700 | 39.3748429 | 55.1170879 | 37.6373991 | 52.5185674 | 54.5439388 | 34.1223146 |
| ENSMUSG00 | Prss16 | 2683 | 0 | 0 | 0 | 0 | 0 | 0 |
| ENSMUSG00 | Cdkal1 | 12791 | 3.965913 | 3.78605069 | 4.85180108 | 4.29057851 | 4.47193865 | 3.6441528 |
| ENSMUSG00 | Rhox6 | 1006 | 0 | 0 | 0 | 0 | 0 | 0 |
| ENSMUSG00 | 5430419D17 | 3123 | 0 | 0 | 0 | 0 | 0 | 0 |
| ENSMUSG00 | Htra1 | 3408 | 0.79324316 | 0.12725293 | 0.02594005 | 0 | 0.94326927 | 0 |
| ENSMUSG00 | Zbtb17 | 3846 | 15.9187311 | 12.566563 | 19.377095 | 14.7668066 | 20.5246499 | 13.9422623 |
| ENSMUSG00 | Clcnkb | 2899 | 683.289607 | 646.635791 | 709.348859 | 598.334642 | 678.761687 | 567.893929 |
| ENSMUSG00 | Fam131c | 1426 | 1.00364467 | 0.84478335 | 2.38677553 | 1.39166371 | 0.93035468 | 1.93179423 |
| ENSMUSG00 | Fblim1 | 4441 | 1.19060547 | 0 | 1.72189121 | 0.2437427 | 0.7927989 | 1.14847051 |
| ENSMUSG00 | Hspb7 | 2766 | 6.8127613 | 5.26113721 | 8.08609276 | 6.39197463 | 5.47897094 | 6.9912217 |
| ENSMUSG00 | Epor | 1821 | 15.6969745 | 18.8141463 | 10.0006366 | 13.2558417 | 9.89143914 | 10.7540862 |
| ENSMUSG00 | Ccdc159 | 1789 | 2.28888457 | 2.07398326 | 1.80365262 | 2.09755877 | 2.48143918 | 2.62226828 |
| ENSMUSG00 | Mob1b | 8972 | 10.506046 | 10.3655632 | 15.2873739 | 13.0903985 | 9.22478618 | 13.7893183 |
| ENSMUSG00 | Atp6v1b1 | 1912 | 576.974283 | 508.351843 | 801.320231 | 559.101848 | 416.724298 | 639.955474 |
| ENSMUSG00 | Vax1 | 2272 | 0 | 0 | 0 | 0 | 0 | 0 |
| ENSMUSG00 | Atp6v1b2 | 5245 | 79.0864983 | 61.8109405 | 81.9229964 | 61.6662435 | 53.8184768 | 51.4292054 |
| ENSMUSG00 | Eps15l1 | 3156 | 8.30128696 | 7.69517327 | 9.46782189 | 10.9526612 | 8.26187517 | 11.9434574 |
| ENSMUSG00 | Tep1 | 8113 | 9.88374266 | 13.0845226 | 7.76923803 | 11.1586158 | 12.2141282 | 7.48682628 |
| ENSMUSG00 | Ttc5 | 1943 | 21.8931365 | 24.4776214 | 28.5730905 | 21.9129246 | 24.764731 | 19.1469933 |
| ENSMUSG00 | Osgep | 4876 | 11.0395653 | 10.8903731 | 12.0385659 | 10.8408932 | 9.64854768 | 8.77643745 |
| ENSMUSG00 | Aamp | 1741 | 140.913883 | 131.910731 | 162.716607 | 122.256164 | 117.586341 | 116.900263 |
| ENSMUSG00 | Tmbim1 | 5505 | 30.0206235 | 22.0843585 | 36.9352367 | 23.4385815 | 23.0151664 | 24.4307441 |
| ENSMUSG00 | Arpc2 | 1669 | 172.551648 | 168.118426 | 191.135231 | 160.71535 | 128.22326 | 230.535252 |
| ENSMUSG00 | Wbp7 | 10101 | 1.02330709 | 1.91295548 | 1.55347533 | 2.9148548 | 1.6165172 | 1.08277699 |
| ENSMUSG00 | Zbtb32 | 2873 | 0.59501763 | 0.58702592 | 0.40001668 | 0.02511803 | 0.39073452 | 0.23733588 |
| ENSMUSG00 | Etv2 | 1290 | 0 | 0 | 0 | 0 | 0 | 0 |
| ENSMUSG00 | Upk1a | 1300 | 0.5504605 | 5.78237306 | 0.88403686 | 0.41633128 | 0.11775318 | 0.16784393 |
| ENSMUSG00 | Tmem147 | 861 | 87.5913432 | 78.1840414 | 94.8209106 | 77.1091312 | 65.0126995 | 94.3366733 |
| ENSMUSG00 | Rps9 | 2549 | 10.0129535 | 10.3594234 | 13.9073669 | 9.88045001 | 6.40582199 | 12.0055493 |
| ENSMUSG00 | Tfpt | 1338 | 6.62591342 | 8.39121152 | 8.85358291 | 7.90137447 | 8.23744184 | 8.21500719 |
| ENSMUSG00 | Susd2 | 3240 | 0.23313399 | 0.22308538 | 0 | 0 | 0 | 0.39565063 |
| ENSMUSG00 | Ggt5 | 4678 | 0 | 0 | 0 | 0.00771314 | 0 | 0 |
| ENSMUSG00 | Ggt1 | 3089 | 7.14285908 | 9.88999816 | 2.14641517 | 3.27062883 | 4.06360908 | 2.74600609 |
| ENSMUSG00 | Crip2 | 1430 | 104.865505 | 94.4184692 | 77.3377701 | 67.9503118 | 85.4602588 | 82.7775765 |
| ENSMUSG00 | Crip1 | 440 | 9.84851676 | 13.2512716 | 26.7220233 | 10.966618 | 11.0170584 | 39.4242513 |
| ENSMUSG00 | Cbfa2t3 | 6273 | 0.08238821 | 0.10754188 | 0.04932455 | 0.18406272 | 0.20335715 | 0.09565472 |
| ENSMUSG00 | Fbln1 | 4556 | 0 | 0 | 0 | 0 | 0 | 0 |
| ENSMUSG00 | Pgrmc1 | 1946 | 2567.39504 | 2148.20904 | 3511.11639 | 2544.15497 | 1653.0081 | 3583.1812 |
| ENSMUSG00 | Gcat | 3840 | 10.3115778 | 12.4481642 | 12.3166594 | 9.9883368 | 6.21883955 | 10.3416341 |
| ENSMUSG00 | Tek | 5081 | 3.2783992 | 4.85563046 | 3.92344641 | 4.85733487 | 6.84904316 | 4.35878903 |
| ENSMUSG00 | Mpl | 4209 | 0 | 0 | 0 | 0 | 0 | 0 |
| ENSMUSG00 | Elovl1 | 2317 | 40.9393943 | 42.3425102 | 43.1716749 | 37.1098455 | 34.7516707 | 37.5865011 |
| ENSMUSG00 | Med8 | 2925 | 17.0710713 | 19.3075249 | 14.9304003 | 17.529089 | 14.2001607 | 23.768566 |
| ENSMUSG00 | Hyi | 1452 | 0.84877403 | 1.52656773 | 1.27856571 | 0.84489636 | 1.01912183 | 1.95355543 |
| ENSMUSG00 | Cdc20 | 1819 | 0.93979421 | 0.87419054 | 0.26730087 | 1.28935288 | 1.0098678 | 0.02998861 |
| ENSMUSG00 | Adamts4 | 5847 | 0 | 0 | 0 | 0 | 0 | 0 |
| ENSMUSG00 | Pvrl4 | 3736 | 7.77870885 | 6.86166682 | 9.16927953 | 5.58228633 | 11.1722632 | 5.56297514 |
| ENSMUSG00 | Pfdn2 | 1989 | 16.7496695 | 14.1240301 | 15.5784273 | 15.4378179 | 17.6501491 | 20.4731204 |
| ENSMUSG00 | Rnf114 | 3348 | 40.8835482 | 40.1553685 | 32.2800198 | 34.0020461 | 29.6739408 | 23.1199003 |
| ENSMUSG00 | C330007P06l | 7441 | 6.31514285 | 5.57566546 | 7.45508843 | 8.85927904 | 6.81631737 | 5.09498032 |
| ENSMUSG00 | Neurl1a | 4404 | 0.85757735 | 1.08321021 | 0.44161696 | 1.02412706 | 1.66843736 | 0.40255485 |
| ENSMUSG00 | Srm | 1591 | 11.0945527 | 7.45057496 | 10.3628457 | 7.59741352 | 9.87814189 | 6.13722241 |
| ENSMUSG00 | Epha2 | 4654 | 1.47780364 | 0.27955177 | 1.28217637 | 0.59697409 | 2.59846392 | 0.42195402 |
| ENSMUSG00 | Rbm14 | 5516 | 17.5497816 | 18.9740668 | 17.0845412 | 19.9183875 | 16.7528583 | 13.0983538 |
| ENSMUSG00 | Actn3 | 2873 | 5.09224394 | 1.49272306 | 5.35406939 | 5.43805266 | 8.79152664 | 5.58213988 |
| ENSMUSG00 | A530013C23 | 6038 | 0 | 0 | 0 | 0 | 0 | 0 |
| ENSMUSG00 | Zdhhc24 | 2939 | 33.6683875 | 28.7741429 | 44.9688706 | 29.8821696 | 36.2167472 | 38.0675639 |
| ENSMUSG00 | Bbs1 | 5564 | 1.80771685 | 5.81112557 | 3.18564685 | 4.57186222 | 7.86855328 | 2.13236306 |
| ENSMUSG00 | Slc34a3 | 2344 | 0 | 0.53449125 | 0 | 0.61573455 | 0 | 0 |
| ENSMUSG00 | Ndor1 | 5895 | 13.1843875 | 12.8497179 | 10.3625016 | 12.3395084 | 14.0744507 | 10.7525465 |
| ENSMUSG00 | Nelf | 3467 | 27.9446513 | 30.9104838 | 26.646049 | 28.2661762 | 34.3070326 | 24.3088074 |
| ENSMUSG00 | Prl7a1 | 1600 | 0 | 0 | 0 | 0 | 0 | 0 |
| ENSMUSG00 | Prl8a9 | 920 | 0 | 0 | 0 | 0 | 0 | 0 |
| ENSMUSG00 | Pdk1 | 6591 | 33.6573481 | 30.0187424 | 24.9813177 | 31.8119801 | 35.3414361 | 24.0427331 |
| ENSMUSG00 | Ptbp1 | 4717 | 27.5094103 | 22.9746255 | 27.0627353 | 21.0433972 | 29.8131634 | 15.0452854 |
| ENSMUSG00 | Mvd | 1751 | 5.17661311 | 2.86201598 | 3.78656565 | 2.74066929 | 2.85584513 | 1.61996715 |
| ENSMUSG00 | Cyba | 1060 | 4.57563078 | 4.81864421 | 4.46188415 | 4.11879942 | 4.23615196 | 6.30404396 |
| ENSMUSG00 | Itih3 | 3852 | 0.89790416 | 0.02501892 | 0.56227682 | 0.05620256 | 1.07298454 | 0.04956451 |
| ENSMUSG00 | Tmem110 | 4345 | 10.4947149 | 8.35083796 | 12.3704122 | 7.88075035 | 9.66506497 | 10.2695765 |
| ENSMUSG00 | Sfmbt1 | 8604 | 3.09116881 | 2.66022315 | 2.36832283 | 1.91229804 | 2.54420214 | 2.72302593 |
| ENSMUSG00 | Itih1 | 3271 | 2.37001489 | 4.43415441 | 2.8512959 | 3.79462651 | 2.69873525 | 1.64264871 |
| ENSMUSG00 | Ihh | 3891 | 2.20693492 | 1.96906818 | 4.40768828 | 1.0849651 | 2.59656192 | 3.22444977 |
| ENSMUSG00 | Prkag3 | 3091 | 0 | 0 | 0 | 0 | 0 | 0 |
| ENSMUSG00 | Cryba2 | 1187 | 0 | 0 | 0 | 0 | 0 | 0 |
| ENSMUSG00 | Atp7b | 6757 | 0.16474078 | 0 | 0 | 0.20291811 | 0 | 0 |
| ENSMUSG00 | Defb2 | 393 | 3.41136136 | 7.31543346 | 1.4588067 | 2.73890107 | 1.34723105 | 3.96067369 |
| ENSMUSG00 | Slc4a1 | 4866 | 122.975646 | 36.669625 | 134.122526 | 93.8756029 | 226.388908 | 107.10873 |
| ENSMUSG00 | Rundc3a | 2853 | 1.37952772 | 0.86137699 | 1.62677656 | 0.60705858 | 1.37715771 | 1.74947739 |
| ENSMUSG00 | Slc4a3 | 5432 | 7.06259917 | 8.01037505 | 5.28111121 | 7.81157665 | 11.864196 | 7.33081246 |
| ENSMUSG00 | Cdt1 | 2179 | 1.44134142 | 0.95090271 | 1.25769356 | 0.41397481 | 0.60885074 | 0 |
| ENSMUSG00 | Runx1t1 | 7944 | 0 | 0.00606577 | 0 | 0 | 0 | 0 |
| ENSMUSG00 | Snai3 | 1586 | 0 | 0 | 0 | 0.81901236 | 0.096519 | 0 |
| ENSMUSG00 | Aprt | 861 | 31.5827511 | 29.4939083 | 29.7245843 | 25.7309818 | 19.6164116 | 24.9938332 |
| ENSMUSG00 | Gtf2h1 | 3226 | 19.7298587 | 16.3708433 | 18.8810104 | 12.9295857 | 21.1318159 | 12.8341297 |
| ENSMUSG00 | Hfe | 4152 | 13.338002 | 19.7411219 | 9.1448418 | 11.4972411 | 13.8012091 | 7.52811575 |
| ENSMUSG00 | Abhd1 | 1669 | 0.28583929 | 0.34645734 | 0.47671251 | 0.34590336 | 0.18343814 | 0.75172763 |
| ENSMUSG00 | Slc5a6 | 4888 | 4.92876866 | 3.9038116 | 4.38582117 | 2.39169035 | 3.10041604 | 3.64368646 |
| ENSMUSG00 | Tcf23 | 4098 | 0 | 0 | 0 | 0 | 0 | 0 |
| ENSMUSG00 | Nphs1 | 5306 | 0 | 0 | 0.00833054 | 0 | 0 | 0 |
| ENSMUSG00 | Aplp1 | 2350 | 3.4172796 | 2.19402098 | 6.39516027 | 4.16095067 | 3.47413623 | 5.06031605 |
| ENSMUSG00 | Qrich1 | 3351 | 45.1653579 | 51.8964099 | 40.4688912 | 45.9343485 | 52.061826 | 27.299057 |
| ENSMUSG00 | P4htm | 1801 | 1.39066922 | 5.05676711 | 1.64437728 | 4.18720006 | 3.85318551 | 1.54470472 |
| ENSMUSG00 | Usp19 | 5174 | 33.7314572 | 29.0478379 | 35.2145336 | 31.5770193 | 36.4206428 | 25.5034219 |
| ENSMUSG00 | Pola1 | 6395 | 1.36144648 | 1.25834806 | 0.89655193 | 1.51211696 | 1.36442694 | 2.18794213 |
| ENSMUSG00 | Cdc42 | 4447 | 187.933373 | 175.73369 | 233.761849 | 176.629194 | 155.511687 | 197.166141 |
| ENSMUSG00 | Pknox1 | 7004 | 5.97126863 | 5.71715212 | 5.13079646 | 7.68108629 | 5.44941878 | 3.18541618 |
| ENSMUSG00 | D130043K22 | 5253 | 0 | 0.03669251 | 0 | 0 | 0 | 0 |
| ENSMUSG00 | Gmnn | 1596 | 15.0951261 | 13.4052508 | 18.2512623 | 8.83964875 | 13.6198221 | 20.2850857 |
| ENSMUSG00 | Acot13 | 718 | 50.3311028 | 48.0522181 | 67.6571385 | 40.2530887 | 37.0261028 | 93.5239023 |
| ENSMUSG00 | Zfp184 | 3405 | 1.86809894 | 1.4010155 | 1.63566291 | 2.31009859 | 3.76141566 | 2.53922486 |
| ENSMUSG00 | Cyp27b1 | 2564 | 0.17055783 | 0 | 0 | 0 | 1.57218583 | 0 |
| ENSMUSG00 | Cdk4 | 2710 | 34.41563 | 49.680044 | 47.9696016 | 35.6826123 | 34.7017013 | 34.5512312 |
| ENSMUSG00 | B4galnt1 | 2417 | 13.8329992 | 12.4602922 | 7.55290077 | 6.21023191 | 9.64793287 | 6.77070068 |
| ENSMUSG00 | Mettl1 | 2900 | 2.54983427 | 1.49544131 | 2.78928871 | 3.19761567 | 2.25219866 | 4.31691704 |
| ENSMUSG00 | Tspan31 | 1510 | 60.5547054 | 58.3662269 | 67.6493107 | 55.9830121 | 50.2829452 | 54.2060877 |
| ENSMUSG00 | Kif5b | 7579 | 38.8165399 | 42.000216 | 55.3704047 | 54.4253771 | 43.7955634 | 52.0157852 |
| ENSMUSG00 | Saal1 | 6895 | 1.64903081 | 1.70522 | 2.59633741 | 2.3862817 | 1.76131653 | 2.18355343 |
| ENSMUSG00 | Tph2 | 2866 | 0 | 0 | 0 | 0 | 0 | 0 |
| ENSMUSG00 | Krt23 | 1557 | 26.2993866 | 18.4761117 | 51.8385134 | 27.0441528 | 30.871449 | 38.170481 |
| ENSMUSG00 | Cnp | 3633 | 4.59600931 | 3.6474736 | 6.22938167 | 3.05897873 | 5.39337416 | 4.1816609 |
| ENSMUSG00 | Ttc25 | 6350 | 0.30677457 | 0.1897104 | 0.09745288 | 0.20455962 | 0.20892689 | 0.39945535 |
| ENSMUSG00 | Sulf2 | 6692 | 5.06151665 | 8.18708678 | 2.44391541 | 4.54530238 | 6.35160954 | 4.42214339 |
| ENSMUSG00 | Sod2 | 3819 | 101.975566 | 93.1049376 | 108.832137 | 98.7319626 | 73.2593922 | 94.4506688 |
| ENSMUSG00 | Tmco6 | 1805 | 6.21110554 | 3.76414867 | 3.69777191 | 5.59721466 | 4.46657473 | 3.62654483 |
| ENSMUSG00 | Stambp | 2147 | 37.9963885 | 33.3287688 | 47.7017561 | 38.8213893 | 37.3131859 | 41.2994188 |
| ENSMUSG00 | Ezh1 | 8686 | 16.8935619 | 18.9505971 | 17.8720784 | 18.4315026 | 22.8285167 | 14.4191968 |
| ENSMUSG00 | Hap1 | 4919 | 2.32762317 | 5.67187437 | 1.4736943 | 3.53558558 | 6.65967336 | 2.98307703 |
| ENSMUSG00 | Leprel4 | 2027 | 8.02170278 | 8.20144181 | 7.04351026 | 3.47113897 | 8.55893822 | 9.01529763 |
| ENSMUSG00 | Ctnnb1 | 4606 | 190.646777 | 145.96119 | 201.096314 | 164.374629 | 181.893631 | 148.832128 |
| ENSMUSG00 | Eif1b | 913 | 56.4327522 | 49.6114519 | 53.9814403 | 59.59663 | 34.3715456 | 58.1938634 |
| ENSMUSG00 | Klk4 | 1243 | 0 | 0 | 0 | 0 | 0 | 0 |
| ENSMUSG00 | Chrd | 4921 | 0 | 0.009792 | 0 | 0 | 0 | 0 |
| ENSMUSG00 | Psmd2 | 3755 | 45.1338518 | 46.4027097 | 51.8886083 | 50.5340805 | 45.9441216 | 37.8866893 |
| ENSMUSG00 | Syngr3 | 2066 | 0 | 0 | 0.32092335 | 0 | 0.44456668 | 0 |
| ENSMUSG00 | Vars | 5706 | 12.0561907 | 16.9175946 | 14.2331793 | 15.4239994 | 17.2531369 | 10.4071766 |
| ENSMUSG00 | D17H6S56E- | 4425 | 0.03593716 | 0.07622714 | 0.03995647 | 0 | 0.02306277 | 0.11094769 |
| ENSMUSG00 | Hspa1l | 2469 | 0.08050928 | 0.3317819 | 0.12531912 | 0.21921048 | 0.51667047 | 0.61862284 |
| ENSMUSG00 | Slc44a4 | 2780 | 31.5326743 | 23.1225589 | 24.1679142 | 16.1330868 | 37.774202 | 14.5693667 |
| ENSMUSG00 | Msh5 | 5510 | 0.00721515 | 0.26235812 | 0.05615479 | 0.15716317 | 0.07408548 | 0.26730137 |
| ENSMUSG00 | Abhd16a | 3344 | 43.2507291 | 44.1228726 | 37.658807 | 28.1729122 | 42.7864706 | 30.1456539 |
| ENSMUSG00 | Neu1 | 3961 | 28.9659977 | 29.8899491 | 28.7574222 | 28.4119911 | 28.7144135 | 22.0483198 |
| ENSMUSG00 | Ddah2 | 1757 | 24.9348546 | 19.417189 | 28.2771039 | 26.1630761 | 16.6118879 | 19.3576979 |
| ENSMUSG00 | Clic1 | 1793 | 55.5423752 | 69.3906267 | 63.0856198 | 55.2600637 | 49.005811 | 60.4057961 |
| ENSMUSG00 | Ly6g6c | 594 | 0 | 0 | 0 | 0 | 0 | 0 |
| ENSMUSG00 | Lsm2 | 1805 | 2.73112442 | 4.93877682 | 4.26100869 | 2.99850786 | 5.31465853 | 5.04694156 |
| ENSMUSG00 | Pole | 9649 | 0 | 0.36455698 | 0.11452441 | 0.00373946 | 0.07932383 | 0.13568066 |
| ENSMUSG00 | Atp1a2 | 6946 | 0 | 0 | 0.01909092 | 0 | 0.01469231 | 0 |
| ENSMUSG00 | Atp1a4 | 3447 | 0 | 0.29356405 | 0 | 0 | 0.01480313 | 0 |
| ENSMUSG00 | Casq1 | 2575 | 0 | 0.07485273 | 0 | 0.44839822 | 0 | 0 |
| ENSMUSG00 | Stx1a | 2705 | 0.55848734 | 0 | 0 | 0.68028993 | 0.3772745 | 0.06049828 |
| ENSMUSG00 | Ceacam9 | 1371 | 0 | 0 | 0 | 0 | 0 | 0 |
| ENSMUSG00 | Zfp775 | 4081 | 4.52010364 | 1.72389624 | 2.7836005 | 2.48445345 | 2.30062562 | 2.47282934 |
| ENSMUSG00 | Scube2 | 4843 | 0 | 0 | 0 | 0 | 0 | 0 |
| ENSMUSG00 | Mrpl49 | 2025 | 40.2266566 | 40.619386 | 52.9767274 | 36.6878665 | 38.9060367 | 37.4167644 |
| ENSMUSG00 | Dennd2c | 6916 | 0.1667017 | 0.97543405 | 0.70303683 | 0.51128403 | 0.00737802 | 0 |
| ENSMUSG00 | Mark3 | 3397 | 68.0185024 | 54.3711018 | 55.366159 | 62.6257682 | 70.7339428 | 50.3019178 |
| ENSMUSG00 | Gatad1 | 4078 | 4.2309658 | 3.95842524 | 7.20800039 | 2.8667441 | 3.70372909 | 5.75857391 |
| ENSMUSG00 | Pcdha4 | 24675 | 0.04350144 | 0 | 0.0232877 | 0 | 0.05790227 | 0 |
| ENSMUSG00 | 2310003L06l | 1600 | 0 | 0 | 0 | 0 | 0 | 0 |
| ENSMUSG00 | M6pr | 4330 | 63.7373085 | 53.6616684 | 71.6008607 | 64.214373 | 60.3949598 | 58.2782823 |
| ENSMUSG00 | Lrrc8a | 5762 | 4.73312385 | 4.35701776 | 4.38029371 | 4.60262109 | 4.66693857 | 4.24598254 |
| ENSMUSG00 | Mc5r | 1301 | 0 | 0 | 0 | 0 | 0 | 0 |
| ENSMUSG00 | Ppp2r1a | 3075 | 115.206858 | 98.6607609 | 106.573159 | 94.3882817 | 106.118268 | 75.5263588 |
| ENSMUSG00 | Fance | 3084 | 7.97945604 | 10.156027 | 7.99760973 | 7.30064714 | 10.5064036 | 6.53565448 |
| ENSMUSG00 | Tinf2 | 2786 | 10.4739852 | 6.14005275 | 10.1699144 | 7.61530588 | 15.7511426 | 8.46825662 |
| ENSMUSG00 | Tssk4 | 1685 | 0.25953133 | 0.14298648 | 0.10493019 | 0.02141368 | 0.03028272 | 1.08451088 |
| ENSMUSG00 | Hapln4 | 3571 | 0 | 0 | 0 | 0 | 0 | 0 |
| ENSMUSG00 | Dus3l | 2290 | 36.4743513 | 33.2254988 | 24.5136859 | 29.1177372 | 34.5597855 | 28.4418509 |
| ENSMUSG00 | Gtpbp3 | 3919 | 3.30703921 | 3.54113175 | 2.33472353 | 1.2245246 | 2.26552421 | 2.67248315 |
| ENSMUSG00 | Tgfbr1 | 6610 | 25.6756651 | 30.5958393 | 20.6498171 | 42.6270326 | 33.7731307 | 27.8234331 |
| ENSMUSG00 | Homer1 | 12527 | 1.58679175 | 0.79624759 | 2.01831677 | 1.29903425 | 1.20162696 | 1.33248816 |
| ENSMUSG00 | Rad51c | 5269 | 0.30180665 | 0.42982782 | 0.40267384 | 0.71219067 | 0.37768621 | 0.11388158 |
| ENSMUSG00 | Gabrb2 | 10181 | 0 | 0.00473298 | 0.02604961 | 0.06733708 | 0 | 0.01607385 |
| ENSMUSG00 | Cav1 | 2380 | 255.988546 | 416.488782 | 131.881213 | 365.398923 | 298.740135 | 246.193729 |
| ENSMUSG00 | Arpp19 | 1879 | 94.765725 | 94.4495298 | 105.246964 | 100.4498 | 59.9335878 | 114.178985 |
| ENSMUSG00 | Bcl2l1 | 3376 | 53.4861945 | 45.3032486 | 57.8447104 | 51.1304802 | 49.8021056 | 58.3544785 |
| ENSMUSG00 | Khsrp | 4574 | 18.686988 | 23.2609673 | 13.8770981 | 5.67183866 | 6.53726634 | 12.6057253 |
| ENSMUSG00 | Dio2 | 5811 | 0 | 0 | 0 | 0 | 0 | 0 |
| ENSMUSG00 | Ccdc124 | 906 | 27.1179768 | 37.1237711 | 38.8839612 | 34.6084952 | 25.4005469 | 37.4499462 |
| ENSMUSG00 | Cct4 | 4377 | 52.7439057 | 56.883561 | 52.0687006 | 58.5951959 | 42.656045 | 57.7646575 |
| ENSMUSG00 | 0610009B22l | 995 | 51.3025497 | 48.8160138 | 54.2416586 | 48.5566408 | 32.7184199 | 72.3120586 |
| ENSMUSG00 | Cpt1c | 2737 | 0.18882764 | 0.88027845 | 1.13048192 | 0 | 1.84567454 | 0.47832762 |
| ENSMUSG00 | Twist2 | 1327 | 0 | 0 | 0 | 0 | 0 | 0 |
| ENSMUSG00 | Zfp655 | 4583 | 32.0871749 | 21.5855915 | 28.4616275 | 38.821855 | 29.5046685 | 35.1422092 |
| ENSMUSG00 | Rhoa | 2120 | 60.2895612 | 55.7780797 | 63.905023 | 56.6079623 | 37.9568843 | 51.7832184 |
| ENSMUSG00 | Zmiz1 | 10261 | 10.7631542 | 8.22752623 | 8.86535356 | 11.1927831 | 11.2087956 | 8.87269719 |
| ENSMUSG00 | Ankrd26 | 5670 | 2.08944148 | 3.69684345 | 1.42662033 | 3.86911518 | 3.26676798 | 2.27529178 |
| ENSMUSG00 | Aldh16a1 | 6665 | 6.13182805 | 7.81538544 | 6.31360158 | 6.71835218 | 6.91325092 | 5.3362535 |
| ENSMUSG00 | Hnrnpa0 | 2677 | 4.39582452 | 5.25604823 | 5.0030476 | 4.78488076 | 2.95445956 | 6.62253101 |
| ENSMUSG00 | Prrg2 | 2152 | 7.64812683 | 7.0980958 | 7.39436036 | 6.65639948 | 6.94736437 | 5.89345133 |
| ENSMUSG00 | Hnrnph1 | 3739 | 78.5114919 | 68.7420386 | 79.3126945 | 88.3860513 | 73.4757977 | 85.0116731 |
| ENSMUSG00 | 1700019E19l | 854 | 29.5140218 | 23.1340062 | 34.6782609 | 18.886035 | 16.5507097 | 29.8935156 |
| ENSMUSG00 | Id3 | 1786 | 104.753243 | 136.222478 | 124.636328 | 115.29688 | 133.708532 | 120.06339 |
| ENSMUSG00 | Tcap | 946 | 0 | 0.10187409 | 0 | 0 | 0.21575635 | 0.89377782 |
| ENSMUSG00 | Arid1a | 13569 | 5.39683065 | 7.24447947 | 6.04277536 | 6.50162863 | 5.13685823 | 6.60508988 |
| ENSMUSG00 | Crlf1 | 1674 | 0.21373914 | 0.48934858 | 0.23764432 | 0.38797897 | 0.15240853 | 0.35844807 |
| ENSMUSG00 | Ctsd | 2509 | 981.607022 | 1311.61613 | 970.396938 | 1126.06467 | 857.015336 | 609.7931 |
| ENSMUSG00 | Rplp1 | 492 | 21.5746205 | 21.5467831 | 42.944067 | 16.794285 | 17.2162162 | 44.6261882 |
| ENSMUSG00 | 4931407G18l | 1490 | 0 | 0 | 0 | 0 | 0 | 0 |
| ENSMUSG00 | Hmgcll1 | 6675 | 1.00058738 | 1.45822641 | 1.55616975 | 0.16216649 | 0.76444009 | 0.33914533 |
| ENSMUSG00 | Ttc9b | 911 | 0 | 0 | 0 | 0 | 0 | 0 |
| ENSMUSG00 | Phox2a | 2046 | 0 | 0 | 0 | 0 | 0 | 0.15996856 |
| ENSMUSG00 | Abhd8 | 1980 | 7.44913296 | 7.52000537 | 4.33088765 | 3.06150681 | 5.82422269 | 3.05806561 |
| ENSMUSG00 | Rabl5 | 906 | 10.2240916 | 11.4881584 | 15.5633421 | 12.1866508 | 12.2215492 | 7.61642796 |
| ENSMUSG00 | Fzd3 | 4821 | 3.77681188 | 2.7386611 | 2.06293605 | 4.10142302 | 4.20192309 | 2.3987652 |
| ENSMUSG00 | 1700008O03l | 3999 | 0.16900304 | 0.4217368 | 0.28738383 | 0.0541366 | 0.42107287 | 0.35465898 |
| ENSMUSG00 | Mid1ip1 | 1968 | 21.5746205 | 21.1305384 | 17.5190232 | 14.3924822 | 15.1419733 | 14.2471185 |
| ENSMUSG00 | Ap2s1 | 1404 | 14.2428823 | 18.7048511 | 21.5971968 | 15.6252728 | 12.8292811 | 19.0378536 |
| ENSMUSG00 | Fgfrl1 | 2649 | 5.64290701 | 3.92913232 | 5.58988955 | 7.13740707 | 7.95541458 | 4.29351626 |
| ENSMUSG00 | 4930432K21l | 2396 | 2.05746226 | 0.62344729 | 0.20293 | 0 | 0.59630155 | 0.07968384 |
| ENSMUSG00 | Fhl2 | 1557 | 1.73627018 | 3.43525695 | 1.16395348 | 1.34409671 | 2.7856403 | 0.52552291 |
| ENSMUSG00 | 2310044H10l | 3942 | 11.5978698 | 11.3681865 | 12.9622858 | 9.4552897 | 8.28434311 | 12.3780643 |
| ENSMUSG00 | Clstn3 | 5134 | 0.10066639 | 0.26280101 | 0 | 0.06325251 | 0.34786193 | 0.04781296 |
| ENSMUSG00 | Fbxw9 | 3632 | 6.10780785 | 8.39813268 | 6.69356103 | 6.7951868 | 6.44854254 | 6.11276331 |
| ENSMUSG00 | Spib | 2834 | 0 | 0 | 0 | 0 | 0 | 0 |
| ENSMUSG00 | Fnbp4 | 6549 | 11.5824487 | 10.0875878 | 9.92162303 | 16.1264536 | 14.6557662 | 16.2256825 |
| ENSMUSG00 | Lass4 | 5747 | 9.49095515 | 15.7630957 | 10.3370936 | 12.5317141 | 17.6155089 | 13.008489 |
| ENSMUSG00 | Scrn3 | 3362 | 24.5485953 | 21.4416768 | 16.2239959 | 25.5214461 | 24.1168683 | 15.8277279 |
| ENSMUSG00 | Phax | 2259 | 21.6992065 | 21.3095421 | 23.7347656 | 23.9588609 | 19.4483 | 25.5480906 |
| ENSMUSG00 | Tle1 | 9188 | 10.6744417 | 5.35987635 | 10.146026 | 7.41433392 | 8.16377584 | 7.02051827 |
| ENSMUSG00 | 1700109H08l | 1011 | 0.58984392 | 0.23831079 | 1.18046465 | 0.46396299 | 0.55518312 | 0.70142495 |
| ENSMUSG00 | Relt | 4013 | 0 | 0 | 0 | 0 | 0 | 0 |
| ENSMUSG00 | Snrpb2 | 2886 | 17.5772672 | 9.40019644 | 12.4671865 | 14.2277778 | 11.7929982 | 24.949774 |
| ENSMUSG00 | Ubc | 4948 | 124.6818 | 116.278521 | 93.7457843 | 123.268165 | 201.578935 | 81.1844961 |
| ENSMUSG00 | Prpf31 | 5489 | 9.37212457 | 10.2184403 | 8.40712773 | 9.80769 | 11.7316973 | 9.83854722 |
| ENSMUSG00 | Sertad1 | 1341 | 10.3465046 | 13.2952898 | 15.7887269 | 10.0631504 | 8.44731949 | 13.8508794 |
| ENSMUSG00 | Carhsp1 | 2845 | 14.0436759 | 13.9901586 | 23.6312841 | 12.872856 | 12.6624328 | 12.7026 |
| ENSMUSG00 | Elk3 | 5202 | 2.89644889 | 4.26100796 | 2.24323079 | 3.89120088 | 2.91327059 | 6.32318626 |
| ENSMUSG00 | Herpud2 | 2736 | 38.6366311 | 37.4783439 | 37.5618732 | 41.7264579 | 40.3026309 | 34.0932991 |
| ENSMUSG00 | Rdh13 | 6586 | 4.35221704 | 4.47037901 | 5.36918834 | 3.51178113 | 5.45438333 | 4.48917536 |
| ENSMUSG00 | Adam21 | 2909 | 0.15033011 | 0.26503371 | 0 | 0 | 0 | 0.71257222 |
| ENSMUSG00 | Nutf2 | 1961 | 10.278444 | 11.9421779 | 9.2415888 | 11.16869 | 9.21128865 | 8.27557897 |
| ENSMUSG00 | Fut1 | 2578 | 0 | 0 | 0 | 0 | 0 | 0 |
| ENSMUSG00 | Arpc5 | 2272 | 87.1750897 | 112.088621 | 116.535669 | 102.084235 | 92.0585891 | 94.9330313 |
| ENSMUSG00 | Rnf151 | 1185 | 0 | 0 | 0.1119034 | 0 | 0 | 0.3452486 |
| ENSMUSG00 | Elavl2 | 11068 | 0 | 0 | 0 | 0 | 0 | 0 |
| ENSMUSG00 | Pou2f2 | 3568 | 0 | 0 | 0 | 0 | 0 | 0 |
| ENSMUSG00 | Mgst1 | 2084 | 77.1263953 | 169.623675 | 57.8399357 | 118.738321 | 78.1310774 | 65.0063019 |
| ENSMUSG00 | Nfib | 20790 | 4.17250883 | 5.49775088 | 4.07575436 | 6.31913055 | 6.26110076 | 5.55856886 |
| ENSMUSG00 | Htr3b | 2402 | 0 | 0 | 0 | 0 | 0 | 0 |
| ENSMUSG00 | Rab25 | 1728 | 36.626577 | 36.8090878 | 33.6885574 | 27.5000305 | 22.4716852 | 24.5124507 |
| ENSMUSG00 | Ubqln4 | 3361 | 26.3538264 | 24.5735084 | 29.6827015 | 25.4109489 | 30.6522621 | 30.4476187 |
| ENSMUSG00 | Rbfox1 | 4213 | 1.92502208 | 1.63557114 | 2.33966556 | 1.81566424 | 0.75092222 | 2.64136074 |
| ENSMUSG00 | Rps18 | 2514 | 2.40367265 | 1.93589127 | 2.05712635 | 1.57876885 | 1.33959459 | 3.40661763 |
| ENSMUSG00 | Rpl10 | 1731 | 3.74358366 | 2.81157173 | 3.75370937 | 4.46074958 | 4.21535052 | 2.61559221 |
| ENSMUSG00 | Rps15a | 9620 | 0.57029691 | 0.57603335 | 0.74435536 | 0.33381517 | 0.48798613 | 0.77968044 |
| ENSMUSG00 | Zfp955a | 1634 | 13.6978798 | 13.8897272 | 12.6329625 | 12.5205136 | 14.3648304 | 19.3793489 |
| ENSMUSG00 | Ncaph2 | 4139 | 25.0116625 | 25.6590767 | 24.7547408 | 23.0405518 | 26.5672481 | 17.429674 |
| ENSMUSG00 | Hipk1 | 13193 | 7.09047568 | 7.52764779 | 9.94399076 | 9.47929708 | 7.05852618 | 8.77799729 |
| ENSMUSG00 | Gprc5b | 4521 | 1.55645212 | 5.54234681 | 1.54476691 | 3.85481697 | 2.78777147 | 1.70127146 |
| ENSMUSG00 | Man1a2 | 13145 | 10.987574 | 10.0148619 | 12.6031577 | 14.314786 | 10.7875465 | 13.1964021 |
| ENSMUSG00 | Ceacam5 | 3244 | 0 | 0 | 0 | 0 | 0 | 0 |
| ENSMUSG00 | Tppp2 | 737 | 0 | 0 | 0 | 0 | 0 | 0 |
| ENSMUSG00 | Acyp1 | 1793 | 8.46993506 | 10.0780345 | 11.6852613 | 11.168731 | 5.37868658 | 13.4319612 |
| ENSMUSG00 | Cldn13 | 1065 | 0 | 0 | 0 | 0 | 0 | 0 |
| ENSMUSG00 | Cd163 | 4475 | 0.15102641 | 0.08614336 | 0.02963252 | 0.17738659 | 0 | 0 |
| ENSMUSG00 | Hdac5 | 7756 | 10.2464164 | 8.7538289 | 7.38597067 | 5.52209733 | 10.5855388 | 6.77646078 |
| ENSMUSG00 | Rala | 2679 | 59.3290075 | 54.4638099 | 81.7544353 | 59.3016954 | 49.3121639 | 106.349713 |
| ENSMUSG00 | Clec3a | 588 | 0 | 0 | 0 | 0 | 0 | 0 |
| ENSMUSG00 | Vdac3 | 1330 | 67.9729043 | 59.3091773 | 61.8493458 | 47.069434 | 45.19479 | 73.7030477 |
| ENSMUSG00 | Slc1a7 | 3349 | 0.01187085 | 0.24460123 | 0 | 0 | 0 | 0 |
| ENSMUSG00 | Vps72 | 3650 | 15.1070826 | 15.6440915 | 15.7552323 | 13.4146122 | 15.2100539 | 14.2425924 |
| ENSMUSG00 | Gabpa | 4457 | 16.7870349 | 15.6873519 | 15.5405627 | 16.8145404 | 19.2107379 | 9.93807811 |
| ENSMUSG00 | Bmp7 | 4012 | 46.3747878 | 37.9414184 | 47.2646826 | 37.7728283 | 40.101237 | 32.2033814 |
| ENSMUSG00 | Dynll1 | 3324 | 27.0657799 | 27.4709109 | 37.4997585 | 30.5134257 | 13.9846656 | 40.4606637 |
| ENSMUSG00 | Pdcl | 5540 | 20.3370093 | 19.3267643 | 25.8429187 | 13.3451461 | 18.0434423 | 26.4376918 |
| ENSMUSG00 | Tmem184b | 3352 | 12.4769587 | 13.0960169 | 7.41092953 | 10.8719764 | 15.5423418 | 5.86665122 |
| ENSMUSG00 | Gm5965 | 586 | 0 | 0 | 0 | 0 | 0 | 0 |
| ENSMUSG00 | Rtdr1 | 3004 | 0 | 0 | 0 | 0 | 0 | 0.06355608 |
| ENSMUSG00 | Nf2 | 9019 | 10.980253 | 9.87343886 | 8.84624977 | 9.50159172 | 12.1017206 | 8.72159437 |
| ENSMUSG00 | Cabp7 | 2623 | 0 | 0 | 0.03370327 | 0.09629215 | 0.03890688 | 0.25995653 |
| ENSMUSG00 | Zmat5 | 1255 | 11.5940286 | 8.10146557 | 15.9197076 | 11.1839963 | 7.92840102 | 17.1906292 |
| ENSMUSG00 | Ewsr1 | 6125 | 44.740331 | 40.2169947 | 35.9676711 | 45.7785417 | 48.410493 | 49.5708219 |
| ENSMUSG00 | Ap1b1 | 5027 | 49.5460683 | 42.013363 | 46.0131777 | 41.5083476 | 50.5087022 | 32.358454 |
| ENSMUSG00 | Derl3 | 1329 | 1.82474365 | 1.08773007 | 1.92904958 | 0.10859908 | 0.84468041 | 0.41045356 |
| ENSMUSG00 | Gstt4 | 1266 | 0 | 0 | 0 | 0 | 0 | 0 |
| ENSMUSG00 | Tbx1 | 1803 | 0 | 0 | 0 | 0 | 0 | 0 |
| ENSMUSG00 | Gnat2 | 3178 | 0 | 0 | 0 | 0 | 0 | 0 |
| ENSMUSG00 | Bcl2l13 | 6926 | 7.7031266 | 5.64239165 | 7.40952061 | 6.93925546 | 7.34526374 | 4.26880002 |
| ENSMUSG00 | 2610028H24 | 1887 | 1.62224271 | 0.30643207 | 1.31176641 | 0.78397659 | 1.1627632 | 2.26927311 |
| ENSMUSG00 | 1700027D21l | 1586 | 0.05013302 | 0 | 0.02787002 | 0 | 0 | 0.08598562 |
| ENSMUSG00 | Dqx1 | 3137 | 0.31682723 | 0.55298435 | 0.32408109 | 0.40257302 | 1.28501233 | 0.24344594 |
| ENSMUSG00 | Ccl8 | 508 | 0 | 0 | 0 | 0 | 0 | 0 |
| ENSMUSG00 | Lnp | 6881 | 3.10833433 | 3.41737884 | 3.87994669 | 5.34335175 | 4.93133846 | 3.72593531 |
| ENSMUSG00 | 2310007L24l | 1269 | 0 | 0 | 0 | 0 | 0 | 0 |
| ENSMUSG00 | Tmem8c | 2204 | 0 | 0 | 0 | 0.03274233 | 0 | 0 |
| ENSMUSG00 | Fam163b | 2956 | 0 | 0 | 0 | 0 | 0 | 0 |
| ENSMUSG00 | Trpm5 | 6371 | 0 | 0.01512681 | 0 | 0 | 0.02402749 | 0.03852955 |
| ENSMUSG00 | Ascl2 | 2119 | 0 | 0.50028397 | 0.25031719 | 0.06811146 | 0.14448242 | 0 |
| ENSMUSG00 | Rarres2 | 702 | 0.33979043 | 0.27456662 | 0.06296559 | 0 | 0 | 0.07770553 |
| ENSMUSG00 | Pttg1ip | 3588 | 150.855872 | 121.755375 | 184.66712 | 123.63229 | 135.544144 | 155.72694 |
| ENSMUSG00 | Trpm2 | 12123 | 0.64603066 | 0.15899181 | 0.41930314 | 0.33930158 | 0.28621575 | 0.49946135 |
| ENSMUSG00 | Ube2g2 | 2521 | 17.5201661 | 17.986292 | 11.8175495 | 12.280204 | 22.2848234 | 9.55315607 |
| ENSMUSG00 | Mpo | 3689 | 0 | 0 | 0 | 0 | 0 | 0 |
| ENSMUSG00 | Lpo | 3121 | 0 | 2.45486841 | 0 | 0 | 0.24524051 | 0.41073632 |
| ENSMUSG00 | Met | 7964 | 59.8129291 | 43.8724124 | 70.2267604 | 60.0355565 | 82.8313645 | 75.6045877 |
| ENSMUSG00 | Slc16a12 | 4044 | 5.26927239 | 2.69291195 | 3.81464966 | 3.94368538 | 7.96183065 | 6.40050263 |
| ENSMUSG00 | Syn2 | 5683 | 70.6336595 | 41.3777649 | 108.455129 | 45.6184216 | 50.0117743 | 60.3132837 |
| ENSMUSG00 | Elk1 | 5375 | 1.03549159 | 1.45231695 | 1.2582106 | 0.80555262 | 1.61385747 | 1.1924726 |
| ENSMUSG00 | Nav1 | 12737 | 4.30421666 | 3.88911537 | 6.91640678 | 6.22094448 | 3.74575343 | 7.84169969 |
| ENSMUSG00 | Tnpo1 | 11095 | 9.28045917 | 9.07703146 | 13.4537741 | 15.2068175 | 10.1638838 | 10.7967747 |
| ENSMUSG00 | Myod1 | 1851 | 0 | 0 | 0 | 0 | 0 | 0 |
| ENSMUSG00 | Otog | 9988 | 0.00398032 | 0 | 0 | 0 | 0.00510877 | 0.00273074 |
| ENSMUSG00 | Rnmt | 7359 | 6.0721783 | 5.55921856 | 7.08767153 | 5.68761675 | 6.28208948 | 5.50385097 |
| ENSMUSG00 | Kcnq1 | 3260 | 91.3888254 | 82.0795439 | 88.4308037 | 79.4026342 | 93.1935711 | 75.0136244 |
| ENSMUSG00 | Srp14 | 1608 | 73.8492663 | 70.3917616 | 75.4839931 | 48.9396388 | 46.044323 | 87.2686685 |
| ENSMUSG00 | 6330409D20 | 1846 | 0 | 0 | 0 | 0 | 0 | 0 |
| ENSMUSG00 | Cdk9 | 5316 | 10.3277499 | 12.7898928 | 10.7012363 | 10.0250526 | 10.3953283 | 9.74314145 |
| ENSMUSG00 | Tor2a | 1909 | 14.452752 | 11.2073234 | 10.5121198 | 9.4505093 | 13.5250635 | 10.244063 |
| ENSMUSG00 | Fpgs | 3786 | 11.0571899 | 11.0220441 | 8.96645944 | 8.28190615 | 8.66612775 | 8.65209766 |
| ENSMUSG00 | Mkl2 | 9018 | 3.56644308 | 2.11062815 | 1.90178699 | 1.97655023 | 3.58735 | 2.23508077 |
| ENSMUSG00 | Cbx5 | 12103 | 12.1667603 | 18.3660297 | 12.4903167 | 21.9062103 | 17.8379407 | 15.3669144 |
| ENSMUSG00 | Odam | 1364 | 0.02914625 | 0 | 0 | 0 | 0 | 0.31993712 |
| ENSMUSG00 | Apobec3 | 6192 | 4.1540368 | 2.80153733 | 2.77689227 | 4.62679964 | 9.08948523 | 4.88494427 |
| ENSMUSG00 | St6galnac1 | 2642 | 0 | 0 | 0 | 0 | 0.01931354 | 0.07226437 |
| ENSMUSG00 | Taf7l | 2338 | 0.25506082 | 0 | 0 | 0 | 0 | 0 |
| ENSMUSG00 | Sardh | 5435 | 8.79228842 | 10.6036771 | 5.73363373 | 6.77824608 | 11.6041583 | 8.47094592 |
| ENSMUSG00 | Vav2 | 6883 | 3.96803939 | 3.1503558 | 3.28800576 | 2.45334837 | 4.18856637 | 2.75401341 |
| ENSMUSG00 | Tex15 | 5814 | 0.06837888 | 1.64102435 | 1.22402764 | 0.01241212 | 0.41249392 | 1.74981776 |
| ENSMUSG00 | Ppp2cb | 1819 | 51.2734235 | 54.6236634 | 69.7655257 | 60.2822063 | 40.2264007 | 86.4421634 |
| ENSMUSG00 | G0s2 | 840 | 4.82745121 | 4.01553685 | 3.84135064 | 3.47934001 | 3.21952133 | 9.80588221 |
| ENSMUSG00 | Dmap1 | 2736 | 12.0893859 | 12.6101947 | 13.8130759 | 11.935033 | 15.2183928 | 12.0323427 |
| ENSMUSG00 | Pla2g12b | 1178 | 0 | 0 | 0 | 0 | 0 | 0 |
| ENSMUSG00 | Ccdc109a | 7183 | 5.58447443 | 8.49954367 | 4.46756759 | 5.32465087 | 5.84640224 | 4.34009643 |
| ENSMUSG00 | Oit3 | 2779 | 0 | 0 | 0 | 0 | 0 | 0 |
| ENSMUSG00 | Tex11 | 3593 | 0 | 0 | 0 | 0 | 0 | 0 |
| ENSMUSG00 | Bcr | 7332 | 1.44230194 | 1.84675262 | 0.88017854 | 2.14563167 | 1.70505484 | 1.59213661 |
| ENSMUSG00 | Fxyd5 | 2277 | 0.29681299 | 0.52905624 | 0.03882463 | 0 | 0.8291506 | 0.53902449 |
| ENSMUSG00 | Kcnd1 | 6030 | 0.04615064 | 0.19277795 | 0.01466064 | 0 | 0.18616588 | 0.28948208 |
| ENSMUSG00 | Tfcp2 | 3332 | 7.80314653 | 6.10284471 | 9.97592615 | 13.0055629 | 13.0628747 | 7.30161411 |
| ENSMUSG00 | Pou6f2 | 7404 | 0 | 0 | 0 | 0 | 0 | 0 |
| ENSMUSG00 | Pou6f1 | 10168 | 1.32153348 | 1.4785769 | 1.06070512 | 1.31652621 | 1.68615876 | 1.70332375 |
| ENSMUSG00 | Ubp1 | 3451 | 25.2633349 | 23.9605722 | 29.7667584 | 24.4241252 | 24.8995702 | 24.318767 |
| ENSMUSG00 | Nuak2 | 4218 | 10.1509371 | 38.3618001 | 5.36542049 | 24.328434 | 35.3240914 | 7.64957284 |
| ENSMUSG00 | Ick | 11726 | 1.11204141 | 1.91496521 | 1.38719753 | 2.36628793 | 1.8276546 | 1.60493784 |
| ENSMUSG00 | Sdhb | 2306 | 204.173528 | 188.838195 | 257.409605 | 172.930944 | 139.422746 | 248.866299 |
| ENSMUSG00 | Cox4i2 | 783 | 0 | 0 | 0 | 0 | 0.06516779 | 0.06966702 |
| ENSMUSG00 | Snap47 | 2731 | 39.0712961 | 43.5283606 | 37.7601244 | 34.5230254 | 27.7459422 | 28.7726971 |
| ENSMUSG00 | Wnt3a | 2757 | 0 | 0 | 0 | 0 | 0 | 0 |
| ENSMUSG00 | Kdsr | 5572 | 7.6557126 | 11.8563554 | 7.29028961 | 9.22773034 | 12.1613473 | 7.8172288 |
| ENSMUSG00 | Vps4b | 9167 | 21.3110539 | 18.8761333 | 21.6452573 | 22.0971526 | 18.5358167 | 25.8107882 |
| ENSMUSG00 | Rps25 | 678 | 4.45636655 | 3.62464388 | 5.34594562 | 3.93815825 | 4.3650882 | 9.45359915 |
| ENSMUSG00 | Nxf2 | 2805 | 0 | 0 | 0 | 0 | 0 | 0 |
| ENSMUSG00 | Taz | 3381 | 50.0324074 | 50.0392187 | 54.2292945 | 49.4433932 | 53.5014795 | 55.9045401 |
| ENSMUSG00 | Kif19a | 4211 | 0.48148409 | 0.21741686 | 0.34639296 | 0 | 0.0363522 | 0.16192495 |

| | | | | | | | | |
|---|---|---|---|---|---|---|---|---|
| ENSMUSG00 | Aldh3a2 | 5606 | 30.3094762 | 23.4313666 | 38.1305945 | 23.3316824 | 24.6939989 | 21.0957609 |
| ENSMUSG00 | Zmynd10 | 1743 | 0 | 0.58055954 | 0.0760789 | 0.08280446 | 0.32202532 | 0.23472151 |
| ENSMUSG00 | Tmem115 | 2175 | 20.0331065 | 26.8957889 | 22.1720509 | 22.5782357 | 21.0439812 | 15.8506409 |
| ENSMUSG00 | Hyal2 | 1889 | 15.4055118 | 16.9634643 | 20.2172538 | 17.0000104 | 18.3143901 | 13.4568363 |
| ENSMUSG00 | Ifrd2 | 1968 | 3.13114811 | 3.23201746 | 4.33483521 | 3.9602244 | 1.65939434 | 3.38161178 |
| ENSMUSG00 | Hyal1 | 3939 | 1.3524332 | 1.40681413 | 1.07727264 | 1.06258369 | 1.95607584 | 2.02880663 |
| ENSMUSG00 | Tusc2 | 1658 | 39.5396787 | 34.1490166 | 45.4815104 | 39.5205023 | 31.422153 | 37.8028474 |
| ENSMUSG00 | Nprl2 | 1445 | 56.8407078 | 45.3519456 | 39.3687003 | 44.9215212 | 40.7151635 | 43.2996695 |
| ENSMUSG00 | Slc38a3 | 4246 | 0.88012604 | 2.09950349 | 2.32148163 | 2.60885249 | 0.27640288 | 0.34687483 |
| ENSMUSG00 | Cacna2d2 | 6399 | 0 | 0 | 0 | 0 | 0 | 0.00852466 |
| ENSMUSG00 | Rassf1 | 5579 | 2.30167059 | 2.99707751 | 2.26594858 | 2.23127897 | 3.60358345 | 2.28796042 |
| ENSMUSG00 | Epn3 | 5052 | 41.6204941 | 33.2593277 | 33.5976004 | 33.3894612 | 37.2496584 | 29.2451941 |
| ENSMUSG00 | Rnf112 | 4258 | 0 | 0 | 0 | 0 | 0 | 0 |
| ENSMUSG00 | Slc3a2 | 2948 | 55.2504083 | 58.3533238 | 55.417236 | 56.7667987 | 53.1034332 | 48.3782526 |
| ENSMUSG00 | Nxf1 | 2262 | 87.6659314 | 97.4380134 | 93.8556375 | 103.747842 | 106.338787 | 96.0761828 |
| ENSMUSG00 | Stx5a | 4420 | 25.5082676 | 26.4589355 | 27.5311479 | 30.8003516 | 36.0763404 | 37.7278607 |
| ENSMUSG00 | Slc47a1 | 4491 | 0.30982895 | 1.65235184 | 0.20668865 | 0 | 0.65899127 | 1.09317191 |
| ENSMUSG00 | 1700027A23l | 1079 | 0 | 0 | 0 | 0 | 0 | 0 |
| ENSMUSG00 | Tnfrsf13b | 2885 | 0 | 0 | 0 | 0 | 0 | 0 |
| ENSMUSG00 | Spire2 | 2315 | 18.1690274 | 24.8321492 | 27.3421336 | 24.2209491 | 22.0636727 | 17.5076086 |
| ENSMUSG00 | Prox1 | 9269 | 0 | 0.202748 | 0.10968199 | 0.3853838 | 0 | 0 |
| ENSMUSG00 | Raver1 | 3560 | 8.21911058 | 6.93018494 | 6.6426927 | 8.53401165 | 9.35961332 | 7.00253379 |
| ENSMUSG00 | 2610507B11l | 8888 | 25.0842411 | 29.1406533 | 30.3813073 | 41.0023233 | 42.8740971 | 25.3137375 |
| ENSMUSG00 | AI597479 | 3454 | 17.6563136 | 15.0669825 | 13.2196016 | 14.0713706 | 15.762925 | 25.6084699 |
| ENSMUSG00 | Tmem86a | 1788 | 7.40412473 | 6.36017916 | 5.98257607 | 4.33872459 | 8.10486059 | 3.3864485 |
| ENSMUSG00 | Optc | 2971 | 0 | 0 | 0.01487777 | 0.0242895 | 0 | 0.10098318 |
| ENSMUSG00 | Tex14 | 4982 | 0 | 0 | 0 | 0.48524628 | 0 | 0 |
| ENSMUSG00 | Ifi35 | 1901 | 10.5192039 | 10.2912654 | 9.39376359 | 10.9328026 | 11.0856882 | 8.70894583 |
| ENSMUSG00 | Rdm1 | 1567 | 9.9198423 | 11.4085323 | 9.39324424 | 13.5624277 | 11.4622108 | 6.56192661 |
| ENSMUSG00 | Nedd8 | 733 | 63.7823262 | 73.2986125 | 87.6800542 | 61.8268048 | 44.9004262 | 127.666149 |
| ENSMUSG00 | Gosr1 | 4280 | 26.6770421 | 28.1125108 | 31.075548 | 28.3513822 | 28.6367651 | 21.4564744 |
| ENSMUSG00 | Mrpl52 | 407 | 12.6983107 | 14.9176701 | 14.0099208 | 11.8795921 | 10.2804983 | 29.7541519 |
| ENSMUSG00 | Spaca7 | 754 | 0 | 0 | 0 | 0 | 0 | 0 |
| ENSMUSG00 | 4632411B12l | 4797 | 24.1085455 | 23.3448596 | 22.7874 | 19.9703624 | 31.2944752 | 19.2008457 |
| ENSMUSG00 | Eya4 | 4172 | 0 | 0 | 0 | 0 | 0 | 0 |
| ENSMUSG00 | Ebf3 | 6951 | 0 | 0 | 0 | 0 | 0 | 0 |
| ENSMUSG00 | Gm16119 | 1673 | 0.78417864 | 0.51844349 | 1.03040758 | 0.94895993 | 0.76249814 | 0.52169065 |
| ENSMUSG00 | Myt1 | 6879 | 0 | 0 | 0 | 0 | 0 | 0 |
| ENSMUSG00 | Faf1 | 5148 | 29.0598044 | 21.5565999 | 27.8880315 | 25.2532258 | 21.9250861 | 23.8096793 |
| ENSMUSG00 | Gm266 | 1212 | 0.951245 | 0 | 0.29176134 | 0.05954133 | 0.16840388 | 0 |
| ENSMUSG00 | 1700021C14 | 1139 | 1.78009614 | 1.64993086 | 1.82395665 | 0.60189539 | 1.11998191 | 2.20304373 |
| ENSMUSG00 | Mettl16 | 13643 | 4.74396557 | 4.11825783 | 5.3749804 | 4.89803903 | 4.48813685 | 4.5221164 |
| ENSMUSG00 | Dazl | 4918 | 0 | 0 | 0 | 0 | 0.04150173 | 0 |
| ENSMUSG00 | Apol7a | 4815 | 0 | 0 | 0.00918003 | 0 | 0 | 0 |
| ENSMUSG00 | Pigyl | 821 | 8.4740671 | 12.5601688 | 18.6821432 | 12.0419978 | 8.39045157 | 16.2451871 |
| ENSMUSG00 | Rbm25 | 4225 | 39.4731931 | 39.8721424 | 38.7302303 | 40.3435688 | 41.6906626 | 51.9283309 |
| ENSMUSG00 | Psen2 | 4534 | 6.78666562 | 1.31784712 | 5.58616146 | 4.28146007 | 3.69136552 | 3.30255336 |
| ENSMUSG00 | Acaa1b | 1950 | 0.59123535 | 1.38381577 | 1.45072716 | 0.77715173 | 1.04669489 | 0.85320666 |
| ENSMUSG00 | Plcd1 | 2636 | 8.17430593 | 6.36148781 | 4.27597495 | 4.01063697 | 7.0461308 | 7.50156049 |
| ENSMUSG00 | Fads1 | 3444 | 8.16118604 | 15.8522761 | 4.0171826 | 10.0891431 | 10.637903 | 2.93020224 |
| ENSMUSG00 | Lmbr1 | 4935 | 8.45861287 | 6.85448478 | 5.59800443 | 7.55273597 | 6.33823067 | 7.56062948 |
| ENSMUSG00 | Tnfrsf22 | 5991 | 0 | 0.61932166 | 0.361524 | 0 | 0.10220606 | 0.32323475 |
| ENSMUSG00 | Cars | 4925 | 4.31861565 | 5.3323068 | 4.48749675 | 4.56428705 | 6.67228143 | 5.12264798 |
| ENSMUSG00 | Phlda2 | 755 | 0.10531253 | 0 | 0 | 0 | 0.0675846 | 0 |
| ENSMUSG00 | Asz1 | 3237 | 0 | 0 | 0 | 0 | 0 | 0 |
| ENSMUSG00 | Wnt2 | 3078 | 0 | 0 | 0 | 0 | 0 | 0 |
| ENSMUSG00 | Gabra1 | 5148 | 0 | 0 | 0 | 0 | 0 | 0 |
| ENSMUSG00 | Grid2ip | 4762 | 0 | 0 | 0 | 0 | 0 | 0 |
| ENSMUSG00 | Kdelr3 | 1409 | 0 | 0 | 0 | 0 | 0 | 0 |
| ENSMUSG00 | 1700006E09l | 885 | 0 | 0 | 0 | 0 | 0 | 0.0616376 |
| ENSMUSG00 | Apip | 1139 | 30.0173075 | 16.4993086 | 29.028076 | 26.6101119 | 15.9933417 | 33.9077166 |
| ENSMUSG00 | Pdhx | 3919 | 26.780931 | 27.8003434 | 35.6524689 | 31.6627076 | 26.4311158 | 25.5277818 |
| ENSMUSG00 | Vac14 | 3681 | 14.1482422 | 9.16340926 | 13.1368694 | 7.69475821 | 14.4720284 | 8.08384445 |
| ENSMUSG00 | Mcoln2 | 3300 | 0 | 0 | 0 | 0 | 0 | 0 |
| ENSMUSG00 | Slc5a1 | 4102 | 0.50397001 | 0.37590594 | 0.64654085 | 0 | 1.81615087 | 0.1795259 |
| ENSMUSG00 | 1700049I03R | 416 | 0 | 0 | 0 | 0 | 0 | 0 |
| ENSMUSG00 | Akt1s1 | 3301 | 6.86477549 | 7.92645807 | 9.21262284 | 5.24670747 | 5.54936956 | 7.57674771 |
| ENSMUSG00 | Tbrg1 | 3809 | 21.8034127 | 27.2622165 | 23.9634565 | 18.5288735 | 18.1787325 | 20.8301196 |
| ENSMUSG00 | Panx3 | 2654 | 0 | 0 | 0 | 0 | 0.07690486 | 0 |
| ENSMUSG00 | Adssl1 | 1777 | 8.94889826 | 4.71831229 | 3.30829776 | 2.70056946 | 2.44076643 | 4.1441489 |
| ENSMUSG00 | 1700026D08 | 3636 | 0 | 0.03975779 | 0 | 0 | 0 | 0 |
| ENSMUSG00 | Brf1 | 2737 | 23.2693752 | 26.0386364 | 29.2310325 | 20.9742539 | 24.5717075 | 26.5870433 |
| ENSMUSG00 | Vipr2 | 4611 | 0 | 0.38666197 | 0.22048198 | 0 | 0 | 0 |
| ENSMUSG00 | Odc1 | 2288 | 13.9178933 | 12.3203971 | 15.3392759 | 11.2283285 | 12.0875418 | 9.92997137 |
| ENSMUSG00 | Thg1l | 3458 | 3.28804726 | 0.98936882 | 1.29103127 | 1.99296429 | 0.98865448 | 1.18311044 |
| ENSMUSG00 | Adam19 | 5766 | 0.00689481 | 0 | 0.04599567 | 0 | 0.07964575 | 0.15609835 |
| ENSMUSG00 | Pabpc4 | 3647 | 1.29720378 | 1.71764121 | 1.57560724 | 1.50383076 | 1.27321091 | 2.22115927 |
| ENSMUSG00 | Exoc3l2 | 2197 | 1.62858136 | 0.43865673 | 1.16691256 | 1.39598261 | 1.55610705 | 0.34760578 |
| ENSMUSG00 | Zfp296 | 1832 | 0.2170059 | 1.42034273 | 0.50668052 | 0.86659932 | 0.30638108 | 0.08932742 |
| ENSMUSG00 | Plin5 | 3012 | 0 | 0.4959428 | 0 | 0 | 0.10164617 | 0 |
| ENSMUSG00 | Sugp1 | 2273 | 17.2804288 | 12.7620933 | 18.1630098 | 14.8741204 | 15.8040337 | 17.5431248 |
| ENSMUSG00 | Dmrtc2 | 1610 | 0 | 0 | 0 | 0 | 0 | 0 |
| ENSMUSG00 | Gm5893 | 2077 | 0 | 0 | 0 | 0 | 0 | 0 |
| ENSMUSG00 | Dhdh | 3670 | 0.6391208 | 1.44428028 | 1.79457074 | 1.05198331 | 1.01496606 | 1.91739971 |
| ENSMUSG00 | Zfp790 | 2103 | 31.1351766 | 20.324001 | 28.6061381 | 27.3660775 | 24.7003569 | 21.8664012 |
| ENSMUSG00 | Cpb1 | 3484 | 0 | 0 | 0 | 0.0103565 | 0 | 0 |
| ENSMUSG00 | Slc25a41 | 1581 | 0.4023325 | 0 | 0.5591631 | 0 | 0 | 0.01725151 |
| ENSMUSG00 | Fsd1 | 1735 | 0.04582764 | 0 | 0.10190627 | 0 | 0.14705007 | 0 |
| ENSMUSG00 | 2310033E01l | 750 | 0 | 0.12849718 | 0 | 0 | 0 | 0 |
| ENSMUSG00 | Fuz | 3956 | 12.51152 | 10.4022299 | 13.0840137 | 9.76842004 | 11.6731219 | 12.5204107 |
| ENSMUSG00 | Spnb4 | 10752 | 0 | 0 | 0.00411104 | 0 | 0 | 0 |
| ENSMUSG00 | Pgam1 | 1827 | 8.07295198 | 7.85963862 | 8.97585318 | 6.81352237 | 5.30651965 | 5.76245799 |
| ENSMUSG00 | Evi5 | 11690 | 8.82510453 | 9.78568124 | 10.4398023 | 12.7382889 | 8.96999164 | 11.3461566 |
| ENSMUSG00 | Evi5l | 7711 | 8.80590853 | 6.574003 | 6.8271813 | 6.65863693 | 10.1510129 | 6.80185855 |
| ENSMUSG00 | Snapc2 | 1508 | 11.2042966 | 11.5033947 | 12.252235 | 8.2548444 | 12.8581053 | 10.3274662 |
| ENSMUSG00 | Git1 | 4721 | 17.2546028 | 21.9957176 | 14.1565742 | 17.6856283 | 18.9038618 | 12.6580671 |
| ENSMUSG00 | Gltp | 2258 | 35.3362486 | 51.1527466 | 41.5199774 | 35.8103905 | 37.241571 | 33.4712247 |
| ENSMUSG00 | Bnip2 | 3628 | 24.6992429 | 20.4672843 | 32.1645164 | 25.8581356 | 21.448518 | 25.891361 |
| ENSMUSG00 | Ccnt1 | 11225 | 7.08338183 | 7.61109683 | 7.59996054 | 8.03609009 | 7.96875161 | 6.91524484 |
| ENSMUSG00 | Scarf2 | 3291 | 1.24424628 | 0.70281046 | 0.99390349 | 0.78939751 | 1.02331839 | 1.19342086 |
| ENSMUSG00 | 4930579F01l | 1707 | 0 | 0 | 0 | 0 | 0 | 0 |
| ENSMUSG00 | Brms1l | 2556 | 28.4323233 | 26.4874614 | 22.5678424 | 30.2942361 | 24.5749095 | 29.3127676 |
| ENSMUSG00 | Med15 | 4570 | 20.1387172 | 16.3116906 | 19.2573019 | 13.1300744 | 20.4105503 | 17.5106765 |
| ENSMUSG00 | Dhdds | 5588 | 7.03617935 | 6.20870407 | 5.82977063 | 8.03258114 | 8.35524944 | 6.28175747 |
| ENSMUSG00 | Aim1l | 6922 | 0.31588434 | 0.04176808 | 0.06385704 | 0 | 0.01474325 | 0.24429755 |
| ENSMUSG00 | Ubxn11 | 3280 | 1.53931281 | 3.65805613 | 0.67380858 | 3.69620944 | 3.03358027 | 1.20573863 |
| ENSMUSG00 | Mogat1 | 1549 | 0.48763985 | 0.43551336 | 0.34242874 | 0.18635013 | 0.23059047 | 0.45780544 |
| ENSMUSG00 | Tex22 | 1504 | 0 | 0.12815543 | 0 | 0 | 0 | 0.12694314 |
| ENSMUSG00 | Wnt8a | 1740 | 0 | 0 | 0 | 0 | 0 | 0.0783754 |
| ENSMUSG00 | Tjap1 | 4152 | 4.06938324 | 3.30759538 | 2.55501983 | 3.16326209 | 4.48570019 | 2.73928819 |
| ENSMUSG00 | Ehf | 7022 | 38.0287044 | 53.4360332 | 30.3092957 | 50.9577949 | 48.8318493 | 36.8063916 |
| ENSMUSG00 | Nanog | 1159 | 0.30871383 | 0 | 0 | 0 | 0.66039313 | 0 |
| ENSMUSG00 | Rpl15 | 2762 | 23.7064723 | 28.9956071 | 14.2271682 | 30.1772349 | 13.9666691 | 26.6623917 |
| ENSMUSG00 | Tmem167 | 10849 | 10.9749898 | 11.4281239 | 12.5976679 | 12.9741041 | 9.7358833 | 17.9425846 |
| ENSMUSG00 | Steap4 | 3135 | 0 | 0 | 0 | 0 | 0.30925076 | 0 |
| ENSMUSG00 | 2810021B07l | 3070 | 18.8158675 | 10.4691716 | 13.1597669 | 15.2555354 | 8.72600889 | 27.0169961 |
| ENSMUSG00 | Kif11 | 4797 | 0 | 0.73329378 | 0 | 0 | 0.07446 | 0 |
| ENSMUSG00 | Rpa3 | 1132 | 9.62279299 | 9.83309906 | 15.150455 | 15.6185528 | 7.79820055 | 22.5039868 |
| ENSMUSG00 | Mlkl | 2567 | 0.43363983 | 0.22525801 | 0 | 1.06826466 | 0.37767867 | 0 |
| ENSMUSG00 | Phox2b | 3644 | 0 | 0 | 0 | 0 | 0 | 0 |
| ENSMUSG00 | Tnpo3 | 10705 | 16.1770082 | 11.4693185 | 13.5557824 | 14.2204714 | 17.2884321 | 11.7786695 |
| ENSMUSG00 | Ttll5 | 13635 | 3.12270771 | 2.28298068 | 4.78488598 | 2.81828951 | 4.38972782 | 3.60261278 |
| ENSMUSG00 | Zfp715 | 5168 | 16.7545349 | 14.6013871 | 14.5571859 | 13.9566577 | 15.1558605 | 18.9149201 |
| ENSMUSG00 | Retn | 1628 | 0 | 0 | 0 | 0 | 0 | 0.03350693 |
| ENSMUSG00 | Acpt | 1889 | 0 | 0.22958072 | 0 | 0 | 0 | 0.12994799 |
| ENSMUSG00 | Cdh23 | 17146 | 0 | 0 | 0 | 0 | 0 | 0 |
| ENSMUSG00 | Rps5 | 1750 | 13.607733 | 17.4297245 | 21.0653355 | 17.8554575 | 8.80569457 | 24.3133927 |
| ENSMUSG00 | Podnl1 | 3466 | 0 | 0 | 0.11477686 | 0.12492341 | 0 | 0.28329112 |
| ENSMUSG00 | Lphn1 | 8863 | 1.06756227 | 1.09823551 | 2.63325884 | 1.48594677 | 1.21477663 | 2.15107445 |
| ENSMUSG00 | Amotl1 | 10303 | 0.37428726 | 1.01021756 | 0.25741149 | 0.49729694 | 0.51506776 | 0.0555923 |
| ENSMUSG00 | 2200002J24R | 331 | 0.36032158 | 0 | 0.13354031 | 0 | 0 | 1.40081229 |
| ENSMUSG00 | Etv5 | 6467 | 0.73769254 | 0.34275187 | 0.23922445 | 0.34592342 | 0.83636862 | 0.49766622 |
| ENSMUSG00 | Tmem190 | 581 | 3.28444589 | 1.16111909 | 0.45647342 | 0 | 2.72257771 | 0.18777721 |
| ENSMUSG00 | Gfod2 | 4836 | 3.30473597 | 4.09524974 | 2.39472351 | 3.05906498 | 3.80904088 | 3.79566423 |
| ENSMUSG00 | E130303B06l | 1265 | 2.3570443 | 2.2474309 | 1.85193493 | 3.76508291 | 3.18662742 | 4.85122042 |
| ENSMUSG00 | 4933405L10F | 1120 | 0.6034314 | 0 | 0.19732966 | 0 | 0 | 0 |
| ENSMUSG00 | Atp6v0d1 | 1785 | 74.9674775 | 79.5548712 | 50.6897326 | 71.5980962 | 56.4577548 | 56.5967864 |
| ENSMUSG00 | Ptprs | 7912 | 3.6730607 | 6.01721497 | 3.15088972 | 7.71623096 | 5.6301853 | 5.09848224 |
| ENSMUSG00 | Slc41a1 | 4900 | 27.6341156 | 26.8172302 | 29.064967 | 25.5298874 | 31.9904136 | 21.0348697 |
| ENSMUSG00 | Fer1l4 | 6808 | 0 | 0.02123374 | 0 | 0 | 0 | 0 |
| ENSMUSG00 | 4931406B18l | 5695 | 0 | 0 | 0 | 0 | 0 | 0 |
| ENSMUSG00 | Iglon5 | 2611 | 0 | 0 | 0 | 0 | 0 | 0 |
| ENSMUSG00 | Igf2bp1 | 9821 | 0.04048007 | 0 | 0 | 0 | 0 | 0.00277718 |
| ENSMUSG00 | B4galnt2 | 1711 | 0.25558754 | 0.22530189 | 0.02583392 | 0 | 0 | 0.0797038 |
| ENSMUSG00 | Zfp651 | 6347 | 6.59563904 | 5.21570596 | 5.88475774 | 4.97997022 | 6.06174372 | 3.06393852 |
| ENSMUSG00 | Cobra1 | 4174 | 26.3544357 | 32.509109 | 24.9601687 | 26.6422763 | 32.8603014 | 20.7859673 |
| ENSMUSG00 | Card14 | 4583 | 0 | 0 | 0.03857896 | 0 | 0 | 0.113074 |
| ENSMUSG00 | Tmem175 | 3931 | 12.3382565 | 16.1683839 | 13.9093564 | 13.3185059 | 13.1881958 | 12.8220639 |
| ENSMUSG00 | Bcas1 | 4762 | 0 | 0 | 0 | 0 | 0.10715325 | 0 |
| ENSMUSG00 | D16H22S680 | 7285 | 13.5556093 | 10.384724 | 16.758475 | 11.1094064 | 10.8846929 | 10.8087692 |
| ENSMUSG00 | 4833413E03l | 1855 | 0.27860984 | 0 | 0.04765697 | 0 | 0.1375374 | 0.23525295 |
| ENSMUSG00 | Aldh1a2 | 2250 | 0 | 0 | 0 | 0 | 0 | 0 |
| ENSMUSG00 | Ndufs2 | 2128 | 228.239053 | 182.963559 | 260.807491 | 148.092376 | 162.095066 | 158.828632 |
| ENSMUSG00 | Snx31 | 3978 | 0.20987056 | 1.0175116 | 0.07778102 | 0 | 0 | 0 |
| ENSMUSG00 | 0610007C21l | 1581 | 61.8083309 | 66.4430385 | 78.3946665 | 54.2257671 | 52.4464647 | 58.8966593 |
| ENSMUSG00 | Cad | 8254 | 1.83990714 | 1.31937678 | 1.54229838 | 2.18573083 | 1.94733564 | 1.61255439 |
| ENSMUSG00 | 2210415F13l | 1277 | 0 | 0 | 0 | 0 | 0 | 0 |
| ENSMUSG00 | Sh3bp5l | 3303 | 9.91780682 | 9.67230128 | 8.4576339 | 11.6122353 | 13.8109537 | 8.84382036 |
| ENSMUSG00 | 1810065E05l | 920 | 0 | 0 | 0 | 0 | 0 | 0 |
| ENSMUSG00 | Atad1 | 2830 | 86.478706 | 72.6202034 | 89.2002563 | 99.104499 | 78.1442801 | 84.0309469 |
| ENSMUSG00 | Pten | 12426 | 11.4313803 | 10.1445141 | 11.3617163 | 13.3775937 | 8.61116289 | 14.6382238 |
| ENSMUSG00 | 4933402N03l | 2157 | 0 | 0 | 0 | 0 | 0 | 0 |
| ENSMUSG00 | Pea15a | 3622 | 47.8558517 | 57.1133065 | 34.3290404 | 46.9405284 | 54.6328784 | 32.0638912 |
| ENSMUSG00 | Timm23 | 2230 | 2.99503172 | 3.06837434 | 3.82554067 | 3.36550012 | 2.08224224 | 2.60515613 |
| ENSMUSG00 | Tnfaip8l2 | 1120 | 0 | 0 | 0.15786373 | 0 | 0 | 0 |
| ENSMUSG00 | Trnt1 | 3724 | 13.8994725 | 13.8840098 | 18.575694 | 17.1399401 | 12.1125983 | 18.3686346 |
| ENSMUSG00 | Ly6g6e | 1299 | 5.47823788 | 4.45140343 | 6.49927022 | 5.11092856 | 2.08190756 | 6.82390897 |
| ENSMUSG00 | Ehmt2 | 5392 | 12.2761267 | 10.947402 | 12.5998206 | 8.22419004 | 16.1444728 | 7.33967017 |
| ENSMUSG00 | Elof1 | 1778 | 13.2816397 | 15.3665426 | 13.1760274 | 11.8717638 | 8.38003472 | 18.745562 |
| ENSMUSG00 | Med16 | 5366 | 14.7730951 | 16.3165795 | 10.9474934 | 13.8249503 | 15.7472379 | 9.15931791 |
| ENSMUSG00 | St3gal1 | 4790 | 0.2406908 | 0.201196 | 0.00922794 | 0 | 0.13848495 | 0 |
| ENSMUSG00 | ORF61 | 6618 | 2.22866172 | 2.44645581 | 1.91688259 | 2.02273172 | 1.98153197 | 2.28318981 |
| ENSMUSG00 | Rnf170 | 5241 | 6.75865925 | 5.56244944 | 5.70972099 | 5.35619741 | 5.86107198 | 6.89021606 |
| ENSMUSG00 | Clip3 | 3783 | 0 | 0 | 0 | 0 | 0 | 0.00720979 |
| ENSMUSG00 | Thap8 | 1181 | 0 | 0 | 0.07485494 | 0 | 0 | 0.27713435 |
| ENSMUSG00 | Myl2 | 1901 | 0 | 0 | 0 | 0 | 0 | 0 |
| ENSMUSG00 | Dedd | 5978 | 1.12390034 | 1.54764083 | 1.28657087 | 1.85299225 | 1.03281892 | 1.20450063 |
| ENSMUSG00 | 1810033B17l | 1125 | 0 | 0 | 0 | 0 | 0 | 0 |
| ENSMUSG00 | D430018E03l | 3830 | 0 | 0 | 0 | 0 | 0.02664563 | 0.00712132 |
| ENSMUSG00 | Nit1 | 4599 | 14.591705 | 13.379884 | 17.3001343 | 15.4323509 | 13.9021633 | 14.0317018 |
| ENSMUSG00 | Pax5 | 3496 | 0 | 0 | 0 | 0 | 0 | 0 |
| ENSMUSG00 | Prdm15 | 13654 | 1.04527739 | 0.98462116 | 0.78989671 | 0.75049807 | 0.92680103 | 0.83897382 |
| ENSMUSG00 | Rnf168 | 5410 | 4.52668688 | 5.24617642 | 6.25035303 | 5.29559027 | 4.4707028 | 4.42141564 |
| ENSMUSG00 | Tctex1d2 | 2562 | 6.68798287 | 5.54839986 | 6.59059486 | 5.29541325 | 5.93515222 | 8.78281709 |
| ENSMUSG00 | 1500003O03 | 3688 | 44.4769828 | 40.3992622 | 39.4077169 | 35.9157227 | 41.1752968 | 23.976242 |
| ENSMUSG00 | Sval2 | 1203 | 0 | 0 | 0 | 0 | 0 | 0 |
| ENSMUSG00 | Trpv4 | 4716 | 40.0926771 | 44.6920055 | 24.3316337 | 44.7200064 | 41.6347528 | 21.0401055 |
| ENSMUSG00 | Klhl3 | 3237 | 3.7950088 | 4.74867935 | 2.82682388 | 3.34402637 | 1.45023991 | 1.75258726 |
| ENSMUSG00 | Fam18b | 2332 | 66.4523426 | 78.0240161 | 85.6742412 | 69.4409159 | 57.984518 | 95.4963248 |
| ENSMUSG00 | Dnajc7 | 4835 | 32.6101832 | 32.0412433 | 29.1265919 | 23.7686271 | 36.6630051 | 33.5757296 |
| ENSMUSG00 | Zfp385c | 5945 | 0.03343606 | 0 | 0 | 0 | 0 | 0.04587828 |
| ENSMUSG00 | Cacybp | 1436 | 82.5562973 | 102.278744 | 98.807741 | 97.3411145 | 68.9353545 | 115.366406 |
| ENSMUSG00 | Cluap1 | 2157 | 7.83313826 | 8.28797869 | 8.60675664 | 6.00531015 | 9.39150264 | 9.7111372 |
| ENSMUSG00 | Zswim7 | 727 | 10.2259627 | 6.09787163 | 13.1936725 | 6.50171641 | 5.40444421 | 10.8423256 |
| ENSMUSG00 | Pigl | 4932 | 3.82078219 | 6.26267425 | 3.74622271 | 6.3136262 | 7.428414 | 6.29882687 |
| ENSMUSG00 | Ndufa2 | 526 | 74.44705 | 78.6923076 | 79.2439885 | 60.4339947 | 61.0182327 | 162.040395 |
| ENSMUSG00 | Pam16 | 883 | 1.89097416 | 1.80085231 | 1.45170266 | 2.65609614 | 2.02256303 | 2.0077594 |
| ENSMUSG00 | Glis2 | 4203 | 10.9154948 | 12.1411949 | 4.98493305 | 11.6410308 | 9.56668669 | 8.83197336 |
| ENSMUSG00 | Cox6c | 1590 | 149.370592 | 146.892884 | 196.517502 | 131.620037 | 101.603463 | 337.176295 |
| ENSMUSG00 | Bicc1 | 7694 | 19.3765339 | 27.5691082 | 9.85263332 | 19.9778411 | 30.8850835 | 9.21325545 |
| ENSMUSG00 | Ube2z | 3990 | 13.1222977 | 12.9825627 | 15.6977473 | 10.8607858 | 12.9292396 | 12.5982858 |
| ENSMUSG00 | Gip | 680 | 0 | 0 | 0 | 0 | 0 | 0 |
| ENSMUSG00 | Tmem87b | 7302 | 18.8324031 | 17.5535382 | 19.4676975 | 16.5042493 | 20.2512239 | 20.7417244 |
| ENSMUSG00 | Anapc1 | 15679 | 6.26543736 | 6.13125531 | 7.38341902 | 6.70598111 | 10.0497129 | 6.59121169 |
| ENSMUSG00 | Mertk | 4317 | 4.56769014 | 8.02549268 | 2.23210604 | 7.69783716 | 6.20543139 | 1.58580831 |
| ENSMUSG00 | Cyp2d10 | 1633 | 0 | 0 | 0 | 0 | 0 | 0 |
| ENSMUSG00 | Tsg101 | 3395 | 34.3922964 | 35.4833889 | 39.3715385 | 36.4646524 | 34.0726934 | 39.252986 |
| ENSMUSG00 | Hps5 | 6930 | 12.5232632 | 8.61515178 | 8.29821036 | 10.1113031 | 13.5923073 | 7.8872117 |
| ENSMUSG00 | Map3k4 | 5364 | 3.87623347 | 3.62027147 | 3.48571581 | 5.11902235 | 3.48166546 | 3.34577044 |
| ENSMUSG00 | Fam38a | 9248 | 6.02264578 | 4.47579518 | 3.87148495 | 8.30261578 | 9.83228825 | 3.51845206 |
| ENSMUSG00 | Blk | 2462 | 0 | 0 | 0 | 0 | 0 | 0 |
| ENSMUSG00 | Rnf166 | 2192 | 15.3979941 | 12.1125591 | 15.0229804 | 13.9258256 | 13.4316692 | 11.8953262 |
| ENSMUSG00 | Ankrd28 | 5201 | 15.7691898 | 13.5915229 | 18.943647 | 22.7411925 | 21.1620636 | 22.9272684 |
| ENSMUSG00 | Ankrd52 | 8502 | 0.38810926 | 2.08003108 | 1.67927491 | 1.33259962 | 1.30836861 | 1.10997709 |
| ENSMUSG00 | Pkd2l2 | 2699 | 1.9590511 | 1.39256854 | 1.99800847 | 1.60424059 | 2.3064905 | 1.80887752 |
| ENSMUSG00 | Srp19 | 2394 | 20.4921733 | 18.5177639 | 23.2087371 | 18.402747 | 15.0691929 | 35.3066445 |
| ENSMUSG00 | 4930511M11 | 2303 | 0 | 0 | 0 | 0 | 0 | 0 |
| ENSMUSG00 | Clec4f | 2549 | 0 | 0 | 0 | 0 | 0 | 0 |
| ENSMUSG00 | Klra17 | 1521 | 0 | 0 | 0 | 0 | 0 | 0 |
| ENSMUSG00 | Wdfy2 | 1999 | 2.5058482 | 3.56758051 | 3.3389086 | 1.80500473 | 3.31837362 | 3.30188229 |
| ENSMUSG00 | Zfyve20 | 6360 | 7.83232973 | 5.37172759 | 4.71903324 | 5.73001022 | 6.74735567 | 5.43777399 |
| ENSMUSG00 | Mrps25 | 6942 | 4.82196983 | 6.24022061 | 5.22119148 | 4.93256414 | 3.46938193 | 6.98956831 |
| ENSMUSG00 | Dguok | 1724 | 17.9868183 | 19.956566 | 23.4085166 | 21.159482 | 16.5450952 | 22.0696762 |
| ENSMUSG00 | Camta1 | 17895 | 0.90641163 | 0.56816649 | 0.73607987 | 0.93355611 | 0.76988664 | 1.54701083 |
| ENSMUSG00 | Csf1 | 5381 | 0.85702207 | 1.68352558 | 0.40250703 | 1.66295243 | 3.2620467 | 1.4597837 |
| ENSMUSG00 | Fam40a | 3213 | 15.0459584 | 19.1066067 | 18.283302 | 24.4140569 | 23.011895 | 12.9964123 |
| ENSMUSG00 | Kif1a | 9009 | 0.18534024 | 0.0427896 | 0 | 0.036046 | 0.33417207 | 0 |
| ENSMUSG00 | Alx3 | 1863 | 0 | 0 | 0 | 0 | 0 | 0 |
| ENSMUSG00 | Slc25a11 | 2172 | 207.709573 | 199.268243 | 245.694683 | 159.329139 | 160.408883 | 170.240462 |
| ENSMUSG00 | Chrne | 1920 | 0.04141196 | 0 | 0 | 0 | 0 | 0 |
| ENSMUSG00 | 2310061C15l | 2618 | 4.00895602 | 5.88986306 | 8.00293109 | 5.05810173 | 4.05404361 | 8.38658694 |
| ENSMUSG00 | Chfr | 3386 | 12.1638151 | 10.2748409 | 10.2476216 | 11.2743069 | 11.5736139 | 7.87790821 |
| ENSMUSG00 | Ceacam16 | 1629 | 0 | 0 | 0 | 0 | 0 | 0 |
| ENSMUSG00 | Hoxa2 | 2421 | 1.23158242 | 0.23884234 | 0.23734984 | 0.08942266 | 0.61122053 | 0.46190013 |
| ENSMUSG00 | Adam28 | 3656 | 0 | 0.23724178 | 0 | 0 | 0 | 0 |
| ENSMUSG00 | Ankrd53 | 1897 | 0.88019514 | 0.07620418 | 0.90873583 | 0.11412349 | 1.1028368 | 0.60386655 |
| ENSMUSG00 | Tex261 | 4137 | 29.482672 | 26.8595445 | 24.3820657 | 18.7867353 | 20.0923293 | 21.4729273 |
| ENSMUSG00 | Fam120b | 4507 | 23.9926574 | 22.4841553 | 26.1758862 | 17.8448807 | 26.6623286 | 18.1366971 |
| ENSMUSG00 | Tbp | 13512 | 4.89587919 | 5.12105765 | 6.00938319 | 6.83881853 | 5.30958292 | 6.30594827 |
| ENSMUSG00 | Psmb1 | 1030 | 80.7846803 | 87.0162906 | 92.3088974 | 73.9156446 | 55.3363707 | 94.852192 |
| ENSMUSG00 | Pdcd2 | 2574 | 0 | 0 | 0 | 0 | 0 | 0 |
| ENSMUSG00 | Dll1 | 4576 | 43.7866306 | 25.9360155 | 55.9960848 | 28.3941089 | 31.5569588 | 49.4829666 |
| ENSMUSG00 | Nol3 | 2865 | 2.3728402 | 3.27970549 | 1.45025244 | 1.75057738 | 1.67416381 | 1.4755913 |
| ENSMUSG00 | Fhod1 | 4546 | 0.7608286 | 0.6783837 | 1.14734216 | 0.64290489 | 0.41530486 | 0.8879557 |
| ENSMUSG00 | Plekhg4 | 4816 | 0 | 0 | 0 | 0 | 0 | 0 |
| ENSMUSG00 | Slc9a5 | 5206 | 0.02290942 | 0 | 0 | 0 | 0.18622765 | 0 |
| ENSMUSG00 | Elmo3 | 3165 | 25.8504831 | 19.5790725 | 22.6805033 | 17.8985181 | 25.8275684 | 17.8556248 |
| ENSMUSG00 | Stc1 | 4193 | 13.8049081 | 37.7056324 | 5.96666901 | 42.6909189 | 36.9341881 | 15.8131763 |
| ENSMUSG00 | 4931428F04l | 3544 | 1.49195229 | 2.03949388 | 1.08509039 | 0.73304379 | 1.16623489 | 1.23905669 |
| ENSMUSG00 | Tppp3 | 2198 | 1.28418522 | 0.54807145 | 0.88484126 | 1.90423893 | 1.32324996 | 1.40219938 |
| ENSMUSG00 | Msh3 | 7430 | 4.75674592 | 3.33349008 | 3.17682156 | 7.04158339 | 7.73293394 | 4.82720735 |
| ENSMUSG00 | Adamts13 | 5210 | 0.06867549 | 0.02774651 | 0.03393616 | 0 | 0.06855751 | 0.20940222 |
| ENSMUSG00 | Tmem208 | 1607 | 23.9967742 | 21.7393718 | 32.5943895 | 20.9262386 | 16.5748402 | 29.022796 |
| ENSMUSG00 | E2f4 | 1982 | 17.4506902 | 15.2679544 | 12.9795523 | 17.4948763 | 14.0824559 | 18.9353701 |
| ENSMUSG00 | Surf4 | 3337 | 79.6777506 | 75.9113354 | 85.5028159 | 64.4654329 | 62.4335309 | 56.3392533 |
| ENSMUSG00 | Surf2 | 2473 | 14.6611396 | 17.5559985 | 20.7514516 | 21.5354216 | 21.3968264 | 20.9991561 |
| ENSMUSG00 | Dnajb9 | 1862 | 18.8528944 | 17.9599306 | 15.2166388 | 15.2118179 | 14.9900256 | 32.6944118 |
| ENSMUSG00 | Naf1 | 3338 | 1.94132514 | 1.50131515 | 0.86073092 | 1.05932905 | 1.34521302 | 1.73224192 |
| ENSMUSG00 | Yes1 | 4564 | 16.3324991 | 18.3285854 | 21.2680209 | 25.9784396 | 18.5591113 | 21.2866488 |
| ENSMUSG00 | Ppp1cb | 7249 | 85.3681625 | 63.6148781 | 102.178271 | 88.6446587 | 54.0602244 | 113.639941 |
| ENSMUSG00 | Gorasp2 | 4301 | 34.3204137 | 35.9073581 | 32.8559154 | 33.984739 | 27.9274794 | 30.6165911 |
| ENSMUSG00 | Tsen15 | 1891 | 8.05200901 | 9.37737214 | 6.68520735 | 10.6662416 | 7.82530354 | 8.45210925 |
| ENSMUSG00 | Oc90 | 2428 | 0.75319279 | 0.17861531 | 0.74640674 | 1.41177687 | 0.3782845 | 1.28060497 |
| ENSMUSG00 | Efr3a | 11134 | 21.673771 | 20.8819457 | 31.1683734 | 24.7363253 | 23.5516926 | 25.6554066 |
| ENSMUSG00 | Trappc2l | 602 | 47.548083 | 53.4693411 | 67.0370144 | 34.4637469 | 31.955056 | 85.8562149 |
| ENSMUSG00 | Acsf3 | 2196 | 7.16902108 | 11.9149536 | 7.76954071 | 6.86807587 | 7.6214259 | 8.16003551 |
| ENSMUSG00 | Ddx19a | 2946 | 1.24151535 | 1.53751716 | 1.14030552 | 2.18010995 | 1.88794126 | 2.434905 |
| ENSMUSG00 | Galns | 3296 | 4.4749039 | 2.88007558 | 3.84868621 | 3.07616945 | 2.97241024 | 1.95291713 |
| ENSMUSG00 | Gata2 | 3344 | 76.1940414 | 99.2404387 | 70.2285861 | 84.3460951 | 87.7702493 | 74.7931944 |
| ENSMUSG00 | C8g | 1580 | 5.66138172 | 5.15411944 | 6.9380108 | 4.59018415 | 5.5870652 | 5.76565159 |
| ENSMUSG00 | Entpd2 | 2295 | 1.66297435 | 1.80567931 | 2.06082667 | 2.48407975 | 1.42295776 | 0.9745187 |
| ENSMUSG00 | B230208H17 | 4171 | 12.2097268 | 11.7953386 | 9.9403809 | 12.2147799 | 10.9980129 | 8.34390786 |
| ENSMUSG00 | Ptgds | 1306 | 0.18264386 | 0.36896204 | 0.13538084 | 0.11051162 | 0 | 0.39679797 |
| ENSMUSG00 | Edf1 | 1470 | 47.6524875 | 52.1200293 | 53.8841515 | 43.0039061 | 28.776099 | 53.4174737 |
| ENSMUSG00 | Clic3 | 1071 | 2.1158379 | 2.33958449 | 1.98103498 | 3.87435586 | 1.38166657 | 2.95411592 |
| ENSMUSG00 | Npdc1 | 3441 | 13.9796953 | 18.7928525 | 15.9799746 | 10.8738972 | 11.2996508 | 11.0810653 |
| ENSMUSG00 | Fbxw5 | 3256 | 39.0350956 | 35.5774591 | 46.6861604 | 40.4260745 | 39.7899165 | 46.7421643 |
| ENSMUSG00 | Slc25a13 | 3117 | 23.9655271 | 22.1994645 | 22.1647355 | 22.1562506 | 23.8024865 | 14.9717381 |
| ENSMUSG00 | Ube2i | 4847 | 9.77687905 | 7.29705973 | 9.25621429 | 13.4665605 | 16.8964995 | 9.67864236 |
| ENSMUSG00 | 0610007P22l | 1373 | 14.3907311 | 11.4412091 | 10.5273144 | 10.2490877 | 10.0714842 | 10.5085828 |
| ENSMUSG00 | Unkl | 5654 | 1.09689688 | 1.33803881 | 1.20394125 | 1.30824534 | 1.46202209 | 0.74288901 |
| ENSMUSG00 | Lrrk1 | 9063 | 2.48718498 | 3.13161364 | 1.77529194 | 3.36415399 | 4.48725825 | 2.2209736 |
| ENSMUSG00 | Aldh1a3 | 4600 | 2.48039628 | 2.45122336 | 2.47914685 | 0.47063536 | 0.98724945 | 0.02371708 |
| ENSMUSG00 | Actn1 | 3941 | 2.15875992 | 2.5432073 | 2.25439494 | 3.67137779 | 3.23689266 | 2.04853926 |
| ENSMUSG00 | Sirt2 | 3662 | 19.5411974 | 20.6062175 | 19.3609383 | 16.7798257 | 15.3134864 | 18.2699864 |
| ENSMUSG00 | Hnrnpl | 5874 | 8.93381585 | 9.565099 | 12.687148 | 12.4020511 | 10.3025675 | 19.2928373 |
| ENSMUSG00 | Nolc1 | 2690 | 19.8038453 | 19.9373644 | 15.478861 | 23.1783543 | 23.2179494 | 13.3230022 |
| ENSMUSG00 | Casd1 | 4198 | 37.5110668 | 38.0969006 | 56.5737262 | 42.3564353 | 39.4913518 | 32.3553367 |
| ENSMUSG00 | Cnksr3 | 3824 | 4.77190522 | 4.90181119 | 3.87228489 | 3.63273722 | 5.07061266 | 3.5448472 |
| ENSMUSG00 | Mtmr1 | 5972 | 18.2134954 | 15.3305827 | 19.214331 | 15.3161391 | 19.4638453 | 16.4415106 |
| ENSMUSG00 | Hmgb3 | 2308 | 3.72061689 | 5.03159989 | 4.25165041 | 3.17359404 | 3.8689843 | 5.2823933 |
| ENSMUSG00 | Mtap2 | 9478 | 1.37579633 | 2.88772939 | 1.69289608 | 3.20162476 | 3.02561967 | 2.43739391 |
| ENSMUSG00 | Cyp2j9 | 3625 | 3.16947141 | 4.5727272 | 4.24337693 | 2.35901919 | 6.01055517 | 4.56709132 |
| ENSMUSG00 | Nipsnap3a | 1594 | 0 | 0 | 0 | 0 | 0 | 0 |
| ENSMUSG00 | Abca1 | 10313 | 10.6125122 | 6.00870402 | 7.75343101 | 6.68950539 | 6.9862545 | 10.4438622 |
| ENSMUSG00 | Nipsnap3b | 1778 | 54.4681387 | 41.0858528 | 63.8167216 | 45.559162 | 39.8912612 | 58.1388542 |
| ENSMUSG00 | Lage3 | 2513 | 3.543664 | 4.12259652 | 5.80446009 | 4.17822322 | 3.75641844 | 6.44693025 |
| ENSMUSG00 | Ubl4 | 4482 | 44.2171063 | 47.8101536 | 53.2552326 | 37.1044496 | 41.4973538 | 48.5551867 |
| ENSMUSG00 | Gdi1 | 4038 | 132.213063 | 97.0530792 | 134.039517 | 110.587267 | 110.228597 | 114.319779 |
| ENSMUSG00 | Sash1 | 7363 | 0.09435197 | 0.01345426 | 0.27768853 | 0.17630484 | 0.04986523 | 0.14850076 |
| ENSMUSG00 | Gadd45b | 1262 | 15.3729592 | 22.8713779 | 29.9465735 | 16.6972377 | 15.6879824 | 24.8756813 |
| ENSMUSG00 | Slamf6 | 3047 | 0 | 0 | 0 | 0 | 0 | 0 |
| ENSMUSG00 | Slamf1 | 2957 | 0 | 0 | 0 | 0 | 0 | 0 |
| ENSMUSG00 | Zdhhc12 | 2396 | 6.63697504 | 5.75180403 | 4.98100902 | 4.81897089 | 3.93984955 | 4.2346268 |
| ENSMUSG00 | Endog | 1060 | 9.63882877 | 8.72811028 | 9.63266579 | 8.30567817 | 6.4986422 | 9.54612374 |
| ENSMUSG00 | Cybb | 5793 | 0 | 0 | 0 | 0 | 0 | 0 |
| ENSMUSG00 | Golga7 | 2743 | 31.0739155 | 25.8411434 | 27.0561044 | 28.7946028 | 19.8859627 | 30.9337961 |
| ENSMUSG00 | Xk | 5073 | 0.94040167 | 0.58891374 | 0.3398131 | 0.66858115 | 0.78455693 | 0 |
| ENSMUSG00 | Pcolce2 | 4615 | 0 | 0 | 0 | 0 | 0 | 0 |
| ENSMUSG00 | Cd48 | 1341 | 0 | 0 | 0 | 0 | 0 | 0 |
| ENSMUSG00 | Clpx | 3750 | 48.2048453 | 30.1839874 | 44.720478 | 35.3796474 | 38.3446206 | 44.3667465 |
| ENSMUSG00 | Trabd | 3582 | 23.1962463 | 19.0485768 | 20.1018432 | 21.445749 | 26.9804455 | 19.4623054 |
| ENSMUSG00 | Mov10l1 | 4223 | 0 | 0 | 0 | 0 | 0 | 0 |
| ENSMUSG00 | Fam116b | 4586 | 22.053629 | 12.7768674 | 11.7588854 | 9.21327391 | 12.6508917 | 13.1080037 |
| ENSMUSG00 | Cd83 | 2299 | 0 | 0 | 0.34607794 | 0 | 0.66585092 | 0 |
| ENSMUSG00 | Tmem27 | 1536 | 15.9953691 | 22.3050523 | 9.72670764 | 7.14123798 | 9.50100487 | 30.488643 |
| ENSMUSG00 | Ace2 | 3743 | 0.38236636 | 1.1328899 | 1.11006499 | 0 | 0.87247877 | 0 |
| ENSMUSG00 | Gzmb | 1413 | 0 | 0 | 0 | 0 | 0 | 0 |
| ENSMUSG00 | Gzmf | 901 | 0 | 0 | 0 | 0 | 0 | 0 |
| ENSMUSG00 | Gzmn | 1233 | 0 | 0 | 0 | 0 | 0 | 0 |
| ENSMUSG00 | C4a | 6491 | 0.0734965 | 0 | 0.04766799 | 0 | 0 | 0.13025941 |
| ENSMUSG00 | Ager | 1814 | 0.32873881 | 0.45158187 | 0.02436706 | 0.03978175 | 0.53445487 | 0.37589084 |
| ENSMUSG00 | Atf6b | 2854 | 11.7009824 | 12.2407746 | 11.1511307 | 14.2608781 | 11.1564326 | 11.3055004 |
| ENSMUSG00 | Egfl8 | 1820 | 0.13106202 | 0.52952134 | 0.02428673 | 0.19825299 | 0.05607294 | 0 |
| ENSMUSG00 | Notch4 | 8237 | 0.61295933 | 0.71369988 | 0.49906172 | 0.90237965 | 0.97877472 | 1.0629063 |
| ENSMUSG00 | Ppt2 | 3328 | 1.48127391 | 2.54832146 | 1.87273433 | 2.00576269 | 1.19593068 | 2.73729853 |
| ENSMUSG00 | Prrt1 | 2302 | 0 | 0 | 0 | 0.06269686 | 0 | 0 |
| ENSMUSG00 | Rnf5 | 1735 | 32.5605405 | 37.7437319 | 36.1002948 | 27.8466038 | 31.4981259 | 29.5697962 |
| ENSMUSG00 | Fam163a | 3877 | 0 | 0 | 0 | 0 | 0 | 0 |
| ENSMUSG00 | 5930434B04l | 4214 | 92.0112011 | 83.7030746 | 68.6208963 | 88.2614416 | 81.9158596 | 41.4038603 |
| ENSMUSG00 | Hivep2 | 9757 | 1.95171417 | 2.00015467 | 1.13709774 | 2.70328734 | 1.46432154 | 1.80302781 |
| ENSMUSG00 | 2310057J18R | 2166 | 0 | 0 | 0 | 0 | 0 | 0 |
| ENSMUSG00 | Arnt | 6715 | 12.3558731 | 12.0914602 | 12.5858412 | 13.4333747 | 17.1810329 | 10.9139175 |
| ENSMUSG00 | Itga2 | 4184 | 0.19953755 | 0 | 0 | 0 | 0 | 0.09126313 |
| ENSMUSG00 | Mocs2 | 2266 | 52.3172299 | 51.0997443 | 58.8707689 | 41.113786 | 44.9914823 | 74.3011245 |
| ENSMUSG00 | Nat9 | 2025 | 8.18668413 | 8.13815467 | 7.70530893 | 9.15860291 | 6.75312036 | 8.26994 |
| ENSMUSG00 | Lpl | 7082 | 16.262585 | 22.1268441 | 5.62353298 | 18.3772853 | 16.7518108 | 10.3406391 |
| ENSMUSG00 | Atp6v0e | 3423 | 120.183965 | 127.286535 | 191.386362 | 120.378892 | 71.4786655 | 189.241216 |
| ENSMUSG00 | Nkx2-5 | 1523 | 0 | 0 | 0 | 0 | 0 | 0 |
| ENSMUSG00 | Zfp318 | 9365 | 3.12318014 | 3.27026372 | 2.29087566 | 1.8542982 | 3.14565219 | 2.13971256 |
| ENSMUSG00 | Ttbk1 | 6944 | 0.04007609 | 0.15266442 | 0.03819284 | 0.11431523 | 0.08083095 | 0.19638997 |
| ENSMUSG00 | Srf | 4086 | 5.80862013 | 6.20314943 | 4.83559076 | 3.32915585 | 5.03270422 | 2.3095999 |
| ENSMUSG00 | Gata3 | 4748 | 28.5690184 | 37.5403643 | 26.5508964 | 42.2451739 | 41.0854749 | 33.3924764 |
| ENSMUSG00 | Gata5 | 3428 | 0.09277825 | 0 | 0 | 0 | 0.47632556 | 0 |
| ENSMUSG00 | Lama5 | 11476 | 11.972372 | 13.8437347 | 11.1236426 | 11.7370401 | 13.7481312 | 9.85841788 |
| ENSMUSG00 | Steap1 | 1719 | 1.38762585 | 1.31748853 | 0.56570131 | 0 | 0 | 0.98372754 |
| ENSMUSG00 | Steap2 | 19419 | 5.62378624 | 4.59804713 | 5.82028491 | 4.92948474 | 3.92833987 | 7.62521327 |
| ENSMUSG00 | Hspa8 | 3064 | 32.1521151 | 40.4489325 | 23.8032118 | 48.7531542 | 32.4243974 | 18.7112571 |
| ENSMUSG00 | Serac1 | 4905 | 3.22582696 | 4.06711255 | 1.13546019 | 1.21376908 | 3.55780235 | 1.40682645 |
| ENSMUSG00 | Awat1 | 1383 | 0 | 0 | 0 | 0 | 0 | 0 |
| ENSMUSG00 | Pdzd11 | 2365 | 33.3508978 | 27.3430044 | 32.0533449 | 23.9987552 | 23.9058031 | 41.4828234 |
| ENSMUSG00 | Psma2 | 2660 | 53.8641924 | 49.4363912 | 64.70751 | 48.208867 | 35.2580747 | 67.6841706 |
| ENSMUSG00 | Mrpl32 | 765 | 39.3397369 | 44.7220574 | 58.5313294 | 41.1759802 | 27.3474694 | 58.7206939 |
| ENSMUSG00 | Setdb1 | 6832 | 8.53648857 | 10.2480826 | 7.0973978 | 12.9022885 | 10.4487558 | 6.15196415 |
| ENSMUSG00 | Anxa9 | 2553 | 10.8225848 | 7.58752438 | 5.99053572 | 11.5468196 | 14.0907148 | 3.93147953 |
| ENSMUSG00 | Arnt2 | 5969 | 1.20551884 | 1.54997435 | 0.41469312 | 0.77374798 | 0.70953077 | 0.63971344 |
| ENSMUSG00 | Prune | 3609 | 20.7534844 | 14.1528481 | 32.2113735 | 15.1866516 | 18.0974677 | 31.113796 |
| ENSMUSG00 | Lass2 | 3036 | 182.670933 | 193.729813 | 198.005621 | 165.720695 | 174.642645 | 169.873568 |
| ENSMUSG00 | Nlrp5 | 3662 | 0 | 0 | 0 | 0 | 0 | 0 |
| ENSMUSG00 | Capza2 | 3522 | 172.002843 | 158.281928 | 200.953978 | 160.176822 | 138.79406 | 197.977329 |
| ENSMUSG00 | Plekho1 | 2980 | 0.25347454 | 0.16169947 | 0.10382983 | 0.01210807 | 0.05136884 | 0.58576406 |
| ENSMUSG00 | Vps45 | 3976 | 11.3587087 | 10.4226208 | 10.1722048 | 12.3691717 | 10.4208796 | 7.00387492 |
| ENSMUSG00 | Prpf3 | 6021 | 6.45093912 | 6.76258821 | 6.67322294 | 8.69540718 | 6.95775694 | 4.69752548 |
| ENSMUSG00 | Anp32e | 3929 | 38.8650549 | 31.3843486 | 35.1454715 | 34.3463595 | 34.597675 | 36.2157783 |
| ENSMUSG00 | Aph1a | 2242 | 0.40783945 | 1.78388702 | 1.4786522 | 1.52890019 | 1.32004005 | 1.24086227 |
| ENSMUSG00 | Tab2 | 6375 | 6.41076611 | 6.99931693 | 8.098471 | 7.83900105 | 6.55538852 | 5.89560045 |
| ENSMUSG00 | Ppil4 | 6549 | 20.8156272 | 24.2587723 | 19.7015086 | 26.7378473 | 19.3150688 | 28.1450622 |
| ENSMUSG00 | Cnih | 1564 | 66.6234747 | 67.2268649 | 69.524638 | 75.3939396 | 43.9466293 | 76.1213557 |
| ENSMUSG00 | Eps8 | 7160 | 146.684397 | 72.979578 | 206.952288 | 125.944477 | 125.734406 | 144.04895 |
| ENSMUSG00 | Med22 | 4224 | 42.9084365 | 37.9080368 | 23.618267 | 26.924859 | 31.6619628 | 16.0199763 |
| ENSMUSG00 | Abo | 1913 | 0 | 0 | 0 | 0 | 0 | 0 |
| ENSMUSG00 | Surf1 | 1745 | 27.9085751 | 32.5016862 | 34.0442848 | 23.9237396 | 21.6386923 | 29.040848 |
| ENSMUSG00 | Med28 | 5182 | 12.213571 | 12.2930291 | 13.9548852 | 9.32339977 | 8.84247115 | 13.2530956 |
| ENSMUSG00 | Qdpr | 2256 | 30.1161863 | 30.0738079 | 32.4264407 | 26.5657252 | 28.3856834 | 28.0000286 |
| ENSMUSG00 | Gnrh1 | 517 | 2.69137682 | 0.83883555 | 3.07788462 | 1.67496853 | 1.67784968 | 0.84408942 |
| ENSMUSG00 | Tnr | 4927 | 0 | 0 | 0 | 0 | 0 | 0 |
| ENSMUSG00 | Sqstm1 | 4024 | 136.703922 | 131.207467 | 88.7222381 | 112.397721 | 124.395812 | 65.3330636 |
| ENSMUSG00 | Nfe2l2 | 3790 | 152.570835 | 147.343715 | 204.308149 | 160.598419 | 137.178824 | 146.484243 |
| ENSMUSG00 | Rxrg | 2583 | 0 | 0 | 0 | 0.05587618 | 0 | 0 |
| ENSMUSG00 | Rxra | 5359 | 12.871013 | 15.6365409 | 12.207264 | 10.6044449 | 12.3781095 | 10.5454474 |
| ENSMUSG00 | Adamtsl4 | 4944 | 0.27339934 | 0.86743393 | 0.59007315 | 0.19705 | 0.36123041 | 0.40823691 |
| ENSMUSG00 | Fcrls | 2325 | 0 | 0 | 0 | 0 | 0 | 0 |
| ENSMUSG00 | Cd5l | 2031 | 0 | 0 | 0 | 0 | 0 | 0 |
| ENSMUSG00 | Prpsap1 | 3653 | 28.4807261 | 31.6054518 | 26.9712313 | 32.1113405 | 31.1494164 | 26.5130698 |
| ENSMUSG00 | Fam184b | 4106 | 0.33888013 | 0.62198769 | 0.22606885 | 0.28999208 | 0.68349992 | 0.31220362 |
| ENSMUSG00 | Ncapg | 6244 | 0.0573029 | 0.40901368 | 0.01415818 | 0 | 0 | 0.14414848 |
| ENSMUSG00 | Lcorl | 12290 | 5.89053946 | 4.39519948 | 6.02425444 | 6.567578 | 5.16906736 | 7.59207005 |
| ENSMUSG00 | Lta4h | 2047 | 30.4526592 | 31.1905402 | 25.5666742 | 27.3391554 | 34.7986422 | 23.677105 |
| ENSMUSG00 | Amdhd1 | 2412 | 1.13728365 | 1.21864551 | 1.94253539 | 0.19447205 | 2.39053917 | 0.27138945 |
| ENSMUSG00 | Dstn | 1907 | 359.467095 | 367.273171 | 359.293534 | 390.791058 | 305.221746 | 410.721455 |
| ENSMUSG00 | H2afy | 7022 | 21.7970094 | 26.2067819 | 26.0540342 | 21.2936475 | 21.2476677 | 20.4734582 |
| ENSMUSG00 | Gtf2ird2 | 4504 | 2.81571898 | 3.66961582 | 2.28664064 | 2.59560001 | 4.46367497 | 2.1134212 |
| ENSMUSG00 | Bola1 | 1493 | 5.96465347 | 6.09995818 | 9.44433216 | 6.64605643 | 3.75947846 | 9.77356463 |
| ENSMUSG00 | Gatsl2 | 7551 | 5.30178372 | 5.90285512 | 3.88690554 | 9.63812526 | 5.223598 | 6.74009758 |
| ENSMUSG00 | Fcgr1 | 2953 | 0 | 0 | 0 | 0 | 0 | 0 |
| ENSMUSG00 | Ncf1 | 3200 | 0.01242359 | 0.04517479 | 0 | 0.02255128 | 0.01594574 | 0 |
| ENSMUSG00 | Wnt11 | 3843 | 0.27931251 | 0.48901151 | 0.06901147 | 0 | 0.15933295 | 0 |
| ENSMUSG00 | Adss | 5639 | 49.0052039 | 43.6489354 | 41.9991975 | 43.7156461 | 27.6984814 | 47.5117541 |
| ENSMUSG00 | 1700016C15l | 762 | 3.86076845 | 2.02357762 | 0.81210735 | 0.09470353 | 1.5401662 | 0.10738047 |
| ENSMUSG00 | Il17rb | 3277 | 4.17329426 | 2.66150321 | 2.96747191 | 3.7216146 | 4.85817188 | 3.18772866 |
| ENSMUSG00 | Cacna1d | 9297 | 2.22788054 | 1.22319031 | 1.50239673 | 0.69082542 | 1.2239305 | 1.54899532 |
| ENSMUSG00 | Chdh | 2784 | 57.1627832 | 49.6230351 | 44.8210499 | 48.8999835 | 71.6275419 | 34.4851761 |
| ENSMUSG00 | Actr8 | 2108 | 34.059961 | 42.7231785 | 45.187368 | 48.7484169 | 48.5815637 | 36.5128235 |
| ENSMUSG00 | Lrrc27 | 3781 | 4.47919458 | 1.40188009 | 3.68251271 | 1.98494188 | 2.6990942 | 2.92150989 |
| ENSMUSG00 | Stk32c | 2289 | 0 | 0 | 0.09655274 | 0 | 0 | 0 |
| ENSMUSG00 | Fnta | 1865 | 148.619416 | 128.566078 | 188.562929 | 128.69585 | 119.672583 | 166.163245 |
| ENSMUSG00 | Skiv2l2 | 3282 | 40.0946498 | 32.1977049 | 40.0402435 | 41.6889436 | 35.3391079 | 35.9174256 |
| ENSMUSG00 | Lbp | 4963 | 0 | 0 | 0 | 0 | 0.49350515 | 0 |
| ENSMUSG00 | Celsr1 | 11039 | 3.90748223 | 4.39130001 | 2.40649585 | 4.95846196 | 3.91514995 | 2.2261482 |
| ENSMUSG00 | Fli1 | 3099 | 1.79598815 | 2.39455053 | 0.68463648 | 1.94440188 | 2.14050625 | 5.2982681 |
| ENSMUSG00 | Stard13 | 8028 | 2.08978734 | 1.79468687 | 1.59672826 | 0.8359816 | 3.49582794 | 1.06679581 |
| ENSMUSG00 | Odz1 | 18543 | 0 | 0 | 0 | 0 | 0 | 0 |
| ENSMUSG00 | Camk1g | 2907 | 0.02735155 | 0 | 0 | 0 | 0 | 0 |
| ENSMUSG00 | Diexf | 5536 | 1.03410209 | 0.92264503 | 1.38130759 | 1.18622329 | 2.05543385 | 1.98549126 |
| ENSMUSG00 | Hsd11b1 | 3290 | 0 | 0.01464634 | 0.32244506 | 0 | 0 | 0.7129541 |
| ENSMUSG00 | Syt14 | 2226 | 0.25003447 | 0 | 0 | 0 | 0.0458458 | 0.11027482 |
| ENSMUSG00 | H2-M3 | 1430 | 2.02947558 | 2.59465458 | 1.11277367 | 1.86718272 | 6.17312099 | 2.63209805 |
| ENSMUSG00 | Lonrf3 | 3983 | 3.45352655 | 2.00827251 | 3.08513993 | 2.52746433 | 2.24193215 | 4.5469145 |
| ENSMUSG00 | Atp5e | 727 | 48.7237045 | 53.0912519 | 74.5412098 | 49.9292115 | 41.9721771 | 130.107908 |
| ENSMUSG00 | Th1l | 2975 | 25.6840449 | 21.0562604 | 26.0753729 | 21.3702731 | 26.0363155 | 14.8979122 |
| ENSMUSG00 | Tubb1 | 1402 | 0 | 0.20621873 | 0 | 0 | 0 | 0 |
| ENSMUSG00 | Ctsz | 1555 | 38.5794341 | 36.9687624 | 70.3534157 | 43.4144729 | 31.2392989 | 54.777298 |
| ENSMUSG00 | Slmo2 | 2204 | 58.8215163 | 60.276779 | 92.8559588 | 67.8093595 | 51.7903824 | 85.4993115 |
| ENSMUSG00 | Sertad4 | 6621 | 0.04203117 | 0.35661315 | 0.02670403 | 0.02179855 | 0.0154135 | 0.15653773 |
| ENSMUSG00 | H2-M2 | 2176 | 0 | 0 | 0 | 0 | 0 | 0 |
| ENSMUSG00 | Ube2a | 863 | 179.613695 | 152.26469 | 208.665479 | 154.906112 | 115.060634 | 222.71655 |
| ENSMUSG00 | Slc25a5 | 1830 | 816.703571 | 771.66769 | 845.317978 | 671.323198 | 562.098749 | 894.459126 |
| ENSMUSG00 | Atp1b4 | 4193 | 0.00948139 | 0.16088962 | 0 | 0 | 0.01216942 | 0.05203843 |
| ENSMUSG00 | Ppdpf | 1490 | 9.04503885 | 10.7691847 | 11.7772696 | 9.49272581 | 7.0546533 | 10.9647711 |
| ENSMUSG00 | Kcnq2 | 14716 | 0 | 0 | 0.00300366 | 0 | 0 | 0 |
| ENSMUSG00 | Eef1a2 | 2459 | 0 | 0 | 0 | 0 | 0 | 0 |

| | | | | | | | | |
|---|---|---|---|---|---|---|---|---|
| ENSMUSG00 | Col20a1 | 4157 | 0.57381016 | 0.57958194 | 0.58482111 | 1.00686004 | 0.20867173 | 0.61018558 |
| ENSMUSG00 | Pls3 | 5453 | 69.9894761 | 93.774899 | 70.465179 | 100.716083 | 94.6229071 | 72.0454619 |
| ENSMUSG00 | Mpped2 | 3413 | 11.5783615 | 17.0833856 | 7.30437722 | 11.111113 | 12.6631527 | 12.5385025 |
| ENSMUSG00 | Nkap | 4317 | 8.24210217 | 7.72411813 | 10.5461891 | 11.8769019 | 11.8553289 | 10.3867285 |
| ENSMUSG00 | Ndufa1 | 416 | 387.711501 | 319.466845 | 310.475445 | 226.033192 | 211.587736 | 839.481925 |
| ENSMUSG00 | Wt1 | 4845 | 0.00820547 | 0 | 0 | 0 | 0.01053176 | 0 |
| ENSMUSG00 | E2f3 | 4786 | 3.91241774 | 1.75186814 | 8.9955276 | 4.1389558 | 2.51613554 | 5.03207406 |
| ENSMUSG00 | Cr1l | 1776 | 40.9194923 | 43.7096612 | 43.1814176 | 44.1273653 | 33.2993072 | 35.4447452 |
| ENSMUSG00 | Ppfibp1 | 7148 | 12.0912723 | 13.3004827 | 10.4568154 | 12.1804664 | 15.9475271 | 13.7365279 |
| ENSMUSG00 | Cd46 | 3358 | 0.76953729 | 1.14797956 | 0.90825705 | 2.65403961 | 1.29161464 | 1.67319109 |
| ENSMUSG00 | Cd34 | 2613 | 7.9571819 | 2.36045335 | 3.24789662 | 3.13456721 | 4.99913977 | 3.1731689 |
| ENSMUSG00 | 5033414D02 | 3045 | 10.9670291 | 11.2039412 | 8.23068801 | 11.0082822 | 12.886464 | 11.6891311 |
| ENSMUSG00 | Cd274 | 3615 | 2.28745227 | 0.79977497 | 1.56509984 | 0.81845855 | 4.88385229 | 3.84787442 |
| ENSMUSG00 | Pdcd1lg2 | 1639 | 0 | 0 | 0 | 0 | 0 | 0 |
| ENSMUSG00 | Gtf3a | 2818 | 2.22901559 | 4.46297424 | 3.27827721 | 1.56210413 | 3.83874793 | 3.71662934 |
| ENSMUSG00 | Mtif3 | 6417 | 4.04555536 | 3.19891294 | 4.23626826 | 3.17692928 | 3.66575647 | 3.79983286 |
| ENSMUSG00 | Lnx2 | 5236 | 4.19118129 | 4.04928085 | 3.09818113 | 3.07345146 | 2.7189379 | 2.35970427 |
| ENSMUSG00 | Il19 | 1459 | 0 | 0 | 0 | 0 | 0.41968232 | 0 |
| ENSMUSG00 | Dyrk3 | 2069 | 0.82623763 | 0.32605616 | 0.0640916 | 0.85452885 | 1.80035062 | 0 |
| ENSMUSG00 | Mapkapk2 | 2581 | 79.0950765 | 57.4279334 | 51.2234454 | 40.7093815 | 54.1895762 | 40.7058932 |
| ENSMUSG00 | Il10 | 1305 | 0 | 0 | 0 | 0 | 0 | 0 |
| ENSMUSG00 | Lamp2 | 9540 | 92.1001965 | 100.94403 | 108.271349 | 108.223441 | 84.6481576 | 132.536449 |
| ENSMUSG00 | Atxn10 | 2666 | 70.5488291 | 76.5271553 | 75.8030106 | 66.5609509 | 78.1664362 | 68.1559065 |
| ENSMUSG00 | Foxred2 | 8797 | 0.24855649 | 1.06813189 | 0.36679943 | 1.18947288 | 1.32249786 | 0.54567881 |
| ENSMUSG00 | Eif3d | 1895 | 15.9021922 | 20.3171331 | 18.4971301 | 16.8128999 | 19.8720134 | 14.7815591 |
| ENSMUSG00 | H3f3b | 4546 | 152.7429 | 135.379947 | 153.316027 | 147.518891 | 116.420055 | 258.305113 |
| ENSMUSG00 | Nup50 | 5088 | 9.73571712 | 8.90236942 | 9.0957802 | 7.20506235 | 8.22359044 | 4.1115661 |
| ENSMUSG00 | Phf21b | 6107 | 0.14321608 | 0.04734217 | 0.06514108 | 0.33677363 | 0.04177696 | 0.01786451 |
| ENSMUSG00 | Nlrp14 | 3634 | 0 | 0 | 0 | 0 | 0 | 0 |
| ENSMUSG00 | Ift27 | 1005 | 7.87198072 | 11.3633699 | 9.3681518 | 7.86265448 | 9.39291495 | 10.28566 |
| ENSMUSG00 | Pacsin2 | 5187 | 82.6763771 | 73.7428143 | 66.8524115 | 61.9801459 | 80.626986 | 66.617005 |
| ENSMUSG00 | Cmah | 9953 | 0.03195457 | 0 | 0.02664634 | 0 | 0.30760399 | 0 |
| ENSMUSG00 | Ttll12 | 5401 | 2.13462125 | 1.6416044 | 2.8480358 | 2.93279347 | 3.07046327 | 2.16136745 |
| ENSMUSG00 | Bik | 938 | 8.60379802 | 3.44188873 | 15.833496 | 5.77004976 | 6.03830247 | 13.0266934 |
| ENSMUSG00 | Scube1 | 10802 | 0 | 0.01784353 | 0.01636802 | 0.13361246 | 0 | 0 |
| ENSMUSG00 | Tox4 | 6057 | 14.0591447 | 11.9809777 | 13.6101102 | 10.9074168 | 10.9600982 | 8.80335475 |
| ENSMUSG00 | Mrps18c | 1194 | 24.0397462 | 26.6760789 | 27.6539169 | 18.2525585 | 18.4190687 | 65.6510161 |
| ENSMUSG00 | Sulf1 | 4586 | 0 | 0 | 0 | 0 | 0 | 0.32115799 |
| ENSMUSG00 | Snx6 | 4003 | 44.194826 | 42.9380562 | 44.2791383 | 49.124942 | 39.9874447 | 55.2715148 |
| ENSMUSG00 | Plcg1 | 7278 | 5.82840034 | 7.70665274 | 6.4195312 | 9.56337784 | 7.51583881 | 5.62131889 |
| ENSMUSG00 | Kctd2 | 2620 | 32.7906845 | 34.742057 | 27.7864257 | 29.5129089 | 38.8735291 | 24.2661008 |
| ENSMUSG00 | Tmprss6 | 3173 | 0 | 0 | 0 | 0 | 0 | 0 |
| ENSMUSG00 | Kctd5 | 2744 | 13.1117747 | 10.4485908 | 9.55236622 | 9.13885603 | 10.9714146 | 7.22618904 |
| ENSMUSG00 | Pom121l2 | 4900 | 0 | 0 | 0 | 0 | 0 | 0 |
| ENSMUSG00 | Etaa1 | 4179 | 6.52602528 | 5.68459344 | 4.31547068 | 6.11296662 | 7.61915734 | 12.7138065 |
| ENSMUSG00 | Matn4 | 5221 | 0.34265402 | 0.60913717 | 1.4646464 | 1.47203131 | 0.92846307 | 0.32388961 |
| ENSMUSG00 | Svs4 | 728 | 0 | 0 | 0 | 0 | 0 | 0 |
| ENSMUSG00 | Svs6 | 531 | 0 | 0 | 0 | 0 | 0 | 0 |
| ENSMUSG00 | Slpi | 1325 | 0.03000414 | 0 | 0 | 0 | 0 | 0 |
| ENSMUSG00 | Svs3a | 1288 | 0 | 0 | 0 | 0 | 0 | 0 |
| ENSMUSG00 | Svs5 | 643 | 0 | 0 | 0 | 0 | 0 | 0 |
| ENSMUSG00 | Rbpjl | 4128 | 0 | 0.14007687 | 0.01070781 | 0.03496322 | 0 | 0.30393251 |
| ENSMUSG00 | Sdc4 | 2878 | 193.459175 | 220.472227 | 321.484704 | 319.710986 | 194.566869 | 258.77738 |
| ENSMUSG00 | Ccdc70 | 1112 | 0 | 0 | 0 | 0 | 0 | 0 |
| ENSMUSG00 | Il13ra1 | 3815 | 64.9009522 | 74.0037653 | 38.9879952 | 80.818105 | 62.4889195 | 67.0748795 |
| ENSMUSG00 | Prl5a1 | 1056 | 0 | 0 | 0 | 0 | 0 | 0 |
| ENSMUSG00 | Nbr1 | 7212 | 45.9128393 | 38.6921362 | 39.568372 | 39.4390955 | 46.9935092 | 35.2770154 |
| ENSMUSG00 | Cyth1 | 3950 | 17.2206651 | 17.566703 | 21.3847398 | 15.4284995 | 14.1452865 | 16.8688465 |
| ENSMUSG00 | Rnd3 | 5138 | 5.12225148 | 5.49576783 | 2.40881978 | 4.74726783 | 6.69361178 | 3.58848893 |
| ENSMUSG00 | Brca1 | 8453 | 0 | 0 | 0.13072827 | 0 | 0.15694851 | 0 |
| ENSMUSG00 | Gast | 457 | 0 | 0 | 0 | 0 | 0 | 0 |
| ENSMUSG00 | Cntnap1 | 5850 | 0.0747539 | 0.09884398 | 0.16622915 | 0.28986993 | 0.34017584 | 0.36832419 |
| ENSMUSG00 | Nt5c3l | 3406 | 8.15886109 | 8.50265758 | 8.03315937 | 9.82033332 | 7.41575339 | 9.25704139 |
| ENSMUSG00 | Ccdc56 | 944 | 87.9337323 | 76.7205747 | 120.290821 | 65.6280407 | 54.1614709 | 148.450314 |
| ENSMUSG00 | Zpbp2 | 1747 | 0 | 0.93780139 | 0.07590471 | 1.63164369 | 0.847032 | 0.01561227 |
| ENSMUSG00 | Gsdma | 2714 | 0 | 0 | 0.03257321 | 0 | 0.11280702 | 0 |
| ENSMUSG00 | Med24 | 6749 | 8.85942253 | 5.90460626 | 7.27637392 | 8.91224896 | 11.3030718 | 6.45797504 |
| ENSMUSG00 | Gsdma2 | 3163 | 0 | 0 | 0 | 0 | 0 | 0 |
| ENSMUSG00 | Psmd3 | 3525 | 60.0787077 | 62.9909576 | 52.9544349 | 52.4801822 | 62.7515856 | 43.4227732 |
| ENSMUSG00 | Exosc10 | 3692 | 22.5482059 | 22.6967018 | 22.8671506 | 22.3216115 | 29.7008889 | 20.4781419 |
| ENSMUSG00 | Glod4 | 5898 | 21.9133718 | 17.8513693 | 19.7776642 | 19.5337349 | 18.6006626 | 16.5044401 |
| ENSMUSG00 | Vps53 | 5462 | 21.2825018 | 13.3302296 | 21.5425313 | 20.0558563 | 20.6459704 | 15.9243546 |
| ENSMUSG00 | Taok1 | 12401 | 9.39628364 | 9.27902781 | 10.7180826 | 10.1865364 | 5.91693641 | 8.49644416 |
| ENSMUSG00 | Dnttip1 | 2251 | 11.1795732 | 11.9235221 | 10.584093 | 12.7433254 | 10.450093 | 9.19655762 |
| ENSMUSG00 | Tnnc2 | 793 | 0 | 0 | 0 | 0 | 0 | 0 |
| ENSMUSG00 | Acot8 | 1308 | 7.05143079 | 7.22059836 | 11.219428 | 6.06884541 | 5.18846176 | 10.0924506 |
| ENSMUSG00 | Cd300lg | 2729 | 2.06862522 | 1.51851742 | 1.73308802 | 0.44953811 | 4.99232039 | 1.65906563 |
| ENSMUSG00 | Spint4 | 747 | 0 | 0 | 0 | 0 | 0 | 0 |
| ENSMUSG00 | Pyy | 813 | 0 | 0 | 0 | 0 | 0 | 0 |
| ENSMUSG00 | Mpp2 | 6761 | 1.54059102 | 2.28067763 | 0.85644748 | 1.4355968 | 2.1811304 | 1.7750098 |
| ENSMUSG00 | Ppy | 496 | 0 | 0 | 0 | 0 | 0 | 0 |
| ENSMUSG00 | Vtn | 2436 | 0 | 0 | 0 | 0 | 0 | 0 |
| ENSMUSG00 | Nlk | 6137 | 2.1766077 | 4.00441348 | 3.53643554 | 4.59771205 | 3.32581886 | 3.20433679 |
| ENSMUSG00 | Traf4 | 4590 | 14.3777991 | 17.0594703 | 14.6472774 | 12.1374023 | 14.5186159 | 13.6729728 |
| ENSMUSG00 | Aldoc | 3308 | 6.92235695 | 8.98761633 | 4.60992619 | 2.21422462 | 5.24454891 | 6.4971027 |
| ENSMUSG00 | Stac2 | 3127 | 0 | 0 | 0 | 0 | 0 | 0 |
| ENSMUSG00 | Rpl19 | 1015 | 3.05510097 | 1.80402444 | 2.17743069 | 1.81298943 | 1.60871333 | 1.61229395 |
| ENSMUSG00 | Nek8 | 4506 | 12.8548014 | 16.0835351 | 10.4569828 | 11.2346002 | 19.9757051 | 12.214874 |
| ENSMUSG00 | Cacnb4 | 8920 | 1.43957626 | 2.8090751 | 1.45687689 | 3.38572548 | 2.98607267 | 1.40959739 |
| ENSMUSG00 | Plxdc1 | 5773 | 0.0275458 | 0 | 0 | 0 | 0 | 0 |
| ENSMUSG00 | Arl5b | 7223 | 5.43796411 | 6.62455172 | 5.8809319 | 7.93275463 | 6.91607669 | 3.34560853 |
| ENSMUSG00 | Zfp207 | 5191 | 49.5660701 | 55.4919189 | 50.1538924 | 54.8980905 | 51.0755247 | 65.346694 |
| ENSMUSG00 | Psmd11 | 6518 | 9.67354896 | 12.9965887 | 10.8910954 | 12.2063376 | 13.5746756 | 11.189381 |
| ENSMUSG00 | C1qtnf1 | 3193 | 0 | 0 | 0 | 0 | 0 | 0.11958815 |
| ENSMUSG00 | Pipox | 2656 | 3.66720359 | 3.42892981 | 1.41459211 | 3.72231935 | 5.05268706 | 4.4362365 |
| ENSMUSG00 | Timp2 | 4085 | 12.7684677 | 16.1958348 | 5.4968265 | 5.53817428 | 17.1253761 | 4.67374479 |
| ENSMUSG00 | Zc3h18 | 6400 | 4.65263358 | 5.08969295 | 4.85529619 | 4.41441264 | 4.83155996 | 5.04580826 |
| ENSMUSG00 | Top2b | 6319 | 30.4756366 | 27.1320401 | 20.1108243 | 28.7331616 | 28.6261279 | 27.8184918 |
| ENSMUSG00 | Rarb | 3479 | 9.81602695 | 8.25499268 | 22.7679513 | 13.534656 | 12.8335955 | 21.0655233 |
| ENSMUSG00 | Igfbp4 | 4493 | 0.6370787 | 1.42639591 | 0.29513806 | 0 | 0.7836234 | 0.73452734 |
| ENSMUSG00 | Cdc6 | 5347 | 0.1115265 | 0.09011865 | 0.04959997 | 0.08772519 | 0.14314487 | 0.4947896 |
| ENSMUSG00 | Suz12 | 4928 | 6.89751133 | 8.10603907 | 5.01396718 | 8.4420855 | 7.27912788 | 8.28533583 |
| ENSMUSG00 | Atad5 | 7603 | 0.35556658 | 0.10774293 | 0.68020724 | 0 | 1.63085747 | 0.36232258 |
| ENSMUSG00 | Crlf3 | 2722 | 7.66775432 | 5.73563823 | 9.2236028 | 6.87971385 | 8.60437417 | 7.86575746 |
| ENSMUSG00 | Krt27 | 1542 | 0 | 0 | 0 | 0 | 0 | 0 |
| ENSMUSG00 | Tns4 | 5301 | 0.30748438 | 0.12726093 | 0 | 0 | 0 | 0 |
| ENSMUSG00 | Tnfaip1 | 6409 | 18.5347754 | 19.8565133 | 20.9457009 | 16.9854156 | 19.4105639 | 15.3544856 |
| ENSMUSG00 | Abr | 12142 | 17.2059834 | 24.5019843 | 12.5120849 | 14.8018995 | 18.5496972 | 10.903566 |
| ENSMUSG00 | Rab11fip4 | 7435 | 11.9025823 | 9.71506076 | 8.92954515 | 10.9823358 | 7.60419965 | 6.74620869 |
| ENSMUSG00 | Cd40 | 3519 | 0.23724498 | 0.10954576 | 0.18841365 | 0 | 0.5945102 | 0 |
| ENSMUSG00 | Slc35c2 | 3025 | 10.3955653 | 8.25143042 | 8.9864904 | 7.21640891 | 8.33290237 | 7.42952157 |
| ENSMUSG00 | Zfp334 | 8200 | 1.47386172 | 1.20466105 | 1.26676014 | 1.40807979 | 2.61976881 | 0.52553573 |
| ENSMUSG00 | Elmo2 | 8032 | 15.3438732 | 10.8347503 | 10.4341004 | 11.5182224 | 12.642242 | 8.84252498 |
| ENSMUSG00 | Wsb1 | 4976 | 155.330798 | 160.343872 | 105.72555 | 176.617837 | 172.99336 | 230.930029 |
| ENSMUSG00 | Ttpal | 6037 | 7.54017313 | 3.76743427 | 4.5688173 | 9.70635089 | 7.80144855 | 4.22876633 |
| ENSMUSG00 | Rhot1 | 7325 | 21.6552037 | 24.7411889 | 26.2133524 | 27.8459792 | 23.482582 | 29.4677809 |
| ENSMUSG00 | Hn1l | 4342 | 0 | 1.02099325 | 0.49682319 | 0.1495801 | 0.38780986 | 0.39573981 |
| ENSMUSG00 | Rhbdl3 | 4725 | 1.8762904 | 4.27304111 | 0.57064813 | 3.39820314 | 3.18577373 | 0.62342033 |
| ENSMUSG00 | Ada | 1990 | 3.11651003 | 1.76764336 | 5.24202761 | 3.00985899 | 2.4872153 | 5.53716294 |
| ENSMUSG00 | Serinc3 | 3987 | 1130.20434 | 1146.8905 | 1460.20184 | 1072.50642 | 928.278177 | 1259.55201 |
| ENSMUSG00 | Tha1 | 2684 | 7.05052486 | 7.00175575 | 9.23890981 | 9.67923698 | 8.86916072 | 7.99744452 |
| ENSMUSG00 | Pgs1 | 5450 | 6.3973498 | 6.06530263 | 5.76651567 | 5.91877941 | 7.19986844 | 5.75019459 |
| ENSMUSG00 | Birc5 | 3553 | 0.01118927 | 0.16274622 | 0 | 0.55854558 | 0.01436149 | 0.10747114 |
| ENSMUSG00 | Afmid | 3560 | 0.72587254 | 0.66324036 | 1.01813234 | 0.71961381 | 0.84566185 | 0.55928333 |
| ENSMUSG00 | Trp53tg5 | 929 | 0 | 0.25934576 | 0 | 0 | 0 | 0.17615483 |
| ENSMUSG00 | Pigt | 3953 | 10.1173826 | 17.6874595 | 7.54774704 | 7.23831553 | 11.3721824 | 9.1973423 |
| ENSMUSG00 | Wfdc2 | 1133 | 133.758069 | 253.350958 | 82.3956685 | 161.33419 | 118.941446 | 145.544986 |
| ENSMUSG00 | Etv4 | 3747 | 0.62598702 | 0.28292013 | 0.17694893 | 0.3274058 | 0.08170757 | 0.45130174 |
| ENSMUSG00 | Spinlw1 | 774 | 0 | 0 | 0 | 0 | 0 | 0 |
| ENSMUSG00 | Dbndd2 | 1831 | 2.93117961 | 5.0791826 | 2.63134948 | 4.94625515 | 5.71294759 | 5.31788433 |
| ENSMUSG00 | Mmp9 | 3524 | 0 | 0 | 0 | 0 | 0 | 0 |
| ENSMUSG00 | Slc12a5 | 7038 | 0.23159629 | 0.41079661 | 0.07536546 | 0 | 0.21025361 | 0.32940386 |
| ENSMUSG00 | Ghdc | 2897 | 10.4843587 | 9.24807105 | 5.90476812 | 4.7079782 | 8.89482906 | 5.88424216 |
| ENSMUSG00 | Pltp | 2547 | 1.45161354 | 0.43513474 | 0.52063419 | 0.90665522 | 0.72122086 | 0.59967797 |
| ENSMUSG00 | Slc12a7 | 5718 | 14.3712099 | 9.51425204 | 15.468326 | 10.7085046 | 20.1142797 | 8.34266248 |
| ENSMUSG00 | Ctsa | 4871 | 41.8693523 | 43.1808294 | 44.1746195 | 38.2302262 | 41.0536577 | 32.4148813 |
| ENSMUSG00 | Zswim1 | 2965 | 13.7836877 | 12.0425476 | 11.3448912 | 7.80053644 | 11.5648312 | 6.74276916 |
| ENSMUSG00 | Slc12a4 | 4805 | 4.21962436 | 2.73775207 | 4.85714321 | 1.94490105 | 5.44776916 | 4.48427992 |
| ENSMUSG00 | 1700020C07 | 1134 | 0.28046194 | 0 | 0 | 0.09545514 | 0 | 0 |
| ENSMUSG00 | Myo1c | 7137 | 35.2602202 | 32.8602233 | 30.5331492 | 35.621982 | 37.7782499 | 23.4759856 |
| ENSMUSG00 | Crk | 3453 | 121.580618 | 92.0468498 | 111.791108 | 122.854503 | 104.786571 | 97.6689587 |
| ENSMUSG00 | Cox7c | 1215 | 9.58712411 | 8.36818049 | 16.2255325 | 12.5321998 | 7.93743624 | 15.8260252 |
| ENSMUSG00 | Pitpna | 4289 | 75.2007958 | 77.2061624 | 73.6250796 | 79.9878945 | 71.2632295 | 76.7492705 |
| ENSMUSG00 | Mlx | 2068 | 28.7785079 | 30.7340025 | 34.8185698 | 29.8706287 | 22.2561852 | 25.6260271 |
| ENSMUSG00 | Fam134c | 5685 | 7.9790683 | 5.48401549 | 6.51559269 | 3.3003805 | 8.21268846 | 4.36586134 |
| ENSMUSG00 | Jph2 | 5924 | 0 | 0 | 0 | 0 | 0 | 0 |
| ENSMUSG00 | Dhx58 | 3832 | 0.34236192 | 0.6790365 | 0.28837319 | 0.51787903 | 0.81226747 | 0.6761719 |
| ENSMUSG00 | Rab5a | 2351 | 82.4533062 | 71.6750269 | 101.846612 | 86.8670234 | 75.3566894 | 116.175346 |
| ENSMUSG00 | Hspb9 | 1376 | 0 | 0 | 0 | 0 | 0 | 0 |
| ENSMUSG00 | Nkiras2 | 3570 | 2.59468542 | 2.21360687 | 3.07060422 | 2.13258022 | 2.47270673 | 2.68926415 |
| ENSMUSG00 | Ppp2r5c | 4570 | 32.6308112 | 31.400268 | 34.8875378 | 33.3739173 | 34.0994643 | 35.2720171 |
| ENSMUSG00 | Ift52 | 2708 | 30.2863945 | 25.3388086 | 32.1883436 | 26.7817243 | 27.2844137 | 27.455933 |
| ENSMUSG00 | Mybl2 | 4617 | 0.41331234 | 0.13567766 | 0.46911204 | 0.67990857 | 0.56364418 | 0.68526276 |
| ENSMUSG00 | Sgk2 | 3511 | 67.5877138 | 42.0104527 | 62.721043 | 43.2039064 | 56.9704908 | 50.1834718 |
| ENSMUSG00 | Eya2 | 2930 | 5.72587467 | 0.93741543 | 5.1292241 | 2.43830881 | 5.50318592 | 4.7288453 |
| ENSMUSG00 | BC034902 | 2314 | 0 | 0 | 0 | 0 | 0 | 0 |
| ENSMUSG00 | Prl3c1 | 924 | 0 | 0 | 0 | 0 | 0 | 0 |
| ENSMUSG00 | B4galt5 | 4581 | 1.29307282 | 1.43055144 | 1.87189643 | 1.33899751 | 1.30322767 | 1.91118952 |
| ENSMUSG00 | Gdap1l1 | 2821 | 0 | 0.102488 | 0.04700657 | 0 | 0.43411309 | 0 |
| ENSMUSG00 | Hnf4a | 5054 | 0 | 0.15254909 | 0.00874591 | 0 | 0 | 0 |
| ENSMUSG00 | Ptgis | 2556 | 0 | 0 | 0 | 0 | 0 | 0 |
| ENSMUSG00 | Cadps2 | 10481 | 8.10585458 | 6.19743574 | 5.28009803 | 6.67178726 | 8.80218371 | 6.84663447 |
| ENSMUSG00 | Ddx27 | 4916 | 10.8041745 | 7.61612399 | 5.55670036 | 6.54702679 | 8.63587158 | 4.89345643 |
| ENSMUSG00 | Cyth3 | 4804 | 13.6297411 | 15.3867615 | 12.1085815 | 12.1900829 | 16.5060341 | 10.9689016 |
| ENSMUSG00 | Cyth4 | 2778 | 0.41501492 | 0 | 0.14320252 | 0 | 0.51430472 | 0 |
| ENSMUSG00 | Rac3 | 1334 | 5.03648891 | 3.35932468 | 7.98549038 | 5.22026581 | 3.02180187 | 6.23595575 |
| ENSMUSG00 | Rrp7a | 5820 | 3.39492677 | 3.39457754 | 3.09109108 | 4.38316245 | 3.71738546 | 3.05082305 |
| ENSMUSG00 | Cyb5r3 | 5513 | 54.1851407 | 57.4251632 | 59.138907 | 45.1794583 | 57.949599 | 53.109605 |
| ENSMUSG00 | Ints2 | 9573 | 6.55323899 | 6.11076369 | 7.63708851 | 8.27327773 | 5.97519767 | 8.08295744 |
| ENSMUSG00 | Med13l | 9289 | 2.79045896 | 2.09055186 | 2.95027911 | 2.98320704 | 2.71364298 | 2.48405047 |
| ENSMUSG00 | Hist1h2bc | 734 | 216.759446 | 192.089278 | 185.298462 | 181.737491 | 118.806643 | 227.115252 |
| ENSMUSG00 | Baiap2l2 | 3171 | 5.7545776 | 5.36417978 | 4.58606318 | 6.05349974 | 5.06493685 | 8.97112858 |
| ENSMUSG00 | Mafk | 4075 | 15.0046452 | 13.1492819 | 12.4307514 | 9.41232229 | 10.5684077 | 7.61012634 |
| ENSMUSG00 | Med1 | 8223 | 8.02071533 | 5.49664146 | 5.41839447 | 6.19138573 | 9.1032097 | 6.5740405 |
| ENSMUSG00 | Erbb3 | 5988 | 12.116525 | 12.7708557 | 13.2502185 | 12.491329 | 20.1958101 | 8.38553956 |
| ENSMUSG00 | Stard3 | 3609 | 14.441517 | 10.1206215 | 17.991828 | 10.2477406 | 13.8134578 | 10.9808952 |
| ENSMUSG00 | Ikzf3 | 9172 | 0 | 0.0682974 | 0 | 0 | 0 | 0.07731581 |
| ENSMUSG00 | Mfng | 2823 | 0.30981954 | 0.66570005 | 0.20355082 | 0 | 0.32535415 | 0 |
| ENSMUSG00 | Vmp1 | 4225 | 45.0718939 | 55.0979176 | 54.0779471 | 50.3867604 | 44.1181896 | 51.4183436 |
| ENSMUSG00 | Uchl5 | 1840 | 47.8578203 | 46.143756 | 51.0002787 | 46.2595343 | 32.9730222 | 79.2446856 |
| ENSMUSG00 | Glrx2 | 2340 | 20.3874259 | 17.5448071 | 19.3430287 | 26.5989431 | 18.535222 | 32.6130088 |
| ENSMUSG00 | Trove2 | 11579 | 14.6950045 | 12.6344277 | 12.0659345 | 13.7111146 | 13.0006698 | 13.3676119 |
| ENSMUSG00 | Stk4 | 5846 | 10.6563185 | 9.09162602 | 10.252771 | 13.3625772 | 12.2983516 | 11.9344042 |
| ENSMUSG00 | Wfdc15b | 706 | 25.114652 | 40.7468923 | 39.3183533 | 26.6782256 | 24.7181594 | 53.235769 |
| ENSMUSG00 | Pmp22 | 2346 | 246.294183 | 204.104295 | 554.48254 | 268.615903 | 141.312176 | 336.236079 |
| ENSMUSG00 | Gdf9 | 2005 | 0 | 0 | 0 | 0 | 0.25449564 | 0 |
| ENSMUSG00 | Zcchc10 | 1386 | 25.9299815 | 26.2139808 | 23.2171297 | 29.0530748 | 24.5191676 | 34.5951052 |
| ENSMUSG00 | Prl8a2 | 998 | 0 | 0 | 0 | 0 | 0 | 0 |
| ENSMUSG00 | Tbx5 | 3932 | 0 | 0 | 0 | 0 | 0 | 0 |
| ENSMUSG00 | Psmb6 | 1977 | 65.4346654 | 81.0175689 | 85.9666598 | 70.5032565 | 47.6452654 | 90.99824 |
| ENSMUSG00 | Spag7 | 1590 | 136.318792 | 117.496123 | 135.05192 | 120.613878 | 104.331288 | 124.34834 |
| ENSMUSG00 | Pfn1 | 1023 | 149.733985 | 167.781141 | 198.108945 | 133.958558 | 96.5161654 | 216.517444 |
| ENSMUSG00 | Tomm34 | 3033 | 16.6073834 | 17.4761248 | 18.5959617 | 14.0735439 | 13.3748661 | 14.388204 |
| ENSMUSG00 | Ywhab | 3375 | 138.59644 | 132.380649 | 126.738144 | 118.840891 | 104.910229 | 143.525213 |
| ENSMUSG00 | Ksr1 | 9548 | 0.4621762 | 0.34317952 | 0.31480156 | 0.11337048 | 0.65733601 | 0.37992533 |
| ENSMUSG00 | Gpx3 | 1806 | 913.341441 | 665.805358 | 1147.41229 | 708.234992 | 642.915383 | 876.760344 |
| ENSMUSG00 | Anxa6 | 2904 | 106.863389 | 117.014735 | 100.732713 | 102.070935 | 109.397457 | 72.9671737 |
| ENSMUSG00 | Il12rb2 | 4495 | 0.29186448 | 0.18224016 | 0.41300943 | 0.38530326 | 0.18162892 | 0.0687455 |
| ENSMUSG00 | Zkscan6 | 2299 | 10.5311386 | 7.7131843 | 9.11338564 | 6.85857045 | 11.7189763 | 7.3554921 |
| ENSMUSG00 | Kpna2 | 3208 | 1.35079407 | 1.74240252 | 1.66721415 | 1.03477185 | 1.68603382 | 0.61214901 |
| ENSMUSG00 | Smurf2 | 6525 | 5.53225691 | 6.91226205 | 7.30261866 | 7.73621154 | 5.99022282 | 8.83656512 |
| ENSMUSG00 | Ccdc45 | 4280 | 4.32851727 | 5.89946161 | 4.89525084 | 5.96871204 | 3.86274434 | 6.14316641 |
| ENSMUSG00 | Vezf1 | 4595 | 27.945637 | 22.7876254 | 39.1804367 | 30.3026354 | 28.9279019 | 32.2012879 |
| ENSMUSG00 | Cuedc1 | 4512 | 4.41433859 | 4.40000314 | 9.66915316 | 7.94892559 | 5.22477518 | 9.81089085 |
| ENSMUSG00 | Srsf1 | 3019 | 137.346692 | 95.3354153 | 114.670035 | 121.954681 | 108.948665 | 199.613499 |
| ENSMUSG00 | Abi3 | 6949 | 0.02860518 | 0.0138686 | 0.05088714 | 0.04153927 | 0.02202894 | 0.03139979 |
| ENSMUSG00 | Shroom1 | 4127 | 0.25645856 | 1.54121889 | 0 | 0.44588909 | 0.346193 | 0.01982649 |
| ENSMUSG00 | Kif3a | 5982 | 10.4672153 | 11.4867714 | 7.26352586 | 8.97527286 | 9.04178508 | 8.04287256 |
| ENSMUSG00 | 8-Sep | 3877 | 5.59878575 | 4.59865453 | 3.86495352 | 7.27783307 | 6.51484551 | 3.5245278 |
| ENSMUSG00 | Mtmr4 | 6178 | 13.8931827 | 10.4671747 | 15.5543553 | 11.2778291 | 11.5631153 | 9.18278563 |
| ENSMUSG00 | Mrm1 | 2513 | 12.8774219 | 8.43694172 | 11.1164205 | 9.60560596 | 7.59405673 | 7.90128825 |
| ENSMUSG00 | Mapt | 4731 | 5.13434762 | 9.19728541 | 3.50363372 | 8.82412292 | 4.88584829 | 3.63777159 |
| ENSMUSG00 | 1700081L11 | 8804 | 10.4626816 | 9.26072923 | 8.52507149 | 11.2746143 | 13.1565054 | 10.4092217 |
| ENSMUSG00 | 4933439F18 | 5438 | 9.4819526 | 4.64319523 | 9.39634618 | 6.21715258 | 8.28545234 | 6.735903 |
| ENSMUSG00 | Myo1b | 11552 | 1.47637648 | 2.8531446 | 1.07137431 | 1.93653632 | 1.61224266 | 1.04121502 |
| ENSMUSG00 | Dhx40 | 5576 | 14.1882005 | 16.41934 | 13.5078803 | 12.2236338 | 13.9736865 | 11.5780257 |
| ENSMUSG00 | Ypel2 | 4842 | 2.70127077 | 4.35887271 | 1.99008711 | 4.41151804 | 3.12987064 | 3.13754111 |
| ENSMUSG00 | Akap1 | 4448 | 5.03199989 | 7.09580027 | 5.27679376 | 5.75949901 | 7.00924362 | 4.60504813 |
| ENSMUSG00 | Nol11 | 5505 | 10.3414801 | 9.56726272 | 9.08927998 | 10.8737715 | 9.90866408 | 6.77778501 |
| ENSMUSG00 | Derl2 | 6959 | 12.699588 | 11.4113029 | 14.5645676 | 11.4224377 | 10.8373306 | 14.8150792 |
| ENSMUSG00 | C1qbp | 1652 | 50.2959772 | 47.165543 | 56.2155401 | 47.0027602 | 36.5091866 | 64.3892815 |
| ENSMUSG00 | Rpain | 3194 | 6.88314988 | 6.50230325 | 5.36954136 | 8.90189452 | 9.13809861 | 12.2197899 |
| ENSMUSG00 | 6330403K07 | 1484 | 0 | 0 | 0 | 0 | 0 | 0 |
| ENSMUSG00 | Slc13a3 | 6763 | 0.2880406 | 0.12112517 | 0 | 0.11737468 | 0.07544932 | 0.35086406 |
| ENSMUSG00 | Kcnab3 | 2851 | 0.11155519 | 1.53804498 | 0.49612732 | 1.55667888 | 1.19914668 | 0.49746799 |
| ENSMUSG00 | Chd3 | 8424 | 5.35641861 | 7.60206334 | 5.89777677 | 9.21754034 | 6.4388696 | 6.23910611 |
| ENSMUSG00 | Kdm6b | 6931 | 1.95593981 | 1.71721991 | 1.51144666 | 1.32750272 | 1.00123894 | 1.57800178 |
| ENSMUSG00 | 1700125H20 | 988 | 0 | 0 | 0 | 0 | 0 | 0 |
| ENSMUSG00 | Appbp2 | 6450 | 19.1381034 | 22.0985265 | 24.5131771 | 27.6405243 | 21.8661865 | 16.698845 |
| ENSMUSG00 | Wnt9b | 4514 | 3.47001757 | 4.41940342 | 1.00659323 | 3.26129246 | 4.2164019 | 0.12688689 |
| ENSMUSG00 | Adora2b | 1843 | 0.81970063 | 2.30081764 | 4.12518557 | 1.74243188 | 2.27029996 | 2.97460797 |
| ENSMUSG00 | Ncor1 | 13827 | 22.6508769 | 26.6250393 | 21.5750516 | 27.4288255 | 23.9318758 | 25.6729175 |
| ENSMUSG00 | Trpv2 | 3728 | 0 | 0 | 0 | 0.01935732 | 0.24637199 | 0 |
| ENSMUSG00 | Cenpv | 1629 | 3.19703373 | 3.7567085 | 3.14758366 | 3.16742318 | 1.94207201 | 4.45368572 |
| ENSMUSG00 | Pcgf2 | 3994 | 0.98542628 | 1.06169427 | 1.6047239 | 2.13203869 | 0.7537697 | 1.91892179 |
| ENSMUSG00 | Cwc25 | 6897 | 5.91980259 | 4.88361941 | 4.26188569 | 4.03352999 | 5.24542561 | 4.04156609 |
| ENSMUSG00 | 1700001P01 | 621 | 0 | 0 | 0 | 0 | 0 | 0 |
| ENSMUSG00 | Pip4k2b | 5193 | 0.35981275 | 0.15774495 | 0.22981894 | 0.36130682 | 0.12773789 | 0.26260966 |
| ENSMUSG00 | Trim37 | 7379 | 3.7444178 | 4.48625637 | 5.27738497 | 5.86290438 | 6.13367602 | 4.99733192 |
| ENSMUSG00 | Ybx2 | 1933 | 21.2659942 | 17.5495371 | 27.0973533 | 27.1595317 | 18.9798056 | 32.6364411 |
| ENSMUSG00 | Ctdnep1 | 2308 | 1.5847072 | 2.17131282 | 2.81528203 | 0.75040647 | 1.50337676 | 2.41075668 |
| ENSMUSG00 | Rai12 | 2356 | 22.0545047 | 26.3021072 | 30.0369909 | 29.0371632 | 22.0045831 | 22.7250072 |
| ENSMUSG00 | Slc2a4 | 2783 | 57.5404511 | 41.5203336 | 63.9918166 | 38.1953659 | 43.5640205 | 37.4279006 |
| ENSMUSG00 | Gabarap | 2003 | 64.6845287 | 71.1128922 | 81.9598827 | 62.0222063 | 43.4603081 | 102.548831 |
| ENSMUSG00 | Cldn7 | 1593 | 88.9195085 | 96.0401468 | 125.80738 | 91.1451018 | 68.5156421 | 85.6077817 |
| ENSMUSG00 | 2810408A11 | 2447 | 1.51093571 | 1.16183085 | 2.90825367 | 0.66354393 | 0.62557878 | 3.02060778 |
| ENSMUSG00 | Phf23 | 3443 | 6.31607547 | 5.71015637 | 7.81844972 | 7.29395963 | 5.05373051 | 5.67198423 |
| ENSMUSG00 | Acadvl | 2827 | 62.6918755 | 60.1179984 | 81.6960134 | 60.1154686 | 65.105107 | 58.2733714 |
| ENSMUSG00 | Dnahc11 | 14374 | 0.23232645 | 0.19778768 | 0.05842737 | 0.10542966 | 0 | 0.12713238 |
| ENSMUSG00 | G3bp1 | 7135 | 11.1883679 | 13.9663017 | 12.0741965 | 10.7715143 | 11.4281918 | 11.5214804 |
| ENSMUSG00 | Atox1 | 677 | 28.7742769 | 34.378215 | 38.2603841 | 26.541888 | 23.3651056 | 69.0124919 |
| ENSMUSG00 | Glra2 | 3141 | 0 | 0 | 0 | 0 | 0 | 0 |
| ENSMUSG00 | Sparc | 3335 | 5.17357677 | 5.72168848 | 3.59181393 | 2.12056394 | 5.90590137 | 2.1836068 |
| ENSMUSG00 | Glra4 | 2001 | 0 | 0.07224354 | 0.04417975 | 0.01803201 | 0 | 0.05452202 |
| ENSMUSG00 | Smcr7 | 2837 | 7.72127944 | 3.85559477 | 3.739317 | 3.87910595 | 4.13678761 | 3.8359468 |
| ENSMUSG00 | Tbx3 | 6735 | 2.402447 | 1.88882268 | 3.18305774 | 2.70012627 | 3.46994508 | 2.13013515 |
| ENSMUSG00 | Mmp20 | 3266 | 0 | 0 | 0 | 0 | 0 | 0 |
| ENSMUSG00 | Mmp7 | 1218 | 0 | 0 | 0 | 0 | 0 | 0 |
| ENSMUSG00 | Crhr1 | 2449 | 0 | 0 | 0 | 0 | 0 | 0 |
| ENSMUSG00 | Dusp14 | 3203 | 0.75712904 | 1.2937977 | 1.17301176 | 0.14644601 | 1.35411862 | 0.32358298 |
| ENSMUSG00 | Tada2a | 3902 | 4.68670452 | 4.00112949 | 4.22534276 | 4.21666128 | 4.48540413 | 4.07511909 |
| ENSMUSG00 | Ikzf1 | 2661 | 0 | 0 | 0 | 0 | 0 | 0 |
| ENSMUSG00 | Fam115e | 3564 | 0 | 0 | 0 | 0 | 0 | 0 |
| ENSMUSG00 | Pnpo | 2919 | 25.1280786 | 19.8259394 | 22.4719202 | 17.2066482 | 28.3712943 | 13.3616766 |
| ENSMUSG00 | Cog1 | 5304 | 11.5953485 | 10.8928251 | 10.0754201 | 9.92528336 | 12.4006407 | 9.63662782 |
| ENSMUSG00 | Cbx1 | 4483 | 8.93899718 | 10.7916993 | 12.8474239 | 14.3587703 | 8.63908527 | 15.7697669 |
| ENSMUSG00 | Cdk5rap3 | 3641 | 23.977763 | 24.1527759 | 21.0993692 | 19.0567898 | 21.6241961 | 16.9970218 |
| ENSMUSG00 | Copz2 | 2314 | 17.1632347 | 23.1977945 | 19.8086911 | 16.2166492 | 15.2594002 | 21.534471 |
| ENSMUSG00 | Slc25a39 | 2990 | 165.151446 | 170.312482 | 177.694366 | 142.143947 | 131.115601 | 128.601292 |
| ENSMUSG00 | Sp2 | 4634 | 5.59356351 | 3.59786556 | 3.52927965 | 4.49273623 | 5.30744781 | 6.0270243 |
| ENSMUSG00 | Aatf | 3018 | 14.3846868 | 11.0327474 | 11.5996884 | 18.9257377 | 15.9943511 | 14.5500892 |
| ENSMUSG00 | Lhx1 | 4931 | 7.27224567 | 8.25746169 | 4.52685677 | 14.7079516 | 10.6067806 | 7.20723077 |
| ENSMUSG00 | Dync1h1 | 15582 | 6.99841375 | 8.85675589 | 7.54568749 | 7.89396033 | 9.66036508 | 7.25188233 |
| ENSMUSG00 | Cpsf4l | 2237 | 33.713074 | 11.7181156 | 74.4337696 | 25.500989 | 34.9908183 | 61.12104 |
| ENSMUSG00 | Pex12 | 4566 | 6.80875728 | 9.55075124 | 11.5006109 | 8.68466205 | 5.57646989 | 9.30061614 |
| ENSMUSG00 | Ndel1 | 3795 | 18.8877817 | 21.9537177 | 20.1499838 | 26.2890259 | 21.9568041 | 19.7139224 |
| ENSMUSG00 | Slc25a35 | 4562 | 18.5705675 | 18.0408687 | 15.0181624 | 15.7394276 | 18.8245047 | 15.2635147 |
| ENSMUSG00 | Zbtb4 | 4079 | 6.42286385 | 9.41519843 | 7.37961635 | 8.38582452 | 5.91700803 | 9.16064128 |
| ENSMUSG00 | BC096441 | 2515 | 0.09484409 | 0 | 0.123027 | 0.27258801 | 0.12173291 | 0.22774053 |
| ENSMUSG00 | Mpdu1 | 3001 | 6.54422105 | 13.4877079 | 13.6243601 | 9.79902242 | 8.48455899 | 10.5426796 |
| ENSMUSG00 | Fxr2 | 3748 | 11.9224013 | 9.69385736 | 10.1187784 | 11.8315989 | 11.3134787 | 12.2037273 |
| ENSMUSG00 | Atp5g3 | 2177 | 365.195268 | 309.349433 | 417.389659 | 284.380211 | 204.949302 | 607.847346 |
| ENSMUSG00 | Cd68 | 1449 | 0.10974598 | 0.83137409 | 1.40323311 | 1.07075771 | 0.95080203 | 0.77174618 |
| ENSMUSG00 | Amac1 | 2878 | 0 | 0 | 0 | 0 | 0.0177298 | 0 |
| ENSMUSG00 | Acsl1 | 9557 | 23.0828881 | 13.9260179 | 26.9225623 | 16.9253747 | 20.1072859 | 16.8236893 |
| ENSMUSG00 | Abca5 | 10710 | 2.03046198 | 3.08195265 | 2.05945095 | 2.17974629 | 2.55846534 | 4.20452189 |
| ENSMUSG00 | Smyd4 | 4021 | 4.93359482 | 6.92657636 | 7.33216346 | 10.6514267 | 12.5503819 | 6.62025564 |
| ENSMUSG00 | Lsp1 | 5278 | 0 | 0 | 0 | 0 | 0 | 0 |
| ENSMUSG00 | Zfyve27 | 8412 | 9.11181246 | 8.47215263 | 8.91708602 | 8.38139741 | 10.3484305 | 7.19477229 |
| ENSMUSG00 | Avpi1 | 1232 | 10.8424038 | 15.6449488 | 12.01917 | 10.7777536 | 8.65626018 | 13.5709041 |
| ENSMUSG00 | Sfrp5 | 1897 | 0.02095703 | 0 | 0.06990276 | 0 | 0 | 0.1150222 |
| ENSMUSG00 | Myh11 | 6590 | 0 | 0 | 0.03353706 | 0 | 0.32520604 | 0.00413879 |
| ENSMUSG00 | Rad51l3 | 4586 | 8.92894569 | 7.93300563 | 7.04569281 | 5.7042046 | 8.22252327 | 8.29658128 |
| ENSMUSG00 | Fndc8 | 2162 | 0.36776578 | 0 | 0.04088977 | 0.10013518 | 0.18881175 | 0 |
| ENSMUSG00 | Unc45b | 4472 | 0.27558584 | 0.18317744 | 0 | 0.03227374 | 0.04564077 | 0.21346431 |
| ENSMUSG00 | Pank3 | 7369 | 10.5039925 | 9.87400294 | 10.8150255 | 11.1541454 | 12.1732079 | 10.493093 |
| ENSMUSG00 | Rara | 2934 | 24.5118147 | 21.1534211 | 28.6091683 | 30.3020306 | 28.5045092 | 20.8696541 |
| ENSMUSG00 | Wwc1 | 4937 | 101.969546 | 101.711751 | 78.6536745 | 102.596866 | 116.398429 | 77.8849229 |
| ENSMUSG00 | Ict1 | 3534 | 9.3370257 | 11.3716731 | 9.59332586 | 6.93257166 | 7.95572526 | 10.3649803 |
| ENSMUSG00 | Fdxr | 2900 | 9.4590626 | 4.51955595 | 5.50236735 | 5.58649586 | 4.66275503 | 3.7996394 |
| ENSMUSG00 | Otop3 | 2587 | 0 | 0 | 0 | 0 | 0 | 0 |
| ENSMUSG00 | Sult4a1 | 2476 | 0 | 0.07784563 | 0.24992965 | 0 | 0 | 0 |
| ENSMUSG00 | Pnpla5 | 1684 | 0 | 0 | 0 | 0 | 0 | 0 |
| ENSMUSG00 | Mrpl45 | 2679 | 16.6946307 | 18.130621 | 21.5151935 | 18.3979369 | 16.3992944 | 15.3120782 |
| ENSMUSG00 | Mb | 1188 | 0 | 0 | 0 | 0.09111627 | 0.25776897 | 0 |
| ENSMUSG00 | Irf1 | 4007 | 3.27409747 | 8.83879086 | 2.58129056 | 6.78958362 | 18.3119362 | 12.8715605 |
| ENSMUSG00 | Slc22a5 | 3492 | 18.6937282 | 19.9120951 | 18.4427507 | 18.1133517 | 15.2260835 | 14.9729334 |
| ENSMUSG00 | P4ha2 | 4936 | 0.34633016 | 0 | 0 | 0.38742876 | 0.39282866 | 0 |
| ENSMUSG00 | Alox12e | 2815 | 0.01412273 | 0 | 0 | 0 | 0 | 0 |
| ENSMUSG00 | Arrb1 | 8623 | 2.51727848 | 2.46436518 | 1.8710046 | 2.51063745 | 3.09483875 | 1.73016673 |
| ENSMUSG00 | Il3 | 624 | 0 | 0 | 0 | 0 | 0 | 0 |
| ENSMUSG00 | Csf2 | 796 | 0 | 0 | 0 | 0 | 0 | 0 |
| ENSMUSG00 | Tm4sf5 | 1004 | 0 | 0 | 0 | 0 | 0 | 0 |
| ENSMUSG00 | Cxcl16 | 3619 | 6.07482197 | 13.7542401 | 6.02141713 | 8.14562873 | 11.0540698 | 10.0612444 |
| ENSMUSG00 | Pelp1 | 3543 | 1.72801129 | 1.19684079 | 1.13529995 | 1.35447698 | 1.41139848 | 1.97073318 |
| ENSMUSG00 | Med11 | 1188 | 13.3187553 | 10.3430495 | 14.6595338 | 13.9104178 | 7.34470561 | 12.6041894 |
| ENSMUSG00 | Alox15 | 2556 | 0 | 0 | 0 | 0 | 0 | 0 |
| ENSMUSG00 | Gm11435 | 1860 | 0.02137391 | 0 | 0 | 0 | 0 | 0 |
| ENSMUSG00 | Ccl6 | 2335 | 0 | 0.45400502 | 0 | 0 | 0 | 0 |
| ENSMUSG00 | Ccl4 | 657 | 0 | 0.22002942 | 0 | 0 | 0.31066287 | 0.74725039 |
| ENSMUSG00 | Gm16515 | 4410 | 9.85322907 | 5.44146217 | 10.5843869 | 5.06457723 | 8.94407903 | 4.60762046 |
| ENSMUSG00 | Map2k3 | 3519 | 31.2259586 | 36.4650456 | 30.208989 | 28.1663473 | 29.363004 | 26.2360483 |
| ENSMUSG00 | Ywhah | 3150 | 22.565968 | 24.2920858 | 24.2899652 | 22.6686877 | 18.4342907 | 21.3954392 |
| ENSMUSG00 | Hoxb1 | 2453 | 0 | 0 | 0 | 0 | 0.04160324 | 0 |
| ENSMUSG00 | Sart3 | 3567 | 12.5385241 | 11.8608624 | 10.2480864 | 13.7672169 | 21.0428368 | 5.55127224 |
| ENSMUSG00 | E2f2 | 4874 | 0.46492868 | 0.80080054 | 0.24486044 | 0.28131262 | 0.52345482 | 0.16787837 |
| ENSMUSG00 | Slfn3 | 1795 | 0 | 0 | 0.02462498 | 0 | 0.71067376 | 0 |
| ENSMUSG00 | Nars2 | 4621 | 9.09360374 | 8.4568718 | 8.14017927 | 6.17634651 | 7.85106107 | 7.23624924 |
| ENSMUSG00 | Slc35b4 | 11070 | 12.7382737 | 16.7847264 | 12.2303745 | 12.3435142 | 17.3037954 | 10.4688295 |
| ENSMUSG00 | Dnahc1 | 12789 | 0.39789675 | 0.31649552 | 0.13479333 | 0.25392009 | 0.18353376 | 0.19407242 |
| ENSMUSG00 | Dalrd3 | 1725 | 59.6447441 | 51.957555 | 52.4528537 | 63.776321 | 69.7804177 | 55.5612071 |
| ENSMUSG00 | Fis1 | 1890 | 40.0289309 | 49.257252 | 56.1059372 | 40.4538901 | 27.9700134 | 58.2291901 |
| ENSMUSG00 | Plod1 | 4588 | 11.7758714 | 12.319681 | 12.2836421 | 8.21047395 | 13.6018434 | 6.00422529 |
| ENSMUSG00 | Rab3d | 5561 | 3.51014942 | 4.61848115 | 3.59274106 | 4.04227904 | 5.88165922 | 2.69264106 |
| ENSMUSG00 | Mfsd3 | 2164 | 4.96024942 | 4.31985664 | 3.18645449 | 2.13424293 | 4.52729397 | 1.33600359 |
| ENSMUSG00 | Slc25a22 | 3753 | 5.61428315 | 4.59652179 | 4.91131589 | 4.59558148 | 5.16653952 | 5.12353335 |
| ENSMUSG00 | Atp6ap1 | 3134 | 592.729605 | 436.876058 | 914.51554 | 460.017285 | 457.934858 | 580.756613 |
| ENSMUSG00 | Dnase1l1 | 2536 | 10.1896934 | 12.5976385 | 17.0985836 | 10.6567237 | 14.4467421 | 15.3258521 |
| ENSMUSG00 | Aldh3a1 | 1930 | 0 | 0 | 0 | 0 | 0 | 0 |
| ENSMUSG00 | Ccl9 | 3196 | 0 | 0 | 0 | 0.07902826 | 0 | 0 |
| ENSMUSG00 | Scrn1 | 8615 | 2.15505623 | 0.93967757 | 1.68290244 | 0.07957735 | 1.705815 | 0.47489215 |
| ENSMUSG00 | BC005537 | 3800 | 20.8297794 | 19.7437606 | 22.9849584 | 23.7191966 | 14.4082372 | 30.3896902 |
| ENSMUSG00 | Isyna1 | 1825 | 64.458338 | 65.5335613 | 44.0807421 | 39.5221956 | 50.0757475 | 36.286479 |
| ENSMUSG00 | Hars2 | 4529 | 18.4249842 | 12.8312761 | 18.6898939 | 14.2766668 | 18.2180724 | 13.6945307 |
| ENSMUSG00 | Cacng2 | 5506 | 0.00722039 | 0 | 0 | 0 | 0 | 0 |
| ENSMUSG00 | Tmem160 | 811 | 12.9413648 | 19.1319783 | 16.9503985 | 14.3705307 | 7.04679913 | 15.2011553 |
| ENSMUSG00 | Rab5c | 3337 | 2.39462139 | 2.5270085 | 3.28500362 | 2.72480528 | 2.17133514 | 3.45734864 |
| ENSMUSG00 | Styxl1 | 2030 | 0.03916796 | 0 | 0.08709723 | 0 | 0.17595302 | 0.17466518 |
| ENSMUSG00 | Mdh2 | 2219 | 189.855262 | 193.853253 | 271.386169 | 175.141821 | 145.789645 | 211.338507 |
| ENSMUSG00 | H13 | 9255 | 6.1340709 | 7.77335042 | 8.35321701 | 7.92206531 | 6.26871846 | 7.13178033 |
| ENSMUSG00 | Rnf145 | 8063 | 11.3897011 | 14.0322171 | 11.1834007 | 14.185797 | 16.4539969 | 7.2389592 |
| ENSMUSG00 | Scn1b | 1508 | 0.39544576 | 0.51126199 | 0.67416604 | 1.57918763 | 0.13534848 | 1.68205667 |
| ENSMUSG00 | Atp6v1e1 | 1194 | 1175.1174 | 960.338842 | 1592.37695 | 954.119256 | 855.396933 | 1222.33329 |
| ENSMUSG00 | Chtf18 | 4152 | 0.47875097 | 0.22050636 | 0.03193775 | 0 | 0.34410851 | 0 |
| ENSMUSG00 | Lhx9 | 7513 | 0 | 0 | 0 | 0.00480262 | 0 | 0 |
| ENSMUSG00 | Agxt2l1 | 2854 | 0 | 0 | 0 | 0 | 0 | 0 |
| ENSMUSG00 | Rps6kl1 | 2507 | 0.30129802 | 0.30753214 | 0.28210191 | 0.28785038 | 0.12212136 | 0.46781392 |
| ENSMUSG00 | Ppp1r12c | 5151 | 6.1666917 | 9.01800237 | 6.71909204 | 6.79471621 | 12.35292 | 6.47051236 |
| ENSMUSG00 | Ahr | 6392 | 4.86992197 | 5.30714256 | 7.87639224 | 5.41343565 | 8.22233528 | 5.41481809 |
| ENSMUSG00 | Mtap1s | 3264 | 13.056947 | 9.12353592 | 13.1359644 | 10.3360023 | 13.9603412 | 8.03866512 |
| ENSMUSG00 | Dpep1 | 2230 | 0.37437896 | 0 | 0 | 0 | 0 | 0.07338468 |
| ENSMUSG00 | Tmem129 | 4131 | 43.8166794 | 36.8367912 | 46.042249 | 35.156192 | 24.2594534 | 26.1588285 |
| ENSMUSG00 | 2610027L16 | 3684 | 6.62591342 | 6.42224297 | 5.19527634 | 4.1233824 | 5.58187553 | 4.22001747 |
| ENSMUSG00 | Hsd17b1 | 1333 | 0.47718506 | 1.51825249 | 0.26527738 | 0.0541366 | 0.30623481 | 0 |
| ENSMUSG00 | Atp6v0a1 | 6364 | 15.0738489 | 13.9546846 | 14.7871973 | 11.9801006 | 12.1953359 | 12.2058725 |
| ENSMUSG00 | Psmc3ip | 2136 | 0.65142407 | 2.23336038 | 1.01399359 | 0.37163155 | 1.7199902 | 1.92812291 |
| ENSMUSG00 | Grb7 | 3171 | 67.4250944 | 38.8713084 | 52.2030596 | 34.739351 | 47.3735889 | 48.4165703 |
| ENSMUSG00 | Noxo1 | 2672 | 0.92246998 | 0.90169241 | 0.67824684 | 0.51314285 | 0.87644809 | 0.48996358 |
| ENSMUSG00 | Aoc3 | 5375 | 0.01479274 | 0 | 0 | 0 | 0.05695968 | 0 |
| ENSMUSG00 | Zfp687 | 7226 | 7.64740076 | 7.47536696 | 7.73195815 | 8.55363165 | 11.6867772 | 8.41716656 |
| ENSMUSG00 | Gdpd2 | 3489 | 0 | 0 | 0 | 0 | 1.00912007 | 0 |
| ENSMUSG00 | D8Ertd738e | 656 | 46.9066188 | 57.8824945 | 55.9261124 | 48.3477395 | 39.3587594 | 74.6310632 |
| ENSMUSG00 | Sec14l4 | 3526 | 0 | 0 | 0 | 0 | 0 | 0 |
| ENSMUSG00 | Calm3 | 3368 | 11.6976488 | 17.1971804 | 14.9614314 | 11.7416629 | 6.96916059 | 13.6940068 |
| ENSMUSG00 | Cops3 | 3292 | 46.6631764 | 36.3447557 | 39.6097925 | 34.711979 | 27.419702 | 38.7495406 |
| ENSMUSG00 | Fkbp8 | 4230 | 43.167121 | 42.160289 | 40.3145887 | 35.6299527 | 38.3120023 | 35.0701567 |
| ENSMUSG00 | Ffar3 | 1623 | 0 | 0 | 0 | 0 | 0 | 0 |
| ENSMUSG00 | Ddx39b | 3524 | 15.1395729 | 18.186143 | 15.8168343 | 17.0376056 | 15.7394071 | 17.1124368 |
| ENSMUSG00 | Gipc1 | 1608 | 42.4009012 | 47.4672415 | 44.1193769 | 34.9376513 | 36.7783393 | 37.9605987 |
| ENSMUSG00 | Tlcd1 | 3161 | 32.9262411 | 32.4698266 | 40.4682486 | 34.278513 | 35.3681713 | 44.0570414 |
| ENSMUSG00 | Plscr3 | 3140 | 4.25408963 | 3.20731414 | 5.67303909 | 3.55074897 | 4.16011217 | 4.31703685 |
| ENSMUSG00 | Ptger1 | 5010 | 20.9013844 | 31.5472116 | 10.4902178 | 22.4990633 | 21.5818144 | 6.33141825 |
| ENSMUSG00 | Arhgef25 | 2370 | 0.05032339 | 0.48796397 | 0.26110793 | 0.21314288 | 0.06459035 | 0.46033146 |
| ENSMUSG00 | Xab2 | 4620 | 26.3659724 | 23.3214036 | 24.6650111 | 26.6710351 | 23.1827625 | 19.3461023 |
| ENSMUSG00 | Cdc37 | 2448 | 38.277642 | 32.1046108 | 33.0069318 | 30.9379949 | 29.4527243 | 21.1913251 |
| ENSMUSG00 | Rab4a | 1847 | 16.509179 | 13.1488721 | 17.49407 | 12.1315374 | 14.946004 | 15.3724415 |
| ENSMUSG00 | Trip10 | 2238 | 15.4900711 | 16.7296093 | 13.3514057 | 14.3328586 | 11.3771946 | 18.4755823 |
| ENSMUSG00 | Cd70 | 839 | 0 | 0 | 0 | 0 | 0 | 0 |
| ENSMUSG00 | Cops6 | 2601 | 81.2075617 | 82.5338715 | 97.9883994 | 75.0495428 | 59.4228729 | 90.8315745 |
| ENSMUSG00 | Ubb | 1529 | 135.517047 | 136.01876 | 154.952177 | 117.119154 | 144.30219 | 136.391035 |
| ENSMUSG00 | Ap4m1 | 3871 | 3.08102406 | 1.31949441 | 3.18582129 | 1.87354455 | 2.95270168 | 2.3392354 |
| ENSMUSG00 | Gyg | 3066 | 11.6958393 | 11.6030065 | 11.7929249 | 11.5801474 | 13.2641949 | 9.55412999 |
| ENSMUSG00 | Rcn3 | 1590 | 0.15002068 | 0 | 0 | 0 | 0.06418412 | 0 |
| ENSMUSG00 | Slc6a8 | 5343 | 7.50019172 | 5.88915342 | 3.57387364 | 5.55783692 | 4.03015733 | 4.98222869 |
| ENSMUSG00 | Arid3a | 5667 | 4.8685907 | 4.40454862 | 3.81413468 | 3.97303614 | 4.3399882 | 3.95138148 |
| ENSMUSG00 | Pdk4 | 3500 | 7.32636712 | 12.2255888 | 9.5223399 | 12.9689177 | 26.7378209 | 20.2689533 |
| ENSMUSG00 | Ubxn6 | 4701 | 10.5287329 | 9.97349675 | 10.0232216 | 9.14907404 | 11.7010068 | 10.1881018 |
| ENSMUSG00 | D17Wsu104e | 2215 | 24.2302026 | 26.7581598 | 21.3924947 | 19.727023 | 16.9320028 | 19.9480432 |
| ENSMUSG00 | Cyb561 | 3455 | 27.59295 | 31.8546552 | 26.0861239 | 26.0876837 | 25.948869 | 20.6276794 |
| ENSMUSG00 | Sema6a | 8597 | 0.12948162 | 0.13452072 | 0.23136942 | 0.22244369 | 0.2374148 | 0.03172576 |
| ENSMUSG00 | Ccdc12 | 811 | 34.3142249 | 29.4703765 | 33.7372883 | 31.0990741 | 29.0051286 | 43.9219222 |
| ENSMUSG00 | 1110001J03 | 686 | 29.3819659 | 26.9731688 | 38.7249383 | 25.8780844 | 18.0005582 | 72.4010468 |
| ENSMUSG00 | Akt3 | 4965 | 0.18416436 | 0.21351495 | 0.71221499 | 0.26888936 | 0.98661271 | 0.78006018 |
| ENSMUSG00 | Mrpl24 | 2296 | 14.0252349 | 13.5576836 | 13.5531784 | 14.7879808 | 11.9565288 | 14.8371187 |
| ENSMUSG00 | Gle1 | 6243 | 8.61591625 | 9.94139636 | 10.145962 | 8.10877919 | 12.096594 | 9.22261148 |
| ENSMUSG00 | 2810055F11 | 2592 | 0.27607973 | 0.50194211 | 0.68212721 | 1.1971666 | 1.33865492 | 0.11574886 |
| ENSMUSG00 | Lyst | 11757 | 1.5757467 | 3.08209616 | 1.20683777 | 2.56260161 | 2.28288455 | 1.55430878 |
| ENSMUSG00 | Slc35e1 | 10461 | 7.7869209 | 5.73022981 | 5.20568498 | 6.74662834 | 7.19959189 | 3.46766755 |
| ENSMUSG00 | Calr3 | 2522 | 1.02462579 | 2.35009214 | 1.7526504 | 0.61519743 | 2.12441295 | 0.82191617 |
| ENSMUSG00 | Tmc4 | 4062 | 53.8587911 | 39.9774274 | 64.017588 | 40.7632945 | 52.3453737 | 44.846705 |
| ENSMUSG00 | AI428936 | 2801 | 12.1210926 | 11.4746008 | 18.5265847 | 12.4953885 | 18.6179779 | 12.6684419 |
| ENSMUSG00 | Polr2i | 1503 | 5.58110871 | 6.86087732 | 6.17590621 | 4.60928313 | 4.14186151 | 11.1965751 |
| ENSMUSG00 | Prl8a1 | 1945 | 0 | 0 | 0 | 0 | 0 | 0 |
| ENSMUSG00 | Krt10 | 2083 | 7.92055901 | 6.03776351 | 4.03185318 | 4.20928316 | 3.2825417 | 15.8305515 |
| ENSMUSG00 | Iyd | 1618 | 3.85760843 | 2.85902253 | 4.86274911 | 3.9917713 | 3.05905961 | 3.92768291 |
| ENSMUSG00 | Rmnd1 | 5958 | 9.06809299 | 8.19282744 | 9.52587553 | 12.2938151 | 9.71196881 | 8.36367337 |
| ENSMUSG00 | Gm221 | 3731 | 0 | 0 | 0 | 0.22243019 | 0.27352654 | 0 |
| ENSMUSG00 | Esr1 | 16230 | 1.05818654 | 0.40081144 | 0.18791911 | 0.36904617 | 0.64765455 | 0.4386125 |
| ENSMUSG00 | Syne1 | 30771 | 0.71058836 | 0.90826221 | 0.40221364 | 0.87358627 | 0.8390805 | 0.38645942 |
| ENSMUSG00 | Vip | 1520 | 0 | 0 | 0 | 0 | 0 | 0 |
| ENSMUSG00 | Fbxo5 | 1997 | 5.27551444 | 3.8124476 | 3.34225253 | 4.17373675 | 2.17187879 | 3.70126552 |
| ENSMUSG00 | Mtrf1l | 3273 | 14.3693044 | 10.8356925 | 15.3416724 | 9.16106905 | 8.90194335 | 17.0414413 |
| ENSMUSG00 | Rgs17 | 8450 | 0.19760121 | 0 | 0 | 0 | 0 | 0 |
| ENSMUSG00 | Hdac2 | 4700 | 61.1473125 | 57.6084295 | 53.5876812 | 51.175087 | 44.9900641 | 66.056855 |
| ENSMUSG00 | Frk | 4979 | 49.3689769 | 56.8288388 | 47.8683144 | 58.5545129 | 43.975533 | 51.7829212 |
| ENSMUSG00 | Rwdd1 | 1880 | 10.3618008 | 9.63728843 | 10.791833 | 8.84777794 | 6.97541414 | 11.7513073 |
| ENSMUSG00 | Clvs2 | 7252 | 0 | 0 | 0 | 0.25374852 | 0 | 0.09778552 |
| ENSMUSG00 | Trdn | 4562 | 0 | 0 | 0 | 0 | 0 | 0 |
| ENSMUSG00 | Hey2 | 2545 | 4.35826289 | 5.71799824 | 11.2545361 | 7.42907321 | 3.32824298 | 7.85552478 |
| ENSMUSG00 | Stxbp5 | 11334 | 3.49009203 | 4.37478816 | 4.60582112 | 3.89026455 | 5.38896875 | 4.84898519 |
| ENSMUSG00 | Hint3 | 1040 | 12.1177763 | 12.3709424 | 15.3856415 | 7.42457455 | 8.38991372 | 16.7057165 |

| | | | | | | | | |
|---|---|---|---|---|---|---|---|---|
| ENSMUSG00 | Trmt11 | 2870 | 12.7854734 | 14.0529798 | 18.7126269 | 12.132116 | 8.30289809 | 21.2495099 |
| ENSMUSG00 | Katna1 | 1889 | 42.7439814 | 37.8553098 | 46.6588009 | 42.8438464 | 34.170654 | 37.2373185 |
| ENSMUSG00 | Pcmt1 | 6301 | 22.8841657 | 24.655307 | 27.8497567 | 21.3022069 | 24.0271794 | 24.469693 |
| ENSMUSG00 | Lrp11 | 4207 | 2.1734634 | 1.31719535 | 3.48823672 | 4.04818755 | 1.91636971 | 1.32256392 |
| ENSMUSG00 | 1700021F05I | 1608 | 20.3227643 | 18.8790165 | 29.220497 | 23.7181101 | 15.3269524 | 30.5652363 |
| ENSMUSG00 | Sec63 | 8030 | 25.6950117 | 23.4751384 | 27.0440417 | 28.658939 | 27.8834044 | 40.2122608 |
| ENSMUSG00 | Nr2e1 | 3953 | 0 | 0 | 0 | 0 | 0 | 0 |
| ENSMUSG00 | Snx3 | 1804 | 92.3367313 | 107.110661 | 111.288676 | 95.4454082 | 75.945576 | 106.558568 |
| ENSMUSG00 | Aig1 | 9222 | 24.2404106 | 23.6752081 | 24.511844 | 22.6852846 | 22.0826573 | 20.8863048 |
| ENSMUSG00 | Adat2 | 1460 | 1.22534015 | 0.99013237 | 0.48440376 | 0.61784323 | 0.94363846 | 1.04615055 |
| ENSMUSG00 | Pex3 | 4633 | 23.4946051 | 25.5857215 | 31.3410435 | 22.655443 | 22.2806731 | 26.5093245 |
| ENSMUSG00 | Fuca2 | 2704 | 149.685853 | 157.995931 | 166.754068 | 163.209869 | 125.96193 | 160.60163 |
| ENSMUSG00 | Cep57l1 | 2966 | 3.19008913 | 2.74561993 | 3.05508355 | 3.78338363 | 4.33534953 | 4.17487735 |
| ENSMUSG00 | Ltv1 | 1755 | 55.7709362 | 44.3425094 | 66.995386 | 50.5559815 | 51.4334238 | 66.1118606 |
| ENSMUSG00 | Fam164b | 1151 | 0 | 0 | 0 | 0 | 0 | 0 |
| ENSMUSG00 | Plagl1 | 5750 | 206.140809 | 126.432844 | 129.853484 | 151.419084 | 92.0693303 | 198.417071 |
| ENSMUSG00 | Cd164 | 2881 | 238.864064 | 268.12803 | 265.778732 | 230.882503 | 190.99911 | 278.246762 |
| ENSMUSG00 | Utrn | 13154 | 6.52516971 | 5.06261693 | 5.55799364 | 7.46383177 | 6.00492852 | 8.12184597 |
| ENSMUSG00 | Smpd2 | 4245 | 10.4890785 | 10.6816118 | 10.277319 | 15.3593061 | 11.4073099 | 9.9974885 |
| ENSMUSG00 | Mical1 | 5962 | 0 | 0 | 0.06672536 | 0 | 0.0171172 | 0 |
| ENSMUSG00 | Zbtb24 | 2858 | 13.2842841 | 7.94115264 | 12.2645422 | 18.44502 | 15.9613646 | 8.59847794 |
| ENSMUSG00 | Grm1 | 8849 | 0 | 0 | 0 | 0 | 0 | 0 |
| ENSMUSG00 | Wasf1 | 2844 | 0.26559569 | 0 | 0 | 0.32986398 | 0.10765058 | 0.56582409 |
| ENSMUSG00 | Rab32 | 2105 | 46.4412003 | 22.2275702 | 45.6927365 | 26.6715731 | 30.5431038 | 45.9457818 |
| ENSMUSG00 | Slc22a16 | 2407 | 0 | 0 | 0 | 0 | 0 | 0 |
| ENSMUSG00 | Amd-ps4 | 1679 | 0 | 0 | 0 | 0 | 0 | 0 |
| ENSMUSG00 | Gtf3c6 | 1014 | 32.1101958 | 24.6634748 | 29.1191629 | 29.4990285 | 23.5003131 | 40.4815898 |
| ENSMUSG00 | Slc16a10 | 6139 | 3.45164866 | 2.33122224 | 3.81609005 | 2.99753098 | 4.21410206 | 4.74496086 |
| ENSMUSG00 | Rev3l | 13681 | 4.73950652 | 4.98031059 | 4.00307605 | 6.5301617 | 6.31442543 | 7.09527382 |
| ENSMUSG00 | Traf3ip2 | 3677 | 4.42208092 | 4.23285853 | 3.26975831 | 2.98312253 | 4.92639745 | 3.76815794 |
| ENSMUSG00 | Fyn | 4343 | 5.0529646 | 9.37544177 | 7.72489036 | 6.66308996 | 8.94107115 | 7.86901289 |
| ENSMUSG00 | Tube1 | 2917 | 2.12610729 | 1.22241917 | 1.25771442 | 0.79165267 | 2.58892822 | 0.69191749 |
| ENSMUSG00 | Lama4 | 11964 | 0.1960526 | 0.00402762 | 0.14778283 | 0 | 0 | 0 |
| ENSMUSG00 | Popdc3 | 2077 | 0 | 0 | 0 | 0 | 0 | 0 |
| ENSMUSG00 | Prep | 2692 | 27.8376972 | 21.676739 | 28.7180622 | 23.9385333 | 21.5326757 | 16.1702542 |
| ENSMUSG00 | Tnfaip3 | 8636 | 1.03117504 | 0.52449347 | 0.61931716 | 1.6043892 | 3.63376808 | 2.3023636 |
| ENSMUSG00 | Perp | 1898 | 72.1379741 | 77.3318771 | 87.8680473 | 73.3997755 | 54.9783659 | 75.0411834 |
| ENSMUSG00 | D10Bwg1379 | 8182 | 3.00279723 | 2.2615001 | 0.52942809 | 2.05502722 | 3.5485221 | 1.15338856 |
| ENSMUSG00 | Hebp2 | 1906 | 17.7710752 | 17.8234217 | 26.3216641 | 16.7915916 | 19.3289839 | 20.8351914 |
| ENSMUSG00 | Reps1 | 5490 | 22.1805167 | 18.8444975 | 26.9880834 | 21.1825919 | 23.9703139 | 21.1987041 |
| ENSMUSG00 | Fam184a | 4682 | 2.44544605 | 2.98463653 | 0.2549017 | 0.77065452 | 2.77909565 | 1.22916465 |
| ENSMUSG00 | Asf1a | 2176 | 22.5451576 | 26.7284171 | 23.2790956 | 23.2477144 | 15.4533004 | 35.008303 |
| ENSMUSG00 | Gopc | 4198 | 29.1016178 | 31.2098466 | 27.2076137 | 29.7904636 | 31.5298759 | 25.7023518 |
| ENSMUSG00 | Qrsl1 | 3240 | 21.2029229 | 12.4779089 | 15.3205771 | 19.3773943 | 16.8198054 | 17.4843907 |
| ENSMUSG00 | Rtn4ip1 | 3125 | 36.422381 | 27.3698991 | 43.8482283 | 40.2848811 | 29.5708053 | 20.7636374 |
| ENSMUSG00 | Nmbr | 1265 | 0 | 0.03809205 | 0.31447952 | 0.54194375 | 0 | 0.2156098 |
| ENSMUSG00 | Aim1 | 7523 | 5.96094404 | 9.6718766 | 1.89193054 | 10.5325229 | 9.99094132 | 3.15781082 |
| ENSMUSG00 | Gje1 | 1477 | 0 | 0 | 0 | 0 | 0 | 0 |
| ENSMUSG00 | Vta1 | 2768 | 33.6657682 | 41.4145758 | 44.8246291 | 32.22356 | 31.7255754 | 44.1045106 |
| ENSMUSG00 | Smpdl3a | 2092 | 583.846858 | 494.855413 | 748.008435 | 446.385944 | 446.529571 | 572.285033 |
| ENSMUSG00 | Reep3 | 3015 | 18.7371601 | 25.7154181 | 26.0959471 | 25.1676748 | 25.9278301 | 26.9851572 |
| ENSMUSG00 | Fabp7 | 1404 | 0 | 0.17160414 | 0 | 0 | 0.21806144 | 0 |
| ENSMUSG00 | Pkib | 7816 | 11.8411923 | 9.19833312 | 22.9605275 | 10.6778108 | 6.54803671 | 15.4937818 |
| ENSMUSG00 | Serinc1 | 2971 | 240.754159 | 189.112728 | 257.370475 | 225.370078 | 216.763342 | 251.677629 |
| ENSMUSG00 | Hsf2 | 2654 | 10.5155793 | 7.40771228 | 9.37665321 | 8.56506709 | 8.07501049 | 11.2942082 |
| ENSMUSG00 | Rspo3 | 2389 | 0 | 0 | 0 | 0 | 0 | 0 |
| ENSMUSG00 | Echdc1 | 3325 | 8.14240067 | 6.49248905 | 10.129866 | 11.7632891 | 9.05430428 | 10.483305 |
| ENSMUSG00 | Mgat4c | 7138 | 3.35844141 | 5.29929351 | 2.13640177 | 1.75405848 | 2.13741754 | 1.82646085 |
| ENSMUSG00 | Ptprk | 7373 | 18.6025238 | 11.0188989 | 17.104009 | 16.3697869 | 16.9418918 | 10.9461084 |
| ENSMUSG00 | Nts | 1172 | 0 | 0 | 0 | 0 | 0 | 0 |
| ENSMUSG00 | Dcbld1 | 5328 | 7.52130712 | 11.0517703 | 4.78687377 | 8.9798782 | 6.17717951 | 6.72651576 |
| ENSMUSG00 | Lrriq1 | 5523 | 0.09357618 | 0.85501925 | 0.16806784 | 0.38545005 | 0.35107773 | 0.93829105 |
| ENSMUSG00 | Ros1 | 8131 | 0.05867246 | 0.41483839 | 0.19570365 | 0.33281925 | 1.07939204 | 0.45284421 |
| ENSMUSG00 | Slc6a15 | 3594 | 0.68582075 | 0.2145195 | 0 | 0.5320947 | 0.09938359 | 0 |
| ENSMUSG00 | Ccdc59 | 4380 | 11.6543463 | 10.7594385 | 9.53669901 | 17.9009778 | 10.7994179 | 16.0472021 |
| ENSMUSG00 | Lama2 | 9727 | 0 | 0 | 0.01363273 | 0 | 0 | 0 |
| ENSMUSG00 | Rfx6 | 4206 | 0 | 0 | 0 | 0 | 0 | 0 |
| ENSMUSG00 | Gprc6a | 2850 | 0 | 0 | 0 | 0 | 0 | 0 |
| ENSMUSG00 | Lin7a | 5945 | 0.00668721 | 0.21884507 | 0 | 0 | 0.11157996 | 0.11010788 |
| ENSMUSG00 | Ppp1r12a | 3502 | 10.6029751 | 10.8839165 | 10.0470209 | 9.72628499 | 14.0315248 | 12.1809069 |
| ENSMUSG00 | Fam162b | 601 | 0 | 0 | 0 | 0 | 0 | 0 |
| ENSMUSG00 | Sim1 | 7344 | 8.3256984 | 9.3564633 | 8.78740343 | 13.9729486 | 11.5615318 | 10.4842465 |
| ENSMUSG00 | P4ha1 | 4210 | 8.56489806 | 14.5703898 | 7.20248558 | 9.25620145 | 9.86590734 | 5.75941907 |
| ENSMUSG00 | 10-Sep | 2633 | 14.7667527 | 12.1335401 | 15.3438984 | 13.6763693 | 10.1742679 | 15.0512916 |
| ENSMUSG00 | Lims1 | 5169 | 15.8821953 | 15.5214599 | 20.0272231 | 21.6464345 | 19.7037427 | 14.4631043 |
| ENSMUSG00 | Zwint | 3616 | 23.5828279 | 21.7078856 | 20.389567 | 16.1451184 | 18.0483226 | 17.0994489 |
| ENSMUSG00 | Ube2d1 | 4580 | 8.88856158 | 11.4363892 | 8.58944111 | 9.40654174 | 8.77921051 | 13.83384 |
| ENSMUSG00 | Dcn | 1928 | 32.8477596 | 27.7421944 | 69.9021885 | 23.4308712 | 18.4732419 | 62.8957708 |
| ENSMUSG00 | Kera | 2137 | 0 | 0 | 0 | 0 | 0 | 0 |
| ENSMUSG00 | 2310015B20I | 422 | 0 | 0 | 0.52371852 | 0 | 0.36274675 | 0 |
| ENSMUSG00 | Slc17a8 | 5351 | 0.11887268 | 0 | 0.06608386 | 0 | 0 | 0.08155377 |
| ENSMUSG00 | Epyc | 2553 | 0.12457652 | 0.20761883 | 0.17313687 | 0 | 0 | 0.29913431 |
| ENSMUSG00 | Cdk1 | 3703 | 0.21472039 | 1.22320431 | 0 | 0.31180811 | 0.16535688 | 0.03682776 |
| ENSMUSG00 | Atp2b1 | 7123 | 7.50682596 | 10.8238534 | 8.34641006 | 9.44728528 | 6.28248373 | 8.89498624 |
| ENSMUSG00 | Rhobtb1 | 6058 | 1.38468247 | 0.68405976 | 1.28417372 | 1.11379041 | 1.60036504 | 2.06203116 |
| ENSMUSG00 | 1700040L02I | 1254 | 53.673069 | 53.8350921 | 49.4538961 | 43.4193024 | 47.282814 | 38.5629469 |
| ENSMUSG00 | Arid5b | 6839 | 3.35413251 | 3.23403669 | 4.10413369 | 3.92001156 | 4.275203 | 2.31708808 |
| ENSMUSG00 | Actr6 | 1692 | 16.5177913 | 18.0271973 | 16.7193732 | 23.9267458 | 19.7832757 | 29.6281247 |
| ENSMUSG00 | Uhrf1bp1l | 6475 | 20.0281664 | 19.4680668 | 27.3402905 | 19.8270138 | 24.5951072 | 23.1676472 |
| ENSMUSG00 | Poc1b | 3876 | 5.90793518 | 7.22299352 | 7.16170212 | 6.18123776 | 9.95251293 | 4.63021482 |
| ENSMUSG00 | Dusp6 | 2793 | 5.4658448 | 2.86392746 | 12.1068421 | 1.9378112 | 1.75386039 | 11.9527957 |
| ENSMUSG00 | Tmpo | 7798 | 9.25316711 | 12.7788616 | 10.2370516 | 10.7718645 | 11.2025078 | 9.09737583 |
| ENSMUSG00 | Kitl | 5922 | 7.08240999 | 11.0254355 | 6.06823681 | 14.3731076 | 8.4354647 | 13.5406011 |
| ENSMUSG00 | Psen1 | 3102 | 38.5764011 | 35.9611907 | 37.8750802 | 31.2897162 | 42.9825661 | 34.5373253 |
| ENSMUSG00 | Sgk1 | 5223 | 39.9001013 | 152.834884 | 33.2677474 | 154.165596 | 89.0397069 | 20.1465744 |
| ENSMUSG00 | Cep290 | 77413 | 0.18333751 | 0.24960618 | 0.17472213 | 0.39012403 | 0.29002371 | 0.38051245 |
| ENSMUSG00 | Ikbip | 2943 | 13.9542682 | 7.49894343 | 11.1893894 | 11.9047452 | 14.4427357 | 14.0126587 |
| ENSMUSG00 | Hbs1l | 4874 | 15.399743 | 13.6037227 | 12.3699864 | 16.4419821 | 17.9440312 | 11.9641319 |
| ENSMUSG00 | Epb4.1l2 | 4993 | 39.6121601 | 23.1522681 | 35.1189087 | 28.2340372 | 33.2545217 | 26.6355179 |
| ENSMUSG00 | Apaf1 | 11277 | 0.715648 | 0.87596175 | 0.05487504 | 0.43834709 | 1.77825359 | 0.29265153 |
| ENSMUSG00 | Myb | 3761 | 0.13741591 | 0 | 0 | 0 | 0 | 0 |
| ENSMUSG00 | Med23 | 6770 | 6.91757106 | 6.02152586 | 7.12322167 | 8.5541627 | 7.26579413 | 5.20917408 |
| ENSMUSG00 | Ahi1 | 5161 | 6.44745925 | 4.95776028 | 6.9887045 | 5.23647575 | 5.12142272 | 5.34289097 |
| ENSMUSG00 | Arg1 | 1468 | 0.13540695 | 0.13129821 | 0 | 0.02457905 | 0.27807289 | 0.18579455 |
| ENSMUSG00 | Nedd1 | 3519 | 11.8735465 | 12.9127067 | 13.289443 | 13.5858792 | 11.9917058 | 13.0521434 |
| ENSMUSG00 | Enpp3 | 2790 | 0 | 0 | 0 | 0 | 0 | 0 |
| ENSMUSG00 | Pde7b | 8920 | 1.27912813 | 1.69084713 | 0.82754572 | 2.97717318 | 1.93351065 | 0.83169304 |
| ENSMUSG00 | Fam54a | 2448 | 0.04871995 | 0.19684004 | 0.01805631 | 0.39796373 | 0.62532323 | 1.1141601 |
| ENSMUSG00 | Mtap7 | 4330 | 19.9603729 | 24.3380483 | 19.7428093 | 23.9074794 | 20.5401785 | 20.408737 |
| ENSMUSG00 | Ctgf | 2915 | 6.30086861 | 15.0758268 | 5.2920903 | 16.0048314 | 12.5334083 | 7.65374096 |
| ENSMUSG00 | Stx7 | 2490 | 273.546445 | 310.0575 | 260.986143 | 307.523353 | 264.087111 | 293.810301 |
| ENSMUSG00 | Moxd1 | 3507 | 0.03400811 | 0.30228164 | 0.7688373 | 0.27779162 | 0 | 0.65328477 |
| ENSMUSG00 | Pex7 | 3238 | 39.3503135 | 38.4836749 | 46.072026 | 30.4769586 | 35.5671804 | 36.3970711 |
| ENSMUSG00 | Il20ra | 2081 | 0 | 0 | 0 | 0 | 0 | 0 |
| ENSMUSG00 | Ifngr1 | 3078 | 25.9224332 | 24.9699075 | 22.6609838 | 28.849224 | 20.0259461 | 21.0186629 |
| ENSMUSG00 | Vnn3 | 1804 | 0 | 0 | 0 | 0 | 0 | 0 |
| ENSMUSG00 | Gm872 | 14383 | 0.43119342 | 0.19766392 | 0.15673323 | 0.40891144 | 0.23414731 | 0.40012159 |
| ENSMUSG00 | Cdk17 | 5431 | 10.8044723 | 9.68874882 | 7.878362 | 6.58392674 | 8.9538089 | 10.1696086 |
| ENSMUSG00 | Hal | 6102 | 0 | 0 | 0 | 0 | 0 | 0.00446979 |
| ENSMUSG00 | Snrpf | 857 | 0.46389125 | 0.84340331 | 1.23785791 | 0.92626019 | 0.83356973 | 3.18257167 |
| ENSMUSG00 | Ntn4 | 3730 | 42.2495241 | 98.7757471 | 12.7865385 | 69.7070276 | 104.77507 | 11.6191426 |
| ENSMUSG00 | Usp44 | 2928 | 0 | 0 | 0 | 0 | 0.08713521 | 0 |
| ENSMUSG00 | Fgd6 | 8219 | 0.3144064 | 0.20519838 | 0.64536089 | 0.97459714 | 0.51529252 | 0.74997791 |
| ENSMUSG00 | Ndufa12 | 1422 | 60.919263 | 58.3184718 | 66.1784274 | 49.8094609 | 33.0487286 | 80.9224353 |
| ENSMUSG00 | Tmcc3 | 7269 | 13.3410542 | 9.84411426 | 12.4597023 | 14.6532116 | 14.6010265 | 11.3766276 |
| ENSMUSG00 | Ccdc41 | 2970 | 6.97394119 | 6.87914191 | 8.61712697 | 9.98634364 | 7.69690787 | 10.6068042 |
| ENSMUSG00 | Socs2 | 10122 | 1.08009851 | 0.92354967 | 0.5982664 | 1.63620415 | 0.9880626 | 0.88921468 |
| ENSMUSG00 | Nudt4 | 3226 | 809.293912 | 839.931809 | 677.702219 | 807.662901 | 725.204062 | 802.124651 |
| ENSMUSG00 | Nuak1 | 5709 | 0.04874555 | 0.19413 | 0.05419739 | 0.04424143 | 0.01787577 | 0.0764397 |
| ENSMUSG00 | 4930463O16I | 2531 | 0 | 0 | 0 | 0 | 0 | 0 |
| ENSMUSG00 | Tcp11l2 | 3028 | 31.4052541 | 32.3364764 | 41.5742566 | 31.9113987 | 45.1789014 | 41.7046168 |
| ENSMUSG00 | Rfx4 | 4702 | 0 | 0 | 0.01880129 | 0 | 0 | 0 |
| ENSMUSG00 | Cry1 | 3001 | 4.04046037 | 4.6083002 | 5.52338925 | 5.60287662 | 4.40380918 | 2.74473211 |
| ENSMUSG00 | Btbd11 | 11664 | 3.30954832 | 2.56548187 | 3.30831695 | 4.2937138 | 2.26171436 | 2.45527874 |
| ENSMUSG00 | Timp3 | 8556 | 36.2101986 | 50.8446937 | 16.9089098 | 45.0350811 | 47.2811019 | 18.2213462 |
| ENSMUSG00 | Hsp90b1 | 4304 | 193.983468 | 174.127262 | 196.895384 | 222.276796 | 185.776791 | 226.777472 |
| ENSMUSG00 | Pah | 2333 | 142.901612 | 77.8253382 | 121.806108 | 93.7545452 | 92.0793151 | 130.633013 |
| ENSMUSG00 | Ascl1 | 2479 | 0 | 0 | 0 | 0 | 0 | 0 |
| ENSMUSG00 | Igf1 | 9913 | 0.13635492 | 0.35970914 | 0.1382283 | 0.50230224 | 0.0514742 | 0.35768215 |
| ENSMUSG00 | Ccdc53 | 1452 | 23.5740143 | 24.3255249 | 24.7798211 | 26.1669373 | 13.1431574 | 27.2182867 |
| ENSMUSG00 | Dram1 | 3036 | 1.61064694 | 0.17458856 | 3.2030321 | 1.23601207 | 7.36151272 | 3.37788681 |
| ENSMUSG00 | Sycp3 | 1304 | 0.39633531 | 0.81296145 | 0.27117695 | 0.30437308 | 0.19565328 | 0.71114857 |
| ENSMUSG00 | Mybpc1 | 4782 | 0 | 0 | 0 | 0 | 0.02134102 | 0 |
| ENSMUSG00 | Slc5a8 | 5306 | 0.29220953 | 0 | 0 | 0 | 0 | 0.43178848 |
| ENSMUSG00 | Sirt1 | 7273 | 8.20472656 | 8.25523263 | 5.70071893 | 7.37714839 | 5.99856336 | 12.1653966 |
| ENSMUSG00 | Herc4 | 3896 | 14.9389177 | 19.2448932 | 24.1090648 | 19.5598763 | 24.2951558 | 19.8679225 |
| ENSMUSG00 | Mypn | 5226 | 0 | 0 | 0 | 0 | 0 | 0 |
| ENSMUSG00 | Pbld2 | 2469 | 6.56955693 | 5.17189436 | 4.4935855 | 8.51998054 | 6.86138387 | 8.25198685 |
| ENSMUSG00 | Hnrnph3 | 3429 | 5.91288862 | 6.16910123 | 5.27225249 | 5.18764886 | 6.01185647 | 6.32351654 |
| ENSMUSG00 | Rufy2 | 10654 | 4.16436233 | 4.22434175 | 5.0159591 | 4.70753162 | 6.49922958 | 5.77544454 |
| ENSMUSG00 | Pbld1 | 2395 | 25.3140325 | 21.0853409 | 25.6536792 | 27.9767669 | 29.1031438 | 28.2084682 |
| ENSMUSG00 | Ccar1 | 4453 | 48.5315051 | 50.6536572 | 49.9193956 | 61.9382772 | 52.1149691 | 51.9094021 |
| ENSMUSG00 | Ddx21 | 4727 | 6.07223544 | 8.51188475 | 7.1067063 | 6.98436022 | 7.76136328 | 7.73752723 |
| ENSMUSG00 | Ddx50 | 3237 | 40.2221806 | 38.1829546 | 41.6483229 | 34.7109937 | 39.7870166 | 34.0827667 |
| ENSMUSG00 | Srgn | 2867 | 1.76105547 | 1.98325782 | 1.38757094 | 1.52282085 | 0.88989145 | 3.26306288 |
| ENSMUSG00 | Vps26a | 4823 | 32.9303638 | 35.0982731 | 41.9656312 | 43.8775018 | 29.4012644 | 35.4858721 |
| ENSMUSG00 | Supv3l1 | 3467 | 10.320142 | 9.11748083 | 12.5963141 | 11.1253838 | 12.3334592 | 10.5338165 |
| ENSMUSG00 | Hkdc1 | 5116 | 9.3949132 | 6.83802912 | 4.48411973 | 5.48001341 | 10.9413476 | 3.90246989 |
| ENSMUSG00 | Tacr2 | 2253 | 0 | 0 | 0 | 0 | 0 | 0 |
| ENSMUSG00 | 2010107G23I | 4407 | 2.58000169 | 3.1052756 | 2.92873568 | 2.28429554 | 2.18833334 | 3.99805241 |
| ENSMUSG00 | Aifm2 | 4174 | 8.06729564 | 6.71885705 | 8.55656185 | 8.86923279 | 9.74317718 | 7.51457478 |
| ENSMUSG00 | H2afy2 | 2337 | 1.36090648 | 1.81446594 | 0.6809013 | 1.76009973 | 3.25328626 | 2.99909336 |
| ENSMUSG00 | Tysnd1 | 2282 | 22.2470415 | 17.6317613 | 16.812971 | 15.1791248 | 15.8311455 | 10.8763899 |
| ENSMUSG00 | Sar1a | 2604 | 101.755099 | 97.5756949 | 102.984862 | 84.5517034 | 103.208879 | 71.1403033 |
| ENSMUSG00 | Ppa1 | 3056 | 20.2029651 | 15.4366907 | 20.8425575 | 19.8356789 | 20.2869917 | 18.9208885 |
| ENSMUSG00 | Npffr1 | 1303 | 0 | 0 | 0.16961567 | 0 | 0.11748206 | 0 |
| ENSMUSG00 | Eif4ebp2 | 2780 | 1.41575272 | 2.11465681 | 2.63939063 | 1.14216544 | 1.54180416 | 1.87390507 |
| ENSMUSG00 | X99384 | 4254 | 5.71006079 | 4.15712841 | 4.51993458 | 2.96018654 | 4.10226153 | 1.76958167 |
| ENSMUSG00 | Tbata | 2477 | 0.04814955 | 0.0389071 | 0 | 0.0437005 | 0.04120014 | 0.03303348 |
| ENSMUSG00 | Sgpl1 | 8479 | 50.4035164 | 31.2510019 | 53.2830567 | 31.9202047 | 44.4908594 | 43.2070945 |
| ENSMUSG00 | Pcbd1 | 805 | 166.03469 | 142.823417 | 157.53427 | 119.67585 | 88.1710419 | 228.869799 |
| ENSMUSG00 | Unc5b | 5835 | 0.03406639 | 0.27251972 | 0 | 0.07420472 | 0 | 0.04674317 |
| ENSMUSG00 | Slc29a3 | 8291 | 1.82210621 | 2.52817541 | 1.80731212 | 1.04011683 | 2.17251365 | 2.111967 |
| ENSMUSG00 | 4632428N05I | 5097 | 0 | 0.04726942 | 0.06070491 | 0 | 0.30033182 | 0 |
| ENSMUSG00 | Slc16a7 | 4678 | 135.022461 | 107.724201 | 114.529829 | 133.614634 | 114.193059 | 121.878807 |
| ENSMUSG00 | Lrig3 | 4031 | 1.23280453 | 1.87677286 | 0.89916922 | 2.11246899 | 1.93674907 | 1.10289412 |
| ENSMUSG00 | Anapc16 | 1325 | 33.1845747 | 30.1483476 | 35.6283535 | 35.5918734 | 24.3386185 | 28.2832863 |
| ENSMUSG00 | Ddit4 | 1682 | 2.15086131 | 2.72158859 | 1.47164281 | 1.480179 | 0.91010183 | 1.378326 |
| ENSMUSG00 | Dnajb12 | 3606 | 5.77700271 | 6.97540843 | 6.827628 | 7.47456663 | 6.05637517 | 5.34752356 |
| ENSMUSG00 | Micu1 | 5268 | 11.3802704 | 12.7051952 | 11.5538986 | 10.0068085 | 10.3060106 | 8.45472397 |
| ENSMUSG00 | Cand1 | 8341 | 17.86399 | 15.4594077 | 17.3129626 | 14.7382239 | 19.117303 | 15.4276164 |
| ENSMUSG00 | Tbk1 | 3010 | 24.9495358 | 31.8094553 | 32.3657349 | 29.5369295 | 38.6851128 | 35.2758042 |
| ENSMUSG00 | Pno1 | 1711 | 35.5034332 | 48.4962322 | 46.0877294 | 36.5038101 | 33.2819611 | 33.6350022 |
| ENSMUSG00 | Plek | 7163 | 0 | 0.31617518 | 0.21597997 | 0 | 0 | 0 |
| ENSMUSG00 | Srgap1 | 9643 | 0.39990476 | 0.17489635 | 0.238359 | 0.14592966 | 0.19578719 | 0.52043281 |
| ENSMUSG00 | Egfr | 11655 | 2.50369135 | 3.49356874 | 1.47529103 | 1.95657247 | 2.97270778 | 1.1420012 |
| ENSMUSG00 | Avpr1a | 2669 | 315.556334 | 164.34664 | 509.91935 | 221.575388 | 217.392626 | 407.985274 |
| ENSMUSG00 | Usp15 | 3688 | 44.9620687 | 52.9946338 | 48.8042041 | 56.5983264 | 44.7726009 | 53.8541006 |
| ENSMUSG00 | Elane | 915 | 0 | 0 | 0 | 0 | 0 | 0 |
| ENSMUSG00 | Vps54 | 6209 | 13.4652562 | 14.365132 | 10.0164267 | 21.2575485 | 21.9506281 | 18.1816286 |
| ENSMUSG00 | Tbc1d15 | 3756 | 27.5197682 | 28.6989007 | 30.0327751 | 23.4879124 | 29.49368 | 33.2219048 |
| ENSMUSG00 | Pcsk4 | 3508 | 1.41660065 | 1.27746269 | 1.10882617 | 0.99770762 | 0.71274014 | 1.90487076 |
| ENSMUSG00 | Rab21 | 1816 | 40.3246669 | 43.2245123 | 44.8833692 | 46.3344316 | 29.6998234 | 69.7335628 |
| ENSMUSG00 | 2310011J03I | 1647 | 14.2897659 | 16.1499187 | 15.1365146 | 9.90229759 | 11.8968599 | 12.9169512 |
| ENSMUSG00 | Peli1 | 5625 | 12.4955893 | 10.3397397 | 13.5395157 | 11.1549645 | 18.4964943 | 17.126049 |
| ENSMUSG00 | Apc2 | 9834 | 1.33003387 | 0.47039846 | 0.47644858 | 1.42728445 | 0.61746377 | 0.2024658 |
| ENSMUSG00 | Thap2 | 3495 | 5.48754063 | 6.14182552 | 7.15277988 | 6.78194082 | 4.15104438 | 6.61640294 |
| ENSMUSG00 | Lgr5 | 8283 | 0.38877145 | 0.06981013 | 0.14408424 | 0.02178078 | 0.28337719 | 0.02304992 |
| ENSMUSG00 | Slc1a4 | 6213 | 0.01279752 | 0.77557448 | 0.1138306 | 0.6155958 | 0.14783112 | 0.09657848 |
| ENSMUSG00 | Dock2 | 11370 | 0.44405858 | 0.28818629 | 0.30323164 | 0.25070198 | 0.15258547 | 0.39100846 |
| ENSMUSG00 | Rab1 | 4023 | 157.223389 | 140.391073 | 157.381854 | 150.920349 | 113.087538 | 172.305483 |
| ENSMUSG00 | Gamt | 1731 | 1.2631724 | 0.64025891 | 0.61285051 | 1.16729896 | 1.47389878 | 1.6229277 |
| ENSMUSG00 | Ptprr | 6461 | 0.23997272 | 0.82038518 | 0.48573454 | 0.32949089 | 0.38698227 | 0.41369984 |
| ENSMUSG00 | Actr2 | 5195 | 16.8740779 | 13.8483846 | 14.3028485 | 19.4197106 | 14.2913141 | 23.4735153 |
| ENSMUSG00 | Ndufs7 | 1549 | 109.898623 | 104.336557 | 132.46285 | 89.4713577 | 72.1418741 | 140.475837 |
| ENSMUSG00 | Ptprb | 13165 | 1.04182611 | 0.65151437 | 0.25181453 | 0.49059521 | 1.01548883 | 0.43299655 |
| ENSMUSG00 | Kcnmb1 | 4187 | 1.30081224 | 0.2186613 | 1.58353868 | 0.71526384 | 0.93838809 | 2.05846334 |
| ENSMUSG00 | Mum1 | 7104 | 6.1950052 | 5.77911722 | 4.156367 | 4.4594644 | 9.68236974 | 5.70909186 |
| ENSMUSG00 | Gabrp | 3366 | 0 | 0 | 0 | 0 | 0 | 0 |
| ENSMUSG00 | Meis1 | 10398 | 0.13764159 | 0 | 0 | 0 | 0 | 0 |
| ENSMUSG00 | Uqcr11 | 795 | 115.26589 | 132.315727 | 164.130617 | 103.026718 | 86.9052992 | 307.054115 |
| ENSMUSG00 | 1700058G18I | 822 | 0 | 0 | 0 | 0 | 0 | 0 |
| ENSMUSG00 | Cnot2 | 4834 | 50.0849972 | 35.9454511 | 58.8138714 | 35.6192628 | 45.716846 | 46.3849419 |
| ENSMUSG00 | Tcf3 | 4745 | 6.13298456 | 5.35179242 | 4.5179966 | 4.88951626 | 6.30167676 | 4.97783722 |
| ENSMUSG00 | Olfr299 | 992 | 0 | 0 | 0 | 0 | 0 | 0 |
| ENSMUSG00 | Best3 | 2709 | 0.29350669 | 0.30238816 | 0.96268318 | 2.45075533 | 0.37671743 | 0.35238552 |
| ENSMUSG00 | Frs2 | 5672 | 10.01597 | 9.60840375 | 11.8063809 | 10.9098566 | 16.4900118 | 12.1322135 |
| ENSMUSG00 | Yeats4 | 1983 | 14.1539936 | 17.1799368 | 14.0875264 | 13.0826979 | 12.5829036 | 11.3059775 |
| ENSMUSG00 | Cobl | 9507 | 4.36988294 | 2.55453527 | 3.80785836 | 3.62072899 | 3.83757849 | 3.72383315 |
| ENSMUSG00 | Rab36 | 6066 | 1.85473145 | 1.7158377 | 0.7359687 | 1.31456179 | 1.94314092 | 0.51707608 |
| ENSMUSG00 | Grb10 | 9656 | 21.4010965 | 14.3920567 | 25.5341633 | 19.2592023 | 15.3036852 | 21.4530743 |
| ENSMUSG00 | 9530003J23I | 1682 | 0 | 0 | 0 | 0 | 0 | 0 |
| ENSMUSG00 | Adora2a | 2531 | 0 | 0.0571155 | 0 | 0 | 0 | 0 |
| ENSMUSG00 | Snrpd3 | 600 | 79.510961 | 89.6267824 | 92.2345124 | 83.5900698 | 51.3665517 | 137.191435 |
| ENSMUSG00 | Nav3 | 9264 | 0 | 0 | 0 | 0 | 0 | 0.21492322 |
| ENSMUSG00 | Ddc | 4746 | 1.16435141 | 0.57872465 | 0.43773422 | 0.342118 | 0.37630071 | 0.01149374 |
| ENSMUSG00 | Cpm | 5981 | 2.47931688 | 2.82786176 | 0.82033181 | 1.95462003 | 1.31383747 | 1.35894374 |
| ENSMUSG00 | Mdm2 | 5734 | 20.2382713 | 22.9839413 | 22.0546691 | 29.0720345 | 22.6477374 | 17.3189678 |
| ENSMUSG00 | E2f7 | 7642 | 0 | 0.03152738 | 0 | 0 | 0 | 0.02498331 |
| ENSMUSG00 | Csrp2 | 1066 | 53.2186404 | 53.6108071 | 44.9067505 | 48.6396792 | 37.5757083 | 48.1016152 |
| ENSMUSG00 | Osbpl8 | 7128 | 3.66433091 | 6.59115896 | 5.23377602 | 8.63579797 | 4.89647041 | 6.24469855 |
| ENSMUSG00 | Mknk2 | 4448 | 19.2967811 | 21.6232326 | 18.8712455 | 23.0785559 | 17.3911838 | 17.7824761 |
| ENSMUSG00 | 4930415F15I | 1843 | 1.59625912 | 0.60135007 | 1.17519821 | 0.15662309 | 0.22149268 | 1.15432548 |
| ENSMUSG00 | Zpbp | 4240 | 3.33796016 | 0.96600179 | 1.56374445 | 0.69781313 | 1.40803914 | 4.70873489 |
| ENSMUSG00 | Cabin1 | 8447 | 5.05473968 | 7.24479478 | 5.61484285 | 7.74010473 | 5.52730364 | 4.78202211 |
| ENSMUSG00 | Ap3d1 | 4699 | 29.9667827 | 24.375223 | 23.2344227 | 25.5853101 | 33.4673953 | 18.4055929 |
| ENSMUSG00 | Phlda1 | 1951 | 11.085075 | 13.2630033 | 16.924027 | 12.5760073 | 7.8984959 | 32.6009527 |
| ENSMUSG00 | Sf3a2 | 2721 | 1.85554797 | 2.12509116 | 1.78691758 | 2.41342598 | 2.32534752 | 3.87919345 |
| ENSMUSG00 | Mdm1 | 3134 | 2.1818579 | 2.38318396 | 2.5669226 | 1.70393829 | 3.10977594 | 4.28178766 |
| ENSMUSG00 | Glipr1l1 | 891 | 0 | 0 | 0 | 0 | 0 | 0 |
| ENSMUSG00 | Glipr1l2 | 1896 | 0 | 0 | 0 | 0 | 0 | 0 |
| ENSMUSG00 | Jsrp1 | 1318 | 0 | 0 | 0 | 0 | 0 | 0 |
| ENSMUSG00 | Wif1 | 2883 | 0 | 0 | 0 | 0 | 0 | 0.02838152 |
| ENSMUSG00 | Timm13 | 1222 | 18.3486833 | 27.5632758 | 35.4844583 | 23.562579 | 13.8213833 | 39.7066966 |
| ENSMUSG00 | Vps13d | 22805 | 1.75722536 | 1.24031754 | 2.14176876 | 1.07273081 | 1.30446731 | 1.99132972 |
| ENSMUSG00 | Liph | 1513 | 11.4037532 | 9.17230359 | 9.08577209 | 8.13217263 | 7.82426913 | 12.618802 |
| ENSMUSG00 | Tmbim4 | 2901 | 91.5294224 | 77.071731 | 133.30642 | 70.4601111 | 65.6254423 | 124.047429 |
| ENSMUSG00 | Slc5a4b | 3187 | 0 | 0 | 0 | 0 | 0 | 0.01711618 |
| ENSMUSG00 | Irak3 | 5188 | 0.25287796 | 0.02786417 | 0.6390012 | 0 | 0.25572201 | 0.47841021 |
| ENSMUSG00 | Hells | 5263 | 0.53631752 | 0.95219266 | 1.15060849 | 0.61016568 | 2.75346584 | 0.89654429 |
| ENSMUSG00 | Slc5a4a | 2040 | 0 | 0 | 0 | 0 | 0 | 0 |
| ENSMUSG00 | Prmt2 | 3639 | 15.6006667 | 16.2872558 | 16.5559527 | 15.0614525 | 16.5881293 | 12.4418525 |
| ENSMUSG00 | Dip2a | 6866 | 6.47019277 | 5.18103344 | 3.4216174 | 6.89523207 | 6.73805991 | 3.03274117 |
| ENSMUSG00 | Hmg20b | 2867 | 2.45436439 | 2.62193407 | 3.3931268 | 2.56740045 | 3.09682225 | 5.74603492 |
| ENSMUSG00 | 4930404N11I | 1987 | 0.04001558 | 0.09700341 | 0.42266483 | 0.19974962 | 0.10272043 | 0.27453084 |
| ENSMUSG00 | Fzr1 | 4634 | 24.0042802 | 20.9008953 | 27.8240777 | 22.6739132 | 21.9675485 | 18.128159 |
| ENSMUSG00 | Ncln | 4600 | 19.0048482 | 15.3672849 | 11.9825431 | 9.91471832 | 13.9546045 | 13.9871465 |
| ENSMUSG00 | Col6a2 | 8297 | 0 | 0 | 0 | 0 | 0 | 0 |
| ENSMUSG00 | Hcfc2 | 5470 | 10.1896131 | 9.02064292 | 11.4262168 | 13.7336045 | 11.4179678 | 9.3641266 |
| ENSMUSG00 | Nfyb | 5682 | 6.69588083 | 6.47065388 | 4.17747091 | 7.29642189 | 4.86735229 | 5.37620485 |
| ENSMUSG00 | Txnrd1 | 4002 | 66.487614 | 90.1478991 | 55.1031971 | 93.2074404 | 70.5469609 | 47.1888061 |
| ENSMUSG00 | Glt8d2 | 4383 | 0 | 0 | 0 | 0 | 0 | 0 |
| ENSMUSG00 | Ppm1m | 3996 | 3.45224017 | 1.94144574 | 1.736659 | 1.68852411 | 2.08140622 | 1.67906938 |
| ENSMUSG00 | D10Wsu102e | 3505 | 6.81684559 | 9.32108638 | 8.76470211 | 7.9679037 | 11.5446265 | 4.54447627 |
| ENSMUSG00 | Aldh1l2 | 8549 | 22.9399679 | 11.9719269 | 30.6036623 | 16.5194903 | 23.7255637 | 22.3040125 |
| ENSMUSG00 | Wdr82 | 5380 | 10.4709137 | 10.4344247 | 10.4835598 | 7.2968893 | 14.5870941 | 3.91375864 |
| ENSMUSG00 | Glyctk | 7339 | 0.74213119 | 0.28232961 | 0.13852601 | 0.23599103 | 0.36154402 | 0.44968406 |
| ENSMUSG00 | Polut2 | 2210 | 24.4829 | 30.9178167 | 25.2410524 | 24.5717091 | 32.4629341 | 15.5872712 |
| ENSMUSG00 | Slc36a1 | 3247 | 1.17540072 | 0.50457008 | 1.47021837 | 3.30039028 | 1.25719435 | 1.6631902 |
| ENSMUSG00 | Adarb1 | 15466 | 0.13623694 | 0.1682444 | 0.04287002 | 0.21230222 | 0.32332761 | 0.14813589 |
| ENSMUSG00 | Appl2 | 5299 | 14.2396494 | 14.6314371 | 12.4038763 | 12.0523153 | 18.6907332 | 11.4832459 |
| ENSMUSG00 | Slc36a2 | 3245 | 0 | 0 | 0 | 0 | 0.14152154 | 0 |
| ENSMUSG00 | Sumo3 | 2500 | 84.3293253 | 81.6085584 | 79.7224441 | 54.902439 | 67.7834376 | 65.2082075 |
| ENSMUSG00 | Hint1 | 989 | 46.3478959 | 37.3213495 | 55.7327586 | 37.7237958 | 26.570863 | 73.0541146 |
| ENSMUSG00 | Lyrm7 | 4112 | 2.21400901 | 2.47260197 | 2.75186571 | 2.43062411 | 1.38982347 | 3.8139391 |
| ENSMUSG00 | 1700008A04I | 685 | 0 | 0 | 0 | 0 | 0 | 0 |
| ENSMUSG00 | Fbxw11 | 4506 | 39.3143411 | 33.1829849 | 40.2289743 | 35.6816668 | 44.5376689 | 28.684979 |
| ENSMUSG00 | Stk10 | 5168 | 1.03850423 | 1.12820424 | 0.25658965 | 0.32116371 | 0.23696459 | 1.35106572 |
| ENSMUSG00 | Papolg | 5029 | 9.37562137 | 9.30384477 | 8.61360234 | 5.87615718 | 11.6480969 | 10.2449381 |
| ENSMUSG00 | Rel | 2789 | 0.34210525 | 0.24188246 | 0.38036724 | 0.06468635 | 0.49398069 | 0 |
| ENSMUSG00 | Pfdn | 4372 | 48.3121839 | 46.3017483 | 43.1402768 | 42.2387703 | 52.6253268 | 25.9832737 |
| ENSMUSG00 | Il9r | 6988 | 0 | 0 | 0 | 0 | 0 | 0 |
| ENSMUSG00 | Pus10 | 4286 | 11.2049978 | 10.5457029 | 10.8184166 | 8.1239321 | 9.73858503 | 8.03730037 |
| ENSMUSG00 | Rhbdf1 | 5288 | 8.7961256 | 9.49513478 | 4.17944809 | 8.35181969 | 12.2451723 | 9.13968622 |
| ENSMUSG00 | Pex13 | 4772 | 22.427023 | 19.2765964 | 22.082396 | 18.6912854 | 17.4614568 | 22.35926 |
| ENSMUSG00 | 1810043G02I | 2833 | 11.5070575 | 14.5937054 | 9.84516871 | 17.2067922 | 12.8421482 | 11.995835 |
| ENSMUSG00 | 1700093K21I | 2000 | 0.25841062 | 0.53005086 | 0.06630277 | 0.01804102 | 0 | 0.39548227 |
| ENSMUSG00 | Mpg | 2762 | 3.44010132 | 3.24499574 | 2.89664503 | 2.15551678 | 3.06675539 | 4.35485732 |
| ENSMUSG00 | Ahsa2 | 5622 | 5.4167019 | 7.10540031 | 5.09476953 | 7.72086438 | 5.40033682 | 5.82654602 |
| ENSMUSG00 | Nprl3 | 7124 | 4.60949283 | 4.84299447 | 5.14364512 | 3.53020565 | 6.38187828 | 4.22289824 |
| ENSMUSG00 | Xpo1 | 7275 | 23.1264871 | 23.5004119 | 22.9485032 | 29.4360047 | 26.2321162 | 34.6603491 |
| ENSMUSG00 | Hbq1a | 601 | 0 | 0 | 0 | 0 | 0 | 0 |
| ENSMUSG00 | Nsg2 | 4245 | 0.19667022 | 0.76053984 | 0.43733272 | 1.09648616 | 0.24040695 | 0.71961357 |
| ENSMUSG00 | 4930524B15I | 1995 | 0 | 0 | 0 | 0 | 0 | 0 |
| ENSMUSG00 | Cpeb4 | 7568 | 10.4274087 | 9.38515007 | 7.58113005 | 14.8180489 | 9.9315343 | 13.7742695 |
| ENSMUSG00 | Stc2 | 7070 | 0.05060811 | 0.04089373 | 0 | 0 | 0.13712887 | 0 |
| ENSMUSG00 | Asb3 | 5427 | 5.12052347 | 5.17646873 | 5.2615424 | 4.50111372 | 4.29685899 | 6.9606367 |
| ENSMUSG00 | Cdc34 | 1467 | 8.94295063 | 13.4015463 | 9.52132406 | 7.55091184 | 6.88699551 | 6.50724181 |
| ENSMUSG00 | Gm16517 | 1099 | 13.7823822 | 17.3190579 | 13.99658 | 11.0642848 | 13.0467804 | 15.039519 |
| ENSMUSG00 | Chac2 | 2106 | 23.4455398 | 23.0635962 | 25.1232698 | 23.4379093 | 12.9625409 | 26.2256147 |
| ENSMUSG00 | Madcam1 | 1415 | 0 | 0 | 0.12495221 | 0 | 0 | 0.11565218 |
| ENSMUSG00 | Erlec1 | 5095 | 21.0286595 | 19.1516281 | 18.3487533 | 19.843354 | 11.8277036 | 33.9019216 |
| ENSMUSG00 | Shc2 | 7573 | 0 | 0 | 0 | 0 | 0 | 0 |
| ENSMUSG00 | Sptbn2 | 18773 | 16.0478896 | 17.1307897 | 18.0334478 | 18.7415978 | 23.6716938 | 20.0408232 |
| ENSMUSG00 | Theg | 1845 | 0 | 0 | 0 | 0 | 0 | 0 |
| ENSMUSG00 | Wdpcp | 3716 | 4.46125817 | 5.88714874 | 3.23544168 | 4.90350713 | 7.55234303 | 4.72682123 |
| ENSMUSG00 | Mdh1 | 2558 | 893.254981 | 780.062771 | 948.317883 | 726.055934 | 746.525054 | 810.551525 |
| ENSMUSG00 | Prss57 | 4795 | 0 | 0 | 0 | 0 | 0 | 0 |
| ENSMUSG00 | Fstl3 | 2560 | 2.22071629 | 1.67523178 | 1.95109698 | 0.70472743 | 1.35538811 | 1.43831105 |
| ENSMUSG00 | Ccng1 | 3992 | 128.587867 | 107.176207 | 145.505115 | 160.471097 | 99.0617266 | 140.424765 |
| ENSMUSG00 | Fgf22 | 769 | 0 | 0 | 0 | 0 | 0 | 0 |
| ENSMUSG00 | Nudcd2 | 2460 | 17.9707701 | 15.6703877 | 20.3580033 | 21.9719117 | 13.4410941 | 23.7045442 |
| ENSMUSG00 | Polrmt | 4482 | 20.081751 | 17.6748128 | 15.9371214 | 17.8558623 | 23.0996244 | 14.1850031 |
| ENSMUSG00 | Hmmr | 6893 | 0.00576752 | 1.02762324 | 0.71179524 | 0.24602584 | 0.61441886 | 0.24136849 |
| ENSMUSG00 | Hcn2 | 3187 | 0.19958823 | 0.30239374 | 0.94312059 | 0.38493552 | 0.17611865 | 0.52204361 |
| ENSMUSG00 | 4930404A10I | 1705 | 0 | 0 | 0 | 0 | 0 | 0 |
| ENSMUSG00 | Acsl6 | 5812 | 0.76610699 | 0.15752622 | 0.41068471 | 0.89398046 | 0.28094357 | 0.93387013 |
| ENSMUSG00 | Slc22a4 | 2542 | 0.64121743 | 0.41703451 | 0 | 0.99360469 | 0.94344597 | 0.65450551 |
| ENSMUSG00 | Zfp354b | 4431 | 1.65087078 | 0.59811652 | 1.05741263 | 2.9722289 | 2.57953243 | 1.66811718 |
| ENSMUSG00 | Cyfip2 | 10944 | 28.9520449 | 31.2260826 | 25.6066963 | 29.0858732 | 35.4536332 | 22.7911711 |
| ENSMUSG00 | Mgat1 | 4814 | 18.7711274 | 17.7971579 | 17.1518577 | 16.9542137 | 18.4220692 | 17.0424003 |
| ENSMUSG00 | Ppp2ca | 7483 | 85.6312755 | 82.2189621 | 94.5882817 | 81.1858057 | 69.5398878 | 100.226918 |
| ENSMUSG00 | Sgcd | 8248 | 2.10634639 | 2.15577075 | 3.34407736 | 2.24856582 | 2.01680389 | 2.23871615 |
| ENSMUSG00 | Flt4 | 8203 | 0.25201572 | 0.1644789 | 0.08621596 | 0.17154696 | 0.22393631 | 0 |
| ENSMUSG00 | Hnrnpab | 2724 | 72.2282941 | 66.9374072 | 76.330936 | 63.4217435 | 62.2843978 | 50.3739758 |
| ENSMUSG00 | Agpat2l2 | 8274 | 5.31418437 | 5.73065737 | 5.4491032 | 6.54134237 | 4.87198899 | 5.439105 |
| ENSMUSG00 | Hspa4 | 5509 | 44.1791708 | 40.6379035 | 34.7661256 | 41.6361512 | 39.4854674 | 34.8346999 |
| ENSMUSG00 | Cnot6 | 7306 | 20.4871864 | 24.2185348 | 24.0308952 | 27.1035122 | 20.7360128 | 25.0384623 |
| ENSMUSG00 | Gfpt2 | 5629 | 0.67094877 | 0.393778 | 0.55752902 | 0.22435096 | 0.50763493 | 0.00969076 |
| ENSMUSG00 | Zfp354a | 4425 | 3.18942273 | 4.85675778 | 3.81584272 | 5.5774279 | 4.65867928 | 4.69062158 |
| ENSMUSG00 | Mapk9 | 6990 | 19.2748675 | 16.399935 | 16.9914667 | 19.0527649 | 23.2648154 | 14.5776899 |
| ENSMUSG00 | Canx | 4308 | 192.594447 | 173.137869 | 191.336112 | 178.592719 | 161.962549 | 230.409923 |
| ENSMUSG00 | Gnb2l1 | 6622 | 10.998496 | 11.5554319 | 12.508948 | 11.202761 | 7.6978782 | 12.1627922 |
| ENSMUSG00 | Rasgef1c | 3659 | 0 | 0 | 0 | 0 | 0 | 0 |
| ENSMUSG00 | Rufy1 | 3119 | 28.2583842 | 31.2539828 | 30.625259 | 23.3104584 | 31.5745128 | 26.3826825 |
| ENSMUSG00 | Rnf130 | 1848 | 71.7878996 | 63.5445668 | 91.5609605 | 64.6079466 | 60.6076271 | 74.5329698 |
| ENSMUSG00 | Ltc4s | 860 | 7.07277735 | 11.3182108 | 5.0369542 | 8.30726141 | 6.1112985 | 7.16752148 |
| ENSMUSG00 | Rad50 | 5388 | 10.3299318 | 9.71241206 | 9.37689431 | 11.1366815 | 13.7604536 | 8.9599316 |
| ENSMUSG00 | 3010026O09I | 2029 | 1.11683706 | 1.49617834 | 1.4160275 | 0.62241082 | 1.0310899 | 1.02162276 |
| ENSMUSG00 | Il13 | 1201 | 0 | 0 | 0 | 0 | 0 | 0 |
| ENSMUSG00 | Clk4 | 4285 | 43.8562792 | 39.7748999 | 60.0257934 | 45.7151505 | 52.2886386 | 67.6041875 |
| ENSMUSG00 | Sar1b | 1344 | 93.2064874 | 100.424274 | 125.501661 | 90.9836673 | 73.5402455 | 120.807982 |
| ENSMUSG00 | Phf15 | 15011 | 0.28338128 | 0.2214952 | 0.43580526 | 0.41343759 | 0.1971574 | 0.53964212 |
| ENSMUSG00 | Pdlim4 | 2842 | 0.06994279 | 0 | 0 | 0 | 0 | 0 |
| ENSMUSG00 | Cdkl3 | 3761 | 4.01677283 | 4.2792533 | 3.52580508 | 3.03162086 | 3.51391421 | 5.60576872 |
| ENSMUSG00 | Ube2b | 2262 | 96.9105745 | 90.1525301 | 120.529163 | 92.9487504 | 76.5621218 | 128.149808 |
| ENSMUSG00 | Cdkn2aipnl | 2019 | 22.5655179 | 27.9715256 | 21.9148315 | 23.3219753 | 15.2649485 | 15.7244577 |
| ENSMUSG00 | Kremen1 | 7748 | 6.95259113 | 6.2192104 | 4.70086715 | 5.30426545 | 5.24884119 | 5.64642764 |
| ENSMUSG00 | Itk | 5336 | 0 | 0 | 0 | 0 | 0.12431463 | 0 |
| ENSMUSG00 | Nefh | 5897 | 0 | 0 | 0 | 0 | 0 | 0 |
| ENSMUSG00 | Med7 | 2928 | 21.5477963 | 16.9672889 | 20.4101406 | 22.3910775 | 18.1415496 | 26.2034017 |
| ENSMUSG00 | Havcr2 | 2803 | 0 | 0 | 0.41000639 | 0 | 0 | 0 |
| ENSMUSG00 | Tnip1 | 3694 | 1.6573752 | 1.74796515 | 2.72821338 | 3.08660695 | 4.32356677 | 2.47347162 |
| ENSMUSG00 | Fam71b | 2489 | 4.48826437 | 1.37454295 | 7.06803275 | 3.36321186 | 2.33708592 | 8.81028926 |
| ENSMUSG00 | Vdac1 | 2627 | 192.224634 | 180.712915 | 209.281509 | 167.842628 | 135.830775 | 152.652844 |
| ENSMUSG00 | Fabp6 | 475 | 0 | 0 | 0 | 0 | 0 | 0 |
| ENSMUSG00 | Upp1 | 2044 | 12.4479 | 7.68531318 | 13.9266081 | 6.40791691 | 7.51415808 | 5.32415903 |
| ENSMUSG00 | Stu7 | 4090 | 28.3440052 | 30.6437496 | 32.9514473 | 28.1598744 | 31.3394269 | 29.6353293 |
| ENSMUSG00 | Nipal4 | 3438 | 0 | 0 | 0 | 0 | 0 | 0 |
| ENSMUSG00 | Ascc2 | 2745 | 20.971197 | 23.1716225 | 17.6485319 | 20.0850142 | 23.7937199 | 17.8452649 |
| ENSMUSG00 | Hus1 | 9653 | 5.77407891 | 3.66402658 | 4.33638717 | 4.64995995 | 4.87375101 | 3.82573931 |
| ENSMUSG00 | Pttg1 | 1519 | 39.1535213 | 76.8635611 | 64.2803629 | 40.3577312 | 60.4321594 | 57.1708041 |
| ENSMUSG00 | Hormad2 | 3001 | 0.15896893 | 0.12845436 | 0 | 0.26451349 | 0 | 0.03635407 |
| ENSMUSG00 | Zfp607 | 2057 | 8.87105763 | 3.65439231 | 4.25472286 | 6.22709082 | 9.02946077 | 4.7468745 |
| ENSMUSG00 | Tns3 | 9539 | 7.2392567 | 6.3245021 | 6.16295746 | 7.33064287 | 6.39233873 | 5.43262549 |
| ENSMUSG00 | Btg2 | 2741 | 72.4039981 | 81.1311312 | 62.1018962 | 67.5830779 | 70.5545291 | 51.4446862 |
| ENSMUSG00 | Gatsl3 | 1565 | 1.44796319 | 1.7242433 | 0.96029563 | 0 | 1.79325921 | 0 |
| ENSMUSG00 | Igfbp3 | 3125 | 9.23599323 | 3.46942383 | 2.74405041 | 7.52815777 | 16.4427393 | 7.18304899 |
| ENSMUSG00 | Gabra6 | 3577 | 0 | 0 | 0 | 0 | 0 | 0 |
| ENSMUSG00 | Igfbp1 | 1521 | 0.44434133 | 0 | 0.61028186 | 0 | 0.33547913 | 0 |
| ENSMUSG00 | Pes1 | 3060 | 10.3546137 | 11.8576441 | 12.7116411 | 9.78695979 | 9.77171772 | 8.42305036 |
| ENSMUSG00 | Adcy1 | 12500 | 0.62336593 | 0.4009112 | 0.67186801 | 0.47050986 | 0.72661559 | 0.7789637 |
| ENSMUSG00 | Tcn2 | 2166 | 143.916308 | 153.48027 | 154.920929 | 106.663588 | 128.34335 | 106.177174 |
| ENSMUSG00 | 4921536K21I | 2587 | 0 | 0.01862638 | 0.01708614 | 0.06973723 | 0.0394483 | 0.0527148 |
| ENSMUSG00 | Osbp2 | 7843 | 0.24837671 | 0.51608583 | 0.22543334 | 0.21622544 | 0.61806779 | 0.10432732 |
| ENSMUSG00 | Gabrg2 | 8302 | 0 | 0 | 0 | 0 | 0 | 0 |
| ENSMUSG00 | Myo1g | 5831 | 0 | 0 | 0 | 0.01237594 | 0 | 0 |
| ENSMUSG00 | Smtn | 4353 | 30.0563488 | 32.002528 | 16.2672531 | 19.9516382 | 24.0303401 | 18.6655526 |
| ENSMUSG00 | Arf5 | 1212 | 38.771434 | 45.4033968 | 24.2891316 | 29.7111225 | 26.776217 | 27.5446852 |
| ENSMUSG00 | 2310033P09I | 1278 | 14.6827753 | 14.6293737 | 13.8692794 | 11.8297157 | 12.6168504 | 16.6891768 |
| ENSMUSG00 | Guk1 | 2157 | 39.2947077 | 44.3887161 | 41.7222774 | 30.4949315 | 26.2110452 | 43.3587102 |
| ENSMUSG00 | Npc1l1 | 4650 | 0 | 0.22797887 | 0.02851732 | 0.75267921 | 0 | 0.03519308 |
| ENSMUSG00 | Rnf185 | 5798 | 20.6114133 | 18.2091229 | 26.7818342 | 16.6594745 | 16.9061173 | 15.998801 |
| ENSMUSG00 | Limk2 | 5716 | 25.880011 | 18.2680231 | 23.8485801 | 15.7874726 | 24.5490786 | 24.9842567 |
| ENSMUSG00 | Patz1 | 6074 | 8.45638473 | 7.60796806 | 5.90910381 | 8.23340693 | 7.21627541 | 5.70728786 |
| ENSMUSG00 | Eif4enif1 | 5250 | 19.7868706 | 24.5062477 | 18.9689052 | 20.563329 | 21.2075337 | 20.3027219 |
| ENSMUSG00 | Trim11 | 4103 | 10.7358207 | 12.6015239 | 10.7838277 | 8.96994526 | 9.5635591 | 9.3197768 |
| ENSMUSG00 | Ogdh | 5411 | 278.310405 | 196.156489 | 258.66778 | 210.424136 | 268.541509 | 168.239814 |
| ENSMUSG00 | Drg1 | 3449 | 59.7657775 | 62.7583352 | 73.8064407 | 59.788021 | 44.6351336 | 63.2243377 |
| ENSMUSG00 | Rtn4 | 7603 | 8.48130861 | 9.48137807 | 6.0055904 | 9.71457913 | 5.44290291 | 19.1385243 |
| ENSMUSG00 | Mtif2 | 3569 | 20.5962128 | 18.1593625 | 21.2773231 | 20.0478269 | 18.5147539 | 22.0932984 |
| ENSMUSG00 | Rps27a | 1678 | 2.5587556 | 3.87673998 | 1.68588674 | 2.0427856 | 1.55086124 | 1.07278081 |
| ENSMUSG00 | 1700034F02I | 2205 | 0.28847514 | 0.02185326 | 1.52351024 | 0.14727365 | 0.09256485 | 2.44915128 |
| ENSMUSG00 | Ccdc104 | 2733 | 26.8527688 | 22.6210045 | 17.7260227 | 20.4504526 | 19.4546226 | 21.8157195 |
| ENSMUSG00 | Smek2 | 6219 | 28.3575031 | 29.9082918 | 33.533413 | 44.320749 | 34.9365345 | 39.8089565 |
| ENSMUSG00 | Pnpt1 | 3659 | 38.5929125 | 30.500082 | 30.1282855 | 34.6916187 | 30.1779386 | 39.2683246 |
| ENSMUSG00 | Efemp1 | 2140 | 86.4403041 | 134.156459 | 76.1758865 | 92.6499227 | 131.309463 | 52.3188758 |
| ENSMUSG00 | Myl7 | 583 | 0 | 0 | 0 | 0 | 0 | 0 |
| ENSMUSG00 | Pold2 | 2115 | 5.92102901 | 8.06524847 | 5.26680257 | 4.43561777 | 5.50071569 | 5.04226191 |
| ENSMUSG00 | Zkscan17 | 4560 | 6.22487129 | 4.64956898 | 4.77883961 | 1.67749856 | 5.40476744 | 3.97157028 |
| ENSMUSG00 | Aebp1 | 4038 | 11.1744602 | 6.12175454 | 3.73274603 | 7.29146814 | 7.24074129 | 6.56536636 |
| ENSMUSG00 | Polm | 4606 | 4.34825568 | 1.36988366 | 1.96644484 | 1.44077608 | 3.31095624 | 2.37350919 |
| ENSMUSG00 | Pgam2 | 837 | 227.18096 | 308.577455 | 168.569036 | 227.355678 | 169.600212 | 336.224287 |
| ENSMUSG00 | Dbnl | 2453 | 34.9906573 | 38.3448573 | 40.0753766 | 31.5074355 | 31.5560588 | 35.7138774 |
| ENSMUSG00 | Mrps24 | 1868 | 16.813078 | 19.1662348 | 26.5968263 | 15.9742242 | 11.7185842 | 29.1143633 |
| ENSMUSG00 | Ankrd36 | 5619 | 0 | 0 | 0 | 0 | 0 | 0 |
| ENSMUSG00 | Ccdc117 | 3367 | 31.6201891 | 35.7927805 | 28.9996648 | 30.5416771 | 42.2517184 | 29.4860965 |
| ENSMUSG00 | Dynll2 | 3233 | 6.68944677 | 6.52819723 | 6.74033672 | 6.42847437 | 3.53538762 | 10.0054816 |
| ENSMUSG00 | Xbp1 | 3467 | 83.8454207 | 97.7905414 | 94.5870993 | 86.6718393 | 77.4299865 | 87.9365208 |
| ENSMUSG00 | Supt4h1 | 1051 | 7.75439915 | 9.94905609 | 9.84132375 | 8.99476276 | 4.70938007 | 9.86142999 |
| ENSMUSG00 | 4-Sep | 2700 | 8.64313595 | 11.7432144 | 4.55115273 | 9.58178738 | 7.21928725 | 4.7074007 |
| ENSMUSG00 | Btnl10 | 4867 | 0 | 0 | 0 | 0 | 0 | 0 |
| ENSMUSG00 | 2810021J22I | 1828 | 30.4473046 | 24.0141405 | 27.4447986 | 39.5757655 | 29.5606849 | 28.6473235 |
| ENSMUSG00 | Fam33a | 3320 | 8.64561956 | 7.8956098 | 7.76195014 | 9.14006007 | 7.25435223 | 7.58267832 |
| ENSMUSG00 | Prr11 | 3814 | 0 | 0 | 0.05794683 | 0 | 0 | 0.17162856 |
| ENSMUSG00 | 1200011M11 | 3628 | 6.2350795 | 8.10191006 | 6.05521389 | 5.70868069 | 7.9886939 | 4.03706761 |
| ENSMUSG00 | Rnf187 | 1945 | 72.8066949 | 65.9745478 | 51.6786586 | 43.6323747 | 54.9615719 | 38.3808163 |
| ENSMUSG00 | Tubd1 | 2299 | 3.02618925 | 6.01544537 | 4.65282558 | 3.68824727 | 2.9519391 | 1.22196078 |
| ENSMUSG00 | Mrpl22 | 1012 | 15.8314809 | 14.2845184 | 16.6848854 | 15.1530325 | 8.62204572 | 16.8175641 |
| ENSMUSG00 | Cnot8 | 2626 | 26.5389784 | 29.9100917 | 30.298293 | 29.2668526 | 24.4833334 | 25.2804539 |
| ENSMUSG00 | Rps6kb1 | 9568 | 6.98878744 | 5.51464822 | 6.96659482 | 8.66225505 | 8.76749183 | 7.91044355 |
| ENSMUSG00 | Sap30l | 1147 | 7.52130712 | 7.60396341 | 8.24690009 | 5.15907175 | 6.71750894 | 10.581704 |
| ENSMUSG00 | Galnt10 | 6205 | 1.78755505 | 0.43488167 | 1.38197543 | 0.88969425 | 2.05585721 | 0.56703118 |
| ENSMUSG00 | Rnft1 | 4823 | 45.7645507 | 38.335347 | 39.4728049 | 34.7054955 | 25.3809404 | 53.2655665 |
| ENSMUSG00 | Mfap3 | 5823 | 16.3514298 | 16.045597 | 16.0016289 | 17.8272441 | 16.5531211 | 15.5460291 |
| ENSMUSG00 | Fam114a2 | 4242 | 20.5993744 | 24.2976897 | 25.0185349 | 21.1797008 | 24.1298703 | 21.3464881 |
| ENSMUSG00 | Gria1 | 6022 | 0 | 0 | 0 | 0 | 0 | 0 |
| ENSMUSG00 | Ppm1d | 3166 | 12.0421686 | 15.6004748 | 13.2493838 | 9.77839362 | 11.2174219 | 12.9912053 |
| ENSMUSG00 | Znhit3 | 2242 | 3.52869787 | 3.33135528 | 3.13474266 | 5.55232175 | 2.2304125 | 3.69825617 |
| ENSMUSG00 | Myo19 | 7428 | 12.3740806 | 14.3171079 | 10.3185239 | 17.1958047 | 15.1196894 | 14.1880999 |
| ENSMUSG00 | Prpsap2 | 2597 | 14.864294 | 11.0029111 | 21.8371061 | 9.55889358 | 12.5355517 | 18.1585873 |

| | | | | | | | | |
|---|---|---|---|---|---|---|---|---|
| ENSMUSG00 | Ggnbp2 | 5077 | 47.9461901 | 50.6446435 | 46.3784159 | 49.4075977 | 55.1470798 | 49.5638806 |
| ENSMUSG00 | Acaca | 10128 | 0.36505329 | 0.29022245 | 0.30113815 | 0.99752888 | 0.33251785 | 0.32585223 |
| ENSMUSG00 | Shmt1 | 4275 | 19.7428971 | 14.5630136 | 18.993868 | 14.1796105 | 19.5750307 | 13.117347 |
| ENSMUSG00 | Llgl1 | 4937 | 8.6403951 | 10.677733 | 10.1887173 | 12.702166 | 13.3948112 | 6.5521009 |
| ENSMUSG00 | Drg2 | 2822 | 28.8938662 | 27.2179974 | 34.4436048 | 24.8048074 | 30.0697601 | 25.5735987 |
| ENSMUSG00 | Srebf1 | 5159 | 19.2188735 | 21.370533 | 11.875122 | 10.1273116 | 12.5711424 | 10.4202973 |
| ENSMUSG00 | Tom1l1 | 7978 | 7.74878067 | 6.468749 | 6.80369306 | 5.28702808 | 7.16978783 | 6.45456492 |
| ENSMUSG00 | Myocd | 9063 | 0 | 0 | 0 | 0 | 0 | 0 |
| ENSMUSG00 | Cox11 | 1419 | 11.0105031 | 12.021142 | 13.0518479 | 7.34863345 | 8.55833506 | 14.0890138 |
| ENSMUSG00 | 1700108M19 | 1956 | 0 | 0 | 0 | 0 | 0 | 0 |
| ENSMUSG00 | Stxbp4 | 11814 | 3.63432529 | 3.75653574 | 3.67039174 | 4.97525398 | 5.33414374 | 4.8251224 |
| ENSMUSG00 | Bzw2 | 1848 | 43.9289455 | 44.4838043 | 40.5901123 | 41.6856954 | 42.5219799 | 30.034573 |
| ENSMUSG00 | 1700086D15 | 1089 | 0.40157051 | 0.39823506 | 0 | 0.76206338 | 0.18742471 | 0 |
| ENSMUSG00 | Elac2 | 3715 | 9.22455564 | 4.63056793 | 11.0534353 | 10.6643566 | 13.2132903 | 8.06125273 |
| ENSMUSG00 | Pctp | 2717 | 9.81815511 | 9.98489765 | 6.32849353 | 5.3518822 | 8.33850234 | 5.52118203 |
| ENSMUSG00 | Twistnb | 2320 | 12.0294601 | 10.0734588 | 12.8413975 | 10.9490342 | 13.0425176 | 15.5888671 |
| ENSMUSG00 | Efcab10 | 671 | 1.42195459 | 1.72350911 | 0.39524748 | 0.91415016 | 0.91254323 | 0.28453424 |
| ENSMUSG00 | Atxn7l1 | 12068 | 2.64860841 | 3.06655515 | 2.75803677 | 2.54141016 | 3.42910099 | 4.40489492 |
| ENSMUSG00 | Atp6v1c2 | 2189 | 398.535525 | 257.772151 | 552.411978 | 287.584939 | 275.761547 | 375.602558 |
| ENSMUSG00 | Sypl | 6631 | 31.60174 | 38.6232755 | 40.5755721 | 42.9599972 | 33.535356 | 50.5718126 |
| ENSMUSG00 | Pdia6 | 3433 | 96.3372684 | 105.987715 | 91.1719343 | 91.5872228 | 94.2940076 | 79.5755277 |
| ENSMUSG00 | Nampt | 4536 | 43.3488992 | 51.267145 | 33.2098503 | 56.143534 | 43.1294367 | 39.246378 |
| ENSMUSG00 | Pik3cg | 6801 | 0.30981333 | 0 | 0.38995892 | 0 | 0.16506106 | 0 |
| ENSMUSG00 | Nbas | 8166 | 1.81105055 | 2.21282339 | 2.02442925 | 1.32557107 | 4.33030596 | 1.14896839 |
| ENSMUSG00 | Tspan13 | 2466 | 56.7474823 | 52.1140475 | 135.383828 | 48.0801291 | 32.8381421 | 116.243495 |
| ENSMUSG00 | Rock2 | 8161 | 9.41641267 | 16.6565315 | 12.4898236 | 15.8635432 | 12.6049717 | 10.4773918 |
| ENSMUSG00 | Agr2 | 1283 | 0 | 0 | 0 | 0 | 0 | 0 |
| ENSMUSG00 | Matn3 | 3006 | 0 | 0 | 0 | 0 | 0 | 0 |
| ENSMUSG00 | Laptm4a | 2780 | 172.407221 | 190.475113 | 209.815655 | 157.618831 | 128.740648 | 207.797431 |
| ENSMUSG00 | Fam49a | 4833 | 0.03290336 | 2.87144534 | 1.97550136 | 4.33760974 | 0.03167373 | 2.44924342 |
| ENSMUSG00 | Snx13 | 7618 | 42.3126065 | 44.3975633 | 37.9237656 | 56.3397112 | 37.7439784 | 37.9331587 |
| ENSMUSG00 | Ntsr2 | 2192 | 0 | 0 | 0 | 0 | 0 | 0 |
| ENSMUSG00 | Sdc1 | 3526 | 13.9922154 | 11.5478002 | 9.75298748 | 11.6862436 | 10.7957108 | 9.70779134 |
| ENSMUSG00 | Lpin1 | 5703 | 10.0521485 | 7.22416413 | 13.4550937 | 9.86356434 | 7.08625103 | 10.0671779 |
| ENSMUSG00 | Pum2 | 7257 | 6.39308885 | 6.51383488 | 5.53665086 | 5.82722284 | 4.72505506 | 11.5796009 |
| ENSMUSG00 | Nrcam | 8372 | 0 | 0.241738 | 0.00527972 | 0 | 0.63996291 | 0.40723004 |
| ENSMUSG00 | Rgs9 | 6784 | 0.02930091 | 0.22729454 | 0.07167162 | 0 | 0.33094937 | 0.08442916 |
| ENSMUSG00 | Slc7a15 | 3211 | 0 | 0 | 0 | 0 | 0 | 0 |
| ENSMUSG00 | Trib2 | 4102 | 2.44231621 | 1.72681789 | 1.77798735 | 1.0115639 | 1.75395392 | 3.96286811 |
| ENSMUSG00 | Arsg | 3891 | 5.09842837 | 3.88860006 | 4.35088817 | 1.61353784 | 4.10466606 | 3.7712043 |
| ENSMUSG00 | Hs1bp3 | 2951 | 6.52038379 | 2.85754909 | 3.9243927 | 5.74671668 | 4.97986996 | 3.04078493 |
| ENSMUSG00 | Fam84a | 3377 | 20.3074338 | 28.167023 | 8.01052056 | 26.0812173 | 28.0592182 | 9.83728475 |
| ENSMUSG00 | Smc6 | 5576 | 11.07963 | 13.9996478 | 11.9620841 | 14.5984743 | 12.4820618 | 13.2655706 |
| ENSMUSG00 | Apob | 13986 | 0 | 0 | 0 | 0 | 0 | 0 |
| ENSMUSG00 | Amz2 | 5626 | 7.5892972 | 7.97397398 | 9.16092232 | 6.07353291 | 6.69347056 | 7.01500231 |
| ENSMUSG00 | Gna13 | 6416 | 18.5331425 | 16.0496301 | 21.5980015 | 14.2056179 | 21.2583389 | 17.1359213 |
| ENSMUSG00 | Prkar1a | 4191 | 163.224184 | 162.829013 | 167.884738 | 161.57283 | 145.116529 | 147.254417 |
| ENSMUSG00 | Fam20a | 3622 | 0.17561781 | 0.05321529 | 0.90307465 | 0.55786706 | 0 | 0.41416487 |
| ENSMUSG00 | 1700012B07 | 2198 | 0 | 0 | 0 | 0 | 0 | 0 |
| ENSMUSG00 | Abca8b | 7722 | 0.49424063 | 1.2105892 | 0.22896578 | 0.8410733 | 0.09251091 | 0.51215005 |
| ENSMUSG00 | Rdh14 | 1470 | 73.4800276 | 79.2945602 | 87.2309841 | 73.3422783 | 61.5093455 | 87.2788456 |
| ENSMUSG00 | Nt5c1b | 2224 | 0.07150266 | 0.0649997 | 0 | 0.0811197 | 0.20649163 | 0.02452755 |
| ENSMUSG00 | Map2k6 | 3595 | 3.12956912 | 2.88180391 | 2.01644013 | 3.18164343 | 3.74713859 | 1.25182628 |
| ENSMUSG00 | 4933434M16 | 2145 | 0 | 0 | 0 | 0 | 0 | 0 |
| ENSMUSG00 | Klhl29 | 7027 | 0.38471221 | 0.10285992 | 0.2327406 | 0.32349065 | 0.03630737 | 0.17854467 |
| ENSMUSG00 | Ttc15 | 5426 | 15.9139152 | 17.761313 | 12.9607689 | 16.0926185 | 19.710889 | 14.1651186 |
| ENSMUSG00 | Adi1 | 1881 | 79.5743669 | 73.9319894 | 92.539531 | 78.4175428 | 51.6231755 | 85.4634363 |
| ENSMUSG00 | Rnaseh1 | 2460 | 5.94716131 | 9.32388068 | 11.46373 | 7.86177881 | 10.7445763 | 10.1670098 |
| ENSMUSG00 | Dcdc2c | 1380 | 0 | 0 | 0 | 0 | 0 | 0 |
| ENSMUSG00 | Ubxn2a | 4234 | 46.1497843 | 38.4672117 | 51.4261405 | 60.2674112 | 39.3122433 | 48.48109 |
| ENSMUSG00 | Fkbp1b | 975 | 1.0193713 | 3.80549338 | 2.40276685 | 0 | 1.09902963 | 1.28680349 |
| ENSMUSG00 | Allc | 1488 | 0 | 0 | 0 | 0 | 0 | 0 |
| ENSMUSG00 | Cmpk2 | 3193 | 3.52358315 | 5.10085106 | 2.39490098 | 3.77431972 | 6.8557204 | 4.68102171 |
| ENSMUSG00 | Pfn4 | 1302 | 0.61068326 | 0.62916246 | 0.33949188 | 0.63739403 | 0.39190765 | 0 |
| ENSMUSG00 | Itsn2 | 6427 | 14.8518607 | 10.8788747 | 15.6807534 | 21.7828431 | 13.6557284 | 12.32812 |
| ENSMUSG00 | Rsad2 | 7136 | 2.45129084 | 1.13443418 | 1.07779158 | 1.07700049 | 3.31785852 | 1.88430453 |
| ENSMUSG00 | Rnf144a | 5153 | 0.77921668 | 0.12156487 | 0.07720097 | 0.01400429 | 0.63374501 | 0.91038967 |
| ENSMUSG00 | Id2 | 1267 | 382.117002 | 539.749003 | 437.58778 | 544.135458 | 427.30059 | 641.201582 |
| ENSMUSG00 | Mboat2 | 2369 | 31.4486157 | 23.4728384 | 28.3235111 | 22.2980638 | 22.2715376 | 20.8503046 |
| ENSMUSG00 | Ncoa1 | 7305 | 14.1497946 | 16.9196748 | 15.2240708 | 14.353788 | 19.2370485 | 12.690828 |
| ENSMUSG00 | Dus4l | 1761 | 6.16311538 | 3.28357357 | 4.61847763 | 4.91748477 | 5.24461898 | 5.93196299 |
| ENSMUSG00 | Rrm2 | 2374 | 5.59323104 | 13.3761017 | 1.99224819 | 6.58109742 | 6.16873204 | 6.14655939 |
| ENSMUSG00 | Bcap29 | 1947 | 120.287384 | 100.679223 | 171.33606 | 100.221724 | 77.8892599 | 209.791986 |
| ENSMUSG00 | Slc26a4 | 3054 | 2063.39275 | 1119.08042 | 3196.99165 | 1347.39381 | 1557.65781 | 2018.05538 |
| ENSMUSG00 | Cenpo | 6983 | 1.49161333 | 2.11846452 | 1.49386151 | 2.51122349 | 1.86334324 | 1.92949618 |
| ENSMUSG00 | Kif11 | 4222 | 9.77408545 | 7.03051832 | 12.1654528 | 8.13597972 | 9.36651854 | 11.1760128 |
| ENSMUSG00 | Adcy3 | 6838 | 0.11627809 | 0.06342176 | 0.05817733 | 0.242728 | 0.08208396 | 0.12763797 |
| ENSMUSG00 | Grhl1 | 3483 | 1.52949595 | 2.36574264 | 1.02794984 | 2.04081619 | 2.59307164 | 1.19811077 |
| ENSMUSG00 | Dnajc27 | 3302 | 5.68279431 | 3.02077333 | 4.91280327 | 3.91198423 | 4.18780977 | 4.00611752 |
| ENSMUSG00 | Efr3b | 6524 | 0.02437491 | 0.42838958 | 0 | 0.26547182 | 0.32849598 | 0.14214251 |
| ENSMUSG00 | Cbll1 | 4723 | 10.2692539 | 10.8452653 | 8.07668654 | 13.4152171 | 8.69706384 | 11.1743441 |
| ENSMUSG00 | Pomc | 1004 | 0.35637383 | 0.52793911 | 0.35220592 | 0.21562975 | 0.45740775 | 0.62481743 |
| ENSMUSG00 | Dnmt3a | 12336 | 5.97814659 | 5.39050666 | 5.92654413 | 7.45859383 | 7.81362062 | 4.85088834 |
| ENSMUSG00 | Dld | 2757 | 323.6099 | 251.908303 | 283.327882 | 272.297069 | 222.965089 | 398.425398 |
| ENSMUSG00 | Kif3c | 6914 | 1.92624906 | 2.96199565 | 1.86678307 | 3.50696181 | 3.32106872 | 2.44185735 |
| ENSMUSG00 | Sh3yl1 | 3080 | 19.8777403 | 25.5325564 | 22.6893227 | 19.9857039 | 22.1500859 | 23.4136838 |
| ENSMUSG00 | Rab10 | 3505 | 104.850118 | 97.074028 | 91.7330117 | 92.9176987 | 74.7561883 | 116.42886 |
| ENSMUSG00 | Sntg2 | 4041 | 0 | 0 | 0 | 0 | 0.07576299 | 0.08099373 |
| ENSMUSG00 | Tpo | 3898 | 0 | 0 | 0 | 0 | 0 | 0 |
| ENSMUSG00 | Pxdn | 9123 | 0.09586984 | 0.47536773 | 0.25194518 | 0.21752849 | 0.38033471 | 0 |
| ENSMUSG00 | Ccl11 | 1079 | 0 | 0 | 0 | 0 | 0 | 0 |
| ENSMUSG00 | Ddx52 | 3317 | 22.3047782 | 16.0815169 | 20.0021002 | 23.2243488 | 24.5055824 | 20.6142661 |
| ENSMUSG00 | Hnf1b | 3284 | 114.61781 | 107.803225 | 76.4110423 | 54.7273641 | 83.9045157 | 97.271795 |
| ENSMUSG00 | Taf15 | 2847 | 1.84324673 | 1.28632582 | 1.55257615 | 3.76409105 | 1.50552005 | 3.25724529 |
| ENSMUSG00 | Ace | 5262 | 0.06044163 | 0.02747232 | 0.0084002 | 0.13714194 | 0.12606288 | 0 |
| ENSMUSG00 | Mmp28 | 4934 | 1.12804363 | 1.03521744 | 0.75252428 | 0.78979749 | 0.45503862 | 0.79601704 |
| ENSMUSG00 | Rasl10b | 4004 | 0 | 0 | 0 | 0 | 0 | 0 |
| ENSMUSG00 | Gas2l2 | 2604 | 0 | 0 | 0 | 0 | 0 | 0 |
| ENSMUSG00 | Cdc27 | 6972 | 15.6239267 | 10.7541744 | 14.7719871 | 18.3774154 | 14.2496204 | 18.2417732 |
| ENSMUSG00 | Itgb3 | 5885 | 0.46612203 | 0 | 0 | 0 | 0 | 0.06024984 |
| ENSMUSG00 | Efcab3 | 2867 | 0 | 0 | 0.10792218 | 0 | 0.01779783 | 0 |
| ENSMUSG00 | Mettl2 | 2943 | 7.56475334 | 7.95739412 | 8.57602867 | 5.84816012 | 6.55384644 | 7.33996408 |
| ENSMUSG00 | Nle1 | 3501 | 0.84030436 | 1.348835 | 1.55293536 | 1.4119509 | 2.41941685 | 1.45682878 |
| ENSMUSG00 | Tlk2 | 8043 | 6.25272695 | 6.17682729 | 5.58911779 | 5.6615119 | 6.50914604 | 9.28823038 |
| ENSMUSG00 | Mrc2 | 6899 | 0 | 0 | 0 | 0 | 0.0295848 | 0 |
| ENSMUSG00 | Rffl | 8277 | 6.36894613 | 5.30359415 | 3.57267524 | 6.54769009 | 5.62233354 | 4.86705838 |
| ENSMUSG00 | Lig3 | 7271 | 5.41300037 | 4.71194614 | 6.06703883 | 5.37434386 | 7.79676847 | 3.02718111 |
| ENSMUSG00 | Cct6b | 1842 | 0.34532448 | 1.22951291 | 0.86387967 | 0.27423921 | 2.07762117 | 1.08091676 |
| ENSMUSG00 | Map3k3 | 3993 | 15.4222989 | 17.1603107 | 14.5900398 | 12.5333824 | 16.702097 | 14.0300799 |
| ENSMUSG00 | Tmem132e | 4534 | 0 | 0 | 0 | 0 | 0 | 0 |
| ENSMUSG00 | Ccl1 | 2508 | 0 | 0 | 0 | 0 | 0 | 0 |
| ENSMUSG00 | 5530401A14 | 5200 | 0 | 0 | 0 | 0 | 0 | 0 |
| ENSMUSG00 | Accn1 | 4253 | 0 | 0 | 0 | 0 | 0 | 0 |
| ENSMUSG00 | Ddx42 | 4210 | 38.8961578 | 29.6558365 | 29.7758734 | 36.6391307 | 38.2637217 | 26.0372388 |
| ENSMUSG00 | Ftsj3 | 4206 | 11.6733282 | 11.5024222 | 9.70002404 | 12.6450151 | 8.78342751 | 8.71543394 |
| ENSMUSG00 | Rnf135 | 1735 | 12.3963775 | 4.7492113 | 11.7192206 | 8.65137206 | 8.52890432 | 7.57716203 |
| ENSMUSG00 | Psmc5 | 1384 | 101.342005 | 110.787037 | 112.16537 | 95.4192796 | 88.2269634 | 104.014123 |
| ENSMUSG00 | Adap2 | 2784 | 0 | 0.06923339 | 0 | 0.12960505 | 0 | 0.06857848 |
| ENSMUSG00 | Tcam1 | 3789 | 0 | 0 | 0 | 0 | 0 | 0 |
| ENSMUSG00 | Gh | 857 | 0 | 0 | 0 | 0 | 0 | 0 |
| ENSMUSG00 | Ern1 | 9454 | 1.93436863 | 2.28342776 | 2.01980074 | 3.74407507 | 3.98323095 | 2.17239299 |
| ENSMUSG00 | Nf1 | 15083 | 0.97787465 | 0.77312995 | 0.78539375 | 1.53820557 | 0.86267492 | 1.01445817 |
| ENSMUSG00 | Pecam1 | 5241 | 2.11634785 | 2.06867954 | 1.29038008 | 0.69534182 | 3.85545599 | 1.59265175 |
| ENSMUSG00 | Polg2 | 4312 | 0.69147983 | 2.91666545 | 2.73698796 | 1.95807013 | 3.53823895 | 2.56806668 |
| ENSMUSG00 | Ddx5 | 5758 | 126.695566 | 112.20629 | 130.433052 | 145.744389 | 132.386771 | 188.577746 |
| ENSMUSG00 | Psmd12 | 2793 | 52.8649676 | 44.0285716 | 44.8190546 | 51.9204214 | 43.9743955 | 45.8288156 |
| ENSMUSG00 | Helz | 14191 | 3.90803293 | 3.89131581 | 3.9246228 | 5.63440284 | 6.27447155 | 3.3653649 |
| ENSMUSG00 | Cacng1 | 1245 | 0 | 0 | 0 | 0 | 0 | 0 |
| ENSMUSG00 | Cacng4 | 3692 | 0 | 0 | 0 | 0 | 0 | 0 |
| ENSMUSG00 | Ccdc46 | 4970 | 0.59193271 | 0.6495959 | 0.44468655 | 0.80585653 | 1.4373627 | 0.42805269 |
| ENSMUSG00 | Rab37 | 5319 | 0.00747424 | 0.34425358 | 0 | 0.1424559 | 0.12471195 | 0 |
| ENSMUSG00 | Slc9a3r1 | 1919 | 95.0692549 | 95.7953501 | 141.565673 | 89.8572646 | 70.3839587 | 98.1403773 |
| ENSMUSG00 | Grin2c | 5021 | 0 | 0.03838792 | 0 | 0 | 0 | 0.09777803 |
| ENSMUSG00 | Nt5c | 1430 | 16.2358046 | 14.3548422 | 17.6498268 | 12.9188859 | 10.1339096 | 16.4601494 |
| ENSMUSG00 | Hn1 | 2410 | 27.3339549 | 26.4325629 | 41.1022117 | 28.3865379 | 20.5799325 | 31.6431084 |
| ENSMUSG00 | Sumo2 | 1730 | 1.99926405 | 1.22555113 | 1.25195972 | 1.12626035 | 2.1826311 | 2.20719624 |
| ENSMUSG00 | Nup85 | 3156 | 11.6646309 | 10.5961315 | 13.4314219 | 14.4282447 | 13.2416356 | 10.0335412 |
| ENSMUSG00 | Gga3 | 5654 | 10.7791213 | 10.5679498 | 8.41977095 | 8.54507563 | 13.5191919 | 7.31311516 |
| ENSMUSG00 | 1300001I01R | 6656 | 32.3550838 | 22.5656761 | 23.7810697 | 24.6058293 | 34.9809725 | 18.4398853 |
| ENSMUSG00 | Mif4gd | 2228 | 71.9274425 | 76.0428773 | 78.8610081 | 58.6414198 | 61.4698351 | 64.7099385 |
| ENSMUSG00 | Slc25a19 | 3916 | 26.5781021 | 27.920081 | 25.2501335 | 29.6875249 | 24.9007672 | 24.7324423 |
| ENSMUSG00 | Pafah1b1 | 13402 | 5.39881917 | 6.97160956 | 8.1926114 | 8.46993822 | 4.34420943 | 8.34805031 |
| ENSMUSG00 | 2310067B10 | 5888 | 13.99004 | 14.9355056 | 9.43643286 | 14.0087557 | 12.0286362 | 8.27318371 |
| ENSMUSG00 | Recql5 | 6800 | 6.43688001 | 6.250065 | 4.96620707 | 5.52904271 | 6.31826595 | 5.25036806 |
| ENSMUSG00 | Sap30bp | 3904 | 19.6435251 | 19.9089988 | 17.1984118 | 17.5326943 | 20.115162 | 17.1234992 |
| ENSMUSG00 | Itgb4 | 7058 | 9.63752156 | 6.00794405 | 15.2871492 | 8.37381538 | 9.40568361 | 10.1593974 |
| ENSMUSG00 | Galk1 | 1398 | 24.086475 | 29.5736533 | 19.9825213 | 16.4666269 | 25.8051843 | 16.3881953 |
| ENSMUSG00 | Unk | 4480 | 13.080263 | 9.45443809 | 7.52812639 | 11.3883953 | 8.78154816 | 6.92215733 |
| ENSMUSG00 | Trim47 | 3048 | 38.1642178 | 39.5071913 | 26.2194659 | 37.8577357 | 24.3748042 | 24.7869917 |
| ENSMUSG00 | Aspa | 5612 | 0.38962071 | 1.93192257 | 0.99241486 | 1.06728785 | 0.19093975 | 0.40824478 |
| ENSMUSG00 | Mrpl38 | 2852 | 15.0825491 | 14.3444212 | 16.8469156 | 16.8517824 | 15.0645892 | 18.5433119 |
| ENSMUSG00 | Fbf1 | 8490 | 5.14152151 | 4.67675245 | 2.94158319 | 2.77946491 | 8.00555154 | 2.57647358 |
| ENSMUSG00 | Acox1 | 5067 | 57.5187345 | 54.9764796 | 47.8308082 | 53.9984495 | 51.0565868 | 36.7214504 |
| ENSMUSG00 | 2310004N24 | 1727 | 15.9528361 | 16.0435462 | 15.5871004 | 12.5566351 | 8.62750535 | 21.7944425 |
| ENSMUSG00 | Srp68 | 2720 | 26.6449415 | 30.2936824 | 22.7346979 | 27.2207189 | 34.611641 | 18.6410126 |
| ENSMUSG00 | Tsen54 | 2426 | 13.5686471 | 14.3208676 | 17.4548086 | 16.3008742 | 19.266348 | 18.0556763 |
| ENSMUSG00 | Llgl2 | 4576 | 42.8483457 | 45.1747895 | 37.1504126 | 41.1915648 | 43.0981081 | 26.2911787 |
| ENSMUSG00 | 1200014J11R | 8516 | 2.89903163 | 3.406322 | 2.93779276 | 2.60573713 | 3.90666945 | 1.8159606 |
| ENSMUSG00 | Camkk1 | 3797 | 0 | 0.48224514 | 0.05820627 | 0 | 0.22845625 | 0 |
| ENSMUSG00 | P2rx1 | 2749 | 0 | 0 | 0.33766413 | 0 | 0 | 0 |
| ENSMUSG00 | Atp2a3 | 5440 | 0.94273106 | 1.22237666 | 0.22750949 | 0.41122918 | 2.31682258 | 0.23063114 |
| ENSMUSG00 | Ankfy1 | 11307 | 7.64731318 | 6.76323372 | 7.29074354 | 9.49040421 | 11.4490064 | 9.44372625 |
| ENSMUSG00 | Exoc7 | 4568 | 15.1519903 | 13.7344019 | 16.6047204 | 13.8941148 | 15.9736684 | 10.1682817 |
| ENSMUSG00 | Galr2 | 2818 | 0.04232308 | 0 | 0 | 0 | 0 | 0 |
| ENSMUSG00 | Ube2g1 | 4888 | 30.1582082 | 30.8756008 | 37.7632767 | 37.9348664 | 31.2546991 | 36.9725368 |
| ENSMUSG00 | Spns3 | 5814 | 0 | 0.016576 | 0 | 0.07447274 | 0.01755293 | 0.07036801 |
| ENSMUSG00 | Tekt1 | 1857 | 0.06422533 | 0 | 0.14281694 | 0.27202403 | 0.54955709 | 0 |
| ENSMUSG00 | Med31 | 1726 | 15.3862462 | 14.0148285 | 16.8253829 | 15.6996613 | 10.9680101 | 28.175366 |
| ENSMUSG00 | Ube2o | 5270 | 2.08961444 | 3.41053945 | 2.53300884 | 2.87560886 | 3.74709819 | 2.7636921 |
| ENSMUSG00 | Txndc17 | 1473 | 72.4397214 | 78.8059329 | 77.7808398 | 69.6655361 | 50.7146057 | 101.229092 |
| ENSMUSG00 | Aanat | 1656 | 0 | 0 | 0 | 0.23967542 | 0 | 0 |
| ENSMUSG00 | Slc13a5 | 3531 | 0 | 0 | 0 | 0 | 0 | 0 |
| ENSMUSG00 | Rhbdf2 | 6543 | 2.68560636 | 3.49817515 | 1.90507714 | 1.83084805 | 3.27542073 | 2.07592394 |
| ENSMUSG00 | 4933427D14 | 5712 | 3.47999654 | 2.8429326 | 2.6388005 | 3.227928 | 5.55644359 | 3.82952743 |
| ENSMUSG00 | Fam64a | 2522 | 0 | 0.34391592 | 0 | 0.0858415 | 0 | 0 |
| ENSMUSG00 | Cygb | 2327 | 0.25626653 | 0 | 0 | 0 | 0 | 0 |
| ENSMUSG00 | Wscd1 | 4274 | 3.99043078 | 1.01468877 | 0.61017986 | 1.63779051 | 2.2086756 | 1.39755405 |
| ENSMUSG00 | 1810032O08 | 2858 | 1.64140892 | 1.85462164 | 0.72690225 | 2.03261343 | 1.67826429 | 1.26925368 |
| ENSMUSG00 | Mxra7 | 2385 | 4.70064801 | 7.63709649 | 3.05798914 | 4.13014598 | 5.77657085 | 4.26841394 |
| ENSMUSG00 | Rabep1 | 6059 | 26.4817824 | 18.1802619 | 26.4379401 | 23.1951747 | 21.4918817 | 29.840499 |
| ENSMUSG00 | Mfsd11 | 6877 | 5.89655229 | 7.04894006 | 8.45857575 | 5.58781117 | 6.65561424 | 6.24655472 |
| ENSMUSG00 | Kif1c | 7178 | 1.94401974 | 2.32272346 | 2.56171172 | 2.22685229 | 2.33876813 | 1.99867097 |
| ENSMUSG00 | Sec14l1 | 6140 | 12.3410335 | 14.047839 | 11.3960126 | 12.3172582 | 14.6929369 | 8.64881476 |
| ENSMUSG00 | Nos2 | 4165 | 0 | 0.15040186 | 0 | 0 | 0 | 0 |
| ENSMUSG00 | Mink1 | 6396 | 2.20656591 | 1.70264789 | 1.80373374 | 2.5780948 | 2.67258222 | 3.18545271 |
| ENSMUSG00 | Pld2 | 6142 | 2.1748358 | 1.85151422 | 1.59765698 | 1.84463725 | 4.40312263 | 2.36688094 |
| ENSMUSG00 | Slc46a1 | 2046 | 3.07007132 | 9.23220201 | 5.7682757 | 2.66294659 | 5.63634454 | 3.19937117 |
| ENSMUSG00 | Vmo1 | 669 | 0 | 0 | 0 | 0 | 0 | 0 |
| ENSMUSG00 | 0610010K14 | 1919 | 2.32027818 | 2.38546743 | 1.91180457 | 1.57941206 | 1.96766639 | 2.62939461 |
| ENSMUSG00 | Eral1 | 3056 | 14.2708646 | 13.6864633 | 16.6914028 | 10.1893994 | 15.578406 | 10.0673407 |
| ENSMUSG00 | Dhrs13 | 2024 | 5.55869614 | 5.45192453 | 7.03211139 | 4.84897041 | 5.72281982 | 7.7889039 |
| ENSMUSG00 | Coro6 | 3107 | 0.05118182 | 0.07754497 | 0.24185109 | 0.48775213 | 0.27919163 | 0.44770087 |
| ENSMUSG00 | Slc6a4 | 3815 | 0 | 0.06315392 | 0.10427696 | 0 | 0 | 0 |
| ENSMUSG00 | Tmigd1 | 1490 | 0.18677071 | 0.35573884 | 0.444985 | 0.50853888 | 0.44519657 | 2.25153062 |
| ENSMUSG00 | Blmh | 5078 | 17.3254979 | 21.9106922 | 18.384067 | 17.6786386 | 20.6396565 | 17.7193846 |
| ENSMUSG00 | Cpd | 8311 | 13.9964548 | 11.8567289 | 15.5724938 | 13.0201001 | 21.0834526 | 13.4748729 |
| ENSMUSG00 | Timm22 | 3179 | 15.0568099 | 16.4158279 | 16.0594931 | 14.8800127 | 15.8745158 | 14.0534316 |
| ENSMUSG00 | Nxn | 3230 | 1.16927884 | 0.4773889 | 0.79371731 | 0.24576006 | 0.7582867 | 0.48976132 |
| ENSMUSG00 | Fam101b | 3509 | 1.99400529 | 0.76900563 | 0.86917275 | 1.16194672 | 1.90494592 | 0.71509456 |
| ENSMUSG00 | Rph3al | 3878 | 2.59364017 | 2.62179971 | 2.42779592 | 2.04694425 | 1.77631788 | 3.2141594 |
| ENSMUSG00 | Doc2b | 4352 | 0 | 0 | 0 | 0 | 0 | 0 |
| ENSMUSG00 | Ywhae | 2739 | 87.827825 | 82.5977897 | 101.991839 | 90.0931371 | 59.4283092 | 134.421451 |
| ENSMUSG00 | Prpf8 | 7833 | 15.5052972 | 14.6595547 | 20.320546 | 24.773297 | 24.2917599 | 16.9365307 |
| ENSMUSG00 | Nme2 | 1395 | 1.90940301 | 1.34714785 | 1.29912227 | 1.9916254 | 0.6949829 | 1.3881716 |
| ENSMUSG00 | Spag9 | 11634 | 8.02695751 | 8.73933238 | 8.02046507 | 12.774793 | 8.07018493 | 8.7750537 |
| ENSMUSG00 | Luc7l3 | 6613 | 21.3355815 | 19.8560494 | 22.1978407 | 19.5169323 | 20.8179589 | 42.6627655 |
| ENSMUSG00 | Ankrd40 | 2705 | 50.8958333 | 48.2398837 | 59.1700087 | 51.1017791 | 45.7822603 | 51.4941156 |
| ENSMUSG00 | Abcc3 | 5782 | 1.2582589 | 2.45015808 | 1.84238052 | 3.55079269 | 3.96244254 | 0.41511039 |
| ENSMUSG00 | Cacna1g | 11174 | 0 | 0 | 0.00791155 | 0 | 0.00456653 | 0 |
| ENSMUSG00 | Spata20 | 2858 | 0 | 0 | 0 | 0.59337162 | 0 | 0 |
| ENSMUSG00 | Xylt2 | 4289 | 8.52763863 | 6.22412892 | 8.04887839 | 7.33586916 | 12.2301502 | 4.99197757 |
| ENSMUSG00 | Lrrc59 | 3361 | 38.9038308 | 35.8280033 | 33.5623634 | 33.1083086 | 34.9639225 | 22.3325817 |
| ENSMUSG00 | Cdc34-ps | 1251 | 0 | 0 | 0 | 0 | 0 | 0 |
| ENSMUSG00 | Dlx4 | 3256 | 0.61049571 | 0.62156959 | 0.10860404 | 0.34353298 | 0.81491755 | 0.29318562 |
| ENSMUSG00 | Tac4 | 1247 | 0 | 0 | 0 | 0 | 0 | 0 |
| ENSMUSG00 | Slc35b1 | 2570 | 20.8368219 | 22.3870085 | 21.6365442 | 23.3901503 | 16.4396261 | 21.5756193 |
| ENSMUSG00 | Hoxb9 | 2868 | 31.3414022 | 28.932027 | 31.8875918 | 23.7527546 | 24.1788162 | 23.1853454 |
| ENSMUSG00 | Snx11 | 5667 | 3.55673701 | 3.63077657 | 3.03414804 | 3.92846682 | 4.13289333 | 2.28130916 |
| ENSMUSG00 | Scrn2 | 2146 | 12.541687 | 11.8108428 | 9.63954452 | 7.95284579 | 11.6984981 | 7.2571384 |
| ENSMUSG00 | Lrrc46 | 1328 | 0.59872712 | 0.18142486 | 0.13313808 | 0 | 0.61477561 | 0.26699572 |
| ENSMUSG00 | Cacnb1 | 4972 | 0.79159143 | 0.62025992 | 0.79122366 | 0.52250748 | 1.0365374 | 0.9709596 |
| ENSMUSG00 | Fbxl20 | 17848 | 3.65969602 | 3.03460113 | 3.84116195 | 2.44415015 | 3.77666083 | 2.70943161 |
| ENSMUSG00 | Asgr1 | 1580 | 0 | 0.12199099 | 0.02797585 | 0 | 0.35524692 | 0 |
| ENSMUSG00 | Dlg4 | 4632 | 0.37764273 | 0.5513561 | 0.1526834 | 0.1869536 | 0.27540142 | 0.28263875 |
| ENSMUSG00 | A230052G05 | 2301 | 0.05183244 | 0 | 0 | 0 | 0.04435148 | 0 |
| ENSMUSG00 | Dvl2 | 6693 | 2.37000399 | 1.92227403 | 1.90861088 | 2.17257791 | 2.2032904 | 1.57706341 |
| ENSMUSG00 | Nr1d1 | 3064 | 41.1308333 | 33.324762 | 34.5362964 | 31.5011322 | 38.9858831 | 27.0788031 |
| ENSMUSG00 | Gucy2e | 10472 | 0.06074176 | 0 | 0.06753528 | 0 | 0 | 0.01823171 |
| ENSMUSG00 | Alox8 | 3726 | 0.29875294 | 0 | 0.49824944 | 0.51324433 | 1.32838391 | 1.58845859 |
| ENSMUSG00 | Aloxe3 | 6718 | 0.02367102 | 0 | 0 | 0 | 0.00759547 | 0.16645731 |
| ENSMUSG00 | Per1 | 5607 | 1.19826578 | 2.26881055 | 1.28498313 | 2.13647918 | 1.68358829 | 1.07016598 |
| ENSMUSG00 | Vamp2 | 3924 | 79.4197787 | 66.2993377 | 86.2972272 | 64.6148132 | 65.8635559 | 92.8199624 |
| ENSMUSG00 | Tmem107 | 1555 | 4.24399342 | 6.91033865 | 5.23031454 | 8.74786546 | 4.49557137 | 6.45470562 |
| ENSMUSG00 | Aurkb | 2825 | 0 | 0.57994302 | 0 | 0.42148937 | 0.25287408 | 0 |
| ENSMUSG00 | 1500010J02R | 6926 | 7.22670372 | 5.88589607 | 6.03100515 | 4.32401053 | 6.66746609 | 5.30843398 |
| ENSMUSG00 | Plxs | 10183 | 2.96321415 | 1.982728 | 1.90558864 | 1.63703276 | 2.78107027 | 2.79630005 |
| ENSMUSG00 | Myh10 | 10412 | 4.7651594 | 6.40974091 | 2.97594045 | 6.81303299 | 6.18961897 | 4.0786221 |
| ENSMUSG00 | Pik3r5 | 5576 | 0 | 0 | 0 | 0 | 0 | 0 |
| ENSMUSG00 | Ntn1 | 6268 | 1.09092895 | 1.18390429 | 0.53595087 | 1.05920488 | 1.43277635 | 0.97471589 |
| ENSMUSG00 | Stx8 | 7049 | 4.22990642 | 5.5029204 | 6.47758602 | 4.1666597 | 3.94515177 | 5.26997569 |
| ENSMUSG00 | Wdr16 | 2731 | 2.43103817 | 1.72913634 | 0.74452024 | 0.51526904 | 2.57841079 | 1.00869226 |
| ENSMUSG00 | Usp43 | 5915 | 1.62651332 | 0.83908767 | 1.27038264 | 0.73811114 | 1.56141573 | 1.22194073 |
| ENSMUSG00 | Rcvrn | 1037 | 0 | 0 | 0 | 0 | 0 | 0 |
| ENSMUSG00 | Myh3 | 6132 | 0 | 0 | 0 | 0 | 0 | 0 |
| ENSMUSG00 | 2310004I24R | 2375 | 19.0156741 | 13.9588515 | 17.0479529 | 13.3997319 | 18.1761322 | 20.4760765 |
| ENSMUSG00 | Krt19 | 2107 | 4.13215483 | 9.33083455 | 3.39852803 | 5.22307716 | 3.705285 | 4.01288 |
| ENSMUSG00 | Krt12 | 1853 | 0 | 0 | 0 | 0 | 0 | 0 |
| ENSMUSG00 | Krt24 | 1839 | 0 | 0 | 0 | 0 | 0 | 0 |
| ENSMUSG00 | Top2a | 9357 | 0.01699497 | 0.54587612 | 0 | 0 | 0 | 0 |
| ENSMUSG00 | Krt36 | 1664 | 0 | 0 | 0 | 0 | 0 | 0 |
| ENSMUSG00 | Acly | 6232 | 28.2728321 | 27.7195756 | 35.1799007 | 21.873871 | 35.6988124 | 24.6968532 |
| ENSMUSG00 | Kat2a | 4085 | 5.19692205 | 3.23208939 | 3.91702991 | 5.68833211 | 7.59462338 | 4.90075525 |
| ENSMUSG00 | Stat5b | 5519 | 5.40974196 | 4.40042885 | 4.4209852 | 4.2626369 | 5.80622649 | 3.66198726 |
| ENSMUSG00 | Tmem101 | 2017 | 4.71073864 | 7.45372828 | 5.85121075 | 5.86758087 | 4.78135103 | 4.23250475 |
| ENSMUSG00 | Lsm12 | 2530 | 6.22259695 | 5.48525507 | 5.59074694 | 7.51590414 | 5.28415433 | 5.89692793 |
| ENSMUSG00 | Ubtf | 7371 | 11.2454452 | 10.093592 | 14.727951 | 16.1294718 | 11.0692138 | 8.07767433 |
| ENSMUSG00 | Ccdc43 | 5144 | 9.26648933 | 6.70713308 | 9.76152676 | 9.74299375 | 6.15014639 | 11.2884345 |
| ENSMUSG00 | Adam11 | 5643 | 0.27475877 | 0.96492432 | 0.28982247 | 0.16624724 | 0.75956328 | 0.23683454 |
| ENSMUSG00 | Higd1b | 754 | 0 | 0 | 0 | 0 | 0 | 0 |
| ENSMUSG00 | Eftud2 | 8075 | 13.1943886 | 14.3992427 | 14.9492549 | 13.369347 | 17.8260565 | 9.63647626 |
| ENSMUSG00 | Ccdc103 | 1955 | 1.40313461 | 2.0950627 | 1.33396866 | 0.46140722 | 0.6003103 | 0.502244 |
| ENSMUSG00 | Gfap | 4207 | 0 | 0.09163098 | 0 | 0.11149669 | 0.04851569 | 0.13614629 |
| ENSMUSG00 | Dcakd | 3671 | 7.52657558 | 6.73376323 | 7.67000109 | 5.20933904 | 6.4634336 | 6.81308749 |
| ENSMUSG00 | Nmt1 | 5088 | 17.5024128 | 16.744031 | 18.2263103 | 13.9420793 | 20.0274514 | 17.4058085 |
| ENSMUSG00 | Plcd3 | 3720 | 7.78010479 | 12.927438 | 3.27949158 | 6.5762436 | 10.9459806 | 3.29201963 |
| ENSMUSG00 | 1700023F06 | 1612 | 0 | 0 | 0 | 0 | 0 | 0 |
| ENSMUSG00 | Map3k14 | 5633 | 0.65635713 | 1.54833056 | 0.51789839 | 1.58855797 | 2.28273508 | 0.97807157 |
| ENSMUSG00 | Lyzl6 | 1294 | 0 | 0 | 0 | 0 | 0 | 0 |
| ENSMUSG00 | Gosr2 | 3931 | 43.1535577 | 48.4683776 | 41.5144249 | 41.0294426 | 40.2395739 | 32.0829132 |
| ENSMUSG00 | Klhl28 | 2849 | 5.47004154 | 5.42922005 | 8.47111488 | 6.21842185 | 7.50440557 | 3.68576206 |
| ENSMUSG00 | Fkbp3 | 940 | 97.4009273 | 87.4020041 | 79.8454569 | 81.7219923 | 65.9543049 | 154.014665 |
| ENSMUSG00 | Foxg1 | 3565 | 0 | 0 | 0 | 0 | 0 | 0 |
| ENSMUSG00 | Scfd1 | 2153 | 84.8104607 | 77.0132594 | 84.5029196 | 93.6490779 | 85.9129644 | 101.738332 |
| ENSMUSG00 | Coch | 2589 | 2.840774 | 1.24700333 | 5.08773628 | 0.93375704 | 0 | 6.58426015 |
| ENSMUSG00 | Strn3 | 4294 | 141.366479 | 122.265647 | 140.648919 | 146.988843 | 111.155935 | 150.179198 |
| ENSMUSG00 | Ap4s1 | 984 | 19.069702 | 28.2066978 | 28.7042456 | 21.1945343 | 13.3788868 | 29.353499 |
| ENSMUSG00 | 6530401N04 | 3104 | 10.668916 | 12.2949943 | 10.3669271 | 13.7167566 | 9.43593419 | 10.8079273 |
| ENSMUSG00 | Ston2 | 6901 | 1.37683812 | 1.91321332 | 0.91593444 | 1.47967231 | 1.14607858 | 0.41103644 |
| ENSMUSG00 | Gtf2a1 | 6740 | 6.33492375 | 7.38525144 | 9.16171731 | 9.94129921 | 8.60786192 | 7.12216099 |
| ENSMUSG00 | Tshr | 4863 | 8.07699254 | 10.7609451 | 8.27137099 | 8.54008498 | 12.8641449 | 5.22721854 |
| ENSMUSG00 | Sel1l | 12255 | 21.056942 | 19.1802093 | 21.5184166 | 21.4107407 | 22.4341177 | 15.0383303 |
| ENSMUSG00 | 1110034A24 | 3263 | 7.59045797 | 6.99980802 | 4.8496046 | 5.40733061 | 4.95720537 | 7.53124424 |
| ENSMUSG00 | Pole2 | 1676 | 0.35580681 | 0.46001377 | 0.81757586 | 0 | 0.15222666 | 0.39056763 |
| ENSMUSG00 | Klhdc2 | 4848 | 25.6508133 | 23.2583075 | 27.5167414 | 22.8564271 | 20.6610511 | 34.0370395 |
| ENSMUSG00 | Nemf | 3678 | 70.4638329 | 76.1052318 | 84.0772414 | 83.7794074 | 63.5263118 | 96.4697613 |
| ENSMUSG00 | Sec23a | 5202 | 12.4188112 | 15.0524738 | 16.2804174 | 19.6224729 | 16.3221625 | 10.5648593 |
| ENSMUSG00 | L2hgdh | 3328 | 22.5894272 | 19.271681 | 33.3373275 | 23.7655774 | 25.3905284 | 18.5874044 |
| ENSMUSG00 | Cdkl1 | 1679 | 125.517452 | 196.01751 | 109.438393 | 157.308258 | 172.438152 | 103.916538 |
| ENSMUSG00 | 4930512B01 | 1855 | 9.47273444 | 14.7286861 | 6.36220598 | 10.425863 | 15.0465917 | 7.836864 |
| ENSMUSG00 | Trappc6b | 1248 | 116.39946 | 104.944511 | 122.617613 | 118.249649 | 85.779917 | 139.542125 |
| ENSMUSG00 | Pnn | 3460 | 37.3770457 | 36.4044392 | 33.8795629 | 43.6947302 | 38.7122071 | 66.5312009 |
| ENSMUSG00 | Ctage5 | 9445 | 16.5461931 | 13.0758974 | 15.9819263 | 14.9370878 | 16.8341119 | 17.1098187 |
| ENSMUSG00 | Galc | 3843 | 10.1483545 | 8.90251728 | 15.5850891 | 14.740778 | 13.4105229 | 16.146189 |
| ENSMUSG00 | Spata7 | 4802 | 4.08979745 | 4.17441898 | 2.92715245 | 4.16273483 | 5.50430293 | 3.79981959 |
| ENSMUSG00 | Ptpn21 | 5527 | 3.79068902 | 4.19353694 | 3.63083711 | 7.12892937 | 7.19188471 | 3.26190276 |
| ENSMUSG00 | Npas3 | 5873 | 0 | 0 | 0 | 0 | 0 | 0 |
| ENSMUSG00 | Zc3h14 | 36640 | 3.68150506 | 3.32596922 | 3.27170847 | 3.69191007 | 4.49266071 | 2.94259413 |
| ENSMUSG00 | Ttc8 | 2902 | 13.8637306 | 14.744852 | 14.5003979 | 13.9006636 | 20.5723155 | 13.5621311 |
| ENSMUSG00 | Polr2h | 1442 | 5.81720277 | 8.35409885 | 7.87785968 | 5.78013335 | 4.98940291 | 9.00327898 |
| ENSMUSG00 | Srp54a | 2500 | 0 | 0 | 0 | 0 | 0 | 0 |
| ENSMUSG00 | Ppp2r3c | 2381 | 11.7045745 | 15.6034216 | 13.6819649 | 10.7291422 | 10.2438503 | 18.6832997 |
| ENSMUSG00 | 1110008L16 | 3110 | 12.7831127 | 10.8148349 | 13.0615736 | 7.33242834 | 10.3201255 | 11.0238655 |
| ENSMUSG00 | Psma6 | 1226 | 139.987283 | 140.23591 | 189.606438 | 111.660095 | 97.3912883 | 147.919312 |
| ENSMUSG00 | Nfkbia | 1571 | 37.0224494 | 34.9972823 | 54.0494847 | 30.8454397 | 37.0598949 | 50.3304769 |
| ENSMUSG00 | Ralgapa1 | 8004 | 22.5797619 | 17.9766012 | 16.7938287 | 34.1706519 | 20.8019821 | 17.5935735 |
| ENSMUSG00 | Mbip | 1500 | 12.2181843 | 16.3191417 | 16.6788288 | 19.5083587 | 11.9401719 | 23.7471192 |
| ENSMUSG00 | Ngb | 2239 | 0.14204727 | 0.92542073 | 0.01974178 | 0 | 0.02278981 | 0.0730897 |
| ENSMUSG00 | Gatz1 | 1647 | 36.8830445 | 42.3935365 | 44.4433831 | 40.4197767 | 42.5064892 | 44.4475603 |
| ENSMUSG00 | Sptlc2 | 10052 | 27.5701805 | 28.9923997 | 24.4446921 | 24.4160433 | 28.63 | 23.1069268 |
| ENSMUSG00 | Ahsa1 | 1338 | 38.864102 | 43.720733 | 48.9259563 | 45.8980865 | 29.1742732 | 42.7058066 |
| ENSMUSG00 | Vipar | 2680 | 36.5216392 | 34.9711306 | 43.2617292 | 46.8931944 | 37.9651468 | 29.5237793 |
| ENSMUSG00 | Snw1 | 2217 | 38.320912 | 39.6228615 | 36.3663336 | 39.8904335 | 42.5565038 | 32.8230661 |
| ENSMUSG00 | 1810035L17 | 2811 | 8.68369443 | 7.97107634 | 8.20823979 | 8.13803494 | 7.40617621 | 15.7379811 |
| ENSMUSG00 | 2700073G19 | 1334 | 0.53643077 | 0.97528781 | 0.26507852 | 0.67620024 | 0.2677546 | 0.79738451 |
| ENSMUSG00 | Adck1 | 2318 | 20.6152233 | 18.2310223 | 12.3566843 | 9.66650114 | 12.4153908 | 8.07178711 |
| ENSMUSG00 | Nova1 | 6995 | 0.17050242 | 0 | 0 | 0.12379829 | 0.34285056 | 0.0311933 |
| ENSMUSG00 | Mthfd1 | 3523 | 225.679777 | 150.563825 | 256.754617 | 160.018695 | 183.799236 | 146.569251 |
| ENSMUSG00 | Ppp2r5e | 6144 | 13.1288851 | 7.90558816 | 8.77705867 | 8.95591112 | 9.25185265 | 9.84177657 |
| ENSMUSG00 | Sgpp1 | 5696 | 48.47991 | 36.7658493 | 70.6717318 | 40.9280354 | 43.3491279 | 51.3841562 |
| ENSMUSG00 | Esr2 | 4836 | 0 | 0 | 0 | 0 | 0 | 0 |
| ENSMUSG00 | Tex21 | 2315 | 0 | 0 | 0.05728101 | 0 | 0 | 0 |
| ENSMUSG00 | Akap5 | 6906 | 0.17269974 | 0.46051335 | 0.4416344 | 0.22466376 | 0.96053126 | 1.12953183 |
| ENSMUSG00 | Spnb1 | 12278 | 1.54773739 | 1.68758512 | 3.42368079 | 2.19818939 | 2.55173438 | 2.09035963 |
| ENSMUSG00 | Rab15 | 4707 | 12.1876266 | 14.1273189 | 8.94930027 | 12.4182945 | 13.4964601 | 6.37393312 |
| ENSMUSG00 | Fut8 | 3375 | 7.47992004 | 14.4773488 | 7.08539172 | 12.3480775 | 16.1621321 | 9.66532401 |
| ENSMUSG00 | Atl1 | 2803 | 1.29067026 | 0.92831533 | 0.7411654 | 1.30013789 | 2.16629994 | 1.39146394 |
| ENSMUSG00 | Sav1 | 2519 | 1.57822471 | 0.7651678 | 3.03569625 | 3.40910147 | 1.55975821 | 1.37510091 |
| ENSMUSG00 | Nin | 10313 | 0.5396846 | 0.37846425 | 0.35574062 | 0.69274167 | 1.54370496 | 1.00233572 |
| ENSMUSG00 | Pygl | 4143 | 17.7426704 | 19.0977886 | 14.4245454 | 17.9408669 | 26.3568536 | 14.1606925 |
| ENSMUSG00 | Bdkrb2 | 1677 | 4.76496815 | 3.56298081 | 5.08703381 | 1.46307634 | 3.52955253 | 3.30158126 |
| ENSMUSG00 | Trim9 | 7559 | 0 | 0 | 0 | 0 | 0 | 0 |
| ENSMUSG00 | Tmx1 | 3291 | 50.3859341 | 54.0285541 | 64.0530505 | 60.2354156 | 45.5531729 | 60.1683017 |
| ENSMUSG00 | Actr10 | 4099 | 43.4543566 | 41.8521306 | 47.0354358 | 39.8801545 | 37.9929488 | 42.0923405 |
| ENSMUSG00 | Tomm20l | 591 | 0.20180447 | 0 | 0.07479161 | 0 | 0 | 0 |
| ENSMUSG00 | Timm9 | 1402 | 18.9986961 | 23.3370858 | 23.3620297 | 18.2211751 | 11.4645566 | 28.675334 |
| ENSMUSG00 | Serpina1f | 3466 | 0 | 0.33366261 | 0 | 0 | 0 | 0 |
| ENSMUSG00 | 4930403N07 | 2609 | 0 | 0 | 0 | 0 | 0 | 0 |
| ENSMUSG00 | Rtn1 | 4816 | 0.96582044 | 0.80043924 | 0.06424687 | 0.1198739 | 0.15892767 | 0.01132668 |
| ENSMUSG00 | Lrrc9 | 7006 | 0 | 0 | 0 | 0 | 0.05098268 | 0 |
| ENSMUSG00 | Serpina3n | 2059 | 0 | 0.14041702 | 0 | 0 | 0 | 0 |
| ENSMUSG00 | Dhrs7 | 1294 | 16.5289401 | 22.8643551 | 10.2135634 | 18.9611981 | 15.6549237 | 9.96978699 |
| ENSMUSG00 | Gsc | 1262 | 0 | 0 | 0 | 0 | 0 | 0 |
| ENSMUSG00 | Ppm1a | 5394 | 8.31371561 | 7.44147318 | 9.70244385 | 8.6425661 | 7.78544815 | 10.86637 |
| ENSMUSG00 | Clmn | 11974 | 6.14227818 | 6.77684722 | 5.59629147 | 7.52437491 | 6.65635536 | 6.93368982 |
| ENSMUSG00 | 4930447C04 | 5752 | 0 | 0 | 0 | 0 | 0 | 0 |
| ENSMUSG00 | Six6 | 5370 | 0 | 0 | 0 | 0 | 0 | 0 |
| ENSMUSG00 | 4930408O17 | 619 | 0.06422533 | 0 | 0 | 0 | 0 | 0 |
| ENSMUSG00 | Glrx5 | 928 | 56.8057836 | 50.9903946 | 67.6364408 | 47.2799204 | 38.0498406 | 61.1915937 |
| ENSMUSG00 | Mnat1 | 2497 | 11.7976816 | 9.30150786 | 10.4972739 | 8.6700948 | 10.4627571 | 10.2457395 |
| ENSMUSG00 | Prkch | 5329 | 5.28183153 | 6.56471327 | 3.87357116 | 7.21776308 | 7.00906495 | 2.56419483 |
| ENSMUSG00 | Hif1a | 5647 | 97.1605076 | 112.474853 | 91.3233403 | 120.009379 | 87.4505517 | 94.6377336 |
| ENSMUSG00 | Papola | 6574 | 43.1057294 | 48.1717824 | 47.0728785 | 45.3742413 | 45.1506584 | 52.0143219 |
| ENSMUSG00 | Mpp5 | 5497 | 31.0912881 | 30.3038986 | 38.8303984 | 34.4935681 | 34.0021129 | 29.6116149 |
| ENSMUSG00 | Snapc1 | 2115 | 11.0901813 | 19.8441566 | 9.40464745 | 17.8789516 | 12.3766103 | 13.3729555 |
| ENSMUSG00 | Atp6v1d | 4287 | 165.735934 | 123.068196 | 228.731907 | 137.704299 | 120.501756 | 191.02427 |
| ENSMUSG00 | Vrk1 | 3781 | 5.91968673 | 7.09861102 | 6.19597376 | 5.39178925 | 6.5587989 | 4.05404582 |
| ENSMUSG00 | Eif2s1 | 3397 | 22.961511 | 22.7527387 | 24.059231 | 26.8517541 | 21.4950674 | 26.7366938 |
| ENSMUSG00 | Plek2 | 1574 | 3.05617099 | 1.77561223 | 2.10618693 | 2.68208336 | 2.43137115 | 5.16381353 |
| ENSMUSG00 | Pigh | 4713 | 9.8945849 | 8.17932394 | 10.3634705 | 10.4961772 | 8.06591343 | 8.95844294 |
| ENSMUSG00 | Rdh12 | 3138 | 0.40540962 | 1.21310673 | 1.02827742 | 1.1613405 | 0.06504318 | 0 |
| ENSMUSG00 | Vti1b | 2004 | 66.9733045 | 66.4607416 | 70.4494444 | 57.1118988 | 49.4222532 | 76.7609608 |
| ENSMUSG00 | Arg2 | 1420 | 3.94755123 | 6.6850205 | 1.99219574 | 0.38114836 | 4.45582437 | 3.49576362 |
| ENSMUSG00 | Zfp36l1 | 4734 | 2.56974589 | 2.22915734 | 1.99813993 | 2.18748348 | 2.22041264 | 3.93506097 |
| ENSMUSG00 | Galntl1 | 4082 | 0 | 0 | 0.12994172 | 0 | 0 | 0 |
| ENSMUSG00 | Erh | 1515 | 2.36171171 | 3.84855413 | 2.7425566 | 2.61981841 | 1.44827337 | 2.25038278 |
| ENSMUSG00 | 4933426M11 | 5030 | 17.775363 | 17.9908824 | 13.9987149 | 19.0739874 | 21.44528 | 10.0097384 |
| ENSMUSG00 | Srsf5 | 3600 | 117.808741 | 98.2200312 | 119.234472 | 134.656186 | 122.718434 | 169.102763 |
| ENSMUSG00 | Slc10a1 | 1574 | 0 | 0.21429803 | 0.14041246 | 0 | 0.16209141 | 0.19061057 |
| ENSMUSG00 | Smoc1 | 3813 | 0.61515168 | 0.12637409 | 0.13910887 | 0 | 0 | 0 |
| ENSMUSG00 | Gm20498 | 3233 | 0.29512265 | 0.35770944 | 0.38281831 | 0.56918784 | 0.26831067 | 0.35432566 |
| ENSMUSG00 | Pcnx | 12132 | 2.25123765 | 1.52121722 | 2.43379749 | 2.30494432 | 3.48671823 | 2.30660896 |
| ENSMUSG00 | Pacs2 | 5832 | 3.22433853 | 3.6519903 | 2.87251346 | 3.39661222 | 2.84354804 | 3.00713187 |
| ENSMUSG00 | Mta1 | 3000 | 2.62386171 | 3.0839323 | 1.81227556 | 4.58241966 | 3.11260893 | 3.427513 |
| ENSMUSG00 | Wdr37 | 11312 | 6.01674175 | 5.30340528 | 7.95570654 | 7.30763473 | 5.76031488 | 9.33828481 |
| ENSMUSG00 | Gm9745 | 735 | 0 | 0 | 0 | 0 | 0 | 0 |
| ENSMUSG00 | Gtpbp4 | 2960 | 30.3807084 | 23.4583321 | 23.0715701 | 38.3493622 | 33.2878148 | 25.6344751 |
| ENSMUSG00 | Zmynd11 | 8077 | 18.3592847 | 19.2161112 | 19.6191169 | 21.2864854 | 22.5408083 | 20.1258945 |
| ENSMUSG00 | Esyt2 | 5517 | 17.6835144 | 16.1320208 | 19.4850249 | 17.1286704 | 16.7960664 | 20.2545218 |
| ENSMUSG00 | Cdca7l | 3245 | 1.26188428 | 2.3907603 | 0.5857255 | 1.05633104 | 1.83977996 | 0.57995381 |
| ENSMUSG00 | 2610021K21 | 2072 | 0.4796752 | 1.13954424 | 0.72531982 | 0 | 0 | 0 |
| ENSMUSG00 | Tdp1 | 5464 | 1.54976525 | 2.19590485 | 2.07903983 | 2.2320152 | 2.93233565 | 2.0066627 |
| ENSMUSG00 | Psmc1 | 1507 | 79.6428637 | 91.1289715 | 88.286362 | 80.7835556 | 73.035098 | 75.2541141 |
| ENSMUSG00 | BC002230 | 3709 | 8.39270457 | 7.09350159 | 4.63589025 | 4.59172958 | 8.21319665 | 5.59611768 |
| ENSMUSG00 | Rps6ka5 | 4388 | 3.07135549 | 2.43787378 | 7.3736894 | 2.05572268 | 4.53516102 | 4.40073942 |
| ENSMUSG00 | Ccdc88c | 9317 | 0.71685314 | 1.24642402 | 0.30837392 | 0.26334432 | 0.60791325 | 0.24590208 |
| ENSMUSG00 | 9030617O03 | 4645 | 5.53752334 | 8.13308302 | 6.67071947 | 6.59497434 | 3.8118735 | 4.33340878 |
| ENSMUSG00 | Fbln5 | 5818 | 0.75848373 | 0.14908146 | 0 | 0.34730053 | 0.61393027 | 0.02812785 |
| ENSMUSG00 | Tc2n | 6457 | 1.1698221 | 1.21641475 | 2.58077975 | 0.00558805 | 1.59630291 | 3.37078552 |
| ENSMUSG00 | Trip11 | 10354 | 4.31189923 | 6.09660413 | 5.04602844 | 6.28665331 | 6.00252325 | 5.59243376 |
| ENSMUSG00 | Atxn3 | 5519 | 0.01440677 | 0 | 0.00800903 | 0 | 0 | 0 |
| ENSMUSG00 | Lgmn | 2823 | 21.2508041 | 20.1416939 | 21.044023 | 18.328605 | 21.4914491 | 23.8544401 |
| ENSMUSG00 | Golga5 | 2654 | 53.0123005 | 38.9268018 | 43.6522346 | 38.4476345 | 49.0653018 | 48.5887319 |
| ENSMUSG00 | Pitrm1 | 3497 | 21.6796401 | 20.3383439 | 20.7926885 | 21.7503431 | 20.2529624 | 16.9169839 |
| ENSMUSG00 | Chga | 1874 | 0 | 0 | 0 | 0 | 0 | 0 |
| ENSMUSG00 | Pfkp | 7118 | 34.7846491 | 35.4459414 | 36.6941122 | 31.6921807 | 38.2446916 | 25.8875334 |
| ENSMUSG00 | Unc79 | 8938 | 0.14678126 | 0 | 0.11868922 | 0.14532934 | 0.00570893 | 0 |
| ENSMUSG00 | Asb2 | 3479 | 0 | 0 | 0 | 0 | 0 | 0 |
| ENSMUSG00 | Otub2 | 3858 | 3.73029651 | 6.65717306 | 5.19010754 | 4.41439218 | 5.06560444 | 4.55991247 |
| ENSMUSG00 | D830046C22 | 1039 | 0 | 0 | 0 | 0 | 0 | 0 |
| ENSMUSG00 | Akr1c21 | 1409 | 5.02233892 | 13.4744203 | 3.10573631 | 5.81307602 | 5.14247365 | 18.4089297 |
| ENSMUSG00 | Ifi27l2b | 1300 | 0 | 0 | 0 | 0 | 0.07850212 | 0 |
| ENSMUSG00 | Ppp4r4 | 3740 | 0 | 0.39940634 | 0.15364277 | 0.28942817 | 0.43658931 | 0.14585369 |
| ENSMUSG00 | Akr1c6 | 1881 | 0 | 0 | 0 | 0 | 0 | 0 |
| ENSMUSG00 | Akr1c12 | 1256 | 7.2167592 | 5.40946524 | 4.22310602 | 6.26264786 | 8.28772346 | 7.47012413 |
| ENSMUSG00 | Akr1c13 | 1512 | 7.15177956 | 12.3971733 | 7.01616556 | 12.719398 | 10.6979902 | 7.5582499 |
| ENSMUSG00 | Akr1c18 | 1198 | 0 | 0 | 0 | 0 | 0 | 0 |
| ENSMUSG00 | Net1 | 3721 | 11.0152917 | 11.6419273 | 10.536693 | 10.9574604 | 17.7173011 | 11.0828419 |
| ENSMUSG00 | Tuba l3 | 3091 | 0 | 0 | 0 | 0 | 0 | 0 |
| ENSMUSG00 | Tshz3 | 5517 | 0 | 0.05240505 | 0 | 0 | 0.24047232 | 0 |
| ENSMUSG00 | Gdi2 | 2864 | 157.147973 | 143.919981 | 166.297088 | 160.781792 | 130.951069 | 154.991185 |
| ENSMUSG00 | Rgs6 | 3373 | 4.40810842 | 5.81437354 | 4.8749142 | 4.93146236 | 5.18886656 | 2.92719224 |
| ENSMUSG00 | Dpf3 | 3938 | 0.16152557 | 0.23248918 | 0.22448879 | 0.1191129 | 0.12957434 | 0.20085438 |
| ENSMUSG00 | Dcaf4 | 2067 | 7.05866538 | 7.52986009 | 7.42043518 | 5.46380258 | 8.76360107 | 7.79840919 |
| ENSMUSG00 | Papln | 5231 | 2.7283918 | 1.40939125 | 1.8589955 | 3.41437814 | 1.83386707 | 1.43907483 |
| ENSMUSG00 | Numb | 5928 | 12.4604728 | 11.9978388 | 16.3370847 | 12.4716783 | 16.4751153 | 16.282886 |
| ENSMUSG00 | Acot2 | 2193 | 6.23615381 | 5.58110183 | 8.8081192 | 7.8317616 | 6.98035237 | 9.05423501 |
| ENSMUSG00 | Acot3 | 1923 | 158.835859 | 85.2972694 | 171.405691 | 92.1283612 | 87.272881 | 137.848268 |
| ENSMUSG00 | Fam161b | 5742 | 2.47173573 | 1.52733063 | 2.22471833 | 1.20022127 | 2.0350122 | 2.3892622 |
| ENSMUSG00 | Coq6 | 2895 | 14.8035952 | 15.0135305 | 14.9935094 | 14.3580364 | 14.8232062 | 9.13865288 |
| ENSMUSG00 | Entpd5 | 9526 | 43.1651202 | 39.3392778 | 44.2946455 | 42.7408976 | 46.9554074 | 36.8519716 |
| ENSMUSG00 | Aldh6a1 | 5304 | 139.758803 | 121.76525 | 168.823706 | 139.42336 | 134.165505 | 161.004705 |
| ENSMUSG00 | Vax2 | 3798 | 0 | 0 | 0 | 0 | 0 | 0 |
| ENSMUSG00 | Abcd4 | 3196 | 6.10761606 | 5.26191124 | 2.84905496 | 4.16591816 | 10.2180477 | 5.38494911 |
| ENSMUSG00 | Isca2 | 864 | 141.35282 | 118.737194 | 144.372223 | 118.060116 | 88.2331082 | 172.991925 |
| ENSMUSG00 | Npc2 | 3257 | 31.0402785 | 27.932454 | 35.2176182 | 28.0502723 | 20.8993507 | 30.2558402 |
| ENSMUSG00 | Fcf1 | 1108 | 15.4644514 | 18.352598 | 18.2312656 | 11.821103 | 12.9408047 | 25.4038156 |
| ENSMUSG00 | Ylpm1 | 10173 | 3.75553099 | 3.66620527 | 2.81122505 | 7.01920438 | 3.73681805 | 5.42918947 |
| ENSMUSG00 | Mlh3 | 5877 | 6.3113601 | 4.95229046 | 4.71576579 | 4.98530205 | 10.3494028 | 5.75008985 |
| ENSMUSG00 | Tmed10 | 3521 | 18.4155606 | 21.8419658 | 22.5591343 | 16.8881594 | 14.7238846 | 21.0079016 |
| ENSMUSG00 | Fos | 2348 | 102.978208 | 149.669388 | 74.6800303 | 158.296908 | 193.848072 | 71.3229493 |
| ENSMUSG00 | 0610007P14 | 2245 | 21.1970201 | 21.1848634 | 37.9210466 | 23.8511154 | 16.2738909 | 25.7803049 |

| | | | | | | | |
|---|---|---|---|---|---|---|---|
| ENSMUSG00 | Tgfb3 | 3377 | 6.45128911 | 5.02269104 | 11.7016428 | 5.86586984 | 3.36952378 | 2.7298869 |
| ENSMUSG00 | 1700020O03I | 4018 | 4.75918022 | 3.39391815 | 4.38938162 | 3.9332841 | 7.8482579 | 4.71095063 |
| ENSMUSG00 | Esrrb | 6037 | 0.72438345 | 0.48689299 | 0.47591847 | 0.69331078 | 0.1605932 | 0.15360903 |
| ENSMUSG00 | Vash1 | 5074 | 0.03134054 | 0.07597389 | 0 | 0 | 0 | 0.09138133 |
| ENSMUSG00 | Angel1 | 4695 | 2.93826448 | 2.82242206 | 1.39337013 | 2.47463649 | 2.26059343 | 1.69631409 |
| ENSMUSG00 | Ccnk | 2626 | 1.09002079 | 1.00923622 | 0.97627833 | 1.0854842 | 1.0298545 | 2.17075385 |
| ENSMUSG00 | Cyp46a1 | 2104 | 1.41713928 | 1.94669562 | 0.46218657 | 0.66882117 | 2.37670382 | 0.33704402 |
| ENSMUSG00 | Hhipl1 | 3093 | 0.42416129 | 0 | 0 | 0 | 0 | 0.16754547 |
| ENSMUSG00 | Evl | 2678 | 1.2469979 | 2.8429641 | 1.25442124 | 2.39828377 | 1.98160683 | 2.3628515 |
| ENSMUSG00 | Degs2 | 3570 | 18.4300617 | 45.5679072 | 16.6778383 | 31.109393 | 22.182895 | 18.6949698 |
| ENSMUSG00 | Yy1 | 3289 | 8.29196097 | 6.5342515 | 6.08800088 | 8.43633087 | 5.89541587 | 6.44341584 |
| ENSMUSG00 | Slc25a29 | 2866 | 8.48930707 | 4.35460172 | 3.85570857 | 1.48558313 | 4.75367842 | 4.30151324 |
| ENSMUSG00 | Wars | 3891 | 20.2098022 | 21.3501481 | 20.6638788 | 23.0902829 | 22.0445484 | 13.2482828 |
| ENSMUSG00 | Meg3 | 16216 | 0 | 0 | 0 | 0 | 0 | 0.00840979 |
| ENSMUSG00 | Hsp90aa1 | 4440 | 24.8650832 | 26.3071927 | 20.0799814 | 26.224495 | 19.3877243 | 23.8530235 |
| ENSMUSG00 | Zfp839 | 2821 | 8.59654134 | 9.25808282 | 4.27759771 | 8.28825411 | 10.6357706 | 7.79275407 |
| ENSMUSG00 | Fdft1 | 2852 | 46.8088722 | 27.1175455 | 58.5689919 | 24.3667664 | 27.445463 | 35.0783229 |
| ENSMUSG00 | Tecpr2 | 7809 | 1.69529709 | 2.11035513 | 1.83962082 | 3.45156707 | 2.56144696 | 1.52981073 |
| ENSMUSG00 | Cirp | 3139 | 10.7146023 | 7.78290098 | 9.30787456 | 8.25323605 | 7.36379364 | 8.0112193 |
| ENSMUSG00 | Traf3 | 7546 | 2.0441461 | 3.64623091 | 1.90373694 | 2.67770275 | 5.46373067 | 1.38071988 |
| ENSMUSG00 | Amn | 1673 | 0.16634092 | 0.63365315 | 0.07926212 | 0 | 0.1219997 | 0.21193683 |
| ENSMUSG00 | Cdc42bpb | 6681 | 5.24836608 | 4.16880161 | 6.60943589 | 5.82194941 | 5.84271479 | 3.58844603 |
| ENSMUSG00 | 1200009I06R | 4578 | 56.1596095 | 33.2610654 | 68.5331327 | 46.3360288 | 42.7226078 | 44.379377 |
| ENSMUSG00 | Tnfaip2 | 4997 | 16.9857817 | 25.7373652 | 5.2366402 | 20.6368788 | 26.2433032 | 9.18615527 |
| ENSMUSG00 | Eif5 | 4117 | 171.11176 | 139.479677 | 149.719383 | 157.509474 | 140.040569 | 236.826282 |
| ENSMUSG00 | Ppp1r13b | 3826 | 7.02423022 | 10.2267096 | 10.4092683 | 9.36473346 | 12.6032214 | 10.1727418 |
| ENSMUSG00 | Zfyve21 | 1300 | 55.2906991 | 45.5547211 | 58.8904555 | 36.1097999 | 45.2957213 | 62.4589239 |
| ENSMUSG00 | Xrcc3 | 4182 | 2.30053235 | 3.46822551 | 0.53904687 | 3.7445259 | 4.73415443 | 0.97828692 |
| ENSMUSG00 | Klc1 | 4138 | 8.96371757 | 9.95635767 | 7.80649378 | 11.8238889 | 10.5554804 | 7.85678347 |
| ENSMUSG00 | 2010107E04f | 361 | 269.037781 | 247.606233 | 302.31122 | 195.602662 | 182.055324 | 716.847028 |
| ENSMUSG00 | Kif26a | 6874 | 0 | 0 | 0.08359383 | 0.01574718 | 0.08907718 | 0.02777458 |
| ENSMUSG00 | Gpr132 | 2436 | 0 | 0 | 0 | 0 | 0 | 0 |
| ENSMUSG00 | Hecw1 | 9482 | 0 | 0 | 0 | 0 | 0 | 0 |
| ENSMUSG00 | Ggps1 | 2386 | 25.3595312 | 25.527101 | 37.4400355 | 37.4733053 | 23.5243141 | 31.4012716 |
| ENSMUSG00 | Gng4 | 3224 | 52.1729026 | 32.5378054 | 81.2880668 | 45.7292909 | 29.8497968 | 54.5323588 |
| ENSMUSG00 | Gpr137b | 4203 | 0.82291858 | 0.87132277 | 0.70462134 | 1.07310387 | 1.11692281 | 0.69435794 |
| ENSMUSG00 | Mtr | 4271 | 6.67400598 | 7.17550391 | 8.81760015 | 6.90213776 | 11.3378672 | 7.42692705 |
| ENSMUSG00 | Ryr2 | 18086 | 0 | 0.02397866 | 0 | 0.01795524 | 0 | 0 |
| ENSMUSG00 | Amph | 3172 | 1.20319235 | 1.59507454 | 2.46649628 | 0.64838479 | 0.88475746 | 1.43595989 |
| ENSMUSG00 | Gli3 | 9730 | 0 | 0.03466651 | 0 | 0 | 0.00524423 | 0 |
| ENSMUSG00 | Sfrp4 | 4573 | 0.21733807 | 0 | 0 | 0 | 0 | 0 |
| ENSMUSG00 | Aoah | 3541 | 0 | 1.30638194 | 0 | 0 | 0 | 0.23877826 |
| ENSMUSG00 | Trim27 | 4692 | 11.108149 | 9.83857877 | 8.97791057 | 10.612366 | 10.8099355 | 10.719189 |
| ENSMUSG00 | Zkscan3 | 9114 | 16.1482103 | 19.3824333 | 16.3683586 | 18.9317903 | 16.2529701 | 17.0877979 |
| ENSMUSG00 | Slc17a1 | 2358 | 4.45099527 | 4.29141342 | 0.03749096 | 1.69851864 | 2.57512244 | 4.67300859 |
| ENSMUSG00 | Slc17a4 | 3659 | 0.93440047 | 0.67163393 | 0.33824859 | 0.24652941 | 0.79489025 | 0 |
| ENSMUSG00 | Scgn | 1501 | 0 | 0 | 0 | 0 | 0 | 0 |
| ENSMUSG00 | Lrrc16a | 9916 | 5.03157806 | 5.9722809 | 5.29120489 | 7.05557527 | 6.00007606 | 5.86696304 |
| ENSMUSG00 | Mrs2 | 2024 | 15.5172083 | 10.4276985 | 11.1159773 | 15.4382661 | 15.27766 | 9.20384319 |
| ENSMUSG00 | Gpld1 | 3214 | 5.14570003 | 3.83812358 | 3.57575581 | 4.33343783 | 6.03298781 | 2.8937934 |
| ENSMUSG00 | Prl | 970 | 0 | 0 | 0 | 0 | 0 | 0 |
| ENSMUSG00 | Prl8a6 | 1422 | 0 | 0 | 0 | 0 | 0 | 0 |
| ENSMUSG00 | Prl8a8 | 1542 | 0 | 0 | 0 | 0 | 0 | 0 |
| ENSMUSG00 | Prl7b1 | 919 | 0 | 0 | 0 | 0 | 0 | 0 |
| ENSMUSG00 | Prl7d1 | 1100 | 0 | 0 | 0 | 0 | 0 | 0 |
| ENSMUSG00 | Irf4 | 4706 | 0 | 0 | 0 | 0 | 0.21685668 | 0 |
| ENSMUSG00 | Exoc2 | 4397 | 20.5694151 | 11.0356487 | 21.9652097 | 19.7109554 | 18.5793104 | 17.6971903 |
| ENSMUSG00 | Tfap2a | 3235 | 1.26578501 | 1.81723213 | 2.15885354 | 2.29765106 | 1.29340427 | 2.925595 |
| ENSMUSG00 | Gcnt2 | 6261 | 2.10810087 | 2.69369985 | 1.14369886 | 1.2448046 | 2.32271476 | 0.92353035 |
| ENSMUSG00 | Tmem14c | 939 | 52.3298977 | 58.6037454 | 72.4458322 | 56.6015459 | 35.8108431 | 56.3792064 |
| ENSMUSG00 | Gcm2 | 2874 | 0 | 0 | 0 | 0 | 0 | 0 |
| ENSMUSG00 | Mak | 3550 | 0 | 0.02714729 | 0 | 0 | 0.08624176 | 0 |
| ENSMUSG00 | Elovl2 | 6751 | 0 | 0 | 0.13094902 | 0 | 0 | 0 |
| ENSMUSG00 | Nedd9 | 4714 | 36.2049805 | 34.3049851 | 25.1764846 | 33.7475041 | 51.7949105 | 25.2727247 |
| ENSMUSG00 | Hivep1 | 8744 | 1.44581917 | 1.21788698 | 1.23850086 | 1.06876138 | 1.24298011 | 2.07117572 |
| ENSMUSG00 | Edn1 | 2339 | 1.59769781 | 7.95210204 | 1.17166065 | 5.35291554 | 8.20261535 | 2.22721509 |
| ENSMUSG00 | Tbc1d7 | 1089 | 19.1658653 | 24.2923386 | 16.3575232 | 12.3586801 | 17.6647784 | 16.7054035 |
| ENSMUSG00 | Ccdc90a | 2266 | 43.7380463 | 32.3440329 | 42.855891 | 31.4324607 | 39.0691801 | 27.3950039 |
| ENSMUSG00 | Cap2 | 3394 | 0 | 0 | 0 | 0 | 0 | 0 |
| ENSMUSG00 | Nup153 | 5999 | 11.6569245 | 9.59873285 | 9.12920212 | 9.6114531 | 10.2750228 | 7.8109402 |
| ENSMUSG00 | Kif13a | 6961 | 2.81560866 | 3.95958123 | 3.77185674 | 4.73767974 | 5.94489165 | 3.85551573 |
| ENSMUSG00 | Tpmt | 2201 | 20.0132087 | 18.3463146 | 13.7164283 | 14.3934735 | 14.7445593 | 21.6858786 |
| ENSMUSG00 | Dek | 4418 | 41.7621514 | 41.8277514 | 48.0337366 | 63.4988448 | 38.1831595 | 69.2916593 |
| ENSMUSG00 | Id4 | 1657 | 58.9014512 | 59.5278498 | 85.4159936 | 61.2545511 | 78.125477 | 85.0336671 |
| ENSMUSG00 | Barx1 | 1395 | 0 | 0 | 0 | 0 | 0 | 0 |
| ENSMUSG00 | Susd3 | 3331 | 0.035805 | 0 | 0 | 0 | 0 | 0.04094062 |
| ENSMUSG00 | Ippk | 3035 | 5.85525528 | 7.74793534 | 9.08795718 | 6.88352678 | 7.27987503 | 6.20981078 |
| ENSMUSG00 | Aspn | 2338 | 0 | 0 | 0 | 0 | 0.08729919 | 0 |
| ENSMUSG00 | Ogn | 2533 | 0 | 0.47558668 | 0.01745039 | 0 | 0 | 0 |
| ENSMUSG00 | Cenpp | 1260 | 2.08242993 | 4.97161705 | 1.68387973 | 2.52001581 | 2.67281016 | 5.73633286 |
| ENSMUSG00 | Nol8 | 4224 | 4.61178633 | 5.33883876 | 6.94839578 | 6.98747927 | 4.15555712 | 4.19063468 |
| ENSMUSG00 | Spin1 | 1468 | 69.5991723 | 53.175774 | 81.0868959 | 61.7671512 | 63.539656 | 85.1124812 |
| ENSMUSG00 | Nxnl2 | 1161 | 0.85606117 | 0.87158939 | 1.44674422 | 0.49725471 | 0.08790073 | 0.86921762 |
| ENSMUSG00 | Wrnip1 | 2626 | 22.8298799 | 18.9919907 | 13.2807518 | 18.3295687 | 25.2605821 | 10.7914509 |
| ENSMUSG00 | Serpinb9b | 3246 | 0.01224753 | 0.17813842 | 0 | 0.24454867 | 0 | 0.30249138 |
| ENSMUSG00 | Serpinb9c | 2792 | 0 | 0 | 0 | 0 | 0 | 0 |
| ENSMUSG00 | Ripk1 | 6432 | 8.12168865 | 8.22585719 | 9.71038622 | 8.58855879 | 6.79875685 | 7.86181299 |
| ENSMUSG00 | 1300014I06R | 3143 | 0 | 0 | 0.2531445 | 0 | 0.37340332 | 0.39050549 |
| ENSMUSG00 | Prpf4b | 4450 | 24.5411921 | 19.5561156 | 29.1434174 | 31.6305743 | 30.9368903 | 28.9907963 |
| ENSMUSG00 | 1700026J04R | 2388 | 0 | 0 | 0 | 0 | 0.55556356 | 0 |
| ENSMUSG00 | 4933417A18f | 1166 | 0 | 0 | 0 | 0 | 0 | 0 |
| ENSMUSG00 | Eci3 | 1408 | 0.05647085 | 1.19781639 | 0.47090032 | 0.07687936 | 0.1449613 | 0.44553743 |
| ENSMUSG00 | Eci2 | 2920 | 29.2583999 | 27.047116 | 30.6234001 | 31.114585 | 44.4908058 | 27.3680456 |
| ENSMUSG00 | Rpp40 | 6775 | 0.45770147 | 0.73257617 | 0.77638662 | 0.96396311 | 0.76068841 | 1.29227442 |
| ENSMUSG00 | Fars2 | 2118 | 22.2427972 | 21.636122 | 28.3618058 | 21.3800594 | 19.1770515 | 20.8873772 |
| ENSMUSG00 | Ly86 | 992 | 0 | 0 | 0 | 0 | 0 | 0 |
| ENSMUSG00 | Sar1 | 10921 | 15.311011 | 17.9226562 | 23.4183558 | 20.9996772 | 16.778291 | 28.9828943 |
| ENSMUSG00 | Riok1 | 2472 | 12.544205 | 10.9745012 | 15.4849499 | 12.3776593 | 19.83671 | 20.0477425 |
| ENSMUSG00 | Snrnp48 | 1780 | 8.68813591 | 12.1278236 | 9.36185102 | 11.1084047 | 10.6639392 | 12.2582648 |
| ENSMUSG00 | Slc35b3 | 2203 | 26.2570241 | 31.8909817 | 24.4785513 | 38.7845127 | 37.4764758 | 42.3914502 |
| ENSMUSG00 | Ctsq | 2015 | 0 | 0 | 0 | 0 | 0.02532326 | 0 |
| ENSMUSG00 | Cts7 | 1485 | 0 | 0 | 0 | 0 | 0 | 0 |
| ENSMUSG00 | Cts6 | 1348 | 0 | 0 | 0 | 0 | 0 | 0 |
| ENSMUSG00 | Shc3 | 1525 | 0 | 0 | 0.08695445 | 0 | 0 | 0 |
| ENSMUSG00 | Sema4d | 8707 | 1.34694691 | 2.03105826 | 1.39098484 | 2.38695965 | 3.72134474 | 1.37203308 |
| ENSMUSG00 | Gadd45g | 1258 | 37.1641058 | 75.4588959 | 35.4528296 | 49.2184328 | 76.0122641 | 56.4788833 |
| ENSMUSG00 | Fbp2 | 1318 | 0 | 0.62152467 | 0.13414823 | 0 | 0 | 0 |
| ENSMUSG00 | Syk | 8286 | 0 | 0 | 0 | 0 | 0 | 0 |
| ENSMUSG00 | 2010111I01R | 6603 | 18.3394205 | 21.6083682 | 19.1454295 | 20.8579682 | 24.4042549 | 23.0489909 |
| ENSMUSG00 | Auh | 9564 | 13.3224922 | 14.263469 | 12.4277244 | 14.59278 | 12.3564499 | 11.4956159 |
| ENSMUSG00 | Fancc | 7791 | 4.84760704 | 3.29654392 | 5.11745121 | 2.76022315 | 4.23091244 | 2.62209021 |
| ENSMUSG00 | Ror2 | 6582 | 0.48924247 | 0.11713508 | 0.12759572 | 0 | 0 | 0.0745888 |
| ENSMUSG00 | Ptch1 | 6550 | 10.4396071 | 10.1228312 | 14.2863056 | 9.53557543 | 14.5210908 | 12.5921388 |
| ENSMUSG00 | Sptlc1 | 3831 | 22.5395207 | 24.0114639 | 31.3370414 | 30.3179591 | 26.172495 | 21.052234 |
| ENSMUSG00 | Msx2 | 2454 | 3.17525435 | 1.66904954 | 2.59375118 | 0.70576126 | 1.0812435 | 1.62269655 |
| ENSMUSG00 | 0610007P08f | 7582 | 5.00220633 | 5.14149719 | 5.67243382 | 5.42515573 | 6.79050557 | 4.44984552 |
| ENSMUSG00 | Stxn1 | 2800 | 15.4620422 | 16.2801337 | 13.1184756 | 9.13648913 | 11.2804738 | 15.5075806 |
| ENSMUSG00 | Habp4 | 2558 | 22.9860656 | 15.220737 | 16.7441696 | 17.885861 | 19.5887025 | 17.2199149 |
| ENSMUSG00 | Ctsl | 1966 | 281.099153 | 220.172335 | 363.394908 | 229.009029 | 202.9116 | 271.886256 |
| ENSMUSG00 | Drd1a | 3524 | 0 | 0 | 0 | 0 | 0 | 0 |
| ENSMUSG00 | Zfp346 | 4373 | 7.11834924 | 5.89520845 | 5.20556807 | 5.20644182 | 7.78289336 | 3.43037997 |
| ENSMUSG00 | 1110018J18R | 4954 | 36.5695851 | 41.1734375 | 55.4798264 | 45.6524536 | 44.5579535 | 45.05766 |
| ENSMUSG00 | Cdk20 | 1946 | 2.55366653 | 3.91236272 | 1.88527902 | 2.29916419 | 4.82469329 | 3.7281881 |
| ENSMUSG00 | Lman2 | 4307 | 40.8631326 | 43.286126 | 39.1936008 | 34.9175671 | 41.524831 | 30.345965 |
| ENSMUSG00 | Mxd3 | 1949 | 0.10198943 | 0 | 0.34018863 | 0 | 0 | 0 |
| ENSMUSG00 | Prelid1 | 1083 | 47.6478428 | 66.1173158 | 47.344541 | 44.9443011 | 35.0541308 | 33.2433276 |
| ENSMUSG00 | Nsd1 | 13098 | 6.00672591 | 7.5270622 | 7.08687359 | 8.99158599 | 7.87718216 | 8.00247661 |
| ENSMUSG00 | Slc34a1 | 3071 | 2.38196301 | 8.22197841 | 0.56133894 | 1.292421 | 4.15388927 | 5.64854138 |
| ENSMUSG00 | F12 | 2148 | 0 | 0 | 0 | 0 | 0 | 0 |
| ENSMUSG00 | Pdlim7 | 4269 | 2.35608727 | 3.04763162 | 2.17437035 | 4.95293467 | 2.34274295 | 5.16231167 |
| ENSMUSG00 | Ddx41 | 2188 | 21.1859645 | 20.5915555 | 19.6362849 | 25.4124454 | 25.5131879 | 16.32988 |
| ENSMUSG00 | Fam193b | 4194 | 13.3087016 | 13.7987404 | 9.12703768 | 13.515711 | 16.047637 | 13.7803912 |
| ENSMUSG00 | Pcbd2 | 542 | 16.5036589 | 20.0925017 | 20.5514104 | 15.844145 | 11.8622202 | 29.9417165 |
| ENSMUSG00 | Txndc15 | 1516 | 67.2643849 | 67.2893786 | 67.0900005 | 58.0026006 | 66.7785816 | 65.4339465 |
| ENSMUSG00 | Catsper3 | 1533 | 0.33713062 | 0.1257311 | 0.28833557 | 0.04707377 | 0.96527391 | 0.9607505 |
| ENSMUSG00 | Ddx46 | 5830 | 18.8344146 | 18.5059081 | 17.1348482 | 22.1319711 | 27.7275401 | 20.2805422 |
| ENSMUSG00 | Caml | 1809 | 25.5806409 | 25.4650297 | 23.4570311 | 25.7700395 | 14.9778914 | 32.2651896 |
| ENSMUSG00 | B4galt7 | 3017 | 3.5578322 | 5.79770583 | 5.24503142 | 4.19781161 | 4.70180062 | 4.41167516 |
| ENSMUSG00 | Pitx1 | 2649 | 0 | 0 | 0 | 0 | 0 | 0 |
| ENSMUSG00 | Cxcl14 | 1819 | 0 | 0.10596249 | 0 | 0 | 0.1683113 | 0 |
| ENSMUSG00 | Slc25a48 | 3366 | 0.30708333 | 0.95914784 | 0.02626372 | 0.33230641 | 0.57605534 | 0.01620597 |
| ENSMUSG00 | A530054K11f | 6990 | 2.95748925 | 5.94919879 | 5.70387159 | 9.41540046 | 3.87625258 | 4.95938004 |
| ENSMUSG00 | Zfp369 | 6022 | 2.6736914 | 4.08888607 | 3.1782461 | 3.95452497 | 2.74535798 | 3.19306222 |
| ENSMUSG00 | Ptdss1 | 3411 | 37.4827398 | 31.7711253 | 31.9041154 | 37.9543758 | 32.9853881 | 24.4999984 |
| ENSMUSG00 | Mterfd1 | 4567 | 14.6678311 | 14.5181836 | 16.7341771 | 16.7808764 | 12.345992 | 21.3084985 |
| ENSMUSG00 | Uqcrb | 872 | 238.350519 | 145.554001 | 194.447557 | 146.562617 | 118.378851 | 546.83775 |
| ENSMUSG00 | Fastkd3 | 1825 | 8.8224491 | 14.8387838 | 14.2899109 | 10.696102 | 9.19872748 | 13.301057 |
| ENSMUSG00 | 1700001L19f | 1805 | 0.37442835 | 0.93436314 | 0.36732834 | 0.73963194 | 0.36750298 | 1.61683457 |
| ENSMUSG00 | Adcy2 | 4186 | 0 | 0 | 0 | 0 | 0 | 0 |
| ENSMUSG00 | Cetn3 | 994 | 87.6700335 | 68.1590922 | 82.5782922 | 82.7636434 | 56.2111484 | 190.016979 |
| ENSMUSG00 | Il9 | 531 | 0 | 0.0907466 | 0 | 0 | 0 | 0 |
| ENSMUSG00 | Lect2 | 954 | 0 | 0.45458908 | 0 | 0 | 0 | 0 |
| ENSMUSG00 | Smad5 | 5680 | 13.0745137 | 15.7878466 | 12.6302097 | 12.9018719 | 14.9575556 | 7.11637595 |
| ENSMUSG00 | Trpc7 | 4325 | 0 | 0 | 0 | 0 | 0 | 0.04414393 |
| ENSMUSG00 | 1700067P10f | 791 | 0 | 0 | 0 | 0 | 0 | 0 |
| ENSMUSG00 | Hnrnpk | 6657 | 98.2211789 | 94.5705072 | 104.200018 | 95.2648362 | 81.6866587 | 126.540068 |
| ENSMUSG00 | Ccnh | 3311 | 21.876921 | 23.4746998 | 25.9657459 | 28.6498626 | 22.3616041 | 32.8020578 |
| ENSMUSG00 | Rasa1 | 9216 | 19.610288 | 18.1954013 | 17.0505156 | 23.635775 | 15.9291323 | 25.1408486 |
| ENSMUSG00 | 2210016F16f | 2818 | 35.128157 | 31.6170091 | 35.5591122 | 28.2459156 | 25.3321149 | 34.2046044 |
| ENSMUSG00 | Gkap1 | 1506 | 6.81069985 | 6.46325452 | 6.1929541 | 5.60637345 | 5.42112891 | 7.31670269 |
| ENSMUSG00 | Slc28a3 | 7858 | 0 | 0 | 0 | 0 | 0 | 0 |
| ENSMUSG00 | Naa35 | 3719 | 11.8443325 | 16.2089914 | 16.401867 | 18.1525962 | 15.9843312 | 13.4649738 |
| ENSMUSG00 | Golm1 | 3682 | 6.56472899 | 16.3980478 | 3.0012115 | 6.09533723 | 10.8510734 | 4.62972557 |
| ENSMUSG00 | Agtpbp1 | 11598 | 6.27970687 | 5.29726795 | 6.68477605 | 5.57812604 | 6.96894114 | 6.62464725 |
| ENSMUSG00 | Dapk1 | 5907 | 6.97252037 | 8.21461778 | 5.52242427 | 7.41554123 | 9.07020393 | 7.627849 |
| ENSMUSG00 | Slc6a19 | 6606 | 3.4413037 | 2.552248 | 2.23417306 | 3.45640651 | 1.27411501 | 2.05963151 |
| ENSMUSG00 | 5430425J12R | 1653 | 0 | 0 | 0 | 0 | 0 | 0 |
| ENSMUSG00 | Nkd2 | 6561 | 0 | 0.04406625 | 0 | 0 | 0 | 0.04157086 |
| ENSMUSG00 | Trip13 | 2254 | 0.37039268 | 0.32067286 | 0.1176624 | 0 | 0.58859174 | 0 |
| ENSMUSG00 | Cep72 | 3620 | 1.47161171 | 1.81031937 | 1.31872902 | 2.78090896 | 1.79015186 | 1.08495803 |
| ENSMUSG00 | Tppp | 3050 | 3.40202637 | 1.24810785 | 9.27514083 | 2.99303517 | 1.37185666 | 4.48019484 |
| ENSMUSG00 | Ahrr | 12855 | 0 | 0 | 0.12722429 | 0 | 0 | 0 |
| ENSMUSG00 | Pdcd6 | 1518 | 75.3730388 | 74.723903 | 79.3769592 | 80.2219369 | 58.7576449 | 90.4662771 |
| ENSMUSG00 | Sdha | 4308 | 676.876737 | 564.423412 | 722.06604 | 582.488827 | 608.218291 | 532.786146 |
| ENSMUSG00 | Ccdc127 | 5786 | 13.7488276 | 16.1648607 | 18.0825721 | 14.5488092 | 13.5458889 | 18.2333745 |
| ENSMUSG00 | Lrrc14b | 2753 | 0 | 0 | 0 | 0 | 0 | 0 |
| ENSMUSG00 | Erap1 | 4675 | 12.4836461 | 9.2559198 | 11.2135585 | 13.0667169 | 20.8471396 | 16.7731739 |
| ENSMUSG00 | Cast | 3883 | 48.7345061 | 38.2835885 | 45.2149679 | 46.6195255 | 42.4321832 | 42.5380415 |
| ENSMUSG00 | Pcsk1 | 3552 | 0 | 0 | 0 | 0 | 0 | 0 |
| ENSMUSG00 | Rhobtb3 | 6119 | 8.64108336 | 6.82752825 | 9.49194668 | 10.9855939 | 7.6218512 | 10.3009791 |
| ENSMUSG00 | Spata9 | 1098 | 0.43448613 | 0.30719954 | 0.88564713 | 0 | 0 | 0.57132669 |
| ENSMUSG00 | Glrx | 1342 | 43.7843519 | 44.9548626 | 32.4761678 | 33.8504427 | 37.680431 | 37.6198912 |
| ENSMUSG00 | Ank | 5671 | 10.8589736 | 12.3376325 | 9.2674998 | 7.4505509 | 8.58387631 | 10.3356021 |
| ENSMUSG00 | Srd5a1 | 3013 | 10.4501628 | 8.92400754 | 16.8122509 | 11.7119945 | 10.3136618 | 10.3558537 |
| ENSMUSG00 | Nsun2 | 5857 | 29.7911566 | 28.0134601 | 33.9834214 | 22.0422666 | 26.7459405 | 24.3175971 |
| ENSMUSG00 | Mctp1 | 8173 | 0.02918547 | 0 | 0.14061519 | 0 | 0.09989258 | 0 |
| ENSMUSG00 | Ankrd32 | 5919 | 8.05990482 | 5.38932669 | 7.54993467 | 10.6069872 | 7.68111182 | 14.5566118 |
| ENSMUSG00 | Med10 | 858 | 0.04633506 | 1.12322709 | 0.82427679 | 0.50464398 | 0.5352417 | 0.34967486 |
| ENSMUSG00 | 8030423J24R | 866 | 0 | 0 | 0 | 0 | 0.17676574 | 0 |
| ENSMUSG00 | Irx4 | 2672 | 0 | 0 | 0 | 0 | 0.11458019 | 0 |
| ENSMUSG00 | Ndufs6 | 1203 | 25.3800574 | 27.7182194 | 40.4173128 | 24.4746038 | 20.1899874 | 54.4585898 |
| ENSMUSG00 | Mrpl36 | 897 | 33.4176503 | 40.5582651 | 45.7294429 | 38.4151087 | 23.3230926 | 58.2284662 |
| ENSMUSG00 | Lpcat1 | 4403 | 0.57786753 | 1.19289625 | 0.76296618 | 1.49146766 | 1.52974826 | 0.34689525 |
| ENSMUSG00 | Slc6a3 | 3441 | 0 | 0 | 0 | 0 | 0 | 0 |
| ENSMUSG00 | Cliptm1l | 2221 | 114.379793 | 105.463438 | 118.276251 | 112.632515 | 112.161534 | 94.8287998 |
| ENSMUSG00 | Tert | 3454 | 1.28911807 | 2.008931 | 0.8574185 | 0 | 2.51143135 | 0.45799915 |
| ENSMUSG00 | Slc6a18 | 4392 | 2.29010396 | 1.03131274 | 0 | 0.0164308 | 0.33692279 | 0.2049324 |
| ENSMUSG00 | Hapln1 | 5050 | 0 | 0 | 0 | 0 | 0 | 0 |
| ENSMUSG00 | Vcan | 16350 | 0.08267195 | 0.08546831 | 0.02703477 | 0.21406473 | 0.03745055 | 0.01334539 |
| ENSMUSG00 | Xrcc4 | 4201 | 12.283412 | 7.69652527 | 7.9228726 | 5.78034182 | 7.90720558 | 14.8741249 |
| ENSMUSG00 | Atg10 | 1601 | 42.462131 | 32.9269005 | 62.2580612 | 48.1169051 | 48.4129075 | 61.6362553 |
| ENSMUSG00 | Acot12 | 3404 | 0.66570576 | 0.75025894 | 0 | 1.52638496 | 0 | 1.08970357 |
| ENSMUSG00 | Zcchc9 | 2246 | 22.4443229 | 18.8369084 | 17.7122256 | 19.3904843 | 19.0156174 | 20.4984822 |
| ENSMUSG00 | Ckmt2 | 1476 | 0.08080382 | 0.03264664 | 0 | 0 | 0 | 0 |
| ENSMUSG00 | Cd180 | 2840 | 0 | 0 | 0 | 0 | 0.01796703 | 0 |
| ENSMUSG00 | Slc30a5 | 3776 | 44.7880864 | 40.3765633 | 44.389139 | 47.0710147 | 42.7291844 | 38.196052 |
| ENSMUSG00 | Mrps36-ps1 | 624 | 0 | 0 | 0 | 0.05782379 | 0 | 0.21854679 |
| ENSMUSG00 | Rad17 | 2829 | 25.9836986 | 21.1890894 | 20.3275354 | 25.9550939 | 16.4316113 | 20.275209 |
| ENSMUSG00 | Marveld2 | 3695 | 13.6750246 | 17.5662077 | 13.792889 | 9.93110671 | 14.7624346 | 7.80224457 |
| ENSMUSG00 | Ocln | 5326 | 20.3629273 | 16.8372078 | 31.3296953 | 20.5612101 | 23.0892914 | 18.2206471 |
| ENSMUSG00 | Gtf2h2 | 4354 | 10.0347435 | 11.0118305 | 10.2738321 | 9.66276158 | 9.24662621 | 12.9607783 |
| ENSMUSG00 | Naip1 | 5338 | 0 | 0 | 0 | 0 | 0 | 0.12262858 |
| ENSMUSG00 | Serf1 | 1323 | 4.38722612 | 4.15211973 | 4.77767464 | 4.30911794 | 4.01114368 | 10.3284915 |
| ENSMUSG00 | Smn1 | 2479 | 3.35171256 | 3.94588453 | 3.20949243 | 3.76976585 | 1.31734088 | 7.27250365 |
| ENSMUSG00 | Mccc2 | 2213 | 55.0792152 | 53.3033938 | 62.2181387 | 48.3267872 | 46.7837852 | 43.1242489 |
| ENSMUSG00 | Cartpt | 871 | 0 | 0 | 0 | 0 | 0 | 0 |
| ENSMUSG00 | Ptcd2 | 1665 | 38.2750963 | 24.4260403 | 35.9720704 | 35.4969303 | 27.7044107 | 54.4346104 |
| ENSMUSG00 | Btf3 | 2037 | 5.99162127 | 6.97839982 | 8.05050749 | 6.99676367 | 5.58609809 | 8.70324767 |
| ENSMUSG00 | Ankra2 | 3721 | 10.9511872 | 10.813136 | 14.6824719 | 8.96960258 | 14.1381869 | 17.3425951 |
| ENSMUSG00 | Rgnef | 6395 | 26.9180971 | 24.5641597 | 23.6872113 | 26.3492022 | 30.807324 | 20.1563636 |
| ENSMUSG00 | Hexb | 1890 | 780.385358 | 486.734057 | 1340.27472 | 556.026215 | 595.874678 | 1014.18365 |
| ENSMUSG00 | Gfm2 | 10405 | 9.37243572 | 9.79474533 | 12.66369 | 10.2298925 | 11.764755 | 11.3423704 |
| ENSMUSG00 | Polk | 4510 | 13.5838571 | 13.8148713 | 16.6908511 | 19.4331011 | 14.4706729 | 15.1189796 |
| ENSMUSG00 | Col4a3bp | 5345 | 16.3782167 | 19.0311655 | 20.4841843 | 12.0701021 | 10.3007408 | 20.8858931 |
| ENSMUSG00 | Hmgcr | 7397 | 12.9633931 | 7.79764381 | 13.2838579 | 8.59003385 | 10.8233586 | 8.20414659 |
| ENSMUSG00 | Poc5 | 3020 | 8.37234623 | 6.09510626 | 6.99618575 | 9.06830192 | 6.97811034 | 5.37364581 |
| ENSMUSG00 | F2rl2 | 2500 | 0.6042833 | 0 | 0 | 0 | 0 | 0 |
| ENSMUSG00 | Iqgap2 | 6683 | 32.908472 | 32.5977712 | 30.3652044 | 45.1039054 | 45.8573116 | 30.5151096 |
| ENSMUSG00 | F2rl1 | 2752 | 9.15878439 | 11.3812454 | 9.12305481 | 10.2922983 | 8.49203493 | 10.3171873 |
| ENSMUSG00 | S100z | 298 | 0.80044592 | 0 | 0.14832833 | 0 | 0.17122945 | 0.36610254 |
| ENSMUSG00 | Crhbp | 1679 | 0 | 0 | 0 | 0 | 0.45586399 | 0 |
| ENSMUSG00 | Aggf1 | 3683 | 25.6257157 | 29.3593202 | 26.4154919 | 27.8428375 | 29.025321 | 25.6972572 |
| ENSMUSG00 | Pde8b | 7602 | 9.81074473 | 9.09596744 | 6.80296716 | 12.4592695 | 12.2229716 | 9.45391666 |
| ENSMUSG00 | Otp | 1179 | 0 | 0 | 0 | 0 | 0 | 0 |
| ENSMUSG00 | Ap3b1 | 3993 | 23.2977271 | 21.2874741 | 20.9219843 | 22.7353929 | 35.4360481 | 17.7186111 |
| ENSMUSG00 | Scamp1 | 5360 | 54.6637857 | 46.6311708 | 61.5279759 | 49.8484216 | 49.4746409 | 51.4147304 |
| ENSMUSG00 | Jmy | 4547 | 0.93552593 | 1.4200535 | 1.65258705 | 1.20617347 | 1.15586468 | 1.60756616 |
| ENSMUSG00 | Dimt1 | 2533 | 6.15244704 | 4.85098411 | 5.07806408 | 9.10241865 | 4.67355673 | 6.28834952 |
| ENSMUSG00 | Kif2a | 8659 | 2.71341829 | 1.85311066 | 1.75602656 | 2.45436242 | 1.87393319 | 4.23341207 |
| ENSMUSG00 | Ercc8 | 6434 | 1.53238408 | 2.43403694 | 1.78621063 | 1.49173513 | 2.32370658 | 1.39043856 |
| ENSMUSG00 | Elovl7 | 5427 | 110.519796 | 101.39841 | 141.458929 | 111.011958 | 103.293858 | 106.374611 |
| ENSMUSG00 | Depdc1b | 3437 | 1.73503697 | 1.87866839 | 0.72019296 | 2.34108116 | 1.46977341 | 1.68234609 |
| ENSMUSG00 | Pde4d | 12032 | 3.40987832 | 4.34126523 | 2.18951948 | 4.14439707 | 3.21459382 | 5.58323109 |
| ENSMUSG00 | Rab3c | 10722 | 0 | 0 | 0 | 0 | 0 | 0.0025438 |
| ENSMUSG00 | Plk2 | 2788 | 4.71989385 | 3.87150755 | 2.07691587 | 2.7695687 | 3.25778153 | 2.17179696 |
| ENSMUSG00 | Thbs4 | 3267 | 0 | 0 | 0.10823837 | 0 | 0 | 0 |
| ENSMUSG00 | Serinc5 | 5304 | 4.06249443 | 7.33153447 | 7.18363286 | 8.14975289 | 6.64766699 | 6.97807041 |
| ENSMUSG00 | Mtx3 | 6266 | 2.4934414 | 2.68386025 | 3.32254517 | 3.21893758 | 2.71174324 | 5.10583336 |
| ENSMUSG00 | Zfyve16 | 7840 | 13.4580415 | 9.48977891 | 14.6756884 | 19.0995516 | 13.2772713 | 13.4320641 |
| ENSMUSG00 | Dhfr | 5284 | 1.04580087 | 2.12479959 | 1.42208806 | 0.64188346 | 1.16846924 | 1.68272755 |
| ENSMUSG00 | Rasgrf2 | 8867 | 0.50663915 | 1.11404315 | 0.25423413 | 0.97662013 | 0.71357512 | 0.23992578 |
| ENSMUSG00 | Erbb2ip | 7763 | 34.4960604 | 33.7547176 | 42.5050571 | 50.0630815 | 43.6514443 | 36.8732885 |
| ENSMUSG00 | Nln | 5552 | 8.67144937 | 11.6039757 | 13.4070432 | 7.36328466 | 9.3284891 | 9.12265942 |
| ENSMUSG00 | 2410002O22I | 7223 | 18.8127139 | 20.2272314 | 21.002454 | 17.0544234 | 17.3573038 | 23.2229034 |
| ENSMUSG00 | Trim23 | 5858 | 12.405773 | 9.32800024 | 11.8465165 | 11.9678069 | 11.6024467 | 15.4112421 |
| ENSMUSG00 | Ppwd1 | 2386 | 6.18159399 | 4.32183513 | 5.6132265 | 6.21530608 | 6.05216444 | 4.77820587 |
| ENSMUSG00 | Cenpk | 1865 | 0.70344818 | 0.8267915 | 0.78212376 | 0 | 0.27359987 | 3.58299551 |
| ENSMUSG00 | Cwc27 | 2856 | 8.51903154 | 10.4775142 | 14.7494245 | 7.84557061 | 14.3288352 | 14.7928628 |
| ENSMUSG00 | Srek1ip1 | 3248 | 8.88623117 | 8.8717649 | 7.37604646 | 9.15381918 | 6.83388962 | 15.1656399 |
| ENSMUSG00 | 4933425L06f | 2149 | 0 | 0 | 0 | 0 | 0 | 0 |
| ENSMUSG00 | Rgs7bp | 5822 | 0.04097095 | 0 | 0 | 0 | 0.00876441 | 0 |
| ENSMUSG00 | Rnf180 | 1978 | 2.55255107 | 1.24242091 | 0.11173368 | 1.02153412 | 0.74811168 | 0.60671594 |
| ENSMUSG00 | Htr1a | 4440 | 0 | 0 | 0 | 0 | 0 | 0 |
| ENSMUSG00 | Parp8 | 3306 | 6.89046895 | 8.78899716 | 2.40683392 | 8.66580259 | 10.9276086 | 7.60653884 |
| ENSMUSG00 | Emb | 2244 | 459.189973 | 254.052494 | 690.703042 | 316.714809 | 285.397524 | 648.368251 |
| ENSMUSG00 | Hcn1 | 3517 | 0 | 0 | 0 | 0 | 0 | 0 |
| ENSMUSG00 | Mrps30 | 1585 | 26.4116852 | 30.7967608 | 27.5529469 | 28.7518121 | 19.1228185 | 28.2899097 |
| ENSMUSG00 | Fgf10 | 4725 | 0 | 0 | 0 | 0 | 0 | 0 |
| ENSMUSG00 | Slc4a7 | 7756 | 0.37930706 | 0.71447149 | 0.58130325 | 0.30704164 | 0.63157969 | 0 |
| ENSMUSG00 | Psmd6 | 2318 | 45.8783047 | 48.4774733 | 41.7038096 | 49.795712 | 37.5323429 | 46.3598269 |
| ENSMUSG00 | Atxn7 | 6835 | 2.64648773 | 5.5412646 | 4.52042257 | 5.47433507 | 4.31503222 | 3.97846603 |
| ENSMUSG00 | Gm5457 | 335 | 1.89876922 | 1.29456113 | 1.8472412 | 2.26185951 | 1.21854032 | 1.70975351 |
| ENSMUSG00 | Fezf2 | 2299 | 0 | 0.02095974 | 0 | 0.09416802 | 0.06658509 | 0 |
| ENSMUSG00 | Ptprg | 11663 | 2.19519244 | 2.03686153 | 1.57660694 | 2.93593932 | 3.61380318 | 2.82732049 |
| ENSMUSG00 | 4930452B06I | 3262 | 1.08468356 | 1.35902902 | 0.70462779 | 0.3429011 | 1.47041059 | 1.26255994 |
| ENSMUSG00 | Pdhb | 1475 | 261.981878 | 210.092888 | 293.829879 | 222.192007 | 174.00859 | 252.794301 |
| ENSMUSG00 | Oit1 | 1241 | 2.08227738 | 3.10629764 | 0.14247169 | 0.90132424 | 0.94569431 | 0.04395591 |
| ENSMUSG00 | Fam107a | 5105 | 0.56849169 | 0 | 0.29439034 | 1.61149974 | 0.1799167 | 0.36330568 |
| ENSMUSG00 | Acox2 | 3686 | 0 | 1.47722951 | 0 | 0.75374862 | 0 | 0 |
| ENSMUSG00 | Kctd6 | 3249 | 18.366567 | 20.9415994 | 24.2028559 | 17.4024511 | 17.1030235 | 21.6501369 |
| ENSMUSG00 | Map3k1 | 8315 | 4.06399981 | 7.05845899 | 5.55513241 | 7.20773012 | 5.73164582 | 9.16152344 |
| ENSMUSG00 | Il6st | 5192 | 18.4611833 | 19.1929049 | 16.422449 | 14.2674186 | 23.636062 | 11.5255313 |
| ENSMUSG00 | Ddx4 | 2827 | 0.04218834 | 0 | 0 | 0 | 0 | 0 |
| ENSMUSG00 | Ppap2a | 1971 | 18.6171022 | 13.5929284 | 13.6126427 | 17.519288 | 23.1961607 | 17.3528146 |
| ENSMUSG00 | Gpx8 | 1013 | 100.742664 | 106.171904 | 96.9560663 | 84.0608342 | 77.6729235 | 158.612932 |
| ENSMUSG00 | BC067074 | 8397 | 6.03646989 | 10.2547543 | 2.23720177 | 6.30372268 | 8.53781803 | 3.9140098 |
| ENSMUSG00 | Ndufs4 | 1529 | 98.8296805 | 91.4565436 | 126.534642 | 80.2819591 | 62.8401999 | 161.203999 |
| ENSMUSG00 | Fst | 2680 | 3.67886536 | 2.19356192 | 2.75436858 | 0.01346345 | 3.96025603 | 0.88540807 |
| ENSMUSG00 | Myo14 | 8404 | 2.5166487 | 1.6914565 | 2.03547278 | 3.00111246 | 2.14330208 | 2.61257551 |
| ENSMUSG00 | Dusp13 | 2758 | 0 | 0 | 0 | 0 | 0 | 0 |
| ENSMUSG00 | Samd8 | 7863 | 3.74651037 | 2.67804594 | 2.96252973 | 2.28524204 | 2.6476868 | 3.00045313 |
| ENSMUSG00 | Vdac2 | 4828 | 76.8018572 | 75.4235591 | 95.901886 | 71.3195839 | 67.3129635 | 72.0337459 |
| ENSMUSG00 | Nkiras1 | 5316 | 6.33425357 | 5.21203992 | 4.34867643 | 6.4141332 | 4.7225314 | 4.48933584 |
| ENSMUSG00 | Comtd1 | 2706 | 1.76299248 | 2.546438 | 2.15618746 | 3.3601904 | 2.50794826 | 2.30816422 |
| ENSMUSG00 | Ube2e1 | 4450 | 5.0744074 | 4.24473827 | 3.89373538 | 5.22986938 | 4.28850889 | 5.86557972 |
| ENSMUSG00 | Nr1d2 | 5458 | 17.6343016 | 20.1115272 | 28.8713027 | 16.5734125 | 20.3525871 | 25.9853654 |
| ENSMUSG00 | Thrb | 6608 | 10.7570822 | 6.26394579 | 6.88981512 | 10.696841 | 9.15818428 | 5.84043804 |
| ENSMUSG00 | Dlg5 | 10193 | 3.3542346 | 4.87395486 | 4.31914408 | 4.13812519 | 7.53406216 | 4.31877443 |
| ENSMUSG00 | Ngly1 | 4019 | 35.1359708 | 36.8322345 | 30.1461189 | 27.4812486 | 34.0387444 | 39.0151533 |
| ENSMUSG00 | Oxsm | 5229 | 2.22004213 | 1.44679124 | 3.76167912 | 2.01490859 | 2.80064446 | 6.55655413 |
| ENSMUSG00 | Sftpa1 | 2898 | 0 | 0 | 0 | 0 | 0 | 0 |
| ENSMUSG00 | Dydc1 | 1081 | 0 | 0 | 0 | 0 | 0 | 0.10092374 |
| ENSMUSG00 | Dydc2 | 749 | 0 | 0 | 0 | 0 | 0 | 0 |
| ENSMUSG00 | 5730469M10 | 3049 | 50.6690053 | 51.6791295 | 48.1740651 | 48.3421292 | 44.8175252 | 56.6961835 |
| ENSMUSG00 | Glud1 | 4074 | 61.5848889 | 62.710976 | 54.2595525 | 67.3903478 | 51.4522219 | 39.2784262 |
| ENSMUSG00 | Sftpd | 1725 | 0 | 0.08380251 | 0 | 0 | 0 | 0 |
| ENSMUSG00 | Bmpr1a | 6520 | 32.4690082 | 39.399068 | 34.4530316 | 42.8778652 | 41.736758 | 41.9577656 |
| ENSMUSG00 | 9230112D13I | 777 | 0.15349606 | 0 | 0 | 0.18575055 | 0.13134202 | 0 |
| ENSMUSG00 | Ldb3 | 5999 | 0 | 0 | 0.0147364 | 0 | 0.02551744 | 0 |
| ENSMUSG00 | Opn4 | 2859 | 0 | 0 | 0 | 0 | 0 | 0 |
| ENSMUSG00 | Cdhr1 | 4286 | 0 | 0 | 0 | 0 | 0 | 0 |
| ENSMUSG00 | Rgr | 1652 | 0 | 0 | 0 | 0 | 0 | 0 |
| ENSMUSG00 | Nid2 | 4863 | 2.1991002 | 3.1311774 | 1.04528315 | 0 | 2.05658434 | 0.07291185 |
| ENSMUSG00 | 2700060E02f | 1502 | 34.461808 | 33.846003 | 34.0195276 | 30.220514 | 22.1839037 | 31.3240697 |
| ENSMUSG00 | Nudt13 | 2570 | 10.9366244 | 10.3497728 | 13.4153454 | 13.0148075 | 20.4701142 | 19.3681777 |
| ENSMUSG00 | Ecd | 3111 | 25.8519245 | 33.6267329 | 29.0842728 | 24.8317767 | 37.101145 | 26.3716216 |
| ENSMUSG00 | Dnajc9 | 1626 | 28.9486402 | 28.4495599 | 29.5494486 | 26.8063406 | 26.5801601 | 32.172668 |
| ENSMUSG00 | Anxa7 | 2835 | 55.5875572 | 55.0362256 | 62.4126905 | 49.636675 | 48.2905701 | 42.080872 |
| ENSMUSG00 | Zmynd17 | 1822 | 0.67641048 | 0.29091705 | 0 | 0.73273087 | 0.75615376 | 0.14969615 |
| ENSMUSG00 | Ppp3cb | 7654 | 4.20720397 | 3.79624048 | 5.37074915 | 4.20501486 | 2.59998518 | 7.54025825 |
| ENSMUSG00 | 2310021P13f | 6046 | 4.65545488 | 6.24845694 | 4.37193221 | 5.19269043 | 5.84026663 | 6.91113916 |
| ENSMUSG00 | Camk2g | 4804 | 1.63027262 | 1.32402381 | 1.03971862 | 1.32941754 | 1.68884133 | 2.24260668 |
| ENSMUSG00 | Plau | 2332 | 1.53430242 | 1.61172491 | 1.25099556 | 0.04641773 | 0.89711896 | 2.21050893 |
| ENSMUSG00 | Vcl | 5190 | 6.15865247 | 3.7137913 | 3.75587915 | 4.5328503 | 6.82318012 | 3.46319601 |
| ENSMUSG00 | Ap3m1 | 8568 | 6.55167349 | 6.6700654 | 5.69547557 | 6.96120677 | 5.34204705 | 5.66311662 |
| ENSMUSG00 | Txndc16 | 11246 | 6.32778856 | 6.03723075 | 6.23762448 | 4.65543719 | 5.85764282 | 4.56194171 |
| ENSMUSG00 | Ero1l | 4275 | 16.9716379 | 12.3988506 | 17.1947754 | 20.1215425 | 11.4585546 | 17.232468 |
| ENSMUSG00 | Psmc6 | 2166 | 73.8761815 | 72.3908969 | 71.3229184 | 81.3928299 | 56.5624785 | 92.9302113 |
| ENSMUSG00 | Bmp4 | 3225 | 0.49309123 | 0 | 0 | 0.02237646 | 0.03164426 | 0.12685879 |
| ENSMUSG00 | Samd4 | 8714 | 0 | 0 | 0.00507251 | 0.0041407 | 0.01756703 | 0 |
| ENSMUSG00 | Mapk1ip1l | 4726 | 15.1922128 | 12.3575895 | 14.2818905 | 15.8880099 | 17.1023655 | 11.6404882 |
| ENSMUSG00 | Ktn1 | 4442 | 30.2506808 | 22.4985774 | 30.3104038 | 30.4366098 | 34.2320126 | 33.371829 |
| ENSMUSG00 | Peli2 | 7390 | 6.14891938 | 9.77421878 | 4.65345519 | 10.3314758 | 9.70122571 | 5.40325736 |
| ENSMUSG00 | Otx2 | 4521 | 0 | 0 | 0 | 0 | 0 | 0 |
| ENSMUSG00 | 1700011H14I | 1371 | 274.721679 | 385.772713 | 181.256573 | 297.157086 | 314.90457 | 251.479734 |
| ENSMUSG00 | Slc35f4 | 6477 | 0 | 0 | 0 | 0 | 0 | 0 |
| ENSMUSG00 | Anxa11 | 4095 | 15.960442 | 18.5214798 | 13.8488314 | 21.4201344 | 20.3233258 | 23.2317318 |
| ENSMUSG00 | Gm9746 | 1597 | 0 | 0 | 0 | 0 | 0 | 0 |
| ENSMUSG00 | Ppif | 3293 | 20.8616673 | 14.7061568 | 21.0203724 | 16.5344079 | 13.8993802 | 19.7871586 |
| ENSMUSG00 | Slmap | 9653 | 10.7985984 | 12.4247441 | 12.9038427 | 13.0228782 | 13.2098739 | 14.5033382 |
| ENSMUSG00 | Pnp | 2936 | 14.2989739 | 14.869522 | 12.5258629 | 13.18666 | 12.3047255 | 11.7236358 |
| ENSMUSG00 | Rnase10 | 1662 | 0 | 0 | 0 | 0 | 0 | 0 |
| ENSMUSG00 | 4933413J09R | 4822 | 0.05771223 | 0.08993737 | 0 | 0 | 0.12698393 | 0 |
| ENSMUSG00 | Rnase4 | 1993 | 15.1003004 | 12.4757673 | 16.4343029 | 8.76252361 | 10.1131051 | 17.3665415 |
| ENSMUSG00 | Arf4 | 2693 | 237.617607 | 238.409267 | 203.348174 | 223.513059 | 158.61188 | 260.167446 |
| ENSMUSG00 | Dnahc12 | 11036 | 0.16931022 | 0.44972848 | 0.20026207 | 0.61139383 | 0.85074784 | 0.20759965 |
| ENSMUSG00 | Rnase6 | 1070 | 0.55731982 | 0.40530652 | 0.57834187 | 0 | 0 | 0 |
| ENSMUSG00 | Hacl1 | 4930 | 4.20940382 | 4.59383931 | 4.86847886 | 4.06197459 | 5.07158705 | 1.3830953 |
| ENSMUSG00 | Gpr65 | 2047 | 0 | 0 | 0 | 0 | 0 | 0 |
| ENSMUSG00 | Eaf1 | 4232 | 30.4554036 | 22.8748966 | 28.1483854 | 21.6816255 | 29.5885459 | 18.2453931 |
| ENSMUSG00 | Mettl6 | 2430 | 13.202746 | 11.2236734 | 18.9540413 | 13.1409915 | 16.2528456 | 18.4973685 |
| ENSMUSG00 | Sh3bp5 | 3181 | 20.1089494 | 20.2834474 | 18.0781508 | 16.6515062 | 20.6126668 | 21.152636 |
| ENSMUSG00 | Capn7 | 5021 | 49.3914932 | 56.8141282 | 55.3381369 | 52.4522691 | 42.2865465 | 59.0742286 |
| ENSMUSG00 | Arhgef3 | 3758 | 13.075512 | 14.5149155 | 11.5503486 | 8.09397646 | 16.3887269 | 10.0810202 |
| ENSMUSG00 | Asb14 | 2609 | 0.01523782 | 0.05540794 | 0.03388413 | 0.41489511 | 0.1369048 | 0.21953523 |
| ENSMUSG00 | Btd | 2427 | 10.4179998 | 10.999295 | 13.003756 | 11.9232797 | 15.6211773 | 8.94545235 |
| ENSMUSG00 | Bap1 | 3389 | 44.7527761 | 42.3284268 | 35.4371223 | 44.4397916 | 52.4717969 | 32.7392247 |
| ENSMUSG00 | Phf7 | 2314 | 8.59020754 | 7.57988978 | 6.32273554 | 4.72465838 | 7.85021166 | 9.09940428 |
| ENSMUSG00 | Galntl2 | 6937 | 0.01146187 | 0 | 0.11469413 | 0 | 0.07355683 | 0 |
| ENSMUSG00 | Sema3g | 4522 | 0.45716165 | 0 | 0.3812189 | 0.15160523 | 0.12412431 | 0.1387255 |
| ENSMUSG00 | Dph3 | 4252 | 10.939302 | 8.30683492 | 11.1856028 | 7.70519671 | 13.7646409 | 12.6558945 |
| ENSMUSG00 | Oxnad1 | 5157 | 10.1373777 | 9.24110748 | 15.0596545 | 10.1452326 | 10.4882603 | 12.1273508 |
| ENSMUSG00 | Msmb | 581 | 0 | 0 | 0 | 0 | 0 | 0 |
| ENSMUSG00 | Gm6768 | 1849 | 0.27951393 | 0.26060812 | 0.35858715 | 0.40980148 | 0.27596742 | 0.38352657 |
| ENSMUSG00 | Tnnc1 | 1081 | 0 | 0 | 0 | 0 | 0 | 0 |
| ENSMUSG00 | Nisch | 10589 | 32.6145868 | 30.2661278 | 26.9994825 | 36.75673 | 36.3000934 | 28.7917573 |
| ENSMUSG00 | Parg | 5990 | 14.0305652 | 15.5017152 | 12.6775904 | 16.5411176 | 15.0267992 | 12.64924 |
| ENSMUSG00 | Ogdhl | 3425 | 1.57861178 | 1.43504149 | 1.96165843 | 0.94814132 | 1.47492298 | 0.86801042 |
| ENSMUSG00 | Glt8d1 | 2510 | 18.3888896 | 13.5344389 | 20.4103331 | 15.6115938 | 19.5160641 | 22.8737512 |
| ENSMUSG00 | Spcs1 | 1765 | 52.6168853 | 46.4391717 | 57.1743952 | 38.8009748 | 28.7655773 | 72.8302407 |
| ENSMUSG00 | Nek4 | 4375 | 8.36000962 | 8.96543175 | 9.46677186 | 9.27824002 | 9.15558972 | 4.94995739 |
| ENSMUSG00 | Chat | 3059 | 0 | 0 | 0 | 0 | 0 | 0 |
| ENSMUSG00 | Itih4 | 3411 | 0.11655081 | 0 | 0 | 0.51832899 | 0 | 0 |
| ENSMUSG00 | Gm5803 | 962 | 0 | 0 | 0 | 0 | 0.15912591 | 0 |
| ENSMUSG00 | Ebpl | 2073 | 3.81637271 | 4.37001984 | 3.43294584 | 2.57604563 | 2.85531095 | 3.93396871 |
| ENSMUSG00 | Kpna3 | 4152 | 20.7682171 | 30.5111167 | 30.2131095 | 24.3501659 | 22.4899489 | 22.9259371 |
| ENSMUSG00 | 6330409N04 | 4933 | 13.4103322 | 15.1797549 | 19.1036548 | 21.5629165 | 9.15433663 | 22.0497185 |
| ENSMUSG00 | Rnaseh2b | 4460 | 2.03234295 | 2.00956911 | 2.75518214 | 1.14880052 | 1.78477907 | 2.15873266 |
| ENSMUSG00 | Gucy1b2 | 2902 | 0 | 0 | 0 | 0 | 0 | 0 |
| ENSMUSG00 | Mapk8 | 10517 | 2.06016325 | 2.13968513 | 1.28608576 | 2.25062482 | 1.70783344 | 2.9564599 |
| ENSMUSG00 | Pspc1 | 3994 | 6.84821497 | 7.10611278 | 7.7026747 | 7.39889696 | 7.06499395 | 8.8366007 |
| ENSMUSG00 | Ctsb | 4728 | 166.547547 | 124.94027 | 197.33767 | 122.738478 | 135.217737 | 148.620248 |
| ENSMUSG00 | Ptpn20 | 3118 | 0 | 0 | 0 | 0 | 0 | 0 |
| ENSMUSG00 | Gdf10 | 4344 | 0 | 0 | 0 | 0 | 0 | 0 |
| ENSMUSG00 | Gata4 | 5513 | 0 | 0 | 0 | 0 | 0 | 0 |
| ENSMUSG00 | Zmym2 | 4979 | 36.0745654 | 37.7342715 | 26.9259269 | 49.3075379 | 40.849796 | 36.340622 |
| ENSMUSG00 | Cryl1 | 1989 | 10.7333801 | 20.3260055 | 15.8006588 | 14.9480164 | 10.4412946 | 13.5481868 |
| ENSMUSG00 | Prkcd | 5632 | 101.478123 | 70.6369436 | 117.379752 | 78.852593 | 83.9235297 | 82.8699622 |
| ENSMUSG00 | Anxa8 | 1919 | 0 | 0 | 0 | 0 | 0 | 0 |
| ENSMUSG00 | N6amt2 | 831 | 22.3415276 | 18.323605 | 28.1381167 | 17.9324722 | 11.1754517 | 22.3514192 |
| ENSMUSG00 | Xpo4 | 7083 | 5.2704216 | 6.16361945 | 4.48071767 | 6.64789893 | 5.76325013 | 6.41529705 |
| ENSMUSG00 | Tdh | 2799 | 0 | 0 | 0 | 0 | 0 | 0 |
| ENSMUSG00 | Tkt | 6518 | 29.7525673 | 39.8621197 | 24.2370953 | 35.0191798 | 29.5370537 | 18.4913518 |
| ENSMUSG00 | Pinx1 | 1344 | 5.91599412 | 7.1347485 | 4.11103451 | 2.65782917 | 2.23999715 | 3.53109169 |
| ENSMUSG00 | Lats2 | 6618 | 12.5790233 | 14.0962454 | 11.0604793 | 14.5843864 | 14.7034298 | 11.7662579 |
| ENSMUSG00 | 4930578I06R | 1029 | 0 | 0.23414209 | 0 | 0 | 0.04958832 | 0 |
| ENSMUSG00 | Dcp1a | 5606 | 8.79357757 | 7.67579251 | 5.88989952 | 8.32216974 | 7.82780649 | 4.87012378 |
| ENSMUSG00 | Sap18 | 3842 | 14.3935121 | 16.1290382 | 18.9715875 | 20.8960305 | 18.6202444 | 15.5966638 |
| ENSMUSG00 | F630043A04f | 2251 | 0 | 0.04281337 | 0.05890961 | 0 | 0.18134652 | 0 |
| ENSMUSG00 | Prss52 | 1059 | 0 | 0 | 0 | 0 | 0 | 0 |
| ENSMUSG00 | Mrp63 | 2319 | 14.5204364 | 12.7790694 | 13.3043926 | 9.17999408 | 7.76727497 | 18.806446 |
| ENSMUSG00 | Zdhhc20 | 5195 | 17.4327208 | 19.7105273 | 21.09266 | 16.0858547 | 18.2005533 | 23.8882788 |
| ENSMUSG00 | Hmbox1 | 9685 | 1.78561012 | 2.13940838 | 1.77537604 | 2.88358311 | 1.91776983 | 2.27828427 |
| ENSMUSG00 | Efha1 | 2298 | 85.1161724 | 83.4350972 | 94.9050885 | 85.5103631 | 80.8028641 | 84.3638537 |
| ENSMUSG00 | Fgf9 | 4260 | 20.3070248 | 18.7881879 | 16.0413261 | 16.8298175 | 21.7760449 | 22.6392311 |
| ENSMUSG00 | Ints9 | 4178 | 9.19190861 | 6.23955606 | 7.5433016 | 6.75350857 | 7.60876786 | 7.56613377 |
| ENSMUSG00 | 1700129C05I | 1350 | 0 | 0 | 0 | 0 | 0 | 0 |
| ENSMUSG00 | Extl3 | 6441 | 2.82071955 | 2.7830083 | 5.29104502 | 4.55437078 | 3.77885131 | 5.90294164 |
| ENSMUSG00 | Cab39l | 3112 | 68.320151 | 87.0359869 | 53.8603434 | 81.1961947 | 81.9832516 | 61.8675059 |
| ENSMUSG00 | Cdadc1 | 1889 | 52.9301395 | 52.0382964 | 54.0296747 | 46.4730621 | 42.7876015 | 50.9396121 |
| ENSMUSG00 | Atp8a2 | 6221 | 0.1981064 | 0.13167811 | 0.07105263 | 0.33640228 | 0.16404557 | 0 |
| ENSMUSG00 | Fam123a | 3479 | 0.28568181 | 0 | 0.11434797 | 0 | 0 | 0.23519379 |
| ENSMUSG00 | Mtmr6 | 3799 | 43.8053281 | 36.2127645 | 47.6574754 | 41.0968694 | 29.723972 | 46.6231922 |
| ENSMUSG00 | Spata13 | 9054 | 15.0081989 | 19.6598981 | 15.5785378 | 16.8813277 | 21.0834628 | 12.6552639 |
| ENSMUSG00 | Cacna2d3 | 3653 | 0 | 0 | 0 | 0 | 0 | 0 |
| ENSMUSG00 | Mipep | 3847 | 15.5528979 | 10.62181 | 14.431379 | 14.6597961 | 15.7442964 | 10.804926 |
| ENSMUSG00 | Wnt5a | 7797 | 0.07648226 | 0.46968957 | 0.12471983 | 0.14345817 | 0.47773829 | 0 |
| ENSMUSG00 | Esd | 2384 | 45.9089085 | 55.4427059 | 41.068407 | 42.3933753 | 34.4599266 | 40.557297 |
| ENSMUSG00 | Lrrc63 | 2509 | 0 | 0 | 0 | 0 | 0 | 0 |
| ENSMUSG00 | Lcp1 | 9526 | 0.3756029 | 0.25292065 | 0.02320063 | 0.55679829 | 0.32139225 | 0.09448489 |
| ENSMUSG00 | Cpb2 | 1462 | 0 | 0 | 0 | 0 | 0 | 0.27983556 |
| ENSMUSG00 | Zc3h13 | 6396 | 6.6010507 | 8.21188586 | 5.50104238 | 7.6835123 | 9.64522958 | 5.58200481 |
| ENSMUSG00 | 4930564B18I | 864 | 0 | 0 | 0 | 0 | 0 | 0 |
| ENSMUSG00 | Slc25a30 | 1696 | 31.3402579 | 38.2138943 | 24.0034773 | 26.5569384 | 31.8613986 | 12.1417174 |
| ENSMUSG00 | 1200011I18R | 5440 | 9.12037494 | 9.17668273 | 13.8293266 | 17.4175458 | 13.5163617 | 9.85196069 |
| ENSMUSG00 | Nufip1 | 3964 | 4.69363393 | 5.16630623 | 5.09592388 | 5.94388877 | 6.14015673 | 6.19252657 |
| ENSMUSG00 | Tsc22d1 | 8272 | 44.6720492 | 48.8621708 | 55.7011619 | 50.0053867 | 30.3863549 | 53.9557781 |
| ENSMUSG00 | Enox1 | 3126 | 0 | 0.44702713 | 0 | 0 | 0.52234294 | 0 |
| ENSMUSG00 | Dnajc15 | 642 | 91.0908284 | 64.1735328 | 116.632277 | 62.8906664 | 44.6679489 | 168.108641 |
| ENSMUSG00 | Epsti1 | 1881 | 3.48732285 | 3.45835709 | 4.93481501 | 2.30188482 | 3.87919816 | 3.10301584 |
| ENSMUSG00 | Tnfsf11 | 2077 | 0 | 0 | 0 | 0 | 0 | 0 |

| | | | | | | | | |
|---|---|---|---|---|---|---|---|---|
| ENSMUSG00 | Akap11 | 9407 | 5.40103158 | 5.45023647 | 5.05593527 | 9.16722509 | 6.87258618 | 7.76748789 |
| ENSMUSG00 | 1190002H23 | 913 | 8.70875805 | 6.49171126 | 6.72952484 | 3.51730774 | 6.20364482 | 13.6522565 |
| ENSMUSG00 | Tdrd3 | 6027 | 17.1831801 | 17.8370586 | 15.5480185 | 18.283485 | 17.7368935 | 16.4000817 |
| ENSMUSG00 | Naa16 | 6807 | 4.13499048 | 5.11100503 | 4.09745305 | 7.84507505 | 4.66261286 | 4.89637273 |
| ENSMUSG00 | Diap3 | 5069 | 0 | 0.01901221 | 0.00872003 | 0.01423636 | 0 | 0 |
| ENSMUSG00 | Mtrf1 | 1851 | 12.1994127 | 11.6105636 | 12.2743097 | 10.7992721 | 12.0743126 | 8.133766 |
| ENSMUSG00 | Wbp4 | 3442 | 18.2145243 | 16.08548 | 21.1377785 | 23.376804 | 22.1183477 | 29.2556561 |
| ENSMUSG00 | Sugt1 | 1938 | 72.2696377 | 54.8748596 | 65.4588697 | 63.0784143 | 45.2865668 | 88.3540687 |
| ENSMUSG00 | Lect1 | 2183 | 0.14569118 | 0 | 0 | 0 | 0.30386756 | 0 |
| ENSMUSG00 | Olfm4 | 2563 | 1.31846112 | 4.38058565 | 0.01724614 | 2.33695645 | 7.04773197 | 1.50047762 |
| ENSMUSG00 | Elp3 | 2755 | 20.0003978 | 17.630466 | 23.4406144 | 19.3965546 | 21.7996531 | 12.1869622 |
| ENSMUSG00 | Scara5 | 6390 | 0.28618969 | 0.3167184 | 0.33894997 | 0 | 0.63882787 | 1.0243996 |
| ENSMUSG00 | Pbk | 1979 | 0 | 6.47680324 | 0.06700633 | 1.40389966 | 0.30940703 | 0.06891016 |
| ENSMUSG00 | Esco2 | 3999 | 0 | 0.18074434 | 0 | 0.0270683 | 0 | 0 |
| ENSMUSG00 | Ccdc25 | 2218 | 39.1639878 | 35.1947864 | 31.9855537 | 39.4169495 | 40.5588371 | 27.8648929 |
| ENSMUSG00 | Clu | 3470 | 511.95206 | 1027.9404 | 257.160463 | 629.314529 | 827.921329 | 290.164433 |
| ENSMUSG00 | Adam2 | 2530 | 0 | 0 | 0 | 0 | 0 | 0 |
| ENSMUSG00 | Ephx2 | 2147 | 35.4040423 | 38.6927929 | 27.3404972 | 38.9894473 | 20.4866027 | 24.0860345 |
| ENSMUSG00 | Chrna2 | 3520 | 0.21458924 | 0.67077718 | 0 | 0.01025058 | 0.07248065 | 0 |
| ENSMUSG00 | Trim35 | 5092 | 19.4561385 | 14.6868339 | 25.338802 | 13.5413957 | 23.759532 | 14.5800409 |
| ENSMUSG00 | Stmn4 | 2013 | 0 | 0 | 0 | 0 | 0 | 0 |
| ENSMUSG00 | Dpysl2 | 4506 | 0.69700021 | 0.89828254 | 0.88285971 | 0.96090665 | 0.64547346 | 0.54476643 |
| ENSMUSG00 | Bnip3l | 4351 | 10.507654 | 11.2852182 | 11.2968167 | 14.1641306 | 13.7094538 | 13.7094084 |
| ENSMUSG00 | Ppp2r2a | 2960 | 43.0594833 | 38.6142705 | 39.6173951 | 37.959286 | 39.5454413 | 40.9948885 |
| ENSMUSG00 | Ebf2 | 7281 | 0 | 0 | 0 | 0 | 0 | 0 |
| ENSMUSG00 | Nefm | 5777 | 0.01376337 | 0.00834108 | 0.00765135 | 0 | 0.08832677 | 0.04249122 |
| ENSMUSG00 | Nefl | 3376 | 1.04805621 | 1.39877705 | 1.38785408 | 1.53904455 | 1.22427027 | 2.30250953 |
| ENSMUSG00 | Adam7 | 4268 | 0 | 0 | 0 | 0 | 0.07173342 | 0 |
| ENSMUSG00 | Adamdec1 | 2411 | 0 | 0 | 0 | 0 | 0 | 0 |
| ENSMUSG00 | Nkx3-1 | 3193 | 0 | 0 | 0 | 0 | 0 | 0 |
| ENSMUSG00 | Pibf1 | 3274 | 6.54496151 | 6.74080345 | 6.89894373 | 6.32592961 | 7.88617781 | 7.75586107 |
| ENSMUSG00 | Entpd4 | 4302 | 11.8471702 | 12.5450524 | 15.0832881 | 10.3750556 | 13.7588554 | 10.8984437 |
| ENSMUSG00 | 6720463M24 | 2661 | 1.28484454 | 3.44059527 | 1.21260223 | 1.45087515 | 1.72580753 | 5.8321194 |
| ENSMUSG00 | Tnfrsf10b | 3163 | 0.05027566 | 0.25898499 | 0.62885961 | 0.31941108 | 0.37104225 | 0.40528234 |
| ENSMUSG00 | Rhobtb2 | 5498 | 2.74774147 | 2.32255496 | 2.27521309 | 2.36915571 | 2.34806713 | 3.10051829 |
| ENSMUSG00 | Klhl1 | 4111 | 0 | 0 | 0 | 0 | 0 | 0 |
| ENSMUSG00 | Pebp4 | 1040 | 0 | 0 | 0 | 0 | 0 | 0 |
| ENSMUSG00 | Bin3 | 1703 | 30.6978608 | 29.9078505 | 24.3460533 | 36.2303559 | 27.7753672 | 34.7859756 |
| ENSMUSG00 | Pdlim2 | 4901 | 0 | 0.09831961 | 0.07215155 | 0 | 0.21863985 | 0 |
| ENSMUSG00 | Sorbs3 | 2893 | 10.2514996 | 9.06098324 | 11.5202868 | 7.14656464 | 9.94776217 | 9.59750527 |
| ENSMUSG00 | Ppp3cc | 3291 | 15.1846366 | 7.81876636 | 9.57639443 | 7.43349322 | 8.15553743 | 9.09154642 |
| ENSMUSG00 | Slc39a14 | 10975 | 1.14104568 | 1.03178259 | 0.73300551 | 0.94027014 | 0.69274989 | 0.44235861 |
| ENSMUSG00 | Fam160b2 | 4016 | 42.131306 | 26.0369969 | 39.6011532 | 33.6292561 | 33.0858274 | 36.6740667 |
| ENSMUSG00 | Hr | 6577 | 0.09066933 | 0 | 0.05376536 | 0 | 0.1318912 | 0.08708643 |
| ENSMUSG00 | Sftpc | 798 | 0 | 0 | 0 | 0 | 0 | 0 |
| ENSMUSG00 | Bmp1 | 4045 | 0.0098283 | 0.88153195 | 0.34968084 | 0.66901195 | 0.37844037 | 0.31691175 |
| ENSMUSG00 | Epb4.9 | 6685 | 36.3835497 | 32.5087291 | 24.1738127 | 29.2758429 | 29.5014125 | 22.1053097 |
| ENSMUSG00 | Xpo7 | 15093 | 5.70005034 | 5.7148169 | 5.25396584 | 5.86186796 | 6.60607821 | 4.5936615 |
| ENSMUSG00 | Fgf17 | 1682 | 0.04727168 | 0.14324151 | 0 | 0.2788743 | 0.15168364 | 0.3891744 |
| ENSMUSG00 | Dok2 | 2746 | 0 | 0.66681894 | 0.86922779 | 0.6569928 | 0.05574622 | 0.44696241 |
| ENSMUSG00 | Gfra2 | 3869 | 0 | 0 | 0 | 0 | 0 | 0 |
| ENSMUSG00 | Rb1 | 5984 | 35.4969138 | 30.7929403 | 29.6057799 | 39.0245642 | 27.9434214 | 46.6503898 |
| ENSMUSG00 | Rcbtb2 | 7949 | 4.58120772 | 5.69217124 | 7.75158752 | 7.17646401 | 9.18590311 | 8.01529215 |
| ENSMUSG00 | Itm2b | 1767 | 3652.44196 | 3744.79831 | 4511.21458 | 3511.27097 | 3054.1033 | 4784.25265 |
| ENSMUSG00 | Med4 | 1297 | 27.4334272 | 27.3440412 | 28.4227734 | 34.1624909 | 31.2767685 | 21.1972524 |
| ENSMUSG00 | Sucla2 | 3414 | 189.345083 | 162.230511 | 222.083835 | 178.011094 | 161.762878 | 249.602096 |
| ENSMUSG00 | Uchl3 | 1662 | 30.0438529 | 28.471171 | 31.1434165 | 30.4374407 | 18.4517761 | 41.1252984 |
| ENSMUSG00 | Gpc5 | 4983 | 2.13816351 | 4.15817181 | 0.11531687 | 2.48367274 | 1.99681784 | 1.89384411 |
| ENSMUSG00 | Trim52 | 3082 | 0 | 0 | 0 | 0 | 0 | 0 |
| ENSMUSG00 | Spry2 | 2083 | 3.11096655 | 4.16397484 | 4.45625878 | 3.82819579 | 2.74361694 | 2.73662967 |
| ENSMUSG00 | 4930449E01f | 592 | 0 | 0 | 0 | 0 | 0 | 0 |
| ENSMUSG00 | Rbm26 | 8072 | 12.1797948 | 13.556914 | 13.0546573 | 14.4784121 | 11.4101348 | 16.691863 |
| ENSMUSG00 | Rnf219 | 3450 | 4.85132096 | 3.44986992 | 5.82951843 | 8.4818951 | 3.69756347 | 5.06754892 |
| ENSMUSG00 | Ednrb | 6600 | 4.22251392 | 7.00893704 | 2.81284456 | 7.12347031 | 8.72860277 | 4.17797883 |
| ENSMUSG00 | Scel | 3129 | 8.42374036 | 7.93097402 | 4.35096441 | 9.46735652 | 9.88238535 | 3.69589231 |
| ENSMUSG00 | Fbxl3 | 7229 | 16.0418781 | 17.5708205 | 21.5597868 | 21.2678921 | 14.8159307 | 19.857024 |
| ENSMUSG00 | Cln5 | 2392 | 42.9631761 | 35.1930244 | 50.6325459 | 40.1397661 | 25.2358707 | 44.7431791 |
| ENSMUSG00 | Irg1 | 2618 | 0 | 0 | 0 | 0 | 0 | 0 |
| ENSMUSG00 | Dct | 2475 | 0 | 0 | 0 | 0 | 0 | 0 |
| ENSMUSG00 | Tgds | 2253 | 8.32871141 | 8.64062256 | 8.55392967 | 7.15875451 | 9.71607422 | 12.3722509 |
| ENSMUSG00 | Gpr180 | 2868 | 19.1014826 | 22.9339238 | 21.2532572 | 27.1873425 | 26.7586016 | 17.0514045 |
| ENSMUSG00 | Cldn10 | 2622 | 1.59203869 | 0.49619906 | 0.64060642 | 1.52750074 | 0.48652151 | 0.83217816 |
| ENSMUSG00 | Dnajc3 | 5178 | 36.8609621 | 37.9219297 | 41.1457867 | 48.4508412 | 38.639324 | 37.9464081 |
| ENSMUSG00 | Mbnl2 | 4752 | 56.4373888 | 64.3804127 | 65.4284015 | 77.8816352 | 65.29701 | 67.210864 |
| ENSMUSG00 | Nipbl | 8768 | 12.5641465 | 14.2778714 | 10.0875785 | 14.7982473 | 13.8739599 | 18.0918456 |
| ENSMUSG00 | Nup155 | 6367 | 2.67242746 | 2.91373963 | 2.83941476 | 1.44509523 | 3.63043007 | 3.50839164 |
| ENSMUSG00 | Gdnf | 3506 | 0 | 0 | 0 | 0 | 0 | 0 |
| ENSMUSG00 | Osmr | 12364 | 0.01929253 | 0.09353564 | 0 | 0 | 0.23936669 | 0 |
| ENSMUSG00 | Fyb | 6328 | 0 | 0.03045919 | 0 | 0 | 0 | 0 |
| ENSMUSG00 | C9 | 2673 | 0 | 0 | 0 | 0 | 0 | 0 |
| ENSMUSG00 | Dab2 | 7126 | 118.295707 | 65.6257958 | 150.023263 | 82.1390579 | 94.7274664 | 112.049546 |
| ENSMUSG00 | Ttc33 | 2168 | 32.8606186 | 34.6284487 | 34.0688559 | 33.9517393 | 33.0682925 | 47.6677159 |
| ENSMUSG00 | Heatr7b2 | 5190 | 0 | 0 | 0 | 0 | 0 | 0 |
| ENSMUSG00 | Gzme | 938 | 0 | 0 | 0 | 0 | 0 | 0 |
| ENSMUSG00 | Mcpt8 | 850 | 0 | 0 | 0 | 0 | 0 | 0 |
| ENSMUSG00 | Rab2b | 5219 | 6.28439767 | 8.14340716 | 5.97940241 | 7.12100131 | 8.06605864 | 7.7554253 |
| ENSMUSG00 | Mettl3 | 3506 | 7.72204285 | 9.62079564 | 9.55647376 | 11.5367861 | 6.08357818 | 6.17685783 |
| ENSMUSG00 | Dad1 | 3913 | 21.4677563 | 25.022962 | 27.0768765 | 22.9420206 | 12.1926045 | 28.5362569 |
| ENSMUSG00 | Lrp10 | 4194 | 56.5809402 | 58.974966 | 57.3021985 | 55.7404784 | 55.1751584 | 51.7593834 |
| ENSMUSG00 | Rem2 | 2051 | 0.50397001 | 0.16445885 | 0.2801677 | 0.29907107 | 0.19903023 | 0.79789291 |
| ENSMUSG00 | Haus4 | 1577 | 15.453462 | 14.9112136 | 9.61397093 | 8.58006766 | 15.9194527 | 7.43696568 |
| ENSMUSG00 | Jub | 3247 | 17.8881297 | 28.6566122 | 18.3641165 | 29.8368616 | 21.2151547 | 15.7415072 |
| ENSMUSG00 | 4931414P19f | 2578 | 1.63463186 | 1.73830067 | 1.18305941 | 2.58928559 | 2.79081419 | 0.71942415 |
| ENSMUSG00 | Slc7a8 | 4022 | 47.7915834 | 61.1375968 | 35.1790402 | 54.4729424 | 62.5206315 | 39.7184391 |
| ENSMUSG00 | C6 | 4118 | 0 | 0 | 0 | 0 | 0 | 0 |
| ENSMUSG00 | Fbxo4 | 2039 | 12.9073703 | 12.3597397 | 13.7656549 | 12.5110375 | 11.3864644 | 11.2897477 |
| ENSMUSG00 | Acin1 | 8657 | 3.46718122 | 2.95007674 | 2.91036739 | 4.63060546 | 3.78999187 | 3.99809602 |
| ENSMUSG00 | Oxct1 | 3846 | 400.490467 | 366.48508 | 340.524287 | 348.652372 | 310.722237 | 363.271814 |
| ENSMUSG00 | Gm5546 | 2775 | 0 | 0 | 0 | 0 | 0 | 0 |
| ENSMUSG00 | Drosha | 5849 | 7.48346453 | 8.04892357 | 6.51427401 | 10.1725579 | 8.86353186 | 7.60556277 |
| ENSMUSG00 | Psmb5 | 918 | 117.837323 | 206.078401 | 102.559832 | 36.6716204 | 35.7406968 | 66.6119185 |
| ENSMUSG00 | Pabpn1 | 2115 | 30.1690526 | 30.8940026 | 27.5660666 | 35.0243203 | 32.2322639 | 47.6758115 |
| ENSMUSG00 | 6030458C11l | 3965 | 7.00859543 | 8.17893457 | 9.0298847 | 5.19617842 | 9.291562 | 2.86848035 |
| ENSMUSG00 | Pdzd2 | 16835 | 0.93514525 | 0.36637152 | 0.96884349 | 0.80158507 | 0.65468947 | 1.28637146 |
| ENSMUSG00 | Slc22a17 | 3429 | 18.5038632 | 20.615197 | 14.6437135 | 17.5096302 | 19.0028235 | 16.4729594 |
| ENSMUSG00 | Golph3 | 3343 | 104.341785 | 94.484633 | 124.592871 | 121.942848 | 89.3839237 | 115.119701 |
| ENSMUSG00 | Zfr | 5984 | 30.5075479 | 31.0264642 | 36.4236778 | 40.0375628 | 31.8659035 | 34.3986802 |
| ENSMUSG00 | Efs | 3565 | 1.28243486 | 1.24352109 | 0.89271604 | 1.09308859 | 1.38837544 | 1.40007265 |
| ENSMUSG00 | Ngdn | 1187 | 22.607371 | 27.2393451 | 25.2475565 | 26.5067758 | 23.6862115 | 23.1616145 |
| ENSMUSG00 | Sub1 | 4179 | 48.1841371 | 39.4577662 | 41.197937 | 43.2570096 | 30.3545275 | 78.6258864 |
| ENSMUSG00 | Npr3 | 10083 | 0.2089696 | 0 | 0 | 0.20397466 | 0.20748601 | 0 |
| ENSMUSG00 | Jph4 | 7984 | 0 | 0 | 0 | 0 | 0 | 0 |
| ENSMUSG00 | Dhrs2 | 1257 | 0 | 0 | 0 | 0 | 0 | 0 |
| ENSMUSG00 | Dhrs4 | 1382 | 31.7295912 | 32.0080708 | 34.5426848 | 27.7534105 | 25.6608764 | 26.2286509 |
| ENSMUSG00 | Lrrc16b | 4540 | 0.23643127 | 1.15689916 | 0.04868044 | 0.32585106 | 0.15735006 | 0.26433571 |
| ENSMUSG00 | Cpne6 | 3393 | 0 | 0 | 0 | 0 | 0 | 0 |
| ENSMUSG00 | Dcaf11 | 3810 | 114.873513 | 115.280688 | 107.894263 | 108.302956 | 113.436589 | 79.3040561 |
| ENSMUSG00 | Fitm1 | 967 | 0 | 0 | 0.7770748 | 0 | 0.31660626 | 0.19743793 |
| ENSMUSG00 | Psme1 | 1816 | 46.1259898 | 43.5959936 | 47.0739892 | 38.2874999 | 37.2301476 | 63.9061619 |
| ENSMUSG00 | Fam158a | 2682 | 5.21772153 | 5.08455001 | 6.69125588 | 6.10784795 | 7.32481532 | 6.76272748 |
| ENSMUSG00 | Tgm1 | 2990 | 0 | 0 | 0 | 0.06033787 | 0 | 0 |
| ENSMUSG00 | Cideb | 1752 | 0.8622764 | 0.90762134 | 0.95871577 | 0.78260142 | 1.19411039 | 1.08974015 |
| ENSMUSG00 | Adcy4 | 3645 | 1.54877318 | 0.70065334 | 0.56995518 | 0.21777914 | 1.02192742 | 0.24693089 |
| ENSMUSG00 | Ripk3 | 1954 | 0.40691382 | 1.08505806 | 0.11310605 | 1.458793 | 2.14133205 | 1.35396111 |
| ENSMUSG00 | Sdr39u1 | 1673 | 66.0373463 | 66.936815 | 75.0083875 | 56.8728939 | 56.9433614 | 63.3202025 |
| ENSMUSG00 | Cma1 | 1069 | 0 | 0 | 0 | 0 | 0 | 0 |
| ENSMUSG00 | Mcpt2 | 966 | 0 | 0 | 0 | 0 | 0 | 0 |
| ENSMUSG00 | Mcpt1 | 1023 | 0 | 0 | 0 | 0 | 0 | 0 |
| ENSMUSG00 | Zfp187 | 5660 | 2.78850279 | 2.78391636 | 4.44361284 | 3.50620574 | 3.04715813 | 3.93217414 |
| ENSMUSG00 | Atp12a | 3928 | 0.06072629 | 0.08587197 | 0 | 0.56952311 | 0.80540614 | 0.3055204 |
| ENSMUSG00 | Sema5a | 12502 | 5.21826456 | 5.16090594 | 1.95871229 | 5.40566864 | 4.93448155 | 1.79547497 |
| ENSMUSG00 | Cct5 | 2796 | 61.7518068 | 65.9202937 | 59.2203519 | 67.1828054 | 50.5518816 | 43.5360211 |
| ENSMUSG00 | Cmbl | 1255 | 0.95033021 | 1.95817414 | 0.98617658 | 0.54626203 | 0.4879016 | 0.34772448 |
| ENSMUSG00 | Ropn1l | 1000 | 0.63608769 | 1.34922038 | 0.6188258 | 0.14432818 | 1.37771215 | 0.60004206 |
| ENSMUSG00 | Ankrd33b | 15204 | 0.02876284 | 0.05704788 | 0 | 0 | 0.16109353 | 0.01255739 |
| ENSMUSG00 | Ctnnd2 | 5891 | 0 | 0.13087474 | 0 | 0.20212315 | 0 | 0 |
| ENSMUSG00 | Tars | 2617 | 31.7648872 | 37.0282519 | 29.507306 | 33.90361 | 35.5254325 | 28.9421755 |
| ENSMUSG00 | Slc45a2 | 2976 | 0 | 0 | 0 | 0 | 0 | 0 |
| ENSMUSG00 | Amacr | 2570 | 9.32784231 | 11.5497464 | 12.8477731 | 13.0569266 | 7.40577361 | 11.3874272 |
| ENSMUSG00 | Skor1 | 4575 | 0 | 0 | 0 | 0 | 0 | 0 |
| ENSMUSG00 | Rai14 | 4989 | 6.45458794 | 6.84790288 | 4.52738861 | 6.37890625 | 6.61737125 | 6.8500948 |
| ENSMUSG00 | Brix1 | 2720 | 35.0052852 | 36.1575472 | 34.6139432 | 44.8637774 | 28.9837319 | 46.3869416 |
| ENSMUSG00 | Rad1 | 7320 | 1.89001465 | 2.12625967 | 1.89005148 | 2.29702633 | 1.39416327 | 2.08658442 |
| ENSMUSG00 | Ttc23l | 2171 | 0 | 0 | 0 | 0 | 0 | 0 |
| ENSMUSG00 | 1110020G09l | 3800 | 26.2490791 | 27.5169946 | 25.4974842 | 28.3433955 | 27.5542429 | 26.8726972 |
| ENSMUSG00 | Mtdh | 9938 | 10.8689516 | 11.2296035 | 10.154237 | 12.0902806 | 9.49363744 | 9.86091556 |
| ENSMUSG00 | Laptm4b | 1890 | 219.265148 | 217.017458 | 290.024897 | 218.115005 | 191.90237 | 203.001244 |
| ENSMUSG00 | Sdc2 | 3720 | 39.9157849 | 42.5776439 | 41.2075246 | 45.8591147 | 42.0556098 | 42.9208974 |
| ENSMUSG00 | Dnahc5 | 15551 | 0 | 0 | 0 | 0 | 0 | 0 |
| ENSMUSG00 | Trio | 12310 | 0.43921571 | 0.92771623 | 0.41293355 | 1.10209982 | 1.01555338 | 0.81314319 |
| ENSMUSG00 | Ank | 3596 | 68.9087625 | 42.5584372 | 125.91871 | 46.4972732 | 52.3034542 | 86.0258505 |
| ENSMUSG00 | 11-Mar | 3572 | 0 | 0 | 0 | 0 | 0 | 0 |
| ENSMUSG00 | Fam134b | 3437 | 36.366375 | 28.0538466 | 38.7746746 | 24.2926538 | 20.2502114 | 41.5666454 |
| ENSMUSG00 | Myo10 | 10563 | 9.88335623 | 11.1171409 | 8.64955122 | 12.8642412 | 12.5645748 | 6.88902182 |
| ENSMUSG00 | Rnf19a | 4266 | 41.5820333 | 37.5348212 | 31.478012 | 44.6500499 | 52.5215228 | 45.093303 |
| ENSMUSG00 | Pabpc1 | 5945 | 6.90120368 | 8.97264785 | 7.68048845 | 8.98864726 | 9.27830316 | 10.3914311 |
| ENSMUSG00 | Ywhaz | 5198 | 40.6885641 | 42.9487854 | 47.4417242 | 43.3359377 | 30.2153107 | 47.8276908 |
| ENSMUSG00 | Grhl2 | 5635 | 9.30567503 | 11.108108 | 11.8917469 | 11.493748 | 15.2128325 | 7.5555833 |
| ENSMUSG00 | 4930447A16l | 583 | 0 | 0.4132628 | 0 | 0.4332321 | 0 | 0 |
| ENSMUSG00 | Rrm2b | 5195 | 8.83115005 | 12.2529914 | 13.5030462 | 14.7453668 | 11.4625179 | 12.9049208 |
| ENSMUSG00 | Atp6v1c1 | 2084 | 99.2170125 | 100.881693 | 105.265714 | 89.2182224 | 78.2779863 | 94.8984689 |
| ENSMUSG00 | Baalc | 2492 | 0 | 0 | 0 | 0 | 0 | 0 |
| ENSMUSG00 | Fzd6 | 4074 | 22.0050586 | 16.7954708 | 18.4228595 | 16.1368398 | 19.6139677 | 12.8540273 |
| ENSMUSG00 | Slc25a32 | 2938 | 28.0236896 | 26.3073701 | 35.5811296 | 39.4962067 | 24.436389 | 25.6686105 |
| ENSMUSG00 | Dcaf13 | 1531 | 88.5474778 | 83.5945071 | 75.98906 | 71.0328427 | 51.3263349 | 103.362153 |
| ENSMUSG00 | Dcstamp | 1742 | 0 | 0 | 0.35523869 | 0 | 0 | 0 |
| ENSMUSG00 | Dpys | 2889 | 0.0825659 | 0 | 0.1224004 | 0.69940966 | 0 | 0 |
| ENSMUSG00 | Lrp12 | 4567 | 4.29153753 | 5.99297113 | 4.82958609 | 5.68053208 | 4.5808658 | 4.52686152 |
| ENSMUSG00 | Zfpm2 | 6359 | 0 | 0.26521866 | 0 | 0 | 0.2407283 | 0 |
| ENSMUSG00 | Oxr1 | 5060 | 39.3940671 | 28.0262251 | 38.174319 | 38.1285953 | 30.5452356 | 34.1741527 |
| ENSMUSG00 | Angpt1 | 4247 | 10.9615417 | 32.8806945 | 7.01484394 | 28.8350504 | 18.0100159 | 19.3561956 |
| ENSMUSG00 | Csmd3 | 18781 | 0.00211679 | 0 | 0 | 0 | 0 | 0 |
| ENSMUSG00 | Eif3h | 1484 | 33.8886003 | 34.353946 | 34.0449505 | 32.9698466 | 27.8513237 | 29.0941738 |
| ENSMUSG00 | Utp23 | 1295 | 13.6611497 | 8.37216176 | 13.41415 | 18.55648 | 10.9145221 | 10.4254413 |
| ENSMUSG00 | Rad21 | 3590 | 18.4713486 | 28.2810121 | 18.6164865 | 28.1017818 | 19.5008879 | 28.4674299 |
| ENSMUSG00 | Slc30a8 | 1963 | 0 | 0 | 0 | 0 | 0 | 0 |
| ENSMUSG00 | Cdh10 | 6877 | 0 | 0 | 0 | 0 | 0 | 0 |
| ENSMUSG00 | Shcbp1 | 2119 | 1.2382547 | 1.93291533 | 0 | 0 | 0.04816081 | 0 |
| ENSMUSG00 | Hrsp12 | 2742 | 60.8946091 | 80.3458838 | 54.5833116 | 76.4144539 | 49.7981698 | 67.5897789 |
| ENSMUSG00 | Matn2 | 3813 | 0.0104263 | 0.02527482 | 0.04636962 | 0 | 0 | 0.20028584 |
| ENSMUSG00 | Pop1 | 3869 | 0.67817568 | 0.87181467 | 0.43413544 | 2.09833549 | 0.738557 | 0.43002145 |
| ENSMUSG00 | Stk3 | 4372 | 16.8587971 | 14.2399093 | 16.4796441 | 17.7439149 | 15.7327435 | 18.3411344 |
| ENSMUSG00 | Osr2 | 1719 | 1.66515102 | 0.50457008 | 0.35999174 | 0 | 0.62335887 | 0.36493118 |
| ENSMUSG00 | Khdrbs3 | 1878 | 14.2679414 | 9.26267605 | 16.2638304 | 10.5287329 | 7.82513112 | 20.6229967 |
| ENSMUSG00 | Zfat | 7545 | 0.43206752 | 0.38319238 | 0.57998442 | 0.34910394 | 1.16998847 | 0.4988602 |
| ENSMUSG00 | Eif3e | 1528 | 47.8730917 | 49.7632236 | 46.1978687 | 51.5019235 | 38.3365703 | 53.0855871 |
| ENSMUSG00 | Ttc35 | 1234 | 96.9724444 | 100.785419 | 111.829946 | 110.556086 | 85.9261013 | 135.621707 |
| ENSMUSG00 | Eny2 | 2424 | 23.0594908 | 39.300576 | 32.2578632 | 46.1147582 | 27.4919335 | 57.103463 |
| ENSMUSG00 | Ebag9 | 1554 | 40.7787876 | 40.9305686 | 32.5113942 | 33.458318 | 34.3459389 | 35.9098532 |
| ENSMUSG00 | Sybu | 4753 | 0.2174716 | 0.74008211 | 0.65098443 | 0.12146281 | 0.5904588 | 0.08607608 |
| ENSMUSG00 | Kcnv1 | 4493 | 0 | 0 | 0 | 0 | 0 | 0 |
| ENSMUSG00 | Myc | 3100 | 0.24366262 | 1.69429748 | 0.15684525 | 1.78082349 | 0.55964412 | 0.19356196 |
| ENSMUSG00 | A1bg | 1816 | 0 | 0 | 0 | 0 | 0 | 0 |
| ENSMUSG00 | E430025E21f | 4060 | 21.5130026 | 14.8238587 | 21.6327739 | 17.3833692 | 21.8056064 | 20.0125986 |
| ENSMUSG00 | Sqle | 6583 | 14.9347263 | 11.0675833 | 15.8396093 | 5.80996008 | 10.8129719 | 15.3381003 |
| ENSMUSG00 | Mtss1 | 5883 | 3.34505573 | 3.31727164 | 2.03615493 | 2.33064371 | 3.49543251 | 2.64725803 |
| ENSMUSG00 | Ndufb9 | 647 | 225.014791 | 254.412498 | 360.651514 | 239.691851 | 154.104387 | 402.037457 |
| ENSMUSG00 | Klhl38 | 2208 | 0 | 0 | 0.10009475 | 0 | 0 | 0 |
| ENSMUSG00 | Fbxo32 | 2713 | 3.60480951 | 3.71211442 | 3.77988484 | 4.80118617 | 3.31022563 | 6.91667962 |
| ENSMUSG00 | Wdyhv1 | 1675 | 14.7154614 | 24.6829656 | 16.7571166 | 10.4691783 | 18.5218128 | 12.0496914 |
| ENSMUSG00 | Atad2 | 5671 | 0.63793841 | 1.31703377 | 1.06782798 | 2.1505433 | 1.01674845 | 0.21642722 |
| ENSMUSG00 | Zhx1 | 7757 | 23.7548862 | 23.8167873 | 24.7648846 | 25.9416736 | 31.7064756 | 19.6833093 |
| ENSMUSG00 | 9130401M01 | 1598 | 5.59761146 | 5.09606303 | 5.91939575 | 7.1351227 | 6.35434843 | 6.40049419 |
| ENSMUSG00 | Wdr67 | 6334 | 1.37455798 | 1.89428862 | 1.1095821 | 2.29001925 | 0.89421025 | 0.70619527 |
| ENSMUSG00 | Derl1 | 3137 | 34.128629 | 29.6921877 | 31.1540564 | 20.1516551 | 26.188226 | 22.0927186 |
| ENSMUSG00 | Slc22a22 | 2757 | 0.519114 | 0 | 0 | 0 | 0 | 0 |
| ENSMUSG00 | Has2 | 4233 | 0 | 0 | 0 | 0 | 0 | 0 |
| ENSMUSG00 | Mtbp | 5598 | 2.05950149 | 1.82485276 | 1.60288927 | 1.30199589 | 1.51310797 | 2.2266006 |
| ENSMUSG00 | Mrpl13 | 8171 | 7.76523643 | 7.28900288 | 7.88178745 | 7.29942719 | 5.53290527 | 7.74745291 |
| ENSMUSG00 | Col14a1 | 6961 | 1.76475269 | 0.18690331 | 2.266924 | 0.93820573 | 0.59375613 | 1.63389234 |
| ENSMUSG00 | Sla | 8555 | 0 | 0 | 0 | 0 | 0.17893528 | 0.251864 |
| ENSMUSG00 | Lrrc6 | 2072 | 0 | 0.02325601 | 0.5333234 | 0.57466577 | 0 | 1.13205549 |
| ENSMUSG00 | Adcy8 | 5014 | 0 | 0 | 0 | 0 | 0 | 0 |
| ENSMUSG00 | Asap1 | 8424 | 2.41628751 | 3.77529105 | 2.62356618 | 3.86348577 | 3.69492987 | 3.57121642 |
| ENSMUSG00 | Fam49b | 4591 | 8.91056185 | 12.4795643 | 10.5329593 | 8.77097837 | 10.3141966 | 10.5153804 |
| ENSMUSG00 | Wnt7b | 4407 | 4.33909374 | 3.06153932 | 1.39415843 | 1.80123662 | 3.31144622 | 0.01856681 |
| ENSMUSG00 | Ppara | 3824 | 2.59907692 | 1.05848879 | 1.38708712 | 1.59463011 | 1.24096573 | 1.61907508 |
| ENSMUSG00 | Gtse1 | 2767 | 0 | 0.76624628 | 0 | 0.62592632 | 0.71920082 | 0.05914269 |
| ENSMUSG00 | Trmu | 2418 | 20.6669309 | 14.6073871 | 14.587705 | 17.6828878 | 18.4859823 | 16.671595 |
| ENSMUSG00 | Brd1 | 8315 | 9.59582073 | 11.0570934 | 10.9242077 | 10.9005527 | 10.6900718 | 13.2683002 |
| ENSMUSG00 | Ttll8 | 3415 | 0 | 0 | 0 | 0 | 0 | 0 |
| ENSMUSG00 | Tef | 8068 | 8.09103855 | 12.6916447 | 8.88638936 | 9.68240288 | 10.6884699 | 4.30011665 |
| ENSMUSG00 | Zc3h7b | 5121 | 1.97185941 | 1.8160483 | 1.47598421 | 1.35985835 | 1.44480072 | 1.4114 |
| ENSMUSG00 | Rangap1 | 4097 | 9.05342038 | 11.9986246 | 9.82862558 | 10.5771383 | 10.0757476 | 7.20977162 |
| ENSMUSG00 | L3mbtl2 | 6009 | 3.6321782 | 5.13218888 | 3.67796996 | 5.39219104 | 6.06304416 | 2.46465787 |
| ENSMUSG00 | Rbx1 | 1746 | 71.5188799 | 73.54918 | 92.4289398 | 71.1514773 | 58.6540299 | 92.3838583 |
| ENSMUSG00 | Xpnpep3 | 5928 | 3.80252319 | 3.01571694 | 2.78871277 | 4.41895485 | 4.27802105 | 1.3526896 |
| ENSMUSG00 | St13 | 3572 | 54.6248316 | 63.012002 | 61.6870626 | 61.7899962 | 52.1691838 | 69.2250502 |
| ENSMUSG00 | Slc25a17 | 1658 | 76.3218905 | 76.5519232 | 76.1402073 | 56.7998409 | 65.2448231 | 78.550303 |
| ENSMUSG00 | Adsl | 4573 | 17.8304156 | 17.2598715 | 19.9019451 | 20.2700136 | 17.1054962 | 12.0001256 |
| ENSMUSG00 | Fam83f | 5305 | 3.16245293 | 3.4970368 | 3.31617974 | 3.15590361 | 2.14493531 | 2.68890412 |
| ENSMUSG00 | Smcr7l | 5607 | 7.91280832 | 7.97521283 | 8.2065487 | 9.00924957 | 7.26218083 | 5.3994738 |
| ENSMUSG00 | Tab1 | 2920 | 14.2820202 | 12.2941436 | 11.0353231 | 13.0364921 | 14.469123 | 9.71425508 |
| ENSMUSG00 | Syngr1 | 4949 | 4.15307808 | 3.48570343 | 3.75121723 | 1.95392765 | 5.16553128 | 3.8522879 |
| ENSMUSG00 | Cacna1i | 10068 | 0 | 0 | 0 | 0 | 0 | 0 |
| ENSMUSG00 | Deptor | 9058 | 41.0414549 | 35.7062707 | 51.5021253 | 38.9341911 | 42.8468331 | 52.3481567 |
| ENSMUSG00 | Dnalc4 | 2547 | 11.4724296 | 12.6945829 | 13.0679183 | 10.6956983 | 10.117126 | 12.4326094 |
| ENSMUSG00 | Nptxr | 5639 | 0 | 0 | 0 | 0.00639866 | 0.07239067 | 0 |
| ENSMUSG00 | Dscc1 | 2017 | 0.13797143 | 0.71670464 | 0 | 0.59033588 | 0.83483907 | 0.31101473 |
| ENSMUSG00 | Enpp2 | 4081 | 103.114864 | 48.3989773 | 156.66147 | 65.4003635 | 49.3509202 | 89.8773214 |
| ENSMUSG00 | Josd1 | 3390 | 11.7976441 | 14.6833613 | 9.81828548 | 10.5478779 | 7.9324189 | 7.06405111 |
| ENSMUSG00 | Tomm22 | 1109 | 60.7265861 | 49.2292506 | 69.4315695 | 51.9269099 | 45.1369474 | 89.6447791 |
| ENSMUSG00 | Cby1 | 1114 | 14.1677969 | 20.459773 | 17.2204667 | 23.45009 | 9.71058499 | 18.3381551 |
| ENSMUSG00 | Dmc1 | 3166 | 0.07534204 | 0 | 0.05584566 | 0 | 0 | 0 |
| ENSMUSG00 | Ribc2 | 1645 | 0 | 0 | 0 | 0 | 0.96159128 | 0.34818688 |
| ENSMUSG00 | Smc1b | 4013 | 0.11888008 | 0 | 0.15420528 | 0 | 0 | 0.16991427 |
| ENSMUSG00 | Csnk1e | 7727 | 3.77129121 | 4.29668159 | 3.44942556 | 4.48282163 | 3.91596232 | 3.99218545 |
| ENSMUSG00 | Fam118a | 2534 | 42.3284477 | 23.0853752 | 58.5229611 | 38.8871601 | 24.52649 | 46.92874 |
| ENSMUSG00 | Upk3a | 1100 | 0.68668557 | 17.5223426 | 1.2456883 | 0.72164089 | 0.3247133 | 1.01660019 |
| ENSMUSG00 | Sh3bp1 | 3706 | 9.1289568 | 14.406524 | 13.9308561 | 14.1271038 | 12.6120238 | 11.4220831 |
| ENSMUSG00 | Samm50 | 1658 | 76.5376923 | 72.9481121 | 87.3106368 | 62.2186763 | 71.5230986 | 61.6722693 |
| ENSMUSG00 | Parvb | 4521 | 0.00879352 | 0.06395016 | 0 | 0.03990494 | 0 | 0.04826302 |
| ENSMUSG00 | Parvg | 3075 | 0 | 0 | 0 | 0 | 0 | 0 |
| ENSMUSG00 | C1qtnf6 | 4029 | 0.029602 | 0.07175941 | 0.01097092 | 0 | 0 | 0.11508287 |
| ENSMUSG00 | Efcab6 | 6828 | 0 | 0 | 0 | 0 | 0 | 0 |
| ENSMUSG00 | Ttll1 | 3389 | 10.9095889 | 8.09031737 | 13.0688246 | 10.5296967 | 12.0752887 | 8.41819789 |
| ENSMUSG00 | Myh9 | 12053 | 5.14878496 | 6.44058395 | 5.25889346 | 7.04697028 | 6.65506204 | 6.11433463 |
| ENSMUSG00 | Cyp2d26 | 1636 | 0 | 2.82756629 | 0.43229186 | 0 | 1.49710638 | 0 |
| ENSMUSG00 | Adamts20 | 8956 | 0.2352658 | 0.39814613 | 0 | 0.04431694 | 0.05127706 | 0 |
| ENSMUSG00 | Ndufa6 | 557 | 118.053079 | 101.303992 | 147.207215 | 96.5210886 | 72.2797316 | 268.045557 |
| ENSMUSG00 | Twf1 | 3369 | 23.5063571 | 21.2540971 | 19.8508129 | 24.1725065 | 21.7948812 | 25.4692832 |
| ENSMUSG00 | 1500032L24f | 1022 | 94.3317419 | 98.5417457 | 130.918766 | 95.783353 | 55.0705406 | 163.06071 |
| ENSMUSG00 | Naga | 1903 | 74.1210956 | 89.8399878 | 84.5943837 | 78.7813426 | 82.3983462 | 58.9350061 |
| ENSMUSG00 | Nell2 | 4002 | 0.09933903 | 0.39733948 | 0.05522469 | 0.18933607 | 0.36975635 | 0.08859828 |
| ENSMUSG00 | Wbp2nl | 1312 | 0 | 0 | 0 | 0 | 0 | 0 |
| ENSMUSG00 | 3-Sep | 4960 | 0.7614457 | 0.61204554 | 0.69510963 | 1.23668298 | 0.88473152 | 0.35742975 |
| ENSMUSG00 | Slc38a2 | 4610 | 52.4841333 | 60.9490551 | 64.231702 | 54.6005082 | 72.720887 | 64.8438278 |
| ENSMUSG00 | Srebf2 | 3719 | 36.1423177 | 24.1774405 | 30.402881 | 20.4131814 | 26.3981573 | 24.3117583 |
| ENSMUSG00 | Slc38a4 | 4338 | 0 | 0.56650727 | 0.10189452 | 0 | 0.38816745 | 0 |
| ENSMUSG00 | Rpap3 | 2243 | 50.1418878 | 36.0055627 | 69.485376 | 40.1036723 | 46.4992921 | 52.1051401 |
| ENSMUSG00 | Endou | 2707 | 0.14686177 | 0.21360817 | 0.32657439 | 0 | 0.28274682 | 0.50377982 |
| ENSMUSG00 | Rapgef3 | 6622 | 29.7775873 | 36.8638653 | 25.9190209 | 33.4012282 | 33.4961727 | 20.0173281 |
| ENSMUSG00 | Xrcc6 | 3918 | 11.0803943 | 12.0404612 | 11.5863432 | 14.127069 | 14.3129114 | 9.18900557 |
| ENSMUSG00 | Pppde2 | 4622 | 16.1275478 | 16.9830624 | 12.9870409 | 18.6187097 | 16.1513604 | 12.6164385 |
| ENSMUSG00 | Pmm1 | 2978 | 8.57052333 | 6.01926007 | 7.43624021 | 6.02175155 | 6.37401337 | 6.31035139 |
| ENSMUSG00 | Hdac7 | 6577 | 7.81569656 | 10.9531292 | 2.93693255 | 10.5881627 | 8.4720697 | 7.92486477 |
| ENSMUSG00 | Polr3h | 2508 | 4.85054906 | 3.8234059 | 3.08425938 | 3.66862893 | 3.03147129 | 2.28376166 |
| ENSMUSG00 | Aco2 | 3938 | 276.138054 | 228.365559 | 229.146934 | 201.62148 | 261.325533 | 156.46556 |
| ENSMUSG00 | Vdr | 5730 | 92.5407677 | 81.4460196 | 51.7153849 | 74.0532013 | 74.1173693 | 39.6743662 |
| ENSMUSG00 | Col2a1 | 6556 | 0 | 0 | 0 | 0 | 0 | 0 |
| ENSMUSG00 | Hoxc10 | 1907 | 1.9596304 | 2.88057389 | 1.22847283 | 4.86265624 | 1.87301852 | 4.69118074 |
| ENSMUSG00 | Hoxc5 | 2921 | 0.02722046 | 0.03299311 | 0.03026487 | 0.20999478 | 0.06987522 | 0 |
| ENSMUSG00 | Gtsf1 | 1472 | 0 | 0 | 0 | 0 | 0 | 0 |
| ENSMUSG00 | Nckap1l | 5414 | 1.60813636 | 1.4151541 | 2.81670407 | 1.89940574 | 1.48913324 | 2.41814312 |
| ENSMUSG00 | Pde1b | 3779 | 0.62068625 | 1.41537314 | 0.45617144 | 1.20305309 | 1.75533973 | 1.47957159 |
| ENSMUSG00 | Ppp1r1a | 1288 | 34.7551794 | 47.3633818 | 21.0370573 | 30.7873967 | 29.8710306 | 26.7240642 |
| ENSMUSG00 | Glycam1 | 621 | 0 | 0 | 0 | 0 | 0 | 0 |
| ENSMUSG00 | Shisa9 | 1270 | 0 | 0 | 0 | 0 | 0 | 0 |
| ENSMUSG00 | Tnfrsf17 | 812 | 0.14687985 | 0 | 0 | 0 | 0 | 0 |
| ENSMUSG00 | Txndc11 | 9040 | 8.35128955 | 5.31969793 | 8.13626845 | 8.36592537 | 8.51756649 | 4.92995139 |
| ENSMUSG00 | Litaf | 2863 | 159.410729 | 111.200777 | 169.288581 | 84.7669687 | 119.322943 | 111.680264 |
| ENSMUSG00 | Prm1 | 467 | 0 | 0 | 0 | 0 | 0 | 0 |
| ENSMUSG00 | Nubp1 | 3151 | 8.07474057 | 4.32775726 | 6.60713045 | 5.25600077 | 7.07665066 | 3.86051701 |
| ENSMUSG00 | Ciita | 6091 | 0.00652692 | 0.06328871 | 0.52249757 | 0.37320125 | 0.36860295 | 0.28210512 |
| ENSMUSG00 | Emp2 | 3615 | 14.7364713 | 12.8097292 | 18.8178801 | 7.34616453 | 14.3833685 | 14.1918662 |
| ENSMUSG00 | 1810013L24f | 4062 | 26.6308869 | 30.9617464 | 27.3350639 | 24.5521347 | 33.4020515 | 24.0784974 |
| ENSMUSG00 | Bcl6 | 5350 | 2.87577027 | 2.43183914 | 1.51195089 | 4.29612381 | 2.1268938 | 1.4631442 |
| ENSMUSG00 | Trp63 | 8961 | 0.04880151 | 0.37641457 | 0.34035567 | 0 | 0.6206757 | 0.44742461 |
| ENSMUSG00 | Cldn1 | 3243 | 0.40454236 | 0 | 0 | 0 | 1.25874501 | 0 |
| ENSMUSG00 | Il1rap | 8400 | 2.18181863 | 2.00203194 | 2.76261519 | 1.51630497 | 2.39338001 | 1.62998439 |
| ENSMUSG00 | Anks3 | 2537 | 33.3149986 | 25.4512544 | 18.224331 | 24.9601845 | 23.9745526 | 18.5235331 |
| ENSMUSG00 | Nudt16l1 | 1065 | 25.1971355 | 31.6718399 | 25.4419998 | 17.8885629 | 27.3578034 | 23.8172906 |
| ENSMUSG00 | Mgrn1 | 3378 | 124.162321 | 115.345048 | 99.6307971 | 92.394815 | 111.871326 | 67.4195494 |
| ENSMUSG00 | 4930562C15l | 4365 | 0 | 0 | 0 | 0 | 0 | 0 |
| ENSMUSG00 | Srl | 8373 | 0.41308095 | 0.04028486 | 0.9396785 | 0.08187733 | 0.56675659 | 0.36809199 |
| ENSMUSG00 | Crebbp | 7477 | 5.23196373 | 7.50798517 | 7.19454901 | 9.41018949 | 8.03237185 | 5.16164431 |
| ENSMUSG00 | Fgf12 | 3074 | 3.18147307 | 1.47349563 | 3.29285037 | 3.40396647 | 2.02511967 | 5.39459358 |
| ENSMUSG00 | Hrasls | 3280 | 0 | 0 | 0 | 0 | 0 | 0 |
| ENSMUSG00 | Zfp251 | 3493 | 9.45800295 | 11.0361162 | 7.9722763 | 7.50977566 | 10.576323 | 6.49656452 |
| ENSMUSG00 | Hes1 | 1901 | 40.9056917 | 54.7515597 | 25.2283007 | 35.2468999 | 36.424404 | 46.1416306 |
| ENSMUSG00 | Zfp263 | 4098 | 10.6907124 | 13.0284475 | 8.40244893 | 11.8336427 | 14.3317127 | 11.4076944 |
| ENSMUSG00 | Atp13a3 | 9494 | 3.84825012 | 5.36983946 | 3.58959564 | 5.79577817 | 4.88550407 | 5.98123014 |
| ENSMUSG00 | Mefv | 4088 | 0 | 0 | 0 | 0 | 0 | 0 |
| ENSMUSG00 | Glyr1 | 3332 | 72.9725447 | 48.6781405 | 49.4418574 | 45.0592399 | 67.3051986 | 37.4576078 |
| ENSMUSG00 | Tmem44 | 9673 | 1.27819233 | 0.84188036 | 0.24675897 | 1.06310169 | 1.13942905 | 0.73029376 |
| ENSMUSG00 | Lsg1 | 6033 | 7.84170758 | 8.12292585 | 8.46231206 | 10.2689989 | 7.44293233 | 6.12130972 |
| ENSMUSG00 | Rogdi | 1449 | 115.617388 | 144.127012 | 122.843907 | 121.966773 | 104.376934 | 137.257881 |
| ENSMUSG00 | 12-Sep | 1612 | 0 | 0 | 0 | 0 | 0 | 0 |
| ENSMUSG00 | 4930451G09l | 2200 | 1.12038172 | 0.74469956 | 0.84385337 | 1.52528643 | 1.925086 | 0.57028791 |
| ENSMUSG00 | Fam86 | 3146 | 11.208872 | 11.931735 | 11.4649408 | 12.6734454 | 8.61252561 | 10.5162547 |
| ENSMUSG00 | Ercc4 | 6943 | 6.72802673 | 8.07849901 | 6.06716929 | 7.15092802 | 7.64329985 | 7.42461014 |
| ENSMUSG00 | Gpt | 1769 | 1.82034704 | 2.88737302 | 2.47370405 | 3.12071951 | 1.81721972 | 2.35897106 |
| ENSMUSG00 | Apod | 2790 | 0 | 0 | 0 | 0 | 0 | 0 |
| ENSMUSG00 | Adck5 | 3486 | 9.38576032 | 5.9438239 | 6.37794809 | 7.93887784 | 11.8271118 | 5.93063011 |
| ENSMUSG00 | Cyc1 | 1187 | 422.138228 | 345.586505 | 429.171222 | 314.311997 | 314.239939 | 373.572944 |
| ENSMUSG00 | Sharpin | 3850 | 10.8010994 | 10.4883736 | 10.0343924 | 10.9277049 | 9.70163821 | 12.2629612 |
| ENSMUSG00 | Maf1 | 2651 | 63.7047458 | 64.7635961 | 72.2469203 | 59.6966606 | 70.7940439 | 62.0289702 |
| ENSMUSG00 | Fam203a | 1953 | 6.31039373 | 5.87218291 | 5.97505712 | 7.8704306 | 5.43445272 | 4.28741129 |
| ENSMUSG00 | Dgat1 | 3110 | 23.6231923 | 18.8097559 | 19.0025287 | 14.1775751 | 20.6894726 | 15.6894951 |
| ENSMUSG00 | Hsf1 | 2433 | 16.8956707 | 19.7459444 | 16.7323869 | 21.1924544 | 18.6027108 | 16.9948019 |
| ENSMUSG00 | Bop1 | 2697 | 18.3226037 | 20.7074848 | 15.4550753 | 14.194679 | 16.8763542 | 10.3657787 |
| ENSMUSG00 | Heatr7a | 14972 | 10.7646753 | 9.8130151 | 11.4431167 | 9.54588421 | 11.7171173 | 6.41059682 |
| ENSMUSG00 | Fbxl6 | 1703 | 15.08047 | 13.6665012 | 16.1182293 | 11.3352283 | 14.8015441 | 12.9566548 |
| ENSMUSG00 | Slc52a2 | 2695 | 7.67081628 | 6.9910571 | 5.88811193 | 5.07424671 | 5.05530328 | 4.51372509 |
| ENSMUSG00 | Gpaa1 | 2742 | 39.0305447 | 42.8968291 | 33.2239236 | 36.0557265 | 39.0978904 | 25.3847117 |
| ENSMUSG00 | Oplah | 5583 | 21.9392147 | 19.3160053 | 18.9379919 | 17.2234728 | 22.6113655 | 14.558199 |
| ENSMUSG00 | Grina | 1686 | 91.8431771 | 99.0308553 | 86.5160628 | 87.893806 | 83.1070154 | 100.961461 |
| ENSMUSG00 | Plec | 25477 | 4.58302965 | 5.63250087 | 6.69004619 | 5.080123 | 6.05859351 | 4.58306436 |
| ENSMUSG00 | Scrib | 6070 | 3.70046894 | 4.92978263 | 5.14110398 | 6.45553548 | 6.0945836 | 6.43897167 |
| ENSMUSG00 | Tsta3 | 1300 | 31.712641 | 27.1697401 | 29.5812334 | 22.6484218 | 23.7076392 | 28.9950396 |
| ENSMUSG00 | Pycrl | 1347 | 31.7276478 | 31.9096558 | 42.7907968 | 29.3585158 | 24.2441578 | 31.4863133 |
| ENSMUSG00 | Naprt1 | 1877 | 12.0092474 | 16.4301135 | 15.9663557 | 9.91919818 | 16.3110418 | 12.4966381 |
| ENSMUSG00 | Gsdmd | 1765 | 1.23883934 | 0.0546022 | 0.12521769 | 0.36797553 | 0.98294435 | 0 |
| ENSMUSG00 | Ly6h | 4896 | 0 | 0 | 0 | 0 | 0 | 0 |
| ENSMUSG00 | Gpihbp1 | 792 | 0.05019631 | 1.15598788 | 0.22324163 | 0.45558137 | 0.51541794 | 1.37750703 |
| ENSMUSG00 | Rhpn1 | 2966 | 0.53614943 | 1.13723903 | 0.29805693 | 0.75424368 | 0.70535449 | 0.13793648 |
| ENSMUSG00 | Ly6g | 334 | 0 | 0 | 0 | 0 | 0 | 0 |
| ENSMUSG00 | Ly6f | 873 | 1.00185632 | 2.53903361 | 0 | 2.43853451 | 0.17534837 | 0 |
| ENSMUSG00 | Ly6c2 | 793 | 0.05013302 | 0.85070642 | 0.11148006 | 0.40950618 | 0.77215197 | 0.58470223 |
| ENSMUSG00 | Ly6i | 875 | 0 | 0 | 0 | 0.45360285 | 0.11663172 | 0.62342033 |
| ENSMUSG00 | Ly6e | 2559 | 278.738895 | 224.644101 | 125.229919 | 154.000817 | 177.385947 | 133.250309 |
| ENSMUSG00 | Cyp11b2 | 1616 | 0 | 0 | 0 | 0 | 0 | 0 |
| ENSMUSG00 | Gm9747 | 2821 | 0 | 0 | 0 | 0 | 0 | 0 |
| ENSMUSG00 | Lynx1 | 3948 | 2.50737453 | 1.68432853 | 1.97049756 | 1.81872514 | 1.05981834 | 0.9188265 |
| ENSMUSG00 | Lypd2 | 472 | 4.29561336 | 1.0208992 | 1.87295945 | 0 | 1.40538747 | 1.04013455 |
| ENSMUSG00 | Slurp1 | 522 | 0 | 0 | 0 | 0.27649076 | 0 | 0 |
| ENSMUSG00 | Psca | 851 | 0 | 5.09609846 | 0 | 0.12719875 | 0 | 0 |
| ENSMUSG00 | Rpl24 | 6107 | 8.79476898 | 8.97134186 | 8.52624383 | 9.19923749 | 12.2657147 | 6.89570051 |
| ENSMUSG00 | Arc | 3054 | 0.82010323 | 0 | 0.49209648 | 0.02362937 | 0 | 0.02679238 |
| ENSMUSG00 | 1700016M24 | 3966 | 0 | 0 | 0 | 0 | 0 | 0 |
| ENSMUSG00 | Cep97 | 9532 | 1.23870937 | 1.97153928 | 1.85488221 | 2.27500092 | 2.47851574 | 0.77543299 |
| ENSMUSG00 | Ptk2 | 5083 | 28.4537553 | 23.7661638 | 23.2444475 | 20.1670448 | 24.1931076 | 22.2092725 |
| ENSMUSG00 | Mapk12 | 1877 | 3.57947587 | 1.41196288 | 0.11774599 | 1.30718116 | 1.7942146 | 2.12152229 |
| ENSMUSG00 | Miox | 1694 | 1.4550412 | 14.7062518 | 1.35684524 | 3.72748394 | 1.98804062 | 4.07348508 |
| ENSMUSG00 | Lmf2 | 2892 | 26.2562129 | 31.8077172 | 21.7646696 | 29.2823508 | 23.6076386 | 19.3242517 |
| ENSMUSG00 | Tymp | 1407 | 1.44103021 | 3.63025079 | 2.57608467 | 1.05143129 | 2.86502039 | 1.957881 |
| ENSMUSG00 | Chkb | 5434 | 12.1739271 | 10.8095828 | 9.67982944 | 11.3013691 | 11.7659273 | 16.1469478 |
| ENSMUSG00 | Mapk8ip2 | 5688 | 0 | 0 | 0 | 0 | 0 | 0 |
| ENSMUSG00 | Arsa | 4599 | 3.38859499 | 5.01876621 | 4.69986981 | 2.95780187 | 4.02752216 | 2.30105676 |
| ENSMUSG00 | Rabl2 | 3118 | 9.0782239 | 10.7098154 | 8.61921762 | 12.2202178 | 10.4900279 | 13.0249961 |
| ENSMUSG00 | Acr | 1498 | 0 | 0 | 0 | 0 | 0 | 0 |
| ENSMUSG00 | Shank3 | 8934 | 0 | 0 | 0 | 0.0848134 | 0.02284593 | 0.15264517 |
| ENSMUSG00 | Kif21a | 6461 | 12.9523737 | 10.8663746 | 15.1193427 | 14.4361686 | 13.2837586 | 13.3565947 |
| ENSMUSG00 | Yaf2 | 3209 | 14.3089997 | 17.5837718 | 22.6725564 | 19.8231987 | 14.0565024 | 29.2975012 |
| ENSMUSG00 | Zcrb1 | 2957 | 27.910848 | 23.8569331 | 29.5526018 | 24.0994379 | 18.1016802 | 45.4085387 |
| ENSMUSG00 | Alcam | 7664 | 20.8166419 | 15.6555638 | 43.6077942 | 38.6008198 | 17.0842485 | 25.6838559 |
| ENSMUSG00 | Cblb | 6632 | 1.00108041 | 1.37322641 | 2.37938148 | 2.55708096 | 2.21586192 | 2.20434358 |
| ENSMUSG00 | 5330426P16f | 2629 | 11.1902075 | 12.5369062 | 9.91977458 | 12.4894106 | 8.98638722 | 10.0736686 |
| ENSMUSG00 | Bbx | 15633 | 7.39009956 | 9.04362702 | 8.28448794 | 9.51616897 | 10.0009477 | 8.93452489 |
| ENSMUSG00 | Retnlb | 691 | 0 | 0 | 0 | 0 | 0 | 0 |
| ENSMUSG00 | Retnlg | 604 | 0 | 0 | 0 | 0 | 0 | 0 |
| ENSMUSG00 | Morc1 | 3023 | 0 | 0 | 0 | 0 | 0 | 0 |
| ENSMUSG00 | Pvrl3 | 12691 | 14.8671902 | 28.9209778 | 6.22050994 | 21.5735997 | 21.3899866 | 10.178291 |
| ENSMUSG00 | Cd96 | 2169 | 0 | 0 | 0 | 0 | 0 | 0 |
| ENSMUSG00 | Tagln3 | 1236 | 0 | 0 | 0.17881005 | 1.05093334 | 0.1651339 | 1.56674708 |
| ENSMUSG00 | Gcet2 | 3403 | 0 | 0 | 0 | 0 | 0 | 0 |
| ENSMUSG00 | Cd200 | 4583 | 14.6339724 | 19.4512149 | 11.4000826 | 17.9347365 | 23.5369318 | 18.4786722 |
| ENSMUSG00 | Atg3 | 3142 | 57.3177361 | 60.1640396 | 50.1948364 | 56.0753098 | 47.2749141 | 58.9677353 |
| ENSMUSG00 | Slc35a5 | 4608 | 22.66442 | 21.458025 | 23.4726367 | 26.6935091 | 18.3265304 | 27.4226136 |
| ENSMUSG00 | Ccdc80 | 4242 | 0.39361862 | 0.85195266 | 1.02116469 | 0.50184833 | 0.82999655 | 0.2764756 |
| ENSMUSG00 | Cd200r1 | 3612 | 0 | 0 | 0 | 0 | 0 | 0 |
| ENSMUSG00 | Gtpbp8 | 3743 | 12.8623796 | 12.7063902 | 14.6552197 | 15.2406392 | 17.6676952 | 14.5809653 |
| ENSMUSG00 | Mzt2 | 1519 | 3.03596823 | 2.91846786 | 3.4919165 | 2.61291962 | 5.60987805 | 5.62011987 |
| ENSMUSG00 | Prkdc | 12588 | 4.06145122 | 4.23373094 | 3.75371546 | 3.64604072 | 4.5643231 | 3.75925477 |
| ENSMUSG00 | Mcm4 | 4237 | 8.62291399 | 5.98207896 | 4.65282558 | 6.49766343 | 6.21421051 | 6.829925 |
| ENSMUSG00 | Ube2v2 | 5308 | 4.30659406 | 5.07464604 | 5.18797065 | 6.95401875 | 6.11393652 | 5.4384296 |
| ENSMUSG00 | Snai2 | 2028 | 0 | 0 | 0.43591561 | 0.78284515 | 1.03159633 | 0.05379613 |
| ENSMUSG00 | 0610037P05f | 2227 | 35.1497715 | 33.4513873 | 49.2035783 | 29.1313498 | 28.6407588 | 32.0388219 |
| ENSMUSG00 | Nde1 | 3177 | 15.8045867 | 18.5950828 | 17.5861283 | 16.320396 | 21.0401487 | 13.2123607 |
| ENSMUSG00 | Mpv17l | 3753 | 13.4636866 | 23.7401363 | 17.1837168 | 19.3822014 | 14.0856183 | 10.5377636 |
| ENSMUSG00 | Pdxdc1 | 8955 | 17.824484 | 17.9401003 | 16.8811059 | 17.3056819 | 22.3479002 | 12.466231 |
| ENSMUSG00 | Ntan1 | 2323 | 15.830314 | 12.1140259 | 11.664111 | 12.5658089 | 9.48919258 | 15.8035577 |
| ENSMUSG00 | Rrn3 | 3579 | 17.750561 | 14.3657261 | 15.894879 | 15.9793352 | 15.0983325 | 15.271969 |
| ENSMUSG00 | Pla2g10 | 1773 | 0 | 0 | 0 | 0 | 0.05755936 | 0.1692166 |
| ENSMUSG00 | Bfar | 4353 | 23.4441347 | 20.6450068 | 23.0909697 | 20.946319 | 25.2494403 | 20.7896285 |
| ENSMUSG00 | Parn | 2878 | 19.5738415 | 16.3579409 | 16.6332856 | 13.0386818 | 16.1163918 | 13.07818 |
| ENSMUSG00 | B3gnt5 | 5433 | 0.31464857 | 0.01773843 | 0 | 0.34534628 | 0 | 0.40161442 |
| ENSMUSG00 | Boc | 5187 | 2.05407148 | 2.19240415 | 0.67321103 | 0.94604937 | 2.45933949 | 0.38911188 |
| ENSMUSG00 | Sidt1 | 6769 | 0 | 0 | 0 | 0 | 0 | 0 |
| ENSMUSG00 | Naa50 | 4904 | 47.2947539 | 46.1524712 | 55.468626 | 58.9644178 | 41.5057531 | 53.6260342 |
| ENSMUSG00 | 2610015P09f | 3947 | 3.27350676 | 4.71243138 | 2.72131945 | 4.24171995 | 3.95593387 | 4.33961831 |
| ENSMUSG00 | Hira | 5396 | 1.78229461 | 2.16026658 | 1.56401482 | 3.37558957 | 1.84330183 | 1.08128433 |
| ENSMUSG00 | Qtrtd1 | 6354 | 2.59030043 | 2.00966433 | 1.28000301 | 1.58433907 | 1.99158659 | 1.83290383 |
| ENSMUSG00 | Drd3 | 1448 | 0 | 0 | 0 | 0 | 0 | 0 |
| ENSMUSG00 | Mrpl40 | 1646 | 26.3748388 | 36.154469 | 30.0497341 | 20.2769691 | 18.0731331 | 37.0345193 |
| ENSMUSG00 | Gbe1 | 5568 | 33.5365467 | 30.3069182 | 37.7557409 | 33.0233655 | 29.3713244 | 32.0849294 |
| ENSMUSG00 | Zbtb20 | 30702 | 13.1650697 | 14.2823472 | 12.3397172 | 14.9901139 | 15.976697 | 12.1395331 |
| ENSMUSG00 | Usp7 | 9474 | 15.2618411 | 14.7549999 | 15.984327 | 19.2635615 | 16.9603185 | 10.4820521 |
| ENSMUSG00 | Pmm2 | 4614 | 26.7621418 | 21.9314106 | 24.7258251 | 18.1192235 | 24.3851666 | 23.1781232 |
| ENSMUSG00 | Cldn26 | 1402 | 0 | 0 | 0 | 0 | 0 | 0 |
| ENSMUSG00 | Dgcr8 | 8918 | 3.54402411 | 3.95520023 | 4.35178495 | 4.28469221 | 4.98934736 | 2.87487788 |
| ENSMUSG00 | Trmt2a | 4306 | 17.0802692 | 17.8377122 | 16.4858709 | 14.672471 | 21.8633682 | 11.7434234 |
| ENSMUSG00 | Arl6 | 2609 | 27.1842764 | 29.10764 | 30.8345551 | 29.2639349 | 23.1369117 | 39.8195097 |
| ENSMUSG00 | Crybg3 | 9064 | 1.38161699 | 0.85059916 | 1.86775219 | 1.89486465 | 2.13360508 | 2.51562207 |
| ENSMUSG00 | Mina | 5419 | 6.88146166 | 5.61093283 | 6.12577673 | 6.59184796 | 5.90395601 | 6.05487932 |
| ENSMUSG00 | Gsc2 | 1162 | 0 | 0 | 0 | 0 | 0 | 0 |
| ENSMUSG00 | Cpox | 3179 | 18.1957295 | 15.0819471 | 18.5761756 | 13.9266023 | 16.7573251 | 16.0095869 |
| ENSMUSG00 | Cldn25 | 2716 | 37.3842037 | 35.3237137 | 48.5797133 | 35.3912248 | 33.2911406 | 50.5926482 |
| ENSMUSG00 | St3gal6 | 6595 | 16.1192047 | 15.6359342 | 19.4702584 | 20.5002913 | 15.884329 | 20.2977907 |
| ENSMUSG00 | 2610528E23f | 1825 | 1.4813001 | 3.32684477 | 1.59853241 | 2.03641128 | 2.20881298 | 2.97405655 |
| ENSMUSG00 | Tbc1d23 | 3630 | 18.2968557 | 21.6507127 | 24.390163 | 16.0927907 | 20.9868813 | 20.9320978 |
| [illegible] | | | | | | | | |

| | | | | | | | | |
|---|---|---|---|---|---|---|---|---|
| ENSMUSG00 | Klhl22 | 5629 | 8.65877049 | 6.74558828 | 8.96757945 | 8.63430698 | 4.38741622 | 7.09848046 |
| ENSMUSG00 | Nit2 | 1270 | 17.9995286 | 27.3182979 | 19.7690133 | 30.6555323 | 22.4194628 | 18.8130583 |
| ENSMUSG00 | Tomm70a | 6354 | 32.2911607 | 29.2500956 | 36.0418238 | 34.7418868 | 26.9426331 | 44.1141749 |
| ENSMUSG00 | Tmem30c | 3941 | 0.01008766 | 0 | 0 | 0.21057778 | 0.03884271 | 0 |
| ENSMUSG00 | Tmem45a | 3334 | 0 | 0 | 0 | 0 | 0 | 0 |
| ENSMUSG00 | Gpr128 | 2682 | 0 | 0 | 0 | 0 | 0 | 0 |
| ENSMUSG00 | Slc7a4 | 4891 | 11.322712 | 8.55159104 | 12.8511594 | 8.0633152 | 7.44895366 | 5.44823606 |
| ENSMUSG00 | Tfg | 3010 | 56.3444784 | 55.0221932 | 56.1994861 | 61.1237027 | 49.7040976 | 55.038591 |
| ENSMUSG00 | P2rx6 | 4230 | 0 | 0.10252435 | 0.02089922 | 0.14501058 | 0.26538541 | 0.01289581 |
| ENSMUSG00 | 4930451C15I | 4142 | 0.00959814 | 0 | 0 | 0 | 0.02463852 | 0 |
| ENSMUSG00 | Thap7 | 2358 | 7.28344681 | 6.74364966 | 9.82263178 | 6.50333712 | 8.72079281 | 8.66357286 |
| ENSMUSG00 | Lztr1 | 5675 | 13.821597 | 15.3687155 | 14.1056454 | 15.2339346 | 15.3663571 | 8.33378405 |
| ENSMUSG00 | Ncam2 | 8380 | 0.19925181 | 0.18400551 | 0.13714176 | 0.54252239 | 0.15222666 | 0 |
| ENSMUSG00 | Aifm3 | 4194 | 0.18010351 | 0.12638313 | 0.53750453 | 0.55060821 | 0.01216652 | 0.30565285 |
| ENSMUSG00 | Snap29 | 3424 | 8.56879224 | 7.43061958 | 6.53216488 | 10.3693726 | 5.2009945 | 7.55150643 |
| ENSMUSG00 | Serpind1 | 2250 | 0 | 0 | 0 | 0.14432818 | 0.65767329 | 0.70307959 |
| ENSMUSG00 | Ccdc116 | 5487 | 0.02173618 | 0.41275064 | 0.23361645 | 0 | 0 | 0.06959084 |
| ENSMUSG00 | Sdf2l1 | 1105 | 22.01842 | 30.5689556 | 21.7209057 | 27.0696968 | 15.9774896 | 26.4107367 |
| ENSMUSG00 | Dlg1 | 8735 | 9.92180281 | 11.2315508 | 9.69556167 | 12.9044427 | 10.1760672 | 10.994162 |
| ENSMUSG00 | Ppil2 | 6433 | 17.9403327 | 17.4379041 | 18.1465984 | 17.752164 | 20.7817686 | 16.4504275 |
| ENSMUSG00 | Senp5 | 4434 | 3.9002332 | 4.54261002 | 5.34329902 | 9.50469734 | 5.7309732 | 3.11253213 |
| ENSMUSG00 | Ypel1 | 4831 | 4.11462228 | 3.52097166 | 2.59849377 | 4.51118917 | 3.18980863 | 2.00424279 |
| ENSMUSG00 | Ncbp2 | 2426 | 46.3923188 | 54.9595571 | 50.1597996 | 42.0907609 | 51.4680716 | 48.3208572 |
| ENSMUSG00 | Top3b | 5444 | 20.4984632 | 24.1374775 | 18.8937709 | 24.7086447 | 26.7316723 | 18.1814228 |
| ENSMUSG00 | Mfi2 | 4117 | 0 | 0 | 0 | 0.30674558 | 0.18591101 | 0 |
| ENSMUSG00 | Pak2 | 6779 | 19.8454855 | 17.7704832 | 21.6021103 | 18.7888505 | 16.0704104 | 18.1495645 |
| ENSMUSG00 | Spag6 | 4547 | 0.74317481 | 1.03854659 | 0.26246971 | 0.96017757 | 0.37032558 | 0.16795467 |
| ENSMUSG00 | Wdr53 | 1534 | 6.21989265 | 8.73261468 | 9.24953821 | 8.27958258 | 7.05188505 | 8.83669864 |
| ENSMUSG00 | Fgd4 | 17746 | 4.4244942 | 3.40775301 | 3.12596151 | 4.97536137 | 3.52808313 | 3.91613777 |
| ENSMUSG00 | Dnm1l | 4164 | 76.618091 | 60.2677691 | 65.3049323 | 65.7691446 | 59.4940093 | 69.9354763 |
| ENSMUSG00 | Igsf11 | 3612 | 2.15727414 | 0.98720839 | 0.33041245 | 0.75920138 | 1.5680863 | 2.34084667 |
| ENSMUSG00 | Tnk2 | 5300 | 0.60758376 | 0.4454973 | 0.60047787 | 0.6399457 | 0.4813809 | 0.77192375 |
| ENSMUSG00 | Yars2 | 3821 | 8.10508226 | 5.85148107 | 5.0205705 | 5.98691867 | 5.04788539 | 5.32501462 |
| ENSMUSG00 | B4galt4 | 3068 | 2.35837596 | 5.8269785 | 0.63392474 | 3.72816431 | 3.59246975 | 1.04013455 |
| ENSMUSG00 | Tfrc | 6219 | 11.3084813 | 16.1861196 | 14.1084834 | 13.2921634 | 12.2089158 | 30.9366949 |
| ENSMUSG00 | 4930435E12f | 1526 | 0 | 0 | 0 | 0 | 0 | 0 |
| ENSMUSG00 | Arhgap31 | 11129 | 0.29649608 | 0.14288369 | 0.07943543 | 0.42796566 | 0.68316381 | 0.19361097 |
| ENSMUSG00 | Fyttd1 | 5726 | 32.9999649 | 35.8326704 | 41.7547623 | 39.2327644 | 29.9332163 | 42.0265704 |
| ENSMUSG00 | Lrch3 | 6868 | 5.84639419 | 8.67187573 | 8.64986561 | 8.96797467 | 9.16810538 | 9.11405024 |
| ENSMUSG00 | Lmln | 3685 | 3.73280767 | 2.53681676 | 3.44258587 | 5.70850257 | 4.84646717 | 2.93840754 |
| ENSMUSG00 | Popdc2 | 3256 | 0 | 0.47357683 | 0 | 0 | 0.12537193 | 0.00837673 |
| ENSMUSG00 | 4932425I24R | 4516 | 0.17606502 | 0.25608384 | 0 | 0 | 0 | 0 |
| ENSMUSG00 | Osbpl11 | 4507 | 7.55945125 | 10.4348719 | 8.09108509 | 13.3056042 | 8.36665004 | 6.0455657 |
| ENSMUSG00 | Snx4 | 2428 | 43.7506771 | 46.2415199 | 46.0587573 | 55.2227667 | 47.2435308 | 45.5738103 |
| ENSMUSG00 | Nr1i2 | 2544 | 0 | 0 | 0 | 0 | 0 | 0 |
| ENSMUSG00 | Zfp148 | 2249 | 54.4803873 | 51.9145173 | 57.3307141 | 55.2701838 | 49.3020519 | 62.4199948 |
| ENSMUSG00 | Gsk3b | 10143 | 7.60774797 | 7.1213129 | 6.15330793 | 8.5411603 | 6.53990817 | 9.15877169 |
| ENSMUSG00 | Umps | 5619 | 16.9804508 | 11.8772419 | 15.4261994 | 16.451717 | 17.3811147 | 14.1251469 |
| ENSMUSG00 | Fstl1 | 3645 | 6.48957775 | 12.6381999 | 4.22009365 | 3.81113502 | 9.77130699 | 6.02361717 |
| ENSMUSG00 | Itgb5 | 3697 | 47.8313166 | 55.4463416 | 44.4648781 | 42.306808 | 64.3454055 | 35.3161207 |
| ENSMUSG00 | Cyp2ab1 | 2712 | 0 | 0.05330359 | 0 | 0 | 0 | 0 |
| ENSMUSG00 | Ndufb4 | 1285 | 9.96207372 | 10.0122802 | 10.9730645 | 9.94011188 | 6.47260643 | 20.0155524 |
| ENSMUSG00 | Hgd | 2686 | 0.42922894 | 0.91493245 | 0.13165106 | 0.17463387 | 0.7408893 | 1.27945069 |
| ENSMUSG00 | Abcc5 | 8382 | 8.84087517 | 8.37025289 | 10.0933342 | 8.8160376 | 11.3168734 | 9.11757805 |
| ENSMUSG00 | Muc13 | 3025 | 0 | 0 | 0 | 0 | 0 | 0.00901641 |
| ENSMUSG00 | Rabl3 | 3087 | 11.551884 | 14.4855906 | 10.8965347 | 10.1221414 | 13.0747856 | 10.2666443 |
| ENSMUSG00 | Gtf2e1 | 8220 | 3.90300155 | 3.34139562 | 3.45763809 | 5.58349886 | 2.77479197 | 1.66567748 |
| ENSMUSG00 | Stxbp5l | 8920 | 0 | 0 | 0.01982145 | 0 | 0.02288178 | 0 |
| ENSMUSG00 | Hcls1 | 2511 | 0 | 0.05757042 | 0 | 0 | 0.02032114 | 0 |
| ENSMUSG00 | Ropn1 | 969 | 0 | 0 | 0 | 0 | 0 | 0 |
| ENSMUSG00 | Ccdc14 | 4094 | 1.66052447 | 2.77772358 | 2.24572138 | 2.14165531 | 1.85709086 | 2.30508676 |
| ENSMUSG00 | Mylk | 10594 | 1.27214536 | 3.09750492 | 0.88870989 | 1.33851647 | 2.50941493 | 0.9036623 |
| ENSMUSG00 | Iqcb1 | 5363 | 8.19127465 | 7.45753253 | 8.08540145 | 11.1482282 | 8.91510612 | 15.0638601 |
| ENSMUSG00 | Eaf2 | 4543 | 0.65631984 | 0.80611261 | 1.41080062 | 0.34152056 | 1.21304173 | 2.79170312 |
| ENSMUSG00 | Adcy5 | 5620 | 10.7806677 | 5.16160822 | 12.6470754 | 7.35765534 | 4.13113897 | 4.61048172 |
| ENSMUSG00 | Ap2m1 | 4126 | 14.5397528 | 15.2524221 | 13.9590406 | 15.2688439 | 14.9641092 | 15.4485778 |
| ENSMUSG00 | Ece2 | 6396 | 0.60913651 | 1.09994067 | 0.89841144 | 0.54156915 | 0.36698144 | 0.37526083 |
| ENSMUSG00 | Clcn2 | 6186 | 3.31617005 | 2.47709159 | 1.31476546 | 2.73560926 | 2.89528903 | 2.79536393 |
| ENSMUSG00 | Pdia5 | 1791 | 2.77467061 | 1.37214324 | 1.01187915 | 2.17580168 | 2.90602475 | 0.67006372 |
| ENSMUSG00 | Thpo | 1830 | 0 | 0 | 0 | 0 | 0 | 0.55145446 |
| ENSMUSG00 | Dirc2 | 5858 | 9.08037435 | 12.6347516 | 10.5034082 | 9.19605539 | 10.1739172 | 7.51938245 |
| ENSMUSG00 | Hspbap1 | 4582 | 2.07367085 | 1.18836053 | 3.98414692 | 1.59857159 | 2.04907315 | 1.91077241 |
| ENSMUSG00 | Ehhadh | 3011 | 31.4505329 | 28.7262251 | 23.3267116 | 18.8019687 | 23.8947824 | 16.0604236 |
| ENSMUSG00 | Senp2 | 4500 | 19.1798107 | 14.6700946 | 16.6886514 | 18.8187908 | 15.3192519 | 16.4132718 |
| ENSMUSG00 | Tmem41a | 1334 | 2.71195557 | 4.91256082 | 5.93113186 | 8.08735481 | 7.0381208 | 5.15233065 |
| ENSMUSG00 | Tmprss15 | 4645 | 0 | 0 | 0 | 0 | 0 | 0 |
| ENSMUSG00 | Tra2b | 5430 | 12.2707524 | 13.5241506 | 10.9405667 | 13.3696268 | 9.10581182 | 20.5288123 |
| ENSMUSG00 | Chodl | 3813 | 0 | 0 | 0 | 0 | 0 | 0 |
| ENSMUSG00 | Dgkg | 5679 | 0.01400087 | 0.00848502 | 0 | 0.02541437 | 0.15274668 | 0 |
| ENSMUSG00 | Btg3 | 1433 | 1.38714168 | 1.34505072 | 2.06665979 | 1.88845313 | 2.31452507 | 1.38942684 |
| ENSMUSG00 | D16Ertd472e | 3043 | 0.23516223 | 0 | 0 | 0 | 0 | 0 |
| ENSMUSG00 | Cxadr | 6419 | 22.9094131 | 20.6738529 | 33.9140173 | 31.3434305 | 27.5442269 | 23.1105722 |
| ENSMUSG00 | Usp25 | 4385 | 46.3557062 | 33.5162254 | 43.607109 | 38.7480839 | 52.1318503 | 38.8189335 |
| ENSMUSG00 | Ahsg | 1467 | 0 | 0 | 0 | 0 | 0.45217647 | 0 |
| ENSMUSG00 | Fetub | 1903 | 23.2516289 | 23.5234917 | 22.8325863 | 30.0146487 | 16.4635811 | 11.093311 |
| ENSMUSG00 | Kng1 | 3146 | 0.63184171 | 0.41355179 | 0.3091038 | 0.52758234 | 0.21085279 | 0.07802662 |
| ENSMUSG00 | Samsn1 | 5447 | 0 | 0 | 0 | 0 | 0 | 0 |
| ENSMUSG00 | Hrg | 1727 | 0 | 0 | 0 | 0 | 0 | 0 |
| ENSMUSG00 | Adipoq | 1377 | 0 | 0 | 0 | 0 | 0.40761811 | 0 |
| ENSMUSG00 | Rfc4 | 1588 | 4.30600922 | 4.15714268 | 4.31441163 | 5.11238036 | 3.82376494 | 8.31292531 |
| ENSMUSG00 | Robo1 | 10506 | 0.01892037 | 0.38527138 | 0.57639944 | 0.24384401 | 0.30112653 | 0.22586081 |
| ENSMUSG00 | Eif4a2 | 4742 | 70.942825 | 66.0810699 | 69.1737404 | 74.6826797 | 66.1664245 | 85.125403 |
| ENSMUSG00 | St6gal1 | 3496 | 26.1549214 | 28.5176627 | 16.7527008 | 28.1039494 | 24.316345 | 11.452852 |
| ENSMUSG00 | Prl2a1 | 879 | 0 | 0 | 0 | 0 | 0 | 0 |
| ENSMUSG00 | Masp1 | 6000 | 0.1987774 | 0.1847147 | 0 | 0 | 0.18709671 | 0 |
| ENSMUSG00 | Mrpl39 | 2523 | 19.3813876 | 21.6008189 | 24.6675367 | 23.511249 | 18.5054038 | 27.0476209 |
| ENSMUSG00 | Atp5j | 2589 | 144.848763 | 112.04418 | 124.239788 | 92.8182374 | 87.0936867 | 341.854787 |
| ENSMUSG00 | App | 4328 | 514.708548 | 450.534327 | 696.720316 | 445.548222 | 481.567317 | 448.834186 |
| ENSMUSG00 | Adamts1 | 5916 | 2.9164771 | 3.55126585 | 0.61266922 | 3.20200699 | 3.40693348 | 0.64083415 |
| ENSMUSG00 | Adamts5 | 8083 | 0 | 0 | 0 | 0 | 0 | 0 |
| ENSMUSG00 | Ets2 | 4889 | 25.695913 | 18.6378732 | 45.9286894 | 21.1444176 | 26.1028771 | 30.5884326 |
| ENSMUSG00 | Dyrk1a | 8298 | 7.79011946 | 6.76514884 | 8.22991654 | 8.27912904 | 6.94863879 | 7.5171246 |
| ENSMUSG00 | Dscr3 | 3662 | 19.8994527 | 14.4875131 | 28.1602675 | 17.1542435 | 21.068236 | 16.1473015 |
| ENSMUSG00 | Slc15a2 | 7306 | 5.39795191 | 9.82723772 | 5.92302276 | 6.19311304 | 6.36258259 | 4.96140098 |
| ENSMUSG00 | Ildr1 | 4502 | 49.0188077 | 58.7185299 | 51.2415412 | 46.4450129 | 44.9512676 | 41.2148758 |
| ENSMUSG00 | Cd86 | 4140 | 0.02880832 | 0.25606322 | 0 | 0.8628315 | 1.13391946 | 0.59951502 |
| ENSMUSG00 | Stfa2 | 778 | 0 | 0 | 0 | 0 | 0 | 0 |
| ENSMUSG00 | Kpna1 | 8145 | 17.2981489 | 17.0146357 | 20.5841118 | 25.98173 | 16.0690797 | 21.2671865 |
| ENSMUSG00 | Parp9 | 4908 | 5.54859498 | 3.63263724 | 2.88194576 | 3.02889208 | 5.71811465 | 5.4071361 |
| ENSMUSG00 | Arl13b | 5806 | 1.98571983 | 2.4151317 | 2.71788804 | 3.22538428 | 2.75081909 | 3.02529585 |
| ENSMUSG00 | Pros1 | 4072 | 64.0363941 | 49.1686314 | 93.7226701 | 65.4119972 | 90.9502543 | 81.5158054 |
| ENSMUSG00 | Psmg1 | 1411 | 13.2142596 | 15.3677526 | 10.5257401 | 6.39299159 | 9.72791998 | 16.1598855 |
| ENSMUSG00 | Brwd1 | 21764 | 3.73369795 | 3.52475721 | 4.50061037 | 4.9670008 | 4.36551699 | 5.37246731 |
| ENSMUSG00 | 1700093J21R | 646 | 0 | 0 | 0 | 0 | 0 | 0 |
| ENSMUSG00 | Krtap15 | 377 | 0 | 0 | 0 | 0 | 0 | 0 |
| ENSMUSG00 | Grik1 | 3312 | 0 | 0 | 0 | 0 | 0 | 0 |
| ENSMUSG00 | Fam3b | 1503 | 0.21160602 | 0 | 0 | 0 | 0 | 0 |
| ENSMUSG00 | Pigp | 1882 | 33.2703942 | 23.5299364 | 23.9328789 | 25.2497623 | 14.071567 | 50.2305524 |
| ENSMUSG00 | Ripply3 | 1527 | 0.91122581 | 0.66268192 | 0 | 0 | 1.06931501 | 1.41106516 |
| ENSMUSG00 | Chaf1b | 3581 | 0.35525702 | 0.90156147 | 0.5184243 | 0.92698914 | 1.082939 | 0.18279568 |
| ENSMUSG00 | Dopey2 | 7276 | 9.39794207 | 17.0334702 | 7.07132288 | 12.4620915 | 13.2755538 | 8.70794214 |
| ENSMUSG00 | Cbr3 | 1167 | 4.90556058 | 21.2235058 | 1.85594714 | 24.6421504 | 4.19754248 | 1.75286885 |
| ENSMUSG00 | Setd4 | 4273 | 1.321151 | 1.82686722 | 1.01375629 | 0.83597529 | 0.76426119 | 1.09149621 |
| ENSMUSG00 | Clic6 | 3907 | 0.09157904 | 0 | 0.54304798 | 0.74805365 | 0 | 0.00698097 |
| ENSMUSG00 | Rcan1 | 2501 | 23.3032925 | 29.2471088 | 29.0378361 | 27.0507131 | 26.1354608 | 21.058508 |
| ENSMUSG00 | Runx1 | 6999 | 0.31240912 | 0.19277331 | 0.04420816 | 0 | 0.36452619 | 0.01948467 |
| ENSMUSG00 | Atp5o | 2249 | 61.038539 | 67.341035 | 81.5640945 | 56.3611483 | 46.239108 | 92.593095 |
| ENSMUSG00 | Itsn1 | 14973 | 1.31164145 | 0.97833957 | 0.72326531 | 1.80735547 | 2.00384085 | 1.3370457 |
| ENSMUSG00 | Donson | 2824 | 1.80194813 | 1.96226659 | 1.78435202 | 2.1465239 | 2.42122322 | 2.30829985 |
| ENSMUSG00 | Son | 14067 | 24.2936028 | 21.087349 | 23.6578998 | 28.5229498 | 20.4693139 | 29.6750092 |
| ENSMUSG00 | Gart | 4325 | 8.80595383 | 11.3864841 | 7.27669647 | 8.85157208 | 12.7064524 | 5.3918651 |
| ENSMUSG00 | Tmem50b | 2838 | 143.977036 | 144.117872 | 158.94285 | 143.133072 | 149.231473 | 105.657993 |
| ENSMUSG00 | Ifngr2 | 3390 | 8.16218715 | 7.93157366 | 9.62270211 | 6.99289182 | 7.60127427 | 7.37783015 |
| ENSMUSG00 | Ifnar1 | 6035 | 11.0867396 | 15.3382196 | 16.0694024 | 17.1053404 | 22.1945711 | 10.064726 |
| ENSMUSG00 | Il10rb | 3937 | 10.1383039 | 9.48552011 | 14.606705 | 13.5731542 | 10.4463447 | 9.93442537 |
| ENSMUSG00 | Ifnar2 | 3553 | 10.0143977 | 9.37146961 | 9.40517684 | 6.09322452 | 9.04773903 | 9.11969362 |
| ENSMUSG00 | 1110004E09f | 1481 | 12.2138715 | 15.9428472 | 15.7885044 | 12.6445517 | 15.1597605 | 14.2911007 |
| ENSMUSG00 | Synj1 | 10987 | 3.83189714 | 3.75421812 | 4.09551981 | 3.39900938 | 3.19059982 | 2.4526571 |
| ENSMUSG00 | Gcfc1 | 5491 | 23.1249326 | 16.1996307 | 14.8761621 | 23.9386065 | 27.9804075 | 37.3182234 |
| ENSMUSG00 | 2610039C10I | 2847 | 2.69505014 | 3.09733717 | 4.02117223 | 4.85402987 | 1.79228577 | 2.77823863 |
| ENSMUSG00 | Sod1 | 641 | 19.1644984 | 31.6481937 | 20.68729 | 20.545938 | 16.1596791 | 32.8912576 |
| ENSMUSG00 | Scaf4 | 5446 | 7.29264139 | 6.83068919 | 5.5110267 | 11.1638349 | 8.69490943 | 4.73775408 |
| ENSMUSG00 | Krt75 | 3257 | 0 | 0.01479473 | 0 | 0 | 0 | 0 |
| ENSMUSG00 | Zfp641 | 7346 | 3.62594227 | 3.79797849 | 3.09882237 | 4.04731891 | 4.36912475 | 5.45047242 |
| ENSMUSG00 | Lalba | 729 | 0.1636028 | 0 | 0 | 0 | 0 | 0 |
| ENSMUSG00 | 2310037I24R | 2856 | 4.64927538 | 4.96036904 | 5.41689253 | 4.85136733 | 5.41351256 | 4.50757343 |
| ENSMUSG00 | 4930415O20I | 536 | 0.22251202 | 0.62930055 | 1.15452575 | 0.20195174 | 0.19039693 | 1.06859594 |
| ENSMUSG00 | Adcy6 | 5995 | 13.5480311 | 15.4084086 | 13.4264481 | 20.5357691 | 19.3721487 | 11.8561651 |
| ENSMUSG00 | Enah | 4272 | 1.92635404 | 1.46634772 | 1.15884982 | 3.23488367 | 1.58860206 | 0.92575438 |
| ENSMUSG00 | Wnt10b | 3057 | 0 | 0 | 0 | 0 | 0 | 0 |
| ENSMUSG00 | Wnt1 | 1454 | 0 | 0 | 0 | 0 | 0 | 0 |
| ENSMUSG00 | Lmbr1l | 4094 | 4.97186273 | 5.22588674 | 3.83284179 | 7.33274574 | 6.15706635 | 6.31567123 |
| ENSMUSG00 | Dhh | 2360 | 0 | 0 | 0 | 0 | 0.32432019 | 0.01155705 |
| ENSMUSG00 | Tuba1b | 1872 | 13.5279066 | 14.4404883 | 19.8813845 | 12.7019591 | 7.87746934 | 12.2823295 |
| ENSMUSG00 | Prpf40b | 4603 | 12.739373 | 15.2525845 | 8.6041389 | 13.0437806 | 17.6923954 | 15.0031255 |
| ENSMUSG00 | Fmnl3 | 5457 | 1.63917594 | 3.80581942 | 1.71720556 | 3.09444692 | 3.22596659 | 2.03422726 |
| ENSMUSG00 | Nckap5l | 4988 | 0.30286854 | 0.25117231 | 0.46966674 | 0.49189636 | 0.25574567 | 0.31714697 |
| ENSMUSG00 | Tmbim6 | 3846 | 42.9185531 | 53.1855036 | 49.9483125 | 51.8902206 | 45.3081314 | 52.6699807 |
| ENSMUSG00 | Faim2 | 4508 | 0.05291324 | 0.17102553 | 0.0196104 | 0 | 0.04527629 | 0 |
| ENSMUSG00 | Aqp2 | 1412 | 1983.35362 | 3687.35487 | 756.126994 | 1437.22552 | 1211.26209 | 613.061261 |
| ENSMUSG00 | Racgap1 | 4369 | 3.43959067 | 2.53670902 | 3.166669 | 0.85889966 | 4.4848085 | 1.82288731 |
| ENSMUSG00 | Accn2 | 4484 | 0.36350908 | 0.25791138 | 0.41402262 | 1.16679225 | 0.6145014 | 0.86962012 |
| ENSMUSG00 | Smarcd1 | 3140 | 5.64679755 | 4.88002822 | 5.49003783 | 5.92940604 | 7.42645024 | 6.55807409 |
| ENSMUSG00 | Gpd1 | 2901 | 16.2941283 | 10.6305836 | 12.4636703 | 11.2562049 | 10.2369358 | 11.6112304 |
| ENSMUSG00 | 2310016M24 | 721 | 70.4681055 | 68.8377745 | 98.8870635 | 75.3168891 | 46.0723588 | 121.43078 |
| ENSMUSG00 | Lass5 | 6461 | 12.9892926 | 15.9825949 | 12.6975113 | 13.4756189 | 12.1622998 | 10.88706 |
| ENSMUSG00 | Lima1 | 9973 | 22.2715201 | 17.848262 | 26.5219922 | 18.4444263 | 23.6533576 | 20.3855571 |
| ENSMUSG00 | Larp4 | 6728 | 14.3292272 | 15.4915687 | 19.4467086 | 22.3474851 | 16.1694758 | 19.4425044 |
| ENSMUSG00 | Dip2b | 171295 | 0.20238057 | 0.20872962 | 0.19714649 | 0.25572026 | 0.19213645 | 0.14951333 |
| ENSMUSG00 | Atf1 | 2580 | 15.0855099 | 14.1010713 | 15.7276325 | 17.5235666 | 13.3697016 | 13.2250286 |
| ENSMUSG00 | Slc11a2 | 6334 | 11.2224186 | 11.5787441 | 11.954177 | 9.95189956 | 10.5130124 | 8.91786831 |
| ENSMUSG00 | Cela1 | 1095 | 0.39937012 | 2.20029416 | 0.8073396 | 1.31806555 | 0.41939487 | 0.79706708 |
| ENSMUSG00 | Slc4a8 | 13099 | 1.82100071 | 2.24396746 | 1.55224428 | 2.32485271 | 2.78913544 | 0.99320581 |
| ENSMUSG00 | Scn8a | 11039 | 0.21968333 | 0.17460437 | 0 | 0.12420669 | 0.00462237 | 0.01482452 |
| ENSMUSG00 | Nr4a1 | 2467 | 5.83359706 | 21.7005015 | 11.5924574 | 9.41906635 | 8.2734294 | 8.21445357 |
| ENSMUSG00 | Pcdhga9 | 28969 | 3.04797964 | 3.48644353 | 2.77854107 | 3.00797879 | 3.38719737 | 2.58633139 |
| ENSMUSG00 | Krt7 | 7163 | 132.59782 | 139.924333 | 144.151201 | 131.916318 | 105.814015 | 123.438399 |
| ENSMUSG00 | Krt6b | 2217 | 0 | 0 | 0 | 0 | 0 | 0 |
| ENSMUSG00 | Krt18 | 1393 | 104.254681 | 138.81708 | 82.6920337 | 106.303454 | 89.2686848 | 55.410789 |
| ENSMUSG00 | Csad | 2250 | 39.7201423 | 33.8804229 | 35.2043123 | 35.1358931 | 33.3372322 | 23.7228751 |
| ENSMUSG00 | Soat2 | 2203 | 0 | 0 | 0 | 0 | 0 | 0.01238068 |
| ENSMUSG00 | Igfbp6 | 1108 | 0 | 0.04348957 | 0 | 0 | 0 | 0.246161 |
| ENSMUSG00 | Amhr2 | 1991 | 0.39935189 | 0.0484026 | 0.39961485 | 0.18122574 | 0.07688555 | 0.13698965 |
| ENSMUSG00 | Prr13 | 1557 | 45.0664246 | 44.1012717 | 67.7931928 | 37.4724894 | 39.1955976 | 72.1542961 |
| ENSMUSG00 | Map3k12 | 9631 | 1.6181236 | 1.50598267 | 1.80368854 | 1.82452037 | 2.60668434 | 2.17211591 |
| ENSMUSG00 | Tarbp2 | 3557 | 6.95190016 | 4.83625523 | 4.26236496 | 4.37204419 | 5.02086914 | 3.43520899 |
| ENSMUSG00 | Npff | 439 | 3.44124888 | 0.65858463 | 1.91306382 | 2.30136047 | 1.85973124 | 6.08864384 |
| ENSMUSG00 | Calcoco1 | 4328 | 14.7521492 | 14.8300233 | 13.5424316 | 12.5886985 | 19.9601789 | 14.3620386 |
| ENSMUSG00 | Fabp2 | 961 | 0 | 0 | 0 | 0 | 0 | 0 |
| ENSMUSG00 | Sncg | 847 | 0.46936813 | 0.62579795 | 0 | 0 | 0.36146193 | 0.25761171 |
| ENSMUSG00 | Rttn | 7371 | 1.03015829 | 1.36629581 | 1.15736748 | 0.93007576 | 1.95216904 | 1.18408419 |
| ENSMUSG00 | Cdkn1a | 2219 | 8.15175468 | 4.88596191 | 6.37430814 | 6.94323254 | 2.78242067 | 4.58469601 |
| ENSMUSG00 | Nus1 | 4871 | 18.8942594 | 16.4116829 | 14.8640154 | 20.437356 | 15.1581125 | 22.7503304 |
| ENSMUSG00 | Rgn | 1600 | 11.7527139 | 12.8597568 | 11.0780869 | 11.0275749 | 9.66311992 | 7.27891935 |
| ENSMUSG00 | Ccdc123 | 5217 | 5.82958455 | 6.7887742 | 5.3716635 | 4.56472099 | 7.21822223 | 3.36162412 |
| ENSMUSG00 | Slc10a2 | 1623 | 0 | 0.02968974 | 0 | 0.0222317 | 0 | 0.65539799 |
| ENSMUSG00 | Mospd1 | 2078 | 34.3986304 | 23.5134997 | 33.417274 | 31.150716 | 25.046633 | 46.2015063 |
| ENSMUSG00 | Akirin1 | 3445 | 14.5173859 | 14.9524838 | 14.7938244 | 16.9255686 | 14.7376615 | 14.1242254 |
| ENSMUSG00 | Cxcl13 | 1170 | 0 | 0 | 0 | 0 | 0 | 0 |
| ENSMUSG00 | Gtf2ird1 | 4041 | 10.3594459 | 12.711395 | 8.2146954 | 13.1256138 | 14.874801 | 6.90471565 |
| ENSMUSG00 | H2-M10.2 | 1108 | 0 | 0 | 0 | 0 | 0 | 0 |
| ENSMUSG00 | 4933430H15I | 5117 | 0 | 0.00941693 | 0.10365881 | 0 | 0.04985966 | 0 |
| ENSMUSG00 | Ccrn4l | 11459 | 3.26120531 | 4.13783568 | 5.66264993 | 5.674129 | 5.19659487 | 2.62059323 |
| ENSMUSG00 | Abcc1 | 11188 | 10.1520744 | 9.78114141 | 7.09568377 | 8.38839809 | 11.4795663 | 5.1901776 |
| ENSMUSG00 | Ndufa5 | 1184 | 104.458868 | 86.4018722 | 91.502295 | 71.2193734 | 64.2570324 | 198.915971 |
| ENSMUSG00 | Fhl1 | 3748 | 1.24103288 | 2.03133881 | 0.44815102 | 1.03009039 | 1.42950092 | 1.87021939 |
| ENSMUSG00 | Gm7257 | 515 | 0 | 0 | 0 | 0 | 0 | 0 |
| ENSMUSG00 | Msrb2 | 1370 | 37.9273088 | 40.4484733 | 44.2986354 | 36.2400681 | 31.1000174 | 38.8016583 |
| ENSMUSG00 | Rfc2 | 1507 | 23.8216317 | 22.4145295 | 21.3529806 | 20.44729 | 21.3992499 | 17.9176462 |
| ENSMUSG00 | Denr | 2932 | 42.0334207 | 40.5771916 | 50.6240753 | 31.8486805 | 34.0059817 | 60.5214198 |
| ENSMUSG00 | Prmt5 | 3701 | 15.1567098 | 16.4050033 | 20.7811961 | 12.6545512 | 16.9582389 | 9.88991242 |
| ENSMUSG00 | Sympk | 7789 | 5.6655005 | 6.09986288 | 4.47182595 | 6.07312368 | 5.86321817 | 3.08498616 |
| ENSMUSG00 | Gm853 | 2574 | 0.0772251 | 0.13104316 | 0 | 0 | 0 | 0 |
| ENSMUSG00 | Sult1c2 | 1838 | 8.45723225 | 8.93992207 | 8.20066837 | 4.75073709 | 3.47023775 | 6.45509688 |
| ENSMUSG00 | Gzma | 933 | 0 | 0 | 0 | 0 | 0 | 0 |
| ENSMUSG00 | Reg2 | 686 | 0 | 0 | 0 | 0 | 0 | 0 |
| ENSMUSG00 | Nagpa | 2945 | 7.87010022 | 11.8134503 | 4.4727162 | 5.03555868 | 9.16568856 | 3.3248202 |
| ENSMUSG00 | Wrb | 4423 | 12.3769606 | 11.4610303 | 10.183513 | 12.8077798 | 9.97915783 | 7.84384832 |
| ENSMUSG00 | Ivns1abp | 4460 | 550.613405 | 685.684428 | 321.622917 | 702.904116 | 694.107443 | 504.501326 |
| ENSMUSG00 | Lrrc69 | 1388 | 0.40099188 | 0.4513194 | 0.15922854 | 0 | 0.88230045 | 0.05895095 |
| ENSMUSG00 | Tmem52 | 1232 | 37.1416867 | 29.3733913 | 48.5789736 | 35.4962971 | 22.1997869 | 52.3575664 |
| ENSMUSG00 | Rpp14 | 2461 | 34.2145907 | 26.0218536 | 44.6329053 | 23.8983066 | 29.9813651 | 25.756303 |
| ENSMUSG00 | Psg29 | 5436 | 0 | 0 | 0 | 0 | 0 | 0 |
| ENSMUSG00 | Ssxb2 | 920 | 0 | 0 | 0 | 0 | 0 | 0 |
| ENSMUSG00 | Slc38a1 | 8373 | 0.83090996 | 0.61578279 | 1.95854338 | 0.59468794 | 1.20664307 | 1.76879602 |
| ENSMUSG00 | Gps2 | 1615 | 47.7804258 | 50.9911019 | 47.4861905 | 52.4809428 | 46.1607028 | 57.0656384 |
| ENSMUSG00 | Bsg | 1744 | 241.678672 | 224.492456 | 334.93541 | 203.230461 | 157.438606 | 333.129066 |
| ENSMUSG00 | Cpn2 | 4581 | 0 | 0 | 0 | 0 | 0 | 0 |
| ENSMUSG00 | Ceacam14 | 1106 | 0 | 0 | 0 | 0 | 0 | 0 |
| ENSMUSG00 | Vwa5a | 4995 | 19.9374332 | 22.0336004 | 16.8400615 | 21.0424416 | 25.7941139 | 15.8678882 |
| ENSMUSG00 | Leprel2 | 4551 | 0.30574419 | 1.34468867 | 0.11655067 | 0.68184044 | 1.26697 | 0.28766923 |
| ENSMUSG00 | Grm2 | 3314 | 0 | 0 | 0 | 0 | 0 | 0 |
| ENSMUSG00 | Il15ra | 2734 | 28.7623776 | 24.5867179 | 23.2973138 | 21.9738859 | 29.8991419 | 25.2893967 |
| ENSMUSG00 | Lcn9 | 832 | 0 | 0 | 0 | 0 | 0 | 0 |
| ENSMUSG00 | Epb4.2 | 6402 | 0 | 0 | 0 | 0 | 0 | 0 |
| ENSMUSG00 | Serping1 | 2638 | 0.30140622 | 0.56625463 | 0.75401173 | 0.4376897 | 1.08319827 | 1.12696576 |
| ENSMUSG00 | Serinc2 | 2872 | 48.6423254 | 35.6532693 | 55.083007 | 33.5819307 | 45.5008874 | 41.9376069 |
| ENSMUSG00 | Ccl25 | 2718 | 4.27100821 | 2.42882361 | 2.32555686 | 2.03110847 | 3.86734121 | 2.66926197 |
| ENSMUSG00 | Scg5 | 1562 | 59.6077691 | 59.1687555 | 131.134021 | 53.5455693 | 47.7923098 | 105.850745 |
| ENSMUSG00 | Kcnk5 | 3359 | 0.82848575 | 0.45905512 | 0.61848366 | 0 | 0.19748225 | 0 |
| ENSMUSG00 | Guca2a | 591 | 91.4174247 | 47.6972397 | 194.532984 | 78.9409197 | 30.6503611 | 193.737624 |
| ENSMUSG00 | Parp3 | 4369 | 0.53686733 | 0.9815961 | 0.97124672 | 1.4039706 | 1.02776861 | 0.91768642 |
| ENSMUSG00 | Cypt1 | 970 | 0 | 0 | 0 | 0.18598992 | 0 | 0 |
| ENSMUSG00 | Slc26a6 | 2063 | 6.51350093 | 4.60141983 | 5.03511057 | 2.92084413 | 4.77367452 | 3.68862063 |
| ENSMUSG00 | Acy1 | 1416 | 4.88520735 | 5.3427058 | 5.83739029 | 4.53573724 | 8.07196906 | 3.00483314 |
| ENSMUSG00 | 9530002B09I | 798 | 0 | 0 | 0 | 0 | 0 | 0 |
| ENSMUSG00 | Frs3 | 3121 | 1.69416178 | 2.42398956 | 1.68536345 | 1.17922734 | 1.42239497 | 1.13607917 |
| ENSMUSG00 | Gabrr2 | 2349 | 1.35395421 | 1.43595187 | 0.71505749 | 0.12288478 | 0.36928412 | 0.90567129 |
| ENSMUSG00 | Creld2 | 1354 | 54.0838959 | 47.0120311 | 53.6363575 | 52.2576731 | 29.5455529 | 44.5378341 |
| ENSMUSG00 | Cd4 | 4055 | 0 | 0 | 0 | 0 | 0 | 0 |
| ENSMUSG00 | Twf2 | 1585 | 9.30554149 | 5.80669429 | 11.935892 | 9.4018198 | 4.44267499 | 7.72638046 |
| ENSMUSG00 | Bmp15 | 3058 | 0 | 0.22060503 | 0 | 0 | 0.58401673 | 0 |
| ENSMUSG00 | Zfp605 | 3576 | 4.10228532 | 2.62761639 | 4.49929275 | 3.87458196 | 2.81101679 | 3.92797517 |
| ENSMUSG00 | Ube2j2 | 4412 | 5.03701578 | 6.65130174 | 4.9391452 | 5.97823539 | 5.70172332 | 5.16190475 |
| ENSMUSG00 | Sva | 647 | 0 | 0 | 0 | 0 | 0 | 0 |
| ENSMUSG00 | 5-Mar | 2127 | 50.1476042 | 48.4809526 | 64.4012748 | 46.0906982 | 45.1968463 | 63.8331708 |
| ENSMUSG00 | Ache | 3962 | 8.76986641 | 17.6107946 | 1.61768482 | 13.4601872 | 12.183481 | 1.96884069 |
| ENSMUSG00 | Dtwd1 | 1345 | 16.5524677 | 15.5844627 | 16.3333204 | 3.67527145 | 10.7364047 | 15.4116921 |
| ENSMUSG00 | Gcm1 | 2028 | 0 | 0 | 0 | 0 | 0 | 0 |
| ENSMUSG00 | Wfdc1 | 1206 | 0 | 0 | 0 | 0 | 0.12693128 | 0 |
| ENSMUSG00 | Mx2 | 2403 | 0.34742617 | 1.00263092 | 0.2207333 | 0.85587871 | 0.998019 | 0.09080196 |
| ENSMUSG00 | Poc1a | 2642 | 2.13674422 | 2.73579346 | 3.34608955 | 1.46130915 | 1.46782913 | 2.28148951 |
| ENSMUSG00 | Trip6 | 2035 | 22.8764952 | 28.9829018 | 25.2830198 | 21.7378804 | 25.0242374 | 21.6186698 |
| ENSMUSG00 | Clec4n | 1561 | 0 | 0 | 0 | 0 | 0 | 0 |
| ENSMUSG00 | 4921501E09f | 3915 | 0 | 0 | 0 | 0 | 0 | 0 |
| ENSMUSG00 | Agap3 | 3499 | 9.25999332 | 13.6888607 | 8.91869139 | 9.39432483 | 12.1477482 | 6.32172976 |
| ENSMUSG00 | Tmem176a | 1111 | 108.352471 | 157.050501 | 134.793739 | 120.522473 | 109.217571 | 114.671323 |
| ENSMUSG00 | Kcnk16 | 874 | 0 | 0 | 0 | 0 | 0 | 0 |
| ENSMUSG00 | Dlx2 | 2909 | 0 | 0 | 0 | 0 | 0 | 0 |
| ENSMUSG00 | Slc17a9 | 3257 | 0.83001924 | 0.11835786 | 0.04071401 | 0 | 0.18800016 | 0 |
| ENSMUSG00 | Stk31 | 3262 | 0 | 0 | 0 | 0 | 0 | 0 |
| ENSMUSG00 | Nfatc4 | 4010 | 0 | 0.37251364 | 0.07716032 | 0.15296627 | 0.41991781 | 0.17684305 |
| ENSMUSG00 | Cela3b | 1740 | 0 | 0 | 0 | 0 | 0 | 0 |
| ENSMUSG00 | Gnb3 | 3158 | 0 | 0 | 0 | 0 | 0 | 0 |
| ENSMUSG00 | Esx1 | 1980 | 0 | 0 | 0 | 0 | 0 | 0 |
| ENSMUSG00 | Pisd | 4935 | 9.60253957 | 7.41104551 | 9.20759769 | 7.12867141 | 7.95122249 | 7.90881637 |
| ENSMUSG00 | Tpi1 | 2915 | 186.44843 | 198.019002 | 254.657205 | 183.207664 | 160.291089 | 182.286285 |
| ENSMUSG00 | Rab12 | 3014 | 28.8471254 | 27.1308535 | 22.4675592 | 26.6485173 | 25.7502049 | 23.5553703 |
| ENSMUSG00 | Tspan18 | 664 | 0 | 0 | 0 | 0 | 0 | 0 |
| ENSMUSG00 | Tulp2 | 3142 | 0.61999317 | 0.36806958 | 0.35170149 | 0.96463773 | 1.21800706 | 0.46875574 |
| ENSMUSG00 | Celsr3 | 11506 | 0 | 0.01675176 | 0.00768327 | 0 | 0.0354781 | 0.01896377 |
| ENSMUSG00 | Prph | 1857 | 0 | 0.25948542 | 0 | 0 | 0 | 0 |
| ENSMUSG00 | Pcbp4 | 2002 | 5.65949648 | 4.33244735 | 5.60802604 | 7.49756769 | 4.07803203 | 6.11703947 |
| ENSMUSG00 | Cdca3 | 2774 | 0.14331464 | 2.11923069 | 0.84451971 | 1.58688155 | 0.55183536 | 0.55060555 |
| ENSMUSG00 | Fam132a | 1307 | 4.8667765 | 10.1755669 | 8.3871898 | 4.03058799 | 7.26159595 | 5.6135256 |
| ENSMUSG00 | Ccndbp1 | 3172 | 28.4003527 | 31.4609463 | 33.8341976 | 29.7460739 | 30.0013212 | 28.7105991 |
| ENSMUSG00 | Iqcf3 | 1114 | 0 | 0 | 0 | 0 | 0 | 0 |
| ENSMUSG00 | Ogfod2 | 2303 | 18.1773864 | 16.3411252 | 12.1684622 | 10.6225038 | 14.3130868 | 11.2627798 |
| ENSMUSG00 | Mrps23 | 1401 | 29.8804575 | 32.3994493 | 33.9795753 | 35.0261103 | 27.680261 | 26.7100678 |
| ENSMUSG00 | 1700082M22 | 2235 | 0.23123993 | 0 | 0.07910844 | 0 | 0.11415297 | 0 |
| ENSMUSG00 | Rhebl1 | 2183 | 3.60585668 | 3.33309792 | 2.53102628 | 3.4214307 | 4.02040158 | 7.0591714 |
| ENSMUSG00 | Sf1 | 6903 | 7.71151505 | 7.78326568 | 6.53774905 | 9.44520563 | 11.1839645 | 8.04449739 |
| ENSMUSG00 | Hes7 | 1805 | 0.15417638 | 0.13348045 | 0.07346567 | 0.01999005 | 0.11307784 | 0.33243328 |
| ENSMUSG00 | Pigx | 2119 | 13.5645174 | 13.6668484 | 14.0386222 | 14.933437 | 9.99336761 | 22.5379393 |
| ENSMUSG00 | Pisd-ps2 | 2130 | 8.24972882 | 5.92715861 | 5.83132295 | 7.18252905 | 7.8815388 | 6.28725252 |
| ENSMUSG00 | Tiam2 | 6852 | 1.125593 | 0.87202552 | 1.18697302 | 0.74775983 | 1.57139258 | 0.81202954 |
| ENSMUSG00 | Nox3 | 1779 | 0 | 0 | 0 | 0 | 0 | 0 |
| ENSMUSG00 | Synj2 | 7492 | 0.583703 | 0.65603538 | 1.39236986 | 0.48642372 | 1.17145444 | 0.69169534 |
| ENSMUSG00 | Rsph3b | 2231 | 4.97166251 | 4.42770983 | 4.16063964 | 4.09177825 | 4.27697547 | 3.70426521 |
| ENSMUSG00 | Rps6ka2 | 5654 | 0.34453812 | 0.39203685 | 0.15635601 | 0.38290107 | 1.31762484 | 0.4679236 |
| ENSMUSG00 | Park2 | 1590 | 2.85039294 | 3.12151166 | 5.44878865 | 4.06206665 | 3.49803457 | 2.55592531 |
| ENSMUSG00 | Agpat4 | 2689 | 18.7762961 | 9.89175012 | 20.5310903 | 13.1097648 | 8.50123331 | 10.5284773 |
| ENSMUSG00 | Slc22a3 | 3490 | 0.07973879 | 0.20710505 | 0.20264455 | 0.52727343 | 0 | 0.24226757 |
| ENSMUSG00 | Slc22a1 | 1983 | 0 | 1.62808453 | 0.49038858 | 0 | 0.30878291 | 0.61894038 |
| ENSMUSG00 | Igf2r | 9388 | 3.15062607 | 4.84020185 | 5.06145944 | 6.50306981 | 6.34296768 | 4.81402612 |
| ENSMUSG00 | Acat2 | 3123 | 6.35222055 | 5.64721032 | 6.58144637 | 6.43538226 | 5.1467526 | 6.89944445 |
| ENSMUSG00 | Lnpep | 5163 | 1.62471555 | 5.70248231 | 4.44329973 | 5.8983625 | 6.42400625 | 4.22088645 |
| ENSMUSG00 | Riok2 | 3461 | 13.7610707 | 14.4517558 | 13.8569777 | 12.8335732 | 13.0182866 | 17.8967656 |
| ENSMUSG00 | Chd1 | 9023 | 5.04488808 | 5.64480432 | 6.62807199 | 6.61417507 | 8.48837306 | 3.80871611 |
| ENSMUSG00 | Brp44l | 1975 | 23.5110892 | 27.76515 | 27.5729978 | 25.8694557 | 19.1187433 | 17.9529271 |
| ENSMUSG00 | Pde10a | 12932 | 0.12604197 | 0.09687964 | 0.01367208 | 0.02790136 | 0.45770643 | 0.00843633 |
| ENSMUSG00 | 1700010I14R | 2178 | 0.40157051 | 1.59294024 | 1.40033387 | 0.89459615 | 2.34280881 | 1.31489308 |
| ENSMUSG00 | Zfp54 | 946 | 96.7831624 | 107.782782 | 89.7119858 | 133.457795 | 75.1910866 | 191.845084 |
| ENSMUSG00 | Phf10 | 5172 | 14.5355015 | 17.3292309 | 14.4775565 | 18.9688861 | 13.5557309 | 21.8165473 |
| ENSMUSG00 | Thbs2 | 5904 | 0 | 0.26117313 | 0 | 0 | 0 | 0 |
| ENSMUSG00 | Smoc2 | 2853 | 0.12541161 | 0.25334617 | 0.26338287 | 0.48058805 | 0.23250715 | 0.21031969 |
| ENSMUSG00 | Zfp51 | 2595 | 90.3269029 | 56.8395759 | 57.9989501 | 102.406265 | 54.0545307 | 89.2337908 |
| ENSMUSG00 | Zscan10 | 2774 | 0 | 0 | 0 | 0 | 0 | 0 |
| ENSMUSG00 | Mmp25 | 3363 | 0 | 0 | 0 | 0 | 0 | 0 |
| ENSMUSG00 | Hcfc1r1 | 1240 | 28.405932 | 29.2227456 | 37.3933333 | 28.2254703 | 18.599937 | 51.8658059 |
| ENSMUSG00 | Tnfrsf12a | 959 | 13.0583695 | 15.7774169 | 19.7732958 | 16.3667251 | 9.15175875 | 17.6616799 |
| ENSMUSG00 | Cldn6 | 1503 | 7.88232415 | 4.80902616 | 8.17572346 | 2.1125881 | 6.21279227 | 4.46411261 |
| ENSMUSG00 | Pkmyt1 | 2031 | 2.23147453 | 0.56941143 | 1.47992384 | 0.05329696 | 1.05519832 | 1.75922095 |
| ENSMUSG00 | Paqr4 | 2504 | 3.88981339 | 3.29068754 | 1.94177425 | 2.91077196 | 5.84847039 | 2.25473256 |
| ENSMUSG00 | Flywch2 | 1132 | 2.10718095 | 4.68242812 | 3.16285272 | 2.7412154 | 1.35228911 | 3.66231905 |
| ENSMUSG00 | Slc25a27 | 2836 | 2.14477733 | 1.52918893 | 1.71445795 | 4.28760543 | 3.07669614 | 2.44279209 |
| ENSMUSG00 | Pla2g7 | 2648 | 15.5688948 | 10.9183781 | 9.28105163 | 9.57918327 | 6.08925029 | 6.22125457 |
| ENSMUSG00 | Mep1a | 4113 | 3.13172275 | 0.73808555 | 0.95647071 | 0.13159024 | 0.11165509 | 2.40053961 |
| ENSMUSG00 | Tnfrsf21 | 3620 | 47.42104 | 22.7754151 | 50.6001208 | 28.0483075 | 34.5203299 | 25.4136349 |
| ENSMUSG00 | Gpr115 | 3736 | 0 | 0.20636592 | 0.22479524 | 0.05794761 | 0.06829012 | 0.22631526 |
| ENSMUSG00 | Cenpq | 2594 | 3.21844676 | 5.44280168 | 4.03848758 | 5.11880971 | 2.91129669 | 4.93130525 |
| ENSMUSG00 | Mut | 3284 | 139.507356 | 114.479483 | 141.071716 | 132.956401 | 131.093037 | 112.354239 |
| ENSMUSG00 | Tbc1d5 | 8068 | 10.751916 | 6.51006717 | 8.3659165 | 7.58044936 | 5.57191833 | 6.83218219 |
| ENSMUSG00 | Rhag | 2329 | 0.29018599 | 0.20689756 | 0.37957787 | 0.48026766 | 0.46009184 | 0.97200303 |
| ENSMUSG00 | Satb1 | 15177 | 1.75503538 | 0.80009115 | 2.77262664 | 1.49063991 | 1.41207603 | 1.89234995 |
| ENSMUSG00 | Crisp2 | 1814 | 0 | 0 | 0.07310117 | 0.01989087 | 0.33755045 | 0 |
| ENSMUSG00 | Efhb | 2648 | 0 | 0 | 0 | 0 | 0 | 0 |
| ENSMUSG00 | Cdc5l | 2973 | 20.5797795 | 16.4349318 | 15.5219388 | 19.1272459 | 16.3394248 | 14.0822641 |
| ENSMUSG00 | Spats1 | 1369 | 0.11615918 | 0.73916383 | 0.80719217 | 0.73798192 | 0.63363652 | 0.21915342 |
| ENSMUSG00 | Aars2 | 3326 | 15.2041405 | 11.0686836 | 8.71810253 | 10.3169045 | 11.8897899 | 5.35488257 |
| ENSMUSG00 | Mrpl14 | 1015 | 12.2595718 | 16.2362199 | 15.9823413 | 14.8594036 | 8.09383892 | 22.5721152 |
| ENSMUSG00 | Sgol1 | 3801 | 0 | 0 | 0 | 0 | 0.09397123 | 0 |
| ENSMUSG00 | Slc29a1 | 3962 | 11.3085377 | 17.2094436 | 6.54883439 | 13.8153613 | 17.7729426 | 8.37101497 |
| ENSMUSG00 | Sult1c1 | 1548 | 0 | 0 | 0 | 0 | 0 | 0 |
| ENSMUSG00 | Hsp90ab1 | 2730 | 424.014771 | 429.847775 | 437.09632 | 420.851451 | 353.315598 | 317.894413 |
| ENSMUSG00 | Slc5a7 | 5167 | 0 | 0 | 0 | 0 | 0 | 0 |
| ENSMUSG00 | Nfkbie | 2416 | 2.71508869 | 4.04877804 | 1.22579614 | 1.07529272 | 3.25250906 | 2.5287745 |
| ENSMUSG00 | Tcte1 | 3069 | 0.89381823 | 0.92636041 | 1.77153037 | 0.88177039 | 0.94770395 | 1.60857273 |
| ENSMUSG00 | Vegfa | 6165 | 29.6634567 | 35.7197146 | 30.4358189 | 41.5075198 | 34.4645302 | 43.9137176 |
| ENSMUSG00 | Gtpbp2 | 5507 | 8.85781271 | 11.4362956 | 6.3088158 | 6.66996937 | 11.9249947 | 9.09815005 |
| ENSMUSG00 | Polh | 3429 | 1.91299338 | 1.41931486 | 0.41249897 | 1.97825149 | 1.59224911 | 0.73177801 |
| ENSMUSG00 | Clic5 | 5900 | 0.13476434 | 0.58803794 | 0.78664297 | 1.17419535 | 2.1015948 | 0 |
| ENSMUSG00 | Enpp5 | 4824 | 257.017863 | 135.789074 | 249.954825 | 124.813656 | 137.868529 | 158.932444 |
| ENSMUSG00 | Enpp4 | 7866 | 23.3246304 | 27.6523773 | 28.1923019 | 30.3344216 | 33.4142972 | 23.8106977 |
| ENSMUSG00 | Cyp39a1 | 3325 | 16.6674105 | 17.9702822 | 21.1504157 | 25.8054442 | 21.6535989 | 26.4215378 |
| ENSMUSG00 | Calcr | 5864 | 2.9626782 | 0.320476 | 2.84930025 | 1.85824991 | 1.10510739 | 2.67444024 |
| ENSMUSG00 | Fbxl17 | 8141 | 10.3283186 | 9.97940566 | 11.0708215 | 9.40056706 | 9.59604243 | 8.54656734 |
| ENSMUSG00 | Rsph9 | 1596 | 14.0738387 | 14.431779 | 10.0257313 | 7.16667175 | 10.1988809 | 12.5948679 |
| ENSMUSG00 | Mrps18a | 3522 | 10.2379957 | 8.3731126 | 13.5667781 | 8.01140228 | 8.3450291 | 11.8406938 |
| ENSMUSG00 | Crip3 | 1468 | 0 | 0.03282455 | 0 | 0 | 0.27807289 | 0.55738364 |
| ENSMUSG00 | Rrp36 | 1220 | 15.9999516 | 23.2242852 | 25.5066373 | 19.4606437 | 16.6463095 | 17.6167342 |
| ENSMUSG00 | Ptk7 | 4688 | 3.27338214 | 3.03226999 | 3.65834366 | 3.24799974 | 4.68032031 | 4.56128779 |
| ENSMUSG00 | Cnpy3 | 3804 | 12.3007888 | 15.1374339 | 11.1666591 | 11.856613 | 11.6700702 | 10.2244047 |
| ENSMUSG00 | Ubr2 | 7699 | 9.81106806 | 8.61209825 | 10.5868552 | 11.7258443 | 13.9180919 | 8.0382071 |
| ENSMUSG00 | Prph2 | 3798 | 0 | 0 | 0 | 0 | 0 | 0 |
| ENSMUSG00 | Guca1b | 1716 | 0.34751294 | 0.08424203 | 0.12879325 | 0.88312696 | 0.2973565 | 0 |
| ENSMUSG00 | Taf8 | 2829 | 12.6897133 | 9.48739776 | 12.1090238 | 6.9766272 | 14.429516 | 10.6051973 |
| ENSMUSG00 | Guca1a | 867 | 0.22927036 | 0.05557837 | 0.10196504 | 0.33293697 | 0 | 0 |
| ENSMUSG00 | Gm20517 | 6579 | 0 | 0.02197284 | 0.22171467 | 0.39487874 | 0.05429163 | 0.11193422 |
| ENSMUSG00 | Pgc | 1500 | 0 | 0 | 0 | 0 | 0.23812369 | 0 |
| ENSMUSG00 | Bysl | 3847 | 8.7013555 | 6.72631126 | 5.61858623 | 7.52217305 | 5.8891892 | 2.09859455 |
| ENSMUSG00 | Tfeb | 5648 | 7.3485697 | 7.00443856 | 9.25045652 | 8.60526097 | 8.63690072 | 7.53819261 |
| ENSMUSG00 | Foxp4 | 5498 | 0.41939212 | 0.77126353 | 0.56277356 | 0.60377375 | 0.29698873 | 0.99712668 |
| ENSMUSG00 | Trem2 | 3234 | 0 | 0 | 0 | 0 | 0 | 0 |
| ENSMUSG00 | Treml1 | 1242 | 0 | 0 | 0 | 0 | 0 | 0 |
| ENSMUSG00 | Nfya | 8289 | 0.04796173 | 0.0465064 | 0.21863621 | 0.18282614 | 0.12311829 | 0.15136125 |
| ENSMUSG00 | Tspo2 | 4104 | 0 | 0 | 0 | 0 | 0 | 0 |
| ENSMUSG00 | Kif6 | 6157 | 0.2518213 | 0.45392458 | 0.32306041 | 0.50984861 | 0.21547601 | 0.25250194 |
| ENSMUSG00 | Brd4 | 9921 | 7.36924994 | 8.24235383 | 7.46721993 | 9.81609094 | 6.87140793 | 9.07505132 |
| ENSMUSG00 | Stk38 | 8421 | 16.2638203 | 18.0591867 | 18.87541 | 19.2686087 | 19.2689556 | 16.4438123 |
| ENSMUSG00 | Ppil1 | 2657 | 24.2991721 | 23.0322851 | 33.4549892 | 21.048991 | 18.7628036 | 23.1685588 |
| ENSMUSG00 | Cpne5 | 13296 | 0 | 0 | 0 | 0 | 0 | 0 |
| ENSMUSG00 | Pi16 | 4287 | 0.5656833 | 0.05620066 | 0.34025212 | 0.10941604 | 0.11902564 | 0.19086522 |
| ENSMUSG00 | Mtch1 | 3041 | 87.2371988 | 98.5278847 | 87.2699327 | 80.9443301 | 71.665127 | 56.9170867 |
| ENSMUSG00 | Fgd2 | 4344 | 0.62232336 | 1.09817168 | 0.61052269 | 0.76416853 | 1.04542989 | 0.40183631 |
| ENSMUSG00 | Pim1 | 1966 | 0.64708819 | 0.26960678 | 0.29228075 | 0.49553164 | 0.5969515 | 0.23584378 |
| ENSMUSG00 | 1110021J02R | 4889 | 1.3091905 | 1.11373169 | 0.8860259 | 2.21407514 | 1.96215149 | 1.13807045 |
| ENSMUSG00 | Ftsjd2 | 8058 | 0.89792721 | 0.58005521 | 0.82281912 | 0.95376961 | 0.98152001 | 0.92066293 |
| ENSMUSG00 | Glo1 | 2137 | 44.0714709 | 51.6138353 | 49.434911 | 48.2557245 | 40.1620796 | 62.3475025 |
| ENSMUSG00 | Glp1r | 1579 | 0 | 0 | 0 | 0 | 0 | 0 |
| ENSMUSG00 | Tff2 | 580 | 0 | 0.24924022 | 0 | 0.31105211 | 0 | 0 |
| ENSMUSG00 | Tff3 | 490 | 0 | 0 | 0 | 0 | 0 | 0 |
| ENSMUSG00 | Abcg1 | 4133 | 1.52942691 | 1.91206787 | 1.24060339 | 0.26190693 | 0.38272868 | 0.60712964 |
| ENSMUSG00 | Tff1 | 475 | 0 | 0 | 0 | 0 | 0 | 0 |
| ENSMUSG00 | Rsph1 | 1350 | 3.68106301 | 3.06965483 | 1.99726846 | 4.0358435 | 6.27435436 | 3.67702544 |
| ENSMUSG00 | Tmprss3 | 2866 | 0 | 0 | 0 | 0 | 0 | 0 |
| ENSMUSG00 | Slc37a1 | 3605 | 5.1389886 | 8.75509559 | 4.32822485 | 4.63411557 | 6.77992622 | 3.88124547 |
| ENSMUSG00 | Wdr4 | 7758 | 2.46998474 | 3.08075217 | 2.62088783 | 2.66034151 | 3.56487441 | 2.0285469 |
| ENSMUSG00 | Ndufv3 | 2087 | 30.7833524 | 26.5752731 | 35.2640482 | 25.6913839 | 31.4666726 | 40.4068091 |
| ENSMUSG00 | Cbs | 3975 | 0.1900262 | 1.26072704 | 0.07783972 | 0.20877661 | 0.60333073 | 0 |
| ENSMUSG00 | Cryaa | 1182 | 0 | 0 | 0 | 0 | 0 | 0 |
| ENSMUSG00 | Sik1 | 4497 | 2.73169746 | 2.2073398 | 2.90943863 | 2.7280176 | 1.87221526 | 2.15916646 |
| ENSMUSG00 | Arhgap28 | 5904 | 2.32984354 | 0.84065101 | 2.44817118 | 1.36896646 | 2.04831488 | 0.8223045 |
| ENSMUSG00 | Epb4.1l3 | 5813 | 5.19768883 | 15.4017563 | 3.29251643 | 7.65339083 | 15.2034548 | 8.15001692 |
| ENSMUSG00 | Akap8 | 3681 | 46.9160031 | 40.1095514 | 36.9969786 | 46.3351873 | 46.4657462 | 59.0172512 |
| ENSMUSG00 | Myl12a | 3570 | 36.8712594 | 43.5432668 | 35.5719594 | 33.3834715 | 30.3156703 | 38.6581722 |
| ENSMUSG00 | Myom1 | 5694 | 0.50270365 | 0.27080544 | 0.13196897 | 0.22178988 | 0.50184002 | 0.15328213 |
| ENSMUSG00 | Wiz | 10318 | 2.28484202 | 3.20838203 | 2.261928 | 2.95496488 | 3.86233762 | 3.38091332 |
| ENSMUSG00 | Lpin2 | 11448 | 11.6196574 | 18.1330532 | 11.3052932 | 10.060612 | 20.5032607 | 10.5400947 |
| ENSMUSG00 | Emilin2 | 5060 | 1.30423118 | 2.95213381 | 0.45424819 | 2.4601394 | 0.1411797 | 1.07265874 |
| ENSMUSG00 | Smchd1 | 8402 | 9.06114558 | 6.76743653 | 8.64361201 | 8.83370216 | 9.4194131 | 5.27508793 |
| ENSMUSG00 | Cyp4f13 | 3534 | 7.32337799 | 10.5808373 | 10.7440247 | 8.70910696 | 9.89051144 | 10.0794224 |
| ENSMUSG00 | Ndc80 | 2184 | 0.38226424 | 0.04412678 | 0 | 0.72692764 | 0 | 0.36216325 |
| ENSMUSG00 | Clip4 | 4445 | 0 | 0.31437724 | 0 | 0 | 0 | 0 |
| ENSMUSG00 | Lbh | 3203 | 0 | 0 | 0 | 0 | 0 | 0 |
| ENSMUSG00 | Galnt14 | 5332 | 0.61139336 | 0.36148869 | 0.42278582 | 0 | 0.60289979 | 0 |
| ENSMUSG00 | Ehd3 | 3652 | 7.88142604 | 5.87156811 | 13.5800843 | 7.23221703 | 17.0469511 | 4.20471574 |
| ENSMUSG00 | Xdh | 4655 | 31.7018998 | 12.9601344 | 26.8534505 | 14.9986587 | 20.8161305 | 17.9291936 |
| ENSMUSG00 | Dpy30 | 968 | 46.2855646 | 44.652106 | 53.2887921 | 44.8789066 | 29.0449722 | 81.1195107 |
| ENSMUSG00 | Spast | 4679 | 6.81425419 | 8.46532495 | 7.93535972 | 8.7525369 | 9.4222673 | 7.36805813 |
| ENSMUSG00 | Slc30a6 | 2118 | 33.185878 | 25.9360454 | 37.3357399 | 22.2829623 | 29.3919634 | 23.5530289 |
| ENSMUSG00 | Prkd3 | 12405 | 2.96443527 | 2.68803047 | 3.74138937 | 1.71320631 | 3.31537758 | 3.91145371 |
| ENSMUSG00 | Yipf4 | 2313 | 112.202238 | 102.247755 | 131.822012 | 106.826563 | 83.2349831 | 146.691099 |
| ENSMUSG00 | Birc6 | 16139 | 2.62096978 | 2.54979996 | 3.94664203 | 4.41998897 | 2.61787218 | 2.96423058 |
| ENSMUSG00 | Crim1 | 5978 | 22.8570737 | 18.6764782 | 26.5669491 | 17.3408688 | 23.2683005 | 24.3774502 |
| ENSMUSG00 | Vit | 2617 | 0 | 0 | 0 | 0 | 0 | 0 |
| ENSMUSG00 | Strn | 9901 | 5.15564458 | 3.10016803 | 4.85278289 | 3.83378556 | 4.90112952 | 4.4847099 |
| ENSMUSG00 | Ttc27 | 2800 | 4.79905443 | 3.52793594 | 3.67822479 | 5.78601357 | 5.32132205 | 2.83461429 |
| ENSMUSG00 | Eif2ak2 | 4837 | 9.59161583 | 7.60104514 | 13.8901802 | 12.42022 | 9.86348178 | 16.2283258 |
| ENSMUSG00 | Cebpz | 4058 | 14.8323799 | 11.5538216 | 13.6810042 | 14.9734261 | 13.9700107 | 11.2311292 |
| ENSMUSG00 | 2410091C18I | 2293 | 23.0245434 | 21.2667595 | 25.9852091 | 17.8600613 | 20.4951121 | 20.3756899 |

| | | | | | | | | |
|---|---|---|---|---|---|---|---|---|
| ENSMUSG00 | Pja2 | 6551 | 62.4702971 | 43.7289572 | 61.0971895 | 56.3234601 | 52.1713731 | 62.1641946 |
| ENSMUSG00 | Qpct | 1917 | 10.7010057 | 8.94855159 | 16.9013829 | 6.28659513 | 6.01562907 | 21.6119859 |
| ENSMUSG00 | Man2a1 | 6929 | 6.52933134 | 7.83055764 | 6.02839395 | 8.96192793 | 7.50409539 | 6.72714078 |
| ENSMUSG00 | Cyp1b1 | 6752 | 0 | 0 | 0 | 0 | 0.41564732 | 0 |
| ENSMUSG00 | 4930583I09R | 626 | 0 | 0 | 0 | 0 | 0 | 0 |
| ENSMUSG00 | Vapa | 1634 | 172.038557 | 158.625993 | 163.606332 | 149.075816 | 129.62698 | 177.869373 |
| ENSMUSG00 | Hnrpll | 2753 | 33.1271603 | 41.0626202 | 38.1006079 | 50.7874764 | 41.6292191 | 35.8047752 |
| ENSMUSG00 | Ralbp1 | 5189 | 18.1883813 | 18.0896491 | 22.5396178 | 21.3404885 | 19.1262866 | 22.9908021 |
| ENSMUSG00 | Snf7 | 2656 | 112.919934 | 133.71012 | 133.953552 | 177.842637 | 93.7148574 | 164.233172 |
| ENSMUSG00 | Twsg1 | 4041 | 167.984368 | 148.983274 | 238.980912 | 154.525084 | 175.92167 | 173.765303 |
| ENSMUSG00 | Ndufv2 | 2126 | 233.558773 | 194.921638 | 259.451928 | 180.970292 | 133.326208 | 394.494429 |
| ENSMUSG00 | ORF19 | 2979 | 1.41459582 | 1.43960838 | 1.3205653 | 1.42923171 | 0.83930595 | 1.87690535 |
| ENSMUSG00 | D6Wsu116e | 4268 | 39.0289277 | 33.5205124 | 40.1317108 | 47.8585881 | 47.4755713 | 36.1190865 |
| ENSMUSG00 | 9130404H23I | 1793 | 0 | 0.16124855 | 0 | 0 | 0 | 0 |
| ENSMUSG00 | Lhcgr | 2542 | 0 | 0 | 0 | 0 | 0 | 0 |
| ENSMUSG00 | Nrxn1 | 15676 | 0 | 0 | 0 | 0 | 0 | 0 |
| ENSMUSG00 | Cacna1h | 11518 | 0 | 0 | 0 | 0 | 0 | 0 |
| ENSMUSG00 | Prss41 | 2564 | 0 | 0.07517386 | 0 | 0 | 1.09455954 | 0.17020056 |
| ENSMUSG00 | Prss21 | 1088 | 0 | 0 | 0 | 0 | 0 | 0 |
| ENSMUSG00 | 1600002H07I | 3309 | 0.86503312 | 1.12129225 | 2.07050035 | 1.83190797 | 1.46494582 | 1.15395875 |
| ENSMUSG00 | Lrpprc | 8123 | 17.330316 | 11.5794577 | 17.9898182 | 12.7528684 | 21.7724263 | 13.2024968 |
| ENSMUSG00 | Atp6v0c | 3833 | 52.772211 | 42.5418524 | 69.479808 | 46.0133143 | 35.7703814 | 27.08963 |
| ENSMUSG00 | Pdpk1 | 14050 | 6.37785431 | 7.30856286 | 8.22689107 | 9.32740112 | 8.28406856 | 6.72450892 |
| ENSMUSG00 | Prss30 | 1920 | 0 | 0 | 0 | 0 | 0 | 0 |
| ENSMUSG00 | Sbpl | 867 | 0 | 0 | 0 | 0 | 0 | 0.28312774 |
| ENSMUSG00 | Prepl | 3815 | 25.5310425 | 24.0490133 | 20.8438049 | 27.8820098 | 25.078494 | 16.8151904 |
| ENSMUSG00 | Sbp | 553 | 0.35945281 | 0 | 0.079931 | 0 | 0.09227193 | 1.57827931 |
| ENSMUSG00 | Abca3 | 8993 | 29.5745763 | 26.3517094 | 27.7705341 | 31.1550179 | 35.5420014 | 28.3846713 |
| ENSMUSG00 | Slc3a1 | 2297 | 0.95191616 | 0.31466984 | 0.59654207 | 0.89537507 | 0.42207277 | 1.73360789 |
| ENSMUSG00 | Eci1 | 1084 | 64.3275949 | 74.2800229 | 77.067789 | 67.5706184 | 58.9815949 | 64.5130766 |
| ENSMUSG00 | Six2 | 3074 | 0 | 0 | 0 | 0 | 0 | 0 |
| ENSMUSG00 | Srbd1 | 4218 | 4.09996065 | 1.9078084 | 1.70813191 | 3.61846961 | 2.98803102 | 1.86874603 |
| ENSMUSG00 | Dnase1l2 | 3197 | 2.15129751 | 2.04984552 | 1.96329738 | 2.26853017 | 2.17065596 | 1.26263579 |
| ENSMUSG00 | E4f1 | 2547 | 22.6795105 | 17.291876 | 15.8272795 | 20.3147436 | 25.503171 | 19.7144134 |
| ENSMUSG00 | Epas1 | 5293 | 11.6720228 | 7.49243187 | 11.85842 | 8.37119794 | 9.08121028 | 13.2585768 |
| ENSMUSG00 | Mlst8 | 3350 | 13.4812615 | 10.1982649 | 8.0486938 | 10.1352848 | 9.26090642 | 8.66275109 |
| ENSMUSG00 | Rhoq | 4369 | 18.6811631 | 19.8745637 | 16.7843574 | 16.7980953 | 20.0998839 | 22.954646 |
| ENSMUSG00 | Pigf | 982 | 17.7320779 | 18.3520666 | 17.0145587 | 13.6318111 | 12.8345365 | 36.5791241 |
| ENSMUSG00 | Cript | 1272 | 173.680193 | 143.309207 | 129.44329 | 141.009311 | 87.2502888 | 218.282883 |
| ENSMUSG00 | Mcfd2 | 4003 | 51.4844894 | 51.159725 | 50.2087885 | 48.6381994 | 41.7847764 | 41.242416 |
| ENSMUSG00 | Msh2 | 5669 | 11.6973261 | 9.4944886 | 11.7346576 | 10.3237037 | 14.9055704 | 11.5853469 |
| ENSMUSG00 | Gtf2a1l | 1647 | 0 | 0 | 0 | 0 | 0 | 0 |
| ENSMUSG00 | 4930528F23I | 1689 | 0 | 0 | 0 | 0 | 0 | 0.6459358 |
| ENSMUSG00 | Hagh | 1896 | 177.159312 | 173.76092 | 199.677629 | 148.800372 | 120.541739 | 176.033629 |
| ENSMUSG00 | Spsb3 | 2798 | 5.41345178 | 3.99544481 | 4.42334383 | 4.02344456 | 4.65036663 | 4.25008675 |
| ENSMUSG00 | Mapk8ip3 | 11743 | 9.24908224 | 7.78008127 | 6.01879818 | 8.29920823 | 12.2405945 | 7.84352012 |
| ENSMUSG00 | C3 | 5199 | 0 | 0.17609971 | 0 | 0 | 0 | 0 |
| ENSMUSG00 | Hn1l | 2928 | 110.087922 | 93.5258028 | 109.025174 | 96.1325237 | 96.8594934 | 77.5296525 |
| ENSMUSG00 | Tmem204 | 2587 | 3.27325758 | 1.62049496 | 0.61510102 | 2.18974913 | 3.35310549 | 0.8856087 |
| ENSMUSG00 | Ift140 | 7231 | 6.84491401 | 2.93876656 | 4.93305038 | 7.38506789 | 6.37917905 | 4.80917785 |
| ENSMUSG00 | Telo2 | 5961 | 4.37503694 | 5.57769587 | 6.20649935 | 4.55792309 | 4.75938013 | 3.87088489 |
| ENSMUSG00 | Prss28 | 1184 | 0 | 0 | 0 | 0 | 0.56025582 | 0.29946817 |
| ENSMUSG00 | St6gal2 | 7788 | 0 | 0 | 0 | 0 | 0 | 0 |
| ENSMUSG00 | Tpsab1 | 1597 | 0 | 0 | 0 | 0 | 0 | 0 |
| ENSMUSG00 | Pot1b | 3482 | 1.47284807 | 1.39771127 | 1.19327204 | 0.44558527 | 1.96368017 | 1.75460057 |
| ENSMUSG00 | Tekt4 | 1526 | 0 | 0.28419265 | 0 | 0 | 0 | 0 |
| ENSMUSG00 | Sox8 | 3208 | 0 | 0 | 0 | 0 | 0.01590598 | 0 |
| ENSMUSG00 | Nme4 | 1493 | 4.50011802 | 3.09839146 | 5.9804235 | 4.49515089 | 1.46961431 | 4.40267117 |
| ENSMUSG00 | Tmem8 | 3451 | 45.9877943 | 41.6377776 | 43.894441 | 31.6907207 | 40.764914 | 28.0728828 |
| ENSMUSG00 | Mrpl28 | 1395 | 65.8601545 | 70.0171457 | 74.8104311 | 52.9979278 | 45.9420273 | 63.3084458 |
| ENSMUSG00 | Axin1 | 4124 | 20.3308216 | 18.9520876 | 15.7879037 | 14.7950381 | 23.8304558 | 11.8847058 |
| ENSMUSG00 | Pdia2 | 1830 | 0.04344861 | 0.02633139 | 0 | 0.15773571 | 0 | 0.10432922 |
| ENSMUSG00 | Rgs11 | 3770 | 48.6556464 | 52.9923048 | 39.019558 | 47.8063163 | 59.1743541 | 43.9649822 |
| ENSMUSG00 | Itfg3 | 4972 | 67.1653331 | 60.3299683 | 65.8049159 | 48.0706884 | 71.5723944 | 41.4989232 |
| ENSMUSG00 | Luc7l | 7214 | 21.3711884 | 17.7743447 | 18.7615807 | 26.6689023 | 25.0817201 | 28.6735326 |
| ENSMUSG00 | Dusp1 | 3754 | 17.2407891 | 14.9924785 | 25.2800631 | 18.9637384 | 26.4374803 | 20.9899928 |
| ENSMUSG00 | Bnip1 | 5140 | 5.22080727 | 3.59992097 | 4.88456164 | 6.31786772 | 5.39052959 | 7.80033457 |
| ENSMUSG00 | Phf1 | 2414 | 10.9187587 | 13.7732581 | 9.72294061 | 12.0173835 | 9.74447359 | 12.1572129 |
| ENSMUSG00 | Cuta | 806 | 31.9128981 | 36.1099393 | 31.6432549 | 33.3512694 | 21.4614658 | 45.2772547 |
| ENSMUSG00 | Plin3 | 2151 | 101.578857 | 87.5017401 | 103.117084 | 76.877736 | 70.0041074 | 71.4897332 |
| ENSMUSG00 | Kdm4b | 5025 | 13.077773 | 11.3250126 | 14.3820922 | 12.8602869 | 16.1558137 | 12.5707591 |
| ENSMUSG00 | Rpl36-ps2 | 310 | 0 | 0 | 0 | 0.11639369 | 0.16460121 | 0.43991354 |
| ENSMUSG00 | Rfx2 | 3335 | 1.12054428 | 0.76578154 | 1.20610726 | 2.47759766 | 1.00981733 | 2.21632001 |
| ENSMUSG00 | Acsbg2 | 2878 | 0 | 0 | 0 | 0 | 0 | 0 |
| ENSMUSG00 | 2900010M23 | 1003 | 16.5680866 | 19.4090953 | 20.6245888 | 14.3177006 | 12.4131961 | 35.9220324 |
| ENSMUSG00 | 1700061G19I | 3089 | 0 | 0 | 0.02861887 | 0.01168082 | 0.01651874 | 0 |
| ENSMUSG00 | Ip6k3 | 2301 | 0 | 0.27223979 | 0.53787553 | 0 | 0.42133904 | 0 |
| ENSMUSG00 | Grm8 | 8399 | 0 | 0 | 0 | 0 | 0 | 0 |
| ENSMUSG00 | Mllt1 | 3621 | 7.31210992 | 13.7200281 | 7.70266858 | 9.06784328 | 8.51144187 | 8.01442037 |
| ENSMUSG00 | Nudt3 | 3357 | 17.5033066 | 16.521476 | 16.6036712 | 16.1761921 | 14.1055933 | 12.8532855 |
| ENSMUSG00 | Spdef | 2754 | 0 | 0 | 0 | 0 | 0 | 0 |
| ENSMUSG00 | Snrpc | 6244 | 1.12695709 | 0.30097233 | 0.34687545 | 0.52585939 | 0.90709925 | 0.99593494 |
| ENSMUSG00 | Taf11 | 2945 | 6.5336681 | 6.36486451 | 6.87417457 | 6.67732233 | 5.94297009 | 5.76055199 |
| ENSMUSG00 | Anks1 | 7392 | 4.46388648 | 3.44840745 | 2.63106208 | 4.65181121 | 3.55500319 | 3.0182163 |
| ENSMUSG00 | Zfp523 | 3378 | 24.7500816 | 23.9791028 | 18.2669547 | 17.902163 | 24.7125965 | 14.7111879 |
| ENSMUSG00 | Fkbp5 | 8353 | 4.73562829 | 5.01882014 | 3.87354832 | 6.88984329 | 3.04215673 | 2.41955078 |
| ENSMUSG00 | 4930511I11R | 1077 | 0 | 0 | 0 | 0 | 0 | 0 |
| ENSMUSG00 | Gm749 | 439 | 0 | 0 | 0 | 0 | 0 | 0 |
| ENSMUSG00 | Clps | 774 | 0 | 0 | 0 | 0 | 0 | 0 |
| ENSMUSG00 | 2610034M16 | 4627 | 0 | 0.08331349 | 0.06687117 | 0.17155932 | 0 | 0 |
| ENSMUSG00 | Nudt12 | 6574 | 9.61533526 | 10.7455619 | 9.41323094 | 13.5129288 | 11.3245334 | 7.54678564 |
| ENSMUSG00 | Cul2 | 4439 | 30.3159048 | 34.7910672 | 33.7861801 | 37.6427007 | 29.4157458 | 40.6815469 |
| ENSMUSG00 | Bambi | 5037 | 1.16811815 | 0.5261573 | 0.73713616 | 0.52292818 | 0.94211891 | 1.33205504 |
| ENSMUSG00 | Lyzl1 | 926 | 0 | 0 | 0 | 0 | 0 | 0 |
| ENSMUSG00 | Mtpap | 2642 | 21.0815398 | 22.7618016 | 22.2180346 | 22.9439193 | 19.6804985 | 30.3716838 |
| ENSMUSG00 | Map3k8 | 5591 | 1.2016949 | 0.4481658 | 2.15040267 | 0.63245222 | 1.44199023 | 3.39530476 |
| ENSMUSG00 | Svil | 10894 | 5.75494701 | 4.02954368 | 4.94602962 | 3.83872679 | 5.40053344 | 2.80657891 |
| ENSMUSG00 | Zeb1 | 9794 | 0 | 0 | 0 | 0 | 0 | 0 |
| ENSMUSG00 | Epc1 | 5812 | 6.81972025 | 6.95602632 | 8.57874723 | 5.87295494 | 5.57497396 | 5.23718125 |
| ENSMUSG00 | Sos1 | 8897 | 8.03420861 | 8.47609104 | 10.1400429 | 12.5356412 | 9.29681412 | 13.7829356 |
| ENSMUSG00 | Map4k3 | 4196 | 39.0733083 | 38.4366115 | 32.6668053 | 48.8001913 | 49.9440719 | 38.6109049 |
| ENSMUSG00 | Tmem178 | 1692 | 43.0919334 | 89.2816171 | 18.1039463 | 59.9661402 | 50.8454312 | 28.8704958 |
| ENSMUSG00 | Thumpd2 | 2011 | 6.60284957 | 5.5590525 | 2.79146398 | 4.25233444 | 6.41952913 | 4.82832997 |
| ENSMUSG00 | Pkdcc | 3383 | 2.40906695 | 4.45827857 | 0.95380861 | 2.05847904 | 2.80547026 | 2.08812639 |
| ENSMUSG00 | Cox7a2l | 1143 | 32.0687253 | 28.1614553 | 34.9592879 | 26.1697418 | 23.0801717 | 28.7063788 |
| ENSMUSG00 | Thada | 7849 | 4.17359102 | 4.11939135 | 2.55671254 | 2.3352884 | 4.52469838 | 3.42279522 |
| ENSMUSG00 | Dync2li1 | 1311 | 32.4472648 | 29.2206114 | 23.0618311 | 28.8436176 | 28.3350127 | 31.2690944 |
| ENSMUSG00 | Abcg8 | 3492 | 0 | 0 | 0 | 0 | 0 | 0 |
| ENSMUSG00 | Adcyap1 | 2083 | 0 | 0 | 0 | 0 | 0 | 0 |
| ENSMUSG00 | Polr2d | 952 | 24.7218954 | 26.3709416 | 36.5872398 | 22.1722648 | 18.1165074 | 29.4233766 |
| ENSMUSG00 | Slc25a46 | 6638 | 36.1620355 | 28.0131787 | 39.3741334 | 27.6241263 | 25.1365244 | 33.4214998 |
| ENSMUSG00 | Sap130 | 4009 | 9.64881081 | 14.639832 | 9.50411292 | 16.5694797 | 16.8645418 | 12.2732475 |
| ENSMUSG00 | Syt4 | 3897 | 0 | 0 | 0 | 0 | 0 | 0 |
| ENSMUSG00 | Adad2 | 1918 | 0 | 0 | 0 | 0 | 0 | 0 |
| ENSMUSG00 | Celf4 | 3985 | 0.28931215 | 0.08464369 | 0.08873645 | 0.25352503 | 0.17926456 | 0.17795248 |
| ENSMUSG00 | 5730494M16 | 6832 | 5.35930878 | 6.20668459 | 5.39583388 | 4.71094609 | 4.66795739 | 3.69676756 |
| ENSMUSG00 | Slc39a6 | 4063 | 6.02741226 | 6.71265721 | 5.21109594 | 6.07435847 | 5.96542686 | 8.00181393 |
| ENSMUSG00 | Elp2 | 3444 | 37.8392756 | 29.5078997 | 28.6336561 | 29.942649 | 32.1063172 | 24.2098061 |
| ENSMUSG00 | 2700062C07I | 1035 | 15.6717257 | 11.872022 | 12.8548355 | 15.8621548 | 15.6776691 | 15.9167943 |
| ENSMUSG00 | Zscan30 | 1686 | 0.6366536 | 1.65765934 | 2.88386876 | 3.46695802 | 1.96720903 | 2.15155814 |
| ENSMUSG00 | Zfp397 | 5483 | 25.5368963 | 19.0882642 | 23.4270574 | 33.5747932 | 25.0897518 | 36.2136374 |
| ENSMUSG00 | Mapre2 | 10272 | 7.33030375 | 7.67455406 | 5.61129316 | 9.06617571 | 8.77760963 | 7.38157098 |
| ENSMUSG00 | Wac | 6146 | 23.8752813 | 21.2158335 | 23.0286855 | 25.256257 | 23.6202472 | 23.8684188 |
| ENSMUSG00 | Ccny | 4007 | 12.1736897 | 9.50019698 | 12.4762377 | 12.5886444 | 8.79940746 | 12.1432384 |
| ENSMUSG00 | Thoc1 | 3834 | 25.871133 | 28.2407239 | 31.450868 | 27.8567689 | 27.9487192 | 37.0491187 |
| ENSMUSG00 | Rock1 | 6154 | 21.0728595 | 20.992501 | 23.1998624 | 26.4136514 | 24.7752374 | 28.2674113 |
| ENSMUSG00 | Cyp4f14 | 2186 | 0 | 0 | 0 | 0 | 0 | 0 |
| ENSMUSG00 | Esco1 | 5694 | 17.1128702 | 18.0085615 | 13.5772784 | 23.7315168 | 12.7789975 | 24.6401026 |
| ENSMUSG00 | Mib1 | 11016 | 6.38050922 | 8.24978485 | 5.58943058 | 11.513107 | 8.10604191 | 4.90478035 |
| ENSMUSG00 | Cyp4f41-ps | 2164 | 0.29394071 | 0.55668256 | 0.10212995 | 0.35014923 | 0.28295587 | 0 |
| ENSMUSG00 | Zfp871 | 8127 | 14.7193603 | 12.5224272 | 17.0291584 | 15.570288 | 14.1959685 | 15.3807889 |
| ENSMUSG00 | Adamts10 | 4622 | 6.36500553 | 5.73399896 | 7.56461658 | 10.4764396 | 9.1189499 | 6.78620405 |
| ENSMUSG00 | Myo1f | 4997 | 0 | 0 | 0 | 0 | 0 | 0 |
| ENSMUSG00 | Kifc5b | 3279 | 0.1212427 | 0 | 0.35048732 | 0.03301194 | 0.03112313 | 0.1663595 |
| ENSMUSG00 | Dtna | 9378 | 6.2910144 | 5.6623437 | 7.18786635 | 6.27915298 | 8.21601914 | 5.74111089 |
| ENSMUSG00 | Cdh2 | 9379 | 0.35605719 | 0.21578319 | 0.42415672 | 0.01923555 | 0.53316823 | 0.18611546 |
| ENSMUSG00 | 4921528I01R | 3723 | 0 | 0 | 0 | 0 | 0 | 0 |
| ENSMUSG00 | Tapbp | 3290 | 55.5851703 | 66.8165803 | 51.3762455 | 43.0900769 | 70.2271823 | 43.3243966 |
| ENSMUSG00 | H2-Ke2 | 1369 | 16.1170867 | 15.592837 | 16.8541724 | 18.6340435 | 13.9027306 | 24.5850291 |
| ENSMUSG00 | Wdr46 | 2403 | 8.53675736 | 6.25641696 | 6.84273225 | 6.15632054 | 7.62316643 | 6.45828953 |
| ENSMUSG00 | Mep1b | 2376 | 0 | 0 | 0 | 0 | 0 | 0 |
| ENSMUSG00 | Rnf138 | 3102 | 23.2099211 | 21.9961322 | 33.2155649 | 27.9513711 | 12.6496722 | 36.3310153 |
| ENSMUSG00 | Vps52 | 3664 | 19.7909379 | 17.7542841 | 17.7096904 | 13.6883302 | 21.2377792 | 15.1856616 |
| ENSMUSG00 | Ring1 | 3343 | 4.79253923 | 4.03595687 | 7.10032596 | 5.11603024 | 3.83117569 | 9.17041415 |
| ENSMUSG00 | Slc39a7 | 3180 | 3.83802909 | 3.06089978 | 2.52979102 | 2.22392476 | 3.57826472 | 1.8783478 |
| ENSMUSG00 | Col11a2 | 6764 | 0.19395784 | 0.12823122 | 0.08495328 | 0.03734097 | 0.08298198 | 0.27823035 |
| ENSMUSG00 | Daxx | 5707 | 17.6242099 | 15.1727767 | 21.864693 | 21.5594484 | 13.8228101 | 25.7214138 |
| ENSMUSG00 | H2-Oa | 1416 | 0 | 0 | 0.12486396 | 0 | 0 | 0 |
| ENSMUSG00 | Brd2 | 7171 | 30.3807047 | 35.2108432 | 23.9038833 | 27.3873335 | 31.5152445 | 17.2449051 |
| ENSMUSG00 | Psmb9 | 3965 | 0 | 0 | 0 | 0 | 0 | 0 |
| ENSMUSG00 | Psmb8 | 2730 | 1.58730673 | 1.80037256 | 0.5990726 | 1.70497573 | 3.75688701 | 6.20423113 |
| ENSMUSG00 | Tap2 | 4062 | 5.50038898 | 6.71430976 | 3.2427743 | 5.23198528 | 9.69777502 | 6.0364108 |
| ENSMUSG00 | Btnl2 | 2555 | 0 | 0 | 0 | 0 | 0 | 0 |
| ENSMUSG00 | Pfdn1 | 1073 | 23.1937845 | 23.7115018 | 26.4881501 | 26.5319041 | 16.7393143 | 35.0782872 |
| ENSMUSG00 | Psd2 | 4926 | 0 | 0.27389776 | 0 | 0.24171893 | 0.24860597 | 0.33221242 |
| ENSMUSG00 | Tmem173 | 2956 | 3.2277792 | 1.98875032 | 4.69532433 | 3.39337225 | 3.45239349 | 1.56856856 |
| ENSMUSG00 | Dnajc18 | 5161 | 6.56300511 | 5.02311681 | 5.96951842 | 6.95633294 | 7.35586584 | 4.06926414 |
| ENSMUSG00 | Spata24 | 961 | 2.06844332 | 2.45695699 | 1.74783562 | 1.83977126 | 2.23008094 | 5.05191029 |
| ENSMUSG00 | 2010001M09 | 901 | 0 | 0 | 0 | 0 | 0.22653219 | 0 |
| ENSMUSG00 | Slc23a1 | 3222 | 0.07403255 | 0.41875245 | 0.41156278 | 0.26876756 | 0.04751059 | 0.26241894 |
| ENSMUSG00 | Sil1 | 1685 | 40.4632932 | 41.9236345 | 43.7821223 | 29.3153228 | 32.7356156 | 32.5029529 |
| ENSMUSG00 | Hspa9 | 4011 | 182.482711 | 154.530592 | 204.600204 | 190.710382 | 159.961518 | 167.428613 |
| ENSMUSG00 | Etf1 | 3687 | 33.6633063 | 31.8237013 | 35.6899611 | 31.3552673 | 29.312141 | 34.4058441 |
| ENSMUSG00 | Cyp21a1 | 2131 | 0 | 0 | 0 | 0 | 0 | 0 |
| ENSMUSG00 | Gfra3 | 1984 | 0 | 0 | 0 | 0 | 0 | 0 |
| ENSMUSG00 | Rdbp | 2186 | 21.2417206 | 17.4802601 | 15.225978 | 11.3395995 | 19.2107536 | 18.7154432 |
| ENSMUSG00 | Cdc23 | 9483 | 11.7342181 | 12.1647519 | 11.6575777 | 13.5607304 | 12.8763173 | 13.4978258 |
| ENSMUSG00 | C2 | 4746 | 0.00837663 | 0.03045919 | 0.00931349 | 0.01520524 | 0 | 0.04022808 |
| ENSMUSG00 | Epb4.1l4a | 3829 | 10.0816327 | 13.5410321 | 9.639211 | 10.5164695 | 15.2852581 | 11.2902333 |
| ENSMUSG00 | Stard4 | 5989 | 6.87041615 | 9.80785991 | 5.70513018 | 7.16339137 | 8.7330164 | 9.15378609 |
| ENSMUSG00 | Tslp | 1128 | 7.04884406 | 1.75145756 | 6.34813703 | 1.08757936 | 1.7642098 | 8.82557033 |
| ENSMUSG00 | Gypc | 314 | 0 | 0 | 0 | 0 | 0 | 0 |
| ENSMUSG00 | Bin1 | 2432 | 3.43283343 | 4.63635998 | 2.45363849 | 2.71505516 | 4.95157265 | 1.88410337 |
| ENSMUSG00 | Ercc3 | 2888 | 6.59379334 | 6.45711673 | 5.23442878 | 5.67217736 | 8.18047507 | 5.57203503 |
| ENSMUSG00 | Map3k2 | 10774 | 3.67149648 | 5.63979057 | 4.89855212 | 8.20838047 | 4.05880862 | 5.81997361 |
| ENSMUSG00 | Iws1 | 9609 | 5.75914548 | 5.68167748 | 4.6598467 | 5.75645481 | 5.54392094 | 5.46968778 |
| ENSMUSG00 | Proc | 1633 | 0.97380234 | 0.64917436 | 0.51428966 | 0 | 0.71868135 | 1.31947122 |
| ENSMUSG00 | Csnk2b | 2309 | 11.6735452 | 13.5439588 | 14.9700482 | 13.1889327 | 9.14894742 | 14.5882111 |
| ENSMUSG00 | Myo7b | 6574 | 0 | 0.14659703 | 0 | 0 | 0 | 0 |
| ENSMUSG00 | Apom | 1160 | 0.17135983 | 0.16616015 | 0.34294533 | 0.03110521 | 0.26392953 | 0.54079026 |
| ENSMUSG00 | Bag6 | 6264 | 4.07455595 | 4.10784804 | 5.2711968 | 6.43993068 | 4.34994966 | 7.07555696 |
| ENSMUSG00 | Prrc2a | 7808 | 0.64154592 | 1.12319832 | 0.92275876 | 1.08135226 | 0.745006 | 1.58589731 |
| ENSMUSG00 | Lims2 | 1678 | 2.8193696 | 4.36492205 | 2.71322398 | 1.48370743 | 3.0409044 | 2.1130531 |
| ENSMUSG00 | Aif1 | 1726 | 0 | 0 | 0 | 0 | 0 | 0.09481335 |
| ENSMUSG00 | Ltb | 1335 | 0.02977939 | 0.03609471 | 0 | 0 | 0 | 0 |
| ENSMUSG00 | Wdr33 | 6390 | 24.5812056 | 19.0031039 | 28.4372107 | 25.6018764 | 25.6808802 | 23.4672874 |
| ENSMUSG00 | Tnf | 2691 | 0 | 0 | 0 | 0 | 0 | 0 |
| ENSMUSG00 | Lta | 1379 | 0.49009657 | 0 | 0 | 0 | 0 | 0 |
| ENSMUSG00 | Atp6v1g2 | 2504 | 4.33436349 | 4.58002126 | 1.78290182 | 5.3748413 | 4.01445529 | 4.09555286 |
| ENSMUSG00 | Riok3 | 3648 | 40.9542477 | 37.923047 | 48.7819682 | 43.8464099 | 35.5701956 | 50.0109819 |
| ENSMUSG00 | Pou5f1 | 2250 | 0 | 0 | 0 | 0 | 0 | 0 |
| ENSMUSG00 | Psors1c2 | 841 | 0 | 0.63026262 | 0 | 0 | 0.06067346 | 0.162156 |
| ENSMUSG00 | 3110002H16I | 3151 | 12.6672493 | 16.668747 | 17.6470703 | 16.9474535 | 20.2907169 | 11.7460125 |
| ENSMUSG00 | Aqp4 | 5307 | 33.6352191 | 54.097385 | 23.8791566 | 23.3000126 | 23.3450237 | 10.9211623 |
| ENSMUSG00 | Npc1 | 7241 | 4.89188415 | 5.27715076 | 8.16766551 | 6.14904612 | 5.58111996 | 7.53339021 |
| ENSMUSG00 | Mrpl27 | 1045 | 21.4565464 | 20.3351397 | 22.9257406 | 15.0197984 | 12.5490704 | 30.9286579 |
| ENSMUSG00 | Zfp521 | 6173 | 1.21720166 | 0.031224 | 1.12420045 | 0 | 0.37197261 | 1.96175925 |
| ENSMUSG00 | Lama3 | 10456 | 0.0988564 | 0.1382549 | 0.05072897 | 0 | 0 | 0.09129805 |
| ENSMUSG00 | Dhx16 | 4339 | 10.280168 | 10.4279947 | 9.95280033 | 10.328163 | 11.0661027 | 7.09052618 |
| ENSMUSG00 | Impact | 3421 | 7.90228785 | 9.11330841 | 14.3420186 | 7.84713275 | 9.93381067 | 11.8873099 |
| ENSMUSG00 | Ttc39c | 5182 | 0 | 0.03719525 | 0 | 0.17407393 | 0.02954055 | 0.01579003 |
| ENSMUSG00 | Ndfip1 | 922 | 329.987736 | 371.014699 | 562.063349 | 412.438902 | 341.909924 | 396.547223 |
| ENSMUSG00 | Atat1 | 4613 | 8.67846713 | 11.0725404 | 11.8625367 | 16.9577005 | 9.6013211 | 12.1503108 |
| ENSMUSG00 | Spry4 | 4585 | 0.68499083 | 0.66210379 | 0.41454291 | 0.76334969 | 0.73451272 | 0 |
| ENSMUSG00 | Gnl1 | 3357 | 11.4162297 | 16.5645381 | 15.589807 | 9.84544319 | 12.3271942 | 11.1145983 |
| ENSMUSG00 | Cabyr | 2208 | 1.47642636 | 1.33123776 | 0.74070117 | 1.25829594 | 1.59457424 | 1.67995966 |
| ENSMUSG00 | Nr3c1 | 11456 | 5.70860383 | 5.42181599 | 9.06724259 | 6.38742897 | 5.55873927 | 7.81146049 |
| ENSMUSG00 | Mrps18b | 2651 | 5.08378268 | 4.70776632 | 4.48521153 | 4.31460133 | 3.98433036 | 2.59268544 |
| ENSMUSG00 | Gm8615 | 635 | 0.06260706 | 0 | 0 | 0.62504329 | 0.160713 | 0.12885656 |
| ENSMUSG00 | Pcdh12 | 5368 | 0.25180446 | 0 | 0.01646865 | 0.41002323 | 0.62737347 | 0 |
| ENSMUSG00 | 0610009O20I | 2391 | 3.80761398 | 3.10360188 | 3.84524607 | 4.07450942 | 3.39322198 | 3.67312164 |
| ENSMUSG00 | Rpp21 | 1346 | 13.3798163 | 14.6420913 | 18.7184625 | 12.6260349 | 10.53888 | 25.4306629 |
| ENSMUSG00 | H2-M10.1 | 1435 | 0 | 0 | 0 | 0 | 0 | 0 |
| ENSMUSG00 | Arap3 | 6216 | 0.19826575 | 0.06201601 | 0.14933055 | 0.09287528 | 0.44327933 | 0.24571747 |
| ENSMUSG00 | Hdac3 | 4726 | 19.5160209 | 21.8297022 | 22.4750378 | 25.561296 | 25.6103604 | 21.6765859 |
| ENSMUSG00 | Diap1 | 8175 | 2.89351815 | 2.35185204 | 3.51992658 | 4.03854076 | 3.28940674 | 4.31056072 |
| ENSMUSG00 | Trim26 | 2217 | 101.441928 | 80.245532 | 85.3930959 | 73.7427802 | 85.9185661 | 70.9239041 |
| ENSMUSG00 | H2-M5 | 2427 | 0 | 0.01985432 | 0 | 0 | 0 | 0 |
| ENSMUSG00 | Gabbr1 | 7178 | 1.98832788 | 1.6782684 | 1.25006605 | 2.2922001 | 1.49993944 | 1.8960776 |
| ENSMUSG00 | Myot | 2199 | 0 | 0 | 0 | 0 | 0 | 0 |
| ENSMUSG00 | Dcp2 | 8632 | 1.04546966 | 1.26718285 | 1.53108794 | 0.70224554 | 0.63842083 | 1.14381597 |
| ENSMUSG00 | Ik | 1976 | 66.4536195 | 55.6485126 | 56.0576005 | 67.2338502 | 63.6022083 | 60.5949728 |
| ENSMUSG00 | Pggt1b | 2727 | 16.6923451 | 15.4436929 | 22.7735935 | 23.7239112 | 28.0486019 | 27.8347345 |
| ENSMUSG00 | Mal2 | 2771 | 444.125101 | 467.796276 | 592.074994 | 474.131868 | 319.858588 | 607.97543 |
| ENSMUSG00 | Ap3s1 | 1473 | 6.20757673 | 9.84665247 | 5.55148741 | 9.92072507 | 5.022963 | 11.8690046 |
| ENSMUSG00 | 4833403I15R | 3199 | 0.11184724 | 0 | 0 | 0 | 0 | 0 |
| ENSMUSG00 | Ankhd1 | 9714 | 17.0865896 | 14.3557302 | 14.3881231 | 20.5742685 | 18.7632504 | 14.1595641 |
| ENSMUSG00 | Slc4a9 | 3419 | 1279.82648 | 726.165032 | 1623.53538 | 909.723429 | 1146.93682 | 1405.98811 |
| ENSMUSG00 | Hbegf | 2367 | 15.5192497 | 20.1743828 | 23.5294982 | 25.5333437 | 12.9559999 | 22.7231046 |
| ENSMUSG00 | Yipf5 | 2323 | 31.5237172 | 27.7751382 | 31.4341131 | 28.6885649 | 22.4270038 | 33.790965 |
| ENSMUSG00 | Rbm27 | 6410 | 5.60670115 | 6.11915194 | 4.11677071 | 7.68361791 | 5.68374919 | 6.50591629 |
| ENSMUSG00 | Lars | 3829 | 15.5740979 | 13.100571 | 17.5699151 | 15.0019887 | 14.965427 | 15.8989279 |
| ENSMUSG00 | Pou4f3 | 2358 | 0 | 0 | 0 | 0 | 0 | 0 |
| ENSMUSG00 | Tcerg1 | 4555 | 18.9569492 | 20.977764 | 21.213003 | 26.5367397 | 20.1977071 | 20.8736317 |
| ENSMUSG00 | Ppp2r2b | 3970 | 7.56055108 | 7.986569 | 5.45564309 | 10.7064606 | 7.84032475 | 7.83888246 |
| ENSMUSG00 | Dpysl3 | 7211 | 0.07167123 | 0.02004706 | 0.23906142 | 0.17513127 | 0.00707619 | 0 |
| ENSMUSG00 | Jakmip2 | 3516 | 0 | 0 | 0 | 0 | 0 | 0 |
| ENSMUSG00 | Spink3 | 418 | 0.19021761 | 0.57639285 | 0.31723811 | 0.34528272 | 0.36621801 | 2.28376166 |
| ENSMUSG00 | Dtwd2 | 5197 | 2.2872597 | 1.45569972 | 4.38020957 | 3.03402991 | 2.04223324 | 5.42133969 |
| ENSMUSG00 | Hsd17b4 | 3278 | 114.948885 | 87.1266146 | 151.578071 | 94.8391994 | 93.3978813 | 104.455711 |
| ENSMUSG00 | Ftmt | 1450 | 0 | 0 | 0 | 0 | 0 | 0 |
| ENSMUSG00 | Rab27b | 12391 | 1.69404356 | 3.09550544 | 1.0059656 | 2.1024321 | 2.41727727 | 1.9348243 |
| ENSMUSG00 | 2310002L13I | 1038 | 0 | 0 | 0 | 0 | 0 | 0 |
| ENSMUSG00 | Mbd2 | 3880 | 28.4231193 | 29.2471833 | 31.7615305 | 30.6046414 | 31.3654398 | 31.5907806 |
| ENSMUSG00 | Smad4 | 4166 | 38.2858828 | 35.4284859 | 48.3185773 | 38.7323339 | 40.1009012 | 34.1162134 |
| ENSMUSG00 | Sec11c | 2463 | 32.4274301 | 27.8201871 | 32.2316322 | 31.48206 | 25.3163749 | 52.8660689 |
| ENSMUSG00 | Grp | 1054 | 0 | 0 | 0 | 0 | 0 | 0 |
| ENSMUSG00 | Rax | 1666 | 0.04772567 | 0 | 0 | 0.17326312 | 0 | 0 |
| ENSMUSG00 | Cplx4 | 1748 | 0 | 0.05513323 | 0 | 0 | 0 | 0 |
| ENSMUSG00 | Pmaip1 | 2643 | 2.88802583 | 3.08116108 | 2.80965177 | 5.9931962 | 2.08507325 | 4.99467476 |
| ENSMUSG00 | Gnal | 7906 | 2.04660771 | 2.13322221 | 2.0742327 | 2.25912118 | 2.29767136 | 3.65340791 |
| ENSMUSG00 | Impa2 | 1653 | 18.4226849 | 20.8137445 | 13.3167077 | 12.4857408 | 14.7862275 | 12.540064 |
| ENSMUSG00 | Cidea | 1159 | 0 | 0 | 0 | 0 | 0 | 0 |
| ENSMUSG00 | Afg3l2 | 3601 | 45.9269089 | 41.0942971 | 50.7811787 | 44.4687902 | 44.5364896 | 34.5761533 |
| ENSMUSG00 | Srfbp1 | 1455 | 11.3118687 | 11.4256512 | 11.0580556 | 12.1017441 | 9.32853331 | 11.3784921 |
| ENSMUSG00 | Lox | 4688 | 0.25440794 | 0.28780298 | 0.21686058 | 0 | 0.39184077 | 0 |
| ENSMUSG00 | Simo1 | 2160 | 0 | 0.20077684 | 0.22510198 | 0 | 0 | 0 |
| ENSMUSG00 | 1700034E13I | 576 | 0 | 0 | 0 | 0 | 0 | 0 |
| ENSMUSG00 | Spire1 | 8945 | 23.5155112 | 15.9723646 | 22.4591813 | 18.6642392 | 27.5924628 | 18.9047248 |
| ENSMUSG00 | Sncaip | 4000 | 0.44724916 | 0.60233053 | 0 | 0.51416913 | 0.40821101 | 0 |
| ENSMUSG00 | Snx24 | 1956 | 19.5118923 | 23.1324485 | 21.0840079 | 22.6527355 | 20.7653349 | 16.4400816 |
| ENSMUSG00 | Psmg2 | 1429 | 50.2438053 | 41.3749227 | 34.9222399 | 36.4860422 | 29.9230951 | 46.3420743 |
| ENSMUSG00 | Ppic | 1281 | 51.8900573 | 81.0630623 | 29.1573438 | 60.1930751 | 60.5465193 | 38.3463117 |
| ENSMUSG00 | Ptpn2 | 3475 | 24.3223458 | 22.6580278 | 26.1649471 | 27.2562207 | 26.8420763 | 33.0199733 |
| ENSMUSG00 | Cep192 | 8050 | 1.15068658 | 1.83168339 | 1.81200102 | 3.15549806 | 2.96016367 | 1.7211822 |
| ENSMUSG00 | D18Ertd653e | 2637 | 3.7690065 | 1.88214014 | 3.00042848 | 1.01254126 | 1.58671324 | 2.16169875 |
| ENSMUSG00 | 4930546C10I | 360 | 0.11043189 | 0 | 0 | 0 | 0 | 0 |
| ENSMUSG00 | Setbp1 | 9947 | 1.60668575 | 2.46575843 | 2.56403573 | 2.62263177 | 1.47738979 | 1.77132974 |
| ENSMUSG00 | Slc14a2 | 4870 | 33.2003161 | 19.1855259 | 9.72077492 | 1.87448815 | 5.84655394 | 0 |
| ENSMUSG00 | Galr1 | 2868 | 0 | 0 | 0 | 0 | 0 | 0 |
| ENSMUSG00 | Me2 | 2439 | 32.0945228 | 22.6806214 | 32.6212864 | 28.1821627 | 35.5448186 | 22.1417736 |
| ENSMUSG00 | Mapk4 | 7290 | 1.32518268 | 1.58638493 | 0.824616 | 1.13839098 | 1.32990554 | 0.85677536 |
| ENSMUSG00 | Cxxc1 | 2603 | 29.5226062 | 32.6179451 | 23.281109 | 29.0957401 | 27.2872513 | 23.8168479 |
| ENSMUSG00 | Mbd1 | 2880 | 9.23486683 | 10.9256065 | 7.58184391 | 9.69704947 | 14.6523656 | 9.9912307 |
| ENSMUSG00 | Smad2 | 3814 | 24.5787685 | 21.9327907 | 29.7614926 | 20.6237171 | 23.5732759 | 23.6489858 |
| ENSMUSG00 | Sall3 | 6882 | 3.55846789 | 4.27110284 | 1.46440282 | 5.11188512 | 5.27168748 | 4.20890248 |
| ENSMUSG00 | Atp9b | 4543 | 22.1573578 | 18.3284552 | 12.8528802 | 19.7764233 | 20.0039567 | 16.4200173 |
| ENSMUSG00 | Rbfa | 1371 | 18.819334 | 20.7366892 | 24.9542133 | 19.3174476 | 18.6091816 | 16.9098785 |
| ENSMUSG00 | Gm16286 | 2740 | 37.5645763 | 27.7335837 | 34.5226073 | 29.1290082 | 29.070136 | 25.7317308 |
| ENSMUSG00 | Pde6a | 3884 | 0.02047141 | 0 | 0 | 0 | 0 | 0.02808923 |
| ENSMUSG00 | Csnk1a1 | 10038 | 42.1238584 | 42.9059992 | 52.2645621 | 47.7786945 | 32.4315877 | 56.0246849 |
| ENSMUSG00 | Il17b | 689 | 0 | 0 | 0 | 0 | 0 | 0 |
| ENSMUSG00 | Pcyox1l | 2639 | 1.37087864 | 2.00852923 | 0.41873667 | 1.9004942 | 0.73474888 | 0.89916393 |
| ENSMUSG00 | Grpel2 | 4026 | 9.38095044 | 8.5816391 | 10.3313299 | 6.69480558 | 8.04812435 | 7.76371997 |
| ENSMUSG00 | Napg | 5288 | 12.735587 | 13.3496857 | 12.6553688 | 16.6012886 | 11.5118129 | 10.4652502 |
| ENSMUSG00 | Txnl1 | 2550 | 45.4147901 | 42.9709684 | 42.641778 | 48.2791906 | 32.3967461 | 56.2071487 |
| ENSMUSG00 | Nars | 2617 | 121.165347 | 113.515253 | 120.680997 | 136.311856 | 112.269726 | 109.848948 |
| ENSMUSG00 | Fech | 3024 | 102.188939 | 80.836581 | 126.802575 | 87.5800948 | 104.280093 | 87.8849487 |
| ENSMUSG00 | Nedd4l | 9178 | 71.9306504 | 56.1090115 | 75.5014483 | 58.9389858 | 66.7101202 | 61.336735 |
| ENSMUSG00 | Lmnb1 | 2951 | 2.30368931 | 3.51070317 | 1.87232476 | 0.1711788 | 2.2824404 | 1.81153145 |
| ENSMUSG00 | C330018D20 | 3531 | 12.0583743 | 10.630767 | 13.006433 | 11.220075 | 9.33532676 | 16.452841 |
| ENSMUSG00 | Megf10 | 7692 | 0 | 0 | 0 | 0 | 0 | 0 |
| ENSMUSG00 | Prrc1 | 4792 | 11.5400404 | 10.5885483 | 13.8822566 | 16.279086 | 14.2579961 | 7.88869992 |
| ENSMUSG00 | Slc12a2 | 6635 | 3.16366145 | 4.2412784 | 3.9172093 | 5.59040569 | 5.2295306 | 2.55275825 |
| ENSMUSG00 | Fbn2 | 10415 | 0.0305371 | 0.09253277 | 0.05092867 | 0 | 0.00979863 | 0 |
| ENSMUSG00 | Slc27a6 | 2475 | 0 | 0.07787708 | 0 | 0 | 0 | 0 |
| ENSMUSG00 | Isoc1 | 2558 | 93.3585502 | 109.879405 | 69.0157002 | 113.394666 | 111.667573 | 80.1818949 |
| ENSMUSG00 | Dctn4 | 5248 | 14.650743 | 14.1217127 | 17.2747675 | 17.3053556 | 10.7925452 | 18.7669059 |
| ENSMUSG00 | Rbm22 | 4656 | 5.63544183 | 6.39588085 | 5.74358141 | 7.32335311 | 5.84129259 | 9.58361465 |
| ENSMUSG00 | Rps14 | 1397 | 9.76100917 | 13.2797282 | 13.7952781 | 9.09153878 | 8.87574039 | 18.5475356 |
| ENSMUSG00 | Cd74 | 1555 | 2.37765896 | 1.30149876 | 1.3928555 | 0.44087386 | 3.90491236 | 24.9593644 |
| ENSMUSG00 | Tcof1 | 5668 | 6.23546616 | 6.1210724 | 5.20159303 | 7.49268991 | 8.51639935 | 4.13835387 |
| ENSMUSG00 | Tns3 | 4595 | 35.5679609 | 51.0073677 | 44.7790162 | 52.1324687 | 36.9788534 | 56.1459875 |
| ENSMUSG00 | Camk2a | 7653 | 0.05194758 | 0.09444618 | 0.19059987 | 0.08486565 | 0.0200025 | 0.31718841 |
| ENSMUSG00 | Cdx1 | 1747 | 0 | 0 | 0 | 0 | 0 | 0 |
| ENSMUSG00 | Pdgfrb | 5826 | 0 | 0 | 0 | 0 | 0.42040268 | 0 |
| ENSMUSG00 | Csf1r | 5206 | 0.66437319 | 0.16660698 | 0.24622617 | 0 | 1.05855716 | 0.01047816 |
| ENSMUSG00 | Hmgxb3 | 5177 | 5.85158898 | 6.15245089 | 5.88276412 | 6.10544157 | 9.94506726 | 3.02934471 |
| ENSMUSG00 | Gnaq | 6342 | 13.7909267 | 11.0018871 | 14.3924323 | 12.7783463 | 7.66763421 | 13.349177 |
| ENSMUSG00 | Psat1 | 5266 | 2.95939012 | 2.43402841 | 0.51202287 | 1.14426537 | 2.00578424 | 1.52791845 |
| ENSMUSG00 | Tle4 | 6191 | 6.26095841 | 5.12919809 | 6.3543273 | 4.45270264 | 4.61553391 | 3.73588985 |
| ENSMUSG00 | Cndp2 | 2102 | 32.6252159 | 25.2394257 | 34.2974339 | 20.2038851 | 24.3722556 | 29.5194121 |
| ENSMUSG00 | 1700034H14I | 1333 | 35.3116946 | 31.5579625 | 35.2155719 | 41.2250216 | 32.3460522 | 35.8887601 |
| ENSMUSG00 | Cyb5 | 2366 | 203.330968 | 186.554442 | 249.798328 | 176.872169 | 95.4318314 | 254.175199 |
| ENSMUSG00 | Cbln2 | 2888 | 0 | 0 | 0 | 0 | 0 | 0 |
| ENSMUSG00 | Slc22a6 | 4016 | 1.27700622 | 0.19197786 | 0 | 0 | 0.0889404 | 2.24798446 |
| ENSMUSG00 | Scgb1a1 | 473 | 0 | 0 | 0 | 0 | 0 | 0.57663085 |
| ENSMUSG00 | Asrgl1 | 4289 | 8.76863711 | 7.6734297 | 4.31815627 | 5.12333064 | 10.374213 | 4.87115264 |
| ENSMUSG00 | Gm9750 | 2011 | 0 | 0 | 0 | 0 | 0 | 0.0542509 |
| ENSMUSG00 | Anxa1 | 2379 | 11.7979694 | 14.2189501 | 7.32052381 | 11.7240103 | 16.2151913 | 7.08521524 |
| ENSMUSG00 | Incenp | 3680 | 0.97228077 | 1.66295602 | 0.57654578 | 1.59819926 | 1.20633008 | 0.85974406 |
| ENSMUSG00 | Fth1 | 862 | 1203.13367 | 1371.02167 | 1371.02631 | 1174.17335 | 743.55256 | 1396.13246 |
| ENSMUSG00 | Rab3il1 | 4526 | 0.57973083 | 0.68138142 | 0.6055047 | 0.17538776 | 0.77790984 | 0.0482097 |
| ENSMUSG00 | Fads3 | 3638 | 0.16391759 | 0.95366241 | 0.31590102 | 0.06942669 | 1.5148017 | 0.22491456 |
| ENSMUSG00 | Fads2 | 1496 | 0.31889423 | 2.06145207 | 0.29546687 | 1.25418871 | 0.78449642 | 0.03646342 |
| ENSMUSG00 | Tmem138 | 1585 | 8.37749558 | 9.63728843 | 7.64120188 | 8.21805557 | 8.7243835 | 8.24261969 |
| ENSMUSG00 | Tmem216 | 1969 | 7.36960406 | 10.3518868 | 9.76526242 | 8.7227289 | 8.08543893 | 8.83759261 |
| ENSMUSG00 | Sdhaf2 | 3092 | 28.1708466 | 22.2698661 | 32.2936492 | 27.7617024 | 22.9882735 | 24.7606444 |
| ENSMUSG00 | Cd5 | 2058 | 0 | 0.18731367 | 0 | 0 | 0 | 0 |
| ENSMUSG00 | Cd6 | 4077 | 0 | 0 | 0 | 0 | 0 | 0 |
| ENSMUSG00 | Ms4a14 | 3498 | 0 | 0 | 0 | 0 | 0 | 0 |
| ENSMUSG00 | Ms4a7 | 2619 | 0 | 0 | 0 | 0 | 0 | 0 |
| ENSMUSG00 | Ms4a1 | 3797 | 0 | 0 | 0 | 0 | 0 | 0 |
| ENSMUSG00 | Ms4a4c | 1743 | 0 | 0 | 0 | 0 | 0 | 0 |
| ENSMUSG00 | Ms4a6b | 2829 | 2.16413715 | 0.83461848 | 0.74997825 | 1.04585636 | 1.22650886 | 1.2533415 |
| ENSMUSG00 | Ms4a4d | 1506 | 0 | 0 | 0 | 0 | 0 | 1.19530292 |
| ENSMUSG00 | Ms4a6d | 1927 | 0 | 0 | 0 | 0 | 0.26479697 | 0 |
| ENSMUSG00 | Ms4a2 | 3363 | 0 | 0 | 0 | 0 | 0 | 0 |
| ENSMUSG00 | Ms4a3 | 1876 | 0 | 0 | 0 | 0 | 0 | 0 |
| ENSMUSG00 | Gif | 1528 | 0 | 0 | 0 | 0 | 0 | 0 |
| ENSMUSG00 | Mrpl16 | 1251 | 22.0863781 | 25.7687688 | 26.8179048 | 20.5647464 | 13.7865028 | 23.7208693 |
| ENSMUSG00 | Osbp | 4767 | 28.0131443 | 20.8838241 | 27.3074109 | 25.197635 | 31.2238176 | 19.1614783 |
| ENSMUSG00 | Fam111a | 4118 | 4.35398779 | 1.57969152 | 3.28454686 | 0.87620312 | 2.73842376 | 3.25865044 |
| ENSMUSG00 | Keg1 | 4928 | 1.59731841 | 2.11206808 | 0.6906538 | 0.98112702 | 1.0457922 | 1.48881452 |
| ENSMUSG00 | Zfp91 | 10034 | 4.72675785 | 4.38931713 | 4.62546693 | 4.55250831 | 4.54630058 | 6.73575404 |
| ENSMUSG00 | Lpxn | 1666 | 0.26249117 | 1.01232021 | 1.40618108 | 0 | 0.58193346 | 1.06413659 |
| ENSMUSG00 | Gna14 | 3376 | 0 | 0.28546471 | 0.26185926 | 0 | 0.25694561 | 0 |
| ENSMUSG00 | Rfk | 2597 | 167.104669 | 138.083751 | 150.76824 | 141.243768 | 132.782537 | 182.982688 |
| ENSMUSG00 | Pcsk5 | 8746 | 0.20455027 | 0.57850176 | 0.03537765 | 0.21040296 | 0.29171265 | 0.25571923 |
| ENSMUSG00 | Ostf1 | 5111 | 44.9203483 | 52.1461967 | 44.2450849 | 46.9963159 | 40.6833265 | 39.7619521 |
| ENSMUSG00 | 2410127L17I | 2082 | 2.31047701 | 3.72623304 | 3.20580129 | 2.70355377 | 2.81845976 | 2.84274578 |
| ENSMUSG00 | Trpm6 | 6469 | 4.67060832 | 1.68343421 | 2.19335162 | 4.03824397 | 4.22787717 | 4.52398948 |
| ENSMUSG00 | 1700025F22I | 643 | 0 | 0 | 0 | 0 | 0 | 0 |
| ENSMUSG00 | 1700017D01I | 1187 | 0 | 0 | 0 | 0 | 0 | 0 |
| ENSMUSG00 | Ms4a8a | 1074 | 0 | 0 | 0 | 0 | 0 | 0 |
| ENSMUSG00 | Ms4a10 | 998 | 0 | 0 | 0 | 0 | 0 | 0 |
| ENSMUSG00 | Ccdc86 | 3362 | 5.28575247 | 5.17409447 | 4.49643971 | 3.86363356 | 4.11307194 | 2.97733272 |
| ENSMUSG00 | Zp1 | 2133 | 0 | 0 | 0 | 0.08458051 | 0 | 0 |
| ENSMUSG00 | Prpf19 | 2160 | 41.9273077 | 37.8798976 | 50.6274813 | 41.0934396 | 29.0566862 | 39.9396685 |
| ENSMUSG00 | Tmem132a | 5376 | 4.31867571 | 3.16402792 | 5.27856831 | 4.50354388 | 4.65084155 | 5.23068323 |
| ENSMUSG00 | Slc15a3 | 3387 | 0 | 0.34144512 | 0.01305044 | 0.27698056 | 0.13558824 | 0 |
| ENSMUSG00 | Pga5 | 1407 | 1.72358515 | 0.41097179 | 3.42430767 | 1.02578663 | 2.17596485 | 2.55881477 |
| ENSMUSG00 | Ddb1 | 4329 | 51.0879506 | 41.841727 | 42.0167669 | 54.4439628 | 66.5617797 | 31.2313206 |
| ENSMUSG00 | Fen1 | 4250 | 0.17773038 | 0.22675973 | 0.17680737 | 0.20375743 | 0.24012412 | 0.07701075 |
| ENSMUSG00 | Syt7 | 6984 | 8.27669941 | 3.44287437 | 11.158054 | 4.06077993 | 3.00284085 | 10.6536678 |
| ENSMUSG00 | Aldh1a7 | 2053 | 1.27806221 | 2.27670964 | 3.81087492 | 1.95085579 | 3.10681782 | 6.85519428 |
| ENSMUSG00 | Tmc1 | 4483 | 0 | 0 | 0.1478982 | 0.13682685 | 0 | 0 |
| ENSMUSG00 | Zfand5 | 3418 | 64.5299608 | 65.0615654 | 71.3592071 | 81.9921708 | 54.8182715 | 84.5529776 |
| ENSMUSG00 | Tmem2 | 6642 | 9.16973745 | 7.10971293 | 9.15048692 | 5.66600007 | 8.98838598 | 8.20456628 |
| ENSMUSG00 | Slc22a19 | 5421 | 0 | 0 | 0 | 0 | 0 | 0 |
| ENSMUSG00 | Rtn3 | 5894 | 132.216988 | 124.145084 | 157.181274 | 130.15615 | 114.666499 | 144.517642 |
| ENSMUSG00 | Atl3 | 6684 | 3.30700885 | 2.73229527 | 4.31172975 | 5.20933168 | 3.16052059 | 3.57459365 |
| ENSMUSG00 | Gm16437 | 506 | 0.86424958 | 1.14276147 | 0.78619879 | 0.49915872 | 0.5042132 | 0.3234147 |
| ENSMUSG00 | Naa40 | 3267 | 18.6426067 | 21.4899437 | 17.9946285 | 18.6539863 | 16.3684242 | 14.9522127 |
| ENSMUSG00 | Lipo1 | 3501 | 7.13123158 | 6.12481198 | 8.14344152 | 8.32741845 | 6.93760494 | 13.5944717 |
| ENSMUSG00 | Otub1 | 3567 | 30.4825453 | 22.776098 | 25.7627226 | 18.1371197 | 15.5496693 | 19.6664907 |
| ENSMUSG00 | Lipf | 1401 | 0 | 0 | 0 | 0 | 0 | 0 |
| ENSMUSG00 | Cdc42bpg | 6062 | 2.93149122 | 1.86005072 | 1.90311465 | 1.94635905 | 2.9460956 | 3.18099141 |
| ENSMUSG00 | Lipn | 2017 | 0 | 0 | 0 | 0 | 0 | 0 |
| ENSMUSG00 | Lipk | 2088 | 0 | 0.39232257 | 0.27520305 | 0 | 0.34213087 | 0 |
| ENSMUSG00 | Ehd1 | 3714 | 9.13070137 | 8.92629838 | 7.80732607 | 7.65553341 | 6.49851259 | 4.69999169 |
| ENSMUSG00 | Atg2a | 7654 | 8.08198689 | 6.6670293 | 6.49687397 | 7.27392144 | 8.66661727 | 6.70284772 |
| ENSMUSG00 | Ankrd22 | 2572 | 0 | 0 | 0 | 0 | 0 | 0 |
| ENSMUSG00 | Stambpl1 | 3088 | 1.90537925 | 5.21187554 | 3.10615283 | 3.10810358 | 4.29626221 | 2.26994245 |
| ENSMUSG00 | Ppp2r5b | 2839 | 13.779286 | 11.13431 | 16.0365968 | 16.0774168 | 12.8329807 | 13.3731236 |
| ENSMUSG00 | Fas | 1791 | 2.59709169 | 1.91023863 | 1.77695851 | 1.47068077 | 2.56413949 | 2.22339326 |
| ENSMUSG00 | Cdc37l1 | 5404 | 19.5687598 | 20.9634207 | 23.4423542 | 23.9167068 | 22.3216788 | 22.2477072 |
| ENSMUSG00 | Lipa | 2959 | 5.01142083 | 5.29252913 | 3.97353506 | 6.95057972 | 5.17333989 | 4.67328222 |
| ENSMUSG00 | Ak3 | 3115 | 258.123465 | 260.129442 | 194.715149 | 269.62496 | 205.006451 | 248.221107 |
| ENSMUSG00 | Gpha2 | 803 | 0 | 0 | 0 | 0 | 0 | 0 |
| ENSMUSG00 | Rcl1 | 1794 | 10.8585203 | 7.0372619 | 6.20895454 | 5.18905657 | 11.3202328 | 6.8414647 |
| ENSMUSG00 | 1700123I01R | 1745 | 0 | 0 | 0 | 0 | 0 | 0 |
| ENSMUSG00 | Snx15 | 2158 | 9.69016809 | 8.7753808 | 10.610081 | 9.73111673 | 9.45808634 | 15.2424536 |
| ENSMUSG00 | Jak2 | 5005 | 13.1141455 | 13.1995229 | 11.3661882 | 17.0569665 | 14.2017466 | 5.03531802 |
| ENSMUSG00 | Sac3d1 | 1372 | 7.18612184 | 8.74520707 | 5.18695097 | 8.28414288 | 4.35137461 | 6.34155242 |
| ENSMUSG00 | Cdca5 | 2122 | 0.48710768 | 0.22708031 | 0 | 0 | 0 | 0.03855981 |
| ENSMUSG00 | Zfpl1 | 2196 | 11.3871572 | 9.76455681 | 14.1904824 | 10.1049442 | 9.61972659 | 11.7618777 |
| ENSMUSG00 | Tnfrsf25 | 2169 | 0.05498683 | 0.13329583 | 0.06113671 | 0.43251875 | 0.0470506 | 0 |
| ENSMUSG00 | Kif20b | 5535 | 0.22984198 | 1.51480414 | 0.47116689 | 0.91916319 | 0.78360288 | 1.00524416 |
| ENSMUSG00 | 1110014N23I | 5122 | 11.0759607 | 13.8199695 | 13.2467452 | 9.08038957 | 11.546187 | 12.8598699 |
| ENSMUSG00 | Htr7 | 5047 | 0 | 0 | 0 | 0 | 0 | 0 |
| ENSMUSG00 | Tm7sf2 | 3027 | 5.05644532 | 4.42544794 | 4.55598778 | 5.26866986 | 2.37684803 | 3.33386737 |
| ENSMUSG00 | Rpp30 | 1144 | 14.2827819 | 14.6159925 | 12.2868759 | 11.1021676 | 6.55671088 | 17.4996374 |
| ENSMUSG00 | Ankrd1 | 1756 | 0.27167754 | 0.08232308 | 0.15103135 | 0 | 1.39479843 | 0 |
| ENSMUSG00 | Pcgf5 | 1708 | 14.4776984 | 13.5418573 | 15.4758209 | 13.9638357 | 13.144968 | 7.01028491 |
| ENSMUSG00 | Mlana | 618 | 0.64329257 | 0 | 0.21457205 | 0 | 0.1651339 | 0.04413372 |
| ENSMUSG00 | Syvn1 | 4954 | 9.58978587 | 8.18021757 | 6.02265725 | 11.2383114 | 10.1249349 | 9.70083051 |
| ENSMUSG00 | Il33 | 2922 | 0 | 0.19789093 | 0.16639982 | 1.17309864 | 0.68105019 | 1.12944263 |
| ENSMUSG00 | Tnks2 | 9688 | 13.9480675 | 18.5026388 | 19.0303352 | 17.5456763 | 23.2062564 | 18.2684903 |
| ENSMUSG00 | Tjp2 | 5077 | 18.033656 | 15.6034097 | 16.2024089 | 22.9128445 | 22.0105895 | 11.6254322 |
| ENSMUSG00 | Trpd52l3 | 2230 | 0 | 0 | 0 | 0 | 0 | 0 |
| ENSMUSG00 | Frmd8 | 5071 | 7.62025775 | 6.82269088 | 5.36942128 | 8.38190662 | 8.35178307 | 6.1369283 |
| ENSMUSG00 | Uhrf2 | 8874 | 6.70655334 | 6.27173098 | 7.92983255 | 8.5874778 | 9.01615475 | 8.50757285 |
| ENSMUSG00 | Slc25a45 | 2411 | 10.5860715 | 19.7662345 | 9.88170609 | 12.9751691 | 13.1216728 | 10.7356419 |
| ENSMUSG00 | Rad9 | 2035 | 9.37721408 | 11.4132015 | 10.4259875 | 7.89017681 | 10.1551264 | 6.27249689 |
| ENSMUSG00 | Dpf2 | 7731 | 10.5263832 | 9.28700023 | 11.3377642 | 9.57240633 | 9.38552664 | 10.3898348 |
| ENSMUSG00 | Gldc | 4209 | 0.25502447 | 0.54952464 | 0 | 0 | 0.40006424 | 0 |
| ENSMUSG00 | Mrpl21 | 2120 | 7.85733318 | 9.47818225 | 11.3632096 | 10.1948796 | 6.73933266 | 13.096973 |
| ENSMUSG00 | Rps6kb2 | 2843 | 13.9556699 | 16.3728819 | 13.4020361 | 11.0162556 | 14.6635769 | 12.2990107 |
| ENSMUSG00 | Ighmbp2 | 5994 | 2.28159581 | 2.83780682 | 1.92470488 | 2.52827139 | 4.46076425 | 1.82467974 |
| ENSMUSG00 | Pola2 | 2853 | 1.42133159 | 1.57074627 | 3.70285331 | 3.41470453 | 2.8616264 | 1.99803702 |
| ENSMUSG00 | Coro1b | 2465 | 60.9799074 | 58.7424984 | 54.1181185 | 45.5819418 | 57.4641863 | 44.5577981 |
| ENSMUSG00 | Dmrt1 | 3082 | 0 | 0 | 0 | 0 | 0 | 0 |
| ENSMUSG00 | Eif1ad | 2463 | 31.604235 | 30.1678822 | 26.7400512 | 28.2151919 | 26.9944652 | 21.7266919 |
| ENSMUSG00 | Cabp4 | 1425 | 0 | 0 | 0 | 0.10128293 | 0.03580798 | 0.13398068 |
| ENSMUSG00 | Chka | 12362 | 28.7022863 | 27.0439683 | 17.978229 | 31.006274 | 32.9512686 | 32.0976745 |
| ENSMUSG00 | Banf1 | 816 | 28.4037318 | 28.640226 | 38.0265855 | 31.7928799 | 20.5731344 | 48.4325396 |
| ENSMUSG00 | Tmem134 | 4176 | 10.03407 | 9.1157302 | 10.7752577 | 9.8931851 | 7.72238255 | 10.6459919 |
| ENSMUSG00 | Cst6 | 3948 | 0.07048844 | 0.01220528 | 0.1231561 | 0.03655729 | 0 | 0.08981011 |
| ENSMUSG00 | Aip | 2251 | 17.0960929 | 22.7553034 | 25.193676 | 21.1426996 | 15.8224835 | 18.9989491 |
| ENSMUSG00 | Pitpnm1 | 6745 | 8.15149437 | 7.29404854 | 6.84161959 | 8.32374519 | 9.07051514 | 8.48364613 |

| | | | | | | | | |
|---|---|---|---|---|---|---|---|---|
| ENSMUSG00 | Sf3b2 | 3210 | 57.0076252 | 45.7846257 | 43.1415496 | 41.7360222 | 56.1449095 | 41.4982362 |
| ENSMUSG00 | Pold4 | 906 | 8.46888271 | 8.19063145 | 11.2212184 | 7.7261773 | 6.25157585 | 13.6674241 |
| ENSMUSG00 | Pacs1 | 4353 | 13.0783019 | 8.92218524 | 10.1035685 | 10.2452118 | 15.309085 | 8.92863794 |
| ENSMUSG00 | Cdk2ap2 | 1512 | 29.4485041 | 38.6893788 | 35.3147 | 29.113819 | 22.8133797 | 31.279249 |
| ENSMUSG00 | Cabp2 | 1338 | 0 | 0 | 0 | 0 | 0 | 0 |
| ENSMUSG00 | Adrbk1 | 4504 | 9.33865417 | 9.24358038 | 10.1966507 | 7.67464446 | 9.00665381 | 6.64910166 |
| ENSMUSG00 | Klc2 | 4529 | 1.59759273 | 1.4469764 | 1.90314846 | 1.07553014 | 1.52098934 | 1.74042189 |
| ENSMUSG00 | Mbl2 | 1095 | 0.10891913 | 0 | 0.32293584 | 0 | 0 | 0 |
| ENSMUSG00 | Acy3 | 1499 | 7.16076033 | 10.8331094 | 3.45004378 | 4.01981417 | 5.99108882 | 8.42438824 |
| ENSMUSG00 | Pip5k1b | 4175 | 1.24741747 | 0.98104134 | 0.45525252 | 1.27043366 | 1.5277358 | 1.85532875 |
| ENSMUSG00 | Dkk1 | 2423 | 0 | 0 | 0 | 0.55098458 | 1.03189947 | 0.78795904 |
| ENSMUSG00 | Nudt8 | 4511 | 2.70559355 | 1.91207563 | 2.05772269 | 1.79970295 | 3.72149804 | 2.79336806 |
| ENSMUSG00 | Rab1b | 1852 | 95.3101153 | 95.1239916 | 109.764728 | 91.5689035 | 80.0386727 | 94.3567029 |
| ENSMUSG00 | Doc2g | 2817 | 0.50805726 | 0.68422353 | 0.47073315 | 0.70447726 | 0.94191393 | 0.95853365 |
| ENSMUSG00 | Cnih2 | 1231 | 0.96885818 | 1.29175678 | 1.97489957 | 1.14313545 | 0.33160927 | 0.64253821 |
| ENSMUSG00 | Yif1a | 1447 | 62.2570275 | 53.1149794 | 71.0528589 | 54.6342499 | 41.4699502 | 61.4291979 |
| ENSMUSG00 | Cbwd1 | 1360 | 21.163947 | 15.0582632 | 22.7184473 | 20.6410519 | 14.6325637 | 22.6018518 |
| ENSMUSG00 | Rin1 | 4166 | 2.63382444 | 2.22078658 | 2.52521331 | 2.17393019 | 2.64563062 | 2.19977887 |
| ENSMUSG00 | Aldh3b1 | 1855 | 0.70724035 | 0.62343645 | 0.52422671 | 0.79750072 | 0.79771693 | 0.11762648 |
| ENSMUSG00 | Asah2 | 4910 | 2.2509213 | 2.17869453 | 3.63697446 | 2.99826358 | 1.75630499 | 1.51093725 |
| ENSMUSG00 | Rce1 | 1438 | 21.3153515 | 19.7705152 | 26.0047004 | 17.2882675 | 22.3196039 | 19.8205828 |
| ENSMUSG00 | Slc29a2 | 2460 | 5.3168915 | 4.46606049 | 4.4201843 | 5.19229421 | 3.81660697 | 4.8229545 |
| ENSMUSG00 | Pcx | 4304 | 12.0541129 | 9.71789772 | 14.2649535 | 12.3822444 | 12.756594 | 5.73502521 |
| ENSMUSG00 | 2700046G09I | 1639 | 3.95371771 | 3.38098892 | 4.23409967 | 3.4783179 | 3.86044576 | 5.07551249 |
| ENSMUSG00 | Minpp1 | 2614 | 21.0792257 | 24.7568446 | 26.2268778 | 18.5379441 | 23.7563502 | 16.4962527 |
| ENSMUSG00 | Apba1 | 6883 | 0.08086252 | 0.10501186 | 0.00642189 | 0 | 0.08154731 | 0.01188783 |
| ENSMUSG00 | Papss2 | 3622 | 2.86476544 | 0.9179637 | 1.1715563 | 0.27893353 | 1.35243845 | 2.55276165 |
| ENSMUSG00 | Cpt1a | 4281 | 100.20127 | 100.841471 | 90.530661 | 101.334833 | 135.116794 | 86.9590169 |
| ENSMUSG00 | Peli3 | 4755 | 2.04002886 | 1.39847088 | 1.74762282 | 0.11382348 | 1.97452222 | 0.7342069 |
| ENSMUSG00 | Mrpl11 | 816 | 19.146941 | 24.4475332 | 33.4763958 | 28.8302611 | 23.0744273 | 36.7672833 |
| ENSMUSG00 | Lao1 | 2497 | 0 | 0 | 0 | 0 | 0 | 0 |
| ENSMUSG00 | Mll5 | 4349 | 0 | 0.07755923 | 0 | 0.07466967 | 0.0117329 | 0.0250859 |
| ENSMUSG00 | Mus81 | 3532 | 8.70073229 | 11.787397 | 10.5874177 | 10.7367579 | 10.5028809 | 8.02331545 |
| ENSMUSG00 | Gal | 706 | 0.50679791 | 0 | 0 | 0 | 0 | 0 |
| ENSMUSG00 | Ppp6r3 | 4928 | 36.052606 | 35.7389298 | 41.5468622 | 35.5841592 | 39.450181 | 33.4180746 |
| ENSMUSG00 | Efemp2 | 2392 | 3.70630107 | 7.47373329 | 3.49253693 | 4.4951711 | 4.77838971 | 4.54957402 |
| ENSMUSG00 | Ctsw | 1222 | 0.13013251 | 0 | 0.43406065 | 1.44682503 | 0.79337246 | 0.04463934 |
| ENSMUSG00 | Fibp | 1199 | 45.4253614 | 38.3401717 | 46.5612408 | 39.001109 | 43.6639379 | 45.381906 |
| ENSMUSG00 | Fosl1 | 1703 | 0 | 0 | 0 | 0 | 0 | 0 |
| ENSMUSG00 | Lrp5 | 10175 | 0.35945987 | 0.69142217 | 1.07300788 | 1.2482437 | 0.78232085 | 1.18480497 |
| ENSMUSG00 | Drap1 | 1907 | 12.9043746 | 12.6846324 | 16.9204748 | 10.5199878 | 8.3750685 | 25.1006774 |
| ENSMUSG00 | Smarca2 | 7924 | 15.3021473 | 19.6479549 | 15.0277341 | 19.8168927 | 22.0229941 | 21.6813418 |
| ENSMUSG00 | Ovol1 | 2896 | 3.7613956 | 5.4409415 | 5.2810213 | 4.68468534 | 5.26826187 | 3.15504287 |
| ENSMUSG00 | Vldlr | 9715 | 2.84815383 | 3.23888283 | 0.5687319 | 2.67040556 | 2.25850145 | 1.55534227 |
| ENSMUSG00 | Rnaseh2c | 1610 | 2.00012045 | 1.97534483 | 3.7887294 | 1.86013025 | 1.77483046 | 2.93075316 |
| ENSMUSG00 | Kat5 | 2476 | 9.95492646 | 13.7981371 | 13.0320458 | 10.4049191 | 12.4474681 | 8.12951687 |
| ENSMUSG00 | Rela | 2787 | 31.3821518 | 27.092987 | 26.0104135 | 29.4016229 | 27.8109311 | 22.6652546 |
| ENSMUSG00 | Slc1a1 | 6416 | 1.22067171 | 0.60833881 | 0.09645041 | 0 | 1.15318337 | 0.66316143 |
| ENSMUSG00 | Kcnk7 | 1223 | 0 | 0 | 0.03614215 | 0 | 0 | 0 |
| ENSMUSG00 | Ehbp1l1 | 6420 | 0.65020646 | 0.89317549 | 0.32359605 | 0.37093691 | 0.24638904 | 0.48431602 |
| ENSMUSG00 | Fam89b | 1949 | 0 | 0.04944735 | 0 | 0.05553932 | 0.0523616 | 0.02798834 |
| ENSMUSG00 | Ltbp3 | 5165 | 12.3923182 | 15.6920805 | 9.79885968 | 10.9049509 | 20.4500674 | 7.83650816 |
| ENSMUSG00 | Scyl1 | 2620 | 41.8494715 | 36.4157082 | 38.1283073 | 30.1877258 | 47.8518341 | 28.3969374 |
| ENSMUSG00 | Capn1 | 3628 | 7.11171633 | 6.65419171 | 9.8077408 | 6.9021331 | 5.59771157 | 10.0738745 |
| ENSMUSG00 | Smc5 | 3437 | 6.44277063 | 5.5659056 | 10.3013315 | 6.897266 | 6.96286595 | 10.2765952 |
| ENSMUSG00 | Arl2 | 3305 | 9.79151623 | 13.6467503 | 15.4071174 | 12.7078668 | 10.7919627 | 12.3870601 |
| ENSMUSG00 | Men1 | 3201 | 19.3871619 | 21.1050896 | 17.1504808 | 16.9307188 | 20.6432854 | 13.0195716 |
| ENSMUSG00 | Map4k2 | 4502 | 8.23897453 | 6.36848802 | 4.10403607 | 8.76804903 | 10.1780732 | 3.64711969 |
| ENSMUSG00 | Sf1 | 5763 | 19.8053071 | 22.5923593 | 18.3771674 | 20.9617708 | 21.2764847 | 24.117918 |
| ENSMUSG00 | Rps6ka4 | 3764 | 10.61484 | 7.83477752 | 9.48859967 | 10.7747657 | 9.31326254 | 5.75347066 |
| ENSMUSG00 | Prdx5 | 1645 | 152.665879 | 167.407609 | 159.368469 | 144.635259 | 112.971466 | 175.867537 |
| ENSMUSG00 | Esrra | 2531 | 14.2937524 | 11.7848312 | 17.8658575 | 12.8304386 | 10.523812 | 17.4036122 |
| ENSMUSG00 | Kcnk4 | 1708 | 0 | 0.39497084 | 0.23291369 | 0.7816368 | 0.29874927 | 0.35131268 |
| ENSMUSG00 | Gpr137 | 3048 | 13.1083523 | 11.0506309 | 14.4004036 | 11.4709649 | 10.881609 | 15.3912006 |
| ENSMUSG00 | Bad | 2112 | 9.01651286 | 11.6359306 | 11.9713325 | 11.224386 | 7.68294863 | 12.8237295 |
| ENSMUSG00 | Plcb3 | 4200 | 22.4713121 | 19.1254284 | 16.5546426 | 25.1972944 | 28.5869196 | 14.6179079 |
| ENSMUSG00 | Vegfb | 1598 | 44.7560134 | 34.2551929 | 46.9955765 | 36.7142706 | 31.9633305 | 47.9951725 |
| ENSMUSG00 | Dnajc4 | 1080 | 25.4361454 | 27.9749067 | 24.9658558 | 17.0721526 | 18.7096711 | 27.7797252 |
| ENSMUSG00 | Fermt3 | 2572 | 2.82863644 | 1.91097088 | 2.97314107 | 0.70143944 | 3.88847965 | 3.23435652 |
| ENSMUSG00 | Stip1 | 2159 | 93.2845133 | 95.6363616 | 98.3946538 | 86.3194812 | 85.9578179 | 71.7175206 |
| ENSMUSG00 | Rcor2 | 3142 | 0.25305844 | 0 | 0.18288477 | 0 | 0 | 0 |
| ENSMUSG00 | Mark2 | 6874 | 10.2829858 | 12.7931709 | 10.7964641 | 15.175035 | 13.7624237 | 12.0859109 |
| ENSMUSG00 | AI846148 | 3177 | 4.35470797 | 4.17100145 | 4.27131439 | 3.13460632 | 2.65009506 | 3.88043341 |
| ENSMUSG00 | Lgals12 | 4265 | 0 | 1.06202241 | 0 | 1.62432651 | 0.55034309 | 0.01918498 |
| ENSMUSG00 | Hrasls5 | 1676 | 0 | 0 | 0 | 0 | 0 | 0 |
| ENSMUSG00 | Smc3 | 5672 | 17.5857723 | 15.1389704 | 14.9391631 | 18.3654553 | 17.3266572 | 16.9264334 |
| ENSMUSG00 | Pdcd4 | 2543 | 24.7944129 | 43.2976879 | 23.8999348 | 30.0943832 | 31.9040258 | 39.8233329 |
| ENSMUSG00 | Shoc2 | 4193 | 24.6516216 | 32.809991 | 47.7122684 | 36.039018 | 31.2023915 | 31.7499438 |
| ENSMUSG00 | Gpam | 6322 | 24.6066427 | 15.4574667 | 27.5754617 | 19.7189914 | 28.9111797 | 17.2699116 |
| ENSMUSG00 | Tectb | 3888 | 0 | 0 | 0 | 0 | 0 | 0 |
| ENSMUSG00 | Acsl5 | 3510 | 132.45031 | 120.370007 | 130.892865 | 128.517869 | 130.749637 | 108.391437 |
| ENSMUSG00 | Zdhhc6 | 2518 | 41.4922172 | 48.7796827 | 53.7689615 | 50.0821071 | 54.1876604 | 46.49037 |
| ENSMUSG00 | Vti1a | 8152 | 9.02203618 | 7.48332136 | 11.5709928 | 10.4722911 | 9.02601005 | 8.81273305 |
| ENSMUSG00 | Tcf7l2 | 8767 | 2.68452657 | 4.22668804 | 4.54774295 | 5.67138786 | 4.29536505 | 5.90478673 |
| ENSMUSG00 | Hhex | 3833 | 0.11409087 | 0.52800171 | 0 | 0.33888693 | 0 | 0.37713433 |
| ENSMUSG00 | Cyp26a1 | 1766 | 0 | 0 | 0 | 0.34733565 | 0 | 0.03088861 |
| ENSMUSG00 | Cep55 | 1882 | 0 | 0.48647311 | 0 | 0 | 0 | 0 |
| ENSMUSG00 | Rbp4 | 1290 | 0 | 0 | 0 | 0 | 0 | 0 |
| ENSMUSG00 | Eif3a | 5124 | 49.4926249 | 42.5345975 | 48.523686 | 42.8632719 | 43.428186 | 43.1369392 |
| ENSMUSG00 | Pde6c | 2937 | 0 | 0 | 0 | 0 | 0 | 0 |
| ENSMUSG00 | Fam45a | 5486 | 14.9137402 | 16.8292423 | 22.6649261 | 15.3115202 | 18.5279877 | 30.5559484 |
| ENSMUSG00 | Prdx3 | 1471 | 237.695072 | 211.646909 | 254.393476 | 189.829329 | 166.780976 | 211.355277 |
| ENSMUSG00 | Plce1 | 10759 | 0.02217054 | 0.23737164 | 0.08627556 | 0.18780505 | 0.66397366 | 0.2991363 |
| ENSMUSG00 | Noc3l | 3956 | 6.11004352 | 2.04634031 | 5.16209592 | 5.48162507 | 5.15039088 | 3.88160311 |
| ENSMUSG00 | Hells | 7057 | 0.00563348 | 0.30043977 | 0.17537928 | 1.06860526 | 0.60014017 | 0.01545962 |
| ENSMUSG00 | Cyp2c55 | 1986 | 0 | 0 | 0 | 0 | 0 | 0 |
| ENSMUSG00 | Cyp2c39 | 1808 | 0 | 0 | 0 | 0 | 0 | 0 |
| ENSMUSG00 | Cyp2c40 | 2327 | 0 | 0 | 0 | 0 | 0 | 0 |
| ENSMUSG00 | Sorbs1 | 4886 | 0.58583598 | 1.40042464 | 1.12178234 | 1.15941076 | 2.05734671 | 0.35726093 |
| ENSMUSG00 | Aldh18a1 | 4231 | 3.1383433 | 2.56250283 | 3.94901836 | 3.41120724 | 3.37683414 | 3.04913835 |
| ENSMUSG00 | Tctn3 | 3259 | 11.7839197 | 12.4347339 | 14.3225364 | 7.83862765 | 16.064149 | 10.8462511 |
| ENSMUSG00 | Ccnj | 4484 | 1.04619686 | 1.93433532 | 3.26289252 | 2.75202035 | 3.45941531 | 2.0072772 |
| ENSMUSG00 | Tll2 | 3421 | 0.02324202 | 0 | 0 | 0 | 0 | 0 |
| ENSMUSG00 | Dntt | 2116 | 0 | 0.38713115 | 0 | 0 | 0.02411454 | 0 |
| ENSMUSG00 | Tm9sf3 | 10424 | 49.3205942 | 49.240405 | 55.5745735 | 57.1552877 | 47.164153 | 63.5501188 |
| ENSMUSG00 | Pik3ap1 | 4604 | 0.00863499 | 0 | 0 | 0 | 0 | 0 |
| ENSMUSG00 | Lcor | 7620 | 2.75992772 | 3.64243972 | 2.73216116 | 5.78638562 | 3.9039865 | 4.24510789 |
| ENSMUSG00 | Slit1 | 6526 | 0 | 0.01476753 | 0.00677319 | 0 | 0 | 0.12538143 |
| ENSMUSG00 | Smndc1 | 2119 | 22.1947775 | 19.3064131 | 22.4868272 | 17.6068117 | 19.8422528 | 23.4003276 |
| ENSMUSG00 | Mxi1 | 5611 | 14.0855276 | 11.3273779 | 16.968592 | 10.6361971 | 13.0044052 | 10.9613815 |
| ENSMUSG00 | Add3 | 4920 | 16.742552 | 14.7693404 | 15.0843281 | 18.2096985 | 20.0786715 | 11.1704061 |
| ENSMUSG00 | Xpnpep1 | 2715 | 46.3594296 | 39.6850404 | 54.2306958 | 33.8759232 | 41.9863439 | 35.241044 |
| ENSMUSG00 | Trim8 | 3224 | 57.5985885 | 54.8972711 | 53.8949891 | 48.8853507 | 53.5428751 | 48.8473223 |
| ENSMUSG00 | Arl3 | 816 | 27.4293328 | 26.6324576 | 31.6888213 | 32.7656801 | 20.8232637 | 38.8396211 |
| ENSMUSG00 | Sfxn2 | 3198 | 19.591819 | 20.8838051 | 22.1700301 | 13.0089423 | 19.5617063 | 19.7438367 |
| ENSMUSG00 | Maoa | 4152 | 11.4325732 | 12.3019337 | 15.702726 | 10.6021422 | 11.2941328 | 16.1532606 |
| ENSMUSG00 | Efhc2 | 2539 | 0.95513364 | 2.96064788 | 0 | 1.22215669 | 1.647957 | 0.65527885 |
| ENSMUSG00 | Fundc1 | 2816 | 49.3131724 | 50.8558615 | 52.5838686 | 49.3949864 | 37.943619 | 51.633914 |
| ENSMUSG00 | Nt5c2 | 5677 | 10.3082733 | 15.4736452 | 14.9026471 | 15.7433899 | 13.6891264 | 12.9286691 |
| ENSMUSG00 | Dusp21 | 820 | 0 | 0 | 0 | 0 | 0 | 0 |
| ENSMUSG00 | Msr1 | 4002 | 0 | 0 | 0 | 0 | 0 | 0 |
| ENSMUSG00 | Pdcd11 | 6005 | 2.36348152 | 3.47455944 | 1.72979735 | 2.79403009 | 3.24597428 | 2.39362779 |
| ENSMUSG00 | Taf5 | 3247 | 4.16287754 | 4.65985304 | 6.547917 | 6.30074507 | 3.37870983 | 4.28397475 |
| ENSMUSG00 | Pcgf6 | 4075 | 6.62428743 | 9.3061914 | 6.11775201 | 8.94303435 | 12.1336339 | 9.49759831 |
| ENSMUSG00 | 4930542N07I | 960 | 0 | 0 | 0 | 0 | 0 | 0 |
| ENSMUSG00 | Nr0b1 | 1806 | 0 | 0 | 0 | 0 | 0 | 0 |
| ENSMUSG00 | 5430427O19I | 3752 | 0 | 0.10274295 | 0 | 0 | 0 | 0.00726936 |
| ENSMUSG00 | Gyk | 7721 | 2.3942881 | 3.27650332 | 1.9292865 | 2.38334966 | 3.68108941 | 2.55048433 |
| ENSMUSG00 | Slk | 8590 | 16.6935993 | 18.5453292 | 19.2759143 | 21.3761961 | 20.0125565 | 18.139699 |
| ENSMUSG00 | Col17a1 | 5711 | 0.16706908 | 0 | 0.31733069 | 0 | 0 | 0 |
| ENSMUSG00 | Sfr1 | 1619 | 98.0013111 | 89.7952415 | 102.409582 | 91.1968861 | 83.5521447 | 122.441061 |
| ENSMUSG00 | Gsta1 | 2001 | 48.8152002 | 65.0914308 | 30.8816474 | 47.5684325 | 28.0759819 | 35.698291 |
| ENSMUSG00 | Gsta2 | 1777 | 2.37145804 | 2.73879047 | 3.58191637 | 2.07111342 | 2.61305582 | 5.31065007 |
| ENSMUSG00 | Habp2 | 3033 | 83.9806672 | 47.9163566 | 61.4133091 | 50.3697252 | 78.8864742 | 46.4918843 |
| ENSMUSG00 | Casp7 | 2343 | 8.61962616 | 11.2291069 | 6.22561169 | 10.1639562 | 14.9834172 | 6.37921567 |
| ENSMUSG00 | Dclre1a | 4083 | 6.15367712 | 3.89456917 | 4.88244702 | 4.4539188 | 5.02390475 | 4.30863148 |
| ENSMUSG00 | Nhlrc2 | 4369 | 10.9011366 | 10.2350694 | 7.2438818 | 9.18361948 | 9.4835013 | 8.15305078 |
| ENSMUSG00 | Tdrd1 | 5416 | 0.21287093 | 0.10676464 | 0.12242017 | 0 | 0.24495675 | 0.01007188 |
| ENSMUSG00 | Vwa2 | 4201 | 1.55198734 | 2.38580813 | 1.21000046 | 2.31041894 | 2.18632413 | 1.64907364 |
| ENSMUSG00 | Afap1l2 | 6613 | 5.14001752 | 3.46114623 | 6.53034941 | 3.8520979 | 3.5339604 | 3.89755544 |
| ENSMUSG00 | Ablim1 | 15884 | 12.0237553 | 10.5449555 | 12.3778518 | 10.4856911 | 9.80115038 | 10.5619684 |
| ENSMUSG00 | Trub1 | 5700 | 3.87790301 | 2.20643182 | 3.334525 | 3.96269471 | 2.18428697 | 3.89500989 |
| ENSMUSG00 | Gfra1 | 8272 | 0.28355577 | 0.66990339 | 1.08474059 | 0 | 0.6662051 | 1.00895063 |
| ENSMUSG00 | 1700011F14I | 1081 | 0 | 0 | 0 | 0 | 0 | 0 |
| ENSMUSG00 | Pnliprp2 | 1555 | 0 | 0 | 0 | 0 | 0 | 0 |
| ENSMUSG00 | Hspa12a | 5721 | 0.47253499 | 1.23814141 | 0.54083709 | 1.04064628 | 0.19622099 | 0.52442061 |
| ENSMUSG00 | Slc18a2 | 3842 | 1.89361086 | 1.78096689 | 1.95583376 | 2.29151975 | 2.29764784 | 3.03130426 |
| ENSMUSG00 | 3110040N11I | 1216 | 4.6751922 | 6.61771039 | 6.39763517 | 4.89600111 | 4.40606041 | 7.04295783 |
| ENSMUSG00 | Btbd1 | 2992 | 40.0478002 | 33.5147012 | 41.7199214 | 43.2695106 | 37.8263708 | 38.3595194 |
| ENSMUSG00 | Hdgfrp3 | 9569 | 2.48030326 | 2.10491512 | 1.03009834 | 1.101051 | 2.36761531 | 1.73869056 |
| ENSMUSG00 | Bnc1 | 5282 | 0 | 0 | 0 | 0 | 0 | 0 |
| ENSMUSG00 | Gcgr | 2510 | 102.667341 | 85.7949044 | 103.812695 | 108.907398 | 100.914315 | 98.9928142 |
| ENSMUSG00 | Bhlhe22 | 3184 | 0.27469239 | 0.55995552 | 0.1249424 | 0.33996901 | 0.76924185 | 0.09422771 |
| ENSMUSG00 | Ppp1r27 | 768 | 0.1035299 | 0 | 0.17266345 | 0.84567292 | 0 | 0.49719395 |
| ENSMUSG00 | P4hb | 3386 | 163.131319 | 161.408927 | 183.70004 | 165.83248 | 162.874504 | 125.909594 |
| ENSMUSG00 | Arhgdia | 3572 | 49.8390374 | 44.7599706 | 54.0890974 | 37.6376534 | 34.555656 | 50.7161685 |
| ENSMUSG00 | Ints4 | 3220 | 22.3099855 | 31.2164343 | 28.1821067 | 28.5854955 | 32.8026702 | 21.7773595 |
| ENSMUSG00 | Alyref | 4191 | 16.0217148 | 13.371709 | 16.2632407 | 16.0221152 | 11.3107858 | 18.9249943 |
| ENSMUSG00 | Anapc11 | 5217 | 1.75268555 | 2.08743261 | 1.9741287 | 1.78439093 | 1.51602228 | 4.45951069 |
| ENSMUSG00 | Pcyt2 | 4396 | 82.1245493 | 65.9878029 | 72.215113 | 70.8097812 | 73.9627085 | 73.9505062 |
| ENSMUSG00 | Sirt7 | 4358 | 27.1847939 | 21.1631139 | 24.900304 | 21.6674416 | 24.4828251 | 20.8283615 |
| ENSMUSG00 | Tollip | 4324 | 46.0994402 | 42.5253153 | 56.9390068 | 55.9088109 | 47.0967312 | 34.3834548 |
| ENSMUSG00 | Pycr1 | 3899 | 0.12235593 | 0.09886934 | 0.12470384 | 0.3979297 | 0.14395746 | 0.24483518 |
| ENSMUSG00 | Myadml2 | 1398 | 0.19906178 | 0 | 0 | 0 | 0.07299911 | 0.13656829 |
| ENSMUSG00 | Aspscr1 | 4641 | 12.2067571 | 12.2309047 | 11.7147742 | 12.8514587 | 12.1491371 | 11.0132889 |
| ENSMUSG00 | Stra13 | 1347 | 9.06082592 | 10.266896 | 11.7149651 | 7.68785052 | 9.01579617 | 16.1177527 |
| ENSMUSG00 | Lrrc45 | 2981 | 19.6176826 | 24.6670616 | 16.8296182 | 22.828157 | 30.0920391 | 18.8937035 |
| ENSMUSG00 | Mob2 | 3221 | 13.3546818 | 16.0372139 | 13.8328028 | 13.1288905 | 10.4714171 | 9.99195104 |
| ENSMUSG00 | Cbr2 | 1640 | 15.4173693 | 4.75988026 | 13.0449342 | 13.0027368 | 5.04041029 | 7.63357281 |
| ENSMUSG00 | Maged1 | 4425 | 67.0587359 | 74.2125657 | 61.9924605 | 62.9580714 | 66.1555521 | 48.0711664 |
| ENSMUSG00 | Fasn | 10251 | 2.95131408 | 3.49258868 | 2.03524241 | 2.99187766 | 4.35548521 | 1.47667786 |
| ENSMUSG00 | Arhgap19 | 8733 | 1.09711105 | 0.58488067 | 0.17715155 | 0.76436256 | 1.61265083 | 1.20554343 |
| ENSMUSG00 | Dus1l | 4099 | 13.9275116 | 14.3301471 | 15.7979263 | 17.5612781 | 18.7474316 | 11.2917938 |
| ENSMUSG00 | Gps1 | 2935 | 12.3262308 | 11.8372827 | 11.2349489 | 11.2487464 | 10.709454 | 6.88603328 |
| ENSMUSG00 | Zdhhc16 | 1819 | 19.4734102 | 19.9474387 | 19.5858634 | 21.4032579 | 19.1313845 | 18.2030852 |
| ENSMUSG00 | Rfng | 3348 | 17.8472184 | 15.6159766 | 14.6679354 | 17.9224731 | 17.206923 | 14.6637845 |
| ENSMUSG00 | Mms19 | 10357 | 7.66551073 | 7.21144978 | 6.38466324 | 6.93979267 | 9.35590303 | 7.28412206 |
| ENSMUSG00 | Slc16a3 | 5309 | 0.12730141 | 0.19968011 | 0.00832583 | 0 | 0 | 0.06678665 |
| ENSMUSG00 | Csnk1d | 5838 | 28.7100216 | 27.4855862 | 22.5097772 | 29.1907325 | 34.4546032 | 16.6086575 |
| ENSMUSG00 | Cd7 | 1129 | 0 | 0 | 0 | 0 | 0 | 0 |
| ENSMUSG00 | Sectm1a | 2047 | 0 | 0 | 0 | 0 | 0 | 0 |
| ENSMUSG00 | 1110031I02R | 1299 | 22.9535107 | 23.7779133 | 31.0671922 | 27.082366 | 16.3802915 | 17.2382439 |
| ENSMUSG00 | Rab40b | 3189 | 2.63641906 | 4.91081646 | 3.29885188 | 3.85825562 | 2.03209462 | 7.69745228 |
| ENSMUSG00 | Ubtd1 | 1524 | 12.3388073 | 9.73846731 | 6.29383198 | 14.9394817 | 8.4709141 | 4.88581136 |
| ENSMUSG00 | Ankrd2 | 1091 | 0 | 0.22083612 | 0 | 0.23150716 | 0 | 0 |
| ENSMUSG00 | Wdr45l | 4783 | 18.4605733 | 20.7938149 | 21.3107778 | 18.7388247 | 17.357289 | 16.1891492 |
| ENSMUSG00 | Fn3k | 4031 | 5.83856226 | 5.116298 | 9.45224229 | 6.17628646 | 5.70897928 | 6.08959944 |
| ENSMUSG00 | Hoga1 | 2569 | 10.8170809 | 9.69731047 | 13.3517439 | 6.05346874 | 13.069426 | 8.05817485 |
| ENSMUSG00 | Pi4k2a | 3762 | 54.7404011 | 34.673232 | 54.8939422 | 39.237545 | 45.777251 | 40.6727077 |
| ENSMUSG00 | D19Ertd386e | 6615 | 8.55809588 | 9.99422504 | 8.59313229 | 8.72732747 | 13.0439307 | 12.5302538 |
| ENSMUSG00 | Loxl4 | 4492 | 0 | 0 | 0 | 0.2008128 | 0.28398473 | 0.03035913 |
| ENSMUSG00 | Hps1 | 6435 | 2.41560107 | 1.58749429 | 2.09503685 | 2.42789826 | 1.92687014 | 1.49194608 |
| ENSMUSG00 | Cnnm1 | 6447 | 1.43062998 | 2.90000614 | 0.53478265 | 3.06140505 | 1.58294946 | 1.10418302 |
| ENSMUSG00 | Got1 | 3066 | 123.15641 | 88.4518253 | 150.107498 | 100.79671 | 86.1257322 | 106.990243 |
| ENSMUSG00 | Entpd7 | 9045 | 0.58457478 | 1.68215257 | 0.76724039 | 1.79512659 | 0.9928847 | 0.78401395 |
| ENSMUSG00 | Cutc | 1535 | 10.333835 | 10.7673939 | 9.93461618 | 11.8941463 | 12.0003399 | 14.4457861 |
| ENSMUSG00 | Abcc2 | 5818 | 0 | 0.60460816 | 0 | 0 | 0 | 0.00468798 |
| ENSMUSG00 | Dnmbp | 4884 | 11.3389403 | 10.5469506 | 9.6205076 | 12.6479239 | 17.5416227 | 9.06920847 |
| ENSMUSG00 | Cpn1 | 3679 | 0.16209084 | 0.44532184 | 0 | 0.09807569 | 0.29126227 | 0 |
| ENSMUSG00 | Cyp2c44 | 1955 | 0.02033528 | 0.59154712 | 0 | 0 | 0.02610045 | 0 |
| ENSMUSG00 | Erlin1 | 4725 | 3.43285419 | 6.85318288 | 5.04228431 | 3.58911342 | 7.25708456 | 2.73035013 |
| ENSMUSG00 | Chuk | 6271 | 54.9197461 | 41.2400949 | 50.6019823 | 66.4793402 | 50.6683531 | 55.8540771 |
| ENSMUSG00 | Cwf19l1 | 3560 | 4.29939888 | 3.55984109 | 5.20240793 | 3.71969392 | 4.68697328 | 4.35168402 |
| ENSMUSG00 | Gm13540 | 869 | 0.04574854 | 0.05545045 | 0.10173036 | 0.04152134 | 0.1761555 | 0.06277247 |
| ENSMUSG00 | Scd3 | 1990 | 0.45948545 | 0.26635722 | 0.55529954 | 0.12692177 | 0 | 0.37005792 |
| ENSMUSG00 | Scd2 | 5447 | 178.896013 | 170.80665 | 99.2127286 | 125.084863 | 127.242751 | 85.0636261 |
| ENSMUSG00 | Ndufb8 | 3113 | 54.6589967 | 56.1272211 | 76.7888105 | 55.0561232 | 38.4050108 | 85.2321525 |
| ENSMUSG00 | Sema4g | 4921 | 1.28451969 | 0.56793612 | 1.2036267 | 1.39312913 | 1.49315142 | 2.45532721 |
| ENSMUSG00 | Mrpl43 | 1715 | 19.3098048 | 15.9591249 | 19.0725154 | 16.5998444 | 16.5426618 | 19.9748962 |
| ENSMUSG00 | Peo1 | 3597 | 9.30612026 | 8.01097926 | 7.3731181 | 4.94535667 | 8.38381654 | 3.31359949 |
| ENSMUSG00 | Sfxn3 | 3016 | 62.849513 | 54.577217 | 49.5365482 | 51.6466135 | 60.2977465 | 43.4260006 |
| ENSMUSG00 | Kazald1 | 1675 | 1.13926153 | 1.32332916 | 1.47779296 | 1.61561393 | 0.67019718 | 0.37451743 |
| ENSMUSG00 | Tlx1 | 3581 | 0 | 0 | 0 | 0 | 0 | 0 |
| ENSMUSG00 | Lbx1 | 2477 | 0 | 0 | 0 | 0 | 0 | 0 |
| ENSMUSG00 | Btrc | 6308 | 10.4871781 | 7.64658031 | 12.7462195 | 10.015799 | 10.8556431 | 8.97192081 |
| ENSMUSG00 | Poll | 2315 | 6.0277208 | 4.4127541 | 2.90223764 | 6.56178866 | 4.36424294 | 3.6169824 |
| ENSMUSG00 | Fgf8 | 1811 | 0 | 0 | 0.09762969 | 0 | 0 | 0 |
| ENSMUSG00 | Mgea5 | 5059 | 35.8105801 | 35.0706622 | 32.467691 | 44.3411881 | 49.2411083 | 40.8283936 |
| ENSMUSG00 | Kcnip2 | 3422 | 0.5227927 | 0.22530189 | 0 | 0 | 0.49207201 | 0.09564456 |
| ENSMUSG00 | Ldb1 | 4110 | 10.4466956 | 10.5283273 | 8.44244447 | 10.0257165 | 7.68499431 | 10.5515028 |
| ENSMUSG00 | Gbf1 | 8684 | 8.60666782 | 9.27217237 | 8.2305827 | 12.4068384 | 14.4429792 | 6.77468872 |
| ENSMUSG00 | Nfkb2 | 3266 | 4.17517753 | 7.46550512 | 3.09927191 | 6.96010044 | 9.06163441 | 5.19437404 |
| ENSMUSG00 | Fbxl15 | 1344 | 4.022876 | 4.62503797 | 3.8479283 | 4.64448936 | 2.46779347 | 2.86140189 |
| ENSMUSG00 | Tmem180 | 2794 | 5.81961043 | 4.44957125 | 2.83182888 | 3.0864741 | 5.02228109 | 3.71926899 |
| ENSMUSG00 | Actr1a | 2731 | 40.963721 | 35.2532081 | 39.1520536 | 40.4155892 | 36.1164352 | 35.6937242 |
| ENSMUSG00 | Pitx3 | 1576 | 0 | 0 | 0 | 0 | 0 | 0 |
| ENSMUSG00 | Sufu | 7297 | 4.84344555 | 3.60556358 | 3.65875198 | 3.584964 | 4.26560083 | 4.7469908 |
| ENSMUSG00 | Hexa | 2144 | 61.2835182 | 36.3421068 | 84.1772971 | 39.8013224 | 39.4121634 | 44.830144 |
| ENSMUSG00 | Arih1 | 7687 | 13.436287 | 12.1923546 | 11.9604311 | 12.4200715 | 12.3267829 | 8.75328932 |
| ENSMUSG00 | Bbs4 | 2526 | 26.4564381 | 30.5600476 | 31.2352691 | 33.1109181 | 28.8260643 | 23.0311978 |
| ENSMUSG00 | Adpgk | 2803 | 6.85048059 | 5.58708302 | 7.20665082 | 7.23443062 | 6.91759644 | 6.13022573 |
| ENSMUSG00 | Parp6 | 2679 | 12.4356449 | 11.0258638 | 10.0976215 | 11.1788343 | 12.9708705 | 13.1435458 |
| ENSMUSG00 | Limd1 | 4841 | 28.299398 | 23.2222618 | 26.0864833 | 28.6509769 | 27.2997962 | 16.598035 |
| ENSMUSG00 | Sacm1l | 3413 | 102.784166 | 113.583336 | 129.859912 | 118.268336 | 105.880104 | 108.155572 |
| ENSMUSG00 | Fyco1 | 8041 | 15.1091591 | 14.7477719 | 17.5740943 | 20.632414 | 22.8447896 | 15.7657659 |
| ENSMUSG00 | Slc6a20b | 5737 | 0.19403058 | 0.21838025 | 0 | 0.01886807 | 0.04447131 | 0.11409994 |
| ENSMUSG00 | Lztfl1 | 12326 | 3.18340575 | 2.16186131 | 1.9006147 | 3.35177194 | 2.59147422 | 3.17532917 |
| ENSMUSG00 | Tbl1x | 5599 | 41.2607782 | 36.1721051 | 50.3991009 | 48.8161256 | 50.1514817 | 33.8996633 |
| ENSMUSG00 | Zfhx4 | 16081 | 0.00988881 | 0 | 0.0027487 | 0.1795015 | 0 | 0 |
| ENSMUSG00 | Ribc1 | 1539 | 1.39492914 | 2.37957739 | 2.09684363 | 3.25887212 | 2.42035441 | 1.61273046 |
| ENSMUSG00 | Hsd17b10 | 1213 | 68.5315002 | 63.9175477 | 96.2747479 | 69.5761765 | 46.6935508 | 99.4748591 |
| ENSMUSG00 | Huwe1 | 22312 | 6.6728341 | 6.59345679 | 5.29740625 | 10.4937427 | 6.90886883 | 6.99835558 |
| ENSMUSG00 | Fam120c | 5808 | 3.16236777 | 2.67149352 | 1.89501703 | 3.74612136 | 4.72661677 | 2.1320052 |
| ENSMUSG00 | Tsr2 | 7119 | 4.47870422 | 5.11037559 | 4.00480246 | 4.18144216 | 3.93503025 | 5.12237571 |
| ENSMUSG00 | Fgd1 | 4797 | 1.30114873 | 1.64739973 | 0.58051201 | 1.00792036 | 1.09562575 | 0.38663237 |
| ENSMUSG00 | Gnl3l | 5553 | 22.0505817 | 19.4811026 | 14.6145478 | 19.4802754 | 17.8174217 | 16.6015272 |
| ENSMUSG00 | Maged2 | 3178 | 33.6633096 | 37.7698009 | 38.3740985 | 32.6532285 | 35.7409417 | 28.9567756 |
| ENSMUSG00 | Apex2 | 9756 | 1.49551674 | 0.85941379 | 0.89255464 | 0.6990064 | 0.29289433 | 0.46967399 |
| ENSMUSG00 | Alas2 | 2418 | 0 | 0.45834912 | 0 | 0 | 0 | 0.16919751 |
| ENSMUSG00 | Pfkfb1 | 4037 | 0.1378689 | 0.26259642 | 0.0547459 | 0 | 0 | 0.31753977 |
| ENSMUSG00 | Tro | 6920 | 0 | 0 | 0 | 0 | 0 | 0 |
| ENSMUSG00 | Abhd6 | 2162 | 36.5926948 | 37.1539311 | 43.1795987 | 41.4058985 | 38.2815826 | 35.0709977 |
| ENSMUSG00 | Flnb | 9054 | 31.06583 | 28.8831259 | 27.6615466 | 40.3940362 | 37.9400886 | 28.0307619 |
| ENSMUSG00 | Dnase1l3 | 5142 | 0 | 0 | 0 | 0 | 0 | 0 |
| ENSMUSG00 | Polr3a | 4652 | 7.8194894 | 8.35908401 | 4.76984634 | 6.59280693 | 7.53549445 | 4.91905037 |
| ENSMUSG00 | Sat1 | 2906 | 34.9536314 | 45.8650031 | 27.8961391 | 38.7515695 | 38.8931254 | 45.3138191 |
| ENSMUSG00 | Acot9 | 2425 | 35.165157 | 44.8084235 | 20.7976507 | 44.6822184 | 41.8311073 | 38.4881713 |
| ENSMUSG00 | 4933436I01R | 1424 | 0 | 0 | 0 | 0 | 0 | 0 |
| ENSMUSG00 | Prdx4 | 2097 | 6.90080825 | 9.62809693 | 7.2931987 | 3.88867051 | 6.10759196 | 7.30965535 |
| ENSMUSG00 | Rps24 | 933 | 5.36890734 | 6.40420024 | 7.67491808 | 9.12686228 | 9.51617298 | 9.17924622 |
| ENSMUSG00 | Ptprj | 8639 | 11.9096173 | 10.2240709 | 10.4121716 | 13.0394887 | 12.8702944 | 10.7816753 |
| ENSMUSG00 | Banp | 6614 | 2.92726323 | 2.86321012 | 3.04748116 | 4.14065192 | 2.16788806 | 2.15260987 |
| ENSMUSG00 | Car5a | 1331 | 0 | 0 | 0 | 0 | 0 | 0 |
| ENSMUSG00 | Jph3 | 4450 | 0 | 0 | 0.03973199 | 0 | 0 | 0 |
| ENSMUSG00 | Itgb8 | 11046 | 0 | 0.0305364 | 0 | 0 | 0 | 0.12099016 |
| ENSMUSG00 | Sp4 | 5764 | 4.89011722 | 4.10470907 | 6.01985544 | 6.04080031 | 4.78925561 | 6.52528236 |
| ENSMUSG00 | Atp10a | 5448 | 1.32080433 | 0.55722207 | 0.51114466 | 0.9735794 | 0.44957159 | 0.80602366 |
| ENSMUSG00 | Ube3a | 5084 | 64.113923 | 56.7451277 | 65.4159455 | 71.2698448 | 52.7827912 | 87.2530946 |
| ENSMUSG00 | Padi3 | 4696 | 0 | 0 | 0 | 0 | 0.01086592 | 0 |
| ENSMUSG00 | Padi1 | 3738 | 0.0106355 | 0.06445485 | 0.03547499 | 0.0289583 | 0.0819043 | 0.13863514 |
| ENSMUSG00 | Padi4 | 4814 | 0 | 0.08007718 | 0 | 0 | 0 | 0 |
| ENSMUSG00 | Kdm5c | 6418 | 7.87304701 | 7.5605714 | 8.69849295 | 9.95095338 | 10.5662283 | 8.83514724 |
| ENSMUSG00 | Gpr143 | 2292 | 0.72850357 | 0.73583136 | 1.11854577 | 2.29841994 | 0.55657042 | 0 |
| ENSMUSG00 | Sbds | 2084 | 54.3871761 | 50.3367968 | 53.2797647 | 51.7364429 | 47.5985003 | 45.3617561 |
| ENSMUSG00 | Rabgef1 | 3477 | 29.1563058 | 32.4984777 | 29.340769 | 32.3980854 | 32.0951061 | 30.7575695 |
| ENSMUSG00 | Mettl7b | 1225 | 0.12981381 | 1.37675549 | 0.14433255 | 0.50073041 | 0.41654184 | 0 |
| ENSMUSG00 | Itga7 | 4231 | 0 | 0 | 0 | 0 | 0.37386378 | 0 |
| ENSMUSG00 | Rdh5 | 2476 | 1.34873197 | 1.8488336 | 1.14253552 | 1.98188936 | 3.4003845 | 1.17866979 |
| ENSMUSG00 | Cd63 | 1022 | 148.09111 | 162.711753 | 138.314573 | 137.726082 | 112.986975 | 150.224016 |
| ENSMUSG00 | Gdf11 | 4059 | 0.20568246 | 0 | 0 | 0.08889393 | 0.12571169 | 0.3897337 |
| ENSMUSG00 | Ormdl2 | 2571 | 10.3911641 | 12.5948227 | 12.5676963 | 11.157225 | 9.08987947 | 12.2316838 |
| ENSMUSG00 | Dnajc14 | 4084 | 18.3007599 | 15.3974797 | 18.2045788 | 16.6009211 | 18.6913463 | 12.495311 |
| ENSMUSG00 | Mmp19 | 3401 | 0 | 0 | 0.02599344 | 0 | 0 | 0 |
| ENSMUSG00 | Dgka | 2784 | 13.9801061 | 10.9388762 | 9.00231501 | 9.5000498 | 14.1128985 | 8.43515244 |
| ENSMUSG00 | Cdk2 | 2424 | 11.7265547 | 10.7942401 | 14.4239513 | 10.9556043 | 12.0829784 | 10.2504936 |
| ENSMUSG00 | Pmel | 1971 | 0.80680833 | 1.39351964 | 1.4576965 | 0 | 0.18122001 | 0 |
| ENSMUSG00 | Rps26 | 443 | 6.9998363 | 5.32987735 | 8.18183099 | 4.39826276 | 4.03142924 | 12.6214471 |
| ENSMUSG00 | Pa2g4 | 2492 | 5.31242978 | 4.67942175 | 5.7114741 | 6.25499327 | 4.75044911 | 4.67346347 |
| ENSMUSG00 | Esyt1 | 3795 | 10.0043436 | 11.0339969 | 7.31456058 | 7.21640891 | 11.9935513 | 4.29782194 |
| ENSMUSG00 | Smarcc2 | 4998 | 21.3254188 | 19.494395 | 22.0920776 | 23.8164596 | 29.157929 | 25.3810218 |
| ENSMUSG00 | Cdh9 | 3208 | 0 | 0 | 0 | 0 | 0 | 0 |
| ENSMUSG00 | Chmp6 | 2423 | 30.5836631 | 23.2082451 | 25.8315352 | 21.0416545 | 18.3214804 | 23.3686137 |
| ENSMUSG00 | Baiap2 | 4685 | 17.7351023 | 22.3498695 | 17.1995645 | 17.4980587 | 20.5957047 | 18.7575 |
| ENSMUSG00 | Rnf41 | 4367 | 8.73946904 | 11.5859318 | 9.41326174 | 10.5759141 | 10.7264055 | 6.83895809 |
| ENSMUSG00 | Obfc2b | 2812 | 6.20649216 | 5.98046526 | 7.32505649 | 5.73565028 | 4.93569496 | 9.28230077 |
| ENSMUSG00 | Aatk | 6998 | 0 | 0.07574319 | 0.01894906 | 0.04640446 | 0 | 0.03117993 |
| ENSMUSG00 | 2410002I01R | 3317 | 8.06615568 | 5.52030389 | 6.43638534 | 8.29984925 | 6.15331635 | 7.0303939 |
| ENSMUSG00 | Fscn2 | 2106 | 0.01887725 | 1.00674428 | 0.29383941 | 0.11993082 | 0 | 0.09065645 |
| ENSMUSG00 | Cnpy2 | 1011 | 66.888301 | 77.8323047 | 87.7478723 | 68.0954906 | 51.9348573 | 95.8254388 |
| ENSMUSG00 | Il23a | 1355 | 0 | 0 | 0 | 0 | 0 | 0 |
| ENSMUSG00 | 2310003H01I | 4987 | 17.0118699 | 14.8897749 | 17.3968807 | 14.405324 | 15.6650053 | 10.6429613 |
| ENSMUSG00 | Pde6g | 2864 | 0 | 0.18507363 | 0.38584011 | 0.61732548 | 0.07126589 | 0.04761634 |
| ENSMUSG00 | Mip | 2140 | 0 | 0 | 0 | 0 | 0 | 0 |
| ENSMUSG00 | Atp5b | 3852 | 1432.40483 | 1160.12737 | 1680.63394 | 1171.87024 | 1173.1695 | 1152.86332 |
| ENSMUSG00 | Prim1 | 3397 | 3.92054341 | 2.93629483 | 2.42023633 | 5.31086908 | 8.35168235 | 7.56335301 |
| ENSMUSG00 | Hsd17b6 | 1387 | 0 | 0 | 0 | 0 | 0 | 0 |
| ENSMUSG00 | Tac2 | 673 | 0 | 0 | 0 | 0 | 0 | 0 |
| ENSMUSG00 | Myo1a | 3359 | 0 | 0.1578002 | 0 | 0 | 0 | 0 |
| ENSMUSG00 | Nab2 | 3470 | 4.41091066 | 7.26268278 | 3.47755134 | 5.15754873 | 3.88212196 | 3.38771456 |
| ENSMUSG00 | Shmt2 | 2268 | 11.8495171 | 15.6159766 | 10.6996525 | 12.8705353 | 12.7790924 | 10.7992178 |
| ENSMUSG00 | R3hdm2 | 5919 | 15.3272523 | 13.741969 | 12.1948994 | 15.2277323 | 13.0604763 | 10.9808186 |
| ENSMUSG00 | Inhbc | 1964 | 0 | 0 | 0 | 0 | 0 | 0 |
| ENSMUSG00 | Gli1 | 3945 | 0 | 0 | 0 | 0 | 0 | 0 |
| ENSMUSG00 | Ddit3 | 1807 | 12.540467 | 14.2932667 | 16.0956351 | 20.2874141 | 9.09269121 | 16.8598628 |
| ENSMUSG00 | Mbd6 | 4466 | 0.89018093 | 1.43501944 | 1.07881793 | 1.4865867 | 0.99402031 | 3.47498203 |
| ENSMUSG00 | Dctn2 | 1627 | 137.030568 | 123.353738 | 115.163868 | 93.3653396 | 103.965849 | 90.5243097 |
| ENSMUSG00 | Ttc4 | 4031 | 15.7700356 | 18.0743985 | 18.0491773 | 22.3868006 | 22.9245266 | 21.5030522 |
| ENSMUSG00 | Pip4k2c | 3364 | 35.2173995 | 28.0466866 | 33.834645 | 37.9269053 | 33.688936 | 29.9745367 |
| ENSMUSG00 | Bsnd | 2752 | 216.285994 | 171.576652 | 160.50473 | 131.990531 | 142.77002 | 104.995468 |
| ENSMUSG00 | Katnal2 | 4277 | 0.18590358 | 0.24786105 | 0.23769988 | 0.0084363 | 0.26246908 | 0.00637705 |
| ENSMUSG00 | Hdhd2 | 4805 | 22.1323435 | 16.9680458 | 23.0438328 | 19.3213529 | 21.5680685 | 18.8737098 |
| ENSMUSG00 | Agap2 | 5614 | 0 | 0 | 0 | 0 | 0.00908913 | 0 |
| ENSMUSG00 | Pias2 | 5439 | 34.3100249 | 38.7245704 | 37.4484451 | 37.3225929 | 40.3782904 | 35.689209 |
| ENSMUSG00 | St8sia5 | 2463 | 0.20983404 | 0 | 0 | 0 | 0 | 0 |
| ENSMUSG00 | Rnf165 | 7289 | 0 | 0 | 0 | 0 | 0 | 0 |
| ENSMUSG00 | Atp5a1 | 2845 | 872.859872 | 706.147327 | 860.125916 | 640.332635 | 725.058794 | 669.642725 |
| ENSMUSG00 | Pstpip2 | 4670 | 0 | 0 | 0.0757205 | 0 | 0 | 0.07008473 |
| ENSMUSG00 | Crisp1 | 1408 | 0 | 0 | 0.3453269 | 0 | 0 | 0 |
| ENSMUSG00 | Avil | 3470 | 0 | 0.13886583 | 0.24202738 | 0.5095159 | 0.30880516 | 0.13362215 |
| ENSMUSG00 | Crisp3 | 2171 | 0 | 0 | 0.16288104 | 0 | 0 | 0 |
| ENSMUSG00 | Xrcc6bp1 | 1355 | 4.25427651 | 4.80086324 | 5.41513354 | 3.80792057 | 4.82020377 | 2.85830168 |
| ENSMUSG00 | Usp33 | 6085 | 35.10373 | 29.9096454 | 31.6640647 | 39.2247699 | 31.4795423 | 27.6914908 |
| ENSMUSG00 | Clns1a | 1682 | 46.2553361 | 40.1076213 | 37.9736404 | 32.2421599 | 34.6748796 | 36.3391597 |
| ENSMUSG00 | Gm9752 | 2657 | 0.91271521 | 1.50525968 | 1.29760774 | 1.72465926 | 1.72840565 | 2.20701823 |
| ENSMUSG00 | Paip1 | 3192 | 5.8039016 | 5.60061718 | 4.5835871 | 6.11540918 | 4.85965484 | 5.51132278 |
| ENSMUSG00 | Nnt | 8686 | 32.7847694 | 27.2608999 | 28.2024653 | 22.8887941 | 29.4138918 | 23.4468952 |
| ENSMUSG00 | Cd163l1 | 3821 | 0 | 0 | 0 | 0 | 0 | 0 |
| ENSMUSG00 | Paox | 1932 | 0.55558901 | 0.84800157 | 0.84651563 | 0.50425218 | 1.26773605 | 0.79056925 |
| ENSMUSG00 | Echs1 | 3429 | 67.6504021 | 83.6693137 | 85.9286922 | 68.5443097 | 67.5143386 | 67.4031184 |
| ENSMUSG00 | 1810014F10I | 3265 | 7.13528685 | 5.12119309 | 7.83855042 | 5.65819504 | 5.18859319 | 10.8430266 |
| ENSMUSG00 | Prap1 | 615 | 0.25857223 | 0 | 0 | 0 | 0.16593943 | 0 |
| ENSMUSG00 | Caly | 1442 | 0.11027873 | 0 | 0.09195945 | 0 | 0 | 0 |
| ENSMUSG00 | Msx3 | 4287 | 0 | 0 | 0 | 0 | 0 | 0 |
| ENSMUSG00 | Zfp511 | 1291 | 19.7699601 | 17.5053767 | 16.6056498 | 17.1326826 | 15.1774813 | 18.971825 |
| ENSMUSG00 | Adam8 | 4746 | 0.20103909 | 0.05076532 | 0.16764289 | 0.09883409 | 0.05375724 | 0.10344364 |
| ENSMUSG00 | Tubgcp2 | 3141 | 5.30326213 | 10.8921916 | 6.95183395 | 8.76491562 | 13.9221917 | 6.97279061 |
| ENSMUSG00 | Gpr123 | 5151 | 0 | 0.31806232 | 0.00858122 | 0.04202917 | 0.00990611 | 0 |
| ENSMUSG00 | Inpp5a | 3865 | 15.799332 | 14.5369706 | 15.3248304 | 13.2845406 | 12.0271742 | 15.4544528 |
| ENSMUSG00 | Dpysl4 | 6129 | 0 | 0.14151672 | 0.01442384 | 0.26491956 | 0.041627 | 0.07565163 |
| ENSMUSG00 | Cyp2e1 | 2000 | 2.02752951 | 0.9155424 | 0.35361474 | 0 | 0 | 4.67760063 |
| ENSMUSG00 | Syce1 | 1449 | 0.60360288 | 0 | 0 | 0 | 0 | 0 |
| ENSMUSG00 | 1190003J15R | 1574 | 0 | 0 | 0 | 0 | 0.03241828 | 0 |
| ENSMUSG00 | Odf3 | 1383 | 0 | 0 | 0 | 0 | 0 | 0 |
| ENSMUSG00 | Bet1l | 1548 | 13.072054 | 14.1633276 | 14.791056 | 9.23029046 | 12.1303012 | 14.8530497 |
| ENSMUSG00 | Ric8 | 4314 | 22.5502227 | 21.9151135 | 18.1459119 | 17.212992 | 24.3895195 | 13.7700891 |
| ENSMUSG00 | Sirt3 | 1562 | 45.1766824 | 39.3018741 | 44.1452465 | 34.3264521 | 35.8687328 | 56.6970254 |
| ENSMUSG00 | Psmd13 | 3060 | 31.7654085 | 42.3284825 | 37.7449071 | 25.0805584 | 34.2510379 | 36.375099 |
| ENSMUSG00 | Cox8b | 360 | 0 | 0 | 0 | 0 | 0 | 0 |
| ENSMUSG00 | Ifitm5 | 706 | 0.16893264 | 0.1365055 | 0.12521769 | 0.05110771 | 0.21682596 | 0.2317958 |
| ENSMUSG00 | Ifitm1 | 1091 | 0 | 0 | 0 | 0 | 0 | 0 |
| ENSMUSG00 | Ifitm3 | 676 | 8.52743295 | 12.2604557 | 3.98862785 | 9.44751758 | 10.2656614 | 26.8308212 |
| ENSMUSG00 | Sigirr | 1980 | 14.8179518 | 10.8297812 | 9.86728011 | 9.94989713 | 9.1228975 | 7.76913967 |
| ENSMUSG00 | Ptdss2 | 4859 | 20.3891042 | 20.7165009 | 19.0307174 | 16.5669976 | 19.6271448 | 18.4001374 |
| ENSMUSG00 | Drd4 | 2770 | 0 | 0 | 0.01595734 | 0 | 0 | 0 |
| ENSMUSG00 | Cdhr5 | 2625 | 6.80008029 | 5.87415676 | 1.11136062 | 5.56694401 | 5.67607685 | 1.08059523 |
| ENSMUSG00 | Irf7 | 3035 | 3.22235526 | 4.01685992 | 0.94666221 | 5.0288978 | 4.35447491 | 3.22622586 |
| ENSMUSG00 | Hras1 | 4894 | 19.2116288 | 19.977583 | 25.8581685 | 19.3017557 | 12.3656072 | 23.7301622 |
| ENSMUSG00 | 1600016N20I | 2066 | 8.71695676 | 9.04953512 | 9.19980276 | 8.52276754 | 10.6449022 | 7.74937717 |
| ENSMUSG00 | Taldo1 | 2066 | 91.5953956 | 114.541925 | 108.365119 | 91.0259516 | 78.4907175 | 95.6064557 |
| ENSMUSG00 | Eps8l2 | 4451 | 55.046737 | 49.2690403 | 48.8196496 | 41.8944071 | 62.8572497 | 41.0314501 |
| ENSMUSG00 | Tmem80 | 3611 | 9.6003265 | 10.1283605 | 10.5761264 | 11.2712673 | 10.4568037 | 9.08651094 |
| ENSMUSG00 | Lrdd | 4743 | 0 | 0.8737158 | 1.10900681 | 0.4184087 | 1.40933064 | 1.529634 |
| ENSMUSG00 | Rplp2 | 2142 | 2.02303799 | 2.31708849 | 2.33184326 | 2.34145854 | 1.59606311 | 4.17650872 |
| ENSMUSG00 | Pnpla2 | 3276 | 87.0591628 | 62.4246827 | 84.9090959 | 48.8253063 | 71.0101488 | 66.6713404 |
| ENSMUSG00 | Cd151 | 2001 | 10.8279545 | 13.9911659 | 11.9506232 | 10.5306917 | 6.29860811 | 19.1644892 |
| ENSMUSG00 | Tspan4 | 2068 | 8.28559579 | 10.2524345 | 8.72067309 | 10.9921122 | 8.61131778 | 7.29346011 |
| ENSMUSG00 | Chid1 | 12684 | 3.44146307 | 3.38870281 | 4.16788113 | 2.24730489 | 2.97291799 | 2.88787768 |
| ENSMUSG00 | Muc2 | 6990 | 0 | 0 | 0 | 0 | 0 | 0 |
| ENSMUSG00 | Tktl2 | 2038 | 0 | 0 | 0 | 0 | 0 | 0 |
| ENSMUSG00 | Tmem192 | 1117 | 72.9265708 | 63.0263133 | 109.812099 | 67.9324437 | 55.3204486 | 85.9261016 |
| ENSMUSG00 | Apool | 3599 | 30.2225603 | 29.0805647 | 29.316421 | 20.8832728 | 26.1724617 | 32.3900551 |
| ENSMUSG00 | Satl1 | 2732 | 0 | 0 | 0 | 0 | 0 | 0 |
| ENSMUSG00 | 2010106E10I | 1424 | 0 | 0 | 0 | 0 | 0 | 0 |
| ENSMUSG00 | Zfp711 | 5526 | 3.07914326 | 3.56645944 | 4.13542397 | 5.84390696 | 5.9373796 | 4.30885995 |
| ENSMUSG00 | Chm | 7219 | 11.2123782 | 11.0136625 | 11.7255201 | 11.8407485 | 8.51028626 | 14.9086752 |
| ENSMUSG00 | Crcp | 1637 | 16.149905 | 18.1324669 | 19.4682522 | 13.5996466 | 15.8658676 | 17.5110848 |
| ENSMUSG00 | Asl | 3821 | 13.4737889 | 14.6413136 | 16.7737954 | 13.6547072 | 11.831816 | 15.7109346 |
| ENSMUSG00 | Gusb | 4093 | 11.6459372 | 14.8338424 | 14.2767741 | 11.2398257 | 9.72405892 | 11.4882673 |
| ENSMUSG00 | 4930579G22I | 618 | 2.12286546 | 1.87131814 | 4.93515723 | 2.91925927 | 2.22930768 | 4.41337205 |
| ENSMUSG00 | Phkg1 | 3221 | 3.65340647 | 3.56050085 | 3.71893805 | 2.41965899 | 4.73669245 | 2.76895593 |
| ENSMUSG00 | Sumf2 | 2745 | 6.37246318 | 9.23354046 | 5.41049882 | 6.03339105 | 7.52848168 | 2.76224033 |
| ENSMUSG00 | Tm9sf2 | 3443 | 241.165543 | 188.057282 | 240.369186 | 214.993652 | 213.205182 | 286.142098 |
| ENSMUSG00 | Clybl | 1231 | 77.2825872 | 81.7721183 | 76.733825 | 76.6780085 | 67.8140949 | 87.1636319 |
| ENSMUSG00 | Fgf14 | 6367 | 0 | 0 | 0 | 0 | 0 | 0 |
| ENSMUSG00 | Farp1 | 5496 | 7.48670348 | 7.89079293 | 9.07199398 | 8.8038613 | 6.54541393 | 10.9177959 |
| ENSMUSG00 | Slc15a1 | 3191 | 0.52562873 | 0.12431201 | 0.12828655 | 0.11635616 | 0.13163851 | 0.1055452 |
| ENSMUSG00 | Dock9 | 10450 | 3.16141668 | 5.09992392 | 2.67325979 | 3.79810995 | 6.22082323 | 3.57571826 |
| ENSMUSG00 | Tnrc6c | 12934 | 2.8923695 | 4.03106003 | 3.30813237 | 4.75923674 | 4.41460604 | 4.73837285 |
| ENSMUSG00 | Tmc6 | 4338 | 10.3925115 | 12.1188124 | 11.2287762 | 12.0772542 | 9.65713567 | 7.19275871 |
| ENSMUSG00 | 6030468B19I | 1065 | 0.67292361 | 0 | 0 | 0 | 0 | 0 |
| ENSMUSG00 | Tk1 | 5706 | 1.12870449 | 0.52358209 | 0.24788976 | 1.39749945 | 0.54549753 | 0.54491919 |
| ENSMUSG00 | Cant1 | 6122 | 6.79908332 | 7.82380325 | 9.53061627 | 9.83891314 | 8.79333984 | 7.60499987 |
| ENSMUSG00 | Rbfox3 | 6525 | 0.04874235 | 0 | 0 | 0.11612612 | 0 | 0 |
| ENSMUSG00 | Cbx2 | 4252 | 2.00086379 | 1.45057963 | 1.60091341 | 3.37738799 | 2.40011175 | 1.48817411 |
| ENSMUSG00 | Cbx8 | 4255 | 1.56032086 | 1.04186902 | 2.39967702 | 1.3907063 | 0.87542313 | 1.57045514 |
| ENSMUSG00 | Gaa | 4958 | 34.6958379 | 34.7743223 | 28.2078724 | 34.3136022 | 32.4703945 | 19.6280583 |
| ENSMUSG00 | Eif4a3 | 2240 | 43.4470608 | 44.3573409 | 40.0579202 | 39.6741409 | 36.5385298 | 30.6230883 |
| ENSMUSG00 | Nptx1 | 5212 | 0 | 0 | 0 | 0 | 0.12727223 | 0 |
| ENSMUSG00 | Rptor | 12980 | 3.66007698 | 2.65062555 | 3.26235482 | 4.28925999 | 4.13557376 | 2.86194597 |
| ENSMUSG00 | Pde8a | 4251 | 23.0995147 | 23.4754462 | 18.3524472 | 24.0122569 | 25.8672877 | 19.2930699 |
| ENSMUSG00 | Cpeb1 | 5590 | 12.3960291 | 9.68900912 | 15.2611014 | 7.84251952 | 9.72148305 | 8.27509627 |
| ENSMUSG00 | Nat1 | 1346 | 0.67932842 | 1.03819229 | 0.91950342 | 0.21445495 | 1.55429525 | 1.56028768 |
| ENSMUSG00 | 1810029B16I | 3797 | 0.79573782 | 0.98987161 | 1.327103 | 2.09061096 | 1.08852685 | 1.46537435 |
| ENSMUSG00 | Dach2 | 10047 | 0 | 0 | 0.10118865 | 0 | 0 | 0.00271471 |
| ENSMUSG00 | Klhl4 | 3898 | 0 | 0 | 0 | 0 | 0 | 0 |
| ENSMUSG00 | Zfp202 | 3990 | 7.18388507 | 3.69550158 | 3.67794784 | 2.64059073 | 5.83158581 | 5.41391336 |
| ENSMUSG00 | Copg2 | 4632 | 25.0016655 | 23.2401796 | 22.2631476 | 15.1977696 | 19.3662281 | 20.7621714 |
| ENSMUSG00 | Podxl | 5706 | 2.72421887 | 0.85293212 | 0.96057282 | 1.59352878 | 3.11201871 | 2.82019579 |
| ENSMUSG00 | Mkln1 | 22326 | 22.3332991 | 16.0966804 | 21.7821677 | 20.5929146 | 17.1802055 | 26.2044707 |
| ENSMUSG00 | ORF63 | 1647 | 0 | 0 | 0 | 0 | 0.09294422 | 0 |
| ENSMUSG00 | Bach1 | 5520 | 1.56285132 | 1.16101391 | 1.26519768 | 1.34000347 | 0.9613665 | 0.72139444 |
| ENSMUSG00 | Cct8 | 3135 | 130.844353 | 129.357926 | 114.938891 | 131.886491 | 116.685196 | 116.380494 |
| ENSMUSG00 | Usp16 | 7419 | 13.7019495 | 15.4061519 | 15.2939926 | 14.3569478 | 13.4529709 | 14.6538229 |
| ENSMUSG00 | Phf6 | 6963 | 11.0879137 | 10.851119 | 14.3911501 | 17.9037001 | 18.4377943 | 13.4512583 |
| ENSMUSG00 | Hprt | 1280 | 206.138378 | 191.766982 | 192.519746 | 162.67928 | 139.604975 | 237.694219 |
| ENSMUSG00 | Gm7628 | 2008 | 0.01979855 | 0.11998616 | 0.02201287 | 0.16172231 | 0.12705771 | 0.13582988 |
| ENSMUSG00 | Ccdc51 | 2395 | 8.18348722 | 6.90102282 | 5.99816241 | 6.67404832 | 6.86033111 | 6.77595422 |
| ENSMUSG00 | Atrip | 6784 | 1.64671138 | 2.48603401 | 2.12408621 | 2.26044648 | 1.85782943 | 1.9097072 |
| ENSMUSG00 | Shisa5 | 3054 | 109.841763 | 84.8232852 | 128.00298 | 69.3876383 | 90.474075 | 94.0948262 |
| ENSMUSG00 | Pfkfb4 | 4325 | 1.92113189 | 2.15028516 | 1.59433237 | 1.47665246 | 2.02925703 | 1.20449852 |
| ENSMUSG00 | Col7a1 | 9920 | 0.20839566 | 0.53918297 | 0.08911662 | 0.82566775 | 0.79214333 | 0.12097622 |

| | | | | | | | | |
|---|---|---|---|---|---|---|---|---|
| ENSMUSG00 | Uqcrc1 | 1718 | 511.938589 | 514.456183 | 495.585507 | 298.842498 | 316.316008 | 373.049752 |
| ENSMUSG00 | Tmem89 | 536 | 0 | 0 | 0.3298645 | 0 | 0 | 0 |
| ENSMUSG00 | Arhgef9 | 9209 | 1.69227368 | 1.33952972 | 2.10713531 | 1.91596479 | 2.24961544 | 2.1887239 |
| ENSMUSG00 | Cnksr2 | 9521 | 0 | 0.20750385 | 0 | 0 | 0.16613987 | 0 |
| ENSMUSG00 | Rps6ka6 | 6702 | 0.72962162 | 1.02096013 | 0.29019414 | 0.6299014 | 0.23602173 | 0.04883552 |
| ENSMUSG00 | Tmem47 | 1993 | 1.15695829 | 0.58026825 | 1.68556953 | 0 | 0.43524756 | 0 |
| ENSMUSG00 | Alox5 | 3878 | 0 | 0.02485118 | 0.02279621 | 0 | 0 | 0 |
| ENSMUSG00 | 8-Mar | 6538 | 36.3259775 | 29.5619179 | 28.3681452 | 27.5885807 | 33.0290031 | 23.0737274 |
| ENSMUSG00 | Myo3a | 8046 | 0 | 0 | 0 | 0 | 0 | 0 |
| ENSMUSG00 | Wdr73 | 3712 | 8.13959192 | 9.86575863 | 10.1335583 | 9.82730254 | 12.0555311 | 9.14053104 |
| ENSMUSG00 | Nmb | 1060 | 0.18752585 | 0.72734252 | 0.41699852 | 0.71483296 | 1.05903799 | 1.08069325 |
| ENSMUSG00 | Sec11a | 3888 | 20.0720186 | 24.0560402 | 22.7375736 | 23.7298837 | 14.8170726 | 27.5833029 |
| ENSMUSG00 | Slc28a1 | 2959 | 0 | 0 | 0 | 0 | 0 | 0 |
| ENSMUSG00 | A930017K11I | 2536 | 18.0906248 | 34.8667671 | 9.39463462 | 25.382637 | 24.1449728 | 14.056764 |
| ENSMUSG00 | Pigq | 5356 | 65.7271807 | 61.1327249 | 40.0672046 | 64.1406172 | 67.5270634 | 41.8540441 |
| ENSMUSG00 | Rab40c | 4876 | 7.7130198 | 9.85272412 | 7.60569038 | 6.23073862 | 9.93109735 | 5.82299005 |
| ENSMUSG00 | 0610011F06I | 1329 | 43.2853125 | 52.3198164 | 49.4235807 | 47.0234019 | 32.4818013 | 42.8308293 |
| ENSMUSG00 | Fam195a | 811 | 53.2850892 | 45.156222 | 61.1522415 | 46.2259485 | 25.9221539 | 72.1718568 |
| ENSMUSG00 | Rhot2 | 4186 | 12.8687712 | 12.0638775 | 11.44644 | 16.1877878 | 11.4218142 | 10.4576675 |
| ENSMUSG00 | Rhbdl1 | 1561 | 0.45842322 | 0.2160827 | 0.16989818 | 0.34672048 | 0.09806478 | 0.08736271 |
| ENSMUSG00 | Jmjd8 | 2537 | 30.572701 | 28.8320927 | 33.730693 | 32.0855705 | 28.6809665 | 26.1028081 |
| ENSMUSG00 | Wdr24 | 4461 | 5.75701421 | 11.7198587 | 6.77741776 | 7.10155461 | 12.3762696 | 4.5427162 |
| ENSMUSG00 | Fbxl16 | 3472 | 0.08015218 | 0.12490725 | 0.25461891 | 0.47804552 | 0.07348268 | 0.33779075 |
| ENSMUSG00 | Gng13 | 769 | 0 | 0 | 0 | 0 | 0 | 0 |
| ENSMUSG00 | Prps2 | 3865 | 99.3218421 | 92.7448753 | 85.8190504 | 96.6886766 | 70.8956373 | 80.8359619 |
| ENSMUSG00 | Sdc3 | 6080 | 0.3007816 | 0 | 0.10178056 | 0.05341092 | 1.0574545 | 0.07626133 |
| ENSMUSG00 | Hadha | 5857 | 25.1348035 | 20.4444782 | 20.4594838 | 20.8163272 | 28.5667554 | 19.153251 |
| ENSMUSG00 | Il6 | 1082 | 0 | 0 | 0 | 0 | 0 | 0 |
| ENSMUSG00 | Tyms | 8387 | 2.60707217 | 3.28635327 | 3.94744014 | 2.73616076 | 3.25493157 | 3.2422577 |
| ENSMUSG00 | Agbl1 | 4573 | 0 | 0 | 0 | 0 | 0 | 0 |
| ENSMUSG00 | Hspa4l | 11771 | 11.0914109 | 10.7417572 | 9.17758086 | 13.1901315 | 11.2751341 | 9.21975954 |
| ENSMUSG00 | Plk4 | 6100 | 1.30997567 | 1.06642126 | 0.52897288 | 1.27174419 | 1.00379756 | 0.98814677 |
| ENSMUSG00 | Mfsd8 | 3172 | 14.4633747 | 17.5306287 | 20.4147856 | 15.970741 | 16.6334403 | 23.5514618 |
| ENSMUSG00 | Larp1b | 4635 | 1.14077215 | 1.18516816 | 0.44821718 | 1.6270005 | 0.95777663 | 2.05368913 |
| ENSMUSG00 | Phf17 | 5936 | 20.6948172 | 26.2687545 | 23.1955426 | 20.533212 | 24.5590222 | 28.2671127 |
| ENSMUSG00 | D3Ertd751e | 3783 | 32.5778455 | 27.3858976 | 25.7990138 | 28.5661442 | 31.5087534 | 42.1772772 |
| ENSMUSG00 | Crisp4 | 3281 | 0 | 0 | 0 | 0 | 0 | 0 |
| ENSMUSG00 | Crispld1 | 4575 | 0.04344861 | 0 | 0.65698914 | 0.56784857 | 0.03345992 | 0.57828197 |
| ENSMUSG00 | Gdap1 | 5402 | 0 | 0 | 0 | 0 | 0 | 0 |
| ENSMUSG00 | Ly96 | 598 | 15.8224153 | 10.2335755 | 29.7142824 | 18.161698 | 11.9459743 | 35.9861043 |
| ENSMUSG00 | Itih5 | 7886 | 0.26214621 | 0.68436235 | 1.03133897 | 0 | 0.26529057 | 0.39774075 |
| ENSMUSG00 | Atp5c1 | 22595 | 31.5369433 | 27.3934432 | 35.607964 | 23.6341783 | 17.1721426 | 37.4819572 |
| ENSMUSG00 | Taf3 | 8292 | 4.37732196 | 7.13614942 | 4.93619231 | 7.74989403 | 5.83984933 | 5.58847227 |
| ENSMUSG00 | 4930412O13I | 1376 | 0.08667619 | 0.31517295 | 0.1927406 | 0.28844658 | 0.22249873 | 0.79286742 |
| ENSMUSG00 | Clec3b | 987 | 0 | 0 | 0 | 0 | 0.05169846 | 0.05526776 |
| ENSMUSG00 | Exosc7 | 1898 | 15.981787 | 12.2877966 | 11.2717029 | 14.5430791 | 13.0119947 | 15.5198158 |
| ENSMUSG00 | Zdhhc3 | 11248 | 7.88535535 | 7.33849355 | 8.99127105 | 8.13834877 | 7.64397611 | 5.23038735 |
| ENSMUSG00 | Tgm4 | 4004 | 0 | 0.16848406 | 0.03311826 | 0.36947149 | 0.08920695 | 0.00681185 |
| ENSMUSG00 | St8sia2 | 5370 | 0 | 0 | 0 | 0 | 0 | 0 |
| ENSMUSG00 | Slco3a1 | 8140 | 3.92670839 | 8.32311273 | 1.1783538 | 4.57009679 | 6.56322058 | 3.23676923 |
| ENSMUSG00 | Pgm2 | 3193 | 31.8492074 | 27.0888392 | 30.8983757 | 19.4027155 | 22.069347 | 23.003634 |
| ENSMUSG00 | Slc25a10 | 3633 | 26.3723391 | 20.9298667 | 35.9162787 | 25.176984 | 21.4470894 | 19.1815864 |
| ENSMUSG00 | Hgs | 3358 | 25.2645013 | 25.7864909 | 20.9162384 | 23.6069839 | 31.5305926 | 18.3726129 |
| ENSMUSG00 | Rpl14 | 732 | 8.09230409 | 14.1531217 | 12.9827818 | 9.02051113 | 11.0138899 | 13.413757 |
| ENSMUSG00 | Rassf3 | 3420 | 8.57881422 | 6.43894856 | 14.6305516 | 9.67463007 | 8.89231579 | 12.0104113 |
| ENSMUSG00 | Ccr1 | 2829 | 0 | 0 | 0 | 0 | 0 | 0 |
| ENSMUSG00 | Ccdc7 | 2082 | 0 | 0 | 0 | 0 | 0 | 0 |
| ENSMUSG00 | Itgb1 | 4266 | 133.114694 | 102.280411 | 148.510318 | 122.354162 | 113.679015 | 122.070953 |
| ENSMUSG00 | Nrp1 | 5890 | 4.85300348 | 8.03380071 | 5.77850919 | 6.59768454 | 12.8822107 | 4.94555428 |
| ENSMUSG00 | Pard3 | 13443 | 2.58766982 | 2.99664254 | 3.8339172 | 4.54414204 | 4.64600789 | 2.83032967 |
| ENSMUSG00 | Homer2 | 11139 | 0.00356904 | 0 | 0.07936411 | 0.06154582 | 0 | 0.05631715 |
| ENSMUSG00 | Dhtkd1 | 6159 | 1.01986782 | 1.18138541 | 0.58849669 | 1.00179081 | 0.90304838 | 1.3905239 |
| ENSMUSG00 | Sec61a2 | 4778 | 4.82590596 | 4.2458125 | 3.64494059 | 6.7512239 | 6.07660273 | 6.15934193 |
| ENSMUSG00 | Nudt5 | 6500 | 6.50766635 | 6.32354387 | 5.84144356 | 6.82228196 | 6.1231651 | 5.29128001 |
| ENSMUSG00 | Zfp282 | 6312 | 1.71316393 | 2.31313244 | 2.19888763 | 2.92680716 | 4.13902161 | 2.48461939 |
| ENSMUSG00 | Pdia4 | 2470 | 113.504311 | 104.312918 | 146.599797 | 101.058941 | 96.1857513 | 118.628117 |
| ENSMUSG00 | Iscu | 2311 | 41.8370094 | 44.3915774 | 47.3960048 | 38.7362841 | 35.0847733 | 49.5569927 |
| ENSMUSG00 | Pramel6 | 2124 | 0 | 0 | 0 | 0 | 0 | 0 |
| ENSMUSG00 | Pramel7 | 2130 | 0 | 0 | 0 | 0 | 0 | 0 |
| ENSMUSG00 | Fam20c | 5352 | 10.9119583 | 9.44461469 | 7.60648307 | 9.77559129 | 8.59973673 | 6.95115993 |
| ENSMUSG00 | Prkar1b | 5311 | 11.1369357 | 7.88439432 | 13.9155492 | 6.20276909 | 8.6276945 | 6.66587869 |
| ENSMUSG00 | Pdgfa | 3818 | 2.00964058 | 1.95623325 | 1.85235592 | 1.80504728 | 1.20282185 | 1.89307899 |
| ENSMUSG00 | Heatr2 | 3500 | 13.9371356 | 9.33440222 | 12.363887 | 7.91743148 | 14.4185959 | 12.5073703 |
| ENSMUSG00 | Get4 | 2588 | 14.7930942 | 12.3817558 | 10.9992221 | 10.7353842 | 12.4805626 | 9.52715374 |
| ENSMUSG00 | Xiap | 7291 | 5.12006531 | 5.86881918 | 8.40871708 | 6.71558557 | 4.91997561 | 6.75974121 |
| ENSMUSG00 | Stag2 | 8506 | 22.8963158 | 19.4829996 | 23.6595196 | 24.1946479 | 25.2732896 | 22.9148004 |
| ENSMUSG00 | Cpbx2 | 4921 | 0.51703937 | 0.27417606 | 0.27845095 | 0.57191617 | 0.16590571 | 0.31592246 |
| ENSMUSG00 | Higd2a | 646 | 80.5571579 | 91.4498113 | 113.583683 | 79.0341997 | 54.1859037 | 159.67908 |
| ENSMUSG00 | Nop16 | 2210 | 11.0991545 | 12.6462156 | 13.9805829 | 9.12663479 | 8.81994369 | 9.74975792 |
| ENSMUSG00 | Arl10 | 1905 | 5.21725466 | 4.27480773 | 7.30896617 | 6.38301785 | 5.11603033 | 6.29965421 |
| ENSMUSG00 | 4833439L19I | 4804 | 194.896195 | 148.320758 | 204.180495 | 150.92269 | 155.022888 | 141.068477 |
| ENSMUSG00 | Thoc3 | 2422 | 10.8006219 | 15.180122 | 15.2388683 | 14.7784427 | 8.76423301 | 11.5089518 |
| ENSMUSG00 | Faf2 | 5091 | 19.9284986 | 16.6489789 | 21.6624628 | 18.5335978 | 21.3687357 | 15.1079322 |
| ENSMUSG00 | Tspan17 | 1910 | 6.95200549 | 7.16489506 | 5.50787363 | 5.45953449 | 3.90044548 | 4.79805172 |
| ENSMUSG00 | Unc5a | 3620 | 0 | 0 | 0.1587359 | 0 | 0 | 0 |
| ENSMUSG00 | Hk3 | 3989 | 0 | 0 | 0 | 0 | 0 | 0 |
| ENSMUSG00 | Uimc1 | 3797 | 19.2233506 | 15.6222045 | 14.551568 | 18.9580403 | 17.2148505 | 14.2658503 |
| ENSMUSG00 | Smad7 | 4744 | 2.33806473 | 2.89484317 | 2.45979902 | 2.92063767 | 3.48493798 | 3.73703953 |
| ENSMUSG00 | Myo6b | 9817 | 6.98970759 | 8.32476376 | 8.20370357 | 8.97548668 | 15.2294266 | 5.38158055 |
| ENSMUSG00 | Casp12 | 3878 | 0 | 0 | 0.14817534 | 0 | 0.19736865 | 0 |
| ENSMUSG00 | Casp1 | 1487 | 0 | 0 | 0 | 0 | 1.09807937 | 1.87088985 |
| ENSMUSG00 | Snca | 1456 | 1.6382753 | 1.35689644 | 0.57680977 | 0.99126496 | 1.85741615 | 0.31845389 |
| ENSMUSG00 | Gria4 | 6446 | 0 | 0 | 0 | 0.01119517 | 0 | 0 |
| ENSMUSG00 | Kbtbd3 | 2307 | 6.65176224 | 7.10159962 | 8.58362621 | 7.80448211 | 6.3700027 | 15.6885333 |
| ENSMUSG00 | Aasdhppt | 3083 | 34.5974551 | 27.3676485 | 34.5671889 | 33.2731925 | 20.3741384 | 47.5426245 |
| ENSMUSG00 | Cwf19l2 | 3935 | 10.7294334 | 11.0822694 | 11.188065 | 11.3885386 | 13.8490901 | 15.5607535 |
| ENSMUSG00 | Alkbh8 | 3645 | 10.6778094 | 12.3341428 | 10.2228131 | 14.5120102 | 16.8268049 | 13.5587507 |
| ENSMUSG00 | Rp1 | 12436 | 1.45454677 | 0.84857115 | 2.2712269 | 1.18377888 | 0.74676748 | 2.28531474 |
| ENSMUSG00 | Sox17 | 4104 | 1.28837205 | 1.77294171 | 0.09693387 | 0.70335369 | 0.78329963 | 0.14620908 |
| ENSMUSG00 | Lypla1 | 3478 | 97.8570065 | 90.35997 | 102.72671 | 110.538868 | 72.3876188 | 110.227812 |
| ENSMUSG00 | Oprk1 | 6943 | 0 | 0 | 0 | 0 | 0 | 0 |
| ENSMUSG00 | Rb1cc1 | 14517 | 9.67254647 | 10.6848631 | 9.47247597 | 13.2850126 | 8.84710257 | 11.3592663 |
| ENSMUSG00 | Sntg1 | 10053 | 0 | 0 | 0 | 0 | 0 | 0 |
| ENSMUSG00 | Adhfe1 | 3407 | 43.2211036 | 36.1221524 | 62.3392591 | 39.8205129 | 40.5874607 | 41.7805525 |
| ENSMUSG00 | Mybl1 | 6531 | 0.21395188 | 0.41317421 | 0 | 0.01657421 | 0.27345325 | 0.19628052 |
| ENSMUSG00 | Sgk3 | 5835 | 9.61353608 | 9.02618359 | 11.1053816 | 11.0750543 | 9.45321547 | 11.2277093 |
| ENSMUSG00 | Ppp1r42 | 1620 | 1.96323361 | 0.80310737 | 0.81855265 | 1.33637202 | 0 | 2.35706759 |
| ENSMUSG00 | Cops5 | 3326 | 82.8338785 | 54.763906 | 93.0152433 | 52.0293101 | 53.6804838 | 99.8074667 |
| ENSMUSG00 | Stau2 | 7792 | 14.5052401 | 10.6242695 | 14.6526387 | 11.914669 | 11.77431 | 14.942946 |
| ENSMUSG00 | Rdh10 | 4154 | 13.7622487 | 23.1536203 | 10.6620719 | 12.5687815 | 14.1016561 | 15.022719 |
| ENSMUSG00 | Gm5045 | 880 | 0 | 0 | 0.05022937 | 0 | 0 | 0 |
| ENSMUSG00 | Terf1 | 1957 | 14.1185789 | 22.0372334 | 17.8433602 | 20.944917 | 20.9111668 | 20.2364723 |
| ENSMUSG00 | Tfap2b | 6020 | 11.1539878 | 21.4917936 | 6.46139898 | 20.5703616 | 17.172684 | 12.4593452 |
| ENSMUSG00 | Il17a | 1168 | 0 | 0 | 0 | 0 | 0 | 0 |
| ENSMUSG00 | Msc | 1715 | 0 | 0 | 0 | 0 | 0 | 0 |
| ENSMUSG00 | Paqr8 | 4652 | 2.80305194 | 1.92662903 | 3.69615583 | 0.75235562 | 2.47892539 | 4.58485982 |
| ENSMUSG00 | Eya1 | 5682 | 0 | 0 | 0 | 0 | 0 | 0 |
| ENSMUSG00 | Tmem14a | 1573 | 9.57872035 | 7.99533465 | 10.3690274 | 7.29439934 | 8.95313396 | 11.5479401 |
| ENSMUSG00 | Gsta3 | 1435 | 43.5232473 | 83.7470291 | 15.8633792 | 81.3668962 | 29.3712797 | 20.9454024 |
| ENSMUSG00 | Tram1 | 2877 | 80.2433352 | 86.1895833 | 100.54114 | 94.4253435 | 72.2208559 | 90.185486 |
| ENSMUSG00 | Gm4956 | 1213 | 0 | 0.03972501 | 0 | 0 | 0 | 0 |
| ENSMUSG00 | Lactb2 | 3044 | 21.5886364 | 20.2148773 | 30.0584149 | 25.7932092 | 16.7964614 | 28.1348118 |
| ENSMUSG00 | Slco5a1 | 13692 | 0 | 0 | 0 | 0 | 0 | 0 |
| ENSMUSG00 | Ube2w | 2790 | 38.4302978 | 41.6752276 | 58.9833196 | 54.4075847 | 33.3957571 | 46.347335 |
| ENSMUSG00 | Tmem70 | 2111 | 56.9307521 | 53.6647681 | 68.4489933 | 48.5424 | 43.1947577 | 67.5082852 |
| ENSMUSG00 | Crygf | 649 | 0 | 0 | 0 | 0 | 0 | 0 |
| ENSMUSG00 | Pth2r | 2568 | 0 | 0 | 0 | 0 | 0 | 0 |
| ENSMUSG00 | Pikfyve | 6631 | 3.37540877 | 3.12474289 | 2.62638006 | 3.16690543 | 4.48625805 | 3.45508927 |
| ENSMUSG00 | Idh1 | 3453 | 27.2980725 | 32.5429433 | 27.3941338 | 24.2845849 | 23.6881785 | 27.7011758 |
| ENSMUSG00 | Crygc | 1120 | 0 | 0 | 0 | 0 | 0 | 0 |
| ENSMUSG00 | Akr1cl | 3887 | 0 | 0 | 0 | 0 | 0 | 0.04210132 |
| ENSMUSG00 | Mettl21a | 4657 | 6.62449063 | 6.67387915 | 9.15928248 | 6.09760984 | 9.34625265 | 7.51414262 |
| ENSMUSG00 | Creb1 | 8640 | 6.72714265 | 10.3678931 | 9.00919509 | 12.2987987 | 6.98069169 | 7.49105543 |
| ENSMUSG00 | Klf7 | 8208 | 1.38524213 | 1.67901102 | 2.08946334 | 0.74291734 | 1.47956597 | 1.1364433 |
| ENSMUSG00 | 4933402D24 | 1755 | 0 | 0 | 0.07555871 | 0 | 0 | 0 |
| ENSMUSG00 | Fastkd2 | 5449 | 10.980363 | 13.3001301 | 15.7371216 | 14.3891141 | 9.41117778 | 14.0302466 |
| ENSMUSG00 | Mdh1b | 1918 | 0 | 0 | 0.25350379 | 0 | 0 | 0 |
| ENSMUSG00 | Adam23 | 10350 | 0 | 0 | 0 | 0 | 0 | 0 |
| ENSMUSG00 | Eef1b2 | 2383 | 17.6672488 | 20.7264386 | 18.6786638 | 17.9274615 | 13.3186764 | 24.619282 |
| ENSMUSG00 | Ndufs1 | 4073 | 199.890127 | 134.893644 | 194.214629 | 161.416875 | 150.410672 | 181.446899 |
| ENSMUSG00 | Nrp2 | 10068 | 0.28831215 | 0.29201003 | 0.08782405 | 0.13262822 | 0.40553448 | 0.02709581 |
| ENSMUSG00 | 9430016H08I | 3535 | 7.14136637 | 5.58880941 | 8.96540918 | 6.93061053 | 5.42741649 | 10.0534243 |
| ENSMUSG00 | Boll | 3501 | 0 | 0 | 0.18938236 | 0.05153106 | 0 | 0 |
| ENSMUSG00 | Rftn2 | 6193 | 0 | 0.02334238 | 0 | 0.02913131 | 0.34605325 | 0 |
| ENSMUSG00 | Mob4 | 3836 | 48.3885137 | 56.7409174 | 75.6363132 | 61.4410623 | 36.7134508 | 72.0545345 |
| ENSMUSG00 | Hspd1 | 2630 | 50.3367871 | 43.019683 | 49.4119462 | 47.7847 | 39.2301642 | 52.7655378 |
| ENSMUSG00 | Coq10b | 1727 | 44.2443738 | 55.2735043 | 40.2090882 | 53.1306539 | 24.6711198 | 32.2652507 |
| ENSMUSG00 | Sf3b1 | 6167 | 130.876563 | 114.937987 | 139.414634 | 141.531483 | 140.022597 | 131.893026 |
| ENSMUSG00 | Ccdc150 | 5069 | 0 | 0.20913429 | 0 | 0.0640636 | 0 | 0 |
| ENSMUSG00 | Slc39a10 | 5949 | 6.79632269 | 9.99530503 | 5.37941408 | 4.77940008 | 8.22563362 | 3.98872687 |
| ENSMUSG00 | Cps1 | 5517 | 0 | 0 | 0 | 0 | 0 | 0 |
| ENSMUSG00 | Slc40a1 | 3372 | 39.1188269 | 82.5971636 | 34.1214108 | 67.0171545 | 63.389528 | 34.7646499 |
| ENSMUSG00 | Wdr75 | 4853 | 9.4453058 | 8.91642799 | 8.16090075 | 12.4090114 | 10.4092545 | 9.08779964 |
| ENSMUSG00 | Ikzf2 | 9409 | 2.01122422 | 2.2892273 | 1.75228903 | 2.19353061 | 2.01198697 | 2.83210995 |
| ENSMUSG00 | Lancl1 | 4596 | 15.319181 | 19.1236011 | 14.6858604 | 16.8241561 | 21.0611477 | 21.2630619 |
| ENSMUSG00 | Acadl | 1993 | 140.909541 | 115.908582 | 169.51063 | 133.501754 | 149.264311 | 122.373218 |
| ENSMUSG00 | 1110028C15I | 8136 | 9.46000618 | 10.5185744 | 9.5890183 | 14.1472126 | 10.4862464 | 14.9950174 |
| ENSMUSG00 | Rpe | 3292 | 32.7753263 | 32.3780103 | 45.9205052 | 45.1902399 | 28.241208 | 33.0328028 |
| ENSMUSG00 | Icos | 3425 | 0 | 0 | 0 | 0 | 0 | 0 |
| ENSMUSG00 | Ctla4 | 2323 | 0 | 0 | 0 | 0 | 0 | 0 |
| ENSMUSG00 | Cd28 | 6115 | 0 | 0 | 0 | 0 | 0 | 0 |
| ENSMUSG00 | Raph1 | 3026 | 3.8100097 | 3.18482764 | 6.48586368 | 3.88722621 | 2.68116119 | 3.60537201 |
| ENSMUSG00 | Carf | 3339 | 1.17873392 | 3.46353582 | 1.07228191 | 1.30755537 | 2.96469509 | 1.90326174 |
| ENSMUSG00 | Ica1l | 3637 | 0.04372338 | 0.23848115 | 0.0607669 | 0 | 0.05611919 | 0 |
| ENSMUSG00 | Wdr12 | 4858 | 5.16379337 | 4.9098989 | 5.87780786 | 3.95134782 | 4.84214888 | 6.93375452 |
| ENSMUSG00 | Nop58 | 2572 | 19.1821739 | 32.1118048 | 16.4811693 | 26.9072167 | 16.5260385 | 19.3531169 |
| ENSMUSG00 | Sumo1 | 1731 | 27.4682581 | 40.1692871 | 33.3237465 | 32.1424106 | 25.6163608 | 36.2558896 |
| ENSMUSG00 | Cdk15 | 2682 | 0 | 0 | 0 | 0 | 0 | 0 |
| ENSMUSG00 | Als2 | 6990 | 2.08730491 | 2.26800279 | 1.08765622 | 1.57439536 | 4.5040449 | 1.84172099 |
| ENSMUSG00 | Stradb | 5966 | 10.1554395 | 8.15761089 | 9.12783617 | 9.70090503 | 6.81663116 | 7.15010657 |
| ENSMUSG00 | Trak2 | 7749 | 11.9948785 | 7.59889435 | 10.2675594 | 12.3812307 | 11.9120808 | 8.78884685 |
| ENSMUSG00 | Casp8 | 2767 | 16.1349493 | 18.7904485 | 21.9811117 | 19.0907529 | 18.6992212 | 18.3638066 |
| ENSMUSG00 | Cflar | 9545 | 11.8370954 | 8.39539584 | 10.2018502 | 11.616566 | 11.5524356 | 10.8984258 |
| ENSMUSG00 | Ndufb3 | 760 | 65.4396133 | 71.1384054 | 89.4505718 | 67.9861681 | 45.4537585 | 152.163777 |
| ENSMUSG00 | Clk1 | 6474 | 73.9289821 | 60.7503461 | 88.0691339 | 88.5275248 | 70.9277658 | 126.476976 |
| ENSMUSG00 | Ppil3 | 2458 | 13.4728703 | 15.6439304 | 11.6708691 | 10.2756026 | 5.66729072 | 18.5307761 |
| ENSMUSG00 | Nif3l1 | 4179 | 9.0279854 | 7.01061422 | 10.3761685 | 5.85394262 | 8.05872411 | 6.65712659 |
| ENSMUSG00 | Orc2 | 3925 | 25.7777574 | 20.8460073 | 22.0952907 | 17.0987523 | 13.4423624 | 19.8809536 |
| ENSMUSG00 | Sgol2 | 4930 | 0.04838395 | 0.07819301 | 0.0986248 | 0.64406084 | 0.14490249 | 0.55323812 |
| ENSMUSG00 | Col5a2 | 8545 | 0.03721988 | 0 | 0 | 0 | 0.48966212 | 0.07660519 |
| ENSMUSG00 | Col3a1 | 7122 | 0 | 0 | 0 | 0 | 0 | 0 |
| ENSMUSG00 | Kdelc1 | 3550 | 13.5056647 | 15.1753359 | 11.5671865 | 14.0974073 | 13.2093632 | 15.189285 |
| ENSMUSG00 | Ercc5 | 4771 | 10.1492717 | 11.3421452 | 9.26469147 | 9.718178 | 14.2779735 | 10.0271887 |
| ENSMUSG00 | 1700029F09I | 2364 | 10.4601983 | 12.1892946 | 11.0504608 | 9.9057728 | 4.66230845 | 14.6641567 |
| ENSMUSG00 | 1500015O10I | 898 | 0.75260932 | 0.96587523 | 0.24611271 | 0.44198496 | 0.28411122 | 0.69857094 |
| ENSMUSG00 | Khdrbs2 | 1350 | 0 | 0 | 0 | 0 | 0 | 0 |
| ENSMUSG00 | Slc9a2 | 3481 | 17.6678335 | 13.3582065 | 13.2821396 | 14.1073435 | 16.9159545 | 11.3454977 |
| ENSMUSG00 | 4930521A18I | 1448 | 0 | 0 | 0 | 0 | 0 | 0 |
| ENSMUSG00 | Ptp4a1 | 7131 | 3.95269046 | 5.58154554 | 4.2707993 | 5.65189226 | 3.95703069 | 8.81998571 |
| ENSMUSG00 | Slc9a4 | 3810 | 15.5474189 | 11.2940926 | 8.16747965 | 12.5766286 | 16.151656 | 10.2727316 |
| ENSMUSG00 | Il18rap | 6810 | 0 | 0 | 0 | 0.50334717 | 0 | 0 |
| ENSMUSG00 | Il1rl1 | 6881 | 0 | 0 | 0.08350879 | 0 | 0 | 0 |
| ENSMUSG00 | Il18r1 | 4906 | 0.28362043 | 0.3830557 | 1.05414098 | 0.29418707 | 1.92414992 | 1.50660971 |
| ENSMUSG00 | Il1r1 | 5044 | 21.8166475 | 14.4349156 | 19.5245254 | 17.0323846 | 18.0170451 | 20.4559794 |
| ENSMUSG00 | Il1r2 | 1355 | 0 | 0 | 0 | 0 | 0 | 0 |
| ENSMUSG00 | Map4k4 | 6152 | 1.28597865 | 1.31588464 | 1.10648307 | 0.82697794 | 1.28561253 | 1.07289706 |
| ENSMUSG00 | Npas2 | 4766 | 1.72668579 | 2.32540531 | 1.0109108 | 1.4308658 | 1.65948138 | 1.16744155 |
| ENSMUSG00 | Pdcl3 | 1700 | 27.9925354 | 34.184029 | 39.7816587 | 36.7824607 | 23.0519157 | 45.2438133 |
| ENSMUSG00 | Chst10 | 4695 | 0 | 0 | 0 | 0.26129702 | 0.23910123 | 0 |
| ENSMUSG00 | Rev1 | 4238 | 5.22505961 | 5.42353301 | 6.65426518 | 10.6764709 | 9.3191163 | 9.95607997 |
| ENSMUSG00 | Eif5b | 7773 | 11.1548569 | 11.0903827 | 8.84832983 | 11.9670025 | 11.100682 | 15.007543 |
| ENSMUSG00 | Lyg1 | 1146 | 0 | 0 | 0 | 0 | 0 | 0 |
| ENSMUSG00 | Mrpl30 | 2013 | 19.3346326 | 18.9586002 | 23.9344306 | 19.394323 | 11.4321388 | 28.9411969 |
| ENSMUSG00 | Mitd1 | 2920 | 11.6815761 | 11.3700201 | 10.5206441 | 15.1124454 | 11.8129555 | 27.0971673 |
| ENSMUSG00 | 2010300C02I | 5783 | 0.02749817 | 0.74158626 | 0 | 0.30572716 | 0.55588132 | 0.22166835 |
| ENSMUSG00 | Stk17b | 3285 | 19.4844212 | 24.4379338 | 22.3902182 | 24.8016002 | 18.2359103 | 27.091978 |
| ENSMUSG00 | Asnsd1 | 4607 | 15.9988411 | 18.38762 | 19.0066966 | 17.7551541 | 15.1074401 | 21.2359737 |
| ENSMUSG00 | Osgepl1 | 4704 | 11.7981826 | 7.04767691 | 11.7270196 | 10.7923973 | 8.34168431 | 12.5124727 |
| ENSMUSG00 | Ormdl1 | 4707 | 12.8717553 | 13.0012283 | 14.5743064 | 14.3883573 | 8.82419161 | 16.3172688 |
| ENSMUSG00 | Pms1 | 5772 | 2.01119202 | 2.40431312 | 3.23932425 | 2.53799551 | 3.55381204 | 2.37211866 |
| ENSMUSG00 | Mstn | 2679 | 0 | 0 | 0 | 0 | 0 | 0 |
| ENSMUSG00 | Inpp1 | 2518 | 7.51533309 | 8.3245045 | 13.8503789 | 10.2600254 | 11.0037022 | 10.7993707 |
| ENSMUSG00 | Gls | 13846 | 47.6830323 | 49.4462639 | 34.8353684 | 64.166415 | 56.1783961 | 40.2165994 |
| ENSMUSG00 | Stat1 | 5219 | 5.3017464 | 5.70592475 | 6.82634326 | 6.75458086 | 20.4340152 | 25.5239199 |
| ENSMUSG00 | Obfc2a | 2822 | 21.5119131 | 19.1413897 | 12.7029393 | 26.5053431 | 16.526615 | 27.7772194 |
| ENSMUSG00 | Tmeff2 | 5636 | 0.02116154 | 0 | 0 | 0 | 0 | 0 |
| ENSMUSG00 | Mgat4a | 8324 | 1.64772234 | 1.37195901 | 0.95583034 | 0.39012302 | 1.66123833 | 1.34014025 |
| ENSMUSG00 | Unc50 | 1792 | 64.8910605 | 67.0630506 | 73.7272928 | 55.633197 | 64.1246643 | 73.1179503 |
| ENSMUSG00 | 6330578E17I | 3860 | 121.954571 | 130.51535 | 141.308483 | 86.5221254 | 118.193478 | 111.331403 |
| ENSMUSG00 | Inpp4a | 8036 | 1.57320095 | 0.47370942 | 0.89657795 | 0.97883097 | 1.1746988 | 0.77384492 |
| ENSMUSG00 | Cnga3 | 3709 | 0 | 0 | 0 | 0 | 0 | 0 |
| ENSMUSG00 | Vwa3b | 1688 | 0 | 0 | 0 | 0 | 0 | 0 |
| ENSMUSG00 | Tmem131 | 6505 | 8.20166869 | 5.28902686 | 6.18350148 | 8.880454 | 9.18553207 | 3.75681426 |
| ENSMUSG00 | Zap70 | 2315 | 0 | 0 | 0 | 0 | 0 | 0 |
| ENSMUSG00 | Sema4c | 4574 | 0.22598218 | 0.47406863 | 0.30923895 | 1.06494884 | 0.68050042 | 1.26415031 |
| ENSMUSG00 | Plekhb2 | 6040 | 104.963683 | 92.5993434 | 124.335979 | 83.3112902 | 93.0133109 | 69.1303729 |
| ENSMUSG00 | Cfc1 | 1032 | 0 | 0 | 0 | 0 | 0 | 0 |
| ENSMUSG00 | Prss39 | 1241 | 0 | 0 | 0 | 0 | 0 | 0 |
| ENSMUSG00 | Ptpn18 | 1564 | 40.6451492 | 38.6662308 | 38.2688129 | 32.5753497 | 35.2029792 | 40.3015929 |
| ENSMUSG00 | Imp4 | 2518 | 26.5404935 | 27.8440323 | 25.2080407 | 21.9243559 | 21.8047579 | 18.305854 |
| ENSMUSG00 | Dst | 29746 | 7.56725377 | 6.4748608 | 7.563618 | 7.91243113 | 8.43635165 | 8.08079727 |
| ENSMUSG00 | Prim2 | 2154 | 0.71980675 | 1.20801666 | 1.78531121 | 3.01521263 | 4.5009338 | 0.36720731 |
| ENSMUSG00 | Zfp142 | 11572 | 2.87550442 | 2.10284767 | 2.20780032 | 2.33229946 | 2.62804355 | 1.15254914 |
| ENSMUSG00 | Col19a1 | 7657 | 0 | 0 | 0 | 0 | 0 | 0 |
| ENSMUSG00 | Rhbdd1 | 3876 | 17.2827618 | 11.1390743 | 13.5479333 | 14.0660395 | 21.8402367 | 9.08450962 |
| ENSMUSG00 | Col9a1 | 5229 | 0 | 0 | 0 | 0 | 0 | 0 |
| ENSMUSG00 | Tm4sf20 | 1813 | 0 | 0 | 0 | 0 | 0 | 0.03008785 |
| ENSMUSG00 | Mff | 3024 | 82.8765043 | 73.9368689 | 93.6950442 | 77.4618496 | 75.7464951 | 100.061127 |
| ENSMUSG00 | Fam135a | 6446 | 19.3165009 | 17.1784741 | 24.6929626 | 21.6178802 | 19.0300043 | 24.8374391 |
| ENSMUSG00 | 1110058L19I | 871 | 18.2117528 | 24.8954064 | 28.1653535 | 24.109931 | 16.5791784 | 44.8105726 |
| ENSMUSG00 | Smap1 | 3560 | 7.9510961 | 9.01465463 | 8.41821617 | 6.16232671 | 6.46429649 | 11.6070445 |
| ENSMUSG00 | B3gat2 | 6532 | 0.35300335 | 0.79671399 | 0.37218331 | 0.88382228 | 0.75774012 | 0.50524056 |
| ENSMUSG00 | Ogfrl1 | 4837 | 4.11773739 | 4.9511395 | 3.30805606 | 4.47575495 | 4.71548274 | 3.7272145 |
| ENSMUSG00 | Arfgef1 | 3130 | 26.3427689 | 24.2625664 | 25.3631023 | 21.1650587 | 23.7525334 | 23.1006609 |
| ENSMUSG00 | Nhej1 | 1122 | 3.25980767 | 1.46019522 | 3.03345982 | 3.56961403 | 3.91111259 | 5.17780585 |
| ENSMUSG00 | Sphkap | 11441 | 0 | 0 | 0 | 0 | 0 | 0 |
| ENSMUSG00 | Ccl20 | 1546 | 0 | 0.09350539 | 0 | 0 | 3.66360137 | 0 |
| ENSMUSG00 | Wnt10a | 2464 | 0.08067265 | 0.05866856 | 0 | 0.62967854 | 0 | 0 |
| ENSMUSG00 | Cyp27a1 | 1881 | 4.29046387 | 2.07501426 | 3.75985906 | 1.592137 | 1.84465367 | 0.37700193 |
| ENSMUSG00 | Rnf25 | 2767 | 8.82177992 | 9.89154288 | 11.7094149 | 8.67268761 | 8.70417397 | 8.43769107 |
| ENSMUSG00 | Bcs1l | 2277 | 14.6311342 | 11.9778332 | 11.0261954 | 9.82471129 | 8.11223015 | 13.8229836 |
| ENSMUSG00 | Plcd4 | 5268 | 4.98830156 | 5.49735226 | 5.067941 | 5.41093682 | 5.1723775 | 3.06503159 |
| ENSMUSG00 | Rqcd1 | 3973 | 7.64489985 | 8.453549 | 9.2787156 | 8.84568622 | 8.12980012 | 10.5789628 |
| ENSMUSG00 | Vil1 | 3531 | 0.68679816 | 0.27293369 | 0 | 0.20437295 | 0.08670582 | 0.06951905 |
| ENSMUSG00 | Ctdsp1 | 3496 | 1.91044643 | 2.8531446 | 2.59192729 | 1.97130163 | 1.79526437 | 2.7539896 |
| ENSMUSG00 | Slc11a1 | 3780 | 0 | 1.10905303 | 0.01169361 | 1.15500725 | 0 | 0 |
| ENSMUSG00 | Pnkd | 4442 | 31.4231184 | 28.7252811 | 29.1262482 | 27.1955617 | 26.8916582 | 33.1078016 |
| ENSMUSG00 | Cxcr2 | 2535 | 0 | 0 | 0 | 0.01423355 | 0 | 0 |
| ENSMUSG00 | Ppm1f | 4967 | 4.82636496 | 5.99541398 | 3.6664303 | 4.7944734 | 6.44121958 | 1.96034754 |
| ENSMUSG00 | Tnp1 | 416 | 0 | 0 | 0 | 0 | 0 | 0 |
| ENSMUSG00 | Igfbp5 | 9460 | 34.136762 | 19.3306076 | 45.3466053 | 22.6103975 | 25.6102783 | 40.03548 |
| ENSMUSG00 | Xrcc5 | 2534 | 28.914503 | 28.9993387 | 29.1480977 | 21.4726611 | 36.4474113 | 23.259864 |
| ENSMUSG00 | Tmem169 | 3121 | 0 | 0 | 0 | 0 | 0.01634937 | 0 |
| ENSMUSG00 | Pecr | 1767 | 6.05219256 | 6.65392863 | 7.30443586 | 4.67616763 | 5.89099076 | 13.5832952 |
| ENSMUSG00 | Atic | 5591 | 0.44796911 | 1.11179593 | 0.68781262 | 0.61309144 | 0.50195863 | 0.98541891 |
| ENSMUSG00 | Fn1 | 9058 | 0.03072294 | 0 | 0 | 0 | 0 | 0 |
| ENSMUSG00 | Bard1 | 5648 | 0.00703886 | 0.33273216 | 0 | 0 | 0.06324091 | 0 |
| ENSMUSG00 | Zfand2b | 1660 | 18.8000315 | 20.8711156 | 20.3701265 | 13.2808007 | 16.4759864 | 20.2752439 |
| ENSMUSG00 | Abcb6 | 3731 | 6.56375663 | 6.78045621 | 7.46372584 | 8.40399268 | 6.8518398 | 6.68159214 |
| ENSMUSG00 | Ankzf1 | 4052 | 13.6181162 | 10.8930851 | 8.70510136 | 12.2707447 | 15.7537009 | 13.9940708 |
| ENSMUSG00 | Glb1l | 4953 | 12.6899686 | 17.3074259 | 7.8979671 | 14.3293724 | 14.2684293 | 9.28428059 |
| ENSMUSG00 | Stk16 | 3261 | 24.6262099 | 19.2686662 | 25.7945744 | 21.7864292 | 23.2834245 | 22.4235841 |
| ENSMUSG00 | Tuba4a | 2049 | 154.617581 | 130.143373 | 163.863933 | 134.642905 | 95.1795063 | 119.215066 |
| ENSMUSG00 | Dnajb2 | 1400 | 15.4478439 | 16.9340925 | 10.8610243 | 16.7781507 | 16.0733084 | 5.72767424 |
| ENSMUSG00 | Ptprn | 4077 | 0.0487558 | 0.04727637 | 0.10841757 | 0.07965131 | 0.20025068 | 0.14717735 |
| ENSMUSG00 | Slc23a3 | 2519 | 0.2998627 | 0.49735907 | 0 | 0 | 0.0202566 | 0.04331027 |
| ENSMUSG00 | Speg | 12490 | 0 | 0.003858 | 0 | 0.05777749 | 0.02042689 | 0 |
| ENSMUSG00 | Des | 3651 | 0 | 0 | 0 | 0 | 0 | 0 |
| ENSMUSG00 | Dnpep | 3342 | 17.2131 | 16.9993463 | 19.5085932 | 15.9896792 | 14.6116821 | 14.5758545 |
| ENSMUSG00 | Obsl1 | 8213 | 0.24686832 | 0.33442435 | 0.11302066 | 0.09665226 | 0.23608941 | 0.31548651 |
| ENSMUSG00 | Stk11ip | 5809 | 7.44602561 | 6.79371598 | 4.85466962 | 7.13069153 | 8.90699692 | 3.5308192 |
| ENSMUSG00 | Trip12 | 9591 | 13.5585629 | 13.1230306 | 14.5311658 | 17.2415817 | 17.7323439 | 17.4038987 |
| ENSMUSG00 | Slc16a14 | 3193 | 0 | 0.1810953 | 0 | 0.51981649 | 0.1917684 | 0.01708402 |
| ENSMUSG00 | Sp100 | 5591 | 1.31546484 | 0.76705301 | 2.24527337 | 1.27135804 | 1.91656929 | 5.21491493 |
| ENSMUSG00 | Itm2c | 2485 | 48.7784548 | 46.2473499 | 61.6335558 | 37.8679968 | 36.9402214 | 42.1906291 |
| ENSMUSG00 | 4933407L21I | 1578 | 0 | 0 | 0 | 0 | 0 | 0 |
| ENSMUSG00 | Spata3 | 2032 | 0 | 0 | 0 | 0 | 0 | 0 |
| ENSMUSG00 | 2810459M11 | 3632 | 0 | 0 | 0 | 0 | 0 | 0 |
| ENSMUSG00 | Htr2b | 2258 | 0.08803251 | 0 | 0 | 0.52732837 | 0.18078433 | 0 |
| ENSMUSG00 | Psmd1 | 5173 | 54.4188203 | 54.5951551 | 52.0288077 | 57.321137 | 53.0189001 | 44.9902371 |
| ENSMUSG00 | Ncl | 8243 | 52.3337036 | 51.4892858 | 47.1135985 | 52.724521 | 53.6386688 | 53.6095724 |
| ENSMUSG00 | Epha4 | 6310 | 0 | 0.35127993 | 0.06304542 | 0.13151934 | 0.12938542 | 0.03025713 |
| ENSMUSG00 | Nmur1 | 1320 | 0 | 0 | 0 | 0 | 0 | 0 |
| ENSMUSG00 | Ptma | 1187 | 15.6409515 | 16.8875821 | 19.1032397 | 15.8371906 | 14.7447741 | 22.9548143 |
| ENSMUSG00 | Pde6d | 2298 | 15.293231 | 12.8329428 | 20.9083566 | 10.5514073 | 10.6804555 | 15.5600749 |
| ENSMUSG00 | Cops7b | 3267 | 5.99922005 | 6.32751261 | 8.84848648 | 5.58846481 | 5.65397859 | 9.75108009 |
| ENSMUSG00 | Nppc | 1040 | 0.26758497 | 0.50966429 | 0 | 0 | 0.34344676 | 0 |
| ENSMUSG00 | Farsb | 2426 | 31.8732109 | 27.0924596 | 36.7680623 | 34.6096115 | 26.0390162 | 31.130864 |
| ENSMUSG00 | Alppl2 | 2102 | 0 | 0 | 0 | 0 | 0 | 0 |
| ENSMUSG00 | Ecel1 | 3908 | 0 | 0 | 0 | 0 | 0 | 0 |
| ENSMUSG00 | Mrpl44 | 1420 | 41.3233023 | 40.41553 | 47.376905 | 37.0964401 | 29.7174738 | 37.4354027 |
| ENSMUSG00 | Serpine2 | 3821 | 0.41617881 | 0.59271468 | 0.10411321 | 0 | 0.32050066 | 0.5995995 |
| ENSMUSG00 | Chrnd | 3027 | 0.01313362 | 0.04775663 | 0.01460253 | 0.01192007 | 0 | 0.02703136 |
| ENSMUSG00 | Chrng | 1974 | 0 | 0.07323168 | 0 | 0 | 0 | 0 |
| ENSMUSG00 | Eif4e2 | 7300 | 9.84084292 | 10.9904693 | 11.3108278 | 11.8279908 | 10.4079827 | 7.98063417 |
| ENSMUSG00 | Efhd1 | 2117 | 201.51916 | 123.186124 | 216.394852 | 131.732698 | 146.161522 | 179.842555 |
| ENSMUSG00 | 3110079O15I | 459 | 0 | 0 | 0 | 0 | 0.11116858 | 0 |
| ENSMUSG00 | Ngef | 3889 | 21.651352 | 16.5164637 | 26.5051936 | 15.4849402 | 21.0587126 | 20.212268 |
| ENSMUSG00 | Ndufa10 | 2327 | 162.951342 | 155.88635 | 182.16402 | 147.910023 | 128.739086 | 164.32765 |
| ENSMUSG00 | Rnpepl1 | 2701 | 33.1614578 | 42.3526892 | 25.5130223 | 34.8129611 | 39.5402461 | 22.740647 |
| ENSMUSG00 | Capn10 | 4582 | 9.01482851 | 13.5767562 | 8.15158388 | 10.7883896 | 10.7130889 | 7.50618073 |
| ENSMUSG00 | Gpr35 | 4494 | 0 | 0.39672861 | 0 | 0.23283918 | 0.21573234 | 0 |
| ENSMUSG00 | Agxt | 1523 | 0 | 0 | 0 | 0 | 0 | 0 |
| ENSMUSG00 | Mterfd2 | 5511 | 5.25167661 | 3.50621726 | 4.74822919 | 3.18197671 | 3.53693986 | 6.60708463 |
| ENSMUSG00 | Pask | 10512 | 0.35549992 | 0.42630699 | 0 | 0.32951639 | 0.16503965 | 0.04151391 |
| ENSMUSG00 | Ppp1r7 | 10489 | 5.66635936 | 6.13758096 | 5.4909907 | 5.80670142 | 5.45338615 | 7.36526285 |
| ENSMUSG00 | 2-Sep | 5052 | 8.26271863 | 10.7208157 | 13.902757 | 11.0774448 | 9.1609111 | 12.6385452 |
| ENSMUSG00 | Stk25 | 4008 | 68.7488112 | 69.4543603 | 64.1742826 | 69.5443598 | 79.5059174 | 55.5360107 |
| ENSMUSG00 | Bok | 1427 | 4.26250071 | 2.5663417 | 1.6726696 | 0.58156063 | 4.04063102 | 2.63763155 |
| ENSMUSG00 | Thap4 | 2165 | 19.2993118 | 21.0328812 | 25.8065264 | 19.5826339 | 16.6159792 | 22.3992188 |
| ENSMUSG00 | Atg4b | 6024 | 19.772148 | 16.534093 | 24.2875333 | 14.8724629 | 14.7895173 | 20.2839283 |
| ENSMUSG00 | Dtymk | 2845 | 10.494329 | 12.736803 | 9.83471586 | 9.80366271 | 12.8776583 | 13.1435959 |
| ENSMUSG00 | Ing5 | 4531 | 9.8796449 | 7.79533483 | 9.40423233 | 9.26936655 | 13.6152921 | 4.7313764 |
| ENSMUSG00 | Pdcd1 | 1923 | 0 | 0 | 0 | 0 | 0 | 0 |
| ENSMUSG00 | Inpp5d | 7848 | 0.05572251 | 0 | 0.10701262 | 0.02758566 | 0.09102565 | 0 |
| ENSMUSG00 | Atg16l1 | 5129 | 9.92918123 | 10.3719696 | 8.13541427 | 8.69514662 | 11.1225362 | 6.61525662 |
| ENSMUSG00 | Spp2 | 958 | 0.16599366 | 0.704186 | 0.92279422 | 0 | 0.58589784 | 0.99646386 |
| ENSMUSG00 | Iqca | 3532 | 0.30390656 | 0 | 0.26280824 | 0 | 0 | 0.45560694 |
| ENSMUSG00 | Mlph | 5341 | 4.5107323 | 2.32767311 | 3.39313904 | 3.37108036 | 3.99345162 | 3.5338046 |
| ENSMUSG00 | Rab17 | 2832 | 33.2699466 | 26.3902443 | 39.7379563 | 29.0108811 | 28.4681052 | 36.9151456 |
| ENSMUSG00 | Lrrfip1 | 5821 | 44.6113724 | 41.6799066 | 44.6422668 | 46.2043566 | 40.9280448 | 43.2992512 |
| ENSMUSG00 | Scly | 3980 | 8.70025717 | 8.09968085 | 6.63027645 | 10.8336792 | 12.3591523 | 10.0943577 |
| ENSMUSG00 | Klhl30 | 3081 | 0 | 0.1094791 | 0 | 0 | 0 | 0 |
| ENSMUSG00 | Ilkap | 1487 | 34.3282024 | 43.4228867 | 37.6622294 | 34.55335 | 31.9129318 | 43.0671506 |
| ENSMUSG00 | Asb1 | 6221 | 3.13135918 | 5.19741242 | 6.51552645 | 3.39302299 | 6.14350675 | 4.53335107 |
| ENSMUSG00 | Cdh7 | 8320 | 0 | 0 | 0 | 0 | 0 | 0 |
| ENSMUSG00 | Hdac4 | 4492 | 3.85872429 | 5.18122252 | 5.55967082 | 5.50227082 | 6.74747712 | 4.26242136 |
| ENSMUSG00 | Serpinb8 | 1656 | 10.5870573 | 6.43067857 | 12.7587445 | 3.24651246 | 4.22138496 | 5.58339535 |
| ENSMUSG00 | Cln8 | 7599 | 9.79369121 | 7.55231644 | 13.1692292 | 9.74818231 | 8.91736111 | 8.71467615 |
| ENSMUSG00 | 2310035C23I | 7374 | 8.94417442 | 10.1025549 | 9.4290072 | 12.903221 | 12.3241084 | 10.3454253 |
| ENSMUSG00 | Tnfrsf11a | 5012 | 0.1745053 | 0 | 0 | 0 | 0.28506355 | 0.68023342 |
| ENSMUSG00 | Htr4 | 4650 | 0 | 0 | 0.38023091 | 0.01551916 | 0.0768139 | 0 |
| ENSMUSG00 | Serpinb11 | 2076 | 0 | 0 | 0 | 0 | 0 | 0 |
| ENSMUSG00 | Slco6c1 | 2780 | 0 | 0 | 0 | 0 | 0 | 0 |
| ENSMUSG00 | Gin1 | 3935 | 8.10264177 | 7.61676417 | 9.9749013 | 5.46503139 | 7.35246635 | 7.15309472 |
| ENSMUSG00 | Pam | 7944 | 28.4403821 | 15.5162285 | 35.4382601 | 24.2590886 | 25.5324577 | 21.8258915 |
| ENSMUSG00 | Slco6d1 | 3524 | 0 | 0 | 0 | 0 | 0 | 0 |
| ENSMUSG00 | Ccdc93 | 11602 | 3.23128927 | 2.3341476 | 1.60775537 | 2.79898639 | 2.53328672 | 1.21069119 |
| ENSMUSG00 | Actr3 | 2496 | 67.6289144 | 88.8244472 | 86.5088154 | 83.0638733 | 63.8647428 | 60.6030246 |
| ENSMUSG00 | Slc35f5 | 2581 | 74.0736559 | 90.7346412 | 80.4401614 | 81.3628435 | 91.7921205 | 53.682746 |
| ENSMUSG00 | Gpr39 | 3012 | 4.75165106 | 5.63135048 | 3.80088889 | 3.96519148 | 5.40418788 | 3.98434305 |
| ENSMUSG00 | Lypd1 | 8319 | 0 | 0.1448064 | 0.00531336 | 0.3946948 | 0 | 0.26228767 |
| ENSMUSG00 | Tmem163 | 2830 | 41.7783742 | 21.1134941 | 69.6764741 | 32.1423443 | 27.0998738 | 45.4609446 |
| ENSMUSG00 | Acmsd | 2293 | 0.01733776 | 0.84058338 | 0 | 0 | 0.71209944 | 1.87936895 |
| ENSMUSG00 | Ccnt2 | 8542 | 20.0545968 | 17.290031 | 23.534299 | 31.5496126 | 23.9361611 | 26.0581048 |
| ENSMUSG00 | Ubxn4 | 4080 | 31.5510407 | 35.4902595 | 39.2074681 | 42.0161259 | 30.3281768 | 32.2749898 |
| ENSMUSG00 | Lct | 6389 | 0 | 0 | 0 | 0 | 0 | 0.11953199 |
| ENSMUSG00 | Mcm6 | 4808 | 4.6883023 | 7.29611686 | 9.15682139 | 7.18188781 | 9.81684643 | 6.89240572 |
| ENSMUSG00 | Dars | 2490 | 145.227249 | 109.609642 | 137.114472 | 132.402265 | 99.2452763 | 120.052227 |
| ENSMUSG00 | Rgs18 | 2026 | 0 | 0 | 0 | 0 | 0 | 0.09423617 |
| ENSMUSG00 | Rgs1 | 4720 | 0 | 0 | 0 | 0 | 0 | 0 |
| ENSMUSG00 | Rgs2 | 3817 | 131.025398 | 108.93393 | 198.196108 | 135.338389 | 92.3340785 | 165.219861 |
| ENSMUSG00 | Cdc73 | 14343 | 7.01257517 | 6.2353787 | 6.35461203 | 8.33185048 | 5.79529849 | 6.79251282 |
| ENSMUSG00 | Cfh | 7473 | 3.61220009 | 1.17999718 | 1.01735809 | 2.23551273 | 3.89201582 | 0.50366656 |
| ENSMUSG00 | F13b | 2518 | 0.06315406 | 1.81799524 | 0 | 1.30399764 | 0 | 0.9640361 |
| ENSMUSG00 | Tsn | 3034 | 36.3093067 | 35.1789616 | 33.7705577 | 31.8363986 | 31.4668257 | 31.8144194 |
| ENSMUSG00 | Mki67ip | 2100 | 50.0919054 | 49.7008732 | 41.234005 | 46.2021989 | 36.7875871 | 37.5740626 |
| ENSMUSG00 | Tfcp2l1 | 9254 | 246.020867 | 210.309295 | 292.112374 | 287.159111 | 281.714779 | 231.159494 |
| ENSMUSG00 | Epb4.1l5 | 7744 | 12.0693614 | 16.8192088 | 14.1783804 | 15.0450571 | 16.5189985 | 14.7432386 |
| ENSMUSG00 | Ptpn4 | 5869 | 4.47748724 | 4.2529523 | 5.00838739 | 4.46952571 | 3.72982028 | 4.96325008 |
| ENSMUSG00 | Dbi | 1660 | 28.5233598 | 28.4474177 | 36.8792485 | 20.671099 | 18.6277011 | 52.7419229 |
| ENSMUSG00 | Sctr | 2201 | 13.9803462 | 7.09332451 | 16.6083254 | 9.09838015 | 7.51137927 | 8.27780964 |
| ENSMUSG00 | 3110009E18I | 2209 | 0.48592031 | 0.32720536 | 0.38018788 | 0.75136897 | 0.48508551 | 1.23470526 |
| ENSMUSG00 | Steap3 | 5542 | 0.05738792 | 0 | 0.07975793 | 0.44272447 | 0.09207213 | 0.16732908 |
| ENSMUSG00 | Marco | 1968 | 0 | 0 | 0 | 0 | 0 | 0 |
| ENSMUSG00 | Nek7 | 4759 | 49.9219902 | 42.5567591 | 62.4900189 | 58.1983135 | 52.7847968 | 55.2026319 |
| ENSMUSG00 | Atp6v1g3 | 1882 | 739.722325 | 545.720419 | 1211.60493 | 651.585741 | 417.808964 | 1489.27936 |
| ENSMUSG00 | Ptprc | 7127 | 2.68866867 | 6.02415041 | 2.63586127 | 6.72329944 | 4.88283826 | 3.78102197 |
| ENSMUSG00 | Nr5a2 | 3826 | 0 | 0 | 0 | 0.00943075 | 0 | 0.02851504 |
| ENSMUSG00 | Cd55 | 5068 | 20.9288915 | 10.8010652 | 17.0423049 | 15.043678 | 6.25244266 | 24.9012936 |
| ENSMUSG00 | Daf2 | 1867 | 0 | 0 | 0 | 0 | 0 | 0 |
| ENSMUSG00 | Ddx59 | 2944 | 3.01136962 | 3.91187489 | 3.78358167 | 4.75537815 | 5.21703095 | 4.78973794 |
| ENSMUSG00 | C4bp | 3051 | 0.1172728 | 0.85285738 | 1.20247557 | 1.19445641 | 0.75260141 | 1.20684245 |
| ENSMUSG00 | Cacna1s | 6481 | 0 | 0 | 0 | 0 | 0 | 0 |
| ENSMUSG00 | Pfkfb2 | 12172 | 8.22741172 | 6.23510075 | 6.40948512 | 7.98595366 | 7.78475024 | 7.81356121 |
| ENSMUSG00 | Tmem9 | 1931 | 58.0582367 | 46.8638728 | 53.197868 | 40.7908354 | 34.7750961 | 55.4532541 |
| ENSMUSG00 | Pkp1 | 5705 | 0 | 0 | 0 | 0 | 0 | 0 |
| ENSMUSG00 | Tnnt2 | 1090 | 0.07294584 | 0.08841549 | 0 | 0.4634391 | 1.2639561 | 0.05004521 |
| ENSMUSG00 | Fcamr | 1904 | 1.12751888 | 0.70862415 | 0.02321525 | 0.34111177 | 0.80398701 | 0.55867171 |
| ENSMUSG00 | Il20 | 896 | 0 | 0 | 0 | 0 | 0 | 0 |
| ENSMUSG00 | Pigr | 4012 | 1.91246454 | 3.12275049 | 2.42382988 | 0.93532718 | 5.48164705 | 2.3521997 |
| ENSMUSG00 | Tnni1 | 1695 | 0 | 0 | 0 | 0 | 0 | 0 |
| ENSMUSG00 | Il24 | 1842 | 0 | 0 | 0 | 0 | 0 | 0 |
| ENSMUSG00 | Csrp1 | 1949 | 63.2334477 | 44.774575 | 68.5593491 | 40.7473474 | 44.2717299 | 46.7545253 |
| ENSMUSG00 | Gpr37l1 | 2267 | 0 | 0 | 0.07799178 | 0 | 0.22508326 | 0 |
| ENSMUSG00 | Sngap2 | 8655 | 1.74547459 | 1.4085696 | 0.43410244 | 0.85879852 | 2.91831959 | 1.13762505 |
| ENSMUSG00 | Arl8a | 2292 | 11.5693305 | 11.1005417 | 14.4639539 | 10.1697211 | 7.70293457 | 16.4457031 |
| ENSMUSG00 | Eif2d | 7169 | 9.58257363 | 8.84549559 | 6.68977538 | 5.6722645 | 8.71911151 | 6.92423538 |
| ENSMUSG00 | Ube2t | 1777 | 0.11186123 | 0.40675106 | 0.47261397 | 0.62945604 | 0.48815329 | 0 |
| ENSMUSG00 | Rassf5 | 3831 | 3.44526743 | 7.62228764 | 2.18067041 | 5.51920598 | 4.63512889 | 1.5022054 |
| ENSMUSG00 | Avpr1b | 4021 | 0 | 0 | 0 | 0 | 0 | 0 |
| ENSMUSG00 | Rab7l1 | 1779 | 14.2127519 | 17.0372525 | 9.61557799 | 8.61993756 | 9.32185056 | 12.7404301 |
| ENSMUSG00 | Nucks1 | 6089 | 14.1158399 | 11.8151352 | 15.5058526 | 14.0500127 | 13.123223 | 19.4313327 |
| ENSMUSG00 | Slc45a3 | 3334 | 7.04723725 | 9.85697467 | 1.08714791 | 2.74890201 | 6.38212319 | 3.44409812 |
| ENSMUSG00 | Elk4 | 5538 | 4.63742153 | 5.01180763 | 3.81518255 | 4.4369578 | 5.23347444 | 6.47144745 |
| ENSMUSG00 | Cdk18 | 3153 | 9.94832671 | 12.3942926 | 6.16835107 | 8.17081503 | 7.54148149 | 8.60712529 |
| ENSMUSG00 | Rbbp5 | 3618 | 16.7900426 | 14.6370647 | 14.9660745 | 13.2041534 | 13.3841986 | 7.65921323 |
| ENSMUSG00 | Nfasc | 10344 | 0 | 0 | 0 | 0 | 0 | 0 |
| ENSMUSG00 | Lrrn2 | 3913 | 0 | 0 | 0 | 0 | 0 | 0 |
| ENSMUSG00 | Pik3c2b | 9203 | 0.57885846 | 0.42934785 | 0.21133121 | 0.51753014 | 0.7152453 | 0.45047758 |
| ENSMUSG00 | Chit1 | 1803 | 0 | 0 | 0 | 0 | 0 | 0 |
| ENSMUSG00 | Syt2 | 6870 | 0.01736047 | 0 | 0.07077442 | 0 | 0.01485484 | 0 |
| ENSMUSG00 | Klhl12 | 6653 | 15.4886826 | 18.0201215 | 17.127965 | 23.5647803 | 15.5464398 | 16.9600455 |
| ENSMUSG00 | Cyb5r1 | 3957 | 18.3757326 | 24.1480199 | 10.5449936 | 23.571414 | 22.8632207 | 15.4190973 |
| ENSMUSG00 | Adipor1 | 4990 | 54.398882 | 57.7561344 | 52.767611 | 47.9262106 | 48.326784 | 43.2841419 |
| ENSMUSG00 | Ppfia4 | 6043 | 0.09868148 | 0 | 0 | 0.08359236 | 0 | 0.14894309 |
| ENSMUSG00 | Myog | 1476 | 0 | 0 | 0 | 0 | 0 | 0 |
| ENSMUSG00 | Atp2b4 | 11734 | 1.94135762 | 2.16005357 | 0.60271816 | 2.77672458 | 1.84380291 | 1.63638538 |
| ENSMUSG00 | Zc3h11a | 4378 | 29.9573619 | 31.2474439 | 26.634185 | 39.2138803 | 33.2056064 | 25.031864 |
| ENSMUSG00 | Tor1aip1 | 6829 | 8.14437212 | 8.60850189 | 7.88370842 | 8.71802218 | 7.69617325 | 11.2229809 |
| ENSMUSG00 | Lhx4 | 5596 | 0 | 0 | 0 | 0 | 0 | 0.00487395 |
| ENSMUSG00 | Xpr1 | 14767 | 4.24018973 | 4.67930914 | 2.46048046 | 4.81843244 | 4.07741068 | 5.14019916 |
| ENSMUSG00 | Stx6 | 2527 | 25.1559214 | 26.009619 | 26.3426892 | 23.945227 | 30.5915945 | 20.9389791 |
| ENSMUSG00 | Mr1 | 2503 | 28.7325866 | 30.1670615 | 25.7122986 | 28.6868832 | 35.7368106 | 23.1992437 |
| ENSMUSG00 | Glul | 3319 | 1.47331247 | 1.4373178 | 1.8378591 | 1.58721859 | 1.5374021 | 1.78324698 |
| ENSMUSG00 | Rgs16 | 2334 | 0 | 0.0619363 | 0 | 0 | 0.0655866 | 0 |
| ENSMUSG00 | Lamc1 | 9472 | 5.28841907 | 4.46660644 | 3.99459223 | 4.5140649 | 5.37091392 | 2.98316367 |
| ENSMUSG00 | Lamc2 | 5140 | 38.6185047 | 39.2522633 | 34.0715374 | 28.2338489 | 55.6525025 | 24.8655563 |
| ENSMUSG00 | Ncf2 | 2581 | 0.06161252 | 0.20536647 | 0 | 0 | 0.05931001 | 0 |
| ENSMUSG00 | Rgl1 | 8004 | 0.99835727 | 1.04151106 | 0.84493778 | 1.42002049 | 1.12839437 | 1.48572498 |
| ENSMUSG00 | Fam129a | 7363 | 13.941145 | 15.2419155 | 16.959148 | 16.0930917 | 15.3155359 | 10.9906091 |

| | | | | | | | | |
|---|---|---|---|---|---|---|---|---|
| ENSMUSG00 | Rnf2 | 3021 | 30.7805591 | 50.5471152 | 39.0076509 | 47.4883181 | 33.5784294 | 39.5622209 |
| ENSMUSG00 | Adck3 | 5706 | 28.5629337 | 28.8122507 | 34.0728993 | 18.5720124 | 33.3709637 | 23.7673538 |
| ENSMUSG00 | Cdc42bpa | 19461 | 2.45547955 | 2.41167436 | 1.78978739 | 2.48630706 | 3.03887619 | 1.73506004 |
| ENSMUSG00 | Ahctf1 | 10810 | 4.63752645 | 3.89592511 | 3.89270625 | 7.59358476 | 7.04739862 | 3.70894724 |
| ENSMUSG00 | Tfb2m | 3215 | 2.2876404 | 3.55215763 | 3.45090594 | 4.01784508 | 3.14252641 | 3.13043293 |
| ENSMUSG00 | Kif26b | 7804 | 1.14111067 | 1.77210519 | 0.70799979 | 1.07728298 | 1.81770022 | 0.46133424 |
| ENSMUSG00 | Efcab2 | 2529 | 31.8169603 | 20.4444652 | 38.8011433 | 31.9445226 | 29.5585767 | 49.9441892 |
| ENSMUSG00 | Parp1 | 3823 | 22.1603267 | 13.1715491 | 13.4004541 | 14.3271105 | 18.0454251 | 10.3234117 |
| ENSMUSG00 | Mixl1 | 2292 | 0 | 0 | 0 | 0 | 0 | 0 |
| ENSMUSG00 | Acbd3 | 3462 | 44.5096021 | 37.8031129 | 51.198553 | 43.8154001 | 41.8439864 | 46.8049196 |
| ENSMUSG00 | Fam36a | 550 | 2.74674229 | 3.94252708 | 5.30422116 | 4.98588252 | 0.92775229 | 7.33935747 |
| ENSMUSG00 | Pppde1 | 4869 | 11.0554366 | 12.6379311 | 13.9259863 | 11.5456614 | 10.699924 | 12.5982057 |
| ENSMUSG00 | Sdccag8 | 4282 | 6.58258657 | 8.58623432 | 6.88524738 | 7.27202345 | 8.05554182 | 5.08931265 |
| ENSMUSG00 | Capn2 | 4178 | 37.3195296 | 30.944046 | 26.1317741 | 29.0262689 | 31.265563 | 21.1251155 |
| ENSMUSG00 | Trp53bp2 | 4343 | 17.4565281 | 17.2308415 | 14.0350775 | 21.9499797 | 21.9590828 | 12.2023081 |
| ENSMUSG00 | Srp9 | 1602 | 18.9843587 | 22.4088011 | 22.3768381 | 20.901459 | 17.8369354 | 31.8204125 |
| ENSMUSG00 | Cnih3 | 3597 | 0 | 0 | 0 | 0 | 0 | 0 |
| ENSMUSG00 | Nvl | 3037 | 12.0038774 | 10.3131997 | 10.5083078 | 13.7342257 | 12.382759 | 10.5703821 |
| ENSMUSG00 | Tmem63a | 4524 | 24.2891574 | 15.2100441 | 25.139554 | 11.0064592 | 22.7498232 | 13.5529154 |
| ENSMUSG00 | Pycr2 | 1591 | 29.3355966 | 23.4723398 | 88.7648576 | 28.8021349 | 26.4913805 | 19.1831058 |
| ENSMUSG00 | Wdr64 | 3967 | 0 | 0 | 0 | 0 | 0 | 0 |
| ENSMUSG00 | Opn3 | 2208 | 3.13290471 | 6.98354234 | 5.52523038 | 0.39219614 | 4.99171068 | 3.58226693 |
| ENSMUSG00 | Fh1 | 3571 | 74.0447216 | 59.1704141 | 76.1123306 | 59.4025034 | 61.9718264 | 60.2241754 |
| ENSMUSG00 | Rgs7 | 1613 | 0 | 0 | 0.0274035 | 0 | 0 | 0 |
| ENSMUSG00 | 1810030J14R | 797 | 0 | 0 | 0 | 0 | 0 | 0 |
| ENSMUSG00 | Spna1 | 9640 | 0 | 0 | 0 | 0 | 0 | 0 |
| ENSMUSG00 | Ifi202b | 1789 | 0 | 0 | 0.12353785 | 0 | 0 | 0 |
| ENSMUSG00 | Mnda | 1644 | 0.04836433 | 0 | 0 | 0 | 0 | 0.14931372 |
| ENSMUSG00 | Apcs | 981 | 0 | 0 | 0 | 0 | 0 | 0 |
| ENSMUSG00 | Dusp23 | 1415 | 5.28200024 | 4.97188732 | 4.37332722 | 7.16540955 | 4.39944724 | 4.27913068 |
| ENSMUSG00 | Ccdc19 | 2610 | 0 | 0.03692448 | 0 | 0.11059631 | 0.07820134 | 0.08360043 |
| ENSMUSG00 | Tagln2 | 2077 | 56.0060356 | 58.9280515 | 79.4867037 | 65.5105392 | 43.4350662 | 63.2031 |
| ENSMUSG00 | Slamf9 | 1309 | 0 | 0.07362329 | 0 | 0 | 0 | 0 |
| ENSMUSG00 | Copa | 7807 | 80.5192337 | 70.7026636 | 71.6503552 | 68.2864363 | 75.9153779 | 55.6357922 |
| ENSMUSG00 | Dcaf8 | 3901 | 76.5249688 | 56.697198 | 63.4870357 | 52.4072967 | 90.1888904 | 52.4867257 |
| ENSMUSG00 | Vangl2 | 7411 | 0.63836219 | 0.58518146 | 0.61432868 | 0.14606144 | 0.70917781 | 1.21817644 |
| ENSMUSG00 | Uck2 | 1324 | 10.9597813 | 9.71735654 | 13.5877267 | 7.14010247 | 7.82353043 | 12.2777077 |
| ENSMUSG00 | Fmo9 | 4319 | 0 | 0 | 0 | 0 | 0 | 0 |
| ENSMUSG00 | Tada1 | 4260 | 16.4061349 | 20.8355461 | 18.7598432 | 14.7800863 | 15.1881325 | 15.109894 |
| ENSMUSG00 | Dusp27 | 4014 | 0 | 0 | 0 | 0 | 0 | 0 |
| ENSMUSG00 | Pou2f1 | 17172 | 1.53493382 | 2.13544622 | 1.57532774 | 1.79233543 | 2.00872525 | 1.18965204 |
| ENSMUSG00 | Mpzl1 | 2350 | 74.2835383 | 75.8680153 | 75.839077 | 67.1433109 | 74.6722155 | 61.1184044 |
| ENSMUSG00 | Adcy10 | 6165 | 0.19345733 | 0.0781613 | 0 | 0 | 0.07449106 | 0.0884822 |
| ENSMUSG00 | Brp44 | 4086 | 87.2557879 | 74.4849654 | 67.6333633 | 69.7179984 | 46.7429576 | 147.073452 |
| ENSMUSG00 | Dcaf6 | 3278 | 9.45981537 | 7.87917419 | 7.64565131 | 9.99466029 | 9.93130805 | 12.4225248 |
| ENSMUSG00 | Tbx19 | 3549 | 0 | 0 | 0 | 0 | 0 | 0 |
| ENSMUSG00 | Xcl1 | 520 | 0 | 0 | 0 | 0 | 0 | 0 |
| ENSMUSG00 | Dpt | 1632 | 1.09619891 | 1.15151424 | 0.81253388 | 1.96770954 | 1.65710657 | 0.46794724 |
| ENSMUSG00 | Nme7 | 1202 | 11.6752784 | 23.5720699 | 16.106828 | 11.0467491 | 20.8011016 | 15.8837333 |
| ENSMUSG00 | Atp1b1 | 2416 | 1167.01094 | 1257.9334 | 1112.2179 | 1240.48456 | 1139.96218 | 1393.62569 |
| ENSMUSG00 | Blzf1 | 6350 | 12.0393369 | 13.6136185 | 17.2491602 | 18.8024386 | 19.0766325 | 17.0176568 |
| ENSMUSG00 | 4930455F23f | 1903 | 32.7361208 | 32.4365909 | 27.7103514 | 26.8292659 | 21.6386155 | 27.4752935 |
| ENSMUSG00 | F5 | 7405 | 0 | 0 | 0 | 0 | 0.0551264 | 0.04051601 |
| ENSMUSG00 | Selp | 4396 | 0 | 0 | 0 | 0 | 0 | 0 |
| ENSMUSG00 | Sell | 2365 | 0 | 0 | 0 | 0 | 0 | 0 |
| ENSMUSG00 | Sele | 2898 | 0 | 0 | 0 | 0 | 0 | 0 |
| ENSMUSG00 | Scyl3 | 5445 | 6.39592304 | 7.06203505 | 6.38065172 | 6.46097216 | 9.28689411 | 6.66212492 |
| ENSMUSG00 | Kifap3 | 3991 | 16.9740263 | 17.9416319 | 12.7034613 | 16.8702318 | 15.8922037 | 12.8821586 |
| ENSMUSG00 | Prrx1 | 8165 | 0 | 0 | 0 | 0 | 0 | 0 |
| ENSMUSG00 | Astn1 | 7342 | 0 | 0 | 0.13846941 | 0 | 0 | 0.1745993 |
| ENSMUSG00 | Sec16b | 5431 | 0.08052114 | 0.22181202 | 0 | 0 | 0.01879079 | 0.06528638 |
| ENSMUSG00 | 1700057K13f | 1321 | 0 | 0 | 0 | 0 | 0 | 0 |
| ENSMUSG00 | Ralgps2 | 7007 | 0 | 0.76333596 | 0 | 0 | 0.2111838 | 0.29972131 |
| ENSMUSG00 | Abl2 | 10870 | 0.79730403 | 1.53380947 | 0.91494155 | 1.02237992 | 1.33316382 | 0.85060743 |
| ENSMUSG00 | Soat1 | 4769 | 5.71865373 | 4.73882184 | 16.6278269 | 7.56595608 | 2.84609268 | 7.1832558 |
| ENSMUSG00 | 9430070O13l | 2143 | 0 | 0 | 0.14438306 | 0 | 0 | 0 |
| ENSMUSG00 | Nphs2 | 3108 | 0.11512205 | 0 | 0 | 0 | 0 | 0 |
| ENSMUSG00 | Smyd2 | 3076 | 5.50579802 | 6.64208435 | 5.86292326 | 4.72726396 | 6.55247674 | 4.18518522 |
| ENSMUSG00 | Ptpn14 | 10969 | 3.49862342 | 3.21565098 | 0.97518881 | 3.06906259 | 4.3867146 | 2.32738284 |
| ENSMUSG00 | Cenpf | 13304 | 0 | 0.2535386 | 0 | 0.33630288 | 0 | 0.11890628 |
| ENSMUSG00 | Kctd3 | 4701 | 7.89866386 | 8.2617044 | 9.94800041 | 7.06136587 | 10.550444 | 5.79607841 |
| ENSMUSG00 | Ush2a | 17277 | 0.09204256 | 0 | 0 | 0 | 0.01181371 | 0 |
| ENSMUSG00 | Esrrg | 6196 | 56.8485406 | 48.6375096 | 69.6485787 | 66.6900539 | 55.8523048 | 71.8974633 |
| ENSMUSG00 | Spata17 | 2857 | 1.47500208 | 1.7540742 | 2.05769868 | 3.82669216 | 3.00050092 | 4.75420733 |
| ENSMUSG00 | Slc30a10 | 5832 | 0 | 0 | 0 | 0 | 0 | 0 |
| ENSMUSG00 | Eprs | 7273 | 26.1775053 | 22.6256978 | 23.592954 | 24.0315446 | 24.4643163 | 21.0681869 |
| ENSMUSG00 | Cr2 | 7309 | 6.49990412 | 7.25862263 | 2.39484606 | 6.16588776 | 8.16812966 | 2.94800452 |
| ENSMUSG00 | Bpnt1 | 2286 | 89.5628717 | 90.6815722 | 94.5524988 | 91.0416592 | 80.2225699 | 89.0422715 |
| ENSMUSG00 | Iars2 | 9583 | 14.9015638 | 12.9781078 | 13.5469908 | 12.0185627 | 18.0506537 | 10.3742106 |
| ENSMUSG00 | Mark1 | 4528 | 0.65849405 | 0.23412141 | 0.14642837 | 0.613586 | 0.57472287 | 0.34936596 |
| ENSMUSG00 | Mosc1 | 2714 | 0.27831766 | 0 | 0.06514642 | 0 | 0.03760234 | 0 |
| ENSMUSG00 | Nek2 | 4667 | 0 | 0.42332208 | 0 | 0.44068492 | 0 | 0 |
| ENSMUSG00 | Lpgat1 | 7779 | 7.02198615 | 8.31292009 | 10.9155085 | 8.46970219 | 9.70806482 | 9.23530014 |
| ENSMUSG00 | Ppp2r5a | 3068 | 74.7682929 | 67.0966365 | 101.644069 | 67.2833692 | 70.4855869 | 71.3870121 |
| ENSMUSG00 | Tmem206 | 2362 | 11.1591378 | 7.36465098 | 6.99893704 | 10.9529322 | 11.2767859 | 5.21936365 |
| ENSMUSG00 | Atf3 | 2234 | 1.86854318 | 3.04131081 | 5.02563479 | 13.8901335 | 5.50463589 | 4.4684503 |
| ENSMUSG00 | Batf3 | 709 | 0.95323437 | 0.06796395 | 0.06234393 | 1.17050356 | 0.287878 | 0.61550667 |
| ENSMUSG00 | Tatdn3 | 3228 | 6.05938551 | 4.26930683 | 5.25821181 | 5.48831594 | 6.21231899 | 3.422004 |
| ENSMUSG00 | Angel2 | 5579 | 18.4489943 | 17.8874568 | 25.4324997 | 17.2810939 | 23.212931 | 21.6378479 |
| ENSMUSG00 | Traf5 | 1967 | 6.00273397 | 8.67208974 | 2.31458558 | 4.82439131 | 7.15977617 | 1.8580588 |
| ENSMUSG00 | Irf6 | 4427 | 50.3341921 | 44.1373442 | 78.3991126 | 46.1560014 | 51.5911584 | 54.9435809 |
| ENSMUSG00 | Lamb3 | 4339 | 0.32984496 | 0 | 0.48898098 | 0 | 0.34103823 | 0 |
| ENSMUSG00 | Plxna2 | 15059 | 1.15367189 | 1.34713408 | 0.95982487 | 2.69075875 | 2.20247987 | 1.0957671 |
| ENSMUSG00 | Usf1 | 2911 | 6.91043392 | 5.39635182 | 5.25380889 | 6.5817814 | 5.78450842 | 4.69412376 |
| ENSMUSG00 | Nmt2 | 5046 | 7.16165909 | 5.29038624 | 6.10556571 | 7.20782816 | 4.52017241 | 9.12397762 |
| ENSMUSG00 | Acbd7 | 527 | 0.22631203 | 0 | 0 | 0.34233439 | 0.58094546 | 1.91491775 |
| ENSMUSG00 | Olah | 2093 | 0.18994496 | 0 | 0.08447557 | 0 | 0 | 0 |
| ENSMUSG00 | Suv39h2 | 5552 | 0.22913822 | 0.41659748 | 0.91556411 | 1.28028868 | 0.51467526 | 0.74671203 |
| ENSMUSG00 | Dclre1c | 7507 | 3.20395174 | 1.46350191 | 2.68496609 | 3.74422708 | 2.88879842 | 2.68495516 |
| ENSMUSG00 | 1700009P17f | 1871 | 0.21248253 | 0.23178941 | 0.49611903 | 0.07713959 | 0.19090574 | 0.14577573 |
| ENSMUSG00 | Meig1 | 1373 | 0.17373116 | 0.03509573 | 0.12877449 | 0 | 0.11149245 | 0 |
| ENSMUSG00 | Fam107b | 3751 | 223.694794 | 171.549386 | 256.514455 | 179.015422 | 145.774098 | 207.748259 |
| ENSMUSG00 | Fcgr2b | 4205 | 0.24581272 | 0.19480845 | 0.18921122 | 0.13729197 | 0 | 0.38268816 |
| ENSMUSG00 | Frmd4a | 20281 | 0.36264306 | 0.20670679 | 0.22884441 | 0 | 0.15095817 | 0.00134484 |
| ENSMUSG00 | Dusp12 | 2896 | 3.58293523 | 3.96007363 | 2.67103678 | 3.72532159 | 3.2243877 | 4.32287963 |
| ENSMUSG00 | Sephs1 | 5915 | 14.2353521 | 8.14648219 | 13.3464906 | 11.1509683 | 11.9047166 | 13.0540158 |
| ENSMUSG00 | Atl6 | 7447 | 9.39034379 | 8.24352455 | 14.0197064 | 9.74849921 | 12.7103434 | 8.47868804 |
| ENSMUSG00 | Phyh | 1948 | 335.227681 | 338.393496 | 322.98205 | 294.157571 | 269.407739 | 331.174046 |
| ENSMUSG00 | Uhmk1 | 8092 | 22.1720815 | 18.8767946 | 19.7466217 | 30.1872765 | 23.9052139 | 18.2718511 |
| ENSMUSG00 | Ucma | 1063 | 0 | 0 | 0 | 0 | 0 | 0 |
| ENSMUSG00 | Mcm10 | 7218 | 0.07710955 | 0.3538212 | 0.04286685 | 0.05498788 | 0.23328767 | 0.01511479 |
| ENSMUSG00 | Uap1 | 4337 | 35.0988552 | 34.5982432 | 34.9782627 | 30.7325507 | 36.7432261 | 30.1108999 |
| ENSMUSG00 | Optn | 4250 | 17.7917468 | 18.9004233 | 15.1014297 | 18.380618 | 18.4415325 | 16.0824108 |
| ENSMUSG00 | Sh2d1b1 | 2484 | 0 | 0 | 0.07117849 | 0 | 0 | 0 |
| ENSMUSG00 | Ddr2 | 8591 | 0 | 0 | 0 | 0 | 0.00593952 | 0 |
| ENSMUSG00 | Hsd17b7 | 5352 | 16.7504874 | 14.5855823 | 23.257173 | 14.0835932 | 12.4419694 | 15.742033 |
| ENSMUSG00 | Ccdc3 | 2667 | 0 | 0 | 0.03314724 | 0.09470353 | 0.114795 | 0 |
| ENSMUSG00 | Rgs5 | 4642 | 0 | 0 | 0.54276283 | 0.52856076 | 2.3523577 | 0 |
| ENSMUSG00 | Enkur | 1556 | 0.4598963 | 1.02194897 | 1.16470152 | 0.32464565 | 0.81983252 | 0.85890574 |
| ENSMUSG00 | Nuf2 | 3128 | 0.27961016 | 0.41593157 | 0 | 0.17302771 | 0 | 0 |
| ENSMUSG00 | Lmx1a | 3384 | 0.48167101 | 0.34174782 | 0.18286815 | 1.62070649 | 0.10555101 | 0.68508993 |
| ENSMUSG00 | Aldh9a1 | 2822 | 40.2485499 | 31.7941727 | 33.8014094 | 36.2610483 | 32.4926993 | 26.414454 |
| ENSMUSG00 | Mgst3 | 1091 | 43.9460215 | 63.1591313 | 33.5464033 | 46.3675769 | 31.9441015 | 71.1490589 |
| ENSMUSG00 | Fmo3 | 2284 | 0.29590331 | 0.21097392 | 0.67734873 | 0 | 0 | 0 |
| ENSMUSG00 | Fmo4 | 2976 | 0 | 0.09715009 | 0.01485277 | 0 | 0 | 0.62321084 |
| ENSMUSG00 | Mettl13 | 7405 | 2.50719911 | 1.99122907 | 2.03549338 | 2.41196651 | 3.26623932 | 1.33702823 |
| ENSMUSG00 | Vamp4 | 4743 | 11.625733 | 13.0041421 | 13.8765643 | 16.0973237 | 12.6194263 | 23.3412947 |
| ENSMUSG00 | Myoc | 2076 | 0 | 0.0232112 | 0 | 0 | 0.1228959 | 0 |
| ENSMUSG00 | Pigc | 4088 | 13.0897448 | 13.6968312 | 12.8237245 | 12.7628758 | 16.463745 | 16.6596806 |
| ENSMUSG00 | Tnfsl4 | 1594 | 0 | 0 | 0 | 0 | 0 | 0 |
| ENSMUSG00 | Prdx6 | 3948 | 88.1508299 | 97.2394591 | 86.9258129 | 80.2798073 | 72.0288735 | 69.008706 |
| ENSMUSG00 | Klhl20 | 10239 | 1.23083185 | 2.47073758 | 2.01604265 | 2.57250635 | 2.54658444 | 1.46242572 |
| ENSMUSG00 | Nuf6 | 5024 | 2.54010932 | 3.3089814 | 3.41367737 | 5.19253945 | 3.61572249 | 2.61671499 |
| ENSMUSG00 | Cenpl | 5497 | 0.73768583 | 1.9285096 | 1.52780611 | 1.43094155 | 1.53162671 | 1.27516505 |
| ENSMUSG00 | Dars2 | 5868 | 7.65570773 | 5.46901337 | 9.78496832 | 5.98292933 | 8.59135298 | 5.08029664 |
| ENSMUSG00 | Mrc1 | 5917 | 0.1746903 | 0.21988067 | 0.26146096 | 0.01219606 | 0.11210798 | 0.02765723 |
| ENSMUSG00 | Serpinc1 | 2149 | 0.57348529 | 1.12113639 | 0.24682276 | 0.52049483 | 0.49862908 | 0.59651375 |
| ENSMUSG00 | Stam | 5563 | 19.8241422 | 16.5703153 | 21.0401726 | 17.4605184 | 17.198356 | 17.6208976 |
| ENSMUSG00 | Rabgap1l | 10583 | 6.43119934 | 5.50025722 | 9.20958744 | 5.9324159 | 7.54089122 | 9.72639975 |
| ENSMUSG00 | Trdmt1 | 3971 | 5.31608163 | 5.55764052 | 5.36521993 | 5.47909163 | 4.65446095 | 7.23248429 |
| ENSMUSG00 | Tnn | 5810 | 0.51319467 | 0 | 0 | 0 | 0.02634753 | 0.2018605 |
| ENSMUSG00 | Cubn | 12165 | 0.11111273 | 0.21389789 | 0.02906821 | 0.01186422 | 0.22650426 | 0.04035705 |
| ENSMUSG00 | Rsu1 | 8979 | 7.44280685 | 5.96762709 | 7.18729155 | 6.86358526 | 6.08634018 | 4.41971268 |
| ENSMUSG00 | Vim | 2179 | 1.07644486 | 0.3980523 | 0.12171228 | 0.97698055 | 1.24111882 | 0.16272157 |
| ENSMUSG00 | 4930562F07f | 1251 | 0 | 0 | 0 | 0 | 0.20394235 | 1.04650894 |
| ENSMUSG00 | Pter | 3835 | 4.08439617 | 3.43022131 | 1.75193745 | 0.94086166 | 1.63656955 | 3.02261322 |
| ENSMUSG00 | 4921504E06l | 2276 | 0 | 0 | 0 | 0 | 0 | 0 |
| ENSMUSG00 | Ptf1a | 1506 | 0 | 0 | 0 | 0 | 0 | 0 |
| ENSMUSG00 | 4930426L09f | 1458 | 0 | 0.33049686 | 0 | 0.27222393 | 0.06999503 | 0 |
| ENSMUSG00 | Pip4k2a | 3556 | 4.22597628 | 3.02181569 | 2.63520549 | 4.48488855 | 4.59179985 | 3.95006727 |
| ENSMUSG00 | Bmi1 | 4149 | 35.4531882 | 27.443857 | 29.4039737 | 26.2984099 | 21.2025722 | 45.0896001 |
| ENSMUSG00 | Dnajc1 | 10556 | 6.51167353 | 6.46837709 | 6.68722464 | 7.51994107 | 6.98978206 | 6.23213621 |
| ENSMUSG00 | Mllt10 | 5846 | 11.0167428 | 8.25087728 | 8.34738876 | 11.2146895 | 11.3818869 | 12.8488465 |
| ENSMUSG00 | Plxdc2 | 7283 | 18.5430959 | 17.1494244 | 29.5872559 | 21.4619548 | 21.9505198 | 26.0163283 |
| ENSMUSG00 | Nek6 | 4261 | 7.27746416 | 5.90315015 | 12.0852258 | 4.58117486 | 4.95773752 | 8.21887745 |
| ENSMUSG00 | Psmb7 | 1206 | 30.8879645 | 34.3618078 | 35.9185789 | 31.863497 | 26.7825007 | 51.3152209 |
| ENSMUSG00 | Nr5a1 | 5837 | 0 | 0 | 0 | 0 | 0 | 0 |
| ENSMUSG00 | Ppp6c | 4303 | 95.4667395 | 67.380391 | 79.9290124 | 73.5812086 | 46.9945451 | 99.2485014 |
| ENSMUSG00 | Golga1 | 6316 | 12.5069886 | 12.4585909 | 15.5364299 | 15.9615631 | 15.4307121 | 14.2418873 |
| ENSMUSG00 | Arpc5l | 4619 | 17.4376713 | 19.1326986 | 17.8568173 | 20.1618872 | 12.8477322 | 25.5031753 |
| ENSMUSG00 | Orc4 | 8159 | 11.7332022 | 12.4201603 | 14.9632909 | 15.4561522 | 8.64928027 | 22.4742493 |
| ENSMUSG00 | Kif5c | 7113 | 0.8048347 | 1.93748383 | 0.63385182 | 0.23334374 | 2.55382958 | 2.25850731 |
| ENSMUSG00 | Lypd6b | 2633 | 14.7214559 | 30.1416902 | 7.89018846 | 13.5256278 | 17.3640838 | 12.3165766 |
| ENSMUSG00 | Mmadhc | 2056 | 56.6167543 | 49.2411065 | 59.7240856 | 65.0389382 | 42.9356178 | 64.3527602 |
| ENSMUSG00 | Fam188a | 4678 | 15.9939748 | 22.0845943 | 20.8914654 | 20.3781021 | 18.7285779 | 23.1117294 |
| ENSMUSG00 | Itga8 | 8962 | 0 | 0.03763726 | 0 | 0.24961914 | 0 | 0.10956115 |
| ENSMUSG00 | Il2ra | 5296 | 0 | 0 | 0 | 0 | 0 | 0 |
| ENSMUSG00 | Spopl | 7496 | 4.55045388 | 5.10406017 | 4.6466185 | 8.44289035 | 5.68396796 | 8.66340929 |
| ENSMUSG00 | Pfkfb3 | 7251 | 4.44651699 | 7.21035578 | 1.64591058 | 6.52372919 | 8.53606315 | 2.91140129 |
| ENSMUSG00 | 4931423N10l | 2599 | 0 | 0 | 0.068029 | 0 | 0 | 0 |
| ENSMUSG00 | Yme1l1 | 8202 | 17.6820279 | 15.815399 | 16.4315313 | 19.7127093 | 14.3709983 | 14.9308864 |
| ENSMUSG00 | Prkcq | 6732 | 3.49602562 | 4.0370103 | 3.3551904 | 2.50837743 | 6.77622994 | 3.72737197 |
| ENSMUSG00 | Mastl | 5765 | 1.87571391 | 0.50150677 | 0.44470198 | 0.32545816 | 1.33651045 | 1.99651306 |
| ENSMUSG00 | Acbd5 | 4225 | 50.6235468 | 43.6244121 | 53.2723751 | 49.3277844 | 42.7655377 | 52.0897193 |
| ENSMUSG00 | Abi2 | 6000 | 14.0734401 | 15.146605 | 12.1702408 | 16.3932756 | 14.4999951 | 9.72340889 |
| ENSMUSG00 | Pdss1 | 4401 | 4.36307591 | 2.75914188 | 5.6645852 | 10.0022936 | 3.9072685 | 8.403638 |
| ENSMUSG00 | Pkn3 | 4099 | 0.16488001 | 0.11755658 | 0.10783568 | 0.03521058 | 0.88384306 | 0.37927652 |
| ENSMUSG00 | Apbb1ip | 7531 | 0 | 0 | 0 | 0 | 0.00677551 | 0 |
| ENSMUSG00 | Gad2 | 5860 | 0 | 0 | 0 | 0 | 0 | 0 |
| ENSMUSG00 | Zbtb43 | 6822 | 7.75645625 | 6.55482532 | 6.1294259 | 4.52215598 | 12.1470001 | 4.4098398 |
| ENSMUSG00 | Odf2 | 8427 | 5.06673621 | 5.91823515 | 4.22243783 | 6.17852026 | 6.01877508 | 4.66714487 |
| ENSMUSG00 | Slc2a8 | 2218 | 27.7105378 | 31.327705 | 20.7856277 | 23.1816562 | 26.3183832 | 15.8876618 |
| ENSMUSG00 | Lrsam1 | 6096 | 3.15643907 | 4.61628645 | 5.43821888 | 3.2968666 | 4.64561 | 3.89254202 |
| ENSMUSG00 | Fam129b | 4653 | 7.06593217 | 5.19870921 | 5.5192931 | 5.09475677 | 6.79913024 | 3.88046662 |
| ENSMUSG00 | Stxbp1 | 6315 | 1.60532819 | 0.98433112 | 0.91693451 | 1.21130537 | 1.97964562 | 0.9372283 |
| ENSMUSG00 | Coq4 | 3529 | 8.6968634 | 8.54766585 | 6.25013309 | 7.93416451 | 5.59569494 | 6.2757175 |
| ENSMUSG00 | Med27 | 3508 | 5.55307453 | 3.37909486 | 6.26234777 | 4.55654097 | 3.88370648 | 4.81271428 |
| ENSMUSG00 | Ttf1 | 6049 | 4.85031321 | 5.14604755 | 3.79248744 | 5.19548037 | 4.82510943 | 2.59715527 |
| ENSMUSG00 | Barhl1 | 6138 | 0 | 0 | 0 | 0 | 0 | 0 |
| ENSMUSG00 | Ddx31 | 3913 | 3.99281979 | 3.7559072 | 2.34959963 | 5.07158817 | 3.42957752 | 2.45353259 |
| ENSMUSG00 | Ak8 | 1925 | 0.18586978 | 1.15146823 | 0 | 0.61854934 | 0.23856487 | 0.32587881 |
| ENSMUSG00 | 1700026L06f | 759 | 0.31427257 | 0.88881448 | 0 | 0.33277248 | 0.06722843 | 0 |
| ENSMUSG00 | Dpm2 | 2014 | 13.1465492 | 11.0058408 | 10.3810684 | 11.9676295 | 12.4652318 | 15.3572199 |
| ENSMUSG00 | St6galnac6 | 4786 | 0.31565154 | 0.1308867 | 0.21242006 | 0 | 0.29852456 | 0.24505004 |
| ENSMUSG00 | Tsc1 | 5873 | 14.6485373 | 15.1213371 | 13.193569 | 12.2137076 | 16.3426891 | 15.5947963 |
| ENSMUSG00 | Eng | 4137 | 1.17238787 | 0.80368975 | 0.77996967 | 1.91879376 | 2.44216157 | 1.5954708 |
| ENSMUSG00 | Gfi1b | 2571 | 0 | 0 | 0 | 0 | 0 | 0 |
| ENSMUSG00 | Gtf3c5 | 4545 | 6.9714231 | 6.20221533 | 5.96165671 | 5.02528805 | 8.8917249 | 4.32073493 |
| ENSMUSG00 | Ak1 | 2783 | 15.2850752 | 15.7735713 | 18.1858104 | 15.1173783 | 9.2225178 | 14.4066546 |
| ENSMUSG00 | Cel | 2093 | 0 | 0 | 0 | 0.34478781 | 0 | 0.14334498 |
| ENSMUSG00 | Slc25a25 | 6166 | 9.12325737 | 10.5188049 | 8.91778993 | 8.29196515 | 12.3469596 | 4.49416696 |
| ENSMUSG00 | Ptges2 | 2128 | 34.8794559 | 31.1807946 | 38.8843281 | 26.2476527 | 25.7529735 | 25.3649018 |
| ENSMUSG00 | Ralgds | 4745 | 6.74460734 | 6.44855443 | 4.94650762 | 4.72982754 | 7.5060928 | 2.67860525 |
| ENSMUSG00 | Lcn2 | 2185 | 4.14839797 | 18.5909248 | 2.26572376 | 8.27327428 | 9.94839181 | 2.90846268 |
| ENSMUSG00 | Kcnj3 | 5211 | 0.03814573 | 0 | 0.2375075 | 0 | 0.52877073 | 0 |
| ENSMUSG00 | Dnm1 | 5017 | 0.07924154 | 0.63390576 | 0.02643124 | 0.31644608 | 0.3864864 | 0.26638575 |
| ENSMUSG00 | Nr4a2 | 4745 | 0.99702891 | 0.15232806 | 0.29809462 | 1.93907721 | 0.19356686 | 0.2586636 |
| ENSMUSG00 | Gpd2 | 6837 | 5.73334851 | 6.51225282 | 5.72807268 | 6.65488638 | 5.97061586 | 4.54377857 |
| ENSMUSG00 | Galnt5 | 4451 | 0 | 0 | 0 | 0 | 0 | 0 |
| ENSMUSG00 | Gbgt1 | 1330 | 0 | 0 | 0 | 0 | 0 | 0 |
| ENSMUSG00 | Ermn | 3632 | 0 | 0 | 0 | 0 | 0 | 0 |
| ENSMUSG00 | 1700007K13f | 1388 | 0.65877237 | 0.90262788 | 0.89167983 | 0.64989273 | 0.36762519 | 0.96286551 |
| ENSMUSG00 | Cytip | 6848 | 26.7397405 | 17.7321604 | 26.6192174 | 15.1957676 | 16.5343937 | 21.3800034 |
| ENSMUSG00 | Olfm1 | 3914 | 56.4438439 | 83.1751669 | 31.8695966 | 68.8453522 | 64.6108121 | 48.7515013 |
| ENSMUSG00 | Acvr1c | 12502 | 0 | 0 | 0 | 0.0028861 | 0 | 0 |
| ENSMUSG00 | Fcnb | 1330 | 0 | 0 | 0 | 0 | 0 | 0 |
| ENSMUSG00 | Acvr1 | 4812 | 3.27990976 | 1.70234729 | 2.05760865 | 2.67690979 | 2.42831257 | 1.81377485 |
| ENSMUSG00 | Col5a1 | 8424 | 0.26900076 | 0 | 0.01049427 | 0 | 0.09085893 | 0.08418099 |
| ENSMUSG00 | Upp2 | 7250 | 0.99251613 | 0.53171247 | 1.00597298 | 0.83610807 | 0.52785906 | 0.96683894 |
| ENSMUSG00 | Lamc3 | 6461 | 0 | 0.00745805 | 0.20523995 | 0 | 0 | 0 |
| ENSMUSG00 | Fibcd1 | 4744 | 0 | 0 | 0 | 0 | 0 | 0 |
| ENSMUSG00 | Abl1 | 12870 | 3.33303524 | 2.03678512 | 2.88496878 | 3.54652574 | 3.84977552 | 3.60906843 |
| ENSMUSG00 | Fubp3 | 4563 | 11.3699106 | 9.89495864 | 10.3747916 | 14.2256406 | 10.6906016 | 10.9983205 |
| ENSMUSG00 | Tor1b | 3436 | 18.3851742 | 23.5182373 | 24.5966018 | 19.2801612 | 24.6518579 | 17.4237 |
| ENSMUSG00 | Tor1a | 2235 | 11.2240305 | 15.8034282 | 18.9069181 | 10.3645068 | 13.1275911 | 14.2657957 |
| ENSMUSG00 | BC005624 | 2615 | 17.7873469 | 18.924465 | 20.7064083 | 18.2824891 | 19.0251306 | 19.8588578 |
| ENSMUSG00 | Crat | 7628 | 27.0126711 | 29.6838086 | 29.5065266 | 25.9026319 | 26.4229398 | 23.1519781 |
| ENSMUSG00 | Usp20 | 7440 | 13.2571703 | 10.278479 | 9.47606715 | 7.5025434 | 12.2970822 | 9.60111293 |
| ENSMUSG00 | Dolpp1 | 3319 | 7.96546988 | 9.48049012 | 6.2061039 | 5.89227723 | 7.24116391 | 4.97172544 |
| ENSMUSG00 | Mettl11a | 2931 | 10.2677921 | 7.72692863 | 12.9694933 | 10.537779 | 8.96573987 | 9.90113141 |
| ENSMUSG00 | Fam73b | 4450 | 17.5996172 | 14.3801338 | 11.2342212 | 11.4084127 | 15.0327143 | 9.44499305 |
| ENSMUSG00 | Sh3glb2 | 4467 | 14.5245006 | 15.5012128 | 13.7642184 | 13.8690106 | 14.8955438 | 12.7489247 |
| ENSMUSG00 | Hspa5 | 3478 | 279.831489 | 240.807519 | 253.429831 | 309.757816 | 194.950685 | 302.77359 |
| ENSMUSG00 | Kynu | 7523 | 0.01585358 | 0 | 0 | 0 | 0 | 0 |
| ENSMUSG00 | Gapvd1 | 9538 | 13.050368 | 11.2458608 | 13.7313966 | 14.5833803 | 17.7185318 | 12.6250296 |
| ENSMUSG00 | Psmd5 | 4149 | 36.8521519 | 39.2436703 | 46.6947886 | 36.1342239 | 35.3089238 | 36.3661857 |
| ENSMUSG00 | D730039F16l | 2405 | 18.0346068 | 11.520667 | 25.4551154 | 11.2822027 | 13.8757795 | 12.9738825 |
| ENSMUSG00 | Zeb2 | 24976 | 0.00159175 | 0.20643615 | 0.0584025 | 0 | 0.05720446 | 0.04477339 |
| ENSMUSG00 | Phf19 | 7299 | 0 | 0 | 0.03633526 | 0.03460396 | 0.00699087 | 0.14947055 |
| ENSMUSG00 | Hc | 6328 | 0 | 0 | 0 | 0 | 0 | 0 |
| ENSMUSG00 | Traf1 | 5897 | 0.15505784 | 0.75176406 | 0.59215628 | 0.17132394 | 1.53157004 | 0.07862792 |
| ENSMUSG00 | Rab14 | 6602 | 114.298512 | 96.4093781 | 140.54606 | 111.093565 | 92.91716 | 120.20754 |
| ENSMUSG00 | Gsn | 3170 | 156.250325 | 110.540002 | 171.132246 | 89.1920823 | 112.24193 | 86.2033472 |
| ENSMUSG00 | Stom | 3096 | 3.36432038 | 2.41243493 | 4.19746184 | 1.32860242 | 3.32924028 | 1.96454927 |
| ENSMUSG00 | 4930568D16l | 1221 | 0 | 0 | 0 | 0 | 0 | 0 |
| ENSMUSG00 | Dab2ip | 8306 | 2.8766005 | 3.03413307 | 2.91095691 | 2.7367792 | 2.69077205 | 2.96192206 |
| ENSMUSG00 | Ttll11 | 4841 | 0.89513476 | 0.51759864 | 0.91307257 | 0.71552908 | 1.12782942 | 0.53523874 |
| ENSMUSG00 | Mrrf | 2909 | 5.90387335 | 6.60927825 | 7.20236287 | 7.1196609 | 5.66569934 | 6.62879682 |
| ENSMUSG00 | Grb14 | 5445 | 34.1626985 | 38.9739378 | 24.0695068 | 40.8002109 | 31.2249558 | 24.3743608 |
| ENSMUSG00 | Rbm18 | 6083 | 26.7955729 | 36.0031859 | 27.5107969 | 28.1989215 | 31.7918732 | 32.7672306 |
| ENSMUSG00 | Lhx6 | 6971 | 0 | 0.11059863 | 0 | 0 | 0 | 0 |
| ENSMUSG00 | Gca | 3401 | 51.2578012 | 54.3213229 | 62.085329 | 63.9949699 | 51.9265765 | 67.9660593 |
| ENSMUSG00 | Morn5 | 709 | 0 | 0.1359279 | 0 | 0 | 0.3598475 | 0.5770375 |
| ENSMUSG00 | Ndufa8 | 857 | 169.22753 | 177.283375 | 216.367248 | 159.653574 | 111.102937 | 238.533747 |
| ENSMUSG00 | Ifih1 | 8789 | 0.75991816 | 1.20616877 | 0.42245475 | 1.11665902 | 1.70107272 | 1.91161427 |
| ENSMUSG00 | Slc4a10 | 6502 | 0.12228693 | 0.13339833 | 0.06798192 | 0 | 0 | 0.15940269 |
| ENSMUSG00 | Psmd14 | 2853 | 48.729378 | 47.6797498 | 45.1934021 | 47.3252754 | 35.6093636 | 74.319329 |
| ENSMUSG00 | Strbp | 16113 | 19.9061141 | 16.9174395 | 21.2381722 | 21.3092991 | 19.0323663 | 22.8668219 |
| ENSMUSG00 | Wdr5 | 3582 | 30.4436859 | 29.2454844 | 26.259498 | 26.9255458 | 26.424882 | 17.5511009 |
| ENSMUSG00 | Brd3 | 9407 | 10.2949241 | 10.8236369 | 10.7086213 | 8.59187623 | 10.485169 | 12.3804305 |
| ENSMUSG00 | Lcn4 | 767 | 0 | 0 | 0 | 0 | 0 | 0 |
| ENSMUSG00 | Egfl7 | 2904 | 1.23209134 | 1.19470518 | 1.73519632 | 1.14309507 | 1.862533 | 2.01930008 |
| ENSMUSG00 | Agpat2 | 1707 | 24.8035072 | 25.7728305 | 24.7809981 | 19.5735051 | 19.6393257 | 20.1484008 |
| ENSMUSG00 | Notch1 | 17230 | 1.16520706 | 1.45146625 | 1.65211764 | 2.53600441 | 1.70877649 | 1.04317976 |
| ENSMUSG00 | Sec16a | 8477 | 13.5394682 | 9.46448345 | 9.84464782 | 13.4504259 | 15.7888621 | 12.1427968 |
| ENSMUSG00 | Inpp5e | 5514 | 9.88479575 | 9.37686841 | 6.98219174 | 7.53835968 | 12.6131575 | 6.46004332 |
| ENSMUSG00 | Pmpca | 4354 | 38.1393298 | 28.9849086 | 31.1057526 | 29.3529173 | 34.8652889 | 23.9420467 |
| ENSMUSG00 | Sdccag3 | 3780 | 28.1128041 | 32.7744293 | 31.1868559 | 28.1879041 | 26.0666486 | 27.1519755 |
| ENSMUSG00 | Card9 | 2504 | 0.31753579 | 0.30790059 | 0.01765249 | 0.04322929 | 0.18340151 | 0.22874098 |
| ENSMUSG00 | Gpsm1 | 4405 | 2.83387534 | 1.76118438 | 3.96361589 | 3.25188914 | 1.47113501 | 1.5355529 |
| ENSMUSG00 | 1700019N19l | 1380 | 0 | 0 | 0 | 0 | 0 | 0 |
| ENSMUSG00 | Nacc2 | 7724 | 12.5638436 | 16.076704 | 11.3652072 | 12.4166331 | 13.7409194 | 9.12097271 |
| ENSMUSG00 | Camsap1 | 11307 | 2.18695577 | 2.45897029 | 2.69737965 | 2.22740489 | 3.3846097 | 1.65958714 |
| ENSMUSG00 | Lhx3 | 3968 | 0 | 0 | 0 | 0 | 0 | 0 |
| ENSMUSG00 | Lcn3 | 677 | 0 | 0 | 0 | 0 | 0 | 0 |
| ENSMUSG00 | Lcn5 | 1011 | 0 | 0 | 0 | 0 | 0 | 0 |
| ENSMUSG00 | Fcna | 1798 | 0 | 0 | 0 | 0 | 0 | 0 |
| ENSMUSG00 | Tmem141 | 1543 | 15.2271478 | 11.1800041 | 19.5656894 | 16.3222729 | 12.4341655 | 17.3051665 |
| ENSMUSG00 | 4921530D09l | 2904 | 0 | 0.03318626 | 0.03044204 | 0.03727484 | 0.0527132 | 0.02817628 |
| ENSMUSG00 | Mamdc4 | 4716 | 0.4383556 | 0.29631188 | 0.63734634 | 0.1530197 | 0.55181195 | 0.37013929 |
| ENSMUSG00 | Traf2 | 5853 | 7.79758955 | 7.80467233 | 6.58534207 | 8.60593442 | 7.01797925 | 7.10175128 |
| ENSMUSG00 | Lcn12 | 889 | 0 | 0 | 0 | 0 | 0.34438499 | 0 |
| ENSMUSG00 | Abca2 | 8248 | 6.66126019 | 7.41374819 | 6.4148407 | 7.98372105 | 7.64653256 | 5.30744374 |
| ENSMUSG00 | Nmi | 2267 | 9.59252221 | 12.1369468 | 6.8632769 | 6.55747788 | 11.0966049 | 10.16633 |
| ENSMUSG00 | Neb | 26653 | 0.05071423 | 0.02531085 | 0.0082921 | 0.08393377 | 0.04977623 | 0.0828892 |
| ENSMUSG00 | 2010317E24l | 4084 | 0 | 0 | 0 | 0 | 0 | 0 |
| ENSMUSG00 | Uap1l1 | 3999 | 36.0871203 | 20.8940462 | 42.9528287 | 24.4246298 | 20.4411738 | 30.8757919 |
| ENSMUSG00 | Dpp7 | 2454 | 9.59056417 | 5.73367608 | 7.54709545 | 12.1302717 | 10.0430886 | 5.57940868 |
| ENSMUSG00 | Grin1 | 7778 | 0 | 0.00619522 | 0.01136586 | 0 | 0 | 0 |
| ENSMUSG00 | Arl6ip6 | 2698 | 8.28116384 | 7.57266548 | 10.1739602 | 9.9098573 | 8.18918485 | 10.7460866 |
| ENSMUSG00 | Lrrc26 | 1188 | 0.23424946 | 0 | 0.85575959 | 0.6985581 | 0.12885448 | 0.1377507 |
| ENSMUSG00 | Tmem210 | 575 | 0 | 0 | 0 | 0 | 0 | 0 |
| ENSMUSG00 | Anapc2 | 3252 | 38.2762637 | 40.0960985 | 43.8348535 | 42.7392483 | 46.8526317 | 28.4320501 |
| ENSMUSG00 | Ssna1 | 1741 | 11.3032526 | 10.2960118 | 16.2995883 | 11.5852171 | 9.02706706 | 13.9898064 |
| ENSMUSG00 | Fam166a | 1118 | 0 | 0 | 0 | 0 | 0 | 0.07318776 |
| ENSMUSG00 | Rbms1 | 6493 | 11.014956 | 12.1783385 | 11.7567508 | 12.6478875 | 13.2418671 | 12.1734028 |
| ENSMUSG00 | Itgb6 | 7365 | 16.5607351 | 14.7797926 | 19.2891682 | 21.0221389 | 23.0987016 | 21.1012756 |
| ENSMUSG00 | Arrdc1 | 2727 | 51.3016267 | 55.7139172 | 59.3734327 | 53.7988973 | 54.0763572 | 50.3185589 |
| ENSMUSG00 | Zmynd19 | 4161 | 2.67520657 | 4.18055891 | 5.10960983 | 4.69127279 | 3.09027799 | 3.46094918 |
| ENSMUSG00 | Wdr85 | 1863 | 16.0046218 | 14.8206287 | 14.4017814 | 9.62575208 | 13.3386178 | 12.2099029 |
| ENSMUSG00 | Pax8 | 6290 | 32.7650892 | 40.0123668 | 31.2504922 | 34.6249954 | 38.192715 | 29.4427346 |
| ENSMUSG00 | 7-Mar | 10175 | 12.3271293 | 12.4882209 | 12.5546267 | 9.42211227 | 11.2082506 | 20.0317817 |
| ENSMUSG00 | Pad4 | 6185 | 0.45636849 | 0.88036669 | 0.98623352 | 0.25085334 | 1.08900269 | 0.32191571 |
| ENSMUSG00 | Ly75 | 8375 | 1.00634769 | 0.66166458 | 0.51722754 | 1.39589044 | 2.15681636 | 1.35477611 |
| ENSMUSG00 | Il1rn | 3336 | 0 | 0 | 0 | 0 | 0 | 0 |
| ENSMUSG00 | Il1f5 | 1832 | 3.75420204 | 0.86798722 | 6.05603854 | 3.68304712 | 5.79338765 | 6.29758319 |
| ENSMUSG00 | Il1f6 | 907 | 0 | 0 | 0 | 0 | 0 | 0 |
| ENSMUSG00 | Il1f8 | 1212 | 0 | 0 | 0 | 0 | 0 | 0 |
| ENSMUSG00 | Hnmt | 2775 | 34.8845388 | 30.9782388 | 56.1801443 | 36.5111283 | 30.1929042 | 50.2934519 |
| ENSMUSG00 | Baz2b | 18996 | 5.57740343 | 6.42029296 | 4.96561029 | 7.34519194 | 6.71541059 | 5.30244487 |
| ENSMUSG00 | Wdsub1 | 3425 | 11.9904851 | 12.9435115 | 15.0996077 | 10.5559733 | 12.8571569 | 10.575393 |
| ENSMUSG00 | Dapl1 | 771 | 0.25781764 | 0.18749588 | 0.22932214 | 0.51478922 | 1.19127726 | 2.19329135 |
| ENSMUSG00 | Pkp4 | 5912 | 25.4120367 | 17.6868369 | 29.3009173 | 22.7770958 | 17.0289309 | 22.5643201 |
| ENSMUSG00 | Galnt3 | 4569 | 6.98701047 | 16.3468998 | 4.66300905 | 14.0016338 | 11.0116014 | 6.95446591 |
| ENSMUSG00 | Nup35 | 3354 | 2.29951199 | 2.126295 | 1.21245365 | 2.15158286 | 2.14511598 | 1.54507497 |
| ENSMUSG00 | Dusp19 | 3493 | 2.50392377 | 1.35192423 | 1.75896255 | 2.39651713 | 1.49003445 | 2.23319405 |
| ENSMUSG00 | Nckap1 | 5826 | 89.1461719 | 85.9763244 | 84.7543406 | 96.608962 | 88.1006376 | 71.7258407 |
| ENSMUSG00 | Frzb | 3589 | 6.43575764 | 2.45698493 | 0.50495279 | 0 | 3.04253119 | 0 |
| ENSMUSG00 | Dnajc10 | 4982 | 24.1150265 | 27.9040316 | 24.2302757 | 22.2344193 | 34.0551384 | 27.5757443 |
| ENSMUSG00 | Safa2 | 6076 | 10.665147 | 10.9839076 | 9.7773662 | 14.3829348 | 14.2682377 | 8.62769201 |
| ENSMUSG00 | Itga4 | 14127 | 0.33488371 | 0.15690354 | 0.38485359 | 0.13792245 | 0.34313766 | 0.3108386 |
| ENSMUSG00 | Slc25a12 | 7983 | 38.5801964 | 36.0960697 | 37.7124831 | 29.306774 | 35.7050401 | 29.1947629 |
| ENSMUSG00 | Ube2e3 | 4546 | 29.5586283 | 33.8555865 | 26.3888698 | 28.6052988 | 24.0764576 | 50.4574826 |
| ENSMUSG00 | Dync1i2 | 8366 | 7.5984955 | 7.20550312 | 6.88969678 | 7.84523536 | 7.88633803 | 11.1954472 |
| ENSMUSG00 | Cwc22 | 6797 | 3.17014425 | 2.82156892 | 1.76235096 | 2.63833693 | 3.21307766 | 2.07459004 |
| ENSMUSG00 | Cybrd1 | 5335 | 0.46201308 | 1.39094885 | 0.30655449 | 1.1429926 | 0.38257826 | 0.57258849 |
| ENSMUSG00 | Zfp385b | 9245 | 2.81664032 | 1.46461766 | 2.5435782 | 2.5915065 | 1.78274953 | 2.63158228 |
| ENSMUSG00 | Hat1 | 2602 | 41.2528045 | 38.3898903 | 33.3977031 | 30.3826901 | 35.4164622 | 43.2914143 |
| ENSMUSG00 | Xirp2 | 11970 | 0 | 0 | 0 | 0 | 0 | 0 |
| ENSMUSG00 | Stk39 | 6479 | 29.6923553 | 28.0833471 | 27.2824773 | 32.4175924 | 30.6914318 | 38.0430491 |
| ENSMUSG00 | Lass6 | 8299 | 4.71853817 | 2.94379156 | 5.64573486 | 5.79990931 | 4.07645345 | 5.94855971 |
| ENSMUSG00 | Abcb11 | 5684 | 0 | 0 | 0 | 0 | 0 | 0 |
| ENSMUSG00 | Ssrp1 | 3758 | 32.7945688 | 30.6069817 | 31.6164327 | 32.7119547 | 38.7382252 | 26.4109665 |
| ENSMUSG00 | Dhrs9 | 4276 | 0 | 0 | 0 | 0 | 0 | 0 |
| ENSMUSG00 | Lrp2 | 16621 | 0.37791745 | 1.45536333 | 0.08510072 | 0.9682084 | 0.57715894 | 0.35609149 |
| ENSMUSG00 | P2rx3 | 3258 | 0.09761935 | 0.14790191 | 0.25777625 | 0.19934831 | 0.57948923 | 0 |
| ENSMUSG00 | Prg3 | 1049 | 0 | 0 | 0 | 0 | 0 | 0 |
| ENSMUSG00 | Prg2 | 832 | 0 | 0 | 0 | 0 | 0 | 0 |
| ENSMUSG00 | Slc43a3 | 4477 | 5.32796254 | 7.27586216 | 12.3018755 | 8.76059468 | 6.95244343 | 10.0520784 |
| ENSMUSG00 | Slc43a1 | 5098 | 0 | 0 | 0 | 0 | 0 | 0 |
| ENSMUSG00 | Timm10 | 811 | 20.9316772 | 24.1229291 | 33.5192768 | 26.8279567 | 16.2328051 | 35.2115256 |
| ENSMUSG00 | Smtnl1 | 1753 | 0.15874978 | 0 | 0 | 0 | 0.14554015 | 0 |
| ENSMUSG00 | Ube2l6 | 2089 | 4.0535746 | 4.05974812 | 7.1730133 | 2.05541565 | 2.83344164 | 5.75783433 |
| ENSMUSG00 | Clp1 | 2668 | 29.6974035 | 19.4335127 | 34.5761793 | 18.4196944 | 23.6962817 | 20.7524429 |
| ENSMUSG00 | Med19 | 2054 | 28.0069038 | 23.7178642 | 23.9946714 | 30.4606939 | 24.61886 | 31.2051618 |
| ENSMUSG00 | Tfpi | 5497 | 0.44839727 | 0 | 0.29752014 | 0 | 0.47341189 | 0.27289524 |
| ENSMUSG00 | Fastkd1 | 6042 | 11.1528533 | 7.79181628 | 9.19591471 | 9.35194997 | 8.07368674 | 6.55458063 |
| ENSMUSG00 | Itgav | 8531 | 38.5951107 | 29.3942381 | 52.6214884 | 50.1874974 | 47.4734902 | 53.7084357 |
| ENSMUSG00 | Phospho2 | 3268 | 26.556369 | 27.0274628 | 34.5851018 | 25.2397655 | 24.0767049 | 34.719247 |
| ENSMUSG00 | Zc3h15 | 3025 | 51.3601381 | 41.4164461 | 43.7196411 | 39.6246453 | 45.3417845 | 61.8976524 |
| ENSMUSG00 | Mtx2 | 3878 | 66.8810611 | 68.4525812 | 83.2631416 | 60.7663313 | 46.3026861 | 93.3934849 |
| ENSMUSG00 | Hoxd8 | 2115 | 207.611954 | 256.037464 | 231.855909 | 270.009702 | 200.824375 | 250.230307 |
| ENSMUSG00 | Atf2 | 10919 | 13.6353397 | 13.7423373 | 14.5450336 | 16.1392715 | 15.9495394 | 14.9899359 |
| ENSMUSG00 | Chrna1 | 4377 | 0 | 0 | 0 | 0 | 0.01165784 | 0 |
| ENSMUSG00 | Ola1 | 4071 | 10.6248988 | 7.82393472 | 9.31593744 | 9.54565511 | 9.40058508 | 11.905437 |
| ENSMUSG00 | Sp3 | 6610 | 19.1499924 | 16.8980594 | 21.6930074 | 18.0683158 | 24.0464691 | 23.6806588 |
| ENSMUSG00 | Itga6 | 8994 | 10.8560663 | 9.66514806 | 8.98878929 | 11.3533674 | 11.8687101 | 10.4197977 |
| ENSMUSG00 | Kif18a | 7225 | 0.56675633 | 0.44685005 | 0.73414826 | 0.67419737 | 1.15118329 | 0.34730337 |
| ENSMUSG00 | Fshb | 1597 | 0 | 0 | 0 | 0 | 0 | 0 |
| ENSMUSG00 | 2700007P21f | 5748 | 5.40171021 | 4.18320018 | 6.97478629 | 3.22026598 | 6.03652324 | 5.9550578 |
| ENSMUSG00 | Slc12a6 | 11024 | 4.46455777 | 3.57988898 | 3.99356274 | 4.22877372 | 4.57311859 | 4.54000027 |
| ENSMUSG00 | Tmem85 | 1047 | 47.007913 | 49.5671425 | 56.1916457 | 46.7309001 | 34.5049418 | 52.5694575 |
| ENSMUSG00 | 2410042D21l | 4172 | 10.8441363 | 10.8569644 | 12.4913646 | 13.7513065 | 9.4665424 | 15.5789706 |
| ENSMUSG00 | Nop10 | 1107 | 35.3382049 | 37.5218727 | 46.1983129 | 36.0820445 | 21.986975 | 67.5829589 |
| ENSMUSG00 | Lpcat4 | 2811 | 11.7385446 | 18.9934464 | 6.4313603 | 10.5897142 | 10.0564255 | 5.09398278 |
| ENSMUSG00 | 4930430A15f | 2415 | 0 | 0 | 0 | 0 | 0 | 0 |
| ENSMUSG00 | Ccdc34 | 4504 | 1.18277851 | 1.92574591 | 0.71641531 | 1.36188889 | 1.42746966 | 2.65237389 |
| ENSMUSG00 | Lin7c | 4567 | 47.6856847 | 53.5358019 | 59.9681671 | 62.6122625 | 44.9706947 | 62.9580134 |
| ENSMUSG00 | Commd9 | 1661 | 10.4833838 | 9.54445483 | 12.0816597 | 7.3424028 | 7.52646723 | 5.87857968 |
| ENSMUSG00 | Traf6 | 8055 | 9.97968803 | 7.99817171 | 10.0201819 | 7.09994296 | 9.71749975 | 8.55316434 |
| ENSMUSG00 | B230118H07 | 4804 | 7.65483336 | 7.38243577 | 8.85140986 | 6.31661104 | 7.49888038 | 10.0434715 |
| ENSMUSG00 | Dnajc24 | 1673 | 5.18033159 | 6.33653154 | 7.21285304 | 6.42704678 | 5.0934876 | 9.9936365 |
| ENSMUSG00 | Elp4 | 7013 | 2.64734199 | 2.31553273 | 3.37832423 | 3.20020415 | 1.91358004 | 2.90908743 |
| ENSMUSG00 | Pax6 | 17502 | 0 | 0.00825959 | 0.01767872 | 0 | 0 | 0 |
| ENSMUSG00 | Eif3m | 2330 | 31.0706996 | 24.6309239 | 36.0254506 | 25.0716009 | 23.2356158 | 34.7780915 |
| ENSMUSG00 | Prrg4 | 3159 | 0.28945111 | 1.15928129 | 0.61566353 | 0.79953882 | 1.63142259 | 0.37125973 |
| ENSMUSG00 | Depdc7 | 2087 | 40.7841321 | 94.179443 | 26.7922048 | 105.721947 | 42.9579982 | 52.1054081 |
| ENSMUSG00 | Tcp11l1 | 6862 | 1.92925896 | 2.40861989 | 1.33339865 | 1.74573576 | 2.20851554 | 2.08276173 |
| ENSMUSG00 | Cstf3 | 6020 | 11.6822998 | 12.2147028 | 11.4396132 | 16.0151533 | 11.9937412 | 17.859905 |
| ENSMUSG00 | Hipk3 | 8519 | 20.3187419 | 17.6817488 | 21.0450376 | 29.1824494 | 24.3362091 | 18.6238498 |
| ENSMUSG00 | Fbxo3 | 7315 | 26.4293782 | 27.0805829 | 32.9746353 | 29.2010531 | 28.0906651 | 28.8928475 |
| ENSMUSG00 | Caprin1 | 11660 | 53.9119775 | 41.2849534 | 51.1998363 | 58.8762418 | 44.628386 | 50.219488 |
| ENSMUSG00 | Nat10 | 7839 | 3.65147927 | 5.13890384 | 3.94145787 | 4.4003616 | 3.43039929 | 3.67071622 |
| ENSMUSG00 | Elf5 | 5134 | 0.34071701 | 0.47867327 | 0.51657783 | 0.30220645 | 0.70566277 | 0.86594589 |
| ENSMUSG00 | Cat | 2826 | 126.848962 | 160.655576 | 90.7967794 | 134.790568 | 110.340475 | 95.6832178 |
| ENSMUSG00 | Pamr1 | 6227 | 15.9609284 | 9.3246608 | 20.0459298 | 12.4812468 | 11.3164325 | 10.2624827 |
| ENSMUSG00 | Trim44 | 5649 | 71.7766234 | 63.6600138 | 66.0875847 | 92.8207242 | 77.0589506 | 69.8306085 |
| ENSMUSG00 | Api5 | 4103 | 51.0048378 | 42.1029479 | 45.3006946 | 49.8007843 | 43.8630338 | 40.6493831 |
| ENSMUSG00 | Ttc17 | 8957 | 3.6839398 | 4.30380191 | 3.08430857 | 3.73026384 | 5.57718231 | 2.98720789 |
| ENSMUSG00 | Hsd17b12 | 3354 | 58.8627657 | 58.6455149 | 85.7152018 | 65.2037185 | 47.877163 | 77.7741949 |
| ENSMUSG00 | 4921507L20f | 1546 | 0.23143553 | 0.40519001 | 0.22872881 | 1.49369395 | 0.16502709 | 1.0938083 |
| ENSMUSG00 | Ext2 | 6813 | 13.0475935 | 14.3364037 | 12.1128491 | 11.9479073 | 13.241543 | 8.64717757 |
| ENSMUSG00 | Gatm | 2834 | 1.29058017 | 5.78101282 | 1.34134033 | 0.96762011 | 4.78934932 | 4.00361677 |
| ENSMUSG00 | Sema6d | 8279 | 0.79232447 | 0.5936728 | 0.70475218 | 0.08716522 | 0.82588892 | 0.05600542 |
| ENSMUSG00 | Myef2 | 6362 | 0.43742277 | 0.21964898 | 0.91016055 | 0.55580641 | 1.12286901 | 0.89172037 |
| ENSMUSG00 | Slc12a1 | 5411 | 11.0133922 | 4.68417456 | 7.76044185 | 3.24745069 | 8.51539778 | 2.71183809 |
| ENSMUSG00 | Dut | 2401 | 9.2558574 | 11.0782658 | 12.7211468 | 5.75569473 | 7.56576001 | 13.0522951 |
| ENSMUSG00 | Fbn1 | 11803 | 0.06399679 | 0.06532094 | 0 | 0 | 0 | 0 |
| ENSMUSG00 | Cops2 | 4273 | 28.5350009 | 27.6398244 | 30.4644109 | 30.5257643 | 20.0737977 | 37.1747014 |
| ENSMUSG00 | Galk2 | 6317 | 10.9316558 | 8.88868753 | 11.8953828 | 9.43605154 | 9.03886569 | 10.0040113 |
| ENSMUSG00 | Fgf7 | 3244 | 0 | 0 | 0 | 0 | 0.11010624 | 0 |
| ENSMUSG00 | 4930525F21f | 2655 | 0 | 0 | 0.06659411 | 0.04077067 | 0 | 0.05136467 |
| ENSMUSG00 | Meis2 | 13113 | 0.27285848 | 0.30867545 | 0.0573043 | 0.1375812 | 0.40080201 | 0.36607462 |
| ENSMUSG00 | Cd82 | 3438 | 106.916572 | 134.355798 | 94.5621162 | 93.2801663 | 107.484879 | 88.4640786 |
| ENSMUSG00 | Tspan18 | 10339 | 5.82162661 | 7.54092885 | 1.93241445 | 4.37982067 | 6.67256602 | 1.5195079 |
| ENSMUSG00 | Slc28a2 | 3549 | 0 | 0 | 0 | 0 | 0 | 0 |
| ENSMUSG00 | Syt13 | 3755 | 0 | 0 | 0 | 0 | 0 | 0 |
| ENSMUSG00 | Chst1 | 2677 | 0 | 0 | 0 | 0 | 0 | 0 |
| ENSMUSG00 | Pex16 | 1764 | 18.4128274 | 20.4601163 | 19.4698594 | 19.186484 | 18.7733095 | 20.2549759 |
| ENSMUSG00 | Mapk8ip1 | 5372 | 1.73171676 | 1.74913586 | 1.70323557 | 1.42393772 | 1.71924312 | 0.95958802 |
| ENSMUSG00 | Duoxa1 | 3547 | 0 | 0 | 0.19938807 | 0 | 0 | 0 |
| ENSMUSG00 | Duoxa2 | 1586 | 0 | 0 | 0 | 0 | 0 | 0 |
| ENSMUSG00 | Sord | 2343 | 74.2848885 | 68.8142703 | 57.5774754 | 108.061334 | 42.7506512 | 37.1228445 |
| ENSMUSG00 | 4933406I08R | 2108 | 0 | 0 | 0 | 0 | 0 | 0 |
| ENSMUSG00 | Creb3l1 | 2591 | 0.56771624 | 0.27896435 | 0.2217769 | 0.16711097 | 0.92560388 | 0.56844096 |
| ENSMUSG00 | Patl2 | 2416 | 0 | 0 | 0 | 0 | 0 | 0 |
| ENSMUSG00 | Eif3j | 5894 | 1.79419033 | 1.88854227 | 2.3998286 | 3.948578 | 1.57563631 | 3.34570142 |
| ENSMUSG00 | Frmd5 | 9079 | 0 | 0.11145669 | 0 | 0 | 0 | 0 |
| ENSMUSG00 | Mdk | 1612 | 0 | 0.26903101 | 0.0274205 | 0 | 0 | 0.33839503 |
| ENSMUSG00 | Wdr76 | 5807 | 0.92422763 | 0.68873337 | 0.83730028 | 0.11184378 | 0.68538958 | 0.61998491 |
| ENSMUSG00 | Harbi1 | 3141 | 6.8347531 | 8.51431881 | 7.57102564 | 4.05506581 | 9.0161218 | 5.81789503 |
| ENSMUSG00 | Atg13 | 5926 | 13.9741252 | 11.2212775 | 13.7170417 | 8.88351383 | 10.7632416 | 9.24189599 |
| ENSMUSG00 | 2310003F16f | 4934 | 4.46382979 | 2.76383525 | 3.54761448 | 3.15918995 | 2.65783919 | 3.0956218 |
| ENSMUSG00 | Ell3 | 2701 | 0.0883128 | 0.21408268 | 0.14728493 | 0.1469465 | 0.07556664 | 0.18176361 |
| ENSMUSG00 | Arhgap1 | 4394 | 23.3520472 | 21.3187172 | 22.1210407 | 22.2289701 | 28.4163727 | 19.1245252 |
| ENSMUSG00 | Pdia3 | 3448 | 382.68109 | 326.683954 | 370.844233 | 351.475531 | 282.539318 | 499.462876 |
| ENSMUSG00 | F2 | 4243 | 0.03747865 | 0 | 0.09375833 | 0 | 0.24052027 | 0 |
| ENSMUSG00 | Lrp4 | 9176 | 0.05632315 | 0.20480288 | 0.01445134 | 0.00393222 | 0.02780426 | 0.00891717 |
| ENSMUSG00 | Mtap1a | 12260 | 0 | 0 | 0.00360537 | 0.00294307 | 0.05826829 | 0.00687406 |
| ENSMUSG00 | Arfgap2 | 4222 | 6.90212393 | 6.83649428 | 6.53291095 | 7.11898226 | 8.53259624 | 3.90837441 |
| ENSMUSG00 | Pacsin3 | 4189 | 12.4894276 | 13.8152106 | 10.9634077 | 12.1709069 | 11.937419 | 15.3204168 |

| | | | | | | | | |
|---|---|---|---|---|---|---|---|---|
| ENSMUSG00 | Adal | 5539 | 8.1750302 | 6.68120375 | 9.67987644 | 5.83019134 | 7.5263674 | 7.2926053 |
| ENSMUSG00 | Hao1 | 2275 | 0 | 0 | 0 | 0 | 0 | 0 |
| ENSMUSG00 | Tubgcp4 | 8376 | 5.02164498 | 4.29167692 | 5.74687286 | 8.83527618 | 7.63324366 | 5.08306016 |
| ENSMUSG00 | 6330527O06I | 2927 | 0 | 0 | 0 | 0 | 0 | 0 |
| ENSMUSG00 | Ubr1 | 10058 | 10.1661459 | 9.67274077 | 9.03549306 | 9.03306702 | 12.1960039 | 9.72699652 |
| ENSMUSG00 | Snap25 | 6928 | 0 | 0 | 0 | 0 | 0 | 0 |
| ENSMUSG00 | Mkks | 4136 | 33.5845379 | 26.6446792 | 48.391186 | 34.8519748 | 30.8428287 | 37.7136513 |
| ENSMUSG00 | Jag1 | 6390 | 37.3975263 | 23.5879798 | 48.0548049 | 31.4348579 | 31.9014666 | 42.0174568 |
| ENSMUSG00 | 2210009G21I | 5892 | 1.26850481 | 0.85871969 | 2.46065929 | 2.27809242 | 1.57617115 | 0.41661873 |
| ENSMUSG00 | Mtch2 | 6694 | 55.5590858 | 49.3597899 | 60.3730879 | 47.6817696 | 42.1001903 | 53.058015 |
| ENSMUSG00 | Cdan1 | 13193 | 0.59062186 | 0.53325404 | 0.70693465 | 0.74663823 | 1.29180698 | 0.43827966 |
| ENSMUSG00 | Haus2 | 3211 | 12.9505552 | 14.4514306 | 12.9122793 | 16.0689267 | 19.7049847 | 18.7720189 |
| ENSMUSG00 | Lrrc57 | 2932 | 17.11167 | 15.6622372 | 14.8193764 | 18.2256166 | 15.3496806 | 19.1536433 |
| ENSMUSG00 | Snap23 | 4395 | 36.6166519 | 30.5893455 | 34.416088 | 28.2581109 | 29.6754076 | 44.675797 |
| ENSMUSG00 | Zfp106 | 11534 | 8.03450885 | 6.95181548 | 8.30077961 | 8.19619877 | 15.1433401 | 6.7228888 |
| ENSMUSG00 | Vps39 | 8115 | 9.60696208 | 9.20975623 | 10.6705373 | 10.0531735 | 11.0918702 | 7.42784384 |
| ENSMUSG00 | Ehd4 | 3998 | 7.45788154 | 7.55700331 | 6.31296958 | 7.01244337 | 9.96788382 | 6.97897848 |
| ENSMUSG00 | Itpka | 1811 | 0.54880564 | 0.47893758 | 0.78103754 | 0 | 1.18338365 | 1.34038812 |
| ENSMUSG00 | Ltk | 4058 | 0.77394849 | 0.54622384 | 0.15249527 | 0.60462766 | 1.16940684 | 0.08737563 |
| ENSMUSG00 | Tyro3 | 5849 | 0.27187882 | 0.48606601 | 0.20404339 | 0 | 0 | 0 |
| ENSMUSG00 | Ubox5 | 5578 | 5.9013155 | 5.53738068 | 4.5723312 | 3.74533951 | 6.48578352 | 4.0144279 |
| ENSMUSG00 | Oxt | 534 | 0 | 0 | 0 | 0.40541623 | 0 | 0 |
| ENSMUSG00 | Ptpra | 3604 | 70.1015757 | 58.5082882 | 76.0531711 | 63.8243712 | 70.5930937 | 61.9279614 |
| ENSMUSG00 | Rtf1 | 5884 | 3.16881719 | 3.0710258 | 2.2010775 | 3.05998304 | 2.06394926 | 4.32018419 |
| ENSMUSG00 | Ndufaf1 | 1831 | 27.7267879 | 26.3696423 | 33.5798818 | 31.5102071 | 24.6910808 | 30.611351 |
| ENSMUSG00 | Nusap1 | 4249 | 0.17777221 | 1.1794281 | 0.27047492 | 0.11039458 | 0.25218967 | 0.48784951 |
| ENSMUSG00 | 4930402H24I | 10803 | 16.5454633 | 12.8550705 | 15.0899192 | 14.2284097 | 15.9743778 | 13.8733817 |
| ENSMUSG00 | Atrn | 13645 | 4.10228777 | 5.41015972 | 3.86786373 | 4.59057745 | 6.51806325 | 4.23761343 |
| ENSMUSG00 | Chac1 | 1634 | 1.07052702 | 0.76673653 | 0.13525656 | 0.66246104 | 0 | 0.10015167 |
| ENSMUSG00 | Dll4 | 3841 | 17.5747998 | 14.4270785 | 18.4241475 | 8.14452814 | 11.7170694 | 7.31394908 |
| ENSMUSG00 | Spint1 | 2756 | 194.348907 | 170.593293 | 227.29627 | 143.608108 | 173.59336 | 141.845938 |
| ENSMUSG00 | Gfra4 | 2165 | 12.6336123 | 13.8438647 | 14.0261737 | 8.09971069 | 10.0874313 | 12.5349959 |
| ENSMUSG00 | Ppp1r14d | 1130 | 2.32200152 | 2.81442936 | 0.89968353 | 0.41510317 | 1.39983863 | 0.86892656 |
| ENSMUSG00 | Adam33 | 5926 | 0.01341731 | 0.04878816 | 0.14917936 | 0.29226091 | 0.42191907 | 0.12887106 |
| ENSMUSG00 | Siglec1 | 7776 | 0 | 0 | 0 | 0.1670465 | 0 | 0.10171869 |
| ENSMUSG00 | Rad51 | 4882 | 0 | 0.93767145 | 0.11770257 | 0.79820992 | 0.4076257 | 0.02793388 |
| ENSMUSG00 | Rpusd2 | 2703 | 1.02955369 | 1.47964288 | 1.04658452 | 0.69414218 | 1.32143778 | 2.6134412 |
| ENSMUSG00 | Casc5 | 8594 | 0.11564894 | 0.38127508 | 0.03600336 | 0.75153432 | 0.10687395 | 0.03173684 |
| ENSMUSG00 | 1700037H04I | 2194 | 9.0419256 | 7.24773287 | 12.249189 | 9.71945684 | 5.46545046 | 10.2435291 |
| ENSMUSG00 | Spef1 | 4213 | 0.86814721 | 0.52612778 | 0.66098175 | 0.25693362 | 0.41179606 | 0.72507942 |
| ENSMUSG00 | Cdc25b | 4089 | 9.39197705 | 8.9915029 | 4.4212653 | 10.3154586 | 9.07218763 | 7.21053681 |
| ENSMUSG00 | D2Ertd750e | 1885 | 0.1054522 | 0.38344649 | 0.09379701 | 0.63167505 | 0.48725452 | 0.99838202 |
| ENSMUSG00 | Ivd | 5629 | 45.469139 | 47.2533595 | 42.3093854 | 47.626504 | 45.2701583 | 30.2739289 |
| ENSMUSG00 | Smox | 3724 | 1.67605006 | 3.01488749 | 1.57863725 | 3.55588355 | 2.91853331 | 1.45015536 |
| ENSMUSG00 | Adra1d | 2486 | 0 | 0 | 0 | 0 | 0 | 0 |
| ENSMUSG00 | Prnd | 3453 | 0.46053264 | 0 | 0.21761695 | 0 | 0 | 0.30805414 |
| ENSMUSG00 | Rassf2 | 13121 | 0.0242393 | 0 | 0.25602775 | 0.00274995 | 0.01944455 | 0.15382389 |
| ENSMUSG00 | Slc23a2 | 8331 | 11.0471657 | 7.93563781 | 14.7976162 | 9.51100345 | 9.48743922 | 12.0871406 |
| ENSMUSG00 | 5730494N06I | 3225 | 19.3784854 | 21.7548712 | 18.3797431 | 18.8745455 | 14.9044484 | 23.3081878 |
| ENSMUSG00 | Pcna | 2382 | 9.6301059 | 9.62919667 | 11.6906638 | 11.3911408 | 9.61832189 | 13.3052606 |
| ENSMUSG00 | Fsip1 | 1963 | 0 | 0 | 0.06755249 | 0.38600251 | 0 | 0.0416831 |
| ENSMUSG00 | 4921508D12I | 1071 | 0 | 0 | 0 | 0 | 0 | 0 |
| ENSMUSG00 | Gpcpd1 | 7258 | 14.6138912 | 10.2640176 | 15.5053585 | 14.8792449 | 15.3191613 | 20.5330158 |
| ENSMUSG00 | Rasgrp1 | 5715 | 2.35819911 | 3.17027161 | 5.15881527 | 1.93195199 | 2.38605242 | 3.56025912 |
| ENSMUSG00 | Fam98b | 3382 | 2.26871902 | 0.74089148 | 1.2416248 | 1.67500916 | 0.70911877 | 1.32260226 |
| ENSMUSG00 | Chgb | 2732 | 0 | 0 | 0 | 0 | 0 | 0.22961812 |
| ENSMUSG00 | Spred1 | 6895 | 4.07645029 | 5.69571434 | 3.15406915 | 7.94380618 | 4.79551726 | 3.9478013 |
| ENSMUSG00 | Mcm8 | 3940 | 1.12001481 | 0.51366258 | 0.45996842 | 2.26199619 | 1.83902167 | 0.69224973 |
| ENSMUSG00 | Tmco5 | 1295 | 0 | 0 | 0 | 0 | 0.39402607 | 0 |
| ENSMUSG00 | Fermt1 | 8486 | 4.41779615 | 4.28147271 | 1.63556195 | 3.26549731 | 3.39133584 | 1.42704924 |
| ENSMUSG00 | Crls1 | 2523 | 25.621249 | 25.611581 | 24.9828887 | 29.2460488 | 20.5278523 | 36.0418737 |
| ENSMUSG00 | Bmp2 | 3552 | 0 | 0.06783001 | 0.16177476 | 0 | 0.30167621 | 0.15357342 |
| ENSMUSG00 | Slc27a2 | 3782 | 3.50041645 | 4.95624696 | 1.04018086 | 1.15439645 | 2.69838053 | 4.75250853 |
| ENSMUSG00 | Hdc | 4494 | 0 | 0.38600621 | 0 | 0 | 0.13625201 | 0.13958983 |
| ENSMUSG00 | Gabpb1 | 9209 | 2.2362188 | 2.29185163 | 3.29749877 | 2.85239748 | 3.71241957 | 3.19275286 |
| ENSMUSG00 | Usp8 | 4345 | 76.1804674 | 49.5837935 | 80.3059491 | 57.6150115 | 74.1732082 | 69.5142359 |
| ENSMUSG00 | Usp50 | 3103 | 0.1537434 | 0.03105797 | 0.21367311 | 0.24419044 | 0.23021891 | 0.35159058 |
| ENSMUSG00 | Trpm7 | 10385 | 18.9876922 | 17.4742553 | 19.5535163 | 20.4991876 | 22.9753812 | 22.6128689 |
| ENSMUSG00 | 2010106G01I | 8672 | 12.7490765 | 16.8530303 | 15.5358876 | 16.2019697 | 17.9816195 | 15.3702908 |
| ENSMUSG00 | Stard7 | 3763 | 75.1582483 | 42.3087975 | 69.1277827 | 49.6307899 | 62.6066376 | 58.2675591 |
| ENSMUSG00 | Dusp2 | 1991 | 0.31948151 | 0 | 0 | 0 | 0 | 0.02739793 |
| ENSMUSG00 | Fahd2a | 1403 | 36.1284659 | 38.0545815 | 37.8377858 | 29.2153974 | 32.041509 | 33.1261477 |
| ENSMUSG00 | Mrps5 | 6472 | 6.63410367 | 4.30342453 | 5.64816505 | 4.85033864 | 6.97749422 | 5.46167065 |
| ENSMUSG00 | Mal | 2954 | 345.767707 | 373.273648 | 367.844924 | 341.655676 | 337.216286 | 264.842841 |
| ENSMUSG00 | Prom2 | 6812 | 5.35753539 | 3.7915345 | 2.0894001 | 5.66231732 | 7.56556659 | 2.80273745 |
| ENSMUSG00 | Mall | 2002 | 0.13900518 | 0 | 0 | 0 | 0 | 0 |
| ENSMUSG00 | Nphp1 | 3120 | 11.4806692 | 12.6798298 | 12.0705033 | 8.02594196 | 18.2517421 | 11.6266892 |
| ENSMUSG00 | Bub1 | 4329 | 0.17448698 | 0.72352015 | 0.02042127 | 0 | 0 | 0 |
| ENSMUSG00 | Acoxl | 3158 | 0.03776645 | 0.15258531 | 0.01399678 | 0 | 0.04847344 | 0 |
| ENSMUSG00 | Bcl2l11 | 6027 | 5.9695885 | 5.70849837 | 6.8719308 | 6.52553983 | 4.07228916 | 4.11359666 |
| ENSMUSG00 | 2310003L22F | 3627 | 7.63980422 | 6.04489418 | 7.50712305 | 5.45160199 | 4.69887222 | 8.91106907 |
| ENSMUSG00 | Fbln7 | 3219 | 1.35852838 | 1.93105034 | 1.38688603 | 0.65012693 | 0.44384546 | 0.17793334 |
| ENSMUSG00 | Zc3h8 | 2986 | 12.8479701 | 10.699133 | 11.5611652 | 10.972035 | 10.8170448 | 15.1809948 |
| ENSMUSG00 | Ttl | 6255 | 2.01478614 | 1.45599322 | 2.04225941 | 0.77874916 | 1.07681561 | 1.20784574 |
| ENSMUSG00 | Polr1b | 6024 | 3.51094217 | 2.28773613 | 2.50212956 | 3.23444622 | 4.6249006 | 1.97406088 |
| ENSMUSG00 | Slc20a1 | 4703 | 4.82678702 | 3.04303069 | 3.77825662 | 5.16334594 | 4.99088515 | 1.94860276 |
| ENSMUSG00 | Il1b | 2017 | 0 | 0 | 0 | 0 | 0 | 0 |
| ENSMUSG00 | Il1a | 1967 | 0.14147858 | 0 | 0 | 0 | 0.12970609 | 0 |
| ENSMUSG00 | Pdyn | 3001 | 0 | 0 | 0 | 0 | 0 | 0 |
| ENSMUSG00 | Tgm3 | 2284 | 0.01740608 | 0 | 0 | 0 | 0 | 0 |
| ENSMUSG00 | Tgm6 | 3531 | 0 | 0 | 0 | 0 | 0 | 0 |
| ENSMUSG00 | Snrpb | 2277 | 46.4948813 | 50.2180181 | 50.1420116 | 40.8993223 | 36.3033504 | 47.314372 |
| ENSMUSG00 | Nop56 | 3319 | 15.8351146 | 17.1171483 | 11.9993554 | 13.1760886 | 12.5144531 | 16.0081342 |
| ENSMUSG00 | Idh3b | 3597 | 135.447154 | 128.256046 | 148.764946 | 112.569559 | 112.521883 | 119.031773 |
| ENSMUSG00 | Cpxm1 | 2784 | 0 | 0.0346167 | 0 | 0 | 0 | 0 |
| ENSMUSG00 | 1700020A23I | 730 | 0 | 0 | 0 | 0 | 0 | 0 |
| ENSMUSG00 | Vps16 | 3659 | 20.6002709 | 18.9111044 | 20.6331642 | 24.6529411 | 29.3969938 | 19.8428778 |
| ENSMUSG00 | Lpin3 | 3673 | 10.2067569 | 9.47198781 | 10.1328483 | 6.63092297 | 11.1555077 | 6.16334075 |
| ENSMUSG00 | Otor | 945 | 0 | 0 | 0 | 0 | 0 | 0 |
| ENSMUSG00 | Pcsk2 | 8291 | 0.11508039 | 0.19760452 | 0.24523999 | 0.20018985 | 0.1046253 | 0.15790408 |
| ENSMUSG00 | Bfsp1 | 2382 | 0 | 0 | 0.05566983 | 0.36354705 | 0 | 0 |
| ENSMUSG00 | Rrbp1 | 8760 | 4.63813939 | 5.42372511 | 4.15779893 | 4.55144511 | 4.94536451 | 5.34284029 |
| ENSMUSG00 | Snx5 | 4242 | 30.6835085 | 40.7119775 | 38.6479575 | 39.3227939 | 42.0167682 | 37.5106661 |
| ENSMUSG00 | 8430408I07R | 3457 | 8.164996 | 11.2486141 | 9.02704691 | 6.81561327 | 7.05542599 | 4.56023763 |
| ENSMUSG00 | Csrp2bp | 4612 | 10.6284708 | 14.2093585 | 7.97396648 | 12.2750928 | 14.5821256 | 8.75839998 |
| ENSMUSG00 | Polr3f | 7401 | 11.6725658 | 9.46670543 | 13.6768302 | 10.145485 | 10.8106144 | 10.2118667 |
| ENSMUSG00 | Rbbp9 | 2544 | 62.1491926 | 74.5147262 | 61.4030319 | 71.0578 | 69.1583899 | 101.025521 |
| ENSMUSG00 | Sec23b | 6346 | 49.0772824 | 34.0251256 | 66.3515217 | 41.8701821 | 39.2547694 | 38.8360606 |
| ENSMUSG00 | Dtd1 | 2342 | 10.0492077 | 16.5216537 | 11.2297594 | 12.8028091 | 13.1160881 | 12.5309615 |
| ENSMUSG00 | 1700010M22 | 772 | 0 | 0 | 0 | 0 | 0 | 0 |
| ENSMUSG00 | Xrn2 | 3952 | 54.7744411 | 34.1889635 | 46.8078727 | 43.4500126 | 47.8114043 | 57.8412833 |
| ENSMUSG00 | Nkx2-2 | 2827 | 0 | 0 | 0 | 0 | 0 | 0 |
| ENSMUSG00 | Cd93 | 6675 | 0.54198483 | 0.26710088 | 0.27812396 | 0.0216222 | 0.28284283 | 0.38000621 |
| ENSMUSG00 | Napb | 7041 | 3.59103616 | 1.67670478 | 2.94427842 | 3.36683756 | 4.30473899 | 4.00926738 |
| ENSMUSG00 | Gzf1 | 4561 | 14.5563807 | 10.4381067 | 15.4478706 | 18.2269306 | 17.7658156 | 12.4981355 |
| ENSMUSG00 | Cst8 | 945 | 0.37862362 | 0 | 0.23387219 | 0 | 0 | 0.25975847 |
| ENSMUSG00 | Cst12 | 874 | 0 | 0 | 0 | 0 | 0 | 0 |
| ENSMUSG00 | 8030411F24I | 599 | 0 | 0 | 0 | 0 | 0 | 0 |
| ENSMUSG00 | Cst9 | 684 | 0 | 0 | 0 | 0 | 0 | 0 |
| ENSMUSG00 | 9230104L09F | 994 | 0 | 0 | 0 | 0 | 0 | 0 |
| ENSMUSG00 | Cst3 | 1262 | 268.700522 | 295.151504 | 375.925817 | 233.618372 | 145.194703 | 371.557526 |
| ENSMUSG00 | Acss1 | 4037 | 246.263389 | 164.552463 | 226.680891 | 167.325231 | 230.080039 | 147.210085 |
| ENSMUSG00 | Gins1 | 1507 | 0.39570817 | 0 | 0.90926154 | 0.88588961 | 0.54175316 | 0.76014257 |
| ENSMUSG00 | Nsfl1c | 4150 | 11.8116886 | 15.9189427 | 13.8463605 | 11.8070883 | 11.8651693 | 12.3754568 |
| ENSMUSG00 | Sdcbp2 | 1390 | 0.31461172 | 0.76266311 | 0.92219673 | 0 | 0 | 0.25508655 |
| ENSMUSG00 | Snph | 11759 | 0 | 0 | 0 | 0 | 0 | 0 |
| ENSMUSG00 | Fam110a | 3725 | 3.05236709 | 4.42409751 | 3.80907157 | 2.363495 | 1.7396912 | 2.44556497 |
| ENSMUSG00 | Angpt4 | 2785 | 0 | 0.22492774 | 0.0158714 | 0 | 0 | 0.14690111 |
| ENSMUSG00 | Slc52a3 | 2877 | 4.87788829 | 4.68967807 | 2.181669 | 4.10108744 | 3.56492928 | 3.50768736 |
| ENSMUSG00 | Tbc1d20 | 4352 | 14.5063656 | 15.0582632 | 13.8841727 | 10.4548387 | 17.3761693 | 11.8010445 |
| ENSMUSG00 | Rbck1 | 3819 | 31.5004148 | 26.5851882 | 32.2804242 | 32.1422141 | 37.7988 | 27.117519 |
| ENSMUSG00 | Defb22 | 626 | 0 | 0 | 0 | 0 | 0 | 0 |
| ENSMUSG00 | Tpx2 | 4977 | 0 | 0.26140927 | 0 | 0.31173959 | 0 | 0.06576164 |
| ENSMUSG00 | Mylk2 | 2941 | 0 | 0 | 0 | 0 | 0 | 0 |
| ENSMUSG00 | Pdrg1 | 2668 | 29.1162702 | 32.1844977 | 32.6212252 | 31.632647 | 27.559598 | 32.1202836 |
| ENSMUSG00 | BC020535 | 2256 | 0 | 0.68349563 | 0.82290665 | 0 | 0.9725772 | 0.90673668 |
| ENSMUSG00 | Kif3b | 5676 | 8.7341586 | 9.39788433 | 7.15671137 | 8.15596602 | 7.81217769 | 5.55007294 |
| ENSMUSG00 | Dnmt3b | 5369 | 0.31839927 | 0.41284715 | 0.9550035 | 1.20967927 | 0.59874496 | 1.07188469 |
| ENSMUSG00 | Mapre1 | 7802 | 23.6433517 | 22.672703 | 26.6446126 | 26.8927312 | 25.042296 | 27.6521913 |
| ENSMUSG00 | Sun5 | 1362 | 0 | 0 | 0 | 0 | 0 | 0 |
| ENSMUSG00 | Bpifb2 | 2120 | 0 | 0 | 0 | 0 | 0 | 0 |
| ENSMUSG00 | Bpifa3 | 1179 | 0 | 0 | 0 | 0 | 0 | 0 |
| ENSMUSG00 | Bpifa1 | 1550 | 0 | 0 | 0 | 0 | 0 | 0 |
| ENSMUSG00 | Bpifa5 | 1080 | 0 | 0 | 0 | 0 | 0 | 0 |
| ENSMUSG00 | Bpifb1 | 2702 | 0.02942671 | 0 | 0 | 0 | 0 | 0.01009424 |
| ENSMUSG00 | Cdk5rap1 | 3649 | 12.104231 | 9.08530342 | 6.12938684 | 3.38176466 | 4.89428105 | 2.10034904 |
| ENSMUSG00 | Snta1 | 3085 | 14.2398074 | 9.93406068 | 8.33888902 | 9.36846602 | 10.8668813 | 7.3734357 |
| ENSMUSG00 | Necab3 | 2656 | 0 | 0 | 0.06656904 | 0 | 0 | 0 |
| ENSMUSG00 | E2f1 | 4449 | 0 | 0 | 0 | 0.06488118 | 0.1605685 | 0 |
| ENSMUSG00 | 2010011I20R | 3614 | 9.25134452 | 13.226605 | 16.3769418 | 16.2339249 | 13.342536 | 12.2109481 |
| ENSMUSG00 | Aurka | 3307 | 0 | 0.80140741 | 0.81533487 | 1.4947808 | 0.63262214 | 0.35464454 |
| ENSMUSG00 | Cstf1 | 3620 | 10.6197651 | 10.6356263 | 10.1102558 | 10.7847437 | 11.2483558 | 14.5565202 |
| ENSMUSG00 | Pkia | 3680 | 14.9623208 | 19.4971773 | 19.6145678 | 26.7477765 | 12.1880936 | 27.4302826 |
| ENSMUSG00 | Stmn2 | 1293 | 0 | 0 | 0 | 0 | 0 | 0 |
| ENSMUSG00 | 2410001C21I | 2771 | 41.0036677 | 33.8052846 | 33.8014094 | 32.9699519 | 35.7792307 | 26.2805077 |
| ENSMUSG00 | 1700029I11R | 696 | 1.14239887 | 0 | 0.44455877 | 0 | 0 | 0.1175631 |
| ENSMUSG00 | Tpd52 | 6016 | 27.3913675 | 32.8718681 | 36.8177253 | 28.6209303 | 22.7311648 | 33.0732203 |
| ENSMUSG00 | Pag1 | 11801 | 1.2868904 | 0.46140734 | 1.38212694 | 1.40341144 | 0.6442615 | 1.37055004 |
| ENSMUSG00 | Rae1 | 2595 | 36.3237936 | 34.3711385 | 35.1230059 | 29.1298202 | 29.3573715 | 24.1845645 |
| ENSMUSG00 | Rbm38 | 2530 | 0.20427717 | 0.28569037 | 0.06988434 | 0.31375691 | 0.10084264 | 0.23717078 |
| ENSMUSG00 | Pck1 | 3176 | 0.93881015 | 1.71444205 | 0.22267931 | 1.19288875 | 3.24538033 | 1.36544951 |
| ENSMUSG00 | Zbp1 | 4132 | 0 | 0.10495595 | 0 | 0 | 0.23463242 | 0.2442308 |
| ENSMUSG00 | Ankrd60 | 1281 | 0 | 0 | 0 | 0 | 0 | 0.06387503 |
| ENSMUSG00 | 1700021F07I | 642 | 0 | 0 | 0 | 0 | 0 | 0 |
| ENSMUSG00 | Rab22a | 7484 | 17.6626099 | 16.2832058 | 15.6631865 | 16.0498565 | 20.5632749 | 14.0272697 |
| ENSMUSG00 | Zdbf2 | 12855 | 0 | 0 | 0 | 0 | 0 | 0 |
| ENSMUSG00 | Stx16 | 4020 | 13.8451922 | 20.0177508 | 15.9764373 | 17.7897045 | 21.2990693 | 14.2140222 |
| ENSMUSG00 | Gnas | 10102 | 0.46831342 | 0.22895954 | 0.48568646 | 0.33931838 | 0.32832255 | 0.39688893 |
| ENSMUSG00 | Edn3 | 4326 | 34.5631905 | 34.708496 | 93.2468838 | 35.197926 | 36.1053482 | 60.9423632 |
| ENSMUSG00 | Phactr3 | 6514 | 0 | 0 | 0 | 0 | 0 | 0 |
| ENSMUSG00 | Fabp9 | 579 | 0 | 0 | 0 | 0 | 0 | 0 |
| ENSMUSG00 | Fabp12 | 2945 | 0.01349931 | 0 | 0.24014584 | 0 | 0.74503707 | 0.51863491 |
| ENSMUSG00 | Impa1 | 4793 | 32.9954729 | 25.3147217 | 36.3722238 | 37.3844008 | 29.6065828 | 35.5884819 |
| ENSMUSG00 | Fabp5 | 3349 | 0.77160533 | 0.04316492 | 0.07919112 | 0.06464385 | 0.31996235 | 0.26061166 |
| ENSMUSG00 | Snx16 | 2930 | 12.0080547 | 10.9036216 | 10.3942218 | 14.6175382 | 8.2199486 | 10.3699482 |
| ENSMUSG00 | Chmp4c | 1851 | 2.64177423 | 3.87886541 | 3.4625971 | 2.45615214 | 1.90211774 | 3.66903575 |
| ENSMUSG00 | Ptpn1 | 5522 | 13.9021791 | 13.8398583 | 10.4621168 | 8.61865569 | 11.1441161 | 9.7056621 |
| ENSMUSG00 | Nfatc2 | 14020 | 1.08888049 | 1.01734571 | 1.14445571 | 1.71917302 | 0.94264132 | 1.35984114 |
| ENSMUSG00 | Atp9a | 5878 | 11.3355198 | 10.4931347 | 7.87331229 | 5.81315008 | 11.3372655 | 5.93935663 |
| ENSMUSG00 | Sall4 | 4892 | 0 | 0 | 0 | 0 | 0 | 0 |
| ENSMUSG00 | Lrrcc1 | 7671 | 7.37997709 | 9.65487701 | 4.25826646 | 10.1035369 | 8.99330728 | 8.62221356 |
| ENSMUSG00 | Zfp64 | 6481 | 4.06081285 | 2.49073571 | 3.59425571 | 2.40509848 | 3.55869802 | 2.3609123 |
| ENSMUSG00 | E2f5 | 1953 | 39.3280022 | 41.7467794 | 36.1672018 | 40.2574373 | 38.1979321 | 60.9594472 |
| ENSMUSG00 | Car13 | 2292 | 0 | 0.37842756 | 0.52070234 | 0.11019822 | 0 | 0 |
| ENSMUSG00 | Car1 | 1173 | 0 | 0 | 0 | 0 | 0 | 0 |
| ENSMUSG00 | Car3 | 1500 | 0 | 0.35336724 | 0.58935791 | 0 | 0.34017584 | 0 |
| ENSMUSG00 | Dok5 | 1760 | 0 | 0 | 0 | 0 | 0 | 0 |
| ENSMUSG00 | Car2 | 1782 | 2508.34326 | 2144.43188 | 2839.55914 | 2322.8778 | 1865.26888 | 1993.98735 |
| ENSMUSG00 | Cypt12 | 508 | 0 | 0 | 0 | 0 | 0 | 0 |
| ENSMUSG00 | Psma7 | 3167 | 47.262199 | 44.0631312 | 52.8970587 | 39.8645639 | 35.76841 | 64.8063342 |
| ENSMUSG00 | Ntsr1 | 4056 | 0 | 0 | 0 | 0 | 0 | 0 |
| ENSMUSG00 | 1600027N09I | 3050 | 10.0105603 | 13.4922038 | 15.8546939 | 15.1307983 | 14.5717945 | 18.4126171 |
| ENSMUSG00 | Col9a3 | 3820 | 0 | 0.05045701 | 0 | 0 | 0.36065763 | 0.23561861 |
| ENSMUSG00 | 2310003C23I | 2458 | 12.227479 | 17.5847187 | 12.3382345 | 12.2426465 | 9.96446721 | 9.94226069 |
| ENSMUSG00 | Nkain4 | 1230 | 0 | 0 | 0 | 0 | 0 | 0.39914106 |
| ENSMUSG00 | Arfgap1 | 4038 | 17.8200643 | 17.9476001 | 12.0411162 | 17.8176317 | 17.6658924 | 16.2040266 |
| ENSMUSG00 | Chrna4 | 4930 | 1.87891013 | 1.21199165 | 0.7083054 | 1.13442534 | 1.33517292 | 0.7358067 |
| ENSMUSG00 | Srms | 3371 | 0 | 0 | 0 | 0 | 0.06054747 | 0 |
| ENSMUSG00 | BC006779 | 11012 | 0.91698983 | 0.63886856 | 0.5298441 | 0.65859889 | 0.5745796 | 0.96347936 |
| ENSMUSG00 | Stmn3 | 1150 | 0 | 0 | 0 | 0 | 0 | 0 |
| ENSMUSG00 | Zgpat | 3968 | 19.8176261 | 16.2726392 | 10.0813175 | 14.3127868 | 22.3240394 | 9.28629981 |
| ENSMUSG00 | Zbtb46 | 8502 | 2.09485477 | 1.84198938 | 3.24937096 | 1.72304282 | 2.58072708 | 1.26396235 |
| ENSMUSG00 | Oprl1 | 3933 | 0 | 0 | 0 | 0 | 0.01297391 | 0.10402227 |
| ENSMUSG00 | Pcmtd2 | 4245 | 30.2965794 | 40.3540169 | 44.0144147 | 48.2793906 | 45.328731 | 47.0832877 |
| ENSMUSG00 | Raly | 3720 | 3.66562629 | 4.8445509 | 6.30945662 | 4.98552981 | 3.20972364 | 5.86551382 |
| ENSMUSG00 | a | 1162 | 0 | 0 | 0 | 0 | 0 | 0 |
| ENSMUSG00 | Ahcy | 2795 | 1.02411256 | 0.6378885 | 0.56932606 | 0.54219888 | 0.49292027 | 0.185409 |
| ENSMUSG00 | Itch | 9757 | 18.6288877 | 13.24053 | 14.4107884 | 13.6902459 | 20.4482043 | 14.5304474 |
| ENSMUSG00 | Armc1 | 2922 | 21.2927197 | 18.733675 | 29.6040407 | 31.4019984 | 25.5830392 | 28.5627639 |
| ENSMUSG00 | Mtfr1 | 4319 | 31.1582083 | 24.6120147 | 45.2048022 | 31.0611349 | 25.2001064 | 39.4121379 |
| ENSMUSG00 | Map1lc3a | 1280 | 22.3935168 | 21.3074424 | 23.5512945 | 20.4934737 | 14.3910326 | 27.2959476 |
| ENSMUSG00 | Ggt7 | 3247 | 0.53872533 | 0.50457008 | 0.10890506 | 0.85565674 | 0.58145239 | 0.39479767 |
| ENSMUSG00 | Acss2 | 7912 | 6.9793178 | 4.42764705 | 3.80453174 | 7.08689297 | 6.37829698 | 1.71673033 |
| ENSMUSG00 | Dnajc5b | 1336 | 0 | 0 | 0 | 0 | 0 | 0 |
| ENSMUSG00 | Gss | 6146 | 7.47760096 | 11.0548134 | 8.4433719 | 7.94321612 | 6.33470953 | 6.75874969 |
| ENSMUSG00 | Procr | 2184 | 0 | 0.06619017 | 0.5666903 | 0 | 0 | 0.33718647 |
| ENSMUSG00 | Mmp24 | 4674 | 0 | 0 | 0 | 0 | 0 | 0.0058354 |
| ENSMUSG00 | Eif6 | 2811 | 18.7533857 | 20.2276776 | 20.5048749 | 17.1489191 | 15.9559532 | 17.4262725 |
| ENSMUSG00 | Hps3 | 10164 | 3.34033652 | 4.30947225 | 4.07053572 | 3.38313542 | 2.4097462 | 3.31943416 |
| ENSMUSG00 | Nfs1 | 4170 | 36.0850105 | 33.9153975 | 37.8418656 | 33.4775612 | 37.5661808 | 36.7390045 |
| ENSMUSG00 | Rbm39 | 7105 | 79.6900146 | 84.2601488 | 88.3539162 | 102.517696 | 95.8908051 | 145.789761 |
| ENSMUSG00 | Epb4.1l1 | 11759 | 2.96839118 | 5.72877338 | 5.87904435 | 5.43731464 | 4.55197451 | 5.56672627 |
| ENSMUSG00 | 0610011L14F | 3897 | 12.8335629 | 10.0032927 | 12.8851151 | 6.80531248 | 11.1166007 | 10.2183663 |
| ENSMUSG00 | Tbl1xr1 | 8111 | 7.90110154 | 8.03800471 | 8.621294 | 6.10783469 | 8.06507383 | 8.29235118 |
| ENSMUSG00 | Ndrg3 | 6066 | 57.0968919 | 49.4653767 | 69.3559416 | 51.8092001 | 56.2080849 | 51.7345861 |
| ENSMUSG00 | Dsn1 | 3580 | 1.47694942 | 0.8883534 | 0.65438483 | 0.93732686 | 1.53934318 | 0.93708956 |
| ENSMUSG00 | Sla2 | 4706 | 0 | 0 | 0 | 0.08433967 | 0.08674267 | 0 |
| ENSMUSG00 | 1110008F13I | 3144 | 5.0453043 | 6.29918185 | 6.48124988 | 5.08407943 | 3.94383248 | 4.90145394 |
| ENSMUSG00 | Samhd1 | 11107 | 2.85271612 | 2.17786927 | 2.73799118 | 3.4532469 | 4.03359665 | 3.26843836 |
| ENSMUSG00 | Rbl1 | 5971 | 2.97616811 | 2.16278119 | 2.26524267 | 1.91559339 | 3.76010808 | 3.12441019 |
| ENSMUSG00 | Rpn2 | 4560 | 52.571392 | 49.539044 | 45.1130214 | 40.6318637 | 42.4995999 | 29.3262185 |
| ENSMUSG00 | Ghrh | 661 | 0 | 0 | 0 | 0 | 0 | 0 |
| ENSMUSG00 | Src | 4353 | 3.80841612 | 4.28397728 | 4.27497724 | 3.58085073 | 4.32546122 | 6.11533377 |
| ENSMUSG00 | Ctnnbl1 | 2686 | 14.652988 | 15.4103328 | 14.7284622 | 12.8288729 | 15.3687037 | 12.8046612 |
| ENSMUSG00 | Tti1 | 4764 | 12.3672591 | 11.0755991 | 12.0525177 | 11.5426188 | 15.5842521 | 5.77095642 |
| ENSMUSG00 | Rprd1b | 6344 | 7.99621582 | 11.2946468 | 11.7472111 | 7.78630501 | 11.7190148 | 8.33200676 |
| ENSMUSG00 | Ralgapb | 13129 | 3.95768246 | 3.9785287 | 3.46436869 | 5.62845816 | 5.37508399 | 4.55581414 |
| ENSMUSG00 | Fam83d | 3987 | 0 | 0.2296319 | 0 | 0 | 0 | 0 |
| ENSMUSG00 | Dhx35 | 4791 | 5.94963046 | 2.60494438 | 4.12402919 | 2.3723323 | 3.64245889 | 2.90337425 |
| ENSMUSG00 | Wisp2 | 2138 | 0 | 0 | 0 | 0 | 0 | 0 |
| ENSMUSG00 | Skil | 6807 | 4.92928244 | 5.21718934 | 7.49359877 | 7.71785762 | 7.59361227 | 4.96448921 |
| ENSMUSG00 | Slc2a10 | 3564 | 0 | 0 | 0.23564394 | 0.71880616 | 0 | 0 |
| ENSMUSG00 | Zmat3 | 13706 | 0.81506567 | 0.73478523 | 1.16099982 | 0.7002644 | 0.6291739 | 1.27358596 |
| ENSMUSG00 | Pik3ca | 8905 | 14.7860923 | 11.6231867 | 14.2408858 | 12.8323229 | 16.3364624 | 11.360094 |
| ENSMUSG00 | Zfp639 | 1454 | 28.7365956 | 29.0643121 | 30.6129683 | 27.0739413 | 27.7942845 | 41.4934677 |
| ENSMUSG00 | Mfn1 | 5916 | 27.1017331 | 21.1935636 | 20.5468337 | 18.1934988 | 23.8830349 | 21.7422581 |
| ENSMUSG00 | Gnb4 | 10840 | 15.8728524 | 7.97476727 | 18.3576289 | 7.10323644 | 11.3491321 | 13.0510659 |
| ENSMUSG00 | Ocstamp | 3654 | 0.02175998 | 0 | 0.29032409 | 0.02962401 | 0 | 0.05225027 |
| ENSMUSG00 | Actl6a | 2750 | 23.4051356 | 27.107064 | 28.3856199 | 26.8581619 | 23.3979127 | 23.4363537 |
| ENSMUSG00 | Ndufb5 | 1952 | 83.8897665 | 68.2805835 | 97.6652402 | 66.6926315 | 43.3410508 | 115.847354 |
| ENSMUSG00 | Pex5l | 6260 | 0 | 0 | 0 | 0 | 0 | 0 |
| ENSMUSG00 | Ccdc39 | 7024 | 0.40185637 | 1.26914747 | 0.38387136 | 0.63184674 | 0.60295974 | 0.92805222 |
| ENSMUSG00 | Ttc14 | 6558 | 26.2308576 | 23.872789 | 27.917663 | 34.3709259 | 26.7347709 | 44.0395174 |
| ENSMUSG00 | Ncoa3 | 7992 | 11.5207323 | 11.4738238 | 10.3314618 | 11.7248586 | 11.0965767 | 9.94814482 |
| ENSMUSG00 | Dnajc19 | 3992 | 19.2304191 | 21.9687687 | 18.2808725 | 26.103443 | 13.5874342 | 32.7200019 |
| ENSMUSG00 | Fxr1 | 2446 | 18.1223879 | 22.5172132 | 17.6192956 | 22.9090005 | 19.588621 | 24.0743852 |
| ENSMUSG00 | Mecom | 12100 | 24.63197 | 24.5392443 | 25.0927653 | 29.71252 | 31.6911747 | 23.2984026 |
| ENSMUSG00 | Slc2a2 | 2755 | 0 | 0.29733921 | 0.27275184 | 0 | 0 | 0.9009046 |
| ENSMUSG00 | Tnik | 15796 | 0 | 0 | 0.01119317 | 0 | 0 | 0.0345336 |
| ENSMUSG00 | Gm8325 | 1707 | 0.02328968 | 0.08468619 | 0 | 0.08455078 | 0 | 0.01597811 |
| ENSMUSG00 | Pld1 | 8573 | 0.38489501 | 0.28103606 | 0.48981396 | 0 | 0.42259101 | 0.38177496 |
| ENSMUSG00 | Nceh1 | 5455 | 25.6096716 | 18.1880631 | 27.3719204 | 17.6805325 | 22.0755723 | 25.5046627 |
| ENSMUSG00 | Ect2 | 6205 | 0.33957139 | 0.10095467 | 0.30631414 | 0.62220447 | 0.83056631 | 0.29890016 |
| ENSMUSG00 | Lrrc34 | 1880 | 1.1842058 | 2.94757492 | 0 | 1.90006511 | 2.90416075 | 4.33782826 |
| ENSMUSG00 | Lrriq4 | 2621 | 1.39546135 | 1.3420871 | 0.03372899 | 0.46806162 | 1.03181912 | 0.43706023 |
| ENSMUSG00 | Sec62 | 3958 | 71.7667782 | 61.7609452 | 69.496733 | 61.2154949 | 54.8939637 | 87.1989098 |
| ENSMUSG00 | Dcun1d1 | 6567 | 6.66526329 | 8.70980765 | 8.68959637 | 7.86805052 | 6.81439494 | 7.23502689 |
| ENSMUSG00 | Mccc1 | 10075 | 19.4377665 | 19.9824273 | 17.9922341 | 21.8175499 | 18.1517152 | 15.5336854 |
| ENSMUSG00 | Acad9 | 8505 | 9.83953986 | 7.75628916 | 8.79359417 | 9.06611748 | 9.68931181 | 4.50248013 |
| ENSMUSG00 | Anxa5 | 1665 | 505.933259 | 419.381221 | 602.047685 | 425.854809 | 416.516201 | 427.84326 |
| ENSMUSG00 | 1810062G17I | 387 | 1.95181946 | 1.74317879 | 0 | 1.21205834 | 1.97776651 | 0.98667946 |
| ENSMUSG00 | Exosc9 | 3387 | 13.1461878 | 8.59303545 | 14.1988796 | 9.71562581 | 8.55712473 | 11.7570042 |
| ENSMUSG00 | Ccna2 | 4148 | 0.08625828 | 1.46371546 | 0.33034164 | 0.31315179 | 0.20912449 | 0.13808279 |
| ENSMUSG00 | Trpc3 | 5822 | 0.32093913 | 0.00827661 | 0 | 0 | 0.14899491 | 0 |
| ENSMUSG00 | Il21 | 3179 | 0 | 0 | 0 | 0 | 0 | 0 |
| ENSMUSG00 | Adad1 | 3460 | 0 | 0 | 0 | 0 | 0 | 0 |
| ENSMUSG00 | Il2 | 1197 | 0.0996378 | 0.04025601 | 0.18463594 | 0.09043119 | 0.0852571 | 0.63800326 |
| ENSMUSG00 | Spata5 | 3937 | 10.1383039 | 8.95922673 | 11.2272906 | 9.53145194 | 6.6877343 | 6.38047822 |
| ENSMUSG00 | Slc7a11 | 11233 | 0 | 0.04289722 | 0.003935 | 0.02248503 | 0 | 0 |
| ENSMUSG00 | Rab33b | 3163 | 24.019198 | 21.4195819 | 29.9756413 | 29.864936 | 22.6497097 | 22.652696 |
| ENSMUSG00 | Cog6 | 3134 | 43.1678048 | 40.4064997 | 44.9211455 | 39.5705128 | 37.1382142 | 28.7019018 |
| ENSMUSG00 | Stoml3 | 2069 | 0 | 0 | 0 | 0 | 0.04932468 | 0.10546018 |
| ENSMUSG00 | Ufm1 | 4289 | 21.5600951 | 18.4252192 | 29.5159894 | 21.5869728 | 16.2037594 | 24.5147434 |
| ENSMUSG00 | Trpc4 | 3294 | 0 | 0 | 0 | 0 | 0 | 0 |
| ENSMUSG00 | Postn | 5358 | 0 | 0 | 0 | 0 | 0 | 0 |
| ENSMUSG00 | Fam48a | 3832 | 42.9819822 | 33.2350644 | 32.6323105 | 43.9820538 | 45.0342388 | 44.8622263 |
| ENSMUSG00 | Exosc8 | 3604 | 6.05598191 | 7.62105217 | 4.88133561 | 4.00466643 | 5.57835518 | 4.92668983 |
| ENSMUSG00 | Aadac | 1433 | 0.74905651 | 1.78219221 | 0 | 0 | 0 | 1.10392818 |
| ENSMUSG00 | Sucnr1 | 1569 | 1.57096226 | 0.67565438 | 0 | 0 | 0.29269432 | 0 |
| ENSMUSG00 | Mbnl1 | 5633 | 22.0197229 | 27.459343 | 28.4844115 | 32.9689833 | 26.4688568 | 28.707853 |
| ENSMUSG00 | P2ry1 | 3884 | 3.42896137 | 2.61774956 | 1.37703991 | 1.06834066 | 1.61592282 | 2.12073663 |
| ENSMUSG00 | Dhx36 | 4467 | 37.9488178 | 23.958381 | 27.0237818 | 28.8365567 | 22.0348957 | 49.2372265 |
| ENSMUSG00 | Gfm1 | 4606 | 44.9773793 | 32.410247 | 44.6144959 | 44.2839335 | 34.3093109 | 30.2472552 |
| ENSMUSG00 | Mfsd1 | 2962 | 74.16907 | 76.0864247 | 81.6585027 | 69.2404933 | 76.1433427 | 87.7263633 |
| ENSMUSG00 | Il12a | 1362 | 0 | 0 | 0 | 0 | 0 | 0 |
| ENSMUSG00 | Schip1 | 2547 | 9.89594607 | 9.42161295 | 12.5646385 | 11.4465222 | 10.337499 | 15.2917883 |
| ENSMUSG00 | Ifi80 | 11708 | 5.44991 | 3.42425007 | 2.39357435 | 4.86928855 | 4.57616114 | 4.01385407 |
| ENSMUSG00 | Kpna4 | 8922 | 5.98426478 | 7.46938461 | 7.36201958 | 12.1567615 | 7.69799393 | 8.37009216 |
| ENSMUSG00 | Arl14 | 1195 | 0 | 0 | 0 | 0 | 0 | 0 |
| ENSMUSG00 | Ppm1l | 9987 | 5.01571097 | 7.45932107 | 7.02838958 | 8.80464028 | 8.32301654 | 4.85301231 |
| ENSMUSG00 | Nmd3 | 2876 | 37.8340578 | 40.0436706 | 46.0307788 | 46.8589834 | 33.6036008 | 51.4576469 |
| ENSMUSG00 | Otol1 | 2156 | 0 | 0 | 0 | 0 | 0 | 0 |
| ENSMUSG00 | Sis | 6890 | 0 | 0 | 0 | 0 | 0 | 0 |
| ENSMUSG00 | Bche | 6735 | 0.07673664 | 0 | 0.26908323 | 0 | 0 | 0 |
| ENSMUSG00 | Ccna1 | 1557 | 0 | 0.37137913 | 0 | 0 | 0 | 0 |
| ENSMUSG00 | Sohlh2 | 2353 | 0 | 0 | 0 | 0 | 0 | 0 |
| ENSMUSG00 | Smad9 | 5362 | 0 | 0 | 0.04121768 | 0 | 0.29500516 | 0.0559532 |
| ENSMUSG00 | Dclk1 | 11213 | 0.06736414 | 0.15040805 | 0 | 0 | 0.00455064 | 0.04135101 |
| ENSMUSG00 | Nbea | 11562 | 17.5774102 | 12.2695802 | 25.3504948 | 21.7703116 | 18.4166292 | 16.2251314 |
| ENSMUSG00 | Tm4sf1 | 4054 | 3.81472174 | 2.11573426 | 2.60588073 | 1.99367982 | 2.20266793 | 2.77186763 |
| ENSMUSG00 | Tm4sf4 | 1491 | 0 | 0 | 0 | 0 | 0 | 0 |
| ENSMUSG00 | Wwtr1 | 8100 | 10.0026752 | 12.0763553 | 7.91813262 | 13.5285394 | 11.4084897 | 11.8627845 |
| ENSMUSG00 | Ppid | 3333 | 16.8063222 | 12.0140814 | 14.6278526 | 23.7948797 | 15.4472287 | 22.0373848 |
| ENSMUSG00 | Pfn2 | 4421 | 58.153968 | 72.3614091 | 35.363474 | 58.518041 | 51.3726304 | 66.8262593 |
| ENSMUSG00 | Tsc22d2 | 9489 | 9.06218825 | 12.0758098 | 9.5307167 | 14.1111252 | 12.1153362 | 12.8713073 |
| ENSMUSG00 | Serp1 | 2437 | 462.987399 | 488.824466 | 302.520533 | 551.313504 | 258.209793 | 527.910324 |
| ENSMUSG00 | Etfdh | 7119 | 41.5369102 | 37.6407929 | 49.5229526 | 42.3770156 | 46.4175881 | 45.1550637 |
| ENSMUSG00 | Eif2a | 6147 | 11.1434347 | 7.44706687 | 11.2248376 | 9.32721144 | 9.41335776 | 12.9828525 |
| ENSMUSG00 | 4930579G24I | 1965 | 1.45668936 | 1.83917718 | 1.70958782 | 3.01142764 | 2.4409564 | 5.55208942 |
| ENSMUSG00 | Mme | 4355 | 62.0021179 | 30.4277189 | 98.3808184 | 64.0943046 | 108.532105 | 101.576894 |
| ENSMUSG00 | Slc33a1 | 2848 | 37.3824357 | 41.8416683 | 32.9340979 | 39.9841757 | 39.8464395 | 30.6839692 |
| ENSMUSG00 | Gmps | 10283 | 19.4196031 | 20.8059713 | 17.2199342 | 16.9479991 | 16.2462661 | 21.9618801 |
| ENSMUSG00 | Vmn2r1 | 2733 | 0.13091816 | 0.91682949 | 0 | 0.05280943 | 0.63479575 | 0 |
| ENSMUSG00 | Kcnab1 | 8098 | 0.00981859 | 0.00595041 | 0.03820856 | 0.03564539 | 0.09451663 | 0.22902883 |
| ENSMUSG00 | Ssr3 | 4522 | 78.8867595 | 65.3958858 | 92.5188952 | 75.8425107 | 54.6259808 | 96.8605554 |
| ENSMUSG00 | Ccnl1 | 6034 | 28.4956005 | 30.6974948 | 31.8364617 | 37.3378001 | 39.2972437 | 38.2541054 |
| ENSMUSG00 | Veph1 | 9103 | 4.59876096 | 8.77127701 | 3.28733909 | 5.36692171 | 5.35320102 | 4.36849654 |
| ENSMUSG00 | Ptx3 | 1919 | 0 | 0 | 0.11516895 | 0.1504202 | 0.15954052 | 0.17055533 |
| ENSMUSG00 | Shox2 | 4414 | 0 | 0 | 0 | 0 | 0 | 0 |
| ENSMUSG00 | Serpini1 | 5294 | 0.37547677 | 0.34587926 | 1.185618 | 0.57933014 | 1.47469503 | 0.91190237 |
| ENSMUSG00 | Pdcd10 | 3255 | 10.7602391 | 10.7919866 | 12.3303452 | 11.5396032 | 7.72841883 | 16.6999558 |
| ENSMUSG00 | Wnt2b | 3568 | 0.01114223 | 0.22958787 | 0.07433045 | 0.03033804 | 0.08580669 | 0.22168289 |
| ENSMUSG00 | Ptpn22 | 3041 | 0 | 0 | 0 | 0 | 0 | 0.26906912 |
| ENSMUSG00 | Dclre1b | 11082 | 2.46095106 | 1.07834666 | 1.97037633 | 1.41957872 | 1.51946436 | 1.69328206 |
| ENSMUSG00 | Olfml3 | 6963 | 0 | 0 | 0 | 0 | 0.03664109 | 0.07834163 |
| ENSMUSG00 | Syt6 | 6014 | 0 | 0 | 0 | 0 | 0 | 0 |
| ENSMUSG00 | Nras | 3011 | 14.1540602 | 17.1877246 | 15.707729 | 19.6767576 | 9.42233974 | 16.4227569 |
| ENSMUSG00 | Sike1 | 3797 | 16.4906194 | 18.4649127 | 16.2279087 | 17.5136181 | 15.0512354 | 13.4182318 |
| ENSMUSG00 | Sycp1 | 3433 | 0 | 0 | 0.09012901 | 0 | 0 | 0.01588968 |
| ENSMUSG00 | Tshb | 640 | 0 | 0 | 0 | 0 | 0 | 0 |
| ENSMUSG00 | Tspan2 | 4576 | 0 | 0.06318152 | 0 | 0 | 0.1784139 | 0.44702752 |
| ENSMUSG00 | Ngf | 1183 | 0.13442259 | 2.07735296 | 1.6066604 | 2.25703406 | 0.51759637 | 1.72916141 |
| ENSMUSG00 | Vangl1 | 4344 | 2.69063335 | 4.17083385 | 0.98701169 | 0.93029212 | 4.24045158 | 1.60106653 |
| ENSMUSG00 | Casq2 | 3085 | 0 | 0 | 0 | 0 | 0 | 0 |
| ENSMUSG00 | Cd2 | 2110 | 0 | 0 | 0 | 0 | 0 | 0 |
| ENSMUSG00 | Ptgfrn | 6849 | 0.84166224 | 0.44323928 | 0.58083894 | 1.50144294 | 0.42466103 | 1.13893223 |
| ENSMUSG00 | Gdap2 | 41108 | 1.27076728 | 1.63637913 | 1.78170803 | 1.72212152 | 1.29340964 | 1.53862724 |
| ENSMUSG00 | Spag17 | 8598 | 0 | 0 | 0 | 0 | 0 | 0 |
| ENSMUSG00 | Tbx15 | 3737 | 0 | 0.95418697 | 0.1892506 | 0 | 0 | 0.56928602 |
| ENSMUSG00 | Hsd3b6 | 2964 | 0 | 0 | 0 | 0 | 0 | 0 |
| ENSMUSG00 | Hao2 | 1984 | 3.20608714 | 6.09616783 | 1.69321576 | 1.50948069 | 4.60369016 | 9.29317345 |
| ENSMUSG00 | Hsd3b1 | 3016 | 0 | 0 | 0 | 0 | 0 | 0 |
| ENSMUSG00 | Hmgcs2 | 3274 | 2.99926808 | 1.36876577 | 13.3388579 | 4.52954194 | 14.8216504 | 8.53061411 |
| ENSMUSG00 | Reg4 | 1015 | 0 | 0 | 0 | 0 | 0 | 0.53743132 |
| ENSMUSG00 | Notch2 | 10467 | 3.77538432 | 5.35404913 | 1.10641854 | 4.28144638 | 6.11809512 | 1.15696377 |
| ENSMUSG00 | Sec22b | 3574 | 40.0891863 | 36.6454252 | 45.475707 | 40.1809002 | 30.5101749 | 37.3557244 |
| ENSMUSG00 | 4930443G12I | 1998 | 0 | 0 | 0 | 0 | 0 | 0 |
| ENSMUSG00 | Prpf38b | 6690 | 14.4819292 | 14.6792181 | 12.5073376 | 20.3332299 | 14.7434954 | 27.4417932 |
| ENSMUSG00 | Stxbp3a | 5869 | 18.9463416 | 20.5586729 | 19.5440079 | 21.8865358 | 20.2662263 | 19.5881078 |
| ENSMUSG00 | Gpsm2 | 3772 | 1.22259696 | 1.27747726 | 0 | 1.02353626 | 0.48699616 | 0.00723082 |
| ENSMUSG00 | Clcc1 | 3949 | 19.8022361 | 17.7785886 | 20.9648144 | 16.9948349 | 22.0696505 | 21.3693932 |
| ENSMUSG00 | 1700013F07I | 1446 | 0 | 0.19994374 | 0 | 0 | 0.03528795 | 0 |
| ENSMUSG00 | Sypl2 | 4879 | 0 | 0 | 0 | 0 | 0 | 0 |
| ENSMUSG00 | Ampd2 | 6110 | 4.68477021 | 5.05523886 | 4.23209135 | 4.0688263 | 4.16729321 | 3.70506556 |
| ENSMUSG00 | Gstm4 | 1859 | 13.344497 | 11.6124401 | 14.6705418 | 12.7131464 | 7.16400435 | 5.3404888 |
| ENSMUSG00 | Ahcyl1 | 7685 | 54.5971817 | 50.7509516 | 61.9343456 | 59.3323093 | 59.8173871 | 53.1544921 |
| ENSMUSG00 | Slc6a17 | 12212 | 0.1855603 | 0.51295754 | 0.0434345 | 0.01181855 | 0.2507028 | 0.06030259 |
| ENSMUSG00 | Kcnc4 | 3009 | 0 | 0 | 0 | 0 | 0 | 0.02719306 |
| ENSMUSG00 | Slc16a4 | 3989 | 7.68399987 | 6.184873 | 8.00043386 | 6.82926639 | 6.45984452 | 6.9536997 |
| ENSMUSG00 | Dram2 | 2929 | 56.2195972 | 70.0340335 | 62.8846295 | 72.2995969 | 54.4583308 | 75.0355217 |
| ENSMUSG00 | Dennd2d | 4532 | 8.24584106 | 12.5250726 | 9.2265983 | 6.44094749 | 10.2232898 | 12.8850403 |
| ENSMUSG00 | BC051070 | 1653 | 0 | 0 | 0 | 0 | 0 | 0 |
| ENSMUSG00 | Ddx20 | 4994 | 8.8920448 | 11.8681065 | 8.24912248 | 9.67438078 | 9.75774708 | 7.32384686 |
| ENSMUSG00 | S100a11 | 508 | 119.970771 | 107.376088 | 193.078523 | 92.7621066 | 75.3342163 | 236.881316 |
| ENSMUSG00 | Tchhl1 | 2063 | 0 | 0 | 0 | 0.66462322 | 0 | 0 |
| ENSMUSG00 | Lce1m | 875 | 0 | 0 | 0 | 0 | 0 | 0 |
| ENSMUSG00 | Crct1 | 1120 | 0 | 0 | 0 | 0 | 0 | 0 |
| ENSMUSG00 | Lce1g | 923 | 0 | 0 | 0 | 0 | 0 | 0 |
| ENSMUSG00 | Lce1b | 645 | 0 | 0 | 0 | 0 | 0.55377462 | 0 |
| ENSMUSG00 | Sprr2j-ps | 671 | 0 | 0 | 0 | 0 | 0 | 0 |
| ENSMUSG00 | Lelp1 | 868 | 0 | 0 | 0 | 0 | 0 | 0 |
| ENSMUSG00 | Npr1 | 5233 | 1.55739987 | 0.75507133 | 1.40216051 | 0.67571954 | 1.31636934 | 1.42810074 |
| ENSMUSG00 | Slc27a3 | 2752 | 5.22946364 | 4.60502699 | 6.56924193 | 4.20869778 | 3.98643561 | 3.28048895 |
| ENSMUSG00 | Ints3 | 4945 | 8.02345188 | 8.01970513 | 7.74090921 | 8.09200958 | 11.206197 | 5.81895741 |
| ENSMUSG00 | Rab13 | 1642 | 6.17396317 | 9.00927999 | 8.15661293 | 6.04297335 | 7.02311872 | 6.39508898 |
| ENSMUSG00 | Crtc2 | 4180 | 11.9076224 | 13.6835662 | 10.6274766 | 13.6731958 | 11.889878 | 10.5379288 |
| ENSMUSG00 | Jtb | 1536 | 66.0520744 | 63.9976185 | 70.2740239 | 71.1539797 | 54.7470491 | 88.305196 |
| ENSMUSG00 | Creb3l4 | 2019 | 3.50494083 | 3.93797076 | 6.98384741 | 0.62549359 | 4.82716086 | 4.5525277 |
| ENSMUSG00 | Nup210l | 5641 | 0.09161873 | 0.5808683 | 0.28992523 | 0.0767567 | 0.01809125 | 0.00483507 |
| ENSMUSG00 | Tpm3 | 4925 | 12.1082682 | 10.8896467 | 11.1648919 | 11.7294118 | 10.4539316 | 10.8046337 |
| ENSMUSG00 | 4933434E20I | 3764 | 16.0542854 | 14.2613434 | 18.6014132 | 16.0854598 | 12.5532476 | 19.6226682 |
| ENSMUSG00 | Hax1 | 1364 | 8.59814278 | 7.41873376 | 9.39775255 | 5.3699817 | 6.06031736 | 6.37874628 |
| ENSMUSG00 | Il6ra | 3333 | 0.5486805 | 0.44817873 | 0.03978564 | 0.05412848 | 0.26026054 | 0.19639704 |
| ENSMUSG00 | Chrnb2 | 5378 | 0.09609915 | 0.10751902 | 0.25478935 | 0 | 0.27515152 | 0.19271779 |
| ENSMUSG00 | Adar | 6769 | 7.92290489 | 9.03362315 | 6.07293751 | 7.22813597 | 10.5233891 | 6.48724615 |
| ENSMUSG00 | Pmvk | 1632 | 24.5304956 | 25.5990474 | 34.0451695 | 19.2570225 | 17.6966475 | 25.8707975 |
| ENSMUSG00 | Slc50a1 | 1566 | 16.7297967 | 26.4317713 | 20.3226864 | 19.4234761 | 18.1818121 | 31.0192417 |
| ENSMUSG00 | Efna1 | 2692 | 47.7449734 | 42.7448826 | 50.4577502 | 41.7248903 | 46.1928967 | 44.3060912 |
| ENSMUSG00 | Fam198b | 7170 | 0.08317046 | 0.26882255 | 0 | 0 | 0.32024922 | 0.45647932 |
| ENSMUSG00 | Tmem144 | 3460 | 0.22980047 | 0.5570687 | 1.25195972 | 1.06369033 | 1.75495339 | 0.89076134 |
| ENSMUSG00 | Slc35a3 | 10727 | 9.94350277 | 11.1672877 | 9.56394869 | 11.9712871 | 11.697013 | 10.5798242 |
| ENSMUSG00 | Sass6 | 4392 | 1.52070144 | 1.29462663 | 0.69442786 | 1.19123326 | 1.48710749 | 0.72657851 |
| ENSMUSG00 | Lrrc39 | 5366 | 0.22967199 | 0.35919823 | 0.3871574 | 0.20845013 | 0.41840487 | 0.58452917 |
| ENSMUSG00 | Vcam1 | 4907 | 0 | 0 | 0 | 0 | 1.84056827 | 1.2950868 |
| ENSMUSG00 | Extl2 | 3794 | 12.322732 | 14.3136848 | 18.8387929 | 9.38665734 | 19.1517025 | 15.0679082 |
| ENSMUSG00 | Olfm3 | 4930 | 0.20966379 | 0.08796714 | 0 | 0 | 0 | 0 |
| ENSMUSG00 | Col11a1 | 8860 | 0 | 0.0543865 | 0.13470088 | 0.12217397 | 0.15549799 | 0.13852808 |
| ENSMUSG00 | Neurog2 | 2224 | 0 | 0 | 0 | 0 | 0 | 0 |
| ENSMUSG00 | Larp7 | 1867 | 9.73125581 | 12.6208732 | 11.3404836 | 14.1081374 | 8.41784882 | 13.8199297 |
| ENSMUSG00 | Ndst4 | 8184 | 0 | 0 | 0 | 0 | 0 | 0 |
| ENSMUSG00 | 1700006A11I | 2336 | 0 | 0 | 0 | 0 | 0 | 0 |
| ENSMUSG00 | Ndst3 | 8874 | 0 | 0 | 0 | 0 | 0 | 0 |
| ENSMUSG00 | Prss12 | 2596 | 1.10261733 | 0.79815756 | 0.17026904 | 0.05559637 | 0.21621346 | 0.55683971 |
| ENSMUSG00 | Rnpc3 | 7115 | 23.1269057 | 17.4595429 | 25.6265077 | 23.1198933 | 15.9928065 | 37.858656 |
| ENSMUSG00 | Cyp2u1 | 2326 | 7.60584214 | 4.72334859 | 6.70819028 | 7.84931837 | 3.071235 | 3.50606928 |
| ENSMUSG00 | Hadh | 1840 | 117.213849 | 114.835624 | 106.036378 | 102.245533 | 91.0155247 | 90.2286571 |
| ENSMUSG00 | Lef1 | 8641 | 0 | 0.09480031 | 0.01534609 | 0.00835136 | 0.1003875 | 0.1893853 |
| ENSMUSG00 | Trim2 | 18905 | 14.5710513 | 11.571883 | 14.9077467 | 14.232415 | 16.2431489 | 18.1234604 |
| ENSMUSG00 | Ccdc109b | 1514 | 0 | 0.57289033 | 0 | 0.81029691 | 1.04479369 | 0.18014953 |
| ENSMUSG00 | Tlr2 | 3010 | 0.46227303 | 0.19210542 | 0 | 0 | 0.28818883 | 0 |
| ENSMUSG00 | Sfrp2 | 2626 | 0 | 0 | 0 | 0 | 0 | 0 |
| ENSMUSG00 | Casp6 | 1716 | 50.9453972 | 37.3753814 | 59.9146194 | 48.9294392 | 33.0660431 | 50.957164 |
| ENSMUSG00 | Plrg1 | 2923 | 12.9752749 | 14.4575812 | 14.5625641 | 19.3680219 | 18.4518916 | 11.7571144 |
| ENSMUSG00 | Pla2g12a | 1794 | 23.3125783 | 18.1840699 | 21.977728 | 19.971834 | 17.0941203 | 19.505776 |
| ENSMUSG00 | Fga | 2820 | 0.5921029 | 0 | 0 | 0 | 0.03618892 | 0 |
| ENSMUSG00 | Lrat | 5349 | 0 | 0 | 0 | 0 | 0 | 0 |
| ENSMUSG00 | Npy2r | 4443 | 0.05368735 | 0 | 0 | 0 | 0 | 0 |
| ENSMUSG00 | Gucy1b3 | 3237 | 0 | 0 | 0 | 0 | 0 | 0.1011108 |
| ENSMUSG00 | Snx7 | 2667 | 17.4852563 | 18.7000254 | 19.3579875 | 19.1165838 | 21.5431943 | 17.0274745 |
| ENSMUSG00 | Accn5 | 2294 | 0 | 0 | 0 | 0 | 0.9342231 | 0 |
| ENSMUSG00 | 1700061I17R | 1953 | 0 | 0 | 0 | 0 | 0 | 0 |

| | | | | | | | | |
|---|---|---|---|---|---|---|---|---|
| ENSMUSG00 | Gar1 | 1545 | 4.01414561 | 3.52431583 | 3.23288561 | 4.15702519 | 2.57608888 | 4.89001623 |
| ENSMUSG00 | Tdo2 | 4066 | 0.03911016 | 0 | 0 | 0 | 0 | 0.50980634 |
| ENSMUSG00 | Rrh | 2119 | 0 | 0 | 0 | 0 | 0.0240804 | 0 |
| ENSMUSG00 | Ppa2 | 2094 | 59.5952499 | 59.4621617 | 48.6769102 | 61.6358516 | 43.7403747 | 66.4672799 |
| ENSMUSG00 | Ctso | 4697 | 48.2872081 | 58.8352683 | 35.4122919 | 47.5204444 | 53.3837792 | 46.9249222 |
| ENSMUSG00 | Ints12 | 4866 | 14.3629541 | 9.35803285 | 15.2880604 | 12.5834833 | 8.38904658 | 11.0645577 |
| ENSMUSG00 | Egf | 5725 | 25.8323821 | 17.4228708 | 8.87897283 | 13.1975199 | 24.4035314 | 8.55161178 |
| ENSMUSG00 | Gstcd | 5163 | 0.57750553 | 0.74664253 | 0.69346296 | 0.41232629 | 0.13836321 | 1.46331107 |
| ENSMUSG00 | Pdgfc | 5464 | 0.12369019 | 0.44976364 | 0.69570983 | 0.2839546 | 0.82180108 | 1.1530823 |
| ENSMUSG00 | Glrb | 3622 | 0 | 0 | 0 | 0 | 0 | 0 |
| ENSMUSG00 | Pitx2 | 4478 | 0.81677182 | 0.09684636 | 0.13819245 | 0 | 0.29626748 | 0 |
| ENSMUSG00 | Enpep | 5342 | 2.3368066 | 1.01027359 | 0.86053757 | 1.25631978 | 1.91993664 | 0.5258869 |
| ENSMUSG00 | Alpk1 | 4169 | 3.51877484 | 1.10959425 | 4.65449966 | 2.94264695 | 4.07574554 | 6.96095375 |
| ENSMUSG00 | Aimp1 | 1115 | 22.3914276 | 32.6717043 | 23.1514586 | 24.9500057 | 20.3647868 | 24.3392521 |
| ENSMUSG00 | Tbck | 4081 | 16.6873654 | 13.8384 | 6.55283387 | 23.4652649 | 24.34412 | 7.62567101 |
| ENSMUSG00 | Dkk2 | 3690 | 1.48679033 | 0.54846357 | 0.40727985 | 0 | 2.06041463 | 1.04220166 |
| ENSMUSG00 | Papss1 | 2587 | 84.766614 | 96.7640382 | 72.5135762 | 91.3557757 | 80.6914973 | 55.6246605 |
| ENSMUSG00 | Kcnq5 | 12414 | 0 | 0 | 0 | 0 | 0 | 0 |
| ENSMUSG00 | Fubp1 | 5751 | 29.4485041 | 25.1782749 | 21.1132782 | 39.8213766 | 26.7509169 | 30.8505527 |
| ENSMUSG00 | Dnajb4 | 4309 | 20.8326468 | 30.0144838 | 22.9779829 | 27.0720005 | 17.0995792 | 28.1924445 |
| ENSMUSG00 | Ptgfr | 4940 | 41.1316318 | 46.576976 | 43.4144418 | 54.5759184 | 44.6635727 | 41.9444199 |
| ENSMUSG00 | Ifi44 | 3116 | 4.64409336 | 2.95366189 | 1.92922036 | 4.36551052 | 3.2587448 | 4.71791738 |
| ENSMUSG00 | Efna3 | 2428 | 1.30990051 | 2.1632299 | 0.52794623 | 2.30342541 | 0.73555319 | 1.07840419 |
| ENSMUSG00 | Efna4 | 1595 | 11.8892566 | 21.4497642 | 10.6139849 | 14.9531232 | 10.7171385 | 7.11363632 |
| ENSMUSG00 | Adam15 | 5858 | 7.44482112 | 8.62881151 | 5.78743831 | 8.21054376 | 11.0188401 | 5.63837285 |
| ENSMUSG00 | Zbtb7b | 5835 | 9.78386804 | 9.6455466 | 9.95393688 | 4.50793667 | 8.00156536 | 4.15546775 |
| ENSMUSG00 | Cks1b | 1528 | 1.66515102 | 1.26142519 | 1.07033259 | 1.98356789 | 1.40255745 | 2.26693664 |
| ENSMUSG00 | Thbs3 | 4835 | 0.01644487 | 0.21925579 | 0.01828411 | 0.73880505 | 0.17941022 | 0.06205192 |
| ENSMUSG00 | Gba | 2084 | 28.6529423 | 17.0872268 | 24.4764524 | 23.5641375 | 21.4242221 | 24.4346216 |
| ENSMUSG00 | Scamp3 | 2328 | 23.8567037 | 25.0660573 | 25.6894732 | 21.0633585 | 21.1513972 | 16.6131609 |
| ENSMUSG00 | Hcn3 | 4146 | 0.81505447 | 0.30218222 | 0.4264529 | 0.17405714 | 0.76305724 | 0.1841992 |
| ENSMUSG00 | Ash1l | 11241 | 11.5719182 | 12.5042124 | 14.3367956 | 17.6253453 | 14.044623 | 12.5830691 |
| ENSMUSG00 | Rit1 | 3294 | 13.3242412 | 10.8251267 | 18.2094417 | 15.9597872 | 11.1223248 | 10.938008 |
| ENSMUSG00 | Arhgef2 | 9201 | 8.08849685 | 6.73489453 | 9.76178079 | 8.3332621 | 7.96923183 | 8.30007498 |
| ENSMUSG00 | 2810403A07R | 3823 | 45.1421765 | 32.620066 | 31.0788789 | 49.5880361 | 38.1596674 | 52.3947033 |
| ENSMUSG00 | Lamtor2 | 1398 | 30.0014535 | 29.8493984 | 38.0363499 | 29.6554143 | 21.826733 | 55.953981 |
| ENSMUSG00 | Lmna | 4442 | 11.1605322 | 12.5293428 | 8.06021901 | 11.0959191 | 7.94917876 | 5.88228375 |
| ENSMUSG00 | Sema4a | 6291 | 16.013414 | 11.8110783 | 10.9819553 | 12.7213439 | 13.8698303 | 8.86610034 |
| ENSMUSG00 | Pmf1 | 1418 | 4.23348206 | 4.65553073 | 7.48127103 | 3.96953381 | 3.88635302 | 5.75114043 |
| ENSMUSG00 | Iqgap3 | 5766 | 0 | 0.31756587 | 0 | 0.31288627 | 0 | 0 |
| ENSMUSG00 | Gpatch4 | 1557 | 10.0601537 | 7.45853087 | 7.26761195 | 12.7920929 | 9.27454358 | 6.1486181 |
| ENSMUSG00 | Apoa1bp | 899 | 158.623925 | 176.665755 | 148.437557 | 103.309436 | 103.642005 | 133.157277 |
| ENSMUSG00 | Sh2d2a | 4751 | 0 | 0.01014238 | 0 | 0.00759462 | 0 | 0 |
| ENSMUSG00 | Ntrk1 | 2571 | 49.6054382 | 31.693222 | 80.2028773 | 27.5211549 | 25.6818866 | 44.0255748 |
| ENSMUSG00 | Pear1 | 7750 | 0.9233531 | 2.01450416 | 0.68441563 | 0.96373977 | 1.3299778 | 1.01356079 |
| ENSMUSG00 | Cd1d1 | 2515 | 0.64810127 | 0 | 0.05272586 | 0 | 0.3449099 | 0 |
| ENSMUSG00 | Dclk2 | 5477 | 0.04355174 | 0 | 0 | 0 | 0 | 0.02489925 |
| ENSMUSG00 | Lrba | 9951 | 6.36423279 | 5.3992446 | 4.33090111 | 6.39984008 | 5.08674152 | 5.70380105 |
| ENSMUSG00 | Rps3a | 971 | 6.75556569 | 11.2650076 | 4.96189587 | 8.24944787 | 5.57033557 | 6.62905753 |
| ENSMUSG00 | Sh3d19 | 6416 | 2.0076022 | 2.23057564 | 1.32963773 | 3.49797876 | 3.26072539 | 2.98422643 |
| ENSMUSG00 | Pet112l | 4477 | 9.10193601 | 7.90012252 | 8.7771808 | 7.93047394 | 9.15215095 | 6.52471267 |
| ENSMUSG00 | Fbxw7 | 7321 | 4.11619372 | 5.00228057 | 3.21204487 | 3.68164011 | 4.35616513 | 4.37750323 |
| ENSMUSG00 | Fmo5 | 5611 | 7.08527544 | 4.50862273 | 4.77389358 | 2.66869515 | 3.64668984 | 2.80961353 |
| ENSMUSG00 | Chd1l | 2985 | 12.6125427 | 9.91171708 | 16.2739784 | 8.50980213 | 15.0942345 | 12.6642044 |
| ENSMUSG00 | Acp6 | 5122 | 58.4689642 | 46.4742335 | 71.7826884 | 52.777563 | 51.3551283 | 56.9668276 |
| ENSMUSG00 | Gpr89 | 8353 | 3.45058343 | 4.43040675 | 4.79960153 | 4.75161606 | 3.92181651 | 3.48728777 |
| ENSMUSG00 | Rnf115 | 2256 | 87.3175558 | 66.3204357 | 91.6365336 | 98.8097833 | 67.8994593 | 86.1037149 |
| ENSMUSG00 | Polr3c | 3909 | 21.0422331 | 20.2533447 | 25.3518885 | 18.4517797 | 22.7393059 | 20.5414525 |
| ENSMUSG00 | Nudt17 | 1353 | 1.49854361 | 0.39175969 | 1.24144127 | 1.200068 | 2.9416536 | 0.08063456 |
| ENSMUSG00 | Pias3 | 3848 | 4.10159193 | 6.26123209 | 4.59478618 | 4.99785076 | 3.34164415 | 5.10336285 |
| ENSMUSG00 | Pex11b | 5029 | 9.82622038 | 10.2524345 | 10.7318454 | 9.85099366 | 8.7766584 | 9.75682561 |
| ENSMUSG00 | Polr3gl | 1718 | 0.78677901 | 0.6731517 | 0.92623187 | 0.90310123 | 0.50491757 | 0.90492109 |
| ENSMUSG00 | Rprd2 | 9812 | 3.59792771 | 4.76855232 | 4.41928333 | 6.14483249 | 4.67516427 | 6.21823971 |
| ENSMUSG00 | Tars2 | 4420 | 10.0378091 | 10.7056756 | 11.2304683 | 9.59194623 | 11.4405291 | 9.87934331 |
| ENSMUSG00 | Ecm1 | 2602 | 0.79449846 | 2.03709018 | 0.64553038 | 0.74882029 | 1.66688776 | 0.79664588 |
| ENSMUSG00 | Hormad1 | 1758 | 0 | 0.02740981 | 0 | 0 | 0 | 0 |
| ENSMUSG00 | Ctsk | 1474 | 0.7551923 | 0.42498219 | 1.70929786 | 0 | 0.90005819 | 0.46259565 |
| ENSMUSG00 | Mettl14 | 4425 | 9.08311656 | 6.81688424 | 6.91246901 | 5.46327002 | 3.50554085 | 8.14849109 |
| ENSMUSG00 | Bnipl | 2854 | 0.16715689 | 0 | 0.07743841 | 0.24020983 | 0.0178789 | 0.48738844 |
| ENSMUSG00 | Myoz2 | 3392 | 0.01172037 | 0.05682364 | 0 | 0.38294623 | 0 | 0 |
| ENSMUSG00 | Bcar3 | 3286 | 41.0621123 | 28.829746 | 40.9734674 | 35.0388935 | 44.5665547 | 34.5373627 |
| ENSMUSG00 | Gclm | 5934 | 20.8826816 | 43.6796212 | 24.4324309 | 46.0663245 | 23.2860509 | 20.5272226 |
| ENSMUSG00 | Abca4 | 11003 | 0.06503669 | 0.03065574 | 0.01606902 | 0.02295504 | 0.06028746 | 0.00991535 |
| ENSMUSG00 | Pip5k1a | 5908 | 4.421014 | 5.44014166 | 5.5514163 | 5.08128995 | 5.24255418 | 3.1761936 |
| ENSMUSG00 | Abcd3 | 4284 | 119.804681 | 133.176348 | 118.90337 | 141.439256 | 125.743569 | 122.137414 |
| ENSMUSG00 | F3 | 2063 | 32.5289632 | 46.9017818 | 19.3905322 | 39.072849 | 43.5566882 | 33.1975856 |
| ENSMUSG00 | Tmem56 | 7661 | 0 | 0 | 0.07500639 | 0 | 0 | 0 |
| ENSMUSG00 | Rwdd3 | 5583 | 2.42819611 | 2.684217 | 2.77894446 | 2.06810931 | 1.95587398 | 4.74851323 |
| ENSMUSG00 | Ptbp2 | 3350 | 8.80554225 | 5.88306115 | 8.58967158 | 4.11443015 | 10.9211676 | 13.1732437 |
| ENSMUSG00 | Snx27 | 9023 | 6.89982073 | 7.28964796 | 7.20613001 | 8.44966863 | 8.018996 | 9.34042284 |
| ENSMUSG00 | Celf3 | 2731 | 0.18924249 | 0.0352885 | 0.145667 | 0.05284811 | 0.29894618 | 0.05992231 |
| ENSMUSG00 | Adh5 | 2874 | 69.9663258 | 78.8520659 | 65.1800316 | 56.3201293 | 56.8675998 | 69.7145094 |
| ENSMUSG00 | Riiad1 | 1154 | 0.03445016 | 0.04175602 | 0.30642526 | 0.28140243 | 0.53060356 | 0.11817435 |
| ENSMUSG00 | Mrpl9 | 2864 | 18.7117276 | 13.5103747 | 16.4830895 | 9.54964726 | 15.9991919 | 10.9708046 |
| ENSMUSG00 | Oaz3 | 1126 | 0.77675006 | 0 | 0 | 0 | 0.36253198 | 0.46022926 |
| ENSMUSG00 | Them4 | 4177 | 8.937131 | 5.62963461 | 7.98955985 | 6.68601926 | 5.27732687 | 6.03346102 |
| ENSMUSG00 | Them5 | 1535 | 0.23309402 | 0.28252637 | 0.08638797 | 0 | 0.4321452 | 0.28429591 |
| ENSMUSG00 | Rap1gds1 | 3648 | 17.2840439 | 14.9790092 | 11.7047644 | 11.1767298 | 18.4774787 | 12.0223739 |
| ENSMUSG00 | Rorc | 3552 | 7.31984354 | 4.38181892 | 6.8318727 | 10.4832967 | 5.68875136 | 4.47666517 |
| ENSMUSG00 | Tspan5 | 4258 | 1.77396461 | 1.8106695 | 0.86161413 | 1.11856033 | 1.06654473 | 0.5316569 |
| ENSMUSG00 | Eif4e | 2874 | 35.6194719 | 38.7972955 | 39.6186317 | 34.412277 | 30.6797416 | 37.4385706 |
| ENSMUSG00 | Mttp | 4389 | 1.6576106 | 0.55992448 | 1.34952084 | 1.68530853 | 1.32536041 | 0.93214762 |
| ENSMUSG00 | Dapp1 | 5454 | 4.27149094 | 4.98297642 | 5.79470439 | 5.25286824 | 5.18309721 | 3.69062775 |
| ENSMUSG00 | Ppp3ca | 5625 | 15.6194866 | 21.1849016 | 18.7258652 | 20.7768431 | 22.2338928 | 23.3131494 |
| ENSMUSG00 | Nfkb1 | 5626 | 2.49443381 | 4.07691903 | 3.51194878 | 3.91861522 | 3.55533937 | 2.87968996 |
| ENSMUSG00 | Manba | 6433 | 25.5725444 | 23.2430483 | 30.246621 | 20.2986039 | 26.6514259 | 19.3674106 |
| ENSMUSG00 | Cisd2 | 4357 | 37.8119604 | 36.8393479 | 44.2221331 | 50.003072 | 29.0207389 | 51.5444984 |
| ENSMUSG00 | Bdh2 | 1267 | 4.20460488 | 4.22154308 | 2.65141284 | 0.34173996 | 2.05394252 | 2.45407172 |
| ENSMUSG00 | Tacr3 | 3759 | 0 | 0 | 0 | 0 | 0 | 0 |
| ENSMUSG00 | Wls | 3292 | 413.326572 | 448.52034 | 354.138399 | 459.67341 | 389.455868 | 397.644687 |
| ENSMUSG00 | Rpe65 | 1588 | 0 | 0 | 0 | 0 | 0 | 0 |
| ENSMUSG00 | Depdc1a | 3047 | 0 | 0 | 0 | 0.16578557 | 0 | 0 |
| ENSMUSG00 | Lrrc7 | 8314 | 0.14345254 | 0 | 0.21797878 | 0 | 0 | 0.07545306 |
| ENSMUSG00 | 1810013D15 | 649 | 0 | 0.59397772 | 0 | 0 | 0 | 0 |
| ENSMUSG00 | Cth | 2654 | 14.7847247 | 25.7454314 | 7.26149343 | 19.0198871 | 10.1898942 | 7.63566577 |
| ENSMUSG00 | Zranb2 | 4405 | 45.1705289 | 35.9237857 | 35.9936461 | 50.547627 | 32.5734776 | 63.8678556 |
| ENSMUSG00 | Lrriq3 | 4049 | 1.24696123 | 0.24991733 | 1.20064039 | 1.16738647 | 0.51669089 | 3.2333482 |
| ENSMUSG00 | Lphn2 | 10475 | 3.96985514 | 6.70700073 | 1.70055778 | 7.81232239 | 6.66874544 | 1.4971759 |
| ENSMUSG00 | Dnase2b | 2573 | 0 | 0 | 0.17179107 | 0.64507347 | 0 | 0 |
| ENSMUSG00 | Uox | 3149 | 0 | 0 | 0 | 0 | 0 | 0 |
| ENSMUSG00 | Rpf1 | 1254 | 91.1459382 | 75.9685775 | 84.8788182 | 96.5928417 | 53.549211 | 110.468814 |
| ENSMUSG00 | Spata1 | 4004 | 1.00282306 | 3.00864399 | 3.84171862 | 4.60487631 | 3.63199728 | 3.22881594 |
| ENSMUSG00 | Ctbs | 3022 | 15.5890948 | 17.6672991 | 21.428094 | 16.2500538 | 12.2753723 | 21.2998506 |
| ENSMUSG00 | Bcl10 | 1913 | 18.6204446 | 25.6171519 | 30.3844347 | 24.6519771 | 22.4857474 | 18.9910191 |
| ENSMUSG00 | Ddah1 | 3975 | 23.3032125 | 17.9411156 | 16.2462623 | 16.6930515 | 18.1897798 | 15.0542286 |
| ENSMUSG00 | Cyr61 | 2023 | 23.5034872 | 31.8702222 | 40.5529514 | 38.8822823 | 51.9091851 | 25.5893551 |
| ENSMUSG00 | Col24a1 | 7766 | 3.14317088 | 1.36505502 | 8.15053299 | 3.45209362 | 2.64133442 | 5.86864823 |
| ENSMUSG00 | Cryz | 2975 | 38.2454404 | 35.2935319 | 30.9338612 | 27.252556 | 27.923005 | 23.2957507 |
| ENSMUSG00 | Lhx8 | 2535 | 0 | 0 | 0 | 0 | 0 | 0 |
| ENSMUSG00 | Asph | 16079 | 31.7692748 | 27.6849526 | 39.6714221 | 18.6009122 | 20.8180251 | 30.2702866 |
| ENSMUSG00 | Trp53inp1 | 6576 | 21.7337215 | 20.7298426 | 22.4033976 | 30.798132 | 24.7372394 | 25.2589041 |
| ENSMUSG00 | Ccne2 | 3664 | 0.56421534 | 0.81538194 | 0.6635102 | 0.24619299 | 0.59883574 | 0.76672703 |
| ENSMUSG00 | Gem | 2099 | 3.21983406 | 4.88981047 | 3.22195426 | 6.06810944 | 5.56695572 | 3.36547478 |
| ENSMUSG00 | Cdh17 | 4544 | 0 | 0 | 0 | 0 | 0 | 0 |
| ENSMUSG00 | Fam92a | 2513 | 10.0614746 | 11.5049205 | 10.4656174 | 12.7644002 | 8.30473049 | 15.1947851 |
| ENSMUSG00 | Tmem55a | 2332 | 81.2157415 | 63.6011445 | 120.360936 | 82.4842965 | 53.4113994 | 112.712564 |
| ENSMUSG00 | Calb1 | 5047 | 932.595673 | 1450.1188 | 426.700613 | 1345.70226 | 940.090456 | 974.196937 |
| ENSMUSG00 | Decr1 | 3866 | 24.3304363 | 25.9877734 | 30.1043592 | 18.6663448 | 25.2228154 | 30.4635003 |
| ENSMUSG00 | Nbn | 2660 | 34.4796592 | 31.103805 | 32.1044965 | 36.4482908 | 36.6584226 | 30.853154 |
| ENSMUSG00 | Mmp16 | 6075 | 0.63477887 | 0.24588966 | 0.1236924 | 0.52860938 | 0.2435827 | 0.00448965 |
| ENSMUSG00 | Cpne3 | 7115 | 26.529729 | 22.0716254 | 29.4658245 | 22.5062704 | 23.2218419 | 18.5689884 |
| ENSMUSG00 | Fam82b | 2072 | 51.3636203 | 37.3491439 | 52.4576892 | 35.0197836 | 21.6714338 | 69.7662104 |
| ENSMUSG00 | Tmem68 | 3702 | 21.0590214 | 26.6053722 | 30.2798147 | 20.6823605 | 20.7441101 | 39.7331522 |
| ENSMUSG00 | Tgs1 | 5479 | 8.85228805 | 9.32243633 | 7.03486167 | 9.64119606 | 10.0115631 | 8.90072184 |
| ENSMUSG00 | Rps20 | 4291 | 2.5941586 | 2.31330857 | 2.11171704 | 2.74125997 | 1.54589347 | 2.28824753 |
| ENSMUSG00 | Sdr16c5 | 1561 | 0 | 0 | 0 | 0 | 0 | 0 |
| ENSMUSG00 | Atp6v0d2 | 2707 | 1129.05858 | 738.817245 | 1686.5281 | 984.27872 | 837.703424 | 1360.27603 |
| ENSMUSG00 | Cyp7a1 | 4168 | 0 | 0 | 0 | 0 | 0 | 0 |
| ENSMUSG00 | Ubxn2b | 6634 | 8.26391432 | 10.1617173 | 7.54243086 | 8.44126215 | 8.83770061 | 8.18156951 |
| ENSMUSG00 | 1700025O08 | 454 | 0 | 0 | 0 | 0 | 0 | 0 |
| ENSMUSG00 | Nsmaf | 8624 | 15.2678747 | 12.5606589 | 13.8335777 | 12.8487893 | 12.4311706 | 13.3305432 |
| ENSMUSG00 | 6230409E13 | 8853 | 0 | 0 | 0 | 0 | 0 | 0 |
| ENSMUSG00 | Coq3 | 2990 | 5.83700868 | 5.67278516 | 7.80554285 | 4.51327246 | 4.53947022 | 9.14019683 |
| ENSMUSG00 | Sfrs18 | 8234 | 28.2112306 | 27.4933089 | 25.3862662 | 33.3432447 | 38.8058253 | 43.2074805 |
| ENSMUSG00 | Sdcbp | 6810 | 83.8134264 | 83.1552229 | 81.7961271 | 77.6267304 | 70.11069 | 80.2298933 |
| ENSMUSG00 | 2610029I01R | 1584 | 13.9294771 | 9.49126891 | 14.5665165 | 11.5489877 | 8.89095942 | 15.6691425 |
| ENSMUSG00 | Ccnc | 5289 | 8.59903001 | 11.5068021 | 13.3382759 | 14.1012641 | 7.60234149 | 12.89731 |
| ENSMUSG00 | Clca3 | 2921 | 0 | 0 | 0 | 0 | 0 | 0 |
| ENSMUSG00 | Odf2l | 3402 | 10.6108102 | 9.19253408 | 7.9386614 | 10.9137048 | 8.27941195 | 8.92318445 |
| ENSMUSG00 | Fhl5 | 1364 | 0 | 0 | 0 | 0 | 0 | 0 |
| ENSMUSG00 | Ndufaf4 | 2322 | 61.2939794 | 63.2939916 | 67.2926421 | 73.9510981 | 45.2469026 | 101.592745 |
| ENSMUSG00 | Clca2 | 3853 | 0.03095418 | 0.47523613 | 0 | 0.0749173 | 1.9864927 | 0 |
| ENSMUSG00 | Spaca1 | 1858 | 0 | 0 | 0 | 0 | 0 | 0 |
| ENSMUSG00 | Lmo4 | 3948 | 2.80946785 | 4.77226415 | 2.66465011 | 3.38154926 | 2.63662124 | 3.0328183 |
| ENSMUSG00 | Gbp3 | 3365 | 0.24810255 | 0.32935755 | 0.22330797 | 0.88998802 | 1.65286032 | 3.24215623 |
| ENSMUSG00 | Gbp1 | 2812 | 0.0141378 | 0 | 0 | 0 | 0 | 0 |
| ENSMUSG00 | Gbp2 | 2678 | 0.41566597 | 1.88931159 | 1.61754317 | 0.04042051 | 4.47766928 | 5.97842167 |
| ENSMUSG00 | Gtf2b | 1264 | 64.6026558 | 64.2358821 | 89.8724182 | 65.0276087 | 68.8290908 | 74.6168533 |
| ENSMUSG00 | Pdlim5 | 5350 | 23.0804715 | 23.8860644 | 36.6586126 | 31.9073183 | 27.1155115 | 23.7620736 |
| ENSMUSG00 | Rngtt | 4600 | 13.5082209 | 8.10789266 | 10.9639789 | 14.7544186 | 9.10709882 | 17.3253252 |
| ENSMUSG00 | Ube2j1 | 3610 | 26.2430222 | 25.5882021 | 24.855884 | 20.9695649 | 22.3894126 | 18.9940286 |
| ENSMUSG00 | Rragd | 5617 | 145.956475 | 93.0786714 | 111.043654 | 127.542993 | 116.75111 | 100.241704 |
| ENSMUSG00 | Gabrr1 | 2495 | 0 | 0.07725281 | 0 | 0 | 0.20451453 | 0 |
| ENSMUSG00 | Casp8ap2 | 6837 | 6.27993549 | 5.45506892 | 6.6784414 | 8.33839847 | 6.64977342 | 5.73657032 |
| ENSMUSG00 | Map3k7 | 6408 | 15.2185071 | 12.6481891 | 11.7540481 | 12.0104558 | 15.3684309 | 9.5767694 |
| ENSMUSG00 | 1700009N14 | 1256 | 0 | 0 | 0 | 0 | 0 | 0 |
| ENSMUSG00 | Epha7 | 7704 | 0.29930139 | 0.42532166 | 0.62538952 | 0 | 0.8742837 | 0.23720156 |
| ENSMUSG00 | Akirin2 | 2028 | 23.2299036 | 20.0776842 | 19.3982447 | 19.9981351 | 23.0725772 | 26.4811463 |
| ENSMUSG00 | Rars2 | 2816 | 19.7365631 | 25.7359407 | 24.2199729 | 23.7813475 | 24.3534976 | 21.715107 |
| ENSMUSG00 | Slc35a1 | 2620 | 66.4311045 | 76.8959979 | 72.5956223 | 71.4892722 | 60.6084281 | 84.3431783 |
| ENSMUSG00 | 1700003M02 | 2512 | 0 | 0 | 0.22875158 | 0.18673033 | 0 | 0.02171548 |
| ENSMUSG00 | 1810030N24 | 1977 | 20.3905196 | 16.8664734 | 17.8417151 | 22.0470965 | 14.0664516 | 35.0555682 |
| ENSMUSG00 | Cga | 1022 | 0 | 0 | 0 | 0 | 0 | 0 |
| ENSMUSG00 | 3110043O21 | 6481 | 4.81531433 | 4.30488351 | 7.14759011 | 3.95282389 | 4.82628736 | 4.831243 |
| ENSMUSG00 | Aldob | 2382 | 13.585626 | 24.2348269 | 5.51131292 | 10.497421 | 15.402168 | 25.4654902 |
| ENSMUSG00 | Rnf20 | 7267 | 9.06492792 | 8.34161886 | 11.1493021 | 14.4238805 | 9.99883847 | 7.72787701 |
| ENSMUSG00 | Ppp3r2 | 3024 | 0 | 0 | 0 | 0 | 0 | 0 |
| ENSMUSG00 | Smc2 | 5649 | 1.80866675 | 2.73815683 | 2.12832383 | 1.14333262 | 2.05948198 | 3.74187386 |
| ENSMUSG00 | 4930547C10 | 2771 | 0 | 0 | 0 | 0 | 0 | 0 |
| ENSMUSG00 | Polr1e | 4011 | 1.75435554 | 2.67902683 | 2.33627293 | 2.64475719 | 2.50615708 | 1.68639006 |
| ENSMUSG00 | Exosc3 | 1553 | 13.9259378 | 17.5928607 | 21.4889836 | 16.8677169 | 15.1140585 | 21.3033725 |
| ENSMUSG00 | 1300002K09 | 7972 | 0 | 0.25386736 | 0.060991 | 0.10862633 | 0 | 0 |
| ENSMUSG00 | Tmod1 | 4032 | 8.3514117 | 5.86794224 | 5.3059752 | 5.63781946 | 5.78349547 | 4.94488127 |
| ENSMUSG00 | Xpa | 1885 | 6.74894099 | 5.59831874 | 5.51057459 | 5.11082541 | 4.81840578 | 8.31985017 |
| ENSMUSG00 | Ncbp1 | 3363 | 21.739616 | 25.977407 | 19.5444962 | 22.7135558 | 22.2130878 | 17.5099453 |
| ENSMUSG00 | 5830415F09 | 2515 | 1.65977155 | 3.83192383 | 2.56599168 | 2.51067904 | 6.04606759 | 2.25571569 |
| ENSMUSG00 | Hemgn | 2506 | 0.01586412 | 0 | 0.45859853 | 0 | 0.08144673 | 0.57683794 |
| ENSMUSG00 | Anp32b | 1797 | 21.5259224 | 23.0876609 | 28.7299681 | 24.6169096 | 20.8137638 | 31.9797801 |
| ENSMUSG00 | Nans | 4744 | 2.31292425 | 3.28082226 | 3.04679652 | 2.27414235 | 1.5058374 | 2.2537223 |
| ENSMUSG00 | Coro2a | 8632 | 0.45134813 | 0.56939494 | 0.85515614 | 0.7231457 | 1.19999471 | 1.04902459 |
| ENSMUSG00 | Col15a1 | 5937 | 0.32141874 | 0.28407032 | 0 | 0.12154975 | 0.06875712 | 0.03675208 |
| ENSMUSG00 | Nr4a3 | 6463 | 0 | 0 | 0 | 0 | 0 | 0 |
| ENSMUSG00 | Erp44 | 3623 | 37.3414022 | 50.8331719 | 49.9848056 | 50.7020946 | 39.8155022 | 60.7976778 |
| ENSMUSG00 | Invs | 11702 | 0.95804525 | 1.41652163 | 1.57890706 | 2.08438405 | 1.46076191 | 1.19801441 |
| ENSMUSG00 | Tex10 | 4493 | 7.71573092 | 11.3039194 | 7.27023414 | 14.0377062 | 8.85835147 | 11.7706489 |
| ENSMUSG00 | Tmeff1 | 2986 | 0.13313959 | 0.83914769 | 1.19904531 | 2.01798441 | 0.27341662 | 0 |
| ENSMUSG00 | Murc | 2052 | 0.1162441 | 0 | 0 | 0 | 0 | 0 |
| ENSMUSG00 | Dbc1 | 6173 | 0 | 0 | 0 | 0 | 0 | 0 |
| ENSMUSG00 | Fmn2 | 8157 | 0 | 0 | 0 | 0 | 0 | 0 |
| ENSMUSG00 | Ambp | 1398 | 0.39812355 | 0.34468127 | 0.28456122 | 0.92915136 | 2.043975 | 0.13656829 |
| ENSMUSG00 | Kif12 | 3193 | 61.0961926 | 84.9940627 | 19.0207743 | 67.8021507 | 69.5799688 | 44.3586601 |
| ENSMUSG00 | Zfp618 | 9465 | 6.85902796 | 9.46419397 | 8.05113337 | 8.23044206 | 9.12706331 | 9.05408521 |
| ENSMUSG00 | Orm3 | 767 | 0 | 0 | 0 | 0 | 0 | 0 |
| ENSMUSG00 | Slc44a5 | 7119 | 0 | 0 | 0 | 0 | 0 | 0 |
| ENSMUSG00 | Tnfsf8 | 3803 | 0 | 0 | 0 | 0 | 0 | 0 |
| ENSMUSG00 | Tnc | 7483 | 0 | 0 | 0.05906968 | 0 | 0 | 0 |
| ENSMUSG00 | Txn1 | 1046 | 82.6275474 | 94.4841232 | 59.4569724 | 82.4781725 | 50.2458576 | 104.561476 |
| ENSMUSG00 | Svep1 | 14892 | 0.02402628 | 0.02588581 | 0.0148408 | 0.02422915 | 0.01713214 | 0.01098898 |
| ENSMUSG00 | Pappa | 11091 | 4.35872909 | 7.84642634 | 2.66621882 | 6.84163192 | 5.11597962 | 1.60583711 |
| ENSMUSG00 | Astn2 | 5201 | 0.78731292 | 0.20382652 | 2.46462882 | 0.42318876 | 0.69657233 | 1.13797284 |
| ENSMUSG00 | Ptgr1 | 2286 | 4.03467694 | 5.03786512 | 5.9747898 | 4.00911606 | 3.48211489 | 6.99166168 |
| ENSMUSG00 | Ugcg | 5386 | 4.46566389 | 6.69206438 | 6.45875426 | 8.22665256 | 4.92642321 | 3.11435261 |
| ENSMUSG00 | Rod1 | 12431 | 27.9992947 | 26.1534812 | 31.0276955 | 35.7598575 | 27.5881483 | 30.8685463 |
| ENSMUSG00 | Hadl2 | 2967 | 69.0461716 | 69.3969219 | 90.1169358 | 75.5692028 | 65.8854216 | 91.4854094 |
| ENSMUSG00 | Snx30 | 8129 | 1.93177695 | 1.86723204 | 1.21801117 | 2.28592114 | 1.65714888 | 1.30182803 |
| ENSMUSG00 | Slc46a2 | 2829 | 0 | 0.47692484 | 0.03124909 | 0.01275435 | 0 | 0 |
| ENSMUSG00 | Zfp37 | 3820 | 7.69091626 | 2.20749408 | 5.95914899 | 4.68499845 | 7.41351795 | 7.03999851 |
| ENSMUSG00 | Wdr31 | 4717 | 3.30382624 | 3.18723128 | 1.1432319 | 1.05561207 | 4.29456672 | 1.20269768 |
| ENSMUSG00 | Bspry | 2320 | 24.8129034 | 35.6828912 | 14.4227565 | 21.9291736 | 27.6906065 | 15.1773962 |
| ENSMUSG00 | Alad | 1860 | 79.767448 | 48.1346288 | 99.0976803 | 46.86786 | 39.2848227 | 78.6125489 |
| ENSMUSG00 | Pole3 | 2037 | 27.2843209 | 26.5415749 | 30.5745689 | 19.2543851 | 25.400464 | 21.2359243 |
| ENSMUSG00 | 2310002L09 | 1177 | 0 | 0 | 0 | 0 | 0 | 0 |
| ENSMUSG00 | Kdm4c | 5797 | 11.2607381 | 9.07703896 | 14.3272836 | 9.51067173 | 15.3158347 | 19.751412 |
| ENSMUSG00 | 3110001D03 | 739 | 16.138896 | 14.605904 | 20.4560627 | 10.839261 | 10.2881326 | 31.4820938 |
| ENSMUSG00 | Ptprd | 26192 | 0.76954905 | 0.87571573 | 0.49109409 | 2.06777447 | 0.71302892 | 0.86118382 |
| ENSMUSG00 | Mpdz | 12517 | 0 | 0.40421638 | 0.11300303 | 0 | 0.40765659 | 0.20918474 |
| ENSMUSG00 | Zdhhc21 | 15374 | 2.22903761 | 1.94325446 | 3.83251312 | 3.76451148 | 3.88323531 | 4.18149634 |
| ENSMUSG00 | Aco1 | 5450 | 38.2673855 | 34.6500306 | 30.8115232 | 33.1160343 | 36.2334082 | 22.2951409 |
| ENSMUSG00 | 2010003O02 | 1052 | 4.34589378 | 3.66436822 | 4.87396748 | 1.74922459 | 3.54080365 | 7.15570383 |
| ENSMUSG00 | Smu1 | 3106 | 46.4112596 | 45.9679421 | 50.7766181 | 42.8430715 | 44.2086212 | 38.8571406 |
| ENSMUSG00 | Dnaja1 | 8533 | 15.5052431 | 16.1675593 | 18.3893757 | 22.0179545 | 13.0660531 | 18.4047079 |
| ENSMUSG00 | Aptx | 7664 | 5.29623507 | 5.28139502 | 4.84466964 | 3.30500982 | 4.80036756 | 4.8292909 |
| ENSMUSG00 | Slc44a1 | 6978 | 32.3433452 | 36.7578721 | 28.8154462 | 37.1317228 | 37.6811285 | 22.1972881 |
| ENSMUSG00 | B4galt1 | 4227 | 3.70562085 | 4.06968532 | 2.122776 | 3.45711569 | 6.22891174 | 3.49079249 |
| ENSMUSG00 | Fktn | 6518 | 17.6331841 | 13.8985135 | 18.716951 | 18.8492423 | 16.4007759 | 16.1019963 |
| ENSMUSG00 | Spink4 | 574 | 0 | 0 | 0 | 0 | 0 | 0.19006717 |
| ENSMUSG00 | Bag1 | 1604 | 188.912888 | 225.130422 | 176.862487 | 184.5943 | 162.55909 | 211.633516 |
| ENSMUSG00 | Tal2 | 2969 | 0.26780384 | 0.30836726 | 0.08932673 | 0 | 0 | 0.04593237 |
| ENSMUSG00 | Chmp5 | 1511 | 245.031628 | 228.463052 | 211.706643 | 233.231852 | 198.364614 | 348.55942 |
| ENSMUSG00 | Tmem38b | 3962 | 5.97034601 | 8.93918096 | 6.70502465 | 8.24186025 | 5.85991948 | 5.70688439 |
| ENSMUSG00 | Nfx1 | 8539 | 13.2642422 | 13.2387157 | 10.5392847 | 15.1697552 | 13.4512674 | 14.2202476 |
| ENSMUSG00 | Rad23b | 5473 | 2.90557139 | 3.35447368 | 2.6894233 | 4.27209298 | 2.6011964 | 6.26923007 |
| ENSMUSG00 | Aqp7 | 2818 | 0 | 0 | 0 | 0 | 0 | 0 |
| ENSMUSG00 | Nol6 | 5217 | 9.12158524 | 8.88544323 | 9.42159276 | 7.65628202 | 9.29174948 | 7.75317041 |
| ENSMUSG00 | Ikbkap | 8069 | 2.71967099 | 4.26386413 | 3.99344944 | 4.71763006 | 5.88741553 | 4.55309692 |
| ENSMUSG00 | Ubap2 | 5701 | 12.3429575 | 9.48345695 | 9.80798657 | 9.27842085 | 12.2978847 | 7.17149346 |
| ENSMUSG00 | Epb4.1l4b | 8953 | 13.4457271 | 15.635163 | 9.54341143 | 13.8436081 | 16.4540542 | 12.2862303 |
| ENSMUSG00 | Aqp3 | 1823 | 842.214294 | 1151.95465 | 603.744318 | 701.531534 | 595.158436 | 463.953134 |
| ENSMUSG00 | Dcaf12 | 3433 | 27.7929372 | 13.7555238 | 14.2790108 | 21.2204043 | 17.3011073 | 15.4527158 |
| ENSMUSG00 | Ubap1 | 3758 | 31.7366795 | 32.8124283 | 32.2751084 | 35.7555971 | 25.3502511 | 26.7303077 |
| ENSMUSG00 | Kif24 | 9499 | 0.16740912 | 0.12174699 | 0.07910636 | 0.37225396 | 0.22024228 | 0.21822009 |
| ENSMUSG00 | 2310028H24 | 6420 | 0.60066692 | 0.19514759 | 0.39933129 | 0.64632946 | 0.8742837 | 0.15719029 |
| ENSMUSG00 | 1110017D15 | 1354 | 0.26425356 | 0 | 0.4570353 | 0.10659393 | 0 | 0.08057501 |
| ENSMUSG00 | Nudt2 | 842 | 18.6029208 | 22.4908216 | 28.8729378 | 21.8120673 | 16.4229784 | 28.6027393 |
| ENSMUSG00 | Cntfr | 2769 | 0.73222445 | 1.2355498 | 0.20752039 | 0.56032066 | 0.16584954 | 0.35459986 |
| ENSMUSG00 | Enho | 2035 | 0.19535863 | 0.35518262 | 0 | 0 | 0.05014877 | 0.14743048 |
| ENSMUSG00 | Dctn3 | 1475 | 24.0958641 | 32.7667807 | 31.22598 | 31.7032744 | 19.8224497 | 36.9640704 |
| ENSMUSG00 | 1700022I11R | 4475 | 0 | 0 | 0 | 0 | 0.02280508 | 0 |
| ENSMUSG00 | Vcp | 4760 | 13.7724343 | 13.7371853 | 13.9291522 | 13.9401008 | 14.1394516 | 10.4743783 |
| ENSMUSG00 | Fancg | 3277 | 2.92373232 | 6.57288363 | 3.22375358 | 6.08891383 | 6.07271485 | 2.77989915 |
| ENSMUSG00 | Pigo | 3682 | 28.6667031 | 18.0208394 | 27.4790925 | 15.728322 | 23.4205799 | 19.1189147 |
| ENSMUSG00 | Stoml2 | 3284 | 21.6209769 | 17.9305214 | 21.0510604 | 18.5903835 | 15.0095856 | 18.3132094 |
| ENSMUSG00 | Unc13b | 11170 | 5.16429742 | 5.81947274 | 4.55472886 | 6.42176406 | 8.78457661 | 4.15834473 |
| ENSMUSG00 | Atp8b5 | 6515 | 0.34782232 | 0.86535898 | 0.78023207 | 0.63136655 | 0.03132855 | 0.33072855 |
| ENSMUSG00 | Tesk1 | 4737 | 5.07748907 | 7.4054739 | 5.82266203 | 6.17744102 | 6.21537236 | 5.91324773 |
| ENSMUSG00 | Cd72 | 3727 | 0.02133377 | 0.05171606 | 0.11859899 | 0.01936251 | 0.06845503 | 0.10977182 |
| ENSMUSG00 | Sit1 | 1918 | 0 | 0 | 0 | 0 | 0 | 0 |
| ENSMUSG00 | Ccdc107 | 1542 | 19.9550856 | 20.9057911 | 25.9420674 | 22.065738 | 13.6665974 | 26.1956814 |
| ENSMUSG00 | Car9 | 2647 | 0.34543863 | 0.14563337 | 0.1836873 | 0.12268168 | 0 | 0.31942343 |
| ENSMUSG00 | Tpm2 | 2886 | 0.37193277 | 0.30053914 | 0.32163503 | 0.08751709 | 0.03536131 | 0.36857621 |
| ENSMUSG00 | Tln1 | 9742 | 11.181484 | 11.9353197 | 10.6126166 | 13.9076244 | 14.4614888 | 10.3224795 |
| ENSMUSG00 | Creb3 | 2343 | 30.1517238 | 31.527877 | 32.6938941 | 29.2290741 | 25.8289721 | 31.0462923 |
| ENSMUSG00 | Gba2 | 4639 | 12.4005562 | 9.78478735 | 12.5869012 | 6.59572618 | 18.2920593 | 6.57907336 |
| ENSMUSG00 | Rgp1 | 6452 | 2.66186726 | 1.65052754 | 1.60310466 | 1.0513677 | 1.59753997 | 1.24705806 |
| ENSMUSG00 | Npr2 | 5096 | 0.65531012 | 0.1229246 | 0.44236538 | 0.49563248 | 0.34044285 | 0 |
| ENSMUSG00 | Hint2 | 1721 | 13.7215313 | 14.9515166 | 19.0830734 | 11.2376385 | 11.0888231 | 24.0733161 |
| ENSMUSG00 | 5430416O09 | 4084 | 0 | 0 | 0 | 0.32689414 | 0 | 0.04007048 |
| ENSMUSG00 | Reck | 4756 | 1.12010816 | 1.62107459 | 1.62643451 | 0.53106458 | 0.52571329 | 1.32473542 |
| ENSMUSG00 | Cilta | 1788 | 28.5937069 | 22.9343749 | 31.1489498 | 20.9671389 | 17.8649392 | 25.8712462 |
| ENSMUSG00 | Gne | 6708 | 15.4979996 | 12.1256283 | 15.6960033 | 11.1989888 | 13.2282152 | 14.5846945 |
| ENSMUSG00 | Glipr2 | 2396 | 1.72561351 | 0.16088962 | 6.82582718 | 0 | 0.38333671 | 6.42024063 |
| ENSMUSG00 | Snapc3 | 7069 | 11.1128631 | 10.7633883 | 12.6809079 | 10.157486 | 9.03734935 | 19.3958992 |
| ENSMUSG00 | Psip1 | 5029 | 17.763087 | 13.3856551 | 17.1568896 | 19.0347314 | 16.8633598 | 30.0026455 |
| ENSMUSG00 | Bnc2 | 12061 | 0 | 0 | 0 | 0.00299163 | 0 | 0.05879617 |
| ENSMUSG00 | Sh3gl2 | 4942 | 18.3412577 | 27.3206012 | 18.3086145 | 28.7152329 | 19.886038 | 14.6914386 |
| ENSMUSG00 | Fam154a | 2189 | 0 | 0 | 0 | 0 | 0 | 0 |
| ENSMUSG00 | Plin2 | 3499 | 16.6452641 | 14.597779 | 14.6539405 | 18.2627325 | 15.7643648 | 10.0945007 |
| ENSMUSG00 | Rps6 | 1601 | 0.74494967 | 0.63205202 | 0.93870248 | 0.92402488 | 1.33860574 | 1.03919612 |
| ENSMUSG00 | Mllt3 | 13461 | 2.15596915 | 1.31733235 | 2.66307813 | 2.49553402 | 2.65726836 | 2.65431821 |
| ENSMUSG00 | Ptplad2 | 6875 | 4.88052735 | 8.73313555 | 3.03465744 | 8.29755817 | 6.59817427 | 4.24889289 |
| ENSMUSG00 | Usp24 | 12307 | 4.51920998 | 7.30216256 | 5.91895972 | 7.00707617 | 6.96134598 | 7.6857438 |
| ENSMUSG00 | Ppap2b | 3153 | 24.8140773 | 30.1528228 | 15.5610675 | 24.2492396 | 31.9137371 | 16.9547393 |
| ENSMUSG00 | Prkaa2 | 7960 | 19.543115 | 22.9551493 | 22.2675114 | 21.3863173 | 22.372117 | 19.4314672 |
| ENSMUSG00 | Dab1 | 8006 | 2.03097571 | 3.04550833 | 1.40787784 | 1.46022763 | 1.47228239 | 0.32705038 |
| ENSMUSG00 | 4921539E11 | 2744 | 0 | 0 | 0 | 0 | 0 | 0 |
| ENSMUSG00 | Slc35d1 | 11382 | 1.35522109 | 1.98977577 | 1.54174413 | 2.66605161 | 1.23732909 | 1.6726145 |
| ENSMUSG00 | Mier1 | 10032 | 3.81227794 | 4.36137255 | 3.50283744 | 4.46709523 | 3.66218008 | 5.05690082 |
| ENSMUSG00 | Tctex1d1 | 2536 | 0 | 0 | 0 | 0 | 0 | 0 |
| ENSMUSG00 | Sgip1 | 11673 | 0.04427493 | 0.00412803 | 0 | 0 | 0 | 0 |
| ENSMUSG00 | Pde4b | 10597 | 4.53190719 | 6.08867095 | 4.73844425 | 4.3174514 | 4.3577305 | 3.719152 |
| ENSMUSG00 | Ak4 | 5344 | 0.23805677 | 0.2254231 | 0.07444173 | 0 | 0 | 0.1990477 |
| ENSMUSG00 | Dnajc6 | 5966 | 0.63304905 | 0.25038212 | 0.41490164 | 0.70760966 | 0.47040742 | 0.28344412 |
| ENSMUSG00 | Jak1 | 7174 | 21.0470191 | 25.5642039 | 22.8033205 | 20.0981112 | 30.0155152 | 22.3017889 |
| ENSMUSG00 | Cachd1 | 5991 | 1.49970599 | 1.44776491 | 0.83371862 | 0.92749706 | 2.1378101 | 0.53720705 |
| ENSMUSG00 | Izumo3 | 1670 | 0 | 0 | 0 | 0 | 0 | 0 |
| ENSMUSG00 | 2610528J11R | 1200 | 27.5969294 | 29.1527975 | 39.8553285 | 29.70755 | 17.8167096 | 46.2305135 |
| ENSMUSG00 | St3gal3 | 3732 | 8.31967585 | 6.02976112 | 5.40086828 | 5.04684546 | 4.97684909 | 7.14022576 |
| ENSMUSG00 | Artn | 3305 | 4.4506892 | 3.65954523 | 3.1563192 | 3.22063635 | 3.53556432 | 3.43305595 |
| ENSMUSG00 | Dph2 | 3593 | 11.0978978 | 6.73242248 | 5.58520365 | 8.25478447 | 7.82508574 | 10.2251431 |
| ENSMUSG00 | B4galt2 | 4981 | 3.40008727 | 1.93480996 | 2.74209386 | 2.24562012 | 1.47516525 | 2.29433324 |
| ENSMUSG00 | Slc6a9 | 5119 | 3.55694668 | 4.24537711 | 4.42104779 | 3.90495266 | 3.97724633 | 2.8025904 |
| ENSMUSG00 | Slc5a9 | 7485 | 0 | 0.35407539 | 0 | 0.4675963 | 0.39539476 | 0.08016594 |
| ENSMUSG00 | Bend5 | 2450 | 6.08502253 | 2.41915608 | 8.60582821 | 5.59639874 | 5.91489418 | 9.44036493 |
| ENSMUSG00 | Elavl4 | 8251 | 0 | 0 | 0.0964287 | 0 | 0 | 0.01322247 |
| ENSMUSG00 | Itgb3bp | 6295 | 3.45452706 | 2.31172447 | 2.54186929 | 3.11239876 | 2.59387455 | 3.63084157 |
| ENSMUSG00 | Atg4c | 3340 | 5.46340286 | 4.37140478 | 3.69230964 | 6.10369915 | 4.4762659 | 8.99083168 |
| ENSMUSG00 | Cdkn2c | 2683 | 1.55584251 | 2.8556259 | 1.74632701 | 1.84243015 | 2.16809797 | 2.29745377 |
| ENSMUSG00 | Eps15 | 7048 | 17.170216 | 15.3761788 | 20.5330355 | 21.1229449 | 16.5575087 | 18.7725986 |
| ENSMUSG00 | Angptl3 | 3571 | 2.53829447 | 3.30598665 | 2.53749029 | 2.53615043 | 3.35793848 | 4.01749097 |
| ENSMUSG00 | Ttc39a | 4883 | 0.02442483 | 0.28617793 | 0.12673066 | 0.857161 | 0.25079521 | 0 |
| ENSMUSG00 | Dock7 | 8687 | 11.7110941 | 14.5385824 | 9.40313179 | 14.9777659 | 17.439543 | 11.0800279 |
| ENSMUSG00 | Rnf11 | 5781 | 58.3850596 | 56.5634313 | 65.534336 | 63.8254605 | 44.0622156 | 66.372535 |
| ENSMUSG00 | Calr4 | 2503 | 0 | 0 | 0 | 0 | 0 | 0 |
| ENSMUSG00 | Osbpl9 | 6363 | 30.8084668 | 26.2931522 | 27.0226574 | 23.8448841 | 29.5749291 | 27.1031815 |
| ENSMUSG00 | Usp1 | 4088 | 12.2047769 | 10.0663458 | 12.1209066 | 12.047943 | 12.4445442 | 11.4622872 |
| ENSMUSG00 | Tm2d1 | 1191 | 78.1423844 | 87.5528638 | 88.2181199 | 78.0717286 | 54.0682505 | 116.518358 |
| ENSMUSG00 | Nfia | 10900 | 1.3786763 | 0.96814962 | 2.33580381 | 1.68824245 | 1.22182423 | 1.52888115 |
| ENSMUSG00 | Txndc12 | 2467 | 6.13978033 | 5.82065657 | 6.27103569 | 5.6163377 | 4.96405764 | 8.94413585 |
| ENSMUSG00 | Btf3l4 | 4706 | 5.56711892 | 6.85013383 | 8.50974708 | 5.45907686 | 6.47317177 | 8.69936964 |
| ENSMUSG00 | Cyp2j13 | 4680 | 0.00849476 | 0.17503622 | 0 | 0 | 0 | 0.46040524 |
| ENSMUSG00 | Hook1 | 5559 | 15.5903845 | 18.4198938 | 17.8111402 | 21.5103248 | 22.7457023 | 17.623764 |
| ENSMUSG00 | Fggy | 5898 | 8.56043748 | 6.89544883 | 5.725705 | 6.44802898 | 6.15119586 | 5.59088485 |
| ENSMUSG00 | 4930579C15 | 1472 | 0.27007799 | 0.22914748 | 0 | 0 | 0.20798795 | 0.64851384 |
| ENSMUSG00 | Ift74 | 2635 | 26.2521959 | 21.7250449 | 15.6677501 | 21.7724671 | 14.8141091 | 28.1027118 |
| ENSMUSG00 | Plaa | 7169 | 15.0670443 | 13.960558 | 17.1714511 | 20.5751706 | 16.5129295 | 11.9614264 |
| ENSMUSG00 | 5830433M19 | 2827 | 7.17201806 | 9.51115483 | 11.4608953 | 13.3504841 | 8.8623808 | 15.7646836 |
| ENSMUSG00 | Pum1 | 6096 | 16.4865247 | 19.4927438 | 16.0608731 | 21.2491043 | 23.6633144 | 17.3106265 |
| ENSMUSG00 | Laptm5 | 3898 | 0.18358098 | 0 | 0 | 0 | 0 | 0 |
| ENSMUSG00 | Cc2d1b | 4011 | 11.9137591 | 6.49934311 | 9.47733358 | 9.1127178 | 11.0423571 | 7.43915616 |
| ENSMUSG00 | Pdpn | 1891 | 0.69377623 | 0.43319382 | 0.32724791 | 0.24805213 | 0.9984008 | 0.33173808 |
| ENSMUSG00 | Lrrc38 | 2322 | 0 | 0 | 0 | 0 | 0 | 0 |
| ENSMUSG00 | Orc1 | 4797 | 0.058013 | 0.41184993 | 0.04607238 | 0.08273973 | 0.75523717 | 0.18194464 |
| ENSMUSG00 | 1700012P22 | 970 | 0 | 0 | 0 | 0 | 0 | 0 |
| ENSMUSG00 | D530049N12 | 765 | 0 | 0 | 0.34668112 | 0 | 0 | 0 |
| ENSMUSG00 | Pramef12 | 2568 | 2.84852353 | 0.30022705 | 1.27372912 | 0.78683586 | 1.4902563 | 1.25327392 |
| ENSMUSG00 | 9430007A20 | 1402 | 0 | 0 | 0 | 0 | 0 | 0 |
| ENSMUSG00 | Gpx7 | 1867 | 0.38328798 | 0.41295291 | 0.3077793 | 0.52180782 | 0.4919522 | 0.23374088 |
| ENSMUSG00 | Tnfrsf1b | 3808 | 0 | 0 | 0 | 0 | 0 | 0 |
| ENSMUSG00 | Podn | 5100 | 1.37195384 | 0.32124295 | 0.2166757 | 0 | 0.58029996 | 0.07487156 |
| ENSMUSG00 | Echdc2 | 2334 | 28.2069733 | 34.6430377 | 21.1161332 | 35.4482126 | 40.0952756 | 29.2378524 |
| ENSMUSG00 | Tnfrsf8 | 5541 | 0 | 0 | 0 | 0 | 0 | 0 |
| ENSMUSG00 | Scp2 | 4037 | 29.0017067 | 26.8325791 | 27.789021 | 24.0785306 | 25.9366171 | 25.2004965 |
| ENSMUSG00 | Cpt2 | 2966 | 25.6413467 | 39.2509927 | 35.4836778 | 29.7926254 | 28.5926624 | 25.168809 |
| ENSMUSG00 | 0610037L13 | 4789 | 17.3333563 | 12.4868187 | 14.1216997 | 12.4618295 | 13.9259706 | 18.8399471 |
| ENSMUSG00 | Magoh | 765 | 21.2029229 | 22.4870063 | 28.0233907 | 17.6401107 | 13.0067233 | 43.6394228 |
| ENSMUSG00 | Dmrtb1 | 2856 | 0 | 0 | 0 | 0 | 0 | 0 |
| ENSMUSG00 | 4930430J20R | 792 | 0 | 0 | 0 | 0 | 0 | 0 |
| ENSMUSG00 | Lrp8 | 8058 | 0.77951923 | 0.59799506 | 0.24684574 | 0.48807928 | 0.33561652 | 0.08461975 |
| ENSMUSG00 | Tmem48 | 5173 | 5.07990965 | 6.34350804 | 4.67396252 | 5.10575229 | 4.24151201 | 6.20045926 |
| ENSMUSG00 | Lrrc42 | 2596 | 6.99855724 | 9.67069968 | 7.13427281 | 8.31165741 | 6.30950179 | 7.25992901 |
| ENSMUSG00 | Tmem59 | 2437 | 296.820258 | 343.217178 | 403.947167 | 298.45788 | 278.247644 | 377.524879 |
| ENSMUSG00 | 2210012G02 | 7628 | 1.60001737 | 1.49082333 | 1.66307406 | 0.69534092 | 1.86632916 | 2.07384515 |
| ENSMUSG00 | Cyb5rl | 4262 | 9.33722102 | 6.44445613 | 3.930666 | 6.69659719 | 10.3680998 | 5.76594321 |
| ENSMUSG00 | Mrpl37 | 4672 | 26.446925 | 27.7959036 | 27.9194432 | 27.8106683 | 21.1117261 | 27.9343212 |
| ENSMUSG00 | Col9a2 | 3102 | 0 | 0 | 0 | 0 | 0 | 0 |
| ENSMUSG00 | Dem1 | 1956 | 13.4550757 | 8.81939994 | 6.57604106 | 6.56707968 | 8.0870023 | 8.89632916 |
| ENSMUSG00 | Dyrk2 | 2134 | 4.22891006 | 4.51606768 | 6.66963142 | 6.54346356 | 8.01023238 | 5.94316179 |
| ENSMUSG00 | Kcnq4 | 4009 | 0 | 0 | 0 | 0 | 0 | 0 |
| ENSMUSG00 | Ctps | 2775 | 3.18043844 | 8.2828587 | 2.85121796 | 4.81093927 | 3.65918875 | 2.86015136 |
| ENSMUSG00 | Hivep3 | 11885 | 0.23415092 | 0.10946857 | 0.27521551 | 0.22769485 | 0.05152011 | 0.18359034 |
| ENSMUSG00 | Edn2 | 1457 | 0 | 0 | 0 | 0 | 0 | 0 |
| ENSMUSG00 | Ppcs | 1559 | 15.4023799 | 19.0087633 | 19.0813601 | 15.7613036 | 11.6192196 | 13.8909965 |
| ENSMUSG00 | Ccdc30 | 6765 | 0.64643058 | 1.17527908 | 0.26788996 | 1.02939166 | 0.55061721 | 0.87891668 |
| ENSMUSG00 | Ybx1 | 2900 | 43.3060562 | 58.4385231 | 40.5285174 | 50.3531152 | 53.9120054 | 50.721424 |
| ENSMUSG00 | Tfap2c | 3670 | 0 | 0 | 0 | 0 | 0 | 0 |
| ENSMUSG00 | Lepre1 | 4327 | 2.57257558 | 1.22498466 | 1.00110483 | 0.25850321 | 2.52360629 | 1.64517699 |
| ENSMUSG00 | 4930538K18 | 1445 | 0.08253733 | 0 | 0.09176853 | 0 | 0 | 0 |
| ENSMUSG00 | Ccdc23 | 1139 | 8.72596148 | 9.1803845 | 10.3228185 | 7.38113817 | 7.2126835 | 17.9835418 |
| ENSMUSG00 | Ermap | 7424 | 0.01606498 | 0 | 0 | 0 | 0 | 0 |
| ENSMUSG00 | Slc2a1 | 3009 | 118.777591 | 126.031007 | 93.104447 | 103.785342 | 90.8266098 | 58.3200495 |
| ENSMUSG00 | Rragc | 3625 | 45.0525556 | 40.3835617 | 55.1517065 | 44.9507622 | 36.6545331 | 44.9034613 |
| ENSMUSG00 | Mycbp | 3102 | 22.8638869 | 27.2310874 | 31.4913195 | 30.8709691 | 24.8552076 | 36.1727485 |
| ENSMUSG00 | Ndufs5 | 914 | 14.3102331 | 15.4997965 | 24.7607698 | 14.5275628 | 11.0538538 | 33.750128 |
| ENSMUSG00 | Macf1 | 28023 | 4.00207724 | 4.81812836 | 4.3818549 | 5.63576737 | 5.50632554 | 4.34576113 |
| ENSMUSG00 | Ppie | 1835 | 12.3491138 | 14.3377643 | 11.8995697 | 7.96361201 | 13.7924155 | 10.9841735 |
| ENSMUSG00 | Trit1 | 2076 | 8.80901764 | 4.8047175 | 12.3705543 | 9.42026403 | 12.6582772 | 8.98644183 |
| ENSMUSG00 | Mycl1 | 4033 | 0 | 0 | 0 | 0 | 0 | 0 |
| ENSMUSG00 | Mfsd2a | 2235 | 0.53363061 | 0 | 0 | 0 | 0.09132237 | 0 |
| ENSMUSG00 | Cap1 | 3470 | 25.7665924 | 7.44320835 | 29.5018641 | 28.9696185 | 7.08761359 | 28.4457981 |
| ENSMUSG00 | Ppt1 | 3797 | 117.905311 | 100.065867 | 117.541746 | 111.714647 | 96.4085384 | 118.781521 |
| ENSMUSG00 | Epha8 | 4696 | 0 | 0 | 0 | 0 | 0 | 0 |
| ENSMUSG00 | Ephb2 | 14278 | 0.3925986 | 1.05633537 | 0.22599345 | 0.83141845 | 0.76121432 | 0.48902561 |
| ENSMUSG00 | Tceb3 | 4697 | 16.0816293 | 18.1276226 | 15.9040057 | 16.7005248 | 17.0558676 | 13.4079502 |
| ENSMUSG00 | 1110049F12 | 2868 | 16.1212078 | 15.5077009 | 15.7973811 | 12.6186509 | 15.4075458 | 13.1903503 |
| ENSMUSG00 | Lypla2 | 2334 | 18.5150845 | 20.7486608 | 19.089742 | 20.1744079 | 16.6808589 | 19.4568442 |
| ENSMUSG00 | Gale | 2281 | 8.62734014 | 7.54167464 | 6.56923489 | 4.08117821 | 4.07137238 | 3.65893889 |
| ENSMUSG00 | Hmgcl | 2497 | 19.7424092 | 17.4451517 | 19.2597538 | 18.1782988 | 17.4515518 | 16.6793649 |
| ENSMUSG00 | Fuca1 | 3268 | 34.3785153 | 37.0982523 | 43.2009445 | 31.0804637 | 27.9489635 | 33.2670478 |
| ENSMUSG00 | Pnrc2 | 2339 | 99.210235 | 97.4647532 | 100.592736 | 111.531929 | 94.9190941 | 115.89681 |
| ENSMUSG00 | Srsf10 | 7385 | 15.3154153 | 12.6909452 | 17.3934492 | 19.8219167 | 14.136759 | 25.6680762 |
| ENSMUSG00 | Rnf220 | 9127 | 10.3406936 | 9.40815601 | 6.19416645 | 5.07607595 | 7.37414153 | 3.59796928 |
| ENSMUSG00 | Kif2c | 4615 | 0.19813132 | 0 | 0.02873359 | 0.00781843 | 0 | 0 |
| ENSMUSG00 | Plk3 | 3038 | 0.15703284 | 0.04758372 | 0.08729791 | 0.01187691 | 1.10853878 | 0 |
| ENSMUSG00 | Ptch2 | 6194 | 0.2246435 | 0 | 0 | 0 | 0 | 0 |
| ENSMUSG00 | Eif2b3 | 6606 | 4.57374587 | 5.01121492 | 3.39911236 | 5.82796571 | 5.29110921 | 3.64157309 |
| ENSMUSG00 | Urod | 2688 | 32.1090581 | 33.2895175 | 39.5645961 | 28.5112584 | 26.6142035 | 26.3918663 |
| ENSMUSG00 | Mutyh | 2071 | 0.40312172 | 1.83811151 | 1.49402656 | 0.94081623 | 1.62614235 | 2.00180839 |
| ENSMUSG00 | Toe1 | 2151 | 11.1263595 | 15.3452873 | 10.6035105 | 8.9240575 | 10.4851967 | 9.00278655 |
| ENSMUSG00 | Ccdc163 | 2857 | 2.64387165 | 1.24809126 | 1.48525619 | 2.17224769 | 1.51811059 | 2.42483667 |
| ENSMUSG00 | Mmachc | 2102 | 27.253876 | 20.2878217 | 26.7271848 | 25.9028569 | 20.6824321 | 29.4675099 |
| ENSMUSG00 | Prdx1 | 3093 | 224.985429 | 285.940498 | 208.433198 | 264.006709 | 154.497902 | 234.801752 |
| ENSMUSG00 | Akr1a1 | 2900 | 185.863726 | 205.589948 | 208.023018 | 176.690039 | 146.234555 | 158.597325 |
| ENSMUSG00 | Nasp | 7697 | 3.10936719 | 4.85808485 | 3.15276235 | 4.2893427 | 3.6063854 | 4.76606337 |
| ENSMUSG00 | Ipp | 2222 | 13.1146567 | 17.6741451 | 13.1889388 | 24.0817606 | 13.043646 | 21.7018732 |
| ENSMUSG00 | Pik3r3 | 5702 | 17.8767191 | 29.8228612 | 23.6203114 | 34.0064727 | 29.3432982 | 22.185159 |
| ENSMUSG00 | Tspan1 | 2320 | 169.52628 | 161.653051 | 160.860414 | 124.311975 | 119.802013 | 127.073955 |
| ENSMUSG00 | Pomgnt1 | 5063 | 44.0192028 | 46.3400725 | 37.1215162 | 46.1875826 | 49.4944759 | 36.1309511 |
| ENSMUSG00 | 1520402A15 | 6197 | 10.1425552 | 13.9497301 | 9.2797479 | 14.7542199 | 19.9099204 | 12.0858737 |
| ENSMUSG00 | Rad54l | 7657 | 0.32190673 | 0.79293349 | 1.06218351 | 0.71155658 | 0.6797297 | 0.40607403 |
| ENSMUSG00 | Lrrc41 | 3841 | 14.1695529 | 15.3930655 | 10.1499676 | 12.0523986 | 10.2557569 | 5.75174637 |
| ENSMUSG00 | Nsun4 | 5781 | 3.93359883 | 3.29244848 | 3.38720229 | 3.1269857 | 4.02491392 | 3.25068785 |
| ENSMUSG00 | Dmbx1 | 9711 | 0 | 0 | 0 | 0 | 0 | 0 |
| ENSMUSG00 | Mknk1 | 4546 | 9.99571364 | 10.7057428 | 8.1966902 | 10.3975975 | 13.2448578 | 10.343484 |
| ENSMUSG00 | Mob3c | 3114 | 8.37495029 | 7.72159108 | 10.8162506 | 9.04947873 | 9.50394925 | 7.82153271 |
| ENSMUSG00 | Atpaf1 | 4086 | 2.67566254 | 3.87991866 | 2.76937636 | 3.75302715 | 2.13546507 | 3.20406922 |
| ENSMUSG00 | Cyp4a31 | 2716 | 0 | 1.11772675 | 0 | 0 | 0.54483244 | 0.54227928 |
| ENSMUSG00 | Cyp4b1 | 2771 | 7.33130658 | 6.29501698 | 3.87623524 | 3.78920063 | 6.26090501 | 9.94131564 |
| ENSMUSG00 | Cyp4a14 | 2589 | 0 | 1.47034721 | 0 | 0 | 0 | 0 |
| ENSMUSG00 | Pdzk1ip1 | 2262 | 146.420384 | 212.195027 | 162.620574 | 160.821916 | 130.881977 | 214.230113 |
| ENSMUSG00 | Tal1 | 4372 | 0 | 0 | 0.04044084 | 0 | 0 | 0 |
| ENSMUSG00 | Stil | 6002 | 0.33118528 | 0.0160568 | 0.63334863 | 0.51099197 | 0.13602499 | 0 |

| | | | | | | | | |
|---|---|---|---|---|---|---|---|---|
| ENSMUSG00 | Cmpk1 | 3411 | 141.166338 | 126.378162 | 115.318734 | 153.954288 | 78.4917602 | 186.900445 |
| ENSMUSG00 | Ebna1bp2 | 3694 | 13.2374773 | 15.849087 | 15.3880807 | 11.7603632 | 16.8108011 | 13.8071401 |
| ENSMUSG00 | Wdr65 | 4211 | 0 | 0.33184679 | 0 | 0 | 0.0121174 | 0.012954 |
| ENSMUSG00 | Pax7 | 6079 | 0 | 0 | 0 | 0 | 0 | 0 |
| ENSMUSG00 | Aldh4a1 | 5194 | 26.1158451 | 29.7152819 | 21.2584143 | 16.6308076 | 22.3301795 | 15.291442 |
| ENSMUSG00 | Tas1r2 | 3251 | 0 | 0 | 0 | 0 | 0 | 0 |
| ENSMUSG00 | Mrto4 | 2750 | 2.74674229 | 2.75100779 | 3.08609231 | 3.17521992 | 3.24713301 | 2.80680833 |
| ENSMUSG00 | Akr7a5 | 1786 | 29.6941831 | 35.8835207 | 30.4661079 | 20.7885912 | 22.9704402 | 26.5874283 |
| ENSMUSG00 | Pqlc2 | 2249 | 7.22987618 | 7.7561103 | 5.9551616 | 8.88815148 | 4.76457933 | 2.74080412 |
| ENSMUSG00 | Capzb | 3389 | 56.659478 | 53.0491637 | 60.2704977 | 44.7698428 | 37.9423037 | 50.4045635 |
| ENSMUSG00 | Htr6 | 2791 | 0 | 0 | 0 | 0 | 0 | 0 |
| ENSMUSG00 | Pla2g2f | 2460 | 0 | 0 | 0 | 0 | 0 | 0 |
| ENSMUSG00 | Pla2g2c | 3605 | 0 | 0 | 0 | 0 | 0 | 0 |
| ENSMUSG00 | Pla2g2e | 938 | 0 | 0 | 0 | 0 | 0 | 0 |
| ENSMUSG00 | Vwa5b1 | 7781 | 0.01532791 | 0 | 0.03408444 | 0 | 0 | 0.28743355 |
| ENSMUSG00 | Cda | 1503 | 0.44966279 | 0.19236105 | 0.14704539 | 0.36010025 | 0.40739621 | 1.43359714 |
| ENSMUSG00 | Pink1 | 3399 | 52.0248242 | 47.6051994 | 35.1117906 | 39.8505429 | 54.8544799 | 28.4622376 |
| ENSMUSG00 | Ddost | 2109 | 91.7448666 | 113.988696 | 99.5746592 | 95.1922692 | 88.1166718 | 78.3838732 |
| ENSMUSG00 | Kif17 | 4566 | 0.31344663 | 0.4537926 | 0.50339374 | 0.35560491 | 0.21233052 | 0.10154815 |
| ENSMUSG00 | Hp1bp3 | 8500 | 22.3706427 | 16.9729656 | 20.9360729 | 17.8839592 | 22.0854161 | 18.9831489 |
| ENSMUSG00 | Eif4g3 | 11152 | 9.55029881 | 9.53617986 | 11.3516928 | 11.9292054 | 11.5944079 | 9.66547474 |
| ENSMUSG00 | Hspg2 | 14408 | 1.77696623 | 3.27753424 | 1.49711961 | 3.30317995 | 2.38699176 | 1.58067211 |
| ENSMUSG00 | Alpl | 4015 | 0.38616781 | 0.28803851 | 1.0238534 | 0 | 0 | 0.36003882 |
| ENSMUSG00 | Ptpn12 | 4242 | 3.98304555 | 2.52177986 | 2.55291173 | 4.06582208 | 2.50200051 | 8.60289187 |
| ENSMUSG00 | Zcchc17 | 4004 | 9.0154786 | 11.0718099 | 8.22436889 | 9.25481012 | 6.85619136 | 9.543399 |
| ENSMUSG00 | Fabp3 | 819 | 7.62101397 | 5.88357047 | 6.63837203 | 4.58184692 | 4.61044177 | 8.45880143 |
| ENSMUSG00 | Tinagl1 | 4345 | 44.2479871 | 42.8077484 | 47.6911945 | 32.8932056 | 32.8941033 | 29.4967848 |
| ENSMUSG00 | Gnat3 | 1281 | 0 | 0 | 0 | 0 | 0 | 0 |
| ENSMUSG00 | Hcrtr1 | 3345 | 0 | 0 | 0 | 0 | 0 | 0 |
| ENSMUSG00 | Pef1 | 1909 | 44.295394 | 51.4173822 | 49.3189762 | 38.4635729 | 38.7308636 | 31.2893976 |
| ENSMUSG00 | Sema3c | 7438 | 15.5002546 | 9.95732207 | 16.7465439 | 12.4962822 | 14.5299629 | 15.9108174 |
| ENSMUSG00 | Bai2 | 6602 | 0 | 0.22626169 | 0 | 0 | 0.20868103 | 0 |
| ENSMUSG00 | Spocd1 | 2037 | 0 | 0 | 0 | 0 | 0 | 0 |
| ENSMUSG00 | Hpca | 3654 | 0 | 0 | 0.07258102 | 0 | 0 | 0.01492865 |
| ENSMUSG00 | Tmem54 | 1372 | 0.26078668 | 0 | 0 | 0 | 0 | 0 |
| ENSMUSG00 | Ptp4a2 | 4502 | 32.2141255 | 35.2246959 | 35.4832209 | 37.2281423 | 30.8175735 | 36.3500301 |
| ENSMUSG00 | Adc | 4012 | 3.70601937 | 3.72327943 | 3.29420515 | 1.66380315 | 4.45145352 | 2.45417367 |
| ENSMUSG00 | Khdrbs1 | 6642 | 10.9174942 | 12.5725842 | 13.1367718 | 17.2370261 | 10.4480384 | 12.565552 |
| ENSMUSG00 | Ak2 | 2131 | 35.4273381 | 33.17199 | 31.652751 | 27.9885592 | 31.0324651 | 39.8176456 |
| ENSMUSG00 | Rnf19b | 2929 | 5.06274982 | 5.67576734 | 4.04441582 | 2.16812558 | 3.3274284 | 3.97619357 |
| ENSMUSG00 | A3galt2 | 3582 | 0 | 0.14797623 | 0.34551971 | 0 | 0 | 0.05330052 |
| ENSMUSG00 | Ccdc28b | 2161 | 6.91719605 | 4.61573046 | 6.85220611 | 6.74555576 | 6.61146933 | 8.68345757 |
| ENSMUSG00 | Phc2 | 6206 | 12.3827496 | 12.6483587 | 11.6380618 | 10.4304202 | 11.3136148 | 11.9892388 |
| ENSMUSG00 | 2510006D16l | 3897 | 24.1674964 | 28.1798567 | 25.9857384 | 27.6101249 | 24.8912344 | 28.0235195 |
| ENSMUSG00 | Eif3i | 2326 | 34.4057533 | 39.6305519 | 38.7478753 | 34.5928458 | 24.1750069 | 33.5949447 |
| ENSMUSG00 | Zfp362 | 4575 | 2.41574286 | 3.19136437 | 2.17386113 | 2.72094106 | 2.50949389 | 3.85123868 |
| ENSMUSG00 | Hdac1 | 3759 | 7.07539677 | 7.02478593 | 9.12493487 | 8.83093402 | 7.1265888 | 7.72019584 |
| ENSMUSG00 | 4930555I21R | 3045 | 0.18278382 | 0.14242298 | 0.15967825 | 0.03554881 | 0.28487632 | 0.05374313 |
| ENSMUSG00 | Nipal3 | 7489 | 3.73188714 | 2.39999238 | 3.16950056 | 3.46896409 | 2.8821147 | 4.21738981 |
| ENSMUSG00 | Csmd2 | 17103 | 0 | 0 | 0 | 0 | 0.00298348 | 0.00159473 |
| ENSMUSG00 | Zbtb8a | 2316 | 3.98241428 | 2.24703616 | 3.77891404 | 2.99125758 | 3.61326668 | 3.238569 |
| ENSMUSG00 | Srrm1 | 7188 | 18.1244727 | 17.2420046 | 17.5995652 | 23.0708231 | 18.2581857 | 22.7060992 |
| ENSMUSG00 | Yars | 5103 | 9.01373524 | 9.78270705 | 9.21629649 | 6.56872807 | 8.94936437 | 5.38758306 |
| ENSMUSG00 | CK137956 | 2312 | 0 | 0 | 0 | 0 | 0 | 0 |
| ENSMUSG00 | Sfpq | 4731 | 13.798034 | 13.9130585 | 10.3614128 | 18.3651582 | 9.50205815 | 14.7298041 |
| ENSMUSG00 | Syf2 | 2154 | 25.5808245 | 29.462184 | 32.4844557 | 31.3414602 | 30.4168368 | 42.140204 |
| ENSMUSG00 | Tmem50a | 1348 | 52.2896639 | 75.210886 | 71.7132275 | 62.2602638 | 59.0512955 | 67.0332936 |
| ENSMUSG00 | Rhd | 1584 | 0 | 0 | 0 | 0 | 0 | 0 |
| ENSMUSG00 | Tmem57 | 8537 | 3.75807342 | 3.93416308 | 4.88255101 | 5.61285641 | 5.36742245 | 4.1149977 |
| ENSMUSG00 | AU040320 | 6175 | 14.0286951 | 12.3607003 | 14.6170305 | 18.2776737 | 14.064274 | 12.1156819 |
| ENSMUSG00 | Stmn1 | 1948 | 0.16326686 | 0.32157277 | 0.20421796 | 0.22227132 | 0.15716543 | 0.14001355 |
| ENSMUSG00 | Ncdn | 4243 | 11.6277519 | 10.6071499 | 13.074078 | 12.0755369 | 10.594918 | 12.4320416 |
| ENSMUSG00 | Trim63 | 3092 | 0 | 0 | 0 | 0 | 0.01650271 | 0 |
| ENSMUSG00 | Slc30a2 | 3685 | 0.23734615 | 0.66689513 | 0.51578813 | 0.0685412 | 0.59542311 | 1.62093514 |
| ENSMUSG00 | Psmb2 | 1364 | 71.3500119 | 66.4153308 | 83.3483433 | 63.3022966 | 47.3228485 | 90.3022514 |
| ENSMUSG00 | Extl1 | 4320 | 0.12883721 | 0 | 0.19440625 | 0.05011395 | 0.24804488 | 0 |
| ENSMUSG00 | Zfp593 | 3709 | 0.33227821 | 0.42872812 | 0.38135858 | 0.25293426 | 0.16508938 | 0.37503548 |
| ENSMUSG00 | Cnksr1 | 3176 | 16.2476743 | 17.2506249 | 10.0066515 | 11.3608453 | 15.712782 | 10.0304719 |
| ENSMUSG00 | Eif2c3 | 16549 | 1.01376595 | 0.87061129 | 1.74414185 | 1.70500688 | 1.64342609 | 1.18334588 |
| ENSMUSG00 | Sh3bgrl3 | 1211 | 59.6496351 | 101.665037 | 60.8824725 | 63.7618293 | 45.3380515 | 106.238211 |
| ENSMUSG00 | Tekt2 | 2025 | 0.29448504 | 0.07138732 | 0.45838948 | 0 | 0.50396421 | 0 |
| ENSMUSG00 | Trappc3 | 1869 | 27.3757643 | 27.9218389 | 31.809245 | 23.321086 | 19.984648 | 37.0228784 |
| ENSMUSG00 | Gpn2 | 1927 | 5.83850596 | 10.9275948 | 8.71650251 | 7.63958182 | 7.38783542 | 7.10527706 |
| ENSMUSG00 | Mtap7d1 | 4218 | 2.99721261 | 3.27868869 | 2.09586738 | 2.5406276 | 2.06863686 | 2.37957972 |
| ENSMUSG00 | Gpatch3 | 2079 | 3.40378813 | 3.36076677 | 2.35998296 | 6.16119567 | 5.39961649 | 3.17482573 |
| ENSMUSG00 | Nudc | 1417 | 32.7695139 | 30.2312965 | 28.3553107 | 30.2763239 | 33.3453945 | 27.6211061 |
| ENSMUSG00 | Slc9a1 | 6325 | 9.26475546 | 10.9400365 | 10.5106043 | 10.8845124 | 12.0527123 | 8.52952355 |
| ENSMUSG00 | Tmem222 | 2087 | 23.5637419 | 25.7440743 | 28.2112386 | 24.1353781 | 19.0951603 | 25.418866 |
| ENSMUSG00 | Csf3r | 4263 | 0.02797712 | 0 | 0 | 0 | 0 | 0 |
| ENSMUSG00 | Sytl1 | 2357 | 4.72275543 | 2.94393198 | 2.5692204 | 0.94912463 | 3.31227641 | 2.67307665 |
| ENSMUSG00 | Mrps15 | 1907 | 17.6783678 | 24.0047824 | 23.8972733 | 18.1450869 | 16.5093203 | 27.9182464 |
| ENSMUSG00 | Map3k6 | 4497 | 0.14144712 | 0.49290112 | 0.5111176 | 0.49746203 | 0.22693518 | 0.35783939 |
| ENSMUSG00 | Meaf6 | 1815 | 13.8432307 | 11.7346597 | 18.9714797 | 13.6376212 | 13.6070336 | 22.661243 |
| ENSMUSG00 | Hgf | 2736 | 0 | 0 | 0 | 0 | 0 | 0 |
| ENSMUSG00 | Cd164l2 | 1652 | 12.658222 | 9.7422952 | 9.79290227 | 10.0688998 | 7.3203699 | 12.1679232 |
| ENSMUSG00 | Wasf2 | 6463 | 3.99215641 | 3.78751549 | 5.19780299 | 4.84592521 | 3.20543224 | 6.65935026 |
| ENSMUSG00 | Gnl2 | 3542 | 24.0642999 | 18.4202266 | 17.8080265 | 18.9374706 | 20.2405585 | 21.0681573 |
| ENSMUSG00 | Rspo1 | 1932 | 0 | 0 | 0 | 0 | 0 | 0 |
| ENSMUSG00 | Cdca8 | 5164 | 0.25405323 | 0.29859918 | 0 | 0.28647634 | 0.2173858 | 0 |
| ENSMUSG00 | Fgr | 4017 | 0 | 0.31188636 | 0 | 0 | 0 | 0.25801252 |
| ENSMUSG00 | Epha10 | 6469 | 0 | 0 | 0 | 0 | 0 | 0 |
| ENSMUSG00 | Fam76a | 4135 | 3.91305455 | 6.33940134 | 6.48863814 | 5.68935744 | 5.68879305 | 5.21087425 |
| ENSMUSG00 | Stx12 | 3179 | 39.0926807 | 41.7292467 | 42.1300989 | 34.8221808 | 34.6382256 | 39.7065211 |
| ENSMUSG00 | Ppp1r8 | 3293 | 5.78283485 | 5.67760084 | 5.58395587 | 5.22658222 | 3.59493446 | 7.52061113 |
| ENSMUSG00 | Sema3a | 9870 | 0 | 0 | 0 | 0 | 0 | 0 |
| ENSMUSG00 | Rpa2 | 3115 | 2.3738425 | 4.5324647 | 5.13677919 | 4.35532865 | 6.56872446 | 3.21341657 |
| ENSMUSG00 | Smpdl3b | 1998 | 0 | 0 | 0 | 0 | 0 | 0 |
| ENSMUSG00 | Eya3 | 5736 | 3.46543589 | 4.79680238 | 3.77595939 | 3.7113679 | 2.06382831 | 2.1445018 |
| ENSMUSG00 | Yrdc | 2383 | 20.5533999 | 21.6970426 | 24.502994 | 16.973551 | 19.7210626 | 22.7994467 |
| ENSMUSG00 | Mtf1 | 8894 | 2.71771218 | 4.98443071 | 3.51865357 | 4.8033664 | 3.79226832 | 3.31503092 |
| ENSMUSG00 | Sesn2 | 3465 | 13.5386629 | 25.8106888 | 7.02892222 | 21.2535218 | 20.4400028 | 7.99741226 |
| ENSMUSG00 | Inpp5b | 4673 | 6.68688373 | 5.56830275 | 6.90506 | 5.6288916 | 9.95849663 | 6.25688279 |
| ENSMUSG00 | Rcc1 | 4911 | 1.65141886 | 1.95257626 | 2.61016788 | 2.24823979 | 2.30662908 | 1.90494833 |
| ENSMUSG00 | Trnau1ap | 2485 | 14.8463122 | 15.0473558 | 19.0859467 | 13.5761415 | 13.5317431 | 16.9684476 |
| ENSMUSG00 | Taf12 | 4104 | 9.81293902 | 12.3284026 | 12.9999085 | 10.8316469 | 9.95909528 | 10.6865546 |
| ENSMUSG00 | Gmeb1 | 11341 | 0.59242361 | 0.6373306 | 0.51837097 | 0.48359675 | 1.21480658 | 0.46415707 |
| ENSMUSG00 | Sf3a3 | 2875 | 22.3322091 | 23.0959712 | 21.9702378 | 24.3726367 | 25.1493477 | 18.8882806 |
| ENSMUSG00 | Epb4.1 | 11566 | 14.5980915 | 16.1899113 | 14.0141932 | 14.0260135 | 14.9867369 | 11.9677325 |
| ENSMUSG00 | Utp11l | 4894 | 7.22974615 | 7.02021521 | 7.38804814 | 7.43169205 | 6.77710345 | 9.6302772 |
| ENSMUSG00 | Ptpru | 7604 | 0.35551982 | 0.28516437 | 0.4010951 | 0.17082504 | 0.55025813 | 0.18651779 |
| ENSMUSG00 | Mecr | 2942 | 10.9861338 | 11.8254967 | 14.9042244 | 11.1606596 | 11.7593076 | 10.1236934 |
| ENSMUSG00 | Srsf4 | 3008 | 8.04889881 | 7.32087901 | 6.86245369 | 6.34554573 | 9.90671658 | 7.44430812 |
| ENSMUSG00 | Casp9 | 3844 | 12.1624467 | 9.80275696 | 10.2225386 | 11.6018229 | 10.8318217 | 9.23818426 |
| ENSMUSG00 | Plekhm2 | 5166 | 3.9940175 | 2.03341936 | 2.96047961 | 3.61798278 | 3.64474113 | 2.94604117 |
| ENSMUSG00 | Arhgef19 | 4191 | 4.02202905 | 4.13913605 | 3.3538979 | 3.77953175 | 3.44558921 | 2.68125778 |
| ENSMUSG00 | Fbxo42 | 6181 | 6.69559217 | 9.00426156 | 7.32287449 | 6.3629556 | 5.78700687 | 3.53453908 |
| ENSMUSG00 | Necap2 | 2792 | 30.4288904 | 34.5520262 | 36.5235143 | 31.3003982 | 31.3249313 | 26.1805277 |
| ENSMUSG00 | Cdk14 | 18653 | 2.26772269 | 2.94755431 | 0.66351343 | 1.53783442 | 2.15835579 | 1.49584281 |
| ENSMUSG00 | Padi2 | 4996 | 0.77187382 | 0.97414545 | 0.46891467 | 1.81276885 | 1.12347905 | 0.86802795 |
| ENSMUSG00 | Kcnab2 | 7933 | 0.01503422 | 0.13363188 | 0.08357843 | 0 | 0.0578895 | 0.1272106 |
| ENSMUSG00 | Psmc2 | 3466 | 32.2540136 | 39.0246229 | 36.5883115 | 27.5560219 | 35.7596846 | 27.6444915 |
| ENSMUSG00 | Xrcc2 | 5988 | 1.67307633 | 0.93347149 | 0.53148509 | 0.31936345 | 0.9203154 | 1.31636118 |
| ENSMUSG00 | Rpl22 | 2801 | 7.50826462 | 9.10054548 | 9.23173087 | 6.31210347 | 6.59462622 | 7.84839673 |
| ENSMUSG00 | Acot7 | 2112 | 6.32473554 | 5.22476101 | 4.2694962 | 3.7243777 | 4.03475604 | 2.23414422 |
| ENSMUSG00 | Galntl5 | 1760 | 0 | 0 | 0 | 0 | 0 | 0 |
| ENSMUSG00 | Hes2 | 2633 | 0 | 0 | 0 | 0 | 0 | 0 |
| ENSMUSG00 | Espn | 6421 | 0.37148868 | 0.90054089 | 0.08949135 | 0.71928075 | 0.35760581 | 0.42477245 |
| ENSMUSG00 | Prkag2 | 7504 | 9.74284764 | 6.5177557 | 9.01236938 | 7.53472332 | 6.69109193 | 9.50468838 |
| ENSMUSG00 | Rheb | 5621 | 29.7759425 | 34.0674117 | 38.7365733 | 38.5919323 | 22.9305507 | 39.8614413 |
| ENSMUSG00 | Hes3 | 1883 | 0 | 0 | 0 | 0 | 0 | 0.14484673 |
| ENSMUSG00 | Nol9 | 4913 | 4.08640701 | 5.13936407 | 4.72338033 | 7.68937525 | 8.17377524 | 2.02075487 |
| ENSMUSG00 | Smarcd3 | 4816 | 0.81723268 | 1.31071925 | 1.22986856 | 0.77168825 | 1.16546955 | 1.70466495 |
| ENSMUSG00 | Tas1r1 | 5437 | 0.48259366 | 0.3367822 | 0.19511573 | 0.28536471 | 0.47863623 | 0.34112111 |
| ENSMUSG00 | Zbtb48 | 3422 | 6.2618948 | 5.04112983 | 6.25180947 | 4.74486266 | 7.41090263 | 6.3523926 |
| ENSMUSG00 | Abcf2 | 3147 | 20.5283304 | 18.2670561 | 24.7064003 | 24.9031461 | 16.846649 | 15.1583553 |
| ENSMUSG00 | Nub1 | 3440 | 11.9613146 | 12.6209257 | 11.5515863 | 14.0132592 | 14.9222483 | 16.0555653 |
| ENSMUSG00 | Vamp3 | 2884 | 26.1911973 | 31.3111625 | 29.6569231 | 22.0320667 | 23.549274 | 25.922256 |
| ENSMUSG00 | Per3 | 11758 | 1.0278675 | 1.9917172 | 1.49620118 | 1.41775001 | 1.51890045 | 0.68198197 |
| ENSMUSG00 | Tmub1 | 2644 | 3.78910026 | 9.95075545 | 6.62024578 | 4.14861631 | 5.01772228 | 3.91995572 |
| ENSMUSG00 | Fastk | 4934 | 40.9399263 | 35.2950551 | 36.4257588 | 32.1696218 | 37.3235084 | 33.991033 |
| ENSMUSG00 | Ube4b | 7432 | 5.78251809 | 7.7674055 | 4.56768238 | 6.15123122 | 8.63713412 | 3.73595032 |
| ENSMUSG00 | Pgd | 6435 | 10.0083727 | 22.8763917 | 11.1483436 | 18.7559346 | 11.1330275 | 10.973433 |
| ENSMUSG00 | Slc4a2 | 6461 | 15.0505967 | 12.7458025 | 13.1558805 | 12.3196085 | 15.8346825 | 8.86499651 |
| ENSMUSG00 | Uts2 | 534 | 0.29779386 | 0.18047357 | 0 | 0 | 0 | 0.0510761 |
| ENSMUSG00 | Park7 | 2769 | 44.6800489 | 48.6214949 | 52.7421052 | 43.5356124 | 29.0420976 | 49.5848805 |
| ENSMUSG00 | Tnfrsf9 | 2708 | 0.39638035 | 0 | 0.35909917 | 0.02664848 | 0.01884283 | 0.02014375 |
| ENSMUSG00 | Errfi1 | 3576 | 22.5014241 | 24.6187444 | 11.2482319 | 24.2464074 | 18.1360525 | 15.4830866 |
| ENSMUSG00 | Cdk5 | 2038 | 19.7021763 | 21.5633146 | 21.6671448 | 17.7754528 | 18.4025448 | 28.8136891 |
| ENSMUSG00 | Abcb1b | 4316 | 0 | 0 | 0 | 0 | 0 | 0 |
| ENSMUSG00 | Cort | 675 | 0 | 0 | 0 | 0 | 0 | 0 |
| ENSMUSG00 | Car6 | 2103 | 0.1512334 | 0.48117703 | 0.04203694 | 0.03431483 | 0.26689973 | 0 |
| ENSMUSG00 | Abcb8 | 4070 | 31.8434561 | 27.8581569 | 22.3615712 | 20.6119788 | 26.2654195 | 23.3476273 |
| ENSMUSG00 | Dffa | 3533 | 9.66599427 | 8.06062477 | 8.35743876 | 10.4477587 | 11.1353908 | 6.20685252 |
| ENSMUSG00 | Pex14 | 6262 | 4.14569287 | 2.56245373 | 3.72701583 | 2.52954608 | 3.93576166 | 3.17086193 |
| ENSMUSG00 | Slc2a5 | 3388 | 0 | 0.45512578 | 0 | 0 | 0 | 0 |
| ENSMUSG00 | Casz1 | 12264 | 4.08446718 | 4.76206522 | 3.02752349 | 4.43964491 | 3.61977715 | 4.46348671 |
| ENSMUSG00 | Nos3 | 3959 | 0.21087777 | 0.02434273 | 0.30145233 | 0.12759501 | 0.67021256 | 0.2480139 |
| ENSMUSG00 | Masp2 | 3322 | 0.01196733 | 0.04351575 | 0.02661159 | 0.52135405 | 0.15360137 | 0 |
| ENSMUSG00 | H6pd | 5228 | 11.3836944 | 16.7196263 | 5.61400608 | 10.8218527 | 15.6651364 | 3.70409217 |
| ENSMUSG00 | Slc25a33 | 1880 | 15.098624 | 8.3813652 | 12.1555068 | 10.8246134 | 8.0882234 | 11.9254066 |
| ENSMUSG00 | Klhl7 | 3408 | 18.5945528 | 19.8373176 | 20.7131289 | 20.761998 | 17.6825557 | 17.6868993 |
| ENSMUSG00 | Ctnnbip1 | 2786 | 1.65528921 | 2.54250072 | 2.39572085 | 2.6679473 | 2.41761723 | 2.05587733 |
| ENSMUSG00 | Angptl7 | 2152 | 0.59115956 | 0.47022086 | 0.34917813 | 0.31856824 | 0 | 0 |
| ENSMUSG00 | Lzic | 1783 | 9.98903829 | 15.2964253 | 11.1310306 | 16.9178852 | 13.9370976 | 8.03095099 |
| ENSMUSG00 | Mtor | 9487 | 3.87204216 | 4.58652443 | 3.76463467 | 6.14995952 | 5.93792758 | 3.26019194 |
| ENSMUSG00 | Nmnat1 | 3826 | 8.18800801 | 8.36272807 | 6.56211365 | 9.11010324 | 8.73556617 | 7.82025072 |
| ENSMUSG00 | Fam126a | 7349 | 0.64915739 | 0.41963972 | 0.24058698 | 1.01632647 | 0.61795448 | 0.40824742 |
| ENSMUSG00 | Rbp7 | 933 | 0.21305188 | 0 | 0 | 0.15469258 | 0 | 0 |
| ENSMUSG00 | Tomm7 | 1122 | 14.7754326 | 17.2646611 | 16.3491665 | 10.7731595 | 13.0522013 | 43.4400894 |
| ENSMUSG00 | Rint1 | 7059 | 4.17322724 | 5.24938008 | 3.34378583 | 4.87636641 | 3.83113461 | 2.43033689 |
| ENSMUSG00 | Fbxo44 | 3855 | 24.7608064 | 20.1870568 | 21.5677475 | 15.6495923 | 19.1266182 | 30.9041827 |
| ENSMUSG00 | Mad2l2 | 2611 | 3.82176393 | 4.04168167 | 3.97833516 | 3.57918404 | 4.2017123 | 3.62477971 |
| ENSMUSG00 | Mll5 | 7800 | 57.7473839 | 59.5411448 | 58.2897638 | 71.0816277 | 54.8206448 | 70.1482741 |
| ENSMUSG00 | 2610109H07l | 5193 | 0 | 0 | 0 | 0 | 0.18669385 | 0 |
| ENSMUSG00 | Agtrap | 4317 | 9.90894071 | 13.0149158 | 10.4540379 | 10.2972154 | 10.7206215 | 9.83706586 |
| ENSMUSG00 | Mthfr | 6794 | 1.92516236 | 2.37598736 | 2.29682211 | 4.16372136 | 3.71770033 | 1.30873306 |
| ENSMUSG00 | Orc5 | 3537 | 26.4137346 | 30.7891884 | 22.5320676 | 25.0442237 | 19.3458779 | 22.3625824 |
| ENSMUSG00 | Dnajc2 | 4795 | 4.86683359 | 7.07471434 | 3.80716604 | 7.97642694 | 8.13016708 | 6.80870556 |
| ENSMUSG00 | Slc26a5 | 3375 | 0 | 0 | 0 | 0 | 0 | 0 |
| ENSMUSG00 | Clcn6 | 6636 | 1.8392002 | 2.25102427 | 1.52534992 | 2.88177872 | 2.88349773 | 1.27002163 |
| ENSMUSG00 | Pmpcb | 5128 | 32.4059884 | 31.5730198 | 35.5131812 | 29.48905 | 30.1401896 | 28.5298684 |
| ENSMUSG00 | Nppb | 778 | 0 | 0 | 0 | 0 | 0 | 0 |
| ENSMUSG00 | Mfn2 | 6616 | 34.9723242 | 31.2672908 | 32.6569844 | 24.9127538 | 31.313042 | 19.4500828 |
| ENSMUSG00 | Miip | 3648 | 8.36957484 | 8.14995472 | 5.62216424 | 6.85440155 | 9.56744547 | 9.24855503 |
| ENSMUSG00 | Trp73 | 8594 | 0 | 0.24110042 | 0 | 0.04198516 | 0.08906163 | 0 |
| ENSMUSG00 | Dffb | 2343 | 3.20690816 | 3.7224695 | 0.67915764 | 2.47938932 | 4.74765255 | 1.83926291 |
| ENSMUSG00 | Lrrc47 | 2924 | 20.2856282 | 21.5388782 | 16.175093 | 16.5849069 | 18.7945988 | 12.5646166 |
| ENSMUSG00 | Wdr8 | 5306 | 2.61490062 | 1.89803362 | 1.9576768 | 1.86326427 | 1.92334624 | 1.70145224 |
| ENSMUSG00 | Tprgl | 2663 | 87.6770323 | 93.2424846 | 84.6192248 | 70.4162168 | 79.5957811 | 60.6535538 |
| ENSMUSG00 | Arhgef16 | 3972 | 25.8931088 | 32.8885812 | 25.1167069 | 24.7905084 | 30.8444935 | 21.0396512 |
| ENSMUSG00 | Acap3 | 5505 | 1.4443408 | 2.87105414 | 3.08328932 | 2.16951076 | 1.2884044 | 2.34348853 |
| ENSMUSG00 | Cpsf3l | 3136 | 36.3580096 | 44.4218764 | 29.5007836 | 13.703353 | 15.0670995 | 28.9445151 |
| ENSMUSG00 | Atad3a | 3932 | 29.7357244 | 25.2329309 | 29.8013952 | 24.2351678 | 23.5276753 | 21.3646716 |
| ENSMUSG00 | Ssu72 | 3021 | 22.1609497 | 23.670533 | 22.4447624 | 16.6137451 | 17.5661804 | 20.5394917 |
| ENSMUSG00 | 4930584F24f | 758 | 0 | 0 | 0 | 0 | 0 | 0 |
| ENSMUSG00 | Pex10 | 3335 | 9.38157816 | 9.7528781 | 7.17037393 | 4.77126886 | 6.64031398 | 8.30915547 |
| ENSMUSG00 | Rer1 | 1806 | 147.288992 | 143.331986 | 180.821271 | 120.033734 | 98.77531 | 165.188264 |
| ENSMUSG00 | Morn1 | 4168 | 7.0773912 | 4.8903227 | 7.75229061 | 4.4410002 | 9.76944527 | 5.97450711 |
| ENSMUSG00 | Ski | 7523 | 6.6109406 | 7.79514822 | 6.72751698 | 8.56606826 | 6.43001519 | 8.9332329 |
| ENSMUSG00 | Prkcz | 6887 | 7.32535281 | 5.78629122 | 7.16908068 | 5.867851 | 7.37204065 | 5.67512096 |
| ENSMUSG00 | Gabrd | 2271 | 0.3326086 | 0 | 0 | 0 | 0 | 0.72059813 |
| ENSMUSG00 | Plch2 | 8882 | 0.24170186 | 0.46114024 | 0.32347667 | 0.03656141 | 0.5285326 | 0.10133563 |
| ENSMUSG00 | Pank4 | 2934 | 19.8777403 | 15.3066681 | 14.2819861 | 18.3484698 | 20.0175046 | 11.9919171 |
| ENSMUSG00 | 2810405K02f | 1019 | 4.01846368 | 5.34354069 | 4.03412306 | 2.72651367 | 3.0044971 | 5.43351498 |
| ENSMUSG00 | Mib2 | 4964 | 13.7750658 | 14.1627758 | 11.0326518 | 13.2291138 | 16.6113262 | 7.65382201 |
| ENSMUSG00 | Mmp23 | 2473 | 0 | 0.03897003 | 0 | 0.01459039 | 0.02063339 | 0.08823175 |
| ENSMUSG00 | Cdk11b | 3370 | 22.9921162 | 25.7375655 | 21.694317 | 26.5101312 | 30.1010193 | 28.7638184 |
| ENSMUSG00 | Nadk | 4267 | 98.321909 | 75.3004913 | 131.093115 | 83.2415389 | 90.6205475 | 81.6129819 |
| ENSMUSG00 | Gnb1 | 4470 | 37.7988349 | 39.1420518 | 45.1511443 | 37.7691021 | 32.3623659 | 35.6705908 |
| ENSMUSG00 | Mrpl20 | 1168 | 41.0148579 | 41.1730045 | 52.5275326 | 38.5225253 | 23.7220223 | 57.3047643 |
| ENSMUSG00 | Ccnl2 | 5381 | 89.8764952 | 71.899079 | 98.9099409 | 106.837988 | 110.359592 | 103.497651 |
| ENSMUSG00 | Mxra8 | 2680 | 1.27573557 | 1.78002156 | 0.31337128 | 0 | 3.33194618 | 0.55974073 |
| ENSMUSG00 | Dvl1 | 4819 | 22.0515458 | 21.3284252 | 16.9506134 | 26.1387046 | 25.1267047 | 14.7381533 |
| ENSMUSG00 | Tas1r3 | 3517 | 1.06255763 | 0.68505036 | 1.10598868 | 0.68737475 | 0.52230581 | 0.64367218 |
| ENSMUSG00 | Gltpd1 | 3437 | 15.7425688 | 18.8988432 | 20.1911241 | 20.34536 | 21.7199848 | 17.9344442 |
| ENSMUSG00 | Ttll10 | 3635 | 4.21069051 | 5.68692811 | 4.35330393 | 5.32048855 | 3.90242984 | 6.58793212 |
| ENSMUSG00 | Tnfrsf4 | 1508 | 0.47453491 | 1.46987821 | 0.0879347 | 0.33497919 | 0 | 0.14469305 |
| ENSMUSG00 | Sdf4 | 6201 | 26.9524335 | 29.5288631 | 31.9556301 | 24.246715 | 26.2249734 | 28.0443638 |
| ENSMUSG00 | Bst1 | 2634 | 9.14647729 | 2.085518 | 1.47675102 | 0.02739715 | 5.2111219 | 0.07248386 |
| ENSMUSG00 | Cd38 | 3006 | 1.30931223 | 2.5006936 | 3.11736218 | 2.2566282 | 2.91967287 | 3.50233225 |
| ENSMUSG00 | Prom1 | 4677 | 14.6968625 | 19.9875343 | 5.99186839 | 8.53255105 | 18.0779784 | 5.03854785 |
| ENSMUSG00 | Kcnip4 | 5101 | 0.17146061 | 0 | 0.21683322 | 0.48807314 | 0.55017657 | 0.55607969 |
| ENSMUSG00 | Pacrgl | 1455 | 7.13139547 | 10.1340559 | 11.1188141 | 9.54748257 | 9.18825462 | 12.072074 |
| ENSMUSG00 | Gpr125 | 4428 | 12.991459 | 10.0442832 | 6.65822703 | 9.21607777 | 15.7066283 | 6.05487136 |
| ENSMUSG00 | 4930433H24 | 404 | 0 | 0 | 0 | 0 | 0.63151455 | 0.13502297 |
| ENSMUSG00 | Sorcs2 | 8772 | 0 | 0 | 0 | 0 | 0 | 0 |
| ENSMUSG00 | Afap1 | 8599 | 1.18355658 | 1.0871229 | 0.84815724 | 1.16650871 | 0.66460683 | 0.34890224 |
| ENSMUSG00 | Ablim2 | 6779 | 1.84145462 | 1.24393382 | 1.16063255 | 1.35726823 | 1.76887422 | 0.97366318 |
| ENSMUSG00 | Htra3 | 4935 | 0 | 0 | 0 | 0 | 0 | 0 |
| ENSMUSG00 | 2310079F23f | 2952 | 1.34673037 | 3.06878426 | 2.35084328 | 1.49119561 | 5.41031695 | 4.32402817 |
| ENSMUSG00 | Acox3 | 10373 | 6.88716846 | 8.20372668 | 5.84216011 | 7.76478894 | 6.72448238 | 5.40602124 |
| ENSMUSG00 | Rgs12 | 7964 | 0.21964354 | 0.58085114 | 0.24975928 | 0.70224974 | 0.44209191 | 0.11986594 |
| ENSMUSG00 | Hgfac | 2089 | 20.8007373 | 22.1440806 | 16.4831191 | 11.1234259 | 13.7519625 | 12.1032028 |
| ENSMUSG00 | Lrpap1 | 4435 | 38.8321853 | 42.1237511 | 40.5839695 | 33.9911572 | 45.2391679 | 34.611425 |
| ENSMUSG00 | Htt | 15106 | 1.52905694 | 1.89479324 | 2.35844601 | 3.58766258 | 2.09091266 | 2.89790785 |
| ENSMUSG00 | Add1 | 6013 | 34.9686906 | 37.3358779 | 31.6903618 | 31.7495589 | 37.2111522 | 27.3426784 |
| ENSMUSG00 | Pcdh7 | 9373 | 0.09755426 | 0.49867544 | 0.14147608 | 0.27331966 | 0.46273786 | 0 |
| ENSMUSG00 | Rnf4 | 2632 | 79.510961 | 77.955118 | 93.425096 | 79.2516335 | 81.5026155 | 75.5130305 |
| ENSMUSG00 | Whsc2 | 4916 | 8.80669615 | 11.174236 | 8.32605912 | 8.22048207 | 11.1581273 | 9.47066908 |
| ENSMUSG00 | Nkx1-1 | 1386 | 0 | 0 | 0 | 0 | 0 | 0 |
| ENSMUSG00 | Man2b2 | 3509 | 5.69877649 | 6.6738703 | 3.91757571 | 4.53467701 | 6.15108492 | 4.01852052 |
| ENSMUSG00 | Ppp2r2c | 4440 | 0.01790787 | 0.98760501 | 0 | 0.81265866 | 0.22984854 | 0.22728866 |
| ENSMUSG00 | Crmp1 | 3750 | 0.01060146 | 0 | 0 | 0 | 0 | 0 |
| ENSMUSG00 | Evc | 7677 | 4.56227411 | 4.63850212 | 3.50643772 | 4.97261992 | 5.63636404 | 2.95590724 |
| ENSMUSG00 | Stk32b | 3425 | 0.94020261 | 1.646077 | 1.0582631 | 0.63209421 | 1.77288722 | 1.05116858 |
| ENSMUSG00 | Stx18 | 3500 | 8.57582508 | 6.37437792 | 8.57515754 | 8.29887024 | 8.16422013 | 7.23946853 |
| ENSMUSG00 | Nsg1 | 2135 | 0.03724167 | 0 | 0.1035172 | 0.65910995 | 0.62139849 | 0 |
| ENSMUSG00 | Zbtb49 | 7736 | 2.84187962 | 2.84035905 | 2.8340375 | 2.36940003 | 3.16606262 | 1.67117102 |
| ENSMUSG00 | Rab28 | 2167 | 40.6360818 | 46.2296323 | 47.4858747 | 39.6119907 | 33.2719285 | 41.7111927 |
| ENSMUSG00 | Rnf32 | 2848 | 5.63947125 | 7.19074365 | 3.50759007 | 4.09217008 | 4.4970577 | 6.52177996 |
| ENSMUSG00 | Dnajb6 | 5883 | 2.38546398 | 2.2524684 | 2.89269243 | 2.23864461 | 2.23777069 | 2.99033526 |
| ENSMUSG00 | Plb1 | 8879 | 0.20148627 | 0.14110232 | 0.24891228 | 0.04063751 | 0.11493721 | 0.04914903 |
| ENSMUSG00 | Fosl2 | 5851 | 0.92407201 | 1.51534872 | 1.83576275 | 3.027907 | 1.2296563 | 1.14207599 |
| ENSMUSG00 | Rbks | 1479 | 8.44031162 | 5.01738478 | 12.0441804 | 8.48989283 | 4.8300829 | 4.22305097 |
| ENSMUSG00 | 4930548H24 | 1639 | 0 | 0 | 0 | 0 | 0.21792839 | 0 |
| ENSMUSG00 | Slc4a1ap | 5432 | 4.90356626 | 6.43136424 | 4.40228223 | 6.61592716 | 6.07769977 | 4.94076676 |
| ENSMUSG00 | Mrpl33 | 5572 | 7.19194622 | 6.79729783 | 6.90951279 | 5.99640582 | 6.62097447 | 14.7484724 |
| ENSMUSG00 | Gtf3c2 | 6832 | 21.2859408 | 17.0049052 | 21.7968397 | 21.6376078 | 25.4160944 | 20.3362138 |
| ENSMUSG00 | Eif2b4 | 2823 | 21.0958944 | 17.427686 | 15.1097338 | 14.4174801 | 18.2921333 | 12.3474988 |
| ENSMUSG00 | Snx17 | 2467 | 42.2694063 | 44.9831278 | 43.9330843 | 39.9871543 | 40.0020311 | 28.2143816 |
| ENSMUSG00 | Ppm1g | 2155 | 38.4641192 | 44.1839488 | 41.8635553 | 35.1611571 | 40.347538 | 28.2998113 |
| ENSMUSG00 | Nrbp1 | 3379 | 37.3435617 | 35.2093299 | 38.367566 | 40.1398063 | 38.1300974 | 34.6280563 |
| ENSMUSG00 | Krtcap3 | 1621 | 9.1724551 | 11.0284824 | 14.0158836 | 8.19135866 | 8.53062791 | 19.3160308 |
| ENSMUSG00 | Slc30a3 | 2081 | 0 | 0 | 0 | 0 | 0 | 0 |
| ENSMUSG00 | Ociad1 | 4123 | 60.8819073 | 61.8137503 | 83.7615812 | 56.9278922 | 45.3457776 | 69.4732625 |
| ENSMUSG00 | Ociad2 | 2502 | 155.430899 | 120.003885 | 291.0043 | 123.691325 | 108.88482 | 214.043782 |
| ENSMUSG00 | Cwh43 | 3153 | 17.0218518 | 20.9679032 | 10.1777793 | 13.7095748 | 9.20837547 | 10.4064038 |
| ENSMUSG00 | Spata18 | 1983 | 0 | 0 | 0 | 0 | 0 | 0 |
| ENSMUSG00 | Sgcb | 4089 | 10.9184164 | 10.4174162 | 8.11826464 | 9.54775548 | 14.5878781 | 12.7401714 |
| ENSMUSG00 | Yipf7 | 1208 | 0 | 0 | 0 | 0 | 0 | 0 |
| ENSMUSG00 | Cgref1 | 1322 | 0 | 1.27573788 | 0.4346626 | 0.68233821 | 0.88775087 | 1.44419421 |
| ENSMUSG00 | Khk | 1836 | 11.4112953 | 13.0439334 | 12.663491 | 12.9510171 | 10.7833518 | 11.1713119 |
| ENSMUSG00 | Emilin1 | 4873 | 0.06526654 | 0.09888455 | 0.22676915 | 0.12587621 | 0.23036739 | 0.36381111 |
| ENSMUSG00 | Agbl5 | 5270 | 4.5564156 | 5.13866802 | 2.67559543 | 4.66944106 | 4.89930667 | 4.84422435 |
| ENSMUSG00 | Mapre3 | 3566 | 5.85295212 | 4.13489941 | 5.06970101 | 5.47402863 | 5.99552761 | 7.99270836 |
| ENSMUSG00 | Ppargc1a | 6928 | 60.9358325 | 49.3966675 | 59.89069 | 63.3050305 | 55.8285117 | 45.2858221 |
| ENSMUSG00 | Dpysl5 | 6793 | 0 | 0 | 0 | 0 | 0 | 0.00401511 |
| ENSMUSG00 | Dhx15 | 2995 | 73.0730952 | 68.7159581 | 77.5856724 | 69.3570385 | 74.7421405 | 78.0355204 |
| ENSMUSG00 | Pgm1 | 2594 | 19.8624143 | 27.9756291 | 15.6768294 | 22.6173494 | 20.1036839 | 15.3722138 |
| ENSMUSG00 | Sepsecs | 9354 | 6.32415598 | 6.84624563 | 5.67998881 | 4.79473822 | 7.58249544 | 6.07366619 |
| ENSMUSG00 | Tbc1d1 | 6634 | 34.0264725 | 30.7539035 | 16.8105592 | 25.2639579 | 29.5666851 | 19.7056596 |
| ENSMUSG00 | 4930471M23 | 2335 | 8.27458823 | 10.5246619 | 11.1119837 | 11.527711 | 11.7568266 | 7.47570412 |
| ENSMUSG00 | Anapc4 | 6138 | 19.8518325 | 19.414316 | 22.1225255 | 21.2859373 | 20.07636 | 19.9471906 |
| ENSMUSG00 | Cenpa | 2723 | 0 | 0.23004912 | 0.16232774 | 0.17226095 | 0.01873903 | 0.26042623 |
| ENSMUSG00 | Klf3 | 2874 | 23.3497742 | 21.4273741 | 19.3786786 | 21.1671284 | 19.4766647 | 22.0455765 |
| ENSMUSG00 | Zcchc4 | 8851 | 2.37158442 | 2.18856059 | 2.90650465 | 1.6795619 | 3.13041714 | 2.84733552 |
| ENSMUSG00 | 1700001C02l | 1159 | 0 | 0 | 0 | 0 | 0 | 0 |
| ENSMUSG00 | Olfr109 | 944 | 0 | 0 | 0 | 0 | 0 | 0 |
| ENSMUSG00 | Fam114a1 | 2949 | 4.39480727 | 9.23205826 | 2.83287499 | 4.52707917 | 5.26009435 | 3.56077881 |
| ENSMUSG00 | Pi4k2b | 4102 | 8.67409923 | 11.253684 | 9.50415055 | 10.8193356 | 8.69513328 | 8.89650593 |
| ENSMUSG00 | Slc34a2 | 2920 | 1.38871884 | 0.34654633 | 0 | 0.2100667 | 1.43293247 | 0.72856913 |
| ENSMUSG00 | Sel1l3 | 4487 | 0.85943428 | 0.53695612 | 0.00985109 | 0.09649756 | 0.09097638 | 0 |
| ENSMUSG00 | D5Ertd579e | 8199 | 14.0809752 | 10.5259078 | 13.892932 | 14.1177215 | 19.2616945 | 14.2410941 |
| ENSMUSG00 | Rfc1 | 7265 | 8.85400791 | 8.56279378 | 8.60308411 | 9.38182823 | 13.3939847 | 9.42317199 |
| ENSMUSG00 | Tbc1d14 | 12277 | 16.8192527 | 13.8707247 | 10.3798903 | 14.6744032 | 14.6009333 | 10.6814748 |
| ENSMUSG00 | Cckar | 5503 | 0.09391627 | 0.15761511 | 0 | 0.44586208 | 0 | 0 |
| ENSMUSG00 | Klb | 3459 | 0 | 0 | 0 | 0 | 0 | 0 |
| ENSMUSG00 | Tada2b | 4836 | 7.79325797 | 4.5037783 | 5.52066029 | 9.74424114 | 7.51256815 | 8.93362874 |
| ENSMUSG00 | Grpel1 | 3524 | 28.6997566 | 31.2035928 | 29.9152655 | 35.1810173 | 24.2389765 | 35.4632228 |
| ENSMUSG00 | Lias | 4074 | 19.8582236 | 24.6254719 | 20.7664034 | 19.5643683 | 22.1815689 | 24.0477428 |
| ENSMUSG00 | Ugdh | 5815 | 28.5979665 | 60.7157458 | 18.8817503 | 63.8680111 | 26.5530203 | 18.0908484 |
| ENSMUSG00 | Pds5a | 4128 | 13.7622533 | 11.2645147 | 12.2711512 | 15.715968 | 13.7454772 | 13.7694642 |
| ENSMUSG00 | Ube2k | 5435 | 14.8927614 | 16.730049 | 13.6631272 | 13.443632 | 9.70768585 | 18.3068784 |
| ENSMUSG00 | Rhoh | 4957 | 0 | 0.12637155 | 0.18725816 | 0.18925422 | 0 | 0 |
| ENSMUSG00 | Chrna9 | 3114 | 0 | 0 | 0 | 0.01158704 | 0 | 0 |
| ENSMUSG00 | Nsun7 | 5653 | 0.9705035 | 1.78152599 | 0.66463058 | 0.22339847 | 1.16440872 | 0.95531197 |
| ENSMUSG00 | Apbb2 | 11371 | 2.38791603 | 3.5553975 | 3.43632303 | 3.63961877 | 3.22645011 | 2.61209059 |
| ENSMUSG00 | Guf1 | 7620 | 21.2029229 | 16.2834762 | 16.6714038 | 21.3793217 | 23.0556184 | 31.1510742 |
| ENSMUSG00 | Gnpda2 | 2979 | 13.2785173 | 15.2857294 | 12.2411952 | 15.2733999 | 14.7306758 | 12.8362015 |
| ENSMUSG00 | Gabra4 | 4117 | 0 | 0 | 0 | 0 | 0 | 0 |
| ENSMUSG00 | Gabrb1 | 3080 | 0 | 0 | 0 | 0 | 0 | 0 |
| ENSMUSG00 | Commd8 | 3476 | 21.7534304 | 21.6949891 | 23.4361325 | 23.8332492 | 17.2338795 | 21.8291273 |
| ENSMUSG00 | Tec | 4334 | 5.35699138 | 5.0143252 | 6.40488173 | 5.73616259 | 4.48570586 | 4.43669143 |
| ENSMUSG00 | Slc10a4 | 3810 | 0 | 0 | 0 | 0 | 0 | 0 |
| ENSMUSG00 | Slc30a9 | 10065 | 22.8974188 | 19.7916296 | 26.5123225 | 19.8173415 | 18.6665788 | 23.1746364 |
| ENSMUSG00 | Uchl1 | 1686 | 0.04715953 | 0.54302634 | 0.07865097 | 0 | 0 | 0 |
| ENSMUSG00 | Fip1l1 | 9663 | 7.8087449 | 8.70677098 | 8.83304966 | 9.41351902 | 8.6443317 | 10.2798547 |
| ENSMUSG00 | Lnx1 | 4499 | 6.81295298 | 5.8907631 | 7.04439241 | 4.45110796 | 6.18123468 | 4.89640154 |
| ENSMUSG00 | Chic2 | 6706 | 6.27811719 | 8.37119074 | 6.61116143 | 5.13305554 | 5.94267813 | 7.78461967 |
| ENSMUSG00 | Pdgfra | 6713 | 0 | 0 | 0 | 0 | 0 | 0 |
| ENSMUSG00 | Srd5a3 | 3193 | 10.3590854 | 10.4582538 | 10.548639 | 7.52603873 | 7.92642731 | 9.75497589 |
| ENSMUSG00 | Tmem165 | 2982 | 46.0214349 | 38.814163 | 49.4343214 | 45.3385046 | 33.6924661 | 51.6498618 |
| ENSMUSG00 | Pdcl2 | 1658 | 0.19182379 | 0 | 0 | 0 | 0 | 0 |
| ENSMUSG00 | Nmu | 948 | 0 | 0 | 0 | 0 | 0 | 0 |
| ENSMUSG00 | Clock | 13508 | 9.3208178 | 7.40918273 | 10.3501947 | 16.0403226 | 8.4540294 | 12.9851351 |
| ENSMUSG00 | Epha5 | 8633 | 0 | 0 | 0 | 0 | 0 | 0 |
| ENSMUSG00 | Ppat | 6539 | 28.5869811 | 30.3164128 | 35.7724657 | 34.6473708 | 38.5955735 | 24.6885747 |
| ENSMUSG00 | Paics | 3350 | 111.445886 | 105.39166 | 124.068636 | 93.1024456 | 105.586519 | 108.797315 |
| ENSMUSG00 | 1700023E05l | 2233 | 0 | 0 | 0 | 0 | 0 | 0 |
| ENSMUSG00 | Rest | 7542 | 8.21781943 | 9.30888971 | 8.59187243 | 9.34344112 | 10.1552095 | 9.21088653 |
| ENSMUSG00 | Polr2b | 4782 | 34.1770766 | 33.2629539 | 34.9677064 | 28.0763881 | 37.5495225 | 21.4056297 |
| ENSMUSG00 | Cenpc1 | 3154 | 7.75194056 | 9.97645743 | 9.72608721 | 7.31021764 | 6.45514393 | 9.33079764 |
| ENSMUSG00 | Stap1 | 4281 | 33.4313781 | 7.57521036 | 49.9322852 | 16.0392737 | 16.1863626 | 49.24216 |
| ENSMUSG00 | Gnrhr | 3620 | 0 | 0 | 0 | 0 | 0 | 0 |
| ENSMUSG00 | Ugt2b34 | 6021 | 0 | 1.12843188 | 0 | 0.0719124 | 0 | 0 |
| ENSMUSG00 | Pigg | 5002 | 15.697136 | 18.4094944 | 14.5365917 | 14.2972036 | 19.0456305 | 12.3886406 |
| ENSMUSG00 | Dr1 | 3474 | 30.3716308 | 27.8382813 | 35.7660854 | 40.5273857 | 25.0725456 | 34.2149908 |
| ENSMUSG00 | Mtf2 | 5494 | 15.1887065 | 14.7611544 | 11.062529 | 14.008555 | 11.7581711 | 8.87642064 |
| ENSMUSG00 | Ugt2a1 | 3643 | 0 | 0 | 0 | 0 | 0 | 0 |
| ENSMUSG00 | Sult1b1 | 4466 | 0.07121448 | 0.03236886 | 0 | 0 | 0 | 0.23817979 |
| ENSMUSG00 | Fam69a | 3399 | 12.3863059 | 11.9225648 | 14.213773 | 11.1568781 | 11.3041682 | 15.4628508 |
| ENSMUSG00 | Sult1e1 | 1252 | 0 | 0 | 0 | 0 | 0 | 0 |
| ENSMUSG00 | Sult1d1 | 2600 | 38.0276462 | 40.9214093 | 45.255887 | 32.5848618 | 20.3124227 | 50.7413193 |
| ENSMUSG00 | Gfi1 | 4115 | 0 | 0 | 0 | 0 | 0 | 0 |
| ENSMUSG00 | Glmn | 2850 | 4.50562112 | 4.12543575 | 4.85444802 | 3.79810995 | 3.34804641 | 5.58890853 |
| ENSMUSG00 | Brdt | 5228 | 0.73762081 | 0.69127451 | 1.6064174 | 1.21469775 | 0.35136755 | 0.45388172 |
| ENSMUSG00 | Smr3a | 544 | 0 | 0 | 0 | 0 | 0 | 0 |
| ENSMUSG00 | Smr2 | 1382 | 0 | 0 | 0 | 0 | 0 | 0 |
| ENSMUSG00 | Amtn | 1013 | 0 | 0 | 0 | 0 | 0 | 0 |
| ENSMUSG00 | Cdc7 | 3994 | 0.72662746 | 0.20510003 | 0.38734715 | 1.60806308 | 0.56213333 | 0.16389368 |
| ENSMUSG00 | Enam | 5473 | 0 | 0 | 0 | 0 | 0 | 0 |
| ENSMUSG00 | Tgfbr3 | 7003 | 10.3603815 | 8.55286985 | 13.2359367 | 10.4438532 | 13.0644426 | 12.1359097 |
| ENSMUSG00 | Ambn | 1839 | 0 | 0 | 0 | 0 | 0 | 0 |
| ENSMUSG00 | Zfp326 | 7362 | 3.13205361 | 2.87992863 | 2.3836093 | 3.53860855 | 1.36541647 | 3.3009649 |
| ENSMUSG00 | Rufy3 | 6721 | 7.05673088 | 6.80388835 | 7.22776752 | 7.23144085 | 7.22766103 | 7.41013112 |
| ENSMUSG00 | Gbp9 | 3622 | 1.18542018 | 0.31929172 | 0.64679671 | 0.73718147 | 2.6062616 | 2.15365732 |
| ENSMUSG00 | Abcg3 | 3958 | 0 | 0.02434888 | 0 | 0.00911623 | 0 | 0 |
| ENSMUSG00 | Spp1 | 1715 | 3100.16251 | 4592.35057 | 1661.42228 | 2879.17884 | 3442.42081 | 2889.0443 |
| ENSMUSG00 | Ibsp | 2061 | 0 | 0 | 0 | 0 | 0 | 0 |
| ENSMUSG00 | Dmp1 | 2758 | 0 | 0 | 0 | 0.01308269 | 0 | 0.01977856 |
| ENSMUSG00 | Sparcl1 | 3103 | 2.01147613 | 0.88515218 | 0.07122437 | 0.74419944 | 3.60128144 | 0.57133469 |
| ENSMUSG00 | Nudt9 | 2380 | 29.0315232 | 27.9197915 | 33.7456928 | 32.8225321 | 24.8699983 | 28.9936291 |
| ENSMUSG00 | Hsd17b11 | 3258 | 4.63691915 | 3.94898099 | 4.24652451 | 2.90162543 | 2.67817995 | 5.65919464 |
| ENSMUSG00 | Klhl8 | 4622 | 2.81264434 | 5.79655167 | 4.54259529 | 5.15234734 | 5.85114219 | 3.04494025 |
| ENSMUSG00 | Aff1 | 9428 | 15.9772503 | 21.8034962 | 15.4153224 | 21.7227073 | 18.4448334 | 17.2129936 |
| ENSMUSG00 | Agpat9 | 3390 | 1.23136444 | 1.6630719 | 1.56466701 | 0.4044595 | 0.57197707 | 0.65974054 |
| ENSMUSG00 | Coq2 | 2832 | 38.3516853 | 38.2667049 | 36.8192612 | 38.4900623 | 28.8284609 | 28.0354784 |
| ENSMUSG00 | 1700016H13l | 2271 | 0 | 0 | 0.0194636 | 0.20654627 | 0 | 0.01200997 |
| ENSMUSG00 | Slc10a6 | 2118 | 0 | 0 | 0 | 0 | 0.43365192 | 0 |
| ENSMUSG00 | Plac8 | 1849 | 1.16105784 | 1.40728387 | 18.9334017 | 0.74154553 | 1.68340126 | 17.4062057 |
| ENSMUSG00 | Enoph1 | 1820 | 17.6933732 | 16.4416377 | 17.5835903 | 20.6183112 | 17.6910127 | 17.3089057 |
| ENSMUSG00 | Hnrpdl | 5694 | 33.6113248 | 36.3048537 | 28.5285868 | 29.9416334 | 36.5178225 | 53.7014367 |
| ENSMUSG00 | Cds1 | 3926 | 81.4146875 | 63.3935236 | 77.6515821 | 70.877414 | 66.011962 | 74.758633 |
| ENSMUSG00 | 4930522N08 | 914 | 0 | 0 | 0 | 0 | 0 | 0 |
| ENSMUSG00 | Rasgef1b | 3271 | 1.71370307 | 1.84142625 | 2.94588865 | 0.29783406 | 1.96555284 | 1.70935526 |
| ENSMUSG00 | Prkg2 | 6182 | 0 | 0 | 0 | 0 | 0 | 0 |
| ENSMUSG00 | Bmp3 | 3109 | 5.81818708 | 4.43271871 | 4.49269295 | 2.39076911 | 2.65882048 | 4.46535586 |
| ENSMUSG00 | Fgf5 | 2498 | 0 | 0 | 0 | 0 | 0 | 0 |
| ENSMUSG00 | Antxr2 | 3734 | 0 | 1.56147817 | 0 | 0.66675444 | 0.34163368 | 0.40174214 |
| ENSMUSG00 | Crybb1 | 959 | 0 | 0 | 0 | 0 | 0 | 0 |
| ENSMUSG00 | Tpst2 | 2370 | 39.8225784 | 31.8193173 | 40.7701387 | 36.8128201 | 24.5443327 | 26.7107332 |
| ENSMUSG00 | Tfip11 | 4072 | 10.9835254 | 12.4844539 | 9.38963512 | 11.0585441 | 15.6637942 | 6.10866184 |
| ENSMUSG00 | Srrd | 1361 | 6.98130775 | 13.6663973 | 12.2764854 | 12.327811 | 7.12344703 | 11.2425221 |
| ENSMUSG00 | Asphd2 | 1581 | 0 | 0 | 0 | 0 | 0 | 0 |
| ENSMUSG00 | Crybb3 | 2531 | 0.12565936 | 0.152308 | 0.40167617 | 0.71280214 | 0.18144504 | 0.3879443 |
| ENSMUSG00 | Tesc | 1803 | 34.2430733 | 60.6406197 | 34.7387751 | 42.7060915 | 29.0649406 | 43.0676084 |
| ENSMUSG00 | Gm9754 | 3001 | 0.15896893 | 0.11239757 | 0.3093098 | 0.26451349 | 0.22104062 | 0.09997369 |
| ENSMUSG00 | Nos1 | 10917 | 0 | 0 | 0.0890758 | 0.00330513 | 0.00467403 | 0 |
| ENSMUSG00 | Rfc5 | 5497 | 4.52736598 | 4.04987016 | 5.122171 | 3.3279237 | 4.05649013 | 4.55983145 |
| ENSMUSG00 | Wsb2 | 6015 | 21.5730488 | 20.5643553 | 21.8620919 | 19.0458007 | 23.3118006 | 15.14502 |
| ENSMUSG00 | Dck | 3071 | 1.39810873 | 3.85993643 | 2.61958171 | 2.31460852 | 2.11017575 | 1.57199972 |
| ENSMUSG00 | Alb | 2028 | 0 | 0.04752115 | 0 | 0 | 0 | 0 |
| ENSMUSG00 | Afm | 2971 | 10.1563143 | 9.05016315 | 7.39424974 | 10.5052065 | 6.54360457 | 7.78488525 |
| ENSMUSG00 | Rassf6 | 1947 | 42.1853224 | 40.7864698 | 37.0732458 | 40.6780112 | 40.0191453 | 36.7724335 |
| ENSMUSG00 | Cxcl5 | 1824 | 0 | 2.19269446 | 0 | 0 | 0 | 1.76447776 |
| ENSMUSG00 | Ppbp | 1077 | 0.18456583 | 0 | 0.24624982 | 0 | 0 | 0.40519427 |
| ENSMUSG00 | Pf4 | 583 | 0 | 0 | 0 | 0 | 0 | 0 |
| ENSMUSG00 | Cxcl15 | 2123 | 0 | 0.02269734 | 0 | 0 | 0 | 0 |
| ENSMUSG00 | Mthfd2l | 2227 | 7.60477535 | 9.99646891 | 11.1745117 | 8.57090326 | 8.59222763 | 8.32813412 |
| ENSMUSG00 | Ereg | 4131 | 0 | 0 | 0 | 0 | 0 | 0 |
| ENSMUSG00 | Areg | 1193 | 0 | 0 | 0 | 0.0604896 | 0 | 0 |
| ENSMUSG00 | Cxcl3 | 1005 | 0 | 0 | 0 | 0 | 0 | 0 |
| ENSMUSG00 | Cxcl1 | 949 | 0 | 1.87871271 | 0.65208198 | 0.76042244 | 1.07537146 | 0 |
| ENSMUSG00 | Shroom3 | 7679 | 7.17037902 | 9.45029065 | 5.34175157 | 6.568915 | 11.3960457 | 6.9332069 |
| ENSMUSG00 | 2010109A12l | 546 | 0.72812235 | 0.17650711 | 0.48573454 | 0 | 0.56072941 | 0.0999071 |
| ENSMUSG00 | Ccng2 | 3699 | 52.1367494 | 50.7006307 | 77.3980366 | 47.7387202 | 61.9793205 | 78.8228963 |
| ENSMUSG00 | Tcfcp2 | 7547 | 3.18697041 | 2.64332676 | 1.97962408 | 2.72515773 | 5.48329015 | 1.31187148 |
| ENSMUSG00 | Gtf2h3 | 3862 | 6.05288 | 9.68972881 | 9.76286175 | 9.7072098 | 6.51372431 | 6.92812561 |
| ENSMUSG00 | Eif2b1 | 2559 | 20.3981813 | 26.0044848 | 24.4759717 | 26.212005 | 24.2868799 | 20.35739 |
| ENSMUSG00 | Ddx55 | 5278 | 4.85080134 | 3.93489135 | 3.85237738 | 5.25028613 | 6.70941736 | 3.1677442 |
| ENSMUSG00 | Tmed2 | 2443 | 16.5824129 | 17.5940673 | 21.9290355 | 16.2317507 | 14.537191 | 19.0688022 |
| ENSMUSG00 | Rilpl1 | 2238 | 13.0386607 | 12.035845 | 9.87530005 | 11.0600011 | 8.93759576 | 9.87151823 |
| ENSMUSG00 | Cdk2ap1 | 2488 | 1.21439571 | 1.18142 | 1.43904714 | 1.00066763 | 0.82035974 | 1.11817251 |
| ENSMUSG00 | Rchy1 | 3746 | 33.5151648 | 37.162475 | 37.7945817 | 32.7300553 | 24.0829238 | 41.7638363 |
| ENSMUSG00 | Rilpl2 | 1387 | 32.5897486 | 24.2148163 | 47.0381545 | 26.7428564 | 18.0266216 | 29.6147128 |
| ENSMUSG00 | Snrnp35 | 1388 | 4.06720334 | 4.23540774 | 6.08253027 | 5.69306034 | 3.89682697 | 4.30341931 |
| ENSMUSG00 | Cdkl2 | 5282 | 7.50401629 | 4.87155625 | 8.3014442 | 8.2383464 | 3.38114948 | 5.984723 |
| ENSMUSG00 | Arl6ip4 | 1578 | 19.52503 | 20.2456344 | 18.5995081 | 16.4404246 | 16.2650615 | 20.2226413 |
| ENSMUSG00 | G3bp2 | 7556 | 14.6426022 | 14.1893639 | 15.5373339 | 13.561806 | 13.2833353 | 18.7233396 |
| ENSMUSG00 | Pitpnm2 | 15707 | 0.05062135 | 0.05828881 | 0.09849523 | 0.22971952 | 0.07147006 | 0.01041878 |
| ENSMUSG00 | Uso1 | 3873 | 36.5631349 | 36.6779322 | 28.988557 | 40.9637877 | 35.8752444 | 37.6337909 |
| ENSMUSG00 | Abcb9 | 5774 | 2.90557894 | 2.9125508 | 3.65158973 | 3.01829365 | 3.71165186 | 2.3854681 |
| ENSMUSG00 | U90926 | 514 | 0 | 0 | 0 | 0 | 0 | 0 |
| ENSMUSG00 | Ppef2 | 2258 | 0.05281951 | 0.23474352 | 0 | 0.30361331 | 0 | 0 |
| ENSMUSG00 | Naaa | 4989 | 4.2074351 | 5.76614672 | 3.12753068 | 5.47486625 | 5.17525479 | 3.05602793 |
| ENSMUSG00 | Kntc1 | 6897 | 0 | 0 | 0.10895046 | 0.11509424 | 0 | 0.03163653 |
| ENSMUSG00 | Sdad1 | 4998 | 6.45091931 | 6.17033273 | 4.59883121 | 7.13988436 | 8.08581226 | 5.91550799 |
| ENSMUSG00 | Slc15a4 | 4038 | 9.13647497 | 7.29121252 | 10.5086105 | 10.2044811 | 7.48083568 | 6.50457593 |
| ENSMUSG00 | Cxcl9 | 5200 | 0 | 0 | 0 | 0 | 0 | 0.5035318 |
| ENSMUSG00 | Gm996 | 4683 | 0 | 0 | 0 | 0 | 0 | 0 |
| ENSMUSG00 | Rimbp2 | 4242 | 0 | 0 | 0 | 0 | 0 | 0 |

| | | | | | | | | |
|---|---|---|---|---|---|---|---|---|
| ENSMUSG00 | Rsrc2 | 2538 | 55.2629374 | 49.2686436 | 58.6745505 | 61.57263 | 58.1837398 | 77.0417214 |
| ENSMUSG00 | Piwil1 | 3922 | 0 | 0 | 0 | 0 | 0 | 0 |
| ENSMUSG00 | Scarb2 | 2098 | 8.98193411 | 8.03872962 | 15.6750101 | 9.16669673 | 6.56678812 | 10.6732502 |
| ENSMUSG00 | Zcchc8 | 4272 | 10.711273 | 13.885185 | 11.6091919 | 15.6676481 | 11.8488214 | 12.3731861 |
| ENSMUSG00 | Stx2 | 5773 | 1.17069664 | 1.96985975 | 1.26334732 | 2.71255973 | 2.43950801 | 1.43152879 |
| ENSMUSG00 | Ran | 2716 | 21.9269918 | 22.5674354 | 20.6687558 | 14.1750889 | 16.6079957 | 20.5965704 |
| ENSMUSG00 | B3gnt4 | 1422 | 0 | 0 | 0 | 0 | 0 | 0.01918048 |
| ENSMUSG00 | Gbas | 2785 | 76.1278196 | 77.7038821 | 93.4190478 | 73.5244534 | 62.6057904 | 85.0263619 |
| ENSMUSG00 | Diablo | 5015 | 14.3801479 | 14.4030861 | 14.45484 | 12.1736429 | 15.8624367 | 15.5054829 |
| ENSMUSG00 | Vps33a | 4094 | 23.2473425 | 22.3159488 | 25.1563982 | 23.1263519 | 25.1268133 | 15.3961141 |
| ENSMUSG00 | Mmp17 | 3479 | 0.03428182 | 0.40166911 | 0.02541066 | 0 | 0 | 0 |
| ENSMUSG00 | Il31 | 1627 | 0 | 0 | 0 | 0 | 0 | 0 |
| ENSMUSG00 | Bcl7a | 8572 | 3.00531397 | 3.87312863 | 2.3255986 | 5.69938034 | 4.41688881 | 3.45546643 |
| ENSMUSG00 | Sfswap | 3949 | 5.19468927 | 4.62462942 | 3.99596302 | 6.43245362 | 6.65449064 | 5.95359307 |
| ENSMUSG00 | Psmd9 | 4555 | 7.94236823 | 6.88679996 | 8.61717598 | 5.98066819 | 6.32928701 | 7.49079554 |
| ENSMUSG00 | Wdr66 | 7560 | 0.02629331 | 0.00637387 | 0 | 0.03818206 | 0.02024856 | 0 |
| ENSMUSG00 | Hpd | 2328 | 0.08538548 | 0.28978101 | 0 | 0 | 0 | 0.08201137 |
| ENSMUSG00 | Psph | 1808 | 5.91494705 | 11.0338424 | 5.86750129 | 5.5280566 | 7.73297952 | 6.06438328 |
| ENSMUSG00 | Cct6a | 1584 | 135.956716 | 117.241508 | 135.228618 | 117.562773 | 101.85947 | 102.193803 |
| ENSMUSG00 | Rhof | 4595 | 0.3114684 | 0.10486712 | 0.29820612 | 0.02355737 | 0.45529519 | 0.15432875 |
| ENSMUSG00 | Gm13103 | 2037 | 0 | 0 | 0 | 0 | 0 | 0 |
| ENSMUSG00 | Tmem116 | 3714 | 42.5385782 | 34.1223325 | 57.5790298 | 29.5826132 | 30.8576306 | 45.7514816 |
| ENSMUSG00 | Mapkapk5 | 5866 | 5.95722253 | 3.67189561 | 4.14439374 | 5.28990083 | 4.8625544 | 2.49684304 |
| ENSMUSG00 | Aldh2 | 4691 | 47.0607937 | 75.1507164 | 30.9441169 | 56.8728709 | 53.1583668 | 29.6061529 |
| ENSMUSG00 | Acad10 | 6397 | 5.03391265 | 3.77386392 | 3.4134298 | 3.55349196 | 5.08110073 | 2.65626079 |
| ENSMUSG00 | Brap | 5142 | 10.7854717 | 9.06189995 | 11.5791292 | 8.51878686 | 10.6379376 | 11.8656881 |
| ENSMUSG00 | Fam168a | 6858 | 5.91868556 | 6.93496915 | 6.3357264 | 8.13398087 | 7.2023231 | 5.92978815 |
| ENSMUSG00 | Vps29 | 3147 | 47.8531173 | 34.6507526 | 44.4827572 | 38.1229101 | 35.1201557 | 53.4659198 |
| ENSMUSG00 | 1500011H22I | 1381 | 8.89532475 | 12.1425647 | 10.0822452 | 12.2015314 | 7.94400493 | 9.14421287 |
| ENSMUSG00 | Gpn3 | 2614 | 13.2467573 | 10.8944864 | 12.7498813 | 10.8632628 | 12.805395 | 10.4966668 |
| ENSMUSG00 | Arpc3 | 2647 | 23.6400175 | 24.6848567 | 27.9872644 | 21.8918638 | 13.9373138 | 30.2524899 |
| ENSMUSG00 | Anapc7 | 3272 | 25.1873811 | 29.9398035 | 26.3022581 | 24.5582864 | 24.4683324 | 22.1314692 |
| ENSMUSG00 | Atp2a2 | 5653 | 36.7244152 | 30.7803366 | 33.2315289 | 34.7225053 | 36.2320666 | 25.7499999 |
| ENSMUSG00 | P2rx7 | 5767 | 0.42051055 | 0 | 0 | 0 | 0.1327199 | 0 |
| ENSMUSG00 | Ift81 | 4427 | 5.79224869 | 4.00555923 | 4.74268702 | 7.71847666 | 6.9963881 | 5.98230737 |
| ENSMUSG00 | P2rx4 | 3493 | 26.6781696 | 27.5489049 | 37.5203162 | 27.2087333 | 26.6745383 | 27.1497053 |
| ENSMUSG00 | Camkk2 | 3402 | 8.00485131 | 10.4248152 | 6.11965551 | 7.21216645 | 9.2093459 | 8.69870182 |
| ENSMUSG00 | Anapc5 | 2733 | 77.9108502 | 81.6507185 | 74.9959554 | 73.1542659 | 87.0790402 | 66.7145859 |
| ENSMUSG00 | Rnf34 | 1948 | 41.0207987 | 44.8965054 | 48.6265655 | 39.2494109 | 34.2882577 | 30.4809495 |
| ENSMUSG00 | Kdm2b | 12990 | 0.37643758 | 0.72335306 | 0.54444148 | 0.24721339 | 0.76205673 | 0.52071675 |
| ENSMUSG00 | Morn3 | 1546 | 0 | 0.31168462 | 0.17154661 | 0 | 0.13202167 | 0.26463104 |
| ENSMUSG00 | Ncor2 | 11609 | 3.21564271 | 2.97611155 | 3.32018323 | 3.91621812 | 2.62845833 | 4.91032785 |
| ENSMUSG00 | Dhx37 | 5491 | 2.27339663 | 2.64143491 | 2.09296652 | 3.39070023 | 3.15023519 | 1.50504748 |
| ENSMUSG00 | Aacs | 3651 | 14.460498 | 11.8783341 | 10.4481486 | 11.029187 | 8.66511997 | 9.30071922 |
| ENSMUSG00 | Tmem132b | 1805 | 0 | 0 | 0 | 0 | 0 | 0 |
| ENSMUSG00 | Anxa3 | 1584 | 28.310721 | 21.020727 | 19.8685052 | 17.60822 | 17.6530644 | 13.6889761 |
| ENSMUSG00 | Mrpl1 | 4285 | 15.8465253 | 15.7210376 | 16.3603554 | 13.7254919 | 13.3966564 | 26.0461666 |
| ENSMUSG00 | Mfsd7a | 2498 | 1.00264022 | 2.41125111 | 3.43280927 | 2.8166528 | 1.10305216 | 1.55043986 |
| ENSMUSG00 | Pde6b | 5275 | 0.15826826 | 0.04567435 | 0.11731295 | 0 | 0.05803948 | 0.11375205 |
| ENSMUSG00 | Pxmp2 | 1527 | 3.30644795 | 3.34496586 | 3.93677187 | 1.13420965 | 3.14111285 | 5.28702896 |
| ENSMUSG00 | Pgam5 | 3649 | 4.30348446 | 3.56545338 | 4.7605712 | 3.64874608 | 4.53070588 | 2.5861949 |
| ENSMUSG00 | Ankle2 | 5205 | 13.3893098 | 10.6556378 | 9.96982972 | 15.83312 | 12.5090597 | 10.9622565 |
| ENSMUSG00 | Golga3 | 12161 | 3.23640537 | 3.35613161 | 4.69605963 | 5.74120189 | 6.15958554 | 3.83069516 |
| ENSMUSG00 | P2rx2 | 2455 | 0 | 0 | 0.21605789 | 0 | 0 | 0 |
| ENSMUSG00 | Ddx51 | 5478 | 5.28331322 | 5.6296683 | 4.75262606 | 4.51190225 | 5.43983269 | 3.27614323 |
| ENSMUSG00 | Ep400 | 15551 | 4.78313318 | 2.66480228 | 4.07597215 | 3.99777267 | 3.67169407 | 3.91466753 |
| ENSMUSG00 | Pus1 | 2440 | 7.86962995 | 6.24053923 | 7.66286049 | 7.12768257 | 4.22431472 | 5.56670916 |
| ENSMUSG00 | Gpc2 | 7394 | 0.09678099 | 0.28022951 | 0.16738594 | 0.48311096 | 0.34505258 | 0.28403398 |
| ENSMUSG00 | Ulk1 | 5185 | 12.582207 | 10.1298403 | 13.5546635 | 10.6332428 | 14.1220539 | 10.9466778 |
| ENSMUSG00 | Prkab1 | 3618 | 73.4894012 | 71.294092 | 63.5294593 | 51.7495105 | 71.984141 | 58.6351974 |
| ENSMUSG00 | Cit | 14608 | 0.30752802 | 0.35295381 | 0.41151771 | 0.27417216 | 0.37724867 | 0.5134533 |
| ENSMUSG00 | Ankrd7 | 1187 | 0 | 0 | 0 | 0 | 0 | 0 |
| ENSMUSG00 | Rab35 | 2850 | 2.53877104 | 4.12543575 | 2.40395988 | 3.06380869 | 1.77249516 | 3.30166686 |
| ENSMUSG00 | Chek2 | 2232 | 1.8524059 | 3.82123668 | 2.47546164 | 2.27937647 | 2.76621482 | 2.94497673 |
| ENSMUSG00 | Pla2g1b | 762 | 0.20869019 | 0.75884161 | 0 | 0.18940706 | 0.1339275 | 0.89483725 |
| ENSMUSG00 | Sirt4 | 2787 | 6.13378422 | 5.44626095 | 6.26470326 | 4.01343158 | 5.78555248 | 6.49815589 |
| ENSMUSG00 | 1700123K08I | 1041 | 0 | 0 | 0 | 0 | 0 | 0 |
| ENSMUSG00 | Pxn | 8338 | 10.8519398 | 11.2115253 | 8.15332628 | 9.51167353 | 9.43051633 | 7.20955924 |
| ENSMUSG00 | Ccr9 | 3170 | 0 | 0.03040154 | 0 | 0.0227647 | 0.01609665 | 0 |
| ENSMUSG00 | St7 | 9084 | 4.08758463 | 3.02358785 | 2.56432973 | 2.39514232 | 3.85337889 | 2.69623801 |
| ENSMUSG00 | Triap1 | 1086 | 25.9911521 | 21.7859513 | 24.5023108 | 20.5328762 | 19.9219 | 21.975423 |
| ENSMUSG00 | Gatc | 1791 | 46.1483216 | 37.7742964 | 37.8097284 | 49.4793419 | 39.1458629 | 45.4120459 |
| ENSMUSG00 | Srsf9 | 1353 | 39.3734988 | 45.8714985 | 58.9031212 | 54.2430736 | 39.3351885 | 66.1606549 |
| ENSMUSG00 | Cyp2w1 | 1503 | 0 | 0 | 0 | 0 | 0 | 0 |
| ENSMUSG00 | Cabp1 | 2550 | 0 | 0.07558658 | 0 | 0 | 0.04002069 | 0.07487156 |
| ENSMUSG00 | Acads | 2253 | 12.9694976 | 14.2014193 | 13.9491835 | 11.8191517 | 10.7352429 | 12.0938147 |
| ENSMUSG00 | Uncx | 3633 | 0.12037167 | 0.05305416 | 0.07300057 | 0 | 0 | 0 |
| ENSMUSG00 | Ints1 | 7066 | 4.10720362 | 3.45065663 | 4.19748608 | 6.20942063 | 6.81699671 | 4.78251883 |
| ENSMUSG00 | Sppl3 | 4102 | 12.4635661 | 13.4738784 | 15.3445696 | 15.5165106 | 12.8623288 | 10.7117123 |
| ENSMUSG00 | Psmg3 | 836 | 11.6508286 | 14.5827391 | 13.9584767 | 10.6606041 | 9.52166822 | 12.4954388 |
| ENSMUSG00 | Tes | 3320 | 25.4459024 | 28.8392953 | 30.1025202 | 24.6270822 | 20.241487 | 25.5165752 |
| ENSMUSG00 | Tfec | 1772 | 0.40383671 | 0.02719325 | 0 | 0 | 0.51832662 | 0.47715227 |
| ENSMUSG00 | Mad1l1 | 4207 | 2.37191006 | 3.65378537 | 2.15388111 | 3.99672754 | 3.14139086 | 1.73748593 |
| ENSMUSG00 | Hnf1a | 3924 | 2.65441791 | 1.41219186 | 2.28669066 | 0.5884941 | 1.80750924 | 2.18947792 |
| ENSMUSG00 | Ftsj2 | 1440 | 7.17807287 | 11.6450569 | 10.1969805 | 7.51709261 | 5.84677223 | 6.30726033 |
| ENSMUSG00 | 2210016L21F | 3448 | 8.97034914 | 9.74070481 | 10.2812872 | 8.1414822 | 9.42685655 | 14.3096933 |
| ENSMUSG00 | Snx8 | 2837 | 6.41805078 | 3.94051977 | 2.9914536 | 6.11754086 | 7.03253893 | 4.71081186 |
| ENSMUSG00 | Oasl2 | 4365 | 1.36616772 | 1.12600621 | 0 | 1.00021246 | 5.51762873 | 4.16773984 |
| ENSMUSG00 | Foxp2 | 11588 | 0 | 0 | 0 | 0 | 0 | 0 |
| ENSMUSG00 | 4930519G04 | 3729 | 0 | 0 | 0 | 0 | 0 | 0 |
| ENSMUSG00 | Tmem168 | 5613 | 17.0481813 | 12.3621017 | 20.5692524 | 19.8891583 | 20.6450917 | 21.0160012 |
| ENSMUSG00 | Lfng | 2274 | 0.27972194 | 0 | 0 | 0 | 0.06731712 | 0.11994125 |
| ENSMUSG00 | Tmem106b | 6877 | 82.3783041 | 72.88576 | 103.276896 | 89.3472642 | 89.2015545 | 81.4392093 |
| ENSMUSG00 | Mmab | 3879 | 12.4319149 | 10.4472281 | 8.84264764 | 11.4506308 | 10.2210606 | 8.59935133 |
| ENSMUSG00 | Radil | 5191 | 0.00765854 | 0.04641345 | 0.28951313 | 0.06255797 | 0.37353155 | 0.15237229 |
| ENSMUSG00 | Ube3b | 5384 | 22.1963177 | 21.4977404 | 21.0254309 | 21.0634967 | 22.2150492 | 12.0517026 |
| ENSMUSG00 | Wipi2 | 8272 | 17.1479152 | 14.9941855 | 12.1191707 | 13.2603258 | 20.3254241 | 13.5647807 |
| ENSMUSG00 | Actb | 2810 | 96.233729 | 116.916428 | 103.992308 | 109.774163 | 69.9842179 | 84.0175364 |
| ENSMUSG00 | Fscn1 | 3077 | 0 | 0.07830101 | 0.01436524 | 0.03517911 | 0.01658316 | 0.13296054 |
| ENSMUSG00 | 4933411G11 | 2311 | 0 | 0 | 0 | 0 | 0 | 0 |
| ENSMUSG00 | Zfp12 | 5973 | 7.1683664 | 11.6331575 | 7.87389268 | 5.52134417 | 10.9348336 | 6.26955963 |
| ENSMUSG00 | Ung | 2464 | 0.38722871 | 0 | 0.0896953 | 0 | 0.41417513 | 0.25459282 |
| ENSMUSG00 | Usp30 | 3359 | 4.5211651 | 5.82426184 | 4.81627703 | 5.8758197 | 5.39278458 | 4.03557479 |
| ENSMUSG00 | Rbm19 | 3061 | 3.44175183 | 1.10194412 | 1.03970359 | 1.5088212 | 3.11725981 | 1.18507907 |
| ENSMUSG00 | Lhx5 | 2845 | 0 | 0 | 0 | 0 | 0 | 0 |
| ENSMUSG00 | Sdsl | 1699 | 5.03085833 | 5.84249975 | 5.87970366 | 2.27238303 | 1.29142682 | 4.07755054 |
| ENSMUSG00 | Sds | 1605 | 0 | 0.09006812 | 0 | 0 | 0 | 0.1529419 |
| ENSMUSG00 | Plbd2 | 6859 | 15.4292301 | 17.8934055 | 16.9679914 | 15.213482 | 14.9084206 | 13.4921783 |
| ENSMUSG00 | Ddx54 | 3132 | 25.488188 | 21.6931301 | 18.177514 | 19.8727736 | 22.1244631 | 13.9246961 |
| ENSMUSG00 | 1110008J03R | 2827 | 1.78597313 | 3.80105292 | 3.62745935 | 1.5698944 | 2.70744831 | 2.66282293 |
| ENSMUSG00 | Iqcd | 1742 | 0.27386095 | 0.16596938 | 0.17761935 | 0.72495497 | 0.6737122 | 0.29748458 |
| ENSMUSG00 | Rasal1 | 3604 | 0 | 0.0802216 | 0 | 0.71082829 | 0.01415826 | 0 |
| ENSMUSG00 | Dtx1 | 5528 | 0 | 0 | 0 | 0 | 0 | 0 |
| ENSMUSG00 | Oas1b | 2097 | 0.66353926 | 0.73532005 | 0.73775131 | 1.75506368 | 0.77865714 | 0.84542277 |
| ENSMUSG00 | Ankrd61 | 3427 | 0.01160067 | 0.14060824 | 0.09028681 | 0.01052876 | 0.28290083 | 0.00795875 |
| ENSMUSG00 | Rph3a | 4105 | 0 | 0 | 0.01076781 | 0.0439489 | 0 | 0.07973098 |
| ENSMUSG00 | Aimp2 | 1650 | 7.6378711 | 10.1921626 | 8.17064371 | 10.0155008 | 7.4220183 | 6.61203376 |
| ENSMUSG00 | Eif2ak1 | 5837 | 26.5081943 | 23.1232182 | 21.4307377 | 18.7982692 | 24.6521124 | 16.6208484 |
| ENSMUSG00 | Rpl6 | 2367 | 2.11626132 | 2.23933614 | 2.35294982 | 1.7987669 | 1.35811647 | 2.96137824 |
| ENSMUSG00 | Erp29 | 9054 | 5.88823717 | 7.13695813 | 5.755906 | 4.0449829 | 5.66396153 | 6.24478984 |
| ENSMUSG00 | Ccz1 | 1747 | 52.9086962 | 51.744571 | 46.5801906 | 44.6326836 | 52.6912318 | 48.7102888 |
| ENSMUSG00 | Ocm | 753 | 0 | 0 | 0 | 0 | 0 | 0 |
| ENSMUSG00 | 1700018F24I | 1114 | 0 | 0 | 0 | 0 | 0 | 0 |
| ENSMUSG00 | Arpc1a | 2462 | 46.9411787 | 51.4549782 | 60.2164831 | 36.9906907 | 43.6687952 | 40.5131421 |
| ENSMUSG00 | Arpc1b | 4835 | 37.0091867 | 33.9746807 | 40.5084522 | 29.4850378 | 29.2649721 | 34.2131721 |
| ENSMUSG00 | Pdap1 | 3486 | 3.13618966 | 3.06867188 | 2.02877076 | 2.58763945 | 2.32736482 | 3.70859983 |
| ENSMUSG00 | Ptcd1 | 3170 | 7.56232011 | 12.3886279 | 9.04952559 | 11.2343779 | 15.6942323 | 8.04472798 |
| ENSMUSG00 | Cpsf4 | 2683 | 1.77810573 | 2.2090691 | 1.25208351 | 1.10276841 | 3.51840459 | 1.34187565 |
| ENSMUSG00 | Zkscan14 | 2668 | 5.60272139 | 5.72530066 | 6.19621038 | 6.57266628 | 3.99719361 | 4.8456443 |
| ENSMUSG00 | Phf14 | 4460 | 7.60345849 | 5.82342877 | 5.08420302 | 7.13550746 | 7.18487982 | 9.49720062 |
| ENSMUSG00 | Cyp3a25 | 2576 | 0 | 0 | 0 | 0 | 0 | 0 |
| ENSMUSG00 | Ndufa4 | 488 | 552.177555 | 477.026029 | 702.066568 | 450.212232 | 351.224583 | 1367.02951 |
| ENSMUSG00 | Gm5578 | 1235 | 10.8482566 | 12.1343996 | 9.44881503 | 11.2774649 | 12.3950711 | 5.71994459 |
| ENSMUSG00 | Rnf6 | 5847 | 31.4467415 | 31.4073082 | 34.1246997 | 32.1881211 | 30.1166449 | 36.5621041 |
| ENSMUSG00 | Cdk8 | 4416 | 3.41198531 | 6.06695241 | 4.38415019 | 2.83525123 | 3.32780712 | 6.12691172 |
| ENSMUSG00 | Wasf3 | 3986 | 0.38897736 | 0.6648907 | 0.18851765 | 0.8780628 | 0.96010491 | 0.58846437 |
| ENSMUSG00 | Glcci1 | 7763 | 0.29702665 | 0.21725177 | 0.10818434 | 0.05112746 | 0.17089859 | 0.63241467 |
| ENSMUSG00 | Usp12 | 7570 | 4.57949525 | 4.87593325 | 3.96473599 | 4.71878786 | 6.96978502 | 3.28953047 |
| ENSMUSG00 | Rasl11a | 1122 | 10.1337499 | 19.6267416 | 6.65785336 | 14.8573125 | 12.7338549 | 7.60870061 |
| ENSMUSG00 | Polr1d | 2328 | 9.01670692 | 11.4049526 | 12.8352431 | 10.8804103 | 6.59748244 | 14.6917516 |
| ENSMUSG00 | Pdx1 | 4316 | 0 | 0 | 0 | 0 | 0 | 0 |
| ENSMUSG00 | Cdx2 | 2258 | 0 | 0 | 0 | 0 | 0 | 0 |
| ENSMUSG00 | Pan3 | 7840 | 3.64087181 | 3.48491233 | 2.5821995 | 3.41951009 | 5.10263758 | 5.23227773 |
| ENSMUSG00 | Flt1 | 11550 | 0.83641401 | 0.75930151 | 0.52047192 | 0.26553886 | 0.4417868 | 0.40852923 |
| ENSMUSG00 | Pomp | 728 | 74.8145718 | 76.3834536 | 83.3034734 | 67.4555805 | 45.1387171 | 96.5851923 |
| ENSMUSG00 | Slc46a3 | 3225 | 0.76429141 | 1.89757462 | 1.93254569 | 0.96218785 | 1.88283371 | 1.27704513 |
| ENSMUSG00 | Mtus2 | 10159 | 0.29349946 | 0.08063486 | 0.48296137 | 0.23441431 | 0.43698146 | 0.2684776 |
| ENSMUSG00 | N4bp2l2 | 12254 | 6.36204482 | 6.80288436 | 6.47841603 | 9.18687603 | 6.15864288 | 7.91040215 |
| ENSMUSG00 | C8b | 2623 | 0 | 0.42252694 | 0 | 0 | 0 | 0 |
| ENSMUSG00 | Hsph1 | 7024 | 15.8421968 | 14.1389888 | 13.8760057 | 16.1300712 | 16.6504062 | 9.68824387 |
| ENSMUSG00 | Wdr95 | 3342 | 0 | 0 | 0 | 0 | 0 | 0 |
| ENSMUSG00 | 6330406I15R | 5488 | 0 | 0 | 0 | 0 | 0 | 0 |
| ENSMUSG00 | 4930588N13 | 1352 | 0 | 0 | 0 | 0.10675161 | 0 | 0 |
| ENSMUSG00 | Col1a2 | 11111 | 0 | 0 | 0 | 0 | 0 | 0 |
| ENSMUSG00 | Gngt1 | 3508 | 0 | 0 | 0 | 0 | 0 | 0 |
| ENSMUSG00 | Tfpi2 | 1525 | 0.70386752 | 0.094793 | 1.27533186 | 1.11203678 | 0.86995788 | 1.28772067 |
| ENSMUSG00 | Tspan12 | 2615 | 53.1644036 | 91.6740152 | 46.6697088 | 79.4494885 | 62.0511951 | 75.3155526 |
| ENSMUSG00 | Ing3 | 4859 | 5.26091253 | 6.43609816 | 6.03124551 | 8.45057968 | 6.24834197 | 4.9677003 |
| ENSMUSG00 | Wnt16 | 4726 | 0 | 0 | 0 | 0 | 0 | 0 |
| ENSMUSG00 | Fam3c | 4355 | 41.5173193 | 58.6867713 | 50.6570375 | 51.1611079 | 43.844018 | 51.4366503 |
| ENSMUSG00 | Auts2 | 19123 | 0.01871042 | 0.05795577 | 0.06240913 | 0 | 0.04536152 | 0.00142627 |
| ENSMUSG00 | Limk1 | 8482 | 0.25310021 | 0.47152496 | 0.15112632 | 0.19142797 | 0.09625348 | 0.06431181 |
| ENSMUSG00 | Eln | 2546 | 0.29668269 | 0 | 0 | 0.07086026 | 0.20041782 | 0 |
| ENSMUSG00 | Pot1a | 8851 | 4.16823929 | 5.0413112 | 4.43466689 | 4.99384302 | 7.69632942 | 5.3248872 |
| ENSMUSG00 | Hyal5 | 2026 | 0 | 0 | 0 | 0 | 0 | 0 |
| ENSMUSG00 | Hyal6 | 3672 | 0.29231971 | 0 | 0 | 0 | 0 | 0 |
| ENSMUSG00 | Hyal4 | 3264 | 0 | 0 | 0 | 0 | 0 | 0 |
| ENSMUSG00 | Bcl7b | 2606 | 17.3300943 | 14.1637815 | 16.1982963 | 12.8904004 | 15.7621767 | 16.9655373 |
| ENSMUSG00 | Spam1 | 2010 | 0 | 0 | 0 | 0 | 0 | 0 |
| ENSMUSG00 | Lmod2 | 1964 | 0 | 0 | 0.02250603 | 0 | 0 | 0 |
| ENSMUSG00 | Wasl | 4910 | 37.8122391 | 37.5481319 | 29.0417812 | 43.8128614 | 41.2367941 | 51.3607565 |
| ENSMUSG00 | Asb15 | 4487 | 39.3213333 | 26.0316327 | 29.6714845 | 35.969464 | 34.923557 | 29.8154904 |
| ENSMUSG00 | Cul1 | 4089 | 70.8676198 | 73.3343677 | 82.2095906 | 83.9443604 | 81.8368727 | 78.9557116 |
| ENSMUSG00 | Ezh2 | 5116 | 1.67072485 | 3.65448388 | 3.27453059 | 4.9440018 | 5.31607864 | 4.01442599 |
| ENSMUSG00 | Aass | 3792 | 14.3945872 | 16.7356393 | 19.1401441 | 15.1769148 | 14.7481297 | 15.2053237 |
| ENSMUSG00 | Fezf1 | 2304 | 0 | 0 | 0 | 0 | 0 | 0 |
| ENSMUSG00 | Srcrb4d | 3931 | 0.7686127 | 0.56387086 | 0.52848808 | 0.44058462 | 0.44133726 | 0.38160905 |
| ENSMUSG00 | Slc13a1 | 8264 | 0.08659229 | 0.58891949 | 0.1658104 | 0 | 0 | 0.42245327 |
| ENSMUSG00 | Rbm28 | 6030 | 9.38835891 | 8.51054078 | 8.54715571 | 7.8327357 | 13.1670051 | 9.89666844 |
| ENSMUSG00 | Lrwd1 | 6113 | 1.54781684 | 2.64855956 | 2.37893119 | 2.60890949 | 1.61935497 | 1.59730425 |
| ENSMUSG00 | Cux1 | 13871 | 8.58392792 | 8.85844045 | 6.29679488 | 8.4592898 | 12.0217862 | 6.26072031 |
| ENSMUSG00 | Pax4 | 2012 | 0 | 0 | 0 | 0 | 0 | 0 |
| ENSMUSG00 | Fscn3 | 2075 | 0 | 0 | 0 | 0 | 0 | 0 |
| ENSMUSG00 | Gcc1 | 6411 | 5.55621752 | 6.07310018 | 4.78491328 | 7.14774552 | 8.12633438 | 3.05036911 |
| ENSMUSG00 | Ephb4 | 5109 | 16.83908 | 17.4297407 | 8.44411798 | 12.2815963 | 15.3009215 | 8.72318664 |
| ENSMUSG00 | Epo | 2711 | 0 | 0 | 0 | 0 | 0 | 0 |
| ENSMUSG00 | Actl6b | 1945 | 0 | 0 | 0 | 0 | 0 | 0 |
| ENSMUSG00 | Gnb2 | 3087 | 16.7032259 | 18.1069883 | 17.2826772 | 16.5975067 | 13.0086679 | 21.8409162 |
| ENSMUSG00 | Gigyf1 | 5413 | 1.46888899 | 1.94063262 | 2.27827715 | 2.30637122 | 2.21525925 | 2.83176561 |
| ENSMUSG00 | Pop7 | 1866 | 8.88426333 | 8.39260112 | 10.4227283 | 6.61310784 | 5.55110091 | 8.81383037 |
| ENSMUSG00 | Tfr2 | 2907 | 0 | 0 | 0 | 0.09929699 | 0 | 0 |
| ENSMUSG00 | Pcolce | 3737 | 58.9576861 | 57.9604114 | 70.9334901 | 44.2021942 | 45.9879138 | 50.6153661 |
| ENSMUSG00 | Lrch4 | 4442 | 1.79893102 | 1.33429365 | 1.14435208 | 1.73018359 | 2.75694061 | 1.93415384 |
| ENSMUSG00 | Agfg2 | 5126 | 10.3925759 | 9.84221711 | 11.4945487 | 12.3534897 | 7.79431375 | 11.8122706 |
| ENSMUSG00 | Tsc22d4 | 6597 | 4.42330191 | 6.82951696 | 5.755553 | 4.51777608 | 4.86517969 | 6.6646534 |
| ENSMUSG00 | Ppp1r35 | 1081 | 2.46403071 | 4.45758022 | 5.96990664 | 3.63824501 | 2.59616158 | 7.26650888 |
| ENSMUSG00 | Mepce | 3344 | 7.15693758 | 7.50751685 | 8.43324636 | 9.06367147 | 7.76687359 | 6.59843994 |
| ENSMUSG00 | Cyp3a13 | 2943 | 4.48481805 | 2.89806329 | 5.61722368 | 1.49575584 | 2.06324795 | 2.28910496 |
| ENSMUSG00 | Zkscan1 | 8646 | 23.2481737 | 20.2866816 | 34.9432798 | 32.4554777 | 28.7827475 | 26.0222645 |
| ENSMUSG00 | Mcm7 | 4644 | 4.27174521 | 2.94680223 | 3.66445081 | 4.79384614 | 5.52675862 | 3.71179414 |
| ENSMUSG00 | Tpk1 | 5945 | 15.6480781 | 14.930097 | 16.7067353 | 14.5602733 | 14.7199722 | 15.8371833 |
| ENSMUSG00 | Nobox | 2259 | 0 | 0 | 0 | 0 | 0 | 0 |
| ENSMUSG00 | Asns | 2776 | 4.53980091 | 7.82856103 | 8.21619272 | 8.00667847 | 4.85265246 | 6.63198181 |
| ENSMUSG00 | Dlx6 | 1804 | 0 | 0 | 0 | 0 | 0 | 0 |
| ENSMUSG00 | Dlx5 | 1420 | 0 | 0 | 0 | 0 | 0 | 0 |
| ENSMUSG00 | Dync1i1 | 3123 | 0 | 0 | 0 | 0 | 0 | 0 |
| ENSMUSG00 | Pon3 | 3095 | 27.2957661 | 21.9524664 | 42.9593356 | 28.6674467 | 23.5595125 | 30.2003841 |
| ENSMUSG00 | Cald1 | 7452 | 2.99819915 | 1.00226765 | 0.72957953 | 0.59071517 | 1.25306317 | 0.92599084 |
| ENSMUSG00 | Akr1b8 | 1489 | 1.44177028 | 6.18106813 | 0.77182533 | 3.48946569 | 1.30221778 | 0.73269682 |
| ENSMUSG00 | Exoc4 | 12725 | 7.37311859 | 9.46309775 | 7.6211272 | 8.48672371 | 8.96219646 | 6.16224659 |
| ENSMUSG00 | Plxna4 | 17206 | 9.2676527 | 5.52830622 | 14.0908467 | 6.01017898 | 5.98757717 | 8.60278201 |
| ENSMUSG00 | 1700012A03I | 1107 | 0 | 0 | 0 | 0 | 0 | 0 |
| ENSMUSG00 | Calu | 4143 | 99.5470323 | 88.1266406 | 115.225659 | 104.387976 | 95.7344966 | 82.3834988 |
| ENSMUSG00 | Ccdc136 | 4132 | 1.77033118 | 1.45772151 | 0.85579561 | 2.61097079 | 2.16108804 | 1.22115398 |
| ENSMUSG00 | Irf5 | 2890 | 0 | 0.30012317 | 0 | 0 | 0 | 0.00943759 |
| ENSMUSG00 | Ahcyl2 | 11642 | 19.2972187 | 16.9534158 | 14.6668716 | 17.8364685 | 21.800824 | 16.6946469 |
| ENSMUSG00 | Klhdc10 | 8687 | 10.8415717 | 10.006716 | 12.2373009 | 11.679833 | 12.2999 | 6.25743709 |
| ENSMUSG00 | Hibadh | 3040 | 159.231161 | 155.68659 | 180.849514 | 139.544933 | 125.451031 | 175.885536 |
| ENSMUSG00 | Gars | 2373 | 47.327953 | 65.5481817 | 48.8027091 | 47.2426938 | 57.0901929 | 40.9751744 |
| ENSMUSG00 | Adcyap1r1 | 6923 | 0 | 0 | 0 | 0 | 0 | 0 |
| ENSMUSG00 | Nt5c3 | 4792 | 22.1923853 | 19.7089788 | 25.098751 | 18.3798562 | 16.1853279 | 26.5631764 |
| ENSMUSG00 | Fkbp9 | 3304 | 35.207184 | 37.598259 | 24.8568476 | 29.6825052 | 43.0573656 | 23.097591 |
| ENSMUSG00 | Tmem209 | 10157 | 6.18427284 | 7.82786547 | 6.86288337 | 8.18480167 | 7.75669236 | 7.01938633 |
| ENSMUSG00 | 1700025E21I | 1802 | 0 | 0 | 0 | 0 | 0 | 0 |
| ENSMUSG00 | Avl9 | 7723 | 11.721252 | 12.5410784 | 17.1988267 | 13.203141 | 15.4275007 | 14.8221754 |
| ENSMUSG00 | Cpa5 | 2828 | 0 | 0 | 0 | 0 | 0 | 0 |
| ENSMUSG00 | Tsga14 | 7020 | 0.66259134 | 1.30419145 | 0.34001418 | 0.92004074 | 1.04669489 | 0.26031351 |
| ENSMUSG00 | Sspo | 15614 | 0.12221488 | 0.08949704 | 0 | 0.00924351 | 0.1732034 | 0.13799773 |
| ENSMUSG00 | Herc6 | 5381 | 2.31248196 | 3.15213299 | 1.1007335 | 2.79617405 | 3.39480441 | 2.42790414 |
| ENSMUSG00 | Abcg2 | 4967 | 1.76886676 | 2.8327846 | 1.21917707 | 2.0993982 | 1.89024626 | 1.59243918 |
| ENSMUSG00 | Herc3 | 7189 | 6.18811833 | 4.64504165 | 5.675101 | 4.79320525 | 6.88490535 | 5.96786862 |
| ENSMUSG00 | Tmem176b | 2363 | 159.442526 | 191.17558 | 201.536461 | 147.488984 | 135.048224 | 146.772879 |
| ENSMUSG00 | Abp1 | 3127 | 1.89432894 | 4.43802217 | 1.73867179 | 3.04626151 | 4.89539902 | 1.46534678 |
| ENSMUSG00 | 1600015I10R | 4240 | 0 | 0 | 0 | 0 | 0 | 0 |
| ENSMUSG00 | Igf2bp3 | 4329 | 0.1102023 | 0.08904863 | 0.04084254 | 0.20003906 | 0.07072263 | 0.08820627 |
| ENSMUSG00 | 2410003K15I | 3330 | 2.80556694 | 3.29925189 | 4.39363665 | 2.95807752 | 2.92673807 | 4.77510951 |
| ENSMUSG00 | Gpnmb | 4714 | 0 | 0 | 0 | 0 | 0.05412216 | 0 |
| ENSMUSG00 | Tra2a | 3862 | 7.23669156 | 6.31339713 | 6.94731194 | 8.5113264 | 4.62434789 | 7.23180492 |
| ENSMUSG00 | Npy | 562 | 0 | 0 | 0 | 0 | 0 | 0 |
| ENSMUSG00 | Dfna5 | 7363 | 0.6587218 | 1.23034784 | 0.84045335 | 0.55865178 | 1.10188697 | 1.31131635 |
| ENSMUSG00 | Osbpl3 | 12834 | 6.55156937 | 4.41540096 | 7.45996509 | 4.43364378 | 6.59995199 | 5.85701307 |
| ENSMUSG00 | Luc7l2 | 12318 | 4.05042442 | 5.52356359 | 3.90058478 | 6.18649765 | 3.45063899 | 6.43892279 |
| ENSMUSG00 | Zc3hav1 | 6505 | 2.09625362 | 2.91118705 | 1.31144592 | 2.26864815 | 2.57289028 | 2.48218085 |
| ENSMUSG00 | 4921507P07I | 2368 | 2.03142447 | 0.75291316 | 1.32530864 | 0.1523735 | 0.1508381 | 1.84288103 |
| ENSMUSG00 | Tmem213 | 527 | 248.56605 | 300.913816 | 371.060633 | 256.408457 | 182.80417 | 505.486531 |
| ENSMUSG00 | Svopl | 4276 | 13.2394304 | 11.2239702 | 13.986224 | 12.8430477 | 16.3007552 | 20.1179168 |
| ENSMUSG00 | Npvf | 809 | 0 | 0 | 0 | 0 | 0 | 0 |
| ENSMUSG00 | Nfe2l3 | 3583 | 0.62135275 | 1.49279796 | 1.93683767 | 0.22154758 | 0.96840457 | 3.67670967 |
| ENSMUSG00 | Trim24 | 7654 | 4.88243424 | 5.14349663 | 2.66804958 | 6.85436278 | 5.87996648 | 4.71800658 |
| ENSMUSG00 | Cbx3 | 2279 | 1.56998387 | 1.50120114 | 0.79520648 | 1.64656983 | 1.50011724 | 2.1302702 |
| ENSMUSG00 | 1700111E14I | 896 | 0 | 0 | 0 | 0 | 0 | 0 |
| ENSMUSG00 | Ptn | 3844 | 0.07239552 | 0.78973618 | 0 | 0.24405129 | 0.11946862 | 0 |
| ENSMUSG00 | Mtpn | 3956 | 113.136299 | 123.645241 | 142.806814 | 133.830091 | 96.1839445 | 174.796241 |
| ENSMUSG00 | Slc13a4 | 4512 | 0 | 0.01067962 | 0 | 0 | 0 | 0 |
| ENSMUSG00 | Hoxa1 | 3966 | 0.1202889 | 0.01214989 | 0 | 0.16376117 | 0 | 0 |
| ENSMUSG00 | 2010107G12I | 3384 | 0 | 0 | 0.07837206 | 0 | 0 | 0.08059882 |
| ENSMUSG00 | Stra8 | 1791 | 0 | 0 | 0 | 0 | 0 | 0 |
| ENSMUSG00 | Fam115c | 4768 | 6.14509 | 2.3951734 | 5.01535173 | 2.8756663 | 3.29616689 | 5.56590268 |
| ENSMUSG00 | Epha1 | 3255 | 18.0884383 | 11.3989433 | 21.8089585 | 12.7811359 | 12.196166 | 11.6723725 |
| ENSMUSG00 | Zyx | 3977 | 10.4761739 | 7.22130237 | 9.8028729 | 9.48094959 | 7.86501594 | 9.49160189 |
| ENSMUSG00 | Fam131b | 7364 | 0 | 0 | 0 | 0 | 0 | 0 |
| ENSMUSG00 | Clcn1 | 5582 | 0 | 0 | 0.02375592 | 0 | 0 | 0.11238207 |
| ENSMUSG00 | Casp2 | 4528 | 13.0996416 | 11.3016768 | 10.3964141 | 10.2556518 | 20.8252523 | 8.41490085 |
| ENSMUSG00 | Gstk1 | 922 | 50.7507598 | 61.513495 | 51.9683274 | 47.0788499 | 36.9692181 | 61.4714755 |
| ENSMUSG00 | Sval1 | 611 | 0 | 0 | 0 | 0 | 0 | 0 |
| ENSMUSG00 | Kel | 3007 | 0 | 1.08968343 | 0 | 1.15193757 | 0.13575358 | 0.05442229 |
| ENSMUSG00 | 1700034O15I | 739 | 4.35750192 | 4.43393514 | 2.69158719 | 2.245973 | 5.59287746 | 3.72765706 |
| ENSMUSG00 | Trpv6 | 2917 | 4.27947236 | 8.27610816 | 1.84868867 | 5.78896018 | 8.20410362 | 3.87099782 |
| ENSMUSG00 | Ephb6 | 3705 | 6.36302293 | 15.3078117 | 3.16153533 | 7.05085027 | 9.32384789 | 5.75675247 |
| ENSMUSG00 | AI836003 | 2328 | 0 | 0 | 0 | 0 | 0 | 0 |
| ENSMUSG00 | Dbpht2 | 2994 | 0 | 0 | 0 | 0 | 0 | 0 |
| ENSMUSG00 | 2210010C04 | 2137 | 0 | 0 | 0 | 0 | 0 | 0 |
| ENSMUSG00 | 1700074P13I | 2518 | 0 | 0 | 0 | 0 | 0 | 0 |
| ENSMUSG00 | Moxd2 | 1847 | 0 | 0 | 0 | 0 | 0 | 0 |
| ENSMUSG00 | Prss37 | 1117 | 0 | 0 | 0 | 0 | 0 | 0 |
| ENSMUSG00 | Mad2l1 | 2126 | 4.99280964 | 4.26107767 | 2.14148158 | 3.1567546 | 4.82422462 | 4.15662423 |
| ENSMUSG00 | Ssbp1 | 3116 | 19.8777403 | 18.0312553 | 25.0373084 | 26.7604637 | 21.5830434 | 46.7940377 |
| ENSMUSG00 | Prdm5 | 3867 | 0.72992995 | 0.5108984 | 0.36577682 | 0.39189187 | 1.42509661 | 0.21159534 |
| ENSMUSG00 | Clec5a | 5196 | 0 | 0 | 0 | 0 | 0 | 0 |
| ENSMUSG00 | Agk | 2209 | 5.34512346 | 8.92179908 | 5.58275881 | 6.43563854 | 4.27337235 | 6.63036726 |
| ENSMUSG00 | C130060K24I | 2629 | 0 | 0 | 0 | 0 | 0 | 0 |
| ENSMUSG00 | Mrps33 | 1585 | 6.62173303 | 3.98260184 | 5.18709325 | 5.0537627 | 3.83100235 | 10.3247846 |
| ENSMUSG00 | Hpgds | 3580 | 0 | 0 | 0 | 0.02015757 | 0 | 0 |
| ENSMUSG00 | Smarcad1 | 6578 | 6.41236923 | 7.50120352 | 7.30425712 | 11.6726347 | 5.94195863 | 8.53731867 |
| ENSMUSG00 | Mkrn1 | 4494 | 32.8907157 | 30.3336548 | 32.5661554 | 28.4946987 | 26.7848733 | 34.3330294 |
| ENSMUSG00 | Rab19 | 1571 | 7.66767065 | 11.9622613 | 11.2263115 | 12.195777 | 16.5973762 | 10.9549951 |
| ENSMUSG00 | Slc37a3 | 5348 | 7.02485585 | 9.60485178 | 7.29809787 | 7.56319594 | 7.737919 | 6.62486096 |
| ENSMUSG00 | Tbxas1 | 9044 | 0.72090876 | 1.51315232 | 0.73311327 | 2.30998494 | 1.01556254 | 0.19602516 |
| ENSMUSG00 | Gfpt1 | 7553 | 5.80041567 | 6.7689415 | 6.95244135 | 7.17532515 | 5.33706301 | 8.05636438 |
| ENSMUSG00 | Nfu1 | 2067 | 22.6954364 | 24.3613121 | 22.3254592 | 23.6357466 | 18.6380812 | 21.4159359 |
| ENSMUSG00 | Anxa4 | 5606 | 206.76963 | 169.408953 | 288.34488 | 168.741163 | 169.699563 | 220.537303 |
| ENSMUSG00 | Pcyox1 | 5173 | 91.207818 | 105.501574 | 86.5323921 | 98.2717814 | 115.171847 | 61.4878877 |
| ENSMUSG00 | Tgfa | 3756 | 4.5619308 | 4.02836604 | 2.81263059 | 1.40255019 | 4.90429225 | 1.51041666 |
| ENSMUSG00 | Add2 | 3103 | 0 | 0 | 0 | 0 | 0 | 0.15821576 |
| ENSMUSG00 | Figla | 756 | 0 | 0 | 0 | 0 | 0 | 0 |
| ENSMUSG00 | Dusp11 | 7628 | 24.7403338 | 19.7723602 | 29.1124881 | 25.7181537 | 25.4462944 | 26.6810905 |
| ENSMUSG00 | Nat8 | 1121 | 0.8156789 | 0 | 0 | 0 | 0 | 1.58149113 |
| ENSMUSG00 | Cct7 | 2440 | 109.897834 | 103.719342 | 110.214759 | 99.3882874 | 91.7220018 | 83.8359813 |
| ENSMUSG00 | 1700040I03R | 1948 | 6.2041407 | 7.17354632 | 10.5058795 | 7.96472235 | 8.251185 | 9.81494975 |
| ENSMUSG00 | Zfml | 6565 | 19.3660361 | 21.9096009 | 16.1792885 | 30.3935577 | 19.6877091 | 21.6743021 |
| ENSMUSG00 | Reg3g | 1351 | 0 | 0 | 0 | 0 | 0 | 0 |
| ENSMUSG00 | Fbxl14 | 4218 | 3.45904726 | 4.2497289 | 3.76208194 | 3.37894917 | 3.32675518 | 5.32172312 |
| ENSMUSG00 | Prickle2 | 14537 | 0.13673894 | 0.20219942 | 0 | 0.15388916 | 0.07722228 | 0.07317266 |
| ENSMUSG00 | Adamts9 | 14360 | 0.29622816 | 0.09731245 | 0.13235928 | 0.07286764 | 0.25584255 | 0.16144459 |
| ENSMUSG00 | Lrig1 | 5806 | 1.04763839 | 0.35687513 | 0.83744449 | 1.04405503 | 2.79476189 | 0.96771886 |
| ENSMUSG00 | 1700003E16I | 2046 | 3.16722548 | 2.73197815 | 3.43504059 | 3.06856097 | 3.89057411 | 1.34640204 |
| ENSMUSG00 | Kbtbd8 | 5015 | 3.33739926 | 4.66971304 | 4.08966205 | 0.51802736 | 4.84318143 | 3.86141453 |
| ENSMUSG00 | Wdr54 | 5798 | 1.43306234 | 1.36298319 | 0.9453309 | 1.51223471 | 1.21449463 | 0.82322557 |
| ENSMUSG00 | Ino80b | 2034 | 5.90273113 | 6.06476361 | 4.30283428 | 5.80080067 | 5.19294975 | 6.30239943 |
| ENSMUSG00 | Wbp1 | 2751 | 35.0732647 | 35.259654 | 42.3701418 | 35.1245784 | 29.0651875 | 40.4707661 |
| ENSMUSG00 | Mogs | 2888 | 20.5797934 | 20.2890283 | 18.243974 | 20.2524218 | 18.9052016 | 11.0307405 |
| ENSMUSG00 | Mrpl53 | 816 | 0 | 0 | 0.27084463 | 0.04421819 | 0.18759697 | 0.33424803 |
| ENSMUSG00 | D6Mm5e | 2076 | 0 | 0 | 0 | 0 | 0.24579179 | 0.19707109 |
| ENSMUSG00 | Pole4 | 2501 | 28.5012301 | 31.5591332 | 43.9720853 | 35.2597029 | 26.5843134 | 42.2151654 |
| ENSMUSG00 | Tacr1 | 6468 | 0 | 0 | 0 | 0 | 0 | 0 |
| ENSMUSG00 | Mrpl19 | 5303 | 6.07240038 | 7.81450881 | 7.07662921 | 5.13708158 | 5.57123734 | 6.04331532 |
| ENSMUSG00 | Bmp10 | 1315 | 0 | 0 | 0 | 0 | 0 | 0 |
| ENSMUSG00 | Arhgap25 | 4986 | 0 | 0 | 0 | 0 | 0.15350895 | 0 |
| ENSMUSG00 | Gkn3 | 838 | 0 | 0 | 0 | 0 | 0 | 0 |
| ENSMUSG00 | Gkn2 | 771 | 0 | 0 | 0 | 0 | 0 | 0 |
| ENSMUSG00 | Gkn1 | 1222 | 0 | 0 | 0 | 0 | 0 | 0 |
| ENSMUSG00 | Aplf | 4760 | 3.61641241 | 2.06513324 | 4.18803176 | 3.94173596 | 3.54826269 | 3.16867133 |
| ENSMUSG00 | Gp9 | 809 | 0 | 0 | 0 | 0.75821354 | 0 | 0 |
| ENSMUSG00 | Rab43 | 6798 | 4.71358007 | 6.96073642 | 3.86879915 | 5.14320405 | 7.58114733 | 3.63902586 |
| ENSMUSG00 | Isy1 | 1609 | 12.5270532 | 12.8177733 | 15.3566378 | 12.1992742 | 10.2116178 | 9.54358104 |
| ENSMUSG00 | Cnbp | 3507 | 196.895621 | 194.490758 | 231.256178 | 181.819758 | 163.802378 | 225.748775 |
| ENSMUSG00 | Copg | 7944 | 35.221434 | 31.8634668 | 33.7133644 | 27.6837942 | 38.0578143 | 23.1340029 |
| ENSMUSG00 | Tmf1 | 3448 | 10.9650412 | 13.5000299 | 10.8453478 | 11.7831735 | 9.36766122 | 7.28536623 |
| ENSMUSG00 | 8430410A17I | 2353 | 15.3919434 | 14.7037252 | 13.1121489 | 15.3191511 | 15.4401953 | 16.1932304 |
| ENSMUSG00 | Uba3 | 3620 | 54.3617758 | 48.5059103 | 65.0328773 | 62.0072925 | 46.5721397 | 69.2640217 |
| ENSMUSG00 | Rpn1 | 3632 | 167.187558 | 140.327092 | 209.922244 | 145.252085 | 136.346635 | 159.104566 |
| ENSMUSG00 | Frmd4b | 12839 | 3.35346876 | 1.58757419 | 2.73356674 | 1.51196666 | 3.40599767 | 3.52431081 |
| ENSMUSG00 | Foxp1 | 15253 | 13.4073423 | 8.90247125 | 17.6801576 | 10.7633451 | 13.5820551 | 15.1348946 |
| ENSMUSG00 | Eif4e3 | 3988 | 0.54828271 | 0.857883 | 1.57388709 | 1.51095823 | 1.03639329 | 0.95064565 |
| ENSMUSG00 | Prok2 | 1509 | 0 | 0 | 0 | 0 | 0 | 0 |
| ENSMUSG00 | Gxylt2 | 3039 | 0 | 0 | 0 | 0 | 0 | 0 |
| ENSMUSG00 | Cntn3 | 5192 | 0 | 0.22274164 | 0.4256726 | 0.09729365 | 0.00982788 | 0.436016 |
| ENSMUSG00 | Chl1 | 7920 | 4.92425838 | 2.10511477 | 5.58104078 | 4.97950438 | 3.76255094 | 4.80749954 |
| ENSMUSG00 | Ruvbl1 | 1887 | 6.36256233 | 8.12044971 | 5.90294883 | 5.22013681 | 8.22046542 | 4.97216529 |
| ENSMUSG00 | Sec61a1 | 6890 | 26.5998208 | 25.6877797 | 21.0038946 | 17.6639675 | 23.869087 | 14.7061371 |
| ENSMUSG00 | Abtb1 | 2407 | 19.869482 | 21.2805102 | 18.400601 | 16.2946333 | 24.8030161 | 12.9177768 |
| ENSMUSG00 | Plxna1 | 9140 | 3.11867391 | 6.9590923 | 2.03599299 | 4.00297518 | 6.48716069 | 2.67673429 |
| ENSMUSG00 | Chchd6 | 2519 | 16.0663276 | 14.9399013 | 15.5996183 | 11.4161927 | 10.5942019 | 12.5491492 |
| ENSMUSG00 | Klf15 | 2523 | 23.9037114 | 19.289856 | 26.8049226 | 24.4837337 | 12.0335686 | 12.7887032 |
| ENSMUSG00 | Aldh1l1 | 4414 | 522.792675 | 299.882548 | 482.324823 | 290.184979 | 417.378183 | 259.696089 |
| ENSMUSG00 | Slc41a3 | 2800 | 21.3117772 | 6.22981859 | 19.7803248 | 9.75503847 | 8.21889125 | 14.26074 |
| ENSMUSG00 | Nup210 | 8506 | 2.50983929 | 1.94309307 | 3.48168761 | 2.79544643 | 4.9970575 | 2.3407579 |
| ENSMUSG00 | Cntn6 | 4367 | 0 | 0 | 0 | 0 | 0 | 0 |
| ENSMUSG00 | Wnt7a | 3448 | 0 | 0 | 0 | 0 | 0 | 0 |
| ENSMUSG00 | Xpc | 5475 | 24.6738124 | 18.931331 | 28.2407392 | 19.3294313 | 26.5337154 | 18.3425063 |
| ENSMUSG00 | Tmem43 | 3899 | 14.9580123 | 13.2114149 | 17.639925 | 15.4544792 | 14.6705738 | 13.4869209 |
| ENSMUSG00 | Slc6a6 | 6154 | 23.398497 | 23.5294538 | 29.8366032 | 27.4455721 | 29.2195236 | 28.5510605 |
| ENSMUSG00 | Grip2 | 5980 | 0.03324037 | 0.12892694 | 0.04434968 | 0.04223651 | 0.04266419 | 0.02736586 |
| ENSMUSG00 | Sumf1 | 3634 | 31.4849403 | 24.6766563 | 35.2617344 | 24.9317594 | 27.6755605 | 31.3951062 |
| ENSMUSG00 | Itpr1 | 10369 | 14.7074957 | 19.0719605 | 15.3293305 | 18.0427622 | 18.2029669 | 18.6653332 |
| ENSMUSG00 | Bhlhe40 | 3460 | 23.3477273 | 39.7607781 | 29.8426316 | 45.2589807 | 35.4087654 | 18.2645489 |
| ENSMUSG00 | Edem1 | 5846 | 5.88919708 | 8.26736255 | 8.99007721 | 10.2024666 | 11.2334837 | 6.90030644 |
| ENSMUSG00 | Arl8b | 2921 | 116.966313 | 110.840365 | 125.659741 | 135.088408 | 77.0374246 | 150.211271 |
| ENSMUSG00 | Usp18 | 1759 | 2.48613011 | 2.9311821 | 4.84988954 | 3.34358912 | 4.2932937 | 9.56705652 |
| ENSMUSG00 | Slc6a13 | 2928 | 0 | 0.27977101 | 0.01509626 | 0 | 0.40082194 | 0 |
| ENSMUSG00 | Slc6a12 | 3035 | 1.99104878 | 0.65095358 | 0.36410085 | 0 | 0.31944024 | 0.78184304 |
| ENSMUSG00 | Ret | 9588 | 0 | 0 | 0.15674412 | 0 | 0 | 0 |
| ENSMUSG00 | A2m | 4634 | 0 | 0 | 0 | 0 | 0 | 0 |
| ENSMUSG00 | Klrg1 | 1409 | 0 | 0 | 0 | 0 | 0 | 0 |
| ENSMUSG00 | Mfap5 | 3578 | 0 | 0 | 0.11118407 | 0 | 0 | 0.09147448 |
| ENSMUSG00 | Gdf3 | 2123 | 0 | 0 | 0 | 0 | 0 | 0 |
| ENSMUSG00 | Mlf2 | 1524 | 108.805846 | 112.751216 | 125.267559 | 111.868544 | 83.5707573 | 116.507809 |
| ENSMUSG00 | Ptms | 1313 | 9.29545508 | 9.61526873 | 7.47357894 | 8.13426137 | 6.179127 | 17.8438044 |
| ENSMUSG00 | Plxnd1 | 7221 | 0.05505537 | 0.12011577 | 0.04897033 | 0.27982198 | 0.26852268 | 0.10198245 |
| ENSMUSG00 | Lag3 | 2100 | 0 | 0 | 0 | 0 | 0 | 0 |
| ENSMUSG00 | Lrrc23 | 2020 | 0 | 0 | 0.0875284 | 0.0178624 | 0.32838757 | 0 |
| ENSMUSG00 | Tmcc1 | 8240 | 2.02637158 | 1.30407483 | 1.46981857 | 1.60267334 | 2.26646281 | 1.76093545 |
| ENSMUSG00 | Cops7a | 2702 | 48.6129192 | 51.3964938 | 58.6958596 | 45.7902778 | 45.9841692 | 38.8830164 |
| ENSMUSG00 | Mug2 | 4509 | 0 | 0 | 0 | 0 | 0 | 0 |
| ENSMUSG00 | Rasgef1a | 3325 | 0 | 0.15941379 | 0 | 0 | 0 | 0 |
| ENSMUSG00 | Tuba8 | 2220 | 1.82660316 | 0.60775693 | 1.71232365 | 0 | 0.25283339 | 0.51600669 |
| ENSMUSG00 | Brms1 | 4263 | 10.0344586 | 11.8346716 | 9.89175656 | 12.9245325 | 14.2198767 | 8.52212514 |
| ENSMUSG00 | Vmn2r26 | 2562 | 0 | 0 | 0 | 0 | 0 | 0 |
| ENSMUSG00 | Clec4e | 5766 | 0 | 0 | 0 | 0 | 0 | 0 |
| ENSMUSG00 | Gm8882 | 1237 | 0 | 0 | 0 | 0 | 0 | 0 |
| ENSMUSG00 | Clec4d | 1330 | 0 | 0 | 0 | 0 | 0 | 0 |
| ENSMUSG00 | Zfp248 | 6319 | 1.73643181 | 2.40207775 | 0.9583245 | 1.81009782 | 1.52618848 | 0.65607615 |
| ENSMUSG00 | Clec4b1 | 902 | 0 | 0 | 0 | 0 | 0 | 0 |
| ENSMUSG00 | Clec4a2 | 2958 | 0 | 0 | 0 | 0 | 0 | 0 |
| ENSMUSG00 | Klrk1 | 4089 | 0 | 0 | 0 | 0 | 0 | 0 |
| ENSMUSG00 | Klrb1f | 1814 | 1.0519642 | 0.1328182 | 0.02436706 | 0.43759922 | 1.01265134 | 0 |
| ENSMUSG00 | Clec2e | 2263 | 0 | 0 | 0 | 0 | 0 | 0.01205243 |
| ENSMUSG00 | Cd69 | 1616 | 0 | 0 | 0 | 0 | 0 | 0 |
| ENSMUSG00 | Clec2d | 1208 | 0.72402365 | 0.99723597 | 0.14636372 | 0.95581575 | 0.84480755 | 2.14494266 |
| ENSMUSG00 | Clec12b | 1505 | 0 | 0 | 0.46991993 | 0 | 0.03390457 | 2.10223133 |
| ENSMUSG00 | Clec1b | 5168 | 0.55386892 | 0 | 0 | 0.11869094 | 0.12835582 | 0.40637524 |
| ENSMUSG00 | D630042F21I | 2870 | 1285.00241 | 1811.59196 | 617.824715 | 1499.84384 | 1582.5466 | 1060.09015 |
| ENSMUSG00 | Gabarapl1 | 1761 | 177.15006 | 159.362771 | 189.508186 | 151.704405 | 135.172086 | 176.921184 |
| ENSMUSG00 | Olr1 | 3562 | 0.078127 | 0.12175126 | 0.08686494 | 0.08103772 | 0.17190245 | 0.0535998 |
| ENSMUSG00 | Klrd1 | 854 | 0 | 0 | 0 | 0 | 0 | 0 |
| ENSMUSG00 | Rad52 | 3398 | 4.63306954 | 6.9060616 | 3.05692558 | 7.42240998 | 6.57726681 | 6.04408574 |
| ENSMUSG00 | Klrc1 | 1977 | 0 | 0 | 0 | 0 | 0 | 0 |
| ENSMUSG00 | Adipor2 | 6322 | 4.0183094 | 5.41163776 | 5.62136694 | 3.26462029 | 2.62315282 | 4.40915554 |
| ENSMUSG00 | Wnt5b | 3941 | 0.28245457 | 0.04890783 | 0 | 0 | 0.62148339 | 0.00692074 |
| ENSMUSG00 | Erc1 | 9826 | 6.53825122 | 9.95016193 | 3.77870427 | 9.72739018 | 8.93714561 | 5.14348733 |
| ENSMUSG00 | Klra5 | 1367 | 0 | 0 | 0 | 0 | 0 | 0 |
| ENSMUSG00 | Ccdc77 | 7130 | 1.85116683 | 2.02748003 | 1.87842334 | 1.68517824 | 1.84639621 | 1.62194208 |
| ENSMUSG00 | Klra13-ps | 948 | 0 | 0 | 0 | 0 | 0 | 0 |
| ENSMUSG00 | Kdm5a | 14219 | 6.75219674 | 7.1776415 | 5.84425521 | 6.2627812 | 7.64372885 | 6.80187571 |
| ENSMUSG00 | Klra2 | 1928 | 0 | 0 | 0 | 0 | 0 | 0 |
| ENSMUSG00 | Magohb | 2052 | 3.40982679 | 3.87464082 | 3.55424176 | 2.95408552 | 1.84013246 | 5.1970682 |
| ENSMUSG00 | Csda | 1609 | 51.3189142 | 63.3102167 | 52.2235572 | 56.4440684 | 55.0856525 | 46.7008273 |
| ENSMUSG00 | Tas2r116 | 917 | 0 | 0 | 0 | 0 | 0 | 0 |
| ENSMUSG00 | Tas2r103 | 938 | 0 | 0 | 0 | 0 | 0 | 0 |
| ENSMUSG00 | Etv6 | 6142 | 7.30770718 | 7.49235628 | 7.72920537 | 9.51691829 | 8.23300854 | 6.19474467 |
| ENSMUSG00 | Bcl2l14 | 4158 | 6.09048607 | 6.18844639 | 4.43294097 | 4.45168082 | 6.24637453 | 4.10628287 |
| ENSMUSG00 | Lrp6 | 10396 | 7.67117102 | 9.46022782 | 8.05718474 | 10.2665148 | 8.90851021 | 9.05394191 |
| ENSMUSG00 | Dusp16 | 10213 | 1.56094661 | 1.26918172 | 1.4715193 | 1.75941038 | 1.05919825 | 1.7038304 |
| ENSMUSG00 | Ddx47 | 3080 | 32.023814 | 32.4789136 | 30.2237277 | 39.7488237 | 36.1326382 | 29.6124655 |
| ENSMUSG00 | Gprc5d | 1381 | 0 | 0 | 0.73616393 | 0 | 0 | 0 |
| ENSMUSG00 | Gsg1 | 1973 | 0 | 0 | 0.0672101 | 0 | 0 | 0 |
| ENSMUSG00 | 8430419L09I | 5800 | 40.7288043 | 32.2849162 | 43.3330482 | 35.0742357 | 45.2023309 | 28.0646633 |
| ENSMUSG00 | Emp1 | 3039 | 24.5806344 | 31.8230304 | 9.59961053 | 18.7593387 | 21.475069 | 14.5931436 |
| ENSMUSG00 | Grin2b | 10477 | 0 | 0 | 0 | 0 | 0 | 0 |
| ENSMUSG00 | Atf7ip | 4510 | 11.9178292 | 10.406562 | 8.08570299 | 11.0886283 | 17.955401 | 6.95473063 |
| ENSMUSG00 | Plbd1 | 2623 | 0 | 0 | 0 | 0 | 0 | 0 |
| ENSMUSG00 | Wbp11 | 5523 | 10.1782092 | 9.76292391 | 8.7155182 | 11.2303159 | 9.73778746 | 10.2323108 |
| ENSMUSG00 | Art4 | 2514 | 0.3953409 | 0.1916724 | 0.17582277 | 0 | 0.87276617 | 1.13915558 |
| ENSMUSG00 | Mgp | 594 | 1.07085469 | 0.89234152 | 0.52089714 | 0.12148837 | 0.08590299 | 0 |
| ENSMUSG00 | Erp27 | 2974 | 0.10694144 | 0 | 0.37156896 | 0.07279498 | 0.08578745 | 0 |
| ENSMUSG00 | Arhgdib | 2499 | 27.8081552 | 46.9909402 | 14.3978792 | 25.7440118 | 23.6857127 | 21.4901019 |
| ENSMUSG00 | Rerg | 4265 | 0 | 0 | 0 | 0 | 0 | 0 |
| ENSMUSG00 | Ptpro | 6821 | 0 | 0 | 0 | 0 | 0 | 0 |
| ENSMUSG00 | Strap | 3079 | 46.5987428 | 48.2959924 | 47.3170752 | 60.1758034 | 41.3315301 | 48.073877 |
| ENSMUSG00 | Dera | 1731 | 18.2585829 | 14.3919068 | 15.3212627 | 12.4442407 | 15.2695914 | 20.2156916 |
| ENSMUSG00 | Lmo3 | 6809 | 0 | 0 | 0 | 0 | 0 | 0.00400568 |

| | | | | | | | | |
|---|---|---|---|---|---|---|---|---|
| ENSMUSG00 | Pik3c2g | 4544 | 0.96239059 | 0.74230875 | 0.84629409 | 0.73053435 | 2.82980781 | 0.89434887 |
| ENSMUSG00 | Plcz1 | 2643 | 0 | 0 | 0 | 0 | 0 | 0 |
| ENSMUSG00 | Plekha5 | 5139 | 7.03979126 | 7.323139 | 7.70672336 | 8.20078381 | 9.0753274 | 6.00264974 |
| ENSMUSG00 | Aebp2 | 6883 | 9.94609 | 9.78710535 | 6.22280777 | 14.3007144 | 11.9059072 | 7.66764884 |
| ENSMUSG00 | Slco1c1 | 2898 | 0 | 0 | 0 | 0 | 0 | 0 |
| ENSMUSG00 | Slco1b2 | 4914 | 0 | 0 | 0 | 0 | 0 | 0 |
| ENSMUSG00 | Slco1a4 | 3836 | 0 | 0.60295861 | 0 | 0 | 0.9577422 | 0 |
| ENSMUSG00 | Recql | 5420 | 3.98288301 | 2.99609428 | 5.53746336 | 1.98384673 | 5.85579442 | 4.19184031 |
| ENSMUSG00 | Gys2 | 2840 | 0.13998409 | 0 | 0 | 0 | 0 | 0 |
| ENSMUSG00 | Golt1b | 2929 | 11.6321088 | 10.4796052 | 10.7448659 | 12.2573009 | 9.84291647 | 9.33986452 |
| ENSMUSG00 | Ldhb | 1914 | 1132.47038 | 1186.0058 | 1163.45071 | 1008.93993 | 946.307333 | 924.658721 |
| ENSMUSG00 | Kcnj8 | 2185 | 0 | 0 | 0 | 0 | 0 | 0.3619975 |
| ENSMUSG00 | Abcc9 | 7630 | 0 | 0 | 0 | 0 | 0 | 0 |
| ENSMUSG00 | Rad18 | 6703 | 0.20758494 | 1.43775749 | 0.14507542 | 0 | 0.52526032 | 0.13834667 |
| ENSMUSG00 | Sspn | 4990 | 1.3464281 | 1.10085259 | 0.87695039 | 1.75710157 | 3.06771799 | 1.55230412 |
| ENSMUSG00 | Bhlhe41 | 6490 | 0.99848125 | 0.90581602 | 1.4030169 | 0.73387209 | 0.55036153 | 1.52132811 |
| ENSMUSG00 | Srgap3 | 15055 | 1.61345723 | 2.04524321 | 1.12449724 | 2.23370744 | 1.76584137 | 1.1413499 |
| ENSMUSG00 | Rassf8 | 5553 | 0.88775069 | 2.16938782 | 0.78007935 | 2.4626499 | 1.35996824 | 0.50590453 |
| ENSMUSG00 | Lrmp | 3610 | 0 | 0 | 0 | 0 | 0.04240419 | 0 |
| ENSMUSG00 | Thumpd3 | 3897 | 21.9843625 | 19.1533998 | 12.5334967 | 17.4438214 | 13.4211022 | 19.0789496 |
| ENSMUSG00 | Kras | 6083 | 21.8089822 | 23.2178961 | 27.8523203 | 31.5562185 | 21.5329125 | 26.7186545 |
| ENSMUSG00 | Bcat1 | 10448 | 1.6742354 | 0.07840443 | 0.32999079 | 0.28318603 | 0.46884878 | 0.14357821 |
| ENSMUSG00 | Mtmr14 | 3129 | 11.9685723 | 10.2563684 | 6.8513563 | 13.8147297 | 16.405412 | 7.63585298 |
| ENSMUSG00 | Cpne9 | 5270 | 0.10561228 | 0.03657415 | 0.05032468 | 0.00684669 | 0 | 0.03105272 |
| ENSMUSG00 | Ogg1 | 2403 | 1.55514572 | 1.82478828 | 2.09696634 | 1.08110995 | 1.31653571 | 1.91819144 |
| ENSMUSG00 | Camk1 | 6009 | 41.2043822 | 26.9119154 | 54.3971758 | 34.5088218 | 31.9201442 | 36.8563939 |
| ENSMUSG00 | Etnk1 | 6421 | 132.584311 | 88.027872 | 164.712272 | 130.279726 | 117.302653 | 143.81521 |
| ENSMUSG00 | Ttll3 | 4645 | 0.34235075 | 0.9232709 | 0.26644814 | 1.12635015 | 0.69206925 | 0.2759759 |
| ENSMUSG00 | Cidec | 2933 | 0 | 0 | 0 | 0 | 0 | 0 |
| ENSMUSG00 | 5730419I09R | 4447 | 27.793971 | 21.8881523 | 24.5709368 | 21.0715245 | 24.9796313 | 29.1759521 |
| ENSMUSG00 | Il17rc | 2854 | 9.83439707 | 16.7994078 | 6.75262914 | 14.3114486 | 8.17065653 | 5.55240554 |
| ENSMUSG00 | Cmas | 3047 | 16.9225002 | 19.6098419 | 13.5927558 | 21.3034454 | 14.4354237 | 13.8924319 |
| ENSMUSG00 | St8sia1 | 11347 | 0.20671308 | 0.14863184 | 0.02337279 | 0.06359751 | 0 | 0.4518932 |
| ENSMUSG00 | Creld1 | 4459 | 4.69861813 | 8.51556771 | 6.16585476 | 6.6273143 | 5.72172862 | 2.88099464 |
| ENSMUSG00 | Tmem111 | 2414 | 78.2264012 | 80.2042769 | 83.5879923 | 73.1804847 | 79.3086007 | 70.3333179 |
| ENSMUSG00 | Itpr2 | 13711 | 43.214622 | 35.3833491 | 45.2173474 | 45.3874278 | 49.8689692 | 40.8751577 |
| ENSMUSG00 | Med21 | 2170 | 30.613552 | 37.2834269 | 33.996717 | 20.5019175 | 20.2224347 | 52.3120039 |
| ENSMUSG00 | 1700023A16 | 1657 | 0 | 0 | 0 | 0 | 0 | 0 |
| ENSMUSG00 | Sec13 | 1655 | 58.3720953 | 64.6076829 | 65.5415847 | 64.0974084 | 55.8361128 | 62.4597479 |
| ENSMUSG00 | Ccdc91 | 2910 | 31.3262257 | 20.3674652 | 26.8856571 | 27.216525 | 18.0959518 | 21.9790478 |
| ENSMUSG00 | Atp2b2 | 11839 | 2.43119925 | 5.00220774 | 0.76538372 | 4.35825016 | 5.37029008 | 2.42589705 |
| ENSMUSG00 | Far2 | 5336 | 0.57368291 | 0.7404963 | 0.61299408 | 0.66943823 | 0.95626641 | 0.56225831 |
| ENSMUSG00 | Ergic2 | 7060 | 20.7787143 | 23.6222771 | 18.231695 | 24.9098987 | 21.7187336 | 21.0354689 |
| ENSMUSG00 | Tmtc1 | 8556 | 3.83336506 | 3.59877706 | 3.10487467 | 3.5550682 | 3.76912921 | 1.06790605 |
| ENSMUSG00 | Slc6a11 | 4052 | 0 | 0 | 0 | 0 | 0 | 0 |
| ENSMUSG00 | Caprin2 | 7128 | 3.23487355 | 1.9198863 | 1.24643244 | 1.94381385 | 2.62719977 | 0.66579507 |
| ENSMUSG00 | Slc6a1 | 4305 | 0 | 0 | 0 | 0 | 0 | 0 |
| ENSMUSG00 | Dennd5b | 10599 | 0.37883796 | 1.07293144 | 0.27941537 | 0.64000607 | 1.33836517 | 0.32938521 |
| ENSMUSG00 | Atg7 | 3652 | 6.54245449 | 6.14865335 | 4.38145322 | 5.65140456 | 7.58688994 | 4.06281592 |
| ENSMUSG00 | Vgll4 | 5157 | 7.11543698 | 6.83038379 | 5.73415416 | 5.67433355 | 5.85759443 | 5.55858946 |
| ENSMUSG00 | 1500001M20 | 5032 | 3.97396794 | 4.46241694 | 4.11098222 | 4.18758227 | 3.55927224 | 4.20610494 |
| ENSMUSG00 | Timp4 | 1411 | 2.47943465 | 1.40017302 | 3.53990663 | 1.81560961 | 2.45910245 | 4.56188155 |
| ENSMUSG00 | Cand2 | 4795 | 3.40761262 | 4.27095681 | 4.20355379 | 3.37869405 | 2.66039499 | 1.04661807 |
| ENSMUSG00 | BC060267 | 2990 | 6.30237383 | 3.98061913 | 4.80454815 | 2.77554189 | 5.58047655 | 3.67614703 |
| ENSMUSG00 | Mbd4 | 6414 | 1.59914468 | 1.77299682 | 1.99163278 | 1.84516848 | 3.49245073 | 1.67968234 |
| ENSMUSG00 | Ift122 | 8120 | 9.30239076 | 9.9814773 | 6.17301601 | 5.21678821 | 11.8454087 | 5.8109761 |
| ENSMUSG00 | Rho | 3230 | 0.12308198 | 0.14918403 | 0 | 0 | 0 | 0 |
| ENSMUSG00 | Klrb1c | 4891 | 0 | 0 | 0 | 0 | 0 | 0 |
| ENSMUSG00 | Necap1 | 2391 | 43.5132131 | 43.34966 | 60.6550594 | 48.3055728 | 45.8405083 | 48.3779779 |
| ENSMUSG00 | C530028O21 | 2737 | 0 | 0 | 0 | 0 | 0 | 0 |
| ENSMUSG00 | Ing4 | 6340 | 17.6642253 | 16.7740501 | 15.3590947 | 13.2718183 | 20.5715168 | 20.098914 |
| ENSMUSG00 | Mrpl51 | 3397 | 16.1151889 | 18.4121289 | 20.7671891 | 18.0357114 | 11.7013679 | 19.0930504 |
| ENSMUSG00 | Cd27 | 2601 | 0.35154789 | 0.38904855 | 0.30589511 | 0.2635751 | 0.45121363 | 0.62917276 |
| ENSMUSG00 | Vamp1 | 8885 | 7.37836661 | 5.04371313 | 5.03458246 | 3.65896704 | 4.51398215 | 3.88936049 |
| ENSMUSG00 | Ltbr | 2344 | 21.9299643 | 22.818665 | 27.8335837 | 25.4144435 | 18.1117511 | 17.5702668 |
| ENSMUSG00 | Scnn1a | 4555 | 94.2436177 | 81.7952954 | 83.357592 | 99.9603337 | 74.4839457 | 95.7396722 |
| ENSMUSG00 | Tnfrsf1a | 2109 | 17.4177639 | 20.7688364 | 15.4675013 | 16.5611281 | 17.468489 | 19.2306252 |
| ENSMUSG00 | Cd9 | 2171 | 192.825983 | 259.99815 | 309.18894 | 233.245239 | 202.624775 | 247.657745 |
| ENSMUSG00 | Akap3 | 3146 | 0 | 0 | 0 | 0.3096679 | 0 | 0 |
| ENSMUSG00 | Dyrk4 | 2814 | 0 | 0 | 0 | 0 | 0 | 0 |
| ENSMUSG00 | Rad51ap1 | 3960 | 0 | 0.13385123 | 0.03348625 | 0.37357672 | 0.48964704 | 0.16530084 |
| ENSMUSG00 | D6Wsu163e | 2224 | 10.3857618 | 7.49663174 | 18.1259356 | 13.8065737 | 10.9211128 | 14.0297605 |
| ENSMUSG00 | Prmt8 | 2714 | 0.07324149 | 0 | 0 | 0 | 0 | 0 |
| ENSMUSG00 | Tspan11 | 4478 | 0 | 0 | 0 | 0 | 0 | 0 |
| ENSMUSG00 | Tspan9 | 6415 | 13.7951208 | 21.6632423 | 8.64054733 | 14.1178382 | 17.4595316 | 8.3928508 |
| ENSMUSG00 | Tead4 | 3582 | 0.01109868 | 0 | 0 | 0 | 0 | 0 |
| ENSMUSG00 | Fkbp4 | 3561 | 74.9001738 | 80.161888 | 82.2221309 | 72.7315124 | 70.6144285 | 63.3267951 |
| ENSMUSG00 | Pzp | 4818 | 0.21453767 | 0 | 0 | 0 | 0 | 0 |
| ENSMUSG00 | Klrb1a | 1890 | 0 | 0 | 0 | 0 | 0 | 0 |
| ENSMUSG00 | Clec2j | 510 | 0 | 0 | 0 | 0 | 0 | 0 |
| ENSMUSG00 | Clec2h | 2151 | 0.99804554 | 1.05288832 | 0.3698899 | 1.20776718 | 0.28466597 | 0.57059915 |
| ENSMUSG00 | Clec2i | 3181 | 0 | 0 | 0 | 0 | 0 | 0 |
| ENSMUSG00 | Ceacam12 | 2752 | 0 | 0 | 0 | 0 | 0 | 0 |
| ENSMUSG00 | Ceacam11 | 1043 | 0 | 0 | 0 | 0 | 0 | 0 |
| ENSMUSG00 | Psg28 | 1956 | 0 | 0 | 0 | 0 | 0 | 0 |
| ENSMUSG00 | Strn4 | 7830 | 19.5324819 | 16.3883136 | 15.4113706 | 13.4144102 | 18.7292215 | 12.331063 |
| ENSMUSG00 | Slc8a2 | 4484 | 0 | 0 | 0 | 0 | 0 | 0 |
| ENSMUSG00 | 2810007J24R | 3482 | 0 | 0 | 0 | 0 | 0 | 0 |
| ENSMUSG00 | Mzf1 | 5022 | 0.38789696 | 0.24947182 | 0.32566073 | 0.5101205 | 0.40642275 | 0.86353398 |
| ENSMUSG00 | Slc27a5 | 2539 | 0 | 0.03795702 | 0 | 0 | 0 | 0 |
| ENSMUSG00 | 2900092C05 | 881 | 0.09025081 | 0 | 0 | 0 | 0 | 0 |
| ENSMUSG00 | Zfp606 | 3111 | 25.1363003 | 16.9605124 | 29.5247283 | 30.5264999 | 22.6838566 | 22.3474945 |
| ENSMUSG00 | Zik1 | 4195 | 1.42153089 | 2.01016147 | 0.9377745 | 0 | 1.76372455 | 0.72168891 |
| ENSMUSG00 | Mark4 | 3031 | 2.58390949 | 2.7026378 | 2.58123597 | 3.1784579 | 1.86866635 | 2.13265902 |
| ENSMUSG00 | Ckm | 1407 | 0 | 0 | 0 | 0 | 0 | 0 |
| ENSMUSG00 | Ercc2 | 4121 | 0.57882282 | 1.15759713 | 0.69718995 | 0.15760175 | 0.89150669 | 0.7809773 |
| ENSMUSG00 | Rtn2 | 2023 | 0.17686571 | 1.02422986 | 0.83028672 | 0.01783591 | 1.51338732 | 1.76617783 |
| ENSMUSG00 | Ppm1n | 1626 | 0 | 0 | 0 | 0 | 0 | 0 |
| ENSMUSG00 | Vasp | 2595 | 3.60020729 | 3.73236026 | 5.91061253 | 2.96364759 | 3.5000751 | 7.01047568 |
| ENSMUSG00 | Gipr | 1778 | 0 | 0 | 0.24860429 | 0 | 0 | 0 |
| ENSMUSG00 | Qpctl | 2093 | 8.24361134 | 11.6724922 | 8.48979498 | 8.34386505 | 11.165829 | 4.39156877 |
| ENSMUSG00 | Dmpk | 4523 | 2.25014438 | 1.23582297 | 1.94476382 | 1.8427929 | 2.7301311 | 2.03218073 |
| ENSMUSG00 | Dmwd | 3643 | 19.3593803 | 19.3777485 | 15.5428386 | 14.2723651 | 16.415842 | 14.3448281 |
| ENSMUSG00 | Nova2 | 9009 | 0.03089004 | 0.05883571 | 0.04415769 | 0.19224533 | 0.084959 | 0.00605498 |
| ENSMUSG00 | Pglyrp1 | 678 | 0 | 0 | 0 | 0 | 0.30104057 | 0 |
| ENSMUSG00 | Pdcd5 | 2653 | 5.61941394 | 4.5589133 | 6.36453224 | 4.6649609 | 4.48139675 | 8.58436629 |
| ENSMUSG00 | C80913 | 3439 | 22.9122891 | 19.9667578 | 24.4851733 | 18.4029968 | 23.8587997 | 28.6942846 |
| ENSMUSG00 | Pop4 | 1207 | 24.7689489 | 23.9135699 | 20.3247911 | 15.1263584 | 22.1945711 | 16.9251904 |
| ENSMUSG00 | Zfp939 | 925 | 8.0370539 | 11.1479985 | 13.7623036 | 8.81572115 | 20.9621868 | 11.4995776 |
| ENSMUSG00 | Lilra6 | 2918 | 0 | 0 | 0 | 0 | 0 | 0 |
| ENSMUSG00 | Ttyh1 | 4035 | 0.39410637 | 0.15524752 | 0.99686932 | 0.95682249 | 1.1760726 | 1.18291496 |
| ENSMUSG00 | 2210411K11 | 1218 | 7.70303235 | 8.50581696 | 4.7177665 | 5.12495378 | 6.28403643 | 5.06081155 |
| ENSMUSG00 | Rpl28 | 1142 | 0.69624309 | 0.59072696 | 0.69670155 | 0.78988714 | 0.67022385 | 1.07474498 |
| ENSMUSG00 | Sbk2 | 1135 | 0 | 0 | 0.23366613 | 0 | 0.26974296 | 0.26433571 |
| ENSMUSG00 | U2af2 | 2240 | 2.67994534 | 3.7860776 | 2.5258196 | 3.47934001 | 2.46020026 | 2.39870711 |
| ENSMUSG00 | Zfp583 | 4293 | 0.65749805 | 2.18876222 | 1.03992223 | 1.27753803 | 1.74723439 | 2.67473169 |
| ENSMUSG00 | Zfp273 | 543 | 15.2286187 | 22.0078042 | 33.2124345 | 67.6455273 | 31.3863896 | 71.9789283 |
| ENSMUSG00 | Cyfip1 | 7103 | 18.1566632 | 18.9815099 | 22.9909195 | 16.9107597 | 22.6791875 | 17.3523997 |
| ENSMUSG00 | Oca2 | 4053 | 0 | 0 | 0 | 0 | 0 | 0 |
| ENSMUSG00 | Herc2 | 15346 | 2.70718605 | 3.95954018 | 2.9869224 | 5.09982761 | 4.70163531 | 3.66126397 |
| ENSMUSG00 | Nipa2 | 6298 | 14.8972585 | 14.9042854 | 20.6060037 | 18.9691409 | 18.2781048 | 19.0160275 |
| ENSMUSG00 | 4933421I07R | 1409 | 0 | 0 | 0 | 0.64020661 | 0 | 0 |
| ENSMUSG00 | Psd3 | 23399 | 2.0371307 | 1.79574245 | 2.30086093 | 2.02006403 | 2.17852684 | 2.23684913 |
| ENSMUSG00 | Siglecg | 3988 | 0 | 0 | 0 | 0 | 0 | 0 |
| ENSMUSG00 | Zfp719 | 4617 | 7.58600354 | 9.57049327 | 9.5162729 | 12.1054986 | 13.615875 | 7.94550353 |
| ENSMUSG00 | Csrp3 | 1224 | 0 | 0 | 0 | 0 | 0 | 0 |
| ENSMUSG00 | Zdhhc13 | 2504 | 33.7223006 | 29.9818198 | 34.7930641 | 27.0615334 | 30.8114538 | 30.6077222 |
| ENSMUSG00 | Ceacam18 | 1924 | 0 | 0 | 0 | 0 | 0 | 0 |
| ENSMUSG00 | Siglece | 1842 | 0.06474834 | 0.13079925 | 0 | 0 | 0 | 0.31094866 |
| ENSMUSG00 | Cyp2b10 | 2281 | 0 | 0 | 0 | 0 | 0 | 0 |
| ENSMUSG00 | Lypd5 | 930 | 0 | 0 | 0 | 0 | 0 | 0 |
| ENSMUSG00 | Zfp108 | 1926 | 2.35312813 | 5.35404913 | 4.81951559 | 5.07696473 | 3.86804303 | 3.75274112 |
| ENSMUSG00 | Tdrd12 | 4047 | 0 | 0.05953353 | 0.07645488 | 0 | 0.41607868 | 0.10109207 |
| ENSMUSG00 | Slc7a9 | 2551 | 0.09350564 | 0.11333542 | 0.19059987 | 0 | 0.92011497 | 0 |
| ENSMUSG00 | C230052I12 | 2498 | 5.2041802 | 5.20830239 | 5.23768836 | 3.79887018 | 5.04544229 | 6.12532931 |
| ENSMUSG00 | Rhpn2 | 3939 | 15.4621468 | 10.7529532 | 11.6031241 | 11.1388084 | 13.3816314 | 15.7319067 |
| ENSMUSG00 | Slc7a10 | 2680 | 0 | 0 | 0 | 0 | 0 | 0 |
| ENSMUSG00 | Gas2 | 4994 | 1.81502795 | 2.81746918 | 2.21274745 | 2.39150115 | 3.82135854 | 4.32003197 |
| ENSMUSG00 | Kctd15 | 4260 | 0.6439268 | 0.76917326 | 0.36316068 | 0.1863392 | 1.01813191 | 0.65945724 |
| ENSMUSG00 | Slc17a6 | 4307 | 0 | 0 | 0 | 0 | 0 | 0 |
| ENSMUSG00 | Prmt3 | 3212 | 10.359694 | 11.5665464 | 13.3486263 | 11.6042192 | 9.8335388 | 7.95651836 |
| ENSMUSG00 | Dbx1 | 2087 | 0 | 0 | 0 | 0 | 0 | 0 |
| ENSMUSG00 | Asb7 | 9544 | 1.34545476 | 1.52475959 | 1.8340245 | 3.41388162 | 1.47026963 | 1.40316931 |
| ENSMUSG00 | Lass3 | 3241 | 0.02453285 | 0.01486777 | 0.02727667 | 0 | 0 | 0 |
| ENSMUSG00 | Snrpa1 | 2929 | 11.2249172 | 9.49251523 | 14.0649087 | 11.9862852 | 7.87433318 | 19.331564 |
| ENSMUSG00 | Pcsk6 | 5121 | 0.07763226 | 0.05645746 | 0.31936501 | 0.0140918 | 0 | 0 |
| ENSMUSG00 | Tarsl2 | 4077 | 7.86918636 | 6.47686296 | 7.55670458 | 5.96499829 | 8.44807547 | 7.44583603 |
| ENSMUSG00 | Tjp1 | 8290 | 8.98214897 | 10.2359861 | 7.12348158 | 8.64837424 | 11.9041026 | 8.61997043 |
| ENSMUSG00 | Fam189a1 | 8586 | 0 | 0 | 0 | 0 | 0 | 0 |
| ENSMUSG00 | Apba2 | 3729 | 0 | 0 | 0.02370708 | 0 | 0 | 0 |
| ENSMUSG00 | Mphosph10 | 3476 | 10.773781 | 8.80275914 | 7.42631655 | 7.51536255 | 13.6667307 | 8.8901514 |
| ENSMUSG00 | Mtmr10 | 5138 | 10.3528285 | 7.41834872 | 10.3235134 | 9.76841649 | 9.53392775 | 12.4747766 |
| ENSMUSG00 | Trpm1 | 9997 | 0.23462772 | 0.52538984 | 0.13706683 | 0.20933866 | 0.49000021 | 0.33830702 |
| ENSMUSG00 | Chrna7 | 4534 | 0.25428075 | 0.35071738 | 0.02924692 | 0.29444986 | 0 | 0.0601558 |
| ENSMUSG00 | Crtc3 | 9835 | 1.16416659 | 1.36205703 | 1.00673237 | 1.7756695 | 1.38526104 | 1.75267636 |
| ENSMUSG00 | Blm | 4923 | 0.58950845 | 0.94943833 | 0.05387184 | 0.29317119 | 0.27985215 | 0.9418421 |
| ENSMUSG00 | Furin | 10284 | 10.0161853 | 8.91664716 | 7.00593194 | 8.70123205 | 10.9157941 | 5.7153683 |
| ENSMUSG00 | Hddc3 | 856 | 32.974756 | 18.970597 | 33.1513823 | 30.1386236 | 28.2552595 | 36.1962502 |
| ENSMUSG00 | Unc45a | 4001 | 16.6831922 | 19.0529746 | 12.5170428 | 14.7087765 | 19.3979299 | 9.96638905 |
| ENSMUSG00 | Vps33b | 3312 | 21.3781736 | 19.6557619 | 18.1905532 | 19.2829767 | 21.831031 | 23.0994454 |
| ENSMUSG00 | Iqgap1 | 7321 | 41.0316092 | 47.7520336 | 44.3044835 | 55.3773873 | 65.6282413 | 41.4763774 |
| ENSMUSG00 | Cib1 | 2274 | 15.0700194 | 19.4525743 | 19.1852327 | 18.834383 | 10.9502513 | 16.851745 |
| ENSMUSG00 | Sema4b | 5419 | 5.82503258 | 5.79766752 | 3.31167158 | 5.94598003 | 3.59698756 | 3.34201153 |
| ENSMUSG00 | Idh2 | 2932 | 415.412229 | 469.324771 | 407.615768 | 328.639495 | 327.094384 | 279.918544 |
| ENSMUSG00 | Mesp2 | 2169 | 0.07331578 | 0.48875137 | 0.30568356 | 0.28280072 | 0.7292843 | 0.06287377 |
| ENSMUSG00 | Mesp1 | 1273 | 0 | 0 | 0.20833547 | 0.0283441 | 0 | 0.10712741 |
| ENSMUSG00 | Pex11a | 3183 | 5.59549333 | 11.1269353 | 8.70705484 | 10.3383049 | 13.6262706 | 8.91160063 |
| ENSMUSG00 | Plin1 | 1907 | 0 | 0 | 0 | 0 | 0 | 0 |
| ENSMUSG00 | Rhcg | 2530 | 748.722978 | 815.360312 | 601.5469 | 787.51558 | 664.452155 | 573.446595 |
| ENSMUSG00 | Nr2f2 | 5351 | 4.40571855 | 6.31259502 | 4.27066956 | 5.738333 | 4.24345678 | 5.84638595 |
| ENSMUSG00 | Pgpep1l | 951 | 0 | 0 | 0 | 0 | 0.0536555 | 0 |
| ENSMUSG00 | Synm | 10053 | 0.04350048 | 0.0718986 | 0.14509707 | 0.03230264 | 0.24871107 | 0.32828323 |
| ENSMUSG00 | Ttc23 | 4355 | 5.64153547 | 3.17554739 | 8.576477 | 6.18904415 | 6.2684526 | 3.52597518 |
| ENSMUSG00 | Lrrc28 | 4297 | 2.35923843 | 4.12674208 | 2.54080876 | 1.99849351 | 3.63371445 | 3.64338909 |
| ENSMUSG00 | Mef2a | 8067 | 16.5684797 | 19.5445691 | 32.4048221 | 23.6074168 | 21.1392275 | 29.4553932 |
| ENSMUSG00 | Rab38 | 1468 | 2.30191815 | 2.46184139 | 2.28837879 | 2.90032783 | 3.61494761 | 4.14321836 |
| ENSMUSG00 | Ctsc | 5777 | 11.7401506 | 12.1779826 | 17.7434783 | 13.9906145 | 13.8761358 | 10.3772991 |
| ENSMUSG00 | Nox4 | 6992 | 0.17626143 | 0.67538206 | 0 | 0.13417236 | 0.29921073 | 0.62413362 |
| ENSMUSG00 | Cd22 | 5557 | 0.00715413 | 0 | 0 | 0 | 0.00918236 | 0 |
| ENSMUSG00 | Tyrobp | 559 | 0 | 0 | 0 | 0 | 0 | 0.14637552 |
| ENSMUSG00 | Sipa1l3 | 7785 | 3.21720652 | 3.72617061 | 3.89498578 | 4.93144449 | 4.60122233 | 4.93991539 |
| ENSMUSG00 | Dpf1 | 3653 | 0 | 0 | 0 | 0 | 0 | 0 |
| ENSMUSG00 | 2200002D01 | 1688 | 0.2826219 | 0.99912647 | 0.28804519 | 0.89777599 | 0.78595129 | 0.09694777 |
| ENSMUSG00 | Yif1b | 1395 | 41.7218806 | 45.5266887 | 55.4186548 | 38.6168405 | 26.2996159 | 51.8706939 |
| ENSMUSG00 | Rasgrp4 | 5232 | 0.04559115 | 0 | 0.00844836 | 0.03448208 | 0.0195055 | 0 |
| ENSMUSG00 | Fam98c | 2096 | 9.61642587 | 7.88547216 | 7.27558962 | 6.93753051 | 7.44946136 | 8.92671876 |
| ENSMUSG00 | Psmd8 | 1505 | 84.7940814 | 85.6469985 | 88.814866 | 69.6707119 | 70.9622622 | 82.385721 |
| ENSMUSG00 | Ryr1 | 15256 | 0 | 0 | 0 | 0 | 0 | 0 |
| ENSMUSG00 | Nfkbib | 2158 | 10.1138827 | 11.8568122 | 10.2209081 | 9.78127713 | 7.94479253 | 12.7273224 |
| ENSMUSG00 | Fbxo17 | 2696 | 1.66630909 | 1.46561137 | 1.32802273 | 2.08783344 | 2.17657019 | 2.22567531 |
| ENSMUSG00 | Lrfn1 | 2724 | 0.74432067 | 1.30902963 | 0.37321674 | 0.76826673 | 1.19885758 | 0.64081384 |
| ENSMUSG00 | Pak4 | 3524 | 22.5175764 | 27.9492304 | 19.5045022 | 19.8328378 | 19.7502771 | 15.8818274 |
| ENSMUSG00 | Psmc4 | 2364 | 168.658086 | 167.612992 | 124.65892 | 36.6925698 | 54.9116329 | 126.070213 |
| ENSMUSG00 | Zfp626 | 1070 | 0.63162913 | 0.49537464 | 0.74358241 | 3.37215369 | 2.7659157 | 2.62550266 |
| ENSMUSG00 | Mfge8 | 2115 | 6.20298277 | 8.3842131 | 4.26344018 | 8.8371154 | 9.86751192 | 3.30132749 |
| ENSMUSG00 | Hapln3 | 3306 | 0 | 0 | 0 | 0.01091411 | 0 | 0 |
| ENSMUSG00 | Acan | 7338 | 0 | 0 | 0 | 0 | 0 | 0 |
| ENSMUSG00 | Aen | 5768 | 2.41923954 | 1.90473454 | 1.81619917 | 1.2386 | 1.59236263 | 1.28618271 |
| ENSMUSG00 | Det1 | 2871 | 3.08793875 | 2.7189842 | 5.04987966 | 3.48127006 | 3.85674802 | 5.59552857 |
| ENSMUSG00 | Mrps11 | 2276 | 5.86899888 | 8.27807508 | 7.69065458 | 7.03885227 | 5.87386224 | 8.16082132 |
| ENSMUSG00 | Mrpl46 | 1512 | 26.7402934 | 24.8580852 | 34.6715515 | 22.193321 | 18.6286769 | 31.5678695 |
| ENSMUSG00 | Ccdc90b | 1791 | 26.5036537 | 27.900246 | 36.1068098 | 27.5601546 | 22.5359371 | 38.3306907 |
| ENSMUSG00 | Tmem126b | 2340 | 32.7727871 | 38.7344862 | 36.0037234 | 44.7016441 | 29.2638446 | 34.7227138 |
| ENSMUSG00 | Tmem126a | 1243 | 33.2948151 | 27.6790967 | 32.8580072 | 29.4345882 | 18.6781987 | 49.6780235 |
| ENSMUSG00 | Sytl2 | 6354 | 0.99482553 | 0.84936757 | 0.42434882 | 1.56162453 | 2.6179727 | 0.53656435 |
| ENSMUSG00 | Ccdc83 | 2734 | 0 | 0 | 0 | 0 | 0 | 0 |
| ENSMUSG00 | Eed | 2206 | 37.9893712 | 45.4778661 | 42.6389492 | 41.4632742 | 35.1586996 | 56.7005873 |
| ENSMUSG00 | Me3 | 14835 | 13.4899704 | 11.8564659 | 10.679721 | 10.2267387 | 12.823711 | 11.1521313 |
| ENSMUSG00 | 1700019G06 | 331 | 0 | 0 | 0 | 0 | 0 | 0 |
| ENSMUSG00 | Zfand6 | 1650 | 163.214318 | 167.279964 | 163.118195 | 160.18241 | 142.193501 | 228.296996 |
| ENSMUSG00 | Fah | 2088 | 18.0689421 | 16.2006142 | 19.7299414 | 15.4834827 | 16.5933473 | 11.2991202 |
| ENSMUSG00 | 1700010L04 | 325 | 0 | 0 | 0 | 0 | 0 | 0 |
| ENSMUSG00 | Sh3gl3 | 1586 | 0 | 0 | 0 | 0.31850481 | 0 | 0 |
| ENSMUSG00 | 4632434I11R | 5331 | 0.26846695 | 1.25640883 | 1.14422329 | 0.67683445 | 0.22014756 | 0.60883175 |
| ENSMUSG00 | Rab30 | 2111 | 1.48777023 | 1.27827606 | 1.54947247 | 2.05108733 | 0.87017979 | 1.4599878 |
| ENSMUSG00 | Ndufc2 | 1005 | 143.396634 | 123.223041 | 147.119567 | 111.190141 | 71.5892437 | 339.372501 |
| ENSMUSG00 | 3200002M19 | 4287 | 1.23337507 | 2.11314465 | 1.6600179 | 1.22040972 | 1.82109529 | 1.9213765 |
| ENSMUSG00 | Tmc5 | 4446 | 0 | 0 | 0 | 0.00811562 | 0 | 0 |
| ENSMUSG00 | Art2b | 867 | 0 | 0 | 0 | 0 | 0 | 0 |
| ENSMUSG00 | Coq7 | 2028 | 55.183273 | 57.9520377 | 70.3785755 | 57.7882054 | 38.3955865 | 62.7666378 |
| ENSMUSG00 | Pde2a | 4886 | 1.17980857 | 1.2820789 | 1.12178234 | 0.14031086 | 2.6004027 | 0.95455655 |
| ENSMUSG00 | Arl6ip1 | 2128 | 223.008069 | 257.598198 | 277.715527 | 220.426024 | 230.649769 | 223.516181 |
| ENSMUSG00 | Smg1 | 15484 | 2.13104164 | 2.33712336 | 1.92405336 | 3.8822453 | 2.67259047 | 2.30224448 |
| ENSMUSG00 | Xylt1 | 3810 | 1.15823054 | 1.4923885 | 2.35511132 | 1.17432376 | 1.28570396 | 1.82546798 |
| ENSMUSG00 | Nucb2 | 1666 | 21.4049616 | 23.1676712 | 17.1660219 | 19.7736535 | 22.1134714 | 16.2076188 |
| ENSMUSG00 | Pik3c2a | 7948 | 8.61835593 | 8.82124727 | 6.92947866 | 11.2313762 | 11.2478876 | 8.09522823 |
| ENSMUSG00 | Ipo5 | 4562 | 13.2111592 | 9.62250083 | 13.6423049 | 14.1180293 | 14.4735051 | 9.57781063 |
| ENSMUSG00 | 1110004F10 | 1679 | 50.3632561 | 53.1227543 | 68.2640732 | 51.125184 | 44.188416 | 62.0055378 |
| ENSMUSG00 | 1110006G14 | 418 | 0 | 0 | 0.42298414 | 0.34528272 | 0 | 0.26100133 |
| ENSMUSG00 | Calcb | 957 | 0 | 0 | 0 | 0 | 0 | 0 |
| ENSMUSG00 | Calca | 1474 | 1.07884615 | 1.4710922 | 0.38983986 | 0.19583199 | 0.48464672 | 0.79566451 |
| ENSMUSG00 | Cyp2r1 | 1851 | 2.62029639 | 2.05657965 | 1.86263844 | 0.15594617 | 1.26807849 | 2.16605725 |
| ENSMUSG00 | Pde3b | 6883 | 12.9033479 | 17.1729395 | 8.11726421 | 16.334687 | 17.784727 | 9.7559439 |
| ENSMUSG00 | Mylpf | 877 | 0.2266561 | 2.19778527 | 1.20962854 | 0.90513681 | 0.17454861 | 2.05259543 |
| ENSMUSG00 | Qprt | 2639 | 0 | 0.34692778 | 0 | 0 | 0 | 0.62011306 |
| ENSMUSG00 | Kif22 | 2049 | 0 | 1.50509141 | 0.17257918 | 0.58111638 | 0.69728576 | 0.07986717 |
| ENSMUSG00 | Maz | 2657 | 5.20696546 | 6.1117166 | 4.72462304 | 5.32335772 | 4.43624118 | 7.19590595 |
| ENSMUSG00 | 2900092E17 | 4744 | 2.00285831 | 3.1284621 | 2.29208545 | 2.8445794 | 1.83927282 | 2.55268546 |
| ENSMUSG00 | Mvp | 2886 | 29.4791158 | 31.8738455 | 30.8769631 | 32.4563367 | 32.5500894 | 22.1712764 |
| ENSMUSG00 | Cdipt | 1870 | 72.8779611 | 76.9436985 | 52.0987177 | 50.5920432 | 61.2589378 | 37.119763 |
| ENSMUSG00 | Sez6l2 | 4596 | 0 | 0.03145329 | 0.00961746 | 0 | 0.05551172 | 0.13649189 |
| ENSMUSG00 | Kctd13 | 2481 | 6.05706233 | 8.17674008 | 9.74542851 | 4.7411312 | 7.32180161 | 6.80491806 |
| ENSMUSG00 | Stard10 | 2152 | 25.1427551 | 26.7801974 | 26.8867159 | 26.5752977 | 25.7265882 | 28.2885664 |
| ENSMUSG00 | Ino80e | 6048 | 1.08459892 | 1.01185155 | 1.86366898 | 1.34830391 | 1.22335061 | 1.8625043 |
| ENSMUSG00 | Fchsd2 | 9440 | 2.04252204 | 2.59818846 | 2.25223374 | 1.80792448 | 3.69725011 | 2.629229 |
| ENSMUSG00 | Klk10 | 1348 | 0 | 0 | 0 | 0 | 0 | 0 |
| ENSMUSG00 | Aldoa | 2927 | 924.128165 | 1032.42786 | 997.418287 | 884.077891 | 831.431822 | 757.269596 |
| ENSMUSG00 | Ppp4c | 1284 | 32.7580205 | 34.4885828 | 44.0297175 | 25.1787476 | 29.089803 | 34.1357829 |
| ENSMUSG00 | Tbx6 | 1796 | 0.28776239 | 0.42927788 | 0.29533525 | 0.30135338 | 0.02841112 | 0.25816752 |
| ENSMUSG00 | Plekhb1 | 4682 | 6.02870379 | 5.41351315 | 8.60057219 | 4.70099256 | 3.83624184 | 6.41379278 |
| ENSMUSG00 | Gdpd3 | 1056 | 3.0870733 | 5.88641197 | 4.81364768 | 3.79271491 | 3.33410978 | 1.85963449 |
| ENSMUSG00 | Rab6a | 3718 | 42.4393497 | 41.5118812 | 55.5435747 | 45.9031927 | 39.9509898 | 46.9860313 |
| ENSMUSG00 | Mrpl48 | 6244 | 4.19584588 | 3.86633689 | 3.97136994 | 4.53048093 | 3.27699819 | 3.90511331 |
| ENSMUSG00 | Coro1a | 2844 | 0 | 0 | 0 | 0 | 0.07176706 | 0 |
| ENSMUSG00 | Dnajb13 | 1710 | 0.46497638 | 0.19725444 | 0.28433934 | 1.60364642 | 0.2088799 | 0.51040261 |
| ENSMUSG00 | Sult1a1 | 1677 | 9.83811831 | 6.7236896 | 6.19405671 | 12.7158547 | 11.4710457 | 8.76626748 |
| ENSMUSG00 | Klk7 | 1739 | 0 | 0 | 0 | 0 | 0 | 0 |
| ENSMUSG00 | Ccdc101 | 1888 | 10.4863503 | 14.4457237 | 12.2912964 | 14.4863293 | 8.29719141 | 9.40454987 |
| ENSMUSG00 | Nupr1 | 1493 | 8.30791019 | 5.8740338 | 11.8128167 | 7.49191815 | 6.32275923 | 14.8521646 |
| ENSMUSG00 | Ppme1 | 2347 | 17.599465 | 15.8499929 | 15.34917 | 14.5588821 | 16.2406062 | 13.794204 |
| ENSMUSG00 | Clrn3 | 6575 | 17.9640354 | 13.2137118 | 17.7546871 | 12.1608838 | 14.7840678 | 14.5976358 |
| ENSMUSG00 | Nfatc2ip | 3642 | 0.96059371 | 0.17199993 | 0.78888517 | 0.44582427 | 0.2662002 | 0.78633639 |
| ENSMUSG00 | Cd19 | 2695 | 0 | 0 | 0 | 0 | 0 | 0 |
| ENSMUSG00 | Lipt2 | 1661 | 3.92528525 | 2.37886108 | 2.31520791 | 1.95507767 | 1.59745427 | 3.08707536 |
| ENSMUSG00 | Pold3 | 4582 | 11.0884994 | 9.14932446 | 14.8079068 | 8.40234428 | 13.0294325 | 11.0598603 |
| ENSMUSG00 | Rabep2 | 5396 | 45.9147803 | 29.3887289 | 58.5535163 | 34.7848028 | 46.5913554 | 47.3464688 |
| ENSMUSG00 | Pgm2l1 | 7367 | 0.23744281 | 1.15118961 | 0.15599945 | 1.1264925 | 0.40172795 | 1.29579527 |
| ENSMUSG00 | Atp2a1 | 3726 | 1.4937647 | 0.46556949 | 1.61337913 | 0.91028239 | 1.54749878 | 1.76414065 |
| ENSMUSG00 | Syt3 | 3155 | 3.50301857 | 8.55290257 | 6.73885467 | 8.05127079 | 8.24832066 | 6.77759657 |
| ENSMUSG00 | Chrdl2 | 1716 | 0 | 0.19656474 | 0.1030346 | 0.50464398 | 0 | 0 |
| ENSMUSG00 | Sh2b1 | 4208 | 10.9119724 | 9.30978552 | 10.5882742 | 9.37207098 | 8.92476535 | 10.8502248 |
| ENSMUSG00 | Gm9755 | 1358 | 0.38057529 | 0.3193505 | 0.65098443 | 0.53139977 | 0.67634372 | 0.3414351 |
| ENSMUSG00 | Slco2b1 | 7145 | 0 | 0 | 0 | 0 | 0 | 0 |
| ENSMUSG00 | Eif3c | 2856 | 73.4279271 | 70.2550228 | 62.7430923 | 71.1786551 | 69.9647366 | 43.1466457 |
| ENSMUSG00 | Myh14 | 7755 | 3.41933017 | 2.49165226 | 3.921453 | 2.838175 | 3.83602542 | 2.44785931 |
| ENSMUSG00 | Spns1 | 2984 | 21.5431005 | 18.0699158 | 18.4273099 | 18.2586754 | 15.8687925 | 11.3796669 |
| ENSMUSG00 | Lat | 1227 | 0 | 0 | 0 | 0.02940672 | 0 | 0 |
| ENSMUSG00 | Rps3 | 3075 | 17.0786959 | 21.6408054 | 23.8618079 | 21.2502708 | 13.9886943 | 25.2966735 |
| ENSMUSG00 | Il21r | 2956 | 0 | 0 | 0.01495326 | 0 | 0 | 0 |
| ENSMUSG00 | Dgat2 | 2243 | 16.5367202 | 21.4830326 | 14.8981691 | 16.2956358 | 10.8741006 | 10.8587842 |
| ENSMUSG00 | Il4ra | 3653 | 10.4585318 | 2.98114862 | 5.57816853 | 4.18800626 | 5.57337092 | 5.28618795 |
| ENSMUSG00 | Nsmce1 | 1319 | 16.1855141 | 20.8600898 | 21.3469098 | 21.4194396 | 15.4355754 | 19.1067223 |
| ENSMUSG00 | Psma1 | 2149 | 164.109292 | 163.663491 | 155.7863 | 144.796303 | 102.978777 | 180.172536 |
| ENSMUSG00 | Jmjd5 | 2485 | 3.77557079 | 5.3712855 | 5.28287621 | 4.23982173 | 4.47635812 | 4.09393982 |
| ENSMUSG00 | Prkrir | 4269 | 23.0952428 | 19.3581045 | 28.3289394 | 25.6690488 | 27.9216711 | 20.6909697 |
| ENSMUSG00 | Copb1 | 3699 | 82.7352498 | 83.4762696 | 83.4923701 | 94.8824134 | 85.2094954 | 91.8150332 |
| ENSMUSG00 | Zkscan2 | 4957 | 0 | 0 | 0 | 0 | 0 | 0 |
| ENSMUSG00 | Far1 | 8025 | 40.4440801 | 35.0965426 | 55.8347767 | 55.3707637 | 40.4904625 | 51.9662597 |
| ENSMUSG00 | Acer3 | 4306 | 11.9284907 | 32.4525504 | 16.3421584 | 28.2975068 | 21.1642144 | 15.5248818 |
| ENSMUSG00 | Myo7a | 9121 | 1.25529858 | 0.41735873 | 1.33754069 | 1.34106053 | 2.01397822 | 1.22602806 |
| ENSMUSG00 | Aqp8 | 1570 | 0.40515139 | 0 | 0 | 0 | 0 | 0 |
| ENSMUSG00 | Lcmt1 | 1358 | 31.207174 | 41.6574986 | 29.4895948 | 33.5313256 | 34.6062534 | 31.6329579 |
| ENSMUSG00 | Arhgap17 | 3576 | 6.53697498 | 11.898386 | 7.73779467 | 13.7729281 | 10.2166905 | 7.36018649 |
| ENSMUSG00 | Slc5a11 | 2938 | 0.36534989 | 0 | 0 | 0 | 0.26051587 | 0.31563572 |
| ENSMUSG00 | Parva | 10140 | 30.1420251 | 29.2207527 | 32.1836496 | 24.7272315 | 28.8092704 | 22.5916859 |
| ENSMUSG00 | Micalcl | 3483 | 0.75333383 | 0.98226741 | 1.8020849 | 0.62156838 | 0.70320587 | 0.97884867 |
| ENSMUSG00 | Dkk3 | 3349 | 30.7929879 | 17.3666873 | 49.4680524 | 22.3883215 | 22.2602374 | 22.3393059 |
| ENSMUSG00 | Pak1 | 3746 | 0.63676691 | 0.82326069 | 1.8289604 | 0.82836514 | 2.98312235 | 1.8566559 |
| ENSMUSG00 | Trat1 | 3981 | 0 | 0 | 0 | 0 | 0 | 0 |
| ENSMUSG00 | Rbbp6 | 8319 | 10.9770812 | 11.5499177 | 10.9561486 | 11.9492767 | 13.6965858 | 17.1863997 |
| ENSMUSG00 | BC017158 | 6025 | 10.1813621 | 8.36564622 | 7.56383405 | 8.10273963 | 13.95709 | 7.77723323 |
| ENSMUSG00 | Slc5a2 | 3023 | 0.13151003 | 1.17955565 | 0.4678991 | 0.10742256 | 0.55701965 | 0.216537 |
| ENSMUSG00 | Tgfb1i1 | 3370 | 0.15335942 | 0.14298648 | 0.22297666 | 0.18201625 | 0.04542407 | 0.2751744 |
| ENSMUSG00 | Cox6a2 | 655 | 0 | 0 | 0 | 0 | 0 | 0 |
| ENSMUSG00 | Itgam | 6327 | 0 | 0 | 0 | 0 | 0.02419458 | 0.0129325 |
| ENSMUSG00 | Lyve1 | 2601 | 0 | 0 | 0 | 0 | 0 | 0 |
| ENSMUSG00 | Rnf141 | 5604 | 10.2510117 | 14.1274669 | 17.1554262 | 14.9762376 | 14.5503477 | 14.0266792 |
| ENSMUSG00 | Itgax | 4104 | 0 | 0 | 0 | 0 | 0 | 0 |
| ENSMUSG00 | Adm | 1383 | 58.3827772 | 87.6623922 | 56.2830409 | 71.8771024 | 86.8518356 | 72.4168296 |
| ENSMUSG00 | Dkkl1 | 898 | 0 | 0 | 0.3445578 | 0.20090225 | 0 | 0 |
| ENSMUSG00 | Pycard | 1402 | 17.155539 | 21.1374193 | 18.6328739 | 19.7653425 | 14.9221213 | 32.4299741 |
| ENSMUSG00 | Fus | 4516 | 15.1591978 | 14.7888416 | 9.92486023 | 26.5421948 | 13.0164714 | 25.5714842 |
| ENSMUSG00 | Tead2 | 4167 | 0.24805435 | 0.10407439 | 0.03182278 | 0.008659 | 0.13469886 | 0.11781702 |
| ENSMUSG00 | Cd37 | 2916 | 0 | 0 | 0 | 0 | 0 | 0 |
| ENSMUSG00 | Prss8 | 2654 | 49.1775594 | 57.5187071 | 55.0108091 | 47.6652781 | 41.7785662 | 43.9127858 |
| ENSMUSG00 | Myst1 | 1562 | 14.1765702 | 15.6713909 | 13.2718722 | 16.1006306 | 14.8963043 | 12.7467904 |
| ENSMUSG00 | Bckdk | 3118 | 48.6807006 | 44.2919638 | 52.7076499 | 37.6674326 | 56.0995562 | 38.1127656 |
| ENSMUSG00 | Vkorc1 | 2389 | 8.2539633 | 11.8600368 | 9.71367417 | 12.0827273 | 7.94550515 | 10.4805855 |
| ENSMUSG00 | Stx4a | 2053 | 20.5458182 | 18.4953319 | 18.1931599 | 16.3274327 | 16.8265253 | 16.181447 |
| ENSMUSG00 | Stx1b | 4810 | 0 | 0 | 0 | 0 | 0 | 0 |
| ENSMUSG00 | Fbxl19 | 3398 | 0.85407595 | 0.36870144 | 0.40325401 | 0.53093061 | 0.97607841 | 0.76253406 |
| ENSMUSG00 | Bcl7c | 4995 | 1.03467717 | 0.65599161 | 1.8848834 | 1.60364642 | 1.52210611 | 1.10845881 |
| ENSMUSG00 | Phkg2 | 2750 | 20.4560018 | 18.7313842 | 21.0722241 | 25.165586 | 21.1341971 | 25.0430779 |
| ENSMUSG00 | Rnf40 | 4134 | 15.5502206 | 13.4977987 | 14.1244883 | 17.508607 | 20.6253203 | 14.5909773 |
| ENSMUSG00 | Prr14 | 3283 | 19.0361302 | 19.0955106 | 21.3806052 | 24.3221336 | 19.5525985 | 19.5981949 |
| ENSMUSG00 | 9130019O22 | 6176 | 0.65658339 | 1.51362853 | 1.65327489 | 1.53652489 | 1.28887866 | 0.58735865 |
| ENSMUSG00 | Nucb1 | 5776 | 60.0461932 | 57.0378644 | 55.2140404 | 54.0293634 | 55.9028577 | 38.8248068 |
| ENSMUSG00 | Hsd17b14 | 2356 | 3.54356999 | 6.62665843 | 3.47085779 | 2.51165335 | 3.46528868 | 1.19239722 |
| ENSMUSG00 | Bcat2 | 3926 | 61.3141708 | 51.9667336 | 64.4563299 | 47.8183591 | 44.5018622 | 43.6769376 |
| ENSMUSG00 | Fgf21 | 927 | 0 | 0 | 0 | 0 | 0 | 0 |
| ENSMUSG00 | Itgal | 5171 | 0.11532242 | 0.05591156 | 0.11112434 | 0 | 0.66114237 | 0 |
| ENSMUSG00 | Abcc6 | 5477 | 0 | 0.01759593 | 0 | 0 | 0 | 0.20915371 |
| ENSMUSG00 | Nomo1 | 4228 | 15.2044967 | 15.03262 | 15.7550089 | 11.9220947 | 20.1064196 | 9.23776807 |
| ENSMUSG00 | Ush1c | 3906 | 19.1758139 | 16.7320885 | 11.3445365 | 14.0616566 | 17.2742823 | 14.0351329 |
| ENSMUSG00 | Sergef | 2669 | 5.24313568 | 7.11332267 | 5.56456323 | 6.69187413 | 5.12366756 | 6.36646694 |
| ENSMUSG00 | 2400001E08 | 1179 | 16.7923913 | 13.4872993 | 21.107411 | 13.4045255 | 10.213931 | 21.7456835 |
| ENSMUSG00 | Rgs10 | 1860 | 0 | 0 | 0 | 0 | 0 | 0.20529298 |
| ENSMUSG00 | Tial1 | 6376 | 4.63897075 | 5.64543166 | 5.43510742 | 7.99822474 | 4.61766293 | 9.22272933 |
| ENSMUSG00 | Bag3 | 2587 | 12.4629666 | 16.6333563 | 14.4719602 | 21.1861715 | 14.8128366 | 15.4138085 |
| ENSMUSG00 | Fgfr2 | 8713 | 14.7332086 | 17.2050983 | 10.4353485 | 16.0263414 | 22.4532451 | 10.7714947 |
| ENSMUSG00 | Ate1 | 5039 | 30.6430415 | 29.0514807 | 31.3421681 | 34.3133871 | 30.682659 | 44.1568778 |
| ENSMUSG00 | Ldhc | 1570 | 0 | 0 | 0.11261616 | 0 | 0 | 0 |
| ENSMUSG00 | Tacc2 | 9720 | 11.4685563 | 10.7675877 | 12.7421362 | 17.0238947 | 13.276307 | 11.6422303 |
| ENSMUSG00 | Ptpre | 6347 | 0 | 0 | 0 | 0 | 0 | 0.00429725 |
| ENSMUSG00 | 1700007K09 | 420 | 0 | 0 | 0 | 0 | 0 | 0 |
| ENSMUSG00 | Fam24a | 376 | 0 | 0 | 0 | 0 | 0 | 0 |
| ENSMUSG00 | Acadsb | 4393 | 64.6784474 | 83.0127462 | 72.9184512 | 80.3036989 | 67.3924499 | 69.4500602 |
| ENSMUSG00 | Cpxm2 | 4435 | 0 | 0 | 0 | 0 | 0 | 0 |
| ENSMUSG00 | 2010110P09 | 1595 | 0.87237732 | 0.69485152 | 3.07611572 | 0 | 0.70381208 | 0.73530375 |
| ENSMUSG00 | Emi2 | 2951 | 0 | 0 | 0 | 0 | 0 | 0 |
| ENSMUSG00 | Plk1 | 2193 | 0 | 1.4721804 | 0.20155879 | 0 | 0 | 0.32336554 |
| ENSMUSG00 | Dctn5 | 1936 | 68.5453481 | 61.6766548 | 76.394301 | 76.3947833 | 69.8977008 | 73.0610946 |
| ENSMUSG00 | Ndufab1 | 6848 | 22.8907505 | 17.2114541 | 20.4162426 | 20.7598212 | 21.3032139 | 26.1156264 |
| ENSMUSG00 | Ubfd1 | 7481 | 22.069845 | 15.7744415 | 24.9399785 | 16.9823391 | 21.9083972 | 17.6094784 |
| ENSMUSG00 | Ears2 | 5479 | 2.58312668 | 2.90226792 | 2.43638558 | 5.03792007 | 1.80673789 | 1.83689394 |
| ENSMUSG00 | Gga2 | 10290 | 27.4888478 | 22.5525681 | 30.4515904 | 21.2214318 | 23.7528027 | 21.1570622 |
| ENSMUSG00 | Scnn1b | 2605 | 266.750881 | 350.271965 | 121.983512 | 280.581181 | 334.913234 | 149.073441 |
| ENSMUSG00 | Mettl9 | 1817 | 80.9550236 | 79.5063036 | 90.3741589 | 76.8903007 | 70.4034806 | 102.253628 |
| ENSMUSG00 | 4933427G17 | 2105 | 0 | 0 | 0 | 0 | 0 | 0 |
| ENSMUSG00 | Cdr2 | 4503 | 1.39492914 | 2.88926036 | 0.49080439 | 2.33976394 | 2.73092529 | 0.34227972 |
| ENSMUSG00 | Mrpl17 | 4393 | 14.1447339 | 12.9762261 | 13.5030442 | 11.4907308 | 9.8846906 | 19.2778864 |
| ENSMUSG00 | Polr3e | 4213 | 9.54018296 | 10.2938044 | 7.41768408 | 11.9302844 | 11.4212847 | 11.8472796 |
| ENSMUSG00 | Arfip2 | 6467 | 7.02652145 | 5.39461638 | 6.14465082 | 5.67425998 | 6.45424083 | 4.56756368 |
| ENSMUSG00 | Dnhd1 | 15491 | 0.45424569 | 0.93007206 | 0.37950069 | 0.74302319 | 0.64890556 | 0.57045918 |
| ENSMUSG00 | Uqcrc2 | 2573 | 234.283852 | 175.872086 | 242.070801 | 174.211908 | 177.333017 | 204.374289 |
| ENSMUSG00 | Pdzd9 | 2619 | 0.13661677 | 0.29438071 | 0.13501899 | 0.34442578 | 0.07793261 | 0.09372728 |
| ENSMUSG00 | Rrp8 | 3353 | 11.7025527 | 11.3244606 | 10.8494234 | 13.1070475 | 9.45045612 | 7.41043733 |
| ENSMUSG00 | Vwa3a | 5582 | 0 | 0 | 0 | 0 | 0.10969482 | 0.18078854 |
| ENSMUSG00 | Ilk | 2640 | 0.01505889 | 0 | 0.03348625 | 0.01366744 | 0.01932817 | 0.04132521 |
| ENSMUSG00 | Tpp1 | 3525 | 102.890567 | 88.4990146 | 104.31635 | 93.0865569 | 94.2214696 | 83.0541781 |
| ENSMUSG00 | Hpx | 1468 | 0.56870919 | 0.03282455 | 0 | 0.27036954 | 1.00801424 | 0.42732745 |
| ENSMUSG00 | Cnga4 | 2474 | 0.83560428 | 0.25320281 | 0.30373134 | 0 | 1.1343778 | 0 |
| ENSMUSG00 | Cckbr | 3124 | 0.59811382 | 0 | 0 | 0 | 0 | 0 |
| ENSMUSG00 | Crym | 1887 | 0 | 0 | 0 | 0.17209242 | 0.37857407 | 0 |
| ENSMUSG00 | Ankrd4b | 2028 | 0 | 0 | 0.41411983 | 0 | 0.37741402 | 0 |
| ENSMUSG00 | Zp2 | 2401 | 0 | 0 | 0 | 0 | 0 | 0 |
| ENSMUSG00 | Tmem159 | 2040 | 10.4455576 | 12.6843723 | 8.4070172 | 12.7525265 | 10.5304432 | 9.74667256 |
| ENSMUSG00 | Trim30a | 8197 | 1.59080122 | 0.81123936 | 1.81186035 | 0.40937296 | 2.46510246 | 2.08627532 |
| ENSMUSG00 | Lyrm1 | 1530 | 5.89836214 | 4.56668896 | 3.81349234 | 5.21185088 | 4.90253415 | 4.97361069 |
| ENSMUSG00 | 2610020H08 | 5210 | 2.09841788 | 2.4324442 | 2.03616935 | 1.49591586 | 2.77168222 | 2.12543254 |
| ENSMUSG00 | Eri2 | 7043 | 3.38680794 | 1.39571691 | 2.51667458 | 1.75210269 | 2.34737268 | 1.29344449 |
| ENSMUSG00 | Chst15 | 5378 | 1.13101311 | 2.1683003 | 2.76158781 | 0.30191372 | 2.13479631 | 2.41911546 |
| ENSMUSG00 | Oat | 2145 | 267.316222 | 311.965092 | 267.560247 | 233.078233 | 277.921281 | 192.041434 |
| ENSMUSG00 | Acsm3 | 4976 | 0.91079678 | 0.48418853 | 0 | 0.21753644 | 0.17432645 | 0.59745492 |
| ENSMUSG00 | Thumpd1 | 2649 | 29.4151536 | 34.4526695 | 45.6368594 | 37.5394922 | 41.8188984 | 35.2233072 |
| ENSMUSG00 | Acsm2 | 3158 | 3.91512174 | 5.90505165 | 1.52564943 | 2.11373598 | 2.5852502 | 8.96487509 |
| ENSMUSG00 | Lhpp | 2267 | 3.26180828 | 3.31587339 | 5.84938373 | 6.414232 | 3.26370732 | 4.76433928 |
| ENSMUSG00 | Gp2 | 2261 | 0.77365818 | 0.72460815 | 0.09774844 | 0.57450403 | 0.94785837 | 0.10856778 |
| ENSMUSG00 | Fam53b | 10817 | 10.4047116 | 12.9186172 | 11.1229931 | 13.4194384 | 13.6988625 | 8.9007559 |
| ENSMUSG00 | Mettl10 | 2030 | 46.1986101 | 37.291084 | 35.8840578 | 35.228873 | 23.1252541 | 58.5665776 |
| ENSMUSG00 | Umod | 2461 | 182.542434 | 93.2988203 | 159.34935 | 59.9070435 | 128.239795 | 78.1333632 |
| ENSMUSG00 | Fam175b | 5089 | 15.0381804 | 14.7049606 | 19.2910775 | 18.8103093 | 18.0081295 | 14.6583098 |
| ENSMUSG00 | Trim21 | 2767 | 6.07754545 | 2.17683602 | 3.24285296 | 2.73842767 | 9.12831805 | 10.0049725 |
| ENSMUSG00 | Zranb1 | 5421 | 17.0799694 | 12.471053 | 15.6879443 | 17.2056973 | 15.2297871 | 16.9906764 |
| ENSMUSG00 | Pdilt | 2127 | 6.01846014 | 2.78652204 | 4.21860561 | 3.13830665 | 4.5100887 | 2.70566473 |
| ENSMUSG00 | Ctbp2 | 6855 | 11.001626 | 10.3542858 | 10.7941481 | 9.09551757 | 10.3764796 | 13.5676907 |
| ENSMUSG00 | Acsm5 | 2871 | 0 | 0 | 0 | 0 | 0.01777303 | 0.0190001 |
| ENSMUSG00 | 4930404H21 | 911 | 0 | 0 | 0 | 0 | 0 | 0 |
| ENSMUSG00 | Rrm1 | 3978 | 14.3111735 | 14.7175785 | 11.7782689 | 13.6509495 | 11.0441703 | 15.2211411 |
| ENSMUSG00 | Uros | 4324 | 16.4575185 | 17.9974801 | 17.633714 | 18.566732 | 15.7657812 | 15.4097927 |
| ENSMUSG00 | 2310008H09 | 9720 | 8.31511233 | 6.79171047 | 7.08502792 | 5.81321828 | 9.5543214 | 7.70256016 |
| ENSMUSG00 | Mmp21 | 3008 | 0 | 0 | 0 | 0 | 0 | 0 |
| ENSMUSG00 | 9030624J02R | 9437 | 6.18006675 | 7.24047631 | 6.74011361 | 6.93576653 | 8.91623331 | 4.04914375 |
| ENSMUSG00 | Bccip | 2210 | 28.080681 | 23.8315752 | 24.1410066 | 24.4737488 | 22.3731032 | 26.818005 |
| ENSMUSG00 | Dhx32 | 4590 | 20.6399368 | 14.8548618 | 23.7187667 | 14.7551193 | 16.3862479 | 13.2926728 |
| ENSMUSG00 | Stim1 | 3585 | 8.73844314 | 7.99747087 | 9.54315941 | 9.55142545 | 11.3439391 | 10.1338409 |
| ENSMUSG00 | Pgap2 | 3718 | 22.0269741 | 21.2678729 | 26.8444759 | 23.1748043 | 20.7097367 | 16.5569825 |
| ENSMUSG00 | D7Ertd443e | 3842 | 0 | 0 | 0 | 0 | 0 | 0 |
| ENSMUSG00 | Art1 | 1002 | 0 | 0 | 0.04411362 | 0 | 0 | 0.24498179 |
| ENSMUSG00 | Mki67 | 10061 | 0.09878611 | 0.56994202 | 0 | 0.24387029 | 0.27894351 | 0.05692947 |
| ENSMUSG00 | Atp6ap2 | 2609 | 921.233072 | 671.470407 | 1374.47571 | 859.372236 | 634.866687 | 1213.44441 |
| ENSMUSG00 | Usp9x | 17535 | 9.2502059 | 8.35121743 | 9.82599282 | 13.5109612 | 14.7593828 | 8.78045843 |
| ENSMUSG00 | Cask | 9785 | 5.956212 | 5.35295479 | 4.15140457 | 8.4037792 | 8.26538637 | 4.35669506 |
| ENSMUSG00 | Swap70 | 4020 | 8.98948552 | 6.35293889 | 3.94737852 | 7.08177441 | 9.54523249 | 5.49563627 |
| ENSMUSG00 | Wee1 | 1930 | 10.587729 | 12.0840611 | 10.3748367 | 12.6193679 | 9.41211906 | 5.00270585 |
| ENSMUSG00 | Tmem9b | 3151 | 66.4399747 | 68.6018342 | 84.7564378 | 62.625421 | 57.9410894 | 86.9914708 |
| ENSMUSG00 | BC051019 | 2413 | 0 | 0 | 0.03663642 | 0 | 0.04229289 | 0.04521283 |
| ENSMUSG00 | D930014E17 | 1324 | 24.3517332 | 22.6010427 | 28.7779371 | 18.3953022 | 20.8884409 | 31.9714805 |
| ENSMUSG00 | St5 | 5265 | 21.5351625 | 14.2591599 | 22.5248898 | 15.632126 | 15.2739921 | 14.359981 |
| ENSMUSG00 | Trim66 | 10646 | 0.11949797 | 0 | 0 | 0 | 0.0527231 | 0 |
| ENSMUSG00 | Stk33 | 2900 | 0 | 0 | 0 | 0.43547295 | 0 | 0 |
| ENSMUSG00 | Tub | 6905 | 0 | 0.02791394 | 0 | 0 | 0 | 0 |
| ENSMUSG00 | Eif3f | 2884 | 12.461496 | 13.0658106 | 12.2152805 | 9.79620002 | 6.70561596 | 14.157467 |
| ENSMUSG00 | Ndufb11 | 905 | 55.2183856 | 56.9185709 | 69.4530616 | 48.2821612 | 36.3104818 | 93.5776297 |
| ENSMUSG00 | Rbm10 | 4135 | 7.90302418 | 7.60961227 | 6.3176032 | 7.32112101 | 8.21851664 | 6.13432032 |
| ENSMUSG00 | Cdk16 | 4261 | 39.1023748 | 38.449637 | 40.4051111 | 44.2113012 | 37.5901886 | 32.4274401 |
| ENSMUSG00 | Usp11 | 4897 | 7.39580207 | 5.74655549 | 6.78778557 | 7.34608911 | 8.69021798 | 6.70030448 |
| ENSMUSG00 | Glrx3 | 1640 | 26.1561972 | 27.4721484 | 24.337966 | 24.4873875 | 17.5480951 | 28.538586 |
| ENSMUSG00 | Mrgprf | 2303 | 0 | 0 | 0 | 0 | 0 | 0 |
| ENSMUSG00 | Oraov1 | 4375 | 8.54174896 | 8.4257436 | 9.59811448 | 9.92977865 | 9.20224241 | 9.25155763 |
| ENSMUSG00 | Fgf15 | 1806 | 0 | 0 | 0 | 0 | 0 | 0 |

| | | | | | | | | |
|---|---|---|---|---|---|---|---|---|
| ENSMUSG00 | Fgf3 | 2322 | 0 | 0 | 0 | 0 | 0 | 0 |
| ENSMUSG00 | Ano1 | 6731 | 0 | 0.39373857 | 0.21014099 | 0 | 0.83388818 | 0 |
| ENSMUSG00 | Fadd | 3901 | 1.83439797 | 1.7293263 | 2.45880542 | 0.81395025 | 1.34727421 | 1.37736579 |
| ENSMUSG00 | Cttn | 5537 | 55.3503701 | 47.0638033 | 51.4104875 | 50.1447233 | 48.6487697 | 38.3972933 |
| ENSMUSG00 | Zfp300 | 5320 | 1.98777403 | 1.47638911 | 3.23205206 | 1.21403872 | 1.2372937 | 2.01483707 |
| ENSMUSG00 | Gm498 | 1779 | 0 | 0 | 0 | 0 | 0 | 0 |
| ENSMUSG00 | Slc6a14 | 3420 | 0 | 0 | 0 | 0 | 0 | 0 |
| ENSMUSG00 | Nadsyn1 | 3491 | 6.95806319 | 6.12855351 | 5.25458747 | 7.94817881 | 9.42767471 | 4.49238544 |
| ENSMUSG00 | Dock11 | 8119 | 0 | 0 | 0 | 0 | 0.12569621 | 0 |
| ENSMUSG00 | Cul4b | 5958 | 23.4943013 | 21.4000211 | 17.8795639 | 20.8994857 | 15.6641896 | 24.4821703 |
| ENSMUSG00 | Tnni2 | 1343 | 0 | 0.17939852 | 0 | 0 | 0 | 0 |
| ENSMUSG00 | Syt8 | 1811 | 0 | 0.31929172 | 0 | 0 | 0 | 0 |
| ENSMUSG00 | Smarca1 | 6842 | 0.91224941 | 1.40854844 | 0.46514655 | 0.89651382 | 1.02917858 | 0.03587719 |
| ENSMUSG00 | Sash3 | 2669 | 0 | 0 | 0 | 0 | 0 | 0 |
| ENSMUSG00 | Elf4 | 10535 | 0.50944375 | 0.39335871 | 0.81816416 | 0.13014881 | 1.02198057 | 0.95791329 |
| ENSMUSG00 | Rab33a | 1156 | 0 | 0.54188906 | 0 | 0.06242568 | 0.08828093 | 0 |
| ENSMUSG00 | Slc25a14 | 2350 | 4.9905816 | 4.77763448 | 4.66470514 | 7.76915512 | 5.16777764 | 3.79523704 |
| ENSMUSG00 | Rbmx2 | 1866 | 1.66180465 | 1.67852023 | 2.53461801 | 1.50825267 | 2.07824467 | 2.60176087 |
| ENSMUSG00 | Enox2 | 5498 | 5.1700925 | 5.14467834 | 6.77740154 | 4.54142867 | 6.92354972 | 5.86886101 |
| ENSMUSG00 | Igsf1 | 6160 | 0 | 0 | 0 | 0 | 0.25678858 | 0 |
| ENSMUSG00 | 2610018G03I | 5363 | 3.64715577 | 4.34872982 | 1.61543189 | 1.9241963 | 4.06270044 | 3.58542246 |
| ENSMUSG00 | 1700080O16I | 2515 | 0 | 0 | 0 | 0 | 0 | 0 |
| ENSMUSG00 | Gpc4 | 2642 | 32.863728 | 23.272483 | 33.5278173 | 26.358193 | 29.6655993 | 18.0557699 |
| ENSMUSG00 | 3830403N18I | 1014 | 0.07841318 | 0.09504229 | 1.13338059 | 0 | 0 | 0.53796133 |
| ENSMUSG00 | Slc9a9 | 5964 | 2.89967036 | 1.43008053 | 8.40457578 | 2.35343986 | 2.89183686 | 4.2851001 |
| ENSMUSG00 | Brs3 | 2769 | 0 | 0 | 0 | 0 | 0 | 0 |
| ENSMUSG00 | Vgll1 | 2445 | 0 | 0 | 0 | 0 | 0 | 0 |
| ENSMUSG00 | Cd40lg | 1245 | 0 | 0 | 0 | 0 | 0 | 0 |
| ENSMUSG00 | Arhgef6 | 10690 | 2.29458667 | 2.50623591 | 2.16667593 | 3.45294028 | 4.14798135 | 1.68138328 |
| ENSMUSG00 | Rbmx | 3651 | 47.9548442 | 39.581249 | 46.4658103 | 48.8209532 | 55.7642398 | 66.33766 |
| ENSMUSG00 | Fgf13 | 4186 | 0 | 0.28778334 | 0 | 0 | 0 | 0 |
| ENSMUSG00 | F9 | 2726 | 0 | 0 | 0 | 0 | 0 | 0 |
| ENSMUSG00 | Mcf2 | 5206 | 0 | 0 | 0 | 0 | 0 | 0 |
| ENSMUSG00 | Cacna1f | 8854 | 0.00898023 | 0 | 0.0049923 | 0 | 0.10373557 | 0 |
| ENSMUSG00 | Ccdc22 | 3165 | 0 | 0.01522478 | 0.01396583 | 0 | 0 | 0.03447032 |
| ENSMUSG00 | Syp | 5346 | 1.01879926 | 0.39659623 | 0.97564861 | 0.81667179 | 0.90675378 | 0.67344788 |
| ENSMUSG00 | Prickle3 | 4921 | 4.1201575 | 2.44800052 | 2.12880244 | 3.51948412 | 3.84693872 | 2.06735225 |
| ENSMUSG00 | Plp2 | 1876 | 6.63297729 | 6.96083466 | 6.9271545 | 6.53938973 | 4.35192971 | 6.20803356 |
| ENSMUSG00 | Magix | 3468 | 1.82269937 | 2.15366163 | 1.54222117 | 1.58145063 | 1.44192181 | 2.6425256 |
| ENSMUSG00 | Gpkow | 3726 | 20.5072554 | 24.3518708 | 24.5091271 | 25.2942298 | 27.6084739 | 21.272169 |
| ENSMUSG00 | Praf2 | 1336 | 11.0101256 | 11.4695274 | 10.3556713 | 11.8833081 | 9.66292895 | 9.94217763 |
| ENSMUSG00 | Ccdc120 | 4128 | 6.11548695 | 3.32682558 | 4.358079 | 4.7287757 | 7.24356982 | 4.79684789 |
| ENSMUSG00 | Gripap1 | 5376 | 12.3274528 | 14.0005995 | 12.0206649 | 14.0609902 | 13.848118 | 11.1259829 |
| ENSMUSG00 | Otud5 | 5061 | 18.2399182 | 15.1671611 | 11.6858459 | 13.6100816 | 17.0793678 | 14.6370125 |
| ENSMUSG00 | Pim2 | 3043 | 3.50130423 | 5.24144343 | 2.93420056 | 3.69950637 | 2.90094085 | 0.79771373 |
| ENSMUSG00 | Slc35a2 | 2462 | 10.9480974 | 14.600766 | 12.6932181 | 13.5417584 | 9.82392451 | 12.2746955 |
| ENSMUSG00 | Pqbp1 | 1675 | 24.6365306 | 26.5816553 | 25.8349877 | 20.7660244 | 20.8370394 | 24.4250501 |
| ENSMUSG00 | Timm17b | 2551 | 8.78952999 | 12.5046745 | 11.6612467 | 10.2828876 | 7.16089476 | 12.9156269 |
| ENSMUSG00 | Eras | 1075 | 0 | 0 | 0 | 0 | 0 | 0 |
| ENSMUSG00 | Hdac6 | 6047 | 24.9038796 | 29.7868233 | 27.1117307 | 31.6963487 | 35.2045709 | 19.4580443 |
| ENSMUSG00 | Gata1 | 2428 | 0 | 0 | 0 | 0 | 0 | 0 |
| ENSMUSG00 | Glod5 | 669 | 34.8826115 | 27.2264202 | 66.6661579 | 28.4773087 | 15.9409754 | 93.0762353 |
| ENSMUSG00 | Was | 2370 | 0 | 0 | 0 | 0 | 0 | 0 |
| ENSMUSG00 | Wdr13 | 7694 | 22.5749538 | 18.2311844 | 18.3839222 | 20.2967363 | 21.4212625 | 22.9817372 |
| ENSMUSG00 | Rbm3 | 6617 | 4.53610213 | 5.65100183 | 3.92118511 | 5.40385464 | 6.93255431 | 5.11116105 |
| ENSMUSG00 | Ebp | 2332 | 21.3608993 | 21.7582863 | 31.3886158 | 17.5458999 | 15.0322128 | 21.9764353 |
| ENSMUSG00 | Porcn | 3414 | 8.18632185 | 7.02895378 | 8.37685777 | 8.20142547 | 7.60762311 | 8.09291434 |
| ENSMUSG00 | Slc38a5 | 2254 | 0 | 0 | 0 | 0 | 0 | 0 |
| ENSMUSG00 | Ftsj1 | 5067 | 5.14694991 | 4.74541832 | 5.87089804 | 4.00199507 | 4.69277504 | 5.02215086 |
| ENSMUSG00 | Otc | 4376 | 0 | 0 | 0 | 0 | 0 | 0 |
| ENSMUSG00 | Rpgr | 7194 | 3.5367678 | 3.6437899 | 2.84479473 | 4.8199673 | 4.61748272 | 4.99314705 |
| ENSMUSG00 | Dynlt3 | 2624 | 152.264702 | 143.255501 | 142.123075 | 163.868035 | 124.143439 | 172.742847 |
| ENSMUSG00 | 3830417A13I | 1762 | 0 | 0 | 0 | 0 | 0 | 0 |
| ENSMUSG00 | Ctag2 | 851 | 0 | 0 | 0 | 0 | 0 | 0 |
| ENSMUSG00 | 4930447F04I | 701 | 0 | 0 | 0 | 0 | 0 | 0 |
| ENSMUSG00 | Aff2 | 7580 | 0.28846325 | 0.34328072 | 0 | 0.23324805 | 0 | 0.07916122 |
| ENSMUSG00 | 4931400O07I | 645 | 0 | 0 | 0 | 0 | 0 | 0 |
| ENSMUSG00 | F8 | 10144 | 1.73224787 | 0.4085207 | 1.21572498 | 1.48326228 | 0.80986263 | 0.40062315 |
| ENSMUSG00 | Vbp1 | 2236 | 135.517117 | 128.913818 | 110.623217 | 131.999608 | 74.645472 | 188.190132 |
| ENSMUSG00 | Fundc2 | 2809 | 27.9236608 | 29.3338612 | 36.9476424 | 24.4443328 | 16.1126363 | 40.6643607 |
| ENSMUSG00 | Mtcp1 | 4954 | 2.67230016 | 4.77584641 | 4.21139885 | 6.44582346 | 5.38631849 | 7.04163577 |
| ENSMUSG00 | Brcc3 | 6741 | 12.7977144 | 15.7905134 | 16.2879956 | 14.3771505 | 10.4005697 | 28.4115883 |
| ENSMUSG00 | Rab39b | 3399 | 2.26906832 | 1.57360844 | 1.53451529 | 2.40971583 | 2.73221548 | 1.55671669 |
| ENSMUSG00 | Asb12 | 1255 | 0 | 0 | 0 | 0 | 0 | 0 |
| ENSMUSG00 | Msn | 6500 | 4.84405239 | 8.44374732 | 3.89656247 | 6.11174323 | 6.55492674 | 3.66738995 |
| ENSMUSG00 | Heph | 7639 | 11.3088963 | 7.92278719 | 9.91199857 | 14.7086643 | 14.9826104 | 9.24388546 |
| ENSMUSG00 | Gpr165 | 5504 | 0 | 0 | 0 | 0 | 0 | 0 |
| ENSMUSG00 | Pgr15l | 5374 | 0 | 0 | 0 | 0 | 0 | 0 |
| ENSMUSG00 | Ophn1 | 12985 | 2.58402969 | 2.12265267 | 2.77772075 | 3.24002033 | 2.94723002 | 1.98494679 |
| ENSMUSG00 | Stard8 | 5482 | 38.1962999 | 45.1802832 | 38.6463761 | 41.0974619 | 55.1311974 | 29.6677625 |
| ENSMUSG00 | Efnb1 | 3446 | 7.91417865 | 8.94931021 | 9.35088321 | 8.08338316 | 9.06214219 | 8.65095203 |
| ENSMUSG00 | Awat2 | 2831 | 0.87066047 | 0 | 0.1717486 | 0 | 0 | 0.00963428 |
| ENSMUSG00 | Igbp1 | 5912 | 7.79374187 | 6.33302868 | 6.39251958 | 8.73976451 | 6.68901239 | 7.6905994 |
| ENSMUSG00 | Magee2 | 2312 | 0 | 0.02084189 | 0 | 0.76471461 | 0 | 0 |
| ENSMUSG00 | 2610029G23I | 4842 | 6.73265056 | 7.37425726 | 7.2665566 | 5.73795421 | 4.16262256 | 5.9934358 |
| ENSMUSG00 | Magee1 | 3848 | 2.0869561 | 4.83367117 | 2.98661101 | 4.38835677 | 3.59359352 | 2.92734564 |
| ENSMUSG00 | Atrx | 15029 | 22.9501995 | 22.5237711 | 18.6818888 | 26.0417817 | 23.4777689 | 30.2363939 |
| ENSMUSG00 | Fgf16 | 1642 | 6.36765614 | 3.99108169 | 6.72987866 | 3.49393732 | 2.08207502 | 8.38836347 |
| ENSMUSG00 | Cox7b | 1364 | 256.10807 | 212.281768 | 187.015276 | 159.618077 | 144.66202 | 573.567267 |
| ENSMUSG00 | Magt1 | 5253 | 36.8720162 | 37.5547866 | 50.0920562 | 53.4464855 | 39.3698651 | 49.611494 |
| ENSMUSG00 | Pgk2 | 1591 | 0 | 0 | 0 | 0 | 0 | 0 |
| ENSMUSG00 | Itm2a | 1738 | 8.60072536 | 16.468784 | 6.63790623 | 4.46354406 | 7.5159679 | 5.99477111 |
| ENSMUSG00 | Tbx22 | 5539 | 0 | 0 | 0 | 0 | 0 | 0 |
| ENSMUSG00 | 2610002M06 | 5834 | 8.69523365 | 7.83834995 | 6.90985273 | 8.91227737 | 8.30031379 | 8.3731366 |
| ENSMUSG00 | Hmgn5 | 1928 | 32.1054373 | 28.791899 | 21.7570275 | 27.8101235 | 23.3694449 | 40.8412259 |
| ENSMUSG00 | Sh3bgrl | 3418 | 59.4354902 | 73.6189587 | 75.0060531 | 74.6131921 | 62.5961363 | 101.262485 |
| ENSMUSG00 | Tnmd | 1346 | 0 | 0 | 0 | 0 | 0 | 0 |
| ENSMUSG00 | Srpx2 | 3240 | 0.17178294 | 0 | 0 | 0 | 0 | 0 |
| ENSMUSG00 | Sytl4 | 5107 | 3.02817347 | 4.13269271 | 3.53995571 | 5.21412744 | 5.40537876 | 3.15631717 |
| ENSMUSG00 | Cstf2 | 8819 | 7.7356168 | 6.31630878 | 8.3351474 | 7.69182943 | 6.82164612 | 7.261691 |
| ENSMUSG00 | Nox1 | 4034 | 0.12811632 | 0.03583523 | 0.13148788 | 0.07155586 | 0.02529815 | 0.07437309 |
| ENSMUSG00 | Xkrx | 3207 | 0.52065176 | 0.57096501 | 1.21289747 | 0.93383527 | 0.71599218 | 0.91000511 |
| ENSMUSG00 | Cenpi | 5731 | 0.00693692 | 0.0252241 | 0.00771276 | 0 | 0.07122858 | 0 |
| ENSMUSG00 | Btk | 3564 | 0 | 0 | 0 | 0 | 0 | 0 |
| ENSMUSG00 | Gla | 3257 | 3.25904615 | 6.36173476 | 4.70925377 | 3.25702213 | 2.75733563 | 2.16890745 |
| ENSMUSG00 | 4930513O06I | 800 | 0 | 0 | 0 | 0 | 0 | 0 |
| ENSMUSG00 | Serpina7 | 2442 | 0.56979599 | 1.12474496 | 0.3982148 | 0.02955122 | 0.75223159 | 0 |
| ENSMUSG00 | Col4a6 | 8405 | 0.65273722 | 0.73956586 | 0.52064039 | 0 | 0.3763992 | 0.32774998 |
| ENSMUSG00 | Col4a5 | 6898 | 5.87860106 | 6.28701043 | 4.35097875 | 8.8348178 | 11.7172774 | 6.40151362 |
| ENSMUSG00 | Acsl4 | 5551 | 84.1160365 | 64.8359821 | 118.551079 | 76.3306519 | 64.272459 | 116.188686 |
| ENSMUSG00 | Chrdl1 | 7842 | 0.32952133 | 0.946278 | 0.45092418 | 0.17484286 | 0.48801048 | 0.13216479 |
| ENSMUSG00 | Pak3 | 9586 | 0.22395117 | 0.7188255 | 0 | 0.31994302 | 0.48439393 | 0.12519133 |
| ENSMUSG00 | Dcx | 20403 | 0 | 0 | 0 | 0 | 0 | 0 |
| ENSMUSG00 | Gtl28d2 | 4639 | 3.97640504 | 6.66861303 | 6.40302619 | 5.35902752 | 3.77280597 | 6.09108132 |
| ENSMUSG00 | Il13ra2 | 2466 | 0.49976476 | 0 | 0 | 0 | 0 | 0 |
| ENSMUSG00 | Lrch2 | 9094 | 0 | 0 | 0.12151375 | 0 | 0 | 0.82177822 |
| ENSMUSG00 | Cdkl5 | 3463 | 1.04468632 | 2.43506422 | 1.44233562 | 1.16696188 | 1.4882079 | 1.13414613 |
| ENSMUSG00 | Rs1 | 1517 | 0 | 0 | 0.02913767 | 2.68771986 | 0 | 0 |
| ENSMUSG00 | D630029K05I | 2848 | 0.39085444 | 0.25379095 | 0 | 0.20270812 | 0.10749939 | 0 |
| ENSMUSG00 | Phka2 | 6153 | 3.12719853 | 2.85845139 | 1.94680635 | 3.93483697 | 3.64059467 | 2.45130432 |
| ENSMUSG00 | Slc7a3 | 4051 | 0.16683366 | 0.6661172 | 0.19640414 | 0.82834612 | 0.44085983 | 0.1211907 |
| ENSMUSG00 | Gpr64 | 5273 | 5.96369905 | 2.75063893 | 6.97438524 | 4.7967975 | 5.45778038 | 6.73460654 |
| ENSMUSG00 | Pdha1 | 3762 | 197.794612 | 170.919693 | 203.126385 | 173.389474 | 177.371589 | 166.3956 |
| ENSMUSG00 | Nlgn3 | 10994 | 0.01446443 | 0 | 0.04824651 | 0.03610174 | 0.11603233 | 0 |
| ENSMUSG00 | Map3k15 | 4319 | 0.54308251 | 0.40164666 | 0.07163994 | 0.42606721 | 0.42531825 | 0.03789021 |
| ENSMUSG00 | Il2rg | 2188 | 0.43607474 | 0.37439192 | 0.26262521 | 0 | 0.58302532 | 0.28670782 |
| ENSMUSG00 | Rps6ka3 | 8294 | 4.19891499 | 3.71826899 | 3.5919993 | 3.37589421 | 2.8054048 | 4.30132965 |
| ENSMUSG00 | Zmym3 | 6980 | 4.39702449 | 6.30289709 | 5.28775629 | 3.97005877 | 7.08374759 | 5.74408592 |
| ENSMUSG00 | Nono | 3885 | 30.7606112 | 23.4172465 | 31.7092758 | 27.1010054 | 26.8331752 | 27.2535778 |
| ENSMUSG00 | Itgb1bp2 | 1548 | 0.15409101 | 0.15564096 | 0.54252908 | 0.74588206 | 0 | 0 |
| ENSMUSG00 | Taf1 | 7433 | 6.40216739 | 7.38387698 | 6.33918534 | 10.5386948 | 7.87397458 | 9.50374218 |
| ENSMUSG00 | Rps4x | 1521 | 4.07748518 | 4.0551378 | 4.4172782 | 4.00911606 | 1.77803939 | 3.47881658 |
| ENSMUSG00 | Dmrtc1a | 1922 | 0.64121743 | 0.50141979 | 1.10389618 | 1.23902963 | 2.09733803 | 3.59026209 |
| ENSMUSG00 | 4930519F16I | 3272 | 0 | 0 | 0 | 0 | 0 | 0 |
| ENSMUSG00 | Cdx4 | 1960 | 0 | 0 | 0 | 0 | 0 | 0 |
| ENSMUSG00 | Chic1 | 2263 | 40.9325098 | 40.0737446 | 71.4690869 | 39.3984675 | 34.5211584 | 53.8020281 |
| ENSMUSG00 | Flna | 9633 | 7.16036112 | 12.6406249 | 7.95660706 | 12.4206436 | 13.7352219 | 5.04834234 |
| ENSMUSG00 | Tsx | 805 | 0.19754276 | 0.05985893 | 0.21963649 | 0.22411208 | 0.25354721 | 0.57598617 |
| ENSMUSG00 | Zcchc13 | 1078 | 0 | 0 | 0 | 0 | 0 | 0 |
| ENSMUSG00 | Abcb7 | 5889 | 13.6433734 | 13.9101633 | 13.0901765 | 19.5819688 | 10.0943671 | 12.384511 |
| ENSMUSG00 | Mtm1 | 13244 | 1.46186341 | 3.01620058 | 2.36962463 | 3.62890995 | 2.95509138 | 3.33622135 |
| ENSMUSG00 | Gabre | 4726 | 0.26077442 | 0.43842933 | 0 | 0.1832351 | 0.30231454 | 0.1038814 |
| ENSMUSG00 | Gpm6b | 5754 | 1.86548136 | 5.82859988 | 3.79487495 | 5.08559925 | 3.16587003 | 3.74943338 |
| ENSMUSG00 | Gabra3 | 4322 | 0 | 0 | 0 | 0 | 0 | 0 |
| ENSMUSG00 | Gabrq | 5510 | 0 | 0 | 0 | 0 | 0 | 0 |
| ENSMUSG00 | Cetn2 | 2228 | 34.0455372 | 38.84329 | 35.0757893 | 33.782381 | 30.1394572 | 42.7727063 |
| ENSMUSG00 | Nsdhl | 2845 | 18.9345083 | 17.5469786 | 17.4942971 | 13.03773 | 16.3392015 | 12.7409475 |
| ENSMUSG00 | Zfp185 | 3846 | 6.06772414 | 2.66867191 | 9.41271696 | 2.89894742 | 12.604019 | 5.82936908 |
| ENSMUSG00 | Hccs | 5140 | 9.54440913 | 6.09361622 | 9.46813797 | 9.45574202 | 5.54936655 | 8.70772043 |
| ENSMUSG00 | Rbbp7 | 3183 | 115.33211 | 141.380202 | 117.635505 | 145.382413 | 111.04609 | 112.029103 |
| ENSMUSG00 | Amelx | 990 | 0 | 0 | 0 | 0.03644651 | 0 | 0 |
| ENSMUSG00 | Arhgap6 | 9898 | 1.84358108 | 3.33965441 | 2.68837235 | 3.36469257 | 4.01076181 | 1.80765441 |
| ENSMUSG00 | Syap1 | 2441 | 55.6348715 | 48.9168389 | 60.2818252 | 57.7519656 | 42.3303609 | 64.5384336 |
| ENSMUSG00 | Msl3 | 3413 | 20.2213637 | 13.765538 | 19.7762128 | 14.504707 | 20.0188448 | 13.8810572 |
| ENSMUSG00 | Ctps2 | 4822 | 19.3665748 | 17.3479186 | 21.7984203 | 17.3451222 | 22.1904417 | 20.4701202 |
| ENSMUSG00 | Xlr4c | 2216 | 0 | 0 | 0 | 0 | 0 | 0 |
| ENSMUSG00 | Grpr | 2520 | 0.03155197 | 1.01344501 | 0.05262124 | 0.12886445 | 0.48596548 | 0.2489352 |
| ENSMUSG00 | Zfp275 | 6870 | 1.23838032 | 0.41382825 | 1.16456093 | 0.96113743 | 1.12154043 | 2.02078477 |
| ENSMUSG00 | Ap1s2 | 5088 | 2.00808933 | 4.17653714 | 2.99717686 | 5.23359844 | 2.4470196 | 1.29726075 |
| ENSMUSG00 | Znrf2 | 7583 | 6.17067131 | 6.96457083 | 6.45278125 | 9.06452523 | 6.41278335 | 8.65753088 |
| ENSMUSG00 | Haus7 | 1675 | 22.1681306 | 26.0062947 | 26.8113866 | 15.4021862 | 18.3999588 | 22.3407791 |
| ENSMUSG00 | Trex2 | 1015 | 0 | 0 | 0 | 0 | 0 | 0 |
| ENSMUSG00 | Car5b | 6346 | 4.01564182 | 6.85665888 | 3.17618034 | 4.38943246 | 4.24549739 | 4.28933028 |
| ENSMUSG00 | Zfp92 | 6737 | 0.00590107 | 0.00715251 | 0 | 0 | 0 | 0 |
| ENSMUSG00 | Bgn | 3458 | 0.26442338 | 0.66886906 | 0.08947742 | 0.27129357 | 0.14756037 | 0.32338352 |
| ENSMUSG00 | Atp2b3 | 4797 | 0 | 0 | 0 | 0 | 0 | 0.0568577 |
| ENSMUSG00 | Bmx | 3527 | 0.47341372 | 0 | 0 | 0.85933988 | 0.34721662 | 1.14449861 |
| ENSMUSG00 | Abcd1 | 4182 | 7.88074925 | 7.18994259 | 6.69052287 | 6.37604756 | 7.32085736 | 6.24147053 |
| ENSMUSG00 | Pir | 5585 | 0.65487989 | 1.99302921 | 0.28491788 | 1.34378966 | 0.71263336 | 0.32231445 |
| ENSMUSG00 | Figf | 2550 | 3.18043844 | 2.49435701 | 0.9880412 | 3.59405463 | 3.16163427 | 5.35866442 |
| ENSMUSG00 | Piga | 6394 | 3.6186565 | 3.08984068 | 1.39642982 | 4.37905326 | 5.05156332 | 3.34427857 |
| ENSMUSG00 | Asb11 | 3054 | 0.23431521 | 0.18933769 | 0.94077269 | 0 | 0.46782532 | 1.01811031 |
| ENSMUSG00 | Dusp9 | 2718 | 0.67283006 | 0.5850451 | 0.19515163 | 0.31860525 | 0.39424352 | 0.30104458 |
| ENSMUSG00 | Asb9 | 1132 | 2.52861714 | 2.17094395 | 4.1780894 | 1.97622505 | 2.4341204 | 2.240761 |
| ENSMUSG00 | Plxnb3 | 6366 | 0 | 0 | 0 | 0 | 0.21641724 | 0.09425731 |
| ENSMUSG00 | Hcfc1 | 10017 | 7.06049714 | 5.2145456 | 6.10715291 | 11.0619905 | 8.68523218 | 7.19917602 |
| ENSMUSG00 | Renbp | 1919 | 6.54649914 | 2.8876711 | 10.6876786 | 3.49726956 | 3.77579227 | 3.21212531 |
| ENSMUSG00 | Naa10 | 4443 | 4.57237239 | 3.64407935 | 4.17843216 | 2.87486918 | 3.7325168 | 2.75017744 |
| ENSMUSG00 | Arhgap4 | 8213 | 2.86560873 | 3.43224993 | 1.48541443 | 3.14996489 | 4.01973276 | 1.93609092 |
| ENSMUSG00 | Avpr2 | 3048 | 0.36520783 | 0.2687564 | 0 | 0.16573118 | 0.20089124 | 0.05369024 |
| ENSMUSG00 | L1cam | 8730 | 24.3633243 | 43.3346801 | 9.1694774 | 28.8945674 | 44.6086255 | 12.2282861 |
| ENSMUSG00 | Irak1 | 5383 | 22.6140222 | 27.1143848 | 20.9718572 | 25.7326712 | 24.712198 | 25.3948527 |
| ENSMUSG00 | Mecp2 | 20111 | 2.44728183 | 2.04860067 | 3.07265559 | 2.83474866 | 1.63144347 | 3.50172137 |
| ENSMUSG00 | Opn1mw | 1691 | 0 | 0 | 0 | 0 | 0 | 0 |
| ENSMUSG00 | Tktl1 | 2593 | 0.18398217 | 0 | 0 | 0 | 0 | 0 |
| ENSMUSG00 | Plxna3 | 7852 | 1.35184836 | 1.05553592 | 1.51993092 | 1.88405989 | 2.5539182 | 0.96565839 |
| ENSMUSG00 | Fam3a | 2650 | 12.8867765 | 11.9647845 | 10.4916827 | 11.1105465 | 10.2822961 | 11.6303179 |
| ENSMUSG00 | G6pdx | 3067 | 53.0159489 | 75.649729 | 52.2870187 | 73.1640805 | 53.7049694 | 32.8505111 |
| ENSMUSG00 | Mpp1 | 4495 | 3.37855252 | 3.63408318 | 3.45157884 | 4.54336756 | 2.57686036 | 3.34334288 |
| ENSMUSG00 | Dkc1 | 3641 | 13.3100606 | 22.260257 | 15.854877 | 19.5621961 | 14.9533488 | 20.5702168 |
| ENSMUSG00 | Tceal6 | 1111 | 0 | 0 | 0 | 0 | 0 | 0 |
| ENSMUSG00 | Nxf7 | 2724 | 0 | 0 | 0 | 0 | 0 | 0 |
| ENSMUSG00 | Prame | 2771 | 0 | 0 | 0 | 0 | 0 | 0 |
| ENSMUSG00 | Morf4l2 | 4675 | 30.5798306 | 27.0050221 | 20.3375753 | 25.8710189 | 20.9017133 | 27.9689028 |
| ENSMUSG00 | Kir3dl1 | 1599 | 0 | 0 | 0 | 0 | 0 | 0 |
| ENSMUSG00 | Plp1 | 5672 | 0.83408007 | 2.18334197 | 1.09101869 | 1.61580383 | 1.38541288 | 1.60608771 |
| ENSMUSG00 | Zcchc18 | 4897 | 0.33285169 | 1.3972789 | 0.73113116 | 0.25051859 | 1.23997115 | 0.52911798 |
| ENSMUSG00 | Psmd10 | 1464 | 36.1981254 | 29.9190409 | 33.6948475 | 31.7443124 | 25.966291 | 33.0686371 |
| ENSMUSG00 | Vsig1 | 2941 | 5.9748121 | 4.07970897 | 7.12399646 | 5.27551144 | 5.48260277 | 7.72518684 |
| ENSMUSG00 | Tsc22d3 | 4907 | 12.7198093 | 18.6775246 | 13.0164384 | 12.3974581 | 9.24443613 | 10.4162776 |
| ENSMUSG00 | Prps1 | 1986 | 21.999634 | 28.3635201 | 21.6112737 | 18.8040867 | 23.5605169 | 23.8412758 |
| ENSMUSG00 | Rbm41 | 6460 | 1.12620014 | 2.02144363 | 2.40167909 | 2.5916515 | 2.30645538 | 2.51635991 |
| ENSMUSG00 | Morc4 | 7320 | 1.17854362 | 1.30998661 | 2.35501623 | 1.513277 | 2.35613593 | 1.12153913 |
| ENSMUSG00 | Rnf128 | 2051 | 260.087293 | 284.184888 | 257.775838 | 306.02007 | 280.906294 | 277.493855 |
| ENSMUSG00 | Atp11a | 11182 | 6.26090155 | 8.7435424 | 4.76330002 | 7.40549921 | 7.7894852 | 6.46864097 |
| ENSMUSG00 | Mcf2l | 12512 | 0.04766082 | 0.03851218 | 0.04945858 | 0.0028838 | 0.26508268 | 0 |
| ENSMUSG00 | F7 | 1928 | 0 | 0 | 0 | 0 | 0 | 0 |
| ENSMUSG00 | F10 | 2552 | 0 | 0 | 0 | 0 | 0.03998932 | 0 |
| ENSMUSG00 | Proz | 2733 | 0.08727877 | 0 | 0 | 0 | 0 | 0 |
| ENSMUSG00 | Cul4a | 6550 | 24.7212324 | 24.0049406 | 25.8800104 | 20.1398404 | 25.1003027 | 16.2856663 |
| ENSMUSG00 | Lamp1 | 2246 | 568.47151 | 538.546781 | 657.418774 | 505.261079 | 457.488036 | 494.586602 |
| ENSMUSG00 | Adprhl1 | 1308 | 0 | 0 | 0.27034766 | 0 | 0 | 0 |
| ENSMUSG00 | Atp4b | 1897 | 0 | 0 | 0 | 0 | 0 | 0 |
| ENSMUSG00 | Grk1 | 7418 | 0 | 0 | 0 | 0 | 0 | 0.19119456 |
| ENSMUSG00 | Gas6 | 2578 | 144.541551 | 122.484777 | 86.3976288 | 118.701249 | 145.142131 | 70.5247266 |
| ENSMUSG00 | 1700029H14I | 1519 | 0 | 0 | 0 | 0 | 0 | 0 |
| ENSMUSG00 | Rasa3 | 5068 | 0 | 0.48490698 | 0 | 0 | 0.49334894 | 0.28523202 |
| ENSMUSG00 | 2410022L05I | 798 | 13.5507402 | 17.1490596 | 15.4540278 | 14.830715 | 9.20776707 | 20.6781413 |
| ENSMUSG00 | Myom2 | 8813 | 0.20299519 | 0 | 0 | 0 | 0 | 0 |
| ENSMUSG00 | Angpt2 | 3801 | 13.4087151 | 10.3827141 | 2.8258479 | 1.77514926 | 11.2362738 | 0.33007982 |
| ENSMUSG00 | Agpat5 | 11274 | 5.51512964 | 4.0775116 | 5.79085703 | 5.1111429 | 4.42192382 | 5.5400855 |
| ENSMUSG00 | Defb8 | 285 | 0 | 0 | 0 | 0 | 0 | 0 |
| ENSMUSG00 | Nek3 | 2241 | 0.79830282 | 3.18232639 | 0.31558656 | 0.85334599 | 1.5255543 | 0.03651224 |
| ENSMUSG00 | Vps36 | 3816 | 39.8909159 | 39.6755244 | 32.2131356 | 34.4651605 | 40.6231996 | 30.2694175 |
| ENSMUSG00 | Thsd1 | 4501 | 0.53878789 | 0 | 0.08838405 | 0.23247707 | 0.7368839 | 0.72110244 |
| ENSMUSG00 | Tpte | 2727 | 0 | 0 | 0.03241793 | 0 | 0 | 0 |
| ENSMUSG00 | Slc25a15 | 3444 | 8.09192562 | 11.0392285 | 6.75092027 | 11.5663697 | 9.57114947 | 6.28013616 |
| ENSMUSG00 | Erlin2 | 3862 | 79.3359628 | 63.0341548 | 85.2218858 | 70.5384351 | 76.9095116 | 65.7924752 |
| ENSMUSG00 | Prosc | 4526 | 18.6040892 | 14.0534917 | 18.1163099 | 16.1037848 | 20.8006327 | 13.4866643 |
| ENSMUSG00 | Gpr124 | 5480 | 0.03627325 | 0.05275888 | 0.11292442 | 0.03292157 | 0.20485041 | 0.09954248 |
| ENSMUSG00 | Brf2 | 1379 | 2.3063368 | 2.65567049 | 3.17330128 | 1.85774123 | 3.51523256 | 3.59969858 |
| ENSMUSG00 | Rab11fip1 | 8772 | 6.15004412 | 6.59734576 | 7.45767529 | 6.22756674 | 8.37642284 | 7.21664816 |
| ENSMUSG00 | Adrb3 | 3356 | 0 | 0 | 0 | 0 | 0 | 0 |
| ENSMUSG00 | Eif4ebp1 | 978 | 21.381782 | 20.3486714 | 22.1913138 | 16.4545929 | 13.0435521 | 24.9599204 |
| ENSMUSG00 | Chrna6 | 2898 | 0 | 0 | 0 | 0 | 0 | 0 |
| ENSMUSG00 | Chrnb3 | 4582 | 0 | 0 | 0 | 0 | 0 | 0 |
| ENSMUSG00 | Ggn | 1774 | 0.13446048 | 0.32595113 | 0 | 0 | 0.05752692 | 0.16912121 |
| ENSMUSG00 | Cd209a | 1477 | 0 | 0 | 0 | 0 | 0 | 0 |
| ENSMUSG00 | Cd209d | 909 | 0 | 0 | 0 | 0 | 0 | 0 |
| ENSMUSG00 | Tnfsf13b | 969 | 0 | 0 | 0.31931156 | 0 | 0.1053176 | 0.14073601 |
| ENSMUSG00 | Col4a1 | 6766 | 15.5590471 | 22.1916869 | 9.18530761 | 19.6088794 | 25.0229848 | 13.0890857 |
| ENSMUSG00 | Col4a2 | 6571 | 9.72862771 | 8.81450365 | 5.43525935 | 12.4153862 | 13.4036627 | 4.2711313 |
| ENSMUSG00 | Rab20 | 1444 | 53.989264 | 72.8803252 | 47.7526836 | 61.3444732 | 51.9097965 | 52.9815534 |
| ENSMUSG00 | Carkd | 1312 | 79.9048797 | 69.0843732 | 101.138668 | 67.6263319 | 62.6939922 | 75.8368018 |
| ENSMUSG00 | Ptpn7 | 3373 | 0 | 0 | 0 | 0 | 0 | 0 |
| ENSMUSG00 | Ankrd10 | 3944 | 24.8975753 | 24.8751512 | 32.8936129 | 23.4935827 | 24.905115 | 35.0476348 |
| ENSMUSG00 | 1700016D06I | 1260 | 0 | 0 | 0 | 0 | 0 | 0 |
| ENSMUSG00 | 1700128E19I | 2437 | 0 | 0 | 0 | 0 | 0 | 0 |
| ENSMUSG00 | Arhgef7 | 10957 | 1.45132721 | 1.82067966 | 1.10938275 | 0.74093822 | 1.13164272 | 1.15749815 |
| ENSMUSG00 | 1700018L24I | 1132 | 0 | 0 | 0.07809513 | 0.12749839 | 0 | 0 |
| ENSMUSG00 | Leprotl1 | 2875 | 22.1109612 | 27.5375042 | 37.8982758 | 26.7446389 | 30.3850975 | 42.6907397 |
| ENSMUSG00 | Dctn6 | 3195 | 26.229907 | 30.5557846 | 25.4973386 | 23.9756434 | 24.8663747 | 30.7234512 |
| ENSMUSG00 | Gpm6a | 3330 | 2.78168978 | 0.86822418 | 2.99988484 | 1.45195014 | 2.91141484 | 4.61948845 |
| ENSMUSG00 | Spata4 | 1051 | 0 | 0 | 0 | 0 | 0 | 0 |
| ENSMUSG00 | Asb5 | 1572 | 0.22760771 | 1.22611812 | 0 | 0 | 0.09737858 | 0.20820335 |
| ENSMUSG00 | Vegfc | 2207 | 0.64848088 | 0.34933533 | 0.32044834 | 1.91282248 | 0.36992389 | 0.34603077 |
| ENSMUSG00 | Aga | 1257 | 76.4114884 | 53.7066074 | 65.6873848 | 70.7288446 | 52.9342833 | 97.7503976 |
| ENSMUSG00 | Dlc1 | 14388 | 0.0386834 | 0.14735915 | 0.22119354 | 0.15046724 | 0.20924077 | 0.19525214 |
| ENSMUSG00 | Eri1 | 5060 | 8.28893912 | 6.55183245 | 8.21140957 | 11.1526319 | 7.22033302 | 7.67570875 |
| ENSMUSG00 | Tnks | 6417 | 3.79774187 | 2.90605472 | 4.73222162 | 5.57227772 | 7.17247795 | 3.54056015 |
| ENSMUSG00 | Dusp4 | 2736 | 0.43591536 | 1.02149622 | 0.45235804 | 1.18690936 | 0.67139968 | 0.46853364 |
| ENSMUSG00 | Tmem66 | 1891 | 163.100485 | 152.917419 | 154.975261 | 165.145476 | 159.042548 | 119.108393 |
| ENSMUSG00 | Mrps31 | 1529 | 22.5428395 | 23.0059534 | 23.0693726 | 19.7047137 | 13.3155814 | 18.516073 |
| ENSMUSG00 | AI316807 | 1453 | 30.4801138 | 29.3827858 | 31.5466698 | 25.0314528 | 20.6493524 | 55.0748738 |
| ENSMUSG00 | Dkk4 | 1300 | 0 | 0 | 0 | 0 | 0 | 0.04196098 |
| ENSMUSG00 | Polb | 1213 | 54.2418139 | 63.2422225 | 60.2719277 | 54.2271782 | 48.0817375 | 58.5966281 |
| ENSMUSG00 | Ikbkb | 14252 | 7.40324483 | 7.35374064 | 6.47272287 | 8.47619177 | 11.8221368 | 6.77847125 |
| ENSMUSG00 | Plat | 2517 | 8.3080583 | 5.30299741 | 14.1895467 | 6.07818311 | 16.5830653 | 6.81595077 |
| ENSMUSG00 | Ap3m2 | 3458 | 4.6331575 | 4.6542139 | 2.13467547 | 4.30939398 | 4.70717579 | 2.90256427 |
| ENSMUSG00 | Myst3 | 9244 | 3.32442519 | 3.63327025 | 3.8253434 | 6.28039914 | 3.57692466 | 3.40785464 |
| ENSMUSG00 | Ank1 | 9802 | 0.00405585 | 0.00983196 | 0.04058525 | 0 | 0.1457599 | 0.00278256 |
| ENSMUSG00 | Agpat6 | 4856 | 27.0658193 | 30.1760648 | 21.1360543 | 23.4577872 | 31.4396451 | 18.8608399 |
| ENSMUSG00 | Gins4 | 1356 | 26.2691081 | 29.9210799 | 37.8779805 | 26.555959 | 30.329837 | 30.1911752 |
| ENSMUSG00 | Sfrp1 | 4369 | 44.4234964 | 39.3631064 | 8.76145481 | 40.3682842 | 36.1120517 | 18.8968489 |
| ENSMUSG00 | Ido2 | 5683 | 0 | 0 | 0 | 0.02539648 | 0 | 0 |
| ENSMUSG00 | Ido1 | 1814 | 0 | 0 | 0 | 0 | 0 | 0 |
| ENSMUSG00 | Adam18 | 2476 | 0 | 0 | 0 | 0 | 0 | 0 |
| ENSMUSG00 | Adam3 | 3075 | 0 | 0 | 0 | 0 | 0 | 0 |
| ENSMUSG00 | Adam5 | 3335 | 0 | 0 | 0.27833245 | 0.02163841 | 0 | 0 |
| ENSMUSG00 | Adam9 | 5345 | 38.334845 | 36.6739844 | 37.1560114 | 41.8065672 | 31.8091458 | 43.1342424 |
| ENSMUSG00 | Tm2d2 | 1276 | 87.0819499 | 118.086994 | 116.220363 | 89.0457353 | 62.0234396 | 95.1192356 |
| ENSMUSG00 | Plekha2 | 7581 | 1.95080316 | 2.34544218 | 1.25941144 | 0.82815931 | 1.19808665 | 0.93182055 |
| ENSMUSG00 | Slit2 | 15687 | 7.38493467 | 9.54083568 | 6.66113067 | 14.1158607 | 15.2067513 | 9.62184316 |
| ENSMUSG00 | 4930555F03I | 4553 | 0 | 0 | 0 | 0 | 0 | 0 |
| ENSMUSG00 | Odz3 | 12318 | 0.34210756 | 0.10953242 | 0.12200541 | 0.29877972 | 0.3479636 | 0.1549947 |
| ENSMUSG00 | Dctd | 3133 | 0 | 0.53831008 | 0.02821694 | 0.21881866 | 0.48860239 | 0.04352799 |
| ENSMUSG00 | Wwc2 | 6523 | 27.0298262 | 28.1007552 | 29.510812 | 33.2886316 | 37.8923447 | 25.8028206 |
| ENSMUSG00 | Fgfr1 | 8042 | 4.63698591 | 4.82343769 | 5.68325114 | 5.90001101 | 5.19019839 | 4.48020374 |
| ENSMUSG00 | Rwdd4a | 3100 | 34.8309307 | 30.0621223 | 26.3072259 | 29.4243253 | 27.965746 | 34.2692645 |
| ENSMUSG00 | Ppapdc1b | 2555 | 15.0775188 | 21.2359819 | 16.9541316 | 18.4152587 | 13.3207799 | 19.17231 |
| ENSMUSG00 | Star | 3497 | 1.34147747 | 0.99211434 | 1.54207173 | 1.03179996 | 3.64786787 | 3.58774208 |
| ENSMUSG00 | Ash2l | 5248 | 15.3022238 | 19.7777432 | 20.4164001 | 14.8508415 | 20.3113756 | 14.1778232 |
| ENSMUSG00 | Kcnu1 | 3630 | 0.18618269 | 0 | 0 | 0.0695797 | 0 | 0.05259572 |
| ENSMUSG00 | Tti2 | 5068 | 2.4631454 | 1.29308527 | 3.27937904 | 3.22517091 | 2.85941017 | 2.07735019 |
| ENSMUSG00 | Mak16 | 1847 | 25.5709317 | 23.3235946 | 22.615453 | 26.0994107 | 27.7095046 | 27.6881151 |
| ENSMUSG00 | Wrn | 6372 | 6.52608798 | 5.21036702 | 3.97483617 | 8.22774807 | 7.55946308 | 3.75390123 |
| ENSMUSG00 | Gsr | 2734 | 50.0796909 | 89.5168031 | 77.4906487 | 81.5475322 | 41.6758951 | 54.7587765 |
| ENSMUSG00 | Gtf2e2 | 1597 | 27.5325995 | 34.8499315 | 32.4386725 | 35.6979526 | 19.6820586 | 44.7290421 |
| ENSMUSG00 | Rbpms | 4305 | 47.7527502 | 44.9516264 | 49.0892013 | 50.0874095 | 41.769326 | 49.9053039 |
| ENSMUSG00 | Frg1 | 1076 | 67.2444001 | 52.8886507 | 65.8508869 | 61.0309675 | 39.4079166 | 84.5868691 |
| ENSMUSG00 | Asah1 | 4313 | 91.6781844 | 61.9619773 | 86.6614386 | 87.5907735 | 75.0547944 | 109.623434 |
| ENSMUSG00 | Pcm1 | 8458 | 10.2514428 | 5.69144664 | 6.83565981 | 9.52176914 | 8.98300705 | 10.7318514 |
| ENSMUSG00 | Fgl1 | 3949 | 0.28188236 | 0.23184158 | 0.01119317 | 0.3380693 | 0.06460671 | 0.18648145 |
| ENSMUSG00 | Pdgfrl | 3312 | 0 | 0 | 0.0400379 | 0 | 0 | 0 |
| ENSMUSG00 | Slc7a2 | 9265 | 0.09440049 | 0.07801367 | 0 | 0.22198344 | 0 | 0.32970962 |
| ENSMUSG00 | Vps37a | 6431 | 17.9582757 | 20.4704338 | 18.2209743 | 24.0023304 | 14.5676296 | 23.5466952 |
| ENSMUSG00 | Cnot7 | 6589 | 25.9023035 | 27.841218 | 32.8310547 | 26.7288601 | 23.0311795 | 30.2219064 |
| ENSMUSG00 | Fgf20 | 1954 | 0 | 0 | 0 | 0 | 0 | 0 |
| ENSMUSG00 | Sc4mol | 7108 | 26.1251899 | 14.4871169 | 27.8158193 | 16.2694081 | 14.3359134 | 27.0866177 |
| ENSMUSG00 | Klhl2 | 3010 | 23.1136515 | 16.3769868 | 28.8560204 | 24.02273 | 21.3259737 | 22.1912264 |
| ENSMUSG00 | Galnt7 | 6650 | 2.21793733 | 2.71003449 | 5.98220432 | 2.58814064 | 2.78534954 | 4.5977249 |
| ENSMUSG00 | Sap30 | 1174 | 4.43608854 | 7.55221921 | 6.96536709 | 6.45419877 | 6.21530813 | 8.0848164 |
| ENSMUSG00 | Scrg1 | 697 | 0 | 0 | 0 | 0 | 0 | 0 |
| ENSMUSG00 | Hpgd | 1712 | 9.66021022 | 7.37432701 | 10.2242581 | 9.80032167 | 8.28582505 | 4.54046273 |
| ENSMUSG00 | Ednra | 5468 | 0.33444625 | 0 | 0 | 0.13197529 | 0.61589993 | 0 |
| ENSMUSG00 | Tmem184c | 3238 | 65.7598374 | 62.1602127 | 79.3530925 | 64.7314999 | 48.5049984 | 75.1105492 |
| ENSMUSG00 | Nr3c2 | 6461 | 22.446679 | 23.5525127 | 21.1055078 | 26.5938239 | 23.7164845 | 22.7999267 |
| ENSMUSG00 | 1700007B14I | 2191 | 0 | 0 | 0 | 0 | 0 | 0 |
| ENSMUSG00 | Isx | 2560 | 0 | 0 | 0 | 0 | 0 | 0 |
| ENSMUSG00 | Sin3b | 4278 | 21.4389653 | 26.8753742 | 24.2500527 | 26.8295894 | 16.913371 | 20.5739273 |
| ENSMUSG00 | Sorbs2 | 16542 | 6.56583078 | 10.262413 | 7.33223656 | 10.9192791 | 13.1468029 | 9.45923137 |
| ENSMUSG00 | Irf2 | 2475 | 5.38104484 | 5.56821109 | 5.35779915 | 5.61276248 | 7.87558609 | 12.0559416 |
| ENSMUSG00 | Casp3 | 1448 | 17.9558593 | 22.6622701 | 22.5282874 | 22.4266851 | 17.7253225 | 21.2094225 |
| ENSMUSG00 | Mlf1ip | 3670 | 0.9857626 | 0.27572624 | 0.24088198 | 0.42275965 | 0.97325513 | 0.54252007 |
| ENSMUSG00 | 4933411K20I | 5003 | 13.4610802 | 16.4987758 | 13.7915405 | 15.5997326 | 14.7275808 | 10.20005 |
| ENSMUSG00 | Slc25a4 | 2349 | 451.205241 | 355.151925 | 465.615327 | 351.404399 | 264.059866 | 477.010102 |
| ENSMUSG00 | Ufsp2 | 2609 | 63.1150634 | 54.1520308 | 54.7398064 | 46.5374012 | 56.7177041 | 70.115372 |
| ENSMUSG00 | Anxa10 | 2062 | 0 | 0 | 0 | 0 | 0 | 0 |
| ENSMUSG00 | Pdlim3 | 1571 | 20.0675341 | 9.50846408 | 22.3119424 | 14.3776957 | 23.9054186 | 17.6391047 |
| ENSMUSG00 | Lrp2bp | 4523 | 0.36037469 | 0.11719011 | 0.20522633 | 0.07179713 | 0.2039676 | 0.1809063 |
| ENSMUSG00 | Tlr3 | 5619 | 7.03273672 | 7.91530274 | 9.75465548 | 7.17916458 | 10.2888208 | 13.2562804 |
| ENSMUSG00 | Klkb1 | 2481 | 10.9603985 | 5.14687915 | 15.3218803 | 7.37347706 | 9.6047229 | 13.1261263 |
| ENSMUSG00 | Cbr4 | 2581 | 22.6734085 | 17.7361953 | 18.393173 | 11.8828895 | 15.7962318 | 24.8758236 |
| ENSMUSG00 | Sh3rf1 | 5745 | 3.93056796 | 4.36152305 | 2.78521622 | 6.07333978 | 4.36988284 | 3.03842805 |
| ENSMUSG00 | Nek1 | 8414 | 6.01482371 | 5.86438279 | 5.8312391 | 4.50703932 | 5.28214563 | 11.0246076 |
| ENSMUSG00 | F11 | 2229 | 0 | 0 | 0 | 0 | 0 | 0.1101264 |
| ENSMUSG00 | Mfap3l | 7030 | 5.42325829 | 12.1939802 | 6.09268647 | 11.943516 | 13.2392759 | 12.7566165 |
| ENSMUSG00 | Triml1 | 1844 | 0 | 0 | 0 | 0 | 0 | 0 |
| ENSMUSG00 | N4bp1 | 7167 | 3.023125 | 3.65079365 | 3.60793612 | 3.46371517 | 3.53845525 | 2.71338325 |
| ENSMUSG00 | Cbln1 | 3330 | 0 | 0 | 0 | 0 | 0 | 0 |
| ENSMUSG00 | Heatr3 | 4783 | 7.87130254 | 8.3820029 | 9.60186387 | 12.0927279 | 9.76147478 | 7.4929701 |
| ENSMUSG00 | Adcy7 | 6299 | 0 | 0 | 0.18244927 | 0.20621584 | 0.21871919 | 0.21216976 |
| ENSMUSG00 | Brd7 | 5610 | 21.861971 | 15.8645916 | 18.4450115 | 22.890552 | 24.3580454 | 19.1116946 |
| ENSMUSG00 | Nkd1 | 4448 | 0 | 0 | 0.15899944 | 0 | 0 | 0 |
| ENSMUSG00 | Snx20 | 1430 | 0 | 0 | 0 | 0 | 0.24977946 | 0 |
| ENSMUSG00 | Sall1 | 5248 | 6.2345199 | 8.86050241 | 4.19445843 | 11.0350003 | 7.26309122 | 5.17636065 |
| ENSMUSG00 | Rbl2 | 7253 | 21.3275299 | 19.4526269 | 27.9057272 | 26.4807284 | 24.728762 | 27.7522181 |
| ENSMUSG00 | Aktip | 2978 | 153.401688 | 158.377735 | 105.265101 | 140.753899 | 165.073239 | 145.238828 |
| ENSMUSG00 | Eif2ak3 | 5877 | 6.48723938 | 6.19036308 | 4.06894624 | 6.34827872 | 11.1308174 | 3.80554776 |
| ENSMUSG00 | Gins3 | 2409 | 0.89115648 | 2.36031556 | 1.68807371 | 1.5277578 | 3.38904945 | 2.45686871 |
| ENSMUSG00 | Setd6 | 1718 | 17.5636844 | 20.5872227 | 14.4337799 | 17.2219305 | 13.9891869 | 15.2407763 |
| ENSMUSG00 | Got2 | 2575 | 65.2453051 | 62.9511423 | 68.3712391 | 54.6905708 | 49.1834815 | 56.8053943 |
| ENSMUSG00 | Cdh11 | 4203 | 0.03783534 | 0.1719716 | 0.08413389 | 0.12877246 | 0.08498326 | 0.04542529 |
| ENSMUSG00 | Smad1 | 5806 | 13.1399874 | 13.3122723 | 17.3122616 | 14.8467111 | 14.8526654 | 16.3431743 |
| ENSMUSG00 | 1700011L22I | 972 | 0 | 0 | 0.22737574 | 0 | 0 | 0 |
| ENSMUSG00 | Lsm6 | 7487 | 18.2396254 | 13.2453183 | 19.6183684 | 15.6386049 | 15.5730291 | 25.6608199 |
| ENSMUSG00 | Slc10a7 | 3924 | 3.56624086 | 6.42240297 | 3.48072617 | 2.63902823 | 4.61630057 | 4.31639933 |
| ENSMUSG00 | Pou4f2 | 3203 | 0 | 0 | 0 | 0 | 0 | 0 |
| ENSMUSG00 | Tnpo2 | 4964 | 3.19549491 | 2.9315684 | 2.74258011 | 3.83789676 | 3.19685795 | 2.11537794 |
| ENSMUSG00 | Vps35 | 3163 | 125.324659 | 113.922926 | 141.842778 | 115.193325 | 102.649646 | 131.984075 |
| ENSMUSG00 | Orc6 | 1646 | 6.78693197 | 7.02596969 | 6.3644209 | 8.22039289 | 5.27003882 | 8.18570583 |
| ENSMUSG00 | Mylk3 | 5341 | 0 | 0.13532983 | 0 | 0 | 0.19107424 | 0 |
| ENSMUSG00 | Gpt2 | 5713 | 1.34304354 | 0.96153587 | 1.52420148 | 1.29473466 | 2.96529963 | 0.73044282 |
| ENSMUSG00 | Dnaja2 | 2830 | 130.603075 | 123.156373 | 120.500784 | 127.702388 | 108.291312 | 125.097109 |
| ENSMUSG00 | Itfg1 | 3337 | 111.093754 | 110.06205 | 122.790256 | 105.218572 | 100.875338 | 101.05593 |
| ENSMUSG00 | Rfx1 | 4151 | 2.22193965 | 4.26028048 | 2.25747789 | 5.3197329 | 3.82298311 | 2.94363729 |
| ENSMUSG00 | Tecr | 1321 | 174.219892 | 196.904175 | 179.450783 | 162.109716 | 123.606664 | 173.599672 |
| ENSMUSG00 | Tbc1d9 | 6215 | 0.62047974 | 1.27928608 | 0.61164256 | 0.58636951 | 0.70607696 | 0.25014553 |
| ENSMUSG00 | Ucp1 | 1638 | 0.31551969 | 0 | 0 | 0 | 0 | 0 |
| ENSMUSG00 | Zfp330 | 1911 | 20.6994783 | 19.5166438 | 20.4471111 | 25.2253331 | 22.9899056 | 20.15269 |
| ENSMUSG00 | Il15 | 1900 | 9.66685895 | 15.2167712 | 6.25805041 | 12.2489046 | 11.8703464 | 10.0198412 |
| ENSMUSG00 | Gab1 | 4849 | 13.5524457 | 10.1758954 | 15.0134946 | 10.2241141 | 13.5011013 | 17.2231275 |
| ENSMUSG00 | Smarca5 | 7753 | 10.9733946 | 9.34144493 | 10.5644286 | 12.10026 | 9.60899119 | 11.7429054 |
| ENSMUSG00 | Hp | 1338 | 0 | 0 | 0 | 0 | 0 | 0 |
| ENSMUSG00 | Txnl4b | 1886 | 1.93929173 | 1.40522498 | 1.14840419 | 1.68357366 | 1.21749041 | 3.06586613 |
| ENSMUSG00 | Ces1f | 3283 | 0.42383242 | 1.49711151 | 0.21542171 | 1.03311367 | 0 | 0.46523905 |
| ENSMUSG00 | Pmfbp1 | 3377 | 0 | 0 | 0 | 0 | 0 | 0 |
| ENSMUSG00 | Zfp821 | 2387 | 10.7424738 | 7.67107164 | 13.3142544 | 8.22313876 | 8.82861048 | 8.84397603 |
| ENSMUSG00 | 2400003C14I | 4978 | 45.5934187 | 41.4879653 | 41.0674014 | 45.9252379 | 47.4797454 | 42.8953698 |
| ENSMUSG00 | Dhodh | 5567 | 6.24147476 | 5.79068256 | 8.84513259 | 10.0786023 | 8.14845484 | 6.88847544 |
| ENSMUSG00 | Ap1g1 | 4875 | 19.5148441 | 16.6552113 | 17.644469 | 19.1993484 | 20.3896164 | 11.2007852 |
| ENSMUSG00 | Phlpp2 | 8536 | 0.11643475 | 0.24273864 | 0.4556891 | 0.98067413 | 0.62766746 | 0.60390192 |
| ENSMUSG00 | Irx3 | 3205 | 0.1984673 | 0.19545203 | 0.02758305 | 0.40528974 | 0.74828071 | 0.03404011 |
| ENSMUSG00 | 4933436C20I | 1231 | 0 | 0 | 0 | 0 | 0 | 0 |
| ENSMUSG00 | Irx5 | 2398 | 0.31499338 | 0.2009443 | 0.12902957 | 0.04514017 | 0.12767233 | 0.21610431 |
| ENSMUSG00 | Irx6 | 2734 | 0 | 0 | 0 | 0 | 0 | 0 |
| ENSMUSG00 | Mmp2 | 3025 | 0 | 0 | 0.11689744 | 0 | 0 | 0 |
| ENSMUSG00 | Gnao1 | 6529 | 0 | 0.00738037 | 0.07447086 | 0 | 0 | 0 |
| ENSMUSG00 | St3gal2 | 5178 | 3.18627354 | 3.87129392 | 2.31338344 | 4.0486033 | 4.99620945 | 2.10696325 |
| ENSMUSG00 | Il34 | 2558 | 3.24819993 | 3.20238279 | 1.41694727 | 2.27099655 | 2.17430609 | 1.23684838 |
| ENSMUSG00 | Amfr | 4349 | 67.8558133 | 57.5267895 | 70.8815022 | 51.9037182 | 68.5201276 | 38.9772092 |
| ENSMUSG00 | Cog4 | 3650 | 25.5306428 | 24.436467 | 29.8271615 | 20.9572423 | 27.1208682 | 18.8904899 |
| ENSMUSG00 | Nudt21 | 1104 | 1.11632237 | 3.14259405 | 2.16204667 | 2.61464091 | 1.61768402 | 2.91522412 |
| ENSMUSG00 | Bbs2 | 3415 | 9.94178048 | 8.63546196 | 7.90644102 | 11.5589332 | 12.058063 | 9.73580827 |
| ENSMUSG00 | Cenpn | 1756 | 0.18111836 | 1.45437439 | 0.95653191 | 1.10958451 | 1.65632313 | 0.13979029 |
| ENSMUSG00 | Mt4 | 388 | 0 | 0 | 0 | 0 | 0 | 0 |
| ENSMUSG00 | Cdyl2 | 2403 | 2.31617448 | 1.96515661 | 2.02338857 | 0.58560122 | 2.82418143 | 1.21447624 |
| ENSMUSG00 | Mt3 | 535 | 0 | 0.36027247 | 0 | 0 | 0.19075281 | 0.05098063 |
| ENSMUSG00 | Mt2 | 553 | 42.4154313 | 61.3440421 | 80.570448 | 39.3444355 | 41.7069112 | 149.541964 |
| ENSMUSG00 | Mt1 | 878 | 179.533577 | 214.204651 | 254.940926 | 149.136378 | 150.870697 | 475.038478 |
| ENSMUSG00 | Slc12a3 | 3282 | 112.216567 | 182.688574 | 35.5015412 | 171.032409 | 154.913714 | 48.9813296 |
| ENSMUSG00 | Nudt7 | 1573 | 25.0209318 | 18.0431115 | 18.4057261 | 16.3091759 | 16.0248122 | 30.9332209 |
| ENSMUSG00 | Herpud1 | 2566 | 57.8669991 | 44.8813705 | 57.5691969 | 42.8456702 | 40.1688539 | 42.9527737 |
| ENSMUSG00 | Cntnap4 | 6428 | 0 | 0 | 0 | 0 | 0 | 0 |
| ENSMUSG00 | Fam192a | 1789 | 33.6888253 | 35.2577154 | 37.7037521 | 28.6397447 | 31.631219 | 26.1007168 |
| ENSMUSG00 | Pllp | 1885 | 2.08795362 | 1.45709666 | 1.71179551 | 0.8422334 | 1.92194837 | 2.08357987 |
| ENSMUSG00 | Arl2bp | 2013 | 66.515876 | 65.1582193 | 72.8133688 | 55.5122165 | 62.179682 | 59.3186143 |
| ENSMUSG00 | Cx3cl1 | 3832 | 10.3331051 | 22.5213773 | 6.97863127 | 21.0635526 | 18.4557821 | 6.82577741 |
| ENSMUSG00 | Ccl22 | 2427 | 0 | 0 | 0 | 0 | 0 | 0 |
| ENSMUSG00 | Ccl17 | 510 | 0 | 0.18896644 | 0 | 0 | 0 | 0 |
| ENSMUSG00 | Ciapin1 | 5866 | 9.39329969 | 10.1038738 | 8.84639681 | 6.81536061 | 9.24668215 | 7.71371062 |
| ENSMUSG00 | Coq9 | 3137 | 84.5928698 | 82.6558002 | 97.1256946 | 78.1336718 | 85.7217088 | 79.2764299 |
| ENSMUSG00 | Polr2c | 4846 | 3.68349376 | 4.26578285 | 4.91638328 | 5.1450047 | 3.32734931 | 4.02421937 |
| ENSMUSG00 | Gpr56 | 3319 | 257.865618 | 338.031013 | 176.034939 | 296.190209 | 356.323686 | 186.271241 |
| ENSMUSG00 | Ccdc135 | 2814 | 0 | 0 | 0 | 0 | 0 | 0 |
| ENSMUSG00 | Katnb1 | 3405 | 7.250559 | 4.00492309 | 6.5037073 | 4.33408406 | 6.90841681 | 4.49370708 |
| ENSMUSG00 | Kifc3 | 3734 | 24.6262524 | 24.8029839 | 14.1104973 | 21.0752381 | 29.2711561 | 12.9653146 |
| ENSMUSG00 | Cngb1 | 5483 | 0 | 0 | 0 | 0 | 0.16751318 | 0 |
| ENSMUSG00 | Mmp15 | 4080 | 0.23385577 | 0.02362081 | 0.32501355 | 0.08843638 | 0.05002586 | 0.36767283 |
| ENSMUSG00 | Tmem38a | 2088 | 13.1566269 | 12.9697225 | 12.5111538 | 10.8868238 | 11.2658809 | 12.0698116 |
| ENSMUSG00 | AA960436 | 2923 | 14.7161922 | 11.1605273 | 12.9747456 | 11.0850756 | 17.3870237 | 12.0463768 |
| ENSMUSG00 | Gtl3 | 1286 | 29.0283019 | 31.6246945 | 38.805506 | 32.3223292 | 30.433305 | 37.3700734 |
| ENSMUSG00 | Tpm4 | 2194 | 109.789634 | 91.6508765 | 85.2809487 | 94.3494482 | 88.6100692 | 81.1401871 |
| ENSMUSG00 | Rhsr4 | 1375 | 4.51043997 | 4.13527285 | 3.05394552 | 6.2192324 | 3.52545869 | 4.26476177 |
| ENSMUSG00 | B3gnt3 | 3052 | 0.18236459 | 0 | 0.69517971 | 0.56747645 | 0.13375197 | 0.31278256 |
| ENSMUSG00 | Jak3 | 5050 | 5.96725826 | 4.33200886 | 5.1816814 | 4.8299925 | 6.17368626 | 3.98587797 |
| ENSMUSG00 | Pgls | 2786 | 16.4244645 | 13.7329631 | 17.08736 | 11.9280556 | 15.9159801 | 15.693197 |
| ENSMUSG00 | Slc27a1 | 2781 | 69.9759357 | 74.9220337 | 65.9928271 | 55.1026405 | 90.3465928 | 56.8148095 |
| ENSMUSG00 | 1700018B08I | 1073 | 0 | 0 | 0 | 0 | 0 | 0 |
| ENSMUSG00 | Fbxo31 | 1810 | 7.75341692 | 7.40101156 | 9.91488854 | 7.93406283 | 8.14730531 | 7.76046365 |
| ENSMUSG00 | Map1lc3b | 1706 | 143.776114 | 151.414892 | 169.405893 | 121.787463 | 130.374183 | 124.396628 |

| | | | | | | | | |
|---|---|---|---|---|---|---|---|---|
| ENSMUSG00 | Fam125a | 1051 | 41.382013 | 32.0937293 | 42.3513377 | 36.83733 | 33.5968145 | 44.4542883 |
| ENSMUSG00 | Mthfsd | 4866 | 9.32203109 | 12.2496155 | 7.54865013 | 8.4013474 | 9.42719109 | 4.39443427 |
| ENSMUSG00 | Cox4i1 | 674 | 411.652075 | 475.001071 | 602.364878 | 404.772016 | 257.554496 | 628.449774 |
| ENSMUSG00 | Cox4nb | 4928 | 12.9963635 | 7.53890968 | 12.9968487 | 9.56964939 | 16.8983452 | 12.5968099 |
| ENSMUSG00 | 5430437P03l | 1583 | 20.8195157 | 24.3824006 | 21.7239633 | 19.44282 | 17.4708122 | 20.813496 |
| ENSMUSG00 | Gins2 | 4359 | 0.20976739 | 0.56377806 | 0.69968506 | 0.55459899 | 0.43312134 | 0.48179565 |
| ENSMUSG00 | Gse1 | 7308 | 2.48063754 | 2.16931301 | 3.32663022 | 2.1971141 | 2.66722435 | 5.24741965 |
| ENSMUSG00 | Zdhhc7 | 3776 | 12.7815554 | 11.0895175 | 11.8698805 | 10.1862975 | 12.3511937 | 9.06506151 |
| ENSMUSG00 | 6430548M08 | 6773 | 2.43005595 | 2.21260645 | 1.23997493 | 0.81508236 | 2.21493496 | 0.75706957 |
| ENSMUSG00 | Crispld2 | 4852 | 0.33593873 | 0.05958752 | 0.02733008 | 0 | 0 | 0 |
| ENSMUSG00 | Usp10 | 5259 | 27.4561524 | 26.5717213 | 27.9802121 | 24.9946545 | 31.0194568 | 22.4254687 |
| ENSMUSG00 | Cotl1 | 2448 | 0.77951923 | 0.86609618 | 1.82368715 | 0.63379408 | 0.37519394 | 1.96092178 |
| ENSMUSG00 | Klhl36 | 2136 | 6.40256802 | 6.49704838 | 6.29089901 | 3.00683704 | 4.32386424 | 4.12439536 |
| ENSMUSG00 | Lrrc50 | 2809 | 0 | 0.25731457 | 0.2203011 | 0 | 0.19981849 | 0 |
| ENSMUSG00 | Taf1c | 3643 | 5.57646186 | 5.35698849 | 3.80987612 | 5.09090609 | 5.01439543 | 3.05463981 |
| ENSMUSG00 | Mast3 | 7307 | 1.76279946 | 1.82669283 | 1.75428144 | 2.23198086 | 3.9385351 | 2.06043532 |
| ENSMUSG00 | Pik3r2 | 4184 | 18.0724006 | 14.9718869 | 11.58925 | 13.4617762 | 21.3179027 | 10.6125986 |
| ENSMUSG00 | Mbtps1 | 6776 | 19.8894745 | 16.0005158 | 16.7387735 | 13.7650657 | 22.9227107 | 16.8453904 |
| ENSMUSG00 | Necab2 | 2220 | 0 | 0 | 0 | 0 | 0 | 0 |
| ENSMUSG00 | Ifi30 | 984 | 19.7161326 | 23.4566116 | 19.5404489 | 19.4344346 | 14.6234126 | 15.7161793 |
| ENSMUSG00 | Hsbp1 | 1159 | 147.908224 | 169.005943 | 181.078819 | 136.825354 | 123.053253 | 183.227214 |
| ENSMUSG00 | Rab3a | 2381 | 10.8363322 | 9.43086184 | 13.8304801 | 10.0168969 | 7.92934022 | 10.0461397 |
| ENSMUSG00 | Cdh13 | 11275 | 0.1868772 | 0 | 0.22737977 | 0.11840671 | 0.26248601 | 0 |
| ENSMUSG00 | Pde4c | 4454 | 38.1756152 | 33.8084041 | 23.3612794 | 25.9152358 | 42.6059927 | 22.1981516 |
| ENSMUSG00 | Mphosph6 | 1049 | 18.5323451 | 19.6145003 | 21.742756 | 19.3308952 | 15.1279341 | 22.2565216 |
| ENSMUSG00 | Hsd17b2 | 1275 | 0.12472308 | 0 | 0 | 0 | 0.08004137 | 0 |
| ENSMUSG00 | Bcmo1 | 2760 | 0 | 0.17458856 | 0 | 0 | 0 | 0 |
| ENSMUSG00 | 1700030J22R | 2968 | 0.7367087 | 0.21105921 | 0.22339206 | 0.91177673 | 1.35818184 | 0.33082447 |
| ENSMUSG00 | Lsm4 | 1058 | 27.5432582 | 23.6833175 | 36.3892299 | 26.7375453 | 21.8477772 | 36.4005582 |
| ENSMUSG00 | Comp | 2490 | 0 | 0.42574367 | 0.10651047 | 0.08694469 | 0.2049252 | 0.16430506 |
| ENSMUSG00 | Ntpcr | 2698 | 7.79490333 | 5.76879941 | 6.52050909 | 6.21873636 | 6.56269548 | 5.98465027 |
| ENSMUSG00 | BC021891 | 5682 | 8.48704644 | 5.7752494 | 5.92002861 | 7.16306692 | 6.31318941 | 6.13943393 |
| ENSMUSG00 | Mau2 | 5353 | 17.898507 | 16.014138 | 15.9945769 | 15.8132779 | 25.0699362 | 9.70636798 |
| ENSMUSG00 | Pbx4 | 3160 | 0.05032339 | 0.10674212 | 0.04196378 | 0.20553063 | 0.24221381 | 0.21578037 |
| ENSMUSG00 | Lpar2 | 5974 | 0.46583255 | 0.29844298 | 0.3255576 | 1.0207341 | 0.11103831 | 0.07761447 |
| ENSMUSG00 | Atp13a1 | 4291 | 28.3597124 | 30.5558865 | 21.2098799 | 34.2152969 | 40.799696 | 12.1849181 |
| ENSMUSG00 | Ints10 | 6901 | 8.51450517 | 8.03689247 | 8.42275374 | 7.2362773 | 5.59729988 | 5.13400324 |
| ENSMUSG00 | Dctn1 | 6268 | 17.9432442 | 17.3049906 | 19.1743477 | 19.025392 | 21.6544607 | 14.490196 |
| ENSMUSG00 | Pgr | 6382 | 0.3613002 | 0.30956505 | 0.1592984 | 0.29399347 | 0.59965186 | 0.05983155 |
| ENSMUSG00 | Cdh5 | 3974 | 0.58022594 | 0.65477299 | 0.27806897 | 0.02723858 | 0.57780244 | 0.16471851 |
| ENSMUSG00 | Bean1 | 3379 | 0 | 0 | 0 | 0 | 0 | 0.00807181 |
| ENSMUSG00 | Cmtm3 | 1611 | 0.49355035 | 0.38884156 | 0.30181271 | 0 | 0.28506355 | 0.2200933 |
| ENSMUSG00 | Cmtm1 | 2269 | 0 | 0 | 0 | 0 | 0 | 0 |
| ENSMUSG00 | Ces2g | 2847 | 0 | 0 | 0.03105152 | 0 | 0.30468858 | 0 |
| ENSMUSG00 | Nae1 | 4554 | 24.8624525 | 30.6217751 | 23.8189115 | 25.0371675 | 27.8997971 | 45.6433961 |
| ENSMUSG00 | Fam96b | 1316 | 12.3556167 | 12.3029214 | 19.1115871 | 13.3525499 | 9.77100814 | 24.8704917 |
| ENSMUSG00 | Rrad | 1570 | 12.4077614 | 18.8755808 | 5.51819187 | 10.3190051 | 18.7855065 | 8.13027463 |
| ENSMUSG00 | Cdh16 | 3298 | 972.514524 | 1165.7817 | 791.451557 | 909.764225 | 1090.87831 | 609.536086 |
| ENSMUSG00 | Car7 | 2450 | 1.21700451 | 1.18007613 | 0.99228627 | 0.38291149 | 1.52037773 | 0.50096276 |
| ENSMUSG00 | Ces2d-ps | 1867 | 0 | 0 | 0 | 0 | 0 | 0 |
| ENSMUSG00 | Cbfb | 2905 | 11.3313383 | 23.2223818 | 21.028209 | 19.3514029 | 13.7358437 | 21.2563798 |
| ENSMUSG00 | Ces2e | 2687 | 0.36988724 | 0 | 0.87186367 | 0.9802714 | 2.2408308 | 0 |
| ENSMUSG00 | Tradd | 2320 | 7.77973627 | 7.8926069 | 7.8115326 | 9.59595753 | 7.1920797 | 6.17206276 |
| ENSMUSG00 | D230025D16 | 4969 | 8.58475158 | 5.69238107 | 9.36698344 | 6.73860683 | 8.03031312 | 6.37268186 |
| ENSMUSG00 | Hsd11b2 | 1840 | 372.318718 | 717.847047 | 175.245894 | 505.638869 | 481.117714 | 416.249535 |
| ENSMUSG00 | Tsnaxip1 | 2293 | 0 | 0.02101458 | 0 | 0 | 0 | 0.09515792 |
| ENSMUSG00 | Ctrl | 881 | 0.54150484 | 0.05469517 | 0.50172353 | 0.45051361 | 0.86878052 | 0.58821583 |
| ENSMUSG00 | Psmb10 | 1262 | 8.66304052 | 6.37649433 | 6.26951656 | 8.66312163 | 11.7659868 | 15.5391962 |
| ENSMUSG00 | Dpep3 | 1671 | 0 | 0 | 0 | 0 | 0 | 0 |
| ENSMUSG00 | Dus2l | 3349 | 7.88224517 | 5.45316858 | 7.81352376 | 8.55453668 | 7.64862367 | 8.00566449 |
| ENSMUSG00 | Nfatc3 | 6185 | 11.1135369 | 9.86322324 | 13.3355924 | 15.0512005 | 13.6042835 | 13.1764951 |
| ENSMUSG00 | Pla2g15 | 2698 | 16.1939485 | 14.3594883 | 16.9566003 | 15.4331651 | 12.614749 | 16.0837476 |
| ENSMUSG00 | Slc7a6 | 3720 | 6.89308735 | 8.48444075 | 8.06802457 | 9.43758853 | 9.65660446 | 8.4463399 |
| ENSMUSG00 | Smpd3 | 1961 | 0.38518824 | 0 | 0 | 0 | 0 | 0 |
| ENSMUSG00 | Zfp90 | 2467 | 0.87020509 | 0.66410176 | 0.60918632 | 0.58503518 | 0.76529222 | 0.38695273 |
| ENSMUSG00 | Has3 | 6374 | 0 | 0.14363703 | 0 | 0 | 0.28018876 | 0 |
| ENSMUSG00 | Vps4a | 2512 | 34.9443077 | 31.8429514 | 34.4886992 | 32.9938122 | 30.2867541 | 24.2996189 |
| ENSMUSG00 | Cog8 | 5955 | 6.08849676 | 5.63995805 | 6.41316413 | 4.51404251 | 7.09485125 | 3.76944217 |
| ENSMUSG00 | Nip7 | 3026 | 9.38050664 | 10.5099312 | 10.122894 | 11.4708945 | 7.79054383 | 8.76105398 |
| ENSMUSG00 | Mtmr2 | 5234 | 21.4424382 | 17.2344325 | 16.8733822 | 16.6622663 | 17.0607867 | 15.7738504 |
| ENSMUSG00 | Tmed6 | 859 | 3.14711604 | 6.8437089 | 7.51276959 | 3.82242847 | 3.62352611 | 3.74669084 |
| ENSMUSG00 | Terf2 | 5436 | 3.26176312 | 4.64490355 | 4.01687094 | 4.17505629 | 2.88173241 | 4.47051206 |
| ENSMUSG00 | Cep57 | 5215 | 13.7143067 | 11.891841 | 12.8494715 | 14.4397367 | 14.207152 | 15.4286071 |
| ENSMUSG00 | Cyb5b | 4289 | 158.076466 | 154.625788 | 147.466582 | 162.844518 | 169.901533 | 112.036511 |
| ENSMUSG00 | Maml2 | 6017 | 0.72679123 | 2.32243115 | 0.93296228 | 2.47663028 | 2.27273869 | 2.61550056 |
| ENSMUSG00 | 1700012B09l | 675 | 0 | 0 | 0 | 0 | 0 | 0 |
| ENSMUSG00 | Mre11a | 5962 | 3.7341612 | 3.23290454 | 3.432649 | 5.18656695 | 5.04101566 | 2.68537626 |
| ENSMUSG00 | Wwp2 | 4577 | 26.1185777 | 21.2348818 | 22.0767781 | 24.7458024 | 27.4585894 | 17.674586 |
| ENSMUSG00 | Ankrd49 | 1751 | 18.4360081 | 16.676747 | 17.0647892 | 16.340983 | 15.5905831 | 18.5361626 |
| ENSMUSG00 | Gpr83 | 4689 | 0 | 0 | 0 | 0 | 0 | 0 |
| ENSMUSG00 | Folr4 | 2197 | 0 | 0 | 0 | 0 | 0 | 0 |
| ENSMUSG00 | Panx1 | 3171 | 0.43880221 | 0.41029137 | 0.04181821 | 0.35274153 | 1.49651623 | 0 |
| ENSMUSG00 | Med17 | 3887 | 6.79126294 | 9.16125051 | 10.1435668 | 7.6304195 | 11.9984841 | 4.93287147 |
| ENSMUSG00 | Hephl1 | 5241 | 0 | 0 | 0 | 0 | 0 | 0 |
| ENSMUSG00 | 2200002K05l | 1964 | 0.95137861 | 0 | 0 | 0 | 0.3637318 | 0.02777458 |
| ENSMUSG00 | 4931406C07l | 2554 | 8.70137573 | 9.09391743 | 7.82272241 | 7.96800044 | 7.19244138 | 10.9034482 |
| ENSMUSG00 | Taf1d | 1709 | 12.212772 | 6.8515538 | 8.40585078 | 12.8366782 | 5.52362758 | 10.820483 |
| ENSMUSG00 | Kars | 2495 | 27.1357047 | 28.2359032 | 24.3243008 | 24.425881 | 30.186345 | 20.1362267 |
| ENSMUSG00 | Adat1 | 4389 | 2.93478598 | 2.2177401 | 1.33944979 | 3.73234181 | 4.02258511 | 2.87722898 |
| ENSMUSG00 | Gabarapl2 | 2415 | 19.9847426 | 20.5116615 | 20.4444963 | 15.7625495 | 16.670729 | 22.7796884 |
| ENSMUSG00 | Tmem231 | 2880 | 11.1950329 | 11.0092635 | 4.71179368 | 11.7266645 | 15.5913926 | 3.87337759 |
| ENSMUSG00 | Chst5 | 1933 | 0 | 0 | 0 | 0 | 0 | 0 |
| ENSMUSG00 | Tmem170 | 1940 | 3.31978755 | 4.66961398 | 4.46575321 | 2.62245788 | 2.26199398 | 5.59551877 |
| ENSMUSG00 | Cfdp1 | 1208 | 64.5368355 | 63.4242078 | 63.9243541 | 54.3022822 | 54.3211253 | 56.287811 |
| ENSMUSG00 | Bcar1 | 4195 | 39.3669287 | 44.5107183 | 39.1125724 | 42.0082731 | 42.5361707 | 34.19245 |
| ENSMUSG00 | Ctrb1 | 880 | 0.22588341 | 0 | 0 | 0 | 0 | 0 |
| ENSMUSG00 | Ldhd | 6705 | 3.90143269 | 5.73494121 | 3.42803257 | 4.26742152 | 5.80658087 | 2.7132266 |
| ENSMUSG00 | Wdr59 | 5194 | 2.01303261 | 3.39550208 | 2.23817572 | 3.08441043 | 1.67009701 | 3.41326829 |
| ENSMUSG00 | Aars | 8373 | 17.4633533 | 16.1254522 | 15.4043924 | 18.1207449 | 16.9173796 | 9.48895547 |
| ENSMUSG00 | Cdh15 | 3237 | 0 | 0 | 0 | 0 | 0 | 0 |
| ENSMUSG00 | Bmper | 3778 | 0 | 0.02550897 | 0.1520974 | 0 | 0.05402475 | 0 |
| ENSMUSG00 | Tbx20 | 4485 | 0 | 0 | 0 | 0 | 0 | 0 |
| ENSMUSG00 | Glb1l3 | 2448 | 0 | 0 | 0 | 0 | 0 | 0 |
| ENSMUSG00 | Afg3l1 | 4847 | 17.1587508 | 17.0297866 | 16.2508117 | 16.40702 | 19.6441546 | 16.1723361 |
| ENSMUSG00 | Acad8 | 5419 | 19.0817503 | 15.6056848 | 21.4932379 | 17.944475 | 17.13748 | 18.3860966 |
| ENSMUSG00 | Dbndd1 | 2640 | 0.27106009 | 0.47456345 | 0.1506881 | 0.42369067 | 0.2126099 | 0.19629475 |
| ENSMUSG00 | 1700054N08l | 4437 | 0.77951923 | 0.30408393 | 1.41461837 | 0 | 1.10401895 | 1.15565296 |
| ENSMUSG00 | Acta1 | 1439 | 0 | 0.03348606 | 0 | 0 | 0 | 0 |
| ENSMUSG00 | Abcb10 | 4293 | 12.4554208 | 10.1917749 | 10.1315196 | 10.3884014 | 8.17753239 | 9.07883985 |
| ENSMUSG00 | Urb2 | 6143 | 3.07404416 | 4.0789435 | 2.40329083 | 1.85608434 | 4.01200383 | 1.91806026 |
| ENSMUSG00 | Cog2 | 4670 | 16.1916325 | 14.5591156 | 17.4157155 | 12.5475889 | 17.0452133 | 12.188902 |
| ENSMUSG00 | Agt | 1886 | 3.41483979 | 0.281045 | 0.72654143 | 0 | 1.24454575 | 0.33261755 |
| ENSMUSG00 | Capn9 | 2501 | 0 | 0 | 0 | 0 | 0 | 0 |
| ENSMUSG00 | Arv1 | 1159 | 7.16902108 | 5.03068119 | 4.53841184 | 5.38584444 | 3.83028015 | 6.77747723 |
| ENSMUSG00 | 2310022B05l | 3293 | 1.85919951 | 1.94618792 | 1.50337274 | 1.53400736 | 1.25512798 | 1.18441343 |
| ENSMUSG00 | 2810004N23l | 1375 | 33.915039 | 21.6926601 | 26.3925186 | 20.4421183 | 22.9711466 | 19.578232 |
| ENSMUSG00 | Gnpat | 3812 | 45.7938916 | 43.2692003 | 52.3998239 | 48.4343709 | 49.1523746 | 45.6628404 |
| ENSMUSG00 | Gm505 | 2111 | 1.67609558 | 1.43806057 | 0.98412441 | 1.96562535 | 1.69201625 | 1.21450312 |
| ENSMUSG00 | Egln1 | 3510 | 18.371336 | 23.9010244 | 15.6028728 | 16.3140184 | 14.6101162 | 15.0671013 |
| ENSMUSG00 | Vps26b | 6866 | 6.34026379 | 7.32692188 | 8.78757875 | 7.12601986 | 8.39787426 | 5.22373298 |
| ENSMUSG00 | Jam3 | 2528 | 0.70767272 | 0.20967202 | 0.24478869 | 0.22836737 | 0.80737936 | 0.67970818 |
| ENSMUSG00 | Spata19 | 816 | 0 | 0 | 0 | 0 | 0 | 0 |
| ENSMUSG00 | Snx19 | 7678 | 8.09296901 | 10.047733 | 10.5985404 | 10.7240461 | 13.6438069 | 6.49362341 |
| ENSMUSG00 | Adamts8 | 4747 | 0 | 0 | 0 | 0 | 0 | 0.04596527 |
| ENSMUSG00 | St14 | 4131 | 123.125543 | 102.520126 | 152.742994 | 111.381804 | 127.189202 | 83.3886937 |
| ENSMUSG00 | Aplp2 | 4195 | 410.433875 | 414.17367 | 400.166292 | 373.171772 | 504.802294 | 323.056561 |
| ENSMUSG00 | Trpc6 | 2780 | 0 | 0 | 0 | 0 | 0 | 0 |
| ENSMUSG00 | 1700128F08l | 1107 | 0 | 0 | 0 | 0 | 0 | 0 |
| ENSMUSG00 | Birc3 | 5624 | 7.49303153 | 9.06494591 | 6.42907318 | 6.1783444 | 8.16567181 | 4.26772449 |
| ENSMUSG00 | Dcun1d5 | 1193 | 118.866554 | 85.50771 | 96.221447 | 104.072351 | 84.8158451 | 174.530257 |
| ENSMUSG00 | Pdgfd | 3369 | 3.65811783 | 2.34567424 | 4.53957782 | 6.15825931 | 1.51458521 | 4.50934226 |
| ENSMUSG00 | Sesn3 | 2152 | 1.51484638 | 1.74653461 | 2.42370701 | 2.95094788 | 1.70720217 | 2.71225502 |
| ENSMUSG00 | Usp2 | 4931 | 13.3754496 | 24.3326386 | 12.2449235 | 23.6132139 | 15.522118 | 10.5259863 |
| ENSMUSG00 | Thy1 | 1731 | 0 | 0 | 0 | 0 | 0 | 0 |
| ENSMUSG00 | Pvrl1 | 5048 | 0.29926867 | 0.16227605 | 0.27144555 | 0.50034531 | 0.05054118 | 0.19991316 |
| ENSMUSG00 | Trim29 | 2821 | 0.43687341 | 0.03416267 | 0.34471483 | 0.12790516 | 0.68734572 | 0 |
| ENSMUSG00 | Oaf | 2384 | 8.7215253 | 9.23708224 | 7.11975995 | 10.1556426 | 5.88601231 | 6.28094671 |
| ENSMUSG00 | Pou2f3 | 5132 | 0.0542261 | 0 | 0 | 0 | 0 | 0 |
| ENSMUSG00 | Grik4 | 6643 | 0 | 0 | 0 | 0 | 0 | 0 |
| ENSMUSG00 | Sc5d | 3385 | 44.1949404 | 35.1469204 | 22.9039978 | 38.1393073 | 24.5861208 | 35.7833598 |
| ENSMUSG00 | Ubash3b | 7622 | 0.19820366 | 0.10747435 | 0.35955317 | 0.06627508 | 0.09372465 | 0.23617505 |
| ENSMUSG00 | Crtam | 1866 | 0 | 0.30988066 | 0 | 0 | 0 | 0 |
| ENSMUSG00 | 4931429I11R | 4218 | 0 | 0 | 0 | 0 | 0 | 0 |
| ENSMUSG00 | 9030425E11l | 4929 | 0 | 0.00977611 | 0 | 0 | 0 | 0 |
| ENSMUSG00 | Rexo2 | 1068 | 59.3354269 | 62.7145642 | 66.0130521 | 57.9407363 | 45.4364077 | 69.131506 |
| ENSMUSG00 | Fam55b | 4876 | 0.62780394 | 0.02964711 | 0.01813037 | 0 | 0.05232401 | 0 |
| ENSMUSG00 | Cul5 | 9555 | 9.19098445 | 8.81022652 | 7.8874037 | 12.4351829 | 7.23607946 | 11.717676 |
| ENSMUSG00 | Barx2 | 1911 | 0 | 0 | 0 | 0 | 0.0534028 | 0 |
| ENSMUSG00 | Kcnj5 | 4492 | 0 | 0.70799314 | 0.36408464 | 0.04819507 | 0.05679695 | 0 |
| ENSMUSG00 | Ets1 | 6411 | 2.30682245 | 2.08199103 | 0.73773159 | 1.67155939 | 3.70898317 | 0.78280044 |
| ENSMUSG00 | Kirrel3 | 4398 | 0 | 0 | 0 | 0 | 0 | 0 |
| ENSMUSG00 | St3gal4 | 2018 | 0.05910131 | 1.43269897 | 0.04380758 | 0 | 1.18842402 | 0.01351568 |
| ENSMUSG00 | Dcps | 1876 | 7.67136671 | 13.0226685 | 13.4066358 | 12.0786375 | 10.8526247 | 11.747286 |
| ENSMUSG00 | Tirap | 6066 | 4.33207593 | 3.67792989 | 3.79643261 | 3.29532685 | 3.73486826 | 2.66181774 |
| ENSMUSG00 | Srpr | 3348 | 44.8614711 | 39.9394794 | 46.2350222 | 43.1087748 | 57.8390371 | 45.6695423 |
| ENSMUSG00 | Rpusd4 | 2047 | 12.9928757 | 9.67495248 | 10.278494 | 9.60660199 | 9.09849886 | 7.46155251 |
| ENSMUSG00 | Abhd12 | 3430 | 28.5474485 | 23.9667843 | 36.5986105 | 25.215353 | 24.2486859 | 20.8257377 |
| ENSMUSG00 | Acat1 | 3924 | 188.068549 | 170.506817 | 188.815314 | 162.599081 | 160.894329 | 160.672926 |
| ENSMUSG00 | 4930510E17l | 1637 | 0 | 0.50040899 | 0 | 0 | 0 | 0 |
| ENSMUSG00 | Rdx | 4825 | 62.990808 | 58.2731378 | 59.4549142 | 65.1720245 | 56.7582506 | 71.1967008 |
| ENSMUSG00 | Fdx1 | 3097 | 19.6916071 | 16.4148197 | 22.3221448 | 18.9322965 | 11.5497221 | 23.6550472 |
| ENSMUSG00 | Pou2af1 | 2561 | 0 | 0 | 0 | 0 | 0 | 0 |
| ENSMUSG00 | Btg4 | 1481 | 0 | 0 | 0 | 0 | 0 | 0 |
| ENSMUSG00 | 4833427G06l | 1184 | 0 | 0 | 0 | 0 | 0 | 0 |
| ENSMUSG00 | Ppp2r1b | 6090 | 20.9874992 | 15.7535643 | 27.1307864 | 21.9276924 | 19.3632109 | 18.1203925 |
| ENSMUSG00 | Alg9 | 4082 | 7.52841412 | 7.23622955 | 7.56910541 | 4.83510004 | 7.6252056 | 6.9355893 |
| ENSMUSG00 | Cryab | 1216 | 55.1868841 | 34.3566162 | 28.3895061 | 34.1829896 | 46.8301278 | 47.2820227 |
| ENSMUSG00 | 2310030G06l | 1238 | 46.049563 | 45.5007681 | 49.2004359 | 31.7976661 | 42.4532852 | 48.9313197 |
| ENSMUSG00 | Dixdc1 | 12937 | 6.36726873 | 3.88113726 | 7.40740478 | 3.15721376 | 6.86294303 | 9.37124306 |
| ENSMUSG00 | Tex12 | 612 | 0 | 0 | 0 | 0 | 0 | 0.44566404 |
| ENSMUSG00 | Bco2 | 2106 | 1.18926651 | 0.663536 | 1.95193324 | 1.67903151 | 0.7995586 | 0.07770553 |
| ENSMUSG00 | Pts | 1029 | 195.184342 | 199.863688 | 221.39582 | 170.837435 | 136.913337 | 280.592159 |
| ENSMUSG00 | 1600029D21l | 1981 | 472.169079 | 348.761841 | 875.803402 | 377.23131 | 274.424537 | 603.910904 |
| ENSMUSG00 | Cadm1 | 7281 | 0.39313207 | 1.54201909 | 0.63136817 | 0.68387888 | 0.97413353 | 1.82430289 |
| ENSMUSG00 | Bud13 | 3970 | 3.32463968 | 2.68241907 | 3.09524241 | 3.34463284 | 4.19007519 | 2.92669933 |
| ENSMUSG00 | Zfp259 | 3457 | 18.5264909 | 17.6465247 | 20.2661039 | 20.154593 | 20.4873039 | 19.629535 |
| ENSMUSG00 | Apoa5 | 2514 | 0 | 0 | 0 | 0 | 0 | 0 |
| ENSMUSG00 | Apoa4 | 1812 | 0 | 0 | 0 | 0 | 0 | 0 |
| ENSMUSG00 | Apoc3 | 1300 | 0 | 0 | 0 | 0 | 0 | 0 |
| ENSMUSG00 | Apoa1 | 1248 | 0 | 0 | 0 | 0 | 0 | 0.34967486 |
| ENSMUSG00 | Tagln | 1581 | 2.3134119 | 1.40200907 | 2.46031764 | 0.89006941 | 3.16292526 | 0.75906649 |
| ENSMUSG00 | Bace1 | 6386 | 16.3479317 | 17.6341552 | 13.9956352 | 21.7588402 | 18.4577088 | 14.1840083 |
| ENSMUSG00 | Dscaml1 | 7279 | 0.20754338 | 0.41705535 | 0.4979463 | 0.90217504 | 0.87626006 | 0.11241093 |
| ENSMUSG00 | Il10ra | 6092 | 0 | 0 | 0 | 0 | 0 | 0 |
| ENSMUSG00 | Tmprss4 | 3257 | 0.32956646 | 0.71014714 | 0.35285475 | 0.62038517 | 0.70500059 | 0.02512248 |
| ENSMUSG00 | Mpzl2 | 3101 | 13.8330098 | 17.2638301 | 13.9974879 | 14.7074183 | 15.4675244 | 10.6952471 |
| ENSMUSG00 | Cd3e | 1428 | 0 | 0 | 0 | 0 | 0 | 0 |
| ENSMUSG00 | Cd3d | 1318 | 0 | 0 | 0 | 0 | 0 | 0 |
| ENSMUSG00 | Arcn1 | 5037 | 86.4091723 | 73.4802585 | 89.1934748 | 86.3404572 | 84.8616141 | 71.9363872 |
| ENSMUSG00 | Ddx6 | 6111 | 7.70258369 | 10.2034456 | 8.93295306 | 8.25440979 | 7.59007947 | 6.61893144 |
| ENSMUSG00 | Treh | 2138 | 0 | 0 | 0 | 0 | 0 | 0.15308497 |
| ENSMUSG00 | Pate4 | 1063 | 0 | 0 | 0.04158217 | 0 | 0 | 0 |
| ENSMUSG00 | Ddx25 | 1663 | 0 | 0 | 0 | 0 | 0 | 0 |
| ENSMUSG00 | Pus3 | 2751 | 0.04335385 | 0.03503195 | 0.03213511 | 0.02623195 | 0.0370966 | 0.00991445 |
| ENSMUSG00 | Pdzd3 | 2170 | 0.32976896 | 0.17764587 | 0.08147805 | 0 | 0 | 0.47762041 |
| ENSMUSG00 | D730048I06l | 1407 | 0 | 0 | 0 | 0 | 0 | 0 |
| ENSMUSG00 | Nlrx1 | 3968 | 5.36017693 | 3.91029092 | 2.35045083 | 1.79137167 | 4.97661478 | 3.07252111 |
| ENSMUSG00 | Acrv1 | 1091 | 0 | 0 | 0 | 0 | 0 | 0 |
| ENSMUSG00 | Trappc4 | 1734 | 42.5296519 | 45.3241564 | 63.9065862 | 42.0332929 | 39.6087093 | 44.5926195 |
| ENSMUSG00 | Chek1 | 5613 | 0.4178823 | 0.11160231 | 0.23624716 | 0.43712436 | 0.50908196 | 0.09232463 |
| ENSMUSG00 | Slc37a4 | 2422 | 25.7704808 | 19.4576137 | 28.6344722 | 20.0224062 | 25.3235771 | 18.0404509 |
| ENSMUSG00 | Hyou1 | 5921 | 6.02945812 | 7.73131566 | 9.09269461 | 6.06344102 | 6.60128422 | 6.85896603 |
| ENSMUSG00 | Stt3a | 5688 | 68.7962719 | 61.6223945 | 69.939625 | 72.9697184 | 57.8532169 | 51.9695043 |
| ENSMUSG00 | Fez1 | 4015 | 0 | 0.40805455 | 0 | 0 | 0 | 0 |
| ENSMUSG00 | Hinfp | 2090 | 6.37228994 | 6.4555999 | 7.00038758 | 7.44084268 | 10.791224 | 4.20212145 |
| ENSMUSG00 | C2cd2l | 3866 | 4.0619283 | 6.16975915 | 4.8134961 | 4.330592 | 4.85714079 | 3.2311911 |
| ENSMUSG00 | Tmem218 | 1015 | 22.7174174 | 16.7584375 | 20.5984943 | 13.9351344 | 16.3384947 | 22.3302712 |
| ENSMUSG00 | Slc37a2 | 6401 | 0.1863247 | 0.19572684 | 0.0345273 | 0.12401265 | 0.83702069 | 0.21731083 |
| ENSMUSG00 | Dpagt1 | 1896 | 18.7035278 | 21.3738385 | 21.3782121 | 19.6966857 | 17.4124816 | 16.3705377 |
| ENSMUSG00 | Robo4 | 4212 | 0.87779195 | 1.08682621 | 0.27285088 | 1.07081091 | 1.06607813 | 1.71599701 |
| ENSMUSG00 | Hmbs | 1740 | 31.210337 | 27.3056506 | 30.9412901 | 24.2828012 | 19.4134832 | 24.7039261 |
| ENSMUSG00 | Vps11 | 3163 | 34.7781899 | 23.2934145 | 31.946068 | 22.8150772 | 37.330077 | 21.8852464 |
| ENSMUSG00 | Robo3 | 6562 | 0.50285049 | 0.18358139 | 0.00673603 | 0.06048499 | 0.5598749 | 0.36992425 |
| ENSMUSG00 | Abcg4 | 6512 | 0 | 0 | 0.22399583 | 0 | 0 | 0 |
| ENSMUSG00 | Muc16 | 1026 | 0 | 0 | 0 | 0 | 0 | 0 |
| ENSMUSG00 | Mcam | 3528 | 0.81133634 | 0.09560802 | 0.88954956 | 0.61364021 | 0 | 0.70351252 |
| ENSMUSG00 | Pin1 | 3804 | 10.6599869 | 10.8432162 | 14.292394 | 10.0354372 | 8.41049885 | 12.3825715 |
| ENSMUSG00 | Olfm2 | 1820 | 0.28396772 | 0.0794282 | 0.04857345 | 0.87231316 | 0.16821882 | 0.25476311 |
| ENSMUSG00 | Icam5 | 2971 | 0 | 0 | 0 | 0 | 0 | 0 |
| ENSMUSG00 | Tyk2 | 4588 | 10.441446 | 7.35189832 | 9.98106133 | 10.8922475 | 14.2691456 | 8.05517354 |
| ENSMUSG00 | Pde4a | 6642 | 2.04703016 | 4.18602486 | 2.86826172 | 3.25401154 | 3.6337683 | 3.03051547 |
| ENSMUSG00 | Ilf3 | 4916 | 5.24843508 | 6.14582979 | 5.91635734 | 7.28099841 | 6.26931062 | 5.58142536 |
| ENSMUSG00 | Bmp5 | 4017 | 0 | 0 | 0 | 0 | 0 | 0 |
| ENSMUSG00 | Tmed1 | 2318 | 14.0121776 | 15.6533179 | 17.9438888 | 14.3518745 | 14.4846226 | 14.6727675 |
| ENSMUSG00 | Scg3 | 2237 | 0 | 0 | 0 | 0 | 0 | 0 |
| ENSMUSG00 | Yipf2 | 2417 | 20.6261368 | 16.3877764 | 13.0941331 | 14.2118769 | 19.1269741 | 9.9190765 |
| ENSMUSG00 | Lysmd2 | 1197 | 0 | 0.64409614 | 0.25849031 | 0 | 0.04262855 | 0.91143323 |
| ENSMUSG00 | Carm1 | 4627 | 11.1181309 | 12.1116992 | 10.6325159 | 11.7596117 | 14.3914892 | 9.90304602 |
| ENSMUSG00 | Tmod2 | 10686 | 0.16741499 | 0.04509306 | 0 | 0 | 0 | 0.07657114 |
| ENSMUSG00 | Smarca4 | 6576 | 17.362795 | 19.3742325 | 16.4547007 | 20.8338691 | 21.8507108 | 13.0193267 |
| ENSMUSG00 | Bcl2l10 | 1218 | 0 | 0.31649552 | 0 | 0 | 0 | 0.40307349 |
| ENSMUSG00 | Gnb5 | 2332 | 41.5284522 | 47.0499695 | 42.0978959 | 33.6528503 | 35.4909012 | 31.0172999 |
| ENSMUSG00 | Ldlr | 4707 | 19.6961505 | 11.783003 | 16.7341585 | 6.94504617 | 9.61555032 | 8.98145121 |
| ENSMUSG00 | Kank2 | 7056 | 1.09305034 | 1.68679864 | 1.09627587 | 1.02273369 | 1.59819006 | 1.09392332 |
| ENSMUSG00 | Dock6 | 6641 | 0.97577825 | 1.28429457 | 0.59903115 | 1.22247553 | 1.89015035 | 0.28338354 |
| ENSMUSG00 | Polr2m | 3488 | 118.103867 | 111.596927 | 127.054954 | 98.553222 | 96.2303613 | 122.352718 |
| ENSMUSG00 | Rab27a | 2965 | 13.3009904 | 9.31225341 | 4.12948078 | 7.48413402 | 4.87030838 | 5.41813238 |
| ENSMUSG00 | Pigb | 2864 | 5.33034376 | 5.98965552 | 5.52523038 | 6.80317879 | 7.58981707 | 4.54259879 |
| ENSMUSG00 | Aqp9 | 5711 | 0 | 0 | 0 | 0 | 0 | 0 |
| ENSMUSG00 | Lipc | 2328 | 0 | 0 | 0 | 0 | 0 | 0.05857955 |
| ENSMUSG00 | Sltm | 3847 | 24.8329659 | 21.1935171 | 19.886126 | 21.197146 | 21.7926528 | 21.468055 |
| ENSMUSG00 | Rsl24d1 | 1741 | 8.40322621 | 9.10588137 | 9.31767742 | 7.08791455 | 8.85121511 | 6.6737486 |
| ENSMUSG00 | Nedd4 | 5249 | 91.1217729 | 85.4944438 | 94.3657559 | 91.6246088 | 101.109799 | 75.0896955 |
| ENSMUSG00 | Rnf111 | 6664 | 5.39896607 | 5.99438183 | 6.62629669 | 11.3812211 | 6.88366024 | 8.02604555 |
| ENSMUSG00 | Ccnb2 | 1532 | 0 | 1.16377178 | 0 | 1.01274668 | 1.26566729 | 0 |
| ENSMUSG00 | Tex9 | 1993 | 0.09973778 | 0.50773471 | 0.17742837 | 0.38019214 | 0.58886435 | 0.27370436 |
| ENSMUSG00 | Myo1e | 4934 | 26.7024042 | 25.5483852 | 32.4839649 | 24.505661 | 29.8877637 | 27.6892315 |
| ENSMUSG00 | Mns1 | 1795 | 0 | 1.15432703 | 0.02462498 | 0.22111559 | 0.3979773 | 0.03038957 |
| ENSMUSG00 | Fam81a | 3586 | 6.04203482 | 5.06589199 | 4.74559664 | 6.63080517 | 5.86248378 | 5.11114266 |
| ENSMUSG00 | Gcnt3 | 4298 | 0.46248814 | 0.07847955 | 1.01814389 | 0.15950648 | 0 | 0 |
| ENSMUSG00 | Tcf12 | 5424 | 9.13994915 | 8.80397569 | 10.2925752 | 13.2912841 | 10.0284138 | 13.0238414 |
| ENSMUSG00 | Anxa2 | 3738 | 16.6658208 | 28.9660074 | 10.7607483 | 14.9424839 | 14.4834095 | 10.8573203 |
| ENSMUSG00 | Cgnl1 | 8357 | 25.4317098 | 19.2814961 | 18.5862482 | 25.3355794 | 23.17771 | 26.8634146 |
| ENSMUSG00 | Narg2 | 6302 | 5.05932964 | 4.5647899 | 6.64220015 | 3.79600056 | 7.66772103 | 7.26660097 |
| ENSMUSG00 | Rora | 13160 | 6.11436874 | 4.82596738 | 4.38323747 | 2.91452989 | 5.90525611 | 5.19171515 |
| ENSMUSG00 | Rp9 | 2015 | 22.1959879 | 18.4615054 | 25.1171763 | 17.7455614 | 14.3836136 | 23.7959384 |
| ENSMUSG00 | Itga11 | 5237 | 0 | 0 | 0.0084403 | 0.06889831 | 0.00974344 | 0.0156242 |
| ENSMUSG00 | Fem1b | 6486 | 14.5696542 | 10.8987836 | 11.2310572 | 11.2429559 | 14.6643794 | 7.49779244 |
| ENSMUSG00 | Cln6 | 2587 | 17.2114952 | 11.8836297 | 11.0718184 | 8.38241545 | 9.76345421 | 10.3742733 |
| ENSMUSG00 | Calml4 | 856 | 0.32510323 | 1.51989946 | 0.67128967 | 0.88519035 | 0.29805126 | 1.81618509 |
| ENSMUSG00 | Anp32a | 2833 | 3.97133815 | 3.99711045 | 4.21267124 | 4.87801732 | 2.59364565 | 5.28548427 |
| ENSMUSG00 | Irak1bp1 | 1889 | 5.93491027 | 8.62203147 | 3.43974109 | 7.83146546 | 5.51052444 | 5.94873021 |
| ENSMUSG00 | Glce | 4623 | 8.17812286 | 8.55744517 | 8.28963831 | 9.74831789 | 14.2163037 | 6.20655862 |
| ENSMUSG00 | Phip | 10702 | 12.2104527 | 13.462667 | 15.7031776 | 17.7274706 | 11.3142954 | 15.8851454 |
| ENSMUSG00 | Kif23 | 3644 | 6.09860417 | 5.65965896 | 7.60553117 | 5.65390983 | 6.9174066 | 7.72432154 |
| ENSMUSG00 | Ankk1 | 2678 | 0 | 0 | 0 | 0 | 0 | 0 |
| ENSMUSG00 | Lca5 | 5619 | 1.59899245 | 1.8180327 | 1.68343022 | 1.4769301 | 1.77080322 | 2.14546904 |
| ENSMUSG00 | Drd2 | 2539 | 0 | 0 | 0 | 0 | 0 | 0 |
| ENSMUSG00 | Sh3bgrl2 | 6383 | 85.3780553 | 41.248742 | 99.1165563 | 59.609143 | 70.3721113 | 70.017584 |
| ENSMUSG00 | Elovl4 | 3047 | 0 | 0 | 0 | 0 | 0 | 0.06265917 |
| ENSMUSG00 | Bckdhb | 1475 | 34.4727184 | 44.2661892 | 59.5750942 | 33.8559658 | 38.2265392 | 68.0848961 |
| ENSMUSG00 | Zw10 | 2828 | 16.4898086 | 22.8152921 | 17.6932413 | 19.393461 | 13.153546 | 20.8610483 |
| ENSMUSG00 | Fam46a | 5476 | 0.07985944 | 0.18479096 | 0 | 0.00658912 | 0.26090915 | 0.11455747 |
| ENSMUSG00 | Usp28 | 4471 | 11.8706255 | 9.04236747 | 13.57395 | 6.15759337 | 12.0518571 | 11.9566748 |
| ENSMUSG00 | Tmprss5 | 3044 | 0 | 0 | 0 | 0 | 0 | 0 |
| ENSMUSG00 | Htr3a | 2073 | 0 | 0.8600571 | 0 | 1.25321139 | 1.72303247 | 0.40786967 |
| ENSMUSG00 | Nnmt | 1877 | 0 | 0 | 0 | 0 | 0 | 0 |
| ENSMUSG00 | Cyp19a1 | 2404 | 0 | 0 | 0 | 0 | 0 | 0 |
| ENSMUSG00 | Paqr5 | 6494 | 91.0813118 | 80.3527844 | 90.0032292 | 75.0255385 | 85.7720848 | 78.3127384 |
| ENSMUSG00 | Idh3a | 2548 | 300.896563 | 253.659634 | 359.825207 | 284.323112 | 229.3183 | 292.131939 |
| ENSMUSG00 | Tle3 | 8256 | 2.00318313 | 1.85601848 | 2.44673477 | 2.25512778 | 2.37331981 | 2.47110346 |
| ENSMUSG00 | Acsbg1 | 3586 | 0.13303563 | 0 | 0 | 0 | 0 | 0 |
| ENSMUSG00 | Dnaja4 | 3581 | 7.19395459 | 5.62466708 | 7.23325328 | 6.52922783 | 5.54293778 | 4.01388855 |
| ENSMUSG00 | Imp3 | 897 | 17.329312 | 22.5622137 | 20.9921796 | 15.9291969 | 14.1076268 | 25.6935007 |
| ENSMUSG00 | Thsd4 | 16755 | 0.20880229 | 0.71898601 | 0.30866103 | 0.00215351 | 0.72481513 | 0.06674188 |
| ENSMUSG00 | Ptpn9 | 4041 | 15.1505667 | 17.7434858 | 11.6712117 | 11.2505261 | 16.9456561 | 11.6495985 |
| ENSMUSG00 | Crabp1 | 798 | 0 | 0 | 0 | 0 | 0 | 0 |
| ENSMUSG00 | Nr2e3 | 3402 | 0.64272529 | 0 | 0 | 0 | 0.13499041 | 0 |
| ENSMUSG00 | Ireb2 | 5786 | 16.5727997 | 16.1148921 | 17.6089264 | 20.5042797 | 20.2571008 | 18.4360723 |
| ENSMUSG00 | Pkm2 | 2420 | 218.573003 | 222.175339 | 207.127645 | 201.567422 | 202.73496 | 160.942913 |
| ENSMUSG00 | Man2c1 | 6070 | 7.6956655 | 10.2803036 | 5.8765877 | 5.43905614 | 9.14607857 | 4.39000371 |
| ENSMUSG00 | Celf6 | 2863 | 0 | 0 | 0 | 0 | 0 | 0 |
| ENSMUSG00 | Neil1 | 1748 | 8.21037098 | 9.26238247 | 7.83899962 | 6.68797622 | 10.8883514 | 8.87830076 |
| ENSMUSG00 | Commd4 | 816 | 50.9123494 | 62.7132372 | 51.0812965 | 48.7726656 | 36.8315384 | 60.2983446 |
| ENSMUSG00 | 1700017B05l | 4321 | 4.02983117 | 2.33070271 | 1.6060378 | 3.25665294 | 3.6489586 | 1.62221298 |
| ENSMUSG00 | Psma4 | 2466 | 33.2101743 | 35.5829323 | 38.1075094 | 38.0719707 | 29.5274283 | 36.8085967 |
| ENSMUSG00 | Chrna3 | 3117 | 0 | 0 | 0 | 0 | 0 | 0 |
| ENSMUSG00 | 2310046O06l | 3050 | 8.58979071 | 7.47284824 | 4.89843375 | 4.8503732 | 6.574874 | 2.72746393 |
| ENSMUSG00 | Mpi | 2766 | 23.6290708 | 23.2395929 | 25.2490643 | 20.5717224 | 18.7059816 | 21.930151 |
| ENSMUSG00 | Ube2q2 | 4334 | 26.3538291 | 26.8171893 | 18.3171458 | 25.9084731 | 22.6051319 | 29.8863087 |
| ENSMUSG00 | Ulk3 | 2810 | 22.990269 | 22.0354371 | 18.7661205 | 18.1694281 | 27.4380263 | 15.995945 |
| ENSMUSG00 | Fbxo22 | 2588 | 20.5075605 | 23.2553579 | 20.6320813 | 18.9890822 | 21.6881814 | 15.5975526 |
| ENSMUSG00 | Cyp1a2 | 1885 | 0 | 0 | 0 | 0 | 0 | 0 |
| ENSMUSG00 | Nrg4 | 7244 | 0.01646417 | 0.14634204 | 0.09762969 | 0.09961912 | 0.0422637 | 0.24473379 |
| ENSMUSG00 | Csk | 2735 | 1.97687216 | 2.67800337 | 1.82625531 | 2.26914503 | 2.70523748 | 2.71250526 |
| ENSMUSG00 | AI118078 | 3766 | 0.08445137 | 0.1535415 | 0.28168892 | 0 | 0.37937826 | 0.07242337 |
| ENSMUSG00 | Etfa | 1813 | 238.883731 | 214.858907 | 302.927689 | 233.88648 | 205.45645 | 313.560566 |
| ENSMUSG00 | Cyp1a1 | 2612 | 69.4046674 | 43.9802749 | 113.787593 | 56.4714388 | 22.153105 | 49.7563767 |
| ENSMUSG00 | Clk3 | 2481 | 28.3784587 | 22.5880017 | 25.0494927 | 24.7382337 | 31.7963632 | 20.7665431 |
| ENSMUSG00 | Isl2 | 2186 | 0 | 0 | 0 | 0 | 0 | 0 |
| ENSMUSG00 | Rcn2 | 3538 | 45.4524926 | 52.4766426 | 48.9368624 | 48.4221445 | 46.2669909 | 65.5809373 |
| ENSMUSG00 | Pstpip1 | 4284 | 0 | 0 | 0 | 0 | 0 | 0.03183315 |
| ENSMUSG00 | Cyp11a1 | 1766 | 0 | 0 | 0 | 0 | 0 | 0 |
| ENSMUSG00 | Tspan3 | 3224 | 146.135918 | 134.978833 | 129.644115 | 126.734824 | 124.62211 | 142.870381 |
| ENSMUSG00 | Stra6 | 4789 | 0 | 0 | 0 | 0 | 0.18113351 | 0 |
| ENSMUSG00 | Tmem30a | 5639 | 111.433787 | 109.746139 | 137.998483 | 136.413036 | 101.699847 | 158.322534 |
| ENSMUSG00 | Hmg20a | 3681 | 23.6956056 | 23.3666936 | 22.3711039 | 22.0844462 | 25.4508085 | 18.4942949 |
| ENSMUSG00 | Cox7a2 | 750 | 156.583586 | 136.142761 | 184.82264 | 133.07058 | 95.5213755 | 334.496176 |
| ENSMUSG00 | Col12a1 | 15966 | 0 | 0 | 0 | 0.09265714 | 0 | 0.01195807 |
| ENSMUSG00 | Stoml1 | 1980 | 11.4648381 | 13.2877765 | 10.1574942 | 9.56720878 | 10.9526312 | 6.46050798 |
| ENSMUSG00 | Loxl1 | 4481 | 0.05323207 | 0 | 0 | 0 | 0.05693637 | 0 |
| ENSMUSG00 | Nptn | 3801 | 89.6354822 | 75.6328108 | 115.196911 | 89.1276812 | 71.2301895 | 116.396191 |
| ENSMUSG00 | Hcn4 | 3790 | 0.10489573 | 0 | 0 | 0 | 0 | 0 |
| ENSMUSG00 | Neo1 | 7358 | 6.26751256 | 6.54230167 | 6.1034347 | 10.3175621 | 12.170602 | 4.88927007 |

| | | | | | | | | |
|---|---|---|---|---|---|---|---|---|
| ENSMUSG00 | Myd88 | 3094 | 12.3352493 | 12.1322684 | 12.1290772 | 9.27124286 | 8.2625127 | 8.40982736 |
| ENSMUSG00 | Scn5a | 11226 | 0 | 0 | 0 | 0 | 0 | 0 |
| ENSMUSG00 | Wdr48 | 3817 | 18.5601955 | 18.5575242 | 26.2408636 | 23.2637966 | 31.4820841 | 15.7273987 |
| ENSMUSG00 | Gorasp1 | 3474 | 13.206052 | 12.899652 | 11.5784908 | 12.4324143 | 14.9965255 | 11.2191881 |
| ENSMUSG00 | Ttc21a | 5473 | 0 | 0 | 0.00807635 | 0.01318547 | 0 | 0.09468631 |
| ENSMUSG00 | Carnp1 | 2866 | 1.94199835 | 1.86625788 | 2.35969364 | 2.15283657 | 1.0504383 | 2.30302257 |
| ENSMUSG00 | Mobp | 6489 | 0 | 0 | 0 | 0 | 0 | 0 |
| ENSMUSG00 | Rpsa | 1854 | 0.81483725 | 0.49382007 | 0.92981223 | 0.48654321 | 0.52292402 | 0.51489341 |
| ENSMUSG00 | Slc25a38 | 2905 | 11.0986901 | 14.331527 | 14.9418967 | 12.395828 | 9.57293453 | 11.8205752 |
| ENSMUSG00 | Hhat1 | 2116 | 90.1074123 | 63.8310952 | 109.418362 | 65.2409746 | 48.0602869 | 66.4851556 |
| ENSMUSG00 | Nktr | 8841 | 9.07437616 | 7.49966569 | 8.8543676 | 9.66840442 | 8.54768268 | 10.5476747 |
| ENSMUSG00 | Deb1 | 709 | 33.363203 | 28.408932 | 38.9026094 | 31.7562705 | 26.2688677 | 58.9347635 |
| ENSMUSG00 | Pccb | 3522 | 47.5327452 | 41.5645688 | 57.5929181 | 36.358653 | 39.0448844 | 42.6079117 |
| ENSMUSG00 | Vipr1 | 5551 | 0 | 0 | 0 | 0 | 0 | 0 |
| ENSMUSG00 | Lyzl4 | 1524 | 0 | 0 | 0 | 0 | 0 | 0 |
| ENSMUSG00 | Amotl2 | 4625 | 25.3833371 | 28.9535401 | 20.3567407 | 15.9307103 | 33.0761243 | 17.290647 |
| ENSMUSG00 | Cck | 844 | 0 | 0 | 0 | 0 | 0 | 0 |
| ENSMUSG00 | Cep63 | 7862 | 8.24741652 | 6.09225687 | 7.89921005 | 8.30685584 | 7.39239914 | 6.24798083 |
| ENSMUSG00 | Trak1 | 5496 | 25.2956542 | 17.1493233 | 28.3338961 | 25.7813299 | 39.318905 | 24.3119465 |
| ENSMUSG00 | Ephb1 | 7494 | 0 | 0.02572001 | 0 | 0.00962958 | 0.2451227 | 0 |
| ENSMUSG00 | Abhd5 | 3717 | 12.6207875 | 13.3656771 | 19.4074274 | 11.7167145 | 11.8883082 | 11.5864018 |
| ENSMUSG00 | Ryk | 4662 | 10.1307402 | 12.2688346 | 9.82263178 | 10.9902409 | 14.6884162 | 7.09655457 |
| ENSMUSG00 | Slco2a1 | 4247 | 2.56486971 | 0.55595377 | 0.69732128 | 0 | 2.13861429 | 0.49450135 |
| ENSMUSG00 | Rab6b | 4726 | 0.46266429 | 1.18273959 | 0.25252852 | 0.28248744 | 0.17275117 | 1.58130579 |
| ENSMUSG00 | 1110059G10 | 3931 | 6.78604106 | 6.80322447 | 7.23016664 | 6.5720539 | 5.77632593 | 19.8228553 |
| ENSMUSG00 | Srprb | 5556 | 9.78145102 | 15.116805 | 12.4108847 | 13.1183819 | 9.20238095 | 9.24863577 |
| ENSMUSG00 | Trf | 2828 | 0 | 0 | 0 | 0 | 0 | 0 |
| ENSMUSG00 | Topbp1 | 5200 | 5.41286158 | 4.7352446 | 4.9982084 | 4.96821998 | 6.57455227 | 3.16805425 |
| ENSMUSG00 | Bfsp2 | 1822 | 0 | 0 | 0 | 0 | 0 | 0 |
| ENSMUSG00 | Uba5 | 2565 | 37.3686018 | 33.7023303 | 41.996059 | 33.4233676 | 22.8773225 | 32.2191645 |
| ENSMUSG00 | Nphp3 | 6804 | 1.07510412 | 0.84276699 | 0.91599939 | 0.5674278 | 1.4473972 | 1.7918524 |
| ENSMUSG00 | Dnajc13 | 7761 | 9.53291447 | 7.61818896 | 9.22081998 | 7.90820226 | 11.5254783 | 7.4890164 |
| ENSMUSG00 | Acpp | 7778 | 1.51293677 | 0.7372315 | 0.72741526 | 0.43606482 | 0.03936208 | 0.52599587 |
| ENSMUSG00 | Gnai2 | 3118 | 22.8103126 | 32.3303518 | 30.961137 | 29.3586103 | 24.531282 | 25.0265372 |
| ENSMUSG00 | Mrpl3 | 3769 | 12.9846104 | 11.2123931 | 17.2163188 | 15.3652644 | 8.66460083 | 13.5323893 |
| ENSMUSG00 | Cpne4 | 5034 | 0.02369218 | 0.66048163 | 0.12292924 | 0.21503006 | 0 | 0 |
| ENSMUSG00 | Nudt16 | 1603 | 10.9370973 | 5.08016764 | 10.3955676 | 3.35385192 | 7.03482786 | 9.44318452 |
| ENSMUSG00 | 1700080E11 | 868 | 0 | 0 | 0 | 0 | 0 | 0 |
| ENSMUSG00 | Aste1 | 2765 | 11.7469178 | 11.5194352 | 11.030478 | 7.26860716 | 12.2906207 | 8.59175797 |
| ENSMUSG00 | Atp2c1 | 6892 | 44.8720085 | 49.9763042 | 58.6578723 | 46.6888673 | 51.2558909 | 61.0275989 |
| ENSMUSG00 | Pik3r4 | 4864 | 10.4047547 | 11.6503405 | 13.0133715 | 8.70152924 | 15.1064928 | 8.47846516 |
| ENSMUSG00 | Col6a4 | 8712 | 0 | 0 | 0 | 0 | 0 | 0 |
| ENSMUSG00 | Manf | 3517 | 5.48234519 | 5.65851595 | 6.69877234 | 8.33057155 | 3.03227539 | 8.00324928 |
| ENSMUSG00 | Mapkapk3 | 6697 | 14.7636076 | 13.0593389 | 15.7679861 | 14.8110408 | 17.1814958 | 14.1240031 |
| ENSMUSG00 | Cish | 1821 | 25.2155848 | 27.3347582 | 18.2293158 | 19.279423 | 21.0158055 | 19.815677 |
| ENSMUSG00 | Hemk1 | 3039 | 4.12075563 | 4.70923768 | 4.47981825 | 5.35472263 | 3.60996078 | 4.89130582 |
| ENSMUSG00 | Rbm5 | 3062 | 74.6681151 | 86.4428892 | 66.5914768 | 112.771119 | 108.001941 | 74.2882075 |
| ENSMUSG00 | Rbm6 | 4202 | 26.1220566 | 23.473976 | 28.8437538 | 29.9252559 | 29.9212256 | 27.6835641 |
| ENSMUSG00 | Mon1a | 2075 | 45.6756942 | 45.0514206 | 24.4121986 | 15.5457098 | 21.1482811 | 22.29291 |
| ENSMUSG00 | Mst1r | 4687 | 0 | 0.81218881 | 0.24519904 | 0.85451396 | 0.35926401 | 0.17457631 |
| ENSMUSG00 | Traip | 2682 | 0.08893844 | 0.48509841 | 0.14832833 | 0.09417387 | 0.19025494 | 0 |
| ENSMUSG00 | Bsn | 9812 | 0.01215516 | 0.12277426 | 0.08559264 | 0.01470935 | 0.04160324 | 0.07783224 |
| ENSMUSG00 | Apeh | 2366 | 29.1697017 | 35.3964651 | 31.7035197 | 37.3935643 | 41.1057787 | 18.8939703 |
| ENSMUSG00 | Mst1 | 2344 | 0.03392106 | 0 | 0.4525786 | 0 | 0 | 0 |
| ENSMUSG00 | Amigo3 | 2542 | 1.25115596 | 1.64918193 | 1.06070512 | 2.48401172 | 2.83033792 | 0.98712306 |
| ENSMUSG00 | Ip6k1 | 4572 | 20.0429533 | 26.327588 | 28.3657497 | 27.2667244 | 31.1381427 | 21.0644688 |
| ENSMUSG00 | Cdhr4 | 3105 | 0 | 0 | 0 | 0 | 0.03286723 | 0 |
| ENSMUSG00 | Uba7 | 5028 | 3.28132944 | 1.84963869 | 0.58021512 | 0.76785576 | 5.73387079 | 1.04151367 |
| ENSMUSG00 | Nckipsd | 2924 | 7.12444316 | 7.99262122 | 4.76182645 | 6.37975958 | 8.07975787 | 5.07435149 |
| ENSMUSG00 | Ip6k2 | 2835 | 12.2141176 | 14.0225096 | 25.0087323 | 17.181926 | 13.3730502 | 36.8568406 |
| ENSMUSG00 | Prkar2a | 4178 | 22.1043517 | 19.6182714 | 20.281219 | 18.3173807 | 24.0476146 | 16.6683844 |
| ENSMUSG00 | Slc25a20 | 1774 | 26.2197927 | 31.1011704 | 46.9426563 | 40.7600999 | 33.854591 | 41.0657054 |
| ENSMUSG00 | Qars | 3751 | 19.6180737 | 17.6122666 | 16.3090778 | 10.0714211 | 14.8005057 | 14.2299304 |
| ENSMUSG00 | Nicn1 | 2108 | 27.6855054 | 25.7390578 | 19.9201854 | 26.4966817 | 28.1516485 | 13.8055219 |
| ENSMUSG00 | Amt | 1617 | 3.54037674 | 4.38058565 | 2.733571 | 5.84631767 | 2.80850182 | 1.87228507 |
| ENSMUSG00 | Klhdc8b | 1891 | 13.4130072 | 9.50478208 | 11.2666781 | 10.9142937 | 12.9522265 | 12.8945148 |
| ENSMUSG00 | 1700102P08 | 1090 | 0.32825626 | 0.44207745 | 1.01380374 | 0.43033631 | 2.34065944 | 0.27524865 |
| ENSMUSG00 | Usp4 | 3751 | 47.3758992 | 39.0141895 | 58.0127095 | 50.5975884 | 52.4955437 | 41.0392066 |
| ENSMUSG00 | Nt5m | 3354 | 5.99769623 | 9.03675376 | 8.09181026 | 8.1329832 | 5.46167827 | 5.82249305 |
| ENSMUSG00 | Srek1 | 4268 | 36.9983057 | 36.2075726 | 28.7084135 | 36.1412231 | 37.2057361 | 44.311703 |
| ENSMUSG00 | Oas1d | 2065 | 0 | 0 | 0 | 0 | 0 | 0 |
| ENSMUSG00 | Eml4 | 5457 | 13.6088029 | 15.7177693 | 11.6397377 | 10.8437884 | 23.3111151 | 9.15152363 |
| ENSMUSG00 | Thsd7a | 14070 | 2.54299449 | 1.44182602 | 5.74592544 | 4.0877597 | 2.83600753 | 3.82659118 |
| ENSMUSG00 | Flcn | 5557 | 16.6333439 | 13.501222 | 18.3425319 | 11.8693499 | 16.6843485 | 11.6028877 |
| ENSMUSG00 | Atxn2l | 4908 | 13.4786307 | 11.6637109 | 13.3380052 | 17.2617442 | 16.010721 | 13.9540789 |
| ENSMUSG00 | Chsy1 | 4208 | 0.17005671 | 0.44659488 | 0.07352968 | 0.54020171 | 0.25464684 | 0.11666908 |
| ENSMUSG00 | Gpr19 | 3952 | 1.51899736 | 1.35341475 | 1.30860719 | 2.24599771 | 1.73014533 | 1.51832506 |
| ENSMUSG00 | Fhl3 | 1809 | 0 | 0.02663706 | 0 | 0 | 0 | 0 |
| ENSMUSG00 | Pygm | 3504 | 27.2297812 | 29.9652561 | 8.41399238 | 33.2090735 | 17.9990298 | 4.7715051 |
| ENSMUSG00 | Glt25d2 | 4231 | 6.35185649 | 12.9150143 | 2.39239472 | 9.67077251 | 11.2038533 | 7.12325133 |
| ENSMUSG00 | Crebl2 | 4134 | 14.011547 | 14.1855104 | 15.1937153 | 10.552296 | 13.2071171 | 15.2009601 |
| ENSMUSG00 | 3-Mar | 6000 | 1.06014615 | 1.87927124 | 1.9154132 | 2.17093635 | 1.91348909 | 1.0864398 |
| ENSMUSG00 | Fam189b | 3880 | 2.27466925 | 2.62044827 | 3.12147036 | 2.07378761 | 2.35405187 | 2.67122756 |
| ENSMUSG00 | Oas3 | 5371 | 0 | 0 | 0 | 0 | 0 | 0 |
| ENSMUSG00 | 1700025G04 | 12436 | 1.54086053 | 1.37553771 | 2.05085746 | 2.01648608 | 1.19400735 | 2.98713884 |
| ENSMUSG00 | Pon2 | 4642 | 15.8268264 | 12.249031 | 18.8919553 | 12.9341926 | 11.4320187 | 15.5116432 |
| ENSMUSG00 | A930018P22 | 820 | 0.19392917 | 0 | 0 | 0 | 0 | 0.06652351 |
| ENSMUSG00 | Prorsd1 | 2925 | 9.85392252 | 14.0193717 | 16.4566862 | 11.5215828 | 9.45514383 | 10.0053634 |
| ENSMUSG00 | Cd59a | 1893 | 86.5464 | 86.419953 | 81.6554913 | 85.392266 | 68.331676 | 71.8822901 |
| ENSMUSG00 | 6820408C15 | 1775 | 0.69432107 | 1.68313206 | 1.49414681 | 1.09770727 | 1.20738467 | 0.98342361 |
| ENSMUSG00 | Malt1 | 5490 | 4.57658719 | 4.9327469 | 2.3590419 | 3.22700981 | 4.10813445 | 5.87224474 |
| ENSMUSG00 | Oas2 | 6063 | 0.39342386 | 0.18279534 | 0.13851806 | 0.32136408 | 2.6762968 | 0.00899708 |
| ENSMUSG00 | Nlrp3 | 4046 | 0 | 0 | 0 | 0 | 0.03783468 | 0 |
| ENSMUSG00 | Lmo2 | 2248 | 1.27330721 | 4.07269751 | 1.49436836 | 0.78648585 | 3.15510064 | 2.17177955 |
| ENSMUSG00 | Kank1 | 5583 | 21.6472615 | 19.1433868 | 19.0171641 | 18.7293149 | 24.2656328 | 19.0331353 |
| ENSMUSG00 | Exd2 | 3314 | 14.7313609 | 15.3399808 | 14.8050832 | 20.6105342 | 24.7895187 | 14.764847 |
| ENSMUSG00 | 2810474O19 | 7064 | 15.7299785 | 13.1857861 | 15.0363857 | 33.3288987 | 22.6165887 | 23.7108663 |
| ENSMUSG00 | Syde1 | 3280 | 3.50284569 | 5.70010352 | 4.33932727 | 4.2902431 | 3.50028493 | 4.58180678 |
| ENSMUSG00 | Trib3 | 3376 | 1.15403942 | 0.22837176 | 0.48443963 | 0.64126856 | 0.77083684 | 0.11310573 |
| ENSMUSG00 | Mdfi | 3914 | 0 | 0 | 0 | 0 | 0 | 0 |
| ENSMUSG00 | Mansc1 | 2369 | 2.24872705 | 3.25446633 | 2.48157244 | 2.37602319 | 3.31703752 | 3.3388119 |
| ENSMUSG00 | Gm106 | 2743 | 6.17420149 | 7.6241035 | 2.70722188 | 4.52505407 | 5.58071919 | 3.48017635 |
| ENSMUSG00 | Abtb2 | 6016 | 3.58169389 | 3.38810921 | 5.59870419 | 2.32709995 | 5.13995741 | 3.91710245 |
| ENSMUSG00 | Folr2 | 1052 | 2.64532665 | 0.27482762 | 0.08403392 | 0 | 1.01858735 | 1.81485242 |
| ENSMUSG00 | Bmp8a | 2398 | 17.1920072 | 13.3828901 | 27.1699402 | 16.2956023 | 17.3634373 | 21.6104314 |
| ENSMUSG00 | Mier3 | 5861 | 4.80240236 | 6.19081913 | 5.61855879 | 5.00506777 | 6.99969394 | 6.5243208 |
| ENSMUSG00 | Snx33 | 4562 | 3.18950151 | 2.28151502 | 3.85627653 | 3.89135596 | 3.88122587 | 4.61552422 |
| ENSMUSG00 | Ablim3 | 4537 | 0.31545014 | 0.01062077 | 0 | 0 | 0.30366148 | 0 |
| ENSMUSG00 | Inppl1 | 5065 | 10.8160024 | 8.22927393 | 9.10217616 | 10.0730525 | 11.0716658 | 8.27682794 |
| ENSMUSG00 | Pram1 | 2417 | 0.0657931 | 0 | 0 | 0 | 0.27444886 | 0.04513801 |
| ENSMUSG00 | Ccdc88a | 11497 | 4.24630165 | 4.90791717 | 5.65162297 | 6.03198135 | 4.93087793 | 4.20139046 |
| ENSMUSG00 | Tpcn1 | 4617 | 2.02350832 | 1.28371938 | 2.02962762 | 1.57082325 | 1.73513992 | 1.22874701 |
| ENSMUSG00 | D430042O09 | 7344 | 1.39663861 | 1.62721101 | 1.84776223 | 2.05860249 | 2.320644 | 1.54125481 |
| ENSMUSG00 | Heyl | 5200 | 1.57492865 | 3.96611485 | 0.73103048 | 2.04002329 | 3.71903778 | 0.84970992 |
| ENSMUSG00 | Gpbp1 | 4611 | 42.4282628 | 47.2563633 | 38.9294479 | 57.8439543 | 40.3806648 | 55.1053002 |
| ENSMUSG00 | Gab3 | 4193 | 0 | 0.40222406 | 0.04216727 | 0 | 0 | 0 |
| ENSMUSG00 | Slc24a6 | 3494 | 4.66506783 | 4.49593021 | 4.12415593 | 0.86745614 | 4.64407198 | 3.38786303 |
| ENSMUSG00 | Bet1 | 2829 | 33.8532883 | 32.2264929 | 38.9363707 | 37.0131118 | 22.7805983 | 46.7496379 |
| ENSMUSG00 | Kap | 604 | 32.2519627 | 41.3254587 | 6.36682176 | 13.202205 | 2.70338415 | 36.3511334 |
| ENSMUSG00 | Ilvbl | 2368 | 29.9005535 | 33.270622 | 38.1352894 | 35.8534843 | 33.8954768 | 30.9143293 |
| ENSMUSG00 | Gng11 | 878 | 33.1446603 | 16.2999696 | 47.7762517 | 19.6848512 | 10.635338 | 62.1290188 |
| ENSMUSG00 | Trpa1 | 4236 | 0 | 0 | 0 | 0 | 0 | 0 |
| ENSMUSG00 | Chrm1 | 6936 | 0.67634756 | 0.32652289 | 1.31917265 | 0.35374554 | 0.38990743 | 0.79039828 |
| ENSMUSG00 | Mctp2 | 5764 | 1.24149662 | 0.65207191 | 0.19938375 | 0.24413597 | 0.71706045 | 2.33282393 |
| ENSMUSG00 | Gtf3c1 | 6923 | 12.4325604 | 12.6399796 | 12.5333263 | 14.859152 | 17.8957158 | 11.9570317 |
| ENSMUSG00 | Cntrob | 6952 | 0.85206654 | 1.9476971 | 1.33521102 | 1.58300109 | 2.81114815 | 1.45945998 |
| ENSMUSG00 | Troap | 2568 | 0.03096221 | 0 | 0.03442511 | 0 | 0 | 0 |
| ENSMUSG00 | Alas1 | 4660 | 41.4446619 | 46.9869513 | 39.0512849 | 41.1381765 | 31.634163 | 33.0222134 |
| ENSMUSG00 | Pdxk | 5329 | 28.3338929 | 24.4051806 | 19.442507 | 23.6371585 | 27.8351801 | 19.0855939 |
| ENSMUSG00 | Lama1 | 9455 | 0.26910108 | 0.44338662 | 0.15427402 | 0 | 0.11872874 | 0.19904285 |
| ENSMUSG00 | Snn1 | 2858 | 72.4862872 | 83.5085542 | 48.9344406 | 77.441309 | 53.4902106 | 40.2343874 |
| ENSMUSG00 | Cdv3 | 7472 | 2.19740544 | 1.65737629 | 1.7687836 | 1.5259537 | 1.61164677 | 1.87622652 |
| ENSMUSG00 | Slc10a3 | 3140 | 11.4075439 | 8.80860439 | 9.40344941 | 11.606008 | 9.40900369 | 6.49727075 |
| ENSMUSG00 | Alox12b | 2416 | 0 | 0 | 0 | 0 | 0 | 0 |
| ENSMUSG00 | Cyp2c38 | 2860 | 0 | 0 | 0 | 0 | 0 | 0 |
| ENSMUSG00 | Arap1 | 7510 | 7.29468071 | 9.0854863 | 7.13350649 | 5.8855532 | 8.62896102 | 5.58567179 |
| ENSMUSG00 | Fanca | 8447 | 4.04755691 | 7.01661227 | 2.16116505 | 3.74190494 | 6.03472824 | 2.42168574 |
| ENSMUSG00 | Igdcc4 | 6407 | 0 | 0 | 0 | 0 | 0.00796416 | 0 |
| ENSMUSG00 | Loxhd1 | 8174 | 0 | 0 | 0 | 0 | 0 | 0 |
| ENSMUSG00 | Ank2 | 17181 | 0.1087543 | 0.3085099 | 0.12863583 | 0.28141517 | 0.07424826 | 0.5635584 |
| ENSMUSG00 | Ppp1r9a | 14561 | 1.06480581 | 1.73075402 | 1.96101851 | 2.17815515 | 1.87130586 | 1.7626149 |
| ENSMUSG00 | Pwp2 | 3845 | 5.60402352 | 5.4640543 | 6.40323188 | 4.22286607 | 4.44573105 | 3.65317015 |
| ENSMUSG00 | Trpc1 | 3001 | 0 | 0.93129412 | 0.33876787 | 0.01202334 | 0.35706561 | 0 |
| ENSMUSG00 | 2410131K14 | 5644 | 6.0365781 | 5.61776735 | 4.58934798 | 3.86775991 | 3.91467412 | 2.58055588 |
| ENSMUSG00 | Prr5l | 6152 | 4.95004845 | 7.01021875 | 1.19270253 | 3.46626923 | 4.08078298 | 3.97237919 |
| ENSMUSG00 | Abcc10 | 6175 | 4.5903899 | 3.34768967 | 2.50536762 | 4.67459686 | 5.98268763 | 2.8754316 |
| ENSMUSG00 | Alpk2 | 8238 | 0 | 0 | 0 | 0 | 0 | 0 |
| ENSMUSG00 | Zswim6 | 5443 | 2.95080179 | 2.54964089 | 3.61378287 | 4.2028323 | 2.4561658 | 3.37738506 |
| ENSMUSG00 | Abcc4 | 5698 | 10.9540373 | 20.2792363 | 7.21441102 | 21.460521 | 11.2386981 | 5.70575114 |
| ENSMUSG00 | Rnft2 | 8500 | 0.18708461 | 0.07369691 | 0.33281388 | 0.16979786 | 0.42622032 | 0.34975714 |
| ENSMUSG00 | Rspo4 | 2366 | 0 | 0 | 0 | 0 | 0 | 0 |
| ENSMUSG00 | Ugt8a | 3577 | 11.4253939 | 14.8587212 | 3.46002685 | 15.2519014 | 9.88573622 | 3.29400172 |
| ENSMUSG00 | Pkd1 | 14142 | 7.93591582 | 8.39906519 | 6.66997122 | 10.5322217 | 9.02396874 | 4.75599352 |
| ENSMUSG00 | P2ry2 | 2997 | 3.80708138 | 5.45051848 | 1.17989571 | 1.31229324 | 5.56744241 | 1.67451906 |
| ENSMUSG00 | Rag2 | 3545 | 0 | 0 | 0 | 0.04071317 | 0 | 0 |
| ENSMUSG00 | Fbxw8 | 4967 | 11.5416555 | 9.94385005 | 4.50294596 | 6.08752835 | 9.38959281 | 4.06346548 |
| ENSMUSG00 | Psmf1 | 9056 | 6.87028788 | 8.21553298 | 6.3842768 | 6.74546173 | 6.37829698 | 5.79164436 |
| ENSMUSG00 | Smap2 | 3150 | 3.28140474 | 2.30988977 | 2.9748542 | 2.40546964 | 3.67713883 | 3.01319824 |
| ENSMUSG00 | Cyb5r4 | 6987 | 14.0313461 | 13.6621357 | 13.0891102 | 15.1000284 | 13.0432385 | 13.8891035 |
| ENSMUSG00 | Arhgef17 | 10372 | 0.92374381 | 2.16959781 | 1.05688942 | 2.40036741 | 1.83994066 | 1.02293045 |
| ENSMUSG00 | Ccdc85a | 10219 | 0 | 0.13674595 | 0.03460365 | 0.00706176 | 0.05991942 | 0.01067605 |
| ENSMUSG00 | Acsl3 | 3804 | 63.6045884 | 45.0576169 | 72.9376889 | 54.9293165 | 38.7526813 | 53.9900193 |
| ENSMUSG00 | Gm6685 | 686 | 0 | 0 | 0 | 0 | 0 | 0.0795179 |
| ENSMUSG00 | Rims3 | 8265 | 0.30303633 | 0 | 0.08556921 | 0 | 0.03086895 | 0.01650008 |
| ENSMUSG00 | Rangrf | 1300 | 0 | 0.07413299 | 0.06800284 | 0 | 0 | 0 |
| ENSMUSG00 | 1700031F05 | 1498 | 0 | 0.12866874 | 0 | 0 | 0 | 0 |
| ENSMUSG00 | Nfyc | 2845 | 18.5711893 | 16.6831794 | 16.7640733 | 15.3586488 | 15.8370087 | 17.055038 |
| ENSMUSG00 | Fbxo21 | 3897 | 40.4593882 | 34.5107416 | 36.7838278 | 32.9432678 | 34.1616152 | 23.8101932 |
| ENSMUSG00 | Styk1 | 4800 | 5.20962443 | 5.37078053 | 2.77182391 | 4.63052905 | 4.06084908 | 2.73882836 |
| ENSMUSG00 | 1700065I17R | 1006 | 0 | 0 | 0 | 0 | 0 | 0 |
| ENSMUSG00 | Slc16a1 | 4421 | 0.35070431 | 0.7411622 | 0.26995018 | 0.27749141 | 0 | 0.64161106 |
| ENSMUSG00 | Atg12 | 2488 | 46.6264036 | 39.1224329 | 53.8310226 | 49.6418161 | 46.3913433 | 53.2337814 |
| ENSMUSG00 | Sgpp2 | 3993 | 74.4432076 | 66.4569839 | 68.92078 | 70.9261126 | 79.6895767 | 48.2036387 |
| ENSMUSG00 | Cspg4 | 8040 | 0.13350721 | 0.20377351 | 0.36834869 | 0.09424415 | 0.09519846 | 0.50885521 |
| ENSMUSG00 | Lrig2 | 7734 | 6.07075543 | 9.38308532 | 6.14390758 | 6.02300485 | 8.64294703 | 6.99675256 |
| ENSMUSG00 | Emr4 | 3474 | 0 | 0 | 0 | 0 | 0 | 0 |
| ENSMUSG00 | Odf4 | 1500 | 0.07951096 | 0 | 0 | 0.31271105 | 0.17008792 | 0 |
| ENSMUSG00 | Itgbl1 | 4498 | 0 | 0 | 0 | 0 | 0.18150778 | 0 |
| ENSMUSG00 | Hspa13 | 6183 | 9.97905641 | 9.73392629 | 12.7751405 | 12.4241748 | 10.2003236 | 9.69147379 |
| ENSMUSG00 | Camkv | 3020 | 0 | 0 | 0 | 0 | 0 | 0 |
| ENSMUSG00 | Fshr | 2350 | 0 | 0 | 0 | 0 | 0 | 0 |
| ENSMUSG00 | Nup93 | 2860 | 14.1368265 | 14.0347225 | 12.4414278 | 13.5496909 | 16.9850034 | 8.4589528 |
| ENSMUSG00 | Rbm11 | 1774 | 0.6498923 | 0.9506908 | 1.39532312 | 2.52208203 | 0.69032301 | 0 |
| ENSMUSG00 | Ucp3 | 2739 | 0 | 0 | 0.01613795 | 0.210775 | 0.05588869 | 0.05974729 |
| ENSMUSG00 | Rasgrp2 | 4506 | 0.10587345 | 0.24595832 | 0 | 0 | 0.0113241 | 0.12711217 |
| ENSMUSG00 | Lipi | 2068 | 0 | 0 | 0 | 0 | 0 | 0 |
| ENSMUSG00 | Ap4b1 | 2978 | 16.3801124 | 18.6888317 | 19.1917337 | 21.0094913 | 13.3820012 | 10.5966278 |
| ENSMUSG00 | Pebp1 | 2092 | 22.5382409 | 22.273561 | 28.291715 | 19.0413849 | 15.4884076 | 23.1808358 |
| ENSMUSG00 | Ift57 | 3034 | 18.7377512 | 22.917942 | 15.3992578 | 19.88437 | 23.5454602 | 20.8919781 |
| ENSMUSG00 | Fkbp1a | 3474 | 39.6982619 | 41.8337103 | 61.7349863 | 41.7841293 | 31.7703083 | 48.6688223 |
| ENSMUSG00 | Inha | 1473 | 0.21591571 | 0.91596767 | 0.36009648 | 0.5633992 | 1.0045926 | 0.51845886 |
| ENSMUSG00 | 1810008A18 | 4310 | 3.66193173 | 5.88075837 | 3.35359891 | 2.68731701 | 3.74114496 | 1.55041453 |
| ENSMUSG00 | Guca2b | 867 | 2.84295247 | 2.16755622 | 1.37652798 | 1.70630199 | 1.23593298 | 4.46712661 |
| ENSMUSG00 | 5730522E02 | 6140 | 0 | 0 | 0.02159699 | 0 | 0.01662097 | 0.19101132 |
| ENSMUSG00 | Olfr281 | 1857 | 0 | 0 | 0 | 0 | 0 | 0 |
| ENSMUSG00 | Slc16a8 | 1800 | 0.04417276 | 0 | 0.02455658 | 0.04009116 | 0.02834799 | 0.01515258 |
| ENSMUSG00 | Chpf | 4603 | 6.31354687 | 6.28109174 | 8.70977007 | 3.92724404 | 5.78661051 | 2.98640412 |
| ENSMUSG00 | Foxj3 | 10955 | 5.77007887 | 4.3633912 | 6.84311508 | 6.96609075 | 6.89356789 | 5.25326233 |
| ENSMUSG00 | Nlrp4f | 4283 | 0 | 0 | 0.0103263 | 0 | 0 | 0 |
| ENSMUSG00 | Mycbp2 | 19987 | 5.35854628 | 5.60290647 | 4.59777013 | 5.58191233 | 6.2267139 | 4.3244775 |
| ENSMUSG00 | Sox10 | 4267 | 0 | 0 | 0 | 0 | 0 | 0 |
| ENSMUSG00 | Accn4 | 3431 | 0 | 0 | 0 | 0 | 0 | 0 |
| ENSMUSG00 | Ogfod1 | 5695 | 2.71551921 | 3.05448738 | 2.46816261 | 2.60398952 | 3.05531065 | 2.73943699 |
| ENSMUSG00 | Trim33 | 8848 | 3.65743232 | 3.7250821 | 4.37622225 | 5.46042695 | 2.61244894 | 5.41608738 |
| ENSMUSG00 | 4933427E13 | 2092 | 0 | 0 | 0 | 0 | 0 | 0 |
| ENSMUSG00 | Nfatc1 | 7422 | 1.71406006 | 1.44130829 | 1.98318697 | 1.30774319 | 3.14188275 | 1.35601481 |
| ENSMUSG00 | Polr2f | 538 | 27.4889196 | 32.6019794 | 35.246451 | 30.3813498 | 18.5895347 | 52.622817 |
| ENSMUSG00 | Gmppa | 2313 | 22.7051404 | 22.853665 | 20.1421282 | 16.5668531 | 22.6122072 | 17.3930363 |
| ENSMUSG00 | Cdo1 | 1525 | 322.605948 | 480.695308 | 176.546509 | 387.438347 | 392.183706 | 275.572224 |
| ENSMUSG00 | Klra9 | 1998 | 0 | 0 | 0 | 0 | 0 | 0 |
| ENSMUSG00 | Klrc3 | 823 | 0 | 0 | 0 | 0 | 0 | 0 |
| ENSMUSG00 | 1700088E04 | 1975 | 0.36232843 | 0.73194596 | 0.35809088 | 0.43846535 | 0.41337823 | 0.3452486 |
| ENSMUSG00 | C330027C09 | 5584 | 0.34173766 | 0.62994453 | 0.2770531 | 0 | 0.48431195 | 0.90850339 |
| ENSMUSG00 | Afap1l1 | 4033 | 1.32091108 | 0.6690902 | 0.50416186 | 0.69784267 | 2.54309485 | 1.10911005 |
| ENSMUSG00 | Calhm2 | 4356 | 0.32855769 | 0.24336587 | 0.20294694 | 0.76206338 | 0.6911286 | 0.10644373 |
| ENSMUSG00 | Gm7879 | 533 | 0 | 0 | 0 | 0.27078457 | 0 | 0 |
| ENSMUSG00 | Micall1 | 6675 | 15.7592512 | 13.3333871 | 10.8004803 | 14.719312 | 13.0566367 | 9.3775726 |
| ENSMUSG00 | Dhrs7c | 1716 | 0 | 0 | 0 | 0 | 0 | 0 |
| ENSMUSG00 | Eif3l | 1880 | 130.833595 | 120.10727 | 103.357076 | 105.520788 | 98.5786154 | 79.9233977 |
| ENSMUSG00 | 1700028P14 | 862 | 0 | 0 | 0 | 0.25115112 | 0 | 0 |
| ENSMUSG00 | Npat | 6230 | 4.22441864 | 5.34459575 | 3.83129939 | 6.38820146 | 6.92910336 | 3.6731015 |
| ENSMUSG00 | Ankrd54 | 5164 | 11.9251044 | 11.9626295 | 9.23582274 | 11.4590536 | 11.6004968 | 10.7904895 |
| ENSMUSG00 | Pygb | 4788 | 29.9245513 | 35.6265675 | 21.3439142 | 35.0571577 | 40.5717236 | 19.1970623 |
| ENSMUSG00 | Lmo7 | 9817 | 13.3638673 | 10.2685176 | 13.912977 | 13.5551167 | 18.4052559 | 10.9993172 |
| ENSMUSG00 | Resp18 | 901 | 0 | 0 | 0 | 0 | 0 | 0 |
| ENSMUSG00 | Cntnap3 | 4844 | 0.45960093 | 0.91518429 | 0.67525524 | 0.68529069 | 0.11587327 | 1.29503861 |
| ENSMUSG00 | Pfkm | 3027 | 51.8778157 | 38.2689815 | 37.4116689 | 34.7946772 | 47.6718173 | 24.2291064 |
| ENSMUSG00 | Gas7 | 11371 | 0.01048865 | 0.00847532 | 0.01943622 | 0 | 0 | 0 |
| ENSMUSG00 | Entpd6 | 4026 | 9.63769225 | 8.73723367 | 5.66521386 | 9.32074672 | 13.6754743 | 6.84914562 |
| ENSMUSG00 | Senp1 | 7742 | 4.26721833 | 2.99998258 | 3.03166864 | 4.58132908 | 5.29904497 | 2.25116178 |
| ENSMUSG00 | Vax1 | 3789 | 0 | 0 | 0.04666333 | 0 | 0 | 0.00719837 |
| ENSMUSG00 | Clec1a | 5229 | 2.25805655 | 0.57134431 | 2.87409192 | 0.84874574 | 1.39544306 | 3.36434162 |
| ENSMUSG00 | Tbc1d4 | 9921 | 3.90702485 | 2.61793089 | 3.49747473 | 4.69892163 | 3.22482992 | 2.16085742 |
| ENSMUSG00 | Triobp | 10672 | 4.94708378 | 5.90590951 | 4.36137001 | 3.85096034 | 4.27451087 | 3.98947825 |
| ENSMUSG00 | 2310001A20 | 2418 | 49.6861299 | 53.7464162 | 62.4273341 | 40.2005906 | 49.9923426 | 51.1089291 |
| ENSMUSG00 | Nol12 | 1731 | 7.53310087 | 9.2141608 | 7.58402506 | 6.67027975 | 7.42844986 | 9.59575696 |
| ENSMUSG00 | Cdc14b | 7382 | 6.62950373 | 5.8748033 | 5.58061741 | 7.07759421 | 7.17493807 | 7.66291003 |
| ENSMUSG00 | Lss | 10780 | 4.99709426 | 3.2049795 | 5.15414811 | 4.18725767 | 3.94294722 | 2.98300554 |
| ENSMUSG00 | Slc7a6os | 1520 | 16.8176145 | 15.3752792 | 15.7614467 | 15.8333709 | 15.2743428 | 22.8783981 |
| ENSMUSG00 | Rnf125 | 1491 | 4.93277257 | 3.2318204 | 9.51629216 | 5.13037789 | 4.58587147 | 12.1281595 |
| ENSMUSG00 | 3830406C13 | 2546 | 39.7867103 | 32.7614813 | 41.5108431 | 32.3406228 | 26.2146503 | 49.4500137 |
| ENSMUSG00 | Slc35d2 | 2384 | 0.383547 | 3.23398941 | 1.35349603 | 1.25621212 | 3.0607264 | 2.47119215 |
| ENSMUSG00 | Hsd17b3 | 1420 | 0 | 0 | 0 | 0 | 0 | 0 |
| ENSMUSG00 | Atg9a | 4114 | 15.8867307 | 12.4155636 | 14.3220848 | 12.1384418 | 15.404657 | 12.3378959 |
| ENSMUSG00 | Ybey | 4515 | 3.0994306 | 1.53684334 | 1.75240972 | 0.71125182 | 0.90412183 | 2.63383006 |
| ENSMUSG00 | Gja1 | 3017 | 9.57979262 | 7.50667146 | 7.47197214 | 7.98899759 | 6.2577922 | 7.91027821 |
| ENSMUSG00 | Slc22a15 | 8277 | 0.51873769 | 1.04791103 | 1.62345781 | 1.06803201 | 1.41174823 | 0.33611371 |
| ENSMUSG00 | Phldb2 | 9248 | 32.2626385 | 35.0664744 | 20.2894489 | 32.3208972 | 37.6352627 | 22.3022083 |
| ENSMUSG00 | Podxl2 | 4208 | 0.44403697 | 0.36643682 | 1.58614028 | 0.58307486 | 0.47291556 | 1.12780115 |
| ENSMUSG00 | Cst10 | 831 | 0 | 0 | 0 | 0 | 0 | 0 |
| ENSMUSG00 | Abhd10 | 4188 | 6.48352273 | 5.31569634 | 6.69149199 | 6.35829724 | 5.2634657 | 10.5308242 |
| ENSMUSG00 | 1810031K17 | 5383 | 9.03970057 | 6.97329341 | 7.18495498 | 7.63467373 | 8.60708699 | 8.72504715 |
| ENSMUSG00 | Atp1a1 | 4332 | 220.637411 | 280.019736 | 223.049928 | 259.37093 | 222.987147 | 230.317077 |
| ENSMUSG00 | Dis3 | 3896 | 9.83682834 | 8.21247371 | 6.71650181 | 6.770014 | 8.29047636 | 8.04377836 |
| ENSMUSG00 | Card10 | 6531 | 12.8683335 | 7.84293186 | 11.5732892 | 6.53576155 | 6.07066208 | 7.80528262 |
| ENSMUSG00 | Mgll | 2634 | 27.5903639 | 29.3984862 | 31.9179595 | 24.6848308 | 27.2179415 | 31.3233803 |
| ENSMUSG00 | Tmprss7 | 2934 | 0.1083994 | 0.03284693 | 0 | 0.29514965 | 0.26087104 | 0 |
| ENSMUSG00 | Kbtbd12 | 2523 | 0 | 0 | 0 | 1.34431716 | 0.56628558 | 0 |
| ENSMUSG00 | Tmed7 | 4275 | 104.005917 | 105.108438 | 125.067952 | 133.735672 | 94.938899 | 133.999824 |
| ENSMUSG00 | Mzt1 | 2253 | 43.3022411 | 32.8514759 | 46.5168974 | 39.6534142 | 39.226668 | 60.8080395 |
| ENSMUSG00 | BC016579 | 2186 | 1.09118428 | 1.12420336 | 0.24264507 | 1.7166206 | 1.63396446 | 0.67375596 |
| ENSMUSG00 | Tie1 | 4895 | 1.34007238 | 1.25018961 | 0.6411299 | 1.7543466 | 3.16895163 | 0.3176005 |
| ENSMUSG00 | Lpcat2 | 3505 | 0.04537002 | 0 | 0 | 0 | 0 | 0 |
| ENSMUSG00 | Myh2 | 7120 | 0 | 0 | 0 | 0 | 0 | 0 |
| ENSMUSG00 | Tpsg1 | 1510 | 0 | 0 | 0 | 0 | 0 | 0 |
| ENSMUSG00 | Mamdc2 | 3324 | 0 | 0 | 0 | 0 | 0 | 0 |
| ENSMUSG00 | S100b | 1481 | 0.02684367 | 0 | 0 | 0 | 0.034454 | 0.0552491 |
| ENSMUSG00 | Ttc28 | 12456 | 1.22560248 | 1.09479473 | 1.63237378 | 1.68881117 | 1.54029522 | 1.57218939 |
| ENSMUSG00 | Slc9a10 | 4028 | 0 | 0 | 0 | 0 | 0 | 0 |
| ENSMUSG00 | AA467197 | 1197 | 0 | 0 | 0 | 0 | 0 | 0 |
| ENSMUSG00 | Slitrk5 | 2882 | 0.04138322 | 0.05015938 | 0.06134885 | 0 | 0 | 0.0851741 |
| ENSMUSG00 | Eefsec | 3269 | 10.5317364 | 8.9179558 | 11.3986399 | 9.315768 | 11.5976131 | 9.28622621 |
| ENSMUSG00 | Gm9758 | 1279 | 0 | 0.11302528 | 0 | 0 | 0.03989552 | 0 |
| ENSMUSG00 | Rac2 | 3029 | 0 | 0 | 0 | 0.28589273 | 0 | 0 |
| ENSMUSG00 | Tlf2 | 6038 | 2.66002221 | 3.6311413 | 3.11125924 | 3.31658409 | 1.89299572 | 4.30485777 |
| ENSMUSG00 | Wnt6 | 2064 | 0.4044889 | 1.44746096 | 0.34264995 | 1.60830819 | 0.14833249 | 0.54179274 |
| ENSMUSG00 | Scaf11 | 7709 | 15.4504371 | 19.6896216 | 16.1635745 | 21.8860605 | 20.4264361 | 17.9236376 |
| ENSMUSG00 | Trim45 | 6085 | 0.15026722 | 0.23756668 | 0.14528133 | 0.50995166 | 0.1593264 | 0.14343278 |
| ENSMUSG00 | Arid2 | 11451 | 4.07241102 | 4.79297508 | 4.01062919 | 5.41971152 | 6.56378059 | 5.10670042 |
| ENSMUSG00 | Hsf4 | 1907 | 0.20847132 | 0.22741373 | 0 | 0.3216543 | 0.08027222 | 0.18593094 |
| ENSMUSG00 | Szt2 | 12481 | 1.68819844 | 1.49412332 | 2.46844681 | 2.2751838 | 1.59444648 | 1.86186945 |
| ENSMUSG00 | Gm5134 | 3131 | 0 | 0 | 0 | 0 | 0 | 0 |
| ENSMUSG00 | Shf | 2943 | 0.40525464 | 0.81866195 | 1.06637134 | 0.26972646 | 0.41611723 | 1.34380656 |
| ENSMUSG00 | Ttll4 | 9712 | 4.22442915 | 2.8975373 | 4.24632615 | 7.0105867 | 4.90719059 | 5.26845379 |
| ENSMUSG00 | Duox1 | 5234 | 0.03797811 | 0 | 0.02533541 | 0 | 0 | 0 |
| ENSMUSG00 | Slc35a4 | 2940 | 59.8901439 | 46.9080263 | 53.3278698 | 52.1225997 | 41.1682189 | 43.5280977 |
| ENSMUSG00 | Stk36 | 6567 | 1.22287301 | 0.73376644 | 1.21829353 | 2.13184609 | 1.19659843 | 1.30828557 |
| ENSMUSG00 | Ptprm | 6907 | 3.14268023 | 4.40912573 | 1.59989297 | 4.98367895 | 7.80133964 | 0.92797745 |
| ENSMUSG00 | Rpgrip1l | 9680 | 1.32655167 | 1.25444044 | 0.81280249 | 1.31580183 | 2.56186143 | 0.94108776 |
| ENSMUSG00 | Wdr3 | 3776 | 8.15955969 | 9.95376718 | 8.99020536 | 15.2507794 | 9.68906554 | 5.98077365 |
| ENSMUSG00 | Kctd17 | 2302 | 0.08634987 | 0.02093243 | 0 | 0 | 0.39898991 | 0.1303306 |
| ENSMUSG00 | Noc4l | 2627 | 7.40023981 | 8.12584464 | 8.36251084 | 10.6034786 | 11.9068018 | 11.8671156 |
| ENSMUSG00 | Ptprf | 9997 | 22.2419128 | 34.0394773 | 22.0191236 | 28.0441618 | 27.6696992 | 17.8347821 |
| ENSMUSG00 | Lpp | 15978 | 10.5497082 | 12.5819149 | 9.68799939 | 15.1979071 | 14.2847027 | 12.4560673 |
| ENSMUSG00 | Mif | 551 | 75.6869311 | 65.3271729 | 149.211303 | 70.2650341 | 44.4512893 | 114.345584 |
| ENSMUSG00 | Dpyd | 8212 | 2.1543094 | 1.37306715 | 3.15958133 | 1.77510302 | 1.01903624 | 1.9197201 |
| ENSMUSG00 | Fbxl8 | 2177 | 5.09498349 | 3.82923955 | 2.7816502 | 2.56900179 | 4.1955541 | 1.74146755 |
| ENSMUSG00 | Galnt9 | 2985 | 0 | 0 | 0 | 0 | 0 | 0 |
| ENSMUSG00 | Gstt2 | 1365 | 29.4743929 | 30.4651279 | 34.7138284 | 30.4252258 | 27.6626506 | 35.2672075 |
| ENSMUSG00 | Fem1c | 2863 | 27.8552197 | 22.1324385 | 28.2842391 | 28.0666131 | 22.1001418 | 36.2772708 |
| ENSMUSG00 | Ctdp1 | 4736 | 17.8882874 | 9.47246149 | 14.0464049 | 9.23383403 | 14.1356852 | 12.9289622 |
| ENSMUSG00 | Kdm4a | 7169 | 4.19792143 | 6.54002067 | 5.16068386 | 6.26113313 | 5.68699599 | 4.4855349 |
| ENSMUSG00 | Trxb | 12491 | 0 | 0 | 0 | 0 | 0.00408505 | 0 |
| ENSMUSG00 | Dnmt2 | 5617 | 47.8311443 | 43.1507573 | 42.0928713 | 43.0903249 | 45.2851137 | 30.3968741 |
| ENSMUSG00 | 4833424O15 | 4860 | 0 | 0 | 0 | 0 | 0 | 0 |
| ENSMUSG00 | Magea4 | 947 | 0 | 0 | 0 | 0 | 0 | 0 |
| ENSMUSG00 | Chst2 | 6237 | 0.40157051 | 0.0154518 | 0.02126111 | 0.37603542 | 0 | 0.14431026 |
| ENSMUSG00 | Map2k4 | 5122 | 23.6111229 | 18.2133838 | 23.9563288 | 23.8738869 | 21.1895942 | 24.3245904 |
| [illegible] | [illegible] | [illegible] | [illegible] | [illegible] | [illegible] | [illegible] | [illegible] | [illegible] |

| | | | | | | | | |
|---|---|---|---|---|---|---|---|---|
| ENSMUSG00 | Calm4 | 914 | 0 | 0 | 0 | 0 | 0 | 0 |
| ENSMUSG00 | D930015E06I | 4942 | 2.18003546 | 3.10062497 | 1.27900922 | 4.66540398 | 5.6271499 | 2.22965485 |
| ENSMUSG00 | Nrxn2 | 13316 | 0.00597108 | 0 | 0 | 0 | 0.00383196 | 0 |
| ENSMUSG00 | Exoc6b | 14379 | 9.45571621 | 8.44159175 | 11.0204887 | 9.48787888 | 14.3756762 | 10.9637259 |
| ENSMUSG00 | Clcnka | 2921 | 18.319369 | 8.29776802 | 17.2661085 | 9.65975995 | 17.2591781 | 17.7971456 |
| ENSMUSG00 | Rpap2 | 3649 | 6.35172517 | 4.8595809 | 4.89381874 | 4.93421217 | 5.48159477 | 5.27703352 |
| ENSMUSG00 | Npbwr1 | 3691 | 0 | 0 | 0 | 0 | 0 | 0 |
| ENSMUSG00 | Tlr13 | 4174 | 0 | 0 | 0 | 0 | 0 | 0 |
| ENSMUSG00 | Asb13 | 3426 | 12.4627513 | 13.2772917 | 16.9143655 | 12.3222394 | 13.076812 | 23.4374016 |
| ENSMUSG00 | Dysf | 8358 | 0.21404602 | 0.10377554 | 0.15865701 | 0.23743868 | 1.15386277 | 0.2512739 |
| ENSMUSG00 | Tubgcp5 | 3658 | 7.82502624 | 8.74680087 | 11.0323353 | 11.964877 | 13.0704522 | 11.3855588 |
| ENSMUSG00 | Atp7a | 8767 | 12.3025686 | 16.439563 | 12.6751949 | 20.1420698 | 21.7387377 | 11.0318092 |
| ENSMUSG00 | Atp6v1h | 1962 | 840.660232 | 717.221544 | 869.730963 | 787.680963 | 587.063395 | 871.078578 |
| ENSMUSG00 | Lpcat2b | 3210 | 0.12384885 | 0 | 0 | 0 | 0.07948034 | 0 |
| ENSMUSG00 | BC016423 | 8321 | 4.52928681 | 4.26792547 | 5.37582804 | 6.39165168 | 5.55581018 | 5.83122019 |
| ENSMUSG00 | Ephx4 | 1637 | 2.76855515 | 1.94276431 | 2.37615283 | 2.79927896 | 1.40267985 | 1.78276506 |
| ENSMUSG00 | Tmem87a | 8736 | 8.93315113 | 6.90032499 | 9.61349608 | 2.6888062 | 6.46007002 | 8.03939973 |
| ENSMUSG00 | Alg3 | 1671 | 25.6233707 | 20.5607021 | 20.9502452 | 23.5364623 | 22.2915945 | 19.325657 |
| ENSMUSG00 | Tcea1 | 2833 | 5.03784592 | 6.14024201 | 11.2181875 | 5.68040659 | 4.6109256 | 4.48640377 |
| ENSMUSG00 | Ppp1r16a | 3879 | 17.9970672 | 19.3043906 | 14.8365042 | 18.1665978 | 19.2976781 | 12.1853441 |
| ENSMUSG00 | Tpsb2 | 1182 | 0 | 0 | 0 | 0 | 0 | 0 |
| ENSMUSG00 | Dnahc8 | 14717 | 0 | 0 | 0 | 0 | 0 | 0.01297292 |
| ENSMUSG00 | Fgb | 1636 | 0 | 0 | 0 | 0 | 0 | 0 |
| ENSMUSG00 | Tpbpa | 1367 | 0 | 0 | 0 | 0 | 0 | 0 |
| ENSMUSG00 | Foxh1 | 1303 | 0 | 0 | 0 | 0 | 0 | 0 |
| ENSMUSG00 | Mrpl15 | 5106 | 7.83274841 | 10.2582575 | 9.32342047 | 10.7129611 | 7.8448306 | 11.2816369 |
| ENSMUSG00 | Pla2g4c | 3728 | 0 | 0.01292555 | 0.03557015 | 0.17421588 | 0 | 0 |
| ENSMUSG00 | B3galt2 | 5078 | 0.15657929 | 0 | 0.34818309 | 0.12079456 | 0 | 0 |
| ENSMUSG00 | Olfr1015 | 1015 | 0 | 0 | 0 | 0 | 0 | 0 |
| ENSMUSG00 | Pla2g4b | 6613 | 2.2724288 | 3.329987 | 2.65358108 | 2.6790086 | 2.80864975 | 2.7344754 |
| ENSMUSG00 | Kcnk10 | 3880 | 0 | 0 | 0 | 0 | 0 | 0 |
| ENSMUSG00 | Ston1 | 4558 | 0 | 0.06343103 | 0.00969764 | 0 | 0.16792357 | 0 |
| ENSMUSG00 | Engase | 5190 | 3.83000776 | 3.70450682 | 1.9247816 | 0.87598027 | 1.82869093 | 2.18092009 |
| ENSMUSG00 | Fgg | 2075 | 0.72805217 | 3.48335726 | 0.27692721 | 0.06955575 | 1.27873327 | 0 |
| ENSMUSG00 | Cdk10 | 1630 | 9.43887789 | 8.18873894 | 10.1691357 | 8.9873068 | 11.8018059 | 12.114625 |
| ENSMUSG00 | Klf9 | 3261 | 17.6284651 | 20.2586974 | 16.468948 | 17.9248427 | 16.9618496 | 20.4831621 |
| ENSMUSG00 | Ppargc1b | 4249 | 8.32722468 | 7.17863447 | 10.4652987 | 10.8781123 | 9.06681896 | 11.4708826 |
| ENSMUSG00 | Best4-ps | 1371 | 0 | 0 | 0 | 0 | 0 | 0 |
| ENSMUSG00 | Lgals3bp | 3495 | 10.6128364 | 13.1806119 | 9.15654774 | 5.9052731 | 16.7459952 | 15.6546284 |
| ENSMUSG00 | Rbm46 | 1784 | 0 | 0 | 0.12388409 | 0 | 0 | 0 |
| ENSMUSG00 | D3Ertd254e | 2385 | 17.1356956 | 13.9205277 | 12.3246229 | 22.4359212 | 13.6926124 | 16.1703731 |
| ENSMUSG00 | Pxk | 3026 | 52.1314431 | 39.7466489 | 65.8353293 | 49.6992603 | 52.4765636 | 57.0550121 |
| ENSMUSG00 | Cfhr2 | 4227 | 0 | 0 | 0 | 0 | 0 | 0 |
| ENSMUSG00 | Mtap9 | 6204 | 3.15275571 | 4.52039157 | 1.71706063 | 5.02496558 | 2.72239368 | 2.88836847 |
| ENSMUSG00 | Mapkbp1 | 7572 | 1.24432764 | 1.29820843 | 0.9807065 | 1.21988951 | 2.11599076 | 2.10359077 |
| ENSMUSG00 | 6330503K22I | 5956 | 4.13841469 | 3.26043254 | 2.0111987 | 5.62191694 | 5.19173669 | 3.37040539 |
| ENSMUSG00 | Zdhhc15 | 5497 | 5.74237794 | 4.36544446 | 4.2778571 | 2.58619712 | 2.39490722 | 3.08619713 |
| ENSMUSG00 | Usp36 | 9559 | 3.2065567 | 5.05103202 | 3.34322968 | 3.30660854 | 3.93947749 | 3.09011762 |
| ENSMUSG00 | Gucy1a3 | 4644 | 0 | 0 | 0 | 0 | 0.01098759 | 0 |
| ENSMUSG00 | Chmp2a | 921 | 97.0361783 | 111.0224 | 116.575762 | 103.074766 | 75.1821846 | 105.841 |
| ENSMUSG00 | Gde1 | 1685 | 177.448646 | 146.646929 | 204.299082 | 153.000717 | 135.545435 | 165.428376 |
| ENSMUSG00 | Parl | 3963 | 15.539046 | 15.9283975 | 16.027769 | 12.6100509 | 14.2533934 | 16.9511573 |
| ENSMUSG00 | Rbm34 | 3935 | 6.22347548 | 12.7721624 | 8.55954368 | 4.82316529 | 6.78190459 | 9.82857394 |
| ENSMUSG00 | Vhl | 2992 | 20.9805828 | 17.6833935 | 25.2919637 | 13.4584097 | 18.8279141 | 21.3766808 |
| ENSMUSG00 | Ndufb7 | 4692 | 11.6334771 | 11.0606987 | 15.0542509 | 9.25121474 | 6.4381105 | 23.949598 |
| ENSMUSG00 | 6720456B07I | 1089 | 74.1810252 | 85.2223026 | 84.0200322 | 67.8899075 | 66.6294825 | 72.8075081 |
| ENSMUSG00 | Mga | 15430 | 5.66573569 | 5.32767792 | 5.22229163 | 8.16581332 | 6.04842783 | 5.52740097 |
| ENSMUSG00 | Zswim5 | 5568 | 7.26851278 | 9.42439574 | 9.63739896 | 8.93626785 | 7.86290058 | 7.45545994 |
| ENSMUSG00 | Trim36 | 4838 | 1.73385829 | 4.48199648 | 0.95002069 | 4.19888199 | 10.177853 | 1.92805428 |
| ENSMUSG00 | Aspm | 11894 | 0 | 0.00810265 | 0 | 0.02426907 | 0 | 0.03898343 |
| ENSMUSG00 | Ppp3r1 | 3222 | 72.046012 | 75.3455293 | 85.3169651 | 91.3137775 | 53.8928482 | 92.2868147 |
| ENSMUSG00 | Tnks1bp1 | 6247 | 0.59184564 | 1.45014425 | 0.69341774 | 1.80208066 | 1.57645118 | 1.28361517 |
| ENSMUSG00 | 9430020K01I | 6726 | 0.69746457 | 1.84120066 | 0.76889914 | 0.12874949 | 0.86485383 | 0.42984118 |
| ENSMUSG00 | Zfp446 | 3211 | 7.09432897 | 7.90851916 | 5.64396002 | 3.57336972 | 6.84907131 | 5.94588835 |
| ENSMUSG00 | 4931417G12I | 1538 | 0 | 0.12532235 | 0 | 0 | 0 | 0 |
| ENSMUSG00 | Zbtb41 | 8349 | 5.80451919 | 7.3298337 | 4.78601822 | 6.43072012 | 4.9687919 | 6.71983866 |
| ENSMUSG00 | Slc16a2 | 4294 | 2.62012133 | 1.61594031 | 1.10144322 | 0.99994488 | 0.59415901 | 1.23860146 |
| ENSMUSG00 | Cdkl4 | 2921 | 0.35386597 | 0.59387604 | 0 | 0.51881064 | 0.78609617 | 1.21386618 |
| ENSMUSG00 | 2310014L17I | 3237 | 0.15966056 | 0.44658427 | 0.31406932 | 0 | 0 | 0 |
| ENSMUSG00 | Rfc3 | 3576 | 2.81267801 | 6.11762996 | 6.73657842 | 4.28827431 | 6.97760005 | 7.26866085 |
| ENSMUSG00 | Zfp944 | 5021 | 10.6415785 | 12.3801056 | 10.2207408 | 15.4719462 | 11.2499897 | 19.2025194 |
| ENSMUSG00 | Gria2 | 6939 | 0 | 0 | 0 | 0 | 0 | 0 |
| ENSMUSG00 | Coil | 3509 | 3.09297412 | 1.41442107 | 1.86431256 | 1.69664786 | 3.82443341 | 2.47951266 |
| ENSMUSG00 | Tesk2 | 4546 | 24.0229443 | 8.80838835 | 32.3978313 | 12.6755445 | 15.2540354 | 17.6091215 |
| ENSMUSG00 | Dnahc17 | 16723 | 0.01901835 | 0.03169592 | 0.04757718 | 0 | 0.15256346 | 0.05055994 |
| ENSMUSG00 | Ttc37 | 7235 | 6.22020787 | 6.94657348 | 6.86701749 | 3.37131473 | 7.56756064 | 5.53032423 |
| ENSMUSG00 | 1700018A14I | 710 | 0 | 0 | 0 | 0 | 0 | 0 |
| ENSMUSG00 | Kcnk1 | 2298 | 31.5725639 | 65.1502505 | 30.1218826 | 48.5019302 | 48.58386 | 27.0253932 |
| ENSMUSG00 | Neu4 | 4602 | 0 | 0 | 0 | 0 | 0 | 0 |
| ENSMUSG00 | Gal3st2 | 2842 | 0 | 0 | 0 | 0 | 0 | 0 |
| ENSMUSG00 | Pqlc1 | 5761 | 10.8549507 | 6.44883647 | 12.0459805 | 8.0168403 | 5.79261409 | 7.23883413 |
| ENSMUSG00 | Scaper | 6136 | 8.92813757 | 8.20645894 | 7.10283853 | 8.8323388 | 9.72128966 | 7.42762748 |
| ENSMUSG00 | Rxfp1 | 2259 | 0 | 0 | 0 | 0 | 0 | 0 |
| ENSMUSG00 | Pds5b | 9486 | 6.38283753 | 6.11601058 | 4.7808445 | 8.61161172 | 6.75618048 | 10.2531482 |
| ENSMUSG00 | Cpsf1 | 4478 | 18.8745314 | 17.1202835 | 16.5633525 | 22.9722887 | 30.435786 | 14.9651158 |
| ENSMUSG00 | Fancd2 | 7055 | 0.31556441 | 0.59421977 | 0 | 0 | 0.02893062 | 0 |
| ENSMUSG00 | Cct2 | 1946 | 75.4863826 | 65.5939801 | 76.4105858 | 68.9378425 | 51.7605683 | 57.758884 |
| ENSMUSG00 | Cd226 | 2491 | 0 | 0 | 0 | 0 | 0 | 0 |
| ENSMUSG00 | 1700106J16I | 1218 | 0.16319984 | 0 | 0.18145256 | 0.47398417 | 0 | 0.82853994 |
| ENSMUSG00 | Rpap1 | 7087 | 5.0879386 | 6.6292904 | 5.73183205 | 2.81548901 | 8.94239576 | 4.31037053 |
| ENSMUSG00 | Ccdc17 | 2550 | 18.3498826 | 9.63728843 | 18.0967546 | 16.4420925 | 21.1909537 | 19.0815515 |
| ENSMUSG00 | Fgd5 | 6138 | 0.33680107 | 0.54953583 | 0.21604029 | 0.64663162 | 1.32179761 | 0.21773496 |
| ENSMUSG00 | Prss29 | 1172 | 0 | 0 | 0 | 0 | 0 | 0 |
| ENSMUSG00 | Wbscr17 | 8479 | 0.0046887 | 0.00568303 | 0 | 0.00425546 | 0 | 0.00321673 |
| ENSMUSG00 | Lyl1 | 1796 | 0.04427114 | 0 | 0 | 0 | 0.11364449 | 0 |
| ENSMUSG00 | Gpbp1l1 | 4163 | 8.31780531 | 6.43566222 | 8.33496199 | 8.10394226 | 8.26129649 | 13.7781807 |
| ENSMUSG00 | Phka1 | 6225 | 9.57963386 | 7.52405169 | 12.0072798 | 12.4041085 | 10.057729 | 9.30185689 |
| ENSMUSG00 | Gm239 | 3089 | 0.01287002 | 0 | 0.01430943 | 0.12848899 | 0 | 0 |
| ENSMUSG00 | Ypel4 | 2804 | 0.2268501 | 0.30932809 | 0.34680476 | 0.12868062 | 0.27296563 | 0.37935483 |
| ENSMUSG00 | 4930590J08I | 2785 | 0.08564915 | 0 | 0.0158714 | 0 | 0 | 0 |
| ENSMUSG00 | Poglut1 | 6288 | 12.3919755 | 10.6825541 | 12.9976475 | 10.627218 | 12.9188916 | 11.8849414 |
| ENSMUSG00 | Farp2 | 5281 | 7.24948452 | 7.00760984 | 7.23165922 | 8.23307397 | 10.0680711 | 5.88772747 |
| ENSMUSG00 | Zfp551 | 2901 | 3.54935176 | 3.55460139 | 1.81317453 | 0.90795907 | 2.63838551 | 0.11282167 |
| ENSMUSG00 | Zdhhc5 | 5028 | 47.899503 | 37.5294565 | 47.7182983 | 34.0009401 | 45.4954739 | 34.2343374 |
| ENSMUSG00 | C130022K22I | 8890 | 3.25556691 | 3.52861348 | 2.34682451 | 2.6503459 | 3.28313689 | 3.33493058 |
| ENSMUSG00 | Nlrp4b | 4475 | 0 | 0 | 0 | 0 | 0 | 0 |
| ENSMUSG00 | Hdlbp | 6809 | 41.1217285 | 32.0016289 | 34.3799325 | 43.3260091 | 41.6739133 | 29.6459987 |
| ENSMUSG00 | Fstl5 | 6515 | 0 | 0 | 0.25103119 | 0 | 0 | 0 |
| ENSMUSG00 | Ctnnd1 | 10123 | 16.9931803 | 16.6841331 | 20.2386192 | 14.6495311 | 18.0555644 | 15.6755755 |
| ENSMUSG00 | 4632415K11I | 2342 | 9.42113223 | 15.0196852 | 12.3432986 | 6.71723801 | 9.69544716 | 4.099348 |
| ENSMUSG00 | Ano7 | 3193 | 0.11205742 | 0 | 0.27686717 | 0.11300358 | 0 | 0 |
| ENSMUSG00 | Ccs | 1049 | 168.458638 | 140.516994 | 193.53581 | 129.09048 | 129.584619 | 160.709767 |
| ENSMUSG00 | Golim4 | 9918 | 0.58522889 | 1.04457401 | 1.17657658 | 0.31287113 | 1.36337866 | 1.75725897 |
| ENSMUSG00 | Kctd7 | 4666 | 4.34532685 | 5.33913215 | 6.27124305 | 3.90515055 | 7.52382052 | 0.9703365 |
| ENSMUSG00 | Tmed8 | 7173 | 3.17578284 | 2.8147385 | 2.88393455 | 2.79682375 | 3.54260911 | 3.2890789 |
| ENSMUSG00 | Atp2c2 | 3196 | 0.02487827 | 0 | 0.06915182 | 0 | 0.23948549 | 0 |
| ENSMUSG00 | Scn11a | 5810 | 0 | 0 | 0 | 0 | 0 | 0 |
| ENSMUSG00 | Vav1 | 5050 | 0 | 0 | 0 | 0 | 0 | 0 |
| ENSMUSG00 | Gpr44 | 2641 | 0 | 0 | 0 | 0 | 0 | 0 |
| ENSMUSG00 | Tpst1 | 2781 | 0.17154468 | 0.62377271 | 0.41324988 | 0.05189794 | 1.08254447 | 1.32401161 |
| ENSMUSG00 | Snrf2 | 2584 | 3.07704958 | 2.70396057 | 21.0916889 | 4.27287369 | 2.92256332 | 32.3305805 |
| ENSMUSG00 | Mks1 | 4130 | 4.05255625 | 4.52695873 | 2.5258196 | 4.07997937 | 5.0161522 | 2.94539688 |
| ENSMUSG00 | Nlrp4d | 2778 | 0 | 0 | 0 | 0 | 0 | 0 |
| ENSMUSG00 | Pomt2 | 5038 | 7.5675875 | 6.34132813 | 6.36092421 | 10.8647204 | 8.69008147 | 3.73551034 |
| ENSMUSG00 | Tspan8 | 1616 | 154.74132 | 238.576562 | 97.7309314 | 169.469504 | 162.551846 | 137.45338 |
| ENSMUSG00 | Sik3 | 7343 | 6.40483909 | 4.79042663 | 5.31529721 | 5.48871629 | 7.19219651 | 5.02554636 |
| ENSMUSG00 | Serpini2 | 1402 | 0 | 0 | 0 | 0 | 0 | 0 |
| ENSMUSG00 | Tmem63c | 4060 | 0 | 0 | 0 | 0 | 0.10054458 | 0 |
| ENSMUSG00 | Zbtx | 3184 | 0 | 0 | 0 | 0 | 0 | 0 |
| ENSMUSG00 | Exoc3 | 4583 | 35.331458 | 27.547126 | 29.1656935 | 25.3904852 | 31.7425698 | 20.9782028 |
| ENSMUSG00 | Ino80 | 6768 | 1.53312358 | 1.75145756 | 0.89474771 | 1.70600683 | 1.08566757 | 1.57167691 |
| ENSMUSG00 | Bzrap1 | 10515 | 1.66734826 | 1.3381304 | 0.61373933 | 1.33484692 | 1.28111871 | 1.46035396 |
| ENSMUSG00 | 2310044G17I | 4023 | 18.6671893 | 17.5833202 | 22.2492498 | 17.3997431 | 21.6129616 | 23.3966642 |
| ENSMUSG00 | Lrrc58 | 9165 | 7.8469901 | 7.25556903 | 10.0509155 | 10.3344645 | 8.39582921 | 8.7731191 |
| ENSMUSG00 | 2310007B03I | 2878 | 0.04144074 | 0.71995032 | 0 | 0 | 0.31913647 | 0 |
| ENSMUSG00 | Ogt | 7803 | 104.221211 | 89.4935178 | 96.4929122 | 118.747521 | 122.246196 | 124.366483 |
| ENSMUSG00 | Scx | 1134 | 6.13510502 | 4.84413969 | 4.95029459 | 3.05456462 | 4.18470278 | 4.44956637 |
| ENSMUSG00 | Zfc3h1 | 7827 | 6.62337378 | 9.74564174 | 11.1422303 | 13.8667165 | 11.5456384 | 13.4822511 |
| ENSMUSG00 | Emid1 | 3573 | 0.03337992 | 0.09440389 | 0.63092471 | 0.35344852 | 0.39987084 | 0.28244099 |
| ENSMUSG00 | Ccnd3 | 2248 | 51.1975605 | 43.8993921 | 60.1676519 | 45.0383527 | 40.5396384 | 30.1986553 |
| ENSMUSG00 | 6430527G18I | 4111 | 12.2331994 | 11.6627365 | 7.4942069 | 10.1286012 | 10.8234006 | 8.72443459 |
| ENSMUSG00 | Faah | 7400 | 8.05854335 | 12.2484727 | 5.39381949 | 8.25986263 | 9.95014329 | 6.5385419 |
| ENSMUSG00 | 2410018M08I | 2652 | 1.81388128 | 0.34522715 | 1.25005212 | 0.61225189 | 1.61622005 | 0.95646359 |
| ENSMUSG00 | Rhbdd3 | 3267 | 9.44299139 | 9.43964584 | 5.2225012 | 7.28930192 | 6.93471407 | 4.0406873 |
| ENSMUSG00 | Rnf43 | 7413 | 6.5910543 | 5.86973658 | 6.87504721 | 5.40281525 | 9.6022925 | 4.37100653 |
| ENSMUSG00 | 6430628N08I | 1600 | 0 | 0 | 0 | 0 | 0 | 0 |
| ENSMUSG00 | Nsf | 8658 | 11.9982757 | 9.42802414 | 10.5220603 | 12.0398506 | 13.5669574 | 8.30714065 |
| ENSMUSG00 | Hsdl1 | 3163 | 44.9841495 | 28.6559272 | 55.3256707 | 26.7620855 | 34.9747653 | 35.7855684 |
| ENSMUSG00 | Chmp7 | 1346 | 36.2702304 | 31.4679663 | 44.8914706 | 37.2347399 | 46.2497611 | 26.9301602 |
| ENSMUSG00 | Lsm3 | 871 | 2.05395709 | 1.16178563 | 1.52245154 | 1.36705794 | 1.11308971 | 2.88090334 |
| ENSMUSG00 | R3hcc1 | 3281 | 8.62721796 | 9.04689069 | 7.0593617 | 10.6013688 | 9.47121697 | 6.01022986 |
| ENSMUSG00 | Gas2l1 | 7405 | 1.9649569 | 3.26665685 | 2.5548128 | 3.15261078 | 3.52808973 | 2.01475053 |
| ENSMUSG00 | Chchd4 | 1242 | 62.6740989 | 53.773276 | 82.0688003 | 65.278868 | 43.1382404 | 76.1362112 |
| ENSMUSG00 | Loxl2 | 5214 | 0.48035966 | 0.63768019 | 0.05086518 | 0.39445273 | 0.4110295 | 0.08369663 |
| ENSMUSG00 | Polq | 8612 | 0.2492796 | 0 | 0.10778434 | 0.14245117 | 0.02962516 | 0.12351497 |
| ENSMUSG00 | Rasl10a | 1777 | 0 | 0.32540085 | 0 | 0.28427047 | 0.28714899 | 0.0920922 |
| ENSMUSG00 | 4732418C07I | 9249 | 10.513775 | 13.2748464 | 14.0983281 | 11.4772813 | 13.3620805 | 12.6715456 |
| ENSMUSG00 | Mrps17 | 1541 | 42.9544939 | 45.591066 | 55.0443457 | 52.2147692 | 38.7416351 | 62.7263605 |
| ENSMUSG00 | Ankmy1 | 3228 | 0 | 0.23884234 | 0.19170564 | 0.38004632 | 0.58487482 | 0 |
| ENSMUSG00 | Vps18 | 4166 | 6.99490332 | 7.8768524 | 6.19632173 | 8.25400586 | 7.54494659 | 3.7710495 |
| ENSMUSG00 | Atm | 10732 | 2.03370842 | 3.29115375 | 1.55274831 | 2.63588547 | 1.35506123 | 2.39656959 |
| ENSMUSG00 | 14-Sep | 1372 | 0 | 0 | 0 | 0 | 0 | 0 |
| ENSMUSG00 | Gpc1 | 3586 | 7.53868565 | 6.00650854 | 5.9535667 | 5.1718268 | 6.47434495 | 3.21728176 |
| ENSMUSG00 | Slc38a8 | 2889 | 0 | 0 | 0 | 0 | 0 | 0 |
| ENSMUSG00 | Rhov | 1700 | 0 | 0.96372884 | 0.28601293 | 0 | 0.84043443 | 0.38505373 |
| ENSMUSG00 | Foxj1 | 2620 | 0 | 0 | 0 | 0 | 0 | 0 |
| ENSMUSG00 | Usp54 | 7301 | 5.31997048 | 5.81458095 | 3.07554256 | 2.39196131 | 5.26967366 | 3.66476114 |
| ENSMUSG00 | Gm884 | 10963 | 0 | 0 | 0 | 0 | 0 | 0 |
| ENSMUSG00 | Golgb1 | 11623 | 6.03703201 | 7.01051998 | 5.40020796 | 6.7768307 | 7.65198082 | 6.32411191 |
| ENSMUSG00 | Hdac11 | 3867 | 6.98059769 | 5.81925743 | 8.24140905 | 8.36035994 | 6.10944195 | 9.94498095 |
| ENSMUSG00 | Plekhm1 | 5120 | 4.25507877 | 3.95279408 | 4.17845547 | 4.31997916 | 6.75700836 | 4.38418518 |
| ENSMUSG00 | Slc25a37 | 4158 | 5.55505872 | 7.85724093 | 5.57041023 | 4.54713596 | 5.10509195 | 7.36638285 |
| ENSMUSG00 | Senp6 | 7627 | 11.0973408 | 12.8758318 | 9.45236737 | 15.6637799 | 13.6882083 | 16.6966423 |
| ENSMUSG00 | Agpat1 | 2813 | 7.50450059 | 7.69133044 | 7.07103781 | 8.01680691 | 5.40556701 | 7.80522026 |
| ENSMUSG00 | Arhgap27 | 7725 | 0.34480482 | 0.49278044 | 0.26893031 | 0.87344237 | 1.69097116 | 0.38484604 |
| ENSMUSG00 | Mfsd7c | 3425 | 0.46429758 | 0.29544972 | 0.03871694 | 0.20016317 | 0.2532696 | 0.65299866 |
| ENSMUSG00 | Exosc4 | 2014 | 19.7593029 | 22.9447855 | 23.725021 | 14.672887 | 18.6978477 | 17.6865337 |
| ENSMUSG00 | 2010321M09 | 3196 | 17.1908868 | 15.7254253 | 18.090116 | 13.0396625 | 14.0977127 | 14.0128153 |
| ENSMUSG00 | Zdhhc14 | 3106 | 6.36138886 | 3.72335677 | 10.3602517 | 5.34376706 | 6.09490838 | 9.37840139 |
| ENSMUSG00 | Batf | 952 | 0 | 0 | 0 | 0 | 0 | 0.34379797 |
| ENSMUSG00 | Setd5 | 11735 | 3.43181097 | 3.46564612 | 2.57640056 | 4.36305251 | 4.20907813 | 4.75766396 |
| ENSMUSG00 | Jdp2 | 1752 | 3.31295671 | 4.56561039 | 1.84174346 | 4.94274583 | 2.9998383 | 2.3662929 |
| ENSMUSG00 | Thoc5 | 2874 | 11.4535622 | 9.72447336 | 12.0117047 | 8.6627038 | 9.14355725 | 8.90174378 |
| ENSMUSG00 | Igsf9b | 5020 | 0.17422721 | 0.21117564 | 0.04402574 | 0.33781994 | 0.10164617 | 0.15756266 |
| ENSMUSG00 | Dnajc17 | 1297 | 2.78932053 | 3.34370069 | 2.82864531 | 3.42181301 | 3.89484287 | 4.73152955 |
| ENSMUSG00 | Evpl | 6362 | 2.12462486 | 3.39319806 | 1.75779099 | 2.47844286 | 4.45039405 | 1.76200515 |
| ENSMUSG00 | Nipsnap1 | 2741 | 12.3574131 | 13.1673277 | 15.9004076 | 9.25416903 | 10.7227992 | 13.8015048 |
| ENSMUSG00 | Nek9 | 5364 | 11.2951813 | 9.31568615 | 8.6195242 | 10.5945601 | 11.2916309 | 8.37459561 |
| ENSMUSG00 | Traf3ip1 | 3481 | 5.35631151 | 5.27406908 | 4.40621647 | 7.42164432 | 9.57202626 | 4.11352646 |
| ENSMUSG00 | Fhod3 | 5369 | 3.8430051 | 4.82851665 | 3.77061727 | 4.67070608 | 4.71392855 | 5.88774574 |
| ENSMUSG00 | Med13 | 10809 | 5.33309712 | 6.32143352 | 6.54296871 | 10.4918 | 4.78682071 | 8.29165183 |
| ENSMUSG00 | Fam53c | 4509 | 16.0556066 | 12.311107 | 9.01878367 | 12.5554952 | 13.9080541 | 8.28703832 |
| ENSMUSG00 | Ccdc15 | 3509 | 0.57780835 | 0.05492897 | 0.42828802 | 1.18251215 | 0.8288696 | 1.22809718 |
| ENSMUSG00 | Sdr42e1 | 5383 | 3.72222965 | 3.92975072 | 6.3884514 | 2.9828181 | 4.27510598 | 2.72594388 |
| ENSMUSG00 | Tmem132d | 4269 | 0 | 0 | 0 | 0 | 0 | 0 |
| ENSMUSG00 | Kif4 | 5484 | 0.1739846 | 0.69415188 | 0 | 0.57899707 | 0.16748263 | 0 |
| ENSMUSG00 | Iqsec1 | 8821 | 4.79084863 | 3.44696123 | 4.57502354 | 6.06616847 | 3.69060512 | 4.2205966 |
| ENSMUSG00 | Trim59 | 4520 | 0.62447768 | 3.29416164 | 1.25173361 | 1.37303355 | 1.38854961 | 4.33859859 |
| ENSMUSG00 | Slc26a2 | 10875 | 6.22196118 | 6.12798616 | 6.10899955 | 7.5846946 | 7.26334067 | 6.04932689 |
| ENSMUSG00 | Exosc1 | 1614 | 14.2617244 | 15.2560545 | 11.1189271 | 12.0944152 | 13.5627728 | 19.2139187 |
| ENSMUSG00 | Tmem132c | 4725 | 0.00841386 | 0.19376559 | 0.01870978 | 0.48109393 | 0.23758313 | 0 |
| ENSMUSG00 | Kctd9 | 4736 | 9.07425558 | 10.6120057 | 11.3117892 | 9.9423708 | 7.61732426 | 7.96470145 |
| ENSMUSG00 | Brip1 | 11340 | 0.00701155 | 0.22521 | 0.16371053 | 0 | 0.08099425 | 0.2741895 |
| ENSMUSG00 | Plcg2 | 4263 | 182.783818 | 105.619075 | 258.751347 | 159.089399 | 150.6493 | 151.472457 |
| ENSMUSG00 | Zbed4 | 5802 | 0.4042698 | 0.45678289 | 0.55614177 | 0.90174017 | 0.58923943 | 0.61581829 |
| ENSMUSG00 | Fam151b | 1375 | 5.11761458 | 6.76362424 | 4.82201924 | 5.64191969 | 4.23055043 | 7.7757517 |
| ENSMUSG00 | Ina | 3237 | 0 | 0 | 0 | 0 | 0 | 0 |
| ENSMUSG00 | Gm5689 | 311 | 0 | 0 | 0 | 0 | 0 | 0 |
| ENSMUSG00 | Wbp2 | 4139 | 50.0329302 | 45.0081627 | 50.6842104 | 32.1329829 | 40.424133 | 43.8476563 |
| ENSMUSG00 | Cbl | 5099 | 1.11493111 | 3.87457174 | 3.63219695 | 3.26217347 | 2.23159086 | 2.55683027 |
| ENSMUSG00 | Ube2f | 4712 | 11.4322318 | 11.3512205 | 14.0898065 | 12.7956486 | 11.2080431 | 9.41762268 |
| ENSMUSG00 | Gtf2h5 | 3925 | 6.71538435 | 8.16407236 | 8.70522921 | 8.04376789 | 6.55217667 | 11.74373 |
| ENSMUSG00 | Smc4 | 5104 | 8.61472599 | 12.7263566 | 7.19665587 | 15.3970793 | 9.81737873 | 15.7962057 |
| ENSMUSG00 | Ramp1 | 3017 | 0.10541725 | 0.38331909 | 0.80580092 | 0.38270647 | 0.33825904 | 0.40681431 |
| ENSMUSG00 | Mtmr3 | 9632 | 16.3033792 | 13.4023545 | 12.8585511 | 10.0282011 | 16.0252063 | 10.2308214 |
| ENSMUSG00 | Skint2 | 2363 | 0 | 0 | 0 | 0 | 0 | 0 |
| ENSMUSG00 | Cpne2 | 3335 | 0.58411351 | 0.60684575 | 0.10603141 | 0.30293771 | 0.64261103 | 0.3025972 |
| ENSMUSG00 | Csta | 2980 | 0 | 0.22637926 | 0 | 0 | 0 | 0 |
| ENSMUSG00 | Dak | 3329 | 1.50471329 | 1.57774773 | 2.04478337 | 1.50657981 | 1.94663555 | 1.58125725 |
| ENSMUSG00 | Tulp4 | 17602 | 3.07392393 | 2.20920684 | 3.02095314 | 3.18756842 | 2.62640021 | 3.49881465 |
| ENSMUSG00 | Wdr5b | 1783 | 16.9302839 | 11.945265 | 11.4037284 | 9.79456509 | 10.0736311 | 11.120955 |
| ENSMUSG00 | AI661453 | 4701 | 2.6808099 | 2.48056137 | 3.42256347 | 3.00875589 | 2.23599945 | 4.17735381 |
| ENSMUSG00 | Barhl2 | 3745 | 0 | 0 | 0 | 0 | 0 | 0 |
| ENSMUSG00 | D630042P16I | 2530 | 0 | 0 | 0 | 0 | 0.1411797 | 0 |
| ENSMUSG00 | 4933407C03I | 4063 | 11.301398 | 14.955231 | 11.8038395 | 12.6018296 | 13.7518783 | 11.8617493 |
| ENSMUSG00 | Fbxo15 | 1289 | 0 | 0 | 0 | 0 | 0 | 0 |
| ENSMUSG00 | Lif | 4345 | 0 | 0.01109009 | 0 | 0.04152134 | 0 | 0 |
| ENSMUSG00 | Spata6 | 5223 | 7.43655073 | 11.3108516 | 6.27948832 | 7.73729463 | 11.58669 | 13.4623818 |
| ENSMUSG00 | Kcnh5 | 4050 | 1.46260904 | 2.65322679 | 0.12005439 | 2.83310868 | 1.90246488 | 0.37039634 |
| ENSMUSG00 | Pja1 | 5061 | 52.8580574 | 44.1113982 | 35.7649767 | 43.1758272 | 36.7095504 | 42.5906884 |
| ENSMUSG00 | Tbc1d10a | 2645 | 15.7518879 | 18.8920002 | 12.2682203 | 17.2157052 | 13.600603 | 11.3017031 |
| ENSMUSG00 | Neurl1b | 6179 | 6.94225012 | 5.68504877 | 10.859103 | 7.12998485 | 7.10190698 | 6.32538567 |
| ENSMUSG00 | Pkd1l2 | 8258 | 0 | 0 | 0.03746826 | 0 | 0 | 0 |
| ENSMUSG00 | Parp14 | 9290 | 0.53492304 | 1.50939232 | 0.12846607 | 0.37674471 | 2.2190157 | 1.26831476 |
| ENSMUSG00 | Gcsh | 1573 | 44.1530988 | 44.1428246 | 40.3239954 | 43.2388137 | 35.3583914 | 46.1050641 |
| ENSMUSG00 | Myo15b | 7684 | 0.26903761 | 0.50795182 | 0.02300981 | 0.4930524 | 0.29218643 | 0.56792586 |
| ENSMUSG00 | Zfp707 | 3217 | 9.93578064 | 11.8032068 | 8.62877134 | 5.83234789 | 7.88315474 | 5.82458103 |
| ENSMUSG00 | Zxdc | 5872 | 8.24628325 | 7.01624796 | 10.7644134 | 8.41832442 | 10.7405655 | 6.10799568 |
| ENSMUSG00 | Cops8 | 1835 | 39.7554805 | 40.9650407 | 44.5149896 | 35.708443 | 32.9516378 | 33.175474 |
| ENSMUSG00 | Tmem30b | 3281 | 65.9158226 | 56.146531 | 72.695259 | 60.4300014 | 62.8926132 | 63.3775829 |
| ENSMUSG00 | Gm9761 | 341 | 0.34975496 | 0 | 0 | 0.10581245 | 0 | 0.07998428 |
| ENSMUSG00 | Gbp8 | 3555 | 0 | 0 | 0 | 0 | 0.24400799 | 0.01534438 |
| ENSMUSG00 | Trmt5 | 2286 | 7.25198398 | 6.2182854 | 9.74528822 | 9.62819211 | 12.5445421 | 5.47640394 |
| ENSMUSG00 | Cybasc3 | 3369 | 10.4551368 | 9.49712009 | 7.91146077 | 5.87979889 | 9.60247025 | 5.05175865 |
| ENSMUSG00 | Dhrs11 | 1596 | 28.4216812 | 26.2368535 | 26.7814425 | 24.6877147 | 24.1384171 | 27.5138905 |
| ENSMUSG00 | Gulo | 2253 | 0 | 0.02138768 | 0 | 0.25624177 | 0 | 0 |
| ENSMUSG00 | Slc24a1 | 5203 | 0 | 0 | 0 | 0 | 0 | 0 |
| ENSMUSG00 | Polr3b | 4801 | 7.77554597 | 5.71091139 | 7.64164335 | 5.46378804 | 6.79147139 | 5.86850705 |
| ENSMUSG00 | Uroc1 | 3447 | 2.27207127 | 1.24415241 | 0.19234919 | 2.99375246 | 1.3026751 | 1.47173858 |
| ENSMUSG00 | Eda2r | 6195 | 0.0256694 | 0.01555656 | 0.21405251 | 0 | 0 | 0 |
| ENSMUSG00 | Ifit1 | 2654 | 4.10437139 | 0.79887093 | 0.34975083 | 8.89135536 | 6.03703165 | 11.2428242 |
| ENSMUSG00 | Six4 | 10223 | 9.42651714 | 4.43072049 | 5.75925412 | 5.61542921 | 6.88307999 | 9.69539656 |
| ENSMUSG00 | Pkd2 | 8527 | 27.5961873 | 24.0282341 | 24.8664526 | 27.3482179 | 28.8433366 | 25.4418296 |
| ENSMUSG00 | Scara3 | 3399 | 0 | 0.31188636 | 0.06502183 | 0.12738586 | 0.10508521 | 0.20060782 |
| ENSMUSG00 | Dynlrb2 | 492 | 0.24241147 | 1.37115892 | 0.71872916 | 0.80671238 | 0.82969717 | 0.38805381 |
| ENSMUSG00 | Caskin2 | 5233 | 7.37675742 | 10.2302957 | 10.0347391 | 12.5835527 | 10.7942286 | 12.5662441 |
| ENSMUSG00 | Rasd2 | 3172 | 24.0387805 | 14.781024 | 24.4838078 | 17.165134 | 15.1052229 | 19.5359333 |
| ENSMUSG00 | Sec22a | 5258 | 16.7626284 | 13.9390602 | 14.3080139 | 10.9110785 | 14.8382139 | 15.1053156 |
| ENSMUSG00 | Diap2 | 11894 | 5.11733989 | 3.8203981 | 4.56363404 | 6.32816083 | 5.25107483 | 6.95968809 |
| ENSMUSG00 | Ly6g5c | 506 | 0 | 0 | 0 | 0 | 0 | 0 |
| ENSMUSG00 | Snx2 | 1998 | 145.093576 | 127.146608 | 172.537625 | 139.813408 | 122.253884 | 212.723075 |
| ENSMUSG00 | Uaca | 4476 | 4.15673925 | 4.12319199 | 3.15022072 | 3.74846977 | 3.10079853 | 2.61412427 |
| ENSMUSG00 | Gbx2 | 4238 | 0 | 0 | 0 | 0 | 0 | 0 |
| ENSMUSG00 | Kdelc2 | 3636 | 4.51568027 | 6.48051986 | 5.00856967 | 6.83732912 | 6.97472739 | 3.73563541 |
| ENSMUSG00 | Edil3 | 14224 | 0 | 0 | 0 | 0.07356435 | 0 | 0 |
| ENSMUSG00 | 4930556J24I | 1218 | 0.26111974 | 0.23737164 | 0 | 0 | 0.29325503 | 0 |
| ENSMUSG00 | 1810048J11I | 4409 | 4.74288563 | 7.45365243 | 5.65430698 | 5.43399355 | 9.86039288 | 6.996511 |
| ENSMUSG00 | Mad2l1bp | 2161 | 7.94741674 | 9.72202165 | 7.5067452 | 6.19455739 | 7.17816671 | 6.48735057 |
| ENSMUSG00 | Hmgxb4 | 5285 | 5.0249122 | 5.67113851 | 7.17600403 | 8.57503272 | 6.69087577 | 6.01745116 |
| ENSMUSG00 | Gjc1 | 4576 | 1.05991447 | 2.35877689 | 0.02897848 | 0.51252904 | 1.97370379 | 0.26821651 |
| ENSMUSG00 | Zfp395 | 4375 | 4.74339676 | 3.94302772 | 4.42523594 | 3.43088698 | 3.35899343 | 3.38517237 |
| ENSMUSG00 | BC018507 | 7656 | 3.79587987 | 7.97442819 | 4.26083596 | 7.79986725 | 9.28418776 | 5.61096615 |
| ENSMUSG00 | Hsd17b13 | 2149 | 0 | 0 | 0 | 0 | 0 | 0 |
| ENSMUSG00 | Fbxo16 | 3771 | 1.05424239 | 1.85283323 | 1.54724033 | 0.95682961 | 2.5438766 | 1.48271043 |
| ENSMUSG00 | Scn10a | 6344 | 0 | 0 | 0 | 0 | 0 | 0 |
| ENSMUSG00 | Zfp418 | 2721 | 5.21598917 | 8.87225561 | 6.12425388 | 9.0834989 | 7.59488505 | 4.44052377 |
| ENSMUSG00 | Morc2a | 6374 | 8.37023138 | 7.65812141 | 7.80848372 | 10.5800661 | 8.28558189 | 11.536308 |
| ENSMUSG00 | Rsrc1 | 5721 | 10.1456042 | 8.53222617 | 8.73065594 | 6.88718627 | 9.5989901 | 12.0378368 |
| ENSMUSG00 | Hdx | 16987 | 0 | 0 | 0 | 0 | 0 | 0 |
| ENSMUSG00 | Zswim2 | 2214 | 0 | 0 | 0 | 0 | 0 | 0 |
| ENSMUSG00 | Tex16 | 4434 | 0 | 0 | 0 | 0 | 0 | 0 |
| ENSMUSG00 | Zfyve9 | 6601 | 3.95687785 | 4.08792722 | 3.99765191 | 4.32372326 | 4.21290332 | 3.90050893 |
| ENSMUSG00 | A230046K03I | 5821 | 17.3814977 | 13.8160405 | 21.2770253 | 23.4183069 | 18.303225 | 15.6029117 |
| ENSMUSG00 | Ccpg1 | 5272 | 0.53540196 | 0.37474282 | 0.49467161 | 0.22585498 | 0.4549013 | 0.2586745 |
| ENSMUSG00 | Atp5h | 1187 | 64.0374716 | 64.3839067 | 71.9816028 | 54.0470726 | 51.4562526 | 106.9846 |
| ENSMUSG00 | Inpp5j | 4236 | 29.5725966 | 20.8398399 | 20.4000479 | 22.6833061 | 24.1158651 | 20.3529591 |
| ENSMUSG00 | Ptpn13 | 8593 | 6.56499789 | 9.56863218 | 3.96083081 | 4.82465602 | 10.5520621 | 4.592855 |
| ENSMUSG00 | Daam1 | 6096 | 7.12807418 | 4.93247046 | 6.89566153 | 4.39187615 | 7.00608211 | 9.8118904 |
| ENSMUSG00 | Papd7 | 3959 | 3.93638504 | 5.30671603 | 5.12468955 | 3.68202728 | 4.13727372 | 2.97616675 |
| ENSMUSG00 | Pla2g3 | 5314 | 0.1346629 | 0.45339144 | 0.18299596 | 0.62467973 | 0.72016902 | 0.22070182 |
| ENSMUSG00 | Olfr1347 | 963 | 0 | 0 | 0 | 0 | 0 | 0 |
| ENSMUSG00 | Exph5 | 10512 | 4.83706807 | 5.78035612 | 5.71865549 | 7.56171462 | 12.3342866 | 4.88307373 |
| ENSMUSG00 | C630004H02 | 4576 | 1.68543777 | 1.10567667 | 1.31369114 | 0.55983942 | 2.74311374 | 0.98346055 |
| ENSMUSG00 | 8430429K09I | 2569 | 6.53048376 | 6.07723132 | 5.5402855 | 4.91580988 | 7.46824341 | 6.93279075 |
| ENSMUSG00 | Slc41a2 | 5335 | 1.11777359 | 1.30062749 | 0.33141026 | 0.0676327 | 2.02766479 | 0 |
| ENSMUSG00 | Myo5a | 15232 | 9.65960041 | 2.78704409 | 8.53740954 | 3.86119568 | 3.50069346 | 6.29759101 |
| ENSMUSG00 | Ppp1r18 | 4337 | 0.10083244 | 0 | 0 | 0.04991752 | 0 | 0 |
| ENSMUSG00 | 2700049A03I | 5843 | 3.69454491 | 4.32960502 | 3.85054563 | 5.53920827 | 5.20481259 | 2.88010481 |
| ENSMUSG00 | Mon2 | 9267 | 7.95367011 | 6.88452028 | 7.59830969 | 5.2096445 | 5.2915018 | 6.63397398 |
| ENSMUSG00 | Pof1b | 3871 | 4.00533128 | 1.5435595 | 4.93288457 | 5.36896348 | 3.61178687 | 2.64925455 |
| ENSMUSG00 | Zcchc11 | 8564 | 5.40347727 | 2.55448911 | 2.33809375 | 4.85363327 | 4.02777091 | 5.78041455 |
| ENSMUSG00 | Chst11 | 5524 | 2.41814653 | 1.62249787 | 1.4563243 | 1.19533203 | 2.79887615 | 0.82455915 |
| ENSMUSG00 | Ppm1h | 10864 | 3.7435435 | 4.52413209 | 2.6324183 | 3.30796358 | 3.03415305 | 3.76331777 |
| ENSMUSG00 | Pik3ip1 | 3982 | 13.5080776 | 19.1196832 | 10.4898899 | 20.7956535 | 21.0666403 | 11.0482153 |
| ENSMUSG00 | Sah3 | 2726 | 0.48126591 | 0.91918378 | 0.94046475 | 1.74718631 | 0.67386263 | 0.91048869 |
| ENSMUSG00 | Mtrr | 5477 | 3.54220823 | 2.06752125 | 2.88922034 | 2.62858057 | 2.98127447 | 2.90325264 |
| ENSMUSG00 | Tmem5 | 5696 | 16.1087867 | 15.4135705 | 19.2684649 | 13.030331 | 12.5595117 | 18.8231403 |
| ENSMUSG00 | Gpatch8 | 12810 | 6.62280995 | 5.80795212 | 6.09026174 | 8.47829462 | 8.45659609 | 7.02838284 |
| ENSMUSG00 | Prss55 | 1736 | 0 | 0 | 0 | 0 | 0 | 0 |
| ENSMUSG00 | Ly6d | 688 | 0.28892064 | 0.2101153 | 0.83520924 | 0 | 0 | 0.87215416 |
| ENSMUSG00 | Zyg11b | 9031 | 6.62077762 | 7.60333076 | 5.76077613 | 8.35029045 | 9.00066623 | 8.39591375 |
| ENSMUSG00 | Setmar | 1939 | 6.58149007 | 6.36190262 | 5.08355389 | 4.83823186 | 6.00000706 | 4.67002591 |
| ENSMUSG00 | Tiparp | 4151 | 13.7147309 | 9.70461711 | 19.7316346 | 10.3700022 | 12.2802576 | 16.5842422 |
| ENSMUSG00 | Cd300ld | 2641 | 0 | 0 | 0 | 0 | 0 | 0 |
| ENSMUSG00 | Zyg11a | 2281 | 0 | 0 | 0 | 0 | 0 | 0 |
| ENSMUSG00 | Ankrd12 | 11840 | 9.64339021 | 10.2152002 | 7.47399406 | 11.0988857 | 7.41261541 | 10.5113327 |
| ENSMUSG00 | Lrrn1 | 3677 | 0.96226211 | 2.54233608 | 0.28850809 | 2.86536769 | 1.47098065 | 0.41538749 |
| ENSMUSG00 | Cd300a | 4870 | 0.43265718 | 0.16820729 | 0.04538177 | 0.26672559 | 0.37719703 | 0 |
| ENSMUSG00 | Ythdc2 | 6269 | 1.24295329 | 2.8747375 | 1.11403592 | 3.15408525 | 1.9778927 | 1.58366066 |
| ENSMUSG00 | Cacna1a | 11052 | 0.1438852 | 0.0174399 | 0.15197883 | 0.03917703 | 0.00461694 | 0.08143893 |
| ENSMUSG00 | Tmem109 | 3839 | 20.4524809 | 17.9240009 | 23.4653181 | 16.7486854 | 13.2782885 | 20.5891911 |
| ENSMUSG00 | Galp | 644 | 0 | 0 | 0 | 0 | 0 | 0 |
| ENSMUSG00 | Bmp2k | 8464 | 0.84546154 | 0.95074856 | 1.36825176 | 1.1637994 | 1.19366995 | 1.73366681 |
| ENSMUSG00 | Itga2b | 5366 | 1.45211968 | 1.16739424 | 2.25704528 | 0.78673112 | 1.01748457 | 1.62143308 |
| ENSMUSG00 | Xpot | 5967 | 9.1210412 | 10.1508895 | 11.5338142 | 14.5186843 | 7.98703453 | 9.41608126 |
| ENSMUSG00 | Pbx2 | 3630 | 1.14995192 | 1.3407247 | 1.47339477 | 1.82895212 | 1.05426396 | 1.63046742 |
| ENSMUSG00 | Tdg | 3746 | 1.61314283 | 1.6207933 | 1.46316832 | 1.98422348 | 1.98874823 | 1.47076271 |
| ENSMUSG00 | Dbn1 | 5135 | 8.19109998 | 5.7898802 | 6.19772677 | 4.67977442 | 8.1781319 | 7.65389584 |
| ENSMUSG00 | Gpr142 | 1113 | 0.39291131 | 0 | 0 | 0 | 0 | 0.39208826 |
| ENSMUSG00 | Rnps1 | 1900 | 12.0731117 | 10.8799914 | 10.5386499 | 10.1599441 | 11.7897784 | 6.64639893 |
| ENSMUSG00 | Ppp1r1c | 6151 | 0 | 0 | 0 | 0 | 0 | 0 |
| ENSMUSG00 | Sema3f | 3467 | 6.96036247 | 9.08968364 | 2.01439031 | 7.35794793 | 6.57882607 | 1.62058716 |
| ENSMUSG00 | Fam171a2 | 3272 | 4.13107071 | 7.43708841 | 5.17399324 | 5.33731954 | 5.75450265 | 4.85141811 |
| ENSMUSG00 | Prr7 | 1357 | 0 | 0 | 0 | 0 | 0 | 0 |
| ENSMUSG00 | Fras1 | 15774 | 2.61860937 | 3.96818742 | 1.51879035 | 2.90047118 | 4.11148242 | 1.84666633 |
| ENSMUSG00 | 4921530L21I | 1154 | 0 | 0 | 0 | 0 | 0 | 0 |
| ENSMUSG00 | Nlrp4c | 3888 | 0 | 0 | 0 | 0 | 0 | 0 |
| ENSMUSG00 | Neurod1 | 4775 | 0 | 0 | 0 | 0 | 0 | 0 |
| ENSMUSG00 | Dnaic2 | 4287 | 0 | 0.3259638 | 0.09279603 | 0 | 0.10712325 | 0 |
| ENSMUSG00 | Gns | 3841 | 49.1638981 | 42.60379 | 48.1145289 | 35.1614404 | 45.9782053 | 34.9365335 |
| ENSMUSG00 | Grn | 3404 | 127.26659 | 114.945332 | 122.139405 | 107.016546 | 135.660606 | 75.1174333 |
| ENSMUSG00 | Klraq1 | 3111 | 8.02521432 | 7.97686201 | 10.4998914 | 8.65226783 | 7.79091241 | 7.32934697 |
| ENSMUSG00 | Ttyh2 | 3937 | 0.27264363 | 0.01223938 | 0.21331852 | 0.31160516 | 0 | 0.09006104 |
| ENSMUSG00 | Tme4 | 10899 | 17.8551314 | 14.1079858 | 19.1788711 | 16.2781368 | 17.5425113 | 18.2131227 |
| ENSMUSG00 | Cnot6l | 12113 | 3.7907686 | 4.2883666 | 4.11621225 | 5.76182518 | 4.72224612 | 5.57066437 |
| ENSMUSG00 | Mrps10 | 7139 | 5.4406926 | 3.91485312 | 5.45480091 | 3.11339675 | 3.48085797 | 5.37164068 |
| ENSMUSG00 | Bai1 | 6506 | 0 | 0 | 0 | 0 | 0 | 0 |
| ENSMUSG00 | Dgkh | 4200 | 0.4070204 | 0.26387814 | 0.27363046 | 0.77318667 | 1.56723869 | 0.35067393 |
| ENSMUSG00 | Pabpc5 | 2351 | 0 | 0 | 0 | 0 | 0 | 0 |
| ENSMUSG00 | Nostrin | 1872 | 0.16989522 | 0 | 0.5194661 | 0 | 0.95401878 | 0.48080294 |
| ENSMUSG00 | Mfrp | 4487 | 0 | 0 | 0 | 0.13670487 | 0 | 0 |
| ENSMUSG00 | Nagk | 3447 | 13.3094936 | 16.7191717 | 19.3118583 | 16.1516086 | 12.2569885 | 15.8330586 |
| ENSMUSG00 | Sirt6 | 3068 | 1.99554889 | 1.75908785 | 1.81532993 | 2.24630721 | 0.93138105 | 1.75133766 |
| ENSMUSG00 | Mast4 | 18836 | 2.08528428 | 2.49681289 | 1.76234785 | 1.87153098 | 2.61145818 | 0.95278789 |
| ENSMUSG00 | Pcdh11x | 9302 | 0 | 0 | 0 | 0 | 0 | 0 |
| ENSMUSG00 | Tmub2 | 2688 | 25.5275146 | 31.9091767 | 32.3949519 | 25.598385 | 26.8040337 | 21.9678549 |
| ENSMUSG00 | Tle6 | 3191 | 2.13042531 | 3.44296735 | 2.02239708 | 1.70742361 | 2.17473742 | 1.94025168 |
| ENSMUSG00 | Map4k5 | 10174 | 9.02645567 | 9.38721529 | 8.74565657 | 10.7494277 | 10.6727077 | 8.70193425 |
| ENSMUSG00 | Glis1 | 3707 | 0 | 0 | 0 | 0 | 0.17894332 | 0 |
| ENSMUSG00 | 1700006J14I | 2258 | 0.07042601 | 0.04268064 | 0.09787831 | 0 | 0.1129902 | 0 |
| ENSMUSG00 | Dusp5 | 2469 | 0.67627792 | 1.09292862 | 0.30434643 | 0.6868595 | 0.74400548 | 0.22093673 |
| ENSMUSG00 | Asb16 | 2838 | 0.01400827 | 0.03395803 | 0 | 0 | 0 | 0.12493668 |
| ENSMUSG00 | Tle2 | 5491 | 29.7423992 | 39.577646 | 20.567421 | 40.96439 | 39.4662207 | 34.5763547 |
| ENSMUSG00 | BC030867 | 3021 | 0 | 0.11165346 | 0 | 0 | 0.08445279 | 0 |
| ENSMUSG00 | Dsg1c | 4073 | 0 | 0 | 0 | 0 | 0 | 0 |
| ENSMUSG00 | Vax2 | 1230 | 0 | 0 | 0.03593646 | 0.11733998 | 0 | 0 |
| ENSMUSG00 | B3galt1 | 5028 | 0 | 0 | 0 | 0 | 0 | 0.12476466 |
| ENSMUSG00 | Gna11 | 4788 | 34.2670986 | 31.0373825 | 34.5546154 | 29.6991098 | 27.0478158 | 22.6377227 |
| ENSMUSG00 | Cd207 | 1524 | 0 | 0 | 0 | 0 | 0 | 0 |
| ENSMUSG00 | Dio1 | 2169 | 4.08735461 | 1.06636663 | 4.11653863 | 0.49906009 | 3.78757331 | 2.65327288 |
| ENSMUSG00 | Gpsm3 | 1507 | 0.10552218 | 0.06395016 | 0 | 0 | 0.10157872 | 0.14478906 |
| ENSMUSG00 | Rab24 | 1857 | 52.5791385 | 57.2943803 | 60.5305799 | 58.0577001 | 62.5945526 | 68.9283156 |
| ENSMUSG00 | Gna15 | 2920 | 0 | 0 | 0.06055047 | 0 | 0 | 0 |
| ENSMUSG00 | G6pc3 | 2219 | 30.4750213 | 29.5980715 | 21.7722462 | 26.2607039 | 20.9256431 | 27.8892098 |
| ENSMUSG00 | Ccdc122 | 2166 | 0.4772126 | 0 | 0.46936398 | 0.54972644 | 0.37692614 | 0 |
| ENSMUSG00 | Cpne7 | 3663 | 1.26983107 | 0.51304156 | 1.25497998 | 1.06384407 | 0.80795244 | 1.04243776 |
| ENSMUSG00 | Unc13a | 15848 | 0.12793599 | 0.25236463 | 0.15619026 | 0.1798638 | 0.23504073 | 0.14628624 |
| ENSMUSG00 | Zfp661 | 7569 | 1.33936419 | 3.10038279 | 1.74027602 | 1.68277999 | 4.57747512 | 1.62156 |
| ENSMUSG00 | Sos2 | 5592 | 15.8325206 | 15.6312958 | 16.8839658 | 22.6867791 | 16.2423013 | 25.4846191 |
| ENSMUSG00 | Glt25d1 | 3167 | 22.8339814 | 13.8914499 | 19.1211004 | 11.8488558 | 19.1731567 | 17.8185124 |
| ENSMUSG00 | Scn7a | 7313 | 0 | 0 | 0 | 0 | 0 | 0 |
| ENSMUSG00 | Grip1 | 7938 | 0.41568467 | 0.80128626 | 0.35637666 | 1.88637547 | 0.9963578 | 0.16836197 |
| ENSMUSG00 | Celf5 | 9765 | 0 | 0 | 0.02263279 | 0.0295603 | 0 | 0 |
| ENSMUSG00 | Cpsf7 | 3594 | 7.43341205 | 7.3875152 | 5.50985689 | 6.09399027 | 6.94265381 | 7.20190113 |
| ENSMUSG00 | Nrip3 | 7134 | 0.2229071 | 0 | 0.22305388 | 0.38944736 | 0.25749222 | 0.30203203 |
| ENSMUSG00 | Nup54 | 2980 | 9.11174268 | 8.7317714 | 7.78723744 | 8.53618839 | 8.93817724 | 7.92612 |
| ENSMUSG00 | Nxnl1 | 4028 | 0.42440061 | 0.47851482 | 0.32920936 | 0.21498736 | 1.00076556 | 0.06094135 |
| ENSMUSG00 | Tet3 | 11230 | 5.26060944 | 3.53138396 | 4.62092286 | 4.25723143 | 4.2302365 | 5.79252134 |
| ENSMUSG00 | Tespa1 | 1740 | 0 | 0 | 0 | 0 | 0 | 0 |
| ENSMUSG00 | Gnat1 | 2196 | 0 | 0.06582847 | 0 | 0 | 0 | 0 |
| ENSMUSG00 | Larp6 | 2437 | 0 | 0 | 0 | 0 | 0 | 0 |
| ENSMUSG00 | Art3 | 3998 | 0 | 0 | 0 | 0.36100095 | 0.15315571 | 0 |
| ENSMUSG00 | Plvap | 2009 | 9.14237531 | 7.53138021 | 4.04835197 | 9.03398128 | 12.3692608 | 8.20004087 |
| ENSMUSG00 | Ttc21b | 9791 | 2.4809109 | 2.8397076 | 3.34075823 | 2.77497493 | 4.492364 | 2.71604262 |
| ENSMUSG00 | Tmem127 | 4922 | 36.1369402 | 35.3321556 | 35.2213792 | 34.2346908 | 37.1760633 | 29.3914438 |
| ENSMUSG00 | Acot11 | 7309 | 8.18607172 | 5.22146151 | 8.81738776 | 5.08475932 | 6.24128882 | 4.01152513 |
| ENSMUSG00 | Mfsd12 | 4295 | 1.61983914 | 1.35752258 | 2.08916744 | 1.83980623 | 1.73454037 | 1.91144736 |
| ENSMUSG00 | Cxcl10 | 1516 | 5.34968207 | 2.79710218 | 0.81639288 | 6.16441262 | 4.24097847 | 3.54426381 |
| ENSMUSG00 | BC031353 | 6438 | 7.48425635 | 5.83057447 | 7.35324229 | 5.62135611 | 10.8504362 | 8.93903212 |
| ENSMUSG00 | Ano8 | 3736 | 14.3549634 | 8.56998139 | 15.2505824 | 14.8056141 | 14.2589765 | 11.4763739 |
| ENSMUSG00 | Ankrd27 | 4355 | 11.8125354 | 11.7838257 | 10.6165621 | 12.7094963 | 16.7549294 | 6.21899352 |
| ENSMUSG00 | Myl12b | 1932 | 109.636232 | 114.03127 | 121.921129 | 96.8350936 | 84.5157364 | 129.441598 |
| ENSMUSG00 | Fam151a | 1915 | 0.04152008 | 0 | 0.0230819 | 0 | 0 | 0.82607263 |
| ENSMUSG00 | Gipc3 | 1018 | 0.11715761 | 0 | 0 | 0 | 0 | 0.05358475 |
| ENSMUSG00 | Nudt19 | 3279 | 75.2917151 | 61.4564505 | 70.4479511 | 52.7970874 | 47.4627771 | 72.2915187 |
| ENSMUSG00 | Mrpl34 | 643 | 39.8173086 | 35.2218162 | 60.2189961 | 37.8216143 | 23.5689481 | 56.3308257 |
| ENSMUSG00 | Tbxa2r | 1760 | 0.3840018 | 0.43805857 | 0.75344051 | 0.6970395 | 0.60883744 | 0.35642995 |
| ENSMUSG00 | Lrr1 | 1738 | 0 | 0 | 0 | 0.29064938 | 0 | 0 |
| ENSMUSG00 | 2510012J08I | 2715 | 11.7143221 | 13.7016329 | 11.4452654 | 9.58200888 | 15.4488696 | 13.6624344 |
| ENSMUSG00 | Sncb | 2298 | 0 | 0 | 0 | 0 | 0 | 0 |
| ENSMUSG00 | Rps29 | 690 | 0.34569983 | 0.20950627 | 1.79369798 | 1.20273482 | 0.88741523 | 3.99237473 |
| ENSMUSG00 | Cog3 | 4199 | 20.0055562 | 16.6741844 | 15.7901321 | 16.9540066 | 24.2797568 | 15.0111196 |
| ENSMUSG00 | Filip1 | 5720 | 0.26410984 | 0.45490697 | 0.03863798 | 0.08200465 | 0.24085877 | 0.01430488 |
| ENSMUSG00 | Pip5k1c | 5883 | 14.637153 | 19.6333336 | 18.3253944 | 17.2038305 | 18.8822744 | 10.8347496 |
| ENSMUSG00 | Cobll1 | 9480 | 22.7587545 | 20.9821189 | 21.1590679 | 24.8692066 | 22.1329596 | 21.8197114 |
| ENSMUSG00 | Ncaph | 4741 | 0 | 0 | 0.00932332 | 0 | 0.37669757 | 0 |
| ENSMUSG00 | Sidt2 | 6454 | 26.1176383 | 27.8561537 | 25.8129449 | 29.2167283 | 28.7151839 | 18.1337894 |
| ENSMUSG00 | Pygo1 | 7913 | 0.59786455 | 0.0426267 | 0.12847749 | 0.43774501 | 0.73512029 | 0.19302158 |
| ENSMUSG00 | Ushbp1 | 3536 | 1.92256424 | 4.31988183 | 1.0000417 | 2.05104382 | 5.02182658 | 1.7278052 |
| ENSMUSG00 | Mdga2 | 4354 | 0 | 0 | 0 | 0 | 0 | 0 |
| ENSMUSG00 | Spert | 1831 | 0 | 0 | 0 | 0 | 0 | 0 |
| ENSMUSG00 | Tjp3 | 3008 | 35.4600912 | 38.3024545 | 27.185309 | 25.4781458 | 32.1120111 | 28.8432937 |
| ENSMUSG00 | Cdhr2 | 4097 | 0 | 0 | 0 | 0 | 0 | 0 |
| ENSMUSG00 | Ttc22 | 3311 | 2.4494467 | 1.52811127 | 2.62994958 | 3.29108349 | 2.04968529 | 2.07587106 |
| ENSMUSG00 | Ly6g6f | 1135 | 0 | 0 | 0 | 0 | 0 | 0 |
| ENSMUSG00 | Dhcr24 | 4019 | 1.86956602 | 0.41963809 | 3.29946576 | 2.2624223 | 1.25693237 | 1.60638256 |
| ENSMUSG00 | Rnf44 | 7399 | 6.8560607 | 7.07915429 | 8.88935564 | 9.88489042 | 5.27573692 | 13.3368894 |
| ENSMUSG00 | Rtkn | 3927 | 13.9301098 | 10.0127672 | 10.7718777 | 10.1621445 | 11.7853127 | 7.34130219 |
| ENSMUSG00 | Dhx8 | 5247 | 7.48588045 | 9.15606686 | 8.19675877 | 6.16152314 | 14.1399563 | 4.0285583 |
| ENSMUSG00 | Mrpl54 | 822 | 18.378446 | 17.2345669 | 24.5207304 | 17.602068 | 11.5461142 | 34.5080594 |
| ENSMUSG00 | Arl4d | 1380 | 63.4935355 | 67.3213482 | 52.6258174 | 65.6797796 | 52.1356449 | 33.9747137 |

| | | | | | | | | |
|---|---|---|---|---|---|---|---|---|
| ENSMUSG00 | Synrg | 8521 | 4.93152716 | 5.47971628 | 4.44560257 | 4.67910565 | 3.68280966 | 3.12085122 |
| ENSMUSG00 | Tmem106a | 3211 | 2.41430044 | 2.0859282 | 2.80821426 | 1.01133105 | 1.23950711 | 2.42082597 |
| ENSMUSG00 | Zfr2 | 7303 | 3.19546311 | 3.23970192 | 2.11234331 | 3.71047726 | 4.96778765 | 3.36871486 |
| ENSMUSG00 | Cog7 | 4392 | 38.5968508 | 39.7603976 | 21.5071354 | 29.772616 | 43.6373105 | 23.5920661 |
| ENSMUSG00 | 1700047I17R | 3832 | 0 | 0 | 0 | 0 | 0 | 0 |
| ENSMUSG00 | Cebpa | 2633 | 1.69108007 | 1.93990234 | 1.76270168 | 2.09667786 | 1.70540109 | 1.94744861 |
| ENSMUSG00 | Atcay | 3983 | 0 | 0 | 0 | 0 | 0 | 0 |
| ENSMUSG00 | 5031414D18I | 2637 | 0 | 0 | 0 | 0 | 0 | 0 |
| ENSMUSG00 | Lbx2 | 933 | 0.08522075 | 0.05164678 | 0.04737604 | 0.03867315 | 0.05469065 | 0 |
| ENSMUSG00 | Dopey1 | 9334 | 4.33161814 | 3.20589036 | 5.95261589 | 7.78156799 | 7.70807691 | 6.59219907 |
| ENSMUSG00 | Dapk3 | 1475 | 16.8994483 | 18.817214 | 19.3589089 | 18.1511031 | 19.0959725 | 15.8840103 |
| ENSMUSG00 | Parm1 | 5064 | 84.6060058 | 63.9536093 | 107.013151 | 73.8456377 | 69.0831028 | 83.9136808 |
| ENSMUSG00 | Hrh2 | 2786 | 0 | 0.84750024 | 0 | 0 | 0 | 0 |
| ENSMUSG00 | Otoa | 4357 | 0.50184793 | 0 | 0 | 0 | 0 | 0 |
| ENSMUSG00 | Vat1 | 2760 | 0.46093311 | 0.71581309 | 0.76872771 | 0.40526934 | 0.38824416 | 0.71151233 |
| ENSMUSG00 | Eef2 | 3084 | 147.458801 | 139.700058 | 145.963544 | 103.449234 | 120.782277 | 86.5288815 |
| ENSMUSG00 | Htr2a | 2968 | 0 | 0 | 0.23828487 | 0 | 0 | 0 |
| ENSMUSG00 | Foxn2 | 7498 | 1.26191042 | 1.66448233 | 1.72138412 | 1.33779786 | 1.37467697 | 0.7638936 |
| ENSMUSG00 | Dpp4 | 6291 | 16.1208442 | 16.3532116 | 11.6705232 | 17.1319452 | 14.5268223 | 19.2496262 |
| ENSMUSG00 | Igsf6 | 2127 | 0 | 0.11327325 | 0.18703178 | 0.28838494 | 0.67171534 | 0.12823056 |
| ENSMUSG00 | Rundc1 | 3218 | 20.9896089 | 19.5560887 | 15.7412406 | 23.3782047 | 22.5797263 | 10.8657823 |
| ENSMUSG00 | Zbtb7a | 6801 | 5.72277833 | 5.92322682 | 5.92737551 | 4.64753286 | 4.62921245 | 5.34985572 |
| ENSMUSG00 | Epgn | 1755 | 0 | 0 | 0 | 0 | 0 | 0.01554111 |
| ENSMUSG00 | Baz1a | 8079 | 1.12195192 | 1.10341525 | 0.95746039 | 1.28178571 | 1.08002355 | 0.75284621 |
| ENSMUSG00 | Ncapd3 | 5467 | 6.37745682 | 5.45590044 | 5.64347657 | 5.80797497 | 14.4949628 | 7.76780928 |
| ENSMUSG00 | Map2k2 | 3160 | 31.8672887 | 33.8677494 | 34.2844042 | 27.57536 | 26.5627811 | 28.4312221 |
| ENSMUSG00 | C8a | 3855 | 0 | 0.09999781 | 0.28665268 | 0 | 0.15883697 | 0.0424508 |
| ENSMUSG00 | Nek11 | 3796 | 0 | 0.17771607 | 0.45412852 | 0.08554752 | 1.19635075 | 0.4095507 |
| ENSMUSG00 | Tbr1 | 5478 | 0 | 0 | 0 | 0 | 0 | 0 |
| ENSMUSG00 | Creb3l3 | 3004 | 0 | 0 | 0 | 0 | 0 | 0 |
| ENSMUSG00 | Ccl5 | 957 | 0.04154178 | 0 | 0 | 0.45243943 | 0.10663819 | 0 |
| ENSMUSG00 | Zc3h12b | 7717 | 1.20549209 | 1.3674784 | 1.22576058 | 2.06196471 | 1.04472818 | 0.93307046 |
| ENSMUSG00 | Kri1 | 2580 | 3.89850255 | 4.51981357 | 2.00450218 | 3.09074878 | 2.4919858 | 3.3194716 |
| ENSMUSG00 | Anapc13 | 402 | 30.4594229 | 33.8024296 | 35.8452757 | 23.426402 | 27.797951 | 81.5525283 |
| ENSMUSG00 | Rrp12 | 4285 | 2.20812237 | 1.42816293 | 1.39259015 | 1.05256839 | 2.33399525 | 0.45829032 |
| ENSMUSG00 | Dhx57 | 5053 | 6.31776189 | 4.59645065 | 3.87520611 | 7.24782806 | 7.16974606 | 4.45311248 |
| ENSMUSG00 | 4933434I20R | 2104 | 0 | 0 | 0 | 0 | 0 | 0 |
| ENSMUSG00 | Zc4h2 | 3707 | 4.83672018 | 6.92834466 | 6.20042039 | 9.45121803 | 4.06063687 | 9.18964781 |
| ENSMUSG00 | Eef2k | 7361 | 0.35645452 | 0.64807197 | 0.27622399 | 0.14705357 | 0.12477581 | 0.1852645 |
| ENSMUSG00 | Xkr6 | 7859 | 0.76408188 | 0.34948813 | 0.79865908 | 0.68408508 | 0.52591124 | 0.669806 |
| ENSMUSG00 | Oma1 | 2133 | 15.3393157 | 17.1239208 | 14.3816592 | 16.4593668 | 15.310305 | 11.8791092 |
| ENSMUSG00 | Mtmr9 | 2392 | 4.57055064 | 6.36576744 | 6.43070291 | 2.85096422 | 10.0687497 | 5.62140349 |
| ENSMUSG00 | 1700020L24F | 936 | 0 | 0 | 0 | 0 | 0 | 0.203977 |
| ENSMUSG00 | Becn1 | 6977 | 24.9347833 | 25.2638678 | 29.681186 | 27.9471648 | 31.7479572 | 21.4381714 |
| ENSMUSG00 | Secisbp2l | 8215 | 9.67391425 | 11.590555 | 12.7951288 | 8.92937267 | 16.8390146 | 12.3009785 |
| ENSMUSG00 | Fam167a | 4493 | 0.51320229 | 0.25739475 | 0 | 0.27304463 | 0.42020385 | 0.05463427 |
| ENSMUSG00 | Fam176a | 2798 | 27.5787661 | 31.5157931 | 13.7281635 | 29.4923645 | 25.3490573 | 22.7808549 |
| ENSMUSG00 | Egln3 | 2657 | 0.43391379 | 5.73086779 | 0 | 3.31351858 | 1.42113354 | 0.25663003 |
| ENSMUSG00 | Dcbld2 | 8148 | 3.60082776 | 2.8918962 | 2.40321753 | 3.82607836 | 2.5613387 | 0.71969164 |
| ENSMUSG00 | Shc4 | 8955 | 0 | 0.16680957 | 0.11352791 | 0.29413615 | 0 | 0.27107123 |
| ENSMUSG00 | Wnk4 | 4320 | 49.6483374 | 47.2494836 | 29.6316059 | 48.1845636 | 47.2112093 | 42.4777252 |
| ENSMUSG00 | Neil2 | 2239 | 0.71023637 | 0.1721713 | 0.76992938 | 0.322305 | 0.29626748 | 0.75526022 |
| ENSMUSG00 | AW146020 | 7325 | 1.78560452 | 1.13147687 | 1.20687626 | 1.07876693 | 2.18734225 | 1.73142761 |
| ENSMUSG00 | Wdr78 | 8561 | 0.97983955 | 0.74297516 | 0.78996402 | 0.83451055 | 1.82982098 | 1.32215574 |
| ENSMUSG00 | Gm6781 | 1097 | 0.14496073 | 0.04392565 | 0.08058677 | 0.09867469 | 0.18605789 | 0.19890348 |
| ENSMUSG00 | Fam5c | 6123 | 0 | 0 | 0 | 0 | 0 | 0 |
| ENSMUSG00 | Arhgap5 | 10223 | 32.2383775 | 32.0048853 | 37.5908073 | 54.5695481 | 36.0973051 | 40.6651718 |
| ENSMUSG00 | Secisbp2 | 4689 | 4.49358172 | 5.33349616 | 4.80762208 | 4.94791099 | 3.83051488 | 5.36302354 |
| ENSMUSG00 | Nubpl | 5084 | 3.16698851 | 2.92871964 | 3.15603246 | 3.22211806 | 3.37231752 | 2.71458841 |
| ENSMUSG00 | Gpr33 | 1386 | 0 | 0 | 0 | 0 | 0 | 0 |
| ENSMUSG00 | Eif2s3x | 3799 | 1.25576669 | 0.71030291 | 1.24495846 | 0.89279078 | 1.06109073 | 0.59589236 |
| ENSMUSG00 | Elmod2 | 11772 | 2.84353675 | 3.95004389 | 2.94003084 | 3.6351771 | 5.11477434 | 4.09629309 |
| ENSMUSG00 | Ap2b1 | 8730 | 5.83353614 | 5.20501889 | 4.6682817 | 5.34824348 | 5.06172296 | 4.82695506 |
| ENSMUSG00 | Mitf | 5342 | 4.93408528 | 4.54623116 | 8.11718607 | 9.57099534 | 6.09412725 | 8.46524745 |
| ENSMUSG00 | Ints6 | 4974 | 7.920724 | 15.0255673 | 7.85573567 | 16.9746651 | 14.0030163 | 9.27249999 |
| ENSMUSG00 | Zc3h12c | 6150 | 3.94969083 | 4.08213599 | 3.12647182 | 2.78682458 | 6.32229242 | 3.2330426 |
| ENSMUSG00 | Kcne3 | 1059 | 0 | 0 | 0 | 0 | 0 | 0 |
| ENSMUSG00 | Tanc1 | 10032 | 33.6209626 | 32.9504579 | 24.7710089 | 41.0131134 | 38.2443501 | 30.4256866 |
| ENSMUSG00 | 1110059E24I | 1424 | 36.0144451 | 39.8620989 | 44.0466399 | 46.1414348 | 31.0314898 | 43.4785331 |
| ENSMUSG00 | Plekhh3 | 3951 | 4.84994726 | 5.70773347 | 4.97844094 | 2.43834621 | 5.68251211 | 3.21690253 |
| ENSMUSG00 | A630007B06I | 7108 | 15.4200703 | 12.1821942 | 17.262847 | 18.3862078 | 17.6022332 | 10.6213002 |
| ENSMUSG00 | Nlrp2 | 3385 | 0 | 0 | 0 | 0 | 0 | 0 |
| ENSMUSG00 | Ppp1r32 | 1444 | 0 | 0.43381146 | 0 | 0 | 0.35336825 | 0 |
| ENSMUSG00 | Heatr5a | 7725 | 2.08941425 | 2.59489449 | 1.48769957 | 1.97575467 | 2.40434962 | 0.98153394 |
| ENSMUSG00 | Slc24a5 | 2320 | 0 | 0.02077002 | 0 | 0 | 0.02199413 | 0.02351262 |
| ENSMUSG00 | Fam124a | 3902 | 0 | 0.30872913 | 0.12460796 | 0 | 0 | 0.53822328 |
| ENSMUSG00 | Ubd | 929 | 0 | 0 | 0 | 0 | 0 | 0 |
| ENSMUSG00 | Nkx6-1 | 3365 | 0 | 0 | 0 | 0 | 0 | 0 |
| ENSMUSG00 | Ano4 | 5114 | 0 | 0 | 0 | 0 | 0 | 0 |
| ENSMUSG00 | Rfpl4 | 1674 | 0 | 0 | 0 | 0 | 0 | 0 |
| ENSMUSG00 | Tubg1 | 2255 | 14.4741683 | 12.9708073 | 11.2513782 | 10.976622 | 13.5994909 | 10.8130942 |
| ENSMUSG00 | Arl6ip5 | 1439 | 41.7170087 | 43.5988518 | 53.8777155 | 41.0969222 | 36.2751789 | 59.3635651 |
| ENSMUSG00 | Chrnb4 | 3480 | 0.05711994 | 1.27389445 | 0.01270168 | 1.33752406 | 0.82111409 | 0 |
| ENSMUSG00 | B020004J07F | 2038 | 0 | 0 | 0 | 0 | 0 | 0 |
| ENSMUSG00 | Lars2 | 3860 | 24.4197524 | 26.6273785 | 24.013281 | 21.7333558 | 19.8817796 | 18.8732024 |
| ENSMUSG00 | Epn1 | 2874 | 5.57462027 | 5.16403068 | 3.93725533 | 4.59499941 | 5.77020603 | 7.13657923 |
| ENSMUSG00 | 3110056O03I | 3638 | 22.6315558 | 25.7621303 | 15.7221509 | 21.8991628 | 24.0124122 | 14.2520861 |
| ENSMUSG00 | Slfn8 | 5450 | 0 | 0 | 0 | 0 | 0.36514287 | 0 |
| ENSMUSG00 | Xrra1 | 2612 | 0 | 0.0368962 | 0.21999386 | 0 | 0.09767683 | 0.13574667 |
| ENSMUSG00 | Leprot | 1926 | 77.2403988 | 77.2334096 | 79.1548061 | 71.658266 | 98.6350972 | 63.6833088 |
| ENSMUSG00 | Lsm7 | 464 | 3.76991626 | 3.63475318 | 2.47682741 | 3.88815135 | 4.50879614 | 5.81937346 |
| ENSMUSG00 | Rims4 | 4956 | 0 | 0 | 0 | 0 | 0 | 0 |
| ENSMUSG00 | Spcs2 | 1386 | 92.8775223 | 101.622634 | 111.875949 | 79.219092 | 83.1663658 | 120.276043 |
| ENSMUSG00 | Ccdc106 | 2361 | 0.8755972 | 1.14292281 | 0.67397982 | 0.91695158 | 2.07477005 | 0.53139916 |
| ENSMUSG00 | Pdk3 | 2785 | 30.7480449 | 35.607791 | 32.1713234 | 36.7816605 | 30.9639408 | 32.9156418 |
| ENSMUSG00 | Fam175a | 2813 | 13.1434756 | 8.32513718 | 14.1106488 | 11.0439532 | 8.10835051 | 15.6977039 |
| ENSMUSG00 | Trim13 | 1695 | 6.5203679 | 5.71414447 | 9.20545757 | 8.85553423 | 6.71320461 | 8.43180588 |
| ENSMUSG00 | Scai | 12160 | 2.6187615 | 2.55990474 | 2.61721439 | 3.78030631 | 2.56810378 | 4.97493009 |
| ENSMUSG00 | Lcat | 1336 | 0.53562773 | 0 | 0.23159648 | 0.54015037 | 0.15277358 | 0.44913328 |
| ENSMUSG00 | Kcnk15 | 2357 | 0 | 0 | 0 | 0 | 0.15154206 | 0 |
| ENSMUSG00 | Neu3 | 3338 | 1.17908705 | 2.28084418 | 1.29771738 | 2.81046482 | 2.46112837 | 1.55248096 |
| ENSMUSG00 | Oaz1 | 1065 | 30.4605372 | 33.7078868 | 33.6183031 | 25.4437704 | 26.5912099 | 24.7136403 |
| ENSMUSG00 | A130022J15R | 5115 | 10.7879975 | 11.04096 | 6.71453216 | 9.07870798 | 13.5072752 | 3.32201374 |
| ENSMUSG00 | Pcyt1b | 5348 | 0.01486742 | 0.3604072 | 0 | 0 | 0 | 0 |
| ENSMUSG00 | Hectd1 | 8945 | 17.377745 | 19.1991593 | 16.8851535 | 23.0368425 | 24.1869015 | 12.8674095 |
| ENSMUSG00 | Zcchc6 | 5523 | 18.7440288 | 17.2836035 | 21.2965968 | 21.8922562 | 26.6634294 | 17.234925 |
| ENSMUSG00 | Abi3bp | 5072 | 0 | 0.23751204 | 0 | 0 | 0 | 0 |
| ENSMUSG00 | Amh | 1660 | 0 | 0 | 0 | 0 | 0 | 0 |
| ENSMUSG00 | Helq | 5518 | 2.68014475 | 3.53669972 | 4.54995412 | 4.42688368 | 5.08599252 | 4.14210723 |
| ENSMUSG00 | Pkig | 4076 | 8.65140118 | 9.78860994 | 7.33082578 | 7.60414039 | 5.7085445 | 10.8536469 |
| ENSMUSG00 | Impg2 | 9537 | 0.30847285 | 0.32336503 | 0.57007725 | 0.02270025 | 0.29426976 | 0.66349128 |
| ENSMUSG00 | Hpse | 3504 | 0.13614891 | 0 | 0 | 0 | 0.39318269 | 0.17124488 |
| ENSMUSG00 | Tpbg | 3719 | 1.89210004 | 2.37109947 | 0.46353102 | 1.58143944 | 1.12507739 | 1.04874252 |
| ENSMUSG00 | Raver2 | 6327 | 0.84826772 | 0.68544014 | 0 | 0.55317818 | 0.22581611 | 0.13794665 |
| ENSMUSG00 | Arx | 3159 | 0 | 0 | 0 | 0 | 0 | 0 |
| ENSMUSG00 | Plekhj1 | 1253 | 11.6125346 | 16.8441035 | 16.6506544 | 14.1678898 | 11.5654355 | 20.0260719 |
| ENSMUSG00 | A430110N23I | 4270 | 0 | 0 | 0 | 0.10140153 | 0 | 0 |
| ENSMUSG00 | Adrb1 | 2490 | 2.90582227 | 0.94824726 | 2.39648547 | 2.05769089 | 1.88531188 | 2.3659928 |
| ENSMUSG00 | Vps13c | 11643 | 1.31459761 | 1.24573727 | 0.66817181 | 1.12185005 | 1.06496688 | 1.11740985 |
| ENSMUSG00 | Nat14 | 1609 | 2.74260928 | 1.73698797 | 3.24165163 | 2.91526774 | 1.96621212 | 2.39013309 |
| ENSMUSG00 | G2e3 | 8027 | 4.96757779 | 5.28267959 | 5.37448596 | 11.9569252 | 6.83360583 | 7.63839405 |
| ENSMUSG00 | Wdr38 | 2859 | 0.83432278 | 0.89327787 | 1.23684765 | 1.2368102 | 1.07085784 | 1.7171861 |
| ENSMUSG00 | Sgcg | 2200 | 0 | 0 | 0 | 0 | 0 | 0 |
| ENSMUSG00 | Cops4 | 5100 | 26.0983037 | 23.6963916 | 29.9099138 | 23.4108795 | 22.3515536 | 24.6594827 |
| ENSMUSG00 | Klhl35 | 2564 | 0 | 0 | 0 | 0.1407256 | 0 | 0 |
| ENSMUSG00 | Mid1 | 6874 | 0.97162071 | 0.75006536 | 0.23792089 | 0.53540421 | 0.83138698 | 0.89275441 |
| ENSMUSG00 | Ror1 | 343221 | 0.07679566 | 0.06430082 | 0.14179269 | 0.07453556 | 0.08905282 | 0.13890779 |
| ENSMUSG00 | Lin54 | 8154 | 3.80295251 | 3.78210976 | 3.97350392 | 3.01347465 | 3.74844237 | 5.4522519 |
| ENSMUSG00 | Gnptab | 5857 | 14.1998404 | 9.82322211 | 9.13922347 | 14.3416424 | 15.9865801 | 8.98754546 |
| ENSMUSG00 | Gdpd5 | 3564 | 1.18240206 | 1.41963424 | 0.62011564 | 0.12148837 | 2.09030608 | 1.91320421 |
| ENSMUSG00 | Sec31a | 4433 | 70.2020982 | 57.8606884 | 67.6139629 | 55.632929 | 69.0979775 | 53.6017724 |
| ENSMUSG00 | Fbxo33 | 5790 | 10.1620227 | 13.4156381 | 13.5201147 | 12.5196593 | 12.4878022 | 9.43542356 |
| ENSMUSG00 | Uchl4 | 1161 | 1.26697052 | 0.78858088 | 2.20818854 | 3.263234 | 1.14270954 | 3.50036283 |
| ENSMUSG00 | Lzts2 | 2820 | 20.3288663 | 16.83108 | 19.5146435 | 12.0529383 | 19.3610717 | 15.2235043 |
| ENSMUSG00 | Mia2 | 1625 | 8.63611361 | 12.8398335 | 9.13958108 | 7.77151728 | 13.2511573 | 5.90810647 |
| ENSMUSG00 | Nup37 | 1353 | 12.5760131 | 13.9608909 | 13.6885235 | 15.147525 | 10.9746307 | 13.6473989 |
| ENSMUSG00 | Ccl12 | 638 | 0 | 0 | 0 | 0 | 0 | 0 |
| ENSMUSG00 | Uvrag | 3423 | 12.2878464 | 10.839486 | 9.62032517 | 11.5635416 | 13.4758527 | 11.0915273 |
| ENSMUSG00 | Kcnh4 | 3937 | 0 | 0 | 0 | 0 | 0 | 0 |
| ENSMUSG00 | Nfkbiz | 6408 | 3.90854443 | 4.17345122 | 2.51084127 | 5.02265664 | 4.62645511 | 4.56705492 |
| ENSMUSG00 | Pdzrn3 | 9695 | 0.3895586 | 0.15904757 | 0.07294786 | 0.56197924 | 0.33157934 | 0 |
| ENSMUSG00 | 4930524J08R | 602 | 0 | 0.40021962 | 1.83562471 | 0.17981085 | 0.67809137 | 0.58898723 |
| ENSMUSG00 | 4930547N16I | 3575 | 0 | 0.71437243 | 0.69239251 | 0.71659445 | 0 | 0 |
| ENSMUSG00 | Rmi1 | 3519 | 4.43987037 | 3.62870337 | 4.3083922 | 3.79379269 | 3.24805575 | 5.89051601 |
| ENSMUSG00 | Adat3 | 1409 | 3.21655414 | 2.97531616 | 2.32145946 | 2.17670247 | 3.36795809 | 1.64538278 |
| ENSMUSG00 | Six9 | 595 | 0 | 0 | 0 | 0 | 0 | 0.04583973 |
| ENSMUSG00 | 1810055G02I | 1878 | 10.1399761 | 14.4456693 | 9.17929647 | 12.2194783 | 12.9060322 | 7.8715945 |
| ENSMUSG00 | Ccl7 | 798 | 0 | 0 | 0 | 0 | 0 | 0 |
| ENSMUSG00 | Ptplb | 3596 | 10.0162584 | 11.1220097 | 12.9311287 | 9.79312443 | 11.3659975 | 14.5702397 |
| ENSMUSG00 | Shq1 | 2876 | 2.44670377 | 3.18338804 | 2.29001203 | 2.22062652 | 2.87422562 | 2.03895947 |
| ENSMUSG00 | Pcsk7 | 3766 | 26.4227211 | 17.0175167 | 16.1971703 | 13.6050194 | 24.063421 | 14.0139211 |
| ENSMUSG00 | Pmch | 746 | 0 | 0 | 0 | 0 | 0 | 0 |
| ENSMUSG00 | Ccl2 | 1263 | 0 | 0 | 0 | 0 | 0 | 0 |
| ENSMUSG00 | 4930415L06F | 2615 | 0 | 0 | 0 | 0 | 0 | 0 |
| ENSMUSG00 | Brsk1 | 4077 | 0.0975116 | 0.02363819 | 0.06505054 | 0 | 0.075094 | 0.07358868 |
| ENSMUSG00 | Dennd1a | 7437 | 3.04166578 | 3.77742312 | 2.64485951 | 4.52177834 | 5.02236212 | 3.55740219 |
| ENSMUSG00 | Ccdc11 | 2754 | 0 | 0.6998757 | 0 | 0.37994891 | 0.38909001 | 0.22778384 |
| ENSMUSG00 | Pet2 | 4171 | 0 | 0 | 0 | 0 | 0 | 0 |
| ENSMUSG00 | Klf16 | 2672 | 0.65465612 | 0.9377601 | 0.99255636 | 1.40439096 | 1.31767213 | 0.67369992 |
| ENSMUSG00 | 3230401D17I | 3828 | 2.32633951 | 2.2154686 | 2.89829221 | 1.91344176 | 2.73260372 | 2.78588922 |
| ENSMUSG00 | 2210018M11 | 4026 | 12.2841077 | 13.1537258 | 13.4384143 | 15.0745154 | 15.6146287 | 10.9545682 |
| ENSMUSG00 | Crb2 | 6684 | 0 | 0 | 0.07935699 | 0 | 0.16031626 | 0 |
| ENSMUSG00 | Kank4 | 5061 | 0.36919731 | 0.23802827 | 0.00873382 | 0.77710786 | 0.3528795 | 0 |
| ENSMUSG00 | Tmem98 | 1626 | 18.3862985 | 24.2710308 | 22.8349005 | 17.508446 | 20.2724716 | 20.1456591 |
| ENSMUSG00 | Fam170a | 1341 | 0 | 0 | 0 | 0 | 0 | 0 |
| ENSMUSG00 | Mageb4 | 4797 | 0 | 0 | 0 | 0 | 0 | 0 |
| ENSMUSG00 | Ptprh | 4295 | 0.48132363 | 0.16828792 | 0.14408051 | 0 | 0 | 0.01270065 |
| ENSMUSG00 | Sstr1 | 2120 | 0 | 0 | 0 | 0 | 0 | 0 |
| ENSMUSG00 | Abca17 | 5549 | 0.08597329 | 0.06078665 | 0.11948597 | 0 | 0.03678239 | 0.27525316 |
| ENSMUSG00 | Rabgap1 | 7750 | 18.682511 | 16.2528206 | 16.2833886 | 17.7523659 | 20.9570264 | 17.8182579 |
| ENSMUSG00 | Haus8 | 5685 | 4.66436332 | 1.41550323 | 4.50959876 | 2.90052671 | 2.71063597 | 2.26928837 |
| ENSMUSG00 | Myo1d | 6442 | 14.6321398 | 15.1022084 | 11.0881991 | 17.5201235 | 20.2457958 | 12.6635278 |
| ENSMUSG00 | Thyn1 | 1006 | 7.5875271 | 9.005021 | 11.5996884 | 8.64390928 | 5.37653662 | 7.97091843 |
| ENSMUSG00 | Ccr3 | 3471 | 0 | 0 | 0 | 0 | 0 | 0 |
| ENSMUSG00 | Foxa1 | 3186 | 0 | 0.52935514 | 0 | 0 | 0 | 0 |
| ENSMUSG00 | 1700072E05I | 1045 | 0 | 0 | 0 | 0 | 0 | 0 |
| ENSMUSG00 | Fignl1 | 4204 | 0.25532778 | 0.33239934 | 0 | 0.06866231 | 0.71611707 | 0.38277919 |
| ENSMUSG00 | Prdm8 | 3138 | 0 | 0 | 0 | 0 | 0 | 0 |
| ENSMUSG00 | Tnni3 | 1593 | 0.29947631 | 0 | 0.49945585 | 0.15855261 | 0 | 0 |
| ENSMUSG00 | Stab2 | 9022 | 0 | 0 | 0.0244967 | 0 | 0 | 0.03930063 |
| ENSMUSG00 | Rcbtb1 | 6063 | 20.1695296 | 14.6315751 | 17.0377218 | 20.2875952 | 19.5756804 | 12.2945059 |
| ENSMUSG00 | Slc25a21 | 4851 | 1.30305533 | 2.09592647 | 0.30980471 | 1.48017457 | 1.42002901 | 1.90601994 |
| ENSMUSG00 | Galm | 2469 | 35.1986555 | 21.1559754 | 25.8873491 | 24.9753804 | 26.1228591 | 15.9626787 |
| ENSMUSG00 | Tab3 | 6603 | 4.44938666 | 5.15944489 | 5.7971825 | 7.48089035 | 7.24863555 | 6.92295856 |
| ENSMUSG00 | Mbd3 | 2383 | 37.1863055 | 43.8187244 | 45.8155905 | 40.0188182 | 31.6265032 | 35.9045503 |
| ENSMUSG00 | Plk5 | 2641 | 0 | 0 | 0.08368391 | 0 | 0 | 0 |
| ENSMUSG00 | 1600014K23I | 833 | 0 | 0 | 0 | 0 | 0 | 0 |
| ENSMUSG00 | Tgfbi | 3127 | 0 | 0 | 0 | 0 | 0 | 0 |
| ENSMUSG00 | Tstd2 | 6721 | 10.5111574 | 12.8119584 | 5.16269108 | 11.8967134 | 11.5703313 | 4.08652905 |
| ENSMUSG00 | Cdcp1 | 7618 | 1.83173716 | 2.84640312 | 3.98617303 | 3.55705112 | 2.72614006 | 2.64941363 |
| ENSMUSG00 | Reep6 | 2647 | 0.75095354 | 2.31192979 | 0.90173764 | 3.77586941 | 1.6578271 | 0.76249464 |
| ENSMUSG00 | Cox18 | 2195 | 9.2008128 | 5.33453551 | 6.9474423 | 7.3807918 | 7.8108712 | 5.95196 |
| ENSMUSG00 | Slc12a8 | 4427 | 0 | 0.25034745 | 0 | 0 | 0.0922094 | 0.01848293 |
| ENSMUSG00 | Fbxl21 | 2741 | 0.94276039 | 1.17785174 | 1.49973419 | 1.18474426 | 2.75516367 | 1.09456779 |
| ENSMUSG00 | Ntng2 | 3886 | 0.04092175 | 0 | 0 | 0 | 0 | 0 |
| ENSMUSG00 | Tdrd7 | 3944 | 18.8193413 | 20.8066711 | 15.5446137 | 21.4077039 | 27.9325419 | 12.496266 |
| ENSMUSG00 | Gnptg | 1703 | 37.5844531 | 43.7441219 | 47.7577165 | 35.9337331 | 28.3446574 | 47.7746617 |
| ENSMUSG00 | Tsga8 | 1068 | 0 | 0 | 0 | 0 | 0 | 0 |
| ENSMUSG00 | Vmn1r172 | 1367 | 0 | 0 | 0 | 0 | 0 | 0 |
| ENSMUSG00 | Npffr2 | 1329 | 1.22646704 | 0 | 0.46563266 | 1.35748851 | 0 | 0 |
| ENSMUSG00 | Prdm4 | 4501 | 12.1536417 | 8.98209841 | 12.432689 | 9.78808628 | 14.4315877 | 9.92576296 |
| ENSMUSG00 | Eif1 | 2094 | 72.0496889 | 75.7544258 | 92.4354683 | 81.5550701 | 70.6181648 | 100.593142 |
| ENSMUSG00 | E230008N13I | 5141 | 0 | 0.00937297 | 0.03439163 | 0.03509244 | 0 | 0.00530532 |
| ENSMUSG00 | Gc | 1653 | 0 | 0 | 0 | 0 | 0 | 0 |
| ENSMUSG00 | Leng8 | 6446 | 39.1880737 | 37.4069123 | 30.0004752 | 49.9024863 | 53.8207151 | 56.8383849 |
| ENSMUSG00 | Capn5 | 5762 | 15.4826967 | 16.8343124 | 15.948565 | 16.6007463 | 17.1003005 | 14.1390745 |
| ENSMUSG00 | Igfbpl1 | 3245 | 0 | 0 | 0 | 0 | 0 | 0 |
| ENSMUSG00 | Krt17 | 1841 | 0 | 1.25635482 | 0 | 0 | 0 | 0 |
| ENSMUSG00 | Mpv17l2 | 950 | 32.5576461 | 34.3391804 | 34.7566071 | 25.6372422 | 24.8686442 | 34.3660455 |
| ENSMUSG00 | Wdr20b | 2226 | 0 | 0 | 0 | 0 | 0.0229229 | 0 |
| ENSMUSG00 | Aldh1b1 | 598 | 25.6615643 | 57.6948036 | 31.1926049 | 32.3410968 | 36.1792364 | 16.3282957 |
| ENSMUSG00 | Pcdh17 | 9505 | 0.07528655 | 0.30924492 | 0.00930076 | 0 | 0.29526046 | 0 |
| ENSMUSG00 | Ankrd11 | 12685 | 12.4108556 | 17.9450337 | 13.4539469 | 21.2424682 | 18.0935466 | 13.6620463 |
| ENSMUSG00 | Dcaf10 | 7904 | 5.98042337 | 6.32203195 | 6.57659 | 5.55108377 | 6.47513347 | 6.3631623 |
| ENSMUSG00 | Utp6 | 4497 | 19.07768 | 20.8732909 | 16.3066172 | 14.081385 | 20.9347706 | 14.8654972 |
| ENSMUSG00 | L3mbtl1 | 4245 | 0.90842912 | 0 | 0.33320588 | 0.32299592 | 0 | 0.30840582 |
| ENSMUSG00 | Iqcg | 3563 | 0.62484056 | 0.6491578 | 0.62028969 | 2.01524751 | 0.67309561 | 0.42102306 |
| ENSMUSG00 | Kcnh8 | 1979 | 0 | 0 | 0 | 0 | 0 | 0 |
| ENSMUSG00 | Gdpd4 | 3035 | 0 | 0 | 0 | 0 | 0 | 0 |
| ENSMUSG00 | Tsen34 | 2674 | 10.4666635 | 12.5241501 | 15.3896469 | 10.1067507 | 11.8310969 | 9.40434457 |
| ENSMUSG00 | Krt33a | 2055 | 0 | 0 | 0 | 0 | 0 | 0 |
| ENSMUSG00 | Chrna5 | 2987 | 0 | 0 | 0 | 0 | 0 | 0 |
| ENSMUSG00 | 1600002K03I | 675 | 2.23808631 | 4.06907734 | 4.3874422 | 8.12514188 | 3.70413691 | 6.10143782 |
| ENSMUSG00 | Mboat7 | 6527 | 3.31955521 | 3.80943836 | 3.09487395 | 3.73148155 | 3.28344995 | 3.61878927 |
| ENSMUSG00 | Prpf39 | 4518 | 33.3055542 | 29.9698766 | 28.0297212 | 36.8487281 | 27.2185847 | 50.8365952 |
| ENSMUSG00 | Rg9mtd3 | 2514 | 6.72079523 | 5.29015833 | 5.43292343 | 7.13316473 | 6.08906633 | 3.88397806 |
| ENSMUSG00 | Ky | 5972 | 0 | 0 | 0 | 0 | 0 | 0 |
| ENSMUSG00 | Fam179b | 6971 | 13.0598265 | 11.9861269 | 9.2449128 | 11.9514332 | 13.7099988 | 9.02633664 |
| ENSMUSG00 | Frmpd1 | 5358 | 0.34131245 | 0.87235627 | 0 | 0 | 0.39998279 | 0.3461507 |
| ENSMUSG00 | Ric8b | 3058 | 26.9630041 | 16.7659825 | 28.1862635 | 22.135355 | 22.6097905 | 17.793625 |
| ENSMUSG00 | Midn | 4296 | 0.91615283 | 1.03192567 | 0.9465944 | 1.2766459 | 0.36820709 | 0.70472182 |
| ENSMUSG00 | Raf1 | 11895 | 5.32412614 | 4.85712855 | 8.08602021 | 7.1011405 | 7.50703301 | 9.14657721 |
| ENSMUSG00 | Olfr1509 | 1009 | 0 | 0 | 0 | 0 | 0 | 0 |
| ENSMUSG00 | 1700021K19I | 9855 | 2.96905466 | 3.05596411 | 3.52538291 | 3.81506752 | 6.64818534 | 2.38566606 |
| ENSMUSG00 | Cnot3 | 3384 | 3.43043744 | 4.11521329 | 3.44837073 | 7.28251667 | 5.20215711 | 5.28728231 |
| ENSMUSG00 | Grhpr | 1724 | 9.36236954 | 10.5372904 | 12.5375297 | 8.37170407 | 7.57700244 | 7.78371375 |
| ENSMUSG00 | Muc20 | 4022 | 71.5242807 | 52.1999822 | 69.1381886 | 54.715164 | 97.3459427 | 61.988433 |
| ENSMUSG00 | Dos | 6432 | 0.16070312 | 0.32963362 | 0.30924797 | 0.34780578 | 0.05553244 | 0.14417564 |
| ENSMUSG00 | 1810020D17I | 2002 | 25.3982316 | 27.1018651 | 49.1475038 | 29.9362018 | 17.943341 | 42.9282659 |
| ENSMUSG00 | Zcchc7 | 9594 | 5.52781019 | 5.60517714 | 5.9986241 | 6.67559194 | 5.21751873 | 10.2770295 |
| ENSMUSG00 | 4930480E11I | 1485 | 0 | 0 | 0 | 0 | 0 | 0 |
| ENSMUSG00 | Lrfn5 | 9516 | 0 | 0 | 0 | 0 | 0 | 0 |
| ENSMUSG00 | Gtf3c4 | 11715 | 2.6843009 | 2.64069107 | 2.90528546 | 3.41878527 | 3.21882132 | 1.24790496 |
| ENSMUSG00 | Zswim4 | 4611 | 2.48310093 | 3.06194482 | 0.89151408 | 1.87805046 | 4.66994808 | 2.34830444 |
| ENSMUSG00 | Sbno2 | 4498 | 8.40539395 | 8.28104041 | 8.04831246 | 3.93068071 | 9.52915867 | 5.45128961 |
| ENSMUSG00 | Ndufa3 | 881 | 62.2730569 | 75.971587 | 59.8556172 | 45.2561398 | 55.254441 | 140.614542 |
| ENSMUSG00 | Tnfsf9 | 1227 | 1.84683161 | 2.9061098 | 1.08072966 | 1.82321659 | 1.91296927 | 0.60017544 |
| ENSMUSG00 | Kcnc2 | 6191 | 4.21250125 | 7.12172421 | 0.52119763 | 3.18216707 | 3.82429953 | 2.11906017 |
| ENSMUSG00 | Melk | 2953 | 0 | 0.48953378 | 0 | 0.76978287 | 0.17279504 | 0.46181238 |
| ENSMUSG00 | Thrsp | 1275 | 0.71715769 | 0.07558658 | 0.48535357 | 0.65089178 | 0.3201655 | 0.55618872 |
| ENSMUSG00 | Isg15 | 754 | 0.68543932 | 0.89470648 | 0.11724627 | 0.62210422 | 0.33837119 | 4.5216577 |
| ENSMUSG00 | Caps2 | 3398 | 1.82514861 | 1.10610432 | 0.98862274 | 1.01938678 | 1.14126091 | 1.59730818 |
| ENSMUSG00 | Rnf38 | 11759 | 6.79551967 | 5.26981585 | 7.26610788 | 6.28727862 | 7.89327133 | 5.35565457 |
| ENSMUSG00 | Hmha1 | 4635 | 0 | 0 | 0.20026725 | 0 | 0.01100893 | 0.58256511 |
| ENSMUSG00 | Osta | 1591 | 0.19990185 | 0.33315579 | 0.30560671 | 0 | 0 | 0.42857698 |
| ENSMUSG00 | Alg8 | 2890 | 5.75010064 | 3.53478399 | 5.65905949 | 7.04161699 | 5.34982418 | 5.43605267 |
| ENSMUSG00 | Dok3 | 1541 | 0.77395484 | 0.34396552 | 0.31552257 | 0.60878206 | 1.19205031 | 1.25665113 |
| ENSMUSG00 | Usp35 | 4889 | 1.25226918 | 0.91661672 | 0.19890377 | 0 | 1.32549596 | 0.31799027 |
| ENSMUSG00 | Abca7 | 6705 | 4.5773648 | 4.26886602 | 5.23434204 | 4.03064151 | 4.90857754 | 3.06305792 |
| ENSMUSG00 | Prkx | 5798 | 17.6287238 | 13.2142882 | 16.9778379 | 16.8710629 | 12.3737641 | 14.9074391 |
| ENSMUSG00 | Supt16h | 4465 | 19.980134 | 16.0909261 | 20.3932355 | 23.0634166 | 25.3589088 | 17.3788015 |
| ENSMUSG00 | Dagla | 9083 | 2.40292401 | 2.05838815 | 4.58904965 | 3.26935183 | 2.21902658 | 4.25800269 |
| ENSMUSG00 | Grin3b | 3287 | 0.03628428 | 0.0146597 | 0.10757978 | 0.03293159 | 0 | 0 |
| ENSMUSG00 | Wdr18 | 2944 | 12.4100838 | 14.1580409 | 15.0592556 | 12.9792409 | 8.99547862 | 10.0519645 |
| ENSMUSG00 | 1300018J18R | 3807 | 12.5939347 | 9.40439362 | 9.06793785 | 10.6341093 | 11.8485202 | 7.50106312 |
| ENSMUSG00 | Bbs10 | 2751 | 5.79496463 | 5.42995168 | 4.65959087 | 7.59414896 | 9.19995725 | 5.90900945 |
| ENSMUSG00 | Tmem161b | 2696 | 29.2415126 | 30.9208253 | 39.6111472 | 28.4400388 | 29.8663283 | 38.4839495 |
| ENSMUSG00 | Fbxo45 | 4176 | 7.19711285 | 10.3619313 | 12.8922042 | 8.10463548 | 7.96676174 | 8.22941702 |
| ENSMUSG00 | Dym | 2467 | 22.8187112 | 24.3178437 | 18.9206106 | 15.3571734 | 22.6071458 | 15.6328901 |
| ENSMUSG00 | Xylb | 3604 | 0.95969112 | 1.17658349 | 1.80290536 | 1.49173825 | 1.10434443 | 1.62709418 |
| ENSMUSG00 | Dync1li2 | 4620 | 31.0557422 | 24.5834295 | 26.7028883 | 23.4220891 | 27.8657026 | 17.5336967 |
| ENSMUSG00 | Mrps2 | 2976 | 35.4539803 | 26.3762481 | 30.6858225 | 29.6682671 | 29.8339697 | 25.0384121 |
| ENSMUSG00 | Kiss1r | 3401 | 1.63650905 | 1.10512864 | 1.32566539 | 0.98665985 | 1.21527093 | 0.87413575 |
| ENSMUSG00 | Krt20 | 1919 | 0 | 0 | 0 | 0 | 0.05318017 | 0 |
| ENSMUSG00 | Cd99l2 | 5016 | 13.7590738 | 13.4203468 | 15.3155509 | 13.3725121 | 11.4036219 | 11.1523694 |
| ENSMUSG00 | Ggta1 | 7316 | 0.10868092 | 0.2832172 | 0.12083609 | 0 | 0.00697463 | 0 |
| ENSMUSG00 | Ugt2a3 | 2111 | 0 | 0 | 0 | 0 | 0 | 0 |
| ENSMUSG00 | C030046I01F | 2620 | 68.1002277 | 68.9323608 | 72.7643317 | 59.1359921 | 59.4788366 | 59.60862 |
| ENSMUSG00 | Acta2 | 1772 | 0.11217686 | 0.89737731 | 0.32427989 | 0.3665219 | 0.54712254 | 0.96969654 |
| ENSMUSG00 | Cmtm2b | 861 | 0 | 0 | 0 | 0 | 0 | 0 |
| ENSMUSG00 | 1500031L02F | 2042 | 20.6759649 | 19.4444801 | 20.0878111 | 21.6633627 | 23.664044 | 18.1919974 |
| ENSMUSG00 | Zdhhc17 | 4930 | 13.7007224 | 10.301929 | 10.947353 | 11.0222229 | 11.5714986 | 13.9526654 |
| ENSMUSG00 | Twist1 | 1614 | 1.77347868 | 1.13450112 | 5.75116917 | 2.21321091 | 0.88521594 | 3.31216189 |
| ENSMUSG00 | Ins1 | 1014 | 0 | 0 | 0 | 0 | 0 | 0 |
| ENSMUSG00 | Mlc1 | 2789 | 0 | 0 | 0 | 0 | 0 | 0 |
| ENSMUSG00 | Ugt2b35 | 4045 | 0 | 3.99071894 | 0 | 8.92015934 | 0.20183486 | 0 |
| ENSMUSG00 | 9930023K05I | 2372 | 0 | 0 | 0.57768007 | 0 | 0.04302393 | 0 |
| ENSMUSG00 | Tk2 | 2390 | 22.8219746 | 17.3592162 | 17.4588033 | 11.1869435 | 24.63784 | 17.3119781 |
| ENSMUSG00 | Pim3 | 2715 | 54.0323105 | 62.7577383 | 60.7103767 | 56.814269 | 53.3568622 | 39.8018861 |
| ENSMUSG00 | Ppp1r26 | 967 | 8.83911924 | 21.42727 | 10.3762341 | 22.313405 | 9.7620264 | 6.17698655 |
| ENSMUSG00 | Krt25 | 1981 | 0 | 0 | 0 | 0 | 0 | 0 |
| ENSMUSG00 | Polr3g | 6169 | 2.80975677 | 2.39799607 | 3.21715473 | 1.98863594 | 3.19276723 | 4.00564486 |
| ENSMUSG00 | BC005764 | 3979 | 1.57862928 | 1.33212079 | 0.98868159 | 1.00656118 | 1.29521587 | 0.98706913 |
| ENSMUSG00 | Ugt2b1 | 2567 | 0 | 0 | 0 | 1.01204021 | 0 | 0 |
| ENSMUSG00 | Lysmd3 | 3121 | 17.1581648 | 21.815906 | 24.912221 | 18.6826607 | 19.4720966 | 29.4943632 |
| ENSMUSG00 | Ddx11 | 4130 | 0.79896002 | 1.72677807 | 0.89902054 | 1.79973394 | 1.77912787 | 0.6405908 |
| ENSMUSG00 | Alg12 | 3152 | 5.58746125 | 5.64111585 | 5.0764807 | 4.08670365 | 4.03095418 | 3.99774217 |
| ENSMUSG00 | Ids | 8681 | 7.57923284 | 8.66478933 | 8.17741733 | 10.3412195 | 6.23649173 | 11.2730568 |
| ENSMUSG00 | Krt222 | 4525 | 0 | 0 | 0.05861018 | 0 | 0 | 0 |
| ENSMUSG00 | Ythdc1 | 8591 | 12.4389165 | 13.6914335 | 15.0752415 | 17.2829255 | 12.645228 | 17.702641 |
| ENSMUSG00 | 9130017N09I | 2481 | 7.88379541 | 5.32167882 | 3.1712729 | 3.12681966 | 3.88713625 | 3.68279087 |
| ENSMUSG00 | Cdhr3 | 2933 | 0 | 0 | 0 | 0 | 0 | 0 |
| ENSMUSG00 | Tmprss11bnl | 4161 | 0 | 0 | 0 | 0 | 0 | 0 |
| ENSMUSG00 | Palm | 2642 | 2.64835904 | 4.52317852 | 1.12094 | 1.13353887 | 1.91204058 | 0.90846641 |
| ENSMUSG00 | Syt1 | 11419 | 0.10792713 | 0 | 0 | 0.12639301 | 0 | 0.10748391 |
| ENSMUSG00 | 3110052M02 | 1380 | 24.4294547 | 22.5219241 | 23.0298008 | 24.9698207 | 22.4072346 | 34.8641041 |
| ENSMUSG00 | Pawr | 1821 | 13.0335101 | 21.1427607 | 21.3848565 | 23.7574802 | 17.4571291 | 14.8280576 |
| ENSMUSG00 | AI182371 | 2136 | 0.44669079 | 0 | 0.72428114 | 0 | 0 | 0 |
| ENSMUSG00 | Zhx3 | 12703 | 8.36858655 | 8.11768765 | 8.57034869 | 8.8877208 | 10.8174471 | 9.39357213 |
| ENSMUSG00 | Agphd1 | 7855 | 0.87052103 | 1.07967076 | 0.81594745 | 1.95224302 | 1.42263225 | 1.62501988 |
| ENSMUSG00 | Cox8a | 543 | 238.240025 | 256.018206 | 328.868224 | 214.299436 | 172.531171 | 513.144962 |
| ENSMUSG00 | Rnf126 | 1599 | 4.37583776 | 4.30935661 | 5.77747667 | 5.03208001 | 4.27613156 | 8.30691014 |
| ENSMUSG00 | Cerk | 7281 | 8.69804703 | 7.87554816 | 11.6985238 | 7.0271033 | 9.25777262 | 8.96043635 |
| ENSMUSG00 | Rnase1 | 968 | 28.8309373 | 14.0875652 | 63.9282854 | 13.9035151 | 22.2449696 | 43.2787664 |
| ENSMUSG00 | Uba6 | 7138 | 10.5208886 | 8.97167016 | 12.3168206 | 16.9643235 | 10.3082143 | 10.546092 |
| ENSMUSG00 | Gramd4 | 5546 | 3.28308873 | 2.20687997 | 1.47445744 | 3.03177655 | 4.49908002 | 2.14911961 |
| ENSMUSG00 | Dennd5a | 8218 | 6.07603839 | 4.06342229 | 6.15856781 | 5.55851404 | 4.58231508 | 4.54024173 |
| ENSMUSG00 | Dcdc2a | 8581 | 1.05168326 | 0.92093888 | 0.21119631 | 0 | 1.00494785 | 0.33056316 |
| ENSMUSG00 | Cd276 | 5348 | 0.20071017 | 0.1802036 | 0.2479535 | 0 | 0.2671538 | 0.59669648 |
| ENSMUSG00 | Ptprq | 7116 | 0 | 0 | 0 | 0 | 0 | 0 |
| ENSMUSG00 | Bbs9 | 6542 | 4.96487734 | 4.1174291 | 4.99314613 | 4.62194334 | 5.82646022 | 5.21978727 |
| ENSMUSG00 | Myf6 | 1245 | 0 | 0 | 0 | 0 | 0 | 0 |
| ENSMUSG00 | H2-Q4 | 2474 | 0.03213863 | 0.01947714 | 0 | 0.02916899 | 0.18562546 | 0.62839711 |
| ENSMUSG00 | Chst4 | 1998 | 0 | 0 | 0 | 0 | 0 | 0 |
| ENSMUSG00 | Olfr750 | 1131 | 0 | 0 | 0 | 0 | 0 | 0 |
| ENSMUSG00 | Cog5 | 2974 | 11.282322 | 10.8719242 | 13.3467569 | 9.34202229 | 16.3682456 | 12.8577822 |
| ENSMUSG00 | Pknox2 | 9679 | 0 | 0 | 0 | 0 | 0 | 0.08735549 |
| ENSMUSG00 | Aldh5a1 | 5637 | 2.60946031 | 3.38510397 | 2.32104764 | 2.41955168 | 3.20442381 | 3.19341173 |
| ENSMUSG00 | Ibtk | 6332 | 15.3509397 | 12.4194999 | 14.1080109 | 12.9466843 | 18.5587087 | 11.3414601 |
| ENSMUSG00 | Ttc38 | 3942 | 10.1254446 | 9.25345934 | 9.19470098 | 10.6818229 | 10.6013703 | 5.67356778 |
| ENSMUSG00 | Gsx2 | 2322 | 0 | 0 | 0 | 0 | 0 | 0 |
| ENSMUSG00 | Acss3 | 4531 | 6.36122785 | 8.16755409 | 5.41426239 | 10.8700046 | 8.50251903 | 7.48833618 |
| ENSMUSG00 | Fbxw2 | 7357 | 19.291432 | 17.880792 | 20.6619734 | 15.4588289 | 16.978737 | 23.0705559 |
| ENSMUSG00 | Ascl3 | 704 | 0.50823768 | 1.30048637 | 1.38130759 | 0.20501162 | 0.79728712 | 0.77484771 |
| ENSMUSG00 | Tmem55b | 4041 | 24.4376673 | 20.8199772 | 28.1771709 | 23.6796788 | 24.8628889 | 26.0799817 |
| ENSMUSG00 | Dock4 | 8103 | 1.16278525 | 1.72455488 | 0.95462453 | 0.56106845 | 1.02014968 | 0.58904873 |
| ENSMUSG00 | Tdp2 | 2194 | 29.336428 | 23.6759274 | 40.4747049 | 29.9971054 | 25.0945577 | 56.9485517 |
| ENSMUSG00 | Apex1 | 1478 | 7.34319769 | 5.02077949 | 9.92896609 | 5.19991575 | 6.80121518 | 5.81293056 |
| ENSMUSG00 | Odf3l2 | 2015 | 0 | 0 | 0 | 0 | 0 | 0 |
| ENSMUSG00 | Tmem59l | 1474 | 0 | 0 | 0 | 0 | 0 | 0 |
| ENSMUSG00 | Ddx26b | 6800 | 20.4506869 | 21.3083282 | 25.7735746 | 29.6774816 | 33.0695939 | 29.6130385 |
| ENSMUSG00 | Rusc2 | 7606 | 2.29459058 | 2.91424709 | 2.19672583 | 4.72491669 | 4.51495542 | 2.25196864 |
| ENSMUSG00 | Gm7008 | 1710 | 0.65096693 | 1.69075236 | 0.72377287 | 2.38436902 | 1.04439951 | 2.6636636 |
| ENSMUSG00 | Nme5 | 1316 | 3.98763178 | 3.03911451 | 2.82139424 | 3.18048417 | 1.55095367 | 2.5492254 |
| ENSMUSG00 | Fnip1 | 7100 | 3.56119516 | 2.23965154 | 4.20228789 | 4.177386 | 3.27000014 | 5.92359067 |
| ENSMUSG00 | B230312A22I | 3943 | 11.3932775 | 9.31221631 | 8.84485269 | 9.44374079 | 9.73163445 | 7.11783003 |
| ENSMUSG00 | Fam65b | 10731 | 3.17125071 | 1.95332237 | 5.19827844 | 2.12504446 | 3.67564892 | 4.12512716 |
| ENSMUSG00 | BC067068 | 2949 | 4.32740225 | 4.19936101 | 2.72795369 | 6.05649781 | 3.5471031 | 4.86485597 |
| ENSMUSG00 | Fam122c | 985 | 0 | 0 | 0 | 0 | 0.05180343 | 0 |
| ENSMUSG00 | Tmtc2 | 8535 | 0.1304222 | 0.85815339 | 0.01553668 | 0.00845508 | 0.1673976 | 0 |
| ENSMUSG00 | Fam122b | 6345 | 1.86716047 | 1.71633348 | 0.96136396 | 3.2300396 | 3.24091875 | 0.71356818 |
| ENSMUSG00 | Parp2 | 1772 | 37.7587323 | 42.5030525 | 39.2628109 | 35.7155226 | 36.0236999 | 33.908595 |
| ENSMUSG00 | Tmem63b | 5072 | 13.1917338 | 10.8115479 | 13.4034768 | 10.0093527 | 10.1610094 | 9.18475628 |
| ENSMUSG00 | 1810046K07F | 1313 | 0 | 0.073399 | 0.10099431 | 0 | 0 | 0.12463659 |
| ENSMUSG00 | Prtg | 8728 | 0 | 0 | 0 | 0 | 0.02923143 | 0.00312496 |
| ENSMUSG00 | Zfp57 | 3727 | 0 | 0.34908343 | 0.0237198 | 0.01936251 | 0.08214603 | 0.03659061 |
| ENSMUSG00 | Adamtsl2 | 4804 | 1.07581442 | 1.06323124 | 0.23002624 | 0.07510834 | 0.88159642 | 0.12490468 |
| ENSMUSG00 | Gm12221 | 1130 | 0.07036368 | 0.08528574 | 0.19558338 | 0.22351709 | 0.27093651 | 0.12068424 |
| ENSMUSG00 | 5031439G07I | 12145 | 2.88386812 | 1.71003348 | 3.37018581 | 1.83901075 | 1.63435654 | 2.25473346 |
| ENSMUSG00 | Dnajb5 | 3863 | 2.00681302 | 1.59665146 | 0.9840431 | 1.13953131 | 0.84537615 | 1.20028182 |
| ENSMUSG00 | Fmnl2 | 6404 | 0.30418778 | 0.86530931 | 0.64190684 | 0.37749797 | 1.01989009 | 2.30837828 |
| ENSMUSG00 | Sugp2 | 7588 | 3.13831482 | 4.50239687 | 4.62523239 | 6.55734573 | 4.82154869 | 4.00421035 |
| ENSMUSG00 | Ptpn23 | 5311 | 0.89825977 | 1.95068444 | 1.72279825 | 2.60883169 | 0.84547563 | 3.02994486 |
| ENSMUSG00 | Smug1 | 3229 | 11.9795858 | 7.9539714 | 14.3871592 | 11.0402787 | 10.9353521 | 10.5415763 |
| ENSMUSG00 | N28178 | 7256 | 0.00547898 | 0 | 0 | 0.01989087 | 0.03516151 | 0.31950721 |
| ENSMUSG00 | Slc2a6 | 2954 | 0.10766549 | 0.4730558 | 0.02992677 | 0.20764887 | 1.07096659 | 0 |
| ENSMUSG00 | Galt | 2709 | 24.0528729 | 19.5662925 | 24.5728961 | 18.1009592 | 20.4557564 | 24.4555551 |
| ENSMUSG00 | Sigmar1 | 3402 | 12.8077621 | 12.0961851 | 16.0722163 | 8.82429778 | 10.109282 | 15.9783527 |
| ENSMUSG00 | Slc17a3 | 4567 | 0.30467305 | 0.01055101 | 0 | 0 | 0.79327188 | 0.72262565 |
| ENSMUSG00 | Zranb3 | 6709 | 0.60442078 | 0.99116545 | 0.82355498 | 0.40873981 | 0.44112831 | 1.15863351 |
| ENSMUSG00 | Slain2 | 5661 | 23.1468051 | 20.5479723 | 23.3619351 | 24.1503033 | 24.2287402 | 26.927744 |
| ENSMUSG00 | Hyal3 | 1943 | 3.0486704 | 0.94240082 | 4.32236242 | 2.95267374 | 2.86251928 | 2.79343964 |
| ENSMUSG00 | Arl5a | 5448 | 24.0882601 | 21.3601795 | 25.5410062 | 19.2662752 | 21.3452846 | 24.5661995 |
| ENSMUSG00 | Dgkb | 7467 | 0 | 0 | 0 | 0 | 0 | 0 |
| ENSMUSG00 | Fam178a | 7382 | 5.70320426 | 7.74168523 | 6.41291979 | 10.4355412 | 9.89145811 | 6.05569408 |
| ENSMUSG00 | Gm98 | 6411 | 0.11782158 | 0.19542154 | 0.00689469 | 0 | 0.02387757 | 0.17442836 |
| ENSMUSG00 | Vezt | 17735 | 1.77313702 | 1.67097051 | 1.89917138 | 2.01823446 | 2.36790005 | 1.06883982 |
| ENSMUSG00 | Colec12 | 3312 | 0 | 0.30552998 | 0 | 0 | 0.26191075 | 0 |
| ENSMUSG00 | Rab3gap1 | 6052 | 19.9302921 | 20.9641279 | 20.7789563 | 20.7537338 | 27.4692552 | 20.5235802 |
| ENSMUSG00 | Prr5 | 1908 | 1.93776713 | 2.82855426 | 1.1814958 | 2.19366728 | 0.58835444 | 0.68615445 |
| ENSMUSG00 | Mbnl3 | 8438 | 0.64076148 | 0.83375451 | 0.46621996 | 0.80818991 | 0.33259667 | 1.43516708 |
| ENSMUSG00 | Slc17a2 | 5098 | 0 | 0 | 0 | 0 | 0 | 0 |
| ENSMUSG00 | Lmo1 | 1067 | 0 | 0 | 0 | 0 | 0 | 0 |
| ENSMUSG00 | Metap2 | 4978 | 88.7031181 | 64.9810338 | 66.8710486 | 83.3627148 | 55.5057905 | 99.3020413 |
| ENSMUSG00 | 2810432D09I | 854 | 18.5277298 | 15.1217406 | 17.6496821 | 12.0836824 | 10.4562246 | 13.7650697 |
| ENSMUSG00 | Il5 | 1533 | 0.23339812 | 0 | 0 | 0.25890573 | 0.83213268 | 0.26687514 |
| ENSMUSG00 | Rfxank | 3687 | 4.61844415 | 3.86332533 | 3.4112685 | 3.40452343 | 4.91200182 | 4.2842084 |
| ENSMUSG00 | Slc9a3 | 2474 | 0.64277252 | 0.29215709 | 0 | 0 | 0.0825002 | 0 |
| ENSMUSG00 | Frmd7 | 3570 | 0 | 0 | 0 | 0 | 0 | 0 |
| ENSMUSG00 | Fam110c | 2665 | 15.5740044 | 14.609623 | 17.2494997 | 13.3090618 | 13.6517096 | 23.5697914 |
| ENSMUSG00 | Acaa1a | 4401 | 28.3374102 | 24.9089198 | 25.9626822 | 28.3425649 | 25.6349277 | 20.2530155 |
| ENSMUSG00 | Hoxc9 | 2426 | 2.34337746 | 3.41635122 | 1.67624467 | 1.63603664 | 1.95608118 | 4.35090082 |
| ENSMUSG00 | Meox2 | 4608 | 0 | 0 | 0 | 0 | 0 | 0 |
| ENSMUSG00 | Tm6sf2 | 1937 | 0.16419403 | 0.67167472 | 0.45639487 | 0.33530036 | 1.92303843 | 0.32385994 |
| ENSMUSG00 | Mgat5 | 3818 | 1.17662894 | 0.63104298 | 2.01443706 | 1.2191157 | 2.75312557 | 0.87153117 |
| ENSMUSG00 | Prickle1 | 4336 | 2.09963216 | 3.07833129 | 1.69222923 | 4.72661469 | 4.31888375 | 2.46578842 |
| ENSMUSG00 | Surf6 | 3493 | 6.74921269 | 8.49780944 | 7.76980579 | 6.3838258 | 11.2190829 | 6.04367901 |
| ENSMUSG00 | 1700066J24R | 705 | 0 | 0 | 0 | 0.05118021 | 0 | 0 |
| ENSMUSG00 | Pphln1 | 4253 | 24.1168915 | 21.141759 | 38.7142876 | 29.0828236 | 26.514999 | 34.0083263 |
| ENSMUSG00 | Ccdc38 | 4286 | 0.05565396 | 0.01124275 | 0.01031308 | 0.21888315 | 0.04762144 | 0.00636366 |
| ENSMUSG00 | Sostdc1 | 1749 | 115.425003 | 163.431661 | 51.9858154 | 130.134669 | 122.27075 | 97.7926031 |
| ENSMUSG00 | Cd200r3 | 6092 | 0 | 0 | 0 | 0 | 0 | 0 |
| ENSMUSG00 | Gatad2a | 5488 | 17.7262502 | 18.6142962 | 14.6909916 | 18.7313675 | 18.8280632 | 14.1094572 |
| ENSMUSG00 | Hist1h1c | 1559 | 40.1379899 | 45.2192206 | 35.2423931 | 30.8977097 | 43.7602723 | 33.4328644 |
| ENSMUSG00 | Ng23 | 1194 | 0 | 0 | 0 | 0 | 0.08547132 | 0.13705849 |
| ENSMUSG00 | Fam69b | 2676 | 1.20336096 | 0.79230323 | 0.90848332 | 1.40229172 | 0.99154393 | 0.49942351 |
| ENSMUSG00 | Ankmy2 | 2436 | 30.0287702 | 28.1285389 | 34.9114721 | 31.0755868 | 30.5404171 | 28.9765051 |
| ENSMUSG00 | Rorb | 12536 | 5.83203005 | 3.8015628 | 12.7253072 | 6.60276086 | 5.71889423 | 10.7088206 |
| ENSMUSG00 | Slc26a8 | 4159 | 1.0419205 | 0.13903277 | 0.72270385 | 0.32967485 | 0.51529401 | 0.04590586 |
| ENSMUSG00 | Gxylt1 | 9988 | 4.74056641 | 4.55426526 | 4.49630281 | 6.90717552 | 4.36288796 | 5.00544791 |
| ENSMUSG00 | Arhgap36 | 3405 | 0 | 0.25473009 | 0 | 0 | 0 | 0 |
| ENSMUSG00 | Ndufa13 | 1238 | 54.6236449 | 56.8273066 | 75.5501614 | 50.5381787 | 38.8674252 | 110.178086 |
| ENSMUSG00 | Rif1 | 10515 | 8.85471568 | 11.4703438 | 11.8754357 | 15.613248 | 11.3359595 | 13.8150004 |
| ENSMUSG00 | Sh3bp4 | 5249 | 21.0024667 | 27.9810013 | 15.1493838 | 25.7090582 | 30.4661196 | 12.507155 |
| ENSMUSG00 | BC027231 | 5843 | 0.52390416 | 1.12982074 | 0.87753103 | 1.75377386 | 1.10907919 | 0.77487423 |
| ENSMUSG00 | Hist1h1t | 908 | 0 | 0.53068769 | 0.14604133 | 0 | 0 | 0 |
| ENSMUSG00 | Znrd1as | 1149 | 8.75381773 | 9.10048559 | 6.38599299 | 8.03916745 | 7.10550055 | 12.5572534 |
| ENSMUSG00 | Leap2 | 528 | 0 | 0 | 0 | 0 | 0 | 0 |
| ENSMUSG00 | Pdzrn4 | 4179 | 0 | 0.01153062 | 0 | 0 | 0 | 0 |
| ENSMUSG00 | Ska1 | 2662 | 0 | 0.19911753 | 0 | 0.25753525 | 0 | 0 |
| ENSMUSG00 | Kctd1 | 3608 | 5.06860339 | 6.42396859 | 4.47163878 | 9.2705253 | 4.45490809 | 4.46009899 |
| ENSMUSG00 | Agr3 | 1068 | 0 | 0 | 0 | 0 | 0 | 0 |
| ENSMUSG00 | Ube2r2 | 4184 | 32.6956521 | 38.0746601 | 42.469266 | 33.3741664 | 33.2208049 | 41.4660565 |
| ENSMUSG00 | 3632451O06I | 4399 | 0.47898169 | 0.15335535 | 0 | 0.43472343 | 0 | 0 |
| ENSMUSG00 | Tbc1d21 | 1086 | 0 | 0 | 0 | 0 | 0 | 0 |
| ENSMUSG00 | Gmip | 4804 | 2.2178328 | 3.31005951 | 1.89541625 | 3.07193094 | 1.64635476 | 2.22557423 |
| ENSMUSG00 | Rbm43 | 3955 | 7.43844642 | 7.26147143 | 12.4726313 | 7.40799498 | 6.57988664 | 7.22036594 |
| ENSMUSG00 | Trpm8 | 4701 | 0 | 0 | 0 | 0 | 0 | 0 |
| ENSMUSG00 | Igfbp7 | 1462 | 353.040632 | 546.200902 | 206.79932 | 306.37654 | 339.105518 | 249.389451 |
| ENSMUSG00 | Pnpla8 | 3932 | 100.794096 | 87.4145198 | 98.6447539 | 100.381863 | 102.169547 | 104.215204 |
| ENSMUSG00 | Fat4 | 4589 | 0 | 0 | 0 | 0 | 0 | 0 |

| | | | | | | | |
|---|---|---|---|---|---|---|---|
| ENSMUSG00 Edc4 | 5424 | 13.9114864 | 10.2431725 | 11.4253289 | 14.6949182 | 15.0143982 | 6.91923002 |
| ENSMUSG00 Lrrk2 | 11780 | 1.89327883 | 2.53612853 | 0.78797853 | 2.32481085 | 2.31308019 | 2.00044892 |
| ENSMUSG00 9530068E07R | 3425 | 100.752575 | 105.953897 | 126.514063 | 102.736379 | 105.449544 | 94.4777577 |
| ENSMUSG00 Macrod1 | 1263 | 13.7869362 | 11.4838631 | 15.9938577 | 12.3130334 | 10.4638411 | 19.457205 |
| ENSMUSG00 Zbtb20 | 3078 | 0 | 0 | 0 | 0 | 0 | 0 |
| ENSMUSG00 Snapc4 | 6442 | 3.727462 | 5.29587101 | 2.00356072 | 6.10515811 | 4.31688526 | 2.70121388 |
| ENSMUSG00 Naa30 | 6941 | 11.3464352 | 12.3850472 | 12.1633079 | 18.1943749 | 12.0710718 | 20.5159043 |
| ENSMUSG00 Noa1 | 3625 | 6.44861311 | 7.74970918 | 5.1335106 | 5.12613874 | 6.30615624 | 4.66490382 |
| ENSMUSG00 Mudeng | 3157 | 32.5146185 | 24.864021 | 24.1941035 | 31.6817952 | 25.9576685 | 26.419337 |
| ENSMUSG00 Gramd1c | 3606 | 10.7712436 | 11.7860349 | 13.0055894 | 14.6289376 | 13.2447831 | 10.1882804 |
| ENSMUSG00 Lrrn3 | 3178 | 0 | 0 | 0 | 0 | 0.14450516 | 0 |
| ENSMUSG00 Slc2a13 | 12952 | 0.02455558 | 0 | 0.11603325 | 0.15043471 | 0 | 0 |
| ENSMUSG00 BC031181 | 1044 | 82.9000777 | 104.542425 | 117.490531 | 97.4629938 | 73.9002684 | 103.76903 |
| ENSMUSG00 Zdhhc23 | 1652 | 4.86114229 | 3.29604598 | 7.06373278 | 3.38542185 | 3.39764004 | 3.28550437 |
| ENSMUSG00 Gm15430 | 563 | 0 | 0 | 0.07851127 | 0 | 0.09063299 | 0 |
| ENSMUSG00 Lzts1 | 1910 | 0 | 0 | 0 | 0 | 0 | 0 |
| ENSMUSG00 Skp1a | 2310 | 190.516524 | 193.162967 | 202.276053 | 198.576204 | 134.877511 | 236.285749 |
| ENSMUSG00 Znrd1 | 954 | 6.04249966 | 8.23311328 | 12.6489551 | 9.9093246 | 7.91604153 | 17.5827077 |
| ENSMUSG00 H2-Ea-ps | 1038 | 0 | 0 | 0 | 0 | 0 | 0 |
| ENSMUSG00 Srp72 | 5591 | 29.6868416 | 24.7697791 | 26.318715 | 21.5937258 | 24.3769361 | 26.6502154 |
| ENSMUSG00 Qsox2 | 6966 | 11.3057145 | 6.94504564 | 6.83396281 | 4.04019448 | 5.26671895 | 5.24271346 |
| ENSMUSG00 Slc18a1 | 5180 | 0.85957796 | 0.31628167 | 1.57863725 | 1.49761382 | 1.17222755 | 0.93197126 |
| ENSMUSG00 Kidins220 | 7429 | 53.4978515 | 45.9616542 | 50.6515399 | 58.3073017 | 64.907693 | 34.492561 |
| ENSMUSG00 Igsf10 | 10077 | 0 | 0 | 0 | 0 | 0 | 0 |
| ENSMUSG00 6720456H20R | 8171 | 6.6364552 | 5.87955976 | 9.01780349 | 9.09226896 | 9.49836221 | 6.39223279 |
| ENSMUSG00 Ubac1 | 2558 | 31.5961266 | 27.5593295 | 34.7497679 | 26.0670908 | 31.1983002 | 30.5266975 |
| ENSMUSG00 P2ry12 | 4270 | 0 | 0 | 0 | 0 | 0.01194997 | 0 |
| ENSMUSG00 Csgalnact1 | 5507 | 0 | 0.63875255 | 0.39329768 | 0.4193301 | 0.41695786 | 0.39621775 |
| ENSMUSG00 Gpr101 | 5408 | 0 | 0 | 0 | 0 | 0 | 0 |
| ENSMUSG00 P2ry13 | 2522 | 0 | 0 | 0 | 0 | 0 | 0 |
| ENSMUSG00 Fam82a1 | 1791 | 11.6536166 | 8.20595469 | 12.0438299 | 12.0273482 | 5.92601126 | 18.7160981 |
| ENSMUSG00 Serbp1 | 6677 | 26.5493017 | 25.3669828 | 28.9824275 | 24.0907226 | 20.0452574 | 29.5458238 |
| ENSMUSG00 1810006K21R | 1653 | 10.2214635 | 13.3511134 | 10.5624489 | 7.3561095 | 8.89025786 | 19.6516003 |
| ENSMUSG00 Abt1 | 1275 | 12.4723076 | 21.5043809 | 19.3794747 | 18.2532696 | 13.6470542 | 17.9477822 |
| ENSMUSG00 C530008M17 | 9682 | 0 | 0 | 0.00913073 | 0.03354043 | 0 | 0.1408523 |
| ENSMUSG00 P2ry14 | 3230 | 14.4129002 | 6.32540293 | 20.2261065 | 12.1986355 | 32.0060178 | 21.3046176 |
| ENSMUSG00 Gadd45a | 2075 | 43.4340599 | 29.9568725 | 45.1391351 | 40.8987801 | 36.1979881 | 38.0267621 |
| ENSMUSG00 Sec24a | 13655 | 1.19659484 | 1.52093421 | 2.19795323 | 2.0372945 | 2.06272863 | 2.69850147 |
| ENSMUSG00 Glb1l2 | 6787 | 28.4854725 | 36.8125391 | 19.9093906 | 31.3292306 | 45.6433074 | 25.1367126 |
| ENSMUSG00 Ppp1r11 | 2235 | 13.8210328 | 15.9974676 | 17.4434119 | 16.8382874 | 11.4609578 | 17.7559732 |
| ENSMUSG00 Glt6d1 | 1397 | 0 | 0 | 0 | 0 | 0 | 0 |
| ENSMUSG00 Gng12 | 4780 | 185.386958 | 200.608828 | 182.808836 | 186.532093 | 219.103842 | 170.689029 |
| ENSMUSG00 Cep135 | 6731 | 1.94318126 | 2.09755275 | 1.22144448 | 2.25680296 | 2.47892214 | 2.18002614 |
| ENSMUSG00 9530077C05R | 1994 | 1.87412997 | 0.74913727 | 0.9310318 | 1.68286366 | 1.48421755 | 0.53345583 |
| ENSMUSG00 Arsi | 2973 | 0 | 0 | 0 | 0 | 0 | 0 |
| ENSMUSG00 Pcdh8 | 3992 | 0 | 0 | 0 | 0 | 0 | 0 |
| ENSMUSG00 Gpi1 | 4868 | 76.5709501 | 83.6432695 | 69.6623952 | 66.2788912 | 67.7242018 | 43.6405434 |
| ENSMUSG00 Tbcc | 1842 | 15.6475154 | 19.1490096 | 20.9730786 | 18.33485 | 14.266332 | 18.3163565 |
| ENSMUSG00 Siah2 | 3226 | 6.86416697 | 5.95982344 | 5.72733118 | 4.06006887 | 4.55536151 | 5.94360552 |
| ENSMUSG00 Exoc1 | 8937 | 5.42706571 | 7.43528071 | 5.43558526 | 7.59430746 | 8.35880208 | 5.03865161 |
| ENSMUSG00 Npy1r | 3004 | 0 | 0 | 0 | 0 | 0 | 0 |
| ENSMUSG00 Calm2 | 1797 | 270.36824 | 262.116011 | 325.942472 | 237.595344 | 169.803966 | 386.519735 |
| ENSMUSG00 Thap11 | 1847 | 18.8122848 | 14.8968373 | 8.47182048 | 11.5845438 | 14.724991 | 9.03740077 |
| ENSMUSG00 Lum | 2072 | 0.46048819 | 0 | 0.78931863 | 0 | 0 | 0 |
| ENSMUSG00 Lcn8 | 644 | 0 | 0 | 0 | 0 | 0 | 0 |
| ENSMUSG00 Hif1an | 6598 | 8.90551685 | 11.1008475 | 14.8255045 | 12.4192669 | 12.2113743 | 9.75983984 |
| ENSMUSG00 Arhgap26 | 17044 | 0.31022524 | 0.33360715 | 0.49274526 | 0.42974977 | 0.37721916 | 0.39046069 |
| ENSMUSG00 Wtip | 2227 | 3.71312975 | 7.33507135 | 1.32982644 | 2.77055663 | 4.07844405 | 0.90629695 |
| ENSMUSG00 Elf1 | 4317 | 25.9326924 | 30.4499917 | 32.4474497 | 38.3638138 | 36.1688001 | 28.2476053 |
| ENSMUSG00 4930544G11R | 1441 | 0 | 0 | 0 | 0 | 0 | 0 |
| ENSMUSG00 Megf11 | 10251 | 0.15125 | 0.16922368 | 0.21559771 | 0.29918777 | 0.06471007 | 0.16496221 |
| ENSMUSG00 1-Mar | 5914 | 0 | 0 | 0 | 0.1037191 | 0 | 0 |
| ENSMUSG00 Tbc1d24 | 13296 | 7.63654462 | 3.17836265 | 6.90487575 | 5.19412103 | 6.20943713 | 3.75395531 |
| ENSMUSG00 Btg1 | 5006 | 26.3634087 | 25.728943 | 24.5281393 | 23.9562296 | 24.0357869 | 27.2909779 |
| ENSMUSG00 Prss56 | 924 | 0 | 0 | 0 | 0 | 0 | 0 |
| ENSMUSG00 Rnf39 | 2186 | 0.18186405 | 0.13225922 | 0 | 0 | 0.1867388 | 0 |
| ENSMUSG00 Eea1 | 7954 | 14.6796387 | 11.1712094 | 12.1535618 | 18.2995936 | 14.1454814 | 9.7042028 |
| ENSMUSG00 Akp3 | 1669 | 0 | 0 | 0 | 0 | 0 | 0 |
| ENSMUSG00 Fam13b | 3322 | 12.3861897 | 11.7202424 | 15.3016615 | 16.0207753 | 15.4983785 | 13.1611218 |
| ENSMUSG00 Fam70a | 4480 | 0 | 0 | 0 | 0 | 0 | 0 |
| ENSMUSG00 Rnf13 | 4163 | 124.480588 | 117.751784 | 118.101633 | 102.482367 | 76.2147501 | 168.188149 |
| ENSMUSG00 Phpt1 | 1268 | 7.49334372 | 9.00645646 | 10.0395353 | 6.28874751 | 5.23141077 | 12.7984309 |
| ENSMUSG00 Cdh8 | 7801 | 0 | 0 | 0 | 0 | 0 | 0 |
| ENSMUSG00 Commd2 | 5269 | 3.28214728 | 5.97186311 | 4.81530801 | 5.36197397 | 3.43791297 | 4.54491047 |
| ENSMUSG00 Abpe | 517 | 0 | 0 | 0 | 0 | 0 | 0 |
| ENSMUSG00 Greb1 | 9533 | 0 | 0 | 0.06955079 | 0 | 0.06423125 | 0.23460825 |
| ENSMUSG00 Card11 | 4087 | 0 | 0 | 0 | 0 | 0 | 0 |
| ENSMUSG00 Ppfibp2 | 4701 | 46.4786473 | 31.3452755 | 73.0115531 | 29.9954542 | 48.8120851 | 46.3976367 |
| ENSMUSG00 Sbf1 | 7212 | 4.68554609 | 6.76828423 | 5.87151492 | 6.35388194 | 8.90768263 | 5.01472153 |
| ENSMUSG00 Cdc42ep3 | 1965 | 2.81221974 | 3.92357799 | 2.02451189 | 1.83623636 | 3.92111081 | 3.17857119 |
| ENSMUSG00 Slc38a7 | 4132 | 4.72409025 | 4.45479693 | 3.23062841 | 3.08251736 | 3.7294205 | 3.14859703 |
| ENSMUSG00 Rnf113a1 | 1217 | 2.15600798 | 4.47417252 | 1.92497755 | 1.60101101 | 2.34796799 | 2.33078265 |
| ENSMUSG00 Adamts2 | 9312 | 0 | 0 | 0 | 0 | 0 | 0.03807671 |
| ENSMUSG00 Cnot1 | 9372 | 12.1658897 | 9.77405319 | 11.7767821 | 11.1033522 | 16.2465541 | 11.6700068 |
| ENSMUSG00 Akap14 | 1786 | 0 | 0 | 0 | 0 | 0 | 0 |
| ENSMUSG00 2210404J11R | 4272 | 4.00160501 | 5.00814146 | 4.13874935 | 3.90213122 | 5.42274687 | 5.36299086 |
| ENSMUSG00 Sh3tc1 | 5946 | 37.5356992 | 37.8376217 | 38.7751115 | 37.3260437 | 36.9525016 | 27.4030768 |
| ENSMUSG00 Iqce | 11797 | 1.43897518 | 1.28666011 | 1.07909899 | 1.57210909 | 1.76042511 | 0.91786317 |
| ENSMUSG00 1700011E24R | 840 | 0 | 0 | 0 | 0 | 0 | 0 |
| ENSMUSG00 Lgi4 | 4988 | 0 | 0.09660474 | 0.15950946 | 0 | 0 | 0.41010384 |
| ENSMUSG00 Ppp6r2 | 3066 | 4.6290628 | 4.6677669 | 2.92660604 | 3.271627 | 5.1259373 | 3.19360584 |
| ENSMUSG00 Ndrg4 | 4867 | 2.89977308 | 1.14847489 | 1.87088138 | 1.60875358 | 4.7283533 | 3.33998048 |
| ENSMUSG00 Ttyh3 | 4636 | 0.60885227 | 0.96663915 | 0.3146378 | 0.78608423 | 0.74844555 | 0.62950526 |
| ENSMUSG00 BC032203 | 6568 | 3.62569014 | 4.90081354 | 3.29764054 | 5.36725906 | 5.60140331 | 5.0039495 |
| ENSMUSG00 Fxyd1 | 700 | 0 | 0.13767555 | 0.31572745 | 0 | 0.07289482 | 0 |
| ENSMUSG00 Upf3b | 4512 | 15.8687102 | 15.8805939 | 12.559123 | 17.3852759 | 11.9423433 | 22.6442373 |
| ENSMUSG00 1700019O17R | 1501 | 0 | 0 | 0 | 0 | 0 | 0 |
| ENSMUSG00 Fxyd7 | 678 | 0.29318201 | 0 | 0 | 0 | 0 | 0 |
| ENSMUSG00 Spg20 | 5439 | 19.1723893 | 23.9647593 | 34.8884928 | 13.9511932 | 12.327387 | 23.5387308 |
| ENSMUSG00 Fgf1 | 7972 | 3.03701551 | 1.34791478 | 0.92040968 | 0.38924433 | 2.71389656 | 0.82111307 |
| ENSMUSG00 Grifin | 608 | 0 | 0 | 0 | 0 | 0 | 0 |
| ENSMUSG00 Fut7 | 2177 | 0 | 0 | 0 | 0 | 0 | 0 |
| ENSMUSG00 Arhgap21 | 11456 | 4.91044037 | 3.50798645 | 4.95418701 | 5.83309589 | 7.4962806 | 4.48307836 |
| ENSMUSG00 H2-Aa | 1978 | 0 | 0 | 0.11173368 | 0 | 0.30956345 | 2.52338675 |
| ENSMUSG00 Cpz | 2237 | 0.21326141 | 0 | 0 | 0 | 0 | 0 |
| ENSMUSG00 Ccdc113 | 1262 | 0.72454521 | 0.1145478 | 0.66547941 | 0 | 0.727793 | 0.15128564 |
| ENSMUSG00 Chst12 | 1564 | 0.83883047 | 2.18749194 | 2.28922589 | 1.75334743 | 3.49093492 | 1.70902471 |
| ENSMUSG00 Alx1 | 4157 | 0 | 0 | 0 | 0 | 0 | 0 |
| ENSMUSG00 Plxnb2 | 7338 | 63.1493433 | 58.4042268 | 51.9844515 | 59.6745288 | 77.958122 | 38.6595152 |
| ENSMUSG00 Eepd1 | 2926 | 1.10054474 | 0.18115204 | 0.58915649 | 1.10983732 | 0.05231686 | 0.47539528 |
| ENSMUSG00 Tssc1 | 1683 | 17.149423 | 10.9085172 | 6.56593034 | 9.24026215 | 12.8854484 | 9.73978502 |
| ENSMUSG00 Rfxap | 2158 | 12.5272135 | 16.0993627 | 15.7307764 | 10.2828811 | 13.6432896 | 14.2313456 |
| ENSMUSG00 Etl4 | 15352 | 11.2958185 | 6.41565189 | 13.022729 | 7.66202874 | 8.96750664 | 19.2834116 |
| ENSMUSG00 Mgat4b | 2704 | 30.7575685 | 34.4112499 | 24.6019873 | 27.1282532 | 26.3434988 | 20.0827688 |
| ENSMUSG00 Atp13a2 | 6151 | 10.1925529 | 10.7951418 | 7.41607901 | 5.43782398 | 12.5263904 | 7.32083066 |
| ENSMUSG00 Alg5 | 3617 | 11.6397716 | 15.5070552 | 17.2187992 | 13.4172933 | 12.8518187 | 17.1776688 |
| ENSMUSG00 Mag | 2414 | 0.34584304 | 0.4591086 | 0.18310623 | 0 | 0.33820299 | 0 |
| ENSMUSG00 Clcn7 | 5415 | 17.8918017 | 25.2011088 | 14.0727567 | 19.2171037 | 20.6838218 | 9.21243124 |
| ENSMUSG00 Nudt1 | 1430 | 4.50376772 | 5.76215497 | 6.12025519 | 3.68390105 | 3.24713301 | 9.38400176 |
| ENSMUSG00 Ccdc148 | 9528 | 0.90960272 | 0.68779976 | 0.81185165 | 1.08306725 | 1.0657272 | 1.82059935 |
| ENSMUSG00 Tbc1d9b | 6558 | 15.5918109 | 16.7969208 | 15.6977878 | 16.8856046 | 20.7435679 | 10.9256599 |
| ENSMUSG00 Man1b1 | 5231 | 4.62078611 | 3.51887228 | 2.56879378 | 2.15209289 | 3.97012712 | 2.69565829 |
| ENSMUSG00 Olfr6 | 950 | 0 | 0 | 0 | 0 | 0 | 0 |
| ENSMUSG00 Gm3448 | 727 | 0 | 0 | 0 | 0 | 0 | 0 |
| ENSMUSG00 Colec11 | 1376 | 0 | 0 | 0 | 0 | 0 | 0 |
| ENSMUSG00 Olfr11 | 941 | 0 | 0 | 0 | 0 | 0 | 0 |
| ENSMUSG00 Dennd3 | 5854 | 9.01866726 | 5.88542981 | 8.26802496 | 5.60276366 | 6.63325453 | 5.80063647 |
| ENSMUSG00 Fam115a | 9989 | 7.56981919 | 5.49930364 | 5.64194112 | 6.98233978 | 7.29459052 | 5.83500892 |
| ENSMUSG00 Cenpt | 1754 | 3.10518861 | 2.44503612 | 2.41925709 | 2.81826688 | 1.9491261 | 3.04779321 |
| ENSMUSG00 Tmtc3 | 7895 | 5.99731188 | 6.69544358 | 7.87738988 | 8.28127482 | 8.36974752 | 5.1854634 |
| ENSMUSG00 Aaas | 1800 | 11.2861392 | 8.21846541 | 9.9208581 | 12.7890802 | 12.614854 | 6.06103094 |
| ENSMUSG00 Cc2d1a | 4017 | 10.708347 | 9.17665627 | 6.86630571 | 7.90445586 | 8.90452812 | 6.51142122 |
| ENSMUSG00 Tmem184a | 5922 | 18.5753824 | 14.036071 | 17.2866377 | 17.1088114 | 12.442095 | 14.0978843 |
| ENSMUSG00 Nop14 | 2716 | 10.9342209 | 7.30957812 | 11.0504608 | 10.5482855 | 11.948739 | 8.68651066 |
| ENSMUSG00 Eif2c2 | 10938 | 3.22754312 | 4.1675237 | 4.45332154 | 4.4104675 | 4.24986546 | 3.63063401 |
| ENSMUSG00 Zcchc12 | 2342 | 0.93362572 | 0.08229965 | 0 | 0 | 0.74077574 | 0 |
| ENSMUSG00 Cab39 | 4413 | 145.725052 | 111.474142 | 179.88237 | 119.521262 | 131.433676 | 189.000333 |
| ENSMUSG00 Cyld | 8051 | 14.4682099 | 10.7194035 | 13.000865 | 16.1161386 | 16.4341563 | 13.3713826 |
| ENSMUSG00 Micall2 | 4343 | 0.44854215 | 0.31066553 | 0.59030783 | 0.64803119 | 0.64620094 | 0.60917338 |
| ENSMUSG00 Zscan12 | 1846 | 11.5217671 | 5.53387093 | 11.349769 | 10.105318 | 13.2679634 | 8.24444675 |
| ENSMUSG00 2310002J15R | 1020 | 0 | 0 | 0.04333514 | 0 | 0 | 0.16043905 |
| ENSMUSG00 Rbm42 | 2404 | 0.13229777 | 0.02004428 | 0.07354716 | 0.12007336 | 0.12735368 | 0.03403657 |
| ENSMUSG00 Oxsr1 | 5303 | 8.45637979 | 7.74181571 | 7.80179616 | 7.71582849 | 6.46609929 | 6.33133715 |
| ENSMUSG00 Psma8 | 1846 | 0 | 0 | 0 | 0.76229672 | 0 | 0.97514962 |
| ENSMUSG00 Olfr701 | 1597 | 0 | 0.0905193 | 0 | 0 | 0 | 0 |
| ENSMUSG00 Ttll7 | 8143 | 1.55741106 | 1.80484648 | 1.09106846 | 1.09003843 | 0.8271499 | 2.6293248 |
| ENSMUSG00 Cuedc2 | 1931 | 23.5732911 | 26.8506534 | 31.4517516 | 23.0394412 | 21.2720003 | 28.9837182 |
| ENSMUSG00 Pramel5 | 2927 | 0 | 0 | 0 | 0 | 0 | 0 |
| ENSMUSG00 Cox6b1 | 501 | 271.940183 | 307.200591 | 428.872573 | 260.352477 | 198.809353 | 664.281733 |
| ENSMUSG00 Tubb4b | 2005 | 28.9887843 | 34.8720836 | 35.0748789 | 34.2824413 | 27.9690708 | 23.6697617 |
| ENSMUSG00 Kcnk9 | 1207 | 0 | 0 | 0 | 0 | 0 | 0 |
| ENSMUSG00 Dnajc12 | 2735 | 8.0673827 | 10.852961 | 6.4807821 | 9.84175693 | 8.11571243 | 8.85553187 |
| ENSMUSG00 Dner | 3617 | 0 | 0 | 0 | 0 | 0 | 0 |
| ENSMUSG00 Kif15 | 4780 | 0.03326819 | 0 | 0.0184945 | 0.02264564 | 0.27754933 | 0.38800742 |
| ENSMUSG00 Wdr44 | 8795 | 2.02506598 | 3.67630502 | 2.31689024 | 3.29025579 | 2.465743 | 2.69180067 |
| ENSMUSG00 4933433C11R | 2904 | 0 | 0 | 0.04566306 | 0 | 0 | 0 |
| ENSMUSG00 Decr2 | 2128 | 30.5638939 | 34.1018712 | 53.4036365 | 29.1810144 | 28.8701863 | 24.147284 |
| ENSMUSG00 Anln | 5397 | 0.45670554 | 0.68748491 | 0.09828092 | 0.04679902 | 0.06618207 | 0.01010733 |
| ENSMUSG00 Papd5 | 5023 | 8.57952237 | 8.97919766 | 11.2550582 | 9.74064352 | 11.8143888 | 9.19833544 |
| ENSMUSG00 Rps27l | 899 | 35.1564038 | 36.8768323 | 41.5467824 | 35.2793294 | 24.1793505 | 84.2813658 |
| ENSMUSG00 Klhl13 | 5898 | 2.45354271 | 2.06700066 | 3.46989714 | 2.58165866 | 1.71299125 | 3.17233003 |
| ENSMUSG00 Slitrk2 | 8077 | 0.00984412 | 0.01789765 | 0.04378046 | 0.04467258 | 0 | 0.01350731 |
| ENSMUSG00 Mbd5 | 12403 | 3.32390819 | 4.10651208 | 3.16109286 | 3.35132753 | 4.04409638 | 3.45908309 |
| ENSMUSG00 Fam135b | 4603 | 0 | 0 | 0 | 0 | 0 | 0 |
| ENSMUSG00 Noxa1 | 1605 | 0 | 0 | 0 | 0 | 0 | 0 |
| ENSMUSG00 Tmem188 | 2500 | 26.0159864 | 32.0150722 | 34.4420761 | 36.0964773 | 29.3503714 | 43.7812509 |
| ENSMUSG00 Entpd8 | 2712 | 0 | 0 | 0 | 0 | 0.2634105 | 0.06034212 |
| ENSMUSG00 Slc6a20a | 2870 | 0 | 0 | 0.15401339 | 0 | 0.53337675 | 0 |
| ENSMUSG00 Dpp10 | 7118 | 0 | 0 | 0 | 0 | 0 | 0 |
| ENSMUSG00 Atoh7 | 449 | 0.17708455 | 0.32195841 | 0 | 0 | 0 | 0.2429812 |
| ENSMUSG00 Sun1 | 6863 | 15.7967646 | 18.1427607 | 22.0075328 | 11.8450891 | 17.2491901 | 13.7585314 |
| ENSMUSG00 Jmjd4 | 7501 | 4.07571851 | 2.70450502 | 3.52978322 | 4.15128709 | 5.47610086 | 3.15980023 |
| ENSMUSG00 Amdhd2 | 2887 | 12.26953 | 14.3708094 | 18.3880892 | 13.9853855 | 7.86516704 | 10.9495348 |
| ENSMUSG00 Topors | 3853 | 42.3762674 | 31.5531777 | 38.1331238 | 41.6540187 | 37.9552539 | 33.3270183 |
| ENSMUSG00 Ssx2ip | 4299 | 18.2085464 | 20.7137811 | 16.4098957 | 21.6710488 | 17.9702101 | 18.8302231 |
| ENSMUSG00 Tmem149 | 1265 | 0 | 0 | 0 | 0 | 0 | 0 |
| ENSMUSG00 Lpar3 | 2491 | 0 | 0.4836054 | 0 | 0 | 0 | 0 |
| ENSMUSG00 Pnpla7 | 7581 | 3.81245671 | 4.61460981 | 3.84820161 | 1.71343306 | 6.48178339 | 3.06529385 |
| ENSMUSG00 Ptch1 | 11611 | 0.00342395 | 0.08300136 | 0.21318605 | 0.20820747 | 0.36915129 | 0.03523552 |
| ENSMUSG00 Psenen | 644 | 5.12376535 | 5.76142243 | 5.69682138 | 7.17158649 | 3.48627412 | 7.62334637 |
| ENSMUSG00 Siah1a | 2105 | 14.6557021 | 16.0468864 | 17.2607672 | 19.6437164 | 12.7747744 | 16.3906977 |
| ENSMUSG00 Lin37 | 1511 | 6.68291995 | 8.83364955 | 9.15630501 | 7.33103089 | 6.4500581 | 8.84485323 |
| ENSMUSG00 Mrpl41 | 774 | 32.2563847 | 30.0698341 | 40.147152 | 27.5044009 | 18.6569307 | 35.1328364 |
| ENSMUSG00 Mcoln3 | 3445 | 126.121233 | 154.490349 | 87.197598 | 112.341367 | 132.446401 | 65.0949445 |
| ENSMUSG00 Hspb6 | 2497 | 3.45492161 | 2.62449184 | 2.49597912 | 1.87852054 | 1.7778513 | 1.94428746 |
| ENSMUSG00 Gjd4 | 2659 | 0 | 0 | 0 | 0 | 0 | 0 |
| ENSMUSG00 Wnt4 | 4189 | 2.7332486 | 3.62347321 | 2.49024468 | 2.99750573 | 3.24015659 | 1.29569186 |
| ENSMUSG00 Ptcra | 596 | 0 | 0.32339894 | 0 | 0 | 0 | 0 |
| ENSMUSG00 Mrpl55 | 1193 | 34.3903235 | 31.2626205 | 39.348162 | 38.0479564 | 18.5200509 | 45.5644225 |
| ENSMUSG00 Dchs1 | 14129 | 0.01406875 | 0 | 0 | 0 | 0.28530566 | 0.07528565 |
| ENSMUSG00 Syde2 | 5491 | 3.17117109 | 2.6677615 | 2.73695622 | 3.03585951 | 4.32111906 | 1.71366792 |
| ENSMUSG00 BC053749 | 4009 | 1.78498042 | 1.89909151 | 1.25692445 | 2.02505862 | 1.60372246 | 1.26542352 |
| ENSMUSG00 Smad6 | 3246 | 5.47464565 | 7.12553673 | 10.0495872 | 7.66993553 | 6.79094096 | 10.6964312 |
| ENSMUSG00 Abcc12 | 6617 | 0 | 0 | 0 | 0 | 0 | 0 |
| ENSMUSG00 2410004B18R | 2220 | 13.1981032 | 16.908666 | 12.5039448 | 17.0008192 | 11.492427 | 19.3748226 |
| ENSMUSG00 Dna2 | 5899 | 0.28979245 | 0.55546331 | 0.44958647 | 0.3425317 | 0.32005016 | 0.33752308 |
| ENSMUSG00 Phkb | 7492 | 17.6490561 | 15.3074923 | 20.1244643 | 18.2384814 | 23.2928731 | 17.6891981 |
| ENSMUSG00 Acaa2 | 1505 | 158.308701 | 154.740914 | 169.141803 | 121.072641 | 174.066056 | 136.626914 |
| ENSMUSG00 Arhgap33 | 4393 | 0.81447604 | 0.5265079 | 0.422599 | 1.54414395 | 1.32415362 | 0.79470836 |
| ENSMUSG00 Arhgef26 | 8933 | 0.44059024 | 0.65809313 | 1.00942304 | 0.09290127 | 0.66831814 | 0.97093197 |
| ENSMUSG00 C1qa | 1037 | 0 | 0 | 0 | 0 | 0 | 0 |
| ENSMUSG00 Gtdc1 | 9053 | 0.14052528 | 0.21290817 | 0.3564271 | 0.37066499 | 0.44527601 | 0.17775364 |
| ENSMUSG00 Prodh2 | 2647 | 0 | 1.51094624 | 0.10019307 | 0.47709541 | 0 | 0 |
| ENSMUSG00 Ehmt1 | 7631 | 5.36981633 | 6.13144219 | 7.26946835 | 5.81587142 | 7.18822618 | 7.43788813 |
| ENSMUSG00 Rap2b | 4196 | 0.37898456 | 1.02206705 | 0.03160284 | 0.28377204 | 0.68100025 | 0.31200731 |
| ENSMUSG00 C1qc | 1928 | 0 | 0 | 0 | 0 | 0 | 0 |
| ENSMUSG00 Zfp157 | 5303 | 19.7315528 | 12.1942684 | 21.8633668 | 18.8813263 | 18.2628816 | 19.6266309 |
| ENSMUSG00 Trpv5 | 3228 | 42.6496992 | 59.9494274 | 17.650612 | 49.5513331 | 60.574062 | 29.5644246 |
| ENSMUSG00 Neto2 | 5086 | 0 | 0 | 0 | 0 | 0 | 0 |
| ENSMUSG00 Fzd8 | 3345 | 3.30404292 | 7.17394565 | 3.46214735 | 4.00192482 | 5.44586433 | 3.70184939 |
| ENSMUSG00 C1qb | 1085 | 0 | 0 | 0 | 0 | 0 | 0 |
| ENSMUSG00 C1ql2 | 1995 | 0 | 0 | 0 | 0 | 0 | 0 |
| ENSMUSG00 Unc93b1 | 2957 | 10.1102879 | 15.9535093 | 4.85816672 | 7.33355048 | 13.563318 | 3.99388529 |
| ENSMUSG00 Piwil4 | 5362 | 0.31140063 | 0.20669306 | 0 | 0.08075057 | 0.31403775 | 0.15768628 |
| ENSMUSG00 Trim67 | 10620 | 0 | 0 | 0.19145808 | 0 | 0.00480474 | 0.10529757 |
| ENSMUSG00 Kirrel2 | 4225 | 0 | 0 | 0 | 0 | 0 | 0 |
| ENSMUSG00 Zfp280c | 6850 | 3.09918636 | 3.77050117 | 3.74909063 | 3.96639117 | 3.32975036 | 4.43163117 |
| ENSMUSG00 Ttc7 | 6809 | 1.64650397 | 2.19383126 | 2.42139631 | 2.30513869 | 3.18493314 | 2.45547861 |
| ENSMUSG00 4930549C01R | 681 | 0 | 0 | 0 | 0 | 0 | 0 |
| ENSMUSG00 Stox1 | 3675 | 0.01081782 | 0.07867174 | 0.21649882 | 0.27491082 | 0 | 0.26718014 |
| ENSMUSG00 Cst13 | 779 | 0 | 0 | 0 | 0 | 0 | 0 |
| ENSMUSG00 Stag3 | 5812 | 0 | 0.26530732 | 0 | 0 | 0 | 0 |
| ENSMUSG00 Nfkbid | 2334 | 3.16817454 | 2.64261551 | 2.42409422 | 4.52957971 | 2.38297985 | 3.6459672 |
| ENSMUSG00 Aifm1 | 4233 | 61.619586 | 48.0043058 | 66.1931214 | 54.9115358 | 48.6395999 | 64.7812953 |
| ENSMUSG00 4921524J17R | 1591 | 14.31797 | 20.5345115 | 19.0309632 | 14.3557097 | 13.5022654 | 15.9087776 |
| ENSMUSG00 Try5 | 1157 | 0 | 0 | 0 | 0 | 0 | 0 |
| ENSMUSG00 Kdm1a | 5186 | 24.0633346 | 15.2661636 | 21.5724768 | 17.8322947 | 23.5748547 | 18.3864518 |
| ENSMUSG00 Elac1 | 4967 | 8.50016515 | 5.28721783 | 9.74451744 | 10.1991525 | 8.96839664 | 5.94693664 |
| ENSMUSG00 Rab8b | 6281 | 5.99402007 | 4.14276051 | 6.28438876 | 7.31291875 | 6.79160169 | 5.66683716 |
| ENSMUSG00 Tmem71 | 2921 | 0 | 0 | 0 | 1.03762127 | 0.68128335 | 0 |
| ENSMUSG00 BC037034 | 3693 | 0 | 0 | 0 | 0 | 0 | 0 |
| ENSMUSG00 Slc39a12 | 3082 | 0 | 0 | 0 | 0 | 0 | 0 |
| ENSMUSG00 C130079G13 | 2783 | 0 | 0 | 0 | 0 | 0 | 0 |
| ENSMUSG00 2510003E04R | 6489 | 17.7487482 | 18.9879384 | 16.8387973 | 14.2960297 | 18.9982623 | 16.9599583 |
| ENSMUSG00 Lrfn3 | 2858 | 2.07262652 | 1.29823515 | 1.70126058 | 1.37611716 | 1.51757941 | 3.989083 |
| ENSMUSG00 Cst11 | 612 | 0 | 0 | 0 | 0 | 0 | 0 |
| ENSMUSG00 Bcorl1 | 7738 | 0.57542179 | 0.44213458 | 0.38272467 | 0.78337858 | 0.90341348 | 0.60625975 |
| ENSMUSG00 Clca5 | 3970 | 0 | 0 | 0 | 0 | 0 | 0 |
| ENSMUSG00 Wnt8b | 3370 | 0.51906265 | 0 | 0 | 0 | 0 | 0 |
| ENSMUSG00 Gm101 | 2999 | 0 | 0 | 0 | 0 | 0 | 0 |
| ENSMUSG00 Trim17 | 3728 | 0.25593657 | 0.42654308 | 0.28456122 | 0.483933 | 0.17793532 | 0 |
| ENSMUSG00 Spryd3 | 2352 | 22.7343203 | 23.7039599 | 27.5509787 | 14.313158 | 19.6555682 | 23.6797676 |
| ENSMUSG00 Cnpy4 | 2393 | 9.55261233 | 10.3098447 | 8.88469975 | 12.6656571 | 11.6424577 | 10.7708041 |
| ENSMUSG00 Zic4 | 10231 | 0 | 0 | 0 | 0 | 0 | 0 |
| ENSMUSG00 Tmem177 | 3339 | 7.16765477 | 5.03655834 | 4.8054115 | 6.7755142 | 8.3592176 | 4.88476618 |
| ENSMUSG00 Anapc10 | 2797 | 11.5983097 | 17.4863206 | 14.0175312 | 10.2041106 | 6.43986795 | 20.3901576 |
| ENSMUSG00 Taf6 | 4290 | 16.0041293 | 15.0737075 | 12.8278076 | 10.7909704 | 17.5202451 | 10.1278556 |
| ENSMUSG00 Tfb1m | 1320 | 15.1191297 | 11.0974837 | 11.5527544 | 10.9886227 | 13.5297209 | 9.07088381 |
| ENSMUSG00 Zdhhc9 | 6366 | 9.96897249 | 7.39524882 | 9.97075818 | 4.60235943 | 6.14785269 | 6.28953353 |
| ENSMUSG00 Pml | 8699 | 4.80319692 | 4.89674846 | 2.92680326 | 4.24323848 | 6.14146632 | 4.14496759 |
| ENSMUSG00 Trim3 | 4598 | 10.3149822 | 10.1235544 | 6.69084009 | 6.64669241 | 14.2270148 | 8.90370455 |
| ENSMUSG00 Otud4 | 8386 | 10.2967927 | 11.1588446 | 11.1427017 | 14.4353994 | 12.1755042 | 13.3283415 |
| ENSMUSG00 Nxt1 | 1322 | 9.4126062 | 11.5180906 | 11.836197 | 8.87039673 | 9.37928088 | 11.2647149 |
| ENSMUSG00 Asap3 | 4200 | 1.1074741 | 0.91783699 | 0.28415471 | 1.22850771 | 1.65228265 | 0.81823918 |
| ENSMUSG00 1810009J06R | 865 | 0 | 0 | 0 | 0 | 0 | 0 |
| ENSMUSG00 Zfp39 | 5535 | 3.13159702 | 4.56182397 | 2.76311431 | 2.49673407 | 3.39253953 | 2.20266734 |
| ENSMUSG00 Tenc1 | 4826 | 10.2642621 | 14.4479448 | 10.0566978 | 15.207393 | 19.3807184 | 10.2124478 |
| ENSMUSG00 Xpnpep2 | 5129 | 0 | 0 | 0.04309012 | 0 | 0 | 0 |
| ENSMUSG00 Zfp113 | 7788 | 9.10169771 | 5.71703551 | 5.90833572 | 6.6159681 | 5.1694672 | 5.24620051 |
| ENSMUSG00 Apln | 3146 | 0 | 0 | 0 | 0 | 0 | 0 |
| ENSMUSG00 Hk1 | 9211 | 46.9331338 | 44.0902545 | 36.4469653 | 44.2221475 | 54.5385593 | 30.1498618 |
| ENSMUSG00 Ss18 | 6192 | 18.3689325 | 21.5951836 | 19.2597824 | 23.0057997 | 25.0599498 | 17.3417724 |
| ENSMUSG00 Sstr4 | 3559 | 0 | 0 | 0 | 0 | 0 | 0 |
| ENSMUSG00 Tmem185b | 1775 | 14.2447806 | 18.9759567 | 18.6021277 | 19.9823379 | 20.0943305 | 19.3918844 |
| ENSMUSG00 Frem2 | 12353 | 1.58983303 | 1.88408092 | 0.61903334 | 1.80220363 | 2.1190424 | 0.88096665 |
| ENSMUSG00 Zscan21 | 3264 | 30.0358502 | 31.2385146 | 41.2090099 | 27.2605153 | 32.4230096 | 22.8458529 |
| ENSMUSG00 Wdr62 | 6053 | 0.28241957 | 1.066741 | 0.51847528 | 0.47688147 | 0.42149658 | 0.68040154 |
| ENSMUSG00 Mmaa | 2836 | 13.5835898 | 11.5538719 | 17.1289935 | 10.8526037 | 9.69789019 | 11.9158244 |
| ENSMUSG00 Foxa2 | 3648 | 0 | 0 | 0 | 0 | 0 | 0 |
| ENSMUSG00 Gm12800 | 1452 | 0 | 0 | 0 | 0 | 0 | 0 |
| ENSMUSG00 Zfp146 | 2572 | 10.7117216 | 13.095771 | 11.1192039 | 9.27302933 | 13.2327343 | 6.25662409 |
| ENSMUSG00 Tspan15 | 3605 | 51.6324992 | 56.9014381 | 30.1259163 | 49.8042312 | 46.5111431 | 29.6351628 |
| ENSMUSG00 Apbb1 | 2517 | 23.7869502 | 19.7953044 | 16.2442212 | 20.513868 | 30.1860443 | 8.23548901 |
| ENSMUSG00 Clca4 | 3733 | 0 | 0 | 0 | 0 | 0 | 0 |
| ENSMUSG00 Pax1 | 4485 | 0 | 0 | 0 | 0 | 0 | 0 |
| ENSMUSG00 Inhbb | 4253 | 0 | 0 | 0 | 0 | 0 | 0.25652141 |
| ENSMUSG00 Smpd1 | 2405 | 90.6524138 | 75.9161868 | 132.752562 | 86.3118512 | 67.2147853 | 80.9053124 |
| ENSMUSG00 Azgp1 | 1311 | 0 | 0.55133229 | 0 | 0.52292818 | 0 | 0.2704579 |
| ENSMUSG00 Paip2 | 2915 | 109.665118 | 108.704646 | 124.265559 | 101.648628 | 99.3045043 | 166.595181 |
| ENSMUSG00 Prkcdbp | 1033 | 2.30912762 | 2.89211947 | 1.71159121 | 2.68956189 | 2.81558899 | 1.58419976 |
| ENSMUSG00 Sh3glb1 | 4132 | 36.0127695 | 33.2360504 | 30.8407341 | 40.5268075 | 27.1556148 | 46.6282797 |
| ENSMUSG00 Gm5229 | 352 | 0 | 0 | 0 | 0 | 0 | 0 |
| ENSMUSG00 Rbmxl1 | 1721 | 3.41883272 | 4.36785879 | 3.44163101 | 2.4110605 | 4.50668746 | 4.40578136 |
| ENSMUSG00 Scd1 | 4830 | 0.70786156 | 0.52875392 | 0.26539409 | 0.62751382 | 0 | 0.17505462 |
| ENSMUSG00 15-Sep | 2098 | 313.628674 | 303.496495 | 338.951025 | 312.923165 | 228.743119 | 439.085291 |
| ENSMUSG00 Rnf139 | 4346 | 61.12428 | 50.0713237 | 52.6943738 | 71.5663193 | 56.4154937 | 68.9647977 |
| ENSMUSG00 Trmt12 | 4132 | 7.12943152 | 4.71135591 | 4.35386014 | 6.97714269 | 5.40889463 | 5.83513579 |
| ENSMUSG00 1110059M19 | 1108 | 0 | 0 | 0 | 0 | 0 | 0 |
| ENSMUSG00 Slc35b2 | 2310 | 28.104199 | 35.3158643 | 38.040374 | 27.6316609 | 29.2462864 | 29.2700566 |
| ENSMUSG00 Lrg1 | 1349 | 0.088411 | 0 | 0.58979479 | 0.4012088 | 0 | 0.18196572 |
| ENSMUSG00 Gm9762 | 443 | 0.80767342 | 1.41404909 | 1.99556853 | 1.30318897 | 2.07330647 | 0.43097624 |
| ENSMUSG00 Rab11fip3 | 8619 | 28.9206245 | 23.2238636 | 35.7040528 | 26.8009339 | 30.9686706 | 26.2493483 |
| ENSMUSG00 Ttc29 | 3667 | 0 | 0 | 0 | 0 | 0 | 0 |
| ENSMUSG00 Dcaf15 | 2299 | 2.28261132 | 4.35962591 | 1.71116313 | 2.05600167 | 2.41925836 | 3.22692557 |
| ENSMUSG00 Socs5 | 4745 | 11.051102 | 8.00230061 | 10.7966146 | 8.23537497 | 9.22668713 | 9.51882036 |
| ENSMUSG00 Fer1l6 | 8295 | 0.1198176 | 0.34854569 | 0 | 0.23489215 | 0.47366256 | 0 |
| ENSMUSG00 Zcwpw1 | 2524 | 0.85055307 | 0.68728681 | 1.57613545 | 1.91560775 | 1.17255539 | 0.97255053 |
| ENSMUSG00 Ralgapa2 | 19373 | 16.5646125 | 11.7897969 | 20.1170521 | 20.7593284 | 17.8498812 | 16.7663077 |
| ENSMUSG00 Setd7 | 7781 | 2.12036042 | 1.09613164 | 1.97689775 | 1.63229401 | 1.5607605 | 2.12420401 |
| ENSMUSG00 Sik2 | 8847 | 3.37474464 | 3.51853076 | 3.50237278 | 4.02542986 | 4.15847372 | 4.83094379 |
| ENSMUSG00 D15Ertd621e | 5597 | 52.4698472 | 42.0393777 | 46.3341456 | 46.7835264 | 47.1244125 | 40.5830259 |
| ENSMUSG00 Trim58 | 2186 | 0 | 0 | 0 | 0 | 0 | 0 |
| ENSMUSG00 Psd | 4079 | 0.48731896 | 1.01594362 | 0.49847629 | 0.47767355 | 1.16338636 | 0.23403098 |
| ENSMUSG00 Tmprss13 | 3232 | 0 | 0.1043642 | 1.8189496 | 0.16745998 | 0 | 0.6666759 |
| ENSMUSG00 H2-M10.6 | 1401 | 0 | 0 | 0 | 0 | 0 | 0 |
| ENSMUSG00 Prmt10 | 4852 | 6.60406375 | 6.84263368 | 4.71899313 | 5.89716845 | 6.86731727 | 6.07664572 |
| ENSMUSG00 Aff3 | 8919 | 0 | 0 | 0.00991184 | 0 | 0 | 0 |
| ENSMUSG00 Myom3 | 6001 | 0 | 0 | 0 | 0 | 0 | 0 |
| ENSMUSG00 Tas2r108 | 893 | 0 | 0 | 0 | 0 | 0 | 0 |
| ENSMUSG00 4930529M08 | 10511 | 0 | 0.03209068 | 0 | 0 | 0 | 0.01297433 |
| ENSMUSG00 2210407C18R | 1517 | 0 | 0 | 0 | 0 | 0 | 0 |
| ENSMUSG00 Arhgap10 | 3015 | 0.13185897 | 0.20776907 | 0.01466064 | 0.05983755 | 0.15231754 | 0.09046315 |
| ENSMUSG00 Ddx1 | 2481 | 81.2575742 | 77.2031872 | 74.382384 | 77.1815439 | 72.5804435 | 71.5121033 |
| ENSMUSG00 Lrrc20 | 2614 | 3.6348737 | 2.65449414 | 4.53178804 | 2.40178873 | 3.37703252 | 2.34766405 |
| ENSMUSG00 Ndufc1 | 1631 | 48.3354493 | 35.8370045 | 42.7925874 | 30.8832214 | 27.0930972 | 102.493111 |
| ENSMUSG00 Il22ra1 | 2562 | 0 | 0 | 0 | 0 | 0 | 0 |
| ENSMUSG00 Wee2 | 3262 | 0 | 0 | 0 | 0 | 0 | 0 |
| ENSMUSG00 4930583H14R | 2610 | 0 | 0 | 0 | 0 | 0.01955034 | 0 |
| ENSMUSG00 Ppp1r14a | 527 | 0 | 0 | 0.08387447 | 0.20540063 | 0.09682424 | 0.05175453 |
| ENSMUSG00 Spaca5 | 636 | 0 | 0 | 0 | 0 | 0 | 0 |
| ENSMUSG00 Mycn | 3288 | 0.30227707 | 0.07327622 | 0.86037651 | 0.65843147 | 1.17944178 | 0.30692264 |
| ENSMUSG00 Nodal | 2092 | 0 | 0 | 0 | 0 | 0 | 0 |
| ENSMUSG00 E330009J07R | 7502 | 1.60569323 | 1.58009394 | 0.99574933 | 1.58237705 | 2.14933814 | 1.09796602 |
| ENSMUSG00 Elf2 | 5658 | 9.68943224 | 10.0665208 | 9.23410718 | 7.66536188 | 12.5266235 | 10.3111441 |
| ENSMUSG00 Krt80 | 4804 | 29.8248859 | 17.5332849 | 38.1383512 | 22.5700549 | 20.7015834 | 24.3904767 |
| ENSMUSG00 Grhl3 | 2773 | 0 | 0.31278614 | 0 | 0.02602383 | 0.01840115 | 0 |
| ENSMUSG00 Cyb561d2 | 1779 | 8.64832544 | 12.6221934 | 20.0262425 | 11.6014219 | 9.55131149 | 18.4897217 |
| ENSMUSG00 Pacrg | 1566 | 1.11701222 | 4.06169244 | 1.12903814 | 1.15204484 | 1.23818792 | 2.24676147 |
| ENSMUSG00 Rbm17 | 2547 | 22.4453793 | 23.1378166 | 18.1527789 | 22.4397167 | 21.8369649 | 20.8388096 |
| ENSMUSG00 Prf1 | 2051 | 0 | 0 | 0 | 0 | 0 | 0 |
| ENSMUSG00 9430023L20R | 1227 | 26.698057 | 27.6473148 | 23.9921984 | 26.2602003 | 24.951773 | 20.6060233 |
| ENSMUSG00 Islr | 4139 | 0 | 0.17463074 | 0 | 0 | 0.13561008 | 0 |
| ENSMUSG00 Fam193a | 5034 | 4.79371805 | 4.87224852 | 6.17280406 | 5.64812298 | 5.76758201 | 3.87934877 |
| ENSMUSG00 Spry1 | 3002 | 13.0973252 | 16.5650927 | 11.5878916 | 10.5169184 | 12.0511994 | 9.35805411 |
| ENSMUSG00 Thap1 | 1612 | 2.39223425 | 3.07891039 | 4.4421207 | 1.81305559 | 2.43736411 | 4.1791786 |
| ENSMUSG00 Lipt1 | 3206 | 2.82727684 | 3.36673832 | 4.59114589 | 4.28797847 | 5.29999474 | 2.96907333 |
| ENSMUSG00 Syn1 | 4342 | 0.05493618 | 0 | 0.05090033 | 0.08310006 | 0 | 0.05025268 |
| ENSMUSG00 Mospd3 | 1645 | 8.24110569 | 10.3988979 | 13.0590247 | 9.65112437 | 8.71635964 | 12.6839508 |
| ENSMUSG00 Zfyve28 | 7027 | 0 | 0 | 0 | 0 | 0 | 0 |
| ENSMUSG00 Fgf2 | 1795 | 0.0664437 | 0.10737926 | 0.14774989 | 0.36182552 | 0 | 0 |
| ENSMUSG00 Hook3 | 23710 | 6.84277166 | 6.07055684 | 8.93171784 | 9.40172379 | 6.12058257 | 10.8224296 |
| ENSMUSG00 Mxd4 | 4446 | 18.7600086 | 20.2565138 | 24.1489601 | 17.4972758 | 18.7877142 | 21.5755525 |
| ENSMUSG00 Matr3 | 4457 | 73.3117106 | 72.0147837 | 88.9590603 | 77.0295386 | 68.5198248 | 94.9196611 |
| ENSMUSG00 Spred3 | 5991 | 0.11944561 | 0.04825883 | 0.02951216 | 0.05420437 | 0.10220606 | 0.16844628 |
| ENSMUSG00 Clic4 | 6567 | 10.6971119 | 4.21915703 | 4.36835635 | 4.01644199 | 8.32179816 | 2.42551992 |
| ENSMUSG00 Zfp692 | 2029 | 35.3665068 | 36.4307552 | 32.8954081 | 39.5141956 | 46.3487485 | 35.8240087 |
| ENSMUSG00 H2-M10.5 | 1377 | 0 | 0 | 0 | 0 | 0 | 0 |
| ENSMUSG00 Pldi | 850 | 0 | 0 | 0 | 0 | 0 | 0 |
| ENSMUSG00 4930444A02R | 3596 | 13.2776229 | 15.2224133 | 18.5239648 | 11.7397086 | 13.0687687 | 14.456469 |
| ENSMUSG00 Mex3c | 3736 | 16.9514134 | 14.3295335 | 14.5052087 | 19.113053 | 12.7429359 | 14.7907974 |
| ENSMUSG00 Itih2 | 3621 | 0 | 0 | 0.04882833 | 0 | 0.11273433 | 0 |
| ENSMUSG00 Aagab | 2590 | 32.0192789 | 25.525791 | 33.8767021 | 25.8564767 | 33.2163975 | 23.5572849 |
| ENSMUSG00 6330439K17R | 8779 | 2.12838317 | 1.65762678 | 1.78237298 | 6.28014171 | 3.55714113 | 3.33670834 |
| ENSMUSG00 Hgsnat | 2661 | 49.4366347 | 50.9027016 | 58.371017 | 45.4652744 | 63.1837311 | 33.2400057 |
| ENSMUSG00 Kin | 1820 | 13.3246391 | 16.1504009 | 15.9563796 | 15.1267033 | 20.3544774 | 12.5433368 |
| ENSMUSG00 1700055N04 | 891 | 0 | 0 | 0 | 0 | 0 | 0 |
| ENSMUSG00 D4Wsu53e | 3263 | 443.828576 | 316.97021 | 473.852427 | 325.335529 | 519.443189 | 742.709407 |
| ENSMUSG00 4932438A13R | 25325 | 10.1472626 | 9.15209424 | 7.79313836 | 12.6760889 | 12.8004172 | 11.102636 |
| ENSMUSG00 Gemin6 | 7084 | 2.46367249 | 4.1629168 | 2.73921642 | 1.58915837 | 4.27860915 | 1.62092365 |
| ENSMUSG00 Tmem97 | 1372 | 2.98455867 | 3.40676741 | 3.54387954 | 1.78832291 | 4.53733079 | 1.82891167 |
| ENSMUSG00 Ovol2 | 2109 | 3.58157482 | 5.07225707 | 2.99709035 | 3.95208738 | 3.82274413 | 2.23732223 |
| ENSMUSG00 Galnt6 | 5509 | 0 | 0 | 0 | 0 | 0 | 0 |
| ENSMUSG00 Stag1 | 10229 | 8.53096879 | 7.58433589 | 6.04107699 | 8.28592459 | 6.82912391 | 5.52745402 |
| ENSMUSG00 Tbcel | 8275 | 3.44467426 | 3.86656164 | 2.62273171 | 3.02173497 | 3.72446296 | 3.37183758 |
| ENSMUSG00 Ldlrap1 | 4443 | 5.87876451 | 4.19719854 | 4.39730241 | 4.84829633 | 4.5708975 | 4.38923409 |
| ENSMUSG00 Lsm1 | 2356 | 8.16708513 | 11.4125784 | 11.4632114 | 10.8429913 | 10.0709952 | 15.1538637 |
| ENSMUSG00 Ttc13 | 7673 | 4.50765907 | 5.23752023 | 5.92775922 | 6.07558993 | 5.96515823 | 4.58546652 |
| ENSMUSG00 Man1c1 | 5727 | 1.05514808 | 0.47117876 | 0.0463089 | 0.2394129 | 1.10481414 | 0.30479779 |
| ENSMUSG00 Banf2 | 633 | 0 | 0 | 0 | 0 | 0 | 0 |
| ENSMUSG00 Tacc3 | 3202 | 1.09259284 | 1.42964148 | 1.04913806 | 1.46491749 | 2.32662426 | 2.18060826 |
| ENSMUSG00 Phf16 | 5113 | 3.90323709 | 5.30588049 | 3.20729195 | 3.23207049 | 8.08358389 | 2.42713883 |
| ENSMUSG00 Bag4 | 4793 | 49.9328172 | 29.7583703 | 80.0391812 | 49.5648198 | 40.8381343 | 71.598062 |
| ENSMUSG00 Traf3ip3 | 4710 | 0 | 0 | 0 | 0 | 0 | 0 |
| ENSMUSG00 Tap1 | 4503 | 2.69274296 | 3.25309314 | 1.5214936 | 3.32534943 | 13.6092999 | 8.02551566 |
| ENSMUSG00 Bbs7 | 3938 | 2.32193005 | 2.93670546 | 2.96325205 | 1.54846763 | 1.63263871 | 2.18862011 |
| ENSMUSG00 Solh | 5098 | 8.08678566 | 8.45956566 | 4.68203123 | 11.324298 | 7.9772502 | 5.97067427 |
| ENSMUSG00 Larp1 | 6697 | 14.941697 | 14.577532 | 12.349059 | 15.2636154 | 13.8747246 | 13.8429892 |
| ENSMUSG00 H2-M1 | 1052 | 0 | 0 | 0 | 0 | 0 | 0 |
| ENSMUSG00 Hand1 | 2802 | 0 | 0 | 0 | 0 | 0 | 0 |
| ENSMUSG00 Mfsd2b | 5675 | 0.81962841 | 0.74720827 | 0.48291 | 1.13173637 | 0.89914319 | 0.61998739 |
| ENSMUSG00 Map4k1 | 2877 | 0.31782275 | 0.46896781 | 0 | 0.73995156 | 0.40792723 | 0.2370059 |
| ENSMUSG00 Fam53a | 9547 | 3.94764801 | 4.83025297 | 4.1437781 | 3.42792651 | 4.46287041 | 3.99106222 |
| ENSMUSG00 Slc9a7 | 3801 | 5.2505265 | 4.19618846 | 5.87264686 | 6.21776879 | 5.30266205 | 5.56112745 |
| ENSMUSG00 Taf2 | 5007 | 13.196247 | 8.67105739 | 15.952213 | 21.5324843 | 12.5247485 | 9.44017372 |
| ENSMUSG00 Slc12a9 | 5005 | 4.67851709 | 5.17968149 | 3.02921721 | 2.02578512 | 6.99382494 | 4.71379879 |
| ENSMUSG00 Hrh4 | 1873 | 0 | 0 | 0 | 0 | 0 | 0 |
| ENSMUSG00 Chst7 | 2204 | 0.05411363 | 1.39924333 | 0 | 0 | 1.94474391 | 0.49500253 |
| ENSMUSG00 Paqr7 | 4315 | 0.39617281 | 0.99388027 | 0.36877552 | 0.20905008 | 0.42571252 | 0.47406673 |
| ENSMUSG00 Nudt22 | 993 | 52.7270572 | 46.4880278 | 47.5403508 | 32.5574138 | 37.5118372 | 48.2318898 |
| ENSMUSG00 Actr1b | 3014 | 42.1693003 | 42.3190154 | 45.1697666 | 35.6390997 | 40.6822764 | 33.6906045 |
| ENSMUSG00 Letmd1 | 2570 | 19.6921116 | 16.8183808 | 21.292561 | 22.5758473 | 20.9664797 | 20.3657715 |
| ENSMUSG00 4933407H18R | 9056 | 4.83334629 | 4.41638107 | 2.46975846 | 3.37870735 | 5.00346971 | 3.87916689 |
| ENSMUSG00 4930578C19R | 3576 | 0 | 0 | 0.03708208 | 0 | 0 | 0 |
| ENSMUSG00 0610009D07R | 1111 | 49.5601625 | 61.5016876 | 59.559099 | 55.3084805 | 39.4065081 | 77.3313354 |
| ENSMUSG00 Nov | 2864 | 0 | 0 | 0.20063686 | 0 | 0.10689883 | 0 |
| ENSMUSG00 Letm2 | 2087 | 3.82886995 | 5.65676968 | 4.76541192 | 8.5753206 | 3.71634362 | 4.49567604 |
| ENSMUSG00 Srrt | 3004 | 17.9720182 | 16.6182271 | 14.1698984 | 20.0829489 | 19.1943424 | 16.7969649 |
| ENSMUSG00 Pafah2 | 3658 | 31.8869819 | 19.4958815 | 20.9892295 | 26.3365388 | 23.9787698 | 18.9237382 |
| ENSMUSG00 Kdm6a | 8653 | 6.26678324 | 5.65785568 | 7.25374056 | 10.687424 | 8.3972679 | 11.0226989 |
| ENSMUSG00 Enpp1 | 10212 | 6.1626445 | 3.66635973 | 7.72623284 | 2.13057901 | 4.56699055 | 8.47725274 |
| ENSMUSG00 Ctbp1 | 2245 | 51.3545183 | 49.9679454 | 52.8846995 | 51.7524202 | 46.8669875 | 46.6280915 |
| ENSMUSG00 Hhat | 8904 | 0.02232451 | 0.21105921 | 0.43685559 | 0.33634431 | 0.2521519 | 0.00612638 |
| ENSMUSG00 Trmt6 | 6553 | 8.31146167 | 7.81660125 | 7.67613266 | 5.72643466 | 8.05925515 | 7.58345685 |
| ENSMUSG00 Spon2 | 2231 | 0 | 0 | 0 | 0 | 0 | 0 |
| ENSMUSG00 Gm6755 | 836 | 0 | 0 | 0 | 0 | 0 | 0 |
| ENSMUSG00 Rims2 | 5569 | 4.72582656 | 2.31890223 | 6.64337271 | 2.76657084 | 1.37438613 | 4.78004092 |
| ENSMUSG00 Muc3 | 2487 | 0 | 0.42625723 | 0 | 0.10155783 | 0 | 0 |
| ENSMUSG00 Nmur2 | 3298 | 0 | 0 | 0 | 0 | 0 | 0 |
| ENSMUSG00 Rcor3 | 3555 | 11.7309421 | 9.63728843 | 7.34832327 | 9.06364719 | 7.24847251 | 10.2960804 |
| ENSMUSG00 Atp11b | 6582 | 17.1053663 | 18.009518 | 21.78529 | 30.2712017 | 28.1102459 | 22.5050996 |
| ENSMUSG00 Icam1 | 2533 | 4.31612994 | 8.44641939 | 0.85506921 | 4.37314949 | 7.15134758 | 1.6797646 |
| ENSMUSG00 Htra4 | 1786 | 0.40067114 | 0.80940272 | 0.2722398 | 0.06060814 | 0.51426359 | 0.82465315 |
| ENSMUSG00 Cnnm4 | 6040 | 0.88199245 | 1.42804191 | 1.57340998 | 1.19476969 | 0.5153326 | 1.46307668 |
| ENSMUSG00 Tbc1d2b | 8767 | 0.86158792 | 2.09410682 | 0.38822196 | 1.39932647 | 1.85084835 | 1.55241759 |
| ENSMUSG00 Serpine1 | 4412 | 0 | 0 | 0 | 0 | 0 | 0 |
| ENSMUSG00 Ranbp10 | 5294 | 5.82739949 | 11.2410759 | 7.89655374 | 10.4688365 | 13.0023765 | 9.98455815 |
| ENSMUSG00 Dmxl1 | 11800 | 26.1543894 | 22.1657634 | 38.8976219 | 39.6382664 | 21.7856679 | 40.7755857 |
| ENSMUSG00 Best1 | 2140 | 0.6130518 | 0.27020435 | 1.07406348 | 0.01686077 | 0.71532303 | 0.44608055 |
| ENSMUSG00 Endod1 | 5891 | 15.5890613 | 20.9317782 | 13.3183282 | 14.1179957 | 18.1030598 | 10.0236961 |
| ENSMUSG00 Taar9 | 1193 | 0 | 0 | 0 | 0 | 0 | 0 |
| ENSMUSG00 Depdc5 | 11763 | 0.88886265 | 1.6262873 | 1.20622219 | 1.35273159 | 2.04747508 | 0.56112069 |
| ENSMUSG00 Vgf | 2551 | 0 | 0 | 0 | 0 | 0 | 0 |
| ENSMUSG00 Fer1l5 | 1711 | 4.87939854 | 3.21055196 | 4.9601133 | 7.29654436 | 7.06794335 | 4.33588654 |
| ENSMUSG00 Slc30a1 | 5595 | 3.34670801 | 3.88419757 | 4.02912242 | 2.9729799 | 2.61743876 | 5.00644406 |
| ENSMUSG00 Adam32 | 3155 | 0 | 0.27491473 | 0.15411102 | 0.40027625 | 0.25877084 | 0.17289787 |
| ENSMUSG00 Gm9763 | 269 | 0.14778989 | 0 | 0.16431912 | 0 | 0 | 0.30417813 |
| ENSMUSG00 Vnn1 | 2409 | 3.11904766 | 3.68049205 | 1.54128469 | 1.31806555 | 1.14380419 | 2.37761488 |
| ENSMUSG00 Ccdc21 | 6825 | 5.78420398 | 4.43385871 | 3.56852974 | 5.15986454 | 6.04092479 | 3.00520567 |
| ENSMUSG00 Tulp1 | 6380 | 0 | 0 | 0 | 0 | 0 | 0.01710009 |
| ENSMUSG00 Arid5a | 6906 | 0.43174935 | 0.09070718 | 0.0192015 | 0.05224739 | 0.31771419 | 0 |
| ENSMUSG00 Slc22a20 | 1887 | 0 | 0 | 0 | 0 | 0 | 0 |

| | | | | | | | |
|---|---|---|---|---|---|---|---|
| ENSMUSG00 1110021L09f | 3367 | 6.23430167 | 7.49916712 | 4.88360131 | 8.66954975 | 5.22842283 | 4.58492599 |
| ENSMUSG00 Azin1 | 8364 | 28.3336226 | 30.8741205 | 26.3868726 | 37.1432811 | 24.8116058 | 24.0136829 |
| ENSMUSG00 Ints7 | 4184 | 8.41858407 | 9.28256988 | 9.0220779 | 9.30509705 | 12.4151173 | 10.6517114 |
| ENSMUSG00 Fbxo27 | 3053 | 0.26043551 | 0.44193265 | 1.042428 | 0.49637926 | 0 | 0.96484148 |
| ENSMUSG00 Klf10 | 3809 | 21.6886029 | 20.1398834 | 19.2287881 | 14.1145304 | 17.4285416 | 21.8684033 |
| ENSMUSG00 4930427A07f | 3783 | 2.08077852 | 2.39467384 | 3.31835142 | 0.65811818 | 3.47999074 | 1.73755962 |
| ENSMUSG00 C330005M16 | 2217 | 0 | 0 | 0 | 0 | 0 | 0 |
| ENSMUSG00 Uggt1 | 9726 | 10.3946316 | 10.6024045 | 13.0751288 | 10.4692093 | 11.0436481 | 7.55478903 |
| ENSMUSG00 Dtl | 4373 | 0.00909112 | 0.68318304 | 0 | 0.94062499 | 0 | 0 |
| ENSMUSG00 Thoc2 | 12160 | 22.5422729 | 18.676209 | 21.4429829 | 25.4265658 | 17.8172691 | 26.7901555 |
| ENSMUSG00 Tbx10 | 1769 | 0.15731394 | 0 | 0.02498691 | 0 | 0 | 0 |
| ENSMUSG00 Erv3 | 3376 | 0 | 0 | 0 | 0 | 0 | 0 |
| ENSMUSG00 Asxl2 | 14883 | 5.04055578 | 4.04062889 | 5.3548292 | 6.16034907 | 5.57474213 | 9.8502443 |
| ENSMUSG00 Ubr5 | 9731 | 6.39372387 | 6.1501964 | 6.86806974 | 7.51601112 | 7.34117009 | 10.4070225 |
| ENSMUSG00 Slc2a12 | 4231 | 7.06597054 | 4.11139343 | 6.80108717 | 3.89730427 | 6.5486462 | 9.13452229 |
| ENSMUSG00 Zmat4 | 6026 | 0 | 0 | 0 | 0.29339864 | 0 | 0.24441263 |
| ENSMUSG00 Cib2 | 1427 | 2.06160165 | 2.19489751 | 1.54876815 | 1.82053764 | 1.14424949 | 1.68196794 |
| ENSMUSG00 Nenf | 738 | 18.2616638 | 21.2856099 | 22.1608156 | 17.2587557 | 10.7860632 | 26.0919991 |
| ENSMUSG00 Fam168b | 9394 | 13.5297287 | 11.931197 | 10.5963967 | 10.7547078 | 14.3779877 | 7.34853225 |
| ENSMUSG00 Arhgef4 | 8945 | 0.63110992 | 1.15281147 | 0.34096447 | 0.12101301 | 0.83285085 | 0.24698108 |
| ENSMUSG00 Samd4b | 4299 | 8.11940263 | 6.93821998 | 6.6626644 | 6.31163003 | 6.90796714 | 7.112089 |
| ENSMUSG00 Pank2 | 5238 | 9.11537459 | 8.83142124 | 12.3964314 | 9.03773242 | 10.3845202 | 8.32089987 |
| ENSMUSG00 Ppfia1 | 6538 | 10.0209593 | 8.76318136 | 9.30956528 | 9.59722781 | 12.9009788 | 6.34517074 |
| ENSMUSG00 Mavs | 3395 | 105.120751 | 88.3820251 | 105.837049 | 94.1746676 | 96.4466137 | 72.2074986 |
| ENSMUSG00 Bcdin3d | 1269 | 4.76188577 | 5.24013319 | 6.82707741 | 7.36426283 | 5.79022704 | 4.10516635 |
| ENSMUSG00 Atg14 | 3440 | 15.4168055 | 9.52522694 | 11.4230926 | 14.674064 | 16.0050754 | 10.6719955 |
| ENSMUSG00 Prss40 | 2092 | 0 | 0 | 0 | 0 | 0 | 0 |
| ENSMUSG00 Mrpl47 | 3554 | 9.40752929 | 9.13834046 | 9.91245607 | 10.6398376 | 6.17370332 | 11.4194325 |
| ENSMUSG00 Rapgef6 | 10270 | 8.77175451 | 7.67604862 | 7.03700227 | 10.6243527 | 9.01283795 | 7.23959753 |
| ENSMUSG00 1700021A07f | 442 | 0 | 0 | 0 | 0 | 0 | 0 |
| ENSMUSG00 Fbxo34 | 3801 | 6.93446029 | 6.87110014 | 6.09359793 | 4.51856175 | 9.42397154 | 6.87427109 |
| ENSMUSG00 H2-M11 | 1001 | 0 | 0 | 0 | 0 | 0 | 0 |
| ENSMUSG00 Shank2 | 10654 | 12.3960678 | 10.4161232 | 4.16544494 | 4.54496938 | 6.92069768 | 6.19017061 |
| ENSMUSG00 Aldh8a1 | 4560 | 0.42719705 | 0.39098648 | 0 | 0.17408004 | 0.53711975 | 1.32784428 |
| ENSMUSG00 Dlgap5 | 3710 | 0 | 0.40263604 | 0.33359882 | 0.20423799 | 0.19255236 | 0 |
| ENSMUSG00 H2-DMb2 | 1702 | 0.16350668 | 0 | 0 | 0 | 0 | 0.38460126 |
| ENSMUSG00 Plekhg2 | 8117 | 0.36733535 | 0.48085522 | 0.33218091 | 0.28449561 | 0.82351303 | 0.47714658 |
| ENSMUSG00 Zdhhc18 | 5477 | 10.9532628 | 9.23786092 | 11.3309088 | 9.32190853 | 7.7606301 | 9.75054659 |
| ENSMUSG00 Rps16 | 891 | 1.20471153 | 0.75713826 | 0.64492027 | 0.76942631 | 0.17180598 | 1.16322816 |
| ENSMUSG00 Vash2 | 4862 | 0.31071745 | 0.06937579 | 0.00909129 | 0 | 0 | 0.24121956 |
| ENSMUSG00 Mcrs1 | 1900 | 8.07663972 | 6.82218576 | 5.35074942 | 5.45028778 | 8.75505185 | 5.6415438 |
| ENSMUSG00 Wdhd1 | 8035 | 0.24738942 | 0 | 0.87468488 | 0.56132614 | 0.85096881 | 0.70265717 |
| ENSMUSG00 Tob1 | 2286 | 102.84948 | 106.996667 | 99.8505325 | 98.3653987 | 101.159902 | 122.676221 |
| ENSMUSG00 Ephx3 | 2955 | 0 | 0.71749694 | 0 | 0 | 0 | 0.02768999 |
| ENSMUSG00 Pkd2l1 | 4426 | 0 | 0 | 0 | 0 | 0 | 0 |
| ENSMUSG00 Kcnh3 | 3758 | 0 | 0.02564473 | 0 | 0.29764326 | 0.13578067 | 0.00725775 |
| ENSMUSG00 Gch1 | 2769 | 2.74225236 | 1.89682997 | 3.52784663 | 1.44640915 | 1.38207952 | 1.35929946 |
| ENSMUSG00 Nr0b2 | 1130 | 2.67381993 | 1.57778616 | 5.00693443 | 1.94779178 | 1.98686773 | 4.99632772 |
| ENSMUSG00 BC030499 | 4185 | 0.00949952 | 0 | 0.08449576 | 0.06035229 | 0.34139511 | 0 |
| ENSMUSG00 BC022687 | 1910 | 11.1773262 | 6.13052641 | 6.82698622 | 5.9695948 | 4.1408839 | 4.6838124 |
| ENSMUSG00 1810019J16f | 2436 | 10.9180692 | 8.98056024 | 10.7782819 | 8.3539709 | 7.60368408 | 8.59890104 |
| ENSMUSG00 Nme1 | 3378 | 12.628073 | 11.6115346 | 11.2401963 | 11.1407852 | 7.73401552 | 14.7434847 |
| ENSMUSG00 Gm5519 | 511 | 0 | 0 | 0 | 0 | 0 | 0 |
| ENSMUSG00 Lphn3 | 13688 | 0 | 0 | 0 | 0 | 0 | 0 |
| ENSMUSG00 Osbpl5 | 4714 | 8.34072343 | 7.0940583 | 5.9167083 | 4.47772508 | 7.6420495 | 6.32396706 |
| ENSMUSG00 Bclaf1 | 5753 | 59.2358733 | 42.8510061 | 49.3880492 | 62.8064651 | 57.8293013 | 61.3999397 |
| ENSMUSG00 Kcnmb2 | 3331 | 0 | 0.28932118 | 0 | 0 | 0 | 0.28658432 |
| ENSMUSG00 Tnfrsf23 | 4325 | 1.01112205 | 0.05570687 | 0.43946341 | 1.89378592 | 0.04719202 | 0.08198158 |
| ENSMUSG00 Spag1 | 3847 | 1.98415707 | 2.37988687 | 0.73535689 | 1.32247681 | 2.55994035 | 2.08441485 |
| ENSMUSG00 Atoh8 | 2352 | 0 | 0 | 0 | 0 | 0.08677955 | 0.19713812 |
| ENSMUSG00 Wdtc1 | 6684 | 0.1962793 | 0.54069168 | 0.56872509 | 0.51823403 | 0.29009609 | 1.0487107 |
| ENSMUSG00 Kcnk2 | 3926 | 0.43542681 | 0.08591571 | 0.05629374 | 0 | 0 | 0.0416831 |
| ENSMUSG00 Cldn11 | 1841 | 0 | 0 | 0.14405815 | 0.23518986 | 0 | 0 |
| ENSMUSG00 Rgs22 | 4765 | 0 | 0.01011258 | 0 | 0.00757231 | 0 | 0.2346821 |
| ENSMUSG00 Cdkn3 | 1144 | 0 | 0.12636305 | 0 | 0 | 0 | 0 |
| ENSMUSG00 Slc25a43 | 2082 | 5.74755026 | 4.23540774 | 6.53898542 | 6.23897024 | 2.99001818 | 3.19645148 |
| ENSMUSG00 Zbtb42 | 3440 | 1.13256892 | 1.48481479 | 1.47767789 | 0.83911732 | 1.46849163 | 1.4430187 |
| ENSMUSG00 Zfp60 | 8402 | 16.3384521 | 14.2517625 | 16.713789 | 15.7477812 | 15.7961918 | 20.0518266 |
| ENSMUSG00 Prkci | 5312 | 19.6981221 | 19.0949663 | 19.4048754 | 19.3384 | 20.6430124 | 18.3046101 |
| ENSMUSG00 Vps13b | 13418 | 6.54492894 | 5.5699189 | 4.99074319 | 7.68538406 | 7.75777364 | 7.54534862 |
| ENSMUSG00 H2-DMa | 1636 | 1.31222246 | 2.53302813 | 1.26985735 | 0.52932095 | 2.27684929 | 2.23398634 |
| ENSMUSG00 Phc3 | 15646 | 2.66290065 | 2.71637748 | 1.82220304 | 3.09485516 | 3.16346571 | 2.99313279 |
| ENSMUSG00 Kctd8 | 3400 | 1.18097163 | 0.87869395 | 0.67602819 | 0.97633768 | 0.48024824 | 0.68988793 |
| ENSMUSG00 Slc20a2 | 3638 | 13.1571189 | 14.3976533 | 15.5520503 | 11.6140941 | 12.1324396 | 9.73880055 |
| ENSMUSG00 Gdf7 | 1405 | 0 | 0 | 0.09438116 | 0 | 0 | 0 |
| ENSMUSG00 Gpr160 | 4320 | 24.6907301 | 29.2241848 | 21.6814133 | 26.4184044 | 25.8557261 | 25.4752707 |
| ENSMUSG00 Cdkn1c | 1927 | 2.43443005 | 1.1252672 | 4.12886961 | 1.59157955 | 1.6417412 | 3.51017672 |
| ENSMUSG00 1110057K04f | 2829 | 19.4772344 | 22.1259061 | 28.3898016 | 16.631667 | 22.672377 | 22.1648624 |
| ENSMUSG00 Rfx7 | 8432 | 6.24715509 | 4.9889426 | 5.64579992 | 7.36446853 | 6.86241819 | 6.22995199 |
| ENSMUSG00 Inf2 | 4577 | 0 | 0 | 0 | 0 | 0.02229687 | 0 |
| ENSMUSG00 Esyt3 | 4468 | 1.58381279 | 0.37746765 | 0.35614735 | 0.29072373 | 0.44539585 | 0.09767105 |
| ENSMUSG00 Armc3 | 3061 | 0 | 0 | 0 | 0 | 0 | 0 |
| ENSMUSG00 Atp8a1 | 11327 | 5.15940287 | 4.94329 | 2.65359347 | 7.97965229 | 6.92394629 | 5.95963028 |
| ENSMUSG00 Aspg | 2717 | 0.57065283 | 1.50748899 | 0.86223691 | 0.94288744 | 0.8826793 | 0.46177159 |
| ENSMUSG00 1700090G07f | 748 | 0 | 0 | 0 | 0 | 0 | 0 |
| ENSMUSG00 Ahdc1 | 4406 | 1.54293853 | 2.70132801 | 1.74560161 | 3.43131563 | 1.62135556 | 2.99612469 |
| ENSMUSG00 Ddhd1 | 8685 | 2.40318103 | 2.54109275 | 1.84237964 | 3.96757081 | 3.11974733 | 2.18259923 |
| ENSMUSG00 Prosapip1 | 6869 | 3.92403782 | 3.1006562 | 2.54181511 | 2.79453307 | 3.52853814 | 2.62462317 |
| ENSMUSG00 Tecta | 8090 | 0 | 0.07147556 | 0 | 0.08474151 | 0 | 0 |
| ENSMUSG00 Cd81 | 2225 | 184.215283 | 231.92297 | 177.204692 | 168.409902 | 189.955717 | 134.362841 |
| ENSMUSG00 BC061194 | 2614 | 0.13687809 | 0 | 0 | 0.0828203 | 0 | 0 |
| ENSMUSG00 Fam13a | 6435 | 78.3433175 | 44.5771391 | 83.6503565 | 63.2711407 | 73.0386865 | 78.1279604 |
| ENSMUSG00 Cisd1 | 1081 | 94.5525813 | 95.1693376 | 113.959793 | 84.7143654 | 52.9144933 | 121.133712 |
| ENSMUSG00 Fermt2 | 4952 | 30.6113989 | 30.1749106 | 29.4202897 | 30.2821036 | 37.2702749 | 33.0688478 |
| ENSMUSG00 Ccdc33 | 5521 | 0 | 0 | 0 | 0 | 0 | 0 |
| ENSMUSG00 Tmem33 | 4029 | 41.4329269 | 47.361209 | 44.9259238 | 58.8650481 | 35.9046353 | 57.1555669 |
| ENSMUSG00 Gnpnat1 | 2623 | 27.7666947 | 30.4586813 | 28.6814856 | 34.4038099 | 23.6942912 | 36.9866153 |
| ENSMUSG00 Ckap2 | 2592 | 0 | 2.41675829 | 0.01705318 | 0.73778872 | 0 | 0 |
| ENSMUSG00 Avp | 574 | 0 | 0 | 0 | 0 | 0 | 0 |
| ENSMUSG00 Mycn | 4144 | 25.0678259 | 17.6396797 | 28.4474699 | 27.8625826 | 24.1587233 | 24.5563897 |
| ENSMUSG00 BC013712 | 3073 | 0 | 0 | 0 | 0 | 0 | 0 |
| ENSMUSG00 2810032G03f | 3169 | 2.28321157 | 1.00356743 | 0.58582436 | 0.34157821 | 0.32203456 | 0.89509703 |
| ENSMUSG00 Limch1 | 5019 | 20.6500374 | 16.8302118 | 25.1877408 | 16.6571224 | 16.9376255 | 21.7697161 |
| ENSMUSG00 Arpm1 | 1959 | 0 | 0 | 0 | 0 | 0 | 0 |
| ENSMUSG00 Nek5 | 2393 | 0.78082223 | 1.0470936 | 0.24012702 | 0 | 0.66101866 | 0.79783734 |
| ENSMUSG00 Mrps26 | 4355 | 7.67723516 | 7.60139742 | 9.40875051 | 6.58673373 | 6.32703627 | 8.92453753 |
| ENSMUSG00 Eef1a1 | 2400 | 67.9818717 | 59.7311106 | 68.5128567 | 57.4305875 | 55.1722689 | 42.9689213 |
| ENSMUSG00 Phyhipl | 8875 | 1.67980904 | 1.46595373 | 0.7968783 | 1.35383897 | 1.06939785 | 1.49972101 |
| ENSMUSG00 BC017647 | 1154 | 53.294392 | 53.071896 | 54.8501206 | 65.4416977 | 61.4615792 | 59.4653313 |
| ENSMUSG00 Xkr8 | 4355 | 0.29211834 | 0.91836388 | 0.9236206 | 0.50539718 | 0.85532157 | 0.20041067 |
| ENSMUSG00 Ppp1r16b | 8831 | 0.41416648 | 0.01091302 | 0.28029705 | 0.26966538 | 0.13867433 | 0.10809788 |
| ENSMUSG00 Ptger2 | 3682 | 0.23753954 | 0.31408871 | 0 | 0 | 0.05543333 | 0 |
| ENSMUSG00 Actr5 | 4176 | 4.18879584 | 2.96549704 | 2.42390375 | 4.68306229 | 4.82648909 | 4.00367669 |
| ENSMUSG00 Slc16a9 | 3863 | 0.56602419 | 0.94801181 | 0.42336738 | 0.11208505 | 0.76612213 | 0.88256016 |
| ENSMUSG00 Slc32a1 | 3299 | 0 | 0 | 0 | 0 | 0 | 0 |
| ENSMUSG00 Mrpl23 | 1771 | 0.65099319 | 0.8162582 | 0.2995043 | 0.30560738 | 0.54743147 | 0.2464112 |
| ENSMUSG00 Fam113a | 5972 | 1.79072744 | 1.79125756 | 2.19084821 | 2.09652871 | 2.76834323 | 2.13739638 |
| ENSMUSG00 Mbl1 | 918 | 0 | 0 | 0 | 0 | 0 | 0 |
| ENSMUSG00 Dzip1l | 4762 | 2.19565128 | 3.71365482 | 1.50371663 | 4.4401676 | 4.81118075 | 1.96455297 |
| ENSMUSG00 2810002N01f | 4086 | 8.19239221 | 8.43203772 | 8.51366874 | 7.53254625 | 8.09228868 | 10.1328689 |
| ENSMUSG00 Vopp1 | 9651 | 4.01220993 | 5.96651107 | 6.49905867 | 4.12376905 | 3.8596264 | 4.73088241 |
| ENSMUSG00 Defb7 | 304 | 0 | 0 | 0 | 0 | 0 | 0 |
| ENSMUSG00 Phf12 | 5477 | 2.96877698 | 3.0968829 | 3.16361557 | 4.17015413 | 4.91910287 | 4.83543449 |
| ENSMUSG00 N4bp2 | 9849 | 2.5026295 | 2.76427453 | 3.39737995 | 4.27533211 | 3.40383074 | 3.92406984 |
| ENSMUSG00 Adh4 | 1502 | 0 | 0 | 0 | 0 | 0 | 0 |
| ENSMUSG00 Mat1a | 3199 | 0 | 0.31632238 | 0 | 0 | 0 | 0.38366951 |
| ENSMUSG00 Iqch | 7241 | 0.01647099 | 0 | 0.03052192 | 0 | 0.1127499 | 0.01883348 |
| ENSMUSG00 Rpl10a | 1378 | 8.82814008 | 12.1340315 | 10.9702687 | 14.8988994 | 6.59121545 | 10.8267255 |
| ENSMUSG00 Fam76b | 4641 | 13.6201711 | 12.2412875 | 12.2195571 | 11.9573765 | 9.97218765 | 18.6767515 |
| ENSMUSG00 D630003M21 | 5088 | 0.21096658 | 0.06629424 | 0.02606241 | 0 | 0.09025892 | 0.12865396 |
| ENSMUSG00 Larp1b | 1853 | 28.1914201 | 29.2031163 | 35.3757869 | 32.0512927 | 26.8762778 | 41.7288733 |
| ENSMUSG00 Ctnna1 | 3680 | 178.16505 | 228.636812 | 133.362245 | 197.186412 | 216.141072 | 134.372068 |
| ENSMUSG00 Fbxw17 | 3423 | 14.3435637 | 9.84000089 | 15.8703082 | 7.3049538 | 13.3118766 | 12.3345433 |
| ENSMUSG00 3110057O12f | 5574 | 5.73437501 | 4.11495272 | 7.81105407 | 5.71590336 | 4.07368806 | 5.60270218 |
| ENSMUSG00 Tgm2 | 4594 | 1.76537179 | 1.48943726 | 2.61708779 | 1.41375011 | 2.42136481 | 0.93804811 |
| ENSMUSG00 1110003E01f | 7521 | 21.0855753 | 22.1551279 | 25.2775066 | 22.5818619 | 19.9871963 | 24.2139032 |
| ENSMUSG00 Tspan14 | 2466 | 105.321393 | 86.446399 | 101.900843 | 59.9904228 | 86.8855442 | 67.257535 |
| ENSMUSG00 Ppm1k | 5553 | 5.26207062 | 5.21520831 | 7.10827405 | 5.64655082 | 6.79984119 | 4.91660614 |
| ENSMUSG00 Gm5884 | 1478 | 0.2689816 | 0.26081971 | 0.35887829 | 0.097651 | 0.17261968 | 0.62742743 |
| ENSMUSG00 Sh2d4b | 4110 | 33.0521842 | 19.4738882 | 52.7948534 | 25.6612691 | 33.7941107 | 63.4948293 |
| ENSMUSG00 Vstm2l | 1523 | 0.13051701 | 0 | 0.05804576 | 0.04738286 | 0 | 0.21490195 |
| ENSMUSG00 Fdxacb1 | 3151 | 3.36868083 | 3.44079641 | 1.78154048 | 3.43529462 | 2.73673675 | 4.44046015 |
| ENSMUSG00 Rtkn2 | 4480 | 0.37270763 | 0.25814165 | 0 | 0.08054028 | 0 | 0.25569974 |
| ENSMUSG00 BC016495 | 3384 | 11.3251428 | 12.5165138 | 9.14340724 | 12.9336643 | 14.6112761 | 11.1065168 |
| ENSMUSG00 Gm4955 | 3016 | 1.21270033 | 0.54321586 | 0.96728171 | 2.16540121 | 1.26889197 | 1.03998127 |
| ENSMUSG00 Iars | 5311 | 21.9025675 | 21.98376 | 23.0954838 | 24.2675698 | 16.09286 | 10.2401865 |
| ENSMUSG00 Cpe | 2078 | 6.75345747 | 3.57108378 | 1.12738098 | 0 | 0.56477702 | 0.06562714 |
| ENSMUSG00 Zfp365 | 8121 | 0.18602491 | 0.14240544 | 1.60565739 | 0.27102632 | 0.79169109 | 1.77330495 |
| ENSMUSG00 Nufip2 | 6024 | 21.1118496 | 27.8287902 | 25.4835659 | 33.799963 | 31.5526643 | 27.9583163 |
| ENSMUSG00 Aim2 | 3398 | 0 | 0 | 0 | 0.14866057 | 0 | 0.07224007 |
| ENSMUSG00 Egr2 | 3107 | 0.01279546 | 0.01550899 | 0.12803881 | 0.19742348 | 0.34488378 | 0.00877845 |
| ENSMUSG00 Darc | 1156 | 0 | 0 | 0 | 0 | 0 | 0 |
| ENSMUSG00 Jmjd1c | 14137 | 2.99213633 | 2.90407079 | 3.80829347 | 3.05511829 | 2.95250587 | 3.64832304 |
| ENSMUSG00 1700017G19f | 2414 | 0 | 0 | 0 | 0 | 0 | 0 |
| ENSMUSG00 Stk35 | 6191 | 1.13660476 | 1.16749577 | 3.58412618 | 1.80089675 | 1.98632799 | 2.25122608 |
| ENSMUSG00 Dusp8 | 6400 | 0.18014202 | 0.62491792 | 0.33151382 | 0.82312164 | 0.55012811 | 0.31110135 |
| ENSMUSG00 Wdr19 | 4371 | 5.96650543 | 3.1969957 | 2.49779346 | 4.42461127 | 6.74748232 | 3.75642481 |
| ENSMUSG00 Pcdh18 | 5164 | 0 | 0 | 0.61629216 | 0 | 0 | 0 |
| ENSMUSG00 H2afz | 1340 | 28.5112065 | 28.2286247 | 19.8248565 | 21.5145922 | 11.081101 | 33.1366513 |
| ENSMUSG00 Rcor1 | 5942 | 14.2108113 | 11.6938488 | 18.9245185 | 13.7903607 | 15.6634314 | 12.0353592 |
| ENSMUSG00 Sirpa | 3212 | 2.6363379 | 4.17055757 | 0.77064234 | 1.60639239 | 5.09946035 | 1.82566857 |
| ENSMUSG00 Ankrd9 | 4034 | 24.401232 | 21.3936336 | 27.2618209 | 16.0285136 | 19.6313672 | 13.6102749 |
| ENSMUSG00 Bri3bp | 6972 | 4.32794173 | 3.99480258 | 3.81028509 | 3.11034262 | 2.12975837 | 3.85334476 |
| ENSMUSG00 Ankrd13b | 5159 | 0.20035714 | 0.24284697 | 0.23990146 | 0.132886 | 0.20770574 | 0.32249525 |
| ENSMUSG00 1700018B24f | 590 | 0 | 0 | 0 | 0 | 0 | 0 |
| ENSMUSG00 Tmem156 | 882 | 0 | 0 | 0 | 0 | 0 | 0 |
| ENSMUSG00 Ndufv1 | 1991 | 217.526974 | 185.049491 | 223.096093 | 177.619346 | 162.331022 | 163.688932 |
| ENSMUSG00 Ddx60 | 7077 | 0.00561756 | 0.43576831 | 0 | 0.06628043 | 0 | 0.95193386 |
| ENSMUSG00 Bank1 | 4031 | 0 | 0 | 0 | 0 | 0 | 0 |
| ENSMUSG00 Olfr16 | 1049 | 0 | 0 | 0 | 0 | 0 | 0 |
| ENSMUSG00 Ssh2 | 9604 | 2.61200627 | 1.75104835 | 1.95143497 | 2.75762398 | 2.60338652 | 1.93398889 |
| ENSMUSG00 Bicd2 | 6436 | 8.57374253 | 9.74809628 | 7.58216822 | 10.4669324 | 10.1164785 | 6.1278944 |
| ENSMUSG00 Smarce1 | 7864 | 2.39119307 | 2.20589003 | 2.22020956 | 2.62907064 | 2.21910231 | 3.35036896 |
| ENSMUSG00 Scarb1 | 6861 | 8.35554626 | 10.0853711 | 10.4625846 | 9.86589645 | 9.33363965 | 8.43164405 |
| ENSMUSG00 Chchd5 | 1695 | 6.49691333 | 8.50014525 | 8.81429082 | 7.00353549 | 8.85059262 | 10.8293995 |
| ENSMUSG00 Inpp4b | 10384 | 0.23736901 | 0.04176406 | 0 | 0.19111252 | 0.48156634 | 0.07617147 |
| ENSMUSG00 Crp | 1824 | 0 | 0 | 0 | 0 | 0 | 0 |
| ENSMUSG00 Ccr7 | 1975 | 0 | 0 | 0 | 0 | 0 | 0 |
| ENSMUSG00 Fgd3 | 6916 | 0.44837008 | 0.73157554 | 0 | 0.02608592 | 0.71566779 | 0.20507248 |
| ENSMUSG00 Ano10 | 2627 | 24.2588638 | 17.5907111 | 32.204921 | 14.8888223 | 19.5792108 | 20.9102107 |
| ENSMUSG00 A4gnt | 2754 | 0 | 0 | 0 | 0 | 0 | 0 |
| ENSMUSG00 Wdr20a | 3586 | 25.2767695 | 24.3350939 | 30.0883153 | 29.6927254 | 27.889486 | 41.5280341 |
| ENSMUSG00 Ccdc55 | 3620 | 11.4434284 | 12.7521026 | 13.236132 | 13.2765976 | 14.9273293 | 16.9524692 |
| ENSMUSG00 1110007C09f | 926 | 24.643246 | 32.0028746 | 28.067693 | 26.2237753 | 18.5700741 | 27.3040935 |
| ENSMUSG00 Fam101a | 1934 | 1.80893603 | 0.64780119 | 1.82841129 | 1.49253545 | 2.61200166 | 1.01539505 |
| ENSMUSG00 Zc3h7a | 8941 | 25.1311907 | 22.0479516 | 29.3558376 | 30.6017385 | 30.8292677 | 38.1863253 |
| ENSMUSG00 Ninj1 | 1675 | 40.3013767 | 41.5697964 | 43.8323948 | 33.8848096 | 24.5840509 | 34.7812713 |
| ENSMUSG00 1110032A03f | 5598 | 16.7458776 | 15.0722508 | 15.3893162 | 17.1515399 | 13.9278854 | 19.2793404 |
| ENSMUSG00 Snn | 2900 | 7.27936557 | 15.0042611 | 7.04181085 | 12.0190535 | 10.4164188 | 11.154387 |
| ENSMUSG00 Ccdc129 | 3574 | 0 | 0 | 0 | 0 | 0 | 0 |
| ENSMUSG00 Muc5ac | 9347 | 0 | 0 | 0 | 0 | 0 | 0 |
| ENSMUSG00 6430571L13f | 1932 | 1.44041596 | 1.72094436 | 1.34984924 | 0.78439227 | 2.9316396 | 1.93407121 |
| ENSMUSG00 Ccdc92 | 2945 | 5.38622639 | 5.4322237 | 5.55337247 | 6.93461365 | 4.27963152 | 4.99186097 |
| ENSMUSG00 Gm9725 | 2326 | 0.61530408 | 0 | 0 | 0 | 0 | 0.10553386 |
| ENSMUSG00 Neurod6 | 2152 | 0 | 0 | 0 | 0 | 0 | 0 |
| ENSMUSG00 Wnk2 | 7905 | 0.02011663 | 0 | 0.05032468 | 0 | 0 | 0.12421088 |
| ENSMUSG00 Sh3rf3 | 7898 | 0 | 0.01220219 | 0.22386347 | 0 | 0.02584268 | 0.02762688 |
| ENSMUSG00 A630055G03 | 3873 | 0 | 0.01244163 | 0.02282564 | 0.01863261 | 0 | 0 |
| ENSMUSG00 Rara | 4558 | 2.4596414 | 2.00864941 | 2.66685099 | 2.29569831 | 3.99658099 | 3.32705121 |
| ENSMUSG00 Dhx38 | 4170 | 5.87275204 | 7.88085217 | 6.03137858 | 5.44259137 | 8.29637477 | 4.02251874 |
| ENSMUSG00 Nhedc2 | 2912 | 0 | 0 | 0 | 0 | 0 | 0 |
| ENSMUSG00 Igsf9 | 6107 | 0.35153037 | 0.03945181 | 0.23885063 | 0.99259595 | 0.09190931 | 0 |
| ENSMUSG00 Slc24a2 | 22154 | 0 | 0 | 0 | 0 | 0 | 0 |
| ENSMUSG00 Parp11 | 5709 | 2.59744162 | 2.03414478 | 3.37572317 | 4.06389116 | 4.23655669 | 3.9557543 |
| ENSMUSG00 Arap2 | 10890 | 0.89805769 | 1.01771182 | 1.22174056 | 1.57051323 | 1.3822572 | 1.64299021 |
| ENSMUSG00 Acd | 1687 | 11.5708008 | 16.0240629 | 12.864911 | 13.0254684 | 14.8511859 | 18.576503 |
| ENSMUSG00 Cramp1l | 7620 | 0.77215369 | 2.01093026 | 1.3631802 | 1.46316952 | 1.28570396 | 2.06170501 |
| ENSMUSG00 2700029M09 | 1784 | 20.6131275 | 19.8525981 | 19.3506947 | 17.6567404 | 15.7312257 | 29.9501224 |
| ENSMUSG00 Acer2 | 7333 | 5.81721404 | 7.9708379 | 2.32070225 | 5.94396697 | 8.90682682 | 2.48458462 |
| ENSMUSG00 Dnajc22 | 1845 | 0.08619074 | 0.83575401 | 0.40727985 | 0 | 2.21252578 | 0.44349007 |
| ENSMUSG00 Ccdc138 | 2415 | 5.23488315 | 5.7464577 | 3.18472906 | 9.02424633 | 7.88109241 | 8.45909101 |
| ENSMUSG00 Dnahc10 | 14002 | 0 | 0 | 0 | 0.00515384 | 0 | 0 |
| ENSMUSG00 Wipf2 | 7767 | 1.67375333 | 2.20241882 | 1.9235513 | 1.8442863 | 2.24024645 | 2.50728627 |
| ENSMUSG00 Fam120a | 5087 | 29.7208753 | 27.697495 | 23.8952365 | 31.7411341 | 31.1855715 | 29.8589475 |
| ENSMUSG00 Prm2 | 616 | 0 | 0 | 0 | 0.41002323 | 0 | 0 |
| ENSMUSG00 Rapgefl1 | 5662 | 0.29490113 | 0.39999343 | 0.21078237 | 0.87942814 | 0.61282119 | 0.47207959 |
| ENSMUSG00 Fam188b | 4677 | 2.89857149 | 3.21449005 | 1.79567034 | 0.97977756 | 3.53486119 | 1.80781231 |
| ENSMUSG00 Atp6v0a2 | 5283 | 19.8288266 | 17.2296402 | 18.6663471 | 24.9494054 | 21.5579918 | 13.4850194 |
| ENSMUSG00 Dennd4c | 8721 | 3.84289303 | 3.83458214 | 3.47694809 | 5.1303446 | 6.22544019 | 2.64896451 |
| ENSMUSG00 Phf2 | 5238 | 1.99612283 | 1.43510595 | 1.48520893 | 2.7002981 | 1.44175327 | 2.3015255 |
| ENSMUSG00 Kcnj9 | 3266 | 0 | 0 | 0 | 0 | 0 | 0 |
| ENSMUSG00 9630033F20f | 3645 | 1.84325822 | 1.62604455 | 1.03077 | 1.36606918 | 1.20391449 | 1.30199924 |
| ENSMUSG00 Igsf8 | 4359 | 11.1176717 | 11.4192693 | 7.60527237 | 6.96973652 | 10.8163274 | 7.18009218 |
| ENSMUSG00 Socs1 | 1218 | 1.37087864 | 1.1077343 | 1.66906353 | 0.68135224 | 1.17302013 | 2.41844092 |
| ENSMUSG00 Gcc2 | 8409 | 7.95204165 | 8.21730979 | 9.1147575 | 11.7613134 | 9.47225266 | 11.1738771 |
| ENSMUSG00 Ptpdc1 | 6542 | 3.15394289 | 4.58146851 | 3.30398979 | 4.88116928 | 6.52064356 | 2.28886838 |
| ENSMUSG00 Cct8l1 | 1954 | 0.02034569 | 0 | 0 | 0 | 0 | 0 |
| ENSMUSG00 Sult6b1 | 2343 | 0.27148429 | 0.4113226 | 0.13205843 | 0.47739794 | 0.23956045 | 0.83814512 |
| ENSMUSG00 Rnmtl1 | 2280 | 9.13678587 | 9.23573474 | 8.20060509 | 8.2767146 | 7.96727622 | 7.45267555 |
| ENSMUSG00 Haus6 | 7171 | 2.17876221 | 0.57788789 | 3.43333239 | 3.79387276 | 3.91361131 | 3.12264382 |
| ENSMUSG00 Cntnap5c | 3894 | 0 | 0 | 0 | 0 | 0 | 0 |
| ENSMUSG00 Dexi | 1387 | 17.5417117 | 15.0430638 | 24.6663493 | 12.4349079 | 19.9396508 | 22.9681173 |
| ENSMUSG00 Mll3 | 22197 | 5.56293745 | 4.40032068 | 6.65706002 | 6.68258256 | 5.31251766 | 3.99344855 |
| ENSMUSG00 Dbil5 | 579 | 0.54929852 | 0 | 0 | 0 | 0.52877073 | 0.18842583 |
| ENSMUSG00 Nod1 | 4402 | 2.14040184 | 2.16740471 | 3.10276454 | 2.25410319 | 3.744098 | 2.29250716 |
| ENSMUSG00 2010002N04 | 1982 | 3.40990499 | 5.59176675 | 3.74667489 | 3.93225107 | 5.76685579 | 4.37605211 |
| ENSMUSG00 Dlec1 | 7075 | 4.5009385 | 4.51556341 | 2.20540644 | 4.45734373 | 3.75034847 | 3.64689876 |
| ENSMUSG00 Cldn22 | 993 | 0.36032158 | 0.1941045 | 0.17805375 | 0.54504599 | 0.61663294 | 0.65920578 |
| ENSMUSG00 2410066E13f | 5343 | 8.86927433 | 12.8244658 | 5.61726584 | 8.31311937 | 12.2050736 | 4.16546989 |
| ENSMUSG00 Csf3 | 1406 | 0.19764799 | 0.1368933 | 0 | 0 | 0 | 0.19371193 |
| ENSMUSG00 Rnf144b | 4685 | 0.03394278 | 0.37026935 | 0.55665074 | 0.55451595 | 0.96933777 | 0.6462081 |
| ENSMUSG00 Cdkn2aip | 3491 | 13.062027 | 11.9120308 | 11.7626789 | 11.0902417 | 12.6579322 | 15.6335013 |
| ENSMUSG00 Cntln | 7496 | 0.32882068 | 0.02571315 | 0.26535258 | 0.32250493 | 0.60583611 | 0.42571142 |
| ENSMUSG00 Npy6r | 2565 | 20.9239371 | 13.1690822 | 2.75722997 | 16.9226899 | 15.1984995 | 28.4336785 |
| ENSMUSG00 Galnt11 | 2979 | 28.9191428 | 22.6617004 | 36.5900453 | 21.0387752 | 37.3234216 | 33.5645611 |
| ENSMUSG00 Fkbp14 | 6256 | 2.54191052 | 1.56359459 | 1.1799405 | 1.48803828 | 2.52032451 | 2.37170776 |
| ENSMUSG00 Kcna6 | 5795 | 0 | 0 | 0 | 0 | 0 | 0 |
| ENSMUSG00 Als2cr4 | 2661 | 4.54177605 | 6.24739687 | 7.62444417 | 6.14249011 | 7.05663521 | 6.14966229 |
| ENSMUSG00 Kdm1b | 4937 | 31.6385017 | 28.3438177 | 17.7183406 | 32.7055194 | 29.2701431 | 30.5451652 |
| ENSMUSG00 Opa1 | 8074 | 21.5961776 | 14.6039415 | 22.1720912 | 16.9327306 | 23.0547707 | 16.5863858 |
| ENSMUSG00 4921517L17f | 4214 | 1.46229224 | 1.02913616 | 2.695746 | 1.77242127 | 1.54992314 | 1.76049929 |
| ENSMUSG00 Hspb2 | 838 | 2.18228175 | 1.61004819 | 0.15824049 | 1.11949064 | 0.85246929 | 0.81368256 |
| ENSMUSG00 Hsd3b5 | 1601 | 0 | 0 | 0 | 0 | 0 | 0 |
| ENSMUSG00 Atp13a4 | 4447 | 0 | 0 | 0 | 0 | 0 | 0 |
| ENSMUSG00 Sbno1 | 10822 | 15.675165 | 14.0169026 | 18.5352027 | 15.5437527 | 17.5117316 | 15.9383495 |
| ENSMUSG00 D030016E14f | 7171 | 12.2742482 | 9.91816882 | 10.786951 | 12.3829189 | 13.0145365 | 9.47823191 |
| ENSMUSG00 AW551984 | 7458 | 0 | 0 | 0 | 0 | 0 | 0 |
| ENSMUSG00 Ano2 | 5391 | 0 | 0 | 0 | 0 | 0 | 0 |
| ENSMUSG00 Phf20 | 9275 | 4.95068248 | 4.5926485 | 6.20970366 | 4.36874782 | 4.19213999 | 5.80486862 |
| ENSMUSG00 Cdon | 7691 | 0.28946911 | 0.476163 | 0.77012703 | 0.41754024 | 0.84922326 | 0.61705724 |
| ENSMUSG00 4933403F05f | 9806 | 14.3448864 | 12.0957403 | 21.0508573 | 15.5099816 | 11.3571144 | 15.6784682 |
| ENSMUSG00 D630037F22f | 7202 | 5.43174019 | 4.23521492 | 4.54170561 | 5.07513345 | 7.95648709 | 7.51737836 |
| ENSMUSG00 Mphosph9 | 10868 | 0.73160619 | 1.104014 | 1.30555683 | 1.54381217 | 0.82164297 | 0.4718101 |
| ENSMUSG00 Ccdc50 | 8783 | 11.4699291 | 13.2659475 | 10.7598247 | 12.3368302 | 14.605523 | 12.2445523 |
| ENSMUSG00 Camk4 | 12441 | 0.00639104 | 0.08521033 | 0.38371506 | 0.59455182 | 0.2912043 | 0.73442683 |
| ENSMUSG00 Rbm24 | 3211 | 0 | 0 | 0 | 0 | 0 | 0 |
| ENSMUSG00 Crygn | 797 | 0.24940703 | 0 | 0.05546028 | 0 | 0.06402306 | 0 |
| ENSMUSG00 Tmem181a | 1852 | 1.58850192 | 2.15954357 | 1.43202515 | 1.26637845 | 2.17661106 | 0.83944624 |
| ENSMUSG00 Stox2 | 10833 | 3.03863545 | 4.23905468 | 2.1707173 | 3.45732135 | 3.54212449 | 2.54543158 |
| ENSMUSG00 Snrk | 3410 | 13.7803454 | 16.4059998 | 15.1789907 | 17.8611411 | 20.0663842 | 13.3013857 |
| ENSMUSG00 Notch3 | 8011 | 0.63521427 | 0.67969891 | 0.2151881 | 0.78821093 | 0.78982282 | 0.24513469 |
| ENSMUSG00 Cd84 | 6340 | 0 | 0 | 0 | 0 | 0.06438659 | 0.55065518 |
| ENSMUSG00 Cldn16 | 1410 | 4.70862783 | 0.7518452 | 1.15990652 | 1.58658636 | 0.97710081 | 1.23799781 |
| ENSMUSG00 Ormdl3 | 2180 | 21.2454747 | 21.4628602 | 24.5746026 | 23.370572 | 20.3637371 | 19.7928804 |
| ENSMUSG00 Prdm1 | 7432 | 0 | 0 | 0.40443021 | 0 | 0.29522796 | 0.62755158 |
| ENSMUSG00 5033430I15f | 719 | 0.66351289 | 0.67018696 | 0.86067566 | 0.65238745 | 0.56774827 | 1.47943106 |
| ENSMUSG00 Gstp2 | 742 | 0 | 0.12988259 | 0.05957122 | 0 | 0 | 0 |
| ENSMUSG00 Spon1 | 8553 | 3.07241583 | 0.33803186 | 0.45478337 | 0 | 0.65624943 | 0.07653354 |
| ENSMUSG00 Atg5 | 2355 | 33.6613283 | 31.3876867 | 39.8473515 | 44.2943485 | 27.8640846 | 32.3705379 |
| ENSMUSG00 1700022A21f | 1055 | 0 | 0 | 0 | 0 | 0 | 0 |
| ENSMUSG00 Plekhg6 | 3282 | 1.85331765 | 3.27409403 | 1.21211635 | 1.17634941 | 2.2699119 | 1.39614241 |
| ENSMUSG00 Leprel1 | 3704 | 42.0201691 | 57.0197559 | 36.7433787 | 60.1042696 | 94.9029976 | 37.097633 |
| ENSMUSG00 Pde4dip | 17447 | 9.87791643 | 10.9535983 | 7.37499656 | 14.7806713 | 10.9908363 | 8.16503931 |
| ENSMUSG00 Ttc39b | 10980 | 2.93640207 | 2.1108997 | 4.51277468 | 3.77251249 | 4.19178424 | 4.66003855 |
| ENSMUSG00 Enpp6 | 4689 | 0 | 1.05847804 | 0 | 0.02308512 | 0.02176429 | 0 |
| ENSMUSG00 Fam126b | 12566 | 5.53337064 | 7.43925655 | 5.44168798 | 7.44267543 | 4.78752961 | 7.13012812 |
| ENSMUSG00 Mylip | 3055 | 12.05027 | 18.8329335 | 10.5042678 | 9.72030201 | 11.9423433 | 9.90100652 |
| ENSMUSG00 Slc43a2 | 6872 | 99.9267193 | 66.095082 | 135.030456 | 87.69533 | 90.7440831 | 89.8134185 |
| ENSMUSG00 Slamf7 | 4202 | 0 | 0 | 0 | 0 | 0 | 0 |
| ENSMUSG00 Spag4 | 4122 | 0 | 0 | 0 | 0.00875353 | 0 | 0 |
| ENSMUSG00 Chpf2 | 5913 | 3.36170138 | 2.86038241 | 2.94529446 | 2.95954534 | 3.21018295 | 3.12738126 |
| ENSMUSG00 Btbd10 | 5533 | 9.44848308 | 13.6033296 | 11.6556098 | 12.2990667 | 11.4908484 | 7.42867908 |
| ENSMUSG00 Scarf1 | 3025 | 5.75633073 | 3.80712716 | 3.78455449 | 4.90238688 | 3.93029606 | 3.49836695 |
| ENSMUSG00 Cer1 | 1737 | 0 | 0 | 0 | 0 | 0 | 0 |
| ENSMUSG00 Hand2 | 2233 | 0 | 0 | 0 | 0 | 0 | 0 |
| ENSMUSG00 Lhb | 2419 | 0 | 0 | 0.03654555 | 0 | 0 | 0 |
| ENSMUSG00 Rilp | 1576 | 18.2380789 | 17.1526612 | 13.6868651 | 15.4539214 | 14.5373368 | 17.0639558 |
| ENSMUSG00 4931409K22f | 2651 | 0 | 0 | 0 | 0 | 0 | 0 |
| ENSMUSG00 Kcna7 | 3492 | 0 | 0.05519638 | 0 | 0.05166387 | 0.01461236 | 0 |
| ENSMUSG00 Hoxa13 | 2670 | 0 | 0 | 0 | 0 | 0 | 0 |
| ENSMUSG00 Asb10 | 2799 | 0 | 0 | 0 | 0 | 0 | 0 |
| ENSMUSG00 Prkab2 | 5245 | 3.33506414 | 4.40981739 | 3.77548631 | 6.2189072 | 4.0081729 | 3.99889372 |
| ENSMUSG00 Fbxo8 | 3734 | 32.8030625 | 26.545129 | 32.6719568 | 28.8057244 | 24.9865744 | 29.0496091 |
| ENSMUSG00 Pgap3 | 3716 | 1.82943681 | 2.00992964 | 1.54635081 | 2.45660852 | 1.66151547 | 1.15234617 |
| ENSMUSG00 Itln1 | 1133 | 0 | 0 | 0 | 0 | 0 | 0 |
| ENSMUSG00 Hoxa11 | 3750 | 0.11661608 | 0.05139887 | 0.29467895 | 0 | 0.31296177 | 0.12364503 |
| ENSMUSG00 Hiatl1 | 5723 | 21.0760314 | 24.5941984 | 22.4291721 | 27.4004133 | 22.218719 | 28.5232773 |
| ENSMUSG00 Tapbpl | 2345 | 9.39212631 | 9.47289971 | 4.7500488 | 11.6324203 | 9.61776465 | 8.43245776 |
| ENSMUSG00 Bend3 | 6024 | 1.25390792 | 1.33584592 | 1.62161476 | 1.38362422 | 1.2790476 | 1.84275866 |
| ENSMUSG00 BC088983 | 2629 | 5.45900664 | 6.06683619 | 8.89417077 | 6.39567621 | 9.49102236 | 10.2292865 |
| ENSMUSG00 Pnmt | 1243 | 0 | 0 | 0 | 0 | 0 | 0 |
| ENSMUSG00 Tlcd2 | 1928 | 0.80418244 | 2.44931086 | 3.57649767 | 0.71116063 | 0.44992136 | 1.40051312 |
| ENSMUSG00 Gm9765 | 318 | 0 | 0 | 0 | 0 | 0 | 0 |
| ENSMUSG00 Serpinf2 | 3392 | 0.07032219 | 0.48300089 | 0 | 0.27657228 | 0 | 0.16081745 |
| ENSMUSG00 Ccdc111 | 6847 | 2.22960487 | 1.9001518 | 2.54352653 | 1.32797944 | 3.14490004 | 2.74060929 |
| ENSMUSG00 Hoxa9 | 2788 | 36.048011 | 42.9322533 | 23.9558769 | 34.6196087 | 27.5264237 | 46.811029 |
| ENSMUSG00 Fam198a | 3765 | 0.17950682 | 0 | 0.10566178 | 0 | 0 | 0 |
| ENSMUSG00 F11r | 2776 | 107.16508 | 95.3834653 | 102.686486 | 90.2701006 | 89.1674889 | 93.5551567 |
| ENSMUSG00 Hoxa7 | 4402 | 3.82021087 | 5.29810041 | 2.9521449 | 3.51640098 | 3.54704021 | 4.74610942 |
| ENSMUSG00 Hrc | 4315 | 0.00921332 | 0.01116719 | 0 | 0.01672401 | 0.15372952 | 0.06952979 |
| ENSMUSG00 Pdss2 | 6448 | 3.30473597 | 1.9878402 | 4.27074259 | 2.82030869 | 2.65894266 | 3.13861388 |
| ENSMUSG00 Cep250 | 8595 | 1.72990689 | 1.65947491 | 0.96683496 | 2.03184521 | 3.40175839 | 1.06623372 |
| ENSMUSG00 Aox4 | 6488 | 0 | 0 | 0 | 0.11678837 | 0.18088882 | 0 |
| ENSMUSG00 Mical2 | 9184 | 3.70543242 | 2.0252577 | 4.4615754 | 3.09196092 | 4.66704657 | 5.23278684 |
| ENSMUSG00 Fam50b | 1217 | 0 | 0 | 0 | 0 | 0 | 0 |
| ENSMUSG00 Sobp | 4972 | 0.02398762 | 0.07753249 | 0.2489243 | 0.02902819 | 0.06157648 | 0.55953604 |
| ENSMUSG00 Usp38 | 4674 | 8.01233689 | 9.86615857 | 9.49479041 | 8.07484351 | 6.70307975 | 6.35474586 |
| ENSMUSG00 Ncapd2 | 4609 | 2.42379909 | 3.02145406 | 2.19618617 | 2.08240917 | 3.60915568 | 3.3375779 |
| ENSMUSG00 Hoxa5 | 1875 | 1.86585722 | 2.54424415 | 0.54220927 | 2.40546964 | 3.15683178 | 1.74557691 |
| ENSMUSG00 Neurod2 | 3140 | 0 | 0 | 0 | 0 | 0 | 0 |
| ENSMUSG00 Bcl9 | 8737 | 1.94750437 | 1.16371058 | 1.58351572 | 1.75929392 | 2.54051431 | 2.82517438 |
| ENSMUSG00 Glra3 | 5069 | 0 | 0 | 0 | 0 | 0 | 0 |
| ENSMUSG00 Gdf5 | 2594 | 0 | 0 | 0 | 0 | 0 | 0 |
| ENSMUSG00 Trpm4 | 5280 | 4.17884312 | 5.0376735 | 4.24438152 | 3.10934285 | 6.47493784 | 1.91645666 |
| ENSMUSG00 Sema7a | 3282 | 0.15747143 | 0.13213832 | 0.1346796 | 0.20888448 | 0.27985215 | 0.54848452 |
| ENSMUSG00 Slc22a23 | 10042 | 4.83780095 | 2.91748226 | 2.41213005 | 3.79792084 | 2.79471288 | 2.68346381 |
| ENSMUSG00 Ovca2 | 3206 | 2.82727684 | 4.25351314 | 5.01854986 | 2.41972538 | 3.86756373 | 2.01624751 |
| ENSMUSG00 Iffo1 | 5827 | 1.85575608 | 1.77794492 | 2.16950868 | 2.79888178 | 2.74966226 | 2.66333786 |
| ENSMUSG00 Fau | 556 | 1.9305719 | 3.46665051 | 2.30549181 | 2.20645596 | 1.2848368 | 5.34700662 |
| ENSMUSG00 Accn3 | 1931 | 0 | 0.07486242 | 0.16023454 | 0 | 0 | 0.2259939 |
| ENSMUSG00 Nop2 | 2607 | 17.2929478 | 14.8607209 | 13.6318687 | 16.7192596 | 14.6991976 | 11.6861419 |
| ENSMUSG00 Ostm1 | 5567 | 12.8400133 | 14.5329686 | 18.730492 | 15.6655832 | 17.8917704 | 19.5973697 |
| ENSMUSG00 Bphl | 1254 | 47.9982436 | 54.1425016 | 43.0033879 | 49.4329765 | 33.5699841 | 43.1522203 |
| ENSMUSG00 Smg6 | 8431 | 3.29134449 | 3.43798003 | 2.7105151 | 4.18553428 | 4.51496577 | 2.47804219 |
| ENSMUSG00 Snx25 | 4974 | 3.34093101 | 5.93853821 | 3.40355969 | 4.46854432 | 3.82646526 | 4.16741573 |
| ENSMUSG00 Ccdc155 | 2560 | 0 | 0 | 0 | 0 | 0.01993218 | 0 |
| ENSMUSG00 Atg9b | 4383 | 0 | 0 | 0 | 0 | 0 | 0.03733695 |
| ENSMUSG00 Galntl4 | 2564 | 30.7624311 | 41.7214905 | 20.9631206 | 25.5979871 | 32.0805452 | 20.4559794 |
| ENSMUSG00 Pdzk1 | 3213 | 0.68053266 | 1.12480024 | 0.31641531 | 0.57273087 | 0.47643675 | 1.1799486 |
| ENSMUSG00 Wdr36 | 3188 | 18.5932941 | 19.0599447 | 18.1493766 | 20.497046 | 17.6543578 | 14.7686334 |
| ENSMUSG00 Pth2 | 479 | 0 | 0 | 0 | 0 | 0 | 0 |
| ENSMUSG00 Snx10 | 4851 | 20.660393 | 11.5126956 | 22.2694917 | 14.712489 | 8.80417987 | 19.4031705 |
| ENSMUSG00 Lace1 | 3719 | 8.39151713 | 7.33356447 | 10.007516 | 7.49000763 | 6.59953933 | 6.36579373 |
| ENSMUSG00 Cd160 | 3100 | 0.55144699 | 0.17098415 | 0.21387989 | 1.5480361 | 0.16460121 | 0.68626512 |
| ENSMUSG00 Spata2l | 8283 | 0.23998238 | 0.20943039 | 0.39489755 | 0.47482106 | 0.45586766 | 0.10537106 |
| ENSMUSG00 2410017I17f | 1710 | 0 | 0 | 0 | 0 | 0 | 0 |
| ENSMUSG00 Edem2 | 3451 | 19.123181 | 16.6439406 | 22.7605549 | 17.7221285 | 21.2474361 | 17.1503817 |
| ENSMUSG00 Kcnh2 | 4895 | 0 | 0.11812815 | 0 | 0 | 0.27102876 | 0 |
| ENSMUSG00 1700066M21 | 3006 | 12.8947417 | 12.7920096 | 11.7195173 | 15.8564141 | 9.35313808 | 12.2037225 |
| ENSMUSG00 Trpc4ap | 7625 | 24.5415143 | 23.5086642 | 23.819721 | 22.7896559 | 28.9227536 | 21.9377525 |
| ENSMUSG00 Serpinb9f | 1905 | 0 | 0.02529472 | 0 | 0.01894071 | 0 | 0 |
| ENSMUSG00 C87499 | 2069 | 0.03842966 | 0 | 0 | 0 | 0 | 0 |
| ENSMUSG00 Satb2 | 10026 | 0.01189572 | 0.00480615 | 0 | 0.10796542 | 0.32572193 | 0.09793407 |
| ENSMUSG00 Sesn1 | 3316 | 11.017879 | 13.5578862 | 8.83770263 | 14.4828713 | 13.7875853 | 14.9451205 |
| ENSMUSG00 Tsr1 | 3961 | 6.49376316 | 4.08751441 | 4.77616481 | 4.80061537 | 4.7277657 | 4.20722156 |
| ENSMUSG00 Mlxip | 7963 | 9.26612732 | 9.37345208 | 9.84175156 | 8.92195726 | 11.6752552 | 7.44632246 |
| ENSMUSG00 Txlng | 7531 | 4.80908814 | 4.67723931 | 3.4218131 | 3.79937078 | 4.26857214 | 4.3459789 |
| ENSMUSG00 Zfp384 | 5704 | 7.12308925 | 7.30737596 | 6.98209337 | 9.20395771 | 9.36616681 | 11.2799569 |
| ENSMUSG00 Tcte2 | 8286 | 1.32902101 | 1.02932661 | 0.98155191 | 1.35862876 | 1.12694023 | 0.51020626 |
| ENSMUSG00 Pick1 | 3626 | 6.95117889 | 10.3921119 | 5.79036829 | 11.4336098 | 11.2156706 | 6.16048802 |
| ENSMUSG00 Sgsm2 | 8489 | 2.06528059 | 2.05483473 | 1.23404839 | 0.96485147 | 1.64097074 | 2.35829723 |
| ENSMUSG00 Arl5c | 2067 | 0.21156763 | 0.06993678 | 0.1069227 | 0.12219367 | 0.1728034 | 0.22431972 |
| ENSMUSG00 Ankrd35 | 3795 | 0.16761204 | 0 | 0 | 0.00950779 | 0 | 0.02874797 |
| ENSMUSG00 Camp | 778 | 0 | 0 | 0 | 0 | 0 | 0 |
| ENSMUSG00 Fbxo25 | 2084 | 20.774337 | 20.3011978 | 28.4215305 | 20.3691354 | 16.4293177 | 19.4874942 |
| ENSMUSG00 Lasp1 | 6480 | 30.8411729 | 35.8423844 | 25.873085 | 29.6674588 | 32.608059 | 18.877586 |
| ENSMUSG00 BC057079 | 10854 | 0.96696581 | 1.00332927 | 1.17692396 | 2.04444973 | 1.3069221 | 0.71616659 |
| ENSMUSG00 Ncoa6 | 10269 | 6.31426514 | 4.59387738 | 4.52822464 | 6.35977219 | 7.76153462 | 6.11415129 |
| ENSMUSG00 Pcp4l1 | 1574 | 0.63144029 | 1.16333215 | 0.14041246 | 0.16046653 | 0.42143767 | 0.7277858 |
| ENSMUSG00 Sbf2 | 12256 | 7.63904672 | 9.09000116 | 8.03178071 | 10.3629893 | 14.359495 | 10.6218875 |
| ENSMUSG00 Gmds | 1576 | 7.59289317 | 10.3649771 | 10.9663202 | 9.20366872 | 7.80289123 | 11.8028579 |
| ENSMUSG00 Rbm8a | 2598 | 23.7645347 | 24.4641714 | 23.1727907 | 27.5962365 | 21.2904509 | 26.8442081 |
| ENSMUSG00 Trp53inp2 | 5252 | 93.0832338 | 63.7378548 | 83.463381 | 66.9496428 | 73.4014222 | 63.4400215 |
| ENSMUSG00 Ttk | 2951 | 0 | 0.60416752 | 0 | 0 | 0 | 0 |
| ENSMUSG00 Pigu | 6169 | 16.620098 | 14.7941516 | 17.0673999 | 13.8327177 | 16.0051933 | 11.7516601 |
| ENSMUSG00 Setd1b | 10771 | 0.15871188 | 0.13421161 | 0.16415131 | 0.4388402 | 0.16580848 | 0.27094851 |
| ENSMUSG00 Rras | 1096 | 7.61738221 | 14.596623 | 13.4702697 | 6.38678525 | 9.31138245 | 10.2031041 |
| ENSMUSG00 Mpp6 | 3041 | 12.5502339 | 20.0446726 | 13.6050395 | 17.2045263 | 13.423578 | 14.0543768 |
| ENSMUSG00 Gpr162 | 3934 | 0.15158419 | 0.15923329 | 0.10112267 | 0.46776418 | 0.18158853 | 0.81810052 |
| ENSMUSG00 Txnip | 2740 | 325.980431 | 371.86216 | 194.149336 | 267.138816 | 329.101501 | 185.457593 |
| ENSMUSG00 Upf3a | 1393 | 12.2434323 | 16.0506168 | 15.1993416 | 12.9771029 | 14.4690728 | 11.062578 |
| ENSMUSG00 Pmepa1 | 5462 | 15.7289625 | 17.0002333 | 12.2522135 | 18.8007138 | 11.5748223 | 19.7194344 |
| ENSMUSG00 Foxf2 | 2361 | 0 | 0 | 0 | 0 | 0 | 0 |
| ENSMUSG00 Hfe2 | 2017 | 0 | 0 | 0 | 0 | 0 | 0 |
| ENSMUSG00 Scaf1 | 8149 | 1.75140723 | 2.57223001 | 2.0232283 | 3.14372949 | 2.11644558 | 2.36967046 |
| ENSMUSG00 1700018A04f | 2062 | 0.19280058 | 0 | 0.36441869 | 0 | 0.19796848 | 0.27777276 |
| ENSMUSG00 Higd1a | 1869 | 71.9810305 | 61.1805389 | 65.9125396 | 49.8855019 | 39.9965974 | 52.3457095 |
| ENSMUSG00 Foxq1 | 2800 | 17.5350066 | 14.9033782 | 19.0067925 | 14.9353891 | 15.6541632 | 15.0010516 |
| ENSMUSG00 Cdc16 | 3174 | 41.1332697 | 35.4035233 | 41.1798519 | 28.9429381 | 38.6154237 | 38.2223048 |
| ENSMUSG00 Fig4 | 3240 | 20.5157912 | 13.0133139 | 17.926303 | 21.6603632 | 19.7333484 | 13.2753414 |
| ENSMUSG00 Egr1 | 5024 | 30.418007 | 69.0570031 | 16.954011 | 56.0980991 | 48.0606709 | 23.4147132 |
| ENSMUSG00 Fcrla | 2259 | 0 | 0 | 0 | 0 | 0 | 0 |
| ENSMUSG00 Hdhd3 | 1513 | 8.30319355 | 2.8344966 | 8.58912217 | 4.7457547 | 2.46194675 | 6.0029444 |
| ENSMUSG00 Poli | 4794 | 4.44491814 | 4.00045974 | 3.27318612 | 3.56003485 | 5.01322966 | 5.71208548 |
| ENSMUSG00 Usp5 | 5222 | 31.2897405 | 24.4254141 | 20.0525021 | 19.215658 | 31.0438139 | 18.4320571 |
| ENSMUSG00 Mll6 | 7539 | 2.93723341 | 1.91748676 | 3.40059278 | 1.65597392 | 2.78854846 | 3.88190581 |
| ENSMUSG00 Cdc40 | 2307 | 18.3871251 | 17.5451285 | 20.9800685 | 25.2589952 | 14.1555616 | 18.1476252 |
| ENSMUSG00 Spab2 | 1607 | 8.55967409 | 6.02705344 | 10.0396221 | 7.00535028 | 9.04948171 | 10.6586643 |
| ENSMUSG00 Srcin1 | 8350 | 0.19044542 | 0.46166651 | 0.25938806 | 0.7389257 | 0.32999093 | 0.47689788 |
| ENSMUSG00 Dennd2a | 5607 | 0 | 0 | 0 | 0 | 0 | 0 |
| ENSMUSG00 Fam70b | 1464 | 0 | 0 | 0.5736578 | 0 | 0 | 0 |
| ENSMUSG00 Fam108c | 3851 | 56.1800376 | 56.8076986 | 43.6508982 | 52.2819549 | 46.7598235 | 43.380204 |
| ENSMUSG00 Uqcrfs1 | 1361 | 586.575903 | 482.60793 | 679.8185 | 482.295778 | 354.410236 | 587.29653 |
| ENSMUSG00 Olfml2b | 3087 | 0.24468874 | 0 | 0 | 0 | 0 | 0.95421478 |
| ENSMUSG00 Chmp4b | 2021 | 5.82267305 | 5.19774586 | 2.9307506 | 5.26680017 | 3.83770862 | 5.95156651 |
| ENSMUSG00 Nos1ap | 7235 | 4.29699872 | 5.28152711 | 3.65344881 | 4.5383083 | 5.17669106 | 6.14857452 |
| ENSMUSG00 Cdk19 | 5794 | 5.26962531 | 3.75910802 | 4.30269925 | 4.97576002 | 8.31358281 | 4.97100778 |
| ENSMUSG00 Tfdp1 | 2529 | 6.25649713 | 7.94533269 | 6.39694525 | 10.3152701 | 9.82595691 | 5.4355153 |
| ENSMUSG00 Socs7 | 7254 | 0.88235902 | 0.98976839 | 1.52336101 | 1.01968833 | 0.74562942 | 1.06782431 |
| ENSMUSG00 Sv2a | 4623 | 8.4705058 | 9.28706899 | 10.5747866 | 8.95221827 | 11.412778 | 13.1623881 |
| ENSMUSG00 Polr2l | 1648 | 9.7941293 | 7.36831518 | 7.10769927 | 6.59022779 | 3.6845502 | 10.4265915 |
| ENSMUSG00 4930520K10f | 1612 | 0 | 0 | 0.08226149 | 0.24621743 | 0.06330816 | 0 |
| ENSMUSG00 Otud7b | 8753 | 4.15586252 | 3.58384255 | 3.41373768 | 3.99858142 | 2.6641213 | 3.83272093 |
| ENSMUSG00 Slc19a3 | 4540 | 1.49739805 | 2.00599946 | 1.96668993 | 2.06637259 | 1.34871478 | 1.82031183 |
| ENSMUSG00 Tmco3 | 4561 | 25.3298457 | 22.3764272 | 21.6115128 | 23.9307574 | 24.4671528 | 16.8395931 |
| ENSMUSG00 Catsper1 | 2049 | 0 | 0 | 0 | 0 | 0 | 0 |
| ENSMUSG00 Prr3 | 3533 | 4.28724542 | 4.10532666 | 3.315451 | 3.72769495 | 5.22828985 | 2.83322746 |
| ENSMUSG00 Ptov1 | 3635 | 54.6842923 | 40.74969 | 44.372084 | 29.1336453 | 42.7161655 | 37.1865508 |
| ENSMUSG00 Mesdc2 | 4840 | 22.1037186 | 24.1230887 | 26.3110557 | 25.6450895 | 20.9376823 | 24.3499414 |
| ENSMUSG00 Dcun1d2 | 3027 | 20.5672555 | 23.4166688 | 22.560901 | 21.9925247 | 16.3007947 | 18.2011137 |
| ENSMUSG00 Parp12 | 3983 | 8.41422799 | 9.82359605 | 11.652507 | 10.0464442 | 17.1283617 | 8.89524388 |

| | | | | | | | | |
|---|---|---|---|---|---|---|---|---|
| ENSMUSG00 | Gdf15 | 2629 | 9.84435824 | 56.5993661 | 0.20175813 | 26.3650086 | 15.8960068 | 1.64955026 |
| ENSMUSG00 | Rpf2 | 1820 | 11.0310537 | 22.8488459 | 19.5265285 | 17.822944 | 11.4949528 | 21.2502409 |
| ENSMUSG00 | Grtp1 | 3081 | 4.3484573 | 4.87963971 | 4.84914734 | 3.73585596 | 3.41169537 | 2.99215452 |
| ENSMUSG00 | Tbkbp1 | 5463 | 0 | 0 | 0 | 0.01981441 | 0.12142465 | 0.04992612 |
| ENSMUSG00 | Jarid2 | 7988 | 4.98187732 | 8.74691301 | 5.45606124 | 7.38534587 | 11.6834303 | 7.8771398 |
| ENSMUSG00 | Tbc1d17 | 3828 | 21.5394113 | 19.4986413 | 15.7154411 | 14.7231331 | 26.1796766 | 15.8175808 |
| ENSMUSG00 | C1s | 4862 | 0 | 0 | 0 | 0 | 0.20989871 | 0 |
| ENSMUSG00 | AI317395 | 4733 | 13.1034333 | 48.0235469 | 1.81178059 | 37.4085342 | 20.7965094 | 3.59013316 |
| ENSMUSG00 | 1700003F12f | 973 | 0 | 0 | 0 | 0 | 0 | 0 |
| ENSMUSG00 | Fchsd1 | 7519 | 0.56045763 | 1.09587467 | 0.5173244 | 0.48467702 | 0.84829059 | 0.88146527 |
| ENSMUSG00 | Armc10 | 3757 | 9.05794286 | 8.5547929 | 7.6003169 | 7.23177523 | 8.66511253 | 11.7679506 |
| ENSMUSG00 | Car14 | 2530 | 1.22566304 | 1.14276147 | 2.20135661 | 0.41358865 | 3.5294924 | 0.78697576 |
| ENSMUSG00 | C1rl | 3004 | 0.6352407 | 0.46518203 | 0.05885731 | 0.36033999 | 0.25479216 | 0.01815888 |
| ENSMUSG00 | BC021785 | 6014 | 0.73376427 | 1.30601764 | 0.04409895 | 1.26593139 | 1.13693621 | 0.88436226 |
| ENSMUSG00 | Rgs4 | 5338 | 0 | 0 | 0 | 0 | 0.21985887 | 0 |
| ENSMUSG00 | Cbfa2t2 | 8789 | 6.39597786 | 5.4442072 | 6.45249341 | 7.24596753 | 6.84493083 | 6.2499856 |
| ENSMUSG00 | Osbpl7 | 4980 | 2.11550248 | 4.05424082 | 2.71601686 | 2.30403417 | 4.00628774 | 2.38242331 |
| ENSMUSG00 | Zfp280d | 4452 | 13.9572812 | 8.96189894 | 16.6203695 | 27.0696381 | 13.6849714 | 17.7542463 |
| ENSMUSG00 | Mc3r | 2622 | 0 | 0 | 0 | 0 | 0 | 0 |
| ENSMUSG00 | Ubn2 | 19377 | 5.57647706 | 4.78954882 | 5.54775673 | 7.64953496 | 6.61233574 | 7.10686141 |
| ENSMUSG00 | Atf5 | 1988 | 0.41995226 | 0.31510249 | 0.28904626 | 0.07259969 | 0.69301416 | 0.90549607 |
| ENSMUSG00 | Tmc3 | 9253 | 0.42964963 | 0.01562297 | 0.44426363 | 0.2534673 | 0.52939934 | 0.15327799 |
| ENSMUSG00 | Srd5a2 | 849 | 0 | 0 | 0 | 0 | 0 | 0 |
| ENSMUSG00 | Pcid2 | 4078 | 10.9576165 | 9.47658818 | 10.9691675 | 10.1397801 | 10.310381 | 6.04616819 |
| ENSMUSG00 | BC028528 | 1675 | 0.71203846 | 0.77673668 | 0.02638916 | 0.36620583 | 0.3960256 | 0.50478437 |
| ENSMUSG00 | 1110054O05i | 7150 | 10.9146865 | 11.2547354 | 8.53126481 | 8.00365351 | 7.3078334 | 8.73551383 |
| ENSMUSG00 | Cul7 | 6558 | 9.34781198 | 10.1619167 | 10.5146196 | 11.108516 | 13.1650851 | 5.16546233 |
| ENSMUSG00 | Ranbp9 | 3029 | 25.4755324 | 25.7238286 | 28.6020509 | 28.9347264 | 33.3549766 | 25.9419728 |
| ENSMUSG00 | Gm129 | 2818 | 0.67716929 | 0.32489085 | 0.2980252 | 0.89628925 | 0.54321905 | 0.522651 |
| ENSMUSG00 | Fndc4 | 2604 | 1.25190069 | 1.79496348 | 0.64503458 | 1.13622414 | 1.09734142 | 1.30926648 |
| ENSMUSG00 | Reep2 | 1838 | 0.28118675 | 0.70785307 | 0 | 0 | 0 | 0.26710746 |
| ENSMUSG00 | Sp6 | 3838 | 0 | 0 | 0 | 0 | 0 | 0 |
| ENSMUSG00 | Eftud1 | 4397 | 6.86249937 | 7.24385678 | 5.69989652 | 6.19557508 | 5.39623943 | 8.99436591 |
| ENSMUSG00 | Ift172 | 5359 | 7.44070665 | 9.31538609 | 4.68494996 | 16.3948084 | 15.3298125 | 8.37731969 |
| ENSMUSG00 | Cyp24a1 | 3548 | 0 | 0 | 0 | 0 | 0 | 0.07687328 |
| ENSMUSG00 | Rad9b | 2907 | 1.9829875 | 2.3703685 | 1.39888874 | 1.34050939 | 3.44037484 | 1.1165057 |
| ENSMUSG00 | Fam154b | 4881 | 2.76928158 | 1.18466975 | 0.9055899 | 1.60413925 | 1.80855624 | 1.67078819 |
| ENSMUSG00 | Bpifb5 | 1622 | 0 | 0 | 0.13625722 | 0.06673621 | 0 | 0 |
| ENSMUSG00 | Susd4 | 3624 | 8.83089454 | 11.8737563 | 1.9271223 | 10.0061961 | 7.13862377 | 2.95023692 |
| ENSMUSG00 | Susd1 | 5403 | 21.4118912 | 14.0376568 | 16.0838096 | 16.6419498 | 18.1987463 | 10.2677432 |
| ENSMUSG00 | Sct | 697 | 0 | 0.27653625 | 0 | 0 | 0 | 0 |
| ENSMUSG00 | Pptc7 | 4712 | 23.0669533 | 16.9756991 | 22.0633987 | 21.2265338 | 17.976185 | 19.1304505 |
| ENSMUSG00 | Pln | 2477 | 8.79531826 | 11.6137691 | 6.17433899 | 13.4743202 | 6.40662208 | 3.99705048 |
| ENSMUSG00 | Akap12 | 6240 | 1.52268587 | 0.43244243 | 0.45335224 | 0.32381322 | 1.01398567 | 0.62504382 |
| ENSMUSG00 | Colec10 | 4581 | 0 | 0 | 0 | 0 | 0.23391266 | 0 |
| ENSMUSG00 | Tctn1 | 5362 | 0.90878472 | 1.51224402 | 1.63551957 | 1.55176557 | 1.30482015 | 1.79118611 |
| ENSMUSG00 | Gm9766 | 1338 | 0.83195325 | 0.39615162 | 0.9250012 | 1.15958738 | 1.67799741 | 0.16307707 |
| ENSMUSG00 | AI481877 | 5753 | 0 | 0 | 0 | 0 | 0 | 0 |
| ENSMUSG00 | Capn8 | 3683 | 0 | 0 | 0 | 0 | 0 | 0 |
| ENSMUSG00 | Atp6v0a4 | 5065 | 86.7477928 | 72.3415017 | 121.408892 | 91.4695857 | 82.4279974 | 103.368998 |
| ENSMUSG00 | Slc35f1 | 4933 | 1.97447653 | 1.24055912 | 1.85481077 | 2.95502655 | 2.69975352 | 1.54259565 |
| ENSMUSG00 | Fam65a | 4087 | 19.4740573 | 22.0240455 | 9.76615225 | 24.9493168 | 23.6841289 | 16.8839827 |
| ENSMUSG00 | Samd10 | 3537 | 20.7825512 | 20.6805256 | 22.3071218 | 19.7191383 | 16.3596014 | 16.2938402 |
| ENSMUSG00 | Gng10 | 1310 | 0.42486773 | 0.44140252 | 0.43864424 | 0.16526127 | 0.46741718 | 0.29148469 |
| ENSMUSG00 | Dock10 | 8088 | 0.16220708 | 0.35746619 | 0.09837205 | 0.16506376 | 0.7002878 | 0.04046682 |
| ENSMUSG00 | Phrf1 | 6672 | 13.2041584 | 15.8888149 | 11.7659582 | 18.6413081 | 17.1846922 | 8.92394147 |
| ENSMUSG00 | Mcl1 | 3481 | 55.6759458 | 60.7694573 | 46.6652609 | 67.8831685 | 55.2187181 | 57.7460762 |
| ENSMUSG00 | Nfe2l1 | 8248 | 57.7919752 | 50.196158 | 36.7794918 | 41.4322313 | 60.5288629 | 24.6854003 |
| ENSMUSG00 | Rassf7 | 1860 | 46.103533 | 50.5957643 | 45.8891177 | 38.0995352 | 47.4325827 | 45.4723959 |
| ENSMUSG00 | Ensa | 2943 | 40.3363455 | 33.7616187 | 44.1718044 | 36.5724563 | 38.3868148 | 33.3820084 |
| ENSMUSG00 | Med30 | 867 | 25.6782804 | 32.6800784 | 34.2092695 | 33.8763371 | 24.3655359 | 43.2870861 |
| ENSMUSG00 | Tm6sf1 | 6390 | 0.11820878 | 0.22622743 | 0.17293366 | 0.16375263 | 0.33538463 | 0.07256164 |
| ENSMUSG00 | Nepn | 1977 | 0 | 0.02437352 | 0 | 0 | 0 | 0 |
| ENSMUSG00 | Polr3k | 3330 | 23.6145167 | 23.9051058 | 30.264325 | 29.2665472 | 23.9348946 | 38.4629748 |
| ENSMUSG00 | Zkscan16 | 2788 | 0.04277849 | 0 | 0.12683456 | 0.2588382 | 0.07320857 | 0 |
| ENSMUSG00 | Degs1 | 2587 | 18.7789707 | 28.1444585 | 18.7434951 | 20.4190619 | 18.284287 | 16.4153898 |
| ENSMUSG00 | Lrrc56 | 5806 | 2.39655842 | 1.95866351 | 1.1267435 | 1.16213268 | 2.68051061 | 1.13213711 |
| ENSMUSG00 | Akr1d1 | 3019 | 0.30287382 | 0.31922121 | 0 | 0 | 0 | 0.53302541 |
| ENSMUSG00 | Ctss | 1350 | 0 | 0.1784683 | 0 | 0.80182321 | 1.28510872 | 1.89912303 |
| ENSMUSG00 | Pold1 | 4626 | 2.29457702 | 3.03118346 | 2.41743759 | 5.58468307 | 5.38280834 | 1.2853159 |
| ENSMUSG00 | Fam103a1 | 1743 | 23.5385289 | 26.3739907 | 21.4796105 | 27.4289782 | 20.6681706 | 23.8790014 |
| ENSMUSG00 | Creb3l2 | 7270 | 18.0731586 | 17.206603 | 14.8839218 | 17.857386 | 14.8306372 | 13.7498697 |
| ENSMUSG00 | Rnh1 | 1900 | 97.1079921 | 90.0579242 | 100.407976 | 68.7267995 | 71.8934779 | 54.4918582 |
| ENSMUSG00 | Sycp2l | 3005 | 0 | 0 | 0 | 0 | 0 | 0 |
| ENSMUSG00 | Cyp3a16 | 1717 | 0 | 0 | 0 | 0 | 0 | 0 |
| ENSMUSG00 | C030046E11l | 10099 | 2.8815736 | 3.82666864 | 2.39413884 | 2.56886722 | 4.09767213 | 5.06656335 |
| ENSMUSG00 | Fad2 | 3972 | 0.33029478 | 0.30328828 | 0 | 0.51779369 | 0 | 0.13733454 |
| ENSMUSG00 | Herc1 | 25887 | 5.33973831 | 5.23057529 | 4.64437799 | 5.67985211 | 6.92257241 | 4.50626307 |
| ENSMUSG00 | Dgki | 6714 | 0 | 0 | 0 | 0 | 0 | 0 |
| ENSMUSG00 | Lpar1 | 4280 | 3.00023837 | 3.14112555 | 1.18766634 | 1.60177301 | 2.22942343 | 0.72647404 |
| ENSMUSG00 | Mybpc2 | 3584 | 0.24403476 | 0 | 0 | 0 | 0 | 0.068491 |
| ENSMUSG00 | Arfrp1 | 5038 | 21.7400454 | 17.417132 | 23.8732066 | 22.975625 | 17.3092648 | 23.885611 |
| ENSMUSG00 | Ucn | 502 | 0 | 0 | 0 | 0 | 0 | 0 |
| ENSMUSG00 | Scube3 | 8721 | 2.39781937 | 3.80143013 | 1.37861499 | 3.27266337 | 5.13130737 | 1.77952869 |
| ENSMUSG00 | Trps1 | 16609 | 1.90770775 | 2.16141245 | 0.95009079 | 1.37515408 | 2.20584851 | 1.90326371 |
| ENSMUSG00 | Pak1ip1 | 4099 | 15.9545659 | 12.5550427 | 16.8331488 | 15.4310378 | 14.2535253 | 12.8088999 |
| ENSMUSG00 | Rtel1 | 5876 | 4.039146 | 7.40509824 | 2.91870577 | 6.55199821 | 8.68386246 | 4.04292219 |
| ENSMUSG00 | Atp5j2 | 1387 | 39.8128064 | 42.4193553 | 57.2998657 | 39.7760967 | 30.0934214 | 95.0187865 |
| ENSMUSG00 | Mbd3l1 | 841 | 0 | 0 | 0 | 0 | 0 | 0 |
| ENSMUSG00 | Hoxb4 | 4147 | 0.84361762 | 0.72041462 | 0.79940637 | 1.13109857 | 0.81209086 | 2.11120309 |
| ENSMUSG00 | Josd2 | 2024 | 10.8424038 | 13.9750205 | 9.21599691 | 14.6360467 | 14.5213402 | 8.36835522 |
| ENSMUSG00 | Mapkap1 | 4040 | 28.0945289 | 24.6299512 | 24.1797211 | 25.5075047 | 25.5131879 | 25.0265069 |
| ENSMUSG00 | Taf5l | 3027 | 13.2386932 | 12.0187525 | 16.0043673 | 15.5556882 | 13.5868051 | 13.7950025 |
| ENSMUSG00 | Hoxb5 | 2810 | 1.57041222 | 2.12637681 | 3.0359273 | 3.89069733 | 2.7964633 | 4.50371267 |
| ENSMUSG00 | Dsel | 5752 | 1.83157203 | 4.95100614 | 1.42165177 | 5.01209207 | 3.43310282 | 1.39882103 |
| ENSMUSG00 | Aspdh | 1182 | 0 | 0.52996933 | 0.18697903 | 0.09157879 | 0.04316952 | 0.20767492 |
| ENSMUSG00 | Gmeb2 | 5427 | 1.34789173 | 1.28745792 | 1.22986517 | 2.20734085 | 1.80524492 | 2.08065241 |
| ENSMUSG00 | Golga4 | 7857 | 28.057037 | 23.5933872 | 28.0164412 | 26.7825485 | 27.2374469 | 21.2899787 |
| ENSMUSG00 | Txndc8 | 879 | 0 | 0 | 0 | 0 | 0 | 0 |
| ENSMUSG00 | Fam63a | 4665 | 147.056929 | 106.877839 | 198.524548 | 109.011862 | 129.463705 | 138.922337 |
| ENSMUSG00 | Atp5l | 508 | 6.26070559 | 5.78616727 | 8.8751732 | 5.96632232 | 3.51559676 | 7.83877427 |
| ENSMUSG00 | Pbx3 | 7455 | 2.57570719 | 2.12653782 | 2.69183591 | 1.50039354 | 1.58109897 | 1.82928499 |
| ENSMUSG00 | Hoxb7 | 4081 | 12.4984762 | 13.4841747 | 12.1200349 | 9.43384992 | 10.6153867 | 15.3315419 |
| ENSMUSG00 | Bud31 | 1649 | 13.8143665 | 18.3219522 | 20.3987887 | 14.2446398 | 15.4409712 | 19.0376651 |
| ENSMUSG00 | Pkhd1l1 | 12676 | 0 | 0 | 0 | 0 | 0 | 0.01075838 |
| ENSMUSG00 | Akap2 | 8024 | 0.38645657 | 0.25222216 | 0.6500271 | 0.3372574 | 0.31796097 | 0.2719306 |
| ENSMUSG00 | Mboat1 | 3691 | 36.0502827 | 40.7319695 | 45.6508875 | 38.3109002 | 42.9390201 | 33.733061 |
| ENSMUSG00 | Wdr26 | 12857 | 8.46006025 | 8.41398169 | 7.99325543 | 11.3407126 | 6.96517769 | 10.4541823 |
| ENSMUSG00 | Nudcd1 | 5795 | 4.28083172 | 4.60660724 | 6.28512833 | 7.2101825 | 4.24412651 | 5.02682894 |
| ENSMUSG00 | Shank1 | 8267 | 0.01442681 | 0.01748631 | 0.02673391 | 0.03055211 | 0.05555067 | 0.11547265 |
| ENSMUSG00 | Fam125b | 5280 | 0.48188461 | 0.57495187 | 0.67809646 | 0.44419184 | 0.19328173 | 0.53722774 |
| ENSMUSG00 | Angptl6 | 1575 | 0.73200567 | 1.46853919 | 0.58935791 | 1.2600079 | 0.51836318 | 0.64073756 |
| ENSMUSG00 | Nlrp6 | 3260 | 0 | 0.65036916 | 0 | 0 | 1.14261516 | 0 |
| ENSMUSG00 | Omt2b | 865 | 0 | 0 | 0 | 0 | 0 | 0 |
| ENSMUSG00 | Ptk6 | 3681 | 0 | 0 | 0 | 0 | 0 | 0 |
| ENSMUSG00 | Elovl3 | 1877 | 0 | 0 | 0 | 0 | 0 | 0 |
| ENSMUSG00 | Ttll6 | 3842 | 0.15521401 | 0 | 0.02300981 | 0 | 0.15937442 | 0 |
| ENSMUSG00 | Nup205 | 6377 | 5.63571497 | 3.49856088 | 6.14819425 | 5.6638116 | 4.15284445 | 2.70308484 |
| ENSMUSG00 | Trhr | 2378 | 0 | 0 | 0 | 0 | 0 | 0 |
| ENSMUSG00 | Abcf1 | 4562 | 19.7034506 | 20.1533826 | 17.5276488 | 20.2160688 | 21.2069287 | 12.4834386 |
| ENSMUSG00 | Alpk3 | 6428 | 0 | 0 | 0 | 0 | 0 | 0 |
| ENSMUSG00 | Ptprd | 7212 | 2.91606339 | 5.15137921 | 1.66094002 | 3.18694708 | 3.16261647 | 2.68132546 |
| ENSMUSG00 | Lmx1b | 5323 | 0 | 0 | 0 | 0.00677852 | 0 | 0 |
| ENSMUSG00 | Gabpb2 | 21542 | 2.3529959 | 2.51423085 | 1.99238648 | 3.54254592 | 2.61503662 | 2.33598131 |
| ENSMUSG00 | 9130409I23R | 1895 | 0 | 0 | 0 | 0.93299218 | 0 | 0.01439295 |
| ENSMUSG00 | Kpna7 | 3606 | 0 | 0.01336285 | 0 | 0 | 0 | 0.00756368 |
| ENSMUSG00 | Kdm3b | 6604 | 5.27946039 | 7.1433263 | 7.98497861 | 6.63835313 | 8.01246998 | 5.76550521 |
| ENSMUSG00 | Ascc3 | 7435 | 12.8115846 | 10.5640754 | 11.164904 | 16.4031352 | 12.1131881 | 8.77484023 |
| ENSMUSG00 | Vill | 4415 | 2.9895401 | 3.54713334 | 3.05358145 | 3.08107153 | 4.25316111 | 0.88341414 |
| ENSMUSG00 | Ephx1 | 1994 | 62.9029794 | 86.9724199 | 58.012148 | 58.2126064 | 61.3647188 | 35.3311902 |
| ENSMUSG00 | Sema6c | 5275 | 0.09797559 | 0.10961845 | 0.2262464 | 0.11628337 | 0.03869299 | 0.13443424 |
| ENSMUSG00 | Smurf1 | 9272 | 4.92227045 | 5.2645455 | 5.76835958 | 5.52983016 | 6.59292474 | 4.92720241 |
| ENSMUSG00 | Stap2 | 1478 | 35.2365896 | 37.9166659 | 40.4037144 | 34.373152 | 49.3692269 | 38.6052404 |
| ENSMUSG00 | 1700028J19R | 1240 | 16.1907401 | 12.9403913 | 11.9416269 | 14.4910147 | 9.01191637 | 13.0214407 |
| ENSMUSG00 | Cnot4 | 7172 | 8.72492001 | 10.3938129 | 8.67149932 | 11.4756196 | 9.75420542 | 14.1393069 |
| ENSMUSG00 | Scgb3a2 | 981 | 0 | 0 | 0 | 0 | 0 | 0 |
| ENSMUSG00 | 4931403M11 | 984 | 0 | 0 | 0 | 0 | 0 | 0 |
| ENSMUSG00 | Lefty1 | 1754 | 0.40798099 | 0.1648339 | 0.90722141 | 0.74056648 | 0.32000578 | 0.57534872 |
| ENSMUSG00 | Zscan2 | 2832 | 0.23864519 | 0.40835968 | 0.42141588 | 0.10192668 | 0.19819567 | 0.30818801 |
| ENSMUSG00 | Scgb1c1 | 433 | 0 | 0 | 0 | 0 | 0 | 0 |
| ENSMUSG00 | Ost4 | 1518 | 11.2876233 | 10.9514641 | 14.2098151 | 12.8117405 | 7.22705586 | 20.0337436 |
| ENSMUSG00 | Six3 | 7722 | 0 | 0 | 0 | 0 | 0 | 0 |
| ENSMUSG00 | BC031781 | 3166 | 12.6951329 | 14.6720563 | 17.284233 | 13.7786456 | 11.6042295 | 15.6273517 |
| ENSMUSG00 | Rap1gap2 | 10610 | 9.40492517 | 7.59810723 | 12.2148731 | 8.41007503 | 11.9366131 | 9.15924031 |
| ENSMUSG00 | Gngt2 | 1655 | 0.04804288 | 0.05823135 | 0.16024837 | 0.28342392 | 0.03083165 | 0.47792419 |
| ENSMUSG00 | Trmt112 | 1238 | 0.93126731 | 1.05091594 | 1.92802869 | 0.43718148 | 0.86555242 | 0.83718602 |
| ENSMUSG00 | Ctnnal1 | 7558 | 13.6656177 | 17.0354821 | 8.78422443 | 15.6731083 | 26.8769519 | 14.6044542 |
| ENSMUSG00 | Hace1 | 3979 | 5.46525455 | 6.63638359 | 5.48773824 | 8.30639678 | 5.61687676 | 8.07477382 |
| ENSMUSG00 | BC026590 | 6084 | 7.97200628 | 7.92811121 | 6.40069423 | 5.2248983 | 8.36181734 | 7.16833468 |
| ENSMUSG00 | Tmem214 | 6058 | 18.8802439 | 17.5310199 | 14.5709938 | 18.0052857 | 17.7640519 | 8.36968544 |
| ENSMUSG00 | Ralgps1 | 8369 | 3.95702178 | 4.84225091 | 4.61613225 | 3.04815456 | 5.77392495 | 4.35077589 |
| ENSMUSG00 | Agbl3 | 10632 | 2.66980936 | 2.71026076 | 3.16796505 | 1.7240104 | 1.99171802 | 3.05274837 |
| ENSMUSG00 | Vars2 | 6300 | 3.66633876 | 3.03651072 | 2.75735306 | 2.74338085 | 2.94009118 | 1.75336967 |
| ENSMUSG00 | Birc7 | 1526 | 0 | 0 | 0 | 0 | 0 | 0 |
| ENSMUSG00 | Gcnt1 | 9924 | 7.99195724 | 4.03010348 | 17.0990402 | 6.66085304 | 9.52759718 | 14.7256668 |
| ENSMUSG00 | Kif16b | 8180 | 16.4805421 | 12.8006282 | 12.7255917 | 20.3523904 | 20.2670776 | 12.2202387 |
| ENSMUSG00 | Phb | 2745 | 23.5781138 | 19.6958791 | 16.8111928 | 15.2477857 | 17.36198 | 15.3512997 |
| ENSMUSG00 | Ythdf1 | 3608 | 22.3789858 | 17.5624089 | 23.6078025 | 27.4215538 | 21.4966994 | 19.2993775 |
| ENSMUSG00 | Itpkb | 6541 | 1.30674642 | 2.21004933 | 0.90552621 | 1.37355589 | 2.66014282 | 0.62129969 |
| ENSMUSG00 | Baiap2l1 | 3351 | 17.0719596 | 17.5144991 | 14.3118471 | 21.2228319 | 16.8412927 | 13.5518575 |
| ENSMUSG00 | Garnl3 | 10925 | 0.02183367 | 0.25140752 | 0.12137806 | 0.01651352 | 0.18215365 | 0.08238564 |
| ENSMUSG00 | Pi4kb | 4120 | 5.78963308 | 3.68415759 | 4.14124063 | 4.39640445 | 4.9911722 | 4.26993746 |
| ENSMUSG00 | Zcchc2 | 7763 | 3.51310829 | 4.66780942 | 3.1772032 | 3.65328958 | 5.63308052 | 2.61398063 |
| ENSMUSG00 | Bpgm | 3624 | 12.4510128 | 13.3363691 | 9.53803566 | 8.38330063 | 12.8129145 | 9.59579609 |
| ENSMUSG00 | Zfhx3 | 16429 | 2.75619285 | 1.60142415 | 3.54873887 | 3.05277509 | 2.2641809 | 4.47743028 |
| ENSMUSG00 | Rnf146 | 6161 | 13.0603948 | 18.301619 | 17.4339358 | 16.5505369 | 15.7029717 | 14.7418518 |
| ENSMUSG00 | Nipal2 | 3814 | 31.1977329 | 26.4431629 | 45.5577989 | 26.5837822 | 21.7136361 | 31.0290141 |
| ENSMUSG00 | Mrps34 | 2424 | 4.7398242 | 4.87032934 | 5.85346188 | 3.43851167 | 3.26282519 | 4.92833828 |
| ENSMUSG00 | Prl3a1 | 1330 | 0 | 0 | 0 | 0 | 0 | 0 |
| ENSMUSG00 | A230050P20f | 3012 | 1.96665558 | 0.83190405 | 1.23272072 | 0.61095095 | 3.37126454 | 2.1642227 |
| ENSMUSG00 | Man2a2 | 5188 | 3.14181708 | 4.62545262 | 2.75196517 | 2.7889167 | 4.01286841 | 0.97784944 |
| ENSMUSG00 | Ctu1 | 2472 | 2.79832266 | 3.93756526 | 3.6119629 | 3.1819926 | 7.12139964 | 3.34312933 |
| ENSMUSG00 | Prl3b1 | 870 | 0 | 0 | 0 | 0 | 0 | 0 |
| ENSMUSG00 | Fam117a | 3089 | 1.9891125 | 2.49589859 | 2.01763026 | 2.41792917 | 2.6099603 | 2.66653967 |
| ENSMUSG00 | Irs2 | 4022 | 10.8729559 | 9.92003334 | 9.49537354 | 11.0973369 | 14.9704434 | 10.6670829 |
| ENSMUSG00 | Zfp653 | 2099 | 6.36390731 | 4.84389676 | 5.11722146 | 7.95902173 | 7.00123689 | 8.22527234 |
| ENSMUSG00 | Rpl12 | 2471 | 0.80444113 | 1.07254323 | 1.19851213 | 0.68630356 | 0.4749521 | 1.4128506 |
| ENSMUSG00 | Pogz | 7405 | 5.78212728 | 5.14726208 | 4.81116617 | 6.36369347 | 5.4644046 | 8.00743629 |
| ENSMUSG00 | Ccdc68 | 2495 | 4.62087749 | 4.19096511 | 4.73021727 | 4.00590234 | 5.39918365 | 4.83181986 |
| ENSMUSG00 | Myst2 | 6083 | 19.9332921 | 20.3661586 | 24.9893372 | 17.1779716 | 21.3483686 | 16.8185389 |
| ENSMUSG00 | Plcl2 | 4044 | 0.35390636 | 0 | 0 | 0 | 0.25235596 | 0 |
| ENSMUSG00 | Dido1 | 13289 | 4.19722621 | 4.99668277 | 4.84627024 | 5.59327351 | 6.589003 | 5.78167347 |
| ENSMUSG00 | 6330407J23R | 3941 | 0 | 0 | 0 | 0 | 0 | 0 |
| ENSMUSG00 | 3930402G23 | 2844 | 0 | 0 | 0 | 0 | 0.21530116 | 0.00959024 |
| ENSMUSG00 | E330034G19 | 1887 | 0 | 0 | 0 | 0 | 0 | 0 |
| ENSMUSG00 | Rccd1 | 3599 | 3.24759969 | 3.4810989 | 3.23008744 | 2.79713543 | 3.70044015 | 4.25147892 |
| ENSMUSG00 | Tcf15 | 3912 | 0 | 0 | 0 | 0 | 0 | 0 |
| ENSMUSG00 | Sccpdh | 3444 | 17.465169 | 18.8604309 | 16.7489562 | 21.1840575 | 18.1051775 | 16.432891 |
| ENSMUSG00 | Prc1 | 6768 | 0.58740367 | 1.00388421 | 0.56166645 | 0.2612323 | 0.69362095 | 0.51583242 |
| ENSMUSG00 | Cnst | 8964 | 5.71673521 | 5.50456456 | 5.54742006 | 6.56512881 | 7.58792492 | 3.41389394 |
| ENSMUSG00 | Supt3h | 4212 | 2.83158693 | 3.80961188 | 3.04333677 | 3.52082628 | 3.04074557 | 3.45789586 |
| ENSMUSG00 | Edc3 | 5940 | 13.4258407 | 9.39392256 | 14.3097886 | 12.0820179 | 13.9850067 | 8.70125276 |
| ENSMUSG00 | Slco4a1 | 4384 | 20.9478011 | 10.4638442 | 12.3410255 | 9.77770732 | 15.1891926 | 14.4958735 |
| ENSMUSG00 | Ube2l3 | 6816 | 10.6737866 | 8.78752165 | 10.3695345 | 8.88816795 | 8.48942344 | 9.53972124 |
| ENSMUSG00 | Pdk2 | 3519 | 79.5561505 | 78.5716979 | 53.3085029 | 48.6938418 | 78.4318462 | 52.5728554 |
| ENSMUSG00 | Klk1b16 | 1379 | 1.81624023 | 0.76874672 | 0.28848193 | 0.75879572 | 0.55503672 | 0.39557127 |
| ENSMUSG00 | Lmtk2 | 8100 | 5.41852475 | 5.11014245 | 5.78989573 | 7.06940798 | 8.54219329 | 3.7847771 |
| ENSMUSG00 | Cldnd2 | 653 | 0.18264386 | 0.44275445 | 0.27076167 | 0 | 0 | 0 |
| ENSMUSG00 | Rabggtb | 2056 | 29.9306373 | 22.4995061 | 29.0665816 | 29.9396731 | 26.7292834 | 18.4395664 |
| ENSMUSG00 | Ppp1r9b | 5110 | 0.88691287 | 1.18815885 | 1.02935799 | 1.03091556 | 0.83878974 | 1.06216306 |
| ENSMUSG00 | BC066135 | 2306 | 0.91372093 | 0.2716495 | 0.19168189 | 1.32999731 | 1.54893595 | 0.35483052 |
| ENSMUSG00 | Muted | 2470 | 13.6971311 | 16.5823789 | 15.5153838 | 11.6864922 | 13.8205042 | 14.7084289 |
| ENSMUSG00 | Tspyl5 | 4007 | 0.47623236 | 0 | 0 | 0 | 0.39476358 | 0 |
| ENSMUSG00 | 1700019L03f | 1770 | 0.29198941 | 0.08167194 | 0.09989117 | 0.34655071 | 0.05765692 | 0.21573161 |
| ENSMUSG00 | Cables2 | 5164 | 11.8096257 | 11.1134881 | 8.8249613 | 11.2145007 | 11.501685 | 8.40844804 |
| ENSMUSG00 | Txndc5 | 5915 | 40.5284104 | 36.5206797 | 39.1726223 | 29.8233501 | 33.8508028 | 26.1449205 |
| ENSMUSG00 | Hils1 | 795 | 0 | 0 | 0 | 0 | 0 | 0 |
| ENSMUSG00 | Asb17 | 2550 | 0.01559039 | 0 | 0 | 0 | 0 | 0 |
| ENSMUSG00 | Ube3c | 5010 | 18.9731245 | 17.0239526 | 19.3570546 | 17.8105581 | 19.7281617 | 13.1528001 |
| ENSMUSG00 | Rps21 | 740 | 2.14894489 | 2.93025662 | 3.76312988 | 1.17022847 | 2.41340967 | 4.93892116 |
| ENSMUSG00 | Bmp6 | 3671 | 0.37903618 | 0.51192352 | 0.69836745 | 0 | 0.54209443 | 0 |
| ENSMUSG00 | Tlr4 | 5858 | 1.55411489 | 0.55112523 | 0.5810075 | 0 | 0.18292146 | 0.70305062 |
| ENSMUSG00 | Pgcp | 1915 | 163.236211 | 133.085166 | 178.561597 | 120.323727 | 107.115421 | 136.658049 |
| ENSMUSG00 | Siglec5 | 2755 | 0 | 0 | 0 | 0 | 0 | 0 |
| ENSMUSG00 | Timm8b | 525 | 42.935919 | 47.7275237 | 59.6935366 | 42.0613548 | 27.4084533 | 122.190384 |
| ENSMUSG00 | Mtg1 | 2292 | 11.812165 | 11.0374704 | 14.0011073 | 12.2634872 | 11.5098762 | 8.53224971 |
| ENSMUSG00 | Ttc16 | 5664 | 0.21758826 | 0 | 0 | 0 | 0.67566705 | 0 |
| ENSMUSG00 | Arhgap18 | 3866 | 117.723937 | 77.9383918 | 159.428479 | 116.533991 | 102.290329 | 171.443613 |
| ENSMUSG00 | Tsga13 | 1133 | 0 | 0 | 0 | 0 | 0 | 0 |
| ENSMUSG00 | Tasp1 | 3638 | 5.86824987 | 7.89420548 | 7.21712335 | 7.57742771 | 7.77037169 | 8.56924482 |
| ENSMUSG00 | St6galnac5 | 5996 | 0 | 0 | 0 | 0 | 0 | 0 |
| ENSMUSG00 | Adrm1 | 3647 | 6.21349709 | 5.37753824 | 4.60562115 | 5.39202475 | 5.90434072 | 5.43697909 |
| ENSMUSG00 | 2610034B18I | 2271 | 16.3153271 | 14.2161674 | 21.3515728 | 22.5612079 | 10.9422479 | 10.4366629 |
| ENSMUSG00 | Usp6nl | 8225 | 8.2314387 | 5.95812908 | 7.92677428 | 8.9009688 | 8.12079343 | 5.56767408 |
| ENSMUSG00 | Pigk | 9197 | 9.40177994 | 9.43085744 | 8.39147743 | 9.05091625 | 7.67309751 | 7.53855964 |
| ENSMUSG00 | Foxred1 | 4393 | 24.8415192 | 17.7915796 | 16.4511754 | 12.5584899 | 17.5740739 | 14.695896 |
| ENSMUSG00 | Osbpl2 | 4306 | 46.4767972 | 46.7316726 | 47.722798 | 44.5955971 | 52.3061827 | 38.4353254 |
| ENSMUSG00 | Eme1 | 2753 | 0.05776314 | 0.52509744 | 0.01605588 | 0.1441709 | 0.01853483 | 0.01981449 |
| ENSMUSG00 | Myo16 | 6848 | 0 | 0 | 0 | 0 | 0 | 0 |
| ENSMUSG00 | Ak5 | 3089 | 0.16731021 | 0 | 0.10016604 | 0 | 0.51208082 | 0 |
| ENSMUSG00 | Hrh3 | 4338 | 0.08248025 | 0.02221597 | 0 | 0.08317668 | 0 | 0.11317278 |
| ENSMUSG00 | Anpep | 4429 | 4.70351587 | 2.90489502 | 4.87028661 | 4.4318429 | 6.29044958 | 4.65559433 |
| ENSMUSG00 | Echdc3 | 2822 | 7.49465472 | 6.93256397 | 7.94129497 | 6.02219808 | 5.4244907 | 7.0747824 |
| ENSMUSG00 | Fam173b | 5286 | 3.52730238 | 2.80768524 | 2.54206589 | 2.38226136 | 1.77617351 | 3.13715109 |
| ENSMUSG00 | Psmd7 | 1592 | 63.4289702 | 76.396093 | 62.5822576 | 59.0412852 | 64.2316942 | 60.2714704 |
| ENSMUSG00 | Zzz3 | 8499 | 8.76594546 | 8.71426775 | 9.95958694 | 13.2712638 | 12.4158778 | 10.0318299 |
| ENSMUSG00 | Gtpbp5 | 3250 | 10.5199118 | 11.312694 | 11.342873 | 6.93885472 | 8.66663368 | 6.8480325 |
| ENSMUSG00 | Cpa4 | 2728 | 0 | 0 | 0 | 0 | 0 | 0 |
| ENSMUSG00 | Trhr2 | 1337 | 0 | 0 | 0 | 0 | 0 | 0 |
| ENSMUSG00 | Zfp503 | 5701 | 3.60525933 | 5.3756494 | 3.24865326 | 5.42401546 | 3.26690531 | 6.99447862 |
| ENSMUSG00 | Chad | 1676 | 0 | 0 | 0.05274683 | 0 | 0 | 0 |
| ENSMUSG00 | Ss18l1 | 4332 | 18.1707875 | 19.3880352 | 14.703337 | 8.26255498 | 20.6956007 | 13.4484371 |
| ENSMUSG00 | Rreb1 | 13814 | 3.97151897 | 3.89984381 | 6.009198 | 5.78294821 | 4.02995338 | 6.51163749 |
| ENSMUSG00 | L3mbtl3 | 6514 | 1.45253368 | 1.07261807 | 0.88892254 | 1.12998727 | 1.33166778 | 1.14725993 |
| ENSMUSG00 | Spdc3 | 3155 | 0 | 0 | 0 | 1.00640885 | 0 | 0.44088957 |
| ENSMUSG00 | En2 | 4981 | 0 | 0 | 0 | 0 | 0 | 0 |
| ENSMUSG00 | Rsad1 | 4326 | 2.05853621 | 1.92701213 | 2.01289022 | 2.23531852 | 1.25029955 | 3.20284714 |
| ENSMUSG00 | Rln1 | 695 | 1.02963834 | 1.94132429 | 1.39919503 | 1.4536651 | 0.44051548 | 1.80522792 |
| ENSMUSG00 | Gm9767 | 1192 | 0 | 0 | 0 | 0 | 0 | 0 |
| ENSMUSG00 | Wdr93 | 2267 | 1.12234264 | 0.80771275 | 1.26736647 | 1.60753705 | 0.51769151 | 0.32484131 |
| ENSMUSG00 | 6-Mar | 6344 | 7.89594979 | 9.00077773 | 9.6081875 | 9.7940228 | 8.76714213 | 10.4558516 |
| ENSMUSG00 | Nexn | 2549 | 0.0467895 | 1.19095561 | 0.05202257 | 0.5662149 | 0.38034568 | 1.17701464 |
| ENSMUSG00 | Atp6v1g1 | 1132 | 60.4760931 | 55.2526519 | 61.421819 | 53.1987035 | 38.6303923 | 73.4873231 |
| ENSMUSG00 | Htr5a | 5574 | 0 | 0 | 0 | 0 | 0 | 0 |
| ENSMUSG00 | Lsm14b | 2882 | 8.09731681 | 3.76195333 | 2.975419 | 5.39603095 | 6.58633303 | 3.46374669 |
| ENSMUSG00 | F13a1 | 6425 | 0 | 0 | 0 | 0 | 0 | 0 |
| ENSMUSG00 | Mycbpap | 3804 | 12.551612 | 8.14507947 | 14.3853527 | 24.6143285 | 21.6902339 | 16.4981454 |
| ENSMUSG00 | Nrn1 | 2160 | 0 | 0 | 0 | 0 | 0 | 0 |
| ENSMUSG00 | Itga9 | 7659 | 0.02595344 | 0.0629148 | 0.05194106 | 0.01413319 | 0.20653057 | 0.18517838 |
| ENSMUSG00 | Gpr126 | 6480 | 2.53379837 | 5.72585809 | 0.89358664 | 6.5816322 | 3.26001846 | 0.2735882 |
| ENSMUSG00 | Taf4a | 5849 | 4.20052778 | 3.87205126 | 2.68279266 | 5.25592442 | 5.42629608 | 3.64656657 |
| ENSMUSG00 | Prune2 | 13628 | 0.00291719 | 0 | 0 | 0.00264764 | 0 | 0 |
| ENSMUSG00 | Cdc123 | 3664 | 28.991988 | 28.0386175 | 31.9691277 | 27.8985896 | 21.7948357 | 34.4357208 |
| ENSMUSG00 | Zc3hc1 | 4917 | 1.99707214 | 2.45979194 | 2.79576432 | 2.45830484 | 3.2585483 | 2.53498274 |
| ENSMUSG00 | Gipc2 | 4030 | 38.108045 | 46.8592225 | 19.7317965 | 43.1283396 | 35.136028 | 25.0006247 |
| ENSMUSG00 | 9330171B17I | 1986 | 0 | 0 | 0 | 0 | 0 | 0 |
| ENSMUSG00 | Whrn | 7285 | 1.80086597 | 1.24352109 | 0.70383168 | 1.15403107 | 1.91918009 | 1.31038075 |
| ENSMUSG00 | 4930591A17f | 734 | 0 | 0 | 0 | 0 | 0 | 0.03715891 |
| ENSMUSG00 | Camk1d | 7628 | 0.0625414 | 0.23373078 | 0.01158937 | 0.0473021 | 0.00668935 | 0.0035756 |
| ENSMUSG00 | H28 | 483 | 0 | 0 | 0.82363683 | 0 | 0 | 3.89637704 |
| ENSMUSG00 | Sart1 | 5071 | 19.5837488 | 16.2395247 | 14.5567103 | 17.546504 | 20.7788338 | 15.0653253 |
| ENSMUSG00 | Runx2 | 9430 | 0 | 0 | 0.04687364 | 0 | 0 | 0 |
| ENSMUSG00 | Shd | 1542 | 1.8047235 | 3.90616425 | 0.83129277 | 0.28079412 | 2.64728279 | 2.74161419 |
| ENSMUSG00 | Cdh26 | 3228 | 0 | 0 | 0 | 0 | 0.09484457 | 0 |
| ENSMUSG00 | Stim2 | 5712 | 4.56575547 | 5.05316506 | 4.24839142 | 4.90821929 | 6.14603406 | 4.15900049 |
| ENSMUSG00 | Fam102a | 6748 | 31.289472 | 26.8067433 | 30.0334099 | 29.2431389 | 25.9290816 | 18.9726077 |
| ENSMUSG00 | Akna | 5448 | 0.91945495 | 0.3361022 | 0.46246622 | 0.18544369 | 0.3371787 | 0.14518439 |
| ENSMUSG00 | Ube2h | 5046 | 21.2486189 | 24.8571758 | 23.6514023 | 23.9116839 | 30.9535744 | 17.1128632 |
| ENSMUSG00 | Cmc1 | 495 | 23.9336024 | 23.3631235 | 37.4152974 | 19.3895431 | 26.2863148 | 75.5424857 |
| ENSMUSG00 | Naif1 | 4648 | 2.93376287 | 2.68508789 | 2.83393916 | 1.83981165 | 1.92117379 | 2.10662554 |
| ENSMUSG00 | Akap7 | 5486 | 8.04385406 | 11.4185882 | 12.472558 | 9.81305331 | 7.02240498 | 13.7069232 |
| ENSMUSG00 | Eltd1 | 4326 | 2.72020856 | 2.32800888 | 1.35895634 | 7.13133335 | 5.18992265 | 3.39199166 |
| ENSMUSG00 | Dap | 1477 | 98.9445811 | 99.2764681 | 116.355292 | 83.8657135 | 86.679199 | 100.511755 |
| ENSMUSG00 | 1700003H04 | 1034 | 0 | 0 | 0 | 0 | 0 | 0 |
| ENSMUSG00 | Polg | 6971 | 8.54876851 | 6.9608015 | 5.22483412 | 8.40068258 | 9.06924109 | 6.30317656 |
| ENSMUSG00 | Tbc1d19 | 2667 | 23.5670846 | 22.0967449 | 23.0704782 | 19.1977582 | 22.6720118 | 31.2630567 |
| ENSMUSG00 | Tekt5 | 1800 | 0.30920929 | 0 | 1.44883819 | 0.42095719 | 0.70869966 | 0.39396701 |
| ENSMUSG00 | AW209491 | 2008 | 35.8749655 | 37.8196379 | 39.6671918 | 38.8852313 | 44.5210211 | 33.0066601 |
| ENSMUSG00 | Nubp2 | 1423 | 55.7917741 | 52.2838135 | 90.6712437 | 57.6602735 | 57.0147137 | 68.2153486 |
| ENSMUSG00 | Fanci | 5728 | 0.00694055 | 0.05888706 | 0.08488482 | 0.02519696 | 0.00890824 | 0.04761634 |
| ENSMUSG00 | Rbpj | 5804 | 1.51377691 | 1.80989738 | 2.23141627 | 2.34371654 | 1.86381662 | 2.42013081 |
| ENSMUSG00 | Nlrc4 | 3829 | 0 | 0 | 0.01154397 | 0 | 0 | 0 |
| ENSMUSG00 | Rlbp1 | 2293 | 0 | 0 | 0 | 0 | 0 | 0.41042549 |
| ENSMUSG00 | 1110008P14f | 3124 | 5.31939528 | 6.78682284 | 6.53689228 | 4.58533024 | 3.36473541 | 8.31160582 |
| ENSMUSG00 | Orm1 | 769 | 0 | 0 | 0 | 0 | 0 | 0 |
| ENSMUSG00 | Adk | 1690 | 65.7729725 | 73.9048864 | 88.9529397 | 72.0146368 | 62.8318864 | 72.527946 |
| ENSMUSG00 | Ptchd3 | 4555 | 0 | 0 | 0 | 0 | 0 | 0 |
| ENSMUSG00 | Zdhhc1 | 1862 | 55.0000633 | 60.9964791 | 44.201843 | 51.6426685 | 73.8813691 | 38.5682734 |
| ENSMUSG00 | Atf7ip2 | 4696 | 0 | 0 | 0 | 0 | 0 | 0 |
| ENSMUSG00 | Tbc1d25 | 4372 | 5.5013874 | 7.48366748 | 8.02750763 | 5.39745131 | 8.11146642 | 4.57280663 |
| ENSMUSG00 | Abhd2 | 10643 | 1.35220182 | 1.40806009 | 1.33731029 | 0.8645045 | 2.41156319 | 1.18908509 |
| ENSMUSG00 | Ciz1 | 6739 | 13.3501487 | 9.28119927 | 10.2322118 | 10.3389862 | 14.5908631 | 7.45509504 |
| ENSMUSG00 | Dagla | 5192 | 6.03376707 | 6.74721561 | 5.40604205 | 3.75970456 | 3.58717781 | 1.72305117 |
| ENSMUSG00 | Metrnl | 1581 | 25.0703441 | 31.6366394 | 23.2891431 | 30.7872727 | 21.7854546 | 16.3026781 |
| ENSMUSG00 | Rpl39l | 924 | 0 | 0 | 0 | 0 | 0 | 0 |
| ENSMUSG00 | Gpatch2 | 7755 | 4.11652493 | 6.10175193 | 4.17794338 | 5.52280939 | 5.79022704 | 4.66711106 |
| ENSMUSG00 | Svs1 | 2773 | 0 | 0 | 0 | 0 | 0 | 0 |
| ENSMUSG00 | Il18 | 697 | 3.42227953 | 1.86661971 | 3.36111575 | 4.14141114 | 1.02492003 | 7.00453433 |
| ENSMUSG00 | Srrm2 | 9051 | 59.9955926 | 58.6478673 | 46.4727365 | 71.5700967 | 74.3662494 | 56.1102401 |
| ENSMUSG00 | Arid4b | 10707 | 12.3718372 | 14.351972 | 12.3230131 | 14.5581799 | 14.2685131 | 14.0156034 |
| ENSMUSG00 | Ppp1r10 | 5631 | 10.4066024 | 8.12948322 | 8.7289024 | 6.68328404 | 11.5083462 | 8.33109208 |
| ENSMUSG00 | Rpl22l1 | 485 | 18.197354 | 14.3069024 | 18.0452885 | 23.211542 | 9.46881201 | 40.6026588 |
| ENSMUSG00 | D1Pas1 | 3211 | 0 | 0 | 0 | 0 | 0 | 0 |
| ENSMUSG00 | Tbcd | 7934 | 5.16610794 | 5.1198854 | 5.66590299 | 4.28400377 | 6.05833703 | 3.28299477 |
| ENSMUSG00 | Suv39h1 | 6820 | 2.76889344 | 4.45124026 | 2.86469423 | 2.35961757 | 2.7458475 | 1.44371624 |
| ENSMUSG00 | Stx11 | 2393 | 0.56485012 | 0.86586586 | 0.53566797 | 0 | 0.27720137 | 0 |
| ENSMUSG00 | Tbce | 4532 | 19.6935247 | 13.5883215 | 12.7085175 | 15.1111475 | 15.3461938 | 20.058776 |
| ENSMUSG00 | Sec24d | 3833 | 4.14875873 | 6.12230559 | 5.18936326 | 4.32080836 | 6.84256644 | 2.87475979 |
| ENSMUSG00 | Isg20 | 1857 | 0.79211243 | 0.23353688 | 0.19042259 | 0.23316345 | 0.19234498 | 0.07343737 |
| ENSMUSG00 | Zfp750 | 3229 | 17.4338063 | 11.1474984 | 17.9873712 | 16.1246176 | 14.3328965 | 18.3210568 |
| ENSMUSG00 | Tgfb2 | 5330 | 1.245622 | 0.6509238 | 2.15618746 | 0.71080951 | 0.62227288 | 2.56371375 |
| ENSMUSG00 | B3galnt2 | 3669 | 13.2301558 | 17.2441533 | 21.1190599 | 19.6194273 | 21.3618188 | 13.3065152 |
| ENSMUSG00 | E130309D02l | 5429 | 2.80463235 | 2.74260649 | 3.56610927 | 3.43606871 | 2.6504767 | 1.97940834 |
| ENSMUSG00 | Lyplal1 | 1573 | 8.99742598 | 17.3998087 | 14.893183 | 14.6576138 | 11.7428786 | 12.4669203 |
| ENSMUSG00 | Lgi2 | 7275 | 0 | 0 | 0.02430342 | 0 | 0 | 0 |
| ENSMUSG00 | Fn3krp | 2514 | 0.82230906 | 0.72835513 | 0.73845561 | 1.1051382 | 0.14207821 | 1.5188741 |
| ENSMUSG00 | Pomt1 | 6077 | 6.86251424 | 5.39986294 | 5.68797782 | 8.5796535 | 9.14393998 | 3.72069704 |
| ENSMUSG00 | Vstm2b | 1450 | 0.3015933 | 0 | 0 | 0 | 0 | 0 |
| ENSMUSG00 | Prrc2b | 11141 | 5.60594739 | 7.01971058 | 6.85582845 | 6.32512638 | 6.95710124 | 5.54501911 |
| ENSMUSG00 | Npepl1 | 2986 | 27.5598944 | 27.4013995 | 25.6092393 | 21.110292 | 29.2555778 | 17.0900369 |
| ENSMUSG00 | Gimap3 | 904 | 0.43977301 | 0 | 0.19558338 | 0 | 2.08846892 | 0.63359228 |
| ENSMUSG00 | 2300002M23 | 1637 | 0 | 0 | 0 | 0 | 0 | 0 |
| ENSMUSG00 | Megf9 | 8218 | 8.93990362 | 6.73132582 | 6.95998842 | 10.8492008 | 11.033569 | 9.34599466 |
| ENSMUSG00 | Foxk2 | 5580 | 3.63356542 | 4.79273753 | 4.0716393 | 4.27423502 | 4.94718088 | 2.8447742 |
| ENSMUSG00 | Pcsk1n | 2327 | 0 | 0 | 0.01899521 | 0.01550582 | 0 | 0 |
| ENSMUSG00 | Azi2 | 6011 | 17.4141042 | 20.690269 | 20.6338998 | 20.1029391 | 18.4717009 | 20.5002196 |
| ENSMUSG00 | Fndc3b | 11381 | 5.42136067 | 7.11724886 | 4.88973907 | 6.28051403 | 8.8100192 | 5.21479791 |
| ENSMUSG00 | BC017643 | 4357 | 7.45472288 | 8.15088544 | 8.48123497 | 7.02262423 | 10.7158903 | 7.59332968 |
| ENSMUSG00 | Spdyb | 2588 | 0 | 0 | 0 | 0 | 0 | 0 |
| ENSMUSG00 | Cdk5rap2 | 13068 | 2.07782317 | 1.90267862 | 1.72504897 | 1.02989001 | 2.39747435 | 1.61961433 |
| ENSMUSG00 | Tnfsf10 | 5489 | 2.1583409 | 4.57371768 | 0.78112202 | 4.06900812 | 7.98536288 | 3.49815012 |
| ENSMUSG00 | Hexdc | 3224 | 3.84730457 | 3.40772606 | 5.30586639 | 4.00662901 | 3.90927879 | 3.26551202 |
| ENSMUSG00 | Ndst2 | 3518 | 13.911028 | 17.2035734 | 11.1823878 | 7.04615253 | 12.7348374 | 9.23368259 |
| ENSMUSG00 | Edaradd | 3386 | 3.07617717 | 3.78546769 | 3.66825691 | 5.20024741 | 4.27982597 | 7.26571902 |
| ENSMUSG00 | AF529169 | 4185 | 0.00949952 | 0 | 0.07393379 | 0.25003089 | 0 | 0 |
| ENSMUSG00 | Clnk | 1709 | 5.46666935 | 1.38158904 | 5.97462009 | 4.09591377 | 8.09136796 | 6.04861807 |
| ENSMUSG00 | Rftn1 | 5714 | 0.11827847 | 0.00843305 | 0.10056422 | 0 | 0.12502087 | 0.27685143 |
| ENSMUSG00 | Rab3gap2 | 6950 | 7.39051523 | 8.0634291 | 5.92113897 | 7.92766647 | 8.23029745 | 4.54838948 |
| ENSMUSG00 | Uts2r | 1702 | 0 | 0 | 0 | 0 | 0 | 0 |
| ENSMUSG00 | Igfbp2 | 2233 | 0 | 0 | 0.13856377 | 0.67865915 | 0 | 0.08550044 |
| ENSMUSG00 | Rnf122 | 7416 | 4.94263121 | 2.94992513 | 7.9093643 | 3.65880495 | 2.55269491 | 5.83668453 |
| ENSMUSG00 | Tex19.1 | 1935 | 0 | 0 | 0 | 0 | 0 | 0 |
| ENSMUSG00 | Tsga10ip | 483 | 0.4938569 | 0 | 0.18303041 | 0 | 0.10564467 | 0 |
| ENSMUSG00 | Spata16 | 2188 | 0 | 0 | 0 | 0 | 0 | 0 |
| ENSMUSG00 | Tex19.2 | 2043 | 0.03891873 | 0 | 0.28126479 | 0 | 0.1498572 | 0.32040692 |
| ENSMUSG00 | 2010001E11f | 2281 | 0.3485794 | 0 | 0 | 0.22145928 | 0 | 0 |
| ENSMUSG00 | Ankar | 4549 | 0 | 0 | 0 | 0 | 0 | 0 |
| ENSMUSG00 | Mettl22 | 2728 | 3.42468399 | 5.12245903 | 4.90951556 | 3.84892777 | 4.18984904 | 1.81964236 |
| ENSMUSG00 | Atp6v0e2 | 1828 | 2.21830362 | 1.02804773 | 14.9918724 | 0 | 1.3119473 | 13.6074787 |
| ENSMUSG00 | C130074G19 | 3033 | 6.50139081 | 5.68768424 | 8.07379526 | 7.72081995 | 5.23218031 | 8.1383279 |
| ENSMUSG00 | Smarcal1 | 5710 | 3.22360551 | 3.5781176 | 3.9015287 | 4.13900861 | 6.33584947 | 4.9868167 |
| ENSMUSG00 | Exosc2 | 2194 | 10.7995745 | 7.6869894 | 8.90483802 | 8.88072199 | 10.3262128 | 7.78209853 |
| ENSMUSG00 | Fut11 | 3341 | 10.828341 | 12.1006959 | 14.129772 | 17.052843 | 13.6996885 | 11.804588 |
| ENSMUSG00 | Drd5 | 2801 | 0 | 0 | 0 | 0 | 0 | 0 |
| ENSMUSG00 | Picalm | 4295 | 23.3719647 | 25.669518 | 27.1694681 | 25.790891 | 16.1930036 | 36.7112199 |
| ENSMUSG00 | Sectm1b | 3328 | 0.43004727 | 0 | 0 | 0 | 0 | 0.28684266 |
| ENSMUSG00 | Sec24c | 4491 | 60.0448507 | 43.7014872 | 49.4477977 | 50.8892608 | 57.4799455 | 29.7403492 |
| ENSMUSG00 | 4-Mar | 4549 | 0 | 0 | 0 | 0 | 0 | 0 |
| ENSMUSG00 | Wdr17 | 7210 | 0.34737799 | 0.14703214 | 0.13487386 | 0.03503111 | 0.04954017 | 0 |
| ENSMUSG00 | Synpo2l | 8093 | 0 | 0.05954089 | 0.0873878 | 0 | 0 | 0 |
| ENSMUSG00 | Hlx | 2945 | 0.01349931 | 0.01636212 | 0.01500912 | 0.11026771 | 0.13861155 | 0.08335204 |
| ENSMUSG00 | Wdr45 | 2532 | 24.1484712 | 18.7074536 | 21.1756855 | 18.6680404 | 26.3192918 | 19.1741145 |
| ENSMUSG00 | Dusp10 | 3036 | 2.85464254 | 4.04728022 | 6.85740055 | 5.58582376 | 2.13450255 | 1.99439062 |
| ENSMUSG00 | Cdh6 | 2620 | 2.21538174 | 1.25064048 | 0.65796637 | 1.08797005 | 0.19475716 | 0 |
| ENSMUSG00 | Ccdc81 | 3031 | 0 | 0 | 0 | 0.10713903 | 0.01683483 | 0.06298293 |
| ENSMUSG00 | Mreg | 2488 | 6.98277531 | 5.32607379 | 11.0149287 | 10.9348318 | 8.69581324 | 4.264403 |
| ENSMUSG00 | Neil3 | 3970 | 0 | 0.97101143 | 0 | 0 | 0 | 0 |
| ENSMUSG00 | Prss23 | 3224 | 52.8511134 | 68.6626908 | 45.5865782 | 49.5680444 | 56.7715912 | 48.052094 |
| ENSMUSG00 | Prdm16 | 14042 | 3.15676975 | 5.05130628 | 3.00932644 | 4.31946424 | 4.69128694 | 3.58171448 |
| ENSMUSG00 | Heatr5b | 7193 | 7.06346505 | 7.2751948 | 7.31883707 | 9.07945233 | 8.02319353 | 3.97763055 |
| ENSMUSG00 | Cntnap2 | 7157 | 0.01110954 | 0.10099157 | 0.00617603 | 0 | 0 | 0.15624706 |
| ENSMUSG00 | Alg1 | 2612 | 14.6115089 | 11.8990257 | 9.67972979 | 12.9851156 | 12.2877452 | 6.78733366 |
| ENSMUSG00 | Tmem135 | 6093 | 34.7379252 | 33.9037055 | 29.4606408 | 32.5229917 | 32.4180372 | 35.726062 |
| ENSMUSG00 | Mtmr7 | 5250 | 0.35590621 | 0.53234546 | 0.26942076 | 0.6735315 | 0.76782547 | 0.18183093 |
| ENSMUSG00 | Ttc36 | 761 | 7.57496015 | 8.3582265 | 4.70479538 | 6.25864636 | 8.24736429 | 10.1428685 |
| ENSMUSG00 | Prpf18 | 5807 | 15.8487924 | 18.8364429 | 12.8868125 | 15.8321077 | 13.4705414 | 12.5265133 |
| ENSMUSG00 | Dcxr | 1320 | 51.923067 | 41.9076027 | 86.9302913 | 41.7403651 | 20.5265194 | 67.7113582 |
| ENSMUSG00 | Snx22 | 2548 | 1.06097829 | 0.51060987 | 0.55512519 | 0.66556362 | 2.04265712 | 0.96338993 |
| ENSMUSG00 | Morc3 | 4161 | 20.6564165 | 16.5948502 | 20.5234224 | 26.9943294 | 29.504797 | 24.6985919 |
| ENSMUSG00 | Ppl | 6255 | 7.30916109 | 6.70989466 | 2.66412387 | 7.64327882 | 12.7749488 | 3.32266589 |
| ENSMUSG00 | Mtmr12 | 4962 | 21.7765812 | 9.42947104 | 19.1256262 | 14.2524803 | 17.3789954 | 11.4166517 |
| ENSMUSG00 | Tcta | 1625 | 46.77691 | 48.3940145 | 43.7666249 | 50.0707756 | 38.7486448 | 38.6712424 |
| ENSMUSG00 | Col10a1 | 3127 | 0 | 0.077049 | 0 | 0.42693817 | 0.03263599 | 0.02616691 |
| ENSMUSG00 | Slc9a8 | 8639 | 7.95201647 | 5.53315784 | 6.99943526 | 4.16829268 | 9.28504026 | 4.63469967 |
| ENSMUSG00 | Zdhhc2 | 5716 | 0.52858931 | 0.514236 | 1.26821243 | 0.55549684 | 0.50883545 | 1.77027487 |
| ENSMUSG00 | Ubn1 | 6436 | 9.51883087 | 7.876342 | 8.9282778 | 8.84111004 | 11.0361584 | 10.4208107 |
| ENSMUSG00 | Wfs1 | 6076 | 8.16570765 | 10.6032378 | 5.45611953 | 8.95518814 | 10.5983026 | 4.17020077 |
| ENSMUSG00 | Prrx2 | 1946 | 0 | 0 | 0 | 0 | 0 | 0 |
| ENSMUSG00 | Tnrc18 | 7148 | 2.2580757 | 2.91221922 | 1.9602664 | 3.05899818 | 3.28373431 | 1.9116668 |
| ENSMUSG00 | Efha2 | 6497 | 2.09271577 | 0.74908891 | 1.11576147 | 0.87192258 | 1.24876001 | 1.44412435 |
| ENSMUSG00 | Nt5dc1 | 3178 | 27.3584757 | 26.792147 | 32.9636148 | 32.0174215 | 31.5663483 | 35.3591925 |
| ENSMUSG00 | Nrtn | 1020 | 4.71609132 | 7.1334831 | 5.11354655 | 5.624554 | 3.90201697 | 4.27837478 |
| ENSMUSG00 | Asb6 | 5374 | 7.81941624 | 8.90382156 | 7.66582019 | 6.54633298 | 7.32068027 | 6.16140902 |
| ENSMUSG00 | Tspyl4 | 3887 | 11.4551423 | 6.60750544 | 5.65174067 | 7.89033647 | 10.1212595 | 11.0656306 |
| ENSMUSG00 | Cntn5 | 1239378 | 0 | 0 | 0 | 0 | 0 | 0 |
| ENSMUSG00 | Ccdc27 | 2077 | 0.32539392 | 0.09280008 | 0.08512632 | 0.38218824 | 0.54048159 | 0.28889844 |
| ENSMUSG00 | Cdnf | 3137 | 0.83642388 | 0.76803382 | 0.84542894 | 0.2530459 | 0.35785154 | 0.59992034 |
| ENSMUSG00 | Dse | 4218 | 0 | 0.02284801 | 0.10479337 | 0 | 0 | 0 |
| ENSMUSG00 | Znfx1 | 11238 | 8.90058631 | 10.2693121 | 7.20964391 | 8.29328359 | 14.7657745 | 7.17664249 |
| ENSMUSG00 | Fam26d | 3712 | 0 | 0 | 0 | 0 | 0 | 0 |
| ENSMUSG00 | Nup133 | 5782 | 4.64111888 | 4.10026453 | 2.75974841 | 4.03130418 | 4.80964629 | 0.96701851 |
| ENSMUSG00 | Uhrf1bp1 | 8809 | 5.37504567 | 4.90671343 | 5.22353486 | 7.67598495 | 5.50869377 | 5.05303795 |
| ENSMUSG00 | Ppp2r4 | 5366 | 11.0835257 | 13.0837954 | 12.2242891 | 12.6616642 | 11.2874223 | 9.92683013 |
| ENSMUSG00 | Cdsn | 2676 | 0 | 0 | 0 | 0 | 0 | 0 |
| ENSMUSG00 | Cyp7b1 | 2160 | 13.141395 | 36.0952145 | 5.85265144 | 21.5991128 | 26.0092777 | 7.37425432 |
| ENSMUSG00 | Foxp3 | 4091 | 0 | 0 | 0.12965586 | 0 | 0.08730986 | 0.20000958 |
| ENSMUSG00 | BC046331 | 5083 | 7.63355282 | 16.2865055 | 8.59166258 | 10.9601961 | 18.7923226 | 8.36536742 |
| ENSMUSG00 | Atp8b1 | 6412 | 9.06464637 | 8.95792873 | 8.16203714 | 12.5094175 | 13.0351222 | 9.43042346 |
| ENSMUSG00 | Tusc3 | 1865 | 34.9592429 | 47.9022326 | 58.9199902 | 36.7204936 | 31.8470249 | 56.2310927 |
| ENSMUSG00 | Zufsp | 2629 | 34.2359863 | 28.4646423 | 25.0180078 | 29.3981512 | 28.1625224 | 23.4464377 |
| ENSMUSG00 | Mmd2 | 3303 | 21.6771482 | 12.7796922 | 37.3367066 | 21.2144506 | 9.90248468 | 18.1748353 |
| ENSMUSG00 | Stau1 | 7135 | 24.7447917 | 22.8066735 | 25.5856498 | 19.3229842 | 23.1925069 | 19.0712229 |
| ENSMUSG00 | Sgcz | 3916 | 0 | 0 | 0 | 0 | 0 | 0 |
| ENSMUSG00 | 4921524L21F | 1517 | 0 | 0 | 0 | 0 | 0 | 0 |
| ENSMUSG00 | Ncam1 | 11332 | 0.12980522 | 0.24663022 | 0.07411181 | 0.1018907 | 0.14859428 | 0.05535799 |
| ENSMUSG00 | Ttc18 | 4502 | 0.04415313 | 0.43883699 | 0.20618363 | 1.47469929 | 0.52137123 | 0.08481674 |
| ENSMUSG00 | 4930524L23F | 3567 | 0.25634316 | 0.10807164 | 0 | 0.01011552 | 0 | 0 |
| ENSMUSG00 | Ajap1 | 3147 | 0 | 0.88808823 | 0 | 0 | 0.11350004 | 0.5460128 |
| ENSMUSG00 | Rsph4a | 3297 | 0 | 0 | 0 | 0 | 0 | 1.02356665 |
| ENSMUSG00 | Cylc2 | 1352 | 0 | 0 | 0 | 0 | 0 | 0 |
| ENSMUSG00 | Ppp1r3f | 10169 | 0.48477526 | 0.36486931 | 0.47379299 | 0.3796616 | 1.10392395 | 0.45596309 |

| | | | | | | | | |
|---|---|---|---|---|---|---|---|---|
| ENSMUSG00 | 2210406O10I | 2426 | 0.09832353 | 0.25821259 | 0.27330076 | 0.83289138 | 0.82029211 | 0.07869846 |
| ENSMUSG00 | Fam100a | 3152 | 6.45774303 | 5.44237735 | 5.60937094 | 7.39498755 | 6.15165698 | 8.10797493 |
| ENSMUSG00 | Nphp4 | 6019 | 0.02641999 | 0.70450356 | 0.27906131 | 1.07304967 | 1.60225702 | 0.52564515 |
| ENSMUSG00 | Fam190a | 8020 | 4.81824527 | 6.42285619 | 7.22551324 | 6.708021 | 7.73030507 | 6.3289406 |
| ENSMUSG00 | Grin3a | 19722 | 0 | 0 | 0 | 0 | 0.01552369 | 0 |
| ENSMUSG00 | Myo9a | 12989 | 5.12055731 | 8.92946079 | 7.60915551 | 10.9810087 | 8.23006062 | 7.60766941 |
| ENSMUSG00 | Fam149b | 4492 | 11.7443283 | 11.2420729 | 11.7884702 | 11.6873052 | 12.3590153 | 16.3635692 |
| ENSMUSG00 | Rcan2 | 3816 | 281.913863 | 151.479182 | 355.062656 | 204.871504 | 179.875998 | 263.84068 |
| ENSMUSG00 | Rbms3 | 11796 | 2.61531476 | 2.49183873 | 1.92605521 | 2.85389518 | 2.3402229 | 1.58154063 |
| ENSMUSG00 | Olfr303 | 1039 | 0 | 0 | 0 | 0 | 0 | 0 |
| ENSMUSG00 | 2810432L12F | 3796 | 17.343276 | 16.045222 | 16.7678224 | 12.0051692 | 18.8458849 | 10.6626883 |
| ENSMUSG00 | Stub1 | 1665 | 44.0533703 | 45.379184 | 48.343277 | 39.9394643 | 41.8017878 | 39.1018402 |
| ENSMUSG00 | Mocos | 2869 | 1.10855296 | 1.66275977 | 0.40057439 | 0.71686181 | 0.01778542 | 0 |
| ENSMUSG00 | Gm7488 | 683 | 0.17462144 | 0.42330696 | 0.06471719 | 0.31697257 | 0.29883675 | 0.03993359 |
| ENSMUSG00 | 6430573F11F | 4210 | 5.14649332 | 3.90298736 | 8.10542109 | 2.69972542 | 4.26633831 | 4.66454638 |
| ENSMUSG00 | Prex1 | 9011 | 0.12794462 | 0.35828339 | 0.18640218 | 0.94900062 | 0.45867678 | 0.34203021 |
| ENSMUSG00 | C330006K01I | 3906 | 9.67932974 | 12.1267979 | 6.78983764 | 8.01823212 | 9.10532103 | 8.42120198 |
| ENSMUSG00 | Hs3st6 | 1849 | 2.49412425 | 3.38790561 | 1.79293576 | 3.4735554 | 1.04867619 | 4.07128201 |
| ENSMUSG00 | Fam40b | 8329 | 4.16695095 | 5.1779164 | 1.87336422 | 6.45915817 | 5.39731506 | 1.17560277 |
| ENSMUSG00 | Hnrnpu | 5357 | 139.518953 | 121.316137 | 128.958485 | 157.799167 | 145.201806 | 139.590962 |
| ENSMUSG00 | Ccdc151 | 2187 | 0 | 1.74061679 | 0 | 0.29697156 | 0.11665838 | 0.07482754 |
| ENSMUSG00 | Lonrf1 | 3918 | 2.79039233 | 3.03778745 | 1.64713351 | 1.05906971 | 1.02886261 | 0.62652311 |
| ENSMUSG00 | Zfp189 | 3037 | 11.3231777 | 9.63094184 | 13.0262264 | 8.91061356 | 16.2471207 | 5.42439319 |
| ENSMUSG00 | Coro7 | 6960 | 3.83274819 | 3.32320291 | 2.50858161 | 2.92388982 | 5.13196309 | 2.38261216 |
| ENSMUSG00 | Kcne1 | 3752 | 77.7308649 | 115.73993 | 24.7398375 | 77.3186668 | 102.855139 | 39.2981609 |
| ENSMUSG00 | Mrpl12 | 1527 | 45.6914658 | 42.5378677 | 55.2884873 | 36.8618137 | 30.9433032 | 42.367678 |
| ENSMUSG00 | Vasn | 3184 | 13.1602627 | 18.4633981 | 5.6224078 | 12.9981486 | 11.6668347 | 2.63837591 |
| ENSMUSG00 | Ccbl1 | 3253 | 22.4014128 | 22.3230766 | 17.1073226 | 20.0209316 | 23.3093128 | 18.9488733 |
| ENSMUSG00 | Cpeb3 | 11181 | 2.6347206 | 1.94797172 | 4.28537678 | 3.60142757 | 2.6104183 | 3.97130066 |
| ENSMUSG00 | Baat | 2530 | 0 | 0 | 0 | 0.28523355 | 0.02016853 | 0.6360489 |
| ENSMUSG00 | Rxrb | 3779 | 9.85733931 | 8.39022994 | 9.17021565 | 4.15339756 | 9.15477185 | 7.93916464 |
| ENSMUSG00 | D2Wsu81e | 3629 | 9.0268713 | 7.51543848 | 8.45303914 | 6.8107469 | 8.23958562 | 7.57587114 |
| ENSMUSG00 | Dusp26 | 2557 | 2.16113093 | 0.54650247 | 1.64222725 | 0.86077619 | 0.47893352 | 0.60799939 |
| ENSMUSG00 | Icmt | 4187 | 18.4487458 | 22.0734645 | 14.8641497 | 14.5896588 | 24.2640349 | 6.56623749 |
| ENSMUSG00 | 1810049H13I | 956 | 12.3508135 | 11.0889302 | 11.0504608 | 11.2095892 | 7.57923156 | 16.9753246 |
| ENSMUSG00 | Zmynd8 | 12576 | 4.41938309 | 4.57495324 | 4.38995721 | 4.49590997 | 4.9987671 | 3.39848598 |
| ENSMUSG00 | Kcne2 | 1764 | 0.1126856 | 0 | 0.32575054 | 0 | 0.23141214 | 0.27831265 |
| ENSMUSG00 | Capsl | 941 | 1.30969171 | 2.56038481 | 2.53655635 | 2.60741661 | 1.68099644 | 3.56512288 |
| ENSMUSG00 | Tbc1d13 | 4742 | 0.41080104 | 1.43278961 | 1.67784305 | 1.08048299 | 0.36585761 | 1.5069497 |
| ENSMUSG00 | Mrps6 | 1741 | 88.6449026 | 72.3211794 | 154.185981 | 71.3143683 | 53.9572418 | 158.133377 |
| ENSMUSG00 | Lap3 | 3116 | 33.9886393 | 30.7582906 | 44.9962279 | 28.6595187 | 30.5732489 | 31.9925566 |
| ENSMUSG00 | Sdk1 | 11214 | 0.52468442 | 0.99260461 | 0.14976331 | 1.03928129 | 1.697239 | 0.30402443 |
| ENSMUSG00 | Gm5422 | 2693 | 0.1771503 | 0.46522373 | 0.67295788 | 0.56273519 | 0.94738908 | 0.45575892 |
| ENSMUSG00 | Zer1 | 9177 | 6.68439647 | 6.88377745 | 5.78954073 | 6.62113577 | 6.86690358 | 6.7079504 |
| ENSMUSG00 | Tspan10 | 2355 | 0 | 0 | 0.03753872 | 0 | 0 | 0 |
| ENSMUSG00 | 5730528L13F | 1966 | 0.64708819 | 1.54412302 | 1.14663988 | 0.89929816 | 1.76490008 | 1.09597991 |
| ENSMUSG00 | Ncoa7 | 5342 | 33.7199705 | 26.4024179 | 41.8601879 | 33.9273886 | 32.8681691 | 35.4897075 |
| ENSMUSG00 | Batf2 | 2390 | 0.11643865 | 0 | 0 | 0 | 0.27754933 | 0.07988388 |
| ENSMUSG00 | Usp53 | 5043 | 9.86989126 | 10.7303935 | 6.65262718 | 14.5673295 | 13.6090572 | 7.7232173 |
| ENSMUSG00 | Nploc4 | 5067 | 20.321037 | 18.4396115 | 16.5484303 | 18.1442766 | 20.704604 | 9.24226477 |
| ENSMUSG00 | Lmbrd2 | 3845 | 1.84043577 | 1.89236743 | 3.48326617 | 6.39060399 | 2.81341786 | 4.62498434 |
| ENSMUSG00 | Ldb2 | 2619 | 0.89559884 | 0 | 0.32067011 | 0.71640562 | 1.05209022 | 1.4683941 |
| ENSMUSG00 | Slc7a12 | 2003 | 0 | 0.02405714 | 0 | 0 | 0 | 0 |
| ENSMUSG00 | Plekhg5 | 4751 | 3.23834371 | 2.75872706 | 1.66536096 | 3.12898387 | 3.76978697 | 3.22634125 |
| ENSMUSG00 | Cpb3 | 2829 | 0 | 0 | 0 | 0 | 0 | 0 |
| ENSMUSG00 | Wdr34 | 2410 | 5.1797597 | 6.21825042 | 4.58525343 | 6.87205744 | 7.47398783 | 2.0823791 |
| ENSMUSG00 | Dock3 | 17159 | 0.02780266 | 0.00842469 | 0.11076865 | 0.02733648 | 0.32413748 | 0.05563333 |
| ENSMUSG00 | Ralyl | 2276 | 0 | 0 | 0.13594591 | 0 | 1.16580472 | 0 |
| ENSMUSG00 | Got1l1 | 1346 | 0 | 0.42959681 | 0 | 0.40210302 | 0.45491568 | 0.95238339 |
| ENSMUSG00 | 2810408M09 | 2649 | 4.33723438 | 3.5835142 | 4.75557767 | 3.09196833 | 3.62135095 | 3.14034163 |
| ENSMUSG00 | Slc6a5 | 7050 | 0 | 0 | 0 | 0 | 0 | 0 |
| ENSMUSG00 | Fnbp1l | 8992 | 6.69370524 | 5.75536464 | 5.05824027 | 5.94278336 | 4.88586628 | 6.69429813 |
| ENSMUSG00 | Prkrip1 | 4372 | 3.62818772 | 3.53793411 | 3.04317355 | 4.08522691 | 3.24458657 | 2.12732205 |
| ENSMUSG00 | Six4 | 8280 | 4.13399381 | 6.18043497 | 4.17995689 | 7.17718929 | 6.79119156 | 4.63471225 |
| ENSMUSG00 | Alg2 | 2249 | 28.6543503 | 28.4962063 | 33.1366418 | 33.2102411 | 36.55113 | 34.7087672 |
| ENSMUSG00 | Bahcc1 | 10718 | 0.25222734 | 0.39563416 | 0.54025391 | 0.33664904 | 0.47608113 | 0.74306724 |
| ENSMUSG00 | Fam71f1 | 1866 | 0 | 0 | 0 | 0 | 0 | 0 |
| ENSMUSG00 | Htatip2 | 1459 | 14.4144271 | 23.6143291 | 11.9366183 | 23.5683266 | 13.7445961 | 7.36546118 |
| ENSMUSG00 | Orai2 | 4369 | 1.09193355 | 0.70586686 | 0.66773212 | 1.23879759 | 1.47157779 | 0.9925996 |
| ENSMUSG00 | Exo1 | 5491 | 0 | 0.27204147 | 0.21734652 | 0 | 0 | 0 |
| ENSMUSG00 | Fbxl5 | 8162 | 27.2326503 | 23.7567071 | 25.9134794 | 25.1584067 | 26.3134055 | 25.6038086 |
| ENSMUSG00 | Alkbh4 | 2123 | 3.48305199 | 2.20164149 | 5.12183391 | 5.18371342 | 4.75893661 | 3.67430373 |
| ENSMUSG00 | Dnttip2 | 2394 | 39.9713624 | 36.3713037 | 32.7359514 | 36.2026194 | 36.6179255 | 34.4293901 |
| ENSMUSG00 | Thap3 | 1880 | 7.61275159 | 9.66291952 | 7.47669472 | 6.87094252 | 8.19679016 | 8.38550079 |
| ENSMUSG00 | Il22ra2 | 2493 | 0 | 0 | 0 | 0 | 0 | 0 |
| ENSMUSG00 | Dnajc28 | 5213 | 5.84168388 | 7.68136072 | 8.63178135 | 7.87672486 | 5.08012451 | 7.1835948 |
| ENSMUSG00 | Cc2d2a | 6618 | 5.03401219 | 4.82010044 | 2.39777299 | 4.18721822 | 6.67706882 | 4.86723315 |
| ENSMUSG00 | Dnajc11 | 7207 | 16.4714673 | 17.5977127 | 15.5353501 | 13.9431794 | 19.0879852 | 11.7999619 |
| ENSMUSG00 | Ypel5 | 2866 | 62.6294468 | 81.2410637 | 79.4892878 | 72.1137303 | 63.3823789 | 92.8441663 |
| ENSMUSG00 | Polr2j | 1205 | 7.918104 | 10.1571422 | 10.6011059 | 8.89325081 | 4.99677373 | 16.4326872 |
| ENSMUSG00 | Galnt12 | 2683 | 5.21577679 | 7.07620507 | 1.00496177 | 4.85487069 | 6.33312827 | 2.95822588 |
| ENSMUSG00 | Defb3 | 273 | 0 | 0 | 0 | 0 | 0 | 0 |
| ENSMUSG00 | Azi1 | 4082 | 3.70090215 | 2.79769397 | 1.52681525 | 2.90813147 | 3.15008494 | 1.73723817 |
| ENSMUSG00 | Cpeb2 | 7388 | 20.7602387 | 15.9925766 | 29.6214571 | 27.0859528 | 22.0046066 | 29.7185769 |
| ENSMUSG00 | Kmo | 4707 | 0 | 0.72684032 | 0.1220786 | 1.16517331 | 0.31437538 | 0.69533816 |
| ENSMUSG00 | Defb5 | 205 | 0 | 0 | 0 | 0 | 0 | 0 |
| ENSMUSG00 | Cercam | 3370 | 0.87296901 | 0.32886889 | 0.40660449 | 0 | 0.34825123 | 0.09712038 |
| ENSMUSG00 | Zfp597 | 4289 | 2.6231758 | 4.42654656 | 3.81316902 | 4.26523585 | 4.04499132 | 4.28610559 |
| ENSMUSG00 | Zfand1 | 7210 | 3.69985124 | 2.84707689 | 3.78259877 | 4.39390223 | 3.62350963 | 4.02499531 |
| ENSMUSG00 | 2600006K01F | 869 | 0.04574854 | 0 | 0 | 0 | 0 | 0 |
| ENSMUSG00 | 2410089E03F | 12303 | 1.32809091 | 1.82907165 | 0.76885267 | 1.75380498 | 1.30645439 | 0.90449913 |
| ENSMUSG00 | Ncoa5 | 6186 | 9.28656148 | 8.46729108 | 7.00255515 | 11.1757513 | 9.69220685 | 7.70268263 |
| ENSMUSG00 | Gabbr2 | 5830 | 0 | 0 | 0 | 0 | 0 | 0 |
| ENSMUSG00 | Zc3h10 | 5119 | 3.49481661 | 3.68999518 | 3.68708478 | 4.8776665 | 3.11999524 | 2.20584111 |
| ENSMUSG00 | Tbc1d2 | 5411 | 14.0330748 | 11.1494041 | 13.1845823 | 8.8221299 | 11.5990468 | 6.65862104 |
| ENSMUSG00 | Xkr5 | 6006 | 0 | 0 | 0 | 0 | 0 | 0 |
| ENSMUSG00 | Myl6b | 1019 | 3.43325053 | 2.50626245 | 2.64603771 | 0.81441317 | 3.30494682 | 2.35541536 |
| ENSMUSG00 | Trub2 | 6307 | 4.94185773 | 3.56795124 | 4.66057168 | 3.18657029 | 4.89471339 | 2.79362882 |
| ENSMUSG00 | Wdr70 | 2133 | 7.32484943 | 7.38723234 | 8.22697219 | 13.6851261 | 9.04264888 | 8.77187179 |
| ENSMUSG00 | Olig2 | 2435 | 0 | 0 | 0 | 0 | 0 | 0 |
| ENSMUSG00 | Arhgap29 | 5574 | 13.2304299 | 17.6873808 | 7.3035338 | 17.5878929 | 19.9015682 | 8.75880952 |
| ENSMUSG00 | Zfp335 | 4714 | 4.19144544 | 4.64078166 | 3.178707 | 4.35525739 | 5.2606743 | 4.14269022 |
| ENSMUSG00 | Nhsl1 | 11044 | 3.56013856 | 2.29064489 | 2.76961928 | 2.2151056 | 3.62692005 | 3.05246778 |
| ENSMUSG00 | Slc45a1 | 4386 | 0 | 0 | 0 | 0 | 0.10470529 | 0 |
| ENSMUSG00 | 5430411K18F | 9579 | 2.02948429 | 2.23350875 | 2.54256347 | 2.56894815 | 2.98306404 | 1.7482648 |
| ENSMUSG00 | Zfp800 | 6236 | 10.3213796 | 10.3775484 | 8.98780259 | 15.720801 | 6.76696806 | 9.37292366 |
| ENSMUSG00 | Mcph1 | 8003 | 4.74403147 | 3.84142823 | 4.2362631 | 3.72407457 | 5.99972756 | 2.42653294 |
| ENSMUSG00 | Rapgef1 | 7680 | 2.00330351 | 2.69166454 | 2.8949905 | 2.75783335 | 2.6642678 | 2.82335133 |
| ENSMUSG00 | Pcif1 | 6650 | 14.311973 | 11.1444734 | 9.85734334 | 9.34335048 | 13.8960551 | 9.84758028 |
| ENSMUSG00 | A730011L01F | 6213 | 2.59789556 | 1.6985081 | 2.51850192 | 3.25220424 | 2.89913257 | 2.92369383 |
| ENSMUSG00 | 4932438H23 | 3279 | 0.03637281 | 0 | 0 | 0.0990358 | 0 | 0.1330876 |
| ENSMUSG00 | Rere | 8761 | 5.18668123 | 5.14260055 | 5.08061362 | 7.18263733 | 4.75842359 | 5.11808092 |
| ENSMUSG00 | Trim14 | 4828 | 0.2552651 | 0.50901171 | 0.29296996 | 0.45588333 | 0.70811251 | 0.86998642 |
| ENSMUSG00 | Mrgpra2a | 2427 | 0 | 0 | 0 | 0 | 0 | 0 |
| ENSMUSG00 | Srrm3 | 6532 | 0 | 0 | 0 | 0 | 0 | 0 |
| ENSMUSG00 | Slc44a3 | 2544 | 88.3715575 | 102.415131 | 115.630214 | 86.2054507 | 89.2761001 | 76.9350673 |
| ENSMUSG00 | Neurl2 | 1012 | 0.98210179 | 0.90468617 | 1.13562047 | 1.06962583 | 0.55463452 | 0.48512204 |
| ENSMUSG00 | Slc39a5 | 2129 | 0.01867331 | 1.0185016 | 0.02076179 | 0.27116614 | 0.07190189 | 0.3971413 |
| ENSMUSG00 | Heca | 4684 | 11.5854464 | 7.9522032 | 7.14363688 | 10.9001053 | 12.2881622 | 15.2502733 |
| ENSMUSG00 | Lrrc17 | 2182 | 0 | 0 | 0 | 0 | 0 | 0 |
| ENSMUSG00 | Tmem120a | 1843 | 9.14613334 | 15.5305191 | 15.0137568 | 12.9605608 | 7.69687058 | 9.23460385 |
| ENSMUSG00 | Atg14 | 906 | 25.4066481 | 20.902066 | 24.8330443 | 24.6520812 | 24.6683804 | 25.4683718 |
| ENSMUSG00 | Txlnb | 5659 | 0 | 0 | 0.07810893 | 0.17852929 | 0 | 0 |
| ENSMUSG00 | Fgl2 | 3786 | 0 | 0.34364341 | 0.32690217 | 0.17154696 | 0 | 0.30257128 |
| ENSMUSG00 | 9130011E15F | 2919 | 9.45197104 | 10.581538 | 11.0391037 | 10.8530439 | 13.6874451 | 12.1189473 |
| ENSMUSG00 | 4931408A02F | 3793 | 0.46117615 | 0.85117101 | 0.24472415 | 0.55174231 | 0.57846933 | 0.05033548 |
| ENSMUSG00 | Gpr37 | 3937 | 2.17105113 | 3.31687219 | 6.53428312 | 3.59262419 | 2.56622363 | 2.6949034 |
| ENSMUSG00 | Slc26a11 | 3640 | 3.97554805 | 5.52025999 | 3.98302322 | 6.00706565 | 2.94382937 | 3.86640491 |
| ENSMUSG00 | Cited2 | 1014 | 45.2051963 | 89.6248815 | 31.4295156 | 80.5974663 | 59.7823811 | 89.7588473 |
| ENSMUSG00 | Spsb1 | 4665 | 7.84030913 | 4.51392824 | 7.07798001 | 3.85184527 | 5.5675081 | 3.25658606 |
| ENSMUSG00 | Pak7 | 7973 | 0.13961538 | 0.16317998 | 0.08315912 | 0.15839352 | 0.16639731 | 0.08210101 |
| ENSMUSG00 | Coq10a | 2497 | 7.22826918 | 6.94718429 | 9.9662145 | 8.62674433 | 10.6875429 | 8.55268023 |
| ENSMUSG00 | Rhbdd2 | 6266 | 13.4886932 | 9.96642659 | 9.43856782 | 13.3882467 | 9.85348145 | 6.890481 |
| ENSMUSG00 | Urb1 | 8151 | 1.5949015 | 1.32422562 | 1.49671312 | 2.29745812 | 1.09552396 | 0.73615761 |
| ENSMUSG00 | Pion | 3714 | 4.09971703 | 3.60684731 | 1.93993011 | 1.89445307 | 1.63493234 | 1.39531003 |
| ENSMUSG00 | Pik3cd | 7834 | 0.22328838 | 0.01845281 | 0.15234232 | 0 | 0.22797079 | 0 |
| ENSMUSG00 | Ptger4 | 6643 | 4.53630201 | 2.79993477 | 3.53987362 | 3.74779629 | 1.88190006 | 2.97668425 |
| ENSMUSG00 | Plcb4 | 8631 | 8.72400418 | 8.06177992 | 12.2962142 | 12.8550906 | 13.6626063 | 9.23058988 |
| ENSMUSG00 | Wfdc3 | 1617 | 0 | 0 | 0 | 0 | 0 | 0 |
| ENSMUSG00 | Dag1 | 8261 | 23.6097794 | 24.8835931 | 19.7118496 | 18.0606772 | 26.757809 | 13.6950979 |
| ENSMUSG00 | Clstn1 | 4776 | 57.0278051 | 60.0716241 | 55.9649382 | 50.6931572 | 76.9028168 | 45.857486 |
| ENSMUSG00 | Stk32a | 3767 | 1.28754145 | 1.50942399 | 1.31420399 | 0.96742408 | 2.96649225 | 1.74493976 |
| ENSMUSG00 | Mrap | 1459 | 0 | 0 | 0 | 0 | 0 | 0 |
| ENSMUSG00 | 4833442J19R | 2490 | 8.9409916 | 10.5468317 | 6.51489012 | 5.08626411 | 11.6397516 | 11.797103 |
| ENSMUSG00 | Hip1 | 7819 | 0.48810924 | 0.88743416 | 0.2261253 | 0.39686096 | 1.08330712 | 0.73950934 |
| ENSMUSG00 | Rhou | 3830 | 6.8611939 | 3.96311469 | 12.7989149 | 9.55279194 | 7.64729497 | 9.33604492 |
| ENSMUSG00 | Olfr906 | 935 | 0 | 0 | 0 | 0 | 0 | 0 |
| ENSMUSG00 | Ccdc40 | 6292 | 0.08845784 | 0 | 0.14050173 | 0.14909936 | 0.1054264 | 0.22541024 |
| ENSMUSG00 | Zfp292 | 9978 | 10.4189799 | 10.049708 | 9.93633332 | 12.3202571 | 14.2319507 | 15.4359479 |
| ENSMUSG00 | Rsbn1l | 5542 | 4.81341166 | 3.31270921 | 2.48047152 | 5.42337479 | 5.75450828 | 4.83777885 |
| ENSMUSG00 | Tbc1d16 | 7106 | 1.70636386 | 3.23458105 | 1.23785066 | 2.50837743 | 2.39836892 | 1.57368451 |
| ENSMUSG00 | Ccdc67 | 4812 | 0.21480517 | 0 | 0.09185753 | 0.19495702 | 0.31811955 | 0.12469702 |
| ENSMUSG00 | Zc3h12d | 4082 | 0.00973922 | 0 | 0.08862782 | 0 | 0.13750371 | 0 |
| ENSMUSG00 | Dtx4 | 5756 | 0.10360184 | 0.3264891 | 0.23037792 | 0 | 0.37232588 | 0.18480037 |
| ENSMUSG00 | Ccdc32 | 2210 | 7.08762865 | 9.44105405 | 13.2605529 | 12.4736118 | 8.65832169 | 7.14570865 |
| ENSMUSG00 | Fam60a | 5799 | 1.17230336 | 1.56217471 | 1.43299646 | 1.29419991 | 1.24948187 | 1.20405374 |
| ENSMUSG00 | Phtf2 | 8278 | 1.93542627 | 1.73466535 | 2.17324838 | 2.95525806 | 2.56426339 | 1.68695522 |
| ENSMUSG00 | Ankrd13c | 3135 | 24.3098106 | 22.9327501 | 22.3053638 | 22.9267728 | 24.4470866 | 25.4215298 |
| ENSMUSG00 | Cbx4 | 6133 | 4.08380119 | 3.83417312 | 4.0360396 | 4.75955878 | 4.11006517 | 5.58567535 |
| ENSMUSG00 | 2700050L05F | 5155 | 8.86882688 | 12.0302524 | 10.5810037 | 12.7809531 | 16.807524 | 12.825165 |
| ENSMUSG00 | Timeless | 5919 | 0.31567961 | 1.06646797 | 0.95587699 | 0.79857203 | 0.26724407 | 0.20275116 |
| ENSMUSG00 | Ifi203 | 6368 | 0.49944071 | 0.06053573 | 0.26376728 | 0.55528272 | 0.08012936 | 0.02569847 |
| ENSMUSG00 | Magi2 | 7723 | 0.62801614 | 1.7906913 | 0.23465953 | 0.26630539 | 0.25106853 | 0.23308639 |
| ENSMUSG00 | BC013529 | 2510 | 99.4045401 | 115.628263 | 98.4943856 | 80.4442714 | 97.6209788 | 101.92674 |
| ENSMUSG00 | Bahd1 | 4656 | 3.49226622 | 5.5161885 | 3.47462941 | 6.08342031 | 2.67406265 | 3.53820492 |
| ENSMUSG00 | Gnaz | 3781 | 3.55391495 | 4.85560287 | 5.03861263 | 3.99851274 | 5.02031521 | 2.68346094 |
| ENSMUSG00 | Slc7a5 | 4477 | 2.01574583 | 2.27101615 | 1.53032961 | 2.26469835 | 0.72943669 | 1.79718977 |
| ENSMUSG00 | Fkbp6 | 1720 | 0.7627505 | 1.23267643 | 0.07709624 | 0.56640419 | 0.77132894 | 0.0317147 |
| ENSMUSG00 | Ptger3 | 5499 | 32.5765039 | 19.4796256 | 41.4367159 | 24.9601923 | 21.9731693 | 34.8534079 |
| ENSMUSG00 | Saa4 | 2011 | 0 | 0 | 0 | 0 | 0 | 0 |
| ENSMUSG00 | Cox15 | 4979 | 32.138142 | 31.2790884 | 24.7153908 | 28.0018116 | 32.5896516 | 17.2226282 |
| ENSMUSG00 | Lats1 | 4420 | 15.6143697 | 16.5054917 | 22.0509194 | 15.3226239 | 14.696058 | 23.6894435 |
| ENSMUSG00 | Rab11fip2 | 8899 | 1.82717064 | 2.40959285 | 1.79310769 | 2.29896834 | 1.57683485 | 3.5920752 |
| ENSMUSG00 | Ythdf2 | 5444 | 18.8188599 | 19.4108868 | 17.5215976 | 20.1089131 | 19.4582579 | 19.0180989 |
| ENSMUSG00 | Saa3 | 527 | 0 | 0 | 0 | 0 | 0 | 0 |
| ENSMUSG00 | Elavl1 | 6024 | 23.0653062 | 25.5410541 | 25.1093471 | 22.0780903 | 21.625222 | 19.9398259 |
| ENSMUSG00 | Ipo8 | 9275 | 10.8357795 | 8.52549343 | 7.85386925 | 12.8300359 | 9.47907769 | 11.2803791 |
| ENSMUSG00 | Stat2 | 3417 | 8.17913456 | 4.65365113 | 5.05792234 | 7.49729342 | 12.0659384 | 5.82689103 |
| ENSMUSG00 | Nup43 | 2719 | 2.70495178 | 2.7646506 | 3.83657041 | 3.22469173 | 3.99728505 | 6.60048276 |
| ENSMUSG00 | Disp2 | 9351 | 0 | 0 | 0 | 0 | 0.00545678 | 0.15458837 |
| ENSMUSG00 | Negr1 | 7090 | 0.5607261 | 0 | 0.26184449 | 0 | 0.79886146 | 0.36161017 |
| ENSMUSG00 | Ift88 | 4131 | 5.5817426 | 4.80581316 | 4.20511361 | 3.7732857 | 5.90428653 | 5.83654832 |
| ENSMUSG00 | Rbms2 | 5729 | 8.33414769 | 10.3707249 | 7.67687795 | 8.40801701 | 9.75281577 | 6.75081837 |
| ENSMUSG00 | Orc3 | 6067 | 24.8938636 | 17.219088 | 18.4471842 | 17.7644745 | 19.2095339 | 18.9443431 |
| ENSMUSG00 | Tph1 | 3984 | 0.17961814 | 0.29027977 | 0.74335429 | 0.12679433 | 0.47388953 | 0.58875978 |
| ENSMUSG00 | Ndufb10 | 1245 | 104.226416 | 108.642043 | 145.919337 | 96.4216563 | 65.0022747 | 206.673853 |
| ENSMUSG00 | Baz2a | 8457 | 4.31542286 | 3.86312023 | 5.19007096 | 5.24356542 | 4.16923563 | 3.55083101 |
| ENSMUSG00 | Gjb6 | 2799 | 0 | 0 | 0.03158403 | 0 | 0 | 0 |
| ENSMUSG00 | Plcb2 | 7499 | 0 | 0 | 0 | 0 | 0 | 0.105476 |
| ENSMUSG00 | Pfpl | 3117 | 0 | 0 | 0 | 0 | 0 | 0 |
| ENSMUSG00 | Gm9769 | 961 | 1.65475465 | 1.2034075 | 1.28787888 | 1.87731761 | 0.90265181 | 1.87318022 |
| ENSMUSG00 | Bub1b | 4255 | 0.61665375 | 0.48696052 | 0.51941061 | 0.71231298 | 0 | 0 |
| ENSMUSG00 | Tnni3k | 3256 | 0 | 0 | 0 | 0 | 0 | 0 |
| ENSMUSG00 | Bmf | 7586 | 0.82278018 | 0.53356988 | 1.10125868 | 0.50417832 | 0.36995131 | 0.8808709 |
| ENSMUSG00 | Flywch1 | 3567 | 13.8313853 | 14.4410728 | 12.2679632 | 16.4275975 | 15.1777361 | 9.67267134 |
| ENSMUSG00 | Klhdc5 | 6280 | 5.05806193 | 4.19713119 | 5.22257445 | 6.69930954 | 4.00573301 | 6.78825814 |
| ENSMUSG00 | Ppapdc2 | 2878 | 38.1669189 | 39.0614905 | 45.2615814 | 37.5865078 | 32.9597056 | 42.6178084 |
| ENSMUSG00 | Gramd1b | 7438 | 1.35226359 | 1.37342373 | 0.99837451 | 1.30493009 | 0.71346371 | 1.36043172 |
| ENSMUSG00 | Mrps35 | 3195 | 22.061492 | 19.1388404 | 27.2543445 | 18.1821883 | 14.4534805 | 24.7051036 |
| ENSMUSG00 | Mettl11b | 2131 | 0 | 0.11306063 | 0 | 0 | 0 | 0 |
| ENSMUSG00 | Cacna2d1 | 8570 | 0.55666951 | 0.04498151 | 1.70721237 | 1.76410463 | 0.42869301 | 1.20301209 |
| ENSMUSG00 | Rep15 | 981 | 0 | 0 | 0.63081122 | 0.36780881 | 0.46813189 | 0.16681737 |
| ENSMUSG00 | Zmym5 | 7044 | 13.7823514 | 12.1355407 | 16.9616101 | 19.224292 | 14.2416035 | 20.5992449 |
| ENSMUSG00 | Gorab | 2504 | 9.76422544 | 7.21642005 | 9.42643138 | 10.0724238 | 11.4524054 | 21.9809193 |
| ENSMUSG00 | Gpr26 | 11568 | 0 | 0 | 0 | 0 | 0 | 0 |
| ENSMUSG00 | Sdr9c7 | 3178 | 0 | 0 | 0 | 0 | 0 | 0 |
| ENSMUSG00 | Pnrc1 | 2594 | 78.2848861 | 85.3015198 | 70.0515716 | 73.2212345 | 63.6354379 | 151.713867 |
| ENSMUSG00 | Svs2 | 1982 | 0 | 0 | 0 | 0 | 0 | 0 |
| ENSMUSG00 | Gpr176 | 3896 | 0 | 0 | 0 | 0 | 0 | 0 |
| ENSMUSG00 | Rdh7 | 1587 | 0 | 0 | 0 | 0 | 0 | 0 |
| ENSMUSG00 | Abcc8 | 5118 | 0 | 0 | 0 | 0 | 0 | 0 |
| ENSMUSG00 | Ndp | 1936 | 0 | 0 | 0 | 0 | 0 | 0 |
| ENSMUSG00 | 9430038I01R | 5855 | 1.91478147 | 1.65422286 | 2.93672364 | 0.88125233 | 1.35954135 | 1.29967965 |
| ENSMUSG00 | Tdrd6 | 7606 | 0 | 0 | 0 | 0 | 0 | 0 |
| ENSMUSG00 | Rgl3 | 2531 | 63.1438292 | 57.9722309 | 41.6346084 | 54.9855574 | 54.6754371 | 43.6545095 |
| ENSMUSG00 | Maob | 2924 | 7.53233112 | 4.53189863 | 7.81544215 | 1.64121475 | 8.04485611 | 3.68450154 |
| ENSMUSG00 | Hmx3 | 1455 | 1.39349107 | 0.69547442 | 1.33668804 | 1.11593952 | 1.01702055 | 1.68709109 |
| ENSMUSG00 | Hs2st1 | 4908 | 7.2091234 | 6.92164664 | 7.60113193 | 6.78560047 | 8.84748285 | 5.56273714 |
| ENSMUSG00 | Thbs1 | 6051 | 64.5574866 | 67.4180167 | 72.4717377 | 69.546337 | 81.7973633 | 37.3848716 |
| ENSMUSG00 | Wfdc5 | 3016 | 0 | 0 | 0 | 0 | 0 | 0 |
| ENSMUSG00 | Tax1bp3 | 2067 | 4.44292017 | 4.52257851 | 6.94997532 | 3.89274114 | 3.25857843 | 4.51278501 |
| ENSMUSG00 | 1700034J05R | 4199 | 0.12308198 | 0 | 0 | 0 | 0 | 0 |
| ENSMUSG00 | Kcns1 | 2708 | 0 | 0 | 0 | 0 | 0 | 0 |
| ENSMUSG00 | Cd209c | 1776 | 0 | 0 | 0 | 0 | 0 | 0 |
| ENSMUSG00 | Ikzf5 | 6685 | 3.8179534 | 4.22397234 | 5.41530433 | 7.17862666 | 4.2515619 | 3.60261877 |
| ENSMUSG00 | Fmo2 | 5998 | 2.80369594 | 3.03675811 | 1.3117586 | 0.07820383 | 3.11364681 | 0.41835059 |
| ENSMUSG00 | Alkbh3 | 3494 | 12.2088239 | 10.150321 | 10.5507547 | 12.2063471 | 13.844592 | 9.42981231 |
| ENSMUSG00 | 2310057M21 | 2208 | 7.83226179 | 8.83854577 | 10.6901196 | 6.35684571 | 10.7229341 | 9.52388898 |
| ENSMUSG00 | Fmo1 | 4050 | 0.59878625 | 0.95183096 | 0.72032633 | 0 | 0.42836958 | 0.51182039 |
| ENSMUSG00 | Ankrd6 | 6346 | 4.25369859 | 1.95904539 | 8.079757 | 3.37167545 | 6.25567608 | 7.13025946 |
| ENSMUSG00 | Arntl2 | 3097 | 0.03851031 | 0 | 0 | 0.02330129 | 0.01647607 | 0 |
| ENSMUSG00 | Scamp2 | 2332 | 55.2007916 | 62.8779346 | 55.2143948 | 43.9730577 | 54.0897088 | 34.8067437 |
| ENSMUSG00 | Ccdc114 | 5730 | 1.58189347 | 1.25301569 | 0.90255072 | 2.50622229 | 2.46672009 | 1.21855282 |
| ENSMUSG00 | Tmem194 | 7519 | 0.7455144 | 1.86490952 | 1.00525537 | 1.15170777 | 2.7416752 | 1.27322761 |
| ENSMUSG00 | Cd209e | 1796 | 0 | 0 | 0 | 0 | 0 | 0 |
| ENSMUSG00 | 2810417H13I | 4963 | 0.22429044 | 0.80585829 | 0.11578158 | 0.00727021 | 0.02056271 | 0.30775333 |
| ENSMUSG00 | Cuzd1 | 2246 | 0 | 0 | 0 | 0 | 0 | 0 |
| ENSMUSG00 | Zfp704 | 13834 | 3.77036218 | 4.01960057 | 3.58177433 | 4.4365735 | 5.34829008 | 4.45179525 |
| ENSMUSG00 | Emp3 | 966 | 0.32923793 | 0.14964734 | 0.41181841 | 0.14940805 | 0.21128934 | 0.67763079 |
| ENSMUSG00 | Ccbl2 | 2697 | 21.7129488 | 14.7578796 | 13.3408603 | 14.9840155 | 13.0734986 | 18.7999831 |
| ENSMUSG00 | Gm6369 | 428 | 0 | 0 | 0 | 0 | 0 | 0 |
| ENSMUSG00 | Ttc12 | 4524 | 3.19871682 | 6.64640581 | 2.38400745 | 6.59589983 | 7.30881776 | 4.58194647 |
| ENSMUSG00 | Gas8 | 1437 | 17.1250121 | 11.266976 | 7.1977949 | 12.7052989 | 16.3696307 | 8.54111876 |
| ENSMUSG00 | Prrc2c | 8786 | 5.08142552 | 4.41498815 | 3.54681304 | 7.39628525 | 6.13292202 | 3.49237072 |
| ENSMUSG00 | Gpr34 | 2086 | 0 | 0.66989781 | 0 | 0.19026965 | 0.02446135 | 0 |
| ENSMUSG00 | Syngr4 | 995 | 0.15982103 | 0 | 0.17769585 | 0.43516034 | 0 | 0 |
| ENSMUSG00 | Tm7sf3 | 3425 | 6.82517446 | 12.4651643 | 8.23380317 | 8.31203887 | 12.1867374 | 6.33886506 |
| ENSMUSG00 | Trappc5 | 2544 | 17.7524473 | 16.1947359 | 16.6973157 | 21.8421182 | 15.8454548 | 14.5378973 |
| ENSMUSG00 | Fgfr1op2 | 4610 | 43.5326823 | 41.2877324 | 53.3298592 | 40.5981703 | 40.6440026 | 44.8818684 |
| ENSMUSG00 | Tbc1d10c | 1748 | 0.04548682 | 0 | 0.40459353 | 0 | 0.35029549 | 0.62413362 |
| ENSMUSG00 | Lrp1 | 18050 | 0.20483433 | 0.33637073 | 0.00489771 | 0.47576325 | 0.18657844 | 0.0649756 |
| ENSMUSG00 | 4933424B01I | 4004 | 19.9671007 | 18.617489 | 20.7430727 | 23.8894855 | 21.422412 | 15.5105778 |
| ENSMUSG00 | Gbp7 | 5638 | 0.59936429 | 1.29910238 | 0.27439952 | 1.70234549 | 4.89628757 | 2.87356079 |
| ENSMUSG00 | Sema3d | 7081 | 0.03368633 | 0.05444027 | 0 | 1.20766058 | 1.15297559 | 0.48147577 |
| ENSMUSG00 | Nxph4 | 2649 | 0 | 0 | 0 | 0 | 0 | 0 |
| ENSMUSG00 | Daam2 | 6019 | 0.40290485 | 0.52037195 | 0.09546834 | 1.00111338 | 0.33062447 | 0.13594271 |
| ENSMUSG00 | Klhdc4 | 4106 | 6.62268599 | 7.9450125 | 6.0930938 | 10.1233598 | 6.87228101 | 6.48984962 |
| ENSMUSG00 | Gbp5 | 4074 | 0.02927502 | 0 | 0 | 0.21255991 | 0.71391837 | 0 |
| ENSMUSG00 | Dnm3 | 12707 | 1.29212351 | 1.96431707 | 0.44177493 | 1.64977319 | 2.03189944 | 1.12687382 |
| ENSMUSG00 | Plekha1 | 8509 | 28.0984205 | 35.6882076 | 19.1269463 | 31.7822216 | 33.2100211 | 22.5563094 |
| ENSMUSG00 | Mrps28 | 780 | 17.8389977 | 21.2514565 | 20.8542029 | 15.9131068 | 11.5790622 | 34.303104 |
| ENSMUSG00 | Bach2 | 10556 | 0.20337211 | 0.07760227 | 0.10468417 | 0.00341816 | 0.09667748 | 0.4134087 |
| ENSMUSG00 | Accs | 6711 | 0.84119777 | 0.46671416 | 0.65864764 | 0.83336565 | 1.44464482 | 1.25989244 |
| ENSMUSG00 | Cdk6 | 3727 | 3.20006551 | 4.99060013 | 1.76712493 | 2.46872051 | 3.17631317 | 2.24666334 |
| ENSMUSG00 | Pacsin1 | 5045 | 0.59101309 | 0.88827336 | 0.98128968 | 0.42197039 | 1.13279566 | 1.16234835 |
| ENSMUSG00 | Ndufa4l2 | 956 | 0 | 0 | 0 | 0.03774272 | 0.16012461 | 0 |
| ENSMUSG00 | BC052040 | 4590 | 1.14329486 | 1.40675016 | 2.36898767 | 1.68225654 | 2.10108606 | 3.713867 |
| ENSMUSG00 | Btnl9 | 6679 | 0.19047393 | 0 | 0.17206886 | 0.74551911 | 0.27503362 | 0 |
| ENSMUSG00 | Gzmg | 904 | 0 | 0 | 0 | 0 | 0 | 0 |
| ENSMUSG00 | Stac3 | 2413 | 0.08237771 | 0.01996952 | 0.14654569 | 0.19439145 | 0.12687868 | 0.27127698 |
| ENSMUSG00 | Hey1 | 2198 | 0.18087116 | 0 | 0 | 0 | 0 | 0 |
| ENSMUSG00 | Ddx58 | 6301 | 4.48597788 | 4.71081548 | 3.61976686 | 3.10943504 | 7.15065701 | 6.28948593 |
| ENSMUSG00 | AI848100 | 6284 | 15.949008 | 14.8071324 | 16.2626768 | 19.2812708 | 17.7423029 | 20.3561518 |
| ENSMUSG00 | Btbd16 | 3416 | 0.04655209 | 0 | 0.01293965 | 0 | 0 | 0 |
| ENSMUSG00 | C030048B08 | 3891 | 6.4675454 | 5.47376187 | 6.31617186 | 5.61956283 | 6.34715135 | 6.16150293 |
| ENSMUSG00 | Alx4 | 7757 | 0.14350309 | 0 | 0 | 0 | 0.01315621 | 0 |
| ENSMUSG00 | Cchcr1 | 5265 | 3.05056299 | 2.54431736 | 2.98037118 | 1.45972944 | 4.0607885 | 3.60035599 |
| ENSMUSG00 | Ctsg | 997 | 0 | 0 | 0 | 0 | 0 | 0 |
| ENSMUSG00 | Zfp770 | 6790 | 2.41226185 | 1.35546546 | 3.02056777 | 3.15651465 | 2.56259119 | 4.77205765 |
| ENSMUSG00 | Slc25a24 | 3646 | 1.95179128 | 5.04860694 | 5.79497558 | 2.20688314 | 4.70237583 | 5.84990891 |
| ENSMUSG00 | Gm15429 | 795 | 0.10001379 | 0.42428314 | 0.33359882 | 0.18154488 | 0.12836824 | 0.06861545 |
| ENSMUSG00 | Vprbp | 5022 | 10.7627897 | 11.226232 | 13.2552719 | 13.2056547 | 15.311977 | 8.24973656 |
| ENSMUSG00 | Cul9 | 7809 | 5.61535344 | 6.25084721 | 5.08301383 | 5.28131347 | 7.67127228 | 4.07600256 |
| ENSMUSG00 | Olfr56 | 2048 | 0 | 0 | 0 | 0 | 0.04983045 | 0.46611932 |
| ENSMUSG00 | Il7 | 4241 | 1.14363797 | 0.89760173 | 0.40647769 | 1.08050452 | 1.01066154 | 1.18333733 |
| ENSMUSG00 | Nsmce4a | 2343 | 65.8519078 | 58.9836603 | 65.5387122 | 54.5773648 | 66.0097931 | 63.1519069 |
| ENSMUSG00 | Fam102b | 5691 | 0.2864127 | 0.63503482 | 1.14174502 | 0.77350368 | 0.9504122 | 0.61824433 |
| ENSMUSG00 | 1700019B03I | 1682 | 0 | 0 | 0.18395535 | 0.08580748 | 0 | 0 |
| ENSMUSG00 | Arhgap9 | 2307 | 0.18955799 | 0.22975764 | 0.01915988 | 0.75073175 | 0 | 0.80393388 |
| ENSMUSG00 | Trim7 | 6126 | 0.22064746 | 0.64500298 | 0.38241882 | 0.93061754 | 0.59139286 | 0.2893979 |
| ENSMUSG00 | Ankib1 | 6590 | 9.33258397 | 6.92451604 | 7.81413461 | 12.9325932 | 10.0426722 | 8.87357004 |
| ENSMUSG00 | Mars | 3488 | 20.2766828 | 16.4535701 | 18.9581299 | 13.4066312 | 18.2278854 | 12.894461 |
| ENSMUSG00 | Skiv2l | 5342 | 22.3931563 | 22.4785874 | 18.4850089 | 22.7555987 | 25.7233302 | 12.1107159 |
| ENSMUSG00 | 1810074P20F | 5666 | 18.7130015 | 15.0189299 | 18.2909149 | 16.4426119 | 18.5157481 | 15.5339324 |
| ENSMUSG00 | Bcor | 8347 | 3.82932866 | 2.19370409 | 2.69013253 | 4.08500444 | 4.1263692 | 2.64675426 |
| ENSMUSG00 | Sec1 | 2463 | 0 | 0 | 0 | 0.01464963 | 0 | 0 |
| ENSMUSG00 | Trim41 | 2837 | 3.68547458 | 3.3800148 | 3.06935604 | 3.15415828 | 4.94615909 | 4.83579258 |
| ENSMUSG00 | 4932412H11I | 3048 | 0.1434745 | 0.079046 | 0.21752876 | 0.28411059 | 0.05022281 | 0.08948373 |
| ENSMUSG00 | Lyrm5 | 2775 | 51.1305621 | 54.6113011 | 54.8899283 | 57.2501773 | 44.2412469 | 77.6860355 |
| ENSMUSG00 | Gpr63 | 2585 | 0 | 0 | 0 | 0.02791648 | 0.65140054 | 0 |
| ENSMUSG00 | Cacng5 | 6765 | 0 | 0 | 0 | 0 | 0 | 0 |
| ENSMUSG00 | Pxmp3 | 2263 | 82.0758307 | 79.487401 | 85.6300839 | 75.7675102 | 79.5271885 | 91.5381728 |
| ENSMUSG00 | Cbln3 | 5715 | 0.01391268 | 0.05902101 | 0 | 0 | 0.017857 | 0 |
| ENSMUSG00 | Aqr | 10029 | 7.98360133 | 8.09593728 | 7.47908261 | 10.6817819 | 12.2567095 | 5.02849815 |
| ENSMUSG00 | Ppp1ca | 1423 | 187.630223 | 188.716122 | 235.11156 | 191.693785 | 140.385286 | 172.36882 |
| ENSMUSG00 | Klhl32 | 9307 | 0.3972558 | 0.33653366 | 0.05699174 | 0.66294505 | 1.04169028 | 0.31649952 |
| ENSMUSG00 | Wdr47 | 4624 | 10.274178 | 14.4121647 | 11.1078161 | 13.1640158 | 14.5332476 | 8.40535487 |
| ENSMUSG00 | Map3k10 | 4212 | 1.11375753 | 1.51011642 | 1.32227736 | 1.47343582 | 0.83590217 | 0.9972209 |
| ENSMUSG00 | Abhd13 | 5490 | 24.8091578 | 24.9972654 | 29.532306 | 23.3054517 | 23.0966382 | 23.8864236 |
| ENSMUSG00 | C630050I24F | 782 | 0 | 0.55457542 | 0.33914458 | 0 | 0 | 0.24414639 |
| ENSMUSG00 | Havcr1 | 2300 | 0 | 0 | 0 | 0 | 0.95397137 | 0 |
| ENSMUSG00 | Akap9 | 15713 | 6.10007405 | 9.64771508 | 6.47007185 | 10.4092094 | 8.64457535 | 7.21398846 |
| ENSMUSG00 | Fbxl4 | 3866 | 16.0626127 | 16.1410871 | 19.3911862 | 19.7396596 | 18.1746817 | 13.5597146 |
| ENSMUSG00 | 5330417C22I | 6490 | 0.06125652 | 0.04454833 | 0.2656197 | 0.10007347 | 0.04717385 | 0.14708973 |
| ENSMUSG00 | Timd2 | 4471 | 0 | 0.1616633 | 0 | 0 | 0 | 0.71984062 |
| ENSMUSG00 | Slc25a28 | 1510 | 19.03524 | 19.5936924 | 15.9536453 | 17.8259637 | 21.0188118 | 26.4076309 |
| ENSMUSG00 | Dtx3 | 2647 | 12.2705809 | 15.4189333 | 9.919114 | 15.6623609 | 12.028885 | 23.5858138 |
| ENSMUSG00 | Cdh18 | 3329 | 0 | 0 | 0 | 0 | 0 | 0 |
| ENSMUSG00 | Rc3h1 | 11288 | 3.51840229 | 3.81632524 | 2.48654946 | 4.13946205 | 4.06384761 | 4.68027784 |
| ENSMUSG00 | Hipk4 | 4362 | 0.12759668 | 0.04418748 | 0.01013339 | 0 | 0 | 0.15006679 |
| ENSMUSG00 | Plekha4 | 3694 | 0.35515183 | 0.76962645 | 0.31111205 | 1.48469701 | 0.8978653 | 1.21089357 |
| ENSMUSG00 | Mterf | 3753 | 1.21819351 | 1.5792519 | 1.86088228 | 1.70171113 | 2.03942349 | 4.14243122 |
| ENSMUSG00 | Pitpnc1 | 6624 | 1.34438823 | 0.61833448 | 2.9628047 | 1.49252419 | 3.30469735 | 2.67228878 |
| ENSMUSG00 | Ltb4r2 | 1082 | 0 | 0 | 0.24511188 | 0.03334755 | 0.04715931 | 0.20166092 |
| ENSMUSG00 | Zbtb38 | 14545 | 4.61923424 | 4.64802876 | 5.75885132 | 5.92147407 | 5.05527725 | 6.02873601 |
| ENSMUSG00 | Gyltl1b | 5280 | 26.3455353 | 18.6539939 | 22.7539033 | 22.0797511 | 22.952205 | 22.8786699 |
| ENSMUSG00 | Ppp1r15a | 2678 | 31.3976256 | 25.5866769 | 34.4305618 | 42.347224 | 53.4652765 | 38.0500569 |
| ENSMUSG00 | Slc26a10 | 3739 | 0.9250406 | 0.43817572 | 0.10639652 | 0.29915576 | 0.40941195 | 0.0218839 |
| ENSMUSG00 | Rprd1a | 4290 | 7.99743116 | 6.16651672 | 9.3555416 | 7.20799817 | 6.75593974 | 7.5211884 |
| ENSMUSG00 | Spns2 | 5144 | 8.01446992 | 20.4305269 | 2.59505377 | 14.1901198 | 15.8415867 | 3.66383665 |
| ENSMUSG00 | Sgms1 | 12080 | 36.3262412 | 26.2352639 | 58.4649874 | 38.0773099 | 33.3867942 | 40.3700786 |
| ENSMUSG00 | Cdh12 | 5627 | 0 | 0 | 0 | 0 | 0 | 0 |
| ENSMUSG00 | Usp45 | 6159 | 4.55067605 | 3.62239369 | 5.7055472 | 7.65696253 | 4.4821025 | 5.87208502 |
| ENSMUSG00 | 1700054O13I | 553 | 0 | 0 | 0 | 0 | 0 | 0 |
| ENSMUSG00 | Arglu1 | 1592 | 20.4520971 | 18.1001837 | 18.2415897 | 22.2339734 | 19.5195119 | 22.8202384 |
| ENSMUSG00 | Os9 | 4034 | 62.274388 | 61.75605 | 51.7952682 | 52.2538698 | 59.6403971 | 39.6205716 |
| ENSMUSG00 | Mybbp1a | 6000 | 9.35578975 | 8.60127992 | 9.37079072 | 7.92000877 | 12.3738961 | 8.19602909 |
| ENSMUSG00 | Gtpbp10 | 5252 | 3.34575826 | 3.11028254 | 4.72148399 | 4.29384574 | 2.89525133 | 2.90818697 |
| ENSMUSG00 | Blvrb | 1573 | 4.32179731 | 5.94289242 | 5.47956732 | 5.57402214 | 3.89266694 | 5.4098458 |
| ENSMUSG00 | 4921522P10F | 2099 | 0 | 0 | 0.33693639 | 0 | 0 | 0.63671145 |
| ENSMUSG00 | Ggt6 | 2209 | 4.71522675 | 9.75071962 | 7.26358941 | 5.16157812 | 6.6987999 | 5.61790895 |
| ENSMUSG00 | Rabggta | 3274 | 14.5470573 | 17.5731863 | 17.848148 | 15.9911566 | 18.9205926 | 18.4024674 |
| ENSMUSG00 | A330021E22F | 7457 | 3.21477199 | 1.17606711 | 3.34907353 | 2.46288865 | 3.32557579 | 2.37377509 |
| ENSMUSG00 | Prdm13 | 6562 | 0 | 0 | 0 | 0 | 0 | 0 |
| ENSMUSG00 | Dgkz | 6935 | 4.68352236 | 5.59337937 | 5.31569388 | 5.43702041 | 5.14310551 | 6.04092947 |
| ENSMUSG00 | Bptf | 16632 | 1.95526594 | 2.92908207 | 2.35732532 | 2.80074071 | 1.43273915 | 2.64842126 |
| ENSMUSG00 | Dom3z | 1909 | 14.3902761 | 15.8013163 | 13.2674992 | 13.4008222 | 19.2184202 | 12.0585624 |
| ENSMUSG00 | Xaf1 | 4763 | 0.53419079 | 1.16343499 | 0.38049036 | 0.71967126 | 0.86775907 | 2.0328568 |
| ENSMUSG00 | Lrrc52 | 1135 | 0.66551025 | 2.08029574 | 0.85677581 | 0 | 0.67435739 | 0.69688506 |
| ENSMUSG00 | Ltbp4 | 6770 | 0.62246395 | 0.42705857 | 0.4048027 | 1.52962286 | 0.75371308 | 0.44719128 |
| ENSMUSG00 | Sox30 | 2992 | 0 | 0 | 0.04432003 | 0.01205951 | 0.08527135 | 0 |
| ENSMUSG00 | Lrfn2 | 3394 | 0 | 0 | 0 | 0 | 0 | 0 |
| ENSMUSG00 | Chrm4 | 1625 | 0 | 0 | 0.02720113 | 0.02220434 | 0 | 0 |
| ENSMUSG00 | 2210010C17I | 1890 | 0.08413858 | 0.02549547 | 0 | 0 | 0 | 0.20203437 |
| ENSMUSG00 | 9-Mar | 3033 | 0.41944457 | 0.63549545 | 0.72868188 | 0.68999624 | 0.77389162 | 0.56653553 |
| ENSMUSG00 | Abcg5 | 2753 | 0 | 0 | 0 | 0 | 0 | 0 |
| ENSMUSG00 | Ambra1 | 6364 | 9.8389192 | 6.39810553 | 8.67506316 | 6.66191111 | 10.8402986 | 5.99150623 |
| ENSMUSG00 | Pvr | 5521 | 2.92351478 | 4.93123344 | 4.71561049 | 4.24148649 | 4.32536567 | 3.50257548 |
| ENSMUSG00 | 4921511H03I | 3414 | 0 | 0 | 0 | 0 | 0 | 0 |
| ENSMUSG00 | Manea | 5008 | 21.1955138 | 26.806595 | 23.5572522 | 34.900444 | 27.2555023 | 22.100736 |
| ENSMUSG00 | Tsfm | 2897 | 9.71587166 | 12.1588848 | 15.9596575 | 10.761093 | 9.19425895 | 13.2183616 |
| ENSMUSG00 | Tlr8 | 6174 | 0 | 0 | 0 | 0 | 0 | 0 |
| ENSMUSG00 | Zfp609 | 12366 | 0.64296044 | 0.85337464 | 0.68987188 | 1.42974299 | 0.95731192 | 1.31234113 |
| ENSMUSG00 | Cblc | 2527 | 7.12672444 | 8.19951331 | 7.32907488 | 7.06791137 | 9.34911437 | 7.08039705 |
| ENSMUSG00 | Gm885 | 2446 | 0 | 0.11820059 | 0.05421322 | 0 | 0.16688921 | 0.17841138 |
| ENSMUSG00 | Abhd11 | 2065 | 37.8495277 | 33.8121814 | 43.7523327 | 31.1196713 | 33.9764004 | 43.4677181 |
| ENSMUSG00 | Matn1 | 1993 | 0 | 0 | 0 | 0 | 0 | 0 |
| ENSMUSG00 | Necab1 | 6215 | 0 | 0 | 0 | 0 | 0 | 0 |
| ENSMUSG00 | Adam22 | 13079 | 1.41647327 | 2.09265992 | 0.51032023 | 2.81947011 | 2.25500766 | 1.27625042 |
| ENSMUSG00 | Gm9770 | 808 | 0 | 0 | 0 | 0 | 0 | 0 |
| ENSMUSG00 | Tmem225 | 989 | 0 | 0 | 0 | 0 | 0 | 0 |
| ENSMUSG00 | Pitpnm3 | 10852 | 0.82793389 | 1.5097116 | 0.2199502 | 1.84200633 | 1.46703182 | 0.79169843 |
| ENSMUSG00 | Tex2 | 5561 | 8.93622561 | 9.18497189 | 7.98033634 | 11.886766 | 12.2496335 | 6.32206617 |
| ENSMUSG00 | Ckap5 | 9417 | 7.30771337 | 6.86184761 | 9.45807727 | 8.6747105 | 8.56670916 | 9.22477711 |
| ENSMUSG00 | Otud6b | 5620 | 20.5639114 | 15.6991772 | 22.6121528 | 23.0616911 | 16.8060181 | 18.0827946 |
| ENSMUSG00 | C3ar1 | 4303 | 0 | 0 | 0 | 0 | 0 | 0.55145099 |
| ENSMUSG00 | Aipl1 | 3890 | 0 | 0 | 0.21589589 | 0 | 0.11805588 | 0.16827541 |
| ENSMUSG00 | Wbscr27 | 1767 | 48.0350599 | 40.5780565 | 50.6307472 | 38.6345378 | 58.4767465 | 48.2515689 |
| ENSMUSG00 | Wdr7 | 7081 | 3.54267875 | 5.40319659 | 4.5069525 | 4.59624406 | 6.53593036 | 5.40793582 |
| ENSMUSG00 | Gstm2 | 1863 | 102.920388 | 180.925477 | 75.1407605 | 122.597607 | 106.763721 | 80.6819844 |
| ENSMUSG00 | BC018242 | 2612 | 1.50681186 | 1.9186026 | 1.4891892 | 1.38139527 | 1.36747562 | 0.79359594 |
| ENSMUSG00 | Apoc1 | 533 | 0.14917629 | 0.90406083 | 0.08293029 | 0.06769614 | 0.38293715 | 0.35820352 |
| ENSMUSG00 | Btaf1 | 8471 | 6.74402379 | 7.84430454 | 7.88964545 | 9.15361984 | 11.3425411 | 8.22650257 |
| ENSMUSG00 | Slc26a7 | 5624 | 1.92980907 | 0 | 7.9616762 | 1.40504405 | 3.65640638 | 8.82158051 |
| ENSMUSG00 | Rundc3b | 4135 | 0 | 0 | 0 | 0 | 0.03702035 | 0 |
| ENSMUSG00 | Wbscr28 | 929 | 0 | 0 | 0 | 0 | 0 | 0 |
| ENSMUSG00 | Cyp2b13 | 1867 | 0 | 0 | 0 | 0 | 0 | 0 |
| ENSMUSG00 | Abcb1a | 5344 | 0 | 0 | 0 | 0 | 0 | 0 |
| ENSMUSG00 | Ofd1 | 4430 | 4.33451402 | 4.98180373 | 3.85144727 | 5.80733583 | 4.7225314 | 6.27378271 |
| ENSMUSG00 | 1110051M20 | 5349 | 4.19182857 | 4.91863851 | 3.46243639 | 4.49929402 | 6.80160889 | 3.91094565 |
| ENSMUSG00 | Cd79b | 1986 | 0.18016079 | 0.29115675 | 0.44513437 | 0.90840998 | 0.77079118 | 0.28840253 |
| ENSMUSG00 | Ranbp17 | 10510 | 0.13995745 | 0.1237901 | 0.26075302 | 0.26778301 | 0.52919838 | 0.2958429 |
| ENSMUSG00 | Pogk | 8714 | 6.33697067 | 6.8679781 | 7.7609387 | 7.01020213 | 5.41650191 | 5.85306121 |
| ENSMUSG00 | Mis12 | 3112 | 27.121107 | 31.3088001 | 25.0126753 | 31.3283047 | 23.3324334 | 32.3491946 |
| ENSMUSG00 | Eps8l3 | 2250 | 0 | 0 | 0 | 0 | 0 | 0 |
| ENSMUSG00 | Nlrp4a | 4746 | 0 | 0 | 0 | 0 | 0 | 0 |
| ENSMUSG00 | Bace2 | 3727 | 1.04535473 | 0.15514819 | 2.37197977 | 0.5615129 | 0.90360633 | 1.55144178 |
| ENSMUSG00 | Kazn | 10797 | 5.32429608 | 4.07020795 | 3.87282981 | 3.986837 | 3.5444829 | 6.86349864 |
| ENSMUSG00 | Tlx3 | 1529 | 0 | 0 | 0 | 0 | 0 | 0 |
| ENSMUSG00 | Ildr2 | 8514 | 0 | 0 | 0.06229999 | 0 | 0 | 0 |
| ENSMUSG00 | Apobec1 | 3516 | 0.28267549 | 0 | 0.27657581 | 0.05131121 | 0.37732815 | 0.03878646 |
| ENSMUSG00 | Nlrp9c | 5906 | 0 | 0 | 0 | 0 | 0 | 0 |
| ENSMUSG00 | Tmem51 | 1872 | 84.926371 | 45.6896019 | 133.502789 | 57.5924941 | 71.9057582 | 95.1115625 |
| ENSMUSG00 | Pck2 | 2439 | 18.9078956 | 18.8083202 | 10.7287786 | 10.3260628 | 22.4901 | 11.5070126 |
| ENSMUSG00 | Dhx33 | 6318 | 6.6573755 | 6.46756932 | 5.69488766 | 5.39117601 | 6.06533844 | 4.98610452 |
| ENSMUSG00 | Gemin8 | 2390 | 1.77984787 | 2.35891788 | 1.8494495 | 1.22286427 | 1.40909657 | 1.24390614 |
| ENSMUSG00 | Plekhg1 | 11654 | 0.38547875 | 0.77733406 | 0.43617745 | 0.9876585 | 0.4772503 | 0.3065881 |
| ENSMUSG00 | Aicda | 2647 | 0 | 0 | 0 | 0 | 0 | 0 |
| ENSMUSG00 | Mael | 1534 | 0 | 0 | 0 | 0 | 0 | 0 |
| ENSMUSG00 | Dok4 | 2595 | 3.952568 | 4.92077348 | 1.20937605 | 1.44606267 | 4.71920239 | 0.68317979 |
| ENSMUSG00 | Nrl | 3859 | 0 | 0 | 0 | 0 | 0 | 0 |
| ENSMUSG00 | Erc2 | 6396 | 0.89505772 | 0.45956425 | 0.73255087 | 0.62054798 | 0.43878216 | 0.75052166 |
| ENSMUSG00 | Ppip5k2 | 10930 | 4.50659107 | 5.13606634 | 5.4878226 | 6.71792869 | 7.30615537 | 6.3283152 |
| ENSMUSG00 | Rimklb | 5458 | 0.45160128 | 0.30017205 | 0.38873002 | 0.21154735 | 0.22437395 | 0.2098818 |
| ENSMUSG00 | Cyp2b23 | 2180 | 0 | 0 | 0 | 0 | 0 | 0 |
| ENSMUSG00 | D14Abb1e | 7582 | 5.99326947 | 6.43799339 | 5.56166687 | 9.57016507 | 8.04227369 | 10.7199189 |
| ENSMUSG00 | Oaz2 | 2600 | 33.593381 | 33.767576 | 32.9473738 | 27.2974545 | 34.0306676 | 28.8271956 |
| ENSMUSG00 | Ppp1r14c | 2520 | 0.33129567 | 1.8930388 | 0.15786373 | 0.52977605 | 1.19466515 | 0.66021944 |
| ENSMUSG00 | 1700063H04 | 815 | 0 | 0 | 0 | 0 | 0 | 0 |
| ENSMUSG00 | BC048355 | 1102 | 6.89046895 | 7.039943 | 14.8409092 | 6.61395009 | 4.16730637 | 8.6377941 |
| ENSMUSG00 | Efhd2 | 2377 | 11.7075458 | 11.7171912 | 7.68000049 | 9.66944147 | 9.57415381 | 5.62245403 |
| ENSMUSG00 | Cyp2b9 | 1867 | 0 | 0 | 0 | 0 | 0 | 0 |
| ENSMUSG00 | Rad54l2 | 9302 | 1.31635003 | 1.60068917 | 1.88173832 | 2.13342555 | 1.7224556 | 1.51297719 |
| ENSMUSG00 | Clcf1 | 3945 | 4.6255426 | 3.99416136 | 6.77873638 | 3.36582823 | 5.1091048 | 5.37196317 |
| ENSMUSG00 | Sh3bgr | 1634 | 0.29196191 | 0.94367573 | 0.18935918 | 0.41955866 | 0.96806466 | 0.45068253 |
| ENSMUSG00 | Nup88 | 5025 | 30.491069 | 28.4707556 | 30.3651268 | 33.8130045 | 37.3584153 | 30.1459396 |
| ENSMUSG00 | Phc1 | 6047 | 9.45400711 | 5.11587496 | 9.2321693 | 4.85111662 | 8.34547473 | 9.13815431 |
| ENSMUSG00 | Mthfd1l | 4405 | 0 | 0.86418364 | 0.42144777 | 0.84368912 | 0.42859839 | 0.10525968 |
| ENSMUSG00 | Kremen2 | 2106 | 0.28315869 | 0 | 0 | 0 | 0.09691619 | 0.27196934 |
| ENSMUSG00 | Hmgn1 | 3398 | 23.4578394 | 26.929386 | 30.2960837 | 23.9874451 | 18.7106722 | 31.3040297 |
| ENSMUSG00 | Madd | 9298 | 4.57072582 | 4.25997585 | 5.55256535 | 6.98124307 | 5.97630924 | 4.22407835 |

| | | | | | | | | |
|---|---|---|---|---|---|---|---|---|
| ENSMUSG00 | Tbl3 | 5973 | 8.55944215 | 12.8351966 | 7.6888858 | 8.2457711 | 10.328292 | 4.17361799 |
| ENSMUSG00 | Col16a1 | 6609 | 0.36893665 | 0.43017099 | 0.12707445 | 0.14194782 | 0.50184815 | 0.00412689 |
| ENSMUSG00 | Slco4c1 | 2381 | 7.06281741 | 4.65471722 | 9.44928101 | 7.30430301 | 4.2646892 | 7.40000712 |
| ENSMUSG00 | Apobec2 | 1215 | 0.85073456 | 0.63455397 | 0.50932165 | 0.41576018 | 0.20998509 | 0.92037877 |
| ENSMUSG00 | Dnajc16 | 6494 | 3.55068967 | 3.33906867 | 3.64151309 | 3.36706483 | 3.79515545 | 1.96138844 |
| ENSMUSG00 | Limd2 | 4220 | 0.96091446 | 1.68995105 | 1.79111734 | 0.94907747 | 1.02778245 | 0.94362496 |
| ENSMUSG00 | Ap1g2 | 4349 | 13.2183088 | 9.00795067 | 15.5199389 | 13.9466353 | 17.4937518 | 10.9249072 |
| ENSMUSG00 | Cyp2s1 | 5858 | 52.2426919 | 35.864269 | 54.878799 | 32.4664487 | 47.5769998 | 32.014411 |
| ENSMUSG00 | A930016O22I | 1210 | 0 | 0.07964701 | 0.43836539 | 0.02981987 | 0.04217056 | 0 |
| ENSMUSG00 | Agmat | 1400 | 0.39755481 | 0 | 0.15786373 | 0.54123067 | 0.58315858 | 0.50652902 |
| ENSMUSG00 | St8sia4 | 5406 | 0.30886611 | 2.87015064 | 0 | 1.22809771 | 0.45306438 | 0.27244368 |
| ENSMUSG00 | Sh3pxd2b | 9824 | 4.04272445 | 2.17780744 | 6.93353421 | 2.21839932 | 3.72413594 | 4.41158406 |
| ENSMUSG00 | Camta2 | 5786 | 11.0347912 | 10.4267932 | 9.9465606 | 12.10426 | 13.6164404 | 10.2197404 |
| ENSMUSG00 | Creg1 | 2356 | 47.4500896 | 55.8561857 | 38.1419129 | 51.8258229 | 37.121905 | 40.344702 |
| ENSMUSG00 | Klc3 | 2682 | 1.48230725 | 2.78482421 | 0.84052722 | 1.07627277 | 1.84547295 | 2.19661524 |
| ENSMUSG00 | Rsc1a1 | 1748 | 0 | 0 | 0 | 0 | 0 | 0 |
| ENSMUSG00 | Il17rd | 9146 | 0.10432228 | 0.16332601 | 0.15465329 | 0.00789024 | 0.16179367 | 0 |
| ENSMUSG00 | 1110037F02I | 7175 | 7.33606358 | 7.09868562 | 8.88349235 | 10.294069 | 11.9334158 | 9.40452369 |
| ENSMUSG00 | Zfhx2 | 10552 | 0.49732026 | 0.48862294 | 0.41470645 | 0.91983226 | 0.54159914 | 0.63844161 |
| ENSMUSG00 | Scamp5 | 3183 | 16.574151 | 21.2849946 | 12.762015 | 15.133374 | 19.5096134 | 14.2756987 |
| ENSMUSG00 | Rcsd1 | 2663 | 0.53743796 | 0.16285317 | 0.46475839 | 0 | 0.07664495 | 0.61452435 |
| ENSMUSG00 | Kcna2 | 3453 | 0 | 0 | 0 | 0 | 0 | 0 |
| ENSMUSG00 | Hnrnpul1 | 3573 | 13.8304121 | 11.2340628 | 9.95871358 | 13.4108467 | 14.6524102 | 12.8701488 |
| ENSMUSG00 | Hesx1 | 1137 | 0 | 0 | 0 | 0 | 0.35902463 | 0 |
| ENSMUSG00 | Esrp1 | 7723 | 3.60851895 | 4.38625778 | 4.49287152 | 5.33545188 | 3.79906333 | 4.26971887 |
| ENSMUSG00 | AK129341 | 3636 | 7.70836462 | 5.0359868 | 2.321934 | 4.62437644 | 5.01001545 | 5.28839952 |
| ENSMUSG00 | Eif4h | 4189 | 57.9391761 | 56.4571635 | 58.8584102 | 42.6972296 | 47.5060553 | 44.7371798 |
| ENSMUSG00 | Erg | 5606 | 0.10637392 | 0.26646088 | 0 | 0.03218163 | 0.16383781 | 0 |
| ENSMUSG00 | Ppp1r13l | 3617 | 0.27478214 | 0.70607726 | 0.45216151 | 0.44890572 | 0.60661713 | 0.56555099 |
| ENSMUSG00 | Ints8 | 7290 | 10.8263186 | 11.460197 | 13.8445828 | 18.7837403 | 10.0278538 | 6.94953256 |
| ENSMUSG00 | Slc25a34 | 2665 | 0.59670515 | 0 | 0.41465144 | 0.32494149 | 0.21061544 | 0.23539088 |
| ENSMUSG00 | Rnf167 | 2524 | 50.7339155 | 41.5999435 | 46.2858443 | 43.4156772 | 55.9187621 | 36.6651549 |
| ENSMUSG00 | Cd53 | 2794 | 0 | 0 | 0 | 0 | 0 | 0 |
| ENSMUSG00 | Siah1b | 2456 | 3.33453949 | 2.51134552 | 2.12370418 | 2.61506675 | 1.62054451 | 1.12163622 |
| ENSMUSG00 | Lat2 | 1492 | 0.39968647 | 0.4521516 | 0.7998993 | 0.31438779 | 0 | 0.43873414 |
| ENSMUSG00 | Myh6 | 6258 | 0.10799667 | 0 | 0.04944278 | 0 | 0.00815378 | 0.02179182 |
| ENSMUSG00 | Cmtm5 | 873 | 0 | 0 | 0.30379274 | 0 | 0 | 0 |
| ENSMUSG00 | Appl1 | 8697 | 8.37895778 | 9.09209515 | 6.70879991 | 9.84923234 | 8.27264458 | 9.20758202 |
| ENSMUSG00 | Spen | 14377 | 3.14128301 | 3.50245754 | 4.01834937 | 4.92655306 | 4.37967224 | 3.96873794 |
| ENSMUSG00 | Snrnp25 | 1253 | 7.77341798 | 8.9989046 | 8.64281847 | 7.86145104 | 5.94561123 | 11.1667118 |
| ENSMUSG00 | Il25 | 982 | 0 | 0.09813939 | 0 | 0 | 0 | 0 |
| ENSMUSG00 | AI314976 | 1487 | 47.9097654 | 51.1353098 | 45.6583933 | 46.4917265 | 39.4965424 | 43.4890179 |
| ENSMUSG00 | Cept1 | 3842 | 69.7842167 | 54.5577885 | 66.5213577 | 65.1110918 | 54.4927694 | 87.6806531 |
| ENSMUSG00 | Gm9772 | 404 | 0 | 0 | 0 | 0 | 0 | 4.32073493 |
| ENSMUSG00 | Rfwd2 | 5036 | 2.912981 | 5.02340789 | 4.45880386 | 2.78711583 | 4.56967742 | 3.08708189 |
| ENSMUSG00 | Ttc3 | 15207 | 13.6831844 | 14.3383738 | 11.5366026 | 13.8187563 | 15.898137 | 13.3297244 |
| ENSMUSG00 | C1qtnf4 | 1506 | 2.00625267 | 3.26362357 | 3.16985328 | 1.72503799 | 2.23621567 | 3.94812175 |
| ENSMUSG00 | Iqcc | 3583 | 7.87786524 | 4.58598291 | 8.68492813 | 6.67663844 | 9.86918182 | 7.65029652 |
| ENSMUSG00 | Iqsec3 | 17189 | 0 | 0 | 0 | 0 | 0 | 0 |
| ENSMUSG00 | S100g | 477 | 2882.4807 | 4217.3744 | 1526.02925 | 2677.40873 | 3009.80736 | 7968.88338 |
| ENSMUSG00 | Chi3l3 | 2005 | 0 | 0 | 0 | 0 | 0 | 0 |
| ENSMUSG00 | Eml2 | 3517 | 22.5171786 | 20.6611188 | 14.8554389 | 15.8711751 | 18.1356184 | 11.5783442 |
| ENSMUSG00 | Agbl2 | 5788 | 0.8379697 | 1.04897922 | 0.83241204 | 0.01246788 | 1.00501155 | 0.36284506 |
| ENSMUSG00 | Tex264 | 2649 | 29.5352154 | 33.0156257 | 33.3891611 | 23.6052032 | 26.4474194 | 25.1433254 |
| ENSMUSG00 | Fam116a | 5690 | 11.9336311 | 11.0600164 | 12.9498194 | 11.7694683 | 11.2634671 | 12.6642525 |
| ENSMUSG00 | Hlcs | 8367 | 2.03837709 | 2.04448273 | 1.48977169 | 2.97125955 | 3.11634732 | 2.83927463 |
| ENSMUSG00 | 1700123O20I | 1887 | 19.0455508 | 18.4114599 | 20.7071697 | 16.2149305 | 15.5485777 | 21.2039723 |
| ENSMUSG00 | Snrpd2 | 610 | 4.75762308 | 5.05562672 | 6.59404543 | 4.79122231 | 4.09884003 | 10.507443 |
| ENSMUSG00 | Tmem146 | 2543 | 0.90673137 | 0.49266516 | 0.48668958 | 0.79457117 | 0.50163563 | 0.72932578 |
| ENSMUSG00 | Zmynd15 | 3995 | 0.40800368 | 0.49452919 | 0.19915724 | 0.30708123 | 0.01277256 | 0.03413597 |
| ENSMUSG00 | Gpr161 | 3039 | 0 | 0.4122565 | 0.07272432 | 0.16622199 | 0.2182767 | 0 |
| ENSMUSG00 | Gm11639 | 3751 | 0 | 0 | 0 | 0 | 0 | 0 |
| ENSMUSG00 | 4930579G18I | 962 | 0.28928104 | 0 | 0.5973222 | 0.22504394 | 0.15912591 | 0.34022419 |
| ENSMUSG00 | Six5 | 3004 | 0.95286105 | 1.02660862 | 1.97172003 | 3.42322992 | 1.39286379 | 2.11550964 |
| ENSMUSG00 | D4Ertd22e | 3322 | 4.82283523 | 3.13313411 | 3.36636553 | 3.75809374 | 2.16577935 | 3.30875365 |
| ENSMUSG00 | Tiprl | 1879 | 61.6326316 | 69.8818812 | 83.6518115 | 72.5481449 | 62.9207625 | 73.3758922 |
| ENSMUSG00 | Sft2d2 | 5518 | 29.9931253 | 22.8095301 | 25.4573137 | 25.5608394 | 29.0086519 | 17.4680274 |
| ENSMUSG00 | Psme4 | 8332 | 9.56669928 | 9.39785988 | 10.0637177 | 10.8956342 | 9.71289391 | 11.6110352 |
| ENSMUSG00 | Plekhh2 | 6856 | 0.27253611 | 0.45684346 | 0.25788707 | 0.43155304 | 1.02707699 | 0.34212646 |
| ENSMUSG00 | Reps2 | 8278 | 7.5688134 | 10.0994778 | 4.09019229 | 7.91991724 | 9.36942392 | 6.87961425 |
| ENSMUSG00 | Dlk1 | 5889 | 0 | 0 | 0 | 0 | 0 | 0 |
| ENSMUSG00 | Erf | 3688 | 0.59288271 | 0.77087855 | 0.73110424 | 0.53809991 | 0.41507356 | 0.66559586 |
| ENSMUSG00 | Bsdc1 | 5652 | 16.9164775 | 14.476394 | 17.4007609 | 13.9680668 | 15.5281965 | 14.4866361 |
| ENSMUSG00 | Crocc | 8380 | 1.84070721 | 2.35757056 | 1.71954664 | 1.98063729 | 2.69136732 | 2.25227331 |
| ENSMUSG00 | Ino80d | 14395 | 3.4301012 | 1.90134763 | 3.54044634 | 4.64717684 | 2.41041581 | 3.24188314 |
| ENSMUSG00 | Rsph6a | 3164 | 0 | 0 | 0 | 0 | 0 | 0 |
| ENSMUSG00 | Begain | 2644 | 0 | 0 | 0 | 0 | 0 | 0.03094702 |
| ENSMUSG00 | Osbpl10 | 1862 | 6.93906078 | 5.64159205 | 10.0890351 | 2.82920435 | 5.39860152 | 4.5701866 |
| ENSMUSG00 | Wdr25 | 2734 | 1.439573 | 1.8153634 | 1.06705254 | 1.63649361 | 0.37327268 | 0.46887639 |
| ENSMUSG00 | Tmem205 | 781 | 27.6913974 | 48.3098389 | 31.1846549 | 37.1908398 | 21.1684325 | 39.5674344 |
| ENSMUSG00 | Gfer | 1772 | 6.41651661 | 5.13952459 | 9.55378435 | 7.41188725 | 6.82463379 | 5.20249891 |
| ENSMUSG00 | Foxa3 | 3550 | 0.07839109 | 0 | 0.06225612 | 0 | 0 | 0 |
| ENSMUSG00 | Kcnd3 | 4994 | 0.00796065 | 0.40525242 | 0 | 0 | 0.09195783 | 0 |
| ENSMUSG00 | Ccr6 | 1880 | 0 | 0 | 0 | 0 | 0 | 0 |
| ENSMUSG00 | Kcnk18 | 3029 | 0 | 0 | 0 | 0 | 0 | 0 |
| ENSMUSG00 | Rnasek | 2176 | 132.329479 | 119.691048 | 190.965775 | 136.868681 | 83.0116592 | 244.368735 |
| ENSMUSG00 | Atp1a3 | 6359 | 0 | 0 | 0 | 0 | 0 | 0 |
| ENSMUSG00 | Fbxw4 | 3344 | 1.33152327 | 0.99427766 | 0.89884131 | 0.84162664 | 1.90738546 | 1.49260137 |
| ENSMUSG00 | Slc19a2 | 4233 | 19.9951377 | 14.5709062 | 16.4464689 | 13.3911864 | 20.5166293 | 27.1779892 |
| ENSMUSG00 | 4930505A04I | 984 | 0.04040191 | 0.39175969 | 0.35936458 | 1.21006857 | 0.51856073 | 0 |
| ENSMUSG00 | S100pbp | 11032 | 0.79280327 | 0.80368975 | 0.99366 | 1.49142606 | 1.29971099 | 1.08534868 |
| ENSMUSG00 | Rfx3 | 12855 | 1.84010198 | 1.19201 | 2.02527309 | 1.84971352 | 1.85766969 | 2.83036708 |
| ENSMUSG00 | Padi6 | 2514 | 0 | 0.01916724 | 0 | 0 | 0 | 0 |
| ENSMUSG00 | Ulk4 | 7937 | 0.68621656 | 1.32350314 | 1.4479626 | 1.07741521 | 1.08005936 | 0.57731377 |
| ENSMUSG00 | Slc16a11 | 2443 | 9.81275266 | 5.62142285 | 9.80654888 | 5.73059078 | 12.2187597 | 6.48651879 |
| ENSMUSG00 | Arhgef1 | 6479 | 9.85449941 | 10.3825086 | 11.3455263 | 14.3403712 | 15.3181511 | 12.0186904 |
| ENSMUSG00 | Tet2 | 10644 | 6.46157284 | 6.89024473 | 5.01236608 | 5.12891144 | 7.3442698 | 6.20367358 |
| ENSMUSG00 | Rcc2 | 4139 | 6.95408743 | 8.24257092 | 9.32307537 | 7.75864209 | 10.1954126 | 7.15638034 |
| ENSMUSG00 | Mgl2 | 1998 | 0 | 0 | 0 | 0 | 0 | 0 |
| ENSMUSG00 | Rps19 | 2568 | 2.32216592 | 2.23293871 | 2.30648246 | 1.96708966 | 2.14596908 | 2.0604673 |
| ENSMUSG00 | Cables1 | 4646 | 2.43016713 | 4.6464757 | 1.07507711 | 2.50850202 | 3.15208133 | 1.15650106 |
| ENSMUSG00 | Asgr2 | 1544 | 0 | 0 | 0 | 0 | 0 | 0 |
| ENSMUSG00 | Arhgef10I | 5604 | 28.3765029 | 24.136214 | 30.6115904 | 23.3271319 | 29.3647505 | 24.9530827 |
| ENSMUSG00 | Slc22a2 | 2137 | 0.59530902 | 0.92449421 | 0 | 0 | 0 | 1.51880303 |
| ENSMUSG00 | D630013G24 | 8385 | 3.22405805 | 5.11460149 | 4.66003926 | 7.06149494 | 3.46869818 | 6.01440763 |
| ENSMUSG00 | Igsf21 | 1969 | 0.14133487 | 0.14683527 | 0.20203991 | 0 | 0.36280816 | 0 |
| ENSMUSG00 | Gm11992 | 2393 | 1.74439008 | 1.18804851 | 1.75477438 | 2.06570836 | 0.87425048 | 0.94600713 |
| ENSMUSG00 | Sun3 | 1288 | 0 | 0 | 0 | 0 | 0.35655076 | 0.23293558 |
| ENSMUSG00 | Mill2 | 1115 | 0 | 0 | 0 | 0 | 0 | 0 |
| ENSMUSG00 | Sh3kbp1 | 8958 | 1.73525261 | 2.26462292 | 0.83883828 | 1.83270041 | 3.21264523 | 0.76422577 |
| ENSMUSG00 | Abhd4 | 2384 | 145.347638 | 127.601095 | 148.921646 | 126.423372 | 115.023382 | 100.312096 |
| ENSMUSG00 | Npnt | 6841 | 78.1220471 | 98.3099215 | 31.6798182 | 78.5513652 | 74.9545462 | 42.5206881 |
| ENSMUSG00 | Trim62 | 5997 | 1.14022722 | 1.55077261 | 1.71736359 | 1.22138653 | 1.67620411 | 0.85958093 |
| ENSMUSG00 | Iqcf4 | 995 | 0 | 0 | 0 | 0 | 0 | 0 |
| ENSMUSG00 | Cmtm8 | 1060 | 20.5153282 | 17.728974 | 22.3928205 | 11.5394463 | 9.38692763 | 19.5554017 |
| ENSMUSG00 | Nrg3 | 8915 | 0.00445939 | 0 | 0 | 0 | 0 | 0 |
| ENSMUSG00 | Mtap7d2 | 4607 | 0 | 0 | 0 | 0.02349601 | 0.03322751 | 0.16576729 |
| ENSMUSG00 | Iffo2 | 6201 | 0.41031297 | 0.34191315 | 0.38492171 | 0.18038113 | 0.30446314 | 0.47503 |
| ENSMUSG00 | Ghitm | 2843 | 519.436187 | 492.220354 | 566.414823 | 500.084699 | 445.884015 | 506.465971 |
| ENSMUSG00 | Dpcd | 1339 | 10.1838161 | 13.6030434 | 13.6996003 | 9.94344618 | 9.06966202 | 9.36992835 |
| ENSMUSG00 | Irgq | 6120 | 7.82117623 | 10.2435558 | 7.14307561 | 10.0168944 | 12.2063095 | 5.03154701 |
| ENSMUSG00 | Fam117b | 5681 | 7.15892564 | 5.29278998 | 9.02554795 | 10.6766268 | 12.0447756 | 5.06988542 |
| ENSMUSG00 | Lrit1 | 4321 | 0 | 0 | 0 | 0 | 0 | 0 |
| ENSMUSG00 | Ramp3 | 1438 | 0.35940281 | 0.73720565 | 0.30738417 | 0.45165285 | 0.35484267 | 0.26553891 |
| ENSMUSG00 | Slc7a13 | 2137 | 0.59530902 | 2.29996121 | 0 | 0.74291528 | 0.81183752 | 2.10590336 |
| ENSMUSG00 | Wdr43 | 3432 | 20.0514967 | 25.3287709 | 23.3115781 | 28.1339019 | 24.3832332 | 32.8376484 |
| ENSMUSG00 | Wwp1 | 8568 | 11.2148689 | 10.5899942 | 13.2430125 | 14.4488206 | 10.7793815 | 15.5982414 |
| ENSMUSG00 | Mslnl | 2612 | 0.24352515 | 0 | 0 | 0 | 0 | 0 |
| ENSMUSG00 | Pif1 | 4050 | 0 | 0 | 0 | 0 | 0 | 0 |
| ENSMUSG00 | 4930596D02I | 1159 | 0 | 0 | 0 | 0 | 0 | 0 |
| ENSMUSG00 | Nacad | 4901 | 0.12978733 | 0 | 0.17135993 | 0 | 0 | 0.51199078 |
| ENSMUSG00 | Fzd7 | 8930 | 7.54152116 | 15.9182536 | 4.89041667 | 11.2731136 | 10.1309927 | 6.97900904 |
| ENSMUSG00 | Grid1 | 6077 | 0 | 0 | 0 | 0 | 0 | 0 |
| ENSMUSG00 | Rwdd2b | 2004 | 5.55465796 | 6.22768888 | 7.85222361 | 7.41806504 | 6.49287716 | 5.96122355 |
| ENSMUSG00 | Ostc | 1054 | 53.4850772 | 63.3647142 | 68.8189985 | 51.5897923 | 34.7599031 | 62.4935994 |
| ENSMUSG00 | Tspyl2 | 3511 | 8.1186783 | 6.42302903 | 4.88473799 | 8.44757636 | 10.1442354 | 11.5359839 |
| ENSMUSG00 | Elmo1 | 3811 | 0.01043177 | 0.06322021 | 0.01159849 | 0.22722883 | 0.33473088 | 0 |
| ENSMUSG00 | Iqsec2 | 7869 | 0.87402442 | 1.54314188 | 0.84258183 | 1.15550581 | 0.74571524 | 1.34830788 |
| ENSMUSG00 | Ccdc8 | 4807 | 0.92627706 | 0.1704118 | 0.51493722 | 0.43535648 | 0.22291531 | 0.08510913 |
| ENSMUSG00 | Pde9a | 2770 | 7.31960111 | 8.50655964 | 9.31908892 | 5.77052192 | 7.8105355 | 10.3092229 |
| ENSMUSG00 | Nbl1 | 2423 | 5.13556145 | 2.38656484 | 1.09455658 | 0 | 1.81108887 | 1.03560331 |
| ENSMUSG00 | 8430410K20I | 2738 | 60.6496136 | 58.3235461 | 65.2857024 | 74.2198958 | 46.2368292 | 81.8637638 |
| ENSMUSG00 | H2afv | 2837 | 8.73023065 | 13.9276992 | 14.6923997 | 11.6500362 | 13.3636226 | 11.7674157 |
| ENSMUSG00 | Zfp598 | 3274 | 4.21354665 | 4.44480926 | 4.37428135 | 4.63974977 | 3.89633291 | 4.97341467 |
| ENSMUSG00 | N4bp2l1 | 1901 | 3.34606885 | 2.50944649 | 4.62712612 | 2.31562832 | 8.75044635 | 3.51514288 |
| ENSMUSG00 | Smc1a | 5655 | 16.9848348 | 16.726788 | 19.9943969 | 19.4160321 | 21.9896159 | 16.0609282 |
| ENSMUSG00 | Cyyr1 | 8887 | 0.03578754 | 0.24941784 | 0.02984259 | 0.30450696 | 0.09760869 | 0.24859298 |
| ENSMUSG00 | Ripk2 | 3954 | 4.50441459 | 3.58290693 | 4.77344132 | 5.8129141 | 3.44563539 | 2.80748057 |
| ENSMUSG00 | Txndc3 | 2163 | 0 | 0 | 0 | 0 | 0 | 0 |
| ENSMUSG00 | Pnmal1 | 1683 | 0 | 0 | 0 | 0 | 0 | 0 |
| ENSMUSG00 | Tmco4 | 3682 | 37.1425456 | 27.2341081 | 28.8596498 | 20.9416972 | 29.781554 | 21.0893259 |
| ENSMUSG00 | Dnahc7b | 12292 | 0.54658934 | 0.91339416 | 0.467478 | 1.26516118 | 0.8759002 | 0.60353904 |
| ENSMUSG00 | Brca2 | 10697 | 0.26387203 | 0.55857893 | 0.31817724 | 0.81629006 | 0.72506396 | 0.36206402 |
| ENSMUSG00 | Osgin2 | 7238 | 3.12529268 | 7.15673187 | 4.37865729 | 5.62317577 | 3.76458755 | 5.27179059 |
| ENSMUSG00 | Otud3 | 2240 | 0.86965114 | 0.83896038 | 1.61810318 | 1.64302167 | 2.61965769 | 0.52357566 |
| ENSMUSG00 | Zmiz2 | 4637 | 12.6716843 | 15.1511392 | 12.1633711 | 16.6909609 | 14.7235909 | 11.9521387 |
| ENSMUSG00 | Spem1 | 1038 | 0 | 0 | 0 | 0 | 0 | 0 |
| ENSMUSG00 | Lonp1 | 2926 | 50.9239716 | 46.0620228 | 46.8908136 | 43.5672807 | 45.9865185 | 31.1150874 |
| ENSMUSG00 | Hectd2 | 9293 | 0.27379218 | 0.09333433 | 0 | 0.55911056 | 0.11530764 | 0.0088049 |
| ENSMUSG00 | Prkd2 | 3785 | 3.91777919 | 3.51372735 | 3.85379345 | 4.95711048 | 4.44879895 | 1.25384075 |
| ENSMUSG00 | Chrnb1 | 2302 | 6.56259018 | 7.41007842 | 8.92869548 | 10.9876252 | 10.0190799 | 5.91226976 |
| ENSMUSG00 | Pla2g5 | 3145 | 0.18961279 | 0 | 0 | 0 | 0 | 0 |
| ENSMUSG00 | Rpusd1 | 1697 | 2.95179172 | 3.74814989 | 5.28755105 | 5.67702174 | 5.89344117 | 1.65544363 |
| ENSMUSG00 | Pla2g2d | 2238 | 0 | 0 | 0 | 0 | 0 | 0 |
| ENSMUSG00 | 2310036O22I | 888 | 47.7244845 | 53.1787312 | 62.7686082 | 43.8023018 | 37.5227741 | 72.7938007 |
| ENSMUSG00 | Map6d1 | 3270 | 0.30394098 | 0.35366196 | 0 | 0.52964469 | 0.62417585 | 0.28358952 |
| ENSMUSG00 | Yeats2 | 5430 | 4.94930107 | 3.57626817 | 4.94930397 | 6.14657296 | 7.26397578 | 3.85260558 |
| ENSMUSG00 | Clvs1 | 7710 | 0 | 0 | 0 | 0 | 0.2845829 | 0 |
| ENSMUSG00 | Arhgap11a | 7555 | 0.6946027 | 0.54213734 | 0.45050191 | 0.82623345 | 0.82398648 | 0.94585777 |
| ENSMUSG00 | Elovl6 | 5946 | 18.192846 | 14.2630572 | 20.3911294 | 16.6999305 | 21.4712399 | 17.7473224 |
| ENSMUSG00 | Arhgap12 | 6536 | 29.1961913 | 28.6936403 | 26.4358942 | 34.3154818 | 28.2612424 | 39.4722786 |
| ENSMUSG00 | Phf8 | 8856 | 3.89205077 | 4.78273291 | 5.83468321 | 5.90366785 | 7.4730363 | 4.65048614 |
| ENSMUSG00 | Ublcp1 | 3573 | 8.38948567 | 11.0722275 | 10.713349 | 10.7145394 | 10.8536372 | 6.73278249 |
| ENSMUSG00 | Chd7 | 11292 | 3.16860897 | 3.20474832 | 2.53655635 | 3.63312829 | 4.10759614 | 3.47816866 |
| ENSMUSG00 | Vps41 | 3166 | 46.8753028 | 42.8898907 | 43.126814 | 47.592741 | 39.9862408 | 30.6430486 |
| ENSMUSG00 | Pldr | 3325 | 0.31086992 | 0.02898433 | 0.07976272 | 0 | 0 | 0 |
| ENSMUSG00 | Rbbp8 | 3900 | 42.1408093 | 39.970036 | 46.6726127 | 49.0068179 | 50.228271 | 50.9965819 |
| ENSMUSG00 | Mul1 | 4197 | 7.65366411 | 8.56494778 | 11.1215502 | 7.90073837 | 7.29469276 | 6.41412174 |
| ENSMUSG00 | Wnk3-ps | 10505 | 0.00378443 | 0.24311104 | 0.32399257 | 0.07212974 | 0.06800279 | 0.19472613 |
| ENSMUSG00 | Lamp3 | 3337 | 0.01191354 | 0 | 0 | 0 | 0 | 0 |
| ENSMUSG00 | Kcnj1 | 3263 | 358.396327 | 491.758665 | 162.773321 | 372.785132 | 426.241473 | 259.840465 |
| ENSMUSG00 | Tmco5b | 1225 | 0 | 0 | 0 | 0 | 0 | 0 |
| ENSMUSG00 | Zfp236 | 11802 | 1.9807001 | 2.06186691 | 2.29960444 | 3.24071913 | 1.69050271 | 2.08222759 |
| ENSMUSG00 | Car8 | 3839 | 8.01530136 | 7.10433088 | 3.02815439 | 6.6073658 | 4.43938774 | 5.62685967 |
| ENSMUSG00 | Rusc1 | 3828 | 5.56659811 | 4.68269501 | 6.1776348 | 5.74032527 | 4.73206986 | 9.49054845 |
| ENSMUSG00 | Uspl1 | 6884 | 8.95711073 | 8.53972391 | 8.10966411 | 6.0643413 | 10.7404443 | 7.53578789 |
| ENSMUSG00 | Dmxl2 | 13961 | 9.17784641 | 5.49134227 | 7.31367791 | 5.67295235 | 7.0357262 | 8.12710402 |
| ENSMUSG00 | Tox | 8013 | 0.35225747 | 0.31270373 | 0.21513439 | 0.51333497 | 0.28018976 | 0.28591909 |
| ENSMUSG00 | Ttc1 | 2622 | 32.068208 | 26.6293496 | 28.2372567 | 31.5683486 | 27.5176565 | 35.5860186 |
| ENSMUSG00 | Sox15 | 859 | 0 | 0 | 0 | 0.08400942 | 0.3564124 | 0 |
| ENSMUSG00 | Gpr110 | 3837 | 2.27943855 | 1.23071966 | 1.15198965 | 2.10643158 | 2.63310462 | 2.03298067 |
| ENSMUSG00 | Cdk13 | 7067 | 3.61157754 | 3.47062389 | 2.97722899 | 2.43542313 | 2.68597875 | 3.58541671 |
| ENSMUSG00 | Katnal1 | 12098 | 3.61473207 | 3.52894592 | 2.53928591 | 3.54318639 | 3.52604151 | 2.90827283 |
| ENSMUSG00 | Cftr | 7843 | 0 | 0 | 0 | 0 | 0 | 0 |
| ENSMUSG00 | Gtf3c3 | 2987 | 16.2375916 | 7.74338541 | 11.202141 | 7.52564906 | 10.1642764 | 13.6601474 |
| ENSMUSG00 | Sntb2 | 4986 | 1.10830561 | 1.4109949 | 1.10814889 | 1.44009765 | 1.3918145 | 1.61372214 |
| ENSMUSG00 | Nkx6-2 | 1902 | 0.10450968 | 0 | 0 | 0 | 0.02682775 | 0.01433998 |
| ENSMUSG00 | Slc7a1 | 7512 | 4.85829754 | 4.20156012 | 4.48961744 | 2.363201 | 2.62196234 | 2.77393828 |
| ENSMUSG00 | Thoc6 | 2103 | 6.55974881 | 4.39933281 | 7.10424296 | 3.9118907 | 3.63954178 | 5.29151346 |
| ENSMUSG00 | Ak7 | 3075 | 0.06464306 | 0 | 0.15812041 | 0.22294597 | 0.34847281 | 0 |
| ENSMUSG00 | Inhba | 1975 | 0 | 0 | 0 | 0 | 0 | 0 |
| ENSMUSG00 | Pcf11 | 6913 | 6.97575551 | 7.92535581 | 7.3403321 | 6.83225753 | 8.60650285 | 12.297852 |
| ENSMUSG00 | Atp1b2 | 5456 | 0.08743874 | 0.12364556 | 0.02430453 | 0.36373029 | 0.45826474 | 0.16996659 |
| ENSMUSG00 | Mup4 | 915 | 0 | 0 | 0 | 0 | 0 | 0 |
| ENSMUSG00 | Atg2b | 11438 | 2.71802638 | 3.69887185 | 2.59306143 | 3.94952962 | 4.04178147 | 2.68263413 |
| ENSMUSG00 | Ankrd42 | 4732 | 3.427776 | 3.17712805 | 2.18580542 | 1.37252071 | 2.37231671 | 1.71763366 |
| ENSMUSG00 | Wrap53 | 2311 | 5.90053216 | 4.14933015 | 6.06316929 | 6.61998567 | 4.17307877 | 3.10395072 |
| ENSMUSG00 | Bdkrb1 | 1337 | 0.11893936 | 0 | 0 | 0 | 0 | 0 |
| ENSMUSG00 | Rap1gap | 5566 | 25.3060847 | 23.1236053 | 21.3703125 | 24.1865088 | 27.6492184 | 21.2277645 |
| ENSMUSG00 | Tmem29 | 7734 | 0.38038603 | 0.28037108 | 0.43435998 | 0.23326897 | 0.41565318 | 1.36126335 |
| ENSMUSG00 | Rgl2 | 4709 | 6.31494997 | 7.52113718 | 5.60384376 | 6.87313526 | 9.12384974 | 7.40799822 |
| ENSMUSG00 | Sar2 | 1162 | 66.7837332 | 81.2783362 | 80.8338351 | 71.698314 | 52.6072274 | 77.1529597 |
| ENSMUSG00 | BC125332 | 3724 | 0 | 0 | 0 | 0 | 0.15072238 | 0 |
| ENSMUSG00 | Tcl1 | 2785 | 0 | 0 | 0 | 0 | 0 | 0 |
| ENSMUSG00 | D19Bwg1357 | 3444 | 21.0782542 | 14.8448941 | 15.1061467 | 18.7953507 | 16.9495279 | 14.7777227 |
| ENSMUSG00 | Myzap | 5792 | 2.02483887 | 2.29617715 | 2.12919789 | 2.35480194 | 3.31248572 | 2.71239506 |
| ENSMUSG00 | 4930506M07 | 4116 | 4.22088071 | 3.24286795 | 7.19514937 | 3.13833138 | 2.18188585 | 5.81805959 |
| ENSMUSG00 | B4galnt3 | 4601 | 0.52707766 | 0.97399242 | 0.3650663 | 0.36858424 | 1.17556962 | 0.49795038 |
| ENSMUSG00 | Ccdc9 | 5429 | 2.54101156 | 1.5710076 | 1.78305464 | 2.12677367 | 2.41550536 | 2.07988592 |
| ENSMUSG00 | Ninj2 | 932 | 0 | 0 | 0 | 0 | 0 | 0 |
| ENSMUSG00 | Cldn5 | 1413 | 1.09728503 | 0 | 0.25025814 | 0 | 2.02227675 | 1.94956728 |
| ENSMUSG00 | Htr2c | 8402 | 0 | 0 | 0 | 0 | 0 | 0 |
| ENSMUSG00 | Mdfic | 7676 | 4.24693773 | 3.25176876 | 7.02530595 | 8.244907 | 5.09200155 | 6.1755241 |
| ENSMUSG00 | 2810422O20I | 1599 | 7.93120593 | 4.18881517 | 6.46856239 | 4.1971609 | 7.72256595 | 7.48815924 |
| ENSMUSG00 | 1700013G24 | 1046 | 0 | 0 | 0 | 0 | 0 | 0 |
| ENSMUSG00 | BC055324 | 4068 | 0 | 0.37904773 | 0.57588439 | 0.62088081 | 0.77768813 | 0.04022808 |
| ENSMUSG00 | Wapal | 4906 | 36.5222076 | 41.0950008 | 35.1200131 | 51.4680285 | 40.6151646 | 39.5498948 |
| ENSMUSG00 | Dicer1 | 10027 | 3.48113213 | 3.60905735 | 3.53544212 | 3.40057166 | 4.92605284 | 3.54975608 |
| ENSMUSG00 | Pik3r1 | 7378 | 12.7004158 | 10.2734174 | 15.2351975 | 13.5906752 | 10.5815065 | 15.0531758 |
| ENSMUSG00 | Meis3 | 2749 | 0.67970447 | 0.56091893 | 1.27026031 | 0.42001652 | 0.51973027 | 0.88302761 |
| ENSMUSG00 | Paqr6 | 2083 | 0.32445663 | 0.3469979 | 0.25464336 | 0 | 0.22046922 | 0.13093922 |
| ENSMUSG00 | Hibch | 2510 | 33.1190079 | 37.0325286 | 25.939966 | 26.8818419 | 25.3098956 | 31.6863937 |
| ENSMUSG00 | Nthl1 | 1074 | 3.44251368 | 2.19845034 | 4.23909668 | 2.35171612 | 3.32574144 | 4.19023787 |
| ENSMUSG00 | Ccnb1 | 4963 | 0.04005186 | 0.55342075 | 0 | 0.174485 | 0.15422036 | 0 |
| ENSMUSG00 | Cirh1a | 2231 | 16.4831105 | 18.4451912 | 14.2452376 | 22.140887 | 20.7216031 | 13.1177445 |
| ENSMUSG00 | Mfsd6 | 9048 | 29.3991954 | 23.3476307 | 23.2196463 | 24.9120836 | 29.3593405 | 22.0536318 |
| ENSMUSG00 | Glt5 | 6284 | 0.7781547 | 1.66398121 | 0.47127999 | 1.2459904 | 2.47661436 | 0.50347838 |
| ENSMUSG00 | Arhgap32 | 14367 | 4.37761329 | 5.60112469 | 3.53503986 | 5.08820368 | 7.37319943 | 3.4000751 |
| ENSMUSG00 | Mmrn2 | 3928 | 1.62948889 | 3.49621589 | 0.58515678 | 1.36869263 | 4.29982953 | 1.02765952 |
| ENSMUSG00 | Serpina3h | 1429 | 0 | 0 | 0 | 0 | 0 | 0 |
| ENSMUSG00 | Rpl21 | 3419 | 1.2325478 | 1.31071633 | 1.82288969 | 3.45095892 | 1.26857033 | 2.99949235 |
| ENSMUSG00 | Tardbp | 11765 | 12.3878956 | 12.2872356 | 11.8234764 | 15.5645028 | 13.8484673 | 15.8269411 |
| ENSMUSG00 | Cacna2d4 | 6039 | 0.11191309 | 0 | 0 | 0 | 0.05069685 | 0 |
| ENSMUSG00 | Gpr12 | 2466 | 0.20957877 | 0.19540325 | 0.01792451 | 0.35116345 | 0.64145079 | 0.35392881 |
| ENSMUSG00 | Fam35a | 3690 | 5.95793516 | 6.37262433 | 7.10343981 | 5.67143247 | 7.99274938 | 4.19837265 |
| ENSMUSG00 | Smpx | 2452 | 1.45921421 | 1.94553743 | 0.72107411 | 1.04479003 | 1.74804876 | 3.29253394 |
| ENSMUSG00 | Dcp1b | 4494 | 3.49430681 | 5.29686302 | 7.0817372 | 9.65078271 | 5.5522692 | 5.27406798 |
| ENSMUSG00 | Syt15 | 5291 | 0 | 0.76500872 | 0.01670831 | 0.00681951 | 0.24109987 | 0 |
| ENSMUSG00 | Serpina3g | 2365 | 0.48748792 | 0 | 0 | 0 | 1.6181726 | 0.29984804 |
| ENSMUSG00 | Fam38b | 14570 | 0 | 0 | 0.05764139 | 0 | 0.03151938 | 0 |
| ENSMUSG00 | Zfp281 | 4935 | 16.4902672 | 16.4429521 | 17.5374283 | 23.0384037 | 23.4194005 | 20.8359453 |
| ENSMUSG00 | Stx3 | 4964 | 12.9501636 | 8.69763339 | 12.5197001 | 12.1242648 | 10.7932503 | 9.91205665 |
| ENSMUSG00 | Cep78 | 2533 | 1.82062208 | 3.42422408 | 2.07659665 | 3.14809784 | 1.95403018 | 1.27059117 |
| ENSMUSG00 | Kif14 | 8740 | 0 | 0.01653997 | 0 | 0 | 0 | 0 |
| ENSMUSG00 | Gm3402 | 956 | 0 | 0 | 0 | 0 | 0 | 0 |
| ENSMUSG00 | Rrp9 | 2774 | 8.28358606 | 5.73234532 | 4.28633589 | 5.87926609 | 5.73908769 | 3.94272903 |
| ENSMUSG00 | 4930569F06I | 432 | 0 | 0 | 0 | 0 | 0 | 0 |
| ENSMUSG00 | Irf8 | 3851 | 0.09291076 | 0 | 0.16069224 | 0 | 0.22525276 | 0.64450589 |
| ENSMUSG00 | Upk2 | 868 | 0.22900622 | 1.55440136 | 0.15277135 | 0 | 0.17635844 | 0.09426719 |
| ENSMUSG00 | Rnf123 | 6998 | 18.3779622 | 12.304826 | 15.3361067 | 15.1072293 | 15.6987407 | 11.3339027 |
| ENSMUSG00 | Eif2c1 | 9497 | 1.69118818 | 2.24771969 | 2.44350506 | 2.36696996 | 1.76768218 | 1.81218252 |
| ENSMUSG00 | Rbp3 | 5272 | 0 | 0 | 0 | 0 | 0 | 0 |
| ENSMUSG00 | Serpina3a | 1842 | 0 | 0 | 0 | 0 | 0 | 0 |
| ENSMUSG00 | H2-Ob | 3256 | 0 | 0 | 0 | 0 | 0 | 0 |
| ENSMUSG00 | Sox5 | 14713 | 0.10267847 | 0.4617881 | 0.23733743 | 0.38502556 | 0.93985916 | 0.51905791 |
| ENSMUSG00 | Ptchd2 | 5429 | 0 | 0.19526648 | 0 | 0 | 0.02819656 | 0 |
| ENSMUSG00 | Hspb8 | 2496 | 1.8316828 | 7.33607693 | 0.6906538 | 7.99413887 | 4.3953008 | 2.04341248 |
| ENSMUSG00 | Serpina5 | 2157 | 0 | 0 | 0 | 0 | 0 | 0 |
| ENSMUSG00 | Ptchd1 | 4654 | 1.34967037 | 1.84297093 | 1.59559726 | 4.63624036 | 1.32664192 | 1.95739783 |
| ENSMUSG00 | Fbxo2 | 1282 | 0.24808412 | 1.46589021 | 0.34478817 | 1.35096579 | 0.47762598 | 0.91482799 |
| ENSMUSG00 | Fmod | 3389 | 0.42230667 | 0 | 0 | 0 | 0.4968635 | 0 |
| ENSMUSG00 | Gltscr2 | 1542 | 29.2107389 | 33.6242618 | 37.0068608 | 25.1778728 | 29.6495673 | 29.0257348 |
| ENSMUSG00 | L3mbtl4 | 3495 | 0 | 0 | 0 | 0 | 0 | 0 |
| ENSMUSG00 | Tssk1 | 1448 | 0 | 0 | 0.21368294 | 0 | 0 | 0 |
| ENSMUSG00 | Serpina12 | 3685 | 0 | 0 | 0 | 0 | 0 | 0 |
| ENSMUSG00 | Camsap2 | 8404 | 6.49665182 | 7.37933734 | 8.4311702 | 9.11067331 | 9.01644075 | 7.60405518 |
| ENSMUSG00 | Sepw1 | 730 | 12.4167802 | 12.8717209 | 20.102756 | 15.1742297 | 11.0440649 | 30.8614411 |
| ENSMUSG00 | Prelp | 3673 | 0 | 0 | 0 | 0 | 0.80575274 | 0 |
| ENSMUSG00 | Crx | 5633 | 0 | 0 | 0 | 0 | 0 | 0 |
| ENSMUSG00 | Obox6 | 2020 | 0 | 0 | 0 | 0 | 0 | 0 |
| ENSMUSG00 | Sdk2 | 16393 | 0 | 0 | 0 | 0 | 0 | 0 |
| ENSMUSG00 | Tmtc4 | 8481 | 2.83599407 | 2.4317648 | 2.97076412 | 4.23743855 | 3.52569936 | 4.31904734 |
| ENSMUSG00 | Vmn1r90 | 4515 | 0 | 0 | 0 | 0 | 0.03390457 | 0 |
| ENSMUSG00 | Cdc42ep4 | 6288 | 10.8176888 | 6.89691443 | 12.9835884 | 7.08672471 | 10.1517169 | 8.17631918 |
| ENSMUSG00 | 5730559C18I | 3980 | 5.60372477 | 4.83075136 | 5.73069121 | 4.48759097 | 4.24365085 | 5.24248719 |
| ENSMUSG00 | Mbp | 4683 | 13.1244871 | 12.1932381 | 8.50434776 | 13.9766878 | 15.134665 | 7.9674795 |
| ENSMUSG00 | Entpd3 | 3735 | 2.2352479 | 3.35434403 | 2.99412752 | 0.18354989 | 2.11756044 | 2.00087046 |
| ENSMUSG00 | Ccdc64 | 7078 | 2.29164115 | 2.07641493 | 1.76107936 | 2.69162468 | 2.35739261 | 1.3332898 |
| ENSMUSG00 | 5330439B14I | 311 | 0 | 0 | 0 | 0 | 0 | 0 |
| ENSMUSG00 | Nppa | 862 | 0 | 0 | 0 | 0 | 0 | 0 |
| ENSMUSG00 | Ccdc74a | 1270 | 0 | 0.26559456 | 0.20882761 | 1.25008658 | 0.60267373 | 0.90199594 |
| ENSMUSG00 | Mmp1b | 1439 | 0 | 0 | 0 | 0 | 0 | 0 |
| ENSMUSG00 | D11Wsu47e | 2789 | 6.74232423 | 8.32766766 | 13.6139775 | 12.1092842 | 8.17812477 | 10.3465645 |
| ENSMUSG00 | Gucy1a2 | 5273 | 0 | 0.02741501 | 0 | 0 | 0 | 0 |
| ENSMUSG00 | A2ld1 | 1626 | 9.73104628 | 10.9649346 | 14.5708412 | 10.4961913 | 7.97090988 | 6.89414313 |
| ENSMUSG00 | D11Wsu99e | 5113 | 2.43369165 | 2.99692717 | 4.83255641 | 2.73102899 | 2.84422396 | 5.11566185 |
| ENSMUSG00 | Mrps27 | 1837 | 16.1445773 | 17.8108842 | 14.1484397 | 14.9867175 | 18.0272825 | 17.2229622 |
| ENSMUSG00 | Kctd12b | 4903 | 12.1788154 | 5.8771145 | 13.6671413 | 12.061691 | 10.9691618 | 12.7611712 |
| ENSMUSG00 | Gcn1l1 | 13930 | 4.85170975 | 3.93309294 | 4.07431202 | 4.53551041 | 6.28946786 | 3.48324359 |
| ENSMUSG00 | Kif21b | 10871 | 0.03291319 | 0 | 0 | 0 | 0 | 0 |
| ENSMUSG00 | Slc5a12 | 4677 | 0 | 0.9375596 | 0.45364303 | 0 | 0.62187373 | 0.18078123 |
| ENSMUSG00 | Ddx24 | 3148 | 29.9176408 | 28.1495766 | 32.6880211 | 35.3943308 | 34.8658623 | 25.8363959 |
| ENSMUSG00 | Klf8 | 5195 | 0.11478965 | 3.40412402 | 1.43794254 | 1.71554668 | 2.0724861 | 0.68252225 |
| ENSMUSG00 | Pcca | 3783 | 52.2296427 | 51.0906211 | 49.2377918 | 48.4622967 | 45.9277847 | 34.5637379 |
| ENSMUSG00 | Pnpla3 | 4632 | 0 | 0.04161178 | 0.27673865 | 0.2804304 | 0 | 0.00588831 |
| ENSMUSG00 | Slc39a11 | 3333 | 5.92813496 | 6.34678911 | 3.99182561 | 6.19229807 | 6.99641577 | 2.21764994 |
| ENSMUSG00 | Rragb | 2835 | 11.4288242 | 6.95176537 | 8.77800269 | 9.49460501 | 5.16563311 | 10.4095801 |
| ENSMUSG00 | Bbox1 | 4395 | 0 | 0 | 0.26148986 | 0 | 0.12771107 | 0 |
| ENSMUSG00 | Prima1 | 1674 | 0.07124638 | 0.05757042 | 0 | 0 | 0 | 0 |
| ENSMUSG00 | Rims1 | 8739 | 0 | 0 | 0 | 0 | 0 | 0 |
| ENSMUSG00 | Pyroxd1 | 4526 | 4.19865658 | 4.06699534 | 2.80290078 | 3.39614471 | 2.95380258 | 3.30839084 |
| ENSMUSG00 | Lrrc18 | 3271 | 0.27954022 | 0 | 0 | 0.23164871 | 0 | 0 |
| ENSMUSG00 | BC006965 | 5866 | 0.01355455 | 0 | 0 | 0 | 0 | 0 |
| ENSMUSG00 | Lrrc29 | 1903 | 6.58064969 | 2.70937956 | 5.29585928 | 3.3370677 | 4.18292939 | 4.25673561 |
| ENSMUSG00 | Iapp | 1034 | 0 | 0 | 0 | 0 | 0 | 0 |
| ENSMUSG00 | Blvrb | 3709 | 25.5961411 | 27.633476 | 23.2867083 | 19.24246 | 20.3197512 | 18.3546777 |
| ENSMUSG00 | Fcho2 | 5221 | 33.9227477 | 30.0692259 | 39.7317083 | 36.9528236 | 36.7475911 | 47.8573013 |
| ENSMUSG00 | Amot | 8388 | 1.1611937 | 1.50510823 | 0.98015532 | 2.60248414 | 1.53298125 | 1.68759392 |
| ENSMUSG00 | Kcnj2 | 5442 | 0 | 0 | 0 | 0.13923611 | 0.06563481 | 0.01002376 |
| ENSMUSG00 | Rasl12 | 3871 | 0 | 0 | 0.25121171 | 0 | 0 | 0 |
| ENSMUSG00 | Cox6a1 | 969 | 184.787084 | 209.45438 | 254.172001 | 153.935162 | 130.593821 | 405.17898 |
| ENSMUSG00 | Slco1a1 | 6539 | 0.24926972 | 0.27265612 | 0 | 0.40833022 | 0 | 0 |
| ENSMUSG00 | Lhfpl1 | 1897 | 0 | 0 | 0 | 1.55968774 | 0 | 0 |
| ENSMUSG00 | Btbd7 | 4286 | 6.06628191 | 8.13975364 | 9.45709054 | 10.102299 | 9.66715286 | 8.95366715 |
| ENSMUSG00 | Zic5 | 5468 | 0 | 0.02643733 | 0 | 0 | 0 | 0 |
| ENSMUSG00 | 1810011H11I | 1468 | 0 | 0 | 0 | 0 | 0 | 0 |
| ENSMUSG00 | Mpped1 | 4284 | 0 | 0 | 0 | 0 | 0 | 0 |
| ENSMUSG00 | Trpc5 | 5003 | 1.16811026 | 0.76088925 | 2.02323047 | 0.40387657 | 1.0811105 | 2.71492511 |
| ENSMUSG00 | Ubr7 | 3244 | 13.3580375 | 13.9033631 | 16.4326211 | 21.0775198 | 14.7856946 | 11.2411198 |
| ENSMUSG00 | AK010878 | 4786 | 0 | 0 | 0 | 0 | 0 | 0 |
| ENSMUSG00 | Golt1a | 1837 | 7.89915644 | 7.18730819 | 11.3813129 | 7.89601628 | 7.83311812 | 9.26476586 |
| ENSMUSG00 | Alg13 | 6050 | 5.02693266 | 5.27263218 | 3.40463783 | 8.52848325 | 4.72310256 | 5.78853501 |
| ENSMUSG00 | Pi4ka | 11465 | 11.5434797 | 8.87235756 | 10.8991374 | 13.0291901 | 11.9365669 | 9.98206597 |
| ENSMUSG00 | Khdc1c | 1943 | 0 | 0 | 0 | 0 | 0 | 0 |
| ENSMUSG00 | Coro2b | 7823 | 0.19819299 | 0.10471296 | 0.13560581 | 0.44739337 | 0.0652261 | 0.0976211 |
| ENSMUSG00 | Prrxl1 | 3248 | 0 | 0 | 0 | 0 | 0 | 0 |
| ENSMUSG00 | Pgm5 | 3622 | 1.42689466 | 0.05321529 | 0.68340784 | 1.22531515 | 1.69054807 | 1.08435893 |
| ENSMUSG00 | Coq5 | 2030 | 39.4029689 | 27.9861159 | 39.8252073 | 31.727315 | 36.4976836 | 32.2055716 |
| ENSMUSG00 | Kirrel | 12466 | 0 | 0 | 0 | 0 | 0.01637298 | 0 |
| ENSMUSG00 | Gm13178 | 1223 | 0 | 0 | 0 | 0 | 0 | 0 |
| ENSMUSG00 | Tspo | 850 | 123.288761 | 132.087541 | 146.698117 | 116.141734 | 82.4826357 | 125.014111 |
| ENSMUSG00 | Tmem45b | 1566 | 135.513892 | 224.931604 | 58.2865938 | 140.296021 | 149.33198 | 83.8442614 |
| ENSMUSG00 | Rnf10 | 4379 | 40.9538645 | 38.3158693 | 32.785473 | 31.3359249 | 36.7403889 | 27.0379559 |
| ENSMUSG00 | Pde3a | 4740 | 0 | 0 | 0 | 0 | 0.35524692 | 0 |
| ENSMUSG00 | Utp15 | 4058 | 9.00327417 | 5.35536851 | 8.81204805 | 8.10912386 | 7.84634266 | 6.02219733 |
| ENSMUSG00 | Ccrl1 | 2695 | 3.21584221 | 2.16347291 | 1.08249412 | 1.21835475 | 1.19282437 | 1.39682346 |
| ENSMUSG00 | Cd1d2 | 1236 | 0 | 0 | 0 | 0 | 0.33026781 | 0 |
| ENSMUSG00 | Trem3 | 990 | 0 | 0 | 0 | 0 | 0 | 0 |
| ENSMUSG00 | Plekha6 | 7443 | 4.77514437 | 4.33114736 | 2.29828204 | 5.43921187 | 5.58732988 | 3.88067217 |
| ENSMUSG00 | Gpr155 | 7694 | 2.13400227 | 1.08973758 | 2.52204433 | 2.40578228 | 1.37281775 | 1.12728558 |
| ENSMUSG00 | Tpp2 | 4652 | 38.4992347 | 36.0776823 | 56.1169572 | 41.3718026 | 33.9261781 | 44.4649321 |
| ENSMUSG00 | Ubac2 | 2144 | 31.5781639 | 28.0488245 | 34.8419378 | 28.9800749 | 30.0351149 | 19.2601697 |
| ENSMUSG00 | Ppp2r2d | 4313 | 18.3890989 | 20.7135784 | 21.4398923 | 19.0742086 | 18.385112 | 20.0148929 |
| ENSMUSG00 | Slc24a4 | 4509 | 0 | 0 | 0 | 0.17604901 | 0 | 0 |
| ENSMUSG00 | Enc1 | 4739 | 4.72300813 | 3.84352714 | 6.7529298 | 1.41617647 | 1.99195601 | 3.7524931 |
| ENSMUSG00 | Ydjc | 5278 | 0.66284242 | 0.38344649 | 0.49410927 | 0.35548812 | 0.47371967 | 0.48575523 |
| ENSMUSG00 | Mapk1ip1 | 2973 | 16.4076605 | 14.6196336 | 13.1579654 | 15.0250828 | 14.6059354 | 15.3391177 |
| ENSMUSG00 | Cir1 | 3150 | 1.75428946 | 2.32518705 | 1.45936244 | 1.38600869 | 1.76567459 | 3.18637055 |
| ENSMUSG00 | Tram2 | 6484 | 2.57515451 | 1.7538557 | 2.73364267 | 1.73621189 | 1.97526532 | 2.00647793 |
| ENSMUSG00 | Cpsf2 | 5300 | 12.7292548 | 11.319268 | 11.7843782 | 11.5326384 | 12.0730331 | 10.3489244 |
| ENSMUSG00 | Lad1 | 2982 | 92.4828197 | 116.830307 | 50.6201522 | 95.0692904 | 122.244275 | 62.0676398 |
| ENSMUSG00 | 2700046A07I | 2463 | 0 | 0.19564126 | 0 | 0 | 0 | 0 |
| ENSMUSG00 | Capza3 | 1264 | 0 | 0 | 0 | 0 | 0 | 0 |
| ENSMUSG00 | Myrip | 5421 | 0 | 0 | 0 | 0.2462711 | 0 | 0 |
| ENSMUSG00 | Abca9 | 8009 | 0 | 0 | 0 | 0 | 0 | 0 |
| ENSMUSG00 | Gck | 3289 | 0 | 0 | 0 | 0 | 0 | 0 |
| ENSMUSG00 | Phlda3 | 1556 | 2.04398357 | 1.33163047 | 2.55666187 | 2.75948798 | 2.09877124 | 1.48993852 |
| ENSMUSG00 | Pramel1 | 2790 | 0 | 0 | 0 | 0 | 0 | 0 |
| ENSMUSG00 | Efhc1 | 2273 | 0.47223844 | 0 | 0.13612534 | 0.04762258 | 0.56122279 | 0.44397785 |
| ENSMUSG00 | Poldip3 | 5671 | 15.9554706 | 14.9632031 | 14.5520792 | 14.0739698 | 21.2887331 | 13.8128662 |
| ENSMUSG00 | Fam169a | 4575 | 0.23462251 | 0.12639067 | 0.32849457 | 0.14196214 | 0 | 0.33385351 |
| ENSMUSG00 | Oasl1 | 2666 | 0.19385643 | 0 | 0.33159672 | 1.16393692 | 1.56945342 | 1.03328528 |
| ENSMUSG00 | Abca8a | 7016 | 0 | 0 | 0 | 0 | 0 | 0 |
| ENSMUSG00 | Sytl3 | 5020 | 0.03959709 | 0 | 0 | 0 | 0.03049385 | 0 |
| ENSMUSG00 | Ptpre | 6825 | 0.65822261 | 0.00706029 | 0.67355189 | 0.34363852 | 0.17943341 | 0.64340175 |
| ENSMUSG00 | Pdcd7 | 3285 | 6.00265398 | 7.97973349 | 8.40978748 | 5.56882696 | 6.67925163 | 4.29253835 |
| ENSMUSG00 | Haus1 | 995 | 9.34953009 | 10.4605744 | 8.4849769 | 12.94602 | 11.94889 | 11.2682078 |
| ENSMUSG00 | Rpl37 | 863 | 2.5336633 | 1.84258701 | 3.43166105 | 1.71421069 | 2.4241963 | 4.29820503 |
| ENSMUSG00 | Fhdc1 | 5595 | 0.73187033 | 0.42200816 | 0.22120672 | 0.5546123 | 0.50159976 | 0.4094852 |
| ENSMUSG00 | Rhod | 2006 | 12.3666101 | 14.6529061 | 11.568279 | 13.4183476 | 11.6246529 | 6.48554483 |
| ENSMUSG00 | Smek1 | 4579 | 29.5192474 | 37.0737794 | 34.0660197 | 54.6076804 | 36.7626149 | 49.2778097 |
| ENSMUSG00 | Card6 | 6098 | 2.7055632 | 1.65151958 | 1.623682 | 1.57392979 | 2.54378784 | 1.43574274 |
| ENSMUSG00 | Tcf20 | 2807 | 4.24889354 | 4.29163183 | 2.81871389 | 2.22379541 | 4.92630846 | 2.74981222 |
| ENSMUSG00 | Oosp1 | 1133 | 0 | 0 | 0 | 0 | 0 | 0 |
| ENSMUSG00 | Mcm3 | 2974 | 2.32597633 | 2.33317003 | 1.1741579 | 1.61362203 | 1.750064 | 0.63280098 |
| ENSMUSG00 | Ankrd13a | 4462 | 46.3843639 | 45.5298162 | 43.0031826 | 32.2247753 | 48.9221964 | 29.4079537 |
| ENSMUSG00 | Il17f | 1170 | 0 | 0 | 0 | 0 | 0 | 0 |
| ENSMUSG00 | 8430432A02I | 1247 | 1.78533032 | 0.54098652 | 3.43831497 | 0.46296128 | 4.86939753 | 1.74977638 |
| ENSMUSG00 | Ipo9 | 8973 | 11.2935161 | 9.71461873 | 10.6157329 | 9.538238 | 11.9590403 | 8.62952199 |
| ENSMUSG00 | Ndufa7 | 1408 | 33.3178033 | 35.4553651 | 51.0455943 | 35.6207684 | 27.3976847 | 80.6229038 |
| ENSMUSG00 | Olfr680-ps1 | 898 | 0 | 0 | 0 | 0 | 0 | 0 |
| ENSMUSG00 | Macc1 | 3814 | 3.76290731 | 9.43766971 | 5.29634039 | 12.4972157 | 6.70273054 | 4.52670337 |
| ENSMUSG00 | Shisa4 | 3349 | 19.3257457 | 17.5969002 | 22.8202408 | 13.7906889 | 13.7888534 | 20.278845 |
| ENSMUSG00 | Git2 | 9965 | 2.09050394 | 3.32687127 | 2.46625436 | 2.78083394 | 3.79433462 | 3.69227179 |
| ENSMUSG00 | Lman1 | 4893 | 69.0947489 | 66.5234859 | 74.0672207 | 71.9207399 | 73.092963 | 74.4715268 |
| ENSMUSG00 | Wipi1 | 4080 | 3.04986897 | 3.94467443 | 3.42347608 | 2.19322231 | 4.41478203 | 3.39595999 |
| ENSMUSG00 | Gpr45 | 1545 | 0 | 0 | 0 | 0 | 0 | 0 |
| ENSMUSG00 | Dlx1 | 5316 | 0 | 0 | 0 | 0 | 0 | 0 |
| ENSMUSG00 | Tdrkh | 3837 | 2.42449373 | 3.08935751 | 2.07358138 | 1.6644571 | 3.61719422 | 1.27239 |
| ENSMUSG00 | Ammecr1l | 3472 | 14.164035 | 11.9078247 | 11.9798198 | 12.5434988 | 11.9482844 | 10.0865889 |
| ENSMUSG00 | Slc16a6 | 5542 | 0.43040939 | 0.1391164 | 0.66199079 | 0.13021308 | 0.36828853 | 0.68900207 |
| ENSMUSG00 | Metap1d | 4398 | 33.7442494 | 38.6543356 | 41.5887282 | 54.0082081 | 41.4777839 | 44.8561768 |
| ENSMUSG00 | Nol4 | 7655 | 0 | 0 | 0 | 0 | 0 | 0 |
| ENSMUSG00 | Rnpep | 3169 | 21.7281452 | 20.0257319 | 22.8889948 | 20.4036049 | 17.5347817 | 19.8728754 |
| ENSMUSG00 | Gm6363 | 889 | 0 | 0 | 0 | 0 | 0 | 0 |
| ENSMUSG00 | BC057022 | 3545 | 0.59436967 | 0.93790254 | 0.64837682 | 1.09925552 | 0.97878521 | 0.66166967 |
| ENSMUSG00 | AW549877 | 5077 | 89.2051278 | 78.6244804 | 97.4408168 | 105.687627 | 66.3734855 | 99.5736534 |
| ENSMUSG00 | Agrn | 9141 | 6.77160958 | 8.18658185 | 2.57251728 | 7.12877939 | 11.767159 | 4.1623588 |
| ENSMUSG00 | Mvk | 2727 | 9.00949283 | 4.01111931 | 4.92752485 | 2.31549607 | 2.13311582 | 2.03034535 |
| ENSMUSG00 | Mfsd9 | 7624 | 5.06851089 | 7.86885106 | 4.65557187 | 5.8070132 | 9.23614882 | 5.3375868 |
| ENSMUSG00 | Tmco7 | 4391 | 1.94657898 | 1.53634751 | 1.93276107 | 1.95571091 | 2.95165098 | 1.15533658 |
| ENSMUSG00 | Tnfrsf18 | 1664 | 0.9795521 | 1.50582632 | 0.47814494 | 0.75893724 | 1.99321781 | 1.47519083 |
| ENSMUSG00 | Pkp2 | 5597 | 10.5692612 | 10.8477608 | 11.4433573 | 11.8425435 | 12.225544 | 13.4545791 |
| ENSMUSG00 | Pigs | 2548 | 21.6096313 | 23.9230178 | 18.5966938 | 15.2796413 | 24.0112342 | 11.7105398 |
| ENSMUSG00 | S100a10 | 1256 | 53.5559499 | 42.3165969 | 95.5477737 | 43.4363466 | 27.666371 | 79.7826629 |
| ENSMUSG00 | Znrf3 | 9042 | 3.36351942 | 3.43732086 | 3.88635979 | 3.50764401 | 3.25051896 | 2.64541679 |
| ENSMUSG00 | Dcaf17 | 9789 | 7.84225486 | 10.9870404 | 8.05558156 | 13.6712946 | 9.78409515 | 10.6323448 |
| ENSMUSG00 | 2310008H04 | 3327 | 7.16960875 | 9.95461443 | 6.04503714 | 7.41812998 | 8.89549083 | 5.19751166 |
| ENSMUSG00 | Mettl8 | 5845 | 4.04696508 | 3.2893739 | 3.38792569 | 4.38910756 | 2.3570781 | 2.58047485 |
| ENSMUSG00 | Arhgef11 | 9094 | 3.89948192 | 4.22836825 | 5.83266017 | 3.83674258 | 5.77371242 | 5.10462239 |
| ENSMUSG00 | Rptn | 4143 | 0 | 0 | 0 | 0 | 0 | 0.35549856 |
| ENSMUSG00 | Elmod1 | 2648 | 0 | 0 | 0.3004657 | 0.99470893 | 0 | 1.0712094 |
| ENSMUSG00 | Hrnr | 11187 | 0 | 0 | 0.01580472 | 0 | 0 | 0 |
| ENSMUSG00 | Rapgef5 | 3959 | 0.45188094 | 0.14605641 | 0.16747352 | 0.4739243 | 0.95376403 | 0.20667825 |
| ENSMUSG00 | Zbed3 | 1892 | 11.9350491 | 12.4795754 | 12.0316856 | 10.1838329 | 13.5656803 | 13.73823 |
| ENSMUSG00 | Tlk1 | 6491 | 11.1775924 | 9.56156794 | 12.6796844 | 13.0575755 | 13.4896412 | 17.1564246 |
| ENSMUSG00 | Foxn4 | 3530 | 0 | 0 | 0 | 0 | 0 | 0 |
| ENSMUSG00 | Acacb | 9577 | 3.60734182 | 4.60380021 | 4.6384935 | 6.0356516 | 6.99035242 | 4.59086546 |
| ENSMUSG00 | Wdr41 | 5066 | 13.2622902 | 12.6696291 | 13.7160081 | 13.6678728 | 15.773254 | 9.42714041 |
| ENSMUSG00 | Ncapg2 | 6575 | 0.98557313 | 0.37376556 | 0.10756342 | 0.40060673 | 0.69069923 | 0.96239031 |
| ENSMUSG00 | Lce3b | 561 | 0 | 0 | 0 | 0 | 0 | 0 |

| | | | | | | | | |
|---|---|---|---|---|---|---|---|---|
| ENSMUSG00 | Mat2b | 4602 | 67.8400236 | 65.3270779 | 101.773735 | 64.8567302 | 66.4939539 | 112.939054 |
| ENSMUSG00 | Igsf3 | 7319 | 0.85279552 | 1.11923694 | 0.80323065 | 1.3212171 | 1.70111159 | 1.03598165 |
| ENSMUSG00 | 2010003K11f | 1163 | 0 | 0 | 0 | 0 | 0 | 0 |
| ENSMUSG00 | Csgalnact2 | 5489 | 8.87961725 | 11.359675 | 11.8456752 | 12.9103909 | 10.2443189 | 11.7466291 |
| ENSMUSG00 | Tbca | 499 | 116.398311 | 77.3493791 | 75.2937206 | 78.0210943 | 45.6067407 | 241.263041 |
| ENSMUSG00 | Sln | 513 | 0 | 0 | 0 | 0 | 0 | 0 |
| ENSMUSG00 | Dstyk | 3328 | 17.8589073 | 17.664501 | 15.4467378 | 15.8726302 | 19.5641994 | 10.9409986 |
| ENSMUSG00 | Wdr60 | 3703 | 3.62877462 | 6.06397031 | 5.2402401 | 7.71725068 | 9.52180008 | 6.84996341 |
| ENSMUSG00 | Wdr11 | 9639 | 8.25712958 | 9.67828115 | 8.9284146 | 11.1027434 | 14.0866466 | 9.22453823 |
| ENSMUSG00 | Zfp386 | 1212 | 80.2325952 | 88.6598729 | 127.937348 | 107.710262 | 92.0748217 | 218.062091 |
| ENSMUSG00 | Myo3b | 7055 | 2.56396083 | 1.6119065 | 2.21165848 | 3.44198667 | 3.84053941 | 2.46650884 |
| ENSMUSG00 | Tmcc2 | 7185 | 0.03873185 | 0 | 0.02460785 | 0 | 0.56814336 | 0.03036842 |
| ENSMUSG00 | Abhd14b | 1573 | 146.53673 | 158.803888 | 151.236058 | 111.916272 | 138.611382 | 118.201663 |
| ENSMUSG00 | Svop | 4877 | 0 | 0 | 0 | 0 | 0.03138797 | 0 |
| ENSMUSG00 | Hnrnpf | 2807 | 11.6561313 | 11.4157407 | 12.3613989 | 12.5072424 | 9.97986474 | 9.06563535 |
| ENSMUSG00 | Arsb | 4659 | 5.91340373 | 4.35425888 | 3.44392982 | 4.22080152 | 5.13658945 | 2.45290274 |
| ENSMUSG00 | 4933440N22l | 650 | 0 | 0 | 0.20400851 | 0 | 0.07850212 | 0.20980492 |
| ENSMUSG00 | Lce1c | 454 | 0 | 0 | 0 | 0 | 0 | 0 |
| ENSMUSG00 | Dao | 2471 | 0.06435529 | 0.42901729 | 0.0894412 | 0 | 0 | 0.04415158 |
| ENSMUSG00 | Zfp239 | 3832 | 0.28011429 | 0.36466775 | 0 | 0.39547126 | 0.665793 | 0.99646386 |
| ENSMUSG00 | Kank3 | 2991 | 4.58563717 | 4.89758556 | 5.42362968 | 4.47557289 | 8.23996641 | 6.06407058 |
| ENSMUSG00 | Dmgdh | 3395 | 0 | 0 | 0 | 0 | 0.01502986 | 0 |
| ENSMUSG00 | Uggt2 | 13554 | 2.38169176 | 0.97766501 | 1.56862081 | 1.06217617 | 1.45316372 | 1.33012665 |
| ENSMUSG00 | Inpp5f | 7960 | 2.75691272 | 2.9965187 | 1.78251151 | 3.236505 | 2.65388437 | 4.262519 |
| ENSMUSG00 | 6230427J02R | 1580 | 0.07548509 | 0.51846172 | 0.0559517 | 0 | 0 | 0 |
| ENSMUSG00 | Csdc2 | 4623 | 0 | 0 | 0 | 0 | 0 | 0 |
| ENSMUSG00 | Ccdc115 | 1699 | 16.3093407 | 18.6903269 | 18.107406 | 18.2640131 | 17.5994445 | 16.5991624 |
| ENSMUSG00 | Klhdc8a | 4264 | 47.559265 | 33.6762659 | 52.6710986 | 26.2322556 | 35.2661186 | 40.7520466 |
| ENSMUSG00 | Vwa1 | 3619 | 1.12049158 | 1.31817015 | 0.35420101 | 0.77767325 | 0.94467178 | 0.40697168 |
| ENSMUSG00 | Bhmt2 | 2459 | 0.88920351 | 1.31292868 | 0 | 0 | 0 | 1.19791014 |
| ENSMUSG00 | Ssh1 | 12576 | 1.60906008 | 2.203181 | 1.73630013 | 2.00264528 | 2.35331568 | 1.86732382 |
| ENSMUSG00 | Lce1f | 1141 | 0 | 0 | 0 | 0 | 0 | 0 |
| ENSMUSG00 | Rassf4 | 9992 | 2.15647222 | 0.95485544 | 2.20301419 | 1.63943637 | 1.81799337 | 2.0226689 |
| ENSMUSG00 | Ppig | 8059 | 15.1000777 | 15.3785245 | 13.4761048 | 16.812021 | 13.1823942 | 17.453197 |
| ENSMUSG00 | BC024479 | 2285 | 8.09028378 | 5.50400937 | 8.18266065 | 7.84804207 | 8.48578679 | 8.58226067 |
| ENSMUSG00 | Cox10 | 3030 | 20.5600125 | 15.2351853 | 14.8798283 | 9.28844711 | 10.0873925 | 11.2159078 |
| ENSMUSG00 | Klhl23 | 4919 | 1.93160395 | 1.38123365 | 2.7946276 | 0.94624593 | 1.55599845 | 1.86303696 |
| ENSMUSG00 | Dzip1 | 6035 | 0 | 0 | 0 | 0.61581617 | 0.11837105 | 0.06327174 |
| ENSMUSG00 | Sprr2i | 615 | 0 | 0 | 0 | 0 | 0 | 0 |
| ENSMUSG00 | Gm9774 | 930 | 0.17099131 | 0.20725352 | 0.04752886 | 0.15519159 | 0 | 0.11731028 |
| ENSMUSG00 | Papd4 | 1858 | 11.8538946 | 12.0077087 | 13.7506272 | 16.9340919 | 12.6055471 | 13.9896293 |
| ENSMUSG00 | Armc5 | 3828 | 8.0071775 | 6.19324178 | 5.68111462 | 7.26730834 | 8.17115161 | 6.41965778 |
| ENSMUSG00 | Pnliprp1 | 1606 | 0.34656085 | 0.93012435 | 0.24770647 | 0.40440648 | 0 | 0.3736252 |
| ENSMUSG00 | Bend6 | 4709 | 0.35458275 | 0 | 0 | 0 | 0 | 0 |
| ENSMUSG00 | 1700069L16f | 1514 | 0 | 0 | 0 | 0 | 0 | 0 |
| ENSMUSG00 | Nfrkb | 10002 | 4.06219767 | 4.54307288 | 4.17623892 | 4.68251288 | 6.38212319 | 3.23685231 |
| ENSMUSG00 | Tekt3 | 1740 | 0 | 0 | 0 | 0 | 0 | 0 |
| ENSMUSG00 | Cmklr1 | 4303 | 0 | 0 | 0 | 0 | 0 | 0.07606221 |
| ENSMUSG00 | Slc35f2 | 2560 | 13.5727695 | 19.2933997 | 4.4201843 | 12.2058791 | 15.5271667 | 3.52652562 |
| ENSMUSG00 | Zfp451 | 17179 | 4.14934377 | 4.59172279 | 3.89811934 | 5.30970421 | 5.82471174 | 3.86439676 |
| ENSMUSG00 | Chchd7 | 3816 | 7.38643493 | 6.9451109 | 8.68746915 | 7.29016151 | 5.83005761 | 15.4241802 |
| ENSMUSG00 | Cdrt4 | 721 | 0.33083618 | 0 | 0.1226126 | 0 | 0.07077167 | 1.36184052 |
| ENSMUSG00 | Slc35e2 | 2455 | 17.1005244 | 16.6444411 | 25.2967778 | 26.6169379 | 21.6576308 | 25.6748233 |
| ENSMUSG00 | Tbc1d22b | 4126 | 3.23747975 | 2.09049276 | 2.28186926 | 3.45429171 | 2.69601307 | 3.26555303 |
| ENSMUSG00 | Kdm5b | 6609 | 11.2246522 | 12.6718167 | 9.94524748 | 11.5851261 | 13.3337193 | 7.35825114 |
| ENSMUSG00 | 0610010F05f | 5105 | 10.6611661 | 7.58901067 | 9.84475916 | 13.8249714 | 8.89588139 | 12.3951348 |
| ENSMUSG00 | Abhd14a | 1823 | 3.59827443 | 4.07060455 | 2.86111765 | 5.81904613 | 3.21888822 | 5.08687732 |
| ENSMUSG00 | Fbxo38 | 4366 | 22.54571 | 14.9106467 | 21.3517377 | 18.3054807 | 17.9164989 | 18.3163633 |
| ENSMUSG00 | Sprr2d | 679 | 0 | 0 | 0 | 0 | 0 | 0 |
| ENSMUSG00 | Anubl1 | 3653 | 0.96858411 | 1.26632862 | 0.0484006 | 0.97785995 | 1.03365776 | 0.96316136 |
| ENSMUSG00 | Bag2 | 2526 | 43.2494301 | 61.3299333 | 36.4323979 | 50.7519811 | 39.8959193 | 35.9990686 |
| ENSMUSG00 | Sgsm1 | 9303 | 3.70931496 | 4.40271695 | 4.41400754 | 2.63740624 | 3.99303468 | 3.37451465 |
| ENSMUSG00 | Olfr631 | 1387 | 0 | 0 | 0 | 0 | 0 | 0 |
| ENSMUSG00 | Ammecr1 | 5431 | 0.88573249 | 1.54381163 | 0.2278865 | 1.14271988 | 1.18381944 | 0.3816742 |
| ENSMUSG00 | Lyn | 4296 | 10.9660718 | 11.844712 | 13.2317435 | 10.4819347 | 14.2531776 | 12.4818298 |
| ENSMUSG00 | Rabif | 3959 | 21.5597415 | 22.6509141 | 24.7860802 | 18.72001 | 23.0965561 | 18.8421668 |
| ENSMUSG00 | 2010015L04f | 6018 | 0 | 0.10409168 | 0.1689336 | 0 | 0 | 0 |
| ENSMUSG00 | Crybb2 | 2256 | 0 | 0 | 0 | 0 | 0 | 0 |
| ENSMUSG00 | BC051665 | 1700 | 0 | 0 | 0 | 0 | 0 | 0 |
| ENSMUSG00 | Pglyrp3 | 1100 | 0 | 0 | 0 | 0 | 0 | 0 |
| ENSMUSG00 | Tmc7 | 5743 | 0.12460363 | 0 | 0.3771357 | 0 | 0 | 0.11398073 |
| ENSMUSG00 | Cyp2c37 | 2008 | 0 | 0 | 0 | 0 | 0 | 0 |
| ENSMUSG00 | Adrbk2 | 6774 | 6.70807709 | 8.37251881 | 6.64919959 | 7.75545569 | 8.8358339 | 6.7643038 |
| ENSMUSG00 | Pglyrp4 | 2448 | 0 | 0 | 0 | 0 | 0 | 0 |
| ENSMUSG00 | Pm20d1 | 2876 | 44.4691867 | 39.0551449 | 41.8810926 | 45.4162035 | 49.2344203 | 66.6407823 |
| ENSMUSG00 | Cilp | 4313 | 0 | 0 | 0 | 0 | 0 | 0 |
| ENSMUSG00 | 3110082D06 | 6886 | 0 | 0 | 0 | 0 | 0 | 0 |
| ENSMUSG00 | Isl1 | 4284 | 0 | 0 | 0 | 0 | 0 | 0 |
| ENSMUSG00 | Ccr8 | 1154 | 0 | 0 | 0 | 0 | 0 | 0 |
| ENSMUSG00 | Trem1 | 3095 | 0 | 0 | 0 | 0 | 0 | 0 |
| ENSMUSG00 | Slc26a9 | 5204 | 0 | 0 | 0 | 0 | 0 | 0 |
| ENSMUSG00 | Fam92b | 1389 | 0 | 0.45098902 | 0 | 0.10390798 | 0 | 0 |
| ENSMUSG00 | Nxt2 | 3169 | 27.5490802 | 31.5515517 | 34.0057095 | 30.650951 | 35.0856652 | 44.160989 |
| ENSMUSG00 | Sestd1 | 11051 | 18.426167 | 20.7727998 | 14.1593091 | 21.9117366 | 23.6500857 | 20.8477855 |
| ENSMUSG00 | Pelo | 1598 | 3.08490587 | 3.79943161 | 2.26817968 | 4.1094694 | 2.96962012 | 3.43066489 |
| ENSMUSG00 | H1foo | 1191 | 0 | 0 | 0 | 0 | 0 | 0 |
| ENSMUSG00 | Gucy2f | 8102 | 0 | 0 | 0 | 0 | 0 | 0 |
| ENSMUSG00 | Itga1 | 7971 | 9.86031677 | 11.9090818 | 6.09432009 | 10.3705889 | 13.12308 | 8.70489051 |
| ENSMUSG00 | Stab1 | 9761 | 3.16870852 | 1.06137538 | 2.50874101 | 1.31966908 | 1.46372147 | 1.00313446 |
| ENSMUSG00 | Hsd3b7 | 5746 | 8.1365202 | 9.5349782 | 10.8619914 | 12.2952738 | 8.55175773 | 8.53459822 |
| ENSMUSG00 | Mkl1 | 7618 | 1.02285038 | 1.29036941 | 1.22428313 | 1.46829008 | 1.3061359 | 2.10879004 |
| ENSMUSG00 | Gm5617 | 619 | 4.6242239 | 7.70671692 | 8.06915713 | 4.02206958 | 3.3797761 | 13.5271636 |
| ENSMUSG00 | Ttc19 | 4417 | 30.1428807 | 19.6149475 | 26.118816 | 25.8464091 | 20.0316359 | 31.1216169 |
| ENSMUSG00 | Ehbp1 | 5681 | 1.35060865 | 0.89909573 | 1.19043865 | 1.51797371 | 1.28441678 | 2.21327384 |
| ENSMUSG00 | Sgsm3 | 4154 | 9.94365533 | 7.34280642 | 5.3948807 | 6.28873381 | 12.6767501 | 5.2133037 |
| ENSMUSG00 | Tmem183a | 3161 | 22.3994023 | 25.1831707 | 31.0014192 | 30.020809 | 22.9223201 | 25.3246018 |
| ENSMUSG00 | S100a14 | 1082 | 0.66136659 | 1.02429591 | 0.08170396 | 0.366823 | 0.33011519 | 0.75622845 |
| ENSMUSG00 | Setd1a | 6932 | 0.39571958 | 0.6256174 | 0.49736638 | 0.54653991 | 0.52263022 | 0.74757378 |
| ENSMUSG00 | S100a13 | 711 | 8.10765777 | 10.4370072 | 14.9204533 | 7.66299399 | 10.549757 | 24.7811772 |
| ENSMUSG00 | Prox2 | 4527 | 0.16685534 | 0.32997122 | 0.45891023 | 0.67748482 | 0.48467731 | 1.22907586 |
| ENSMUSG00 | Pbrm1 | 8955 | 27.4936562 | 26.4473884 | 29.4382794 | 27.8381737 | 30.0802052 | 32.4067182 |
| ENSMUSG00 | Hps4 | 4005 | 5.0227898 | 4.75247057 | 3.38825613 | 6.33350245 | 3.50368373 | 5.13485094 |
| ENSMUSG00 | Specc1 | 13338 | 3.0610945 | 3.48988627 | 2.64455471 | 4.43383348 | 4.92742331 | 4.30243222 |
| ENSMUSG00 | Tnfrsf14 | 1211 | 0 | 0 | 0 | 0 | 0 | 0 |
| ENSMUSG00 | Ctf1 | 1342 | 27.3726259 | 32.1362636 | 33.3984119 | 14.6264026 | 28.4029082 | 26.8681618 |
| ENSMUSG00 | Ubash3a | 3462 | 0 | 0 | 0 | 0 | 0 | 0 |
| ENSMUSG00 | Arl15 | 3433 | 35.9802732 | 37.5469656 | 45.4636492 | 41.5158974 | 43.3419493 | 46.7633343 |
| ENSMUSG00 | Ikbke | 6211 | 0.89611452 | 0.4422198 | 0.29178483 | 0.27885013 | 0.09037033 | 0.60161417 |
| ENSMUSG00 | 1110018G07 | 8406 | 5.48139447 | 5.97887927 | 6.6781276 | 6.06518307 | 9.90054948 | 4.9351328 |
| ENSMUSG00 | Grap2 | 3151 | 0 | 0.03058486 | 0 | 0 | 0 | 0 |
| ENSMUSG00 | Frem3 | 11576 | 0 | 0.01248785 | 0 | 0 | 0 | 0 |
| ENSMUSG00 | Gnl3 | 1868 | 23.1552263 | 19.8885155 | 21.2490659 | 25.7673701 | 19.7221859 | 27.8586786 |
| ENSMUSG00 | Gjb5 | 1719 | 0.0231271 | 0 | 0 | 0 | 0.11873502 | 0 |
| ENSMUSG00 | Osbpl6 | 10180 | 2.10883688 | 2.76906372 | 1.18971562 | 2.23297525 | 3.1929078 | 1.47625994 |
| ENSMUSG00 | 4930433N12l | 1198 | 0 | 0 | 0 | 0 | 0 | 0 |
| ENSMUSG00 | 1110067D22l | 4275 | 3.98019782 | 3.55057995 | 3.9187131 | 5.12322831 | 2.79302268 | 3.98752036 |
| ENSMUSG00 | Snx18 | 7099 | 8.51783151 | 11.8039474 | 8.66727222 | 8.15264114 | 10.1707736 | 7.81855062 |
| ENSMUSG00 | Gjb3 | 1963 | 0 | 1.2519147 | 0 | 0 | 0.28593486 | 0 |
| ENSMUSG00 | Rbm45 | 1888 | 1.83195276 | 4.31329911 | 2.85626316 | 1.83468023 | 4.97290951 | 1.35795344 |
| ENSMUSG00 | Slc5a10 | 2651 | 0 | 0.16359034 | 0.01667365 | 0 | 0.28871959 | 0.38067207 |
| ENSMUSG00 | Dmrt3 | 2388 | 0 | 0 | 0 | 0 | 0 | 0 |
| ENSMUSG00 | Fam83g | 6653 | 0.79475108 | 1.0284796 | 0.5514434 | 0.0650811 | 1.67199007 | 0.0327968 |
| ENSMUSG00 | Esm1 | 2094 | 5.44881705 | 9.38876237 | 1.83646626 | 6.08259872 | 5.50714467 | 3.73821464 |
| ENSMUSG00 | BC003266 | 1020 | 19.3320768 | 26.3608184 | 28.5578574 | 15.211058 | 14.85768 | 27.1142002 |
| ENSMUSG00 | Gzmk | 1779 | 0 | 0 | 0 | 0 | 0 | 0 |
| ENSMUSG00 | Tex13 | 3235 | 0 | 0 | 0 | 0 | 0 | 0 |
| ENSMUSG00 | Dlgap3 | 4507 | 0 | 0 | 0 | 0 | 0 | 0 |
| ENSMUSG00 | Tsen2 | 4085 | 2.66658547 | 3.47980427 | 2.52118223 | 3.66561774 | 3.95969673 | 3.29832847 |
| ENSMUSG00 | Gatad2b | 1896 | 1.40486139 | 1.98235363 | 4.05649825 | 1.71275528 | 2.71817719 | 2.33042804 |
| ENSMUSG00 | Rbm7 | 1771 | 65.4809354 | 79.4763396 | 73.3036775 | 71.1046501 | 63.6749241 | 109.945596 |
| ENSMUSG00 | Crtac1 | 2601 | 0 | 0 | 0 | 0 | 0 | 0 |
| ENSMUSG00 | Olfr607 | 983 | 0 | 0 | 0 | 0 | 0 | 0 |
| ENSMUSG00 | Dennd4b | 6003 | 1.61591492 | 1.05957194 | 0.99404445 | 0.47484283 | 2.12503647 | 1.8537486 |
| ENSMUSG00 | Atf4 | 1905 | 52.2768917 | 50.3111987 | 58.1468864 | 53.4506717 | 48.6692519 | 46.0733801 |
| ENSMUSG00 | Zmym6 | 7882 | 7.54052821 | 5.78946469 | 6.33698079 | 6.59200001 | 8.07928407 | 4.8929637 |
| ENSMUSG00 | Agps | 8671 | 13.5208064 | 11.7145681 | 15.8945158 | 17.1359543 | 18.4132776 | 11.7673193 |
| ENSMUSG00 | Prdm14 | 2556 | 0 | 0.33934114 | 0 | 0 | 0 | 0 |
| ENSMUSG00 | Ccno | 1873 | 0 | 0 | 0 | 0 | 0 | 0 |
| ENSMUSG00 | Nfkbil1 | 1882 | 1.8377932 | 3.4565195 | 2.11379696 | 1.51460229 | 1.89789921 | 2.55065702 |
| ENSMUSG00 | Fbrs | 2639 | 5.36299775 | 4.60135787 | 4.80709699 | 6.82263745 | 2.62962755 | 7.41035102 |
| ENSMUSG00 | Frmpd3 | 3447 | 0.03460007 | 0.01397924 | 0 | 0.02093533 | 0 | 0.00791257 |
| ENSMUSG00 | Dhx29 | 4378 | 6.42008331 | 5.06298844 | 6.39099283 | 5.93400458 | 7.31945272 | 5.32036133 |
| ENSMUSG00 | Mgat3 | 4663 | 0.74173854 | 0.97137584 | 0.21802325 | 0 | 0.48148414 | 0.59076529 |
| ENSMUSG00 | Adora1 | 8273 | 11.5042452 | 13.7342718 | 9.65462714 | 13.6338053 | 11.2007613 | 8.36404687 |
| ENSMUSG00 | E230019M04 | 1502 | 0 | 0 | 0 | 0 | 0 | 0 |
| ENSMUSG00 | Mfap4 | 1940 | 0.22541767 | 0 | 0 | 0 | 0 | 0 |
| ENSMUSG00 | Zfp532 | 6568 | 2.9840822 | 3.25009042 | 2.64484231 | 4.69154477 | 6.91435361 | 3.77476356 |
| ENSMUSG00 | Fam63b | 8072 | 45.1731005 | 39.6977007 | 40.5110548 | 38.7282995 | 44.1866163 | 43.4191172 |
| ENSMUSG00 | Zmym4 | 10582 | 5.26716912 | 3.93433056 | 5.43437892 | 6.27736193 | 6.23966455 | 4.51054798 |
| ENSMUSG00 | Mios | 6541 | 13.0249189 | 6.59331382 | 11.9678129 | 8.46750319 | 10.0086898 | 13.4434241 |
| ENSMUSG00 | Hoxd1 | 1861 | 0.76904745 | 0 | 0 | 0.3102164 | 0.21935035 | 0 |
| ENSMUSG00 | Mybph | 1817 | 0 | 0 | 0 | 0 | 0.02808276 | 0 |
| ENSMUSG00 | Reln | 20639 | 0.22151656 | 0.04669455 | 0.13492495 | 0.03671316 | 0.19036925 | 0.10043474 |
| ENSMUSG00 | Bpifa2 | 2035 | 0 | 0 | 0 | 0 | 0 | 0 |
| ENSMUSG00 | C1galt1 | 2448 | 20.5273396 | 20.4319963 | 23.7259893 | 26.8404424 | 20.2396286 | 25.0463191 |
| ENSMUSG00 | Dctpp1 | 673 | 4.19411459 | 8.01914045 | 7.15899092 | 5.03969121 | 3.41186763 | 12.1986128 |
| ENSMUSG00 | Hoxd4 | 2554 | 0.29575338 | 0.37734097 | 0.36344507 | 0.72051068 | 0.77918115 | 0.37377149 |
| ENSMUSG00 | Zfp410 | 2357 | 24.3221905 | 27.7220261 | 26.6673826 | 31.9028345 | 25.5673101 | 24.8561413 |
| ENSMUSG00 | Tbc1d8b | 5190 | 7.71363562 | 4.00161013 | 5.71472768 | 5.14464604 | 5.63354785 | 7.2311953 |
| ENSMUSG00 | Faim3 | 1844 | 0 | 0 | 0 | 0 | 0 | 0.0739551 |
| ENSMUSG00 | Abcb4 | 4073 | 0.35138652 | 0 | 0.20619568 | 0.42522419 | 0 | 0.52901952 |
| ENSMUSG00 | Tfap2e | 2078 | 0 | 0 | 0 | 0 | 0 | 0 |
| ENSMUSG00 | Mustn1 | 1174 | 0.67726543 | 0.36940203 | 0.60241013 | 0 | 0.73888278 | 0.11616116 |
| ENSMUSG00 | Leo1 | 2180 | 17.4340548 | 18.6556684 | 13.4430376 | 20.2920122 | 18.7954953 | 12.0233616 |
| ENSMUSG00 | Clspn | 7302 | 0.00544447 | 0 | 0 | 0 | 0 | 0 |
| ENSMUSG00 | Tbc1d10b | 6371 | 6.4709517 | 6.52721701 | 5.07859819 | 5.44260631 | 9.82724268 | 3.51475103 |
| ENSMUSG00 | Prdm10 | 5045 | 2.42709376 | 2.06308652 | 1.78734905 | 3.29709069 | 2.66004695 | 1.2488487 |
| ENSMUSG00 | D330045A20l | 4493 | 3.76938231 | 2.48814925 | 0.75752101 | 2.22451065 | 1.28332528 | 0.06070474 |
| ENSMUSG00 | Hoxd11 | 5423 | 0.19793435 | 0.36430834 | 0.15486539 | 0.16633803 | 0.3763701 | 0.89021043 |
| ENSMUSG00 | Eif2c4 | 11334 | 1.16102559 | 1.61981952 | 1.13098063 | 2.18707999 | 1.9178786 | 1.92034252 |
| ENSMUSG00 | Cpa6 | 7412 | 0 | 0 | 0 | 0 | 0 | 0 |
| ENSMUSG00 | Cd2bp2 | 3643 | 31.8545835 | 20.0787865 | 23.1625908 | 18.5114854 | 24.8478701 | 19.8177244 |
| ENSMUSG00 | Acn9 | 4075 | 4.62431847 | 5.6995988 | 3.67715946 | 2.62092888 | 2.29149123 | 5.76950651 |
| ENSMUSG00 | Usp22 | 5367 | 22.8295865 | 22.0057704 | 26.7418268 | 15.0997339 | 26.07888 | 17.6088364 |
| ENSMUSG00 | C130039O16 | 10880 | 1.72834028 | 2.8344966 | 2.27915753 | 2.54033512 | 1.7212022 | 1.94030981 |
| ENSMUSG00 | Dmtf1 | 5238 | 20.4317972 | 13.7714975 | 19.147381 | 21.3612868 | 21.4412091 | 19.3182344 |
| ENSMUSG00 | AA986860 | 2904 | 49.8860093 | 47.9873246 | 56.3786593 | 39.1137315 | 61.2878784 | 47.5709531 |
| ENSMUSG00 | Klhl14 | 6679 | 2.3094964 | 4.97087268 | 1.71407057 | 5.29966847 | 2.7274167 | 1.35576886 |
| ENSMUSG00 | Mum1l1 | 5023 | 2.62767659 | 2.25439258 | 1.87437638 | 2.74404559 | 1.79806262 | 2.37831814 |
| ENSMUSG00 | Ubap2l | 5415 | 60.7361263 | 49.4767532 | 49.2219969 | 53.4800534 | 58.6120143 | 48.7266593 |
| ENSMUSG00 | Dnalc1 | 6998 | 7.2943751 | 5.47416712 | 9.75245007 | 6.20788534 | 5.62908862 | 8.74986641 |
| ENSMUSG00 | Sun2 | 6440 | 0.63584076 | 0.80061325 | 1.68845549 | 1.31105566 | 0.87156853 | 1.5670212 |
| ENSMUSG00 | 4933428M09 | 1024 | 0 | 0 | 0 | 0 | 0 | 0 |
| ENSMUSG00 | Kcnj12 | 2746 | 0 | 0 | 0 | 0 | 0 | 0 |
| ENSMUSG00 | Golga7b | 2981 | 0 | 0 | 0 | 0 | 0 | 0 |
| ENSMUSG00 | Gtpbp1 | 3391 | 18.0312383 | 16.7394954 | 10.8581938 | 15.4606932 | 18.8094868 | 9.48298428 |
| ENSMUSG00 | Acot5 | 1516 | 1.46854018 | 1.11248382 | 2.01182531 | 0.59502052 | 1.38000093 | 1.43929495 |
| ENSMUSG00 | Shfm1 | 474 | 84.8787896 | 90.9849488 | 92.6000635 | 67.2161294 | 63.4061927 | 192.591176 |
| ENSMUSG00 | Axil1 | 7668 | 8.64272118 | 10.1111092 | 7.93765491 | 6.74303206 | 10.9931629 | 7.51226371 |
| ENSMUSG00 | Gm16516 | 1289 | 3.7935796 | 5.27097622 | 2.77761775 | 2.32335896 | 1.93971483 | 2.47566547 |
| ENSMUSG00 | Zp3r | 2518 | 0 | 0 | 0 | 0 | 0 | 0 |
| ENSMUSG00 | Sin3a | 5375 | 8.63895837 | 7.64707631 | 8.2400459 | 8.39788608 | 9.24645396 | 8.77862808 |
| ENSMUSG00 | Adprhl2 | 3203 | 16.6568389 | 20.1591734 | 19.554796 | 15.2078552 | 20.4551566 | 16.8092844 |
| ENSMUSG00 | 4933432B09l | 6755 | 0.12359217 | 0.25680413 | 0.2028508 | 0.35254107 | 0.21906216 | 0.15343246 |
| ENSMUSG00 | Nek10 | 5680 | 0 | 0 | 0 | 0 | 0 | 0 |
| ENSMUSG00 | Dhrs7b | 3370 | 7.44383033 | 6.90624675 | 8.57804313 | 6.20996612 | 7.67666841 | 7.55110932 |
| ENSMUSG00 | Mier2 | 4760 | 2.87308515 | 3.09769985 | 1.9129369 | 2.10731268 | 3.9127368 | 2.14300737 |
| ENSMUSG00 | Ube2q1 | 3068 | 18.4134739 | 20.0410366 | 19.997444 | 16.5003613 | 18.0288759 | 22.6251489 |
| ENSMUSG00 | 4632404H12l | 2789 | 2.12390341 | 1.67589992 | 2.58332751 | 0.94442067 | 2.72604159 | 2.49373719 |
| ENSMUSG00 | Thsd7b | 9710 | 0 | 0.13895164 | 0 | 0 | 0 | 0.08988553 |
| ENSMUSG00 | Cux2 | 9591 | 0.75440491 | 0.54260617 | 0.45625925 | 0.14672086 | 0.82463646 | 0.76213151 |
| ENSMUSG00 | Ipo11 | 5716 | 12.1297337 | 9.80420438 | 13.1151725 | 11.8863698 | 15.898876 | 13.3701083 |
| ENSMUSG00 | Sh2b3 | 7058 | 3.58238674 | 4.82000966 | 3.25031971 | 2.71970072 | 3.62201959 | 2.13699001 |
| ENSMUSG00 | Fam199x | 8293 | 3.67689058 | 4.10221008 | 6.01759083 | 5.60396399 | 4.33167353 | 6.60406073 |
| ENSMUSG00 | Tfap2d | 1945 | 0 | 0 | 0 | 0 | 0 | 0 |
| ENSMUSG00 | Jhdm1d | 11231 | 11.525585 | 8.30637984 | 12.7044385 | 10.8911166 | 12.5078455 | 17.5727264 |
| ENSMUSG00 | Kcna4 | 4830 | 0 | 0 | 0 | 0 | 0 | 0 |
| ENSMUSG00 | Atxn2 | 6283 | 19.5898405 | 20.0092993 | 14.4290912 | 18.4803469 | 20.0109804 | 15.5321756 |
| ENSMUSG00 | Hirip3 | 3071 | 3.17163553 | 1.31802707 | 1.82794987 | 3.00781615 | 2.59202691 | 2.0249488 |
| ENSMUSG00 | Asb4 | 3924 | 2.32008283 | 2.89806329 | 0.7209271 | 4.13784914 | 2.34065944 | 0.95224913 |
| ENSMUSG00 | Stk40 | 4155 | 17.0312287 | 15.4474948 | 11.1595026 | 16.5777673 | 12.8333484 | 9.48540402 |
| ENSMUSG00 | Pbxip1 | 4731 | 78.1076287 | 88.2043097 | 92.5333023 | 75.6495032 | 82.9731366 | 71.9079635 |
| ENSMUSG00 | Oscp1 | 1677 | 2.5839877 | 8.27531028 | 2.26676118 | 1.78581377 | 4.95962985 | 3.85455546 |
| ENSMUSG00 | Maff | 2115 | 0.24435993 | 0.29618144 | 0.16719373 | 0.25590103 | 0.21713351 | 0.38687432 |
| ENSMUSG00 | Safb2 | 8010 | 9.83216066 | 12.3865024 | 8.70792862 | 13.1174674 | 13.4541455 | 9.45588929 |
| ENSMUSG00 | Shc1 | 4768 | 12.8905144 | 8.65092166 | 8.02827098 | 7.74159638 | 11.6864099 | 6.54960326 |
| ENSMUSG00 | Zfyve1 | 3857 | 6.98838885 | 3.51060157 | 5.90198319 | 4.06004857 | 4.68326084 | 4.51866593 |
| ENSMUSG00 | Xkr7 | 2708 | 0 | 0 | 0 | 0 | 0 | 0 |
| ENSMUSG00 | Pla2g6 | 5623 | 3.8178836 | 3.17929398 | 3.30943907 | 2.29723847 | 2.86756798 | 2.80361755 |
| ENSMUSG00 | Gucy2c | 3931 | 0 | 0 | 0 | 0 | 0 | 0 |
| ENSMUSG00 | Rgsl1 | 4961 | 0 | 0 | 0 | 0 | 0 | 0.29688178 |
| ENSMUSG00 | Fiad1 | 4422 | 18.6460576 | 18.4267919 | 21.6111227 | 21.4354435 | 22.0629649 | 16.7212906 |
| ENSMUSG00 | Itpr3 | 8932 | 0.52075585 | 1.62383778 | 0.58889603 | 0.94931488 | 1.00544583 | 0.55575284 |
| ENSMUSG00 | Acad12 | 4996 | 4.87792425 | 3.73261672 | 3.53897862 | 4.74497663 | 5.21907087 | 2.10728798 |
| ENSMUSG00 | Alkbh5 | 5982 | 11.9027191 | 13.8791775 | 10.6773091 | 10.6521048 | 11.3107613 | 9.08242751 |
| ENSMUSG00 | Fam159b | 1307 | 0 | 0 | 0 | 0 | 0 | 0 |
| ENSMUSG00 | Arrdc4 | 3981 | 41.8825635 | 38.6847197 | 49.7201339 | 45.2453068 | 47.7322842 | 36.2017568 |
| ENSMUSG00 | Wdr55 | 1491 | 15.2515995 | 11.569917 | 12.8069726 | 17.2061259 | 12.4913663 | 14.3415944 |
| ENSMUSG00 | Dusp15 | 3451 | 0.19583981 | 0 | 0.02561683 | 0 | 0.05914387 | 0.09484082 |
| ENSMUSG00 | Immp1l | 948 | 32.0392269 | 34.6149442 | 34.5968012 | 41.3725553 | 23.2525257 | 54.5780492 |
| ENSMUSG00 | Rgs8 | 6974 | 0 | 0 | 0 | 0 | 0 | 0 |
| ENSMUSG00 | Dcst1 | 4042 | 0.10819156 | 0.45301455 | 0.06561382 | 0.21424272 | 0.11361637 | 0.06747808 |
| ENSMUSG00 | Ypel3 | 1697 | 68.9922747 | 77.319789 | 76.5783256 | 69.6338809 | 64.0160013 | 81.8078455 |
| ENSMUSG00 | Zc3h12a | 2926 | 0.38043522 | 0.29643061 | 0.12085261 | 0.51792408 | 0.24414534 | 0.40082348 |
| ENSMUSG00 | Myo15 | 12467 | 0.00956657 | 0.03865119 | 0.09927421 | 0.06077829 | 0.28650408 | 0.11376283 |
| ENSMUSG00 | Fam59a | 4917 | 12.8313906 | 14.2001535 | 11.2549741 | 11.3228787 | 18.4616479 | 13.3793837 |
| ENSMUSG00 | Selk | 1335 | 64.1447978 | 54.9722482 | 58.5384708 | 62.4611273 | 43.458419 | 98.7811841 |
| ENSMUSG00 | Npl | 1391 | 11.689426 | 5.85442755 | 22.2756952 | 8.92323747 | 6.12609976 | 12.2549601 |
| ENSMUSG00 | Jph1 | 2187 | 0.59987694 | 0.28643061 | 0.84886941 | 0.7259348 | 0 | 0.436494 |
| ENSMUSG00 | Mapk6 | 6538 | 5.44220787 | 7.06066252 | 4.88126807 | 9.28265508 | 7.9372628 | 4.82249663 |
| ENSMUSG00 | Obfc1 | 3006 | 4.70823389 | 3.89531119 | 3.9996345 | 4.45323969 | 7.02758469 | 4.37337862 |
| ENSMUSG00 | Dhx9 | 4588 | 6.9494105 | 8.62272646 | 3.33344326 | 5.73317578 | 3.69240557 | 5.5524222 |
| ENSMUSG00 | Sipa1l1 | 8815 | 2.56166908 | 2.0608382 | 1.87036727 | 2.62377658 | 4.35880442 | 1.30881139 |
| ENSMUSG00 | Commd10 | 3507 | 11.75547 | 10.3050561 | 9.06219707 | 9.95934391 | 8.83176793 | 16.2154613 |
| ENSMUSG00 | Dnali1 | 2127 | 0 | 0.43043836 | 0 | 0 | 1.36742051 | 0 |
| ENSMUSG00 | 1700012A16l | 4112 | 0 | 0.24608835 | 0 | 0.0526489 | 0 | 0 |
| ENSMUSG00 | Atpaf2 | 3018 | 26.3719258 | 24.2049855 | 37.7429256 | 27.199023 | 20.170466 | 29.5430072 |
| ENSMUSG00 | Wbp5 | 1034 | 139.990043 | 171.215656 | 182.877645 | 123.216344 | 93.9595934 | 275.938106 |
| ENSMUSG00 | Ppp1r3a | 7116 | 0 | 0 | 0 | 0 | 0 | 0 |
| ENSMUSG00 | Naa25 | 4651 | 8.28274793 | 9.33476336 | 9.70330719 | 12.9634695 | 11.0259101 | 9.01922063 |
| ENSMUSG00 | Map3k9 | 10608 | 2.13243481 | 1.49447016 | 1.74590613 | 1.098652 | 1.130392 | 2.14175853 |
| ENSMUSG00 | Trafd1 | 4682 | 5.62112945 | 4.8886288 | 4.60711222 | 5.88009397 | 4.84979437 | 5.52250278 |
| ENSMUSG00 | Wdr74 | 1192 | 23.5130988 | 33.876039 | 29.3690098 | 19.221559 | 25.170729 | 24.7348029 |
| ENSMUSG00 | Gm9775 | 186 | 0.85495657 | 0.25906689 | 0.23764432 | 0 | 0 | 0 |
| ENSMUSG00 | Ttc9 | 2039 | 0 | 0.51991257 | 0.26013836 | 0 | 0.32532756 | 0.34778844 |
| ENSMUSG00 | Dpm3 | 914 | 4.39311109 | 6.37916794 | 4.78772698 | 3.94770728 | 3.29382514 | 9.01197052 |
| ENSMUSG00 | B630005N14 | 4134 | 16.9350269 | 15.9922106 | 9.66581183 | 15.2829366 | 11.1087901 | 11.1170216 |
| ENSMUSG00 | Sgtb | 3233 | 1.97978112 | 3.91989925 | 2.99418609 | 1.3169444 | 2.99876629 | 2.99489543 |
| ENSMUSG00 | Gm15800 | 16312 | 2.38357405 | 2.89201366 | 1.97000612 | 4.05237283 | 3.03430509 | 2.14023653 |
| ENSMUSG00 | Id1 | 1653 | 5.09870651 | 7.14197116 | 6.68509422 | 7.26879663 | 8.76678205 | 5.90703016 |
| ENSMUSG00 | Krtcap2 | 1332 | 16.1469332 | 13.8554109 | 20.7735388 | 15.2779828 | 9.57702249 | 22.9336306 |
| ENSMUSG00 | Bex2 | 900 | 1.36935544 | 2.19516014 | 0.98226318 | 2.68610776 | 2.26783893 | 2.36380207 |
| ENSMUSG00 | Nmnat2 | 4511 | 0.00881301 | 0 | 0 | 0.22396303 | 0.01131154 | 0 |
| ENSMUSG00 | Tmem108 | 3724 | 0.05337739 | 0 | 0 | 0 | 0.06851017 | 0 |
| ENSMUSG00 | Apobr | 3554 | 0.12304735 | 0 | 0.39799071 | 0 | 0 | 0.0460461 |
| ENSMUSG00 | Mrap2 | 2343 | 0.61083965 | 0 | 0 | 0 | 0 | 0 |
| ENSMUSG00 | Maneal | 2723 | 0.27739777 | 0.07078434 | 0 | 0.39752528 | 0 | 0 |
| ENSMUSG00 | Trim46 | 4890 | 0.27641847 | 0.35474681 | 0.10847078 | 0.3763158 | 0.15652263 | 0.09481981 |
| ENSMUSG00 | Hebp1 | 1057 | 1.95580415 | 2.23380858 | 1.75636462 | 1.67267756 | 1.30341736 | 2.60618596 |
| ENSMUSG00 | Smg7 | 6072 | 10.8947825 | 11.4355506 | 11.1814939 | 16.6921053 | 14.4961295 | 10.8972785 |
| ENSMUSG00 | Olfr1353 | 1263 | 0 | 0 | 0 | 0 | 0 | 0 |
| ENSMUSG00 | Muc1 | 2338 | 148.36038 | 168.673158 | 143.854501 | 115.65391 | 163.424081 | 90.9699046 |
| ENSMUSG00 | Exog | 3880 | 2.03900532 | 3.65124072 | 2.69995794 | 2.63175737 | 2.18308721 | 2.47440026 |
| ENSMUSG00 | Fam166b | 1788 | 0 | 0 | 0 | 0.24216137 | 0 | 0.13728845 |
| ENSMUSG00 | Rnf214 | 5918 | 8.0478313 | 7.96322075 | 8.20101785 | 8.99917163 | 6.94089769 | 6.87626931 |
| ENSMUSG00 | Lgr6 | 5620 | 0 | 0 | 0 | 0 | 0 | 0 |
| ENSMUSG00 | Olfr1032 | 932 | 0 | 0 | 0 | 0 | 0 | 0 |
| ENSMUSG00 | Aqp11 | 1319 | 4.49095269 | 6.10093695 | 3.51872139 | 5.41641002 | 4.29410744 | 2.77088829 |
| ENSMUSG00 | 1700015E13f | 769 | 3.87732255 | 0.56395056 | 4.48341191 | 0.84457321 | 4.44573105 | 1.27683617 |
| ENSMUSG00 | Olfr769 | 1083 | 0 | 0 | 0 | 0 | 0 | 0 |
| ENSMUSG00 | Gpr153 | 3922 | 0 | 0.19657906 | 0 | 0.08279918 | 0 | 0 |
| ENSMUSG00 | Hecw2 | 15067 | 0.11082035 | 0.20787939 | 0.1232148 | 0.0359216 | 0.04402621 | 0.00724089 |
| ENSMUSG00 | Gpx2 | 1065 | 0 | 1.08589165 | 0 | 0.20327912 | 0 | 0 |
| ENSMUSG00 | Krba1 | 9540 | 4.38810492 | 7.35424107 | 4.55455049 | 4.64452313 | 6.2472544 | 4.44285004 |
| ENSMUSG00 | Foxf1a | 2120 | 0 | 0 | 0 | 0 | 0 | 0.0128654 |
| ENSMUSG00 | Mcts2 | 792 | 15.0588941 | 13.8110131 | 18.5290554 | 13.3485342 | 12.1767488 | 24.1408108 |
| ENSMUSG00 | Gpr151 | 1632 | 0 | 0 | 0 | 0 | 0 | 0 |
| ENSMUSG00 | Flt3 | 6690 | 0 | 0 | 0 | 0 | 0 | 0 |
| ENSMUSG00 | Snai1 | 1618 | 0 | 0 | 0 | 0 | 0 | 0 |
| ENSMUSG00 | Fgf11 | 4845 | 1.32928542 | 3.16270147 | 1.42321724 | 0.92346203 | 1.94837555 | 2.67961368 |
| ENSMUSG00 | Trim72 | 2546 | 0.24983806 | 0.37852665 | 0.01736129 | 0.01417205 | 0.18037603 | 0.13926564 |
| ENSMUSG00 | Alkbh6 | 3173 | 17.9544921 | 20.3042147 | 24.963663 | 22.174594 | 18.0755268 | 24.2145158 |
| ENSMUSG00 | D0H4S114 | 2508 | 5.89674591 | 10.6056284 | 4.7056986 | 13.0344228 | 10.5796314 | 6.82953487 |
| ENSMUSG00 | Serpinb6b | 2292 | 102.857766 | 70.4505967 | 135.806884 | 93.4795726 | 73.7567116 | 122.378879 |
| ENSMUSG00 | Wfdc12 | 772 | 0 | 0 | 0 | 0 | 0 | 0 |
| ENSMUSG00 | Lrrtm3 | 4625 | 0 | 0 | 0 | 0 | 0 | 0 |
| ENSMUSG00 | Vmn1r226 | 896 | 0 | 0 | 0 | 0 | 0 | 0 |
| ENSMUSG00 | Olfr1414 | 1187 | 0 | 0 | 0 | 0 | 0 | 0 |
| ENSMUSG00 | Zc3h6 | 6084 | 3.52205851 | 3.74625035 | 3.00781772 | 2.30109028 | 7.44763671 | 4.48749407 |
| ENSMUSG00 | Trp53rk | 5432 | 0.36593778 | 0.41692982 | 0.5940233 | 0.4583323 | 0.54483244 | 0.46696271 |
| ENSMUSG00 | Gm9776 | 2979 | 1.66815544 | 1.7469405 | 1.85472655 | 1.25966184 | 2.67207608 | 1.51983555 |
| ENSMUSG00 | Kcna10 | 2266 | 0 | 0 | 0 | 0 | 0 | 0 |
| ENSMUSG00 | Olfr1026 | 923 | 0 | 0 | 0 | 0 | 0 | 0 |
| ENSMUSG00 | Olfr1370 | 950 | 0 | 0 | 0 | 0 | 0 | 0 |
| ENSMUSG00 | Tom1 | 2342 | 0.84875065 | 0.96702083 | 0.77381536 | 0.49300829 | 1.002226 | 0.59393963 |
| ENSMUSG00 | Lhfpl4 | 6999 | 0 | 0 | 0 | 0 | 0.01458105 | 0 |
| ENSMUSG00 | D930007J09f | 392 | 0 | 0 | 0 | 0 | 0 | 0 |
| ENSMUSG00 | Ccdc85b | 4386 | 1.38681909 | 1.93361008 | 2.06597762 | 1.34094238 | 1.32626695 | 1.26858787 |
| ENSMUSG00 | 1700110M21 | 1858 | 0 | 0 | 0 | 0 | 0 | 0 |
| ENSMUSG00 | Olfr1260 | 900 | 0 | 0 | 0 | 0 | 0 | 0 |
| ENSMUSG00 | Abra | 1760 | 0 | 0 | 0 | 0 | 0.60883744 | 0 |
| ENSMUSG00 | Foxo4 | 4021 | 0.27683498 | 0.11983696 | 0.17588398 | 0.43072324 | 0 | 0.12209488 |
| ENSMUSG00 | Olfr648 | 950 | 0 | 0 | 0 | 0 | 0.10742395 | 0.11484059 |
| ENSMUSG00 | Mamstr | 3381 | 0.04703399 | 0 | 0.0261472 | 0.010672 | 0 | 0 |
| ENSMUSG00 | D130020L05l | 2429 | 0.60557957 | 1.2894684 | 0.98266757 | 0.22282037 | 0.67222891 | 0.60635262 |
| ENSMUSG00 | Gm14117 | 842 | 1.08595731 | 0.68674264 | 0.89243626 | 0.5570862 | 0.54541257 | 0.74503171 |
| ENSMUSG00 | 9230106D20l | 1402 | 0 | 0 | 0 | 0 | 0 | 0 |
| ENSMUSG00 | Greb1l | 9542 | 0.1416565 | 0 | 0 | 0 | 0.52940801 | 0.00285838 |
| ENSMUSG00 | 4922501L14f | 2876 | 0 | 0 | 0 | 0 | 0 | 0 |
| ENSMUSG00 | Olfr957 | 935 | 0 | 0 | 0 | 0 | 0 | 0 |
| ENSMUSG00 | Egflam | 7321 | 0 | 0 | 0 | 0.00492857 | 0.00696986 | 0.0111766 |
| ENSMUSG00 | Gm5436 | 641 | 5.02370347 | 7.96842881 | 11.5848824 | 4.84096073 | 3.74140353 | 7.57392478 |
| ENSMUSG00 | 9930038B18l | 3870 | 0 | 0 | 0 | 0 | 0 | 0 |
| ENSMUSG00 | Sbk1 | 4088 | 7.04084342 | 7.01343764 | 6.16317283 | 4.88978783 | 6.2285131 | 4.4968461 |
| ENSMUSG00 | Upk3b | 2340 | 2.7523025 | 1.29732729 | 0.52891094 | 0 | 0.67599045 | 0.72266138 |
| ENSMUSG00 | Notum | 3076 | 0.06462204 | 0.14098764 | 0 | 0 | 0 | 0 |
| ENSMUSG00 | Loh12cr1 | 1817 | 11.2680641 | 9.01672555 | 13.1364861 | 8.12193517 | 9.04264888 | 9.17160406 |
| ENSMUSG00 | Ifnk | 803 | 0.04950869 | 0 | 0 | 0 | 0.06354468 | 0 |
| ENSMUSG00 | Nhlrc3 | 5594 | 7.6042839 | 5.76273325 | 9.30814642 | 6.13407657 | 9.48649104 | 5.15848558 |
| ENSMUSG00 | Rasef | 5442 | 9.03666472 | 4.88770968 | 12.4596889 | 7.84363445 | 8.99196884 | 6.811142 |
| ENSMUSG00 | Gng2 | 4664 | 0 | 0.75420461 | 0.18954478 | 0.07736287 | 0.40479758 | 1.134494 |
| ENSMUSG00 | Klhl6 | 3405 | 0 | 0.24057842 | 0.3764621 | 0.8477426 | 0.46455731 | 0.36045779 |
| ENSMUSG00 | Onecut1 | 3804 | 0 | 0 | 0 | 0 | 0 | 0 |
| ENSMUSG00 | Tmem194b | 3610 | 1.70695276 | 1.17462795 | 0.71016812 | 1.97901518 | 0.86221854 | 2.66702151 |
| ENSMUSG00 | Ptgir | 3728 | 0 | 0 | 0 | 0 | 0 | 0 |
| ENSMUSG00 | Edem3 | 6865 | 11.6515698 | 11.4131326 | 10.0637262 | 14.41705 | 15.4008231 | 8.90746412 |
| ENSMUSG00 | Wdr63 | 3599 | 0 | 0 | 0 | 0 | 0.12760138 | 0 |
| ENSMUSG00 | Gm16433 | 1978 | 0 | 0.04872239 | 0 | 0 | 0 | 0.13788999 |
| ENSMUSG00 | Trpv3 | 5817 | 0 | 0 | 0 | 0 | 0 | 0 |
| ENSMUSG00 | Ccdc63 | 3069 | 0 | 0 | 0.01440269 | 0 | 0 | 0 |
| ENSMUSG00 | Tnp2 | 547 | 0 | 0 | 0.16161551 | 0.13192704 | 0 | 0 |
| ENSMUSG00 | Disc1 | 3863 | 0.10291349 | 0 | 0.18307779 | 0.0467021 | 0.10567202 | 0.07766529 |
| ENSMUSG00 | Zfp513 | 3487 | 5.68912669 | 5.23735634 | 3.05495961 | 5.08066644 | 5.37042728 | 4.27070635 |
| ENSMUSG00 | Fscb | 3565 | 0 | 0 | 0 | 0 | 0 | 0 |
| ENSMUSG00 | Tmem18 | 3038 | 13.0075535 | 15.2743725 | 23.8468798 | 15.8556713 | 18.3076859 | 16.8244483 |
| ENSMUSG00 | Spice1 | 5460 | 2.73774005 | 3.4860155 | 2.35581251 | 3.83289118 | 4.71947249 | 2.18297021 |
| ENSMUSG00 | Vmn1r66 | 1523 | 0 | 0 | 0 | 0 | 0 | 0 |
| ENSMUSG00 | Dpy19l1 | 8968 | 4.84531001 | 3.86867065 | 4.36202398 | 4.88041035 | 3.41958652 | 2.81931206 |
| ENSMUSG00 | Fam89a | 1313 | 11.7782802 | 9.98226372 | 18.8522712 | 9.89302059 | 7.61703706 | 17.6568495 |
| ENSMUSG00 | Dub3 | 1620 | 0.07362126 | 0 | 0 | 0 | 0 | 0 |
| ENSMUSG00 | Synpo | 10364 | 0.47949007 | 0.33940663 | 0.45208369 | 0.43170335 | 0.41356769 | 0.01579003 |
| ENSMUSG00 | Tmem82 | 4459 | 0.02674735 | 0.21613116 | 0.1784331 | 0.37223011 | 0.30897335 | 0.05505085 |
| ENSMUSG00 | Olfr855 | 991 | 0 | 0 | 0 | 0 | 0 | 0 |
| ENSMUSG00 | Il17re | 3146 | 29.2669081 | 35.7186215 | 20.2041482 | 29.5560803 | 30.4114605 | 18.6917108 |
| ENSMUSG00 | Mmp1a | 2011 | 0 | 0 | 0 | 0 | 0 | 0 |
| ENSMUSG00 | Zfp866 | 5650 | 21.8127415 | 26.0377359 | 28.2031228 | 23.6097909 | 31.3654165 | 24.3782174 |
| ENSMUSG00 | Tuba1c | 1973 | 17.6713413 | 20.1489178 | 17.9002902 | 19.3120319 | 14.1466942 | 12.344781 |
| ENSMUSG00 | Hic1 | 4138 | 0 | 0 | 0 | 0.01743936 | 0.02466234 | 0 |
| ENSMUSG00 | Qrfp | 2925 | 0.16309941 | 0.09884398 | 0.04533522 | 0.04934297 | 0.40123304 | 0.39163585 |
| ENSMUSG00 | Lrrn4 | 3710 | 0 | 0 | 0 | 0 | 0 | 0 |
| ENSMUSG00 | Olfr448 | 932 | 0 | 0 | 0 | 0 | 0 | 0 |
| ENSMUSG00 | A530016L24f | 2631 | 0.01511041 | 0 | 0 | 0 | 0 | 0 |
| ENSMUSG00 | D830039M14 | 1692 | 0.32894606 | 0 | 0.36573629 | 0 | 0.21110203 | 0 |
| ENSMUSG00 | Mob1a | 4071 | 45.4976133 | 47.3578863 | 44.2887049 | 51.5217207 | 58.0956158 | 37.9138255 |
| ENSMUSG00 | Tmem186 | 2734 | 23.7166016 | 18.8410046 | 25.3020791 | 18.8724666 | 25.9611151 | 17.4382112 |
| ENSMUSG00 | Aqp6 | 2055 | 20.5258223 | 9.09797545 | 31.3392142 | 22.8782988 | 32.5029323 | 62.2339579 |
| ENSMUSG00 | Gm11292 | 565 | 0 | 0 | 0 | 0 | 0 | 0 |
| ENSMUSG00 | Ispd | 2680 | 4.91009853 | 6.99422612 | 4.8160217 | 6.34128469 | 4.91224066 | 5.35315681 |
| ENSMUSG00 | Ppp2r3a | 7442 | 15.2568735 | 18.7449274 | 9.02805716 | 23.6215696 | 18.7732084 | 9.6718319 |
| ENSMUSG00 | Hpdl | 1601 | 6.38173547 | 3.31074868 | 2.95415191 | 2.0058101 | 2.77282617 | 1.56731219 |
| ENSMUSG00 | Arl11 | 1480 | 0 | 0 | 0 | 0 | 0 | 0 |
| ENSMUSG00 | Pigy | 3571 | 31.5171564 | 28.0131767 | 36.4655921 | 31.2522441 | 26.877797 | 42.9474367 |
| ENSMUSG00 | Tmem212 | 712 | 0 | 0 | 0 | 0 | 0 | 0 |
| ENSMUSG00 | Lor | 1773 | 0.85206332 | 1.25018406 | 0.32409699 | 0.79368287 | 0.68193268 | 0.06153331 |
| ENSMUSG00 | 4930426D05l | 2571 | 1.0360238 | 0.99334167 | 1.04874073 | 1.37535603 | 1.56790498 | 3.51143741 |
| ENSMUSG00 | Gm11993 | 1100 | 0 | 0.08761171 | 0.08036699 | 0.09840558 | 0.18555046 | 0 |
| ENSMUSG00 | 4732471D19l | 5660 | 4.67091776 | 7.35566891 | 5.13865949 | 12.6478403 | 10.9625571 | 5.36336987 |
| ENSMUSG00 | Dusp8 | 1185 | 0 | 0 | 0 | 0 | 0 | 0 |
| ENSMUSG00 | Rfesd | 2857 | 16.2250229 | 11.3340179 | 13.4601342 | 9.87614939 | 9.10866352 | 19.7614642 |
| ENSMUSG00 | Gm1840 | 1412 | 1.74563725 | 1.43330777 | 1.00174149 | 1.78876991 | 1.30095576 | 0.8112853 |
| ENSMUSG00 | Zmpste24 | 7134 | 23.3940997 | 25.167183 | 28.5199164 | 30.8624384 | 27.1368194 | 30.0999768 |
| ENSMUSG00 | Hoxa6 | 2319 | 0.82288187 | 1.16362258 | 0.07624294 | 1.13582978 | 1.16619143 | 0.30579587 |
| ENSMUSG00 | Gm4835 | 2117 | 0 | 0 | 0.10439736 | 0.01704395 | 0.04820631 | 0.01288363 |
| ENSMUSG00 | Olfr1009 | 944 | 0 | 0 | 0 | 0 | 0 | 0 |
| ENSMUSG00 | Fam124b | 3575 | 0 | 0 | 0 | 0 | 0 | 0 |
| ENSMUSG00 | Upf2 | 5451 | 10.4949801 | 6.3470676 | 8.10894203 | 13.6623262 | 12.7027687 | 14.0200952 |
| ENSMUSG00 | Fam129c | 3015 | 0.80433974 | 1.16670324 | 0.33719482 | 0.88559579 | 0.79543604 | 0.18997261 |
| ENSMUSG00 | Gm9780 | 1327 | 0 | 0 | 0 | 0 | 0 | 0 |
| ENSMUSG00 | Exoc3l | 2483 | 0.54437629 | 0.7956682 | 1.35293599 | 2.2524031 | 1.35631929 | 1.05451686 |
| ENSMUSG00 | Tmem64 | 4667 | 48.5379747 | 51.7175678 | 55.7850558 | 68.9865188 | 57.6301751 | 43.3460466 |
| ENSMUSG00 | Pigv | 4712 | 8.25145584 | 8.67192337 | 8.18934825 | 8.56106235 | 11.9011008 | 6.72604644 |
| ENSMUSG00 | Fam13c | 3524 | 0.49637944 | 0.72471096 | 0.05017235 | 1.04437246 | 0.46334961 | 0.32506664 |
| ENSMUSG00 | Uevld | 6781 | 1.95816701 | 1.39990106 | 1.58399172 | 1.53246259 | 1.45230652 | 2.28461806 |
| ENSMUSG00 | Pyhin1 | 3490 | 0.28478138 | 0.30375408 | 0 | 0 | 0 | 0 |
| ENSMUSG00 | Olfr1031 | 1010 | 0 | 0 | 0 | 0 | 0 | 0 |
| ENSMUSG00 | Olfr1123 | 971 | 0 | 0 | 0 | 0 | 0 | 0 |
| ENSMUSG00 | Trim66 | 12616 | 2.29722141 | 2.53230906 | 3.14976671 | 2.1049766 | 2.54403855 | 2.42349957 |
| ENSMUSG00 | Gm5531 | 2526 | 0 | 0 | 0 | 0 | 0 | 0 |
| ENSMUSG00 | Tmem11 | 2103 | 26.2011869 | 24.2421568 | 33.6505709 | 22.5448438 | 19.3138351 | 28.4678237 |
| ENSMUSG00 | Pnpla1 | 4475 | 0 | 0 | 0 | 0 | 0 | 0 |
| ENSMUSG00 | Mei4 | 4792 | 1.66753998 | 0.52289128 | 2.88714041 | 1.52098769 | 1.24584432 | 0.77407157 |
| ENSMUSG00 | Zfp784 | 2151 | 7.68864709 | 6.72056376 | 6.59636988 | 5.08268689 | 4.12765662 | 6.32731055 |
| ENSMUSG00 | Gm16373 | 607 | 0 | 0 | 0 | 0 | 0.08406322 | 0 |
| ENSMUSG00 | C030030A07l | 3027 | 0.14446987 | 0.31837755 | 0.0584101 | 0.15496088 | 0.5394265 | 0 |
| ENSMUSG00 | B3galnt1 | 1898 | 48.4690105 | 20.3865717 | 76.4566125 | 22.6986097 | 31.8309952 | 68.8332571 |
| ENSMUSG00 | Kcnj6 | 5830 | 0 | 0 | 0 | 0 | 0 | 0 |
| ENSMUSG00 | Vmn1r75 | 917 | 0 | 0 | 0 | 0 | 0 | 0 |
| ENSMUSG00 | Olfr571 | 968 | 0 | 0 | 0 | 0 | 0 | 0 |
| ENSMUSG00 | D17H6S53E | 2030 | 1.78214223 | 1.37675549 | 1.19758688 | 1.83076384 | 0.55299521 | 0.68522493 |
| ENSMUSG00 | Olfr131 | 2384 | 0 | 0 | 0 | 0 | 0 | 0 |
| ENSMUSG00 | Pcdhb19 | 7130 | 0.26206278 | 0.35818814 | 0.37196502 | 0.23784798 | 1.51003721 | 1.48040468 |
| ENSMUSG00 | Olfr30 | 1065 | 0 | 0 | 0 | 0 | 0 | 0 |
| ENSMUSG00 | Cox8c | 474 | 0 | 0 | 0 | 0 | 0 | 0 |
| ENSMUSG00 | Fbrsl1 | 5955 | 3.56497508 | 4.85505714 | 4.70595608 | 5.47744219 | 4.85843075 | 5.56027071 |
| ENSMUSG00 | Gm8849 | 475 | 0 | 0 | 0 | 0 | 0 | 0 |
| ENSMUSG00 | Olfr975 | 932 | 0 | 0 | 0 | 0 | 0 | 0 |
| ENSMUSG00 | Rhbdl2 | 3155 | 5.07811685 | 5.1928337 | 6.22048124 | 6.08419895 | 3.02438424 | 6.22432338 |
| ENSMUSG00 | Filip1l | 5509 | 0.59174976 | 1.12834472 | 0.28884849 | 0.77940884 | 1.21336998 | 1.05454677 |
| ENSMUSG00 | 6530409C15l | 2200 | 0.05421202 | 0.17522343 | 0.58266066 | 0.37722138 | 0.57984518 | 0.55789035 |
| ENSMUSG00 | Hoxd9 | 2181 | 16.2230067 | 21.2762695 | 9.50511892 | 13.764448 | 18.9974402 | 16.3197635 |
| ENSMUSG00 | Gm6741 | 995 | 0 | 0.09685717 | 0 | 0 | 0.05128279 | 0 |
| ENSMUSG00 | Olfr577 | 938 | 0 | 0 | 0 | 0 | 0 | 0 |
| ENSMUSG00 | Olfr187 | 1085 | 0 | 0 | 0 | 0 | 0 | 0 |
| ENSMUSG00 | Vmn1r28 | 1136 | 0 | 0 | 0 | 0 | 0 | 0 |
| ENSMUSG00 | Olfr78 | 3492 | 0 | 0 | 0 | 0 | 0 | 0 |
| ENSMUSG00 | Hexim2 | 2368 | 2.18252216 | 2.13664545 | 3.77059641 | 2.78843503 | 2.75818248 | 2.40726335 |
| ENSMUSG00 | Olfr1342 | 1001 | 0 | 0 | 0 | 0 | 0 | 0 |
| ENSMUSG00 | Gprasp1 | 8255 | 7.39726445 | 7.31406566 | 7.60682119 | 10.7393802 | 11.3549912 | 7.37125623 |
| ENSMUSG00 | Olfr267 | 943 | 0 | 0 | 0 | 0 | 0 | 0 |
| ENSMUSG00 | Tmem130 | 2797 | 0 | 0 | 0 | 0 | 0 | 0.12676808 |
| ENSMUSG00 | 2510009E07f | 5376 | 2.15933786 | 3.50463149 | 1.81707725 | 1.22152755 | 3.549826 | 1.45606798 |

| | | | | | | | | |
|---|---|---|---|---|---|---|---|---|
| ENSMUSG00 | Gpr135 | 1373 | 0 | 0 | 0 | 0 | 0 | 0 |
| ENSMUSG00 | Hfm1 | 7141 | 0 | 0 | 0 | 0 | 0 | 0 |
| ENSMUSG00 | Usp48 | 7848 | 11.6156112 | 11.4571739 | 10.6336748 | 12.0503362 | 14.2715208 | 11.1767635 |
| ENSMUSG00 | Otud1 | 2528 | 6.73075382 | 11.1698003 | 6.11971719 | 5.98037051 | 6.15626765 | 8.46937967 |
| ENSMUSG00 | Lrit2 | 2778 | 0 | 0 | 0 | 0 | 0 | 0 |
| ENSMUSG00 | A030009H04 | 2250 | 3.92254074 | 1.71329572 | 2.65211058 | 1.45931825 | 0.88445718 | 2.40016825 |
| ENSMUSG00 | Hilpda | 1100 | 5.96332208 | 8.19169516 | 5.10330369 | 8.69249254 | 2.50493118 | 6.91784032 |
| ENSMUSG00 | Gm9781 | 2376 | 4.08263352 | 1.70356109 | 3.85091814 | 0.48595346 | 2.53413819 | 4.38506405 |
| ENSMUSG00 | 4922505E12l | 2637 | 0 | 0 | 0 | 0 | 0 | 0 |
| ENSMUSG00 | Psapl1 | 2547 | 0 | 0 | 0 | 0 | 0 | 0 |
| ENSMUSG00 | Leng9 | 1689 | 3.20115178 | 3.56619613 | 5.7836633 | 4.03760001 | 2.96067782 | 3.95635679 |
| ENSMUSG00 | E130012A19l | 2323 | 0.10268312 | 0.10371598 | 0.05708374 | 0.26405284 | 0.17572579 | 0.08218789 |
| ENSMUSG00 | Gpr149 | 4284 | 0 | 0 | 0.01031789 | 0 | 0 | 0 |
| ENSMUSG00 | Pgp | 1025 | 49.8010117 | 54.5799603 | 45.7111742 | 55.7599595 | 41.9163009 | 44.8102366 |
| ENSMUSG00 | Gjc2 | 3917 | 0 | 0.35675436 | 0 | 0 | 0 | 0 |
| ENSMUSG00 | Magea10 | 1817 | 0 | 0 | 0 | 0 | 0 | 0 |
| ENSMUSG00 | Zfp536 | 10749 | 0.18122789 | 0.09862329 | 0.12747764 | 0.06377885 | 0.10443579 | 0.04059859 |
| ENSMUSG00 | Pcdhb12 | 3033 | 1.48116363 | 0.14298648 | 2.02573563 | 0.39258407 | 0 | 0 |
| ENSMUSG00 | Elfn2 | 4844 | 0 | 0 | 0 | 0 | 0 | 0 |
| ENSMUSG00 | 1110032A04l | 1782 | 1.62859151 | 0.48673174 | 0 | 1.17438753 | 0.02863433 | 0 |
| ENSMUSG00 | Rab9b | 3977 | 0 | 0 | 0 | 0 | 0 | 0 |
| ENSMUSG00 | Zbtb37 | 5496 | 1.45393952 | 1.5430884 | 1.22246727 | 2.66544943 | 0.93771178 | 2.58553258 |
| ENSMUSG00 | Adam30 | 2361 | 0 | 0 | 0 | 0 | 0 | 0 |
| ENSMUSG00 | Gm9782 | 373 | 0 | 0 | 0 | 0 | 0 | 0 |
| ENSMUSG00 | Gm6863 | 641 | 0.49616824 | 0.75173857 | 0.4137458 | 0.16887072 | 0.07960433 | 0.63825209 |
| ENSMUSG00 | Gm5867 | 590 | 0.06738217 | 0.49003162 | 0.59934702 | 1.34543217 | 0 | 0.69342303 |
| ENSMUSG00 | Krt34 | 1602 | 0 | 0 | 0 | 0 | 0 | 0.13620294 |
| ENSMUSG00 | Acot6 | 3385 | 11.2160957 | 6.16387871 | 16.1921079 | 9.93455406 | 7.14520004 | 11.7236633 |
| ENSMUSG00 | Gm9783 | 466 | 0 | 0.41361753 | 0.09485374 | 0.30971712 | 0.32849598 | 0.58529269 |
| ENSMUSG00 | Tril | 5354 | 2.1459346 | 1.36801255 | 0.38802514 | 0.68740541 | 1.12460074 | 1.02394518 |
| ENSMUSG00 | 9330132A10l | 3725 | 0 | 0 | 0 | 0 | 0 | 0 |
| ENSMUSG00 | Lgals2 | 586 | 0 | 0.65783539 | 0.60343813 | 1.10832219 | 0 | 0.13963126 |
| ENSMUSG00 | Gimap5 | 2581 | 0.04620939 | 0.37339359 | 0.27401375 | 0.8387922 | 1.7595302 | 0.54950842 |
| ENSMUSG00 | Hscb | 1030 | 8.33707164 | 6.31569873 | 7.25253541 | 7.21640891 | 4.5576957 | 11.7837034 |
| ENSMUSG00 | Rai2 | 2400 | 0.97732223 | 1.04403958 | 0.97612403 | 1.41321341 | 1.59457424 | 1.93195361 |
| ENSMUSG00 | A430060F13l | 457 | 0 | 0 | 0 | 0 | 0 | 0 |
| ENSMUSG00 | Olfr266 | 1026 | 0 | 0 | 0 | 0 | 0 | 0 |
| ENSMUSG00 | Sorcs1 | 7636 | 0 | 0 | 0 | 0 | 0 | 0 |
| ENSMUSG00 | Setx | 11350 | 3.52019894 | 3.57471227 | 3.6958193 | 3.60502542 | 3.71346136 | 4.80850688 |
| ENSMUSG00 | Vmn1r225 | 896 | 0 | 0 | 0 | 0 | 0 | 0 |
| ENSMUSG00 | Casc1 | 4861 | 0.60520583 | 1.30849833 | 0.73654583 | 0.61608922 | 0.82927046 | 1.18390226 |
| ENSMUSG00 | Fam164a | 3352 | 33.1018932 | 26.9251808 | 54.3028608 | 39.2037011 | 28.8621744 | 43.5727008 |
| ENSMUSG00 | 4932442L08l | 4490 | 0.04427114 | 0.1824431 | 0.04922254 | 0.19286616 | 0.27274678 | 0.27942838 |
| ENSMUSG00 | Fbxl7 | 1941 | 0 | 0.04965115 | 0 | 0 | 0 | 0 |
| ENSMUSG00 | Mdga1 | 13813 | 0 | 0 | 0.00960005 | 0 | 0 | 0.01974563 |
| ENSMUSG00 | 4930412D23l | 1542 | 0 | 0 | 0 | 0 | 0 | 0 |
| ENSMUSG00 | Pars2 | 3125 | 3.34582124 | 2.20501159 | 2.60260452 | 0.03463876 | 2.77583485 | 2.18197114 |
| ENSMUSG00 | 4930470P17l | 1393 | 0 | 0 | 0 | 0 | 0 | 0 |
| ENSMUSG00 | Acpl2 | 6416 | 1.17110128 | 1.72738181 | 0.47536271 | 0.35429688 | 1.90076431 | 0.47186486 |
| ENSMUSG00 | Unc5cl | 3213 | 0.02474664 | 0 | 0 | 0.28075042 | 0 | 0 |
| ENSMUSG00 | Olfr706 | 950 | 0 | 0 | 0 | 0 | 0 | 0 |
| ENSMUSG00 | Zfp3 | 2059 | 13.0330011 | 10.9057222 | 11.7427917 | 7.90335215 | 11.4493374 | 11.8689057 |
| ENSMUSG00 | Olfr13 | 1053 | 0 | 0 | 0 | 0 | 0 | 0 |
| ENSMUSG00 | Mmp3 | 1788 | 0 | 0 | 0 | 0 | 0 | 0.41186536 |
| ENSMUSG00 | Vps37d | 2620 | 0.86490931 | 1.32420757 | 1.43402926 | 0.7574475 | 1.36330012 | 2.1549047 |
| ENSMUSG00 | Eif5al3-ps | 446 | 0.08913785 | 0.2160827 | 0 | 0 | 0 | 0.0611539 |
| ENSMUSG00 | Ubxn10 | 4554 | 0 | 0 | 0 | 0 | 0 | 0 |
| ENSMUSG00 | 4933412A08l | 1341 | 0.68186134 | 0.35933216 | 0.0659237 | 0 | 0.11415297 | 0 |
| ENSMUSG00 | 1700019D03l | 3999 | 0.76548437 | 0.86757285 | 1.52534492 | 0.28872854 | 0.34451417 | 1.1682824 |
| ENSMUSG00 | Ecm2 | 3586 | 0.39910689 | 0.08062428 | 0 | 0 | 0 | 0.23578188 |
| ENSMUSG00 | 4930412F15l | 4406 | 0.07218426 | 0.14217516 | 0.01003219 | 0.03275719 | 0.02316222 | 0.03714204 |
| ENSMUSG00 | Adamts3 | 8522 | 0.19126669 | 0.69548608 | 0.29564716 | 1.59621342 | 0.43709522 | 0.41926517 |
| ENSMUSG00 | Rbm20 | 6774 | 3.68562766 | 2.21939326 | 3.13210579 | 2.93492863 | 2.1694119 | 2.39569093 |
| ENSMUSG00 | 0610009L18l | 617 | 6.63665234 | 7.65360021 | 9.38483215 | 7.36845642 | 4.8793455 | 10.1230023 |
| ENSMUSG00 | Pld6 | 2277 | 1.11741359 | 0.78300323 | 0.58236947 | 0.14261678 | 0.56023689 | 0.16769651 |
| ENSMUSG00 | Npsr1 | 3949 | 0 | 0 | 0 | 0.08223307 | 0.20674146 | 0 |
| ENSMUSG00 | Gm9785 | 1597 | 0 | 0 | 0.19374634 | 0 | 0 | 0.66606821 |
| ENSMUSG00 | Tmem221 | 1535 | 16.8345683 | 19.1804014 | 13.8508707 | 16.1957842 | 22.4383086 | 13.4152134 |
| ENSMUSG00 | Tox3 | 3349 | 9.71035624 | 7.55386149 | 8.88260387 | 13.9953944 | 9.43127102 | 18.6663103 |
| ENSMUSG00 | Diras1 | 3411 | 0 | 0 | 0 | 0 | 0 | 0 |
| ENSMUSG00 | Dpy19l3 | 11343 | 4.33549585 | 2.26424876 | 3.87345781 | 1.61912727 | 2.39319686 | 3.00566861 |
| ENSMUSG00 | Kcns3 | 3055 | 22.7081223 | 17.5395495 | 38.9207718 | 25.0743635 | 23.4504195 | 24.1498852 |
| ENSMUSG00 | Fam25c | 405 | 0.19632336 | 0 | 0 | 0.2672744 | 0 | 0 |
| ENSMUSG00 | Fem1a | 6815 | 10.4595123 | 11.991813 | 11.804454 | 8.19059763 | 10.1977878 | 6.71960665 |
| ENSMUSG00 | 1190005I06R | 1061 | 1.34891357 | 0.90832125 | 0.62490824 | 1.22427295 | 0.9618544 | 0.82260929 |
| ENSMUSG00 | Olfr399 | 1002 | 0 | 0 | 0 | 0 | 0 | 0 |
| ENSMUSG00 | Olfr62 | 948 | 0 | 0 | 0 | 0 | 0 | 0 |
| ENSMUSG00 | Pde12 | 3526 | 13.3495431 | 13.4063811 | 13.2755961 | 12.7709562 | 11.4469266 | 13.2118786 |
| ENSMUSG00 | Capn13 | 2591 | 0 | 0.20457386 | 0 | 0 | 0 | 0 |
| ENSMUSG00 | Olfr1326-ps1 | 946 | 0 | 0 | 0 | 0 | 0 | 0 |
| ENSMUSG00 | Rpl7 | 2182 | 11.6788556 | 9.34136802 | 9.78436763 | 9.50832979 | 6.85184607 | 12.3623365 |
| ENSMUSG00 | Col6a6 | 7444 | 0 | 0 | 0 | 0 | 0 | 0 |
| ENSMUSG00 | F830045P16l | 1918 | 0 | 0 | 0 | 0 | 0 | 0 |
| ENSMUSG00 | Ptpn11 | 11089 | 5.70393809 | 4.0108293 | 3.63532156 | 3.45885232 | 4.01713645 | 2.67113881 |
| ENSMUSG00 | B430306N03 | 5753 | 0 | 0 | 0 | 0 | 0 | 0 |
| ENSMUSG00 | 1520401A03l | 3990 | 0 | 0 | 0 | 0.11756055 | 0 | 0 |
| ENSMUSG00 | Dmrta1 | 4191 | 0 | 0 | 0 | 0 | 0 | 0 |
| ENSMUSG00 | Pkhd1 | 14246 | 9.25376761 | 13.8951217 | 5.49808127 | 14.3152966 | 13.1810377 | 9.62059962 |
| ENSMUSG00 | Gm12481 | 475 | 0 | 0 | 0 | 0 | 0 | 0 |
| ENSMUSG00 | 1700048O20l | 1638 | 0.04854149 | 0.11767141 | 0.0539705 | 0 | 0 | 2.06474681 |
| ENSMUSG00 | Ccdc64b | 1903 | 10.2574571 | 11.5211935 | 6.29463976 | 8.19098436 | 9.86742318 | 8.87178229 |
| ENSMUSG00 | Defb12 | 1000 | 0 | 0 | 0 | 0 | 0 | 0 |
| ENSMUSG00 | Vwce | 3520 | 0 | 0 | 0 | 0 | 0.44938001 | 0 |
| ENSMUSG00 | D830025C05 | 974 | 0.40816715 | 0.4452546 | 0.95301715 | 0.62976874 | 0.78582714 | 1.09210568 |
| ENSMUSG00 | Gm14492 | 2375 | 0 | 0 | 0 | 0 | 0 | 0 |
| ENSMUSG00 | Kifc5c-ps | 739 | 0 | 0 | 0 | 0 | 0 | 0 |
| ENSMUSG00 | Gm9786 | 679 | 0.5269504 | 0.85160133 | 1.4972642 | 0.90337961 | 0.90179162 | 1.88793527 |
| ENSMUSG00 | Ly6g5b | 2004 | 0.09919032 | 0.12022565 | 0.17645446 | 0 | 0.25462263 | 0.24498179 |
| ENSMUSG00 | Rtn4r | 1936 | 0 | 0 | 0 | 0 | 0 | 0 |
| ENSMUSG00 | Adamtsl5 | 4625 | 0 | 0 | 0.10512871 | 0 | 0 | 0.02358888 |
| ENSMUSG00 | Olfr94 | 1303 | 0 | 0 | 0 | 0 | 0 | 0 |
| ENSMUSG00 | Lysmd4 | 2895 | 4.21586615 | 4.57729589 | 6.45850763 | 6.55583952 | 5.1643275 | 9.18575933 |
| ENSMUSG00 | Clec4a3 | 1320 | 0 | 0 | 0 | 0 | 0 | 0 |
| ENSMUSG00 | 2900005J15R | 3832 | 9.96999394 | 3.35745826 | 2.89526686 | 3.86055278 | 8.61536147 | 3.4662707 |
| ENSMUSG00 | Tmem145 | 2530 | 0 | 0 | 0 | 0 | 0 | 0 |
| ENSMUSG00 | Clrn1 | 6274 | 0 | 0 | 0 | 0 | 0 | 0 |
| ENSMUSG00 | Olfr479 | 1110 | 0 | 0 | 0 | 0 | 0 | 0 |
| ENSMUSG00 | Mgat5b | 4584 | 0 | 0 | 0 | 0 | 0 | 0 |
| ENSMUSG00 | Nup62 | 2823 | 9.30866901 | 11.6070778 | 6.93638557 | 9.58609047 | 10.0859786 | 5.90322514 |
| ENSMUSG00 | 1700049L16l | 804 | 0 | 0 | 0 | 0 | 0 | 0 |
| ENSMUSG00 | Tas2r118 | 899 | 0 | 0 | 0 | 0 | 0 | 0 |
| ENSMUSG00 | Taf10 | 4747 | 4.3549294 | 5.19727373 | 5.6521 | 4.62902151 | 3.13876169 | 5.06767049 |
| ENSMUSG00 | Foxl1 | 1081 | 0 | 0 | 0 | 0 | 0 | 0 |
| ENSMUSG00 | Gm5809 | 824 | 0 | 0 | 0 | 0 | 0 | 0 |
| ENSMUSG00 | Zmym1 | 6434 | 0.96391902 | 1.13089101 | 1.29843773 | 1.38518262 | 1.18961088 | 1.43706912 |
| ENSMUSG00 | Chi3l7 | 1878 | 0 | 0 | 0 | 0 | 0 | 0 |
| ENSMUSG00 | Olfr323 | 971 | 0 | 0 | 0 | 0 | 0 | 0 |
| ENSMUSG00 | Kbtbd7 | 2948 | 21.2687547 | 16.4762326 | 23.3454103 | 18.5428417 | 20.1301476 | 27.5614486 |
| ENSMUSG00 | Slc36a4 | 5650 | 1.43541912 | 2.49887213 | 1.3847303 | 2.02442622 | 3.84729843 | 2.75160077 |
| ENSMUSG00 | Gm8399 | 1117 | 0.42709558 | 0.86278321 | 0.59357891 | 0.41993427 | 0.22840813 | 0.46393746 |
| ENSMUSG00 | Olfr1018 | 935 | 0 | 0 | 0 | 0 | 0 | 0 |
| ENSMUSG00 | S1pr2 | 2813 | 0.67837293 | 2.84356537 | 1.27278681 | 0.10261513 | 0.87069536 | 0.87263332 |
| ENSMUSG00 | Vmn2r2 | 2721 | 0 | 0 | 0 | 0 | 0 | 0 |
| ENSMUSG00 | Gm22 | 3077 | 0 | 0 | 0 | 0 | 0 | 0 |
| ENSMUSG00 | Trp53bp1 | 14059 | 2.70051098 | 3.80103594 | 2.61582854 | 4.60425264 | 4.30815194 | 2.8304786 |
| ENSMUSG00 | Olfr922 | 1131 | 0 | 0 | 0 | 0 | 0 | 0 |
| ENSMUSG00 | Ccdc60 | 4688 | 0 | 0 | 0 | 0 | 0 | 0 |
| ENSMUSG00 | D630028G08 | 361 | 0 | 0 | 0 | 0 | 0 | 0 |
| ENSMUSG00 | Ccdc84 | 1145 | 22.4991715 | 19.8216718 | 20.6918671 | 21.5862013 | 23.8865829 | 28.9182638 |
| ENSMUSG00 | A330049M08 | 3352 | 0.94881815 | 0.41688748 | 0.39560122 | 0.46286632 | 0.59368397 | 0.71604065 |
| ENSMUSG00 | Olfr544 | 1066 | 0 | 0 | 0 | 0 | 0 | 0 |
| ENSMUSG00 | Klhl15 | 7882 | 2.89515933 | 3.46022916 | 4.00407459 | 2.60933334 | 2.81609661 | 5.21131778 |
| ENSMUSG00 | Gimap7 | 1267 | 0 | 0 | 0 | 0 | 0 | 0 |
| ENSMUSG00 | Klri2 | 3907 | 0 | 0 | 0 | 0 | 0 | 0 |
| ENSMUSG00 | A530064D06 | 2984 | 0 | 0 | 0 | 0 | 0 | 0 |
| ENSMUSG00 | Wdfy3 | 17317 | 18.3843557 | 20.2323509 | 12.6451424 | 21.1862464 | 21.8255105 | 18.4781468 |
| ENSMUSG00 | Naalad2 | 5214 | 19.4202549 | 5.97016525 | 33.7744807 | 6.6088133 | 10.7356989 | 22.8230248 |
| ENSMUSG00 | Adam6a | 2506 | 0 | 0 | 0 | 0 | 0 | 0 |
| ENSMUSG00 | Olfr691 | 968 | 0.04106971 | 0 | 0 | 0 | 0 | 0.16905768 |
| ENSMUSG00 | Ccrl2 | 2108 | 1.75391826 | 0 | 1.61458345 | 0.82160253 | 2.61425455 | 1.56557464 |
| ENSMUSG00 | Thrap3 | 9297 | 4.78930173 | 4.9705064 | 3.9651863 | 6.25623919 | 6.14160638 | 6.09623538 |
| ENSMUSG00 | Orai3 | 1680 | 17.7953103 | 15.9187354 | 24.9424686 | 12.7790574 | 18.16296 | 12.387232 |
| ENSMUSG00 | Emx2 | 2785 | 7.82262238 | 9.25664149 | 12.0146482 | 11.4400163 | 9.10596366 | 14.0241592 |
| ENSMUSG00 | Opn5 | 1817 | 0.04375947 | 0.05303956 | 0.02432683 | 0 | 0 | 0 |
| ENSMUSG00 | Krtap16-4 | 303 | 0 | 0 | 0 | 0 | 0 | 0 |
| ENSMUSG00 | 4932411G14 | 3823 | 0 | 0 | 0 | 0 | 0 | 0 |
| ENSMUSG00 | Cep164 | 7365 | 1.09037435 | 3.04232119 | 1.19431999 | 0.76426327 | 1.79441023 | 1.34799303 |
| ENSMUSG00 | Pura | 14941 | 10.2761305 | 9.91077818 | 10.5674977 | 11.3841616 | 7.80712771 | 15.8105649 |
| ENSMUSG00 | 2900052L18l | 1586 | 2.00532058 | 2.12676605 | 2.17386113 | 1.72902609 | 1.89820692 | 4.1617041 |
| ENSMUSG00 | Mgat2 | 5028 | 23.9497515 | 21.6781488 | 18.3471055 | 20.2871798 | 21.7278183 | 22.0074008 |
| ENSMUSG00 | Gpr75 | 1941 | 0.18433762 | 0 | 0 | 0.03717882 | 0 | 0 |
| ENSMUSG00 | Gls2 | 3700 | 0 | 0 | 0.04778578 | 0.13652666 | 0 | 0.15480201 |
| ENSMUSG00 | Cilp2 | 4306 | 0 | 0.14547695 | 0 | 0.19272806 | 0.47400256 | 0 |
| ENSMUSG00 | Npy5r | 2312 | 0 | 0 | 0 | 0 | 0 | 0 |
| ENSMUSG00 | Gpr133 | 8764 | 0.28578221 | 0 | 0 | 0 | 0 | 0 |
| ENSMUSG00 | Mrpl50 | 2112 | 35.7084027 | 37.5543957 | 49.2247797 | 34.7153004 | 25.8031105 | 49.4481978 |
| ENSMUSG00 | Muc19 | 7688 | 0 | 0 | 0 | 0 | 0 | 0 |
| ENSMUSG00 | Pcdhb21 | 4661 | 0.02558817 | 0.86841046 | 0.42675025 | 0.71993781 | 0.20800282 | 0.02340669 |
| ENSMUSG00 | Retl2 | 4156 | 0 | 0.06956657 | 0.11699237 | 0.03472767 | 0.06138881 | 0.11812885 |
| ENSMUSG00 | Olfr790 | 1069 | 0 | 0 | 0 | 0 | 0 | 0 |
| ENSMUSG00 | Tmem20 | 3468 | 106.897306 | 139.835165 | 70.6617698 | 137.336502 | 135.393515 | 98.2374622 |
| ENSMUSG00 | Olfr178 | 959 | 0 | 0 | 0 | 0 | 0 | 0 |
| ENSMUSG00 | Irf2bp1 | 2696 | 14.9525435 | 12.3504568 | 16.2477843 | 13.1158767 | 14.3464365 | 13.0303173 |
| ENSMUSG00 | Ccdc141 | 11631 | 9.99441364 | 12.366652 | 4.51101261 | 11.3603686 | 11.2748505 | 13.0287934 |
| ENSMUSG00 | Npb | 730 | 0.32675737 | 0.13201765 | 1.99816551 | 0.93912171 | 0.62909231 | 3.10108912 |
| ENSMUSG00 | Als2cl | 5874 | 3.71565523 | 4.68410937 | 2.91217454 | 6.12424215 | 4.06543137 | 3.00884682 |
| ENSMUSG00 | Olfr1288 | 938 | 0 | 0 | 0 | 0 | 0 | 0 |
| ENSMUSG00 | Olfr1497 | 944 | 0 | 0 | 0 | 0 | 0 | 0 |
| ENSMUSG00 | Krt13 | 1812 | 0 | 0 | 0 | 0 | 0 | 0 |
| ENSMUSG00 | Fmn1 | 14995 | 5.15137703 | 6.0060327 | 2.59698724 | 7.46184862 | 4.85592786 | 5.97877554 |
| ENSMUSG00 | Pcdhb14 | 3437 | 0.6130464 | 1.34591168 | 1.37608298 | 0.94483096 | 4.03816533 | 2.5790683 |
| ENSMUSG00 | Ccr10 | 1724 | 0.02306002 | 0 | 0.17947384 | 0.64880707 | 0.11839066 | 0.07910278 |
| ENSMUSG00 | Otos | 3375 | 0 | 0 | 0 | 0 | 0 | 0 |
| ENSMUSG00 | Efcab9 | 1038 | 0 | 1.34624935 | 0 | 0 | 0.24579179 | 0 |
| ENSMUSG00 | Mthfr-ps1 | 598 | 0 | 0 | 0 | 0 | 0 | 0 |
| ENSMUSG00 | A830010M20 | 7917 | 1.57173998 | 2.0511344 | 1.45162046 | 2.22407954 | 2.36537576 | 1.06797249 |
| ENSMUSG00 | Olfr225 | 2175 | 0 | 0 | 0 | 0 | 0 | 0 |
| ENSMUSG00 | 0610009B14l | 942 | 2.53219621 | 6.13840027 | 5.91234843 | 8.04376789 | 2.87091074 | 2.80853504 |
| ENSMUSG00 | Gm9788 | 905 | 0 | 0 | 0 | 0 | 0 | 0 |
| ENSMUSG00 | Cep68 | 5251 | 20.8354756 | 13.2143357 | 12.8539734 | 18.0719438 | 17.7926669 | 14.1333638 |
| ENSMUSG00 | Gpr22 | 7753 | 3.20484655 | 0.60287436 | 1.66476695 | 2.34558886 | 1.08594763 | 0.52417403 |
| ENSMUSG00 | Znrr1 | 3283 | 1.90119112 | 3.05293328 | 0.83475914 | 1.0990571 | 1.42991976 | 7.16966606 |
| ENSMUSG00 | Fam19a2 | 4130 | 0 | 0 | 0 | 0 | 0 | 0 |
| ENSMUSG00 | Eml6 | 9712 | 0.68360433 | 0.89307656 | 0.79191935 | 0.63529659 | 1.25044043 | 0.85935334 |
| ENSMUSG00 | S100a1 | 1018 | 329.603394 | 310.087802 | 488.087345 | 296.631268 | 189.519378 | 677.177315 |
| ENSMUSG00 | 4930441O14 | 3061 | 1.41566396 | 0.99174971 | 0.88085999 | 0.45971896 | 1.35025692 | 1.05142353 |
| ENSMUSG00 | Efcab8 | 4223 | 0.01882808 | 0 | 0 | 0 | 0 | 0.01937578 |
| ENSMUSG00 | 4933402P03l | 1637 | 0 | 0 | 0 | 0 | 0 | 0 |
| ENSMUSG00 | Lmod3 | 2239 | 4.61653637 | 3.29277608 | 3.25739352 | 2.25613499 | 4.4895918 | 2.75304532 |
| ENSMUSG00 | C130050O18 | 3176 | 2.44090639 | 3.50474438 | 1.57267263 | 2.35169497 | 2.1689423 | 1.5200287 |
| ENSMUSG00 | 2410075B13l | 2299 | 10.5830161 | 15.6778855 | 11.4013453 | 12.1162846 | 12.984093 | 12.2196078 |
| ENSMUSG00 | Rsbn1 | 7416 | 3.08780431 | 2.52757902 | 3.69540759 | 4.89949013 | 3.68110991 | 3.8469893 |
| ENSMUSG00 | Il1f9 | 1643 | 0 | 0 | 0 | 0 | 0 | 0 |
| ENSMUSG00 | Olfr868 | 1130 | 0 | 0 | 0 | 0 | 0 | 0 |
| ENSMUSG00 | Olfr964-ps1 | 895 | 0 | 0 | 0 | 0 | 0 | 0 |
| ENSMUSG00 | 2900011O08l | 2625 | 0 | 0 | 0 | 0 | 0 | 0 |
| ENSMUSG00 | Olfr683 | 959 | 0 | 0 | 0 | 0 | 0 | 0 |
| ENSMUSG00 | 5430402E10l | 791 | 0 | 0 | 0 | 0 | 0 | 0 |
| ENSMUSG00 | Proca1 | 2082 | 0.13366396 | 0.41659748 | 0.10615236 | 0.20796567 | 0.98033383 | 1.24452004 |
| ENSMUSG00 | 9530080O11l | 2136 | 0.29779386 | 0.31582874 | 0.80705612 | 0 | 0 | 0.10215221 |
| ENSMUSG00 | Gm5870 | 284 | 0 | 0 | 0 | 0 | 0 | 0.86433716 |
| ENSMUSG00 | Fam109a | 2331 | 25.7191182 | 14.9045151 | 25.6753734 | 17.3831557 | 16.3520818 | 16.3460622 |
| ENSMUSG00 | Sirpb1a | 1490 | 0 | 0 | 0 | 0 | 0 | 0 |
| ENSMUSG00 | Prss53 | 2311 | 0 | 0.04170181 | 0 | 0 | 0.04415956 | 0 |
| ENSMUSG00 | E130201H02l | 1670 | 0 | 0 | 0 | 0 | 0 | 0 |
| ENSMUSG00 | 1810024B03l | 3547 | 0.12329019 | 0.91020345 | 0.99694036 | 1.75985162 | 0.60420293 | 1.13804528 |
| ENSMUSG00 | Arl6 | 3728 | 64.9225765 | 47.876229 | 80.9220973 | 54.345676 | 32.3979162 | 71.4496049 |
| ENSMUSG00 | 1810030O07l | 2692 | 38.972776 | 27.3509969 | 33.7424916 | 29.5545721 | 30.8394924 | 36.7072875 |
| ENSMUSG00 | Nkrf | 3265 | 1.0593344 | 1.07736915 | 0.92058969 | 0.87304181 | 0.54699025 | 2.03828851 |
| ENSMUSG00 | A830080D01l | 4853 | 8.02810033 | 9.6313309 | 10.4652622 | 11.7101216 | 9.54707382 | 16.7087374 |
| ENSMUSG00 | Naa38 | 3364 | 18.8495813 | 17.7619466 | 24.8733915 | 19.7786237 | 13.1358031 | 28.7421511 |
| ENSMUSG00 | Hepacam2 | 2219 | 544.823868 | 364.340751 | 844.954384 | 425.927478 | 452.798723 | 576.368807 |
| ENSMUSG00 | Tnip3 | 4090 | 0 | 0 | 0 | 0 | 0 | 0 |
| ENSMUSG00 | Rnf182 | 3690 | 0 | 0 | 0 | 0 | 0 | 0 |
| ENSMUSG00 | Bcl2l15 | 2379 | 0 | 0 | 0.07432004 | 0 | 0 | 0 |
| ENSMUSG00 | Foxo1 | 5549 | 9.80811909 | 11.7752416 | 13.1593881 | 10.8135592 | 13.0117718 | 12.23402 |
| ENSMUSG00 | Olfr1384 | 1033 | 0 | 0 | 0 | 0 | 0 | 0 |
| ENSMUSG00 | Ptx4 | 1579 | 0 | 0 | 0 | 0 | 0 | 0 |
| ENSMUSG00 | Spink10 | 1696 | 0.04688146 | 0.14205909 | 0.15637445 | 0.23402269 | 0.12034523 | 0.19298094 |
| ENSMUSG00 | Wfikkn2 | 3696 | 0 | 0 | 0 | 0 | 0 | 0 |
| ENSMUSG00 | Nkx2-6 | 1380 | 0 | 0 | 0 | 0 | 0 | 0 |
| ENSMUSG00 | Gpr146 | 6716 | 8.45899072 | 8.70312006 | 9.83286978 | 8.23611886 | 9.16286618 | 8.65429664 |
| ENSMUSG00 | S1pr4 | 1229 | 0.09704348 | 0 | 0 | 0.08807659 | 0 | 0 |
| ENSMUSG00 | Cdc25c | 1926 | 0 | 0 | 0 | 0.50582305 | 0 | 0 |
| ENSMUSG00 | Olfr983 | 1048 | 0 | 0 | 0 | 0 | 0 | 0 |
| ENSMUSG00 | Vsig4 | 1477 | 0.02691637 | 0 | 0 | 0 | 0 | 0 |
| ENSMUSG00 | Gm7887 | 493 | 1.45151856 | 1.36837767 | 1.43454257 | 0.36594366 | 0.72451244 | 2.71086678 |
| ENSMUSG00 | Olfr1094 | 1146 | 0 | 0 | 0 | 0 | 0 | 0 |
| ENSMUSG00 | Kcnj4 | 2146 | 0 | 0 | 0 | 0 | 0 | 0 |
| ENSMUSG00 | Aqp5 | 1654 | 0.14421577 | 0 | 0.05344842 | 0.02181502 | 0.2468023 | 0.11543076 |
| ENSMUSG00 | Nkx2-3 | 2109 | 0 | 0 | 0 | 0 | 0 | 0 |
| ENSMUSG00 | Grsf1 | 3875 | 67.1585485 | 62.9719079 | 58.1182942 | 55.6920538 | 52.2905131 | 65.4169025 |
| ENSMUSG00 | Defb13 | 450 | 0 | 0 | 0 | 0 | 0 | 0 |
| ENSMUSG00 | Dnajc21 | 5226 | 12.308528 | 12.1065439 | 10.6233285 | 15.1412024 | 11.8046093 | 12.7813991 |
| ENSMUSG00 | Gm9789 | 391 | 0 | 0 | 0 | 0 | 0 | 0 |
| ENSMUSG00 | Olfr1010 | 1247 | 0 | 0 | 0 | 0 | 0 | 0 |
| ENSMUSG00 | Fam55d | 4852 | 0.63090932 | 0.19862507 | 0.19131053 | 2.5358568 | 0.49427858 | 0.00562132 |
| ENSMUSG00 | Nhlrc1 | 2292 | 2.72321572 | 2.52285037 | 2.77707914 | 2.97535184 | 2.82737772 | 3.11778162 |
| ENSMUSG00 | Bhlha9 | 1206 | 0 | 0 | 0 | 0 | 0 | 0 |
| ENSMUSG00 | Il20rb | 5718 | 0.47278291 | 0.31180454 | 0.23190894 | 0 | 0.77637193 | 0.27665776 |
| ENSMUSG00 | Vmn1r45 | 3248 | 0 | 0 | 0 | 0 | 0 | 0 |
| ENSMUSG00 | Defb29 | 460 | 3.37057335 | 6.07568184 | 2.21009215 | 3.21600832 | 1.55297666 | 1.54161005 |
| ENSMUSG00 | Fam113b | 2192 | 0.29018599 | 0.54957165 | 0.72594268 | 0.27983337 | 0.60523986 | 1.31893785 |
| ENSMUSG00 | Osbpl1a | 8127 | 40.7191608 | 37.6858652 | 51.6150474 | 39.944648 | 40.6478106 | 44.4207118 |
| ENSMUSG00 | Pcsk9 | 3509 | 0.90636604 | 0.01373224 | 0.30232096 | 0 | 0.15995729 | 0 |
| ENSMUSG00 | Ctla2a | 1368 | 0.4940374 | 0.4931361 | 1.09658382 | 0.15825458 | 0.78329963 | 0.53831525 |
| ENSMUSG00 | E030002O03l | 2738 | 0 | 0 | 0 | 0 | 0 | 0 |
| ENSMUSG00 | Gm4895 | 767 | 0.82931902 | 0.81671936 | 0.57629522 | 0.14112925 | 0.19958165 | 0.14224062 |
| ENSMUSG00 | 4933427E11l | 1213 | 0.22942157 | 0 | 0.54660152 | 0.44619181 | 0.21033131 | 0.47219079 |
| ENSMUSG00 | Crb3 | 1558 | 19.7756723 | 27.124204 | 28.9383054 | 26.8646801 | 17.4563917 | 28.7801713 |
| ENSMUSG00 | Gm1821 | 1150 | 2.2470489 | 2.30456897 | 2.22931034 | 1.44328178 | 2.75098722 | 1.66019543 |
| ENSMUSG00 | Olfr221 | 1061 | 0 | 0 | 0 | 0 | 0 | 0 |
| ENSMUSG00 | Nrn1l | 709 | 0.05607261 | 0 | 0.06234393 | 0 | 0 | 0.15387667 |
| ENSMUSG00 | Cnr1 | 6966 | 0 | 0 | 0 | 0 | 0 | 0 |
| ENSMUSG00 | Olfr978 | 935 | 0 | 0 | 0 | 0 | 0 | 0 |
| ENSMUSG00 | Olfr794 | 962 | 0 | 0 | 0 | 0 | 0 | 0 |
| ENSMUSG00 | Krt84 | 2561 | 0 | 0 | 0 | 0 | 0 | 0 |
| ENSMUSG00 | Zfp879 | 4345 | 0.98816845 | 0.74303605 | 1.39370598 | 0.93838229 | 0.73985309 | 1.13618175 |
| ENSMUSG00 | Cdkn2a | 1399 | 0 | 0 | 0 | 0 | 0 | 0 |
| ENSMUSG00 | 4930500M09 | 696 | 0 | 0 | 0.12701679 | 0 | 0 | 0 |
| ENSMUSG00 | Ubr3 | 10216 | 9.69759763 | 10.6551657 | 9.89955137 | 11.1325964 | 10.9784626 | 8.34845311 |
| ENSMUSG00 | Apol7c | 2039 | 0 | 0 | 0 | 0 | 0 | 0 |
| ENSMUSG00 | Neurog3 | 1538 | 0 | 0 | 0 | 0 | 0 | 0 |
| ENSMUSG00 | Mab21l3 | 5171 | 6.37348547 | 2.85148932 | 4.32530121 | 6.48932535 | 5.84173554 | 4.81037923 |
| ENSMUSG00 | Gpr4 | 2882 | 0.13794407 | 0 | 0.18404654 | 0.28795525 | 0.61968187 | 0.21766714 |
| ENSMUSG00 | 1700001O22l | 2680 | 0.59336538 | 0.28768025 | 0.23090515 | 0.95590491 | 0.2665557 | 0.50685521 |
| ENSMUSG00 | Dsc1 | 3758 | 0 | 0 | 0 | 0 | 0 | 0 |
| ENSMUSG00 | Trp53i13 | 2055 | 6.20998017 | 9.66073682 | 7.29169089 | 8.26989926 | 8.66579327 | 6.84852256 |
| ENSMUSG00 | Gm9790 | 287 | 0 | 0 | 0 | 0.12572141 | 0.17779225 | 0 |
| ENSMUSG00 | Cxcr7 | 3426 | 22.0593019 | 47.4550659 | 13.0308994 | 38.3674513 | 32.0069123 | 9.67270481 |
| ENSMUSG00 | Aplnr | 3562 | 0.66965998 | 0 | 0.18613915 | 0.46596689 | 0 | 0 |
| ENSMUSG00 | Alkbh2 | 1430 | 1.83486833 | 1.95441514 | 2.81284456 | 1.91764712 | 2.10528404 | 3.50946407 |
| ENSMUSG00 | Phlpp1 | 6718 | 1.81083314 | 2.38852117 | 2.71738035 | 3.46963393 | 4.48892351 | 3.47124391 |
| ENSMUSG00 | Gm5601 | 2406 | 1.45406579 | 1.20165691 | 1.28600541 | 1.39469249 | 1.12402241 | 1.02024835 |
| ENSMUSG00 | Marveld1 | 3056 | 5.94510949 | 3.05895608 | 3.52920474 | 2.12525131 | 5.27628755 | 2.84705822 |
| ENSMUSG00 | Mcart6 | 9584 | 0.80473322 | 1.09606056 | 1.61421589 | 1.00144239 | 0.88380409 | 2.90276837 |
| ENSMUSG00 | Snhg11 | 6311 | 0 | 0.43521268 | 0.07003937 | 0 | 0.08085308 | 0 |
| ENSMUSG00 | 9030625A04l | 3636 | 0.29521396 | 0 | 0.29176134 | 0 | 0.57537993 | 0.80263652 |
| ENSMUSG00 | Sowaha | 4311 | 1.99192387 | 4.18040579 | 1.40469786 | 1.93341505 | 3.16029746 | 1.59434217 |
| ENSMUSG00 | P2ry4 | 4431 | 0 | 0 | 0 | 0 | 0 | 0 |
| ENSMUSG00 | BC024139 | 4862 | 0 | 0.42616557 | 0 | 0 | 0 | 0 |
| ENSMUSG00 | Ccdc89 | 2348 | 0.49101743 | 0.67723705 | 0.20707848 | 1.02959837 | 1.06486048 | 0.39494793 |
| ENSMUSG00 | 5430405G05l | 2090 | 0.15217409 | 0.36889142 | 0.08459683 | 0.29349031 | 0.12207267 | 0.40455207 |
| ENSMUSG00 | Cxxc4 | 5208 | 0 | 0 | 0 | 0 | 0.35271688 | 0 |
| ENSMUSG00 | Slc16a13 | 2005 | 12.9676231 | 3.22044052 | 8.68604795 | 8.13420655 | 9.67083432 | 8.42044972 |
| ENSMUSG00 | Olfr1391 | 935 | 0 | 0 | 0 | 0 | 0 | 0 |
| ENSMUSG00 | BC027072 | 5002 | 0 | 0 | 0 | 0 | 0 | 0 |
| ENSMUSG00 | Gm13080 | 419 | 0.56929089 | 0.11500344 | 0.63296195 | 0 | 0 | 0.13018921 |
| ENSMUSG00 | Slc15a5 | 3953 | 0.01005704 | 0.02437968 | 0.05590924 | 0.09127762 | 0 | 0.07589705 |
| ENSMUSG00 | 2410080I02R | 412 | 0.19298777 | 0.35087215 | 0 | 0 | 0.12385043 | 0 |
| ENSMUSG00 | Tigd3 | 2112 | 0 | 0 | 0.43950696 | 0 | 0.02416022 | 0 |
| ENSMUSG00 | Dsg2 | 8442 | 10.0683745 | 8.25368341 | 13.3726021 | 10.3348002 | 11.2847955 | 10.496956 |
| ENSMUSG00 | Sowahd | 1573 | 0 | 0 | 0 | 0 | 0 | 0 |
| ENSMUSG00 | Adig | 919 | 0 | 0.10486712 | 0 | 0.03926229 | 0 | 0 |
| ENSMUSG00 | Qk | 284 | 0.83990452 | 1.187694 | 0.62256117 | 1.27049453 | 1.4373627 | 2.11282417 |
| ENSMUSG00 | 1110002B05l | 1361 | 134.60195 | 147.214715 | 150.013456 | 151.963468 | 90.2053345 | 208.237161 |
| ENSMUSG00 | Gm9493 | 578 | 0 | 0 | 0 | 0 | 0 | 0 |
| ENSMUSG00 | Cryga | 728 | 0 | 0 | 0 | 0 | 0 | 0 |
| ENSMUSG00 | Klk12 | 1180 | 0 | 0 | 0 | 0 | 0 | 0 |
| ENSMUSG00 | Camsap3 | 4281 | 9.38864536 | 7.85660748 | 6.10214651 | 7.10515383 | 9.05864182 | 7.07828176 |
| ENSMUSG00 | Gm9791 | 446 | 0 | 0 | 0 | 0 | 0 | 0.1834617 |
| ENSMUSG00 | Olfr1442 | 1064 | 0 | 0 | 0 | 0 | 0 | 0 |
| ENSMUSG00 | N6amt1 | 6289 | 9.79821828 | 7.67734378 | 11.7023483 | 9.20267124 | 9.29817565 | 10.9506224 |
| ENSMUSG00 | Pfn3 | 788 | 0 | 0.30575154 | 0 | 0 | 0.58278856 | 0 |
| ENSMUSG00 | Dock5 | 10283 | 0.52192841 | 0.74976473 | 0.40406236 | 0.76494075 | 0.66493575 | 0.46151777 |
| ENSMUSG00 | Zfp507 | 7111 | 3.84081214 | 2.03967362 | 2.92151121 | 3.29817592 | 2.4684395 | 2.46626256 |
| ENSMUSG00 | Ftlar1 | 902 | 0 | 0 | 0 | 0 | 0 | 0 |
| ENSMUSG00 | Olfr867 | 995 | 0 | 0 | 0 | 0 | 0 | 0 |
| ENSMUSG00 | Rin3 | 6980 | 0.43856332 | 0.15187704 | 0.12665285 | 0.47557996 | 0.22662144 | 0.28915807 |
| ENSMUSG00 | Gm4776 | 6194 | 0 | 0 | 0 | 0 | 0 | 0 |
| ENSMUSG00 | Shisa2 | 3142 | 5.46606225 | 2.85253922 | 2.23682146 | 1.73405115 | 4.07626363 | 1.5885611 |
| ENSMUSG00 | Fam160a2 | 4674 | 21.0430164 | 17.8456849 | 24.5881027 | 18.9982695 | 21.9542237 | 18.1772574 |
| ENSMUSG00 | Fam46c | 6309 | 14.4742968 | 13.7631905 | 12.7792299 | 14.909794 | 14.7280124 | 9.7400162 |
| ENSMUSG00 | Tnfaip8l1 | 2377 | 0 | 0.24326349 | 0 | 0 | 0 | 0 |
| ENSMUSG00 | AB041803 | 2785 | 4.98192557 | 3.49503099 | 3.7297785 | 9.71688811 | 6.44929418 | 8.42233025 |
| ENSMUSG00 | Ascc1 | 1588 | 20.2532643 | 18.5099054 | 21.8782422 | 16.7686076 | 16.9016837 | 15.0285323 |
| ENSMUSG00 | Zfand3 | 2748 | 4.74519564 | 5.08517766 | 4.43948642 | 4.77942657 | 3.97366973 | 4.78397967 |
| ENSMUSG00 | 4930578I07R | 1371 | 0 | 0 | 0 | 0 | 0 | 0 |
| ENSMUSG00 | Klk1b11 | 858 | 2.03874259 | 5.39149003 | 1.39096709 | 2.94375655 | 3.15197893 | 3.75105761 |
| ENSMUSG00 | Olfr1112 | 956 | 0 | 0 | 0 | 0 | 0 | 0 |
| ENSMUSG00 | 2510039O18l | 2728 | 42.2037652 | 38.7010611 | 34.463827 | 31.1485392 | 33.3878595 | 22.8055176 |
| ENSMUSG00 | Hs3st5 | 5128 | 2.10871504 | 1.90753661 | 7.86117021 | 4.81281561 | 1.69159202 | 3.69654335 |
| ENSMUSG00 | Zfp758 | 2968 | 23.279995 | 20.7649796 | 28.2367569 | 32.8361194 | 19.2208518 | 43.3196258 |
| ENSMUSG00 | Bod1 | 2230 | 23.2828061 | 19.7067423 | 20.4755623 | 18.4617098 | 25.2157247 | 18.7742472 |
| ENSMUSG00 | Lingo4 | 3643 | 0 | 0.02645426 | 0.03640009 | 0.13866281 | 0.01400669 | 0.01497373 |
| ENSMUSG00 | 9930014A18l | 5962 | 0.4734383 | 0.71932126 | 0.35586858 | 0.26628815 | 0.90721165 | 0.2653353 |
| ENSMUSG00 | Gm8385 | 596 | 0 | 0 | 0 | 0 | 0 | 0 |
| ENSMUSG00 | Foxa3 | 976 | 0 | 0 | 0 | 0 | 0 | 0 |
| ENSMUSG00 | Zfp488 | 8540 | 0 | 0 | 0 | 0 | 0 | 0 |
| ENSMUSG00 | A730020M07 | 968 | 1.88920672 | 2.04095468 | 0.86759816 | 2.38558972 | 4.00620306 | 3.94467923 |
| ENSMUSG00 | Znrf4 | 1235 | 0 | 0 | 0 | 0 | 0 | 0 |
| ENSMUSG00 | Tram1l1 | 2223 | 0.57227862 | 0 | 0 | 0 | 0 | 0 |
| ENSMUSG00 | Rps2 | 1826 | 7.75079467 | 8.33894618 | 8.95656184 | 6.65917251 | 6.95814216 | 3.83876358 |
| ENSMUSG00 | Ccbp2 | 3040 | 0.27462668 | 0 | 0 | 0 | 0.48676477 | 0 |
| ENSMUSG00 | Olfr779 | 389 | 0 | 0 | 0 | 0 | 0 | 0 |
| ENSMUSG00 | Gm4830 | 1796 | 0 | 0 | 0 | 0 | 0 | 0 |
| ENSMUSG00 | Prop1 | 1168 | 0 | 0 | 0 | 0 | 0 | 0 |
| ENSMUSG00 | 4921513I03R | 1531 | 0 | 0 | 0 | 0 | 0 | 0 |
| ENSMUSG00 | Dact1 | 4080 | 0 | 0 | 0.03250136 | 0 | 0.02501293 | 0.24734354 |
| ENSMUSG00 | Tceal3 | 2983 | 0.22656492 | 0.14538316 | 0.04445375 | 0.22982194 | 0.11974007 | 0.09143359 |
| ENSMUSG00 | 9930012K11l | 3203 | 0.27306293 | 0.30088319 | 0.59340595 | 0.47313327 | 0.52571664 | 1.66900696 |
| ENSMUSG00 | 4930544O15l | 800 | 0 | 0.30116526 | 0 | 0.04510256 | 0 | 0 |
| ENSMUSG00 | Olfr1302 | 947 | 0 | 0 | 0 | 0 | 0 | 0 |
| ENSMUSG00 | Rasip1 | 3254 | 1.78374313 | 0.79965208 | 1.01878864 | 1.69654358 | 2.22671954 | 1.08964447 |
| ENSMUSG00 | Cage1 | 4497 | 1.14041739 | 0.5143316 | 0.91411417 | 0.95480616 | 0.48791064 | 1.50413843 |
| ENSMUSG00 | Acp1 | 3214 | 8.86891087 | 6.38687752 | 5.92750291 | 8.91385916 | 8.60494577 | 4.98988422 |
| ENSMUSG00 | 5031434C07l | 2892 | 0 | 0 | 0 | 0 | 0.19408373 | 0.33008727 |
| ENSMUSG00 | Fam69b | 4089 | 0 | 0 | 0 | 0 | 0 | 0 |
| ENSMUSG00 | 4932415D10l | 3162 | 0 | 0 | 0 | 0 | 0 | 0 |
| ENSMUSG00 | Tlr7 | 7630 | 0 | 0 | 0 | 0 | 0 | 0 |
| ENSMUSG00 | Serpinb3a | 2472 | 0 | 0 | 0 | 0 | 0 | 0 |
| ENSMUSG00 | Dnd1 | 3224 | 0.07398663 | 0.2241925 | 0.0274205 | 0.0111917 | 0.06330816 | 0.00845988 |
| ENSMUSG00 | Mycs | 2361 | 0 | 0 | 0 | 0 | 0 | 0 |
| ENSMUSG00 | 9130011J15R | 3583 | 28.704278 | 22.6340447 | 36.7752425 | 29.4255468 | 23.8968068 | 20.766178 |
| ENSMUSG00 | Gm9294 | 1579 | 0 | 0 | 0 | 0 | 0 | 0 |
| ENSMUSG00 | Zbtb39 | 5944 | 0.26084518 | 0.44587051 | 0.39412814 | 0.2185319 | 0.62666982 | 0.29825901 |
| ENSMUSG00 | Gm6922 | 1948 | 0.93878445 | 0.09894547 | 0.06807265 | 0.50011047 | 0 | 0 |
| ENSMUSG00 | Liph | 3783 | 0.48341319 | 0.67509422 | 1.56570102 | 0.55320079 | 1.564647 | 1.4347484 |
| ENSMUSG00 | Swi5 | 2434 | 35.9824665 | 34.4867634 | 35.7392059 | 27.113418 | 22.4734079 | 49.6748052 |
| ENSMUSG00 | Rnf208 | 1559 | 7.65018868 | 9.21075033 | 5.75559598 | 8.86428676 | 6.61149963 | 10.0945907 |
| ENSMUSG00 | Cnrip1 | 4021 | 0 | 0 | 0 | 0 | 0 | 0.25775586 |
| ENSMUSG00 | B530045E10l | 2525 | 0.15744745 | 0 | 0.70022722 | 0 | 0 | 0 |
| ENSMUSG00 | Csrnp2 | 4121 | 5.02611146 | 7.7991645 | 3.89353774 | 5.69117422 | 7.96165 | 1.68770517 |
| ENSMUSG00 | Gm11507 | 411 | 0 | 0 | 0 | 0 | 0 | 0 |
| ENSMUSG00 | Pard6b | 3396 | 46.9550154 | 45.1357693 | 38.0062961 | 38.6851367 | 64.5492669 | 36.2698089 |
| ENSMUSG00 | Gm7334 | 1353 | 0.94026266 | 1.21089359 | 1.92750092 | 1.17339982 | 0.71655665 | 1.33047021 |
| ENSMUSG00 | Zbtb7c | 4723 | 1.66664941 | 1.70381873 | 4.53904168 | 2.88778591 | 2.18236882 | 3.86337787 |
| ENSMUSG00 | Csrnp3 | 12369 | 0.01607061 | 0 | 0.22513672 | 0.28004497 | 0.00412534 | 0 |
| ENSMUSG00 | Krtap4-2 | 970 | 0 | 0 | 0 | 0 | 0 | 0 |
| ENSMUSG00 | Prss42 | 1561 | 0 | 0 | 0 | 0 | 0 | 0 |
| ENSMUSG00 | D3Bwg0562e | 9260 | 2.53730983 | 2.12311754 | 2.97861232 | 3.22244609 | 2.45764186 | 3.30182188 |
| ENSMUSG00 | Fzd1 | 4374 | 5.01715255 | 7.22686467 | 3.07209883 | 7.21805875 | 7.02283453 | 2.21988376 |
| ENSMUSG00 | Zfp612 | 2346 | 27.0798201 | 29.8238338 | 34.1782367 | 25.9926067 | 33.8653313 | 29.6928511 |
| ENSMUSG00 | Ly6k | 825 | 0.04818846 | 0 | 0.05357799 | 0 | 0 | 0 |
| ENSMUSG00 | Cnpy1 | 6124 | 0 | 0 | 0 | 0 | 0 | 0 |
| ENSMUSG00 | Gm13749 | 1430 | 0 | 0 | 0 | 0 | 0 | 0 |
| ENSMUSG00 | Gm4798 | 707 | 0 | 0 | 0 | 0 | 0.07217309 | 0 |
| ENSMUSG00 | 2010007H06l | 625 | 14.0575379 | 4.08621029 | 9.19398335 | 12.0081044 | 9.14392655 | 14.8810432 |
| ENSMUSG00 | Gm5479 | 859 | 0 | 0 | 0 | 0 | 0 | 0 |
| ENSMUSG00 | Tmem201 | 10624 | 4.89459417 | 4.62633387 | 4.19801013 | 4.05515448 | 6.23420894 | 3.99466667 |
| ENSMUSG00 | Il27 | 1016 | 0 | 0 | 0 | 0 | 0 | 0 |
| ENSMUSG00 | Palb2 | 5391 | 0.81856026 | 0.49160718 | 1.49225291 | 0.95710116 | 0.68148749 | 1.69486258 |
| ENSMUSG00 | Phf11 | 1346 | 0 | 0 | 0 | 0 | 0 | 0 |
| ENSMUSG00 | Olfr670 | 938 | 0 | 0 | 0 | 0 | 0 | 0 |
| ENSMUSG00 | Ccnjl | 2775 | 0.12893669 | 1.0592335 | 0 | 0 | 0 | 0.46194885 |
| ENSMUSG00 | Kcnj10 | 3503 | 22.9816866 | 35.1459776 | 18.4605471 | 24.370573 | 21.2670593 | 21.3805765 |
| ENSMUSG00 | Gemin7 | 2239 | 22.8696109 | 23.6305107 | 30.6589827 | 19.6122591 | 17.2518832 | 25.5448497 |
| ENSMUSG00 | Slc38a6 | 8004 | 0.41225698 | 0.1264262 | 0.45836494 | 0.50489618 | 0.70126204 | 1.0836251 |
| ENSMUSG00 | 4933433P14l | 2786 | 30.5229622 | 26.9989362 | 44.1225145 | 31.0440275 | 24.6890002 | 31.1612264 |
| ENSMUSG00 | Dok7 | 4344 | 0.10982177 | 0.44370573 | 0.11192916 | 0.06644944 | 0.3641385 | 0 |
| ENSMUSG00 | E230025N22l | 2275 | 0 | 0 | 0.2137232 | 0 | 0 | 0 |
| ENSMUSG00 | Gpr152 | 3970 | 0 | 0 | 0 | 0.02726603 | 0 | 0 |
| ENSMUSG00 | BC030476 | 2373 | 0.05025977 | 0.40612256 | 0.09313494 | 0.53218355 | 0.08601159 | 0.52871193 |
| ENSMUSG00 | 9930104L06l | 3925 | 7.51555835 | 3.64620976 | 2.92802017 | 5.99375619 | 5.48614792 | 4.19021846 |
| ENSMUSG00 | Serpinb1a | 3064 | 66.4061845 | 64.6365135 | 66.4614526 | 43.1948235 | 69.1454022 | 81.1295895 |
| ENSMUSG00 | Klk14 | 1266 | 0 | 0 | 0 | 0 | 0 | 0.19389554 |
| ENSMUSG00 | Defb10 | 296 | 0 | 0 | 0 | 0 | 0 | 0.1842881 |
| ENSMUSG00 | 1700001G17l | 869 | 0.2287427 | 0.05545045 | 0.30519109 | 0.16608536 | 0 | 0.53356602 |
| ENSMUSG00 | Defb1 | 870 | 917.940336 | 1030.80216 | 639.250102 | 1125.05474 | 667.09655 | 1627.16736 |
| ENSMUSG00 | Abca6 | 5359 | 0 | 0 | 0 | 0 | 0 | 0.10687953 |
| ENSMUSG00 | Gm12231 | 1248 | 0.38226424 | 0.46333117 | 0.63752658 | 0.37585463 | 0.40886519 | 0.8304778 |
| ENSMUSG00 | Gm6430 | 2000 | 0 | 0 | 0 | 0.01804102 | 0 | 0 |
| ENSMUSG00 | Rg9mtd1 | 3000 | 20.2885469 | 23.6434809 | 25.1361147 | 28.9498271 | 16.8727216 | 28.3838079 |
| ENSMUSG00 | D1Ertd622e | 4161 | 10.1944479 | 16.9306846 | 13.0980227 | 17.0394659 | 10.815973 | 10.631931 |
| ENSMUSG00 | Scml4 | 7982 | 0.16934181 | 0.31391819 | 0 | 0.09040853 | 0.02557072 | 0.56039098 |
| ENSMUSG00 | Sntn | 1236 | 0 | 0 | 0 | 0 | 0 | 0 |
| ENSMUSG00 | Hjurp | 6844 | 6.70336419 | 4.72429905 | 7.19474768 | 5.72546761 | 5.45007024 | 4.56304236 |
| ENSMUSG00 | Zfp36 | 1790 | 39.4445438 | 62.480856 | 35.9788186 | 63.7382261 | 43.415179 | 38.5044768 |
| ENSMUSG00 | 4933400C05l | 1107 | 0 | 0 | 0 | 0 | 0 | 0 |
| ENSMUSG00 | Fads6 | 6596 | 1.53091147 | 1.63641041 | 0.24794848 | 1.30739973 | 1.10624193 | 0.83527549 |
| ENSMUSG00 | Setd2 | 10505 | 6.31243613 | 6.16034191 | 7.76740621 | 8.89600146 | 8.42263072 | 7.44632702 |
| ENSMUSG00 | Isca1 | 4284 | 12.7599873 | 10.1344501 | 15.4355642 | 11.218787 | 11.5535912 | 9.1743126 |
| ENSMUSG00 | 9330133O14 | 1988 | 6.39927252 | 6.03542459 | 9.16054292 | 2.39578968 | 5.59544765 | 7.72415588 |
| ENSMUSG00 | Cyb5d1 | 1600 | 3.85131217 | 4.93911032 | 4.364932 | 4.26219151 | 4.75183125 | 2.86383712 |
| ENSMUSG00 | Olfr923 | 1067 | 0 | 0 | 0 | 0 | 0.23911141 | 0 |
| ENSMUSG00 | Olfr159 | 959 | 0 | 0 | 0 | 0 | 0 | 0 |
| ENSMUSG00 | Zfp354c | 7019 | 3.52299599 | 3.80328949 | 0.86275146 | 0.46779685 | 1.42487102 | 1.86908413 |
| ENSMUSG00 | AF251705 | 1584 | 0 | 0 | 0 | 0 | 0 | 0 |
| ENSMUSG00 | Shb | 3798 | 5.73617781 | 6.57203187 | 4.37595918 | 5.16814961 | 5.96516874 | 5.93894857 |
| ENSMUSG00 | Olfr543 | 1129 | 0 | 0 | 0 | 1.08661604 | 0 | 0 |
| ENSMUSG00 | D630023F18l | 3006 | 0 | 0 | 0.42643162 | 0 | 0 | 0 |
| ENSMUSG00 | Oxgr1 | 3728 | 248.791674 | 157.459023 | 184.14668 | 210.443105 | 268.791837 | 154.42216 |
| ENSMUSG00 | AY074887 | 688 | 0.17335239 | 0 | 0 | 0.1573345 | 0 | 0.75322405 |
| ENSMUSG00 | Olfr545 | 950 | 0 | 0 | 0 | 0 | 0 | 0 |
| ENSMUSG00 | Tlr1 | 2649 | 1.45574995 | 0.7276171 | 1.98566226 | 1.33485857 | 6.72261426 | 3.54189351 |
| ENSMUSG00 | Ankrd45 | 2343 | 0.52600081 | 1.99491459 | 0.92440901 | 1.5553933 | 1.93826182 | 0.84978603 |
| ENSMUSG00 | Lsm11 | 7875 | 1.0450012 | 1.52972832 | 0.85316573 | 1.75942922 | 2.25487985 | 0.94552083 |
| ENSMUSG00 | 1700056E22l | 542 | 0.80684555 | 0.26871462 | 1.06019181 | 0.5325763 | 0.84730144 | 1.00644425 |
| ENSMUSG00 | Lrrnd2 | 2586 | 26.5651471 | 37.5653006 | 29.1944114 | 33.4449577 | 26.0854095 | 21.3366571 |
| ENSMUSG00 | Gm1123 | 1329 | 0 | 0 | 0 | 0 | 0 | 0 |
| ENSMUSG00 | Defb36 | 518 | 0 | 0 | 0 | 0 | 0 | 0 |

| | | | | | | | | |
|---|---|---|---|---|---|---|---|---|
| ENSMUSG00 | Ankrd50 | 10920 | 2.65400598 | 2.03865717 | 2.35176472 | 3.39673459 | 3.66343211 | 2.28537499 |
| ENSMUSG00 | Gm3345 | 1720 | 0 | 0.33618448 | 0 | 0 | 0 | 0.11100144 |
| ENSMUSG00 | Zfp444 | 3413 | 5.18347168 | 5.9156517 | 6.54026684 | 3.92219119 | 4.39547451 | 4.49116533 |
| ENSMUSG00 | Chchd8 | 1422 | 5.70331788 | 7.31945957 | 8.08191222 | 11.1392529 | 6.78198686 | 7.07759626 |
| ENSMUSG00 | Gm4788 | 941 | 0 | 0 | 0 | 0 | 0 | 0 |
| ENSMUSG00 | Uqcrq | 2021 | 46.2469741 | 45.6114121 | 63.4048208 | 41.3845518 | 26.8892084 | 98.3425513 |
| ENSMUSG00 | Olfr821 | 932 | 0 | 0 | 0 | 0 | 0 | 0 |
| ENSMUSG00 | Olfr649 | 2019 | 0 | 0 | 0 | 0 | 0 | 0 |
| ENSMUSG00 | Psg22 | 1866 | 0 | 0 | 0 | 0 | 0 | 0 |
| ENSMUSG00 | 4930503L19R | 2282 | 0.6445893 | 2.59725346 | 0.52298412 | 1.28073865 | 1.5205055 | 0.57369969 |
| ENSMUSG00 | Syt16 | 3526 | 0 | 0 | 0 | 0 | 0 | 0 |
| ENSMUSG00 | 1700029I15R | 640 | 2.67107135 | 3.53869185 | 2.00289601 | 2.76253153 | 3.02969106 | 2.13083119 |
| ENSMUSG00 | Rassf9 | 2116 | 17.2474155 | 13.1624592 | 10.7371207 | 11.7317801 | 12.9977394 | 16.3183806 |
| ENSMUSG00 | Olfr1030 | 988 | 0 | 0 | 0 | 0 | 0 | 0 |
| ENSMUSG00 | H1fx | 1060 | 0.07501034 | 0.04545891 | 0 | 0.03403967 | 0 | 0 |
| ENSMUSG00 | Sstr3 | 3925 | 0 | 0 | 0 | 0 | 0 | 0 |
| ENSMUSG00 | Zfp367 | 5580 | 5.47172205 | 6.45076564 | 5.09350987 | 4.9014677 | 5.17579368 | 4.35025608 |
| ENSMUSG00 | BC030307 | 4629 | 0.38647583 | 1.04096872 | 1.07902533 | 2.03443803 | 1.84087379 | 1.0075531 |
| ENSMUSG00 | Klhl31 | 6250 | 0.48342664 | 0.67075528 | 0.40312081 | 0.76782591 | 0.22043394 | 0.20510529 |
| ENSMUSG00 | Wdr96 | 6627 | 0 | 0 | 0 | 0.01088941 | 0 | 0 |
| ENSMUSG00 | Ubtd2 | 1009 | 9.77141642 | 8.16638415 | 10.3823952 | 6.07923446 | 7.99025508 | 8.27159525 |
| ENSMUSG00 | Pwwp2a | 4345 | 11.5652307 | 12.7979642 | 11.4344929 | 10.9948509 | 13.2821245 | 14.3874507 |
| ENSMUSG00 | Mylk4 | 6649 | 0 | 0 | 0 | 0 | 0 | 0 |
| ENSMUSG00 | Kctd21 | 2964 | 7.75258696 | 4.43822493 | 4.26509012 | 4.94241231 | 3.42585992 | 2.07964523 |
| ENSMUSG00 | 4921523A10R | 2473 | 0.32151622 | 0.01948502 | 0 | 0 | 0.18570052 | 0.34189803 |
| ENSMUSG00 | Fbxo48 | 2265 | 0 | 0 | 0 | 0.30267499 | 0 | 0 |
| ENSMUSG00 | Napepld | 6390 | 9.19539909 | 3.07669302 | 11.2476051 | 10.5479279 | 8.19296739 | 9.02752144 |
| ENSMUSG00 | Wdr72 | 8548 | 47.5131012 | 23.174364 | 64.9376164 | 39.568681 | 43.8034097 | 38.4678347 |
| ENSMUSG00 | Sft2d3 | 2809 | 8.60495983 | 8.18260338 | 8.89072314 | 6.67948136 | 8.51953373 | 6.94245891 |
| ENSMUSG00 | Olfr124 | 1121 | 0 | 0 | 0 | 0 | 0 | 0 |
| ENSMUSG00 | Tst | 1079 | 1.17903186 | 2.00962919 | 0.98317352 | 0.40128316 | 0.8985182 | 0.93527493 |
| ENSMUSG00 | Ucn3 | 1476 | 0 | 0 | 0 | 0 | 0 | 0 |
| ENSMUSG00 | 1110034G24R | 2111 | 5.08478434 | 4.42831349 | 4.75311149 | 4.76877803 | 5.41445201 | 3.8502333 |
| ENSMUSG00 | Olfr1426 | 1668 | 0 | 0 | 0 | 0 | 0 | 0 |
| ENSMUSG00 | E130304I02R | 434 | 0.82442241 | 0 | 0.40739026 | 0.33255341 | 0.70543377 | 0.62844791 |
| ENSMUSG00 | Spata21 | 2511 | 0.01583253 | 0.01919014 | 0.05280985 | 0.14369592 | 0 | 0 |
| ENSMUSG00 | Fzd5 | 6910 | 0.41423945 | 1.22732472 | 1.79110218 | 0.09921257 | 0.79751789 | 0.90783893 |
| ENSMUSG00 | Tubg2 | 1743 | 0.45617304 | 0.44233108 | 0.53255233 | 0.47612566 | 1.4052014 | 1.62740246 |
| ENSMUSG00 | 9030612E09R | 1858 | 0 | 0 | 0 | 0 | 0 | 0.29359138 |
| ENSMUSG00 | Prrt3 | 4054 | 0.00980648 | 0 | 0.14174247 | 0 | 0.20138678 | 0 |
| ENSMUSG00 | Gm4779 | 1417 | 0 | 0 | 0 | 0 | 0 | 0 |
| ENSMUSG00 | Olfr711 | 1356 | 0 | 0 | 0 | 0 | 0 | 0 |
| ENSMUSG00 | Acer1 | 2423 | 0 | 0 | 0 | 0 | 0 | 0 |
| ENSMUSG00 | 1700024P04R | 475 | 0 | 0 | 0 | 0 | 0 | 0 |
| ENSMUSG00 | Prss22 | 1317 | 0 | 0.03658804 | 0.53700037 | 0 | 0.1549776 | 1.03548365 |
| ENSMUSG00 | Olfr1443 | 4332 | 0 | 0 | 0 | 0 | 0 | 0 |
| ENSMUSG00 | Cetn4 | 2672 | 1.69615448 | 1.64108018 | 0.89330072 | 1.80950373 | 1.24128534 | 0.85743626 |
| ENSMUSG00 | Ankrd34b | 4743 | 0.14249276 | 0.04063794 | 0 | 0 | 0 | 0.08050705 |
| ENSMUSG00 | Tmem232 | 3643 | 0 | 0 | 0 | 0 | 0.26612713 | 0 |
| ENSMUSG00 | Prkce | 8164 | 0.43826473 | 0.49579387 | 0.23822649 | 0.50384041 | 0.53126432 | 0.55457988 |
| ENSMUSG00 | Megf8 | 10118 | 3.80344981 | 3.35276292 | 4.01477503 | 2.32154685 | 2.73841886 | 2.55278547 |
| ENSMUSG00 | Lrfn4 | 4210 | 2.48353714 | 2.33492499 | 1.67988002 | 2.41689704 | 2.76342368 | 1.34753562 |
| ENSMUSG00 | Gm9795 | 698 | 0 | 0 | 0 | 0 | 0 | 0 |
| ENSMUSG00 | Prlhr | 1571 | 0 | 0 | 0 | 0 | 0 | 0 |
| ENSMUSG00 | Kcng3 | 3354 | 0.03555946 | 0.02873372 | 0.02635769 | 0.03227374 | 0.07606794 | 0.02439592 |
| ENSMUSG00 | Rpsa-ps2 | 509 | 0 | 0.18933769 | 0 | 0 | 0 | 0 |
| ENSMUSG00 | Pcdhb7 | 3505 | 0.54444025 | 0 | 1.09716415 | 0 | 0.23293067 | 0.38130024 |
| ENSMUSG00 | Fam164c | 4479 | 6.51496376 | 10.6076874 | 5.04289837 | 6.87161955 | 7.64427287 | 6.30866851 |
| ENSMUSG00 | 9930022D16R | 2704 | 0 | 0 | 0 | 0 | 0 | 0 |
| ENSMUSG00 | Olfr907 | 932 | 0 | 0 | 0 | 0 | 0 | 0 |
| ENSMUSG00 | E130308A19R | 11702 | 3.5875737 | 4.90017656 | 4.37787866 | 4.11635015 | 5.97822263 | 3.09758978 |
| ENSMUSG00 | Gm9796 | 2249 | 0 | 0 | 0 | 0 | 0.02268847 | 0 |
| ENSMUSG00 | Rnf216 | 4628 | 10.1708057 | 8.35037308 | 7.46885074 | 9.05949346 | 10.9704503 | 5.57515314 |
| ENSMUSG00 | Lingo2 | 11470 | 0.0346604 | 0.18484773 | 0 | 0 | 0.02224341 | 0 |
| ENSMUSG00 | S1pr5 | 2497 | 0.22289817 | 0 | 0 | 0 | 0.06130522 | 0.06553778 |
| ENSMUSG00 | Aqp12 | 1085 | 0 | 0 | 0 | 0 | 0 | 0 |
| ENSMUSG00 | S1pr1 | 5095 | 1.04558084 | 1.95772199 | 0.0260266 | 0 | 1.96293811 | 0.78156964 |
| ENSMUSG00 | Arhgef37 | 3584 | 9.58391048 | 5.36450625 | 6.63520969 | 4.59079584 | 5.85151799 | 4.26927249 |
| ENSMUSG00 | Magi1 | 7856 | 6.21938462 | 6.67347875 | 5.01321819 | 8.03303155 | 9.88571079 | 4.68348884 |
| ENSMUSG00 | Suv420h1 | 11245 | 8.62989132 | 9.19591862 | 11.0062393 | 11.8080857 | 11.4077642 | 9.45456148 |
| ENSMUSG00 | Slc25a26 | 1905 | 7.88848905 | 11.2308558 | 7.86583978 | 7.76568937 | 10.4463447 | 8.47589839 |
| ENSMUSG00 | Poln | 2872 | 0 | 0 | 0.04617184 | 0.70354962 | 0 | 0 |
| ENSMUSG00 | Dmd | 23967 | 4.95803109 | 4.52369904 | 5.54021515 | 4.7016408 | 4.45179421 | 5.97112831 |
| ENSMUSG00 | Gm5514 | 305 | 7.03867524 | 5.68758006 | 6.66650747 | 3.54905356 | 4.34978941 | 6.25975327 |
| ENSMUSG00 | Ccdc73 | 4330 | 0.84468919 | 0.32272675 | 0.33687317 | 0.21665893 | 0.76598486 | 0.83776606 |
| ENSMUSG00 | 1810063B07R | 2292 | 3.65986317 | 3.38482425 | 5.38059084 | 3.55782813 | 2.24854448 | 3.64137854 |
| ENSMUSG00 | Krtap4-7 | 977 | 0 | 0 | 0 | 0 | 0 | 0 |
| ENSMUSG00 | Rassf8 | 1130 | 0 | 0 | 0 | 0 | 0 | 0 |
| ENSMUSG00 | Taar6 | 2001 | 0 | 0 | 0 | 0 | 0 | 0 |
| ENSMUSG00 | Prrt2 | 2059 | 0 | 0.04680568 | 0.06440288 | 0 | 0.04956423 | 0 |
| ENSMUSG00 | Olfr1445 | 1000 | 0 | 0 | 0 | 0 | 0 | 0 |
| ENSMUSG00 | Rpl18a | 1286 | 3.70968714 | 3.70953171 | 3.33404259 | 2.52518197 | 2.34102346 | 4.83562811 |
| ENSMUSG00 | Olfr620 | 941 | 0 | 0 | 0 | 0 | 0 | 0 |
| ENSMUSG00 | Tubb2b | 1918 | 5.24407538 | 6.38131194 | 5.13921324 | 4.66545727 | 3.53832533 | 6.75466822 |
| ENSMUSG00 | Pigw | 4399 | 5.16938733 | 6.72572755 | 5.44610114 | 3.53520373 | 2.49390107 | 4.35253393 |
| ENSMUSG00 | Olfr1255 | 1860 | 0 | 0 | 0 | 0 | 0 | 0 |
| ENSMUSG00 | Gm8946 | 1024 | 0.03882371 | 0.04705707 | 0.25899517 | 0.03523637 | 0.04983045 | 0.21308312 |
| ENSMUSG00 | Olfr1161 | 1011 | 0 | 0 | 0 | 0 | 0 | 0 |
| ENSMUSG00 | Bola3 | 3224 | 14.2794189 | 13.0928422 | 17.2749138 | 14.8178123 | 8.89479628 | 16.33602 |
| ENSMUSG00 | AI467606 | 1916 | 3.79710487 | 4.90415252 | 3.99108499 | 2.69297096 | 3.14254298 | 2.91821558 |
| ENSMUSG00 | Fam123c | 4348 | 0 | 0 | 0 | 0 | 0 | 0 |
| ENSMUSG00 | 2310047M10 | 1801 | 1.98667032 | 2.27420743 | 2.69972389 | 2.70465076 | 1.61493805 | 4.87642079 |
| ENSMUSG00 | 4933411O13 | 2224 | 0 | 0 | 0 | 0 | 0 | 0.02452755 |
| ENSMUSG00 | Sox3 | 3187 | 0 | 0 | 0 | 0 | 0 | 0 |
| ENSMUSG00 | Shroom2 | 10971 | 12.0378914 | 4.88408747 | 12.5663612 | 8.37342862 | 7.2881534 | 9.75532626 |
| ENSMUSG00 | Cirbp | 2590 | 18.0664867 | 13.2652252 | 19.7628317 | 12.4267119 | 19.6619008 | 25.6529039 |
| ENSMUSG00 | Lrrc3b | 3073 | 0 | 0 | 0.71919693 | 0.28179924 | 0.16604743 | 0.59466347 |
| ENSMUSG00 | Olfr123 | 940 | 0 | 0 | 0 | 0 | 0 | 0 |
| ENSMUSG00 | Olfr835 | 935 | 0 | 0 | 0 | 0 | 0 | 0 |
| ENSMUSG00 | Dpy19l4 | 8944 | 5.62284021 | 4.73567561 | 8.23348283 | 5.72052092 | 4.58119184 | 8.25801913 |
| ENSMUSG00 | Vcpip1 | 9746 | 11.3522986 | 7.62894318 | 10.0458734 | 13.3724959 | 9.45029842 | 15.2464842 |
| ENSMUSG00 | Nudt18 | 3713 | 11.4566023 | 11.770833 | 10.4522538 | 9.39707435 | 7.72335664 | 9.68899789 |
| ENSMUSG00 | Asxl3 | 15518 | 0 | 0 | 0 | 0 | 0 | 0 |
| ENSMUSG00 | Hs6st1 | 3717 | 22.4500279 | 28.1184808 | 20.6679589 | 23.1713318 | 28.4578093 | 22.1014833 |
| ENSMUSG00 | Ppp1r2-ps9 | 1675 | 0 | 0 | 0 | 0 | 0 | 0 |
| ENSMUSG00 | Olfr1152 | 932 | 0 | 0 | 0 | 0 | 0 | 0 |
| ENSMUSG00 | BC106179 | 4126 | 0 | 0 | 0 | 0.00874504 | 0 | 0 |
| ENSMUSG00 | Rln3 | 473 | 0 | 0 | 0 | 0 | 0 | 0 |
| ENSMUSG00 | Krtap5-4 | 1001 | 0 | 0 | 0 | 0 | 0 | 0 |
| ENSMUSG00 | 1110012L19R | 1091 | 19.5315651 | 21.2002679 | 22.202209 | 16.4039359 | 16.182517 | 24.1246809 |
| ENSMUSG00 | A730035I17R | 682 | 0 | 0 | 0 | 0 | 0 | 0 |
| ENSMUSG00 | Kcng4 | 3990 | 0.03985512 | 0 | 0 | 0 | 0 | 0 |
| ENSMUSG00 | Med26 | 3037 | 2.69661804 | 4.34741032 | 4.75930282 | 5.09687096 | 7.0398589 | 4.25689134 |
| ENSMUSG00 | Zfp688 | 3370 | 4.24687626 | 6.63457245 | 6.55813695 | 7.36830464 | 5.26919252 | 6.23189087 |
| ENSMUSG00 | Zfp574 | 4193 | 9.86064864 | 11.9518006 | 7.77986171 | 9.25930835 | 9.34611415 | 7.33741786 |
| ENSMUSG00 | Morn2 | 639 | 14.3716995 | 18.5506491 | 15.5640292 | 12.4790796 | 7.98534833 | 15.4086773 |
| ENSMUSG00 | Klhdc9 | 1554 | 3.68390553 | 2.13955245 | 0.96709309 | 2.0896937 | 2.82385349 | 2.5273797 |
| ENSMUSG00 | Tas2r119 | 1007 | 0 | 0 | 0 | 0 | 0 | 0 |
| ENSMUSG00 | Zfp691 | 2442 | 5.404922 | 1.20367444 | 3.76493995 | 5.68861062 | 1.69252107 | 3.31718586 |
| ENSMUSG00 | Cenph | 1775 | 0 | 0 | 0.69726851 | 0.14229539 | 0 | 0.01536599 |
| ENSMUSG00 | Lca5l | 6832 | 0.12801823 | 0.12695491 | 0.1552758 | 0.09506394 | 0.43318645 | 0.45510961 |
| ENSMUSG00 | Gpr20 | 2142 | 0 | 0 | 0 | 0 | 0 | 0 |
| ENSMUSG00 | Tmem86b | 1217 | 3.20134519 | 3.16755577 | 2.36082153 | 3.37991214 | 4.57015198 | 4.61674255 |
| ENSMUSG00 | Dcaf12l1 | 3529 | 1.25045575 | 0.3003972 | 0.87677218 | 0.65436408 | 1.40253853 | 1.74668985 |
| ENSMUSG00 | Rtn4rl1 | 3523 | 7.76377252 | 15.6746133 | 4.09020744 | 7.75308195 | 11.0511282 | 5.37286393 |
| ENSMUSG00 | Ush1g | 3303 | 0 | 0 | 0 | 0 | 0 | 0 |
| ENSMUSG00 | E330021D16R | 1115 | 0 | 0 | 0 | 0 | 0 | 0 |
| ENSMUSG00 | Insig1 | 2642 | 15.6794893 | 6.78476778 | 13.1166711 | 3.87861493 | 9.00011019 | 9.62148516 |
| ENSMUSG00 | Preb | 7654 | 10.0141843 | 11.2942876 | 7.5883488 | 11.6722161 | 11.6466003 | 8.78746543 |
| ENSMUSG00 | Olfr734 | 1032 | 0 | 0 | 0 | 0 | 0 | 0 |
| ENSMUSG00 | Lhfpl2 | 8342 | 29.2185149 | 15.3651326 | 44.3925966 | 21.2504297 | 18.7296527 | 33.4312139 |
| ENSMUSG00 | Sowahb | 3899 | 3.54832193 | 2.76834138 | 5.36226513 | 2.60042434 | 3.48115311 | 1.52497342 |
| ENSMUSG00 | Fahd1 | 1406 | 65.146961 | 56.994348 | 65.7054423 | 56.7150202 | 46.054389 | 79.496068 |
| ENSMUSG00 | Adra2c | 3195 | 0.73413876 | 0.27147291 | 0.179851 | 0.83570307 | 0 | 0.37561319 |
| ENSMUSG00 | 5430407P10R | 3638 | 0.78680445 | 0.41060465 | 0.31590102 | 0.08926289 | 0.40675231 | 0.1724345 |
| ENSMUSG00 | Tlr9 | 3469 | 0 | 0 | 0 | 0 | 0 | 0 |
| ENSMUSG00 | Fndc7 | 6412 | 0 | 0 | 0 | 0 | 0 | 0 |
| ENSMUSG00 | 6330549D23 | 5288 | 0.12780695 | 0.42828343 | 0.39286812 | 0.10917411 | 0.64651422 | 0.2682075 |
| ENSMUSG00 | Cenpe | 7767 | 0 | 0.7506836 | 0.03414588 | 0.13936672 | 0.18394986 | 0 |
| ENSMUSG00 | 4933402E13R | 1966 | 0 | 0 | 0 | 0 | 0 | 0 |
| ENSMUSG00 | 2310079G19 | 878 | 0 | 0 | 0 | 0 | 0 | 0 |
| ENSMUSG00 | Zfp423 | 6289 | 0.11378576 | 0.06895818 | 0 | 0.12622118 | 0.25152133 | 0.12143264 |
| ENSMUSG00 | Hsfy2 | 1398 | 0 | 0 | 0 | 0 | 0 | 0 |
| ENSMUSG00 | Defb11 | 315 | 10.3490457 | 19.8864682 | 1.54355642 | 10.7673403 | 8.42340172 | 17.4904036 |
| ENSMUSG00 | A930104D05 | 935 | 0 | 0 | 0 | 0 | 0 | 0 |
| ENSMUSG00 | Vmn1r70 | 896 | 0 | 0 | 0 | 0 | 0 | 0 |
| ENSMUSG00 | Olfr167 | 938 | 0 | 0 | 0 | 0 | 0 | 0 |
| ENSMUSG00 | 9530056K15R | 2313 | 0 | 0.06249863 | 0.15288143 | 0.04679902 | 0.50739587 | 0.51884312 |
| ENSMUSG00 | 6430598A04R | 5273 | 0.14324941 | 0.1553517 | 0.12574012 | 0.31476845 | 0.14515373 | 0.01551753 |
| ENSMUSG00 | Sh2d5 | 3825 | 0 | 0.15117315 | 0.02311208 | 0 | 0 | 0.19252687 |
| ENSMUSG00 | 1700030F18R | 5632 | 0 | 0 | 0 | 0 | 0 | 0 |
| ENSMUSG00 | Tnfrsf26 | 3099 | 0 | 0.01554903 | 0.02852652 | 0.06985875 | 0 | 0.33444217 |
| ENSMUSG00 | Ifne | 663 | 0 | 0 | 0 | 0 | 0 | 0 |
| ENSMUSG00 | Rbm15b | 3015 | 3.88983972 | 3.97957681 | 3.12271727 | 4.20059623 | 4.62029871 | 3.19334914 |
| ENSMUSG00 | Wdr81 | 8168 | 2.09777327 | 1.8524171 | 2.02934514 | 3.49865074 | 2.49259598 | 1.62285439 |
| ENSMUSG00 | Tmem88 | 1580 | 1.15743804 | 0.94543019 | 0.7273721 | 0.41106127 | 1.00115041 | 1.74350542 |
| ENSMUSG00 | Pxt1 | 707 | 0 | 0 | 0 | 0 | 0 | 0 |
| ENSMUSG00 | Olfr433 | 1003 | 0 | 0 | 0 | 0 | 0 | 0 |
| ENSMUSG00 | Cxcr4 | 2608 | 2.24081888 | 2.29107317 | 2.32195264 | 2.32430348 | 1.81957552 | 6.00293057 |
| ENSMUSG00 | 1700120B22R | 451 | 0.17629925 | 0.10684355 | 0.09800852 | 0.2400136 | 0.11314052 | 0 |
| ENSMUSG00 | Olfr1033 | 1103 | 0 | 0 | 0.92170661 | 0 | 0 | 0 |
| ENSMUSG00 | Epcam | 2030 | 438.211149 | 428.360855 | 536.039887 | 378.701498 | 391.872512 | 445.060308 |
| ENSMUSG00 | Olfr1499 | 1051 | 0 | 0 | 0 | 0 | 0 | 0 |
| ENSMUSG00 | Kcnk13 | 3405 | 0 | 0 | 0 | 0 | 0 | 0 |
| ENSMUSG00 | Trim39 | 8694 | 6.79053239 | 5.02704199 | 4.32155125 | 6.63205741 | 7.54185564 | 4.71518091 |
| ENSMUSG00 | Akr1e1 | 4350 | 5.90391734 | 9.17204002 | 10.6795717 | 8.26154399 | 10.9325477 | 20.3587939 |
| ENSMUSG00 | 2410002F23R | 6409 | 21.8968398 | 24.495464 | 21.1801935 | 22.0523277 | 23.869102 | 24.5212157 |
| ENSMUSG00 | 1190002N15R | 4352 | 19.7863904 | 13.4306421 | 18.353109 | 28.2388428 | 12.2407023 | 19.8229972 |
| ENSMUSG00 | Vmn1r230 | 950 | 0 | 0 | 0 | 0 | 0 | 0 |
| ENSMUSG00 | Olfr1390 | 935 | 0 | 0 | 0 | 0 | 0 | 0 |
| ENSMUSG00 | Hnrnph2 | 4474 | 28.8969206 | 28.9937198 | 30.6962732 | 29.081773 | 25.7184795 | 26.1956023 |
| ENSMUSG00 | Tmem60 | 1145 | 51.9425317 | 44.6092827 | 47.058556 | 33.3404394 | 32.3984063 | 69.1513343 |
| ENSMUSG00 | Cox19 | 3767 | 7.52472992 | 7.24011846 | 7.79135221 | 7.7872849 | 4.971245 | 9.02879616 |
| ENSMUSG00 | Insm2 | 3101 | 0.05128085 | 0 | 0 | 0 | 0 | 0 |
| ENSMUSG00 | Gprin3 | 3098 | 4.76090486 | 3.42189066 | 5.87836001 | 4.00652786 | 8.23537376 | 6.56774722 |
| ENSMUSG00 | Gm9797 | 612 | 0.19487981 | 0 | 0 | 0 | 0.33350572 | 0.17826562 |
| ENSMUSG00 | Slco6b1 | 2266 | 0 | 0 | 0 | 0 | 0 | 0 |
| ENSMUSG00 | 2810002D19R | 2038 | 11.7042632 | 11.372757 | 15.8328486 | 13.2961803 | 8.31244199 | 6.82535133 |
| ENSMUSG00 | Zfp956 | 3100 | 5.51446988 | 5.16061251 | 4.54851223 | 5.8196846 | 9.95837335 | 5.3493486 |
| ENSMUSG00 | Ttll13 | 4017 | 0.3463883 | 0.56379457 | 0.50616997 | 0.70062222 | 1.23215296 | 0.57713327 |
| ENSMUSG00 | Hcrt | 746 | 0 | 0.06459309 | 0 | 0 | 0 | 0 |
| ENSMUSG00 | Olfr1368 | 1055 | 0 | 0 | 0 | 0 | 0 | 0 |
| ENSMUSG00 | Lce3c | 562 | 0 | 0 | 0 | 0 | 0 | 0 |
| ENSMUSG00 | Olfr459 | 1108 | 0 | 0 | 0 | 0 | 0 | 0 |
| ENSMUSG00 | Trrap | 15110 | 3.46512031 | 3.68334485 | 3.02772386 | 4.78546772 | 4.01862258 | 3.13902036 |
| ENSMUSG00 | Bhlhe23 | 3242 | 0 | 0 | 0 | 0 | 0 | 0 |
| ENSMUSG00 | Pcdhb3 | 3786 | 0 | 0 | 0.05837539 | 0 | 0 | 0 |
| ENSMUSG00 | Niacr1 | 1937 | 0.53363061 | 0 | 0.66177256 | 1.60199062 | 1.79132347 | 1.09830762 |
| ENSMUSG00 | Sys1 | 5001 | 5.7713415 | 5.48252061 | 4.9938095 | 5.24528022 | 3.66296119 | 5.07752232 |
| ENSMUSG00 | Olfr209 | 917 | 0 | 0 | 0 | 0 | 0 | 0 |
| ENSMUSG00 | A930002H24 | 491 | 1.70033623 | 2.15906665 | 0.9002412 | 2.42506613 | 2.28631419 | 1.38872909 |
| ENSMUSG00 | Olfr1367 | 1054 | 0 | 0 | 0 | 0 | 0 | 0 |
| ENSMUSG00 | Gpr150 | 2129 | 0 | 0 | 0 | 0 | 0 | 0 |
| ENSMUSG00 | Olfr460 | 1060 | 0 | 0 | 0 | 0 | 0 | 0 |
| ENSMUSG00 | Pou3f3 | 3061 | 22.2350156 | 25.1872942 | 15.1767844 | 20.8877435 | 21.0540061 | 17.5623371 |
| ENSMUSG00 | Onecut3 | 4774 | 0 | 0 | 0 | 0 | 0 | 0 |
| ENSMUSG00 | Zfp560 | 4974 | 5.96252281 | 6.77167733 | 4.46994914 | 3.91723041 | 3.63155148 | 8.93252661 |
| ENSMUSG00 | Gm9799 | 1400 | 0 | 0 | 0 | 0 | 0 | 0 |
| ENSMUSG00 | Tssk2 | 1386 | 0 | 0 | 0 | 0 | 0 | 0.01967867 |
| ENSMUSG00 | Olfr891 | 950 | 0 | 0 | 0 | 0 | 0 | 0 |
| ENSMUSG00 | C1ql1 | 1471 | 0.10810464 | 0.32757609 | 0 | 0 | 0.31219401 | 0 |
| ENSMUSG00 | Kcna5 | 3002 | 0 | 0.12841157 | 0 | 0 | 0 | 0 |
| ENSMUSG00 | Ddx28 | 1870 | 7.48338457 | 7.06047334 | 10.4004337 | 7.7373796 | 6.87628166 | 4.50687889 |
| ENSMUSG00 | Sprr3 | 842 | 0 | 0 | 0 | 0 | 0 | 0 |
| ENSMUSG00 | Olfr600 | 947 | 0 | 0 | 0 | 0 | 0 | 0 |
| ENSMUSG00 | Krt14 | 1749 | 0 | 0 | 0 | 0 | 0 | 0 |
| ENSMUSG00 | Rnf113a2 | 1381 | 22.3102805 | 17.8649228 | 28.5503577 | 12.4105512 | 21.7259298 | 20.1251683 |
| ENSMUSG00 | Fpr1 | 1322 | 0 | 0 | 0 | 0 | 0 | 0 |
| ENSMUSG00 | 9030224M15 | 1524 | 0.83476075 | 1.42282802 | 0.66708818 | 1.42055293 | 1.84150306 | 0.62638607 |
| ENSMUSG00 | Olfr827 | 1057 | 0 | 0 | 0 | 0 | 0 | 0 |
| ENSMUSG00 | Sprr4 | 230 | 0 | 0 | 0 | 0 | 0 | 0 |
| ENSMUSG00 | Mc2r | 1559 | 0 | 0 | 0 | 0 | 0 | 0 |
| ENSMUSG00 | Penk | 2026 | 0 | 0 | 0 | 0 | 0 | 0 |
| ENSMUSG00 | Vmn1r233 | 959 | 0 | 0 | 0 | 0 | 0 | 0 |
| ENSMUSG00 | St7l | 4861 | 15.2282873 | 15.2658138 | 12.8122602 | 15.3428484 | 14.3075397 | 14.4200417 |
| ENSMUSG00 | Olfr710 | 1154 | 0 | 0 | 0 | 0 | 0 | 0 |
| ENSMUSG00 | Olfr610 | 947 | 0 | 0 | 0 | 0 | 0 | 0 |
| ENSMUSG00 | BC049730 | 1211 | 0 | 0 | 0 | 0 | 0 | 0 |
| ENSMUSG00 | 6430704M03 | 9836 | 0 | 0 | 0.00898777 | 0 | 0 | 0 |
| ENSMUSG00 | Olig3 | 2046 | 0 | 0 | 0 | 0 | 0 | 0 |
| ENSMUSG00 | Glb1 | 3068 | 22.2361162 | 25.7109537 | 24.1899917 | 23.2510437 | 28.3572265 | 16.233211 |
| ENSMUSG00 | Zfp553 | 3602 | 11.3902432 | 10.3008219 | 8.72501676 | 9.95712167 | 8.1171886 | 5.89865185 |
| ENSMUSG00 | Dbx2 | 3456 | 0 | 0 | 0 | 0.01044041 | 0 | 0 |
| ENSMUSG00 | Gm5455 | 1131 | 0 | 0 | 0 | 0.03190278 | 0.04511616 | 0 |
| ENSMUSG00 | Chrm2 | 6220 | 0 | 0 | 0 | 0 | 0 | 0 |
| ENSMUSG00 | Odf3l1 | 1075 | 0 | 0 | 0 | 0 | 0 | 0 |
| ENSMUSG00 | Esf1 | 4454 | 10.5235096 | 11.0242444 | 11.1844358 | 8.06053756 | 9.54310082 | 19.3506646 |
| ENSMUSG00 | Pigz | 2637 | 0.54273694 | 0.51164975 | 0.23467038 | 1.14937116 | 0.71595598 | 0 |
| ENSMUSG00 | Sh3tc2 | 5736 | 0.99111466 | 0.00840071 | 0.05394228 | 0.21387544 | 1.12087227 | 0 |
| ENSMUSG00 | Tmprss12 | 1824 | 0 | 0 | 0 | 0 | 0 | 0 |
| ENSMUSG00 | Mtus1 | 12746 | 11.0508134 | 5.63674763 | 11.4024525 | 7.91223242 | 12.3822831 | 9.10081914 |
| ENSMUSG00 | Zfp629 | 7665 | 10.3369436 | 7.85819344 | 7.20262256 | 9.67836706 | 9.37979955 | 6.70034517 |
| ENSMUSG00 | Vwc2l | 4991 | 0 | 0 | 0 | 0 | 0 | 0 |
| ENSMUSG00 | AU021034 | 3562 | 0 | 0 | 0 | 0 | 0 | 0 |
| ENSMUSG00 | Pcdhb10 | 2840 | 0 | 0 | 0.35797267 | 0.40655825 | 0 | 0.22088616 |
| ENSMUSG00 | Pid1 | 9251 | 0 | 0.03125269 | 0 | 0 | 0 | 0 |
| ENSMUSG00 | Plekha7 | 6212 | 7.1677621 | 7.84232019 | 5.18012584 | 8.58487795 | 9.68449728 | 6.07225146 |
| ENSMUSG00 | 4921515J06R | 2758 | 0 | 0.22712971 | 0 | 0 | 0 | 0 |
| ENSMUSG00 | Cdc42ep2 | 1434 | 14.0835315 | 14.3820064 | 20.9912518 | 11.9770245 | 11.5645552 | 11.0506035 |
| ENSMUSG00 | Mfsd5 | 1740 | 33.7464625 | 31.0165605 | 40.6453729 | 40.4782477 | 24.6040973 | 27.3373396 |
| ENSMUSG00 | Smtnl2 | 4974 | 34.0966787 | 33.4321294 | 24.5980502 | 29.0020133 | 35.3512045 | 25.2293155 |
| ENSMUSG00 | Spred2 | 5125 | 0.85328836 | 1.06245229 | 0.87972449 | 1.06310024 | 1.20472029 | 1.0537324 |
| ENSMUSG00 | Col27a1 | 12959 | 5.42385134 | 5.67052429 | 2.07382673 | 7.47869215 | 7.3119824 | 7.04859687 |
| ENSMUSG00 | Olfr1489 | 941 | 0 | 0 | 0 | 0 | 0 | 0 |
| ENSMUSG00 | Pqlc3 | 5286 | 0.32339873 | 0.1185062 | 1.25430883 | 0.01365193 | 0.27994039 | 1.82140516 |
| ENSMUSG00 | Tcf21 | 1221 | 0 | 0 | 0 | 0 | 0 | 0 |
| ENSMUSG00 | Lcn6 | 1614 | 0 | 0 | 0 | 0 | 0 | 0 |
| ENSMUSG00 | Pcdhb4 | 3718 | 0.28870306 | 0 | 1.01053165 | 1.75654924 | 0 | 0.05135085 |
| ENSMUSG00 | Wdr89 | 2015 | 1.83486833 | 0.83698535 | 3.29045978 | 1.52207136 | 2.15247739 | 1.13700729 |
| ENSMUSG00 | Thtpa | 2777 | 12.6696436 | 5.55263287 | 9.00909008 | 6.66549832 | 11.1901559 | 7.08138815 |
| ENSMUSG00 | Nlrp4e | 3300 | 0 | 0 | 0 | 0 | 0 | 0 |
| ENSMUSG00 | Gm4997 | 661 | 0 | 0.14579862 | 0 | 0 | 0 | 0 |
| ENSMUSG00 | Gm8935 | 407 | 0 | 0 | 0 | 0 | 0 | 0 |
| ENSMUSG00 | Olfr447 | 932 | 0 | 0 | 0 | 0 | 0 | 0 |
| ENSMUSG00 | 1700023L04R | 1040 | 0.53516993 | 0.83399611 | 0.34001418 | 0.38163701 | 0.49063823 | 0.05245123 |
| ENSMUSG00 | Vmn1r76 | 974 | 0 | 0 | 0 | 0 | 0 | 0 |
| ENSMUSG00 | Prr15 | 1379 | 1.35497287 | 2.30624016 | 1.53857032 | 1.09894552 | 1.70211261 | 0.73180685 |
| ENSMUSG00 | Adrb2 | 2142 | 44.0799562 | 24.9930613 | 65.5598764 | 47.0986912 | 29.5390784 | 68.6959285 |
| ENSMUSG00 | Pnoc | 1890 | 0 | 0 | 0 | 0 | 0 | 0.11544821 |
| ENSMUSG00 | Sprn | 3902 | 0 | 0 | 0 | 0 | 0 | 0 |
| ENSMUSG00 | 9930021D14 | 1131 | 0 | 0 | 0 | 0 | 0 | 0 |
| ENSMUSG00 | B230317F23R | 2471 | 3.41083038 | 3.41263755 | 2.1465889 | 1.81067322 | 2.49866106 | 2.59390539 |
| ENSMUSG00 | D830014E11R | 2972 | 0 | 0 | 0 | 0 | 0 | 0 |
| ENSMUSG00 | Mms22l | 6900 | 0.99676785 | 1.86460581 | 1.76166766 | 1.17135913 | 0.99094701 | 0.92101318 |
| ENSMUSG00 | Tssc4 | 1893 | 19.216199 | 22.2986388 | 18.8435749 | 16.6400554 | 16.4427307 | 15.4023187 |
| ENSMUSG00 | Zfp764 | 4430 | 3.11402974 | 3.26319021 | 2.37472656 | 2.00365304 | 2.48796776 | 3.68176846 |
| ENSMUSG00 | Fam179a | 4340 | 0 | 0 | 0 | 0.12470753 | 0 | 0 |
| ENSMUSG00 | Basp1 | 1845 | 0.17238149 | 0.15670388 | 0.04791528 | 0.01955666 | 0 | 0 |
| ENSMUSG00 | B230219D22 | 4649 | 59.4835712 | 54.5193989 | 62.6660244 | 67.6392811 | 54.2642293 | 59.0373618 |
| ENSMUSG00 | Slc16a5 | 3962 | 43.1069016 | 45.6688769 | 42.0263359 | 44.4331891 | 39.4482052 | 30.0282626 |
| ENSMUSG00 | Lrtm1 | 7315 | 0.00543479 | 0 | 0 | 0.00493261 | 0 | 0 |
| ENSMUSG00 | 6330512M04 | 7362 | 0.19980342 | 0.41889871 | 0.0780527 | 0.20094592 | 0.11089677 | 0.14819147 |
| ENSMUSG00 | Olfr624 | 926 | 0 | 0 | 0 | 0 | 0 | 0 |
| ENSMUSG00 | Ccdc149 | 893 | 6.41073818 | 6.96086342 | 2.6728998 | 7.2729765 | 5.71403985 | 3.45132613 |
| ENSMUSG00 | Olfr578 | 941 | 0 | 0 | 0 | 0 | 0 | 0 |
| ENSMUSG00 | 4930526D03 | 862 | 0 | 0 | 0 | 0 | 0 | 0 |
| ENSMUSG00 | Whamm | 2895 | 5.91869157 | 7.75643594 | 9.09993038 | 5.69585297 | 6.39812588 | 8.658167 |
| ENSMUSG00 | 4930402K13R | 1780 | 0 | 0 | 0 | 0 | 0 | 0 |
| ENSMUSG00 | Gm9800 | 507 | 0.15682635 | 0.19008458 | 0.26154937 | 0.3558387 | 0.20128748 | 0.1613884 |
| ENSMUSG00 | Hsf3 | 4092 | 0.64121743 | 0.3297215 | 0.33486245 | 0.39679668 | 0.66089881 | 1.44638238 |
| ENSMUSG00 | Olfr984 | 944 | 0 | 0 | 0 | 0 | 0 | 0 |
| ENSMUSG00 | Gm9801 | 403 | 0 | 0 | 0 | 0 | 0.7596979 | 0 |
| ENSMUSG00 | 1700101I19R | 389 | 0 | 0 | 0 | 0 | 0 | 0 |
| ENSMUSG00 | Zfp36l2 | 4667 | 2.87070858 | 3.53112561 | 3.87369912 | 4.73929576 | 4.51551172 | 4.9558376 |
| ENSMUSG00 | Zswim3 | 2706 | 6.43492257 | 7.1229035 | 8.26538528 | 3.93355622 | 2.86622658 | 1.25991497 |
| ENSMUSG00 | Olfr574 | 1040 | 0 | 0 | 0 | 0 | 0 | 0 |
| ENSMUSG00 | Ptprcap | 925 | 0.34383118 | 0 | 0.04778578 | 0.23404569 | 0 | 0.05897219 |
| ENSMUSG00 | Serpinb9 | 3486 | 376.388376 | 174.071103 | 487.589693 | 254.530567 | 294.375056 | 576.006581 |
| ENSMUSG00 | Hdgfl1 | 1993 | 0 | 0 | 0 | 0 | 0 | 0 |
| ENSMUSG00 | A430105I19R | 5027 | 5.63077424 | 4.31348696 | 4.00076359 | 3.53858125 | 6.57749981 | 3.90102758 |
| ENSMUSG00 | Gm7148 | 595 | 0 | 0 | 0 | 0 | 0 | 0 |
| ENSMUSG00 | Lyrm2 | 919 | 41.702158 | 37.9618978 | 39.4401646 | 34.4722906 | 33.2587586 | 58.5558904 |
| ENSMUSG00 | Cradd | 2422 | 1.88764668 | 2.28796071 | 4.34353371 | 1.54935286 | 1.39047928 | 3.6035857 |
| ENSMUSG00 | Gvin1 | 8844 | 0 | 0 | 0 | 0.0122395 | 0 | 0.00308397 |
| ENSMUSG00 | Slitrk6 | 4240 | 0.15939697 | 0 | 0.55252304 | 0 | 0 | 0 |
| ENSMUSG00 | Adra1a | 7195 | 0.45861082 | 0.14733867 | 0.49147289 | 0.73217213 | 0.31913647 | 0.20649273 |
| ENSMUSG00 | Pcdhb8 | 2348 | 0 | 0 | 0.43298228 | 0.75298986 | 0 | 0.24393843 |
| ENSMUSG00 | 4933415A04R | 2008 | 0 | 0 | 0 | 0 | 0 | 0.12224689 |
| ENSMUSG00 | Olfr1461 | 938 | 0 | 0 | 0 | 0 | 0 | 0 |
| ENSMUSG00 | Gm9803 | 377 | 0 | 0 | 0.11724627 | 0.09570834 | 0 | 0.72346523 |
| ENSMUSG00 | Paip2b | 6388 | 3.67184307 | 3.37184403 | 3.07918287 | 3.07273516 | 2.82769835 | 2.91191358 |
| ENSMUSG00 | Npas4 | 3269 | 0 | 0 | 0 | 0 | 0 | 0 |
| ENSMUSG00 | C2cd4c | 6742 | 0.00589669 | 0 | 0 | 0.00535183 | 0 | 0 |
| ENSMUSG00 | Ccdc42 | 1640 | 0 | 0 | 0 | 0 | 0 | 0 |
| ENSMUSG00 | 6330416G13 | 4197 | 1.62924533 | 0.84960608 | 3.64399278 | 1.85697441 | 2.10330308 | 0.61736734 |
| ENSMUSG00 | 4933440M02 | 3798 | 0 | 0 | 0 | 0 | 0 | 0 |
| ENSMUSG00 | Gm16532 | 4455 | 0 | 0 | 0 | 0 | 0 | 0 |
| ENSMUSG00 | Clec14a | 4538 | 0.95490246 | 0.99813256 | 0.91559569 | 1.70948426 | 1.49548435 | 0.57097636 |
| ENSMUSG00 | Ifit2 | 4235 | 4.32757415 | 4.66503926 | 2.4527587 | 4.39630106 | 5.94002439 | 1.41686438 |
| ENSMUSG00 | Mtmr11 | 3754 | 2.20275438 | 2.87526986 | 1.40117723 | 3.59474818 | 3.26220836 | 1.55481428 |
| ENSMUSG00 | BC049762 | 931 | 0.42701913 | 0 | 0 | 0 | 0.82212206 | 0 |
| ENSMUSG00 | Gm9804 | 560 | 0 | 0 | 0 | 0 | 0 | 0 |
| ENSMUSG00 | Mrps12 | 1203 | 17.2174608 | 20.6284436 | 21.5314048 | 15.3866075 | 9.58600244 | 23.6924339 |
| ENSMUSG00 | Gm5548 | 395 | 0 | 0 | 0 | 0 | 0 | 0.06904972 |
| ENSMUSG00 | Sdpr | 3052 | 39.9638972 | 63.2486524 | 30.6168729 | 69.9296505 | 65.7056543 | 47.4178361 |
| ENSMUSG00 | Wnk1 | 15450 | 16.8413994 | 20.8433652 | 13.0517028 | 29.5662578 | 18.6204988 | 23.032506 |
| ENSMUSG00 | Gpr158 | 9749 | 0 | 0 | 0 | 0 | 0 | 0 |
| ENSMUSG00 | 5830403L16R | 3500 | 0.05679354 | 0 | 0 | 0.03092747 | 0 | 0 |
| ENSMUSG00 | Ing1 | 2941 | 12.0848009 | 10.7809177 | 11.918416 | 11.3852898 | 11.2948557 | 13.2246296 |
| ENSMUSG00 | Mcart1 | 9613 | 5.36800309 | 6.69186521 | 7.54553463 | 8.22759197 | 6.36436332 | 6.75553064 |
| ENSMUSG00 | C2cd2 | 6541 | 3.324606 | 5.54722381 | 3.83834992 | 3.61316911 | 5.82735098 | 4.03636306 |
| ENSMUSG00 | Tmem104 | 4687 | 6.88744403 | 8.17329241 | 5.46039409 | 3.84146367 | 5.60669587 | 4.14327782 |
| ENSMUSG00 | Eif4g1 | 6905 | 19.6618343 | 18.6185992 | 17.571913 | 22.5009824 | 21.8072172 | 16.2028342 |
| ENSMUSG00 | 4930451I11R | 471 | 0 | 0.30692001 | 0 | 0 | 0 | 0.86861908 |
| ENSMUSG00 | Onecut2 | 14465 | 0.00274839 | 0 | 0 | 0 | 0 | 0 |
| ENSMUSG00 | B3gat1 | 5327 | 0 | 0 | 0 | 0.02709371 | 0 | 0 |
| ENSMUSG00 | Polr2k | 615 | 2.26250702 | 2.35055815 | 2.44367913 | 2.75748958 | 1.57642462 | 2.97138346 |
| ENSMUSG00 | AY036118 | 710 | 109.69153 | 98.8840087 | 116.543451 | 75.2132759 | 80.420443 | 128.882274 |
| ENSMUSG00 | Naa11 | 3347 | 0 | 0 | 0 | 0 | 0 | 0 |
| ENSMUSG00 | D830044D21 | 2058 | 0.11590519 | 0.07024263 | 0.04295612 | 0.08766289 | 0 | 0.06626492 |
| ENSMUSG00 | Gapt | 2052 | 0 | 0 | 0 | 0 | 0 | 0 |
| ENSMUSG00 | Pnlip | 1472 | 0 | 0 | 0 | 0 | 0 | 0 |
| ENSMUSG00 | Zfp830 | 3277 | 7.19407993 | 15.2632063 | 15.0396872 | 17.3088111 | 16.4119012 | 15.8054745 |
| ENSMUSG00 | Olfr1364 | 944 | 0 | 0 | 0 | 0 | 0 | 0 |
| ENSMUSG00 | Pofut1 | 5556 | 4.68679621 | 3.14825027 | 4.08127368 | 6.1760303 | 2.11232297 | 4.09414131 |
| ENSMUSG00 | Stard5 | 5580 | 16.3154212 | 21.899788 | 11.85053 | 18.8040476 | 18.9291394 | 10.484606 |
| ENSMUSG00 | Fam26f | 1078 | 0 | 0 | 0 | 0 | 0 | 0 |
| ENSMUSG00 | Snx12 | 2947 | 19.9654263 | 19.16339 | 18.6586674 | 20.9488898 | 16.2065449 | 20.1112287 |
| ENSMUSG00 | Fam105b | 1484 | 34.0761262 | 26.6584023 | 30.1430358 | 34.2584911 | 28.4014733 | 24.7566975 |
| ENSMUSG00 | Olfr803 | 926 | 0 | 0 | 0 | 0 | 0 | 0 |
| ENSMUSG00 | Rp1l1 | 4847 | 0 | 0 | 0 | 0 | 0 | 0 |
| ENSMUSG00 | Sbsn | 2281 | 1.34203069 | 1.94351279 | 1.20145299 | 0.96492973 | 1.87909495 | 2.55886576 |
| ENSMUSG00 | Gm15428 | 332 | 0 | 0 | 0 | 0 | 0 | 0 |
| ENSMUSG00 | Eid2 | 1284 | 3.68450326 | 2.73957187 | 3.68348692 | 3.70936906 | 3.13947328 | 6.0539503 |
| ENSMUSG00 | Ppp1r15b | 7825 | 13.824238 | 13.2951474 | 15.5172476 | 14.4466512 | 17.1305162 | 10.2441105 |
| ENSMUSG00 | Igfals | 2890 | 1.89835859 | 0.66694038 | 0.5812007 | 0 | 0.98874638 | 0 |
| ENSMUSG00 | Olfr998 | 1011 | 0 | 0 | 0 | 0 | 0 | 0 |
| ENSMUSG00 | Lrrc8d | 10646 | 7.47235736 | 6.21906552 | 6.80092231 | 7.18861699 | 7.47709433 | 4.33996232 |
| ENSMUSG00 | Clec9a | 1432 | 1.08272608 | 0 | 0.64821139 | 0 | 0.39196238 | 0.38093072 |
| ENSMUSG00 | Tmem174 | 2250 | 0 | 1.02797743 | 0 | 0 | 0.09071356 | 1.60011217 |
| ENSMUSG00 | 4931422A03R | 1595 | 0.34895093 | 0.54379684 | 0.60968059 | 0.27146366 | 0.67182062 | 0.59850306 |
| ENSMUSG00 | Gm9805 | 170 | 0 | 0 | 0 | 2.54696785 | 0 | 0 |
| ENSMUSG00 | Hpcal4 | 5718 | 0.67441092 | 2.02251594 | 1.10543259 | 2.40420758 | 2.63252551 | 1.84597506 |
| ENSMUSG00 | Krt32 | 2144 | 0 | 0 | 0 | 0 | 0 | 0 |
| ENSMUSG00 | BC030336 | 4839 | 4.56789567 | 6.95063766 | 8.21191504 | 11.9304136 | 6.63269072 | 10.8219275 |
| ENSMUSG00 | 6030422M02 | 6157 | 0.19370869 | 0.10174172 | 0.24409009 | 0.05274296 | 0.27348878 | 0.1816242 |
| ENSMUSG00 | Il17c | 717 | 0 | 0 | 0 | 0 | 0 | 0 |
| ENSMUSG00 | AB041806 | 2103 | 0 | 0 | 0 | 0 | 0 | 0 |
| ENSMUSG00 | Serinc4 | 4486 | 0 | 0 | 0 | 0 | 0 | 0 |
| ENSMUSG00 | 5830418K08R | 9901 | 1.73460939 | 2.97363056 | 2.5089825 | 4.12897247 | 3.8910124 | 3.51779763 |
| ENSMUSG00 | 1110008E08R | 720 | 0 | 0 | 0 | 0 | 0 | 0 |
| ENSMUSG00 | Vmn1r184 | 944 | 0 | 0 | 0 | 0 | 0 | 0 |
| ENSMUSG00 | C130073F10R | 910 | 0 | 0 | 0 | 0 | 0 | 0 |
| ENSMUSG00 | 9930021J03R | 12418 | 4.33154817 | 4.26840766 | 5.20042621 | 5.57879896 | 6.02389008 | 4.43888275 |
| ENSMUSG00 | Patl1 | 4202 | 4.43724901 | 6.50207346 | 5.61727372 | 9.32534516 | 6.76384848 | 10.2490851 |
| ENSMUSG00 | 1700018M17 | 955 | 1.24886326 | 0.75685511 | 0.41656187 | 0.75564491 | 0.48087684 | 0.85679495 |
| ENSMUSG00 | Olfr918 | 977 | 0 | 0 | 0 | 0 | 0 | 0 |
| ENSMUSG00 | Fut10 | 5130 | 0 | 0 | 0.43943353 | 0 | 0.78578629 | 0 |
| ENSMUSG00 | Tmem229b | 8103 | 0.03434387 | 0.33896424 | 0.07091497 | 0.18702282 | 0.39042766 | 0 |
| ENSMUSG00 | Chrm3 | 1948 | 0 | 0 | 0 | 0 | 0 | 0.07000677 |
| ENSMUSG00 | Olig1 | 2170 | 0 | 0 | 0 | 0 | 0 | 0 |
| ENSMUSG00 | Gldn | 4648 | 0 | 0 | 0 | 0 | 0 | 0 |
| ENSMUSG00 | Kcnrg | 854 | 0.88448961 | 0.22569762 | 0 | 0.76051148 | 0.47799884 | 0.09581255 |
| ENSMUSG00 | Adamts6 | 4996 | 0.24668132 | 0.34722016 | 0.10616936 | 0.96055083 | 0 | 0.2347497 |
| ENSMUSG00 | Pabpc6 | 3207 | 0 | 0 | 0 | 0 | 0 | 0 |
| ENSMUSG00 | Nxph1 | 5451 | 0 | 0 | 0 | 0 | 0 | 0 |
| ENSMUSG00 | E2f8 | 6231 | 0 | 0 | 0 | 0 | 0 | 0 |
| ENSMUSG00 | 4930550L24R | 1681 | 0 | 0 | 0 | 0 | 0 | 0 |
| ENSMUSG00 | Gsg1l | 3917 | 0.02029894 | 0.01230187 | 0 | 0 | 0 | 0 |
| ENSMUSG00 | Zfp84 | 1269 | 25.6891206 | 24.3780109 | 33.7522347 | 42.4511367 | 46.8847551 | 33.7440375 |
| ENSMUSG00 | Cd109 | 7056 | 0 | 0.03414572 | 0.20672631 | 0.32727478 | 0.24587539 | 0 |
| ENSMUSG00 | Pcdhb20 | 3547 | 0.66128372 | 1.10039521 | 0.5732407 | 0.5391453 | 1.0789338 | 2.3683645 |
| ENSMUSG00 | Iqub | 3638 | 0.24041247 | 0 | 0 | 0 | 0 | 0.29988608 |
| ENSMUSG00 | Ttc39d | 2435 | 0.16326686 | 0.07915637 | 0 | 0.10372662 | 0.04191078 | 0.08960867 |
| ENSMUSG00 | Scaf8 | 5422 | 11.5702966 | 9.2515836 | 8.90232618 | 13.0499611 | 10.5779503 | 9.4973292 |
| ENSMUSG00 | Sprr2g | 647 | 0 | 0 | 0 | 0 | 0.39433057 | 0 |
| ENSMUSG00 | Pnma2 | 5088 | 0 | 0 | 0 | 0 | 0 | 0 |
| ENSMUSG00 | Pik3r6 | 3562 | 0.81475297 | 0.17586293 | 1.29056476 | 1.60049496 | 0.48705693 | 0.65085467 |
| ENSMUSG00 | Olfr735 | 1064 | 0 | 0 | 0 | 0 | 0 | 0 |
| ENSMUSG00 | Cym | 1227 | 0 | 0 | 0 | 0 | 0 | 0 |
| ENSMUSG00 | Rprml | 1071 | 0 | 0 | 0 | 0 | 0 | 0 |
| ENSMUSG00 | C230012O17 | 5308 | 0 | 0 | 0 | 0.29229991 | 0 | 0 |
| ENSMUSG00 | Plaur | 1389 | 3.77805862 | 13.3562133 | 1.75025296 | 4.13034203 | 5.10631119 | 1.6690744 |
| ENSMUSG00 | Scand1 | 1307 | 8.88186711 | 8.92204973 | 13.0204358 | 6.29434289 | 6.24653415 | 15.0667862 |
| ENSMUSG00 | Vps13a | 9791 | 3.00064346 | 2.32294972 | 1.92770779 | 2.40276714 | 1.58952555 | 4.11167067 |
| ENSMUSG00 | Hepacam | 3320 | 0.01197454 | 0 | 0 | 0.01086809 | 0 | 0 |
| ENSMUSG00 | 1600029I14R | 1487 | 0.18714752 | 0.22683598 | 0 | 0 | 0 | 0 |
| ENSMUSG00 | Pilra | 3323 | 0 | 0 | 0 | 0 | 0 | 0 |
| ENSMUSG00 | Krtap5-3 | 1071 | 0 | 0 | 0 | 0 | 0 | 0 |
| ENSMUSG00 | Adam29 | 2724 | 0 | 0 | 0 | 0 | 0 | 0 |
| ENSMUSG00 | Sprr2h | 750 | 0 | 0 | 0 | 0 | 0 | 0 |
| ENSMUSG00 | C87977 | 3183 | 0 | 0 | 0 | 0 | 0 | 0 |
| ENSMUSG00 | Usp27x | 4275 | 0.23248819 | 0.22543365 | 0.3308676 | 0.58237686 | 0.53711975 | 0.74646381 |
| ENSMUSG00 | Olfr1444 | 1029 | 0 | 0 | 0 | 0 | 0 | 0 |
| ENSMUSG00 | Tusc5 | 3299 | 0 | 0 | 0 | 0 | 0 | 0 |
| ENSMUSG00 | She | 5975 | 0.55225186 | 0.77420895 | 0.3550943 | 0.6220001 | 1.21267705 | 0.38800742 |
| ENSMUSG00 | 4930529F22R | 2587 | 0 | 0 | 0 | 0 | 0 | 0 |
| ENSMUSG00 | Pnma3 | 3403 | 0 | 0 | 0 | 0 | 0 | 0 |
| ENSMUSG00 | Prps1l1 | 1614 | 0 | 0 | 0 | 0 | 0 | 0 |
| ENSMUSG00 | Ankle1 | 2632 | 0 | 0.36615838 | 0 | 0 | 0.07754768 | 0 |
| ENSMUSG00 | Olfr1412 | 965 | 0 | 0 | 0 | 0 | 0 | 0 |
| ENSMUSG00 | Zfp62 | 5380 | 45.0168006 | 48.9387955 | 51.4483162 | 61.668104 | 47.6878471 | 49.0436543 |
| ENSMUSG00 | AI464131 | 4894 | 17.9118992 | 6.62637425 | 16.9979298 | 13.0791882 | 12.3968862 | 8.73301179 |
| ENSMUSG00 | Stxbp6 | 5464 | 11.0229782 | 12.8579489 | 7.14316021 | 9.87237492 | 12.6538688 | 9.34445913 |
| ENSMUSG00 | BC107364 | 1212 | 0 | 0 | 0.21882101 | 0 | 0 | 0 |
| ENSMUSG00 | Ccbe1 | 9495 | 4.24142199 | 2.23296836 | 4.52492801 | 4.60193322 | 3.56297917 | 4.13643818 |
| ENSMUSG00 | Hs3st2 | 3037 | 0 | 0 | 0 | 0 | 0 | 0 |
| ENSMUSG00 | Dppa3 | 887 | 0 | 0 | 0 | 0 | 0 | 0 |
| ENSMUSG00 | Ermp1 | 11103 | 23.8933911 | 25.2628371 | 22.7159971 | 27.7561688 | 28.9944524 | 31.8781405 |
| ENSMUSG00 | Slc25a23 | 4310 | 24.2591447 | 22.5615407 | 18.6550238 | 22.1096704 | 18.6820467 | 25.0850742 |
| ENSMUSG00 | Rpl37a | 580 | 4.52389951 | 4.90172429 | 6.63027645 | 3.48378361 | 5.54252013 | 10.6747295 |
| ENSMUSG00 | Gm6195 | 359 | 0 | 0 | 0.24624982 | 0.20101418 | 0 | 0 |
| ENSMUSG00 | Fam178b | 3151 | 0.02523356 | 0.04587729 | 0.07013939 | 0.08015688 | 0 | 0 |
| ENSMUSG00 | Gpat2 | 2606 | 0 | 0.20339634 | 0.11873097 | 0.65075061 | 0.15664275 | 0.21978796 |
| ENSMUSG00 | Gm11223 | 1419 | 0.16809928 | 0.50937042 | 0.21804997 | 0.22885018 | 0.28767513 | 0.05766309 |
| ENSMUSG00 | 2310010M20 | 1405 | 0.02829572 | 0 | 0 | 0 | 0 | 0.03882511 |
| ENSMUSG00 | Zfp322a | 5874 | 14.8558529 | 13.7898211 | 21.2355466 | 20.7376545 | 13.064976 | 19.0095971 |
| ENSMUSG00 | Gjb2 | 2404 | 0.13229777 | 0.34075271 | 0.11032074 | 0 | 0.31838421 | 0.45382095 |
| ENSMUSG00 | Defb14 | 202 | 0 | 0 | 0 | 1.0717439 | 0.75781746 | 0 |
| ENSMUSG00 | Rpl27a | 1998 | 2.84536222 | 2.65290723 | 3.42907191 | 2.65468496 | 1.58340105 | 3.20797809 |
| ENSMUSG00 | 4931440L10R | 3048 | 0 | 0 | 0 | 0 | 0 | 0 |

| | | | | | | | | |
|---|---|---|---|---|---|---|---|---|
| ENSMUSG00 | Gm5590 | 344 | 0.69340954 | 0 | 0 | 0 | 0 | 0.15857348 |
| ENSMUSG00 | Asphd1 | 1889 | 0 | 0 | 0 | 0 | 0 | 0 |
| ENSMUSG00 | Jrk | 5740 | 0.47789689 | 1.00738207 | 0.91637967 | 0.92405236 | 1.48456529 | 0.92182579 |
| ENSMUSG00 | Pcdhb17 | 3268 | 3.32106676 | 3.78944787 | 4.74750517 | 2.20820346 | 3.57559365 | 7.77844669 |
| ENSMUSG00 | Gm9806 | 549 | 0 | 0 | 0 | 0 | 0 | 0 |
| ENSMUSG00 | Olfr609 | 964 | 0 | 0 | 0 | 0 | 0 | 0 |
| ENSMUSG00 | Rbp1 | 2624 | 0 | 0 | 0.05053564 | 0.12375701 | 0.03889206 | 0.18709737 |
| ENSMUSG00 | Yod1 | 3823 | 11.0645648 | 4.65100632 | 9.15716968 | 14.166662 | 8.50217144 | 17.0297054 |
| ENSMUSG00 | 1700067K01l | 2152 | 0.83131813 | 0.35826351 | 0.45187758 | 0.30180149 | 0.90102337 | 0.29150404 |
| ENSMUSG00 | Kcnk6 | 4484 | 0 | 0 | 0 | 0 | 0 | 0 |
| ENSMUSG00 | Gm6816 | 767 | 0 | 0 | 0 | 0 | 0 | 0 |
| ENSMUSG00 | D230002A01l | 3310 | 0 | 0 | 0 | 0 | 0 | 0 |
| ENSMUSG00 | B430212C06 | 5708 | 0.4945058 | 0.34611845 | 0.55755653 | 0.36031474 | 0.57212475 | 0.50172339 |
| ENSMUSG00 | BC046404 | 2507 | 6.07353372 | 3.32519126 | 5.62440683 | 5.80018506 | 5.21051145 | 3.50316467 |
| ENSMUSG00 | Olfr518 | 1001 | 0 | 0 | 0 | 0 | 0 | 0 |
| ENSMUSG00 | Ngfrap1 | 1044 | 28.4457317 | 40.893858 | 28.1130496 | 29.3771435 | 22.6783893 | 38.1165696 |
| ENSMUSG00 | Hnrnpa1 | 1116 | 1.74553633 | 3.02244709 | 1.22782897 | 3.39481602 | 1.09734142 | 2.54172265 |
| ENSMUSG00 | Gm13078 | 2102 | 0 | 0 | 0 | 0 | 0 | 0 |
| ENSMUSG00 | Abpz | 520 | 0 | 0 | 0 | 0 | 0 | 0 |
| ENSMUSG00 | Gm5564 | 1108 | 0.17940199 | 0 | 0 | 0 | 0 | 0.1230805 |
| ENSMUSG00 | Ftsjd1 | 3575 | 1.36781094 | 1.76571299 | 4.14199088 | 6.16674943 | 2.76898375 | 4.53941548 |
| ENSMUSG00 | Ppm1e | 6694 | 0.04751178 | 0.00719845 | 0.12546086 | 0 | 0.11434055 | 0.01222347 |
| ENSMUSG00 | Vmn1r79 | 920 | 0 | 0 | 0 | 0 | 0 | 0 |
| ENSMUSG00 | Camk2n1 | 4442 | 4.76134976 | 5.76024331 | 6.56758586 | 4.22392237 | 3.25089247 | 7.03049571 |
| ENSMUSG00 | C77370 | 21892 | 0.41949187 | 0.93106455 | 0.44419904 | 0.33952591 | 0.24939806 | 0.3662865 |
| ENSMUSG00 | Olfr71 | 2433 | 0 | 0 | 0 | 0 | 0 | 0 |
| ENSMUSG00 | Olfr905 | 932 | 0 | 0 | 0 | 0 | 0 | 0 |
| ENSMUSG00 | Tmem150b | 3725 | 0 | 0 | 0 | 0 | 0 | 0 |
| ENSMUSG00 | Sh2d7 | 3039 | 0 | 0 | 0.07272432 | 0 | 0 | 0 |
| ENSMUSG00 | 5930403N24 | 2121 | 0.13120621 | 0 | 0 | 0.05103542 | 0 | 0 |
| ENSMUSG00 | Sox18 | 1618 | 1.7199528 | 0.56584821 | 1.20202787 | 3.83566852 | 2.99598622 | 1.39913168 |
| ENSMUSG00 | Krtap4-16 | 1473 | 0 | 0 | 0 | 0 | 0 | 0 |
| ENSMUSG00 | Scn4b | 4935 | 10.9317502 | 17.3803175 | 4.17387211 | 9.0735192 | 11.8079273 | 5.7423202 |
| ENSMUSG00 | Olfr231 | 923 | 0 | 0 | 0 | 0 | 0 | 0 |
| ENSMUSG00 | Mospd4 | 882 | 0 | 0 | 0 | 0 | 0.23141214 | 0 |
| ENSMUSG00 | Rnf222 | 3460 | 0 | 0 | 0 | 0 | 0 | 0 |
| ENSMUSG00 | C1qtnf2 | 1385 | 0.51667773 | 0 | 0 | 0 | 0 | 0.07877152 |
| ENSMUSG00 | Olfr1352 | 1172 | 0 | 0 | 0 | 0 | 0 | 0 |
| ENSMUSG00 | Fam19a4 | 2176 | 0 | 0 | 0 | 0 | 0 | 0 |
| ENSMUSG00 | Cox17 | 3221 | 13.6262187 | 11.6838284 | 13.9151409 | 8.98410423 | 10.0753726 | 30.9750482 |
| ENSMUSG00 | Ferd3l | 885 | 0 | 0 | 0 | 0 | 0 | 0 |
| ENSMUSG00 | Golph3l | 2637 | 45.1376218 | 36.6560497 | 42.8105829 | 49.8471324 | 44.9117248 | 33.5115022 |
| ENSMUSG00 | Kctd4 | 3013 | 0.56736995 | 0 | 0.32274827 | 0.38321455 | 0.03387081 | 0.43451135 |
| ENSMUSG00 | Pard3b | 1662 | 0 | 0 | 0 | 0 | 0 | 0 |
| ENSMUSG00 | Ar | 9088 | 0.04811953 | 0.56203377 | 0 | 0 | 0.26950551 | 0 |
| ENSMUSG00 | Zfp526 | 3351 | 0.33218545 | 0.25883496 | 0.34295671 | 0.51684218 | 0.42636184 | 0.71625433 |
| ENSMUSG00 | Fam43a | 3074 | 1.25448328 | 0 | 0.04313778 | 0.762958 | 0.84656642 | 0.25730792 |
| ENSMUSG00 | 4731417B20l | 1626 | 0 | 0 | 0 | 0 | 0 | 0 |
| ENSMUSG00 | Spin2 | 1463 | 5.57065858 | 2.83255914 | 8.48989603 | 3.52681638 | 3.69705799 | 10.7196976 |
| ENSMUSG00 | Zfp319 | 4104 | 5.69596066 | 2.43045651 | 2.91878642 | 2.46173793 | 5.30903082 | 2.85107706 |
| ENSMUSG00 | Arsj | 5026 | 0 | 0 | 0 | 0 | 0.07106738 | 0 |
| ENSMUSG00 | Unc119b | 5136 | 24.0111956 | 30.3323145 | 20.6120354 | 29.4150157 | 29.6958406 | 15.6340611 |
| ENSMUSG00 | 4930430F08l | 2412 | 3.74149837 | 4.89455984 | 4.54479978 | 3.75480646 | 3.55407593 | 5.7896415 |
| ENSMUSG00 | Zfp518b | 6806 | 0.41472805 | 0.38939969 | 0.15586897 | 0.2014572 | 0.56979203 | 0.25647619 |
| ENSMUSG00 | Lyrm4 | 1512 | 4.46986223 | 3.69684345 | 3.44961473 | 4.17616256 | 3.64474113 | 3.69795043 |
| ENSMUSG00 | Prr12 | 6865 | 0.34746232 | 0.62470406 | 0.56016902 | 0.64648091 | 0.4757011 | 0.89789781 |
| ENSMUSG00 | Olfr1389 | 1072 | 0 | 0 | 0 | 0 | 0 | 0 |
| ENSMUSG00 | Gm7862 | 434 | 0.18320498 | 0.99925802 | 0 | 0.08313835 | 0.1175723 | 0.06284479 |
| ENSMUSG00 | Ccdc147 | 3327 | 0 | 0 | 0 | 0 | 0 | 0 |
| ENSMUSG00 | Lrrc8e | 3875 | 2.29812326 | 1.7160591 | 1.83651528 | 1.68538066 | 2.51510652 | 1.40772332 |
| ENSMUSG00 | Olfr1165-ps | 1587 | 0 | 0 | 0 | 0 | 0 | 0 |
| ENSMUSG00 | 5730590G19l | 7702 | 0 | 0 | 0 | 0 | 0 | 0 |
| ENSMUSG00 | Tmem215 | 4731 | 0 | 0 | 0 | 0 | 0 | 0 |
| ENSMUSG00 | Bdh1 | 3585 | 16.7560756 | 12.6615421 | 17.5081219 | 10.8296457 | 9.96330909 | 14.8660096 |
| ENSMUSG00 | D9Ertd402e | 3256 | 7.09396013 | 12.6533809 | 11.4713014 | 8.87644891 | 6.77008426 | 7.04483158 |
| ENSMUSG00 | B3gntl1 | 3504 | 2.66624941 | 4.31807729 | 1.65252324 | 2.94505272 | 3.94638923 | 1.78250354 |
| ENSMUSG00 | Hrk | 5364 | 0 | 0 | 0 | 0 | 0 | 0 |
| ENSMUSG00 | 5330437I02R | 3746 | 0.20164285 | 0.05145376 | 0 | 0 | 0.01362156 | 0.37133118 |
| ENSMUSG00 | Vwa5b2 | 7077 | 0 | 0.19064863 | 0 | 0 | 0 | 0.1156195 |
| ENSMUSG00 | Actrt1 | 1338 | 0 | 0 | 0 | 0 | 0 | 0 |
| ENSMUSG00 | Olfml2a | 5571 | 0.46384962 | 0 | 0 | 0 | 0 | 0 |
| ENSMUSG00 | Gm9808 | 476 | 0 | 0 | 0 | 0 | 0 | 0 |
| ENSMUSG00 | Gjb4 | 2839 | 1.07825713 | 0.06789213 | 0.09341707 | 0 | 0.10784017 | 0 |
| ENSMUSG00 | Pkd1l1 | 7581 | 0 | 0.01271242 | 0 | 0 | 0 | 0 |
| ENSMUSG00 | Gm7729 | 961 | 0 | 0 | 0 | 0.03754635 | 0 | 0 |
| ENSMUSG00 | Ttc34 | 3405 | 0.07005371 | 0.31133678 | 0.24664758 | 0.01059678 | 0 | 0.03204069 |
| ENSMUSG00 | Olfr218 | 941 | 0 | 0 | 0 | 0 | 0 | 0 |
| ENSMUSG00 | Olfr1440 | 947 | 0 | 0 | 0 | 0 | 0 | 0 |
| ENSMUSG00 | Tas2r143 | 881 | 0 | 0 | 0 | 0 | 0 | 0 |
| ENSMUSG00 | Zfp316 | 10702 | 0.6946622 | 1.01757951 | 1.04908131 | 1.05528686 | 1.50666555 | 0.50461396 |
| ENSMUSG00 | Rbm12b | 1414 | 2.41794295 | 3.03295145 | 2.34451077 | 3.03660771 | 2.70649094 | 1.71672058 |
| ENSMUSG00 | Cxxc5 | 2268 | 70.9568717 | 81.6917416 | 68.1902314 | 64.6231327 | 65.5828419 | 70.2911228 |
| ENSMUSG00 | Fam54b | 5404 | 43.4558889 | 40.4020669 | 43.8910232 | 36.3290904 | 39.6766897 | 39.7107442 |
| ENSMUSG00 | D230037D09 | 3367 | 10.4141176 | 10.9052774 | 13.0229368 | 12.4845803 | 8.24423773 | 16.9950084 |
| ENSMUSG00 | Lce1l | 668 | 0 | 0 | 0 | 0 | 0 | 0 |
| ENSMUSG00 | Olfr981 | 996 | 0 | 0 | 0 | 0 | 0 | 0 |
| ENSMUSG00 | C87436 | 4165 | 7.9224607 | 8.32994918 | 13.0854436 | 8.83641907 | 10.7443293 | 9.63944033 |
| ENSMUSG00 | 0610025J13R | 1177 | 0.27021567 | 0 | 0 | 0 | 0.08670582 | 0.13903809 |
| ENSMUSG00 | Gm5424 | 1238 | 0 | 0 | 0 | 0 | 0 | 0 |
| ENSMUSG00 | Tifa | 2778 | 11.0193377 | 11.309417 | 14.9407958 | 10.585625 | 10.4697747 | 15.2867579 |
| ENSMUSG00 | Chtf8 | 5667 | 2.27294436 | 3.04407028 | 2.62855498 | 3.19625664 | 2.88131997 | 2.37756693 |
| ENSMUSG00 | Fam46b | 2285 | 2.69676126 | 5.60945016 | 0.54164184 | 1.76857286 | 3.63995591 | 0.7042467 |
| ENSMUSG00 | Enpp7 | 1964 | 0 | 0 | 0 | 0 | 0 | 0 |
| ENSMUSG00 | Slitrk4 | 12071 | 0 | 0 | 0 | 0 | 0 | 0 |
| ENSMUSG00 | Csnk2a2 | 4200 | 3.48333734 | 5.30050864 | 4.16760234 | 5.78171809 | 4.47088245 | 4.66266452 |
| ENSMUSG00 | Mapk10 | 12109 | 0.19042182 | 0.01591756 | 0 | 0.50358126 | 0.08849235 | 0.0247767 |
| ENSMUSG00 | Hmga1 | 2073 | 0.86299885 | 0.97628103 | 0.81026051 | 0.36551999 | 0.54152449 | 0.48681218 |
| ENSMUSG00 | Foxc2 | 2721 | 0 | 0 | 0 | 0.13260582 | 0.35630325 | 0 |
| ENSMUSG00 | Vmn1r67 | 998 | 0 | 0 | 0 | 0 | 0 | 0 |
| ENSMUSG00 | Igbp1b | 1349 | 0 | 0 | 0 | 0 | 0 | 0 |
| ENSMUSG00 | Bst2 | 759 | 5.60452756 | 6.6661086 | 5.4160361 | 5.03912611 | 6.25224368 | 14.805206 |
| ENSMUSG00 | Nxph3 | 2097 | 0 | 0 | 0 | 0 | 0 | 0 |
| ENSMUSG00 | Rpl14-ps1 | 690 | 2.47751545 | 3.56160659 | 1.9858799 | 1.25502764 | 1.922733 | 2.21359391 |
| ENSMUSG00 | Cdc42se1 | 3254 | 63.2251419 | 58.8632783 | 68.9244473 | 61.397136 | 50.038465 | 51.5066559 |
| ENSMUSG00 | Adam24 | 2868 | 0 | 0 | 0 | 0 | 0 | 0 |
| ENSMUSG00 | 0610010O12l | 2834 | 351.823377 | 400.097096 | 441.566118 | 382.286333 | 246.093333 | 507.67978 |
| ENSMUSG00 | Kctd11 | 2720 | 15.2152409 | 6.37761734 | 6.54902306 | 12.3236101 | 10.6930273 | 4.56248561 |
| ENSMUSG00 | Gprc5a | 2079 | 6.44425057 | 3.98656472 | 5.16644918 | 1.59670423 | 4.14788721 | 3.73894766 |
| ENSMUSG00 | Fat4 | 18340 | 0.07153385 | 0.17603553 | 0.25547412 | 0.19673961 | 0.04173368 | 0.16210118 |
| ENSMUSG00 | Gm9812 | 965 | 0.53556606 | 0.04993414 | 0.36644015 | 0 | 0.42301659 | 0.14131937 |
| ENSMUSG00 | 2310005G13l | 3060 | 0 | 0 | 0 | 0.49524375 | 0 | 0 |
| ENSMUSG00 | BC089491 | 1456 | 0 | 0 | 0 | 0 | 0 | 0 |
| ENSMUSG00 | Ccdc66 | 2922 | 9.29260547 | 9.79560118 | 8.45613629 | 8.01411598 | 9.09813203 | 10.062307 |
| ENSMUSG00 | Kif2b | 2294 | 0 | 0 | 0 | 0 | 0 | 0 |
| ENSMUSG00 | Mrps7 | 2068 | 47.2721115 | 43.7592545 | 40.546855 | 34.0581001 | 32.9648153 | 35.2671109 |
| ENSMUSG00 | Fam83h | 4621 | 6.28895396 | 6.55902881 | 5.13663486 | 7.14457276 | 7.47562355 | 6.04988211 |
| ENSMUSG00 | A530053G22l | 1521 | 0 | 0 | 0 | 0 | 0 | 0 |
| ENSMUSG00 | Rhoj | 4668 | 1.36265572 | 0.08258174 | 0.4734559 | 0.89664035 | 1.41011193 | 0.59013251 |
| ENSMUSG00 | 8030474K03l | 2475 | 0 | 0 | 0 | 0 | 0 | 0 |
| ENSMUSG00 | Ttc6 | 6649 | 0 | 0 | 0 | 0 | 0 | 0 |
| ENSMUSG00 | Epm2aip1 | 11487 | 11.3517869 | 8.35617592 | 10.9436791 | 10.9687908 | 15.0187322 | 9.06542811 |
| ENSMUSG00 | Gm9814 | 388 | 0 | 0 | 0 | 0 | 0 | 0 |
| ENSMUSG00 | Olfr1341 | 951 | 0 | 0 | 0 | 0 | 0 | 0 |
| ENSMUSG00 | 2410016O06l | 2345 | 4.89950272 | 5.38373042 | 6.6161394 | 5.81621016 | 6.04918229 | 4.78033123 |
| ENSMUSG00 | Zfp787 | 1896 | 5.59847748 | 3.63431499 | 3.86999258 | 3.76806161 | 3.79468301 | 4.17175389 |
| ENSMUSG00 | Gpr61 | 6390 | 0 | 0.04524549 | 0 | 0 | 0 | 0.10670829 |
| ENSMUSG00 | Ppp1r3b | 4204 | 0 | 0 | 0.70445373 | 0 | 0.44909037 | 0.38926697 |
| ENSMUSG00 | Cldn12 | 7412 | 13.5003433 | 17.3255354 | 12.762 | 18.4888836 | 15.1041377 | 15.9592702 |
| ENSMUSG00 | Phgr1 | 410 | 0 | 0 | 0 | 0 | 0 | 0 |
| ENSMUSG00 | Mpeg1 | 4333 | 0 | 0 | 0 | 0 | 0 | 0 |
| ENSMUSG00 | 3110062M04 | 7120 | 0.27359811 | 0.37899449 | 0.29178183 | 0.1773696 | 0.26516516 | 0.59759041 |
| ENSMUSG00 | AI646023 | 4597 | 0.30268476 | 0.42976814 | 0.10576904 | 0 | 0.38849753 | 0.20172672 |
| ENSMUSG00 | Atp10d | 8020 | 1.29874512 | 0.38453021 | 1.93451956 | 2.39347228 | 2.79308965 | 1.1120707 |
| ENSMUSG00 | Gltpd2 | 1213 | 0 | 0.43697516 | 0 | 0 | 0 | 0 |
| ENSMUSG00 | Gchfr | 642 | 10.651001 | 12.8347066 | 9.57018096 | 9.77924571 | 5.96102521 | 16.5262222 |
| ENSMUSG00 | Ddit4l | 2911 | 18.7100681 | 41.4658322 | 7.18223007 | 34.5202659 | 29.4308777 | 17.5022419 |
| ENSMUSG00 | Slc39a3 | 7172 | 5.28816626 | 4.57542765 | 3.68554129 | 4.34674937 | 4.90200404 | 1.8824521 |
| ENSMUSG00 | Fam187b | 2260 | 1.14340984 | 4.0084297 | 3.30535906 | 2.80992913 | 2.77709922 | 2.19645325 |
| ENSMUSG00 | 4832428D23 | 4285 | 0 | 0 | 0 | 0 | 0 | 0 |
| ENSMUSG00 | Krt1 | 2436 | 0.03263997 | 0.09890485 | 0.01814526 | 0 | 0 | 0.08957189 |
| ENSMUSG00 | 0610008F07l | 949 | 0 | 0 | 0 | 0 | 0 | 0 |
| ENSMUSG00 | Ckap4 | 5092 | 5.70723807 | 9.40638717 | 6.35423195 | 6.33490727 | 8.10690064 | 4.9064355 |
| ENSMUSG00 | Vat1l | 5769 | 0 | 0 | 0 | 0 | 0 | 0 |
| ENSMUSG00 | Il1f10 | 534 | 0 | 0 | 0 | 0.06756937 | 0 | 0 |
| ENSMUSG00 | Spesp1 | 3011 | 0 | 0 | 0 | 0 | 0 | 0 |
| ENSMUSG00 | Gm7094 | 376 | 0 | 0 | 0 | 0 | 0 | 0 |
| ENSMUSG00 | Pip5kl1 | 2482 | 0 | 0 | 0 | 0 | 0 | 0 |
| ENSMUSG00 | Gpr1 | 1937 | 0 | 0 | 0 | 0 | 0 | 0 |
| ENSMUSG00 | C230030N03 | 942 | 0 | 0 | 0 | 0 | 0 | 0 |
| ENSMUSG00 | Hectd3 | 4788 | 12.8034568 | 10.3055382 | 13.9677085 | 12.1931387 | 13.8755934 | 11.8144532 |
| ENSMUSG00 | Pramel6 | 4289 | 9.2413649 | 8.95420713 | 11.5631774 | 13.0396757 | 10.5169774 | 9.01735567 |
| ENSMUSG00 | Fbl | 1152 | 0.51764949 | 0.92022719 | 1.11272001 | 1.22152755 | 0.84158085 | 1.37320232 |
| ENSMUSG00 | Mbtps2 | 9966 | 2.09428329 | 0.85580978 | 1.72531777 | 1.46992877 | 0.97792874 | 0.93871175 |
| ENSMUSG00 | Atxn1 | 6560 | 0.57572723 | 0.65374899 | 0.31669003 | 0.54453086 | 0.8556252 | 0.61534247 |
| ENSMUSG00 | Irgm1 | 5806 | 13.4207271 | 13.2126793 | 14.7542494 | 12.9574686 | 31.6827566 | 31.9863966 |
| ENSMUSG00 | Olfr374 | 1078 | 0 | 0 | 0 | 0 | 0 | 0 |
| ENSMUSG00 | Zfp474 | 1928 | 0 | 0 | 0 | 0 | 0 | 0 |
| ENSMUSG00 | Zfp740 | 4958 | 27.3108444 | 19.7099848 | 26.7279398 | 23.6228956 | 29.2799595 | 25.8773504 |
| ENSMUSG00 | Prl7a2 | 899 | 0 | 0 | 0 | 0 | 0 | 0 |
| ENSMUSG00 | Ltb4r1 | 1387 | 0 | 0 | 0 | 0 | 0.25752317 | 0.15731587 |
| ENSMUSG00 | 1110002N22l | 1552 | 21.3890633 | 14.2199681 | 17.7149139 | 10.7176692 | 18.4115789 | 19.0500702 |
| ENSMUSG00 | Olfr1451 | 932 | 0 | 0 | 0 | 0 | 0 | 0 |
| ENSMUSG00 | Myct1 | 2773 | 0.48744549 | 0.38229417 | 0.36662185 | 0.96288182 | 0.84645268 | 0.03904315 |
| ENSMUSG00 | Gpr6 | 1772 | 0 | 0 | 0 | 0 | 0 | 0 |
| ENSMUSG00 | Vmn1r179 | 971 | 0 | 0 | 0 | 0 | 0 | 0 |
| ENSMUSG00 | Vmn1r193 | 959 | 0 | 0 | 0 | 0 | 0 | 0 |
| ENSMUSG00 | Csl | 1953 | 0.36640996 | 0.22205734 | 0.13579675 | 0.24017746 | 0.31352612 | 0.1536206 |
| ENSMUSG00 | Mageb16 | 3441 | 0 | 0 | 0 | 0 | 0 | 0 |
| ENSMUSG00 | Adck2 | 4326 | 6.46968522 | 11.2501865 | 6.69260453 | 6.50577779 | 9.89623886 | 5.2393031 |
| ENSMUSG00 | Nqo2 | 3966 | 30.0421521 | 25.5876569 | 28.7768933 | 29.3951301 | 29.2443147 | 32.150514 |
| ENSMUSG00 | Gm5815 | 707 | 0 | 0 | 0.12504057 | 0 | 0.14434618 | 0.11573397 |
| ENSMUSG00 | Spz1 | 1601 | 0 | 0 | 0 | 0 | 0 | 0 |
| ENSMUSG00 | 4930432E11l | 4105 | 0 | 0 | 0 | 0 | 0 | 0 |
| ENSMUSG00 | Slc26a1 | 6252 | 0.97290283 | 0.15414729 | 0.73528338 | 0.20199481 | 0.23668664 | 0.25739023 |
| ENSMUSG00 | Gpr156 | 4571 | 0 | 0 | 0 | 0 | 0 | 0 |
| ENSMUSG00 | Zfp295 | 9451 | 3.71853188 | 3.34975055 | 3.99411427 | 3.52383103 | 4.0168896 | 3.99119946 |
| ENSMUSG00 | Pla2g4f | 3846 | 0 | 0 | 0 | 0 | 0 | 0 |
| ENSMUSG00 | BC053393 | 2014 | 0.63166603 | 0 | 0 | 0 | 0 | 0 |
| ENSMUSG00 | Olfr1020 | 1078 | 0 | 0 | 0 | 0 | 0 | 0 |
| ENSMUSG00 | Pcdh15 | 465 | 0 | 0 | 0 | 0 | 0 | 0 |
| ENSMUSG00 | Tshz1 | 9799 | 4.33703528 | 5.79771561 | 4.56047181 | 6.26345114 | 6.22793606 | 3.23710638 |
| ENSMUSG00 | Tapt1 | 3473 | 26.1106395 | 34.8252145 | 34.0072918 | 35.8534799 | 33.7188403 | 31.7353346 |
| ENSMUSG00 | Wdr27 | 2565 | 0.21698898 | 0.63872867 | 1.03396124 | 0 | 1.07423949 | 0.76560391 |
| ENSMUSG00 | Gm5637 | 1118 | 0.21335678 | 0.1724023 | 0.07907306 | 0.12909497 | 0.41076689 | 0.1219796 |
| ENSMUSG00 | Mars2 | 2881 | 3.29800758 | 7.9948349 | 3.85097626 | 5.134897 | 5.66762909 | 3.5596297 |
| ENSMUSG00 | 2610303G11l | 514 | 0 | 0 | 0 | 0 | 0 | 0 |
| ENSMUSG00 | Spsb4 | 5698 | 3.98392056 | 5.90279688 | 1.12482753 | 4.86328715 | 3.98503637 | 2.33112484 |
| ENSMUSG00 | 1110032F04l | 2577 | 0 | 0 | 0 | 0 | 0 | 0 |
| ENSMUSG00 | Msgn1 | 566 | 0 | 0 | 0 | 0 | 0 | 0 |
| ENSMUSG00 | Zfp41 | 4102 | 2.80091026 | 4.3699065 | 1.68100622 | 5.22494745 | 2.87349898 | 3.01869484 |
| ENSMUSG00 | Mtap4 | 3157 | 0 | 0 | 0 | 0 | 0 | 0 |
| ENSMUSG00 | Fbxo41 | 10068 | 0 | 0 | 0 | 0 | 0 | 0 |
| ENSMUSG00 | Catsperb | 3689 | 0 | 0 | 0 | 0 | 0 | 0.00739351 |
| ENSMUSG00 | Ccdc108 | 7579 | 0 | 0 | 0 | 0 | 0 | 0 |
| ENSMUSG00 | Mipol1 | 8944 | 3.62262037 | 4.18614329 | 3.90917462 | 3.93336241 | 4.19895043 | 3.26905336 |
| ENSMUSG00 | 4921510H08 | 1867 | 1.23503903 | 0 | 0 | 0 | 0 | 0 |
| ENSMUSG00 | Acsm4 | 2140 | 0 | 0 | 0 | 0.33721537 | 0 | 0 |
| ENSMUSG00 | Spata2 | 5274 | 6.75405964 | 5.73778644 | 7.4424036 | 5.63739186 | 4.68273908 | 4.47854334 |
| ENSMUSG00 | Vmn1r68 | 959 | 0 | 0 | 0 | 0 | 0 | 0 |
| ENSMUSG00 | Pcdhb15 | 2794 | 0.41263741 | 0 | 0.66445147 | 0.69736235 | 0 | 0.14642791 |
| ENSMUSG00 | Ankrd33 | 1706 | 0 | 0 | 0 | 0 | 0 | 0 |
| ENSMUSG00 | Zfp445 | 2955 | 26.4767464 | 12.6703437 | 21.4053595 | 28.8900566 | 28.7509021 | 29.4436884 |
| ENSMUSG00 | Nipa1 | 3178 | 5.35410499 | 4.67005166 | 4.90977048 | 5.17728518 | 3.9658637 | 5.68150132 |
| ENSMUSG00 | Olfr1151 | 926 | 0 | 0 | 0 | 0 | 0 | 0 |
| ENSMUSG00 | Prr15l | 1732 | 64.5911791 | 73.3090503 | 81.1813295 | 59.3103815 | 57.7140128 | 71.6825392 |
| ENSMUSG00 | D030056L22l | 2098 | 11.2747907 | 11.8284165 | 13.9684566 | 10.7661391 | 11.285147 | 9.4122206 |
| ENSMUSG00 | Tmem164 | 2879 | 37.8222512 | 41.7091399 | 43.1424727 | 46.4843012 | 55.067367 | 30.3725229 |
| ENSMUSG00 | 1700031F10l | 1156 | 0 | 0 | 0 | 0 | 0 | 0 |
| ENSMUSG00 | Olfr432 | 938 | 0 | 0 | 0 | 0 | 0 | 0 |
| ENSMUSG00 | Olfr914 | 950 | 0 | 0 | 0 | 0 | 0 | 0 |
| ENSMUSG00 | Teddm1 | 1414 | 0 | 0 | 0 | 0 | 0 | 0 |
| ENSMUSG00 | Gm9817 | 2530 | 0 | 0 | 0 | 0 | 0 | 0 |
| ENSMUSG00 | Dusp28 | 1489 | 6.96855636 | 7.0871933 | 4.74969435 | 7.12432578 | 6.23693781 | 5.93484427 |
| ENSMUSG00 | 4930524E20l | 741 | 0 | 0 | 0 | 0 | 0 | 0 |
| ENSMUSG00 | Ngrn | 1919 | 37.5180694 | 25.4366159 | 29.4832512 | 27.4328833 | 24.4096993 | 31.1689859 |
| ENSMUSG00 | Lrrc4b | 2924 | 0 | 0 | 0 | 0.03701988 | 0 | 0 |
| ENSMUSG00 | Tmem198b | 4239 | 1.21920558 | 2.13707269 | 1.50154881 | 0.44262003 | 1.39633395 | 0.98443496 |
| ENSMUSG00 | Ofcc1 | 2797 | 0 | 0 | 0 | 0 | 0 | 0 |
| ENSMUSG00 | Rnf31 | 4897 | 11.260129 | 9.30863269 | 6.92318023 | 8.47342275 | 13.1186864 | 5.78687976 |
| ENSMUSG00 | Olfr1437 | 938 | 0 | 0 | 0 | 0 | 0 | 0 |
| ENSMUSG00 | Tas2r139 | 959 | 0 | 0 | 0 | 0 | 0 | 0 |
| ENSMUSG00 | Pbp2 | 789 | 0 | 0 | 0 | 0 | 0 | 0 |
| ENSMUSG00 | Dnajb7 | 1185 | 0.90582108 | 0 | 0.26110793 | 0 | 0.99038535 | 0.20714916 |
| ENSMUSG00 | Cldn14 | 2968 | 0 | 0.04870597 | 0.0148928 | 0.03647107 | 0.03438435 | 0.06432698 |
| ENSMUSG00 | 0610012G03l | 1439 | 26.9364792 | 27.1571957 | 29.1504858 | 24.8988674 | 16.8787734 | 31.5203093 |
| ENSMUSG00 | D330028D13 | 2706 | 29.0012264 | 12.1267432 | 44.9532417 | 29.0549797 | 10.1260768 | 24.9362371 |
| ENSMUSG00 | 9430041J12R | 532 | 0 | 0 | 0 | 0 | 0 | 0 |
| ENSMUSG00 | Ankdd1b | 3301 | 2.61343207 | 2.75893292 | 3.74932325 | 2.85289719 | 2.5505459 | 1.50378199 |
| ENSMUSG00 | Olfr705 | 950 | 0 | 0 | 0 | 0 | 0 | 0 |
| ENSMUSG00 | Ticam1 | 2990 | 6.80762743 | 7.47776226 | 9.23951568 | 4.57361033 | 7.38943837 | 4.98970825 |
| ENSMUSG00 | Cltc | 8757 | 62.7816079 | 45.5011637 | 62.2672074 | 70.1780727 | 64.7313017 | 54.3312558 |
| ENSMUSG00 | 1700113H08 | 1552 | 0 | 0 | 0 | 0 | 0 | 0 |
| ENSMUSG00 | Cd24a | 1811 | 199.743301 | 340.551228 | 115.495926 | 238.607711 | 286.407018 | 164.280378 |
| ENSMUSG00 | Zfp654 | 4739 | 14.2277474 | 9.38512051 | 14.3173305 | 16.0728415 | 12.9423304 | 15.0560153 |
| ENSMUSG00 | Dmrta2 | 2919 | 0 | 0 | 0 | 0 | 0 | 0 |
| ENSMUSG00 | Tet1 | 23532 | 0.26354985 | 0.32558407 | 0.36064737 | 0.67159338 | 0.49222282 | 0.59806705 |
| ENSMUSG00 | Olfr1052 | 939 | 0 | 0 | 0 | 0 | 0.10868238 | 0 |
| ENSMUSG00 | 1700001C19l | 2409 | 0.04950869 | 0.22002942 | 0.25688078 | 0 | 0.02118156 | 0.29437137 |
| ENSMUSG00 | Khnyn | 3868 | 8.31493892 | 5.75546439 | 3.58825716 | 7.86379616 | 14.6430396 | 6.00070269 |
| ENSMUSG00 | Cyp4x1 | 4175 | 0 | 0 | 0.53995066 | 0.05185444 | 0.01222189 | 0.58142345 |
| ENSMUSG00 | Chst9 | 1928 | 3.21672975 | 8.17270051 | 0.80241935 | 0.16843278 | 3.96989438 | 0 |
| ENSMUSG00 | Olfr65 | 1105 | 0 | 0 | 0 | 0 | 0 | 0 |
| ENSMUSG00 | Tigd5 | 1912 | 0.810912 | 0.42843594 | 1.59515019 | 0.8492113 | 3.46936656 | 2.36798646 |
| ENSMUSG00 | Gm6684 | 683 | 0 | 0 | 0 | 0 | 0 | 0 |
| ENSMUSG00 | Helt | 1526 | 0 | 0 | 0 | 0 | 0 | 0 |
| ENSMUSG00 | Neurl3 | 2579 | 4.40871168 | 2.76525531 | 8.03825683 | 3.52565926 | 4.59019744 | 4.60041414 |
| ENSMUSG00 | Samd14 | 4063 | 1.4481445 | 3.15471169 | 2.0017571 | 2.68195359 | 1.7833486 | 2.37637763 |
| ENSMUSG00 | Irs3 | 2522 | 0.898518 | 1.6622603 | 1.36706731 | 0.32905909 | 1.13302024 | 0.30281122 |
| ENSMUSG00 | Rab2a | 2125 | 269.794722 | 229.480845 | 277.96199 | 234.388961 | 193.588067 | 313.29255 |
| ENSMUSG00 | Gm9818 | 253 | 0 | 0 | 0 | 0 | 0 | 0 |
| ENSMUSG00 | Dync2h1 | 15724 | 2.92527925 | 3.23919291 | 2.94041769 | 4.35995196 | 4.42635313 | 2.50127384 |
| ENSMUSG00 | Gjd3 | 836 | 0 | 0 | 0 | 0 | 0 | 0 |
| ENSMUSG00 | Vmn1r54 | 947 | 0 | 0 | 0 | 0 | 0 | 0 |
| ENSMUSG00 | Dusp18 | 2687 | 5.77024094 | 3.19210521 | 4.78702505 | 7.68103069 | 6.43764101 | 4.273399 |
| ENSMUSG00 | Olfr1495 | 1088 | 0 | 0 | 0 | 0 | 0 | 0 |
| ENSMUSG00 | Ythdf3 | 5608 | 68.4094841 | 61.9171723 | 67.2880053 | 85.7334599 | 57.5866499 | 75.749377 |
| ENSMUSG00 | Rpl9 | 2458 | 9.2676527 | 2.39167858 | 1.52854217 | 0.27890921 | 1.26631771 | 2.25254344 |
| ENSMUSG00 | Cdh19 | 3042 | 0 | 0 | 0 | 0 | 0 | 0 |
| ENSMUSG00 | Ccdc36 | 2852 | 0 | 0 | 0 | 0 | 0 | 0 |
| ENSMUSG00 | Fam185a | 4460 | 2.54934247 | 3.00354953 | 2.18035997 | 4.80554584 | 1.99071511 | 3.83434951 |
| ENSMUSG00 | Ear11 | 684 | 0 | 0 | 0 | 0 | 0 | 0 |
| ENSMUSG00 | Olfr684 | 905 | 0 | 0 | 0 | 0 | 0 | 0 |
| ENSMUSG00 | Gm527 | 1458 | 2.56311054 | 4.69305541 | 4.27466383 | 2.79648219 | 4.96964703 | 3.06792924 |
| ENSMUSG00 | BC048546 | 4662 | 0 | 0 | 0 | 0.03869803 | 0 | 0.04680333 |
| ENSMUSG00 | Cldn2 | 3321 | 2.69346074 | 1.81370228 | 0.62556056 | 0 | 1.41355814 | 1.5686408 |
| ENSMUSG00 | Fbxw21 | 2431 | 0 | 0 | 0 | 0 | 0 | 0 |
| ENSMUSG00 | Mageh1 | 1397 | 16.0786303 | 19.2124898 | 13.9851214 | 10.873687 | 10.8481271 | 24.8146503 |
| ENSMUSG00 | Taf9b | 3149 | 22.3458877 | 29.1658808 | 30.614233 | 29.4133402 | 15.6368538 | 24.589616 |
| ENSMUSG00 | Hist1h2be | 4659 | 0.17919405 | 0.51713288 | 0.04743705 | 0.91386161 | 0.35047092 | 0.14635458 |
| ENSMUSG00 | C2cd3 | 8399 | 3.01041619 | 3.04069703 | 3.98390227 | 4.34324884 | 5.80190367 | 3.93905672 |
| ENSMUSG00 | Ptgs1 | 3886 | 46.0471996 | 73.6808644 | 19.132141 | 56.9457486 | 58.2089357 | 30.8962846 |
| ENSMUSG00 | Krtap1-5 | 1039 | 0 | 0 | 0 | 0 | 0 | 0 |
| ENSMUSG00 | Prss45 | 1229 | 0.12939131 | 0 | 0 | 0 | 0 | 0 |
| ENSMUSG00 | Mc4r | 2757 | 0 | 0 | 0 | 0 | 0 | 0 |
| ENSMUSG00 | Tmem93 | 1828 | 45.4534761 | 40.8583071 | 48.6994047 | 45.3788952 | 35.6179735 | 46.8204694 |
| ENSMUSG00 | Gap43 | 2175 | 0 | 0 | 0 | 0 | 0 | 0 |
| ENSMUSG00 | Zfp358 | 1997 | 13.2186475 | 14.2846138 | 10.9785249 | 10.3168991 | 11.8303515 | 7.81226522 |
| ENSMUSG00 | Sfn | 1612 | 2.39223425 | 1.22558569 | 1.5081274 | 3.02175931 | 0.60142751 | 1.23514185 |
| ENSMUSG00 | Neurl4 | 5658 | 4.39853849 | 4.45413737 | 4.95298135 | 6.07106864 | 2.88590319 | 4.80126205 |
| ENSMUSG00 | Olfr370 | 1095 | 0 | 0 | 0 | 0 | 0 | 0 |
| ENSMUSG00 | Gpr15 | 1289 | 0 | 0 | 0 | 0 | 0 | 0 |
| ENSMUSG00 | Smgc | 3499 | 0 | 0 | 0 | 0 | 0 | 0 |
| ENSMUSG00 | Kcnv2 | 5106 | 0 | 0 | 0 | 0 | 0 | 0 |
| ENSMUSG00 | Pcdhb13 | 3692 | 0 | 0 | 0.02394466 | 0 | 0 | 0 |
| ENSMUSG00 | 4931429P17R | 2004 | 0 | 0 | 0 | 0 | 0 | 0 |
| ENSMUSG00 | Kcne4 | 2529 | 0 | 0 | 0 | 0 | 1.49306121 | 0 |
| ENSMUSG00 | Zfp286 | 4543 | 1.22513037 | 0.13788768 | 0.80756174 | 0.39711693 | 0.71883955 | 0.85252009 |
| ENSMUSG00 | A530098C11l | 1421 | 0 | 0 | 0 | 0 | 0 | 0 |
| ENSMUSG00 | Lancl3 | 3986 | 0.69816449 | 0.93084698 | 1.60794462 | 0.66081015 | 0.57606295 | 1.13587309 |
| ENSMUSG00 | Gm5806 | 2906 | 0.01368048 | 0 | 0 | 0 | 0 | 0 |
| ENSMUSG00 | Olfr976 | 3179 | 0 | 0 | 0 | 0 | 0 | 0 |
| ENSMUSG00 | Lcn10 | 1100 | 0 | 0 | 0 | 0 | 0 | 0 |
| ENSMUSG00 | Gm973 | 3574 | 0.57842333 | 0.32357991 | 0.75442429 | 0.31296681 | 1.51337321 | 0.16025949 |
| ENSMUSG00 | Cstad | 1226 | 0.90795551 | 0.66816437 | 1.22582599 | 1.17722821 | 0.58268292 | 2.18019139 |
| ENSMUSG00 | Fam108b | 2471 | 33.7865273 | 32.800322 | 41.5722716 | 36.7391437 | 27.9395737 | 40.3545452 |
| ENSMUSG00 | Dnahc14 | 2379 | 0 | 0 | 0 | 0 | 0 | 0 |
| ENSMUSG00 | Gm7367 | 632 | 0 | 0 | 0 | 0 | 0 | 0 |
| ENSMUSG00 | Zfp768 | 4162 | 17.3560087 | 18.7095845 | 16.8757157 | 16.9400084 | 17.4705876 | 13.9191095 |
| ENSMUSG00 | B3gnt1 | 2033 | 32.6960961 | 30.7890744 | 35.7441367 | 29.6749525 | 35.7661513 | 23.3169321 |
| ENSMUSG00 | Als2cr11 | 780 | 0 | 0 | 0 | 0 | 0 | 0 |
| ENSMUSG00 | A730013G03 | 1550 | 0 | 0 | 0 | 0 | 0 | 0 |
| ENSMUSG00 | Atmin | 4873 | 7.07326115 | 8.16786399 | 5.71458262 | 5.83473242 | 13.2147109 | 5.1661178 |
| ENSMUSG00 | Defb9 | 308 | 0.77445741 | 0.31289698 | 0.57404991 | 0.23429899 | 0.66268021 | 0.61987817 |
| ENSMUSG00 | Odf3b | 1034 | 0.23068944 | 0 | 0.34198718 | 0.06979119 | 0 | 0 |
| ENSMUSG00 | Tgif1 | 3986 | 4.9769154 | 8.17211111 | 5.76642207 | 8.90735871 | 9.80587152 | 8.14270464 |
| ENSMUSG00 | Ctdspl | 5259 | 11.4677817 | 7.68747385 | 18.1547787 | 6.92275773 | 8.9264624 | 13.0123731 |
| ENSMUSG00 | Zbtb44 | 9063 | 6.05346608 | 6.17284115 | 8.72039008 | 8.25312571 | 8.33830548 | 8.74545974 |
| ENSMUSG00 | Flrt2 | 3293 | 0.90545431 | 0.3511918 | 0 | 3.69257486 | 2.78917329 | 0 |
| ENSMUSG00 | Gpr68 | 6228 | 0 | 0 | 0 | 0 | 0 | 0 |
| ENSMUSG00 | Rexo1 | 5208 | 5.32821148 | 5.59769537 | 5.94959523 | 7.23303658 | 8.28884676 | 7.66706449 |
| ENSMUSG00 | Cmya5 | 11837 | 0.02686862 | 0.07327498 | 0 | 0 | 0.11207956 | 0 |
| ENSMUSG00 | Fam180a | 1525 | 0 | 0 | 0 | 0 | 0 | 0 |
| ENSMUSG00 | AI837181 | 1691 | 8.76924555 | 10.0875225 | 10.2205326 | 11.3089791 | 5.85400172 | 9.40337947 |
| ENSMUSG00 | Dlk2 | 1970 | 0 | 0 | 0 | 0 | 0 | 0.09691496 |
| ENSMUSG00 | AI480653 | 3678 | 14.2030183 | 11.9483511 | 11.633329 | 14.8036447 | 13.637555 | 11.4867907 |
| ENSMUSG00 | Antxrl | 2109 | 0 | 0 | 0 | 0 | 0 | 0 |
| ENSMUSG00 | Fam132b | 2806 | 0.08500816 | 0.48083406 | 0.63010468 | 0.20574224 | 0.20003212 | 0 |
| ENSMUSG00 | Olfr139 | 947 | 0 | 0 | 0 | 0 | 0 | 0 |
| ENSMUSG00 | Arl4a | 6156 | 6.41925725 | 6.70038897 | 8.85329312 | 4.9410597 | 8.28888496 | 14.053794 |
| ENSMUSG00 | Gphn | 3372 | 20.3257558 | 15.5191211 | 20.0953219 | 16.6178574 | 15.8435989 | 18.7331152 |
| ENSMUSG00 | Dynlrb1 | 1087 | 54.1288971 | 55.1462135 | 63.5172758 | 55.3678475 | 36.6620051 | 87.2939006 |
| ENSMUSG00 | A530099J19R | 2771 | 0 | 0 | 0 | 0 | 0 | 0 |
| ENSMUSG00 | 8030462N17l | 3670 | 1.59238573 | 1.29985225 | 3.2759949 | 2.41857846 | 1.07058064 | 4.06518465 |
| ENSMUSG00 | Zfp30 | 6951 | 3.78051685 | 2.05889416 | 2.48639341 | 2.53835704 | 1.96735272 | 2.37784943 |
| ENSMUSG00 | Klhl34 | 1934 | 0 | 0.02491543 | 0 | 0 | 0 | 0 |
| ENSMUSG00 | 4932411E22l | 2661 | 0 | 0 | 0.01661099 | 0 | 0 | 0 |
| ENSMUSG00 | Inhbe | 2133 | 0 | 0 | 0 | 0 | 0 | 0 |
| ENSMUSG00 | Dlgap2 | 6402 | 0 | 0 | 0 | 0 | 0 | 0 |
| ENSMUSG00 | Rnf152 | 8526 | 51.3659833 | 29.354959 | 65.4473453 | 54.4785549 | 37.0817997 | 63.0810006 |
| ENSMUSG00 | Adamts12 | 6353 | 0 | 0 | 0 | 0 | 0 | 0 |
| ENSMUSG00 | Cldn4 | 1808 | 34.8520114 | 53.0370685 | 36.5496434 | 55.2007385 | 35.7015295 | 42.9937621 |
| ENSMUSG00 | Heatr8 | 5233 | 0 | 0.24862105 | 0.51525175 | 0 | 0 | 0.11466502 |
| ENSMUSG00 | Baiap3 | 4867 | 0 | 0 | 0 | 0 | 0 | 0 |
| ENSMUSG00 | Mbd3l2 | 2077 | 0 | 0 | 0 | 0 | 0 | 0 |
| ENSMUSG00 | Gm6776 | 899 | 0.57488459 | 0 | 0 | 0.04013576 | 0.11351808 | 0.21237205 |
| ENSMUSG00 | Olfr1396 | 2146 | 0.33345697 | 0 | 0 | 0 | 0 | 0 |
| ENSMUSG00 | Tspyl1 | 2695 | 47.9130986 | 50.0995959 | 46.5144441 | 36.4301459 | 40.6885646 | 47.3536316 |
| ENSMUSG00 | BC049715 | 1088 | 8.00225205 | 6.1561723 | 6.09400409 | 12.2373846 | 12.2407023 | 7.32003186 |
| ENSMUSG00 | Dmbt1 | 6291 | 0 | 0.32170252 | 0 | 0 | 0 | 0 |
| ENSMUSG00 | Slfnl1 | 1757 | 0 | 0 | 0 | 0.67769349 | 0.26137586 | 0 |
| ENSMUSG00 | Als2cr12 | 1564 | 0 | 0 | 0 | 0 | 0 | 0 |
| ENSMUSG00 | Rtp2 | 2419 | 0 | 0 | 0 | 0 | 0 | 0 |
| ENSMUSG00 | Mis18bp1 | 6565 | 0.02422268 | 0.44039399 | 0.11446022 | 0.06045735 | 0.16322222 | 0.03323642 |
| ENSMUSG00 | Olfr67 | 2202 | 0 | 0 | 0 | 0 | 0 | 0 |
| ENSMUSG00 | Fbxo28 | 4908 | 9.07215529 | 8.8754164 | 12.1582087 | 8.7411473 | 11.2075047 | 3.53992364 |
| ENSMUSG00 | Olfr616 | 1890 | 0 | 0 | 0 | 0 | 0 | 0 |
| ENSMUSG00 | Olfr629 | 1121 | 0 | 0 | 0 | 0 | 0 | 0 |
| ENSMUSG00 | Cltb | 3724 | 12.2767998 | 10.9726377 | 14.2433436 | 10.3479118 | 9.56401996 | 14.2818331 |
| ENSMUSG00 | Tmem41b | 6847 | 5.35337418 | 6.41125293 | 5.89400945 | 6.60300888 | 5.14213514 | 5.73217553 |
| ENSMUSG00 | Lxn | 1828 | 10.5043201 | 11.2294444 | 7.10904915 | 9.79031405 | 9.54651014 | 6.71421645 |
| ENSMUSG00 | Mmp10 | 1710 | 0 | 0 | 0 | 0 | 0 | 0 |
| ENSMUSG00 | Krtap3-1 | 603 | 0 | 0 | 0 | 0 | 0 | 0 |
| ENSMUSG00 | Acot10 | 1531 | 0.41547204 | 0.34621219 | 0.05774245 | 0.28281158 | 0.06665758 | 0.28503869 |
| ENSMUSG00 | Tyw3 | 7496 | 1.58576423 | 2.24347229 | 1.82798444 | 1.30446025 | 1.42269378 | 2.08489438 |
| ENSMUSG00 | Nccrp1 | 1525 | 1.09490504 | 0.50556267 | 0.28984815 | 0.52052786 | 0.40151902 | 0.41135522 |
| ENSMUSG00 | Mafa | 1079 | 0 | 0 | 0 | 0 | 0 | 0 |
| ENSMUSG00 | BC055004 | 4136 | 0 | 0.25631086 | 0.11755809 | 0 | 0.02467426 | 0.27037239 |
| ENSMUSG00 | Olfr1122 | 1079 | 0 | 0 | 0 | 0 | 0 | 0 |
| ENSMUSG00 | Zfp235 | 2671 | 3.67637727 | 6.69306254 | 7.71174049 | 12.2930215 | 10.2396621 | 4.07434708 |
| ENSMUSG00 | Frat2 | 2154 | 0.49832775 | 1.63305955 | 0.90291601 | 1.62486459 | 1.27921276 | 1.30421905 |
| ENSMUSG00 | Ankrd34c | 2473 | 0 | 0 | 0 | 0 | 0 | 0 |
| ENSMUSG00 | A430005L14R | 1410 | 12.7443101 | 15.1394283 | 16.8656678 | 14.3304574 | 14.294623 | 13.8501005 |
| ENSMUSG00 | BC029214 | 1839 | 12.5384332 | 12.4724016 | 14.3734106 | 12.9887512 | 10.9599883 | 17.560181 |
| ENSMUSG00 | Ddi1 | 1519 | 0 | 0 | 0 | 0 | 0 | 0 |
| ENSMUSG00 | Olfr791 | 938 | 0 | 0 | 0 | 0 | 0 | 0 |
| ENSMUSG00 | Apof | 1928 | 0 | 0 | 0 | 0 | 0 | 0 |
| ENSMUSG00 | Fgfbp3 | 1770 | 3.99800877 | 3.5663412 | 4.94461295 | 4.79055393 | 3.45941531 | 9.07613702 |
| ENSMUSG00 | 2810006K23l | 3001 | 5.02076878 | 6.23003651 | 4.47762755 | 4.97766292 | 3.46863734 | 6.87091879 |
| ENSMUSG00 | Cdcp2 | 2607 | 0 | 0 | 0 | 0 | 0 | 0 |
| ENSMUSG00 | Nr1h4 | 2867 | 0.47146367 | 0.60506171 | 0.97129966 | 0.16360885 | 0.64072185 | 1.85509405 |
| ENSMUSG00 | Krt83 | 1838 | 0.28118675 | 0.02621678 | 0 | 0.29446718 | 0.0555238 | 0.0148393 |
| ENSMUSG00 | D930020B18 | 3576 | 0.03335191 | 0 | 0 | 0.01009006 | 0 | 0 |
| ENSMUSG00 | Gm5454 | 1454 | 0.02734215 | 0 | 0 | 0 | 0 | 0.01875835 |
| ENSMUSG00 | Fbxo30 | 5193 | 18.7638519 | 18.057157 | 16.5980346 | 18.0861856 | 16.8319241 | 18.7555819 |
| ENSMUSG00 | Cd3eap | 1654 | 3.67750213 | 5.65185597 | 6.17329246 | 4.64659945 | 4.34989056 | 7.55246964 |
| ENSMUSG00 | Pea15b | 383 | 0 | 0 | 0 | 0 | 0 | 0 |
| ENSMUSG00 | Tssk6 | 1332 | 0.77600788 | 0.07235202 | 0.6968759 | 0.8668359 | 0.7661618 | 0.43000557 |
| ENSMUSG00 | Vmn1r74 | 914 | 0 | 0 | 0 | 0 | 0 | 0 |
| ENSMUSG00 | Trpt1 | 1135 | 15.6219774 | 11.8449492 | 15.8114082 | 13.447317 | 13.1724477 | 11.6548018 |
| ENSMUSG00 | Crry-ps | 482 | 0.08248025 | 0 | 0 | 0.07485901 | 0 | 0.05658639 |
| ENSMUSG00 | Gal3st3 | 2418 | 0.14797325 | 0.67755957 | 0.03656066 | 0 | 0.37984895 | 0.28199586 |
| ENSMUSG00 | 1600012P17R | 2855 | 0 | 0 | 0 | 0 | 0 | 0 |
| ENSMUSG00 | Olfr26 | 930 | 0 | 0 | 0 | 0 | 0 | 0 |
| ENSMUSG00 | Msl3l2 | 2138 | 0.18594706 | 0 | 0.28944144 | 1.23198749 | 0.07159922 | 0 |
| ENSMUSG00 | Tctex1d4 | 929 | 0 | 0 | 0.09516005 | 0 | 0.21970453 | 0.44038707 |
| ENSMUSG00 | Pdha2 | 2351 | 0 | 0 | 0 | 0 | 0 | 0 |
| ENSMUSG00 | Rps8 | 1876 | 1.56817994 | 1.00174373 | 1.67288425 | 0.88474096 | 1.16958111 | 1.67195283 |
| ENSMUSG00 | Rpsa-ps10 | 1030 | 1.5053046 | 1.9181011 | 1.75949084 | 0 | 1.04034359 | 1.45641278 |
| ENSMUSG00 | Gpr82 | 2493 | 0 | 0 | 0 | 0.54998704 | 0.26608218 | 0 |
| ENSMUSG00 | Zcchc5 | 2946 | 0 | 0 | 0 | 0 | 0 | 0 |
| ENSMUSG00 | 4930533K18R | 1252 | 1.23838957 | 0 | 0 | 0 | 0.32604713 | 0.23963341 |
| ENSMUSG00 | Yipf6 | 6304 | 31.2355481 | 29.3903664 | 34.2802682 | 26.2831137 | 32.9356477 | 38.7703113 |
| ENSMUSG00 | Ccdc144b | 4949 | 0 | 0 | 0 | 0 | 0 | 0 |
| ENSMUSG00 | Olfr1388 | 935 | 0 | 0 | 0 | 0 | 0 | 0 |
| ENSMUSG00 | Zfp828 | 4145 | 9.06367409 | 7.60287893 | 10.5465917 | 9.81048593 | 11.7809992 | 6.10636073 |
| ENSMUSG00 | Ubl | 4276 | 0.50205705 | 0.27045711 | 0.12404633 | 0 | 0.25059726 | 0.21049184 |
| ENSMUSG00 | Ppp1r2 | 4434 | 48.3987559 | 30.4941716 | 43.2548031 | 35.056709 | 24.2012784 | 51.0738227 |
| ENSMUSG00 | Olfr736 | 938 | 0 | 0 | 0 | 0 | 0 | 0 |
| ENSMUSG00 | Ubiad1 | 3133 | 22.7772383 | 22.3321781 | 17.0994682 | 23.7130321 | 17.0847968 | 7.44328649 |
| ENSMUSG00 | 4922502D21l | 1576 | 0 | 0 | 0 | 0 | 0 | 0 |
| ENSMUSG00 | Bola2 | 2142 | 2.43135759 | 3.12694466 | 2.99218825 | 2.13931823 | 1.64370678 | 5.71723297 |
| ENSMUSG00 | Olfr937 | 935 | 0 | 0 | 0 | 0 | 0 | 0 |
| ENSMUSG00 | BC025446 | 1468 | 0 | 0.29542097 | 0 | 0.02457905 | 0 | 0 |
| ENSMUSG00 | Fcgbp | 9137 | 1.09211181 | 0.4851869 | 0.74016438 | 0.56470749 | 0.79859601 | 0.32238821 |
| ENSMUSG00 | D19Wsu162e | 4636 | 23.3507708 | 17.8152636 | 20.0128707 | 13.1610736 | 24.3464934 | 10.2426978 |
| ENSMUSG00 | BC048609 | 542 | 0 | 0 | 0 | 0 | 0 | 0 |
| ENSMUSG00 | Samd9l | 5753 | 57.4944547 | 53.8317858 | 55.365632 | 86.0374564 | 69.8563768 | 90.5804376 |
| ENSMUSG00 | Fbxo40 | 5867 | 0.04743282 | 0 | 0.01506795 | 0.28904996 | 0 | 0.02789293 |
| ENSMUSG00 | Rnf150 | 10319 | 0 | 0.00466968 | 0.02570124 | 0.05594658 | 0.08406322 | 0.07400813 |
| ENSMUSG00 | Zc3hav1l | 6532 | 0.24345059 | 0.25081737 | 0.2639118 | 0.48610225 | 0.18748209 | 0.02087771 |
| ENSMUSG00 | Utf1 | 1798 | 0 | 0 | 0 | 0 | 0 | 0 |
| ENSMUSG00 | Fancb | 3080 | 1.1487785 | 2.90996047 | 0.50229367 | 0.82004647 | 1.55729848 | 1.46114139 |
| ENSMUSG00 | Hs3st3a1 | 3937 | 0.06058747 | 0.257027 | 0 | 0.54989146 | 0.11664653 | 0 |
| ENSMUSG00 | Lrrc49 | 7717 | 0.53062259 | 0.22479097 | 0.9050008 | 0.92110442 | 0.32399798 | 0.38171064 |
| ENSMUSG00 | Atg16l2 | 4186 | 9.37378387 | 8.05793347 | 4.96293985 | 12.3520234 | 13.3477978 | 7.23240553 |
| ENSMUSG00 | Ankfn1 | 2561 | 0 | 0 | 0 | 0 | 0 | 0 |
| ENSMUSG00 | Phf13 | 3201 | 7.27794801 | 9.58911704 | 11.0470086 | 4.4073975 | 12.6091419 | 7.36185202 |
| ENSMUSG00 | Lix1 | 3765 | 10.0101449 | 10.75076 | 9.89698636 | 11.4810862 | 9.98843001 | 11.1706491 |
| ENSMUSG00 | Flrt1 | 5857 | 7.88729185 | 11.2629741 | 3.01873608 | 8.36595808 | 6.49917643 | 6.48686571 |
| ENSMUSG00 | Slc38a9 | 3034 | 9.7750786 | 15.6597996 | 9.17836556 | 13.557525 | 13.9422761 | 11.4438417 |
| ENSMUSG00 | Sned1 | 9811 | 0.3890048 | 0.23083914 | 0.19372941 | 0.3604159 | 1.50827123 | 0.5977013 |
| ENSMUSG00 | Olfr685 | 1087 | 0 | 0 | 0 | 0 | 0 | 0 |
| ENSMUSG00 | Gjb1 | 1703 | 3.36159084 | 8.1772647 | 1.40158516 | 4.1739065 | 2.42697384 | 2.24218996 |
| ENSMUSG00 | Cd300lf | 5142 | 0 | 0 | 0 | 0 | 0 | 0 |
| ENSMUSG00 | Oog4 | 2844 | 0 | 0 | 0 | 0 | 0 | 0 |
| ENSMUSG00 | Akap10 | 7789 | 10.0702995 | 6.73088318 | 6.36156965 | 8.54684454 | 7.78923621 | 7.63017575 |
| ENSMUSG00 | 9330154K18l | 3720 | 0 | 0.20725352 | 0 | 0.01939895 | 0 | 0.10997838 |
| ENSMUSG00 | Ccdc88b | 4967 | 0.05602745 | 0 | 0 | 0 | 0.13355001 | 0.13727924 |
| ENSMUSG00 | Tigd4 | 3458 | 0 | 0 | 0 | 0 | 0 | 0 |
| ENSMUSG00 | Trim16 | 7682 | 0.7348709 | 1.14162099 | 0.93213988 | 1.29636088 | 1.38824688 | 0.23433041 |
| ENSMUSG00 | Gm6484 | 860 | 1.15568257 | 0.22412299 | 0.51397492 | 1.09085251 | 0.65266295 | 1.07829969 |

| | | | | | | | |
|---|---|---|---|---|---|---|---|
| ENSMUSG00 Pygo2 | 3819 | 10.5036082 | 13.437696 | 12.176051 | 10.9124801 | 13.9624411 | 8.39164732 |
| ENSMUSG00 A730018C14 | 3885 | 0 | 0 | 0 | 0 | 0 | 0 |
| ENSMUSG00 Cdca4 | 3079 | 5.95234703 | 5.68531083 | 8.29771525 | 5.44922076 | 5.23687391 | 4.60630478 |
| ENSMUSG00 BC051628 | 1139 | 0.55846154 | 0.12691776 | 0 | 0.12671482 | 0 | 0 |
| ENSMUSG00 Diras2 | 4346 | 0 | 0 | 0 | 0 | 0 | 0 |
| ENSMUSG00 Bri3 | 1642 | 1.01688805 | 1.0564628 | 0.75374641 | 1.31846691 | 0.62151493 | 1.26240718 |
| ENSMUSG00 Bex4 | 834 | 0.19067377 | 1.21332768 | 1.85499341 | 0.34611074 | 0.42827893 | 2.48545873 |
| ENSMUSG00 Stx19 | 2810 | 0.69324503 | 0.0171482 | 0.4561756 | 1.0400874 | 0 | 0.6406143 |
| ENSMUSG00 Foxi1 | 2355 | 975.6552 | 888.021906 | 1296.56863 | 1034.90352 | 891.650708 | 842.595257 |
| ENSMUSG00 Lonp2 | 5658 | 32.8906688 | 27.6871904 | 31.7334546 | 29.7240031 | 32.7730381 | 29.4969101 |
| ENSMUSG00 Gimap6 | 3254 | 1.99143925 | 1.70296277 | 1.50780718 | 2.19552699 | 3.48120941 | 2.95042195 |
| ENSMUSG00 Olfr770 | 935 | 0 | 0 | 0 | 0 | 0 | 0 |
| ENSMUSG00 Gpr157 | 5529 | 0.1150457 | 0.13944349 | 0 | 0 | 0.13843292 | 0.04439713 |
| ENSMUSG00 A4galt | 2168 | 33.0256552 | 48.8976811 | 30.9494454 | 40.2760829 | 34.4569254 | 31.1494496 |
| ENSMUSG00 Usp14 | 43693 | 4.33467853 | 4.68376673 | 4.59489554 | 5.33059294 | 3.54906177 | 6.55320447 |
| ENSMUSG00 Cxcr5 | 2612 | 0.24352515 | 0.20292912 | 0.06769042 | 0.16576743 | 0.21488903 | 0.44900823 |
| ENSMUSG00 Rell1 | 5351 | 1.62706975 | 5.58317962 | 1.50340785 | 2.98716982 | 4.17670578 | 2.47719579 |
| ENSMUSG00 Klk9 | 1470 | 0 | 0 | 0 | 0 | 0 | 0 |
| ENSMUSG00 Tnrc6b | 17020 | 5.62930129 | 4.71105991 | 5.93426623 | 6.54861548 | 6.0739975 | 5.16968192 |
| ENSMUSG00 Serpinb6d | 1456 | 0 | 0 | 0 | 0 | 0 | 0 |
| ENSMUSG00 Ang2 | 437 | 0 | 0 | 0 | 0 | 0 | 0 |
| ENSMUSG00 Ripply2 | 498 | 0 | 0 | 0 | 0 | 0 | 0 |
| ENSMUSG00 Ccr4 | 4199 | 0 | 0 | 0 | 0 | 0 | 0 |
| ENSMUSG00 Sstr2 | 1886 | 0 | 0 | 0.16405774 | 0 | 0 | 0 |
| ENSMUSG00 Gm8566 | 1120 | 0.07099193 | 0.17209444 | 0.11839779 | 0.22551278 | 0.04555926 | 0.14611414 |
| ENSMUSG00 Tshz2 | 6370 | 0.12482098 | 0.12859804 | 0.08326878 | 0.02832186 | 0.23230218 | 0.04281733 |
| ENSMUSG00 Ankrd16 | 5870 | 13.82976 | 14.1439932 | 18.0948942 | 18.0410223 | 27.1213446 | 16.0859968 |
| ENSMUSG00 Pcdhb16 | 5267 | 1.72849915 | 0 | 0.80565349 | 0.58915053 | 0.7653472 | 0.67837056 |
| ENSMUSG00 Npm2 | 988 | 0 | 0.0487717 | 0.62634191 | 0.36520288 | 0 | 0.35887683 |
| ENSMUSG00 Trappc9 | 6350 | 10.0672146 | 11.0487338 | 9.13272725 | 9.20518301 | 11.6115139 | 8.27688659 |
| ENSMUSG00 Gm9821 | 311 | 0 | 0 | 0 | 0.46407774 | 0.3281439 | 0 |
| ENSMUSG00 Gm5607 | 5052 | 0 | 0 | 0 | 0 | 0 | 0 |
| ENSMUSG00 4930483J18R | 1329 | 0 | 0 | 0 | 0 | 0 | 0 |
| ENSMUSG00 B930007M17 | 2428 | 0 | 0.61523052 | 0.50974119 | 0.57957156 | 0 | 0.11233377 |
| ENSMUSG00 Marcksl1 | 1603 | 4.04250987 | 7.96594334 | 3.47438067 | 5.71730462 | 6.78017346 | 5.90411717 |
| ENSMUSG00 Gp5 | 2194 | 0 | 0 | 0 | 0 | 0 | 0 |
| ENSMUSG00 Kcna3 | 1967 | 0 | 0 | 0 | 0 | 0 | 0 |
| ENSMUSG00 Olfr186 | 1004 | 0 | 0 | 0 | 0 | 0 | 0 |
| ENSMUSG00 Stbd1 | 2337 | 0.34022662 | 1.05156549 | 0.24588103 | 0 | 0.34934618 | 0 |
| ENSMUSG00 Rpl9-ps7 | 578 | 0 | 0 | 0 | 0 | 0 | 0 |
| ENSMUSG00 Olfr1093 | 1629 | 0 | 0 | 0 | 0 | 0 | 0 |
| ENSMUSG00 Kcna1 | 8968 | 0 | 0 | 0 | 0 | 0 | 0 |
| ENSMUSG00 Synb | 4052 | 0 | 0 | 0.03272595 | 0 | 0 | 0.01346231 |
| ENSMUSG00 9230110F15I | 1132 | 0 | 0 | 0 | 0 | 0 | 0 |
| ENSMUSG00 Palm3 | 3606 | 1.43322586 | 0.74831968 | 0.50257226 | 0.63038514 | 2.26406548 | 1.35389918 |
| ENSMUSG00 4933428G20I | 1422 | 0 | 0 | 0 | 0 | 0 | 0 |
| ENSMUSG00 Ino80c | 3105 | 16.9648991 | 13.2221735 | 23.6597305 | 20.1036834 | 15.6612355 | 16.4262723 |
| ENSMUSG00 C2cd4a | 1350 | 0 | 0 | 0 | 0 | 0 | 0 |
| ENSMUSG00 Fam69c | 2774 | 0 | 0 | 0 | 0 | 0 | 0 |
| ENSMUSG00 Cypt4 | 642 | 0 | 0 | 0 | 0 | 0 | 0 |
| ENSMUSG00 Prrg1 | 10315 | 1.42603275 | 1.19590201 | 0.50565366 | 1.80497673 | 1.58792696 | 1.24276107 |
| ENSMUSG00 Gigyf2 | 12332 | 6.5990487 | 6.50978046 | 7.11846093 | 9.58228153 | 9.01195642 | 7.96653021 |
| ENSMUSG00 Hes5 | 1439 | 0 | 0 | 0 | 0 | 0 | 0 |
| ENSMUSG00 Catsper4 | 2389 | 0.41602638 | 0.44374287 | 0.37004473 | 0.09062046 | 0.34174216 | 0.75350615 |
| ENSMUSG00 Tmem196 | 1364 | 0 | 0 | 0 | 0 | 0 | 0 |
| ENSMUSG00 Timm8a1 | 1419 | 2.32537342 | 1.25644705 | 3.02154952 | 1.62737903 | 1.36645686 | 4.59382578 |
| ENSMUSG00 Zfp473 | 13029 | 0 | 0.14053917 | 0.08820692 | 0 | 0 | 0.03977421 |
| ENSMUSG00 Krt35 | 2690 | 0 | 0 | 0 | 0 | 0 | 0 |
| ENSMUSG00 Neurod4 | 3342 | 0 | 0 | 0 | 0 | 0 | 0 |
| ENSMUSG00 Tmem229a | 5156 | 56.3099834 | 98.3822103 | 24.9642682 | 76.2229692 | 79.9637542 | 32.961264 |
| ENSMUSG00 Rgmb | 5744 | 18.3896782 | 21.3500166 | 24.471076 | 20.5411361 | 21.8443346 | 16.0067564 |
| ENSMUSG00 Eno4 | 1813 | 0.08771204 | 0.07973487 | 0.0243805 | 0 | 0.28144719 | 0.25574676 |
| ENSMUSG00 Fcrl5 | 1779 | 0 | 0 | 0 | 0 | 0 | 0 |
| ENSMUSG00 4932418E24I | 3254 | 0 | 0 | 0 | 0 | 0 | 0 |
| ENSMUSG00 Isg20l2 | 2907 | 3.88392035 | 2.66873656 | 2.63051904 | 3.26438862 | 8.26743138 | 3.67790113 |
| ENSMUSG00 Anxa2 | 1539 | 3.69397902 | 2.00385464 | 3.38909518 | 4.43112828 | 2.71875427 | 3.03051547 |
| ENSMUSG00 Zbtb33 | 5390 | 2.51514264 | 1.62707467 | 2.02557611 | 0.67611994 | 1.61883307 | 3.43589611 |
| ENSMUSG00 Ldlrad3 | 6230 | 0.57431673 | 0.04640749 | 0.29798995 | 0.86874907 | 0.49142577 | 1.03757456 |
| ENSMUSG00 Fpr-rs4 | 971 | 0.08188565 | 0 | 0 | 0 | 0 | 0 |
| ENSMUSG00 Cyb5r2 | 2727 | 0 | 0.07068052 | 0.04862689 | 0 | 0 | 0.08001361 |
| ENSMUSG00 Olfr1349 | 1053 | 0 | 0 | 0 | 0 | 0 | 0 |
| ENSMUSG00 Pirt | 4198 | 0 | 0 | 0 | 0 | 0 | 0 |
| ENSMUSG00 Arf1 | 3364 | 79.948224 | 83.7103353 | 93.5150169 | 85.0673886 | 66.4526018 | 85.4805356 |
| ENSMUSG00 H1fnt | 1325 | 0 | 0 | 0 | 0 | 0 | 0 |
| ENSMUSG00 Odz4 | 20988 | 0.32769669 | 0.20433549 | 0.41489247 | 0.17707502 | 0.20179099 | 0.30019257 |
| ENSMUSG00 Olfr731 | 980 | 0 | 0 | 0 | 0 | 0 | 0 |
| ENSMUSG00 Gm4737 | 2080 | 0 | 0 | 0.53127215 | 0 | 0 | 0.09178965 |
| ENSMUSG00 Lmod1 | 4399 | 0 | 0.07667768 | 0.17081867 | 0.09022562 | 0.04639816 | 0.01860057 |
| ENSMUSG00 Taf13 | 2839 | 20.0807887 | 18.3478492 | 19.664293 | 16.5985031 | 15.0976244 | 20.5784703 |
| ENSMUSG00 Olfr19 | 1019 | 0 | 0 | 0 | 0 | 0 | 0 |
| ENSMUSG00 4632415L05F | 1076 | 6.57665012 | 9.0013707 | 13.9671251 | 6.97496772 | 7.39787605 | 11.9389917 |
| ENSMUSG00 Kctd12 | 3803 | 0 | 0 | 0 | 0 | 0 | 0 |
| ENSMUSG00 Tmem72 | 3859 | 66.3552863 | 51.4579757 | 73.650648 | 52.7532768 | 47.6545889 | 47.7853941 |
| ENSMUSG00 Rbm15 | 3817 | 2.69758173 | 4.51945148 | 3.462497 | 4.0269717 | 5.56116645 | 4.33736076 |
| ENSMUSG00 Arid4a | 5722 | 10.0187702 | 11.6718645 | 14.4378267 | 13.2107451 | 14.1076069 | 9.21865647 |
| ENSMUSG00 Entpd1 | 3846 | 0 | 0.01252898 | 0.17239408 | 0 | 0.21227822 | 0.95028644 |
| ENSMUSG00 Col6a3 | 12598 | 0 | 0 | 0.04561232 | 0 | 0 | 0 |
| ENSMUSG00 Dmrt2 | 2672 | 25.2340176 | 10.0628872 | 22.8949666 | 20.2691425 | 37.7923644 | 24.0082154 |
| ENSMUSG00 Nat8l | 6526 | 0 | 0 | 0 | 0 | 0 | 0 |
| ENSMUSG00 Nwd1 | 8977 | 0 | 0 | 0 | 0 | 0 | 0 |
| ENSMUSG00 Gm12663 | 1708 | 4.58538036 | 0.67709286 | 0.93165477 | 3.42230165 | 0.74687318 | 2.09190734 |
| ENSMUSG00 Olfr49 | 1029 | 0 | 0 | 0 | 0 | 0 | 0 |
| ENSMUSG00 Mll2 | 19750 | 3.48036586 | 3.61581303 | 2.92739295 | 3.71782079 | 3.73848941 | 3.7922106 |
| ENSMUSG00 1700014N06 | 1048 | 0 | 0 | 0 | 0 | 0 | 0 |
| ENSMUSG00 Selplg | 4229 | 0 | 0 | 0 | 0 | 0 | 0 |
| ENSMUSG00 Gm7204 | 872 | 6.88426325 | 6.02330527 | 5.27177944 | 6.49642315 | 3.27692322 | 8.75791169 |
| ENSMUSG00 Mcmbp | 5644 | 13.242435 | 13.5577729 | 16.1880244 | 12.0763611 | 12.4672878 | 11.0181037 |
| ENSMUSG00 Olfr1324 | 1151 | 0 | 0 | 0 | 0 | 0 | 0 |
| ENSMUSG00 Tmem81 | 1738 | 1.34958191 | 1.85759012 | 1.06816882 | 0.8304268 | 1.82027348 | 1.28683568 |
| ENSMUSG00 Asb8 | 3128 | 22.4831985 | 19.256091 | 24.9271263 | 20.578762 | 24.3712933 | 20.7350039 |
| ENSMUSG00 Gm7075 | 338 | 0 | 0 | 0 | 0 | 0 | 0 |
| ENSMUSG00 Bend7 | 4643 | 2.23480086 | 1.49447505 | 1.12337228 | 1.75630887 | 2.19799164 | 1.29235853 |
| ENSMUSG00 Gm8181 | 1437 | 0.02766561 | 0 | 0.03075981 | 0.12554643 | 0 | 0.01898026 |
| ENSMUSG00 Muc6 | 8772 | 0 | 0 | 0 | 0 | 0 | 0.03109284 |
| ENSMUSG00 Olfr1153 | 956 | 0 | 0 | 0 | 0 | 0 | 0 |
| ENSMUSG00 Efcab4a | 2752 | 4.60828426 | 3.48441206 | 3.565701 | 2.03223725 | 5.72934234 | 2.46779985 |
| ENSMUSG00 Dnajb8 | 988 | 0 | 0 | 0 | 0 | 0 | 0 |
| ENSMUSG00 A630023P12I | 1434 | 0 | 0 | 0 | 0 | 0 | 0 |
| ENSMUSG00 Gpr85 | 6082 | 0 | 0.0079228 | 0 | 0.0059326 | 0 | 0.00896897 |
| ENSMUSG00 Naga | 2722 | 0 | 0 | 0 | 0 | 0 | 0 |
| ENSMUSG00 Amigo2 | 2869 | 0.96998384 | 2.0826486 | 2.89646096 | 1.54691233 | 3.4681573 | 1.77774749 |
| ENSMUSG00 Mfap1b | 3355 | 7.58375783 | 9.04842282 | 10.1578602 | 9.80829347 | 10.0836027 | 10.1050303 |
| ENSMUSG00 Olfr1212 | 1869 | 0 | 0 | 0 | 0 | 0 | 0 |
| ENSMUSG00 Fbxo43 | 4030 | 0.28608162 | 0 | 0 | 0.03581344 | 0.37984895 | 0 |
| ENSMUSG00 H2-M10.4 | 1404 | 0 | 0 | 0 | 0 | 0 | 0 |
| ENSMUSG00 Fbxo10 | 5752 | 2.10803574 | 1.35712859 | 2.58202699 | 2.40881521 | 2.80325709 | 1.71651937 |
| ENSMUSG00 Rnf149 | 2831 | 34.4332173 | 42.2801562 | 44.4360456 | 39.9821172 | 32.8760542 | 39.3078517 |
| ENSMUSG00 Ovch2 | 1937 | 0 | 0 | 0 | 0 | 0 | 0 |
| ENSMUSG00 Gng7 | 6897 | 1.07213562 | 0.18165108 | 0.21149207 | 0.70102856 | 0.91739461 | 0.53386636 |
| ENSMUSG00 D630032N06 | 445 | 0 | 0 | 0 | 0 | 0 | 0 |
| ENSMUSG00 A930001N09I | 8674 | 12.434473 | 8.94954557 | 15.7769087 | 10.0999774 | 14.3419765 | 15.9641853 |
| ENSMUSG00 Bcl11b | 10299 | 0 | 0.07953874 | 0 | 0.03503451 | 0 | 0 |
| ENSMUSG00 Gm4879 | 2620 | 0.06069539 | 0.05517532 | 0.08435466 | 0.09640241 | 0 | 0.02082034 |
| ENSMUSG00 Dip2c | 8336 | 1.74550214 | 3.56658287 | 1.35744624 | 2.39796697 | 3.31157262 | 2.68950977 |
| ENSMUSG00 Rbm33 | 5591 | 9.15136888 | 13.0485197 | 9.52660004 | 9.98371005 | 13.625895 | 11.9957678 |
| ENSMUSG00 Syngr2 | 1435 | 63.1100938 | 91.3695534 | 74.4500729 | 48.8050512 | 62.974015 | 66.1243693 |
| ENSMUSG00 Sacs | 20700 | 0.00192056 | 0 | 0 | 0 | 0 | 0 |
| ENSMUSG00 Zfp738 | 5555 | 3.35649331 | 4.53672534 | 5.9996741 | 9.4703188 | 6.20032469 | 5.45983778 |
| ENSMUSG00 Dixu7 | 1312 | 0.3636172 | 0.03672747 | 0 | 0 | 0 | 0 |
| ENSMUSG00 Tas2r126 | 929 | 0 | 0.20747661 | 0.23790012 | 0 | 0.05492613 | 0 |
| ENSMUSG00 Frmd6 | 4678 | 2.16709011 | 1.24637869 | 1.85197974 | 0.78673975 | 2.47605543 | 1.9473556 |
| ENSMUSG00 Olfr1417 | 947 | 0 | 0 | 0 | 0 | 0 | 0 |
| ENSMUSG00 Krtap4-13 | 818 | 0 | 0 | 0 | 0 | 0 | 0 |
| ENSMUSG00 Olfr148 | 1016 | 0 | 0 | 0 | 0 | 0 | 0 |
| ENSMUSG00 Slitrk3 | 3729 | 0 | 0 | 0 | 0 | 0.04105099 | 0 |
| ENSMUSG00 Ankrd46 | 2475 | 72.6039482 | 68.4928903 | 71.1337134 | 66.2305973 | 63.5819568 | 59.0895417 |
| ENSMUSG00 Pskh1 | 3252 | 22.9339734 | 19.3516278 | 18.8387929 | 16.0882425 | 23.081119 | 12.4799087 |
| ENSMUSG00 Gm4884 | 1444 | 0 | 0 | 0 | 0 | 0 | 0 |
| ENSMUSG00 Gm9823 | 325 | 0 | 0 | 0 | 0 | 0 | 0 |
| ENSMUSG00 Ckap2l | 3128 | 0 | 1.0783411 | 0 | 0.50754794 | 0 | 0 |
| ENSMUSG00 Mfsd6l | 2058 | 4.21122194 | 5.29161124 | 2.77066946 | 4.48833984 | 5.67786203 | 3.7638472 |
| ENSMUSG00 Ric3 | 9489 | 4.66306774 | 2.36641185 | 6.66591183 | 2.03434438 | 4.90958806 | 5.34340335 |
| ENSMUSG00 Lhfp | 2225 | 0.51816132 | 0.02165683 | 1.8872697 | 0 | 0 | 0 |
| ENSMUSG00 Gm8258 | 622 | 0 | 0.15494033 | 0.28425623 | 0 | 0 | 0 |
| ENSMUSG00 Ppyr1 | 3625 | 0 | 0 | 0 | 0 | 0 | 0 |
| ENSMUSG00 Pcdhb18 | 9970 | 0.29108827 | 0.47848122 | 0.32807787 | 0.29676305 | 0.40432133 | 0.6209973 |
| ENSMUSG00 Pou4f1 | 4587 | 0 | 0.10505002 | 0 | 0.00786615 | 0 | 0 |
| ENSMUSG00 2010305A19I | 3983 | 4.62133755 | 4.92389705 | 7.79053322 | 6.86673104 | 4.90662866 | 9.14176334 |
| ENSMUSG00 Anxa1 | 1579 | 0.30213158 | 1.25119957 | 0.50388421 | 0.6169824 | 0.32315628 | 0.01727336 |
| ENSMUSG00 Olfr1496 | 1066 | 0 | 0 | 0 | 0 | 0 | 0 |
| ENSMUSG00 Gm9825 | 1000 | 0.03975548 | 0.19274577 | 0.04420184 | 0.03608205 | 0 | 5.53675177 |
| ENSMUSG00 Omd | 1553 | 0 | 0 | 0.05692446 | 0 | 0 | 0 |
| ENSMUSG00 Pdp2 | 3483 | 3.78949742 | 2.82228946 | 3.94681975 | 2.53807089 | 2.63702201 | 2.2787597 |
| ENSMUSG00 Fgfbp1 | 1152 | 0.89725911 | 1.08754123 | 0.15347862 | 0.06264244 | 0.97446204 | 1.42055413 |
| ENSMUSG00 F2r | 3305 | 2.02085347 | 0.5686146 | 2.00613508 | 1.40834607 | 5.89775357 | 2.31896328 |
| ENSMUSG00 Foxi2 | 2589 | 0 | 0 | 0 | 0 | 0 | 0 |
| ENSMUSG00 Olfr1394 | 938 | 0 | 0 | 0 | 0 | 0 | 0 |
| ENSMUSG00 Socs4 | 1921 | 21.4402279 | 17.282904 | 21.2840733 | 24.4553993 | 16.2561905 | 24.6195859 |
| ENSMUSG00 Scrt1 | 7172 | 0 | 0 | 0 | 0 | 0 | 0 |
| ENSMUSG00 Osr1 | 2007 | 0 | 0 | 0 | 0 | 0 | 0.01358976 |
| ENSMUSG00 Fam171b | 5736 | 0 | 0 | 0.18494495 | 0 | 0 | 0 |
| ENSMUSG00 Olfr843 | 1106 | 0 | 0 | 0 | 0 | 0 | 0 |
| ENSMUSG00 Gm5065 | 1434 | 0 | 0 | 0 | 0 | 0 | 0 |
| ENSMUSG00 Tpng | 2032 | 0 | 0 | 0 | 0 | 0 | 0 |
| ENSMUSG00 Prss54 | 1432 | 0 | 0 | 0 | 0 | 0 | 0 |
| ENSMUSG00 Gli2 | 6853 | 0 | 0 | 0 | 0 | 0 | 0 |
| ENSMUSG00 B330016D10 | 1696 | 3.34807639 | 5.02296835 | 1.53587087 | 3.22995923 | 1.53259433 | 7.34070091 |
| ENSMUSG00 Zfp407 | 10280 | 2.64907628 | 3.27180317 | 2.32618649 | 1.78304267 | 2.11948079 | 2.48071865 |
| ENSMUSG00 Gm697 | 2858 | 0 | 0 | 0 | 0 | 0 | 0 |
| ENSMUSG00 Mlf1 | 1847 | 0 | 0 | 0 | 0.11721292 | 0 | 0.25103891 |
| ENSMUSG00 Ranbp3l | 3011 | 14.1672636 | 16.8036414 | 9.29251632 | 8.72392176 | 14.2690828 | 12.5186156 |
| ENSMUSG00 Olfr686 | 1017 | 0 | 0 | 0 | 0 | 0 | 0 |
| ENSMUSG00 1810026J23R | 3144 | 39.9957331 | 36.216464 | 47.7025615 | 39.364317 | 33.9851244 | 31.7510113 |
| ENSMUSG00 Nupl2 | 3964 | 3.84115768 | 1.87202626 | 5.93223524 | 6.08042526 | 3.03789725 | 3.6879936 |
| ENSMUSG00 Cyp4f16 | 2922 | 4.53065538 | 5.52465521 | 5.89962997 | 4.44542643 | 5.01183089 | 4.57377592 |
| ENSMUSG00 2210020M01 | 1810 | 97.5656599 | 93.5242935 | 111.261656 | 103.780731 | 77.4416875 | 95.6722014 |
| ENSMUSG00 Ccdc57 | 4103 | 4.9512675 | 3.32360788 | 1.88528456 | 3.40330276 | 7.67323273 | 3.04454906 |
| ENSMUSG00 Msx1 | 1933 | 1.56307114 | 0.2492832 | 1.57782057 | 1.21331241 | 2.19099286 | 0.60673021 |
| ENSMUSG00 Sprr1b | 953 | 0 | 0 | 0 | 0 | 0 | 0 |
| ENSMUSG00 Olfr1448 | 944 | 0 | 0 | 0 | 0 | 0 | 0 |
| ENSMUSG00 6530418L21F | 4778 | 0.58243693 | 0.24204157 | 0 | 0.01510341 | 0.2776655 | 0 |
| ENSMUSG00 Olfr564 | 950 | 0 | 0 | 0 | 0 | 0 | 0 |
| ENSMUSG00 Gm766 | 2286 | 0 | 0 | 0 | 0 | 0 | 0 |
| ENSMUSG00 4732415M23 | 515 | 1.31231683 | 1.40348861 | 1.71657643 | 2.03180445 | 1.88252649 | 3.49539066 |
| ENSMUSG00 Cxcr1 | 1123 | 0 | 0 | 0 | 0 | 0 | 0 |
| ENSMUSG00 Mypop | 2207 | 0.3422538 | 0.58950337 | 0.38053241 | 0.09809346 | 0.20808219 | 0.69206153 |
| ENSMUSG00 Bdnf | 6208 | 0 | 0 | 0 | 0 | 0 | 0 |
| ENSMUSG00 Zdhhc22 | 1016 | 0 | 0 | 0 | 0 | 0 | 0 |
| ENSMUSG00 Gm7461 | 826 | 0.62569158 | 0 | 0.48161814 | 0.65524294 | 0 | 0.26416116 |
| ENSMUSG00 Zbtb8b | 6016 | 0.86568616 | 0.21626229 | 0.8816857 | 0.17993041 | 1.39949335 | 0.46243571 |
| ENSMUSG00 Fitm2 | 3965 | 8.68303811 | 14.4498562 | 12.5526545 | 13.91411 | 12.5346003 | 6.44548221 |
| ENSMUSG00 8430408G22I | 1315 | 1.48138292 | 4.21402346 | 1.00840707 | 0 | 0.54324659 | 2.07411705 |
| ENSMUSG00 Nrip1 | 15604 | 9.26116199 | 9.30130503 | 7.78432867 | 12.4613008 | 11.9554236 | 9.42831352 |
| ENSMUSG00 1110034B05I | 2811 | 6.54812788 | 9.10245492 | 7.68932808 | 6.63622093 | 6.26257547 | 10.6148898 |
| ENSMUSG00 Mmgt2 | 4612 | 8.7320689 | 9.47638834 | 11.1271335 | 8.36333599 | 8.19829672 | 11.1771614 |
| ENSMUSG00 Cd300e | 2084 | 0 | 0 | 0 | 0 | 0 | 0 |
| ENSMUSG00 Defb15 | 343 | 0 | 0.84291152 | 0 | 0 | 1.48764944 | 0 |
| ENSMUSG00 Olfr938 | 1118 | 0 | 0 | 0 | 0 | 0 | 0 |
| ENSMUSG00 Duxbl | 4649 | 0 | 0 | 0 | 0 | 0 | 0 |
| ENSMUSG00 Tmem136 | 3497 | 0.9208447 | 0.12401429 | 0.26543858 | 0.42303798 | 1.31323244 | 0.37437309 |
| ENSMUSG00 Olfr453 | 953 | 0 | 0 | 0 | 0 | 0 | 0 |
| ENSMUSG00 Adam26a | 2487 | 0 | 0 | 0 | 0 | 0 | 0 |
| ENSMUSG00 Fbxl13 | 3737 | 0.03191502 | 0 | 0 | 0 | 0 | 0 |
| ENSMUSG00 Cxcr6 | 1891 | 0 | 0 | 0 | 0 | 0 | 0 |
| ENSMUSG00 Nkx1-2 | 3263 | 0 | 0 | 0 | 0 | 0 | 0 |
| ENSMUSG00 Amica1 | 2446 | 0 | 0 | 0 | 0 | 0 | 0 |
| ENSMUSG00 Phldb1 | 8540 | 0.53069377 | 1.4613921 | 0.24844127 | 0.74783629 | 0.7827231 | 0.74733789 |
| ENSMUSG00 Gm9826 | 1298 | 0 | 0 | 0 | 0 | 0 | 0 |
| ENSMUSG00 Nhlh2 | 3350 | 0 | 0 | 0 | 0 | 0 | 0 |
| ENSMUSG00 5930422O12I | 839 | 0.47384363 | 0 | 0 | 0.25803608 | 0 | 1.33284888 |
| ENSMUSG00 Tob2 | 3992 | 11.213595 | 9.82559216 | 8.08300235 | 11.3343897 | 13.6257807 | 11.6969395 |
| ENSMUSG00 Thnsl1 | 3444 | 8.83070342 | 8.46480764 | 8.41939867 | 8.59096298 | 10.5934549 | 9.82013724 |
| ENSMUSG00 4930555K19I | 170 | 0.23385577 | 0 | 0 | 0 | 0 | 0 |
| ENSMUSG00 Sp8 | 1921 | 0 | 0 | 0 | 0 | 0 | 0 |
| ENSMUSG00 E030010A14I | 2460 | 3.34527824 | 3.11448955 | 2.01244163 | 1.77476723 | 3.94106155 | 5.72102189 |
| ENSMUSG00 Cypt3 | 673 | 0 | 0 | 0 | 0 | 0 | 0 |
| ENSMUSG00 Gm5593 | 2038 | 0 | 0.04728797 | 0 | 0.03540927 | 0 | 0 |
| ENSMUSG00 Mlec | 11165 | 30.9889787 | 32.0796977 | 24.6524744 | 34.0880793 | 37.7224994 | 18.9713254 |
| ENSMUSG00 E130311K13I | 2392 | 3.92236346 | 2.86056638 | 4.85998525 | 2.70011955 | 3.13581825 | 4.53817158 |
| ENSMUSG00 Gja3 | 1546 | 0.33429576 | 1.02855924 | 0.31450212 | 0 | 0.42907043 | 0.6351145 |
| ENSMUSG00 Igf2 | 7654 | 0 | 0 | 0 | 0 | 0 | 0 |
| ENSMUSG00 Gm5946 | 970 | 0.04098503 | 0.19870698 | 0 | 0.03719798 | 0 | 0 |
| ENSMUSG00 Gm5763 | 3050 | 0 | 0 | 0.04347722 | 0 | 0 | 0 |
| ENSMUSG00 Morc2b | 6790 | 0 | 0 | 0 | 0 | 0 | 0 |
| ENSMUSG00 Gm9828 | 2788 | 0.07129749 | 0.06913406 | 0.36464935 | 0.71180504 | 0.82359645 | 0.58697215 |
| ENSMUSG00 Myof | 7091 | 5.83633553 | 3.60158149 | 3.28506152 | 3.3176552 | 4.21681797 | 1.89241609 |
| ENSMUSG00 Nog | 1921 | 0.0206952 | 0.07525212 | 0 | 0.28174423 | 0 | 0.02839629 |
| ENSMUSG00 Rtbdn | 3387 | 0.05868834 | 0.02845376 | 0 | 0 | 0 | 0 |
| ENSMUSG00 Olfr1336 | 944 | 0 | 0 | 0 | 0 | 0 | 0 |
| ENSMUSG00 Gm6377 | 1060 | 0.3750517 | 0 | 0 | 0 | 0 | 3.13915659 |
| ENSMUSG00 Klf17 | 2397 | 0 | 0 | 0 | 0 | 0 | 0 |
| ENSMUSG00 4930542C16 | 1249 | 0 | 0 | 0 | 0 | 0 | 0 |
| ENSMUSG00 A730049H05I | 3733 | 0.0532487 | 0 | 0 | 0 | 0 | 0 |
| ENSMUSG00 Krtap16-10a | 167 | 0 | 0 | 0 | 0 | 0 | 0 |
| ENSMUSG00 Ctxn1 | 1222 | 2.17971947 | 1.38013541 | 1.59155572 | 1.56493319 | 2.04606581 | 1.20526229 |
| ENSMUSG00 Exd1 | 10243 | 0.15136813 | 0.44220693 | 0.63866765 | 0 | 0.21420816 | 0 |
| ENSMUSG00 Gm9830 | 1375 | 1.85043691 | 0.07008937 | 1.9288077 | 3.88374007 | 2.37504586 | 2.16213504 |
| ENSMUSG00 Samd13 | 2922 | 0 | 0 | 0 | 0.07409044 | 0 | 0 |
| ENSMUSG00 A730037C10I | 3314 | 0 | 0 | 0 | 0 | 0.15397217 | 0 |
| ENSMUSG00 Lemd3 | 6499 | 6.86958065 | 5.21976539 | 5.65870648 | 7.43959681 | 5.38251957 | 7.68004152 |
| ENSMUSG00 A430010J10R | 405 | 3.14117377 | 4.75915478 | 4.03819307 | 4.09820753 | 3.52774944 | 2.49175717 |
| ENSMUSG00 5930416I19R | 3351 | 2.17106921 | 3.50865171 | 1.91264316 | 1.52899144 | 4.01998306 | 4.17543716 |
| ENSMUSG00 Tpcn2 | 2985 | 5.18086496 | 7.86157364 | 3.71680489 | 5.98345462 | 4.68382813 | 2.71375808 |
| ENSMUSG00 Hmgb4 | 682 | 0 | 0 | 0 | 0 | 0 | 0 |
| ENSMUSG00 Olfr435 | 941 | 0 | 0 | 0 | 0 | 0 | 0 |
| ENSMUSG00 Mex3d | 4655 | 0.43555951 | 0.96269369 | 0.40830918 | 0.20153237 | 0.559043 | 0.39842652 |
| ENSMUSG00 Vmn1r26 | 1019 | 0 | 0 | 0 | 0 | 0 | 0 |
| ENSMUSG00 4732456N10 | 2525 | 0 | 0 | 0 | 0 | 0 | 0 |
| ENSMUSG00 Ccdc6 | 6362 | 2.6057899 | 2.46915752 | 3.34188722 | 2.08143828 | 2.38604541 | 2.05352911 |
| ENSMUSG00 Speer4b | 2646 | 0 | 0 | 0 | 0 | 0 | 0 |
| ENSMUSG00 D4Bwg0951e | 3064 | 2.01112907 | 0.8964188 | 0.40393329 | 1.37780653 | 1.13243915 | 0.38277072 |
| ENSMUSG00 Tpm | 4562 | 3.39864915 | 2.74626808 | 3.89503308 | 2.61005583 | 3.23249071 | 3.09694501 |
| ENSMUSG00 Gm8686 | 291 | 0 | 0 | 0 | 0 | 0 | 0 |
| ENSMUSG00 Tbc1d12 | 4965 | 3.6112229 | 2.4457167 | 3.72132334 | 4.76006831 | 3.03177862 | 4.18046334 |
| ENSMUSG00 C030019I05F | 2288 | 0 | 0.02106051 | 0.03863798 | 0 | 0 | 0.05960367 |
| ENSMUSG00 Zfp454 | 2541 | 3.69236261 | 3.39447979 | 5.09686738 | 6.29057289 | 3.79535027 | 4.50820484 |
| ENSMUSG00 Ggnbp1 | 6932 | 0.83158463 | 0.81330262 | 0.80981449 | 1.35854207 | 1.39122692 | 2.60470444 |
| ENSMUSG00 Klhl11 | 2391 | 2.44418889 | 3.60743335 | 1.97808331 | 3.78778468 | 1.83532761 | 1.00383449 |
| ENSMUSG00 Olfr820 | 1060 | 0 | 0 | 0 | 0 | 0 | 0 |
| ENSMUSG00 E130114P18I | 2035 | 0 | 0 | 0 | 0 | 0 | 0 |
| ENSMUSG00 Prss50 | 1450 | 0 | 0 | 0.15242015 | 0.02488417 | 0 | 0 |
| ENSMUSG00 Mcat | 1889 | 17.8678682 | 22.8050182 | 21.8552257 | 16.4078752 | 24.8783971 | 16.4889561 |
| ENSMUSG00 Foxo3 | 8073 | 1.74327265 | 2.94860677 | 2.06417624 | 2.38669785 | 2.36391237 | 2.29738074 |
| ENSMUSG00 Rpl29 | 829 | 3.16509254 | 3.6619371 | 4.26555783 | 3.48198258 | 2.33896536 | 4.21128326 |
| ENSMUSG00 Hoxb3 | 2999 | 0.18558744 | 0.11247252 | 0.20634405 | 0.5534425 | 0.20417356 | 0.26374276 |
| ENSMUSG00 Tmprss11f | 6795 | 0 | 0 | 0 | 0 | 0 | 0 |
| ENSMUSG00 Skint10 | 3727 | 0 | 0 | 0 | 0 | 0 | 0 |
| ENSMUSG00 H1f0 | 531 | 171.000975 | 155.085932 | 141.678977 | 134.067559 | 125.211615 | 152.655796 |
| ENSMUSG00 Tmem53 | 1687 | 6.03284114 | 7.62642564 | 8.96089526 | 5.28290753 | 8.13639306 | 10.2825552 |
| ENSMUSG00 Serpinb13 | 1751 | 0.04540889 | 0 | 0 | 0 | 0 | 0 |
| ENSMUSG00 Pthlh | 1599 | 0 | 0 | 0 | 0 | 0 | 0 |
| ENSMUSG00 P2ry6 | 1935 | 0.12327281 | 0 | 0.20558997 | 0 | 0.18459154 | 0 |
| ENSMUSG00 Insc | 3795 | 0 | 0 | 0 | 0 | 0 | 0 |
| ENSMUSG00 Dcun1d3 | 8440 | 2.27039593 | 3.39132069 | 2.36197052 | 2.83868218 | 2.26716717 | 2.50771565 |
| ENSMUSG00 E030011O05I | 1796 | 0 | 0 | 0 | 0 | 0 | 0 |
| ENSMUSG00 Ccdc37 | 3752 | 0 | 0.03852861 | 0 | 0 | 0 | 0.02180808 |
| ENSMUSG00 Cyb561d1 | 8649 | 2.69816939 | 2.91380613 | 2.98971883 | 2.51977742 | 1.75810614 | 4.32975832 |
| ENSMUSG00 Cep120 | 5457 | 8.98268416 | 6.89639203 | 8.13242631 | 8.58246175 | 9.69660101 | 4.86815075 |
| ENSMUSG00 Ifnb1 | 769 | 0 | 0 | 0 | 0 | 0 | 0 |
| ENSMUSG00 Slc35e4 | 4023 | 2.1246404 | 4.25209718 | 1.15366481 | 2.98665693 | 2.81577316 | 1.06440924 |
| ENSMUSG00 Olfr202 | 923 | 0 | 0 | 0 | 0 | 0 | 0 |
| ENSMUSG00 Lonrf2 | 9989 | 0 | 0 | 0 | 0 | 0 | 0 |
| ENSMUSG00 Gm568 | 504 | 0 | 0 | 0 | 0 | 0 | 0 |
| ENSMUSG00 Dact2 | 2804 | 8.52105699 | 10.2250118 | 13.3519833 | 9.62531002 | 12.4836283 | 10.4176671 |
| ENSMUSG00 Pld1l3 | 6724 | 0.41387324 | 0.07166336 | 0.03286871 | 0.06976005 | 0.05312086 | 0.00811262 |
| ENSMUSG00 2310057N15I | 1025 | 0 | 0 | 0 | 0 | 0 | 0 |
| ENSMUSG00 Vps37c | 2410 | 49.2572053 | 40.0687262 | 47.7599997 | 31.036547 | 42.3667129 | 33.2727964 |
| ENSMUSG00 Slc39a9 | 5222 | 26.7295083 | 21.4818149 | 28.7625168 | 21.5925678 | 27.0961586 | 22.8350676 |
| ENSMUSG00 Vstm2a | 4810 | 3.60361528 | 5.93063903 | 2.18711822 | 6.9838635 | 8.31698101 | 3.68009166 |
| ENSMUSG00 Gm12185 | 9403 | 0 | 0.10249164 | 0.01410247 | 0.20337641 | 0.20078442 | 0 |
| ENSMUSG00 Slc25a47 | 1573 | 3.15920856 | 2.45067729 | 1.60171968 | 1.97269919 | 1.16780008 | 1.62988944 |
| ENSMUSG00 Arhgap30 | 5526 | 0 | 0 | 0 | 0 | 0 | 0 |
| ENSMUSG00 Phf3 | 9536 | 5.42802387 | 6.47303192 | 6.63305761 | 7.21943592 | 7.93541478 | 6.59842625 |
| ENSMUSG00 Hdhd1a | 1043 | 0 | 0 | 0 | 0 | 0 | 0 |
| ENSMUSG00 Hexim1 | 3400 | 9.65824321 | 11.9190582 | 18.22676 | 12.6711651 | 8.2092434 | 19.4532354 |
| ENSMUSG00 Cdk5r1 | 7396 | 0 | 0.05863683 | 0.01195291 | 0 | 0.04139511 | 0.00368776 |
| ENSMUSG00 Zfp710 | 4796 | 11.0745042 | 13.5838344 | 10.2025522 | 8.13275441 | 10.3308196 | 8.90008558 |
| ENSMUSG00 Rimklа | 4152 | 0.65110132 | 2.29790837 | 0 | 1.85102974 | 1.40101321 | 0 |
| ENSMUSG00 Neurog1 | 1658 | 0 | 0 | 0 | 0 | 0 | 0 |
| ENSMUSG00 4930539E08I | 4580 | 0 | 0 | 0 | 0 | 0 | 0 |
| ENSMUSG00 2810453I06R | 2328 | 0 | 0 | 0 | 0 | 0 | 0 |
| ENSMUSG00 Rnf24 | 12308 | 0.45220729 | 0.50504152 | 0.10055668 | 0.31954362 | 0.64259736 | 0.35012943 |
| ENSMUSG00 Efna5 | 4171 | 0.22875366 | 0.41589833 | 0.10597421 | 0.18166457 | 0.14680329 | 0.21579072 |
| ENSMUSG00 Gm5540 | 1847 | 0 | 0 | 0.07179509 | 0.03907097 | 0 | 0.014767 |
| ENSMUSG00 Olfr223 | 1101 | 0 | 0 | 0 | 0 | 0 | 0 |
| ENSMUSG00 Fkrp | 2771 | 16.5277205 | 13.8594711 | 14.9466355 | 15.3521222 | 10.8276828 | 11.6539779 |
| ENSMUSG00 Zfp689 | 2963 | 3.56900365 | 1.7075857 | 3.31178169 | 3.31229028 | 1.94599408 | 1.79498976 |
| ENSMUSG00 Cdca2 | 7365 | 0.21051782 | 0.01962788 | 0 | 0 | 0.04849757 | 0.03703278 |
| ENSMUSG00 Ccdc125 | 2692 | 11.6519592 | 9.71962782 | 10.4921908 | 6.32642088 | 11.695124 | 10.719379 |
| ENSMUSG00 Tada3 | 4275 | 7.96039563 | 8.69046711 | 6.82414418 | 7.98447114 | 9.99042727 | 6.12483127 |
| ENSMUSG00 Olfr722 | 1578 | 0 | 0 | 0 | 0 | 0 | 0 |
| ENSMUSG00 Nr1h5 | 2824 | 0 | 0 | 0 | 0 | 0 | 0 |
| ENSMUSG00 Atp13a5 | 5579 | 0 | 0 | 0 | 0 | 0 | 0 |
| ENSMUSG00 Gm6206 | 3780 | 0.54690079 | 0.25495472 | 0.86532709 | 0.10500066 | 0.26998082 | 0.14431026 |
| ENSMUSG00 Prex2 | 11014 | 0 | 0.01312505 | 0.12441049 | 0.65847929 | 0.39379353 | 0.07181447 |
| ENSMUSG00 Rassf10 | 1237 | 5.07790293 | 4.79137622 | 2.7157155 | 3.64612414 | 3.96000976 | 5.31381411 |
| ENSMUSG00 Mrgpre | 3757 | 1.25922337 | 0.66694038 | 4.20017513 | 0.91237536 | 1.4396582 | 2.42473503 |
| ENSMUSG00 Yjefn3 | 1006 | 0.3951837 | 0.47899048 | 0.79088785 | 0.50213581 | 0.55794248 | 0.29823164 |
| ENSMUSG00 C1galt1c1 | 2397 | 35.2939769 | 37.7530823 | 49.7155481 | 46.3180855 | 31.3353464 | 54.3444627 |
| ENSMUSG00 Nrsn1 | 4226 | 0 | 0 | 0 | 0 | 0 | 0 |
| ENSMUSG00 Krt31 | 1574 | 0 | 0 | 0 | 0 | 0 | 0 |
| ENSMUSG00 Gphb5 | 1611 | 0 | 0 | 0 | 0 | 0 | 0.4063261 |
| ENSMUSG00 Elfn1 | 3819 | 0.03122976 | 0 | 0 | 0 | 0.06680594 | 0.04999279 |
| ENSMUSG00 Prss32 | 1827 | 0 | 0 | 0 | 0 | 0 | 0 |
| ENSMUSG00 4930557A04I | 551 | 0 | 0 | 0 | 0 | 0 | 0 |
| ENSMUSG00 Olfr1366 | 2159 | 0 | 0 | 0 | 0 | 0 | 0 |
| ENSMUSG00 Atxn7l2 | 4508 | 1.7461369 | 1.56060793 | 1.95123486 | 1.7288646 | 1.70917985 | 2.32330556 |
| ENSMUSG00 A930038C07I | 4714 | 0 | 0 | 0 | 0 | 0 | 0 |
| ENSMUSG00 BB014433 | 2360 | 0 | 0 | 0 | 0 | 0 | 0 |
| ENSMUSG00 Olfr982 | 965 | 0 | 0 | 0 | 0 | 0 | 0 |
| ENSMUSG00 Olfr729 | 971 | 0 | 0 | 0 | 0 | 0 | 0 |
| ENSMUSG00 Prss48 | 943 | 0 | 0 | 0 | 0 | 0.05411069 | 0.02892327 |
| ENSMUSG00 Olfr1467 | 926 | 0 | 0 | 0 | 0 | 0 | 0 |
| ENSMUSG00 Olfr368 | 984 | 0 | 0 | 0 | 0 | 0 | 0 |
| ENSMUSG00 Olfr873 | 1061 | 0 | 0 | 0 | 0 | 0 | 0 |
| ENSMUSG00 Tmem121 | 1596 | 0 | 0 | 0 | 0 | 0 | 0 |
| ENSMUSG00 Clec4a1 | 1552 | 0 | 0 | 0 | 0.02324874 | 0 | 0 |
| ENSMUSG00 Mterfd3 | 1535 | 28.2820747 | 26.526087 | 28.9687649 | 18.1938127 | 28.7875189 | 37.4026812 |
| ENSMUSG00 Olfr398 | 2246 | 0 | 0 | 0 | 0 | 0 | 0 |
| ENSMUSG00 Rapgef4 | 6796 | 3.79654309 | 2.51000596 | 2.81627399 | 5.84023675 | 3.58896522 | 2.63676251 |
| ENSMUSG00 Armcx3 | 3658 | 3.14087859 | 3.67523711 | 2.98465151 | 4.62615606 | 2.85959733 | 2.64693738 |
| ENSMUSG00 Olfr812 | 1042 | 0 | 0 | 0 | 0 | 0 | 0 |
| ENSMUSG00 Olfr1257 | 1482 | 0 | 0 | 0 | 0 | 0 | 0 |
| ENSMUSG00 Gm9244 | 2127 | 0 | 0 | 0 | 0 | 0 | 0 |
| ENSMUSG00 Olfr985 | 1115 | 0 | 0 | 0 | 0 | 0 | 0 |
| ENSMUSG00 Acap2 | 6590 | 4.95887784 | 3.45128994 | 5.56044429 | 5.82021447 | 5.82273666 | 4.2422618 |
| ENSMUSG00 Bmyc | 838 | 6.97381341 | 8.62525814 | 6.48786001 | 7.10445984 | 6.21084765 | 9.27598113 |
| ENSMUSG00 Zadh2 | 3319 | 15.1643382 | 19.4110495 | 18.3519553 | 12.7847196 | 13.2677802 | 16.829907 |
| ENSMUSG00 Sephs2 | 1358 | 85.0439403 | 51.5218807 | 103.376328 | 64.9901921 | 61.3969795 | 59.9921557 |
| ENSMUSG00 Gpr137c | 4095 | 0.29124894 | 0.01176714 | 0.12952921 | 0.95161436 | 0.09968523 | 0.29306084 |
| ENSMUSG00 Il23r | 2477 | 0 | 0.44743164 | 0.48181258 | 0 | 0.47380163 | 0.37437938 |
| ENSMUSG00 Olfr367-ps | 736 | 0 | 0 | 0 | 0 | 0 | 0 |
| ENSMUSG00 Ankrd34a | 925 | 0 | 0.52093451 | 0 | 0 | 0 | 0 |
| ENSMUSG00 Olfr147 | 1164 | 0 | 0 | 0 | 0 | 0 | 0 |
| ENSMUSG00 Pcdh10 | 14114 | 0 | 0 | 0 | 0 | 0 | 0 |
| ENSMUSG00 Ccr2 | 4749 | 0 | 0 | 0 | 0 | 0 | 0 |
| ENSMUSG00 Dcaf5 | 5698 | 10.8144954 | 11.0275747 | 9.26237286 | 7.46590567 | 13.4058415 | 8.08474301 |
| ENSMUSG00 Ntf3 | 1424 | 0.72587254 | 7.13998546 | 0.83809674 | 1.97640412 | 1.46915829 | 1.8770468 |
| ENSMUSG00 Themis | 9376 | 0 | 0 | 0 | 0 | 0 | 0 |
| ENSMUSG00 Oxtr | 6547 | 0 | 0 | 0 | 0 | 0 | 0 |
| ENSMUSG00 Agtr1a | 2463 | 0.22597512 | 0 | 0 | 0 | 0.41434329 | 0 |
| ENSMUSG00 Fam110b | 8443 | 1.66687671 | 3.85811144 | 0.97377032 | 1.88893328 | 2.0910963 | 1.16942075 |
| ENSMUSG00 Frmd3 | 6532 | 0 | 0 | 0 | 0 | 0 | 0 |
| ENSMUSG00 Catsperg2 | 4623 | 0 | 0 | 0 | 0 | 0 | 0 |
| ENSMUSG00 Gm8186 | 230 | 0 | 0.83802508 | 0 | 0.31375691 | 0 | 2.37170776 |
| ENSMUSG00 Ivl | 3593 | 0.26555289 | 0.134112 | 0 | 0 | 0 | 0 |
| ENSMUSG00 C5ar1 | 4912 | 0 | 0 | 0 | 0 | 0 | 0 |
| ENSMUSG00 Flg2 | 7005 | 0 | 0 | 0 | 0 | 0 | 0 |
| ENSMUSG00 Nrap | 5726 | 0 | 0 | 0 | 0 | 0 | 0 |
| ENSMUSG00 Plcxd3 | 2266 | 0.59650765 | 0 | 0 | 0 | 0.02251826 | 0 |
| ENSMUSG00 Olfr1258 | 1444 | 0 | 0 | 0 | 0 | 0 | 0 |
| ENSMUSG00 Ugt3a2 | 2521 | 2.25507089 | 2.33191033 | 0.17533456 | 0.80150515 | 1.25491285 | 1.22254432 |
| ENSMUSG00 Fam183b | 751 | 0 | 0 | 0 | 0 | 0 | 0 |
| ENSMUSG00 4930469G21I | 1189 | 0 | 0.08105373 | 0 | 0 | 0.04291537 | 0 |
| ENSMUSG00 Zfp518a | 6454 | 9.54772154 | 6.61499655 | 7.41034926 | 11.6732738 | 9.4794894 | 5.7262374 |
| ENSMUSG00 Olfr449 | 1036 | 0 | 0 | 0 | 0 | 0 | 0 |
| ENSMUSG00 Myoz3 | 863 | 0 | 0 | 0 | 0 | 0 | 0 |
| ENSMUSG00 Frmpd4 | 13397 | 0.92882477 | 1.12939784 | 1.10859291 | 0.37436771 | 0.57512747 | 0.68609043 |
| ENSMUSG00 Purg | 3000 | 2.3190697 | 1.15647461 | 1.31132134 | 1.68382874 | 1.41172973 | 2.24561197 |
| ENSMUSG00 Rgag4 | 4922 | 0.09692519 | 0.09790013 | 0.08980464 | 0.1172923 | 0.031101 | 0.04433099 |
| ENSMUSG00 4930515G13I | 489 | 0 | 0 | 0 | 0 | 0 | 0 |
| ENSMUSG00 Skint7 | 4009 | 0 | 0 | 0 | 0 | 0 | 0 |
| ENSMUSG00 Olfr788 | 935 | 0 | 0 | 0 | 0 | 0 | 0 |
| ENSMUSG00 Pdp1 | 6267 | 10.2512935 | 6.24340051 | 9.42295552 | 8.31378208 | 7.75939623 | 11.6941049 |
| ENSMUSG00 Olfr844 | 929 | 0 | 0 | 0 | 0 | 0 | 0 |
| ENSMUSG00 Gm9833 | 1764 | 0.06761136 | 0.08194973 | 0.05011547 | 0.22500141 | 0.14463259 | 0.03092363 |
| ENSMUSG00 Gm12497 | 236 | 0 | 0 | 0.56188784 | 0 | 0 | 0 |
| ENSMUSG00 Tigd2 | 3005 | 16.5372215 | 12.4755581 | 17.0776505 | 16.5581163 | 13.346666 | 19.0423272 |
| ENSMUSG00 Apoo-ps | 596 | 5.80323289 | 5.65948146 | 4.44984997 | 6.71997809 | 3.16774481 | 10.1135827 |
| ENSMUSG00 Gm7324 | 2775 | 0.11461039 | 0.31256071 | 0.07964296 | 0.14302792 | 0.18387883 | 0.12777309 |
| ENSMUSG00 Gpr81 | 4315 | 0.1750531 | 0 | 0.18438776 | 0 | 0 | 0.39821605 |
| ENSMUSG00 Hspe1-ps2 | 311 | 0 | 1.0845823 | 0.14212811 | 0 | 0 | 0 |
| ENSMUSG00 Rpl37-ps1 | 269 | 0 | 0 | 0 | 0 | 0 | 0 |
| ENSMUSG00 Gm5316 | 269 | 0 | 0 | 0 | 0 | 0 | 0 |
| ENSMUSG00 Lrp1b | 23227 | 0.02909731 | 0.01244753 | 0.03235163 | 0.4691427 | 0 | 0.24894405 |
| ENSMUSG00 Kcnk3 | 3808 | 0.01043999 | 0 | 0.01160763 | 0 | 0 | 0.00716246 |
| ENSMUSG00 Olfr509 | 1074 | 0 | 0 | 0 | 0 | 0 | 0 |
| ENSMUSG00 Scn3b | 3429 | 0 | 0 | 0 | 0 | 0 | 0 |
| ENSMUSG00 Mblac1 | 1086 | 4.75894334 | 5.2357276 | 3.58173314 | 6.91074149 | 3.14803608 | 4.69646182 |
| ENSMUSG00 A230051G13I | 4880 | 2.59062353 | 2.65617888 | 3.44194679 | 2.00373649 | 2.23763205 | 1.29666318 |
| ENSMUSG00 Lix1l | 2503 | 0.06353253 | 0 | 0.194255 | 0.01441552 | 0 | 0 |
| ENSMUSG00 Prss38 | 1238 | 0 | 0 | 0 | 0 | 0 | 0 |
| ENSMUSG00 Zfp219 | 4172 | 8.0235174 | 11.295863 | 8.73018184 | 7.55024565 | 7.58301846 | 7.1912999 |
| ENSMUSG00 Trappc1 | 841 | 42.8754112 | 41.8265193 | 54.0828733 | 41.7882418 | 34.8265633 | 68.0406577 |
| ENSMUSG00 Prmt6 | 5938 | 8.63667394 | 12.3428054 | 10.3097933 | 10.4636714 | 7.37295899 | 13.0172327 |
| ENSMUSG00 Syt12 | 5673 | 0.31535283 | 0.26331389 | 0.08570779 | 0.34981711 | 0.29682186 | 0.18750413 |
| ENSMUSG00 Ccdc71 | 3106 | 15.1674966 | 12.3336193 | 13.4484036 | 10.501664 | 12.1076751 | 9.36962012 |
| ENSMUSG00 Fut4 | 3663 | 1.17215176 | 1.94692696 | 0.27754365 | 1.7238214 | 1.99202068 | 0.22337952 |
| ENSMUSG00 Sorl1 | 11237 | 19.0870177 | 19.2882991 | 19.5106471 | 16.4274931 | 22.1687965 | 17.9759703 |
| ENSMUSG00 Fbxw13 | 1502 | 0 | 0 | 0 | 0 | 0 | 0 |
| ENSMUSG00 Olfr348 | 1008 | 0 | 0 | 0 | 0 | 0 | 0 |
| ENSMUSG00 Zfp2 | 5574 | 7.77421488 | 8.82640069 | 12.1487663 | 7.0688182 | 10.8387565 | 10.2120869 |
| ENSMUSG00 Smcr8 | 10931 | 3.62603733 | 3.31058623 | 2.6526767 | 4.96123986 | 5.28889249 | 1.92626672 |
| ENSMUSG00 Setd8 | 4012 | 24.3566728 | 21.4268726 | 27.5545387 | 22.1780463 | 18.7342601 | 19.368257 |
| ENSMUSG00 Olfr902 | 1055 | 0 | 0 | 0 | 0 | 0 | 0 |
| ENSMUSG00 Odz2 | 12573 | 0 | 0 | 0.01054685 | 0.01147922 | 0 | 0.0021693 |
| ENSMUSG00 Fam134a | 1694 | 65.1248279 | 67.8706322 | 53.0213371 | 46.2421007 | 58.3761017 | 39.044274 |
| ENSMUSG00 Gm5105 | 1661 | 0 | 0 | 0 | 0 | 0 | 0 |
| ENSMUSG00 Zg16 | 633 | 0 | 0 | 0 | 0 | 0 | 0 |
| ENSMUSG00 Rd3 | 2357 | 0 | 0 | 0 | 0 | 0 | 0 |
| ENSMUSG00 Dcaf7 | 6487 | 15.9769597 | 16.6984927 | 14.9292798 | 14.5841099 | 26.7048784 | 10.0361591 |
| ENSMUSG00 4933408B17I | 2726 | 0.07291908 | 0 | 0.06485964 | 0.48974162 | 0.26205769 | 0.14007518 |
| ENSMUSG00 Olfr173 | 1163 | 0 | 0 | 0 | 0 | 0 | 0 |
| ENSMUSG00 Olfr1183 | 923 | 0 | 0 | 0 | 0 | 0 | 0 |
| ENSMUSG00 Krt8 | 1796 | 40.8622589 | 53.9816935 | 36.5231264 | 41.5465858 | 37.8152039 | 28.9147623 |
| ENSMUSG00 Cox7b2 | 539 | 0 | 0.08939971 | 0 | 0 | 0 | 0 |
| ENSMUSG00 Gemin4 | 3731 | 0.01065545 | 0 | 0.08293029 | 0 | 0.08205796 | 0.12427469 |
| ENSMUSG00 Ogfr | 2691 | 14.3598391 | 13.9312716 | 14.8325025 | 14.5883554 | 19.7772241 | 12.8822246 |
| ENSMUSG00 Rarres1 | 1319 | 0 | 0 | 0 | 0 | 0 | 0 |
| ENSMUSG00 Prokr1 | 4052 | 0 | 0 | 0 | 0 | 0 | 0 |
| ENSMUSG00 Gm13060 | 1466 | 0 | 0 | 0 | 0 | 0 | 0 |

| | | | | | | | | |
|---|---|---|---|---|---|---|---|---|
| ENSMUSG00 | 6030446N20 | 3319 | 13.4514627 | 7.91250707 | 7.64442841 | 12.1433094 | 9.36277881 | 6.15507828 |
| ENSMUSG00 | Gm5417 | 382 | 0.20814388 | 0 | 0.23142326 | 0 | 0.5343076 | 0 |
| ENSMUSG00 | Tmem200a | 1475 | 0 | 0 | 0 | 0 | 0 | 0 |
| ENSMUSG00 | Zfp260 | 4501 | 80.9241752 | 69.9511693 | 64.6774801 | 101.664628 | 58.6786317 | 95.0279699 |
| ENSMUSG00 | Chchd10 | 935 | 303.587306 | 294.942255 | 446.7459 | 282.752022 | 193.900228 | 619.936506 |
| ENSMUSG00 | Gm9837 | 730 | 0 | 0 | 0 | 0 | 0 | 0 |
| ENSMUSG00 | Upk1b | 1761 | 0 | 7.27858808 | 0 | 0 | 0 | 0 |
| ENSMUSG00 | Cyp20a1 | 2273 | 14.604411 | 16.9595925 | 14.8182157 | 13.8105494 | 11.920372 | 10.5354744 |
| ENSMUSG00 | Gm7103 | 311 | 0 | 0 | 0 | 0 | 0 | 0 |
| ENSMUSG00 | Olfr1404 | 1033 | 0 | 0 | 0 | 0 | 0 | 0 |
| ENSMUSG00 | Gm8396 | 822 | 0 | 0 | 0 | 0 | 0 | 0 |
| ENSMUSG00 | Aff4 | 10704 | 17.9204217 | 16.1116863 | 15.9686591 | 21.1894368 | 15.1210449 | 17.3931883 |
| ENSMUSG00 | 1700104B16I | 553 | 0 | 0.08713642 | 0 | 0 | 0 | 0 |
| ENSMUSG00 | Gm8508 | 406 | 2.74175728 | 2.01765891 | 2.72178836 | 1.77744062 | 1.13112656 | 1.27639937 |
| ENSMUSG00 | Ctu2 | 2574 | 6.99659389 | 8.1059555 | 5.23759212 | 7.6818028 | 6.83919956 | 5.14975706 |
| ENSMUSG00 | Tmem67 | 8243 | 2.89858593 | 1.71280208 | 2.57928988 | 2.81897812 | 3.62130655 | 3.00441252 |
| ENSMUSG00 | Fam58b | 1222 | 24.5625105 | 30.441844 | 35.4121148 | 26.5152831 | 22.7572625 | 31.5153769 |
| ENSMUSG00 | Slc36a3 | 2681 | 0 | 0 | 0.31325439 | 0 | 0 | 0 |
| ENSMUSG00 | Pls1 | 3873 | 12.9849426 | 14.1834609 | 17.8496469 | 20.579715 | 11.8310575 | 21.3943594 |
| ENSMUSG00 | Gm12669 | 1650 | 0.21684808 | 0.29203904 | 0.10715598 | 0.06560372 | 0.03092508 | 0.18183093 |
| ENSMUSG00 | Olfr1449 | 947 | 0 | 0 | 0 | 0 | 0 | 0 |
| ENSMUSG00 | Dtx3l | 5719 | 1.86299507 | 3.42925021 | 1.26754717 | 4.20820488 | 5.21059687 | 2.74701735 |
| ENSMUSG00 | 2810046L04F | 5458 | 6.81043867 | 3.92872238 | 3.49857021 | 3.67563517 | 6.88080114 | 3.89280762 |
| ENSMUSG00 | 4933407P14F | 4439 | 0 | 0 | 0 | 0 | 0 | 0 |
| ENSMUSG00 | 4930429B21I | 1661 | 0.43082399 | 0.84130453 | 1.80958779 | 1.47716979 | 1.13665015 | 1.67490259 |
| ENSMUSG00 | Htr1b | 1272 | 0.06250862 | 0.15152969 | 0.55599803 | 0.62406052 | 0.04011508 | 0.04288465 |
| ENSMUSG00 | Espnl | 5729 | 0 | 0 | 0 | 0 | 0 | 0.0047608 |
| ENSMUSG00 | Spty2d1 | 5217 | 17.3439666 | 13.4574748 | 17.3350529 | 24.9053944 | 14.3484174 | 17.3884321 |
| ENSMUSG00 | Rps23 | 569 | 0.13973807 | 0.76217571 | 0.62146704 | 1.07802242 | 0.627741 | 1.48596453 |
| ENSMUSG00 | Cdc42ep1 | 2539 | 42.3860125 | 55.7019329 | 36.9422256 | 46.4987986 | 42.2841649 | 31.2277969 |
| ENSMUSG00 | Tmem202 | 1250 | 0.22263069 | 0.46258985 | 0.81331391 | 0.46185017 | 0 | 0.63277163 |
| ENSMUSG00 | Olfr429 | 942 | 0 | 0 | 0 | 0 | 0 | 0 |
| ENSMUSG00 | Clrn2 | 941 | 0 | 0 | 0 | 0 | 0 | 0 |
| ENSMUSG00 | Sall2 | 9915 | 0.12830816 | 0.30617709 | 0.75341518 | 0.40758336 | 0.25217271 | 0.68220984 |
| ENSMUSG00 | Tceal1 | 2801 | 9.82177526 | 13.3841671 | 11.5357184 | 9.68714655 | 8.67138697 | 18.598558 |
| ENSMUSG00 | Tecrl | 2338 | 0 | 0 | 0 | 0 | 0 | 0 |
| ENSMUSG00 | Adamts16 | 7966 | 1.7517165 | 6.62971888 | 0.40506334 | 5.60752839 | 4.16999381 | 2.31111995 |
| ENSMUSG00 | Hist1h1a | 746 | 0 | 0 | 0 | 0.0967347 | 0 | 0 |
| ENSMUSG00 | 4930474N05I | 2268 | 0 | 0 | 0 | 0 | 0 | 0 |
| ENSMUSG00 | Krt82 | 2086 | 0 | 0.02309992 | 0 | 0 | 0 | 0 |
| ENSMUSG00 | Clip1 | 11466 | 8.03014939 | 8.8001404 | 8.64299076 | 11.6434297 | 9.45229568 | 7.83795014 |
| ENSMUSG00 | Fzd9 | 2292 | 0 | 0 | 0 | 0 | 0.02226282 | 0 |
| ENSMUSG00 | Polr1a | 6542 | 2.82578698 | 3.16725315 | 1.96617798 | 7.78780906 | 4.21969877 | 2.25134595 |
| ENSMUSG00 | Tmie | 2796 | 5.78701022 | 7.32447708 | 5.91254982 | 5.22647641 | 5.4201837 | 6.76988542 |
| ENSMUSG00 | Lingo1 | 6146 | 0 | 0.06272235 | 0 | 0 | 0 | 0 |
| ENSMUSG00 | Defb20 | 688 | 0 | 0 | 0 | 0 | 0 | 0 |
| ENSMUSG00 | Olfr95 | 973 | 0 | 0 | 0 | 0 | 0 | 0 |
| ENSMUSG00 | Gm962 | 3090 | 2.17432887 | 1.99607268 | 3.53328648 | 3.09441482 | 2.84030312 | 2.1625523 |
| ENSMUSG00 | Aknad1 | 5979 | 0 | 0 | 0 | 0 | 0 | 0 |
| ENSMUSG00 | 9330101J02R | 5244 | 0 | 0.00918887 | 0.0252871 | 0 | 0 | 0 |
| ENSMUSG00 | Olfr780 | 1024 | 0 | 0 | 0 | 0 | 0 | 0 |
| ENSMUSG00 | Zfa | 3421 | 0 | 0 | 0 | 0 | 0 | 0 |
| ENSMUSG00 | Zfpm1 | 2002 | 3.97157647 | 5.07859106 | 2.45075154 | 7.02896971 | 3.69571653 | 1.83919895 |
| ENSMUSG00 | Olfr491 | 932 | 0 | 0 | 0 | 0 | 0 | 0 |
| ENSMUSG00 | Tsku | 5198 | 0 | 0.18540378 | 0 | 0.52061434 | 0.00981654 | 0 |
| ENSMUSG00 | Grm5 | 4706 | 0 | 0 | 0 | 0 | 0 | 0 |
| ENSMUSG00 | Ccdc69 | 1841 | 1.3388592 | 1.43957323 | 0.14405815 | 0.21559071 | 0.2217333 | 0.37037805 |
| ENSMUSG00 | Lce1h | 986 | 0 | 0 | 0 | 0 | 0 | 0 |
| ENSMUSG00 | Vsig8 | 1871 | 0 | 0 | 0.07087415 | 0 | 0 | 0 |
| ENSMUSG00 | Zbtb45 | 2696 | 1.78427787 | 1.94819073 | 2.80360354 | 2.23505246 | 2.0819367 | 3.3992132 |
| ENSMUSG00 | Olfr738 | 935 | 0 | 0 | 0 | 0 | 0 | 0 |
| ENSMUSG00 | Hoxb13 | 2204 | 0 | 0 | 0 | 0 | 0 | 0 |
| ENSMUSG00 | Olfr418-ps1 | 990 | 0 | 0 | 0 | 0 | 0 | 0 |
| ENSMUSG00 | Zfp644 | 9768 | 8.74229854 | 7.50816594 | 9.56620558 | 8.03793293 | 10.22826 | 8.86258241 |
| ENSMUSG00 | Gpr55 | 2548 | 0 | 0 | 0 | 0 | 0 | 0 |
| ENSMUSG00 | Omg | 2226 | 0 | 0 | 0 | 0 | 0 | 0.09802206 |
| ENSMUSG00 | Olfr103 | 2012 | 0 | 0 | 0 | 0 | 0 | 0 |
| ENSMUSG00 | Prss33 | 1948 | 0 | 0 | 0 | 0 | 0 | 0 |
| ENSMUSG00 | Slc17a5 | 5137 | 11.7788284 | 14.3424314 | 13.9480606 | 10.4165217 | 11.3734087 | 6.0740096 |
| ENSMUSG00 | Tifab | 3495 | 0 | 0.15165976 | 0 | 0 | 0 | 0 |
| ENSMUSG00 | C1ql3 | 2474 | 0 | 0 | 0 | 0.02916899 | 0 | 0 |
| ENSMUSG00 | Vgll2 | 1614 | 0 | 0 | 0 | 0 | 0 | 0 |
| ENSMUSG00 | 2310022A10F | 2700 | 2.63564112 | 2.89118653 | 3.09412901 | 3.8353877 | 2.62691342 | 4.40434915 |
| ENSMUSG00 | Purb | 1532 | 19.3846893 | 22.7407296 | 22.44715 | 20.7024263 | 17.752649 | 25.4052939 |
| ENSMUSG00 | Olfr273 | 2328 | 0 | 0 | 0 | 0 | 0 | 0 |
| ENSMUSG00 | Gpr3 | 3728 | 0.01066402 | 0.0258511 | 0 | 0 | 0.50643131 | 0 |
| ENSMUSG00 | Spatc1 | 1600 | 0 | 0 | 0 | 0 | 0 | 0 |
| ENSMUSG00 | Zbtb5 | 7912 | 3.80370308 | 4.42764705 | 4.69281448 | 2.06130993 | 4.28874367 | 3.55411439 |
| ENSMUSG00 | Bdp1 | 13378 | 1.39373003 | 2.55015705 | 2.26328767 | 3.66808047 | 2.55932861 | 1.90013184 |
| ENSMUSG00 | Aftph | 8076 | 16.5450929 | 17.7686014 | 18.6637302 | 17.5450952 | 17.0656564 | 15.9034517 |
| ENSMUSG00 | Mtag2 | 683 | 0 | 0 | 0 | 0 | 0 | 0 |
| ENSMUSG00 | Morn4 | 1811 | 29.3720778 | 24.6652854 | 23.8216448 | 16.5168498 | 20.117522 | 17.575651 |
| ENSMUSG00 | Zfp161 | 4226 | 4.61901111 | 3.80839368 | 4.58126059 | 3.03956646 | 5.36102955 | 3.89822104 |
| ENSMUSG00 | Olfr714 | 953 | 0 | 0 | 0 | 0 | 0 | 0 |
| ENSMUSG00 | Catsperg1 | 6461 | 0.03691888 | 0.17153508 | 0.15735063 | 0.67015096 | 0.25272311 | 0.57833549 |
| ENSMUSG00 | Urgcp | 9155 | 5.48022025 | 5.81605883 | 5.3640904 | 3.46040798 | 6.91127319 | 3.9683036 |
| ENSMUSG00 | Cyp2g1 | 1844 | 0 | 0 | 0 | 0 | 0 | 0 |
| ENSMUSG00 | Orai1 | 2221 | 5.44154258 | 5.55413291 | 8.41845097 | 4.37013506 | 5.35305969 | 7.44188716 |
| ENSMUSG00 | Fam109b | 3306 | 0 | 0.01457545 | 0 | 0 | 0.64824797 | 0.01650008 |
| ENSMUSG00 | Nckap5 | 13462 | 7.13484184 | 6.86178945 | 6.33706411 | 7.79963969 | 7.67556166 | 4.16757784 |
| ENSMUSG00 | Nkx3-2 | 1997 | 0 | 0 | 0 | 0 | 0 | 0 |
| ENSMUSG00 | 4933425O20I | 1351 | 0 | 0 | 0 | 0 | 0 | 0 |
| ENSMUSG00 | BC048671 | 1457 | 0 | 0 | 0 | 0 | 0 | 0 |
| ENSMUSG00 | Ucn2 | 1001 | 0 | 0 | 0 | 0.82905797 | 0 | 0.59944262 |
| ENSMUSG00 | Olfr27 | 982 | 0 | 0 | 0 | 0 | 0 | 0 |
| ENSMUSG00 | Nlrp10 | 4473 | 0 | 0 | 0 | 0 | 0 | 0 |
| ENSMUSG00 | Lig4 | 7182 | 2.17542521 | 3.77064613 | 4.0250634 | 6.41559048 | 4.15628375 | 4.57995195 |
| ENSMUSG00 | Prss46 | 1341 | 0 | 0 | 0 | 0 | 0 | 0 |
| ENSMUSG00 | Gal3st1 | 2163 | 15.053046 | 19.0919006 | 11.7708098 | 14.1792593 | 11.4414204 | 11.4747673 |
| ENSMUSG00 | Mmp12 | 6590 | 0 | 0 | 0 | 0 | 0 | 0 |
| ENSMUSG00 | Zfp668 | 6291 | 1.04270454 | 0.32170252 | 1.62305289 | 0.94635787 | 1.6384244 | 1.17925638 |
| ENSMUSG00 | Trex1 | 2299 | 0 | 0 | 0 | 0 | 0 | 0 |
| ENSMUSG00 | Olfr1361 | 953 | 0 | 0 | 0 | 0 | 0 | 0 |
| ENSMUSG00 | Zfp646 | 3594 | 4.51314303 | 6.36854758 | 10.7491405 | 6.34497834 | 10.0519405 | 5.92695973 |
| ENSMUSG00 | 5330417H12I | 1190 | 0 | 0.12147843 | 0 | 0 | 0.42879307 | 0.18335892 |
| ENSMUSG00 | Rxfp4 | 1244 | 0 | 0 | 0 | 0 | 0 | 0 |
| ENSMUSG00 | C330006D17 | 377 | 2.10904406 | 3.19538741 | 4.92434325 | 2.29700018 | 3.11301497 | 3.25559354 |
| ENSMUSG00 | Arhgap15 | 9149 | 0 | 0 | 0 | 0 | 0 | 0 |
| ENSMUSG00 | Rpl36al | 1279 | 8.92089359 | 7.57269341 | 9.29655651 | 9.33788643 | 5.58537343 | 13.8185819 |
| ENSMUSG00 | Zfp672 | 5954 | 28.3443088 | 24.3117353 | 25.0110865 | 20.5196175 | 23.9277194 | 21.7775672 |
| ENSMUSG00 | Olfr1318 | 1102 | 0 | 0 | 0 | 0 | 0 | 0 |
| ENSMUSG00 | 2410015M20 | 751 | 27.527097 | 30.8624217 | 44.9669881 | 25.3679354 | 23.984435 | 51.2806799 |
| ENSMUSG00 | 4930479M11 | 683 | 0 | 0 | 0 | 0 | 0 | 0 |
| ENSMUSG00 | Zfp280b | 1891 | 17.4495234 | 17.4551628 | 13.160041 | 17.0583542 | 28.1980765 | 20.8418052 |
| ENSMUSG00 | Tmsb4x | 1807 | 120.8285 | 162.852575 | 94.1032042 | 119.707724 | 98.9183146 | 237.894626 |
| ENSMUSG00 | Fzd4 | 3683 | 24.6434325 | 44.967907 | 17.1862718 | 48.7494579 | 49.2529911 | 20.3504504 |
| ENSMUSG00 | Bag5 | 4013 | 16.623398 | 17.7111892 | 17.359109 | 17.8746834 | 17.0638914 | 17.1953245 |
| ENSMUSG00 | Crh | 1318 | 0 | 0 | 0.4024447 | 0 | 0 | 0 |
| ENSMUSG00 | Olfr642 | 1171 | 0 | 0 | 0 | 0 | 0 | 0 |
| ENSMUSG00 | Lrrc19 | 3073 | 0.45279591 | 0 | 0.43151816 | 0.96281408 | 0.23246641 | 0.90530856 |
| ENSMUSG00 | Sertad2 | 6772 | 4.54381895 | 5.58569951 | 4.6146933 | 7.3208401 | 4.54354764 | 4.66391498 |
| ENSMUSG00 | Armcx4 | 7533 | 2.34849181 | 2.61625579 | 2.78131868 | 3.9372681 | 3.61716244 | 4.68154899 |
| ENSMUSG00 | Olfr466 | 926 | 0 | 0 | 0 | 0 | 0 | 0 |
| ENSMUSG00 | Arhgap23 | 6609 | 0.45115162 | 0.10207447 | 0.18057948 | 0.25113846 | 0.34743334 | 0.48284654 |
| ENSMUSG00 | Krtap9-3 | 758 | 0 | 0 | 0 | 0 | 0 | 0 |
| ENSMUSG00 | Fam161a | 4388 | 3.36127695 | 2.7014277 | 2.56870327 | 3.49472856 | 4.80261924 | 2.90274761 |
| ENSMUSG00 | 4921511C20I | 1605 | 0 | 0 | 0 | 0 | 0 | 0 |
| ENSMUSG00 | Zbtb12 | 1991 | 0.03993519 | 0.04840426 | 0.04440165 | 0.01812257 | 0 | 0.08219379 |
| ENSMUSG00 | Gm9839 | 1436 | 0 | 0 | 0 | 0 | 0 | 0 |
| ENSMUSG00 | Gm9840 | 661 | 0.18043335 | 0.36449654 | 0.20061351 | 0.65504468 | 0.23158718 | 0.45388961 |
| ENSMUSG00 | Olfr1102 | 1062 | 0 | 0 | 0 | 0 | 0 | 0 |
| ENSMUSG00 | Ceacam19 | 3354 | 0.02370631 | 0 | 0.25039804 | 0 | 0 | 0.21956329 |
| ENSMUSG00 | Suox | 2345 | 38.9078157 | 23.1993574 | 36.2662456 | 32.4507599 | 33.0529061 | 26.0184938 |
| ENSMUSG00 | Olfr250 | 995 | 0 | 0 | 0 | 0 | 0 | 0 |
| ENSMUSG00 | Arl4c | 5144 | 1.34476159 | 2.59479869 | 1.27174821 | 2.12536149 | 1.80536555 | 1.62248048 |
| ENSMUSG00 | Nlrc3 | 4096 | 0 | 0 | 0 | 0.27308188 | 0.21177939 | 0 |
| ENSMUSG00 | Fam26e | 1784 | 0.8245251 | 0.37814473 | 1.06540317 | 0.52585939 | 0.51484012 | 1.16192409 |
| ENSMUSG00 | Rif | 12119 | 2.82772706 | 2.52085191 | 2.73184095 | 2.35505382 | 5.14095263 | 3.19580722 |
| ENSMUSG00 | 2810025M15 | 1190 | 3.00671701 | 2.99646783 | 1.96865351 | 1.7889417 | 2.35836191 | 1.55855081 |
| ENSMUSG00 | Mettl21d | 1021 | 2.95927181 | 0.7079301 | 2.12134212 | 7.28002074 | 2.44886622 | 7.07911793 |
| ENSMUSG00 | Gm7984 | 532 | 2.61549214 | 2.3549765 | 4.90208409 | 1.49211462 | 1.4387136 | 3.69130456 |
| ENSMUSG00 | Rasd1 | 1609 | 41.0650147 | 95.0851173 | 15.05445 | 50.8377843 | 58.161823 | 27.6645191 |
| ENSMUSG00 | Olfr808 | 938 | 0 | 0 | 0 | 0 | 0 | 0 |
| ENSMUSG00 | Gm711 | 2215 | 1.47176045 | 0.36982822 | 0.01995569 | 0.50498573 | 0.8984328 | 0.1723965 |
| ENSMUSG00 | 4921517D22I | 3137 | 0.01267309 | 0.01536068 | 0 | 0 | 0 | 0 |
| ENSMUSG00 | Tmem17 | 1564 | 6.55812914 | 5.54575421 | 8.76123487 | 6.57505287 | 6.29673308 | 7.16744037 |
| ENSMUSG00 | Rasl11b | 1795 | 13.3994795 | 17.6370432 | 6.91961999 | 9.06573932 | 10.6885333 | 6.03232968 |
| ENSMUSG00 | Gja8 | 1641 | 0 | 0 | 0 | 0 | 0 | 0 |
| ENSMUSG00 | 2610318N02I | 1257 | 0 | 0 | 0 | 0 | 0 | 0 |
| ENSMUSG00 | Slc35c1 | 4775 | 6.41915717 | 5.17688897 | 4.99874246 | 4.43563563 | 4.14622698 | 6.11751595 |
| ENSMUSG00 | Olfr921 | 1517 | 0 | 0 | 0 | 0 | 0 | 0 |
| ENSMUSG00 | Glp2r | 5800 | 0 | 0 | 0 | 0 | 0.25513188 | 0 |
| ENSMUSG00 | Lpar4 | 4235 | 1.10770878 | 0 | 2.0561424 | 0 | 0.57833909 | 1.33958087 |
| ENSMUSG00 | H2afx | 1356 | 4.16318454 | 6.89393051 | 8.47527963 | 7.39735131 | 3.98878749 | 9.2323447 |
| ENSMUSG00 | Lrrc4 | 2597 | 0.0918494 | 1.02050609 | 0.27232556 | 0.750262 | 0.72698341 | 0.27306146 |
| ENSMUSG00 | Pgrmc2 | 3443 | 14.4103513 | 11.5602676 | 13.7753638 | 13.4665778 | 13.6791591 | 19.3924824 |
| ENSMUSG00 | BC030500 | 660 | 0 | 0 | 0 | 0 | 0 | 0 |
| ENSMUSG00 | Rpp38 | 1040 | 7.68351114 | 10.656617 | 7.69282076 | 11.7613587 | 8.29178607 | 5.97944014 |
| ENSMUSG00 | Fam71e2 | 2927 | 0 | 0 | 0 | 0 | 0 | 0 |
| ENSMUSG00 | Ccdc137 | 2271 | 29.3045682 | 19.1599988 | 24.9912666 | 23.0855177 | 28.5352256 | 26.205752 |
| ENSMUSG00 | Mrps16 | 563 | 19.136297 | 18.0592172 | 33.2102657 | 22.8797334 | 15.2263431 | 40.6939555 |
| ENSMUSG00 | Plekhf2 | 3641 | 29.1096158 | 33.0992287 | 41.3125163 | 29.6405919 | 24.497145 | 35.0277982 |
| ENSMUSG00 | Glt1d1 | 3745 | 0.96602102 | 0.65621056 | 1.18028953 | 0.44319734 | 0.77663643 | 0.71372888 |
| ENSMUSG00 | Skint9 | 1365 | 0 | 0 | 0 | 0 | 0 | 0 |
| ENSMUSG00 | Ankrd55 | 6549 | 0 | 0 | 0 | 0 | 0 | 0 |
| ENSMUSG00 | Lrrc25 | 1270 | 0 | 0 | 0 | 0 | 0 | 0 |
| ENSMUSG00 | Ppp1r3d | 3266 | 0.31648576 | 0.59015851 | 0.33834846 | 0.41981558 | 0.24997612 | 0.60127802 |
| ENSMUSG00 | 5133401N09I | 1499 | 10.5820125 | 9.5472804 | 11.7360464 | 12.3964329 | 10.0418818 | 11.7541141 |
| ENSMUSG00 | Shisa3 | 5464 | 0 | 0 | 0 | 0 | 0 | 0.03993359 |
| ENSMUSG00 | Apol10b | 1982 | 0 | 0 | 0 | 0 | 0 | 0 |
| ENSMUSG00 | Olfr350 | 938 | 0 | 0 | 0 | 0 | 0 | 0 |
| ENSMUSG00 | Pitpnb | 2749 | 95.7804829 | 92.0783487 | 141.931491 | 117.013979 | 85.4956301 | 107.005087 |
| ENSMUSG00 | Amz1 | 6713 | 0.07696812 | 0 | 0.00658451 | 0 | 0.30404514 | 0 |
| ENSMUSG00 | Olfr1176 | 948 | 0 | 0 | 0 | 0 | 0 | 0 |
| ENSMUSG00 | Olfr745 | 1174 | 0 | 0 | 0 | 0 | 0 | 0 |
| ENSMUSG00 | Rap2c | 3542 | 3.22129388 | 3.40107582 | 4.3802504 | 3.60616707 | 2.83800001 | 3.88867669 |
| ENSMUSG00 | Olfr728 | 973 | 0 | 0 | 0 | 0 | 0 | 0 |
| ENSMUSG00 | Fhl4 | 2668 | 0 | 0.14448708 | 0 | 0.04057201 | 0 | 0 |
| ENSMUSG00 | Tmx2 | 3535 | 50.1582583 | 42.9793075 | 61.8700762 | 42.7983063 | 38.4393886 | 51.2547181 |
| ENSMUSG00 | Ccdc158 | 6408 | 0.09926462 | 0.01503946 | 0 | 0 | 0.02388875 | 0 |
| ENSMUSG00 | 2610019F03F | 4468 | 123.804332 | 82.2879484 | 212.184675 | 107.349735 | 92.5623939 | 118.102606 |
| ENSMUSG00 | Prm3 | 406 | 0 | 0 | 0 | 0 | 0 | 0 |
| ENSMUSG00 | Klk6 | 1477 | 0 | 0 | 0 | 0 | 0 | 0 |
| ENSMUSG00 | Zfp697 | 5160 | 0.59325039 | 1.00855344 | 0.25698746 | 1.01393342 | 0.81088427 | 0.67129442 |
| ENSMUSG00 | A130023I24R | 1893 | 0.04200262 | 0 | 0 | 0 | 0 | 0 |
| ENSMUSG00 | Grem2 | 3743 | 1.01964364 | 2.04692608 | 0.17713803 | 0.3566753 | 0.0681624 | 0.0364342 |
| ENSMUSG00 | Bex1 | 1794 | 0.53184589 | 1.8533247 | 0.81307738 | 0.50281556 | 1.13771184 | 0.38008137 |
| ENSMUSG00 | Spink8 | 788 | 635.784981 | 545.33844 | 888.636544 | 572.550615 | 288.674598 | 848.836615 |
| ENSMUSG00 | Gpr171 | 2223 | 0 | 0 | 0 | 0 | 0.04590767 | 0.17177011 |
| ENSMUSG00 | Rspry1 | 6862 | 7.50267375 | 8.53901394 | 7.70408106 | 10.2693432 | 7.63685339 | 7.42083233 |
| ENSMUSG00 | Olfr622 | 1104 | 0 | 0 | 0 | 0 | 0 | 0 |
| ENSMUSG00 | Cby3 | 728 | 0 | 0.33095084 | 0.54645136 | 0 | 0 | 0.44958197 |
| ENSMUSG00 | 1600012H06 | 3375 | 35.6562487 | 34.6656834 | 37.142645 | 38.8510074 | 31.326415 | 41.7807066 |
| ENSMUSG00 | Akap4 | 3156 | 0 | 0 | 0 | 0 | 0 | 0 |
| ENSMUSG00 | Sprr2b | 697 | 0 | 0 | 0 | 0 | 0 | 0 |
| ENSMUSG00 | Ces2b | 3542 | 0 | 0 | 0 | 0 | 0 | 0 |
| ENSMUSG00 | Hmx2 | 1667 | 199.94598 | 153.953804 | 288.465417 | 163.397333 | 129.876972 | 247.418773 |
| ENSMUSG00 | Vmn1r235 | 1777 | 0 | 0 | 0 | 0 | 0 | 0 |
| ENSMUSG00 | Agmo | 6796 | 0.25154292 | 0.34033979 | 0 | 0 | 0.01501659 | 0 |
| ENSMUSG00 | Grrp1 | 1824 | 0.28334498 | 0.15850803 | 0 | 0.35607281 | 0.11189995 | 0.22429802 |
| ENSMUSG00 | Tmc8 | 3038 | 0.05234428 | 0 | 0.10184757 | 0.03563072 | 0.16796042 | 0.06284479 |
| ENSMUSG00 | Gsg2 | 2809 | 0 | 0 | 0 | 0 | 0.16348785 | 0.71852022 |
| ENSMUSG00 | Bpifc | 2831 | 0.58980225 | 0.32339894 | 0 | 0 | 0 | 0 |
| ENSMUSG00 | Prdx6b | 2884 | 0 | 0 | 0 | 0.01251111 | 0 | 0.01891445 |
| ENSMUSG00 | Opalin | 1832 | 0 | 0 | 0 | 0 | 0 | 0 |
| ENSMUSG00 | Vwa3b | 1923 | 0 | 0 | 0.06895763 | 0 | 0.02653478 | 0.35458449 |
| ENSMUSG00 | Olfr1023 | 1878 | 0 | 0 | 0 | 0 | 0 | 0 |
| ENSMUSG00 | Sarm1 | 7371 | 0.04854149 | 0.0196119 | 0.11993445 | 0.20070056 | 0 | 0 |
| ENSMUSG00 | Olfr430 | 953 | 0 | 0 | 0 | 0 | 0 | 0 |
| ENSMUSG00 | Kcnk12 | 1934 | 0 | 0 | 0 | 0 | 0 | 0 |
| ENSMUSG00 | BC049635 | 2425 | 0 | 0 | 0 | 0 | 0 | 0 |
| ENSMUSG00 | Slc25a44 | 3614 | 17.5016518 | 20.7732365 | 18.5173189 | 15.1955026 | 25.7249742 | 13.8939156 |
| ENSMUSG00 | F2rl3 | 1749 | 0 | 0 | 0.65708858 | 0 | 0 | 0 |
| ENSMUSG00 | Ubqln2 | 3305 | 3.13954022 | 3.19298966 | 2.52773021 | 3.89751585 | 1.71374515 | 3.51558134 |
| ENSMUSG00 | Nhedc1 | 3763 | 0.06338902 | 0 | 0 | 0 | 0.12204023 | 0.21744331 |
| ENSMUSG00 | Gm867 | 1216 | 0 | 0 | 0 | 0 | 0 | 0 |
| ENSMUSG00 | Olfr165 | 1039 | 0 | 0 | 0 | 0 | 0 | 0 |
| ENSMUSG00 | Mchr1 | 2366 | 0 | 0 | 0 | 0 | 0 | 0 |
| ENSMUSG00 | Nudt6 | 2110 | 2.41170687 | 2.85464705 | 3.20515734 | 2.4111698 | 2.6359597 | 3.08940227 |
| ENSMUSG00 | A930002I21R | 1428 | 0 | 0 | 0 | 0 | 0 | 0 |
| ENSMUSG00 | Lsm10 | 988 | 8.16838314 | 9.02276498 | 9.75303824 | 8.36314595 | 7.17881198 | 11.3184232 |
| ENSMUSG00 | Etos1 | 2014 | 0 | 0 | 0 | 0 | 0 | 0 |
| ENSMUSG00 | Gm5346 | 2319 | 0 | 0 | 0 | 0 | 0 | 0 |
| ENSMUSG00 | Eif5a2 | 4987 | 4.20912246 | 4.22247347 | 3.23514592 | 5.14424176 | 6.01634429 | 4.59408401 |
| ENSMUSG00 | Scd4 | 3762 | 0.62349106 | 0.7685238 | 0 | 0 | 0.09494541 | 0.06525033 |
| ENSMUSG00 | Rhox13 | 872 | 0 | 0 | 0 | 0 | 0 | 0 |
| ENSMUSG00 | Olfr768 | 938 | 0 | 0 | 0 | 0 | 0 | 0 |
| ENSMUSG00 | Lgr4 | 5864 | 9.24735256 | 10.4442306 | 8.82680051 | 14.7921615 | 15.6542377 | 9.9814761 |
| ENSMUSG00 | Otop2 | 2638 | 0 | 0 | 0 | 0 | 0 | 0 |
| ENSMUSG00 | Pla2g4e | 6939 | 0 | 0 | 0 | 0 | 0 | 0 |
| ENSMUSG00 | Fam176b | 2568 | 4.13345533 | 8.29377236 | 5.00885371 | 5.39544591 | 4.98739109 | 5.44855527 |
| ENSMUSG00 | Snip1 | 2674 | 11.7155268 | 9.2804853 | 11.703405 | 13.817507 | 9.77019612 | 9.99594109 |
| ENSMUSG00 | Olfr70 | 3289 | 0 | 0 | 0 | 0 | 0 | 0 |
| ENSMUSG00 | Lgsn | 1858 | 0 | 0 | 0 | 0 | 0 | 0 |
| ENSMUSG00 | Il17d | 1209 | 0.95360541 | 0.59784668 | 1.38930524 | 1.3131596 | 0.46425983 | 0.94750608 |
| ENSMUSG00 | Krtap13 | 881 | 0 | 0 | 0 | 0 | 0 | 0 |
| ENSMUSG00 | Pigm | 7568 | 5.47898602 | 4.62890373 | 6.03336467 | 5.78800239 | 4.19376398 | 5.83838737 |
| ENSMUSG00 | Cxcr3 | 1729 | 0 | 0 | 0 | 0 | 0 | 0 |
| ENSMUSG00 | Gja4 | 1683 | 1.58266025 | 0.9448322 | 0.26263721 | 0 | 0.96956107 | 0.25929544 |
| ENSMUSG00 | Krtap24-1 | 1570 | 0 | 0 | 0 | 0 | 0 | 0 |
| ENSMUSG00 | Hic2 | 4582 | 1.81337744 | 2.1874247 | 1.32161774 | 1.96868423 | 2.06020941 | 2.00601341 |
| ENSMUSG00 | Klre1 | 1296 | 0 | 0 | 0 | 0.02784108 | 0 | 0 |
| ENSMUSG00 | Gm5446 | 782 | 0.15251463 | 0 | 0 | 0.04614072 | 0.3262556 | 0 |
| ENSMUSG00 | Heatr1 | 6492 | 13.8397083 | 8.53579921 | 12.6573053 | 4.82983621 | 11.0195593 | 5.36502069 |
| ENSMUSG00 | Evc2 | 6036 | 3.80034994 | 5.90754924 | 3.36859639 | 3.99317524 | 7.79428736 | 2.59371155 |
| ENSMUSG00 | Gm17291 | 2500 | 1.39939291 | 0.69388477 | 1.29069382 | 0.59174553 | 0.83683256 | 0.0981887 |
| ENSMUSG00 | Olfr809 | 1099 | 0 | 0 | 0 | 0 | 0 | 0 |
| ENSMUSG00 | Olfr690 | 1061 | 0 | 0 | 0.1666422 | 0 | 0 | 0 |
| ENSMUSG00 | Tmem220 | 1662 | 7.05647819 | 8.84287897 | 7.31378269 | 2.36639161 | 6.44737601 | 3.65959359 |
| ENSMUSG00 | D8Ertd82e | 9728 | 0 | 0.28234243 | 0.0363502 | 0.24109096 | 0 | 0.01401863 |
| ENSMUSG00 | Dscam | 7465 | 0.02662792 | 0 | 0.04144848 | 0 | 0 | 0.18999079 |
| ENSMUSG00 | Mrgprg | 1818 | 0 | 0 | 0 | 0 | 0 | 0 |
| ENSMUSG00 | Olfr630 | 2608 | 0 | 0 | 0 | 0 | 0 | 0 |
| ENSMUSG00 | Fzd2 | 3627 | 0.60285399 | 0.45170638 | 0.63371818 | 0.57699437 | 0.77376638 | 0.53391216 |
| ENSMUSG00 | Foxc1 | 3989 | 1.30558234 | 0.55567218 | 0.2326996 | 0.0633177 | 0.46050377 | 0.20512389 |
| ENSMUSG00 | Abca12 | 8270 | 0 | 0 | 0 | 0 | 0 | 0 |
| ENSMUSG00 | Gm9843 | 503 | 1.42266133 | 1.34117334 | 1.75752855 | 1.21947268 | 0.71016861 | 2.54852494 |
| ENSMUSG00 | Slc25a2 | 1345 | 0 | 0 | 0 | 0.34874839 | 0 | 0 |
| ENSMUSG00 | Rictor | 9341 | 18.2413007 | 18.1273052 | 17.1488576 | 18.4833083 | 21.0420297 | 19.4522692 |
| ENSMUSG00 | Nsun3 | 5064 | 11.2106687 | 9.99126466 | 8.21365211 | 8.40774339 | 6.38837328 | 11.2190114 |
| ENSMUSG00 | Synpo2 | 7580 | 0 | 0 | 0 | 0 | 0 | 0 |
| ENSMUSG00 | Neto1 | 3520 | 0 | 0 | 0 | 0 | 0 | 0 |
| ENSMUSG00 | 2310030N02I | 1401 | 13.3653329 | 12.4163495 | 12.8409351 | 13.7786537 | 10.9992616 | 13.6665216 |
| ENSMUSG00 | 1700034J04R | 749 | 0 | 0 | 0 | 0 | 0 | 0 |
| ENSMUSG00 | Hoxc12 | 1029 | 0 | 0 | 0 | 0 | 0 | 0 |
| ENSMUSG00 | Fam123b | 8494 | 1.46965162 | 3.26764665 | 2.83612014 | 3.64473678 | 4.33730202 | 3.79224735 |
| ENSMUSG00 | C130071C03 | 3411 | 1.94639849 | 0.25428202 | 0.86822735 | 2.2319881 | 0.53853695 | 0.75962789 |
| ENSMUSG00 | Lgals3 | 1828 | 157.69529 | 208.720049 | 188.704148 | 160.928287 | 103.420527 | 181.328606 |
| ENSMUSG00 | Olfr1378 | 947 | 0 | 0 | 0 | 0 | 0 | 0 |
| ENSMUSG00 | 4930486L24F | 1356 | 0.17590921 | 0.53303586 | 0 | 0 | 0 | 0 |
| ENSMUSG00 | Gm9844 | 609 | 0 | 0 | 0 | 0 | 0 | 0 |
| ENSMUSG00 | Gpr18 | 1307 | 0.39542559 | 0.33181177 | 0.06763863 | 0 | 0.58561258 | 0 |
| ENSMUSG00 | Rltpr | 5472 | 0.22522293 | 0.29940406 | 0.09693387 | 0.0329697 | 0.009325 | 0.049844 |
| ENSMUSG00 | Sprr1a | 804 | 0 | 12.1065439 | 0.27488708 | 0 | 0.12693128 | 0 |
| ENSMUSG00 | Olfr524 | 1057 | 0 | 0 | 0 | 0 | 0 | 0 |
| ENSMUSG00 | Hoxd10 | 2649 | 29.0699757 | 44.1481673 | 23.7278297 | 36.7767158 | 48.0406875 | 34.0701326 |
| ENSMUSG00 | Ch25h | 1349 | 0.44205501 | 0 | 0 | 0 | 0 | 0 |
| ENSMUSG00 | Snx21 | 3347 | 8.67089954 | 8.07666389 | 7.34276209 | 5.26084187 | 9.48264289 | 9.27354032 |
| ENSMUSG00 | Il31ra | 3933 | 0 | 0 | 0 | 0 | 0 | 0 |
| ENSMUSG00 | 6-Sep | 8671 | 1.22874741 | 1.45042802 | 0.97875145 | 1.81429725 | 1.77129963 | 0.95937781 |
| ENSMUSG00 | Kif7 | 4503 | 0 | 0.19261736 | 0 | 0 | 0.21530116 | 0 |
| ENSMUSG00 | Svs3b | 1246 | 0 | 0 | 0 | 0 | 0 | 0 |
| ENSMUSG00 | C77080 | 5759 | 19.8121599 | 19.1105806 | 16.6015951 | 21.0703101 | 21.1760789 | 20.7531636 |
| ENSMUSG00 | Armcx6 | 4270 | 2.07622299 | 0.5980987 | 2.20491629 | 0.79431199 | 1.64909599 | 1.83960096 |
| ENSMUSG00 | Tnfsf15 | 8783 | 0.50695819 | 0.44439278 | 1.0165971 | 0.15611041 | 0.1394322 | 0.87882533 |
| ENSMUSG00 | Foxl2 | 2519 | 0 | 0 | 0 | 0 | 0 | 0 |
| ENSMUSG00 | Tmem151b | 1683 | 0.47243589 | 0.31494407 | 0.23637349 | 0.02143912 | 0.18191221 | 0.03241193 |
| ENSMUSG00 | Tcf19 | 2221 | 4.5107524 | 5.42395792 | 5.41328289 | 4.76003559 | 6.04229484 | 5.40335041 |
| ENSMUSG00 | Ppp1r3g | 2516 | 0 | 0 | 0 | 0 | 0 | 0 |
| ENSMUSG00 | Pnma5 | 3123 | 0 | 0 | 0 | 0.01155365 | 0 | 0 |
| ENSMUSG00 | Mrgprb2 | 2072 | 0 | 0 | 0 | 0 | 0 | 0 |
| ENSMUSG00 | Fbxo46 | 3022 | 3.80189738 | 3.44416661 | 3.74443144 | 4.28638451 | 5.38630506 | 3.37548481 |
| ENSMUSG00 | Olfr730 | 956 | 0 | 0 | 0 | 0 | 0 | 0 |
| ENSMUSG00 | Ube2dnl | 741 | 0 | 0 | 0 | 0 | 0 | 0 |
| ENSMUSG00 | Enthd1 | 2104 | 0 | 0 | 0 | 0 | 0 | 0 |
| ENSMUSG00 | Hamp | 389 | 0 | 0 | 0 | 0 | 0 | 0 |
| ENSMUSG00 | Cyp8b1 | 1949 | 0 | 0 | 0 | 0 | 0 | 0 |
| ENSMUSG00 | Lypd6 | 3805 | 12.2662113 | 28.6205938 | 13.4754633 | 19.7431844 | 20.0618813 | 13.2108174 |
| ENSMUSG00 | Vmn1r89 | 950 | 0 | 0 | 0 | 0 | 0 | 0 |
| ENSMUSG00 | Gm17646 | 7230 | 0 | 0 | 0 | 0 | 0 | 0 |
| ENSMUSG00 | Krt78 | 4538 | 0 | 0 | 0 | 0 | 0 | 0 |
| ENSMUSG00 | Astl | 2549 | 0 | 0 | 0 | 0 | 0 | 0 |
| ENSMUSG00 | Fam118b | 6697 | 5.87694874 | 8.77098297 | 6.29003381 | 8.85753042 | 5.80589792 | 5.44922612 |
| ENSMUSG00 | Slc35d3 | 2627 | 0 | 0.20177041 | 0 | 0.37084705 | 0 | 0.02076486 |
| ENSMUSG00 | Olfr807 | 996 | 0 | 0 | 0 | 0 | 0 | 0 |
| ENSMUSG00 | Gm8394 | 964 | 0.04124013 | 0.09997187 | 0 | 0.03742951 | 0.05293193 | 0.05658639 |
| ENSMUSG00 | Fam167b | 956 | 0.83170461 | 0.45363805 | 0.23118119 | 0.37742724 | 1.12087227 | 1.1411983 |
| ENSMUSG00 | Fbxl22 | 3933 | 0.30324547 | 0.02450366 | 0.30344515 | 0.04587089 | 0.41516502 | 0.36061054 |
| ENSMUSG00 | Olfr1323 | 929 | 0 | 0 | 0 | 0 | 0 | 0 |
| ENSMUSG00 | Pcdh20 | 5234 | 0 | 0 | 0 | 0 | 0 | 0 |
| ENSMUSG00 | Oprd1 | 4018 | 0.24735864 | 0 | 0 | 0 | 0 | 0 |
| ENSMUSG00 | Cldn8 | 2355 | 1189.23751 | 1060.10173 | 1021.99166 | 1115.50973 | 964.496006 | 1198.13842 |
| ENSMUSG00 | 4933406M09 | 1766 | 0 | 0.16371385 | 0 | 0 | 0 | 0 |
| ENSMUSG00 | Fam171a1 | 4432 | 46.5368757 | 33.5522023 | 70.8705543 | 38.64654 | 40.0658366 | 52.96154 |
| ENSMUSG00 | Gm9845 | 647 | 0 | 0.07447673 | 0.20495445 | 0 | 0 | 0.04215555 |
| ENSMUSG00 | Htr5b | 1933 | 0 | 0 | 0 | 0 | 0 | 0 |
| ENSMUSG00 | B230217C12 | 2447 | 0 | 0 | 0 | 0 | 0 | 0 |
| ENSMUSG00 | Olfr462 | 935 | 0 | 0 | 0 | 0 | 0 | 0 |
| ENSMUSG00 | Adra1b | 3813 | 0.36492049 | 0.03791223 | 0.46369623 | 0.26496125 | 0 | 0 |
| ENSMUSG00 | A830093I24R | 2879 | 0.2209405 | 0.36821873 | 0.18423832 | 0.45118222 | 0.88618228 | 0.38841966 |
| ENSMUSG00 | 5730508B09I | 1601 | 57.6839358 | 49.6612302 | 88.2932504 | 61.3462369 | 32.4771249 | 99.3028558 |
| ENSMUSG00 | Gm11868 | 2703 | 0.1470791 | 0.07130809 | 0.37611631 | 0.14683777 | 0.24540987 | 0.32289621 |
| ENSMUSG00 | 0910001L09F | 1172 | 9.46397531 | 10.7309398 | 11.4653245 | 8.22005622 | 7.53205036 | 22.0151952 |
| ENSMUSG00 | Gk2 | 1847 | 0 | 0 | 0 | 0 | 0 | 0.04430096 |
| ENSMUSG00 | Hyls1 | 5671 | 1.18474276 | 0.98565108 | 0.68590411 | 1.17707252 | 0.97175958 | 0.93304179 |
| ENSMUSG00 | Kcnb1 | 4046 | 0 | 0.25010264 | 0 | 0 | 0.13872717 | 0 |
| ENSMUSG00 | Prokr2 | 7888 | 0 | 0 | 0 | 0 | 0 | 0 |
| ENSMUSG00 | Tor1aip2 | 6922 | 9.12618586 | 8.35361609 | 12.2350088 | 14.439073 | 11.3891579 | 12.8295616 |
| ENSMUSG00 | Maml1 | 8739 | 0.82795485 | 1.05316517 | 1.01665756 | 1.50703127 | 0.71234899 | 0.96751781 |
| ENSMUSG00 | Olfr1038-ps | 1208 | 0 | 0 | 0 | 0 | 0 | 0 |
| ENSMUSG00 | Mmp13 | 2665 | 0 | 0 | 0 | 0 | 0 | 0 |
| ENSMUSG00 | Lrrc4c | 4984 | 0 | 0 | 0.15963748 | 0 | 0 | 0 |
| ENSMUSG00 | Fam78a | 5790 | 0 | 0 | 0 | 0 | 0 | 0.08479163 |
| ENSMUSG00 | Zfp831 | 9802 | 0 | 0 | 0 | 0 | 0 | 0 |
| ENSMUSG00 | Olfr1008 | 941 | 0 | 0 | 0 | 0 | 0 | 0 |
| ENSMUSG00 | 3110021A11F | 820 | 2.13322091 | 2.76190583 | 2.91085308 | 1.49608477 | 3.36027353 | 4.35728993 |
| ENSMUSG00 | Zfp61 | 3111 | 7.96131931 | 5.54508077 | 7.64403457 | 5.70632141 | 8.67661614 | 9.39839707 |
| ENSMUSG00 | Minos1 | 2982 | 36.0892306 | 25.3697901 | 37.8724711 | 29.8747713 | 25.9580859 | 54.2931786 |
| ENSMUSG00 | Txndc2 | 1896 | 0 | 0 | 0 | 0 | 0 | 0 |
| ENSMUSG00 | Olfr125 | 1117 | 0 | 0 | 0 | 0 | 0 | 0 |
| ENSMUSG00 | Zscan29 | 7799 | 10.7557461 | 9.93509411 | 10.082713 | 11.7559271 | 12.3983821 | 8.48769707 |
| ENSMUSG00 | Gm9846 | 343 | 29.2081081 | 36.9476218 | 33.1191652 | 39.6586903 | 27.5215147 | 23.7758517 |
| ENSMUSG00 | 1700019N12I | 776 | 3.48372767 | 1.36611047 | 3.01894031 | 2.69685384 | 1.70964661 | 3.09300033 |
| ENSMUSG00 | Ccdc121 | 1818 | 0 | 0.29155713 | 0 | 0 | 0.25260582 | 0 |
| ENSMUSG00 | Gpd1l | 7131 | 13.2518268 | 17.6501174 | 23.2197594 | 10.2563882 | 10.5687782 | 14.1976526 |
| ENSMUSG00 | Fam100b | 1529 | 0.75402808 | 0.59878509 | 0.9829056 | 0.92033992 | 0.26697908 | 1.32002832 |
| ENSMUSG00 | 1110035H17I | 1228 | 0.3237417 | 0.07847955 | 0.14397995 | 0.23506218 | 0.29086696 | 0.44421237 |
| ENSMUSG00 | Sprr2f | 607 | 0 | 0 | 0 | 0 | 0 | 0 |
| ENSMUSG00 | Tmem150c | 5544 | 1.68516196 | 4.20675289 | 1.29958521 | 2.53823906 | 2.76116752 | 1.71204446 |
| ENSMUSG00 | BC048562 | 685 | 0 | 0 | 0 | 0 | 0 | 0 |
| ENSMUSG00 | Defb19 | 392 | 0 | 0 | 0 | 0 | 0 | 0 |
| ENSMUSG00 | Mrgpra1 | 1086 | 0 | 0 | 0 | 0 | 0 | 0 |
| ENSMUSG00 | Olfr215 | 1093 | 0 | 0 | 0 | 0 | 0 | 0 |
| ENSMUSG00 | Trhde | 8046 | 0 | 0 | 0 | 0 | 0 | 0 |
| ENSMUSG00 | E130203B14I | 2771 | 0 | 0 | 0 | 0 | 0 | 0 |
| ENSMUSG00 | Gm12612 | 2192 | 0 | 0 | 0 | 0 | 0 | 0 |
| ENSMUSG00 | Ccdc75 | 4237 | 11.6442179 | 9.56450268 | 7.82425826 | 8.7629039 | 10.7183088 | 11.747986 |
| ENSMUSG00 | Ism2 | 3113 | 0 | 0 | 0 | 0 | 0 | 0 |
| ENSMUSG00 | Gp1ba | 3560 | 0.13400724 | 0.02707104 | 0.06208124 | 0.05067703 | 0.10033276 | 0.17621256 |
| ENSMUSG00 | Ccdc96 | 3583 | 0.72121302 | 0.59173973 | 0.08635582 | 0.57400964 | 0.24210114 | 0.06089788 |
| ENSMUSG00 | Ccdc54 | 1295 | 0 | 0 | 0 | 0 | 0 | 0 |
| ENSMUSG00 | Prkaa1 | 4646 | 34.4074015 | 25.099266 | 34.7069142 | 26.7004023 | 26.0513481 | 30.3801675 |
| ENSMUSG00 | Emilin3 | 6281 | 0 | 0 | 0 | 0 | 0 | 0 |
| ENSMUSG00 | Rfpl3s | 766 | 0 | 0 | 0 | 0 | 0 | 0 |
| ENSMUSG00 | 2310061N02I | 791 | 0 | 0 | 0 | 0 | 0 | 0 |
| ENSMUSG00 | 2310061I04R | 1901 | 65.3947331 | 55.1824748 | 73.7549953 | 62.0094895 | 51.3484781 | 73.5884401 |
| ENSMUSG00 | Ftl1 | 1632 | 58.8049816 | 99.3845369 | 41.4121434 | 105.438278 | 74.2571339 | 40.1933256 |
| ENSMUSG00 | 4932442E05F | 2716 | 0.14637511 | 0 | 0.292943 | 0.05313998 | 0.05636198 | 0.42177277 |
| ENSMUSG00 | Scg2 | 4640 | 0 | 0 | 0 | 0 | 0 | 0 |
| ENSMUSG00 | Zbtb26 | 7245 | 1.54741829 | 2.30124348 | 2.25127399 | 2.80389111 | 2.57068698 | 2.71805239 |
| ENSMUSG00 | Plekho2 | 3265 | 0.06088129 | 0.76744104 | 0.69044227 | 0.50835346 | 0.51573366 | 0.27567017 |
| ENSMUSG00 | Arhgap42 | 6289 | 2.30100094 | 1.06502074 | 6.76838525 | 2.21460792 | 1.53346876 | 4.43229151 |
| ENSMUSG00 | Vamp8 | 2733 | 34.795137 | 39.4589307 | 47.4041719 | 31.8308852 | 29.5180023 | 60.796598 |
| ENSMUSG00 | Ptges | 7265 | 26.2391644 | 37.9920083 | 21.3799417 | 37.6663766 | 36.1925554 | 27.6688357 |
| ENSMUSG00 | Olfr164 | 1094 | 0 | 0 | 0 | 0 | 0 | 0 |
| ENSMUSG00 | Trim15 | 2755 | 0.11544241 | 0.24486758 | 0.20857494 | 0.379811 | 0.64824797 | 0.35640182 |
| ENSMUSG00 | Pgbd5 | 5718 | 0 | 0.09269865 | 0.02319089 | 0 | 0 | 0 |
| ENSMUSG00 | Defb6 | 321 | 0 | 0 | 0 | 0 | 0 | 0 |
| ENSMUSG00 | Gp1bb | 2489 | 0 | 0.21295736 | 0.01775888 | 0 | 0 | 0 |
| ENSMUSG00 | Prss27 | 1247 | 0.0956427 | 0.30913515 | 0 | 0 | 0 | 0 |
| ENSMUSG00 | Olfr1395 | 953 | 0 | 0 | 0 | 0 | 0 | 0 |
| ENSMUSG00 | Gm5084 | 1001 | 0 | 0 | 0 | 0 | 0 | 0 |
| ENSMUSG00 | Ear14 | 467 | 0 | 0 | 0 | 0 | 0 | 0 |
| ENSMUSG00 | Olfr1124 | 956 | 0 | 0 | 0 | 0 | 0 | 0 |
| ENSMUSG00 | Olfr1317 | 950 | 0 | 0 | 0 | 0 | 0 | 0 |
| ENSMUSG00 | Tmem37 | 1309 | 52.936442 | 38.9099078 | 78.6786052 | 44.4065498 | 22.0633527 | 63.4880258 |
| ENSMUSG00 | Olfr1284 | 1063 | 0 | 0 | 0 | 0 | 0 | 0 |
| ENSMUSG00 | Htr1f | 2506 | 0 | 0 | 0 | 0 | 0 | 0 |
| ENSMUSG00 | Ccdc126 | 2590 | 33.9379797 | 23.2569049 | 85.8949328 | 32.7942598 | 21.100096 | 68.0286369 |
| ENSMUSG00 | Olfr419 | 1087 | 0 | 0 | 0 | 0 | 0 | 0 |
| ENSMUSG00 | B3galt6 | 3212 | 4.83947474 | 2.59534698 | 6.94954257 | 5.2572842 | 4.67053378 | 6.27520498 |
| ENSMUSG00 | Hist1h2ba | 388 | 0 | 0 | 0 | 0 | 0 | 0 |
| ENSMUSG00 | Olfr866 | 1076 | 0 | 0 | 0 | 0 | 0 | 0 |
| ENSMUSG00 | Taar8a | 1108 | 0 | 0 | 0 | 0 | 0 | 0 |
| ENSMUSG00 | Muc15 | 3660 | 0 | 0 | 0 | 0 | 0 | 0 |
| ENSMUSG00 | Oog3 | 1861 | 0 | 0 | 0 | 0 | 0 | 0 |
| ENSMUSG00 | AI314180 | 10982 | 15.457649 | 12.5928874 | 13.3104621 | 15.4519992 | 18.1394073 | 13.4808543 |
| ENSMUSG00 | Olfr332 | 1034 | 0 | 0 | 0 | 0 | 0 | 0 |
| ENSMUSG00 | Olfr1441 | 1087 | 0 | 0 | 0 | 0 | 0 | 0 |
| ENSMUSG00 | Olfr330 | 2027 | 0 | 0 | 0 | 0 | 0 | 0 |
| ENSMUSG00 | Fam131a | 3034 | 1.49377877 | 1.04822188 | 0.80128588 | 0.52327289 | 0.60545469 | 0.16181394 |
| ENSMUSG00 | Slc29a4 | 2778 | 0 | 0 | 0 | 0 | 0 | 0.3239968 |
| ENSMUSG00 | Sstr5 | 2695 | 0 | 0 | 0 | 0 | 0 | 0 |
| ENSMUSG00 | Vwc2 | 6034 | 0 | 0.01597164 | 0 | 0.00597979 | 0 | 0.10396366 |
| ENSMUSG00 | Ntng1 | 326 | 0 | 0 | 0 | 0 | 0 | 0 |
| ENSMUSG00 | Cdh20 | 4000 | 0.2981661 | 0.07227966 | 0.03315138 | 0.04510256 | 0.17859232 | 0.37502629 |
| ENSMUSG00 | 1700020N01I | 446 | 0 | 0 | 0 | 0 | 0 | 0 |
| ENSMUSG00 | Zfp623 | 2461 | 12.4064238 | 13.3144172 | 16.2546396 | 19.5145068 | 18.7850047 | 14.5405635 |
| ENSMUSG00 | Gm12115 | 712 | 7.48207077 | 4.39904317 | 7.2635051 | 8.15900164 | 6.30663072 | 16.7018858 |
| ENSMUSG00 | Olfr958 | 941 | 0 | 0 | 0 | 0 | 0 | 0 |
| ENSMUSG00 | Tmem125 | 2566 | 9.00153319 | 7.09839249 | 12.14431 | 5.68088308 | 8.45136778 | 10.6398729 |
| ENSMUSG00 | Zfp940 | 5738 | 1.93996768 | 2.04905749 | 2.30330273 | 1.72927191 | 1.30722852 | 2.75693461 |
| ENSMUSG00 | Atp5k | 704 | 20.9506865 | 24.5723476 | 32.9002354 | 18.1435281 | 21.9616362 | 70.588626 |
| ENSMUSG00 | Phospho1 | 1450 | 0 | 0 | 0 | 0 | 0 | 0 |
| ENSMUSG00 | Olfr1494 | 947 | 0 | 0 | 0 | 0 | 0 | 0 |
| ENSMUSG00 | Clrn3 | 3302 | 0 | 0.64209675 | 0 | 0 | 0 | 0 |
| ENSMUSG00 | Mrgprb8 | 1122 | 0 | 0 | 0 | 0 | 0 | 0 |
| ENSMUSG00 | A730017C20I | 5623 | 0 | 0 | 0 | 0 | 0 | 0 |
| ENSMUSG00 | Fam75d3 | 4273 | 0 | 0 | 0 | 0 | 0 | 0 |
| ENSMUSG00 | 4930523C07I | 354 | 0 | 0 | 0 | 0 | 0 | 0 |
| ENSMUSG00 | Olfr832 | 938 | 0 | 0 | 0 | 0 | 0 | 0 |
| ENSMUSG00 | Pdik1l | 9917 | 7.49248695 | 5.73358896 | 8.70902754 | 8.56116273 | 7.5070568 | 7.57155207 |
| ENSMUSG00 | Tatdn1 | 1450 | 7.48499737 | 9.80344857 | 10.7303784 | 12.3425476 | 9.36070067 | 17.1736177 |
| ENSMUSG00 | Rtn4rl2 | 2251 | 0 | 0 | 0.01963654 | 0.06411736 | 0.04533663 | 0 |
| ENSMUSG00 | Gm7327 | 1043 | 0.49551414 | 0.46199849 | 1.18662666 | 0.3805393 | 0.9295313 | 0.96755671 |
| ENSMUSG00 | Mtnr1b | 1242 | 0 | 0 | 0 | 0 | 0 | 0 |
| ENSMUSG00 | Gm5512 | 632 | 0.12580848 | 0.38122185 | 0.69939625 | 0.3996429 | 0.40368968 | 0.3452486 |
| ENSMUSG00 | Fam18a | 3927 | 0.03037088 | 0.03681164 | 0.43897934 | 0.23889309 | 0.67567394 | 0 |
| ENSMUSG00 | Cdr2l | 3695 | 0.74238922 | 1.57795927 | 0.960934 | 0.6737919 | 1.17381379 | 0.98173938 |
| ENSMUSG00 | Tmem123 | 3265 | 65.8248477 | 56.5692596 | 73.0650373 | 54.4601272 | 61.2003944 | 56.3870796 |
| ENSMUSG00 | Ankrd37 | 1612 | 2.17027437 | 2.1223557 | 1.64522989 | 4.96911531 | 1.55104989 | 3.11323426 |

| | | | | | | | | |
|---|---|---|---|---|---|---|---|---|
| ENSMUSG00 | Fgl4 | 3027 | 0 | 0 | 0 | 0 | 0 | 0 |
| ENSMUSG00 | Zfp366 | 59456 | 0 | 0 | 0 | 0 | 0.02574662 | 0.00458737 |
| ENSMUSG00 | P2ry10 | 2978 | 0 | 0 | 0 | 0 | 0 | 0 |
| ENSMUSG00 | Dcaf12l2 | 4735 | 0 | 0 | 0 | 0 | 0 | 0 |
| ENSMUSG00 | 4933403G14 | 7022 | 3.78758423 | 1.173438 | 2.42977947 | 4.12101961 | 3.56792232 | 2.6878454 |
| ENSMUSG00 | Sgms2 | 5866 | 29.7928899 | 30.5580574 | 15.1835516 | 31.3764932 | 26.4004519 | 19.9050001 |
| ENSMUSG00 | Vmn1r231 | 1560 | 0 | 0 | 0 | 0 | 0 | 0 |
| ENSMUSG00 | Hist2h2bb | 1889 | 0 | 0 | 0 | 0.03820227 | 0 | 0 |
| ENSMUSG00 | Efcab5 | 5524 | 0 | 0 | 0 | 0.19595607 | 0.60041898 | 0.01974992 |
| ENSMUSG00 | Zfp438 | 3204 | 8.94622392 | 4.87278628 | 9.9192026 | 9.5047583 | 7.30995833 | 5.51621917 |
| ENSMUSG00 | Amigo1 | 2386 | 15.6289358 | 16.9036262 | 12.5047125 | 12.5667138 | 12.7886726 | 16.8608939 |
| ENSMUSG00 | Gja1 | 1531 | 0 | 1.10158424 | 1.81888707 | 0 | 3.63283799 | 2.93946145 |
| ENSMUSG00 | Zfp169 | 9532 | 4.37093407 | 3.599323 | 4.19203379 | 4.77333803 | 3.00847915 | 2.96724726 |
| ENSMUSG00 | Insl6 | 703 | 2.03584253 | 5.07225707 | 5.28158579 | 5.64583912 | 5.51636495 | 7.99228405 |
| ENSMUSG00 | Olfr1392 | 1017 | 0.35181841 | 0 | 0 | 0 | 0 | 0 |
| ENSMUSG00 | AY358078 | 1452 | 0.05475962 | 0 | 0 | 0.04969979 | 0 | 0 |
| ENSMUSG00 | Kcns2 | 4858 | 0 | 0 | 0 | 0 | 0 | 0 |
| ENSMUSG00 | Prkca | 10419 | 13.2518268 | 5.81345213 | 14.9672811 | 13.7069519 | 8.97699845 | 16.9265589 |
| ENSMUSG00 | Lin28a | 4066 | 0 | 0 | 0 | 0 | 0 | 0 |
| ENSMUSG00 | Creg2 | 5795 | 0 | 0 | 0 | 0 | 0 | 0 |
| ENSMUSG00 | D330012F22I | 2246 | 4.4428431 | 3.86178076 | 6.53384322 | 2.76318418 | 6.88378979 | 4.77245469 |
| ENSMUSG00 | Gm9847 | 596 | 0 | 0 | 0 | 0 | 0 | 0 |
| ENSMUSG00 | Gm5215 | 404 | 0 | 0 | 0 | 0 | 0 | 0 |
| ENSMUSG00 | Apol10a | 3344 | 0 | 0 | 0 | 0 | 0 | 0 |
| ENSMUSG00 | Sepn1 | 5412 | 2.60775972 | 2.94710132 | 4.75341327 | 1.35341002 | 3.21507653 | 3.10947014 |
| ENSMUSG00 | 9130014G24 | 6059 | 0.72175324 | 0.36583204 | 0.26262854 | 0.44067854 | 0.46318711 | 0.18456184 |
| ENSMUSG00 | Cetn1 | 1245 | 0 | 0 | 0 | 0 | 0 | 0 |
| ENSMUSG00 | Fam160a1 | 5229 | 8.63687624 | 8.22920115 | 6.74566279 | 9.30860166 | 8.50927514 | 6.74433134 |
| ENSMUSG00 | Gm9848 | 238 | 0 | 0 | 0 | 0 | 0 | 0 |
| ENSMUSG00 | Olfr161 | 941 | 0 | 0 | 0 | 0 | 0 | 0 |
| ENSMUSG00 | Pddc1 | 3646 | 8.72308953 | 7.1632067 | 9.06828814 | 5.53205236 | 7.57138489 | 5.89479312 |
| ENSMUSG00 | 4930412M03 | 1109 | 2.22257871 | 0.69520566 | 0.19928694 | 0 | 1.33432362 | 0.61484759 |
| ENSMUSG00 | Hs3st1 | 3690 | 0 | 0.24811447 | 0.11978819 | 0.32268495 | 0.74672745 | 0.39914106 |
| ENSMUSG00 | Serpinb1b | 1689 | 0.02353788 | 0 | 0.18319296 | 0 | 0 | 0 |
| ENSMUSG00 | Gm9726 | 680 | 0 | 0 | 0.13000542 | 0 | 0 | 0 |
| ENSMUSG00 | Zfp11 | 3926 | 12.3742224 | 10.1380543 | 11.641545 | 11.1021676 | 10.6575721 | 8.71871428 |
| ENSMUSG00 | Ttc24 | 2635 | 0 | 0 | 0 | 0 | 0 | 0 |
| ENSMUSG00 | Zfp455 | 2767 | 1.39366881 | 3.29137606 | 1.03835193 | 0.07824079 | 0.5163493 | 0.01971423 |
| ENSMUSG00 | Rhox11 | 806 | 0 | 0 | 0 | 0 | 0 | 0 |
| ENSMUSG00 | Olfml1 | 1682 | 0 | 0 | 0 | 0 | 0 | 0 |
| ENSMUSG00 | Gprc5c | 5023 | 154.937943 | 134.457729 | 92.5044344 | 124.69604 | 150.854406 | 69.3676123 |
| ENSMUSG00 | Olfr214 | 1007 | 0 | 0 | 0 | 0 | 0 | 0 |
| ENSMUSG00 | P4ha3 | 3194 | 0 | 0 | 0 | 0 | 0 | 0.01707867 |
| ENSMUSG00 | Spink14 | 563 | 0 | 0.08558871 | 0 | 0 | 0 | 0 |
| ENSMUSG00 | Olfr523 | 1055 | 0 | 0 | 0 | 0 | 0 | 0 |
| ENSMUSG00 | Olfr335-ps | 955 | 0 | 0 | 0 | 0 | 0 | 0 |
| ENSMUSG00 | 1700014D04 | 794 | 0.2002795 | 0 | 0 | 0 | 0 | 0 |
| ENSMUSG00 | Gja10 | 1776 | 0 | 0 | 0 | 0.4875952 | 0 | 0 |
| ENSMUSG00 | Fbll1 | 1503 | 0 | 0 | 0 | 0 | 0 | 0 |
| ENSMUSG00 | 1600021P15I | 4551 | 1.03953025 | 1.48233397 | 2.24360047 | 2.64807798 | 0.72878805 | 2.46916093 |
| ENSMUSG00 | Lingo3 | 3431 | 0 | 0 | 0 | 0 | 0 | 0 |
| ENSMUSG00 | 4930579K19I | 651 | 2.0763231 | 2.3686116 | 2.17274804 | 3.7689386 | 1.09734142 | 4.73430758 |
| ENSMUSG00 | Vtcn1 | 1373 | 0.26059674 | 0.07019147 | 0.06438724 | 0 | 0.18582074 | 0.41716491 |
| ENSMUSG00 | Rgs13 | 1636 | 0.36450624 | 2.17958235 | 0.13509121 | 0 | 0 | 0 |
| ENSMUSG00 | Gm4847 | 3003 | 0 | 0 | 0 | 0 | 0 | 0 |
| ENSMUSG00 | Olfr986 | 1015 | 0 | 0 | 0 | 0 | 0 | 0 |
| ENSMUSG00 | Mblac2 | 3879 | 4.5607602 | 5.3788939 | 8.95659928 | 6.01832503 | 3.32809309 | 7.63605523 |
| ENSMUSG00 | Gm15440 | 1227 | 0.48600832 | 2.51339226 | 0.6484378 | 1.64677628 | 0.99807092 | 0.1111436 |
| ENSMUSG00 | Sv2c | 4369 | 0.32758006 | 0.07720419 | 1.24440986 | 0.14039706 | 0.15182945 | 0.41826524 |
| ENSMUSG00 | Fam71e1 | 1169 | 0.81619464 | 1.77246964 | 1.2855968 | 0.37038882 | 0.7420431 | 1.60968033 |
| ENSMUSG00 | Gm8121 | 1275 | 0.12472308 | 0.07558658 | 0.2773449 | 0.11319857 | 0.08004137 | 0.0855675 |
| ENSMUSG00 | Olfr77 | 2668 | 0 | 0 | 0 | 0 | 0 | 0 |
| ENSMUSG00 | Gimap9 | 855 | 5.99819531 | 5.35404913 | 8.89206666 | 4.76873805 | 2.68559873 | 13.5575692 |
| ENSMUSG00 | Gm6293 | 194 | 0.20492516 | 0 | 0 | 0.18598992 | 0 | 0.5623637 |
| ENSMUSG00 | Ghsr | 4430 | 0 | 0 | 0 | 0 | 0 | 0 |
| ENSMUSG00 | Camk2n2 | 1289 | 0.12336844 | 0.11214843 | 0 | 0.02799228 | 0.11875805 | 0.27507394 |
| ENSMUSG00 | Nat2 | 1506 | 10.5328265 | 6.3672656 | 10.4781261 | 6.32513928 | 9.11427297 | 21.280014 |
| ENSMUSG00 | Adnp | 9528 | 0.10013975 | 0.03034411 | 0.13917457 | 0.03408254 | 0.1231745 | 0.06583929 |
| ENSMUSG00 | Tas2r105 | 956 | 0 | 0 | 0 | 0 | 0 | 0 |
| ENSMUSG00 | Commd3 | 2153 | 50.6129921 | 50.3574059 | 60.1333944 | 49.3048653 | 36.071595 | 97.1524424 |
| ENSMUSG00 | Olfr1491 | 959 | 0 | 0 | 0 | 0 | 0 | 0 |
| ENSMUSG00 | Cited1 | 2352 | 0.10141704 | 0 | 0.0375866 | 0 | 0 | 0 |
| ENSMUSG00 | Olfr853 | 917 | 0 | 0 | 0 | 0 | 0 | 0 |
| ENSMUSG00 | Eml5 | 12406 | 2.86439391 | 1.82912752 | 0.99390024 | 2.41942787 | 3.84918502 | 2.14539393 |
| ENSMUSG00 | Rpusd3 | 2467 | 2.1432829 | 4.59011508 | 2.00673142 | 3.6272181 | 1.44785015 | 2.86345017 |
| ENSMUSG00 | Olfr672 | 938 | 0 | 0 | 0 | 0 | 0 | 0 |
| ENSMUSG00 | Zfp42 | 1796 | 0 | 0 | 0 | 0 | 0 | 0 |
| ENSMUSG00 | Plcb1 | 9109 | 2.29131927 | 1.55525458 | 3.94997258 | 2.23012307 | 1.26599637 | 3.72484919 |
| ENSMUSG00 | Olfr522 | 1075 | 0 | 0 | 0 | 0 | 0 | 0 |
| ENSMUSG00 | Olfr655 | 926 | 0 | 0 | 0 | 0 | 0 | 0 |
| ENSMUSG00 | Zfp524 | 1108 | 8.68305621 | 8.52395547 | 9.61429979 | 7.6527802 | 7.69079852 | 10.3633783 |
| ENSMUSG00 | Fam174a | 2104 | 49.6943506 | 49.468971 | 35.5043321 | 48.412363 | 34.7532303 | 53.0585069 |
| ENSMUSG00 | Olfr1356 | 962 | 0 | 0 | 0 | 0 | 0 | 0 |
| ENSMUSG00 | 4933414I15R | 394 | 0 | 0 | 0 | 0 | 0 | 0 |
| ENSMUSG00 | 4930548G14 | 317 | 0 | 0 | 0 | 0 | 0 | 0 |
| ENSMUSG00 | Olfr513 | 929 | 0 | 0 | 0 | 0 | 0 | 0 |
| ENSMUSG00 | Mrgprd | 3115 | 0 | 0 | 0 | 0 | 0 | 0 |
| ENSMUSG00 | Gpr119 | 2267 | 0 | 0 | 0 | 0 | 0 | 0 |
| ENSMUSG00 | Gpr183 | 2940 | 0.79781407 | 0.01638995 | 1.56360261 | 0.66273143 | 0.95457506 | 0 |
| ENSMUSG00 | 3100002H09I | 872 | 0 | 0 | 0 | 0 | 0 | 0 |
| ENSMUSG00 | Ercc6l | 5300 | 0 | 0 | 0 | 0 | 0.1059038 | 0 |
| ENSMUSG00 | Bzw1 | 2821 | 83.0764118 | 77.2417906 | 90.7070079 | 87.7557276 | 86.3523283 | 81.5628702 |
| ENSMUSG00 | Tceanc | 7789 | 3.3431557 | 2.3817923 | 1.74219615 | 1.41752544 | 2.64663703 | 2.80835289 |
| ENSMUSG00 | Fam83a | 3797 | 0 | 0 | 0 | 0 | 0 | 0 |
| ENSMUSG00 | Gm9851 | 2314 | 0 | 0 | 0 | 0 | 0 | 0 |
| ENSMUSG00 | Nyx | 5180 | 0.12279685 | 0.26046726 | 0.46079141 | 0 | 0 | 0.2264111 |
| ENSMUSG00 | Tmem199 | 2327 | 32.3579201 | 28.3486202 | 28.853717 | 26.2048368 | 26.708262 | 33.9672989 |
| ENSMUSG00 | Rnf7 | 1345 | 39.134763 | 38.2625429 | 40.9154606 | 31.0922599 | 21.7383742 | 44.7141855 |
| ENSMUSG00 | Gen1 | 6104 | 0.00651302 | 0.07104816 | 0.00724146 | 0 | 0 | 0.00446832 |
| ENSMUSG00 | Msrb3 | 4525 | 1.28271827 | 0.53244688 | 0.99637304 | 0 | 1.83807718 | 0.35562513 |
| ENSMUSG00 | 9230109A22I | 1938 | 0 | 0 | 0 | 0 | 0 | 0 |
| ENSMUSG00 | 2310047B19I | 1844 | 12.0301291 | 12.5953715 | 12.1531098 | 8.59003121 | 9.04860352 | 10.856608 |
| ENSMUSG00 | Pcdhb9 | 3054 | 0.01301751 | 0.18933769 | 0 | 0 | 2.05508979 | 0.44653961 |
| ENSMUSG00 | Islr2 | 4927 | 0 | 0 | 0 | 0 | 0 | 0.022143 |
| ENSMUSG00 | A930005I04R | 2790 | 0.64121743 | 0.18998239 | 0.06337182 | 0.06466316 | 0.09144512 | 0.30305155 |
| ENSMUSG00 | Nhlh1 | 2462 | 0 | 0 | 0 | 0 | 0 | 0 |
| ENSMUSG00 | Gm6563 | 686 | 1.91243565 | 0.91315415 | 1.48198599 | 0.94675919 | 0.66944225 | 2.62409066 |
| ENSMUSG00 | Jagn1 | 1199 | 34.6161148 | 49.6332411 | 43.8700527 | 46.6448449 | 28.6837175 | 27.8888305 |
| ENSMUSG00 | Trap1a | 1175 | 0.0676689 | 0 | 0 | 0 | 0 | 0 |
| ENSMUSG00 | Olfr1362 | 984 | 0 | 0 | 0 | 0 | 0 | 0 |
| ENSMUSG00 | Cml3 | 3115 | 0 | 0 | 0 | 0 | 0 | 0 |
| ENSMUSG00 | Actrt2 | 1439 | 0 | 0 | 0 | 0 | 0 | 0 |
| ENSMUSG00 | 4930422G04 | 7020 | 0.17555839 | 0.05491332 | 0.03777935 | 0.0668186 | 0.56695973 | 0.03885276 |
| ENSMUSG00 | Gdf6 | 3530 | 0 | 0 | 0 | 0 | 0 | 0 |
| ENSMUSG00 | Pcmtd1 | 1542 | 235.774234 | 193.714497 | 273.151337 | 296.658989 | 204.00623 | 313.622976 |
| ENSMUSG00 | 9630028B13I | 3404 | 0.04671619 | 0.04246749 | 0 | 0 | 0.07495061 | 0 |
| ENSMUSG00 | 2410124H12I | 701 | 0 | 0 | 0 | 0 | 0 | 0 |
| ENSMUSG00 | Usp42 | 5430 | 9.29092169 | 10.0454977 | 8.07518016 | 10.0272201 | 13.8983444 | 9.03629392 |
| ENSMUSG00 | Olfr1262 | 1327 | 0 | 0 | 0 | 0 | 0 | 0 |
| ENSMUSG00 | Ffar2 | 4286 | 0 | 0 | 0 | 0 | 0.10714825 | 0 |
| ENSMUSG00 | Taf7 | 3832 | 4.68932077 | 4.47661101 | 4.56783138 | 3.56865733 | 4.15454834 | 5.39514002 |
| ENSMUSG00 | 1500011K16I | 798 | 22.7672363 | 23.0666929 | 20.3284165 | 20.3470176 | 20.9093044 | 31.6836975 |
| ENSMUSG00 | Pcdh19 | 15143 | 0.09451214 | 0.29911679 | 0.13719122 | 0.22636163 | 0.24598332 | 0.00540342 |
| ENSMUSG00 | Nup160 | 6429 | 6.46822719 | 7.6900435 | 11.1449973 | 12.6839976 | 11.7863071 | 10.6018546 |
| ENSMUSG00 | Cacna1c | 10639 | 0 | 0.00452923 | 0.0041547 | 0.05087233 | 0 | 0.01281823 |
| ENSMUSG00 | Glod1 | 6652 | 0.84865878 | 0.16660977 | 0.63790994 | 1.02518136 | 0.61366658 | 0 |
| ENSMUSG00 | 2900026A02I | 5868 | 1.85633975 | 2.88231786 | 2.84735799 | 2.92075173 | 2.73045024 | 2.31007084 |
| ENSMUSG00 | Olfr645 | 1132 | 0 | 0 | 0 | 0 | 0 | 0 |
| ENSMUSG00 | Zfp52 | 3159 | 43.7197022 | 46.5237887 | 27.1731495 | 56.001983 | 35.8751443 | 66.2741788 |
| ENSMUSG00 | Rab11fip5 | 6163 | 5.77980051 | 5.43397327 | 5.53688509 | 5.5092007 | 5.39821467 | 5.89040156 |
| ENSMUSG00 | Plekhm3 | 10815 | 2.63566154 | 0.96239219 | 1.65035721 | 1.46129778 | 3.35929538 | 1.62916477 |
| ENSMUSG00 | Spryd4 | 1568 | 13.9701976 | 12.3846532 | 14.7151544 | 15.6938485 | 13.0494749 | 12.8197762 |
| ENSMUSG00 | Zfp46 | 5434 | 6.64298423 | 7.03199275 | 8.1668477 | 6.99197514 | 7.95350392 | 3.35788253 |
| ENSMUSG00 | Samd3 | 3561 | 0.01116413 | 0 | 0 | 0 | 0 | 0.00765926 |
| ENSMUSG00 | Commd1 | 8380 | 7.26320294 | 8.07324162 | 10.5704646 | 7.39294397 | 5.69936608 | 8.95050811 |
| ENSMUSG00 | Ncald | 9677 | 33.7081448 | 15.4114951 | 48.9796799 | 22.103375 | 26.8024252 | 31.8039712 |
| ENSMUSG00 | 6030498E09I | 1298 | 0 | 0 | 0 | 0 | 0 | 0 |
| ENSMUSG00 | Olfr589 | 953 | 0 | 0 | 0 | 0 | 0 | 0 |
| ENSMUSG00 | Six1 | 9015 | 0 | 0 | 0 | 0 | 0 | 0 |
| ENSMUSG00 | 1700065D16I | 463 | 0 | 0.10407439 | 0.38187337 | 0 | 0.44083262 | 0.05890851 |
| ENSMUSG00 | Ppapdc3 | 2297 | 0.12115297 | 0.65031768 | 0.94292134 | 0.48695837 | 0.79971682 | 0 |
| ENSMUSG00 | Pcdh1 | 9501 | 0.615099 | 1.19185495 | 0.95838119 | 1.38996193 | 1.94416883 | 1.01336151 |
| ENSMUSG00 | Kif18b | 3555 | 0 | 0.60995496 | 0 | 0.01014966 | 0 | 0 |
| ENSMUSG00 | Flrt3 | 4860 | 1.17794016 | 4.37247345 | 2.12823689 | 1.56652498 | 1.34390455 | 1.96983506 |
| ENSMUSG00 | Zbtb22 | 2669 | 6.25601417 | 7.67300034 | 5.21677803 | 6.502609 | 5.20014021 | 5.4876288 |
| ENSMUSG00 | Ywhag | 3929 | 21.8963756 | 17.7955021 | 18.258995 | 17.283382 | 14.2208912 | 14.2794434 |
| ENSMUSG00 | Olfr1402 | 962 | 0 | 0 | 0 | 0 | 0 | 0 |
| ENSMUSG00 | Hspa14 | 8084 | 5.86693323 | 6.12761783 | 6.72541649 | 6.69954835 | 6.23627651 | 5.59055879 |
| ENSMUSG00 | Tacstd2 | 1763 | 90.6506135 | 147.046545 | 38.5355829 | 97.6446026 | 98.5506578 | 62.0525116 |
| ENSMUSG00 | Kctd16 | 1282 | 2.32578864 | 1.09002092 | 1.31019504 | 0.42217681 | 1.23386712 | 0.51060167 |
| ENSMUSG00 | Ppp1r37 | 2958 | 6.81407323 | 7.34688486 | 6.84396352 | 4.92804124 | 5.08883735 | 7.81909874 |
| ENSMUSG00 | Plagl2 | 8744 | 3.77367896 | 1.66426184 | 3.20999203 | 2.01372801 | 2.61434313 | 5.52417499 |
| ENSMUSG00 | Olfr829 | 965 | 0 | 0 | 0 | 0 | 0 | 0 |
| ENSMUSG00 | Olfr1184 | 935 | 0 | 0 | 0 | 0 | 0 | 0 |
| ENSMUSG00 | Ccdc157 | 5575 | 4.30714085 | 4.81432256 | 2.36271735 | 4.70522805 | 5.69298758 | 2.36298668 |
| ENSMUSG00 | Gm5644 | 464 | 0 | 0 | 0.09526259 | 0 | 0 | 0 |
| ENSMUSG00 | Gpr87 | 2575 | 0 | 0 | 0 | 0 | 0 | 0 |
| ENSMUSG00 | Fhad1 | 6215 | 0.06396698 | 0.12405198 | 0.14224246 | 0.01741692 | 0.06568158 | 0.14043258 |
| ENSMUSG00 | Ubqlnl | 2297 | 0 | 0 | 0 | 0 | 0 | 0 |
| ENSMUSG00 | Cd14 | 1570 | 0.2025757 | 2.3019001 | 1.43585605 | 0 | 3.12008413 | 0.76438479 |
| ENSMUSG00 | Bbs12 | 2463 | 3.05066416 | 3.01287539 | 0.89731715 | 1.86050331 | 3.93626123 | 4.25231891 |
| ENSMUSG00 | Crebzf | 5752 | 16.3666512 | 15.8834309 | 17.5132128 | 18.8627795 | 18.5671427 | 20.0007699 |
| ENSMUSG00 | Gm11437 | 1232 | 0 | 0 | 0 | 0 | 0 | 0 |
| ENSMUSG00 | Gm1564 | 4547 | 0.1049188 | 0 | 0.0388844 | 0.37296153 | 0.02244397 | 0.13196439 |
| ENSMUSG00 | Hspb3 | 780 | 0 | 0 | 0.17000709 | 0 | 0 | 0 |
| ENSMUSG00 | Spn | 4603 | 0 | 0 | 0 | 0 | 0 | 0 |
| ENSMUSG00 | Zfp191 | 6976 | 23.0121316 | 21.0539386 | 23.3998576 | 27.1391181 | 20.6051176 | 27.6695272 |
| ENSMUSG00 | A130050O07 | 2103 | 0 | 0 | 0.10509235 | 0.01715742 | 0 | 0 |
| ENSMUSG00 | Krtap31-2 | 934 | 0 | 0 | 0 | 0 | 0 | 0 |
| ENSMUSG00 | Cbr1 | 1234 | 60.2775559 | 116.639305 | 68.3445028 | 98.5969645 | 54.8719617 | 63.7439705 |
| ENSMUSG00 | Pcdhb11 | 3614 | 0 | 0.26666542 | 0.46476758 | 0.81868502 | 0 | 0 |
| ENSMUSG00 | Foxd4 | 2344 | 0 | 0 | 0.01885744 | 0 | 0 | 0.01163594 |
| ENSMUSG00 | Olfr974 | 1079 | 0 | 0 | 0 | 0 | 0 | 0 |
| ENSMUSG00 | Irf2bp2 | 4528 | 22.7487743 | 19.1234621 | 12.1242688 | 24.6231267 | 20.3970274 | 18.3236424 |
| ENSMUSG00 | Kcnj16 | 3962 | 113.065309 | 204.810622 | 56.0053639 | 160.739042 | 153.645801 | 83.4829758 |
| ENSMUSG00 | Tlr6 | 2601 | 0.53496417 | 0 | 1.08204516 | 0 | 0.1177079 | 2.44328756 |
| ENSMUSG00 | Zfp786 | 2931 | 1.23430526 | 1.15081916 | 0.49766661 | 1.03408111 | 1.65387434 | 1.69361458 |
| ENSMUSG00 | Ufsp1 | 995 | 9.30957483 | 15.7392902 | 7.32995387 | 17.8778371 | 10.1027096 | 9.92303458 |
| ENSMUSG00 | Gm6583 | 2232 | 0.0178116 | 0 | 0.01980369 | 0.08082895 | 0 | 0 |
| ENSMUSG00 | Siglech | 2568 | 0 | 0 | 0 | 0 | 0 | 0 |
| ENSMUSG00 | Wdfy4 | 10883 | 0 | 0 | 0 | 0 | 0 | 0 |
| ENSMUSG00 | Olfr414 | 954 | 0 | 0 | 0 | 0 | 0 | 0 |
| ENSMUSG00 | Mafg | 6336 | 9.04161102 | 12.2443453 | 6.44610211 | 12.5797406 | 9.29362969 | 5.48419987 |
| ENSMUSG00 | Fam181b | 1834 | 0.65030775 | 0.05254792 | 0 | 0 | 0.13911226 | 0.29743336 |
| ENSMUSG00 | E130306D19I | 2837 | 0.95289837 | 0.45859497 | 0.18696585 | 0.85213147 | 0.1618743 | 0.39416997 |
| ENSMUSG00 | Rps19bp1 | 826 | 12.273181 | 18.9012194 | 14.8766493 | 14.4153447 | 7.41303281 | 20.0432276 |
| ENSMUSG00 | Usp29 | 2609 | 0 | 0 | 0 | 0 | 0 | 0 |
| ENSMUSG00 | Olfr424 | 947 | 0 | 0 | 0 | 0 | 0 | 0 |
| ENSMUSG00 | Gm5124 | 3568 | 0 | 0 | 0 | 0 | 0 | 0 |
| ENSMUSG00 | Gm6365 | 743 | 0 | 0.06485389 | 0.05949104 | 0.04856264 | 0 | 0 |
| ENSMUSG00 | Zfp579 | 3301 | 4.34769114 | 3.48880035 | 3.96357029 | 4.29574174 | 3.9262949 | 6.20516634 |
| ENSMUSG00 | Gm9853 | 1528 | 0.18212589 | 0.72531948 | 0.95462096 | 1.25153689 | 0.20036535 | 0.71399579 |
| ENSMUSG00 | Pusl1 | 2318 | 5.31673812 | 5.8414108 | 6.59785922 | 3.58018561 | 6.69198371 | 3.80056449 |
| ENSMUSG00 | Cmtm4 | 1294 | 1.87409916 | 1.75020308 | 4.33820252 | 3.45763023 | 2.44484954 | 1.64406635 |
| ENSMUSG00 | Tceal8 | 3528 | 19.8664717 | 22.2766687 | 23.6294433 | 23.3899195 | 17.515006 | 22.9607933 |
| ENSMUSG00 | Otud6a | 872 | 0 | 0 | 0 | 0 | 0 | 0 |
| ENSMUSG00 | Mical3 | 10519 | 2.75518065 | 2.65691952 | 3.36167643 | 2.27763833 | 2.94933325 | 3.28260227 |
| ENSMUSG00 | Ysk4 | 3926 | 0 | 0 | 0 | 0 | 0 | 0.02084155 |
| ENSMUSG00 | Olfr697 | 1046 | 0 | 0 | 0 | 0 | 0 | 0 |
| ENSMUSG00 | Ccnb3 | 4552 | 0 | 0 | 0 | 0.19023925 | 0 | 0 |
| ENSMUSG00 | Olfr272 | 2268 | 0 | 0 | 0 | 0 | 0 | 0 |
| ENSMUSG00 | Otop1 | 3231 | 0 | 0 | 0 | 0 | 0 | 0 |
| ENSMUSG00 | Pcdhb2 | 2773 | 0 | 0 | 0 | 0 | 0 | 0 |
| ENSMUSG00 | 2010001K21I | 517 | 0 | 0 | 0 | 0 | 0 | 0 |
| ENSMUSG00 | Olfr112 | 1699 | 0 | 0 | 0 | 0 | 0 | 0 |
| ENSMUSG00 | 4930533L02I | 1159 | 0 | 0 | 0 | 0 | 0 | 0 |
| ENSMUSG00 | Rap2a | 4189 | 10.8570708 | 9.09894384 | 14.0973173 | 5.60740296 | 6.51685631 | 9.2326184 |
| ENSMUSG00 | C230029F24I | 524 | 0 | 0 | 0 | 0 | 0 | 0.05205084 |
| ENSMUSG00 | Krt9 | 2572 | 0 | 0 | 0 | 0 | 0 | 0 |
| ENSMUSG00 | Ubqln3 | 2374 | 0 | 0 | 0 | 0 | 0 | 0 |
| ENSMUSG00 | Olfr904 | 932 | 0 | 0 | 0 | 0 | 0 | 0 |
| ENSMUSG00 | Hist1h1e | 747 | 0.37254132 | 0.90309262 | 0.2958624 | 0.67623644 | 0.1366168 | 0.1095367 |
| ENSMUSG00 | Gm9857 | 392 | 0 | 0 | 0 | 0 | 0 | 0 |
| ENSMUSG00 | Gm5812 | 2087 | 0.0190491 | 0.09235542 | 0.06353883 | 0.03457791 | 0 | 0 |
| ENSMUSG00 | Kctd19 | 3138 | 0.27871911 | 0 | 0 | 0 | 0 | 0 |
| ENSMUSG00 | B3gnt2 | 5676 | 0.51130199 | 0.69613958 | 0.71644989 | 0.45770035 | 0.73716752 | 0.50935725 |
| ENSMUSG00 | Lrrc3 | 4294 | 20.2203003 | 17.853896 | 18.6113023 | 15.6377934 | 15.1510548 | 16.0573563 |
| ENSMUSG00 | Pcdhb1 | 2456 | 0 | 0.23543864 | 0.32395488 | 0 | 0 | 0.14436902 |
| ENSMUSG00 | AU021092 | 1355 | 23.8239485 | 11.4865098 | 13.7009403 | 8.70762255 | 11.2220369 | 10.1449581 |
| ENSMUSG00 | 1810063B05I | 760 | 32.4844124 | 33.5402999 | 39.3745365 | 32.7112219 | 21.9547696 | 51.6782638 |
| ENSMUSG00 | Dcun1d4 | 5784 | 11.8909027 | 9.75558848 | 10.1104838 | 10.0123931 | 7.94861076 | 10.1053859 |
| ENSMUSG00 | Trim32 | 3226 | 9.10703661 | 8.7828977 | 8.16624254 | 7.58326912 | 5.86819139 | 7.58380391 |
| ENSMUSG00 | Pcdhb6 | 5336 | 0.17881026 | 0 | 0.05798593 | 0 | 0 | 0 |
| ENSMUSG00 | Olfr693 | 951 | 0 | 0 | 0 | 0 | 0 | 0 |
| ENSMUSG00 | Treml4 | 2138 | 0 | 0 | 0 | 0 | 0 | 0 |
| ENSMUSG00 | Vmn1r73 | 911 | 0 | 0 | 0 | 0 | 0 | 0 |
| ENSMUSG00 | Pcbp1 | 1672 | 135.981814 | 134.61655 | 155.92253 | 131.83326 | 103.548142 | 142.996105 |
| ENSMUSG00 | Tmem198 | 2809 | 0.28305789 | 0.60040067 | 0.18882952 | 0.24405797 | 0.25431444 | 0.5340353 |
| ENSMUSG00 | Senp8 | 3789 | 6.86199109 | 7.52873416 | 6.62619341 | 7.65636416 | 4.47103636 | 9.81138382 |
| ENSMUSG00 | Olfr1325 | 947 | 0 | 0 | 0 | 0 | 0 | 0 |
| ENSMUSG00 | Apon | 1616 | 0 | 0 | 0 | 0 | 0 | 0 |
| ENSMUSG00 | BC068281 | 2156 | 2.58152471 | 2.30204246 | 3.60831372 | 5.80726783 | 4.63876144 | 3.76987129 |
| ENSMUSG00 | Gm9858 | 415 | 0 | 0 | 0 | 0 | 0 | 0 |
| ENSMUSG00 | Kcnf1 | 6267 | 0 | 0 | 0 | 0 | 0 | 0 |
| ENSMUSG00 | Kctd14 | 2549 | 0 | 0 | 0.08670428 | 0 | 0 | 0.0963012 |
| ENSMUSG00 | 4930563D23I | 2678 | 8.89228261 | 4.78625602 | 12.6432456 | 5.47024275 | 9.22209331 | 10.1337812 |
| ENSMUSG00 | Gm5087 | 3454 | 0 | 0 | 0 | 0 | 0 | 0 |
| ENSMUSG00 | Mettl5 | 2532 | 6.233383 | 7.19371056 | 6.35445137 | 7.9659806 | 5.92486354 | 17.0735795 |
| ENSMUSG00 | Pabpc2 | 2585 | 0 | 0 | 0 | 0 | 0.05921823 | 0 |
| ENSMUSG00 | Rinl | 2859 | 0.41716139 | 0.65731768 | 1.56152016 | 1.4387384 | 0.60681944 | 0.97307212 |
| ENSMUSG00 | 1700025K23I | 1870 | 6.16528842 | 7.11200964 | 2.36373492 | 3.37666192 | 4.50232728 | 7.78858682 |
| ENSMUSG00 | Ttn | 116800 | 0.01497638 | 0.09323747 | 0.02081422 | 0.2372518 | 0.08999515 | 0.1274996 |
| ENSMUSG00 | 1100001G20I | 431 | 0 | 0 | 0 | 0 | 0 | 0 |
| ENSMUSG00 | 4930544M13 | 995 | 0 | 0 | 0 | 0 | 0 | 0 |
| ENSMUSG00 | Xrcc1 | 2138 | 15.0245221 | 15.2582887 | 12.4873308 | 13.2818377 | 16.8258162 | 9.31266915 |
| ENSMUSG00 | Wfdc15a | 485 | 0 | 0 | 0 | 0 | 0 | 0 |
| ENSMUSG00 | Iqcj | 1575 | 0 | 0 | 0 | 0 | 0 | 0 |
| ENSMUSG00 | Tubgcp6 | 9050 | 2.68843691 | 3.18403231 | 2.21253424 | 3.37297344 | 3.76636675 | 2.45622442 |
| ENSMUSG00 | 4930564D02I | 732 | 0 | 0 | 0 | 0 | 0 | 0 |
| ENSMUSG00 | Nlgn2 | 5777 | 0.06881683 | 0.01668217 | 0.09181619 | 0.02498324 | 0.15898819 | 0.09914617 |
| ENSMUSG00 | Olfr284 | 917 | 0 | 0 | 0 | 0 | 0 | 0 |
| ENSMUSG00 | Krtap8-2 | 482 | 0 | 0 | 0 | 0 | 0 | 0 |
| ENSMUSG00 | Adam6b | 2473 | 0 | 0 | 0 | 0 | 0 | 0 |
| ENSMUSG00 | A530095I07R | 845 | 0 | 0 | 0 | 0 | 0 | 0 |
| ENSMUSG00 | Cox6b2 | 547 | 2.03501546 | 1.40947546 | 1.29292411 | 0.92348926 | 2.33210127 | 4.78677398 |
| ENSMUSG00 | Sox12 | 4170 | 1.8876703 | 1.44443771 | 2.40619145 | 1.36713742 | 1.63969649 | 2.66205712 |
| ENSMUSG00 | Rhox2a | 929 | 0 | 0 | 0 | 0 | 0 | 0 |
| ENSMUSG00 | E230016K23I | 2856 | 1.3084787 | 6.1582813 | 1.036948 | 3.07000589 | 4.55592641 | 1.45159145 |
| ENSMUSG00 | Gypa | 1819 | 0 | 0 | 0 | 0 | 0 | 0 |
| ENSMUSG00 | B230319C09 | 2508 | 0.28532642 | 0.05763929 | 0.14099472 | 0.2877356 | 0.38656345 | 0.03262517 |
| ENSMUSG00 | Cxx1c | 1531 | 0.80497707 | 1.22747958 | 0.40419713 | 1.60259897 | 0.53326062 | 0.6947818 |
| ENSMUSG00 | Arl3 | 3832 | 83.4844342 | 84.4771708 | 88.2883368 | 61.2509655 | 88.124362 | 48.4139082 |
| ENSMUSG00 | Mest | 3559 | 0.12287448 | 0.81235924 | 0.17387631 | 0.16221206 | 0.24373374 | 0.26056132 |
| ENSMUSG00 | Samd7 | 3323 | 0.11963732 | 0.01450089 | 0.01330179 | 0.06514964 | 0.03071103 | 0.10670187 |
| ENSMUSG00 | Tbc1d22a | 3302 | 19.7091828 | 20.8827373 | 23.6269694 | 17.3307458 | 15.762236 | 12.580035 |
| ENSMUSG00 | Vmn2r72-ps | 2565 | 0 | 0 | 0 | 0 | 0 | 0 |
| ENSMUSG00 | Krt71 | 2176 | 0 | 0 | 0 | 0 | 0 | 0 |
| ENSMUSG00 | Olfr664 | 920 | 0 | 0 | 0 | 0 | 0 | 0 |
| ENSMUSG00 | Klhdc1 | 3121 | 7.37533586 | 3.55106751 | 4.87197501 | 5.20247358 | 3.58051147 | 6.76404062 |
| ENSMUSG00 | 1700011F03I | 810 | 0 | 0 | 0 | 0.22272867 | 0 | 0 |
| ENSMUSG00 | Abca16 | 7117 | 0 | 0 | 0 | 0 | 0 | 0 |
| ENSMUSG00 | Cd209f | 1986 | 0 | 0 | 0 | 0 | 0 | 0 |
| ENSMUSG00 | Sos6 | 11577 | 4.80073955 | 5.84797022 | 4.98258661 | 7.65461703 | 7.63830952 | 3.57866261 |
| ENSMUSG00 | Zfp664 | 506 | 0 | 0 | 0 | 0 | 0 | 0 |
| ENSMUSG00 | Tas2r144 | 959 | 0 | 0 | 0 | 0 | 0 | 0 |
| ENSMUSG00 | Rspo2 | 3318 | 0 | 0 | 0.319724 | 0 | 0 | 0 |
| ENSMUSG00 | Spats2 | 3052 | 19.0831517 | 15.6621725 | 23.1436911 | 21.5759277 | 17.1871279 | 15.6301914 |
| ENSMUSG00 | Prss58 | 981 | 0 | 0 | 0 | 0 | 0 | 0 |
| ENSMUSG00 | 5031410I06R | 2990 | 0 | 0 | 0 | 0 | 0 | 0 |
| ENSMUSG00 | 2010005H15I | 344 | 0 | 0 | 0 | 0 | 0 | 0 |
| ENSMUSG00 | B3galtl | 4782 | 4.11521599 | 4.92747181 | 3.06880215 | 3.07851823 | 4.0974756 | 3.91267305 |
| ENSMUSG00 | Xkr4 | 6884 | 0 | 0 | 0 | 0 | 0 | 0 |
| ENSMUSG00 | Olfr371 | 938 | 0 | 0 | 0 | 0 | 0 | 0 |
| ENSMUSG00 | Rnf133 | 2012 | 0.03951837 | 0.16764667 | 0 | 0 | 0 | 0 |
| ENSMUSG00 | Nanos2 | 1438 | 0 | 0 | 0 | 0 | 0 | 0 |
| ENSMUSG00 | Tlr11 | 3160 | 0 | 0 | 0 | 0 | 0 | 0 |
| ENSMUSG00 | 6330444E15I | 959 | 0 | 0 | 0 | 0 | 0 | 0 |
| ENSMUSG00 | Prdm9 | 5892 | 0.42506406 | 1.46391262 | 1.02777537 | 0.81447928 | 0.6841622 | 2.16178828 |
| ENSMUSG00 | Erich1 | 1573 | 9.12379432 | 9.77207568 | 14.0501726 | 10.9874757 | 8.20703947 | 12.1721531 |
| ENSMUSG00 | Casr | 6217 | 48.714372 | 28.8793121 | 51.3756663 | 39.7442241 | 56.6075139 | 38.4924698 |
| ENSMUSG00 | Sec31b | 4205 | 1.41815032 | 0.58442534 | 1.01963823 | 1.45014638 | 2.00222402 | 1.21941312 |
| ENSMUSG00 | Igfn1 | 9436 | 0.32441416 | 0.42895943 | 0.75418575 | 1.86222506 | 0.70299161 | 1.22845715 |
| ENSMUSG00 | A530006G24 | 1328 | 0 | 0 | 0 | 0 | 0 | 0 |
| ENSMUSG00 | Fam165b | 1015 | 9.63531843 | 13.8625036 | 15.2855635 | 12.7264748 | 6.58567018 | 24.5606111 |
| ENSMUSG00 | Lax1 | 2273 | 0.05247094 | 0.93277759 | 0.01944648 | 2.15889048 | 0.08979565 | 0.26398683 |
| ENSMUSG00 | Gm9864 | 1648 | 0 | 0 | 0 | 0 | 0 | 0 |
| ENSMUSG00 | Olfr796 | 932 | 0 | 0 | 0 | 0 | 0 | 0 |
| ENSMUSG00 | Btla | 3378 | 0 | 0.09985349 | 0 | 0 | 0 | 0.07266778 |
| ENSMUSG00 | A330070K13I | 703 | 0 | 0 | 0 | 0 | 0 | 0 |
| ENSMUSG00 | Ppp1r2-ps7 | 1469 | 0.02706296 | 0 | 0 | 0 | 0 | 0 |
| ENSMUSG00 | Slc6a7 | 3328 | 0 | 0 | 0 | 0 | 0.15332445 | 0.31142917 |
| ENSMUSG00 | Tagap1 | 2354 | 8.91711712 | 8.88399146 | 9.68910408 | 10.6529401 | 12.5506676 | 13.6952522 |
| ENSMUSG00 | Pldn4 | 2511 | 3.40399375 | 4.20264071 | 5.52743079 | 3.5923979 | 2.51982103 | 5.13775562 |
| ENSMUSG00 | Klf13 | 6991 | 0.73926655 | 1.48191747 | 1.09382332 | 0.82579418 | 0.72988665 | 1.4747266 |
| ENSMUSG00 | Zfp217 | 8138 | 0.62530124 | 0.74606681 | 0.66807897 | 0.60299313 | 0.28842631 | 0.54629715 |
| ENSMUSG00 | Olfr901 | 935 | 0 | 0 | 0 | 0 | 0 | 0 |
| ENSMUSG00 | Pard3b | 13017 | 1.9973945 | 3.46859463 | 2.12231327 | 2.81349583 | 3.35158265 | 2.20007461 |
| ENSMUSG00 | 1700029F12I | 1125 | 0 | 0 | 0 | 0 | 0 | 0 |
| ENSMUSG00 | Gm9866 | 1582 | 0 | 0 | 0 | 0 | 0 | 0 |
| ENSMUSG00 | Dock8 | 7762 | 10.8633566 | 13.3099371 | 9.94285209 | 13.1925847 | 12.6809945 | 8.66519716 |
| ENSMUSG00 | Rgs14 | 2640 | 0 | 0 | 0 | 0 | 0 | 0 |
| ENSMUSG00 | Gm9867 | 2646 | 0 | 0 | 0 | 0 | 0 | 0 |
| ENSMUSG00 | Prss51 | 1264 | 0 | 0 | 0 | 0 | 0 | 0 |
| ENSMUSG00 | Gnpda1 | 2675 | 25.7556066 | 21.706416 | 29.7598203 | 21.6762039 | 23.0620143 | 17.364004 |
| ENSMUSG00 | 1110012J17R | 9929 | 0.95294636 | 0.66002177 | 0.7256421 | 1.31551013 | 1.78841563 | 1.62071076 |
| ENSMUSG00 | D630039A03I | 2828 | 5.94645271 | 4.75389581 | 4.51709075 | 4.84836525 | 6.62188116 | 6.93439378 |
| ENSMUSG00 | F730043M19 | 4484 | 0.01773215 | 0 | 0 | 0.02414053 | 0.01137966 | 0 |
| ENSMUSG00 | Gm1604A | 715 | 0 | 0 | 0 | 0 | 0 | 0 |
| ENSMUSG00 | Akr1b7 | 1271 | 0 | 0.18956114 | 0 | 0.45421929 | 0 | 0.25751036 |
| ENSMUSG00 | Sema5b | 6737 | 0 | 0 | 0 | 0 | 0 | 0 |
| ENSMUSG00 | Foxo6 | 4707 | 0.03378414 | 0.13308344 | 0 | 0 | 0 | 0.08112279 |
| ENSMUSG00 | C430048L16I | 3611 | 2.97257816 | 3.88320539 | 3.97828828 | 6.07529301 | 4.08380576 | 6.4051216 |
| ENSMUSG00 | Bre | 4818 | 7.8388764 | 6.80090923 | 9.6330293 | 6.80750902 | 6.15324291 | 11.3332976 |
| ENSMUSG00 | Rasal3 | 5068 | 0 | 0 | 0 | 0 | 0 | 0 |
| ENSMUSG00 | Gm9869 | 372 | 0.42747829 | 1.42486791 | 0.23764432 | 0 | 0.41150303 | 0.07331892 |
| ENSMUSG00 | Ppp4r2 | 3632 | 2.65985181 | 3.43620279 | 3.5536724 | 5.74213154 | 2.99246092 | 4.69346077 |
| ENSMUSG00 | Rps10 | 2001 | 2.16559089 | 1.75792618 | 2.20898766 | 1.91139266 | 1.7595302 | 2.23540272 |
| ENSMUSG00 | Ppap2c | 2407 | 19.1097179 | 20.1394104 | 28.6109146 | 16.3845761 | 16.7473356 | 19.5126418 |
| ENSMUSG00 | Gm9870 | 643 | 0 | 0 | 0 | 0 | 0 | 0 |
| ENSMUSG00 | Acvr2a | 5675 | 8.61660635 | 12.3204454 | 13.5993688 | 13.3328718 | 16.4003717 | 14.2356796 |
| ENSMUSG00 | Gm650 | 1550 | 0 | 0.18652816 | 0 | 0 | 0.06584049 | 0 |
| ENSMUSG00 | Pld4 | 1967 | 0 | 0 | 0 | 0 | 0 | 0 |
| ENSMUSG00 | Serpinb6c | 1919 | 0.02071677 | 0 | 0.18427032 | 0 | 0.18613061 | 0.07106472 |
| ENSMUSG00 | Olfr849 | 938 | 0 | 0 | 0 | 0 | 0 | 0 |
| ENSMUSG00 | Hbb-y | 658 | 0 | 0 | 0 | 0 | 0 | 0 |
| ENSMUSG00 | Gm14964 | 1231 | 0 | 0.07828829 | 0 | 0 | 0.04145116 | 0.11078245 |
| ENSMUSG00 | Gm5963 | 474 | 0 | 0 | 0 | 0 | 0 | 0 |
| ENSMUSG00 | Cd177 | 2695 | 0 | 0 | 0 | 0 | 0 | 0 |
| ENSMUSG00 | Opa3 | 4752 | 21.2999691 | 17.0558914 | 19.8219965 | 20.0303943 | 16.8155102 | 14.624533 |
| ENSMUSG00 | Hbb-bh1 | 573 | 0 | 0 | 0 | 0 | 0 | 0 |
| ENSMUSG00 | Ppp1r36 | 1424 | 0.47460897 | 2.36871556 | 1.27266542 | 1.41895681 | 3.83414481 | 0.86190925 |
| ENSMUSG00 | 1700084E18I | 727 | 0.16405288 | 0.19884364 | 0.24320134 | 0.14889427 | 0.63168828 | 0.0750334 |
| ENSMUSG00 | Gpr17 | 5193 | 0.05358913 | 0 | 0 | 0 | 0 | 0 |
| ENSMUSG00 | Epx | 2723 | 0 | 0 | 0 | 0 | 0 | 0 |
| ENSMUSG00 | Gm9871 | 324 | 11.4112953 | 1.93340663 | 19.3724127 | 7.35004611 | 13.3865492 | 18.1830928 |
| ENSMUSG00 | A930035D04 | 541 | 0.22045553 | 0.35627684 | 0.08170396 | 1.33390183 | 0.84886762 | 0.25207615 |
| ENSMUSG00 | Gm13476 | 1384 | 0 | 0 | 0 | 0 | 0 | 0 |
| ENSMUSG00 | Zfp622 | 2797 | 8.82665477 | 11.8011129 | 9.86126208 | 12.1262502 | 9.32230177 | 9.72213634 |
| ENSMUSG00 | Prpmp5 | 1295 | 0 | 0 | 0 | 0 | 0 | 0 |
| ENSMUSG00 | Fpr2 | 1305 | 0 | 0 | 0 | 0 | 0 | 0 |
| ENSMUSG00 | Bhlha15 | 3492 | 0.34154193 | 0 | 0 | 0 | 0.14612364 | 0 |
| ENSMUSG00 | Dnahc3 | 12191 | 0 | 0.00395262 | 0.03263199 | 0 | 0.03348462 | 0 |
| ENSMUSG00 | Ostn | 1418 | 0 | 0 | 0 | 0 | 0 | 0 |
| ENSMUSG00 | Olfr1463 | 932 | 0 | 0 | 0 | 0 | 0 | 0 |
| ENSMUSG00 | Gm15024 | 970 | 0 | 0 | 0.36455128 | 0.26038589 | 0.4734406 | 0.5623637 |
| ENSMUSG00 | 5330438D12I | 2225 | 7.28999373 | 3.40012199 | 5.92006706 | 6.95694252 | 6.58182915 | 14.4524942 |
| ENSMUSG00 | Taf9 | 1329 | 23.5421845 | 21.2469941 | 23.4146708 | 22.9958553 | 20.1955408 | 31.6870151 |
| ENSMUSG00 | 8030423F21I | 2847 | 0 | 0 | 0 | 0 | 0 | 0.00958013 |
| ENSMUSG00 | Ppp6r1 | 3763 | 44.2455361 | 33.690813 | 23.0582561 | 20.7402238 | 38.7681128 | 21.4978953 |
| ENSMUSG00 | Cdc42se2 | 3652 | 93.0639657 | 85.3950311 | 107.708708 | 87.7943723 | 83.7771493 | 65.7369591 |
| ENSMUSG00 | Ltn1 | 7807 | 11.0349848 | 9.86319131 | 10.0553956 | 7.71833154 | 13.4183617 | 8.9122076 |
| ENSMUSG00 | Doc2a | 2581 | 0 | 0 | 0 | 0 | 0.09885001 | 0 |
| ENSMUSG00 | Tbc1d30 | 5679 | 0.32902053 | 0.21212556 | 0.30355201 | 1.42955802 | 0.56606122 | 0.76363227 |
| ENSMUSG00 | Mrgpra6 | 1014 | 0 | 0 | 0 | 0 | 0 | 0 |
| ENSMUSG00 | Hbb-b1 | 1051 | 0 | 0 | 0 | 0 | 0 | 0 |
| ENSMUSG00 | Slc39a1 | 2360 | 1.36448895 | 2.2868142 | 1.51709715 | 1.31485417 | 1.08106728 | 1.04013455 |
| ENSMUSG00 | Lrrc15 | 7250 | 0 | 0 | 0 | 0 | 0 | 0 |
| ENSMUSG00 | Ankrd44 | 6166 | 12.6758473 | 8.20560562 | 14.1078863 | 16.3557111 | 9.76502165 | 20.2945256 |
| ENSMUSG00 | 1700024B05I | 1569 | 0 | 0 | 0 | 0 | 0 | 0 |
| ENSMUSG00 | Cx3cr1 | 3727 | 0 | 0 | 0 | 0 | 0 | 0 |
| ENSMUSG00 | Immt | 2716 | 316.41908 | 252.198187 | 299.501663 | 254.620201 | 268.489668 | 231.332323 |
| ENSMUSG00 | 9930013L23I | 10626 | 0 | 0 | 0 | 0 | 0 | 0 |
| ENSMUSG00 | Zdhhc19 | 1836 | 0 | 0 | 0 | 0 | 0 | 0 |
| ENSMUSG00 | B630019K06I | 2624 | 3.30285623 | 2.46074057 | 1.09493895 | 2.00761376 | 1.96404876 | 3.72115886 |
| ENSMUSG00 | Gm9873 | 1096 | 0 | 0 | 0 | 0 | 0 | 0 |
| ENSMUSG00 | Tmem106c | 1565 | 38.9425889 | 40.4273473 | 48.2124895 | 39.5403875 | 30.9744773 | 37.4699517 |
| ENSMUSG00 | 1700109F18I | 2867 | 5.4772985 | 3.76482841 | 5.85863284 | 4.01470952 | 3.93332022 | 8.75223862 |
| ENSMUSG00 | Il1rapl1 | 10048 | 0 | 0.03356008 | 0 | 0.0107729 | 0.01015652 | 0 |
| ENSMUSG00 | Mpp3 | 10249 | 0.33359072 | 0.38552915 | 0.38815161 | 0.77451945 | 0.22404009 | 0.91811492 |
| ENSMUSG00 | Actrt2 | 3854 | 0.69113056 | 0.52512469 | 1.12397006 | 0.4587494 | 0.62227288 | 1.15354598 |
| ENSMUSG00 | 4921508M14 | 1098 | 0 | 0 | 0 | 0 | 0 | 0 |
| ENSMUSG00 | Rnase9 | 1832 | 0 | 0 | 0 | 0 | 0 | 0 |
| ENSMUSG00 | Lrrc33 | 6381 | 0.36758711 | 0.11327325 | 0.28401122 | 0.15267438 | 0.46380345 | 0.29065593 |
| ENSMUSG00 | Trpm3 | 8200 | 1.04236931 | 1.88632292 | 0.90020827 | 0.81404613 | 1.85437317 | 0.50890485 |
| ENSMUSG00 | Gm9874 | 452 | 0 | 0.10660717 | 0.19558338 | 1.9956883 | 0 | 0 |
| ENSMUSG00 | Gm9875 | 2476 | 0 | 0 | 0 | 0 | 0 | 0 |
| ENSMUSG00 | Acot4 | 2765 | 6.12507584 | 3.38089323 | 8.5206446 | 4.69784305 | 4.94577531 | 7.37845575 |
| ENSMUSG00 | Rft1 | 3273 | 8.17459162 | 7.22870244 | 12.5461388 | 8.72010303 | 12.9397776 | 5.52492692 |
| ENSMUSG00 | Mogat2 | 1828 | 0.23922882 | 0 | 0 | 0 | 0 | 0 |
| ENSMUSG00 | Ezr | 3277 | 307.270922 | 315.263143 | 286.280109 | 234.359896 | 288.438384 | 214.077264 |
| ENSMUSG00 | Gm13520 | 1606 | 0.49508693 | 0.30004011 | 0.66055058 | 0.53920864 | 0.73076379 | 0.40759112 |
| ENSMUSG00 | Rexo4 | 4460 | 15.5902097 | 16.6167596 | 12.8542131 | 16.3663624 | 11.9328498 | 12.8790113 |
| ENSMUSG00 | 4930473A06I | 6673 | 1.68006077 | 1.81971878 | 2.51711379 | 3.93101249 | 1.68227224 | 2.01504528 |
| ENSMUSG00 | Atf7 | 3633 | 4.62883795 | 3.54136528 | 4.68420302 | 3.51583919 | 3.83435194 | 4.31679537 |
| ENSMUSG00 | Tchh | 5821 | 3.40117322 | 1.61421684 | 3.45504872 | 3.5827902 | 1.67428926 | 4.34351324 |
| ENSMUSG00 | Olfr720 | 1022 | 0 | 0 | 0 | 0 | 0 | 0 |
| ENSMUSG00 | 2610001J05R | 4131 | 12.2605863 | 14.9656755 | 14.2631462 | 15.7482271 | 11.0056889 | 19.4045424 |
| ENSMUSG00 | B4galt3 | 3370 | 13.7551603 | 18.2307756 | 17.956179 | 13.4906161 | 15.7621535 | 13.6292263 |
| ENSMUSG00 | Gm9877 | 2625 | 0 | 0 | 0 | 0 | 0 | 0 |
| ENSMUSG00 | Tmco1 | 4467 | 84.9665486 | 77.5384254 | 81.4671532 | 68.165743 | 59.5250603 | 101.472404 |
| ENSMUSG00 | Prmt1 | 2765 | 10.6110469 | 10.4912254 | 11.6059812 | 11.2356746 | 7.54784366 | 11.6989954 |
| ENSMUSG00 | Bmpr1b | 8585 | 22.8067259 | 16.5860147 | 30.3311657 | 26.4237407 | 24.2382715 | 21.1684242 |
| ENSMUSG00 | Cebpe | 1236 | 0 | 0 | 0 | 0 | 0 | 0 |
| ENSMUSG00 | Thsd4 | 731 | 0 | 0 | 0 | 0 | 0 | 0 |
| ENSMUSG00 | Zfp961 | 3998 | 16.3775079 | 13.2940585 | 15.5447202 | 18.708874 | 15.4049114 | 17.0756433 |
| ENSMUSG00 | 2810055G20I | 2563 | 0.6359636 | 0.33841434 | 0.86230673 | 0.61943424 | 0.91580698 | 1.06416852 |
| ENSMUSG00 | Asna1 | 1243 | 13.6889346 | 18.4914987 | 15.6111095 | 15.297858 | 12.4384488 | 22.5569824 |
| ENSMUSG00 | Atp6v1a | 7612 | 518.424627 | 383.79501 | 705.887442 | 455.012025 | 443.611728 | 591.994397 |
| ENSMUSG00 | D630004N19 | 1864 | 0 | 0 | 0 | 0 | 0 | 0 |
| ENSMUSG00 | Pmp2 | 1360 | 0 | 0 | 0 | 0 | 0 | 0 |
| ENSMUSG00 | Tcp10c | 2613 | 0 | 0 | 0 | 0 | 0 | 0 |
| ENSMUSG00 | Gm9881 | 3204 | 0 | 0 | 0 | 0 | 0 | 0 |
| ENSMUSG00 | C130026I21F | 3813 | 0 | 0 | 0 | 0 | 0 | 0.17167358 |
| ENSMUSG00 | A330008L17F | 5826 | 0 | 0 | 0 | 0 | 0 | 0 |
| ENSMUSG00 | 1700036A12F | 1823 | 0 | 0 | 0 | 0 | 0 | 0 |
| ENSMUSG00 | Gm8174 | 1000 | 0.07951096 | 0.14455933 | 0.08840369 | 0.07216409 | 0.15307913 | 0.08182392 |
| ENSMUSG00 | Tmem171 | 1578 | 58.7766388 | 49.5911166 | 76.4428578 | 57.187068 | 62.2146686 | 60.9963257 |
| ENSMUSG00 | Cherp | 3776 | 6.81191628 | 5.67875179 | 6.07541222 | 8.65739734 | 8.40529814 | 9.13729308 |
| ENSMUSG00 | Pkdrej | 7057 | 0 | 0 | 0 | 0 | 0 | 0 |
| ENSMUSG00 | Epha3 | 5642 | 0 | 0 | 0 | 0 | 0 | 0 |
| ENSMUSG00 | Olfr536 | 3846 | 0 | 0 | 0 | 0 | 0 | 0 |
| ENSMUSG00 | Nav2 | 15825 | 1.16063393 | 1.79043463 | 0.8407428 | 2.07486006 | 2.16680724 | 0.86003444 |
| ENSMUSG00 | Robo2 | 15555 | 0 | 0.02168467 | 0 | 0 | 0.06560768 | 0 |
| ENSMUSG00 | Cyp2j5 | 2336 | 2.3145314 | 2.02152026 | 1.94896825 | 0 | 1.59457424 | 3.42100569 |
| ENSMUSG00 | Spdya | 2260 | 0.38700025 | 0.6183216 | 0 | 0.47896519 | 1.15148016 | 2.02749531 |
| ENSMUSG00 | Nup188 | 6139 | 2.63568669 | 3.13184402 | 2.27525369 | 2.22757695 | 2.47692784 | 2.13256668 |
| ENSMUSG00 | Pbx1 | 13338 | 5.40684073 | 4.27384623 | 7.6784878 | 4.08486184 | 3.75677771 | 6.02217818 |
| ENSMUSG00 | Olfr198 | 1053 | 0 | 0 | 0 | 0 | 0 | 0 |
| ENSMUSG00 | Magi3 | 9194 | 8.66110806 | 5.39306602 | 7.13941015 | 6.51470458 | 8.34714697 | 8.85224315 |
| ENSMUSG00 | St6galnac3 | 913 | 0.30480653 | 0.63333768 | 0.14524154 | 0.79040623 | 2.51499114 | 0 |
| ENSMUSG00 | Arl13a | 1730 | 0 | 0.02785344 | 0 | 0 | 0 | 0.20495394 |

| | | | | | | | | |
|---|---|---|---|---|---|---|---|---|
| ENSMUSG00 | Adarb2 | 10071 | 0 | 0 | 0 | 0 | 0 | 0 |
| ENSMUSG00 | Defb34 | 244 | 0 | 0 | 0 | 0 | 0 | 0 |
| ENSMUSG00 | Gan | 3453 | 0.03453995 | 0.51633315 | 0.15361197 | 0.08359582 | 0.17732885 | 0.30015531 |
| ENSMUSG00 | Gm9884 | 1221 | 0 | 0 | 0 | 0 | 0 | 0 |
| ENSMUSG00 | Cpne8 | 5834 | 4.46346242 | 4.26194792 | 7.46294401 | 4.29224184 | 5.51021885 | 8.79389723 |
| ENSMUSG00 | Slc22a30 | 3804 | 0.01045097 | 0 | 0 | 0.01897058 | 0 | 0.12905981 |
| ENSMUSG00 | D930048N14 | 3682 | 9.03729962 | 6.77908121 | 9.01563935 | 10.4659488 | 9.65925691 | 6.66680481 |
| ENSMUSG00 | Hist1h1d | 3820 | 0.04162878 | 0.23967079 | 0.38184838 | 0.01889112 | 1.48270359 | 0.15707907 |
| ENSMUSG00 | Hook2 | 2744 | 33.2647898 | 24.6376014 | 26.0152976 | 25.5887969 | 26.3127995 | 21.420134 |
| ENSMUSG00 | Dlg2 | 10958 | 0.08707169 | 0.03517901 | 0.14924879 | 0.43464408 | 0.00931308 | 0.01244508 |
| ENSMUSG00 | Lrrtm4 | 6441 | 0 | 0 | 0.00686258 | 0 | 0.00792212 | 0 |
| ENSMUSG00 | Serp2 | 1431 | 0 | 0.16836633 | 0 | 0 | 0.03565785 | 0 |
| ENSMUSG00 | Adam17 | 7187 | 12.296707 | 11.297364 | 13.4136941 | 13.8112849 | 16.6064691 | 15.4304554 |
| ENSMUSG00 | A1cf | 3816 | 0 | 0 | 0 | 0 | 0 | 0 |
| ENSMUSG00 | Isoc2b | 1694 | 6.17219089 | 7.16823106 | 5.45347414 | 3.91918311 | 6.5665584 | 5.31324141 |
| ENSMUSG00 | Plekhg3 | 5023 | 10.2099482 | 7.97191587 | 7.85830098 | 7.88733524 | 7.1109821 | 10.2463158 |
| ENSMUSG00 | Pcdh15 | 19601 | 0.00405648 | 0 | 0 | 0.00368165 | 0.00520651 | 0.00139149 |
| ENSMUSG00 | Ccdc79 | 4005 | 2.03492472 | 1.84083037 | 1.67757307 | 0 | 0.15288802 | 0.09534206 |
| ENSMUSG00 | Pou3f2 | 1689 | 0 | 0.02852957 | 0 | 0 | 0 | 0 |
| ENSMUSG00 | Olfr851 | 1878 | 0 | 0 | 0 | 0 | 0 | 0 |
| ENSMUSG00 | Gm9885 | 2204 | 0 | 0 | 0 | 0 | 0 | 0 |
| ENSMUSG00 | Sh2d6 | 1922 | 0 | 0 | 0 | 0 | 0 | 0 |
| ENSMUSG00 | Asap2 | 6935 | 0.98600471 | 1.11867587 | 0.90507018 | 2.99686484 | 1.59664363 | 1.4748363 |
| ENSMUSG00 | 4930504O13I | 545 | 0 | 0 | 0 | 0 | 0 | 0 |
| ENSMUSG00 | Rnf103 | 5073 | 26.8798143 | 25.3422881 | 23.1857095 | 23.9835707 | 30.9899988 | 24.882127 |
| ENSMUSG00 | 5830454E08I | 726 | 4.05221151 | 3.11950796 | 5.41868323 | 2.48498929 | 2.67080204 | 6.31148677 |
| ENSMUSG00 | Gm9887 | 623 | 1.0848205 | 1.16018721 | 1.13519982 | 1.73749813 | 0.98285154 | 2.62677104 |
| ENSMUSG00 | Zfp112 | 4200 | 2.16762025 | 1.2505529 | 2.61001359 | 0.8590963 | 2.85504722 | 1.41568366 |
| ENSMUSG00 | Zmat1 | 8297 | 4.52322208 | 3.26392437 | 4.19270224 | 3.68778761 | 4.41568493 | 1.8112964 |
| ENSMUSG00 | Rap1b | 2016 | 176.079705 | 163.896049 | 220.241822 | 165.447629 | 138.474852 | 202.313966 |
| ENSMUSG00 | Jun | 5034 | 43.9174071 | 41.1604492 | 42.6740082 | 59.3267943 | 64.8320818 | 33.8901209 |
| ENSMUSG00 | 5430435G22I | 2847 | 0 | 0 | 0 | 0 | 0 | 0 |
| ENSMUSG00 | Tln2 | 13054 | 2.80182642 | 3.16714933 | 5.2077857 | 3.43019896 | 2.12642473 | 4.33544327 |
| ENSMUSG00 | Tnrc6a | 6665 | 12.1622543 | 10.3674956 | 9.17859726 | 13.7506966 | 16.5443358 | 15.1780401 |
| ENSMUSG00 | BC004004 | 3487 | 52.4220532 | 64.6723686 | 61.6949727 | 55.3906465 | 50.2215434 | 55.7381931 |
| ENSMUSG00 | Zfp608 | 8084 | 1.42616147 | 2.08029049 | 2.26914459 | 1.34794377 | 1.04779545 | 1.67345634 |
| ENSMUSG00 | Gm9888 | 962 | 0.33060691 | 0.20035943 | 0.22973931 | 0.22504394 | 0.05304197 | 0.45363225 |
| ENSMUSG00 | Kcnt2 | 6379 | 0 | 0 | 0 | 0 | 0 | 0 |
| ENSMUSG00 | Mtap1b | 12059 | 0.10549593 | 0.03196712 | 0 | 0.31716533 | 0.03385115 | 0 |
| ENSMUSG00 | Gm5111 | 962 | 0 | 0 | 0 | 0 | 0 | 0 |
| ENSMUSG00 | Klrc2 | 4885 | 0 | 0 | 0 | 0 | 0 | 0 |
| ENSMUSG00 | Suclg1 | 4854 | 84.4742534 | 75.6747525 | 88.7679369 | 65.258399 | 67.3308482 | 79.5987927 |
| ENSMUSG00 | 1200016B10I | 9299 | 5.68179735 | 5.5549915 | 8.37548633 | 8.5830178 | 5.82202223 | 10.303883 |
| ENSMUSG00 | Trim30b | 5323 | 0.07468623 | 0.08147247 | 0.17438262 | 0.12879182 | 0 | 0.03586746 |
| ENSMUSG00 | Repin1 | 1900 | 37.3910756 | 34.7956835 | 27.7075763 | 26.3398925 | 26.2114436 | 25.3079942 |
| ENSMUSG00 | Traf7 | 5095 | 16.4717997 | 14.867338 | 17.4291467 | 15.6084055 | 15.1827254 | 12.8530734 |
| ENSMUSG00 | Gpr25 | 1959 | 0 | 0 | 0 | 0 | 0 | 0 |
| ENSMUSG00 | A630001G21I | 3387 | 6.92522395 | 2.91651038 | 9.14835901 | 6.33859359 | 5.04689575 | 10.8470443 |
| ENSMUSG00 | Zfp212 | 2997 | 3.54177955 | 2.79761126 | 4.17388107 | 2.37175935 | 4.71615152 | 4.177197 |
| ENSMUSG00 | Gm12703 | 2037 | 0.03903336 | 0 | 0.28209325 | 0 | 0.02504977 | 0.02677922 |
| ENSMUSG00 | Gm9889 | 1575 | 0 | 0 | 0 | 0 | 0 | 0 |
| ENSMUSG00 | Oas1a | 2292 | 0.19079855 | 0.58866509 | 0 | 0.72415971 | 1.04635238 | 0.89249474 |
| ENSMUSG00 | 4833418N17I | 613 | 0 | 0 | 0 | 0 | 0 | 0 |
| ENSMUSG00 | Grk4 | 5401 | 1.80338691 | 2.68545067 | 0.64653686 | 2.29145367 | 1.19984257 | 1.97957019 |
| ENSMUSG00 | Olfr573-ps1 | 887 | 0 | 0 | 0 | 0 | 0 | 0 |
| ENSMUSG00 | Gm9890 | 695 | 0 | 0 | 0 | 0 | 0 | 0 |
| ENSMUSG00 | 1700030K09I | 3019 | 3.97686489 | 3.8146935 | 2.41580129 | 6.69292644 | 6.42266407 | 4.49909584 |
| ENSMUSG00 | Nup107 | 3407 | 4.67917455 | 6.85953168 | 6.49988945 | 3.46311375 | 10.6186382 | 6.04413051 |
| ENSMUSG00 | Atad2b | 4657 | 2.723212 | 3.41454282 | 2.71456455 | 5.43903699 | 5.13879542 | 1.9385668 |
| ENSMUSG00 | Olfr813 | 932 | 0 | 0 | 0 | 0 | 0 | 0 |
| ENSMUSG00 | Best2 | 2079 | 0 | 0 | 0 | 0 | 0 | 0 |
| ENSMUSG00 | Cysltr1 | 6183 | 0 | 0.43642904 | 0 | 1.21382262 | 0.94905923 | 0 |
| ENSMUSG00 | Gm9892 | 995 | 1.7580313 | 1.40442897 | 1.37714285 | 0.61647714 | 1.38463532 | 1.69952526 |
| ENSMUSG00 | Rbmy1a1 | 1721 | 0 | 0 | 0 | 0 | 0 | 0 |
| ENSMUSG00 | Sae1 | 2521 | 36.3492196 | 40.1585541 | 43.0797018 | 36.3396712 | 31.474024 | 32.846412 |
| ENSMUSG00 | Junb | 1806 | 55.2966595 | 91.9704685 | 32.5762199 | 69.78659 | 74.8443353 | 35.7318917 |
| ENSMUSG00 | C130026L21I | 2468 | 0 | 0 | 0 | 0 | 0 | 0 |
| ENSMUSG00 | Tas2r137 | 1001 | 0 | 0 | 0 | 0.072092 | 0 | 0 |
| ENSMUSG00 | Reep1 | 3917 | 0.27403574 | 0.4797731 | 2.70830797 | 1.40017125 | 0.53410299 | 1.67811796 |
| ENSMUSG00 | Nrk | 9557 | 0 | 0 | 0 | 0 | 0 | 0 |
| ENSMUSG00 | Dnahc6 | 13405 | 0 | 0.01078399 | 0.00989224 | 0.04306697 | 0 | 0 |
| ENSMUSG00 | Gm13619 | 2136 | 0 | 0 | 0 | 0 | 0 | 0 |
| ENSMUSG00 | Prkcb | 4948 | 0.40173283 | 0.18503282 | 0.16079894 | 0.41565815 | 0.37125092 | 0.23702698 |
| ENSMUSG00 | Ubxn8 | 2821 | 11.7392114 | 9.61679083 | 12.958144 | 11.7033218 | 9.11637484 | 16.3589825 |
| ENSMUSG00 | Olfr699 | 950 | 0 | 0 | 0 | 0 | 0 | 0 |
| ENSMUSG00 | Gm9894 | 1279 | 0 | 0 | 0 | 0 | 0 | 0 |
| ENSMUSG00 | Lamb2 | 5591 | 41.9668836 | 32.6126806 | 40.9446155 | 35.7593648 | 39.508707 | 21.9182533 |
| ENSMUSG00 | Smarca5-ps | 3454 | 0.05754991 | 0 | 0 | 0 | 0.01477313 | 0.00789654 |
| ENSMUSG00 | Cyp2j6 | 3596 | 56.5708548 | 82.9594726 | 63.8074992 | 67.3879341 | 73.275919 | 76.1429654 |
| ENSMUSG00 | Msl1 | 5565 | 6.67949223 | 9.00519314 | 7.91900045 | 8.72712164 | 7.99550759 | 10.4099432 |
| ENSMUSG00 | Senp7 | 5505 | 7.19281718 | 8.23677422 | 10.1090137 | 12.9908469 | 9.04663811 | 7.92723393 |
| ENSMUSG00 | Prkg1 | 2308 | 0 | 0.48019418 | 0 | 0 | 0 | 0 |
| ENSMUSG00 | Arhgef15 | 8138 | 0.64484191 | 0.31382175 | 0.26071375 | 0.64289708 | 0.61447344 | 0.23795765 |
| ENSMUSG00 | Bpi | 1911 | 0 | 0 | 0 | 0 | 0 | 0.07136222 |
| ENSMUSG00 | Rnaseh2a | 3568 | 5.08085737 | 4.87536592 | 4.42266198 | 3.95405813 | 3.01753512 | 5.39683165 |
| ENSMUSG00 | Ctif | 7073 | 0.5452116 | 0.2248201 | 0.23122695 | 0.35709644 | 0.31021266 | 0.29306837 |
| ENSMUSG00 | Fbxo31 | 1515 | 4.61845846 | 6.17040909 | 5.68934613 | 6.57336257 | 6.33198591 | 4.44675637 |
| ENSMUSG00 | Glis3 | 9595 | 7.51604394 | 10.4860126 | 5.89665024 | 12.1201073 | 11.4762813 | 4.77554913 |
| ENSMUSG00 | Rnf157 | 9923 | 0.28846061 | 0.09226468 | 0.14254348 | 0.2254446 | 0.45765872 | 0.28036009 |
| ENSMUSG00 | C130021I20I | 4145 | 0 | 0 | 0 | 0 | 0 | 0 |
| ENSMUSG00 | Cpvl | 1584 | 0.37647235 | 0 | 0 | 0 | 0 | 0 |
| ENSMUSG00 | Gas1 | 5723 | 8.34983183 | 9.49751996 | 3.00445866 | 6.05885808 | 7.40921166 | 6.3099113 |
| ENSMUSG00 | Lingo2 | 334 | 0 | 0 | 0 | 0 | 0 | 0 |
| ENSMUSG00 | Mrpl35 | 3711 | 23.2469396 | 20.6587522 | 30.2182904 | 22.7518147 | 14.9462869 | 25.1652829 |
| ENSMUSG00 | Cyp2f2 | 2068 | 0 | 0 | 0 | 0 | 0 | 0 |
| ENSMUSG00 | 6530403G13I | 299 | 0 | 0 | 0 | 0 | 0 | 0 |
| ENSMUSG00 | Ube2ql1 | 2677 | 0.10395531 | 0 | 0 | 0.01347854 | 0.28591544 | 0.30565528 |
| ENSMUSG00 | Uba2 | 4456 | 29.16532 | 26.4614506 | 27.070653 | 29.9118206 | 24.9864345 | 25.8545952 |
| ENSMUSG00 | Hrh1 | 4443 | 0 | 0 | 0 | 0 | 0 | 0 |
| ENSMUSG00 | Creb5 | 3031 | 0.01311629 | 0.19077443 | 0.26249857 | 0.46426913 | 0.43770563 | 0 |
| ENSMUSG00 | Krcc1 | 1865 | 105.858293 | 102.961379 | 97.8602734 | 91.76254 | 78.6599627 | 139.561331 |
| ENSMUSG00 | Cntn2 | 9764 | 0 | 0 | 0 | 0 | 0 | 0 |
| ENSMUSG00 | Sv2b | 5689 | 0 | 0 | 0 | 0 | 0 | 0 |
| ENSMUSG00 | Spink2 | 1096 | 0 | 0 | 0 | 0 | 0 | 0 |
| ENSMUSG00 | Gm6180 | 616 | 0.06453812 | 0.15644949 | 0 | 0.41002323 | 0 | 0 |
| ENSMUSG00 | Aph1c | 5296 | 4.4422885 | 6.15745967 | 7.52549309 | 5.31219229 | 8.14803963 | 4.97306559 |
| ENSMUSG00 | Cd8b1 | 1412 | 0 | 0 | 0 | 0 | 0 | 0 |
| ENSMUSG00 | Brsk2 | 5492 | 0.04343279 | 0.03509573 | 0.00804841 | 0.05912935 | 0.08361933 | 0.09435873 |
| ENSMUSG00 | Gm15413 | 1931 | 0 | 0 | 0 | 0 | 0.42279752 | 0 |
| ENSMUSG00 | Adh6a | 2533 | 1.09865126 | 1.65504164 | 0.06980157 | 0 | 0.76549636 | 1.64746143 |
| ENSMUSG00 | Jam2 | 4529 | 2.65972855 | 1.24482529 | 0.40014916 | 2.38210009 | 1.91531991 | 0.95753315 |
| ENSMUSG00 | Clec12a | 5131 | 0.24019098 | 0.55408304 | 0.09476131 | 0.72431311 | 0.04972362 | 0.3508334 |
| ENSMUSG00 | 9230110C19I | 1815 | 3.41699998 | 4.14164461 | 4.91943377 | 10.1586362 | 9.41809141 | 7.94946786 |
| ENSMUSG00 | 2700081O15I | 5211 | 0.77054376 | 1.07265924 | 1.41656261 | 1.18403946 | 0.89107661 | 1.26140627 |
| ENSMUSG00 | 1700069B07I | 822 | 0.38691465 | 0.17586293 | 0.1613206 | 0.04389543 | 0.24830353 | 0.06636165 |
| ENSMUSG00 | Vmn2r54 | 2416 | 0 | 0 | 0 | 0 | 0 | 0 |
| ENSMUSG00 | Lins | 5894 | 2.73175596 | 3.8424886 | 2.632312 | 3.1833497 | 2.6578041 | 1.64739932 |
| ENSMUSG00 | Myh7 | 7984 | 0.01493818 | 0.22934429 | 0.58684812 | 0.01355788 | 0.22368777 | 0.6968971 |
| ENSMUSG00 | 0610007L01F | 3775 | 29.1715711 | 30.2904443 | 29.7176894 | 34.5527393 | 32.2513729 | 24.1678592 |
| ENSMUSG00 | Gpr141 | 3428 | 0 | 0 | 0 | 0 | 0 | 0 |
| ENSMUSG00 | Yap1 | 9480 | 5.92977315 | 8.29538751 | 5.75369982 | 8.33921514 | 7.8261973 | 7.08910455 |
| ENSMUSG00 | Fank1 | 2205 | 0 | 0 | 0 | 0 | 0 | 0 |
| ENSMUSG00 | Socs3 | 2803 | 1.48923491 | 5.89652146 | 1.27732761 | 4.55691893 | 4.6784797 | 0.94386015 |
| ENSMUSG00 | E330013P04I | 1720 | 0.13868191 | 0.02801537 | 0.53967367 | 0 | 0.20766548 | 1.18930113 |
| ENSMUSG00 | Chmp3 | 3422 | 70.2168686 | 60.5780462 | 61.1618138 | 55.6414228 | 53.9191628 | 70.5856824 |
| ENSMUSG00 | Gm5129 | 299 | 0 | 0 | 0.2956645 | 0 | 0 | 0.63853671 |
| ENSMUSG00 | Rnf26 | 4058 | 2.09651869 | 2.77861692 | 1.74280308 | 1.78720822 | 2.59029902 | 1.02834396 |
| ENSMUSG00 | Gsx1 | 1771 | 0 | 0 | 0 | 0 | 0 | 0 |
| ENSMUSG00 | Supt7l | 6696 | 3.97198573 | 3.58375869 | 4.34361002 | 3.2601011 | 3.49015534 | 4.63538523 |
| ENSMUSG00 | Mapk11 | 2364 | 0 | 1.22300615 | 0.48614548 | 2.50315368 | 0.47486475 | 0.59994976 |
| ENSMUSG00 | Ptprt | 15797 | 0.00754994 | 0 | 0 | 0 | 0 | 0.00172657 |
| ENSMUSG00 | Olfr352 | 947 | 0 | 0 | 0 | 0 | 0 | 0 |
| ENSMUSG00 | Spag16 | 5438 | 0 | 0 | 0 | 0.22559572 | 0 | 0 |
| ENSMUSG00 | Fes | 2812 | 0.622063 | 0.4626721 | 0.03143801 | 0 | 0.09072969 | 0 |
| ENSMUSG00 | Wdr69 | 2203 | 0.23459884 | 0.0656193 | 0.04012877 | 1.11374445 | 0 | 1.07711921 |
| ENSMUSG00 | Gpr21 | 3730 | 0 | 0.38755852 | 0 | 0.60942863 | 0.28727986 | 0.03656118 |
| ENSMUSG00 | Cdh22 | 4342 | 0 | 0 | 0.06108039 | 0 | 0 | 0 |
| ENSMUSG00 | 9030619P08I | 852 | 0 | 1.07458028 | 0 | 0.33879854 | 0.4791209 | 0.12804995 |
| ENSMUSG00 | Rpl18-ps2 | 1181 | 0.06732512 | 0.04080139 | 0.14970988 | 0.15276056 | 0 | 0.06928359 |
| ENSMUSG00 | Bcl3 | 2752 | 1.84909212 | 2.17119143 | 1.79891222 | 2.33379503 | 2.59581854 | 2.16056372 |
| ENSMUSG00 | Gm9897 | 1425 | 1.08804473 | 0.94682132 | 1.45768535 | 0.70898052 | 1.6829752 | 0.8613044 |
| ENSMUSG00 | A830005F24I | 784 | 0.10141704 | 0 | 0.39465931 | 0 | 0 | 0 |
| ENSMUSG00 | Gm609 | 2575 | 0 | 0 | 0 | 0 | 0 | 0 |
| ENSMUSG00 | Spaca3 | 1096 | 0 | 0 | 0 | 0 | 0 | 0 |
| ENSMUSG00 | Gm9898 | 1164 | 0 | 0 | 0 | 0 | 0 | 0 |
| ENSMUSG00 | Mllt11 | 1921 | 8.65059389 | 13.0938692 | 12.6784047 | 8.99703245 | 5.6312294 | 11.8412541 |
| ENSMUSG00 | Cib4 | 762 | 0 | 0 | 0 | 0 | 0 | 0 |
| ENSMUSG00 | Prx | 5573 | 0.27820989 | 0.25074589 | 0.21414853 | 0.00647444 | 0.4303319 | 0.23980932 |
| ENSMUSG00 | Arhgap20 | 12934 | 0 | 0 | 0 | 0 | 0 | 0 |
| ENSMUSG00 | Styx | 1246 | 0.09571946 | 0 | 0.31927495 | 0.14479151 | 0.20476074 | 0.54724397 |
| ENSMUSG00 | 1700047G07I | 376 | 0 | 0 | 0 | 0 | 0 | 0 |
| ENSMUSG00 | Kat2b-ps | 2489 | 0 | 0 | 0.08879438 | 0 | 0 | 0.01095807 |
| ENSMUSG00 | Zfy1 | 69426 | 0 | 0 | 0 | 0 | 0 | 0 |
| ENSMUSG00 | Gm9899 | 552 | 0 | 0 | 0 | 0 | 0 | 0 |
| ENSMUSG00 | Btn2a2 | 2973 | 0.02674435 | 0 | 0 | 0 | 0 | 0 |
| ENSMUSG00 | Tas2r136 | 983 | 0 | 0 | 0 | 0 | 0 | 0 |
| ENSMUSG00 | Gm9900 | 2618 | 3.21931317 | 2.72406166 | 3.66378913 | 3.94173596 | 4.32691193 | 3.51090658 |
| ENSMUSG00 | Raet1c | 1267 | 0 | 0 | 0 | 0 | 0 | 0 |
| ENSMUSG00 | Dand5 | 1653 | 8.34552434 | 7.87074373 | 9.70675681 | 9.73538527 | 5.49467326 | 9.1080465 |
| ENSMUSG00 | Ceacam3 | 3318 | 0 | 0.01452274 | 0 | 0 | 0 | 0 |
| ENSMUSG00 | Gm9901 | 993 | 0.60053596 | 0.04852613 | 0.22256719 | 0.0726728 | 0 | 0.27466908 |
| ENSMUSG00 | Olfr212 | 2670 | 0 | 0 | 0 | 0 | 0 | 0 |
| ENSMUSG00 | Ndfip2 | 8389 | 22.3586074 | 22.4762842 | 24.8381795 | 21.6819152 | 20.0723615 | 21.8255636 |
| ENSMUSG00 | Gm12592 | 2952 | 0.09427113 | 0.45705297 | 0.35936458 | 0.9044957 | 0.1209975 | 0.11087252 |
| ENSMUSG00 | Dspp | 2909 | 0 | 0 | 0 | 0 | 0 | 0 |
| ENSMUSG00 | Gm14496 | 1383 | 0 | 0 | 0 | 0 | 0 | 0 |
| ENSMUSG00 | Aldh1a1 | 2040 | 1.55903845 | 5.81071802 | 1.86341103 | 4.24494642 | 4.20217213 | 1.10970346 |
| ENSMUSG00 | 1190005F20I | 2649 | 28.7398057 | 25.3938366 | 24.8624938 | 24.6948836 | 39.3725603 | 33.0096238 |
| ENSMUSG00 | Olfr1013 | 917 | 0 | 0 | 0 | 0 | 0 | 0 |
| ENSMUSG00 | Ddx10 | 3355 | 11.5296818 | 8.12921796 | 9.28831575 | 10.12018 | 7.71099033 | 13.0479273 |
| ENSMUSG00 | Rab4b | 2231 | 0.42766989 | 0.75595046 | 0.8519405 | 0.80865183 | 0.18297221 | 0.84354554 |
| ENSMUSG00 | Pom121 | 5568 | 3.00593701 | 4.04149937 | 2.44507321 | 4.42601229 | 4.33467596 | 1.81732959 |
| ENSMUSG00 | AI854703 | 4145 | 2.69512425 | 0.62776065 | 1.63157587 | 2.05436972 | 3.00372393 | 1.17126316 |
| ENSMUSG00 | Slc22a26 | 3129 | 0 | 0 | 0 | 0 | 0 | 0 |
| ENSMUSG00 | Nrgn | 1289 | 0.06168422 | 0.03738281 | 0 | 0 | 0.03958802 | 0.02115953 |
| ENSMUSG00 | Sec61b | 1267 | 18.6069455 | 18.7877683 | 17.0597484 | 15.606125 | 8.98096433 | 37.8874231 |
| ENSMUSG00 | Slamf8 | 2035 | 0 | 0 | 0 | 0 | 0 | 0 |
| ENSMUSG00 | D10Jhu81e | 1356 | 36.911615 | 40.4396543 | 37.8779805 | 27.1147518 | 20.771799 | 32.0215529 |
| ENSMUSG00 | Gas5 | 2859 | 19.9820306 | 17.1913854 | 14.5329599 | 19.4229684 | 16.6339917 | 33.3801898 |
| ENSMUSG00 | Dis3l2 | 3251 | 10.272102 | 8.52267125 | 13.2020885 | 12.3640103 | 14.0632522 | 11.2756429 |
| ENSMUSG00 | Ficd | 3151 | 6.97706052 | 6.7133761 | 9.01992544 | 8.66839343 | 7.30336258 | 4.40583668 |
| ENSMUSG00 | Gm9903 | 1745 | 1.89094836 | 1.1874023 | 1.84913154 | 0.02067739 | 0.29241476 | 0.01563017 |
| ENSMUSG00 | Tarm1 | 861 | 2.07781257 | 0.83948506 | 0.15401339 | 0 | 2.4890915 | 0.06335572 |
| ENSMUSG00 | Zfp943 | 5529 | 15.6534059 | 18.0579324 | 17.0363768 | 24.5637214 | 12.8834908 | 30.0469936 |
| ENSMUSG00 | 2310001K24I | 320 | 0 | 0 | 0 | 0 | 0 | 0 |
| ENSMUSG00 | Gm9905 | 1844 | 0.68989988 | 2.71767353 | 0.91088397 | 1.017498 | 1.02384811 | 3.22444217 |
| ENSMUSG00 | F730035P03I | 864 | 0 | 0 | 0 | 0 | 0 | 0 |
| ENSMUSG00 | Gm6792 | 1097 | 0 | 0 | 0 | 0.09867469 | 0 | 0 |
| ENSMUSG00 | Rxfp2 | 5875 | 0.01353378 | 0 | 0 | 0 | 0 | 0 |
| ENSMUSG00 | Atp6v1e2 | 1003 | 0 | 0 | 0 | 0 | 0 | 0 |
| ENSMUSG00 | Trim50 | 2199 | 0 | 0 | 0 | 0 | 0 | 0 |
| ENSMUSG00 | Tas2r107 | 1031 | 0 | 0 | 0 | 0 | 0 | 0 |
| ENSMUSG00 | Zfp952 | 4422 | 14.6363461 | 9.26243234 | 15.1037958 | 15.1035424 | 12.5200402 | 14.451488 |
| ENSMUSG00 | Olfr211 | 929 | 0 | 0 | 0 | 0 | 0 | 0 |
| ENSMUSG00 | Cacng8 | 658 | 0 | 0 | 0 | 0 | 0 | 0 |
| ENSMUSG00 | Phgdh | 2473 | 2.31491678 | 0.79888562 | 0.32172793 | 0.27721749 | 0.55710156 | 0.40807184 |
| ENSMUSG00 | Adamts18 | 5937 | 0 | 0 | 0 | 0 | 0 | 0 |
| ENSMUSG00 | Cbx7 | 6176 | 16.0541076 | 12.4679298 | 8.40235811 | 10.732305 | 20.9525403 | 15.7659427 |
| ENSMUSG00 | Hunk | 5071 | 0 | 0 | 0 | 0 | 0 | 0.06992118 |
| ENSMUSG00 | Gm4792 | 768 | 0 | 0 | 0 | 0 | 0 | 0.24859697 |
| ENSMUSG00 | Mapk14 | 4644 | 19.4668309 | 16.7262155 | 19.7975524 | 24.8316568 | 25.7659025 | 17.366733 |
| ENSMUSG00 | Adamts19 | 4643 | 0 | 0 | 0 | 0 | 0 | 0 |
| ENSMUSG00 | 4930597O21I | 982 | 0 | 0 | 0.09002412 | 0 | 0.05196169 | 0.13887291 |
| ENSMUSG00 | Thoc7 | 875 | 37.9380871 | 43.7257544 | 35.9676711 | 35.4634952 | 28.0499277 | 47.9721941 |
| ENSMUSG00 | Ggcx | 3228 | 10.6162405 | 12.0615382 | 6.47691194 | 5.52184944 | 10.3854798 | 4.6725141 |
| ENSMUSG00 | Hs6st3 | 1411 | 0.08452618 | 0 | 0.12530643 | 0 | 0 | 0 |
| ENSMUSG00 | A530088E08I | 1692 | 0 | 0 | 0 | 0 | 0 | 0 |
| ENSMUSG00 | Tg | 8831 | 0.17557057 | 0.62749868 | 0.67571609 | 0.38406887 | 0.29468295 | 0.45092258 |
| ENSMUSG00 | Kdm3a | 4936 | 16.8413107 | 15.5512566 | 17.3279105 | 22.0615094 | 21.9673923 | 19.5995027 |
| ENSMUSG00 | Gm13534 | 3025 | 0.13142308 | 0.0159294 | 0 | 0 | 0.18555046 | 0.01803282 |
| ENSMUSG00 | Tnfaip6 | 1704 | 0 | 0 | 0 | 0 | 0 | 0 |
| ENSMUSG00 | Tcf4 | 12754 | 3.52855606 | 3.35876957 | 3.67713309 | 3.15159147 | 4.54892499 | 3.55849143 |
| ENSMUSG00 | F930015N05I | 676 | 0 | 0 | 0 | 0 | 0 | 0 |
| ENSMUSG00 | Usp21 | 3183 | 13.1643973 | 15.3809065 | 12.484278 | 14.8839851 | 17.0568847 | 15.5695946 |
| ENSMUSG00 | Trim60 | 3622 | 0.0878089 | 1.02439427 | 0.08542598 | 0.05977147 | 0 | 0 |
| ENSMUSG00 | Gm5444 | 2467 | 0 | 0 | 0 | 0 | 0 | 0 |
| ENSMUSG00 | BC048502 | 711 | 0 | 0 | 0 | 0 | 0 | 0 |
| ENSMUSG00 | Nrd1 | 7502 | 23.0202356 | 23.7014092 | 20.6279195 | 21.9849408 | 26.4586246 | 17.5201928 |
| ENSMUSG00 | Kcnip1 | 7000 | 0.03407613 | 0.28223488 | 0.05683094 | 0 | 0 | 0.04286015 |
| ENSMUSG00 | Lgals7 | 922 | 0 | 0 | 0 | 0.03913454 | 0 | 0 |
| ENSMUSG00 | A530021J07R | 410 | 0 | 0 | 0 | 0 | 0 | 0 |
| ENSMUSG00 | Cstf2t | 3749 | 60.3702189 | 44.728946 | 65.2121616 | 50.5283366 | 46.9431769 | 48.6927609 |
| ENSMUSG00 | Gm4759 | 3088 | 0 | 0 | 0 | 0 | 0 | 0 |
| ENSMUSG00 | 6430503K07I | 473 | 0 | 0 | 0 | 0 | 0 | 0 |
| ENSMUSG00 | Shisa7 | 11355 | 0.67572063 | 0.52196674 | 0.34645214 | 0.71814549 | 0.72349155 | 0.18735551 |
| ENSMUSG00 | Ebf4 | 3219 | 0.04940103 | 0.58380592 | 0.02746309 | 0 | 0.06340649 | 0.09320318 |
| ENSMUSG00 | 3110082I17R | 1424 | 4.02021713 | 5.61723974 | 5.05962108 | 4.66228665 | 4.44330801 | 2.73895605 |
| ENSMUSG00 | Smagp | 1138 | 32.6637735 | 43.2746431 | 37.9871726 | 29.0431923 | 22.7780307 | 33.1945302 |
| ENSMUSG00 | Ier2 | 1521 | 23.3671266 | 33.8033753 | 26.9105238 | 27.0911866 | 25.7983452 | 21.895026 |
| ENSMUSG00 | Eif3k | 1065 | 28.0341464 | 26.8758184 | 35.2369622 | 26.9683638 | 20.3147262 | 30.8344279 |
| ENSMUSG00 | 4930563E22I | 4737 | 0.63783334 | 0.81378834 | 2.23015421 | 1.61481812 | 2.06820016 | 0.7820591 |
| ENSMUSG00 | Tanc2 | 13503 | 12.6747644 | 9.46385578 | 11.2836964 | 13.9646571 | 13.2525735 | 14.2604572 |
| ENSMUSG00 | Zfand2a | 5704 | 4.09124598 | 4.95887825 | 7.70277559 | 8.41956544 | 5.59107379 | 6.51263301 |
| ENSMUSG00 | Gm9912 | 344 | 0 | 0 | 0 | 0 | 0 | 0 |
| ENSMUSG00 | A730061H03I | 464 | 0.94247906 | 0.20770018 | 0.19052519 | 0 | 0 | 0.3526893 |
| ENSMUSG00 | Fate1 | 1494 | 0 | 0 | 0 | 0 | 0 | 0 |
| ENSMUSG00 | Zfp472 | 2372 | 13.0730501 | 13.8546179 | 11.9822028 | 19.6686693 | 13.8537041 | 23.6065912 |
| ENSMUSG00 | 4930442H23I | 1244 | 0.31957782 | 0.30988066 | 0.92383273 | 0.4930826 | 1.31257558 | 1.27164717 |
| ENSMUSG00 | Rpia | 3797 | 1.06796392 | 0.96449028 | 0.15133631 | 0.83624438 | 0.59129854 | 0.74705359 |
| ENSMUSG00 | 4732444A12I | 2658 | 0 | 0 | 0 | 0 | 0 | 0 |
| ENSMUSG00 | Gm9913 | 2792 | 0 | 0 | 0 | 0 | 0 | 0 |
| ENSMUSG00 | Sh3pxd2a | 10449 | 0.97400737 | 0.63178702 | 0.68529989 | 0.88400836 | 0.78133985 | 0.8561663 |
| ENSMUSG00 | Gykl1 | 1989 | 0 | 0 | 0 | 0 | 0 | 0 |
| ENSMUSG00 | Tll1 | 4907 | 0 | 0 | 0 | 0.02941271 | 0 | 0.27235731 |
| ENSMUSG00 | Gm9915 | 1382 | 0 | 0 | 0 | 0 | 0 | 0 |
| ENSMUSG00 | Dennd4a | 8857 | 2.09617358 | 1.70287416 | 1.82157307 | 3.04723601 | 3.80234903 | 4.00943664 |
| ENSMUSG00 | Aldh7a1 | 4832 | 11.5843784 | 13.4427409 | 11.828018 | 10.9694792 | 9.95816896 | 8.52332477 |
| ENSMUSG00 | Plxnb1 | 8912 | 12.5618754 | 11.1328416 | 10.8619879 | 11.2715902 | 16.4152401 | 12.0856767 |
| ENSMUSG00 | Gpr30 | 2608 | 11.1736071 | 9.86639575 | 2.35584976 | 5.43720993 | 5.81090246 | 1.43275521 |
| ENSMUSG00 | Krt42 | 2036 | 0 | 0 | 0 | 0 | 0.22555864 | 0 |
| ENSMUSG00 | 2700023E23I | 2334 | 3.59400445 | 2.89036071 | 3.42782073 | 3.84936979 | 3.45422767 | 10.2250683 |
| ENSMUSG00 | Uck2 | 92 | 4.75337267 | 5.23765676 | 20.1791022 | 5.49074591 | 7.21024876 | 4.74341552 |
| ENSMUSG00 | Gm9917 | 5402 | 0.83897164 | 1.21313516 | 0.45003728 | 0.42080133 | 0.97292053 | 0.84318118 |
| ENSMUSG00 | Tgm5 | 3142 | 0 | 0 | 0 | 0 | 0 | 0 |
| ENSMUSG00 | Defb40 | 387 | 0 | 0 | 0 | 0 | 0 | 0 |
| ENSMUSG00 | BC048403 | 2096 | 1.53635206 | 1.26443431 | 2.23539855 | 1.514895 | 1.82584837 | 2.22517334 |
| ENSMUSG00 | Dpep2 | 2927 | 0.02716466 | 0 | 0.01510142 | 0.07396388 | 0 | 0.27023045 |
| ENSMUSG00 | Mast1 | 6450 | 0.44994575 | 0.5080121 | 0.08223599 | 0.95099962 | 0.18195452 | 0.35943323 |
| ENSMUSG00 | Defb37 | 350 | 0 | 0 | 0 | 0 | 0 | 0 |
| ENSMUSG00 | Nebl | 18638 | 0.21330336 | 0.16546477 | 0.73282377 | 0.23424871 | 0.00547552 | 0.00878033 |
| ENSMUSG00 | B430305J03I | 3701 | 0 | 0.02603969 | 0 | 0.1559883 | 0 | 0 |
| ENSMUSG00 | 4732471J01R | 435 | 0 | 0 | 0.10161343 | 0 | 0.35190604 | 0.06270032 |
| ENSMUSG00 | Dusp7 | 3265 | 7.41534078 | 9.78487324 | 6.66073714 | 5.0835346 | 8.39239321 | 5.71389073 |
| ENSMUSG00 | Klk1b26 | 868 | 0.13740373 | 0.11102867 | 0 | 0.04156918 | 0 | 0.0314224 |
| ENSMUSG00 | Vmn2r43 | 2843 | 0 | 0 | 0 | 0 | 0 | 0 |
| ENSMUSG00 | Gm9918 | 2190 | 0.03630638 | 0 | 0.02018349 | 0.65903278 | 0 | 0 |
| ENSMUSG00 | Spink1 | 588 | 0 | 0 | 0 | 0 | 0 | 0 |
| ENSMUSG00 | Tmem39b | 2485 | 7.10319249 | 8.68713323 | 8.92930591 | 10.0913565 | 4.80489817 | 7.48543419 |
| ENSMUSG00 | Gm6457 | 1522 | 0.0522411 | 0.03165995 | 0.14520973 | 0.07112098 | 0.06705174 | 0.08960131 |
| ENSMUSG00 | Gm5114 | 4027 | 0 | 0 | 0 | 0 | 0 | 0 |
| ENSMUSG00 | Ptrh1 | 1744 | 0.93461852 | 2.34853646 | 0.60828224 | 1.34480097 | 0.84848905 | 2.50226048 |
| ENSMUSG00 | Sox14 | 1761 | 0 | 0 | 0 | 0 | 0 | 0 |
| ENSMUSG00 | Gm9920 | 3125 | 1.31034064 | 0.66304544 | 0.60821736 | 0.43875766 | 1.02869174 | 1.09971346 |
| ENSMUSG00 | Chd8 | 12562 | 1.49375791 | 2.48182042 | 1.56933784 | 2.37540605 | 2.38843409 | 2.5185943 |
| ENSMUSG00 | Oas1f | 1474 | 0 | 0 | 0 | 0 | 0 | 0 |
| ENSMUSG00 | Chchd3 | 2909 | 65.7489229 | 46.5299814 | 70.1090765 | 51.0159674 | 44.5888784 | 64.8815757 |
| ENSMUSG00 | Lysmd1 | 2391 | 1.1306452 | 1.02781621 | 1.14617912 | 1.44871446 | 1.15241501 | 1.18634985 |
| ENSMUSG00 | Rdh8 | 1444 | 0 | 0 | 0 | 0 | 0 | 0 |
| ENSMUSG00 | Ubxn7 | 10410 | 5.5107741 | 3.69845987 | 4.33101632 | 4.85253241 | 4.31346885 | 4.38071055 |
| ENSMUSG00 | 1700016K19I | 919 | 0.3893355 | 0 | 0 | 0.03926229 | 0.11104761 | 0.29678606 |
| ENSMUSG00 | Defb38 | 284 | 0 | 0 | 0 | 0 | 0 | 0 |
| ENSMUSG00 | Krt16 | 1854 | 0 | 0 | 0 | 0 | 0 | 0 |
| ENSMUSG00 | Exoc6 | 2707 | 16.1254221 | 18.8509207 | 24.0522035 | 17.7277869 | 17.9261483 | 14.9421094 |
| ENSMUSG00 | Grwd1 | 2909 | 7.9948285 | 5.54914339 | 6.65534797 | 6.40025266 | 6.7356921 | 3.32846234 |
| ENSMUSG00 | Olfr744 | 1104 | 0 | 0 | 0 | 0 | 0 | 0 |
| ENSMUSG00 | Camk2d | 13943 | 5.1893405 | 5.19085104 | 4.89793068 | 6.26771225 | 6.00913068 | 5.81368628 |
| ENSMUSG00 | Bcl2a1c | 443 | 0 | 0 | 0 | 0 | 0 | 0 |
| ENSMUSG00 | Gm9922 | 434 | 3.11448465 | 2.88674538 | 3.66651232 | 2.16159714 | 3.17445195 | 4.02206662 |
| ENSMUSG00 | Ppfia2 | 7963 | 0.14478324 | 0.10287197 | 0 | 0.01812485 | 0.00640793 | 0 |
| ENSMUSG00 | Gm9923 | 1838 | 0 | 0.02621678 | 0 | 0 | 0 | 0 |
| ENSMUSG00 | H2-T24 | 2700 | 0.95707638 | 1.51698059 | 0.65484212 | 1.47000922 | 0.73704765 | 1.12129073 |
| ENSMUSG00 | Nudcd3 | 8858 | 16.5520673 | 11.5053426 | 16.8813316 | 13.1977675 | 13.6696308 | 7.66695097 |
| ENSMUSG00 | Gm9924 | 2310 | 0 | 0 | 0 | 0 | 0 | 0 |
| ENSMUSG00 | Txlna | 6327 | 18.5613544 | 15.2396192 | 16.019413 | 16.4869908 | 16.5652246 | 12.7557543 |
| ENSMUSG00 | Lipg | 3777 | 0 | 0.05103145 | 0 | 0 | 0 | 0 |
| ENSMUSG00 | Gpr112 | 4706 | 0 | 0 | 0 | 0 | 0 | 0 |
| ENSMUSG00 | Dnajc5g | 1557 | 0 | 0 | 0 | 0 | 0 | 0 |
| ENSMUSG00 | Gm9925 | 482 | 0 | 0 | 0 | 0.29943605 | 0 | 0.28293194 |
| ENSMUSG00 | Ostb | 484 | 0.41069711 | 0 | 0 | 0 | 0 | 0 |
| ENSMUSG00 | Mepe | 1679 | 0 | 0 | 0.34224179 | 0.1289412 | 0 | 0 |
| ENSMUSG00 | Gm5142 | 624 | 0 | 0 | 0 | 0 | 0 | 0 |
| ENSMUSG00 | Fpgt | 5632 | 10.6306381 | 10.7204567 | 11.2780626 | 10.2505808 | 10.4825136 | 10.7994399 |
| ENSMUSG00 | Aym1 | 527 | 0 | 0 | 0 | 0 | 0 | 0 |
| ENSMUSG00 | Srcap | 4895 | 5.40901941 | 5.54217915 | 5.19224918 | 8.32208953 | 10.4137588 | 6.36872577 |
| ENSMUSG00 | Sh2d4a | 2709 | 57.6300376 | 54.8389816 | 47.8078258 | 55.4350201 | 59.2199799 | 59.8048568 |
| ENSMUSG00 | A130066N16I | 3795 | 0 | 0 | 0.24459518 | 0 | 0 | 0 |
| ENSMUSG00 | 4933409G03I | 1343 | 0 | 0 | 0 | 0 | 0 | 0 |
| ENSMUSG00 | Slc39a8 | 7655 | 1.32431712 | 0.48469707 | 0.56010173 | 0.22624927 | 0.61991547 | 0.05344475 |
| ENSMUSG00 | Ech1 | 1608 | 136.820167 | 130.894515 | 170.26506 | 115.9203 | 117.823964 | 126.535329 |
| ENSMUSG00 | Mafg | 491 | 0 | 0 | 0 | 0 | 0 | 0 |
| ENSMUSG00 | Mat2a | 4798 | 150.031781 | 164.775939 | 147.769396 | 176.056012 | 145.134838 | 174.596346 |
| ENSMUSG00 | Rhox10 | 1338 | 0 | 0 | 0 | 0 | 0 | 0 |
| ENSMUSG00 | Kdm4d | 4515 | 0 | 0.23479551 | 0.02937 | 0 | 0 | 0.02416358 |
| ENSMUSG00 | Nanp | 1644 | 0.02418217 | 0.20517342 | 0.34952796 | 0.06584315 | 0.06207588 | 0.11613289 |
| ENSMUSG00 | Gm9929 | 1669 | 0.42875893 | 0.20210012 | 0.21187223 | 0.36752232 | 0.36687628 | 0.34318 |
| ENSMUSG00 | Cyhr1 | 7006 | 23.3959244 | 24.6159401 | 24.8264705 | 18.1440255 | 19.8104114 | 21.5168329 |
| ENSMUSG00 | Shisa6 | 7993 | 0 | 0 | 0 | 0 | 0 | 0 |
| ENSMUSG00 | Cnn3 | 1996 | 325.612523 | 362.532967 | 278.830363 | 313.639014 | 342.382892 | 269.890481 |
| ENSMUSG00 | Atf7ip | 3023 | 5.14204197 | 6.12093741 | 5.74638581 | 8.62961237 | 7.41004928 | 8.37276389 |
| ENSMUSG00 | Adnp2 | 4981 | 6.45697325 | 5.25300905 | 3.25679108 | 3.99140866 | 7.60119873 | 3.55375243 |
| ENSMUSG00 | Gm12474 | 1407 | 0.67813186 | 1.33565831 | 0.62831333 | 0.15386799 | 1.55944148 | 1.1243277 |
| ENSMUSG00 | Ang5 | 544 | 0 | 0 | 0 | 0 | 0 | 0 |
| ENSMUSG00 | 6330403A02I | 6502 | 0 | 0 | 0 | 0 | 0 | 0 |
| ENSMUSG00 | Lgals4 | 2391 | 5.0879034 | 6.56996242 | 4.91747814 | 6.45885197 | 10.6064863 | 9.02310316 |
| ENSMUSG00 | Pde5a | 8773 | 0 | 0.34603281 | 0.03526877 | 0 | 0.11632595 | 0.01243572 |
| ENSMUSG00 | Cd8a | 3125 | 6.53898143 | 3.00683399 | 0.70722949 | 3.92572644 | 4.14742383 | 0.86406057 |
| ENSMUSG00 | A730045E13I | 852 | 0 | 0 | 0 | 0 | 0 | 0 |
| ENSMUSG00 | Gm9930 | 3150 | 0 | 0 | 0 | 0 | 0 | 0 |
| ENSMUSG00 | Zfp14 | 2425 | 3.72144086 | 6.2791405 | 5.26776603 | 5.69873116 | 6.45983397 | 3.68910585 |
| ENSMUSG00 | Gm9931 | 1184 | 0 | 0 | 0 | 0 | 0 | 0 |
| ENSMUSG00 | Tusc1 | 1363 | 2.68342202 | 1.23736278 | 2.39980659 | 0.68828552 | 0.97335713 | 0.6403437 |
| ENSMUSG00 | Tdrd9 | 4773 | 0 | 0 | 0 | 0 | 0 | 0 |
| ENSMUSG00 | Mill1 | 2484 | 0.24006933 | 0.09699364 | 0 | 0 | 0 | 0 |
| ENSMUSG00 | D630008O14 | 2050 | 0.67875211 | 0.30557256 | 0.10780937 | 0 | 0.17423641 | 0.33261755 |
| ENSMUSG00 | Ndst1 | 7335 | 14.6230793 | 11.1876834 | 15.8969955 | 13.7834886 | 17.5235775 | 12.5757096 |
| ENSMUSG00 | Tmem179 | 2369 | 0 | 0 | 0 | 0 | 0 | 0 |
| ENSMUSG00 | Sirt5 | 2130 | 11.4600305 | 10.9267848 | 11.3513653 | 10.5296947 | 8.60022015 | 9.6293562 |
| ENSMUSG00 | Nt5dc3 | 5962 | 1.72704956 | 2.07714117 | 1.96469111 | 2.55999746 | 1.02703205 | 2.20045312 |
| ENSMUSG00 | Rdh18-ps | 949 | 0 | 0.20310408 | 0 | 0 | 0 | 0 |
| ENSMUSG00 | Gm9932 | 2114 | 0 | 0 | 0 | 0 | 0 | 0 |
| ENSMUSG00 | Adam39 | 2439 | 0 | 0 | 0 | 0 | 0 | 0 |
| ENSMUSG00 | Tceal5 | 1453 | 0.46513639 | 0.46428781 | 0.33463198 | 0.12416395 | 0.14047179 | 0.43173896 |
| ENSMUSG00 | Olfr279 | 932 | 0.08531219 | 0 | 0 | 0.61943424 | 0 | 0 |
| ENSMUSG00 | Mrgpra2b | 2495 | 0 | 0 | 0 | 0 | 0 | 0 |
| ENSMUSG00 | Gm9933 | 1717 | 0.60200495 | 0.14032161 | 0.51487295 | 0.54637924 | 0.17830999 | 0.55597692 |
| ENSMUSG00 | Klk13 | 1027 | 0 | 0 | 0 | 0 | 0 | 0 |
| ENSMUSG00 | Ercc6 | 8401 | 3.88516242 | 6.7223557 | 4.56171859 | 3.80104861 | 5.47253477 | 5.5321968 |
| ENSMUSG00 | Rdh19 | 1903 | 0 | 0 | 0 | 0 | 0 | 0 |
| ENSMUSG00 | Olfr309 | 2306 | 0 | 0 | 0 | 0 | 0 | 0 |
| ENSMUSG00 | A930004D18I | 3687 | 0.54991308 | 0.30059348 | 0.43158838 | 0.51867327 | 1.21787933 | 0.42905589 |
| ENSMUSG00 | Gm9934 | 2391 | 0 | 0 | 0 | 0 | 0 | 0 |
| ENSMUSG00 | Pkp3 | 3601 | 5.67462288 | 6.35616774 | 8.46966723 | 6.71340456 | 7.66600092 | 6.77132116 |
| ENSMUSG00 | 6030452D12I | 666 | 0 | 0 | 0 | 0 | 0 | 0 |
| ENSMUSG00 | Iigp1 | 3118 | 5.16387736 | 5.26990916 | 9.35638755 | 4.81402518 | 47.0987523 | 67.3643351 |
| ENSMUSG00 | 2810030E01F | 3979 | 3.21720652 | 2.10717289 | 2.37727932 | 1.44183088 | 2.19289024 | 3.17370143 |
| ENSMUSG00 | Utp18 | 4970 | 10.0068624 | 10.8783075 | 9.67637932 | 11.8845688 | 5.62624828 | 10.3994852 |
| ENSMUSG00 | Capn12 | 2807 | 0 | 0 | 0.11022904 | 0.11568878 | 0.38174346 | 0 |
| ENSMUSG00 | Fbxw14 | 2230 | 0 | 0 | 0 | 0 | 0 | 0 |
| ENSMUSG00 | 1810037I17R | 1008 | 21.5736586 | 24.4278491 | 27.4507477 | 21.4774075 | 16.6544421 | 44.7542196 |
| ENSMUSG00 | Slc25a40 | 2816 | 10.6024027 | 9.47986113 | 15.1315968 | 9.00769791 | 7.59234782 | 11.9132835 |
| ENSMUSG00 | Nlrp9a | 3512 | 0 | 0 | 0 | 0 | 0 | 0 |
| ENSMUSG00 | Try4 | 796 | 0 | 0 | 0 | 0 | 0 | 0 |
| ENSMUSG00 | Skp2 | 8069 | 0.64542917 | 0.45982849 | 0.46015055 | 0.59920609 | 0.13279885 | 0.9667303 |
| ENSMUSG00 | E130116L18F | 305 | 0.13034584 | 0 | 0 | 0.35490536 | 0.33459919 | 0.08942505 |
| ENSMUSG00 | Zdhhc25 | 1224 | 0 | 0 | 0 | 0 | 0 | 0 |
| ENSMUSG00 | Gm9936 | 2924 | 0 | 0 | 0 | 0 | 0 | 0 |
| ENSMUSG00 | H2-T3 | 3710 | 0.03214729 | 0 | 0 | 0 | 0 | 0.01470331 |
| ENSMUSG00 | Umodl1 | 5166 | 0 | 0 | 0 | 0 | 0 | 0.459329 |
| ENSMUSG00 | A430110L20F | 2414 | 0.23056202 | 0.43914736 | 0.25634872 | 0 | 0.19023918 | 0.0451941 |
| ENSMUSG00 | Adm2 | 1040 | 0 | 0 | 0 | 0 | 0 | 0 |
| ENSMUSG00 | Olfr24 | 1053 | 0 | 0 | 0 | 0 | 0 | 0 |
| ENSMUSG00 | Vmn1r236 | 1120 | 0 | 0 | 0 | 0 | 0 | 0 |
| ENSMUSG00 | Krt15 | 1886 | 0 | 0 | 0 | 0 | 0 | 0 |
| ENSMUSG00 | 4831426I19R | 5512 | 0 | 0 | 0 | 0 | 0 | 0.31668667 |
| ENSMUSG00 | Gm4894 | 354 | 0 | 0 | 0 | 0 | 0 | 0 |
| ENSMUSG00 | Nkx2-4 | 1723 | 0 | 0 | 0 | 0 | 0 | 0 |
| ENSMUSG00 | Fam83e | 5271 | 0 | 0 | 0 | 0 | 0 | 0.02069789 |
| ENSMUSG00 | Spock3 | 6100 | 6.58246481 | 4.51056696 | 2.04342946 | 7.19866364 | 6.86764829 | 7.1674175 |
| ENSMUSG00 | 4922502B01I | 1391 | 0 | 0 | 0 | 0 | 0 | 0 |
| ENSMUSG00 | Ceacam10 | 1039 | 0 | 0 | 0 | 0 | 0 | 0 |
| ENSMUSG00 | Gm9938 | 1755 | 1.74425755 | 3.07514617 | 1.23412553 | 3.00169715 | 4.68105214 | 2.20683691 |
| ENSMUSG00 | A930012O16I | 1539 | 0.64580053 | 0.90930686 | 0.45953833 | 1.33637202 | 1.65777699 | 1.98489902 |
| ENSMUSG00 | Klf1 | 1531 | 0.18176902 | 1.03863657 | 0.57742447 | 0.35351448 | 0.39994547 | 0.28503869 |
| ENSMUSG00 | Cthrc1 | 1678 | 0.07107654 | 0.63176503 | 0.21073584 | 1.0321443 | 0 | 0 |
| ENSMUSG00 | Gon4l | 7605 | 1.47939526 | 2.19864503 | 1.89477986 | 2.3058348 | 2.67712347 | 1.68561216 |
| ENSMUSG00 | O3far1 | 1390 | 0 | 0 | 0 | 0 | 0 | 0 |
| ENSMUSG00 | Ifi205 | 1606 | 0 | 0 | 0 | 0 | 0 | 0 |
| ENSMUSG00 | Fam150b | 1504 | 0 | 0.12815543 | 0 | 0 | 0 | 0 |
| ENSMUSG00 | Gzmm | 1466 | 1.00337843 | 0.69025599 | 0.21105928 | 0 | 0.55690451 | 0.11162881 |
| ENSMUSG00 | Sprr2k | 635 | 0 | 0 | 0 | 0 | 0 | 0 |
| ENSMUSG00 | Tprkb | 3517 | 8.88479036 | 9.9743332 | 6.64850013 | 8.15616304 | 6.57234809 | 9.23630803 |
| ENSMUSG00 | Olfr481 | 938 | 0 | 0 | 0 | 0 | 0 | 0 |
| ENSMUSG00 | 5730455O13I | 1388 | 25.4629843 | 22.7045628 | 19.1711164 | 29.7650872 | 27.571889 | 26.390375 |
| ENSMUSG00 | Gm9939 | 1152 | 0 | 0 | 0 | 0 | 0.04429373 | 0.0236759 |
| ENSMUSG00 | Fgfr3 | 6178 | 0.74002594 | 0.16379335 | 0.45074719 | 1.70539932 | 2.378698 | 0.75934574 |
| ENSMUSG00 | Msi1 | 5784 | 0.02062006 | 0 | 0 | 0 | 0.01764398 | 0.03772426 |
| ENSMUSG00 | Gm5082 | 1297 | 0 | 0.44582676 | 0 | 0 | 0 | 0 |
| ENSMUSG00 | Lifr | 10562 | 8.95081733 | 6.03585144 | 8.59596531 | 9.0119706 | 8.11146421 | 10.5436804 |
| ENSMUSG00 | Serpinb9d | 1243 | 0 | 0 | 0 | 0 | 0 | 0 |
| ENSMUSG00 | Zscan4c | 2403 | 0 | 0 | 0 | 0 | 0 | 0 |
| ENSMUSG00 | Arfgap3 | 2602 | 28.9075208 | 26.6303243 | 26.3138566 | 25.2102832 | 21.9636975 | 16.18449 |
| ENSMUSG00 | C330019G07 | 15550 | 2.44157453 | 2.82611159 | 2.5753614 | 3.49914618 | 2.85157046 | 3.12386704 |
| ENSMUSG00 | Gm6543 | 671 | 0.11849622 | 0.07181288 | 0.13174916 | 0 | 0 | 0 |
| ENSMUSG00 | A630033H20I | 3461 | 0 | 0 | 0 | 0 | 0 | 0.0709251 |
| ENSMUSG00 | Eapp | 6359 | 8.54001987 | 10.4193046 | 11.4414583 | 9.50423409 | 8.16871373 | 10.9373062 |
| ENSMUSG00 | D130007C19 | 1503 | 1.69284814 | 1.18622645 | 1.38222663 | 1.94454132 | 1.5277358 | 1.83282672 |
| ENSMUSG00 | Cpsf3 | 2542 | 42.4454658 | 47.0680314 | 42.8976974 | 43.2785813 | 57.5702777 | 32.4033873 |
| ENSMUSG00 | Obox1 | 1786 | 0 | 0 | 0 | 0 | 0 | 0 |
| ENSMUSG00 | Mrps21 | 1555 | 9.79186433 | 8.95555098 | 11.3986746 | 7.51805944 | 7.05509376 | 20.2761948 |
| ENSMUSG00 | Lrrc36 | 2536 | 1.66170384 | 1.21606163 | 2.68418132 | 0.01422794 | 1.14688621 | 1.63475756 |
| ENSMUSG00 | Taf4b | 5142 | 3.57969418 | 3.52355158 | 0.71348755 | 4.54007834 | 0.97249802 | 1.835283 |
| ENSMUSG00 | Lce3a | 543 | 0 | 0 | 0 | 0 | 0 | 0 |
| ENSMUSG00 | Kcnn4 | 2640 | 0 | 0 | 0 | 0 | 0 | 0 |
| ENSMUSG00 | Gm9944 | 2421 | 0 | 0 | 0 | 0 | 0 | 0 |
| ENSMUSG00 | 4930553M12 | 497 | 0 | 0 | 0 | 0 | 0 | 0 |
| ENSMUSG00 | Bax | 872 | 0 | 0 | 0 | 0 | 0 | 0 |
| ENSMUSG00 | BC055111 | 1662 | 0 | 0 | 0.18616901 | 0 | 0 | 0.0984644 |
| ENSMUSG00 | Rhob | 2222 | 1.30609815 | 1.45296653 | 2.6656377 | 3.37761713 | 2.04381073 | 2.4304134 |
| ENSMUSG00 | 2410024N13I | 664 | 0 | 0 | 0 | 0 | 0 | 0 |
| ENSMUSG00 | Zfp747 | 2654 | 4.55375512 | 4.26669703 | 4.28028397 | 3.03176184 | 8.26727265 | 5.24117032 |
| ENSMUSG00 | Pnma1 | 2358 | 7.19914766 | 5.31317852 | 7.51693768 | 7.17662378 | 7.33585301 | 4.82337768 |
| ENSMUSG00 | Ceacam2 | 4046 | 0.00982587 | 0 | 0.02184965 | 0 | 0.07566937 | 0.02022341 |
| ENSMUSG00 | Mdm4 | 4369 | 9.135844 | 7.88585629 | 7.99255115 | 15.9804889 | 12.0178853 | 8.75859896 |
| ENSMUSG00 | 4930517O19I | 1022 | 0.03889969 | 0 | 0 | 0 | 0 | 0 |
| ENSMUSG00 | Cklf | 2291 | 1.1973497 | 2.12432591 | 1.8714879 | 2.25217476 | 1.15817178 | 2.19054283 |
| ENSMUSG00 | Slfn5 | 4419 | 0.26089815 | 1.02501144 | 0 | 0.67771208 | 1.39719201 | 0.03086065 |
| ENSMUSG00 | Dnajc8 | 2641 | 29.7150013 | 27.5142578 | 31.2308365 | 27.2835452 | 29.2904149 | 32.5932455 |
| ENSMUSG00 | Olfr411 | 1365 | 0 | 0 | 0 | 0 | 0 | 0 |
| ENSMUSG00 | Spcs3 | 6422 | 18.5591608 | 28.497681 | 19.3133559 | 22.614486 | 17.4087184 | 30.1329127 |
| ENSMUSG00 | Tmem74 | 1470 | 0 | 0 | 0 | 0 | 0 | 0 |
| ENSMUSG00 | Gm4793 | 2612 | 0 | 0 | 0.18614865 | 0 | 0.03907073 | 0 |

| | | | | | | | | |
|---|---|---|---|---|---|---|---|---|
| ENSMUSG00 | Slc30a7 | 9009 | 5.01301208 | 6.45053271 | 6.18207595 | 6.27600885 | 7.49338386 | 5.99442621 |
| ENSMUSG00 | Cyp3a44 | 1949 | 0 | 0 | 0 | 0 | 0 | 0 |
| ENSMUSG00 | 2900041M22 | 1782 | 0.29002315 | 0.02704065 | 0.12402313 | 0.16198449 | 0.34361196 | 0.0918338 |
| ENSMUSG00 | Fabp1 | 549 | 0.07241435 | 0 | 0 | 0 | 0 | 0 |
| ENSMUSG00 | Cadps | 6733 | 0 | 0 | 0 | 0 | 0.03789275 | 0 |
| ENSMUSG00 | A930005H10 | 308 | 8.26087907 | 15.1756003 | 13.0596354 | 10.4263051 | 3.64474113 | 10.0951587 |
| ENSMUSG00 | Atpif1 | 1439 | 35.998187 | 41.5896881 | 33.2665712 | 33.850424 | 26.5237867 | 63.9693589 |
| ENSMUSG00 | Olfr450 | 932 | 0 | 0 | 0 | 0 | 0 | 0 |
| ENSMUSG00 | Tmem120b | 2424 | 4.24780093 | 2.38546743 | 3.79289742 | 2.02440514 | 2.08399802 | 2.08160407 |
| ENSMUSG00 | Gimap4 | 1881 | 4.18478742 | 3.07409519 | 3.71286082 | 1.82232548 | 6.48341511 | 6.56853354 |
| ENSMUSG00 | Cpa1 | 1740 | 0 | 0 | 0 | 0 | 0 | 0 |
| ENSMUSG00 | Gm9945 | 931 | 0 | 0.36230408 | 0 | 0.77512448 | 0 | 0.02929607 |
| ENSMUSG00 | Aes | 1445 | 90.2683263 | 102.875553 | 111.89643 | 88.0951232 | 75.2153498 | 154.984127 |
| ENSMUSG00 | Sytl5 | 2284 | 4.9781381 | 1.41352523 | 3.01904007 | 3.75986278 | 8.22141257 | 0.94338726 |
| ENSMUSG00 | Vapb | 7124 | 3.76683736 | 4.46421278 | 2.3019208 | 3.24657363 | 2.72178872 | 4.18078412 |
| ENSMUSG00 | 9430021M05 | 1198 | 0 | 0 | 0 | 0 | 0 | 0 |
| ENSMUSG00 | Vsnl1 | 1930 | 0 | 0 | 0 | 0 | 0 | 0 |
| ENSMUSG00 | Lclat1 | 4838 | 7.06691055 | 7.21103434 | 10.1231174 | 7.54008826 | 5.81139584 | 7.59946542 |
| ENSMUSG00 | Thrsl2 | 2515 | 5.29546162 | 13.986522 | 4.65745066 | 10.7026661 | 7.87206116 | 4.67410319 |
| ENSMUSG00 | Kcnn2 | 3778 | 0.64189632 | 0.20407175 | 0.28079519 | 0.98370847 | 0.6881556 | 0.61364329 |
| ENSMUSG00 | Tmem82 | 3333 | 18.1541678 | 15.0501309 | 11.8163343 | 12.070651 | 16.4270331 | 10.8591198 |
| ENSMUSG00 | Gm9946 | 2098 | 0 | 0 | 0 | 0 | 0 | 0 |
| ENSMUSG00 | Gm9947 | 2919 | 0 | 0 | 0 | 0 | 0 | 0 |
| ENSMUSG00 | Tas2r130 | 1089 | 0 | 0 | 0 | 0 | 0 | 0 |
| ENSMUSG00 | Olfr308 | 1055 | 0 | 0 | 0 | 0 | 0 | 0 |
| ENSMUSG00 | Dedd2 | 1823 | 12.6920953 | 15.7537683 | 13.1417438 | 10.6682512 | 14.3590405 | 11.2958599 |
| ENSMUSG00 | Parp4 | 6499 | 1.34577715 | 0.96387713 | 2.32605483 | 1.32691316 | 1.75871799 | 2.43411152 |
| ENSMUSG00 | Gm14461 | 1544 | 0 | 0 | 0 | 0 | 0 | 0 |
| ENSMUSG00 | 2610204G22 | 1638 | 0.89801757 | 0.76486416 | 0.83654282 | 0.8590963 | 0.62303267 | 0.71600091 |
| ENSMUSG00 | Trim65 | 5877 | 1.21762574 | 0.60673757 | 1.06048322 | 3.11274402 | 1.06793334 | 1.65216464 |
| ENSMUSG00 | Zfp867 | 3414 | 17.5371276 | 14.2413943 | 14.0995334 | 15.3671039 | 13.3320232 | 10.1141457 |
| ENSMUSG00 | Sh3bp2 | 3945 | 0.06046461 | 0.48858243 | 0.24649951 | 1.26218559 | 0.37509883 | 0.54618416 |
| ENSMUSG00 | Ms4a5 | 940 | 0 | 0 | 0 | 0 | 0 | 0 |
| ENSMUSG00 | Olfr1500 | 1024 | 0 | 0 | 0 | 0 | 0 | 0 |
| ENSMUSG00 | Tmprss11e | 3406 | 0 | 0 | 0 | 0 | 0 | 0 |
| ENSMUSG00 | Ugt1a6a | 4657 | 3.10736416 | 11.9095115 | 0.79728469 | 12.2494551 | 5.46750301 | 1.09520239 |
| ENSMUSG00 | Adam12 | 10047 | 0 | 0 | 0 | 0 | 0 | 0 |
| ENSMUSG00 | Gm4876 | 3027 | 0.19700436 | 0.15918878 | 0.01460253 | 0.07152041 | 0.74171144 | 2.18052947 |
| ENSMUSG00 | Dub1 | 2653 | 0 | 0 | 0 | 0 | 0 | 0 |
| ENSMUSG00 | 9130219A07 | 3204 | 0 | 0 | 0 | 0 | 0 | 0 |
| ENSMUSG00 | Pla2r1 | 10852 | 0.12455643 | 0 | 0 | 0.17622082 | 0.45609643 | 0.06534654 |
| ENSMUSG00 | Pabpc1l | 2709 | 0.01467533 | 0.01778754 | 0 | 0 | 0.11301523 | 0.01006816 |
| ENSMUSG00 | Gbp10 | 6100 | 0 | 0 | 0 | 0.11830179 | 0.71102327 | 0.56784905 |
| ENSMUSG00 | Gm9949 | 960 | 0.57976742 | 0.85330158 | 0.13813076 | 0.97722204 | 0.2126099 | 1.16485439 |
| ENSMUSG00 | Oscar | 1842 | 0 | 0.26159849 | 0.02399666 | 1.7042008 | 0.41552423 | 0 |
| ENSMUSG00 | 9130230L23 | 5661 | 0.35815748 | 0.28089606 | 0.21862773 | 1.08354488 | 0.55884743 | 0.23608149 |
| ENSMUSG00 | Cggbp1 | 4519 | 68.9364783 | 58.1884078 | 68.5280177 | 75.8529372 | 66.6312555 | 90.4426796 |
| ENSMUSG00 | Kdm2a | 15609 | 3.1684168 | 3.29084165 | 3.55959489 | 4.30885585 | 4.07975636 | 3.6205428 |
| ENSMUSG00 | Mgmt | 813 | 2.05378866 | 0.29634958 | 0.81553216 | 1.73087299 | 0.37657842 | 2.14708107 |
| ENSMUSG00 | Gm9951 | 1851 | 1.35310388 | 1.11940411 | 0 | 0.21442598 | 1.21294464 | 0 |
| ENSMUSG00 | Mettl7a1 | 2385 | 21.2862678 | 29.7806355 | 12.1022237 | 26.0819475 | 18.0571326 | 13.8717557 |
| ENSMUSG00 | D730045B01 | 308 | 1.16168612 | 0 | 0 | 0 | 0.16567005 | 0.35421609 |
| ENSMUSG00 | Xlr | 2314 | 0.01718042 | 0 | 0.11461152 | 0 | 0 | 0.77792835 |
| ENSMUSG00 | Ugt2b5 | 2410 | 0 | 0 | 0 | 0.04491541 | 0 | 0.23766283 |
| ENSMUSG00 | Slc8a1 | 3054 | 34.1319155 | 35.8952704 | 15.8918218 | 51.2639133 | 41.3858327 | 22.8628279 |
| ENSMUSG00 | Mmrn1 | 4877 | 0 | 0 | 0 | 0 | 0 | 0 |
| ENSMUSG00 | Zfp869 | 1586 | 56.2743088 | 58.6075958 | 64.7420437 | 55.397086 | 53.5036967 | 42.4940943 |
| ENSMUSG00 | Pm20d2 | 1827 | 4.37375566 | 5.53867151 | 3.53227645 | 3.75237464 | 2.23432406 | 3.35894572 |
| ENSMUSG00 | Ano9 | 2988 | 5.85422069 | 2.93505103 | 1.49410514 | 4.33515863 | 3.24464907 | 3.72424793 |
| ENSMUSG00 | Olfr63 | 954 | 0 | 0 | 0 | 0 | 0 | 0 |
| ENSMUSG00 | Irs4 | 6292 | 0 | 0 | 0 | 0 | 0 | 0 |
| ENSMUSG00 | 5830411N06 | 3610 | 0 | 0 | 0 | 0 | 0 | 0 |
| ENSMUSG00 | Tmem119 | 2427 | 0 | 0 | 0 | 0 | 0 | 0 |
| ENSMUSG00 | 1600014C10 | 3443 | 5.57708309 | 6.06004341 | 4.26227473 | 4.08713255 | 3.73472167 | 3.08948862 |
| ENSMUSG00 | Sraf12 | 3250 | 0 | 0 | 0 | 0 | 0 | 0.0083922 |
| ENSMUSG00 | Emcn | 5619 | 4.52104503 | 2.29826775 | 4.05124562 | 2.59425983 | 5.603003 | 3.19393355 |
| ENSMUSG00 | Adam10 | 5170 | 20.8620152 | 19.6380723 | 24.169944 | 27.7001228 | 22.3055337 | 27.0530658 |
| ENSMUSG00 | Ap1s3 | 3230 | 0.56617712 | 1.4023299 | 1.28636943 | 0.75962199 | 0.42653627 | 4.50918185 |
| ENSMUSG00 | Ankrd24 | 6276 | 10.9207215 | 10.5110324 | 11.3180946 | 12.3148087 | 16.0819266 | 11.8989742 |
| ENSMUSG00 | Gm9954 | 314 | 0 | 0 | 0 | 0 | 0 | 0 |
| ENSMUSG00 | Zscan22 | 3568 | 10.2954215 | 10.6015575 | 8.98159646 | 12.9644566 | 10.2825012 | 6.74986168 |
| ENSMUSG00 | Zfp771 | 1367 | 14.7156351 | 17.5543879 | 14.7123911 | 11.7458009 | 12.5046349 | 12.5299733 |
| ENSMUSG00 | Hmgb2 | 2687 | 3.43255358 | 2.72584265 | 3.60260648 | 3.49137759 | 3.55114711 | 4.5170132 |
| ENSMUSG00 | Lrrc8c | 7862 | 0.79389601 | 0.01225806 | 0.53411029 | 0.119325 | 0.49974955 | 0.76868727 |
| ENSMUSG00 | Vmac | 2273 | 16.0735973 | 16.0904134 | 18.046331 | 13.6041849 | 13.9856719 | 14.5072762 |
| ENSMUSG00 | 1700013H16 | 1215 | 0 | 0 | 0 | 0 | 0 | 0 |
| ENSMUSG00 | Phactr1 | 6844 | 20.6677381 | 27.6487686 | 10.7339952 | 26.4025247 | 25.2671519 | 20.9979653 |
| ENSMUSG00 | Msra | 1845 | 28.5937792 | 30.0349097 | 29.2522766 | 19.2437571 | 21.0189949 | 25.0128399 |
| ENSMUSG00 | Zfp182 | 8435 | 4.11458619 | 2.35362349 | 3.39570768 | 5.85611369 | 3.76875307 | 4.52044406 |
| ENSMUSG00 | Gm9955 | 308 | 0.25815247 | 0 | 0 | 0 | 0 | 0.44277012 |
| ENSMUSG00 | Abca15 | 5443 | 0 | 0 | 0 | 0.01325815 | 0 | 0 |
| ENSMUSG00 | Fsd1l | 9831 | 0.45695955 | 1.73022216 | 0.69241011 | 0.90654715 | 0.89274099 | 0.70191066 |
| ENSMUSG00 | AU018091 | 3635 | 0 | 0 | 0 | 0 | 0 | 0 |
| ENSMUSG00 | Akr1c20 | 1726 | 0 | 0 | 0 | 0 | 0 | 0 |
| ENSMUSG00 | Gm9956 | 3120 | 0 | 0 | 0 | 0.06938855 | 0 | 0 |
| ENSMUSG00 | Krtap5-2 | 1414 | 0 | 0 | 0 | 0 | 0 | 0 |
| ENSMUSG00 | Defb42 | 2429 | 0 | 0 | 0 | 0 | 0 | 0.08963002 |
| ENSMUSG00 | Mtnr1a | 1539 | 0 | 0 | 0 | 0 | 0 | 0 |
| ENSMUSG00 | Set | 4057 | 7.08484407 | 6.61568851 | 6.34100878 | 6.28789881 | 6.47734374 | 8.28928523 |
| ENSMUSG00 | Kctd18 | 11698 | 1.29482288 | 1.55705891 | 1.02777409 | 1.09806872 | 1.94107858 | 2.16136011 |
| ENSMUSG00 | 4930566N20 | 1198 | 0 | 0 | 0 | 0 | 0 | 0 |
| ENSMUSG00 | Gm12541 | 1437 | 0 | 0 | 0 | 0 | 0 | 0 |
| ENSMUSG00 | Klhl18 | 8175 | 1.33247727 | 0.89594363 | 1.05435589 | 1.6904493 | 1.05485719 | 1.31785719 |
| ENSMUSG00 | Cadm4 | 2152 | 2.84495539 | 1.58979433 | 1.12969394 | 0.82157072 | 2.15771385 | 1.12799391 |
| ENSMUSG00 | Actn4 | 5734 | 87.2069121 | 85.8011117 | 66.2025511 | 78.8154182 | 93.269523 | 59.3487398 |
| ENSMUSG00 | Usp46 | 9012 | 13.1900673 | 11.6509385 | 13.9491835 | 9.21269246 | 11.6355085 | 13.8491732 |
| ENSMUSG00 | 1700041G16 | 457 | 0 | 0 | 0 | 0 | 0.33496527 | 0 |
| ENSMUSG00 | Whsc1l1 | 16486 | 3.59791198 | 2.96379063 | 3.08871304 | 3.86077439 | 3.72344596 | 4.13106722 |
| ENSMUSG00 | Cyp2c50 | 3081 | 0 | 0 | 0 | 0 | 0 | 0 |
| ENSMUSG00 | 2610002I17R | 2082 | 3.89534968 | 3.0087596 | 2.35658241 | 1.35177688 | 2.47534292 | 1.32312131 |
| ENSMUSG00 | Atrnl1 | 6570 | 11.2428741 | 6.04347463 | 11.2018369 | 11.7802109 | 13.001241 | 11.6114408 |
| ENSMUSG00 | Ncrna00086 | 2514 | 1.78694085 | 2.08922919 | 1.59998716 | 1.92322751 | 2.47622031 | 2.45189677 |
| ENSMUSG00 | Rnd1 | 2961 | 0.4430702 | 0.35802152 | 1.71672136 | 0.5605451 | 0.9305722 | 1.14220036 |
| ENSMUSG00 | Fam19a5 | 2700 | 0.30920929 | 1.42774643 | 0.93315002 | 1.08246134 | 0.37797315 | 1.23240963 |
| ENSMUSG00 | Tmem158 | 1731 | 1.95217553 | 4.00857751 | 1.07248839 | 0.54196023 | 2.97727554 | 0.6145066 |
| ENSMUSG00 | Pcnxl3 | 9011 | 5.31190751 | 5.97851984 | 6.02863889 | 7.55196271 | 10.4136617 | 5.44221522 |
| ENSMUSG00 | 4930578G10 | 1013 | 0 | 0 | 0 | 0 | 0 | 0 |
| ENSMUSG00 | Dsp | 10047 | 13.1489461 | 16.1532733 | 9.44574071 | 17.4466579 | 8.65421961 | 17.5397078 |
| ENSMUSG00 | Olfr1535 | 980 | 0 | 0 | 0 | 0 | 0 | 0 |
| ENSMUSG00 | Txk | 3068 | 0.32395274 | 1.4763773 | 0 | 0.35282312 | 0.84822202 | 0.54229237 |
| ENSMUSG00 | Zfp667 | 2530 | 5.65690632 | 4.91387434 | 2.30618311 | 6.33218489 | 8.75314115 | 4.86200091 |
| ENSMUSG00 | Atp5a | 1480 | 48.2438128 | 31.8095635 | 53.7589983 | 48.2475447 | 37.2354634 | 78.9674522 |
| ENSMUSG00 | Arhgef33 | 3242 | 0.93196068 | 0.68370646 | 0.57263338 | 1.17973372 | 1.03878495 | 1.00954865 |
| ENSMUSG00 | Stfa3 | 409 | 0 | 0 | 0 | 0 | 0 | 0 |
| ENSMUSG00 | Wbscr25 | 2113 | 0 | 0 | 0 | 0 | 0.33808294 | 0 |
| ENSMUSG00 | 4931415C17 | 3555 | 0 | 0.21687288 | 0.07460227 | 0 | 0.07176706 | 0 |
| ENSMUSG00 | Klhl5 | 4853 | 3.89117108 | 7.07952468 | 4.18064001 | 3.50931898 | 10.5459417 | 5.40097431 |
| ENSMUSG00 | Zkscan4 | 1613 | 4.28856392 | 5.76564373 | 5.59031369 | 2.66197353 | 4.93497497 | 6.78061397 |
| ENSMUSG00 | Afp | 2071 | 1.91962726 | 6.60789453 | 1.15253478 | 2.16039282 | 2.02035868 | 1.34331879 |
| ENSMUSG00 | Kcnmb4 | 3318 | 0 | 0 | 0.02664367 | 0 | 0 | 0.00822021 |
| ENSMUSG00 | Olfr1346 | 1065 | 0 | 0 | 0 | 0 | 0 | 0 |
| ENSMUSG00 | Zfp174 | 5894 | 1.47717174 | 1.25902818 | 0.69745019 | 3.72207042 | 1.45443352 | 0.42573241 |
| ENSMUSG00 | Olfr137 | 941 | 0 | 0 | 0 | 0 | 0 | 0 |
| ENSMUSG00 | Fam73a | 5339 | 13.9170244 | 12.6445412 | 12.6752241 | 21.3153715 | 19.1814827 | 12.659029 |
| ENSMUSG00 | 5330416C01 | 3668 | 0 | 0 | 0 | 0 | 0 | 0 |
| ENSMUSG00 | Gm9958 | 1283 | 9.48182154 | 3.60553269 | 8.95750521 | 5.42777443 | 5.56795995 | 8.82227224 |
| ENSMUSG00 | 9130008F23 | 2157 | 41.1009372 | 23.6799391 | 56.8250861 | 41.7360691 | 34.7272692 | 63.9316532 |
| ENSMUSG00 | Gm9959 | 388 | 0.10246258 | 0.37257558 | 0.34176683 | 0 | 0.26302256 | 0.42177277 |
| ENSMUSG00 | Nt5c1a | 1214 | 0.42571767 | 0.23815375 | 0.32769076 | 0.44582427 | 0.84063222 | 0 |
| ENSMUSG00 | Ifltd1 | 7730 | 0 | 0 | 0 | 0 | 0 | 0 |
| ENSMUSG00 | Zfp647 | 1604 | 1.53668316 | 0.81111842 | 1.87389359 | 0.80982145 | 0.85892279 | 0.35708692 |
| ENSMUSG00 | 9430031J16R | 12501 | 0 | 0.00770921 | 0 | 0 | 0 | 0 |
| ENSMUSG00 | Kbtbd13 | 1394 | 0 | 0 | 0 | 0 | 0 | 0 |
| ENSMUSG00 | Sec14l3 | 2886 | 0 | 0 | 0 | 0 | 0 | 0 |
| ENSMUSG00 | Agtr1b | 3684 | 0 | 0 | 0 | 0 | 0 | 0 |
| ENSMUSG00 | AV320801 | 3693 | 0 | 0 | 0 | 0 | 0 | 0 |
| ENSMUSG00 | Naaladl1 | 2473 | 0.08037906 | 0 | 0 | 0 | 0 | 0 |
| ENSMUSG00 | Lrtm2 | 6472 | 0.57741273 | 1.55608257 | 0.23220993 | 0.07805139 | 0.68227064 | 0.34556867 |
| ENSMUSG00 | A830031A19 | 754 | 0 | 0 | 0 | 0 | 0 | 0 |
| ENSMUSG00 | Agap1 | 10349 | 9.45388323 | 5.96917759 | 11.1220021 | 8.7232849 | 6.72035465 | 9.56154133 |
| ENSMUSG00 | Gm9961 | 544 | 0 | 0 | 0.08125339 | 0 | 0 | 0 |
| ENSMUSG00 | Cntn1 | 9407 | 0 | 0 | 0 | 0 | 0 | 0 |
| ENSMUSG00 | Ep300 | 11680 | 4.27847937 | 5.64375453 | 5.65011572 | 6.78082943 | 5.85405339 | 7.03115914 |
| ENSMUSG00 | Gabrg3 | 11923 | 0 | 0 | 0 | 0 | 0 | 0 |
| ENSMUSG00 | Smyd1 | 6479 | 0 | 0 | 0 | 0 | 0.00787566 | 0 |
| ENSMUSG00 | Sprr2e | 645 | 0 | 0 | 0 | 0 | 0 | 0 |
| ENSMUSG00 | Olfr420 | 1045 | 0 | 0 | 0 | 0 | 0 | 0 |
| ENSMUSG00 | Commd5 | 1114 | 13.703864 | 19.3783897 | 20.0773183 | 20.146348 | 23.2687603 | 15.7429022 |
| ENSMUSG00 | Pdlim1 | 1496 | 0.61121394 | 0.12884075 | 0.02954669 | 0.55473732 | 0.81860496 | 0 |
| ENSMUSG00 | Gm11190 | 3411 | 1.02564711 | 0.53681759 | 0.63497224 | 0.52890713 | 1.13691134 | 1.13544379 |
| ENSMUSG00 | Gm9962 | 317 | 0 | 0 | 0 | 0 | 0 | 0 |
| ENSMUSG00 | Nfic | 10304 | 10.4983174 | 13.075436 | 10.5742957 | 10.1655838 | 10.0131339 | 11.8585388 |
| ENSMUSG00 | 4931431C16 | 2731 | 0 | 0.12350974 | 0 | 0.02642405 | 0 | 0 |
| ENSMUSG00 | Ddx17 | 4748 | 19.5428162 | 13.7313092 | 13.8619512 | 16.2323604 | 16.3675591 | 21.1338243 |
| ENSMUSG00 | Smyd3 | 10857 | 2.49364302 | 3.13786632 | 3.00867293 | 2.15688007 | 3.60479232 | 3.03972677 |
| ENSMUSG00 | Rab39 | 2969 | 0.49543711 | 0 | 0 | 0 | 0 | 0 |
| ENSMUSG00 | Gm9964 | 727 | 0 | 0 | 0 | 0 | 0 | 0 |
| ENSMUSG00 | Gabra5 | 2643 | 0 | 0 | 0 | 0 | 0 | 0 |
| ENSMUSG00 | Olfr354 | 1083 | 0 | 0 | 0 | 0 | 0 | 0 |
| ENSMUSG00 | Gm8430 | 995 | 0.07991051 | 0.14528576 | 0 | 0.03626336 | 0.10256558 | 0 |
| ENSMUSG00 | Spink6 | 582 | 0.75139224 | 0 | 0 | 0.24798656 | 0 | 0 |
| ENSMUSG00 | Gm9966 | 3398 | 0 | 0 | 0 | 0 | 0 | 0 |
| ENSMUSG00 | Zfp819 | 2483 | 0 | 0 | 0 | 0 | 0.43155614 | 0 |
| ENSMUSG00 | Ptar2 | 804 | 0 | 0.29966693 | 0.21990967 | 0.31414715 | 0.95198462 | 0 |
| ENSMUSG00 | Gm15155 | 1408 | 0 | 0.20533995 | 0 | 0 | 0 | 0 |
| ENSMUSG00 | A630012P03 | 1969 | 0 | 0 | 0 | 0 | 0 | 0 |
| ENSMUSG00 | Anxa13 | 938 | 0 | 0.25685737 | 0 | 0 | 0 | 0 |
| ENSMUSG00 | Arntl | 2904 | 23.327596 | 19.0986897 | 18.8892862 | 24.2162207 | 22.9478123 | 24.334914 |
| ENSMUSG00 | Olfr623 | 1074 | 0 | 0 | 0 | 0 | 0 | 0 |
| ENSMUSG00 | MSC1000I18 | 1495 | 0 | 0 | 0 | 0 | 0 | 0 |
| ENSMUSG00 | Cgrrf1 | 1975 | 20.9345315 | 23.2514832 | 23.7235208 | 21.5396104 | 16.7418185 | 22.8692671 |
| ENSMUSG00 | 9130017K11 | 4752 | 0.61908787 | 0.46645125 | 0.66972489 | 0.87319763 | 0.77312691 | 1.83667604 |
| ENSMUSG00 | 5033411D12 | 1297 | 23.5712911 | 15.5667843 | 24.2309255 | 17.1368847 | 15.97279 | 25.1296792 |
| ENSMUSG00 | Gm4861 | 954 | 0 | 0 | 0 | 0 | 0.05348677 | 0 |
| ENSMUSG00 | Klf2 | 1844 | 0.94861233 | 0.67941838 | 2.32515118 | 2.07413054 | 3.12688746 | 3.091323 |
| ENSMUSG00 | Zfp78 | 1570 | 2.15236678 | 5.95424827 | 8.67144436 | 6.36606773 | 4.35511743 | 6.25405741 |
| ENSMUSG00 | 4930583K01 | 923 | 0.43072027 | 0 | 0 | 0 | 0.2764159 | 0 |
| ENSMUSG00 | Ifng | 1203 | 0 | 0 | 0 | 0 | 0 | 0 |
| ENSMUSG00 | C1ra | 2681 | 0.11862881 | 0.12581317 | 0.03297415 | 0 | 0.01903259 | 0.14242632 |
| ENSMUSG00 | Csfl1 | 760 | 0.05230984 | 0 | 0 | 0 | 0 | 0.03588768 |
| ENSMUSG00 | Fam72a | 2187 | 0.29084942 | 0 | 0.16168941 | 0.09899052 | 0.34997514 | 0.54873531 |
| ENSMUSG00 | 2900002K06 | 967 | 1.68559948 | 2.49154303 | 1.09704678 | 1.641789 | 2.37454696 | 1.1000113 |
| ENSMUSG00 | Klk15 | 760 | 0 | 0.38041928 | 0.17448096 | 0.33233462 | 0.93995956 | 0.25121378 |
| ENSMUSG00 | Actbl2 | 2736 | 0 | 0 | 0 | 0 | 0 | 0 |
| ENSMUSG00 | Fev | 2270 | 0 | 0 | 0 | 0 | 0 | 0 |
| ENSMUSG00 | Gm12830 | 2302 | 0 | 0 | 0 | 0 | 0 | 0 |
| ENSMUSG00 | Sertad3 | 1322 | 12.6604064 | 11.7732381 | 11.7358902 | 7.31466561 | 17.0988536 | 9.86178333 |
| ENSMUSG00 | Zfp811 | 1653 | 0.52911105 | 2.30292131 | 0 | 0.87312872 | 1.08041328 | 1.51800775 |
| ENSMUSG00 | Ankrd17 | 10019 | 9.84066191 | 8.55611145 | 9.26478394 | 10.6780379 | 8.68312659 | 11.0579484 |
| ENSMUSG00 | Foxd2 | 2618 | 0 | 0 | 0 | 0 | 0 | 0 |
| ENSMUSG00 | Pld5 | 11759 | 0 | 0 | 0 | 0.00306846 | 0 | 0 |
| ENSMUSG00 | 9430025C20 | 2113 | 0.39510889 | 0.15963327 | 0.20918998 | 1.60516431 | 0 | 1.48442192 |
| ENSMUSG00 | Zfp935 | 1798 | 18.6837492 | 23.9324209 | 9.73522238 | 27.6535358 | 8.57061485 | 21.055172 |
| ENSMUSG00 | Wdr86 | 3896 | 0 | 0 | 0.14749075 | 0 | 0.15716543 | 0 |
| ENSMUSG00 | Kcmf1 | 3244 | 38.9711554 | 26.6629666 | 35.0044805 | 31.477246 | 27.5894769 | 38.5914285 |
| ENSMUSG00 | Zfp101 | 5018 | 2.82835922 | 5.08944088 | 2.14931241 | 3.55211837 | 3.29464842 | 6.34522927 |
| ENSMUSG00 | Ntrk2 | 15073 | 0 | 0 | 0 | 0 | 0 | 0 |
| ENSMUSG00 | 9330161L09 | 998 | 0 | 0.04828301 | 0.22145212 | 0 | 0.20451453 | 0 |
| ENSMUSG00 | D730040F13 | 21423 | 3.96014542 | 4.26464521 | 3.44362956 | 5.31554556 | 6.36192176 | 4.80103929 |
| ENSMUSG00 | Ctsj | 1268 | 0 | 0 | 0 | 0 | 0 | 0 |
| ENSMUSG00 | Adh7 | 3360 | 0 | 1.00388421 | 0 | 0 | 0.25816916 | 0 |
| ENSMUSG00 | Mrfap1 | 1646 | 91.7081163 | 89.9616869 | 98.1517231 | 91.3888479 | 68.1075016 | 102.255042 |
| ENSMUSG00 | Zfp93 | 2730 | 3.94642316 | 2.73586027 | 6.62218088 | 4.44086702 | 7.04649952 | 2.98722239 |
| ENSMUSG00 | 0610012H03 | 1590 | 2.27531366 | 6.90975397 | 3.50278756 | 3.69897689 | 2.8241013 | 3.07054115 |
| ENSMUSG00 | Pgbd1 | 5925 | 3.53605709 | 4.81457785 | 3.40186336 | 1.4189226 | 1.81714182 | 0.72732371 |
| ENSMUSG00 | Sec23ip | 4559 | 11.1618809 | 9.49142905 | 11.6540064 | 13.3912743 | 13.1958976 | 10.1345117 |
| ENSMUSG00 | Tead1 | 11624 | 0.43777542 | 0.33163415 | 0.21294763 | 0.38801235 | 0.24582562 | 0.4129677 |
| ENSMUSG00 | Tns1 | 11558 | 44.8564822 | 46.9607271 | 32.5987636 | 47.3830483 | 49.0485409 | 35.6189137 |
| ENSMUSG00 | Gm9967 | 1853 | 0 | 1.11819591 | 0.02385421 | 0.99308379 | 1.92760189 | 1.79573988 |
| ENSMUSG00 | Fat2 | 16435 | 0 | 0 | 0 | 0 | 0 | 0 |
| ENSMUSG00 | Snupn | 1382 | 21.6325046 | 21.1295108 | 22.4527452 | 19.8425136 | 24.959356 | 26.7220416 |
| ENSMUSG00 | Olfr115 | 958 | 0 | 0 | 0 | 0 | 0 | 0 |
| ENSMUSG00 | Zfp457 | 3814 | 0.19804775 | 1.04862997 | 0.31291289 | 0.78521492 | 1.05691759 | 0.63645592 |
| ENSMUSG00 | 4933400A11 | 2364 | 0 | 0 | 0 | 0 | 0 | 0 |
| ENSMUSG00 | Prl2c5 | 1267 | 0 | 0 | 0 | 0 | 0.08054677 | 0 |
| ENSMUSG00 | Slc6a2 | 6161 | 0 | 0 | 0 | 0 | 0 | 0 |
| ENSMUSG00 | Gm9968 | 2987 | 1.0115221 | 1.00018728 | 0.50313447 | 0.84557855 | 1.58870203 | 1.38792942 |
| ENSMUSG00 | Stam2 | 5991 | 18.4808902 | 16.2310533 | 23.0563778 | 24.8316257 | 21.1225859 | 17.5593863 |
| ENSMUSG00 | Fut9 | 15734 | 1.49076735 | 0.58801302 | 1.49174899 | 2.42396854 | 1.63126141 | 1.51159816 |
| ENSMUSG00 | Rnf212 | 294 | 0 | 0 | 0 | 0 | 0 | 0 |
| ENSMUSG00 | D830050J10 | 493 | 0.16127984 | 0.19548252 | 0.53795347 | 0.07318873 | 0 | 0.49791431 |
| ENSMUSG00 | Fbxo6 | 2545 | 8.950841 | 8.99354028 | 10.6466522 | 6.69183694 | 5.21291069 | 9.54880298 |
| ENSMUSG00 | 4933427D06 | 3002 | 0 | 0 | 0 | 0 | 0 | 0 |
| ENSMUSG00 | Gm9969 | 490 | 0 | 0 | 0 | 0 | 0 | 0 |
| ENSMUSG00 | Mtap6 | 8776 | 0.53907272 | 0.06785131 | 0.22161362 | 0 | 0.25001529 | 0.20201134 |
| ENSMUSG00 | Gm15053 | 2957 | 0 | 0 | 0 | 0 | 0 | 0 |
| ENSMUSG00 | Nell1 | 6271 | 0 | 0 | 0 | 0 | 0 | 0 |
| ENSMUSG00 | H2-Q5 | 1083 | 0.18354331 | 0.08898697 | 0 | 0 | 1.93174645 | 1.93919411 |
| ENSMUSG00 | Atp10b | 9791 | 2.68799184 | 3.45981706 | 1.6207192 | 3.95424714 | 5.74834976 | 1.12541663 |
| ENSMUSG00 | Fam136b-ps | 416 | 0 | 0 | 0 | 0 | 0 | 0 |
| ENSMUSG00 | Pcdh9 | 3710 | 0 | 0 | 0 | 0 | 0 | 0 |
| ENSMUSG00 | Gm9970 | 557 | 0 | 0 | 0 | 0.06477925 | 0 | 0.39173629 |
| ENSMUSG00 | Nap1l5 | 1892 | 1.05062052 | 0.15281113 | 0.74759988 | 0 | 0.94393401 | 0.98027245 |
| ENSMUSG00 | Maf | 7540 | 0.31635661 | 0.10864317 | 0.09379701 | 0.55510838 | 0.11504621 | 0.18086631 |
| ENSMUSG00 | Srsf11 | 8189 | 38.3281864 | 35.4175351 | 42.0751455 | 47.6438085 | 49.2692091 | 65.4105068 |
| ENSMUSG00 | A630091E08 | 766 | 0 | 0 | 0 | 0 | 0 | 0 |
| ENSMUSG00 | Cd47 | 5391 | 39.3351388 | 58.4744397 | 47.6537028 | 46.2018834 | 58.3902268 | 48.7462714 |
| ENSMUSG00 | Gm7353 | 3335 | 0.03576205 | 0.07224354 | 0.03976178 | 0.08655363 | 0.22950394 | 0.07360472 |
| ENSMUSG00 | Gm9972 | 1532 | 0.83040168 | 0.62906582 | 0.05770476 | 0.35328373 | 0.03330703 | 1.1750171 |
| ENSMUSG00 | Alk | 4837 | 0 | 0 | 0 | 0 | 0 | 0 |
| ENSMUSG00 | Zfp458 | 3484 | 3.94816196 | 2.80764861 | 2.91803212 | 4.14259983 | 4.75992312 | 9.33945024 |
| ENSMUSG00 | 9830001H06 | 7632 | 9.99616969 | 8.49829024 | 13.4887438 | 9.81476997 | 10.3964904 | 10.9820449 |
| ENSMUSG00 | Ano5 | 7778 | 0 | 0.00619522 | 0.02273173 | 0 | 0 | 0 |
| ENSMUSG00 | Pprc1 | 6527 | 1.96736942 | 2.06713709 | 1.53727874 | 3.46613175 | 2.04042961 | 1.13661742 |
| ENSMUSG00 | Epm2a | 3598 | 0.4419731 | 0.64284303 | 1.9901886 | 0.89252417 | 0.68072986 | 4.73023204 |
| ENSMUSG00 | Gm14168 | 2195 | 0.07244735 | 0.48296206 | 0 | 0 | 0.20921976 | 0.13668384 |
| ENSMUSG00 | Gm9974 | 692 | 0.05745012 | 0.20890076 | 0.51100397 | 0 | 0.14747508 | 0.39414219 |
| ENSMUSG00 | Vmn2r81 | 3270 | 0 | 0 | 0 | 0 | 0 | 0 |
| ENSMUSG00 | Gucy2g | 3669 | 0.639295 | 0 | 0.33732668 | 0.10817729 | 0.18079664 | 0.47576367 |
| ENSMUSG00 | Cpsf6 | 9780 | 13.9591329 | 13.1749025 | 17.0344321 | 20.2509553 | 12.1200686 | 18.8468315 |
| ENSMUSG00 | Zcchc24 | 4371 | 0.33652546 | 0.17638597 | 0 | 0.30543026 | 0 | 0 |
| ENSMUSG00 | Epha6 | 5918 | 0 | 0 | 0.00746905 | 0 | 0 | 0 |
| ENSMUSG00 | Lair1 | 5540 | 0 | 0 | 0 | 0 | 0.68157975 | 0 |
| ENSMUSG00 | Timd4 | 2164 | 0 | 0 | 0 | 0 | 0 | 0 |
| ENSMUSG00 | Apobec4 | 1564 | 0 | 0 | 0 | 0 | 0.09787668 | 0 |
| ENSMUSG00 | 2810422J05R | 3243 | 20.0187171 | 17.9640483 | 21.3989804 | 17.1008642 | 15.7185783 | 18.998893 |
| ENSMUSG00 | 4930502E18 | 1422 | 0 | 0 | 0 | 0 | 0 | 0 |
| ENSMUSG00 | Zfp459 | 3238 | 1.47333467 | 1.88995619 | 2.77114705 | 1.78292993 | 2.85230822 | 1.35615099 |
| ENSMUSG00 | Spink5 | 4875 | 0 | 0 | 0 | 0.06661301 | 0 | 0 |
| ENSMUSG00 | Unc80 | 14748 | 0.00808696 | 0 | 0.09291139 | 0 | 0 | 0 |
| ENSMUSG00 | Olfr305 | 959 | 0 | 0 | 0 | 0 | 0 | 0 |
| ENSMUSG00 | Tead1 | 3847 | 0 | 0 | 0 | 0 | 0 | 0 |
| ENSMUSG00 | 5530400C23 | 643 | 0 | 0 | 0 | 0 | 0 | 0 |
| ENSMUSG00 | Tcp10b | 2485 | 0.11198727 | 0 | 0.10672477 | 0.07259969 | 0.26693879 | 0.07682997 |
| ENSMUSG00 | Hba-x | 934 | 0 | 0 | 0 | 0 | 0 | 0 |
| ENSMUSG00 | Olfr307 | 1011 | 0 | 0 | 0 | 0 | 0 | 0 |
| ENSMUSG00 | Cdca7 | 3249 | 0 | 0.04449348 | 0 | 0.2887452 | 0 | 0 |
| ENSMUSG00 | Olfr635-ps1 | 916 | 0 | 0 | 0 | 0 | 0 | 0 |
| ENSMUSG00 | B4galnt4 | 3085 | 0 | 0 | 0 | 0 | 0.01654015 | 0 |
| ENSMUSG00 | Hmcn2 | 15950 | 0.09222275 | 0.05740078 | 0 | 0 | 0 | 0.01026005 |
| ENSMUSG00 | Zfp580 | 1099 | 3.00246122 | 4.99841165 | 2.25232321 | 3.02051692 | 3.29651746 | 4.71536074 |
| ENSMUSG00 | Olfr965 | 938 | 0 | 0 | 0 | 0 | 0 | 0 |
| ENSMUSG00 | Dach1 | 6613 | 5.54881423 | 6.33207595 | 4.77911958 | 7.15311665 | 3.84260323 | 4.88741078 |
| ENSMUSG00 | 4931431F19 | 1910 | 0 | 0 | 0 | 0 | 0 | 0 |
| ENSMUSG00 | Klhl25 | 3728 | 3.19920712 | 3.1926103 | 3.75857946 | 5.85558931 | 3.25758515 | 3.01425734 |
| ENSMUSG00 | Gpc3 | 2682 | 6.25533661 | 11.3189629 | 1.0877411 | 7.07649345 | 8.61854894 | 1.2203418 |
| ENSMUSG00 | E030030I06R | 427 | 0.46552085 | 1.24133692 | 1.34572356 | 0.76051148 | 0.71699826 | 1.21362563 |
| ENSMUSG00 | Mettl4 | 3197 | 9.40104575 | 9.23937724 | 11.2405688 | 10.9476332 | 12.3855076 | 10.3740886 |
| ENSMUSG00 | Zzef1 | 15645 | 2.34543359 | 2.15599294 | 3.08805461 | 2.59689243 | 4.31498212 | 2.85908651 |
| ENSMUSG00 | Kbtbd11 | 6968 | 1.58611131 | 1.47297821 | 2.06165313 | 1.68293118 | 0.57119077 | 2.07847782 |
| ENSMUSG00 | Gm5069 | 1325 | 3.27045085 | 2.69116734 | 2.40191147 | 3.73074725 | 5.27593471 | 2.90243331 |
| ENSMUSG00 | Ctsr | 1396 | 0 | 0 | 0 | 0 | 0 | 0 |
| ENSMUSG00 | Cope | 2502 | 78.1921341 | 86.7432995 | 83.987035 | 68.5010837 | 68.9121199 | 72.1655123 |
| ENSMUSG00 | Gja6 | 2015 | 1.14432649 | 0.66958828 | 1.07488353 | 0 | 0 | 0 |
| ENSMUSG00 | Tmem191c | 2462 | 11.141869 | 13.798311 | 7.98936643 | 13.7029698 | 14.922417 | 13.2274246 |
| ENSMUSG00 | Gdf1 | 1399 | 0 | 0 | 0 | 0 | 0 | 0 |
| ENSMUSG00 | 4933401B06 | 1282 | 0 | 0 | 0 | 0 | 0 | 0 |
| ENSMUSG00 | Gm9978 | 3272 | 0 | 0 | 0 | 0 | 0 | 0 |
| ENSMUSG00 | Klhl26 | 3066 | 9.49152372 | 11.1114855 | 8.65006713 | 6.68447531 | 8.95374762 | 6.71635768 |
| ENSMUSG00 | Slain1 | 3536 | 2.81076644 | 3.67939462 | 2.16259017 | 2.80615448 | 6.3061443 | 4.25009226 |
| ENSMUSG00 | Ubl7 | 1374 | 16.2899094 | 16.4128493 | 16.3102871 | 16.5704295 | 12.4409286 | 16.3568433 |
| ENSMUSG00 | Rras2 | 2266 | 11.7371653 | 14.1837409 | 12.8353101 | 12.5156606 | 12.0022323 | 13.288262 |
| ENSMUSG00 | Paqr3 | 7558 | 0.47340477 | 0.31240218 | 0.02339341 | 0.472628 | 0.22279312 | 0.10465263 |
| ENSMUSG00 | Ces2a | 2451 | 0 | 0.23591893 | 0 | 0 | 0 | 0 |
| ENSMUSG00 | Gm9979 | 3003 | 0 | 0 | 0 | 0 | 0 | 0 |
| ENSMUSG00 | Nap1l3 | 2820 | 6.07610358 | 6.54447069 | 2.71167335 | 2.48223994 | 8.86628516 | 13.6760071 |
| ENSMUSG00 | Ghr | 10747 | 16.4948067 | 17.4506032 | 20.8773197 | 19.8422707 | 14.903861 | 25.5996358 |
| ENSMUSG00 | Ldoc1l | 4094 | 3.9036891 | 6.4381983 | 6.20812402 | 5.41142534 | 6.26923963 | 1.71215982 |
| ENSMUSG00 | Magea2 | 2848 | 0 | 0 | 0 | 0 | 0 | 0 |
| ENSMUSG00 | Gsdmc4 | 1977 | 0 | 0 | 0 | 0 | 0 | 0 |
| ENSMUSG00 | Gemin6 | 1424 | 5.16486229 | 6.83543632 | 4.81129611 | 6.86673741 | 4.69413991 | 6.37812842 |
| ENSMUSG00 | Nkain3 | 6935 | 0.22357084 | 0.13201765 | 0.17209081 | 0.13527515 | 0 | 0.03146317 |
| ENSMUSG00 | Eef1d | 6491 | 5.87971981 | 5.85720272 | 6.12874114 | 5.15854835 | 4.10349225 | 5.33223189 |
| ENSMUSG00 | Gm6910 | 686 | 0 | 0.07024263 | 0.12886835 | 0.10519547 | 0 | 0 |
| ENSMUSG00 | Gm7936 | 374 | 0.10629808 | 0.12884075 | 0 | 0 | 0 | 0.29170737 |
| ENSMUSG00 | Myh8 | 9663 | 0 | 0 | 0 | 0 | 0 | 0 |
| ENSMUSG00 | Usp26 | 4124 | 0 | 0 | 0 | 0 | 0 | 0 |
| ENSMUSG00 | Abcd2 | 6363 | 0 | 0 | 0 | 0 | 0 | 0 |
| ENSMUSG00 | E330017A01 | 1104 | 0 | 0 | 0 | 0 | 0 | 0 |
| ENSMUSG00 | Gm5160 | 505 | 0 | 0 | 0 | 0 | 0 | 0 |
| ENSMUSG00 | Tcf7l1 | 4723 | 1.81816299 | 2.24455161 | 0.34627741 | 0.77160417 | 1.29645672 | 0.53128664 |
| ENSMUSG00 | Gm1045 | 2829 | 0 | 0 | 0 | 0 | 0 | 0 |
| ENSMUSG00 | Fmnl1 | 4968 | 0.08802542 | 0.23278475 | 0.04448656 | 0.03631446 | 0.15406514 | 0.04941058 |
| ENSMUSG00 | 6030429G01 | 2212 | 0.19769904 | 0.26140927 | 0.05994825 | 0.11418369 | 0.04613596 | 0.13563338 |
| ENSMUSG00 | Cldn17 | 1171 | 0.44135034 | 0.82299645 | 0.52845927 | 0.70869942 | 1.08937609 | 0.652169 |
| ENSMUSG00 | Magea3 | 2258 | 0 | 0 | 0 | 0 | 0 | 0 |
| ENSMUSG00 | Mta3 | 5399 | 12.5179324 | 11.2723609 | 12.0267656 | 11.7221534 | 13.8741841 | 12.6597974 |
| ENSMUSG00 | A230083G16 | 989 | 0.12059296 | 0.04872239 | 0.08938694 | 0.03648336 | 0.10318782 | 0.27577997 |
| ENSMUSG00 | Olfr967 | 932 | 0 | 0 | 0 | 0 | 0 | 0 |
| ENSMUSG00 | 1700008P20 | 885 | 0 | 0.10889591 | 0 | 0 | 0 | 0 |
| ENSMUSG00 | Gsdmc3 | 1432 | 0 | 0 | 0 | 0 | 0 | 0 |
| ENSMUSG00 | Gm9982 | 1192 | 0.23346339 | 0 | 3.85653664 | 2.84539613 | 1.9263313 | 2.31102229 |
| ENSMUSG00 | 1700034H15 | 6266 | 0.27281929 | 0.17687331 | 0 | 0.26488574 | 0.10586385 | 0.38739912 |
| ENSMUSG00 | Zfp1 | 1881 | 11.0537567 | 9.11981574 | 14.7104486 | 8.76634468 | 10.8780312 | 9.48304842 |
| ENSMUSG00 | Olfr357 | 1091 | 0 | 0 | 0 | 0 | 0 | 0 |
| ENSMUSG00 | Tceb2 | 491 | 27.6911901 | 30.5213513 | 31.1483456 | 21.2377003 | 20.1611342 | 51.4385254 |
| ENSMUSG00 | Rnf181 | 2795 | 76.8084418 | 70.236696 | 82.2518016 | 61.7590344 | 55.1888136 | 61.9266049 |
| ENSMUSG00 | Izumo4 | 928 | 7.49699255 | 9.08688295 | 5.2870739 | 6.95979092 | 9.34750419 | 11.1684945 |
| ENSMUSG00 | Fam19a3 | 4476 | 0 | 0 | 0 | 0 | 0 | 0.00609353 |
| ENSMUSG00 | Per2 | 5818 | 0.67648549 | 1.79725987 | 0.3114946 | 2.97065991 | 2.26277156 | 0.15470318 |
| ENSMUSG00 | Gm9985 | 508 | 0 | 0.0948552 | 0 | 0 | 0 | 0 |
| ENSMUSG00 | Foxi3 | 1873 | 0 | 0 | 0 | 0 | 0 | 0 |
| ENSMUSG00 | Abhd16b | 1774 | 0 | 0.13581297 | 0 | 0.85425359 | 0 | 0 |
| ENSMUSG00 | Fancm | 7752 | 1.46159855 | 1.93317641 | 1.08337851 | 2.18298231 | 2.59344583 | 2.12159539 |
| ENSMUSG00 | Ubl4b | 1375 | 0 | 0 | 0 | 0 | 0 | 0 |
| ENSMUSG00 | Plac1l | 956 | 0 | 0 | 0 | 0 | 0 | 0 |
| ENSMUSG00 | Ppp4r1l-ps | 10511 | 4.22101762 | 5.67088088 | 3.35582444 | 4.53814698 | 5.00020617 | 7.14107195 |
| ENSMUSG00 | Tmem69 | 2282 | 11.63745 | 9.73442149 | 16.7161221 | 9.91386587 | 6.99879738 | 10.8285816 |
| ENSMUSG00 | Tmem150a | 1538 | 2.19714944 | 3.69700922 | 4.39719245 | 4.10556423 | 2.65416779 | 4.59306344 |
| ENSMUSG00 | Zfp277 | 3172 | 15.0775041 | 14.0366559 | 15.370313 | 13.2862005 | 11.083598 | 14.7723298 |
| ENSMUSG00 | Aasdh | 6222 | 3.15641391 | 4.49182521 | 2.51485896 | 5.06262052 | 4.73195417 | 3.55070038 |
| ENSMUSG00 | Gm14137 | 3066 | 0.1037325 | 0.3614769 | 0 | 0 | 0.48263695 | 0.02668751 |
| ENSMUSG00 | Fto | 13098 | 6.92336624 | 5.83843974 | 6.72578051 | 6.59217686 | 7.25386408 | 7.19452425 |
| ENSMUSG00 | Gm97 | 860 | 0 | 0 | 0 | 0 | 0 | 0 |
| ENSMUSG00 | AU015836 | 3897 | 0 | 0 | 0 | 0 | 0 | 0 |
| ENSMUSG00 | Krt28 | 1616 | 0 | 0.23854674 | 0 | 0 | 0.03157573 | 0 |
| ENSMUSG00 | Gm4745 | 1259 | 0 | 0 | 0 | 0 | 0 | 0 |
| ENSMUSG00 | 2900064A13 | 5821 | 35.9035494 | 29.9582089 | 34.0113477 | 28.0052701 | 26.6396077 | 33.9984165 |
| ENSMUSG00 | E130018O15 | 3774 | 0 | 0 | 0 | 0 | 0 | 0 |
| ENSMUSG00 | Prr18 | 3549 | 0 | 0.29870435 | 0 | 0.2541705 | 0 | 0 |
| ENSMUSG00 | Ccl25 | 1107 | 0 | 0 | 0 | 0 | 0 | 0 |
| ENSMUSG00 | Gm9987 | 565 | 0 | 0 | 0 | 0 | 0.09031217 | 0 |
| ENSMUSG00 | Skint4 | 4356 | 0 | 0 | 0 | 0 | 0 | 0 |
| ENSMUSG00 | BC051076 | 2864 | 0 | 0 | 0 | 0 | 0 | 0 |
| ENSMUSG00 | Gm11818 | 1561 | 8.27708595 | 10.2484938 | 6.68266172 | 9.17653535 | 8.72776576 | 11.9337467 |
| ENSMUSG00 | Gm9988 | 432 | 0.09202658 | 0 | 0 | 0 | 0 | 0 |
| ENSMUSG00 | Olfr378 | 2022 | 0 | 0 | 0 | 0 | 0 | 0 |
| ENSMUSG00 | 2810407A14 | 1695 | 0.02345456 | 0 | 0.05215557 | 0 | 0.09031217 | 0.01609123 |
| ENSMUSG00 | Cldn23 | 1847 | 0 | 2.97414965 | 0 | 0 | 0 | 0 |
| ENSMUSG00 | Fut2 | 2949 | 0 | 0 | 0 | 0 | 0 | 0 |
| ENSMUSG00 | Irs1 | 9142 | 5.8054656 | 4.79123561 | 9.62170943 | 8.48176698 | 6.26248017 | 9.29939297 |
| ENSMUSG00 | Zkscan5 | 5146 | 5.35377925 | 4.2980134 | 6.29614282 | 5.37093784 | 6.55430129 | 3.15359704 |
| ENSMUSG00 | Nod2 | 5025 | 0 | 0 | 0 | 0 | 0.04061801 | 0 |
| ENSMUSG00 | 9330182L06 | 8225 | 0 | 0.18747309 | 0 | 0 | 0 | 0.05968918 |
| ENSMUSG00 | Gm11541 | 1679 | 0 | 0 | 1.76386151 | 4.25505945 | 0 | 4.6946818 |
| ENSMUSG00 | A430033K04 | 3292 | 16.0012186 | 10.9195279 | 12.2051869 | 13.7444969 | 16.4611212 | 19.453479 |
| ENSMUSG00 | 1700008F21 | 2383 | 0 | 0.08088366 | 0.01854882 | 0 | 0 | 0.2174646 |
| ENSMUSG00 | Zfp709 | 2178 | 14.8581089 | 6.81424434 | 11.9129852 | 4.2244818 | 4.31076821 | 26.3604755 |
| ENSMUSG00 | Gm9989 | 3195 | 0 | 0 | 0 | 0 | 0 | 0 |
| ENSMUSG00 | Clca1 | 4924 | 0 | 0.01957207 | 0 | 0 | 0 | 0 |
| ENSMUSG00 | Ovol3 | 813 | 0 | 0 | 0 | 0 | 0 | 0 |
| ENSMUSG00 | 9330154J02R | 1482 | 0 | 0 | 0 | 0.09738744 | 0 | 0 |
| ENSMUSG00 | BC018473 | 5883 | 0 | 0 | 0 | 0 | 0 | 0 |
| ENSMUSG00 | Cyp3a11 | 2331 | 0 | 0 | 0 | 0 | 0 | 0 |
| ENSMUSG00 | Rgs9bp | 6584 | 0 | 0.21224283 | 0.26182744 | 0.87684191 | 0.02325017 | 0.25269638 |
| ENSMUSG00 | S100a8 | 389 | 0 | 0 | 0 | 0 | 0 | 0 |
| ENSMUSG00 | Sag | 6304 | 0.27117476 | 0.03057515 | 0.31552712 | 0.1488154 | 0 | 0.21632804 |
| ENSMUSG00 | Gm14318 | 1673 | 0 | 0 | 0 | 0 | 0.21349948 | 0 |
| ENSMUSG00 | Fam105a | 3737 | 2.15958323 | 2.87545534 | 1.14733176 | 0.57932103 | 0.58713785 | 0.86852611 |
| ENSMUSG00 | S100a9 | 519 | 0 | 0 | 0 | 0 | 0 | 0 |
| ENSMUSG00 | Grik2 | 9682 | 0 | 0 | 0 | 0 | 0 | 0 |
| ENSMUSG00 | Eif3b | 3430 | 44.6003175 | 42.9463422 | 35.6836453 | 29.8965512 | 46.5931805 | 27.9982521 |
| ENSMUSG00 | Lipm | 2691 | 0 | 0 | 0 | 0 | 0 | 0 |
| ENSMUSG00 | Gm5468 | 652 | 0 | 0 | 0 | 0 | 0 | 0 |
| ENSMUSG00 | St3gal5 | 2610 | 0.31987168 | 0.07384895 | 0 | 0 | 0 | 0 |
| ENSMUSG00 | Gm9990 | 2085 | 0.20974114 | 0.27733204 | 0.40279857 | 0.41533289 | 0.02447308 | 0.49709175 |
| ENSMUSG00 | Tas2r134 | 896 | 0 | 0 | 0 | 0 | 0 | 0 |
| ENSMUSG00 | H2-T22 | 3891 | 3.82126695 | 3.57899814 | 2.43104457 | 3.14361683 | 3.96041262 | 2.86695643 |
| ENSMUSG00 | Fez2 | 4362 | 11.1920518 | 15.6202726 | 13.3152732 | 13.0447924 | 11.0896388 | 13.856167 |
| ENSMUSG00 | B4galt6 | 5799 | 4.31215679 | 4.48709756 | 5.5719171 | 5.56257075 | 4.61956325 | 6.55174556 |
| ENSMUSG00 | Gm9991 | 1205 | 0 | 0 | 0 | 0 | 0 | 0 |
| ENSMUSG00 | Ticam2 | 3925 | 0.11141663 | 0.15959841 | 0 | 0 | 0.03900105 | 0 |
| ENSMUSG00 | Pgm3 | 4569 | 5.59482906 | 3.31156551 | 2.65075618 | 5.3384684 | 4.26615793 | 4.72187342 |
| ENSMUSG00 | Gm9992 | 1583 | 0 | 0 | 0 | 0 | 0 | 0 |
| ENSMUSG00 | Trim34a | 4361 | 1.45858218 | 1.01654498 | 1.45954263 | 0.42196383 | 1.41577424 | 1.0069289 |
| ENSMUSG00 | AI504432 | 2372 | 0 | 0 | 0 | 0 | 0 | 0 |
| ENSMUSG00 | Rdh9 | 4048 | 0 | 1.70223845 | 0.85171535 | 0.65068904 | 0.6302665 | 0.88909041 |
| ENSMUSG00 | Socs6 | 8175 | 4.45455904 | 6.48969698 | 5.08794303 | 5.91436571 | 5.33670353 | 5.07458426 |
| ENSMUSG00 | Nanos3 | 728 | 0.16382753 | 0.59571151 | 0.06071682 | 1.3382077 | 0.91118528 | 1.01155942 |
| ENSMUSG00 | Car10 | 5100 | 0 | 0 | 0 | 0 | 0 | 0 |
| ENSMUSG00 | Cndp1 | 5544 | 0.25815247 | 0 | 0.05581041 | 0 | 0.00920389 | 0 |
| ENSMUSG00 | Cnot10 | 2833 | 24.8384047 | 26.9592343 | 25.6504871 | 23.3202342 | 30.8715878 | 21.4885262 |
| ENSMUSG00 | Col8a2 | 2588 | 0 | 0 | 0.03415908 | 0 | 0.72951156 | 0.12646664 |
| ENSMUSG00 | Olfr283 | 1101 | 0 | 0 | 0 | 0 | 0 | 0 |
| ENSMUSG00 | Snx32 | 1637 | 25.0141386 | 24.8438346 | 20.1162939 | 18.0740846 | 24.0014107 | 22.5428142 |
| ENSMUSG00 | 4931417E11 | 1063 | 0 | 0 | 0 | 0 | 0 | 0 |
| ENSMUSG00 | Cfl1 | 1142 | 36.8660717 | 43.7981847 | 54.8458945 | 41.5480635 | 28.9983519 | 81.2746037 |
| ENSMUSG00 | Tas2r135 | 1092 | 0 | 0 | 0 | 0 | 0 | 0 |
| ENSMUSG00 | Pgpep1 | 2145 | 30.9888874 | 34.7301816 | 30.6424898 | 22.5912288 | 19.387644 | 19.4546378 |
| ENSMUSG00 | Npm3 | 1182 | 0.33634078 | 0.36690184 | 0.07479161 | 0.06105253 | 0 | 0.09229996 |
| ENSMUSG00 | R3hdm1 | 6890 | 10.9341997 | 8.9449143 | 7.11463627 | 10.8560346 | 8.58339181 | 9.01765286 |
| ENSMUSG00 | Pard6g | 3157 | 0.02518561 | 0.01526337 | 0.15401339 | 0.65146549 | 0.30709571 | 0.01727883 |
| ENSMUSG00 | Lrguk | 3727 | 0.02133377 | 0.37494146 | 0.28463757 | 0.09681257 | 0.64347724 | 0.57813161 |
| ENSMUSG00 | Cebpg | 5525 | 11.9374375 | 13.8497865 | 10.888454 | 15.8826303 | 13.483893 | 13.7730758 |
| ENSMUSG00 | Gm5423 | 503 | 0 | 0 | 0 | 0 | 0 | 0 |
| ENSMUSG00 | Pla2g4a | 6924 | 6.24122001 | 11.6499284 | 4.68575286 | 9.08825616 | 8.07696532 | 6.96440817 |
| ENSMUSG00 | Spock1 | 4633 | 0 | 0 | 0.05724392 | 0 | 0 | 0.04709629 |
| ENSMUSG00 | 4930458L03 | 3230 | 0 | 0 | 0 | 0 | 0 | 0 |
| ENSMUSG00 | Cars2 | 2606 | 16.1706866 | 16.6784999 | 12.0257508 | 10.7719995 | 19.5020224 | 11.9418125 |
| ENSMUSG00 | Ncoa4 | 4510 | 5.13912309 | 5.57723344 | 5.8119053 | 4.90427789 | 6.38112549 | 3.16289054 |
| ENSMUSG00 | Gm5447 | 1772 | 0 | 0 | 0 | 0 | 0 | 0 |
| ENSMUSG00 | Kcnq3 | 2607 | 0.10674659 | 0 | 0.06782024 | 0.76122457 | 0.2935925 | 0.31386236 |
| ENSMUSG00 | 4933421E11 | 3347 | 19.6579983 | 20.3284303 | 22.9791476 | 20.6983942 | 18.9805312 | 40.5737762 |
| ENSMUSG00 | Cep70 | 2559 | 10.6263184 | 14.5369024 | 16.3921645 | 15.1293606 | 15.5531743 | 14.1542481 |
| ENSMUSG00 | Dennd1b | 13442 | 4.31507559 | 5.8001535 | 4.13015287 | 6.57647739 | 6.5026173 | 5.92282934 |
| ENSMUSG00 | Prr9 | 1181 | 0 | 0 | 0 | 0 | 0 | 0 |
| ENSMUSG00 | Lman1l | 1672 | 0 | 0 | 0.02643651 | 0 | 0 | 0 |
| ENSMUSG00 | Olfr1322 | 932 | 0 | 0 | 0 | 0 | 0 | 0 |
| ENSMUSG00 | Gm11961 | 1556 | 0.0255498 | 0.15484075 | 0 | 0.04637795 | 0 | 0 |
| ENSMUSG00 | Ms4a4b | 5129 | 0 | 0 | 0 | 0 | 0 | 0 |
| ENSMUSG00 | Gsdmc2 | 1970 | 0.58523296 | 0 | 0 | 0 | 0 | 0.04153498 |

| | | | | | | | | |
|---|---|---|---|---|---|---|---|---|
| ENSMUSG00 | Synpr | 8364 | 0 | 0 | 0 | 0 | 0 | 0.00326096 |
| ENSMUSG00 | Gm13247 | 3398 | 0.18719473 | 0.04254247 | 0.19512291 | 0.19113502 | 0.21023227 | 0 |
| ENSMUSG00 | Usp39 | 2412 | 9.59274032 | 14.1043235 | 10.6289672 | 9.49921156 | 9.39291495 | 8.14168336 |
| ENSMUSG00 | 6030405A18I | 5977 | 0 | 0 | 0 | 0 | 0 | 0 |
| ENSMUSG00 | Gm9994 | 2842 | 0 | 0 | 0 | 0 | 0 | 0 |
| ENSMUSG00 | Tyw1 | 7370 | 3.92699997 | 5.11286265 | 3.70047994 | 4.41601142 | 4.23719702 | 2.29817517 |
| ENSMUSG00 | 1810011O10I | 1330 | 6.30707247 | 3.55057995 | 4.28724643 | 4.44921451 | 5.02590619 | 5.59847858 |
| ENSMUSG00 | Gm9996 | 912 | 0 | 0 | 0 | 0 | 0 | 0 |
| ENSMUSG00 | Myh1 | 6334 | 0.01882956 | 0.01521517 | 0.04884953 | 0.03417939 | 0.04833569 | 0.00861214 |
| ENSMUSG00 | 1810010K12I | 2529 | 0 | 0 | 0 | 0 | 0 | 0 |
| ENSMUSG00 | Gm16294 | 1456 | 0 | 0 | 0 | 0 | 0 | 0 |
| ENSMUSG00 | Usp34 | 17049 | 4.0760502 | 3.62620713 | 4.09636413 | 6.13324917 | 4.13322922 | 4.88412656 |
| ENSMUSG00 | Krtap13-1 | 747 | 0 | 0 | 0 | 0 | 0 | 0 |
| ENSMUSG00 | Hsd3b4 | 1568 | 0 | 0 | 0 | 0 | 0 | 0 |
| ENSMUSG00 | D830013O20 | 679 | 0 | 0 | 0 | 0 | 0 | 0 |
| ENSMUSG00 | Six3os1 | 7534 | 0 | 0 | 0 | 0 | 0 | 0 |
| ENSMUSG00 | Fabp3-ps1 | 401 | 0.1982817 | 0 | 0.33068711 | 0 | 0 | 1.02024835 |
| ENSMUSG00 | Actr3b | 1831 | 0.23883686 | 1.13163136 | 0.43453478 | 0.55177348 | 0.41802056 | 0 |
| ENSMUSG00 | Sftpb | 1120 | 0 | 0 | 0 | 0 | 0 | 0 |
| ENSMUSG00 | Taar1 | 1113 | 0 | 0 | 0 | 0 | 0 | 0 |
| ENSMUSG00 | Gpr50 | 1238 | 0 | 0 | 0 | 0 | 0 | 0 |
| ENSMUSG00 | AI987944 | 1818 | 137.001147 | 115.297593 | 90.421702 | 165.04856 | 96.9725679 | 199.548942 |
| ENSMUSG00 | Lig1 | 4837 | 2.687625 | 1.8828277 | 0.97779558 | 0.89515099 | 1.3186473 | 0.63154013 |
| ENSMUSG00 | Prss34 | 1211 | 0 | 0 | 0 | 0 | 0 | 0 |
| ENSMUSG00 | Gm12500 | 2033 | 0 | 0.49774485 | 0.21742176 | 0 | 0 | 0.45614252 |
| ENSMUSG00 | Gm12396 | 305 | 0 | 0 | 0 | 0 | 0 | 0 |
| ENSMUSG00 | Adap1 | 3444 | 1.91620493 | 1.39914176 | 0.14117894 | 1.14196947 | 1.91126669 | 0.17422824 |
| ENSMUSG00 | BC043934 | 1650 | 0 | 0 | 0 | 0 | 0.40202599 | 0 |
| ENSMUSG00 | Uts2d | 830 | 0 | 0 | 0 | 0 | 0 | 0 |
| ENSMUSG00 | Nat3 | 1392 | 0 | 0 | 0 | 0 | 0 | 0 |
| ENSMUSG00 | Slit3 | 6030 | 0 | 0 | 0 | 0 | 0.22001422 | 0 |
| ENSMUSG00 | Tgoln1 | 5009 | 123.830107 | 119.499298 | 132.420215 | 131.333167 | 124.800186 | 141.399048 |
| ENSMUSG00 | Cyct | 651 | 0.18320498 | 0 | 0 | 0 | 0 | 0 |
| ENSMUSG00 | 5730446D14 | 1677 | 0 | 0.11493487 | 0 | 0 | 0.06085435 | 0.14637552 |
| ENSMUSG00 | Prl2c3 | 845 | 0 | 0 | 0 | 0 | 0 | 0 |
| ENSMUSG00 | Stk30 | 2130 | 8.24972882 | 8.89073792 | 3.50709459 | 13.2978427 | 14.4454951 | 7.23482215 |
| ENSMUSG00 | Zbtb25 | 4499 | 2.17378266 | 2.39914715 | 1.61126967 | 2.96740531 | 2.76737846 | 2.15820662 |
| ENSMUSG00 | 5730596B20I | 3353 | 0.01185669 | 0.04311343 | 0 | 0 | 0.27392626 | 0.0244032 |
| ENSMUSG00 | Med12l | 10362 | 0.23019966 | 0.16276061 | 0.16209902 | 0.43526883 | 0.62539565 | 0.12371242 |
| ENSMUSG00 | Cd248 | 2559 | 0 | 0.09415092 | 0 | 0 | 0.1794597 | 0 |
| ENSMUSG00 | Chn1 | 7551 | 1.04772091 | 1.58260332 | 0.99514148 | 1.50521044 | 2.31108734 | 0.85966947 |
| ENSMUSG00 | Mettl7a2 | 2309 | 0.03443524 | 0.27129656 | 0 | 0 | 0.08839563 | 0.01181232 |
| ENSMUSG00 | Gpr116 | 8633 | 65.6773616 | 24.1015936 | 66.7866142 | 60.0977318 | 110.617239 | 73.4484899 |
| ENSMUSG00 | Foxk1 | 7954 | 1.12458928 | 1.15104652 | 1.01140752 | 0.66684191 | 0.93020172 | 1.60136491 |
| ENSMUSG00 | Cngb3 | 4690 | 0 | 0 | 0 | 0 | 0 | 0 |
| ENSMUSG00 | Tmem154 | 4826 | 0 | 0 | 0 | 0 | 0 | 0 |
| ENSMUSG00 | Cebpb | 1503 | 5.81916548 | 8.88066831 | 4.6466342 | 6.84190465 | 6.48438974 | 4.02858943 |
| ENSMUSG00 | 1700001K19I | 948 | 0.62904241 | 0.05082958 | 0 | 0.30448983 | 0.32295175 | 0.28770717 |
| ENSMUSG00 | Gm9999 | 441 | 0 | 0 | 0 | 0 | 0 | 0 |
| ENSMUSG00 | Rab31 | 3805 | 6.54058628 | 5.62280692 | 8.60803303 | 3.01552961 | 5.86032227 | 5.54811321 |
| ENSMUSG00 | Ptafr | 3635 | 0 | 0 | 0 | 0.13896798 | 0 | 0 |
| ENSMUSG00 | Ccdc18 | 4628 | 0.30924747 | 0.63512813 | 0.27697784 | 0.21830105 | 0.27563946 | 1.64425764 |
| ENSMUSG00 | Pigh | 4412 | 7.50596448 | 6.86973529 | 8.78626843 | 11.1059421 | 8.5468023 | 7.67176503 |
| ENSMUSG00 | Rlim | 8912 | 16.5365312 | 12.9063103 | 14.0610665 | 20.8305789 | 19.3181095 | 13.6740449 |
| ENSMUSG00 | Defb21 | 424 | 0 | 0 | 0 | 0 | 0 | 0 |
| ENSMUSG00 | Zfp953 | 4964 | 2.02621607 | 1.14544726 | 1.05963322 | 2.22423562 | 2.19976721 | 2.75823306 |
| ENSMUSG00 | Ptprn2 | 5423 | 0 | 0.07997012 | 0.17931782 | 0 | 0 | 0 |
| ENSMUSG00 | Olfr324 | 1121 | 0 | 0 | 0 | 0 | 0 | 0 |
| ENSMUSG00 | Mpz | 1203 | 0 | 0.04005523 | 0 | 0.14996694 | 0.04241594 | 0.20404967 |
| ENSMUSG00 | Tug1 | 6668 | 37.7879777 | 36.2915886 | 36.8702236 | 56.4444821 | 45.1416603 | 54.5860919 |
| ENSMUSG00 | 9930038K12I | 445 | 0 | 0 | 0 | 0 | 0 | 0 |
| ENSMUSG00 | Zar1l | 891 | 0 | 0 | 0 | 0 | 0 | 0 |
| ENSMUSG00 | H2afy3 | 2069 | 0 | 0 | 0 | 0.01743936 | 0 | 0 |
| ENSMUSG00 | Zfp658 | 2957 | 4.82658691 | 2.75397657 | 1.77883643 | 0.08541573 | 2.9507982 | 1.09762667 |
| ENSMUSG00 | Trnp1 | 2245 | 0.21250146 | 0.53659735 | 0.31502427 | 0.38573233 | 0.13637339 | 0.26727932 |
| ENSMUSG00 | Lrrc48 | 4609 | 0.7504289 | 0.52274292 | 0.26852931 | 0.71240314 | 1.17352915 | 0.88173601 |
| ENSMUSG00 | Olfr90 | 1045 | 0 | 0 | 0 | 0 | 0 | 0 |
| ENSMUSG00 | Fry | 14148 | 0.98911006 | 1.0217651 | 0.60297962 | 1.48939172 | 1.34887366 | 0.43375699 |
| ENSMUSG00 | Krt72-ps | 2399 | 0 | 0 | 0 | 0 | 0 | 0 |
| ENSMUSG00 | Gm10000 | 3001 | 0 | 0 | 0 | 0 | 0 | 0 |
| ENSMUSG00 | Chd9 | 13012 | 5.20316503 | 5.76593071 | 5.30272648 | 7.170932 | 6.18418342 | 5.21513237 |
| ENSMUSG00 | Ppp1r14b | 915 | 6.03935715 | 8.7420212 | 9.85483713 | 6.03339105 | 2.95562614 | 8.76365458 |
| ENSMUSG00 | 4931429L15I | 2111 | 0 | 0.02282636 | 0 | 0 | 0 | 0 |
| ENSMUSG00 | Fkbp2 | 636 | 65.8840825 | 76.1436704 | 94.4501646 | 71.0861663 | 47.0950984 | 151.725902 |
| ENSMUSG00 | Dsg3 | 4255 | 0 | 0 | 0 | 0 | 0.07195259 | 0 |
| ENSMUSG00 | Chst13 | 1648 | 0 | 0.43859019 | 0.05364301 | 0.45978333 | 0.24770085 | 0.34755305 |
| ENSMUSG00 | Hoxb8 | 2292 | 0.08672662 | 0.04204751 | 0.07714109 | 0.18891123 | 0.33394225 | 0.29749825 |
| ENSMUSG00 | Apol8 | 2679 | 0 | 0 | 0 | 0 | 0 | 0 |
| ENSMUSG00 | 4930572J05I | 1486 | 0.08026006 | 0.25941557 | 0 | 0.07284397 | 0.27470458 | 0.18354401 |
| ENSMUSG00 | Retsat | 3693 | 75.1618643 | 63.6875235 | 83.8674016 | 55.9745554 | 64.4565511 | 64.6600235 |
| ENSMUSG00 | Olfr1471 | 944 | 0 | 0 | 0 | 0 | 0 | 0 |
| ENSMUSG00 | Prelid2 | 717 | 12.2537813 | 13.4411275 | 16.8916388 | 11.4737882 | 10.4614745 | 19.3242911 |
| ENSMUSG00 | Kdm5d | 5462 | 7.35134297 | 6.3166409 | 4.53995495 | 10.9725881 | 10.4818004 | 3.54540349 |
| ENSMUSG00 | Gpr173 | 6226 | 0.44059238 | 0.71977821 | 0 | 0.08693072 | 0.10654399 | 0.24532281 |
| ENSMUSG00 | 4930465K10I | 1235 | 0 | 0 | 0 | 0 | 0.0413169 | 0.13250837 |
| ENSMUSG00 | Gm11696 | 2763 | 0.73381452 | 0.1046394 | 0.51192869 | 0.05223604 | 0.18467744 | 0.38498405 |
| ENSMUSG00 | D17Wsu92e | 6994 | 17.461944 | 14.4407753 | 11.6792974 | 10.6017446 | 16.9625856 | 7.17938388 |
| ENSMUSG00 | Olfr1350 | 926 | 0 | 0 | 0 | 0 | 0 | 0 |
| ENSMUSG00 | Elmod3 | 4506 | 6.67885014 | 6.95099587 | 7.46506936 | 8.92041669 | 7.73435745 | 5.43555838 |
| ENSMUSG00 | Gm5533 | 1307 | 0 | 0 | 0 | 0 | 0 | 0 |
| ENSMUSG00 | Krtap7-1 | 619 | 0 | 0 | 0 | 0 | 0 | 0 |
| ENSMUSG00 | Ier5 | 3269 | 1.71475153 | 1.78359116 | 1.70371129 | 2.20752796 | 2.74721387 | 1.49347214 |
| ENSMUSG00 | Gm5420 | 2203 | 0 | 0 | 0 | 0 | 0 | 0 |
| ENSMUSG00 | Gm13199 | 1143 | 0.41738037 | 0.3794208 | 0.5414049 | 1.04173882 | 0.62499498 | 0.66814514 |
| ENSMUSG00 | Nbeal2 | 9853 | 8.05358156 | 6.13273099 | 6.18188772 | 8.52888275 | 7.94940494 | 5.90447635 |
| ENSMUSG00 | Ctsll3 | 1324 | 0 | 0 | 0 | 0 | 0 | 0 |
| ENSMUSG00 | Gm10006 | 967 | 0 | 0.09966172 | 0.7770748 | 0 | 0.47490939 | 0 |
| ENSMUSG00 | A930024E05I | 1893 | 0.08400524 | 0 | 0.84060557 | 0.40027625 | 0 | 0.51869362 |
| ENSMUSG00 | Capg | 2514 | 10.5002542 | 18.5155541 | 6.45269546 | 7.24798428 | 11.3256634 | 6.34672393 |
| ENSMUSG00 | A730036I17R | 3978 | 0 | 0 | 0 | 0 | 0 | 0 |
| ENSMUSG00 | Nfil3 | 2024 | 0.92317568 | 5.30907935 | 1.11378162 | 3.6902091 | 4.2858122 | 0.30993908 |
| ENSMUSG00 | Dnahc9 | 16379 | 0.00970889 | 0 | 0.00539738 | 0 | 0.01557677 | 0.00166522 |
| ENSMUSG00 | C330011M18 | 1847 | 0.83944978 | 0.10435613 | 1.2683799 | 0.52745815 | 0.69066562 | 0.31010689 |
| ENSMUSG00 | Grm7 | 5485 | 0 | 0 | 0 | 0 | 0 | 0 |
| ENSMUSG00 | Hmga2 | 6766 | 0.14689433 | 0.14243702 | 0.01306587 | 0 | 0 | 0.11690283 |
| ENSMUSG00 | Cspp1 | 8307 | 7.43231747 | 7.58151919 | 10.7325289 | 9.73828624 | 8.72859998 | 12.6211284 |
| ENSMUSG00 | Setd3 | 7033 | 12.0798325 | 11.2227204 | 14.5055956 | 9.57331083 | 11.1513635 | 10.6880287 |
| ENSMUSG00 | Gm10010 | 2316 | 0.05149674 | 0 | 0 | 0 | 0 | 0.12954276 |
| ENSMUSG00 | Rps6-ps2 | 749 | 0 | 0 | 0 | 0 | 0 | 0.03641474 |
| ENSMUSG00 | Rpl21-ps13 | 482 | 0 | 0 | 0 | 0 | 0 | 0 |
| ENSMUSG00 | Olfr667 | 1908 | 0 | 0 | 0 | 0 | 0 | 0 |
| ENSMUSG00 | Gm4889 | 443 | 0 | 0 | 0 | 0 | 0 | 0 |
| ENSMUSG00 | St8sia3 | 6657 | 0 | 0 | 0 | 0 | 0 | 0 |
| ENSMUSG00 | Gm6812 | 523 | 0 | 0 | 0 | 0 | 0 | 0 |
| ENSMUSG00 | Tsnax | 2385 | 4.41727561 | 3.53569282 | 4.46651747 | 4.78068179 | 4.12917842 | 3.05338729 |
| ENSMUSG00 | 1700028I16R | 1043 | 0 | 0 | 0 | 0 | 0.3913816 | 0 |
| ENSMUSG00 | Olfr166 | 1037 | 0 | 0 | 0 | 0 | 0 | 0 |
| ENSMUSG00 | Zfp663 | 5493 | 0 | 0.00877234 | 0 | 0 | 0 | 0 |
| ENSMUSG00 | Foxb2 | 1902 | 0 | 0 | 0 | 0 | 0 | 0 |
| ENSMUSG00 | Ttc26 | 4715 | 3.08600337 | 6.53046374 | 4.34049911 | 8.1270692 | 6.80712416 | 3.1931285 |
| ENSMUSG00 | Gm6851 | 832 | 0.04778303 | 0.0579164 | 0 | 0.08673568 | 0.06132978 | 0.09834606 |
| ENSMUSG00 | Gm16415 | 488 | 0 | 0 | 0 | 0 | 0 | 0 |
| ENSMUSG00 | Pcbp2 | 2853 | 8.93209359 | 10.9445547 | 11.8832154 | 8.94146705 | 7.11829568 | 8.68046703 |
| ENSMUSG00 | Olfr765 | 929 | 0 | 0 | 0 | 0 | 0 | 0 |
| ENSMUSG00 | Pou3f4 | 1181 | 0 | 0 | 0 | 0 | 0 | 0 |
| ENSMUSG00 | Jakmip3 | 5147 | 0 | 0 | 0.02576365 | 0 | 0 | 0 |
| ENSMUSG00 | Olfr1502 | 950 | 0 | 0 | 0 | 0 | 0 | 0 |
| ENSMUSG00 | Olfr702 | 1346 | 0 | 0 | 0 | 0 | 0 | 0 |
| ENSMUSG00 | Olfr1360 | 990 | 0 | 0 | 0 | 0 | 0 | 0 |
| ENSMUSG00 | Gulp1 | 3917 | 22.6536208 | 18.8587735 | 30.9424186 | 32.2499969 | 14.1472158 | 35.7070079 |
| ENSMUSG00 | Gm10011 | 343 | 0.34771557 | 0.28097051 | 0.12886835 | 0 | 0.59505978 | 0.79517899 |
| ENSMUSG00 | Gm3386 | 387 | 0.41090936 | 0.12451277 | 0 | 0.09323526 | 0.3955533 | 0.35238552 |
| ENSMUSG00 | Rps15a-ps1 | 392 | 0 | 0 | 0 | 0 | 0 | 0 |
| ENSMUSG00 | Gadl1 | 4012 | 0 | 0.02402116 | 0 | 0 | 0 | 0 |
| ENSMUSG00 | Vmn1r215 | 1047 | 0 | 0 | 0 | 0 | 0 | 0 |
| ENSMUSG00 | Olfr533 | 1051 | 0 | 0 | 0 | 0 | 0 | 0 |
| ENSMUSG00 | Gm4559 | 599 | 0 | 0 | 0 | 0 | 0 | 0 |
| ENSMUSG00 | Glipr1 | 1762 | 0 | 0 | 0.02508618 | 0.04095578 | 0 | 0 |
| ENSMUSG00 | Hist3h2ba | 619 | 1.09183064 | 0.93414751 | 0.49985929 | 0.40803605 | 0.41216782 | 0.96937329 |
| ENSMUSG00 | Immp2l | 1405 | 2.49002298 | 2.09208041 | 2.67413285 | 0.84747863 | 2.65119248 | 1.99949313 |
| ENSMUSG00 | Usp13 | 3083 | 0 | 0.3751134 | 0.20072196 | 0 | 0 | 0 |
| ENSMUSG00 | Tas2r102 | 1043 | 0 | 0 | 0 | 0 | 0 | 0 |
| ENSMUSG00 | Gm5620 | 1352 | 0.02940494 | 0.03564086 | 0.03269367 | 0 | 0 | 0.02017355 |
| ENSMUSG00 | Vmn2r107 | 2580 | 0 | 0 | 0 | 0 | 0 | 0 |
| ENSMUSG00 | 1700017N19I | 1677 | 0 | 0 | 0 | 0 | 0 | 0 |
| ENSMUSG00 | Sipa1 | 3850 | 3.17011234 | 2.09016515 | 4.23648833 | 3.04588688 | 4.37368936 | 3.65551009 |
| ENSMUSG00 | 4922501C03I | 9124 | 3.49015124 | 2.55614182 | 2.93096394 | 3.39702721 | 4.30625925 | 1.71587494 |
| ENSMUSG00 | Olfr394 | 932 | 0 | 0 | 0 | 0 | 0 | 0 |
| ENSMUSG00 | Tas2r113 | 929 | 0 | 0 | 0 | 0 | 0 | 0 |
| ENSMUSG00 | Acbd4 | 2927 | 19.3276559 | 12.9067887 | 11.5827857 | 12.4012767 | 17.9908507 | 12.3001448 |
| ENSMUSG00 | Gm3629 | 2719 | 0 | 0 | 0 | 0 | 0 | 0 |
| ENSMUSG00 | Commd7 | 5707 | 9.71769674 | 10.0814109 | 11.4396569 | 10.6090189 | 9.54006361 | 8.69328348 |
| ENSMUSG00 | Gm10015 | 464 | 0.34271966 | 0 | 0 | 0 | 0 | 0 |
| ENSMUSG00 | Olfr512 | 1055 | 0 | 0 | 0 | 0 | 0 | 0 |
| ENSMUSG00 | Mab21l1 | 2775 | 0.35815748 | 0.1041869 | 0.07964296 | 0.27305331 | 0.09193942 | 0.04914349 |
| ENSMUSG00 | Tatdn2 | 3540 | 0.39306266 | 0.72143543 | 0.6742654 | 0.30578004 | 0.79278268 | 0.53162432 |
| ENSMUSG00 | Olfr315 | 997 | 0 | 0 | 0 | 0 | 0 | 0 |
| ENSMUSG00 | Olfr919 | 947 | 0 | 0 | 0 | 0 | 0 | 0 |
| ENSMUSG00 | Jmjd6 | 2564 | 31.2430941 | 26.7806865 | 29.5483459 | 33.8726526 | 28.1998341 | 32.9018952 |
| ENSMUSG00 | Gjc3 | 3512 | 0 | 0 | 0 | 0 | 0 | 0 |
| ENSMUSG00 | Magel2 | 4661 | 0 | 0 | 0 | 0 | 0 | 0 |
| ENSMUSG00 | Ces1d | 2490 | 28.579241 | 59.855689 | 12.071186 | 44.1679003 | 29.6936621 | 19.8809117 |
| ENSMUSG00 | Hamp2 | 434 | 0 | 0 | 0 | 0 | 0 | 0 |
| ENSMUSG00 | Fam71d | 1814 | 0 | 0 | 0 | 0 | 0 | 0 |
| ENSMUSG00 | Olfr1178 | 971 | 0 | 0 | 0 | 0 | 0 | 0 |
| ENSMUSG00 | Ide | 6203 | 16.3367274 | 19.8867148 | 24.3705148 | 24.4832058 | 21.6345911 | 22.60062 |
| ENSMUSG00 | Nxf3 | 3028 | 0 | 0 | 0 | 0 | 0.43813929 | 0 |
| ENSMUSG00 | Olfr799 | 935 | 0 | 0 | 0 | 0 | 0 | 0 |
| ENSMUSG00 | Myh4 | 5998 | 0 | 0 | 0 | 0 | 0 | 0 |
| ENSMUSG00 | Olfr885 | 929 | 0 | 0 | 0 | 0 | 0 | 0 |
| ENSMUSG00 | Gm7536 | 505 | 0 | 0.0954187 | 0 | 0.28579837 | 0.40416931 | 0 |
| ENSMUSG00 | Cfhr1 | 1902 | 0 | 0 | 0 | 0 | 0 | 0 |
| ENSMUSG00 | 1700010B08I | 896 | 0 | 0 | 0 | 0 | 0 | 0 |
| ENSMUSG00 | Olfr397 | 1090 | 0 | 0 | 0 | 0 | 0 | 0 |
| ENSMUSG00 | Inca1 | 1744 | 1.8236459 | 0.96704442 | 2.50916425 | 0.51723114 | 1.87252755 | 2.04872577 |
| ENSMUSG00 | Skap1 | 7975 | 0.22931061 | 0.45920622 | 0.26604244 | 0.321232 | 0.18555046 | 0.01710009 |
| ENSMUSG00 | Slc35f3 | 2302 | 0.13815979 | 0.43958092 | 1.17129124 | 0.03134843 | 0.02216611 | 0.22511648 |
| ENSMUSG00 | Olfr297 | 932 | 0 | 0 | 0 | 0 | 0 | 0 |
| ENSMUSG00 | Fam47e | 2145 | 0 | 0 | 0 | 0 | 0 | 0 |
| ENSMUSG00 | Ero1lb | 4221 | 4.56797158 | 3.85856845 | 2.69127545 | 4.86393825 | 7.08397447 | 6.358267 |
| ENSMUSG00 | 1700022P22I | 426 | 0 | 0.56556857 | 1.66016312 | 1.10109526 | 0.23956045 | 1.21647452 |
| ENSMUSG00 | Ces1g | 2374 | 0 | 0 | 0 | 0.01519884 | 0 | 0 |
| ENSMUSG00 | Fxyd3 | 4451 | 0.03572724 | 0.07578187 | 0.09930767 | 0.04863902 | 0.11464025 | 0.20834368 |
| ENSMUSG00 | C030039L03I | 5502 | 2.28330277 | 1.83041919 | 1.81563368 | 1.28536545 | 1.87337839 | 1.59622571 |
| ENSMUSG00 | Ebf1 | 9935 | 0.11604519 | 0.08730307 | 0.11122759 | 0.00726362 | 0.28761722 | 0.10981234 |
| ENSMUSG00 | Zfp180 | 3881 | 14.2488723 | 10.727412 | 10.8198276 | 13.3692296 | 11.7935221 | 7.49859317 |
| ENSMUSG00 | Cml1 | 1853 | 2.6818322 | 4.62881096 | 4.26990281 | 4.06969633 | 2.61603114 | 4.87204835 |
| ENSMUSG00 | Olfr452 | 953 | 0 | 0 | 0 | 0 | 0 | 0 |
| ENSMUSG00 | Cep110 | 10019 | 4.36084171 | 3.02517937 | 2.63384572 | 3.10077257 | 4.6040367 | 4.30122068 |
| ENSMUSG00 | Npm1 | 2994 | 23.3433984 | 23.1597496 | 17.2585018 | 20.0295117 | 19.4800092 | 30.0622276 |
| ENSMUSG00 | AF366264 | 2856 | 0 | 0 | 0.03095367 | 0 | 0 | 0 |
| ENSMUSG00 | Gja5 | 5740 | 0.76879065 | 0.31900432 | 0.26952343 | 0.39602244 | 0.73783784 | 0.37538267 |
| ENSMUSG00 | Txnl4a | 2103 | 17.6754038 | 18.170161 | 17.5924596 | 12.4562836 | 17.0330556 | 17.9496437 |
| ENSMUSG00 | Rpgrip1 | 5233 | 0.26589754 | 0.26703742 | 0.6588465 | 0.86878227 | 0.22427033 | 0.5159926 |
| ENSMUSG00 | Chd6 | 17413 | 4.05248825 | 5.40170764 | 3.62235284 | 4.80734757 | 4.69736866 | 4.80552422 |
| ENSMUSG00 | Ado | 4443 | 5.46716151 | 8.41608803 | 6.26764823 | 8.16170487 | 5.76530287 | 14.2419903 |
| ENSMUSG00 | A430084P05I | 1349 | 0 | 0 | 0 | 0 | 0 | 0 |
| ENSMUSG00 | Tmem140 | 5684 | 0.17485697 | 0.18650629 | 0.38105037 | 0.10791604 | 0.61044925 | 0.33589457 |
| ENSMUSG00 | Cyp2d12 | 1826 | 0.17417516 | 0.39583605 | 0 | 0 | 0 | 0 |
| ENSMUSG00 | Trim12c | 4289 | 5.44100421 | 4.280493 | 5.30751904 | 4.22317238 | 4.54466672 | 6.32105186 |
| ENSMUSG00 | Vmn1r30 | 1052 | 0 | 0 | 0 | 0 | 0 | 0 |
| ENSMUSG00 | Olfr469 | 944 | 0 | 0 | 0 | 0 | 0 | 0 |
| ENSMUSG00 | Atpbd4 | 9269 | 8.52669061 | 7.57445746 | 9.4231138 | 8.46287246 | 8.19703027 | 10.8521814 |
| ENSMUSG00 | Olfr1301 | 938 | 0 | 0 | 0 | 0 | 0 | 0 |
| ENSMUSG00 | Olfr782 | 944 | 0 | 0 | 0 | 0 | 0 | 0 |
| ENSMUSG00 | Homez | 9339 | 4.65282581 | 4.93267359 | 3.37465618 | 4.21131048 | 5.6495634 | 3.54549867 |
| ENSMUSG00 | Gm6054 | 377 | 0.52726102 | 0 | 0.46898507 | 0 | 0 | 0.72346523 |
| ENSMUSG00 | Gm16372 | 449 | 0 | 0 | 0 | 0.0803609 | 0.11364449 | 0.0607453 |
| ENSMUSG00 | Vmn1r80 | 926 | 0 | 0 | 0 | 0 | 0 | 0 |
| ENSMUSG00 | Prss2 | 810 | 0 | 0 | 0 | 0 | 0 | 0 |
| ENSMUSG00 | Olfr10 | 941 | 0 | 0 | 0 | 0 | 0 | 0 |
| ENSMUSG00 | Prl3d1 | 845 | 0 | 0 | 0 | 0 | 0 | 0 |
| ENSMUSG00 | Rfx8 | 2118 | 0 | 0 | 0.31304421 | 0 | 0 | 0 |
| ENSMUSG00 | Krtap16-10b | 460 | 0 | 0 | 0 | 0 | 0 | 0 |
| ENSMUSG00 | Gm166 | 1822 | 6.50226849 | 6.95556218 | 7.08393971 | 7.05005919 | 9.0738451 | 9.41588808 |
| ENSMUSG00 | Gsk3a | 2909 | 7.77616652 | 8.08352828 | 8.26600296 | 9.60037898 | 7.27945891 | 11.5699226 |
| ENSMUSG00 | Olfr747 | 954 | 0 | 0 | 0 | 0 | 0 | 0 |
| ENSMUSG00 | 5730455P16I | 4868 | 14.0140519 | 12.6900203 | 17.7606419 | 11.5406211 | 15.6601079 | 17.0718623 |
| ENSMUSG00 | Scn3a | 16023 | 0.04466072 | 0.06014659 | 0.23724387 | 0.00675567 | 0.14649025 | 0.15830628 |
| ENSMUSG00 | AB124611 | 1518 | 1.02138586 | 0.60312411 | 1.36856826 | 0.59423657 | 2.82359392 | 1.13195143 |
| ENSMUSG00 | Slc44a2 | 3418 | 61.4127961 | 67.3031231 | 53.6681242 | 58.2084241 | 76.3902765 | 45.0294878 |
| ENSMUSG00 | Ceacam13 | 1082 | 0 | 0 | 0 | 0 | 0 | 0 |
| ENSMUSG00 | Vmn1r234 | 989 | 0 | 0 | 0 | 0 | 0 | 0 |
| ENSMUSG00 | Olfr1028 | 974 | 0 | 0 | 0 | 0 | 0 | 0 |
| ENSMUSG00 | Vmn1r191 | 896 | 0 | 0 | 0 | 0 | 0 | 0 |
| ENSMUSG00 | Olfr1484 | 947 | 0 | 0 | 0 | 0 | 0 | 0 |
| ENSMUSG00 | Gm15514 | 1288 | 0 | 0 | 0 | 0 | 0 | 0 |
| ENSMUSG00 | 4933403O03I | 1103 | 0 | 0 | 0 | 0 | 0 | 0 |
| ENSMUSG00 | Armc7 | 3394 | 4.60338946 | 4.34444646 | 5.86058614 | 3.93351693 | 6.94584138 | 4.16271748 |
| ENSMUSG00 | Gm6578 | 980 | 0 | 0.09833968 | 0.04510392 | 0 | 0 | 0 |
| ENSMUSG00 | Aadat | 1873 | 3.16260897 | 1.59506642 | 1.74636219 | 1.07880112 | 0.24518814 | 2.94152543 |
| ENSMUSG00 | Atp5sl | 2907 | 36.9929738 | 29.0743101 | 35.8693318 | 37.5218509 | 38.3531583 | 44.3224616 |
| ENSMUSG00 | Aak1 | 8837 | 1.39011469 | 1.00331621 | 1.29049174 | 1.15550737 | 1.03935132 | 0.97839319 |
| ENSMUSG00 | Mettl15 | 3218 | 6.00409059 | 7.77152376 | 5.61794711 | 7.99456114 | 6.64389418 | 8.16205022 |
| ENSMUSG00 | Rbbp4 | 9027 | 19.4571522 | 21.5122811 | 17.3634358 | 17.023754 | 16.4548333 | 10.8560739 |
| ENSMUSG00 | Ms4a13 | 1942 | 0 | 0 | 0 | 0 | 0 | 0 |
| ENSMUSG00 | Gm6139 | 933 | 0 | 0 | 0 | 0 | 0 | 0.11693307 |
| ENSMUSG00 | BC051142 | 3397 | 0.11703115 | 0 | 0.19518035 | 0 | 0 | 0.04817422 |
| ENSMUSG00 | Zfp870 | 1680 | 20.2090359 | 10.8419495 | 17.8912222 | 18.2343189 | 20.9572615 | 10.5526878 |
| ENSMUSG00 | Gm10020 | 614 | 0 | 0 | 0 | 0 | 0 | 0.04442124 |
| ENSMUSG00 | Olfr824 | 947 | 0 | 0 | 0 | 0 | 0 | 0 |
| ENSMUSG00 | 2900006K08I | 2222 | 5.95795455 | 2.77581665 | 3.22263662 | 5.5860591 | 4.89136726 | 6.01465942 |
| ENSMUSG00 | Olfr1501 | 947 | 0 | 0 | 0 | 0 | 0 | 0 |
| ENSMUSG00 | Snrpg | 811 | 2.89219895 | 5.70394383 | 4.74175135 | 3.73722779 | 4.08966021 | 7.76873202 |
| ENSMUSG00 | Musk | 3800 | 0.29293512 | 0 | 0.55833907 | 0 | 1.28908739 | 0.05024276 |
| ENSMUSG00 | St6galnac2 | 8873 | 12.1959222 | 16.6015951 | 4.827182 | 11.2438694 | 13.1059518 | 8.53619669 |
| ENSMUSG00 | Gm15732 | 1036 | 0 | 0 | 0 | 0 | 0 | 0 |
| ENSMUSG00 | 1700054O19I | 569 | 0 | 0 | 0 | 0 | 0 | 0 |
| ENSMUSG00 | Vmn1r192 | 1065 | 0 | 0 | 0 | 0 | 0 | 0 |
| ENSMUSG00 | Olfr792 | 935 | 0 | 0 | 0 | 0 | 0 | 0.02917074 |
| ENSMUSG00 | Arhgap24 | 9109 | 91.6614998 | 106.931688 | 61.2633953 | 93.2334045 | 119.311755 | 62.0947929 |
| ENSMUSG00 | Dub2a | 1985 | 0 | 0 | 0 | 0 | 0 | 0 |
| ENSMUSG00 | Rpl38 | 503 | 2.29206548 | 5.07729907 | 5.88772064 | 3.01281485 | 3.04332261 | 5.58506529 |
| ENSMUSG00 | Bcl2 | 7189 | 8.80935881 | 11.6293611 | 4.67903777 | 11.3832351 | 9.34785602 | 9.94771235 |
| ENSMUSG00 | Cep170 | 9413 | 1.28815697 | 1.31561836 | 0.83116182 | 1.52561922 | 1.96776526 | 0.64615368 |
| ENSMUSG00 | Chst3 | 11842 | 0 | 0 | 0 | 0 | 0 | 0 |
| ENSMUSG00 | Sphk2 | 5695 | 13.4100576 | 9.86574039 | 12.0303524 | 6.17734739 | 9.74832254 | 10.2441534 |
| ENSMUSG00 | Apol9b | 1618 | 0.09828302 | 0.11912594 | 0 | 0 | 0.47305046 | 0 |
| ENSMUSG00 | Olfr948 | 965 | 0 | 0 | 0 | 0 | 0 | 0 |
| ENSMUSG00 | Olfr347 | 942 | 0 | 0 | 0 | 0 | 0 | 0 |
| ENSMUSG00 | 4930430O22I | 538 | 0 | 0 | 0 | 0 | 0 | 0 |
| ENSMUSG00 | Gm17494 | 665 | 0 | 0.14492163 | 0.06646894 | 0.05425871 | 0.15346278 | 0.12304349 |
| ENSMUSG00 | Uxs1 | 5891 | 3.61045359 | 5.20227079 | 3.42149727 | 5.22457715 | 4.59938984 | 3.93540033 |
| ENSMUSG00 | Birc2 | 3123 | 33.3905301 | 33.6826779 | 52.7223392 | 43.037341 | 33.6581281 | 58.6889452 |
| ENSMUSG00 | Rpl21-ps12 | 482 | 0 | 0 | 0 | 0 | 0 | 0 |
| ENSMUSG00 | Yipf1 | 4528 | 12.8450239 | 15.9628232 | 11.9876023 | 9.77753283 | 9.95059405 | 9.5413049 |
| ENSMUSG00 | Ryr3 | 18434 | 0.00215664 | 0.002614 | 0.02158059 | 0 | 0.01107223 | 0.02811208 |
| ENSMUSG00 | Tas2r123 | 1001 | 0 | 0 | 0 | 0 | 0 | 0 |
| ENSMUSG00 | 4922502B01I | 1247 | 0 | 0 | 0 | 0 | 0 | 0 |
| ENSMUSG00 | Mrpl18 | 1237 | 51.9038812 | 48.4201678 | 49.8119395 | 41.3908013 | 34.2788345 | 88.1960849 |
| ENSMUSG00 | Vmn1r220 | 896 | 0 | 0 | 0 | 0 | 0 | 0 |
| ENSMUSG00 | Zfp759 | 3565 | 8.01800575 | 7.28541159 | 10.6134019 | 6.08283556 | 9.63274921 | 17.8566642 |
| ENSMUSG00 | Ska2l-ps | 362 | 0 | 0 | 0 | 0 | 0 | 0 |
| ENSMUSG00 | Ces1c | 2309 | 0 | 0 | 0 | 0 | 0 | 0 |
| ENSMUSG00 | 1700042B14I | 1075 | 0 | 0 | 0 | 0 | 0 | 0 |
| ENSMUSG00 | Olfr804 | 944 | 0 | 0 | 0 | 0 | 0 | 0 |
| ENSMUSG00 | Whsc1 | 16012 | 3.42634044 | 1.96513532 | 2.39615287 | 4.34237446 | 2.80116065 | 2.54656418 |
| ENSMUSG00 | Zfp53 | 1955 | 38.5150282 | 41.4575937 | 36.2884696 | 41.2128928 | 25.865544 | 52.5403025 |
| ENSMUSG00 | Fam173a | 851 | 61.1047809 | 67.7781096 | 68.9257881 | 63.8113713 | 49.3474821 | 74.6767444 |
| ENSMUSG00 | Dcpp3 | 606 | 0 | 0 | 0 | 0 | 0 | 0 |
| ENSMUSG00 | Las1l | 7549 | 9.7532322 | 10.3981606 | 10.8616265 | 11.4091721 | 13.2010216 | 9.77320164 |
| ENSMUSG00 | Olfr934 | 932 | 0 | 0 | 0 | 0 | 0 | 0 |
| ENSMUSG00 | Ugt2b37 | 1888 | 1.45292805 | 0.84224184 | 0 | 0.63067133 | 0 | 4.57948127 |
| ENSMUSG00 | Kir3dl2 | 2263 | 0 | 0 | 0 | 0 | 0 | 0 |
| ENSMUSG00 | Mpp7 | 5029 | 13.3361495 | 18.9047227 | 11.0658422 | 18.3603006 | 15.0877353 | 14.578292 |
| ENSMUSG00 | Olfr138 | 1311 | 0 | 0 | 0 | 0 | 0 | 0 |
| ENSMUSG00 | Olfr910 | 970 | 0 | 0 | 0 | 0 | 0 | 0 |
| ENSMUSG00 | Cts8 | 2485 | 0 | 0 | 0 | 0 | 0 | 0 |
| ENSMUSG00 | Olfr1205 | 923 | 0 | 0 | 0 | 0 | 0 | 0 |
| ENSMUSG00 | Lypd3 | 1622 | 0.49020321 | 0.56445278 | 0.02725145 | 0 | 0.09437677 | 0 |
| ENSMUSG00 | Rit2 | 5679 | 0 | 0 | 0 | 0 | 0 | 0 |
| ENSMUSG00 | Phex | 6269 | 0 | 0.00768646 | 0 | 0 | 0 | 0 |
| ENSMUSG00 | Olfr591 | 944 | 0 | 0 | 0 | 0 | 0 | 0 |
| ENSMUSG00 | Olfr1415 | 935 | 0 | 0 | 0 | 0 | 0 | 0 |
| ENSMUSG00 | Saa2 | 606 | 0 | 0 | 0 | 0 | 0 | 0 |
| ENSMUSG00 | E2f6 | 2555 | 22.9974952 | 28.704409 | 21.9711705 | 29.5999864 | 17.1352761 | 21.8303864 |
| ENSMUSG00 | Fam136a | 1602 | 36.0330572 | 44.6070498 | 43.3189098 | 42.3209499 | 35.5783157 | 43.3125357 |
| ENSMUSG00 | Gm6177 | 464 | 0 | 0 | 0 | 0 | 0 | 0 |
| ENSMUSG00 | Olfr1459 | 1100 | 0 | 0 | 0 | 0 | 0 | 0 |
| ENSMUSG00 | Bloc1s2 | 873 | 8.7890123 | 8.50024294 | 9.62010329 | 6.44765057 | 7.48153048 | 11.5909406 |
| ENSMUSG00 | Vmn1r177 | 929 | 0 | 0 | 0 | 0 | 0 | 0 |
| ENSMUSG00 | Spop | 3181 | 86.1722849 | 74.4685789 | 71.6317198 | 68.0125555 | 86.6373643 | 54.1720122 |
| ENSMUSG00 | Ece1 | 5361 | 20.5562753 | 15.100396 | 25.8235725 | 18.0107352 | 25.8320433 | 17.6590697 |
| ENSMUSG00 | Dtnbp1 | 1339 | 71.7320694 | 63.9486988 | 72.2942765 | 49.7711249 | 55.2944521 | 63.7358822 |
| ENSMUSG00 | Olfr191 | 929 | 0 | 0 | 0 | 0 | 0 | 0 |
| ENSMUSG00 | Gm15698 | 1169 | 0 | 0 | 0 | 0 | 0 | 0 |
| ENSMUSG00 | Olfr310 | 1081 | 0 | 0 | 0 | 0 | 0 | 0 |
| ENSMUSG00 | Pus7 | 6582 | 6.81919439 | 6.25940566 | 5.19113106 | 5.78343315 | 5.70577524 | 5.28751742 |
| ENSMUSG00 | Zfp317 | 645 | 46.0423937 | 37.7273694 | 29.8105453 | 31.159223 | 71.9907008 | 37.0427659 |
| ENSMUSG00 | Lgals8 | 5253 | 35.4189318 | 29.9686097 | 49.048647 | 36.2262903 | 35.5814991 | 45.71215 |
| ENSMUSG00 | Eif1a | 1914 | 48.0222941 | 68.9564603 | 54.4324681 | 91.6755395 | 70.0079744 | 55.233282 |
| ENSMUSG00 | Olfr1508 | 1863 | 0 | 0 | 0 | 0 | 0 | 0 |
| ENSMUSG00 | Zbtb8os | 4618 | 3.39187079 | 2.89035177 | 3.44579114 | 2.46120486 | 2.28723686 | 5.50453958 |
| ENSMUSG00 | Gm10024 | 359 | 0 | 0 | 0 | 0 | 0 | 0 |
| ENSMUSG00 | Gm10025 | 369 | 0 | 0 | 0 | 0 | 0 | 0 |
| ENSMUSG00 | Gm10012 | 562 | 0 | 0.171482 | 0 | 0 | 0 | 0 |
| ENSMUSG00 | Gm6128 | 410 | 0 | 0 | 0.21561875 | 0 | 0 | 0 |
| ENSMUSG00 | Vmn1r48 | 908 | 0 | 0 | 0 | 0 | 0 | 0 |
| ENSMUSG00 | Arl16 | 3535 | 8.72708709 | 6.93830242 | 8.61529557 | 7.6859348 | 8.67520562 | 3.88094584 |
| ENSMUSG00 | Trim30d | 4541 | 0.9017429 | 0.46690232 | 0.86632108 | 1.62095069 | 2.60694102 | 1.24330551 |
| ENSMUSG00 | Lmcd1 | 1718 | 52.8067558 | 26.393156 | 64.5017581 | 33.0787079 | 33.6809722 | 45.3730611 |
| ENSMUSG00 | Gm6807 | 887 | 0 | 0.16297557 | 0.19933187 | 0 | 0.05752692 | 0.36899174 |
| ENSMUSG00 | Colq | 5176 | 0 | 0 | 0.04269884 | 0 | 0.00985826 | 0 |
| ENSMUSG00 | Lce1a1 | 706 | 0 | 0 | 0 | 0 | 0 | 0 |
| ENSMUSG00 | Ighv1-73 | 182 | 0 | 0 | 0 | 0 | 0 | 0 |
| ENSMUSG00 | Vmn1r37 | 908 | 0 | 0 | 0 | 0 | 0 | 0 |
| ENSMUSG00 | Gnai1 | 3185 | 59.9015859 | 51.6207663 | 44.9928964 | 59.1586928 | 43.09606 | 32.7124402 |
| ENSMUSG00 | Ldoc1 | 1335 | 0 | 0 | 0 | 0 | 0 | 0 |
| ENSMUSG00 | Olfr787 | 938 | 0 | 0 | 0 | 0 | 0 | 0 |
| ENSMUSG00 | Gm5621 | 644 | 0.06173211 | 0 | 0 | 0 | 0 | 0 |
| ENSMUSG00 | Prdm2 | 7683 | 2.00769575 | 3.43696086 | 1.27721061 | 1.4464753 | 3.23439347 | 0.97624962 |
| ENSMUSG00 | Brd9 | 2390 | 16.7172627 | 19.0326366 | 15.8497822 | 16.0180122 | 23.2500934 | 12.747185 |
| ENSMUSG00 | Krtap16-1 | 604 | 0 | 0 | 0 | 0 | 0 | 0 |
| ENSMUSG00 | Olfr328 | 2096 | 0 | 0 | 0 | 0 | 0 | 0 |
| ENSMUSG00 | Gm10260 | 545 | 0 | 0 | 0 | 0 | 0 | 0 |
| ENSMUSG00 | Gapdh | 2138 | 24.3590643 | 26.5949494 | 22.6384556 | 22.0238859 | 19.9284489 | 13.8286758 |
| ENSMUSG00 | Bloc1s3 | 1822 | 6.15315341 | 6.34728107 | 7.71470147 | 5.98066819 | 4.00481435 | 6.82614462 |
| ENSMUSG00 | Pkn1 | 5788 | 7.69283659 | 5.23657086 | 3.35255858 | 5.32378473 | 6.06533286 | 5.69242644 |
| ENSMUSG00 | Gm11938 | 662 | 0 | 0 | 0 | 0 | 0 | 0 |
| ENSMUSG00 | Gm8526 | 470 | 0 | 0 | 0 | 0 | 0 | 0 |
| ENSMUSG00 | Zfp746 | 3644 | 14.1282512 | 17.8913985 | 14.4832603 | 14.2684485 | 18.5397699 | 9.28116155 |
| ENSMUSG00 | Rpl30-ps1 | 347 | 0 | 0 | 0 | 0 | 0 | 0 |
| ENSMUSG00 | Vmn1r53 | 1121 | 0 | 0 | 0 | 0.06437475 | 0 | 0 |
| ENSMUSG00 | Tas2r131 | 1353 | 0 | 0 | 0 | 0 | 0 | 0 |
| ENSMUSG00 | Mex3b | 3401 | 0.43250596 | 0.02833663 | 0.33791471 | 0.01060925 | 0.22505017 | 0 |
| ENSMUSG00 | Rpl29-ps2 | 482 | 0.08248025 | 0 | 0 | 0 | 0 | 0 |
| ENSMUSG00 | 9630041A04I | 971 | 0 | 0 | 0 | 0 | 0 | 0 |
| ENSMUSG00 | A830018L16F | 9634 | 0 | 0 | 0 | 0 | 0 | 0 |
| ENSMUSG00 | Gm5567 | 3089 | 0 | 0 | 0.17171321 | 0 | 0 | 0 |
| ENSMUSG00 | Sh3rf2 | 5781 | 1.67109181 | 0.87520782 | 1.51391886 | 1.36688596 | 0.32658293 | 0.46236199 |
| ENSMUSG00 | Lepr | 7123 | 0.2009262 | 0 | 0 | 0 | 0.65905189 | 0.43268767 |
| ENSMUSG00 | Krt33b | 1574 | 0 | 0 | 0 | 0 | 0 | 0 |
| ENSMUSG00 | Serpinb9g | 1914 | 0 | 0 | 0 | 0 | 0 | 0 |
| ENSMUSG00 | Olfr1359 | 990 | 0 | 0 | 0 | 0 | 0 | 0 |
| ENSMUSG00 | Prtn3 | 1726 | 0 | 0 | 0 | 0 | 0 | 0 |
| ENSMUSG00 | Olfr1186 | 1089 | 0 | 0 | 0 | 0 | 0 | 0 |
| ENSMUSG00 | Spna2 | 9829 | 16.6399478 | 17.9430642 | 14.5345769 | 19.3166872 | 24.0362316 | 12.9588529 |
| ENSMUSG00 | Megf6 | 7431 | 0.19259821 | 0.09078323 | 0.11301777 | 0 | 0.07553359 | 0.11745236 |
| ENSMUSG00 | Tas2r106 | 926 | 0 | 0 | 0 | 0 | 0 | 0 |
| ENSMUSG00 | Olfr1022 | 947 | 0 | 0 | 0 | 0 | 0 | 0 |
| ENSMUSG00 | Gm6169 | 1104 | 0.0720208 | 0.4364714 | 0.16015161 | 0.32683011 | 0.04621954 | 0.17293702 |
| ENSMUSG00 | Rpl35a-ps7 | 332 | 0 | 0.14513989 | 0.13313806 | 0 | 0.61477561 | 0 |
| ENSMUSG00 | Ankrd29 | 5620 | 2.68101906 | 0.1629079 | 1.29774094 | 0.526464 | 3.32307003 | 1.0676905 |
| ENSMUSG00 | Olfr1383 | 939 | 0 | 0 | 0 | 0 | 0 | 0 |
| ENSMUSG00 | Olfr668 | 925 | 0 | 0 | 0 | 0 | 0 | 0 |
| ENSMUSG00 | Mab21l2 | 2702 | 0 | 0 | 0.39261445 | 0 | 0.07553868 | 0.10094241 |
| ENSMUSG00 | Cyb5d2 | 6857 | 3.93090503 | 2.79687968 | 3.04906982 | 4.07284563 | 4.23421436 | 2.18769893 |
| ENSMUSG00 | Ddx49 | 3725 | 6.85181704 | 6.83018563 | 6.56204542 | 6.95487462 | 6.05467332 | 5.37438529 |
| ENSMUSG00 | Bak1 | 3488 | 0.04559115 | 0.08288952 | 0.03801764 | 0.02068925 | 0.10240385 | 0.14075215 |
| ENSMUSG00 | Vmn1r216 | 1058 | 0 | 0 | 0 | 0 | 0 | 0 |
| ENSMUSG00 | Olfr135 | 1078 | 0 | 0 | 0 | 0 | 0 | 0 |
| ENSMUSG00 | Gm10030 | 301 | 0 | 0 | 0 | 0 | 0 | 0 |
| ENSMUSG00 | 1700084M14 | 557 | 0 | 0 | 0 | 0 | 0 | 0 |
| ENSMUSG00 | Gm10031 | 742 | 0 | 0 | 0 | 0.04862809 | 0.0687687 | 0 |
| ENSMUSG00 | 1700007G11I | 696 | 0 | 0 | 0.0635084 | 0 | 0 | 0 |
| ENSMUSG00 | Olfr1446 | 1043 | 0 | 0 | 0 | 0 | 0 | 0 |
| ENSMUSG00 | 1600029O15I | 1804 | 0 | 0 | 0 | 0 | 0 | 0 |
| ENSMUSG00 | Gm5278 | 551 | 0 | 0.17490542 | 0.08022113 | 0 | 0.09260685 | 0.1485076 |
| ENSMUSG00 | Olfr695 | 1198 | 0 | 0 | 0 | 0 | 0 | 0 |
| ENSMUSG00 | Zfp119a | 1634 | 16.78781 | 23.7393427 | 19.9969192 | 18.1293504 | 19.1739258 | 42.6979971 |
| ENSMUSG00 | Xlr3a | 2731 | 1.36836879 | 0.77634693 | 1.26244737 | 1.98180398 | 1.43867848 | 0.85888648 |
| ENSMUSG00 | Rpl32 | 513 | 20.0714804 | 20.0072362 | 28.606261 | 16.8101533 | 13.8258601 | 39.3435342 |
| ENSMUSG00 | Zfp595 | 4441 | 5.82770048 | 3.1900054 | 6.02164265 | 7.66164557 | 6.26196236 | 5.0360739 |
| ENSMUSG00 | Krtap6-3 | 443 | 0 | 0 | 0 | 0 | 0 | 0 |
| ENSMUSG00 | D19Ertd737e | 2599 | 14.9293378 | 11.3096305 | 21.5141714 | 13.6609203 | 14.175084 | 23.9269633 |
| ENSMUSG00 | Rpl36 | 400 | 1.68960792 | 2.77072042 | 2.9836244 | 2.16492267 | 2.55131879 | 3.5457031 |
| ENSMUSG00 | Abat | 7823 | 1.88537432 | 3.07363347 | 1.44081171 | 2.05247473 | 3.04605874 | 1.26210142 |
| ENSMUSG00 | Gm6984 | 1664 | 0 | 0 | 0.02656361 | 0.02168392 | 0 | 0 |
| ENSMUSG00 | Olfr1375-ps1 | 941 | 0 | 0 | 0 | 0 | 0 | 0 |
| ENSMUSG00 | Zfp329 | 5398 | 18.3900398 | 17.0321844 | 17.0485804 | 18.2883427 | 13.6876977 | 20.4130312 |
| ENSMUSG00 | Zfp105 | 2245 | 2.70939355 | 2.64005897 | 3.01241959 | 2.13759997 | 5.70495338 | 1.85880615 |
| ENSMUSG00 | Camk2b | 11257 | 1.35614325 | 1.04017993 | 0.53009228 | 0.91030476 | 1.23747007 | 0.42400834 |
| ENSMUSG00 | Gpr111 | 3278 | 0.03638391 | 0 | 0 | 0 | 0 | 0 |
| ENSMUSG00 | Tdpoz2 | 1090 | 0 | 0 | 0 | 0 | 0 | 0 |
| ENSMUSG00 | Olfr739 | 929 | 0 | 0 | 0 | 0 | 0 | 0 |
| ENSMUSG00 | Gm10032 | 1836 | 0.19487981 | 0 | 1.44450467 | 0 | 1.11168575 | 3.07508188 |
| ENSMUSG00 | Cacnb2 | 9204 | 1.66727678 | 1.0470761 | 2.0602554 | 2.36391491 | 4.32981307 | 2.92778613 |
| ENSMUSG00 | Gm10033 | 652 | 101.522815 | 57.7941683 | 73.1499825 | 91.8653281 | 39.052395 | 114.369423 |
| ENSMUSG00 | Gsta2 | 1168 | 4.69713725 | 20.1739471 | 0.94610109 | 18.1336988 | 4.10657477 | 2.17169645 |
| ENSMUSG00 | Gm17136 | 410 | 0 | 0 | 0 | 0.08800499 | 0 | 0 |
| ENSMUSG00 | Vmn1r181 | 1079 | 0 | 0 | 0 | 0 | 0 | 0 |
| ENSMUSG00 | Unc13d | 5268 | 0.56599488 | 0.71346668 | 0.04195315 | 0.00684929 | 0.66834091 | 0.13979029 |
| ENSMUSG00 | Olfr1487 | 947 | 0 | 0 | 0 | 0 | 0 | 0 |
| ENSMUSG00 | Itpk1 | 2846 | 10.5325482 | 5.01166088 | 9.38085495 | 5.10929865 | 7.74539507 | 8.87432043 |
| ENSMUSG00 | Fgf18 | 1181 | 0.47127581 | 0.04080139 | 0.07485494 | 0 | 0.04320608 | 0 |
| ENSMUSG00 | Sema3b | 4906 | 66.7723521 | 46.4774244 | 74.3034242 | 47.7097886 | 52.3992828 | 45.0871024 |
| ENSMUSG00 | Skint11 | 1455 | 0 | 0 | 0 | 0 | 0 | 0 |
| ENSMUSG00 | Vmn1r12 | 934 | 0 | 0 | 0 | 0 | 0 | 0 |
| ENSMUSG00 | Zfp809 | 1278 | 34.6227307 | 28.8817016 | 33.9987572 | 47.6858945 | 42.6018333 | 38.9485263 |
| ENSMUSG00 | Olfr850 | 947 | 0 | 0 | 0 | 0 | 0 | 0 |
| ENSMUSG00 | E030024N20I | 493 | 0.24191976 | 0 | 0.17931782 | 0.73188731 | 0.10350178 | 0.22129525 |
| ENSMUSG00 | Olfr541 | 1102 | 0 | 0 | 0 | 0 | 0 | 0 |
| ENSMUSG00 | Tdpoz3 | 1097 | 0 | 0 | 0 | 0 | 0 | 0 |
| ENSMUSG00 | Mdn1 | 21884 | 0.50321094 | 1.21765228 | 0.63422495 | 1.06016974 | 0.75779438 | 0.70791423 |
| ENSMUSG00 | 11-Sep | 4655 | 4.79969496 | 3.98534484 | 4.10208296 | 2.9067168 | 4.13253355 | 3.86122175 |
| ENSMUSG00 | Olfr502 | 1058 | 0 | 0 | 0 | 0 | 0 | 0 |
| ENSMUSG00 | Serpinb3d | 1628 | 0 | 0 | 0 | 0 | 0 | 0 |
| ENSMUSG00 | Ces5a | 1273 | 0 | 0 | 0 | 0 | 0 | 0 |
| ENSMUSG00 | 9530008L14F | 3353 | 0.1185669 | 0 | 0 | 0.04304449 | 0.0608725 | 0 |
| ENSMUSG00 | Zscan5b | 2029 | 0 | 0 | 0 | 0 | 0 | 0 |
| ENSMUSG00 | Vmn1r237 | 946 | 0 | 0 | 0 | 0 | 0 | 0 |
| ENSMUSG00 | 4933430I17R | 1768 | 0.24734745 | 0 | 0.05000209 | 0 | 0 | 0.15426832 |
| ENSMUSG00 | Gm9234 | 721 | 0.05513936 | 0 | 0.1839189 | 0 | 0 | 0 |
| ENSMUSG00 | Defb35 | 265 | 0 | 0 | 0 | 0 | 0 | 0 |
| ENSMUSG00 | Palld | 8562 | 0.13001092 | 0.14069856 | 0.06711329 | 0.03792787 | 0.07151559 | 0.12105072 |
| ENSMUSG00 | Mettl7a3 | 1616 | 0 | 0.14909172 | 0 | 0 | 0 | 0 |
| ENSMUSG00 | Trim31 | 2249 | 0 | 0 | 0 | 0 | 0 | 0 |
| ENSMUSG00 | Gm10036 | 582 | 0 | 0 | 0 | 0 | 0 | 0.14059092 |
| ENSMUSG00 | Eml1 | 7611 | 0.33429914 | 2.41850229 | 0.9756812 | 1.29423179 | 4.2639305 | 1.33309234 |
| ENSMUSG00 | Olfr818 | 956 | 0 | 0 | 0 | 0 | 0 | 0 |
| ENSMUSG00 | Gm11189 | 374 | 6.05899035 | 0.77304453 | 0.70912048 | 0.96476055 | 0.54573664 | 0.58341474 |

| | | | | | | | | |
|---|---|---|---|---|---|---|---|---|
| ENSMUSG00 | Olfr463 | 935 | 0 | 0 | 0 | 0 | 0 | 0 |
| ENSMUSG00 | Sdhc | 3742 | 29.6838088 | 26.3853608 | 36.3938745 | 24.8775187 | 17.1133356 | 46.3129497 |
| ENSMUSG00 | Olfr1503-ps1 | 796 | 0 | 0 | 0 | 0 | 0 | 0 |
| ENSMUSG00 | Olfr605 | 1860 | 0 | 0 | 0 | 0 | 0 | 0 |
| ENSMUSG00 | Olfr825 | 970 | 0 | 0 | 0 | 0 | 0 | 0 |
| ENSMUSG00 | AA987161 | 5457 | 9.17210005 | 9.37768033 | 7.37912388 | 8.19896192 | 8.10699429 | 12.325318 |
| ENSMUSG00 | Nlam1 | 5283 | 0 | 0 | 0 | 0 | 0 | 0 |
| ENSMUSG00 | 1700084J12R | 1004 | 0 | 0.38395571 | 0.04402574 | 0 | 0 | 0 |
| ENSMUSG00 | Olfr127 | 1094 | 0 | 0 | 0 | 0 | 0 | 0 |
| ENSMUSG00 | Gm5771 | 819 | 0 | 0 | 0 | 0 | 0 | 0 |
| ENSMUSG00 | H2-M10.3 | 1586 | 0 | 0 | 0 | 0 | 0 | 0 |
| ENSMUSG00 | Tpm3-rs7 | 2105 | 0.15108971 | 0.04578284 | 0.125991 | 0.03428223 | 0.02424056 | 0.06478537 |
| ENSMUSG00 | Olfr884 | 1854 | 0 | 0 | 0 | 0 | 0 | 0 |
| ENSMUSG00 | Vmn1r82 | 914 | 0 | 0 | 0 | 0 | 0 | 0 |
| ENSMUSG00 | Gstm1 | 1773 | 300.509278 | 599.02841 | 220.585396 | 445.011883 | 224.251281 | 139.003745 |
| ENSMUSG00 | Adamts17 | 7290 | 0 | 0 | 0 | 0 | 0.22398409 | 0.01122413 |
| ENSMUSG00 | Xlr3c | 1575 | 0 | 0 | 0 | 0 | 0 | 0 |
| ENSMUSG00 | Chsy3 | 2303 | 0.63871158 | 0.04184667 | 0.24951106 | 0.04700223 | 0.4652861 | 0 |
| ENSMUSG00 | Sez6l | 6266 | 0 | 0 | 0 | 0 | 0 | 0 |
| ENSMUSG00 | T2 | 1758 | 0 | 0 | 0 | 0 | 0.46440388 | 0.12411668 |
| ENSMUSG00 | Gm6431 | 5381 | 0.00738812 | 0 | 0 | 0 | 0.05689616 | 0 |
| ENSMUSG00 | Krtap6-1 | 593 | 0 | 0 | 0 | 0 | 0 | 0 |
| ENSMUSG00 | BC017612 | 951 | 3.13529026 | 3.9522003 | 1.81269388 | 3.56646918 | 4.02416213 | 6.99790955 |
| ENSMUSG00 | Gm5148 | 4833 | 1.25032755 | 1.38587119 | 0.82312557 | 1.45582427 | 1.25639121 | 0.17494596 |
| ENSMUSG00 | Gm5407 | 746 | 0 | 0 | 0 | 0 | 0 | 0 |
| ENSMUSG00 | Mmel1 | 3539 | 0 | 0 | 0 | 0 | 0 | 0 |
| ENSMUSG00 | Gm13242 | 4132 | 0 | 0.01166177 | 0 | 0 | 0 | 0 |
| ENSMUSG00 | Olfr746 | 944 | 0 | 0 | 0 | 0 | 0 | 0 |
| ENSMUSG00 | Hist1h2bm | 523 | 0 | 0 | 0 | 0 | 0 | 0 |
| ENSMUSG00 | Zfp846 | 3089 | 16.9755516 | 12.5886885 | 13.3220835 | 8.39850761 | 11.4309654 | 12.7852631 |
| ENSMUSG00 | Olfr1126 | 944 | 0 | 0 | 0 | 0 | 0 | 0 |
| ENSMUSG00 | Olfr66 | 935 | 0 | 0 | 0 | 0 | 0 | 0 |
| ENSMUSG00 | Olfr344 | 929 | 0 | 0 | 0 | 0 | 0 | 0 |
| ENSMUSG00 | Serpina3k | 1649 | 0 | 0 | 0 | 0 | 0 | 0 |
| ENSMUSG00 | BC021614 | 679 | 0 | 0 | 0 | 0.4782598 | 0 | 0 |
| ENSMUSG00 | Olfr318 | 920 | 0 | 0 | 0 | 0 | 0 | 0 |
| ENSMUSG00 | Grlf1 | 7841 | 11.5195067 | 14.2021257 | 9.38041918 | 16.2026373 | 17.3038048 | 9.5031647 |
| ENSMUSG00 | Usf2 | 3310 | 1.69351141 | 1.96530806 | 1.36211117 | 1.78775085 | 1.572414 | 3.17242783 |
| ENSMUSG00 | Cryzl1 | 4910 | 14.452858 | 10.844403 | 16.3573826 | 10.1852777 | 10.6105763 | 17.2369054 |
| ENSMUSG00 | Olfr506 | 1007 | 0 | 0 | 0 | 0 | 0 | 0 |
| ENSMUSG00 | Gm10037 | 311 | 0.51132451 | 0.46482099 | 3.69533093 | 0.92815549 | 0.82035974 | 0.70159844 |
| ENSMUSG00 | Kcnh1 | 7309 | 0 | 0 | 0 | 0 | 0 | 0 |
| ENSMUSG00 | Tas2r138 | 995 | 0 | 0 | 0 | 0 | 0 | 0 |
| ENSMUSG00 | Olfr822 | 941 | 0 | 0 | 0 | 0 | 0 | 0 |
| ENSMUSG00 | 1700008I05R | 1766 | 0 | 0 | 0 | 0 | 0 | 0 |
| ENSMUSG00 | BC016201 | 4011 | 0 | 0 | 0 | 0 | 0 | 0 |
| ENSMUSG00 | Tspan7 | 6785 | 1.73435847 | 1.22152808 | 2.01953889 | 0.60092425 | 1.15063161 | 1.74863199 |
| ENSMUSG00 | Idi1 | 2797 | 1.23658449 | 1.44714378 | 1.65934699 | 1.59963301 | 0.83918959 | 1.52121692 |
| ENSMUSG00 | Serpina9 | 3669 | 0.01083551 | 0 | 0 | 0 | 0 | 0 |
| ENSMUSG00 | Mrps14 | 3263 | 26.3290203 | 28.2945826 | 34.7329223 | 27.4568546 | 16.9827288 | 38.0908768 |
| ENSMUSG00 | Olfr25 | 1067 | 0 | 0 | 0 | 0 | 0 | 0 |
| ENSMUSG00 | Olfr412 | 938 | 0 | 0 | 0 | 0 | 0 | 0 |
| ENSMUSG00 | Gm10038 | 438 | 0 | 0 | 0 | 0 | 0 | 0 |
| ENSMUSG00 | Gm12253 | 1349 | 0 | 0 | 0 | 0 | 0 | 0 |
| ENSMUSG00 | Espl1 | 6599 | 0.07229365 | 0.13873957 | 0.04018958 | 0.20230878 | 0 | 0 |
| ENSMUSG00 | Zfp68 | 6663 | 34.3079713 | 30.4609477 | 29.5209667 | 32.9628403 | 25.471068 | 53.6732808 |
| ENSMUSG00 | Prp2 | 1134 | 0 | 0 | 0 | 0 | 0 | 0 |
| ENSMUSG00 | Spock2 | 1988 | 0.01999773 | 0.12119327 | 0.08893731 | 0.03629984 | 0.05133438 | 0.04115891 |
| ENSMUSG00 | Mcm9 | 4883 | 3.47646737 | 4.53937403 | 3.98296353 | 5.97795905 | 5.14130184 | 3.6194893 |
| ENSMUSG00 | Upf1 | 4592 | 18.4059564 | 12.5712887 | 14.3521228 | 22.2919774 | 27.4577907 | 14.7599039 |
| ENSMUSG00 | Olfr798 | 935 | 0 | 0 | 0 | 0 | 0 | 0 |
| ENSMUSG00 | Ube2e2 | 5471 | 11.6338008 | 10.1375608 | 14.5912134 | 9.80038716 | 7.66654742 | 13.8392247 |
| ENSMUSG00 | Phf21a | 4668 | 3.56845466 | 5.72910784 | 6.01288996 | 5.5498946 | 5.96838072 | 5.66760927 |
| ENSMUSG00 | Dock1 | 6755 | 9.60487701 | 12.0555274 | 8.34960055 | 9.54531659 | 11.9351109 | 7.44954988 |
| ENSMUSG00 | Xlr5a | 1840 | 0 | 0 | 0 | 0 | 0 | 0.08893904 |
| ENSMUSG00 | Zfp85-rs1 | 4229 | 3.69446769 | 0.78620584 | 4.71388773 | 5.98096789 | 2.46142839 | 7.71351822 |
| ENSMUSG00 | Ces5a | 731 | 0 | 0 | 0 | 0 | 0 | 0 |
| ENSMUSG00 | Tas2r117 | 1157 | 0 | 0 | 0 | 0 | 0 | 0 |
| ENSMUSG00 | Gm5246 | 692 | 0 | 0 | 0 | 0 | 0 | 0 |
| ENSMUSG00 | 2010107H07I | 588 | 39.6202578 | 31.3867472 | 28.1899509 | 35.4684043 | 26.0338652 | 80.4323545 |
| ENSMUSG00 | Krt6a | 2349 | 0 | 0 | 0 | 0 | 0 | 0 |
| ENSMUSG00 | Abce1 | 3795 | 41.7773007 | 46.8405231 | 45.2734028 | 48.7749377 | 40.0143595 | 46.0326915 |
| ENSMUSG00 | Gm6612 | 394 | 0.10090224 | 1.22300615 | 16.9403002 | 16.9420767 | 0.12950857 | 19.5906673 |
| ENSMUSG00 | Gm10040 | 1653 | 0 | 0 | 0 | 0 | 0 | 0 |
| ENSMUSG00 | Lhfpl3 | 709 | 0.33643566 | 0 | 0.12468785 | 0.30534876 | 0 | 0.03846917 |
| ENSMUSG00 | Krtap16-7 | 747 | 0 | 0 | 0 | 0 | 0 | 0 |
| ENSMUSG00 | Hist1h2bg | 617 | 0.64433518 | 0.54668573 | 1.00295916 | 0 | 0.41350386 | 0.13261575 |
| ENSMUSG00 | Phtf1 | 7275 | 4.46463609 | 7.72970144 | 6.65913676 | 5.46562379 | 6.69830775 | 4.82133142 |
| ENSMUSG00 | Rrp1b | 7887 | 2.10698502 | 2.35219732 | 2.1408779 | 4.47422842 | 3.13779539 | 2.41726548 |
| ENSMUSG00 | Gpr182 | 5424 | 0.0146591 | 0.22209828 | 0.03259723 | 0 | 0.3198556 | 0.11565573 |
| ENSMUSG00 | Prss43 | 1984 | 0 | 0 | 0 | 0 | 0 | 0 |
| ENSMUSG00 | Vmn1r64 | 962 | 0 | 0 | 0 | 0 | 0 | 0 |
| ENSMUSG00 | Qrfpr | 2148 | 0 | 0 | 0 | 0 | 0 | 0 |
| ENSMUSG00 | Zfp420 | 3429 | 11.8025894 | 15.3735689 | 10.0159907 | 7.29217173 | 13.7945321 | 9.20290394 |
| ENSMUSG00 | Txndc9 | 6225 | 14.3439051 | 15.6131813 | 15.4014133 | 17.0875289 | 13.7955647 | 21.6838859 |
| ENSMUSG00 | Syt17 | 1609 | 9.31498829 | 9.07426474 | 2.00542855 | 10.2707125 | 9.45050341 | 3.79709086 |
| ENSMUSG00 | Cxcl2 | 1977 | 0 | 0 | 0.06707412 | 0 | 0 | 0 |
| ENSMUSG00 | Rpl27a-ps2 | 449 | 0 | 0 | 0 | 0 | 0 | 0 |
| ENSMUSG00 | Btnl4 | 1760 | 0 | 0 | 0 | 0 | 0 | 0 |
| ENSMUSG00 | Nrf1 | 9580 | 5.60228588 | 4.00883031 | 3.8618938 | 3.67599953 | 4.43151823 | 3.96877319 |
| ENSMUSG00 | Parw2 | 3140 | 0 | 0 | 0 | 0.03447329 | 0.45501227 | 0.01737238 |
| ENSMUSG00 | Rpl10-ps3 | 727 | 0.49215863 | 0.39768728 | 0.24320134 | 0.14889427 | 1.26337657 | 0.41268368 |
| ENSMUSG00 | Map2k5 | 2706 | 20.0981143 | 15.6347732 | 20.0264078 | 17.3876519 | 17.7064918 | 15.7640561 |
| ENSMUSG00 | Znrf2 | 6389 | 19.8684065 | 20.7181338 | 18.1539578 | 17.5186261 | 18.385149 | 15.96179 |
| ENSMUSG00 | Zfp82 | 3565 | 1.85116683 | 1.36516989 | 2.38057612 | 3.79544648 | 2.24716438 | 0.97163512 |
| ENSMUSG00 | Dhcr7 | 4145 | 2.23474716 | 4.94070878 | 3.80701036 | 3.56032719 | 4.66562037 | 4.36262625 |
| ENSMUSG00 | Gm5471 | 341 | 0 | 0.14130921 | 0.25924835 | 0 | 0 | 0 |
| ENSMUSG00 | Gm8369 | 1369 | 0.08711939 | 0 | 0.19372612 | 0.21085198 | 0 | 0 |
| ENSMUSG00 | Wdr91 | 2973 | 4.35932951 | 4.5544535 | 1.85846969 | 4.68471887 | 6.57352908 | 3.06415389 |
| ENSMUSG00 | Kl | 5105 | 158.562456 | 227.896858 | 71.8918516 | 213.057228 | 180.546411 | 143.137093 |
| ENSMUSG00 | Olfr869 | 1069 | 0 | 0 | 0 | 0 | 0 | 0 |
| ENSMUSG00 | Scp2-ps2 | 431 | 0.73792075 | 1.56522086 | 2.56391201 | 1.75805785 | 1.3022973 | 1.70862009 |
| ENSMUSG00 | Rnf207 | 5012 | 0.07138853 | 0 | 0.0088192 | 0 | 0 | 0.00544187 |
| ENSMUSG00 | Pip | 567 | 0 | 0 | 0 | 0 | 0 | 0 |
| ENSMUSG00 | Fam133b | 1544 | 7.7760072 | 6.9283615 | 10.1343604 | 10.5395091 | 4.26321404 | 11.6058536 |
| ENSMUSG00 | Gm10042 | 76 | 0 | 0 | 0 | 0 | 0 | 0 |
| ENSMUSG00 | Olfr490 | 944 | 0 | 0 | 0 | 0 | 0 | 0 |
| ENSMUSG00 | Olfr801 | 956 | 0 | 0 | 0 | 0 | 0 | 0 |
| ENSMUSG00 | Olfr933 | 1002 | 0 | 0 | 0 | 0 | 0 | 0 |
| ENSMUSG00 | Rps12-ps2 | 398 | 0 | 0 | 0 | 0 | 0 | 0 |
| ENSMUSG00 | Olfr892-ps1 | 922 | 0 | 0 | 0 | 0 | 0 | 0 |
| ENSMUSG00 | Mup5 | 918 | 0 | 0 | 0 | 0 | 0 | 0 |
| ENSMUSG00 | Olfr968 | 944 | 0 | 0 | 0 | 0 | 0 | 0 |
| ENSMUSG00 | Dux | 3138 | 0 | 0 | 0 | 0 | 0 | 0 |
| ENSMUSG00 | Gm15590 | 1469 | 0 | 0 | 0.0601795 | 0.19649854 | 0.03473545 | 0.11140084 |
| ENSMUSG00 | Rpl23a | 905 | 0.17571483 | 0.53244688 | 0.29305089 | 0.11960899 | 0.05638274 | 0.12055089 |
| ENSMUSG00 | Dppa4 | 1600 | 0 | 0 | 0 | 0 | 0 | 0 |
| ENSMUSG00 | Rpl5 | 3987 | 1.08686917 | 1.47447854 | 1.16408164 | 0.89594242 | 1.35660794 | 1.38870122 |
| ENSMUSG00 | Gm10043 | 521 | 4.27314186 | 2.21972094 | 6.44786961 | 4.0168111 | 3.52581484 | 9.37074938 |
| ENSMUSG00 | Defb50 | 371 | 0 | 0.77929556 | 0 | 0.77804948 | 0.96276181 | 0 |
| ENSMUSG00 | Tmed9 | 1497 | 51.1217769 | 53.8194598 | 42.3416452 | 43.2406065 | 38.9259327 | 43.7633156 |
| ENSMUSG00 | Gpc6 | 10198 | 0.08186557 | 0.0378007 | 0.13003092 | 0 | 0.12008561 | 0.13105092 |
| ENSMUSG00 | Cela2a | 1715 | 0 | 0 | 0 | 0 | 0.17851793 | 0 |
| ENSMUSG00 | Gm5801 | 815 | 0.14633919 | 0 | 0.10847078 | 0.22136224 | 0.1252181 | 0.16732908 |
| ENSMUSG00 | Serhl | 5886 | 7.84167735 | 7.48257019 | 8.9214729 | 7.98758138 | 7.3860809 | 7.77090894 |
| ENSMUSG00 | Tmod3 | 3568 | 22.2398932 | 24.7819847 | 28.567671 | 28.4773087 | 29.8893289 | 21.3044898 |
| ENSMUSG00 | Vmn1r21 | 1029 | 0 | 0 | 0 | 0 | 0 | 0 |
| ENSMUSG00 | Anks1b | 10147 | 0 | 0 | 0 | 0 | 0 | 0 |
| ENSMUSG00 | Gm6462 | 305 | 0 | 0 | 0 | 0 | 0 | 0 |
| ENSMUSG00 | Fbxo18 | 4787 | 61.2069963 | 40.8381859 | 57.7384843 | 41.5165869 | 58.7650741 | 37.6386603 |
| ENSMUSG00 | Vmn1r50 | 932 | 0 | 0 | 0 | 0 | 0 | 0 |
| ENSMUSG00 | Rpl30 | 1633 | 6.69489108 | 7.11141002 | 8.76999212 | 11.2687341 | 3.87462989 | 16.167698 |
| ENSMUSG00 | Rpl28-ps1 | 413 | 0 | 0.35002258 | 0.10702625 | 0.3494629 | 0 | 0.66040289 |
| ENSMUSG00 | Gm6635 | 377 | 0 | 0 | 0 | 0 | 0 | 0 |
| ENSMUSG00 | Gm5062 | 1011 | 0 | 0 | 0 | 0 | 0 | 0 |
| ENSMUSG00 | AY761184 | 501 | 0 | 0 | 0 | 0 | 0 | 0 |
| ENSMUSG00 | Adra2b | 7948 | 0 | 0 | 0 | 0 | 0 | 0 |
| ENSMUSG00 | Olfr643 | 1203 | 0 | 0 | 0 | 0 | 0 | 0 |
| ENSMUSG00 | Gm5265 | 341 | 0 | 0 | 0 | 0 | 0 | 0.07998428 |
| ENSMUSG00 | Gda | 7340 | 0 | 0 | 0 | 0 | 0 | 0.00371589 |
| ENSMUSG00 | Gm17383 | 1039 | 0.03826322 | 0 | 0 | 0.17363833 | 0 | 0 |
| ENSMUSG00 | Capn11 | 3102 | 0.03844824 | 0 | 0 | 0 | 0 | 0 |
| ENSMUSG00 | Olfr875 | 1003 | 0 | 0 | 0 | 0 | 0 | 0 |
| ENSMUSG00 | Vmn1r63 | 2170 | 0 | 0 | 0 | 0 | 0 | 0 |
| ENSMUSG00 | Zfp110 | 4455 | 24.5047249 | 21.5892567 | 23.792597 | 23.4553538 | 25.0607655 | 19.7993678 |
| ENSMUSG00 | Gm10045 | 1037 | 0 | 0 | 0 | 0 | 0 | 0.02630148 |
| ENSMUSG00 | Olfr1431 | 938 | 0 | 0 | 0 | 0 | 0 | 0 |
| ENSMUSG00 | Speer4f | 1251 | 0 | 0.07703668 | 0 | 0 | 0 | 0 |
| ENSMUSG00 | Olfr150 | 938 | 0 | 0 | 0 | 0 | 0 | 0 |
| ENSMUSG00 | Eif4b | 2737 | 39.7554805 | 49.295593 | 39.6476159 | 33.1290383 | 53.4127025 | 29.5865561 |
| ENSMUSG00 | Samd12 | 1881 | 3.19142879 | 7.04480148 | 2.72589782 | 5.14087609 | 5.77810635 | 2.79851429 |
| ENSMUSG00 | Olfr58 | 1183 | 0 | 0 | 0 | 0 | 0 | 0 |
| ENSMUSG00 | Gm7452 | 3767 | 0.02110724 | 0 | 0.03520189 | 0 | 0.04063688 | 0.06516373 |
| ENSMUSG00 | Olfr69 | 2304 | 0 | 0 | 0 | 0 | 0 | 0 |
| ENSMUSG00 | En1 | 3926 | 0 | 0 | 0 | 0 | 0 | 0 |
| ENSMUSG00 | Nkx2-9 | 2109 | 0 | 0 | 0 | 0 | 0 | 0 |
| ENSMUSG00 | 4932411N23I | 3622 | 0 | 0 | 0 | 0 | 0 | 0 |
| ENSMUSG00 | Tubb2a | 1580 | 57.2680213 | 80.3920642 | 35.3334986 | 55.1050465 | 61.8129641 | 44.5543316 |
| ENSMUSG00 | Olfr498 | 992 | 0 | 0 | 0 | 0 | 0 | 0 |
| ENSMUSG00 | Vmn2r51 | 2553 | 0 | 0 | 0 | 0 | 0 | 0 |
| ENSMUSG00 | Gcap14 | 7539 | 10.6573586 | 11.2236891 | 13.4068891 | 16.3970135 | 15.8040307 | 10.3324539 |
| ENSMUSG00 | Olfr846 | 941 | 0 | 0 | 0 | 0 | 0 | 0 |
| ENSMUSG00 | Gm8724 | 386 | 0 | 0 | 0 | 0 | 0 | 0.07065969 |
| ENSMUSG00 | Memo1 | 1504 | 65.0255865 | 54.4340194 | 60.3660808 | 51.0762452 | 39.5929392 | 96.2954351 |
| ENSMUSG00 | 0610030E20I | 4628 | 11.2274013 | 10.9117095 | 6.9053441 | 8.88797121 | 10.3640435 | 9.36460712 |
| ENSMUSG00 | Egln2 | 3116 | 17.1219046 | 19.1601418 | 16.682082 | 10.8964069 | 18.307923 | 14.1625052 |
| ENSMUSG00 | Fcer1g | 795 | 0.15002068 | 0 | 0.38919862 | 0.2269311 | 0 | 0.75476989 |
| ENSMUSG00 | Gm4763 | 974 | 0 | 0 | 0 | 0 | 0 | 0 |
| ENSMUSG00 | Gm11937 | 395 | 0 | 0 | 0 | 0 | 0 | 0 |
| ENSMUSG00 | Cd300c | 1210 | 0 | 0 | 0 | 0 | 0 | 0 |
| ENSMUSG00 | Lin9 | 3190 | 4.87285043 | 3.7008396 | 2.06460019 | 5.29354133 | 4.22287248 | 5.24117676 |
| ENSMUSG00 | Vmn1r15 | 899 | 0 | 0 | 0 | 0 | 0 | 0 |
| ENSMUSG00 | Gm10047 | 1329 | 0 | 0 | 0 | 0.43439632 | 0 | 0.57463499 |
| ENSMUSG00 | Kcnt1 | 6991 | 0 | 0 | 0 | 0 | 0 | 0 |
| ENSMUSG00 | Prr19 | 1290 | 0 | 0 | 0.06852999 | 0 | 0 | 0 |
| ENSMUSG00 | Kcnj14 | 2636 | 0 | 0 | 0 | 0 | 0 | 0 |
| ENSMUSG00 | Gm9174 | 257 | 0 | 0 | 0 | 0 | 0 | 0 |
| ENSMUSG00 | Zfp958 | 1398 | 78.2028408 | 60.8017768 | 51.0629302 | 72.2931379 | 65.4071999 | 175.022023 |
| ENSMUSG00 | Gm10048 | 370 | 0 | 0 | 0 | 0 | 0 | 0 |
| ENSMUSG00 | Osm | 2269 | 0 | 0 | 0 | 0 | 0 | 0 |
| ENSMUSG00 | Thra | 3855 | 7.80671822 | 8.88730489 | 7.95747835 | 6.7296991 | 6.52555208 | 7.92414941 |
| ENSMUSG00 | Rnf169 | 7151 | 2.12926151 | 1.99457235 | 1.71219557 | 4.23337091 | 2.79000321 | 3.4822746 |
| ENSMUSG00 | Btnl6 | 1611 | 0 | 0 | 0 | 0 | 0 | 0 |
| ENSMUSG00 | Hist1h1b | 742 | 0 | 0 | 0 | 0 | 0 | 0 |
| ENSMUSG00 | Tomm20 | 1404 | 7.44707363 | 8.20267783 | 10.2948737 | 11.46196 | 8.10461667 | 8.00366907 |
| ENSMUSG00 | Cds2 | 9366 | 11.4393573 | 12.1520795 | 10.495932 | 12.6745597 | 14.1104328 | 13.0199565 |
| ENSMUSG00 | Nfe2 | 4097 | 0.13584982 | 0 | 0 | 0 | 0 | 0 |
| ENSMUSG00 | Nap1l1 | 3024 | 16.6831034 | 16.3489714 | 12.9652893 | 18.625685 | 11.9129039 | 13.9259402 |
| ENSMUSG00 | Olfr1462 | 923 | 0 | 0 | 0 | 0 | 0 | 0 |
| ENSMUSG00 | Olfr97 | 1058 | 0 | 0 | 0 | 0 | 0 | 0 |
| ENSMUSG00 | Olfr340 | 938 | 0 | 0 | 0 | 0 | 0 | 0 |
| ENSMUSG00 | Col13a1 | 4778 | 0 | 0 | 0 | 0 | 0 | 0 |
| ENSMUSG00 | Olfr331 | 2066 | 0 | 0 | 0 | 0 | 0 | 0 |
| ENSMUSG00 | Gm12141 | 3954 | 0.10054497 | 0.04874703 | 0.10061118 | 0.14600726 | 0.06452501 | 0.12416376 |
| ENSMUSG00 | 0610039K10I | 950 | 0.33478299 | 0.15216771 | 0.13958477 | 0.3418299 | 0.05371198 | 0 |
| ENSMUSG00 | Olfr8 | 932 | 0 | 0 | 0 | 0 | 0 | 0 |
| ENSMUSG00 | Ppp1r2-ps3 | 626 | 0 | 0 | 0 | 0 | 0 | 0 |
| ENSMUSG00 | Lilrb3 | 5578 | 0 | 0 | 0 | 0 | 0 | 0.1026833 |
| ENSMUSG00 | Olfr146 | 1060 | 0 | 0 | 0 | 0 | 0 | 0 |
| ENSMUSG00 | Olfr972 | 944 | 0 | 0 | 0 | 0 | 0 | 0 |
| ENSMUSG00 | Opn1sw | 3228 | 0.03694747 | 0 | 0.10954608 | 0.067067 | 0.18968913 | 0 |
| ENSMUSG00 | 2810428I15R | 1943 | 11.151177 | 11.8296104 | 15.8789946 | 10.9936029 | 7.77344686 | 17.71518 |
| ENSMUSG00 | Abi1 | 5486 | 23.1749958 | 20.3163034 | 21.7383472 | 18.7842361 | 16.4166156 | 27.2249224 |
| ENSMUSG00 | Rps27a-ps2 | 525 | 0 | 0 | 0 | 0 | 0 | 0.41561355 |
| ENSMUSG00 | Olfr952 | 1055 | 0 | 0 | 0 | 0 | 0 | 0 |
| ENSMUSG00 | Gm4832 | 290 | 0 | 0 | 0 | 0 | 0 | 0 |
| ENSMUSG00 | Gm5582 | 521 | 0 | 0 | 0 | 0 | 0.1958786 | 0 |
| ENSMUSG00 | Zfp516 | 7810 | 1.41002152 | 1.91265007 | 1.8790028 | 2.17139064 | 2.98579433 | 0.55876342 |
| ENSMUSG00 | Zfp708 | 4034 | 6.96755695 | 4.15688693 | 4.21856955 | 7.37025401 | 6.31188932 | 9.37100892 |
| ENSMUSG00 | Olfr1025-ps1 | 1860 | 0 | 0 | 0 | 0 | 0 | 0 |
| ENSMUSG00 | Deaf1 | 3130 | 13.5270245 | 17.257828 | 16.6357096 | 11.3203092 | 15.2100986 | 8.25210331 |
| ENSMUSG00 | Col25a1 | 8301 | 1.81512193 | 0.57468471 | 2.0074804 | 2.56021253 | 0.62084857 | 2.08642283 |
| ENSMUSG00 | Rai1 | 1644 | 8.27030069 | 10.7862596 | 5.75376789 | 11.654237 | 4.22116004 | 12.6750757 |
| ENSMUSG00 | Olfr1413 | 1081 | 0 | 0 | 0 | 0 | 0 | 0 |
| ENSMUSG00 | Gm10051 | 909 | 0.13120621 | 0 | 0 | 0.03969422 | 0 | 0.09001531 |
| ENSMUSG00 | Zfp353 | 5106 | 0 | 0 | 0 | 0 | 0 | 0 |
| ENSMUSG00 | Pla2g2a | 963 | 0 | 0 | 0 | 0 | 0 | 0 |
| ENSMUSG00 | C1qtnf3 | 2731 | 0.56772748 | 0.0352885 | 0.17803745 | 0.13212027 | 0.28026204 | 1.56796718 |
| ENSMUSG00 | Slc10a5 | 3922 | 0 | 0.30715478 | 0.1127023 | 0.06439937 | 1.69133831 | 0.66760973 |
| ENSMUSG00 | Gm10052 | 1015 | 0 | 0 | 0 | 0 | 0 | 0 |
| ENSMUSG00 | Vmn1r6 | 1057 | 0 | 0 | 0 | 0 | 0 | 0 |
| ENSMUSG00 | 1700011I03R | 1476 | 0 | 0 | 0 | 0 | 0 | 0 |
| ENSMUSG00 | Gm10053 | 475 | 0.50217449 | 0.71011599 | 1.48890419 | 0.83558419 | 0.75196764 | 1.14840586 |
| ENSMUSG00 | Gm2174 | 401 | 0 | 0 | 0 | 0 | 0 | 0 |
| ENSMUSG00 | Gm5185 | 296 | 0 | 0 | 0 | 0 | 0 | 0 |
| ENSMUSG00 | Gm16176 | 2784 | 0 | 0 | 0 | 0 | 0 | 0 |
| ENSMUSG00 | 1700018C11I | 684 | 0 | 0 | 0 | 0 | 0 | 0 |
| ENSMUSG00 | Gm10056 | 2871 | 0.26309792 | 0.70492183 | 0.53885911 | 0 | 0.58651007 | 0.56050286 |
| ENSMUSG00 | Olfr1472 | 944 | 0 | 0 | 0 | 0 | 0 | 0 |
| ENSMUSG00 | Cfi | 2061 | 9.5289701 | 9.91317393 | 27.7735986 | 13.1828139 | 14.7062917 | 16.2377401 |
| ENSMUSG00 | Gm4822 | 479 | 0 | 0 | 0 | 0 | 0 | 0 |
| ENSMUSG00 | Fam57b | 2795 | 0 | 0 | 0 | 0 | 0 | 0 |
| ENSMUSG00 | Kcnc1 | 9722 | 0.00408923 | 0 | 0.00454658 | 0 | 0 | 0 |
| ENSMUSG00 | Usp17l5 | 2450 | 0.01622673 | 0 | 0 | 0.10309156 | 0 | 0 |
| ENSMUSG00 | Cecr5 | 3347 | 10.3100559 | 11.2439831 | 10.0500755 | 12.796351 | 11.5560182 | 13.8125227 |
| ENSMUSG00 | Olfr1406 | 1062 | 0 | 0 | 0 | 0 | 0 | 0 |
| ENSMUSG00 | Gm10057 | 981 | 0 | 0 | 0 | 0 | 0 | 0 |
| ENSMUSG00 | 1300010F03I | 8335 | 12.4298479 | 12.7128958 | 14.8700621 | 12.3635656 | 16.3455817 | 9.53876106 |
| ENSMUSG00 | Zfp799 | 1941 | 32.668723 | 17.7254609 | 22.7954894 | 24.2963587 | 20.7943654 | 25.6446248 |
| ENSMUSG00 | Grin2a | 7081 | 0 | 0 | 0 | 0 | 0 | 0 |
| ENSMUSG00 | Hnrnpa3 | 5509 | 2.29483442 | 2.24794257 | 2.31881151 | 3.34687325 | 1.75984959 | 3.24780602 |
| ENSMUSG00 | Sh2d3c | 5215 | 2.81299565 | 2.18063286 | 3.2462715 | 1.70204851 | 3.13105278 | 3.24785254 |
| ENSMUSG00 | Gm5409 | 736 | 0 | 0 | 0 | 0 | 0 | 0 |
| ENSMUSG00 | Kcp | 7709 | 3.99689184 | 3.66289468 | 4.55836882 | 2.19983927 | 2.84619816 | 5.09121882 |
| ENSMUSG00 | Olfr1201 | 923 | 0 | 0 | 0 | 0 | 0 | 0 |
| ENSMUSG00 | 9630013D21I | 3407 | 1.23688903 | 1.30118952 | 0.53192708 | 1.26027687 | 0.79377691 | 1.17680422 |
| ENSMUSG00 | Olfr128 | 1017 | 0 | 0 | 0 | 0 | 0 | 0 |
| ENSMUSG00 | Olfr482 | 971 | 0 | 0 | 0 | 0 | 0 | 0 |
| ENSMUSG00 | Rpl18a-ps1 | 522 | 0.07615992 | 0 | 0 | 0.48385884 | 0.09775168 | 0.1567508 |
| ENSMUSG00 | Gm5900 | 374 | 0.74408653 | 0.64420377 | 0.47274698 | 1.0612366 | 0.54573664 | 0.58341474 |
| ENSMUSG00 | Gm5506 | 1755 | 5.86704812 | 7.11127551 | 6.62397989 | 5.7361199 | 5.34977388 | 0.55947978 |
| ENSMUSG00 | Olfr192 | 922 | 0 | 0 | 0 | 0 | 0 | 0 |
| ENSMUSG00 | Klk1b9 | 865 | 7.72129564 | 8.80168539 | 3.52592736 | 6.21528859 | 7.19678364 | 8.89184771 |
| ENSMUSG00 | Olfr15 | 938 | 0 | 0 | 0 | 0 | 0 | 0 |
| ENSMUSG00 | Gm10058 | 884 | 0 | 0 | 0 | 0 | 0 | 0 |
| ENSMUSG00 | Frem1 | 16438 | 0 | 0 | 0 | 0 | 0 | 0 |
| ENSMUSG00 | Gm15431 | 194 | 0 | 0 | 0 | 0 | 0 | 0 |
| ENSMUSG00 | Rad51l1 | 2055 | 0.61906344 | 0.28138068 | 0.06452824 | 0.33360528 | 0.72008024 | 0.03981699 |
| ENSMUSG00 | Gm10059 | 337 | 0 | 0 | 0 | 0 | 0 | 0.08093365 |
| ENSMUSG00 | Olfr749 | 1019 | 0 | 0 | 0 | 0 | 0 | 0 |
| ENSMUSG00 | Rpl18 | 3751 | 1.58979528 | 1.88840496 | 2.46285929 | 1.98157856 | 1.74123597 | 2.55222564 |
| ENSMUSG00 | Pth | 714 | 0 | 0 | 0 | 0 | 0 | 0 |
| ENSMUSG00 | Gm10060 | 562 | 3.32474659 | 2.48648901 | 4.95501087 | 4.49420483 | 5.9016271 | 4.46488756 |
| ENSMUSG00 | Olfr698 | 935 | 0 | 0 | 0 | 0 | 0 | 0 |
| ENSMUSG00 | Fcgr4 | 1251 | 0 | 0 | 0 | 0 | 0 | 0 |
| ENSMUSG00 | Olfr157 | 1765 | 0 | 0 | 0 | 0 | 0 | 0 |
| ENSMUSG00 | Olfr1504 | 1084 | 0 | 0 | 0 | 0 | 0 | 0 |
| ENSMUSG00 | Olfr961 | 948 | 0 | 0 | 0 | 0 | 0 | 0 |
| ENSMUSG00 | Ifitm6 | 539 | 0 | 0 | 0 | 0 | 0 | 0 |
| ENSMUSG00 | Olfr1263 | 1028 | 0 | 0 | 0 | 0 | 0 | 0 |
| ENSMUSG00 | Gm10061 | 164 | 0 | 0 | 0 | 0 | 0 | 0 |
| ENSMUSG00 | Gm6772 | 1419 | 0 | 0 | 0 | 0 | 0 | 0 |
| ENSMUSG00 | Nap1l4 | 2643 | 74.1259962 | 65.2513343 | 73.5192213 | 49.2698065 | 64.6565769 | 55.8082668 |
| ENSMUSG00 | Il28a | 697 | 0 | 0 | 0 | 0 | 0 | 0 |
| ENSMUSG00 | Gm9242 | 889 | 0.04471933 | 0 | 0 | 0 | 0 | 0.06136027 |
| ENSMUSG00 | Olfr814 | 932 | 0 | 0 | 0 | 0 | 0 | 0 |
| ENSMUSG00 | Olfr539 | 959 | 0 | 0 | 0 | 0 | 0 | 0 |
| ENSMUSG00 | Zfp945 | 9107 | 9.48159698 | 12.920972 | 10.2750963 | 14.679255 | 10.6512727 | 20.446246 |
| ENSMUSG00 | Ntrk3 | 5221 | 0 | 0 | 0 | 0 | 0 | 0 |
| ENSMUSG00 | Mfsd4 | 6870 | 4.9535213 | 6.06012897 | 3.38430414 | 5.57249625 | 4.93180692 | 2.48131725 |
| ENSMUSG00 | Rpl26-ps4 | 437 | 0 | 0 | 0 | 0 | 0 | 0.18724009 |
| ENSMUSG00 | Gm8129 | 427 | 0 | 0 | 0 | 0 | 0 | 0 |
| ENSMUSG00 | Krt40 | 1313 | 0 | 0 | 0 | 0 | 0 | 0 |
| ENSMUSG00 | Pde1a | 8396 | 92.087254 | 59.8198293 | 138.217626 | 86.2814683 | 86.5916868 | 113.925869 |
| ENSMUSG00 | Gm8991 | 900 | 0 | 0 | 0 | 0 | 0 | 0 |
| ENSMUSG00 | Skap2 | 1625 | 28.4037618 | 39.2904836 | 24.4810208 | 34.5055367 | 38.5602397 | 27.1739329 |
| ENSMUSG00 | Mtfmt | 3866 | 6.519652 | 4.68652412 | 5.27083539 | 8.66118399 | 8.02484131 | 7.04794739 |
| ENSMUSG00 | Fam19a1 | 4551 | 1.22297677 | 0.53999309 | 1.46659598 | 0.04757026 | 0.41484858 | 0.17380016 |
| ENSMUSG00 | Olfr913 | 1273 | 0 | 0 | 0 | 0 | 0 | 0 |
| ENSMUSG00 | Gm12715 | 1964 | 0.95137861 | 1.12860302 | 0.94525326 | 0.75324024 | 0.72746361 | 0.48605518 |
| ENSMUSG00 | Lep | 3350 | 0 | 0 | 0 | 0 | 0 | 0 |
| ENSMUSG00 | Il1rapl2 | 4051 | 0 | 0 | 0 | 0 | 0 | 0 |
| ENSMUSG00 | Olfr1130 | 1123 | 0 | 0 | 0 | 0 | 0 | 0 |
| ENSMUSG00 | Vmn1r71 | 2036 | 0 | 0 | 0 | 0 | 0 | 0 |
| ENSMUSG00 | Hnrnpm | 5031 | 50.5576554 | 55.389027 | 55.2369285 | 58.5660854 | 57.7102124 | 42.0639885 |
| ENSMUSG00 | Ddn | 3738 | 0 | 0 | 0 | 0 | 0 | 0 |
| ENSMUSG00 | Gm13084 | 2244 | 0 | 0 | 0 | 0 | 0 | 0 |
| ENSMUSG00 | Olfr113 | 958 | 0 | 0 | 0 | 0 | 0 | 0 |
| ENSMUSG00 | Olfr494 | 946 | 0 | 0 | 0 | 0 | 0 | 0 |
| ENSMUSG00 | Gm6802 | 1982 | 0 | 0 | 0 | 0 | 0 | 0 |
| ENSMUSG00 | Defb4 | 452 | 0 | 0 | 0 | 0 | 0 | 0 |
| ENSMUSG00 | Ppp1r2-ps4 | 1060 | 0.41255687 | 0.04545891 | 1.91819319 | 0.68079329 | 0.33696863 | 0.56607742 |
| ENSMUSG00 | Gm10062 | 1469 | 0.02706296 | 0 | 0 | 0 | 0 | 0 |
| ENSMUSG00 | Foxb1 | 2982 | 0 | 0 | 0 | 0 | 0 | 0 |
| ENSMUSG00 | 9-Sep | 7548 | 12.1615533 | 8.19705773 | 12.5203419 | 11.6353599 | 9.63335792 | 11.2813227 |
| ENSMUSG00 | Gm12350 | 461 | 0 | 0.31357772 | 0 | 0.15653815 | 0 | 0.29582038 |
| ENSMUSG00 | Olfr345 | 935 | 0 | 0 | 0 | 0 | 0 | 0 |
| ENSMUSG00 | Tpt1-ps5 | 518 | 0 | 0 | 0 | 0.06965646 | 0 | 0.10530749 |
| ENSMUSG00 | Gzmd | 959 | 0 | 0 | 0 | 0 | 0 | 0 |
| ENSMUSG00 | Usp47 | 5786 | 65.9407789 | 50.609922 | 70.2294405 | 66.6887978 | 80.6580078 | 62.8928857 |
| ENSMUSG00 | Zc3h4 | 3529 | 3.09797595 | 4.05536223 | 3.53213934 | 3.43541144 | 5.34988922 | 4.8381763 |
| ENSMUSG00 | Olfr888 | 944 | 0 | 0 | 0 | 0 | 0 | 0 |
| ENSMUSG00 | R74862 | 3883 | 0.46072537 | 0.1365055 | 0.34150278 | 0.17655391 | 0.05256387 | 0.31608518 |
| ENSMUSG00 | Lsmd1 | 921 | 4.87770825 | 6.80156078 | 7.63093707 | 4.11358814 | 3.71201648 | 11.1349233 |
| ENSMUSG00 | Olfr224 | 1273 | 0 | 0 | 0 | 0 | 0 | 0 |
| ENSMUSG00 | Vpreb2 | 764 | 0 | 0 | 0 | 0 | 0 | 0 |
| ENSMUSG00 | Cdyl | 3938 | 2.22097657 | 3.82995338 | 3.14284308 | 2.40974548 | 2.87655039 | 3.01281566 |
| ENSMUSG00 | Rpl11 | 2241 | 1.68530596 | 1.41914555 | 1.06510465 | 1.15926248 | 0.95631762 | 1.98383141 |
| ENSMUSG00 | Gm5434 | 1450 | 0 | 0 | 0 | 0 | 0 | 0.0188101 |
| ENSMUSG00 | Olfr836 | 954 | 0 | 0 | 0 | 0 | 0 | 0 |
| ENSMUSG00 | Vpreb1 | 1684 | 0.14164661 | 0.11445711 | 0.44621813 | 0.19283753 | 0.0606014 | 0.37251586 |
| ENSMUSG00 | Hist1h3f | 1003 | 0.15854628 | 0.76867704 | 0.04406963 | 0.14389649 | 0.10174751 | 0.05438612 |
| ENSMUSG00 | Slc27a4 | 4399 | 9.32658693 | 11.0196774 | 7.11409522 | 5.20027648 | 11.6575376 | 4.38353488 |
| ENSMUSG00 | Olfr295 | 1048 | 0 | 0 | 0 | 0 | 0 | 0 |
| ENSMUSG00 | Tonsl | 5173 | 1.50629696 | 2.68271706 | 0.99118767 | 2.32269879 | 2.74218683 | 1.36030484 |
| ENSMUSG00 | Hopx | 2924 | 0.14955892 | 0.21423521 | 0 | 0.13573957 | 0.13960705 | 0.09327852 |
| ENSMUSG00 | Csf2ra | 1837 | 14.975935 | 12.6171359 | 15.2071665 | 12.1582937 | 10.1941644 | 20.4151491 |
| ENSMUSG00 | Eda | 7374 | 0 | 0.01306928 | 0.34766842 | 0.02935887 | 0.43594544 | 0 |
| ENSMUSG00 | Gm10063 | 383 | 0 | 0 | 0 | 0 | 0.13322814 | 0 |
| ENSMUSG00 | Zfp36l3 | 2613 | 0 | 0 | 0 | 0 | 0 | 0 |
| ENSMUSG00 | Slc14a1 | 4903 | 5.80561372 | 5.47416852 | 0.83841962 | 3.16444608 | 8.356961 | 1.86355377 |
| ENSMUSG00 | Taar7a | 2150 | 0 | 0 | 0 | 0 | 0 | 0 |
| ENSMUSG00 | Aldoart1 | 1913 | 1.3508135 | 0.93199078 | 1.0859836 | 0.90535188 | 0.66683711 | 0.86970883 |
| ENSMUSG00 | Olfr106-ps | 943 | 0 | 0 | 0 | 0 | 0 | 0 |
| ENSMUSG00 | Gm10064 | 388 | 0 | 0 | 0 | 0 | 0 | 0 |
| ENSMUSG00 | BC056474 | 1001 | 7.5459953 | 6.35425611 | 9.14064086 | 5.87549776 | 3.41535183 | 10.3812563 |
| ENSMUSG00 | Nrsn2 | 1452 | 0 | 0 | 0 | 0 | 0 | 0 |
| ENSMUSG00 | Fxn | 2600 | 1.95719289 | 2.29812263 | 3.2641361 | 1.98451245 | 2.04105503 | 3.24148597 |
| ENSMUSG00 | 5430401F13I | 755 | 0 | 0 | 0 | 0 | 0 | 0 |
| ENSMUSG00 | Olfr959 | 1054 | 0 | 0 | 0 | 0 | 0 | 0 |
| ENSMUSG00 | Olfr427 | 947 | 0 | 0 | 0 | 0 | 0 | 0 |
| ENSMUSG00 | Vmn1r33 | 2851 | 0 | 0 | 0 | 0 | 0 | 0 |
| ENSMUSG00 | Olfr1014 | 917 | 0 | 0 | 0 | 0 | 0 | 0 |
| ENSMUSG00 | Tas2r120 | 887 | 0 | 0 | 0 | 0 | 0 | 0 |
| ENSMUSG00 | Gfral | 2092 | 0 | 0 | 0 | 0 | 0 | 0 |
| ENSMUSG00 | Nkapl | 1454 | 0.32810575 | 0.13256243 | 1.64160903 | 0.52112994 | 0.5615007 | 0.30013358 |
| ENSMUSG00 | Olfr54 | 941 | 0 | 0 | 0 | 0 | 0 | 0 |
| ENSMUSG00 | Mamld1 | 3453 | 0 | 1.29781034 | 0.19201496 | 0.40752961 | 0 | 0 |
| ENSMUSG00 | Olfr776 | 938 | 0 | 0 | 0 | 0 | 0 | 0 |
| ENSMUSG00 | Tmprss9 | 3479 | 0.04570909 | 0 | 0.01270533 | 0.08297107 | 0.20533753 | 0.07839793 |
| ENSMUSG00 | Mrgprh | 1808 | 0 | 0 | 0 | 0 | 0 | 0 |
| ENSMUSG00 | Ppp2r5d | 3634 | 24.0239503 | 20.1550336 | 21.8454898 | 17.6339326 | 23.6456843 | 12.9168008 |
| ENSMUSG00 | Tas2r125 | 936 | 0 | 0 | 0 | 0 | 0 | 0 |
| ENSMUSG00 | Olfr434 | 1055 | 0 | 0 | 0 | 0 | 0 | 0 |
| ENSMUSG00 | Fxyd2 | 1002 | 752.576801 | 787.862756 | 697.039243 | 709.073392 | 473.139778 | 1212.36044 |
| ENSMUSG00 | Olfr802 | 938 | 0 | 0 | 0 | 0 | 0 | 0 |
| ENSMUSG00 | Gm8116 | 1202 | 0.06614889 | 0.04008855 | 0.07354716 | 0.21012838 | 0.08490246 | 0.13614629 |
| ENSMUSG00 | Zfp933 | 4464 | 10.9808485 | 8.40887958 | 9.10969883 | 17.6126288 | 10.9276916 | 4.09363985 |
| ENSMUSG00 | Olfr365 | 938 | 0 | 0 | 0 | 0 | 0 | 0 |
| ENSMUSG00 | Actg2 | 1767 | 0 | 0 | 0 | 0 | 0.02887741 | 0 |
| ENSMUSG00 | Gckr | 1993 | 0 | 0 | 0.04435709 | 0 | 0 | 0 |
| ENSMUSG00 | Max | 2564 | 7.44252365 | 8.40067849 | 5.27525896 | 7.6132551 | 4.19912843 | 5.47833043 |
| ENSMUSG00 | Bcas3 | 4800 | 9.09406617 | 7.87045222 | 5.99487496 | 7.65991737 | 9.78005537 | 3.39796547 |
| ENSMUSG00 | Hadhb | 5152 | 18.3112879 | 14.9460277 | 21.1743301 | 15.8979505 | 19.4419208 | 16.8772418 |
| ENSMUSG00 | Olfr156 | 1098 | 0 | 0 | 0 | 0 | 0 | 0 |
| ENSMUSG00 | Plac8l1 | 759 | 0 | 0 | 0 | 0 | 0 | 0 |
| ENSMUSG00 | Ptk2b | 8374 | 6.50406118 | 4.25817616 | 4.4339083 | 4.7310825 | 5.47799282 | 4.89210749 |
| ENSMUSG00 | Gm10065 | 344 | 0 | 0 | 0 | 0 | 0 | 0 |
| ENSMUSG00 | Olfr286 | 968 | 0 | 0 | 0 | 0 | 0 | 0 |
| ENSMUSG00 | Gm7331 | 863 | 0 | 0 | 0.35853175 | 0.12543005 | 0.05912674 | 0.03160445 |
| ENSMUSG00 | Spag11b | 441 | 0 | 0 | 0 | 0 | 0 | 0 |
| ENSMUSG00 | Gm4891 | 617 | 0.19330055 | 0.46858777 | 0.07163994 | 0 | 0.24810231 | 0 |
| ENSMUSG00 | Olfr385 | 938 | 0 | 0 | 0 | 0 | 0 | 0 |
| ENSMUSG00 | Olfr979 | 1044 | 0 | 0 | 0 | 0 | 0 | 0 |
| ENSMUSG00 | Mbtd1 | 11149 | 3.40893707 | 4.65916088 | 3.46113633 | 4.07456221 | 3.26781167 | 5.11292457 |
| ENSMUSG00 | Zfp426 | 7019 | 11.8603756 | 10.9018479 | 12.0596281 | 7.22257765 | 12.0387061 | 4.60470829 |
| ENSMUSG00 | B3gnt8 | 1473 | 3.91347228 | 2.58433736 | 2.67071557 | 2.03313625 | 4.12229377 | 1.62944213 |
| ENSMUSG00 | Plg | 2736 | 0 | 0 | 0 | 0 | 0 | 0 |
| ENSMUSG00 | 2610301B20I | 2627 | 19.0681026 | 22.3414871 | 20.0902172 | 24.5445807 | 17.4231668 | 32.8292384 |
| ENSMUSG00 | Rpl21-ps1 | 482 | 0.08248025 | 0 | 0 | 0 | 0 | 0 |
| ENSMUSG00 | Kbtbd2 | 6965 | 19.6465706 | 16.0990454 | 19.2863462 | 15.826367 | 16.2639705 | 13.2985894 |
| ENSMUSG00 | Olfr727 | 1034 | 0 | 0 | 0 | 0 | 0 | 0 |
| ENSMUSG00 | Nhs | 9402 | 0 | 0.04100101 | 0.00470132 | 0 | 0.08683493 | 0.02610846 |
| ENSMUSG00 | Arhgef12 | 15602 | 6.45434125 | 4.91376935 | 5.36586916 | 5.39079899 | 5.68413288 | 4.92628723 |
| ENSMUSG00 | Fcgr3 | 2303 | 0 | 0 | 0 | 0 | 0 | 0 |
| ENSMUSG00 | Olfr248 | 1009 | 0 | 0 | 0 | 0 | 0 | 0 |
| ENSMUSG00 | Olfr314 | 1130 | 0 | 0 | 0 | 0 | 0 | 0 |
| ENSMUSG00 | Gm12060 | 260 | 0 | 0 | 0 | 0 | 0 | 0 |
| ENSMUSG00 | Znhit1 | 1980 | 5.01963138 | 4.69696128 | 5.64801327 | 3.64465096 | 4.9222413 | 9.43592318 |
| ENSMUSG00 | Olfr700 | 950 | 0 | 0 | 0 | 0 | 0 | 0 |
| ENSMUSG00 | Uqcr10 | 826 | 77.3451055 | 85.5805213 | 90.0090798 | 60.7628619 | 56.8950268 | 170.945287 |
| ENSMUSG00 | Magea1 | 2403 | 0 | 0 | 0 | 0 | 0 | 0 |
| ENSMUSG00 | Tcea2 | 2859 | 6.36866389 | 6.0844021 | 6.36976542 | 5.67923051 | 7.88865272 | 7.72733746 |
| ENSMUSG00 | Vmn1r40 | 932 | 0 | 0 | 0 | 0 | 0 | 0 |
| ENSMUSG00 | Fbxw26 | 1814 | 0 | 0 | 0.07310117 | 0 | 0.08438761 | 0.10524944 |
| ENSMUSG00 | Trp53 | 2782 | 9.18863191 | 13.1811224 | 6.05352343 | 12.3213308 | 13.9396282 | 10.2255387 |
| ENSMUSG00 | Ccdc28a | 3452 | 4.04234463 | 3.44787115 | 5.67248449 | 5.58163725 | 5.23271641 | 4.44042505 |
| ENSMUSG00 | A830007P12I | 4037 | 19.4296664 | 19.8379655 | 22.2158879 | 14.9351242 | 18.1505761 | 12.6002483 |
| ENSMUSG00 | Ccdc154 | 2617 | 0.01519124 | 0 | 0 | 0 | 0.05849413 | 0.0104221 |
| ENSMUSG00 | Gm5292 | 581 | 0 | 0 | 0.0760789 | 0 | 0 | 0 |
| ENSMUSG00 | Gm16218 | 387 | 0.30818202 | 0 | 0 | 0.6526468 | 0 | 0 |
| ENSMUSG00 | Ube2nl | 458 | 0.17360472 | 0 | 0 | 0 | 0 | 0 |
| ENSMUSG00 | Nsmce2 | 3273 | 13.9320306 | 12.8673848 | 13.5049933 | 10.4398705 | 12.8306469 | 21.0830545 |
| ENSMUSG00 | Calcrl | 7657 | 0.88783952 | 0.70482977 | 1.01022888 | 0.74925494 | 0.85299414 | 0.01781027 |
| ENSMUSG00 | Olfr935 | 926 | 0 | 0 | 0 | 0 | 0 | 0 |
| ENSMUSG00 | Syn3 | 12264 | 1.11836601 | 1.22980727 | 0.50458725 | 1.28864445 | 0.91118528 | 0.97631822 |
| ENSMUSG00 | Ear5 | 758 | 0 | 0 | 0 | 0 | 0 | 0 |
| ENSMUSG00 | Olfr222 | 1114 | 0 | 0 | 0 | 0 | 0 | 0 |
| ENSMUSG00 | Olfr39 | 1096 | 0 | 0 | 0 | 0 | 0 | 0 |
| ENSMUSG00 | Sohlh1 | 1281 | 0 | 0 | 0 | 0 | 0 | 0 |
| ENSMUSG00 | Rhox3a | 1047 | 0 | 0 | 0 | 0 | 0 | 0 |
| ENSMUSG00 | 1500035N22I | 863 | 0 | 0.11167194 | 0 | 0.16724007 | 0.35476043 | 0 |
| ENSMUSG00 | Krtap8-1 | 578 | 0 | 0 | 0 | 0 | 0 | 0 |
| ENSMUSG00 | Gm11942 | 852 | 0.09332272 | 0.22622743 | 0.15564029 | 0.04234982 | 0.05989011 | 0.22408741 |
| ENSMUSG00 | Clec4a4 | 770 | 0 | 0 | 0 | 0 | 0 | 0 |
| ENSMUSG00 | Gm7361 | 2990 | 0 | 0 | 0 | 0 | 0 | 0 |
| ENSMUSG00 | Gm10068 | 1709 | 0 | 0.0281957 | 0.05172831 | 0.06333887 | 0 | 0 |
| ENSMUSG00 | Rnase11 | 712 | 0 | 0 | 0 | 0 | 0 | 0 |
| ENSMUSG00 | Reg1 | 767 | 0 | 0 | 0 | 0 | 0 | 0 |
| ENSMUSG00 | Stfa2l1 | 411 | 0 | 0 | 0 | 0 | 0 | 0 |
| ENSMUSG00 | Gm16379 | 874 | 0.04548682 | 0.11026646 | 0 | 0 | 0 | 0.18724009 |
| ENSMUSG00 | Gm10069 | 3691 | 0.15079294 | 0 | 0.14370689 | 0.12708387 | 0.0691227 | 0.16995847 |
| ENSMUSG00 | 4930524N10I | 916 | 0 | 0 | 0 | 0 | 0 | 0 |
| ENSMUSG00 | Krt4 | 2123 | 0 | 0 | 0 | 0 | 0 | 0 |
| ENSMUSG00 | Taf1b | 2004 | 11.9821907 | 14.883936 | 14.5133796 | 11.8653031 | 20.6753579 | 9.94898268 |
| ENSMUSG00 | Cdh24 | 2716 | 0.32202525 | 0 | 0.08137305 | 0 | 0 | 0.33139289 |
| ENSMUSG00 | Tdpoz1 | 1234 | 0 | 0 | 0 | 0 | 0 | 0 |
| ENSMUSG00 | Olfr108 | 1037 | 0 | 0 | 0 | 0 | 0 | 0 |
| ENSMUSG00 | Zfp637 | 2648 | 13.7973137 | 15.9226348 | 17.5772435 | 13.2446176 | 13.0841802 | 15.6046369 |
| ENSMUSG00 | 1700020N15I | 523 | 0 | 0 | 0 | 0 | 0 | 0 |
| ENSMUSG00 | 1700019M22 | 601 | 0 | 0 | 0 | 0 | 0 | 0 |

| | | | | | | | | |
|---|---|---|---|---|---|---|---|---|
| ENSMUSG00 | A830035A12f | 859 | 0 | 0 | 0.05145733 | 0 | 0 | 0 |
| ENSMUSG00 | Akap17b | 6781 | 11.6551977 | 7.76696376 | 13.3954855 | 8.92340196 | 6.48646748 | 12.4567996 |
| ENSMUSG00 | Rcan3 | 5288 | 17.5922512 | 15.892049 | 19.643406 | 13.858289 | 18.9612006 | 10.6560902 |
| ENSMUSG00 | Flot1 | 3006 | 24.0437337 | 34.1440858 | 29.8060997 | 28.0278024 | 33.9666594 | 27.9188506 |
| ENSMUSG00 | Hoxa3 | 2083 | 0 | 0 | 0 | 0 | 0 | 0 |
| ENSMUSG00 | Olfr1385 | 929 | 0 | 0 | 0 | 0 | 0 | 0 |
| ENSMUSG00 | Olfr837 | 943 | 0 | 0 | 0 | 0 | 0 | 0 |
| ENSMUSG00 | Olfr1354 | 932 | 0 | 0 | 0 | 0 | 0 | 0 |
| ENSMUSG00 | Ndufs8 | 956 | 153.324745 | 164.569805 | 189.383629 | 127.268467 | 91.4311525 | 214.48822 |
| ENSMUSG00 | Myl3 | 1298 | 0 | 0 | 0.30648427 | 0 | 0 | 0.02101282 |
| ENSMUSG00 | Kcnh7 | 15210 | 1.80611683 | 2.13211542 | 2.24060624 | 3.20966511 | 2.59996327 | 3.50930105 |
| ENSMUSG00 | Fdps | 1223 | 31.6613557 | 27.9347404 | 41.5273243 | 18.3508027 | 18.9836476 | 34.8571228 |
| ENSMUSG00 | Gm9000 | 861 | 0 | 0 | 10.2162216 | 0.04190714 | 0 | 8.33127773 |
| ENSMUSG00 | Olfr767 | 1085 | 0 | 0 | 0 | 0 | 0 | 0 |
| ENSMUSG00 | Taar2 | 1018 | 0 | 0 | 0 | 0 | 0 | 0 |
| ENSMUSG00 | Olfr682-ps1 | 894 | 0 | 0 | 0 | 0 | 0 | 0 |
| ENSMUSG00 | 4933400F03f | 941 | 0 | 0 | 0 | 0 | 0 | 0 |
| ENSMUSG00 | Six1b | 4494 | 5.38742493 | 5.28614063 | 6.02931237 | 5.25092504 | 5.83612754 | 6.18443645 |
| ENSMUSG00 | Rps26-ps1 | 412 | 0 | 0 | 0 | 0.26273333 | 0 | 0 |
| ENSMUSG00 | Gm10071 | 568 | 0 | 0 | 0 | 0 | 0 | 0.04801873 |
| ENSMUSG00 | Olfr781 | 935 | 0 | 0 | 0 | 0 | 0 | 0 |
| ENSMUSG00 | Nrm | 1641 | 0.79947036 | 0.70473773 | 0.37710287 | 0.41776895 | 0.1865681 | 1.18007275 |
| ENSMUSG00 | Eif4a1 | 3828 | 6.42858998 | 5.68972619 | 7.90968194 | 6.24931962 | 6.6782169 | 9.61167407 |
| ENSMUSG00 | Rgs3 | 9762 | 0.32987031 | 0.16782822 | 0.0090559 | 0 | 0.24044389 | 0.01955772 |
| ENSMUSG00 | Atl2 | 5091 | 21.0998445 | 23.4448669 | 31.2043653 | 27.5558808 | 23.0024617 | 27.7728796 |
| ENSMUSG00 | Gm8355 | 2023 | 0.03930349 | 0 | 0.0873986 | 0.08917955 | 0.05044624 | 0.01348227 |
| ENSMUSG00 | Fam159a | 590 | 0 | 0 | 0 | 0 | 0 | 0 |
| ENSMUSG00 | AU019823 | 6572 | 6.71463533 | 6.07829588 | 10.2030122 | 5.42438527 | 4.30913551 | 7.32497539 |
| ENSMUSG00 | Olfr847 | 938 | 0 | 0 | 0 | 0 | 0 | 0 |
| ENSMUSG00 | Dbp | 3079 | 1.96259599 | 2.73875524 | 2.92860538 | 0.76171903 | 1.65723858 | 2.17913649 |
| ENSMUSG00 | Gm5670 | 371 | 0 | 0 | 0 | 0 | 0 | 0 |
| ENSMUSG00 | Gm16380 | 457 | 0.60894609 | 0.10544079 | 0 | 0 | 0 | 0 |
| ENSMUSG00 | Kprp | 2727 | 0 | 0 | 0 | 0 | 0 | 0 |
| ENSMUSG00 | Sclt1 | 7366 | 4.71711783 | 6.32586065 | 2.13628239 | 5.75079015 | 4.89758997 | 6.46134204 |
| ENSMUSG00 | Rpl13-ps3 | 623 | 0 | 0 | 0 | 0 | 0 | 0.08755904 |
| ENSMUSG00 | Zfp874b | 3767 | 5.61452499 | 7.39361921 | 8.13163717 | 4.59765898 | 7.13854528 | 9.26772982 |
| ENSMUSG00 | Zfp341 | 3330 | 0.97896378 | 0.54987532 | 0.50440542 | 0.0216709 | 0.70486886 | 1.47430482 |
| ENSMUSG00 | Gm11567 | 1042 | 0 | 0 | 0 | 0 | 0 | 0 |
| ENSMUSG00 | Suv420h2 | 3443 | 17.1815728 | 19.6076693 | 14.8152561 | 21.4207665 | 18.2882799 | 23.4801135 |
| ENSMUSG00 | Kcng2 | 704 | 0 | 0 | 0 | 0 | 0 | 0 |
| ENSMUSG00 | Hydin | 16424 | 0 | 0 | 0 | 0 | 0 | 0 |
| ENSMUSG00 | Ntng1 | 6181 | 0 | 0 | 0 | 0 | 0 | 0 |
| ENSMUSG00 | Olfr826 | 941 | 0 | 0 | 0 | 0 | 0 | 0 |
| ENSMUSG00 | Olfr1393 | 935 | 0.51023077 | 0 | 0 | 0 | 0 | 0 |
| ENSMUSG00 | Tnip2 | 3693 | 9.48404504 | 9.69469984 | 10.7482413 | 13.0434686 | 12.7945909 | 13.367749 |
| ENSMUSG00 | Olfr960 | 1066 | 0 | 0 | 0 | 0 | 0 | 0 |
| ENSMUSG00 | Olfr1029 | 1060 | 0 | 0 | 0 | 0 | 0 | 0 |
| ENSMUSG00 | Olfr603 | 938 | 0 | 0 | 0 | 0 | 0 | 0 |
| ENSMUSG00 | Dph3b-ps | 248 | 0 | 0 | 0 | 0 | 0 | 0 |
| ENSMUSG00 | Zfp422 | 5850 | 12.4906963 | 13.5416258 | 15.2250793 | 11.3612181 | 10.6239531 | 15.8192908 |
| ENSMUSG00 | Irak4 | 5187 | 2.53693157 | 2.73121534 | 2.68432245 | 3.55464136 | 2.04617046 | 2.04020822 |
| ENSMUSG00 | Olfr1507 | 1603 | 0 | 0 | 0 | 0 | 0 | 0 |
| ENSMUSG00 | Ube4a | 7175 | 9.53023366 | 9.75817428 | 9.78293055 | 11.4054463 | 9.92080756 | 8.6784671 |
| ENSMUSG00 | Tsks | 2056 | 0.13535426 | 0.02343699 | 0 | 0 | 0.07445483 | 0.35817863 |
| ENSMUSG00 | Ptp4a3 | 4611 | 0.77596904 | 1.48394595 | 0.95861729 | 0.86077313 | 1.08449031 | 0.68615445 |
| ENSMUSG00 | Zfp930 | 1064 | 184.87793 | 145.691527 | 155.246511 | 252.777782 | 84.5004457 | 260.28824 |
| ENSMUSG00 | Dsc3 | 5522 | 0 | 0 | 0 | 0 | 0 | 0 |
| ENSMUSG00 | Tmem40 | 2384 | 0 | 0.40424868 | 0 | 0 | 0 | 0 |
| ENSMUSG00 | Adamts14 | 5666 | 0 | 0 | 0 | 0 | 0 | 0 |
| ENSMUSG00 | Mug1 | 5027 | 0 | 0 | 0 | 0 | 0 | 0 |
| ENSMUSG00 | Olfr1265 | 1851 | 0 | 0 | 0 | 0 | 0 | 0 |
| ENSMUSG00 | Rpl21-ps6 | 467 | 0 | 0.10318296 | 0 | 0 | 0 | 0 |
| ENSMUSG00 | 4930453N24 | 5938 | 13.7048617 | 13.1137236 | 12.230318 | 12.1043167 | 12.9327544 | 18.3040481 |
| ENSMUSG00 | Unc5c | 8029 | 0.01485446 | 0.0120031 | 0.03303164 | 0 | 0.20972355 | 0.52993445 |
| ENSMUSG00 | Grb2 | 4774 | 5.16304942 | 6.71218769 | 7.4070956 | 6.34874684 | 5.6862237 | 8.64972849 |
| ENSMUSG00 | Prh1 | 972 | 0 | 0 | 0 | 0 | 0 | 0 |
| ENSMUSG00 | Gdap10 | 5001 | 2.27355877 | 5.54033278 | 1.44953055 | 3.41267888 | 2.73446685 | 3.55044794 |
| ENSMUSG00 | 9430015G10 | 3723 | 3.94030951 | 3.0286402 | 3.00377821 | 5.00089578 | 3.87871726 | 2.13186141 |
| ENSMUSG00 | Olfr887 | 929 | 0 | 0 | 0 | 0 | 0 | 0 |
| ENSMUSG00 | Serpinb12 | 2489 | 0.25555954 | 0 | 0.26638314 | 0.27543545 | 0 | 0 |
| ENSMUSG00 | Olfr119 | 1233 | 0 | 0 | 0 | 0 | 0 | 0 |
| ENSMUSG00 | Hspa2 | 2518 | 6.31540596 | 6.12377343 | 7.03929271 | 6.79225143 | 10.9226436 | 4.02945425 |
| ENSMUSG00 | Gm14773 | 449 | 0 | 0 | 0 | 0 | 0 | 0 |
| ENSMUSG00 | Ntm | 8123 | 0 | 0 | 0 | 0 | 0 | 0 |
| ENSMUSG00 | Zfp74 | 6145 | 4.52222634 | 4.97939638 | 4.97045948 | 3.5113202 | 6.30252551 | 4.8069055 |
| ENSMUSG00 | Taok2 | 9636 | 12.3482932 | 10.8464501 | 9.76605685 | 11.001354 | 13.3126099 | 9.49630705 |
| ENSMUSG00 | Olfr996 | 1067 | 0 | 0 | 0 | 0 | 0 | 0 |
| ENSMUSG00 | Nptx2 | 2546 | 0 | 0 | 0 | 0 | 0 | 0 |
| ENSMUSG00 | Fcrl1 | 2031 | 0 | 0 | 0 | 0 | 0 | 0 |
| ENSMUSG00 | Atxn7l3 | 6610 | 1.12470119 | 1.32676739 | 1.51128843 | 0.97710832 | 1.46671882 | 1.91871517 |
| ENSMUSG00 | Olfr484 | 1097 | 0 | 0 | 0 | 0 | 0 | 0 |
| ENSMUSG00 | Gm10072 | 802 | 0 | 0.06008285 | 0 | 0 | 0 | 0 |
| ENSMUSG00 | Chpt1 | 5097 | 168.974442 | 214.017006 | 146.758444 | 225.588878 | 163.540686 | 193.565926 |
| ENSMUSG00 | Kif13b | 5492 | 5.36394957 | 4.72037616 | 8.72447156 | 8.86940279 | 11.0842073 | 7.93606583 |
| ENSMUSG00 | Olfr121 | 1240 | 0 | 0 | 0 | 0 | 0 | 0 |
| ENSMUSG00 | Gm10073 | 344 | 0 | 0 | 0 | 0 | 0 | 0 |
| ENSMUSG00 | Vmn1r213 | 1154 | 0 | 0 | 0 | 0 | 0 | 0 |
| ENSMUSG00 | 4930473A02f | 1395 | 0.17099131 | 0.48359153 | 0.63371818 | 0.28451791 | 0.36578047 | 0.13686199 |
| ENSMUSG00 | Olfr317 | 1127 | 0 | 0 | 0 | 0 | 0 | 0 |
| ENSMUSG00 | H2afj | 1826 | 24.6240134 | 24.4890571 | 26.6760301 | 26.5971697 | 23.3335289 | 30.3068144 |
| ENSMUSG00 | Ctf2 | 1751 | 0 | 0.05503877 | 0.42914411 | 0 | 0.17484766 | 0.09345964 |
| ENSMUSG00 | Rpl3 | 1351 | 4.38457927 | 3.10304994 | 4.15516955 | 3.8459026 | 3.55031779 | 4.70391632 |
| ENSMUSG00 | Dhps | 2573 | 10.8157157 | 8.82075952 | 13.2107335 | 10.2510589 | 7.19882411 | 9.83710269 |
| ENSMUSG00 | Tmem26 | 5393 | 0 | 0 | 0.44259216 | 0.13381066 | 0.13246231 | 0 |
| ENSMUSG00 | Olfr235 | 1078 | 0 | 0 | 0 | 0 | 0 | 0 |
| ENSMUSG00 | Olfr193 | 1006 | 0 | 0 | 0 | 0 | 0 | 0 |
| ENSMUSG00 | Olfr497 | 944 | 0 | 0 | 0 | 0 | 0 | 0 |
| ENSMUSG00 | Alox5ap | 877 | 0.13599366 | 0.1648339 | 0.35280833 | 0 | 0 | 0 |
| ENSMUSG00 | Gm12222 | 519 | 0 | 0 | 0 | 0 | 0 | 0 |
| ENSMUSG00 | Defa26 | 370 | 0 | 0 | 0 | 0 | 0 | 0 |
| ENSMUSG00 | Psma3 | 3888 | 19.100627 | 22.5935916 | 19.5884196 | 20.2311875 | 17.4287621 | 21.185548 |
| ENSMUSG00 | Hist1h2aa | 389 | 0 | 0 | 0 | 0 | 0 | 0 |
| ENSMUSG00 | Olfr748 | 923 | 0 | 0 | 0 | 0 | 0 | 0 |
| ENSMUSG00 | Gm10074 | 271 | 0 | 0 | 0 | 0.13314408 | 0 | 0 |
| ENSMUSG00 | Olfr605 | 959 | 0 | 0 | 0 | 0 | 0 | 0 |
| ENSMUSG00 | Rp2h | 5407 | 12.9258618 | 14.2411568 | 16.1127858 | 17.0967631 | 14.0801466 | 14.1594068 |
| ENSMUSG00 | Hist1h4a | 538 | 0 | 0 | 0 | 0 | 0 | 0 |
| ENSMUSG00 | Amd-ps3 | 1335 | 0 | 0 | 0.09932998 | 0 | 0 | 0 |
| ENSMUSG00 | Prmt7 | 6063 | 7.83569177 | 6.91443257 | 11.424095 | 8.11146737 | 5.52933019 | 6.33394228 |
| ENSMUSG00 | Olfr898 | 950 | 0 | 0 | 0 | 0 | 0 | 0 |
| ENSMUSG00 | Olfr504 | 956 | 0 | 0 | 0 | 0 | 0 | 0 |
| ENSMUSG00 | Olfr229 | 923 | 0 | 0 | 0 | 0 | 0 | 0 |
| ENSMUSG00 | Olfr741 | 935 | 0 | 0 | 0 | 0 | 0 | 0 |
| ENSMUSG00 | Olfr60 | 935 | 0 | 0 | 0 | 0 | 0 | 0 |
| ENSMUSG00 | Olfr912 | 993 | 0 | 0 | 0 | 0 | 0 | 0 |
| ENSMUSG00 | Sip1 | 1732 | 6.97786725 | 9.09755576 | 5.71663559 | 10.145471 | 9.72211549 | 6.94463967 |
| ENSMUSG00 | Gm5528 | 482 | 0 | 0 | 0 | 0 | 0 | 0 |
| ENSMUSG00 | Tpt1 | 2530 | 5.18549746 | 6.49469438 | 4.96178791 | 5.10568061 | 3.50932387 | 5.55195226 |
| ENSMUSG00 | Gm10075 | 663 | 45.6318562 | 55.8904585 | 49.1353821 | 51.2658913 | 34.1715096 | 66.3559474 |
| ENSMUSG00 | Olfr951 | 944 | 0 | 0 | 0 | 0 | 0 | 0 |
| ENSMUSG00 | Atp8b4 | 7184 | 0 | 0 | 0.01230564 | 0 | 0 | 0 |
| ENSMUSG00 | Olfr105-ps | 952 | 0 | 0 | 0 | 0 | 0 | 0 |
| ENSMUSG00 | Gm10076 | 376 | 0.42293064 | 0 | 0.11755809 | 0 | 0 | 0.14507787 |
| ENSMUSG00 | Serpinb6a | 5883 | 8.59577957 | 7.15056332 | 9.78255985 | 8.2185857 | 6.47045323 | 9.11472731 |
| ENSMUSG00 | BC002059 | 1483 | 16.8887072 | 24.1419599 | 18.7179753 | 22.9192758 | 20.7477442 | 32.1851778 |
| ENSMUSG00 | Vmn1r49 | 932 | 0 | 0 | 0 | 0 | 0 | 0 |
| ENSMUSG00 | Pop5 | 2026 | 14.8150976 | 17.1245007 | 20.6173453 | 16.3669294 | 10.7795108 | 18.9280073 |
| ENSMUSG00 | Klk1b7-ps | 445 | 2.85882107 | 1.0828414 | 0.09932998 | 0.89191571 | 3.32531438 | 2.14519635 |
| ENSMUSG00 | Zdhhc8 | 4856 | 2.30051277 | 2.73876813 | 1.54742861 | 1.77586669 | 2.95272068 | 1.75802349 |
| ENSMUSG00 | Olfr1371 | 937 | 0 | 0 | 0 | 0 | 0 | 0 |
| ENSMUSG00 | Kif27 | 4445 | 0.10732638 | 0.5745515 | 0.62648281 | 0.13799657 | 1.14794996 | 0.80381951 |
| ENSMUSG00 | Klk1b22 | 850 | 5.42545381 | 3.17463619 | 0.26001084 | 2.71676571 | 3.24167564 | 2.11779552 |
| ENSMUSG00 | Myh13 | 6022 | 0 | 0 | 0 | 0 | 0 | 0 |
| ENSMUSG00 | Slc35e3 | 3477 | 1.77224604 | 2.88259418 | 1.89418309 | 1.07924436 | 2.27468745 | 1.21586686 |
| ENSMUSG00 | Cxcl11 | 2841 | 0 | 0 | 0 | 0 | 0.12572497 | 0 |
| ENSMUSG00 | Lrrc10 | 1427 | 0 | 0 | 0 | 0 | 0 | 0 |
| ENSMUSG00 | Cxcl17 | 757 | 2.67837451 | 1.40039858 | 1.28459782 | 0 | 0.53924836 | 0.75662804 |
| ENSMUSG00 | Gm11353 | 668 | 0 | 0 | 0 | 0 | 0 | 0 |
| ENSMUSG00 | Spink7 | 254 | 0 | 0 | 0 | 0 | 0 | 0 |
| ENSMUSG00 | Gm5868 | 1377 | 1.27032763 | 2.55454632 | 0.38520125 | 1.78182936 | 0.59289907 | 0.83190621 |
| ENSMUSG00 | Olfr57 | 1079 | 0 | 0 | 0 | 0 | 0 | 0 |
| ENSMUSG00 | Zfp462 | 10688 | 0.00371964 | 0.23444003 | 0 | 0.01350376 | 0.03341922 | 0.02807083 |
| ENSMUSG00 | Defa17 | 395 | 0 | 0 | 0 | 0 | 0 | 0 |
| ENSMUSG00 | Gm6570 | 463 | 0 | 0 | 0.28640503 | 0 | 0 | 0 |
| ENSMUSG00 | Pcnxl2 | 8540 | 0.01862083 | 0 | 0.08798962 | 0.06337596 | 0 | 0.37047519 |
| ENSMUSG00 | Gm8203 | 1054 | 0.07543734 | 0.32002381 | 0.08387447 | 0.03423344 | 0.29047273 | 0.2328954 |
| ENSMUSG00 | Arrb2 | 2465 | 1.17734283 | 0.74283359 | 0.77106663 | 1.39058589 | 0.7866135 | 2.51170106 |
| ENSMUSG00 | Pyroxd2 | 3103 | 0 | 1.10255797 | 0.07122437 | 0 | 0.64132409 | 0 |
| ENSMUSG00 | Casc4 | 4893 | 8.73434326 | 9.98590892 | 7.66056875 | 11.8429928 | 11.9614251 | 8.04916801 |
| ENSMUSG00 | Cend1 | 3325 | 0 | 0.52171787 | 0.27916954 | 0.2278866 | 0.13811651 | 0 |
| ENSMUSG00 | Alyref2 | 1272 | 2.46909038 | 2.53812235 | 3.33598815 | 3.00683704 | 0.96276181 | 3.13057966 |
| ENSMUSG00 | Vmn1r228 | 1439 | 0 | 0 | 0 | 0 | 0 | 0 |
| ENSMUSG00 | Olfr980 | 936 | 0 | 0 | 0 | 0 | 0 | 0 |
| ENSMUSG00 | Tdpoz4 | 1112 | 0 | 0 | 0 | 0 | 0 | 0 |
| ENSMUSG00 | Scrt2 | 3393 | 0.03515073 | 0.02840344 | 0.01302736 | 0.01063426 | 0.01503872 | 0.04823102 |
| ENSMUSG00 | Pwwp2b | 2637 | 9.79941689 | 9.02696338 | 10.8451242 | 8.41503883 | 8.10771766 | 6.57818376 |
| ENSMUSG00 | Gtf2i | 8067 | 14.1536804 | 14.6166139 | 14.7339477 | 14.1608718 | 14.0358904 | 16.1815326 |
| ENSMUSG00 | Hist1h2ao | 518 | 0 | 0 | 0 | 0 | 0 | 0 |
| ENSMUSG00 | Gm1661 | 2376 | 0 | 0.24336587 | 0.20463816 | 0.48595346 | 0 | 0.01147923 |
| ENSMUSG00 | Nrg2 | 3930 | 0 | 0 | 0 | 0.00918118 | 0.01298381 | 0.02776045 |
| ENSMUSG00 | Ap2a1 | 4492 | 12.4258002 | 8.05610375 | 10.2534106 | 12.5869465 | 10.3370441 | 11.2814514 |
| ENSMUSG00 | Sp7 | 4305 | 0 | 0 | 0 | 0 | 0.16593943 | 0 |
| ENSMUSG00 | Ppih | 2054 | 1.23872968 | 2.48673947 | 1.31271296 | 0.73780227 | 0.91917035 | 2.20427951 |
| ENSMUSG00 | 2610008E11f | 4223 | 21.8876373 | 20.0824386 | 18.7044029 | 25.7948597 | 20.7585398 | 23.0571779 |
| ENSMUSG00 | Olfr1447 | 1102 | 0 | 0 | 0 | 0 | 0 | 0 |
| ENSMUSG00 | Zfp941 | 4711 | 1.40929001 | 2.75146528 | 0.91950342 | 1.14120667 | 3.09775918 | 2.06687459 |
| ENSMUSG00 | Acnat2 | 2525 | 0 | 0.1908374 | 0 | 0 | 0.08083386 | 0 |
| ENSMUSG00 | Gm2228 | 437 | 0 | 0 | 0 | 0 | 0 | 0 |
| ENSMUSG00 | Tmc2 | 3201 | 0 | 0 | 0 | 0 | 0 | 0 |
| ENSMUSG00 | Olfr384 | 938 | 0 | 0 | 0 | 0 | 0 | 0 |
| ENSMUSG00 | Zfp937 | 2793 | 7.84291792 | 8.21222573 | 6.45698244 | 7.86751347 | 11.0712437 | 10.1755009 |
| ENSMUSG00 | Gm15454 | 1133 | 0 | 0 | 0.0390131 | 0 | 0 | 0 |
| ENSMUSG00 | Calm5 | 586 | 0 | 0 | 0 | 0 | 0 | 0 |
| ENSMUSG00 | Caln1 | 3555 | 0 | 0 | 0 | 0.13194559 | 0 | 0 |
| ENSMUSG00 | Hnrnpc | 3193 | 43.4160227 | 38.6638474 | 40.0073055 | 38.0257062 | 33.0960302 | 50.85913 |
| ENSMUSG00 | 3110018I06R | 1004 | 0 | 0 | 0 | 0 | 0 | 0 |
| ENSMUSG00 | Bckdha | 1764 | 115.795725 | 101.508401 | 73.1435259 | 59.5844647 | 85.9406817 | 46.5864445 |
| ENSMUSG00 | Rpl36a-ps1 | 320 | 0 | 0 | 0 | 0 | 0 | 0 |
| ENSMUSG00 | C030014I23F | 3077 | 0.86565395 | 0.67338869 | 0.96247107 | 0.62149768 | 0.38141262 | 1.75507916 |
| ENSMUSG00 | Gm5580 | 2697 | 0 | 0 | 0 | 0 | 0 | 0 |
| ENSMUSG00 | Zfp128 | 4084 | 3.21236743 | 6.50115809 | 8.03079518 | 5.25681109 | 4.07311423 | 9.78387525 |
| ENSMUSG00 | Chst8 | 3870 | 6.18418586 | 3.48635757 | 3.89478772 | 5.04402741 | 7.50232761 | 8.47839561 |
| ENSMUSG00 | Olfr1369-ps1 | 953 | 0 | 0 | 0 | 0 | 0 | 0 |
| ENSMUSG00 | Cyp2a12 | 1647 | 0 | 0 | 0 | 0 | 0 | 0 |
| ENSMUSG00 | Npn2 | 74 | 0 | 0 | 0 | 0 | 0 | 0 |
| ENSMUSG00 | Tas2r124 | 929 | 0 | 0 | 0 | 0 | 0 | 0 |
| ENSMUSG00 | Gm839 | 1840 | 0.08642496 | 0 | 0 | 0 | 0 | 0 |
| ENSMUSG00 | Gm8603 | 1392 | 0.17135983 | 0 | 0.0635084 | 0 | 0 | 0 |
| ENSMUSG00 | Rps16-ps2 | 934 | 0.04256475 | 0.10318296 | 0.47325314 | 0.27042218 | 0 | 0 |
| ENSMUSG00 | Vmn1r62 | 1985 | 0 | 0 | 0 | 0 | 0 | 0 |
| ENSMUSG00 | Olfr1034 | 1015 | 0 | 0 | 0 | 0 | 0 | 0 |
| ENSMUSG00 | 2610017I09R | 1173 | 7.72740798 | 9.53048131 | 8.47861439 | 10.3047612 | 5.35059184 | 16.1369136 |
| ENSMUSG00 | Zfp868 | 3929 | 14.9753401 | 19.4879757 | 17.9327406 | 28.8270648 | 10.9091768 | 30.3012981 |
| ENSMUSG00 | Sntb1 | 2861 | 0.81984388 | 1.17897622 | 0.8960877 | 0.93326505 | 0.53505462 | 0.02859976 |
| ENSMUSG00 | Rps10-ps1 | 463 | 0.08586497 | 0.10407439 | 0.38187337 | 0.07793098 | 0.11020816 | 0.17672552 |
| ENSMUSG00 | Trim5 | 4267 | 1.51866495 | 1.10669588 | 0.97374578 | 1.22612994 | 1.6143803 | 3.44528487 |
| ENSMUSG00 | Sycp2 | 6210 | 0 | 0.03879746 | 0.07117849 | 0 | 0.11503531 | 0.3074436 |
| ENSMUSG00 | Rnf14 | 3591 | 68.7943625 | 66.9591608 | 73.4481752 | 63.8544675 | 67.6230918 | 66.0485277 |
| ENSMUSG00 | Kng2 | 1878 | 284.046346 | 321.550848 | 136.183101 | 261.143317 | 290.236287 | 217.412859 |
| ENSMUSG00 | Dppa5a | 616 | 0 | 0 | 0 | 0 | 0 | 0 |
| ENSMUSG00 | Thap6 | 2995 | 9.98201047 | 6.72518625 | 5.51969592 | 5.22858341 | 7.82006895 | 1.93062555 |
| ENSMUSG00 | Gm10080 | 451 | 0 | 0.10684355 | 0.19601704 | 0 | 0.11314052 | 0.24190367 |
| ENSMUSG00 | Krtap16-5 | 525 | 0 | 0 | 0 | 0 | 0 | 0 |
| ENSMUSG00 | Gpr97 | 2424 | 5.57626377 | 6.46064096 | 2.88114323 | 9.45218576 | 9.13591052 | 5.45717823 |
| ENSMUSG00 | Wtap | 6706 | 2.54918828 | 1.68142372 | 2.44540468 | 2.74946686 | 2.16097386 | 3.71742026 |
| ENSMUSG00 | Irak2 | 5674 | 2.00388922 | 0.66241496 | 1.67490241 | 0.90300499 | 2.36516335 | 1.52861038 |
| ENSMUSG00 | Olfr171 | 1010 | 0 | 0 | 0 | 0 | 0 | 0 |
| ENSMUSG00 | Samd5 | 2493 | 0 | 0.79247659 | 0.01773038 | 1.09997408 | 0.24561432 | 0.0984644 |
| ENSMUSG00 | Vmn1r218 | 896 | 0 | 0 | 0 | 0 | 0 | 0 |
| ENSMUSG00 | 4930522H14 | 1417 | 0 | 0 | 0 | 0 | 0 | 0 |
| ENSMUSG00 | Rpl10l | 1020 | 0 | 0 | 0 | 0 | 0 | 0 |
| ENSMUSG00 | Olfr715 | 1130 | 0 | 0 | 0 | 0 | 0 | 0 |
| ENSMUSG00 | Nlrp9b | 3530 | 0.40543833 | 0 | 0 | 0 | 0 | 0 |
| ENSMUSG00 | Xcr1 | 4451 | 0 | 0 | 0 | 0 | 0 | 0 |
| ENSMUSG00 | Zfp266 | 7677 | 13.9353912 | 15.3026633 | 19.1558593 | 16.0176642 | 15.6329342 | 16.5736866 |
| ENSMUSG00 | 0610040I01R | 1866 | 84.5389854 | 94.487777 | 67.7003576 | 71.1392507 | 67.1874627 | 68.2012148 |
| ENSMUSG00 | Tor3a | 4715 | 9.11467114 | 7.02101501 | 5.05297845 | 4.12475546 | 12.1857262 | 5.55905161 |
| ENSMUSG00 | Olfr726 | 965 | 0 | 0 | 0 | 0 | 0 | 0 |
| ENSMUSG00 | A930017M01 | 1672 | 0 | 0 | 0 | 0 | 0.51880885 | 0.24468875 |
| ENSMUSG00 | Dcc | 8127 | 0 | 0 | 0 | 0 | 0 | 0.03691658 |
| ENSMUSG00 | Tmem219 | 2138 | 4.74164992 | 4.19208524 | 4.34162162 | 4.16850561 | 2.93556793 | 6.46784008 |
| ENSMUSG00 | Tnfrsf19 | 5059 | 0.27504286 | 0 | 0.31454168 | 0 | 0.52448538 | 0 |
| ENSMUSG00 | Olfr488 | 944 | 0 | 0 | 0 | 0 | 0 | 0 |
| ENSMUSG00 | H2-Q7 | 3685 | 0.23734615 | 0.33998575 | 0.10795566 | 0 | 1.16315212 | 2.52392184 |
| ENSMUSG00 | Olfr1418 | 1071 | 0 | 0 | 0 | 0 | 0 | 0 |
| ENSMUSG00 | Gm10083 | 2230 | 0.55265466 | 0.47538194 | 0.01982145 | 0.27506491 | 0.22881783 | 0.1834617 |
| ENSMUSG00 | Ces4a | 2419 | 0 | 0 | 0.01827278 | 0 | 0 | 0 |
| ENSMUSG00 | Olfr823 | 947 | 0 | 0 | 0 | 0 | 0 | 0 |
| ENSMUSG00 | Gm5591 | 3438 | 0 | 0 | 0 | 0 | 0 | 0 |
| ENSMUSG00 | Rpl21-ps3 | 482 | 0 | 0 | 0 | 0.07485901 | 0.10586385 | 0.11317278 |
| ENSMUSG00 | Fam78b | 5379 | 0.10347216 | 0.06270777 | 0.13147973 | 0.05366357 | 0.07588976 | 0.05070578 |
| ENSMUSG00 | Mfap2 | 1201 | 0.06620396 | 0.56170707 | 0.73608398 | 0.03004333 | 0.46735232 | 0.29522923 |
| ENSMUSG00 | Fhit | 1775 | 2.93406645 | 2.30751977 | 3.11280585 | 1.34164222 | 1.95481327 | 3.34978668 |
| ENSMUSG00 | Olfr881 | 947 | 0 | 0 | 0 | 0 | 0 | 0 |
| ENSMUSG00 | H2-Eb1 | 1287 | 3.52146447 | 0.89858167 | 2.23241631 | 0.95321641 | 1.38766368 | 3.79344244 |
| ENSMUSG00 | Ifitm2 | 649 | 37.4889893 | 49.6713864 | 47.5391162 | 42.4756271 | 32.4713301 | 74.3013285 |
| ENSMUSG00 | Olfr1454 | 923 | 0 | 0 | 0 | 0 | 0 | 0 |
| ENSMUSG00 | Layn | 2236 | 0 | 0.43100574 | 0 | 0 | 0 | 0 |
| ENSMUSG00 | Eno3 | 1729 | 5.56438767 | 6.1591693 | 5.70098959 | 5.48847757 | 3.92510583 | 8.80234164 |
| ENSMUSG00 | Nr1h2 | 3640 | 25.1966741 | 21.5779947 | 22.0766348 | 21.5897508 | 23.1020515 | 17.7884598 |
| ENSMUSG00 | Cyp2c70 | 1972 | 0 | 0 | 0 | 0 | 0 | 0 |
| ENSMUSG00 | Ang4 | 707 | 0 | 0 | 0 | 0 | 0 | 0 |
| ENSMUSG00 | Nkpd1 | 2749 | 0.0144618 | 0 | 0 | 0 | 0 | 0.05952995 |
| ENSMUSG00 | Zfp735 | 4073 | 0 | 0 | 0 | 0 | 0 | 0 |
| ENSMUSG00 | Rpl35a | 1977 | 5.89193515 | 3.04668956 | 6.43911522 | 6.00454863 | 2.99396034 | 8.89840279 |
| ENSMUSG00 | Hist1h4i | 536 | 0 | 0 | 0.41233063 | 0.40390348 | 0.19039693 | 0.05088552 |
| ENSMUSG00 | Olfr1512 | 941 | 0 | 0 | 0 | 0 | 0 | 0 |
| ENSMUSG00 | Gm7099 | 550 | 0.43369615 | 0.61328199 | 0 | 0.45922602 | 0.55665137 | 0.44631228 |
| ENSMUSG00 | 4921513D23 | 6447 | 1.35663188 | 0.42603183 | 0.65133785 | 1.20329449 | 0.63317978 | 1.34955947 |
| ENSMUSG00 | Olfr175-ps1 | 1027 | 0 | 0 | 0 | 0 | 0 | 0 |
| ENSMUSG00 | Atp8b2 | 10316 | 1.1291543 | 1.779666 | 0.8612418 | 1.14024297 | 2.84908806 | 0.8328336 |
| ENSMUSG00 | 4930595M18 | 2640 | 0 | 0 | 0 | 0 | 0 | 0 |
| ENSMUSG00 | Pla2g16 | 6349 | 6.07384094 | 4.43992261 | 5.61138533 | 6.77994631 | 4.26760207 | 5.42571576 |
| ENSMUSG00 | Hist1h4c | 394 | 1.10992458 | 0.8561043 | 2.13156096 | 0 | 1.29508568 | 2.2844241 |
| ENSMUSG00 | Mrps9 | 2496 | 17.1063246 | 12.5485526 | 17.2486359 | 14.1090046 | 13.8196435 | 16.3910092 |
| ENSMUSG00 | Gm8894 | 455 | 0 | 0 | 0.09714691 | 0.0793012 | 0 | 0 |
| ENSMUSG00 | Slc9a6 | 4969 | 12.2090689 | 8.62099961 | 11.3684756 | 16.0985898 | 9.63226817 | 12.7728085 |
| ENSMUSG00 | Gm14511 | 733 | 0 | 0 | 0 | 0 | 0 | 0 |
| ENSMUSG00 | Olfr292 | 1076 | 0 | 0 | 0 | 0 | 0 | 0 |
| ENSMUSG00 | Krtap16-2 | 492 | 0 | 0 | 0 | 0 | 0 | 0 |
| ENSMUSG00 | Vmn1r35 | 890 | 0 | 0 | 0 | 0 | 0 | 0 |
| ENSMUSG00 | Fpr-rs3 | 1031 | 0 | 0 | 0 | 0 | 0 | 0 |
| ENSMUSG00 | Cd302 | 1348 | 0.35390636 | 0.28597295 | 0.59023233 | 1.28482058 | 0.37853395 | 0.93073695 |
| ENSMUSG00 | Cno | 1305 | 18.4307017 | 22.8193266 | 17.1049278 | 12.331488 | 14.3108456 | 19.2489983 |
| ENSMUSG00 | 1700019A02f | 618 | 0 | 0 | 0 | 0 | 0 | 0 |
| ENSMUSG00 | Plekhh1 | 6429 | 5.86221738 | 6.6707005 | 10.2374466 | 11.4268226 | 10.5719603 | 6.31274898 |
| ENSMUSG00 | Vmn1r47 | 932 | 0 | 0 | 0 | 0 | 0 | 0 |
| ENSMUSG00 | Olfr487 | 944 | 0 | 0 | 0 | 0 | 0 | 0 |
| ENSMUSG00 | Tmsb15a | 560 | 0 | 0 | 0 | 0 | 0 | 0 |
| ENSMUSG00 | Gm12571 | 242 | 1.64278845 | 0 | 0.18265224 | 0 | 0.84341117 | 0.11270512 |
| ENSMUSG00 | Gm10086 | 760 | 0.05230984 | 0.12680643 | 0.11632064 | 0.09495275 | 0 | 0.03588768 |
| ENSMUSG00 | Ipmk | 3365 | 15.0633693 | 17.0693132 | 21.9104529 | 13.2104246 | 15.1941838 | 15.2300289 |
| ENSMUSG00 | Rxfp3 | 4251 | 0 | 0 | 0 | 0 | 0 | 0 |
| ENSMUSG00 | Prl7c1 | 1009 | 0 | 0 | 0 | 0 | 0 | 0 |
| ENSMUSG00 | Nsa2 | 2680 | 1.94327162 | 1.61820142 | 2.42450408 | 1.89834637 | 1.99916771 | 2.4119737 |
| ENSMUSG00 | Olfr1076 | 941 | 0 | 0 | 0 | 0 | 0 | 0 |
| ENSMUSG00 | H3f3a | 4006 | 6.81777711 | 10.2844254 | 10.7359943 | 9.51139267 | 6.99288076 | 8.17694502 |
| ENSMUSG00 | Il28b | 697 | 0 | 0 | 0 | 0 | 0 | 0 |
| ENSMUSG00 | Krtap2-4 | 829 | 0 | 0 | 0 | 0 | 0 | 0 |
| ENSMUSG00 | Olfr503 | 971 | 0 | 0 | 0 | 0 | 0 | 0 |
| ENSMUSG00 | Olfr325 | 2044 | 0 | 0 | 0 | 0 | 0 | 0 |
| ENSMUSG00 | Tsga10 | 8336 | 1.5118147 | 1.39310611 | 2.08389207 | 1.76601178 | 1.51193796 | 1.59996384 |
| ENSMUSG00 | Gm10087 | 661 | 0 | 0 | 0 | 0 | 0 | 0 |
| ENSMUSG00 | Lrrtm1 | 3416 | 1.16380212 | 0.0423183 | 0.40112914 | 0.21125319 | 0.55268616 | 0.08782817 |
| ENSMUSG00 | Gm8994 | 1462 | 0 | 0 | 0 | 0 | 0 | 0 |
| ENSMUSG00 | Olfr464 | 1083 | 0 | 0 | 0 | 0 | 0 | 0 |
| ENSMUSG00 | Gmfg | 2893 | 0.60464609 | 0.83281096 | 0.56531911 | 0.79822014 | 0.79370443 | 1.15961999 |
| ENSMUSG00 | Tssk5 | 1506 | 0 | 0 | 0 | 0 | 0.06776411 | 0.0362213 |
| ENSMUSG00 | Gm13363 | 521 | 0 | 0 | 0 | 0 | 0 | 0.05235056 |
| ENSMUSG00 | Gm6280 | 683 | 0 | 0 | 0 | 0 | 0 | 0 |
| ENSMUSG00 | Intu | 7412 | 2.31710302 | 3.40009569 | 2.68956168 | 4.3617798 | 3.61425355 | 3.45900714 |
| ENSMUSG00 | B2m | 856 | 128.787322 | 126.433118 | 214.451231 | 125.739181 | 239.990875 | 833.278465 |
| ENSMUSG00 | Gstp1 | 746 | 56.435729 | 148.047353 | 65.1769803 | 119.370625 | 40.2191809 | 53.5621267 |
| ENSMUSG00 | Serpina6 | 1566 | 0 | 0 | 0 | 0 | 0 | 0 |
| ENSMUSG00 | 3110043A19f | 673 | 0.17721611 | 0.07159947 | 0 | 0 | 0 | 0.52685039 |
| ENSMUSG00 | Gm10088 | 422 | 0 | 0 | 0.20948741 | 0.3420099 | 0 | 0 |
| ENSMUSG00 | Gm4980 | 680 | 0.17539183 | 0 | 0 | 0 | 0 | 0 |
| ENSMUSG00 | Vmn1r52 | 929 | 0 | 0 | 0 | 0 | 0 | 0 |
| ENSMUSG00 | Taar7f | 1076 | 0 | 0 | 0 | 0 | 0 | 0 |
| ENSMUSG00 | Olfr1193 | 956 | 0 | 0 | 0 | 0 | 0 | 0 |
| ENSMUSG00 | Ctnna3 | 7204 | 0 | 0 | 0 | 0 | 0 | 0 |
| ENSMUSG00 | Olfr671 | 941 | 0 | 0 | 0 | 0 | 0 | 0 |
| ENSMUSG00 | Ube2s | 984 | 1.01004778 | 1.42012888 | 1.1230143 | 1.3200748 | 0.67412895 | 1.77396028 |
| ENSMUSG00 | Zbtb40 | 10078 | 1.46745771 | 1.00886478 | 1.03947577 | 1.94409111 | 1.39742803 | 1.49390761 |
| ENSMUSG00 | Tcl1b2 | 1252 | 0 | 0 | 0 | 0 | 0 | 0 |
| ENSMUSG00 | Olfr1420 | 1051 | 0 | 0 | 0 | 0 | 0 | 0 |
| ENSMUSG00 | Gm6564 | 644 | 0 | 0 | 0 | 0 | 0 | 0 |
| ENSMUSG00 | Kcnd2 | 5146 | 0 | 0 | 0 | 0 | 0 | 0 |
| ENSMUSG00 | Gm12960 | 1307 | 0 | 0.18433987 | 0 | 0 | 0 | 0 |
| ENSMUSG00 | Olfr566 | 950 | 0 | 0 | 0 | 0 | 0 | 0 |
| ENSMUSG00 | Arr3 | 2307 | 0 | 0 | 0 | 0 | 0.15482646 | 0.0827579 |
| ENSMUSG00 | Rpl21-ps11 | 467 | 0 | 0 | 0 | 0 | 0 | 0 |
| ENSMUSG00 | Arl1 | 2316 | 46.5187186 | 64.539872 | 66.4172771 | 72.5535757 | 43.1829433 | 54.8908003 |
| ENSMUSG00 | Olfr944 | 935 | 0 | 0 | 0 | 0 | 0 | 0 |
| ENSMUSG00 | Trim55 | 1777 | 0 | 0 | 0 | 0 | 0 | 0 |
| ENSMUSG00 | Olfr51 | 923 | 0 | 0 | 0 | 0 | 0 | 0 |
| ENSMUSG00 | Acyp2 | 859 | 17.8645116 | 27.7675074 | 22.7955954 | 23.9846885 | 11.2863928 | 32.0691335 |
| ENSMUSG00 | Csmd1 | 22294 | 0 | 0 | 0 | 0 | 0 | 0.00122341 |
| ENSMUSG00 | AI597468 | 3775 | 5.46572037 | 2.2465732 | 4.6602208 | 9.41478513 | 3.67660244 | 10.2595994 |
| ENSMUSG00 | Rpl26 | 1398 | 3.4124876 | 4.75660159 | 7.61991714 | 6.06529361 | 3.32145937 | 3.99950003 |
| ENSMUSG00 | Trmt61a | 2625 | 1.55992933 | 0.6424859 | 0.82510107 | 1.29208083 | 0.73866754 | 1.21566964 |
| ENSMUSG00 | Gm10092 | 177 | 0 | 0 | 0 | 0 | 0 | 0 |
| ENSMUSG00 | Slc4a4 | 10849 | 1.13231113 | 1.93651846 | 0.49298765 | 1.9422909 | 1.82018688 | 1.08103004 |
| ENSMUSG00 | Dmkn | 2707 | 1.05740473 | 0.19580749 | 0.29391695 | 0.01332916 | 0.05654936 | 0 |
| ENSMUSG00 | Etd | 476 | 0 | 0 | 0 | 0 | 0 | 0 |
| ENSMUSG00 | Irx1 | 2369 | 0.13425236 | 0.10170207 | 0.01865844 | 0 | 0 | 0.02302629 |
| ENSMUSG00 | Olfr530 | 1079 | 0 | 0 | 0 | 0 | 0 | 0 |
| ENSMUSG00 | Hist1h4h | 469 | 0.5085989 | 0.41097179 | 0.94247 | 1.30787795 | 0.32639473 | 0.58154881 |
| ENSMUSG00 | Tdrd5 | 7409 | 0 | 0 | 0.0536937 | 0 | 0 | 0 |
| ENSMUSG00 | Galnt13 | 8139 | 0 | 0 | 0.11404825 | 0.00886646 | 0 | 0 |
| ENSMUSG00 | Gm11847 | 5253 | 0 | 0.00917313 | 0.01682918 | 0 | 0.00971376 | 0 |
| ENSMUSG00 | Copz1 | 2457 | 104.267121 | 110.964139 | 102.849791 | 102.753791 | 90.547415 | 77.5057107 |
| ENSMUSG00 | Olfr507 | 950 | 0 | 0 | 0 | 0 | 0 | 0 |
| ENSMUSG00 | Mkx | 3497 | 0.28421133 | 0.01377937 | 0.16431912 | 0 | 0.01459147 | 0 |
| ENSMUSG00 | Vmn1r222 | 1053 | 0 | 0 | 0 | 0 | 0 | 0 |
| ENSMUSG00 | Rrs1 | 2046 | 20.7521277 | 17.3574819 | 19.2059816 | 24.1605088 | 17.8816772 | 21.195834 |
| ENSMUSG00 | Vmn2r50 | 2532 | 0 | 0 | 0 | 0 | 0 | 0 |
| ENSMUSG00 | Clasrp | 2132 | 8.93193019 | 11.3007604 | 10.4284836 | 9.52823221 | 11.7274504 | 15.3387863 |
| ENSMUSG00 | Rrp1 | 2486 | 96.4941953 | 98.0592159 | 78.8556612 | 84.2253035 | 94.3967427 | 69.2946989 |
| ENSMUSG00 | Olfr920 | 1958 | 0.24364952 | 0 | 0.58694991 | 0 | 0.70363235 | 0 |
| ENSMUSG00 | Haghl | 2023 | 14.6012467 | 20.698971 | 12.8912938 | 14.6432815 | 12.3088835 | 16.1247991 |
| ENSMUSG00 | Cdh3 | 4072 | 2.5481779 | 4.98194797 | 0.29308688 | 3.03046641 | 3.07010365 | 0.78367701 |
| ENSMUSG00 | Gm10093 | 1976 | 0.58345594 | 0.80473309 | 1.2079451 | 0.63910504 | 1.03292259 | 1.32508369 |
| ENSMUSG00 | Gm5382 | 508 | 0 | 0 | 0 | 0 | 0 | 0 |
| ENSMUSG00 | Mcpt4 | 985 | 0 | 0 | 0 | 0 | 0 | 0 |
| ENSMUSG00 | Olfr1337 | 947 | 0 | 0 | 0 | 0 | 0 | 0 |
| ENSMUSG00 | Zfp143 | 3001 | 16.0691096 | 16.1370791 | 13.9778571 | 11.1817066 | 8.85862773 | 10.1246078 |
| ENSMUSG00 | Lsamp | 11077 | 1.3710024 | 0.47851482 | 2.72545444 | 0.47883547 | 0.58042099 | 1.9353495 |
| ENSMUSG00 | Plac1 | 2282 | 0 | 0 | 0 | 0 | 0 | 0 |
| ENSMUSG00 | Myl4 | 4314 | 0.4054801 | 0.30158413 | 0.09221525 | 0.58547592 | 0.20107751 | 0.12012474 |
| ENSMUSG00 | Olfr38 | 953 | 0 | 0 | 0 | 0 | 0 | 0 |
| ENSMUSG00 | Olfr890 | 944 | 0 | 0 | 0 | 0 | 0 | 0 |
| ENSMUSG00 | Gapdhs | 1614 | 1.69958374 | 2.50784457 | 2.02660247 | 1.6990306 | 1.45428333 | 1.43639674 |
| ENSMUSG00 | Retnla | 594 | 0 | 0 | 0 | 0.12148837 | 0 | 0 |
| ENSMUSG00 | Gm10094 | 779 | 1.68412177 | 1.11342228 | 0.73764308 | 0.88004987 | 1.44105298 | 1.50553208 |
| ENSMUSG00 | Fam195b | 1278 | 58.886639 | 60.1010867 | 74.0156057 | 48.5328909 | 45.0772913 | 56.3206361 |
| ENSMUSG00 | Gm17113 | 169 | 0 | 0 | 0 | 0 | 0 | 0.3227768 |
| ENSMUSG00 | Dnajc30 | 2287 | 7.85722658 | 8.95462961 | 6.95787647 | 5.85327439 | 7.56359484 | 6.78586346 |
| ENSMUSG00 | Prcp | 2947 | 38.4064652 | 33.9283568 | 36.7773733 | 13.6639164 | 43.2520824 | 16.5572886 |
| ENSMUSG00 | Cyp4f39 | 2431 | 1.04662721 | 1.88305718 | 0.50911214 | 0.80149338 | 0 | 0.16829271 |
| ENSMUSG00 | Ppm1b | 4961 | 18.1027274 | 15.09408 | 21.20548 | 19.6956615 | 15.3253981 | 20.6332838 |
| ENSMUSG00 | Blnk | 5974 | 66.6007447 | 60.3661421 | 66.6727163 | 51.6769289 | 61.8141751 | 71.2363904 |
| ENSMUSG00 | Prpf40a | 10284 | 16.2709858 | 15.813812 | 16.2382889 | 19.5672464 | 17.1377968 | 20.596543 |
| ENSMUSG00 | Maml3 | 9769 | 3.28006114 | 2.31831588 | 3.95459216 | 3.92252342 | 3.285453 | 2.97064348 |
| ENSMUSG00 | Spink12 | 1003 | 0.75309485 | 1.15301557 | 0 | 0.57558596 | 0.35611628 | 1.0877224 |
| ENSMUSG00 | Vmn1r229 | 920 | 0 | 0 | 0 | 0 | 0 | 0 |
| ENSMUSG00 | Olfr854 | 947 | 0 | 0 | 0 | 0 | 0 | 0 |
| ENSMUSG00 | Olfr151 | 2250 | 0 | 0 | 0 | 0 | 0 | 0 |
| ENSMUSG00 | Rpl15-ps3 | 1296 | 0 | 0 | 0 | 0 | 0 | 0 |
| ENSMUSG00 | Slc38a11 | 4669 | 0 | 0 | 0 | 0 | 0.03278628 | 0.00584165 |
| ENSMUSG00 | Fnip2 | 12115 | 1.55215372 | 0.54490653 | 0.94496718 | 0.70287763 | 0.84236691 | 0.54932002 |
| ENSMUSG00 | Tmprss11c | 1605 | 0 | 0 | 0 | 0 | 0 | 0 |
| ENSMUSG00 | Sfmbt2 | 9143 | 0.26089126 | 0.34257013 | 0.14503503 | 0.12233877 | 0.0502283 | 0.02088182 |
| ENSMUSG00 | Olfr1289 | 1013 | 0 | 0 | 0 | 0 | 0 | 0 |
| ENSMUSG00 | Olfr601 | 989 | 0 | 0 | 0 | 0 | 0 | 0 |
| ENSMUSG00 | Stk19 | 2797 | 2.82850934 | 3.06656657 | 2.93941466 | 2.65745483 | 2.60878504 | 2.89616298 |
| ENSMUSG00 | Vmn1r8 | 2140 | 0 | 0 | 0 | 0 | 0 | 0 |
| ENSMUSG00 | Olfr47 | 1058 | 0 | 0 | 0 | 0 | 0 | 0 |
| ENSMUSG00 | Vmn1r189 | 1136 | 0 | 0 | 0 | 0 | 0 | 0 |
| ENSMUSG00 | Gm10096 | 898 | 0 | 0 | 0 | 0 | 0 | 0 |
| ENSMUSG00 | H2-K1 | 2339 | 82.9103181 | 122.618941 | 91.843077 | 80.6639636 | 199.371547 | 211.457165 |
| ENSMUSG00 | Olfr611 | 971 | 0 | 0 | 0 | 0 | 0 | 0 |
| ENSMUSG00 | Rpl21-ps7 | 482 | 0 | 0 | 0 | 0 | 0.10586385 | 0 |
| ENSMUSG00 | Exoc5 | 10131 | 12.7613091 | 11.2867858 | 11.6841907 | 16.9066692 | 12.9693927 | 22.5417584 |
| ENSMUSG00 | Olfr387-ps1 | 1144 | 0 | 0 | 0 | 0 | 0 | 0 |
| ENSMUSG00 | Tmprss11d | 2786 | 0 | 0 | 0 | 0 | 0 | 0 |
| ENSMUSG00 | Gm10097 | 317 | 0 | 0 | 0 | 0 | 0 | 0 |
| ENSMUSG00 | Gm14173 | 303 | 0.52482483 | 0 | 0 | 0 | 0 | 0 |
| ENSMUSG00 | Mmgt1 | 4230 | 33.4209193 | 33.2064962 | 38.7784963 | 36.4317759 | 37.9259871 | 39.4934197 |
| ENSMUSG00 | Tgif2lx1 | 1031 | 0 | 0 | 0 | 0 | 0 | 0 |
| ENSMUSG00 | Exosc5 | 1024 | 11.1424052 | 11.3878115 | 13.0792563 | 7.89294724 | 9.56744547 | 12.1990086 |
| ENSMUSG00 | Taok3 | 10929 | 9.1376857 | 6.4592495 | 10.6854487 | 8.26364328 | 10.5937274 | 7.11501478 |
| ENSMUSG00 | Cyp3a59 | 1986 | 0 | 0 | 0 | 0 | 0 | 0 |
| ENSMUSG00 | Olfr1261 | 1116 | 0 | 0 | 0 | 0 | 0 | 0 |
| ENSMUSG00 | Vmn1r210 | 920 | 0 | 0 | 0 | 0 | 0 | 0 |
| ENSMUSG00 | Agbl4 | 9508 | 0.03345013 | 0.08615582 | 0.06043584 | 0 | 0.01073336 | 0 |
| ENSMUSG00 | Olfr342 | 1053 | 0 | 0 | 0 | 0 | 0 | 0 |
| ENSMUSG00 | Slc38a10 | 6789 | 25.0104076 | 24.6645878 | 24.7996494 | 24.7987656 | 26.8923807 | 20.4047566 |
| ENSMUSG00 | Rag1 | 6606 | 0 | 0 | 0 | 0 | 0 | 0 |
| ENSMUSG00 | Ddhd2 | 4566 | 12.4507966 | 10.7327227 | 13.2528084 | 10.6049286 | 12.4716241 | 12.0065758 |
| ENSMUSG00 | Naca | 7025 | 2.16745182 | 2.05778401 | 2.39098937 | 1.7925457 | 1.67061444 | 2.58963479 |
| ENSMUSG00 | Dnaic1 | 2761 | 0 | 0 | 0 | 0.07841082 | 0 | 0 |
| ENSMUSG00 | Gm11361 | 968 | 0 | 0.14933815 | 0 | 0 | 0 | 0.02817628 |
| ENSMUSG00 | Gm17132 | 1157 | 0 | 0 | 0 | 0 | 0 | 0 |
| ENSMUSG00 | Olfr116 | 965 | 0 | 0 | 0 | 0 | 0 | 0 |
| ENSMUSG00 | Olfr1168 | 938 | 0 | 0 | 0 | 0 | 0 | 0 |
| ENSMUSG00 | Olfr320 | 927 | 0 | 0 | 0 | 0 | 0 | 0 |
| ENSMUSG00 | Cxcl12 | 8314 | 0.21039706 | 0.25501605 | 0 | 0 | 0 | 0 |
| ENSMUSG00 | Gm600 | 2726 | 0 | 0 | 0.03242982 | 0 | 0 | 0 |
| ENSMUSG00 | Phf5a | 1138 | 19.388657 | 25.1094554 | 29.4031592 | 26.2530166 | 18.7425528 | 27.3705079 |
| ENSMUSG00 | Olfr168 | 938 | 0 | 0 | 0 | 0 | 0 | 0 |
| ENSMUSG00 | Olfr771 | 953 | 0 | 0 | 0 | 0 | 0 | 0 |
| ENSMUSG00 | Zfp873 | 1891 | 19.4257345 | 33.5852622 | 25.1045898 | 77.2205363 | 17.1347164 | 40.2701178 |
| ENSMUSG00 | Fiz1 | 2829 | 15.1489601 | 9.64069504 | 15.343305 | 10.4075463 | 11.9764983 | 8.32990043 |
| ENSMUSG00 | Vmn1r219 | 938 | 0 | 0 | 0 | 0 | 0 | 0 |
| ENSMUSG00 | Olfr1490 | 950 | 0 | 0 | 0 | 0 | 0 | 0 |
| ENSMUSG00 | Csn1s2b | 2252 | 0 | 0 | 0 | 0 | 0 | 0 |
| ENSMUSG00 | Gm7866 | 395 | 0 | 0 | 0 | 0 | 0 | 0 |
| ENSMUSG00 | Mageb5 | 1355 | 0 | 0 | 0 | 0 | 0 | 0 |
| ENSMUSG00 | Acvr2b | 1839 | 7.47982396 | 8.98746583 | 9.90275113 | 11.0267042 | 10.9044947 | 7.43044821 |
| ENSMUSG00 | Gm13287 | 967 | 0 | 0 | 0 | 0 | 0.05276771 | 0 |
| ENSMUSG00 | Krt79 | 2088 | 0 | 0 | 0 | 0 | 0 | 0 |
| ENSMUSG00 | Zcchc14 | 9917 | 1.65564318 | 1.71521772 | 1.29703906 | 1.93926891 | 2.1661898 | 2.5412692 |
| ENSMUSG00 | 8430427H17 | 13426 | 3.06471938 | 2.85328627 | 3.28896478 | 5.45288756 | 4.72790195 | 3.20567412 |
| ENSMUSG00 | Efcab4b | 1087 | 0.07314716 | 0.22164877 | 0.20332035 | 0.19916492 | 0.18776955 | 0.02509166 |
| ENSMUSG00 | Hipk2 | 9601 | 16.1614041 | 17.9827125 | 10.3034814 | 16.1563076 | 16.8688383 | 13.3432956 |
| ENSMUSG00 | Tmem151a | 4876 | 18.2715406 | 18.8753291 | 20.5235793 | 14.7554546 | 13.4995949 | 16.5683924 |
| ENSMUSG00 | Gm10100 | 344 | 0 | 0 | 0 | 0 | 0 | 0 |
| ENSMUSG00 | Stx17 | 2428 | 16.7667265 | 20.6399917 | 20.3896475 | 27.2547239 | 21.2049478 | 14.8505243 |
| ENSMUSG00 | Olfr871 | 1146 | 0 | 0 | 0 | 0 | 0 | 0 |
| ENSMUSG00 | Nol10 | 3459 | 3.11469651 | 1.74134295 | 3.32248603 | 4.39159894 | 2.67007055 | 3.26444078 |

| | | | | | | | | |
|---|---|---|---|---|---|---|---|---|
| ENSMUSG00 | 1810013D10I | 6728 | 18.749129 | 14.6249576 | 24.8733915 | 13.6594779 | 16.3742487 | 19.9208646 |
| ENSMUSG00 | Obscn | 29009 | 0 | 0 | 0 | 0 | 0 | 0 |
| ENSMUSG00 | Gm3724 | 353 | 0.11262176 | 0.68252751 | 0.62608843 | 0.40886169 | 0 | 0.30906107 |
| ENSMUSG00 | Gm12569 | 793 | 0 | 0 | 0 | 0 | 0 | 0 |
| ENSMUSG00 | Mrps36 | 1464 | 3.55735516 | 4.27885074 | 4.86099503 | 3.62299218 | 2.16095307 | 5.27235171 |
| ENSMUSG00 | Rps7 | 954 | 2.41699986 | 4.04079179 | 4.95764906 | 3.74436311 | 2.03249715 | 2.85897686 |
| ENSMUSG00 | Snrpa | 2081 | 10.3734867 | 9.88736703 | 10.8752252 | 14.3218495 | 8.41040217 | 13.9977485 |
| ENSMUSG00 | Hist1h4d | 392 | 0.7099193 | 3.19603953 | 0.56379902 | 1.28864445 | 0.39050798 | 1.46114139 |
| ENSMUSG00 | Gm5161 | 857 | 0.13916738 | 0.16868066 | 0.41261931 | 0 | 0.11908139 | 0.12730287 |
| ENSMUSG00 | Rpl27a-ps1 | 496 | 0 | 0 | 0 | 0 | 0 | 0.16496758 |
| ENSMUSG00 | Olfr525 | 1049 | 0 | 0 | 0 | 0 | 0 | 0 |
| ENSMUSG00 | Gm5959 | 415 | 0 | 0 | 0 | 0 | 0 | 0 |
| ENSMUSG00 | Olfr197 | 922 | 0 | 0 | 0 | 0 | 0 | 0 |
| ENSMUSG00 | Gm10101 | 2913 | 0.10918086 | 0.84363493 | 1.50222536 | 2.2915133 | 1.82174497 | 1.31083059 |
| ENSMUSG00 | Sox21 | 6609 | 0 | 0 | 0 | 0 | 0 | 0 |
| ENSMUSG00 | Cox5b | 671 | 77.0817886 | 74.3981432 | 100.129361 | 64.5282465 | 56.5016352 | 181.004424 |
| ENSMUSG00 | Olfr153 | 1257 | 0 | 0 | 0 | 0 | 0 | 0 |
| ENSMUSG00 | Zic2 | 5491 | 0 | 0 | 0 | 0 | 0 | 0 |
| ENSMUSG00 | 4921509C19I | 2680 | 0 | 0 | 0 | 0 | 0 | 0 |
| ENSMUSG00 | Krt5 | 2415 | 0 | 0 | 0 | 0 | 0 | 0 |
| ENSMUSG00 | Fam23a | 3632 | 0.07662125 | 0.43781734 | 0.30425277 | 0 | 0.61816094 | 0 |
| ENSMUSG00 | Zfp955b | 1634 | 16.8851306 | 7.81481467 | 17.5563012 | 19.763421 | 21.8282966 | 34.6190954 |
| ENSMUSG00 | 4930534B04I | 11877 | 0.74309309 | 0.75056764 | 0.81131614 | 0.71696241 | 0.75613725 | 0.71648459 |
| ENSMUSG00 | C1qtnf7 | 4529 | 0 | 0 | 0 | 0 | 0.41686375 | 0 |
| ENSMUSG00 | Sec22c | 8129 | 0.36679309 | 0.67576017 | 0.29906524 | 0.55927391 | 0.77835781 | 0.44624517 |
| ENSMUSG00 | Orm2 | 769 | 0 | 0 | 0 | 0 | 0 | 0.10640301 |
| ENSMUSG00 | Zfp229 | 4519 | 9.1668977 | 10.7590441 | 8.80319954 | 3.51318867 | 7.57660947 | 9.05332128 |
| ENSMUSG00 | Olfr301 | 1206 | 0 | 0 | 0 | 0 | 0 | 0 |
| ENSMUSG00 | Wdr61 | 4664 | 22.2388612 | 18.8964414 | 25.6359317 | 17.2055032 | 15.7433437 | 22.2922223 |
| ENSMUSG00 | Olfr373 | 944 | 0 | 0 | 0 | 0 | 0 | 0 |
| ENSMUSG00 | Dpp6 | 10758 | 0.08129985 | 0.01791651 | 0.07395735 | 0.09726523 | 0.02845866 | 0.26620535 |
| ENSMUSG00 | Gpr114 | 3192 | 0 | 0 | 0 | 0 | 0 | 0 |
| ENSMUSG00 | Kir2 | 3870 | 0.0308182 | 0.09961022 | 0.05710832 | 0.12120583 | 0.17140643 | 0.07752481 |
| ENSMUSG00 | Lyg2 | 2560 | 0 | 0 | 0 | 0 | 0 | 0 |
| ENSMUSG00 | Dot1l | 8986 | 2.26074455 | 3.87164603 | 3.22192379 | 3.56564161 | 2.32815648 | 4.7228287 |
| ENSMUSG00 | Pclo | 20404 | 1.9211382 | 2.21283554 | 1.98002767 | 2.25822245 | 2.45328733 | 2.05990096 |
| ENSMUSG00 | Vmn1r78 | 941 | 0 | 0 | 0 | 0 | 0 | 0 |
| ENSMUSG00 | Akap6 | 10252 | 0 | 0 | 0.00862307 | 0 | 0 | 0.07183138 |
| ENSMUSG00 | Mdc1 | 7408 | 2.27005511 | 2.92709219 | 1.88549978 | 1.88982631 | 2.07329092 | 2.43365774 |
| ENSMUSG00 | U2af1 | 965 | 5.02608147 | 4.69380887 | 4.80952696 | 4.29993277 | 3.33125562 | 4.6070116 |
| ENSMUSG00 | Olfr845 | 941 | 0 | 0 | 0 | 0 | 0 | 0 |
| ENSMUSG00 | Hist1h2ab | 505 | 0.07872372 | 0 | 0.26258521 | 0 | 0 | 0 |
| ENSMUSG00 | Olfr12 | 1120 | 0 | 0 | 0 | 0 | 0 | 0 |
| ENSMUSG00 | Vma21-ps | 475 | 0 | 0 | 0 | 0 | 0 | 0 |
| ENSMUSG00 | Olfr68 | 947 | 0 | 0 | 0 | 0 | 0 | 0 |
| ENSMUSG00 | 1700029P11I | 1018 | 0 | 0 | 0 | 0 | 0 | 0 |
| ENSMUSG00 | Olfr1457 | 953 | 0 | 0 | 0 | 0 | 0 | 0 |
| ENSMUSG00 | Med9 | 1918 | 21.4530356 | 24.4198237 | 13.7122506 | 15.7082936 | 15.8825581 | 16.1969834 |
| ENSMUSG00 | Vmn1r46 | 929 | 0 | 0 | 0 | 0 | 0 | 0 |
| ENSMUSG00 | Cd2ap | 5381 | 31.2886935 | 28.5035208 | 34.2541694 | 33.647965 | 31.0558225 | 49.9925231 |
| ENSMUSG00 | Gdpd1 | 2326 | 6.47778466 | 11.8498043 | 6.10008238 | 8.08200567 | 6.4715309 | 4.77247557 |
| ENSMUSG00 | Gm6404 | 407 | 0 | 0 | 0 | 0 | 0.12537193 | 0 |
| ENSMUSG00 | Rpl21-ps8 | 1017 | 0 | 0 | 0 | 0 | 0 | 0.02681872 |
| ENSMUSG00 | Olfr971 | 923 | 0 | 0 | 0 | 0 | 0 | 0 |
| ENSMUSG00 | Dlgap4 | 7912 | 7.38631905 | 7.38144185 | 7.05039508 | 6.63997179 | 7.28119038 | 9.05247759 |
| ENSMUSG00 | Fbxw20 | 2809 | 0 | 0 | 0 | 0.01284516 | 0 | 0 |
| ENSMUSG00 | Tmem91 | 3751 | 0.15897953 | 0.20554068 | 0.30638441 | 0.65411331 | 0.20405109 | 1.13432251 |
| ENSMUSG00 | Olfr1036 | 1003 | 0 | 0 | 0 | 0 | 0 | 0 |
| ENSMUSG00 | Gm4871 | 1124 | 0 | 0 | 0 | 0 | 0 | 0 |
| ENSMUSG00 | Ppp1r1b | 2919 | 2.72391096 | 1.53523094 | 0.48456971 | 1.29791527 | 2.22005815 | 0.63538043 |
| ENSMUSG00 | Tnnt3 | 1382 | 0 | 0 | 0 | 0 | 0 | 0 |
| ENSMUSG00 | Gm2423 | 737 | 0 | 0 | 0 | 0.048958 | 0 | 0 |
| ENSMUSG00 | Btnl7 | 1754 | 0 | 0 | 0 | 0 | 0 | 0 |
| ENSMUSG00 | Ext1 | 7750 | 6.89436978 | 8.61760114 | 4.7338748 | 5.42394605 | 5.34624738 | 3.40669043 |
| ENSMUSG00 | Cyp2d22 | 2761 | 7.84742372 | 3.78719955 | 11.1104959 | 2.32618759 | 6.28358123 | 8.07076431 |
| ENSMUSG00 | Slc22a12 | 2366 | 0.285648 | 1.58856403 | 0.11209259 | 0.36600552 | 0.02156652 | 0.55333165 |
| ENSMUSG00 | Kalrn | 19219 | 3.94265809 | 3.24435486 | 1.89282048 | 3.10149007 | 4.34357411 | 3.34067854 |
| ENSMUSG00 | Bod1l | 10362 | 9.44969586 | 9.09134283 | 8.77041008 | 10.4290411 | 11.5328867 | 12.1606672 |
| ENSMUSG00 | Akr1b10 | 2783 | 14.9708026 | 11.9643663 | 14.7074763 | 15.4674377 | 14.7780305 | 21.1689618 |
| ENSMUSG00 | 1700052N19I | 7242 | 3.32668064 | 4.05213246 | 2.92359608 | 1.44487613 | 2.76903696 | 2.65138719 |
| ENSMUSG00 | Tac1 | 1027 | 0 | 0 | 0 | 0 | 0 | 0 |
| ENSMUSG00 | Klra6 | 1108 | 0 | 0 | 0 | 0 | 0 | 0 |
| ENSMUSG00 | Olfr126 | 959 | 0 | 0 | 0 | 0 | 0 | 0 |
| ENSMUSG00 | Mospd2 | 6426 | 16.2337972 | 12.9502001 | 15.0641202 | 20.3600208 | 20.2326806 | 22.6482221 |
| ENSMUSG00 | Cfd | 918 | 0 | 0 | 0 | 0 | 0 | 0 |
| ENSMUSG00 | Olfr510 | 944 | 0 | 0 | 0 | 0 | 0 | 0 |
| ENSMUSG00 | Rps17 | 1386 | 3.01177883 | 1.66879453 | 1.94539136 | 2.83762111 | 1.14128258 | 2.83372875 |
| ENSMUSG00 | Abpd | 517 | 0 | 0 | 0 | 0 | 0 | 0 |
| ENSMUSG00 | Olfr1204 | 929 | 0 | 0 | 0 | 0 | 0 | 0 |
| ENSMUSG00 | Armc4 | 3456 | 0 | 0 | 0 | 0 | 0 | 0 |
| ENSMUSG00 | Ttr | 1215 | 0 | 0 | 0.03638012 | 0 | 0.33597614 | 0.89793051 |
| ENSMUSG00 | Tbpl2 | 1520 | 0.49694351 | 0.15850803 | 0.02908016 | 0 | 0.13427994 | 0.01794384 |
| ENSMUSG00 | Rufy4 | 2352 | 0.84514202 | 0.51218582 | 0.88328513 | 0.46023016 | 0.41220287 | 0.75376342 |
| ENSMUSG00 | Myl1 | 2020 | 0.03936186 | 0 | 0 | 0 | 0 | 0.0135023 |
| ENSMUSG00 | Ces2c | 2041 | 0.19478432 | 0.18887386 | 0 | 0 | 0 | 0.12027034 |
| ENSMUSG00 | Vmn1r214 | 1226 | 0 | 0 | 0 | 0 | 0 | 0 |
| ENSMUSG00 | Gm6311 | 661 | 0 | 0 | 0 | 0 | 0 | 0 |
| ENSMUSG00 | Olfr316 | 1010 | 0 | 0 | 0 | 0 | 0 | 0 |
| ENSMUSG00 | Suclg2 | 3380 | 79.4874371 | 65.6504633 | 79.9948739 | 65.2786101 | 79.5588759 | 73.0040418 |
| ENSMUSG00 | Vmn1r-ps32 | 1311 | 0 | 0 | 0 | 0 | 0.31137377 | 0 |
| ENSMUSG00 | Gm10104 | 396 | 0 | 0 | 0 | 0 | 0 | 0 |
| ENSMUSG00 | Defb39 | 250 | 0 | 0 | 0 | 0 | 0 | 0 |
| ENSMUSG00 | Gm5805 | 518 | 0.76748032 | 0.27907206 | 0.76798569 | 0.4875952 | 0 | 0.15796123 |
| ENSMUSG00 | Gm7582 | 599 | 0 | 0 | 0.14758545 | 0 | 0 | 0 |
| ENSMUSG00 | Inadl | 14515 | 6.80622591 | 6.32083953 | 6.52293129 | 7.39041808 | 10.4865091 | 6.33433062 |
| ENSMUSG00 | Gm6822 | 617 | 0.12886704 | 0 | 0.14327988 | 0 | 0 | 0 |
| ENSMUSG00 | Galntl6 | 3015 | 0.0131859 | 0 | 0 | 0 | 0 | 0 |
| ENSMUSG00 | Gm9354 | 347 | 0 | 0 | 0 | 0 | 0 | 0.07860127 |
| ENSMUSG00 | Olfr1148 | 944 | 0 | 0 | 0 | 0 | 0 | 0 |
| ENSMUSG00 | BC048679 | 634 | 0 | 0 | 0 | 0 | 0 | 0 |
| ENSMUSG00 | Sphk1 | 4027 | 1.22415684 | 3.78120579 | 0.71346407 | 4.345615 | 2.6229103 | 1.55100382 |
| ENSMUSG00 | Ccdc62 | 3306 | 2.97023705 | 5.26173794 | 2.68740788 | 3.07777875 | 4.05926704 | 4.69427397 |
| ENSMUSG00 | Sabp3 | 6298 | 8.04199463 | 7.50570018 | 10.6539215 | 8.21556873 | 5.16097196 | 9.14207105 |
| ENSMUSG00 | Zscan20 | 11434 | 0.88662301 | 0.93136293 | 0.83115237 | 1.60624109 | 1.29864224 | 1.02333569 |
| ENSMUSG00 | Gm14292 | 455 | 0.43687341 | 0.31771281 | 0.58288145 | 0.31720479 | 0.22429176 | 0.17983279 |
| ENSMUSG00 | Rbak | 6489 | 3.9884139 | 3.97283812 | 2.49312291 | 2.45773828 | 3.61721881 | 3.518088 |
| ENSMUSG00 | Olfr50 | 1875 | 0 | 0 | 0 | 0 | 0 | 0 |
| ENSMUSG00 | Slc25a3 | 3454 | 541.24543 | 429.54851 | 543.654515 | 408.121979 | 407.162115 | 540.462691 |
| ENSMUSG00 | Ugt2b38 | 1885 | 0.99125071 | 1.30371806 | 0 | 0.38283336 | 0.73088178 | 2.99514606 |
| ENSMUSG00 | Myt1l | 8742 | 0 | 0 | 0 | 0 | 0 | 0 |
| ENSMUSG00 | Odf1 | 1103 | 0 | 0 | 0 | 0 | 0 | 0 |
| ENSMUSG00 | Dsg1b | 3950 | 0 | 0 | 0 | 0 | 0 | 0 |
| ENSMUSG00 | Gm10105 | 272 | 0 | 0 | 0 | 0 | 0 | 0 |
| ENSMUSG00 | Csn1s2a | 906 | 0 | 0 | 0 | 0 | 0 | 0 |
| ENSMUSG00 | Serpina10 | 1604 | 0.02478521 | 0 | 0 | 0 | 0.41355542 | 0 |
| ENSMUSG00 | Ppp4r1 | 4995 | 12.5275528 | 11.942906 | 13.326922 | 12.0490191 | 12.5650535 | 10.6095344 |
| ENSMUSG00 | Olfr1377 | 1062 | 0 | 0 | 0 | 0 | 0 | 0 |
| ENSMUSG00 | Gm14851 | 349 | 0 | 0 | 0 | 0 | 0 | 0 |
| ENSMUSG00 | Ces1e | 2346 | 11.6588962 | 21.8338397 | 6.65100195 | 21.5938579 | 13.6809648 | 4.58065126 |
| ENSMUSG00 | Olfr815 | 950 | 0 | 0 | 0 | 0 | 0 | 0 |
| ENSMUSG00 | Gm5581 | 2531 | 0 | 0.0190385 | 0 | 0 | 0 | 0.01077623 |
| ENSMUSG00 | Gm10106 | 666 | 0 | 0.07235202 | 0 | 0 | 0 | 0 |
| ENSMUSG00 | Olfr99 | 917 | 0 | 0 | 0 | 0 | 0 | 0 |
| ENSMUSG00 | Cldn24 | 662 | 0 | 0 | 0 | 0 | 0 | 0 |
| ENSMUSG00 | Olfr492 | 928 | 0 | 0 | 0 | 0 | 0 | 0 |
| ENSMUSG00 | Tas2r104 | 908 | 0 | 0 | 0 | 0 | 0 | 0 |
| ENSMUSG00 | Wbscr16 | 2863 | 12.2612956 | 9.50937472 | 8.04371646 | 9.51517416 | 12.4758865 | 6.82104146 |
| ENSMUSG00 | Dcpp1 | 613 | 0 | 0 | 0 | 0 | 0 | 0 |
| ENSMUSG00 | Flot2 | 2981 | 40.0355426 | 31.359174 | 39.3827894 | 40.0763668 | 41.0812821 | 41.8497819 |
| ENSMUSG00 | Rps12 | 511 | 0.70019437 | 0.84868489 | 0.77850604 | 0.91793851 | 0.69899145 | 0.85400045 |
| ENSMUSG00 | Olfr392 | 1690 | 0 | 0 | 0 | 0 | 0 | 0 |
| ENSMUSG00 | Rpl10a-ps2 | 644 | 0 | 0 | 0 | 0 | 0 | 0 |
| ENSMUSG00 | Hist1h2af | 398 | 0 | 0 | 0 | 0 | 0 | 0 |
| ENSMUSG00 | Gm4953 | 485 | 0.40985032 | 0.19870698 | 0.82024039 | 0.14879194 | 0.42083609 | 0.28118185 |
| ENSMUSG00 | Rpl34 | 1295 | 3.49971025 | 2.75351098 | 2.62821769 | 2.81412085 | 2.44296162 | 6.00252679 |
| ENSMUSG00 | Hsh2d | 1256 | 0 | 0 | 0 | 0 | 0.97502629 | 0 |
| ENSMUSG00 | Zfp13 | 3659 | 3.28126677 | 4.31952802 | 1.08722762 | 2.07084705 | 1.38059885 | 0.87213249 |
| ENSMUSG00 | Gmfb | 4437 | 67.6748128 | 58.1451907 | 65.9989204 | 63.6335358 | 49.0598421 | 86.6432365 |
| ENSMUSG00 | Abca14 | 5959 | 0 | 0 | 0 | 0 | 0 | 0 |
| ENSMUSG00 | Irgc1 | 2961 | 0 | 0 | 0 | 0 | 0 | 0 |
| ENSMUSG00 | Athl1 | 3477 | 12.4628915 | 14.260526 | 6.03850314 | 12.3386687 | 12.9437053 | 7.16184804 |
| ENSMUSG00 | 4932415M13 | 1216 | 0 | 0 | 0 | 0 | 0 | 0 |
| ENSMUSG00 | Olfr285 | 959 | 0 | 0 | 0 | 0 | 0 | 0 |
| ENSMUSG00 | Gm10108 | 475 | 0 | 0 | 0.18611302 | 0 | 0.10742395 | 0.05742029 |
| ENSMUSG00 | Zfp27 | 6889 | 3.46251826 | 2.66497815 | 2.91941335 | 3.16876716 | 7.45878363 | 4.51344009 |
| ENSMUSG00 | Olfr294 | 1007 | 0 | 0 | 0 | 0 | 0 | 0 |
| ENSMUSG00 | Lmtk3 | 5597 | 0 | 0.00860933 | 0 | 0.07736011 | 0.00911674 | 0.02923849 |
| ENSMUSG00 | 5730460C07I | 1831 | 0 | 0 | 0 | 0 | 0 | 0 |
| ENSMUSG00 | Iah1 | 918 | 13.0352937 | 8.81843386 | 12.6153408 | 10.7302812 | 10.2830932 | 15.5091086 |
| ENSMUSG00 | Lcn13 | 737 | 0 | 0 | 0 | 0 | 0 | 0 |
| ENSMUSG00 | Slc2a7 | 1855 | 0 | 0.12988259 | 0 | 0 | 0 | 0 |
| ENSMUSG00 | Pgk1 | 2385 | 7.55104095 | 8.00076775 | 7.19090779 | 7.92745968 | 5.99051792 | 8.06231475 |
| ENSMUSG00 | Gm10109 | 798 | 16.3904174 | 10.3860502 | 4.70821636 | 10.8517427 | 9.97508099 | 9.43333388 |
| ENSMUSG00 | Wisp3 | 1413 | 0 | 0 | 0.1876936 | 0 | 0.54168127 | 0.03860529 |
| ENSMUSG00 | Lrmb2 | 4374 | 3.73559728 | 7.12771561 | 5.5883903 | 6.50862669 | 3.63974149 | 4.30881932 |
| ENSMUSG00 | Trim54 | 1770 | 0 | 0 | 0 | 0 | 0 | 0 |
| ENSMUSG00 | Qk | 7172 | 22.4553056 | 23.2399475 | 22.1379002 | 27.8966727 | 21.4507143 | 23.1446534 |
| ENSMUSG00 | Gm6055 | 677 | 2.99487372 | 4.12823286 | 5.61500517 | 2.29176945 | 2.86410972 | 2.53811266 |
| ENSMUSG00 | Cd200r4 | 2125 | 0 | 0 | 0 | 0 | 0 | 0 |
| ENSMUSG00 | Rpl13a-ps1 | 611 | 0.06506625 | 0 | 0 | 0.05905408 | 0 | 0.08927869 |
| ENSMUSG00 | 4930423O20I | 566 | 0 | 0 | 0 | 0 | 0 | 0 |
| ENSMUSG00 | Gm10110 | 1846 | 0.04307203 | 0 | 0 | 0 | 0 | 0 |
| ENSMUSG00 | Btbd3 | 5857 | 5.87813661 | 4.77174944 | 5.36580338 | 7.62053766 | 4.78290598 | 5.34816774 |
| ENSMUSG00 | Zfp119b | 1628 | 39.8043202 | 23.4124544 | 28.0469925 | 58.9768553 | 22.5982905 | 55.7052675 |
| ENSMUSG00 | Olfr924 | 926 | 0 | 0 | 0 | 0 | 0 | 0 |
| ENSMUSG00 | Olfr183 | 1194 | 0 | 0 | 0 | 0 | 0 | 0 |
| ENSMUSG00 | Sfrd2 | 8419 | 2.18161682 | 2.70150858 | 2.46761685 | 2.17718002 | 2.12736167 | 1.87251948 |
| ENSMUSG00 | Rai1 | 9800 | 7.11947636 | 4.21877218 | 7.23917939 | 7.90491322 | 4.86312602 | 7.39198386 |
| ENSMUSG00 | Zfp954 | 3506 | 6.23659848 | 8.24639626 | 13.4647942 | 7.81125836 | 8.38311251 | 5.36001895 |
| ENSMUSG00 | Olfr149 | 1029 | 0 | 0 | 0 | 0 | 0 | 0 |
| ENSMUSG00 | Defb45 | 732 | 0 | 0 | 0 | 0 | 0 | 0 |
| ENSMUSG00 | Cttnbp2nl | 4993 | 8.90975019 | 7.01613127 | 9.33081164 | 9.1849146 | 7.83841984 | 7.21606218 |
| ENSMUSG00 | Olfr20 | 1080 | 0 | 0 | 0 | 0 | 0 | 0 |
| ENSMUSG00 | Gm1082 | 2636 | 0 | 0 | 0 | 0 | 0 | 0 |
| ENSMUSG00 | Olfr569 | 944 | 0 | 0 | 0 | 0 | 0 | 0 |
| ENSMUSG00 | Olfr495 | 992 | 0 | 0 | 0 | 0 | 0 | 0 |
| ENSMUSG00 | Ear6 | 1603 | 0 | 0 | 0 | 0 | 0 | 0 |
| ENSMUSG00 | Olfr943 | 944 | 0 | 0 | 0 | 0 | 0 | 0 |
| ENSMUSG00 | Unc13c | 6601 | 0.10238497 | 0 | 0.01339247 | 0 | 0 | 0 |
| ENSMUSG00 | Rpl21-ps14 | 482 | 0 | 0 | 0 | 0 | 0 | 0 |
| ENSMUSG00 | 1700061J05R | 1391 | 0 | 0 | 0 | 0 | 0 | 0 |
| ENSMUSG00 | Il28ra | 4175 | 4.86588037 | 2.13520762 | 3.69495646 | 2.83470913 | 2.77436822 | 2.92018293 |
| ENSMUSG00 | Mageb1 | 2134 | 0 | 0 | 0 | 0 | 0 | 0 |
| ENSMUSG00 | Vmn1r232 | 1158 | 0 | 0 | 0 | 0 | 0 | 0 |
| ENSMUSG00 | Ppef1 | 2873 | 2.85054959 | 0 | 2.80011675 | 0 | 1.15444289 | 1.96514108 |
| ENSMUSG00 | Cnih4 | 3568 | 19.2203486 | 17.6107401 | 15.9314939 | 22.6018412 | 17.0326271 | 30.6075268 |
| ENSMUSG00 | Fmr1nb | 1281 | 0 | 0 | 0 | 0 | 0 | 0 |
| ENSMUSG00 | Olfr644 | 1209 | 0 | 0 | 0 | 0 | 0 | 0 |
| ENSMUSG00 | Tgif2 | 3948 | 0.05034889 | 0.10984751 | 0.05598004 | 0.07311458 | 0.16801998 | 0.08981011 |
| ENSMUSG00 | Ces3b | 2663 | 0 | 0 | 0 | 0 | 0 | 0 |
| ENSMUSG00 | Hs6st2 | 7800 | 0.48929822 | 0.63630815 | 0 | 0 | 0 | 0 |
| ENSMUSG00 | Cyp2d34 | 1613 | 0 | 0 | 0 | 0 | 0 | 0 |
| ENSMUSG00 | Olfr395 | 938 | 0 | 0 | 0 | 0 | 0 | 0 |
| ENSMUSG00 | Lancl2 | 3575 | 10.2641422 | 13.5191613 | 14.9853521 | 12.2123843 | 11.2614855 | 13.3664806 |
| ENSMUSG00 | Olfr1475 | 944 | 0 | 0 | 0 | 0 | 0 | 0 |
| ENSMUSG00 | Fabp5l2 | 407 | 0 | 0 | 0 | 0 | 0 | 0 |
| ENSMUSG00 | 2700097O09I | 5199 | 6.63738355 | 7.24789339 | 6.63155175 | 7.93957664 | 6.37952381 | 6.98784756 |
| ENSMUSG00 | Olfr1465 | 923 | 0 | 0 | 0 | 0 | 0 | 0 |
| ENSMUSG00 | Vmn2r7 | 3629 | 0 | 0 | 0 | 0 | 0 | 0 |
| ENSMUSG00 | Prl3d3 | 848 | 0 | 0 | 0 | 0 | 0 | 0 |
| ENSMUSG00 | Btbd9 | 14642 | 5.05564163 | 4.83773186 | 5.49126844 | 6.40713808 | 5.47482833 | 6.3576249 |
| ENSMUSG00 | Gspt1 | 8973 | 16.1405567 | 17.7859686 | 15.4876401 | 15.6222827 | 16.0420602 | 16.5508092 |
| ENSMUSG00 | Olfr1373 | 935 | 0 | 0 | 0 | 0 | 0 | 0 |
| ENSMUSG00 | Erbb4 | 12771 | 0.73154318 | 0.90932054 | 0.38072216 | 0.99733472 | 0.61530514 | 0.21143131 |
| ENSMUSG00 | Tnfaip8 | 3804 | 108.01075 | 63.5645549 | 145.224667 | 70.0204134 | 67.6729932 | 150.799215 |
| ENSMUSG00 | Olfr889 | 943 | 0 | 0 | 0 | 0 | 0 | 0 |
| ENSMUSG00 | 4933411G06I | 1020 | 0 | 0 | 0 | 0 | 0 | 0 |
| ENSMUSG00 | Gm10112 | 353 | 0 | 0 | 0 | 0 | 0 | 0 |
| ENSMUSG00 | Rapgef2 | 12636 | 1.70209837 | 1.2508035 | 1.29079456 | 1.50770177 | 2.33810317 | 1.31667695 |
| ENSMUSG00 | Gak | 6452 | 21.9604049 | 19.9034204 | 21.3404744 | 25.5236285 | 23.1406038 | 18.037955 |
| ENSMUSG00 | Gm5341 | 491 | 0 | 0 | 0 | 0 | 0 | 0 |
| ENSMUSG00 | Olfr743 | 935 | 0 | 0 | 0 | 0 | 0 | 0 |
| ENSMUSG00 | Olfr784 | 959 | 0 | 0 | 0 | 0 | 0 | 0 |
| ENSMUSG00 | Olfr170 | 1007 | 0 | 0 | 0 | 0 | 0 | 0 |
| ENSMUSG00 | Cks2 | 689 | 1.15400524 | 2.16804021 | 2.63029835 | 3.40396647 | 0.5184102 | 1.58343334 |
| ENSMUSG00 | Lhfpl5 | 1948 | 0 | 0 | 0 | 0 | 0 | 0 |
| ENSMUSG00 | Opcml | 13035 | 0.05489825 | 0 | 0 | 0 | 0 | 0.00209242 |
| ENSMUSG00 | Morf4l1 | 1968 | 5.59566469 | 6.73336971 | 4.64927922 | 6.16034907 | 4.35591013 | 5.07241767 |
| ENSMUSG00 | Olfr1129 | 1123 | 0 | 0 | 0 | 0 | 0 | 0 |
| ENSMUSG00 | Fbxw24 | 1477 | 0 | 0 | 0 | 0 | 0 | 0 |
| ENSMUSG00 | Gm11562 | 816 | 0 | 0 | 0 | 0 | 0 | 0 |
| ENSMUSG00 | Gm10237 | 332 | 0 | 0 | 0 | 0 | 0 | 0 |
| ENSMUSG00 | Olfr1285 | 881 | 0 | 0 | 0 | 0 | 0 | 0 |
| ENSMUSG00 | Gm10113 | 3150 | 0 | 0 | 0 | 0 | 0 | 0 |
| ENSMUSG00 | Gm6030 | 401 | 0.1982817 | 0 | 0.88183228 | 0 | 0.12724782 | 0.61214901 |
| ENSMUSG00 | Olfr1386 | 1127 | 0 | 0 | 0 | 0 | 0 | 0 |
| ENSMUSG00 | Trank1 | 10497 | 0.03787318 | 0.18361986 | 0.11369437 | 0 | 0.37916141 | 0 |
| ENSMUSG00 | Pvrl2 | 3430 | 21.4540509 | 21.5925836 | 23.312284 | 17.3782908 | 16.5724148 | 24.3801877 |
| ENSMUSG00 | Rpp25 | 1345 | 0 | 0.32243716 | 0 | 0 | 0.1517513 | 0.1419498 |
| ENSMUSG00 | Glrp1 | 522 | 0.07615992 | 0 | 0 | 0 | 0 | 0 |
| ENSMUSG00 | Erbb2 | 5426 | 16.7271954 | 18.7026626 | 15.7386584 | 18.2471674 | 19.6638688 | 11.244631 |
| ENSMUSG00 | Olfr1505 | 1102 | 0 | 0 | 0 | 0 | 0 | 0 |
| ENSMUSG00 | Gm10115 | 1227 | 0 | 0 | 0 | 0 | 0 | 0 |
| ENSMUSG00 | T | 2307 | 0 | 0 | 0 | 0 | 0 | 0 |
| ENSMUSG00 | Rpl17 | 768 | 1.7082433 | 1.94502566 | 1.49641656 | 2.4430551 | 1.12949009 | 3.51587147 |
| ENSMUSG00 | Cytl1 | 970 | 0 | 0 | 0 | 0 | 0 | 0 |
| ENSMUSG00 | Olfr117 | 953 | 0 | 0 | 0 | 0 | 0 | 0 |
| ENSMUSG00 | Serpinb9e | 1912 | 0 | 0 | 0 | 0 | 0 | 0 |
| ENSMUSG00 | Serpinb2 | 2337 | 0 | 0 | 0 | 0 | 0 | 0 |
| ENSMUSG00 | Itgb1bp1 | 3310 | 7.3865923 | 7.51184415 | 8.90713876 | 7.35812086 | 6.96795223 | 8.53671488 |
| ENSMUSG00 | Gm15772 | 955 | 0 | 0 | 0 | 0.11334674 | 0 | 1.22807276 |
| ENSMUSG00 | Gm5619 | 854 | 0 | 0 | 0 | 0 | 0 | 0 |
| ENSMUSG00 | Gm4968 | 863 | 0 | 0 | 0 | 0 | 0 | 0 |
| ENSMUSG00 | Dub1a | 2934 | 0 | 0 | 0 | 0 | 0 | 0 |
| ENSMUSG00 | Otof | 7852 | 0 | 0 | 0 | 0 | 0 | 0 |
| ENSMUSG00 | Tmem65 | 3740 | 43.3164661 | 44.1794947 | 29.5112305 | 49.4729209 | 43.9045127 | 30.6803228 |
| ENSMUSG00 | 2010012O05I | 2171 | 8.67391606 | 4.83862017 | 8.79557632 | 5.90010401 | 6.88656293 | 8.09068064 |
| ENSMUSG00 | Tubb3 | 1864 | 0 | 0 | 0 | 0 | 0 | 0 |
| ENSMUSG00 | Vps28 | 1214 | 87.6323442 | 97.1667301 | 87.7483045 | 85.4496524 | 72.2103078 | 95.4163039 |
| ENSMUSG00 | Gm10116 | 921 | 0.25899336 | 0.62783638 | 0.43193983 | 0.23506218 | 0.49862908 | 0.26652742 |
| ENSMUSG00 | B130024G19 | 306 | 0.12391987 | 0.4724161 | 0 | 0 | 0 | 0 |
| ENSMUSG00 | Olfr251 | 950 | 0 | 0 | 0 | 0 | 0 | 0 |
| ENSMUSG00 | 4932435O22I | 6200 | 0.0256487 | 0 | 0 | 0 | 0 | 0 |
| ENSMUSG00 | Dgkk | 7130 | 0 | 0 | 0 | 0 | 0 | 0 |
| ENSMUSG00 | Zfp706 | 4409 | 58.5827528 | 70.8534145 | 71.3806129 | 73.6453433 | 58.248072 | 61.8674682 |
| ENSMUSG00 | Krtap16-8 | 599 | 0 | 0 | 0 | 0 | 0 | 0 |
| ENSMUSG00 | Hsd3b3 | 2989 | 0.22611013 | 0.01612126 | 0.02957634 | 0 | 0.18778526 | 0.14600008 |
| ENSMUSG00 | Hist1h3g | 529 | 0 | 0 | 0 | 0 | 0 | 0 |
| ENSMUSG00 | Arf2 | 2991 | 10.2080271 | 10.9712361 | 8.03938569 | 15.7308546 | 12.0614001 | 7.3042414 |
| ENSMUSG00 | Olfr304 | 1001 | 0 | 0 | 0 | 0 | 0 | 0 |
| ENSMUSG00 | Cyp26c1 | 1741 | 0 | 0 | 0 | 0 | 0.20516062 | 0 |
| ENSMUSG00 | Krtap6-2 | 394 | 0 | 0 | 0 | 0 | 0 | 0 |
| ENSMUSG00 | Olfr516 | 944 | 0 | 0 | 0 | 0 | 0 | 0 |
| ENSMUSG00 | Adam1b | 2883 | 0.55158488 | 0.63513174 | 0.38329729 | 0.08760816 | 0.7079622 | 0.10406557 |
| ENSMUSG00 | Ap3b2 | 6665 | 0.16701477 | 0.26027186 | 0.04642354 | 0.37354255 | 0 | 0.2168876 |
| ENSMUSG00 | Rpl9-ps6 | 672 | 0 | 0.35853008 | 0.32888276 | 0.26846759 | 0.07593211 | 0.08117452 |
| ENSMUSG00 | Gm8623 | 435 | 0 | 0 | 0.10161343 | 0.08294723 | 0.23460403 | 0.06270032 |
| ENSMUSG00 | Gm5453 | 492 | 0.16160764 | 0 | 0 | 0 | 0.20742429 | 0 |
| ENSMUSG00 | Gm9705 | 2629 | 0 | 0 | 0.01681318 | 0 | 0 | 0 |
| ENSMUSG00 | 5730577I03R | 2673 | 3.83722932 | 2.79420072 | 1.27330412 | 3.17219621 | 1.67988069 | 5.02024785 |
| ENSMUSG00 | Gm4945 | 455 | 0 | 0 | 0 | 0.0793012 | 0 | 0 |
| ENSMUSG00 | Hmgb1l | 578 | 0 | 0.08336755 | 0 | 0 | 0 | 0 |
| ENSMUSG00 | Ctrc | 1417 | 0 | 0 | 0 | 0 | 0 | 0 |
| ENSMUSG00 | Acat3 | 2414 | 0.85637323 | 0.69864353 | 1.11694798 | 0.61282677 | 0.82436978 | 0.2259705 |
| ENSMUSG00 | Vmn1r171 | 1580 | 0 | 0 | 0 | 0 | 0 | 0 |
| ENSMUSG00 | I830012O16F | 1975 | 3.26095587 | 3.24496041 | 2.26044868 | 2.26540431 | 3.95292937 | 3.21771693 |
| ENSMUSG00 | 4930431F12F | 426 | 0 | 0 | 0 | 0 | 0 | 0 |
| ENSMUSG00 | Olfr1411 | 971 | 0 | 0 | 0 | 0 | 0 | 0 |
| ENSMUSG00 | 4930425F17F | 769 | 0 | 0 | 0 | 0 | 0 | 0 |
| ENSMUSG00 | Nsl1 | 3627 | 0 | 0.49156282 | 0.42654108 | 0 | 0 | 0 |
| ENSMUSG00 | 4930512M02 | 935 | 0 | 0 | 0 | 0 | 0 | 0 |
| ENSMUSG00 | Fabp4 | 633 | 8.72987645 | 4.03456783 | 1.53624099 | 6.27018151 | 2.57953243 | 1.68042801 |
| ENSMUSG00 | Zfp534 | 5341 | 0.00744345 | 0.0811979 | 0.00827595 | 0 | 0.04776856 | 0.00510665 |
| ENSMUSG00 | Zfp398 | 6728 | 5.07579634 | 4.14684156 | 5.51209071 | 6.90213902 | 4.62635096 | 6.77406691 |
| ENSMUSG00 | Ncr1 | 1547 | 0 | 0 | 0 | 0 | 0 | 0 |
| ENSMUSG00 | Mppe1 | 1996 | 15.9738955 | 19.4821938 | 15.7452457 | 16.377922 | 20.0168599 | 14.867138 |
| ENSMUSG00 | Olfr1408 | 1036 | 0 | 0 | 0 | 0 | 0 | 0 |
| ENSMUSG00 | Tas2r109 | 950 | 0 | 0 | 0 | 0 | 0 | 0 |
| ENSMUSG00 | Gm6003 | 467 | 0 | 0 | 0 | 0 | 0 | 0 |
| ENSMUSG00 | Zfp781 | 729 | 0 | 0 | 0 | 0 | 0 | 0 |
| ENSMUSG00 | Syt9 | 4329 | 0 | 0.24488374 | 0.36758289 | 0 | 0 | 0 |
| ENSMUSG00 | Gsdmcl1 | 1128 | 0 | 0 | 0 | 0 | 0 | 0 |
| ENSMUSG00 | Tlr12 | 3591 | 0.19927559 | 0.36230408 | 0 | 0 | 0 | 0 |
| ENSMUSG00 | V1ra8 | 889 | 0 | 0 | 0 | 0 | 0 | 0 |
| ENSMUSG00 | Prl2c1 | 903 | 0 | 0 | 0 | 0 | 0 | 0 |
| ENSMUSG00 | Olfr709-ps1 | 923 | 0 | 0 | 0 | 0 | 0 | 0 |
| ENSMUSG00 | Gm12751 | 380 | 0 | 0.12680643 | 0 | 0 | 0 | 0 |
| ENSMUSG00 | Abph | 415 | 0 | 0 | 0 | 0 | 0 | 0 |
| ENSMUSG00 | Gm10118 | 2568 | 0 | 0 | 0.01721256 | 0 | 0 | 0 |
| ENSMUSG00 | Cys1 | 2645 | 9.27377371 | 11.2222489 | 7.48673938 | 11.1451911 | 10.8611908 | 10.3220847 |
| ENSMUSG00 | Tex28 | 1420 | 0 | 0 | 0 | 0 | 0 | 0 |
| ENSMUSG00 | Timm17a | 1753 | 70.5302592 | 61.6005801 | 92.8163058 | 65.4540226 | 49.0179218 | 75.4603539 |
| ENSMUSG00 | Rpl30-ps8 | 463 | 0 | 0 | 0 | 0 | 0 | 0 |
| ENSMUSG00 | Cnr2 | 3979 | 0 | 0 | 0.39991615 | 0 | 0 | 0.01370929 |
| ENSMUSG00 | Gm6104 | 1467 | 0.02709985 | 0 | 0.06026155 | 0 | 0.03478281 | 0.31606603 |
| ENSMUSG00 | Armc9 | 12475 | 1.95032898 | 2.03947427 | 1.86728427 | 1.13090817 | 2.16376375 | 0.9926001 |
| ENSMUSG00 | Tubb4a | 2224 | 0 | 0 | 0 | 0 | 0.04588703 | 0 |
| ENSMUSG00 | Lilrb4 | 1481 | 0 | 0 | 0 | 0 | 0 | 0 |
| ENSMUSG00 | Vmn1r178 | 914 | 0 | 0 | 0 | 0 | 0 | 0 |
| ENSMUSG00 | Srpk2 | 8987 | 7.91836098 | 5.72639593 | 6.98415679 | 7.27502667 | 6.09795567 | 9.40818757 |
| ENSMUSG00 | Olfr195 | 994 | 0 | 0 | 0 | 0.65339719 | 0.25667191 | 0 |
| ENSMUSG00 | Kcnj15 | 6180 | 0.53393283 | 1.4814602 | 0 | 0.91080889 | 0.21467407 | 0.44575058 |
| ENSMUSG00 | Gm10119 | 794 | 0 | 0 | 0 | 0 | 0.25705983 | 0 |
| ENSMUSG00 | 2310039H08I | 772 | 10.7113212 | 10.7358394 | 15.1729124 | 11.4041695 | 8.32813906 | 19.3254244 |
| ENSMUSG00 | Olfr911-ps1 | 939 | 0 | 0 | 0 | 0 | 0 | 0 |
| ENSMUSG00 | Cyp2c67 | 1723 | 0 | 0 | 0 | 0 | 0 | 0 |
| ENSMUSG00 | Mysm1 | 13985 | 3.21511966 | 1.86750459 | 2.64547319 | 5.1420461 | 3.14878529 | 4.91860137 |
| ENSMUSG00 | Olfr114 | 999 | 0 | 0 | 0 | 0 | 0 | 0 |
| ENSMUSG00 | Gm12588 | 1340 | 0.83071153 | 0.75516066 | 1.1215393 | 1.26556425 | 0.60927016 | 2.86994338 |
| ENSMUSG00 | Btnl1 | 1885 | 0 | 0 | 0 | 0 | 0 | 0 |
| ENSMUSG00 | Ganc | 8909 | 11.2050748 | 10.4226371 | 10.3943048 | 7.07951665 | 9.76540249 | 8.71600606 |
| ENSMUSG00 | Rpl7a | 1440 | 2.26385375 | 2.00776842 | 3.4072254 | 2.50569754 | 1.41739933 | 2.74640465 |
| ENSMUSG00 | Olfr44 | 1069 | 0 | 0 | 0 | 0 | 0 | 0 |
| ENSMUSG00 | H2afb1 | 524 | 0 | 0 | 0 | 0 | 0 | 0 |
| ENSMUSG00 | Ncs1 | 4721 | 0.02526296 | 0.29599806 | 0.05617688 | 0 | 0 | 0.03466381 |
| ENSMUSG00 | Gm10120 | 356 | 0 | 0 | 0 | 0 | 0 | 0 |
| ENSMUSG00 | Atp5g2 | 642 | 1.36233734 | 0.60045411 | 0.48195156 | 1.34886148 | 1.43064605 | 1.10458041 |
| ENSMUSG00 | Gm6531 | 2300 | 0 | 0 | 0 | 0 | 0 | 0 |
| ENSMUSG00 | 1110001A16I | 1497 | 11.0210584 | 11.0085259 | 12.8737499 | 12.9914643 | 9.20315396 | 17.5454092 |
| ENSMUSG00 | Cav3 | 1158 | 0 | 0 | 0 | 0 | 0 | 0 |
| ENSMUSG00 | Olfr136 | 938 | 0 | 0 | 0 | 0 | 0 | 0 |
| ENSMUSG00 | 9430002A10I | 468 | 0 | 0 | 0.18889677 | 0 | 0 | 0 |
| ENSMUSG00 | Tpbpb | 703 | 0 | 0 | 0 | 0 | 0 | 0 |
| ENSMUSG00 | Olfr493 | 944 | 0 | 0 | 0 | 0 | 0 | 0 |
| ENSMUSG00 | Olfr531 | 914 | 0 | 0 | 0 | 0 | 0 | 0 |
| ENSMUSG00 | Sim2 | 1963 | 1.17463977 | 0.04909469 | 0 | 0.38600251 | 0 | 0 |
| ENSMUSG00 | Olfr857 | 929 | 0 | 0 | 0 | 0 | 0 | 0 |
| ENSMUSG00 | Hist1h2bk | 478 | 0 | 0 | 0 | 0 | 0 | 0 |
| ENSMUSG00 | Ppox | 4503 | 21.5684297 | 26.8059155 | 17.2861305 | 23.3976394 | 32.8277617 | 31.1813997 |
| ENSMUSG00 | Lypd4 | 1380 | 0 | 0 | 0 | 0 | 0 | 0 |
| ENSMUSG00 | Prl3d2 | 932 | 0 | 0 | 0 | 0 | 0 | 0 |
| ENSMUSG00 | Gm5239 | 368 | 0.1080312 | 0 | 0 | 0 | 0 | 0 |
| ENSMUSG00 | Zfp677 | 2913 | 23.8423702 | 23.7541129 | 14.3090759 | 20.8218046 | 20.1793289 | 35.3643366 |
| ENSMUSG00 | Prss1 | 800 | 0 | 0 | 0 | 0 | 0 | 0 |
| ENSMUSG00 | AI413582 | 1639 | 11.8854091 | 7.55577525 | 12.8641117 | 10.5009977 | 8.56147245 | 17.4730758 |
| ENSMUSG00 | Olfr1206 | 923 | 0 | 0 | 0 | 0 | 0 | 0 |
| ENSMUSG00 | Gm16477 | 851 | 0 | 0 | 0 | 0.04239958 | 0 | 0.06410021 |
| ENSMUSG00 | 1810041L15F | 5910 | 0.255619 | 0.26090798 | 0 | 0 | 0 | 0.004615 |
| ENSMUSG00 | Zfp512 | 3279 | 25.5094636 | 21.910944 | 20.0721391 | 20.9405705 | 18.7983721 | 19.8550058 |
| ENSMUSG00 | Ei24 | 3347 | 27.6756109 | 24.791471 | 28.578664 | 25.096803 | 23.096791 | 32.7914994 |
| ENSMUSG00 | Tex101 | 1048 | 0 | 0 | 0.08435466 | 0 | 0 | 0 |
| ENSMUSG00 | Chia | 1911 | 0 | 0 | 0 | 0 | 0 | 0.34253864 |
| ENSMUSG00 | Olfr527 | 1894 | 0 | 0 | 0 | 0 | 0 | 0 |
| ENSMUSG00 | Kcnc3 | 6207 | 0.43553612 | 0.21737077 | 0.71925023 | 0.47086283 | 0.43570129 | 1.23476295 |
| ENSMUSG00 | Gm382 | 4067 | 0 | 0 | 0 | 0 | 0 | 0 |
| ENSMUSG00 | Olfr1158 | 941 | 0 | 0 | 0 | 0 | 0 | 0 |
| ENSMUSG00 | Zfp599 | 3984 | 4.30085645 | 3.91877692 | 1.76407958 | 5.48838328 | 1.0630495 | 1.33497858 |
| ENSMUSG00 | I7Rn6 | 3230 | 13.157464 | 11.9198041 | 12.877379 | 11.4278426 | 9.24161915 | 11.8049367 |
| ENSMUSG00 | Hist1h3d | 1248 | 0 | 0 | 0.31876329 | 0 | 0 | 0 |
| ENSMUSG00 | Gm10121 | 475 | 0 | 0.81156113 | 0.09305651 | 0 | 0.32227185 | 0.28710147 |
| ENSMUSG00 | Saa1 | 433 | 0 | 0 | 0 | 0 | 0 | 0 |
| ENSMUSG00 | Vmn1r51 | 1804 | 0 | 0 | 0 | 0.26001473 | 0.14142565 | 0 |
| ENSMUSG00 | 4833420G17I | 2561 | 20.4443451 | 21.8447713 | 21.4882056 | 22.0070885 | 28.9302217 | 26.4013396 |
| ENSMUSG00 | Actg1 | 3308 | 7.34298627 | 8.97304969 | 5.58535985 | 9.16230877 | 7.94394908 | 4.09779193 |
| ENSMUSG00 | Ces2f | 1810 | 0 | 0 | 0 | 0 | 0 | 0 |
| ENSMUSG00 | Sval3 | 546 | 0 | 0 | 0 | 0 | 0 | 0 |
| ENSMUSG00 | Olfr883 | 929 | 0 | 0 | 0 | 0 | 0 | 0 |
| ENSMUSG00 | Olfr1458 | 905 | 0 | 0 | 0 | 0 | 0 | 0 |
| ENSMUSG00 | Gm14176 | 476 | 0.08351992 | 0.10123202 | 0 | 0 | 0 | 0 |
| ENSMUSG00 | Tcp11 | 4677 | 0.75651866 | 1.1333138 | 0.51979931 | 0.77919318 | 0.67642405 | 0.76977814 |
| ENSMUSG00 | Zfp28 | 2702 | 4.29629915 | 3.94123009 | 6.64172771 | 4.38005574 | 5.41990002 | 4.47174877 |
| ENSMUSG00 | Phactr2 | 6868 | 15.1022203 | 16.0598087 | 15.1179571 | 15.8555053 | 15.4609622 | 14.4355436 |
| ENSMUSG00 | Impdh2 | 1667 | 3.00491334 | 2.86170232 | 2.86370669 | 2.03462039 | 2.75487332 | 2.56875727 |
| ENSMUSG00 | Olfr830 | 947 | 0 | 0 | 0 | 0 | 0 | 0 |
| ENSMUSG00 | Olfr1355 | 1146 | 0 | 0 | 0 | 0 | 0 | 0 |
| ENSMUSG00 | Gm4978 | 815 | 0 | 0 | 0 | 0 | 0 | 0.03346582 |
| ENSMUSG00 | Olfr298 | 998 | 0 | 0 | 0 | 0 | 0 | 0 |
| ENSMUSG00 | Olfr1450 | 1917 | 0 | 0 | 0 | 0 | 0 | 0 |
| ENSMUSG00 | Gm9154 | 376 | 0 | 0.38446629 | 0 | 0 | 0 | 0.2176168 |
| ENSMUSG00 | Klhl24 | 6675 | 21.732996 | 19.6788377 | 29.6202013 | 19.7843121 | 30.8680907 | 23.0577962 |
| ENSMUSG00 | Vmn1r32 | 2439 | 0 | 0 | 0 | 0 | 0 | 0 |
| ENSMUSG00 | Hdac10 | 3276 | 9.21074777 | 13.0468175 | 6.24708587 | 9.09760952 | 11.2145881 | 8.81680188 |
| ENSMUSG00 | Acadm | 2919 | 84.0735119 | 80.3272448 | 86.9954053 | 72.4236721 | 81.5128436 | 69.1163092 |
| ENSMUSG00 | Olfr777 | 935 | 0 | 0 | 0 | 0 | 0 | 0 |
| ENSMUSG00 | Cfl2 | 2936 | 88.3665756 | 71.3113389 | 100.899439 | 98.4022243 | 60.5851315 | 117.217779 |
| ENSMUSG00 | Zfp938 | 3863 | 14.0065258 | 24.1867749 | 19.3147065 | 24.742774 | 14.9261726 | 41.8827751 |
| ENSMUSG00 | Gm10123 | 721 | 0.16541809 | 0.46782954 | 0 | 0 | 0.21231502 | 0.0378289 |
| ENSMUSG00 | Olfr1466 | 932 | 0 | 0 | 0 | 0 | 0 | 0 |
| ENSMUSG00 | Mtap | 3218 | 16.5421344 | 16.0821128 | 13.1451721 | 14.464213 | 14.0171896 | 11.9506654 |
| ENSMUSG00 | Stat4 | 3120 | 0 | 0 | 0 | 0.08095331 | 0 | 0 |
| ENSMUSG00 | Olfr204 | 923 | 0 | 0 | 0 | 0 | 0 | 0 |
| ENSMUSG00 | 9130023H24 | 3580 | 5.48581212 | 5.20898132 | 9.22312199 | 7.51877241 | 9.53537582 | 9.25661639 |
| ENSMUSG00 | Atp11c | 9801 | 1.13169871 | 2.42874221 | 1.21317169 | 2.38558972 | 1.46295394 | 2.07878334 |
| ENSMUSG00 | Tas2r110 | 1001 | 0 | 0 | 0 | 0 | 0 | 0 |
| ENSMUSG00 | Kdr | 6641 | 3.35236698 | 1.56001883 | 2.84872592 | 1.73319865 | 5.27090706 | 3.01865078 |
| ENSMUSG00 | Gm1568 | 5624 | 0 | 0 | 0.09431403 | 0 | 0 | 0.26673278 |
| ENSMUSG00 | Gm6378 | 443 | 0 | 0 | 0 | 0 | 0 | 0.1847041 |
| ENSMUSG00 | Ufc1 | 2047 | 17.828789 | 18.8791044 | 23.601668 | 14.5421039 | 12.2144231 | 22.1181735 |
| ENSMUSG00 | A430107O13I | 9368 | 0 | 0 | 0.29725834 | 0 | 0 | 0.21836016 |
| ENSMUSG00 | Mrpl42 | 660 | 191.72984 | 185.517802 | 231.456923 | 175.599283 | 124.164181 | 308.864627 |
| ENSMUSG00 | Gm10081 | 947 | 0 | 0 | 0 | 0 | 0 | 0 |
| ENSMUSG00 | Nrg1 | 3205 | 0.14885047 | 0.99229491 | 0.38616275 | 0.39403169 | 0.44578425 | 0.11063036 |
| ENSMUSG00 | Ica1 | 3488 | 23.7187944 | 21.0401237 | 25.2817307 | 24.4546884 | 25.527817 | 21.0424467 |
| ENSMUSG00 | Krtap16-3 | 496 | 0 | 0 | 0 | 0 | 0 | 0 |
| ENSMUSG00 | Rpl35 | 1383 | 0.48867908 | 0.17420984 | 0.31960841 | 0.18262785 | 0.59032684 | 0.61136212 |
| ENSMUSG00 | Gm15450 | 842 | 0 | 0 | 0 | 0 | 0 | 0 |
| ENSMUSG00 | Msln | 2098 | 0 | 0 | 0 | 0 | 0 | 0 |
| ENSMUSG00 | Ccni | 3294 | 38.005467 | 39.1021129 | 48.8850377 | 38.4371264 | 35.0089889 | 37.4425983 |
| ENSMUSG00 | 2010204K13I | 2021 | 0 | 0 | 0 | 0 | 0 | 0.09446931 |
| ENSMUSG00 | Manbal | 1104 | 14.7282532 | 15.4510874 | 19.2982684 | 14.0210119 | 9.7061041 | 18.5042616 |
| ENSMUSG00 | Olfr194 | 920 | 0 | 0 | 0 | 0 | 0 | 0 |
| ENSMUSG00 | Hist1h2ak | 512 | 0 | 0 | 0.08633173 | 0 | 0 | 0 |
| ENSMUSG00 | Zfp780b | 2417 | 1.1184827 | 1.51517154 | 1.62762268 | 4.50838951 | 3.08227177 | 2.15533972 |
| ENSMUSG00 | Ing2 | 3596 | 6.90967056 | 7.67820672 | 7.26454094 | 8.1876942 | 6.28606354 | 12.0824528 |
| ENSMUSG00 | Lrrc40 | 4735 | 7.74119388 | 8.89439291 | 11.2488323 | 7.47549856 | 7.78057938 | 9.87301619 |
| ENSMUSG00 | Gm6406 | 761 | 0 | 0 | 0 | 0 | 0 | 0 |
| ENSMUSG00 | Sox7 | 3296 | 0.45834595 | 0 | 0.46937637 | 0 | 0.77406517 | 0 |
| ENSMUSG00 | Ctnna2 | 8122 | 0 | 0 | 0 | 0 | 0 | 0 |
| ENSMUSG00 | Mapk3 | 2154 | 27.7956145 | 24.7643414 | 25.7331064 | 24.4232223 | 24.4708664 | 29.0726888 |
| ENSMUSG00 | Kif1b | 19460 | 2.71914412 | 3.13484254 | 4.06129986 | 5.00809878 | 4.47595188 | 2.52423563 |
| ENSMUSG00 | Gm14501 | 538 | 0 | 0 | 0 | 0 | 0 | 0 |
| ENSMUSG00 | Gm10125 | 3868 | 0 | 0 | 0 | 0 | 0 | 0 |
| ENSMUSG00 | Klk1b8 | 854 | 14.0121776 | 8.29438758 | 3.00199871 | 11.7879279 | 5.07873764 | 4.0241271 |
| ENSMUSG00 | Olfr1510 | 1038 | 0 | 0 | 0 | 0 | 0 | 0 |
| ENSMUSG00 | Zfp26 | 10509 | 5.62531159 | 4.97958686 | 2.96109025 | 3.16906719 | 4.23884538 | 4.47180545 |
| ENSMUSG00 | A930013B10I | 1968 | 0 | 0 | 0 | 0 | 0 | 0 |
| ENSMUSG00 | Olfr410 | 1000 | 0 | 0 | 0 | 0 | 0 | 0 |
| ENSMUSG00 | Olfr675 | 941 | 0 | 0 | 0 | 0 | 0 | 0 |
| ENSMUSG00 | Olfr480 | 1028 | 0 | 0 | 0 | 0 | 0 | 0 |
| ENSMUSG00 | Aldoart2 | 1657 | 0.14395467 | 0.23264426 | 0.24008243 | 0.02177553 | 0.12317773 | 0.04938076 |
| ENSMUSG00 | Calml3 | 1425 | 0.50217449 | 0.60867085 | 0.31018837 | 0.20256586 | 0.68035168 | 0 |

| | | | | | | | | |
|---|---|---|---|---|---|---|---|---|
| ENSMUSG00 | Klk1b1 | 862 | 0.92240094 | 2.29193054 | 0.10255648 | 1.92549194 | 1.47988329 | 0.15820556 |
| ENSMUSG00 | Olfr177 | 929 | 0 | 0 | 0 | 0 | 0 | 0 |
| ENSMUSG00 | Kcnma1 | 5370 | 0.00740326 | 0 | 0 | 0 | 0 | 0 |
| ENSMUSG00 | Bbs5 | 2568 | 5.20165166 | 8.85669809 | 7.17763572 | 10.004056 | 6.31868673 | 3.63237018 |
| ENSMUSG00 | Clip2 | 4978 | 0.27951824 | 0.3387958 | 0.53276629 | 0.59436072 | 0.64577374 | 0.11505975 |
| ENSMUSG00 | Csn2 | 1113 | 0 | 0 | 0 | 0 | 0 | 0 |
| ENSMUSG00 | Numbl | 5769 | 1.24042061 | 0.77679652 | 0.40608384 | 1.09453247 | 0.44224628 | 0.88882513 |
| ENSMUSG00 | Gm16416 | 482 | 0 | 0 | 0.09170507 | 0 | 0 | 0 |
| ENSMUSG00 | Speer2 | 1251 | 0 | 0 | 0 | 0 | 0 | 0 |
| ENSMUSG00 | Gm5449 | 356 | 0 | 0 | 0 | 0 | 0 | 0 |
| ENSMUSG00 | Rps4y2 | 942 | 21.9457005 | 22.6609277 | 27.1217253 | 17.964415 | 18.8505083 | 28.9829235 |
| ENSMUSG00 | Hspb11 | 2063 | 2.4088391 | 2.80289533 | 2.97821434 | 1.66155804 | 0.86569227 | 4.56119755 |
| ENSMUSG00 | Olfr811 | 959 | 0 | 0 | 0 | 0 | 0 | 0 |
| ENSMUSG00 | Olfr955 | 944 | 0 | 0 | 0 | 0 | 0 | 0 |
| ENSMUSG00 | Klk1b27 | 877 | 2.81053568 | 4.9999615 | 1.10882617 | 1.81027361 | 2.85096057 | 2.45689453 |
| ENSMUSG00 | Pstk | 2181 | 7.67402902 | 8.21887046 | 7.94457701 | 7.24618776 | 5.98934077 | 12.0553655 |
| ENSMUSG00 | Gm10126 | 170 | 0 | 0 | 0 | 0 | 0 | 0 |
| ENSMUSG00 | Olfr107 | 1032 | 0 | 0 | 0 | 0 | 0 | 0 |
| ENSMUSG00 | Cd300lb | 2913 | 0 | 0 | 0 | 0 | 0 | 0 |
| ENSMUSG00 | Nol7 | 1803 | 58.5417641 | 47.7053795 | 54.5474768 | 44.0268985 | 47.2340661 | 91.7474692 |
| ENSMUSG00 | Gm15315 | 373 | 0 | 0 | 0 | 0 | 0 | 0 |
| ENSMUSG00 | Olfr1351 | 959 | 0 | 0 | 0 | 0 | 0 | 0 |
| ENSMUSG00 | Olfr930 | 2024 | 0 | 0 | 0 | 0 | 0 | 0 |
| ENSMUSG00 | Olfr917 | 929 | 0 | 0 | 0 | 0 | 0 | 0 |
| ENSMUSG00 | Ldha | 2895 | 200.384101 | 212.103569 | 241.422985 | 198.220869 | 164.870715 | 153.727215 |
| ENSMUSG00 | Olfr535 | 990 | 0 | 0 | 0 | 0 | 0 | 0 |
| ENSMUSG00 | Serpina11 | 1624 | 0 | 0 | 0 | 0 | 0 | 0 |
| ENSMUSG00 | Gpr84 | 1548 | 0 | 0 | 0 | 0 | 0 | 0 |
| ENSMUSG00 | Ptpmt1 | 2004 | 20.2149873 | 13.2007768 | 17.4248782 | 13.161664 | 11.2797827 | 19.3943917 |
| ENSMUSG00 | 1110038F14f | 1005 | 21.7171729 | 18.8430565 | 20.4955809 | 13.7865722 | 16.6533844 | 15.9034215 |
| ENSMUSG00 | Grm4 | 5599 | 0 | 0 | 0 | 0 | 0 | 0.07307012 |
| ENSMUSG00 | Olfr133 | 620 | 0 | 0 | 0 | 0 | 0 | 0 |
| ENSMUSG00 | Tgif2lx2 | 1031 | 0 | 0 | 0 | 0 | 0 | 0 |
| ENSMUSG00 | Gm13248 | 1498 | 0 | 0 | 0.32457962 | 0.26495493 | 0.13625201 | 0.30952528 |
| ENSMUSG00 | Krtap4-1 | 1010 | 0 | 0 | 0 | 0 | 0 | 0 |
| ENSMUSG00 | Gm4744 | 735 | 0 | 0 | 0 | 0 | 0 | 0 |
| ENSMUSG00 | Scoc | 2057 | 23.9460575 | 21.2938629 | 26.1729921 | 25.7678772 | 18.5302395 | 19.6504693 |
| ENSMUSG00 | B230325K18l | 2622 | 0.19710955 | 0 | 0.01685806 | 0.02752254 | 0.03892172 | 0 |
| ENSMUSG00 | Syt10 | 1838 | 0 | 0 | 0 | 0 | 0 | 0 |
| ENSMUSG00 | Parp10 | 4264 | 5.30508171 | 5.44696649 | 2.80926348 | 2.26782081 | 5.09785087 | 4.93809135 |
| ENSMUSG00 | Vmn1r202 | 908 | 0 | 0 | 0 | 0 | 0 | 0 |
| ENSMUSG00 | Naa15 | 6070 | 24.7505702 | 21.529099 | 26.2589532 | 37.7048449 | 25.2189666 | 22.3768869 |
| ENSMUSG00 | Ptpla | 1120 | 4.82745121 | 7.44308437 | 4.22285465 | 5.25122612 | 4.829282 | 7.30570694 |
| ENSMUSG00 | Gm10128 | 610 | 0 | 0 | 0 | 0 | 0 | 0 |
| ENSMUSG00 | Zfp35 | 923 | 119.524874 | 146.543167 | 139.741038 | 117.198234 | 118.250724 | 111.137506 |
| ENSMUSG00 | Gm8995 | 1129 | 0 | 0 | 0 | 0.47938943 | 0.18078433 | 0.53148101 |
| ENSMUSG00 | Olfr339 | 929 | 0 | 0 | 0 | 0 | 0 | 0 |
| ENSMUSG00 | Tmem117 | 2773 | 385.411678 | 218.515871 | 629.043394 | 304.478847 | 265.473323 | 553.912219 |
| ENSMUSG00 | Luzp2 | 1026 | 0.19374016 | 0 | 0.90471609 | 0 | 0 | 0.82408754 |
| ENSMUSG00 | Gm10129 | 359 | 0 | 0 | 0 | 0 | 0 | 0 |
| ENSMUSG00 | Psg20 | 2157 | 0 | 0 | 0 | 0 | 0 | 0 |
| ENSMUSG00 | Gm12657 | 452 | 0 | 0 | 0.19558338 | 0.07982753 | 0 | 0.12068424 |
| ENSMUSG00 | Rpl27 | 1284 | 1.64099725 | 3.30249759 | 1.96223135 | 2.44481143 | 2.06648874 | 2.69772522 |
| ENSMUSG00 | Usp31 | 10182 | 2.85417956 | 3.05247055 | 4.24566907 | 3.0086089 | 3.90390363 | 3.12069876 |
| ENSMUSG00 | 1190007I07R | 1178 | 4.15103914 | 6.46303723 | 5.44080411 | 8.17818836 | 5.45782967 | 6.3671696 |
| ENSMUSG00 | Gm5299 | 275 | 0 | 0 | 0 | 0 | 0 | 0 |
| ENSMUSG00 | Gm9396 | 566 | 0 | 0.17027011 | 0 | 0.12749839 | 0 | 0.09637682 |
| ENSMUSG00 | Krr1 | 6066 | 8.45442958 | 10.6922109 | 10.1286782 | 12.5567418 | 10.50642 | 7.93599378 |
| ENSMUSG00 | Olfr874 | 1857 | 0 | 0 | 0 | 0 | 0 | 0 |
| ENSMUSG00 | Slc39a4 | 2252 | 0 | 0.2781633 | 0 | 0.5768 | 0 | 0 |
| ENSMUSG00 | Mapk1 | 6724 | 45.6679553 | 46.3661928 | 53.181575 | 50.9087383 | 41.8592339 | 46.3393038 |
| ENSMUSG00 | Olfr969 | 935 | 0 | 0 | 0 | 0 | 0 | 0 |
| ENSMUSG00 | Alg11 | 3127 | 6.81449874 | 12.589806 | 7.40702454 | 14.1928093 | 9.74184406 | 15.0198045 |
| ENSMUSG00 | 3300002I08R | 2392 | 0.9473505 | 1.16840035 | 0.77611932 | 2.53419042 | 1.10926904 | 1.42530515 |
| ENSMUSG00 | Olfr763 | 1043 | 0 | 0 | 0 | 0 | 0 | 0 |
| ENSMUSG00 | Ifna13 | 818 | 0 | 0 | 0 | 0 | 0 | 0 |
| ENSMUSG00 | Olfr1287 | 917 | 0 | 0 | 0 | 0 | 0 | 0 |
| ENSMUSG00 | Olfr945 | 962 | 0 | 0 | 0 | 0 | 0 | 0 |
| ENSMUSG00 | Bcl9l | 5295 | 1.09618511 | 0.72802934 | 0.90991518 | 1.26065689 | 1.0311279 | 1.24654631 |
| ENSMUSG00 | Zfp947 | 2867 | 8.22288104 | 11.7987033 | 10.6842962 | 6.26747756 | 4.43165943 | 10.2172873 |
| ENSMUSG00 | Olfr1387 | 935 | 0 | 0 | 0 | 0 | 0 | 0 |
| ENSMUSG00 | BC023105 | 2038 | 0 | 0.11821993 | 5.57403025 | 4.95729758 | 18.7530694 | 34.876207 |
| ENSMUSG00 | Gm10130 | 1064 | 0.07472835 | 0.36230408 | 0.2908016 | 0.33911696 | 0.2397856 | 0.46141307 |
| ENSMUSG00 | Olfr293 | 1010 | 0 | 0 | 0 | 0 | 0 | 0 |
| ENSMUSG00 | Tmed5 | 7682 | 13.0051448 | 12.9969159 | 14.7301117 | 16.8151158 | 11.265391 | 18.5369554 |
| ENSMUSG00 | Lrrc43 | 2180 | 0 | 0 | 0 | 0 | 0 | 0 |
| ENSMUSG00 | Stk24 | 2665 | 1.79011545 | 2.13358356 | 2.37180621 | 1.96318816 | 1.95297949 | 4.64641134 |
| ENSMUSG00 | Gm10131 | 314 | 0.12660981 | 0 | 0.2815404 | 0 | 0 | 0 |
| ENSMUSG00 | Cyp26b1 | 4595 | 0 | 0 | 0.17315194 | 0 | 0.03331428 | 0.73602944 |
| ENSMUSG00 | Gm8017 | 557 | 0 | 0.34604267 | 0 | 0.12955851 | 0.09160929 | 0.14690111 |
| ENSMUSG00 | Ddo | 8576 | 4.61712437 | 6.25929297 | 3.49450205 | 5.70934403 | 5.97370351 | 4.66556621 |
| ENSMUSG00 | Wscd2 | 4481 | 2.76806738 | 3.73146517 | 4.75458343 | 4.03416744 | 2.55074943 | 3.52421694 |
| ENSMUSG00 | Sorcs3 | 5607 | 0.48923277 | 1.22034507 | 0.17343331 | 0 | 0.06370334 | 0 |
| ENSMUSG00 | Olfr46 | 956 | 0 | 0 | 0 | 0 | 0 | 0 |
| ENSMUSG00 | B9d2 | 1018 | 12.4577586 | 15.2890185 | 14.5457931 | 12.5471943 | 8.22035917 | 10.9848744 |
| ENSMUSG00 | Gm7117 | 775 | 0 | 0 | 0.45627709 | 0 | 0 | 0 |
| ENSMUSG00 | Nmral1 | 4170 | 0.62922343 | 1.15555017 | 0.82679706 | 1.79112307 | 1.17470793 | 0.74563762 |
| ENSMUSG00 | E130309F12l | 7597 | 2.89910968 | 3.53295357 | 2.16441827 | 3.59538077 | 3.5732568 | 1.9063885 |
| ENSMUSG00 | 1700013N18l | 1089 | 0 | 0 | 0 | 0 | 0 | 0 |
| ENSMUSG00 | Syne2 | 25144 | 8.65658828 | 11.4046101 | 6.63625348 | 12.6482318 | 13.6738673 | 6.5463907 |
| ENSMUSG00 | Olfr800 | 939 | 0 | 0 | 0 | 0 | 0 | 0 |
| ENSMUSG00 | D630045J12l | 18778 | 4.46079441 | 5.49147866 | 2.25269804 | 6.51390622 | 6.23088081 | 3.3813697 |
| ENSMUSG00 | Rps15 | 644 | 6.66706816 | 7.25789579 | 11.8740976 | 7.05953045 | 6.25944672 | 18.0842717 |
| ENSMUSG00 | 1700112E06l | 2792 | 1.93651338 | 2.50251938 | 3.10299471 | 3.36007578 | 2.02862741 | 1.57278543 |
| ENSMUSG00 | Tas2r114 | 929 | 0 | 0 | 0 | 0 | 0 | 0 |
| ENSMUSG00 | Nhp2l1 | 2846 | 2.61218372 | 3.36932607 | 3.52558621 | 2.38349416 | 2.27700272 | 3.0954703 |
| ENSMUSG00 | Olfr1480 | 947 | 0 | 0 | 0 | 0 | 0 | 0 |
| ENSMUSG00 | Zfp167 | 2223 | 3.16541613 | 6.22110162 | 2.08780635 | 4.09027225 | 5.853228 | 2.08577988 |
| ENSMUSG00 | Arhgap22 | 3862 | 0 | 0 | 0 | 0 | 0 | 0 |
| ENSMUSG00 | Gm1330 | 554 | 0 | 0 | 0 | 0 | 0 | 0 |
| ENSMUSG00 | Snrnp70 | 2395 | 14.0098645 | 17.6448893 | 16.5549283 | 19.2839319 | 16.2133912 | 25.3272642 |
| ENSMUSG00 | Gm6756 | 1578 | 0 | 0 | 0 | 0 | 0 | 0 |
| ENSMUSG00 | 2010109I03R | 1884 | 0.14771145 | 0 | 0 | 0 | 0 | 0 |
| ENSMUSG00 | Eno1 | 4523 | 42.708795 | 42.0712492 | 39.3057291 | 37.7094461 | 35.4127335 | 44.3883749 |
| ENSMUSG00 | Gm1574 | 1358 | 0 | 0 | 0 | 0 | 0 | 0 |
| ENSMUSG00 | Sema3e | 8272 | 0 | 0 | 0.15496294 | 0.06106729 | 0 | 0 |
| ENSMUSG00 | Zfp773 | 3021 | 4.23742626 | 1.03678211 | 4.3894581 | 3.69061627 | 2.24644422 | 3.44882893 |
| ENSMUSG00 | Gm5616 | 386 | 0 | 0 | 0 | 0 | 0 | 0 |
| ENSMUSG00 | Gm4825 | 410 | 0 | 0 | 0 | 0 | 0 | 0 |
| ENSMUSG00 | Olfr179 | 1127 | 0 | 0 | 0 | 0 | 0 | 0 |
| ENSMUSG00 | Nup98 | 6907 | 7.94303795 | 6.56485335 | 7.26991366 | 10.5733398 | 9.13850107 | 6.66564226 |
| ENSMUSG00 | Gm10132 | 527 | 0.30174938 | 0 | 0 | 0 | 0 | 0.05175453 |
| ENSMUSG00 | Aox1 | 6009 | 0.80053472 | 0.09622854 | 0.1471188 | 0.01200933 | 0.26324141 | 0 |
| ENSMUSG00 | Col23a1 | 5706 | 0 | 0 | 0 | 0 | 0 | 0.02389996 |
| ENSMUSG00 | Jazf1 | 2529 | 1.39906594 | 1.56239156 | 0.48938379 | 0.32814829 | 1.35182569 | 1.31573981 |
| ENSMUSG00 | Klhdc3 | 2713 | 28.0764838 | 26.6242082 | 34.8498866 | 19.7367321 | 27.3093615 | 26.2492013 |
| ENSMUSG00 | Olfr666 | 956 | 0 | 0 | 0 | 0 | 0 | 0 |
| ENSMUSG00 | Olfr1410 | 967 | 0 | 0 | 0 | 0 | 0 | 0 |
| ENSMUSG00 | Gm5513 | 1217 | 0.42466824 | 0.31675558 | 0.72640662 | 0.50402199 | 0.335424 | 0.4706388 |
| ENSMUSG00 | Slc22a28 | 1994 | 0.19937553 | 1.03912588 | 0.15517297 | 0 | 0 | 0.93012812 |
| ENSMUSG00 | Gng8 | 916 | 0 | 0 | 0 | 0 | 0 | 0 |
| ENSMUSG00 | Eglem1 | 4153 | 2.19215147 | 1.46195322 | 1.12819537 | 2.34581074 | 4.49690671 | 1.58275658 |
| ENSMUSG00 | Olfr196 | 929 | 0 | 0 | 0 | 0 | 0 | 0 |
| ENSMUSG00 | Ccdc102a | 2494 | 0.78108202 | 1.79684808 | 0.17723273 | 0.46296128 | 0.79792649 | 0.21872205 |
| ENSMUSG00 | Olfr486 | 944 | 0 | 0 | 0 | 0 | 0 | 0 |
| ENSMUSG00 | Gm10134 | 380 | 0 | 0 | 0 | 0.09495275 | 0 | 0.4306522 |
| ENSMUSG00 | Olfr64 | 1661 | 0 | 0 | 0 | 0 | 0 | 0 |
| ENSMUSG00 | C230062I16f | 2371 | 0.41918474 | 0.54872794 | 2.44219377 | 0.82177579 | 0.73171522 | 4.75091776 |
| ENSMUSG00 | Unc5d | 13987 | 0 | 0 | 0 | 0 | 0 | 0 |
| ENSMUSG00 | Gm7665 | 296 | 0 | 0 | 0.14933055 | 0.1218988 | 0 | 0.46072026 |
| ENSMUSG00 | Sox11 | 2012 | 0 | 0 | 0 | 0 | 0 | 0 |
| ENSMUSG00 | Jakmip1 | 4462 | 0 | 0 | 0 | 0 | 0 | 0 |
| ENSMUSG00 | Cnfn | 718 | 0 | 0 | 0 | 0 | 0 | 0 |
| ENSMUSG00 | Slc22a21 | 4427 | 1.31111366 | 0.75104236 | 0.65898388 | 0.7579919 | 1.10651278 | 0.9611122 |
| ENSMUSG00 | Gm10135 | 244 | 0 | 0 | 0 | 0 | 0 | 0.33534393 |
| ENSMUSG00 | Olfr970 | 935 | 0 | 0 | 0 | 0 | 0 | 0 |
| ENSMUSG00 | Zfp238 | 5100 | 4.51341632 | 3.04235968 | 4.83620165 | 6.56551712 | 4.19216696 | 5.13939771 |
| ENSMUSG00 | Olfr98 | 1038 | 0 | 0 | 0 | 0 | 0 | 0 |
| ENSMUSG00 | Krt73 | 2156 | 0 | 0 | 0 | 0 | 0 | 0 |
| ENSMUSG00 | Brwd3 | 13852 | 0.61418372 | 0.6331167 | 0.64139261 | 1.12267984 | 0.50098088 | 1.50234982 |
| ENSMUSG00 | Olfr1537-ps1 | 944 | 0 | 0 | 0 | 0 | 0 | 0 |
| ENSMUSG00 | Nkx6-3 | 2233 | 0 | 0.19421316 | 0 | 0.19390261 | 0 | 0 |
| ENSMUSG00 | Crb1 | 10915 | 0 | 0 | 0 | 0 | 0 | 0 |
| ENSMUSG00 | Glyat | 4863 | 0.93196068 | 0.60443615 | 0.31812965 | 0.22259127 | 0.12591333 | 0.32529901 |
| ENSMUSG00 | Gm13910 | 1427 | 1.39297409 | 1.3169385 | 1.51779279 | 1.28954749 | 1.64485865 | 1.07034324 |
| ENSMUSG00 | Pcdhb5 | 3532 | 0 | 0.34107051 | 0.01251468 | 0 | 0 | 0.00772215 |
| ENSMUSG00 | Hist2h2ab | 443 | 0 | 0 | 0 | 0 | 0 | 0 |
| ENSMUSG00 | Gm5576 | 1441 | 0 | 0.10031876 | 0 | 0 | 0 | 0.01892758 |
| ENSMUSG00 | Cycs | 3549 | 16.0859031 | 13.9169071 | 16.5149743 | 14.1420468 | 13.2849736 | 20.2119756 |
| ENSMUSG00 | Gm8730 | 1098 | 0.07241435 | 0.0877713 | 0.08051338 | 0 | 0 | 0.04968058 |
| ENSMUSG00 | Sfxn4 | 3262 | 7.77559674 | 5.21453528 | 3.71284641 | 3.52856291 | 4.2078771 | 6.48002619 |
| ENSMUSG00 | Mapk15 | 2377 | 33.3163303 | 48.4297056 | 26.6661518 | 44.6737304 | 57.9171906 | 51.3823452 |
| ENSMUSG00 | Rhox2e | 791 | 0 | 0 | 0 | 0 | 0 | 0 |
| ENSMUSG00 | Klk1b24 | 870 | 6.30604174 | 5.31712465 | 5.08067161 | 5.14272819 | 3.98826846 | 6.92838537 |
| ENSMUSG00 | 1700011J10R | 968 | 3.20343748 | 3.73345368 | 4.24666467 | 5.21847751 | 4.79690103 | 12.2003293 |
| ENSMUSG00 | Olfr816 | 938 | 0 | 0 | 0 | 0 | 0 | 0 |
| ENSMUSG00 | Tnfrsf11b | 2813 | 0 | 0 | 0.7228172 | 0.35915295 | 0 | 0.69810666 |
| ENSMUSG00 | Magea6 | 1215 | 0 | 0 | 0 | 0 | 0 | 0 |
| ENSMUSG00 | Hsd3b2 | 6796 | 0.25154292 | 0.04963289 | 0.06504097 | 0 | 0 | 0.04013337 |
| ENSMUSG00 | Olfr908 | 928 | 0 | 0 | 0 | 0 | 0 | 0 |
| ENSMUSG00 | Gm4963 | 377 | 0 | 0 | 0 | 0 | 0 | 0 |
| ENSMUSG00 | Gm13939 | 1344 | 0.02957997 | 0 | 0.09866483 | 0 | 0.22779632 | 0.2029363 |
| ENSMUSG00 | Gm10136 | 437 | 0.54584184 | 0.22053292 | 0.10114838 | 0 | 0.35029549 | 0.56172026 |
| ENSMUSG00 | 4632433K11f | 1128 | 0.1762211 | 0 | 0.23511619 | 0 | 0 | 0.2176368 |
| ENSMUSG00 | Rnf217 | 1887 | 11.9877416 | 9.52493001 | 16.6313241 | 11.3007357 | 13.2771333 | 14.6707784 |
| ENSMUSG00 | Tas2r129 | 1353 | 0 | 0 | 0 | 0 | 0 | 0 |
| ENSMUSG00 | Olfr508 | 932 | 0 | 0 | 0 | 0 | 0 | 0 |
| ENSMUSG00 | Chadl | 4957 | 0 | 0.13609243 | 0.18725816 | 0.62599472 | 0.10293802 | 0 |
| ENSMUSG00 | S100a7a | 738 | 0 | 0 | 0 | 0 | 0 | 0 |
| ENSMUSG00 | Olfr1469 | 929 | 0 | 0 | 0 | 0 | 0 | 0 |
| ENSMUSG00 | Chi3l4 | 1621 | 0 | 0 | 0 | 0 | 0 | 0 |
| ENSMUSG00 | Olfr282 | 927 | 0 | 0 | 0 | 0 | 0 | 0 |
| ENSMUSG00 | Utp14a | 4184 | 2.3564434 | 2.37246823 | 1.83822196 | 2.29393495 | 2.62205325 | 2.7313752 |
| ENSMUSG00 | Chchd1 | 584 | 25.5279199 | 19.5551144 | 20.8899121 | 22.7366308 | 17.2126645 | 41.1454746 |
| ENSMUSG00 | Slc22a8 | 3466 | 0.34410399 | 1.39026088 | 0 | 0.42682165 | 0.22082967 | 0.40132908 |
| ENSMUSG00 | Prpf38a | 5686 | 9.5508242 | 8.94067824 | 10.2070031 | 9.70902711 | 8.6958421 | 9.88142048 |
| ENSMUSG00 | Ap3s2 | 5791 | 6.65909447 | 5.87456884 | 7.47256161 | 5.31479606 | 5.50707734 | 4.97829281 |
| ENSMUSG00 | Hspbp1 | 1480 | 23.5040848 | 14.162907 | 24.4304781 | 19.893884 | 18.7556408 | 16.1436378 |
| ENSMUSG00 | Lin28b | 9056 | 0 | 0 | 0 | 0 | 0 | 0.00301178 |
| ENSMUSG00 | Gpatch1 | 3541 | 13.3266748 | 10.6143533 | 11.4467919 | 10.7502279 | 18.5170553 | 10.7758318 |
| ENSMUSG00 | Alms1 | 9963 | 0.14764155 | 0.17895196 | 0.24401299 | 0.659132 | 0.46606446 | 0.70356141 |
| ENSMUSG00 | Gm4987 | 505 | 0.23617117 | 1.43128046 | 1.57551124 | 0.92884471 | 0.40416931 | 0.75612861 |
| ENSMUSG00 | Arl9 | 592 | 0 | 0.16279203 | 0.44799165 | 0.0609494 | 0.51715921 | 1.01358457 |
| ENSMUSG00 | Dupd1 | 1089 | 0 | 0 | 0 | 0 | 0 | 0 |
| ENSMUSG00 | Olfr532 | 929 | 0 | 0 | 0 | 0 | 0 | 0 |
| ENSMUSG00 | Olfr1382 | 935 | 0 | 0 | 0 | 0 | 0 | 0 |
| ENSMUSG00 | Gm6065 | 491 | 0.08096839 | 0 | 0.18004824 | 0.29394741 | 0.10392337 | 0.49994247 |
| ENSMUSG00 | Cdc42ep5 | 1521 | 4.36500016 | 6.33615281 | 7.17807707 | 3.24999349 | 4.56251618 | 7.81837128 |
| ENSMUSG00 | Olfr862 | 1034 | 0 | 0 | 0 | 0 | 0 | 0 |
| ENSMUSG00 | Olfr1276 | 1845 | 0 | 0 | 0 | 0 | 0 | 0 |
| ENSMUSG00 | Ppcdc | 2740 | 18.7895428 | 16.8124959 | 15.728758 | 13.6690373 | 21.5651618 | 12.8310255 |
| ENSMUSG00 | Rnf183 | 2580 | 2.81986548 | 3.10036798 | 2.22722465 | 1.51041117 | 1.95798884 | 1.09944282 |
| ENSMUSG00 | Gpx1 | 845 | 197.083678 | 231.294922 | 193.389602 | 173.919725 | 134.902868 | 184.72516 |
| ENSMUSG00 | Olfr1511 | 956 | 0 | 0 | 0 | 0 | 0 | 0 |
| ENSMUSG00 | Chd4 | 8079 | 6.52011532 | 7.83126606 | 7.05237971 | 9.29406296 | 9.38546781 | 7.80191748 |
| ENSMUSG00 | Slc24a3 | 4824 | 0.09889423 | 0.08990008 | 0 | 0.01495939 | 0.01057761 | 0 |
| ENSMUSG00 | Rps6-ps1 | 797 | 0.19952562 | 0.7859771 | 0.22184112 | 0 | 0.19206917 | 0 |
| ENSMUSG00 | Olfr376 | 1311 | 0 | 0 | 0 | 0 | 0 | 0 |
| ENSMUSG00 | Uqcrh | 1439 | 51.027361 | 46.2442506 | 69.4819798 | 47.741635 | 36.9134519 | 103.374484 |
| ENSMUSG00 | Ptcd3 | 2561 | 62.6835729 | 53.4735918 | 57.5262565 | 48.2831576 | 52.0624444 | 62.5793753 |
| ENSMUSG00 | Gm6498 | 1596 | 0.22418504 | 0.24153605 | 0.52621242 | 1.22082106 | 0.79928534 | 0 |
| ENSMUSG00 | Gm10138 | 364 | 0.43687341 | 0.79428201 | 0.24286727 | 0 | 0.14018235 | 0.07493033 |
| ENSMUSG00 | Nlgn1 | 5175 | 0 | 0 | 0 | 0 | 0 | 0 |
| ENSMUSG00 | Rpl7l1 | 2599 | 23.2506081 | 21.5624595 | 19.6263666 | 18.0618468 | 19.2011526 | 13.5586125 |
| ENSMUSG00 | Crem | 7297 | 6.06384128 | 7.27716311 | 8.77131269 | 8.41601203 | 6.68510556 | 10.4957088 |
| ENSMUSG00 | Zfp192 | 15176 | 3.48672539 | 3.40696181 | 4.76795052 | 5.25919099 | 4.64670871 | 3.39674935 |
| ENSMUSG00 | Nupl1 | 2854 | 13.4700594 | 18.437139 | 15.0695141 | 21.8717369 | 20.4355808 | 17.4313041 |
| ENSMUSG00 | Adam26b | 2403 | 0 | 0 | 0 | 0 | 0 | 0 |
| ENSMUSG00 | Gm7964 | 1718 | 5.41489083 | 5.32911758 | 4.73407399 | 4.57851322 | 4.81156745 | 4.84212164 |
| ENSMUSG00 | Klk1 | 871 | 10510.4179 | 12210.0351 | 3543.09997 | 9434.43971 | 8156.78001 | 8266.94003 |
| ENSMUSG00 | Dpp3 | 2665 | 28.6418471 | 27.0675812 | 28.9426702 | 20.3359215 | 32.6645393 | 30.130033 |
| ENSMUSG00 | Gm10139 | 170 | 0 | 0 | 0 | 0 | 0 | 0 |
| ENSMUSG00 | Gm17687 | 67 | 0.59336538 | 0.71920063 | 0 | 0.53853798 | 0.7615877 | 0 |
| ENSMUSG00 | Tdgf1-ps2 | 503 | 0.15807348 | 0 | 0 | 0 | 0 | 0 |
| ENSMUSG00 | Olfr346 | 929 | 0 | 0 | 0 | 0 | 0 | 0 |
| ENSMUSG00 | Srrm4 | 13482 | 0 | 0 | 0 | 0.05620256 | 0 | 0 |
| ENSMUSG00 | Cyp4a32 | 1712 | 0.6269848 | 0.90068116 | 1.98805018 | 0.14753172 | 0.59610252 | 2.70834619 |
| ENSMUSG00 | Pepd | 3063 | 53.8120216 | 56.6501398 | 44.3605829 | 41.0649713 | 47.361406 | 35.3955896 |
| ENSMUSG00 | Poteg | 1692 | 0 | 0 | 0 | 0 | 0 | 0 |
| ENSMUSG00 | Zar1 | 1255 | 0.38013209 | 0.15358229 | 0 | 0 | 0 | 0 |
| ENSMUSG00 | Olfr205 | 935 | 0 | 0 | 0 | 0 | 0 | 0 |
| ENSMUSG00 | Brpf3 | 7177 | 2.83057692 | 2.00748867 | 1.88459857 | 2.06125655 | 2.28221634 | 1.12868439 |
| ENSMUSG00 | Amd2 | 3200 | 0.01242359 | 0.01505826 | 0.01381308 | 0.03382692 | 0.04783723 | 0.0340933 |
| ENSMUSG00 | Hist2h2aa1 | 577 | 0 | 0 | 0 | 0 | 0 | 0 |
| ENSMUSG00 | 1700011A15f | 898 | 0.08854227 | 0 | 0.09644509 | 0 | 0.28411122 | 0.3037265 |
| ENSMUSG00 | Nr6a1 | 12028 | 0.92546845 | 0.46872412 | 1.6059366 | 0.74695952 | 0.86118676 | 0.82540478 |
| ENSMUSG00 | Slco1a5 | 3712 | 0 | 0 | 0 | 0 | 0 | 0 |
| ENSMUSG00 | Gm6613 | 557 | 0 | 0 | 0 | 0 | 0 | 0 |
| ENSMUSG00 | Olfr120 | 992 | 0 | 0 | 0 | 0 | 0 | 0 |
| ENSMUSG00 | Vmn1r211 | 984 | 0 | 0 | 0 | 0 | 0 | 0 |
| ENSMUSG00 | Olfr190 | 923 | 0 | 0 | 0 | 0 | 0 | 0 |
| ENSMUSG00 | Dppa1 | 1824 | 0.15257038 | 0.39627008 | 0 | 0 | 0.02797499 | 0 |
| ENSMUSG00 | Gm648 | 1035 | 0 | 0 | 0 | 0 | 0 | 0.34258001 |
| ENSMUSG00 | Mageb16-ps2 | 619 | 0 | 0 | 0 | 0 | 0 | 0 |
| ENSMUSG00 | Klk8 | 1317 | 0.12074558 | 0.40246839 | 0 | 0.32876578 | 0.1549776 | 0.04141935 |
| ENSMUSG00 | Gm10142 | 344 | 0 | 0 | 0 | 0 | 0 | 0 |
| ENSMUSG00 | Wibg | 1207 | 2.20680795 | 1.71666695 | 2.01416849 | 2.03279124 | 1.47963807 | 4.36123063 |
| ENSMUSG00 | Gm10143 | 3331 | 0 | 0 | 0 | 0.05416098 | 0.04595591 | 0 |
| ENSMUSG00 | Mro | 2890 | 0 | 0 | 0 | 0 | 0 | 0 |
| ENSMUSG00 | Gpn1 | 6899 | 11.524998 | 9.66662356 | 8.47645141 | 8.02288104 | 7.90653656 | 10.7651606 |
| ENSMUSG00 | Ccr1l1 | 1459 | 0 | 0 | 0 | 0 | 0 | 0 |
| ENSMUSG00 | Trerf1 | 4524 | 2.08268101 | 4.18595751 | 1.96387499 | 4.17128853 | 3.44010712 | 2.21862671 |
| ENSMUSG00 | Olfr319 | 920 | 0 | 0 | 0 | 0 | 0 | 0 |
| ENSMUSG00 | Gm5089 | 2005 | 0 | 0 | 0 | 0 | 0 | 0 |
| ENSMUSG00 | Scgb3a1 | 594 | 0 | 0 | 0 | 0 | 0 | 0 |
| ENSMUSG00 | Dzip3 | 10069 | 1.68592613 | 2.0243187 | 2.49345981 | 2.49051752 | 2.51863232 | 2.56249963 |
| ENSMUSG00 | BC048507 | 582 | 0 | 0.08279457 | 0 | 0 | 0.35069674 | 0 |
| ENSMUSG00 | Ipcef1 | 8988 | 21.1693068 | 19.1180299 | 19.0764296 | 32.7299632 | 27.182275 | 24.2764924 |
| ENSMUSG00 | Mtx1 | 3746 | 16.9273869 | 16.4008846 | 17.4636219 | 18.840491 | 14.5478295 | 20.3795289 |
| ENSMUSG00 | Fbln2 | 5934 | 0 | 0.47098309 | 0 | 0 | 0 | 0 |
| ENSMUSG00 | Gm8973 | 314 | 0 | 0 | 0 | 0 | 0 | 0 |
| ENSMUSG00 | Olfr1202 | 929 | 0 | 0 | 0 | 0 | 0 | 0 |
| ENSMUSG00 | Vrk2 | 5724 | 2.41005446 | 2.76962604 | 1.83788236 | 2.17475635 | 2.35341775 | 3.59278092 |
| ENSMUSG00 | Cnnm2 | 5334 | 4.18125695 | 5.57387229 | 2.45289568 | 3.44991427 | 2.80291116 | 3.02710657 |
| ENSMUSG00 | Hcst | 566 | 0.2107181 | 0.25540517 | 0.70285616 | 0.12749839 | 0.36061043 | 2.60217406 |
| ENSMUSG00 | Olfr963 | 935 | 0 | 0 | 0 | 0 | 0 | 0 |
| ENSMUSG00 | Olfr916 | 932 | 0 | 0 | 0 | 0 | 0 | 0 |
| ENSMUSG00 | Cadm2 | 11263 | 0.25767114 | 0.43638614 | 0.1098865 | 0.16017955 | 0.54818356 | 0.73617068 |
| ENSMUSG00 | Mocs1 | 3297 | 7.06603324 | 3.27381348 | 3.17738453 | 2.73597547 | 3.62152622 | 1.86959917 |
| ENSMUSG00 | Olfr96 | 1050 | 0 | 0 | 0 | 0 | 0 | 0 |
| ENSMUSG00 | BC068157 | 6376 | 0 | 0.07557472 | 0.07625789 | 0.02263616 | 0.04801729 | 0.02994393 |
| ENSMUSG00 | Med14 | 8757 | 8.98436633 | 7.39001848 | 7.80359133 | 10.2020261 | 10.1097136 | 9.51824798 |
| ENSMUSG00 | Cenpj | 4346 | 2.13139139 | 2.52795877 | 1.71884755 | 3.06356982 | 2.81783944 | 1.73839739 |
| ENSMUSG00 | Gm14781 | 1358 | 0 | 0 | 0 | 0 | 0 | 0 |
| ENSMUSG00 | Gm6421 | 410 | 0.38785835 | 0 | 1.18590311 | 0.35201995 | 0.24890915 | 0.13304702 |
| ENSMUSG00 | Rhox8 | 1198 | 0 | 0 | 0 | 0 | 0 | 0 |
| ENSMUSG00 | Fam172a | 7421 | 9.37502909 | 7.95423685 | 9.11912433 | 8.09062419 | 7.68042876 | 9.61466868 |
| ENSMUSG00 | Trim38 | 1410 | 0 | 0 | 0 | 0.10236041 | 0 | 0 |
| ENSMUSG00 | Zfp69 | 3928 | 0.74895763 | 0.55203409 | 0.52889171 | 0.95532908 | 1.1431571 | 0.49299883 |
| ENSMUSG00 | Arih2 | 3908 | 24.1605082 | 21.7258217 | 18.1987629 | 15.834367 | 24.5469771 | 16.156804 |
| ENSMUSG00 | Rab44 | 4125 | 0.00963769 | 0 | 0 | 0 | 0 | 0 |
| ENSMUSG00 | Olfr134 | 938 | 0 | 0 | 0 | 0 | 0 | 0 |
| ENSMUSG00 | Prol1 | 1156 | 0 | 0 | 0 | 0 | 0 | 0 |
| ENSMUSG00 | Izumo1 | 2981 | 0 | 0 | 0.05931143 | 0 | 0.0684688 | 0.39342821 |
| ENSMUSG00 | Krt39 | 1625 | 0 | 0 | 0 | 0 | 0 | 0.18462833 |
| ENSMUSG00 | Hist1h2bh | 435 | 0.36556764 | 0.33232029 | 0.3048403 | 0.41473614 | 0.58651007 | 0.06270032 |
| ENSMUSG00 | Olfr104-ps | 937 | 0 | 0 | 0 | 0 | 0 | 0 |
| ENSMUSG00 | Olfr954 | 944 | 0 | 0 | 0 | 0 | 0 | 0 |
| ENSMUSG00 | Ghrl | 659 | 0 | 0 | 0.26829647 | 1.14980719 | 0 | 0 |
| ENSMUSG00 | Tnnt1 | 1960 | 0 | 0.27043411 | 0 | 0.03681841 | 0 | 0 |
| ENSMUSG00 | Rab3ip | 4721 | 50.0206639 | 60.9347722 | 42.9565888 | 51.8263808 | 47.729819 | 39.3549762 |
| ENSMUSG00 | Gm13699 | 2264 | 0.17559841 | 0.04256753 | 0.19523782 | 0.23905948 | 0.0450763 | 0.06023551 |
| ENSMUSG00 | Zfp936 | 1365 | 0.05824979 | 0 | 0 | 0 | 0 | 0.01998142 |
| ENSMUSG00 | Krt2 | 3138 | 0 | 0 | 0 | 0 | 0 | 0 |
| ENSMUSG00 | 4430402I18R | 3392 | 2.63708229 | 3.33838853 | 3.33598815 | 2.35086434 | 3.36966633 | 1.51972489 |
| ENSMUSG00 | Gm10145 | 464 | 0.51407949 | 0.51925045 | 0.47631296 | 0.07776303 | 0 | 0.17634465 |
| ENSMUSG00 | Ano6 | 4934 | 36.8467394 | 41.9946699 | 47.4269471 | 43.7167536 | 53.5084046 | 42.023066 |
| ENSMUSG00 | Defa24 | 417 | 0 | 0 | 0 | 0 | 0 | 0 |
| ENSMUSG00 | Ifi27l1 | 1500 | 9.48830802 | 9.05905112 | 10.431635 | 5.77312712 | 9.93313449 | 18.2921914 |
| ENSMUSG00 | Hist2h2aa2 | 577 | 0 | 0 | 0 | 0 | 0 | 0 |
| ENSMUSG00 | Olfr694 | 947 | 0 | 0 | 0 | 0 | 0 | 0 |
| ENSMUSG00 | Gsdma3 | 1603 | 0 | 0 | 0 | 0 | 0 | 0 |
| ENSMUSG00 | Paqr9 | 2304 | 0 | 0 | 0 | 0 | 0.04429373 | 0.24859697 |
| ENSMUSG00 | Olfr393 | 938 | 0 | 0 | 0 | 0 | 0 | 0 |
| ENSMUSG00 | Gm5321 | 446 | 0 | 0 | 0.09910727 | 0 | 0 | 0.0611539 |
| ENSMUSG00 | Gm5414 | 1650 | 0 | 0 | 0 | 0 | 0 | 0 |
| ENSMUSG00 | Vmn1r205 | 1100 | 0 | 0 | 0 | 0 | 0 | 0 |
| ENSMUSG00 | Olfr786 | 938 | 0 | 0 | 0 | 0 | 0 | 0 |
| ENSMUSG00 | Chi3l1 | 2793 | 0.25621148 | 0.32779893 | 0.45895218 | 0.43923721 | 0.20096317 | 0.71287099 |
| ENSMUSG00 | Plcxd1 | 3801 | 0.34515413 | 0.31693266 | 0.10466103 | 0.05695666 | 0.59067628 | 0.20091815 |
| ENSMUSG00 | Olfr329-ps | 1304 | 0 | 0 | 0 | 0 | 0 | 0 |
| ENSMUSG00 | Ethe1 | 1476 | 14.1676035 | 21.0244368 | 14.0451656 | 11.6362149 | 12.2380332 | 14.6721298 |
| ENSMUSG00 | Vmn1r13 | 902 | 0 | 0 | 0 | 0 | 0 | 0 |
| ENSMUSG00 | Gimap8 | 3694 | 0.43048707 | 0.30002387 | 0.01196585 | 0 | 0.64922568 | 0.33964088 |
| ENSMUSG00 | Gm1631 | 637 | 0 | 0 | 0 | 0 | 0 | 0 |
| ENSMUSG00 | Zfp428 | 1651 | 2.04676914 | 1.92629024 | 1.90086666 | 2.382158 | 1.05081574 | 1.25552549 |
| ENSMUSG00 | Olfr1283 | 1025 | 0 | 0 | 0 | 0 | 0 | 0 |
| ENSMUSG00 | Hvcn1 | 3555 | 0.34667226 | 0.50151853 | 0.28597536 | 0.86272118 | 0.30142163 | 0.41429832 |
| ENSMUSG00 | Gpbar1 | 1074 | 0.7773418 | 0 | 0.37040651 | 0 | 0.23755296 | 2.31097968 |
| ENSMUSG00 | Ccdc146 | 4346 | 0 | 0 | 0.02034139 | 0 | 0.023482 | 0.08158544 |
| ENSMUSG00 | Rpl19-ps1 | 1272 | 0.03125431 | 0.11364727 | 0 | 0.02836639 | 0 | 0.02144233 |
| ENSMUSG00 | 2210021J22R | 1548 | 13.688418 | 11.7664568 | 11.1075691 | 7.73852635 | 7.38366162 | 14.4478063 |
| ENSMUSG00 | Hist1h4k | 311 | 0 | 0 | 0.14212811 | 0 | 0 | 0 |
| ENSMUSG00 | Tank | 5922 | 22.3884714 | 17.8278444 | 29.4753594 | 24.9930001 | 18.6803753 | 25.7962267 |
| ENSMUSG00 | Gm5852 | 1073 | 0 | 0 | 0 | 0 | 0 | 0 |
| ENSMUSG00 | Cntn4 | 8678 | 0.01832472 | 0.03331628 | 0.00509355 | 0 | 0.01175994 | 0 |
| ENSMUSG00 | Aox3 | 7174 | 0 | 0.00671682 | 0.14787346 | 0 | 0 | 0 |
| ENSMUSG00 | 4921528I07R | 2686 | 0 | 0 | 0 | 0 | 0 | 0 |
| ENSMUSG00 | Clasp1 | 7889 | 7.30706639 | 8.52073606 | 8.20835341 | 8.7312236 | 9.19755437 | 5.72183141 |
| ENSMUSG00 | Lrrc51 | 1427 | 11.0323548 | 12.1225878 | 9.75723935 | 12.8954749 | 9.5830895 | 12.8632321 |
| ENSMUSG00 | 2410137M14 | 1145 | 0 | 0 | 0 | 0 | 0 | 0 |
| ENSMUSG00 | Zpld1 | 2684 | 0 | 0 | 0 | 0 | 0 | 0 |
| ENSMUSG00 | Rpl21-ps10 | 547 | 0 | 0 | 0 | 0.06596352 | 0 | 0.04986223 |
| ENSMUSG00 | Gm10146 | 438 | 0 | 0 | 0.10091745 | 0 | 0 | 0 |
| ENSMUSG00 | Hhip | 4882 | 0 | 0 | 0 | 0 | 0 | 0 |
| ENSMUSG00 | Siva1 | 778 | 10.0155195 | 11.2724068 | 11.5902005 | 8.53354266 | 6.82100654 | 18.7206393 |
| ENSMUSG00 | Scn1a | 14331 | 0 | 0.00336239 | 0 | 0 | 0 | 0 |
| ENSMUSG00 | Pde6h | 5982 | 0.03322926 | 0.31415434 | 0.22167424 | 0.09650831 | 0.07676987 | 0.19149697 |
| ENSMUSG00 | Olfr926 | 1050 | 0 | 0 | 0 | 0 | 0 | 0 |
| ENSMUSG00 | Gm10147 | 855 | 0 | 0 | 0 | 0 | 0 | 0 |
| ENSMUSG00 | Sepp1 | 2837 | 174.506521 | 151.557146 | 158.344494 | 110.688063 | 125.902232 | 187.163439 |
| ENSMUSG00 | U6 | 106 | 0 | 0 | 0 | 0 | 0 | 0 |
| ENSMUSG00 | SNORD115 | 78 | 0 | 0 | 0 | 0 | 0 | 0 |
| ENSMUSG00 | U6 | 106 | 0 | 0 | 0 | 0 | 0 | 0 |
| ENSMUSG00 | SNORA73 | 202 | 0.39361862 | 1.43128046 | 1.53174704 | 0 | 1.2630291 | 1.48525263 |
| ENSMUSG00 | SNORA63 | 128 | 0 | 0 | 0 | 0 | 0 | 0 |
| ENSMUSG00 | U6 | 103 | 0 | 0 | 0 | 0 | 0 | 0 |
| ENSMUSG00 | Snora73a | 204 | 0 | 0 | 0 | 0 | 0 | 0 |
| ENSMUSG00 | U6 | 102 | 0 | 0 | 0 | 0 | 0 | 1.0695937 |
| ENSMUSG00 | Rnu73b | 70 | 0 | 2.06513324 | 0 | 0 | 0 | 0 |
| ENSMUSG00 | U6 | 99 | 0 | 0 | 0 | 0 | 0 | 0 |
| ENSMUSG00 | SNORA9 | 132 | 0 | 0 | 0 | 0 | 0 | 0 |
| ENSMUSG00 | U6 | 106 | 0 | 0 | 0 | 0 | 0 | 0 |
| ENSMUSG00 | U6 | 106 | 0 | 0 | 0 | 0 | 0 | 0 |
| ENSMUSG00 | U6 | 106 | 0 | 0 | 0 | 0 | 0 | 0 |
| ENSMUSG00 | 7SK | 320 | 0 | 0 | 0 | 0 | 0 | 0 |
| ENSMUSG00 | U2 | 172 | 0 | 1.40076867 | 4.62577427 | 0 | 0 | 0.9514409 |
| ENSMUSG00 | U6 | 106 | 0 | 0 | 0 | 0 | 0 | 0 |
| ENSMUSG00 | U3 | 213 | 0 | 0 | 0 | 0.16939927 | 0 | 0 |
| ENSMUSG00 | U6 | 106 | 1.12515511 | 0 | 1.25099556 | 1.36158659 | 0 | 2.31577126 |
| ENSMUSG00 | U1 | 144 | 0 | 0 | 0 | 0 | 0 | 0 |
| ENSMUSG00 | SNORA71 | 131 | 0 | 0 | 0 | 0 | 0 | 0 |
| ENSMUSG00 | U6 | 89 | 0 | 0 | 0 | 0 | 0 | 0 |
| ENSMUSG00 | U6 | 107 | 0 | 0 | 0 | 0 | 0 | 0 |
| ENSMUSG00 | n-R5s40 | 123 | 0 | 0.39175969 | 0 | 0.58669991 | 0 | 0 |
| ENSMUSG00 | SNORA36 | 130 | 0 | 0 | 0 | 0 | 0 | 0 |
| ENSMUSG00 | U6 | 106 | 0 | 0 | 0 | 0 | 0 | 0 |
| ENSMUSG00 | U6 | 106 | 0 | 0 | 0 | 0 | 0 | 0 |
| ENSMUSG00 | U6 | 102 | 0 | 0 | 0 | 0 | 0 | 0 |
| ENSMUSG00 | n-R5s215 | 103 | 0 | 0 | 0 | 0 | 0 | 0 |
| ENSMUSG00 | U1 | 149 | 0 | 0 | 0 | 0 | 0 | 0 |
| ENSMUSG00 | SNORD113 | 71 | 0 | 0 | 0 | 0 | 0 | 0 |
| ENSMUSG00 | U4 | 135 | 2.35588033 | 2.49855626 | 2.29194742 | 2.13819523 | 1.51189262 | 0 |
| ENSMUSG00 | U6 | 106 | 0 | 0 | 0 | 0 | 0 | 0 |
| ENSMUSG00 | U6 | 106 | 0 | 0 | 0 | 0.34039665 | 0 | 0 |
| ENSMUSG00 | U6 | 106 | 0 | 0 | 0 | 0 | 0 | 0 |
| ENSMUSG00 | U6 | 106 | 0 | 0.45458908 | 0 | 0 | 0.4813809 | 0 |
| ENSMUSG00 | SNORA42 | 135 | 2.06139529 | 3.92630269 | 0.65484212 | 1.33637202 | 1.13391946 | 0.40406873 |
| ENSMUSG00 | U6 | 106 | 0 | 0 | 0 | 0 | 0 | 0 |
| ENSMUSG00 | U6 | 106 | 0 | 0 | 0 | 0 | 0 | 0 |
| ENSMUSG00 | U5 | 115 | 1.03709949 | 0 | 0 | 0 | 0 | 0 |
| ENSMUSG00 | U1 | 165 | 0 | 0 | 0 | 0 | 0 | 0 |
| ENSMUSG00 | Y_RNA | 111 | 0 | 0 | 0 | 0 | 0 | 0 |
| ENSMUSG00 | SNORA4 | 139 | 0 | 0 | 0 | 0 | 0 | 0 |
| ENSMUSG00 | U1 | 165 | 0 | 0 | 0 | 0 | 0 | 0 |
| ENSMUSG00 | Snord49b | 71 | 1.11987269 | 2.03604685 | 0 | 0 | 0 | 0 |
| ENSMUSG00 | SNORD74 | 79 | 0 | 0 | 0 | 0 | 0 | 0 |
| ENSMUSG00 | Snord37 | 63 | 6.94143311 | 13.0026907 | 2.80646622 | 1.14546173 | 5.66959731 | 9.95740798 |
| ENSMUSG00 | SNORA36 | 134 | 0 | 0 | 0 | 0 | 0 | 0 |
| ENSMUSG00 | U6 | 115 | 0 | 0 | 0 | 0 | 0 | 0 |
| ENSMUSG00 | U6 | 100 | 0 | 0 | 0 | 0 | 0 | 0 |
| ENSMUSG00 | U6 | 106 | 0 | 0 | 0 | 0 | 0 | 0 |
| ENSMUSG00 | U1 | 164 | 0 | 0 | 0 | 0 | 0 | 0 |
| ENSMUSG00 | U6 | 106 | 0 | 0 | 0 | 0 | 0 | 0 |
| ENSMUSG00 | U6 | 106 | 0 | 0 | 0 | 0 | 0 | 0 |
| ENSMUSG00 | Snord68 | 85 | 0 | 0 | 0 | 0 | 0 | 0 |
| ENSMUSG00 | Snora23 | 183 | 1.08621531 | 1.05325557 | 0.48308025 | 1.18301785 | 0.55766531 | 1.49041745 |
| ENSMUSG00 | SNORD113 | 74 | 0 | 0 | 0 | 0 | 0 | 0 |
| ENSMUSG00 | Snord21 | 93 | 0 | 0 | 0 | 0 | 0 | 0 |
| ENSMUSG00 | n-R5s56 | 104 | 0 | 0 | 0 | 0 | 0 | 0 |
| ENSMUSG00 | U6 | 106 | 0 | 0 | 0 | 0 | 0 | 1.02923167 |
| ENSMUSG00 | U6 | 106 | 0 | 0 | 0 | 0 | 0 | 0 |
| ENSMUSG00 | SNORA73 | 202 | 0 | 0 | 0 | 0 | 0 | 0 |
| ENSMUSG00 | U6 | 106 | 0 | 0 | 0 | 0 | 0 | 0 |
| ENSMUSG00 | U6 | 106 | 0 | 0 | 0 | 0 | 0 | 0 |
| ENSMUSG00 | n-R5s37 | 117 | 0 | 0 | 0 | 0 | 0 | 0 |
| ENSMUSG00 | U6 | 107 | 0 | 0 | 0 | 0 | 0 | 0 |
| ENSMUSG00 | U6 | 106 | 0 | 0.45458908 | 0.41699852 | 5.44634634 | 1.92552362 | 0.77192375 |
| ENSMUSG00 | SNORA32 | 121 | 0 | 0 | 0 | 0 | 0 | 0 |
| ENSMUSG00 | U6 | 90 | 0.44172756 | 0 | 0 | 0 | 0 | 0 |
| ENSMUSG00 | U1 | 159 | 0 | 0 | 0 | 0 | 0 | 0 |
| ENSMUSG00 | SNORA32 | 96 | 0 | 0 | 0 | 0 | 0 | 0 |
| ENSMUSG00 | U6 | 106 | 0 | 0 | 0 | 0 | 0 | 0 |
| ENSMUSG00 | U1 | 158 | 0 | 0 | 0 | 0 | 0 | 0 |
| ENSMUSG00 | SNORD103 | 82 | 0 | 0 | 0 | 0 | 0 | 0 |
| ENSMUSG00 | U1 | 133 | 0 | 0 | 0 | 0 | 0 | 0 |
| ENSMUSG00 | U6 | 103 | 0 | 0 | 0 | 0 | 0 | 0 |
| ENSMUSG00 | SNORD113 | 74 | 0 | 0 | 0 | 0 | 0 | 0 |
| ENSMUSG00 | U6 | 106 | 0 | 0 | 0.41699852 | 0 | 0 | 0 |
| ENSMUSG00 | U1 | 160 | 0 | 0 | 0 | 0 | 0 | 0 |
| ENSMUSG00 | U6 | 106 | 0 | 0 | 0 | 0 | 0 | 0 |
| ENSMUSG00 | U6 | 106 | 0 | 0 | 0 | 0 | 2.40690452 | 0 |
| ENSMUSG00 | Snora28 | 126 | 31.5519687 | 34.4188873 | 35.7824443 | 38.6593334 | 56.2910019 | 84.6379678 |
| ENSMUSG00 | U6 | 106 | 0 | 0 | 0 | 1.02118994 | 0 | 0.51461584 |
| ENSMUSG00 | SNORA5 | 132 | 0 | 0.3650488 | 0 | 0.54669764 | 0 | 0 |
| ENSMUSG00 | SNORD113 | 72 | 0 | 0 | 0 | 0 | 0 | 0 |
| ENSMUSG00 | U6 | 106 | 0 | 0 | 0 | 0 | 0 | 0 |
| ENSMUSG00 | U6 | 106 | 0 | 0 | 0 | 0 | 0 | 0 |
| ENSMUSG00 | SNORD95 | 66 | 0 | 0 | 0 | 0 | 0 | 0 |
| ENSMUSG00 | U6 | 103 | 0 | 0 | 0 | 0 | 0 | 0 |
| ENSMUSG00 | n-R5s94 | 118 | 0 | 0 | 0 | 0 | 0 | 0 |
| ENSMUSG00 | n-R5s96 | 123 | 0 | 0 | 0 | 0 | 0 | 0 |
| ENSMUSG00 | U6 | 106 | 0 | 0 | 0 | 0 | 0 | 0 |
| ENSMUSG00 | SNORA32 | 107 | 0 | 0 | 0 | 0 | 0 | 0 |
| ENSMUSG00 | U1 | 124 | 0 | 0 | 0 | 0 | 0 | 0 |
| ENSMUSG00 | SNORA9 | 132 | 0 | 0 | 0 | 0 | 0 | 0 |
| ENSMUSG00 | SNORD18 | 70 | 2.27174174 | 1.37675549 | 5.0516392 | 10.3091556 | 7.28948226 | 10.1305803 |
| ENSMUSG00 | n-R5s162 | 118 | 0 | 0 | 0 | 0 | 0 | 0 |
| ENSMUSG00 | U6 | 106 | 0 | 0 | 0 | 0 | 0 | 0 |
| ENSMUSG00 | U6 | 106 | 0 | 0 | 0 | 0 | 0 | 0 |
| ENSMUSG00 | n-R5s97 | 112 | 1.06487894 | 0 | 0.39465931 | 0 | 4.10033377 | 0 |

| | | | | | | | | |
|---|---|---|---|---|---|---|---|---|
| ENSMUSG00 | U1 | 152 | 0 | 0.31701607 | 0 | 0 | 0.33569984 | 0 |
| ENSMUSG00 | U6 | 106 | 0 | 0 | 0 | 0 | 0 | 0 |
| ENSMUSG00 | U6 | 103 | 0 | 0 | 0 | 0 | 0 | 0 |
| ENSMUSG00 | SNORA65 | 128 | 0.93176907 | 1.8822829 | 1.0359807 | 0 | 0 | 0.21308312 |
| ENSMUSG00 | U6 | 105 | 0 | 0 | 0 | 0 | 0 | 0 |
| ENSMUSG00 | Y_RNA | 106 | 0 | 0 | 0 | 0 | 0 | 0 |
| ENSMUSG00 | U6 | 104 | 0 | 0 | 0 | 0 | 0 | 0 |
| ENSMUSG00 | SNORA36 | 86 | 0 | 0 | 0 | 1.25867597 | 0 | 0.63429394 |
| ENSMUSG00 | U6 | 106 | 0 | 0 | 0 | 0 | 0 | 0 |
| ENSMUSG00 | U6 | 106 | 0 | 0 | 0 | 0 | 0 | 0 |
| ENSMUSG00 | U6 | 106 | 0 | 0 | 0 | 0 | 0 | 0 |
| ENSMUSG00 | U6 | 106 | 0 | 0 | 0 | 0 | 0 | 0 |
| ENSMUSG00 | SNORD38 | 64 | 0 | 0 | 0 | 0 | 3.18914849 | 0 |
| ENSMUSG00 | Snord42a | 64 | 0 | 0 | 0 | 0 | 0 | 0 |
| ENSMUSG00 | U1 | 159 | 0 | 0 | 0 | 0 | 0 | 0 |
| ENSMUSG00 | SNORD38 | 69 | 0 | 0 | 0 | 1.04585636 | 0 | 0 |
| ENSMUSG00 | U1 | 158 | 0 | 0 | 0 | 0 | 0 | 0 |
| ENSMUSG00 | SNORD113 | 74 | 0 | 0 | 0 | 0 | 0 | 0 |
| ENSMUSG00 | U6 | 106 | 0 | 0 | 0 | 0 | 0 | 0 |
| ENSMUSG00 | 7SK | 318 | 0 | 0 | 0 | 0 | 0 | 0 |
| ENSMUSG00 | n-R5s65 | 116 | 0 | 0 | 0 | 0 | 0 | 0 |
| ENSMUSG00 | U6 | 106 | 0 | 0 | 0 | 0 | 0 | 0 |
| ENSMUSG00 | U2 | 193 | 0 | 0 | 0 | 0 | 0 | 0 |
| ENSMUSG00 | U1 | 163 | 0 | 0 | 0 | 0 | 0 | 0 |
| ENSMUSG00 | U1 | 161 | 0 | 0 | 0 | 0 | 0 | 0 |
| ENSMUSG00 | n-R5s173 | 118 | 0 | 0 | 0 | 0 | 0 | 0 |
| ENSMUSG00 | U1 | 160 | 0 | 0 | 0 | 0 | 0 | 0 |
| ENSMUSG00 | SNORA19 | 129 | 0 | 0 | 0 | 0 | 0 | 0 |
| ENSMUSG00 | U6 | 106 | 0 | 0 | 0 | 0 | 0 | 0 |
| ENSMUSG00 | SNORA65 | 129 | 0 | 1.12061493 | 0 | 0 | 0.3955533 | 0 |
| ENSMUSG00 | U6 | 106 | 0 | 0 | 0 | 0 | 0 | 0 |
| ENSMUSG00 | Snora7a | 139 | 1.71606391 | 0 | 0.31799887 | 0.77874916 | 0.73419246 | 3.33574725 |
| ENSMUSG00 | U6 | 106 | 0 | 0 | 0 | 0 | 0 | 0 |
| ENSMUSG00 | U1 | 162 | 0 | 0 | 1.09140353 | 0 | 2.51982103 | 0.33672394 |
| ENSMUSG00 | U6 | 110 | 0 | 0 | 0 | 0 | 0 | 0 |
| ENSMUSG00 | SNORA38 | 144 | 0 | 0 | 0 | 0 | 0 | 0 |
| ENSMUSG00 | U6 | 106 | 0 | 0 | 0 | 0 | 0 | 0 |
| ENSMUSG00 | SNORA71 | 130 | 0 | 0 | 0 | 0 | 0 | 0 |
| ENSMUSG00 | SNORA9 | 133 | 0 | 0 | 0 | 0 | 0 | 0 |
| ENSMUSG00 | U6 | 106 | 0 | 0 | 0 | 0 | 0 | 0 |
| ENSMUSG00 | U6 | 105 | 0 | 0 | 0 | 0 | 0 | 0 |
| ENSMUSG00 | SNORD81 | 76 | 0 | 0 | 0 | 0 | 0 | 0 |
| ENSMUSG00 | U6 | 106 | 0 | 0 | 0 | 0 | 0 | 0 |
| ENSMUSG00 | U6 | 106 | 0 | 0 | 0 | 0 | 0 | 0 |
| ENSMUSG00 | U6 | 106 | 0 | 0 | 0 | 0 | 0 | 0 |
| ENSMUSG00 | U1 | 162 | 0 | 0 | 0 | 0 | 0 | 0 |
| ENSMUSG00 | U1 | 159 | 0.25003447 | 0 | 0 | 0 | 0 | 0.85769306 |
| ENSMUSG00 | SNORA30 | 129 | 0 | 0 | 0 | 0 | 0 | 0 |
| ENSMUSG00 | U4 | 142 | 1.11987269 | 0 | 0.93384175 | 2.54098905 | 1.79670337 | 2.11282417 |
| ENSMUSG00 | U1 | 155 | 0 | 0 | 0 | 0 | 0 | 0 |
| ENSMUSG00 | SNORA71 | 132 | 0 | 0 | 0 | 0 | 0 | 0 |
| ENSMUSG00 | U6 | 106 | 0 | 0 | 0 | 0 | 0 | 0 |
| ENSMUSG00 | U4 | 150 | 0 | 0 | 0 | 0 | 0 | 0 |
| ENSMUSG00 | n-R5s74 | 115 | 0 | 0 | 0 | 0 | 0 | 0 |
| ENSMUSG00 | U6 | 106 | 0 | 0 | 0 | 0 | 0 | 0 |
| ENSMUSG00 | U6 | 106 | 0.3750517 | 0 | 0 | 0 | 0 | 2.05846334 |
| ENSMUSG00 | U6 | 105 | 0 | 0 | 0 | 0 | 0 | 0 |
| ENSMUSG00 | U1 | 163 | 0 | 0 | 0 | 0 | 0 | 0 |
| ENSMUSG00 | SNORA44 | 125 | 0 | 0 | 0 | 0 | 0 | 0 |
| ENSMUSG00 | SNORA51 | 132 | 0 | 0 | 0 | 0 | 0 | 0 |
| ENSMUSG00 | SNORA2 | 135 | 0.29448504 | 0 | 0 | 0 | 1.88986577 | 0 |
| ENSMUSG00 | SNORA13 | 132 | 0 | 0 | 2.34403713 | 0 | 0 | 0 |
| ENSMUSG00 | U6 | 104 | 0 | 0 | 0 | 0 | 0 | 0 |
| ENSMUSG00 | Snora41 | 131 | 0.30347695 | 2.20701262 | 0.33741865 | 4.95783818 | 5.45320047 | 12.2839978 |
| ENSMUSG00 | Snora44 | 128 | 0 | 0 | 0 | 0 | 0 | 0 |
| ENSMUSG00 | U1 | 148 | 0 | 0 | 0 | 0 | 0 | 0 |
| ENSMUSG00 | U4 | 140 | 0 | 0 | 0 | 0 | 0 | 0 |
| ENSMUSG00 | U6 | 106 | 0 | 0 | 0 | 0 | 0 | 0 |
| ENSMUSG00 | SNORA36 | 130 | 0 | 0 | 0 | 0 | 0 | 0 |
| ENSMUSG00 | U6 | 106 | 0 | 0 | 0 | 0 | 0 | 0 |
| ENSMUSG00 | SNORD51 | 72 | 2.76079726 | 0 | 0 | 5.01139507 | 2.12609899 | 2.2728866 |
| ENSMUSG00 | U1 | 139 | 0 | 0 | 0 | 0 | 0 | 0 |
| ENSMUSG00 | U6 | 106 | 0 | 0 | 1.25099556 | 4.08475976 | 0 | 0 |
| ENSMUSG00 | U6 | 106 | 0 | 0 | 0 | 0 | 0 | 0 |
| ENSMUSG00 | n-R5s176 | 108 | 0 | 0 | 0 | 0 | 0 | 0 |
| ENSMUSG00 | n-R5s95 | 117 | 0 | 0 | 0 | 0 | 0 | 0 |
| ENSMUSG00 | U6 | 105 | 0 | 0 | 0 | 0 | 0 | 0 |
| ENSMUSG00 | SNORA32 | 121 | 0 | 0 | 0 | 0 | 0 | 0 |
| ENSMUSG00 | AF357355 | 71 | 0 | 0 | 0 | 0 | 0 | 0 |
| ENSMUSG00 | SNORA61 | 128 | 0 | 0 | 0 | 0 | 0.79728712 | 0.85233248 |
| ENSMUSG00 | U6 | 106 | 0 | 0 | 0 | 0 | 0 | 0 |
| ENSMUSG00 | U6 | 107 | 0 | 0 | 0 | 0 | 0 | 0 |
| ENSMUSG00 | U6 | 106 | 0 | 0 | 0 | 0 | 0 | 0 |
| ENSMUSG00 | U6 | 106 | 0 | 0 | 0 | 0 | 0 | 0 |
| ENSMUSG00 | U6 | 108 | 0 | 0 | 0 | 0 | 0 | 0 |
| ENSMUSG00 | n-R5s42 | 103 | 0 | 0 | 0 | 0 | 0 | 0 |
| ENSMUSG00 | U6 | 106 | 0 | 0 | 0 | 0 | 0 | 0 |
| ENSMUSG00 | SNORA1 | 132 | 0.30117788 | 0 | 0 | 1.36674411 | 0 | 3.30601688 |
| ENSMUSG00 | Snora20 | 127 | 4.06945864 | 0 | 0 | 3.97754822 | 4.01782487 | 0 |
| ENSMUSG00 | SNORA32 | 78 | 0 | 0 | 0 | 0 | 0 | 0 |
| ENSMUSG00 | U6 | 106 | 0 | 0 | 1.66799408 | 0 | 0 | 0 |
| ENSMUSG00 | U6 | 99 | 0 | 0 | 1.78593305 | 0 | 2.06167175 | 0 |
| ENSMUSG00 | SNORA42 | 124 | 0 | 0 | 0 | 0 | 0 | 0 |
| ENSMUSG00 | SNORD58 | 64 | 1.86353815 | 4.51747895 | 4.14392278 | 9.58429308 | 6.37829698 | 7.24482605 |
| ENSMUSG00 | U1 | 151 | 6.5820332 | 6.70142573 | 0.87818231 | 0 | 6.08261434 | 3.07065475 |
| ENSMUSG00 | U6 | 106 | 0 | 0 | 0 | 0 | 0 | 0 |
| ENSMUSG00 | U6 | 106 | 0 | 0 | 0 | 0 | 0 | 0 |
| ENSMUSG00 | SNORA31 | 127 | 0 | 0 | 0 | 0 | 0 | 0 |
| ENSMUSG00 | U6 | 106 | 0 | 0 | 0 | 0.34039665 | 0 | 0 |
| ENSMUSG00 | SNORA55 | 130 | 0 | 0 | 0 | 0 | 0.39251058 | 0 |
| ENSMUSG00 | U6 | 106 | 0 | 0 | 0 | 0 | 0 | 0 |
| ENSMUSG00 | SNORA2 | 134 | 0 | 0 | 0.9895935 | 0 | 1.14238155 | 0 |
| ENSMUSG00 | U6 | 109 | 0 | 0 | 0 | 0 | 0 | 0 |
| ENSMUSG00 | U6 | 106 | 0 | 0 | 0 | 0 | 0.4813809 | 0 |
| ENSMUSG00 | U4 | 85 | 0 | 0 | 0 | 0 | 0 | 0 |
| ENSMUSG00 | SNORA4 | 105 | 0 | 0 | 0 | 0 | 0 | 0 |
| ENSMUSG00 | Snora52 | 136 | 0 | 0 | 0 | 0 | 0 | 0 |
| ENSMUSG00 | U6 | 106 | 0 | 0 | 0 | 0 | 0 | 0 |
| ENSMUSG00 | U6 | 102 | 0 | 1.8896644 | 0 | 0 | 0 | 0 |
| ENSMUSG00 | U6 | 106 | 0 | 0 | 0 | 0 | 0 | 0 |
| ENSMUSG00 | SNORA65 | 128 | 0 | 0 | 0 | 0 | 0 | 0 |
| ENSMUSG00 | SNORA2 | 135 | 2.06139529 | 0 | 0 | 0 | 0 | 0 |
| ENSMUSG00 | U6 | 106 | 0 | 0 | 0 | 0 | 0 | 0 |
| ENSMUSG00 | U6 | 106 | 0 | 0 | 0 | 0 | 0 | 0 |
| ENSMUSG00 | U1 | 157 | 0 | 0 | 0 | 0 | 0 | 0 |
| ENSMUSG00 | U1 | 155 | 0 | 0 | 0 | 0 | 0 | 0 |
| ENSMUSG00 | U6 | 106 | 0 | 0 | 0 | 0 | 0 | 0 |
| ENSMUSG00 | SNORD113 | 71 | 0 | 0 | 0 | 0 | 0 | 0 |
| ENSMUSG00 | U6 | 106 | 0 | 0 | 0 | 0 | 0 | 0 |
| ENSMUSG00 | U2 | 190 | 0 | 0 | 0 | 0 | 0 | 0 |
| ENSMUSG00 | U6 | 106 | 0 | 0 | 0 | 0 | 0 | 0 |
| ENSMUSG00 | U6 | 102 | 0 | 0 | 0 | 0 | 0 | 0 |
| ENSMUSG00 | U6 | 107 | 0 | 0 | 0 | 0 | 0 | 0 |
| ENSMUSG00 | U6 | 100 | 0 | 0 | 0 | 0 | 0 | 0 |
| ENSMUSG00 | U6 | 106 | 0 | 0 | 0 | 0 | 0 | 0 |
| ENSMUSG00 | U6 | 106 | 0 | 0 | 0 | 0 | 0 | 0 |
| ENSMUSG00 | U6 | 106 | 0 | 0 | 0 | 0 | 0 | 0 |
| ENSMUSG00 | U6 | 106 | 0 | 0 | 0 | 0 | 0 | 0 |
| ENSMUSG00 | SNORA67 | 140 | 0 | 0 | 0 | 0 | 0 | 0 |
| ENSMUSG00 | U6 | 103 | 0 | 0 | 0 | 0 | 0 | 0 |
| ENSMUSG00 | 7SK | 320 | 0 | 0 | 0 | 0 | 0.15945742 | 0 |
| ENSMUSG00 | SNORD20 | 79 | 2.01293572 | 0 | 1.119034 | 0 | 0.64590349 | 3.79773458 |
| ENSMUSG00 | SNORA67 | 143 | 0 | 0 | 0 | 0 | 0 | 0 |
| ENSMUSG00 | U1 | 162 | 0 | 0 | 0 | 0 | 0 | 0 |
| ENSMUSG00 | U6 | 106 | 0 | 0.45458908 | 0 | 0 | 0 | 2.05846334 |
| ENSMUSG00 | U2 | 190 | 0 | 0 | 0 | 0 | 0 | 0 |
| ENSMUSG00 | U6 | 106 | 0 | 0 | 0 | 0 | 0 | 0 |
| ENSMUSG00 | n-R5s180 | 105 | 0 | 0 | 0 | 0 | 0 | 0 |
| ENSMUSG00 | Y_RNA | 109 | 0 | 0 | 0 | 0 | 0 | 0 |
| ENSMUSG00 | U1 | 159 | 0 | 0 | 0 | 0 | 0 | 0 |
| ENSMUSG00 | U6 | 106 | 0 | 0 | 0 | 0 | 0 | 0 |
| ENSMUSG00 | U6 | 106 | 0 | 0 | 0 | 0 | 0 | 0 |
| ENSMUSG00 | U6 | 106 | 0 | 0 | 0 | 0 | 0 | 0 |
| ENSMUSG00 | n-R5s62 | 95 | 0 | 0 | 0 | 0 | 0 | 0 |
| ENSMUSG00 | U6 | 107 | 0 | 0 | 0 | 0 | 0 | 0 |
| ENSMUSG00 | U8 | 134 | 0 | 0 | 0 | 0 | 0 | 0 |
| ENSMUSG00 | U6 | 106 | 0 | 0 | 0 | 0 | 0 | 0 |
| ENSMUSG00 | U6 | 106 | 0 | 1.36376723 | 0 | 0 | 0 | 0.25730792 |
| ENSMUSG00 | SNORD113 | 74 | 0 | 0 | 0 | 0 | 0 | 0 |
| ENSMUSG00 | SNORD25 | 71 | 0 | 0 | 0 | 0 | 0 | 0 |
| ENSMUSG00 | SNORA44 | 128 | 0.62117938 | 0 | 0 | 0 | 0 | 0.21308312 |
| ENSMUSG00 | U6 | 106 | 0 | 0 | 0 | 0 | 0.96276181 | 0.25730792 |
| ENSMUSG00 | SNORA63 | 129 | 0 | 0 | 0 | 0 | 0 | 0 |
| ENSMUSG00 | Snord96a | 80 | 0 | 0 | 0 | 0 | 0 | 0 |
| ENSMUSG00 | SNORD113 | 71 | 0 | 0 | 0 | 0 | 0 | 0 |
| ENSMUSG00 | SNORA71 | 131 | 0 | 0 | 0 | 0 | 0 | 0 |
| ENSMUSG00 | U6 | 106 | 0 | 0 | 0 | 0 | 0 | 0 |
| ENSMUSG00 | Snord45b | 70 | 10.7907733 | 8.94891068 | 11.3661882 | 6.18549335 | 26.2421361 | 20.6507983 |
| ENSMUSG00 | U1 | 162 | 0 | 0 | 0 | 0 | 0 | 0 |
| ENSMUSG00 | U6 | 106 | 0 | 0 | 0 | 0 | 0 | 0 |
| ENSMUSG00 | U6 | 106 | 0 | 0 | 0 | 0 | 0 | 0 |
| ENSMUSG00 | U6 | 110 | 0 | 0 | 0 | 0 | 0 | 0 |
| ENSMUSG00 | U1 | 161 | 0.24692845 | 0 | 0 | 0 | 0 | 1.01644618 |
| ENSMUSG00 | U6 | 93 | 0 | 0 | 0 | 0 | 0 | 0 |
| ENSMUSG00 | SNORA70 | 136 | 0 | 0 | 0 | 0 | 0 | 0 |
| ENSMUSG00 | Snord14a | 86 | 1.84909212 | 0.56030747 | 2.56987459 | 1.67823463 | 2.37331981 | 3.80576362 |
| ENSMUSG00 | U6 | 106 | 0 | 0 | 0 | 0 | 0 | 0 |
| ENSMUSG00 | U6 | 106 | 0 | 0 | 0 | 0 | 0 | 0 |
| ENSMUSG00 | n-R5s86 | 118 | 0 | 0 | 0 | 0 | 0 | 0 |
| ENSMUSG00 | U6 | 102 | 0 | 0 | 0 | 0 | 0 | 0 |
| ENSMUSG00 | U6 | 106 | 0 | 0 | 0 | 0 | 0 | 0 |
| ENSMUSG00 | U6 | 106 | 0 | 0 | 0 | 0 | 0 | 0 |
| ENSMUSG00 | n-R5s28 | 118 | 0 | 0 | 0 | 0 | 0 | 0 |
| ENSMUSG00 | U6 | 106 | 0 | 0 | 0 | 0 | 0 | 0 |
| ENSMUSG00 | SNORD46 | 103 | 0 | 0 | 0 | 0 | 0 | 0 |
| ENSMUSG00 | U6 | 106 | 0 | 0 | 0 | 0.34039665 | 0 | 0 |
| ENSMUSG00 | U6 | 106 | 0 | 0 | 0 | 0 | 0 | 0 |
| ENSMUSG00 | U6 | 106 | 0 | 0 | 0 | 0 | 0 | 0 |
| ENSMUSG00 | U6 | 106 | 0 | 0 | 0 | 0 | 0 | 0 |
| ENSMUSG00 | U2 | 179 | 0 | 0 | 0 | 0 | 0 | 0 |
| ENSMUSG00 | SNORA61 | 129 | 0 | 0 | 0 | 0 | 0 | 0 |
| ENSMUSG00 | U1 | 170 | 0 | 0 | 0 | 0 | 0 | 0 |
| ENSMUSG00 | U6 | 106 | 0 | 0 | 0 | 0 | 0 | 0 |
| ENSMUSG00 | U2 | 191 | 0 | 0 | 0 | 0 | 0 | 0 |
| ENSMUSG00 | Y_RNA | 110 | 0 | 5.69476134 | 1.60733975 | 3.60820445 | 0 | 2.23156139 |
| ENSMUSG00 | SNORA24 | 127 | 0 | 0 | 0 | 0 | 0 | 0 |
| ENSMUSG00 | Snord35b | 86 | 0.92454606 | 2.80153733 | 6.68167394 | 1.25867597 | 2.37331981 | 8.24582117 |
| ENSMUSG00 | SNORD60 | 77 | 0 | 0 | 0 | 0 | 0 | 0 |
| ENSMUSG00 | U1 | 152 | 0 | 0 | 0 | 0 | 0 | 0 |
| ENSMUSG00 | n-R5s29 | 118 | 0 | 0 | 0 | 0 | 0 | 0 |
| ENSMUSG00 | U2 | 190 | 0 | 0 | 0 | 0 | 0 | 0 |
| ENSMUSG00 | U6 | 102 | 0 | 0 | 0 | 0 | 0 | 0 |
| ENSMUSG00 | U3 | 213 | 0 | 0 | 0 | 0 | 0 | 0 |
| ENSMUSG00 | U6 | 105 | 0 | 0 | 0 | 0 | 0 | 0 |
| ENSMUSG00 | SNORD59 | 73 | 5.99055186 | 2.64035299 | 9.68807518 | 8.4026679 | 5.5919316 | 14.1977574 |
| ENSMUSG00 | U6 | 106 | 0 | 0 | 0 | 0 | 1.44414271 | 0 |
| ENSMUSG00 | Snord95 | 67 | 0 | 0 | 0 | 0 | 0 | 0 |
| ENSMUSG00 | U6 | 103 | 0 | 0 | 0 | 0 | 0 | 0 |
| ENSMUSG00 | U6 | 105 | 0 | 0 | 0.42096993 | 0 | 0 | 0 |
| ENSMUSG00 | SNORA71 | 131 | 0 | 0 | 0 | 0 | 0 | 0 |
| ENSMUSG00 | SNORA2 | 136 | 0 | 0 | 0 | 0 | 0 | 0 |
| ENSMUSG00 | Snord14e | 89 | 0 | 0 | 0 | 0 | 0 | 0 |
| ENSMUSG00 | SNORA32 | 121 | 0 | 0 | 0 | 0.29819872 | 2.9519391 | 0 |
| ENSMUSG00 | U6 | 105 | 0 | 0 | 0 | 0 | 0 | 0 |
| ENSMUSG00 | U2 | 183 | 0 | 0 | 0 | 0 | 0.27883266 | 0 |
| ENSMUSG00 | Telomerase-v | 422 | 0.0942073 | 0 | 0 | 0.08550248 | 0.24183117 | 0 |
| ENSMUSG00 | snoZ40 | 73 | 8.16893435 | 5.94079424 | 15.7431222 | 6.91984416 | 7.68890595 | 13.450507 |
| ENSMUSG00 | n-R5s193 | 108 | 0 | 0 | 0 | 0 | 0 | 0 |
| ENSMUSG00 | SNORA73 | 206 | 0 | 0 | 0 | 0 | 0 | 0 |
| ENSMUSG00 | U6 | 106 | 0 | 0 | 0 | 0 | 0 | 0 |
| ENSMUSG00 | U1 | 155 | 0.25648697 | 1.55440136 | 1.4258659 | 0 | 1.3168097 | 0 |
| ENSMUSG00 | SNORA44 | 128 | 0 | 0 | 0 | 0 | 0 | 0 |
| ENSMUSG00 | U2 | 179 | 0 | 0 | 0 | 0 | 0 | 0 |
| ENSMUSG00 | U1 | 158 | 0 | 0 | 0 | 0 | 0 | 0 |
| ENSMUSG00 | U3 | 213 | 0.74658179 | 0 | 0 | 0.16939927 | 0.23956045 | 0.5121998 |
| ENSMUSG00 | SNORA32 | 121 | 0 | 1.19470518 | 0 | 0 | 0 | 0 |
| ENSMUSG00 | U6 | 106 | 0 | 0 | 0 | 0 | 0 | 0 |
| ENSMUSG00 | SNORA32 | 132 | 0 | 0 | 0 | 0 | 0 | 0 |
| ENSMUSG00 | SNORD24 | 82 | 0 | 0 | 0 | 0 | 0 | 0 |
| ENSMUSG00 | U1 | 164 | 0 | 0 | 0 | 0 | 0 | 0 |
| ENSMUSG00 | U6 | 107 | 0 | 0 | 0 | 0 | 0 | 0 |
| ENSMUSG00 | U6 | 106 | 0 | 0 | 0 | 0 | 0 | 0 |
| ENSMUSG00 | U6 | 106 | 0 | 0 | 0 | 0 | 0 | 0 |
| ENSMUSG00 | U6 | 106 | 0 | 0 | 0 | 0 | 0 | 0 |
| ENSMUSG00 | Snord82 | 69 | 0 | 0 | 0 | 4.18342545 | 3.69756347 | 0 |
| ENSMUSG00 | U6 | 106 | 0 | 0 | 0 | 0 | 0 | 0 |
| ENSMUSG00 | n-R5s195 | 118 | 0 | 0 | 0 | 0 | 0 | 0 |
| ENSMUSG00 | U6 | 106 | 0 | 0 | 0 | 0 | 0 | 0 |
| ENSMUSG00 | SNORD36 | 56 | 0 | 0 | 0 | 0 | 0 | 0 |
| ENSMUSG00 | U1 | 166 | 0 | 0 | 0 | 0 | 0 | 0 |
| ENSMUSG00 | U6 | 106 | 0 | 0 | 0 | 0 | 0 | 0 |
| ENSMUSG00 | n-R5s166 | 120 | 0 | 0 | 0 | 0 | 0 | 0 |
| ENSMUSG00 | U1 | 159 | 0 | 0 | 0 | 0 | 1.28368241 | 0 |
| ENSMUSG00 | U6 | 104 | 0 | 0 | 0 | 0 | 0 | 0 |
| ENSMUSG00 | U6 | 106 | 0 | 0 | 0 | 0 | 0 | 0 |
| ENSMUSG00 | SNORA67 | 97 | 0 | 0 | 0 | 0 | 0 | 0 |
| ENSMUSG00 | Snora75 | 132 | 4.51768824 | 4.74563445 | 1.00458734 | 1.09339529 | 2.70594417 | 5.37227743 |
| ENSMUSG00 | U6 | 106 | 0 | 0 | 0 | 0 | 0 | 0 |
| ENSMUSG00 | SNORA42 | 133 | 12.8532757 | 14.129859 | 17.2819236 | 14.6498527 | 9.20776707 | 24.1985521 |
| ENSMUSG00 | U6 | 109 | 0 | 0 | 0 | 0 | 0 | 0 |
| ENSMUSG00 | U6 | 103 | 0 | 0 | 0 | 0 | 0 | 0 |
| ENSMUSG00 | SNORD58 | 64 | 1.86353815 | 0 | 11.0504608 | 3.94647362 | 6.37829698 | 4.68782862 |
| ENSMUSG00 | U6 | 113 | 0 | 0.42642869 | 0 | 0 | 0 | 0 |
| ENSMUSG00 | U6 | 106 | 0 | 0 | 0 | 0 | 0 | 0 |
| ENSMUSG00 | SNORA61 | 127 | 0 | 0 | 0 | 0 | 0 | 0 |
| ENSMUSG00 | U12 | 150 | 0 | 0 | 0 | 0 | 0 | 0 |
| ENSMUSG00 | SNORA38 | 133 | 0.59782678 | 0 | 0 | 0 | 0 | 0 |
| ENSMUSG00 | U6 | 106 | 0 | 0 | 0 | 0 | 0 | 0 |
| ENSMUSG00 | U1 | 157 | 0 | 0 | 0 | 0 | 0 | 0 |
| ENSMUSG00 | U2 | 190 | 0 | 0 | 0 | 0 | 0 | 0 |
| ENSMUSG00 | n-R5s25 | 118 | 0 | 1.63343872 | 0 | 0 | 0 | 0 |
| ENSMUSG00 | Snora43 | 126 | 0 | 0 | 0 | 0 | 0 | 0 |
| ENSMUSG00 | U1 | 169 | 0 | 0 | 0 | 0 | 0.90579365 | 0 |
| ENSMUSG00 | n-R5s220 | 119 | 0 | 0 | 0 | 0 | 0 | 0 |
| ENSMUSG00 | SNORA75 | 129 | 0 | 0 | 0 | 0 | 0 | 0 |
| ENSMUSG00 | U6 | 103 | 0 | 0 | 0 | 0 | 0 | 0 |
| ENSMUSG00 | U6 | 106 | 0 | 0 | 0 | 0 | 0 | 0 |
| ENSMUSG00 | Snord58b | 65 | 3.66973666 | 5.18930915 | 10.8804537 | 6.66130053 | 4.710127 | 9.23141636 |
| ENSMUSG00 | U1 | 159 | 0 | 0 | 0 | 0 | 0 | 0 |
| ENSMUSG00 | U6 | 106 | 0 | 0 | 0 | 0 | 0 | 0 |
| ENSMUSG00 | U6 | 106 | 0 | 0 | 0 | 0 | 0 | 0 |
| ENSMUSG00 | SNORA17 | 137 | 0 | 0 | 0 | 0 | 0 | 0 |
| ENSMUSG00 | SNORA63 | 124 | 0 | 0 | 0 | 0 | 0 | 0 |
| ENSMUSG00 | U5 | 115 | 0 | 0 | 0 | 0 | 0 | 0 |
| ENSMUSG00 | U6 | 102 | 0 | 0 | 0 | 0 | 0 | 0 |
| ENSMUSG00 | Y_RNA | 113 | 0 | 0 | 0 | 0 | 0 | 0 |
| ENSMUSG00 | U6 | 106 | 0 | 0 | 0 | 0 | 0 | 0 |
| ENSMUSG00 | U6 | 106 | 0 | 0 | 0 | 0 | 0 | 0 |
| ENSMUSG00 | U1 | 156 | 0 | 0 | 0 | 0 | 0 | 0 |
| ENSMUSG00 | U6 | 107 | 0 | 0 | 0 | 0 | 0 | 0 |
| ENSMUSG00 | U6 | 106 | 0 | 0 | 0 | 0 | 0 | 0 |
| ENSMUSG00 | SNORA19 | 131 | 0 | 0 | 0 | 0 | 0 | 0 |
| ENSMUSG00 | SNORA43 | 138 | 0 | 0 | 0 | 0 | 0 | 0 |
| ENSMUSG00 | Snord118 | 134 | 1.18673076 | 0 | 0.659729 | 1.61561393 | 0.7615877 | 0.40708417 |
| ENSMUSG00 | SNORA71 | 130 | 0 | 0 | 0 | 0 | 0 | 0 |
| ENSMUSG00 | Snora21 | 136 | 0 | 0 | 0 | 0.26530915 | 0.75038788 | 1.00274409 |
| ENSMUSG00 | U6 | 106 | 0 | 0 | 0 | 0 | 0 | 0 |
| ENSMUSG00 | Vault | 142 | 0 | 0 | 0 | 0 | 0 | 0 |
| ENSMUSG00 | n-R5s204 | 118 | 1.34764341 | 0 | 0 | 0 | 0 | 0 |
| ENSMUSG00 | U6 | 106 | 0 | 0 | 0 | 0 | 0 | 0 |
| ENSMUSG00 | U6 | 105 | 0 | 0 | 0 | 0 | 0 | 0 |
| ENSMUSG00 | U6 | 106 | 0 | 0 | 0 | 0 | 0 | 0 |
| ENSMUSG00 | U6 | 106 | 0 | 0 | 0 | 0 | 0 | 0 |
| ENSMUSG00 | U6 | 109 | 0 | 0 | 0 | 0 | 0 | 0 |
| ENSMUSG00 | U6 | 101 | 0 | 0 | 0 | 0 | 3.03126985 | 0.27004593 |
| ENSMUSG00 | snoZ39 | 59 | 14.1502558 | 26.1350195 | 52.4428646 | 27.5202035 | 7.78368445 | 93.8432503 |
| ENSMUSG00 | SNORD18 | 70 | 3.40761262 | 2.06513324 | 1.2629098 | 4.12366223 | 2.18684468 | 7.79275407 |
| ENSMUSG00 | U6 | 104 | 0 | 0 | 0 | 0.69388547 | 0 | 0 |
| ENSMUSG00 | U6 | 106 | 0 | 0 | 0 | 0 | 0 | 0 |
| ENSMUSG00 | U6 | 107 | 0 | 0 | 0 | 0 | 0 | 0 |
| ENSMUSG00 | U6 | 106 | 0 | 0 | 0 | 0 | 0 | 0 |
| ENSMUSG00 | U1 | 164 | 0 | 0 | 0 | 0 | 0 | 0 |
| ENSMUSG00 | U6 | 109 | 0 | 0 | 0 | 0 | 0 | 0 |
| ENSMUSG00 | Snora62 | 148 | 0 | 0.65116814 | 0.5973222 | 1.46278559 | 0.34477281 | 2.02716913 |
| ENSMUSG00 | U6 | 106 | 0 | 0 | 0 | 0 | 0 | 0 |
| ENSMUSG00 | U1 | 157 | 5.82405129 | 3.37612015 | 1.97078281 | 13.0998506 | 7.80021032 | 8.51246703 |
| ENSMUSG00 | U6 | 107 | 0 | 0 | 0 | 0 | 0 | 0 |
| ENSMUSG00 | SNORA41 | 131 | 0 | 0 | 0 | 0 | 0 | 0 |
| ENSMUSG00 | n-R5s36 | 118 | 0 | 0 | 0 | 0 | 0 | 0 |
| ENSMUSG00 | SNORA33 | 129 | 1.5409101 | 2.61476818 | 0.68529989 | 4.19558657 | 3.95553301 | 4.22862624 |
| ENSMUSG00 | SNORA58 | 135 | 0 | 0.71387322 | 2.29194742 | 0 | 1.51189262 | 0 |
| ENSMUSG00 | U6 | 106 | 0 | 0 | 0 | 0 | 0 | 0 |
| ENSMUSG00 | U1 | 180 | 0 | 0 | 0 | 0 | 0 | 0 |
| ENSMUSG00 | U1 | 163 | 0 | 0.29562234 | 0 | 0 | 0 | 0 |
| ENSMUSG00 | U1 | 120 | 0 | 0 | 0 | 0 | 0 | 0 |
| ENSMUSG00 | Y_RNA | 101 | 6.2978979 | 0.95418697 | 2.62585206 | 9.28844711 | 2.02084657 | 21.0635828 |
| ENSMUSG00 | U6 | 106 | 0 | 0 | 0 | 0 | 0 | 0 |
| ENSMUSG00 | U6 | 105 | 0 | 0 | 0 | 0 | 0 | 0 |
| ENSMUSG00 | Snora61 | 129 | 0 | 0 | 0 | 0 | 0 | 0 |
| ENSMUSG00 | U6 | 106 | 0 | 0 | 0 | 0 | 0 | 0 |
| ENSMUSG00 | SNORD62 | 84 | 0 | 0 | 0 | 0.8590963 | 1.21491371 | 0.97409426 |
| ENSMUSG00 | U6 | 105 | 0 | 0 | 0 | 0 | 0 | 0 |
| ENSMUSG00 | SNORA24 | 129 | 0 | 0 | 0 | 0 | 0 | 1.6914505 |
| ENSMUSG00 | U6 | 106 | 0 | 0 | 0 | 0 | 0 | 0 |
| ENSMUSG00 | n-R5s41 | 118 | 0 | 0 | 0 | 0 | 0 | 0 |
| ENSMUSG00 | SNORA2 | 127 | 0 | 0 | 0 | 0 | 0 | 0 |
| ENSMUSG00 | SNORA2 | 136 | 0 | 0 | 0 | 0 | 0 | 0 |
| ENSMUSG00 | n-R5s93 | 118 | 0 | 0 | 0 | 0 | 0 | 0 |
| ENSMUSG00 | U6 | 103 | 0 | 0 | 0 | 0.35031111 | 0 | 0 |
| ENSMUSG00 | U6 | 104 | 0 | 0 | 0 | 1.04082821 | 0 | 0 |
| ENSMUSG00 | U6 | 106 | 0 | 0 | 0 | 0 | 0 | 0 |
| ENSMUSG00 | Snord15b | 144 | 0 | 0 | 0 | 0 | 0 | 0 |
| ENSMUSG00 | U1 | 166 | 0 | 0 | 0 | 3.91251085 | 1.22955123 | 1.15013539 |
| ENSMUSG00 | Snord47 | 76 | 0 | 0 | 0 | 0 | 0 | 0 |
| ENSMUSG00 | U6 | 106 | 0 | 0 | 0 | 0 | 0 | 0 |
| ENSMUSG00 | U6 | 93 | 0 | 0 | 0 | 0 | 0 | 0 |
| ENSMUSG00 | U6 | 106 | 0 | 0 | 0 | 0 | 0 | 0 |
| ENSMUSG00 | U6 | 106 | 0 | 0 | 0 | 0 | 0 | 0 |
| ENSMUSG00 | U6 | 106 | 0 | 0 | 0 | 0 | 0 | 0 |
| ENSMUSG00 | U6 | 106 | 0 | 0 | 0 | 0 | 0 | 0 |
| ENSMUSG00 | U6 | 106 | 0 | 0 | 0 | 0 | 0 | 0 |
| ENSMUSG00 | U6 | 103 | 0 | 0 | 0 | 0 | 0 | 0 |
| ENSMUSG00 | Snora70 | 136 | 0 | 0 | 0 | 0 | 0 | 0 |
| ENSMUSG00 | U2 | 192 | 0 | 0 | 1.38130759 | 0 | 0 | 0 |
| ENSMUSG00 | U6 | 106 | 0 | 0 | 0 | 0 | 0 | 0 |
| ENSMUSG00 | Snord73a | 67 | 2.37346152 | 6.47280566 | 0 | 2.69268989 | 3.0463508 | 0 |
| ENSMUSG00 | U1 | 133 | 0 | 0 | 0 | 0 | 0 | 0 |
| ENSMUSG00 | U6 | 106 | 0 | 0 | 0 | 0 | 0 | 0 |
| ENSMUSG00 | U6 | 106 | 0 | 0 | 0 | 0 | 0 | 0 |
| ENSMUSG00 | SNORA70 | 134 | 0 | 0 | 0 | 0 | 2.66555695 | 0.81416834 |
| ENSMUSG00 | SNORA71 | 112 | 0 | 0 | 0 | 0 | 0 | 0 |
| ENSMUSG00 | SNORA25 | 132 | 0 | 0 | 0 | 0 | 0 | 0 |
| ENSMUSG00 | U6 | 106 | 0 | 0 | 0 | 0 | 0 | 0 |
| ENSMUSG00 | 7SK | 307 | 0 | 0 | 0 | 0 | 0 | 0 |
| ENSMUSG00 | U6 | 106 | 0 | 0 | 0 | 0 | 0 | 0 |
| ENSMUSG00 | SNORA33 | 128 | 0 | 0 | 0 | 0 | 0 | 0 |
| ENSMUSG00 | n-R5s79 | 119 | 0 | 0 | 0 | 0 | 0 | 0 |
| ENSMUSG00 | U6 | 106 | 0 | 0 | 0 | 0 | 0 | 0 |
| ENSMUSG00 | U6 | 106 | 0 | 0 | 0 | 0 | 0 | 0 |
| ENSMUSG00 | U6 | 106 | 0 | 0 | 0 | 0 | 0 | 0 |
| ENSMUSG00 | U6 | 106 | 0 | 0 | 0 | 0 | 0 | 0 |
| ENSMUSG00 | U6 | 106 | 0 | 0 | 0 | 0 | 0 | 0 |
| ENSMUSG00 | SNORD113 | 71 | 0 | 0 | 0 | 0 | 0 | 0 |
| ENSMUSG00 | U1 | 159 | 0 | 0 | 0 | 0 | 0 | 0 |
| ENSMUSG00 | Snora3 | 131 | 0 | 0 | 0 | 0 | 0 | 0 |
| ENSMUSG00 | U1 | 162 | 0 | 0 | 0 | 0 | 0 | 0 |
| ENSMUSG00 | U1 | 163 | 0 | 0 | 0 | 0 | 0 | 0 |
| ENSMUSG00 | U1 | 164 | 0 | 0 | 0 | 0 | 0 | 0 |
| ENSMUSG00 | AF357341 | 74 | 0 | 0 | 0 | 0 | 0 | 0 |
| ENSMUSG00 | U6 | 106 | 0 | 0 | 0 | 0 | 0 | 0 |
| ENSMUSG00 | U6 | 106 | 0 | 0 | 0 | 0 | 0 | 0 |
| ENSMUSG00 | U1 | 169 | 0 | 0 | 0 | 0 | 0 | 0 |
| ENSMUSG00 | SNORA51 | 129 | 0 | 0 | 0 | 0 | 0 | 0 |
| ENSMUSG00 | U6 | 103 | 0 | 0 | 0 | 0 | 0 | 0 |
| ENSMUSG00 | n-R5s160 | 119 | 0 | 0 | 0.37144406 | 0 | 0 | 0 |
| ENSMUSG00 | U6 | 97 | 0 | 0 | 0 | 0 | 0 | 0 |
| ENSMUSG00 | U6 | 106 | 0 | 0 | 0 | 0 | 0 | 0 |
| ENSMUSG00 | U5 | 115 | 0 | 0 | 0 | 0 | 0 | 0 |
| ENSMUSG00 | U6 | 107 | 0 | 0.45034058 | 0 | 0 | 0 | 0 |
| ENSMUSG00 | U6 | 106 | 0 | 0 | 0 | 0 | 0 | 0 |
| ENSMUSG00 | SNORA70 | 133 | 0 | 1.08691223 | 0 | 1.62776141 | 2.68559873 | 0 |
| ENSMUSG00 | 7SK | 330 | 0.12047115 | 0.73009761 | 1.60733975 | 0.7653767 | 0.30925076 | 0.99180506 |
| ENSMUSG00 | U1 | 162 | 0 | 0 | 0 | 0 | 0 | 0 |
| ENSMUSG00 | SNORA62 | 155 | 0.76946091 | 0.31088027 | 1.71103908 | 1.16393692 | 0 | 0 |
| ENSMUSG00 | U5 | 115 | 0 | 0 | 0 | 0 | 0 | 0 |
| ENSMUSG00 | n-R5s169 | 119 | 0 | 0 | 0 | 0.30321046 | 0 | 0 |
| ENSMUSG00 | U1 | 164 | 0 | 0 | 0 | 0 | 0 | 0 |
| ENSMUSG00 | SNORD62 | 84 | 0 | 0 | 0 | 0.42954815 | 0 | 0 |
| ENSMUSG00 | Y_RNA | 97 | 3.27880252 | 1.98706978 | 1.36706731 | 1.8598992 | 0 | 6.18600066 |
| ENSMUSG00 | U6 | 102 | 0 | 0 | 0 | 0 | 0 | 0 |
| ENSMUSG00 | U1 | 155 | 0 | 0 | 0 | 0 | 0 | 0 |
| ENSMUSG00 | U1 | 167 | 0 | 0 | 0 | 0 | 0 | 0 |
| ENSMUSG00 | U6 | 106 | 0 | 0 | 0 | 0 | 0 | 0 |
| ENSMUSG00 | U6 | 103 | 0 | 0 | 0 | 0 | 0 | 0 |
| ENSMUSG00 | n-R5s213 | 97 | 0.40985032 | 0.49676745 | 0 | 1.48791936 | 0 | 0 |
| ENSMUSG00 | U12 | 150 | 0 | 0 | 0 | 0 | 0 | 0 |
| ENSMUSG00 | U6 | 106 | 0 | 0 | 0 | 0 | 0 | 0 |
| ENSMUSG00 | SNORA29 | 139 | 0 | 0 | 0 | 0 | 0 | 0 |
| ENSMUSG00 | U6 | 106 | 0 | 0 | 0 | 0 | 0 | 0 |
| ENSMUSG00 | U6 | 106 | 0 | 0 | 0 | 0 | 0 | 0 |
| ENSMUSG00 | n-R5s5 | 118 | 0 | 0 | 0 | 0 | 0 | 0 |
| ENSMUSG00 | U6 | 106 | 0 | 0 | 0 | 0 | 0 | 0 |
| ENSMUSG00 | n-R5s196 | 116 | 0.34271966 | 0 | 0 | 0 | 0 | 0 |
| ENSMUSG00 | U6 | 106 | 0 | 0 | 0 | 0 | 0 | 0 |
| ENSMUSG00 | SNORA73 | 195 | 0 | 0 | 0 | 0 | 0 | 0 |
| ENSMUSG00 | U6 | 106 | 0 | 0 | 0 | 0 | 0 | 0 |
| ENSMUSG00 | U6 | 106 | 0 | 0 | 0 | 0 | 0 | 0 |
| ENSMUSG00 | U1 | 146 | 0 | 0 | 0 | 0 | 0 | 0 |
| ENSMUSG00 | U6 | 106 | 3.00041362 | 0 | 0.83399704 | 0 | 0 | 2.57307917 |
| ENSMUSG00 | U6 | 106 | 0 | 0 | 0 | 0 | 0 | 0 |
| ENSMUSG00 | n-R5s205 | 124 | 0 | 0 | 0 | 0 | 0 | 0 |
| ENSMUSG00 | U6 | 106 | 0 | 0 | 0 | 0 | 0 | 0 |
| ENSMUSG00 | U6 | 103 | 0 | 0 | 0 | 0 | 0 | 0 |
| ENSMUSG00 | SNORA4 | 136 | 0 | 0 | 0 | 0 | 0 | 0 |
| ENSMUSG00 | Snord22 | 125 | 0 | 0 | 0 | 0 | 0 | 0 |
| ENSMUSG00 | SNORA67 | 142 | 0 | 0 | 0 | 0 | 0 | 0 |
| ENSMUSG00 | SNORA46 | 135 | 0.58897008 | 0 | 0 | 0.80182321 | 0.75594631 | 0.60610309 |
| ENSMUSG00 | U6 | 103 | 0 | 0 | 0 | 0 | 0 | 0 |
| ENSMUSG00 | U6 | 106 | 0 | 0 | 0 | 0 | 0 | 0 |
| ENSMUSG00 | U6 | 106 | 0 | 0 | 0 | 0 | 0 | 0 |
| ENSMUSG00 | Snord1a | 73 | 0 | 0 | 0 | 0 | 0 | 0 |
| ENSMUSG00 | SNORA63 | 118 | 0 | 0 | 0 | 0 | 0 | 0 |
| ENSMUSG00 | U1 | 163 | 0 | 0 | 0 | 0 | 0 | 0 |
| ENSMUSG00 | Snora16a | 135 | 0 | 0 | 0 | 0 | 0 | 0 |
| ENSMUSG00 | SNORA57 | 147 | 0 | 0 | 0 | 0 | 0 | 0 |
| ENSMUSG00 | n-R5s50 | 110 | 0 | 0 | 0 | 0 | 0 | 0 |
| ENSMUSG00 | U6 | 109 | 0 | 0 | 0 | 0 | 0 | 0 |
| ENSMUSG00 | SNORA29 | 137 | 0 | 0 | 0 | 0 | 0 | 0 |
| ENSMUSG00 | n-R5s88 | 117 | 0 | 0 | 0 | 0 | 0 | 0 |
| ENSMUSG00 | Snord61 | 70 | 0 | 0 | 0 | 0 | 0 | 0 |
| ENSMUSG00 | U6 | 106 | 0 | 0 | 0 | 0 | 0 | 0 |
| ENSMUSG00 | U6 | 106 | 0 | 0 | 0 | 0 | 0 | 0 |
| ENSMUSG00 | U6 | 106 | 0 | 0 | 0 | 0 | 0 | 0 |
| ENSMUSG00 | SNORA54 | 126 | 3.78623624 | 3.44188873 | 0 | 0.28636543 | 0.80994247 | 0.21646539 |
| ENSMUSG00 | U6 | 106 | 0 | 0 | 0 | 0 | 0 | 0.25730792 |
| ENSMUSG00 | U1 | 163 | 0 | 0 | 0 | 0 | 0 | 0 |
| ENSMUSG00 | Snora65 | 129 | 0 | 0 | 0 | 0 | 0 | 0 |
| ENSMUSG00 | n-R5s209 | 118 | 0 | 0 | 0 | 0 | 0 | 0 |
| ENSMUSG00 | Snord104 | 69 | 1.72849915 | 0.69835423 | 1.28121284 | 2.09171273 | 0.73951269 | 4.74341552 |
| ENSMUSG00 | U6 | 106 | 0 | 0 | 0 | 0 | 0 | 0 |
| ENSMUSG00 | U6 | 104 | 0 | 0 | 0 | 0 | 0 | 0 |
| ENSMUSG00 | U6 | 106 | 0 | 0 | 0 | 0 | 0 | 0 |

| | | | | | | | | |
|---|---|---|---|---|---|---|---|---|
| ENSMUSG00 | U6 | 106 | 0 | 0 | 0 | 0 | 0 | 0 |
| ENSMUSG00 | U6 | 106 | 0 | 0 | 0 | 0 | 0 | 0 |
| ENSMUSG00 | U6 | 103 | 0 | 0 | 0 | 0 | 0 | 0 |
| ENSMUSG00 | SNORA46 | 130 | 0 | 0 | 0 | 0 | 0 | 0 |
| ENSMUSG00 | U6 | 106 | 0 | 0 | 0 | 0 | 0 | 0 |
| ENSMUSG00 | Vault | 142 | 0 | 0.67868228 | 0.93384175 | 0 | 1.07802202 | 1.15244955 |
| ENSMUSG00 | SNORD36 | 67 | 0 | 0 | 0 | 0 | 0 | 0 |
| ENSMUSG00 | Snora31 | 129 | 4.31454827 | 5.97661298 | 5.48239913 | 6.43323275 | 3.95553301 | 11.2058595 |
| ENSMUSG00 | n-R5s92 | 118 | 0 | 0 | 0 | 0 | 0 | 0 |
| ENSMUSG00 | U6 | 106 | 0 | 0 | 0 | 0 | 0 | 0 |
| ENSMUSG00 | SNORA67 | 132 | 0 | 0 | 0 | 0 | 0 | 0 |
| ENSMUSG00 | U6 | 106 | 0 | 0 | 0 | 0 | 0 | 0 |
| ENSMUSG00 | U1 | 157 | 0 | 0 | 0 | 0 | 0 | 0 |
| ENSMUSG00 | U6 | 106 | 0 | 0 | 0 | 0 | 0.4813809 | 0 |
| ENSMUSG00 | U6 | 106 | 0 | 0 | 0 | 0 | 0 | 0 |
| ENSMUSG00 | SNORA73 | 226 | 0 | 0 | 0 | 0 | 0 | 0 |
| ENSMUSG00 | U6 | 106 | 0 | 0 | 0 | 0 | 0 | 0 |
| ENSMUSG00 | SNORD79 | 83 | 0 | 0 | 0 | 0 | 0 | 0 |
| ENSMUSG00 | n-R5s167 | 120 | 0 | 0 | 0 | 0 | 0 | 0 |
| ENSMUSG00 | U6 | 107 | 0 | 0 | 0 | 0 | 0 | 0.25490317 |
| ENSMUSG00 | U6 | 106 | 0 | 0 | 0 | 0 | 0 | 0.25730792 |
| ENSMUSG00 | U6 | 101 | 0 | 0 | 0 | 0 | 0 | 0 |
| ENSMUSG00 | U6 | 106 | 0 | 0 | 0 | 0 | 0 | 0 |
| ENSMUSG00 | U6 | 106 | 0 | 0 | 0 | 0 | 0 | 0 |
| ENSMUSG00 | Snora19 | 131 | 0.91043085 | 0 | 0 | 0.27543545 | 1.55805728 | 2.29023688 |
| ENSMUSG00 | U6 | 106 | 0 | 0 | 0 | 0 | 0 | 0 |
| ENSMUSG00 | U6 | 106 | 0 | 0 | 0 | 0 | 0 | 0 |
| ENSMUSG00 | U6 | 106 | 0 | 0 | 0 | 0 | 0 | 0 |
| ENSMUSG00 | U6 | 105 | 0 | 0 | 0 | 0 | 0 | 0 |
| ENSMUSG00 | U1 | 163 | 0 | 0 | 0 | 0 | 0 | 0 |
| ENSMUSG00 | U12 | 150 | 0 | 0.6424859 | 0 | 1.68382874 | 0.34017584 | 0 |
| ENSMUSG00 | U1 | 141 | 0 | 0 | 0 | 0 | 0 | 0 |
| ENSMUSG00 | SNORA73 | 209 | 0 | 0 | 0 | 0 | 0 | 0 |
| ENSMUSG00 | SNORA42 | 135 | 0 | 0 | 0 | 0 | 0 | 0 |
| ENSMUSG00 | U6 | 109 | 0 | 0 | 0 | 0 | 0 | 0 |
| ENSMUSG00 | SNORA36 | 114 | 0 | 0 | 0 | 0 | 0 | 0 |
| ENSMUSG00 | U1 | 139 | 0 | 0 | 0 | 0 | 0 | 0 |
| ENSMUSG00 | U1 | 168 | 1.18319883 | 1.72094436 | 0.26310621 | 3.86593334 | 0.30372843 | 0.32469809 |
| ENSMUSG00 | U6 | 108 | 0 | 0 | 0 | 0 | 0 | 0 |
| ENSMUSG00 | U4 | 130 | 0 | 0 | 0 | 0 | 0 | 0 |
| ENSMUSG00 | U6 | 106 | 0 | 0 | 0 | 0 | 0 | 0 |
| ENSMUSG00 | U6 | 106 | 0 | 0 | 0 | 0 | 0 | 0 |
| ENSMUSG00 | Snord85 | 72 | 0 | 0 | 0 | 0 | 0.70869966 | 0 |
| ENSMUSG00 | U6 | 103 | 0 | 0 | 0 | 0 | 0 | 0 |
| ENSMUSG00 | U6 | 106 | 0 | 0 | 0 | 0 | 0 | 0 |
| ENSMUSG00 | SNORA65 | 129 | 0 | 0 | 0 | 1.9579404 | 1.1866599 | 3.17146968 |
| ENSMUSG00 | U6 | 106 | 0 | 0 | 0 | 0 | 0 | 0 |
| ENSMUSG00 | SNORA20 | 112 | 2.48471753 | 0 | 0 | 5.15457779 | 0 | 0 |
| ENSMUSG00 | U6 | 106 | 0 | 0 | 0 | 0 | 0 | 0 |
| ENSMUSG00 | SNORA32 | 108 | 0 | 0 | 0 | 1.33637202 | 0 | 0 |
| ENSMUSG00 | U6 | 106 | 0 | 0 | 0 | 0.34039665 | 0 | 0 |
| ENSMUSG00 | SNORA4 | 138 | 27.0798201 | 11.5228449 | 17.2963734 | 23.5317682 | 14.7902539 | 26.48407 |
| ENSMUSG00 | U6 | 106 | 0 | 0 | 0 | 0 | 0 | 0 |
| ENSMUSG00 | SNORA76 | 134 | 0 | 0 | 0 | 0 | 0 | 0.20354208 |
| ENSMUSG00 | U1 | 158 | 0 | 0 | 0 | 0 | 0 | 0 |
| ENSMUSG00 | U6 | 106 | 0 | 0 | 0 | 0 | 0 | 0 |
| ENSMUSG00 | U1 | 163 | 0 | 0 | 0 | 0 | 0 | 0 |
| ENSMUSG00 | U1 | 163 | 0 | 0 | 0 | 0 | 0 | 0 |
| ENSMUSG00 | Snord32a | 82 | 0 | 0 | 0 | 0 | 0 | 0 |
| ENSMUSG00 | U6 | 106 | 0 | 0 | 0.41699852 | 0 | 0 | 0 |
| ENSMUSG00 | U6 | 106 | 0 | 0 | 0 | 0 | 0 | 0 |
| ENSMUSG00 | U6 | 99 | 0 | 0 | 0 | 0 | 0 | 0 |
| ENSMUSG00 | SNORA8 | 136 | 6.43103361 | 16.6526675 | 3.57514907 | 1.59185491 | 2.62635758 | 0.80219527 |
| ENSMUSG00 | U6 | 106 | 0 | 0 | 0 | 0 | 0 | 0 |
| ENSMUSG00 | SNORD81 | 75 | 0 | 0 | 0 | 0 | 0 | 0 |
| ENSMUSG00 | SNORA75 | 130 | 0 | 0 | 0 | 0 | 0 | 0 |
| ENSMUSG00 | U2 | 190 | 0 | 0 | 0.93056512 | 0.56971649 | 0 | 0.8613044 |
| ENSMUSG00 | n-R5s60 | 118 | 0 | 0 | 0 | 0 | 0 | 0 |
| ENSMUSG00 | U6 | 106 | 0 | 0 | 0 | 0 | 0 | 0 |
| ENSMUSG00 | U6 | 102 | 0 | 0 | 0 | 0 | 0 | 0 |
| ENSMUSG00 | n-R5s197 | 118 | 0 | 0 | 1.12377567 | 0 | 0 | 0 |
| ENSMUSG00 | U6 | 106 | 0 | 0 | 0 | 0 | 0 | 0 |
| ENSMUSG00 | U2 | 185 | 0 | 0 | 0 | 0 | 0 | 0 |
| ENSMUSG00 | U6 | 96 | 0 | 0 | 0 | 0 | 0 | 0 |
| ENSMUSG00 | U6 | 106 | 0 | 0 | 0 | 0 | 0 | 0 |
| ENSMUSG00 | U6 | 103 | 0 | 0 | 0 | 0 | 0 | 0 |
| ENSMUSG00 | U6 | 102 | 0 | 0 | 0 | 0 | 0 | 0 |
| ENSMUSG00 | U6 | 106 | 0 | 0 | 0 | 0 | 0 | 0 |
| ENSMUSG00 | U6 | 105 | 0.75724725 | 0 | 0 | 0 | 0 | 0 |
| ENSMUSG00 | SNORA32 | 108 | 0 | 0 | 0 | 0 | 0 | 0 |
| ENSMUSG00 | U6 | 106 | 0 | 0 | 0 | 0 | 0 | 0 |
| ENSMUSG00 | U2 | 190 | 0 | 0 | 0 | 0 | 0 | 0 |
| ENSMUSG00 | U6 | 106 | 0 | 0 | 0 | 0 | 0 | 0 |
| ENSMUSG00 | U6 | 106 | 0 | 0 | 0 | 0 | 0 | 0 |
| ENSMUSG00 | 7SK | 320 | 0 | 0 | 0 | 0 | 0 | 0 |
| ENSMUSG00 | SNORA55 | 130 | 0 | 0 | 0 | 0 | 0 | 0 |
| ENSMUSG00 | U1 | 164 | 0 | 0 | 0 | 0 | 0 | 0 |
| ENSMUSG00 | SNORD24 | 73 | 0 | 1.3201765 | 0 | 0 | 1.3979829 | 1.86812598 |
| ENSMUSG00 | Snora30 | 129 | 0.30818202 | 0 | 3.7691494 | 0 | 0 | 0 |
| ENSMUSG00 | U6 | 105 | 0 | 0 | 0 | 0 | 0 | 0 |
| ENSMUSG00 | U6 | 106 | 0 | 0 | 0.41699852 | 0 | 0 | 0 |
| ENSMUSG00 | SNORA32 | 121 | 0 | 0 | 0 | 0 | 0 | 0 |
| ENSMUSG00 | U6 | 106 | 0 | 0 | 0 | 0 | 0 | 0 |
| ENSMUSG00 | SNORA32 | 99 | 7.62983969 | 3.89385391 | 5.80428242 | 5.10251135 | 3.09250763 | 5.78552954 |
| ENSMUSG00 | SNORA16 | 132 | 0 | 0 | 0 | 0 | 0 | 0 |
| ENSMUSG00 | U6 | 106 | 0 | 0 | 0 | 0 | 0 | 0 |
| ENSMUSG00 | SNORA62 | 136 | 0 | 0 | 0 | 0 | 0 | 0 |
| ENSMUSG00 | U6 | 106 | 0 | 0 | 0 | 0 | 0 | 0 |
| ENSMUSG00 | Snord57 | 71 | 0 | 0 | 0 | 0 | 0 | 0 |
| ENSMUSG00 | SNORD83 | 75 | 1.06014615 | 0.6424859 | 3.53614744 | 3.36765749 | 0.68035168 | 0 |
| ENSMUSG00 | SNORA42 | 135 | 0 | 0 | 0 | 0 | 0 | 0 |
| ENSMUSG00 | n-R5s217 | 115 | 0 | 0 | 0 | 0 | 0 | 0 |
| ENSMUSG00 | U6 | 101 | 0 | 0 | 0 | 0 | 0 | 0 |
| ENSMUSG00 | U1 | 163 | 0 | 0 | 0 | 0 | 0 | 0 |
| ENSMUSG00 | SNORD28 | 75 | 0 | 0 | 0 | 0 | 0 | 0 |
| ENSMUSG00 | SNORD27 | 71 | 0 | 0 | 0 | 0 | 0 | 0 |
| ENSMUSG00 | SNORA18 | 130 | 4.89298222 | 1.11199482 | 1.36005671 | 3.88575864 | 1.17753175 | 0.62941475 |
| ENSMUSG00 | SNORA51 | 131 | 0 | 0 | 0 | 0 | 0 | 0 |
| ENSMUSG00 | U1 | 163 | 0 | 0 | 0 | 0 | 0 | 0 |
| ENSMUSG00 | SNORA61 | 132 | 0 | 0 | 0 | 0 | 0 | 0 |
| ENSMUSG00 | SNORD56 | 69 | 0 | 0 | 0 | 0 | 0 | 0 |
| ENSMUSG00 | U6 | 106 | 0 | 0 | 0 | 0 | 0 | 0 |
| ENSMUSG00 | U6 | 106 | 0 | 0 | 0 | 0 | 0 | 0 |
| ENSMUSG00 | Y_RNA | 107 | 0 | 0 | 0 | 0 | 0 | 0 |
| ENSMUSG00 | U1 | 165 | 0 | 0 | 0 | 0 | 0 | 0 |
| ENSMUSG00 | SNORD36 | 70 | 0 | 0 | 0 | 0 | 0 | 0 |
| ENSMUSG00 | U6 | 81 | 0 | 0 | 0 | 0 | 0 | 0 |
| ENSMUSG00 | SNORA42 | 134 | 0 | 0 | 0 | 0 | 0 | 0 |
| ENSMUSG00 | U1 | 161 | 0 | 0 | 0.27454561 | 0 | 0 | 0 |
| ENSMUSG00 | U6 | 102 | 0 | 0 | 0 | 0 | 0 | 0 |
| ENSMUSG00 | U6 | 106 | 0 | 0 | 0 | 0 | 0 | 0 |
| ENSMUSG00 | U6 | 107 | 0 | 0 | 0 | 0 | 0 | 0 |
| ENSMUSG00 | U1 | 160 | 0 | 0 | 0 | 0 | 0 | 0 |
| ENSMUSG00 | SNORA22 | 134 | 7.71374995 | 2.15760189 | 0.9895935 | 4.03903484 | 4.95032004 | 2.44250501 |
| ENSMUSG00 | SNORD31 | 66 | 0 | 0 | 0 | 0 | 0 | 0 |
| ENSMUSG00 | U6 | 106 | 1.50020681 | 0 | 0.83399704 | 1.36158659 | 8.18347537 | 0 |
| ENSMUSG00 | U6 | 106 | 0 | 0 | 0 | 0 | 0 | 0 |
| ENSMUSG00 | U1 | 164 | 0 | 0 | 0 | 0 | 0 | 0 |
| ENSMUSG00 | U6 | 91 | 0 | 0 | 0 | 0 | 0 | 0 |
| ENSMUSG00 | n-R5s156 | 104 | 0 | 0 | 0 | 0 | 0 | 0 |
| ENSMUSG00 | U6 | 106 | 0 | 0 | 0 | 0 | 0 | 0 |
| ENSMUSG00 | U6 | 106 | 0 | 0 | 0 | 0 | 0 | 0 |
| ENSMUSG00 | U6 | 104 | 0 | 0 | 0 | 0 | 0 | 0 |
| ENSMUSG00 | U6 | 108 | 0 | 0 | 1.22782897 | 0 | 0 | 0 |
| ENSMUSG00 | U6 | 106 | 0 | 0 | 0 | 0 | 0 | 0 |
| ENSMUSG00 | U6 | 106 | 0 | 0 | 0 | 0 | 0 | 0 |
| ENSMUSG00 | U6 | 108 | 0 | 0 | 0 | 0 | 0 | 0 |
| ENSMUSG00 | U6 | 80 | 0 | 0 | 0 | 0 | 0 | 0 |
| ENSMUSG00 | U6 | 106 | 0 | 0 | 0 | 0 | 0 | 0 |
| ENSMUSG00 | U6 | 106 | 0 | 0 | 0 | 0 | 0 | 0 |
| ENSMUSG00 | U6 | 106 | 0 | 0 | 0 | 0 | 0 | 0 |
| ENSMUSG00 | U6 | 106 | 0 | 0 | 0 | 0 | 0 | 0 |
| ENSMUSG00 | 7SK | 309 | 0 | 0 | 0 | 0 | 0 | 0 |
| ENSMUSG00 | SNORA73 | 199 | 0 | 0 | 0 | 0 | 0 | 0 |
| ENSMUSG00 | U1 | 154 | 0 | 0 | 0 | 0 | 0 | 0 |
| ENSMUSG00 | U6 | 106 | 0 | 0 | 0 | 0 | 0 | 0 |
| ENSMUSG00 | U6 | 106 | 0 | 0 | 0 | 0 | 0 | 0 |
| ENSMUSG00 | U6 | 106 | 0 | 0 | 0 | 0 | 0 | 0 |
| ENSMUSG00 | U6 | 106 | 0 | 0 | 0 | 0.68079329 | 1.92552362 | 0 |
| ENSMUSG00 | Snora34 | 137 | 6.09390577 | 2.81380684 | 1.93584714 | 3.42384364 | 1.11736589 | 3.7826142 |
| ENSMUSG00 | U1 | 138 | 0 | 0 | 0 | 0 | 0 | 0 |
| ENSMUSG00 | U6 | 106 | 0 | 0 | 0 | 0 | 0 | 0 |
| ENSMUSG00 | U6 | 103 | 0 | 0 | 0 | 0 | 0 | 0 |
| ENSMUSG00 | U6 | 105 | 0 | 0 | 0 | 0 | 0 | 0 |
| ENSMUSG00 | U6 | 106 | 0 | 0 | 0 | 0 | 0 | 0 |
| ENSMUSG00 | 7SK | 326 | 0 | 0 | 0 | 0 | 0 | 0 |
| ENSMUSG00 | U1 | 163 | 0 | 0 | 0 | 0 | 0 | 0 |
| ENSMUSG00 | SNORA4 | 109 | 0 | 0 | 0 | 0 | 0 | 0 |
| ENSMUSG00 | U6 | 106 | 0 | 0 | 0 | 0 | 0 | 0 |
| ENSMUSG00 | U6 | 106 | 0 | 0 | 0 | 0 | 0 | 0 |
| ENSMUSG00 | Snora73b | 204 | 0 | 0 | 0 | 0 | 0 | 0 |
| ENSMUSG00 | SNORD16 | 98 | 2.43400901 | 0 | 2.25519607 | 2.20910477 | 2.08270922 | 1.11325058 |
| ENSMUSG00 | SNORA32 | 108 | 0 | 0 | 0 | 0 | 0 | 0 |
| ENSMUSG00 | Telomerase-v | 411 | 0.38691465 | 0.2344839 | 0.10754706 | 0.08779086 | 0.7449106 | 0.53089322 |
| ENSMUSG00 | SNORA71 | 130 | 0 | 0 | 0.68002835 | 0 | 0 | 0 |
| ENSMUSG00 | U6 | 106 | 0 | 0 | 0 | 0 | 0 | 0 |
| ENSMUSG00 | U6 | 106 | 0 | 1.36376723 | 0 | 0 | 0 | 0 |
| ENSMUSG00 | U1 | 163 | 0.24389865 | 0 | 0 | 1.54953565 | 0.31304525 | 0.16732908 |
| ENSMUSG00 | U1 | 159 | 0 | 0 | 0 | 0 | 0 | 0 |
| ENSMUSG00 | U1 | 164 | 0 | 0 | 0 | 0 | 0 | 0 |
| ENSMUSG00 | U6 | 108 | 0 | 0 | 0 | 0 | 0 | 0 |
| ENSMUSG00 | U1 | 146 | 0 | 0 | 0 | 2.22423562 | 0.69899145 | 0.56043779 |
| ENSMUSG00 | SNORD29 | 64 | 0 | 0 | 0 | 0 | 0 | 0 |
| ENSMUSG00 | SNORA25 | 132 | 0 | 0 | 0 | 0 | 0 | 0 |
| ENSMUSG00 | U6 | 106 | 0 | 0 | 0 | 0 | 0 | 0 |
| ENSMUSG00 | U6 | 106 | 0 | 0 | 0 | 0 | 0 | 0 |
| ENSMUSG00 | U6 | 104 | 0 | 0 | 0 | 0 | 0 | 0 |
| ENSMUSG00 | U6 | 101 | 0 | 0 | 0 | 0 | 0 | 0 |
| ENSMUSG00 | n-R5s153 | 115 | 0 | 0 | 0 | 0 | 0 | 0 |
| ENSMUSG00 | n-R5s85 | 112 | 0 | 0 | 0 | 0 | 0.45569264 | 0 |
| ENSMUSG00 | U1 | 138 | 0 | 0 | 0 | 0 | 0 | 0 |
| ENSMUSG00 | U1 | 140 | 0 | 0 | 1.2629098 | 0 | 0 | 0 |
| ENSMUSG00 | U6 | 106 | 0 | 0 | 0 | 0 | 0 | 0 |
| ENSMUSG00 | U6 | 106 | 0 | 0 | 0 | 0 | 0 | 0 |
| ENSMUSG00 | SNORD26 | 74 | 0 | 0 | 0 | 0 | 0 | 0 |
| ENSMUSG00 | U6 | 106 | 0 | 0 | 0 | 0 | 0 | 0 |
| ENSMUSG00 | Mir25 | 83 | 0 | 0 | 0 | 0 | 0 | 0 |
| ENSMUSG00 | Mir193 | 65 | 0 | 0 | 0 | 0 | 0 | 0 |
| ENSMUSG00 | Mir99b | 69 | 0 | 0 | 0 | 0 | 0 | 0 |
| ENSMUSG00 | Mir155 | 64 | 0 | 0 | 0 | 0 | 0 | 0 |
| ENSMUSG00 | Mir188 | 67 | 0 | 0 | 0 | 0 | 0 | 0 |
| ENSMUSG00 | Mir133a-1 | 86 | 0 | 0 | 0 | 0 | 0 | 0 |
| ENSMUSG00 | Mir200a | 89 | 1.34007238 | 0 | 0 | 0 | 0 | 0 |
| ENSMUSG00 | Mir144 | 65 | 0 | 0 | 0 | 0 | 0 | 0 |
| ENSMUSG00 | Mir122a | 65 | 0 | 0 | 0 | 0 | 0 | 0 |
| ENSMUSG00 | Mir18 | 95 | 0 | 0 | 0 | 0 | 0 | 0 |
| ENSMUSG00 | Mir201 | 65 | 0 | 0 | 0 | 0 | 0 | 0 |
| ENSMUSG00 | Mir30a | 70 | 0 | 0 | 0 | 0 | 0 | 0 |
| ENSMUSG00 | Mirlet7i | 84 | 1.41983859 | 1.72094436 | 0 | 1.7181926 | 0.60745686 | 0.64939617 |
| ENSMUSG00 | Mir135a-1 | 89 | 0 | 0 | 0 | 0 | 0 | 0 |
| ENSMUSG00 | Mir31 | 105 | 0 | 0 | 1.2629098 | 0 | 0 | 0 |
| ENSMUSG00 | Mir30e | 91 | 0 | 0 | 0 | 0.79301197 | 1.12145881 | 0 |
| ENSMUSG00 | Mir298 | 81 | 0 | 0 | 0 | 0 | 0 | 0 |
| ENSMUSG00 | Mir195 | 93 | 0 | 0 | 0 | 0 | 0 | 0 |
| ENSMUSG00 | Mir29b-2 | 80 | 0 | 0 | 0 | 0 | 0 | 0 |
| ENSMUSG00 | Mir367 | 74 | 0 | 0 | 0 | 0 | 0 | 0.36857621 |
| ENSMUSG00 | Mir138-1 | 98 | 0 | 0 | 0 | 0 | 0 | 0 |
| ENSMUSG00 | Mir217 | 107 | 0 | 0 | 0 | 0 | 0 | 0 |
| ENSMUSG00 | Mir19a | 81 | 0 | 0 | 0 | 0 | 0 | 0 |
| ENSMUSG00 | Mir340 | 97 | 0 | 0 | 0 | 0 | 0 | 0 |
| ENSMUSG00 | Mir322 | 94 | 0 | 0 | 0 | 0 | 0 | 0 |
| ENSMUSG00 | Mir300 | 78 | 0 | 0 | 0 | 0 | 0 | 0 |
| ENSMUSG00 | Mir142 | 63 | 0 | 0 | 0 | 0 | 0 | 0 |
| ENSMUSG00 | Mirlet7a-1 | 93 | 0 | 4.14507029 | 1.90115454 | 0 | 0 | 0 |
| ENSMUSG00 | Mir221 | 94 | 0 | 0 | 0 | 0 | 0 | 0 |
| ENSMUSG00 | Mir181a-2 | 75 | 0 | 0 | 0 | 1.44328178 | 0 | 0 |
| ENSMUSG00 | Mir9-2 | 86 | 0 | 0 | 0 | 0 | 0 | 0 |
| ENSMUSG00 | Mir325 | 97 | 0 | 0 | 0 | 0 | 0 | 0 |
| ENSMUSG00 | Mir134 | 70 | 0 | 0 | 0 | 0 | 0 | 0 |
| ENSMUSG00 | Mir429 | 82 | 0 | 0 | 0 | 0 | 0 | 0 |
| ENSMUSG00 | Mir382 | 79 | 0 | 0 | 0 | 0 | 0 | 0 |
| ENSMUSG00 | Mir345 | 95 | 0 | 0.50722571 | 0 | 0 | 0 | 0 |
| ENSMUSG00 | Mir26a-2 | 83 | 0 | 0 | 0 | 0 | 0 | 0 |
| ENSMUSG00 | Mir186 | 70 | 0 | 0.68837775 | 0 | 2.06183112 | 3.64474113 | 1.16891311 |
| ENSMUSG00 | Mir208a | 82 | 0 | 0 | 0 | 0 | 0 | 0 |
| ENSMUSG00 | Mir370 | 78 | 0 | 0 | 0 | 0 | 0 | 0 |
| ENSMUSG00 | Mir7-1 | 107 | 3.34391892 | 9.00681162 | 0.82620267 | 2.02329222 | 3.33817412 | 4.58825707 |
| ENSMUSG00 | Mir135b | 96 | 0 | 0 | 0 | 0 | 0 | 0 |
| ENSMUSG00 | Mir342 | 98 | 0 | 0 | 1.35311764 | 0 | 1.04135461 | 0 |
| ENSMUSG00 | Mir30d | 81 | 0 | 0 | 0 | 0 | 0 | 0 |
| ENSMUSG00 | Mir377 | 67 | 0 | 0 | 0 | 0 | 0 | 0 |
| ENSMUSG00 | Mir140 | 69 | 0.57616638 | 0 | 2.56242568 | 0 | 0 | 1.18585388 |
| ENSMUSG00 | Mirlet7g | 87 | 0 | 0 | 0 | 0 | 0 | 0 |
| ENSMUSG00 | Mir128-2 | 83 | 0 | 0 | 0 | 0 | 0 | 0 |
| ENSMUSG00 | Mir20a | 106 | 0 | 0 | 0 | 0 | 0 | 0 |
| ENSMUSG00 | Mir196b | 84 | 0 | 0 | 0 | 0 | 0 | 0 |
| ENSMUSG00 | Mir27a | 86 | 0 | 0.56030747 | 0 | 3.35646926 | 0 | 0 |
| ENSMUSG00 | Mir143 | 62 | 0 | 0 | 0 | 0 | 0 | 0 |
| ENSMUSG00 | Mir139 | 67 | 0 | 0 | 0 | 0 | 0 | 0 |
| ENSMUSG00 | Mir302d | 65 | 0 | 0 | 0 | 0 | 0 | 9.6510262 |
| ENSMUSG00 | Mir154 | 83 | 0 | 0 | 0 | 0 | 0 | 0 |
| ENSMUSG00 | Mir365-1 | 86 | 0 | 0 | 0 | 0 | 0 | 0 |
| ENSMUSG00 | Mir448 | 111 | 0 | 0 | 0 | 0 | 0 | 0 |
| ENSMUSG00 | Mir101a | 82 | 0 | 0 | 0 | 0 | 0 | 0 |
| ENSMUSG00 | Mir207 | 78 | 0 | 0 | 0 | 0 | 0.65418431 | 0 |
| ENSMUSG00 | Mirlet7d | 102 | 0 | 1.4172483 | 2.16675701 | 1.0612366 | 0 | 1.0695937 |
| ENSMUSG00 | Mir124a-3 | 86 | 0 | 0 | 0 | 0 | 0 | 0 |
| ENSMUSG00 | Mir21 | 91 | 0 | 0 | 0 | 0 | 0 | 0 |
| ENSMUSG00 | Mir106a | 64 | 0 | 0 | 0 | 0 | 0 | 0 |
| ENSMUSG00 | Mir383 | 73 | 0 | 0 | 0 | 0 | 0 | 0 |
| ENSMUSG00 | Mir181b-1 | 79 | 0 | 0 | 0 | 0 | 0 | 0 |
| ENSMUSG00 | Mir133a-2 | 103 | 0 | 0 | 0 | 0 | 0 | 0.52960465 |
| ENSMUSG00 | Mir200c | 68 | 0 | 0 | 0 | 0 | 0 | 0 |
| ENSMUSG00 | Snord45c | 79 | 0 | 0 | 0 | 0 | 0 | 0 |
| ENSMUSG00 | Mir185 | 79 | 0 | 0 | 0 | 0 | 0 | 0 |
| ENSMUSG00 | Mir33 | 68 | 0 | 0 | 0 | 0 | 0 | 0 |
| ENSMUSG00 | n-R5s89 | 103 | 0 | 0 | 0 | 0 | 0.99080341 | 0 |
| ENSMUSG00 | Mir26b | 84 | 0.94655906 | 0.57364812 | 0 | 0 | 0 | 0 |
| ENSMUSG00 | Mir129-1 | 72 | 0 | 0 | 0 | 0 | 0 | 0 |
| ENSMUSG00 | Mir149 | 65 | 0 | 0 | 0 | 0 | 0 | 0 |
| ENSMUSG00 | Mir222 | 78 | 0 | 0 | 0 | 0 | 0 | 0 |
| ENSMUSG00 | Mir125b-2 | 70 | 0 | 0 | 0 | 0 | 0 | 0 |
| ENSMUSG00 | Mir19b-2 | 83 | 0 | 0 | 0 | 0 | 0 | 0 |
| ENSMUSG00 | Mir141 | 71 | 0 | 0 | 0.62256117 | 0 | 0.71868135 | 0 |
| ENSMUSG00 | Mir27b | 96 | 0 | 2.50971053 | 0 | 3.38269167 | 3.72067324 | 0 |
| ENSMUSG00 | Mir30b | 95 | 0.83695748 | 1.01445141 | 0 | 0.379811 | 0 | 0 |
| ENSMUSG00 | Mir411 | 95 | 0 | 0 | 0 | 0 | 0 | 0 |
| ENSMUSG00 | Mir125a | 84 | 0 | 0 | 0 | 0 | 0 | 0 |
| ENSMUSG00 | Mir133b | 118 | 0 | 0 | 0 | 0 | 0 | 0 |
| ENSMUSG00 | Mir346 | 97 | 0 | 0 | 0 | 0 | 0 | 0 |
| ENSMUSG00 | Mir302c | 67 | 0 | 0 | 0 | 0 | 0 | 7.73459919 |
| ENSMUSG00 | Mir181c | 105 | 0 | 0 | 0 | 0 | 0 | 0 |
| ENSMUSG00 | Mir130a | 63 | 0 | 0 | 0 | 0 | 0 | 0 |
| ENSMUSG00 | Mir219-2 | 96 | 0 | 0 | 0 | 0 | 0 | 0 |
| ENSMUSG00 | Mir450-1 | 90 | 0 | 0 | 0 | 0 | 0 | 0 |
| ENSMUSG00 | Mir196a-2 | 109 | 0 | 0 | 0 | 0 | 0 | 0 |
| ENSMUSG00 | Mir365-2 | 111 | 0 | 0 | 0 | 0 | 0 | 0 |
| ENSMUSG00 | Mir30c-1 | 88 | 0 | 0 | 0 | 0 | 0 | 0 |
| ENSMUSG00 | Mir7b | 110 | 0 | 0 | 0 | 0 | 0 | 0 |
| ENSMUSG00 | Mir34b | 83 | 0 | 0 | 0 | 0 | 0 | 0 |
| ENSMUSG00 | Mir34a | 101 | 0 | 0 | 0 | 0 | 0 | 0 |
| ENSMUSG00 | Mir28 | 85 | 0 | 0 | 0 | 0 | 0 | 0 |
| ENSMUSG00 | Mir150 | 64 | 0 | 0 | 1.38130759 | 0 | 0 | 0 |
| ENSMUSG00 | Mir410 | 80 | 0 | 0 | 0 | 0 | 0 | 0 |
| ENSMUSG00 | Mir379 | 122 | 0 | 0 | 0 | 0 | 0 | 0 |
| ENSMUSG00 | Mir384 | 87 | 0 | 0 | 0 | 0 | 0 | 0 |
| ENSMUSG00 | Mir10b | 67 | 0 | 0 | 0 | 0 | 0 | 0.40708417 |
| ENSMUSG00 | Mir335 | 97 | 0 | 0 | 0 | 0 | 0 | 0 |
| ENSMUSG00 | Mir202 | 71 | 0 | 0 | 0 | 0 | 0 | 0 |
| ENSMUSG00 | Mir351 | 98 | 0 | 0 | 0 | 0 | 0 | 0 |
| ENSMUSG00 | Mir148a | 98 | 0 | 0 | 0 | 0 | 0 | 0 |
| ENSMUSG00 | Mir204 | 67 | 0 | 0 | 0 | 0 | 0 | 0 |
| ENSMUSG00 | Mir17 | 83 | 0 | 0 | 0 | 0 | 0 | 0 |
| ENSMUSG00 | Mir421 | 84 | 0 | 0 | 0 | 0.8590963 | 1.21491371 | 0.32469809 |
| ENSMUSG00 | Mir361 | 69 | 0 | 0 | 0 | 0 | 0 | 0 |
| ENSMUSG00 | Mir129-2 | 89 | 0 | 0 | 0 | 0 | 0 | 0 |
| ENSMUSG00 | Mir138-2 | 70 | 0 | 0 | 0 | 0 | 0 | 0 |
| ENSMUSG00 | Mir26a-1 | 89 | 1.34007238 | 0.5414207 | 3.97319937 | 1.62166492 | 0.57333007 | 2.75810959 |
| ENSMUSG00 | Mir106b | 81 | 0.9816168 | 0 | 0 | 1.33637202 | 0 | 0 |
| ENSMUSG00 | Mir152 | 72 | 0 | 0 | 0 | 0 | 0 | 0 |
| ENSMUSG00 | Mir214 | 109 | 0 | 0 | 0 | 0 | 0 | 0 |
| ENSMUSG00 | Mir423 | 108 | 0 | 0 | 0 | 0 | 1.41739933 | 0 |
| ENSMUSG00 | Mir10a | 109 | 0 | 0 | 0 | 0 | 0 | 0 |
| ENSMUSG00 | Mir128-1 | 81 | 0 | 0 | 0 | 0 | 0 | 0 |
| ENSMUSG00 | Mir296 | 78 | 0 | 0 | 0 | 0 | 0 | 0 |
| ENSMUSG00 | Mir192 | 88 | 0 | 0 | 0 | 0 | 0 | 0 |
| ENSMUSG00 | Mir135a-2 | 99 | 0 | 0 | 0 | 0 | 0 | 0 |
| ENSMUSG00 | Mir337 | 96 | 0 | 0 | 0 | 0 | 0 | 0 |
| ENSMUSG00 | Mir93 | 87 | 0 | 0 | 0 | 0 | 0 | 0 |
| ENSMUSG00 | Mir320 | 81 | 0 | 0 | 0 | 1.78182936 | 0.62995526 | 0 |
| ENSMUSG00 | Mir22 | 94 | 0 | 0 | 0 | 0 | 0 | 0 |
| ENSMUSG00 | Mir99a | 64 | 0 | 0 | 0 | 0 | 0 | 0 |
| ENSMUSG00 | Mir187 | 60 | 0 | 0 | 0 | 0 | 0 | 0 |
| ENSMUSG00 | Mir205 | 67 | 0 | 0 | 0 | 0 | 0 | 0 |
| ENSMUSG00 | Mir324 | 88 | 0 | 0 | 0 | 0 | 0 | 0 |
| ENSMUSG00 | Mir98 | 107 | 0 | 0 | 0 | 0 | 0 | 0 |
| ENSMUSG00 | Mir132 | 100 | 0 | 0 | 0 | 0 | 0 | 0 |
| ENSMUSG00 | Mir153 | 68 | 0 | 0 | 0 | 0 | 0 | 0 |
| ENSMUSG00 | Mir216a | 71 | 0 | 0 | 0 | 0 | 0 | 0 |
| ENSMUSG00 | Mir126 | 72 | 0 | 0 | 0 | 0 | 0 | 0 |
| ENSMUSG00 | Mir24-2 | 106 | 1.12515511 | 5.00047985 | 2.0849926 | 1.02118994 | 1.92552362 | 0.51461584 |
| ENSMUSG00 | Mir224 | 81 | 0 | 0 | 0 | 0.89091468 | 0 | 0 |
| ENSMUSG00 | Mir330 | 97 | 0 | 0 | 0 | 0 | 0 | 0 |
| ENSMUSG00 | Mir32 | 69 | 0 | 0 | 0 | 0 | 0 | 0 |
| ENSMUSG00 | Mir196a-1 | 101 | 0 | 0 | 0 | 0 | 0 | 0 |
| ENSMUSG00 | Mir199a-1 | 69 | 0 | 1.39670847 | 0 | 0 | 0 | 0 |
| ENSMUSG00 | Mir29c | 87 | 0 | 0 | 0 | 0 | 0 | 0 |
| ENSMUSG00 | Mir200b | 69 | 0 | 0 | 0 | 0 | 0 | 0 |
| ENSMUSG00 | Mir210 | 109 | 0 | 0 | 0 | 0 | 0 | 0 |
| ENSMUSG00 | Mir302a | 68 | 0 | 0 | 0 | 0 | 0 | 0 |
| ENSMUSG00 | Mir103-1 | 85 | 0 | 0 | 0 | 0 | 0 | 0 |
| ENSMUSG00 | Mir219-1 | 109 | 0 | 0 | 0 | 0 | 0 | 0 |
| ENSMUSG00 | Mir101b | 96 | 0 | 0 | 0 | 0 | 0 | 0 |
| ENSMUSG00 | Mirlet7c-1 | 93 | 0 | 0 | 0 | 0 | 0 | 0 |
| ENSMUSG00 | Mir206 | 72 | 0 | 0 | 0 | 0 | 0 | 0 |
| ENSMUSG00 | Mir148b | 96 | 0 | 0 | 0 | 0 | 0 | 0 |
| ENSMUSG00 | Mir369 | 78 | 0.50968565 | 0 | 0 | 0 | 0 | 0 |
| ENSMUSG00 | Mir215 | 111 | 0.35815748 | 0 | 0.3982148 | 0 | 0.45969708 | 0 |
| ENSMUSG00 | Mir103-2 | 85 | 0 | 0 | 0 | 0 | 0 | 0 |
| ENSMUSG00 | Mirlet7b | 84 | 0 | 0 | 0 | 0 | 0 | 1.29879235 |
| ENSMUSG00 | Mir181a-1 | 86 | 0 | 0 | 0 | 0 | 0 | 0 |
| ENSMUSG00 | Mir381 | 74 | 0 | 0 | 0 | 0 | 0 | 0 |
| ENSMUSG00 | Mir30c-2 | 83 | 0 | 0 | 0 | 0 | 0 | 0 |
| ENSMUSG00 | Mir344 | 94 | 0 | 0 | 0 | 0 | 0 | 0 |
| ENSMUSG00 | Mir137 | 101 | 0 | 0 | 0 | 0 | 0 | 0 |
| ENSMUSG00 | Mir412 | 79 | 0 | 0 | 0 | 0 | 0 | 0 |
| ENSMUSG00 | Mir326 | 94 | 0 | 0 | 0.47023237 | 0 | 0 | 0.58031147 |
| ENSMUSG00 | Mir130b | 81 | 0 | 0 | 0 | 0 | 0 | 0 |
| ENSMUSG00 | Mir350 | 98 | 0 | 0 | 0 | 0 | 0 | 0 |
| ENSMUSG00 | Mir203 | 75 | 0 | 0 | 0 | 0 | 0 | 0 |
| ENSMUSG00 | Mir449a | 90 | 0 | 0 | 0 | 0 | 0 | 0 |
| ENSMUSG00 | Mir329 | 96 | 0 | 0 | 0 | 0 | 0 | 0 |
| ENSMUSG00 | Mir181b-2 | 88 | 0 | 0 | 0 | 1.2300697 | 1.15969036 | 0.30993908 |
| ENSMUSG00 | Mir425 | 84 | 3.78623624 | 3.44188873 | 1.57863725 | 2.14774075 | 1.82237057 | 3.24698086 |
| ENSMUSG00 | Mir15b | 97 | 0 | 0 | 0 | 0 | 0 | 0 |
| ENSMUSG00 | Mir194-1 | 66 | 0 | 0 | 0 | 0 | 0 | 0 |
| ENSMUSG00 | Mir194-2 | 85 | 0 | 0 | 0 | 0 | 0 | 0 |
| ENSMUSG00 | Mir218-2 | 109 | 0 | 0 | 0 | 0 | 0 | 0 |
| ENSMUSG00 | Mir211 | 105 | 0 | 0 | 0 | 0 | 0 | 0 |
| ENSMUSG00 | Mir96 | 105 | 0 | 0 | 0 | 0 | 0 | 0 |
| ENSMUSG00 | Mir34c | 76 | 0 | 0 | 0 | 0 | 0 | 0 |
| ENSMUSG00 | Mir301 | 85 | 0 | 0 | 0 | 0 | 0 | 0 |
| ENSMUSG00 | Mir212 | 90 | 0 | 0 | 0 | 0 | 0 | 0 |
| ENSMUSG00 | U6 | 106 | 1.87525852 | 0 | 0.83399704 | 0 | 0.96276181 | 2.05846334 |
| ENSMUSG00 | Mir145 | 87 | 0 | 0 | 0 | 0 | 0 | 0 |
| ENSMUSG00 | Mir339 | 95 | 0 | 2.02890283 | 0 | 0 | 0 | 0 |
| ENSMUSG00 | Mir107 | 86 | 0 | 0 | 0 | 0 | 0 | 0 |
| ENSMUSG00 | Mir184 | 68 | 0 | 0 | 0 | 0 | 0 | 0 |
| ENSMUSG00 | Mir124a-1 | 84 | 0 | 0 | 0 | 0 | 0 | 0 |
| ENSMUSG00 | Mir23b | 96 | 0 | 0.50194211 | 0.46043587 | 0 | 0 | 0 |
| ENSMUSG00 | Mir338 | 97 | 0 | 0 | 0 | 0 | 0 | 0 |
| ENSMUSG00 | Mir146 | 64 | 0 | 0 | 0 | 0 | 0 | 0 |
| ENSMUSG00 | Mirlet7f-2 | 82 | 0 | 0 | 0 | 0 | 0 | 0 |
| ENSMUSG00 | Mir218-1 | 109 | 0 | 0 | 0 | 0 | 0 | 0 |
| ENSMUSG00 | Mir29b-1 | 80 | 0 | 0.60233053 | 0 | 0 | 1.2756594 | 0 |
| ENSMUSG00 | Mir16-2 | 94 | 0 | 0 | 1.41069712 | 0 | 0 | 0 |
| ENSMUSG00 | Mir331 | 95 | 0 | 0 | 0 | 0 | 0 | 0 |
| ENSMUSG00 | Mirlet7c-2 | 94 | 0 | 0 | 0 | 0 | 0 | 1.16062295 |
| ENSMUSG00 | Mir7-2 | 96 | 0 | 0 | 0 | 0 | 0 | 0 |
| ENSMUSG00 | Mir29a | 87 | 0 | 0 | 0 | 0 | 1.17302013 | 0 |
| ENSMUSG00 | Mir23a | 74 | 0 | 0 | 0 | 0 | 0 | 0 |
| ENSMUSG00 | Mir151 | 67 | 0 | 0 | 0 | 0 | 0 | 0 |
| ENSMUSG00 | Mir92-2 | 90 | 0 | 0 | 0 | 0 | 0 | 0 |
| ENSMUSG00 | Mir375 | 74 | 0 | 0 | 0 | 0 | 0 | 0 |
| ENSMUSG00 | Mir323 | 85 | 0 | 0 | 0 | 0 | 0 | 0 |
| ENSMUSG00 | Mir183 | 69 | 0 | 0 | 0 | 0 | 0 | 0 |
| ENSMUSG00 | U2 | 198 | 0 | 0 | 0 | 0 | 0 | 0 |
| ENSMUSG00 | U2 | 189 | 0 | 0 | 0 | 0 | 0 | 0 |
| ENSMUSG00 | U6 | 103 | 1.15792662 | 8.88876117 | 1.28743232 | 0 | 0 | 0 |
| ENSMUSG00 | U6 | 106 | 0 | 0 | 0 | 0 | 0 | 0 |
| ENSMUSG00 | U6 | 102 | 0 | 0 | 0 | 0 | 0 | 0 |
| ENSMUSG00 | U6 | 106 | 0 | 0.45458908 | 0 | 0 | 0 | 0 |
| ENSMUSG00 | Snord33 | 83 | 0 | 0 | 0 | 0 | 0 | 0 |
| ENSMUSG00 | U3 | 213 | 0 | 0 | 0 | 0 | 0 | 0 |
| ENSMUSG00 | U4 | 83 | 0 | 0 | 0 | 0 | 0 | 0 |
| ENSMUSG00 | SNORA24 | 131 | 0 | 0 | 0 | 0 | 0.77902864 | 0 |
| ENSMUSG00 | snoR38 | 75 | 0 | 0 | 0 | 0 | 0 | 0 |
| ENSMUSG00 | U6 | 103 | 0 | 0 | 0 | 0 | 0 | 0 |
| ENSMUSG00 | SNORA14 | 133 | 0.59782678 | 0 | 0.66468937 | 0.27129357 | 0 | 0.41014495 |
| ENSMUSG00 | U6 | 106 | 0 | 0 | 0 | 0 | 0 | 0 |
| ENSMUSG00 | Snord1c | 75 | 0 | 0 | 0 | 0 | 0 | 0 |
| ENSMUSG00 | Snora69 | 130 | 0.30581139 | 0 | 0 | 0 | 0 | 0 |
| ENSMUSG00 | U6 | 106 | 0 | 0 | 0.41699852 | 0 | 0 | 0 |
| ENSMUSG00 | SNORA32 | 90 | 0 | 0 | 0 | 0 | 0 | 0 |
| ENSMUSG00 | 7SK | 317 | 0 | 0 | 0 | 0 | 0 | 0 |
| ENSMUSG00 | SNORA2 | 135 | 0 | 0 | 0.98226318 | 1.06909762 | 0 | 0 |
| ENSMUSG00 | U6 | 99 | 0 | 0 | 0 | 0 | 0 | 0 |
| ENSMUSG00 | Snora74a | 195 | 3.87361092 | 2.96531952 | 4.9868746 | 10.7320953 | 1.30836861 | 11.0497256 |
| ENSMUSG00 | SNORA51 | 126 | 0 | 0 | 0 | 0 | 0 | 0 |
| ENSMUSG00 | SNORA38 | 133 | 0 | 0 | 0 | 0 | 0 | 0 |
| ENSMUSG00 | U6 | 106 | 0 | 1.81835631 | 0 | 0 | 0 | 0 |
| ENSMUSG00 | U6 | 103 | 0 | 0 | 0 | 0 | 0 | 0 |
| ENSMUSG00 | U6 | 103 | 2.70182877 | 0 | 0 | 0 | 0 | 0 |
| ENSMUSG00 | U5 | 115 | 0 | 0 | 0 | 0 | 0 | 0 |
| ENSMUSG00 | U6 | 106 | 0 | 0 | 0 | 0 | 0 | 0 |
| ENSMUSG00 | n-R5s170 | 112 | 0 | 0 | 0 | 0 | 0 | 0 |
| ENSMUSG00 | U6 | 106 | 0 | 0 | 0 | 0 | 0 | 0 |
| ENSMUSG00 | n-R5s161 | 113 | 0 | 0 | 0 | 0 | 0 | 0 |
| ENSMUSG00 | SNORA25 | 132 | 8.43298071 | 6.20582967 | 1.67431224 | 3.00683704 | 1.93281727 | 5.99215559 |
| ENSMUSG00 | U6 | 106 | 0 | 0 | 0 | 0 | 0 | 0 |
| ENSMUSG00 | U6 | 108 | 0 | 0 | 0 | 0 | 0 | 0 |
| ENSMUSG00 | U6 | 106 | 0 | 0 | 0 | 0 | 0 | 0 |
| ENSMUSG00 | U6 | 103 | 0 | 0 | 0 | 0 | 0 | 8.73847665 |
| ENSMUSG00 | SNORA5 | 135 | 0 | 0 | 0 | 0 | 0 | 0 |
| ENSMUSG00 | SNORA71 | 132 | 0 | 0 | 0 | 0 | 0 | 0.41325211 |
| ENSMUSG00 | U6 | 106 | 0 | 0 | 0.41699852 | 0 | 0.4813809 | 0 |
| ENSMUSG00 | U6 | 106 | 0 | 0 | 0 | 0 | 0 | 0 |
| ENSMUSG00 | Snord42b | 68 | 0 | 3.54312075 | 0 | 0 | 0 | 2.80768345 |
| ENSMUSG00 | U6 | 103 | 0 | 0.93565907 | 0 | 0 | 0 | 0 |
| ENSMUSG00 | Snord38a | 70 | 0 | 0 | 0 | 0 | 0 | 0 |
| ENSMUSG00 | U6 | 106 | 0 | 0 | 0 | 0 | 0 | 0 |

| | | | | | | | | |
|---|---|---|---|---|---|---|---|---|
| ENSMUSG00 | U6 | 105 | 0 | 0 | 0 | 0 | 0 | 0 |
| ENSMUSG00 | U6 | 106 | 0 | 0 | 0 | 0 | 0 | 0 |
| ENSMUSG00 | Snora5c | 135 | 0 | 0 | 0 | 0 | 0 | 0 |
| ENSMUSG00 | snoU6-53 | 108 | 2.94485041 | 0 | 0 | 0 | 0 | 0 |
| ENSMUSG00 | n-R5s7 | 120 | 0 | 0 | 0 | 0 | 0 | 0 |
| ENSMUSG00 | U6 | 106 | 0 | 0 | 0 | 0 | 0 | 0 |
| ENSMUSG00 | n-R5s77 | 118 | 0 | 0.40835968 | 0 | 0 | 0 | 0 |
| ENSMUSG00 | SNORA36 | 136 | 0 | 0 | 0 | 0 | 0 | 0 |
| ENSMUSG00 | U1 | 150 | 0 | 0 | 0 | 0 | 0 | 0 |
| ENSMUSG00 | U6 | 106 | 0 | 2.27294538 | 0 | 0 | 0 | 0 |
| ENSMUSG00 | SNORA41 | 130 | 0 | 0 | 0 | 0 | 0 | 0 |
| ENSMUSG00 | U6 | 106 | 0 | 0 | 0 | 0 | 0 | 0 |
| ENSMUSG00 | Y_RNA | 111 | 0 | 0 | 0 | 1.95038079 | 0.91939416 | 1.22858735 |
| ENSMUSG00 | U6 | 106 | 0 | 0 | 0 | 0 | 0 | 0 |
| ENSMUSG00 | U6 | 106 | 0 | 0 | 0 | 0 | 0 | 0 |
| ENSMUSG00 | U6 | 105 | 0 | 0 | 0 | 0 | 0 | 0 |
| ENSMUSG00 | SNORA61 | 129 | 0 | 0 | 0 | 0 | 0 | 0 |
| ENSMUSG00 | U1 | 164 | 0 | 0 | 0 | 0 | 0 | 0 |
| ENSMUSG00 | SNORA44 | 128 | 0 | 0 | 0 | 0 | 0 | 0 |
| ENSMUSG00 | U6 | 106 | 1.50020681 | 0 | 0 | 0 | 0 | 0 |
| ENSMUSG00 | U6 | 102 | 0 | 0.4724161 | 0 | 0 | 0 | 0 |
| ENSMUSG00 | Snord7 | 96 | 1.24235877 | 2.00776842 | 11.0504608 | 5.63781946 | 1.59457424 | 4.26166238 |
| ENSMUSG00 | SNORA51 | 118 | 0 | 0 | 0 | 0 | 0 | 0 |
| ENSMUSG00 | U1 | 161 | 0 | 0 | 0 | 0 | 0 | 0 |
| ENSMUSG00 | n-R5s12 | 118 | 0 | 0 | 0 | 0 | 0 | 0 |
| ENSMUSG00 | Y_RNA | 111 | 0 | 0 | 0 | 0 | 0 | 0 |
| ENSMUSG00 | U6 | 106 | 0 | 0 | 0 | 0 | 0 | 0 |
| ENSMUSG00 | U6 | 106 | 0 | 0 | 0 | 0 | 0 | 0 |
| ENSMUSG00 | SNORA70 | 137 | 1.16074396 | 0 | 0.32264119 | 0 | 0 | 0.19908496 |
| ENSMUSG00 | SNORA73 | 177 | 0 | 0 | 0 | 0 | 0 | 0 |
| ENSMUSG00 | U6 | 103 | 0 | 0 | 0 | 0 | 0 | 0 |
| ENSMUSG00 | SNORA36 | 130 | 0 | 0 | 0 | 0 | 0 | 0 |
| ENSMUSG00 | U6 | 104 | 0 | 0 | 0 | 0 | 0 | 0 |
| ENSMUSG00 | U6 | 106 | 0 | 0.45458908 | 0 | 0 | 0 | 0 |
| ENSMUSG00 | U6 | 106 | 0 | 0 | 0 | 0 | 0 | 0 |
| ENSMUSG00 | Snord49a | 71 | 3.35961807 | 3.39341142 | 0.62256117 | 0 | 0.71868135 | 3.07319879 |
| ENSMUSG00 | SNORD77 | 80 | 0 | 0 | 0 | 0 | 0 | 0 |
| ENSMUSG00 | U6 | 97 | 0 | 0 | 0 | 0 | 0 | 0 |
| ENSMUSG00 | SNORD45 | 80 | 4.47249156 | 0.60233053 | 3.86766126 | 0.90205111 | 7.65395637 | 1.02279897 |
| ENSMUSG00 | SNORA67 | 142 | 0 | 0 | 0 | 0 | 0 | 0 |
| ENSMUSG00 | U2 | 132 | 0 | 0 | 0 | 0 | 0 | 0 |
| ENSMUSG00 | U6 | 80 | 0 | 0 | 0 | 0 | 0 | 0 |
| ENSMUSG00 | U1 | 160 | 0 | 0 | 0 | 0 | 0 | 0 |
| ENSMUSG00 | U1 | 155 | 0 | 0 | 0 | 0 | 0 | 0 |
| ENSMUSG00 | U6 | 106 | 0 | 0 | 0 | 0 | 0 | 0 |
| ENSMUSG00 | AF357428 | 68 | 0 | 0 | 0 | 0 | 0 | 0 |
| ENSMUSG00 | SNORA5 | 135 | 0.58897008 | 2.14161965 | 0 | 0 | 0 | 3.23254984 |
| ENSMUSG00 | U2 | 149 | 0 | 0 | 0 | 0 | 0 | 0 |
| ENSMUSG00 | SNORD18 | 67 | 1.18673076 | 0 | 1.979187 | 0 | 0 | 0 |
| ENSMUSG00 | SNORA44 | 128 | 0 | 0 | 0 | 0 | 0 | 0 |
| ENSMUSG00 | SNORD113 | 81 | 0 | 0 | 0 | 0 | 0 | 0 |
| ENSMUSG00 | n-R5s10 | 118 | 0 | 0 | 0 | 0 | 0 | 0 |
| ENSMUSG00 | U6 | 105 | 0 | 0 | 0 | 0 | 0 | 0 |
| ENSMUSG00 | U6 | 109 | 0 | 0 | 0 | 0 | 0 | 0 |
| ENSMUSG00 | U1 | 168 | 0 | 1.4341203 | 2.36795588 | 0.42954815 | 0 | 1.1364433 |
| ENSMUSG00 | U6 | 102 | 0.77951923 | 0 | 0.4333514 | 0 | 1.00051717 | 1.60439054 |
| ENSMUSG00 | U2 | 190 | 0.41847874 | 0.25361285 | 0.23264128 | 0.1899055 | 0.80567962 | 3.30166686 |
| ENSMUSG00 | U1 | 151 | 0 | 0 | 0 | 0 | 0 | 0 |
| ENSMUSG00 | U6 | 106 | 0 | 0 | 0 | 0 | 0 | 0 |
| ENSMUSG00 | U1 | 161 | 0 | 0 | 0 | 0 | 0 | 0 |
| ENSMUSG00 | n-R5s54 | 111 | 0 | 0 | 0 | 0 | 0 | 0 |
| ENSMUSG00 | U1 | 165 | 0 | 0 | 0 | 0 | 0 | 0 |
| ENSMUSG00 | U6 | 106 | 8.62618917 | 16.3652068 | 5.42098075 | 14.6370558 | 25.5131879 | 11.3215484 |
| ENSMUSG00 | U6 | 106 | 0.75010341 | 0.45458908 | 2.0849926 | 1.02118994 | 0 | 1.80115542 |
| ENSMUSG00 | SNORA55 | 130 | 0 | 0 | 1.70007089 | 0 | 0 | 0 |
| ENSMUSG00 | SNORD30 | 68 | 0 | 0 | 0 | 0 | 0 | 0 |
| ENSMUSG00 | U6 | 106 | 0 | 0 | 0 | 0 | 0 | 0 |
| ENSMUSG00 | U6 | 105 | 0 | 0 | 0 | 0 | 0 | 0 |
| ENSMUSG00 | U6 | 106 | 0 | 0 | 0 | 0 | 0 | 0 |
| ENSMUSG00 | U6 | 106 | 0 | 0 | 0 | 0 | 0 | 0 |
| ENSMUSG00 | SNORD103 | 82 | 0 | 0 | 0 | 0 | 0 | 0 |
| ENSMUSG00 | U6 | 106 | 0 | 0 | 0 | 0 | 0 | 0 |
| ENSMUSG00 | U1 | 156 | 0 | 0 | 0 | 0 | 0 | 0 |
| ENSMUSG00 | U1 | 134 | 0 | 0 | 0 | 0 | 0 | 0 |
| ENSMUSG00 | U1 | 161 | 0 | 0 | 0 | 0 | 0 | 0 |
| ENSMUSG00 | U6 | 106 | 0 | 0 | 0 | 0 | 0 | 0 |
| ENSMUSG00 | U6 | 106 | 0 | 3.63671262 | 0 | 0 | 0 | 0 |
| ENSMUSG00 | SNORA5 | 133 | 0 | 0 | 0 | 0 | 0 | 0 |
| ENSMUSG00 | U6 | 100 | 0 | 0 | 0 | 0 | 0 | 0 |
| ENSMUSG00 | U6 | 105 | 0 | 0 | 0 | 0 | 0 | 0 |
| ENSMUSG00 | U1 | 162 | 0 | 0 | 0 | 0 | 0 | 0 |
| ENSMUSG00 | U1 | 160 | 0 | 0 | 0 | 0 | 0 | 0 |
| ENSMUSG00 | n-R5s69 | 118 | 0 | 0 | 0 | 0 | 0 | 0 |
| ENSMUSG00 | U6 | 103 | 0 | 0 | 0 | 0 | 0 | 0 |
| ENSMUSG00 | 7SK | 323 | 0 | 0 | 0 | 0 | 0 | 0 |
| ENSMUSG00 | U4 | 82 | 0 | 0 | 0 | 0 | 0 | 0 |
| ENSMUSG00 | SNORA42 | 133 | 0 | 0 | 0 | 0 | 0 | 0 |
| ENSMUSG00 | n-R5s3 | 111 | 0 | 0 | 0 | 0 | 0 | 0 |
| ENSMUSG00 | U6 | 106 | 0 | 0 | 0 | 0 | 0 | 0 |
| ENSMUSG00 | U6 | 106 | 0 | 0 | 0 | 0 | 0 | 0 |
| ENSMUSG00 | SNORA3 | 129 | 0 | 0 | 0 | 1.11882309 | 0.7911066 | 2.74860706 |
| ENSMUSG00 | Snord35a | 89 | 3.12683555 | 5.41420698 | 0 | 0 | 0 | 2.45165297 |
| ENSMUSG00 | U6 | 106 | 0 | 0 | 0 | 0 | 0 | 0 |
| ENSMUSG00 | U2 | 190 | 0 | 0 | 0 | 0 | 0 | 0 |
| ENSMUSG00 | U6 | 103 | 0 | 0 | 0 | 0 | 0 | 0 |
| ENSMUSG00 | Snord15a | 147 | 0 | 0 | 0 | 0 | 0 | 0 |
| ENSMUSG00 | n-R5s182 | 106 | 0 | 0 | 0 | 0 | 0 | 0 |
| ENSMUSG00 | 7SK | 320 | 0 | 0 | 0 | 0 | 0 | 0 |
| ENSMUSG00 | U6 | 106 | 0 | 0 | 0 | 0 | 0 | 0 |
| ENSMUSG00 | U6 | 106 | 0 | 0 | 0 | 0 | 0 | 0 |
| ENSMUSG00 | U6 | 106 | 0 | 0 | 0 | 0 | 0 | 0 |
| ENSMUSG00 | Snord16a | 98 | 3.24534535 | 2.45849195 | 0.90207843 | 0 | 4.68609574 | 2.50481381 |
| ENSMUSG00 | U6 | 106 | 0 | 0 | 0 | 0 | 0 | 0 |
| ENSMUSG00 | U12 | 144 | 0 | 0.33462807 | 0 | 0 | 0 | 0 |
| ENSMUSG00 | U6 | 106 | 0 | 0 | 0 | 0 | 0 | 0 |
| ENSMUSG00 | SNORA71 | 132 | 0 | 0 | 0 | 0 | 0 | 0 |
| ENSMUSG00 | U6 | 109 | 0 | 0 | 0 | 0 | 0 | 0 |
| ENSMUSG00 | U6 | 107 | 0 | 0 | 0 | 0 | 0 | 0 |
| ENSMUSG00 | U1 | 163 | 0 | 0 | 0 | 0 | 0 | 0 |
| ENSMUSG00 | U6 | 106 | 0 | 0 | 0 | 0 | 0 | 0 |
| ENSMUSG00 | SNORA32 | 121 | 0 | 0 | 0.73060898 | 0 | 0.84341117 | 0 |
| ENSMUSG00 | U6 | 106 | 0 | 0 | 0 | 0 | 0 | 0 |
| ENSMUSG00 | U6 | 106 | 0 | 0 | 0 | 0 | 0 | 0 |
| ENSMUSG00 | SNORA2 | 135 | 0.58897008 | 0.71387322 | 4.25647377 | 2.67274404 | 1.51189262 | 2.02034365 |
| ENSMUSG00 | SNORD58 | 69 | 0 | 0 | 0 | 0 | 0 | 0 |
| ENSMUSG00 | U6 | 106 | 3.75051703 | 0.45458908 | 11.6759585 | 9.19070945 | 3.36966633 | 13.3800117 |
| ENSMUSG00 | U1 | 155 | 0 | 0 | 0 | 0 | 0 | 0 |
| ENSMUSG00 | U6 | 104 | 0 | 0 | 0 | 0 | 0 | 0 |
| ENSMUSG00 | SNORA67 | 142 | 0 | 0 | 0 | 0 | 0 | 0 |
| ENSMUSG00 | SNORA71 | 130 | 0 | 0 | 0 | 0 | 0 | 0 |
| ENSMUSG00 | U1 | 163 | 0 | 0 | 0 | 0 | 0 | 0 |
| ENSMUSG00 | U1 | 158 | 0 | 0 | 0 | 0 | 0 | 0 |
| ENSMUSG00 | U6 | 103 | 0 | 0 | 0 | 0 | 0 | 0 |
| ENSMUSG00 | U3 | 211 | 0.7536584 | 0.22837176 | 1.25692445 | 0.6840198 | 0.7254935 | 0.64631847 |
| ENSMUSG00 | U6 | 105 | 0 | 0 | 0 | 0 | 0 | 0 |
| ENSMUSG00 | n-R5s183 | 114 | 0 | 0 | 0 | 0 | 0 | 0.47850244 |
| ENSMUSG00 | SNORA71 | 130 | 0 | 0 | 0 | 0 | 0 | 0 |
| ENSMUSG00 | U6 | 106 | 0 | 0 | 0 | 0 | 0 | 0 |
| ENSMUSG00 | U1 | 163 | 0 | 0 | 0 | 0 | 0 | 0 |
| ENSMUSG00 | Snord34 | 69 | 0 | 0 | 0 | 0 | 0 | 0 |
| ENSMUSG00 | U1 | 176 | 0 | 0 | 0 | 0 | 0 | 0 |
| ENSMUSG00 | n-R5s168 | 103 | 0 | 0 | 0 | 0 | 0 | 0 |
| ENSMUSG00 | U3 | 212 | 0 | 0 | 0.83399704 | 0 | 0.4813809 | 0 |
| ENSMUSG00 | SNORA63 | 129 | 0 | 0 | 0 | 0 | 0 | 0 |
| ENSMUSG00 | U6 | 106 | 0 | 0 | 0 | 0 | 0 | 0 |
| ENSMUSG00 | n-R5s185 | 118 | 0 | 0 | 0 | 0 | 0 | 0 |
| ENSMUSG00 | U1 | 163 | 0 | 0 | 0 | 0 | 0 | 0 |
| ENSMUSG00 | U6 | 106 | 0 | 0 | 0 | 0 | 0 | 0 |
| ENSMUSG00 | n-R5s151 | 118 | 0 | 0 | 0 | 0 | 0 | 0 |
| ENSMUSG00 | n-R5s46 | 115 | 0 | 0 | 0 | 0 | 0 | 0 |
| ENSMUSG00 | U6 | 107 | 0 | 0 | 0 | 0 | 0 | 0 |
| ENSMUSG00 | SNORA72 | 131 | 0 | 0 | 0 | 0 | 0 | 0 |
| ENSMUSG00 | U6 | 105 | 0 | 0 | 0 | 0 | 0 | 0 |
| ENSMUSG00 | SNORD63 | 69 | 5.76166384 | 0 | 3.2030321 | 4.18342545 | 2.21853808 | 7.11512328 |
| ENSMUSG00 | U1 | 164 | 0 | 0 | 0 | 0 | 0 | 0 |
| ENSMUSG00 | U6 | 106 | 0 | 0 | 0 | 0 | 0 | 0 |
| ENSMUSG00 | U2 | 190 | 0 | 0 | 0 | 0 | 0 | 0 |
| ENSMUSG00 | U6 | 106 | 0 | 0 | 0 | 0 | 0 | 0 |
| ENSMUSG00 | 7SK | 337 | 0 | 0 | 0 | 0 | 0 | 0 |
| ENSMUSG00 | U6 | 106 | 0 | 0 | 0 | 0 | 0 | 0 |
| ENSMUSG00 | U6 | 106 | 0 | 0 | 0 | 0 | 0 | 0 |
| ENSMUSG00 | U6 | 106 | 0 | 0 | 0 | 0 | 0 | 0 |
| ENSMUSG00 | U1 | 155 | 0 | 0 | 0 | 0 | 0 | 0 |
| ENSMUSG00 | U6 | 105 | 0 | 0 | 0 | 0 | 0 | 0 |
| ENSMUSG00 | U6 | 106 | 0 | 0 | 0 | 0 | 0 | 0.25730792 |
| ENSMUSG00 | n-R5-8s1 | 147 | 4.86801802 | 4.26138604 | 5.41247057 | 5.40003387 | 3.12406383 | 5.00962762 |
| ENSMUSG00 | SNORA71 | 130 | 0 | 0 | 0 | 0 | 0 | 0 |
| ENSMUSG00 | U6 | 106 | 0 | 0 | 0 | 0 | 0 | 0 |
| ENSMUSG00 | U6 | 106 | 0 | 0 | 0 | 0 | 0 | 0 |
| ENSMUSG00 | U6 | 106 | 0 | 0 | 0 | 0 | 0 | 0 |
| ENSMUSG00 | U6 | 106 | 0 | 0 | 0 | 0 | 0 | 0 |
| ENSMUSG00 | U1 | 161 | 0 | 0 | 0 | 0 | 0 | 0 |
| ENSMUSG00 | U6 | 106 | 0 | 0 | 0 | 0 | 0 | 0 |
| ENSMUSG00 | U6 | 104 | 0 | 0 | 0 | 0 | 0 | 0 |
| ENSMUSG00 | U1 | 152 | 0.78464764 | 0 | 0 | 0 | 0.67139968 | 0 |
| ENSMUSG00 | U6 | 106 | 0 | 0 | 0 | 0 | 0 | 0 |
| ENSMUSG00 | Snora2b | 136 | 0.29231971 | 0 | 0 | 0 | 0.37519394 | 0 |
| ENSMUSG00 | SNORA21 | 136 | 0 | 0 | 0 | 0 | 0 | 0 |
| ENSMUSG00 | U1 | 163 | 0 | 0 | 0.27117695 | 0 | 0 | 1.50596168 |
| ENSMUSG00 | U6 | 106 | 0 | 0 | 0 | 0 | 0 | 0 |
| ENSMUSG00 | U6 | 103 | 0 | 0 | 0 | 0 | 0 | 0 |
| ENSMUSG00 | U2 | 190 | 0.62771811 | 0.25361285 | 0 | 0.56971649 | 0.80567962 | 0 |
| ENSMUSG00 | U1 | 159 | 0 | 0 | 0 | 0 | 0 | 0 |
| ENSMUSG00 | U6 | 103 | 0 | 0 | 0 | 0 | 0 | 0 |
| ENSMUSG00 | Gm5908 | 446 | 0 | 0 | 0 | 0 | 0 | 0 |
| ENSMUSG00 | C330021F23I | 2868 | 0 | 0.15121269 | 0.61648317 | 0 | 0 | 0.03803994 |
| ENSMUSG00 | Tacc1 | 7682 | 15.9239278 | 14.7407106 | 16.2031229 | 16.3876924 | 18.0007132 | 19.0837264 |
| ENSMUSG00 | Gm14850 | 401 | 0 | 0 | 0 | 0 | 0 | 0 |
| ENSMUSG00 | Gm15308 | 748 | 0 | 0 | 0 | 0 | 0 | 0 |
| ENSMUSG00 | Defa20 | 423 | 0 | 0 | 0 | 0 | 0 | 0 |
| ENSMUSG00 | Ifitm7 | 871 | 0 | 0 | 0 | 0 | 0 | 0 |
| ENSMUSG00 | Cpped1 | 7055 | 12.092879 | 12.2463913 | 16.8349188 | 12.7910976 | 12.9826144 | 11.89182 |
| ENSMUSG00 | Cd209b | 2285 | 0 | 0 | 0 | 0 | 0 | 0 |
| ENSMUSG00 | Aurkaip1 | 1742 | 20.9503623 | 26.5551002 | 25.1711988 | 16.8396683 | 15.231754 | 26.1786434 |
| ENSMUSG00 | Gm13154 | 2952 | 0.75416901 | 0.21220317 | 0.85349087 | 0.77004363 | 0.96798003 | 0 |
| ENSMUSG00 | 2610305D13 | 5687 | 6.77388089 | 5.10079799 | 7.72580129 | 8.43838917 | 3.88507486 | 0 |
| ENSMUSG00 | Gm13237 | 1139 | 1.18673076 | 1.05764798 | 1.12542006 | 1.64729264 | 0.9407848 | 1.46071378 |
| ENSMUSG00 | Zfp600 | 4929 | 0 | 0.01955222 | 0.04483855 | 0.06588322 | 0.04140911 | 0 |
| ENSMUSG00 | Gm13051 | 1925 | 0 | 0 | 0 | 0 | 0 | 0 |
| ENSMUSG00 | Dhrs3 | 2920 | 42.3695395 | 34.489611 | 61.3073508 | 30.200177 | 33.2545182 | 50.9251141 |
| ENSMUSG00 | Gm2042 | 1889 | 0 | 0 | 0 | 0 | 0 | 0 |
| ENSMUSG00 | Oog2 | 2633 | 0 | 0 | 0 | 0 | 0 | 0 |
| ENSMUSG00 | Gm13023 | 1919 | 0 | 0 | 0 | 0 | 0 | 0 |
| ENSMUSG00 | Gm17372 | 2729 | 0 | 0 | 0.06478834 | 0.03966513 | 0 | 0.08994934 |
| ENSMUSG00 | Ubr4 | 17888 | 8.03643882 | 7.72039038 | 7.93942988 | 8.96604919 | 8.60043175 | 6.51828504 |
| ENSMUSG00 | Hnrnpr | 3264 | 23.117617 | 22.6316838 | 23.712447 | 23.4798599 | 21.6361839 | 31.7117819 |
| ENSMUSG00 | Gm10151 | 3199 | 4.92127861 | 3.8109315 | 3.39907889 | 3.33863244 | 2.67972214 | 4.19478666 |
| ENSMUSG00 | Med18 | 2153 | 2.60358697 | 2.61858511 | 4.0855396 | 2.19542398 | 3.15211704 | 4.1298339 |
| ENSMUSG00 | Phactr4 | 7298 | 3.32839115 | 2.55524159 | 2.88904893 | 3.79212499 | 4.4188366 | 4.28665542 |
| ENSMUSG00 | 2610044O15I | 387 | 0.5136367 | 1.74317879 | 2.85541622 | 2.70382246 | 1.7140643 | 0 |
| ENSMUSG00 | Cldn19 | 1962 | 2.08706141 | 2.23494711 | 0.36046355 | 1.02986468 | 1.84652023 | 0.75067814 |
| ENSMUSG00 | Gm12886 | 1669 | 0.50021875 | 1.15485781 | 0.74155279 | 2.31322874 | 1.62037023 | 0.01634191 |
| ENSMUSG00 | Olfr1335 | 2106 | 0 | 0 | 0 | 0 | 0 | 0 |
| ENSMUSG00 | Olfr1338 | 941 | 0 | 0 | 0 | 0 | 0 | 0 |
| ENSMUSG00 | Gm13611 | 700 | 0.22717417 | 0.06883777 | 0 | 0 | 0.14578965 | 0.15585508 |
| ENSMUSG00 | Cyp4a12a | 2415 | 35.4589255 | 39.5468026 | 97.7565398 | 54.4293533 | 29.6438945 | 25.6257377 |
| ENSMUSG00 | Cyp4a10 | 2318 | 0.15435691 | 2.68164411 | 0.51486185 | 0.04669807 | 0.22013104 | 2.27092553 |
| ENSMUSG00 | Insl5 | 953 | 0 | 0 | 0 | 0 | 0 | 0.08585931 |
| ENSMUSG00 | Gm10152 | 1343 | 0 | 0 | 0 | 0 | 0 | 0 |
| ENSMUSG00 | Cyp2j11 | 1908 | 43.7977044 | 75.0829625 | 25.4137431 | 57.3190131 | 55.4657775 | 48.559722 |
| ENSMUSG00 | Krtap5-1 | 564 | 0 | 0 | 0 | 0 | 0 | 0 |
| ENSMUSG00 | Gm10153 | 559 | 0 | 0 | 0 | 0 | 0 | 0 |
| ENSMUSG00 | Gm12666 | 760 | 0 | 0.06340321 | 0 | 0 | 0 | 0 |
| ENSMUSG00 | Muc5b | 15048 | 0 | 0.11527857 | 0.13805733 | 0 | 0.07799087 | 0.03443768 |
| ENSMUSG00 | Adamtsl1 | 10912 | 0.10929843 | 0 | 0 | 0.03306639 | 0.01402851 | 0 |
| ENSMUSG00 | Gm10154 | 407 | 0 | 0 | 0 | 0 | 0 | 0 |
| ENSMUSG00 | Olfr45 | 1034 | 0 | 0 | 0 | 0 | 0 | 0 |
| ENSMUSG00 | Olfr53 | 1001 | 0 | 0 | 0 | 0 | 0 | 0 |
| ENSMUSG00 | Olfr61 | 1095 | 0 | 0 | 0 | 0 | 0 | 0 |
| ENSMUSG00 | Olfr538 | 956 | 0 | 0 | 0 | 0 | 0 | 0 |
| ENSMUSG00 | Kndc1 | 8150 | 0 | 0.01182489 | 0 | 0 | 0 | 0 |
| ENSMUSG00 | Gm11237 | 1858 | 0 | 0 | 0 | 0 | 0 | 0 |
| ENSMUSG00 | Gm11487 | 1317 | 0 | 0 | 0 | 0 | 0 | 0 |
| ENSMUSG00 | Gm11232 | 1677 | 0 | 0 | 0 | 0 | 0 | 0 |
| ENSMUSG00 | Prpf4 | 5543 | 6.46932048 | 6.16348322 | 7.54373868 | 7.42731604 | 7.63140277 | 6.25894662 |
| ENSMUSG00 | Cdc26 | 7175 | 2.65405926 | 2.96170327 | 2.61822764 | 1.88079229 | 2.19040052 | 2.27320338 |
| ENSMUSG00 | Slc31a1 | 4276 | 100.104597 | 123.350981 | 70.4376423 | 94.1710984 | 104.845121 | 108.671195 |
| ENSMUSG00 | Fkbp15 | 7485 | 10.0862602 | 7.9119756 | 8.42699131 | 9.39531126 | 9.91895482 | 11.1795048 |
| ENSMUSG00 | Slc31a2 | 2026 | 51.254351 | 51.4686381 | 64.3610251 | 47.3732668 | 39.692778 | 42.6755214 |
| ENSMUSG00 | Mup21 | 671 | 0 | 0 | 0 | 0 | 0 | 0 |
| ENSMUSG00 | Mup3 | 962 | 0 | 0 | 0 | 0 | 0 | 0 |
| ENSMUSG00 | AY512931 | 2776 | 0 | 0 | 0 | 0 | 0 | 0 |
| ENSMUSG00 | E230001N04 | 2054 | 0.01935515 | 0 | 0 | 0.19323393 | 0 | 0.14606672 |
| ENSMUSG00 | 2510046G10 | 347 | 0.91655287 | 0.55546331 | 1.78335966 | 0.72787986 | 1.91165097 | 3.93006329 |
| ENSMUSG00 | Gm12511 | 1601 | 0 | 0 | 0 | 0 | 0 | 0 |
| ENSMUSG00 | 6030445D17 | 1727 | 0 | 0.02790182 | 0 | 0.0208929 | 0 | 0.04737922 |
| ENSMUSG00 | Gm10155 | 540 | 0 | 0 | 0 | 0 | 0 | 0 |
| ENSMUSG00 | Cacng3 | 1519 | 0 | 0 | 0 | 0 | 0 | 0 |
| ENSMUSG00 | Anks6 | 6347 | 3.14435973 | 3.7048974 | 1.05855997 | 1.72252395 | 4.13227622 | 0.92390853 |
| ENSMUSG00 | Spag8 | 1655 | 0.52847164 | 0.90258593 | 0.96149024 | 0.2180184 | 1.20243423 | 3.83987368 |
| ENSMUSG00 | Gpr139 | 1036 | 0 | 0 | 0 | 0.03482823 | 0 | 0 |
| ENSMUSG00 | Il11ra2 | 2666 | 0 | 0 | 0 | 0 | 0 | 0 |
| ENSMUSG00 | Ccl27b | 927 | 0 | 0 | 0 | 0 | 0 | 0 |
| ENSMUSG00 | Arid3c | 1937 | 0.08209702 | 0 | 0 | 0 | 0.21074394 | 0.1689704 |
| ENSMUSG00 | Ipo7 | 3092 | 38.2254345 | 28.87758 | 31.4502117 | 41.4966851 | 36.3389649 | 23.7815095 |
| ENSMUSG00 | Tmem42 | 1041 | 13.5573445 | 18.3765778 | 20.7209408 | 14.696241 | 12.4012229 | 20.48873 |
| ENSMUSG00 | C85492 | 4176 | 9.4628706 | 11.4581267 | 6.58370363 | 8.29472288 | 7.60019295 | 12.1873747 |
| ENSMUSG00 | Olfr519 | 959 | 0 | 0 | 0 | 0 | 0 | 0 |
| ENSMUSG00 | Olfr517 | 1054 | 0 | 0 | 0 | 0 | 0 | 0 |
| ENSMUSG00 | Olfr514 | 932 | 0 | 0 | 0 | 0 | 0 | 0 |
| ENSMUSG00 | Olfr467 | 926 | 0 | 0 | 0 | 0 | 0 | 0 |
| ENSMUSG00 | Olfr470 | 944 | 0 | 0 | 0 | 0 | 0 | 0 |
| ENSMUSG00 | Gm10156 | 1591 | 0 | 0 | 0 | 0 | 0 | 0 |
| ENSMUSG00 | Bhmt-ps1 | 1871 | 0 | 0 | 0 | 0 | 0 | 0 |
| ENSMUSG00 | Olfr206 | 1004 | 0 | 0 | 0 | 0 | 0 | 0 |
| ENSMUSG00 | Trim12a | 6000 | 4.39960651 | 4.46527697 | 4.67066141 | 6.7293013 | 6.25923543 | 5.27309692 |
| ENSMUSG00 | Olfr640 | 944 | 0 | 0 | 0 | 0 | 0 | 0 |
| ENSMUSG00 | Olfr639 | 920 | 0 | 0 | 0 | 0 | 0 | 0 |
| ENSMUSG00 | Olfr612 | 971 | 0 | 0 | 0 | 0 | 0 | 0 |
| ENSMUSG00 | Olfr586 | 953 | 0 | 0 | 0 | 0 | 0 | 0 |
| ENSMUSG00 | Olfr575 | 954 | 0 | 0 | 0 | 0 | 0 | 0 |
| ENSMUSG00 | Gm10157 | 680 | 0.05846394 | 0.07086242 | 0.13000542 | 0 | 0 | 0.16043905 |
| ENSMUSG00 | Olfr559 | 1021 | 0 | 0 | 0 | 0 | 0 | 0 |
| ENSMUSG00 | Olfr33 | 917 | 0 | 0 | 0 | 0 | 0 | 0 |
| ENSMUSG00 | Vps37b | 2508 | 18.8473949 | 20.2890283 | 20.2856146 | 21.0910197 | 18.7178093 | 13.6155695 |
| ENSMUSG00 | Chrna10 | 2508 | 0 | 0 | 0 | 0 | 0.48829068 | 0 |
| ENSMUSG00 | Numa1 | 7308 | 14.2364659 | 13.7016183 | 16.2944397 | 18.6186905 | 21.8265532 | 13.6671769 |
| ENSMUSG00 | Rad54b | 1601 | 0.64562304 | 0.96312689 | 0.93870248 | 0.63104138 | 1.75293608 | 0.63033208 |
| ENSMUSG00 | Gm12918 | 1323 | 0 | 0 | 0 | 0 | 0 | 0 |
| ENSMUSG00 | Rtp3 | 2450 | 0.08113363 | 0 | 0.10824941 | 0 | 0.5206773 | 0 |
| ENSMUSG00 | Impad1 | 6538 | 28.0684151 | 39.0031587 | 29.3484553 | 33.5489062 | 26.0517043 | 33.5572343 |
| ENSMUSG00 | Wdr6 | 5246 | 33.2078756 | 35.988273 | 30.7458111 | 35.9857524 | 41.8540784 | 16.9387676 |
| ENSMUSG00 | Tcl1b1 | 1033 | 0 | 0 | 0 | 0 | 0 | 0 |
| ENSMUSG00 | Serpina3c | 2208 | 0 | 0 | 0 | 0 | 0 | 0 |
| ENSMUSG00 | Rps13-ps1 | 455 | 0 | 0 | 0.09714691 | 0 | 0.11214588 | 0 |
| ENSMUSG00 | Serpina3f | 2283 | 0 | 0.14774643 | 0 | 0 | 0 | 0 |
| ENSMUSG00 | Serpina3b | 2065 | 0 | 0 | 0 | 0 | 0 | 0 |
| ENSMUSG00 | Serpina1a | 2531 | 0 | 0 | 0 | 0 | 0 | 0 |
| ENSMUSG00 | 4921517D21 | 3992 | 0 | 0 | 0 | 0 | 0 | 0 |
| ENSMUSG00 | Vmn2r65 | 2532 | 0 | 0 | 0 | 0 | 0 | 0 |
| ENSMUSG00 | Gm10160 | 204 | 0 | 0 | 0 | 0 | 0 | 0 |
| ENSMUSG00 | Iqcf5 | 542 | 0 | 0 | 0 | 0 | 0 | 0 |
| ENSMUSG00 | Iqcf1 | 631 | 0 | 0 | 0 | 0.22872928 | 0 | 0 |
| ENSMUSG00 | Rpl31-ps1 | 377 | 0 | 0 | 0 | 0 | 0 | 0 |
| ENSMUSG00 | Nrxn3 | 11016 | 0.01804443 | 0.10498136 | 0 | 0.18342361 | 0.00926405 | 0.21045246 |
| ENSMUSG00 | Akap13 | 15343 | 2.0625277 | 1.94089951 | 2.24135005 | 4.05196915 | 3.10621358 | 2.28784858 |
| ENSMUSG00 | Gm10263 | 209 | 0 | 0 | 0 | 0 | 0 | 0.65250333 |
| ENSMUSG00 | Msi2 | 4906 | 15.7449855 | 11.1577249 | 11.4874337 | 18.0557316 | 15.798831 | 11.7526676 |
| ENSMUSG00 | Gm17627 | 1872 | 0.57339635 | 0 | 0.14167257 | 0.82880765 | 0.05451536 | 0 |
| ENSMUSG00 | Vmn2r68-ps | 2550 | 0 | 0 | 0 | 0 | 0 | 0 |
| ENSMUSG00 | 3830431G21 | 3925 | 58.3418007 | 35.2466944 | 50.5308712 | 46.5527321 | 37.6750158 | 35.4049139 |
| ENSMUSG00 | Zfyve26 | 9326 | 2.41278168 | 2.67128357 | 2.46461059 | 3.99278039 | 3.51264564 | 2.74033208 |
| ENSMUSG00 | Rdh11 | 4147 | 10.6698456 | 15.0357502 | 10.1044966 | 10.3713039 | 9.37595813 | 11.2992115 |
| ENSMUSG00 | Mthfs | 3056 | 2.97905924 | 3.07472389 | 3.08082217 | 1.81827057 | 2.20401885 | 3.56105401 |
| ENSMUSG00 | Gm10163 | 1029 | 0 | 0 | 0 | 0 | 0.04958832 | 0 |
| ENSMUSG00 | Gm8226 | 851 | 0 | 0 | 0 | 0 | 0 | 0 |
| ENSMUSG00 | Hmgn3 | 2043 | 28.8971554 | 18.8924817 | 69.4724023 | 20.7343712 | 19.6063167 | 80.5689906 |
| ENSMUSG00 | Gm5068 | 305 | 0 | 0 | 0 | 0 | 0 | 0 |
| ENSMUSG00 | Omt2a | 1751 | 0 | 0 | 0 | 0 | 0 | 0 |
| ENSMUSG00 | Gm10268 | 603 | 0 | 0 | 0.14660644 | 0 | 0 | 0 |
| ENSMUSG00 | Gm16551 | 4217 | 0 | 0 | 0 | 0 | 0 | 0 |
| ENSMUSG00 | Gm5745 | 314 | 0 | 0 | 0 | 0 | 0 | 0 |
| ENSMUSG00 | Gm5786 | 706 | 0.05631088 | 0 | 0 | 0 | 0 | 0 |
| ENSMUSG00 | Rpl26-ps2 | 437 | 0 | 0 | 0 | 0 | 0 | 0 |
| ENSMUSG00 | Gm6265 | 230 | 0.86424958 | 0.20950627 | 0.19218193 | 0.15687845 | 0.66556142 | 1.30443927 |
| ENSMUSG00 | Izumo2 | 713 | 0 | 0 | 0 | 0 | 0 | 0 |
| ENSMUSG00 | Ankdd1a | 1429 | 0 | 0 | 0 | 0 | 0 | 0 |
| ENSMUSG00 | Klk1b5 | 857 | 30.5240445 | 30.1376114 | 12.8427759 | 29.0087849 | 20.6010806 | 26.7972535 |
| ENSMUSG00 | Klk1b4 | 834 | 1.43005326 | 3.75553806 | 1.16599586 | 3.46110739 | 3.36504877 | 2.02761107 |
| ENSMUSG00 | Klk1b3 | 883 | 2.92650763 | 4.31113129 | 1.75205493 | 4.57665797 | 3.64061345 | 3.21241504 |
| ENSMUSG00 | Klk1b21 | 893 | 0.35615212 | 0.64752218 | 0.19799258 | 0.4848651 | 0.1142808 | 0.12217084 |
| ENSMUSG00 | Hmg1l1 | 1280 | 0.21741278 | 0.26351961 | 0.10359807 | 0.19732368 | 0.15945742 | 0.25569974 |
| ENSMUSG00 | Vmn2r57 | 2619 | 0 | 0 | 0 | 0 | 0 | 0 |
| ENSMUSG00 | Gm6254 | 1523 | 0.0261034 | 0.06327832 | 0 | 0.02369143 | 0 | 0 |
| ENSMUSG00 | Gm10166 | 1157 | 0 | 0 | 0 | 0 | 0 | 0.02357359 |
| ENSMUSG00 | Hmgb1 | 5623 | 8.43469469 | 10.077762 | 9.16580988 | 10.5044117 | 7.21429286 | 12.0002592 |
| ENSMUSG00 | Gm6969 | 1372 | 0.0579526 | 0.59706233 | 1.2242493 | 0.13149433 | 1.00416337 | 1.09337111 |
| ENSMUSG00 | Gm10167 | 1478 | 0.02689816 | 0 | 0.02990652 | 0.1464765 | 0.06904787 | 0.0369075 |
| ENSMUSG00 | Lsm14a | 3092 | 37.1068295 | 35.9528208 | 31.3930295 | 31.6943185 | 36.7185272 | 36.5455467 |
| ENSMUSG00 | 4931406P16 | 9677 | 3.35232738 | 3.31633466 | 3.29789508 | 4.85841728 | 6.00062165 | 4.19111177 |
| ENSMUSG00 | Abpa | 1063 | 0 | 0 | 0 | 0 | 0 | 0 |
| ENSMUSG00 | Abpb | 1805 | 0 | 0 | 0 | 0 | 0 | 0 |
| ENSMUSG00 | Abpg | 576 | 0 | 0 | 0 | 0 | 0 | 0 |
| ENSMUSG00 | Gm7589 | 536 | 0 | 0 | 0 | 0 | 0 | 0 |
| ENSMUSG00 | Mfsd7b | 4077 | 18.9367533 | 19.4542271 | 17.4552287 | 23.6210392 | 19.0613614 | 11.7741882 |
| ENSMUSG00 | 6030419C18 | 877 | 0 | 0 | 0 | 0 | 0 | 0 |
| ENSMUSG00 | Zfp932 | 2985 | 36.7721547 | 25.4572929 | 29.7492471 | 44.1083352 | 20.8720954 | 46.2435341 |
| ENSMUSG00 | Gm4883 | 527 | 0 | 0 | 0 | 0 | 0 | 0 |
| ENSMUSG00 | Tecpr1 | 8858 | 11.9697298 | 9.74826195 | 8.06890722 | 11.1366347 | 9.89651317 | 5.93649859 |
| ENSMUSG00 | Gm5121 | 678 | 0.23454561 | 0.07107145 | 0.06519446 | 0.15965506 | 0 | 0.08045616 |
| ENSMUSG00 | Rpl36-ps3 | 383 | 0 | 0 | 0 | 0 | 0 | 0 |
| ENSMUSG00 | Pgk1-rs7 | 1253 | 0 | 0 | 0 | 0 | 0 | 0 |
| ENSMUSG00 | Ttc32 | 997 | 24.6826905 | 31.3671023 | 35.8668917 | 31.9201236 | 18.6294893 | 49.6797842 |
| ENSMUSG00 | 1700001J11R | 2288 | 0 | 0 | 0 | 0 | 0.02230174 | 0 |
| ENSMUSG00 | Fbxl18 | 11703 | 0.67600962 | 0.72467007 | 0.50611356 | 0.48405375 | 0.72813849 | 0.48941932 |
| ENSMUSG00 | Wdr35 | 7304 | 6.66764288 | 7.58026041 | 5.31341979 | 6.94569477 | 7.97811079 | 8.26379746 |
| ENSMUSG00 | Gm5113 | 2999 | 7.42349753 | 4.980926 | 8.59275574 | 5.90739709 | 9.3069115 | 10.1950219 |
| ENSMUSG00 | Lefty2 | 2607 | 0 | 0 | 0 | 0 | 0 | 0 |
| ENSMUSG00 | Gm16432 | 3403 | 4.08886811 | 2.95943768 | 3.97465882 | 5.54537446 | 2.50408603 | 3.7509642 |
| ENSMUSG00 | Olfr220 | 977 | 0 | 0 | 0 | 0 | 0 | 0 |
| ENSMUSG00 | Olfr417 | 929 | 0 | 0 | 0 | 0 | 0 | 0 |
| ENSMUSG00 | Pydc3 | 2749 | 0 | 0 | 0 | 0 | 0.0185618 | 0 |
| ENSMUSG00 | Olfr1407-ps1 | 238 | 0 | 0 | 0 | 0 | 0 | 0 |
| ENSMUSG00 | Pilrb2 | 1824 | 0 | 0 | 0 | 0 | 0 | 0 |
| ENSMUSG00 | Pilrb1 | 1236 | 0 | 0 | 0 | 0 | 0 | 0 |
| ENSMUSG00 | Zbtb16 | 5093 | 1.32700406 | 2.63970496 | 1.25844635 | 4.17284984 | 1.54291417 | 0.56230849 |
| ENSMUSG00 | Gm13083 | 2461 | 0 | 0 | 0 | 0 | 0 | 0 |
| ENSMUSG00 | F830001A07 | 433 | 0 | 0 | 0.20416556 | 0.08333036 | 0 | 0 |
| ENSMUSG00 | Gm7493 | 625 | 0 | 0 | 0 | 0 | 0.0816422 | 0 |
| ENSMUSG00 | Gm10172 | 556 | 0 | 0 | 0 | 0 | 0 | 0 |
| ENSMUSG00 | Cyp2b19 | 4139 | 0.30736298 | 0 | 0 | 0 | 0 | 0 |
| ENSMUSG00 | Fxyd6 | 1783 | 1.33781763 | 1.32424883 | 1.2395357 | 1.31538581 | 3.80622994 | 0.71696133 |
| ENSMUSG00 | Gm8062 | 1024 | 0 | 0 | 0 | 0 | 0 | 0.07990617 |
| ENSMUSG00 | Cldn9 | 1433 | 5.71502372 | 0.63889909 | 17.7979508 | 4.40639064 | 2.74182201 | 7.17553315 |
| ENSMUSG00 | Zfp575 | 2737 | 0 | 0 | 0 | 0 | 0 | 0 |
| ENSMUSG00 | Vmn1r183 | 917 | 0 | 0 | 0 | 0 | 0 | 0 |
| ENSMUSG00 | Gm10175 | 440 | 0 | 0 | 0 | 0 | 0 | 0 |
| ENSMUSG00 | Vkorc1l1 | 3221 | 8.8619792 | 8.36268897 | 9.22186852 | 7.84148748 | 9.2991253 | 5.43629879 |
| ENSMUSG00 | Olfr878 | 1064 | 0 | 0 | 0 | 0 | 0 | 0 |
| ENSMUSG00 | Olfr145 | 1050 | 0 | 0 | 0 | 0 | 0 | 0 |
| ENSMUSG00 | Olfr877 | 935 | 0 | 0 | 0 | 0 | 0 | 0 |
| ENSMUSG00 | Olfr876 | 1047 | 0 | 0 | 0 | 0 | 0 | 0 |
| ENSMUSG00 | Gm10176 | 2263 | 0 | 0 | 0 | 0 | 0 | 0 |
| ENSMUSG00 | Tnfsf18 | 2063 | 0 | 0 | 0 | 0 | 0 | 0 |
| ENSMUSG00 | Igfl3 | 571 | 0 | 0 | 0 | 0 | 0 | 0 |
| ENSMUSG00 | Psg16 | 5736 | 0 | 0 | 0 | 0 | 0 | 0 |
| ENSMUSG00 | Gm3867 | 877 | 0 | 0 | 0 | 0 | 0 | 0 |
| ENSMUSG00 | Gm5614 | 317 | 0 | 0 | 0 | 0 | 0 | 0.08603987 |
| ENSMUSG00 | Obox3 | 2589 | 0 | 0 | 0 | 0 | 0 | 0 |
| ENSMUSG00 | Zfp648 | 1577 | 0 | 1.61945557 | 0 | 0.32032253 | 0.32356611 | 0 |
| ENSMUSG00 | Zbtb6 | 9497 | 3.93494279 | 3.94238871 | 4.85908435 | 4.4489917 | 3.32582148 | 2.87192158 |
| ENSMUSG00 | Rnasel | 4749 | 3.49921896 | 2.89179533 | 2.56890054 | 5.06014775 | 4.28711812 | 4.14661814 |
| ENSMUSG00 | Vmn1r84 | 956 | 0 | 0 | 0 | 0 | 0.58712357 | 0 |
| ENSMUSG00 | Vmn1r83 | 995 | 0 | 0 | 0 | 0 | 0 | 0 |
| ENSMUSG00 | Vmn1r81 | 920 | 0 | 0 | 0 | 0 | 0 | 0 |
| ENSMUSG00 | Gm10179 | 688 | 0.05778413 | 0.07003843 | 0 | 0 | 0.22249873 | 0 |
| ENSMUSG00 | Gm10180 | 77 | 0 | 0.62579795 | 1.14809982 | 0 | 0 | 0.35421609 |
| ENSMUSG00 | Gm10181 | 68 | 0 | 0 | 0 | 0 | 0 | 0 |
| ENSMUSG00 | Vmn2r28 | 2553 | 0 | 0 | 0 | 0 | 0 | 0 |
| ENSMUSG00 | Gm10182 | 272 | 0 | 0 | 0 | 0 | 0 | 0 |
| ENSMUSG00 | Vmn2r37 | 3075 | 0 | 0 | 0 | 0 | 0 | 0 |
| ENSMUSG00 | Zfp810 | 1381 | 29.67987 | 16.0505166 | 36.9362251 | 26.1536036 | 36.3207295 | 32.6466174 |
| ENSMUSG00 | Vmn2r39 | 3199 | 0 | 0 | 0 | 0 | 0 | 0 |
| ENSMUSG00 | Zfp772 | 2564 | 10.8071646 | 5.31855036 | 6.08551115 | 9.79450195 | 8.3584547 | 11.1694116 |
| ENSMUSG00 | Ecsit | 1580 | 53.9466774 | 35.9263474 | 53.5178011 | 41.6398818 | 34.7496078 | 39.8071634 |
| ENSMUSG00 | Hmcn1 | 20289 | 0.09797299 | 0 | 0.00871445 | 0 | 0 | 0 |
| ENSMUSG00 | Olfr1348 | 955 | 0 | 0 | 0 | 0 | 0 | 0 |
| ENSMUSG00 | Olfr5 | 956 | 0 | 0 | 0 | 0 | 0 | 0 |
| ENSMUSG00 | Vmn1r65 | 2679 | 0 | 0 | 0 | 0 | 0 | 0 |
| ENSMUSG00 | Cdkn2d | 1269 | 0.12531278 | 0.53160772 | 0.24382419 | 0.19903413 | 0.4020991 | 0.21493018 |
| ENSMUSG00 | Oas1g | 2683 | 0.04445264 | 0 | 0.14827305 | 0 | 0.66564411 | 0.26430884 |
| ENSMUSG00 | Oas1e | 1709 | 0 | 0 | 0 | 0 | 0 | 0 |
| ENSMUSG00 | Nck2 | 4186 | 4.86259102 | 9.50836149 | 5.15300989 | 3.81852502 | 7.36262088 | 3.16010512 |
| ENSMUSG00 | Gm10184 | 1324 | 0 | 0 | 0 | 0 | 0 | 0 |
| ENSMUSG00 | Zfp617 | 3019 | 16.2366702 | 17.3975561 | 15.8271588 | 15.6925222 | 14.0960575 | 21.0861239 |
| ENSMUSG00 | Ptprv | 288 | 0 | 0 | 0 | 0 | 0 | 0 |
| ENSMUSG00 | Fbxl12 | 5116 | 2.02818225 | 1.73305421 | 3.23133098 | 3.1032251 | 3.35122406 | 3.69968265 |
| ENSMUSG00 | Vsig10 | 5321 | 4.96102971 | 4.75434733 | 2.32597558 | 2.08178682 | 4.98660316 | 3.40356333 |
| ENSMUSG00 | Olfr18 | 1154 | 0 | 0 | 0 | 0 | 0 | 0 |
| ENSMUSG00 | Olfr872 | 1122 | 0 | 0 | 0 | 0 | 0 | 0 |
| ENSMUSG00 | Olfr870 | 1810 | 0 | 0 | 0 | 0 | 0 | 0 |
| ENSMUSG00 | Suds3 | 3580 | 22.2097657 | 19.799513 | 23.8294852 | 18.2325192 | 21.3797664 | 20.7759612 |
| ENSMUSG00 | Gm10186 | 460 | 0 | 0 | 0 | 0 | 0 | 0 |
| ENSMUSG00 | Olfr859 | 929 | 0 | 0 | 0 | 0 | 0 | 0 |
| ENSMUSG00 | Olfr860 | 1217 | 0 | 0 | 0 | 0 | 0 | 0 |
| ENSMUSG00 | Olfr834 | 938 | 0 | 0 | 0 | 0 | 0 | 0 |
| ENSMUSG00 | Olfr833-ps1 | 265 | 0 | 0 | 0 | 0 | 0 | 0 |
| ENSMUSG00 | Olfr831-ps1 | 328 | 0 | 0 | 0 | 0 | 0 | 0 |
| ENSMUSG00 | Gm17644 | 3197 | 0 | 0 | 0 | 0 | 0 | 0 |
| ENSMUSG00 | Gm10188 | 719 | 0 | 0 | 0 | 0 | 0 | 0 |
| ENSMUSG00 | Gm10190 | 323 | 0 | 0 | 0 | 0 | 0 | 0 |
| ENSMUSG00 | 4921513D11 | 1344 | 0.32537968 | 0 | 0 | 0 | 0 | 0 |
| ENSMUSG00 | Myo1h | 5121 | 0 | 0.03763831 | 0 | 0 | 0 | 0 |
| ENSMUSG00 | Gm6396 | 416 | 0 | 0 | 0.10625443 | 0 | 0 | 0 |
| ENSMUSG00 | Gm10191 | 377 | 0 | 0 | 0 | 0 | 0 | 0 |
| ENSMUSG00 | Gm16391 | 446 | 0 | 0 | 0 | 0 | 0 | 0 |
| ENSMUSG00 | Tmem211 | 1822 | 0.02181969 | 0 | 0 | 0 | 0 | 0.16466577 |
| ENSMUSG00 | Cryba4 | 1165 | 0 | 0 | 0 | 0 | 0 | 0 |
| ENSMUSG00 | Bub3 | 2047 | 48.3785549 | 58.2851152 | 63.0961335 | 57.6043583 | 41.0803456 | 55.6018904 |
| ENSMUSG00 | Gm16519 | 527 | 0 | 0 | 0 | 0 | 0 | 0 |
| ENSMUSG00 | Gm10192 | 326 | 0 | 0 | 0 | 0 | 0 | 0 |
| ENSMUSG00 | Serpinb7 | 1948 | 0 | 0 | 0 | 0 | 0 | 0 |
| ENSMUSG00 | Serpinb6 | 2799 | 0 | 0 | 0 | 0 | 0 | 0 |
| ENSMUSG00 | Vmn2r10 | 5081 | 0 | 0 | 0 | 0 | 0 | 0 |
| ENSMUSG00 | Gm10193 | 230 | 0.17284092 | 0 | 0.19218193 | 0.31375691 | 0 | 0 |
| ENSMUSG00 | Gm3608 | 868 | 0.18320498 | 0.44411467 | 0.20369513 | 0 | 0.05878615 | 0.47133593 |
| ENSMUSG00 | Gm626 | 3147 | 0.11369537 | 0 | 0 | 0 | 0 | 0 |
| ENSMUSG00 | Cntnap5b | 5100 | 0 | 0 | 0.88403686 | 0 | 0 | 0 |
| ENSMUSG00 | Zfp960 | 2444 | 12.3300549 | 13.1310027 | 9.00675852 | 19.2959215 | 5.95029342 | 7.91232374 |
| ENSMUSG00 | Rps12-ps3 | 468 | 0 | 0 | 0 | 0 | 0 | 0 |
| ENSMUSG00 | Olfr719-ps | 242 | 0 | 0 | 0 | 0 | 0 | 0 |

| | | | | | | | | |
|---|---|---|---|---|---|---|---|---|
| ENSMUSG00 | Unc93a | 3173 | 0 | 0 | 0 | 0 | 0 | 0.02578756 |
| ENSMUSG00 | Rps15a-ps5 | 392 | 0 | 0 | 0 | 0 | 0 | 0 |
| ENSMUSG00 | Gm6990 | 347 | 0 | 0 | 0 | 0 | 0.14705007 | 0 |
| ENSMUSG00 | Olfr1416 | 938 | 0 | 0 | 0 | 0 | 0 | 0 |
| ENSMUSG00 | Hes6 | 2566 | 17.9410937 | 17.0323862 | 15.1760809 | 11.1367807 | 14.1187556 | 15.4442911 |
| ENSMUSG00 | Asb18 | 1719 | 0 | 0 | 0 | 0 | 0 | 0 |
| ENSMUSG00 | Gm10197 | 200 | 0 | 0 | 0 | 0 | 0 | 0 |
| ENSMUSG00 | Gm6941 | 314 | 0 | 0 | 0 | 0 | 0 | 0 |
| ENSMUSG00 | 1700010H22I | 376 | 0 | 0 | 0 | 0 | 0 | 0 |
| ENSMUSG00 | AY702103 | 2120 | 0 | 0 | 0 | 0 | 0 | 0 |
| ENSMUSG00 | Gm5529 | 971 | 1.51488443 | 1.29026519 | 1.6387913 | 1.56070635 | 0.47295302 | 1.5729967 |
| ENSMUSG00 | Gm7027 | 861 | 0.04617361 | 0 | 0 | 0 | 0 | 0 |
| ENSMUSG00 | Gm10198 | 71 | 0 | 0 | 0 | 0 | 0 | 0 |
| ENSMUSG00 | Rpl30-ps6 | 347 | 0 | 0 | 0 | 0 | 0 | 0 |
| ENSMUSG00 | Slc22a7 | 2017 | 0.5715959 | 0.16723108 | 0.06574394 | 0 | 0 | 0 |
| ENSMUSG00 | Rpl7a-ps11 | 1651 | 0 | 0 | 0 | 0 | 0 | 0 |
| ENSMUSG00 | Polr1c | 2435 | 8.45722337 | 10.9037904 | 10.1292108 | 10.4022979 | 9.93285509 | 9.39770938 |
| ENSMUSG00 | Igj | 2001 | 0.15894245 | 0 | 0 | 0 | 0.15300263 | 0.42254564 |
| ENSMUSG00 | Xpo5 | 3677 | 8.8009141 | 10.7852711 | 7.90993003 | 10.1759805 | 10.6437939 | 8.91599575 |
| ENSMUSG00 | Gm7337 | 1277 | 0 | 0 | 0 | 0 | 0 | 0 |
| ENSMUSG00 | Col4a4 | 9107 | 8.92719421 | 12.8363137 | 5.35838747 | 13.4985754 | 12.9540991 | 10.1108139 |
| ENSMUSG00 | Gm5560 | 2155 | 0.84860887 | 0.69316924 | 0.90249703 | 0.65299292 | 0.61563145 | 1.21501873 |
| ENSMUSG00 | Mrgpra4 | 1502 | 0 | 0 | 0 | 0 | 0 | 0 |
| ENSMUSG00 | Olfr132 | 941 | 0 | 0 | 0 | 0 | 0 | 0 |
| ENSMUSG00 | Olfr130 | 1848 | 0 | 0 | 0 | 0 | 0 | 0 |
| ENSMUSG00 | Gm7335 | 620 | 1.09006963 | 0.62176054 | 0.49905307 | 0.11639369 | 0.32920243 | 0.92381843 |
| ENSMUSG00 | Olfr91 | 938 | 0 | 0 | 0 | 0 | 0 | 0 |
| ENSMUSG00 | Eif1ax | 5259 | 21.3480656 | 21.8712754 | 20.8359705 | 19.3343224 | 14.728663 | 22.5499394 |
| ENSMUSG00 | Frat1 | 577 | 0.27560125 | 0 | 0.15321263 | 0.12506775 | 0 | 0 |
| ENSMUSG00 | H2-M9 | 1437 | 0 | 0 | 0 | 0 | 0 | 0 |
| ENSMUSG00 | H2-K2 | 1111 | 2.71954682 | 2.08186249 | 1.35271167 | 3.1827546 | 3.53648149 | 2.35676451 |
| ENSMUSG00 | Lrrc66 | 3209 | 2.83702245 | 3.93420001 | 3.27829188 | 0.79832507 | 3.80034398 | 3.06829067 |
| ENSMUSG00 | 4921511M17 | 981 | 0 | 0 | 0 | 0 | 0 | 0 |
| ENSMUSG00 | Gm15144 | 981 | 0 | 0 | 0 | 0 | 0 | 0 |
| ENSMUSG00 | H2-T23 | 3276 | 2.94889553 | 4.89807242 | 4.16922146 | 4.54880476 | 5.40480843 | 10.9897814 |
| ENSMUSG00 | Usp51 | 2202 | 0.43330224 | 0 | 0 | 0 | 0.23172741 | 0.09909042 |
| ENSMUSG00 | Nipal1 | 4199 | 3.84394501 | 6.83951406 | 3.20013343 | 2.9817741 | 2.96509543 | 3.11134846 |
| ENSMUSG00 | Cnga1 | 5965 | 0 | 0 | 0 | 0 | 0 | 0 |
| ENSMUSG00 | Gm3695 | 1008 | 0.31551969 | 0.23902005 | 0.08770207 | 0.93068766 | 0.05062141 | 0.5141053 |
| ENSMUSG00 | Cyp2c54 | 1760 | 0 | 0 | 0 | 0 | 0 | 0 |
| ENSMUSG00 | Cyp2c66 | 2123 | 0 | 0 | 0 | 0 | 0 | 0 |
| ENSMUSG00 | Cyp2c65 | 1919 | 0 | 0 | 0 | 0 | 0 | 0 |
| ENSMUSG00 | H2-Q10 | 1789 | 0 | 0 | 0 | 0 | 0.02852229 | 0 |
| ENSMUSG00 | Lgi1 | 8140 | 0 | 0.00591971 | 0.02715101 | 0.03989415 | 0 | 0.00670139 |
| ENSMUSG00 | Foxd3 | 2372 | 0 | 0 | 0 | 0 | 0 | 0 |
| ENSMUSG00 | Vmn1r239-ps | 1894 | 0 | 0 | 0 | 0 | 0 | 0 |
| ENSMUSG00 | Rplp0 | 1654 | 22.4976601 | 30.3568759 | 30.2250813 | 21.2914604 | 19.7133338 | 21.915354 |
| ENSMUSG00 | Capn6 | 7782 | 1.54281099 | 3.30654846 | 0.94856178 | 2.14674719 | 1.66547153 | 0.81312212 |
| ENSMUSG00 | Ppp1r3c | 2717 | 7.02341945 | 4.45152631 | 12.5431067 | 10.4381623 | 5.4463191 | 3.56367204 |
| ENSMUSG00 | Gm16223 | 2527 | 0 | 0 | 0.0699673 | 0.02855722 | 0.02019247 | 0.06475973 |
| ENSMUSG00 | Rps28 | 1361 | 1.69420857 | 2.79700877 | 1.39653141 | 1.53766244 | 0.59986922 | 1.64329201 |
| ENSMUSG00 | Gm10203 | 195 | 0 | 0 | 0 | 0 | 0 | 0 |
| ENSMUSG00 | 2010002M12 | 6495 | 0.44070741 | 0.27450321 | 0.183749 | 0.01111071 | 0.34567522 | 0.0881859 |
| ENSMUSG00 | Crygd | 1024 | 0 | 0 | 0 | 0 | 0 | 0 |
| ENSMUSG00 | Gm7205 | 327 | 0 | 0 | 0 | 0 | 0 | 0 |
| ENSMUSG00 | Gm7931 | 272 | 0 | 0 | 0 | 0 | 0 | 0 |
| ENSMUSG00 | Gm10205 | 230 | 0 | 0 | 0 | 0 | 0 | 0 |
| ENSMUSG00 | Bmpr2 | 3494 | 2.34391213 | 4.02702951 | 2.77052193 | 3.86224518 | 2.40965999 | 2.77898442 |
| ENSMUSG00 | Tuba3b | 1770 | 0 | 0 | 0 | 0 | 0 | 0 |
| ENSMUSG00 | H2-Eb2 | 2204 | 0 | 0 | 0.02005528 | 0 | 0 | 0 |
| ENSMUSG00 | Rps25-ps1 | 816 | 0.0974399 | 0.05905201 | 0 | 0 | 0 | 0.06684961 |
| ENSMUSG00 | Rps15a-ps2 | 392 | 0 | 0 | 0 | 0 | 0 | 0 |
| ENSMUSG00 | Gm14149 | 1849 | 0 | 0 | 0 | 0 | 0 | 0 |
| ENSMUSG00 | B430203G13 | 1374 | 0 | 0.10521057 | 0.3860423 | 0.02626059 | 0.1485484 | 0.13895377 |
| ENSMUSG00 | Gm7903 | 3672 | 0 | 0 | 0 | 0 | 0 | 0 |
| ENSMUSG00 | Gm5128 | 3672 | 0 | 0 | 0 | 0 | 0 | 0 |
| ENSMUSG00 | Pramel3 | 3819 | 0 | 0 | 0 | 0 | 0 | 0 |
| ENSMUSG00 | Tmem128 | 1623 | 26.5526438 | 37.3199986 | 38.5098004 | 37.7049584 | 29.4902899 | 39.2734639 |
| ENSMUSG00 | Lyar | 2500 | 9.00064079 | 8.03749855 | 8.39835017 | 7.43290117 | 6.85794491 | 10.4734615 |
| ENSMUSG00 | Trmt2b | 6246 | 22.9774711 | 17.1036411 | 18.1662073 | 20.4152971 | 16.371575 | 19.126308 |
| ENSMUSG00 | B3galt4 | 1574 | 2.4247307 | 2.47973432 | 3.53839404 | 0.59601853 | 1.39398613 | 3.98549366 |
| ENSMUSG00 | Tspan6 | 2051 | 14.4406792 | 13.3446607 | 11.0989513 | 7.47677666 | 11.369602 | 9.28215417 |
| ENSMUSG00 | Gm17080 | 809 | 0 | 0.11912594 | 0.05463763 | 0.0446008 | 0 | 0.16857008 |
| ENSMUSG00 | Trim43a | 2265 | 0.01755209 | 0 | 0.03903033 | 0.07965131 | 0.0225282 | 0 |
| ENSMUSG00 | Gm2411 | 3057 | 0 | 0 | 0 | 0 | 0 | 0 |
| ENSMUSG00 | Actl9 | 1367 | 0 | 0 | 0 | 0 | 0 | 0 |
| ENSMUSG00 | Zfp563 | 2247 | 8.93480982 | 15.0971319 | 13.1012138 | 6.18228177 | 6.10863155 | 18.8628346 |
| ENSMUSG00 | Zfp763 | 4998 | 11.8439197 | 13.6518612 | 6.07576354 | 10.7062169 | 9.30072597 | 15.2198817 |
| ENSMUSG00 | Hmx1 | 2600 | 0 | 0 | 0 | 0 | 0 | 0 |
| ENSMUSG00 | Dnahc7a | 607 | 0 | 0 | 0 | 0 | 0 | 0 |
| ENSMUSG00 | Gm14920 | 347 | 0 | 0 | 0 | 0 | 0 | 0 |
| ENSMUSG00 | Hist1h4j | 426 | 0.27996817 | 0 | 0 | 0.16939927 | 0 | 0 |
| ENSMUSG00 | Gm14403 | 4007 | 12.8285092 | 19.1206496 | 6.70694299 | 11.2649458 | 12.0593905 | 7.11985841 |
| ENSMUSG00 | Olfr1456-ps1 | 468 | 0 | 0 | 0 | 0 | 0 | 0 |
| ENSMUSG00 | Olfr1453 | 923 | 0 | 0 | 0 | 0 | 0 | 0 |
| ENSMUSG00 | Olfr1436 | 947 | 0 | 0 | 0 | 0 | 0 | 0 |
| ENSMUSG00 | Olfr1433 | 1072 | 0 | 0 | 0 | 0 | 0 | 0 |
| ENSMUSG00 | Olfr262 | 938 | 0 | 0 | 0 | 0 | 0 | 0 |
| ENSMUSG00 | Olfr1432 | 929 | 0 | 0 | 0 | 0 | 0 | 0 |
| ENSMUSG00 | Olfr76 | 1047 | 0 | 0 | 0 | 0 | 0 | 0 |
| ENSMUSG00 | Olfr1428 | 944 | 0 | 0 | 0 | 0 | 0 | 0 |
| ENSMUSG00 | Olfr1427 | 937 | 0 | 0 | 0 | 0 | 0 | 0 |
| ENSMUSG00 | Olfr1425 | 938 | 0 | 0 | 0 | 0 | 0 | 0 |
| ENSMUSG00 | Olfr1424 | 941 | 0 | 0 | 0 | 0 | 0 | 0 |
| ENSMUSG00 | Olfr1423 | 932 | 0 | 0 | 0 | 0 | 0 | 0 |
| ENSMUSG00 | A630073D07 | 481 | 0 | 0 | 0 | 0 | 0 | 0 |
| ENSMUSG00 | Prb1 | 993 | 0 | 0 | 0 | 0 | 0 | 0 |
| ENSMUSG00 | Olfr1419 | 1159 | 0 | 0 | 0 | 0 | 0 | 0 |
| ENSMUSG00 | Gm7666 | 1467 | 0.32519821 | 0.82117318 | 0.51222313 | 0.6148951 | 0.13913122 | 0.53917146 |
| ENSMUSG00 | Gm17283 | 70 | 0 | 0 | 0 | 0 | 0 | 0 |
| ENSMUSG00 | Gm10212 | 335 | 0 | 0 | 0 | 0 | 0 | 0 |
| ENSMUSG00 | Rpl35a-ps5 | 332 | 0 | 0 | 0 | 0 | 0 | 0 |
| ENSMUSG00 | Dmrtc1c2 | 1627 | 0 | 0 | 0 | 0 | 0 | 0 |
| ENSMUSG00 | Dmrtc1c1 | 5352 | 0 | 0 | 0 | 0 | 0 | 0 |
| ENSMUSG00 | Hdac8 | 4826 | 2.55370886 | 3.39481772 | 4.80853037 | 3.92521205 | 3.27769923 | 4.77560509 |
| ENSMUSG00 | Ms4a15 | 1770 | 0 | 0 | 0 | 0 | 0 | 0 |
| ENSMUSG00 | Rpl35a-ps3 | 332 | 0 | 0 | 0 | 0 | 0 | 0 |
| ENSMUSG00 | A430093F15I | 2111 | 0 | 0 | 0 | 0 | 0 | 0.05168098 |
| ENSMUSG00 | Cbln4 | 2899 | 0 | 0 | 0 | 0 | 0 | 0 |
| ENSMUSG00 | Gm8210 | 482 | 0 | 0 | 0 | 0 | 0 | 0 |
| ENSMUSG00 | AY702102 | 859 | 0 | 0 | 0 | 0 | 0 | 0 |
| ENSMUSG00 | F830016D02 | 2692 | 0 | 0 | 0 | 0 | 0 | 0 |
| ENSMUSG00 | S1pr3 | 4482 | 15.6910855 | 8.49337111 | 17.9588479 | 16.181372 | 12.6939779 | 12.3837329 |
| ENSMUSG00 | Klra3 | 1498 | 0 | 0 | 0 | 0 | 0 | 0.03641474 |
| ENSMUSG00 | Krt77 | 2921 | 0 | 0 | 0 | 0 | 0 | 0 |
| ENSMUSG00 | Krt74 | 1639 | 0 | 0 | 0 | 0 | 0 | 0 |
| ENSMUSG00 | Dgat2l6 | 2163 | 0 | 0 | 0 | 0 | 0 | 0 |
| ENSMUSG00 | Klra7 | 1256 | 0 | 0 | 0 | 0 | 0 | 0 |
| ENSMUSG00 | Nms | 1151 | 0 | 0 | 0 | 0 | 0 | 0 |
| ENSMUSG00 | Pcna-ps2 | 776 | 0.4610816 | 0.18628779 | 7.80367589 | 0.32548236 | 0.06575564 | 7.55676216 |
| ENSMUSG00 | Klri1 | 832 | 0 | 0 | 0 | 0 | 0.36797867 | 0 |
| ENSMUSG00 | 5430421N21I | 1832 | 0 | 0 | 0 | 0 | 0 | 0 |
| ENSMUSG00 | Krt86 | 1619 | 0 | 0 | 0 | 0 | 0 | 0 |
| ENSMUSG00 | Krt81 | 1848 | 0 | 0 | 0 | 0 | 0 | 0 |
| ENSMUSG00 | Klk11 | 1465 | 0 | 0 | 0 | 0 | 0 | 0 |
| ENSMUSG00 | Gm17245 | 6241 | 0.0509604 | 0.00772095 | 0.04957746 | 0.08094033 | 0.24527981 | 0.36272954 |
| ENSMUSG00 | Gm10217 | 133 | 0 | 0 | 0.33234468 | 0 | 0 | 0 |
| ENSMUSG00 | Syngap1 | 2473 | 0.06430324 | 0 | 0 | 0.01459039 | 0.10316696 | 0.02205794 |
| ENSMUSG00 | Gpr113 | 3400 | 0 | 0 | 0 | 0 | 0 | 0 |
| ENSMUSG00 | BC061195 | 575 | 0 | 0 | 0 | 0 | 0 | 0 |
| ENSMUSG00 | 1700003E24I | 575 | 0 | 0 | 0 | 0 | 0 | 0 |
| ENSMUSG00 | Mageb18 | 3728 | 0 | 0 | 0 | 0 | 0 | 0 |
| ENSMUSG00 | Gm5072 | 1034 | 0 | 0 | 0 | 0 | 0 | 0 |
| ENSMUSG00 | Ankrd23 | 2270 | 2.25923215 | 1.52838054 | 2.43402219 | 2.81344576 | 5.25998764 | 1.46586167 |
| ENSMUSG00 | Slc22a27 | 2138 | 0 | 0 | 0 | 0 | 0 | 0 |
| ENSMUSG00 | Gm5474 | 520 | 0 | 0 | 0 | 0.06938855 | 0.09812765 | 0.73431721 |
| ENSMUSG00 | Obp1b | 790 | 0 | 0 | 0 | 0 | 0 | 0 |
| ENSMUSG00 | Obp1a | 750 | 0 | 0 | 0 | 0 | 0 | 0 |
| ENSMUSG00 | Gm10220 | 2022 | 0 | 0 | 0 | 0 | 0 | 0 |
| ENSMUSG00 | Gm5862 | 1967 | 0 | 0 | 0 | 0 | 0 | 0 |
| ENSMUSG00 | Tuba3a | 1504 | 0 | 0 | 0.02938952 | 0 | 0 | 0 |
| ENSMUSG00 | Wfdc13 | 1072 | 0 | 0 | 0 | 0 | 0 | 0 |
| ENSMUSG00 | Prkag1 | 1634 | 15.8875941 | 19.3748424 | 22.1550241 | 13.4700411 | 17.6125312 | 13.4704002 |
| ENSMUSG00 | Lpar5 | 1512 | 0 | 0 | 0.02923402 | 0 | 0 | 0 |
| ENSMUSG00 | Gm7275 | 743 | 0 | 0 | 0 | 0 | 0 | 0 |
| ENSMUSG00 | Gm10221 | 385 | 1.03260988 | 0.50063836 | 0.80366987 | 1.21835475 | 0.79521625 | 1.20433472 |
| ENSMUSG00 | BC003965 | 3020 | 19.232701 | 17.7109109 | 20.8129208 | 19.7375952 | 19.7853927 | 23.8517623 |
| ENSMUSG00 | Gbx1 | 3547 | 0 | 0 | 0 | 0 | 0 | 0 |
| ENSMUSG00 | Gm10222 | 296 | 0.13430906 | 0.81396017 | 1.19464441 | 0.97519039 | 0.34477281 | 0.27643216 |
| ENSMUSG00 | Gm5829 | 353 | 0 | 0.1365055 | 0 | 0 | 0 | 0 |
| ENSMUSG00 | Gm14243 | 2169 | 0 | 0 | 0 | 0 | 0 | 0 |
| ENSMUSG00 | Khdc1a | 1984 | 0 | 0 | 0 | 0 | 0 | 0 |
| ENSMUSG00 | Xlr5c | 1656 | 0 | 0 | 0 | 0 | 0 | 0 |
| ENSMUSG00 | Clec4b2 | 1168 | 0 | 0 | 0 | 0 | 0 | 0 |
| ENSMUSG00 | Xlr4b | 2558 | 0.07770813 | 0 | 0.24191783 | 0 | 0.01994776 | 0 |
| ENSMUSG00 | Gm14685 | 2859 | 0 | 0 | 0 | 0 | 0 | 0 |
| ENSMUSG00 | Defb41 | 465 | 0 | 0 | 0 | 0 | 0 | 0 |
| ENSMUSG00 | Pi15 | 6916 | 0.01149667 | 0 | 0.01917373 | 0.01043437 | 0.01475604 | 0 |
| ENSMUSG00 | Nnat | 1968 | 0.12120573 | 0.17139487 | 0.24706315 | 0.09167186 | 0 | 0 |
| ENSMUSG00 | Blcap | 3237 | 49.3596775 | 32.570879 | 65.2308323 | 42.1235855 | 38.9357888 | 43.5282009 |
| ENSMUSG00 | 4930444P10I | 2057 | 0.03865385 | 0.44508624 | 0.25786199 | 0.21049321 | 0.02480621 | 0.27844795 |
| ENSMUSG00 | Magi2 | 388 | 0 | 0 | 0 | 0 | 0 | 0 |
| ENSMUSG00 | Xlr9 | 1922 | 0 | 0 | 0 | 0 | 0.07964575 | 0 |
| ENSMUSG00 | Myt9 | 1210 | 0.32855769 | 2.50888087 | 1.4246875 | 0.35783846 | 0.63255838 | 0 |
| ENSMUSG00 | Pex26 | 6310 | 6.48310451 | 5.33792758 | 9.77203978 | 7.3765194 | 5.57974633 | 7.50376763 |
| ENSMUSG00 | 2900097C17 | 338 | 0 | 0 | 0 | 0 | 0 | 0 |
| ENSMUSG00 | Gm14651 | 2039 | 0 | 0 | 0 | 0 | 0 | 0 |
| ENSMUSG00 | Dcpp2 | 635 | 0 | 0 | 0 | 0 | 0 | 0 |
| ENSMUSG00 | Romo1 | 627 | 24.5380717 | 27.2825948 | 28.7629218 | 19.0480969 | 22.9496619 | 53.3747725 |
| ENSMUSG00 | Gm6460 | 1120 | 0 | 0 | 0 | 0 | 0 | 0 |
| ENSMUSG00 | Arfgef1 | 8029 | 26.4656923 | 28.7354197 | 24.6030684 | 36.9448858 | 29.1833501 | 29.0988367 |
| ENSMUSG00 | Speer3 | 1187 | 0 | 0 | 0 | 0 | 0 | 0 |
| ENSMUSG00 | Zic3 | 4258 | 0 | 0 | 0 | 0 | 0 | 0 |
| ENSMUSG00 | Tcea1-ps1 | 1326 | 1.94879806 | 1.3809086 | 0.33334723 | 1.6054605 | 0.84659145 | 2.32430938 |
| ENSMUSG00 | Gm8759 | 415 | 0.47898169 | 0.81278336 | 1.27812558 | 1.65294903 | 1.1065961 | 1.57732854 |
| ENSMUSG00 | Ccdc87 | 3162 | 0 | 0.25906689 | 0 | 0 | 0.45184647 | 0 |
| ENSMUSG00 | Htatsf1 | 3353 | 36.2340437 | 26.6584701 | 37.5049936 | 38.8261308 | 34.4081824 | 38.8254856 |
| ENSMUSG00 | Mtap7d3 | 2875 | 0.16593592 | 0.38549154 | 0.1998692 | 1.07932377 | 1.65059233 | 0 |
| ENSMUSG00 | 3110035E14I | 3250 | 0 | 0 | 0 | 0 | 0 | 0.31890347 |
| ENSMUSG00 | Gm5146 | 527 | 0 | 0 | 0.92261911 | 0 | 0 | 0 |
| ENSMUSG00 | Sprrb3 | 8214 | 21.7459671 | 26.9501479 | 20.9493274 | 25.13092 | 35.4525842 | 17.6219273 |
| ENSMUSG00 | Sixl1 | 724 | 0 | 0 | 0 | 0 | 0 | 0 |
| ENSMUSG00 | Gm13139 | 2909 | 0.77898329 | 0.2319045 | 0.85091207 | 0.66979388 | 0.40343989 | 2.48462681 |
| ENSMUSG00 | Rex2 | 1522 | 0 | 0 | 0 | 0 | 0 | 0 |
| ENSMUSG00 | Cxx1b | 1230 | 0.48482293 | 0.70516745 | 1.04215727 | 1.08539484 | 0.82969717 | 0.75393312 |
| ENSMUSG00 | Cxx1a | 1275 | 0.28062692 | 0.49131274 | 0.24267679 | 0.31129607 | 0.36018618 | 0.79149934 |
| ENSMUSG00 | Zfp760 | 1141 | 68.6748923 | 40.2468706 | 51.678579 | 91.1696182 | 53.3518548 | 98.0309339 |
| ENSMUSG00 | Gm10226 | 254 | 31.9295985 | 13.8488594 | 22.6229905 | 39.4913716 | 19.0846681 | 18.8989626 |
| ENSMUSG00 | Zfp948 | 2606 | 23.0508561 | 19.6185016 | 23.9497323 | 22.7070426 | 22.5173963 | 45.5903026 |
| ENSMUSG00 | Gm7168 | 1601 | 0 | 0 | 0 | 0 | 0 | 0 |
| ENSMUSG00 | Zfp160 | 3786 | 16.8115489 | 17.1948979 | 16.4618591 | 21.7102212 | 24.475409 | 18.6225416 |
| ENSMUSG00 | Vmn1r-ps144 | 1186 | 0.16760321 | 0 | 0.22361809 | 0 | 0 | 0 |
| ENSMUSG00 | Vmn1r227 | 1026 | 0 | 0 | 0 | 0 | 0 | 0 |
| ENSMUSG00 | Olfr1321 | 1055 | 0 | 0 | 0 | 0 | 0 | 0 |
| ENSMUSG00 | Gm6994 | 1218 | 0 | 0 | 0 | 0 | 0 | 0 |
| ENSMUSG00 | Vmn2r-ps113 | 2562 | 0 | 0 | 0 | 0 | 0 | 0 |
| ENSMUSG00 | Gtf2f2 | 1459 | 30.0005374 | 27.0491406 | 22.3281414 | 26.8575054 | 24.2016807 | 19.2735799 |
| ENSMUSG00 | Bpifb9b | 2052 | 0 | 0 | 0 | 0 | 0 | 0 |
| ENSMUSG00 | Bpifb9a | 2258 | 0.035213 | 0 | 0 | 0 | 0 | 0 |
| ENSMUSG00 | Bpifb3 | 1856 | 0 | 0 | 0 | 0 | 0 | 0 |
| ENSMUSG00 | Bpifb6 | 1676 | 0 | 0 | 0 | 0 | 0.15222666 | 0 |
| ENSMUSG00 | 2510049J12I | 2985 | 5.59373595 | 5.92443024 | 8.75152068 | 4.66588583 | 3.23081576 | 6.93515953 |
| ENSMUSG00 | Gm14648 | 1022 | 0 | 0 | 0 | 0 | 0 | 0 |
| ENSMUSG00 | Lrch1 | 6777 | 8.47671084 | 7.63645254 | 6.0592463 | 7.56567586 | 8.38769111 | 4.96231816 |
| ENSMUSG00 | Mll4 | 10261 | 10.1432461 | 9.44381007 | 10.6573784 | 14.2872378 | 9.97552967 | 15.1085712 |
| ENSMUSG00 | Mas1 | 4065 | 0 | 0.26078763 | 0 | 0.15977289 | 0 | 0.14761182 |
| ENSMUSG00 | Tcp1 | 4722 | 39.401874 | 36.359232 | 39.334211 | 38.8481818 | 28.398415 | 30.8558075 |
| ENSMUSG00 | Tmtsf4 | 6800 | 13.2303901 | 12.0820417 | 14.7296142 | 10.5805289 | 13.559509 | 10.697274 |
| ENSMUSG00 | 4930430D24I | 1189 | 0 | 0.08105373 | 0 | 0 | 0.08583074 | 0 |
| ENSMUSG00 | Rhox9 | 2158 | 0 | 0 | 0 | 0 | 0 | 0 |
| ENSMUSG00 | Gm14164 | 1819 | 0 | 0 | 0 | 0 | 0 | 0 |
| ENSMUSG00 | 1110057P08I | 457 | 0 | 0 | 0 | 0 | 0 | 0 |
| ENSMUSG00 | Gm7735 | 158 | 0 | 0 | 0 | 0 | 0 | 0 |
| ENSMUSG00 | Gm6358 | 155 | 0 | 0 | 0 | 0 | 0 | 0 |
| ENSMUSG00 | 1110025L11I | 236 | 0 | 0 | 0 | 0 | 0 | 0 |
| ENSMUSG00 | Gm10228 | 531 | 0 | 0 | 0 | 0 | 0 | 0 |
| ENSMUSG00 | Gm10229 | 519 | 0 | 0 | 0 | 0 | 0 | 0 |
| ENSMUSG00 | 2310034C09 | 598 | 0 | 0 | 0 | 0 | 0 | 0 |
| ENSMUSG00 | Tcf15 | 978 | 0 | 0 | 0 | 0 | 0.20869683 | 0 |
| ENSMUSG00 | Grxcr1 | 1067 | 0 | 0 | 0 | 0 | 0 | 0 |
| ENSMUSG00 | Cyp2d40 | 1155 | 0 | 0 | 0 | 0 | 0 | 0 |
| ENSMUSG00 | Cyp2d11 | 1576 | 0 | 0 | 0 | 0 | 0 | 0 |
| ENSMUSG00 | Cyp2d9 | 1642 | 0 | 0 | 0 | 0 | 0 | 1.01324787 |
| ENSMUSG00 | 1500009C09 | 1901 | 0 | 0 | 0 | 0 | 0 | 0 |
| ENSMUSG00 | Cenpm | 1195 | 0.66536369 | 0.64517412 | 0.81375778 | 0.81524285 | 1.28099688 | 1.84874124 |
| ENSMUSG00 | Gm6749 | 410 | 0 | 0 | 0 | 0 | 0 | 0 |
| ENSMUSG00 | Gm10230 | 858 | 0 | 0 | 0 | 0 | 0 | 0 |
| ENSMUSG00 | Tnfrsf13c | 3532 | 0 | 0 | 0 | 0 | 0 | 0 |
| ENSMUSG00 | Gm4907 | 1637 | 0 | 0 | 0 | 0 | 0 | 0 |
| ENSMUSG00 | Ccdc134 | 2036 | 5.97503784 | 6.22447656 | 5.21043336 | 8.18757592 | 10.5511317 | 9.41752033 |
| ENSMUSG00 | Ninl | 6767 | 14.0703969 | 19.5964369 | 10.4054287 | 20.2564929 | 25.4867963 | 10.0279744 |
| ENSMUSG00 | Mei1 | 4099 | 0 | 0 | 0 | 0 | 0 | 0 |
| ENSMUSG00 | Gm10231 | 440 | 0 | 0 | 0 | 0 | 0 | 0 |
| ENSMUSG00 | Agtr2 | 3392 | 0 | 0 | 0 | 0 | 0 | 0 |
| ENSMUSG00 | Cst7 | 982 | 0.28338937 | 0.09813939 | 0 | 0 | 0.05196169 | 0.08332375 |
| ENSMUSG00 | Zfp442 | 4730 | 1.82387722 | 1.50773646 | 3.58847943 | 4.32526834 | 1.51029442 | 2.65250191 |
| ENSMUSG00 | Zfp120 | 5413 | 17.545879 | 13.3707807 | 15.5886419 | 22.9303959 | 10.1242061 | 29.6932291 |
| ENSMUSG00 | Gm10232 | 1565 | 0.02540286 | 0.12316024 | 0.02824399 | 0.09222248 | 0.06520943 | 0.08713942 |
| ENSMUSG00 | BC049702 | 874 | 0 | 0 | 0 | 0 | 0 | 0 |
| ENSMUSG00 | A230006K03I | 896 | 0 | 0 | 0 | 0 | 0 | 0 |
| ENSMUSG00 | Insm1 | 3095 | 0 | 0 | 0 | 0 | 0 | 0 |
| ENSMUSG00 | Gm10233 | 629 | 0 | 0 | 0 | 0 | 0 | 0 |
| ENSMUSG00 | Csnka2ip | 1335 | 0 | 0 | 0 | 0.35136073 | 0 | 0 |
| ENSMUSG00 | 4930525M21 | 1302 | 0 | 0 | 0 | 0 | 0 | 0 |
| ENSMUSG00 | AY512915 | 5726 | 0 | 0 | 0 | 0 | 0 | 0 |
| ENSMUSG00 | Gm10234 | 58 | 0 | 0 | 0 | 0 | 0 | 0 |
| ENSMUSG00 | Olfr203 | 923 | 0 | 0 | 0 | 0 | 0 | 0 |
| ENSMUSG00 | Ndufaf2 | 989 | 16.4006431 | 15.8347762 | 16.1343431 | 11.3463254 | 7.94546196 | 19.525222 |
| ENSMUSG00 | Col8a1 | 5104 | 0.10904716 | 0 | 0.25114684 | 0 | 0.11996797 | 0 |
| ENSMUSG00 | Gm5931 | 476 | 0 | 0 | 0 | 0.22740784 | 0.42879307 | 0 |
| ENSMUSG00 | Macrod2 | 8191 | 1.01924685 | 1.42953308 | 1.57034994 | 0.76648465 | 1.04033754 | 2.31756183 |
| ENSMUSG00 | Pick1 | 7573 | 12.588623 | 13.413049 | 10.8213676 | 11.1919613 | 13.8936204 | 11.0459948 |
| ENSMUSG00 | Ssxb9 | 879 | 0 | 0 | 0 | 0 | 0 | 0 |
| ENSMUSG00 | Ssxb10 | 871 | 0 | 0 | 0 | 0 | 0 | 0 |
| ENSMUSG00 | Lgals1 | 796 | 0.99888142 | 1.69500048 | 2.60990782 | 1.26921765 | 1.21796626 | 2.94675751 |
| ENSMUSG00 | Pdxp | 1999 | 1.21314873 | 0.79547403 | 0.8844791 | 1.91330501 | 1.14866779 | 0.31381526 |
| ENSMUSG00 | Gm6723 | 446 | 0 | 0.10804135 | 0.79285817 | 0.24270434 | 0.8008624 | 0.5503851 |
| ENSMUSG00 | Il2rb | 3632 | 0 | 0 | 0 | 0 | 0 | 0 |
| ENSMUSG00 | Vmn1r-ps34 | 1856 | 0 | 0 | 0 | 0 | 0 | 0 |
| ENSMUSG00 | Vmn1r41 | 935 | 0 | 0 | 0 | 0 | 0 | 0 |
| ENSMUSG00 | Vmn1r43 | 1068 | 0 | 0 | 0 | 0 | 0 | 0 |
| ENSMUSG00 | Vmn1r42 | 1097 | 0 | 0 | 0 | 0 | 0 | 0 |
| ENSMUSG00 | Vmn1r44 | 1107 | 0 | 0 | 0 | 0 | 0 | 0 |
| ENSMUSG00 | Vmn1r-ps25 | 979 | 0 | 0 | 0 | 0 | 0 | 0 |
| ENSMUSG00 | Gm11808 | 503 | 0.3951837 | 0.19159619 | 0.17575286 | 0.07173369 | 0.10144409 | 0.27111967 |
| ENSMUSG00 | Gm7079 | 170 | 0 | 0 | 0.52002168 | 0.42449464 | 0 | 0 |
| ENSMUSG00 | D14Ertd668e | 4829 | 0.37046938 | 0.61867041 | 0.10068757 | 0.17185484 | 0.5600327 | 0.83591771 |
| ENSMUSG00 | Apol9a | 1542 | 0 | 0 | 0 | 0 | 0.26472828 | 0 |
| ENSMUSG00 | Amn1 | 2492 | 22.7493239 | 19.3171171 | 27.5995456 | 25.4833059 | 13.043259 | 37.1140858 |
| ENSMUSG00 | Apol7b | 2411 | 0 | 0 | 0 | 0 | 0 | 0 |
| ENSMUSG00 | Olfr461 | 941 | 0.67596991 | 0.20483079 | 2.01985043 | 0 | 0.81338537 | 0.5217253 |
| ENSMUSG00 | Gm5879 | 1295 | 0.03069921 | 0.07441922 | 0.0682654 | 0.13931291 | 0.19701303 | 0.08424599 |
| ENSMUSG00 | Ccdc48 | 3612 | 0.23113652 | 0.41356027 | 0.37936244 | 0.16982136 | 0.05650761 | 0.23408467 |
| ENSMUSG00 | 2310035K24I | 1713 | 37.7595837 | 34.0933846 | 51.2720736 | 41.0530676 | 33.9877962 | 33.3409776 |
| ENSMUSG00 | Cenpb | 6752 | 5.37570404 | 4.19633116 | 4.63490889 | 2.75211092 | 3.94487088 | 3.78096302 |
| ENSMUSG00 | Shroom4 | 8869 | 7.91164992 | 7.88890675 | 4.56023073 | 11.3058971 | 11.4721607 | 6.10750181 |
| ENSMUSG00 | Gm608 | 7406 | 4.4393441 | 2.82378016 | 2.50672078 | 6.15821014 | 3.96856501 | 4.57400782 |
| ENSMUSG00 | Cma2 | 1091 | 0 | 0 | 0 | 0 | 0 | 0 |
| ENSMUSG00 | Ddrgk1 | 2360 | 27.2055513 | 28.2380718 | 26.2401619 | 22.1231858 | 25.2969745 | 24.8129875 |
| ENSMUSG00 | 1700019G17 | 3438 | 3.87378882 | 1.75197943 | 4.30704404 | 6.61189298 | 8.74186601 | 4.10150916 |
| ENSMUSG00 | Noto | 2610 | 0 | 0 | 0 | 0 | 0 | 0 |
| ENSMUSG00 | Spr-ps1 | 616 | 6.97011671 | 3.98946193 | 0.78931863 | 7.61471719 | 4.30742134 | 0.26566207 |
| ENSMUSG00 | Gm6899 | 621 | 0 | 0 | 0 | 0 | 0 | 0 |
| ENSMUSG00 | Gm5416 | 291 | 0 | 0 | 0 | 0 | 0 | 0 |
| ENSMUSG00 | Slc4a5 | 5811 | 4.62479863 | 5.81288865 | 6.14611756 | 6.28377716 | 3.87241985 | 3.81591494 |
| ENSMUSG00 | Gm10029 | 422 | 0 | 0.22837176 | 0 | 0 | 0 | 0 |
| ENSMUSG00 | Tlx2 | 2236 | 0 | 0 | 0 | 0 | 0 | 0 |
| ENSMUSG00 | Aup1 | 1581 | 69.9807098 | 70.1614104 | 68.0501492 | 56.2341289 | 63.5167031 | 62.0019314 |
| ENSMUSG00 | Htra2 | 2410 | 5.4436965 | 7.15798601 | 4.93373269 | 5.59945421 | 9.46422821 | 6.4961174 |
| ENSMUSG00 | Dok1 | 3093 | 1.0668299 | 0.81011801 | 0.45730972 | 0.79326836 | 1.17131351 | 0.75836372 |
| ENSMUSG00 | Reg3d | 776 | 0 | 0 | 0 | 0 | 0 | 0 |
| ENSMUSG00 | Gm10238 | 353 | 0 | 0 | 0 | 0 | 0 | 0 |
| ENSMUSG00 | Gml | 1748 | 0.79601935 | 0 | 0.12643548 | 0.51604755 | 0.11676516 | 0.51491024 |
| ENSMUSG00 | D430041D05 | 17843 | 0 | 0.17553768 | 0 | 0.01819977 | 0 | 0 |
| ENSMUSG00 | Olfr1513 | 941 | 0 | 0 | 0 | 0 | 0 | 0 |
| ENSMUSG00 | Chrac1 | 1822 | 22.4742837 | 27.0817326 | 22.2464819 | 19.0311991 | 20.6682027 | 33.1277589 |
| ENSMUSG00 | Rnase13 | 613 | 0 | 0 | 0 | 0 | 0 | 0 |
| ENSMUSG00 | Cep152 | 5740 | 1.54450734 | 2.07352809 | 1.20900511 | 0.56574634 | 2.24018235 | 1.03586609 |
| ENSMUSG00 | Rpl34-ps1 | 764 | 0 | 0.06307126 | 0 | 0 | 0.06678845 | 0 |
| ENSMUSG00 | Gm10240 | 482 | 0.57736175 | 0.19994374 | 0.45852534 | 0.44915408 | 0.31759155 | 0 |
| ENSMUSG00 | Gm7247 | 1156 | 0 | 0 | 0 | 0 | 0 | 0 |
| ENSMUSG00 | Rnase12 | 544 | 0 | 0 | 0 | 0 | 0 | 0 |
| ENSMUSG00 | Pnp2 | 1366 | 0.90221076 | 1.5874011 | 0.84132351 | 1.63769163 | 1.08328324 | 0.49916983 |
| ENSMUSG00 | Vmn1r19 | 1117 | 0 | 0 | 0 | 0 | 0 | 0 |
| ENSMUSG00 | Gm606 | 6248 | 0 | 0 | 0 | 0 | 0 | 0 |
| ENSMUSG00 | Olfr742 | 1119 | 0 | 0 | 0 | 0 | 0 | 0 |
| ENSMUSG00 | Olfr740 | 935 | 0 | 0 | 0 | 0 | 0 | 0 |
| ENSMUSG00 | Olfr725 | 1109 | 0 | 0 | 0 | 0 | 0 | 0 |
| ENSMUSG00 | Duox2 | 6858 | 0 | 0 | 0 | 0 | 0 | 0 |
| ENSMUSG00 | Uty | 9082 | 9.12686378 | 7.07781478 | 9.4565273 | 15.5380837 | 9.64680547 | 12.8865313 |
| ENSMUSG00 | B630019A10I | 2404 | 0.09922333 | 0 | 0 | 0 | 0 | 0.36305676 |
| ENSMUSG00 | Gm5518 | 647 | 0 | 0.07447673 | 0 | 0 | 0 | 0 |
| ENSMUSG00 | Mfap1a | 4191 | 11.3356715 | 10.3938305 | 9.35505482 | 9.88360466 | 12.9422662 | 10.5363018 |
| ENSMUSG00 | Gm7551 | 791 | 0.15077932 | 0 | 0 | 0 | 0.12901739 | 0 |
| ENSMUSG00 | Gm4943 | 848 | 0 | 0 | 0 | 0 | 0.30086307 | 0.03216349 |
| ENSMUSG00 | Gm6557 | 782 | 0 | 0 | 0 | 0 | 0 | 0 |
| ENSMUSG00 | Gm5800 | 898 | 0 | 0 | 0 | 0 | 0 | 0 |
| ENSMUSG00 | Aard | 1344 | 0 | 0.75291316 | 0 | 0.69801574 | 0.5694908 | 0 |
| ENSMUSG00 | Gng5 | 2865 | 23.2288567 | 20.536002 | 31.1804275 | 24.923688 | 15.1743359 | 58.3477361 |
| ENSMUSG00 | Olfr169 | 941 | 0 | 0 | 0 | 0 | 0 | 0 |
| ENSMUSG00 | Doxl2 | 3920 | 0 | 0 | 0 | 0 | 0 | 0 |
| ENSMUSG00 | Gm10242 | 338 | 0 | 0 | 0 | 0 | 0 | 0 |
| ENSMUSG00 | Gm10243 | 332 | 0 | 0 | 0 | 0 | 0 | 0 |
| ENSMUSG00 | Clca6 | 3405 | 0 | 0 | 0 | 0 | 0 | 0 |
| ENSMUSG00 | Zfp467 | 4203 | 9.83718766 | 13.0010529 | 11.3054916 | 12.035933 | 10.5622048 | 11.1616416 |
| ENSMUSG00 | Myadm | 3234 | 58.9448142 | 66.1706832 | 59.2911549 | 54.5916648 | 48.5018798 | 36.8384738 |
| ENSMUSG00 | Olfr458 | 941 | 0 | 0 | 0 | 0 | 0 | 0 |
| ENSMUSG00 | Rpl7a-ps3 | 865 | 0 | 0 | 0 | 0 | 0 | 0 |
| ENSMUSG00 | Zfyve19 | 3589 | 11.6087332 | 14.2585683 | 12.8208745 | 15.4723507 | 15.2979605 | 12.485994 |
| ENSMUSG00 | Mgarn | 6449 | 0.7952329 | 0.10460695 | 0.06854062 | 0.58187822 | 0.35605317 | 0.0930442 |
| ENSMUSG00 | Gm5481 | 347 | 0 | 2.36071907 | 0 | 0.10398284 | 0.29410015 | 0.2358038 |
| ENSMUSG00 | Hemt1 | 1048 | 0 | 0 | 0 | 0.30986489 | 0 | 0.65063548 |
| ENSMUSG00 | Gm10244 | 1138 | 0.62882131 | 0.16937238 | 0.81567549 | 0.82437009 | 0.26903186 | 3.35540377 |
| ENSMUSG00 | Gm8225 | 877 | 0 | 0 | 0 | 0 | 0 | 0 |
| ENSMUSG00 | Gm4841 | 2638 | 0 | 0 | 0.56969775 | 0 | 1.33465502 | 0.04135654 |
| ENSMUSG00 | Actc1 | 1610 | 0 | 0 | 0 | 0 | 0 | 0 |
| ENSMUSG00 | Gjd2 | 2877 | 0 | 0 | 0 | 0 | 0 | 0 |
| ENSMUSG00 | Efcab1 | 2531 | 0 | 0 | 0 | 0 | 0 | 0 |
| ENSMUSG00 | Gm6155 | 681 | 0.05837809 | 0 | 0 | 0.10596783 | 0.1498572 | 0.04005087 |
| ENSMUSG00 | Gm7676 | 320 | 0 | 0.15058263 | 0 | 0 | 0 | 0 |
| ENSMUSG00 | 4930403O15I | 1136 | 0.10496807 | 0 | 0.85602161 | 0.03176236 | 0.31442309 | 0 |
| ENSMUSG00 | Olfr1298 | 938 | 0 | 0 | 0 | 0 | 0 | 0 |
| ENSMUSG00 | Olfr1281 | 917 | 0 | 0 | 0 | 0 | 0 | 0 |
| ENSMUSG00 | Olfr1280 | 1002 | 0 | 0 | 0 | 0 | 0 | 0 |
| ENSMUSG00 | Olfr1279 | 941 | 0 | 0 | 0 | 0 | 0 | 0 |
| ENSMUSG00 | Olfr1278 | 941 | 0 | 0 | 0 | 0 | 0 | 0 |
| ENSMUSG00 | Clec16a | 8577 | 4.69074808 | 3.50569429 | 4.11767198 | 3.61788018 | 3.19472587 | 7.77821705 |
| ENSMUSG00 | Gm10247 | 332 | 0 | 0 | 0 | 0 | 0 | 0 |
| ENSMUSG00 | Gm10248 | 2727 | 0 | 0 | 0 | 0 | 0 | 0 |
| ENSMUSG00 | Cd59b | 1320 | 2.37930527 | 2.66485627 | 3.68348692 | 2.67881846 | 2.74460052 | 3.59529336 |
| ENSMUSG00 | Rps19-ps9 | 411 | 0 | 0 | 0 | 0 | 0 | 0 |
| ENSMUSG00 | Gpr88 | 3448 | 0 | 0 | 0 | 0 | 0 | 0 |
| ENSMUSG00 | Myoz1 | 1288 | 0 | 0 | 0 | 0 | 0 | 0 |
| ENSMUSG00 | Flnc | 10052 | 0 | 0.10546177 | 0.04397318 | 0.05025354 | 0.02538121 | 0.05698044 |
| ENSMUSG00 | Gm10250 | 482 | 0.1649605 | 0.19994374 | 0.09170507 | 0.29943605 | 0.31759155 | 1.18831416 |
| ENSMUSG00 | Tmem167b | 2944 | 36.5820641 | 52.4420383 | 45.0726674 | 36.7929 | 50.3330827 | 32.7777424 |
| ENSMUSG00 | Trp53i11 | 3157 | 3.45042815 | 2.82372246 | 0.47604139 | 1.12006347 | 3.79828898 | 0.88122053 |
| ENSMUSG00 | Sars | 4010 | 32.1612915 | 37.0711157 | 41.6114607 | 22.0271434 | 32.4990932 | 26.8801432 |
| ENSMUSG00 | Celsr2 | 12196 | 5.14057009 | 8.92136654 | 3.54818008 | 9.00571856 | 7.29247073 | 4.27368282 |
| ENSMUSG00 | Cry2 | 6307 | 3.92701195 | 2.16216317 | 3.18180383 | 3.54899027 | 4.37692553 | 1.96332428 |
| ENSMUSG00 | Psrc1 | 3060 | 0 | 0.03149441 | 0.07222523 | 0.14149821 | 0 | 0 |
| ENSMUSG00 | Mybphl | 1266 | 0 | 0.03806196 | 0.24440198 | 0 | 0 | 0 |
| ENSMUSG00 | Sort1 | 7179 | 48.5494216 | 54.9522777 | 47.8530051 | 50.5218988 | 58.9585997 | 31.8223121 |
| ENSMUSG00 | Ptprz1 | 7303 | 0 | 0 | 0 | 0 | 0 | 0 |
| ENSMUSG00 | Psma5 | 1311 | 17.4366143 | 21.1711599 | 16.1837412 | 15.3850976 | 16.8141833 | 19.7434269 |
| ENSMUSG00 | Il3ra | 1382 | 1.55339794 | 0.45327333 | 0.25587174 | 0 | 0.11076637 | 0.21709192 |
| ENSMUSG00 | Gstm6 | 2063 | 1.40676203 | 0.88758352 | 0.74991009 | 0.57717279 | 0.4946813 | 0.50239277 |
| ENSMUSG00 | 4930469B13I | 569 | 0 | 0 | 0 | 0 | 0 | 0 |
| ENSMUSG00 | Gm10251 | 1684 | 0 | 0 | 0 | 0 | 0 | 0 |
| ENSMUSG00 | Col28a1 | 4860 | 0.0818014 | 0 | 0.18190059 | 0 | 0.41997017 | 0 |
| ENSMUSG00 | Rap1a | 2393 | 34.7715089 | 29.8623041 | 44.6451544 | 29.0857768 | 22.3466963 | 41.6015183 |
| ENSMUSG00 | Olfr1259 | 929 | 0 | 0 | 0 | 0 | 0 | 0 |
| ENSMUSG00 | Olfr1189 | 920 | 0 | 0 | 0 | 0 | 0 | 0 |
| ENSMUSG00 | Olfr1188 | 967 | 0 | 0 | 0 | 0 | 0 | 0 |
| ENSMUSG00 | Olfr1187-ps1 | 296 | 0 | 0 | 0 | 0 | 0 | 0 |
| ENSMUSG00 | Olfr1145 | 977 | 0 | 0 | 0 | 0 | 0 | 0 |
| ENSMUSG00 | Olfr1143 | 944 | 0 | 0 | 0 | 0 | 0 | 0 |
| ENSMUSG00 | Olfr152 | 1894 | 0 | 0 | 0 | 0 | 0 | 0 |
| ENSMUSG00 | Olfr1140 | 1950 | 0 | 0 | 0 | 0 | 0 | 0 |
| ENSMUSG00 | Olfr1132 | 926 | 0 | 0 | 0 | 0 | 0 | 0 |
| ENSMUSG00 | Olfr1131 | 929 | 0 | 0 | 0 | 0 | 0 | 0 |
| ENSMUSG00 | Cadel | 4181 | 94.0591278 | 92.3506245 | 105.646739 | 101.609661 | 98.9162344 | 114.134678 |
| ENSMUSG00 | Olfr1083-ps | 1703 | 0 | 0 | 0 | 0 | 0 | 0 |
| ENSMUSG00 | Gm13721 | 955 | 0 | 0 | 0 | 0 | 0 | 0 |
| ENSMUSG00 | Hist4h4 | 311 | 0 | 0.15494033 | 1.42128113 | 0 | 0.16407195 | 0.26309941 |
| ENSMUSG00 | Hist2h2be | 2617 | 7.82348967 | 9.24325333 | 7.00946307 | 11.7332136 | 6.64883231 | 6.68056697 |
| ENSMUSG00 | Hist2h2ac | 519 | 0.38300078 | 0.09284478 | 0.08516733 | 0.06952224 | 0.09831672 | 0 |
| ENSMUSG00 | Sf3b4 | 2216 | 4.00066433 | 3.45742071 | 3.78986921 | 3.87523763 | 2.44079235 | 5.48939039 |
| ENSMUSG00 | Sp9 | 5165 | 0 | 0 | 0 | 0 | 0 | 0 |
| ENSMUSG00 | Gm128 | 2079 | 0.13385684 | 0.18542162 | 0.04252222 | 1.66612615 | 0.26998082 | 0.05247646 |
| ENSMUSG00 | Gm12258 | 1159 | 3.8760736 | 1.57988335 | 3.9282052 | 2.67735619 | 5.06301399 | 3.81233094 |
| ENSMUSG00 | Selenbp1 | 2358 | 30.2465361 | 31.7564593 | 28.86804 | 23.8251668 | 30.4253963 | 23.7467491 |
| ENSMUSG00 | Cgn | 6512 | 7.112275 | 8.72417311 | 6.36012391 | 10.5941138 | 10.7193001 | 5.54539656 |
| ENSMUSG00 | Selenbp2 | 2354 | 0.13510764 | 0.02047003 | 0.30043734 | 0.01532797 | 0.32514683 | 0.06951905 |
| ENSMUSG00 | Gm5773 | 1146 | 0 | 0 | 0 | 0 | 0 | 0 |
| ENSMUSG00 | Sab | 2302 | 112.306642 | 108.178772 | 105.91545 | 126.945473 | 93.8734586 | 176.372841 |
| ENSMUSG00 | Lce3f | 608 | 0 | 0 | 0 | 0 | 0 | 0 |
| ENSMUSG00 | Lce1j | 419 | 0 | 0 | 0 | 0 | 0 | 0 |
| ENSMUSG00 | Lce1i | 1261 | 0 | 0 | 0 | 0 | 0 | 0 |
| ENSMUSG00 | Lce1e | 709 | 0 | 0 | 0 | 0 | 0 | 0 |
| ENSMUSG00 | Lce1a2 | 1131 | 0 | 0 | 0 | 0 | 0 | 0 |
| ENSMUSG00 | Sprr2a2 | 3581 | 0 | 0 | 0 | 0 | 0 | 0 |
| ENSMUSG00 | Clk2 | 4429 | 15.6005927 | 16.319635 | 13.163746 | 20.9046119 | 18.7100552 | 21.2396096 |
| ENSMUSG00 | Dap3 | 6032 | 14.8555791 | 17.9341118 | 15.4545237 | 11.7721259 | 14.5668798 | 11.8150916 |
| ENSMUSG00 | Msto1 | 3434 | 10.5813946 | 10.9731502 | 11.687616 | 8.17467404 | 9.06409122 | 10.2617455 |
| ENSMUSG00 | Syt11 | 8541 | 3.68649345 | 2.89987487 | 2.27711169 | 2.26436903 | 3.32767724 | 1.68929682 |
| ENSMUSG00 | Gm10253 | 939 | 0 | 0 | 0 | 0 | 0 | 0 |
| ENSMUSG00 | Tcl1b3 | 1139 | 0 | 0 | 0 | 0 | 0 | 0 |
| ENSMUSG00 | Olfr366 | 929 | 0 | 0 | 0 | 0 | 0 | 0 |
| ENSMUSG00 | Olfr363-ps | 253 | 0 | 0 | 0 | 0 | 0 | 0 |
| ENSMUSG00 | Olfr338 | 920 | 0 | 0 | 0 | 0 | 0 | 0 |
| ENSMUSG00 | 4930589L23F | 1362 | 0.05837809 | 0 | 0 | 0.13245978 | 0 | 0 |
| ENSMUSG00 | Zfp619 | 5965 | 4.67201875 | 3.89369072 | 4.705477 | 0.4052803 | 6.49271069 | 3.75855046 |
| ENSMUSG00 | Zfp114 | 2019 | 1.47680091 | 1.26492394 | 1.62007746 | 0.10722747 | 0.4043695 | 1.16177265 |
| ENSMUSG00 | Gm6361 | 310 | 0 | 0 | 0 | 0 | 0 | 0 |
| ENSMUSG00 | Zbtb34 | 7177 | 0.95275779 | 0.45655261 | 1.38573425 | 1.48812668 | 1.92673093 | 0.58904404 |
| ENSMUSG00 | Rpl32-ps | 407 | 0 | 0 | 0 | 0 | 0 | 0 |
| ENSMUSG00 | Gm5848 | 356 | 0 | 0 | 0 | 0 | 0 | 0 |
| ENSMUSG00 | Vmn2r5 | 2958 | 0 | 0 | 0 | 0 | 0 | 0 |
| ENSMUSG00 | Gm5537 | 1253 | 0 | 0 | 0 | 0 | 0 | 0 |
| ENSMUSG00 | Gm10254 | 686 | 0.17385779 | 0 | 0.06443417 | 0 | 0 | 0.1590358 |
| ENSMUSG00 | Gm5641 | 889 | 0 | 0 | 0 | 0 | 0 | 0 |
| ENSMUSG00 | Urm1 | 4094 | 1.67994581 | 1.28293166 | 1.25242154 | 1.76267926 | 1.58288953 | 2.50494977 |
| ENSMUSG00 | Gm10256 | 1210 | 0 | 0 | 0 | 0 | 0 | 0 |
| ENSMUSG00 | Sry | 1312 | 0 | 0 | 0 | 0 | 0 | 0 |
| ENSMUSG00 | Gm4906 | 329 | 0 | 0 | 0 | 0 | 0 | 0 |
| ENSMUSG00 | Slc25a31 | 1619 | 0 | 0 | 0 | 0 | 0 | 0.05053979 |
| ENSMUSG00 | Usp9y | 8051 | 0 | 0 | 0 | 0 | 0 | 0 |
| ENSMUSG00 | Ddx3y | 4623 | 25.3943184 | 28.4657524 | 29.0758825 | 36.3005817 | 36.0595219 | 28.2127028 |
| ENSMUSG00 | Eif2s3y | 3566 | 45.5192448 | 36.2141517 | 42.4417023 | 39.6032312 | 43.7430261 | 56.1784143 |
| ENSMUSG00 | Ube1y1 | 3955 | 0 | 0 | 0 | 0 | 0 | 0 |

| | | | | | | | | |
|---|---|---|---|---|---|---|---|---|
| ENSMUSG00 | Slc38a9 | 1406 | 0.93309449 | 2.87884861 | 1.06889236 | 0.87253877 | 1.19763187 | 0.79534866 |
| ENSMUSG00 | Gm10257 | 1024 | 0 | 0.04705707 | 0.04316586 | 0.14094549 | 0.09966089 | 0.02663539 |
| ENSMUSG00 | Slc7a14 | 9768 | 0 | 0 | 0 | 0 | 0 | 0 |
| ENSMUSG00 | Gm10258 | 2122 | 0 | 0 | 0 | 0 | 0.19237088 | 0 |
| ENSMUSG00 | Lcn11 | 646 | 0 | 0 | 0 | 0 | 0 | 0 |
| ENSMUSG00 | Gm10259 | 1324 | 0 | 0 | 0 | 0 | 0 | 0 |
| ENSMUSG00 | Dytn | 3374 | 0 | 0 | 0 | 0 | 0 | 0 |
| ENSMUSG00 | Cdk7 | 8373 | 12.3212077 | 9.70289519 | 11.8832376 | 8.91170048 | 10.0614531 | 9.50850018 |
| ENSMUSG00 | Pde7a | 10891 | 2.23763746 | 2.21221385 | 3.75417361 | 4.04850412 | 4.58679845 | 3.75398809 |
| ENSMUSG00 | Gm8910 | 1675 | 0.04746923 | 0.05753605 | 0.18472412 | 0.12924912 | 0 | 0.03256673 |
| ENSMUSG00 | Zbtb10 | 5388 | 1.54211125 | 1.50247258 | 3.1338352 | 4.18546359 | 3.08734197 | 3.41692307 |
| ENSMUSG00 | Gm17352 | 425 | 0 | 0 | 0 | 0 | 0 | 0 |
| ENSMUSG00 | 1700008P02f | 784 | 0 | 0 | 0 | 0 | 0 | 0 |
| ENSMUSG00 | Rps24-ps2 | 398 | 0 | 0 | 0 | 0 | 0 | 0.20558773 |
| ENSMUSG00 | Nxph2 | 2577 | 0 | 0 | 0 | 0 | 0 | 0 |
| ENSMUSG00 | Fgfr1op | 4173 | 16.9387118 | 16.9974701 | 14.0984072 | 16.9731736 | 20.7260261 | 16.6406019 |
| ENSMUSG00 | A130006I12R | 405 | 0.09816168 | 0.35693661 | 1.09140353 | 0.71273174 | 0.75594631 | 0.47141352 |
| ENSMUSG00 | Gpr98 | 25128 | 0.44773961 | 0.22052853 | 0.85138857 | 0.4235993 | 0.40410096 | 0.392925 |
| ENSMUSG00 | Nr2f1 | 4942 | 0 | 0 | 0 | 0 | 0 | 0 |
| ENSMUSG00 | Gm17681 | 1421 | 0 | 0 | 0 | 0 | 0 | 0 |
| ENSMUSG00 | Zfp72 | 1626 | 19.1686942 | 17.75134 | 26.2329511 | 18.0854036 | 24.2579265 | 19.9611444 |
| ENSMUSG00 | Gm13192 | 520 | 0 | 0.92666235 | 0 | 0.62449692 | 0.39251058 | 0.10490246 |
| ENSMUSG00 | Zdhhc11 | 1451 | 0 | 0 | 0 | 0 | 0 | 0 |
| ENSMUSG00 | Gm3511 | 313 | 1.27014315 | 1.38555265 | 2.82439892 | 0.57639049 | 0.81511783 | 2.43990383 |
| ENSMUSG00 | Gm6132 | 1029 | 0 | 0.04682842 | 0.34364893 | 0.03506516 | 0.04958832 | 0.05301193 |
| ENSMUSG00 | Zfp748 | 4599 | 20.2364818 | 16.0935802 | 18.4823101 | 10.6465176 | 17.3860254 | 12.4423121 |
| ENSMUSG00 | Zfp874a | 3435 | 24.3046606 | 27.9158719 | 18.7230514 | 18.298376 | 20.6036633 | 31.7449804 |
| ENSMUSG00 | Zfp825 | 819 | 21.5524217 | 17.7095471 | 29.6298069 | 24.0106401 | 21.0585043 | 54.7490927 |
| ENSMUSG00 | 4930451E10f | 896 | 0.22184978 | 0.10755902 | 0.19732966 | 0.24162083 | 0.2847454 | 0.18264267 |
| ENSMUSG00 | Gprin1 | 8285 | 0 | 0 | 0 | 0 | 0 | 0 |
| ENSMUSG00 | Gm7251 | 1007 | 0.27635389 | 0.14355445 | 0.26336749 | 0.03583123 | 0.15201502 | 0.35210557 |
| ENSMUSG00 | C78339 | 4357 | 9.62635574 | 10.3738542 | 9.5160268 | 13.1094507 | 7.68264919 | 11.4181643 |
| ENSMUSG00 | Serpinb6e | 1486 | 0 | 0 | 0 | 0 | 0 | 0 |
| ENSMUSG00 | Dusp22 | 1820 | 9.74227709 | 9.61081236 | 10.054705 | 9.06016173 | 8.32683166 | 7.35815817 |
| ENSMUSG00 | Prl2b1 | 1184 | 0 | 0 | 0 | 0 | 0 | 0 |
| ENSMUSG00 | Prl6a1 | 1153 | 0 | 0 | 0 | 0 | 0 | 0 |
| ENSMUSG00 | Hist1h3a | 533 | 0 | 0 | 0 | 0 | 0 | 0 |
| ENSMUSG00 | Hist1h4b | 490 | 0 | 0 | 0 | 0 | 0 | 0 |
| ENSMUSG00 | Hist1h3b | 619 | 0.12845066 | 0 | 0 | 0 | 0 | 0 |
| ENSMUSG00 | Hist1h2bf | 460 | 0 | 0 | 0 | 0 | 0 | 0.17787808 |
| ENSMUSG00 | Hist1h2ac | 2481 | 1.55432552 | 0.99053146 | 0.89080699 | 0.91623088 | 0.61700575 | 0.53867687 |
| ENSMUSG00 | Hist1h2bn | 380 | 0 | 0 | 0 | 0.379811 | 0 | 0 |
| ENSMUSG00 | Hist1h2ae | 703 | 0 | 0 | 0 | 0.25662905 | 0.36291875 | 0 |
| ENSMUSG00 | Hist1h3e | 835 | 0.38089083 | 0.34624988 | 0.21174536 | 0.34569624 | 0.36665659 | 0.5879563 |
| ENSMUSG00 | Hist1h4f | 390 | 0 | 0 | 0 | 0 | 0 | 0 |
| ENSMUSG00 | Gm10264 | 428 | 0 | 0 | 0 | 0 | 0 | 0 |
| ENSMUSG00 | Vmn1r223 | 1085 | 0 | 0 | 0 | 0 | 0 | 0 |
| ENSMUSG00 | Vmn1r217 | 896 | 0 | 0 | 0 | 0 | 0 | 0 |
| ENSMUSG00 | Vmn1r204 | 908 | 0 | 0 | 0 | 0 | 0 | 0 |
| ENSMUSG00 | Vmn1r203 | 935 | 0 | 0 | 0 | 0 | 0 | 0 |
| ENSMUSG00 | Vmn1r201 | 902 | 0 | 0 | 0 | 0 | 0 | 0 |
| ENSMUSG00 | Vmn1r200 | 1062 | 0 | 0 | 0 | 0 | 0 | 0 |
| ENSMUSG00 | Vmn1r199 | 1103 | 0 | 0 | 0 | 0 | 0 | 0 |
| ENSMUSG00 | Vmn1r198 | 902 | 0 | 0 | 0 | 0 | 0 | 0 |
| ENSMUSG00 | Vmn1r197 | 896 | 0 | 0 | 0 | 0 | 0 | 0 |
| ENSMUSG00 | V1rh19 | 905 | 0 | 0 | 0 | 0 | 0 | 0 |
| ENSMUSG00 | Vmn1r195 | 1087 | 0 | 0 | 0 | 0 | 0 | 0 |
| ENSMUSG00 | Vmn1r194 | 890 | 0 | 0 | 0 | 0 | 0 | 0 |
| ENSMUSG00 | Vmn1r188 | 926 | 0 | 0 | 0 | 0 | 0 | 0 |
| ENSMUSG00 | Hist1h2bj | 462 | 0.60235577 | 0 | 0 | 0.07809966 | 0 | 0 |
| ENSMUSG00 | Hist1h2ag | 470 | 0 | 0 | 0 | 0 | 0 | 0 |
| ENSMUSG00 | Hist1h2ah | 530 | 0 | 0 | 0 | 0 | 0 | 0 |
| ENSMUSG00 | Hist1h2br | 2603 | 0 | 0 | 0 | 0 | 0 | 0 |
| ENSMUSG00 | Hist1h2ap | 518 | 0 | 0 | 0 | 0 | 0 | 0 |
| ENSMUSG00 | Hist1h4n | 429 | 0 | 0 | 0.1030346 | 0 | 0.1189426 | 0 |
| ENSMUSG00 | Hist1h4m | 401 | 0 | 0 | 0.22045807 | 0.17996032 | 0.25449564 | 0 |
| ENSMUSG00 | Hist1h2bq | 3928 | 0 | 0 | 0 | 0 | 0 | 0 |
| ENSMUSG00 | Hist1h2bp | 1087 | 0.10972074 | 0 | 0 | 0.03319415 | 0 | 0.22582498 |
| ENSMUSG00 | Hist1h2an | 392 | 0 | 0 | 0 | 0 | 0 | 0 |
| ENSMUSG00 | Hist1h3c | 629 | 0 | 0 | 0 | 0 | 0 | 0 |
| ENSMUSG00 | Hist1h3h | 487 | 0 | 0 | 0 | 0 | 0 | 0 |
| ENSMUSG00 | Hist1h2bl | 380 | 0 | 0 | 0 | 0 | 0 | 0 |
| ENSMUSG00 | Gm7731 | 469 | 0 | 0 | 0 | 0 | 0 | 0 |
| ENSMUSG00 | Gm5096 | 1223 | 0 | 0 | 0 | 0 | 0 | 0 |
| ENSMUSG00 | Gm5152 | 923 | 0 | 0 | 0 | 0 | 0 | 0 |
| ENSMUSG00 | Rps2-ps10 | 706 | 0 | 0 | 0 | 0 | 0 | 0 |
| ENSMUSG00 | Ctxn3 | 1470 | 0 | 0 | 0 | 0 | 0 | 0 |
| ENSMUSG00 | Gm5507 | 1007 | 0.5921869 | 0.14355445 | 0.04389458 | 0.32248103 | 0.15201502 | 0.02708504 |
| ENSMUSG00 | Prdm6 | 3389 | 0 | 0 | 0 | 0 | 0 | 0 |
| ENSMUSG00 | Gm4950 | 617 | 0 | 0.15619592 | 0 | 0 | 0 | 0 |
| ENSMUSG00 | Gm10267 | 254 | 0 | 0 | 0 | 0 | 0 | 0 |
| ENSMUSG00 | Olfr707 | 923 | 0 | 0 | 0 | 0 | 0 | 0 |
| ENSMUSG00 | Rps19-ps13 | 470 | 0 | 0 | 0 | 0 | 0 | 0 |
| ENSMUSG00 | Rpl17-ps3 | 581 | 0 | 0 | 0 | 0.12420669 | 0.08782509 | 0.0469443 |
| ENSMUSG00 | Olfr810 | 1005 | 0 | 0 | 0 | 0 | 0 | 0 |
| ENSMUSG00 | Olfr764 | 929 | 0 | 0 | 0 | 0 | 0 | 0 |
| ENSMUSG00 | Olfr9 | 1051 | 0 | 0 | 0 | 0 | 0 | 0 |
| ENSMUSG00 | Gm8444 | 356 | 0 | 0 | 0 | 0.10135406 | 0 | 0 |
| ENSMUSG00 | Dsg1a | 5671 | 0 | 0 | 0 | 0 | 0 | 0 |
| ENSMUSG00 | Rdh16 | 6568 | 0 | 0.388837 | 0.03364939 | 0.02197445 | 0.25637491 | 0 |
| ENSMUSG00 | Gm6020 | 65 | 198.777403 | 171.247202 | 296.492362 | 294.20744 | 196.255292 | 287.432737 |
| ENSMUSG00 | Zfp616 | 5195 | 0 | 0 | 0 | 0 | 0 | 0 |
| ENSMUSG00 | Epc2 | 4683 | 3.5909819 | 3.14863363 | 2.52015633 | 2.89704223 | 3.01821612 | 4.16428936 |
| ENSMUSG00 | Lyz1 | 1260 | 0 | 0 | 0 | 0 | 0 | 0 |
| ENSMUSG00 | Lyz2 | 1309 | 0 | 0 | 0.23637349 | 0 | 0 | 0 |
| ENSMUSG00 | Gm10271 | 117 | 0 | 0 | 0 | 0 | 0 | 0 |
| ENSMUSG00 | Tmem19 | 3003 | 26.9272885 | 30.0222555 | 23.7421155 | 20.9186945 | 27.1698884 | 23.6689011 |
| ENSMUSG00 | Scyl2 | 11527 | 2.56943116 | 3.18121649 | 3.67358078 | 2.35705557 | 2.61174302 | 4.56194192 |
| ENSMUSG00 | Gm4925 | 1956 | 1.48371681 | 0.83759664 | 0.36156927 | 0 | 1.53913915 | 0.87847765 |
| ENSMUSG00 | I830134H01F | 2644 | 0 | 0 | 0 | 0 | 0 | 0.02063135 |
| ENSMUSG00 | Dazap1 | 2931 | 3.6757882 | 3.48533802 | 4.50916106 | 3.57004194 | 2.8899278 | 7.79807154 |
| ENSMUSG00 | C330046G03 | 2630 | 0.39301996 | 0.76951733 | 1.00840707 | 1.55029317 | 0.40743494 | 0.66371746 |
| ENSMUSG00 | Krtap10-4 | 634 | 0 | 0 | 0 | 0 | 0 | 0 |
| ENSMUSG00 | Krtap12-1 | 670 | 0 | 0 | 0 | 0 | 0 | 0 |
| ENSMUSG00 | Gm10272 | 314 | 0 | 0 | 0 | 0 | 0 | 0 |
| ENSMUSG00 | Gm11733 | 2308 | 0 | 0 | 0 | 0 | 0 | 0 |
| ENSMUSG00 | Ank3 | 10574 | 17.749728 | 15.6261878 | 18.1923984 | 16.3928638 | 23.7228734 | 14.9915078 |
| ENSMUSG00 | Cd300lh | 848 | 0 | 0 | 0 | 0 | 0 | 0 |
| ENSMUSG00 | Gm11710 | 848 | 0 | 0 | 0 | 0 | 0 | 0 |
| ENSMUSG00 | Dppa5b | 590 | 0 | 0 | 0 | 0 | 0 | 0 |
| ENSMUSG00 | Gm10273 | 629 | 0 | 0 | 0 | 0 | 0 | 0 |
| ENSMUSG00 | Strada | 3939 | 8.38710442 | 7.98825761 | 9.7627833 | 10.8640022 | 8.45905646 | 9.24388002 |
| ENSMUSG00 | Pex11c | 3739 | 5.17810083 | 4.74260784 | 3.71205635 | 5.61640811 | 8.80235689 | 6.74753712 |
| ENSMUSG00 | Gm3699 | 199 | 0.19977628 | 1.69500048 | 2.44331796 | 0.18131681 | 0.25641395 | 1.50764338 |
| ENSMUSG00 | Marcks | 4184 | 3.20210252 | 3.84662325 | 2.60943003 | 2.99246402 | 3.93917767 | 3.93083353 |
| ENSMUSG00 | Sult3a1 | 1082 | 0 | 0 | 0 | 0 | 0 | 0 |
| ENSMUSG00 | Nkain2 | 557 | 0 | 0 | 0 | 0 | 0 | 0 |
| ENSMUSG00 | Nkain2 | 217 | 0 | 0.22205734 | 0 | 0 | 0 | 0 |
| ENSMUSG00 | Pcgf1 | 1461 | 5.44222868 | 5.83778252 | 8.31999099 | 7.1373791 | 6.42631975 | 10.3610026 |
| ENSMUSG00 | Gm10275 | 490 | 0.08113363 | 0.39335871 | 0.27062353 | 0.22091048 | 0.10413546 | 0.11132506 |
| ENSMUSG00 | Taar7e | 1214 | 0 | 0 | 0 | 0 | 0 | 0 |
| ENSMUSG00 | Taar7d | 1076 | 0 | 0 | 0 | 0 | 0 | 0 |
| ENSMUSG00 | Taar7b | 1076 | 0 | 0 | 0 | 0 | 0 | 0 |
| ENSMUSG00 | Taar5 | 1121 | 0 | 0 | 0 | 0 | 0 | 0 |
| ENSMUSG00 | Taar4 | 1043 | 0 | 0 | 0 | 0 | 0 | 0 |
| ENSMUSG00 | Taar3 | 1106 | 0 | 0 | 0 | 0 | 0 | 0 |
| ENSMUSG00 | 4930444G20l | 1772 | 0 | 0 | 0 | 0 | 0 | 0 |
| ENSMUSG00 | 4933406P04f | 1788 | 2.57921462 | 1.45529523 | 1.28551221 | 0.98882561 | 1.76937097 | 5.55255519 |
| ENSMUSG00 | Gm11568 | 1134 | 0 | 0 | 0 | 0 | 0 | 0 |
| ENSMUSG00 | Gm11563 | 1017 | 0 | 0 | 0 | 0 | 0 | 0 |
| ENSMUSG00 | 4930572O03 | 1164 | 0 | 0 | 0 | 0.03099832 | 0 | 0 |
| ENSMUSG00 | Krtap3-2 | 1030 | 0 | 0 | 0 | 0 | 0 | 0 |
| ENSMUSG00 | Krtap3-3 | 731 | 0 | 0 | 0 | 0 | 0 | 0 |
| ENSMUSG00 | Gm6595 | 3144 | 25.8208305 | 24.6756272 | 23.5208917 | 52.5622659 | 15.8240192 | 45.7700371 |
| ENSMUSG00 | Arid1b | 8111 | 8.65592141 | 10.028198 | 10.3161249 | 10.6764772 | 13.9974955 | 8.98842445 |
| ENSMUSG00 | Ube2u | 1896 | 0 | 0 | 0 | 0 | 0.21530116 | 0 |
| ENSMUSG00 | Zfp820 | 3690 | 5.49465991 | 3.39625067 | 8.42110993 | 11.6655499 | 2.46143493 | 15.1747519 |
| ENSMUSG00 | Psmb3 | 2350 | 6.36087688 | 6.52054834 | 5.26660257 | 6.27981115 | 3.82154985 | 7.19586227 |
| ENSMUSG00 | Zfp125 | 541 | 25.793297 | 52.4617642 | 9.4776595 | 18.4078453 | 21.6932836 | 4.78944682 |
| ENSMUSG00 | Tmem100 | 1764 | 0 | 0 | 0 | 0 | 0.14463259 | 0 |
| ENSMUSG00 | Msi2 | 12668 | 2.31603605 | 1.84483932 | 2.02725535 | 2.8226481 | 1.81661632 | 3.33074415 |
| ENSMUSG00 | Gm11444 | 572 | 0.27801035 | 0 | 0 | 0 | 0 | 0.19073174 |
| ENSMUSG00 | Gm11428 | 530 | 0 | 0 | 0 | 0 | 0 | 0 |
| ENSMUSG00 | Slfn9 | 6294 | 0 | 0.01531187 | 0 | 0.31530226 | 0 | 0 |
| ENSMUSG00 | Gm11426 | 2217 | 0 | 0 | 0 | 0 | 0 | 0 |
| ENSMUSG00 | Gm10277 | 2294 | 1.69835967 | 4.15907391 | 2.4856311 | 1.57288773 | 2.46901819 | 1.6407586 |
| ENSMUSG00 | Fbp1 | 2096 | 3.86933112 | 5.86237727 | 1.68709325 | 2.94371642 | 4.28465751 | 9.68145592 |
| ENSMUSG00 | Cacng7 | 2206 | 0 | 0 | 0 | 0 | 0 | 0 |
| ENSMUSG00 | Fam57a | 6608 | 0.86634219 | 1.87407924 | 1.49167842 | 1.16305622 | 1.72970766 | 0.67691296 |
| ENSMUSG00 | E130309D14 | 1380 | 0.97948285 | 1.29195533 | 0.22421225 | 0 | 0.55463452 | 0.17787808 |
| ENSMUSG00 | Olfr23 | 1917 | 0 | 0 | 0 | 0 | 0 | 0 |
| ENSMUSG00 | Olfr390 | 1029 | 0 | 0 | 0 | 0 | 0 | 0 |
| ENSMUSG00 | Olfr382 | 938 | 0 | 0 | 0 | 0 | 0 | 0 |
| ENSMUSG00 | Olfr1 | 1879 | 0 | 0 | 0 | 0 | 0 | 0 |
| ENSMUSG00 | Olfr377-ps1 | 1326 | 0 | 0 | 0 | 0 | 0 | 0 |
| ENSMUSG00 | Spata22 | 3439 | 0 | 0 | 0 | 0 | 0 | 0 |
| ENSMUSG00 | Nlrp1a | 5565 | 0 | 0.00865884 | 0 | 0 | 0 | 0 |
| ENSMUSG00 | Ahnak | 9814 | 9.5439079 | 16.9737651 | 7.06670997 | 14.0703061 | 17.1370425 | 9.30739422 |
| ENSMUSG00 | Sat2 | 1393 | 6.67823578 | 11.3461256 | 7.7741935 | 4.53292017 | 7.39937395 | 7.16620098 |
| ENSMUSG00 | Sco1 | 3919 | 12.436902 | 6.83635158 | 15.1362269 | 13.1659412 | 8.46316517 | 9.33977185 |
| ENSMUSG00 | Slc47a2 | 2404 | 0 | 0 | 0 | 0 | 0 | 0 |
| ENSMUSG00 | Gm10279 | 866 | 0.04590702 | 0.22257017 | 0.15312417 | 0.33332143 | 0.23568765 | 0.18896979 |
| ENSMUSG00 | Pabpn1l | 1141 | 0.13937066 | 0 | 0.07747913 | 0.03162318 | 0 | 0.14342492 |
| ENSMUSG00 | 4930438A08f | 2055 | 0 | 0 | 0 | 0 | 0 | 0 |
| ENSMUSG00 | Irgm2 | 3472 | 3.97325799 | 3.1226813 | 4.6086023 | 5.73654625 | 20.6339377 | 18.7591701 |
| ENSMUSG00 | Gm10281 | 71 | 0 | 0 | 0 | 0 | 0 | 0 |
| ENSMUSG00 | 1700024J04R | 1819 | 0 | 0 | 0 | 0 | 0 | 0 |
| ENSMUSG00 | 9930111J21R | 7013 | 0.02834413 | 0 | 0.02521138 | 0.09261041 | 0.29831476 | 0 |
| ENSMUSG00 | 9930111J21R | 4891 | 0.03251317 | 0 | 0 | 0 | 0.31298125 | 0.10037692 |
| ENSMUSG00 | Atxn1l | 7522 | 2.26207733 | 1.33246211 | 0.928462 | 1.4294668 | 1.27532022 | 2.05955798 |
| ENSMUSG00 | Gm12166 | 1739 | 0 | 0 | 0 | 0 | 0.02934237 | 0 |
| ENSMUSG00 | Ccdc99 | 3448 | 0.61109062 | 1.00621341 | 0.14101516 | 0 | 0.25158016 | 0.21357751 |
| ENSMUSG00 | Fam196b | 5367 | 3.14814221 | 0.34117474 | 1.44127493 | 2.08411288 | 1.68281508 | 2.63243211 |
| ENSMUSG00 | Anp32-ps | 387 | 0.10272734 | 0 | 0 | 0 | 0 | 0 |
| ENSMUSG00 | Hba-a2 | 754 | 0 | 0 | 0 | 0 | 0 | 0 |
| ENSMUSG00 | Hba-a1 | 930 | 0 | 0 | 0 | 0 | 0 | 0 |
| ENSMUSG00 | B3gnt9-ps | 4838 | 0 | 0 | 0.02740916 | 0 | 0 | 0.06765103 |
| ENSMUSG00 | Ces3a | 2036 | 0 | 0 | 0 | 0 | 0 | 0 |
| ENSMUSG00 | 4932416K20f | 3552 | 0 | 0 | 0 | 0 | 0 | 0 |
| ENSMUSG00 | Gm12070 | 1375 | 0 | 0 | 0 | 0 | 0.03711009 | 0.0198361 |
| ENSMUSG00 | Gm5356 | 617 | 0 | 0 | 0 | 0.05847981 | 0 | 0 |
| ENSMUSG00 | 4933402J07R | 1298 | 0 | 0 | 0 | 0 | 0 | 0 |
| ENSMUSG00 | Gm10159 | 520 | 0.99388701 | 2.40932211 | 1.27505316 | 0.27755419 | 0.49063823 | 1.10147582 |
| ENSMUSG00 | Gm8054 | 467 | 0 | 0 | 0 | 0 | 0 | 0 |
| ENSMUSG00 | Gm7125 | 272 | 0 | 0 | 0.16250678 | 0.13265458 | 0 | 0 |
| ENSMUSG00 | Olfr372 | 1015 | 0 | 0 | 0 | 0 | 0 | 0.05374313 |
| ENSMUSG00 | Fcho1 | 4777 | 1.09853955 | 0.48418447 | 0.40713441 | 1.90342793 | 1.71975016 | 0.37683194 |
| ENSMUSG00 | Ell | 3073 | 9.18528837 | 10.7725629 | 8.08377344 | 8.47746051 | 10.2451266 | 7.31347307 |
| ENSMUSG00 | Ssbp4 | 1457 | 30.3691488 | 33.5023102 | 29.1240695 | 29.4946569 | 33.6907162 | 52.4901088 |
| ENSMUSG00 | Gm10283 | 351 | 0 | 0 | 0 | 0 | 0 | 0 |
| ENSMUSG00 | Gm10284 | 2235 | 0 | 0 | 0.03955422 | 0.04843228 | 0.02283059 | 0.1342376 |
| ENSMUSG00 | Sp140 | 3310 | 1.06895401 | 0.33483026 | 1.53571358 | 0.54504599 | 1.09452347 | 0.73336643 |
| ENSMUSG00 | Sp110 | 6090 | 0.82252718 | 0.11868582 | 0.60968059 | 0.21921768 | 0.80435666 | 0.90915464 |
| ENSMUSG00 | Gm4885 | 1357 | 0.14648298 | 0.14203815 | 0.45602491 | 0.05317914 | 0.30081872 | 0.14069453 |
| ENSMUSG00 | Fam149a | 8586 | 14.983547 | 19.8391653 | 10.2808153 | 19.3605846 | 21.4600796 | 13.9009808 |
| ENSMUSG00 | Fat1 | 14749 | 1.71970958 | 3.0645388 | 1.22574777 | 1.79321572 | 2.13806362 | 1.47570426 |
| ENSMUSG00 | Mfhas1 | 6661 | 2.73352501 | 4.02940223 | 7.39241152 | 3.60224585 | 5.57682054 | 5.52371841 |
| ENSMUSG00 | U6 | 104 | 0 | 0 | 0 | 0 | 0 | 0 |
| ENSMUSG00 | Snora33 | 128 | 2.17412784 | 0.37645658 | 3.10794209 | 4.51025557 | 5.97965342 | 1.70466495 |
| ENSMUSG00 | Mir451 | 71 | 0 | 0 | 0 | 0 | 0 | 0 |
| ENSMUSG00 | Mir465 | 73 | 0 | 0 | 0 | 0 | 0 | 0 |
| ENSMUSG00 | Mir433 | 123 | 0 | 0 | 0 | 0 | 0 | 0 |
| ENSMUSG00 | Mir470 | 74 | 0 | 0 | 0 | 0 | 0 | 0 |
| ENSMUSG00 | Mir484 | 66 | 0 | 0 | 0 | 0 | 1.54625381 | 0 |
| ENSMUSG00 | Mir490 | 83 | 0 | 0 | 0 | 0 | 0 | 0 |
| ENSMUSG00 | Mir127 | 96 | 0 | 0 | 0 | 0 | 0 | 0 |
| ENSMUSG00 | Mir491 | 85 | 0.46771154 | 0 | 0 | 0 | 0 | 0 |
| ENSMUSG00 | Mir463 | 74 | 0 | 0 | 0 | 0 | 0 | 0 |
| ENSMUSG00 | Mir431 | 90 | 0 | 0 | 0 | 0 | 0 | 0 |
| ENSMUSG00 | Mir505 | 89 | 0 | 0 | 0 | 0 | 0 | 0 |
| ENSMUSG00 | Mir486 | 127 | 0 | 0 | 0 | 0 | 0 | 0 |
| ENSMUSG00 | Mir466 | 72 | 0 | 0 | 0 | 0 | 0 | 0 |
| ENSMUSG00 | Mir362 | 64 | 0 | 0 | 0 | 0 | 0 | 0 |
| ENSMUSG00 | Mir341 | 95 | 0 | 0 | 0 | 0 | 0 | 0 |
| ENSMUSG00 | Mir455 | 81 | 0.4908084 | 0 | 0 | 0 | 0 | 2.02034365 |
| ENSMUSG00 | Mir488 | 108 | 0 | 0 | 0 | 0 | 0 | 0 |
| ENSMUSG00 | Mir471 | 66 | 0 | 0 | 0 | 0 | 0 | 0 |
| ENSMUSG00 | Mir495 | 79 | 0 | 0 | 0 | 0 | 0 | 0 |
| ENSMUSG00 | Mir363 | 74 | 0 | 0 | 0 | 0 | 0 | 0 |
| ENSMUSG00 | Mir489 | 106 | 0 | 0 | 0 | 0 | 0 | 0 |
| ENSMUSG00 | Mir500 | 91 | 0 | 0 | 0 | 0 | 0 | 0 |
| ENSMUSG00 | Mir497 | 83 | 0 | 0 | 0 | 0 | 0 | 0 |
| ENSMUSG00 | Mir504 | 78 | 0 | 0 | 0 | 0 | 0 | 0 |
| ENSMUSG00 | Gm10286 | 5075 | 0 | 0 | 0 | 0.2275124 | 0.01005446 | 0.01612294 |
| ENSMUSG00 | Mir199a-2 | 109 | 0 | 0 | 0 | 0 | 0 | 0 |
| ENSMUSG00 | Mir146b | 108 | 0 | 0 | 0 | 0 | 0 | 0 |
| ENSMUSG00 | Mir485 | 78 | 0 | 0 | 0 | 0 | 0 | 0 |
| ENSMUSG00 | Mir136 | 81 | 0 | 0 | 0 | 0 | 0 | 0 |
| ENSMUSG00 | Mir328 | 96 | 0 | 0.50194211 | 0 | 0 | 0 | 0 |
| ENSMUSG00 | Mir434 | 93 | 0 | 0 | 0 | 0 | 0 | 0 |
| ENSMUSG00 | Mir496 | 78 | 0 | 0 | 0 | 0 | 0 | 0 |
| ENSMUSG00 | Mir452 | 84 | 0 | 0 | 0 | 0 | 0 | 0 |
| ENSMUSG00 | Mir532 | 95 | 0 | 0 | 0 | 0 | 0 | 0 |
| ENSMUSG00 | Mir483 | 72 | 0 | 0 | 0 | 0 | 0 | 0 |
| ENSMUSG00 | Mir494 | 84 | 0 | 0 | 0 | 0 | 0 | 0 |
| ENSMUSG00 | Mir1a-1 | 76 | 0 | 0 | 0 | 0 | 0 | 0 |
| ENSMUSG00 | U6 | 106 | 0 | 0 | 0 | 0 | 0 | 0 |
| ENSMUSG00 | U6 | 106 | 0 | 0 | 0 | 0 | 0 | 0 |
| ENSMUSG00 | n-R5s152 | 118 | 0 | 0 | 0 | 0 | 0 | 0 |
| ENSMUSG00 | U6 | 106 | 0 | 0 | 0 | 0 | 0 | 0 |
| ENSMUSG00 | U6 | 106 | 0 | 0 | 0 | 0 | 0 | 0 |
| ENSMUSG00 | U6 | 106 | 0.75010341 | 0 | 0 | 0 | 0 | 0.51461584 |
| ENSMUSG00 | SNORA21 | 132 | 8.13180283 | 9.49126891 | 12.0550481 | 14.2141388 | 11.5969036 | 9.29817247 |
| ENSMUSG00 | SNORA57 | 147 | 0 | 0 | 0 | 0.73636826 | 0.69423641 | 2.96866822 |
| ENSMUSG00 | U3 | 213 | 0 | 0 | 0 | 0 | 0 | 0 |
| ENSMUSG00 | SNORA71 | 131 | 0 | 0 | 0 | 0 | 0 | 0.20820335 |
| ENSMUSG00 | U6 | 106 | 0 | 0 | 0 | 0 | 0 | 0 |
| ENSMUSG00 | U6 | 103 | 0 | 0 | 0 | 0 | 0 | 0 |
| ENSMUSG00 | U6 | 106 | 0 | 0 | 0 | 0 | 0 | 1.80115542 |
| ENSMUSG00 | U6 | 101 | 0.78723724 | 0 | 0 | 0 | 0 | 0 |
| ENSMUSG00 | n-R5s71 | 118 | 0 | 0 | 0 | 0 | 0 | 0 |
| ENSMUSG00 | U6 | 104 | 0 | 0 | 0 | 0 | 0 | 0 |
| ENSMUSG00 | U3 | 213 | 0 | 0 | 0 | 0.16939927 | 0 | 0 |
| ENSMUSG00 | U6 | 106 | 0 | 0 | 0 | 0 | 0 | 0 |
| ENSMUSG00 | n-R5s58 | 118 | 0 | 0 | 0 | 0 | 0 | 0 |
| ENSMUSG00 | U1 | 163 | 0 | 0 | 0 | 0 | 0.31304525 | 0 |
| ENSMUSG00 | n-R5s211 | 118 | 0 | 0 | 0 | 0 | 0 | 0 |
| ENSMUSG00 | n-R5s210 | 118 | 0 | 0 | 0 | 0 | 0 | 0 |
| ENSMUSG00 | U6 | 107 | 0 | 0 | 0 | 0 | 0 | 0 |
| ENSMUSG00 | n-R5s27 | 118 | 0 | 0 | 0 | 0 | 0 | 0 |
| ENSMUSG00 | U6 | 106 | 0 | 0 | 0 | 0 | 0 | 0 |
| ENSMUSG00 | n-R5s13 | 118 | 0 | 0 | 0 | 0 | 0 | 0 |
| ENSMUSG00 | U6 | 106 | 0 | 0 | 0 | 0 | 0 | 0 |
| ENSMUSG00 | n-R5s33 | 118 | 0 | 0 | 0 | 1.52890019 | 0 | 0 |
| ENSMUSG00 | U6 | 106 | 0 | 0 | 0 | 0 | 0 | 0 |
| ENSMUSG00 | U6 | 106 | 0 | 0 | 0 | 0 | 0 | 0 |
| ENSMUSG00 | U5 | 115 | 0 | 0 | 0 | 0 | 0 | 0.9486831 |
| ENSMUSG00 | Y_RNA | 109 | 0 | 0 | 0 | 0 | 0 | 0 |
| ENSMUSG00 | U6 | 110 | 0 | 0 | 0 | 0 | 0 | 0 |
| ENSMUSG00 | U5 | 115 | 0 | 0 | 0 | 0.94127073 | 0 | 0 |
| ENSMUSG00 | U6 | 106 | 0 | 0 | 0 | 0 | 0 | 0 |
| ENSMUSG00 | U6 | 106 | 0 | 0 | 0 | 0 | 0 | 0 |
| ENSMUSG00 | U3 | 213 | 0 | 0 | 0 | 0 | 0 | 0 |
| ENSMUSG00 | U1 | 163 | 0 | 0.29562234 | 0 | 1.10681118 | 0 | 0.16732908 |
| ENSMUSG00 | U6 | 106 | 0 | 0 | 0 | 0 | 0 | 0 |
| ENSMUSG00 | U6 | 106 | 0 | 0 | 0 | 0 | 0 | 0 |
| ENSMUSG00 | U3 | 213 | 0 | 0 | 0 | 0 | 0 | 0 |
| ENSMUSG00 | U6 | 106 | 0 | 0 | 0 | 0 | 0 | 0 |
| ENSMUSG00 | U6 | 106 | 0 | 0 | 0 | 0 | 0 | 0 |
| ENSMUSG00 | U5 | 116 | 0 | 2.49240218 | 0 | 0.62210422 | 0.8797651 | 0.4702524 |
| ENSMUSG00 | U6 | 106 | 0 | 0 | 0 | 0 | 0 | 0 |
| ENSMUSG00 | Gm13268 | 873 | 0 | 0 | 0 | 0 | 0 | 0 |
| ENSMUSG00 | Slc22a14 | 2783 | 0 | 0 | 0 | 0 | 0 | 0 |
| ENSMUSG00 | 3000002C10l | 5186 | 1.12689079 | 0.91058067 | 0.79266707 | 0.92535903 | 0.97408623 | 1.36215418 |
| ENSMUSG00 | Ndufaf3 | 1061 | 31.9617577 | 31.8820758 | 44.8684118 | 30.1307177 | 24.9601216 | 41.8502476 |
| ENSMUSG00 | Gmppb | 2730 | 5.54829234 | 4.46562999 | 6.42788706 | 9.0932039 | 5.68205797 | 5.83457484 |
| ENSMUSG00 | BY080835 | 1140 | 0.13949291 | 0.08453762 | 0.15509419 | 0.12660367 | 0.08951996 | 0.07177537 |
| ENSMUSG00 | Tmem22 | 1578 | 34.7167631 | 33.0098504 | 39.916113 | 38.5058067 | 23.4760132 | 31.0944715 |
| ENSMUSG00 | Rpl7a-ps10 | 865 | 0 | 0.11141374 | 0 | 0 | 0 | 0 |
| ENSMUSG00 | Ddx43 | 2190 | 0 | 0 | 0 | 0 | 0.16309801 | 0 |
| ENSMUSG00 | Trcg1 | 2687 | 0 | 0 | 0 | 0 | 0 | 0 |
| ENSMUSG00 | Scn2b | 4768 | 0 | 0 | 0 | 0 | 0.03210552 | 0.05720352 |
| ENSMUSG00 | Mpzl3 | 6511 | 4.79923325 | 2.87890124 | 4.89472106 | 4.44444781 | 2.10030237 | 3.20878109 |
| ENSMUSG00 | Ccdc153 | 1079 | 0 | 0 | 0 | 0 | 0 | 0 |
| ENSMUSG00 | Gm4893 | 329 | 0 | 0 | 0 | 0 | 0 | 0 |
| ENSMUSG00 | Olfr895 | 950 | 0 | 0 | 0 | 0 | 0 | 0 |
| ENSMUSG00 | Olfr143 | 941 | 0 | 0 | 0 | 0 | 0 | 0 |
| ENSMUSG00 | Olfr894 | 948 | 0 | 0 | 0 | 0 | 0 | 0 |
| ENSMUSG00 | Olfr893 | 941 | 0 | 0 | 0 | 0 | 0 | 0 |
| ENSMUSG00 | A630095E13f | 614 | 0 | 0 | 0 | 0 | 0 | 0 |
| ENSMUSG00 | 4930581F22f | 3403 | 0.43225177 | 0.04247997 | 0.11690173 | 0.07422107 | 0.10496169 | 0.09617857 |
| ENSMUSG00 | Eif3g | 1064 | 73.5700575 | 81.9260093 | 78.3087162 | 77.8273423 | 51.5539041 | 71.570294 |
| ENSMUSG00 | Mmp27 | 2344 | 0 | 0 | 0 | 0 | 0 | 0 |
| ENSMUSG00 | Fbxw22 | 1814 | 0 | 0 | 0 | 0 | 0 | 0 |
| ENSMUSG00 | Rnf213 | 16895 | 1.72952224 | 1.13514081 | 1.10929751 | 2.10362911 | 2.14434607 | 1.94853445 |
| ENSMUSG00 | Cldn27 | 1816 | 0 | 0 | 0 | 0 | 0 | 0 |
| ENSMUSG00 | Qrich2 | 2737 | 0 | 0 | 0 | 0 | 0 | 0 |
| ENSMUSG00 | 4933422H20l | 1926 | 0 | 0.02501892 | 0 | 0.07493675 | 0 | 0.18409673 |
| ENSMUSG00 | Krtap31-1 | 1415 | 0 | 0 | 0 | 0 | 0 | 0 |
| ENSMUSG00 | Krtap9-1 | 611 | 0 | 0 | 0 | 0 | 0 | 0 |
| ENSMUSG00 | Fbxo47 | 2191 | 1.48788197 | 0 | 0.84731967 | 0.60932709 | 0.62880518 | 0.01244849 |
| ENSMUSG00 | Gpr179 | 9531 | 0 | 0 | 0 | 0.05300059 | 0.00535373 | 0.22607245 |
| ENSMUSG00 | Gm10287 | 4052 | 0 | 0 | 0 | 0.00890475 | 0 | 0.02019346 |
| ENSMUSG00 | Gm10288 | 536 | 0 | 0.17980016 | 0 | 0.06731725 | 0 | 0 |
| ENSMUSG00 | Hsf5 | 4152 | 0 | 0 | 0.08516733 | 0.05214168 | 0.15976466 | 0.08539747 |
| ENSMUSG00 | Ccnd1 | 3830 | 5.44951104 | 14.6446524 | 4.20090623 | 11.7478615 | 9.24603259 | 7.07858784 |
| ENSMUSG00 | Evi2b | 3937 | 0.02019582 | 0.12239381 | 0.05613645 | 0.41241859 | 0.50546829 | 0.54036623 |
| ENSMUSG00 | E030019B06l | 1503 | 0 | 0 | 0 | 0 | 0 | 0 |
| ENSMUSG00 | Amy2a1 | 886 | 0 | 0 | 0 | 0 | 0 | 0 |
| ENSMUSG00 | Ppapdc1a | 1081 | 0.69875498 | 0.26745481 | 0 | 0 | 0 | 0 |
| ENSMUSG00 | Prok1 | 434 | 0 | 0 | 0 | 0 | 0 | 0 |
| ENSMUSG00 | Itgad | 4579 | 0 | 0 | 0 | 0 | 0 | 0 |
| ENSMUSG00 | Prss36 | 3418 | 3.76851249 | 2.31204696 | 1.78462678 | 4.81375433 | 3.03053081 | 1.89119061 |
| ENSMUSG00 | Capza1 | 5642 | 4.3335024 | 3.55291739 | 3.82320088 | 4.98190583 | 3.70804929 | 3.96405604 |
| ENSMUSG00 | Olfr59 | 1352 | 0 | 0 | 0 | 0 | 0 | 0 |
| ENSMUSG00 | Olfr406-ps | 1381 | 0 | 0 | 0 | 0 | 0 | 0 |
| ENSMUSG00 | Olfr43 | 1091 | 0 | 0 | 0 | 0 | 0 | 0 |
| ENSMUSG00 | Olfr403 | 941 | 0 | 0 | 0 | 0 | 0 | 0 |
| ENSMUSG00 | Olfr402 | 999 | 0 | 0 | 0 | 0 | 0 | 0 |
| ENSMUSG00 | Olfr401 | 947 | 0 | 0 | 0 | 0 | 0 | 0 |
| ENSMUSG00 | Olfr391-ps | 2056 | 0 | 0 | 0 | 0 | 0 | 0 |
| ENSMUSG00 | Olfr389 | 2080 | 0 | 0 | 0 | 0 | 0 | 0 |
| ENSMUSG00 | Ampd1 | 2598 | 0 | 0 | 0 | 0 | 0 | 0 |
| ENSMUSG00 | Fbxo39 | 1704 | 0 | 0 | 0 | 0 | 0 | 0 |
| ENSMUSG00 | Nlrp1b | 6928 | 0 | 0 | 0 | 0 | 0 | 0 |
| ENSMUSG00 | Hist2h3c1 | 2955 | 0 | 0 | 0 | 0 | 0 | 0 |
| ENSMUSG00 | 1810027O10l | 916 | 14.2355869 | 13.3617427 | 16.1655212 | 10.674928 | 11.5310696 | 26.9173296 |
| ENSMUSG00 | Gm12302 | 248 | 0 | 0 | 0 | 0 | 0 | 0 |
| ENSMUSG00 | Hs3st3b1 | 4909 | 2.44574356 | 3.61226537 | 2.49417611 | 2.59461432 | 3.3158309 | 0.73339833 |
| ENSMUSG00 | Gm10290 | 1248 | 0.22298747 | 0.19305466 | 0.14167257 | 0.08673568 | 0.12265956 | 0.13112807 |
| ENSMUSG00 | Olfr485 | 940 | 0 | 0 | 0 | 0 | 0 | 0 |
| ENSMUSG00 | Olfr478 | 944 | 0 | 0 | 0 | 0 | 0 | 0 |
| ENSMUSG00 | Olfr2 | 1248 | 0 | 0 | 0 | 0 | 0 | 0 |
| ENSMUSG00 | Cyp3a57 | 1697 | 0 | 0 | 0 | 0 | 0 | 0 |
| ENSMUSG00 | Zfp498 | 1991 | 0.09983797 | 0.07260639 | 0.2664099 | 0 | 0 | 0.08219379 |
| ENSMUSG00 | Olfr658 | 2910 | 0 | 0 | 0.03037927 | 0 | 0 | 0 |
| ENSMUSG00 | Olfr558 | 2913 | 0.15012368 | 0 | 0 | 0 | 0.42040268 | 0 |
| ENSMUSG00 | Art5 | 2145 | 2.26115087 | 1.12322709 | 2.26676118 | 0.85789477 | 2.02202422 | 2.59395171 |
| ENSMUSG00 | Trpc2 | 8025 | 2.51165466 | 4.08909247 | 2.9357735 | 3.16982447 | 3.93586625 | 3.34093089 |
| ENSMUSG00 | Rnf121 | 5709 | 8.48172627 | 9.24227608 | 8.23026084 | 10.1881688 | 6.57828212 | 10.0279327 |
| ENSMUSG00 | Il18bp | 1393 | 4.73755188 | 6.50326714 | 6.44147461 | 10.5163748 | 11.0990609 | 10.4360249 |
| ENSMUSG00 | Gm6109 | 347 | 0.91655287 | 0 | 1.27382833 | 0 | 0 | 1.49342405 |
| ENSMUSG00 | Olfr311 | 926 | 0 | 0 | 0 | 0 | 0 | 0 |
| ENSMUSG00 | Serpinh1 | 2429 | 19.8368227 | 25.5909882 | 12.8474686 | 24.4062574 | 19.8097457 | 18.0109186 |
| ENSMUSG00 | Olfr312 | 926 | 0 | 0 | 0 | 0 | 0 | 0 |
| ENSMUSG00 | Olfr313 | 1073 | 0 | 0 | 0 | 0 | 0 | 0 |
| ENSMUSG00 | Gm10291 | 1250 | 0.06360877 | 0.38549154 | 0.17680737 | 0.25979072 | 0.0816422 | 0.06545913 |
| ENSMUSG00 | Vmn2r89 | 3224 | 0 | 0 | 0 | 0 | 0 | 0 |
| ENSMUSG00 | Vmn2r73 | 2550 | 0 | 0 | 0 | 0 | 0 | 0 |
| ENSMUSG00 | Olfr290 | 947 | 0 | 0 | 0 | 0 | 0 | 0 |
| ENSMUSG00 | Olfr291 | 1154 | 0 | 0.50107219 | 0 | 0 | 0 | 0 |
| ENSMUSG00 | 9230112E08f | 1359 | 0.20477437 | 1.06371837 | 0.16262635 | 1.0620175 | 0.075094 | 0.52181061 |
| ENSMUSG00 | Mesdc1 | 3846 | 1.11637855 | 1.40324533 | 1.54005381 | 1.39787432 | 1.15426279 | 2.36862442 |
| ENSMUSG00 | Ccl26 | 279 | 0 | 0 | 0 | 0 | 0 | 0 |
| ENSMUSG00 | Gm9696 | 3045 | 0 | 0 | 0 | 0 | 0 | 0 |
| ENSMUSG00 | Adamtsl3 | 7878 | 0 | 0 | 0 | 0.22900511 | 0 | 0.27350806 |
| ENSMUSG00 | Fam194a | 2181 | 0 | 0 | 0.08106711 | 0.13235046 | 0 | 0.05002226 |
| ENSMUSG00 | Cldn3 | 1856 | 11.031289 | 16.3823518 | 16.2660877 | 12.6753734 | 13.9937636 | 8.89070945 |
| ENSMUSG00 | 9030418K01f | 4681 | 0.72190042 | 0.86470009 | 0.23607051 | 0.43165873 | 0.61044158 | 0.22141344 |
| ENSMUSG00 | 4930527J03R | 541 | 0 | 0 | 0 | 0 | 0 | 0 |
| ENSMUSG00 | Gm10293 | 1231 | 0.03229527 | 0.15657658 | 0.14362906 | 0.46897865 | 0.16580463 | 0.17725192 |
| ENSMUSG00 | Chchd2 | 2503 | 96.9188742 | 88.2487618 | 83.2294391 | 74.4705721 | 59.8127793 | 121.509989 |
| ENSMUSG00 | Ctcfl | 3206 | 0 | 0 | 0 | 0 | 0.0159159 | 0 |
| ENSMUSG00 | Tmem132b | 1219 | 0 | 0 | 0 | 0 | 0 | 0 |
| ENSMUSG00 | BC094916 | 3937 | 0 | 0 | 0 | 0 | 0.06480363 | 0 |
| ENSMUSG00 | Fcrl6 | 1907 | 0 | 0 | 0 | 0 | 0 | 0 |
| ENSMUSG00 | Rgma | 3777 | 0.01052568 | 0.3954937 | 0.15213767 | 0.07642477 | 0.1486074 | 0.22385857 |
| ENSMUSG00 | Gm10295 | 2524 | 0 | 0 | 0 | 0 | 0 | 0 |
| ENSMUSG00 | Rpl35a-ps6 | 332 | 0 | 0 | 0 | 0 | 0 | 0 |
| ENSMUSG00 | Ndnl2 | 5945 | 3.19648775 | 4.30395303 | 3.97779412 | 4.39418002 | 3.35598199 | 3.80789749 |
| ENSMUSG00 | Gm6505 | 1861 | 0 | 0 | 0.09500665 | 0.21327377 | 0 | 0.02931181 |
| ENSMUSG00 | Fcrlb | 1278 | 0.15553787 | 0 | 0 | 0 | 0 | 0 |
| ENSMUSG00 | Peg12 | 2639 | 0 | 0 | 0 | 0 | 0 | 0 |
| ENSMUSG00 | Mkrn3 | 2560 | 2.3138932 | 3.38810921 | 2.24462484 | 1.31079302 | 1.29559157 | 3.98465433 |
| ENSMUSG00 | Gm10297 | 215 | 0.18490921 | 0 | 0 | 0 | 0.71199594 | 0.25371757 |
| ENSMUSG00 | Wfdc10 | 715 | 0 | 0 | 0 | 0 | 0 | 0 |
| ENSMUSG00 | Wfdc16 | 1334 | 0 | 0.10836531 | 0 | 0 | 0.07650131 | 0.10222878 |
| ENSMUSG00 | Wfdc6b | 1153 | 0 | 0 | 0 | 0 | 0 | 0 |
| ENSMUSG00 | 1700084C01l | 1730 | 0 | 0 | 0 | 0 | 0 | 0 |
| ENSMUSG00 | Wfdc8 | 3918 | 0 | 0 | 0 | 0 | 0 | 0 |
| ENSMUSG00 | Gm9343 | 395 | 0 | 0 | 0 | 0 | 0 | 0 |
| ENSMUSG00 | Top1 | 10716 | 14.5465882 | 14.0071638 | 11.1907055 | 15.8355595 | 10.3709823 | 15.1670003 |
| ENSMUSG00 | Mrgprb3 | 938 | 0 | 0 | 0 | 0 | 0 | 0 |
| ENSMUSG00 | Mrgprb1 | 4183 | 0 | 0 | 0 | 0 | 0 | 0 |
| ENSMUSG00 | Mrgprb4 | 965 | 0 | 0 | 0 | 0 | 0 | 0 |
| ENSMUSG00 | Mrgprb5 | 1031 | 0 | 0 | 0 | 0 | 0 | 0 |
| ENSMUSG00 | Mrgprx1 | 1182 | 0 | 0 | 0 | 0 | 0 | 0 |
| ENSMUSG00 | Kcnj11 | 3074 | 0 | 0.43891359 | 0.15817185 | 0.01173782 | 0 | 0 |
| ENSMUSG00 | Spaca4 | 709 | 0 | 0 | 0 | 0 | 0 | 0 |
| ENSMUSG00 | Ntn5 | 1568 | 0 | 0 | 0 | 0 | 0 | 0 |
| ENSMUSG00 | Rasal2 | 14058 | 0.52600081 | 0.78151293 | 0.73575411 | 0.87522956 | 0.45371297 | 0.54518236 |
| ENSMUSG00 | 1700039E15f | 3443 | 0 | 0 | 0 | 0 | 0 | 0 |
| ENSMUSG00 | Slc17a7 | 2927 | 0 | 0 | 0 | 0 | 0 | 0 |
| ENSMUSG00 | Gm13072 | 377 | 0.1054522 | 0.89470848 | 0.3517388 | 0.19141668 | 0.54139391 | 0.43407914 |
| ENSMUSG00 | 2310016G11l | 879 | 0 | 0 | 0 | 0 | 0 | 0 |
| ENSMUSG00 | Mn1 | 3349 | 0 | 0.04316492 | 0 | 0 | 0 | 0.00814411 |
| ENSMUSG00 | Gm572 | 1374 | 0.02893412 | 0 | 0 | 0 | 0 | 0 |
| ENSMUSG00 | Fv1 | 1379 | 7.00549802 | 3.21475901 | 8.68651157 | 5.04991631 | 7.03046512 | 6.05224047 |
| ENSMUSG00 | Gm4788 | 3528 | 0 | 0 | 0 | 0 | 0 | 0 |
| ENSMUSG00 | Vmn2r84 | 3028 | 0 | 0 | 0 | 0 | 0 | 0 |
| ENSMUSG00 | Vsig10l | 3759 | 5.63705004 | 4.33280592 | 6.69083498 | 3.83953653 | 7.03156762 | 5.63051877 |
| ENSMUSG00 | Gm13251 | 4546 | 0.13117734 | 0.2331944 | 0.4278225 | 0.22223873 | 0.31428476 | 0.69596528 |
| ENSMUSG00 | Gm10298 | 263 | 0 | 0 | 0 | 0 | 0 | 0 |
| ENSMUSG00 | Gm13177 | 1220 | 0 | 0 | 0 | 0 | 0 | 0 |
| ENSMUSG00 | Gm13127 | 1388 | 0.11456911 | 0.06943291 | 0.19107425 | 0.07798713 | 0.22057511 | 0.17685285 |
| ENSMUSG00 | Gm13040 | 2146 | 0 | 0 | 0 | 0 | 0 | 0 |
| ENSMUSG00 | Gm13089 | 1803 | 0 | 0 | 0 | 0 | 0 | 0 |
| ENSMUSG00 | BC080695 | 2091 | 0 | 0 | 0 | 0 | 0 | 0 |
| ENSMUSG00 | Gm13119 | 2090 | 0 | 0 | 0 | 0 | 0 | 0 |
| ENSMUSG00 | Vmn2r-ps60 | 1156 | 0 | 0 | 0 | 0 | 0 | 0 |
| ENSMUSG00 | Btbd8 | 1527 | 3.22834288 | 2.42983369 | 3.29994113 | 1.84309068 | 3.67577036 | 2.3934523 |
| ENSMUSG00 | Gm694 | 903 | 0 | 0 | 0 | 0 | 0 | 0 |
| ENSMUSG00 | Lrrc8b | 2799 | 0.92322481 | 1.0845823 | 1.59499326 | 1.16019436 | 1.14850364 | 2.50431664 |
| ENSMUSG00 | Gm16410 | 773 | 0 | 0 | 0 | 0 | 0 | 0 |
| ENSMUSG00 | Sox13 | 4571 | 2.05256911 | 4.5645875 | 1.73094069 | 2.67595999 | 3.5721812 | 3.24598638 |
| ENSMUSG00 | Etnk2 | 4560 | 0.7323378 | 1.38430349 | 0.20356112 | 1.02074205 | 1.34279006 | 0.93307976 |
| ENSMUSG00 | Ren1 | 1878 | 0 | 0 | 0 | 0 | 0 | 0 |
| ENSMUSG00 | Rnf186 | 1253 | 70.658783 | 129.022756 | 35.2415333 | 120.542249 | 80.5915385 | 51.1100183 |
| ENSMUSG00 | Ldlrad2 | 1707 | 0 | 0 | 0 | 0 | 0 | 0 |
| ENSMUSG00 | Gm13004 | 792 | 0.15058894 | 0.24336587 | 0.50229367 | 0.04555814 | 0.06442724 | 0.06887535 |
| ENSMUSG00 | D5Ertd577e | 2458 | 0 | 0 | 0 | 0 | 0 | 0 |

| | | | | | | | | |
|---|---|---|---|---|---|---|---|---|
| ENSMUSG00 | Lactbl1 | 3204 | 0 | 0 | 0 | 0 | 0 | 0 |
| ENSMUSG00 | C87414 | 4644 | 0 | 0 | 0 | 0 | 0 | 0 |
| ENSMUSG00 | Htr1d | 3268 | 0 | 0 | 0 | 0.30914848 | 0 | 0 |
| ENSMUSG00 | 5830473C10 | 1075 | 0 | 0.62754436 | 0 | 0.26851754 | 0.0474664 | 1.04024206 |
| ENSMUSG00 | Runx3 | 4994 | 0 | 0 | 0 | 0 | 0 | 0 |
| ENSMUSG00 | Gm14059 | 892 | 0 | 0 | 0 | 0.04045072 | 0 | 0 |
| ENSMUSG00 | Cntnap5a | 5708 | 0 | 0 | 0 | 0 | 0 | 0 |
| ENSMUSG00 | Utp3 | 1600 | 32.5001053 | 29.0022149 | 29.7257394 | 35.1123396 | 24.7477923 | 27.0871261 |
| ENSMUSG00 | Sars2 | 1844 | 11.6636198 | 7.55199683 | 10.2354593 | 11.0555071 | 13.0056381 | 8.06110542 |
| ENSMUSG00 | Csn1s1 | 1381 | 0 | 0 | 0 | 0 | 0 | 0 |
| ENSMUSG00 | Ugt2b36 | 2078 | 0 | 0 | 0 | 0 | 0 | 0 |
| ENSMUSG00 | Eid2b | 2423 | 7.61310068 | 3.81832311 | 3.55730887 | 4.5716829 | 4.8857281 | 6.64136903 |
| ENSMUSG00 | Gtsf1l | 823 | 0 | 0 | 0 | 0 | 0 | 0 |
| ENSMUSG00 | 1700049G17 | 7777 | 6.00651789 | 4.76473885 | 4.56966462 | 8.61570743 | 5.94443828 | 14.1335729 |
| ENSMUSG00 | Gm10282 | 1150 | 0 | 0 | 0 | 0.09412707 | 0.04437076 | 0 |
| ENSMUSG00 | Gm10300 | 2295 | 0.0692906 | 0.23095898 | 0.44298143 | 0.25155238 | 0.11116858 | 0.24957186 |
| ENSMUSG00 | Pla2g4d | 3668 | 0 | 0 | 0 | 0 | 0 | 0 |
| ENSMUSG00 | Tmem200b | 854 | 9.03110447 | 10.438515 | 17.5461649 | 11.8301785 | 8.90272833 | 6.67494099 |
| ENSMUSG00 | Gm12966 | 809 | 0.93368867 | 0.41694079 | 0.7102892 | 0.35680637 | 1.0722477 | 1.78684287 |
| ENSMUSG00 | Fam82a2 | 3525 | 28.7480056 | 40.9140486 | 24.2514509 | 33.8915318 | 33.7859748 | 19.521678 |
| ENSMUSG00 | Rbm44 | 3835 | 0 | 0 | 0 | 0 | 0 | 0 |
| ENSMUSG00 | Fryl | 15704 | 5.32132069 | 4.46454873 | 6.64829045 | 5.70272762 | 6.17359361 | 5.79742396 |
| ENSMUSG00 | Gm12942 | 1906 | 15.288965 | 22.7533043 | 21.335622 | 22.6033374 | 16.0628675 | 21.5077559 |
| ENSMUSG00 | Dgkd | 5664 | 6.3100595 | 5.16404844 | 5.71252632 | 5.88626574 | 7.4053109 | 6.35156235 |
| ENSMUSG00 | Ceacam20 | 2885 | 0 | 0 | 0 | 0 | 0 | 0 |
| ENSMUSG00 | Rpl7a-ps8 | 812 | 0 | 0 | 0 | 0 | 0 | 0 |
| ENSMUSG00 | Rbm47 | 9723 | 28.1228217 | 26.875972 | 29.4633492 | 31.3208326 | 26.1455728 | 28.7361313 |
| ENSMUSG00 | BC016548 | 1174 | 0 | 0 | 0 | 0 | 0 | 0 |
| ENSMUSG00 | Psg21 | 2154 | 0 | 0 | 0 | 0 | 0 | 0 |
| ENSMUSG00 | Psg27 | 1936 | 0 | 0 | 0 | 0 | 0 | 0 |
| ENSMUSG00 | Psg25 | 3060 | 0 | 0 | 0 | 0 | 0 | 0 |
| ENSMUSG00 | Psg26 | 2052 | 0 | 0 | 0 | 0 | 0 | 0 |
| ENSMUSG00 | Pnmal2 | 2396 | 0.265479 | 1.387673 | 1.1991318 | 1.23486129 | 1.63982927 | 1.87826189 |
| ENSMUSG00 | Cited4 | 1492 | 1.91849504 | 1.67942024 | 0.6221439 | 2.03142878 | 1.81259914 | 1.15167713 |
| ENSMUSG00 | Zmynd12 | 1983 | 0.02004815 | 0.7046933 | 0.13374234 | 0.45489214 | 0.25731909 | 0.48139807 |
| ENSMUSG00 | Gltscr1 | 5346 | 0.63210173 | 0.54081304 | 0.63665206 | 0.98540563 | 0.72540302 | 0.59691971 |
| ENSMUSG00 | Sult2a6 | 982 | 0 | 0 | 0 | 0 | 0 | 0 |
| ENSMUSG00 | Sult2a2 | 736 | 0 | 0 | 0 | 0 | 0 | 0 |
| ENSMUSG00 | 6330408A02 | 13611 | 1.94235505 | 0.49209577 | 1.13338059 | 0.9516901 | 2.61673722 | 0.94582542 |
| ENSMUSG00 | Vmn1r87 | 887 | 0 | 0 | 0 | 0 | 0 | 0 |
| ENSMUSG00 | Vmn1r86 | 950 | 0 | 0 | 0 | 0 | 0 | 0 |
| ENSMUSG00 | Vmn1r85 | 926 | 0 | 0 | 0 | 0 | 0 | 0 |
| ENSMUSG00 | Olfr1339 | 947 | 0 | 0 | 0 | 0 | 0 | 0 |
| ENSMUSG00 | Olfr1340 | 950 | 0 | 0 | 0 | 0 | 0 | 0 |
| ENSMUSG00 | Zscan18 | 3620 | 2.65768682 | 0.79867031 | 0.59831224 | 1.52501459 | 1.62100365 | 1.0548203 |
| ENSMUSG00 | Gm1043 | 1576 | 0 | 0 | 0 | 0 | 0 | 0 |
| ENSMUSG00 | Vmn1r72 | 920 | 0 | 0 | 0 | 0 | 0 | 0 |
| ENSMUSG00 | Zscan4-ps2 | 2273 | 0 | 0 | 0 | 0 | 0 | 0 |
| ENSMUSG00 | Zscan4f | 2487 | 0 | 0 | 0 | 0 | 0 | 0 |
| ENSMUSG00 | Gm13015 | 203 | 0 | 0 | 0 | 0 | 0 | 0 |
| ENSMUSG00 | Aurkc | 941 | 0 | 0 | 0 | 0 | 0 | 0 |
| ENSMUSG00 | Vmn2r34 | 4916 | 0 | 0 | 0 | 0 | 0 | 0 |
| ENSMUSG00 | Vmn2r42 | 4107 | 0 | 0 | 0 | 0 | 0 | 0 |
| ENSMUSG00 | Vmn2r-ps44 | 2556 | 0 | 0 | 0 | 0 | 0 | 0 |
| ENSMUSG00 | Vmn2r30 | 4586 | 0 | 0.02101458 | 0 | 0 | 0 | 0 |
| ENSMUSG00 | Gm10302 | 3450 | 0 | 0 | 0 | 0 | 0 | 0 |
| ENSMUSG00 | Vmn2r32 | 2553 | 0 | 0 | 0 | 0 | 0 | 0 |
| ENSMUSG00 | Gm4073 | 2556 | 0 | 0 | 0 | 0 | 0 | 0 |
| ENSMUSG00 | Olfr1121 | 944 | 0 | 0 | 0 | 0 | 0 | 0 |
| ENSMUSG00 | Olfr1120 | 1010 | 0 | 0 | 0 | 0 | 0 | 0 |
| ENSMUSG00 | Olfr1116-ps | 1918 | 0 | 0 | 0 | 0 | 0 | 0 |
| ENSMUSG00 | Olfr1115 | 1101 | 0 | 0 | 0 | 0 | 0 | 0 |
| ENSMUSG00 | Olfr1113 | 1102 | 0 | 0 | 0 | 0 | 0 | 0 |
| ENSMUSG00 | Gm1673 | 1015 | 0.03916796 | 0.14242298 | 0 | 0 | 0 | 0 |
| ENSMUSG00 | Zfp804a | 5241 | 0 | 0 | 0 | 0 | 0 | 0 |
| ENSMUSG00 | Gm12824 | 6610 | 12.3055542 | 9.17073286 | 7.09503108 | 13.2155264 | 7.81220761 | 11.6154477 |
| ENSMUSG00 | Skint3 | 3833 | 0 | 0 | 0 | 0 | 0 | 0 |
| ENSMUSG00 | Crygs | 1060 | 0 | 0 | 0 | 0 | 0 | 0 |
| ENSMUSG00 | Ccnyl1 | 3270 | 1.83580353 | 2.87350343 | 2.75754617 | 2.16271582 | 1.91934074 | 1.77660494 |
| ENSMUSG00 | Lilra5 | 2103 | 0 | 0 | 0 | 0 | 0 | 0 |
| ENSMUSG00 | Olfr1100 | 968 | 0 | 0 | 0 | 0 | 0 | 0 |
| ENSMUSG00 | Ldlrad1 | 1088 | 0 | 0 | 0 | 0 | 0 | 0 |
| ENSMUSG00 | Gad1 | 7618 | 0.21918222 | 0 | 0 | 0 | 0.46217117 | 0 |
| ENSMUSG00 | 4930578N16 | 2549 | 1.27891307 | 1.07753127 | 1.92483506 | 4.1192134 | 2.20200131 | 3.21003993 |
| ENSMUSG00 | Gm10304 | 2305 | 0.03449499 | 0.02090518 | 0.09588252 | 0.01565382 | 0 | 0.01183282 |
| ENSMUSG00 | Pgap1 | 2844 | 12.4270823 | 11.2163941 | 11.8741941 | 11.9765999 | 12.2362828 | 6.60767456 |
| ENSMUSG00 | Gm12794 | 1449 | 0 | 0 | 0 | 0 | 0 | 0 |
| ENSMUSG00 | Gm12689 | 791 | 2.46272888 | 0.42642869 | 0.55880965 | 1.73339784 | 0 | 0 |
| ENSMUSG00 | Gm10305 | 2474 | 0.17676244 | 0 | 0 | 0 | 0.35062587 | 0 |
| ENSMUSG00 | Rpl23a-ps1 | 467 | 0 | 0 | 0 | 0 | 0 | 0 |
| ENSMUSG00 | Gm10306 | 532 | 0.44837008 | 0.4528801 | 0 | 0 | 0 | 0.05126812 |
| ENSMUSG00 | Zfp352 | 1903 | 0 | 0 | 0 | 0 | 0 | 0 |
| ENSMUSG00 | Ifna1 | 569 | 0 | 0 | 0 | 0 | 0 | 0 |
| ENSMUSG00 | Ifna4 | 643 | 0 | 0 | 0 | 0 | 0 | 0 |
| ENSMUSG00 | Ifna11 | 646 | 0 | 0 | 0 | 0 | 0 | 0 |
| ENSMUSG00 | Ifna7 | 575 | 0 | 0 | 0 | 0 | 0 | 0 |
| ENSMUSG00 | Gm13288 | 2849 | 0 | 0 | 0 | 0 | 0.16119248 | 0 |
| ENSMUSG00 | Klhl9 | 7049 | 12.1257318 | 13.4599382 | 15.3129381 | 19.5484931 | 16.3379955 | 18.6229024 |
| ENSMUSG00 | Speer4d | 1313 | 0 | 0 | 0 | 0 | 0 | 0 |
| ENSMUSG00 | Rraga | 1608 | 47.7659132 | 57.1165166 | 45.6862333 | 40.9513254 | 44.9654071 | 45.8987399 |
| ENSMUSG00 | Tgfbrap1 | 6619 | 3.86803587 | 4.26609081 | 5.24892712 | 4.89525245 | 8.0559847 | 6.17686724 |
| ENSMUSG00 | Gm6455 | 1122 | 0 | 0 | 0 | 0 | 0 | 0 |
| ENSMUSG00 | Il1rl2 | 1939 | 1.27119123 | 1.51592211 | 1.59573441 | 0.96764637 | 3.63158322 | 1.37850162 |
| ENSMUSG00 | Olfr356 | 947 | 0 | 0 | 0 | 0 | 0 | 0 |
| ENSMUSG00 | Gm428 | 1316 | 0 | 0 | 0 | 0 | 0 | 0 |
| ENSMUSG00 | Rabepk | 4007 | 5.74455284 | 6.72229128 | 9.58607476 | 12.9848536 | 6.46902893 | 8.06599638 |
| ENSMUSG00 | Gm10307 | 671 | 0 | 0 | 0.19762374 | 0.05377354 | 0.15209054 | 0 |
| ENSMUSG00 | Dnajc25 | 4535 | 4.64617523 | 3.89954229 | 5.47771461 | 3.58831358 | 3.9043335 | 5.9781679 |
| ENSMUSG00 | Defb44-ps | 217 | 0 | 0 | 0 | 0 | 0 | 0 |
| ENSMUSG00 | Actl7a | 1503 | 0 | 0 | 0 | 0 | 0 | 0 |
| ENSMUSG00 | Actl7b | 1418 | 0 | 0 | 0 | 0 | 0 | 0 |
| ENSMUSG00 | Olfr270 | 1882 | 0 | 0 | 0 | 0 | 0 | 0 |
| ENSMUSG00 | 4930552N02 | 1184 | 0 | 0 | 0 | 0 | 0 | 0 |
| ENSMUSG00 | Acnat1 | 2921 | 0.42191712 | 0 | 0.45397305 | 0.09882107 | 0.08734402 | 0.58825822 |
| ENSMUSG00 | Foxe1 | 2803 | 0 | 0 | 0 | 0 | 0 | 0 |
| ENSMUSG00 | Gm6313 | 2587 | 0 | 0 | 0 | 0 | 0 | 0 |
| ENSMUSG00 | 1700055D18 | 1569 | 0.70946683 | 1.38202033 | 1.63397508 | 0.36794947 | 1.43094999 | 2.34676628 |
| ENSMUSG00 | Ccin | 2029 | 0 | 0 | 0 | 0 | 0 | 0 |
| ENSMUSG00 | Olfr155 | 1346 | 0 | 0 | 0 | 0 | 0 | 0 |
| ENSMUSG00 | Hrct1 | 1441 | 0.11035526 | 0 | 0.21472096 | 0 | 0.07082079 | 0 |
| ENSMUSG00 | Ccl21a | 866 | 0 | 0 | 0 | 0 | 0 | 0 |
| ENSMUSG00 | Ccl19 | 1101 | 0 | 0 | 0 | 0 | 0 | 0 |
| ENSMUSG00 | Ndufb6 | 837 | 154.177168 | 131.260559 | 116.392906 | 110.14292 | 71.2662286 | 353.820828 |
| ENSMUSG00 | Gm136 | 1306 | 0 | 0 | 0 | 0 | 0.27349512 | 0 |
| ENSMUSG00 | Sdr16c6 | 2172 | 0 | 0 | 0 | 0 | 0 | 0 |
| ENSMUSG00 | Gm10308 | 740 | 0 | 0 | 0 | 0 | 0 | 0 |
| ENSMUSG00 | Gm5499 | 887 | 2.95813046 | 3.20518612 | 4.63446607 | 3.82380179 | 4.08441114 | 2.76743803 |
| ENSMUSG00 | Gm10309 | 2003 | 0.23817562 | 0.21651422 | 0 | 0 | 0.02547498 | 0.04085068 |
| ENSMUSG00 | 1700106N22 | 3380 | 3.83440433 | 4.83290056 | 4.43326177 | 4.43019186 | 3.47220901 | 5.10794279 |
| ENSMUSG00 | Gm9386 | 380 | 0 | 0 | 0 | 0 | 0.13427994 | 0 |
| ENSMUSG00 | Gm15210 | 3198 | 0.03729407 | 0.18081217 | 0.05528686 | 0.02256538 | 0.12764572 | 0.05970059 |
| ENSMUSG00 | Rasgrp3 | 4906 | 2.01775676 | 0.62860422 | 1.21631651 | 1.72834906 | 2.26737667 | 0.47811231 |
| ENSMUSG00 | Ces1a | 1966 | 0 | 0 | 0 | 0 | 0 | 0 |
| ENSMUSG00 | Rpl7a-ps5 | 806 | 0 | 0 | 0.16452299 | 0 | 0 | 0 |
| ENSMUSG00 | Safb | 3173 | 25.2465469 | 27.0166028 | 21.884997 | 28.0537043 | 34.671562 | 21.3606929 |
| ENSMUSG00 | Zfp827 | 8497 | 5.13260705 | 4.03207725 | 4.35411823 | 6.77308003 | 5.12245482 | 4.87264945 |
| ENSMUSG00 | Olfr806 | 964 | 0 | 0 | 0 | 0 | 0 | 0 |
| ENSMUSG00 | Treml2 | 3525 | 0 | 0 | 0 | 0 | 0 | 0 |
| ENSMUSG00 | Ptges3 | 2247 | 14.8795546 | 13.4029935 | 15.9339087 | 18.0329933 | 10.241609 | 21.8731197 |
| ENSMUSG00 | Gm88 | 1378 | 0 | 0 | 0 | 0 | 0 | 0 |
| ENSMUSG00 | Yipf3 | 2821 | 32.6104863 | 38.3134313 | 41.2874358 | 37.0029617 | 30.6230607 | 28.821588 |
| ENSMUSG00 | Jund | 4434 | 6.33003366 | 7.06386725 | 7.54641298 | 6.1031875 | 5.63890937 | 9.03003392 |
| ENSMUSG00 | Gm9104 | 515 | 0 | 0 | 0 | 0 | 0 | 0.10592093 |
| ENSMUSG00 | Nr2c2ap | 1170 | 1.46109886 | 2.4299146 | 1.62451218 | 0.83266257 | 1.22114404 | 1.98149089 |
| ENSMUSG00 | Gm10311 | 386 | 20.0837272 | 15.2299118 | 20.0396957 | 13.1802287 | 10.0466439 | 44.2329642 |
| ENSMUSG00 | Olfr761 | 971 | 0 | 0 | 0 | 0 | 0 | 0 |
| ENSMUSG00 | Trim75 | 2504 | 0.09526074 | 0 | 0.01765249 | 0.80694668 | 0 | 0.01089243 |
| ENSMUSG00 | Gm10313 | 1001 | 0.19857882 | 0.14441491 | 0.13247306 | 0.18022999 | 0.1019508 | 0.08174218 |
| ENSMUSG00 | 1700029J07 | 2937 | 13.4007238 | 12.5839296 | 7.55509881 | 11.2288011 | 13.2734597 | 13.1404884 |
| ENSMUSG00 | Ccdc110 | 2658 | 0 | 0 | 0 | 0 | 0 | 0 |
| ENSMUSG00 | 4732465J04 | 1629 | 0.14642903 | 0.11832153 | 0.3256121 | 0.48729587 | 0.09397123 | 0.21766133 |
| ENSMUSG00 | B230216G23 | 1761 | 0.67726543 | 0 | 0 | 0 | 0.60849171 | 0.44915647 |
| ENSMUSG00 | Mboat4 | 1836 | 0 | 0.20996271 | 0.38520125 | 0 | 0 | 0 |
| ENSMUSG00 | Tex24 | 1764 | 0 | 0 | 0 | 0 | 0 | 0 |
| ENSMUSG00 | Rpl36a-ps3 | 320 | 0 | 0 | 0 | 0.11275639 | 0.15945742 | 0 |
| ENSMUSG00 | Tas2r140 | 999 | 0 | 0 | 0 | 0 | 0 | 0 |
| ENSMUSG00 | Tas2r115 | 1025 | 0 | 0 | 0 | 0 | 0 | 0 |
| ENSMUSG00 | Tas2r121 | 917 | 0 | 0 | 0 | 0 | 0 | 0 |
| ENSMUSG00 | Gm4799 | 656 | 0 | 0.07345494 | 0 | 0 | 0 | 0.08315439 |
| ENSMUSG00 | Gm156 | 2126 | 0 | 0 | 0 | 0 | 0 | 0 |
| ENSMUSG00 | Defa25 | 304 | 0 | 0 | 0 | 0 | 0 | 0 |
| ENSMUSG00 | Gm6040 | 854 | 0 | 0 | 0 | 0 | 0 | 0 |
| ENSMUSG00 | Defb46 | 172 | 0 | 0 | 0 | 0 | 0 | 0 |
| ENSMUSG00 | Srsf3 | 4185 | 61.1673928 | 56.372956 | 61.4706634 | 64.0682611 | 52.2700294 | 96.0510474 |
| ENSMUSG00 | Arhgef10 | 9225 | 7.36068951 | 5.28092065 | 8.21747 | 8.77311934 | 9.07688702 | 6.93914128 |
| ENSMUSG00 | Serpina1d | 1389 | 0 | 0 | 0.06364556 | 0 | 0.73472103 | 0 |
| ENSMUSG00 | Serpina1b | 1389 | 0 | 0 | 0 | 0 | 0.40409657 | 0.17672552 |
| ENSMUSG00 | Gm46 | 1789 | 0 | 0 | 0 | 0 | 0 | 0 |
| ENSMUSG00 | 2810008M24 | 1850 | 47.3197665 | 57.4590764 | 72.2043079 | 64.9476801 | 46.9718476 | 63.3803644 |
| ENSMUSG00 | 3830408C21 | 1710 | 0.18599055 | 0.28179206 | 0.07754709 | 0 | 0.17903992 | 0.39875204 |
| ENSMUSG00 | Olfr1357 | 1158 | 0 | 0 | 0 | 0 | 0 | 0 |
| ENSMUSG00 | Olfr1358 | 941 | 0 | 0 | 0 | 0 | 0 | 0 |
| ENSMUSG00 | Wfikkn1 | 2274 | 0.64685698 | 1.90711512 | 0.38875851 | 0.80922791 | 0.13463424 | 1.91905993 |
| ENSMUSG00 | Gm4938 | 407 | 0 | 0 | 0 | 0 | 0 | 0.06701386 |
| ENSMUSG00 | Gm10318 | 637 | 0 | 0 | 0 | 0 | 0 | 0 |
| ENSMUSG00 | Ccdc78 | 1638 | 0.14562447 | 0.29417852 | 0.02698525 | 0 | 0.93454901 | 0 |
| ENSMUSG00 | Naip5 | 6412 | 0 | 0 | 0 | 0 | 0.00795795 | 0 |
| ENSMUSG00 | Gm10319 | 1466 | 0 | 0 | 0 | 0 | 0 | 0 |
| ENSMUSG00 | Gm8587 | 860 | 0 | 0 | 0 | 0 | 0 | 0 |
| ENSMUSG00 | Oog1 | 1858 | 0 | 0 | 0 | 0 | 0 | 0 |
| ENSMUSG00 | Gm2022 | 1717 | 0 | 0 | 0 | 0 | 0 | 0 |
| ENSMUSG00 | Gm10320 | 290 | 0 | 0 | 0 | 0 | 0 | 0 |
| ENSMUSG00 | Cecr2 | 12561 | 0.31649933 | 0.21482691 | 0.00351898 | 0.05457837 | 0.38997947 | 0.35827685 |
| ENSMUSG00 | Timm8a2 | 911 | 0 | 0.26447005 | 0 | 0 | 0 | 0 |
| ENSMUSG00 | Npw | 604 | 0 | 0 | 0 | 0 | 0 | 0 |
| ENSMUSG00 | Tmem90a | 2495 | 0 | 0 | 0 | 0.01446174 | 0 | 0 |
| ENSMUSG00 | Vrtn | 3603 | 0 | 0 | 0 | 0 | 0 | 0 |
| ENSMUSG00 | Gm6253 | 687 | 0 | 0 | 0 | 0 | 0 | 0 |
| ENSMUSG00 | Gm4666 | 683 | 0 | 0 | 0 | 0 | 0 | 0 |
| ENSMUSG00 | Gm6219 | 683 | 0 | 0 | 0 | 0 | 0 | 0 |
| ENSMUSG00 | 2210408I21 | 5065 | 2.30762315 | 4.37626128 | 4.56417846 | 3.56189976 | 4.54351342 | 6.76884927 |
| ENSMUSG00 | Slc25a16 | 5004 | 46.1271622 | 41.0027719 | 63.6527739 | 48.4266609 | 45.1834275 | 45.7030076 |
| ENSMUSG00 | Zfp213 | 2666 | 3.65344813 | 3.25339819 | 3.97916066 | 2.6932959 | 2.16278337 | 2.22002878 |
| ENSMUSG00 | Zfp957 | 5329 | 0 | 0 | 0 | 0 | 0 | 0 |
| ENSMUSG00 | 1700086L19 | 706 | 0 | 0 | 0 | 0 | 0 | 0 |
| ENSMUSG00 | Zfp946 | 3667 | 15.6441665 | 11.3534459 | 17.671094 | 15.9500939 | 8.12637128 | 22.551597 |
| ENSMUSG00 | Zfp942 | 5253 | 13.7361883 | 10.0629216 | 11.6710368 | 14.699329 | 11.2679604 | 20.0367091 |
| ENSMUSG00 | Rpl9-ps4 | 672 | 0 | 0 | 0 | 0 | 0 | 0 |
| ENSMUSG00 | Gm5145 | 836 | 0 | 0 | 0 | 0 | 0 | 0 |
| ENSMUSG00 | Fpr-rs6 | 1019 | 0 | 0 | 0 | 0 | 0 | 0 |
| ENSMUSG00 | Fpr-rs7 | 1016 | 0 | 0 | 0 | 0 | 0 | 0 |
| ENSMUSG00 | Gm10322 | 335 | 0 | 0 | 0 | 0 | 0.15231754 | 0.08141683 |
| ENSMUSG00 | Zfp71-rs1 | 4720 | 4.65779253 | 5.54348265 | 4.7666818 | 5.20590515 | 3.23239118 | 4.24143754 |
| ENSMUSG00 | Zfp738 | 254 | 0 | 0 | 0 | 0 | 0 | 0 |
| ENSMUSG00 | Zfp87 | 2553 | 22.6417817 | 13.9293358 | 14.3184192 | 13.2569361 | 25.2233789 | 12.5422744 |
| ENSMUSG00 | Sowahc | 4475 | 28.4373839 | 25.9506873 | 25.4049475 | 28.5189255 | 24.2418045 | 27.2197852 |
| ENSMUSG00 | Zfp58 | 4654 | 10.3446255 | 6.82313394 | 8.85176573 | 7.59785209 | 8.74925825 | 12.1428991 |
| ENSMUSG00 | Gm5451 | 693 | 0 | 0 | 0 | 0 | 0 | 0.03935734 |
| ENSMUSG00 | Gm5196 | 551 | 0 | 0 | 0 | 0 | 0 | 0 |
| ENSMUSG00 | Gm10323 | 808 | 0 | 0 | 0 | 0 | 0 | 0 |
| ENSMUSG00 | Gm10324 | 463 | 0 | 0 | 0 | 0 | 0 | 0 |
| ENSMUSG00 | Rps8-ps1 | 686 | 0 | 0 | 0 | 0 | 0 | 0 |
| ENSMUSG00 | Gpr31c | 959 | 0 | 0 | 0 | 0 | 0 | 0 |
| ENSMUSG00 | Bves | 1599 | 0 | 0 | 0 | 0 | 0 | 0 |
| ENSMUSG00 | Gm4795 | 1007 | 0.03947913 | 0 | 0.08778916 | 0.03583123 | 0.15201502 | 0.02708504 |
| ENSMUSG00 | Gm9034 | 1007 | 0.19739563 | 0.09570296 | 0.17557832 | 0.17915613 | 0.15201502 | 0.21668035 |
| ENSMUSG00 | Tcp10a | 2802 | 0 | 0 | 0 | 0 | 0.05463209 | 0 |
| ENSMUSG00 | Armc2 | 9394 | 0 | 0 | 0 | 0.02688677 | 0.04345444 | 0 |
| ENSMUSG00 | Gm7555 | 386 | 0 | 0 | 0 | 0 | 0 | 0 |
| ENSMUSG00 | E230015B07 | 2728 | 0 | 0.10598191 | 0.21063928 | 0 | 0 | 0.56988799 |
| ENSMUSG00 | G630090E17 | 1783 | 7.22421519 | 33.6467305 | 2.95009496 | 25.5791948 | 13.0785495 | 3.82426237 |
| ENSMUSG00 | Tia1 | 6874 | 20.2131807 | 17.3356228 | 16.6994743 | 24.4763709 | 18.6765146 | 23.2116147 |
| ENSMUSG00 | Bet3l | 1176 | 0 | 0 | 0 | 0 | 0 | 0 |
| ENSMUSG00 | Egr4 | 2136 | 0 | 0 | 0 | 0 | 0 | 0 |
| ENSMUSG00 | Gm889 | 1209 | 0 | 0 | 0 | 0.02984454 | 0.04220544 | 0.02255967 |
| ENSMUSG00 | Gm10327 | 1001 | 0 | 0.52952134 | 0.17663074 | 0 | 0.2039016 | 0.10898957 |
| ENSMUSG00 | C1qtnf9 | 1899 | 0.5652438 | 1.01498562 | 0.30259292 | 0.20900605 | 0.67175324 | 0.31597792 |
| ENSMUSG00 | Setdb2 | 4437 | 3.06431696 | 2.58471337 | 2.90893355 | 3.602512 | 3.46155941 | 2.7600435 |
| ENSMUSG00 | Reg3b | 753 | 0 | 0 | 0 | 0 | 0 | 0 |
| ENSMUSG00 | Tbpl1 | 3776 | 13.1711086 | 18.4782755 | 19.0105384 | 27.2430903 | 15.2430487 | 14.7858015 |
| ENSMUSG00 | Mcpt9 | 1995 | 0 | 0 | 0 | 0 | 0 | 0 |
| ENSMUSG00 | Gm10330 | 1353 | 0 | 0 | 0 | 0 | 0 | 0 |
| ENSMUSG00 | Map3k5 | 6777 | 12.2662992 | 6.17173259 | 13.9969242 | 8.82218501 | 13.5979983 | 12.2146274 |
| ENSMUSG00 | Hpcal1 | 1557 | 33.9594021 | 37.5402404 | 30.0072884 | 27.5771567 | 30.5764988 | 25.2426173 |
| ENSMUSG00 | Ect2l | 2246 | 0 | 0 | 0 | 0 | 0.70428212 | 0 |
| ENSMUSG00 | 2410004P03 | 1620 | 0 | 0 | 0 | 0 | 0 | 0 |
| ENSMUSG00 | Gm6736 | 1350 | 0 | 0 | 0 | 0 | 0 | 0 |
| ENSMUSG00 | Rpl23 | 1824 | 4.20658319 | 5.09867507 | 5.47676344 | 5.0641466 | 3.3849734 | 6.78875341 |
| ENSMUSG00 | Rps15-ps2 | 817 | 0 | 0 | 0 | 0 | 0 | 0 |
| ENSMUSG00 | Grid2 | 16177 | 0 | 0.0059574 | 0.00546478 | 0 | 0 | 0 |
| ENSMUSG00 | Gm10332 | 1063 | 0.93498308 | 1.40524902 | 0.04158217 | 0.98436434 | 0.72003353 | 0.46184714 |
| ENSMUSG00 | Vmn1r27 | 911 | 0 | 0 | 0 | 0 | 0 | 0 |
| ENSMUSG00 | 9230019H11 | 2795 | 0 | 0 | 0 | 0 | 0 | 0 |
| ENSMUSG00 | Psmg4 | 551 | 6.71009018 | 6.82131123 | 8.34299759 | 6.28652681 | 5.74162487 | 15.7410804 |
| ENSMUSG00 | Gm11397 | 1856 | 0 | 0 | 0 | 0 | 0 | 0 |
| ENSMUSG00 | Dtnb | 8006 | 5.40269333 | 5.67572008 | 4.6542785 | 5.19642735 | 5.91462363 | 4.8444338 |
| ENSMUSG00 | 1110002L01 | 1826 | 1.3498575 | 2.29584911 | 1.16193235 | 1.95625543 | 1.87227118 | 2.1210289 |
| ENSMUSG00 | Vmn2r31 | 2580 | 0 | 0 | 0 | 0 | 0 | 0 |
| ENSMUSG00 | Gm7102 | 451 | 0 | 0 | 0 | 0 | 0 | 0 |
| ENSMUSG00 | Gm11360 | 260 | 0 | 0 | 0 | 0 | 0 | 0 |
| ENSMUSG00 | Ccnb1ip1 | 1504 | 2.14108638 | 0.73689373 | 4.05575421 | 0 | 0.27141689 | 2.6476711 |
| ENSMUSG00 | Krtap26-1 | 972 | 0 | 0 | 0 | 0 | 0 | 0 |
| ENSMUSG00 | Rpl21-ps5 | 482 | 0 | 0.09997187 | 0 | 0 | 0 | 0 |
| ENSMUSG00 | Rps19-ps1 | 455 | 0 | 0 | 0 | 0 | 0 | 0 |
| ENSMUSG00 | Zfp777 | 5192 | 4.05058729 | 5.60566469 | 2.57957597 | 4.18362689 | 4.76652394 | 2.84198379 |
| ENSMUSG00 | Hist1h2ad | 571 | 0 | 0.33755826 | 0.07741128 | 0.25276389 | 0 | 0 |
| ENSMUSG00 | Olfr437 | 1017 | 0 | 0 | 0 | 0 | 0 | 0 |
| ENSMUSG00 | Ptgdr | 3185 | 0 | 0 | 0 | 0 | 0 | 0 |
| ENSMUSG00 | Vmn1r212 | 1163 | 0 | 0 | 0 | 0 | 0 | 0 |
| ENSMUSG00 | Vmn1r209 | 938 | 0 | 0 | 0 | 0 | 0 | 0 |
| ENSMUSG00 | Vmn1r208 | 996 | 0 | 0 | 0 | 0 | 0 | 0 |
| ENSMUSG00 | Olfr455 | 1902 | 0 | 0 | 0 | 0 | 0 | 0 |
| ENSMUSG00 | Vmn1r206 | 1134 | 0 | 0 | 0 | 0 | 0 | 0 |
| ENSMUSG00 | Nutf2-ps1 | 864 | 0.04601329 | 0 | 0 | 0 | 0 | 0 |
| ENSMUSG00 | Tmem139 | 1144 | 13.0317353 | 17.1011324 | 16.1893114 | 15.8962854 | 15.0313712 | 11.9922583 |
| ENSMUSG00 | Olfr172 | 1088 | 0 | 0 | 0 | 0 | 0 | 0 |
| ENSMUSG00 | Hist1h2ai | 1040 | 0 | 0 | 0 | 0.20816564 | 0 | 0.1835793 |
| ENSMUSG00 | Gm10334 | 825 | 0 | 0 | 0 | 0 | 0 | 0 |
| ENSMUSG00 | Prss3 | 836 | 0 | 0 | 0 | 0 | 0 | 0 |
| ENSMUSG00 | Try10 | 885 | 0 | 0 | 0 | 0 | 0 | 0 |
| ENSMUSG00 | Olfr263 | 1086 | 0 | 0 | 0 | 0 | 0 | 0 |
| ENSMUSG00 | Usmg5 | 337 | 44.946107 | 39.7502401 | 61.2529991 | 39.6153011 | 28.7685798 | 95.5017042 |
| ENSMUSG00 | Gprin2 | 1599 | 0.14917629 | 0 | 0 | 0 | 0 | 0 |
| ENSMUSG00 | Gm10335 | 470 | 0.25375839 | 0.20504869 | 0.3761859 | 0 | 0.32570027 | 0.40621803 |
| ENSMUSG00 | Pcnp | 3109 | 15.2551587 | 15.7470007 | 19.1508146 | 18.0468251 | 12.9166156 | 18.9492508 |
| ENSMUSG00 | Klrg2 | 2277 | 2.74115522 | 1.77762896 | 1.18415126 | 2.10755904 | 2.39781388 | 1.66498676 |
| ENSMUSG00 | 3425401B19 | 4238 | 0 | 0 | 0 | 0 | 0 | 0 |
| ENSMUSG00 | Nt5dc2 | 1394 | 15.8850808 | 10.7849139 | 11.7321965 | 10.4570631 | 7.17444022 | 9.91982934 |
| ENSMUSG00 | Wdr52 | 8505 | 0 | 0.28894868 | 0 | 0.17818294 | 0.01199915 | 0 |
| ENSMUSG00 | Akr1c19 | 1719 | 110.917559 | 217.273481 | 65.9556296 | 167.060496 | 130.519473 | 61.0863066 |
| ENSMUSG00 | Tigit | 959 | 0 | 0 | 0 | 0 | 0 | 0 |
| ENSMUSG00 | Cpa2 | 2916 | 0 | 0 | 0.06063353 | 0 | 0 | 0 |
| ENSMUSG00 | BC100530 | 987 | 0 | 0 | 0 | 0 | 0.05169646 | 0 |
| ENSMUSG00 | Stfa1 | 436 | 0 | 0 | 0 | 0 | 0 | 0 |
| ENSMUSG00 | Gm5301 | 2075 | 0 | 0 | 0 | 0 | 0 | 0 |
| ENSMUSG00 | Gm5989 | 2075 | 0 | 0 | 0 | 0 | 0 | 0 |
| ENSMUSG00 | A830039N20 | 2555 | 0 | 0 | 0 | 0 | 0 | 0 |
| ENSMUSG00 | Gm5874 | 506 | 0.549977 | 0.85707111 | 1.74710842 | 0.78439227 | 0.60505584 | 1.07804898 |
| ENSMUSG00 | Rnls | 1620 | 5.00624569 | 4.2832393 | 4.63846501 | 3.60820445 | 4.06321141 | 5.67379841 |
| ENSMUSG00 | Gm6458 | 1328 | 0 | 0 | 0 | 0.08151064 | 0 | 0 |
| ENSMUSG00 | Atf7 | 389 | 18.3958522 | 10.1575534 | 14.6581947 | 19.7570064 | 15.8719575 | 16.4068524 |
| ENSMUSG00 | Gm10337 | 164 | 0.96964587 | 0.58763954 | 0 | 0 | 0 | 0 |
| ENSMUSG00 | Gm10338 | 979 | 0 | 0 | 0 | 0 | 0 | 0 |
| ENSMUSG00 | Gm10339 | 581 | 0 | 0 | 0 | 0 | 0 | 0 |
| ENSMUSG00 | Fam189a2 | 2536 | 0.47029354 | 0.64603274 | 0.41831397 | 0.35569839 | 0.90543648 | 1.02172347 |
| ENSMUSG00 | Gm10340 | 1825 | 0 | 0 | 0 | 0 | 0 | 0 |
| ENSMUSG00 | Gm3488 | 887 | 0 | 0 | 0 | 0 | 0 | 0 |
| ENSMUSG00 | Olfr1477 | 1098 | 0 | 0 | 0 | 0 | 0 | 0 |
| ENSMUSG00 | Olfr1474 | 944 | 0 | 0 | 0 | 0 | 0 | 0 |
| ENSMUSG00 | 2510002D24 | 1468 | 26.9189017 | 23.2397827 | 22.2514727 | 21.5312473 | 26.1388519 | 25.751124 |
| ENSMUSG00 | Gm4952 | 1571 | 0 | 0 | 0 | 0 | 0 | 0 |
| ENSMUSG00 | Rimbp3 | 5836 | 0.14986645 | 0.07431083 | 0.06059197 | 0.00618267 | 0.13115073 | 0.46267808 |
| ENSMUSG00 | Cebpd | 1178 | 19.3377677 | 30.3517318 | 45.4776178 | 26.1885807 | 28.7185799 | 20.9074697 |
| ENSMUSG00 | Stxbp3b | 1005 | 0.47469231 | 0.28768025 | 0.21990967 | 0.17961266 | 0.15231754 | 0.16283367 |
| ENSMUSG00 | Eef1g | 2589 | 4.11528342 | 3.21987427 | 3.75604692 | 2.85701782 | 2.44390522 | 2.1385677 |
| ENSMUSG00 | Tut1 | 2738 | 8.21826222 | 9.08115564 | 6.26381121 | 5.66664688 | 7.56636544 | 7.7799464 |
| ENSMUSG00 | Mta2 | 4073 | 24.8898785 | 29.4939357 | 32.6548848 | 25.5488869 | 29.3279513 | 20.5313145 |
| ENSMUSG00 | Eml3 | 3139 | 9.75206116 | 13.7850988 | 8.80094039 | 7.24169737 | 14.8576322 | 6.62098602 |
| ENSMUSG00 | Rom1 | 1523 | 3.73278642 | 5.82160562 | 0.89970921 | 2.51129135 | 4.32199769 | 1.57594764 |
| ENSMUSG00 | B3gat3 | 1596 | 11.7572599 | 11.3220024 | 11.8536271 | 9.38223589 | 11.8933658 | 10.3732494 |
| ENSMUSG00 | Ganab | 3882 | 68.0717875 | 63.578299 | 72.8499206 | 58.5938199 | 67.4436719 | 48.3734908 |
| ENSMUSG00 | Ints5 | 3292 | 6.35218188 | 8.35797645 | 8.78128959 | 8.52728756 | 7.2230532 | 3.93543549 |
| ENSMUSG00 | 1810009A15 | 611 | 30.4510064 | 29.8897898 | 34.5078218 | 33.2474486 | 21.5463256 | 37.5416884 |
| ENSMUSG00 | AI462493 | 827 | 21.6804374 | 25.0546193 | 22.6086815 | 16.710306 | 9.93379263 | 27.8353029 |
| ENSMUSG00 | Ubxn1 | 1058 | 51.9677028 | 43.6775029 | 54.0198327 | 37.1392689 | 46.0587797 | 56.6889714 |
| ENSMUSG00 | Lrrn4cl | 798 | 0 | 1.02652822 | 0.2769539 | 0 | 0 | 0 |
| ENSMUSG00 | Bscl2 | 4480 | 18.1916819 | 14.0687202 | 14.8983891 | 15.922813 | 15.2851331 | 13.7895219 |
| ENSMUSG00 | Gng3 | 1696 | 0 | 0 | 0 | 0.08509916 | 0 | 0.06432698 |
| ENSMUSG00 | Hnrnpul2 | 5712 | 6.89735315 | 8.31789775 | 6.58539363 | 9.08999686 | 4.78818903 | 12.2955525 |
| ENSMUSG00 | Ttc9c | 4683 | 13.8036326 | 14.7553615 | 16.9898179 | 12.8671822 | 15.6685732 | 14.3391334 |
| ENSMUSG00 | Zbtb3 | 2070 | 0.32649428 | 1.1872022 | 1.13173801 | 1.02842542 | 0.93671608 | 1.37032004 |
| ENSMUSG00 | Polr2g | 860 | 45.164075 | 43.7039824 | 53.6589815 | 39.3965579 | 31.1498225 | 66.0299987 |
| ENSMUSG00 | Foxr2 | 3209 | 0 | 0 | 0 | 0 | 0 | 0 |
| ENSMUSG00 | Snx29 | 5230 | 1.29224315 | 2.85617535 | 1.40296481 | 1.91793659 | 4.56603134 | 1.0951576 |
| ENSMUSG00 | Gm10343 | 347 | 0 | 0 | 0 | 0 | 0 | 0.55020886 |
| ENSMUSG00 | Zcchc16 | 2475 | 24.1263563 | 12.421394 | 31.0573758 | 18.6368771 | 12.1226299 | 20.6956657 |
| ENSMUSG00 | Gm7123 | 212 | 0 | 0 | 0 | 0 | 0 | 0 |
| ENSMUSG00 | Tex13a | 1315 | 0 | 0 | 0 | 0 | 0 | 0 |
| ENSMUSG00 | Gm10344 | 1418 | 0 | 0 | 0 | 0 | 0 | 0 |
| ENSMUSG00 | Gm960 | 2032 | 1.27170582 | 0.87741061 | 1.08764378 | 1.68690661 | 1.90846681 | 1.67781984 |
| ENSMUSG00 | Gm5167 | 659 | 0 | 0 | 0 | 0 | 0 | 0 |
| ENSMUSG00 | Gm6604 | 3052 | 0 | 0 | 0 | 0 | 0 | 0 |
| ENSMUSG00 | Sms | 6164 | 0.64496237 | 1.2664185 | 1.49873219 | 1.80293149 | 1.44039413 | 1.80090496 |
| ENSMUSG00 | Mpst | 1509 | 0.79036741 | 0.73445207 | 1.14239356 | 0.43040212 | 0.06762939 | 0.61453793 |
| ENSMUSG00 | Csf2rb | 5105 | 0.00778756 | 0 | 0 | 0 | 0 | 0 |
| ENSMUSG00 | Csf2rb2 | 6550 | 0 | 0 | 0 | 0 | 0 | 0.41224264 |
| ENSMUSG00 | Ncf4 | 2031 | 0 | 0 | 0.06529076 | 0 | 0 | 0 |
| ENSMUSG00 | Apol7e | 2227 | 0 | 0 | 0 | 0 | 0 | 0 |
| ENSMUSG00 | Tmem28 | 3906 | 0 | 0.16037474 | 0 | 0 | 0 | 0 |
| ENSMUSG00 | 1700010D01 | 554 | 0 | 0 | 0 | 0 | 0 | 0 |
| ENSMUSG00 | Spin4 | 4175 | 1.0093607 | 0.32316656 | 1.00579044 | 3.3791807 | 1.80883919 | 0.77087603 |
| ENSMUSG00 | Gspt2 | 4410 | 4.06569653 | 1.69362779 | 4.63066927 | 3.29729341 | 2.99678715 | 6.42592975 |
| ENSMUSG00 | Gm10345 | 2381 | 0.38403026 | 1.27498776 | 0.6497541 | 0.21215818 | 0.90008727 | 0.19473703 |
| ENSMUSG00 | Gm5941 | 558 | 0 | 0 | 0 | 0 | 0 | 0 |
| ENSMUSG00 | Gm16462 | 455 | 0 | 0.21180854 | 0 | 0 | 0 | 0 |
| ENSMUSG00 | Gm4937 | 1075 | 0 | 0 | 0 | 0 | 0 | 0 |
| ENSMUSG00 | DXBay18 | 2857 | 0 | 0 | 0 | 0 | 0 | 0 |
| ENSMUSG00 | Gm14698 | 4225 | 0.01881916 | 0.02281015 | 0.0418479 | 0.07686116 | 0.0241545 | 0.26467697 |
| ENSMUSG00 | 4933412E24 | 1752 | 0 | 0 | 0 | 0 | 0 | 0 |
| ENSMUSG00 | Gm7713 | 398 | 0.09988814 | 0.12107146 | 0.11105991 | 0 | 0 | 0 |
| ENSMUSG00 | C230004F18 | 2435 | 0 | 0 | 0 | 0 | 0 | 0 |
| ENSMUSG00 | Zfx2 | 4430 | 0 | 0 | 0 | 0 | 0 | 0 |
| ENSMUSG00 | 4933416I08 | 1147 | 0 | 0 | 0 | 0 | 0 | 0 |
| ENSMUSG00 | Olfr1320 | 959 | 0 | 0 | 0 | 0 | 0 | 0 |
| ENSMUSG00 | Rhox12 | 1011 | 0 | 0 | 0 | 0 | 0 | 0 |
| ENSMUSG00 | Rhox7 | 975 | 0 | 0 | 0 | 0 | 0 | 0 |
| ENSMUSG00 | Rhox3h | 1043 | 0 | 0 | 0 | 0 | 0 | 0 |
| ENSMUSG00 | Rhox4e | 973 | 0 | 0 | 0 | 0 | 0 | 0 |
| ENSMUSG00 | Rhox4b | 2307 | 0 | 0 | 0 | 0 | 0 | 0 |
| ENSMUSG00 | Rhox3e | 1047 | 0 | 0 | 0 | 0 | 0 | 0 |
| ENSMUSG00 | Rhox1 | 878 | 0 | 0 | 0 | 0 | 0 | 0 |
| ENSMUSG00 | Gm5169 | 1166 | 0 | 0 | 0 | 0 | 0 | 0 |
| ENSMUSG00 | Gm14525 | 884 | 0 | 0 | 0 | 0 | 0 | 0 |
| ENSMUSG00 | 2610005L07 | 8637 | 2.43955131 | 2.51616133 | 3.70523728 | 3.90606827 | 2.93030941 | 5.32419148 |
| ENSMUSG00 | 6820431F20 | 8208 | 1.08010138 | 1.47940831 | 1.17397683 | 1.94741054 | 2.00176572 | 1.51858067 |
| ENSMUSG00 | Fthl17 | 855 | 0 | 0 | 0 | 0 | 0 | 0 |
| ENSMUSG00 | Ssxb5 | 595 | 0 | 0 | 0 | 0.24256837 | 0 | 0 |
| ENSMUSG00 | 4930417O22 | 1178 | 0 | 0 | 0 | 0 | 0 | 0 |
| ENSMUSG00 | Apcdd1 | 2961 | 0 | 0 | 0 | 0 | 0 | 0 |
| ENSMUSG00 | Ccdc112 | 2791 | 2.53546239 | 3.47025255 | 1.02942307 | 4.53772756 | 2.5595459 | 5.50183515 |
| ENSMUSG00 | Mcc | 8406 | 0.58171831 | 1.29551938 | 0.39963598 | 1.41649711 | 0.58274234 | 0.73004923 |
| ENSMUSG00 | Gm94 | 744 | 0 | 0 | 0 | 0 | 0 | 0.62321084 |
| ENSMUSG00 | 2900055J20 | 722 | 1.59682678 | 0.20022067 | 1.22442778 | 0.59970157 | 0.56538921 | 1.92680194 |
| ENSMUSG00 | Lrrtm2 | 5663 | 0.75116306 | 1.1572234 | 1.90451169 | 3.23037199 | 0.78391218 | 1.78202658 |
| ENSMUSG00 | Ppia | 3616 | 10.4226204 | 5.94334989 | 4.42507389 | 8.02266698 | 5.08005954 | 7.49750869 |
| ENSMUSG00 | Gm6304 | 446 | 0.26741355 | 0.10804135 | 3.17143268 | 1.77983179 | 0.45763566 | 2.20154039 |
| ENSMUSG00 | Gm10349 | 864 | 0 | 0.22308538 | 0.15347862 | 0 | 0 | 0.03156787 |
| ENSMUSG00 | 4921528O07 | 2934 | 0 | 0 | 0 | 0 | 0 | 0 |
| ENSMUSG00 | Vmn1r4 | 1629 | 0.53690643 | 0.20706267 | 0.40701513 | 0.39869663 | 0.56382736 | 0 |
| ENSMUSG00 | Gm806 | 2541 | 0 | 0 | 0 | 0 | 0 | 0 |
| ENSMUSG00 | Adam25 | 2618 | 0 | 0 | 0 | 0 | 0 | 0 |
| ENSMUSG00 | Gm10351 | 122 | 0 | 0 | 0 | 0 | 0 | 0 |
| ENSMUSG00 | Gm10352 | 1721 | 0 | 0 | 0 | 0 | 0 | 0 |
| ENSMUSG00 | Fndc1 | 7722 | 0 | 0 | 0 | 0 | 0 | 0 |
| ENSMUSG00 | Vmn2r121 | 3965 | 0 | 0 | 0 | 0 | 0 | 0 |
| ENSMUSG00 | 6720489N17 | 316 | 3.52263751 | 4.11719601 | 0.8392755 | 4.79571478 | 2.26066222 | 3.79773458 |
| ENSMUSG00 | Gm16261 | 204 | 0 | 0 | 0 | 0 | 0 | 0 |
| ENSMUSG00 | Ccnf | 4834 | 1.23362062 | 0.85726811 | 0.32003817 | 0 | 0.72834504 | 0.5247293 |
| ENSMUSG00 | Gm16390 | 1077 | 0 | 0 | 0 | 0 | 0 | 0 |
| ENSMUSG00 | A530040E14 | 2372 | 0 | 0 | 0 | 0 | 0 | 0.01149858 |
| ENSMUSG00 | Ang | 1270 | 1.12692701 | 0.64501537 | 1.42698863 | 0.68186541 | 0.56249548 | 2.4697508 |
| ENSMUSG00 | BC061237 | 894 | 0 | 0 | 0 | 0 | 0 | 0 |
| ENSMUSG00 | Slx | 1506 | 0 | 0 | 0 | 0 | 0 | 0 |
| ENSMUSG00 | Gm10354 | 1210 | 0 | 0.07964701 | 0.0730609 | 0.32801859 | 0 | 0.0901641 |
| ENSMUSG00 | Gm7736 | 1526 | 0 | 0 | 0 | 0 | 0 | 0 |
| ENSMUSG00 | 5-Sep | 2124 | 0 | 0.0907466 | 0.04162132 | 0.18686558 | 0.09609487 | 0.11557051 |
| ENSMUSG00 | Tuba1a | 2399 | 16.1905396 | 17.1334036 | 22.423361 | 11.942119 | 9.40127474 | 15.3597489 |
| ENSMUSG00 | Vmn2r66 | 2631 | 0 | 0 | 0 | 0 | 0 | 0 |
| ENSMUSG00 | Trim6 | 4972 | 2.32679904 | 1.58941603 | 2.23142852 | 1.39335329 | 1.21100409 | 5.91901363 |
| ENSMUSG00 | Gm5635 | 732 | 0 | 0 | 0 | 0 | 0 | 0 |
| ENSMUSG00 | Gm16382 | 415 | 0.09579634 | 0 | 0 | 0 | 0 | 0 |
| ENSMUSG00 | Gm5592 | 3030 | 0 | 0 | 0 | 0 | 0 | 0 |
| ENSMUSG00 | Klf12 | 5717 | 1.87060071 | 2.07344145 | 1.0669677 | 3.7300137 | 2.45447846 | 0.55341231 |
| ENSMUSG00 | Als2cr11 | 2190 | 0 | 0 | 0 | 0 | 0 | 0.29890016 |
| ENSMUSG00 | Olfr474 | 932 | 0 | 0 | 0 | 0 | 0 | 0 |
| ENSMUSG00 | Gm8420 | 611 | 0.84586129 | 0.78864881 | 0 | 0.59054083 | 0.50107734 | 0 |
| ENSMUSG00 | Gm10355 | 518 | 0 | 0 | 0 | 0 | 0 | 0 |
| ENSMUSG00 | Gm10356 | 1259 | 0 | 0 | 0.03510869 | 0 | 0 | 0 |
| ENSMUSG00 | Olfr775 | 938 | 0 | 0 | 0 | 0 | 0 | 0 |
| ENSMUSG00 | Olfr774 | 938 | 0 | 0 | 0 | 0 | 0 | 0 |
| ENSMUSG00 | Gm6419 | 431 | 1.19912122 | 1.22981639 | 2.25624257 | 1.17203857 | 0.23678133 | 1.70862009 |
| ENSMUSG00 | Dppa2 | 1941 | 0 | 0 | 0 | 0 | 0 | 0 |
| ENSMUSG00 | Gm10357 | 532 | 0 | 0.09057602 | 0 | 0.27129357 | 0 | 0 |
| ENSMUSG00 | Psmb11 | 4246 | 0 | 0 | 0 | 0 | 0 | 0 |
| ENSMUSG00 | Gm10358 | 1228 | 0.09712251 | 0.23543864 | 0.07198997 | 0.08814832 | 0.04155242 | 0.04442124 |
| ENSMUSG00 | Nanos1 | 3926 | 0 | 0 | 0.58545487 | 0.00919054 | 0.02599408 | 0 |
| ENSMUSG00 | Gm10359 | 1179 | 0.50579492 | 0.32696483 | 0 | 0.09181182 | 0.17311748 | 0.25447077 |
| ENSMUSG00 | Atp5l-ps1 | 311 | 0 | 0 | 0 | 0 | 0 | 0.08769981 |
| ENSMUSG00 | Gm16451 | 403 | 0 | 0 | 0 | 0 | 0 | 0 |
| ENSMUSG00 | 1700024G13 | 395 | 0 | 0 | 0 | 0 | 0 | 0 |
| ENSMUSG00 | Gm9008 | 2306 | 0 | 0.12537689 | 0 | 0 | 0 | 0.01182768 |
| ENSMUSG00 | Gm10360 | 249 | 0 | 0 | 0 | 0 | 0 | 0 |
| ENSMUSG00 | Xlr5b | 1735 | 0 | 0 | 0 | 0 | 0 | 0 |
| ENSMUSG00 | Gm8556 | 527 | 0 | 0.09143537 | 0 | 0 | 0 | 0 |
| ENSMUSG00 | Gm628 | 7469 | 0 | 0 | 0 | 0 | 0 | 0 |
| ENSMUSG00 | Gm10362 | 584 | 0 | 0 | 0 | 0 | 0 | 0.04670315 |
| ENSMUSG00 | Gm17515 | 1485 | 0.05354274 | 0 | 0 | 0.38876277 | 0 | 0 |
| ENSMUSG00 | Gm10364 | 1119 | 0.71055372 | 0.34449646 | 0.2370072 | 0.48367352 | 0.18239991 | 0.29248943 |
| ENSMUSG00 | Ppp1r3e | 2189 | 0.12713036 | 0.52831184 | 0.02019271 | 0 | 0.09324144 | 0.17443808 |
| ENSMUSG00 | Phf20l1 | 3351 | 16.8702755 | 18.3053837 | 15.2615734 | 18.9724149 | 21.318092 | 15.9447981 |
| ENSMUSG00 | Hhla1 | 2245 | 1.08021573 | 0.21463894 | 0.49222542 | 1.33399096 | 0.0227289 | 0.17008684 |
| ENSMUSG00 | Rps19-ps4 | 449 | 0 | 0 | 0.19689017 | 0 | 0 | 0 |
| ENSMUSG00 | Gm6755 | 863 | 0 | 0 | 0 | 0 | 0 | 0 |
| ENSMUSG00 | Gm5046 | 311 | 0 | 0 | 0 | 0 | 0 | 0 |
| ENSMUSG00 | Rps19-ps2 | 455 | 0 | 0 | 0 | 0 | 0 | 0 |
| ENSMUSG00 | Gm6740 | 863 | 0 | 0 | 0 | 0 | 0 | 0 |
| ENSMUSG00 | Gm525 | 759 | 0 | 0 | 0 | 0 | 0 | 0 |
| ENSMUSG00 | Pvt1 | 5729 | 0 | 0.00841097 | 0 | 0 | 0 | 0 |
| ENSMUSG00 | Fam84b | 5561 | 15.4989897 | 20.0942923 | 13.3058596 | 16.214535 | 22.1869766 | 17.4653822 |
| ENSMUSG00 | G630016D24 | 1220 | 0.39103751 | 0.11849125 | 0.54346528 | 0.32532991 | 0.0418249 | 0 |
| ENSMUSG00 | Slc39a2 | 2802 | 0.14188251 | 0 | 0 | 0.01287725 | 0.25494977 | 0.30175368 |

| | | | | | | | | |
|---|---|---|---|---|---|---|---|---|
| ENSMUSG00 | Gm10369 | 1146 | 0.20814388 | 0 | 0.03857054 | 0 | 0 | 0.73779565 |
| ENSMUSG00 | Gm17184 | 459 | 0 | 0 | 0 | 0 | 0 | 0 |
| ENSMUSG00 | Eddm3b | 1255 | 0.69690882 | 0.537538 | 2.46544144 | 1.58128482 | 0.4879016 | 3.02085646 |
| ENSMUSG00 | Gm16020 | 518 | 0 | 0 | 0 | 0 | 0 | 0.26326872 |
| ENSMUSG00 | Ptrh2 | 3129 | 7.30565717 | 7.82317437 | 15.8781948 | 8.09510874 | 4.46827324 | 11.7675817 |
| ENSMUSG00 | Gm7489 | 1220 | 0 | 0 | 0 | 0 | 0 | 0 |
| ENSMUSG00 | Gm6616 | 335 | 0 | 0 | 0 | 0 | 0 | 0 |
| ENSMUSG00 | Gm10371 | 1407 | 0 | 0.03424765 | 0 | 0 | 0 | 0 |
| ENSMUSG00 | 5930412G12I | 3544 | 0 | 0 | 0 | 0 | 0 | 0 |
| ENSMUSG00 | Gm10373 | 665 | 0.83695748 | 0 | 0.06646894 | 0.21703485 | 0.15346278 | 0.57420293 |
| ENSMUSG00 | Gm16439 | 563 | 0 | 0 | 0.2355338 | 0.12817778 | 0 | 0.04844519 |
| ENSMUSG00 | 4930503E14I | 1283 | 0 | 0 | 0 | 0 | 0 | 0 |
| ENSMUSG00 | Ear2 | 717 | 0 | 0 | 0 | 0 | 0 | 0 |
| ENSMUSG00 | Ang6 | 1155 | 0 | 0 | 0 | 0 | 0 | 0 |
| ENSMUSG00 | Ear-ps2 | 465 | 0 | 0 | 0 | 0 | 0 | 0 |
| ENSMUSG00 | Ear-ps9 | 778 | 0 | 0 | 0 | 0 | 0 | 0 |
| ENSMUSG00 | Ear1 | 711 | 0 | 0 | 0 | 0 | 0 | 0 |
| ENSMUSG00 | Gm10375 | 1241 | 0 | 0 | 0 | 0 | 0 | 0 |
| ENSMUSG00 | 1700001F09I | 894 | 0 | 0 | 0 | 0 | 0 | 0 |
| ENSMUSG00 | Gm10376 | 895 | 0 | 0 | 0 | 0 | 0 | 0 |
| ENSMUSG00 | Gm10377 | 894 | 0 | 0 | 0 | 0 | 0 | 0 |
| ENSMUSG00 | Gm10378 | 1398 | 0 | 0 | 0 | 0 | 0 | 0 |
| ENSMUSG00 | Gm3633 | 1407 | 0 | 0 | 0 | 0 | 0 | 0 |
| ENSMUSG00 | Gm9611 | 1407 | 0 | 0 | 0 | 0 | 0 | 0 |
| ENSMUSG00 | Gm7395 | 1568 | 0 | 0 | 0 | 0 | 0 | 0 |
| ENSMUSG00 | Gm10382 | 1232 | 2.51698659 | 2.2294052 | 0.35878119 | 3.133749 | 2.73355585 | 1.66038794 |
| ENSMUSG00 | 1700049E17I | 2041 | 0 | 0 | 0 | 0 | 0 | 0 |
| ENSMUSG00 | Gm7853 | 474 | 0 | 0 | 0 | 0 | 0 | 0 |
| ENSMUSG00 | Gm10384 | 2044 | 0.09724922 | 0 | 0 | 0 | 0 | 0 |
| ENSMUSG00 | Slfn2 | 1662 | 4.25780718 | 2.00052016 | 0.74467606 | 1.41115096 | 5.40351513 | 3.8072902 |
| ENSMUSG00 | Slfn10-ps | 3616 | 0 | 0 | 0.17113545 | 0 | 0.4233383 | 0 |
| ENSMUSG00 | Zfp9 | 3962 | 0.06020517 | 0.17027011 | 0 | 0.21856867 | 0 | 0.01376812 |
| ENSMUSG00 | Gm5460 | 1898 | 0 | 0 | 0 | 0 | 0 | 0 |
| ENSMUSG00 | Gdf2 | 2986 | 0 | 0 | 0 | 0 | 0 | 0 |
| ENSMUSG00 | 1810012P15I | 8174 | 4.29946719 | 4.56279743 | 3.92592831 | 4.63937225 | 7.22884062 | 3.37345979 |
| ENSMUSG00 | Cux2 | 412 | 0 | 0 | 0 | 0 | 0 | 0 |
| ENSMUSG00 | Adam1a | 3091 | 2.6752313 | 2.4631051 | 3.40344181 | 3.00002764 | 1.58477259 | 1.81772102 |
| ENSMUSG00 | Zfp783 | 3693 | 0.39830836 | 0.74373875 | 0.19150541 | 0.37127476 | 0.59413327 | 0.39881682 |
| ENSMUSG00 | Gm17524 | 1308 | 0 | 0 | 0 | 0 | 0 | 0 |
| ENSMUSG00 | Gm6288 | 1298 | 1.53141296 | 1.74480954 | 1.94106707 | 0.77834919 | 1.72970766 | 1.82811527 |
| ENSMUSG00 | Gm5887 | 1275 | 0.15590385 | 0.26455302 | 0.48535357 | 0.82068964 | 0.08004137 | 0 |
| ENSMUSG00 | Spef2 | 7201 | 0.03864579 | 0.25428202 | 0.06752122 | 0.0050107 | 0.24801044 | 0.20831901 |
| ENSMUSG00 | Ugt3a1 | 2660 | 3.8559827 | 2.22817007 | 3.88843281 | 1.49211462 | 2.55131879 | 2.33782622 |
| ENSMUSG00 | Gm10390 | 8062 | 0.04438096 | 0.00597698 | 0.05482739 | 0.01342671 | 0.12025566 | 0.09134399 |
| ENSMUSG00 | Gm10391 | 5073 | 0 | 0 | 0 | 0 | 0 | 0 |
| ENSMUSG00 | Gm10392 | 1730 | 0 | 0 | 0 | 0 | 0 | 0 |
| ENSMUSG00 | Gm10393 | 446 | 0 | 0 | 0 | 0 | 0 | 0 |
| ENSMUSG00 | Gm10394 | 4758 | 0 | 0 | 0 | 0 | 0 | 0 |
| ENSMUSG00 | Gm10395 | 1617 | 0 | 0 | 0 | 0 | 0 | 0 |
| ENSMUSG00 | Gm10396 | 1205 | 0 | 0 | 0 | 0 | 0 | 0 |
| ENSMUSG00 | Cphx | 1706 | 0 | 0 | 0 | 0 | 0 | 0 |
| ENSMUSG00 | D6Ertd474e | 2623 | 0.13640844 | 0 | 0.38758764 | 0.06878011 | 0.05836032 | 0.57190437 |
| ENSMUSG00 | D14Ertd449e | 1272 | 0 | 0 | 0 | 0 | 0 | 0 |
| ENSMUSG00 | Plac9 | 1327 | 0 | 0 | 0 | 0 | 0 | 0 |
| ENSMUSG00 | Sah2 | 1511 | 0 | 0 | 0 | 0 | 0 | 0 |
| ENSMUSG00 | Gm7457 | 4108 | 0 | 0 | 0 | 0 | 0 | 0 |
| ENSMUSG00 | Gm10397 | 1698 | 0 | 0 | 0 | 0 | 0 | 0 |
| ENSMUSG00 | Gm10398 | 395 | 0 | 0 | 0 | 0 | 0 | 0 |
| ENSMUSG00 | Gm10399 | 497 | 0.63992725 | 0.7756369 | 0 | 0.29039875 | 0.41067506 | 0.8780568 |
| ENSMUSG00 | Gm13826 | 1011 | 0.78645857 | 1.42986475 | 0.39348822 | 0.46396299 | 1.21130863 | 0.78235859 |
| ENSMUSG00 | Gm10401 | 606 | 0 | 0 | 0 | 0 | 0 | 0 |
| ENSMUSG00 | 1500011B03I | 1331 | 7.52695799 | 5.6476972 | 12.3871431 | 7.86160249 | 7.74404802 | 15.4918312 |
| ENSMUSG00 | 2700089E24I | 2508 | 81.7460179 | 64.74813 | 77.1946062 | 91.5143083 | 62.4401704 | 121.093743 |
| ENSMUSG00 | Olfr31 | 953 | 0 | 0 | 0 | 0 | 0 | 0 |
| ENSMUSG00 | Olfr381 | 1998 | 0 | 0 | 0 | 0 | 0 | 0 |
| ENSMUSG00 | Olfr380 | 2212 | 0 | 0 | 0 | 0 | 0 | 0 |
| ENSMUSG00 | Rpl21-ps4 | 440 | 0 | 0 | 0 | 0 | 0 | 0.06198782 |
| ENSMUSG00 | Klra11 | 838 | 0 | 0 | 0 | 0 | 0 | 0 |
| ENSMUSG00 | Klra10 | 819 | 0 | 0 | 0 | 0 | 0 | 0 |
| ENSMUSG00 | Myo18b | 8236 | 0 | 0 | 0 | 0 | 0 | 0 |
| ENSMUSG00 | Klra14 | 877 | 0 | 0 | 0 | 0 | 0 | 0 |
| ENSMUSG00 | Gm6588 | 2312 | 0 | 0 | 0 | 0 | 0 | 0 |
| ENSMUSG00 | Gm10044 | 2622 | 0.27292092 | 0.07351097 | 0.3708774 | 0.3027479 | 0 | 0.54091581 |
| ENSMUSG00 | Gm5797 | 992 | 0 | 0 | 0 | 0.07274606 | 0 | 0 |
| ENSMUSG00 | Gm16434 | 2061 | 0 | 0 | 0 | 0 | 0 | 0 |
| ENSMUSG00 | Gm6676 | 2064 | 0 | 0 | 0 | 0 | 0 | 0 |
| ENSMUSG00 | Gm3715 | 2172 | 0 | 0 | 0 | 0 | 0.14095684 | 0 |
| ENSMUSG00 | Gm10406 | 2714 | 0 | 0 | 0 | 0 | 0 | 0 |
| ENSMUSG00 | Gm3591 | 1740 | 0 | 0 | 0 | 0 | 0 | 0 |
| ENSMUSG00 | Gm16440 | 1084 | 0 | 0 | 0 | 0 | 0 | 0 |
| ENSMUSG00 | Gm10408 | 1507 | 0 | 0 | 0 | 0 | 0 | 0 |
| ENSMUSG00 | Gm10409 | 2061 | 0 | 0 | 0 | 0 | 0 | 0 |
| ENSMUSG00 | Gm10410 | 1686 | 0 | 0 | 0 | 0 | 0 | 0 |
| ENSMUSG00 | Gm3483 | 606 | 0 | 0 | 0 | 0 | 0 | 0 |
| ENSMUSG00 | Gm10412 | 608 | 0 | 0 | 0 | 0 | 0 | 0 |
| ENSMUSG00 | Gm8165 | 1399 | 0.05683414 | 0.48220886 | 0.06319063 | 0.07737393 | 0.07294693 | 0.31193297 |
| ENSMUSG00 | Gm10413 | 1111 | 0 | 0 | 0 | 0 | 0 | 0 |
| ENSMUSG00 | 9230020A06I | 2053 | 0 | 0 | 0 | 0 | 0 | 0 |
| ENSMUSG00 | Gm1679 | 681 | 0 | 0 | 0 | 0 | 0 | 0 |
| ENSMUSG00 | Gm6712 | 468 | 44.5125466 | 30.2709701 | 39.1960787 | 57.0528054 | 21.5880821 | 37.4734894 |
| ENSMUSG00 | 4930522L14I | 2405 | 20.4149765 | 13.2637941 | 25.6940443 | 28.5955829 | 20.3044664 | 28.6015136 |
| ENSMUSG00 | 5430403G16I | 3337 | 24.7920752 | 19.4074253 | 19.1007065 | 20.3603506 | 16.3920512 | 21.9374084 |
| ENSMUSG00 | Gm10418 | 574 | 0 | 0.08394851 | 0.0770067 | 0 | 0 | 0 |
| ENSMUSG00 | Gm10419 | 1666 | 0.143177 | 0 | 0.18572203 | 1.10455238 | 0.55130538 | 0.13097066 |
| ENSMUSG00 | Acrbp | 3129 | 2.18534441 | 2.57179158 | 1.15837364 | 3.2403498 | 3.71812518 | 2.71961887 |
| ENSMUSG00 | Grcc10 | 1110 | 9.34791028 | 10.0279893 | 11.7075152 | 9.42684046 | 6.80351678 | 11.5732929 |
| ENSMUSG00 | Zfp951 | 1775 | 2.79968173 | 7.05829575 | 1.14551255 | 4.61443612 | 0.83367037 | 1.84391927 |
| ENSMUSG00 | Vmn2r27 | 2568 | 0 | 0 | 0 | 0 | 0 | 0 |
| ENSMUSG00 | Vmn2r25 | 2562 | 0.27931251 | 0 | 0 | 0 | 0 | 0.11710423 |
| ENSMUSG00 | Vmn2r24 | 2559 | 0 | 0.05649055 | 0 | 0 | 0 | 0.02131664 |
| ENSMUSG00 | Gm10420 | 1830 | 0.02172431 | 0 | 0 | 0 | 0 | 0.01490417 |
| ENSMUSG00 | Abcb5 | 4348 | 0.10972074 | 0 | 0 | 0 | 0 | 0 |
| ENSMUSG00 | Cecr6 | 787 | 0 | 0 | 0 | 0 | 0 | 0 |
| ENSMUSG00 | 9330160F10I | 2765 | 2.40114475 | 1.08049165 | 1.5826338 | 0.87432079 | 0.84890173 | 4.92225858 |
| ENSMUSG00 | Ahnak2 | 16977 | 0.09132731 | 0.11353347 | 0 | 0 | 0 | 0.09639385 |
| ENSMUSG00 | E330014E10I | 2119 | 0 | 0 | 0 | 0 | 0 | 0 |
| ENSMUSG00 | Gm7982 | 1586 | 0 | 0.03038237 | 0 | 0 | 0 | 0 |
| ENSMUSG00 | Gm7978 | 1592 | 0 | 0 | 0 | 0 | 0 | 0 |
| ENSMUSG00 | Gm6468 | 1215 | 0 | 0 | 0 | 0 | 0 | 0 |
| ENSMUSG00 | Gm10424 | 1844 | 0 | 0 | 0 | 0 | 0 | 0 |
| ENSMUSG00 | Gm16513 | 1689 | 0 | 0 | 0 | 0 | 0 | 0 |
| ENSMUSG00 | Gm6367 | 2511 | 0 | 0 | 0 | 0 | 0 | 0 |
| ENSMUSG00 | Gm6351 | 1592 | 0 | 0 | 0 | 0 | 0 | 0 |
| ENSMUSG00 | BC061212 | 1893 | 0 | 0 | 0 | 0 | 0 | 0 |
| ENSMUSG00 | Gm16429 | 1844 | 0 | 0 | 0 | 0 | 0 | 0 |
| ENSMUSG00 | Gm7792 | 2063 | 0 | 0 | 0 | 0 | 0 | 0 |
| ENSMUSG00 | AW555464 | 6621 | 11.7927449 | 13.8642459 | 9.83375846 | 13.0137324 | 13.64865 | 7.78157281 |
| ENSMUSG00 | Gm16427 | 1706 | 0 | 0 | 0 | 0 | 0 | 0 |
| ENSMUSG00 | 4932430I15R | 3304 | 0.02406506 | 0.01458427 | 0 | 0.03276215 | 0 | 0.12382554 |
| ENSMUSG00 | Gm7292 | 994 | 0 | 0 | 0.08893731 | 0 | 0 | 0 |
| ENSMUSG00 | Gm10425 | 3362 | 6.25539833 | 3.08152441 | 5.08807652 | 3.87436588 | 4.12824932 | 6.87950449 |
| ENSMUSG00 | Gm12298 | 727 | 0 | 0 | 0 | 0 | 0 | 0 |
| ENSMUSG00 | Mir546 | 120 | 0 | 0 | 0 | 0 | 0 | 0 |
| ENSMUSG00 | Gm10428 | 2510 | 0.04751651 | 0.44154907 | 0.15849266 | 0.01437532 | 0.26428003 | 0.49985395 |
| ENSMUSG00 | G530011O06 | 2378 | 0.40123277 | 1.15501564 | 0.16729041 | 0.2275991 | 0.7295613 | 0.26380013 |
| ENSMUSG00 | Tmprss11a | 2648 | 0 | 0 | 0 | 0 | 0 | 0 |
| ENSMUSG00 | A530017D24 | 4334 | 2.2290221 | 1.73444508 | 1.64201586 | 2.66411035 | 2.62549188 | 4.82057537 |
| ENSMUSG00 | Serpina1e | 1809 | 0 | 0 | 0 | 0 | 0.22565561 | 0 |
| ENSMUSG00 | 2310040G07 | 937 | 2.07899525 | 1.33708377 | 0.80195446 | 5.58366751 | 1.85154405 | 2.38689479 |
| ENSMUSG00 | Gm10430 | 1843 | 0 | 0 | 0 | 0 | 0 | 0 |
| ENSMUSG00 | Rybp | 4453 | 1.09811904 | 1.6664523 | 1.54850382 | 1.87986399 | 1.8792557 | 2.18050114 |
| ENSMUSG00 | Gm6116 | 1428 | 0 | 0 | 0 | 0 | 0 | 0 |
| ENSMUSG00 | Gpr27 | 1139 | 0.06980769 | 0.08461184 | 0.23284553 | 0 | 0.08959855 | 0.23946128 |
| ENSMUSG00 | 1700123L14I | 1702 | 0 | 0 | 0 | 0 | 0 | 0 |
| ENSMUSG00 | Gm10431 | 389 | 0 | 0 | 0 | 0 | 0 | 0 |
| ENSMUSG00 | Gm10433 | 2012 | 0.90892252 | 0.19159619 | 0.57119678 | 0.64560318 | 0.25361022 | 1.69449797 |
| ENSMUSG00 | Gm5647 | 920 | 0 | 0 | 0 | 0 | 0 | 0 |
| ENSMUSG00 | Nfxl1 | 5924 | 4.99292328 | 5.18142533 | 4.66342875 | 4.69602571 | 8.24311979 | 6.10042151 |
| ENSMUSG00 | 4933439C10I | 4836 | 5.39280298 | 3.98564451 | 2.56838666 | 7.52828431 | 4.57929014 | 9.3171431 |
| ENSMUSG00 | Ppp1r2-ps2 | 873 | 0 | 0 | 0 | 0 | 0 | 0 |
| ENSMUSG00 | Gm10434 | 469 | 0.25429945 | 1.13017242 | 0.094247 | 0.23080199 | 0.43519297 | 0.23261953 |
| ENSMUSG00 | Mir540 | 83 | 0 | 0 | 0 | 0 | 0 | 0 |
| ENSMUSG00 | Gm10435 | 4148 | 0 | 0 | 0 | 0 | 0 | 0 |
| ENSMUSG00 | Gm5662 | 2838 | 0 | 0 | 0 | 0 | 0 | 0 |
| ENSMUSG00 | Gm8332 | 434 | 0 | 0 | 0 | 0 | 0 | 0 |
| ENSMUSG00 | Gm10436 | 2731 | 0 | 0 | 0 | 0 | 0 | 0 |
| ENSMUSG00 | Gm16368 | 434 | 0 | 0 | 0 | 0 | 0 | 0 |
| ENSMUSG00 | Gm16381 | 493 | 0 | 0 | 0 | 0 | 0 | 0 |
| ENSMUSG00 | Gm12258 | 2523 | 15.3002662 | 7.50585484 | 16.2756687 | 12.885423 | 19.3952812 | 20.4208456 |
| ENSMUSG00 | Gm1965 | 2402 | 0 | 0 | 0 | 0 | 0 | 0 |
| ENSMUSG00 | 1700022K14I | 930 | 0 | 0.05181338 | 0.04752886 | 0 | 0 | 0.05865514 |
| ENSMUSG00 | 4933437F05I | 1095 | 0.98027212 | 0.61608237 | 0.52477074 | 2.40546964 | 0 | 0.24908346 |
| ENSMUSG00 | Gm15091 | 2019 | 0 | 0 | 0 | 0 | 0 | 0 |
| ENSMUSG00 | Gm15097 | 2107 | 0 | 0 | 0 | 0 | 0 | 0 |
| ENSMUSG00 | Gm10439 | 1970 | 0 | 0 | 0 | 0 | 0 | 0 |
| ENSMUSG00 | Gm15085 | 2036 | 0 | 0 | 0 | 0 | 0 | 0 |
| ENSMUSG00 | Ott | 2288 | 0 | 0 | 0 | 0 | 0 | 0 |
| ENSMUSG00 | Gm15127 | 2019 | 0 | 0 | 0 | 0 | 0 | 0 |
| ENSMUSG00 | Gm15109 | 2020 | 0 | 0 | 0 | 0 | 0 | 0 |
| ENSMUSG00 | Gm15107 | 2021 | 0 | 0 | 0 | 0 | 0 | 0 |
| ENSMUSG00 | Gm15080 | 2022 | 0 | 0 | 0 | 0 | 0 | 0 |
| ENSMUSG00 | Gm15128 | 2105 | 0 | 0 | 0 | 0 | 0 | 0 |
| ENSMUSG00 | Gm15070 | 3581 | 0.55508909 | 0 | 0 | 0 | 0 | 0.88351247 |
| ENSMUSG00 | Gm10443 | 515 | 0.07719511 | 0.09356591 | 0 | 0 | 0 | 0.15888139 |
| ENSMUSG00 | Sod3 | 3568 | 0.86909403 | 2.49845622 | 0.49553636 | 0.86969054 | 0.7007546 | 0.25990407 |
| ENSMUSG00 | Nup62cl | 2316 | 0 | 0 | 0 | 0 | 0 | 0 |
| ENSMUSG00 | Ripply1 | 807 | 0 | 0 | 0 | 0 | 0 | 0 |
| ENSMUSG00 | Ptgr2 | 5971 | 41.2801841 | 36.2507952 | 61.6797448 | 36.0276586 | 37.6865378 | 47.0123241 |
| ENSMUSG00 | Acot1 | 2025 | 1.94360127 | 1.30876756 | 1.89904214 | 0.85527809 | 2.41902819 | 3.20561192 |
| ENSMUSG00 | Gm10444 | 368 | 0 | 0 | 0.26310621 | 0 | 0 | 0 |
| ENSMUSG00 | Gm5876 | 4671 | 0 | 0 | 0 | 0 | 0 | 0 |
| ENSMUSG00 | Tmsb15l | 812 | 0.78335922 | 0.11868582 | 0.76210074 | 0.62210422 | 0.37704219 | 1.8810096 |
| ENSMUSG00 | BC065397 | 1285 | 9.2195589 | 5.09988804 | 7.49883407 | 6.54250302 | 5.44016614 | 8.65996328 |
| ENSMUSG00 | Gm16401 | 2071 | 0.05758882 | 0.09306894 | 0 | 0.12195766 | 0.09855408 | 0 |
| ENSMUSG00 | Gm10447 | 2306 | 0 | 0 | 0 | 0 | 0 | 0 |
| ENSMUSG00 | Bhlhb9 | 3195 | 24.574984 | 24.2364964 | 10.5282011 | 18.3064145 | 24.2275468 | 18.2940695 |
| ENSMUSG00 | Gprasp2 | 4480 | 0.0976139 | 0.4732597 | 0.286128 | 0.93426722 | 2.00460762 | 0.16437841 |
| ENSMUSG00 | Gm17728 | 398 | 0 | 0 | 0 | 0 | 0 | 0 |
| ENSMUSG00 | Armcx5 | 3603 | 3.1446883 | 5.80430638 | 4.37969968 | 3.96572013 | 5.69320097 | 4.36030869 |
| ENSMUSG00 | Adam4 | 3919 | 0.44634885 | 0.75003138 | 0.2932503 | 0 | 0 | 0.20878775 |
| ENSMUSG00 | Gm4787 | 2494 | 0.01594045 | 0.15456758 | 0.23040255 | 0.53529898 | 1.4526354 | 1.20297126 |
| ENSMUSG00 | Gm5830 | 316 | 0 | 0 | 0 | 0 | 0 | 0 |
| ENSMUSG00 | Olp5 | 2166 | 0.14683465 | 0.02224674 | 0.04081426 | 0 | 0.11778942 | 0 |
| ENSMUSG00 | 9230009I02R | 1145 | 0 | 0 | 0 | 0 | 0 | 0 |
| ENSMUSG00 | 4933414I15R | 3399 | 0 | 0 | 0 | 0 | 0 | 0 |
| ENSMUSG00 | Cpxcr1 | 1593 | 0 | 0 | 0 | 0 | 0 | 0 |
| ENSMUSG00 | Gm15401 | 1591 | 0.59970555 | 0.93889359 | 0.44451885 | 0.45357693 | 1.02630046 | 1.01144168 |
| ENSMUSG00 | Gm10451 | 1594 | 5.9109466 | 4.05080505 | 8.70726393 | 2.55788647 | 4.86575227 | 6.79299359 |
| ENSMUSG00 | Cylc1 | 1918 | 0 | 0 | 0 | 0 | 0 | 0 |
| ENSMUSG00 | Vamp5 | 2775 | 1.10312505 | 1.45861663 | 0.49378635 | 0.44208631 | 0.68035168 | 0.99269858 |
| ENSMUSG00 | Gm732 | 2077 | 0 | 0 | 0 | 0 | 0 | 0 |
| ENSMUSG00 | Fam46d | 3153 | 0 | 0 | 0.11215184 | 0 | 0 | 0 |
| ENSMUSG00 | Gpr174 | 5528 | 0 | 0 | 0 | 0 | 0 | 0 |
| ENSMUSG00 | Gm5127 | 2143 | 0 | 0 | 0 | 0 | 0 | 0 |
| ENSMUSG00 | Fnd3c2 | 3642 | 0 | 0 | 0 | 0 | 0 | 0 |
| ENSMUSG00 | Uprt | 2297 | 2.07690799 | 2.03486499 | 1.75113962 | 0.64404172 | 0.37764405 | 1.09241045 |
| ENSMUSG00 | Gm1070 | 2710 | 0 | 0 | 0.19572772 | 0 | 0 | 0.22141774 |
| ENSMUSG00 | Dmrtc1b | 1376 | 0 | 0 | 0 | 0 | 0 | 0 |
| ENSMUSG00 | Igkv4-71 | 287 | 0 | 0 | 0 | 0 | 0 | 0 |
| ENSMUSG00 | Atoh1 | 2087 | 0 | 0 | 0 | 0 | 0 | 0 |
| ENSMUSG00 | Gm5570 | 2521 | 0 | 0 | 0 | 0 | 0 | 0 |
| ENSMUSG00 | Gm14812 | 1787 | 0 | 0 | 0 | 0 | 0.02855421 | 0 |
| ENSMUSG00 | D130052B06 | 2291 | 0 | 0 | 0 | 0 | 0 | 0 |
| ENSMUSG00 | Gm10462 | 319 | 0 | 0 | 0 | 0 | 0 | 0 |
| ENSMUSG00 | Gm10463 | 2885 | 0 | 0 | 0 | 0 | 0.38910928 | 0 |
| ENSMUSG00 | Zxda | 4452 | 0 | 0 | 0 | 0 | 0 | 0 |
| ENSMUSG00 | Zxdb | 5622 | 2.12849513 | 0.80567868 | 0.95133814 | 3.58766683 | 1.11637215 | 1.18859598 |
| ENSMUSG00 | Hbq1b | 772 | 0 | 0 | 0 | 0 | 0 | 0 |
| ENSMUSG00 | 9130019P16I | 5283 | 0 | 0 | 0.01673362 | 0 | 0 | 0.23748503 |
| ENSMUSG00 | Gm8914 | 1034 | 0 | 0 | 0 | 0 | 0 | 0 |
| ENSMUSG00 | Mageb2 | 1142 | 0 | 0 | 0 | 0 | 0 | 0 |
| ENSMUSG00 | G930045G22 | 1742 | 0 | 0 | 0 | 0 | 0 | 0 |
| ENSMUSG00 | Gm8098 | 1497 | 0 | 0.32188672 | 1.18107797 | 0 | 0.57945784 | 0.61946408 |
| ENSMUSG00 | Gm7173 | 2816 | 0.64941481 | 0.42779157 | 0.1412701 | 0.82004647 | 0.01812016 | 0.81359009 |
| ENSMUSG00 | Srp54c | 2889 | 5.71080803 | 6.43820238 | 7.25222346 | 4.04658443 | 5.01609233 | 7.90199829 |
| ENSMUSG00 | Fam177a | 3832 | 0 | 0 | 0 | 0 | 0 | 0 |
| ENSMUSG00 | 4930428D18I | 1100 | 0 | 0 | 0 | 0 | 0 | 0 |
| ENSMUSG00 | Gm7932 | 378 | 3.57588978 | 4.97161705 | 0.81855265 | 3.34093005 | 2.42982742 | 0.93801669 |
| ENSMUSG00 | 4930428E23I | 815 | 0.39023785 | 0 | 0.37964773 | 0 | 0 | 0.6693163 |
| ENSMUSG00 | Lrrc61 | 1847 | 14.0769271 | 16.8274257 | 8.63934235 | 9.12307244 | 13.0950201 | 8.06277912 |
| ENSMUSG00 | 5730522E02I | 83 | 0 | 0 | 0 | 0 | 0 | 0 |
| ENSMUSG00 | Ccdc164 | 2469 | 0 | 0 | 0.19693004 | 0.11691225 | 0 | 0 |
| ENSMUSG00 | Gm10466 | 2463 | 0 | 0 | 0 | 0 | 0 | 0 |
| ENSMUSG00 | Olfr237-ps1 | 999 | 0 | 0 | 0 | 0 | 0 | 0 |
| ENSMUSG00 | Olfr441 | 999 | 0 | 0 | 0 | 0 | 0 | 0 |
| ENSMUSG00 | Olfr444 | 1049 | 0 | 0 | 0 | 0 | 0 | 0 |
| ENSMUSG00 | Olfr446 | 926 | 0 | 0 | 0 | 0 | 0 | 0 |
| ENSMUSG00 | Heatr5b | 1780 | 1.22839968 | 1.57012003 | 1.2664573 | 1.56085249 | 1.80598971 | 2.43633014 |
| ENSMUSG00 | Gm10471 | 5635 | 0 | 0 | 0 | 0 | 0 | 0 |
| ENSMUSG00 | Gm7347 | 2990 | 0 | 0 | 0 | 0 | 0 | 0 |
| ENSMUSG00 | Speer4a | 2923 | 0 | 0 | 0 | 0 | 0 | 0 |
| ENSMUSG00 | Xlr3b | 3811 | 0.11474949 | 0.72071037 | 1.33382628 | 0.59647568 | 0.14728159 | 1.51724574 |
| ENSMUSG00 | Gm10472 | 533 | 0 | 0 | 0 | 0 | 0 | 0 |
| ENSMUSG00 | Gm1141 | 2294 | 0 | 0.3570922 | 0.26975841 | 0 | 0 | 0.14267466 |
| ENSMUSG00 | Vma21 | 7168 | 9.70592786 | 7.93920036 | 12.5365997 | 10.7017894 | 6.84812689 | 10.6275206 |
| ENSMUSG00 | BC023829 | 2839 | 51.3642489 | 32.5373052 | 54.9136669 | 36.4506177 | 45.2928732 | 29.2056722 |
| ENSMUSG00 | 4930567H17I | 839 | 0 | 0 | 0 | 0 | 0 | 0 |
| ENSMUSG00 | 6030469F06I | 1401 | 0 | 0 | 0 | 0 | 0 | 0 |
| ENSMUSG00 | 4930599N23I | 2423 | 2.05094307 | 1.65062926 | 2.35329664 | 2.06991506 | 3.32734931 | 2.08246317 |
| ENSMUSG00 | Gm12000 | 627 | 0 | 0 | 0 | 0 | 0 | 0 |
| ENSMUSG00 | Gm10474 | 2739 | 0 | 0 | 0 | 0 | 0 | 0 |
| ENSMUSG00 | 5031425E22I | 2109 | 6.93694492 | 6.55737758 | 9.24277514 | 7.21983063 | 5.7341162 | 16.0233656 |
| ENSMUSG00 | Gm10475 | 1792 | 0 | 0 | 0 | 0 | 0 | 0 |
| ENSMUSG00 | 9330158H04 | 8495 | 0 | 0 | 0 | 0 | 0 | 0 |
| ENSMUSG00 | 1810058I24R | 2847 | 10.0261451 | 7.44714947 | 11.4890635 | 8.0478041 | 4.62409728 | 14.37978 |
| ENSMUSG00 | 9030624G23 | 464 | 0.85679915 | 1.45390127 | 0.28578778 | 1.01091935 | 0.54985319 | 1.11684945 |
| ENSMUSG00 | 2410018L13I | 4513 | 0.07047282 | 0.07474077 | 0.05876602 | 0.00799514 | 0.04522613 | 0.07856643 |
| ENSMUSG00 | Gm20472 | 1234 | 0 | 0 | 0 | 0 | 0 | 0 |
| ENSMUSG00 | Gm10477 | 163 | 0 | 0 | 0 | 0 | 0 | 0 |
| ENSMUSG00 | Magi2 | 876 | 0 | 0 | 0 | 0 | 0 | 0 |
| ENSMUSG00 | Zfp449 | 3733 | 15.2930271 | 11.9013929 | 6.21643921 | 16.3833553 | 11.2085797 | 12.6546143 |
| ENSMUSG00 | Gm773 | 943 | 5.77571668 | 6.23408902 | 3.60927032 | 8.95355082 | 13.3112285 | 13.4493184 |
| ENSMUSG00 | Gm10478 | 394 | 1.21082682 | 1.7122086 | 0.33656226 | 1.28210311 | 0.3885257 | 0.06922497 |
| ENSMUSG00 | Gm17680 | 407 | 0 | 0 | 0 | 0 | 0 | 0 |
| ENSMUSG00 | Gm10479 | 315 | 0.25241575 | 0 | 1.2629098 | 0 | 0.48596548 | 0 |
| ENSMUSG00 | Gm5784 | 407 | 0 | 0 | 0.21720807 | 0 | 0 | 0.73715241 |
| ENSMUSG00 | 5730507C01I | 1175 | 4.90599547 | 1.10726293 | 1.80569231 | 2.73302295 | 2.25818855 | 0.67316131 |
| ENSMUSG00 | Gm10480 | 656 | 0.30301433 | 0.44072965 | 1.01071287 | 0.3300187 | 0.07778411 | 0.7068123 |
| ENSMUSG00 | Ccdc160 | 1534 | 29.8554847 | 43.5374243 | 35.5574148 | 38.9516726 | 33.9621445 | 39.329532 |
| ENSMUSG00 | Speer4c | 1154 | 0 | 0 | 0 | 0 | 0 | 0 |
| ENSMUSG00 | Klf14 | 2969 | 0 | 0 | 0 | 0 | 0 | 0 |
| ENSMUSG00 | Gm10481 | 1231 | 0 | 0.11743243 | 0.07181453 | 0.02931117 | 0 | 0.04431298 |
| ENSMUSG00 | Gm10482 | 632 | 0 | 0 | 0 | 0 | 0 | 0 |
| ENSMUSG00 | Rpl15-ps2 | 686 | 0 | 0 | 0 | 0 | 0.07438247 | 0 |
| ENSMUSG00 | Gm10483 | 2745 | 0 | 0 | 0 | 0 | 0 | 0 |
| ENSMUSG00 | Gm9144 | 1263 | 0.12590809 | 0.26706658 | 0.1049925 | 0.17141114 | 0.12120279 | 0.17276098 |
| ENSMUSG00 | Gm8773 | 1113 | 0 | 0 | 0 | 0 | 0 | 0 |
| ENSMUSG00 | 2500004C02I | 2492 | 3.41399391 | 6.80643164 | 4.00867437 | 8.74540726 | 4.56616445 | 8.87629712 |
| ENSMUSG00 | 2410017P09I | 581 | 3.83185354 | 5.97146959 | 8.52083721 | 5.34088783 | 2.63475262 | 7.08858955 |
| ENSMUSG00 | Gm10485 | 1381 | 0 | 0 | 0.06401426 | 0 | 0 | 0 |
| ENSMUSG00 | Gm9 | 1659 | 0 | 0 | 0 | 0 | 0 | 0 |
| ENSMUSG00 | Gm14819 | 919 | 0 | 0 | 0 | 0 | 0 | 0 |
| ENSMUSG00 | Gm10486 | 915 | 0 | 0 | 0 | 0 | 0 | 0 |
| ENSMUSG00 | Gm10487 | 898 | 0 | 0 | 0 | 0 | 0 | 0 |
| ENSMUSG00 | Gm14632 | 855 | 0 | 0 | 0 | 0 | 0 | 0 |
| ENSMUSG00 | Gm10488 | 865 | 0 | 0 | 0 | 0 | 0 | 0 |
| ENSMUSG00 | Gm4836 | 898 | 0 | 0 | 0 | 0 | 0 | 0 |
| ENSMUSG00 | Gm5935 | 1399 | 0 | 0 | 0 | 0 | 0 | 0 |
| ENSMUSG00 | Gm14636 | 7006 | 0.57312354 | 1.06607173 | 1.24290081 | 2.10641682 | 2.16312213 | 1.22241461 |
| ENSMUSG00 | Gm10489 | 1464 | 0 | 0 | 0 | 0 | 0 | 0.11178131 |
| ENSMUSG00 | Gm10490 | 358 | 0 | 0.134599 | 0 | 0 | 0 | 0 |
| ENSMUSG00 | Gm10491 | 112 | 0 | 0 | 0 | 0 | 0 | 0 |
| ENSMUSG00 | Nudt10 | 1773 | 1.59201304 | 3.91361967 | 0.54847183 | 4.07016859 | 2.24481518 | 3.56893193 |
| ENSMUSG00 | AU022751 | 3756 | 0 | 0 | 0 | 0 | 0 | 0 |
| ENSMUSG00 | Nudt11 | 3034 | 5.03167585 | 11.0222119 | 1.92308612 | 7.51610156 | 4.17091009 | 7.94686259 |
| ENSMUSG00 | Gm10492 | 1592 | 0.84904921 | 0.54482158 | 0.97177419 | 2.13047248 | 2.01925985 | 1.81602498 |
| ENSMUSG00 | Gm10493 | 610 | 0 | 0 | 0 | 0 | 0 | 0 |
| ENSMUSG00 | Gm6594 | 287 | 0 | 0 | 0 | 0 | 0 | 0 |
| ENSMUSG00 | C030034I22I | 1470 | 1.46040541 | 1.14729624 | 1.41325621 | 1.00636995 | 0.83308369 | 3.82216033 |
| ENSMUSG00 | Lrrc30 | 1758 | 0 | 0 | 0 | 0 | 0 | 0 |
| ENSMUSG00 | Gm10495 | 465 | 0 | 0 | 0 | 0.0775958 | 0 | 0 |
| ENSMUSG00 | AU016765 | 1456 | 0 | 0 | 0 | 0 | 0 | 0 |
| ENSMUSG00 | Arrdc5 | 1051 | 0.18913169 | 0 | 0.25234164 | 0 | 0 | 0 |
| ENSMUSG00 | Kcnh8 | 1603 | 0 | 0 | 0 | 0 | 0 | 0 |
| ENSMUSG00 | 9830107B12I | 3665 | 0 | 0 | 0 | 0 | 0 | 0 |
| ENSMUSG00 | A330017A19I | 1189 | 0.96964587 | 0.89159102 | 1.07809373 | 0.84970332 | 0.17166148 | 1.39928763 |
| ENSMUSG00 | B230354K17I | 4894 | 7.85524104 | 7.81774317 | 8.02930904 | 12.9538521 | 10.3846078 | 10.4383734 |
| ENSMUSG00 | Gm10497 | 790 | 0 | 0 | 0.1119034 | 0 | 0 | 0 |
| ENSMUSG00 | Gm6084 | 1046 | 0 | 0 | 0 | 0 | 0 | 0 |
| ENSMUSG00 | Trim40 | 2669 | 0 | 0 | 0 | 0 | 0 | 0 |
| ENSMUSG00 | Trim10 | 2230 | 0 | 0 | 0 | 0.25888462 | 0 | 0 |
| ENSMUSG00 | Gm8909 | 1767 | 0 | 0 | 0 | 0 | 0 | 0 |
| ENSMUSG00 | Gm10499 | 1704 | 0 | 0 | 0 | 0 | 0.02994506 | 0 |
| ENSMUSG00 | C920025E04I | 1809 | 0 | 0.58601533 | 0 | 0 | 0.25386257 | 0 |
| ENSMUSG00 | H2-Bl | 1226 | 0 | 0 | 0 | 0 | 0.4994425 | 0 |
| ENSMUSG00 | Gm6034 | 1630 | 0.26828852 | 0.17737341 | 0.13558848 | 0.0885449 | 0.21913168 | 0.23426071 |
| ENSMUSG00 | Dpcr1 | 2608 | 0 | 0 | 0 | 0 | 0 | 0 |
| ENSMUSG00 | H2-Q6 | 1650 | 0.60235577 | 0.61328199 | 0.13394498 | 0.15307534 | 7.45294338 | 6.95916553 |
| ENSMUSG00 | H2-D1 | 2664 | 81.600213 | 101.690757 | 76.7724954 | 71.1482657 | 150.742334 | 151.576965 |
| ENSMUSG00 | Lst1 | 402 | 0 | 0 | 0 | 0 | 0 | 0 |
| ENSMUSG00 | Ly6g6d | 1397 | 1.70746516 | 1.89710402 | 1.74023004 | 1.21392705 | 0.40178249 | 1.75713496 |
| ENSMUSG00 | AU023871 | 2480 | 2.22823056 | 2.37046207 | 0.92681284 | 0.74200979 | 2.75707031 | 1.48470819 |
| ENSMUSG00 | Gm10501 | 2198 | 0.4521779 | 0.4165343 | 0.50275072 | 0.42681217 | 0.9982412 | 0.94307215 |
| ENSMUSG00 | C4b | 5655 | 0 | 0 | 0 | 0 | 0 | 0 |
| ENSMUSG00 | Btnl5 | 1919 | 0 | 0 | 0 | 0 | 0 | 0 |
| ENSMUSG00 | H2-Ab1 | 1801 | 0 | 0.16053229 | 0.19634356 | 0 | 0.59497718 | 4.34637505 |
| ENSMUSG00 | H2-Ke6 | 1545 | 1.41524364 | 1.09160225 | 1.40187075 | 1.42459852 | 1.58528546 | 1.8006558 |
| ENSMUSG00 | Zfp414 | 2074 | 8.70250152 | 9.6884023 | 5.86090011 | 6.75016069 | 9.12766896 | 11.8093665 |
| ENSMUSG00 | Cyp4f15 | 2412 | 0 | 0 | 0.14660644 | 0 | 0 | 0 |
| ENSMUSG00 | Gm4924 | 2644 | 4.40558086 | 3.04354608 | 0.38450922 | 5.02200922 | 1.31232737 | 3.19785861 |
| ENSMUSG00 | C130040N14 | 1108 | 1.36345511 | 0.78281224 | 0.67818712 | 0.35821524 | 1.2894752 | 0.39385761 |
| ENSMUSG00 | Gm10505 | 1585 | 0 | 0 | 0 | 0 | 0 | 0 |
| ENSMUSG00 | Arhgdig | 2958 | 0.65855935 | 1.98740566 | 1.46442888 | 1.32959529 | 0.8970154 | 1.21712386 |
| ENSMUSG00 | Wdr90 | 8098 | 0.5105668 | 0.94016521 | 0.76962952 | 1.27432264 | 1.03968289 | 1.29670735 |
| ENSMUSG00 | Nme3 | 927 | 0 | 0 | 0 | 0 | 0 | 0 |
| ENSMUSG00 | Eme2 | 6688 | 4.54144544 | 4.30133163 | 3.09307155 | 5.44359793 | 4.09706397 | 4.32283457 |
| ENSMUSG00 | D330041H03 | 282 | 3.80637579 | 0.34174782 | 1.72418537 | 0 | 0.72377838 | 0.67703005 |
| ENSMUSG00 | Gm10506 | 109 | 1.8236459 | 3.09454216 | 2.02760748 | 5.2964469 | 7.02197832 | 0.75067814 |
| ENSMUSG00 | Gm16386 | 1420 | 0.89589815 | 2.64686091 | 0.31128059 | 0.83852639 | 0.71868135 | 0.99878961 |
| ENSMUSG00 | Vmn2r112 | 2568 | 0 | 0 | 0 | 0 | 0 | 0 |
| ENSMUSG00 | Vmn2r111 | 2568 | 0 | 0 | 0 | 0 | 0 | 0.01062097 |
| ENSMUSG00 | Gm10509 | 258 | 15.409101 | 5.41630551 | 11.6500982 | 11.048378 | 10.8777158 | 7.40009592 |
| ENSMUSG00 | Zfp97 | 2705 | 14.0650628 | 12.9862907 | 7.87626186 | 18.968084 | 11.997329 | 18.5931367 |
| ENSMUSG00 | Gm3435 | 2658 | 0.25426756 | 0.54386504 | 0.34922449 | 0.29864747 | 0.19197282 | 0.31810151 |
| ENSMUSG00 | Smok2b | 1940 | 0 | 0 | 0 | 0 | 0 | 0 |
| ENSMUSG00 | Smok2a | 1784 | 0 | 0 | 0 | 0 | 0 | 0 |
| ENSMUSG00 | Pnldc1 | 2133 | 0.55914881 | 0 | 0.0414457 | 0.50748305 | 0.07176706 | 0.17901779 |
| ENSMUSG00 | Park2 | 306 | 0 | 1.8896644 | 1.58895514 | 0.35374554 | 0.66701145 | 0.53479685 |
| ENSMUSG00 | 6530411M01 | 1030 | 0 | 0 | 0 | 0 | 0 | 0 |
| ENSMUSG00 | Sft2d1 | 3743 | 3.25011409 | 2.65199227 | 3.5545698 | 4.10658589 | 4.06247929 | 3.30822501 |
| ENSMUSG00 | Rnaset2a | 1004 | 17.3831235 | 18.0459186 | 15.585112 | 13.8003039 | 14.2812865 | 16.489747 |
| ENSMUSG00 | Gm1604b | 715 | 0 | 0 | 0 | 0 | 0 | 0 |
| ENSMUSG00 | Rsph3a | 2181 | 0.96608916 | 0.83956204 | 0.68907046 | 0.54594565 | 0.53810483 | 1.30057886 |
| ENSMUSG00 | A330023F24I | 6398 | 1.2738158 | 0.49707802 | 0.35925224 | 2.40246186 | 0.29508845 | 1.66683088 |
| ENSMUSG00 | Gm10516 | 2596 | 3.66007698 | 2.28310184 | 2.84349298 | 4.21142507 | 3.12526724 | 1.28178197 |
| ENSMUSG00 | D730003I15I | 545 | 10.7959837 | 15.2958798 | 14.923191 | 12.3142391 | 8.8008795 | 15.2137437 |
| ENSMUSG00 | Mosc2 | 6716 | 27.4192057 | 26.8053228 | 35.3298829 | 23.569376 | 22.1625876 | 27.5386136 |
| ENSMUSG00 | Gm10517 | 401 | 0.29742255 | 0 | 0.44091614 | 0.08998016 | 0.25449564 | 1.49636425 |
| ENSMUSG00 | Gm16067 | 3131 | 0.33013175 | 0.01539011 | 0.01411748 | 0 | 0.39113159 | 0.2352013 |
| ENSMUSG00 | Gm10518 | 1300 | 0.48929822 | 0 | 0 | 0 | 0 | 0 |
| ENSMUSG00 | Ifi204 | 2676 | 0.87652218 | 0.27010338 | 0 | 0.66069514 | 0.4004312 | 0.66250058 |
| ENSMUSG00 | AI607873 | 3313 | 0 | 0 | 0 | 0 | 0.50826152 | 0 |
| ENSMUSG00 | Pydc4 | 4183 | 0.02851218 | 0.04607836 | 0 | 0.06900702 | 0.18297768 | 0.22821238 |
| ENSMUSG00 | Gm10521 | 2573 | 0 | 0 | 0 | 0 | 0 | 0 |
| ENSMUSG00 | Gm17286 | 1253 | 0 | 0 | 0 | 0 | 0 | 0 |
| ENSMUSG00 | Sh2d1b2 | 2765 | 0 | 0 | 0 | 0 | 0 | 0 |
| ENSMUSG00 | AA792892 | 1878 | 0 | 0 | 0 | 0 | 0 | 0 |
| ENSMUSG00 | Gm6502 | 1215 | 0 | 0 | 0 | 0 | 0 | 0 |
| ENSMUSG00 | Gm16522 | 1844 | 0 | 0 | 0 | 0 | 0 | 0 |
| ENSMUSG00 | Gm3106 | 1433 | 0 | 0 | 0 | 0 | 0 | 0 |
| ENSMUSG00 | Gm3176 | 1840 | 0 | 0 | 0 | 0 | 0 | 0 |
| ENSMUSG00 | Gm7647 | 1211 | 0 | 0 | 0 | 0 | 0 | 0 |
| ENSMUSG00 | Dok6 | 5138 | 0 | 0 | 0 | 0 | 0 | 0 |
| ENSMUSG00 | Gm10524 | 1982 | 2.3267587 | 3.50093222 | 3.18913398 | 2.40304232 | 4.32514185 | 3.77056062 |
| ENSMUSG00 | Gm10525 | 245 | 0 | 0 | 0 | 0 | 0 | 0 |
| ENSMUSG00 | Gm10526 | 245 | 0 | 0 | 0 | 0 | 0 | 0 |
| ENSMUSG00 | Gm10527 | 245 | 0 | 0 | 0 | 0 | 0 | 0 |
| ENSMUSG00 | Gm10528 | 245 | 0 | 0 | 0 | 0 | 0 | 0 |
| ENSMUSG00 | Gm10529 | 376 | 0 | 0 | 0 | 0 | 0 | 0 |
| ENSMUSG00 | Gm10530 | 1166 | 0 | 0 | 0 | 0 | 0 | 0 |
| ENSMUSG00 | F830208F22I | 2763 | 0 | 0 | 0 | 0 | 0.14774195 | 0 |
| ENSMUSG00 | Pappa2 | 9056 | 0 | 0 | 0 | 0 | 0 | 0 |
| ENSMUSG00 | Gm7276 | 1251 | 0 | 0 | 0.03533321 | 0 | 0 | 0 |
| ENSMUSG00 | Gm10532 | 2297 | 0 | 0 | 0.01924329 | 0 | 0 | 0 |
| ENSMUSG00 | Gm5532 | 1749 | 0 | 0 | 0 | 0 | 0 | 0 |
| ENSMUSG00 | E330020D12I | 2120 | 0.01875259 | 0 | 0 | 0 | 0 | 0.03859619 |
| ENSMUSG00 | Cep76 | 3967 | 3.05657211 | 5.68471261 | 3.72155674 | 6.07582701 | 3.0613253 | 3.25205555 |
| ENSMUSG00 | Chmp1b | 2382 | 62.6207233 | 68.4158469 | 82.0387691 | 54.4108749 | 55.9319342 | 78.1712688 |
| ENSMUSG00 | F730048M01 | 2146 | 0 | 0 | 0 | 0 | 0 | 0 |
| ENSMUSG00 | Spink13 | 790 | 0 | 0 | 0 | 0 | 0 | 0 |
| ENSMUSG00 | Gm6960 | 637 | 0 | 0 | 0 | 0 | 0 | 0 |
| ENSMUSG00 | Gm4951 | 2642 | 0.33104488 | 0.56539732 | 0.28441761 | 0.16388514 | 2.08586245 | 4.37715634 |
| ENSMUSG00 | Ppp1r12b | 15492 | 0.75446109 | 0.75271876 | 0.54210884 | 0.68241925 | 1.00129217 | 0.81866171 |
| ENSMUSG00 | Gm8618 | 502 | 0 | 0 | 0 | 0 | 0 | 0 |
| ENSMUSG00 | Csnk1g3 | 4381 | 15.0274083 | 17.2463687 | 18.9681499 | 19.8900109 | 17.9017438 | 14.9166938 |
| ENSMUSG00 | Prr16 | 3817 | 0 | 0 | 0 | 0 | 0 | 0 |
| ENSMUSG00 | Tnfaip8 | 5038 | 0.6707455 | 0 | 0.07018951 | 0.40823274 | 0.2734641 | 0 |
| ENSMUSG00 | Gm949 | 2165 | 0 | 0 | 0 | 0 | 0 | 0 |
| ENSMUSG00 | Gm10540 | 275 | 0 | 0.17522343 | 0 | 0 | 0 | 0 |
| ENSMUSG00 | Gm10542 | 267 | 0 | 0 | 0 | 0 | 0 | 0 |
| ENSMUSG00 | Spink11 | 741 | 0 | 0 | 0 | 0 | 0 | 0 |
| ENSMUSG00 | Grxcr2 | 1117 | 0 | 0 | 0 | 0 | 0 | 0 |
| ENSMUSG00 | 9630014M24 | 3864 | 0 | 0 | 0 | 0 | 0 | 0 |
| ENSMUSG00 | Gm10543 | 1586 | 0 | 0 | 0 | 0 | 0 | 0 |
| ENSMUSG00 | 3222401L13I | 2830 | 0.87096812 | 0.66405344 | 1.10895083 | 2.11647328 | 1.87517424 | 1.44565226 |
| ENSMUSG00 | Pcdhb22 | 5324 | 0 | 0 | 0.46493298 | 0.04066346 | 0.15334749 | 0 |
| ENSMUSG00 | Gm10544 | 4057 | 0 | 0 | 0 | 0 | 0.01257737 | 0 |
| ENSMUSG00 | Gm10545 | 4966 | 0 | 0 | 0 | 0 | 0 | 0 |
| ENSMUSG00 | 2900060B14I | 77 | 2.06521977 | 0 | 2.29619964 | 1.87439192 | 1.32536041 | 3.54216094 |
| ENSMUSG00 | 1700066B19I | 2833 | 0.21049496 | 0.06803592 | 0.07801243 | 0.36935379 | 0.18011428 | 0.31770671 |
| ENSMUSG00 | Ecscr | 1829 | 0.1304171 | 0.1844205 | 0 | 0.11836647 | 0.50217319 | 0.26842182 |
| ENSMUSG00 | Gm1614 | 3020 | 0.05265627 | 0.15955776 | 0.08781823 | 0 | 0 | 0 |
| ENSMUSG00 | Serpinb3c | 1651 | 0 | 0 | 0 | 0 | 0 | 0 |
| ENSMUSG00 | Serpinb3b | 2491 | 0 | 0 | 0 | 0 | 0 | 0 |
| ENSMUSG00 | Gm10548 | 2898 | 0.39782917 | 0.16627482 | 0.45757602 | 0.09960537 | 0.10564467 | 0.74351156 |
| ENSMUSG00 | Gm6086 | 4053 | 0 | 0 | 0 | 0 | 0 | 0 |
| ENSMUSG00 | D2hgdh | 3942 | 8.32021091 | 7.70102957 | 7.42303909 | 9.29968474 | 9.33283028 | 5.4798362 |
| ENSMUSG00 | Gm10549 | 447 | 0 | 0 | 0 | 0 | 0 | 0 |
| ENSMUSG00 | Myeov2 | 1495 | 21.5397587 | 19.016723 | 22.1157047 | 13.636358 | 13.652542 | 38.2574706 |
| ENSMUSG00 | Hdac4 | 2391 | 2.02851051 | 2.92222255 | 4.88050462 | 2.52015953 | 3.77736032 | 2.89743136 |
| ENSMUSG00 | 5830472F04I | 350 | 0.68152252 | 1.51443104 | 0.50516392 | 0 | 0.29157929 | 0 |
| ENSMUSG00 | C130036L24I | 3784 | 0.04202482 | 0.0636713 | 0.0233625 | 0.64840883 | 0.02696954 | 0.19461291 |
| ENSMUSG00 | Gm10552 | 1053 | 0 | 0 | 0.12593118 | 0 | 0 | 0.07770553 |
| ENSMUSG00 | Gm10553 | 371 | 0 | 0 | 0 | 0 | 0 | 0 |
| ENSMUSG00 | Gm15433 | 890 | 0 | 0 | 0 | 0 | 0 | 0 |
| ENSMUSG00 | Fbxo36 | 2256 | 5.30425516 | 5.83107212 | 5.01581197 | 5.16600194 | 3.66412805 | 5.76684527 |
| ENSMUSG00 | Rab18 | 2346 | 195.049267 | 196.381319 | 226.435528 | 221.829211 | 158.407969 | 244.878826 |
| ENSMUSG00 | Rpl27-ps3 | 474 | 0.16774464 | 0 | 0 | 0 | 0 | 0 |
| ENSMUSG00 | Wdfy1 | 9597 | 4.66444421 | 2.53560001 | 4.25575731 | 4.16577111 | 3.76437158 | 9.32459012 |
| ENSMUSG00 | Gm10556 | 3896 | 0 | 0 | 0 | 0 | 0 | 0 |
| ENSMUSG00 | Gm10557 | 748 | 3.45468748 | 2.96333735 | 3.36832226 | 4.43789852 | 4.36589312 | 4.0839032 |
| ENSMUSG00 | Gm216 | 2092 | 0 | 0.04606734 | 0 | 0 | 0 | 0 |
| ENSMUSG00 | Apol7d | 2216 | 0 | 0 | 0 | 0 | 0 | 0 |
| ENSMUSG00 | Gm10558 | 1658 | 0 | 0.23250394 | 0.26659736 | 0.19586152 | 0.18465516 | 0.41125813 |
| ENSMUSG00 | Crygb | 620 | 0 | 0 | 0 | 0 | 0 | 0 |
| ENSMUSG00 | Nbeal1 | 16026 | 3.97654033 | 5.49336265 | 8.3764506 | 10.4220507 | 5.22171823 | 8.80562732 |
| ENSMUSG00 | 9930005F22I | 2899 | 0.02742703 | 0 | 0.15247273 | 0 | 0 | 0 |
| ENSMUSG00 | Gm6644 | 841 | 0.85089019 | 0.85944903 | 0.21023469 | 0.94388226 | 1.88087711 | 0.4540368 |
| ENSMUSG00 | Gm10561 | 1940 | 0 | 0 | 0 | 0 | 0 | 0 |
| ENSMUSG00 | Hspe1 | 1568 | 16.2013725 | 11.0939449 | 16.0400821 | 16.2461246 | 10.3484614 | 26.3701232 |
| ENSMUSG00 | Pgap1 | 2966 | 5.72339521 | 5.73493394 | 7.54084038 | 9.18474161 | 3.52677244 | 12.2947379 |

| | | | | | | | | |
|---|---|---|---|---|---|---|---|---|
| ENSMUSG00 | Gm10562 | 686 | 0 | 0 | 0 | 0 | 0 | 0 |
| ENSMUSG00 | Tmem88b | 3469 | 0.74491388 | 0.51395168 | 0.49693626 | 0.26003203 | 0.57366061 | 0.19655981 |
| ENSMUSG00 | 1500002C15I | 386 | 0.92694126 | 0 | 0 | 1.86953599 | 0 | 0 |
| ENSMUSG00 | Gm10563 | 250 | 0 | 0 | 0 | 0 | 0.61231651 | 0.10909856 |
| ENSMUSG00 | 2610002J02R | 1682 | 27.819382 | 26.6429199 | 28.6444762 | 16.8397176 | 20.0525769 | 26.6746821 |
| ENSMUSG00 | Gm10564 | 3111 | 0 | 0 | 0 | 0 | 0 | 0 |
| ENSMUSG00 | Gm833 | 881 | 0 | 0 | 0 | 0 | 0 | 0 |
| ENSMUSG00 | Klhl21 | 3982 | 59.6931236 | 45.7299259 | 37.9523107 | 45.8591731 | 55.2166382 | 25.5828171 |
| ENSMUSG00 | Rpl31 | 1032 | 0.7319323 | 0.32684602 | 1.19927481 | 0.80415409 | 0.93943909 | 0.66072285 |
| ENSMUSG00 | Aptd1 | 1851 | 0.45103463 | 0.75494696 | 0.97907918 | 0.35087888 | 0.96484233 | 1.70926967 |
| ENSMUSG00 | Gm13241 | 1534 | 0 | 0 | 0 | 0 | 0 | 0.12446054 |
| ENSMUSG00 | Vps13d | 2060 | 0 | 0 | 0 | 0 | 0 | 0 |
| ENSMUSG00 | Gm13125 | 1447 | 0 | 0 | 0 | 0 | 0 | 0 |
| ENSMUSG00 | 4931408C20I | 4316 | 0 | 0 | 0 | 0 | 0 | 0 |
| ENSMUSG00 | Gm13102 | 2289 | 0 | 0 | 0 | 0 | 0 | 0 |
| ENSMUSG00 | Pramel4 | 2527 | 0 | 0 | 0 | 0 | 0 | 0 |
| ENSMUSG00 | Lmbrd1 | 5374 | 51.4069286 | 37.2920381 | 69.0170571 | 49.3492794 | 54.6439883 | 58.1172114 |
| ENSMUSG00 | Gm13043 | 2109 | 0 | 0 | 0 | 0 | 0 | 0 |
| ENSMUSG00 | Gm10565 | 4264 | 0.5687346 | 0.53113574 | 1.25432059 | 0.45694897 | 1.47191469 | 1.91255092 |
| ENSMUSG00 | Gm6420 | 419 | 0 | 0.34501033 | 0.84394927 | 0.86114665 | 0 | 0.97641907 |
| ENSMUSG00 | 4933415F23I | 1780 | 0.08933816 | 0 | 0 | 0 | 0 | 0 |
| ENSMUSG00 | AI507597 | 940 | 0 | 0 | 0 | 0 | 0 | 0 |
| ENSMUSG00 | 6330545A04I | 1871 | 0.8074336 | 0.87564887 | 0.42524488 | 0.61711674 | 0.35453922 | 0.26239632 |
| ENSMUSG00 | Defb18 | 256 | 0 | 0 | 0 | 0 | 0 | 0 |
| ENSMUSG00 | Gm10566 | 998 | 0.11950545 | 0.14484902 | 0 | 0 | 0 | 0 |
| ENSMUSG00 | Gm10567 | 2638 | 2.81814816 | 1.11424298 | 0.97183734 | 0.91641281 | 2.59193873 | 0.55831331 |
| ENSMUSG00 | Gm16287 | 2321 | 0 | 0 | 0 | 0 | 0.0439693 | 0 |
| ENSMUSG00 | 4732440D04I | 680 | 0 | 0 | 0 | 0 | 0 | 0.04010976 |
| ENSMUSG00 | Gm7534 | 2330 | 0 | 0 | 0 | 0 | 0 | 0 |
| ENSMUSG00 | A330069E16I | 885 | 1.07811473 | 0.7622714 | 0.84907495 | 0.16308269 | 0.57656922 | 1.38684606 |
| ENSMUSG00 | Gm10570 | 512 | 0.23294227 | 0 | 0.08633173 | 0 | 0 | 0.15981234 |
| ENSMUSG00 | Gm10571 | 1636 | 0 | 0.08836145 | 0 | 0 | 0 | 0 |
| ENSMUSG00 | 5730409E04I | 3047 | 1.50045299 | 1.20189354 | 1.49418767 | 1.11313167 | 0.92105372 | 2.77490586 |
| ENSMUSG00 | 1700029G01I | 2891 | 6.40818192 | 4.98365486 | 3.94468194 | 5.97831177 | 4.97732203 | 5.99079762 |
| ENSMUSG00 | 4933407E24I | 966 | 0 | 0 | 0 | 0 | 0 | 0 |
| ENSMUSG00 | 4933427I04R | 2181 | 0.0182281 | 0.19884364 | 0.04053356 | 0.03308762 | 0 | 0 |
| ENSMUSG00 | Gm12888 | 1177 | 0 | 0 | 0 | 0 | 0 | 0 |
| ENSMUSG00 | Gm12863 | 534 | 0 | 0 | 0 | 0 | 0 | 0 |
| ENSMUSG00 | Olfr1328 | 941 | 0 | 0 | 0 | 0 | 0 | 0 |
| ENSMUSG00 | Olfr1329 | 941 | 0 | 0 | 0 | 0 | 0 | 0 |
| ENSMUSG00 | Olfr1330 | 947 | 0 | 0 | 0 | 0 | 0 | 0 |
| ENSMUSG00 | Olfr1331 | 1172 | 0 | 0 | 0 | 0 | 0 | 0 |
| ENSMUSG00 | Olfr1333 | 2123 | 0 | 0 | 0 | 0 | 0 | 0 |
| ENSMUSG00 | Btbd19 | 3322 | 0.90951731 | 0.73976778 | 1.22413292 | 0.64083101 | 0.36864329 | 0.94418528 |
| ENSMUSG00 | Kncn | 2517 | 0.06317915 | 0 | 0 | 0 | 0 | 0 |
| ENSMUSG00 | Kti12 | 1555 | 12.8342452 | 9.63728843 | 9.32360419 | 10.2561181 | 9.71305932 | 9.94515785 |
| ENSMUSG00 | Gm10574 | 1051 | 0.03782634 | 0.18339274 | 0 | 0.54929849 | 0.09710062 | 0 |
| ENSMUSG00 | B230314M03 | 2989 | 0 | 0 | 0 | 0 | 0 | 0 |
| ENSMUSG00 | Gm6471 | 1162 | 0 | 0 | 0 | 0 | 0 | 0 |
| ENSMUSG00 | Krtap5-5 | 1176 | 0 | 0 | 0 | 0 | 0 | 0 |
| ENSMUSG00 | Gm7579 | 731 | 0 | 0 | 0 | 0 | 0 | 0 |
| ENSMUSG00 | Gm10575 | 2168 | 0.09168699 | 0.15558353 | 0.12232982 | 0.08321505 | 0.11768076 | 0.01258055 |
| ENSMUSG00 | Gm10576 | 704 | 0 | 0 | 0 | 0 | 0 | 0 |
| ENSMUSG00 | Gm10577 | 398 | 0 | 0 | 0 | 0 | 0 | 0 |
| ENSMUSG00 | Efcab7 | 2135 | 7.96971694 | 3.06948765 | 5.79696302 | 3.75185662 | 4.49318907 | 6.1192282 |
| ENSMUSG00 | Alg6 | 8571 | 1.87390201 | 2.18134868 | 2.08864151 | 2.44167376 | 2.55399781 | 1.67701959 |
| ENSMUSG00 | I0C0044D17I | 2770 | 0 | 0 | 0 | 0 | 0 | 0 |
| ENSMUSG00 | 6430531B16I | 1921 | 0.0620856 | 0 | 0 | 0.15026359 | 0.18593682 | 0 |
| ENSMUSG00 | Gm12650 | 502 | 0 | 0 | 0 | 0 | 0 | 0 |
| ENSMUSG00 | Cdkn2b | 1391 | 3.74404597 | 5.64657805 | 2.00195263 | 1.71201649 | 2.60450948 | 1.54902696 |
| ENSMUSG00 | 4930544L04F | 3202 | 0 | 0 | 0 | 0 | 0 | 0 |
| ENSMUSG00 | Nps | 857 | 0.09277825 | 0 | 0 | 0 | 0.35724417 | 0 |
| ENSMUSG00 | Fam196a | 4485 | 0.85095343 | 0.54793948 | 0.63075094 | 1.19871229 | 0.63711863 | 1.16152867 |
| ENSMUSG00 | Gm10580 | 394 | 0 | 0 | 0 | 0 | 0 | 0 |
| ENSMUSG00 | Gm10581 | 229 | 0 | 0 | 0.57906345 | 0 | 0 | 0 |
| ENSMUSG00 | Gm10582 | 394 | 0 | 0 | 0 | 0 | 0 | 0 |
| ENSMUSG00 | Gm10583 | 394 | 0 | 0 | 0 | 0 | 0 | 0 |
| ENSMUSG00 | Ifnz | 1461 | 0 | 0 | 0 | 0 | 0 | 0.11201084 |
| ENSMUSG00 | Ifna12 | 796 | 0 | 0 | 0 | 0 | 0 | 0 |
| ENSMUSG00 | Gm12597 | 743 | 0 | 0 | 0 | 0 | 0 | 0 |
| ENSMUSG00 | Gm11239 | 1320 | 0 | 0 | 0 | 0 | 0 | 0 |
| ENSMUSG00 | Gm11238 | 1316 | 0 | 0 | 0 | 0 | 0 | 0 |
| ENSMUSG00 | Gm11236 | 1316 | 0 | 0 | 0 | 0 | 0 | 0 |
| ENSMUSG00 | 8030451A03I | 3142 | 0 | 0 | 0 | 0 | 0 | 0 |
| ENSMUSG00 | Gm2529 | 703 | 0.84826772 | 0.06854401 | 1.32039645 | 0.61590972 | 0.87100499 | 1.39670983 |
| ENSMUSG00 | Gm10586 | 375 | 0.63608769 | 1.54196615 | 0.35361474 | 0.19243757 | 0.27214067 | 1.67284454 |
| ENSMUSG00 | Mup14 | 1686 | 0 | 0 | 0 | 0 | 0 | 0 |
| ENSMUSG00 | Mup15 | 1313 | 0 | 0 | 0 | 0 | 0 | 0 |
| ENSMUSG00 | Mup11 | 1569 | 0 | 0 | 0 | 0 | 0 | 0 |
| ENSMUSG00 | Gm16002 | 543 | 0 | 0 | 0 | 0 | 0 | 0 |
| ENSMUSG00 | Tufm | 1767 | 15.1642297 | 14.7259076 | 16.0597528 | 14.1306026 | 13.0525874 | 11.1907829 |
| ENSMUSG00 | Mup12 | 1186 | 0 | 0 | 0 | 0 | 0 | 0 |
| ENSMUSG00 | Mup-ps2 | 538 | 0 | 0 | 0 | 0 | 0 | 0 |
| ENSMUSG00 | Mup7 | 1278 | 0 | 0 | 0 | 0 | 0 | 0 |
| ENSMUSG00 | Gm12542 | 645 | 0 | 0.14941532 | 0 | 0 | 0.15822132 | 0.63429394 |
| ENSMUSG00 | Gm6252 | 651 | 0 | 0.07401911 | 0 | 0 | 0 | 0 |
| ENSMUSG00 | Gm12526 | 613 | 0 | 0 | 0 | 0 | 0 | 0 |
| ENSMUSG00 | Eef2k | 1780 | 0 | 0 | 0 | 0 | 0 | 0 |
| ENSMUSG00 | AI427809 | 654 | 0 | 0 | 0.54069533 | 0 | 0 | 0 |
| ENSMUSG00 | Iqck | 3910 | 0.29486162 | 0.75175779 | 0.06782892 | 0 | 0.33930582 | 0.53014644 |
| ENSMUSG00 | Itpripl2 | 1607 | 0.22265048 | 0 | 0 | 0 | 0 | 0 |
| ENSMUSG00 | Itpripl2 | 1369 | 0 | 0.10559483 | 0 | 0.05271299 | 0 | 1.69345824 |
| ENSMUSG00 | Gm829 | 972 | 0 | 0 | 0 | 0 | 0 | 0 |
| ENSMUSG00 | Gm5600 | 485 | 0 | 0 | 0 | 0 | 0 | 0 |
| ENSMUSG00 | B430319F04I | 2668 | 4.39575216 | 1.86027119 | 2.76675704 | 2.0015527 | 4.03544427 | 3.72112769 |
| ENSMUSG00 | AA474408 | 3338 | 1.6793058 | 1.03937203 | 0.42374445 | 0.34590336 | 1.11591535 | 2.79446574 |
| ENSMUSG00 | Gm10590 | 4967 | 0 | 0 | 0 | 0 | 0 | 0 |
| ENSMUSG00 | Gm10591 | 864 | 0 | 0 | 0 | 0 | 0 | 0 |
| ENSMUSG00 | Gm10592 | 926 | 0 | 0 | 0 | 0 | 0 | 0 |
| ENSMUSG00 | Gm10593 | 3534 | 0 | 0 | 0 | 0 | 0 | 0 |
| ENSMUSG00 | Gm10594 | 425 | 0 | 0 | 0 | 0 | 0 | 0 |
| ENSMUSG00 | 4933409K07I | 1709 | 0 | 0 | 0 | 0 | 0 | 0 |
| ENSMUSG00 | Gm10595 | 425 | 0 | 0 | 0 | 0 | 0 | 0 |
| ENSMUSG00 | Gm13305 | 3327 | 0 | 0 | 0 | 0 | 0 | 0 |
| ENSMUSG00 | Gm13306 | 1139 | 0 | 0 | 0 | 0 | 0 | 0 |
| ENSMUSG00 | Gm13304 | 833 | 0 | 0 | 0 | 0 | 0 | 0 |
| ENSMUSG00 | Gm5859 | 2994 | 0 | 0 | 0 | 0 | 0 | 0.00910977 |
| ENSMUSG00 | Gm10596 | 926 | 0 | 0 | 0 | 0 | 0 | 0 |
| ENSMUSG00 | Gm3892 | 3534 | 0 | 0 | 0 | 0 | 0 | 0 |
| ENSMUSG00 | Gm10597 | 425 | 0 | 0 | 0 | 0 | 0 | 0 |
| ENSMUSG00 | Gm10598 | 1709 | 0 | 0 | 0 | 0 | 0 | 0 |
| ENSMUSG00 | Ccl21b | 866 | 0 | 0 | 0 | 0 | 0 | 0 |
| ENSMUSG00 | Gm10600 | 671 | 0 | 0 | 0 | 0 | 0 | 0 |
| ENSMUSG00 | Ccl27a | 2305 | 3.58747937 | 2.59224244 | 3.52847684 | 3.02118637 | 3.09921589 | 4.98161524 |
| ENSMUSG00 | Il11ra1 | 4330 | 7.39103044 | 7.14450251 | 6.36996537 | 6.59976426 | 6.92921686 | 5.44862886 |
| ENSMUSG00 | Olfr477 | 932 | 0 | 0 | 0 | 0 | 0 | 0 |
| ENSMUSG00 | Olfr476 | 932 | 0 | 0 | 0 | 0 | 0 | 0 |
| ENSMUSG00 | Olfr473 | 932 | 0 | 0 | 0 | 0 | 0 | 0 |
| ENSMUSG00 | Olfr472 | 1082 | 0 | 0 | 0 | 0 | 0 | 0 |
| ENSMUSG00 | Rbmxl2 | 1471 | 0 | 0 | 0 | 0 | 0 | 0 |
| ENSMUSG00 | Olfr716 | 1103 | 0 | 0 | 0 | 0 | 0 | 0 |
| ENSMUSG00 | Olfr17 | 965 | 0 | 0 | 0 | 0 | 0 | 0 |
| ENSMUSG00 | Olfr713 | 974 | 0 | 0 | 0 | 0 | 0 | 0 |
| ENSMUSG00 | Olfr1532-ps1 | 923 | 0 | 0 | 0 | 0 | 0 | 0 |
| ENSMUSG00 | Olfr704 | 1893 | 0 | 0 | 0 | 0 | 0 | 0 |
| ENSMUSG00 | Olfr703 | 959 | 0 | 0 | 0 | 0 | 0 | 0 |
| ENSMUSG00 | Gm1966 | 2792 | 0 | 0 | 0 | 0 | 0 | 0 |
| ENSMUSG00 | Olfr692 | 1125 | 0 | 0 | 0 | 0 | 0 | 0 |
| ENSMUSG00 | Olfr689 | 1075 | 0 | 0 | 0 | 0 | 0 | 0 |
| ENSMUSG00 | Olfr687 | 914 | 0 | 0 | 0 | 0 | 0 | 0 |
| ENSMUSG00 | Olfr688 | 965 | 0 | 0 | 0 | 0 | 0 | 0 |
| ENSMUSG00 | Mob3b | 6196 | 5.02397373 | 7.45038922 | 4.03780554 | 4.58304859 | 4.36474809 | 3.57880596 |
| ENSMUSG00 | Olfr681 | 951 | 0 | 0 | 0 | 0 | 0 | 0 |
| ENSMUSG00 | Olfr679 | 951 | 0 | 0 | 0 | 0 | 0 | 0 |
| ENSMUSG00 | Olfr678 | 941 | 0 | 0 | 0 | 0 | 0 | 0 |
| ENSMUSG00 | Olfr677 | 938 | 0 | 0 | 0 | 0 | 0 | 0 |
| ENSMUSG00 | Olfr676 | 953 | 0 | 0 | 0 | 0 | 0 | 0 |
| ENSMUSG00 | Olfr669 | 953 | 0 | 0 | 0 | 0 | 0 | 0 |
| ENSMUSG00 | Olfr665 | 950 | 0 | 0 | 0 | 0 | 0 | 0 |
| ENSMUSG00 | Olfr663 | 955 | 0 | 0 | 0 | 0 | 0 | 0 |
| ENSMUSG00 | Olfr661 | 963 | 0 | 0 | 0 | 0 | 0 | 0 |
| ENSMUSG00 | Olfr659 | 968 | 0 | 0 | 0 | 0 | 0 | 0 |
| ENSMUSG00 | Olfr657 | 960 | 0 | 0 | 0 | 0 | 0 | 0 |
| ENSMUSG00 | Olfr656 | 990 | 0 | 0 | 0 | 0 | 0 | 0 |
| ENSMUSG00 | Olfr654 | 1154 | 0 | 0 | 0 | 0 | 0 | 0 |
| ENSMUSG00 | Olfr653 | 1045 | 0 | 0 | 0 | 0 | 0 | 0 |
| ENSMUSG00 | Olfr652 | 2007 | 0 | 0 | 0 | 0 | 0 | 0 |
| ENSMUSG00 | Olfr651 | 1068 | 0 | 0 | 0 | 0 | 0 | 0 |
| ENSMUSG00 | Trim30e-ps1 | 2487 | 0 | 0 | 0 | 0 | 0 | 0 |
| ENSMUSG00 | Olfr646 | 938 | 0 | 0 | 0 | 0 | 0 | 0 |
| ENSMUSG00 | Olfr641 | 938 | 0 | 0 | 0 | 0 | 0 | 0 |
| ENSMUSG00 | Olfr638 | 965 | 0 | 0 | 0 | 0 | 0 | 0 |
| ENSMUSG00 | Olfr635 | 965 | 0 | 0 | 0 | 0 | 0 | 0 |
| ENSMUSG00 | Olfr633 | 938 | 0 | 0 | 0 | 0 | 0 | 0 |
| ENSMUSG00 | Olfr632 | 953 | 0 | 0 | 0 | 0 | 0 | 0 |
| ENSMUSG00 | Hbb-b2 | 700 | 0 | 0 | 0 | 0 | 0 | 0 |
| ENSMUSG00 | Olfr628 | 2366 | 0 | 0 | 0 | 0 | 0 | 0 |
| ENSMUSG00 | Olfr243 | 950 | 0 | 0 | 0 | 0 | 0 | 0 |
| ENSMUSG00 | Olfr625-ps1 | 965 | 0 | 0 | 0 | 0 | 0 | 0 |
| ENSMUSG00 | Olfr619 | 1123 | 0 | 0 | 0 | 0 | 0 | 0 |
| ENSMUSG00 | Olfr618 | 956 | 0 | 0 | 0 | 0 | 0 | 0 |
| ENSMUSG00 | Olfr617 | 1905 | 0 | 0 | 0 | 0 | 0 | 0 |
| ENSMUSG00 | Olfr615 | 961 | 0 | 0 | 0 | 0 | 0 | 0 |
| ENSMUSG00 | Olfr608 | 950 | 0 | 0 | 0 | 0 | 0 | 0 |
| ENSMUSG00 | Olfr606 | 959 | 0 | 0 | 0 | 0 | 0 | 0 |
| ENSMUSG00 | Olfr599 | 1049 | 0 | 0 | 0 | 0 | 0 | 0 |
| ENSMUSG00 | Olfr598 | 959 | 0 | 0 | 0 | 0 | 0 | 0 |
| ENSMUSG00 | Olfr597 | 2010 | 0 | 0 | 0 | 0 | 0 | 0 |
| ENSMUSG00 | Olfr596 | 938 | 0 | 0 | 0 | 0 | 0 | 0 |
| ENSMUSG00 | Olfr594 | 1086 | 0 | 0 | 0 | 0 | 0 | 0 |
| ENSMUSG00 | Olfr593 | 983 | 0 | 0 | 0 | 0 | 0 | 0 |
| ENSMUSG00 | Olfr592 | 1138 | 0 | 0 | 0 | 0 | 0 | 0 |
| ENSMUSG00 | Olfr584 | 963 | 0 | 0 | 0 | 0 | 0 | 0 |
| ENSMUSG00 | Olfr583 | 959 | 0 | 0 | 0 | 0 | 0 | 0 |
| ENSMUSG00 | Olfr582 | 959 | 0 | 0 | 0 | 0 | 0 | 0 |
| ENSMUSG00 | Olfr576 | 1817 | 0 | 0 | 0 | 0 | 0 | 0 |
| ENSMUSG00 | Olfr572 | 956 | 0 | 0 | 0 | 0 | 0 | 0 |
| ENSMUSG00 | Olfr570 | 1044 | 0 | 0 | 0 | 0 | 0 | 0 |
| ENSMUSG00 | Olfr568 | 941 | 0 | 0 | 0 | 0 | 0 | 0 |
| ENSMUSG00 | Olfr561 | 1070 | 0 | 0 | 0 | 0 | 0 | 0 |
| ENSMUSG00 | Olfr557 | 1056 | 0 | 0 | 0 | 0 | 0 | 0 |
| ENSMUSG00 | Trim68 | 2382 | 10.5647436 | 10.6406669 | 13.9916833 | 10.8912637 | 14.9308917 | 10.4999346 |
| ENSMUSG00 | Olfr556 | 1022 | 0 | 0 | 0 | 0 | 0 | 0 |
| ENSMUSG00 | Olfr555 | 947 | 0 | 0 | 0 | 0 | 0 | 0 |
| ENSMUSG00 | Olfr554 | 953 | 0 | 0 | 0 | 0 | 0 | 0 |
| ENSMUSG00 | Olfr553 | 980 | 0 | 0 | 0 | 0 | 0 | 0 |
| ENSMUSG00 | Olfr552 | 953 | 0 | 0 | 0 | 0 | 0 | 0 |
| ENSMUSG00 | Olfr551 | 1078 | 0 | 0 | 0 | 0 | 0 | 0 |
| ENSMUSG00 | Olfr550 | 1037 | 0 | 0 | 0 | 0 | 0 | 0 |
| ENSMUSG00 | 1700026J12R | 1378 | 0 | 0 | 0 | 0 | 0 | 0 |
| ENSMUSG00 | Olfr549 | 950 | 0 | 0 | 0 | 0 | 0 | 0 |
| ENSMUSG00 | Olfr548-ps1 | 932 | 0 | 0 | 0 | 0 | 0 | 0 |
| ENSMUSG00 | Olfr547 | 939 | 0 | 0 | 0 | 0 | 0 | 0 |
| ENSMUSG00 | Gm9574 | 872 | 0 | 0 | 0 | 0 | 0 | 0 |
| ENSMUSG00 | Rhog | 1353 | 23.4184168 | 27.0670333 | 28.2591236 | 24.8014053 | 20.6292887 | 26.9117837 |
| ENSMUSG00 | Gm10602 | 606 | 1.37766517 | 1.82885837 | 0.51058235 | 1.36945054 | 1.51563493 | 2.4304134 |
| ENSMUSG00 | Ggh | 1986 | 55.6096298 | 58.3769284 | 53.8167454 | 57.0844833 | 45.8620749 | 65.6871095 |
| ENSMUSG00 | Ttpa | 3476 | 0.02287427 | 0.99810812 | 0.59766589 | 1.74389628 | 0.79269974 | 0 |
| ENSMUSG00 | Cnbd1 | 1321 | 0 | 0 | 0 | 0 | 0 | 0 |
| ENSMUSG00 | Gm10604 | 1705 | 0.09326799 | 0 | 0.05184967 | 0.59254971 | 0.14963747 | 0.5438931 |
| ENSMUSG00 | Olfr521 | 966 | 0 | 0 | 0 | 0 | 0 | 0.05646923 |
| ENSMUSG00 | Olfr520 | 950 | 0 | 0 | 0 | 0 | 0 | 0 |
| ENSMUSG00 | Gm10606 | 3585 | 0.53229095 | 0.43011608 | 0.45619754 | 0.95614902 | 1.75069574 | 0.45647932 |
| ENSMUSG00 | Kbtbd5 | 2449 | 0 | 0 | 0 | 0 | 0 | 0 |
| ENSMUSG00 | Gucy2d | 3333 | 0 | 0 | 0 | 0 | 0 | 0 |
| ENSMUSG00 | B3gnt6 | 2366 | 0.05040847 | 0 | 0.09341049 | 0.71676082 | 0 | 0 |
| ENSMUSG00 | Omp | 2142 | 1.002239 | 0.33744007 | 0.72225234 | 0.40428061 | 0.45261491 | 0.98046089 |
| ENSMUSG00 | Gm6205 | 2511 | 0 | 0 | 0 | 0 | 0 | 0 |
| ENSMUSG00 | Gm6509 | 1215 | 0 | 0 | 0 | 0 | 0 | 0 |
| ENSMUSG00 | Gm3147 | 1844 | 0 | 0 | 0 | 0 | 0 | 0 |
| ENSMUSG00 | Gm7682 | 1440 | 0 | 0 | 0 | 0 | 0 | 0 |
| ENSMUSG00 | 4632427E13I | 5407 | 20.3666878 | 20.1140743 | 21.1648921 | 28.77488 | 21.2429022 | 47.2602358 |
| ENSMUSG00 | Gm6091 | 164 | 0 | 0 | 0 | 0 | 0 | 0 |
| ENSMUSG00 | Slc22a13 | 1970 | 0 | 0 | 0 | 0 | 0 | 0 |
| ENSMUSG00 | Gm10608 | 2508 | 0 | 0 | 0 | 0 | 0 | 0 |
| ENSMUSG00 | Exoc8 | 7405 | 9.652985 | 7.76319048 | 9.59845558 | 10.4323643 | 10.2948556 | 7.61700931 |
| ENSMUSG00 | Gm5921 | 881 | 0 | 0 | 0 | 0 | 0 | 0 |
| ENSMUSG00 | Mc1r | 3660 | 0 | 0 | 0 | 0 | 0 | 0 |
| ENSMUSG00 | Gm10610 | 485 | 0 | 0 | 0 | 0 | 0 | 0 |
| ENSMUSG00 | 4930520O04 | 3902 | 0.05094244 | 0 | 0.09062397 | 0 | 0 | 0.11882852 |
| ENSMUSG00 | Gm16378 | 695 | 0.22880852 | 0.13866602 | 0.19079932 | 0.10383322 | 0 | 0.15697634 |
| ENSMUSG00 | BC048644 | 1584 | 0 | 0 | 0 | 0 | 0 | 0 |
| ENSMUSG00 | Gm10614 | 2536 | 0 | 0 | 0 | 0.25610284 | 0 | 0.2043447 |
| ENSMUSG00 | Gm10615 | 2213 | 3.82644254 | 1.69839245 | 4.57398195 | 2.77177929 | 4.58845404 | 6.08841925 |
| ENSMUSG00 | Fbxw18 | 1509 | 0 | 0 | 0.64442714 | 0 | 0 | 0 |
| ENSMUSG00 | Fbxw15 | 1641 | 0 | 0 | 0.32323103 | 0 | 0 | 0 |
| ENSMUSG00 | Fbxw19 | 1478 | 0 | 0 | 0 | 0 | 0 | 0 |
| ENSMUSG00 | Fbxw16 | 1500 | 0 | 0 | 0 | 0 | 0 | 0 |
| ENSMUSG00 | Osgin1 | 2668 | 5.51331626 | 17.0494758 | 1.01061185 | 8.27669087 | 4.18844689 | 0.96095056 |
| ENSMUSG00 | Mlycd | 3550 | 15.5998266 | 12.0805447 | 12.9990772 | 10.7433017 | 14.3017589 | 9.45008628 |
| ENSMUSG00 | Gm10617 | 1170 | 4.68910796 | 2.75939455 | 1.43561541 | 3.63904381 | 2.31145121 | 4.87213641 |
| ENSMUSG00 | Gm10619 | 2690 | 0 | 0 | 0 | 0 | 0 | 0 |
| ENSMUSG00 | Fam169b | 3061 | 0 | 0.01574206 | 0 | 0 | 0 | 0 |
| ENSMUSG00 | Gm10620 | 1328 | 0 | 0 | 0 | 0 | 0 | 0 |
| ENSMUSG00 | Gm10621 | 2113 | 0.47036773 | 0 | 0.58573195 | 2.69804214 | 0.2656366 | 0.2323443 |
| ENSMUSG00 | Snrnp40 | 2933 | 12.659945 | 11.1224757 | 11.965995 | 11.1949064 | 12.2303247 | 9.42941841 |
| ENSMUSG00 | Gm6181 | 671 | 0 | 0 | 0.06587458 | 0 | 0 | 0 |
| ENSMUSG00 | Svip | 3055 | 24.1135533 | 21.9559828 | 25.4793602 | 30.1648254 | 21.5463256 | 29.8547933 |
| ENSMUSG00 | Rbm15b | 1360 | 0 | 0 | 0 | 0 | 0 | 0 |
| ENSMUSG00 | Rpl10-ps2 | 644 | 0 | 0 | 0 | 0 | 0 | 0 |
| ENSMUSG00 | Mrgprx2 | 4557 | 0 | 0 | 0 | 0 | 0 | 0 |
| ENSMUSG00 | Mrgpra9 | 992 | 0 | 0 | 0 | 0 | 0 | 0 |
| ENSMUSG00 | Gm10629 | 2012 | 0 | 0 | 0 | 0 | 0 | 0 |
| ENSMUSG00 | Saa1 | 601 | 0 | 0 | 0 | 0 | 0 | 0 |
| ENSMUSG00 | Gm5897 | 1425 | 0.13949291 | 0 | 0 | 0 | 0.07161597 | 0.09570049 |
| ENSMUSG00 | Ntf5 | 1959 | 0.6494004 | 0.14758482 | 0.85741196 | 0.73674415 | 1.01583903 | 0.73790499 |
| ENSMUSG00 | Foxl2os | 3724 | 0 | 0 | 0 | 0 | 0 | 0 |
| ENSMUSG00 | 7420426K07I | 1503 | 0 | 0 | 0 | 0 | 0 | 0 |
| ENSMUSG00 | Cmtm2a | 1275 | 0 | 0 | 0 | 0 | 0 | 0 |
| ENSMUSG00 | Gm10631 | 491 | 0 | 0 | 0 | 0 | 0 | 0 |
| ENSMUSG00 | Rpl13a | 3585 | 3.36008664 | 3.4678109 | 3.82219563 | 3.51258955 | 1.92149532 | 5.15821629 |
| ENSMUSG00 | 4930513N10I | 5006 | 0.06353253 | 0 | 0 | 0.08649312 | 0.05096522 | 0.19069364 |
| ENSMUSG00 | 1700057G04I | 1987 | 0 | 0 | 0 | 0 | 0 | 0 |
| ENSMUSG00 | Zfp319 | 2249 | 2.7929595 | 4.69223247 | 1.78851388 | 2.4225828 | 1.79238937 | 3.01973552 |
| ENSMUSG00 | Il4i1 | 3936 | 0.24241147 | 1.02836919 | 0.46043587 | 0.74254208 | 1.76310648 | 0.88698014 |
| ENSMUSG00 | 4930579C12I | 756 | 0 | 0 | 0 | 0 | 0 | 0 |
| ENSMUSG00 | Bcl2a1a | 814 | 0 | 0 | 0 | 0 | 0 | 0 |
| ENSMUSG00 | Bcl2a1d | 814 | 0 | 0 | 0 | 0 | 0 | 0 |
| ENSMUSG00 | Gm10634 | 2438 | 0 | 0.01976474 | 0 | 0 | 0 | 0 |
| ENSMUSG00 | Nlrc5 | 7249 | 0 | 0.12629913 | 0.03658588 | 0.13439305 | 1.56971745 | 2.24999783 |
| ENSMUSG00 | 9330175E14I | 5923 | 0 | 0 | 0 | 0.01827556 | 0.19814396 | 0 |
| ENSMUSG00 | Ap3s1-ps2 | 1278 | 0.3732909 | 0 | 0.03458673 | 0.36703175 | 0.15970697 | 0.23475824 |
| ENSMUSG00 | Klk5 | 1234 | 0 | 0 | 0 | 0 | 0 | 0 |
| ENSMUSG00 | Ces1h | 2209 | 0 | 0 | 0 | 0 | 0 | 0 |
| ENSMUSG00 | 9830147E19I | 9715 | 9.55522872 | 7.35072643 | 6.84298218 | 13.7902863 | 6.8385323 | 10.4438145 |
| ENSMUSG00 | Vmn2r-ps54 | 3206 | 0 | 0 | 0.19302115 | 0 | 0 | 0 |
| ENSMUSG00 | Zfp788 | 4507 | 27.229902 | 26.001648 | 18.9870797 | 20.5348223 | 13.0537855 | 17.9975099 |
| ENSMUSG00 | AW146154 | 4616 | 11.8594664 | 9.33246951 | 13.2337407 | 13.7809022 | 8.58914515 | 24.4620898 |
| ENSMUSG00 | Gm16387 | 2963 | 0 | 0 | 0 | 0 | 0 | 0 |
| ENSMUSG00 | Plekhf1 | 1557 | 2.96187267 | 1.64025783 | 3.49186043 | 1.34409671 | 1.96633433 | 1.90939992 |
| ENSMUSG00 | Gm6658 | 476 | 0.83519917 | 0.80985617 | 1.11433218 | 0.30321046 | 0.21439654 | 0.05729966 |
| ENSMUSG00 | Gm10638 | 904 | 0.21988651 | 0.31982152 | 0.19558338 | 0.03991377 | 0.33867064 | 0.27153955 |
| ENSMUSG00 | Gm10639 | 1140 | 6.62591342 | 11.2435032 | 5.04056105 | 7.94437998 | 3.13319852 | 6.31623225 |
| ENSMUSG00 | Gm3776 | 854 | 0.04655209 | 0.39497084 | 0.0517586 | 0.25350383 | 0.05974986 | 0.1916251 |
| ENSMUSG00 | Znhit6 | 2803 | 6.80793102 | 11.1054019 | 7.9005078 | 15.9105983 | 7.48192667 | 8.07632914 |
| ENSMUSG00 | Gsta1 | 949 | 0.20945986 | 0.45698417 | 0.23288642 | 0.26614785 | 0.05376857 | 0.1149616 |
| ENSMUSG00 | Smt3h2-ps2 | 287 | 0 | 0.16789701 | 1.38612051 | 0 | 0 | 0 |
| ENSMUSG00 | Gm5058 | 467 | 0 | 0 | 0 | 0 | 0 | 0 |
| ENSMUSG00 | Gm5116 | 467 | 0 | 0 | 0 | 0 | 0 | 0 |
| ENSMUSG00 | Zfp791 | 1583 | 3.56618966 | 4.35291297 | 5.91964038 | 2.735215 | 9.79912713 | 4.04898309 |
| ENSMUSG00 | AI747448 | 2983 | 0 | 0 | 0 | 0 | 0 | 0 |
| ENSMUSG00 | Gm6289 | 872 | 0 | 0 | 0 | 0 | 0 | 0 |
| ENSMUSG00 | Krtdap | 450 | 0 | 0 | 0 | 0 | 0 | 0 |
| ENSMUSG00 | G430095P16I | 719 | 0 | 0.20105609 | 0.24590733 | 0.25091825 | 0.92259094 | 0.91041911 |
| ENSMUSG00 | Adh6b | 1830 | 0 | 0.15798834 | 0 | 0 | 0 | 0 |
| ENSMUSG00 | Adh1 | 2222 | 107.779935 | 156.551722 | 69.0678663 | 103.358332 | 93.4411896 | 59.6556016 |
| ENSMUSG00 | E130208F15I | 1732 | 0 | 0 | 0.35728972 | 0.43748437 | 0.64814103 | 0.34644461 |
| ENSMUSG00 | Sdhaf1 | 964 | 4.08277238 | 5.09856546 | 5.59400918 | 2.91950153 | 3.44057513 | 7.3562305 |
| ENSMUSG00 | Dnajb14 | 12714 | 6.33511118 | 4.73753757 | 7.9301875 | 9.62074335 | 6.97930372 | 7.96958352 |
| ENSMUSG00 | Gm10642 | 1609 | 0.32120649 | 0.11979227 | 0.35713111 | 0.29152677 | 0.28541789 | 0.13561039 |
| ENSMUSG00 | Gm10643 | 1576 | 0.10090224 | 0 | 0.67312451 | 0.11447349 | 1.10082283 | 0.01730624 |
| ENSMUSG00 | 2210011C24I | 624 | 8.47352389 | 5.79163968 | 5.52523038 | 3.35377978 | 4.00687887 | 5.55108844 |
| ENSMUSG00 | Cox7a1 | 360 | 25.3993348 | 23.8255186 | 39.9044416 | 16.5376037 | 23.5288289 | 80.4601858 |
| ENSMUSG00 | Gm10644 | 2793 | 2.00698089 | 1.12141738 | 1.36103061 | 3.83686617 | 2.63079059 | 2.53899256 |
| ENSMUSG00 | Zfp382 | 2114 | 0.77103817 | 1.73234135 | 0.87818231 | 0.98995203 | 0.89308226 | 0.42576305 |
| ENSMUSG00 | Zfp568 | 7173 | 3.28108803 | 5.46153318 | 3.42621284 | 2.33403996 | 3.46435871 | 4.48683596 |
| ENSMUSG00 | 4932431P20I | 9642 | 0.00412316 | 0 | 0 | 0 | 0.0052921 | 0 |
| ENSMUSG00 | Spint2 | 1505 | 323.749614 | 376.206442 | 409.887655 | 301.219141 | 263.167262 | 332.859335 |
| ENSMUSG00 | Gm10645 | 1089 | 0 | 0 | 0.12176816 | 0 | 0.65598647 | 0 |
| ENSMUSG00 | Gm10647 | 2035 | 1.87544282 | 0.37886146 | 0.93399472 | 0.23049955 | 1.32894247 | 3.00222073 |
| ENSMUSG00 | Gm10649 | 4030 | 0.00986488 | 0.02391387 | 0 | 0 | 0.0379849 | 0.06091111 |
| ENSMUSG00 | Gm9387 | 1316 | 0.15104666 | 0.03661584 | 0 | 0.21934374 | 0 | 0.02072541 |
| ENSMUSG00 | Gm10650 | 1967 | 0.12126735 | 0 | 0 | 0 | 0 | 0 |
| ENSMUSG00 | Ap1ar | 2381 | 100.933171 | 81.4170755 | 124.604271 | 99.8507314 | 85.9583341 | 96.818669 |
| ENSMUSG00 | Cib3 | 641 | 0 | 0 | 0.4137458 | 0 | 0 | 0 |
| ENSMUSG00 | Gm10651 | 608 | 0.0653873 | 0.23776205 | 0 | 0 | 0 | 0.0448596 |
| ENSMUSG00 | Dda1 | 2960 | 16.7617702 | 21.3583149 | 21.6678629 | 15.834654 | 18.1178111 | 18.5854552 |
| ENSMUSG00 | 4930432M17 | 520 | 0 | 0 | 0 | 0 | 0 | 0 |
| ENSMUSG00 | Gm10653 | 2198 | 0.01808712 | 0 | 0 | 0.19699023 | 0 | 0 |
| ENSMUSG00 | Gm10654 | 1632 | 0 | 0 | 0 | 0 | 0 | 0 |
| ENSMUSG00 | Cyp2a4 | 1708 | 0.88448961 | 0.47960745 | 0 | 0 | 0 | 0 |
| ENSMUSG00 | Gm10655 | 1396 | 0.02847814 | 0 | 0.06332642 | 0 | 0 | 0 |
| ENSMUSG00 | Gramd2 | 4476 | 0 | 1.10884798 | 0.098753 | 0 | 0.41039981 | 0.32905061 |
| ENSMUSG00 | Gm7092 | 908 | 0.48161926 | 0.47761892 | 0.04868044 | 0 | 0.28098225 | 0 |
| ENSMUSG00 | Amy1 | 3652 | 15.6213348 | 7.46810686 | 20.0191261 | 13.1009833 | 13.3713696 | 25.1610788 |
| ENSMUSG00 | Amy2a3 | 2080 | 0 | 0 | 0 | 0 | 0 | 0 |
| ENSMUSG00 | Amy2a4 | 1707 | 0 | 0 | 0 | 0 | 0 | 0 |
| ENSMUSG00 | Amy2a5 | 1965 | 0 | 0 | 0 | 0 | 0 | 0 |
| ENSMUSG00 | 2410076I21R | 1276 | 0.46734499 | 0.4909277 | 0.31176849 | 0.70693661 | 0.15995729 | 0 |
| ENSMUSG00 | Ceacam1 | 3934 | 3.37527465 | 2.80495558 | 3.47187837 | 3.53116094 | 2.99621068 | 2.50283294 |
| ENSMUSG00 | Gm10657 | 1030 | 0 | 0 | 0 | 0 | 0 | 0.02648023 |
| ENSMUSG00 | D930028M14 | 996 | 0 | 0.3870397 | 0 | 0 | 0 | 0 |
| ENSMUSG00 | Phldb3 | 2258 | 0.05281951 | 0.08536128 | 0 | 0.30361331 | 0.22598041 | 0.33821519 |
| ENSMUSG00 | Gm6166 | 669 | 0 | 0 | 0 | 0 | 0 | 0 |
| ENSMUSG00 | Gm12522 | 2436 | 0 | 0.17802873 | 0 | 0 | 0.02094679 | 0 |
| ENSMUSG00 | Zfp94 | 5076 | 1.73088282 | 1.58533015 | 1.78514141 | 1.15866297 | 3.19668785 | 1.54749726 |
| ENSMUSG00 | Zfp109 | 1958 | 2.98470666 | 4.89739632 | 4.62787427 | 4.7544267 | 5.96784477 | 0.86365048 |
| ENSMUSG00 | Gm10658 | 1463 | 0.54347889 | 0.88929182 | 1.38980504 | 0.61657629 | 0.52316858 | 1.84565228 |
| ENSMUSG00 | Vmn1r173 | 1227 | 0 | 0 | 0 | 0 | 0 | 0 |
| ENSMUSG00 | Vmn1r168 | 929 | 0 | 0 | 0 | 0 | 0 | 0 |
| ENSMUSG00 | Gm10660 | 1199 | 0 | 0 | 0 | 0 | 0 | 0 |
| ENSMUSG00 | Gm6902 | 1333 | 0 | 0 | 0 | 0 | 0 | 0 |
| ENSMUSG00 | BC030870 | 655 | 0.24278156 | 0.44140252 | 0.20245119 | 1.10174182 | 0.15580573 | 1.7072675 |
| ENSMUSG00 | Gm10663 | 64 | 0 | 0 | 0 | 0 | 0 | 0 |
| ENSMUSG00 | Gm10664 | 1041 | 246.323582 | 369.75341 | 102.840407 | 270.11275 | 255.327946 | 199.856819 |
| ENSMUSG00 | Vmn1r157 | 893 | 0 | 0 | 0 | 0 | 0 | 0 |
| ENSMUSG00 | C230081A13I | 10970 | 2.34473983 | 1.52422018 | 2.3249283 | 2.60172263 | 3.45602527 | 3.65733768 |
| ENSMUSG00 | Gm4567 | 1333 | 0 | 0 | 0 | 0 | 0 | 0 |
| ENSMUSG00 | Gm10665 | 965 | 0 | 0 | 0 | 0 | 0 | 0 |
| ENSMUSG00 | Vmn1r148 | 923 | 0 | 0 | 0 | 0 | 0 | 0 |
| ENSMUSG00 | Gm10666 | 965 | 0 | 0 | 0 | 0 | 0 | 0 |
| ENSMUSG00 | Vmn1r139 | 1322 | 0 | 0 | 0 | 0 | 0 | 0 |
| ENSMUSG00 | Gm6176 | 1333 | 0 | 0 | 0 | 0 | 0 | 0 |
| ENSMUSG00 | Vmn1r132 | 893 | 0 | 0 | 0 | 0 | 0 | 0 |
| ENSMUSG00 | Vmn1r112 | 2239 | 0 | 0 | 0 | 0 | 0 | 0 |
| ENSMUSG00 | Vmn1r114 | 965 | 0 | 0 | 0 | 0 | 0 | 0 |
| ENSMUSG00 | Vmn1r125 | 923 | 0 | 0 | 0 | 0 | 0 | 0 |
| ENSMUSG00 | Gm10670 | 965 | 0 | 0 | 0 | 0 | 0 | 0 |
| ENSMUSG00 | Vmn1r100 | 923 | 0 | 0 | 0 | 0 | 0 | 0 |
| ENSMUSG00 | Vmn1r93 | 965 | 0 | 0 | 0 | 0 | 0 | 0 |
| ENSMUSG00 | Gm16442 | 923 | 0 | 0 | 0 | 0 | 0 | 0 |
| ENSMUSG00 | Apoc4 | 461 | 0 | 0 | 0 | 0 | 0 | 0 |
| ENSMUSG00 | Ovgp1 | 3405 | 0.33859293 | 0.1981234 | 0.07788871 | 0.26491956 | 0.53948591 | 0.44055952 |
| ENSMUSG00 | I830077J02Ri | 1322 | 0.1202889 | 0 | 0.10030675 | 0.21834823 | 0.11579359 | 0.53641499 |
| ENSMUSG00 | 1700104A03I | 838 | 0 | 0 | 0 | 0 | 0 | 0 |
| ENSMUSG00 | Adora3 | 893 | 0.13355705 | 0 | 0.14849443 | 0 | 0 | 0 |
| ENSMUSG00 | Tnfaip8l3 | 2223 | 0 | 0 | 0 | 0 | 0 | 0 |
| ENSMUSG00 | 1700095B22I | 1292 | 0 | 0.03729601 | 0 | 0 | 0 | 0.18999362 |
| ENSMUSG00 | 9230115E21I | 2642 | 0 | 0 | 0 | 0 | 0 | 0 |
| ENSMUSG00 | Gm10676 | 3220 | 0.01234642 | 0.224471 | 0.04118184 | 0.05602802 | 0.0475401 | 0.04235192 |
| ENSMUSG00 | AA386476 | 1664 | 0.9795521 | 0.43437298 | 0.23907247 | 0.86735684 | 1.47191469 | 1.50797284 |
| ENSMUSG00 | Ccdc61 | 2371 | 8.23278825 | 11.0355285 | 9.43320648 | 11.0331009 | 9.3831716 | 10.7327028 |
| ENSMUSG00 | Psg23 | 2174 | 0 | 0 | 0 | 0 | 0 | 0 |
| ENSMUSG00 | Gpr77 | 3358 | 0 | 0 | 0 | 0 | 0 | 0 |
| ENSMUSG00 | Ehd2 | 6895 | 1.25118771 | 1.62834533 | 0.31412477 | 1.03091556 | 1.76131653 | 0.19778564 |
| ENSMUSG00 | Crxos1 | 3088 | 0 | 0 | 0 | 0 | 0 | 0 |
| ENSMUSG00 | Obox5 | 2100 | 0 | 0 | 0 | 0 | 0.02429827 | 0 |
| ENSMUSG00 | Obox2 | 1642 | 0 | 0 | 0 | 0 | 0 | 0 |
| ENSMUSG00 | Gm10680 | 700 | 0 | 0 | 0 | 0 | 0 | 0 |
| ENSMUSG00 | Sult2a3 | 1006 | 0.51373881 | 0.33529334 | 0.3075675 | 0.14346737 | 0.15216613 | 0 |
| ENSMUSG00 | Sult2a4 | 995 | 0 | 0 | 0 | 0 | 0 | 0 |
| ENSMUSG00 | Bsph1 | 947 | 0 | 0 | 0 | 0 | 0 | 0 |
| ENSMUSG00 | Gm10681 | 1659 | 0 | 0 | 0 | 0 | 0 | 0 |
| ENSMUSG00 | AI429214 | 1956 | 3.27230693 | 7.68618096 | 7.11839496 | 0.88544895 | 5.47829188 | 7.62741701 |
| ENSMUSG00 | Gm10684 | 2200 | 0 | 0 | 0 | 0 | 0 | 0 |
| ENSMUSG00 | Gm5544 | 889 | 0 | 0 | 0 | 0 | 0 | 0 |
| ENSMUSG00 | Vmn2r29 | 2590 | 0.32234173 | 0.11162882 | 0.51199046 | 0.26469454 | 0.37432477 | 0.89511364 |
| ENSMUSG00 | Foxr1 | 1927 | 0 | 0 | 0 | 0 | 0 | 0 |
| ENSMUSG00 | Gm15441 | 1362 | 0.43783569 | 0.42455015 | 0.5517117 | 0.66229891 | 0.78675029 | 1.62206004 |
| ENSMUSG00 | Vmn1r59 | 1438 | 0 | 0 | 0 | 0 | 0 | 0 |
| ENSMUSG00 | Vmn1r55 | 1283 | 0 | 0 | 0 | 0 | 0 | 0 |
| ENSMUSG00 | Hist2h3b | 487 | 0 | 0 | 0 | 0 | 0 | 0 |
| ENSMUSG00 | Gm1698 | 2307 | 0 | 0 | 0 | 0 | 0 | 0 |
| ENSMUSG00 | Zfp865 | 2001 | 1.21193619 | 1.83016972 | 1.54629136 | 2.10974473 | 1.37702364 | 3.81654122 |
| ENSMUSG00 | Zfp628 | 1119 | 1.24346901 | 1.11961349 | 2.44907441 | 2.35387779 | 0.68399968 | 2.41303779 |
| ENSMUSG00 | Gm10687 | 1649 | 0 | 0 | 0 | 0 | 0 | 0 |
| ENSMUSG00 | Gm10689 | 433 | 0 | 0 | 0 | 0 | 0 | 0 |
| ENSMUSG00 | Gm10691 | 782 | 0 | 0 | 0 | 0 | 0 | 0.03487806 |
| ENSMUSG00 | 2610203C20I | 3265 | 0.15829135 | 1.04785219 | 0.04061425 | 0.59676276 | 0 | 0 |
| ENSMUSG00 | Gm14548 | 2383 | 0 | 0 | 0 | 0 | 0 | 0 |
| ENSMUSG00 | Gm15930 | 2027 | 0 | 0 | 0 | 0 | 0 | 0 |
| ENSMUSG00 | Gm15448 | 2383 | 0 | 0 | 0 | 0 | 0 | 0 |
| ENSMUSG00 | Gm10693 | 2027 | 0 | 0 | 0 | 0 | 0 | 0 |
| ENSMUSG00 | Gm17312 | 1112 | 0 | 0 | 0 | 0 | 0 | 0 |
| ENSMUSG00 | Gm10696 | 4449 | 0 | 0.07581594 | 0 | 0 | 0 | 0 |
| ENSMUSG00 | Gm10697 | 1022 | 0 | 0 | 0 | 0 | 0 | 0 |
| ENSMUSG00 | Gm9125 | 1238 | 0 | 0 | 0 | 0 | 0 | 0 |
| ENSMUSG00 | Tdpoz5 | 1022 | 0 | 0 | 0 | 0 | 0 | 0 |
| ENSMUSG00 | Gm9117 | 1238 | 0 | 0 | 0 | 0 | 0 | 0 |
| ENSMUSG00 | Gm9116 | 1073 | 0 | 0 | 0 | 0 | 0 | 0 |
| ENSMUSG00 | Gm4858 | 1686 | 0 | 0 | 0 | 0 | 0 | 0 |
| ENSMUSG00 | 2310001H18I | 302 | 0 | 0 | 0 | 0 | 0 | 0 |
| ENSMUSG00 | Gm6696 | 461 | 0 | 0 | 0 | 0 | 0 | 0 |
| ENSMUSG00 | Smcp | 811 | 0 | 0.59416082 | 0 | 0 | 0 | 0 |
| ENSMUSG00 | Defa-ps18 | 369 | 0 | 0 | 0 | 0 | 0 | 0 |
| ENSMUSG00 | Defa-rs1 | 506 | 0 | 0 | 0 | 0 | 0 | 0 |
| ENSMUSG00 | Defa5 | 374 | 0 | 0 | 0 | 0 | 0 | 0 |
| ENSMUSG00 | Defa3 | 408 | 0 | 0 | 0 | 0 | 0 | 0 |
| ENSMUSG00 | Gm15292 | 383 | 0 | 0 | 0 | 0 | 0 | 0 |
| ENSMUSG00 | Defa-rs7 | 375 | 0 | 0 | 0 | 0 | 0 | 0 |
| ENSMUSG00 | Defa22 | 417 | 0 | 0 | 0 | 0 | 0 | 0 |
| ENSMUSG00 | Gm15284 | 396 | 0 | 0 | 0 | 0 | 0 | 0 |
| ENSMUSG00 | Sprr2a3 | 720 | 0 | 0 | 0 | 0 | 0 | 0 |
| ENSMUSG00 | Defa23 | 409 | 0 | 0 | 0 | 0 | 0 | 0 |
| ENSMUSG00 | Defa21 | 416 | 0 | 0 | 0 | 0 | 0 | 0 |
| ENSMUSG00 | Gm5615 | 2339 | 0 | 0 | 0 | 0 | 0 | 0 |
| ENSMUSG00 | Gm15319 | 2512 | 0 | 0 | 0.05278883 | 0.04309161 | 0 | 0.08686191 |
| ENSMUSG00 | Pate2 | 4132 | 1.55866114 | 0.06997063 | 0.06418467 | 0.75098157 | 0.30872686 | 0.01320167 |
| ENSMUSG00 | 4930467E23I | 1043 | 0 | 0 | 0 | 0 | 0.0489227 | 0 |
| ENSMUSG00 | S100a16 | 1713 | 34.7657384 | 32.8838009 | 35.015704 | 28.2884914 | 23.3833655 | 37.480736 |
| ENSMUSG00 | Gm10699 | 2129 | 1.77396461 | 1.22220191 | 0.97580396 | 2.05069394 | 2.6124354 | 0.21778716 |
| ENSMUSG00 | Gm10701 | 902 | 0 | 0 | 0 | 0 | 0 | 0 |
| ENSMUSG00 | Gm15417 | 1997 | 0.99538008 | 0.50671772 | 0.64188956 | 3.08964928 | 1.04761212 | 1.40675405 |
| ENSMUSG00 | Gm10702 | 1266 | 0 | 0 | 0 | 0 | 0 | 0.19389554 |
| ENSMUSG00 | Gm15348 | 3651 | 0.03266679 | 0 | 0 | 0 | 0 | 0 |
| ENSMUSG00 | Zfp872 | 1392 | 0 | 0 | 0 | 0.02592101 | 0.40322567 | 0 |
| ENSMUSG00 | A230072I06R | 1838 | 0 | 0 | 0 | 0 | 0 | 0 |
| ENSMUSG00 | Spc24 | 1544 | 0.38622552 | 1.59165062 | 0.40079391 | 0.5374916 | 0.56181891 | 0.42395812 |
| ENSMUSG00 | Gm10704 | 734 | 6.4995336 | 5.90841934 | 5.48006501 | 5.55486517 | 4.72723918 | 9.10393272 |
| ENSMUSG00 | Mex3a | 5791 | 0.04805532 | 0.1747393 | 0.27478265 | 0.04361497 | 0.16741515 | 0.43801441 |
| ENSMUSG00 | Gm16589 | 2482 | 0.06407007 | 0.25238668 | 0.10685377 | 0.18898734 | 0.06167572 | 0.14285669 |

| | | | | | | | | |
|---|---|---|---|---|---|---|---|---|
| ENSMUSG00 | Bglap | 614 | 0 | 0 | 0 | 0 | 0 | 0.13326371 |
| ENSMUSG00 | Gm7381 | 299 | 0 | 0 | 0 | 0 | 0 | 0 |
| ENSMUSG00 | Bglap2 | 467 | 0 | 0 | 0 | 0 | 0 | 0 |
| ENSMUSG00 | Bglap-rs1 | 2542 | 83.4208234 | 18.6149041 | 128.710481 | 32.0650427 | 54.880453 | 156.459005 |
| ENSMUSG00 | Clec4g | 2451 | 0 | 0 | 0 | 0 | 0 | 0 |
| ENSMUSG00 | A430078G23 | 3294 | 0 | 0 | 0.09393227 | 0 | 0 | 0 |
| ENSMUSG00 | Zfp558 | 2418 | 4.04460224 | 2.6504536 | 5.08193232 | 4.20807963 | 4.89583093 | 5.25640277 |
| ENSMUSG00 | Gm5612 | 377 | 0 | 0.1278155 | 0 | 0.09570834 | 0 | 0.14469305 |
| ENSMUSG00 | Ubtfl1 | 2869 | 0 | 0 | 0 | 0 | 0 | 0 |
| ENSMUSG00 | Fat3 | 19888 | 0.08595564 | 0 | 0 | 0 | 0 | 0 |
| ENSMUSG00 | Gm10705 | 464 | 0.17135983 | 0 | 0 | 0 | 0 | 0 |
| ENSMUSG00 | Gm14340 | 1911 | 0 | 0 | 0 | 0 | 0 | 0 |
| ENSMUSG00 | Gm10706 | 1955 | 0 | 0.39436474 | 0 | 0 | 0 | 0 |
| ENSMUSG00 | Arfip1 | 5157 | 21.3386019 | 20.7621242 | 26.4765354 | 24.7333774 | 24.568255 | 20.6847225 |
| ENSMUSG00 | Gm10709 | 482 | 0 | 0 | 0 | 0 | 0 | 0 |
| ENSMUSG00 | Gm10710 | 3970 | 0 | 0 | 0 | 0 | 0 | 0 |
| ENSMUSG00 | Gm10711 | 67 | 0 | 0 | 0 | 0 | 0 | 0 |
| ENSMUSG00 | Etohi1 | 2043 | 5.70159363 | 4.38343211 | 6.18782531 | 7.1175056 | 10.1153609 | 7.80991873 |
| ENSMUSG00 | Gm14327 | 1448 | 0 | 0 | 0 | 0 | 0.17619605 | 0.05650823 |
| ENSMUSG00 | Gm14296 | 9862 | 0 | 0.01954429 | 0.00896407 | 0 | 0.01034808 | 0.00276563 |
| ENSMUSG00 | C330013J21 | 859 | 0.60165454 | 0.28047987 | 0.66894524 | 0.46205179 | 0.1782062 | 0.1905097 |
| ENSMUSG00 | Gm10712 | 611 | 0 | 0 | 0 | 0 | 0 | 0 |
| ENSMUSG00 | Gm10713 | 335 | 0 | 0.5753605 | 0.3958374 | 1.18478355 | 0.45695262 | 0.56991784 |
| ENSMUSG00 | Gm10714 | 1003 | 0 | 0 | 0 | 0 | 0 | 0 |
| ENSMUSG00 | Gm10715 | 671 | 0 | 0 | 0 | 0 | 0 | 0 |
| ENSMUSG00 | Gm10717 | 681 | 0 | 1.76895896 | 0 | 0 | 0 | 0 |
| ENSMUSG00 | Gm10718 | 671 | 0 | 0.64631592 | 0 | 0 | 0.15209054 | 0 |
| ENSMUSG00 | Gm10719 | 671 | 0 | 0 | 0 | 0 | 0 | 0 |
| ENSMUSG00 | Gm10720 | 673 | 0 | 0.07159947 | 0 | 0.53613736 | 0 | 0 |
| ENSMUSG00 | Gm10721 | 641 | 0 | 0 | 0 | 0 | 0 | 0 |
| ENSMUSG00 | Gcnt7 | 3189 | 1.28404343 | 0.72528982 | 0.36037878 | 0.73544462 | 0.67203129 | 0.47895259 |
| ENSMUSG00 | Cass4 | 5545 | 0 | 0 | 0 | 0 | 0.16564017 | 0 |
| ENSMUSG00 | 1700101G07 | 1152 | 0 | 0 | 0 | 0 | 0 | 0 |
| ENSMUSG00 | Kcng1 | 5893 | 0 | 0 | 0 | 0 | 0 | 0 |
| ENSMUSG00 | Mocs3 | 1972 | 0.98783902 | 1.17289514 | 1.16556584 | 0.78677886 | 1.47490032 | 0.45642145 |
| ENSMUSG00 | Fam65c | 4976 | 0.13582057 | 0.20335918 | 0.10659608 | 0.1957828 | 0.0102545 | 0.36724293 |
| ENSMUSG00 | 1500012F01 | 1233 | 3.32101743 | 3.20461335 | 4.01509036 | 3.97985244 | 3.26932984 | 5.99466929 |
| ENSMUSG00 | Lekr1 | 3371 | 2.21715525 | 1.68673989 | 4.09106349 | 2.94350704 | 2.34621425 | 3.94839038 |
| ENSMUSG00 | 4931440P22 | 3378 | 0.09415152 | 0.42794354 | 0.99447604 | 0.10681482 | 0.34742648 | 0.54904543 |
| ENSMUSG00 | Arfgef2 | 8731 | 10.9144298 | 9.86246388 | 11.2441064 | 10.5878133 | 12.0625862 | 10.91173 |
| ENSMUSG00 | 4930449A18 | 475 | 0 | 0 | 0 | 0 | 0 | 0 |
| ENSMUSG00 | F630111L10 | 2976 | 0 | 0.0809584 | 0.01485277 | 0 | 0.25718939 | 0.17413244 |
| ENSMUSG00 | Gm410 | 2692 | 0 | 0 | 0 | 0 | 0 | 0 |
| ENSMUSG00 | F830104G03 | 522 | 0 | 0 | 0 | 0 | 0 | 0 |
| ENSMUSG00 | Gm14316 | 439 | 0 | 0 | 0 | 0 | 0 | 0 |
| ENSMUSG00 | Wfdc9 | 572 | 0 | 0 | 0 | 0 | 0 | 0 |
| ENSMUSG00 | Wfdc6a | 862 | 0 | 0 | 0 | 0 | 0 | 0 |
| ENSMUSG00 | Spint3 | 555 | 0 | 0 | 0 | 0 | 0.5516365 | 0.09828699 |
| ENSMUSG00 | Ufm1 | 1607 | 5.86312936 | 5.36737595 | 6.87645349 | 2.55964722 | 3.84206066 | 3.22475511 |
| ENSMUSG00 | Gm10728 | 1827 | 0 | 0.07912388 | 0.04838735 | 0 | 0 | 0 |
| ENSMUSG00 | Gm10729 | 1319 | 0 | 0 | 0 | 0 | 0.03868565 | 0.08271308 |
| ENSMUSG00 | Mgst2 | 3166 | 0.01255701 | 0.30439951 | 0 | 0 | 0.14505287 | 0 |
| ENSMUSG00 | Tox2 | 2322 | 0 | 0 | 0 | 0 | 0 | 0 |
| ENSMUSG00 | Zfp850 | 940 | 0.63439597 | 0.30757304 | 0.3761859 | 1.03639915 | 1.62850136 | 1.21865409 |
| ENSMUSG00 | 1700034I23R | 2344 | 0.15264476 | 0.04111471 | 0.11314465 | 0.35404737 | 0.19592039 | 0.80287974 |
| ENSMUSG00 | Gm10731 | 1805 | 0 | 0 | 0.02448856 | 0 | 0.02826946 | 0.04533181 |
| ENSMUSG00 | Mafb | 3388 | 26.4488941 | 22.9411839 | 33.6210595 | 19.8728144 | 18.7658985 | 22.9515928 |
| ENSMUSG00 | Gm826 | 1811 | 0.19757003 | 0.23946879 | 0.36611135 | 0 | 0.0845274 | 0.03012108 |
| ENSMUSG00 | Arhgap40 | 2191 | 0 | 0 | 0 | 0 | 0 | 0 |
| ENSMUSG00 | 4922505G16 | 4818 | 0.02475435 | 0.02000267 | 0.04587157 | 0.36896143 | 0.1059078 | 0.03396593 |
| ENSMUSG00 | Gm1332 | 635 | 630.703481 | 345.804121 | 807.32752 | 343.091945 | 308.167167 | 548.499438 |
| ENSMUSG00 | 4930518I15R | 1330 | 2.72011182 | 1.52167712 | 1.22967533 | 2.55015954 | 2.18684468 | 4.73717419 |
| ENSMUSG00 | Gm7120 | 1445 | 1.26557239 | 1.36722777 | 1.52947554 | 2.47205703 | 2.01280514 | 1.22688689 |
| ENSMUSG00 | 3110070M22 | 1122 | 0.70865384 | 0.90188528 | 0.94549397 | 0.16079343 | 1.45529771 | 0.72926843 |
| ENSMUSG00 | Sox2 | 4828 | 0 | 0 | 0 | 0 | 0 | 0 |
| ENSMUSG00 | BC089597 | 1627 | 0 | 0.82926883 | 0 | 0.73184233 | 0 | 0.08381881 |
| ENSMUSG00 | Cpne1 | 7739 | 11.9384593 | 11.2698618 | 12.8167639 | 13.1059087 | 13.4967039 | 12.4090974 |
| ENSMUSG00 | 6430550D23 | 4905 | 1.03745189 | 1.34588024 | 0.82906617 | 1.24319379 | 1.69567773 | 0.63946657 |
| ENSMUSG00 | Fam83c | 3968 | 0 | 0.01214376 | 0 | 0.05455954 | 0 | 0 |
| ENSMUSG00 | BC029722 | 1613 | 3.32733408 | 4.74993447 | 5.97396266 | 2.43827827 | 2.72056313 | 6.13806203 |
| ENSMUSG00 | Gm10735 | 2942 | 0.17567004 | 0.14740924 | 0 | 0 | 0 | 0 |
| ENSMUSG00 | Gm6320 | 2347 | 0 | 0 | 0.09416669 | 0 | 0 | 0 |
| ENSMUSG00 | Myh7b | 6592 | 0 | 0 | 0 | 0 | 0 | 0.01655015 |
| ENSMUSG00 | Lrrc31 | 4361 | 3.79231367 | 0.64086532 | 2.13863538 | 1.3238081 | 5.87370802 | 1.98884093 |
| ENSMUSG00 | Gm1527 | 2105 | 0 | 0.06867426 | 0.083994 | 0.34282228 | 0.33936782 | 0.02591415 |
| ENSMUSG00 | Eif2s2 | 7784 | 4.9132544 | 5.32997517 | 4.73591175 | 4.51489098 | 3.80862338 | 7.22161247 |
| ENSMUSG00 | Kif5a | 6690 | 0.18421822 | 0.73468118 | 0.29732181 | 0.23191748 | 0.63306266 | 0.3669234 |
| ENSMUSG00 | Gm10736 | 199 | 0 | 0 | 0 | 0 | 0 | 0 |
| ENSMUSG00 | Pde4d | 1854 | 0 | 0 | 0 | 0 | 0.66053561 | 0 |
| ENSMUSG00 | Gm10737 | 391 | 0 | 0.86267288 | 0 | 0 | 0 | 0 |
| ENSMUSG00 | A830092H15 | 1228 | 0 | 0 | 0 | 0 | 0 | 0 |
| ENSMUSG00 | Bpifb4 | 3166 | 0 | 0 | 0 | 0 | 0 | 0 |
| ENSMUSG00 | Tspyl3 | 4029 | 0.04933666 | 0.17939852 | 0.559517 | 0 | 0.68389781 | 0.21662657 |
| ENSMUSG00 | Ttll9 | 3524 | 0.06768811 | 0.08204275 | 0 | 0 | 0.01447968 | 0.01547936 |
| ENSMUSG00 | Gm10742 | 3384 | 0 | 0 | 0 | 0 | 0 | 0 |
| ENSMUSG00 | Foxs1 | 1310 | 0 | 0 | 0 | 0 | 0 | 0 |
| ENSMUSG00 | Sirpb1c | 2841 | 0 | 0 | 0 | 0 | 0 | 0 |
| ENSMUSG00 | Defb25 | 291 | 0 | 0 | 0 | 0 | 0 | 0.09372728 |
| ENSMUSG00 | Defb28 | 394 | 0 | 0 | 0 | 0 | 0 | 0 |
| ENSMUSG00 | Defb26 | 559 | 0 | 0 | 0 | 0 | 0 | 0 |
| ENSMUSG00 | Defb23 | 622 | 0 | 0 | 0 | 0 | 0 | 0 |
| ENSMUSG00 | Zcchc3 | 3088 | 10.6726986 | 7.13121893 | 12.5391239 | 6.96402155 | 5.56861679 | 9.31824625 |
| ENSMUSG00 | Gm9049 | 2821 | 0 | 0 | 0 | 0 | 0 | 0 |
| ENSMUSG00 | Gm9048 | 1244 | 0 | 0 | 0 | 0 | 0 | 0 |
| ENSMUSG00 | Gm9046 | 1244 | 0 | 0 | 0 | 0 | 0 | 0 |
| ENSMUSG00 | Gm9045 | 1244 | 0 | 0 | 0 | 0 | 0 | 0 |
| ENSMUSG00 | Gm9044 | 1244 | 0 | 0 | 0 | 0 | 0 | 0 |
| ENSMUSG00 | Gm9040 | 1244 | 0 | 0 | 0 | 0 | 0 | 0 |
| ENSMUSG00 | Il22 | 1100 | 0 | 0 | 0 | 0 | 0 | 0 |
| ENSMUSG00 | Csnk2a1 | 7150 | 3.33056403 | 3.67295258 | 2.85611909 | 3.40634686 | 3.26854268 | 5.19171805 |
| ENSMUSG00 | Rad21l | 1637 | 0 | 0 | 0 | 0 | 0 | 0 |
| ENSMUSG00 | Gm10748 | 2371 | 0 | 0 | 0 | 0 | 0 | 0 |
| ENSMUSG00 | Zfp345 | 4224 | 0.02823543 | 0.03422333 | 0.03139335 | 0.02562645 | 0.02416022 | 0.01291413 |
| ENSMUSG00 | Zfp950 | 1387 | 22.930342 | 19.2815252 | 19.4398877 | 28.2516946 | 17.106896 | 26.5470533 |
| ENSMUSG00 | 5830428H23 | 4686 | 10.095822 | 13.1006247 | 10.3665867 | 11.6731497 | 11.6949056 | 16.8443888 |
| ENSMUSG00 | 4933416C03 | 2253 | 0 | 0 | 0 | 0 | 0 | 0 |
| ENSMUSG00 | Gm10749 | 988 | 0 | 0 | 0 | 0.03652029 | 0 | 0 |
| ENSMUSG00 | Tmem90b | 2456 | 0 | 0 | 0 | 0 | 0 | 0 |
| ENSMUSG00 | C530025M09 | 689 | 0 | 0 | 0 | 0 | 0 | 0 |
| ENSMUSG00 | B930041F14 | 2139 | 0 | 0 | 0 | 0 | 0 | 0 |
| ENSMUSG00 | Gm10750 | 1505 | 0 | 0 | 0 | 0 | 0 | 0 |
| ENSMUSG00 | Thbd | 3717 | 0.66312612 | 0.94635735 | 0.47567224 | 1.07751061 | 1.56497359 | 1.37217044 |
| ENSMUSG00 | Gm807 | 1721 | 0 | 0 | 0 | 0 | 0 | 0 |
| ENSMUSG00 | Pdzd8 | 7071 | 8.24795501 | 8.46380443 | 8.70784434 | 13.5990169 | 10.0017758 | 11.0240304 |
| ENSMUSG00 | Atxn7l3b | 4044 | 64.9123239 | 54.5493403 | 63.406254 | 52.0173886 | 51.1147004 | 58.8320174 |
| ENSMUSG00 | Plk1s1 | 2398 | 16.3962345 | 21.2800009 | 16.5342173 | 24.0747587 | 26.2366645 | 17.2201017 |
| ENSMUSG00 | Gm14092 | 614 | 0 | 0 | 0 | 0 | 0 | 0 |
| ENSMUSG00 | Gm561 | 469 | 14.4103021 | 15.1032132 | 17.247201 | 9.38594762 | 7.50707875 | 30.4150028 |
| ENSMUSG00 | Gm5535 | 1827 | 0 | 0 | 0 | 0 | 0 | 0 |
| ENSMUSG00 | Gm14105 | 3564 | 0 | 0 | 0 | 0 | 0.41519778 | 0 |
| ENSMUSG00 | Sel1l2 | 2716 | 0 | 0 | 0 | 0 | 0 | 0 |
| ENSMUSG00 | Ism1 | 2938 | 0.01353148 | 1.08247283 | 0 | 0.36843476 | 0.12157407 | 0.03713361 |
| ENSMUSG00 | Bhmt | 5199 | 0 | 0 | 0 | 0 | 0 | 0 |
| ENSMUSG00 | Ankrd5 | 7593 | 0 | 0.29826983 | 0.12224927 | 0.14731244 | 0.3494497 | 0 |
| ENSMUSG00 | Gm10754 | 314 | 0 | 0 | 0 | 0 | 0 | 0 |
| ENSMUSG00 | Gm6843 | 424 | 0.09376293 | 0 | 0 | 0 | 0 | 0.12865396 |
| ENSMUSG00 | Ube2n | 3939 | 14.8464869 | 12.722493 | 16.3161919 | 15.0960166 | 12.3323457 | 20.5165667 |
| ENSMUSG00 | 4833422C13 | 5911 | 0.05380542 | 0 | 0.01495579 | 0.03662532 | 0.06042711 | 0 |
| ENSMUSG00 | AU019990 | 2430 | 0 | 0 | 0 | 0 | 0 | 0 |
| ENSMUSG00 | Plxnc1 | 7008 | 0.15316752 | 0 | 0 | 0.03604086 | 0.05824929 | 0.00778386 |
| ENSMUSG00 | 5830416P10 | 8653 | 0.42268626 | 0.33969409 | 0.03575788 | 0.57127471 | 0.73711973 | 0.08195315 |
| ENSMUSG00 | Hspa12b | 3744 | 0.12742141 | 0.23166559 | 0 | 0.15419677 | 0.31346331 | 0 |
| ENSMUSG00 | Arrdc3 | 6363 | 42.8856857 | 42.9384138 | 36.7758222 | 47.8825686 | 46.062471 | 36.9019911 |
| ENSMUSG00 | Slc4a11 | 4363 | 10.8979039 | 2.36348811 | 2.36053849 | 0.32253031 | 2.666517 | 1.44406181 |
| ENSMUSG00 | Itpa | 2086 | 7.37553737 | 9.4016692 | 9.09040779 | 6.65943775 | 4.94119267 | 7.93658007 |
| ENSMUSG00 | Gm10760 | 710 | 0 | 0 | 0 | 0 | 0 | 0 |
| ENSMUSG00 | Gm4149 | 360 | 0.55215945 | 0.26770246 | 0.1227829 | 0 | 0.8504396 | 0.37881443 |
| ENSMUSG00 | Gas2l3 | 7784 | 0 | 0.33428416 | 0 | 0 | 0 | 0 |
| ENSMUSG00 | AU015228 | 4300 | 0 | 0 | 0 | 0 | 0 | 0 |
| ENSMUSG00 | Gm14047 | 2997 | 0 | 0.20901693 | 0 | 0.3852604 | 0.187284 | 0.464133 |
| ENSMUSG00 | Gm10762 | 1132 | 4.0036438 | 2.29864653 | 3.20190029 | 3.88870091 | 1.84812845 | 5.46938437 |
| ENSMUSG00 | Gm10764 | 2334 | 0 | 0 | 0 | 0 | 0 | 0 |
| ENSMUSG00 | Hps6 | 2695 | 6.52019384 | 3.55810834 | 3.44429946 | 2.95886154 | 6.17239277 | 2.3175853 |
| ENSMUSG00 | Gm355 | 1508 | 0 | 0 | 0 | 0 | 0 | 0 |
| ENSMUSG00 | Gm14005 | 2036 | 0.07810507 | 0 | 0.17368111 | 0.12405418 | 0.17543449 | 0.17415045 |
| ENSMUSG00 | Papolb | 4238 | 0 | 0 | 0.3024666 | 0.01702786 | 0 | 0 |
| ENSMUSG00 | Pdzd7 | 5174 | 0.35345035 | 0.97788489 | 0.41006735 | 0.30684383 | 0.12820697 | 0.16868737 |
| ENSMUSG00 | Gm10766 | 323 | 4.67711535 | 3.87878482 | 8.75825992 | 5.92058316 | 1.73774035 | 9.45746005 |
| ENSMUSG00 | Zfp708 | 1376 | 2.9180985 | 3.11671028 | 1.22069043 | 5.45426254 | 4.07914342 | 1.03072765 |
| ENSMUSG00 | Itpripl1 | 5052 | 0.01573851 | 0.02861428 | 0.0349975 | 0.00714213 | 0 | 0 |
| ENSMUSG00 | Gm10767 | 2200 | 5.89103938 | 3.63588609 | 13.2002777 | 8.97130834 | 6.42468459 | 6.95503301 |
| ENSMUSG00 | Gm10768 | 1925 | 0 | 0 | 0 | 0 | 0 | 0 |
| ENSMUSG00 | Zfp640 | 807 | 0 | 0 | 0 | 0 | 0.06322971 | 0 |
| ENSMUSG00 | 2410141K09 | 1742 | 0 | 0 | 0 | 0.020713 | 0.02929184 | 0 |
| ENSMUSG00 | Vmn2r-ps104 | 668 | 0 | 0 | 0 | 0 | 0 | 0 |
| ENSMUSG00 | Gm10770 | 712 | 0.05583635 | 0 | 0 | 0 | 0 | 0 |
| ENSMUSG00 | Gm16512 | 283 | 0 | 0 | 0 | 0.12749839 | 0 | 0 |
| ENSMUSG00 | Gm5792 | 548 | 0 | 0 | 0 | 0 | 0 | 0 |
| ENSMUSG00 | Gm17449 | 170 | 0 | 0 | 0 | 0 | 0 | 0 |
| ENSMUSG00 | Gm6917 | 536 | 0 | 0 | 0 | 0 | 0 | 0 |
| ENSMUSG00 | Gm6901 | 548 | 0 | 0 | 0 | 0 | 0 | 0 |
| ENSMUSG00 | Gm10774 | 203 | 0 | 0 | 0 | 0 | 0 | 0 |
| ENSMUSG00 | Gm10775 | 1201 | 0 | 0.16048773 | 0 | 0 | 0 | 0 |
| ENSMUSG00 | 4932441B19 | 3386 | 0 | 0 | 0 | 0 | 0 | 0 |
| ENSMUSG00 | Hpse2 | 1767 | 0 | 0 | 0 | 0 | 0 | 0 |
| ENSMUSG00 | Gm8290 | 467 | 0 | 0 | 0 | 0 | 0 | 0 |
| ENSMUSG00 | Gm8158 | 467 | 0 | 0 | 0 | 0 | 0 | 0 |
| ENSMUSG00 | Gm8137 | 467 | 0 | 0 | 0 | 0 | 0 | 0 |
| ENSMUSG00 | Gm8264 | 467 | 0 | 0 | 0 | 0 | 0 | 0 |
| ENSMUSG00 | Gm8112 | 467 | 0 | 0 | 0 | 0 | 0 | 0 |
| ENSMUSG00 | Gm10778 | 1589 | 0.22517264 | 0.18195007 | 0.36162615 | 0.40873304 | 0.2247858 | 0.05149397 |
| ENSMUSG00 | 1810034E14 | 1753 | 2.17713983 | 2.72131077 | 5.14385395 | 3.7255277 | 1.54272557 | 3.57853225 |
| ENSMUSG00 | BC025920 | 2339 | 0 | 0.0824052 | 0.05669326 | 0.30852539 | 0.1308928 | 0.04664325 |
| ENSMUSG00 | Gm5665 | 3442 | 0 | 0.15399502 | 0.15410288 | 0 | 0.02964926 | 0.0475444 |
| ENSMUSG00 | Zfp934 | 1487 | 2.11209345 | 2.91646254 | 4.99388677 | 10.239827 | 3.22560816 | 14.2517786 |
| ENSMUSG00 | Zfp808 | 2567 | 5.7457278 | 4.11095864 | 5.51016352 | 17.4858058 | 4.85018921 | 4.09066463 |
| ENSMUSG00 | Gm6478 | 743 | 0.21402681 | 0.32426946 | 0.23796416 | 0.72843966 | 0 | 0 |
| ENSMUSG00 | Cts3 | 1372 | 0 | 0 | 0 | 0 | 0.03719124 | 0 |
| ENSMUSG00 | Ctsm | 1196 | 0 | 0 | 0 | 0 | 0 | 0 |
| ENSMUSG00 | Ctsr2 | 1026 | 0 | 0 | 0 | 0 | 0 | 0 |
| ENSMUSG00 | AI606181 | 2795 | 1.74952562 | 1.29301723 | 0.91724755 | 2.45280446 | 1.07712207 | 1.79553976 |
| ENSMUSG00 | Ctla2b | 1511 | 0 | 0.06378086 | 0 | 0.47759159 | 0.16884969 | 0 |
| ENSMUSG00 | Spata5l1 | 2658 | 1.01707023 | 0.92457056 | 0.83148689 | 1.94120857 | 0.57591846 | 0.66698704 |
| ENSMUSG00 | Gm10780 | 283 | 0 | 0 | 0 | 0 | 0 | 0 |
| ENSMUSG00 | Mageb3 | 1608 | 0 | 0 | 0.05497742 | 0 | 0 | 0.1696184 |
| ENSMUSG00 | Cyp2c68 | 1715 | 0 | 0 | 0 | 0 | 0.05950598 | 0.09542148 |
| ENSMUSG00 | Serf2 | 5521 | 9.06577612 | 7.82887857 | 10.5841037 | 8.8489564 | 8.74315369 | 11.7427672 |
| ENSMUSG00 | Gm10782 | 427 | 0 | 0 | 0 | 0 | 0 | 0 |
| ENSMUSG00 | Grk6 | 3224 | 7.09038502 | 8.89296932 | 6.30671457 | 8.66237669 | 8.56242845 | 7.32625235 |
| ENSMUSG00 | EU599041 | 807 | 0.0492633 | 0.05971059 | 0 | 0.04471133 | 0.50583768 | 0.03379757 |
| ENSMUSG00 | Lcmt2 | 3379 | 7.36517632 | 6.51707726 | 6.39677456 | 4.27132815 | 6.81056392 | 5.28703424 |
| ENSMUSG00 | B3galt5 | 6152 | 0.40711887 | 0.37596704 | 0 | 0.00586509 | 0.6137763 | 0.11970339 |
| ENSMUSG00 | Eif4e1b | 2188 | 0 | 0 | 0 | 0 | 0 | 0 |
| ENSMUSG00 | Ifit3 | 1743 | 12.3166721 | 6.52438345 | 4.74225166 | 3.89180974 | 8.48975846 | 7.16683005 |
| ENSMUSG00 | Spnb5 | 10233 | 0 | 0.03767141 | 0.03023677 | 0 | 0.15458005 | 0.01066144 |
| ENSMUSG00 | Gm8765 | 3148 | 0 | 0 | 0 | 0 | 0 | 0 |
| ENSMUSG00 | Gm904 | 645 | 0 | 0 | 0 | 0 | 0 | 0 |
| ENSMUSG00 | Gm2058 | 3268 | 0 | 0 | 0 | 0 | 0 | 0 |
| ENSMUSG00 | Gm906 | 3148 | 0 | 0 | 0 | 0 | 0 | 0 |
| ENSMUSG00 | Ranbp6 | 4866 | 23.3009927 | 16.5275734 | 24.3446238 | 27.8067544 | 26.5618187 | 27.6557892 |
| ENSMUSG00 | Tes3-ps | 3258 | 0.12202419 | 0.10353134 | 0.48841816 | 0.14397378 | 0.07830936 | 0.71158513 |
| ENSMUSG00 | Gm17362 | 625 | 2.09908937 | 0.61678646 | 1.83879667 | 4.44530789 | 2.20433944 | 3.49115382 |
| ENSMUSG00 | Gm14207 | 1372 | 5.36061508 | 4.24967894 | 10.6638557 | 7.81076328 | 3.31002001 | 12.7427433 |
| ENSMUSG00 | Gm815 | 606 | 0 | 0 | 0 | 0 | 0 | 0 |
| ENSMUSG00 | Chst14 | 4077 | 1.9892367 | 2.58838137 | 1.53952948 | 1.56647581 | 1.97747544 | 1.50522292 |
| ENSMUSG00 | 5430417L22 | 3052 | 26.9508811 | 23.4616819 | 29.7334154 | 25.1581228 | 28.1380703 | 17.4711402 |
| ENSMUSG00 | Gm10787 | 1545 | 0 | 0 | 0 | 0 | 0 | 0 |
| ENSMUSG00 | Fam122a | 854 | 25.2777821 | 25.8423776 | 32.2973654 | 25.8151396 | 23.3024433 | 33.9495802 |
| ENSMUSG00 | Pak6 | 5366 | 5.77884361 | 6.19616942 | 3.69035141 | 3.81934426 | 6.79907915 | 4.54407892 |
| ENSMUSG00 | Ptar1 | 2057 | 3.24692306 | 3.79494586 | 4.74895834 | 4.59576843 | 3.54728816 | 2.50603151 |
| ENSMUSG00 | Krtap14 | 199 | 0 | 0 | 0 | 0 | 0 | 0 |
| ENSMUSG00 | Gm13981 | 446 | 0 | 0 | 0 | 0 | 0 | 0 |
| ENSMUSG00 | Gm17719 | 3187 | 0 | 0 | 0 | 0 | 0 | 0 |
| ENSMUSG00 | Grem1 | 4146 | 0 | 0 | 0.15991984 | 0 | 0 | 0 |
| ENSMUSG00 | Chrm5 | 1745 | 0 | 0 | 0 | 0 | 0 | 0 |
| ENSMUSG00 | Olfr1316 | 944 | 0 | 0 | 0 | 0 | 0 | 0 |
| ENSMUSG00 | Olfr1314 | 938 | 0 | 0.10274295 | 0 | 0 | 0 | 0 |
| ENSMUSG00 | Olfr1313 | 937 | 0 | 0 | 0 | 0 | 0 | 0 |
| ENSMUSG00 | Olfr1312 | 953 | 0 | 0 | 0 | 0 | 0 | 0 |
| ENSMUSG00 | Olfr1311 | 942 | 0 | 0 | 0 | 0 | 0 | 0 |
| ENSMUSG00 | Olfr1310 | 1008 | 0 | 0 | 0 | 0 | 0 | 0 |
| ENSMUSG00 | Olfr1309 | 963 | 0 | 0 | 0 | 0 | 0 | 0 |
| ENSMUSG00 | Olfr1308 | 962 | 0 | 0 | 0 | 0 | 0 | 0 |
| ENSMUSG00 | Olfr1307 | 938 | 0 | 0 | 0 | 0 | 0 | 0 |
| ENSMUSG00 | Olfr1306 | 938 | 0 | 0 | 0 | 0 | 0 | 0 |
| ENSMUSG00 | Olfr1305 | 938 | 0 | 0 | 0 | 0 | 0 | 0 |
| ENSMUSG00 | Olfr1303 | 938 | 0 | 0 | 0 | 0 | 0 | 0 |
| ENSMUSG00 | Olfr1299 | 994 | 0 | 0 | 0 | 0 | 0 | 0 |
| ENSMUSG00 | Olfr1297 | 938 | 0 | 0 | 0 | 0 | 0 | 0 |
| ENSMUSG00 | Olfr1295 | 1042 | 0 | 0 | 0 | 0 | 0 | 0 |
| ENSMUSG00 | Olfr1294 | 938 | 0 | 0 | 0 | 0 | 0 | 0 |
| ENSMUSG00 | Olfr1290 | 945 | 0 | 0 | 0 | 0 | 0 | 0 |
| ENSMUSG00 | Olfr1286 | 917 | 0 | 0 | 0 | 0 | 0 | 0 |
| ENSMUSG00 | Olfr1282 | 917 | 0 | 0 | 0 | 0 | 0 | 0 |
| ENSMUSG00 | Olfr1277 | 961 | 0 | 0 | 0 | 0 | 0 | 0 |
| ENSMUSG00 | Olfr1275 | 1019 | 0 | 0 | 0 | 0 | 0 | 0 |
| ENSMUSG00 | Ano3 | 7416 | 0 | 0.11045975 | 0 | 0 | 0 | 0.01838905 |
| ENSMUSG00 | Gm10794 | 205 | 0 | 0 | 0 | 0 | 0 | 0 |
| ENSMUSG00 | Fibin | 2068 | 0 | 1.88737999 | 0.12824519 | 1.46561496 | 0.56750805 | 0.75176714 |
| ENSMUSG00 | Gm11382 | 167 | 4.99919216 | 0.57708314 | 1.32340847 | 2.16060147 | 4.88875457 | 4.08302983 |
| ENSMUSG00 | Actg-ps1 | 407 | 0 | 0.11839421 | 0 | 0 | 0 | 0 |
| ENSMUSG00 | Dcdc5 | 5648 | 0 | 0 | 0 | 0 | 0.04517208 | 0 |
| ENSMUSG00 | Olfr1452-ps1 | 919 | 0 | 0 | 0 | 0 | 0 | 0 |
| ENSMUSG00 | AI314831 | 5437 | 0 | 0 | 0 | 0 | 0 | 0 |
| ENSMUSG00 | Gabrr3 | 1674 | 0 | 0 | 0 | 0 | 0 | 0 |
| ENSMUSG00 | Qser1 | 8961 | 3.11885981 | 3.07584476 | 2.64392231 | 2.26292925 | 3.03504724 | 3.17762787 |
| ENSMUSG00 | Olfr201 | 926 | 0 | 0 | 0 | 0 | 0 | 0 |
| ENSMUSG00 | Olfr199 | 930 | 0 | 0 | 0 | 0 | 0 | 0 |
| ENSMUSG00 | Pin1-ps1 | 1560 | 0 | 0.06177749 | 0 | 0.11564758 | 0 | 0.57696352 |
| ENSMUSG00 | Gm10797 | 2221 | 0 | 0 | 0 | 0 | 0 | 0 |
| ENSMUSG00 | Nrbf2 | 3323 | 3.17038891 | 3.71222666 | 3.76440613 | 5.55943629 | 4.36096622 | 3.65248705 |
| ENSMUSG00 | Gm813 | 698 | 0 | 0 | 0 | 0 | 0 | 0 |
| ENSMUSG00 | Jmjd1c | 593 | 0 | 0 | 0 | 0 | 0 | 0 |
| ENSMUSG00 | Gm10799 | 310 | 0 | 0 | 0 | 0 | 0 | 0 |
| ENSMUSG00 | AW112010 | 771 | 3.71257406 | 2.74993963 | 0 | 2.76114219 | 2.1840083 | 2.54704802 |
| ENSMUSG00 | Fjx1 | 2424 | 1.36126439 | 2.66377197 | 0.47411218 | 1.56295985 | 0.44206019 | 0.61885526 |
| ENSMUSG00 | Gm10800 | 655 | 0 | 1.02993922 | 0 | 0 | 0.31161146 | 0 |
| ENSMUSG00 | Gm10801 | 507 | 2.74446118 | 3.32648023 | 0 | 0.28467096 | 0.70450618 | 0 |
| ENSMUSG00 | E530001K10 | 1227 | 0.12960222 | 0.47126105 | 0 | 0.02940672 | 0 | 0.06668616 |
| ENSMUSG00 | Gm13889 | 745 | 0 | 0 | 0 | 0 | 0 | 0 |
| ENSMUSG00 | Accsl | 3201 | 0.7700218 | 0 | 0.57996795 | 0 | 0 | 0.81798356 |
| ENSMUSG00 | Gm10803 | 2481 | 0 | 0 | 0 | 0 | 0 | 0 |
| ENSMUSG00 | Gm10804 | 2202 | 0.52357354 | 1.33486511 | 0 | 0.36049273 | 0.27807289 | 0.03715891 |
| ENSMUSG00 | 4631405J19 | 4921 | 0.01615748 | 0.03916801 | 0.02694687 | 0.05865807 | 0.15553661 | 0.12193499 |
| ENSMUSG00 | Prdm11 | 17034 | 0.38975961 | 0.46958726 | 0.35290893 | 0.43423853 | 0.47329854 | 1.12243291 |
| ENSMUSG00 | 4930558J22 | 1097 | 0.03624018 | 0 | 0 | 0 | 0 | 0 |
| ENSMUSG00 | Hist1h2bb | 468 | 0 | 0.10296248 | 0.18889677 | 0.30839354 | 0.10903072 | 0 |
| ENSMUSG00 | Hist1h3i | 482 | 0 | 0 | 0 | 0 | 0 | 0 |
| ENSMUSG00 | Fam55c | 6424 | 0.02475435 | 0.16502206 | 0 | 0.22467027 | 0.15886169 | 0.18256686 |
| ENSMUSG00 | Gm10805 | 397 | 0 | 0 | 0 | 0 | 0 | 0 |
| ENSMUSG00 | Zfp408 | 6776 | 4.54113665 | 2.22584953 | 2.97462226 | 3.55708467 | 3.02724367 | 3.09134046 |
| ENSMUSG00 | 4930431P03 | 2980 | 0.50694908 | 0 | 0 | 0 | 0 | 0 |
| ENSMUSG00 | Tmem223 | 746 | 24.5141033 | 30.1003781 | 37.2101306 | 26.0700027 | 16.4843922 | 31.5522972 |
| ENSMUSG00 | Slc22a29 | 2712 | 0 | 0.26651793 | 0 | 0 | 0 | 0 |
| ENSMUSG00 | Gm4981 | 1812 | 0 | 0 | 0 | 0 | 0 | 0 |
| ENSMUSG00 | Gm10807 | 2443 | 0 | 0 | 0 | 0 | 0 | 0 |
| ENSMUSG00 | Vdac3-ps1 | 851 | 0.98104006 | 2.03843939 | 0.6752338 | 0.50879499 | 0.41972342 | 1.08970357 |
| ENSMUSG00 | 1600012F09 | 3433 | 7.4230303 | 5.38991954 | 7.00431186 | 11.1619957 | 6.56966447 | 9.55764376 |
| ENSMUSG00 | Olfr1272 | 926 | 0 | 0 | 0 | 0 | 0 | 0 |
| ENSMUSG00 | Olfr1271 | 925 | 0 | 0 | 0 | 0 | 0 | 0 |
| ENSMUSG00 | Olfr142 | 917 | 0 | 0 | 0 | 0 | 0 | 0 |
| ENSMUSG00 | Olfr1506 | 1082 | 0 | 0 | 0 | 0 | 0 | 0 |
| ENSMUSG00 | Olfr1270 | 1017 | 0 | 0 | 0 | 0 | 0 | 0 |
| ENSMUSG00 | Olfr32 | 1007 | 0 | 0 | 0 | 0 | 0 | 0 |
| ENSMUSG00 | Olfr1269 | 1017 | 0 | 0 | 0 | 0 | 0 | 0 |
| ENSMUSG00 | Olfr140 | 2321 | 0 | 0 | 0 | 0 | 0 | 0 |
| ENSMUSG00 | Olfr1264 | 926 | 0 | 0 | 0 | 0 | 0 | 0 |
| ENSMUSG00 | 4932412D23 | 5084 | 0 | 0 | 0 | 0 | 0 | 0 |
| ENSMUSG00 | Olfr48 | 905 | 0 | 0 | 0 | 0 | 0 | 0 |
| ENSMUSG00 | Olfr1256 | 2063 | 0 | 0 | 0 | 0 | 0 | 0 |
| ENSMUSG00 | Olfr1254 | 2062 | 0 | 0 | 0 | 0 | 0 | 0 |
| ENSMUSG00 | Olfr1253 | 956 | 0 | 0 | 0 | 0 | 0 | 0 |
| ENSMUSG00 | Olfr1252 | 1098 | 0 | 0 | 0 | 0 | 0 | 0 |
| ENSMUSG00 | Olfr1251 | 1039 | 0 | 0 | 0 | 0 | 0 | 0 |
| ENSMUSG00 | Olfr1250 | 2799 | 0 | 0 | 0 | 0 | 0 | 0 |
| ENSMUSG00 | Olfr1249 | 956 | 0 | 0 | 0 | 0 | 0 | 0 |
| ENSMUSG00 | Olfr1248 | 1077 | 0 | 0 | 0 | 0 | 0 | 0 |
| ENSMUSG00 | Olfr1247 | 2717 | 0 | 0 | 0 | 0 | 0 | 0 |
| ENSMUSG00 | Olfr1246 | 1077 | 0 | 0 | 0 | 0 | 0 | 0 |
| ENSMUSG00 | Olfr1245 | 1061 | 0 | 0 | 0 | 0 | 0 | 0 |
| ENSMUSG00 | Olfr1243 | 1352 | 0 | 0 | 0 | 0 | 0 | 0 |
| ENSMUSG00 | Olfr1242 | 2257 | 0 | 0 | 0 | 0 | 0 | 0 |
| ENSMUSG00 | Olfr1241 | 944 | 0 | 0 | 0 | 0 | 0 | 0 |
| ENSMUSG00 | Olfr1240 | 1073 | 0 | 0 | 0 | 0 | 0 | 0 |
| ENSMUSG00 | Olfr1239 | 917 | 0 | 0 | 0 | 0 | 0 | 0 |
| ENSMUSG00 | Olfr1238 | 948 | 0 | 0 | 0 | 0 | 0 | 0 |
| ENSMUSG00 | Olfr1234 | 2332 | 0 | 0 | 0 | 0 | 0 | 0 |
| ENSMUSG00 | Olfr1233 | 1041 | 0 | 0 | 0 | 0 | 0 | 0 |
| ENSMUSG00 | Olfr1232 | 935 | 0 | 0 | 0 | 0 | 0 | 0 |
| ENSMUSG00 | Olfr1231 | 941 | 0 | 0 | 0 | 0 | 0 | 0 |
| ENSMUSG00 | Olfr1230 | 917 | 0 | 0 | 0 | 0 | 0 | 0 |
| ENSMUSG00 | Olfr1229 | 2841 | 0 | 0 | 0 | 0 | 0 | 0 |
| ENSMUSG00 | Olfr1228 | 1093 | 0 | 0 | 0 | 0 | 0 | 0 |
| ENSMUSG00 | Olfr1226 | 932 | 0 | 0 | 0 | 0 | 0 | 0 |
| ENSMUSG00 | Olfr1225 | 1026 | 0 | 0 | 0 | 0 | 0 | 0 |
| ENSMUSG00 | Olfr1224-ps1 | 935 | 0 | 0 | 0 | 0 | 0 | 0 |
| ENSMUSG00 | Olfr1223 | 2464 | 0 | 0 | 0 | 0 | 0 | 0 |
| ENSMUSG00 | Olfr1222 | 1587 | 0 | 0 | 0 | 0 | 0 | 0 |
| ENSMUSG00 | Olfr1221 | 935 | 0 | 0 | 0 | 0 | 0 | 0 |
| ENSMUSG00 | Olfr1220 | 1025 | 0 | 0 | 0 | 0 | 0 | 0 |
| ENSMUSG00 | Olfr1219 | 935 | 0 | 0 | 0 | 0 | 0 | 0 |
| ENSMUSG00 | Olfr1218 | 935 | 0 | 0 | 0 | 0 | 0 | 0 |
| ENSMUSG00 | Olfr1217 | 1104 | 0 | 0 | 0 | 0 | 0 | 0 |
| ENSMUSG00 | Olfr1216 | 935 | 0 | 0 | 0 | 0 | 0 | 0 |
| ENSMUSG00 | Olfr1215 | 942 | 0 | 0 | 0 | 0 | 0 | 0 |
| ENSMUSG00 | Olfr1214 | 935 | 0 | 0 | 0 | 0 | 0 | 0 |
| ENSMUSG00 | Gm13762 | 1443 | 0 | 0 | 0 | 0 | 0 | 0 |
| ENSMUSG00 | Olfr1211 | 935 | 0 | 0 | 0 | 0 | 0 | 0 |
| ENSMUSG00 | Olfr1209 | 932 | 0 | 0 | 0 | 0 | 0 | 0 |
| ENSMUSG00 | Olfr1208 | 1074 | 0 | 0 | 0 | 0 | 0 | 0 |
| ENSMUSG00 | Olfr1200 | 1010 | 0 | 0 | 0 | 0 | 0 | 0 |
| ENSMUSG00 | Gm13760 | 895 | 0 | 0 | 0 | 0 | 0 | 0 |
| ENSMUSG00 | Olfr1198 | 926 | 0 | 0 | 0 | 0 | 0 | 0 |
| ENSMUSG00 | Gpr137b-ps | 2992 | 0.41190505 | 0.37041717 | 0.08864006 | 0.02411901 | 0.61395372 | 0.54695132 |
| ENSMUSG00 | Olfr1197 | 944 | 0 | 0 | 0 | 0 | 0 | 0 |
| ENSMUSG00 | Olfr1196 | 944 | 0 | 0 | 0 | 0 | 0 | 0 |
| ENSMUSG00 | Olfr1195 | 926 | 0 | 0 | 0 | 0 | 0 | 0 |
| ENSMUSG00 | Cd80 | 3090 | 0.56609746 | 0.51461249 | 0.14304804 | 0.60720593 | 0.47888832 | 0.2206686 |
| ENSMUSG00 | Olfr1182 | 917 | 0 | 0 | 0 | 0 | 0 | 0 |
| ENSMUSG00 | Olfr1181 | 935 | 0 | 0 | 0 | 0 | 0 | 0 |
| ENSMUSG00 | Olfr1180 | 1062 | 0 | 0 | 0 | 0 | 0 | 0 |
| ENSMUSG00 | Olfr1179 | 923 | 0 | 0 | 0 | 0 | 0 | 0 |
| ENSMUSG00 | Olfr1177-ps | 957 | 0 | 0 | 0 | 0 | 0 | 0 |
| ENSMUSG00 | Olfr1173 | 938 | 0 | 0 | 0 | 0 | 0 | 0 |
| ENSMUSG00 | Olfr1170 | 950 | 0 | 0 | 0 | 0 | 0 | 0 |
| ENSMUSG00 | Olfr1167 | 1099 | 0 | 0 | 0 | 0 | 0 | 0 |
| ENSMUSG00 | Olfr1166 | 1627 | 0 | 0 | 0 | 0 | 0 | 0 |
| ENSMUSG00 | Olfr1164 | 1111 | 0 | 0 | 0 | 0 | 0 | 0 |
| ENSMUSG00 | Olfr1163 | 1042 | 0 | 0 | 0 | 0 | 0 | 0 |
| ENSMUSG00 | Olfr1162 | 944 | 0 | 0 | 0 | 0 | 0 | 0 |
| ENSMUSG00 | Olfr73 | 941 | 0 | 0 | 0 | 0 | 0 | 0.02898474 |
| ENSMUSG00 | Olfr1160 | 959 | 0 | 0 | 0 | 0 | 0 | 0 |
| ENSMUSG00 | Olfr74 | 956 | 0 | 0 | 0 | 0 | 0 | 0 |
| ENSMUSG00 | Olfr1157 | 2048 | 0 | 0 | 0 | 0 | 0 | 0 |
| ENSMUSG00 | Olfr1156 | 1075 | 0 | 0 | 0 | 0 | 0 | 0 |
| ENSMUSG00 | Olfr1155 | 944 | 0 | 0 | 0 | 0 | 0 | 0 |
| ENSMUSG00 | Olfr1154 | 932 | 0 | 0 | 0 | 0 | 0 | 0 |
| ENSMUSG00 | Olfr1141 | 935 | 0 | 0 | 0 | 0 | 0 | 0 |
| ENSMUSG00 | Olfr1138 | 935 | 0 | 0 | 0 | 0 | 0 | 0 |
| ENSMUSG00 | Olfr1137 | 932 | 0 | 0 | 0 | 0 | 0 | 0 |
| ENSMUSG00 | Olfr1136 | 1056 | 0 | 0 | 0 | 0 | 0 | 0 |
| ENSMUSG00 | Olfr1135 | 932 | 0 | 0 | 0 | 0 | 0 | 0 |
| ENSMUSG00 | Olfr1134 | 2061 | 0 | 0 | 0 | 0 | 0 | 0 |
| ENSMUSG00 | Olfr1133 | 941 | 0 | 0 | 0 | 0 | 0 | 0 |
| ENSMUSG00 | Olfr1128 | 935 | 0 | 0 | 0 | 0 | 0 | 0 |
| ENSMUSG00 | Olfr1111 | 939 | 0 | 0 | 0 | 0 | 0 | 0 |
| ENSMUSG00 | Olfr1110 | 938 | 0 | 0 | 0 | 0 | 0 | 0 |
| ENSMUSG00 | Olfr259 | 938 | 0 | 0 | 0 | 0 | 0 | 0 |
| ENSMUSG00 | Olfr1109 | 938 | 0 | 0 | 0 | 0 | 0 | 0 |
| ENSMUSG00 | Gm10812 | 544 | 0 | 0 | 0 | 0 | 0 | 0 |
| ENSMUSG00 | Olfr1107 | 1074 | 0 | 0 | 0 | 0 | 0 | 0 |
| ENSMUSG00 | Olfr1106 | 938 | 0 | 0 | 0 | 0 | 0 | 0 |
| ENSMUSG00 | Olfr1105 | 938 | 0 | 0 | 0 | 0 | 0 | 0 |
| ENSMUSG00 | Olfr1104 | 932 | 0 | 0 | 0 | 0 | 0 | 0 |
| ENSMUSG00 | Olfr1101 | 932 | 0 | 0 | 0 | 0 | 0 | 0 |
| ENSMUSG00 | Olfr1099 | 1030 | 0 | 0 | 0 | 0 | 0 | 0 |
| ENSMUSG00 | Olfr1098 | 2155 | 0 | 0 | 0 | 0 | 0 | 0 |
| ENSMUSG00 | Olfr1097 | 1221 | 0 | 0 | 0 | 0 | 0 | 0 |
| ENSMUSG00 | Olfr1095 | 926 | 0 | 0 | 0 | 0 | 0 | 0 |
| ENSMUSG00 | Olfr1090 | 941 | 0 | 0 | 0 | 0 | 0 | 0 |
| ENSMUSG00 | Olfr1089 | 935 | 0 | 0 | 0 | 0 | 0 | 0 |
| ENSMUSG00 | Olfr1087 | 941 | 0 | 0 | 0 | 0 | 0 | 0 |
| ENSMUSG00 | Olfr1086 | 932 | 0 | 0 | 0 | 0 | 0 | 0 |
| ENSMUSG00 | Olfr1085 | 941 | 0 | 0 | 0 | 0 | 0 | 0 |
| ENSMUSG00 | Olfr1084 | 1929 | 0 | 0 | 0 | 0 | 0 | 0 |
| ENSMUSG00 | Olfr1080 | 1918 | 0 | 0 | 0 | 0 | 0 | 0 |
| ENSMUSG00 | Olfr1079 | 2080 | 0 | 0 | 0 | 0 | 0 | 0 |
| ENSMUSG00 | Olfr228 | 941 | 0 | 0 | 0 | 0 | 0 | 0 |
| ENSMUSG00 | Olfr1066 | 941 | 0 | 0 | 0 | 0 | 0 | 0 |
| ENSMUSG00 | Olfr1065 | 991 | 0 | 0 | 0 | 0 | 0 | 0 |
| ENSMUSG00 | F930017D23 | 1832 | 0 | 0 | 0 | 0 | 0 | 0 |
| ENSMUSG00 | Olfr1061 | 941 | 0 | 0 | 0 | 0 | 0 | 0 |
| ENSMUSG00 | Olfr1058 | 950 | 0 | 0 | 0 | 0 | 0 | 0 |
| ENSMUSG00 | Olfr1057 | 947 | 0 | 0 | 0 | 0 | 0 | 0 |
| ENSMUSG00 | Olfr1056 | 1078 | 0 | 0 | 0 | 0 | 0 | 0 |
| ENSMUSG00 | Olfr1055 | 1040 | 0 | 0 | 0 | 0 | 0 | 0 |
| ENSMUSG00 | Olfr1054 | 938 | 0 | 0 | 0 | 0 | 0 | 0 |
| ENSMUSG00 | Olfr1053 | 941 | 0 | 0 | 0 | 0 | 0 | 0 |
| ENSMUSG00 | Olfr1051 | 926 | 0 | 0 | 0 | 0 | 0 | 0 |
| ENSMUSG00 | Olfr1049 | 926 | 0 | 0 | 0 | 0 | 0 | 0 |
| ENSMUSG00 | Olfr1048 | 962 | 0 | 0 | 0 | 0 | 0 | 0 |
| ENSMUSG00 | Olfr1047 | 959 | 0 | 0 | 0 | 0 | 0 | 0 |
| ENSMUSG00 | Olfr1046 | 950 | 0 | 0 | 0 | 0 | 0 | 0 |
| ENSMUSG00 | Olfr1045 | 947 | 0 | 0 | 0 | 0 | 0 | 0 |
| ENSMUSG00 | Olfr52 | 959 | 0 | 0 | 0 | 0 | 0 | 0 |
| ENSMUSG00 | Olfr1044 | 944 | 0 | 0 | 0 | 0 | 0 | 0 |
| ENSMUSG00 | Olfr1043 | 944 | 0 | 0 | 0 | 0 | 0 | 0 |
| ENSMUSG00 | Olfr1042 | 1096 | 0 | 0 | 0 | 0 | 0 | 0 |
| ENSMUSG00 | Olfr1040 | 941 | 0 | 0 | 0 | 0 | 0 | 0 |
| ENSMUSG00 | Olfr1039 | 959 | 0 | 0 | 0 | 0 | 0 | 0 |
| ENSMUSG00 | Olfr1037 | 1124 | 0 | 0 | 0 | 0 | 0 | 0 |
| ENSMUSG00 | Olfr1024 | 983 | 0 | 0 | 0 | 0 | 0 | 0 |
| ENSMUSG00 | Olfr1019 | 1863 | 0 | 0 | 0 | 0 | 0 | 0 |
| ENSMUSG00 | Olfr1016 | 929 | 0 | 0 | 0 | 0 | 0 | 0 |
| ENSMUSG00 | Olfr1012 | 935 | 0 | 0 | 0 | 0 | 0 | 0 |
| ENSMUSG00 | Olfr1006 | 1947 | 0 | 0 | 0 | 0 | 0 | 0 |
| ENSMUSG00 | Olfr154 | 1078 | 0 | 0 | 0 | 0 | 0 | 0 |
| ENSMUSG00 | Olfr1002 | 956 | 0 | 0 | 0 | 0 | 0 | 0 |
| ENSMUSG00 | Olfr1000 | 944 | 0 | 0 | 0 | 0 | 0 | 0 |
| ENSMUSG00 | 4833423E24 | 2260 | 0 | 0 | 0 | 0 | 0 | 0 |
| ENSMUSG00 | Olfr995 | 1112 | 0 | 0 | 0 | 0 | 0 | 0 |
| ENSMUSG00 | Olfr994 | 944 | 0 | 0 | 0 | 0 | 0 | 0 |
| ENSMUSG00 | Olfr993 | 944 | 0 | 0 | 0 | 0 | 0 | 0 |
| ENSMUSG00 | Olfr992 | 929 | 0 | 0 | 0 | 0 | 0 | 0 |
| ENSMUSG00 | Olfr988 | 2208 | 0 | 0 | 0 | 0 | 0 | 0 |
| ENSMUSG00 | Olfr987 | 2488 | 0 | 0 | 0 | 0 | 0 | 0 |
| ENSMUSG00 | Lrrc55 | 4076 | 0 | 0 | 0 | 0 | 0 | 0 |
| ENSMUSG00 | Ccdc162 | 5527 | 0 | 0 | 0.27991035 | 0 | 0.0276966 | 0.0542828 |
| ENSMUSG00 | Znhit2-ps | 1280 | 6.2428528 | 9.26083185 | 5.90508997 | 6.79357245 | 5.97965342 | 5.26315304 |
| ENSMUSG00 | Gm10813 | 88 | 0 | 0 | 0 | 0 | 0 | 0 |
| ENSMUSG00 | Ccdc58 | 1219 | 15.5238792 | 17.9859157 | 18.3479348 | 17.4046285 | 12.4321851 | 37.0523401 |

| | | | | | | | |
|---|---|---|---|---|---|---|---|
| ENSMUSG00 Amd1 | 3195 | 9.13318394 | 10.4969527 | 7.63675034 | 10.8754331 | 8.06520181 | 8.17812344 |
| ENSMUSG00 Gm13695 | 2492 | 0 | 0 | 0 | 0 | 0 | 0 |
| ENSMUSG00 Gm13693 | 2492 | 0 | 0 | 0 | 0 | 0 | 0 |
| ENSMUSG00 Gm13698 | 2492 | 0 | 0 | 0 | 0 | 0 | 0 |
| ENSMUSG00 Gm8055 | 1007 | 0.03947913 | 0.09570296 | 0.04389458 | 0.21498736 | 0.15201502 | 0.02708504 |
| ENSMUSG00 4930547M16 | 2398 | 0.0165786 | 0.92434376 | 0.07373118 | 0 | 0.06383617 | 0.01137391 |
| ENSMUSG00 Gm6043 | 494 | 0 | 0 | 0.08947742 | 0 | 0 | 0.16563546 |
| ENSMUSG00 Fsip2 | 19266 | 0 | 0 | 0 | 0 | 0 | 0 |
| ENSMUSG00 Heg1 | 12921 | 0.61228548 | 0.96962116 | 0.45156283 | 0.84333855 | 1.29530619 | 0.66703707 |
| ENSMUSG00 Cerkl | 5266 | 0.4907152 | 0.3202669 | 0.29378361 | 0.02055567 | 0.63952541 | 0.25896923 |
| ENSMUSG00 Nkain2 | 322 | 0 | 0 | 0 | 0 | 0 | 0 |
| ENSMUSG00 Gm10818 | 1389 | 0 | 0 | 0 | 0 | 0 | 0 |
| ENSMUSG00 Cenpw | 1507 | 1.31902722 | 1.63072896 | 0.14665509 | 0.16760074 | 0 | 0.7239453 |
| ENSMUSG00 Dfnb59 | 1478 | 0 | 0 | 0 | 0 | 0 | 0 |
| ENSMUSG00 Gm10819 | 560 | 0.21297579 | 0.17209444 | 0.23679559 | 1.22421223 | 0 | 0.14611414 |
| ENSMUSG00 Bex6 | 698 | 0 | 0 | 0 | 0 | 0 | 0 |
| ENSMUSG00 Pde11a | 6561 | 0 | 0 | 0.00673706 | 0 | 0 | 0 |
| ENSMUSG00 Ttc30a1 | 2861 | 1.93149661 | 3.52008613 | 2.44106648 | 0.52969097 | 1.46248263 | 3.98490011 |
| ENSMUSG00 Ttc30a2 | 2128 | 0 | 0 | 0 | 0 | 0 | 0.05126812 |
| ENSMUSG00 Ttc30b | 2599 | 16.015386 | 15.2958118 | 15.0514163 | 12.4808611 | 13.6646239 | 22.3633152 |
| ENSMUSG00 Gm13653 | 997 | 0.03987511 | 0 | 0 | 0.03619062 | 0 | 0 |
| ENSMUSG00 6720416L17F | 2524 | 4.23701437 | 4.77282509 | 6.05936517 | 5.68964094 | 4.06351091 | 8.00733268 |
| ENSMUSG00 Mrpl23-ps1 | 1122 | 0.17716346 | 0.21473459 | 0.27576907 | 0.12863474 | 0.22739027 | 0.48617895 |
| ENSMUSG00 Gm10822 | 2318 | 0 | 0 | 0 | 0 | 0.08805242 | 0 |
| ENSMUSG00 Wipf1 | 8308 | 0.11484491 | 0.12760011 | 0.17557304 | 0.08686096 | 0.23338978 | 0.72881198 |
| ENSMUSG00 Gm1968 | 345 | 0 | 0 | 0 | 0 | 0 | 0 |
| ENSMUSG00 Gm6913 | 347 | 0.11456911 | 0.13886583 | 0.25476567 | 0.10398284 | 0.14705007 | 0.07860127 |
| ENSMUSG00 Carns1 | 3958 | 0.72319217 | 0.41393103 | 0.75940509 | 0 | 1.00557284 | 0.25496757 |
| ENSMUSG00 Aldh3b2 | 2131 | 0 | 0 | 0 | 0 | 0 | 0 |
| ENSMUSG00 H60b | 1206 | 0.26371795 | 0.59933386 | 0 | 0 | 0.04231043 | 0 |
| ENSMUSG00 4933404M02 | 3669 | 0.3792428 | 0.28893479 | 0.39756359 | 0.0196686 | 0.29205612 | 0.2973523 |
| ENSMUSG00 Sp5 | 2098 | 0 | 0 | 0 | 0 | 0 | 0 |
| ENSMUSG00 Kbtbd10 | 2523 | 0 | 0.03819774 | 0 | 0.62925484 | 0.06067346 | 0.0756728 |
| ENSMUSG00 Gm10827 | 2584 | 0 | 0 | 0 | 0 | 0.0789882 | 0 |
| ENSMUSG00 Gm13597 | 1623 | 1.39621835 | 1.09852025 | 1.25279407 | 1.26720674 | 0.94318624 | 0.87386398 |
| ENSMUSG00 Gm1322 | 702 | 0 | 0 | 0 | 0 | 0 | 0 |
| ENSMUSG00 Scn9a | 13206 | 0 | 0 | 0 | 0.00273225 | 0 | 0 |
| ENSMUSG00 Scn2a1 | 9136 | 0 | 0 | 0.26126308 | 0 | 0 | 0.0029854 |
| ENSMUSG00 Gm10828 | 1571 | 0 | 0 | 0 | 0 | 0 | 0 |
| ENSMUSG00 Fign | 13526 | 0.34976358 | 0.23156282 | 0.30064835 | 0.26942825 | 0.25652769 | 0.3690122 |
| ENSMUSG00 Gm13582 | 2651 | 0.68983482 | 0.19994374 | 0.35014663 | 0 | 0.09623986 | 0.3909605 |
| ENSMUSG00 Zbtb2 | 3334 | 8.89549744 | 7.89136095 | 8.9888571 | 6.45017953 | 8.96864314 | 7.84534464 |
| ENSMUSG00 A930003A15F | 744 | 0 | 0 | 0 | 0 | 0 | 0 |
| ENSMUSG00 Rprm | 1459 | 1.33517378 | 1.05686508 | 1.75716717 | 2.0031841 | 1.53883519 | 0.878621 |
| ENSMUSG00 4930588K23F | 1961 | 0 | 0 | 0 | 0 | 0 | 0 |
| ENSMUSG00 Igll1 | 1327 | 0 | 0 | 0 | 1.11481826 | 0.30761945 | 0 |
| ENSMUSG00 Rc3h2 | 14765 | 4.14383234 | 4.81374888 | 4.93060856 | 5.37381754 | 5.75407489 | 8.58416855 |
| ENSMUSG00 Olfr362 | 953 | 0 | 0 | 0 | 0 | 0 | 0 |
| ENSMUSG00 Olfr361 | 968 | 0 | 0 | 0 | 0 | 0 | 0 |
| ENSMUSG00 Olfr358 | 1058 | 0 | 0 | 0 | 0 | 0 | 0 |
| ENSMUSG00 Olfr355 | 932 | 0 | 0 | 0 | 0 | 0 | 0 |
| ENSMUSG00 Olfr353 | 935 | 0 | 0 | 0 | 0 | 0 | 0 |
| ENSMUSG00 Olfr351 | 932 | 0 | 0 | 0 | 0 | 0 | 0 |
| ENSMUSG00 Olfr3 | 941 | 0 | 0 | 0 | 0 | 0 | 0 |
| ENSMUSG00 Olfr341 | 941 | 0 | 0 | 0 | 0 | 0 | 0 |
| ENSMUSG00 2810410L24F | 3069 | 0.81609491 | 0.54953583 | 0.70573161 | 0.63487468 | 0.89782479 | 1.17310276 |
| ENSMUSG00 Rpl36-ps8 | 314 | 0.37982943 | 0 | 0 | 0 | 0 | 0 |
| ENSMUSG00 Gm13443 | 554 | 0.78936875 | 1.30468706 | 0.3989336 | 0.19539013 | 0.27631612 | 8.56640294 |
| ENSMUSG00 Gm5561 | 651 | 0 | 0 | 0 | 0 | 0 | 0 |
| ENSMUSG00 Hoxc4 | 2553 | 0.17129271 | 0.05662332 | 0.08656844 | 0.26853069 | 0.13990781 | 0.405968 |
| ENSMUSG00 A630010A05F | 1953 | 0 | 0 | 0 | 0 | 0 | 0 |
| ENSMUSG00 Krt76 | 2303 | 0 | 0 | 0 | 0 | 0 | 0 |
| ENSMUSG00 Ciita | 1618 | 0 | 0 | 0 | 0.0223004 | 0.15768349 | 0 |
| ENSMUSG00 9430097D07 | 1500 | 0 | 0 | 0 | 0 | 0 | 0 |
| ENSMUSG00 6030408B16F | 1152 | 0 | 0.08365702 | 0 | 0 | 0 | 0 |
| ENSMUSG00 Gm11744 | 1344 | 0 | 0.35853008 | 0.09886483 | 0 | 0 | 0.04058726 |
| ENSMUSG00 Bin2 | 1465 | 0 | 0.36180946 | 0 | 0 | 0 | 0 |
| ENSMUSG00 Fnbp1 | 8586 | 8.31133095 | 7.05454901 | 7.31034441 | 9.3798187 | 9.81185028 | 6.29610238 |
| ENSMUSG00 Gm14488 | 1389 | 0.08586497 | 0.55506341 | 0.38187337 | 0.23379295 | 0.36736052 | 0.35345105 |
| ENSMUSG00 Dolk | 2125 | 24.6390437 | 22.9254085 | 29.8492446 | 22.2095596 | 24.1324742 | 27.0051016 |
| ENSMUSG00 1110017F19F | 976 | 29.7351442 | 22.3158523 | 19.20244 | 23.2167254 | 26.9770594 | 16.9069229 |
| ENSMUSG00 Gm13547 | 3521 | 0 | 0 | 0.11298398 | 0.21520106 | 0 | 0 |
| ENSMUSG00 Olfr288 | 4628 | 0 | 0 | 0 | 0 | 0 | 0 |
| ENSMUSG00 Gm11691 | 656 | 0 | 0 | 0 | 0 | 0 | 0 |
| ENSMUSG00 Gm11681 | 867 | 0 | 0 | 0 | 0 | 0 | 0 |
| ENSMUSG00 4930594M22 | 4562 | 0.32243595 | 0.09506313 | 0.20347188 | 0.16609446 | 0.26844214 | 0.57992986 |
| ENSMUSG00 Muc19 | 308 | 0 | 0 | 0 | 0 | 0 | 0 |
| ENSMUSG00 Gm10837 | 1535 | 0 | 0 | 0 | 0 | 0 | 0 |
| ENSMUSG00 Dnlz | 2913 | 7.23323195 | 5.78965148 | 7.70838869 | 7.93978391 | 4.60691275 | 9.2507187 |
| ENSMUSG00 Gm17205 | 1206 | 0 | 0.03995559 | 0 | 0 | 0 | 0 |
| ENSMUSG00 Alg10b | 3724 | 3.88587404 | 3.71361678 | 4.5460005 | 5.94908038 | 3.45291265 | 5.22934813 |
| ENSMUSG00 Slitrk1 | 4248 | 0 | 0.02268665 | 0 | 0 | 0 | 0 |
| ENSMUSG00 Gm10840 | 308 | 0 | 0 | 0 | 0 | 0 | 0 |
| ENSMUSG00 Gm10842 | 368 | 0 | 0 | 0 | 1.17658841 | 0 | 0 |
| ENSMUSG00 Commd6 | 1943 | 32.6760177 | 33.0584289 | 45.5895488 | 29.2481833 | 25.2374406 | 44.6809968 |
| ENSMUSG00 Gm6999 | 2653 | 0 | 0 | 0 | 0 | 0 | 0 |
| ENSMUSG00 Gm10845 | 3386 | 0.01174113 | 0 | 0 | 0.02131249 | 0 | 0 |
| ENSMUSG00 Gm5465 | 2981 | 0.06668145 | 0 | 0 | 0 | 0.085586 | 0.01829899 |
| ENSMUSG00 Fam187a | 1562 | 0 | 0 | 0 | 0 | 0 | 0 |
| ENSMUSG00 1700001L05F | 2266 | 0.87721713 | 0.06379494 | 0.0390131 | 0.47769697 | 0.45036519 | 0.06018235 |
| ENSMUSG00 Gm1587 | 369 | 0 | 0 | 0 | 0 | 0 | 0 |
| ENSMUSG00 Gm13375 | 852 | 0.09332272 | 0.16967057 | 0.20752039 | 0.21174909 | 0.29945056 | 0.28811239 |
| ENSMUSG00 Rps2-ps6 | 665 | 0 | 0 | 0 | 0 | 0 | 0 |
| ENSMUSG00 Cyp2d37-ps | 586 | 0 | 0 | 0 | 0 | 0 | 0 |
| ENSMUSG00 Gm13318 | 958 | 67.7669099 | 49.6954122 | 48.4466964 | 42.4095847 | 58.8028381 | 29.9793269 |
| ENSMUSG00 4930407I10F | 4671 | 0 | 0 | 0 | 0 | 0 | 0 |
| ENSMUSG00 Gm10849 | 616 | 0.12907624 | 0.23467423 | 0.21526872 | 0.05857475 | 0.24850508 | 0 |
| ENSMUSG00 Aarsd1 | 4863 | 11.1426527 | 11.8112768 | 9.37119065 | 11.4263518 | 9.43300676 | 10.7797361 |
| ENSMUSG00 BC028471 | 4106 | 0 | 0 | 0 | 0 | 0 | 0 |
| ENSMUSG00 Gm10852 | 315 | 0 | 0 | 0 | 0 | 0 | 0.08658616 |
| ENSMUSG00 Gm13262 | 2020 | 13.2452665 | 10.9015862 | 10.4815261 | 16.6120304 | 16.0657302 | 18.160589 |
| ENSMUSG00 Gm10855 | 2540 | 0 | 0 | 0 | 0 | 0 | 0 |
| ENSMUSG00 Prhoxnb | 762 | 0 | 0 | 0 | 0 | 0 | 0 |
| ENSMUSG00 Gm10858 | 2902 | 0 | 0 | 0 | 0 | 0 | 0 |
| ENSMUSG00 Gm6878 | 2014 | 0 | 0 | 0 | 0 | 0 | 0 |
| ENSMUSG00 Cyp3a41a | 2036 | 0 | 0 | 0 | 0 | 0 | 0 |
| ENSMUSG00 Cyp3a41b | 2040 | 0 | 0 | 0 | 0 | 0 | 0 |
| ENSMUSG00 Gm5464 | 2155 | 0 | 0.0223603 | 0.38971463 | 0 | 0 | 0 |
| ENSMUSG00 Gm10863 | 1949 | 0 | 0 | 0 | 0 | 0 | 0.19591839 |
| ENSMUSG00 Gm17151 | 365 | 1.41594862 | 2.1122824 | 0.24220188 | 0.79083933 | 0.83878974 | 1.12087559 |
| ENSMUSG00 Krtap4-6 | 917 | 0 | 0 | 0 | 0 | 0 | 0 |
| ENSMUSG00 Krtap1-4 | 592 | 0 | 0 | 0 | 0 | 0 | 0 |
| ENSMUSG00 Rsph10b2 | 3135 | 0.78623279 | 0.78389427 | 0.38068573 | 0.93226335 | 0.47201432 | 1.47030751 |
| ENSMUSG00 Krt26 | 4832 | 0 | 0 | 0 | 0 | 0 | 0 |
| ENSMUSG00 Defb30 | 1683 | 0 | 0 | 0 | 0 | 0 | 0 |
| ENSMUSG00 Defb43 | 406 | 0 | 0 | 0 | 0 | 0 | 0 |
| ENSMUSG00 Defb47 | 238 | 0 | 0 | 0 | 0 | 0 | 0 |
| ENSMUSG00 Defb48 | 443 | 0 | 0 | 0 | 0 | 0 | 0 |
| ENSMUSG00 Gm12359 | 1818 | 1.00591425 | 1.08671294 | 0.38901512 | 0.17862398 | 0.78588478 | 0.60010207 |
| ENSMUSG00 Gm16409 | 797 | 0.19952562 | 0 | 0 | 0 | 0 | 0 |
| ENSMUSG00 6330403L08F | 5520 | 0.21606239 | 0.01745886 | 0.01601516 | 0.01960981 | 0.01848782 | 0.17293702 |
| ENSMUSG00 Gm11529 | 636 | 0 | 0 | 0 | 0 | 0 | 0 |
| ENSMUSG00 Gm11531 | 1135 | 0 | 0 | 0 | 0 | 0.04495716 | 0 |
| ENSMUSG00 Hoxb2 | 3302 | 0.73442893 | 1.15285552 | 0.92386056 | 0.81954977 | 0.9580967 | 0.56994249 |
| ENSMUSG00 Gm11536 | 532 | 1.19565355 | 0.90576019 | 1.32937874 | 0.33911696 | 1.24688512 | 1.02536238 |
| ENSMUSG00 Nrbp2 | 2483 | 73.2186115 | 67.8452685 | 68.3410694 | 71.0451533 | 75.3784722 | 67.565971 |
| ENSMUSG00 Gm10874 | 2351 | 0 | 0 | 0.03760259 | 0 | 0 | 0 |
| ENSMUSG00 Nynrin | 7427 | 0.38005105 | 0.49308868 | 0.7558414 | 0.5052555 | 1.13361411 | 0.64266351 |
| ENSMUSG00 Gal3st4 | 4630 | 0 | 0 | 0 | 0 | 0 | 0 |
| ENSMUSG00 K230010J24F | 1115 | 0.92703363 | 0.4321654 | 0.63428654 | 0.22652405 | 1.7390155 | 0.2446156 |
| ENSMUSG00 Zfp652 | 15435 | 1.02769269 | 1.2019294 | 1.05958419 | 1.82806342 | 1.42483758 | 1.50023769 |
| ENSMUSG00 B130006D01 | 3123 | 0.42008673 | 1.55838318 | 0.05661459 | 0.03466095 | 0.21240566 | 0.74234529 |
| ENSMUSG00 Smok3b | 3163 | 0 | 0 | 0 | 0 | 0 | 0 |
| ENSMUSG00 Smok3a | 2984 | 0 | 0 | 0 | 0 | 0 | 0 |
| ENSMUSG00 Zc3h3 | 3348 | 0.33248311 | 0.15831866 | 0.22444186 | 0.29098423 | 0.65535668 | 0.52137901 |
| ENSMUSG00 Ly6a | 816 | 442.7182 | 467.16047 | 119.875832 | 290.248211 | 242.000091 | 271.108577 |
| ENSMUSG00 Cyp11b1 | 3681 | 0 | 0 | 0 | 0 | 0 | 0 |
| ENSMUSG00 2300005B03F | 461 | 0 | 0.20905181 | 0 | 0 | 0 | 0 |
| ENSMUSG00 Gm7935 | 1548 | 0.02568184 | 0 | 0.02855416 | 0.02330881 | 0.06592555 | 0.03523855 |
| ENSMUSG00 Tmem92 | 3166 | 0 | 0 | 0 | 0 | 0 | 0 |
| ENSMUSG00 Gm11545 | 2834 | 0 | 0.01700298 | 0.6394762 | 0 | 0 | 0.00962408 |
| ENSMUSG00 2810407C02 | 3605 | 17.049091 | 17.122561 | 23.419007 | 20.9285867 | 13.7580131 | 19.4818852 |
| ENSMUSG00 H47 | 1192 | 128.104699 | 97.9494541 | 143.099759 | 95.7748229 | 84.6301552 | 151.337638 |
| ENSMUSG00 Selm | 1768 | 5.73396354 | 6.35036257 | 5.97524914 | 6.28578604 | 3.98282798 | 8.731587 |
| ENSMUSG00 Ept1 | 8464 | 4.26958079 | 5.05547739 | 4.9612182 | 2.93294502 | 4.23210253 | 2.15902745 |
| ENSMUSG00 Txnrd2 | 3442 | 10.5914514 | 6.41179271 | 13.0602192 | 7.5686334 | 9.10232271 | 6.85431812 |
| ENSMUSG00 Sepx1 | 1334 | 1.31127522 | 1.40874906 | 2.41884148 | 1.92040867 | 1.45352495 | 4.53895796 |
| ENSMUSG00 Gpx4 | 1666 | 8.5667572 | 11.1933692 | 11.6739561 | 9.13962953 | 5.91121881 | 10.3630532 |
| ENSMUSG00 Dio3 | 3896 | 0 | 0 | 0 | 0 | 0 | 0 |
| ENSMUSG00 U6 | 106 | 0 | 0 | 0 | 0 | 0 | 0 |
| ENSMUSG00 SNORD115 | 78 | 0 | 0 | 0 | 0 | 0 | 0 |
| ENSMUSG00 U6 | 106 | 0 | 0 | 0 | 0 | 0 | 0 |
| ENSMUSG00 SNORD115 | 78 | 0 | 0 | 0 | 0 | 0 | 0 |
| ENSMUSG00 SNORD115 | 78 | 0 | 0 | 0 | 0 | 0 | 0 |
| ENSMUSG00 SNORD115 | 78 | 0 | 0 | 0 | 0 | 0 | 0 |
| ENSMUSG00 SNORD115 | 78 | 0 | 0 | 0 | 0 | 0 | 0 |
| ENSMUSG00 SNORD115 | 78 | 0 | 0 | 0 | 0 | 0 | 0 |
| ENSMUSG00 SNORD115 | 78 | 0 | 0 | 0 | 0 | 0 | 0 |
| ENSMUSG00 n-R5s122 | 118 | 0 | 0 | 0 | 0 | 0 | 0 |
| ENSMUSG00 U1 | 170 | 0 | 0 | 0 | 0 | 0 | 0 |
| ENSMUSG00 n-R5s143 | 118 | 0 | 0 | 0 | 0 | 0 | 0 |
| ENSMUSG00 SNORD115 | 78 | 0 | 0 | 0 | 0 | 0 | 0 |
| ENSMUSG00 SNORD115 | 78 | 0 | 0 | 0 | 0 | 0 | 0 |
| ENSMUSG00 SNORD116 | 93 | 0 | 0 | 0 | 0 | 0 | 0 |
| ENSMUSG00 SNORD50 | 71 | 0 | 0 | 0 | 0 | 0 | 0 |
| ENSMUSG00 SNORD50 | 71 | 1.67980904 | 0 | 0 | 1.52459343 | 0 | 0.7682997 |
| ENSMUSG00 SNORD115 | 78 | 0 | 0 | 0 | 0 | 0 | 0 |
| ENSMUSG00 U1 | 164 | 0 | 0 | 0 | 0 | 0 | 0 |
| ENSMUSG00 U6 | 106 | 0 | 0 | 0 | 0 | 0 | 0 |
| ENSMUSG00 SNORD115 | 78 | 0 | 0 | 0 | 0 | 0 | 0 |
| ENSMUSG00 n-R5s142 | 118 | 0 | 0 | 0 | 0 | 0 | 0 |
| ENSMUSG00 U6 | 106 | 0 | 0 | 0 | 0 | 0 | 0 |
| ENSMUSG00 SNORD115 | 78 | 0 | 0 | 0 | 0 | 0 | 0 |
| ENSMUSG00 SNORD115 | 78 | 0 | 0 | 0 | 0 | 0 | 0 |
| ENSMUSG00 n-R5s104 | 118 | 0 | 0 | 0 | 0 | 0 | 0 |
| ENSMUSG00 U1 | 163 | 0.24389865 | 0 | 0 | 0 | 0 | 0 |
| ENSMUSG00 U6 | 106 | 0 | 0 | 0 | 0 | 0 | 0 |
| ENSMUSG00 SNORD115 | 78 | 0 | 0 | 0 | 0 | 0 | 0 |
| ENSMUSG00 SNORD115 | 77 | 0 | 0 | 0 | 0 | 0 | 0 |
| ENSMUSG00 n-R5s102 | 118 | 0 | 0 | 0 | 0 | 0 | 0 |
| ENSMUSG00 SNORD115 | 78 | 0 | 0 | 0 | 0 | 0 | 0 |
| ENSMUSG00 SNORD115 | 78 | 0 | 0 | 0 | 0 | 0 | 0 |
| ENSMUSG00 SNORD116 | 93 | 0 | 0 | 0 | 0 | 0 | 0 |
| ENSMUSG00 Snord116l1 | 93 | 0 | 0 | 0 | 0 | 0 | 0 |
| ENSMUSG00 n-R5s110 | 118 | 0 | 0 | 0 | 0 | 0 | 0 |
| ENSMUSG00 SNORD115 | 78 | 0 | 0 | 0 | 0 | 0 | 0 |
| ENSMUSG00 SNORD116 | 93 | 0 | 0 | 0 | 0 | 0 | 0 |
| ENSMUSG00 SNORD115 | 78 | 0 | 0 | 0 | 0 | 0 | 0 |
| ENSMUSG00 SNORD115 | 78 | 0 | 0 | 0 | 0 | 0 | 0 |
| ENSMUSG00 U2 | 190 | 0 | 0.25361285 | 0.93056512 | 0.56971649 | 0.53711975 | 0.57420293 |
| ENSMUSG00 SNORD115 | 78 | 0 | 0 | 0 | 0 | 0 | 0 |
| ENSMUSG00 n-R5s120 | 118 | 0 | 0 | 0 | 0 | 0 | 0 |
| ENSMUSG00 SNORD115 | 78 | 0 | 0 | 0 | 0 | 0 | 0 |
| ENSMUSG00 SNORD115 | 78 | 0 | 0 | 0 | 0 | 0 | 0 |
| ENSMUSG00 SNORD115 | 78 | 0 | 0 | 0 | 0 | 0 | 0 |
| ENSMUSG00 SNORD116 | 93 | 0 | 0 | 0 | 0 | 0 | 0 |
| ENSMUSG00 SNORD116 | 93 | 0 | 0 | 0 | 0 | 0 | 0 |
| ENSMUSG00 U6 | 106 | 0 | 0 | 0 | 0 | 0 | 0 |
| ENSMUSG00 SNORD115 | 78 | 0 | 0 | 0 | 0 | 0 | 0 |
| ENSMUSG00 SNORD115 | 78 | 0 | 0 | 0 | 0 | 0 | 0 |
| ENSMUSG00 U5 | 115 | 0 | 1.25703762 | 0 | 0 | 0 | 0.9486831 |
| ENSMUSG00 U6 | 106 | 0 | 0 | 0 | 0 | 0 | 0 |
| ENSMUSG00 n-R5s138 | 118 | 0 | 0 | 0 | 0 | 0 | 0 |
| ENSMUSG00 SNORD115 | 78 | 0 | 0 | 0 | 0 | 0 | 0 |
| ENSMUSG00 U6 | 106 | 0 | 0 | 0 | 0 | 0 | 0 |
| ENSMUSG00 SNORD115 | 78 | 0 | 0 | 0 | 0 | 0 | 0 |
| ENSMUSG00 n-R5s112 | 118 | 0 | 0 | 0 | 0 | 0 | 0 |
| ENSMUSG00 SNORD83 | 76 | 0 | 0 | 0 | 0 | 0 | 0 |
| ENSMUSG00 SNORD115 | 78 | 0 | 0 | 0 | 0 | 0 | 0 |
| ENSMUSG00 U6 | 106 | 0 | 0 | 0 | 0 | 0 | 0 |
| ENSMUSG00 n-R5s121 | 118 | 0 | 0 | 0 | 0 | 0 | 0 |
| ENSMUSG00 U6 | 105 | 0 | 0 | 0 | 0 | 0 | 0 |
| ENSMUSG00 SNORD115 | 78 | 0 | 0 | 0 | 0 | 0 | 0 |
| ENSMUSG00 SNORD115 | 77 | 0 | 0 | 0 | 0 | 0 | 0 |
| ENSMUSG00 U4 | 140 | 0 | 1.03256882 | 0.6314549 | 0.51545778 | 1.09342234 | 0.19481885 |
| ENSMUSG00 U6 | 106 | 0 | 0 | 0 | 0 | 0 | 0 |
| ENSMUSG00 SNORD115 | 78 | 0 | 0 | 0 | 0 | 0 | 0 |
| ENSMUSG00 SNORD115 | 78 | 0 | 0 | 0 | 0.46259031 | 0 | 0 |
| ENSMUSG00 SNORD115 | 78 | 0 | 0 | 0 | 0 | 0 | 0 |
| ENSMUSG00 U4 | 140 | 0 | 0 | 0 | 0 | 0 | 0 |
| ENSMUSG00 U6 | 106 | 0 | 0 | 0 | 0 | 0 | 0 |
| ENSMUSG00 SNORD115 | 78 | 0 | 0 | 0 | 0 | 0 | 0 |
| ENSMUSG00 n-R5s131 | 118 | 0 | 0 | 0 | 0 | 0 | 0 |
| ENSMUSG00 U1 | 163 | 0 | 0 | 0 | 0 | 0 | 0 |
| ENSMUSG00 5S_rRNA | 108 | 0 | 0 | 0 | 0 | 0 | 0 |
| ENSMUSG00 SNORD115 | 78 | 0 | 0 | 0 | 0 | 0 | 0 |
| ENSMUSG00 n-R5s130 | 118 | 0 | 0 | 0 | 0 | 0 | 0 |
| ENSMUSG00 SNORD115 | 78 | 0 | 0 | 0 | 0 | 0 | 0 |
| ENSMUSG00 U6 | 104 | 0 | 0 | 0 | 0 | 0 | 0 |
| ENSMUSG00 SNORD115 | 78 | 0 | 0 | 0 | 0 | 0 | 0 |
| ENSMUSG00 U6 | 106 | 0 | 0 | 0 | 0 | 0 | 0 |
| ENSMUSG00 n-R5s100 | 118 | 0 | 0 | 0 | 0 | 0 | 0 |
| ENSMUSG00 SNORD115 | 78 | 0 | 0 | 0 | 0 | 0 | 0 |
| ENSMUSG00 SNORD115 | 78 | 0 | 0 | 0 | 0 | 0 | 0 |
| ENSMUSG00 n-R5s117 | 118 | 0 | 0 | 0 | 0 | 0 | 0 |
| ENSMUSG00 SNORD115 | 78 | 0 | 0 | 0 | 0 | 0 | 0 |
| ENSMUSG00 n-R5s149 | 118 | 0 | 0 | 0 | 0 | 0 | 0 |
| ENSMUSG00 SNORD115 | 78 | 0 | 0 | 0 | 0 | 0 | 0 |
| ENSMUSG00 SNORD116 | 93 | 0 | 0 | 0 | 0 | 0 | 0 |
| ENSMUSG00 SNORD115 | 78 | 0 | 0 | 0 | 0 | 0 | 0 |
| ENSMUSG00 SNORD115 | 78 | 0 | 0 | 0 | 0 | 0 | 0 |
| ENSMUSG00 SNORD115 | 78 | 0 | 0 | 0 | 0 | 0 | 0 |
| ENSMUSG00 n-R5s139 | 118 | 0 | 0 | 0 | 0 | 0 | 0 |
| ENSMUSG00 SNORD115 | 78 | 0 | 0 | 0 | 0 | 0 | 0 |
| ENSMUSG00 U6 | 106 | 0 | 0 | 0 | 0 | 0 | 0 |
| ENSMUSG00 U6 | 105 | 0 | 0 | 0 | 0 | 0 | 0 |
| ENSMUSG00 SNORD115 | 78 | 0 | 0 | 0 | 0 | 0 | 0 |
| ENSMUSG00 n-R5s109 | 118 | 0 | 0 | 0 | 0 | 0 | 0 |
| ENSMUSG00 SNORD50 | 71 | 0 | 0 | 0 | 0 | 0 | 0 |
| ENSMUSG00 SNORD115 | 78 | 0 | 0 | 0 | 0 | 0 | 0 |
| ENSMUSG00 n-R5s133 | 118 | 0 | 0 | 0 | 0 | 0 | 0 |
| ENSMUSG00 SNORD115 | 78 | 0 | 0 | 0 | 0 | 0 | 0 |
| ENSMUSG00 SNORD115 | 78 | 0 | 0 | 0 | 0 | 0 | 0 |
| ENSMUSG00 SNORD115 | 78 | 0 | 0 | 0 | 0 | 0 | 0 |
| ENSMUSG00 U1 | 163 | 0 | 0 | 0 | 0 | 0 | 0 |
| ENSMUSG00 U6 | 106 | 0 | 0 | 0 | 0 | 0 | 0 |
| ENSMUSG00 SNORD115 | 78 | 0 | 0 | 0 | 0 | 0 | 0 |
| ENSMUSG00 SNORD115 | 78 | 0 | 0 | 0 | 0 | 0 | 0 |
| ENSMUSG00 U1 | 159 | 0 | 0 | 0 | 0 | 0 | 0 |
| ENSMUSG00 SNORD115 | 78 | 0 | 0 | 0 | 0 | 0 | 0 |
| ENSMUSG00 SNORD115 | 78 | 0 | 0 | 0 | 0 | 0 | 0 |
| ENSMUSG00 U6 | 106 | 0 | 0 | 0 | 0 | 0 | 0 |
| ENSMUSG00 U6 | 106 | 6.00082725 | 3.18212354 | 2.91898963 | 2.72317317 | 0.4813809 | 2.05846334 |
| ENSMUSG00 SNORD115 | 78 | 0 | 0 | 0 | 0 | 0 | 0 |
| ENSMUSG00 SNORD115 | 78 | 0 | 0 | 0 | 0 | 0 | 0 |
| ENSMUSG00 n-R5s103 | 118 | 0 | 0 | 0 | 0 | 0 | 0 |
| ENSMUSG00 U4 | 140 | 0 | 0 | 1.8943647 | 0 | 0.36447411 | 0 |
| ENSMUSG00 n-R5s106 | 118 | 0 | 0 | 0 | 0 | 0 | 0 |
| ENSMUSG00 SNORD116 | 93 | 0 | 0 | 0 | 0 | 0 | 0 |
| ENSMUSG00 SNORD116 | 93 | 0 | 0 | 0 | 0 | 0 | 0 |
| ENSMUSG00 SNORD115 | 78 | 0 | 0 | 0 | 0 | 0 | 0 |
| ENSMUSG00 U6 | 104 | 0 | 0 | 0 | 0 | 0 | 0 |
| ENSMUSG00 SNORD115 | 78 | 0 | 0 | 0 | 0 | 0 | 0 |
| ENSMUSG00 SNORD115 | 78 | 0 | 0 | 0 | 0 | 0 | 0 |
| ENSMUSG00 SNORD116 | 93 | 0 | 0 | 0 | 0 | 0 | 0 |
| ENSMUSG00 SNORD115 | 78 | 0 | 0 | 0 | 0 | 0 | 0 |
| ENSMUSG00 SNORD115 | 78 | 0 | 0 | 0 | 0 | 0 | 0 |
| ENSMUSG00 U6 | 106 | 0 | 0 | 0 | 0 | 0 | 0 |
| ENSMUSG00 SNORD116 | 93 | 0 | 0 | 0 | 0 | 0 | 0 |
| ENSMUSG00 U6 | 106 | 0 | 0 | 0 | 0 | 0 | 0 |
| ENSMUSG00 SNORD115 | 78 | 0 | 0 | 0 | 0 | 0 | 0 |
| ENSMUSG00 n-R5s127 | 118 | 0 | 0 | 0 | 0 | 0 | 0 |
| ENSMUSG00 SNORD115 | 78 | 0 | 0 | 0 | 0 | 0 | 0 |
| ENSMUSG00 n-R5s113 | 118 | 0 | 0 | 0 | 0 | 0 | 0 |
| ENSMUSG00 n-R5s146 | 118 | 0 | 0 | 0 | 0 | 0 | 0 |
| ENSMUSG00 SNORD115 | 78 | 0 | 0 | 0 | 0 | 0 | 0 |
| ENSMUSG00 SNORD115 | 78 | 0 | 0 | 0 | 0 | 0 | 0 |
| ENSMUSG00 n-R5s115 | 118 | 0 | 0 | 0 | 0 | 0 | 0 |
| ENSMUSG00 n-R5s118 | 118 | 0 | 0 | 0 | 0 | 0 | 0 |
| ENSMUSG00 U6 | 106 | 0 | 0 | 0 | 0 | 0 | 0 |
| ENSMUSG00 n-R5s125 | 118 | 0 | 0 | 0 | 0 | 0 | 0 |
| ENSMUSG00 SNORD115 | 78 | 0 | 0 | 0 | 0 | 0 | 0 |
| ENSMUSG00 SNORD115 | 78 | 0 | 0 | 0 | 0 | 0 | 0 |
| ENSMUSG00 n-R5s107 | 118 | 0 | 0 | 0 | 0 | 0 | 0 |
| ENSMUSG00 SNORD115 | 78 | 0 | 0 | 0 | 0 | 0 | 0 |
| ENSMUSG00 SNORD115 | 78 | 0 | 0 | 0 | 0 | 0 | 0 |
| ENSMUSG00 n-R5s108 | 118 | 0 | 0 | 0 | 0 | 0 | 0 |
| ENSMUSG00 n-R5s129 | 118 | 0 | 0 | 0 | 0 | 0 | 0 |
| ENSMUSG00 SNORD115 | 78 | 0 | 0 | 0 | 0.46259031 | 0 | 0 |
| ENSMUSG00 n-R5s1 | 118 | 0 | 0 | 1.49836756 | 0 | 0 | 0 |
| ENSMUSG00 Snord14c | 85 | 0 | 0 | 0 | 0 | 0 | 0 |
| ENSMUSG00 SNORD115 | 78 | 0 | 0 | 0 | 0 | 0 | 0 |
| ENSMUSG00 SNORD115 | 78 | 0 | 0 | 0 | 0 | 0 | 0 |
| ENSMUSG00 SNORD115 | 78 | 0 | 0 | 0 | 0 | 0 | 0 |
| ENSMUSG00 n-R5s123 | 118 | 0 | 0 | 0 | 0 | 0 | 0 |
| ENSMUSG00 SNORD115 | 78 | 0 | 0 | 0 | 0 | 0 | 0 |
| ENSMUSG00 SNORD115 | 78 | 0 | 0 | 0 | 0 | 0 | 0 |
| ENSMUSG00 SNORD115 | 78 | 0 | 0 | 0 | 0 | 0 | 0 |
| ENSMUSG00 SNORD115 | 78 | 0 | 0 | 0 | 0 | 0 | 0 |
| ENSMUSG00 SNORD115 | 78 | 0 | 0 | 0 | 0 | 0 | 0 |
| ENSMUSG00 SNORD115 | 78 | 0 | 0 | 0 | 0 | 0 | 0 |
| ENSMUSG00 U6 | 104 | 0 | 0 | 0 | 0 | 0 | 0 |
| ENSMUSG00 SNORD115 | 78 | 0 | 0 | 0 | 0 | 0 | 0 |
| ENSMUSG00 SNORD116 | 93 | 0 | 0 | 0 | 0 | 0 | 0 |
| ENSMUSG00 SNORD116 | 93 | 0 | 0 | 0 | 0 | 0 | 0 |
| ENSMUSG00 SNORD115 | 78 | 0 | 0 | 0 | 0 | 0 | 0 |
| ENSMUSG00 U6 | 106 | 0 | 0 | 0 | 0 | 0 | 0 |
| ENSMUSG00 SNORD115 | 78 | 0 | 0 | 0 | 0 | 0 | 0 |
| ENSMUSG00 U6 | 104 | 0 | 0 | 0 | 0 | 0 | 0 |
| ENSMUSG00 SNORD115 | 78 | 0 | 0 | 0 | 0 | 0 | 0 |
| ENSMUSG00 SNORD116 | 93 | 0 | 0 | 0 | 0 | 0 | 0 |
| ENSMUSG00 n-R5s114 | 118 | 0 | 0 | 0 | 0 | 0 | 0 |
| ENSMUSG00 SNORD115 | 78 | 0 | 0 | 0 | 0 | 0 | 0 |
| ENSMUSG00 U6 | 106 | 0 | 0 | 0 | 0 | 0 | 0 |
| ENSMUSG00 SNORD116 | 93 | 0 | 0 | 0 | 0 | 0 | 0 |
| ENSMUSG00 n-R5s128 | 118 | 0 | 0 | 0 | 0 | 0 | 0 |
| ENSMUSG00 U6 | 106 | 0 | 0 | 0 | 0 | 0 | 0 |
| ENSMUSG00 SNORD116 | 93 | 0 | 0 | 0 | 0 | 0 | 0 |
| ENSMUSG00 U6 | 104 | 0 | 0.92866235 | 0.42501772 | 0 | 0 | 0 |
| ENSMUSG00 SNORD115 | 78 | 0 | 0 | 0 | 0 | 0 | 0 |
| ENSMUSG00 U6 | 106 | 0 | 0 | 0 | 0 | 0.4813809 | 0 |
| ENSMUSG00 n-R5s144 | 118 | 0 | 0 | 0 | 0 | 0 | 0 |
| ENSMUSG00 SNORD116 | 93 | 0 | 0 | 0 | 0 | 0 | 0 |
| ENSMUSG00 U6 | 106 | 0 | 0 | 0 | 0 | 0 | 0 |
| ENSMUSG00 SNORD115 | 78 | 0 | 0 | 0 | 0 | 0 | 0 |
| ENSMUSG00 SNORD115 | 78 | 0 | 0 | 0 | 0 | 0 | 0 |
| ENSMUSG00 SNORD115 | 78 | 0 | 0 | 0 | 0 | 0 | 0 |
| ENSMUSG00 SNORD115 | 78 | 0 | 0 | 0 | 0 | 0 | 0 |
| ENSMUSG00 n-R5s2 | 118 | 0 | 0 | 0 | 0 | 0 | 0 |
| ENSMUSG00 SNORD115 | 78 | 0 | 0 | 0 | 0 | 0 | 0 |
| ENSMUSG00 SNORD115 | 78 | 0 | 0 | 0 | 0 | 0 | 0 |
| ENSMUSG00 n-R5s111 | 118 | 0 | 0 | 0 | 0 | 0 | 0 |
| ENSMUSG00 SNORD115 | 78 | 0 | 0 | 0 | 0 | 0 | 0 |
| ENSMUSG00 U1 | 163 | 0 | 0 | 0 | 0 | 0 | 0 |
| ENSMUSG00 Snord14d | 85 | 0 | 0 | 0 | 0 | 0 | 0 |
| ENSMUSG00 SNORD115 | 78 | 0 | 0 | 0 | 0 | 0 | 0 |
| ENSMUSG00 SNORD115 | 78 | 0 | 0 | 0 | 0 | 0 | 0 |
| ENSMUSG00 SNORD115 | 78 | 0 | 0 | 0 | 0 | 0 | 0 |
| ENSMUSG00 U6 | 106 | 0 | 0 | 0 | 0 | 0 | 0 |
| ENSMUSG00 U5 | 115 | 0 | 0 | 0 | 0 | 0 | 0 |
| ENSMUSG00 SNORD115 | 78 | 0 | 0 | 0 | 0 | 0 | 0 |
| ENSMUSG00 SNORD115 | 78 | 0 | 0 | 0 | 0 | 0 | 0 |
| ENSMUSG00 U6 | 105 | 0 | 0 | 0 | 0 | 0 | 0 |
| ENSMUSG00 SNORD115 | 78 | 0 | 0 | 0 | 0 | 0 | 0 |
| ENSMUSG00 SNORD116 | 93 | 0 | 0 | 0 | 0 | 0 | 0 |
| ENSMUSG00 SNORD116 | 93 | 0 | 0 | 0 | 0 | 0 | 0 |
| ENSMUSG00 U6 | 106 | 0 | 0 | 0 | 0 | 0 | 0 |
| ENSMUSG00 SNORD115 | 78 | 0 | 0 | 0 | 0 | 0 | 0 |
| ENSMUSG00 SNORD115 | 78 | 0 | 0 | 0 | 0 | 0 | 0 |
| ENSMUSG00 SNORD115 | 78 | 0 | 0 | 0 | 0 | 0 | 0 |
| ENSMUSG00 n-R5s136 | 118 | 0 | 0 | 0 | 0 | 0 | 0 |
| ENSMUSG00 U1 | 163 | 0 | 0 | 0 | 0 | 0 | 0 |
| ENSMUSG00 SNORD115 | 77 | 0 | 0 | 0 | 0 | 0 | 0 |
| ENSMUSG00 SNORD50 | 71 | 0 | 0 | 0 | 0 | 0 | 0 |
| ENSMUSG00 SNORA25 | 132 | 0 | 0 | 0 | 0 | 0 | 1.44638238 |
| ENSMUSG00 SNORD115 | 77 | 0 | 0 | 0 | 0 | 0 | 0 |
| ENSMUSG00 n-R5s119 | 118 | 0 | 0 | 0 | 0 | 0 | 0 |
| ENSMUSG00 SNORD115 | 78 | 0 | 0 | 0 | 0 | 0 | 0 |
| ENSMUSG00 Snord116l2 | 93 | 0 | 0 | 0 | 0 | 0 | 0 |
| ENSMUSG00 U6 | 106 | 0 | 0 | 0 | 0 | 0 | 0 |
| ENSMUSG00 SNORD115 | 78 | 0 | 0 | 0 | 0 | 0 | 0 |
| ENSMUSG00 SNORD115 | 78 | 0 | 0 | 0 | 0 | 0 | 0 |
| ENSMUSG00 n-R5s105 | 118 | 0 | 0 | 0 | 0 | 0 | 0 |
| ENSMUSG00 SNORD115 | 78 | 0 | 0 | 0 | 0 | 0 | 0 |
| ENSMUSG00 SNORD116 | 93 | 0 | 0 | 0 | 0 | 0 | 0 |
| ENSMUSG00 U6 | 105 | 0 | 0 | 0 | 0 | 0 | 0 |
| ENSMUSG00 SNORD116 | 93 | 0 | 0 | 0 | 0 | 0 | 0 |
| ENSMUSG00 U1 | 163 | 0 | 0 | 0 | 0 | 0 | 0 |
| ENSMUSG00 SNORD115 | 78 | 0 | 0 | 0 | 0 | 0 | 0 |
| ENSMUSG00 n-R5s101 | 118 | 0 | 0 | 0 | 0 | 0 | 0 |
| ENSMUSG00 SNORD116 | 93 | 0 | 0 | 0 | 0 | 0 | 0 |
| ENSMUSG00 n-R5s134 | 118 | 0 | 0 | 0 | 0 | 0 | 0 |
| ENSMUSG00 U6 | 104 | 0 | 0 | 0 | 0 | 0 | 0 |
| ENSMUSG00 SNORD115 | 78 | 0 | 0 | 0 | 0 | 0 | 0 |
| ENSMUSG00 SNORD115 | 78 | 0 | 0 | 0 | 0 | 0 | 0 |
| ENSMUSG00 SNORD115 | 78 | 0 | 0 | 0 | 0 | 0 | 0 |
| ENSMUSG00 SNORD115 | 78 | 0 | 0 | 0 | 0 | 0 | 0 |
| ENSMUSG00 SNORD50 | 71 | 0 | 0 | 0 | 0 | 0 | 0 |
| ENSMUSG00 n-R5s64 | 118 | 0 | 0 | 0 | 0 | 0 | 0 |
| ENSMUSG00 SNORD115 | 78 | 0 | 0 | 0 | 0 | 0 | 0 |
| ENSMUSG00 SNORD115 | 78 | 0 | 0 | 0 | 0 | 0 | 0 |
| ENSMUSG00 SNORD115 | 78 | 0 | 0 | 0 | 0 | 0 | 0 |
| ENSMUSG00 SNORD115 | 78 | 0 | 0 | 0 | 0 | 0 | 0 |
| ENSMUSG00 U1 | 165 | 0 | 0 | 0 | 0 | 0 | 0 |
| ENSMUSG00 SNORD116 | 93 | 0 | 0 | 0 | 0 | 0 | 0 |
| ENSMUSG00 U6 | 106 | 0 | 0 | 0 | 0 | 0 | 0 |
| ENSMUSG00 SNORD115 | 78 | 0 | 0 | 0 | 0 | 0 | 0 |
| ENSMUSG00 SNORD115 | 78 | 0 | 0 | 0 | 0 | 0 | 0 |
| ENSMUSG00 U6 | 105 | 0 | 0 | 0 | 0 | 0 | 0 |
| ENSMUSG00 SNORD115 | 78 | 0 | 0 | 0 | 0 | 0 | 0 |
| ENSMUSG00 SNORD115 | 78 | 0 | 0 | 0 | 0 | 0 | 0 |
| ENSMUSG00 U6 | 106 | 0 | 0 | 0 | 0 | 0 | 0 |
| ENSMUSG00 SNORD116 | 93 | 0 | 0 | 0 | 0 | 0 | 0 |
| ENSMUSG00 SNORD115 | 78 | 0 | 0 | 0 | 0 | 0 | 0 |
| ENSMUSG00 SNORD115 | 78 | 0 | 0 | 0 | 0 | 0 | 0 |
| ENSMUSG00 SNORD115 | 78 | 0 | 0 | 0 | 0 | 0 | 0 |
| ENSMUSG00 SNORD115 | 78 | 0 | 0 | 0 | 0 | 0 | 0 |
| ENSMUSG00 SNORD115 | 78 | 0 | 0 | 0 | 0 | 0 | 0 |
| ENSMUSG00 SNORD115 | 78 | 0 | 0 | 0 | 0 | 0 | 0 |
| ENSMUSG00 SNORD116 | 93 | 0 | 0 | 0 | 0 | 0 | 0 |
| ENSMUSG00 SNORD115 | 78 | 0 | 0 | 0 | 0 | 0 | 0 |
| ENSMUSG00 U1 | 163 | 0 | 0 | 0 | 0 | 0 | 0 |
| ENSMUSG00 SNORD115 | 78 | 0 | 0 | 0 | 0 | 0 | 0 |
| ENSMUSG00 U4 | 140 | 0.85190315 | 1.03256882 | 0.6314549 | 0.25772889 | 1.45789645 | 0.3896377 |
| ENSMUSG00 n-R5s124 | 118 | 0 | 0 | 0 | 0 | 0 | 0 |
| ENSMUSG00 SNORD115 | 78 | 0 | 0 | 0 | 0 | 0 | 0 |
| ENSMUSG00 n-R5s141 | 118 | 0 | 0 | 0 | 0 | 0 | 0 |
| ENSMUSG00 SNORD115 | 78 | 0 | 0 | 0 | 0 | 0 | 0 |
| ENSMUSG00 Mir547 | 77 | 0 | 0 | 0 | 0 | 0 | 0 |
| ENSMUSG00 Mir376b | 85 | 0 | 0 | 0 | 0 | 0 | 0 |
| ENSMUSG00 Mir688 | 74 | 0 | 3.90700882 | 0 | 0 | 0 | 0 |
| ENSMUSG00 Mir615 | 91 | 0 | 0 | 0 | 0 | 0 | 0 |
| ENSMUSG00 Mir652 | 97 | 0.81970063 | 1.49030233 | 0 | 0 | 0 | 0 |
| ENSMUSG00 Mir705 | 81 | 0 | 0 | 0 | 0 | 0 | 0 |
| ENSMUSG00 Mir669a-2 | 96 | 0 | 0 | 0 | 0 | 0 | 0 |
| ENSMUSG00 Mir669a-11 | 96 | 0 | 0 | 0 | 0 | 0 | 0 |
| ENSMUSG00 mmu-mir-669 | 96 | 0 | 0 | 0 | 0 | 0 | 0 |
| ENSMUSG00 Mir669a-12 | 96 | 0 | 0 | 0 | 0 | 0 | 0 |
| ENSMUSG00 Mir669a-10 | 96 | 0 | 0 | 0 | 0 | 0 | 0 |
| ENSMUSG00 Mir669a-9 | 96 | 0 | 0 | 0 | 0 | 0 | 0 |
| ENSMUSG00 Mir669a-8 | 96 | 0 | 0 | 0 | 0 | 0 | 0 |

| | | | | | | | | |
|---|---|---|---|---|---|---|---|---|
| ENSMUSG00 | Mir669a-7 | 96 | 0 | 0 | 0 | 0 | 0 | 0 |
| ENSMUSG00 | Mir669a-6 | 96 | 0 | 0 | 0 | 0 | 0 | 0 |
| ENSMUSG00 | Mir669a-5 | 96 | 0 | 0 | 0 | 0 | 0 | 0 |
| ENSMUSG00 | Mir669a-4 | 96 | 0 | 0 | 0 | 0 | 0 | 0 |
| ENSMUSG00 | Mir669a-1 | 96 | 0 | 0 | 0 | 0 | 0 | 0 |
| ENSMUSG00 | Mir669a-3 | 108 | 0 | 0 | 0 | 0 | 0 | 0 |
| ENSMUSG00 | AC101886.1 | 108 | 0 | 0 | 0 | 0 | 0 | 0 |
| ENSMUSG00 | AC124732.1 | 102 | 0 | 0 | 0 | 0 | 0 | 0 |
| ENSMUSG00 | AL645948.1 | 108 | 0 | 0 | 0 | 0 | 0 | 0 |
| ENSMUSG00 | AC122459.1 | 108 | 0 | 0 | 0 | 0 | 0 | 0 |
| ENSMUSG00 | Mir692-2b | 108 | 0 | 0 | 0 | 0 | 0 | 0 |
| ENSMUSG00 | Mir692-2a | 108 | 0 | 0 | 0 | 0 | 0 | 0 |
| ENSMUSG00 | AC151602.1 | 108 | 0 | 0 | 0 | 0 | 0 | 0 |
| ENSMUSG00 | AC123859.1 | 108 | 0 | 0 | 0 | 0 | 0 | 0 |
| ENSMUSG00 | AC154650.1 | 108 | 0 | 0 | 0 | 0 | 0 | 0 |
| ENSMUSG00 | AC115005.1 | 107 | 0 | 0 | 0 | 0 | 0 | 0 |
| ENSMUSG00 | Mir20b | 79 | 0 | 0 | 0 | 0 | 0 | 0 |
| ENSMUSG00 | Mir710 | 109 | 0 | 0 | 0 | 0 | 0 | 0 |
| ENSMUSG00 | Mir467b | 72 | 0 | 0 | 0 | 0 | 0 | 0 |
| ENSMUSG00 | Mir720 | 63 | 0 | 0 | 0 | 0 | 0 | 0 |
| ENSMUSG00 | Mir598 | 78 | 0 | 0 | 0 | 0 | 0 | 0 |
| ENSMUSG00 | Mir450b | 81 | 0 | 0 | 0 | 0 | 0 | 0 |
| ENSMUSG00 | Mir707 | 72 | 0 | 0 | 0 | 0 | 0 | 0 |
| ENSMUSG00 | Mir541 | 89 | 0 | 0 | 0 | 0 | 0 | 0 |
| ENSMUSG00 | AC162934.1 | 108 | 0 | 0 | 0 | 0 | 0 | 0 |
| ENSMUSG00 | AC122194.1 | 108 | 0 | 0 | 0 | 0 | 0 | 0 |
| ENSMUSG00 | AC122313.1 | 108 | 0 | 0 | 0 | 0 | 0 | 0 |
| ENSMUSG00 | Mir683-1 | 108 | 0 | 0 | 0 | 0 | 0 | 0 |
| ENSMUSG00 | Mir683-2 | 108 | 0 | 0 | 0 | 0 | 0 | 0 |
| ENSMUSG00 | Mir704 | 76 | 0 | 0 | 3.48961919 | 0.47476374 | 0 | 2.51213783 |
| ENSMUSG00 | Mir92-1 | 79 | 0 | 0 | 0 | 0 | 0 | 0 |
| ENSMUSG00 | Mir539 | 73 | 0 | 0 | 0 | 0 | 0 | 0 |
| ENSMUSG00 | Mir690 | 108 | 0 | 0 | 0 | 0 | 0 | 0 |
| ENSMUSG00 | Mir681 | 109 | 6.9298544 | 5.74700686 | 1.62208598 | 5.62747483 | 15.4483523 | 2.50226048 |
| ENSMUSG00 | Mir223 | 109 | 0 | 0 | 0 | 0 | 0 | 0 |
| ENSMUSG00 | Mir592 | 95 | 0 | 0 | 0 | 0 | 0 | 0 |
| ENSMUSG00 | Mir678 | 83 | 0 | 0 | 0 | 0 | 0 | 0 |
| ENSMUSG00 | Mir680-2 | 109 | 0 | 0 | 0 | 0 | 0 | 0 |
| ENSMUSG00 | Mir669c | 108 | 0 | 0 | 0 | 0 | 0 | 0 |
| ENSMUSG00 | Mir698 | 108 | 0 | 0 | 0.81855265 | 0.66818601 | 0 | 0 |
| ENSMUSG00 | Mir551b | 97 | 0 | 0 | 0 | 0 | 0 | 0 |
| ENSMUSG00 | Mir687 | 88 | 0 | 0 | 0 | 2.4601394 | 0 | 0 |
| ENSMUSG00 | Mir503 | 70 | 0 | 0 | 0 | 0 | 0 | 0 |
| ENSMUSG00 | Mir700 | 78 | 0 | 0 | 0 | 0 | 0 | 0 |
| ENSMUSG00 | Mir669b | 96 | 0 | 0 | 0 | 0 | 0 | 0 |
| ENSMUSG00 | Mir718 | 87 | 0 | 0 | 0 | 0 | 0 | 0 |
| ENSMUSG00 | Mir686 | 108 | 0 | 0 | 0 | 0 | 0 | 0 |
| ENSMUSG00 | AL672269.1 | 108 | 0 | 0 | 0 | 0 | 0 | 0 |
| ENSMUSG00 | AC163389.1 | 109 | 0 | 0 | 0 | 0 | 0.46813189 | 0 |
| ENSMUSG00 | AC122119.1 | 108 | 0 | 0 | 0.40927632 | 0 | 0.47246644 | 1.01017182 |
| ENSMUSG00 | CT009594.1 | 109 | 0 | 0 | 0 | 0 | 0 | 0 |
| ENSMUSG00 | mmu-mir-70 | 107 | 0 | 0 | 0 | 0 | 0 | 0 |
| ENSMUSG00 | Mir542 | 84 | 0 | 0 | 0 | 0 | 0 | 0 |
| ENSMUSG00 | Mir693 | 88 | 0 | 0 | 0 | 0 | 0 | 0 |
| ENSMUSG00 | Mir711 | 81 | 0 | 0 | 0 | 0 | 0 | 0 |
| ENSMUSG00 | Mir708 | 108 | 0 | 0 | 0 | 0 | 0 | 0 |
| ENSMUSG00 | Mir709 | 87 | 0 | 0 | 0 | 0 | 0 | 0 |
| ENSMUSG00 | Mir679 | 73 | 0 | 0 | 0 | 0 | 0 | 0 |
| ENSMUSG00 | Mir449c | 108 | 0 | 0 | 0 | 0 | 0 | 0 |
| ENSMUSG00 | Mir694 | 69 | 0 | 0 | 0 | 0 | 0 | 0 |
| ENSMUSG00 | mmu-mir-46 | 82 | 0 | 0 | 0 | 0 | 0 | 0 |
| ENSMUSG00 | Mir467a-9 | 82 | 0 | 0 | 0 | 0 | 0 | 0 |
| ENSMUSG00 | Mir467a-8 | 82 | 0 | 0 | 0 | 0 | 0 | 0 |
| ENSMUSG00 | Mir467a-7 | 82 | 0 | 0 | 0 | 0 | 0 | 0 |
| ENSMUSG00 | Mir467a-5 | 82 | 0 | 0 | 0 | 0 | 0 | 0 |
| ENSMUSG00 | Mir467a-4 | 82 | 0 | 0 | 0 | 0 | 0 | 0 |
| ENSMUSG00 | Mir467a-2 | 82 | 0 | 0 | 0 | 0 | 0 | 0 |
| ENSMUSG00 | Mir467a-1 | 72 | 0 | 0 | 0 | 0 | 0 | 0 |
| ENSMUSG00 | Mir701 | 108 | 0 | 0 | 0 | 0 | 0 | 0.25254296 |
| ENSMUSG00 | AC102703.1 | 109 | 0 | 0 | 0 | 0 | 0 | 0 |
| ENSMUSG00 | AC153658.1 | 108 | 0 | 0 | 0 | 0 | 0 | 0 |
| ENSMUSG00 | AC102628.1 | 109 | 0 | 0 | 0 | 0 | 0 | 0 |
| ENSMUSG00 | AC102561.1 | 109 | 0 | 0 | 0 | 0 | 0 | 0 |
| ENSMUSG00 | AC138317.1 | 109 | 0 | 0 | 0 | 0 | 0 | 0 |
| ENSMUSG00 | AC112681.1 | 109 | 0 | 0 | 0 | 0 | 0 | 0 |
| ENSMUSG00 | Mir680-1 | 108 | 0 | 0 | 0 | 0 | 0 | 0 |
| ENSMUSG00 | Mir717 | 108 | 0 | 0 | 0 | 0 | 0 | 0 |
| ENSMUSG00 | Mir695 | 108 | 0 | 0 | 0 | 0 | 1.88986577 | 1.01017182 |
| ENSMUSG00 | Mir706 | 83 | 0 | 0 | 0 | 0 | 0 | 0 |
| ENSMUSG00 | Mir692-1 | 108 | 0 | 0 | 0 | 0 | 0.47246644 | 0 |
| ENSMUSG00 | Mir487b | 81 | 0 | 0 | 0 | 0 | 0 | 0 |
| ENSMUSG00 | Mir297b | 108 | 0 | 0 | 0 | 0 | 0 | 0 |
| ENSMUSG00 | Mir297-1 | 75 | 0 | 0 | 0 | 0 | 0 | 0 |
| ENSMUSG00 | AL590969.1 | 95 | 0 | 0 | 0 | 0 | 0 | 0 |
| ENSMUSG00 | AC159474.1 | 96 | 0 | 0 | 0 | 0 | 0 | 0 |
| ENSMUSG00 | AC155261.1 | 81 | 0 | 0 | 0 | 0 | 0 | 0 |
| ENSMUSG00 | AL928935.1 | 95 | 0 | 0 | 0 | 0 | 0 | 0 |
| ENSMUSG00 | AL929404.1 | 95 | 0 | 0 | 0 | 0 | 0 | 0 |
| ENSMUSG00 | AL596117.1 | 92 | 0 | 0 | 0 | 0 | 0 | 0 |
| ENSMUSG00 | AL928544.1 | 95 | 0 | 0 | 0 | 0 | 0 | 0 |
| ENSMUSG00 | AL928544.2 | 95 | 0 | 0 | 0 | 0 | 0 | 0 |
| ENSMUSG00 | Mir682 | 95 | 0 | 0 | 0 | 0 | 0 | 0 |
| ENSMUSG00 | AL590614.1 | 93 | 0 | 0 | 0 | 0 | 0 | 0 |
| ENSMUSG00 | AC122372.1 | 93 | 0 | 0 | 0 | 0 | 0 | 0 |
| ENSMUSG00 | Mir702 | 108 | 0 | 0 | 0 | 0 | 0 | 0 |
| ENSMUSG00 | Mir543 | 79 | 0 | 0 | 0 | 0 | 0 | 0 |
| ENSMUSG00 | CT030240.1 | 107 | 0 | 0 | 0 | 0 | 0 | 0 |
| ENSMUSG00 | AC122357.1 | 106 | 0.3750517 | 0 | 0 | 0 | 0 | 0 |
| ENSMUSG00 | CT573034.1 | 107 | 0 | 0 | 0 | 0 | 0 | 0 |
| ENSMUSG00 | AC159290.1 | 107 | 0 | 0 | 0 | 0 | 0 | 0 |
| ENSMUSG00 | Mir713 | 107 | 0 | 0 | 0 | 0 | 0 | 1.01961268 |
| ENSMUSG00 | Mir501 | 108 | 0 | 0 | 0 | 0 | 0 | 0 |
| ENSMUSG00 | Mir302b | 73 | 0 | 0 | 0 | 0 | 0 | 1.12087559 |
| ENSMUSG00 | Mir680-3 | 86 | 0 | 0 | 0 | 0 | 0 | 0 |
| ENSMUSG00 | Mir92b | 82 | 0 | 0 | 0 | 0 | 0 | 0 |
| ENSMUSG00 | Mir19b-1 | 86 | 0 | 0 | 0 | 0 | 0 | 0 |
| ENSMUSG00 | mmu-mir-68 | 56 | 200.90716 | 216.83899 | 209.169436 | 164.946489 | 148.523201 | 224.528727 |
| ENSMUSG00 | Mir374 | 94 | 0 | 0 | 0 | 1.15155461 | 0 | 0 |
| ENSMUSG00 | Mir719 | 109 | 0 | 0 | 0 | 0 | 0 | 0 |
| ENSMUSG00 | Mir697 | 108 | 1.1043189 | 0 | 2.45565795 | 2.67274404 | 2.36233221 | 0 |
| ENSMUSG00 | Mir666 | 98 | 0 | 0 | 0 | 0 | 0 | 0 |
| ENSMUSG00 | AC134908.1 | 91 | 0 | 0 | 0 | 0 | 0 | 0 |
| ENSMUSG00 | Mir675 | 83 | 0 | 0 | 0 | 0 | 0 | 0 |
| ENSMUSG00 | mmu-mir-21 | 56 | 0.7099193 | 0 | 0 | 0 | 0 | 0.97409426 |
| ENSMUSG00 | AC156273.1 | 110 | 0 | 0 | 0 | 0 | 0 | 0 |
| ENSMUSG00 | AC134452.1 | 97 | 4.50835346 | 16.8900931 | 6.37964745 | 8.55553633 | 7.89067667 | 1.40590924 |
| ENSMUSG00 | Mir672 | 99 | 0 | 0 | 0 | 0 | 0 | 0 |
| ENSMUSG00 | AC109299.1 | 163 | 0 | 0 | 0 | 0 | 0 | 0 |
| ENSMUSG00 | Mir301b | 96 | 0 | 0 | 0 | 0 | 0 | 0 |
| ENSMUSG00 | AC122184.1 | 105 | 0 | 0 | 0 | 0 | 0 | 0 |
| ENSMUSG00 | AL627124.1 | 95 | 0 | 0 | 0 | 0 | 0 | 0 |
| ENSMUSG00 | CR974423.1 | 102 | 0 | 0 | 0 | 0 | 0 | 0 |
| ENSMUSG00 | AC157912.1 | 108 | 0 | 0 | 0 | 0 | 0 | 0 |
| ENSMUSG00 | AC133509.1 | 115 | 0 | 0 | 0 | 0 | 0 | 0 |
| ENSMUSG00 | CT573086.1 | 109 | 0 | 0 | 0 | 0 | 0 | 0 |
| ENSMUSG00 | AC133509.2 | 92 | 0 | 0 | 0 | 0 | 0 | 0 |
| ENSMUSG00 | Mir147 | 78 | 0 | 0 | 0 | 0 | 0 | 0 |
| ENSMUSG00 | Mir499 | 78 | 0 | 0 | 0 | 0 | 0 | 0 |
| ENSMUSG00 | Mir665 | 93 | 0 | 0 | 0 | 0 | 0 | 0 |
| ENSMUSG00 | Mir343 | 74 | 0 | 0 | 0 | 0 | 0 | 0 |
| ENSMUSG00 | Mir673 | 90 | 0 | 0 | 0 | 0 | 0 | 0 |
| ENSMUSG00 | Mir216b | 85 | 0 | 0 | 0 | 0 | 0 | 0 |
| ENSMUSG00 | AC167419.1 | 98 | 0.40566817 | 0 | 0 | 0.36818413 | 0 | 0 |
| ENSMUSG00 | AL603804.1 | 98 | 0 | 0 | 0 | 0 | 0 | 0 |
| ENSMUSG00 | AC144778.1 | 109 | 0.36472918 | 0 | 0 | 0 | 0 | 0 |
| ENSMUSG00 | AC151412.1 | 94 | 0 | 0 | 0 | 0 | 0 | 0 |
| ENSMUSG00 | Mir599 | 87 | 2.28479773 | 0.55386715 | 0 | 0 | 0 | 0 |
| ENSMUSG00 | Mir670 | 99 | 0 | 0 | 0 | 0 | 0 | 0 |
| ENSMUSG00 | BX255277.1 | 100 | 0 | 0 | 0 | 0 | 0 | 0 |
| ENSMUSG00 | Mir181d | 71 | 0 | 0 | 0 | 0 | 0 | 0 |
| ENSMUSG00 | AC137692.1 | 112 | 0 | 0 | 0 | 0 | 0 | 0 |
| ENSMUSG00 | AC151983.1 | 100 | 0 | 0 | 0 | 0 | 0 | 0 |
| ENSMUSG00 | AL607145.1 | 89 | 3.12683555 | 0 | 0 | 0 | 0 | 0 |
| ENSMUSG00 | Mir653 | 84 | 0 | 0 | 0 | 0 | 0 | 0 |
| ENSMUSG00 | AC171202.1 | 103 | 0 | 0 | 0 | 0 | 0 | 0 |
| ENSMUSG00 | Mir182 | 74 | 0 | 0 | 0 | 0 | 0 | 0 |
| ENSMUSG00 | AL731552.1 | 99 | 0 | 0.48673174 | 0 | 0 | 0 | 0 |
| ENSMUSG00 | AC099576.1 | 105 | 0 | 0 | 0 | 0 | 0 | 0 |
| ENSMUSG00 | AC132253.1 | 103 | 0 | 0 | 0 | 0 | 0 | 0 |
| ENSMUSG00 | AL671993.1 | 108 | 0 | 0 | 0 | 0 | 0 | 0 |
| ENSMUSG00 | Mir568 | 82 | 0 | 0 | 0 | 0 | 0 | 0 |
| ENSMUSG00 | Mir674 | 99 | 0 | 0 | 0 | 0 | 0.51541794 | 0 |
| ENSMUSG00 | AC153972.1 | 82 | 0 | 0 | 0 | 0 | 0 | 0 |
| ENSMUSG00 | Mir190 | 84 | 0 | 0 | 0 | 0 | 0 | 0 |
| ENSMUSG00 | AC163625.1 | 81 | 0 | 0 | 0 | 0.44545734 | 0 | 0 |
| ENSMUSG00 | AC114901.1 | 105 | 0 | 0 | 0 | 0 | 0 | 0 |
| ENSMUSG00 | Mir671 | 97 | 0 | 0 | 0 | 0 | 0 | 0 |
| ENSMUSG00 | AC099601.1 | 95 | 0 | 0 | 0 | 0 | 0 | 0 |
| ENSMUSG00 | AC116121.1 | 108 | 0 | 0 | 0 | 0 | 0 | 0 |
| ENSMUSG00 | AL663045.1 | 98 | 0 | 0 | 0 | 0 | 0 | 0 |
| ENSMUSG00 | AC113593.1 | 85 | 0 | 0 | 0 | 0 | 0 | 0 |
| ENSMUSG00 | Mir667 | 91 | 0 | 0 | 0 | 0 | 0 | 0 |
| ENSMUSG00 | Mir676 | 88 | 0 | 0 | 0 | 0 | 0 | 0 |
| ENSMUSG00 | Mir450-2 | 68 | 0 | 0 | 0 | 0 | 0 | 0 |
| ENSMUSG00 | CT010465.1 | 109 | 0 | 0 | 0 | 0 | 0 | 0 |
| ENSMUSG00 | AC103388.1 | 68 | 0 | 0 | 0 | 0 | 0 | 0 |
| ENSMUSG00 | AC154309.1 | 67 | 0 | 0 | 0 | 0 | 0 | 0 |
| ENSMUSG00 | BX255946.1 | 108 | 0 | 0 | 0 | 0 | 0 | 0 |
| ENSMUSG00 | Hus1b | 1186 | 0.36872705 | 0 | 0 | 0.12169324 | 0 | 0.16098016 |
| ENSMUSG00 | Sox4 | 5703 | 9.08318273 | 12.9021037 | 16.0825573 | 10.9707637 | 9.34096727 | 9.48371202 |
| ENSMUSG00 | Ywhaq | 4139 | 42.0991232 | 47.20851 | 39.8873843 | 43.7012054 | 34.0874436 | 44.9349517 |
| ENSMUSG00 | BC100451 | 3542 | 0.04489608 | 0 | 0 | 0 | 0 | 0 |
| ENSMUSG00 | Wfdc3 | 2524 | 0 | 0 | 0 | 0 | 0 | 0 |
| ENSMUSG00 | Acsf2 | 3415 | 0.08149 | 0.23987394 | 0.20709502 | 0.23244655 | 0.1195347 | 0.20765465 |
| ENSMUSG00 | Oxct2a | 1759 | 0.20341065 | 0 | 0.25128961 | 0.26666682 | 0.29008741 | 0.27910376 |
| ENSMUSG00 | 2700094K13 | 996 | 2.35499332 | 5.75721548 | 7.4113532 | 5.3978159 | 4.14973538 | 8.18786861 |
| ENSMUSG00 | Oxct2b | 1685 | 0 | 0 | 0 | 0 | 0 | 0 |
| ENSMUSG00 | Mog | 1706 | 0 | 0 | 0 | 0 | 0 | 0 |
| ENSMUSG00 | Ifna5 | 569 | 0 | 0 | 0 | 0 | 0 | 0 |
| ENSMUSG00 | Ass1 | 2291 | 0.24294052 | 0.27342809 | 0.03858738 | 0.11024632 | 0.0668176 | 0.48811009 |
| ENSMUSG00 | Mir761 | 75 | 0 | 0 | 0 | 0 | 0 | 0 |
| ENSMUSG00 | AC119211.1 | 95 | 0 | 0 | 0 | 0 | 0 | 0 |
| ENSMUSG00 | Mir763 | 119 | 0 | 0 | 0 | 0 | 0 | 0 |
| ENSMUSG00 | Mir804 | 94 | 0 | 0 | 0 | 0 | 0 | 0 |
| ENSMUSG00 | Mir770 | 93 | 0 | 0 | 0 | 0 | 0 | 0 |
| ENSMUSG00 | Mir759 | 97 | 0 | 0 | 0 | 0 | 0 | 0 |
| ENSMUSG00 | Mir762 | 75 | 0 | 0 | 0 | 0 | 0 | 0 |
| ENSMUSG00 | Mir767 | 106 | 0 | 0 | 0 | 0 | 0 | 0 |
| ENSMUSG00 | Mir760 | 118 | 0 | 0 | 0 | 0 | 0 | 0 |
| ENSMUSG00 | Mir802 | 96 | 0 | 0 | 0 | 0 | 0 | 0 |
| ENSMUSG00 | Mir764 | 107 | 0 | 0 | 0 | 0 | 0 | 0 |
| ENSMUSG00 | Mir758 | 80 | 0 | 0 | 0 | 0 | 0 | 0 |
| ENSMUSG00 | Mir744 | 99 | 0 | 0 | 0 | 0 | 0 | 0 |
| ENSMUSG00 | Trbv1 | 348 | 0 | 0 | 0 | 0 | 0 | 0 |
| ENSMUSG00 | Trbv2 | 342 | 0 | 0 | 0 | 0 | 0 | 0 |
| ENSMUSG00 | Trbv3 | 345 | 0 | 0 | 0 | 0 | 0 | 0 |
| ENSMUSG00 | Trbv4 | 337 | 0 | 0 | 0 | 0 | 0 | 0 |
| ENSMUSG00 | Trbv5 | 339 | 0 | 0 | 0 | 0 | 0 | 0 |
| ENSMUSG00 | Trbv12-1 | 370 | 0 | 0 | 0 | 0 | 0 | 0 |
| ENSMUSG00 | Trbv13-1 | 332 | 0 | 0 | 0 | 0 | 0 | 0 |
| ENSMUSG00 | Trbv12-2 | 370 | 0 | 0 | 0 | 0 | 0 | 0 |
| ENSMUSG00 | Trbv13-2 | 317 | 0 | 0 | 0 | 0 | 0 | 0 |
| ENSMUSG00 | Trbv13-3 | 338 | 0 | 0 | 0 | 0 | 0 | 0 |
| ENSMUSG00 | Trbv14 | 337 | 0 | 0 | 0 | 0 | 0 | 0 |
| ENSMUSG00 | Trbv15 | 345 | 0 | 0 | 0 | 0 | 0 | 0 |
| ENSMUSG00 | Trbv16 | 340 | 0 | 0 | 0 | 0 | 0 | 0 |
| ENSMUSG00 | Trbv17 | 337 | 0 | 0 | 0 | 0 | 0 | 0 |
| ENSMUSG00 | Trbv19 | 350 | 0 | 0 | 0 | 0 | 0 | 0 |
| ENSMUSG00 | Trbv20 | 333 | 0 | 0 | 0 | 0 | 0 | 0 |
| ENSMUSG00 | Trbv21 | 289 | 0 | 0 | 0 | 0 | 0 | 0 |
| ENSMUSG00 | Trbv23 | 342 | 0 | 0 | 0 | 0 | 0 | 0 |
| ENSMUSG00 | Trbv26 | 342 | 0 | 0 | 0 | 0 | 0 | 0 |
| ENSMUSG00 | Trbv29 | 342 | 0 | 0 | 0 | 0 | 0 | 0 |
| ENSMUSG00 | Trbv30 | 330 | 0 | 0 | 0 | 0 | 0 | 0 |
| ENSMUSG00 | Trbd1 | 11 | 0 | 0 | 0 | 0 | 0 | 0 |
| ENSMUSG00 | Trbj1-1 | 47 | 0 | 0 | 0 | 0 | 0 | 0 |
| ENSMUSG00 | Trbj1-2 | 47 | 0 | 0 | 0 | 0 | 0 | 0 |
| ENSMUSG00 | Trbj1-3 | 53 | 0 | 0 | 0 | 0 | 0 | 0 |
| ENSMUSG00 | Trbj1-4 | 50 | 0 | 0 | 0 | 0 | 0 | 0 |
| ENSMUSG00 | Trbj1-5 | 49 | 0 | 0 | 0 | 0 | 0 | 0 |
| ENSMUSG00 | Trbj1-6 | 52 | 0 | 0 | 0 | 0 | 0 | 0 |
| ENSMUSG00 | Trbj1-7 | 44 | 0 | 0 | 0 | 0 | 0 | 0 |
| ENSMUSG00 | Trbc1 | 509 | 0 | 0 | 0 | 0 | 0 | 0 |
| ENSMUSG00 | Trbd2 | 13 | 0 | 0 | 0 | 0 | 0 | 0 |
| ENSMUSG00 | Trbj2-1 | 49 | 0 | 0 | 0 | 0 | 0 | 0 |
| ENSMUSG00 | Trbj2-2 | 50 | 0 | 0 | 0 | 0 | 0 | 0 |
| ENSMUSG00 | Trbj2-3 | 48 | 0 | 0 | 0 | 0 | 0 | 0 |
| ENSMUSG00 | Trbj2-4 | 48 | 0 | 0 | 0 | 0 | 0 | 0 |
| ENSMUSG00 | Trbj2-5 | 48 | 0 | 0 | 0 | 0 | 0 | 0 |
| ENSMUSG00 | Trbj2-7 | 46 | 0 | 0 | 0 | 0 | 0 | 0 |
| ENSMUSG00 | Trbc2 | 517 | 0 | 0 | 0 | 0 | 0 | 0 |
| ENSMUSG00 | Trbv31 | 332 | 0 | 0 | 0 | 0 | 0 | 0 |
| ENSMUSG00 | AC158671.1 | 357 | 0 | 0 | 0 | 0 | 0 | 0 |
| ENSMUSG00 | Igkv2-137 | 360 | 0.44172756 | 1.07080983 | 1.22782897 | 0.50113951 | 0.28347987 | 0.30305155 |
| ENSMUSG00 | Igkv1-135 | 360 | 0 | 0 | 0 | 0 | 0 | 0 |
| ENSMUSG00 | Igkv1-133 | 360 | 0 | 0 | 0 | 0 | 0 | 0 |
| ENSMUSG00 | Igkv1-132 | 360 | 0 | 0 | 0 | 0 | 0 | 0 |
| ENSMUSG00 | Igkv1-131 | 360 | 0 | 0 | 0 | 0 | 0 | 0 |
| ENSMUSG00 | Igkv14-130 | 348 | 0 | 0 | 0 | 0 | 0 | 0 |
| ENSMUSG00 | Igkv9-129 | 348 | 0 | 0 | 0 | 0 | 0 | 0 |
| ENSMUSG00 | Igkv17-127 | 339 | 0 | 0 | 0 | 0 | 0 | 0 |
| ENSMUSG00 | Igkv14-126 | 351 | 0 | 0 | 0 | 0 | 0 | 0 |
| ENSMUSG00 | Igkv11-125 | 348 | 0 | 0 | 0 | 0 | 0 | 0 |
| ENSMUSG00 | Igkv9-124 | 286 | 0 | 0 | 0 | 0 | 0 | 0 |
| ENSMUSG00 | Igkv9-123 | 351 | 0 | 0 | 0 | 0 | 0 | 0 |
| ENSMUSG00 | Igkv1-122 | 357 | 0 | 0 | 0 | 0 | 0 | 0 |
| ENSMUSG00 | Igkv17-121 | 348 | 0 | 0 | 0 | 0 | 0 | 0 |
| ENSMUSG00 | Igkv9-120 | 351 | 0 | 0 | 0 | 0 | 0 | 0 |
| ENSMUSG00 | Igkv1-117 | 357 | 0 | 0 | 0 | 0 | 0 | 0 |
| ENSMUSG00 | Igkv2-116 | 362 | 0 | 0 | 0 | 0 | 0 | 0 |
| ENSMUSG00 | Igkv2-112 | 356 | 0 | 0 | 0 | 0 | 0 | 0 |
| ENSMUSG00 | Igkv14-111 | 351 | 0 | 0 | 0 | 0 | 0 | 0 |
| ENSMUSG00 | Igkv1-110 | 357 | 0 | 0 | 0 | 0 | 0 | 0 |
| ENSMUSG00 | Igkv2-109 | 362 | 0 | 0 | 0 | 0 | 0 | 0 |
| ENSMUSG00 | Igkv16-104 | 346 | 0 | 0 | 0 | 0 | 0 | 0 |
| ENSMUSG00 | Igkv15-103 | 345 | 0 | 0 | 0 | 0 | 0 | 0 |
| ENSMUSG00 | Igkv14-100 | 348 | 0 | 0 | 0 | 0 | 0 | 0 |
| ENSMUSG00 | Igkv1-99 | 357 | 0 | 0 | 0 | 0 | 0 | 0 |
| ENSMUSG00 | Igkv12-98 | 342 | 0 | 0 | 0 | 0 | 0 | 0 |
| ENSMUSG00 | Igkv10-96 | 345 | 0 | 0 | 0 | 0 | 0 | 0 |
| ENSMUSG00 | Igkv10-95 | 286 | 0 | 0 | 0 | 0 | 0 | 0 |
| ENSMUSG00 | Igkv10-94 | 345 | 0 | 0 | 0 | 0 | 0 | 0 |
| ENSMUSG00 | Igkv19-93 | 342 | 0 | 0 | 0 | 0 | 0 | 0 |
| ENSMUSG00 | Igkv4-92 | 349 | 0 | 0 | 0 | 0 | 0 | 0 |
| ENSMUSG00 | Igkv4-91 | 343 | 0 | 0 | 0 | 0 | 0 | 0 |
| ENSMUSG00 | Igkv4-90 | 324 | 0 | 0 | 0 | 0 | 0 | 0 |
| ENSMUSG00 | Igkv12-89 | 333 | 0 | 0 | 0 | 0 | 0 | 0 |
| ENSMUSG00 | Igkv1-88 | 360 | 0 | 0 | 0 | 0 | 0 | 0 |
| ENSMUSG00 | Igkv4-86 | 338 | 0 | 0 | 0 | 0 | 0 | 0 |
| ENSMUSG00 | Igkv13-84 | 345 | 0 | 0 | 0 | 0 | 0 | 0 |
| ENSMUSG00 | Igkv4-81 | 289 | 0 | 0 | 0 | 0 | 0 | 0 |
| ENSMUSG00 | Igkv4-80 | 344 | 0 | 0 | 0 | 0 | 0 | 0 |
| ENSMUSG00 | Igkv4-79 | 350 | 0 | 0 | 0 | 0 | 0 | 0 |
| ENSMUSG00 | Igkv4-78 | 354 | 0 | 0 | 0 | 0 | 0 | 0 |
| ENSMUSG00 | Igkv4-74 | 356 | 0 | 0 | 0 | 0 | 0 | 0 |
| ENSMUSG00 | Igkv4-73 | 354 | 0 | 0 | 0 | 0 | 0 | 0 |
| ENSMUSG00 | Igkv4-72 | 350 | 0 | 0 | 0 | 0 | 0 | 0 |
| ENSMUSG00 | Igkv4-70 | 346 | 0 | 0 | 0 | 0 | 0 | 0 |
| ENSMUSG00 | Igkv4-69 | 346 | 0 | 0 | 0 | 0 | 0 | 0 |
| ENSMUSG00 | Igkv4-68 | 350 | 0 | 0 | 0 | 0 | 0 | 0 |
| ENSMUSG00 | Igkv4-63 | 285 | 0 | 0 | 0 | 0 | 0 | 0 |
| ENSMUSG00 | Igkv4-62 | 278 | 0 | 0 | 0 | 0 | 0 | 0 |
| ENSMUSG00 | Igkv4-61 | 283 | 0 | 0 | 0 | 0 | 0 | 0 |
| ENSMUSG00 | Igkv4-59 | 350 | 0 | 0 | 0 | 0 | 0 | 0 |
| ENSMUSG00 | Igkv4-58 | 291 | 0 | 0 | 0 | 0 | 0 | 0 |
| ENSMUSG00 | Igkv4-57-1 | 356 | 0 | 0 | 0 | 0 | 0 | 0 |
| ENSMUSG00 | Igkv4-57 | 348 | 0 | 0 | 0 | 0 | 0 | 0 |
| ENSMUSG00 | Igkv4-56 | 285 | 0 | 0 | 0 | 0 | 0 | 0 |
| ENSMUSG00 | Igkv4-55 | 350 | 0 | 0 | 0 | 0 | 0 | 0 |
| ENSMUSG00 | Igkv4-50 | 350 | 0 | 0 | 0 | 0 | 0 | 0 |
| ENSMUSG00 | Igkv5-48 | 345 | 0 | 0 | 0 | 0 | 0 | 0 |
| ENSMUSG00 | Igkv12-46 | 345 | 0 | 0 | 0 | 0 | 0 | 0 |
| ENSMUSG00 | Igkv5-45 | 345 | 0 | 0 | 0 | 0 | 0 | 0 |
| ENSMUSG00 | Igkv12-44 | 345 | 0 | 0 | 0 | 0 | 0 | 0 |
| ENSMUSG00 | Igkv5-43 | 345 | 0 | 0 | 0 | 0 | 0 | 0 |
| ENSMUSG00 | Igkv12-41 | 345 | 0 | 0 | 0 | 0 | 0 | 0 |
| ENSMUSG00 | Igkv5-39 | 286 | 0 | 0 | 0 | 0 | 0 | 0 |
| ENSMUSG00 | Igkv12-38 | 342 | 0 | 0 | 0 | 0 | 0 | 0 |
| ENSMUSG00 | Igkv5-37 | 345 | 0 | 0 | 0 | 0 | 0 | 0 |
| ENSMUSG00 | Igkv18-36 | 342 | 0 | 0 | 0 | 0 | 0 | 0 |
| ENSMUSG00 | Igkv1-35 | 357 | 0 | 0 | 0 | 0 | 0 | 0 |
| ENSMUSG00 | Igkv7-33 | 360 | 0 | 0 | 0 | 0 | 0 | 0 |
| ENSMUSG00 | Igkv6-32 | 345 | 0 | 0 | 0 | 0 | 0 | 0 |
| ENSMUSG00 | Igkv8-30 | 363 | 0 | 0 | 0 | 0 | 0 | 0 |
| ENSMUSG00 | Igkv6-29 | 342 | 0 | 0 | 0 | 0 | 0 | 0 |
| ENSMUSG00 | Igkv8-28 | 304 | 0 | 0 | 0 | 0 | 0 | 0 |
| ENSMUSG00 | Igkv8-27 | 299 | 0 | 0 | 0 | 0 | 0 | 0 |
| ENSMUSG00 | Igkv8-26 | 369 | 0 | 0 | 0 | 0 | 0 | 0 |
| ENSMUSG00 | Igkv6-25 | 345 | 0 | 0 | 0 | 0 | 0 | 0 |
| ENSMUSG00 | Igkv8-24 | 363 | 0 | 0 | 0 | 0 | 0 | 0 |
| ENSMUSG00 | Igkv6-23 | 345 | 0 | 0 | 0 | 0 | 0 | 0 |
| ENSMUSG00 | Igkv8-21 | 360 | 0 | 0 | 0 | 0 | 0 | 0 |
| ENSMUSG00 | Igkv6-20 | 372 | 0 | 0 | 0 | 0 | 0 | 0 |
| ENSMUSG00 | Igkv8-19 | 304 | 0 | 0 | 0 | 0 | 0 | 0 |
| ENSMUSG00 | Igkv8-18 | 368 | 0 | 0 | 0 | 0 | 0 | 0 |
| ENSMUSG00 | Igkv6-17 | 345 | 0 | 0 | 0 | 0 | 0 | 0 |
| ENSMUSG00 | Igkv8-16 | 360 | 0 | 0 | 0 | 0 | 0 | 0 |
| ENSMUSG00 | Igkv6-15 | 345 | 0 | 0 | 0 | 0 | 0 | 0 |
| ENSMUSG00 | Igkv6-14 | 345 | 0 | 0 | 0 | 0 | 0 | 0 |
| ENSMUSG00 | Igkv6-13 | 286 | 0 | 0 | 0 | 0 | 0 | 0 |
| ENSMUSG00 | Igkv3-12 | 357 | 0 | 0 | 0 | 0 | 0 | 0 |
| ENSMUSG00 | Igkv3-10 | 354 | 0 | 0 | 0 | 0 | 0 | 0 |
| ENSMUSG00 | Igkv3-9 | 357 | 0 | 0 | 0 | 0 | 0 | 0 |
| ENSMUSG00 | Igkv3-7 | 298 | 0 | 0 | 0 | 0 | 0 | 0 |
| ENSMUSG00 | Igkv3-5 | 357 | 0 | 0 | 0 | 0 | 0 | 0 |
| ENSMUSG00 | Igkv3-4 | 357 | 0 | 0 | 0 | 0 | 0 | 0 |
| ENSMUSG00 | Igkv3-3 | 357 | 0 | 0 | 0 | 0 | 0 | 0 |
| ENSMUSG00 | Igkv3-2 | 357 | 0 | 0 | 0 | 0 | 0 | 0 |
| ENSMUSG00 | Igkv3-1 | 357 | 0 | 0 | 0 | 0 | 0 | 0 |
| ENSMUSG00 | Igkj1 | 37 | 0 | 0 | 0 | 0 | 0 | 0 |
| ENSMUSG00 | Igkj2 | 42 | 0 | 0 | 0 | 0 | 0 | 0 |
| ENSMUSG00 | Igkj3 | 35 | 0 | 0 | 0 | 0 | 0 | 0 |
| ENSMUSG00 | Igkj4 | 37 | 0 | 0 | 0 | 0 | 0 | 0 |
| ENSMUSG00 | Igkj5 | 41 | 0 | 0 | 0 | 0 | 0 | 0 |
| ENSMUSG00 | Igkc | 322 | 0 | 0 | 0 | 0 | 0 | 0 |
| ENSMUSG00 | Ighe | 1163 | 0 | 0 | 0 | 0 | 0 | 0 |
| ENSMUSG00 | Ighg2c | 1206 | 0 | 0 | 0 | 0 | 0 | 0 |
| ENSMUSG00 | Ighg2b | 1006 | 0 | 0 | 0 | 0 | 0 | 0 |
| ENSMUSG00 | Ighg1 | 970 | 0 | 0 | 0 | 0 | 0 | 0 |
| ENSMUSG00 | Ighg3 | 988 | 0 | 0 | 0 | 0 | 0 | 0 |
| ENSMUSG00 | Ighm | 1363 | 0 | 0 | 0 | 0 | 0 | 0.22011815 |
| ENSMUSG00 | Ighj4 | 53 | 0 | 0 | 0 | 0 | 0 | 0 |
| ENSMUSG00 | Ighj3 | 47 | 0 | 0 | 0 | 0 | 0 | 0 |
| ENSMUSG00 | Ighj2 | 53 | 0 | 0 | 0 | 0 | 0 | 0 |
| ENSMUSG00 | Ighj1 | 52 | 0 | 0 | 0 | 0 | 0 | 0 |
| ENSMUSG00 | Ighd4-1 | 10 | 0 | 0 | 0 | 0 | 0 | 0 |
| ENSMUSG00 | Ighd3-2 | 15 | 0 | 0 | 0 | 0 | 0 | 0 |
| ENSMUSG00 | Ighd2-8 | 16 | 0 | 0 | 0 | 0 | 0 | 0 |
| ENSMUSG00 | Ighd2-7 | 16 | 0 | 0 | 0 | 0 | 0 | 0 |
| ENSMUSG00 | Ighd2-6 | 16 | 0 | 0 | 0 | 0 | 0 | 0 |
| ENSMUSG00 | Ighd2-5 | 16 | 0 | 0 | 0 | 0 | 0 | 0 |
| ENSMUSG00 | Ighd2-4 | 16 | 0 | 0 | 0 | 0 | 0 | 0 |
| ENSMUSG00 | Ighd2-3 | 16 | 0 | 0 | 0 | 0 | 0 | 0 |
| ENSMUSG00 | Ighd1-1 | 22 | 0 | 0 | 0 | 0 | 0 | 0 |
| ENSMUSG00 | Ighd3-1 | 16 | 0 | 0 | 0 | 0 | 0 | 0 |
| ENSMUSG00 | Gm16968 | 22 | 0 | 0 | 0 | 0 | 0 | 0 |
| ENSMUSG00 | Ighv5-2 | 348 | 0 | 0 | 0 | 0 | 0 | 0 |
| ENSMUSG00 | Ighv2-2 | 348 | 0 | 0 | 0 | 0 | 0 | 0 |
| ENSMUSG00 | Ighv5-4 | 351 | 0 | 0 | 0 | 0 | 0 | 0 |
| ENSMUSG00 | Ighv2-3 | 345 | 0 | 0 | 0 | 0 | 0 | 0 |
| ENSMUSG00 | Ighv5-6 | 351 | 0 | 0 | 0 | 0 | 0 | 0 |
| ENSMUSG00 | Ighv2-4 | 348 | 0 | 0 | 0 | 0 | 0 | 0 |
| ENSMUSG00 | Ighv5-9 | 351 | 0 | 0 | 0 | 0 | 0 | 0 |
| ENSMUSG00 | Ighv2-5 | 345 | 0 | 0 | 0 | 0 | 0 | 0 |
| ENSMUSG00 | Ighv5-12 | 348 | 0 | 0 | 0 | 0 | 0 | 0 |
| ENSMUSG00 | Ighv2-6 | 343 | 0 | 0 | 0 | 0 | 0 | 0 |
| ENSMUSG00 | Ighv5-9-1 | 348 | 0 | 0 | 0 | 0 | 0 | 0 |
| ENSMUSG00 | Ighv2-9-1 | 345 | 0 | 0 | 0 | 0 | 0 | 0 |
| ENSMUSG00 | Ighv2-6-8 | 340 | 0 | 0 | 0 | 0 | 0 | 0 |
| ENSMUSG00 | Ighv2-7 | 348 | 0 | 0 | 0 | 0 | 0 | 0 |
| ENSMUSG00 | Ighv5-15 | 348 | 0 | 0 | 0 | 0 | 0 | 0 |
| ENSMUSG00 | Ighv5-16 | 348 | 0 | 0 | 0 | 0 | 0 | 0 |
| ENSMUSG00 | Ighv5-17 | 346 | 0 | 0 | 0 | 0 | 0 | 0 |
| ENSMUSG00 | Ighv2-9 | 345 | 0 | 0 | 0 | 0 | 0 | 0 |
| ENSMUSG00 | Ighv7-1 | 355 | 0 | 0 | 0 | 0 | 0 | 0 |
| ENSMUSG00 | Ighv7-2 | 352 | 0 | 0 | 0 | 0 | 0 | 0 |
| ENSMUSG00 | Ighv14-1 | 349 | 0 | 0 | 0 | 0 | 0 | 0 |
| ENSMUSG00 | Ighv4-1 | 348 | 0 | 0 | 0 | 0 | 0 | 0 |
| ENSMUSG00 | Ighv11-1 | 351 | 0 | 0 | 0 | 0 | 0 | 0 |
| ENSMUSG00 | Ighv14-2 | 349 | 0 | 0 | 0 | 0 | 0 | 0 |
| ENSMUSG00 | Ighv11-2 | 351 | 0 | 0 | 0 | 0 | 0 | 0 |
| ENSMUSG00 | Ighv14-3 | 349 | 0 | 0 | 0 | 0 | 0 | 0 |
| ENSMUSG00 | Ighv16-1 | 298 | 0 | 0 | 0 | 0 | 0 | 0 |
| ENSMUSG00 | Ighv9-1 | 293 | 0 | 0 | 0 | 0 | 0 | 0 |
| ENSMUSG00 | Ighv9-2 | 346 | 0 | 0 | 0 | 0 | 0 | 0 |
| ENSMUSG00 | Ighv9-3 | 349 | 0 | 0 | 0 | 0 | 0 | 0 |
| ENSMUSG00 | Ighv7-3 | 352 | 0 | 0 | 0 | 0 | 0 | 0 |
| ENSMUSG00 | Ighv14-4 | 331 | 0 | 0 | 0 | 0 | 0 | 0 |
| ENSMUSG00 | Ighv3-3 | 295 | 0 | 0 | 0 | 0 | 0 | 0 |
| ENSMUSG00 | Ighv7-4 | 358 | 0 | 0 | 0 | 0 | 0 | 0 |
| ENSMUSG00 | Ighv3-1 | 295 | 0 | 0 | 0 | 0 | 0 | 0 |
| ENSMUSG00 | Ighv3-5 | 352 | 0 | 0 | 0 | 0 | 0 | 0 |
| ENSMUSG00 | Ighv13-1 | 299 | 0 | 0 | 0 | 0 | 0 | 0 |
| ENSMUSG00 | Ighv3-6 | 348 | 0 | 0 | 0 | 0 | 0 | 0 |
| ENSMUSG00 | Ighv9-4 | 293 | 0 | 0 | 0 | 0 | 0 | 0 |
| ENSMUSG00 | Ighv3-8 | 345 | 0 | 0 | 0 | 0 | 0 | 0 |
| ENSMUSG00 | Ighv12-3 | 346 | 0 | 0 | 0 | 0 | 0 | 0 |
| ENSMUSG00 | Ighv6-3 | 354 | 0 | 0 | 0 | 0 | 0 | 0 |
| ENSMUSG00 | Ighv6-6 | 355 | 0 | 0 | 0 | 0 | 0 | 0 |
| ENSMUSG00 | Ighv10-1 | 357 | 0 | 0 | 0 | 0 | 0 | 0 |
| ENSMUSG00 | Ighv1-4 | 349 | 0 | 0 | 0 | 0 | 0 | 0 |
| ENSMUSG00 | Ighv1-5 | 293 | 0 | 0 | 0 | 0 | 0 | 0 |
| ENSMUSG00 | Ighv10-3 | 357 | 0 | 0 | 0 | 0 | 0 | 0 |
| ENSMUSG00 | Ighv1-7 | 293 | 0 | 0 | 0 | 0 | 0 | 0 |
| ENSMUSG00 | Ighv15-2 | 349 | 0 | 0 | 0 | 0 | 0 | 0 |
| ENSMUSG00 | Ighv1-9 | 349 | 0 | 0 | 0 | 0 | 0 | 0 |
| ENSMUSG00 | Ighv1-12 | 293 | 0 | 0 | 0 | 0 | 0 | 0 |
| ENSMUSG00 | Ighv1-14 | 293 | 0 | 0 | 0 | 0 | 0 | 0 |
| ENSMUSG00 | Ighv1-15 | 337 | 0 | 0 | 0 | 0 | 0 | 0 |
| ENSMUSG00 | Ighv1-30 | 291 | 0 | 0 | 0 | 0 | 0 | 0 |
| ENSMUSG00 | Ighv1-18 | 304 | 0 | 0 | 0 | 0 | 0 | 0 |
| ENSMUSG00 | Ighv1-19 | 346 | 0 | 0 | 0 | 0 | 0 | 0 |
| ENSMUSG00 | Ighv1-20 | 346 | 0 | 0 | 0 | 0 | 0 | 0 |
| ENSMUSG00 | Ighv1-22 | 349 | 0 | 0 | 0 | 0 | 0 | 0 |
| ENSMUSG00 | Ighv1-23 | 337 | 0 | 0 | 0 | 0 | 0 | 0 |
| ENSMUSG00 | Ighv1-24 | 293 | 0 | 0 | 0 | 0 | 0 | 0 |
| ENSMUSG00 | Ighv1-26 | 349 | 0 | 0 | 0 | 0 | 0 | 0 |
| ENSMUSG00 | Ighv1-31 | 293 | 0 | 0 | 0 | 0 | 0 | 0 |
| ENSMUSG00 | Ighv1-34 | 349 | 0 | 0 | 0 | 0 | 0 | 0 |
| ENSMUSG00 | Ighv1-36 | 346 | 0 | 0 | 0 | 0 | 0 | 0 |
| ENSMUSG00 | Ighv1-37 | 346 | 0 | 0 | 0 | 0 | 0 | 0 |
| ENSMUSG00 | Ighv1-39 | 349 | 0 | 0 | 0 | 0 | 0 | 0 |
| ENSMUSG00 | Ighv1-42 | 293 | 0 | 0 | 0 | 0 | 0 | 0 |
| ENSMUSG00 | Ighv1-43 | 293 | 0 | 0 | 0 | 0 | 0 | 0 |
| ENSMUSG00 | Ighv1-47 | 293 | 0 | 0 | 0 | 0 | 0 | 0 |
| ENSMUSG00 | Ighv1-49 | 293 | 0 | 0 | 0 | 0 | 0 | 0 |
| ENSMUSG00 | Ighv8-5 | 334 | 0 | 0 | 0 | 0 | 0 | 0 |
| ENSMUSG00 | Ighv1-50 | 349 | 0 | 0 | 0 | 0 | 0 | 0 |
| ENSMUSG00 | Ighv1-52 | 346 | 0 | 0 | 0 | 0 | 0 | 0 |
| ENSMUSG00 | Ighv1-53 | 349 | 0 | 0 | 0 | 0 | 0 | 0 |
| ENSMUSG00 | Ighv8-6 | 300 | 0 | 0 | 0 | 0 | 0 | 0 |

| | | | | | | | | |
|---|---|---|---|---|---|---|---|---|
| ENSMUSG00 | Ighv1-54 | 349 | 0 | 0 | 0 | 0 | 0 | 0 |
| ENSMUSG00 | Ighv1-55 | 349 | 0 | 0 | 0 | 0 | 0 | 0 |
| ENSMUSG00 | Ighv1-56 | 293 | 0 | 0 | 0 | 0 | 0 | 0 |
| ENSMUSG00 | Ighv8-8 | 334 | 0 | 0 | 0 | 0 | 0 | 0 |
| ENSMUSG00 | Ighv1-58 | 343 | 0 | 0 | 0 | 0 | 0 | 0 |
| ENSMUSG00 | Ighv1-59 | 346 | 0 | 0 | 0 | 0 | 0 | 0 |
| ENSMUSG00 | Ighv1-61 | 346 | 0 | 0 | 0 | 0 | 0 | 0 |
| ENSMUSG00 | Ighv1-62-3 | 349 | 0 | 0 | 0 | 0 | 0 | 0 |
| ENSMUSG00 | Ighv8-9 | 300 | 0 | 0 | 0 | 0 | 0 | 0 |
| ENSMUSG00 | Ighv1-63 | 349 | 0 | 0 | 0 | 0 | 0 | 0 |
| ENSMUSG00 | Ighv1-64 | 349 | 0 | 0 | 0 | 0 | 0 | 0 |
| ENSMUSG00 | Ighv8-11 | 334 | 0 | 0 | 0 | 0 | 0 | 0 |
| ENSMUSG00 | Ighv1-66 | 349 | 0 | 0 | 0 | 0 | 0 | 0 |
| ENSMUSG00 | Ighv1-67 | 293 | 0 | 0 | 0 | 0 | 0 | 0 |
| ENSMUSG00 | Ighv1-69 | 346 | 0 | 0 | 0 | 0 | 0 | 0 |
| ENSMUSG00 | Ighv8-12 | 313 | 0 | 0 | 0 | 0 | 0 | 0 |
| ENSMUSG00 | Ighv1-72 | 349 | 0 | 0 | 0 | 0 | 0 | 0 |
| ENSMUSG00 | Ighv8-13 | 302 | 0 | 0 | 0 | 0 | 0 | 0 |
| ENSMUSG00 | Gm16710 | 346 | 0 | 0 | 0 | 0 | 0 | 0 |
| ENSMUSG00 | Ighv1-75 | 349 | 0 | 0 | 0 | 0 | 0 | 0 |
| ENSMUSG00 | Ighv1-80 | 349 | 0 | 0 | 0 | 0 | 0 | 0 |
| ENSMUSG00 | Ighv1-81 | 349 | 0 | 0 | 0 | 0 | 0 | 0 |
| ENSMUSG00 | Ighv1-82 | 293 | 0 | 0 | 0 | 0 | 0 | 0 |
| ENSMUSG00 | Ighv1-83 | 290 | 0 | 0 | 0 | 0 | 0 | 0 |
| ENSMUSG00 | Ighv1-84 | 349 | 0 | 0 | 0 | 0 | 0 | 0 |
| ENSMUSG00 | Ighv1-85 | 349 | 0 | 0 | 0 | 0 | 0 | 0 |
| ENSMUSG00 | Tcrg-V7 | 348 | 0 | 0 | 0 | 0 | 0 | 0 |
| ENSMUSG00 | Tcrg-V4 | 412 | 0 | 0 | 0 | 0 | 0 | 0 |
| ENSMUSG00 | Tcrg-V6 | 401 | 0 | 0 | 0 | 0 | 0 | 0 |
| ENSMUSG00 | Tcrg-V5 | 292 | 0 | 0 | 0 | 0 | 0 | 0 |
| ENSMUSG00 | Trgj1 | 59 | 0 | 0 | 0 | 0 | 0 | 0 |
| ENSMUSG00 | Gm17004 | 515 | 0 | 0 | 0 | 0 | 0 | 0 |
| ENSMUSG00 | Tcrg-V3 | 348 | 0 | 0 | 0 | 0 | 0 | 0 |
| ENSMUSG00 | Trgj3 | 59 | 0 | 0 | 0 | 0 | 0 | 0 |
| ENSMUSG00 | Gm16803 | 500 | 0 | 0 | 0 | 0 | 0 | 0 |
| ENSMUSG00 | Trgj2 | 59 | 0 | 0 | 0 | 0 | 0 | 0 |
| ENSMUSG00 | Trgv2 | 348 | 0 | 0 | 0 | 0 | 0 | 0 |
| ENSMUSG00 | Gm16802 | 345 | 0 | 0 | 0 | 0 | 0 | 0 |
| ENSMUSG00 | Trgj4 | 60 | 0 | 0 | 0 | 0 | 0 | 0 |
| ENSMUSG00 | Tcrg-C | 638 | 0 | 0 | 0 | 0 | 0 | 0 |
| ENSMUSG00 | Trav1 | 367 | 0 | 0 | 0 | 0 | 0 | 0 |
| ENSMUSG00 | Trav2 | 323 | 0 | 0 | 0 | 0 | 0 | 0 |
| ENSMUSG00 | Trav3-1 | 341 | 0 | 0 | 0.12962417 | 0 | 0 | 0 |
| ENSMUSG00 | Trav5-1 | 341 | 0 | 0 | 0 | 0 | 0 | 0 |
| ENSMUSG00 | Trav6-1 | 336 | 0 | 0 | 0 | 0 | 0 | 0 |
| ENSMUSG00 | Trav6-2 | 336 | 0 | 0 | 0 | 0 | 0 | 0 |
| ENSMUSG00 | Trav6-4 | 374 | 0 | 0 | 0 | 0 | 0 | 0 |
| ENSMUSG00 | Trav7d-3 | 399 | 0 | 0 | 0 | 0 | 0 | 0 |
| ENSMUSG00 | Trav6d-4 | 374 | 0 | 0 | 0 | 0 | 0 | 0 |
| ENSMUSG00 | Trav7d-4 | 447 | 0 | 0 | 0 | 0 | 0 | 0 |
| ENSMUSG00 | Trav8d-1 | 397 | 0 | 0 | 0 | 0 | 0 | 0 |
| ENSMUSG00 | Trav9d-1 | 442 | 0 | 0 | 0 | 0 | 0 | 0 |
| ENSMUSG00 | Trav6d-5 | 471 | 0 | 0 | 0 | 0 | 0 | 0 |
| ENSMUSG00 | Trav10d | 403 | 0 | 0 | 0 | 0 | 0 | 0 |
| ENSMUSG00 | Trav6d-6 | 402 | 0 | 0 | 0 | 0 | 0 | 0 |
| ENSMUSG00 | Trav11d | 357 | 0 | 0 | 0 | 0 | 0 | 0 |
| ENSMUSG00 | Trav7d-5 | 355 | 0 | 0 | 0 | 0 | 0 | 0 |
| ENSMUSG00 | Trav12d-1 | 432 | 0 | 0 | 0 | 0 | 0 | 0 |
| ENSMUSG00 | Trav13d-1 | 393 | 0 | 0 | 0 | 0 | 0 | 0 |
| ENSMUSG00 | Trav14d-1 | 412 | 0 | 0 | 0 | 0 | 0 | 0 |
| ENSMUSG00 | Trav15d-1-dv | 415 | 0 | 0 | 0 | 0 | 0 | 0 |
| ENSMUSG00 | Trav9d-2 | 335 | 0 | 0 | 0 | 0 | 0 | 0 |
| ENSMUSG00 | Trav4d-3 | 329 | 0 | 0 | 0 | 0 | 0 | 0 |
| ENSMUSG00 | Trav12d-2 | 339 | 0 | 0 | 0 | 0 | 0 | 0 |
| ENSMUSG00 | Trav9d-3 | 338 | 0 | 0 | 0 | 0 | 0 | 0 |
| ENSMUSG00 | Trav12d-3 | 345 | 0 | 0 | 0 | 0 | 0 | 0 |
| ENSMUSG00 | Trav13d-2 | 263 | 0 | 0 | 0 | 0 | 0 | 0 |
| ENSMUSG00 | Trav14d-2 | 275 | 0 | 0 | 0 | 0 | 0 | 0 |
| ENSMUSG00 | Trav15d-2-dv | 346 | 0 | 0 | 0 | 0 | 0 | 0 |
| ENSMUSG00 | Trav3d-3 | 275 | 0 | 0 | 0 | 0 | 0 | 0 |
| ENSMUSG00 | Trav4d-4 | 329 | 0 | 0 | 0 | 0 | 0 | 0 |
| ENSMUSG00 | Trav5d-4 | 341 | 0 | 0 | 0 | 0 | 0 | 0 |
| ENSMUSG00 | Trav6d-7 | 315 | 0 | 0 | 0 | 0 | 0 | 0 |
| ENSMUSG00 | C920008G01 | 1408 | 0 | 0 | 0 | 0 | 0 | 0 |
| ENSMUSG00 | Trav8d-2 | 332 | 0 | 0 | 0 | 0 | 0 | 0 |
| ENSMUSG00 | Trav16d-dv11 | 351 | 0 | 0 | 0 | 0 | 0 | 0 |
| ENSMUSG00 | Trav13d-4 | 263 | 0 | 0 | 0 | 0 | 0 | 0 |
| ENSMUSG00 | Trav14d-3-dv | 326 | 0 | 0 | 0 | 0 | 0 | 0 |
| ENSMUSG00 | Trav7n-4 | 339 | 0 | 0 | 0 | 0 | 0 | 0 |
| ENSMUSG00 | Trav6n-5 | 336 | 0 | 0 | 0 | 0 | 0 | 0 |
| ENSMUSG00 | Trav10n | 341 | 0 | 0 | 0 | 0 | 0 | 0 |
| ENSMUSG00 | Trav6n-6 | 286 | 0 | 0 | 0 | 0 | 0 | 0 |
| ENSMUSG00 | Trav14n-1 | 338 | 0 | 0 | 0 | 0 | 0 | 0 |
| ENSMUSG00 | Trav15n-1 | 347 | 0 | 0 | 0 | 0 | 0 | 0 |
| ENSMUSG00 | Trav9n-2 | 335 | 0 | 0 | 0 | 0 | 0 | 0 |
| ENSMUSG00 | Trav4n-3 | 329 | 0 | 0 | 0 | 0 | 0 | 0 |
| ENSMUSG00 | Trav12n-2 | 339 | 0 | 0 | 0 | 0 | 0 | 0 |
| ENSMUSG00 | Trav9n-3 | 338 | 0 | 0 | 0 | 0 | 0 | 0 |
| ENSMUSG00 | Trav12n-3 | 345 | 0 | 0 | 0 | 0 | 0 | 0 |
| ENSMUSG00 | Trav13n-2 | 263 | 0 | 0 | 0 | 0 | 0 | 0 |
| ENSMUSG00 | Trav14n-2 | 275 | 0 | 0 | 0 | 0 | 0 | 0 |
| ENSMUSG00 | Trav15n-2 | 346 | 0 | 0 | 0 | 0 | 0 | 0 |
| ENSMUSG00 | Trav3n-3 | 275 | 0 | 0 | 0 | 0 | 0 | 0 |
| ENSMUSG00 | Trav9n-4 | 338 | 0 | 0 | 0 | 0 | 0 | 0 |
| ENSMUSG00 | Trav4n-4 | 329 | 0 | 0 | 0 | 0 | 0 | 0 |
| ENSMUSG00 | Trav5n-4 | 341 | 0 | 0 | 0 | 0 | 0 | 0 |
| ENSMUSG00 | Trav6n-7 | 315 | 0 | 0 | 0 | 0 | 0 | 0 |
| ENSMUSG00 | Trav7n-6 | 1408 | 0 | 0 | 0 | 0 | 0 | 0 |
| ENSMUSG00 | Trav8n-2 | 331 | 0 | 0 | 0 | 0 | 0 | 0 |
| ENSMUSG00 | Trav16 | 351 | 0 | 0 | 0 | 0 | 0 | 0 |
| ENSMUSG00 | Trav13n-4 | 266 | 0 | 0 | 0 | 0 | 0 | 0 |
| ENSMUSG00 | Trav14n-3 | 279 | 0 | 0 | 0 | 0 | 0 | 0 |
| ENSMUSG00 | Trav7-2 | 399 | 0 | 0 | 0 | 0 | 0 | 0 |
| ENSMUSG00 | Trav4-2 | 367 | 0 | 0 | 0 | 0 | 0 | 0 |
| ENSMUSG00 | Trav6-3 | 384 | 0 | 0 | 0 | 0 | 0 | 0 |
| ENSMUSG00 | Trav7-3 | 399 | 0 | 0 | 0 | 0 | 0 | 0 |
| ENSMUSG00 | Trav6d-3 | 343 | 0 | 0 | 0 | 0 | 0 | 0 |
| ENSMUSG00 | Trav7-4 | 447 | 0 | 0 | 0 | 0 | 0 | 0 |
| ENSMUSG00 | Trav8-1 | 325 | 0 | 0 | 0 | 0 | 0 | 0 |
| ENSMUSG00 | Trav9-1 | 395 | 0 | 0 | 0 | 0 | 0 | 0 |
| ENSMUSG00 | Trav6-5 | 371 | 0 | 0 | 0 | 0 | 0 | 0 |
| ENSMUSG00 | Trav10 | 341 | 0 | 0 | 0 | 0 | 0 | 0 |
| ENSMUSG00 | Trav6-6 | 342 | 0 | 0 | 0 | 0 | 0 | 0 |
| ENSMUSG00 | Trav11 | 341 | 0 | 0 | 0 | 0 | 0 | 0 |
| ENSMUSG00 | Trav12-1 | 342 | 0 | 0 | 0 | 0 | 0 | 0 |
| ENSMUSG00 | Trav13-1 | 325 | 0 | 0 | 0 | 0 | 0 | 0 |
| ENSMUSG00 | Trav14-1 | 355 | 0 | 0 | 0 | 0 | 0 | 0 |
| ENSMUSG00 | Trav15-1-dv6 | 347 | 0 | 0 | 0 | 0 | 0 | 0 |
| ENSMUSG00 | Trav9-2 | 338 | 0 | 0 | 0 | 0 | 0 | 0 |
| ENSMUSG00 | Trav4-3 | 329 | 0 | 0 | 0 | 0 | 0 | 0 |
| ENSMUSG00 | Trav12-2 | 422 | 0 | 0 | 0 | 0 | 0 | 0 |
| ENSMUSG00 | Trav12-3 | 438 | 0 | 0 | 0 | 0 | 0 | 0 |
| ENSMUSG00 | Trav13-2 | 387 | 0 | 0 | 0 | 0 | 0 | 0 |
| ENSMUSG00 | Trav14-2 | 384 | 0 | 0 | 0 | 0 | 0 | 0 |
| ENSMUSG00 | Trav15-2-dv6 | 384 | 0 | 0 | 0 | 0 | 0 | 0 |
| ENSMUSG00 | Trav3-3 | 416 | 0 | 0 | 0 | 0 | 0 | 0 |
| ENSMUSG00 | Trav9-4 | 392 | 0 | 0 | 0 | 0 | 0 | 0 |
| ENSMUSG00 | Trav4-4-dv10 | 358 | 0 | 0 | 0 | 0 | 0 | 0 |
| ENSMUSG00 | Trav5-4 | 340 | 0 | 0 | 0 | 0 | 0 | 0 |
| ENSMUSG00 | Trav6-7-dv9 | 375 | 0 | 0 | 0 | 0 | 0 | 0 |
| ENSMUSG00 | Trav7-6 | 378 | 0 | 0 | 0 | 0 | 0 | 0 |
| ENSMUSG00 | Gm13896 | 419 | 0 | 0 | 0 | 0 | 0 | 0 |
| ENSMUSG00 | Trav13-4-dv7 | 382 | 0 | 0 | 0 | 0 | 0 | 0 |
| ENSMUSG00 | Trav14-3 | 962 | 0 | 0 | 0 | 0 | 0 | 0 |
| ENSMUSG00 | Trav3-4 | 455 | 3.75711135 | 1.05904268 | 7.77175262 | 4.75807181 | 4.82227288 | 5.81459342 |
| ENSMUSG00 | Trav17 | 451 | 4.14303234 | 1.38896618 | 4.11635789 | 2.72015413 | 2.03652941 | 0.36285551 |
| ENSMUSG00 | Trav18 | 325 | 0 | 0.88959586 | 0 | 0 | 0 | 0.5035318 |
| ENSMUSG00 | Trav19 | 443 | 0 | 0 | 0 | 0 | 0 | 0 |
| ENSMUSG00 | Trav21-dv12 | 357 | 0 | 0 | 0 | 0 | 0 | 0 |
| ENSMUSG00 | Trdv1 | 414 | 0 | 0 | 0 | 0 | 0 | 0 |
| ENSMUSG00 | Trdv2-1 | 343 | 0 | 0 | 0 | 0 | 0 | 0 |
| ENSMUSG00 | Trdv2-2 | 343 | 0 | 0 | 0 | 0 | 0 | 0 |
| ENSMUSG00 | Trdv4 | 343 | 0 | 0 | 0 | 0 | 0 | 0 |
| ENSMUSG00 | Trdd1 | 10 | 0 | 0 | 0 | 0 | 0 | 0 |
| ENSMUSG00 | Trdd2 | 15 | 0 | 0 | 0 | 0 | 0 | 0 |
| ENSMUSG00 | Trdj1 | 50 | 0 | 0 | 0 | 0 | 0 | 0 |
| ENSMUSG00 | Trdj2 | 58 | 0 | 0 | 0 | 0 | 0 | 0 |
| ENSMUSG00 | Trdv5 | 343 | 0 | 0 | 0 | 0 | 0 | 0 |
| ENSMUSG00 | Traj59 | 63 | 0 | 0 | 0 | 0 | 0 | 0 |
| ENSMUSG00 | Traj58 | 62 | 0 | 0 | 0 | 0 | 0 | 0 |
| ENSMUSG00 | Traj57 | 64 | 0 | 0 | 0 | 0 | 0 | 0 |
| ENSMUSG00 | Traj56 | 62 | 0 | 0 | 0 | 0 | 0 | 0 |
| ENSMUSG00 | Traj53 | 65 | 0 | 0 | 0 | 0 | 0 | 0 |
| ENSMUSG00 | Traj52 | 65 | 0 | 0 | 0 | 0 | 0 | 0 |
| ENSMUSG00 | Traj51 | 42 | 0 | 0 | 0 | 0 | 0 | 0 |
| ENSMUSG00 | Traj50 | 62 | 0 | 0 | 0 | 0 | 0 | 0 |
| ENSMUSG00 | Traj49 | 58 | 1.37087864 | 0 | 0 | 0 | 0 | 0 |
| ENSMUSG00 | Traj48 | 59 | 0 | 0 | 0 | 0 | 0 | 0 |
| ENSMUSG00 | Traj46 | 62 | 0 | 0 | 0 | 0 | 0 | 0 |
| ENSMUSG00 | Traj45 | 62 | 0 | 0 | 0 | 0 | 0 | 0 |
| ENSMUSG00 | Traj44 | 60 | 0 | 0 | 0 | 0 | 0 | 0 |
| ENSMUSG00 | Traj43 | 56 | 0 | 0 | 0 | 0 | 0 | 0 |
| ENSMUSG00 | Traj42 | 63 | 0 | 0 | 0 | 0 | 0 | 0 |
| ENSMUSG00 | Traj41 | 54 | 0 | 0 | 0 | 0 | 0 | 0 |
| ENSMUSG00 | Traj40 | 60 | 0 | 0 | 0 | 0 | 0 | 0 |
| ENSMUSG00 | Traj39 | 62 | 0 | 0 | 0 | 0 | 0 | 0 |
| ENSMUSG00 | Traj38 | 61 | 0 | 0 | 0 | 0 | 0 | 0 |
| ENSMUSG00 | Traj37 | 59 | 0 | 0 | 0 | 0 | 0 | 0 |
| ENSMUSG00 | Traj35 | 64 | 0 | 0 | 0 | 0 | 0 | 0 |
| ENSMUSG00 | Traj34 | 57 | 0 | 0 | 0 | 0 | 0 | 0 |
| ENSMUSG00 | Traj33 | 56 | 0 | 0 | 0 | 0 | 0 | 0 |
| ENSMUSG00 | Traj32 | 65 | 0 | 0 | 0 | 0 | 0 | 0 |
| ENSMUSG00 | Traj31 | 56 | 0 | 0 | 0 | 0 | 0 | 0 |
| ENSMUSG00 | Traj30 | 58 | 0 | 0 | 0 | 0 | 0 | 0 |
| ENSMUSG00 | Traj29 | 59 | 0 | 0 | 0 | 0 | 0 | 0 |
| ENSMUSG00 | Traj28 | 64 | 0 | 0 | 0 | 0 | 0 | 0 |
| ENSMUSG00 | Traj27 | 58 | 0 | 0 | 0 | 0 | 0.8797651 | 1.4107572 |
| ENSMUSG00 | Traj26 | 59 | 0 | 0 | 0 | 0 | 0 | 0 |
| ENSMUSG00 | Traj25 | 56 | 0 | 0 | 0 | 0 | 0 | 0 |
| ENSMUSG00 | Traj24 | 62 | 0 | 0 | 0 | 0 | 0 | 0 |
| ENSMUSG00 | Traj23 | 59 | 0 | 0 | 0 | 0 | 0 | 0 |
| ENSMUSG00 | Traj22 | 59 | 0 | 0 | 0 | 0 | 0 | 0 |
| ENSMUSG00 | Traj21 | 56 | 0 | 0 | 0 | 0 | 0 | 0 |
| ENSMUSG00 | Traj20 | 57 | 0 | 0 | 0 | 0 | 0 | 0 |
| ENSMUSG00 | Traj19 | 60 | 0 | 0 | 0 | 0 | 0 | 0 |
| ENSMUSG00 | Traj18 | 65 | 0 | 0 | 0 | 0 | 0 | 0 |
| ENSMUSG00 | Traj17 | 62 | 0 | 0 | 0 | 0 | 0 | 0 |
| ENSMUSG00 | Traj16 | 60 | 0 | 0 | 0 | 0 | 0 | 0 |
| ENSMUSG00 | Traj15 | 59 | 0 | 0 | 0 | 0 | 0 | 0 |
| ENSMUSG00 | Traj14 | 32 | 0 | 0 | 0 | 0 | 0 | 0 |
| ENSMUSG00 | Traj13 | 56 | 0 | 0 | 0 | 0 | 0 | 0 |
| ENSMUSG00 | Traj12 | 57 | 9.06703941 | 0 | 9.30565116 | 3.16509163 | 4.47599788 | 4.30652199 |
| ENSMUSG00 | Traj11 | 58 | 0.68543932 | 0 | 0 | 0 | 0 | 0 |
| ENSMUSG00 | Traj9 | 57 | 0.69746457 | 0 | 0 | 0 | 0 | 0 |
| ENSMUSG00 | Traj8 | 36 | 2.20863781 | 0 | 0 | 0 | 0 | 0 |
| ENSMUSG00 | Traj7 | 58 | 0 | 0 | 0 | 0 | 0 | 0 |
| ENSMUSG00 | Traj6 | 61 | 0 | 0 | 0 | 0 | 0 | 0 |
| ENSMUSG00 | Traj5 | 61 | 0 | 0 | 0 | 0 | 0 | 0 |
| ENSMUSG00 | Traj4 | 62 | 0 | 0 | 0 | 0 | 0 | 0 |
| ENSMUSG00 | Traj3 | 65 | 0 | 0 | 0 | 0 | 0 | 0 |
| ENSMUSG00 | Traj2 | 65 | 0 | 0 | 0 | 0 | 0 | 0.41960984 |
| ENSMUSG00 | Traj1 | 26 | 0 | 0 | 0 | 0 | 0 | 0 |
| ENSMUSG00 | Trac | 410 | 2.13322091 | 0.47011163 | 3.98894681 | 3.87221941 | 3.23581895 | 4.65664573 |
| ENSMUSG00 | Iglc3 | 579 | 0 | 0 | 0 | 0 | 0 | 0 |
| ENSMUSG00 | Iglv1 | 349 | 0 | 0 | 0 | 0 | 0 | 0 |
| ENSMUSG00 | Iglc2 | 604 | 0 | 0 | 0 | 0 | 0 | 0 |
| ENSMUSG00 | Iglv3 | 364 | 0 | 0 | 0 | 0 | 0 | 0 |
| ENSMUSG00 | Iglv2 | 346 | 0 | 0 | 0 | 0 | 0 | 0 |
| ENSMUSG00 | AC134447.1 | 109 | 0 | 0 | 0 | 0 | 0 | 0 |
| ENSMUSG00 | AC158299.1 | 65 | 0 | 0 | 0 | 0 | 0 | 0 |
| ENSMUSG00 | Mir582 | 80 | 0 | 0 | 0 | 0 | 0 | 0 |
| ENSMUSG00 | Mir467e | 86 | 0 | 0 | 0 | 0 | 0 | 0 |
| ENSMUSG00 | AC121540.1 | 109 | 0 | 0 | 0 | 0 | 0 | 0 |
| ENSMUSG00 | AL929011.1 | 109 | 0 | 0 | 0 | 0 | 0 | 0 |
| ENSMUSG00 | AC127431.1 | 109 | 0 | 0 | 0 | 0 | 0 | 0 |
| ENSMUSG00 | AC087183.1 | 110 | 0 | 0 | 0 | 0 | 0 | 0 |
| ENSMUSG00 | Mir466b-1 | 89 | 0 | 0 | 0 | 0 | 0 | 0 |
| ENSMUSG00 | Mir297a-3 | 101 | 0 | 0 | 0 | 0 | 0 | 0 |
| ENSMUSG00 | AC159715.1 | 109 | 0 | 0 | 0 | 0 | 0 | 0 |
| ENSMUSG00 | AC154424.1 | 109 | 0 | 0 | 0 | 0 | 0 | 0 |
| ENSMUSG00 | Mir669l | 97 | 0 | 0 | 0 | 0 | 0 | 0 |
| ENSMUSG00 | AC158749.1 | 109 | 0 | 0 | 0 | 0 | 0 | 0 |
| ENSMUSG00 | AC132266.1 | 109 | 0 | 0 | 0 | 0 | 0 | 0 |
| ENSMUSG00 | Mir466b-2 | 81 | 0 | 0 | 0 | 0 | 0 | 0 |
| ENSMUSG00 | Mir193b | 78 | 0 | 0 | 0 | 0 | 0 | 0 |
| ENSMUSG00 | AL691478.1 | 109 | 0 | 0 | 0 | 0 | 0 | 0 |
| ENSMUSG00 | AC098884.1 | 109 | 0 | 0 | 0 | 0 | 0 | 0 |
| ENSMUSG00 | Mir297c | 97 | 0 | 0 | 0 | 0 | 0 | 0 |
| ENSMUSG00 | AL683818.1 | 109 | 0 | 0 | 0 | 0 | 0 | 0 |
| ENSMUSG00 | mmu-mir-466 | 89 | 0 | 0 | 0 | 0 | 0 | 0 |
| ENSMUSG00 | AC117684.1 | 109 | 0 | 0 | 0 | 0 | 0 | 0 |
| ENSMUSG00 | mmu-mir-297 | 89 | 0 | 0 | 0 | 0 | 0 | 0 |
| ENSMUSG00 | AC174081.1 | 109 | 0 | 0 | 0 | 0 | 0 | 0 |
| ENSMUSG00 | Mir467a-6 | 82 | 0 | 0 | 0 | 0 | 0 | 0 |
| ENSMUSG00 | BX813309.1 | 109 | 0 | 0 | 0 | 0 | 0 | 0 |
| ENSMUSG00 | AC155315.1 | 75 | 0 | 0 | 0 | 0 | 0 | 0 |
| ENSMUSG00 | Mir466b-5 | 87 | 0 | 0 | 0 | 0 | 0 | 0 |
| ENSMUSG00 | AC137969.1 | 109 | 0 | 0 | 0 | 0 | 0 | 0 |
| ENSMUSG00 | Mir466b-3 | 80 | 0 | 0 | 0 | 0 | 0 | 0 |
| ENSMUSG00 | AC122327.1 | 109 | 0 | 0 | 0 | 0 | 0 | 0 |
| ENSMUSG00 | AC161167.1 | 109 | 0 | 0 | 0 | 0 | 0 | 0 |
| ENSMUSG00 | AC122386.1 | 109 | 0 | 0 | 0 | 0 | 0 | 0 |
| ENSMUSG00 | AC132082.1 | 95 | 0 | 0 | 0 | 0 | 0 | 0 |
| ENSMUSG00 | Mir544 | 77 | 0 | 0 | 0 | 0 | 0 | 0 |
| ENSMUSG00 | AC137969.2 | 109 | 0 | 0 | 0 | 0 | 0 | 0 |
| ENSMUSG00 | AC122386.2 | 109 | 0 | 0 | 0 | 0 | 0 | 0 |
| ENSMUSG00 | AL731707.1 | 66 | 0 | 0 | 0.66972489 | 1.09339529 | 7.73126906 | 0 |
| ENSMUSG00 | AC134254.1 | 85 | 0 | 0 | 0 | 0 | 0 | 0 |
| ENSMUSG00 | AC118240.1 | 109 | 0 | 0 | 0 | 0 | 0 | 0 |
| ENSMUSG00 | AC102463.1 | 109 | 0 | 0 | 0 | 0 | 0 | 0 |
| ENSMUSG00 | AC124506.1 | 109 | 0 | 0 | 0 | 0 | 0 | 0 |
| ENSMUSG00 | AC154322.1 | 109 | 0 | 0 | 0 | 0 | 0 | 0 |
| ENSMUSG00 | AC115121.1 | 108 | 0 | 0 | 0 | 0 | 0 | 0 |
| ENSMUSG00 | Mir467d | 84 | 0 | 0 | 0 | 0 | 0 | 0 |
| ENSMUSG00 | AC123045.1 | 109 | 0 | 0 | 0 | 0 | 0 | 0 |
| ENSMUSG00 | Mir493 | 82 | 0 | 0 | 0 | 0 | 0 | 0 |
| ENSMUSG00 | AL606907.1 | 109 | 0 | 0 | 0 | 0 | 0 | 0 |
| ENSMUSG00 | mmu-mir-466 | 89 | 0 | 0 | 0 | 0 | 0 | 0 |
| ENSMUSG00 | AC115121.2 | 108 | 0 | 0 | 0 | 0 | 0 | 0 |
| ENSMUSG00 | Mir467a-10 | 82 | 0 | 0 | 0 | 0 | 0 | 0 |
| ENSMUSG00 | AC167229.1 | 109 | 0 | 0 | 0 | 0 | 0 | 0 |
| ENSMUSG00 | Mir466b-4 | 89 | 0 | 0 | 0 | 0 | 0 | 0 |
| ENSMUSG00 | AC126045.1 | 109 | 0 | 0 | 0 | 0 | 0 | 0 |
| ENSMUSG00 | AL833773.1 | 109 | 0 | 0 | 0 | 0 | 0 | 0 |
| ENSMUSG00 | Mir669f | 120 | 0 | 0 | 0 | 0 | 0 | 0 |
| ENSMUSG00 | AC111074.1 | 108 | 0 | 0 | 0 | 0 | 0 | 0 |
| ENSMUSG00 | AL671906.1 | 109 | 0 | 0 | 0 | 0 | 0 | 0 |
| ENSMUSG00 | Mir574 | 77 | 0 | 0 | 0 | 0 | 0 | 0 |
| ENSMUSG00 | AC167229.2 | 109 | 0 | 0 | 0 | 0 | 0 | 0 |
| ENSMUSG00 | BX537299.1 | 109 | 0 | 0 | 0 | 0 | 0 | 0 |
| ENSMUSG00 | AC102227.1 | 109 | 0 | 0 | 0 | 0 | 0 | 0 |
| ENSMUSG00 | Mir467c | 96 | 0 | 0 | 0 | 0 | 0 | 0 |
| ENSMUSG00 | AC131665.1 | 109 | 0 | 0 | 0 | 0 | 0 | 0 |
| ENSMUSG00 | AC122806.1 | 109 | 0 | 0 | 0 | 0 | 0 | 0 |
| ENSMUSG00 | AC154006.1 | 109 | 0 | 0 | 0 | 0 | 0 | 0 |
| ENSMUSG00 | Mir467a-3 | 82 | 0 | 0 | 0 | 0 | 0 | 0 |
| ENSMUSG00 | AC121967.1 | 81 | 0 | 0 | 0 | 0 | 0 | 0 |
| ENSMUSG00 | Mir669p-2 | 86 | 0 | 0 | 0 | 0 | 0 | 0 |
| ENSMUSG00 | AC124426.1 | 109 | 0 | 0 | 0 | 0 | 0 | 0 |
| ENSMUSG00 | AC123824.1 | 109 | 0 | 0 | 0 | 0 | 0 | 0 |
| ENSMUSG00 | AC110903.1 | 95 | 0 | 0 | 0 | 0 | 1.07423949 | 0 |
| ENSMUSG00 | Mir466b-6 | 89 | 0 | 0 | 0 | 0 | 0 | 0 |
| ENSMUSG00 | AC143329.1 | 109 | 0 | 0 | 0 | 0 | 0 | 0 |
| ENSMUSG00 | AL591544.1 | 65 | 0 | 0 | 0 | 0 | 0 | 0 |
| ENSMUSG00 | AC124080.1 | 109 | 0 | 0 | 0 | 0 | 0 | 0 |
| ENSMUSG00 | AC020968.1 | 109 | 0 | 0 | 0 | 0 | 0 | 0 |
| ENSMUSG00 | Mir466e | 85 | 0 | 0 | 0 | 0 | 0 | 0 |
| ENSMUSG00 | Mir669p-1 | 86 | 0 | 0 | 0 | 0 | 0 | 0 |
| ENSMUSG00 | AC126798.1 | 109 | 0 | 0 | 0 | 0 | 0 | 0 |
| ENSMUSG00 | AC087801.1 | 109 | 0 | 0 | 0 | 0 | 0 | 0 |
| ENSMUSG00 | AC108942.1 | 109 | 0 | 0 | 0 | 0 | 0 | 0 |
| ENSMUSG00 | AC154654.1 | 109 | 0 | 0 | 0 | 0 | 0 | 0 |
| ENSMUSG00 | AC164303.1 | 109 | 0 | 0 | 0 | 0 | 0 | 0 |
| ENSMUSG00 | AC159378.1 | 79 | 0 | 0 | 0 | 0 | 0 | 0 |
| ENSMUSG00 | AC138311.1 | 109 | 0 | 0 | 0 | 0 | 0 | 0 |
| ENSMUSG00 | AC102528.1 | 109 | 0 | 0 | 0 | 0 | 0 | 0 |
| ENSMUSG00 | AC163215.1 | 109 | 0 | 0 | 0 | 0 | 0 | 0 |
| ENSMUSG00 | Mir669o | 95 | 0 | 0 | 0 | 0 | 0 | 0 |
| ENSMUSG00 | Mir466b-7 | 87 | 0 | 0 | 0 | 0 | 0 | 0 |
| ENSMUSG00 | AC155640.1 | 66 | 0 | 0 | 0 | 0 | 0 | 0 |
| ENSMUSG00 | AC122806.2 | 109 | 0 | 0 | 0 | 0 | 0 | 0 |
| ENSMUSG00 | Mir742 | 64 | 0 | 0 | 0 | 0 | 0 | 0 |
| ENSMUSG00 | AC122520.1 | 109 | 0 | 0 | 0 | 0 | 0 | 0 |
| ENSMUSG00 | CR974486.1 | 108 | 0 | 0 | 0 | 0 | 0 | 0 |
| ENSMUSG00 | AL772216.1 | 103 | 0 | 0 | 0 | 0 | 0 | 0 |
| ENSMUSG00 | AC154375.1 | 64 | 0 | 0 | 0 | 0 | 0 | 0 |
| ENSMUSG00 | Mir466b-8 | 85 | 0 | 0 | 0 | 0 | 0 | 0 |
| ENSMUSG00 | AL606904.1 | 109 | 0 | 0 | 0 | 0 | 0 | 0 |
| ENSMUSG00 | AL772216.2 | 99 | 0 | 0 | 0 | 0 | 0 | 0 |
| ENSMUSG00 | Mir344c | 91 | 0 | 0 | 0 | 0 | 0 | 0 |
| ENSMUSG00 | AC138311.2 | 109 | 0 | 0 | 0 | 0 | 0 | 0 |
| ENSMUSG00 | AC131111.1 | 109 | 0 | 0 | 0 | 0 | 0 | 0 |
| ENSMUSG00 | Mir18b | 82 | 0 | 0 | 0 | 0 | 0 | 0 |
| ENSMUSG00 | Mir466p | 88 | 0 | 0 | 0 | 0 | 0 | 0 |
| ENSMUSG00 | AC117684.2 | 109 | 0 | 0 | 0 | 0 | 0 | 0 |
| ENSMUSG00 | AC122520.2 | 109 | 0 | 0 | 0 | 0 | 0 | 0 |
| ENSMUSG00 | Mir741 | 70 | 0 | 0 | 0 | 0 | 0 | 0 |
| ENSMUSG00 | AC164603.1 | 109 | 0 | 0 | 0 | 0 | 0 | 0 |
| ENSMUSG00 | Mir743 | 61 | 0 | 0 | 0 | 0 | 0 | 0 |
| ENSMUSG00 | AC161197.1 | 109 | 0 | 0 | 0 | 0 | 0 | 0 |
| ENSMUSG00 | AL772216.3 | 103 | 0 | 0 | 0 | 0 | 0 | 0 |
| ENSMUSG00 | AC125134.1 | 109 | 0 | 0 | 0 | 0 | 0 | 0 |
| ENSMUSG00 | AC122775.1 | 109 | 0 | 0 | 0 | 0 | 0 | 0 |
| ENSMUSG00 | AC134445.1 | 109 | 0 | 0 | 0 | 0 | 0 | 0 |
| ENSMUSG00 | Mir466c-1 | 89 | 0 | 0 | 0 | 0 | 0 | 0 |
| ENSMUSG00 | AC114404.1 | 109 | 0 | 0 | 0 | 0 | 0 | 0 |
| ENSMUSG00 | Mir466c-2 | 89 | 0 | 0 | 0 | 0 | 0 | 0 |
| ENSMUSG00 | SNORA17 | 130 | 0 | 0 | 0 | 0 | 0 | 0 |
| ENSMUSG00 | SNORA17 | 125 | 0 | 0 | 0 | 0 | 0 | 0 |
| ENSMUSG00 | SNORA17 | 119 | 0 | 0 | 0 | 0 | 0 | 0 |
| ENSMUSG00 | SNORA17 | 130 | 0 | 0 | 0 | 0 | 0 | 0 |
| ENSMUSG00 | U6 | 103 | 0 | 0 | 0 | 0 | 0 | 0 |
| ENSMUSG00 | SNORA17 | 131 | 0 | 0 | 0 | 0 | 0 | 0 |
| ENSMUSG00 | SNORA17 | 132 | 0.30117788 | 0 | 0 | 0 | 0 | 0 |
| ENSMUSG00 | SNORA17 | 143 | 0 | 0 | 0 | 0 | 0 | 0 |
| ENSMUSG00 | SNORA17 | 130 | 0 | 0 | 0 | 0 | 0 | 0 |
| ENSMUSG00 | U6 | 102 | 0 | 0 | 0 | 0 | 0 | 0 |
| ENSMUSG00 | n-R5s90 | 117 | 0 | 0 | 0 | 0 | 0 | 0 |
| ENSMUSG00 | SNORA17 | 128 | 0 | 0 | 0 | 0 | 0 | 0 |
| ENSMUSG00 | SNORA17 | 125 | 0 | 0 | 0 | 0 | 0 | 0 |
| ENSMUSG00 | SNORA17 | 132 | 0 | 0 | 0 | 0 | 0 | 0 |
| ENSMUSG00 | U6 | 106 | 0 | 0 | 0 | 0 | 0 | 0.51461584 |
| ENSMUSG00 | SNORA17 | 132 | 0 | 0 | 0 | 0 | 0 | 0 |
| ENSMUSG00 | SNORA17 | 131 | 0 | 0 | 0 | 0 | 0 | 0 |
| ENSMUSG00 | SNORA17 | 128 | 0 | 0 | 0 | 0 | 0 | 0 |
| ENSMUSG00 | SNORA63 | 245 | 15.2531231 | 13.9642343 | 17.8611529 | 8.10005081 | 22.9098014 | 10.5758805 |
| ENSMUSG00 | SNORA17 | 132 | 0 | 0 | 0 | 0 | 0 | 0 |
| ENSMUSG00 | SNORA43 | 131 | 0 | 0 | 0 | 0 | 0 | 0 |
| ENSMUSG00 | SNORA17 | 127 | 0 | 0 | 0 | 0 | 0 | 0 |
| ENSMUSG00 | SNORA17 | 131 | 0 | 0 | 0 | 0 | 0 | 0 |
| ENSMUSG00 | SNORA17 | 122 | 0 | 0 | 0 | 0 | 0 | 0 |
| ENSMUSG00 | SNORA17 | 128 | 0 | 0 | 0 | 1.69134584 | 0 | 0 |
| ENSMUSG00 | SNORA17 | 128 | 0 | 0 | 0 | 0 | 0 | 0 |
| ENSMUSG00 | SNORA17 | 131 | 0 | 0 | 0 | 0 | 0 | 0 |
| ENSMUSG00 | SNORA17 | 135 | 0 | 0 | 0 | 0 | 0 | 0 |
| ENSMUSG00 | SNORA17 | 130 | 0 | 0 | 0 | 0 | 0 | 0 |
| ENSMUSG00 | SNORA17 | 131 | 0 | 0 | 0 | 0 | 0 | 0 |
| ENSMUSG00 | SNORA17 | 131 | 0 | 0 | 0 | 0 | 0 | 0 |
| ENSMUSG00 | SNORA17 | 131 | 0 | 0 | 0 | 0 | 0 | 0 |
| ENSMUSG00 | SNORA17 | 128 | 0 | 0 | 0 | 0 | 0 | 0 |
| ENSMUSG00 | SNORD102 | 70 | 0 | 0 | 0 | 0 | 0 | 0 |
| ENSMUSG00 | SNORA17 | 131 | 0 | 0 | 0 | 0 | 0 | 0 |
| ENSMUSG00 | SNORA17 | 129 | 0 | 0 | 0 | 0 | 0 | 0 |
| ENSMUSG00 | Snord64 | 66 | 0 | 0 | 0 | 2.73348822 | 0 | 0 |
| ENSMUSG00 | Snora17 | 131 | 0 | 0 | 0 | 0 | 0 | 0 |
| ENSMUSG00 | U6 | 106 | 0 | 0 | 0 | 0 | 0 | 0 |
| ENSMUSG00 | U6 | 106 | 0 | 0 | 0 | 0 | 0 | 0 |
| ENSMUSG00 | SNORA17 | 130 | 0 | 0 | 0 | 0 | 0 | 0 |
| ENSMUSG00 | SNORA17 | 130 | 0 | 0 | 0 | 0 | 0 | 0 |
| ENSMUSG00 | SNORA17 | 131 | 0 | 0 | 0 | 0 | 0 | 0 |
| ENSMUSG00 | SNORA17 | 128 | 0 | 0 | 0 | 0.56378195 | 0 | 0 |
| ENSMUSG00 | SNORA17 | 132 | 0 | 0 | 0 | 0 | 0 | 0 |
| ENSMUSG00 | SNORA61 | 125 | 0 | 0 | 0 | 0.86596907 | 0 | 0 |
| ENSMUSG00 | SNORA17 | 128 | 0 | 0 | 0 | 0 | 0 | 0 |
| ENSMUSG00 | SNORA17 | 132 | 0 | 0 | 0 | 0 | 0 | 0 |
| ENSMUSG00 | U6 | 103 | 0 | 0 | 0 | 0 | 0 | 0 |
| ENSMUSG00 | SNORA17 | 126 | 0 | 0 | 0 | 0 | 0 | 0 |
| ENSMUSG00 | SNORA17 | 131 | 0 | 0 | 0 | 0 | 0 | 0 |
| ENSMUSG00 | Snord69 | 79 | 8.55497682 | 10.9791894 | 8.392755 | 13.7020422 | 24.5443327 | 11.3932037 |
| ENSMUSG00 | SNORA17 | 128 | 0 | 0 | 0 | 0 | 0 | 0 |
| ENSMUSG00 | SNORA17 | 126 | 0 | 0 | 0 | 0 | 0 | 0 |
| ENSMUSG00 | SNORA17 | 131 | 0 | 0 | 0 | 0 | 0 | 0 |
| ENSMUSG00 | SNORA17 | 131 | 0 | 0 | 0 | 0 | 0 | 0 |
| ENSMUSG00 | SNORD78 | 65 | 0 | 0 | 0 | 0 | 0 | 0 |
| ENSMUSG00 | Snord67 | 110 | 0 | 0 | 0 | 0 | 0 | 0 |
| ENSMUSG00 | SNORA66 | 131 | 0 | 0.36783544 | 0.33741865 | 0 | 0 | 2.08203353 |
| ENSMUSG00 | SNORA17 | 127 | 0 | 0 | 0 | 0 | 0 | 0 |
| ENSMUSG00 | SNORA17 | 126 | 0 | 0 | 0 | 0 | 0 | 0 |
| ENSMUSG00 | SNORD115 | 78 | 0 | 0 | 0 | 0 | 0 | 0 |
| ENSMUSG00 | SNORA17 | 133 | 0 | 0 | 0 | 2.98422925 | 0 | 0 |
| ENSMUSG00 | SNORA17 | 131 | 0 | 0 | 0 | 0 | 0 | 0 |
| ENSMUSG00 | U11 | 144 | 0 | 0 | 0.30695724 | 0 | 0 | 0 |
| ENSMUSG00 | SNORA17 | 128 | 0 | 0 | 0 | 0 | 0 | 0 |
| ENSMUSG00 | SNORA17 | 130 | 0 | 0 | 0 | 0 | 0 | 0 |
| ENSMUSG00 | SNORA17 | 131 | 0 | 0 | 0 | 0 | 0 | 0 |
| ENSMUSG00 | SNORA17 | 131 | 0 | 0 | 0 | 0 | 0 | 0 |
| ENSMUSG00 | Snord66 | 74 | 0.53723622 | 0 | 2.38928881 | 0.4875952 | 0 | 0.36857621 |
| ENSMUSG00 | SNORA17 | 123 | 0 | 0 | 0 | 0 | 0 | 0 |
| ENSMUSG00 | SNORA17 | 128 | 0 | 0 | 0 | 0 | 0 | 0 |
| ENSMUSG00 | SNORA17 | 130 | 0 | 0 | 0 | 0 | 0 | 0 |
| ENSMUSG00 | SNORA17 | 129 | 0 | 0 | 0 | 0 | 0 | 0 |
| ENSMUSG00 | SNORA17 | 131 | 0 | 0 | 0 | 0 | 0 | 0 |
| ENSMUSG00 | SNORA17 | 134 | 0 | 0 | 0 | 0 | 0 | 0 |
| ENSMUSG00 | U6 | 106 | 0 | 0 | 0 | 0 | 0 | 0 |
| ENSMUSG00 | U6 | 103 | 0 | 0 | 0 | 0 | 0 | 0 |
| ENSMUSG00 | SNORA65 | 120 | 0 | 0 | 0 | 0 | 0 | 0 |
| ENSMUSG00 | SNORA17 | 128 | 0 | 0 | 0 | 0 | 0 | 0 |
| ENSMUSG00 | SNORA17 | 122 | 0 | 0 | 0 | 0 | 0 | 0 |
| ENSMUSG00 | U6 | 104 | 0 | 0 | 0 | 0 | 0 | 0 |
| ENSMUSG00 | SNORA49 | 132 | 0.60235577 | 0 | 0 | 0 | 0 | 0 |
| ENSMUSG00 | SNORA17 | 127 | 0 | 0 | 0 | 0 | 0 | 0 |
| ENSMUSG00 | SNORA17 | 131 | 0 | 0 | 0 | 0 | 0 | 0 |
| ENSMUSG00 | SNORA17 | 130 | 0.61162278 | 1.11199482 | 0 | 1.38777094 | 0 | 0 |
| ENSMUSG00 | SNORA17 | 126 | 0 | 0 | 0 | 0 | 0 | 0 |
| ENSMUSG00 | SNORA17 | 127 | 0 | 0 | 0 | 0 | 0 | 0 |
| ENSMUSG00 | SNORA17 | 127 | 0 | 0 | 0 | 0 | 0 | 0 |
| ENSMUSG00 | snoU2-30 | 65 | 0 | 0 | 0 | 0 | 0 | 0 |
| ENSMUSG00 | n-R5s165 | 99 | 0 | 0 | 0 | 0 | 0 | 0 |
| ENSMUSG00 | SNORA17 | 125 | 0 | 0 | 0 | 0 | 0 | 0 |
| ENSMUSG00 | SNORA17 | 131 | 0 | 0 | 0 | 0 | 0 | 0 |
| ENSMUSG00 | U6 | 103 | 0 | 0.46782954 | 0 | 0 | 0.99080341 | 0 |
| ENSMUSG00 | SNORA17 | 132 | 0 | 0 | 0 | 0 | 0 | 0 |
| ENSMUSG00 | SNORA17 | 131 | 0 | 0 | 0 | 0 | 0 | 0 |
| ENSMUSG00 | SNORA17 | 131 | 0 | 0 | 0 | 0 | 0 | 0 |
| ENSMUSG00 | SNORA17 | 136 | 0 | 0 | 0 | 0 | 1.12558182 | 2.00548818 |
| ENSMUSG00 | SNORA17 | 131 | 0 | 0 | 0 | 0 | 0 | 0 |
| ENSMUSG00 | U6 | 106 | 1.87525852 | 0.45458908 | 1.66799408 | 0 | 0 | 1.80115542 |
| ENSMUSG00 | SNORA26 | 121 | 0 | 0 | 0 | 0 | 0 | 0 |
| ENSMUSG00 | SNORA17 | 131 | 0 | 0 | 0 | 0 | 0 | 0 |
| ENSMUSG00 | SNORA69 | 131 | 0 | 0 | 1.01225595 | 0 | 0.38951432 | 0 |
| ENSMUSG00 | U6 | 106 | 0 | 0 | 0 | 0 | 0 | 0 |

| | | | | | | | | |
|---|---|---|---|---|---|---|---|---|
| ENSMUSG00 | SNORA17 | 127 | 0 | 0 | 0 | 0 | 0 | 0 |
| ENSMUSG00 | SNORA17 | 128 | 0 | 0 | 0 | 0 | 0 | 0 |
| ENSMUSG00 | SNORA17 | 133 | 0 | 0 | 0 | 0 | 0 | 0 |
| ENSMUSG00 | SNORA17 | 131 | 0 | 0 | 0 | 0 | 0 | 0 |
| ENSMUSG00 | SNORA17 | 131 | 0 | 0 | 0 | 0 | 0 | 0 |
| ENSMUSG00 | n-R5s158 | 117 | 0 | 0 | 0 | 0 | 0 | 0 |
| ENSMUSG00 | SNORA42 | 114 | 0 | 0 | 0 | 0 | 0 | 0 |
| ENSMUSG00 | SNORA17 | 131 | 0 | 0 | 0 | 0 | 0 | 0 |
| ENSMUSG00 | SNORA17 | 128 | 0 | 0 | 0 | 0 | 0 | 0 |
| ENSMUSG00 | SNORA17 | 131 | 0 | 0 | 0 | 0 | 0 | 0 |
| ENSMUSG00 | SNORA17 | 130 | 0 | 0 | 0 | 0 | 0 | 0 |
| ENSMUSG00 | SNORA17 | 133 | 0 | 0 | 0 | 0 | 0 | 0 |
| ENSMUSG00 | SNORA17 | 131 | 0 | 0 | 0 | 0 | 0 | 0 |
| ENSMUSG00 | SNORA17 | 132 | 0 | 0 | 0 | 0 | 0 | 0 |
| ENSMUSG00 | SNORA17 | 136 | 0 | 0 | 0 | 0 | 0 | 0 |
| ENSMUSG00 | SNORD12 | 82 | 0 | 0 | 0 | 0 | 0 | 0 |
| ENSMUSG00 | SNORA17 | 131 | 0 | 0 | 0 | 0 | 0 | 0 |
| ENSMUSG00 | SNORA17 | 132 | 0 | 0 | 0 | 0 | 0 | 0 |
| ENSMUSG00 | SNORD115 | 78 | 0 | 0 | 0 | 0 | 0 | 0 |
| ENSMUSG00 | SNORA17 | 129 | 0 | 0 | 0 | 0 | 0 | 0 |
| ENSMUSG00 | U6 | 105 | 0 | 0 | 0 | 0 | 0 | 0 |
| ENSMUSG00 | SNORA17 | 131 | 0 | 0 | 0 | 0 | 0 | 0 |
| ENSMUSG00 | U11 | 133 | 0 | 0 | 0 | 0 | 0 | 0 |
| ENSMUSG00 | SNORA17 | 129 | 0 | 0 | 0 | 0 | 0 | 0 |
| ENSMUSG00 | SNORA17 | 129 | 0 | 0 | 0 | 0 | 0 | 0 |
| ENSMUSG00 | SNORA17 | 132 | 0 | 0 | 0 | 0 | 0 | 0 |
| ENSMUSG00 | SNORA17 | 131 | 0 | 0 | 0 | 0 | 0 | 0 |
| ENSMUSG00 | n-R5s8 | 117 | 0 | 0 | 0 | 0 | 0 | 0 |
| ENSMUSG00 | SNORA17 | 131 | 0 | 0 | 0 | 0 | 0 | 0 |
| ENSMUSG00 | SNORA17 | 132 | 0 | 0 | 0 | 0 | 0 | 0 |
| ENSMUSG00 | SNORA17 | 130 | 0 | 0 | 0 | 0 | 0 | 0 |
| ENSMUSG00 | SNORA24 | 116 | 0 | 0 | 0 | 0 | 0 | 0 |
| ENSMUSG00 | n-R5s171 | 78 | 0 | 0 | 0 | 0 | 0 | 0 |
| ENSMUSG00 | U11 | 124 | 0 | 0 | 0 | 0 | 0 | 0 |
| ENSMUSG00 | SNORA40 | 127 | 0 | 0 | 0 | 0 | 0 | 0 |
| ENSMUSG00 | SNORA17 | 131 | 0 | 0 | 0 | 0 | 0 | 0 |
| ENSMUSG00 | U11 | 133 | 0 | 0 | 0 | 0 | 0.38365696 | 0.20507248 |
| ENSMUSG00 | SNORA17 | 131 | 0 | 0 | 0 | 0 | 0 | 0 |
| ENSMUSG00 | SNORA17 | 132 | 0 | 0 | 0 | 0 | 0 | 0 |
| ENSMUSG00 | U1 | 165 | 0 | 0 | 0 | 0 | 0 | 0 |
| ENSMUSG00 | SNORA17 | 129 | 0 | 0 | 0 | 0 | 0 | 0 |
| ENSMUSG00 | SNORA17 | 131 | 0 | 0 | 0 | 0 | 0 | 0 |
| ENSMUSG00 | SNORA17 | 129 | 0 | 0 | 0 | 0 | 0 | 0 |
| ENSMUSG00 | SNORA17 | 127 | 0 | 0 | 0 | 0 | 0 | 0 |
| ENSMUSG00 | U6 | 105 | 0 | 0 | 0 | 0 | 0 | 0 |
| ENSMUSG00 | SNORA17 | 131 | 0 | 0 | 0 | 0 | 0 | 0 |
| ENSMUSG00 | SNORA24 | 82 | 0 | 0 | 0 | 0 | 0 | 0 |
| ENSMUSG00 | U1 | 163 | 0 | 0 | 0 | 0 | 0 | 0 |
| ENSMUSG00 | SNORA17 | 125 | 0 | 0 | 1.06084423 | 0 | 1.63284403 | 0 |
| ENSMUSG00 | SNORA17 | 133 | 0 | 0 | 0 | 0 | 0 | 0 |
| ENSMUSG00 | SNORA17 | 131 | 0 | 0 | 0 | 0 | 0 | 0 |
| ENSMUSG00 | SNORA17 | 124 | 0 | 0 | 0 | 0 | 0 | 0 |
| ENSMUSG00 | Snord70 | 86 | 0 | 0 | 0 | 0 | 0 | 0.9514409 |
| ENSMUSG00 | U1 | 157 | 0 | 0 | 0 | 0 | 0 | 0 |
| ENSMUSG00 | SNORA17 | 128 | 0 | 0 | 0 | 0 | 0 | 0 |
| ENSMUSG00 | U6 | 106 | 0 | 0 | 0.41699852 | 0 | 0 | 0 |
| ENSMUSG00 | n-R5s31 | 109 | 0 | 0.44207745 | 0 | 0 | 0 | 0 |
| ENSMUSG00 | SNORA17 | 131 | 0 | 0 | 0 | 0 | 0 | 0 |
| ENSMUSG00 | SNORA17 | 131 | 0 | 0 | 0 | 0 | 0 | 0 |
| ENSMUSG00 | SNORA17 | 131 | 0 | 0 | 0 | 0 | 0 | 0 |
| ENSMUSG00 | SNORA42 | 133 | 0 | 0 | 0 | 0 | 0 | 0 |
| ENSMUSG00 | SNORA17 | 131 | 0 | 0 | 0 | 0 | 0 | 0 |
| ENSMUSG00 | SNORA17 | 130 | 0 | 0 | 0 | 0 | 0 | 0 |
| ENSMUSG00 | SNORD70 | 83 | 0 | 0 | 7.98828488 | 0 | 0 | 2.62888089 |
| ENSMUSG00 | U1 | 157 | 0 | 0 | 0 | 0 | 0 | 0 |
| ENSMUSG00 | SNORA24 | 121 | 0 | 0 | 0 | 0.89459615 | 0 | 0 |
| ENSMUSG00 | SNORA17 | 130 | 0 | 0 | 0 | 0 | 0 | 0 |
| ENSMUSG00 | SNORA17 | 130 | 0 | 0 | 0 | 0 | 0 | 0 |
| ENSMUSG00 | U6 | 107 | 0 | 0 | 0 | 1.01164611 | 0 | 0 |
| ENSMUSG00 | SNORA17 | 130 | 0 | 0 | 0 | 0 | 0 | 0 |
| ENSMUSG00 | U1 | 154 | 0 | 0 | 0 | 0 | 0 | 0 |
| ENSMUSG00 | SNORA17 | 119 | 0 | 0 | 0 | 0 | 0 | 0 |
| ENSMUSG00 | SNORA17 | 131 | 0 | 0 | 0 | 0 | 0 | 0 |
| ENSMUSG00 | SNORA17 | 128 | 0 | 0 | 0 | 0 | 0 | 0 |
| ENSMUSG00 | SNORA17 | 128 | 0 | 0 | 0 | 0 | 0 | 0 |
| ENSMUSG00 | SNORA68 | 131 | 0 | 0 | 0 | 0 | 0 | 0 |
| ENSMUSG00 | SNORA17 | 129 | 0 | 0 | 0 | 0 | 0 | 0 |
| ENSMUSG00 | n-R5s200 | 112 | 0 | 0 | 0 | 0 | 0 | 0 |
| ENSMUSG00 | SNORA17 | 129 | 0 | 0 | 0 | 0 | 0 | 0 |
| ENSMUSG00 | U1 | 154 | 0 | 0 | 0 | 0 | 0 | 0 |
| ENSMUSG00 | SNORA17 | 131 | 0 | 0 | 2.36193054 | 0 | 0 | 1.24922012 |
| ENSMUSG00 | U1 | 170 | 0 | 0 | 0.26001084 | 0 | 0 | 0 |
| ENSMUSG00 | SNORA12 | 145 | 0.54835146 | 1.66160145 | 0 | 0.74652506 | 0 | 0 |
| ENSMUSG00 | SNORA17 | 125 | 0 | 0 | 0 | 0 | 0 | 0 |
| ENSMUSG00 | SNORA24 | 126 | 0 | 0 | 0 | 0.57273087 | 0 | 0 |
| ENSMUSG00 | snoU2_19 | 75 | 0 | 0 | 0 | 0 | 0 | 0 |
| ENSMUSG00 | U6 | 96 | 0 | 0 | 0 | 0 | 0 | 0 |
| ENSMUSG00 | SNORA17 | 124 | 0 | 0 | 0 | 0 | 0 | 0 |
| ENSMUSG00 | SNORA17 | 127 | 0 | 0 | 0 | 0 | 0 | 0 |
| ENSMUSG00 | SNORA17 | 131 | 0 | 0 | 0 | 0 | 0 | 0 |
| ENSMUSG00 | SNORA17 | 130 | 0 | 0 | 0 | 0 | 0 | 0 |
| ENSMUSG00 | n-R5s82 | 108 | 0 | 0 | 0 | 0 | 0 | 0 |
| ENSMUSG00 | SNORA17 | 128 | 0 | 0 | 0 | 0 | 0 | 0 |
| ENSMUSG00 | U3 | 189 | 0 | 0 | 0 | 0 | 0 | 0 |
| ENSMUSG00 | SNORA17 | 126 | 0 | 0 | 0 | 0 | 0 | 0 |
| ENSMUSG00 | SNORA17 | 131 | 0 | 0 | 0 | 0 | 0 | 0 |
| ENSMUSG00 | U1 | 163 | 0 | 0 | 0 | 0 | 0 | 0 |
| ENSMUSG00 | SNORA17 | 128 | 0 | 0 | 0 | 0 | 0 | 0 |
| ENSMUSG00 | SNORA17 | 131 | 0 | 0 | 0 | 0 | 0 | 0 |
| ENSMUSG00 | U6 | 106 | 0 | 0 | 0 | 0 | 0 | 0 |
| ENSMUSG00 | SNORA17 | 131 | 0 | 0 | 0 | 0 | 0 | 0 |
| ENSMUSG00 | SNORA17 | 130 | 0 | 0 | 0 | 0 | 0 | 0 |
| ENSMUSG00 | SNORD30 | 68 | 0 | 0 | 0 | 0 | 0 | 0 |
| ENSMUSG00 | SNORA17 | 130 | 0 | 0 | 0 | 0 | 0 | 0 |
| ENSMUSG00 | SNORA17 | 130 | 0 | 0 | 0 | 0.27755419 | 0 | 0 |
| ENSMUSG00 | SNORA17 | 124 | 0 | 0 | 0 | 0 | 0 | 0 |
| ENSMUSG00 | SNORA17 | 124 | 0 | 0 | 0 | 0 | 0 | 0 |
| ENSMUSG00 | U2 | 148 | 0 | 0 | 0 | 0 | 0 | 0 |
| ENSMUSG00 | AC161183.1 | 65 | 7.9510961 | 8.15462867 | 6.80028354 | 13.8777094 | 3.92510583 | 3.77648851 |
| ENSMUSG00 | U6 | 106 | 0 | 0.90917815 | 0 | 0 | 0.4813809 | 0 |
| ENSMUSG00 | SNORA17 | 135 | 0 | 0 | 0 | 0 | 0 | 0 |
| ENSMUSG00 | SNORA17 | 130 | 0 | 0 | 0 | 0 | 0 | 0 |
| ENSMUSG00 | SNORA17 | 133 | 0 | 0 | 0 | 0 | 0 | 0 |
| ENSMUSG00 | SNORA66 | 131 | 4.85563121 | 3.67835436 | 0 | 5.78414454 | 0.77902864 | 6.03789724 |
| ENSMUSG00 | SNORA17 | 126 | 0 | 0 | 0 | 0 | 0 | 0 |
| ENSMUSG00 | SNORA42 | 141 | 0 | 0 | 0 | 0 | 0 | 0 |
| ENSMUSG00 | SNORA17 | 130 | 0 | 0 | 0 | 0 | 0 | 0 |
| ENSMUSG00 | U6 | 105 | 0 | 0 | 0 | 0 | 0 | 0 |
| ENSMUSG00 | U6 | 108 | 0 | 0 | 0 | 0 | 0 | 0 |
| ENSMUSG00 | SNORA17 | 127 | 0 | 0 | 0 | 0 | 0 | 0 |
| ENSMUSG00 | n-R5s178 | 113 | 0 | 0 | 0 | 0 | 0 | 0 |
| ENSMUSG00 | SNORA17 | 129 | 0 | 0 | 0 | 0 | 0 | 0 |
| ENSMUSG00 | SNORA17 | 129 | 0 | 0 | 0 | 0 | 0 | 0 |
| ENSMUSG00 | SNORD101 | 73 | 4.35676499 | 3.30044124 | 3.63302819 | 0.49427458 | 0.69899145 | 0 |
| ENSMUSG00 | SNORA17 | 131 | 0 | 0 | 0 | 0 | 0 | 0 |
| ENSMUSG00 | SNORA17 | 131 | 0 | 0 | 0 | 0 | 0 | 0 |
| ENSMUSG00 | U12 | 134 | 0 | 0 | 0 | 0 | 0 | 0 |
| ENSMUSG00 | SNORA17 | 131 | 0 | 0 | 0 | 0 | 0 | 0 |
| ENSMUSG00 | SNORA17 | 131 | 0 | 0 | 0 | 0 | 0 | 0 |
| ENSMUSG00 | SNORA17 | 131 | 14.8703706 | 2.94268349 | 28.6805852 | 15.14895 | 12.4644582 | 31.855113 |
| ENSMUSG00 | SNORA17 | 133 | 0 | 0 | 0 | 0 | 0 | 0 |
| ENSMUSG00 | SNORA42 | 129 | 0.92454606 | 0 | 0 | 0.27970577 | 1.1866599 | 0 |
| ENSMUSG00 | Rab11b | 3701 | 10.1617629 | 11.4965221 | 11.1788503 | 7.89690788 | 9.6372431 | 11.7101869 |
| ENSMUSG00 | SNORA17 | 130 | 0 | 0 | 0 | 0 | 0 | 0 |
| ENSMUSG00 | SNORA17 | 125 | 0 | 0 | 0 | 0 | 0 | 0 |
| ENSMUSG00 | U6 | 102 | 0 | 0 | 0 | 0 | 0 | 0 |
| ENSMUSG00 | SNORA17 | 129 | 0 | 0 | 0 | 0 | 0 | 0 |
| ENSMUSG00 | SNORA17 | 130 | 0 | 0 | 0 | 0 | 0 | 0 |
| ENSMUSG00 | Snord65 | 71 | 2.23974538 | 0 | 2.49024468 | 2.03279124 | 1.4373627 | 9.21959637 |
| ENSMUSG00 | SNORA17 | 131 | 0 | 0 | 0 | 0 | 0 | 0 |
| ENSMUSG00 | SNORA17 | 132 | 0.30117788 | 0 | 0 | 0.82004647 | 0.38656345 | 0 |
| ENSMUSG00 | U6 | 102 | 0 | 0 | 0 | 0 | 0 | 0 |
| ENSMUSG00 | SNORA17 | 134 | 0 | 0 | 0 | 0 | 0 | 0 |
| ENSMUSG00 | SNORA64 | 136 | 0 | 0 | 0 | 0 | 0 | 0 |
| ENSMUSG00 | SNORA17 | 131 | 0 | 0 | 0 | 0 | 0 | 0 |
| ENSMUSG00 | SNORA17 | 127 | 0 | 0 | 0 | 0 | 0 | 0 |
| ENSMUSG00 | SNORA17 | 131 | 0 | 0 | 0 | 0 | 0 | 0 |
| ENSMUSG00 | SNORA17 | 133 | 0 | 0 | 0 | 0 | 0 | 0 |
| ENSMUSG00 | SNORA17 | 129 | 0 | 0 | 0 | 0 | 0 | 0 |
| ENSMUSG00 | SNORA17 | 130 | 0 | 0 | 0 | 0 | 0 | 0 |
| ENSMUSG00 | U1 | 155 | 0 | 0 | 0 | 0 | 0 | 0 |
| ENSMUSG00 | SNORA17 | 130 | 0 | 0 | 0 | 0 | 0 | 0 |
| ENSMUSG00 | SNORA17 | 131 | 0 | 0 | 0 | 0 | 0 | 0 |
| ENSMUSG00 | SNORA17 | 127 | 0 | 0 | 0 | 0 | 0 | 0 |
| ENSMUSG00 | SNORA17 | 139 | 0 | 0 | 0 | 0 | 0 | 0 |
| ENSMUSG00 | SNORA17 | 131 | 0 | 0 | 0 | 0 | 0 | 0 |
| ENSMUSG00 | SNORA17 | 131 | 0 | 0 | 0 | 0 | 0 | 0 |
| ENSMUSG00 | SNORA17 | 127 | 0 | 0 | 0 | 0 | 0 | 0 |
| ENSMUSG00 | SNORA42 | 114 | 0 | 0 | 0 | 0 | 0 | 0 |
| ENSMUSG00 | SNORA17 | 134 | 0 | 0 | 0 | 0 | 0 | 0 |
| ENSMUSG00 | SNORA17 | 128 | 0 | 0 | 0 | 0 | 0 | 0 |
| ENSMUSG00 | SNORA17 | 129 | 0 | 0 | 0 | 0 | 0 | 0 |
| ENSMUSG00 | SNORA17 | 123 | 0 | 0 | 0 | 0 | 0 | 0 |
| ENSMUSG00 | SNORA17 | 131 | 0 | 0 | 0 | 0 | 0 | 0 |
| ENSMUSG00 | Snord91a | 92 | 32.8414839 | 14.1416732 | 22.1009215 | 8.23611886 | 23.8492844 | 7.70805022 |
| ENSMUSG00 | SNORA17 | 129 | 0 | 0 | 0 | 0 | 0 | 0 |
| ENSMUSG00 | SNORA17 | 133 | 0 | 0 | 0 | 0 | 0 | 0 |
| ENSMUSG00 | SNORA17 | 131 | 0 | 0 | 0 | 0 | 0 | 0 |
| ENSMUSG00 | SNORA26 | 119 | 0 | 0 | 0 | 0 | 0 | 0 |
| ENSMUSG00 | SNORA17 | 131 | 0 | 0 | 0 | 0 | 0 | 0 |
| ENSMUSG00 | SNORA17 | 131 | 0 | 0 | 0 | 0 | 0 | 0 |
| ENSMUSG00 | SNORA17 | 128 | 0 | 0 | 0 | 0 | 0 | 0 |
| ENSMUSG00 | SNORA17 | 131 | 0 | 0 | 0 | 0 | 0 | 0 |
| ENSMUSG00 | SNORA17 | 132 | 0 | 0 | 0 | 0 | 0 | 0 |
| ENSMUSG00 | U6 | 106 | 0 | 0 | 0 | 0 | 0 | 0 |
| ENSMUSG00 | SNORA26 | 121 | 0 | 0 | 0 | 0 | 0.42170559 | 0 |
| ENSMUSG00 | Scarna9 | 69 | 0 | 0 | 0 | 0 | 0 | 0 |
| ENSMUSG00 | SNORA17 | 128 | 0 | 0 | 0 | 0 | 0 | 0 |
| ENSMUSG00 | AC120367.1 | 109 | 0 | 0 | 0 | 0 | 0 | 0 |
| ENSMUSG00 | U6 | 106 | 0 | 0 | 0 | 0 | 0 | 0 |
| ENSMUSG00 | SNORA17 | 131 | 0 | 0 | 0 | 0 | 0 | 0 |
| ENSMUSG00 | SNORA66 | 128 | 0 | 0 | 0 | 0 | 0 | 0 |
| ENSMUSG00 | SNORD12 | 92 | 0 | 0 | 0 | 0 | 0 | 0 |
| ENSMUSG00 | n-R5s219 | 131 | 0 | 0 | 0 | 0 | 0 | 0 |
| ENSMUSG00 | SNORA32 | 118 | 0 | 0 | 0 | 0 | 0 | 0 |
| ENSMUSG00 | U6 | 106 | 0 | 0 | 0 | 0 | 0 | 0 |
| ENSMUSG00 | SNORA17 | 131 | 0 | 0 | 0 | 0 | 0 | 0 |
| ENSMUSG00 | SNORA17 | 131 | 0 | 1.10350631 | 0 | 0 | 0 | 0 |
| ENSMUSG00 | SNORA17 | 128 | 0 | 0 | 0 | 0 | 0 | 0 |
| ENSMUSG00 | SNORA17 | 128 | 0 | 0 | 0 | 0 | 0 | 0 |
| ENSMUSG00 | n-R5s67 | 113 | 0.35181841 | 5.54357299 | 1.17350026 | 2.2351709 | 3.61246678 | 3.86189582 |
| ENSMUSG00 | SNORA65 | 113 | 0 | 0 | 0 | 0 | 0 | 0.72410547 |
| ENSMUSG00 | SNORD41 | 69 | 0 | 0 | 0 | 0 | 0 | 0 |
| ENSMUSG00 | SNORA17 | 132 | 0 | 0 | 0 | 0 | 0 | 0 |
| ENSMUSG00 | SNORA17 | 131 | 0 | 0 | 0 | 0 | 0 | 0 |
| ENSMUSG00 | U6 | 93 | 0.42747829 | 0 | 0 | 4.65574768 | 0 | 0 |
| ENSMUSG00 | SNORA17 | 126 | 0 | 0 | 0 | 0 | 0 | 0 |
| ENSMUSG00 | SNORA17 | 131 | 0 | 0 | 0 | 0 | 0 | 0 |
| ENSMUSG00 | SNORA17 | 129 | 0 | 0 | 0 | 0 | 0 | 0 |
| ENSMUSG00 | SNORA17 | 131 | 0 | 0 | 0 | 0 | 0 | 0 |
| ENSMUSG00 | U6 | 106 | 0 | 0 | 0 | 0 | 0 | 0 |
| ENSMUSG00 | SNORA17 | 129 | 0 | 0 | 0 | 0 | 0 | 0 |
| ENSMUSG00 | SNORA17 | 126 | 0 | 0 | 0 | 0 | 0 | 0 |
| ENSMUSG00 | SNORA17 | 130 | 0 | 0 | 0 | 0 | 1.96255292 | 0 |
| ENSMUSG00 | SNORA17 | 135 | 0 | 0 | 0 | 0 | 0 | 0 |
| ENSMUSG00 | U6 | 95 | 0 | 0 | 0 | 0 | 0 | 0 |
| ENSMUSG00 | Snord71 | 84 | 0 | 0 | 0 | 0 | 0.60745686 | 0.64939617 |
| ENSMUSG00 | CT009748.1 | 109 | 0 | 0 | 0 | 0 | 0 | 0 |
| ENSMUSG00 | SNORA17 | 129 | 0 | 0 | 0 | 0 | 0 | 0 |
| ENSMUSG00 | SNORA17 | 129 | 0 | 0 | 0 | 0 | 0 | 0 |
| ENSMUSG00 | SNORA17 | 124 | 0 | 0 | 0 | 0 | 0 | 0 |
| ENSMUSG00 | SNORA17 | 122 | 0 | 0 | 0 | 0 | 0 | 0 |
| ENSMUSG00 | SNORD66 | 81 | 0 | 0 | 0 | 1.78182936 | 1.25991051 | 2.02034365 |
| ENSMUSG00 | SNORA17 | 129 | 0 | 0 | 0 | 0 | 0 | 0 |
| ENSMUSG00 | SNORA17 | 131 | 0 | 0 | 0 | 0 | 0 | 0 |
| ENSMUSG00 | SNORA17 | 132 | 0 | 0 | 0 | 0 | 0 | 0 |
| ENSMUSG00 | U6 | 106 | 0 | 0 | 0 | 0 | 0 | 0 |
| ENSMUSG00 | Snora68 | 130 | 5.50460499 | 4.81864421 | 5.44022683 | 7.49396309 | 3.53259525 | 11.5392705 |
| ENSMUSG00 | SNORA27 | 128 | 0.31058969 | 0 | 2.76261519 | 1.97323681 | 0 | 0 |
| ENSMUSG00 | SNORD12 | 88 | 1.35530047 | 0 | 0 | 0.41002323 | 2.31938072 | 0.61987817 |
| ENSMUSG00 | SNORA17 | 131 | 0 | 0 | 0 | 0 | 0.77902864 | 0 |
| ENSMUSG00 | SNORA68 | 135 | 0 | 0.35693661 | 1.6371053 | 4.00911606 | 2.64581208 | 3.03051547 |
| ENSMUSG00 | SNORA17 | 131 | 0 | 0 | 0 | 0 | 0 | 0 |
| ENSMUSG00 | Snora36b | 131 | 0 | 0 | 0 | 0 | 0 | 0 |
| ENSMUSG00 | SNORA17 | 122 | 0 | 0 | 0 | 0 | 0 | 0 |
| ENSMUSG00 | SNORA17 | 131 | 0 | 0 | 0 | 0 | 0 | 0 |
| ENSMUSG00 | U6 | 107 | 0 | 0 | 0 | 1.01164611 | 0 | 0 |
| ENSMUSG00 | SNORA3 | 129 | 1.23272808 | 2.24122987 | 3.42649946 | 1.67823463 | 0 | 3.17146968 |
| ENSMUSG00 | SNORA17 | 131 | 0 | 0 | 0 | 0 | 0 | 0 |
| ENSMUSG00 | U11 | 101 | 0 | 0 | 0 | 0 | 0 | 0 |
| ENSMUSG00 | SNORA17 | 129 | 0 | 0 | 0 | 0 | 0 | 0 |
| ENSMUSG00 | SNORA17 | 135 | 0 | 0 | 0 | 0 | 0 | 0 |
| ENSMUSG00 | SNORA17 | 131 | 0 | 0 | 0 | 0 | 0 | 0 |
| ENSMUSG00 | U6 | 105 | 0 | 0 | 0 | 0 | 0 | 0 |
| ENSMUSG00 | SNORA17 | 131 | 0 | 0 | 0.33741865 | 0 | 0 | 0 |
| ENSMUSG00 | SNORA17 | 130 | 0 | 0 | 0 | 0 | 0 | 0 |
| ENSMUSG00 | SNORA17 | 126 | 0 | 0 | 0 | 0 | 0 | 0 |
| ENSMUSG00 | Y_RNA | 109 | 0 | 0 | 0 | 0 | 0 | 0 |
| ENSMUSG00 | SNORA17 | 131 | 0 | 0 | 0 | 0 | 0 | 0 |
| ENSMUSG00 | SNORA17 | 124 | 0 | 0 | 0 | 0 | 0 | 0 |
| ENSMUSG00 | SNORA17 | 128 | 0 | 0 | 0 | 0 | 0 | 0 |
| ENSMUSG00 | SNORA17 | 118 | 0 | 0 | 0 | 0 | 0 | 0 |
| ENSMUSG00 | SNORA17 | 131 | 0 | 0 | 1.3496746 | 0.55087091 | 0 | 0.20820335 |
| ENSMUSG00 | SNORA17 | 127 | 0 | 0 | 0 | 0 | 0 | 0 |
| ENSMUSG00 | SNORA17 | 118 | 0 | 0 | 0 | 0 | 0 | 0 |
| ENSMUSG00 | U6 | 106 | 0 | 0 | 0 | 0 | 0 | 0 |
| ENSMUSG00 | U6 | 73 | 0 | 0 | 0 | 0 | 0 | 0 |
| ENSMUSG00 | U6 | 103 | 0 | 0 | 0 | 0 | 0 | 0 |
| ENSMUSG00 | SNORA17 | 127 | 0 | 0 | 0 | 0 | 0 | 0 |
| ENSMUSG00 | SNORA17 | 130 | 0 | 0 | 0 | 0 | 0 | 0 |
| ENSMUSG00 | SNORA17 | 133 | 1.19565355 | 0 | 0 | 0 | 0.38365696 | 0 |
| ENSMUSG00 | SNORA3 | 130 | 0 | 0 | 0 | 0 | 0 | 0 |
| ENSMUSG00 | Snord1b | 74 | 0 | 0 | 0 | 0 | 0 | 0 |
| ENSMUSG00 | SNORA17 | 131 | 0 | 0 | 0 | 0 | 0 | 0 |
| ENSMUSG00 | SNORA17 | 130 | 0 | 0 | 0 | 0 | 0 | 0 |
| ENSMUSG00 | SNORA17 | 133 | 0 | 0 | 0 | 0 | 0 | 0 |
| ENSMUSG00 | SNORA17 | 131 | 0 | 0 | 0 | 0 | 0 | 0 |
| ENSMUSG00 | SNORA74 | 106 | 1.21700451 | 0 | 0 | 0.92046032 | 0 | 1.2524069 |
| ENSMUSG00 | SNORA17 | 128 | 0 | 0 | 0 | 0 | 0 | 0 |
| ENSMUSG00 | SNORA17 | 131 | 0 | 0 | 0 | 0 | 0 | 0 |
| ENSMUSG00 | SNORA40 | 122 | 5.21383351 | 6.71450423 | 2.17386113 | 4.73207141 | 1.25474695 | 0.44712523 |
| ENSMUSG00 | SNORD115 | 68 | 0 | 0 | 0 | 0 | 0 | 0 |
| ENSMUSG00 | SNORA17 | 130 | 0 | 0 | 0 | 0 | 0 | 0 |
| ENSMUSG00 | SNORA17 | 131 | 0 | 0 | 0 | 0 | 0 | 0 |
| ENSMUSG00 | SNORA17 | 119 | 0 | 0 | 0 | 0 | 0 | 0 |
| ENSMUSG00 | SNORA17 | 130 | 0 | 0 | 0 | 0 | 0 | 0 |
| ENSMUSG00 | n-R5s9 | 116 | 0 | 0 | 0 | 0 | 0 | 0 |
| ENSMUSG00 | SNORA63 | 101 | 0.78723724 | 0.95418697 | 0 | 0 | 0 | 0 |
| ENSMUSG00 | SNORA61 | 128 | 0 | 0 | 0 | 0 | 0 | 0 |
| ENSMUSG00 | SNORA17 | 128 | 0.31058969 | 0 | 0 | 0 | 0 | 0 |
| ENSMUSG00 | Snord4a | 70 | 0 | 0 | 0 | 0 | 0 | 0 |
| ENSMUSG00 | SNORA17 | 131 | 0 | 0 | 0 | 0 | 0 | 0 |
| ENSMUSG00 | SNORA17 | 130 | 0 | 0 | 0 | 0 | 0 | 0 |
| ENSMUSG00 | SNORA17 | 120 | 0 | 0 | 0 | 0 | 0 | 0 |
| ENSMUSG00 | SNORA17 | 135 | 0 | 0 | 0 | 0 | 1.51189262 | 0 |
| ENSMUSG00 | SNORA17 | 133 | 0 | 0 | 0 | 0 | 0 | 0 |
| ENSMUSG00 | U6 | 106 | 0 | 0 | 0 | 0 | 0 | 0 |
| ENSMUSG00 | SNORA17 | 134 | 0 | 0 | 0 | 0 | 0 | 0 |
| ENSMUSG00 | SNORD91 | 90 | 2.20863781 | 6.96026387 | 8.34923701 | 1.20273482 | 6.23655704 | 1.51525774 |
| ENSMUSG00 | SNORA33 | 120 | 0.33129567 | 0 | 0.73669738 | 0 | 0 | 0 |
| ENSMUSG00 | SNORA17 | 131 | 0 | 0 | 0 | 0 | 0 | 0 |
| ENSMUSG00 | SNORA17 | 133 | 0 | 0 | 0 | 0 | 0 | 0 |
| ENSMUSG00 | SNORA17 | 132 | 0 | 0 | 0 | 0 | 0 | 0 |
| ENSMUSG00 | SNORA17 | 128 | 0 | 0 | 0 | 0 | 0 | 0 |
| ENSMUSG00 | SNORA17 | 131 | 0 | 0 | 0 | 0 | 0 | 0 |
| ENSMUSG00 | SNORD115 | 68 | 0 | 0 | 0 | 0 | 0 | 0 |
| ENSMUSG00 | SNORA17 | 131 | 0 | 0 | 0 | 0 | 0 | 0 |
| ENSMUSG00 | SNORA26 | 120 | 0 | 0 | 0 | 0 | 0 | 0 |
| ENSMUSG00 | SNORD66 | 87 | 2.28479773 | 2.76933576 | 1.52420148 | 6.22104216 | 6.45161074 | 6.27003201 |
| ENSMUSG00 | SNORA40 | 126 | 0 | 0 | 0 | 0 | 0 | 0 |
| ENSMUSG00 | U1 | 166 | 0 | 0 | 0 | 0 | 0 | 0.49291517 |
| ENSMUSG00 | SNORA17 | 125 | 0 | 0 | 0 | 0 | 0 | 0 |
| ENSMUSG00 | SNORA17 | 130 | 0 | 0 | 0 | 0 | 0 | 0 |
| ENSMUSG00 | SNORA17 | 132 | 0 | 0 | 0 | 0 | 0 | 0 |
| ENSMUSG00 | SNORA17 | 130 | 0 | 0 | 0 | 0 | 0 | 0 |
| ENSMUSG00 | n-R5s51 | 122 | 0 | 0 | 0 | 0 | 0 | 0 |
| ENSMUSG00 | SNORA17 | 131 | 0 | 0 | 0 | 0 | 0 | 0 |
| ENSMUSG00 | SNORA17 | 126 | 0 | 0 | 0 | 0 | 0 | 0 |
| ENSMUSG00 | U1 | 184 | 0 | 0 | 0 | 0 | 0 | 0 |
| ENSMUSG00 | SNORA24 | 113 | 0 | 0 | 0 | 0 | 0 | 0 |
| ENSMUSG00 | U6 | 107 | 0 | 0 | 0 | 0 | 0 | 0 |
| ENSMUSG00 | SNORA17 | 127 | 0 | 0 | 0 | 0 | 0 | 0 |
| ENSMUSG00 | SNORA17 | 130 | 0 | 0 | 0 | 0 | 0 | 0 |
| ENSMUSG00 | SNORA17 | 131 | 0 | 0 | 0 | 0 | 0 | 0 |
| ENSMUSG00 | SNORA17 | 130 | 0 | 0 | 0 | 0 | 0 | 0 |
| ENSMUSG00 | SNORA17 | 131 | 0 | 0 | 0 | 0 | 0 | 0 |
| ENSMUSG00 | SNORA17 | 130 | 0 | 0 | 0 | 0 | 0 | 0 |
| ENSMUSG00 | U6 | 103 | 0 | 0 | 0 | 0 | 0 | 0 |
| ENSMUSG00 | SNORA17 | 130 | 0 | 0 | 0 | 0 | 0 | 0 |
| ENSMUSG00 | SNORA17 | 125 | 0 | 0 | 0 | 0 | 0 | 0 |
| ENSMUSG00 | SNORD115 | 87 | 0 | 0 | 0 | 1.24420843 | 0 | 0 |
| ENSMUSG00 | SNORA17 | 131 | 0 | 0 | 0 | 0 | 0 | 0 |
| ENSMUSG00 | SNORA17 | 127 | 0 | 0 | 0 | 0 | 0 | 0 |
| ENSMUSG00 | SNORD66 | 85 | 0 | 0 | 0 | 3.39595713 | 1.80093091 | 0 |
| ENSMUSG00 | SNORA17 | 131 | 4.55215426 | 1.47134175 | 8.09804758 | 2.20348364 | 1.55805728 | 7.91172742 |
| ENSMUSG00 | SNORA17 | 130 | 0 | 0 | 0 | 0 | 0 | 0 |
| ENSMUSG00 | SNORA17 | 134 | 0 | 0.71920063 | 1.6493225 | 0 | 1.5231754 | 0 |
| ENSMUSG00 | SNORA17 | 131 | 0 | 0 | 0 | 0 | 0 | 0 |
| ENSMUSG00 | SNORA17 | 131 | 0 | 0 | 0 | 0 | 0 | 0 |
| ENSMUSG00 | SNORA17 | 132 | 0 | 0 | 0 | 0 | 0 | 0 |
| ENSMUSG00 | SNORA17 | 131 | 0 | 0 | 0 | 0 | 0 | 0 |
| ENSMUSG00 | n-R5s172 | 113 | 0 | 0 | 0 | 0 | 0 | 0 |
| ENSMUSG00 | SNORA17 | 127 | 0 | 0 | 0 | 0 | 0 | 0 |
| ENSMUSG00 | Snord12 | 89 | 0 | 0 | 0 | 0 | 0 | 0 |
| ENSMUSG00 | SNORA17 | 126 | 0 | 0 | 0 | 0 | 0 | 0 |
| ENSMUSG00 | SNORA17 | 131 | 0 | 0 | 0 | 0 | 0 | 0 |
| ENSMUSG00 | SNORD105 | 84 | 0 | 0 | 0 | 0 | 0 | 0 |
| ENSMUSG00 | SNORA17 | 130 | 0 | 0 | 0 | 0 | 0 | 0 |
| ENSMUSG00 | Snord89 | 109 | 2.55310425 | 5.74700686 | 0 | 1.65513966 | 0 | 1.00090419 |
| ENSMUSG00 | n-R5s202 | 113 | 0 | 0 | 0 | 0 | 0 | 0 |
| ENSMUSG00 | n-R5s72 | 118 | 0 | 0 | 0 | 0 | 0 | 1.38684606 |
| ENSMUSG00 | SNORA17 | 129 | 0 | 0 | 0 | 0 | 0 | 0 |
| ENSMUSG00 | SNORA17 | 131 | 0 | 0 | 0 | 0 | 0 | 0 |
| ENSMUSG00 | Snora64 | 133 | 0 | 0 | 0 | 0 | 0 | 0 |
| ENSMUSG00 | SNORA17 | 130 | 0 | 0 | 0 | 0 | 0 | 0 |
| ENSMUSG00 | SNORA40 | 124 | 5.77095685 | 4.66320408 | 1.78233238 | 1.16393692 | 0 | 2.63948122 |
| ENSMUSG00 | SNORA17 | 131 | 0 | 0 | 0 | 0 | 0 | 0 |
| ENSMUSG00 | U1 | 158 | 0 | 0 | 0 | 0 | 0 | 0 |
| ENSMUSG00 | Snord17 | 237 | 0 | 0.40663664 | 0 | 0 | 0.64590349 | 0 |
| ENSMUSG00 | SNORA17 | 129 | 0 | 0 | 0 | 0 | 0 | 0 |
| ENSMUSG00 | SNORA17 | 128 | 0 | 0 | 0 | 0 | 0 | 0 |
| ENSMUSG00 | SNORA17 | 125 | 0 | 0 | 0 | 0 | 0 | 0 |
| ENSMUSG00 | SNORA17 | 116 | 0 | 0 | 0 | 0 | 0 | 0 |
| ENSMUSG00 | SNORA17 | 129 | 0 | 0 | 0 | 0 | 0 | 0 |
| ENSMUSG00 | SNORA17 | 130 | 0 | 0 | 0 | 0 | 0 | 0 |
| ENSMUSG00 | SNORA17 | 126 | 0 | 0 | 0 | 0 | 0 | 0 |
| ENSMUSG00 | SNORA17 | 127 | 0 | 0 | 0.34804601 | 0 | 0 | 0 |
| ENSMUSG00 | Snord83b | 93 | 2.992348 | 0.51813379 | 0 | 3.49181076 | 0 | 3.81258398 |
| ENSMUSG00 | SNORA17 | 129 | 0 | 0 | 0 | 0 | 0 | 0 |
| ENSMUSG00 | SNORA17 | 123 | 0 | 0 | 0 | 0 | 0 | 0 |
| ENSMUSG00 | Snord72 | 80 | 7.4541526 | 5.42097474 | 7.73532253 | 7.21640891 | 5.10263758 | 12.2735877 |
| ENSMUSG00 | SNORA17 | 125 | 0 | 0 | 0 | 0 | 0 | 0 |
| ENSMUSG00 | SNORA17 | 131 | 0 | 0 | 0 | 0 | 0 | 0 |
| ENSMUSG00 | n-R5s16 | 118 | 0 | 0 | 0 | 0 | 0 | 0 |
| ENSMUSG00 | SNORA17 | 130 | 0 | 0 | 0 | 0 | 0 | 0 |
| ENSMUSG00 | SNORA17 | 128 | 0 | 0 | 0 | 0 | 0 | 0 |
| ENSMUSG00 | SNORA17 | 133 | 0 | 0 | 0 | 0 | 0 | 0 |
| ENSMUSG00 | SNORA17 | 131 | 0 | 0 | 0 | 0 | 0 | 0 |
| ENSMUSG00 | SNORA17 | 131 | 0 | 0 | 0 | 0 | 0 | 0 |
| ENSMUSG00 | SNORA17 | 136 | 0 | 0 | 0 | 0 | 0 | 0 |
| ENSMUSG00 | SNORD95 | 67 | 0 | 0 | 0 | 0 | 0 | 0 |
| ENSMUSG00 | Snord90 | 110 | 0 | 0 | 0 | 0 | 0 | 2.47951266 |
| ENSMUSG00 | SNORA42 | 111 | 0 | 0 | 0 | 0 | 0 | 0 |
| ENSMUSG00 | SNORA17 | 129 | 2.77363818 | 0.37353831 | 2.39854962 | 1.39852886 | 1.97776651 | 2.32574443 |
| ENSMUSG00 | SNORA17 | 131 | 0 | 0 | 0 | 0 | 0 | 0 |
| ENSMUSG00 | SNORA17 | 128 | 0 | 0 | 0 | 0 | 0 | 0 |
| ENSMUSG00 | SNORA17 | 131 | 0 | 0 | 0 | 0 | 0 | 0 |
| ENSMUSG00 | Snora35 | 127 | 0 | 0 | 0 | 0 | 0 | 0 |
| ENSMUSG00 | SNORA17 | 125 | 0 | 0 | 0 | 0 | 0 | 0 |
| ENSMUSG00 | SNORA29 | 120 | 0 | 0 | 0 | 0 | 0 | 0 |
| ENSMUSG00 | SNORA17 | 131 | 0 | 0 | 0 | 0 | 0 | 0 |
| ENSMUSG00 | SNORD116 | 88 | 0 | 0 | 0 | 0 | 0 | 0 |
| ENSMUSG00 | SNORA17 | 131 | 0 | 0 | 0 | 0 | 0 | 0 |
| ENSMUSG00 | SNORA17 | 129 | 0 | 0 | 0 | 0 | 0 | 0 |
| ENSMUSG00 | SNORA17 | 135 | 0 | 0 | 0 | 0 | 0 | 0 |
| ENSMUSG00 | SNORA17 | 129 | 1.84909212 | 1.86769156 | 1.02794984 | 1.39852886 | 5.93329951 | 2.32574443 |
| ENSMUSG00 | SNORA17 | 125 | 0 | 0 | 0 | 0 | 0 | 0 |
| ENSMUSG00 | U2 | 188 | 0 | 0 | 0 | 0 | 0 | 0 |
| ENSMUSG00 | SNORA17 | 133 | 0 | 0 | 0 | 0 | 0 | 0 |
| ENSMUSG00 | SNORA17 | 131 | 0.91043085 | 0.36783544 | 2.69934919 | 0.55087091 | 0 | 0.62461006 |
| ENSMUSG00 | SNORA17 | 130 | 0 | 0 | 0 | 0 | 0 | 0 |
| ENSMUSG00 | SNORA17 | 131 | 0 | 0 | 0 | 0 | 0 | 0 |
| ENSMUSG00 | n-R5s26 | 121 | 0 | 0 | 0 | 0.29819872 | 0.84341117 | 0 |
| ENSMUSG00 | U6 | 73 | 0 | 0 | 0 | 0 | 0 | 0 |
| ENSMUSG00 | SNORA17 | 131 | 0 | 0 | 0 | 0 | 0 | 0 |
| ENSMUSG00 | U6 | 104 | 0 | 0 | 0 | 0 | 0 | 0 |
| ENSMUSG00 | U2 | 175 | 0 | 0 | 0 | 0 | 0 | 0 |
| ENSMUSG00 | SNORA17 | 124 | 0.96182614 | 0 | 0 | 0 | 2.46901819 | 0 |
| ENSMUSG00 | SNORA17 | 132 | 0 | 0 | 0 | 0 | 0 | 0 |
| ENSMUSG00 | Snord19 | 75 | 4.77065766 | 1.28497179 | 5.30422116 | 5.77312712 | 2.04105503 | 8.72788456 |
| ENSMUSG00 | SNORA17 | 131 | 0 | 0 | 0 | 0 | 0 | 0 |
| ENSMUSG00 | SNORA17 | 122 | 0 | 0 | 0 | 0 | 0 | 0 |
| ENSMUSG00 | n-R5s52 | 112 | 0 | 0 | 0 | 0 | 0 | 0 |
| ENSMUSG00 | SNORA17 | 129 | 0 | 0 | 0 | 0 | 0 | 0 |
| ENSMUSG00 | CR974466.1 | 88 | 0 | 0 | 0 | 0 | 0 | 0 |
| ENSMUSG00 | AC127299.1 | 83 | 0 | 0 | 0 | 0 | 0 | 0 |
| ENSMUSG00 | AC158665.1 | 92 | 0 | 0 | 0.48045482 | 0 | 0 | 0 |
| ENSMUSG00 | Mir290 | 82 | 0 | 0 | 0 | 0 | 0 | 0 |
| ENSMUSG00 | AC166933.1 | 93 | 0 | 0 | 0 | 0 | 0 | 0 |
| ENSMUSG00 | AC123803.1 | 84 | 0 | 0 | 0 | 0 | 0 | 0 |
| ENSMUSG00 | AC100600.1 | 93 | 0 | 0 | 0 | 0 | 0 | 0 |
| ENSMUSG00 | Mir465b-1 | 78 | 0 | 0 | 0 | 0 | 0 | 0 |
| ENSMUSG00 | AL593843.1 | 77 | 0 | 1.2515959 | 0 | 0 | 0 | 0 |
| ENSMUSG00 | AC107454.1 | 91 | 0 | 0 | 0 | 0 | 0 | 0 |
| ENSMUSG00 | CR974568.1 | 62 | 0 | 3.10880272 | 0 | 0 | 1.64601212 | 1.75965415 |
| ENSMUSG00 | AC116151.1 | 92 | 0 | 0 | 0 | 0 | 0 | 0 |
| ENSMUSG00 | Mir669d | 120 | 0 | 0 | 0 | 0 | 0 | 0 |
| ENSMUSG00 | AL596212.1 | 88 | 0 | 0 | 0 | 0 | 0 | 0 |
| ENSMUSG00 | AC157101.1 | 92 | 0 | 0 | 0 | 0 | 0 | 0 |
| ENSMUSG00 | AC154473.1 | 76 | 0 | 0 | 0 | 0 | 0 | 0 |
| ENSMUSG00 | CT030170.1 | 90 | 0 | 0 | 0 | 0 | 0 | 0 |
| ENSMUSG00 | AC155640.2 | 90 | 0 | 0 | 0 | 0 | 0 | 0 |
| ENSMUSG00 | AC126023.1 | 72 | 1.1043189 | 0 | 0 | 0 | 0 | 1.89407217 |
| ENSMUSG00 | AC131591.1 | 93 | 0 | 0 | 0 | 0 | 0 | 0.29327569 |
| ENSMUSG00 | Mir105 | 79 | 0 | 0 | 0 | 0 | 0 | 0 |
| ENSMUSG00 | Mir876 | 80 | 0 | 0 | 0 | 0 | 0 | 0 |
| ENSMUSG00 | AC158626.1 | 92 | 0 | 0 | 0 | 0 | 0 | 0 |
| ENSMUSG00 | AC154495.1 | 87 | 0 | 0 | 0 | 0 | 0 | 0 |
| ENSMUSG00 | Mir465b-2 | 78 | 0 | 0 | 0 | 0 | 0 | 0 |
| ENSMUSG00 | AC156822.1 | 109 | 0 | 0 | 0 | 0 | 0 | 0 |
| ENSMUSG00 | U6 | 106 | 0 | 0 | 0 | 0 | 0 | 0 |
| ENSMUSG00 | AL662925.1 | 93 | 0 | 0 | 0 | 0 | 0 | 0 |
| ENSMUSG00 | AC102463.2 | 90 | 0 | 0 | 0 | 0 | 0 | 0 |
| ENSMUSG00 | AC147251.1 | 93 | 0 | 0 | 0 | 0 | 0 | 0 |
| ENSMUSG00 | AC122209.1 | 88 | 0 | 0 | 0 | 0 | 0 | 0 |
| ENSMUSG00 | AC154247.1 | 81 | 0 | 0 | 0 | 0 | 0 | 0 |
| ENSMUSG00 | Mir344e | 65 | 0 | 0 | 0 | 0 | 0 | 0 |
| ENSMUSG00 | AC124717.1 | 91 | 0 | 0 | 0 | 0 | 0 | 0 |
| ENSMUSG00 | Mir878 | 77 | 0 | 0 | 0 | 0 | 0 | 0 |
| ENSMUSG00 | AL671913.1 | 80 | 0 | 0 | 0 | 0 | 0 | 0 |
| ENSMUSG00 | AC118561.1 | 90 | 0 | 0 | 0 | 0 | 0 | 0 |

| | | | | | | | | |
|---|---|---|---|---|---|---|---|---|
| ENSMUSG00 | AC163663.1 | 96 | 0 | 0 | 0 | 0 | 0 | 0 |
| ENSMUSG00 | AC152063.1 | 86 | 0 | 0 | 0 | 0 | 0 | 0 |
| ENSMUSG00 | Mir882 | 76 | 0 | 0 | 0 | 0 | 0 | 0 |
| ENSMUSG00 | AL683882.1 | 86 | 0 | 0 | 0 | 0 | 0 | 0 |
| ENSMUSG00 | AC121892.1 | 90 | 0 | 0 | 0 | 0 | 0 | 0 |
| ENSMUSG00 | Mir295 | 68 | 0 | 0 | 0 | 0 | 0 | 0 |
| ENSMUSG00 | Mir875 | 77 | 0 | 0 | 0 | 0 | 0 | 0 |
| ENSMUSG00 | AC124987.1 | 83 | 0 | 0 | 0 | 0 | 0 | 0 |
| ENSMUSG00 | Mir465c-2 | 80 | 0 | 0 | 0 | 0 | 0 | 0 |
| ENSMUSG00 | Mir879 | 75 | 0 | 0 | 0 | 0 | 0 | 0 |
| ENSMUSG00 | Mir511 | 78 | 0 | 0 | 0 | 0 | 0 | 0 |
| ENSMUSG00 | AC126435.1 | 91 | 0 | 0 | 0 | 0 | 0 | 0 |
| ENSMUSG00 | Mir294 | 83 | 0 | 0 | 0 | 0 | 0 | 0 |
| ENSMUSG00 | AC153556.1 | 80 | 0 | 0 | 0 | 0 | 0 | 0 |
| ENSMUSG00 | n-R5s194 | 118 | 0 | 0 | 0 | 0 | 0 | 0 |
| ENSMUSG00 | AC104196.1 | 93 | 0.42747829 | 0.51813379 | 0 | 0 | 0 | 0 |
| ENSMUSG00 | Mir872 | 80 | 0 | 0 | 0 | 0 | 0 | 0 |
| ENSMUSG00 | AC125010.1 | 92 | 0 | 0 | 0 | 0 | 0 | 0 |
| ENSMUSG00 | Mir453 | 81 | 0 | 0 | 0 | 0 | 0 | 0 |
| ENSMUSG00 | AL583883.1 | 70 | 0 | 0 | 0 | 0 | 0 | 0 |
| ENSMUSG00 | Mir208b | 76 | 0 | 0 | 0 | 0 | 0 | 0 |
| ENSMUSG00 | Mir877 | 84 | 0 | 0.57364812 | 0 | 0 | 2.42982742 | 0 |
| ENSMUSG00 | AC128668.1 | 90 | 0 | 0 | 0 | 0 | 0 | 0 |
| ENSMUSG00 | AC084389.1 | 74 | 0 | 0 | 0 | 0 | 0 | 0 |
| ENSMUSG00 | Mir466h | 80 | 0 | 0 | 0 | 0 | 0 | 0 |
| ENSMUSG00 | Mir871 | 76 | 0 | 0 | 0 | 0 | 0 | 0 |
| ENSMUSG00 | CT485999.1 | 93 | 0 | 0.51813379 | 0 | 0 | 0 | 0 |
| ENSMUSG00 | Mir465c-1 | 80 | 0 | 0 | 0 | 0 | 0 | 0 |
| ENSMUSG00 | Mir297a-4 | 97 | 0 | 0 | 0 | 0 | 0 | 0 |
| ENSMUSG00 | AC160978.1 | 92 | 0 | 0 | 0 | 0 | 0.55463452 | 0 |
| ENSMUSG00 | Mir743b | 76 | 0 | 0 | 0 | 0 | 0 | 0 |
| ENSMUSG00 | AC157352.1 | 89 | 0 | 0 | 0 | 0 | 0 | 0 |
| ENSMUSG00 | Mir874 | 75 | 0 | 0 | 0 | 0 | 0 | 0 |
| ENSMUSG00 | AC122270.1 | 100 | 0 | 0.96372884 | 0 | 2.16492267 | 0 | 4.09119589 |
| ENSMUSG00 | AC161218.1 | 95 | 0 | 0 | 0 | 0 | 0 | 0 |
| ENSMUSG00 | AL732528.1 | 94 | 0 | 0 | 0 | 0.38385154 | 0 | 0.29015574 |
| ENSMUSG00 | AL845457.1 | 85 | 0 | 0 | 0 | 0 | 1.20062061 | 2.88790298 |
| ENSMUSG00 | AC154816.1 | 96 | 0 | 0 | 0 | 0 | 0 | 0 |
| ENSMUSG00 | Mir466f-1 | 93 | 0 | 0 | 0 | 0 | 0 | 0 |
| ENSMUSG00 | AC162467.1 | 91 | 0 | 0 | 0 | 0 | 2.24291762 | 0 |
| ENSMUSG00 | AC142100.1 | 91 | 0 | 0 | 0 | 0 | 0 | 0 |
| ENSMUSG00 | AC188608.1 | 109 | 0 | 0 | 0 | 0 | 0 | 0 |
| ENSMUSG00 | AL844871.1 | 89 | 0 | 0 | 0 | 0 | 0 | 0 |
| ENSMUSG00 | Mir654 | 83 | 0 | 0 | 0 | 0 | 0 | 0 |
| ENSMUSG00 | Mir466f-3 | 93 | 0 | 0 | 0 | 0 | 0 | 0 |
| ENSMUSG00 | AL672259.1 | 86 | 0 | 0 | 0 | 0 | 0 | 0 |
| ENSMUSG00 | Mir466n | 93 | 0 | 0 | 0 | 0 | 0 | 0 |
| ENSMUSG00 | AC113244.1 | 83 | 0.47898169 | 2.90279772 | 0 | 0 | 1.84432684 | 2.30027078 |
| ENSMUSG00 | AC122735.1 | 92 | 0 | 0 | 0 | 0 | 0 | 0 |
| ENSMUSG00 | Mir190b | 79 | 0 | 0 | 0 | 0 | 0 | 0 |
| ENSMUSG00 | Mir466f-2 | 93 | 0 | 0 | 0 | 0 | 0 | 0 |
| ENSMUSG00 | AL589870.1 | 88 | 0 | 0 | 0 | 0 | 0 | 0.30993908 |
| ENSMUSG00 | Mir880 | 77 | 0 | 0 | 0 | 0 | 0 | 0 |
| ENSMUSG00 | CT025680.1 | 91 | 0 | 0 | 0 | 0 | 0 | 0 |
| ENSMUSG00 | AC141869.1 | 94 | 0 | 0 | 0 | 0 | 0 | 0 |
| ENSMUSG00 | Mir291a | 81 | 0 | 0 | 0 | 0 | 0 | 0 |
| ENSMUSG00 | AL954388.1 | 84 | 0 | 0 | 0 | 0 | 0 | 1.62349043 |
| ENSMUSG00 | AL732472.1 | 91 | 0 | 0 | 0 | 0 | 1.12145881 | 0 |
| ENSMUSG00 | AL603708.1 | 78 | 0 | 0 | 0 | 0 | 0 | 0 |
| ENSMUSG00 | Mir883a | 75 | 0 | 0 | 0 | 0 | 0 | 0 |
| ENSMUSG00 | AC124406.1 | 86 | 0 | 0 | 0 | 0 | 0 | 0 |
| ENSMUSG00 | AL805935.1 | 91 | 0 | 0 | 0 | 0 | 0 | 0 |
| ENSMUSG00 | AC168268.1 | 109 | 0 | 0 | 0 | 0 | 0 | 0 |
| ENSMUSG00 | AC159288.1 | 109 | 0 | 0 | 0 | 0 | 0 | 0 |
| ENSMUSG00 | Mir883b | 77 | 0 | 0 | 0 | 0 | 0 | 0 |
| ENSMUSG00 | Mir466g | 79 | 0 | 0 | 0 | 0 | 0 | 0 |
| ENSMUSG00 | Mir466o | 83 | 0 | 0 | 0 | 0 | 0 | 0 |
| ENSMUSG00 | Mir449b | 79 | 0 | 0 | 0 | 0 | 0 | 0 |
| ENSMUSG00 | Mir466d | 95 | 0 | 0 | 0 | 0 | 0 | 0 |
| ENSMUSG00 | Mir291b | 78 | 0 | 0 | 0 | 0 | 0 | 0 |
| ENSMUSG00 | BX119956.1 | 87 | 0.91391909 | 0 | 0 | 0 | 0 | 0 |
| ENSMUSG00 | Mir293 | 79 | 0 | 0 | 0 | 0 | 0 | 0 |
| ENSMUSG00 | AC114558.1 | 95 | 0 | 0 | 0 | 0 | 0 | 3.44521759 |
| ENSMUSG00 | Mir881 | 77 | 0 | 0 | 0 | 0 | 0 | 0 |
| ENSMUSG00 | Mir292 | 81 | 0 | 0 | 0 | 0 | 0 | 0 |
| ENSMUSG00 | AC153982.1 | 96 | 0 | 0 | 0 | 0 | 0 | 0 |
| ENSMUSG00 | AC157475.1 | 94 | 0 | 0 | 0 | 0 | 0 | 0 |
| ENSMUSG00 | AL626775.1 | 81 | 1.4724252 | 0.59489435 | 0 | 0 | 0.62995526 | 2.02034365 |
| ENSMUSG00 | AL590992.1 | 77 | 0 | 0 | 0 | 0 | 0 | 0 |
| ENSMUSG00 | Mir873 | 76 | 0 | 0 | 0 | 0 | 0 | 0 |
| ENSMUSG00 | Mir466l | 120 | 0 | 0 | 0 | 0 | 0 | 0 |
| ENSMUSG00 | AL731851.1 | 91 | 0 | 0 | 0 | 0 | 0 | 0 |
| ENSMUSG00 | AC167132.1 | 86 | 0 | 3.92215227 | 3.08384951 | 0.41955866 | 1.1866599 | 0 |
| ENSMUSG00 | AL713967.1 | 89 | 0 | 0 | 0 | 0 | 0 | 0 |
| ENSMUSG00 | AC121600.1 | 80 | 0 | 0 | 0 | 0 | 0 | 0 |
| ENSMUSG00 | Olfr585 | 956 | 0 | 0 | 0 | 0 | 0 | 0 |
| ENSMUSG00 | Gm15887 | 494 | 0 | 0 | 0 | 0 | 0 | 0 |
| ENSMUSG00 | Gm7336 | 597 | 0.19977628 | 0.08071431 | 0.22211981 | 0 | 0.08547132 | 0.18274465 |
| ENSMUSG00 | Gm10912 | 1660 | 0 | 0 | 0 | 0 | 0 | 0 |
| ENSMUSG00 | Golgb1 | 497 | 0 | 0 | 0 | 0 | 0 | 0 |
| ENSMUSG00 | Gm10913 | 926 | 0 | 0.0520372 | 0 | 0 | 0 | 0 |
| ENSMUSG00 | Gm6551 | 1248 | 0 | 0 | 0 | 0 | 0 | 0 |
| ENSMUSG00 | Gm4478 | 386 | 0.30898042 | 0.12483534 | 0 | 0 | 0 | 0.28263875 |
| ENSMUSG00 | Gm16496 | 4170 | 0 | 0.03466651 | 0 | 0 | 0 | 0 |
| ENSMUSG00 | Olfr828 | 938 | 0 | 0 | 0 | 0 | 0 | 0 |
| ENSMUSG00 | Gm16485 | 3982 | 0 | 0.82287244 | 0.43291609 | 1.70352194 | 0.02562852 | 0.59590498 |
| ENSMUSG00 | Olfr483 | 1064 | 0 | 0 | 0 | 0 | 0 | 0 |
| ENSMUSG00 | Gm14958 | 398 | 0 | 0 | 0 | 0 | 0 | 0 |
| ENSMUSG00 | Gm7671 | 398 | 0 | 0 | 0 | 0 | 0 | 0 |
| ENSMUSG00 | F630028O10I | 3307 | 0 | 0 | 0 | 0 | 0 | 0 |
| ENSMUSG00 | Gm5071 | 1034 | 0 | 0 | 0 | 0 | 0 | 0 |
| ENSMUSG00 | Gm10916 | 482 | 0 | 0 | 0 | 0.22457704 | 0 | 0 |
| ENSMUSG00 | Rpl23a-ps3 | 470 | 0 | 0 | 0.28213942 | 0 | 0 | 0 |
| ENSMUSG00 | Fam170b | 1692 | 0 | 0 | 0 | 0 | 0 | 0 |
| ENSMUSG00 | Gm2178 | 356 | 0 | 0 | 0 | 0 | 0 | 0 |
| ENSMUSG00 | 1700007I08R | 1500 | 0 | 0 | 0 | 0 | 0 | 0 |
| ENSMUSG00 | Gm11555 | 1127 | 0 | 0 | 0 | 0 | 0 | 0 |
| ENSMUSG00 | Krtap1-3 | 878 | 0 | 0 | 0 | 0 | 0 | 0 |
| ENSMUSG00 | Gm11939 | 299 | 0 | 0 | 0 | 0 | 0 | 0 |
| ENSMUSG00 | Gm12355 | 1310 | 0 | 0.11035063 | 0.1012256 | 0.302979 | 0.07790286 | 0.08328134 |
| ENSMUSG00 | Eid1 | 968 | 0 | 0 | 0 | 0 | 0 | 0 |
| ENSMUSG00 | Gm1337 | 4860 | 0 | 0 | 0 | 0 | 0 | 0 |
| ENSMUSG00 | AK157302 | 1923 | 0.02067368 | 0.07517386 | 0.04597176 | 0 | 0.02653478 | 0.01418338 |
| ENSMUSG00 | Gm2399 | 227 | 0 | 0 | 0 | 0 | 0 | 0 |
| ENSMUSG00 | Gm17344 | 1638 | 16.8924386 | 7.97223799 | 10.1734401 | 12.0053201 | 13.519809 | 6.89359014 |
| ENSMUSG00 | Capns2 | 1000 | 0 | 0 | 0 | 0 | 0 | 0 |
| ENSMUSG00 | Gm12226 | 875 | 0 | 0 | 0.05051639 | 0 | 0 | 0.06234203 |
| ENSMUSG00 | Psme2b-ps | 836 | 0.47554403 | 0.92222856 | 0.74022225 | 0.25896204 | 0.61036335 | 0.32625167 |
| ENSMUSG00 | Gm12184 | 2220 | 0.0716315 | 0.21705605 | 0.0995537 | 0.14627856 | 0.11492427 | 0.14743048 |
| ENSMUSG00 | 4931440F15I | 3273 | 0 | 0.32389298 | 0 | 0.33072451 | 0 | 0.50832661 |
| ENSMUSG00 | Gm16503 | 3282 | 0 | 0 | 0 | 0 | 0 | 0 |
| ENSMUSG00 | BC007180 | 5630 | 0 | 0 | 0.03140451 | 0 | 0 | 0 |
| ENSMUSG00 | Gm2574 | 1231 | 0 | 0 | 0 | 0 | 0 | 0 |
| ENSMUSG00 | Gm4392 | 338 | 0 | 0 | 0 | 0 | 0 | 0 |
| ENSMUSG00 | Gm16499 | 65 | 0 | 0 | 0 | 0 | 0 | 0 |
| ENSMUSG00 | Lenep | 2471 | 0 | 0 | 0 | 0 | 0 | 0.0110379 |
| ENSMUSG00 | Gm6525 | 320 | 0.12423588 | 0 | 0 | 0.78929472 | 0 | 0.1704665 |
| ENSMUSG00 | Gm4877 | 809 | 0 | 0 | 0 | 0 | 0 | 0 |
| ENSMUSG00 | Rnf148 | 1238 | 0 | 0.07784563 | 0 | 0 | 0 | 0 |
| ENSMUSG00 | Gm6083 | 356 | 0 | 0 | 0 | 0 | 0 | 0 |
| ENSMUSG00 | Gm15610 | 642 | 0.18577327 | 0.52539734 | 0.48195156 | 0.44962049 | 0.23844101 | 1.69935447 |
| ENSMUSG00 | B020018G12 | 1337 | 0.02973484 | 0.03604072 | 0 | 0.16192391 | 0 | 0.1019994 |
| ENSMUSG00 | Chml | 1865 | 0.59686512 | 2.19616492 | 0.8295252 | 2.68922477 | 1.96991907 | 0.1023713 |
| ENSMUSG00 | Gm13821 | 513 | 0 | 0 | 0 | 0 | 0 | 0 |
| ENSMUSG00 | Gm13820 | 137 | 0 | 0 | 0 | 0 | 0 | 0 |
| ENSMUSG00 | Dnm3os | 8528 | 0.17714684 | 0 | 0 | 0 | 0.26925269 | 0.25585966 |
| ENSMUSG00 | Gm5559 | 1250 | 0.12721754 | 0.03854915 | 0.1414459 | 0.05773127 | 0.0816422 | 0.10909856 |
| ENSMUSG00 | Gm7290 | 1228 | 0 | 0 | 0 | 0 | 0 | 0 |
| ENSMUSG00 | Gm2000 | 444 | 0.53723622 | 0.21705605 | 0.2986611 | 0 | 0.57462135 | 0.24571747 |
| ENSMUSG00 | Gm17374 | 100 | 0 | 0 | 0 | 0 | 0 | 0 |
| ENSMUSG00 | Olfr364-ps1 | 923 | 0 | 0 | 0 | 0 | 0 | 0 |
| ENSMUSG00 | Tmem203 | 1675 | 2.06491153 | 1.03564891 | 2.08474364 | 2.0249028 | 2.07151854 | 2.86587254 |
| ENSMUSG00 | Nrarp | 2572 | 10.912663 | 18.1542233 | 3.24811366 | 10.6338218 | 9.04666694 | 6.35206412 |
| ENSMUSG00 | Gm14458 | 855 | 0 | 0 | 0 | 0 | 0 | 0 |
| ENSMUSG00 | Gm5634 | 841 | 0 | 0 | 0 | 0 | 0 | 0 |
| ENSMUSG00 | Gm14374 | 1959 | 0 | 0 | 0 | 0 | 0 | 0 |
| ENSMUSG00 | Gm5924 | 869 | 0 | 0 | 0 | 0 | 0 | 0 |
| ENSMUSG00 | Gm3763 | 1959 | 0 | 0 | 0 | 0 | 0 | 0 |
| ENSMUSG00 | Gm3757 | 1451 | 0 | 0 | 0 | 0 | 0 | 0 |
| ENSMUSG00 | Gm3750 | 1490 | 0 | 0 | 0 | 0 | 0 | 0 |
| ENSMUSG00 | Gm5925 | 3455 | 0 | 0 | 0 | 0 | 0 | 0 |
| ENSMUSG00 | Gm10921 | 1959 | 0 | 0 | 0 | 0 | 0 | 0 |
| ENSMUSG00 | Gm10922 | 1959 | 0 | 0 | 0 | 0 | 0 | 0 |
| ENSMUSG00 | Gm14347 | 1959 | 0 | 0 | 0 | 0 | 0 | 0 |
| ENSMUSG00 | Gm3706 | 1942 | 0 | 0 | 0 | 0 | 0 | 0 |
| ENSMUSG00 | Gm3701 | 1959 | 0 | 0 | 0 | 0 | 0 | 0 |
| ENSMUSG00 | Gm14351 | 1959 | 0 | 0 | 0 | 0 | 0 | 0 |
| ENSMUSG00 | Gm14345 | 1959 | 0 | 0 | 0 | 0 | 0 | 0 |
| ENSMUSG00 | Gm14346 | 1959 | 0 | 0 | 0 | 0 | 0 | 0 |
| ENSMUSG00 | Gm6919 | 482 | 0 | 0 | 0 | 0 | 0 | 0 |
| ENSMUSG00 | Gm16508 | 2740 | 0 | 0 | 0 | 0 | 0 | 0 |
| ENSMUSG00 | Klhdc7a | 9877 | 6.50851594 | 7.1813752 | 6.70389398 | 7.28435727 | 6.49389029 | 9.20936741 |
| ENSMUSG00 | Fam43b | 2366 | 0 | 0 | 0 | 0 | 0 | 0 |
| ENSMUSG00 | Pou3f1 | 3849 | 0 | 0 | 0 | 0 | 0 | 0 |
| ENSMUSG00 | Gm12854 | 601 | 0.4630422 | 0.24053133 | 0.14709432 | 0 | 0.33960982 | 0.63534933 |
| ENSMUSG00 | Gm3550 | 644 | 0 | 0.14964734 | 0 | 0 | 0 | 0 |
| ENSMUSG00 | Airn | 3669 | 0.87767619 | 0.51220257 | 0.09637905 | 0.53105217 | 0.87616835 | 0.2973523 |
| ENSMUSG00 | Hmga1-rs1 | 1625 | 0 | 0 | 0 | 0 | 0 | 0 |
| ENSMUSG00 | Gm16487 | 4035 | 0 | 0 | 0 | 0.28615252 | 0 | 0.20954494 |
| ENSMUSG00 | Krtap17-1 | 760 | 0 | 0 | 0 | 0 | 0 | 0 |
| ENSMUSG00 | Gm11570 | 1887 | 0 | 0 | 0 | 0 | 0 | 0 |
| ENSMUSG00 | Krtap29-1 | 1028 | 0 | 0 | 0 | 0 | 0 | 0 |
| ENSMUSG00 | Krtap9-5 | 1076 | 0 | 0 | 0 | 0 | 0 | 0 |
| ENSMUSG00 | Gm11565 | 920 | 0 | 0 | 0 | 0 | 0 | 0 |
| ENSMUSG00 | 2300003K06I | 918 | 0 | 0 | 0 | 0 | 0 | 0 |
| ENSMUSG00 | Gm11564 | 690 | 0 | 0 | 0 | 0 | 0 | 0 |
| ENSMUSG00 | Gm11554 | 889 | 0 | 0 | 0 | 0 | 0 | 0 |
| ENSMUSG00 | Gm11569 | 868 | 0 | 0 | 0 | 0 | 0 | 0 |
| ENSMUSG00 | Gm11596 | 1078 | 0 | 0 | 0 | 0 | 0 | 0 |
| ENSMUSG00 | Krtap4-9 | 1058 | 0 | 0 | 0 | 0 | 0 | 0 |
| ENSMUSG00 | Gm14188 | 167 | 0 | 0 | 0 | 0 | 0 | 0 |
| ENSMUSG00 | Gm14194 | 140 | 0 | 0 | 0 | 0 | 0 | 0 |
| ENSMUSG00 | Gm14181 | 122 | 0 | 0 | 0 | 0 | 0 | 0 |
| ENSMUSG00 | Gm14178 | 122 | 0 | 0 | 0 | 0 | 0 | 0 |
| ENSMUSG00 | Gm14179 | 128 | 0 | 0 | 0 | 0 | 0 | 0 |
| ENSMUSG00 | Gm14191 | 152 | 0 | 0 | 0 | 0 | 0 | 0 |
| ENSMUSG00 | Gm14180 | 77 | 0 | 0 | 0 | 0 | 0 | 0 |
| ENSMUSG00 | Gm14193 | 152 | 0 | 0 | 0 | 0 | 0 | 0 |
| ENSMUSG00 | Gm14185 | 137 | 0 | 0 | 0 | 0 | 0 | 0 |
| ENSMUSG00 | Gm14196 | 149 | 0 | 0 | 0 | 0 | 0 | 0 |
| ENSMUSG00 | Gm14183 | 152 | 0 | 0 | 0 | 0 | 0 | 0 |
| ENSMUSG00 | Gm14184 | 137 | 0 | 0 | 0 | 0 | 0 | 0 |
| ENSMUSG00 | Gm14192 | 149 | 0 | 0 | 0 | 0 | 0 | 0 |
| ENSMUSG00 | Gm14190 | 152 | 0 | 0 | 0 | 0 | 0 | 0 |
| ENSMUSG00 | Gm14177 | 152 | 0 | 0 | 0 | 0 | 0 | 0 |
| ENSMUSG00 | Gm14187 | 137 | 0 | 0 | 0 | 0 | 0 | 0 |
| ENSMUSG00 | Gm14189 | 152 | 0 | 0 | 0 | 0 | 0 | 0 |
| ENSMUSG00 | Tas2r122 | 929 | 0 | 0 | 0 | 0 | 0 | 0 |
| ENSMUSG00 | Gm6134 | 170 | 0 | 0 | 0 | 0 | 0 | 0 |
| ENSMUSG00 | Cdc73 | 1446 | 0 | 0 | 0 | 0 | 0 | 0 |
| ENSMUSG00 | Gm10923 | 461 | 1.29356227 | 1.14978495 | 1.2464728 | 0.54788354 | 0.44274513 | 1.24244561 |
| ENSMUSG00 | Gm3940 | 431 | 0 | 0 | 0 | 0.25115112 | 0 | 0 |
| ENSMUSG00 | Gm3150 | 451 | 0.61704737 | 0.53421776 | 0.09800852 | 0.64003627 | 0.45256209 | 0.36285551 |
| ENSMUSG00 | Gm2606 | 1252 | 0.06350716 | 0 | 0.07060997 | 0.05763905 | 0.04075589 | 0.13070913 |
| ENSMUSG00 | Foxd1 | 2460 | 0 | 0 | 0 | 0 | 0 | 0 |
| ENSMUSG00 | Gm3436 | 1912 | 0.04158523 | 0.07560634 | 0.06935436 | 0.09435681 | 0.02668744 | 0.08558987 |
| ENSMUSG00 | Gm16493 | 3757 | 0.19047076 | 0.19238665 | 0.42354707 | 0.15366322 | 0.47535884 | 0.42106177 |
| ENSMUSG00 | AI593442 | 5613 | 0 | 0 | 0 | 0 | 0 | 0.00485919 |
| ENSMUSG00 | Gm7293 | 3449 | 0.04610668 | 0.08382681 | 0.02563169 | 0.04184638 | 0.01479454 | 0.02372395 |
| ENSMUSG00 | Sox4 | 404 | 0 | 0 | 0 | 0 | 0 | 0 |
| ENSMUSG00 | Gm6344 | 482 | 0 | 0 | 0 | 0 | 0 | 0 |
| ENSMUSG00 | Gm14762 | 3198 | 0 | 0 | 0 | 0 | 0 | 0 |
| ENSMUSG00 | Fam47a-ps | 2638 | 0 | 0 | 0 | 0 | 0 | 0 |
| ENSMUSG00 | F8a | 3693 | 3.60630544 | 2.2181682 | 3.27953019 | 3.40009518 | 1.38170528 | 4.35005755 |
| ENSMUSG00 | Gm362 | 1814 | 0 | 0 | 0 | 0 | 0 | 0 |
| ENSMUSG00 | Gm2964 | 1959 | 0 | 0 | 0 | 0 | 0 | 0 |
| ENSMUSG00 | Gm2933 | 1700 | 0 | 0 | 0 | 0 | 0 | 0 |
| ENSMUSG00 | Gm2927 | 1959 | 0 | 0 | 0 | 0 | 0 | 0 |
| ENSMUSG00 | Gm2913 | 1959 | 0 | 0 | 0 | 0 | 0 | 0 |
| ENSMUSG00 | Gm2882 | 777 | 0 | 0 | 0 | 0 | 0 | 0 |
| ENSMUSG00 | Gm2825 | 1959 | 0 | 0 | 0 | 0 | 0 | 0 |
| ENSMUSG00 | Gm5926 | 777 | 0 | 0 | 0 | 0 | 0 | 0 |
| ENSMUSG00 | Gmcl1l | 1959 | 0 | 0 | 0 | 0 | 0 | 0 |
| ENSMUSG00 | Gm2799 | 1496 | 0 | 0 | 0 | 0 | 0 | 0 |
| ENSMUSG00 | Gm2784 | 1959 | 0 | 0 | 0 | 0 | 0 | 0 |
| ENSMUSG00 | Gm2777 | 1959 | 0 | 0 | 0 | 0 | 0 | 0 |
| ENSMUSG00 | Gm5132 | 508 | 0 | 0 | 0 | 0 | 0 | 0 |
| ENSMUSG00 | 1700012L04I | 499 | 0 | 0 | 0 | 0 | 0 | 0 |
| ENSMUSG00 | Sf3b5 | 732 | 27.4812475 | 29.1620135 | 29.7094355 | 25.4348838 | 17.5664573 | 29.2494424 |
| ENSMUSG00 | AW011738 | 1847 | 0.34438965 | 0 | 0 | 0.19535487 | 0 | 0.22150492 |
| ENSMUSG00 | 1190007F08I | 6103 | 5.05493247 | 4.02672219 | 4.18624697 | 3.74241098 | 5.91949436 | 4.23666361 |
| ENSMUSG00 | Ifnab | 669 | 0 | 0 | 0 | 0 | 0 | 0 |
| ENSMUSG00 | Ifna2 | 1031 | 0 | 0 | 0 | 0 | 0 | 0 |
| ENSMUSG00 | Gm13280 | 1009 | 0 | 0 | 0 | 0 | 0 | 0 |
| ENSMUSG00 | Ptprd | 371 | 0 | 0 | 0 | 0 | 0 | 0 |
| ENSMUSG00 | Gm10929 | 926 | 0 | 0 | 0 | 0 | 0 | 0 |
| ENSMUSG00 | Gm2500 | 490 | 0 | 0 | 0 | 0 | 0 | 0 |
| ENSMUSG00 | Gm10930 | 4967 | 0 | 0 | 0 | 0 | 0 | 0 |
| ENSMUSG00 | Gm10931 | 2384 | 0 | 0 | 0 | 0 | 0 | 0 |
| ENSMUSG00 | Mos | 2619 | 0 | 0 | 0.20252849 | 0 | 0 | 0 |
| ENSMUSG00 | Vmn1r3 | 1010 | 0 | 0 | 0 | 0 | 0 | 0 |
| ENSMUSG00 | Vmn1r2 | 1010 | 0 | 0 | 0 | 0 | 0 | 0 |
| ENSMUSG00 | Gm2001 | 493 | 0 | 0 | 0 | 0 | 0 | 0 |
| ENSMUSG00 | Gm6316 | 1019 | 0.23408526 | 0.04728797 | 0.04337767 | 0.03540927 | 0.05007495 | 0 |
| ENSMUSG00 | 2010109K11I | 809 | 0 | 0 | 0 | 0 | 0 | 0 |
| ENSMUSG00 | Gm10476 | 1482 | 0 | 0 | 0 | 0 | 0 | 0 |
| ENSMUSG00 | Gm9257 | 335 | 0 | 0 | 0 | 0 | 0 | 0 |
| ENSMUSG00 | Gm4294 | 682 | 0 | 0 | 0.06481209 | 0 | 0 | 0.03999214 |
| ENSMUSG00 | Gm6096 | 455 | 0.08737468 | 0.21180854 | 0.97146908 | 0.0793012 | 0.11214588 | 0.23977705 |
| ENSMUSG00 | Gm5329 | 467 | 0 | 0 | 0 | 0 | 0 | 0 |
| ENSMUSG00 | Gm6666 | 467 | 0 | 0 | 0 | 0 | 0 | 0 |
| ENSMUSG00 | Gm10935 | 2827 | 0 | 0 | 0 | 0 | 0 | 0 |
| ENSMUSG00 | Gm8660 | 917 | 0 | 0 | 0 | 0 | 0 | 0 |
| ENSMUSG00 | Gm4179 | 920 | 0 | 0 | 0 | 0 | 0 | 0 |
| ENSMUSG00 | Vmn1r137 | 923 | 0 | 0 | 0 | 0 | 0 | 0 |
| ENSMUSG00 | Gm5725 | 923 | 0 | 0 | 0 | 0 | 0 | 0 |
| ENSMUSG00 | Gm4177 | 917 | 0 | 0 | 0 | 0 | 0 | 0 |
| ENSMUSG00 | Vmn1r121 | 944 | 0 | 0 | 0 | 0 | 0 | 0 |
| ENSMUSG00 | Vmn1r127 | 917 | 0 | 0 | 0 | 0 | 0 | 0 |
| ENSMUSG00 | Vmn1r124 | 923 | 0 | 0 | 0 | 0 | 0 | 0 |
| ENSMUSG00 | Gm5728 | 917 | 0 | 0 | 0 | 0 | 0 | 0 |
| ENSMUSG00 | Olfr141 | 922 | 0 | 0 | 0 | 0 | 0 | 0 |
| ENSMUSG00 | Samp | 1051 | 26.3271308 | 19.4396303 | 16.1919216 | 18.6761486 | 13.0114831 | 27.0929813 |
| ENSMUSG00 | Ctdsp2 | 5057 | 10.6129916 | 11.8060118 | 9.26516781 | 11.4446509 | 13.0063276 | 9.72437702 |
| ENSMUSG00 | Gm10936 | 136 | 0 | 0 | 0 | 0 | 0 | 0 |
| ENSMUSG00 | AU041133 | 1491 | 24.9038355 | 14.9310102 | 36.8793378 | 31.7744631 | 27.9943497 | 37.8844922 |
| ENSMUSG00 | Gm4767 | 1494 | 0 | 0 | 0 | 0 | 0 | 0 |
| ENSMUSG00 | Gm3055 | 729 | 0 | 0 | 0 | 0 | 0 | 0 |
| ENSMUSG00 | 2210404O07I | 1054 | 7.20426639 | 4.75463945 | 13.2102282 | 6.05931867 | 2.71107879 | 12.5245972 |
| ENSMUSG00 | Dohh | 3063 | 11.2919582 | 11.4370041 | 9.553255 | 10.0954333 | 7.81304938 | 7.81819565 |
| ENSMUSG00 | Scamp4 | 1795 | 41.5937562 | 57.3673687 | 57.4008335 | 50.8163836 | 43.8627843 | 31.5139843 |
| ENSMUSG00 | Ccdc105 | 1711 | 0 | 0 | 0 | 0 | 0 | 0 |
| ENSMUSG00 | Gm10941 | 706 | 1.40777197 | 1.43330777 | 1.18956801 | 0.91993881 | 0.21682596 | 2.35659064 |
| ENSMUSG00 | Gm10942 | 70 | 0 | 0 | 0 | 0 | 0 | 0 |
| ENSMUSG00 | Gm10943 | 57 | 0 | 0 | 0 | 0 | 0 | 0 |
| ENSMUSG00 | Ppil6 | 1203 | 10.2115075 | 21.9903215 | 8.5243787 | 18.9258272 | 11.1129763 | 13.4899504 |
| ENSMUSG00 | Raet1d | 1264 | 0 | 0.45746822 | 0 | 0 | 0 | 0 |
| ENSMUSG00 | 3110003A17I | 804 | 19.4821633 | 23.9733543 | 22.9805602 | 16.4254083 | 9.51984624 | 36.9089646 |
| ENSMUSG00 | Mrpl48-ps | 635 | 0.06260706 | 0.15176832 | 0.0696092 | 0.17046635 | 0.0803565 | 0.30066532 |
| ENSMUSG00 | Gm10944 | 49 | 0 | 0 | 0 | 0 | 0 | 0 |
| ENSMUSG00 | Klhl17 | 3002 | 20.4604322 | 13.0016716 | 12.6627532 | 13.1972301 | 24.4933403 | 13.7463455 |
| ENSMUSG00 | Plekhn1 | 2194 | 3.02605526 | 0.83458742 | 1.63188206 | 1.39789142 | 3.65138605 | 0.88263418 |
| ENSMUSG00 | 2310042D19I | 3838 | 0.4764857 | 0.96674207 | 0.39157443 | 1.06234263 | 0.41214634 | 1.12282256 |
| ENSMUSG00 | E230028L10I | 1316 | 0 | 0 | 0.13435211 | 0 | 0 | 0 |
| ENSMUSG00 | Gm10945 | 58 | 0 | 0 | 0 | 0 | 0 | 0 |
| ENSMUSG00 | Gm17106 | 457 | 0.0869923 | 0 | 0 | 0.07895415 | 0.22331018 | 0 |
| ENSMUSG00 | C030017K20I | 3614 | 0.13200492 | 0 | 0.09784581 | 0.06988775 | 0.1976672 | 0.04528164 |
| ENSMUSG00 | Gm13090 | 742 | 0 | 0 | 0 | 0 | 0 | 0 |
| ENSMUSG00 | 1700045H11I | 589 | 0 | 0 | 0 | 0.24503935 | 0 | 0 |
| ENSMUSG00 | A930039A15I | 457 | 0 | 0.31632238 | 0 | 0 | 0.11165509 | 0 |
| ENSMUSG00 | Gm13201 | 597 | 0 | 0.16142862 | 0 | 0 | 0 | 0.13705849 |
| ENSMUSG00 | Gm13157 | 4939 | 6.73726203 | 6.49770611 | 5.11019485 | 10.2350566 | 5.17598993 | 6.3672118 |
| ENSMUSG00 | Gm13152 | 1484 | 0.66973518 | 0 | 0.53614095 | 1.38590063 | 0.51576525 | 0.40434101 |
| ENSMUSG00 | Gm13145 | 1423 | 0 | 0 | 0.03106243 | 0 | 0 | 0 |
| ENSMUSG00 | Gm13151 | 1531 | 0.07790101 | 0 | 0 | 0 | 0.03332879 | 0 |
| ENSMUSG00 | Rex2 | 746 | 0 | 0 | 0 | 0 | 0 | 0 |
| ENSMUSG00 | Gm13235 | 1410 | 0 | 0 | 0 | 0 | 0 | 0 |
| ENSMUSG00 | Gm13212 | 2159 | 0.11048304 | 0.3794208 | 0.06141988 | 0.10027433 | 0.87446777 | 0 |
| ENSMUSG00 | Gm13225 | 3638 | 0.36061871 | 0.3576234 | 0.43740142 | 2.61837816 | 1.43064605 | 0 |
| ENSMUSG00 | Gm438 | 1428 | 0 | 0 | 0 | 0 | 0 | 0 |
| ENSMUSG00 | Gm436 | 1220 | 0 | 0 | 0 | 0 | 0 | 0 |
| ENSMUSG00 | Gm13124 | 1217 | 0 | 0 | 0 | 0 | 0 | 0 |
| ENSMUSG00 | Aadacl3 | 3149 | 0 | 0 | 0 | 0 | 0 | 0 |
| ENSMUSG00 | Gm13128 | 1742 | 0 | 0 | 0 | 0 | 0 | 0 |
| ENSMUSG00 | Pramef17 | 1750 | 0 | 0 | 0 | 0 | 0 | 0 |
| ENSMUSG00 | Gm13101 | 1675 | 0 | 0 | 0 | 0 | 0 | 0 |
| ENSMUSG00 | Gm13109 | 2888 | 0 | 0 | 0 | 0 | 0 | 0 |
| ENSMUSG00 | Pramef6 | 2081 | 0 | 0 | 0 | 0 | 0 | 0 |
| ENSMUSG00 | Gm13088 | 1815 | 0 | 0 | 0 | 0 | 0 | 0 |
| ENSMUSG00 | Gm13057 | 2122 | 0 | 0 | 0 | 0 | 0 | 0 |
| ENSMUSG00 | Ddi2 | 9543 | 2.87032653 | 4.15060835 | 3.71012012 | 3.87173987 | 3.60387481 | 2.97240122 |
| ENSMUSG00 | C230096C10 | 6783 | 11.0597953 | 7.87833766 | 8.36074961 | 8.82502018 | 14.1200807 | 5.48066244 |
| ENSMUSG00 | Gm13030 | 652 | 0 | 0 | 0 | 0 | 0 | 0 |
| ENSMUSG00 | 2310026L22I | 398 | 1.69809841 | 3.39000096 | 0.77741935 | 1.26921765 | 0.64103487 | 3.49499146 |
| ENSMUSG00 | Gm13011 | 950 | 0 | 0 | 0 | 0 | 0 | 0 |
| ENSMUSG00 | 2610002D18I | 1272 | 0 | 0 | 0 | 0 | 0.44126583 | 0 |
| ENSMUSG00 | Gm5776 | 715 | 0.16680621 | 0.60654263 | 0.80366987 | 0.20185759 | 0.28546224 | 0.22887809 |
| ENSMUSG00 | Gm10948 | 260 | 0 | 0 | 0 | 0 | 0 | 0 |
| ENSMUSG00 | Gm17068 | 979 | 0.32486603 | 0.14766019 | 0.54179991 | 1.28996073 | 0.15636275 | 0.05571939 |
| ENSMUSG00 | Gm831 | 4509 | 0 | 0 | 0 | 0 | 0 | 0 |
| ENSMUSG00 | Nkain1 | 2545 | 0 | 0 | 0 | 0 | 0 | 0 |
| ENSMUSG00 | 2210404O09I | 1740 | 39.7097846 | 21.5454322 | 19.8146193 | 32.1835248 | 12.9618725 | 71.0394627 |
| ENSMUSG00 | Dear1 | 518 | 0 | 0.18604804 | 0 | 0 | 0 | 0 |
| ENSMUSG00 | Dcdc2b | 1391 | 1.25754216 | 1.59351282 | 0.76264862 | 1.08946504 | 1.57737898 | 0.35294285 |
| ENSMUSG00 | 1700125D06I | 730 | 0.05445956 | 0.06600883 | 0.18165141 | 0.04942746 | 0.27959658 | 0.48571275 |
| ENSMUSG00 | Ebf1 | 451 | 0 | 0 | 0 | 0 | 0 | 0 |
| ENSMUSG00 | Gm9631 | 2015 | 2.22946367 | 1.38700429 | 3.44401457 | 3.58134437 | 1.87392149 | 3.22152067 |
| ENSMUSG00 | Bnip3 | 2259 | 49.5403619 | 47.9731334 | 49.4457978 | 46.8635319 | 40.7036429 | 51.2893615 |
| ENSMUSG00 | 1110065P20I | 1402 | 1.72973203 | 2.16529662 | 2.77443808 | 2.31625108 | 1.67418922 | 3.9686351 |
| ENSMUSG00 | 1810043H04I | 777 | 15.144945 | 18.4807719 | 22.0155898 | 13.7919784 | 8.79991552 | 33.909011 |
| ENSMUSG00 | Gm10955 | 202 | 0 | 0 | 0 | 0 | 0 | 0 |
| ENSMUSG00 | Gm12887 | 356 | 0 | 0 | 0 | 0 | 0 | 0 |
| ENSMUSG00 | Gm12886 | 1145 | 0 | 0 | 0 | 0 | 0 | 0 |
| ENSMUSG00 | Tmco2 | 717 | 0 | 0 | 0 | 0.25161816 | 0 | 0 |
| ENSMUSG00 | Ube2d3 | 2666 | 157.217566 | 139.80575 | 154.590392 | 145.613922 | 103.373389 | 182.073051 |
| ENSMUSG00 | E430018J23I | 4890 | 1.82110995 | 1.97081563 | 1.37396322 | 0.82641902 | 1.52348689 | 1.13783771 |
| ENSMUSG00 | Hivep3 | 2972 | 0.20065014 | 0.08106737 | 0.19334588 | 0.26709454 | 0.84128278 | 0.55980922 |
| ENSMUSG00 | AU022252 | 9537 | 2.08010745 | 4.52205785 | 2.13663098 | 2.31542532 | 3.24766804 | 2.11058863 |
| ENSMUSG00 | Gm11735 | 1572 | 0 | 0 | 0 | 0 | 0.12983811 | 0 |
| ENSMUSG00 | Gm2464 | 2549 | 0 | 0 | 0.06936343 | 0 | 0.06005458 | 0 |
| ENSMUSG00 | Ccdc24 | 5446 | 0.05839953 | 0.03539217 | 0 | 0.17226095 | 0 | 0.10517213 |
| ENSMUSG00 | Gm10959 | 68 | 0 | 0 | 0 | 2.12247321 | 0 | 4.81317163 |
| ENSMUSG00 | Hs3st4 | 3187 | 0 | 0 | 0 | 0 | 0 | 0 |
| ENSMUSG00 | Gm4609 | 1001 | 0 | 0 | 0.04415769 | 0.036046 | 0.1529262 | 0.08174218 |
| ENSMUSG00 | 1700042G07I | 383 | 0 | 0 | 0 | 0 | 0 | 0 |
| ENSMUSG00 | Dnahc7c | 607 | 0 | 0 | 0 | 0 | 0 | 0 |
| ENSMUSG00 | Cyp4a12b | 2383 | 0.73405 | 0.16176733 | 1.74358928 | 0.66622323 | 0 | 0.60661178 |
| ENSMUSG00 | Skint5 | 4530 | 0.00877604 | 0 | 0 | 0 | 0 | 0 |
| ENSMUSG00 | Skint8 | 4834 | 0 | 0 | 0 | 0 | 0 | 0 |
| ENSMUSG00 | 1700092K14I | 392 | 0 | 0 | 0 | 0 | 0 | 0 |
| ENSMUSG00 | Gm12525 | 552 | 0 | 0 | 0 | 0 | 0 | 0 |
| ENSMUSG00 | Gm10961 | 141 | 1.40976881 | 0.34174782 | 0 | 0 | 0 | 0.19343716 |
| ENSMUSG00 | E030025P04I | 1075 | 0 | 0 | 0 | 0 | 0 | 0 |
| ENSMUSG00 | Gm4070 | 9598 | 0 | 0 | 0 | 0.02255598 | 0 | 0.0056834 |
| ENSMUSG00 | 1810010H24I | 1247 | 2.96492357 | 3.94147322 | 1.94956004 | 2.51735194 | 1.1457406 | 5.07435149 |
| ENSMUSG00 | Gm5901 | 999 | 0.95508662 | 0.91645886 | 0.57519916 | 1.91426262 | 0.61292944 | 1.44700288 |
| ENSMUSG00 | 1700024P16I | 3359 | 2.60381236 | 1.7214567 | 1.48699264 | 1.83686502 | 0.7139743 | 0.73078819 |
| ENSMUSG00 | Trim30c | 1535 | 0 | 0 | 0 | 0 | 0 | 0 |
| ENSMUSG00 | Gm10963 | 61 | 0 | 0 | 0 | 0 | 0 | 0 |
| ENSMUSG00 | 4930456L15I | 1181 | 0 | 0 | 0 | 0 | 0 | 0 |
| ENSMUSG00 | Smarcd2 | 3848 | 9.84588694 | 11.0322909 | 9.45377256 | 11.5897627 | 11.4968472 | 6.7973958 |
| ENSMUSG00 | Gm10964 | 118 | 0 | 0 | 0 | 0 | 0 | 0 |
| ENSMUSG00 | Hbb-bh2 | 441 | 0 | 0 | 0 | 0 | 0 | 0 |
| ENSMUSG00 | Ccdc47 | 6145 | 43.6566286 | 42.3993641 | 46.057673 | 46.5749027 | 43.536418 | 49.0810351 |
| ENSMUSG00 | Olfr613 | 5136 | 0 | 0 | 0 | 0 | 0 | 0 |
| ENSMUSG00 | Gm12789 | 1296 | 0 | 0 | 0 | 0 | 0 | 0 |
| ENSMUSG00 | Gm12790 | 947 | 0 | 0 | 0 | 0 | 0 | 0 |
| ENSMUSG00 | 10-Mar | 4234 | 0 | 0 | 0 | 0 | 0.01205158 | 0 |
| ENSMUSG00 | Tomt | 3654 | 0.04351996 | 0.18462238 | 0.06048419 | 0.1974934 | 0.04189358 | 0.02239297 |
| ENSMUSG00 | Gm10967 | 90 | 0 | 0.53540491 | 0 | 0 | 0 | 0.30305155 |
| ENSMUSG00 | Lrrc37a | 9636 | 0 | 0 | 0 | 0 | 0 | 0 |
| ENSMUSG00 | 1700028N14I | 744 | 0.21373914 | 0.32383362 | 0.05941108 | 0.14549212 | 0 | 0.18329731 |
| ENSMUSG00 | Gm3200 | 1001 | 0.27801035 | 0.57765965 | 0.70652297 | 0.57673598 | 0.2039016 | 0.24522653 |
| ENSMUSG00 | Gm12695 | 1893 | 0 | 0 | 0 | 0 | 0 | 0 |
| ENSMUSG00 | Gm11627 | 1508 | 0 | 0.06390775 | 0 | 0 | 0 | 0 |
| ENSMUSG00 | Ifna6 | 569 | 0 | 0 | 0 | 0 | 0 | 0 |
| ENSMUSG00 | Gm13283 | 3016 | 0 | 0.01597694 | 0 | 0 | 0.01691856 | 0 |
| ENSMUSG00 | Ifna9 | 572 | 0 | 0 | 0 | 0 | 0 | 0 |
| ENSMUSG00 | Fam63a | 1124 | 0 | 0 | 0 | 0 | 0 | 0 |
| ENSMUSG00 | Gm17546 | 730 | 5.11919886 | 7.12895308 | 6.8422031 | 2.32309054 | 3.4250581 | 10.7977681 |
| ENSMUSG00 | G6pc | 2409 | 0 | 0.5400722 | 0 | 0 | 0.88962548 | 0.02264395 |
| ENSMUSG00 | Aoc2 | 4135 | 2.1920797 | 1.32847749 | 0.3313802 | 1.54450348 | 0.98720921 | 0.23745756 |
| ENSMUSG00 | Psme3 | 2937 | 71.186269 | 60.2125443 | 72.6312885 | 68.9944712 | 67.878805 | 64.0587203 |
| ENSMUSG00 | Cntd1 | 4250 | 0.72963 | 0.65760321 | 0.78003252 | 1.00180735 | 1.50077576 | 0.59041572 |
| ENSMUSG00 | Gm10972 | 79 | 0 | 1.21990993 | 0 | 2.74040845 | 0 | 0 |
| ENSMUSG00 | Vps25 | 1714 | 1.1829227 | 0.81528986 | 0.82523861 | 1.2420307 | 0.89311043 | 0.50921147 |
| ENSMUSG00 | Crnn | 1625 | 0 | 0 | 0 | 0 | 0 | 0 |
| ENSMUSG00 | Lce1d | 676 | 0 | 0 | 0 | 0 | 0 | 0 |
| ENSMUSG00 | Gm11756 | 1745 | 0 | 0 | 0 | 0 | 0 | 0 |
| ENSMUSG00 | Gm13871 | 1731 | 0 | 0 | 0 | 0 | 0 | 0 |
| ENSMUSG00 | Gm11757 | 1734 | 0 | 0 | 0 | 0 | 0 | 0 |
| ENSMUSG00 | Gm11758 | 1739 | 0 | 0 | 0 | 0 | 0 | 0 |
| ENSMUSG00 | Sprr2a1 | 3539 | 0 | 0 | 0 | 0 | 0 | 0 |
| ENSMUSG00 | Gm10973 | 76 | 0 | 0 | 0 | 0.47476374 | 0 | 0 |
| ENSMUSG00 | 1700094D03I | 3093 | 1.82517887 | 5.343663 | 3.77280522 | 4.10633032 | 4.80073565 | 5.74945515 |
| ENSMUSG00 | Gm11595 | 1935 | 0 | 0 | 0 | 0 | 0 | 0 |
| ENSMUSG00 | Fam174b | 2675 | 32.6515105 | 57.229281 | 13.2027187 | 40.2635151 | 35.3274198 | 18.251067 |
| ENSMUSG00 | Chd2 | 9502 | 4.08349284 | 6.4911225 | 3.16325545 | 6.52377947 | 7.49661341 | 5.22988768 |
| ENSMUSG00 | Mup20 | 921 | 0 | 0 | 0 | 0 | 0 | 0 |
| ENSMUSG00 | Mup19 | 1354 | 0 | 0 | 0 | 0 | 0 | 0 |
| ENSMUSG00 | Mup18 | 923 | 0 | 0 | 0 | 0.31273711 | 0 | 0 |
| ENSMUSG00 | Mup16 | 1031 | 0 | 0 | 0 | 0 | 0 | 0 |
| ENSMUSG00 | Casc3 | 4015 | 13.0504915 | 12.0976174 | 9.15963472 | 12.3388909 | 12.607264 | 10.3256418 |
| ENSMUSG00 | Gm10974 | 123 | 0 | 0 | 0 | 0.58669991 | 0.41484858 | 0 |
| ENSMUSG00 | Mup-ps8 | 539 | 0 | 0 | 0 | 0 | 0 | 0 |
| ENSMUSG00 | Mup-ps7 | 638 | 0 | 0 | 0 | 0 | 0 | 0 |
| ENSMUSG00 | Mup10 | 1384 | 0 | 0 | 0 | 0 | 0 | 0 |
| ENSMUSG00 | Tm2d3 | 1482 | 25.6184102 | 31.5715488 | 35.492708 | 30.7987762 | 22.9308814 | 28.5445111 |
| ENSMUSG00 | Mup1 | 1694 | 0 | 0 | 0 | 0 | 0 | 0 |
| ENSMUSG00 | 5830417I10R | 9873 | 1.01472512 | 1.47394971 | 0.81482178 | 1.55686731 | 1.91226163 | 1.27077221 |
| ENSMUSG00 | Mup-ps4 | 623 | 0 | 0 | 0 | 0 | 0 | 0 |
| ENSMUSG00 | Mup9 | 1591 | 0 | 0 | 0 | 0 | 0 | 0 |
| ENSMUSG00 | Mup8 | 1229 | 0 | 0 | 0 | 0 | 0 | 0 |
| ENSMUSG00 | Mup2 | 1510 | 0 | 0 | 0 | 0 | 0 | 0 |
| ENSMUSG00 | Mup6 | 992 | 0 | 0 | 0 | 0 | 0 | 0 |
| ENSMUSG00 | A230006K03I | 2292 | 0 | 0 | 0 | 0 | 0 | 0 |
| ENSMUSG00 | Gm6821 | 560 | 0 | 0 | 0 | 0 | 0 | 0 |
| ENSMUSG00 | Cisd3 | 729 | 11.7794016 | 8.65901772 | 12.0054388 | 5.9394312 | 7.13949291 | 12.3091307 |
| ENSMUSG00 | Gm12537 | 2253 | 0.0705823 | 0 | 0.05885731 | 0.09609067 | 0 | 0.03631776 |
| ENSMUSG00 | Mrgpra3 | 1293 | 0 | 0 | 0 | 0 | 0 | 0 |
| ENSMUSG00 | D030028A08I | 6217 | 2.21894028 | 1.18586547 | 1.15179324 | 2.82063272 | 1.18188807 | 1.19767999 |
| ENSMUSG00 | Gm10979 | 46 | 0 | 0 | 0 | 0 | 0 | 0 |
| ENSMUSG00 | Gm53 | 3931 | 9.19301241 | 6.06774074 | 10.9970497 | 11.2991597 | 6.71092249 | 11.9478323 |
| ENSMUSG00 | Gm10982 | 64 | 0 | 0 | 0 | 0 | 0 | 0 |
| ENSMUSG00 | Tomm5 | 1290 | 10.9712799 | 9.07698096 | 12.815108 | 11.5238778 | 6.24974216 | 16.3013542 |
| ENSMUSG00 | Tmem8b | 3405 | 4.93878657 | 1.92462735 | 1.32410807 | 0.68879087 | 1.54352914 | 1.32968873 |
| ENSMUSG00 | Msmp | 740 | 0 | 0 | 0 | 0 | 0 | 0 |
| ENSMUSG00 | Gm12429 | 4142 | 0 | 0 | 0 | 0 | 0 | 0 |
| ENSMUSG00 | Gm12394 | 3344 | 0 | 0 | 0 | 0 | 0 | 0 |
| ENSMUSG00 | Gm2163 | 907 | 0 | 0 | 0 | 0 | 0 | 0 |
| ENSMUSG00 | Gm2542 | 3923 | 0 | 0 | 0 | 0 | 0 | 0 |

| | | | | | | | |
|---|---|---|---|---|---|---|---|
| ENSMUSG00 Gm7819 | 3538 | 0 | 0 | 0 | 0 | 0 | 0 |
| ENSMUSG00 Gm7819 | 907 | 0 | 0 | 0 | 0 | 0 | 0 |
| ENSMUSG00 Ccl21c | 864 | 0 | 0 | 0 | 0 | 0 | 0 |
| ENSMUSG00 Gm2506 | 927 | 0 | 0 | 0 | 0 | 0 | 0 |
| ENSMUSG00 Gm2002 | 1939 | 0 | 0 | 0 | 0 | 0 | 0 |
| ENSMUSG00 Gm2023 | 716 | 0 | 0 | 0 | 0 | 0 | 0 |
| ENSMUSG00 Gm2128 | 1569 | 0 | 0 | 0 | 0 | 0 | 0 |
| ENSMUSG00 Gm1987 | 864 | 0 | 0 | 0 | 0 | 0 | 0 |
| ENSMUSG00 Gm10985 | 79 | 0 | 0 | 0 | 0 | 0 | 0 |
| ENSMUSG00 Gm3893 | 907 | 0 | 0 | 0 | 0 | 0 | 0 |
| ENSMUSG00 Gm13298 | 4126 | 0 | 0 | 0 | 0 | 0 | 0 |
| ENSMUSG00 Gm13308 | 1516 | 0 | 0 | 0 | 0 | 0 | 0 |
| ENSMUSG00 Gm12776 | 1067 | 0 | 0 | 0 | 0 | 0.09564457 | 0 |
| ENSMUSG00 Gm12769 | 404 | 0 | 0 | 0 | 0 | 0 | 0 |
| ENSMUSG00 Gm4362 | 336 | 0 | 0 | 0 | 0 | 0 | 0 |
| ENSMUSG00 Gm5326 | 415 | 0 | 0 | 0 | 0 | 0 | 0 |
| ENSMUSG00 Gm4684 | 517 | 0 | 0 | 0 | 0 | 0 | 0 |
| ENSMUSG00 Gm6662 | 416 | 0 | 0 | 0 | 0 | 0 | 0 |
| ENSMUSG00 Haus5 | 5172 | 5.37298548 | 3.15839209 | 3.45273484 | 4.15096993 | 7.50794122 | 2.61039181 |
| ENSMUSG00 Slfn1 | 1862 | 0 | 0 | 0 | 0 | 0 | 0 |
| ENSMUSG00 U2af1l4 | 2388 | 16.9809841 | 13.1766109 | 16.3443163 | 13.1303588 | 20.5131159 | 16.2300094 |
| ENSMUSG00 Fut9 | 622 | 0 | 0 | 0 | 0 | 0 | 0 |
| ENSMUSG00 Gm10968 | 868 | 0 | 0 | 0 | 0 | 0 | 0 |
| ENSMUSG00 Zfp566 | 1688 | 5.79374894 | 2.76900764 | 3.90170297 | 4.70263614 | 2.26716717 | 4.91202034 |
| ENSMUSG00 Gm10969 | 61 | 0 | 0 | 0 | 0 | 0 | 0 |
| ENSMUSG00 Evi2a | 3133 | 0.30454246 | 0.32298605 | 0.16930167 | 0.57583857 | 0.24430119 | 0 |
| ENSMUSG00 Gm12353 | 549 | 1.7379445 | 4.38856486 | 3.30104839 | 0.920125 | 1.0223864 | 2.98083489 |
| ENSMUSG00 Rad54b | 3587 | 0.1884146 | 0.02686727 | 0.29574693 | 0 | 0.98155002 | 0.47143229 |
| ENSMUSG00 Hps3 | 61 | 0 | 0 | 0 | 0 | 0 | 0 |
| ENSMUSG00 9530053A07f | 10869 | 0.02926156 | 0 | 0.04473459 | 0 | 0.05164138 | 0 |
| ENSMUSG00 Zfp59 | 3603 | 2.79160049 | 2.91552717 | 4.98083493 | 3.31478122 | 5.42411933 | 3.96666971 |
| ENSMUSG00 Gm5150 | 909 | 0 | 0 | 0 | 0 | 0 | 0.0300051 |
| ENSMUSG00 Sirpb1b | 1490 | 0 | 0 | 0 | 0 | 0 | 0 |
| ENSMUSG00 Gm9733 | 615 | 0 | 0 | 0.14374583 | 0 | 0 | 0 |
| ENSMUSG00 1810022K09f | 524 | 34.9757185 | 19.2194015 | 28.6805852 | 23.7563079 | 23.4682377 | 80.6787993 |
| ENSMUSG00 BC024978 | 2589 | 0.32246624 | 0.24195587 | 0.56340704 | 0.33448014 | 0.25621587 | 0.76904159 |
| ENSMUSG00 Cyp2t4 | 1608 | 0.42030048 | 0.14983346 | 0.49479675 | 0.08975633 | 0.03173282 | 0.03392368 |
| ENSMUSG00 Dph1 | 3226 | 7.24619422 | 5.1980415 | 6.30280465 | 6.92336812 | 6.16871871 | 3.41567088 |
| ENSMUSG00 Gm5890 | 1334 | 0 | 0 | 0 | 0 | 0 | 0 |
| ENSMUSG00 Gm5155 | 1833 | 0 | 0 | 0 | 0 | 0 | 0 |
| ENSMUSG00 Dact3 | 2248 | 0.22990269 | 0.1286115 | 0.05898823 | 0 | 0 | 0 |
| ENSMUSG00 Ceacam15 | 1388 | 0 | 0 | 0 | 0 | 0 | 0 |
| ENSMUSG00 Zfp541 | 4545 | 0 | 0 | 0 | 0 | 0 | 0 |
| ENSMUSG00 Sult2a1 | 1009 | 0 | 0 | 0 | 0 | 0 | 0 |
| ENSMUSG00 Sult2a5 | 1162 | 0 | 0 | 0 | 0 | 0 | 0 |
| ENSMUSG00 9230107M04 | 525 | 0 | 0 | 0 | 0 | 0 | 0 |
| ENSMUSG00 Gm10991 | 312 | 0 | 0 | 0 | 0 | 0 | 0 |
| ENSMUSG00 Vmn2r49 | 2553 | 0 | 0 | 0 | 0 | 0 | 0 |
| ENSMUSG00 Vmn2r38 | 2696 | 0 | 0 | 0 | 0 | 0 | 0 |
| ENSMUSG00 Vmn1r58 | 3579 | 0 | 0 | 0 | 0 | 0 | 0 |
| ENSMUSG00 Gp6 | 1346 | 0 | 0 | 0 | 0 | 0 | 0 |
| ENSMUSG00 Eif5a | 2967 | 64.5976311 | 70.5662592 | 84.1727041 | 61.1947584 | 51.3187413 | 67.7039832 |
| ENSMUSG00 Leng1 | 2218 | 2.72445132 | 3.49775347 | 3.34802057 | 3.12342315 | 3.01373094 | 4.64824781 |
| ENSMUSG00 Gm3144 | 554 | 0 | 0 | 0 | 0 | 0.27631612 | 0 |
| ENSMUSG00 Cacng6 | 779 | 0 | 0 | 0 | 0 | 0 | 0 |
| ENSMUSG00 Prkcc | 2735 | 0.02907165 | 0 | 0 | 0 | 0 | 0 |
| ENSMUSG00 Nlrp12 | 4341 | 0 | 0 | 0 | 0 | 0 | 0 |
| ENSMUSG00 Gm10995 | 90 | 0 | 0 | 0 | 0 | 0 | 0 |
| ENSMUSG00 Gm17414 | 3516 | 0.30528953 | 0.37003241 | 0.03771488 | 0.37970297 | 0.23220194 | 0.62058337 |
| ENSMUSG00 Gm17382 | 1512 | 0.02629331 | 0.06373868 | 0.96472276 | 0 | 0.47246644 | 0.23450417 |
| ENSMUSG00 Gm10999 | 200 | 0 | 0 | 0 | 0 | 0 | 0 |
| ENSMUSG00 Gm6746 | 482 | 0 | 0 | 0 | 0 | 0 | 0 |
| ENSMUSG00 Gm2348 | 299 | 0.26592295 | 0 | 0 | 0 | 0 | 0 |
| ENSMUSG00 Hist3h2a | 1989 | 17.0095093 | 12.5735362 | 15.2895264 | 12.6259945 | 12.6988718 | 15.2897048 |
| ENSMUSG00 Igtp | 2196 | 3.89227154 | 4.82742134 | 5.99824646 | 5.38930355 | 36.2714812 | 33.1121076 |
| ENSMUSG00 Gm11003 | 64 | 0 | 0 | 0 | 0 | 0 | 0 |
| ENSMUSG00 Gm17491 | 2307 | 2.27469589 | 2.58999516 | 3.08474067 | 2.26783548 | 3.73795298 | 5.59206951 |
| ENSMUSG00 Gm11006 | 49 | 0 | 0 | 1.80415686 | 0 | 0 | 0 |
| ENSMUSG00 Zfp931 | 2596 | 13.3845493 | 12.5849028 | 9.39885106 | 12.2728988 | 4.87463066 | 20.1407872 |
| ENSMUSG00 Gm14326 | 2857 | 3.18656109 | 2.05766396 | 2.84674103 | 3.90246824 | 3.08980155 | 3.35085698 |
| ENSMUSG00 Gm14325 | 1973 | 11.5055141 | 10.9658959 | 6.02650571 | 10.9361696 | 9.36216323 | 7.89347137 |
| ENSMUSG00 Gm14322 | 1248 | 8.31424713 | 1.58304818 | 4.14392278 | 4.22113662 | 3.35269457 | 3.54045798 |
| ENSMUSG00 Gm14406 | 1616 | 0 | 0.08945503 | 0 | 0.06698399 | 0.03157573 | 0.06751148 |
| ENSMUSG00 Gm14420 | 2169 | 7.78980139 | 13.7072544 | 16.0381975 | 24.8199218 | 10.6099103 | 9.29274246 |
| ENSMUSG00 Gm14418 | 2542 | 0.98528532 | 1.27005965 | 1.02592791 | 1.66073926 | 1.62593881 | 1.43776619 |
| ENSMUSG00 Gm14412 | 1577 | 18.680286 | 18.1806805 | 6.25048256 | 68.1371773 | 27.4060497 | 21.4115431 |
| ENSMUSG00 Gm14409 | 1047 | 0.11391255 | 0.04602335 | 0.08443523 | 0.17231158 | 0.14620738 | 0.10420111 |
| ENSMUSG00 Gm14410 | 2392 | 4.9361947 | 7.91691963 | 4.34257237 | 18.6896543 | 6.9329315 | 7.41158675 |
| ENSMUSG00 Gm14401 | 1324 | 0.18016079 | 0.03639459 | 0.43400601 | 0.29977529 | 0.15415824 | 0.06180054 |
| ENSMUSG00 Gm14419 | 2743 | 0 | 0 | 0 | 0 | 0 | 0 |
| ENSMUSG00 Gm14408 | 404 | 0 | 0 | 0 | 0 | 0 | 0 |
| ENSMUSG00 Gm14295 | 3522 | 7.11128697 | 4.9116788 | 5.08283544 | 14.8856362 | 7.57717052 | 6.98515747 |
| ENSMUSG00 Gm14305 | 2564 | 0 | 0 | 0 | 0 | 0 | 0 |
| ENSMUSG00 Gm14306 | 2798 | 0 | 0 | 0 | 0 | 0.01823673 | 0.01949581 |
| ENSMUSG00 Gm14308 | 3323 | 0 | 0 | 0 | 0 | 0 | 0 |
| ENSMUSG00 Gm14434 | 2844 | 0 | 0 | 0 | 0 | 0 | 0 |
| ENSMUSG00 Gm14431 | 1753 | 0 | 0 | 0 | 0 | 0 | 0 |
| ENSMUSG00 Gm14430 | 1497 | 0 | 0 | 0 | 0 | 0 | 0 |
| ENSMUSG00 Gm14436 | 2035 | 0 | 0 | 0 | 0 | 0 | 0 |
| ENSMUSG00 Gm14432 | 1497 | 0 | 0 | 0 | 0 | 0 | 0 |
| ENSMUSG00 Gm2026 | 3093 | 0 | 0 | 0 | 0 | 0 | 0 |
| ENSMUSG00 Gm6710 | 3151 | 0 | 0 | 0 | 0 | 0 | 0 |
| ENSMUSG00 Gm2007 | 1497 | 0 | 0 | 0 | 0 | 0 | 0 |
| ENSMUSG00 Gm14288 | 3574 | 0 | 0 | 0 | 0 | 0 | 0 |
| ENSMUSG00 Gm11007 | 1497 | 0 | 0 | 0 | 0 | 0 | 0 |
| ENSMUSG00 Gm11008 | 1026 | 0 | 0 | 0 | 0 | 0 | 0 |
| ENSMUSG00 Gm2004 | 1749 | 0 | 0 | 0 | 0.0412602 | 0 | 0 |
| ENSMUSG00 2210418O10 | 5784 | 0.00687335 | 0.02499297 | 0 | 0.02495301 | 0.02646596 | 0 |
| ENSMUSG00 Gm11009 | 414 | 0 | 0 | 0 | 0 | 0 | 0 |
| ENSMUSG00 Gm8923 | 1296 | 0 | 0 | 0 | 0 | 0 | 0 |
| ENSMUSG00 Gm4724 | 2625 | 0 | 0 | 0 | 0.01374554 | 0 | 0 |
| ENSMUSG00 Gm4723 | 3148 | 0 | 0 | 0 | 0 | 0 | 0 |
| ENSMUSG00 Gm4631 | 5702 | 0 | 0 | 0 | 0 | 0 | 0 |
| ENSMUSG00 Gm14440 | 3379 | 0.01176546 | 0 | 0 | 0.01067832 | 0.01510103 | 0 |
| ENSMUSG00 Gm14443 | 5759 | 2.95456601 | 1.38894763 | 1.30479481 | 0.87088109 | 1.62143198 | 2.17382522 |
| ENSMUSG00 Gm14391 | 2792 | 0.01423907 | 0 | 0 | 0 | 0 | 0 |
| ENSMUSG00 Gm14393 | 933 | 0.04261038 | 0.30988066 | 1.46865716 | 0.88948234 | 0.4375252 | 0.32156595 |
| ENSMUSG00 Gm14444 | 2006 | 0.31709257 | 0 | 0.02203482 | 0.10792237 | 0.20349502 | 0.04078959 |
| ENSMUSG00 Fam186b | 3179 | 0.1125509 | 0 | 0.01390432 | 0.06810075 | 0.30497048 | 0 |
| ENSMUSG00 Mon1b | 2104 | 13.8879649 | 9.57316199 | 12.2689526 | 11.0441239 | 15.8608602 | 6.9353289 |
| ENSMUSG00 Gm17407 | 1217 | 0 | 0 | 0 | 0 | 0 | 0 |
| ENSMUSG00 Gm11011 | 252 | 0 | 0 | 0 | 0 | 0 | 0 |
| ENSMUSG00 Hsp25-ps1 | 592 | 0 | 0 | 0 | 0 | 0 | 0 |
| ENSMUSG00 Gm11281 | 1048 | 0 | 0 | 0 | 0 | 0 | 0 |
| ENSMUSG00 Dpm1 | 3646 | 4.1216598 | 4.71820073 | 5.27367024 | 5.64091206 | 3.75070453 | 4.06950185 |
| ENSMUSG00 Ifi47 | 2313 | 2.21723173 | 1.27080544 | 1.56703464 | 3.43192814 | 9.06694356 | 11.155127 |
| ENSMUSG00 Tgtp2 | 2935 | 0.17608901 | 0.34477523 | 0.37650633 | 0.17211197 | 2.95553114 | 3.56847069 |
| ENSMUSG00 Tgtp1 | 3614 | 0.06600246 | 0.06666636 | 0.14676871 | 0.1098236 | 2.62614995 | 2.72444515 |
| ENSMUSG00 Ube2v1 | 3900 | 0.18348683 | 0.18533247 | 0.27201134 | 0.2960578 | 0.28784109 | 0.27274639 |
| ENSMUSG00 Gm12169 | 1058 | 0 | 0 | 0 | 0 | 0 | 0 |
| ENSMUSG00 D030018L15l | 4301 | 0 | 0 | 0 | 0 | 0 | 0 |
| ENSMUSG00 Cdc20b | 1548 | 0 | 0 | 0 | 0 | 0 | 0 |
| ENSMUSG00 Gm17594 | 2195 | 0 | 0 | 0 | 0 | 0 | 0 |
| ENSMUSG00 Pdf | 1837 | 0.80073641 | 0.78693155 | 0.31280564 | 0.41247846 | 0.47220925 | 0.17816857 |
| ENSMUSG00 CN725425 | 1917 | 0 | 0 | 0 | 0 | 0 | 0 |
| ENSMUSG00 Ipo11 | 1613 | 0.49293838 | 0.35848562 | 0 | 0.35791241 | 0.09490337 | 0.86237235 |
| ENSMUSG00 9230113P08f | 2035 | 0 | 0 | 0 | 0 | 0 | 0 |
| ENSMUSG00 1700025C18l | 699 | 0 | 0 | 0 | 0 | 0 | 0 |
| ENSMUSG00 2410002O22l | 1607 | 0 | 0 | 0 | 0 | 0 | 0 |
| ENSMUSG00 Cpt1b | 3353 | 0.45055421 | 0.18682486 | 0.56685931 | 0.17217796 | 0.31958064 | 0.49619833 |
| ENSMUSG00 Syce3 | 549 | 0 | 0 | 0 | 0 | 0 | 0 |
| ENSMUSG00 Gm11016 | 46 | 0 | 0 | 0 | 0 | 0 | 0 |
| ENSMUSG00 Wfdc11 | 632 | 0 | 0 | 0 | 0 | 0 | 0 |
| ENSMUSG00 Taf9 | 983 | 27.1372609 | 27.8922539 | 27.6992221 | 14.5723008 | 14.6901977 | 32.7961583 |
| ENSMUSG00 Naip6 | 7485 | 0 | 0 | 0 | 0 | 0 | 0 |
| ENSMUSG00 Naip2 | 5301 | 0.157492 | 0.24543179 | 0.0083384 | 0 | 0 | 0 |
| ENSMUSG00 R3hdml | 1089 | 0 | 0 | 0 | 0 | 0 | 0 |
| ENSMUSG00 Gm11019 | 2722 | 0 | 0 | 0 | 0 | 0.52488557 | 0 |
| ENSMUSG00 Gm11020 | 183 | 0 | 0 | 0 | 0 | 0 | 0 |
| ENSMUSG00 Arhgap8 | 1998 | 6.14837011 | 11.4316184 | 7.92004995 | 8.99342251 | 6.79330129 | 6.86643821 |
| ENSMUSG00 Gm14222 | 573 | 0 | 0 | 0 | 0 | 0 | 0 |
| ENSMUSG00 Gm14221 | 337 | 0 | 0 | 0 | 0 | 0 | 0 |
| ENSMUSG00 1700060C20l | 594 | 0 | 0 | 0 | 0 | 0 | 0.22958451 |
| ENSMUSG00 Atp6ap1l | 1158 | 0 | 0 | 0 | 0 | 0 | 0 |
| ENSMUSG00 Tob2 | 859 | 0.92562236 | 0.56095975 | 0.51457326 | 0.88209888 | 1.00983514 | 0.38101941 |
| ENSMUSG00 4932414J04R | 3485 | 0 | 0 | 0 | 0 | 0 | 0 |
| ENSMUSG00 Hsbp1l1 | 1246 | 23.9617704 | 12.9940968 | 33.4174447 | 16.1587326 | 11.9170749 | 29.4198356 |
| ENSMUSG00 Ces1b | 1951 | 0 | 0 | 0 | 0 | 0 | 0 |
| ENSMUSG00 Gm12033 | 1250 | 0 | 0 | 0 | 0 | 0 | 0.02181971 |
| ENSMUSG00 Fam81b | 1522 | 0 | 0.0633199 | 0.11616779 | 0 | 0 | 0 |
| ENSMUSG00 Gm3272 | 1228 | 0 | 0.03923977 | 0 | 0.05876555 | 0 | 0.06663186 |
| ENSMUSG00 Wdr92 | 3890 | 7.44010021 | 8.96837638 | 8.94263508 | 8.17179466 | 5.20757614 | 6.07193768 |
| ENSMUSG00 Ighv1-33 | 289 | 0 | 0 | 0 | 0 | 0 | 0 |
| ENSMUSG00 Gm15557 | 1378 | 0 | 0 | 0 | 0 | 0 | 0 |
| ENSMUSG00 Sec61g | 2858 | 1.07108887 | 0.79242924 | 1.12901838 | 1.07311889 | 1.1605019 | 0.88752325 |
| ENSMUSG00 Gm2035 | 434 | 0 | 0 | 0 | 0 | 0 | 0 |
| ENSMUSG00 Gm11025 | 1016 | 0 | 0.0474276 | 0 | 0 | 0 | 0 |
| ENSMUSG00 Gm11027 | 196 | 0 | 0 | 0 | 0 | 0 | 0 |
| ENSMUSG00 Ighv1-2 | 122 | 0 | 0 | 0 | 0 | 0 | 0 |
| ENSMUSG00 Ighv3-2 | 158 | 0 | 0 | 0 | 0 | 0 | 0 |
| ENSMUSG00 Gm11029 | 160 | 0 | 0 | 0 | 0 | 0 | 0 |
| ENSMUSG00 Zfp429 | 2029 | 12.265614 | 10.2832575 | 10.8271641 | 5.7083964 | 10.0342651 | 17.3675869 |
| ENSMUSG00 Zfp456 | 4825 | 5.81707135 | 1.62785286 | 1.77723472 | 2.59491595 | 3.55333933 | 7.33730192 |
| ENSMUSG00 Vmn2r-ps106 | 668 | 0 | 0 | 0 | 0 | 0 | 0 |
| ENSMUSG00 Vmn2r-ps105 | 668 | 0 | 0 | 0 | 0 | 0 | 0 |
| ENSMUSG00 Bpifa6 | 1561 | 0 | 0 | 0 | 0 | 0 | 0 |
| ENSMUSG00 7530422B04l | 716 | 0.99943945 | 1.1440915 | 0.67907859 | 0.80630267 | 0.92645654 | 1.14279215 |
| ENSMUSG00 4930404H24 | 329 | 0 | 0 | 0 | 0 | 0 | 0.33160656 |
| ENSMUSG00 C030006K11l | 1755 | 9.03842548 | 5.87572571 | 9.14260343 | 7.07249192 | 5.2625493 | 10.1949649 |
| ENSMUSG00 Samd1 | 2066 | 1.67411752 | 2.56558985 | 2.07530434 | 1.92111563 | 1.160813 | 4.35654935 |
| ENSMUSG00 Gm11031 | 58 | 0 | 0 | 0 | 0 | 0 | 0 |
| ENSMUSG00 Gm14147 | 2677 | 0 | 0 | 0 | 0 | 0 | 0 |
| ENSMUSG00 Gm14151 | 2712 | 0 | 0 | 0 | 0 | 0 | 0 |
| ENSMUSG00 Tcl1b4 | 1707 | 0 | 0 | 0 | 0 | 0 | 0 |
| ENSMUSG00 Gm14124 | 3014 | 0 | 0.01598754 | 0 | 0 | 0.01692979 | 0.01809863 |
| ENSMUSG00 Gm14139 | 1724 | 0 | 0 | 0 | 0 | 0 | 0.01582056 |
| ENSMUSG00 Gm11032 | 100 | 0 | 0 | 0 | 0 | 0 | 0 |
| ENSMUSG00 Gm11033 | 3710 | 0 | 0 | 0 | 0 | 0 | 0 |
| ENSMUSG00 Serpina3m | 1654 | 0 | 0 | 0 | 0 | 0 | 0 |
| ENSMUSG00 Serpina3j | 2151 | 0 | 0 | 0 | 0 | 0 | 0 |
| ENSMUSG00 Serpina3i | 2131 | 0 | 0 | 0 | 0 | 0.52678567 | 0 |
| ENSMUSG00 Serpina1c | 1356 | 0 | 0 | 0 | 0 | 0 | 0 |
| ENSMUSG00 Gm11034 | 88 | 0 | 0 | 0 | 0 | 0 | 0 |
| ENSMUSG00 Ifi27l2a | 459 | 0.08661325 | 0 | 0 | 0.23583036 | 0 | 0 |
| ENSMUSG00 Ly6c1 | 836 | 46.3179881 | 34.6988495 | 11.3676989 | 20.28536 | 27.8325686 | 14.9097011 |
| ENSMUSG00 Insl3 | 621 | 0.57616638 | 0.15518983 | 0.49824944 | 0.34861879 | 0.65734462 | 0.13176154 |
| ENSMUSG00 Slc45a4 | 10298 | 11.6239806 | 9.06361803 | 12.3231202 | 9.62491516 | 14.8550277 | 12.0005235 |
| ENSMUSG00 Flrt3 | 733 | 0 | 0 | 0 | 0 | 0 | 0 |
| ENSMUSG00 Col22a1 | 9137 | 0 | 0 | 0 | 0 | 0 | 0 |
| ENSMUSG00 Gm11035 | 112 | 0 | 0 | 0 | 0 | 0.45559264 | 0 |
| ENSMUSG00 Gsdmc | 2390 | 0 | 0 | 0 | 0 | 0 | 0 |
| ENSMUSG00 Gm2016 | 1840 | 0 | 0 | 0 | 0 | 0 | 0 |
| ENSMUSG00 Gm2046 | 433 | 0 | 0 | 0 | 0 | 0 | 0 |
| ENSMUSG00 BB287469 | 1849 | 0 | 0 | 0 | 0 | 0 | 0 |
| ENSMUSG00 Mef2b | 1396 | 0 | 0 | 0 | 0 | 0 | 0 |
| ENSMUSG00 Gm8300 | 1858 | 0 | 0 | 0 | 0 | 0 | 0 |
| ENSMUSG00 Alkbh1 | 6126 | 0.35692971 | 0.38542861 | 0.37520337 | 0.45941878 | 0.3248496 | 0.38734796 |
| ENSMUSG00 Prnp | 2399 | 17.234556 | 27.3572048 | 10.5023137 | 23.0118917 | 22.907631 | 16.4057126 |
| ENSMUSG00 D130040H23 | 1216 | 12.8159115 | 4.35897092 | 5.23442878 | 7.00276522 | 2.97933609 | 10.0261215 |
| ENSMUSG00 Gm11037 | 58 | 0 | 0 | 0 | 0 | 0 | 0 |
| ENSMUSG00 Trim61 | 2959 | 0.47024056 | 3.68034333 | 1.13529573 | 1.68277193 | 2.67289228 | 0.45165844 |
| ENSMUSG00 Fastkd5 | 3013 | 6.94038591 | 5.1816818 | 4.10770529 | 5.64043909 | 4.45401156 | 6.42714699 |
| ENSMUSG00 C230021G24 | 3677 | 0 | 0 | 0 | 0 | 0 | 0 |
| ENSMUSG00 Gm17566 | 826 | 0.24065061 | 0 | 0 | 0 | 0 | 0 |
| ENSMUSG00 4933413L06F | 1198 | 0 | 0 | 0 | 0 | 0 | 0 |
| ENSMUSG00 Serpinb1c | 1346 | 1.97891322 | 0.25059814 | 2.36443737 | 1.31353654 | 0.83401209 | 2.39109022 |
| ENSMUSG00 Gm14025 | 1857 | 0 | 0 | 0.11901412 | 0 | 0.10991142 | 0 |
| ENSMUSG00 Gm14010 | 372 | 0.42747829 | 0 | 0 | 0 | 0.13716768 | 0 |
| ENSMUSG00 Slc8a3 | 4893 | 0 | 0 | 0 | 0 | 0 | 0 |
| ENSMUSG00 Kcnip3 | 3793 | 0 | 0 | 0 | 0 | 0 | 0 |
| ENSMUSG00 Cyp4v3 | 6220 | 10.8272965 | 5.60109287 | 12.2230177 | 9.18293834 | 10.3775507 | 10.773921 |
| ENSMUSG00 Adam34 | 2333 | 0 | 0 | 0 | 0 | 0 | 0 |
| ENSMUSG00 Gm11042 | 199 | 0 | 0 | 0.22211981 | 0 | 0 | 0.41117547 |
| ENSMUSG00 Gm11043 | 46 | 0 | 0 | 0 | 0 | 0 | 0 |
| ENSMUSG00 Gm11044 | 318 | 0 | 0 | 0 | 0 | 0 | 0 |
| ENSMUSG00 BC005561 | 5179 | 42.864376 | 46.4744697 | 37.8775399 | 54.6490746 | 48.6913206 | 70.8961563 |
| ENSMUSG00 Hmgn2-ps1 | 1433 | 0 | 0 | 0.12338267 | 0 | 0 | 0.03806649 |
| ENSMUSG00 8430423G03 | 636 | 0 | 0 | 0.13899951 | 0.34039665 | 0.1604603 | 0.12865396 |
| ENSMUSG00 Gm3985 | 2511 | 0 | 0 | 0 | 0 | 0 | 0 |
| ENSMUSG00 Gm14085 | 5151 | 0 | 0 | 0 | 0 | 0 | 0 |
| ENSMUSG00 Gm3086 | 538 | 0 | 0 | 0 | 0.134134 | 0 | 0.60835627 |
| ENSMUSG00 Gm3415 | 918 | 0 | 0 | 0 | 0 | 0 | 0 |
| ENSMUSG00 Gm3409 | 918 | 0 | 0 | 0 | 0 | 0 | 0 |
| ENSMUSG00 Gm3404 | 1125 | 0 | 0 | 0 | 0 | 0 | 0 |
| ENSMUSG00 Prl2c2 | 1042 | 0 | 0 | 0 | 0 | 0.04896965 | 0 |
| ENSMUSG00 4930449I24R | 1134 | 0 | 0 | 0 | 0 | 0 | 0 |
| ENSMUSG00 Gm6370 | 972 | 0 | 0 | 0 | 0 | 0 | 0 |
| ENSMUSG00 Gm6309 | 2458 | 0.12939131 | 0 | 0.14386279 | 0.26422978 | 0.47746405 | 0.36617701 |
| ENSMUSG00 Trav12n-1 | 342 | 0 | 0 | 0 | 0 | 0 | 0 |
| ENSMUSG00 Gm5456 | 1691 | 0 | 0 | 0 | 0 | 0 | 0 |
| ENSMUSG00 Gm2904 | 1694 | 0.02346841 | 0 | 0 | 0.02129991 | 0 | 0.01610073 |
| ENSMUSG00 Dab2 | 1373 | 0 | 0 | 0 | 0 | 0 | 0 |
| ENSMUSG00 Tgm7 | 2139 | 0 | 0 | 0 | 0 | 0 | 0 |
| ENSMUSG00 Prps1l3 | 1718 | 32.2579394 | 35.1441281 | 41.6032481 | 29.6343218 | 32.1959205 | 37.7844246 |
| ENSMUSG00 C7 | 2658 | 0 | 0 | 0 | 0 | 0 | 0 |
| ENSMUSG00 Srp54b | 3510 | 0.05663174 | 0.02745686 | 0.05037247 | 0.12335742 | 0.07268715 | 0.04662332 |
| ENSMUSG00 Prss2 | 7870 | 2.39947309 | 1.68377022 | 2.35331286 | 1.96686113 | 3.47523983 | 1.85758661 |
| ENSMUSG00 Capn3 | 4267 | 0.20497318 | 0 | 0.4557959 | 0.10147282 | 0.10762535 | 0.52414352 |
| ENSMUSG00 Kdelr2 | 1841 | 25.9997819 | 25.8076219 | 20.6963545 | 17.9528261 | 19.9005637 | 18.489272 |
| ENSMUSG00 C86695 | 1882 | 0 | 0 | 0 | 0 | 0.05422569 | 0 |
| ENSMUSG00 Gm7861 | 504 | 0 | 0 | 0 | 0 | 0 | 0 |
| ENSMUSG00 Gm7849 | 486 | 0 | 0 | 0 | 0 | 0 | 0 |
| ENSMUSG00 Gm11048 | 93 | 0 | 0 | 0 | 0 | 0 | 0 |
| ENSMUSG00 Gm15293 | 416 | 0 | 0 | 0 | 0 | 0 | 0 |
| ENSMUSG00 AY761185 | 485 | 0 | 0 | 0 | 0 | 0 | 0 |
| ENSMUSG00 Gm11051 | 64 | 0 | 0 | 0 | 0 | 0 | 0 |
| ENSMUSG00 Gm1977 | 1496 | 0 | 0 | 0 | 0 | 0 | 0 |
| ENSMUSG00 Gm3090 | 475 | 0 | 0 | 0 | 0 | 0 | 0 |
| ENSMUSG00 Rpl27-ps1 | 410 | 0 | 0 | 0 | 0.08800499 | 0.37336373 | 0 |
| ENSMUSG00 Gm8818 | 1046 | 0 | 0 | 0 | 0 | 0 | 0 |
| ENSMUSG00 Gm4204 | 779 | 1.68412177 | 1.05156549 | 0.56741775 | 0.97268669 | 0.19650722 | 0.98034647 |
| ENSMUSG00 Gm6374 | 1572 | 0 | 0 | 0 | 0.04590591 | 0 | 0 |
| ENSMUSG00 Gm11052 | 1831 | 0 | 0 | 0 | 0.15764957 | 0 | 0 |
| ENSMUSG00 A130010J15R | 1508 | 8.96343725 | 5.75169734 | 5.36401676 | 4.49829202 | 7.34265488 | 6.81865981 |
| ENSMUSG00 Gm15353 | 958 | 0.08299683 | 0 | 0 | 0 | 0 | 0 |
| ENSMUSG00 Gm11053 | 102 | 1.55903845 | 0 | 0 | 0 | 0.50025859 | 0 |
| ENSMUSG00 Smok3c | 2328 | 0 | 0 | 0 | 0 | 0 | 0 |
| ENSMUSG00 Fam155a | 1935 | 0 | 0 | 0 | 0 | 0 | 0 |
| ENSMUSG00 Gm17608 | 1534 | 0 | 0.21988598 | 0 | 2.70497726 | 0.66527218 | 0.03556016 |
| ENSMUSG00 Gm17607 | 82 | 0 | 0 | 0 | 0 | 0 | 0 |
| ENSMUSG00 Trim43b | 2255 | 0 | 0 | 0 | 0 | 0 | 0.16933257 |
| ENSMUSG00 Gm15498 | 721 | 0.27569681 | 0 | 0 | 0 | 0 | 0 |
| ENSMUSG00 Tlr5 | 3520 | 0.28235427 | 0 | 0 | 0 | 0.01449613 | 0 |
| ENSMUSG00 Sap25 | 1593 | 0.24956359 | 0.21174206 | 0.38846566 | 0.3397556 | 0.57656922 | 0.47940358 |
| ENSMUSG00 Cd209g | 1080 | 0 | 0 | 0 | 0 | 0 | 0 |
| ENSMUSG00 Gm15130 | 653 | 0 | 0 | 0 | 0 | 0 | 0 |
| ENSMUSG00 Gm13941 | 726 | 0 | 0 | 0 | 0 | 0 | 0 |
| ENSMUSG00 Gm13942 | 322 | 0 | 0 | 0 | 0 | 0 | 0 |
| ENSMUSG00 Gm17290 | 1070 | 0 | 0 | 0 | 0 | 0 | 0 |
| ENSMUSG00 Zan | 16867 | 0.17206083 | 0.1028465 | 0.12316871 | 0.08556838 | 0.04235307 | 0.0194045 |
| ENSMUSG00 Gm3054 | 240 | 0 | 0 | 0 | 0 | 0 | 0 |
| ENSMUSG00 Gm11060 | 70 | 0 | 0 | 0 | 0 | 0 | 0 |
| ENSMUSG00 4930417G10l | 2996 | 0.07961712 | 0.01608359 | 0.02950724 | 0.51786646 | 0 | 0.04551842 |
| ENSMUSG00 1700012B15l | 771 | 9.59081631 | 5.24988475 | 7.56763071 | 4.86709809 | 7.47857389 | 10.4358218 |
| ENSMUSG00 Gm11062 | 658 | 0 | 0 | 0 | 0 | 0 | 0 |
| ENSMUSG00 C030005K15l | 1881 | 0 | 0 | 0 | 0 | 0 | 0 |
| ENSMUSG00 Mphosph8 | 3572 | 10.3284115 | 10.6166657 | 10.9267154 | 12.3539587 | 9.58530184 | 18.5699672 |
| ENSMUSG00 Gzmc | 929 | 0 | 0 | 0 | 0 | 0 | 0 |
| ENSMUSG00 Gm7694 | 811 | 4.1667273 | 4.39679004 | 1.85309823 | 5.07195201 | 6.92096343 | 3.8339197 |
| ENSMUSG00 Psme2 | 1879 | 7.57448751 | 8.28324684 | 10.5152868 | 8.02676669 | 5.94719335 | 13.7171549 |
| ENSMUSG00 Zfp664 | 5840 | 26.569459 | 23.6476615 | 27.4747757 | 28.142759 | 23.7307597 | 18.6579082 |
| ENSMUSG00 Gm8778 | 1723 | 0 | 0 | 0 | 0 | 0 | 0.06331895 |
| ENSMUSG00 Gm6565 | 599 | 0 | 0 | 0 | 0 | 0 | 0 |
| ENSMUSG00 Gm9531 | 439 | 0 | 0 | 0.20137514 | 0 | 0 | 0 |
| ENSMUSG00 Gm336 | 569 | 0 | 0 | 0 | 0 | 0 | 0 |
| ENSMUSG00 Ccr5 | 2310 | 0 | 0 | 0 | 0 | 0 | 0 |
| ENSMUSG00 Gm4694 | 1269 | 0 | 0 | 0 | 0 | 0 | 0 |
| ENSMUSG00 Gm11065 | 226 | 0 | 0 | 0 | 0 | 0 | 0 |
| ENSMUSG00 Gm11067 | 52 | 0 | 0 | 0 | 0 | 0 | 0 |
| ENSMUSG00 Ccdc13 | 4012 | 0.72336742 | 0.14412695 | 0.84834046 | 0.15289002 | 0.11446595 | 0.53026467 |
| ENSMUSG00 Trav9n-1 | 334 | 0 | 0 | 0 | 0 | 0 | 0 |
| ENSMUSG00 Olfr1082 | 1929 | 0 | 0 | 0 | 0 | 0 | 0 |
| ENSMUSG00 C730034F03l | 1279 | 1.0568306 | 2.18515531 | 2.76477517 | 0.59243388 | 0.71811944 | 6.12026698 |
| ENSMUSG00 Xirp1 | 5836 | 0 | 0 | 0 | 0 | 0 | 0 |
| ENSMUSG00 Gm5622 | 1576 | 0 | 0 | 0 | 0 | 0 | 0 |
| ENSMUSG00 Slc35f4 | 1309 | 0 | 0 | 0 | 0 | 0 | 0 |
| ENSMUSG00 Gm13691 | 2492 | 0 | 0 | 0 | 0 | 0 | 0 |
| ENSMUSG00 Gm13694 | 2492 | 0 | 0 | 0 | 0 | 0 | 0 |
| ENSMUSG00 Gm13696 | 2492 | 0 | 0 | 0 | 0 | 0 | 0 |
| ENSMUSG00 Gm13697 | 2492 | 0 | 0 | 0 | 0 | 0 | 0 |
| ENSMUSG00 Tor1aip2 | 3228 | 32.8832506 | 39.5731903 | 32.6858114 | 32.706339 | 30.4292979 | 30.6374975 |
| ENSMUSG00 Itprip | 3762 | 2.26147603 | 0.75571507 | 4.31208835 | 1.20848953 | 5.01854308 | 1.76900903 |
| ENSMUSG00 Gm11074 | 146 | 0 | 0 | 0 | 0 | 0 | 0 |
| ENSMUSG00 Calhm1 | 1045 | 0 | 0 | 0 | 0 | 0 | 0 |
| ENSMUSG00 Trim71 | 5186 | 0 | 0 | 0.0085233 | 0.00695759 | 0.00983926 | 0 |
| ENSMUSG00 Glb1 | 2919 | 9.37025371 | 6.07489233 | 5.4211236 | 4.72193937 | 6.29307821 | 3.55065533 |
| ENSMUSG00 Gm15217 | 1030 | 0 | 0.14034886 | 0 | 0 | 0 | 0.45016395 |
| ENSMUSG00 Slco1a6 | 3790 | 0.01048957 | 0 | 0 | 0.93299218 | 0 | 0.43178848 |
| ENSMUSG00 Gm6614 | 1939 | 0 | 0 | 0 | 0 | 0 | 0 |
| ENSMUSG00 Gm11077 | 82 | 0 | 0 | 0 | 0 | 0 | 0 |
| ENSMUSG00 Gm8257 | 2855 | 0 | 0 | 0 | 0 | 0 | 0 |
| ENSMUSG00 Gm6490 | 458 | 0 | 0 | 0 | 0 | 0 | 0 |
| ENSMUSG00 Gm5930 | 1741 | 0 | 0 | 0 | 0 | 0 | 0 |
| ENSMUSG00 Gm3676 | 894 | 0 | 0 | 0 | 0 | 0 | 0 |
| ENSMUSG00 Gm3543 | 894 | 0 | 0 | 0 | 0 | 0 | 0 |
| ENSMUSG00 1700049E17f | 1776 | 0 | 0 | 0 | 0 | 0 | 0 |
| ENSMUSG00 Gm7152 | 601 | 0 | 0 | 0 | 0 | 0 | 0 |
| ENSMUSG00 Gm17165 | 490 | 0 | 0 | 0 | 0 | 0 | 0 |
| ENSMUSG00 Gm11078 | 69 | 0 | 0 | 0 | 0 | 0 | 0 |
| ENSMUSG00 Gm4736 | 1313 | 0 | 0 | 0 | 0 | 0 | 0 |
| ENSMUSG00 Hoxd3 | 5006 | 9.75224332 | 11.7145226 | 14.3218916 | 16.2607056 | 11.48756 | 16.5249662 |
| ENSMUSG00 Tmem233 | 2474 | 0 | 0 | 0 | 0 | 0 | 0 |
| ENSMUSG00 Gm11081 | 31 | 0 | 0 | 0 | 0 | 0 | 0 |
| ENSMUSG00 Gm11082 | 70 | 0 | 0 | 0 | 0 | 0 | 0 |
| ENSMUSG00 2310009B15l | 568 | 7.76911679 | 2.54505856 | 2.56806482 | 1.2069698 | 1.79670337 | 6.91469728 |
| ENSMUSG00 Fbxw28 | 1826 | 0 | 0 | 0 | 0 | 0 | 0 |
| ENSMUSG00 Gm11084 | 67 | 0.59336538 | 1.43840126 | 2.638916 | 0.53853798 | 1.5231754 | 19.9471242 |
| ENSMUSG00 Clec7a | 2264 | 0 | 0 | 0 | 0 | 0 | 0 |
| ENSMUSG00 Klrb1-ps1 | 544 | 0 | 0 | 0 | 0 | 0 | 0 |
| ENSMUSG00 Gm2223 | 2461 | 0.11307938 | 0.11748015 | 0.08980464 | 0 | 0.020734 | 0.03324824 |
| ENSMUSG00 Klrb1b | 3155 | 0 | 0.04581912 | 0 | 0 | 0 | 0 |
| ENSMUSG00 Klrb1 | 639 | 0 | 0 | 0 | 0 | 0 | 0 |
| ENSMUSG00 4933413G19l | 1871 | 0 | 0 | 0 | 0 | 0 | 0 |
| ENSMUSG00 Gm4461 | 947 | 0 | 0 | 0 | 0 | 0 | 0 |
| ENSMUSG00 Olfr239 | 947 | 0 | 0 | 0 | 0 | 0 | 0 |
| ENSMUSG00 Olfr55 | 947 | 0 | 0 | 0 | 0 | 0 | 0 |
| ENSMUSG00 Gm3222 | 989 | 0.08039531 | 0.29233433 | 0.13408041 | 0.62021715 | 0.20637564 | 0.027578 |
| ENSMUSG00 Rab9 | 2336 | 23.7920213 | 24.7945648 | 33.756887 | 21.5163904 | 21.5595175 | 26.4456584 |
| ENSMUSG00 Trappc2 | 2438 | 6.53894491 | 6.00848171 | 8.10427557 | 9.97510173 | 6.61375503 | 8.21147876 |
| ENSMUSG00 Gm11091 | 40 | 0 | 0 | 0 | 0 | 0 | 0 |
| ENSMUSG00 Gm11092 | 108 | 0 | 0 | 0 | 0 | 0 | 0 |
| ENSMUSG00 Cdhr4 | 2250 | 0 | 0 | 0 | 0 | 0.11339195 | 0 |
| ENSMUSG00 4932414N04 | 4321 | 0 | 0 | 0 | 0 | 0 | 0 |
| ENSMUSG00 Gm15262 | 370 | 0 | 0 | 0 | 0 | 0 | 0 |
| ENSMUSG00 Lemd1 | 978 | 0 | 0 | 0 | 0 | 0 | 0 |
| ENSMUSG00 Gm11096 | 71 | 6.1592998 | 0.67868228 | 2.49024468 | 6.09837372 | 2.15604405 | 7.68299697 |
| ENSMUSG00 Nat6 | 2620 | 9.86300089 | 10.8695371 | 9.6164315 | 7.32658309 | 6.27118054 | 13.5540383 |
| ENSMUSG00 Gm14446 | 4404 | 1.11936412 | 0.49236828 | 0.62227844 | 0.27036954 | 0.27807289 | 1.00329055 |
| ENSMUSG00 Gm5097 | 2215 | 0 | 0 | 0 | 0 | 0 | 0 |
| ENSMUSG00 Gm5077 | 4386 | 0 | 0.10986421 | 0.16124703 | 0.32083897 | 0.20941057 | 0.18655704 |
| ENSMUSG00 Lipo4 | 2706 | 0 | 0.01780726 | 0 | 0 | 0 | 0 |
| ENSMUSG00 1700013D24 | 532 | 0 | 0 | 0 | 0 | 0 | 0.05126812 |
| ENSMUSG00 Magea5 | 2754 | 0 | 0 | 0 | 0 | 0 | 0 |
| ENSMUSG00 Magea8 | 1642 | 0 | 0 | 0 | 0 | 0 | 0 |
| ENSMUSG00 Gm15140 | 981 | 0 | 0 | 0 | 0 | 0 | 0 |
| ENSMUSG00 Gm11099 | 111 | 0 | 0 | 0 | 0 | 0 | 0 |
| ENSMUSG00 Gm2237 | 1963 | 0 | 0 | 0 | 0 | 0.07798224 | 0.15283802 |
| ENSMUSG00 Ccrl1 | 2106 | 1.227021 | 0.73217766 | 2.68653177 | 2.55281322 | 3.10131819 | 2.60313509 |
| ENSMUSG00 Mettl4-ps1 | 2135 | 0 | 0.1805581 | 0 | 0 | 0 | 0 |
| ENSMUSG00 Gm11100 | 2128 | 0.54178051 | 0.24908405 | 0.04154309 | 0.86474825 | 0.26376416 | 0.41014495 |
| ENSMUSG00 Gm9006 | 840 | 0 | 0 | 0 | 0 | 0 | 0 |
| ENSMUSG00 Gm3839 | 1001 | 0 | 0.14441491 | 0.04415769 | 0.108138 | 0 | 0 |
| ENSMUSG00 Gm11102 | 58 | 0 | 0 | 0 | 0.62210422 | 0.8797651 | 6.11328121 |
| ENSMUSG00 Gbp6 | 4303 | 0 | 0.64950352 | 0 | 0.55343131 | 1.52972403 | 2.11706472 |
| ENSMUSG00 Gbp4 | 4527 | 0.34249254 | 0.04257693 | 0.24410119 | 0.08767451 | 7.69847906 | 2.57262447 |
| ENSMUSG00 Gm3558 | 2043 | 0 | 0 | 0 | 0 | 0 | 0 |
| ENSMUSG00 Gm3618 | 382 | 0 | 0 | 0 | 0 | 0 | 0 |
| ENSMUSG00 Gm3358 | 2662 | 0 | 0 | 0 | 0 | 0 | 0.01024592 |
| ENSMUSG00 Gm3476 | 1992 | 0 | 0 | 0 | 0 | 0 | 0 |
| ENSMUSG00 Gm3468 | 1954 | 0 | 0 | 0 | 0 | 0 | 0 |
| ENSMUSG00 Gm10021 | 2059 | 0 | 0 | 0 | 0 | 0 | 0 |
| ENSMUSG00 Gm8334 | 1659 | 0 | 0 | 0 | 0 | 0 | 0 |
| ENSMUSG00 Gm3194 | 2134 | 0 | 0 | 0 | 0 | 0 | 0 |
| ENSMUSG00 Gm3383 | 3201 | 0 | 0 | 0 | 0 | 0 | 0 |
| ENSMUSG00 Gm3029 | 979 | 0 | 0 | 0 | 0 | 0 | 0 |
| ENSMUSG00 Gm8279 | 1950 | 0 | 0 | 0 | 0 | 0 | 0 |
| ENSMUSG00 Gm3242 | 482 | 0 | 0 | 0 | 0 | 0 | 0 |
| ENSMUSG00 Gm8271 | 1467 | 0 | 0 | 0 | 0 | 0 | 0 |
| ENSMUSG00 4930555G01l | 2060 | 0 | 0 | 0 | 0 | 0 | 0 |
| ENSMUSG00 Gm3443 | 475 | 0 | 0 | 0 | 0 | 0 | 0 |
| ENSMUSG00 Gm3298 | 870 | 0 | 0 | 0 | 0 | 0 | 0 |
| ENSMUSG00 Gm3173 | 596 | 0 | 0 | 0 | 0 | 0 | 0 |
| ENSMUSG00 Luzp4 | 2107 | 0 | 0 | 0 | 0 | 0 | 0 |
| ENSMUSG00 2610042L04F | 1879 | 0 | 0 | 0 | 0 | 0 | 0 |
| ENSMUSG00 Gm3149 | 1946 | 0 | 0 | 0 | 0 | 0 | 0 |
| ENSMUSG00 Gm11105 | 477 | 0 | 0 | 0 | 0 | 0 | 0 |
| ENSMUSG00 Gm2974 | 2173 | 0 | 0 | 0 | 0 | 0 | 0 |
| ENSMUSG00 Gm17170 | 315 | 0 | 0 | 0 | 0 | 0 | 0 |
| ENSMUSG00 Gm3099 | 2060 | 0 | 0 | 0 | 0 | 0 | 0 |
| ENSMUSG00 Gm15114 | 1967 | 0 | 0 | 0 | 0 | 0 | 0 |
| ENSMUSG00 Gm5796 | 2067 | 0.01923342 | 0.02331226 | 0.1069227 | 0.08728119 | 0.0246862 | 0.01319528 |
| ENSMUSG00 Gm2916 | 382 | 0 | 0 | 0 | 0 | 0 | 0 |
| ENSMUSG00 Gm3033 | 499 | 0 | 0 | 0 | 0 | 0 | 0 |
| ENSMUSG00 Gm3002 | 1222 | 0 | 0 | 0 | 0 | 0 | 0 |
| ENSMUSG00 Gm3020 | 1932 | 0 | 0 | 0 | 0 | 0 | 0 |
| ENSMUSG00 Gm3012 | 1111 | 0 | 0 | 0 | 0 | 0 | 0 |
| ENSMUSG00 Gm11109 | 1698 | 0 | 0 | 0 | 0 | 0 | 0 |
| ENSMUSG00 AC195488.1 | 561 | 0 | 0 | 0 | 0 | 0 | 0 |
| ENSMUSG00 Gm1973 | 810 | 0 | 0 | 0 | 0 | 0 | 0 |
| ENSMUSG00 1700110I01R | 3678 | 0 | 0 | 0 | 0 | 0 | 0 |
| ENSMUSG00 D830030K20l | 603 | 0 | 0 | 0 | 0 | 0 | 0 |
| ENSMUSG00 Gm5795 | 1934 | 0 | 0 | 0 | 0 | 0 | 0 |
| ENSMUSG00 Gm2897 | 2013 | 0 | 0 | 0 | 0 | 0 | 0 |
| ENSMUSG00 Gm11110 | 1008 | 0 | 1.05168822 | 0 | 0.25056975 | 0.91118528 | 0.54116348 |
| ENSMUSG00 Cntf | 1071 | 0.03711996 | 0.13497603 | 0.16508625 | 0 | 0 | 0.17826562 |
| ENSMUSG00 Gm11111 | 96 | 0 | 0 | 0 | 0 | 0 | 0 |
| ENSMUSG00 Atg4a | 2668 | 5.21529917 | 7.31465857 | 5.31813778 | 5.28788584 | 6.36873431 | 7.40136387 |
| ENSMUSG00 Ms4a6c | 1840 | 0.06481872 | 0 | 0 | 0 | 0 | 0 |
| ENSMUSG00 Gm17418 | 3536 | 0.84322993 | 0.73587892 | 0.08750365 | 0.35714693 | 0.23088858 | 0.68649403 |
| ENSMUSG00 4930402F06l | 2135 | 0 | 0 | 0 | 0 | 0 | 0 |
| ENSMUSG00 Gm7971 | 1844 | 0 | 0 | 0 | 0 | 0 | 0 |
| ENSMUSG00 Gm3286 | 1282 | 0 | 0 | 0 | 0 | 0 | 0 |
| ENSMUSG00 Gm3259 | 2888 | 0 | 0 | 0 | 0 | 0 | 0 |
| ENSMUSG00 Arpc4 | 2409 | 19.0938526 | 20.8827919 | 21.11927 | 18.2432255 | 15.8649878 | 16.3149869 |
| ENSMUSG00 Gm2382 | 1190 | 6.81522523 | 5.99293566 | 8.95180182 | 3.91141491 | 3.08731013 | 9.48882408 |
| ENSMUSG00 Tceal7 | 1212 | 0 | 0 | 0 | 0 | 0 | 0 |
| ENSMUSG00 Heatr7b1 | 8657 | 0.15613796 | 0.12245602 | 0.04084726 | 0.19172624 | 0.18272123 | 0.0787647 |
| ENSMUSG00 Gm15023 | 3693 | 0 | 0 | 0 | 0 | 0 | 0 |
| ENSMUSG00 Gm11114 | 55 | 0 | 0 | 0 | 0 | 0 | 0 |
| ENSMUSG00 Neu2 | 2584 | 8.52342732 | 11.3193384 | 12.8979062 | 8.44800191 | 6.95096141 | 8.43360556 |
| ENSMUSG00 Rpl36a | 1243 | 0.92752127 | 1.43435105 | 0.92457596 | 0.4644511 | 0.41050986 | 1.60180906 |
| ENSMUSG00 Kcnj13 | 1287 | 23.5690999 | 22.4271009 | 29.7769991 | 24.5032688 | 14.2731121 | 25.2613597 |
| ENSMUSG00 Tmem179b | 843 | 57.1101861 | 49.5011375 | 63.7072826 | 57.0978024 | 42.4311856 | 68.5586721 |
| ENSMUSG00 Gm11115 | 52 | 0 | 0 | 0 | 0 | 0 | 0 |
| ENSMUSG00 Gm11116 | 52 | 0 | 0 | 0 | 0 | 0 | 0 |
| ENSMUSG00 Alpi | 2324 | 0 | 0 | 0 | 0 | 0.02195627 | 0 |
| ENSMUSG00 St6galnac4 | 4117 | 0.59869803 | 0 | 0 | 0 | 0 | 0.15237229 |
| ENSMUSG00 Prickle4 | 3549 | 0.31365271 | 0.24439447 | 0.13700205 | 0.2541705 | 0.1869098 | 0.57638714 |
| ENSMUSG00 B3gnt7 | 2591 | 27.2503795 | 22.2427576 | 27.1591409 | 19.6773172 | 19.5952312 | 21.9376103 |
| ENSMUSG00 3110007F17f | 5920 | 0.39621172 | 0.12209403 | 0.37332638 | 0.2133229 | 0.12067048 | 0.79704605 |
| ENSMUSG00 Tmprss11g | 4063 | 0 | 0 | 0 | 0 | 0 | 0 |

| | | | | | | | | |
|---|---|---|---|---|---|---|---|---|
| ENSMUSG00 | Gm16026 | 3308 | 0 | 0 | 0.08017263 | 0 | 0 | 0.00824505 |
| ENSMUSG00 | Gm7609 | 2268 | 0 | 0 | 0 | 0 | 0 | 0.06012928 |
| ENSMUSG00 | 4933403O08l | 1186 | 0 | 0 | 0 | 0 | 0 | 0 |
| ENSMUSG00 | Gm15737 | 4892 | 0 | 0 | 0 | 0 | 0 | 0 |
| ENSMUSG00 | Gm17682 | 1791 | 0.39955257 | 0 | 0.09871992 | 0.3424873 | 0.02849044 | 0 |
| ENSMUSG00 | Col4a3 | 10370 | 14.4875565 | 15.0088918 | 7.22489141 | 15.2852866 | 16.2871074 | 13.5084424 |
| ENSMUSG00 | Prdm12 | 2466 | 0 | 0 | 0 | 0 | 0 | 0 |
| ENSMUSG00 | Gm14966 | 1040 | 1.10856628 | 1.57532599 | 2.42260101 | 1.24899385 | 2.94382937 | 1.9669211 |
| ENSMUSG00 | Pigb | 2181 | 8.16618765 | 8.52818279 | 5.33016264 | 7.57706391 | 7.97798907 | 4.90218184 |
| ENSMUSG00 | Utp14b | 3750 | 3.48788082 | 1.9403074 | 2.93500238 | 2.29000709 | 2.89829815 | 2.02195992 |
| ENSMUSG00 | Gm7325 | 941 | 26.9965484 | 52.7951348 | 11.1796372 | 34.4715813 | 36.8734703 | 48.6943613 |
| ENSMUSG00 | Gm550 | 838 | 0 | 0 | 0 | 0 | 0 | 0 |
| ENSMUSG00 | 1700011M02 | 417 | 0 | 0 | 0 | 0 | 0 | 0 |
| ENSMUSG00 | Rab7 | 4013 | 84.2463509 | 74.4950628 | 92.5672287 | 77.5498714 | 74.3334645 | 98.5094995 |
| ENSMUSG00 | Ssxa1 | 3014 | 3.35032915 | 5.14798752 | 4.03301487 | 3.41187216 | 3.35209768 | 4.59705267 |
| ENSMUSG00 | Gm9112 | 417 | 0 | 0 | 0 | 0 | 0 | 0 |
| ENSMUSG00 | Pin4 | 646 | 0.12308198 | 0.89510419 | 0.41054343 | 0.44683647 | 0.31595279 | 1.47772813 |
| ENSMUSG00 | Nhsl2 | 15927 | 0.04992212 | 0.19060375 | 0.17484248 | 0.15631701 | 0.32358033 | 0.29283376 |
| ENSMUSG00 | Phyhd1 | 3000 | 0.05300731 | 0.08031074 | 0.04420184 | 0 | 0.01700879 | 0.03636619 |
| ENSMUSG00 | Med12 | 7441 | 4.74971404 | 7.25937396 | 5.67893589 | 6.09530036 | 8.37296128 | 4.963024 |
| ENSMUSG00 | Cd47 | 1813 | 0 | 0 | 0 | 0 | 0 | 0 |
| ENSMUSG00 | C030013D06 | 1220 | 0.09775938 | 0 | 0.07246204 | 1.2125933 | 0.0836498 | 0.17885009 |
| ENSMUSG00 | H2-T10 | 3847 | 0.15501227 | 0.02505144 | 0 | 0 | 0.01326394 | 0.19142585 |
| ENSMUSG00 | Gm11127 | 1338 | 0.86166587 | 0.8643308 | 0.85892969 | 0.7820473 | 0.99154393 | 0.81538533 |
| ENSMUSG00 | Cml5 | 980 | 0 | 0 | 0 | 0 | 0 | 0 |
| ENSMUSG00 | Gm11128 | 2445 | 0 | 0.09854078 | 0 | 0 | 0 | 0.02231054 |
| ENSMUSG00 | Gm11129 | 51 | 2.33855768 | 0 | 0 | 0 | 3.00155152 | 1.60439054 |
| ENSMUSG00 | Gm13420 | 605 | 6.50544227 | 8.68152429 | 4.09141026 | 5.18865764 | 10.2052752 | 5.99591243 |
| ENSMUSG00 | 6530402F18l | 5492 | 0.18096996 | 0 | 0 | 0.22994748 | 0 | 0 |
| ENSMUSG00 | Gm5138 | 1228 | 0.12949668 | 0 | 0 | 0 | 0 | 0 |
| ENSMUSG00 | 1700101E01l | 1394 | 0 | 0.13826813 | 0 | 0 | 0 | 0 |
| ENSMUSG00 | Gm11131 | 3120 | 0.17838998 | 0.18533247 | 0.07083629 | 0.33537798 | 0.39251058 | 0.03496749 |
| ENSMUSG00 | H2-Q1 | 2611 | 0 | 0 | 0.01692909 | 0 | 0 | 0.22981312 |
| ENSMUSG00 | Apoo | 1238 | 54.3667435 | 41.9198693 | 33.8476148 | 41.5030948 | 27.3679431 | 103.87716 |
| ENSMUSG00 | Zfx | 13094 | 1.6516711 | 1.75905906 | 3.11242548 | 2.09977989 | 2.0147118 | 3.3181992 |
| ENSMUSG00 | Mdga2 | 422 | 0 | 0 | 0 | 0 | 0 | 0 |
| ENSMUSG00 | Ccdc142 | 3753 | 1.90673767 | 2.37529757 | 0.71844189 | 1.40367133 | 2.21617353 | 1.19912483 |
| ENSMUSG00 | 4932429P05l | 3091 | 0 | 0 | 0 | 0 | 0 | 0 |
| ENSMUSG00 | Gm16309 | 519 | 0 | 0 | 0 | 0 | 0 | 0 |
| ENSMUSG00 | Reg3a | 831 | 0 | 0 | 0 | 0 | 0 | 0 |
| ENSMUSG00 | Gm8787 | 1382 | 0 | 0 | 0 | 0 | 0.22153275 | 0 |
| ENSMUSG00 | Gm14743 | 752 | 0 | 0 | 0 | 0 | 0 | 0 |
| ENSMUSG00 | Gm5938 | 1124 | 0 | 0 | 0 | 0 | 0 | 0 |
| ENSMUSG00 | Gm14744 | 793 | 0 | 0 | 0 | 0 | 0 | 0 |
| ENSMUSG00 | Tmsb10 | 1571 | 0.17714091 | 0.15336232 | 0.25322507 | 0.22967565 | 0.19488113 | 0.57292368 |
| ENSMUSG00 | Gm15402 | 668 | 0 | 0 | 0 | 0 | 0 | 0 |
| ENSMUSG00 | 4930468A15l | 1001 | 0 | 0 | 0 | 0 | 0 | 0 |
| ENSMUSG00 | 1700021O21l | 435 | 0.18278382 | 0.11077343 | 0 | 0 | 0.11730201 | 0.18810096 |
| ENSMUSG00 | Gm5936 | 2853 | 0 | 0 | 0 | 0 | 0 | 0 |
| ENSMUSG00 | Gm6890 | 773 | 0 | 0 | 0 | 0 | 0 | 0 |
| ENSMUSG00 | Gm5640 | 2857 | 0 | 0 | 0 | 0 | 0 | 0 |
| ENSMUSG00 | Gm6880 | 624 | 0 | 0 | 0 | 0 | 0 | 0 |
| ENSMUSG00 | C030048H21 | 659 | 0 | 0 | 0 | 0 | 0 | 0 |
| ENSMUSG00 | Gm11140 | 31 | 0 | 0 | 0 | 0 | 0 | 0 |
| ENSMUSG00 | Lcn14 | 725 | 0 | 0 | 0 | 0 | 0 | 0 |
| ENSMUSG00 | Gm4485 | 1386 | 0.3728869 | 1.00823003 | 1.27566647 | 0 | 0.55223351 | 1.08232695 |
| ENSMUSG00 | Gm11143 | 192 | 0 | 0 | 0 | 0 | 0 | 0 |
| ENSMUSG00 | Igkv12-47 | 328 | 0 | 0 | 0 | 0 | 0 | 0 |
| ENSMUSG00 | Igkv13-87 | 122 | 0 | 0 | 0 | 0 | 0 | 0 |
| ENSMUSG00 | Gm11146 | 93 | 0 | 0 | 0 | 0 | 1.64601212 | 0 |
| ENSMUSG00 | H2-DMb1 | 1471 | 5.94575507 | 7.3049467 | 6.85113542 | 4.04727216 | 6.69482701 | 5.8220508 |
| ENSMUSG00 | Mpp4 | 3062 | 0 | 0.36194911 | 0.25984101 | 0.11783816 | 0.28329471 | 0.21377901 |
| ENSMUSG00 | Kifc1 | 3416 | 0.11638021 | 0 | 0 | 0.29575446 | 0 | 0 |
| ENSMUSG00 | Aox3l1 | 4740 | 0 | 0 | 0.0559517 | 0 | 0.03229518 | 0 |
| ENSMUSG00 | Haus3 | 1945 | 10.1790382 | 8.47288597 | 6.59050616 | 7.84714902 | 7.58181111 | 11.7091639 |
| ENSMUSG00 | Gm11147 | 58 | 0 | 0 | 0 | 0 | 0 | 0 |
| ENSMUSG00 | 2-Mar | 5795 | 6.78484387 | 7.90773192 | 7.20806586 | 5.87773081 | 9.50085583 | 7.15400374 |
| ENSMUSG00 | Gm6981 | 1007 | 0.90801991 | 0.14355445 | 0.83399704 | 0.50163716 | 0.86141846 | 0.08125513 |
| ENSMUSG00 | Gm684 | 463 | 1.11624459 | 0.52037195 | 1.5274935 | 0.93517178 | 0.11020816 | 1.53162121 |
| ENSMUSG00 | Hoxa3 | 7361 | 0.01080165 | 0.1047389 | 0.16213147 | 0.28430357 | 0.29114357 | 0.32606552 |
| ENSMUSG00 | Maea | 2159 | 85.035113 | 64.0105216 | 71.2061186 | 68.5207886 | 65.2542027 | 71.8564836 |
| ENSMUSG00 | Pglyrp2 | 2345 | 0 | 0 | 0 | 0 | 0 | 0 |
| ENSMUSG00 | Gm11149 | 1246 | 0 | 0 | 0 | 0 | 0 | 0 |
| ENSMUSG00 | 4930408F14l | 1013 | 0 | 0 | 0 | 0 | 0 | 0 |
| ENSMUSG00 | Rbpsuh-rs3 | 1462 | 0.29911784 | 0.03295926 | 0 | 0.39487874 | 0.69803524 | 0.72757246 |
| ENSMUSG00 | Gm14692 | 774 | 0 | 0 | 0 | 0 | 0 | 0 |
| ENSMUSG00 | Gm1140 | 774 | 0 | 0 | 0 | 0 | 0 | 0 |
| ENSMUSG00 | Gm6760 | 603 | 0 | 0 | 0 | 0 | 0 | 0 |
| ENSMUSG00 | Tmem217 | 901 | 0 | 0 | 0.04905865 | 0 | 0 | 0 |
| ENSMUSG00 | Gm7073 | 1979 | 0 | 0 | 0 | 0 | 0 | 0 |
| ENSMUSG00 | Gm364 | 2139 | 0 | 0 | 0 | 0 | 0 | 0 |
| ENSMUSG00 | CT030161.1 | 75 | 0 | 0 | 0 | 0 | 0 | 0 |
| ENSMUSG00 | Gpr45 | 2034 | 0 | 0 | 0 | 0 | 0 | 0 |
| ENSMUSG00 | Tmem182 | 1768 | 0 | 0.84489802 | 0 | 0.85715264 | 0 | 0.21597565 |
| ENSMUSG00 | Gm16430 | 758 | 0 | 0 | 0 | 0 | 0 | 0 |
| ENSMUSG00 | Gm16405 | 697 | 0 | 0 | 0 | 0 | 0 | 0 |
| ENSMUSG00 | C1qtnf5 | 1382 | 0 | 0 | 0 | 0 | 0 | 0 |
| ENSMUSG00 | Gm14597 | 822 | 0 | 0 | 0 | 0 | 0 | 0 |
| ENSMUSG00 | Gm1975 | 384 | 0 | 0 | 0 | 0 | 0 | 0 |
| ENSMUSG00 | Gm4758 | 288 | 0 | 0 | 0 | 0 | 0 | 0 |
| ENSMUSG00 | Gm5483 | 411 | 0 | 0 | 0 | 0 | 0 | 0 |
| ENSMUSG00 | Clec2l | 1419 | 0 | 0 | 0.06229999 | 0 | 0 | 0 |
| ENSMUSG00 | Gm17604 | 72 | 0 | 0 | 0 | 1.50341852 | 0 | 0 |
| ENSMUSG00 | 9230102O04 | 1215 | 0.91617568 | 1.30876756 | 0.94588306 | 1.81152652 | 1.17591648 | 1.01017182 |
| ENSMUSG00 | Gm13218 | 263 | 0 | 0 | 0 | 0 | 0 | 0 |
| ENSMUSG00 | Gm13219 | 4968 | 0 | 0 | 0 | 0 | 0 | 0 |
| ENSMUSG00 | Zbtb9 | 2796 | 0.0142187 | 0 | 0 | 0 | 0 | 0 |
| ENSMUSG00 | Gm595 | 2101 | 0 | 0 | 0 | 0 | 0 | 0 |
| ENSMUSG00 | Stard6 | 1637 | 0.60713929 | 0.11774329 | 0.62103995 | 0.17633253 | 0.03117066 | 0.09996813 |
| ENSMUSG00 | Gm17371 | 1743 | 0.02280865 | 0 | 0.15215781 | 0 | 0.35130035 | 0 |
| ENSMUSG00 | Ankrd39 | 1782 | 16.6428667 | 11.1677893 | 13.3696933 | 14.7810845 | 28.7202329 | 13.0710112 |
| ENSMUSG00 | Gm11157 | 145 | 0 | 0 | 0 | 0 | 0 | 0 |
| ENSMUSG00 | Seh1l | 3507 | 53.211356 | 50.357374 | 55.7596104 | 52.3379984 | 41.7726618 | 57.4268424 |
| ENSMUSG00 | Hspa14 | 4309 | 0 | 0 | 0 | 0 | 0 | 0 |
| ENSMUSG00 | Gm11158 | 167 | 0 | 0 | 0 | 0 | 0 | 0 |
| ENSMUSG00 | Cypt15 | 517 | 0 | 0 | 0 | 0 | 0 | 0 |
| ENSMUSG00 | Muc4 | 11982 | 0.0199076 | 0.62334323 | 0.29143262 | 0.06624979 | 0.12775758 | 0.11836766 |
| ENSMUSG00 | Gm8123 | 85 | 0 | 0 | 0 | 0 | 0 | 0 |
| ENSMUSG00 | Gm11160 | 110 | 0 | 0 | 0 | 0 | 0 | 0 |
| ENSMUSG00 | Tm4sf19 | 1252 | 0 | 0 | 0 | 0 | 0 | 0 |
| ENSMUSG00 | Gm14543 | 1199 | 0 | 0 | 0 | 0 | 0 | 0 |
| ENSMUSG00 | Rhox2h | 775 | 0 | 0 | 0 | 0 | 0 | 0 |
| ENSMUSG00 | Rhox4g | 974 | 0 | 0 | 0 | 0 | 0 | 0 |
| ENSMUSG00 | Rhox2g | 791 | 0 | 0 | 0 | 0 | 0 | 0 |
| ENSMUSG00 | Rhox4f | 925 | 0 | 0 | 0 | 0 | 0 | 0 |
| ENSMUSG00 | Rhox3f | 824 | 0 | 0 | 0 | 0 | 0 | 0 |
| ENSMUSG00 | Rhox2f | 775 | 0 | 0 | 0 | 0 | 0 | 0 |
| ENSMUSG00 | Rhox4d | 2333 | 0 | 0 | 0 | 0 | 0 | 0 |
| ENSMUSG00 | Rhox2d | 775 | 0 | 0 | 0 | 0 | 0 | 0 |
| ENSMUSG00 | Rhox4c | 2343 | 0 | 0 | 0 | 0 | 0 | 0 |
| ENSMUSG00 | Rhox3c | 1047 | 0 | 0 | 0 | 0 | 0 | 0 |
| ENSMUSG00 | Rhox2c | 753 | 0 | 0 | 0 | 0 | 0 | 0 |
| ENSMUSG00 | Rhox2b | 1131 | 0 | 0 | 0 | 0 | 0 | 0 |
| ENSMUSG00 | Rhox4a | 2386 | 0 | 0 | 0 | 0 | 0 | 0.02286223 |
| ENSMUSG00 | Gm11161 | 110 | 0 | 0 | 0 | 0 | 0 | 0 |
| ENSMUSG00 | Rpl39 | 1430 | 7.11706504 | 5.69476134 | 3.43105215 | 3.50727566 | 2.96167076 | 10.2422946 |
| ENSMUSG00 | Gm6268 | 298 | 0 | 0 | 0 | 0 | 0 | 0 |
| ENSMUSG00 | Gm1110 | 2373 | 0 | 0 | 0 | 0 | 0 | 0 |
| ENSMUSG00 | Gm17193 | 689 | 0.05770026 | 0.06993678 | 0.38492171 | 0.10473743 | 0.4443516 | 0.3562725 |
| ENSMUSG00 | Gm2940 | 1013 | 0 | 0 | 0 | 0 | 0 | 0 |
| ENSMUSG00 | Gm2892 | 710 | 0 | 0 | 0 | 0 | 0 | 0 |
| ENSMUSG00 | Gm2864 | 1013 | 0 | 0 | 0 | 0 | 0 | 0 |
| ENSMUSG00 | Fam71f2 | 1270 | 0.03130353 | 0 | 0 | 0 | 0 | 0 |
| ENSMUSG00 | Gm2790 | 1013 | 0 | 0 | 0 | 0 | 0 | 0 |
| ENSMUSG00 | Prr14 | 3406 | 0 | 0 | 0 | 0 | 0 | 0 |
| ENSMUSG00 | Gm5168 | 906 | 0 | 0 | 0 | 0 | 0 | 0 |
| ENSMUSG00 | Rab26 | 2843 | 0.46146003 | 0.13559322 | 0.21766648 | 1.12954694 | 0.23332497 | 0.00959361 |
| ENSMUSG00 | Tceb1 | 950 | 65.4082274 | 71.2144892 | 83.7508604 | 65.3654722 | 51.9394794 | 131.521182 |
| ENSMUSG00 | 4930420K17l | 983 | 17.6331531 | 15.49025 | 27.6092895 | 20.6655047 | 12.3543006 | 21.4201643 |
| ENSMUSG00 | Ntn3 | 4888 | 0.81332816 | 0.33517574 | 0.37980307 | 0.4650509 | 0.42800356 | 2.28776639 |
| ENSMUSG00 | 1700015F17l | 1195 | 0.03326819 | 0 | 0 | 0.12077672 | 0 | 0.15976776 |
| ENSMUSG00 | Gm17396 | 1653 | 1.08227261 | 1.80735597 | 1.55094186 | 1.61528814 | 0.43216531 | 0.8415043 |
| ENSMUSG00 | 2610203C22l | 2755 | 0.33189693 | 0.13992433 | 0.09626536 | 0.35361713 | 0.98163264 | 0.44550227 |
| ENSMUSG00 | Fdx1l | 777 | 13.9681418 | 18.914884 | 17.5214513 | 18.6679304 | 8.99692856 | 20.1839351 |
| ENSMUSG00 | Vwde | 3964 | 0 | 0 | 0 | 0 | 0 | 0 |
| ENSMUSG00 | Zglp1 | 1005 | 0.27690384 | 0.5274138 | 0.3958374 | 0.96936836 | 0.55849765 | 0.05427789 |
| ENSMUSG00 | Vmn1r-ps150 | 1174 | 0 | 0 | 0 | 0 | 0 | 0 |
| ENSMUSG00 | Gm14862 | 867 | 0 | 0 | 0 | 0 | 0 | 0 |
| ENSMUSG00 | Gm5751 | 439 | 1.08671017 | 0.98787695 | 0 | 1.15068024 | 0.81363242 | 1.3047094 |
| ENSMUSG00 | Vmn2r93 | 2568 | 0 | 0 | 0 | 0 | 0 | 0 |
| ENSMUSG00 | Gm6592 | 501 | 0 | 0 | 0 | 0 | 0 | 0 |
| ENSMUSG00 | Fpr3 | 1266 | 0 | 0 | 0 | 0 | 0 | 0 |
| ENSMUSG00 | Ssxb3 | 856 | 0 | 0 | 0 | 0 | 0 | 0 |
| ENSMUSG00 | Ssxb6 | 694 | 0 | 0 | 0 | 0 | 0 | 0 |
| ENSMUSG00 | Ssxb8 | 700 | 0 | 0 | 0 | 0 | 0 | 0 |
| ENSMUSG00 | Gm14459 | 534 | 0 | 0 | 0 | 0 | 0 | 0 |
| ENSMUSG00 | Ssxb1 | 867 | 0 | 0 | 0 | 0 | 0 | 0 |
| ENSMUSG00 | Tcte3 | 790 | 0 | 0 | 0 | 0 | 0 | 0 |
| ENSMUSG00 | 9030025P20l | 1354 | 0.73403768 | 1.42352857 | 1.76285046 | 1.06593928 | 2.11039664 | 2.01437513 |
| ENSMUSG00 | Gm3417 | 628 | 0 | 0 | 0 | 0 | 0 | 0 |
| ENSMUSG00 | Smok4a | 2144 | 0 | 0.06742506 | 0 | 0 | 0 | 0 |
| ENSMUSG00 | Gm8597 | 525 | 0 | 0 | 0 | 0 | 0 | 0 |
| ENSMUSG00 | Map3k4 | 2983 | 0 | 0 | 0 | 0 | 0 | 0 |
| ENSMUSG00 | Gm11167 | 676 | 0 | 0 | 0.06538734 | 0.10675161 | 0 | 0 |
| ENSMUSG00 | Ttll2 | 1621 | 0 | 0 | 0 | 0 | 0 | 0 |
| ENSMUSG00 | Rnaset2b | 1004 | 20.7884734 | 20.9735809 | 18.8870425 | 16.531614 | 17.7880792 | 18.6358591 |
| ENSMUSG00 | AC175035.1 | 802 | 0 | 0 | 0 | 0 | 0 | 0 |
| ENSMUSG00 | AC175035.2 | 839 | 0 | 0 | 0 | 0 | 0 | 0 |
| ENSMUSG00 | Gm2833 | 1095 | 0 | 0 | 0 | 0 | 0 | 0 |
| ENSMUSG00 | Tmem181c-p | 839 | 0 | 0 | 0 | 0 | 0 | 0 |
| ENSMUSG00 | Dynlt1f | 802 | 0 | 0 | 0 | 0 | 0 | 0 |
| ENSMUSG00 | Tmem181b-p | 2565 | 0.728463 | 0.54479798 | 0.48251525 | 0.29540855 | 0.85541293 | 0.52103599 |
| ENSMUSG00 | Gm2792 | 1818 | 0 | 0 | 0 | 0 | 0 | 0 |
| ENSMUSG00 | Gm11169 | 261 | 0 | 0 | 0 | 0 | 0.19550336 | 0 |
| ENSMUSG00 | 3110001I22R | 1720 | 5.59350365 | 4.6225386 | 4.60007552 | 3.75504998 | 4.65764012 | 6.80280246 |
| ENSMUSG00 | Snx29 | 1197 | 2.65700789 | 4.91123303 | 1.62479623 | 3.345954 | 7.24685371 | 3.44066042 |
| ENSMUSG00 | Gm11172 | 61 | 0 | 0 | 0 | 0 | 0 | 0 |
| ENSMUSG00 | Spag11a | 434 | 0 | 0 | 0 | 0 | 0 | 0 |
| ENSMUSG00 | Gm15093 | 2106 | 0 | 0 | 0 | 0 | 0 | 0 |
| ENSMUSG00 | Xlr4a | 1153 | 0 | 0 | 0 | 0 | 0 | 0 |
| ENSMUSG00 | Amy2a2 | 2677 | 0 | 0 | 0 | 0 | 0 | 0 |
| ENSMUSG00 | Klra4 | 1698 | 0 | 0 | 0 | 0 | 0 | 0 |
| ENSMUSG00 | Klra1 | 2145 | 0 | 0 | 0 | 0 | 0 | 0 |
| ENSMUSG00 | Gm13275 | 548 | 0 | 0 | 0 | 0 | 0 | 0 |
| ENSMUSG00 | Gm13271 | 548 | 0 | 0 | 0 | 0 | 0 | 0 |
| ENSMUSG00 | Gm4984 | 1297 | 0.09195562 | 0 | 0 | 0 | 0.07868369 | 0 |
| ENSMUSG00 | Olfr22-ps1 | 936 | 0 | 0 | 0 | 0 | 0 | 0 |
| ENSMUSG00 | Gm10698 | 605 | 3.67984613 | 4.46023266 | 5.8448718 | 3.28018587 | 2.61457463 | 5.09427146 |
| ENSMUSG00 | Gm11951 | 430 | 0 | 0 | 0 | 0 | 0 | 0 |
| ENSMUSG00 | Gm12106 | 1194 | 0.03329605 | 0 | 0 | 0.06043894 | 0.08547132 | 0 |
| ENSMUSG00 | Gm12772 | 277 | 0 | 0 | 0 | 0 | 0 | 0 |
| ENSMUSG00 | Gm12775 | 282 | 0 | 0 | 0 | 0 | 0 | 0 |
| ENSMUSG00 | Gm15846 | 319 | 0.87237732 | 0 | 1.10851017 | 1.47042815 | 2.55931686 | 1.96651004 |
| ENSMUSG00 | Gm16073 | 516 | 0 | 0 | 0 | 0.06992644 | 0 | 0 |
| ENSMUSG00 | Gm11273 | 389 | 1.22639014 | 0.49549041 | 0.45451767 | 0.37102359 | 0 | 1.68275409 |
| ENSMUSG00 | Rpl28-ps3 | 882 | 0.04507424 | 0 | 0.05011547 | 0.04090935 | 0 | 0.03092363 |
| ENSMUSG00 | Gm12643 | 373 | 0 | 0 | 0 | 0 | 0 | 0 |
| ENSMUSG00 | Gm7146 | 738 | 0 | 0 | 0 | 0 | 0 | 0 |
| ENSMUSG00 | Gm14853 | 548 | 0.145093 | 0.35172586 | 0.0806603 | 0 | 0 | 0.14931372 |
| ENSMUSG00 | Gm14058 | 734 | 0 | 0 | 0 | 0 | 0 | 0 |
| ENSMUSG00 | Gm11970 | 319 | 0 | 0 | 0 | 0 | 0 | 0 |
| ENSMUSG00 | Gm11799 | 459 | 0 | 0.31494407 | 0 | 0 | 0 | 0 |
| ENSMUSG00 | Gm12947 | 419 | 0 | 0 | 0 | 0 | 0 | 0.32547302 |
| ENSMUSG00 | Gm5566 | 458 | 0 | 0 | 0 | 0 | 0 | 0 |
| ENSMUSG00 | Rps19-ps7 | 926 | 0 | 0 | 0 | 0 | 0 | 0 |
| ENSMUSG00 | Gm5915 | 905 | 0.26357225 | 0.74542562 | 0 | 0.11960899 | 0.28191368 | 0.21096406 |
| ENSMUSG00 | Gm15108 | 801 | 0 | 0 | 0 | 0 | 0 | 0 |
| ENSMUSG00 | Gm14864 | 1086 | 0 | 0 | 0 | 0 | 0 | 0 |
| ENSMUSG00 | Gm14510 | 1091 | 0 | 0 | 0 | 0 | 0 | 0 |
| ENSMUSG00 | Rpl12-ps1 | 624 | 0 | 0 | 0 | 0 | 0 | 0 |
| ENSMUSG00 | Gm12378 | 1460 | 0 | 0 | 0 | 0 | 0 | 0 |
| ENSMUSG00 | Gm11175 | 542 | 0 | 0 | 0 | 0 | 0 | 0 |
| ENSMUSG00 | Rps19-ps3 | 874 | 1.27363096 | 1.37833073 | 0.50574191 | 1.0320951 | 0.81735613 | 0.03120668 |
| ENSMUSG00 | Gm11864 | 1219 | 0 | 0 | 0 | 0 | 0 | 0 |
| ENSMUSG00 | Gm41 | 1022 | 0 | 0 | 0 | 0 | 0 | 0 |
| ENSMUSG00 | Gm15125 | 798 | 0 | 0 | 0 | 0 | 0 | 0 |
| ENSMUSG00 | Gm14757 | 777 | 0 | 0 | 0 | 0 | 0 | 0 |
| ENSMUSG00 | Gm9427 | 650 | 0 | 0 | 0 | 0 | 0 | 0 |
| ENSMUSG00 | Hist1h2aj | 724 | 0 | 0 | 0 | 0 | 0 | 0 |
| ENSMUSG00 | Gm15583 | 3509 | 0 | 0 | 0 | 0 | 0 | 0 |
| ENSMUSG00 | Fam119b | 2538 | 0.46992294 | 0.17087391 | 0.26124021 | 0.14216724 | 0.68356847 | 0 |
| ENSMUSG00 | Gm17516 | 2008 | 0.21778401 | 0 | 0 | 0 | 0 | 0.24449378 |
| ENSMUSG00 | Gm12838 | 324 | 0 | 0 | 0 | 0 | 0 | 0 |
| ENSMUSG00 | Gm12260 | 410 | 0 | 0 | 0 | 0 | 0 | 0 |
| ENSMUSG00 | Gm14777 | 175 | 0 | 0 | 0.25258196 | 0.20618311 | 0.29157929 | 0 |
| ENSMUSG00 | Gm14385 | 875 | 0 | 0 | 0 | 0 | 0 | 0 |
| ENSMUSG00 | Tpm3-rs2 | 322 | 0 | 0 | 0 | 0 | 0 | 0 |
| ENSMUSG00 | Gm17235 | 1113 | 0 | 0.17317679 | 0.99285362 | 0.87530566 | 0.229229 | 0.07351655 |
| ENSMUSG00 | Gm11678 | 446 | 0 | 0 | 0 | 0 | 0 | 0 |
| ENSMUSG00 | Gm14448 | 209 | 0 | 0.69167142 | 0.63447622 | 0.34528272 | 0.48829068 | 0 |
| ENSMUSG00 | Gm14756 | 239 | 0 | 0 | 0 | 0 | 0 | 0 |
| ENSMUSG00 | Gm13573 | 630 | 0 | 0 | 0 | 0 | 0 | 0 |
| ENSMUSG00 | Gm15526 | 203 | 0 | 0 | 0 | 0 | 0 | 0 |
| ENSMUSG00 | Gm11486 | 475 | 0 | 0 | 0 | 0 | 0 | 0 |
| ENSMUSG00 | Gm13768 | 923 | 0 | 0 | 0 | 0 | 0 | 0 |
| ENSMUSG00 | Gm13343 | 104 | 1.52905694 | 0 | 0 | 0.69388547 | 0 | 0 |
| ENSMUSG00 | Gm14239 | 683 | 0 | 0.07055116 | 0 | 0 | 0 | 0 |
| ENSMUSG00 | Gm6269 | 452 | 0 | 0 | 0 | 0 | 0 | 0 |
| ENSMUSG00 | Gm15487 | 931 | 0.85403825 | 0.72460815 | 1.70920123 | 1.66651763 | 0.21923255 | 32.342859 |
| ENSMUSG00 | Gm16400 | 1027 | 0 | 0 | 0 | 0 | 0 | 0 |
| ENSMUSG00 | Gm15380 | 782 | 0 | 0 | 0 | 0 | 0 | 0 |
| ENSMUSG00 | Gm14772 | 513 | 0 | 0 | 0 | 0 | 0 | 0 |
| ENSMUSG00 | Brms1 | 2065 | 22.92919 | 21.8180743 | 25.8789483 | 17.0537896 | 21.7943165 | 25.0953097 |
| ENSMUSG00 | Gm14806 | 2109 | 0 | 0 | 0 | 0 | 0 | 0 |
| ENSMUSG00 | Gm14807 | 982 | 0.0404842 | 0 | 0 | 0 | 0.05196169 | 0 |
| ENSMUSG00 | Gm14350 | 639 | 0 | 0 | 0 | 0 | 0 | 0 |
| ENSMUSG00 | Gm14354 | 873 | 0 | 0 | 0 | 0 | 0 | 0 |
| ENSMUSG00 | Gm12419 | 635 | 0.06260706 | 0.22765248 | 0 | 0 | 0 | 0 |
| ENSMUSG00 | Ncrna00085 | 1939 | 3.38300891 | 3.40461195 | 2.18843576 | 2.64241894 | 2.3157922 | 4.59969419 |
| ENSMUSG00 | Vmn2r-ps94 | 1453 | 0 | 0 | 0 | 0 | 0 | 0 |
| ENSMUSG00 | Gm14644 | 645 | 0 | 0 | 0 | 0 | 0 | 0 |
| ENSMUSG00 | AC107756.1 | 84 | 0 | 0 | 0 | 0 | 0 | 0.32469809 |
| ENSMUSG00 | AC140486.1 | 119 | 0 | 0 | 0 | 0 | 0 | 0 |
| ENSMUSG00 | Mir1298 | 97 | 0 | 0 | 0 | 0 | 0 | 0 |
| ENSMUSG00 | AL772216.4 | 126 | 0 | 0 | 0 | 0 | 0 | 0 |
| ENSMUSG00 | AC159266.1 | 120 | 0 | 0 | 0 | 0 | 0 | 0 |
| ENSMUSG00 | SNORD93 | 70 | 0 | 0 | 0 | 0 | 0 | 0 |
| ENSMUSG00 | AC153902.1 | 146 | 0 | 0 | 0 | 0 | 0 | 0 |
| ENSMUSG00 | AC161816.1 | 120 | 0 | 0 | 0 | 0 | 0 | 0 |
| ENSMUSG00 | AC132367.1 | 93 | 0.42747829 | 0 | 0 | 0 | 0 | 0 |
| ENSMUSG00 | AC164156.1 | 89 | 0 | 0 | 0 | 0 | 0 | 0 |
| ENSMUSG00 | AC156572.1 | 82 | 0 | 0 | 0 | 0 | 0 | 0 |
| ENSMUSG00 | AC152946.1 | 88 | 0 | 0 | 0 | 0 | 0 | 0 |
| ENSMUSG00 | AC165441.1 | 111 | 0 | 0 | 0 | 0 | 0 | 0 |
| ENSMUSG00 | AC159639.1 | 118 | 0 | 0 | 0 | 0 | 0 | 0 |
| ENSMUSG00 | AC151974.1 | 120 | 0 | 0 | 0 | 0 | 0 | 0 |
| ENSMUSG00 | AC111029.1 | 120 | 0 | 0 | 0 | 0 | 0 | 0 |
| ENSMUSG00 | SNORD88 | 90 | 0 | 0 | 0 | 0 | 0 | 0 |
| ENSMUSG00 | CT571271.1 | 120 | 0 | 0 | 0 | 0 | 0 | 0 |
| ENSMUSG00 | AC148019.1 | 120 | 0 | 0 | 0 | 0 | 0 | 0 |
| ENSMUSG00 | Mir1247 | 81 | 0 | 0 | 0 | 0 | 0 | 0 |
| ENSMUSG00 | U6atac | 117 | 0 | 0 | 0 | 0 | 0 | 0 |
| ENSMUSG00 | AL772167.1 | 63 | 0 | 0 | 0 | 0 | 0 | 0 |
| ENSMUSG00 | AL805898.1 | 94 | 0 | 0 | 0 | 0 | 0 | 0 |
| ENSMUSG00 | AL772216.5 | 119 | 0 | 0 | 0 | 0 | 0 | 0 |
| ENSMUSG00 | AC136741.1 | 120 | 0 | 0 | 0 | 0 | 0 | 0 |
| ENSMUSG00 | SNORA77 | 125 | 0 | 0 | 0 | 0.57731271 | 0 | 0.21819711 |
| ENSMUSG00 | SNORA79 | 142 | 0 | 1.69670571 | 0 | 0 | 0 | 0 |
| ENSMUSG00 | AC171332.1 | 120 | 0 | 0 | 0 | 0 | 0 | 0 |
| ENSMUSG00 | AC162877.1 | 120 | 0 | 0 | 0 | 0 | 0 | 0 |
| ENSMUSG00 | AC107456.1 | 126 | 0 | 0 | 0 | 0 | 0 | 0 |
| ENSMUSG00 | SNORA42 | 119 | 0 | 0 | 0 | 0 | 0.85758615 | 1.14599325 |
| ENSMUSG00 | Snord88a | 89 | 3.57352634 | 2.70710349 | 4.96649922 | 4.05416231 | 0.57333007 | 3.37102283 |
| ENSMUSG00 | AL845466.1 | 100 | 0 | 0 | 0 | 0 | 0 | 0 |
| ENSMUSG00 | AC102435.1 | 69 | 0 | 0 | 0 | 0 | 0 | 0 |
| ENSMUSG00 | U6atac | 125 | 0 | 0 | 0 | 0 | 0 | 0 |
| ENSMUSG00 | AC133085.1 | 120 | 0 | 0 | 0 | 0 | 0 | 0 |
| ENSMUSG00 | AL732460.1 | 120 | 0 | 0 | 0 | 0 | 0 | 0 |
| ENSMUSG00 | AL772216.6 | 122 | 0 | 0 | 0 | 0 | 0 | 0 |
| ENSMUSG00 | mmu-mir-11S | 120 | 0 | 0 | 0 | 0 | 0 | 0 |
| ENSMUSG00 | U6atac | 115 | 0 | 0 | 0 | 2.82381218 | 0 | 0 |
| ENSMUSG00 | AC165279.1 | 89 | 0 | 0 | 0 | 0 | 0 | 0 |
| ENSMUSG00 | AL929371.1 | 64 | 0 | 0 | 0 | 0 | 0 | 0 |
| ENSMUSG00 | AC191865.1 | 120 | 0 | 0 | 0 | 0 | 0 | 0 |
| ENSMUSG00 | AC160543.1 | 120 | 0 | 0 | 0 | 0 | 0 | 0 |
| ENSMUSG00 | CT025603.1 | 92 | 0 | 0 | 0 | 0 | 0 | 0 |
| ENSMUSG00 | AC135289.1 | 89 | 0 | 0 | 0.49664992 | 0 | 0 | 0 |
| ENSMUSG00 | AC163659.1 | 95 | 0 | 0 | 0 | 0 | 0 | 0 |
| ENSMUSG00 | AC173478.1 | 120 | 0 | 0 | 0 | 0 | 0 | 0 |
| ENSMUSG00 | SNORA76 | 118 | 0 | 0 | 0 | 0 | 0 | 0 |
| ENSMUSG00 | Snord111 | 88 | 0 | 0 | 1.00458734 | 0 | 1.73953554 | 0 |
| ENSMUSG00 | AL772216.7 | 120 | 0 | 0 | 0 | 0 | 0 | 0 |
| ENSMUSG00 | AC160992.1 | 96 | 0 | 0 | 0 | 0 | 0 | 0 |
| ENSMUSG00 | AC126690.1 | 106 | 0 | 0 | 0 | 0 | 0 | 0.25730792 |
| ENSMUSG00 | AL772216.8 | 119 | 0 | 0 | 0 | 0 | 0 | 0 |
| ENSMUSG00 | Mir1188 | 119 | 0 | 0 | 0 | 0 | 0 | 0 |
| ENSMUSG00 | AC134455.1 | 120 | 0 | 0 | 0 | 0 | 0 | 0 |
| ENSMUSG00 | AC121814.1 | 125 | 1.59021922 | 0.38549154 | 2.82891795 | 0 | 1.22463302 | 0.21819711 |
| ENSMUSG00 | Mir467h | 120 | 0 | 0 | 0 | 0 | 0 | 0 |
| ENSMUSG00 | AC084287.1 | 82 | 3.39376053 | 0.58763954 | 1.6171406 | 0.44002493 | 0 | 0 |
| ENSMUSG00 | Mir1193 | 120 | 0 | 0 | 0 | 0 | 0 | 0 |
| ENSMUSG00 | AC111115.1 | 121 | 0 | 0 | 0 | 0.59639743 | 0 | 0 |
| ENSMUSG00 | U6atac | 126 | 0 | 0 | 0 | 0 | 0 | 0 |
| ENSMUSG00 | AC167245.1 | 65 | 0 | 0 | 0 | 0 | 0 | 0 |
| ENSMUSG00 | AL772258.1 | 120 | 0 | 0 | 0 | 0 | 0 | 0 |
| ENSMUSG00 | AL646096.1 | 86 | 0 | 0 | 0 | 0 | 0.59332995 | 0 |
| ENSMUSG00 | AL772216.9 | 126 | 0 | 0 | 0 | 0 | 0 | 0 |
| ENSMUSG00 | AL807401.1 | 120 | 0 | 0 | 0 | 0 | 0 | 0 |
| ENSMUSG00 | U3 | 221 | 0 | 0 | 0 | 0 | 0 | 0 |
| ENSMUSG00 | AL772216.10 | 123 | 0 | 0 | 0 | 0 | 0 | 0 |
| ENSMUSG00 | BX649539.1 | 58 | 0 | 0 | 0 | 0 | 0 | 0 |
| ENSMUSG00 | Mir466k | 121 | 0 | 0 | 0 | 0 | 0 | 0 |
| ENSMUSG00 | U6atac | 119 | 0 | 0 | 0 | 0 | 0 | 0 |
| ENSMUSG00 | AC125541.1 | 87 | 0 | 0 | 0 | 0 | 0 | 0 |
| ENSMUSG00 | AL929540.1 | 110 | 0 | 0 | 0 | 0 | 0 | 0 |
| ENSMUSG00 | U6atac | 125 | 0 | 0 | 0 | 0 | 0 | 0 |
| ENSMUSG00 | AL672141.1 | 145 | 0 | 0 | 0 | 0 | 0 | 0 |
| ENSMUSG00 | CT010493.1 | 94 | 0 | 0 | 0 | 0 | 0 | 0 |
| ENSMUSG00 | AC134443.1 | 120 | 0 | 0 | 0 | 0 | 0 | 0 |
| ENSMUSG00 | U6atac | 123 | 0 | 0 | 0 | 0 | 0 | 0 |
| ENSMUSG00 | mmu-mir-11S | 120 | 0 | 0 | 0 | 0 | 0 | 0 |
| ENSMUSG00 | SNORA79 | 143 | 0.27801035 | 0 | 0 | 0 | 0.71365561 | 0 |
| ENSMUSG00 | Mir1249 | 97 | 0 | 0 | 0 | 0 | 0 | 0 |
| ENSMUSG00 | AL772216.11 | 120 | 0 | 0 | 0 | 0 | 0 | 0 |
| ENSMUSG00 | AC153147.1 | 95 | 0 | 0 | 0 | 0 | 0 | 0 |
| ENSMUSG00 | AL772216.12 | 120 | 0 | 0 | 0 | 0 | 0 | 0 |
| ENSMUSG00 | AC134855.1 | 97 | 0 | 0 | 0 | 0 | 0 | 0 |
| ENSMUSG00 | AC087898.1 | 78 | 0 | 0 | 0 | 0 | 0 | 0 |
| ENSMUSG00 | AC090648.1 | 91 | 0 | 0 | 0 | 0 | 2.80364702 | 0 |
| ENSMUSG00 | AC159290.2 | 120 | 0 | 0 | 0 | 0 | 0 | 0 |
| ENSMUSG00 | Mir1306 | 78 | 0 | 0 | 0 | 0 | 0 | 0 |
| ENSMUSG00 | AC155262.1 | 83 | 0.47898169 | 0 | 0.53255233 | 1.30417028 | 0 | 1.97166067 |
| ENSMUSG00 | CT010479.1 | 89 | 0 | 0 | 0.99329984 | 5.270411 | 0 | 0 |
| ENSMUSG00 | Mir669g | 122 | 0 | 0 | 0 | 0 | 0 | 0 |
| ENSMUSG00 | mmu-mir-11S | 120 | 0 | 0 | 0 | 0 | 0 | 0 |
| ENSMUSG00 | SNORA11 | 126 | 0 | 0 | 0 | 0 | 0.40497124 | 0 |
| ENSMUSG00 | mmu-mir-11S | 120 | 0 | 0 | 0 | 0 | 0 | 0 |
| ENSMUSG00 | SNORD75 | 61 | 0 | 0 | 0 | 0 | 0 | 0 |
| ENSMUSG00 | SNORD94 | 136 | 2.9231971 | 3.54312075 | 4.22517617 | 1.59185491 | 0.75038788 | 1.40384173 |
| ENSMUSG00 | mmu-mir-66S | 126 | 0 | 0 | 0 | 0 | 0 | 0 |
| ENSMUSG00 | AL808131.1 | 95 | 0 | 0 | 0 | 0 | 0 | 0 |
| ENSMUSG00 | Mir1195 | 97 | 0 | 0 | 0 | 0 | 0 | 0 |
| ENSMUSG00 | Snord98 | 64 | 6.21179383 | 3.76456579 | 1.38130759 | 4.51025557 | 4.78372273 | 0 |
| ENSMUSG00 | AL928542.1 | 120 | 0 | 0 | 0 | 0 | 0 | 0 |
| ENSMUSG00 | AL670236.1 | 120 | 0 | 0 | 0 | 0 | 0 | 0 |
| ENSMUSG00 | AL772216.13 | 123 | 0 | 0 | 0 | 0 | 0 | 0 |
| ENSMUSG00 | AC024607.1 | 120 | 0 | 0 | 0 | 0 | 0 | 0 |
| ENSMUSG00 | AL670720.1 | 128 | 0 | 0 | 0 | 0 | 0 | 0 |
| ENSMUSG00 | AL772216.14 | 117 | 0 | 0 | 0 | 0 | 0 | 0 |
| ENSMUSG00 | AL772216.15 | 120 | 0 | 0 | 0 | 0 | 0 | 0 |
| ENSMUSG00 | Snord23 | 108 | 0 | 0 | 1.6371053 | 1.00227902 | 1.41739933 | 1.26271478 |
| ENSMUSG00 | U6atac | 125 | 0 | 0 | 0 | 0 | 0 | 0 |
| ENSMUSG00 | AC138200.1 | 82 | 0 | 0 | 0 | 0 | 0 | 0 |
| ENSMUSG00 | AC111029.2 | 120 | 0 | 0 | 0 | 0 | 0 | 0 |
| ENSMUSG00 | AC132270.1 | 90 | 0 | 0 | 0 | 0 | 0 | 0 |
| ENSMUSG00 | AC129021.1 | 87 | 0 | 0 | 0 | 0 | 0 | 0 |
| ENSMUSG00 | Snord100 | 73 | 7.62433873 | 1.3201765 | 3.02752349 | 4.44847124 | 0.69899145 | 0 |
| ENSMUSG00 | AL606479.1 | 114 | 0 | 0 | 0 | 0 | 0 | 0 |
| ENSMUSG00 | AL772216.16 | 126 | 0 | 0 | 0 | 0 | 0 | 0 |
| ENSMUSG00 | AL845174.1 | 94 | 0 | 0 | 0 | 0 | 0 | 0 |
| ENSMUSG00 | AL772216.17 | 123 | 0 | 0 | 0 | 0 | 0 | 0 |
| ENSMUSG00 | AC161265.1 | 118 | 0 | 0 | 0 | 0 | 0 | 0 |
| ENSMUSG00 | AC120540.1 | 89 | 0 | 0 | 0 | 0 | 0 | 0 |
| ENSMUSG00 | BX571729.1 | 81 | 0 | 0 | 0 | 0 | 0 | 0 |
| ENSMUSG00 | U6atac | 119 | 0 | 0 | 0 | 0 | 0 | 0 |
| ENSMUSG00 | AL772216.18 | 120 | 0 | 0 | 0 | 0 | 0 | 0 |
| ENSMUSG00 | AL929433.1 | 120 | 0 | 0 | 0 | 0 | 0 | 0 |
| ENSMUSG00 | Snord88c | 89 | 7.59374347 | 7.03846908 | 4.96649922 | 7.70290838 | 5.15997059 | 5.20976255 |
| ENSMUSG00 | Mir669d-2 | 85 | 0 | 0 | 0 | 0 | 0 | 0 |
| ENSMUSG00 | mmu-mir-11S | 120 | 0 | 0 | 0 | 0 | 0 | 0 |
| ENSMUSG00 | AC125040.1 | 120 | 0 | 0 | 0 | 0 | 0 | 0 |
| ENSMUSG00 | AL663067.1 | 85 | 0 | 0 | 0 | 0 | 0 | 0 |
| ENSMUSG00 | AC125028.1 | 64 | 0 | 0 | 0 | 0 | 0 | 0 |
| ENSMUSG00 | AC134426.1 | 120 | 0 | 0 | 0 | 0 | 0 | 0 |
| ENSMUSG00 | AC115802.1 | 107 | 0 | 0 | 0 | 0 | 0 | 0.25490317 |
| ENSMUSG00 | AC133514.1 | 108 | 0 | 0 | 0 | 0 | 0 | 0 |
| ENSMUSG00 | AL672234.1 | 57 | 0 | 0 | 0 | 0 | 0 | 0 |
| ENSMUSG00 | U6atac | 125 | 0 | 0 | 0 | 0 | 0 | 0 |
| ENSMUSG00 | SNORD111 | 75 | 0.53007307 | 2.56994358 | 0 | 3.36765749 | 0 | 0.72732371 |
| ENSMUSG00 | AC153564.1 | 94 | 0 | 0 | 0 | 0 | 0 | 0 |
| ENSMUSG00 | AL772216.19 | 120 | 0 | 0 | 0 | 0 | 0 | 0 |
| ENSMUSG00 | AC171241.1 | 65 | 0 | 0 | 0 | 0 | 0 | 0 |
| ENSMUSG00 | AC117578.1 | 53 | 0 | 0 | 0 | 0 | 0 | 0 |
| ENSMUSG00 | U6atac | 125 | 0 | 0 | 0.70722949 | 0 | 0 | 0 |
| ENSMUSG00 | AC132471.1 | 77 | 1.03260988 | 0 | 0 | 0 | 0 | 0 |
| ENSMUSG00 | Mir466j | 121 | 0 | 0 | 0 | 0 | 0 | 0 |
| ENSMUSG00 | AC160528.1 | 53 | 0 | 0 | 0 | 0 | 0 | 0 |
| ENSMUSG00 | AC141471.1 | 83 | 0 | 0 | 0 | 0 | 0 | 0 |
| ENSMUSG00 | mmu-mir-11S | 120 | 0 | 0 | 0 | 0 | 0 | 0 |
| ENSMUSG00 | Mir1191 | 119 | 4.67711535 | 1.61971234 | 1.48577624 | 4.24494642 | 0.42879307 | 0 |
| ENSMUSG00 | Mir1186 | 121 | 0 | 0 | 0 | 0 | 0 | 0 |
| ENSMUSG00 | SNORA3 | 121 | 0 | 0 | 0 | 0 | 0 | 0 |
| ENSMUSG00 | AC160109.1 | 89 | 0 | 0 | 0 | 0 | 0 | 0 |
| ENSMUSG00 | U6atac | 125 | 0.95413153 | 0 | 0 | 0 | 0 | 0 |
| ENSMUSG00 | U6atac | 93 | 0 | 0 | 0 | 0 | 0 | 0 |
| ENSMUSG00 | SNORA76 | 134 | 0 | 0 | 0.3298645 | 0 | 0 | 0 |
| ENSMUSG00 | AC156561.1 | 120 | 0 | 0 | 0 | 0 | 0 | 0 |
| ENSMUSG00 | U4atac | 124 | 3.20608714 | 0.38860034 | 4.27759771 | 3.20082653 | 1.23450909 | 3.51930829 |
| ENSMUSG00 | AC127028.1 | 110 | 0 | 0 | 0 | 0 | 0 | 0 |
| ENSMUSG00 | CPEB3_riboz | 77 | 20.6521977 | 6.2579795 | 9.18479855 | 15.4637334 | 6.62680205 | 19.1276691 |
| ENSMUSG00 | U6atac | 129 | 0 | 0 | 0 | 0 | 0 | 0 |
| ENSMUSG00 | AC129309.1 | 120 | 0 | 0 | 0 | 0 | 0 | 0 |
| ENSMUSG00 | AL928581.1 | 73 | 2.17838249 | 0 | 0.6055047 | 0 | 1.3979829 | 0 |
| ENSMUSG00 | AL772216.20 | 126 | 0 | 0 | 0 | 0 | 0 | 0 |
| ENSMUSG00 | AC105256.1 | 92 | 0 | 0 | 0.48045482 | 0 | 0 | 0 |
| ENSMUSG00 | AL772216.21 | 126 | 0 | 0 | 0 | 0 | 0 | 0 |
| ENSMUSG00 | AL772216.22 | 120 | 0 | 0 | 0 | 0 | 0 | 0 |
| ENSMUSG00 | AC102572.1 | 92 | 0 | 0 | 0 | 0 | 0 | 0 |
| ENSMUSG00 | Mir467g | 119 | 0 | 0 | 0 | 0 | 0 | 0 |
| ENSMUSG00 | CT010480.1 | 93 | 0 | 0 | 0 | 0.38797897 | 0 | 0 |
| ENSMUSG00 | AL772216.23 | 126 | 0 | 0 | 0 | 0 | 0 | 0 |
| ENSMUSG00 | AC133891.1 | 111 | 0.35815748 | 2.60467255 | 0 | 0 | 0 | 0.49143494 |
| ENSMUSG00 | AC157561.1 | 87 | 0 | 0 | 0 | 0 | 0 | 0 |
| ENSMUSG00 | AC191865.2 | 120 | 0 | 0 | 0 | 0 | 0 | 0 |
| ENSMUSG00 | U6atac | 125 | 0 | 0 | 0 | 0 | 0 | 0 |
| ENSMUSG00 | AL772216.24 | 125 | 0 | 0 | 0 | 0 | 0 | 0 |
| ENSMUSG00 | AC117577.1 | 120 | 0 | 0 | 0 | 0 | 0 | 0 |
| ENSMUSG00 | AL772216.25 | 122 | 0 | 0 | 0 | 0 | 0 | 0 |
| ENSMUSG00 | AC158909.1 | 120 | 0 | 0 | 0 | 0 | 0 | 0 |
| ENSMUSG00 | Mir467f | 119 | 0.33407967 | 4.45420894 | 0.74288812 | 1.21284183 | 9.00465456 | 4.35477433 |
| ENSMUSG00 | Mir466i | 120 | 9.27627879 | 6.02330527 | 0.73669738 | 3.30752075 | 0.4252198 | 3.63661857 |

| | | | | | | | | |
|---|---|---|---|---|---|---|---|---|
| ENSMUSG00 | Mir1190 | 120 | 0 | 0 | 0 | 0 | 0 | 0 |
| ENSMUSG00 | AC108410.1 | 120 | 0 | 0 | 0 | 0 | 0 | 0 |
| ENSMUSG00 | U6atac | 102 | 0 | 0 | 0 | 0 | 0 | 0 |
| ENSMUSG00 | Mir466l-4 | 120 | 0 | 0 | 0 | 0 | 0 | 0 |
| ENSMUSG00 | SNORA11 | 127 | 0 | 0 | 0 | 0.85233176 | 1.20534746 | 0 |
| ENSMUSG00 | AL929465.1 | 120 | 0 | 0 | 0 | 0 | 0 | 0 |
| ENSMUSG00 | AC163028.1 | 89 | 0 | 0 | 0 | 0 | 0 | 0 |
| ENSMUSG00 | Mir1187 | 121 | 0 | 0 | 0 | 0 | 0 | 0 |
| ENSMUSG00 | AC131713.1 | 120 | 0 | 0 | 0 | 0 | 0 | 0 |
| ENSMUSG00 | AC111132.1 | 120 | 0 | 0 | 0 | 0 | 0 | 0 |
| ENSMUSG00 | AL590988.1 | 62 | 0 | 0 | 0 | 0 | 0 | 0 |
| ENSMUSG00 | Mir669h | 124 | 0 | 0 | 0 | 0 | 0 | 0 |
| ENSMUSG00 | U6atac | 125 | 0 | 0 | 0 | 0 | 0 | 0 |
| ENSMUSG00 | AC161111.1 | 115 | 0 | 0 | 0 | 0 | 0 | 0 |
| ENSMUSG00 | AL606976.1 | 92 | 0 | 0 | 0.48045482 | 0 | 3.88244164 | 0 |
| ENSMUSG00 | AL772216.26 | 123 | 0 | 0 | 0 | 0 | 0 | 0 |
| ENSMUSG00 | AL772216.27 | 125 | 0 | 0 | 0 | 0 | 0 | 0 |
| ENSMUSG00 | AL591376.1 | 96 | 0 | 0 | 0 | 0 | 0 | 0 |
| ENSMUSG00 | AC119834.1 | 113 | 0 | 0.42642869 | 0 | 0 | 0 | 0 |
| ENSMUSG00 | AL929160.1 | 89 | 0 | 0 | 2.48324961 | 0 | 0 | 0 |
| ENSMUSG00 | AC138376.1 | 90 | 0 | 0 | 0 | 0 | 0 | 0 |
| ENSMUSG00 | AL772216.28 | 126 | 0 | 0 | 0 | 0 | 0 | 0 |
| ENSMUSG00 | AC141438.1 | 120 | 0 | 0 | 0 | 0 | 0 | 0 |
| ENSMUSG00 | SNORD23 | 108 | 0 | 0 | 0 | 0 | 0 | 0 |
| ENSMUSG00 | AC158296.1 | 120 | 0 | 0 | 0 | 0 | 0 | 0 |
| ENSMUSG00 | Snord110 | 67 | 0 | 2.87680252 | 0 | 0 | 0 | 1.2212525 |
| ENSMUSG00 | Mir669j | 120 | 0 | 0 | 0 | 0 | 0 | 0 |
| ENSMUSG00 | AL772216.29 | 125 | 0 | 0 | 0 | 0 | 0 | 0 |
| ENSMUSG00 | AC121604.1 | 57 | 0 | 0 | 0 | 0 | 0 | 0 |
| ENSMUSG00 | AC102689.1 | 86 | 0 | 0 | 0 | 0 | 0 | 0 |
| ENSMUSG00 | Snord99 | 73 | 0 | 0 | 0 | 0 | 0 | 0 |
| ENSMUSG00 | U3 | 124 | 0 | 0 | 0 | 0 | 0 | 0 |
| ENSMUSG00 | mmu-mir-119 | 120 | 0 | 0 | 0 | 0 | 0 | 0 |
| ENSMUSG00 | SNORA11 | 128 | 0 | 0 | 0 | 0 | 0 | 0 |
| ENSMUSG00 | AL772216.30 | 100 | 0 | 0 | 0 | 0 | 0 | 0 |
| ENSMUSG00 | AC163659.2 | 99 | 0 | 0 | 0 | 0 | 0 | 0 |
| ENSMUSG00 | Mir1192 | 120 | 0 | 0 | 0.73669738 | 0.60136741 | 0 | 0 |
| ENSMUSG00 | AL772216.31 | 107 | 0 | 0 | 0 | 0 | 0 | 0 |
| ENSMUSG00 | AL691457.1 | 143 | 0 | 0 | 0 | 0 | 0 | 0 |
| ENSMUSG00 | AC098726.1 | 88 | 0 | 0 | 1.00458734 | 0 | 1.73953554 | 1.54969541 |
| ENSMUSG00 | AC109145.1 | 120 | 0 | 0 | 0 | 0 | 0 | 0 |
| ENSMUSG00 | AC139348.1 | 105 | 0 | 0 | 0 | 0 | 0 | 0.25975847 |
| ENSMUSG00 | AC161265.2 | 122 | 0 | 0 | 0 | 0 | 0 | 0 |
| ENSMUSG00 | AC122490.1 | 81 | 0 | 0 | 0 | 0 | 0 | 0 |
| ENSMUSG00 | AL772216.32 | 125 | 0 | 0 | 0 | 0 | 0 | 0 |
| ENSMUSG00 | AL772216.33 | 120 | 0 | 0 | 0 | 0 | 0 | 0 |
| ENSMUSG00 | Mir1198 | 120 | 0.99388701 | 0 | 1.10504608 | 0 | 0.4252198 | 0 |
| ENSMUSG00 | AC131332.1 | 90 | 0 | 0 | 0 | 0 | 0 | 0 |
| ENSMUSG00 | AL772216.34 | 122 | 0 | 0 | 0 | 0 | 0 | 0 |
| ENSMUSG00 | AC119939.1 | 93 | 0 | 0 | 0 | 0 | 0 | 0 |
| ENSMUSG00 | AC103351.1 | 87 | 0 | 0 | 0 | 0 | 0 | 0 |
| ENSMUSG00 | Mir669k | 109 | 0 | 0 | 0 | 0 | 0 | 0 |
| ENSMUSG00 | Mir669e | 118 | 0 | 0 | 0 | 0 | 0 | 0 |
| ENSMUSG00 | Mir1196 | 118 | 0 | 0 | 0 | 0 | 0 | 0 |
| ENSMUSG00 | AC155833.1 | 92 | 0 | 0 | 0 | 0 | 0 | 0 |
| ENSMUSG00 | Mir669i | 126 | 0 | 0 | 0 | 0 | 0 | 0 |
| ENSMUSG00 | AL844480.1 | 86 | 0 | 0 | 0 | 0 | 0 | 0 |
| ENSMUSG00 | AC132253.2 | 120 | 0 | 0 | 0 | 0 | 0 | 0 |
| ENSMUSG00 | AC108410.2 | 120 | 0 | 0 | 0 | 0 | 0 | 0 |
| ENSMUSG00 | AC101970.1 | 89 | 0 | 0 | 0 | 0 | 0 | 0 |
| ENSMUSG00 | Mir1b | 121 | 0 | 0 | 0 | 0 | 0 | 0 |
| ENSMUSG00 | AL772302.1 | 65 | 0 | 0 | 0 | 0 | 0 | 0 |
| ENSMUSG00 | U3 | 192 | 0 | 0 | 0 | 0 | 0 | 0 |
| ENSMUSG00 | Mir1199 | 118 | 4.04293022 | 7.7588339 | 7.11724591 | 2.14046027 | 2.16213457 | 2.0802691 |
| ENSMUSG00 | AC123619.1 | 113 | 0 | 0 | 0 | 0 | 0 | 0 |
| ENSMUSG00 | AC122402.1 | 100 | 0 | 0 | 0 | 0 | 0 | 0 |
| ENSMUSG00 | Mir1224 | 84 | 0 | 0 | 0 | 0 | 0 | 0 |
| ENSMUSG00 | AC138299.1 | 88 | 0 | 0 | 0 | 0 | 0 | 0 |
| ENSMUSG00 | U6atac | 122 | 0 | 0 | 0 | 0 | 0 | 0 |
| ENSMUSG00 | AL772216.35 | 126 | 0 | 0 | 0 | 0 | 0 | 0 |
| ENSMUSG00 | AC161513.1 | 120 | 0 | 0 | 0 | 0 | 0 | 0 |
| ENSMUSG00 | AC165239.1 | 123 | 0 | 0 | 0 | 0 | 0 | 0 |
| ENSMUSG00 | AC112790.1 | 97 | 0 | 3.97413956 | 0 | 0 | 0 | 0.28118185 |
| ENSMUSG00 | AC161049.1 | 65 | 0 | 0 | 0 | 0 | 0 | 0 |
| ENSMUSG00 | AL627312.1 | 116 | 0.34271966 | 0 | 0.38105037 | 0 | 0.8797651 | 0 |
| ENSMUSG00 | mmu-mir-119 | 120 | 0 | 0 | 0 | 0 | 0 | 0 |
| ENSMUSG00 | SNORD88 | 92 | 0 | 0 | 0 | 0 | 0 | 0 |
| ENSMUSG00 | AL772216.36 | 121 | 0 | 0 | 0 | 0 | 0 | 0 |
| ENSMUSG00 | AL772216.37 | 119 | 0 | 0 | 0 | 0 | 0 | 0 |
| ENSMUSG00 | AL772216.38 | 89 | 0 | 0 | 0 | 0 | 0 | 0 |
| ENSMUSG00 | Mir1197 | 119 | 0 | 0 | 0 | 0 | 0 | 0 |
| ENSMUSG00 | SNORA79 | 128 | 0 | 0 | 0 | 0 | 0 | 0 |
| ENSMUSG00 | Gm11621 | 588 | 0 | 0 | 0 | 0 | 0 | 0 |
| ENSMUSG00 | Gm5394 | 974 | 0 | 0 | 0 | 0 | 0 | 0 |
| ENSMUSG00 | Gm12825 | 561 | 0 | 0 | 0 | 0 | 0 | 0 |
| ENSMUSG00 | Gm11299 | 580 | 0 | 0 | 0 | 0 | 0 | 0 |
| ENSMUSG00 | Gm11333 | 147 | 0 | 0 | 0 | 0 | 0 | 0 |
| ENSMUSG00 | Olfr1296-ps1 | 938 | 0 | 0 | 0 | 0 | 0 | 0 |
| ENSMUSG00 | Gm15981 | 463 | 0 | 0 | 0 | 0 | 0 | 0 |
| ENSMUSG00 | Gm12340 | 461 | 0.17247497 | 0.10452591 | 0 | 0 | 0 | 0 |
| ENSMUSG00 | Gm7438 | 631 | 0 | 0 | 0 | 0 | 0 | 0 |
| ENSMUSG00 | Gm13183 | 788 | 0 | 0.24460123 | 0 | 0 | 0 | 0 |
| ENSMUSG00 | Gm6031 | 1053 | 0.03775449 | 0 | 0 | 0 | 0 | 0 |
| ENSMUSG00 | Gm15439 | 383 | 0 | 0 | 0 | 0 | 0 | 0 |
| ENSMUSG00 | Gm12890 | 372 | 0 | 0 | 0 | 0 | 0 | 0 |
| ENSMUSG00 | Gm13823 | 344 | 0 | 0 | 0 | 0 | 0 | 0 |
| ENSMUSG00 | Ccnd3-ps | 880 | 0 | 0 | 0.3013762 | 0 | 0 | 0 |
| ENSMUSG00 | Gm14324 | 3048 | 0 | 0 | 0 | 0.02367588 | 0 | 0 |
| ENSMUSG00 | Gm8107 | 2008 | 0 | 0 | 0 | 0 | 0 | 0 |
| ENSMUSG00 | Gm11686 | 332 | 0 | 0 | 0 | 0 | 0 | 0 |
| ENSMUSG00 | Hist3h2bb-ps | 464 | 0.08567992 | 0 | 0 | 0.38881514 | 0 | 0.41147085 |
| ENSMUSG00 | Olfr1174-ps | 938 | 0 | 0 | 0 | 0 | 0 | 0 |
| ENSMUSG00 | Gm14602 | 339 | 0 | 0 | 0 | 0 | 0 | 0.08045616 |
| ENSMUSG00 | Gm5406 | 765 | 0.36377564 | 0.18896644 | 0.05778019 | 0 | 0 | 0.10695937 |
| ENSMUSG00 | Gm13503 | 445 | 0 | 0 | 0.39731994 | 0 | 0 | 0 |
| ENSMUSG00 | B230307C23 | 1295 | 5.28026459 | 5.73027961 | 8.36251064 | 3.00915893 | 4.13727372 | 11.3942701 |
| ENSMUSG00 | Gm8019 | 586 | 0 | 0 | 0 | 0 | 0 | 0 |
| ENSMUSG00 | Gm15532 | 883 | 0 | 0 | 0 | 0 | 0 | 0 |
| ENSMUSG00 | Gm16077 | 749 | 0 | 0 | 0 | 0 | 0 | 0 |
| ENSMUSG00 | Gm14944 | 131 | 0 | 0 | 0 | 0 | 0 | 0 |
| ENSMUSG00 | Defa-ps7 | 172 | 0 | 0 | 0 | 0 | 0 | 0 |
| ENSMUSG00 | Gm12564 | 320 | 0 | 0 | 0 | 0 | 0 | 0 |
| ENSMUSG00 | Gm6121 | 1105 | 0 | 0 | 0 | 0 | 0 | 0 |
| ENSMUSG00 | Gm15366 | 4236 | 0 | 0 | 0 | 0 | 0 | 0 |
| ENSMUSG00 | C920021L13I | 2249 | 1.83840372 | 2.29255194 | 2.57467383 | 1.09096444 | 1.678947 | 3.46845123 |
| ENSMUSG00 | Gm13616 | 221 | 0 | 0 | 0 | 0 | 0 | 0 |
| ENSMUSG00 | Gm8084 | 748 | 0 | 0 | 0 | 0 | 0 | 0 |
| ENSMUSG00 | Gm11313 | 431 | 0 | 0 | 0 | 0 | 0 | 0 |
| ENSMUSG00 | Rps2-ps9 | 816 | 0.58463942 | 0.11810403 | 0 | 0 | 0.12506465 | 0.26739842 |
| ENSMUSG00 | Gm13222 | 137 | 0 | 0 | 0 | 0 | 0 | 0 |
| ENSMUSG00 | Gm14867 | 861 | 0 | 0 | 0 | 0 | 0 | 0 |
| ENSMUSG00 | Gm12006 | 402 | 0 | 0 | 0 | 0 | 0 | 0 |
| ENSMUSG00 | Gm12075 | 396 | 0 | 0 | 0 | 0 | 0 | 0 |
| ENSMUSG00 | Gm15617 | 770 | 0 | 0 | 0 | 0 | 0 | 0 |
| ENSMUSG00 | Gm6982 | 1559 | 0 | 0 | 0 | 0 | 0 | 0 |
| ENSMUSG00 | Mup-ps22 | 242 | 0 | 0 | 0 | 0 | 0 | 0 |
| ENSMUSG00 | Gm13795 | 294 | 0 | 0 | 0 | 0 | 0 | 0 |
| ENSMUSG00 | Gm15013 | 800 | 0 | 0 | 0 | 0 | 0 | 0.71595928 |
| ENSMUSG00 | Gm11398 | 2417 | 2.74686191 | 3.26958068 | 1.82878953 | 4.49346107 | 5.55231148 | 3.15966032 |
| ENSMUSG00 | Gm14035 | 1474 | 0 | 0 | 0 | 0 | 0 | 0 |
| ENSMUSG00 | Gm7150 | 972 | 0.0409007 | 0.24787265 | 0 | 0.07424289 | 0 | 0.02806033 |
| ENSMUSG00 | Gm12710 | 215 | 0 | 0 | 0 | 0 | 0 | 0 |
| ENSMUSG00 | Rpsa-ps12 | 1733 | 0.38998452 | 0.52829914 | 0.66315518 | 0.37477023 | 0.29443956 | 0.42493668 |
| ENSMUSG00 | Gm14016 | 485 | 0 | 0 | 0 | 0 | 0 | 0 |
| ENSMUSG00 | Gm15681 | 555 | 0 | 0 | 0 | 0 | 0 | 0 |
| ENSMUSG00 | Gm12134 | 519 | 0 | 0 | 0 | 0 | 0 | 0 |
| ENSMUSG00 | Gm15553 | 429 | 0 | 0 | 0 | 0 | 0 | 0 |
| ENSMUSG00 | Olfr634-ps1 | 954 | 0 | 0 | 0 | 0 | 0 | 0 |
| ENSMUSG00 | Olfr1069-ps1 | 927 | 0 | 0 | 0 | 0 | 0 | 0 |
| ENSMUSG00 | Gm15613 | 325 | 0.24464911 | 0.29653195 | 0 | 0 | 0.15700423 | 0.7552977 |
| ENSMUSG00 | Rpl36-ps7 | 314 | 0 | 0 | 0 | 0 | 0 | 0 |
| ENSMUSG00 | Gm15062 | 366 | 0 | 0 | 0 | 0 | 0 | 0 |
| ENSMUSG00 | Gm12633 | 1704 | 0 | 0 | 0 | 0 | 0 | 0 |
| ENSMUSG00 | Gm15029 | 198 | 0 | 0 | 0 | 0 | 0 | 0 |
| ENSMUSG00 | Gm5392 | 787 | 0 | 0 | 0 | 0 | 0 | 0 |
| ENSMUSG00 | Gm15573 | 539 | 0 | 0 | 0 | 0.66942569 | 0.1893372 | 0.1518069 |
| ENSMUSG00 | Gm7571 | 923 | 0 | 0 | 0 | 0 | 0 | 0 |
| ENSMUSG00 | Gm7386 | 631 | 0 | 0 | 0 | 0 | 0 | 0 |
| ENSMUSG00 | Gm11879 | 701 | 0 | 0 | 0 | 0 | 0 | 0 |
| ENSMUSG00 | Gm12341 | 502 | 0 | 0.09598893 | 0 | 0 | 0 | 0.05433195 |
| ENSMUSG00 | Gm14606 | 1154 | 0 | 0 | 0 | 0 | 0 | 0 |
| ENSMUSG00 | Gm8908 | 1023 | 0 | 0 | 0 | 0 | 0 | 0 |
| ENSMUSG00 | Gm13593 | 690 | 0 | 0 | 0 | 0 | 0 | 0 |
| ENSMUSG00 | Gm15595 | 960 | 0 | 0 | 0 | 0 | 0 | 0 |
| ENSMUSG00 | Gm12219 | 442 | 0 | 0 | 0 | 0 | 0 | 0 |
| ENSMUSG00 | Gm11278 | 996 | 0 | 0 | 0 | 0 | 0 | 0 |
| ENSMUSG00 | Gm11925 | 534 | 0 | 0 | 0 | 0 | 0 | 0 |
| ENSMUSG00 | Gm12543 | 542 | 0 | 0.97795362 | 0.08155322 | 0.46600427 | 0.18828921 | 0.20128885 |
| ENSMUSG00 | Gm12955 | 956 | 0 | 2.11697758 | 0 | 0.45291269 | 0.58712357 | 0 |
| ENSMUSG00 | Gm9143 | 744 | 0 | 0 | 0 | 0 | 0 | 0 |
| ENSMUSG00 | Gm6368 | 2146 | 0.18525387 | 1.30233627 | 0.06179195 | 0.99200402 | 1.21264919 | 0.71173336 |
| ENSMUSG00 | Gm12174 | 812 | 0.24479976 | 0.11868582 | 0 | 0.17774406 | 0.12568073 | 0.13435783 |
| ENSMUSG00 | Gm15365 | 299 | 0 | 0 | 0 | 0 | 0 | 0 |
| ENSMUSG00 | Gm7199 | 1484 | 0 | 0 | 0 | 0 | 0 | 0 |
| ENSMUSG00 | Gm8731 | 411 | 0 | 0 | 0 | 0 | 0 | 0 |
| ENSMUSG00 | Gm11252 | 938 | 0.5085989 | 0 | 0 | 1.03860896 | 0 | 0 |
| ENSMUSG00 | Olfr388-ps1 | 940 | 0 | 0 | 0 | 0 | 0 | 0 |
| ENSMUSG00 | Gm13495 | 536 | 0 | 0 | 0 | 0 | 0 | 0 |
| ENSMUSG00 | Gm8250 | 773 | 0.66859152 | 0.93505386 | 0.74336864 | 2.24054093 | 0.72611919 | 0.35284139 |
| ENSMUSG00 | Gm13765 | 227 | 0 | 0 | 0 | 0 | 0 | 0 |
| ENSMUSG00 | Gm11591 | 701 | 0 | 0 | 0 | 0 | 0 | 0 |
| ENSMUSG00 | Gm11320 | 434 | 0 | 0 | 0 | 0 | 0 | 0 |
| ENSMUSG00 | Gm7841 | 1628 | 0.02441983 | 0 | 0 | 0.08865367 | 0 | 0 |
| ENSMUSG00 | Gm14007 | 209 | 0 | 0 | 0 | 0 | 0 | 0 |
| ENSMUSG00 | Gm7638 | 966 | 0 | 0 | 0.18303041 | 0.03735201 | 0 | 0.93174234 |
| ENSMUSG00 | Gm11442 | 457 | 0 | 0.63264476 | 0 | 0 | 0.11165509 | 0 |
| ENSMUSG00 | Gm14367 | 1496 | 0 | 0 | 0 | 0 | 0 | 0 |
| ENSMUSG00 | Cbx3-ps3 | 548 | 0 | 0 | 0 | 0 | 0 | 0 |
| ENSMUSG00 | Gm11328 | 891 | 0 | 0 | 0 | 0 | 0 | 0 |
| ENSMUSG00 | Gm6783 | 1436 | 0.02768488 | 0 | 0.03078123 | 0 | 0.03553369 | 0 |
| ENSMUSG00 | Gm9159 | 565 | 0 | 0 | 0 | 0.76634431 | 0 | 0.43446328 |
| ENSMUSG00 | Gm15760 | 282 | 0.28195376 | 0 | 0 | 1.53540615 | 1.08566757 | 0 |
| ENSMUSG00 | Gm5135 | 572 | 0 | 0 | 0 | 0 | 0 | 0 |
| ENSMUSG00 | Gm15166 | 384 | 0 | 0 | 0 | 0 | 0 | 0 |
| ENSMUSG00 | Gm11582 | 235 | 0 | 0 | 0 | 0 | 0 | 0 |
| ENSMUSG00 | Gm13310 | 1391 | 0 | 0.10392475 | 0 | 0 | 0 | 0 |
| ENSMUSG00 | Gm13804 | 171 | 0 | 0 | 0 | 0 | 0 | 0 |
| ENSMUSG00 | Gm12683 | 414 | 0 | 0 | 0 | 0 | 0 | 0 |
| ENSMUSG00 | Gm11219 | 357 | 0 | 0 | 0 | 0 | 0 | 0 |
| ENSMUSG00 | Gm13005 | 583 | 0.20457366 | 0 | 0 | 0 | 0 | 0 |
| ENSMUSG00 | Gm13117 | 690 | 0 | 0 | 0.12812128 | 0.15687845 | 0.36975635 | 0 |
| ENSMUSG00 | Gm12608 | 1712 | 0 | 0 | 0 | 0.02107596 | 0 | 0 |
| ENSMUSG00 | Olfr1273-ps | 924 | 0 | 0 | 0 | 0 | 0 | 0 |
| ENSMUSG00 | Gm11737 | 467 | 1.44720165 | 0.20636592 | 1.98766318 | 1.69979653 | 0.43705675 | 0 |
| ENSMUSG00 | Gm14513 | 122 | 0 | 0 | 0 | 0 | 0 | 0 |
| ENSMUSG00 | Gm13526 | 188 | 0 | 0 | 0 | 0 | 0 | 0 |
| ENSMUSG00 | Gm14828 | 1154 | 0 | 0.08351203 | 0.11490947 | 0 | 0.13265089 | 0 |
| ENSMUSG00 | Gm12314 | 537 | 0 | 0 | 0 | 0 | 0 | 0 |
| ENSMUSG00 | Hmgb1-rs18 | 626 | 0 | 0 | 0 | 0 | 0 | 0.04356971 |
| ENSMUSG00 | Gm12447 | 469 | 0 | 0.10274295 | 0 | 0 | 0 | 0 |
| ENSMUSG00 | Gm13751 | 751 | 0 | 0 | 0 | 0 | 0 | 0 |
| ENSMUSG00 | Gm9434 | 1543 | 0 | 0 | 0 | 0.02338435 | 0.03306959 | 0 |
| ENSMUSG00 | Gm2005 | 535 | 0 | 0 | 0 | 0 | 0 | 0 |
| ENSMUSG00 | Gm13197 | 224 | 0 | 0 | 0 | 0 | 0 | 0 |
| ENSMUSG00 | Gm14694 | 279 | 0 | 0 | 0 | 0 | 0 | 0 |
| ENSMUSG00 | Gm11477 | 313 | 0 | 0 | 0 | 0 | 0 | 0.08713942 |
| ENSMUSG00 | Gm9001 | 398 | 1.29854585 | 0.60535731 | 2.22119814 | 2.44777689 | 0.5128279 | 3.7005792 |
| ENSMUSG00 | Gm14892 | 265 | 0 | 0 | 0 | 0 | 0 | 0 |
| ENSMUSG00 | Gm12019 | 531 | 0 | 0 | 0 | 0 | 0 | 0 |
| ENSMUSG00 | Gm12086 | 227 | 0 | 0 | 0 | 0 | 0 | 0 |
| ENSMUSG00 | Gm14669 | 249 | 0 | 0 | 0 | 0 | 0 | 0 |
| ENSMUSG00 | Gm8545 | 800 | 0.14908305 | 0 | 0 | 0.09020511 | 0 | 0.0340933 |
| ENSMUSG00 | Gm12671 | 2253 | 0 | 0 | 0 | 0 | 0 | 0.01210592 |
| ENSMUSG00 | Gm12529 | 267 | 0 | 0 | 1.32439979 | 0.67569372 | 0.19111002 | 0.51076104 |
| ENSMUSG00 | M6pr-ps | 3450 | 0.04609331 | 0.11173668 | 0.06406064 | 0.01045856 | 0.10353178 | 0.00790569 |
| ENSMUSG00 | Gm11976 | 293 | 0 | 0 | 0 | 0 | 0 | 0 |
| ENSMUSG00 | Gm15216 | 276 | 0 | 0 | 0 | 0 | 0 | 0 |
| ENSMUSG00 | Gm12137 | 345 | 0 | 0 | 0 | 0 | 0 | 0 |
| ENSMUSG00 | Gm6444 | 457 | 0 | 0.52720396 | 0.19344351 | 0 | 0 | 0.05968192 |
| ENSMUSG00 | Gm14361 | 867 | 0 | 0 | 0 | 0 | 0 | 0 |
| ENSMUSG00 | Gm12573 | 285 | 0 | 0 | 0 | 0 | 0 | 0 |
| ENSMUSG00 | Gm11625 | 295 | 0.53905736 | 0.32668774 | 0 | 0 | 0.17297077 | 1.75667368 |
| ENSMUSG00 | Gm11349 | 1115 | 0 | 0 | 0.39642909 | 0 | 0 | 0 |
| ENSMUSG00 | Gm14880 | 291 | 0 | 0 | 0 | 0 | 0 | 0 |
| ENSMUSG00 | Gm15400 | 606 | 0 | 0 | 0 | 0.05954133 | 0 | 0 |
| ENSMUSG00 | Gm14726 | 895 | 0 | 0 | 0 | 0 | 0 | 0 |
| ENSMUSG00 | Gm14107 | 329 | 0 | 0 | 0 | 0 | 0 | 0 |
| ENSMUSG00 | Gm9115 | 1347 | 0.08854227 | 0 | 0 | 0 | 0 | 0.02024843 |
| ENSMUSG00 | Gm13379 | 620 | 0 | 0.07772007 | 0 | 0.23278738 | 0 | 0.04399135 |
| ENSMUSG00 | Gm4773 | 701 | 0 | 0 | 0 | 0 | 0 | 0.15563275 |
| ENSMUSG00 | Gm9385 | 671 | 0 | 0 | 0.39524748 | 0.05377354 | 0 | 0.0812955 |
| ENSMUSG00 | Gm15702 | 443 | 0 | 0 | 0 | 0 | 0 | 0 |
| ENSMUSG00 | Gm12439 | 364 | 0 | 0 | 0 | 0 | 0 | 0 |
| ENSMUSG00 | Gm12094 | 471 | 0 | 0 | 0 | 0 | 0 | 0 |
| ENSMUSG00 | Gm14918 | 267 | 0 | 0 | 0 | 0 | 0 | 0 |
| ENSMUSG00 | Gm15256 | 882 | 0 | 0 | 0 | 0 | 0 | 0 |
| ENSMUSG00 | Gm15084 | 1501 | 0 | 0 | 0 | 0 | 0 | 0 |
| ENSMUSG00 | Rpl35-ps1 | 371 | 0 | 0 | 0 | 0 | 0 | 0 |
| ENSMUSG00 | Gm14433 | 987 | 0 | 0 | 0 | 0 | 0 | 0 |
| ENSMUSG00 | Gm12859 | 425 | 0 | 0 | 0 | 0 | 0 | 0 |
| ENSMUSG00 | Gm14310 | 201 | 0 | 0 | 0 | 0 | 0 | 0 |
| ENSMUSG00 | Rpl10-ps1 | 644 | 0 | 0 | 0 | 0 | 0 | 0 |
| ENSMUSG00 | Gm13815 | 411 | 0.09672866 | 0 | 0 | 0 | 0 | 0 |
| ENSMUSG00 | Gm11325 | 468 | 0 | 0 | 0 | 0 | 0 | 0 |
| ENSMUSG00 | Gm14523 | 557 | 0 | 0 | 0.07935699 | 0 | 0 | 0 |
| ENSMUSG00 | Gm14787 | 387 | 0 | 0 | 0 | 0 | 0 | 0 |
| ENSMUSG00 | Rps12-ps6 | 398 | 0 | 0 | 0 | 0 | 0 | 0 |
| ENSMUSG00 | Gm13637 | 341 | 0 | 0.14130921 | 0 | 0 | 0 | 0 |
| ENSMUSG00 | Gm5687 | 639 | 0 | 0.45245486 | 0.20752039 | 0 | 0.15970697 | 0.17073327 |
| ENSMUSG00 | Gm14079 | 445 | 0 | 0 | 0 | 0 | 0 | 0 |
| ENSMUSG00 | Gm15745 | 114 | 0 | 0 | 0 | 0 | 0 | 0 |
| ENSMUSG00 | Gm11856 | 274 | 0 | 0 | 0 | 0 | 0 | 0 |
| ENSMUSG00 | Gm14111 | 998 | 0.1593406 | 0.24141504 | 0.26574254 | 0.25308047 | 0 | 0.16397579 |
| ENSMUSG00 | Gm12875 | 341 | 0 | 0 | 0 | 0 | 0 | 0 |
| ENSMUSG00 | Gm11506 | 469 | 0 | 0 | 0 | 0 | 0 | 0 |
| ENSMUSG00 | Rpl30-ps3 | 780 | 0 | 0 | 0 | 0 | 0 | 0 |
| ENSMUSG00 | Gm15010 | 406 | 0 | 0 | 0 | 0 | 0 | 0 |
| ENSMUSG00 | Gm7332 | 700 | 0 | 0.06883777 | 0 | 0 | 0 | 0 |
| ENSMUSG00 | Gm14332 | 897 | 0 | 0 | 0 | 0 | 0 | 0 |
| ENSMUSG00 | Rpl22-ps1 | 385 | 0 | 0 | 0.57404991 | 0.0937196 | 0 | 0 |
| ENSMUSG00 | Gm15618 | 451 | 0 | 0 | 0 | 0.2400136 | 0 | 0 |
| ENSMUSG00 | Gm14278 | 866 | 0 | 0.05564254 | 0 | 0 | 0 | 0 |
| ENSMUSG00 | Gm12674 | 205 | 0 | 0 | 0 | 0 | 0 | 0 |
| ENSMUSG00 | Gm4167 | 641 | 0 | 0 | 0 | 0 | 0.15920866 | 0.51060167 |
| ENSMUSG00 | Gm14847 | 518 | 0 | 0 | 0 | 0 | 0 | 0 |
| ENSMUSG00 | Gm14831 | 408 | 0 | 0 | 0 | 0 | 0 | 0 |
| ENSMUSG00 | Gm14387 | 2182 | 0.16397769 | 0.02208361 | 0.10128745 | 0.0661449 | 0.02338514 | 0.26249653 |
| ENSMUSG00 | Gm7974 | 404 | 0 | 0 | 0 | 0 | 0 | 0 |
| ENSMUSG00 | Gm16458 | 497 | 0 | 0 | 0 | 0 | 0 | 0 |
| ENSMUSG00 | Olfr587-ps1 | 1423 | 0 | 0 | 0 | 0 | 0 | 0 |
| ENSMUSG00 | Gm9266 | 483 | 0 | 0.09976489 | 0 | 0 | 0 | 0 |
| ENSMUSG00 | Gm15920 | 401 | 0.89226764 | 0.96132553 | 39.1313074 | 0.26994048 | 0.89073474 | 52.0326659 |
| ENSMUSG00 | Defa-ps15 | 474 | 0 | 0 | 0 | 0 | 0 | 0 |
| ENSMUSG00 | Gm11645 | 648 | 0.06135105 | 0 | 0.06821272 | 0 | 0 | 0 |
| ENSMUSG00 | Gm14567 | 349 | 0 | 0 | 0.12665285 | 0.10338695 | 0.29241476 | 0.15630166 |
| ENSMUSG00 | Gm14500 | 112 | 0 | 0 | 0 | 0 | 0 | 0 |
| ENSMUSG00 | Gm14836 | 470 | 0 | 0 | 0 | 0 | 0 | 0 |
| ENSMUSG00 | Gm9109 | 1404 | 0.05663174 | 0 | 0 | 0.05139892 | 0 | 0.01942638 |
| ENSMUSG00 | Gm3657 | 631 | 0 | 0 | 0 | 0 | 0 | 0 |
| ENSMUSG00 | Gm6215 | 1383 | 0 | 0 | 0 | 0 | 0 | 0 |
| ENSMUSG00 | Gm11226 | 737 | 0.05394231 | 0.2615275 | 0.17992609 | 0.24478999 | 0.20770574 | 0.29606121 |
| ENSMUSG00 | Gm16383 | 230 | 0 | 0 | 0 | 0 | 0 | 0 |
| ENSMUSG00 | Gm11966 | 86 | 6.93409544 | 4.48245973 | 6.16769903 | 3.35646926 | 1.1866599 | 13.3201727 |
| ENSMUSG00 | Gm13295 | 255 | 0 | 0 | 0 | 0 | 0 | 0 |
| ENSMUSG00 | Gm12095 | 815 | 0 | 0 | 0 | 0.57554181 | 0 | 0 |
| ENSMUSG00 | 4930455H04 | 1338 | 0 | 0 | 0 | 0 | 0 | 0 |
| ENSMUSG00 | Gm13044 | 252 | 0 | 0 | 0 | 0 | 0 | 0 |
| ENSMUSG00 | Gm14821 | 434 | 0 | 0 | 0 | 0 | 0 | 0 |
| ENSMUSG00 | Gm11918 | 178 | 0 | 0.5414207 | 0.49664992 | 0.81083246 | 0 | 0.76614155 |
| ENSMUSG00 | Gm12269 | 716 | 0 | 0 | 0 | 0 | 0 | 0 |
| ENSMUSG00 | Gm12719 | 240 | 0 | 0 | 0 | 0 | 0 | 0 |
| ENSMUSG00 | Gm7129 | 996 | 0.27940599 | 0 | 0.04437936 | 0.07245391 | 0 | 0.19168923 |
| ENSMUSG00 | Gm6446 | 817 | 0 | 0 | 0 | 0 | 0 | 0 |
| ENSMUSG00 | Gm12463 | 848 | 0 | 0 | 0.05212482 | 0 | 0 | 0.09649047 |
| ENSMUSG00 | Gm15925 | 1824 | 0 | 0 | 0 | 0 | 0 | 0 |
| ENSMUSG00 | Gm12202 | 806 | 0 | 0 | 0 | 0 | 0 | 0 |
| ENSMUSG00 | Gm11854 | 445 | 0 | 0 | 0 | 0 | 0 | 0 |
| ENSMUSG00 | Gm6228 | 1383 | 0 | 0 | 0 | 0 | 0 | 0 |
| ENSMUSG00 | Rpl38-ps2 | 502 | 0.23758255 | 0.57593357 | 0.26415444 | 0 | 0.20329233 | 0.2173278 |
| ENSMUSG00 | Gm2214 | 329 | 0 | 0 | 0 | 0.65803121 | 0 | 0 |
| ENSMUSG00 | Gm5400 | 942 | 0 | 0 | 0 | 0 | 0.10833625 | 0 |
| ENSMUSG00 | Olfr1078-ps1 | 630 | 0 | 0 | 0 | 0 | 0 | 0 |
| ENSMUSG00 | Gm13682 | 701 | 0 | 0 | 0 | 0 | 0 | 0 |
| ENSMUSG00 | Gm11430 | 215 | 0 | 0 | 0 | 0 | 0 | 0 |
| ENSMUSG00 | Hmgb1-ps6 | 615 | 0 | 0 | 0.14374583 | 0 | 0.08296972 | 0 |
| ENSMUSG00 | Gm12891 | 239 | 0 | 0 | 0 | 0 | 0 | 0 |
| ENSMUSG00 | Gm13367 | 801 | 0 | 0 | 0 | 0 | 0 | 0 |
| ENSMUSG00 | Gm10224 | 1300 | 0.12232456 | 0.11119948 | 0 | 0.02775542 | 0 | 0.02098049 |
| ENSMUSG00 | Rhox3g-ps | 641 | 0 | 0 | 0 | 0 | 0 | 0 |
| ENSMUSG00 | Gm12931 | 323 | 0 | 0 | 0 | 0 | 0 | 0 |
| ENSMUSG00 | Got2-ps1 | 1292 | 0 | 0 | 0.03421195 | 0.02792728 | 0.0394941 | 0.0422208 |
| ENSMUSG00 | Olfr337-ps1 | 452 | 0 | 0 | 0 | 0 | 0 | 0 |
| ENSMUSG00 | Gm13510 | 428 | 0.09288664 | 0 | 0 | 0 | 0 | 0 |
| ENSMUSG00 | Gm6062 | 630 | 0 | 0 | 0 | 0 | 0 | 0.08658616 |
| ENSMUSG00 | Gm9208 | 947 | 0 | 0 | 0 | 0 | 0 | 0 |
| ENSMUSG00 | Gm15145 | 308 | 0 | 0 | 0 | 0 | 0 | 0 |
| ENSMUSG00 | Gm16056 | 134 | 0 | 0 | 0 | 0 | 0 | 0 |
| ENSMUSG00 | Gm14556 | 898 | 0 | 0 | 0 | 0 | 0 | 0 |
| ENSMUSG00 | Gm13130 | 1364 | 0.20492516 | 0 | 0.07594818 | 0 | 0 | 0.89040919 |
| ENSMUSG00 | Gm12967 | 288 | 0 | 0 | 0 | 0 | 0 | 0 |
| ENSMUSG00 | Gm12985 | 182 | 0 | 0 | 0 | 0 | 0.84109411 | 0 |
| ENSMUSG00 | Gm12779 | 191 | 0 | 0 | 0 | 0 | 0 | 0 |
| ENSMUSG00 | Gm14481 | 628 | 0 | 0 | 0 | 0 | 0 | 0 |
| ENSMUSG00 | Gm13683 | 1074 | 0 | 0 | 0 | 0 | 0 | 0 |
| ENSMUSG00 | Gm7278 | 1400 | 0 | 0 | 0.37887294 | 0.02577289 | 0.58315858 | 0 |
| ENSMUSG00 | Gm12208 | 162 | 1.227021 | 0.59489435 | 0 | 0.66818601 | 0 | 0 |
| ENSMUSG00 | Gm5379 | 825 | 0 | 0 | 0 | 0 | 0 | 0 |
| ENSMUSG00 | Gm15025 | 417 | 0 | 0 | 0 | 0 | 0 | 0 |
| ENSMUSG00 | Gm14919 | 389 | 0 | 0 | 0 | 0 | 0 | 0 |
| ENSMUSG00 | Gm14263 | 745 | 0 | 0 | 0 | 0 | 0 | 0 |
| ENSMUSG00 | Gm16179 | 372 | 0 | 0 | 0 | 0 | 0 | 0.58655138 |
| ENSMUSG00 | Gm15739 | 871 | 0.04564349 | 0 | 0 | 0.41425998 | 0 | 0 |
| ENSMUSG00 | Gm11902 | 269 | 0 | 0.35826351 | 0.32863824 | 0.40240198 | 0 | 0.10139271 |
| ENSMUSG00 | Gm2003 | 628 | 0 | 0 | 0 | 0 | 0 | 0 |
| ENSMUSG00 | Olfr343-ps1 | 302 | 0 | 0 | 0 | 0 | 0 | 0 |
| ENSMUSG00 | Gm11798 | 419 | 0 | 0 | 0 | 0 | 0 | 0 |
| ENSMUSG00 | Gm11453 | 1006 | 0.07903674 | 0.19159619 | 0.21969107 | 0.07173369 | 0 | 0.10844787 |
| ENSMUSG00 | Ifna-ps1 | 553 | 0 | 0 | 0 | 0 | 0 | 0 |
| ENSMUSG00 | Gm6461 | 632 | 0.12580848 | 0 | 0 | 0 | 0 | 0 |
| ENSMUSG00 | Gm6263 | 1341 | 0 | 0.14373286 | 0.13184741 | 0.26906819 | 0.19025494 | 0.04067806 |
| ENSMUSG00 | Gm12706 | 697 | 0.17111398 | 0 | 0.12683456 | 0 | 0 | 0 |
| ENSMUSG00 | Gm1862 | 809 | 0 | 0.17868891 | 0 | 0 | 0 | 0.06742803 |
| ENSMUSG00 | Gm12372 | 767 | 0 | 0 | 0 | 0 | 0 | 0 |
| ENSMUSG00 | Gm12173 | 272 | 0 | 0 | 0 | 0 | 0 | 0 |
| ENSMUSG00 | Gm16128 | 625 | 0 | 0 | 0 | 0 | 0 | 0 |
| ENSMUSG00 | Gm16061 | 856 | 0.09288664 | 0 | 0 | 0 | 0 | 0 |
| ENSMUSG00 | Gm9009 | 325 | 0 | 0 | 0 | 0 | 0 | 0 |
| ENSMUSG00 | Gm11490 | 271 | 0 | 0 | 0 | 0 | 0 | 0 |
| ENSMUSG00 | Gm13772 | 316 | 0 | 0 | 0 | 0 | 0 | 0.1726243 |
| ENSMUSG00 | Gm12297 | 973 | 0 | 0 | 0 | 0.03708329 | 0 | 0 |
| ENSMUSG00 | Gm13675 | 872 | 0 | 0 | 0 | 0 | 0 | 0 |
| ENSMUSG00 | 1700071K01I | 818 | 0 | 0 | 0 | 0.04411008 | 0 | 0 |
| ENSMUSG00 | Gm12161 | 360 | 0 | 0.13385123 | 0 | 0 | 0.28347987 | 0 |
| ENSMUSG00 | Gm13549 | 795 | 0 | 0 | 0.27799901 | 0 | 0 | 0 |
| ENSMUSG00 | Gm5419 | 525 | 0 | 0 | 0 | 0 | 0 | 0 |
| ENSMUSG00 | Gm14469 | 201 | 0 | 0 | 0 | 0 | 0 | 0 |
| ENSMUSG00 | Gm13559 | 862 | 0 | 0 | 0 | 0.12557556 | 0 | 0 |
| ENSMUSG00 | Olfr1081-ps1 | 944 | 0 | 0 | 0 | 0 | 0 | 0 |
| ENSMUSG00 | Gm12073 | 523 | 0 | 0 | 0 | 0 | 0 | 0 |
| ENSMUSG00 | Gm14048 | 1267 | 0 | 0 | 0 | 0 | 0 | 0 |
| ENSMUSG00 | Olfr118 | 965 | 0 | 0 | 0 | 0 | 0 | 0 |
| ENSMUSG00 | Psmb5-ps | 797 | 0 | 0.42321844 | 0 | 0 | 0.06402306 | 0 |
| ENSMUSG00 | Gm12903 | 543 | 0 | 0 | 0 | 0 | 0 | 0 |
| ENSMUSG00 | Gm13464 | 932 | 0.12796829 | 0 | 0 | 0.15485856 | 0 | 0.0877939 |
| ENSMUSG00 | Gm15165 | 501 | 0 | 0 | 0 | 0 | 0 | 0 |
| ENSMUSG00 | Gm4986 | 1123 | 0 | 0 | 0 | 0 | 0 | 0 |
| ENSMUSG00 | Gm11885 | 1427 | 0 | 0 | 0 | 0 | 0 | 0 |
| ENSMUSG00 | Gm5384 | 462 | 0 | 0 | 0 | 0 | 0 | 0 |
| ENSMUSG00 | Gm11336 | 333 | 0 | 0 | 0 | 0 | 0 | 0 |
| ENSMUSG00 | Gm11200 | 447 | 0 | 0 | 0 | 0 | 0 | 0 |
| ENSMUSG00 | Gm14678 | 481 | 0 | 0 | 0 | 0 | 0 | 0 |
| ENSMUSG00 | Gm14240 | 545 | 0 | 0 | 0 | 0 | 0 | 0 |
| ENSMUSG00 | Gm14301 | 323 | 0 | 0 | 0 | 0 | 0 | 0 |
| ENSMUSG00 | Gm9157 | 224 | 0.17747982 | 0.21511805 | 0.98664828 | 0 | 0.45559264 | 0.24352357 |
| ENSMUSG00 | Gm13198 | 825 | 0.53007307 | 1.10974837 | 0.48220192 | 0.69977299 | 0.37110092 | 0.09918051 |
| ENSMUSG00 | Gm12832 | 312 | 0 | 0 | 0 | 0 | 0 | 0 |
| ENSMUSG00 | Gm11543 | 367 | 0 | 0 | 0 | 0 | 0 | 0 |
| ENSMUSG00 | Gm13314 | 318 | 0 | 0 | 0 | 0 | 0 | 0 |
| ENSMUSG00 | Gm14065 | 584 | 0 | 0 | 0 | 0 | 0 | 0 |
| ENSMUSG00 | Gm13880 | 949 | 0.50270365 | 0.2538801 | 0.46577285 | 0.11406337 | 0.16130572 | 0.63228879 |
| ENSMUSG00 | Gm11913 | 334 | 0 | 0 | 0 | 0 | 0 | 0 |
| ENSMUSG00 | Gm15224 | 387 | 0 | 0 | 0 | 0 | 0 | 0 |
| ENSMUSG00 | Gm14497 | 579 | 0.41197389 | 0.08322356 | 0 | 0.31158933 | 0 | 0.04710646 |
| ENSMUSG00 | Gm13714 | 220 | 0 | 0 | 0 | 0 | 0 | 0 |
| ENSMUSG00 | Gm13805 | 103 | 0 | 0 | 0 | 0 | 0 | 0 |
| ENSMUSG00 | Olfr1397-ps1 | 961 | 0 | 0 | 0 | 0 | 0.05309717 | 0 |
| ENSMUSG00 | Gm15000 | 1242 | 0 | 0 | 0 | 0 | 0 | 0 |
| ENSMUSG00 | Gm15012 | 105 | 0 | 0 | 0 | 0 | 0 | 0 |
| ENSMUSG00 | Gm12567 | 285 | 0 | 0 | 0 | 0 | 0 | 0 |
| ENSMUSG00 | Gm14509 | 1285 | 0 | 0 | 0 | 0 | 0 | 0 |
| ENSMUSG00 | Gm11964 | 443 | 0.08974149 | 0.10877301 | 0.29933528 | 0.48869586 | 0 | 0.67724838 |
| ENSMUSG00 | Gm15936 | 459 | 0 | 0 | 0 | 0 | 0 | 0 |
| ENSMUSG00 | Gm15766 | 812 | 0 | 0 | 0 | 0 | 0 | 0 |
| ENSMUSG00 | Gm14627 | 566 | 0 | 0 | 0 | 0 | 0.09015261 | 0 |
| ENSMUSG00 | Gm11943 | 326 | 0 | 0 | 0 | 0 | 0 | 0 |
| ENSMUSG00 | Gm11855 | 1867 | 0.31940665 | 0.5936198 | 0 | 0 | 0.02733068 | 0.14608805 |
| ENSMUSG00 | Olfr1077-ps1 | 941 | 0 | 0 | 0 | 0 | 0 | 0 |
| ENSMUSG00 | Gm15077 | 361 | 0 | 0 | 0 | 0 | 0 | 0 |
| ENSMUSG00 | Gm12704 | 463 | 0 | 0 | 0 | 0 | 0.11020816 | 0 |
| ENSMUSG00 | Gm15042 | 882 | 0 | 0 | 0 | 0 | 0 | 0 |
| ENSMUSG00 | Gm14797 | 1479 | 0 | 0 | 0 | 0 | 0 | 0 |
| ENSMUSG00 | Gm6944 | 1000 | 0 | 0.09637288 | 0.22100922 | 0.18041022 | 0.10205275 | 0 |
| ENSMUSG00 | Olfr1300-ps1 | 933 | 0 | 0 | 0 | 0 | 0 | 0 |
| ENSMUSG00 | Gm14628 | 634 | 0 | 0 | 0 | 0 | 0 | 0 |
| ENSMUSG00 | Gm13929 | 2245 | 0 | 0 | 0 | 0 | 0 | 0 |
| ENSMUSG00 | Olfr626-ps1 | 284 | 0 | 0 | 0 | 0 | 0 | 0 |
| ENSMUSG00 | Gm14693 | 446 | 0 | 0 | 0 | 0 | 0 | 0 |
| ENSMUSG00 | Nlk-ps1 | 1228 | 0 | 0 | 0 | 0.02938277 | 0 | 0 |
| ENSMUSG00 | Psmb7-ps2 | 819 | 0 | 0.17650711 | 0 | 0 | 0 | 0 |
| ENSMUSG00 | Gm11512 | 662 | 0.54048236 | 0.50952431 | 0.53416125 | 0.272523 | 0.07707912 | 0.45320398 |
| ENSMUSG00 | AU015836 | 4188 | 0 | 0 | 0 | 0 | 0 | 0 |
| ENSMUSG00 | Gm12797 | 272 | 0 | 0 | 0 | 0 | 0 | 0 |
| ENSMUSG00 | Gm12846 | 1371 | 0 | 0.28117545 | 0.19344351 | 0.65795121 | 0.03721836 | 0.41777347 |
| ENSMUSG00 | Olfr937-ps1 | 921 | 0 | 0 | 0 | 0 | 0 | 0 |
| ENSMUSG00 | Gm11760 | 795 | 0 | 0.06061188 | 0 | 0 | 0 | 0 |
| ENSMUSG00 | Rps24-ps3 | 401 | 0.1982817 | 0 | 0 | 0.35992064 | 0.12724782 | 0.20404967 |
| ENSMUSG00 | Gm12461 | 1238 | 0 | 0 | 0 | 0 | 0 | 0 |
| ENSMUSG00 | Gm15427 | 1193 | 0.03332396 | 0.04039098 | 0.111153 | 0.15122399 | 0 | 0.29720898 |
| ENSMUSG00 | Gm6305 | 1011 | 0.51119807 | 0.38129727 | 0.13116274 | 1.78447302 | 0.90848147 | 0.43164612 |
| ENSMUSG00 | Mup-ps17 | 216 | 0 | 0 | 0 | 0 | 0 | 0 |
| ENSMUSG00 | Gm12025 | 143 | 0 | 0 | 0 | 0 | 0 | 0 |
| ENSMUSG00 | Gm6987 | 426 | 0 | 0 | 0 | 0 | 0 | 0.5121998 |
| ENSMUSG00 | Gm13998 | 334 | 0 | 0 | 0 | 0 | 0 | 0 |
| ENSMUSG00 | Olfr1017-ps1 | 686 | 0 | 0 | 0 | 0 | 0 | 0 |
| ENSMUSG00 | Hist2h3c2 | 1010 | 0.23617117 | 0.14312805 | 0.43764201 | 0.07144959 | 0.30312699 | 0.35105971 |
| ENSMUSG00 | Gm11945 | 475 | 0.41847874 | 0 | 0 | 0 | 0 | 0.05742029 |
| ENSMUSG00 | Gm11318 | 960 | 0 | 0 | 0 | 0 | 0 | 0 |
| ENSMUSG00 | Gm14712 | 771 | 0 | 0 | 0 | 0 | 0 | 0 |
| ENSMUSG00 | Gm13640 | 263 | 0 | 0 | 0 | 0 | 0.19401664 | 0 |
| ENSMUSG00 | Gm11825 | 735 | 0 | 0 | 0 | 0 | 0 | 0 |
| ENSMUSG00 | Gm6071 | 438 | 0 | 0 | 0 | 0 | 0 | 0 |

| | | | | | | | |
|---|---|---|---|---|---|---|---|
| ENSMUSG00 Gm13576 | 777 | 0 | 0 | 0 | 0 | 0 | 0 |
| ENSMUSG00 Gm12817 | 608 | 0 | 0 | 0 | 0 | 0 | 0 |
| ENSMUSG00 Gm12912 | 1380 | 0.02880832 | 0 | 0 | 0 | 0 | 0.03952846 |
| ENSMUSG00 Gm12804 | 998 | 0.27884606 | 0.04828301 | 0.1771617 | 0.07230871 | 0.10225727 | 0.10931719 |
| ENSMUSG00 Gm13928 | 1731 | 0 | 0 | 0 | 0 | 0 | 0 |
| ENSMUSG00 Gm13181 | 1068 | 0.40946656 | 0.49630231 | 0.24832496 | 0.20270812 | 0.38222004 | 0.71506545 |
| ENSMUSG00 Gm9836 | 215 | 0 | 0 | 0 | 0 | 0 | 0 |
| ENSMUSG00 Gm14977 | 810 | 0 | 0 | 0 | 0 | 0 | 0 |
| ENSMUSG00 Gm13758 | 741 | 0 | 0 | 0 | 0 | 0 | 0.03680788 |
| ENSMUSG00 Gm7698 | 398 | 0 | 0 | 0 | 0 | 0 | 0 |
| ENSMUSG00 Gm13160 | 1732 | 0 | 0 | 0 | 0 | 0 | 0 |
| ENSMUSG00 Rpsa-ps4 | 1917 | 0.08295353 | 0 | 0 | 0 | 0 | 0 |
| ENSMUSG00 Gm13899 | 1254 | 0 | 0 | 0.17624339 | 0 | 0 | 0 |
| ENSMUSG00 Gm12568 | 623 | 0.25525188 | 0.15469163 | 0 | 0 | 0.0819043 | 0.17511807 |
| ENSMUSG00 Gm13059 | 396 | 0 | 0 | 0 | 0 | 0 | 0 |
| ENSMUSG00 Gm11297 | 885 | 0 | 0 | 0 | 0 | 0 | 0 |
| ENSMUSG00 Gm14856 | 612 | 0 | 0 | 0 | 0 | 0 | 0 |
| ENSMUSG00 Gm14937 | 207 | 0 | 0 | 0 | 0 | 0 | 0 |
| ENSMUSG00 Gm6079 | 403 | 0 | 0 | 0.54840996 | 0 | 0 | 0 |
| ENSMUSG00 Gm5638 | 1257 | 0 | 0 | 0 | 0 | 0 | 0 |
| ENSMUSG00 Gm13463 | 232 | 0 | 0 | 0 | 0 | 0 | 0 |
| ENSMUSG00 Gm9506 | 319 | 0 | 0 | 0 | 0 | 0 | 0 |
| ENSMUSG00 Rps15a-ps7 | 391 | 0 | 0 | 0 | 0 | 0 | 0 |
| ENSMUSG00 Gm11679 | 778 | 0 | 0 | 0 | 0 | 0 | 0 |
| ENSMUSG00 Gm14624 | 190 | 0 | 0 | 0 | 0 | 0 | 0 |
| ENSMUSG00 Gm14128 | 1009 | 0 | 0 | 0 | 0 | 0 | 0.02703136 |
| ENSMUSG00 Gm12565 | 648 | 0 | 0 | 0 | 0.05568217 | 0 | 0.04209049 |
| ENSMUSG00 Rpl19-ps11 | 717 | 0.16634092 | 0 | 0 | 0 | 0 | 0.03803994 |
| ENSMUSG00 Gm14946 | 203 | 0 | 0 | 0 | 0 | 0 | 0 |
| ENSMUSG00 Gm12691 | 455 | 0 | 0 | 0 | 0 | 0 | 0 |
| ENSMUSG00 Gm12135 | 465 | 0 | 0 | 0 | 0 | 0 | 0 |
| ENSMUSG00 Gm12220 | 171 | 0 | 0 | 0 | 0 | 0 | 0 |
| ENSMUSG00 Gm5270 | 998 | 0.11950545 | 0.24141504 | 0.13287127 | 0.03615435 | 0.10225727 | 0.0546586 |
| ENSMUSG00 Gm14630 | 245 | 0.16226727 | 0 | 0 | 0 | 0 | 0 |
| ENSMUSG00 Gm12024 | 1252 | 0 | 0 | 0 | 0 | 0 | 0 |
| ENSMUSG00 Gm11717 | 332 | 0 | 0 | 0 | 0 | 0 | 0 |
| ENSMUSG00 Gm15989 | 497 | 0.07999091 | 0 | 0.80043579 | 0.72599687 | 0.10266876 | 0.2195142 |
| ENSMUSG00 Gm14087 | 139 | 0 | 0 | 0 | 0 | 0 | 0.39244085 |
| ENSMUSG00 Gm7155 | 1670 | 0 | 0 | 0 | 0 | 0 | 0 |
| ENSMUSG00 Gm12235 | 169 | 0 | 0 | 0 | 0 | 0 | 0.3227768 |
| ENSMUSG00 Gm4234 | 2005 | 0.01982817 | 0.16823197 | 0 | 0 | 0 | 0 |
| ENSMUSG00 Gm12745 | 213 | 0 | 0 | 0 | 0 | 0 | 0 |
| ENSMUSG00 Gm5913 | 2278 | 0.20942308 | 0.40190623 | 0.07761518 | 0.04751806 | 0.33599457 | 0.28735353 |
| ENSMUSG00 Gm12140 | 543 | 0 | 0 | 0 | 0 | 0 | 0 |
| ENSMUSG00 Gm7308 | 446 | 0.26741355 | 0.10804135 | 0.19821454 | 0.24270434 | 0.11440891 | 0.1223078 |
| ENSMUSG00 Gm15390 | 737 | 0 | 0 | 0 | 0 | 0 | 0 |
| ENSMUSG00 Gm15913 | 131 | 0 | 0 | 0 | 0 | 0 | 0 |
| ENSMUSG00 Gm13329 | 264 | 0 | 0 | 0 | 0 | 0 | 0 |
| ENSMUSG00 Gm11233 | 632 | 0 | 0 | 0 | 0 | 0 | 0 |
| ENSMUSG00 Gm6758 | 1054 | 0.07543734 | 0.04571769 | 0 | 0 | 0 | 0 |
| ENSMUSG00 Gm14601 | 235 | 0 | 0 | 0 | 0 | 0 | 0 |
| ENSMUSG00 Gm12458 | 494 | 0 | 0 | 0 | 0 | 0 | 0 |
| ENSMUSG00 Gm12791 | 654 | 0 | 0.14735915 | 0 | 0 | 0 | 0 |
| ENSMUSG00 Gm13678 | 339 | 0 | 0 | 0 | 0 | 0 | 0 |
| ENSMUSG00 Gm11469 | 495 | 0 | 0 | 0 | 0 | 0 | 0 |
| ENSMUSG00 Gm14906 | 389 | 0 | 0 | 0 | 0 | 0 | 0 |
| ENSMUSG00 Rpl26-ps5 | 896 | 0 | 0 | 0 | 0 | 0 | 0 |
| ENSMUSG00 Gm15784 | 1461 | 0.02721114 | 0 | 0 | 0.04939363 | 0 | 0 |
| ENSMUSG00 Gm12677 | 424 | 0 | 0 | 0 | 0 | 0 | 0 |
| ENSMUSG00 Gm13328 | 1019 | 0.62422737 | 0.52016768 | 0.95430868 | 0.17704634 | 0.75112428 | 0.64238601 |
| ENSMUSG00 Gm14605 | 266 | 0 | 0 | 0 | 0 | 0 | 0 |
| ENSMUSG00 Gm8217 | 464 | 0 | 0 | 0 | 0 | 0 | 0 |
| ENSMUSG00 Gm14833 | 1026 | 0 | 0 | 0 | 0 | 0 | 0 |
| ENSMUSG00 Gm13744 | 849 | 0 | 0 | 0.05206342 | 0 | 0 | 0 |
| ENSMUSG00 Gm14544 | 448 | 0 | 0 | 0 | 0 | 0 | 0 |
| ENSMUSG00 Gm14913 | 1092 | 0 | 0 | 0 | 0 | 0 | 0 |
| ENSMUSG00 Gm14800 | 1271 | 0 | 0 | 0 | 0 | 0 | 0 |
| ENSMUSG00 Gm12990 | 467 | 0.42564754 | 0 | 0 | 0.07726348 | 1.31117026 | 0.40882757 |
| ENSMUSG00 BC022960 | 1814 | 0.72322539 | 0.92972738 | 1.21835289 | 5.01250012 | 5.51332396 | 2.84173474 |
| ENSMUSG00 Gm12834 | 2828 | 0 | 0 | 0 | 0 | 0 | 0 |
| ENSMUSG00 Gm13333 | 780 | 0 | 0.80310737 | 0 | 0 | 0 | 0 |
| ENSMUSG00 Gm8178 | 645 | 0 | 0 | 0 | 0 | 0 | 0 |
| ENSMUSG00 Gm11501 | 580 | 0 | 0 | 0 | 0 | 0 | 0 |
| ENSMUSG00 Gm15497 | 550 | 0.14456538 | 0 | 0 | 0.59043346 | 0.09277523 | 0.24795127 |
| ENSMUSG00 Gm15823 | 628 | 0 | 0 | 0 | 0 | 0.32500876 | 0.13029286 |
| ENSMUSG00 Vmn2r-ps2 | 764 | 0 | 0 | 0 | 0 | 0 | 0 |
| ENSMUSG00 Gm14585 | 633 | 0 | 0 | 0 | 0 | 0 | 0.12926369 |
| ENSMUSG00 Gm6207 | 1383 | 0 | 0 | 0 | 0 | 0 | 0 |
| ENSMUSG00 Gm6024 | 1557 | 0 | 0 | 0 | 0 | 0 | 0 |
| ENSMUSG00 Gm11891 | 617 | 0 | 0 | 0 | 0 | 0 | 0 |
| ENSMUSG00 Gm12684 | 998 | 0 | 0 | 0.04429042 | 0.03615435 | 0 | 0.0273293 |
| ENSMUSG00 Gm11448 | 449 | 0 | 0 | 0.29533525 | 0 | 0 | 0 |
| ENSMUSG00 Gm12430 | 1321 | 0.15047495 | 0 | 0 | 0.32777028 | 0 | 0 |
| ENSMUSG00 Gm11401 | 343 | 0 | 0 | 0 | 0 | 0 | 0 |
| ENSMUSG00 Defa-ps13 | 202 | 0 | 0 | 0 | 0 | 0 | 0 |
| ENSMUSG00 Gm12771 | 134 | 0 | 0 | 0 | 0 | 0 | 0 |
| ENSMUSG00 Gm14425 | 605 | 0 | 0 | 0.43836539 | 0.05963974 | 0.16868223 | 0 |
| ENSMUSG00 Gm13552 | 416 | 0 | 0 | 0 | 0 | 0.12265956 | 0.13112807 |
| ENSMUSG00 Gm13521 | 865 | 0 | 0 | 0 | 0 | 0 | 0 |
| ENSMUSG00 Gm6023 | 605 | 0 | 0 | 0 | 0 | 0 | 0 |
| ENSMUSG00 Gm14897 | 206 | 0 | 0 | 0 | 0 | 0 | 0.13240116 |
| ENSMUSG00 Gm12783 | 394 | 0 | 0.24460123 | 0 | 0 | 0 | 0 |
| ENSMUSG00 Olfr400-ps1 | 909 | 0 | 0 | 0 | 0 | 0 | 0 |
| ENSMUSG00 Defa-ps12 | 418 | 0 | 0 | 0 | 0 | 0 | 0 |
| ENSMUSG00 Gm8722 | 768 | 0.36235464 | 0.12548553 | 0.11510897 | 0.28189097 | 0.06644059 | 0.42616624 |
| ENSMUSG00 Gm11783 | 884 | 0 | 0 | 0.25001042 | 0.04081679 | 0.17316643 | 0 |
| ENSMUSG00 Gm12819 | 627 | 0 | 0 | 0 | 0 | 0 | 0 |
| ENSMUSG00 Gm8880 | 453 | 0 | 0 | 0 | 0 | 0 | 0 |
| ENSMUSG00 Gm13730 | 1461 | 0 | 0 | 0 | 0 | 0 | 0 |
| ENSMUSG00 Gm12551 | 723 | 0 | 0 | 0 | 0 | 0 | 0 |
| ENSMUSG00 Gm12922 | 808 | 0 | 0 | 0 | 0 | 0 | 0 |
| ENSMUSG00 Gm7485 | 1108 | 0 | 0 | 0 | 0 | 0 | 0 |
| ENSMUSG00 Gm12849 | 736 | 0 | 0 | 0 | 0 | 0 | 0 |
| ENSMUSG00 Gm13500 | 823 | 0 | 0 | 0 | 0 | 0 | 0 |
| ENSMUSG00 Gm13439 | 408 | 0 | 0 | 0 | 0 | 0 | 0 |
| ENSMUSG00 Gm5793 | 1261 | 0 | 0 | 0 | 0 | 0 | 0 |
| ENSMUSG00 H3f3a-ps2 | 410 | 0 | 0 | 0 | 0 | 0 | 0 |
| ENSMUSG00 Gm12509 | 608 | 0 | 0 | 0 | 0 | 0 | 0 |
| ENSMUSG00 Ppp1r2-ps6 | 620 | 0 | 0 | 0 | 0 | 0 | 0 |
| ENSMUSG00 Gm13136 | 2949 | 0.013481 | 0 | 0 | 0 | 0.05190883 | 0.04624388 |
| ENSMUSG00 Gm7212 | 347 | 0 | 0 | 0 | 0 | 0 | 0 |
| ENSMUSG00 Gm13353 | 72 | 0 | 0 | 0 | 0 | 0 | 0 |
| ENSMUSG00 Gm14935 | 1707 | 0.23289678 | 0 | 0 | 0 | 0.14946214 | 0 |
| ENSMUSG00 Gm12449 | 233 | 0 | 0 | 0 | 0 | 0 | 0 |
| ENSMUSG00 Gm12685 | 213 | 0 | 0 | 0 | 0 | 0 | 0 |
| ENSMUSG00 Gm4852 | 728 | 0 | 0 | 0 | 0.04956325 | 0.07009118 | 0.07493033 |
| ENSMUSG00 Gm12578 | 416 | 0 | 0 | 0 | 0 | 0 | 0 |
| ENSMUSG00 3110067C02i | 2115 | 0.09398459 | 0 | 0.2507906 | 0 | 0.48251892 | 0.06447905 |
| ENSMUSG00 Gm14621 | 688 | 0 | 0 | 0 | 0 | 0 | 0 |
| ENSMUSG00 Gm12725 | 1667 | 0 | 0.05781217 | 0.29167383 | 0 | 0 | 0 |
| ENSMUSG00 Gm11807 | 446 | 0 | 0 | 0 | 0 | 0 | 0 |
| ENSMUSG00 Gm11271 | 439 | 0 | 0 | 0 | 0 | 0 | 0 |
| ENSMUSG00 Gm13243 | 564 | 0 | 0 | 0 | 0 | 0 | 0 |
| ENSMUSG00 Gm8424 | 986 | 0 | 0 | 0 | 0 | 0 | 0 |
| ENSMUSG00 Gm6310 | 740 | 0 | 0 | 0 | 0 | 0 | 0 |
| ENSMUSG00 Gm13858 | 668 | 0 | 0 | 0 | 0 | 0 | 0.0408303 |
| ENSMUSG00 Gm15076 | 473 | 0 | 0 | 0 | 0 | 0 | 0 |
| ENSMUSG00 Gm14536 | 137 | 0 | 0 | 0 | 0 | 0 | 0 |
| ENSMUSG00 Gm16407 | 1057 | 0 | 0 | 0 | 0 | 0 | 0 |
| ENSMUSG00 Gm15615 | 197 | 0 | 0 | 0 | 0 | 0 | 0 |
| ENSMUSG00 Smt3h2-ps4 | 287 | 0 | 0 | 0 | 0 | 0 | 0 |
| ENSMUSG00 Gm13220 | 507 | 0 | 0 | 0 | 0 | 0 | 0 |
| ENSMUSG00 Gm12613 | 381 | 0 | 0 | 0 | 0 | 0 | 0 |
| ENSMUSG00 Gm5940 | 1735 | 0 | 0 | 0 | 0 | 0 | 0 |
| ENSMUSG00 Gm11935 | 398 | 0 | 0 | 0 | 0 | 0.38462092 | 0 |
| ENSMUSG00 Gm13775 | 928 | 0 | 0 | 0 | 0 | 0 | 0 |
| ENSMUSG00 Gm11400 | 1213 | 3.11357844 | 4.40947657 | 2.00420558 | 3.5695345 | 3.36530096 | 3.35030606 |
| ENSMUSG00 Gm14890 | 675 | 0 | 0 | 0.19645264 | 0 | 0 | 0 |
| ENSMUSG00 Gm12933 | 871 | 0 | 0 | 0 | 0 | 0 | 0.03131417 |
| ENSMUSG00 Gm14886 | 2628 | 0 | 0 | 0 | 0 | 0 | 0 |
| ENSMUSG00 Gm6964 | 436 | 0 | 0 | 0 | 0 | 0 | 0 |
| ENSMUSG00 Gm11983 | 2028 | 0.03920659 | 0 | 0 | 0.01779194 | 0 | 0 |
| ENSMUSG00 Rpl35a-ps2 | 754 | 0.1581783 | 0.31953874 | 0.23449254 | 0.38283336 | 0.60906815 | 0.21703957 |
| ENSMUSG00 Gm11446 | 865 | 0 | 0 | 0 | 0 | 0 | 0 |
| ENSMUSG00 Gm6744 | 542 | 0 | 0 | 0 | 0 | 0 | 0 |
| ENSMUSG00 Gm15268 | 350 | 0 | 0 | 0 | 0.10309156 | 0 | 1.32476819 |
| ENSMUSG00 Gm4297 | 2571 | 0 | 0 | 0 | 0 | 0 | 0 |
| ENSMUSG00 Bambi-ps1 | 591 | 0 | 0.24460123 | 0 | 0.12210506 | 0.17267809 | 0 |
| ENSMUSG00 Gm14542 | 867 | 0 | 0 | 0 | 0 | 0 | 0 |
| ENSMUSG00 Gm14760 | 1409 | 0 | 0 | 0 | 0 | 0 | 0 |
| ENSMUSG00 Gm15173 | 432 | 0 | 0 | 0 | 0 | 0 | 0.06313574 |
| ENSMUSG00 Gm12247 | 669 | 0 | 0.07202757 | 0 | 0 | 0.22881783 | 0 |
| ENSMUSG00 Gm2066 | 384 | 0 | 0 | 0 | 0 | 0 | 0 |
| ENSMUSG00 Cyp2j12 | 2183 | 0 | 0 | 0.02024821 | 0.01652865 | 0 | 0 |
| ENSMUSG00 Gm8802 | 739 | 0.16138896 | 0.06520493 | 0.05981305 | 0 | 0 | 0 |
| ENSMUSG00 Gm13082 | 560 | 0.28396772 | 0 | 0 | 0 | 0.09111853 | 0.09740943 |
| ENSMUSG00 Gm16089 | 335 | 0.83071153 | 0.14384013 | 0.659729 | 0.1077076 | 1.37085786 | 1.2212525 |
| ENSMUSG00 Lamr1-ps1 | 873 | 0 | 0 | 0 | 0 | 0 | 0 |
| ENSMUSG00 Gm9097 | 879 | 0 | 0 | 0 | 0 | 0 | 0 |
| ENSMUSG00 Tcp1-ps1 | 1464 | 0 | 0 | 0 | 0.02464621 | 0.03485408 | 0.01863022 |
| ENSMUSG00 Gm14373 | 2561 | 0.03104684 | 0.13170835 | 0.0172596 | 0.05635618 | 0.13947077 | 0.05324998 |
| ENSMUSG00 Gm12857 | 591 | 1.34536313 | 0.24460123 | 0 | 0 | 2.76284945 | 0 |
| ENSMUSG00 Olfr1150-ps1 | 936 | 0 | 0 | 0 | 0 | 0 | 0 |
| ENSMUSG00 Gm14788 | 374 | 0 | 0 | 0 | 0 | 0 | 0 |
| ENSMUSG00 Gm13574 | 393 | 0 | 0.24522362 | 0 | 0 | 0 | 0 |
| ENSMUSG00 Gm11385 | 568 | 0 | 0 | 0 | 0 | 0 | 0 |
| ENSMUSG00 Gm15836 | 792 | 0 | 0 | 0 | 0.68337206 | 0.25770897 | 0 |
| ENSMUSG00 Gm11836 | 440 | 0.09035337 | 0.32854392 | 0.60275241 | 0.08200465 | 0 | 0 |
| ENSMUSG00 Gm16511 | 559 | 0 | 0.25860345 | 0 | 0 | 0 | 0 |
| ENSMUSG00 Gm14737 | 173 | 0 | 0 | 0 | 0 | 0 | 0 |
| ENSMUSG00 Gm15719 | 634 | 0.12541161 | 0.07600385 | 0.069719 | 0 | 0 | 0 |
| ENSMUSG00 Gm14037 | 312 | 0 | 0 | 0 | 0 | 0 | 0 |
| ENSMUSG00 Gm13529 | 792 | 0 | 0 | 0 | 0 | 0 | 0 |
| ENSMUSG00 Gm6587 | 578 | 0 | 0 | 0 | 0.18727705 | 0 | 0.14156387 |
| ENSMUSG00 Gm13787 | 167 | 4.76113539 | 0.28854157 | 0 | 0 | 0 | 1.63321193 |
| ENSMUSG00 Gm11517 | 790 | 0.35226375 | 0.36597298 | 0.1119034 | 0.50240822 | 0.77508419 | 0.31072374 |
| ENSMUSG00 Gm15554 | 554 | 0 | 0 | 0 | 0 | 0 | 0 |
| ENSMUSG00 Gm12164 | 550 | 0.07228269 | 0.17522343 | 0.40183494 | 0.19681115 | 0 | 0.04959025 |
| ENSMUSG00 Gm16032 | 167 | 0 | 0 | 0 | 0 | 0 | 0 |
| ENSMUSG00 Gm8501 | 431 | 0 | 0 | 0 | 0 | 0 | 0 |
| ENSMUSG00 Gm12112 | 230 | 0 | 0 | 0 | 0 | 0.22185381 | 0 |
| ENSMUSG00 Gm5380 | 2135 | 0 | 0 | 0.20703439 | 0 | 0 | 0 |
| ENSMUSG00 Gm15791 | 318 | 0 | 0 | 0 | 0 | 0 | 0 |
| ENSMUSG00 Gm13607 | 538 | 0 | 0 | 0 | 0 | 0 | 0 |
| ENSMUSG00 Gm14013 | 269 | 0 | 0 | 0 | 0 | 0 | 0.10139271 |
| ENSMUSG00 Gm6893 | 518 | 0 | 0 | 0 | 0.27862583 | 0 | 0 |
| ENSMUSG00 Gm13149 | 338 | 0 | 0 | 0 | 0 | 0 | 0 |
| ENSMUSG00 Gm15370 | 252 | 0 | 0 | 0 | 0 | 0 | 0 |
| ENSMUSG00 Gm12261 | 644 | 0 | 0 | 0 | 0.39219614 | 0 | 0.04235192 |
| ENSMUSG00 Gm14859 | 356 | 0 | 0 | 0 | 0 | 0 | 0 |
| ENSMUSG00 Gm14021 | 960 | 0 | 0 | 0 | 0 | 0 | 0 |
| ENSMUSG00 Gm11282 | 1415 | 0 | 0 | 0 | 0 | 0.46879356 | 0 |
| ENSMUSG00 Gm15820 | 689 | 0 | 0 | 0 | 0 | 0 | 0 |
| ENSMUSG00 Gm7421 | 1656 | 0 | 0 | 0 | 0 | 0 | 0 |
| ENSMUSG00 Gm14909 | 350 | 0 | 0 | 0 | 0 | 0 | 0 |
| ENSMUSG00 Gm14659 | 841 | 0 | 0 | 0 | 0 | 0 | 0 |
| ENSMUSG00 Gm11653 | 1008 | 1.34095867 | 0 | 0.65776552 | 0.1789784 | 1.92361337 | 0 |
| ENSMUSG00 Gm14883 | 156 | 0 | 0 | 0 | 0 | 0 | 0 |
| ENSMUSG00 Gm13509 | 1307 | 4.95802856 | 4.35042094 | 5.74928333 | 4.96921807 | 3.3575121 | 4.17362498 |
| ENSMUSG00 Gm15896 | 438 | 0 | 0 | 0 | 0 | 0 | 0 |
| ENSMUSG00 Gm15727 | 437 | 0.09097364 | 0.22053292 | 0 | 0 | 0 | 0.62413362 |
| ENSMUSG00 Gm13875 | 281 | 0 | 0 | 0 | 0 | 0 | 0 |
| ENSMUSG00 Gm13132 | 471 | 0 | 0 | 0 | 0 | 0 | 0 |
| ENSMUSG00 Gm15285 | 472 | 0.42113857 | 0.91880928 | 0.65553581 | 0.15289002 | 0.10810673 | 0.46228202 |
| ENSMUSG00 Gm13186 | 315 | 0 | 0 | 0 | 0 | 0 | 0.25975847 |
| ENSMUSG00 Vmn1r-ps2 | 833 | 0 | 0 | 0 | 0 | 0 | 0 |
| ENSMUSG00 Gm9516 | 423 | 0 | 0 | 0 | 0 | 0 | 0 |
| ENSMUSG00 Gm6274 | 796 | 0 | 0 | 0 | 0 | 0 | 0 |
| ENSMUSG00 Gm13836 | 564 | 0 | 0 | 0 | 0 | 0 | 0 |
| ENSMUSG00 Gm11302 | 770 | 0 | 0 | 0 | 0 | 0 | 0 |
| ENSMUSG00 Gm13673 | 530 | 0 | 0 | 0 | 0 | 0 | 0 |
| ENSMUSG00 Gm12117 | 951 | 0.08360774 | 0.05066924 | 0 | 0.03794116 | 0.0536555 | 0.17207974 |
| ENSMUSG00 Gm14995 | 390 | 0 | 0 | 0 | 0 | 0 | 0 |
| ENSMUSG00 Olfr333-ps1 | 805 | 0 | 0 | 0 | 0 | 0 | 0 |
| ENSMUSG00 Gm11386 | 346 | 0 | 0 | 0 | 0 | 0 | 0 |
| ENSMUSG00 Gm14857 | 3184 | 0 | 0 | 0 | 0.0113323 | 0.04807762 | 0 |
| ENSMUSG00 Gm12520 | 315 | 0 | 0 | 0 | 0 | 0 | 0 |
| ENSMUSG00 Gm14738 | 2479 | 0 | 0 | 0 | 0 | 0 | 0 |
| ENSMUSG00 Gm15018 | 632 | 0 | 0 | 0 | 0 | 0 | 0 |
| ENSMUSG00 Gm11673 | 330 | 0 | 0 | 0 | 0 | 0 | 0 |
| ENSMUSG00 Gm15089 | 1195 | 0 | 0 | 0 | 0 | 0 | 0 |
| ENSMUSG00 Gm6275 | 1121 | 0 | 0 | 0.07886145 | 0 | 0.04551862 | 0 |
| ENSMUSG00 Gm12203 | 597 | 0 | 0 | 0 | 0 | 0 | 0 |
| ENSMUSG00 Gm12209 | 595 | 0 | 0 | 0 | 0 | 0 | 0 |
| ENSMUSG00 Gm14622 | 463 | 0 | 0 | 0 | 0 | 0 | 0 |
| ENSMUSG00 Gm13603 | 82 | 0 | 0 | 0 | 0 | 0 | 0 |
| ENSMUSG00 Gm14587 | 657 | 0 | 0 | 0 | 0 | 0 | 0 |
| ENSMUSG00 Gm15951 | 399 | 0.29891339 | 0 | 0 | 0 | 0 | 0 |
| ENSMUSG00 Gm12020 | 2041 | 0.03895687 | 0.04721846 | 0.02165695 | 0.01767861 | 0.15000405 | 0 |
| ENSMUSG00 Gm16011 | 2661 | 0 | 0 | 0 | 0 | 0 | 0 |
| ENSMUSG00 Gm11509 | 324 | 0 | 0 | 0 | 0 | 0 | 0 |
| ENSMUSG00 Gm12879 | 1181 | 0 | 0 | 0 | 0 | 0 | 0 |
| ENSMUSG00 Gm15266 | 253 | 47.140886 | 58.8522159 | 62.3717706 | 65.8889509 | 55.0600814 | 46.3561062 |
| ENSMUSG00 Gm13627 | 703 | 0 | 0 | 0 | 0 | 0 | 0 |
| ENSMUSG00 Gm14480 | 432 | 0 | 0 | 0.30695724 | 0 | 0 | 0.06313574 |
| ENSMUSG00 Gm12397 | 958 | 0 | 0 | 0 | 0 | 0.05326344 | 0 |
| ENSMUSG00 Gm14054 | 703 | 0 | 0 | 0 | 0 | 0 | 0 |
| ENSMUSG00 Gm12197 | 446 | 0 | 0 | 0 | 0 | 0 | 0 |
| ENSMUSG00 Gm14520 | 386 | 0 | 0 | 0 | 0 | 0 | 0 |
| ENSMUSG00 Gm14213 | 767 | 0 | 0 | 0 | 0 | 0 | 0 |
| ENSMUSG00 Gm14132 | 1718 | 0.04628112 | 0 | 0 | 0 | 0 | 0 |
| ENSMUSG00 Olfr621-ps1 | 934 | 0 | 0 | 0 | 0 | 0 | 0 |
| ENSMUSG00 Gm16205 | 1026 | 0 | 0 | 0 | 0 | 0 | 0 |
| ENSMUSG00 Gm12757 | 799 | 0 | 0 | 0 | 0 | 0 | 0 |
| ENSMUSG00 Rpl36-ps1 | 302 | 0 | 0 | 0 | 0 | 0 | 0 |
| ENSMUSG00 Gm12936 | 300 | 0 | 0 | 0 | 0 | 0 | 0 |
| ENSMUSG00 Gm15034 | 348 | 0 | 0 | 1.52420148 | 0 | 0 | 0 |
| ENSMUSG00 Gm15304 | 2639 | 0 | 0 | 0 | 0 | 0 | 0 |
| ENSMUSG00 Rpl7l1-ps1 | 724 | 0.05491089 | 0 | 0 | 0 | 0 | 0 |
| ENSMUSG00 Gm14550 | 406 | 0 | 0 | 0 | 0 | 0 | 0 |
| ENSMUSG00 Gm12599 | 791 | 0 | 0 | 0 | 0 | 0 | 0 |
| ENSMUSG00 Gm14417 | 1071 | 0 | 0 | 0 | 0 | 0 | 0 |
| ENSMUSG00 Gm11819 | 371 | 0 | 0 | 0 | 0 | 0 | 0 |
| ENSMUSG00 Olfr1001-ps1 | 944 | 0 | 0 | 0 | 0 | 0 | 0 |
| ENSMUSG00 Gm12492 | 562 | 0.07073929 | 0 | 0.07865097 | 0 | 0 | 0 |
| ENSMUSG00 Gm13013 | 522 | 0 | 0 | 0 | 0.55298153 | 0 | 0.20900107 |
| ENSMUSG00 Gm15151 | 176 | 0 | 0.54757321 | 0.25114684 | 0 | 0 | 0 |
| ENSMUSG00 Gm12664 | 912 | 0 | 0 | 0.09693387 | 0.03956365 | 0 | 0.23925122 |
| ENSMUSG00 Gm12307 | 589 | 0 | 0 | 0 | 0 | 0 | 0 |
| ENSMUSG00 Gm11286 | 457 | 0 | 0 | 0 | 0 | 0 | 0 |
| ENSMUSG00 Gm15786 | 449 | 0 | 0 | 0 | 0 | 0 | 0 |
| ENSMUSG00 Gm15073 | 633 | 0 | 0 | 0 | 0 | 0 | 0 |
| ENSMUSG00 Gm7618 | 170 | 0 | 0 | 0.52002168 | 0 | 0 | 0.16043905 |
| ENSMUSG00 Gm16044 | 449 | 0 | 0.10731947 | 0.09844509 | 0.64288721 | 1.25008939 | 0.0607453 |
| ENSMUSG00 Gm12275 | 306 | 0 | 0.15747203 | 0 | 0 | 0 | 0 |
| ENSMUSG00 Gm14891 | 595 | 0 | 0 | 0 | 0 | 0 | 0 |
| ENSMUSG00 Gm11434 | 628 | 0 | 0 | 0 | 0 | 0.16250438 | 0 |
| ENSMUSG00 Gm16071 | 792 | 0.50196314 | 0.24336587 | 0.27905204 | 0.77448833 | 1.61068106 | 0.37881443 |
| ENSMUSG00 Gm14303 | 170 | 0 | 0 | 0 | 0.21224732 | 0.30015515 | 0.16043905 |
| ENSMUSG00 Olfr1159-ps1 | 891 | 0 | 0 | 0 | 0 | 0 | 0 |
| ENSMUSG00 Gm12030 | 222 | 0 | 0 | 0 | 0 | 0 | 0 |
| ENSMUSG00 Gm14702 | 445 | 0 | 0 | 0 | 0 | 0.11466601 | 0 |
| ENSMUSG00 Gm13878 | 575 | 0 | 0 | 0 | 0 | 0 | 0 |
| ENSMUSG00 Gm15041 | 755 | 0 | 0 | 0 | 0 | 0 | 0 |
| ENSMUSG00 Gm5388 | 1116 | 0.07124638 | 0.17271126 | 0.07921477 | 0.32331581 | 0.09144512 | 0.02443964 |
| ENSMUSG00 Gm12389 | 417 | 0 | 0 | 0.10599962 | 0.08652769 | 0 | 0.06540681 |
| ENSMUSG00 Gm13818 | 514 | 2.08832291 | 0 | 0 | 0 | 0.19854621 | 0 |
| ENSMUSG00 Gm14894 | 269 | 0 | 0 | 0 | 0 | 0 | 0 |
| ENSMUSG00 Gm15264 | 496 | 0.56106525 | 0.58290051 | 0.08911662 | 0.07274606 | 0 | 0.05498919 |
| ENSMUSG00 Gm1993 | 1567 | 0 | 0 | 0 | 0 | 0 | 0 |
| ENSMUSG00 Xlr4e-ps | 641 | 0 | 0 | 0 | 0 | 0 | 0 |
| ENSMUSG00 Gm14746 | 739 | 0 | 0.06520493 | 0 | 0 | 0 | 0 |
| ENSMUSG00 Gm15683 | 352 | 0.22588341 | 0 | 0.75344051 | 0 | 0.1449613 | 0.38742385 |
| ENSMUSG00 Gm11718 | 536 | 0 | 0 | 0 | 0 | 0 | 0 |
| ENSMUSG00 Gm11813 | 290 | 0 | 0 | 0 | 0 | 0 | 0 |
| ENSMUSG00 Cyp2j7-ps | 1506 | 2.71900033 | 2.14375274 | 0.23480395 | 4.33655382 | 3.65926201 | 5.50563767 |
| ENSMUSG00 Gm11500 | 646 | 0 | 0 | 0 | 0 | 0 | 0 |
| ENSMUSG00 Gm8595 | 629 | 0 | 0 | 0 | 0 | 0 | 0 |
| ENSMUSG00 Gm15031 | 209 | 0 | 0 | 0 | 0 | 0 | 0 |
| ENSMUSG00 Gm7391 | 631 | 0 | 0 | 0 | 0 | 0 | 0 |
| ENSMUSG00 Gm13265 | 382 | 0 | 0 | 0 | 0 | 0 | 0 |
| ENSMUSG00 Gm11896 | 819 | 0.04854149 | 0 | 0 | 0 | 0 | 0 |
| ENSMUSG00 Gm9105 | 1397 | 0.17074652 | 0 | 0.12656219 | 0.15496941 | 0.14610272 | 0.05857117 |
| ENSMUSG00 Gm13155 | 645 | 0 | 0 | 0 | 0 | 0 | 0 |
| ENSMUSG00 Gm16171 | 323 | 0 | 0 | 0 | 0 | 0 | 0 |
| ENSMUSG00 Gm16436 | 1605 | 0 | 0.03002271 | 0.02754009 | 0 | 0 | 0 |
| ENSMUSG00 Gm15011 | 1479 | 0 | 0 | 0 | 0 | 0 | 0 |
| ENSMUSG00 Gm14686 | 342 | 0 | 0 | 0 | 0 | 0 | 0 |
| ENSMUSG00 Gm11694 | 158 | 0.25161697 | 0 | 0 | 0 | 0 | 0.1726243 |
| ENSMUSG00 Gm12877 | 506 | 0 | 0 | 0 | 0 | 0 | 0 |
| ENSMUSG00 Gm6834 | 455 | 1.31062024 | 2.22398964 | 1.84579125 | 2.77554189 | 1.12145881 | 1.55855081 |
| ENSMUSG00 Gm14771 | 202 | 0 | 0 | 0 | 0 | 0 | 0 |
| ENSMUSG00 Gm12452 | 330 | 0 | 0 | 0 | 0 | 0 | 0 |
| ENSMUSG00 Gm5752 | 593 | 0 | 0 | 0 | 0 | 0 | 0 |
| ENSMUSG00 Rpl18-ps1 | 524 | 0.07586924 | 0 | 0 | 0 | 0 | 0 |
| ENSMUSG00 Gm5742 | 504 | 0 | 0 | 0 | 0 | 0 | 0 |
| ENSMUSG00 Gm12752 | 810 | 0 | 0 | 0 | 0 | 0 | 0 |
| ENSMUSG00 Gm15160 | 906 | 0 | 0 | 0 | 0 | 0 | 0 |
| ENSMUSG00 Gm11801 | 745 | 0 | 0 | 0 | 0 | 0 | 0 |
| ENSMUSG00 Gm13368 | 426 | 0.18664545 | 0 | 0 | 0 | 0 | 0 |
| ENSMUSG00 Gm14538 | 1326 | 0.02998151 | 0.0726794 | 0 | 0.10884478 | 0 | 0.04113822 |
| ENSMUSG00 Gm11765 | 1202 | 0 | 0 | 0 | 0 | 0 | 0 |
| ENSMUSG00 Gm11668 | 699 | 0 | 0 | 0 | 0 | 0 | 0 |
| ENSMUSG00 Gm13215 | 471 | 0 | 0 | 0 | 0 | 0 | 0 |
| ENSMUSG00 Gm14879 | 506 | 0 | 0 | 0 | 0 | 0 | 0 |
| ENSMUSG00 Gm15132 | 651 | 0 | 0 | 0 | 0 | 0 | 0 |
| ENSMUSG00 Gm14815 | 487 | 0 | 0 | 0 | 0.22227132 | 0 | 0 |
| ENSMUSG00 Gm7657 | 665 | 0 | 0 | 0 | 0 | 0 | 0 |
| ENSMUSG00 Gm14921 | 912 | 0.08718307 | 0.15850803 | 0.2908016 | 0.23738187 | 0.05594997 | 0.14953201 |
| ENSMUSG00 Gm13680 | 792 | 0 | 0 | 0 | 0 | 0 | 0 |
| ENSMUSG00 Gm14889 | 227 | 0 | 0 | 0 | 0 | 0 | 0 |
| ENSMUSG00 Gm15455 | 5573 | 0.02140076 | 0.01729282 | 0.00793143 | 0 | 0 | 0 |
| ENSMUSG00 Gm14658 | 195 | 0 | 0 | 0 | 0 | 0 | 0 |
| ENSMUSG00 Gm15694 | 282 | 0.28195376 | 0.68349563 | 3.29162661 | 2.17515871 | 1.08566757 | 0.48359289 |
| ENSMUSG00 Gm13142 | 1451 | 0 | 0 | 0 | 0 | 0 | 0 |
| ENSMUSG00 Gm13654 | 1545 | 0.05146341 | 0.09356591 | 0.08582882 | 0.04670815 | 0 | 0 |
| ENSMUSG00 Gm14568 | 293 | 0 | 0 | 0 | 0 | 0 | 0 |
| ENSMUSG00 Gm15047 | 453 | 0 | 0 | 0 | 0 | 0 | 0 |
| ENSMUSG00 Gm8215 | 989 | 0 | 0 | 0 | 0 | 0 | 0 |
| ENSMUSG00 Gm6797 | 506 | 0 | 0 | 0 | 0 | 0 | 0 |
| ENSMUSG00 Gm15403 | 1226 | 0 | 0 | 0 | 0 | 0 | 0 |
| ENSMUSG00 Gm14088 | 579 | 0.27464926 | 0.91545918 | 0 | 0.12463573 | 0.17625691 | 0.8950227 |
| ENSMUSG00 Gm3943 | 717 | 0 | 0 | 0 | 0 | 0 | 0 |
| ENSMUSG00 Gm15176 | 134 | 0 | 0 | 0 | 0 | 0 | 0 |
| ENSMUSG00 Gm14383 | 385 | 0 | 0 | 0 | 0 | 0 | 0 |
| ENSMUSG00 Gm15603 | 491 | 0 | 0 | 0 | 0 | 0 | 0 |
| ENSMUSG00 Gm14830 | 699 | 0 | 0.06893626 | 0.37941496 | 0.61943424 | 0.07299911 | 0.03901951 |
| ENSMUSG00 Olfr29-ps1 | 947 | 0 | 0 | 0 | 0 | 0 | 0 |
| ENSMUSG00 Gm13445 | 306 | 0 | 0 | 0 | 0 | 0 | 0 |
| ENSMUSG00 Gm14282 | 239 | 0 | 0 | 0 | 0 | 0 | 0 |
| ENSMUSG00 Gm12882 | 516 | 0 | 0 | 0 | 0 | 0 | 0 |
| ENSMUSG00 Gm14774 | 535 | 0 | 0 | 0 | 0 | 0 | 0 |
| ENSMUSG00 Gm15714 | 249 | 0 | 0 | 0 | 0 | 0 | 0 |
| ENSMUSG00 Gm12382 | 921 | 0 | 0 | 0 | 0 | 0 | 0 |
| ENSMUSG00 Tpt1-ps2 | 520 | 0 | 0 | 0 | 0 | 0.09812765 | 0 |
| ENSMUSG00 Gm11603 | 245 | 0.64906907 | 0.59003807 | 1.08249412 | 0.5890946 | 0 | 0.66795035 |
| ENSMUSG00 Gm15483 | 455 | 0 | 0 | 0 | 0 | 0 | 0 |
| ENSMUSG00 Gm12593 | 742 | 0 | 0 | 0 | 0 | 0 | 0 |
| ENSMUSG00 Serpinb1-ps1 | 582 | 0 | 0 | 0 | 0 | 0 | 0 |
| ENSMUSG00 Gm14165 | 951 | 0 | 0 | 0.04647933 | 0 | 0.16096649 | 0 |
| ENSMUSG00 Gm11403 | 368 | 0 | 0 | 0 | 0 | 0 | 0 |
| ENSMUSG00 Gm5912 | 1437 | 0.08299683 | 0.03353267 | 0.03075981 | 0.02510929 | 0 | 0.03796053 |
| ENSMUSG00 Gm12402 | 333 | 0 | 0 | 0 | 0 | 0 | 0.24571747 |
| ENSMUSG00 Gm11254 | 652 | 0 | 0 | 0 | 0 | 0 | 0 |
| ENSMUSG00 Gm7840 | 532 | 0 | 0 | 0 | 0 | 0 | 0 |
| ENSMUSG00 Gm12464 | 1000 | 0.03975548 | 0.24093221 | 0.13260553 | 0.10824613 | 0.10205275 | 0.05454928 |
| ENSMUSG00 Gm6161 | 1990 | 0.03995526 | 0 | 0.02221198 | 0 | 0 | 0 |
| ENSMUSG00 Gm14682 | 523 | 0 | 0.09213469 | 0 | 0 | 0 | 0 |
| ENSMUSG00 Gm3880 | 377 | 0 | 0 | 0 | 0 | 0 | 0 |
| ENSMUSG00 Gm15803 | 393 | 0 | 0 | 0 | 0 | 0 | 0 |
| ENSMUSG00 Rpl23a-ps4 | 454 | 0 | 0 | 0 | 0 | 0 | 0 |
| ENSMUSG00 Gm14068 | 212 | 0 | 0 | 0 | 0 | 0.4813809 | 0 |
| ENSMUSG00 Gm13870 | 1022 | 0 | 0.09429832 | 0.17300134 | 0.10591598 | 0.09985592 | 0.1868126 |
| ENSMUSG00 Gm13086 | 2036 | 0 | 0 | 0 | 0 | 0 | 0 |
| ENSMUSG00 Gm11432 | 229 | 0 | 0 | 0 | 0 | 0 | 0 |
| ENSMUSG00 Gm11700 | 402 | 0 | 0 | 0 | 0 | 0 | 0 |
| ENSMUSG00 Gm12734 | 947 | 0 | 0 | 0 | 0.07620284 | 0.43105703 | 0.23040878 |
| ENSMUSG00 Gm14761 | 261 | 0 | 0 | 0 | 0 | 0 | 0 |
| ENSMUSG00 Gm6767 | 1380 | 0.17284992 | 0 | 0 | 0 | 0 | 0.05929269 |
| ENSMUSG00 Hmgb1-ps3 | 635 | 0.06260706 | 0.15176832 | 0 | 0 | 0 | 0.08590438 |
| ENSMUSG00 Gm7781 | 787 | 0.05051522 | 0 | 0 | 0.04584758 | 0 | 0 |
| ENSMUSG00 Gm13488 | 178 | 0 | 0 | 0 | 0 | 0 | 0 |
| ENSMUSG00 Gm14796 | 1001 | 0 | 0 | 0 | 0 | 0 | 0 |
| ENSMUSG00 Gm12952 | 389 | 0 | 0 | 0 | 0 | 0 | 0 |
| ENSMUSG00 Gm13168 | 631 | 0 | 0 | 0 | 0 | 0 | 0 |
| ENSMUSG00 Rpl30-ps11 | 344 | 0 | 0 | 0 | 0 | 0 | 0 |
| ENSMUSG00 Gm15049 | 462 | 0 | 0 | 0 | 0 | 0.55223351 | 0 |
| ENSMUSG00 Gm14732 | 544 | 0 | 0 | 0 | 0 | 0 | 0 |
| ENSMUSG00 Gm12884 | 324 | 0 | 0 | 0 | 0 | 0 | 0 |
| ENSMUSG00 Gm14912 | 1772 | 0 | 0 | 0 | 0 | 0 | 0 |
| ENSMUSG00 Gm13337 | 1593 | 0 | 0 | 0 | 0 | 0 | 0 |
| ENSMUSG00 Gm15088 | 1507 | 0 | 0 | 0 | 0 | 0 | 0 |
| ENSMUSG00 Gm11288 | 902 | 0 | 0 | 0 | 0 | 0 | 0 |
| ENSMUSG00 Gm14131 | 718 | 0 | 0 | 0 | 0 | 0 | 0 |
| ENSMUSG00 Gm14735 | 3087 | 0 | 0 | 0 | 0 | 0 | 0 |
| ENSMUSG00 Gm12620 | 1985 | 0 | 0 | 0 | 0 | 0 | 0 |
| ENSMUSG00 Gm4853 | 393 | 0 | 0 | 0 | 0 | 0 | 0 |
| ENSMUSG00 Gm12855 | 512 | 0 | 0 | 0 | 0 | 0 | 0 |
| ENSMUSG00 Gm14558 | 380 | 0 | 0 | 0 | 0 | 0 | 0 |
| ENSMUSG00 Itpa-ps1 | 596 | 0.60033444 | 0.32339894 | 1.33495499 | 0.36324206 | 0.42807362 | 0.64067945 |
| ENSMUSG00 Gm7916 | 452 | 0 | 0 | 0 | 0 | 0 | 0 |
| ENSMUSG00 AA623943 | 351 | 0 | 0 | 0 | 0 | 0 | 0 |
| ENSMUSG00 Gm13608 | 1405 | 0.14147858 | 0.9260028 | 0.25168309 | 0 | 0.14527082 | 0.13588788 |
| ENSMUSG00 Gm14293 | 576 | 0 | 0 | 0 | 0 | 0 | 0 |
| ENSMUSG00 Gm16404 | 963 | 0 | 0 | 0 | 0 | 0 | 0 |
| ENSMUSG00 Gm12445 | 882 | 0.09014848 | 0 | 0.20046187 | 0.0818187 | 0.11570607 | 0.12369451 |
| ENSMUSG00 Gm13492 | 664 | 0 | 0 | 0 | 0 | 0 | 0 |
| ENSMUSG00 Gm12338 | 191 | 1.04071939 | 0.25228504 | 0 | 0 | 0.2671538 | 1.28519243 |
| ENSMUSG00 Gm14452 | 1629 | 0 | 0 | 0 | 0 | 0 | 0 |
| ENSMUSG00 Gm13689 | 684 | 1.56929528 | 2.60657655 | 3.55424176 | 1.68804887 | 4.47599788 | 0.67787846 |
| ENSMUSG00 Gm15368 | 857 | 0 | 0 | 0 | 0 | 0 | 0 |
| ENSMUSG00 Gm14672 | 446 | 0 | 0 | 0 | 0 | 0 | 0 |
| ENSMUSG00 Gm11830 | 560 | 0 | 0.17209444 | 0 | 0 | 0 | 0 |
| ENSMUSG00 Gm12778 | 332 | 0 | 0 | 0 | 0 | 0 | 0 |
| ENSMUSG00 Gm9311 | 1267 | 0.15688824 | 0 | 0 | 0 | 0.20136691 | 0.12916167 |
| ENSMUSG00 Gm6113 | 631 | 0 | 0 | 0 | 0 | 0 | 0 |
| ENSMUSG00 Gm14130 | 986 | 0.04031996 | 0 | 0.04482946 | 0 | 0 | 0.02766191 |
| ENSMUSG00 Gm11304 | 364 | 0 | 0 | 0 | 0 | 0 | 0 |
| ENSMUSG00 Gm13428 | 364 | 0 | 0 | 0.60716817 | 0 | 2.80364702 | 0.97409426 |
| ENSMUSG00 Gm15560 | 642 | 0 | 0.52539734 | 0 | 0.61822818 | 0 | 0.33987089 |
| ENSMUSG00 Olfr407-ps1 | 844 | 0 | 0 | 0 | 0 | 0 | 0 |
| ENSMUSG00 Gm12517 | 424 | 0 | 0 | 0 | 0 | 1.32379749 | 0 |
| ENSMUSG00 Gm15233 | 399 | 0 | 0 | 0 | 0 | 0 | 0 |
| ENSMUSG00 Gm13286 | 536 | 0 | 0 | 0 | 0 | 0 | 0 |
| ENSMUSG00 Gm5646 | 712 | 0 | 0 | 0 | 0 | 0 | 0 |
| ENSMUSG00 Gm11214 | 1021 | 0 | 0.09439068 | 0.21646348 | 0.03533991 | 0.19990745 | 0.05342731 |
| ENSMUSG00 Gm12508 | 961 | 0 | 0 | 0 | 0 | 0 | 0 |
| ENSMUSG00 Gm8644 | 595 | 0 | 0 | 0 | 0 | 0 | 0 |
| ENSMUSG00 Amy2-ps1 | 1108 | 0 | 0 | 0 | 0 | 0 | 0 |
| ENSMUSG00 Gm14993 | 163 | 0 | 0 | 0 | 0 | 0 | 0 |
| ENSMUSG00 Gm12654 | 586 | 0 | 0 | 0 | 0 | 0 | 0 |
| ENSMUSG00 Gm14598 | 694 | 0 | 0 | 0 | 0.05199142 | 0 | 0 |
| ENSMUSG00 Gm12768 | 758 | 0 | 0 | 0 | 0 | 0 | 0 |
| ENSMUSG00 Gm11475 | 475 | 0 | 0 | 0 | 0 | 0 | 0 |
| ENSMUSG00 Gm15821 | 1207 | 0.06587486 | 0.39922487 | 0 | 0.11957596 | 0.16910149 | 0.29376165 |
| ENSMUSG00 Gm11623 | 1224 | 0 | 0 | 0 | 0 | 0 | 0 |
| ENSMUSG00 Gm14673 | 902 | 0 | 0 | 0 | 0 | 0 | 0 |
| ENSMUSG00 Gm12311 | 373 | 0 | 0 | 0 | 0 | 0 | 0 |
| ENSMUSG00 Gm12470 | 1048 | 0 | 0.04597943 | 0.04217733 | 0 | 0 | 0.02602542 |
| ENSMUSG00 Gm16204 | 1893 | 0 | 0 | 0 | 0 | 0 | 0 |
| ENSMUSG00 Gm12084 | 610 | 0 | 0 | 0 | 0 | 0 | 0 |

| | | | | | | | | |
|---|---|---|---|---|---|---|---|---|
| ENSMUSG00 | Gm13331 | 447 | 0 | 0 | 0 | 0 | 0 | 0 |
| ENSMUSG00 | Gm16200 | 330 | 0 | 0 | 0 | 0 | 0 | 0 |
| ENSMUSG00 | Mup-ps9 | 539 | 0 | 0 | 0 | 0 | 0 | 0 |
| ENSMUSG00 | Gm13170 | 642 | 0 | 0 | 0 | 0 | 0 | 0 |
| ENSMUSG00 | Gm13058 | 1432 | 0 | 0 | 0 | 0 | 0 | 0 |
| ENSMUSG00 | Gm11658 | 1007 | 0.03947913 | 0.38281185 | 0.13168374 | 0.42997471 | 0.40537339 | 0.02708504 |
| ENSMUSG00 | Gm13325 | 805 | 0.64201397 | 0 | 0 | 0 | 0 | 0 |
| ENSMUSG00 | Gm7219 | 296 | 0 | 0 | 0 | 0 | 0 | 0 |
| ENSMUSG00 | Gm12924 | 317 | 0 | 0.60803082 | 0.41831397 | 0.34147045 | 0 | 0.25811961 |
| ENSMUSG00 | Gm14884 | 195 | 0.61162278 | 0 | 0 | 0 | 0 | 1.11895956 |
| ENSMUSG00 | Gm12646 | 237 | 0 | 0 | 0 | 0 | 0 | 0 |
| ENSMUSG00 | Gm11310 | 1105 | 0 | 0 | 0 | 0 | 0 | 0 |
| ENSMUSG00 | Gm15781 | 322 | 0.12346423 | 0.14964734 | 0.27454561 | 0 | 0 | 2.71052316 |
| ENSMUSG00 | Gm15111 | 876 | 0 | 0 | 0 | 0 | 0 | 0 |
| ENSMUSG00 | Slc48a1 | 2874 | 80.2302669 | 114.446992 | 80.8367734 | 83.2247297 | 69.8816337 | 64.3430947 |
| ENSMUSG00 | Gm15453 | 464 | 0.08567992 | 0 | 0.28578778 | 0 | 0 | 0 |
| ENSMUSG00 | Gm15193 | 863 | 0 | 0 | 0 | 0 | 0 | 0 |
| ENSMUSG00 | Gm15235 | 193 | 0 | 0 | 0 | 0 | 0 | 0 |
| ENSMUSG00 | Gm13636 | 400 | 0 | 0 | 0 | 0 | 0 | 0 |
| ENSMUSG00 | Gm11990 | 896 | 0 | 0 | 0 | 0 | 0 | 0 |
| ENSMUSG00 | Gm12538 | 485 | 0.08197006 | 0.19870698 | 0.18227564 | 0.2231879 | 0.31562707 | 0.05623637 |
| ENSMUSG00 | Gm13940 | 300 | 0.79510961 | 0.80310737 | 1.47339477 | 0.72164089 | 0 | 0.63640825 |
| ENSMUSG00 | Gm12512 | 216 | 0 | 0 | 0 | 0 | 0 | 0 |
| ENSMUSG00 | Gm12733 | 766 | 0 | 0 | 0 | 0 | 0 | 0 |
| ENSMUSG00 | Gm13077 | 483 | 0 | 0 | 0 | 0 | 0 | 0 |
| ENSMUSG00 | Gm8346 | 1768 | 0 | 0 | 0 | 0 | 0 | 0 |
| ENSMUSG00 | Gm14927 | 453 | 0 | 0 | 0 | 0 | 0 | 0 |
| ENSMUSG00 | Gm11553 | 239 | 0 | 0 | 0 | 0 | 0 | 0 |
| ENSMUSG00 | Gm11692 | 900 | 0 | 0 | 0 | 0 | 0 | 0 |
| ENSMUSG00 | Gm15877 | 963 | 0 | 0 | 0 | 0 | 0 | 0 |
| ENSMUSG00 | Gm14654 | 446 | 0 | 0 | 0 | 0 | 0 | 0 |
| ENSMUSG00 | Gm12328 | 196 | 14.6040541 | 7.37547584 | 14.8842941 | 7.54777462 | 10.1532074 | 9.1843173 |
| ENSMUSG00 | Gm12699 | 940 | 0.08458613 | 0.05126217 | 0 | 0.11515546 | 0 | 0.08704672 |
| ENSMUSG00 | Gm7350 | 628 | 0 | 0 | 0 | 0 | 0 | 0 |
| ENSMUSG00 | Gm6427 | 978 | 0 | 0 | 0 | 0 | 0 | 0 |
| ENSMUSG00 | Gm15778 | 607 | 0 | 0.15876917 | 0 | 0.05944324 | 0.16812644 | 0.13480052 |
| ENSMUSG00 | Gm5697 | 2483 | 0.09606641 | 0.07762617 | 0.14241432 | 0.11625306 | 0.06165088 | 0.15378371 |
| ENSMUSG00 | Gm8320 | 1761 | 0 | 0 | 0 | 0 | 0 | 0 |
| ENSMUSG00 | Gm12411 | 482 | 0 | 0 | 0 | 0 | 0 | 0 |
| ENSMUSG00 | Gm9218 | 2692 | 0 | 0 | 0 | 0.01340343 | 0 | 0 |
| ENSMUSG00 | Gm12425 | 362 | 0 | 0 | 0 | 0 | 0 | 0 |
| ENSMUSG00 | Gm11575 | 227 | 0.87567138 | 0.63682523 | 1.94721775 | 2.38427607 | 0.67435739 | 1.44183115 |
| ENSMUSG00 | Gm12635 | 751 | 0 | 0 | 0 | 0 | 0 | 0 |
| ENSMUSG00 | Gm13717 | 364 | 0.10921835 | 0 | 0 | 0.0991265 | 0 | 0 |
| ENSMUSG00 | Rps12-ps5 | 388 | 0 | 0 | 0 | 0 | 0 | 0 |
| ENSMUSG00 | Gm13756 | 220 | 0 | 0 | 0 | 0 | 0 | 0 |
| ENSMUSG00 | Gm14838 | 247 | 0 | 0 | 0 | 0 | 0 | 0 |
| ENSMUSG00 | Gm15032 | 410 | 0 | 0 | 0 | 0 | 0 | 0 |
| ENSMUSG00 | Gm12490 | 559 | 0 | 0 | 0 | 0 | 0 | 0 |
| ENSMUSG00 | Gm12192 | 486 | 0 | 0 | 0 | 0 | 0 | 0 |
| ENSMUSG00 | Gm12125 | 735 | 0 | 0 | 0 | 0 | 0 | 0 |
| ENSMUSG00 | Gm14138 | 744 | 0.10686957 | 0 | 0 | 0 | 0 | 0 |
| ENSMUSG00 | Gm15164 | 270 | 0 | 0 | 0 | 0 | 0 | 0 |
| ENSMUSG00 | Gm14429 | 2021 | 0.01967119 | 2.24122987 | 0.3280691 | 1.37472411 | 1.23715607 | 0.24292108 |
| ENSMUSG00 | Gm12090 | 756 | 0 | 0.06373868 | 0 | 0 | 0.13499041 | 0 |
| ENSMUSG00 | Gm11279 | 1253 | 0 | 0 | 0 | 0 | 0 | 0 |
| ENSMUSG00 | Gm13455 | 516 | 0 | 0 | 0 | 0 | 0 | 0 |
| ENSMUSG00 | Gm12611 | 997 | 0 | 0.09666287 | 0 | 0 | 0 | 0 |
| ENSMUSG00 | Gm11909 | 758 | 0 | 0 | 0 | 0 | 0 | 0 |
| ENSMUSG00 | Gm14129 | 289 | 0 | 0 | 0 | 0 | 0 | 0 |
| ENSMUSG00 | Gm14312 | 145 | 0 | 0 | 0 | 0 | 0 | 0 |
| ENSMUSG00 | Gm14769 | 431 | 0 | 0 | 0 | 0 | 0 | 0 |
| ENSMUSG00 | Gm12716 | 1353 | 0.02938321 | 0 | 0.26135606 | 0.02666818 | 0 | 0 |
| ENSMUSG00 | Gm12926 | 408 | 0.29231971 | 0.11810403 | 0.32501355 | 0.17687277 | 0 | 0 |
| ENSMUSG00 | Gm11544 | 379 | 0 | 0 | 0 | 0 | 0 | 0 |
| ENSMUSG00 | Gm14926 | 506 | 0 | 0 | 0 | 0 | 0 | 0 |
| ENSMUSG00 | Gm11661 | 1326 | 0 | 0 | 0 | 0 | 0 | 0 |
| ENSMUSG00 | Gm14353 | 868 | 0 | 0 | 0 | 0 | 0 | 0 |
| ENSMUSG00 | Gm14413 | 404 | 0 | 0 | 0.21882101 | 0 | 0 | 0.13502297 |
| ENSMUSG00 | Gm13440 | 495 | 0 | 0 | 0 | 0 | 0 | 0 |
| ENSMUSG00 | Gm11936 | 159 | 0 | 0 | 0 | 0 | 0 | 0 |
| ENSMUSG00 | Gm11841 | 1962 | 0 | 0 | 0 | 0.07356176 | 0 | 0.02780289 |
| ENSMUSG00 | Gm15467 | 444 | 0.53723622 | 0.43411209 | 0.6968759 | 0.08126587 | 0 | 0.43000557 |
| ENSMUSG00 | Gm13036 | 747 | 0 | 0 | 0 | 0 | 0 | 0 |
| ENSMUSG00 | Gm15968 | 1167 | 0 | 0.04129087 | 0 | 0 | 0 | 0 |
| ENSMUSG00 | Gm14506 | 535 | 0 | 0 | 0.33048107 | 0 | 0.0953764 | 0 |
| ENSMUSG00 | Gm8312 | 617 | 0 | 0 | 0 | 0 | 0 | 0 |
| ENSMUSG00 | Gm14390 | 1692 | 0 | 0 | 0 | 0 | 0 | 0 |
| ENSMUSG00 | Gm12286 | 1978 | 0.06029648 | 0 | 0.04469347 | 0.36483362 | 0 | 0.05515599 |
| ENSMUSG00 | Gm13065 | 140 | 0 | 0.34418887 | 0 | 0 | 0 | 0 |
| ENSMUSG00 | Gm15369 | 393 | 0 | 0 | 0 | 0 | 0 | 0 |
| ENSMUSG00 | Gm14681 | 410 | 0.58178752 | 0.82269535 | 0.75466561 | 0.26401496 | 0.4978183 | 0.19957053 |
| ENSMUSG00 | Gm11518 | 410 | 0 | 0 | 0 | 0 | 0 | 0 |
| ENSMUSG00 | Gm15953 | 1731 | 0.57416927 | 0.52890953 | 0 | 0 | 0.02947798 | 0 |
| ENSMUSG00 | Gm9059 | 181 | 0 | 0 | 0 | 0 | 0 | 0 |
| ENSMUSG00 | Gm15294 | 982 | 0.0404842 | 0.14720909 | 0.31508442 | 0.03674343 | 0.05196169 | 0.05554916 |
| ENSMUSG00 | Gm16210 | 609 | 0 | 0.23737164 | 0 | 0.05924802 | 0 | 0.17914377 |
| ENSMUSG00 | Gm12939 | 164 | 0 | 0 | 0 | 0 | 0 | 0 |
| ENSMUSG00 | Gm13733 | 534 | 0 | 0 | 0 | 0 | 0 | 0 |
| ENSMUSG00 | Gm13651 | 1109 | 0.03584804 | 0 | 0.51814604 | 0.09760697 | 0.09202232 | 0.19675123 |
| ENSMUSG00 | Xlr5e-ps | 703 | 0 | 0 | 0 | 0 | 0 | 0 |
| ENSMUSG00 | Gm12109 | 696 | 0.05711994 | 0 | 0 | 0 | 0 | 0 |
| ENSMUSG00 | Gm9436 | 1854 | 0 | 0 | 0 | 0 | 0 | 0 |
| ENSMUSG00 | Gm12539 | 1096 | 0 | 0 | 0 | 0 | 0 | 0 |
| ENSMUSG00 | Gm14247 | 236 | 0 | 0 | 0 | 0 | 0 | 0 |
| ENSMUSG00 | Gm11601 | 367 | 0 | 0 | 0 | 0 | 0 | 0 |
| ENSMUSG00 | 5033423K11f | 626 | 0 | 0 | 0 | 0 | 0 | 0 |
| ENSMUSG00 | Gm13351 | 413 | 0.6738217 | 0 | 0.10702625 | 0.17473145 | 0.61775273 | 0 |
| ENSMUSG00 | Gm14633 | 469 | 0 | 0 | 0 | 0.23080199 | 0 | 0 |
| ENSMUSG00 | Gm12793 | 291 | 0 | 0 | 0 | 0 | 0 | 0 |
| ENSMUSG00 | Gm5937 | 642 | 0 | 0.07505676 | 0 | 0 | 0 | 0 |
| ENSMUSG00 | Gm8809 | 742 | 0 | 0 | 0 | 0 | 0 | 0 |
| ENSMUSG00 | Gm6912 | 542 | 0 | 0 | 0 | 0 | 0 | 0 |
| ENSMUSG00 | Rps12-ps7 | 384 | 0 | 0 | 0 | 0 | 0 | 0 |
| ENSMUSG00 | Gm14369 | 284 | 0 | 0 | 0 | 0 | 0 | 0 |
| ENSMUSG00 | Gm13158 | 735 | 0 | 0 | 0 | 0 | 0 | 0 |
| ENSMUSG00 | Gm14804 | 507 | 0 | 0 | 0 | 0 | 0 | 0 |
| ENSMUSG00 | Eif3s6-ps1 | 469 | 0 | 0 | 0 | 0 | 0 | 0 |
| ENSMUSG00 | Gm11498 | 948 | 0 | 0 | 0 | 0 | 0 | 0 |
| ENSMUSG00 | Gm12005 | 378 | 0 | 0 | 0 | 0 | 0.13499041 | 0 |
| ENSMUSG00 | Gm11443 | 478 | 0 | 0 | 0 | 0 | 0 | 0 |
| ENSMUSG00 | Gm8522 | 1928 | 0.08248025 | 0.09997187 | 0 | 0.05614426 | 0 | 0.0141466 |
| ENSMUSG00 | Gm13621 | 343 | 0 | 0 | 0 | 0 | 0 | 0 |
| ENSMUSG00 | Gm15071 | 1365 | 0 | 0 | 0.12952921 | 1.05734929 | 0 | 0.01998142 |
| ENSMUSG00 | Gm15901 | 851 | 0 | 0 | 0 | 0.50879499 | 0 | 0 |
| ENSMUSG00 | Gm14594 | 616 | 0 | 0 | 0 | 0 | 0 | 0 |
| ENSMUSG00 | Gm14973 | 254 | 0 | 0 | 0 | 0 | 0 | 0 |
| ENSMUSG00 | Rplp0-ps1 | 932 | 0 | 0 | 0 | 0 | 0 | 0 |
| ENSMUSG00 | Gm9439 | 969 | 0 | 0 | 0.13684781 | 0.07447274 | 0.1053176 | 0.05629441 |
| ENSMUSG00 | Gm13532 | 239 | 0 | 0 | 0 | 0 | 0 | 0 |
| ENSMUSG00 | Gm11605 | 645 | 0 | 0.07470766 | 0.06852999 | 0 | 0.07911066 | 0.08457253 |
| ENSMUSG00 | Gm15101 | 1501 | 0 | 0 | 0 | 0 | 0 | 0 |
| ENSMUSG00 | Gm15060 | 947 | 0 | 0 | 0 | 0 | 0 | 0 |
| ENSMUSG00 | Gm13022 | 284 | 0 | 0 | 0 | 0 | 0 | 0 |
| ENSMUSG00 | Gm13423 | 886 | 0 | 0.0543865 | 0.64855977 | 0.24434793 | 0.23036739 | 0.58489633 |
| ENSMUSG00 | Olfr213 | 983 | 0 | 0 | 0 | 0 | 0 | 0 |
| ENSMUSG00 | Gm16181 | 565 | 0 | 0 | 0.07823335 | 0.95793039 | 0 | 0.19309479 |
| ENSMUSG00 | Gm15530 | 554 | 0.0717608 | 0 | 0.07978672 | 0.32565022 | 0 | 1.03387622 |
| ENSMUSG00 | Gm14283 | 324 | 0 | 0 | 0 | 0 | 0 | 0 |
| ENSMUSG00 | Gm11354 | 421 | 0 | 0 | 0 | 0 | 0 | 0 |
| ENSMUSG00 | Gm5383 | 616 | 0 | 0 | 0 | 0 | 0 | 0 |
| ENSMUSG00 | Gm16445 | 712 | 0 | 0 | 0 | 0 | 0 | 0 |
| ENSMUSG00 | Gm11246 | 344 | 0 | 0.14007687 | 0.12849373 | 0.20977933 | 0 | 0 |
| ENSMUSG00 | Gm15466 | 332 | 0 | 0 | 0 | 0 | 0 | 0.16430506 |
| ENSMUSG00 | Gm12077 | 362 | 0 | 0 | 0 | 0 | 0 | 0 |
| ENSMUSG00 | Gm14917 | 2104 | 0 | 0 | 0 | 0 | 0 | 0 |
| ENSMUSG00 | Gm11787 | 1041 | 0 | 0 | 0 | 0 | 0.04901669 | 0 |
| ENSMUSG00 | Gm14450 | 239 | 0 | 0 | 0.92472475 | 0 | 0.64049844 | 0.11411983 |
| ENSMUSG00 | Gm14098 | 621 | 0 | 0 | 0 | 0 | 0.16433615 | 0.21960257 |
| ENSMUSG00 | Gm11207 | 413 | 0 | 0 | 0 | 0.78629153 | 0 | 0 |
| ENSMUSG00 | Gm15544 | 1530 | 0 | 0 | 0 | 0.02358304 | 0.06670115 | 0.01782656 |
| ENSMUSG00 | Gm15922 | 2033 | 0 | 0 | 0 | 0 | 0 | 0 |
| ENSMUSG00 | Gm11677 | 976 | 0 | 0 | 0 | 0 | 0 | 0 |
| ENSMUSG00 | Gm12883 | 360 | 0 | 0 | 0 | 0 | 0 | 0 |
| ENSMUSG00 | Gm7869 | 2576 | 0 | 0 | 0 | 0 | 0.01980838 | 0 |
| ENSMUSG00 | Npm3-ps1 | 527 | 0 | 0 | 0 | 0 | 0 | 0.1552636 |
| ENSMUSG00 | Gm15697 | 2898 | 0.89168607 | 0.76486416 | 0 | 0.46067483 | 0.12325212 | 2.85169624 |
| ENSMUSG00 | Gm13167 | 577 | 0 | 0 | 0 | 0 | 0 | 0 |
| ENSMUSG00 | Gm13000 | 495 | 0 | 0 | 0 | 0 | 0 | 0 |
| ENSMUSG00 | Gm14794 | 191 | 1.24886326 | 0.75685511 | 0.69426979 | 0.18891123 | 0 | 1.28519243 |
| ENSMUSG00 | Gm6184 | 1260 | 0 | 0 | 0 | 0 | 0 | 0 |
| ENSMUSG00 | Gm12712 | 800 | 0 | 0 | 0 | 0 | 0 | 0 |
| ENSMUSG00 | Gm11643 | 863 | 0 | 0 | 0 | 0 | 0.23650696 | 0 |
| ENSMUSG00 | Mdk-ps1 | 320 | 0 | 0 | 0 | 0 | 0 | 0 |
| ENSMUSG00 | Gm12959 | 402 | 0 | 0 | 0 | 0 | 0 | 0 |
| ENSMUSG00 | Gm11874 | 129 | 0 | 0 | 0 | 0 | 0 | 0 |
| ENSMUSG00 | Gm14473 | 556 | 0.07150266 | 0 | 0 | 0.19468729 | 0 | 0 |
| ENSMUSG00 | Gm14868 | 215 | 0 | 0 | 0 | 0 | 0 | 0 |
| ENSMUSG00 | Gm13812 | 1346 | 0 | 0 | 0 | 0 | 0 | 0 |
| ENSMUSG00 | Fzd10 | 3249 | 0 | 0 | 0 | 0 | 0 | 0 |
| ENSMUSG00 | Rps2-ps13 | 1255 | 0 | 0 | 0.03522059 | 0 | 0 | 0 |
| ENSMUSG00 | Gm12647 | 264 | 0 | 0 | 0 | 0 | 0 | 0 |
| ENSMUSG00 | Gm11774 | 292 | 0 | 0 | 0 | 0 | 0 | 0 |
| ENSMUSG00 | Gm13996 | 429 | 0 | 0 | 0 | 0 | 0 | 0 |
| ENSMUSG00 | Gm12499 | 541 | 0 | 0 | 0 | 0 | 0 | 0 |
| ENSMUSG00 | Gm9166 | 559 | 0 | 0 | 0 | 0 | 0 | 0 |
| ENSMUSG00 | Gm12317 | 318 | 0 | 0 | 0 | 0 | 0 | 0 |
| ENSMUSG00 | Gm11965 | 276 | 0 | 0 | 0 | 0 | 0 | 0.09882116 |
| ENSMUSG00 | Gm15971 | 210 | 5.49004255 | 1.14729624 | 0 | 0 | 0 | 0.12987924 |
| ENSMUSG00 | Gm13297 | 885 | 0 | 0 | 0 | 0 | 0 | 0 |
| ENSMUSG00 | Olfr660-ps1 | 374 | 0 | 0 | 0 | 0 | 0 | 0 |
| ENSMUSG00 | Rpsa-ps5 | 884 | 0 | 0 | 0 | 0 | 0 | 0 |
| ENSMUSG00 | Gm12717 | 978 | 0 | 0 | 0 | 0 | 0.05217421 | 0 |
| ENSMUSG00 | Gm14714 | 438 | 0 | 0 | 0 | 0 | 0 | 0 |
| ENSMUSG00 | Gm12329 | 1005 | 0 | 0 | 0 | 0 | 0 | 0 |
| ENSMUSG00 | Gm15213 | 215 | 0.36981842 | 0 | 0 | 0.67129385 | 0 | 0.12685879 |
| ENSMUSG00 | Gm14337 | 209 | 0.19021761 | 0 | 1.05746036 | 0 | 0 | 0.13050067 |
| ENSMUSG00 | Gm13890 | 389 | 0 | 0 | 0 | 0 | 0 | 0 |
| ENSMUSG00 | Gm5645 | 712 | 0 | 0 | 0 | 0 | 0 | 0 |
| ENSMUSG00 | Gm6285 | 497 | 0 | 0 | 0 | 0 | 0 | 0 |
| ENSMUSG00 | Gm14603 | 447 | 0 | 0 | 0 | 0 | 0 | 0 |
| ENSMUSG00 | Gm12582 | 696 | 0 | 0 | 0 | 0 | 0 | 0 |
| ENSMUSG00 | Gm7927 | 879 | 0 | 0 | 0 | 0 | 0.17415145 | 0 |
| ENSMUSG00 | Gm11343 | 598 | 0 | 0 | 0 | 0 | 0 | 0 |
| ENSMUSG00 | Gm15022 | 745 | 0 | 0 | 0 | 0 | 0 | 0 |
| ENSMUSG00 | Gm13240 | 564 | 0.21146532 | 0.17087391 | 0 | 0.12795051 | 0.1809446 | 0 |
| ENSMUSG00 | Gm11785 | 600 | 0 | 0 | 0 | 0 | 0 | 0 |
| ENSMUSG00 | Gm12096 | 827 | 0 | 0.75746523 | 0 | 0.17452017 | 0.30850288 | 0.09894065 |
| ENSMUSG00 | Gm12170 | 299 | 0 | 0 | 0 | 0 | 0 | 0 |
| ENSMUSG00 | Gm12782 | 726 | 0 | 0 | 0 | 0 | 0 | 0 |
| ENSMUSG00 | Gm11828 | 485 | 0.32788025 | 0.09935349 | 1.09365385 | 0 | 0 | 0.05623637 |
| ENSMUSG00 | Gm14839 | 709 | 0 | 0.40778371 | 0 | 0.10178292 | 0 | 0 |
| ENSMUSG00 | Gm11303 | 821 | 0 | 0 | 0 | 0 | 0 | 0 |
| ENSMUSG00 | Gm8261 | 1207 | 0.16468716 | 0.07984498 | 0 | 0.05978798 | 0 | 0.0451941 |
| ENSMUSG00 | Gm13726 | 793 | 0 | 0 | 0 | 0 | 0 | 0 |
| ENSMUSG00 | Gm12391 | 280 | 0 | 0 | 0 | 0 | 0 | 0 |
| ENSMUSG00 | Gm13882 | 2232 | 0.03562319 | 0.02158891 | 0.03960739 | 0.04849737 | 0.02286128 | 0.03665946 |
| ENSMUSG00 | Gm11763 | 311 | 0.25566225 | 0 | 0 | 0 | 0 | 0 |
| ENSMUSG00 | Gm15931 | 1844 | 0 | 0 | 0 | 0 | 0 | 0 |
| ENSMUSG00 | Olfr129 | 1618 | 0 | 0 | 0 | 0 | 0 | 0 |
| ENSMUSG00 | Gm13202 | 461 | 2.93207448 | 3.44935486 | 4.31471353 | 2.50461047 | 1.66029422 | 2.07074268 |
| ENSMUSG00 | Gm14930 | 651 | 0 | 0 | 0 | 0 | 0 | 0 |
| ENSMUSG00 | Hspa9-ps1 | 1932 | 0.45270216 | 0.49882445 | 0.57197002 | 0.59763221 | 0.47540102 | 0.32469809 |
| ENSMUSG00 | Gm13785 | 689 | 0 | 0 | 0 | 0 | 0 | 0 |
| ENSMUSG00 | Gm14078 | 484 | 0 | 0 | 0.09132612 | 0.07454968 | 0 | 0.05635256 |
| ENSMUSG00 | Gm15495 | 1311 | 0.15162273 | 0.29404389 | 0.77547093 | 0.57797325 | 0.31137377 | 0.37448017 |
| ENSMUSG00 | Gm12852 | 718 | 0 | 0 | 0 | 0 | 0.21320213 | 0.07597393 |
| ENSMUSG00 | Gm12740 | 307 | 0 | 0 | 0 | 0 | 0 | 0 |
| ENSMUSG00 | Gm14530 | 332 | 0 | 0 | 0 | 0 | 0 | 0 |
| ENSMUSG00 | Gm14971 | 662 | 0 | 0 | 0 | 0 | 0 | 0 |
| ENSMUSG00 | Olfr359-ps1 | 943 | 0 | 0 | 0 | 0 | 0 | 0 |
| ENSMUSG00 | Hmgb1-ps2 | 635 | 0 | 0 | 0 | 0 | 0 | 0.04295219 |
| ENSMUSG00 | Mdm4-ps | 6513 | 0 | 0 | 0 | 0.02216002 | 0.02350363 | 0.03768951 |
| ENSMUSG00 | Gm14279 | 663 | 0 | 0 | 0 | 0 | 0 | 0 |
| ENSMUSG00 | Gm14106 | 437 | 0 | 0 | 0 | 0 | 0 | 0 |
| ENSMUSG00 | Gm9051 | 1033 | 0 | 0 | 0 | 0.03492938 | 0 | 0 |
| ENSMUSG00 | Gm4911 | 944 | 0 | 0 | 0 | 0 | 0 | 0 |
| ENSMUSG00 | Gm5721 | 1695 | 0 | 0 | 0 | 0 | 0 | 0 |
| ENSMUSG00 | Gm15549 | 621 | 0 | 0 | 0 | 0 | 0.08216808 | 0 |
| ENSMUSG00 | Gm13925 | 241 | 0 | 0 | 0 | 0 | 0 | 0 |
| ENSMUSG00 | Rpl7a-ps9 | 776 | 0 | 0 | 0 | 0 | 0 | 0 |
| ENSMUSG00 | Gm14870 | 155 | 1.28243486 | 0 | 0.28517318 | 0 | 0.65840485 | 0 |
| ENSMUSG00 | Gm12241 | 1330 | 0 | 0 | 0 | 0 | 0 | 0 |
| ENSMUSG00 | Gm4995 | 431 | 0.09224009 | 0 | 0 | 0 | 0 | 0 |
| ENSMUSG00 | Gm14251 | 278 | 0.14300533 | 0 | 0 | 0 | 0 | 0 |
| ENSMUSG00 | Gm14680 | 1100 | 0 | 0 | 0 | 0.06560372 | 0 | 0.07438538 |
| ENSMUSG00 | Gm12626 | 1982 | 0 | 0 | 0 | 0 | 0 | 0 |
| ENSMUSG00 | Gm12623 | 1982 | 0 | 0 | 0 | 0 | 0 | 0 |
| ENSMUSG00 | Gm12574 | 628 | 0 | 0 | 0 | 0 | 0 | 0 |
| ENSMUSG00 | Gm12291 | 310 | 0 | 0 | 0 | 0 | 0 | 0 |
| ENSMUSG00 | Gm13798 | 138 | 0 | 0.34917712 | 0.64060642 | 0 | 0.73951269 | 0 |
| ENSMUSG00 | Gm11460 | 360 | 0.66259134 | 0 | 0 | 0.40091161 | 0.28347987 | 0.15152577 |
| ENSMUSG00 | Gm6298 | 975 | 0 | 0.34595394 | 0 | 0 | 0 | 0 |
| ENSMUSG00 | Gm13738 | 468 | 0 | 0 | 0 | 0 | 0 | 0 |
| ENSMUSG00 | Gm9435 | 876 | 0 | 0 | 0 | 0 | 0 | 0 |
| ENSMUSG00 | Gm14677 | 485 | 0 | 0 | 0 | 0 | 0 | 0 |
| ENSMUSG00 | 2210409E12f | 345 | 0 | 0 | 0 | 0 | 0 | 0 |
| ENSMUSG00 | Gm12067 | 1146 | 0 | 0 | 0 | 0 | 0 | 0 |
| ENSMUSG00 | Gm12755 | 287 | 0 | 0 | 0 | 0 | 0 | 0 |
| ENSMUSG00 | Gm12216 | 2636 | 17.9171134 | 13.5090215 | 15.6282692 | 10.5672756 | 13.4341065 | 20.1145291 |
| ENSMUSG00 | Rbpsuh-ps1 | 1005 | 0 | 0 | 0 | 0 | 0 | 0 |
| ENSMUSG00 | Gm15381 | 1155 | 0 | 0 | 0 | 0.65603717 | 0 | 0 |
| ENSMUSG00 | Gm13131 | 320 | 0.12423588 | 2.55990474 | 0 | 0 | 0.31891485 | 1.36373196 |
| ENSMUSG00 | Gm8302 | 851 | 0 | 0 | 0 | 0 | 0 | 0 |
| ENSMUSG00 | Gm13866 | 967 | 0 | 0 | 0 | 0 | 0 | 0 |
| ENSMUSG00 | Gm12420 | 1047 | 0.11391255 | 0.09204669 | 0.71769946 | 0.13784926 | 0.04873579 | 0.26050276 |
| ENSMUSG00 | Gm14159 | 450 | 0 | 0 | 0 | 0 | 0 | 0 |
| ENSMUSG00 | Gm6325 | 3095 | 0 | 0 | 0 | 0.02331635 | 0 | 0 |
| ENSMUSG00 | Gm5126 | 1307 | 0 | 0 | 0 | 0 | 0 | 0 |
| ENSMUSG00 | Gm14753 | 245 | 0.32453454 | 0 | 0 | 0 | 0.20827092 | 0 |
| ENSMUSG00 | Gm15152 | 392 | 0 | 0 | 0 | 0 | 0 | 0 |
| ENSMUSG00 | Gm15599 | 773 | 0 | 0 | 0.0571822 | 0 | 0 | 0 |
| ENSMUSG00 | Gm13281 | 566 | 0 | 0 | 0 | 0 | 0 | 0 |
| ENSMUSG00 | Gm12424 | 446 | 0 | 0.2160827 | 0 | 0 | 0 | 0 |
| ENSMUSG00 | Gm11897 | 827 | 0 | 0 | 0 | 0 | 0 | 0 |
| ENSMUSG00 | Gm13991 | 647 | 0 | 0 | 0.06831815 | 0 | 0.23659834 | 0 |
| ENSMUSG00 | Gm5898 | 2070 | 0.26887765 | 0.27934169 | 0.14947483 | 0.13944752 | 0.14790254 | 0.07905693 |
| ENSMUSG00 | Gm11893 | 490 | 0 | 0 | 0 | 0 | 0 | 0 |
| ENSMUSG00 | Gm14596 | 532 | 0 | 0 | 0 | 0 | 0 | 0 |
| ENSMUSG00 | Gm12190 | 186 | 0 | 0 | 0 | 0 | 0 | 0 |
| ENSMUSG00 | Anp32b-ps1 | 2088 | 0.01903998 | 0 | 0 | 0.01728067 | 0.14662752 | 0.01306257 |
| ENSMUSG00 | 5730416F02f | 811 | 0 | 0 | 0.43602311 | 0 | 0 | 0.10089262 |
| ENSMUSG00 | Gm14590 | 737 | 0 | 0 | 0 | 0 | 0 | 0 |
| ENSMUSG00 | Gm9005 | 606 | 0 | 0 | 0 | 0 | 0 | 0 |
| ENSMUSG00 | Gm8662 | 573 | 0 | 0 | 0 | 0 | 0 | 0 |
| ENSMUSG00 | Gm8152 | 636 | 0 | 0 | 0 | 0 | 0 | 0 |
| ENSMUSG00 | Gm14071 | 508 | 0.07825882 | 0 | 0 | 0 | 0 | 0 |
| ENSMUSG00 | Gm14200 | 639 | 0 | 0 | 0 | 0 | 0 | 0.08536863 |
| ENSMUSG00 | Dnmt3l-ps1 | 1252 | 0.47630368 | 0 | 0 | 0.57639049 | 0.4483148 | 1.11102764 |
| ENSMUSG00 | Gm15282 | 299 | 0 | 0 | 0 | 0 | 0 | 0 |
| ENSMUSG00 | Xlr5d-ps | 773 | 0 | 0 | 0 | 0 | 0 | 0 |
| ENSMUSG00 | Gm14335 | 809 | 0.04914151 | 0 | 0.43710104 | 0.84741514 | 0.31536697 | 0 |
| ENSMUSG00 | Gm14901 | 282 | 0 | 0 | 0 | 0 | 0 | 0 |
| ENSMUSG00 | Gm14849 | 2844 | 0 | 0 | 0.01554214 | 0 | 0 | 0 |
| ENSMUSG00 | Gm12622 | 381 | 0.20869019 | 0.1264736 | 0 | 0.37881412 | 0.1339275 | 0 |
| ENSMUSG00 | Gm15539 | 565 | 0 | 0.17057148 | 0 | 0.19158608 | 0 | 0.0965474 |
| ENSMUSG00 | Gm11878 | 399 | 0 | 0 | 0 | 0 | 0 | 0 |
| ENSMUSG00 | Gm12713 | 762 | 0 | 0 | 0 | 0 | 0 | 0 |
| ENSMUSG00 | Gm4789 | 1215 | 0 | 0 | 0 | 0 | 0 | 0 |
| ENSMUSG00 | Gm12806 | 267 | 0 | 0 | 0 | 0 | 0 | 0 |
| ENSMUSG00 | Gm14695 | 279 | 0 | 0 | 0 | 0 | 0 | 0 |
| ENSMUSG00 | Gm8303 | 1761 | 0 | 0 | 0 | 0 | 0 | 0 |
| ENSMUSG00 | Gm15528 | 863 | 0 | 0 | 0.05121882 | 0.29267012 | 0 | 0.22123114 |
| ENSMUSG00 | Gm13590 | 1587 | 0.72647066 | 0 | 0.58490151 | 1.7279366 | 0.48229089 | 0.73901039 |
| ENSMUSG00 | Gm12652 | 337 | 0 | 0 | 0 | 0 | 0 | 0 |
| ENSMUSG00 | Gm12722 | 492 | 0.16160764 | 0.39175969 | 0 | 0.22001247 | 0 | 0 |
| ENSMUSG00 | Gm12029 | 831 | 0 | 0 | 0 | 0 | 0 | 0 |
| ENSMUSG00 | Gm15158 | 569 | 0 | 0 | 0 | 0 | 0 | 0 |
| ENSMUSG00 | Gm15626 | 867 | 0 | 0 | 0 | 0 | 0 | 0 |
| ENSMUSG00 | Cyp4a28-ps | 1465 | 0 | 0.03289177 | 0 | 0 | 0 | 0 |
| ENSMUSG00 | BC002163 | 638 | 0.49850132 | 0.07552734 | 0.27712754 | 0.22621972 | 0 | 0.38475196 |
| ENSMUSG00 | Rpl23a-ps12 | 467 | 0 | 0 | 0 | 0 | 0 | 0 |
| ENSMUSG00 | Gm15103 | 869 | 0 | 0 | 0 | 0 | 0 | 0 |
| ENSMUSG00 | Gm11420 | 401 | 0 | 0 | 0 | 0 | 0 | 0 |
| ENSMUSG00 | Gm15460 | 390 | 0 | 0 | 0 | 0.37007225 | 0 | 0.13986995 |
| ENSMUSG00 | Gm13475 | 233 | 0 | 0 | 0 | 0 | 0 | 0 |
| ENSMUSG00 | Gm7405 | 1335 | 0 | 0 | 0 | 0 | 0 | 0 |
| ENSMUSG00 | Gm6897 | 137 | 0 | 0 | 0 | 0 | 0 | 0 |
| ENSMUSG00 | Gm13669 | 456 | 0 | 0 | 0 | 0 | 0 | 0 |
| ENSMUSG00 | Gm13770 | 304 | 0 | 0 | 0 | 0 | 0 | 0 |
| ENSMUSG00 | Gm11790 | 454 | 0 | 0 | 0 | 0 | 0 | 0 |
| ENSMUSG00 | Olfr1192-ps1 | 929 | 0 | 0 | 0 | 0 | 0 | 0 |
| ENSMUSG00 | Gm14649 | 323 | 0 | 0 | 0 | 0 | 0 | 0 |
| ENSMUSG00 | Gm13038 | 458 | 0 | 0.42084229 | 0.28953172 | 0 | 0 | 0 |
| ENSMUSG00 | Gm13454 | 482 | 0 | 0 | 0 | 0 | 0 | 0 |
| ENSMUSG00 | Gm15220 | 378 | 0 | 0 | 0 | 0 | 0 | 0 |
| ENSMUSG00 | Gm13444 | 521 | 0 | 0 | 0 | 0 | 0 | 0 |
| ENSMUSG00 | Rap1a-ps2 | 558 | 0 | 0 | 0.15842955 | 0 | 0 | 0 |
| ENSMUSG00 | Gm7851 | 322 | 0 | 0.14964734 | 0 | 0 | 0 | 0.08470385 |
| ENSMUSG00 | Gm11241 | 895 | 0 | 0 | 0 | 0 | 0 | 0 |
| ENSMUSG00 | Prkaca-ps1 | 918 | 0 | 0 | 0 | 0 | 0 | 0 |
| ENSMUSG00 | Gm12142 | 483 | 0 | 0 | 0 | 0 | 0 | 0 |
| ENSMUSG00 | Gm14925 | 437 | 0 | 0 | 0 | 0 | 0 | 0 |
| ENSMUSG00 | Gm15847 | 1230 | 0 | 0 | 0 | 0 | 0 | 0 |
| ENSMUSG00 | Gm15667 | 781 | 0.10180661 | 0.18509517 | 0.22638588 | 0.1847992 | 0 | 0.20953628 |
| ENSMUSG00 | Gm13323 | 1150 | 0 | 0 | 0 | 0 | 0 | 0 |
| ENSMUSG00 | Gm12281 | 297 | 1.4724252 | 0 | 0 | 0 | 0 | 0 |
| ENSMUSG00 | Gm13735 | 669 | 0 | 0 | 0 | 0 | 0 | 0 |
| ENSMUSG00 | Gm11447 | 857 | 0.13916738 | 0 | 0.10315483 | 0.75784924 | 0 | 0 |
| ENSMUSG00 | Rpl17-ps5 | 1157 | 0.20616498 | 0 | 0.07640768 | 0 | 0 | 0 |
| ENSMUSG00 | Gm14580 | 717 | 0 | 0 | 0 | 0 | 0 | 0 |
| ENSMUSG00 | Gm13624 | 461 | 0 | 0.10452591 | 0 | 0 | 0 | 0 |
| ENSMUSG00 | Gm14514 | 938 | 0 | 0 | 0 | 0 | 0 | 0 |
| ENSMUSG00 | Olfr650-ps1 | 995 | 0 | 0 | 0 | 0 | 0 | 0 |
| ENSMUSG00 | Gm11533 | 476 | 0 | 0 | 0 | 0 | 0 | 0.11459933 |
| ENSMUSG00 | Gm14441 | 539 | 0 | 0.08939971 | 0.08200713 | 0 | 0.1893372 | 0 |
| ENSMUSG00 | Gm12413 | 311 | 0 | 0 | 0 | 0 | 0 | 0 |
| ENSMUSG00 | Gm12780 | 644 | 0 | 0 | 0 | 0 | 0 | 0 |
| ENSMUSG00 | Olfr405-ps1 | 1644 | 0 | 0 | 0 | 0 | 0 | 0 |
| ENSMUSG00 | Gm15484 | 827 | 0 | 0.05826656 | 0 | 0.21815021 | 0 | 0 |
| ENSMUSG00 | Gm12401 | 227 | 0 | 0 | 0 | 0 | 0 | 0 |
| ENSMUSG00 | Gm12039 | 179 | 0 | 0.269198 | 0 | 0 | 0.5701271 | 0 |
| ENSMUSG00 | Gm15002 | 450 | 0 | 0 | 0 | 0 | 0 | 0 |
| ENSMUSG00 | Gm15141 | 727 | 0 | 0 | 0 | 0 | 0 | 0 |
| ENSMUSG00 | Gm11488 | 862 | 0.83016085 | 0.05590075 | 0.2563912 | 0.04185852 | 0 | 0.18984668 |
| ENSMUSG00 | Gm15572 | 592 | 0 | 0 | 0 | 0 | 0 | 0 |
| ENSMUSG00 | Gm15197 | 477 | 0 | 0 | 0.92666338 | 0.3782185 | 0 | 1.65820658 |
| ENSMUSG00 | Gm14983 | 322 | 0 | 0 | 0 | 0 | 0 | 0 |
| ENSMUSG00 | Gm15223 | 825 | 0 | 0 | 0 | 0 | 0 | 0 |
| ENSMUSG00 | Gm12644 | 605 | 0 | 0 | 0 | 0 | 0 | 0 |
| ENSMUSG00 | Gm8858 | 341 | 0 | 0 | 0 | 0 | 0 | 0 |
| ENSMUSG00 | Gm13739 | 529 | 0 | 0 | 0 | 0 | 0 | 0 |
| ENSMUSG00 | Gm11933 | 374 | 0 | 0 | 0 | 0 | 0 | 0 |
| ENSMUSG00 | Gm5385 | 1558 | 0 | 0 | 0 | 0 | 0 | 0 |
| ENSMUSG00 | Gm14752 | 2070 | 0 | 0 | 0 | 0 | 0 | 0 |
| ENSMUSG00 | Gm14294 | 750 | 0 | 0 | 0 | 0 | 0 | 0 |
| ENSMUSG00 | Gm15574 | 403 | 0 | 0 | 0 | 0 | 0 | 0 |
| ENSMUSG00 | Gm11821 | 978 | 0.20324888 | 0.34489274 | 0.04519616 | 0.40583077 | 0 | 0 |
| ENSMUSG00 | Gm13231 | 2848 | 0 | 0 | 0 | 0 | 0 | 0 |
| ENSMUSG00 | Gm6175 | 399 | 0 | 0.36230408 | 0.44312625 | 0 | 0 | 0.06835749 |
| ENSMUSG00 | Gm12097 | 1412 | 0.1407772 | 0.71665389 | 0.06260884 | 1.30324665 | 0.83116618 | 0.71470372 |
| ENSMUSG00 | Gm14045 | 680 | 0 | 0.14172483 | 0 | 0 | 0 | 0.20054882 |
| ENSMUSG00 | Gm13188 | 563 | 0 | 0 | 0 | 0 | 0 | 0 |
| ENSMUSG00 | Gm11243 | 393 | 0 | 0 | 0 | 0 | 0 | 0 |
| ENSMUSG00 | Gm14660 | 574 | 0 | 0 | 0 | 0 | 0 | 0 |
| ENSMUSG00 | Gm8864 | 1091 | 0.07287897 | 0.13250167 | 0.08102996 | 0.13228981 | 0.04677028 | 0 |
| ENSMUSG00 | Gm13273 | 156 | 0 | 0 | 0 | 0 | 0 | 0 |
| ENSMUSG00 | Gm14212 | 825 | 0.38550769 | 0 | 0.58935791 | 0 | 0 | 0.03306017 |
| ENSMUSG00 | Gm10294 | 1157 | 0 | 0 | 0 | 0.03118586 | 0 | 0.02357359 |
| ENSMUSG00 | Gm5389 | 1175 | 0 | 0 | 0.03761859 | 0 | 0.0434267 | 0.06963738 |
| ENSMUSG00 | Gm15067 | 155 | 0 | 0 | 0 | 0 | 0 | 0 |
| ENSMUSG00 | Gm5754 | 891 | 0 | 0 | 0 | 0 | 0 | 0 |
| ENSMUSG00 | Gm13606 | 948 | 0 | 0 | 0 | 0 | 0 | 0 |
| ENSMUSG00 | Gm14985 | 785 | 0 | 0 | 0 | 0 | 0 | 0 |
| ENSMUSG00 | Gm15357 | 283 | 0 | 0 | 0 | 0 | 0 | 0 |
| ENSMUSG00 | Gm12453 | 330 | 0 | 0 | 0 | 0 | 0 | 0 |
| ENSMUSG00 | Gm11387 | 768 | 0 | 0 | 0 | 0 | 0 | 0 |
| ENSMUSG00 | Gm13419 | 180 | 0 | 0 | 0 | 0 | 0 | 0 |
| ENSMUSG00 | Gm11370 | 259 | 0 | 0 | 0 | 0 | 0 | 0 |
| ENSMUSG00 | Rpl9-ps1 | 575 | 0 | 0 | 0 | 0 | 0 | 0 |
| ENSMUSG00 | Gm13731 | 195 | 0.20387426 | 0.24710996 | 0.45335224 | 0 | 0.52334744 | 0 |
| ENSMUSG00 | Olfr1147-ps1 | 936 | 0 | 0 | 0 | 0 | 0 | 0 |
| ENSMUSG00 | Gm11341 | 446 | 0 | 0 | 0 | 0 | 0 | 0 |
| ENSMUSG00 | Gm15986 | 399 | 0 | 0 | 0 | 0 | 0 | 0 |
| ENSMUSG00 | Mrps36-ps2 | 308 | 0 | 0 | 0.28702496 | 0 | 0 | 0 |
| ENSMUSG00 | Gm11326 | 477 | 0 | 0 | 0 | 0 | 0 | 0 |
| ENSMUSG00 | Gm11803 | 443 | 0 | 0 | 0 | 0 | 0 | 0 |
| ENSMUSG00 | Gm11305 | 724 | 0 | 0 | 0 | 0 | 0 | 0 |
| ENSMUSG00 | Gm15174 | 1322 | 0 | 0 | 0 | 0 | 0 | 0 |
| ENSMUSG00 | Gm12899 | 550 | 0 | 0 | 0 | 0 | 0 | 0 |
| ENSMUSG00 | Gm15209 | 399 | 0 | 0 | 0 | 0.09043119 | 0 | 0 |
| ENSMUSG00 | Gm14613 | 457 | 0 | 0 | 0 | 0 | 0 | 0 |
| ENSMUSG00 | Gm12399 | 444 | 0 | 0 | 0 | 0 | 0 | 0 |
| ENSMUSG00 | Gm15359 | 590 | 0 | 0 | 0 | 0 | 0 | 0 |
| ENSMUSG00 | Gm12858 | 207 | 1.15233277 | 0 | 0.64060642 | 0 | 0.73951269 | 0.39528463 |
| ENSMUSG00 | Gm14646 | 131 | 0 | 0 | 0 | 0 | 0 | 0 |
| ENSMUSG00 | Gm9062 | 726 | 0 | 0 | 0 | 0 | 0 | 0 |
| ENSMUSG00 | Gm11191 | 607 | 0 | 0 | 0 | 0 | 0 | 0 |
| ENSMUSG00 | Gm15771 | 336 | 0 | 0 | 0 | 0 | 0 | 0 |
| ENSMUSG00 | Gm15536 | 113 | 0.70363682 | 0 | 0.39116675 | 0.95793039 | 0 | 0 |
| ENSMUSG00 | Gm15105 | 203 | 0 | 0 | 0 | 0 | 0 | 0 |
| ENSMUSG00 | Gm11741 | 455 | 0 | 1.05904268 | 0 | 0 | 0 | 0 |
| ENSMUSG00 | Gm12345 | 897 | 0 | 0 | 0 | 0 | 0 | 0 |
| ENSMUSG00 | Gm15211 | 525 | 0 | 0 | 0 | 0 | 0 | 0 |
| ENSMUSG00 | Gm12331 | 621 | 0 | 0 | 0 | 0.23241253 | 0 | 0.21960257 |
| ENSMUSG00 | Gm14767 | 585 | 0 | 0 | 0 | 0 | 0 | 0 |
| ENSMUSG00 | Gm12193 | 378 | 0 | 0 | 0 | 0 | 0 | 0.07215513 |
| ENSMUSG00 | Gm16186 | 214 | 0 | 0 | 0 | 0 | 0 | 0 |
| ENSMUSG00 | Gm11244 | 236 | 0 | 0 | 0 | 0 | 0 | 0 |
| ENSMUSG00 | Gm14599 | 470 | 0 | 0 | 0 | 0 | 0 | 0 |
| ENSMUSG00 | Gm15039 | 874 | 0 | 0 | 0 | 0 | 0 | 0.03120668 |
| ENSMUSG00 | Gm15399 | 675 | 0 | 0 | 0 | 0 | 0 | 0 |
| ENSMUSG00 | Xlr3e-ps | 674 | 0 | 0 | 0 | 0 | 0 | 0 |
| ENSMUSG00 | Gm11218 | 1213 | 0 | 0 | 0 | 0 | 0 | 0 |
| ENSMUSG00 | Gm15149 | 359 | 0 | 0 | 0 | 0 | 0 | 0 |
| ENSMUSG00 | Gm9078 | 615 | 0 | 0 | 0 | 0.17600997 | 0 | 0 |
| ENSMUSG00 | Gm12276 | 300 | 6.22835861 | 3.6942909 | 2.06275267 | 4.93121275 | 3.74193423 | 1.54556289 |
| ENSMUSG00 | Gm15121 | 966 | 0 | 0 | 0 | 0 | 0 | 0 |
| ENSMUSG00 | Gm11472 | 555 | 0 | 0 | 0 | 0 | 0 | 0 |
| ENSMUSG00 | Gm14512 | 175 | 0 | 0 | 0 | 0 | 0 | 0 |
| ENSMUSG00 | Olfr1191-ps1 | 1911 | 0 | 0 | 0 | 0 | 0 | 0 |
| ENSMUSG00 | Gm14392 | 397 | 0 | 0 | 0 | 0 | 0 | 0 |
| ENSMUSG00 | Gm11600 | 238 | 0 | 0 | 0 | 0 | 0 | 0 |
| ENSMUSG00 | Gm15352 | 269 | 0.14778989 | 0.17913176 | 0 | 0.26826799 | 0.37937826 | 0.20278542 |
| ENSMUSG00 | Gm12178 | 329 | 0 | 0 | 0 | 0 | 0.31019073 | 0.91191803 |
| ENSMUSG00 | Gm12435 | 1015 | 0.03916796 | 0 | 0 | 0 | 0.10054458 | 0 |
| ENSMUSG00 | Gm13626 | 399 | 0 | 0 | 0 | 0 | 0 | 0 |
| ENSMUSG00 | Olfr909-ps1 | 930 | 0 | 0 | 0 | 0 | 0 | 0 |
| ENSMUSG00 | Ak3l2-ps | 671 | 0 | 0 | 0 | 0 | 0 | 0 |
| ENSMUSG00 | Gm11594 | 161 | 0 | 0 | 0 | 0 | 0 | 0 |
| ENSMUSG00 | Gm16480 | 831 | 0 | 0 | 0 | 0 | 0 | 0 |
| ENSMUSG00 | Mageb17-ps | 918 | 0 | 0 | 0 | 0 | 0 | 0 |
| ENSMUSG00 | Olfr383-ps1 | 934 | 0 | 0 | 0 | 0 | 0 | 0 |
| ENSMUSG00 | Gm12773 | 586 | 0 | 0 | 0 | 0 | 0 | 0 |
| ENSMUSG00 | Gm8366 | 1768 | 0 | 0 | 0 | 0 | 0 | 0 |
| ENSMUSG00 | Gm11873 | 1303 | 0 | 0 | 0 | 0.38768122 | 0 | 0 |
| ENSMUSG00 | Gm11620 | 1844 | 0 | 0 | 0 | 0 | 0 | 0 |
| ENSMUSG00 | Gm13900 | 497 | 0 | 0 | 0 | 0 | 0 | 0 |
| ENSMUSG00 | Gm14017 | 385 | 0.10326099 | 0 | 0 | 0.18743919 | 0.39760812 | 0.14168644 |
| ENSMUSG00 | Rpl23a-ps2 | 473 | 0.42024821 | 0.81499268 | 1.40174978 | 1.60195124 | 0.43151269 | 0.92260936 |
| ENSMUSG00 | Gm16471 | 617 | 0 | 0 | 0 | 0.05847981 | 0 | 0.13261575 |

| | | | | | | | |
|---|---|---|---|---|---|---|---|
| ENSMUSG00 Gm16057 | 757 | 0 | 0.12730896 | 0 | 0 | 0 | 0 |
| ENSMUSG00 Gm14115 | 481 | 0 | 0 | 0 | 0.07501465 | 0 | 0 |
| ENSMUSG00 Gm16397 | 493 | 0 | 0 | 0.17931782 | 0 | 0 | 0 |
| ENSMUSG00 Gm13600 | 742 | 0 | 0 | 0 | 0 | 0 | 0.03675827 |
| ENSMUSG00 Gm11960 | 904 | 0.17590921 | 0.26651793 | 0.09779169 | 0.2394826 | 0.11289021 | 0.36205273 |
| ENSMUSG00 Gm12482 | 629 | 0 | 0 | 0 | 0.05736414 | 0 | 0.39025716 |
| ENSMUSG00 Gm14823 | 237 | 0 | 0 | 0 | 0 | 0 | 0 |
| ENSMUSG00 Flg | 3161 | 0 | 0 | 0 | 0 | 0 | 0 |
| ENSMUSG00 Gm14895 | 1360 | 0 | 0 | 0 | 0 | 0 | 0 |
| ENSMUSG00 Gm8078 | 649 | 0 | 0 | 0 | 0 | 0 | 0 |
| ENSMUSG00 Gm13702 | 824 | 0 | 0 | 0 | 0 | 0 | 0 |
| ENSMUSG00 Gm6168 | 343 | 0 | 0 | 0 | 0 | 0 | 0 |
| ENSMUSG00 Gm13249 | 344 | 0 | 0 | 0 | 0 | 0 | 0 |
| ENSMUSG00 Dnajb3 | 1016 | 1.5651764 | 0.2845656 | 1.21816103 | 2.20185705 | 0.80356497 | 0 |
| ENSMUSG00 Gng2-ps1 | 216 | 0 | 0 | 0 | 0 | 0 | 0 |
| ENSMUSG00 Gm14841 | 757 | 0 | 0 | 0 | 0 | 0 | 0 |
| ENSMUSG00 Gm12639 | 492 | 0 | 0 | 0 | 0 | 0 | 0 |
| ENSMUSG00 Gm15746 | 1026 | 0 | 0 | 0 | 0 | 0 | 0 |
| ENSMUSG00 Gm13300 | 2393 | 0 | 0 | 0 | 0 | 0 | 0.00911281 |
| ENSMUSG00 Gm13319 | 306 | 0 | 0 | 0 | 0 | 0 | 0 |
| ENSMUSG00 Gm13859 | 1420 | 0 | 0 | 0 | 0 | 0.03593407 | 0.05762248 |
| ENSMUSG00 Gm13408 | 797 | 0.04988141 | 0 | 0 | 0.04527233 | 0 | 0 |
| ENSMUSG00 Gm8475 | 525 | 0 | 0 | 0 | 0 | 0 | 0 |
| ENSMUSG00 Gm14517 | 336 | 0 | 0 | 0 | 0 | 0 | 0.08117452 |
| ENSMUSG00 Gm11719 | 350 | 0 | 0 | 0 | 0 | 0 | 0 |
| ENSMUSG00 Gm7153 | 454 | 0 | 0.31841261 | 0.09736089 | 0.23842761 | 0 | 0.0600763 |
| ENSMUSG00 Gm11886 | 302 | 0 | 0 | 0 | 0 | 0 | 0 |
| ENSMUSG00 Gm13245 | 259 | 0 | 0 | 0 | 0 | 0 | 0.10530749 |
| ENSMUSG00 Gm13461 | 1525 | 0.31283001 | 0.15798834 | 0.31883297 | 0.18928286 | 0.03345992 | 0.33981518 |
| ENSMUSG00 Gm12488 | 498 | 0 | 0 | 0 | 0 | 0 | 0 |
| ENSMUSG00 Gm15607 | 357 | 0 | 0 | 0 | 0 | 0 | 0 |
| ENSMUSG00 Gm6568 | 696 | 0 | 0 | 0.19052519 | 0 | 0 | 0 |
| ENSMUSG00 Gm16422 | 401 | 0 | 0 | 0 | 0 | 0 | 0 |
| ENSMUSG00 Gm13519 | 271 | 0 | 0 | 0 | 0 | 0 | 0 |
| ENSMUSG00 Gm7189 | 428 | 0 | 0 | 0.10327533 | 0.08430384 | 0 | 0.06372579 |
| ENSMUSG00 Gm13649 | 605 | 0 | 0 | 0 | 0.05963974 | 0 | 0 |
| ENSMUSG00 Gm15206 | 453 | 0 | 0 | 0 | 0 | 0 | 0 |
| ENSMUSG00 Gm12390 | 396 | 0 | 0 | 0 | 0 | 0 | 0 |
| ENSMUSG00 Gm12954 | 609 | 0 | 0 | 0 | 0 | 0 | 0 |
| ENSMUSG00 Olfr1171-ps1 | 841 | 0 | 0 | 0 | 0 | 0 | 0 |
| ENSMUSG00 Gm15043 | 1542 | 0 | 0 | 0 | 0 | 0 | 0 |
| ENSMUSG00 Gm6257 | 789 | 0 | 0 | 0 | 0 | 0 | 0 |
| ENSMUSG00 Gm13216 | 590 | 0.13476434 | 0 | 0 | 0 | 0 | 0.0462282 |
| ENSMUSG00 Gm11371 | 595 | 0 | 0 | 0 | 0 | 0.08575862 | 0 |
| ENSMUSG00 Gm13452 | 1096 | 0 | 0 | 0 | 0 | 0 | 0 |
| ENSMUSG00 Pgam1-ps2 | 764 | 0 | 0 | 0 | 0 | 0 | 0 |
| ENSMUSG00 Gm11229 | 436 | 0 | 0 | 0.10138037 | 0 | 0 | 0 |
| ENSMUSG00 Gm6322 | 1011 | 0.19661464 | 0.14298648 | 0.21860457 | 0 | 0 | 0.1618673 |
| ENSMUSG00 Gm15231 | 446 | 0 | 0 | 0 | 0 | 0 | 0 |
| ENSMUSG00 Gm14182 | 209 | 0 | 0 | 0 | 0 | 0 | 0 |
| ENSMUSG00 Gm14991 | 596 | 0 | 0 | 0 | 0 | 0 | 0 |
| ENSMUSG00 Gm12386 | 687 | 0 | 0 | 0 | 0 | 0 | 0 |
| ENSMUSG00 Gm14407 | 446 | 0.1782757 | 0 | 0 | 0 | 0 | 0.2446156 |
| ENSMUSG00 Gm11794 | 236 | 0 | 0 | 0 | 0 | 0 | 0 |
| ENSMUSG00 Gm14359 | 421 | 0 | 0 | 0 | 0 | 0 | 0 |
| ENSMUSG00 Gm5764 | 884 | 0 | 0 | 0 | 0 | 0 | 0 |
| ENSMUSG00 Gm11416 | 861 | 0 | 0 | 0 | 0 | 0 | 0 |
| ENSMUSG00 Gm1826 | 1161 | 0.03424245 | 0.12451277 | 0.03807222 | 0.18647051 | 0.30765257 | 0.28190842 |
| ENSMUSG00 H3f3c | 410 | 0 | 0.23505582 | 0.10780937 | 0.17600997 | 0 | 0 |
| ENSMUSG00 Gm13081 | 5598 | 0 | 0 | 0 | 0 | 0 | 0 |
| ENSMUSG00 Gm8097 | 828 | 0 | 0 | 0 | 0 | 0 | 0 |
| ENSMUSG00 Gm12271 | 413 | 0 | 0 | 0 | 0 | 0.12355055 | 0 |
| ENSMUSG00 Mageb10-ps | 1612 | 0 | 0 | 0 | 0 | 0 | 0 |
| ENSMUSG00 Gm6646 | 398 | 0 | 0 | 0 | 0 | 0 | 0 |
| ENSMUSG00 Gm14934 | 620 | 0 | 0.2331602 | 0.21387989 | 0.05819685 | 0.08230061 | 0.04399135 |
| ENSMUSG00 Gm13724 | 1154 | 0 | 0 | 0 | 0 | 0 | 0 |
| ENSMUSG00 Gm11319 | 291 | 0 | 0 | 0 | 0 | 0 | 0 |
| ENSMUSG00 Gm12071 | 468 | 0 | 0 | 0 | 0 | 0 | 0 |
| ENSMUSG00 Gm15653 | 884 | 0 | 0 | 0 | 0 | 0 | 0 |
| ENSMUSG00 Gm12046 | 330 | 0 | 0 | 0 | 0 | 0 | 0 |
| ENSMUSG00 Gm15923 | 1165 | 0 | 0 | 0 | 0 | 0 | 0 |
| ENSMUSG00 Gm12848 | 840 | 0 | 0 | 0 | 0 | 0 | 0 |
| ENSMUSG00 Gm14284 | 278 | 0.28601065 | 0.34666505 | 0 | 0.90854069 | 0 | 1.27543277 |
| ENSMUSG00 Gm11585 | 1134 | 0 | 0.08498491 | 0 | 0 | 0 | 0.04810342 |
| ENSMUSG00 Gm14464 | 1253 | 0 | 0 | 0 | 0 | 0 | 0 |
| ENSMUSG00 Gm12948 | 681 | 0 | 0 | 0 | 0 | 0 | 0 |
| ENSMUSG00 Gm11481 | 254 | 0 | 0 | 0 | 0 | 0 | 0 |
| ENSMUSG00 Gm12105 | 778 | 0.05109959 | 0 | 0.11362942 | 0 | 0.0655866 | 0.03505738 |
| ENSMUSG00 Gm13989 | 199 | 0 | 0 | 0 | 0 | 0 | 0 |
| ENSMUSG00 Gm16072 | 2376 | 0 | 0 | 0.03720694 | 0 | 0.10737874 | 0.08035458 |
| ENSMUSG00 Gm13819 | 642 | 0 | 0.07505676 | 0.34425111 | 0.16860769 | 0.23844101 | 0.16993545 |
| ENSMUSG00 Gm11220 | 474 | 0 | 0 | 0 | 0 | 0 | 0 |
| ENSMUSG00 Psmb6-ps | 712 | 0 | 0 | 0 | 0.10135406 | 0 | 0.03830708 |
| ENSMUSG00 Gm14311 | 147 | 0 | 0 | 0 | 0 | 0 | 0 |
| ENSMUSG00 Gm13745 | 526 | 0 | 0 | 0 | 0 | 0 | 0 |
| ENSMUSG00 Gm15789 | 1356 | 0 | 0 | 0 | 0 | 0.71497134 | 0 |
| ENSMUSG00 Gm11923 | 507 | 0 | 0 | 0 | 0 | 0 | 0 |
| ENSMUSG00 Gm15183 | 347 | 0 | 0 | 0 | 0 | 0 | 0 |
| ENSMUSG00 Gm15048 | 952 | 0 | 0 | 0 | 0 | 0 | 0 |
| ENSMUSG00 Gm12726 | 600 | 0 | 0.08031074 | 0.14733948 | 0 | 0 | 0 |
| ENSMUSG00 Fit2 | 551 | 0 | 0 | 0.08022113 | 0 | 0 | 0.04950025 |
| ENSMUSG00 Gm12993 | 477 | 0.08334482 | 0 | 0.83399704 | 0.83208069 | 0.42789414 | 1.08641121 |
| ENSMUSG00 Gm14217 | 890 | 0 | 0 | 0 | 0 | 0 | 0 |
| ENSMUSG00 Mup-ps14 | 526 | 0 | 0 | 0 | 0 | 0 | 0 |
| ENSMUSG00 Gm13904 | 836 | 0.47554403 | 0.23055714 | 0.05287302 | 0.25896204 | 0.06103634 | 0.03262517 |
| ENSMUSG00 Gm7950 | 463 | 0 | 0 | 0 | 0 | 0 | 0 |
| ENSMUSG00 Gm2673 | 1607 | 0 | 0 | 0 | 0 | 0 | 0 |
| ENSMUSG00 Gm16242 | 548 | 0 | 0 | 0 | 0 | 0 | 0 |
| ENSMUSG00 Gm1866 | 476 | 0 | 0 | 0 | 0 | 0 | 0 |
| ENSMUSG00 Pramel | 1380 | 0 | 0 | 0 | 0 | 0 | 0 |
| ENSMUSG00 Gm15785 | 693 | 0.05736722 | 0.20859932 | 0.06378332 | 0.05206644 | 0 | 0 |
| ENSMUSG00 Gm13114 | 105 | 0.37862362 | 0 | 0 | 2.06183112 | 0.48596548 | 0 |
| ENSMUSG00 Gm12661 | 3334 | 0 | 0 | 0 | 0 | 0 | 0 |
| ENSMUSG00 Gm13104 | 1445 | 0 | 0 | 0 | 0 | 0 | 0 |
| ENSMUSG00 Gm8659 | 1083 | 0.03670866 | 0.04449348 | 0 | 0 | 0.04711577 | 0.02518434 |
| ENSMUSG00 Olfr260-ps1 | 820 | 0 | 0 | 0 | 0 | 0 | 0 |
| ENSMUSG00 Gm12098 | 506 | 0 | 0 | 0 | 0 | 0 | 0 |
| ENSMUSG00 Gm14490 | 2199 | 0 | 0 | 0 | 0 | 0 | 0.0248064 |
| ENSMUSG00 Gm9246 | 4331 | 0 | 0 | 0 | 0 | 0 | 0 |
| ENSMUSG00 Gm14358 | 401 | 0 | 0 | 0 | 0 | 0 | 0 |
| ENSMUSG00 Gm13230 | 1572 | 0.10115898 | 0.09195886 | 0 | 0.34429432 | 0 | 0 |
| ENSMUSG00 Gm14728 | 492 | 0 | 0 | 0 | 0 | 0 | 0 |
| ENSMUSG00 Usp-ps | 956 | 0 | 0 | 0 | 0 | 0 | 0 |
| ENSMUSG00 Gm6441 | 695 | 0 | 0.20799903 | 0.06359977 | 0.05191661 | 0.22025774 | 0.03924409 |
| ENSMUSG00 Gm12138 | 1018 | 0 | 0 | 0 | 0 | 0.05012414 | 0 |
| ENSMUSG00 Gm15753 | 1060 | 0.15002068 | 0.63642471 | 0.41699852 | 0.30635698 | 0.33696663 | 0.72046217 |
| ENSMUSG00 Gm12431 | 167 | 0 | 0 | 0 | 0 | 0 | 0 |
| ENSMUSG00 Gm16481 | 1226 | 0.03242698 | 0 | 0.10816112 | 0.20601494 | 0 | 0 |
| ENSMUSG00 Gm13922 | 155 | 0 | 0 | 0 | 0 | 0 | 0 |
| ENSMUSG00 Gm14665 | 556 | 0 | 0 | 0 | 0 | 0 | 0 |
| ENSMUSG00 Gm15928 | 286 | 0 | 0 | 0 | 0 | 0 | 0 |
| ENSMUSG00 Olfr1292-ps1 | 938 | 0 | 0 | 0 | 0 | 0 | 0 |
| ENSMUSG00 Gm11991 | 1073 | 0.11115232 | 0 | 0 | 0 | 0 | 0 |
| ENSMUSG00 Gm13505 | 816 | 0 | 0 | 0 | 0 | 0 | 0 |
| ENSMUSG00 Gm14039 | 100 | 0 | 0 | 0 | 0 | 0 | 0 |
| ENSMUSG00 Gm15377 | 244 | 0 | 0 | 0 | 0 | 0 | 0 |
| ENSMUSG00 Gm12013 | 154 | 0 | 0 | 0 | 0 | 0 | 0 |
| ENSMUSG00 Gins-ps1 | 1117 | 0 | 0 | 0 | 0 | 0 | 0 |
| ENSMUSG00 Slfn14-ps | 4521 | 0 | 0 | 0.05866204 | 0 | 0.24830353 | 0 |
| ENSMUSG00 Gm14572 | 575 | 0 | 0 | 0 | 0 | 0 | 0 |
| ENSMUSG00 Gm15320 | 362 | 0 | 0 | 0 | 0 | 0 | 0 |
| ENSMUSG00 Gm15933 | 614 | 0 | 0 | 0 | 0 | 0 | 0 |
| ENSMUSG00 Gm12621 | 381 | 0 | 0 | 0 | 0 | 0 | 0 |
| ENSMUSG00 Mageb16-ps1 | 1643 | 0.04839377 | 0 | 0 | 0 | 0 | 0 |
| ENSMUSG00 Gm16216 | 1110 | 0 | 0 | 0 | 0.03250635 | 0 | 0 |
| ENSMUSG00 Gm13777 | 344 | 0 | 0 | 0 | 1.36356564 | 0 | 0.79286742 |
| ENSMUSG00 Rpl19-ps12 | 590 | 0 | 0 | 0.52442865 | 0 | 0 | 0 |
| ENSMUSG00 Gm11196 | 394 | 0.10090224 | 0 | 0.33656226 | 0.27473638 | 0 | 0 |
| ENSMUSG00 Gm12479 | 1584 | 0 | 0 | 0 | 0 | 0 | 0 |
| ENSMUSG00 Gm14445 | 1481 | 0 | 0 | 0 | 0.0487266 | 0.034454 | 0 |
| ENSMUSG00 Gm13915 | 930 | 0 | 0.05181338 | 0 | 0 | 0.05486707 | 0 |
| ENSMUSG00 Gm13489 | 968 | 0 | 0 | 0 | 0 | 0 | 0.02817628 |
| ENSMUSG00 Gm15490 | 385 | 0 | 0 | 0 | 0 | 0 | 0 |
| ENSMUSG00 Gm12188 | 309 | 0 | 0 | 0 | 0.35031111 | 0.1651339 | 0.17653488 |
| ENSMUSG00 Gm4994 | 2186 | 0 | 0 | 0 | 0 | 0 | 0 |
| ENSMUSG00 Gm13862 | 196 | 0 | 0 | 0.22551961 | 2.76138096 | 0.26033865 | 0.97409426 |
| ENSMUSG00 Gm12659 | 358 | 0 | 0 | 0 | 2.62048368 | 0.85519066 | 0.07618614 |
| ENSMUSG00 Gm15720 | 281 | 1.41478578 | 0.171482 | 3.46064251 | 2.05449364 | 0.7263541 | 4.27076202 |
| ENSMUSG00 Gm8199 | 822 | 0 | 0 | 0 | 0 | 0 | 0 |
| ENSMUSG00 Gm11809 | 949 | 0 | 0.05077602 | 0 | 0 | 0 | 0 |
| ENSMUSG00 Gm16149 | 308 | 0 | 0 | 0 | 0.1171495 | 0 | 0 |
| ENSMUSG00 Gm14679 | 721 | 0 | 0 | 0 | 0 | 0 | 0.0378289 |
| ENSMUSG00 Gm14975 | 663 | 0 | 0 | 0 | 0 | 0 | 0 |
| ENSMUSG00 Gm13577 | 212 | 0 | 0 | 0 | 0 | 0 | 0 |
| ENSMUSG00 Gm11824 | 845 | 0 | 0 | 0 | 0 | 0 | 0 |
| ENSMUSG00 Gm14502 | 112 | 0 | 0 | 0 | 0 | 0 | 0 |
| ENSMUSG00 Gm15799 | 288 | 0 | 0 | 0 | 0 | 0 | 0 |
| ENSMUSG00 Gm12624 | 381 | 0 | 0 | 0 | 0 | 0 | 0 |
| ENSMUSG00 Gm13541 | 282 | 1.69172258 | 1.36699127 | 0 | 3.19876281 | 1.99039055 | 0 |
| ENSMUSG00 Gm7586 | 401 | 0 | 0 | 0 | 0 | 0 | 0 |
| ENSMUSG00 Gm15236 | 378 | 0.52586614 | 0.89234152 | 0.35080828 | 0 | 0.40497124 | 0 |
| ENSMUSG00 Gm14988 | 1546 | 0 | 0 | 0 | 0 | 0 | 0 |
| ENSMUSG00 Gm15660 | 477 | 0 | 0 | 0 | 0 | 0 | 0 |
| ENSMUSG00 Gm11905 | 677 | 0 | 0 | 0 | 0 | 0 | 0 |
| ENSMUSG00 Gm16463 | 719 | 0 | 0 | 0 | 0 | 0 | 0 |
| ENSMUSG00 Gm13094 | 408 | 0 | 0 | 0 | 0 | 0 | 0 |
| ENSMUSG00 Gm11212 | 612 | 0 | 0 | 0 | 0 | 0 | 0 |
| ENSMUSG00 Gm13518 | 232 | 0 | 0 | 0 | 0 | 0 | 0 |
| ENSMUSG00 Gm12864 | 1005 | 0.19778846 | 0 | 0 | 0.1077076 | 0.05077251 | 0.21711156 |
| ENSMUSG00 Gm12788 | 307 | 0.51798672 | 0 | 0 | 0 | 0 | 0 |
| ENSMUSG00 Gm12312 | 643 | 0.06182812 | 0 | 0.06874315 | 0.11223031 | 0 | 0 |
| ENSMUSG00 Rpl23a-ps13 | 440 | 0 | 0 | 0 | 0 | 0 | 0 |
| ENSMUSG00 Gm14570 | 390 | 0 | 0 | 0 | 0 | 0 | 0 |
| ENSMUSG00 Gm12266 | 961 | 0.2482132 | 0.70198771 | 0.09199135 | 0.33791717 | 0.21238866 | 0.02838152 |
| ENSMUSG00 Gm13002 | 681 | 0 | 0 | 0 | 0 | 0 | 0 |
| ENSMUSG00 Gm13144 | 344 | 0 | 0 | 0 | 0 | 0 | 0 |
| ENSMUSG00 Gm11810 | 332 | 0 | 0 | 0 | 0 | 0 | 0 |
| ENSMUSG00 Gm13655 | 744 | 3.63356542 | 4.46890391 | 6.00051901 | 3.2493239 | 3.08627273 | 4.06920021 |
| ENSMUSG00 Gm16464 | 1094 | 0.10901869 | 0.08809222 | 0 | 0 | 0 | 0.29917337 |
| ENSMUSG00 Gm7164 | 449 | 0 | 0 | 0 | 0 | 0 | 0 |
| ENSMUSG00 Gm12008 | 754 | 0 | 0 | 0 | 0 | 0 | 0 |
| ENSMUSG00 Gm12038 | 206 | 0.38597554 | 0 | 0 | 0 | 0 | 0 |
| ENSMUSG00 Gm14116 | 804 | 0 | 0 | 0 | 0 | 0 | 0 |
| ENSMUSG00 Gm7910 | 836 | 0 | 0 | 0 | 0 | 0 | 0 |
| ENSMUSG00 Gm11578 | 458 | 0 | 0 | 0 | 0 | 0 | 0 |
| ENSMUSG00 Gm11350 | 869 | 0 | 0 | 0 | 0 | 0 | 0 |
| ENSMUSG00 Gm14701 | 347 | 0 | 0 | 0 | 0 | 0 | 0 |
| ENSMUSG00 Gm14276 | 656 | 0.78783727 | 0.44072965 | 1.41499802 | 0.82504675 | 0.54448877 | 0.49892633 |
| ENSMUSG00 Gm13360 | 798 | 0 | 0 | 0 | 0 | 0 | 0.06835749 |
| ENSMUSG00 Gm7091 | 585 | 0 | 0 | 0 | 0 | 0 | 0 |
| ENSMUSG00 Gm14360 | 391 | 0 | 0 | 0 | 0 | 0 | 0 |
| ENSMUSG00 Gm14656 | 501 | 0 | 0 | 0 | 0 | 0 | 0 |
| ENSMUSG00 Gm12369 | 941 | 0 | 0 | 0 | 0 | 0 | 0.05796948 |
| ENSMUSG00 Gm8617 | 2221 | 0 | 0 | 0 | 0 | 0 | 0 |
| ENSMUSG00 Rpl27-ps2 | 398 | 0 | 0 | 0 | 0 | 0 | 0 |
| ENSMUSG00 Gm14835 | 314 | 0 | 0 | 0 | 0 | 0 | 0 |
| ENSMUSG00 Mup-ps10 | 550 | 0 | 0 | 0 | 0 | 0 | 0 |
| ENSMUSG00 Gm14877 | 476 | 0 | 0 | 0 | 0 | 0 | 0 |
| ENSMUSG00 Gm14563 | 353 | 0.11262176 | 0 | 0 | 0 | 0 | 0.07726527 |
| ENSMUSG00 Gm13163 | 335 | 0 | 0 | 0 | 0 | 0 | 0 |
| ENSMUSG00 Gm16005 | 268 | 0 | 0 | 0 | 0 | 0 | 0 |
| ENSMUSG00 Gm11307 | 927 | 0 | 0 | 0 | 0 | 0 | 0 |
| ENSMUSG00 Gm11407 | 1057 | 0.11283486 | 1.41322583 | 0.20909103 | 2.69676586 | 1.39996679 | 0.3612535 |
| ENSMUSG00 Gm12230 | 453 | 0 | 0.21274367 | 0 | 0 | 0 | 0.06020892 |
| ENSMUSG00 Gm12252 | 218 | 0 | 0 | 0 | 0 | 0 | 0 |
| ENSMUSG00 Gm6660 | 463 | 0 | 0 | 0 | 0 | 0 | 0 |
| ENSMUSG00 Gm15994 | 401 | 0 | 0 | 0 | 0 | 0.38174346 | 0 |
| ENSMUSG00 Gm12428 | 726 | 0.05475962 | 0 | 0.18265224 | 0 | 0 | 0 |
| ENSMUSG00 Gm12403 | 950 | 1.54837135 | 1.52167712 | 0.18611302 | 0.60769759 | 1.66507121 | 0.08613044 |
| ENSMUSG00 Gm13670 | 459 | 0 | 0 | 0 | 0 | 0 | 0 |
| ENSMUSG00 Gm2768 | 247 | 0 | 0 | 0 | 0 | 0 | 0 |
| ENSMUSG00 Gm4784 | 1822 | 0 | 0 | 0 | 0.01980354 | 0 | 0 |
| ENSMUSG00 Gm13641 | 472 | 0.16845543 | 0 | 0.09364797 | 0 | 0 | 0 |
| ENSMUSG00 Gm5166 | 569 | 0 | 0 | 0 | 0 | 0 | 0 |
| ENSMUSG00 Gm14719 | 230 | 0 | 0.20950627 | 0.19218193 | 0 | 0 | 0.11858539 |
| ENSMUSG00 Rpl5-ps1 | 893 | 0 | 0 | 0 | 0 | 0.1142808 | 0 |
| ENSMUSG00 Gm12444 | 1148 | 0.03463021 | 0.08394851 | 0 | 0 | 0 | 0 |
| ENSMUSG00 Gm13034 | 4359 | 0 | 0 | 0 | 0 | 0 | 0.03128543 |
| ENSMUSG00 Gm14231 | 591 | 0 | 0 | 0 | 0 | 0 | 0 |
| ENSMUSG00 Gm8577 | 985 | 0 | 0 | 0 | 0 | 0.05180343 | 0 |
| ENSMUSG00 Gm14504 | 1496 | 0 | 0 | 0 | 0 | 0 | 0 |
| ENSMUSG00 Gm15763 | 402 | 0 | 0 | 0 | 0 | 0 | 0 |
| ENSMUSG00 Gm12871 | 437 | 0 | 0 | 0 | 0 | 0 | 0 |
| ENSMUSG00 Gm13018 | 446 | 0 | 0 | 0 | 0 | 0 | 0 |
| ENSMUSG00 Gm8910 | 736 | 0 | 0 | 0 | 0 | 0 | 0 |
| ENSMUSG00 Gm15636 | 666 | 0 | 0 | 0 | 0 | 0 | 0 |
| ENSMUSG00 Gm7722 | 744 | 0 | 0 | 0 | 0 | 0 | 0 |
| ENSMUSG00 Gm7058 | 901 | 0 | 0 | 0 | 0 | 0 | 0 |
| ENSMUSG00 Gm12998 | 787 | 0.60618268 | 0 | 0 | 0.04584758 | 0.38901938 | 0 |
| ENSMUSG00 Gm15379 | 647 | 0.67590462 | 0 | 0 | 0.33460938 | 0 | 2.27639957 |
| ENSMUSG00 Eif4a-ps4 | 1217 | 0.35933466 | 0.11878334 | 0.69008629 | 0.50402199 | 0.503136 | 0.60510703 |
| ENSMUSG00 Gm14382 | 436 | 0 | 0 | 0 | 0 | 0.35109892 | 0 |
| ENSMUSG00 Gm15299 | 392 | 0 | 0 | 0 | 0 | 0 | 0 |
| ENSMUSG00 Gm15826 | 1382 | 0 | 0 | 0 | 0 | 0 | 0 |
| ENSMUSG00 Gm12102 | 603 | 1.12080128 | 0 | 0 | 0.59837553 | 0 | 0 |
| ENSMUSG00 Gm15009 | 321 | 0 | 0 | 0 | 0 | 0 | 0 |
| ENSMUSG00 Gm14314 | 1014 | 0 | 0 | 0 | 0 | 0 | 0 |
| ENSMUSG00 Olfr271-ps1 | 951 | 0 | 0 | 0 | 0 | 0 | 0 |
| ENSMUSG00 Gm1401 | 771 | 0 | 0 | 0 | 0 | 0 | 0 |
| ENSMUSG00 Gm13602 | 492 | 0 | 0.09793992 | 0.08984114 | 0 | 0 | 0 |
| ENSMUSG00 Gm14355 | 263 | 0 | 0 | 0 | 0 | 0 | 0 |
| ENSMUSG00 Gm12943 | 209 | 0.38043522 | 0 | 0 | 0 | 0 | 0 |
| ENSMUSG00 Gm13120 | 1445 | 0 | 0 | 0 | 0 | 0 | 0 |
| ENSMUSG00 Gm14865 | 492 | 0 | 0 | 0 | 0 | 0 | 0 |
| ENSMUSG00 Gm15079 | 215 | 0 | 0 | 0 | 0 | 0 | 0 |
| ENSMUSG00 Gm15098 | 1507 | 0 | 0 | 0 | 0 | 0 | 0 |
| ENSMUSG00 Gm13728 | 399 | 0 | 0 | 0 | 0 | 0 | 0 |
| ENSMUSG00 Gm715 | 830 | 0 | 0 | 0 | 0 | 0 | 0 |
| ENSMUSG00 Olfr1144-ps1 | 319 | 0 | 0 | 0 | 0 | 0 | 0 |
| ENSMUSG00 Gm8344 | 855 | 0 | 0 | 0 | 0 | 0 | 0 |
| ENSMUSG00 Nap1l2 | 2464 | 0 | 0 | 0 | 0 | 0 | 0 |
| ENSMUSG00 Gm3681 | 631 | 0 | 0 | 0 | 0 | 0 | 0 |
| ENSMUSG00 Gm14309 | 201 | 0 | 0 | 0 | 0 | 0 | 0 |
| ENSMUSG00 Gm6506 | 1135 | 0 | 0 | 0 | 0 | 0 | 0.1201526 |
| ENSMUSG00 Gm16221 | 392 | 0 | 0 | 0 | 0 | 0 | 0 |
| ENSMUSG00 Gm13326 | 1010 | 0.03936186 | 0.14312805 | 0.0437642 | 0 | 0.05052116 | 0.05400919 |
| ENSMUSG00 Gm11355 | 305 | 0 | 0 | 0 | 0 | 0.33459919 | 0 |
| ENSMUSG00 Olfr1172-ps1 | 203 | 0 | 0 | 0 | 0 | 0 | 0 |
| ENSMUSG00 Gm12383 | 759 | 0 | 0 | 0 | 0 | 0 | 0 |
| ENSMUSG00 Gm14234 | 844 | 0 | 0 | 0 | 0 | 0 | 0 |
| ENSMUSG00 Gm14380 | 430 | 0 | 0 | 0 | 0 | 0 | 0 |
| ENSMUSG00 Gm15776 | 410 | 0.09696459 | 0.35258372 | 0 | 0 | 0 | 0.39914106 |
| ENSMUSG00 Gm12443 | 322 | 0 | 0 | 0 | 0 | 0 | 0 |
| ENSMUSG00 Gm12076 | 475 | 0 | 0 | 0 | 0 | 0 | 0 |
| ENSMUSG00 Gm8503 | 454 | 0 | 0 | 0 | 0 | 0 | 0 |
| ENSMUSG00 Gm6292 | 1094 | 0 | 0 | 0 | 0 | 0 | 0 |
| ENSMUSG00 Gm13047 | 814 | 0 | 0 | 0 | 0.08865367 | 0 | 0 |
| ENSMUSG00 Gm16074 | 443 | 0 | 0 | 0.09977843 | 0.08144931 | 0.23036739 | 0 |
| ENSMUSG00 Gm13161 | 558 | 0.21373914 | 0.17271126 | 0.23764432 | 0 | 0.36578047 | 0.34215497 |
| ENSMUSG00 Gm12997 | 639 | 0 | 0 | 0 | 0 | 0 | 0 |
| ENSMUSG00 Gm12181 | 530 | 0 | 0 | 0.0833997 | 0 | 0.09627618 | 0 |
| ENSMUSG00 Gm13438 | 572 | 0 | 0 | 0 | 0 | 0 | 0 |
| ENSMUSG00 Gm11204 | 146 | 0 | 0 | 0 | 0 | 0 | 0 |
| ENSMUSG00 Gm14639 | 865 | 0.13788028 | 0.05570687 | 0 | 0 | 0.05899003 | 0.03153138 |
| ENSMUSG00 Gm11922 | 474 | 0 | 0 | 0 | 0.07612246 | 0 | 0 |
| ENSMUSG00 Gm11822 | 414 | 0 | 0 | 0 | 0 | 0 | 0 |
| ENSMUSG00 Gm12287 | 705 | 0 | 0 | 0 | 0 | 0 | 0 |
| ENSMUSG00 Gm12234 | 427 | 0 | 0 | 0 | 0 | 0 | 0 |
| ENSMUSG00 Gm15289 | 243 | 0 | 0.19829812 | 0.54570177 | 0 | 0 | 0 |
| ENSMUSG00 Gm13808 | 183 | 0 | 0 | 0 | 0 | 0 | 0 |
| ENSMUSG00 Gm7429 | 780 | 0 | 0 | 0 | 0 | 0.06541843 | 0.06993497 |
| ENSMUSG00 Gm14999 | 623 | 0 | 0 | 0 | 0 | 0 | 0 |
| ENSMUSG00 Gm14730 | 1044 | 0.11423989 | 0 | 0 | 0.10368404 | 0 | 0 |
| ENSMUSG00 Gm11188 | 387 | 0 | 0 | 0 | 0 | 0 | 0 |
| ENSMUSG00 Gm12799 | 2559 | 0.09321332 | 0.03766037 | 0 | 0.02820011 | 0.0598199 | 0.04263328 |
| ENSMUSG00 Gm15547 | 332 | 0 | 0 | 0 | 0 | 1.07585732 | 0 |
| ENSMUSG00 Gm12303 | 672 | 0 | 0.07170602 | 0 | 0 | 0 | 0.16234904 |
| ENSMUSG00 Gm12467 | 744 | 0 | 0 | 0 | 0 | 0 | 0 |
| ENSMUSG00 Gm11317 | 375 | 0 | 0 | 0 | 0 | 0 | 0 |
| ENSMUSG00 Gm14676 | 857 | 0 | 0 | 0 | 0 | 0 | 0 |
| ENSMUSG00 Gm14375 | 854 | 0 | 0 | 0 | 0 | 0 | 0 |
| ENSMUSG00 Gm12217 | 694 | 0 | 0 | 0 | 0 | 0 | 0 |
| ENSMUSG00 Gm11675 | 831 | 0 | 0.11597218 | 0.05319115 | 0 | 0 | 0 |
| ENSMUSG00 Gm14801 | 446 | 0 | 0 | 0 | 0 | 0 | 0 |
| ENSMUSG00 Gm14026 | 1788 | 0.60033444 | 0.05389982 | 0.07416417 | 0.04036023 | 0.05707648 | 0.10677991 |
| ENSMUSG00 Gm14854 | 840 | 0 | 0 | 0 | 0 | 0 | 0 |
| ENSMUSG00 Gm13885 | 186 | 0 | 0 | 0 | 0 | 0 | 0 |
| ENSMUSG00 Gm15807 | 1125 | 0 | 0 | 0 | 0 | 0 | 0 |
| ENSMUSG00 Gm13471 | 432 | 0 | 0 | 0 | 0 | 0 | 0 |
| ENSMUSG00 Gm12870 | 1015 | 0.23500777 | 0.71211491 | 0.17419446 | 1.17311081 | 0 | 0.67178914 |
| ENSMUSG00 Gm7901 | 1596 | 0 | 0.03019201 | 0 | 0.0452156 | 0 | 0 |
| ENSMUSG00 Gm4886 | 1174 | 0 | 0 | 0 | 0 | 0 | 0 |
| ENSMUSG00 Gm14472 | 281 | 0.14147858 | 0.685928 | 0 | 0.89884097 | 0 | 0 |
| ENSMUSG00 H3f3a-ps1 | 409 | 0 | 0 | 0 | 0 | 0 | 0 |
| ENSMUSG00 Gm14069 | 228 | 0 | 0 | 0 | 0 | 0 | 0 |
| ENSMUSG00 Pisd-ps1 | 1655 | 45.2563899 | 36.0452057 | 35.9223437 | 38.4802469 | 47.604063 | 42.3539715 |
| ENSMUSG00 Gm15333 | 363 | 0 | 0 | 0 | 0 | 0 | 0 |
| ENSMUSG00 Gm14755 | 383 | 0.20760042 | 0.12581317 | 0 | 0.18841799 | 0 | 0.07121316 |
| ENSMUSG00 Gm15596 | 783 | 0 | 0 | 0 | 0 | 0 | 0 |
| ENSMUSG00 Gm14307 | 290 | 0 | 0 | 0 | 0 | 0 | 0 |
| ENSMUSG00 Gm5762 | 257 | 0 | 0 | 0 | 0 | 0 | 0 |
| ENSMUSG00 Gm12250 | 2700 | 0 | 0.42832393 | 0.01637105 | 0.33469301 | 2.02215638 | 0.70712028 |
| ENSMUSG00 Gm12072 | 350 | 0 | 0 | 0 | 0 | 0 | 0 |
| ENSMUSG00 Gm12133 | 660 | 0 | 0 | 0 | 0 | 0 | 0 |
| ENSMUSG00 Gm11450 | 332 | 0 | 0 | 0 | 0 | 0 | 0.08215253 |
| ENSMUSG00 Rhox3-ps | 644 | 0 | 0 | 0 | 0 | 0 | 0 |
| ENSMUSG00 Gm8331 | 1761 | 0 | 0 | 0 | 0 | 0 | 0 |
| ENSMUSG00 Gm14032 | 1025 | 0 | 0 | 0.04312375 | 0 | 0 | 0.0266094 |
| ENSMUSG00 Gm12385 | 794 | 0 | 0 | 0 | 0 | 0 | 0 |
| ENSMUSG00 Gm13633 | 897 | 0 | 0 | 0 | 0 | 0 | 0 |
| ENSMUSG00 Gm11198 | 973 | 0.08171733 | 0.19809432 | 0 | 0 | 0 | 0.16818894 |
| ENSMUSG00 Gm2420 | 1538 | 0.10339527 | 0.06266117 | 0.1436991 | 0 | 0.16588549 | 0.08866918 |
| ENSMUSG00 Gm11358 | 489 | 0 | 0 | 0 | 0 | 0 | 0 |
| ENSMUSG00 Gm13418 | 943 | 0 | 0 | 0 | 1.10962809 | 0 | 0 |
| ENSMUSG00 Gm15770 | 583 | 0 | 0 | 0.07581791 | 0 | 0 | 0 |
| ENSMUSG00 Gm12123 | 471 | 0 | 0 | 0 | 0 | 0 | 0 |
| ENSMUSG00 Gm12904 | 1398 | 0 | 0 | 0 | 0 | 0 | 0 |
| ENSMUSG00 Gm14791 | 1365 | 0 | 0 | 0 | 0 | 0 | 0 |
| ENSMUSG00 Gm11996 | 914 | 0 | 0 | 0.04836088 | 0.07895415 | 0 | 0 |
| ENSMUSG00 Gm11701 | 1140 | 0 | 0 | 0 | 0 | 0 | 0 |
| ENSMUSG00 Gm13296 | 1014 | 0.19603294 | 0.28512688 | 0 | 0 | 0.05032187 | 0 |
| ENSMUSG00 Gm16523 | 557 | 1.14198867 | 0.08651067 | 0 | 1.29558508 | 0.73287434 | 0 |
| ENSMUSG00 Gm11329 | 1083 | 0 | 0 | 0 | 0 | 0 | 0 |
| ENSMUSG00 Gm14933 | 479 | 0 | 0 | 0 | 0 | 0 | 0 |
| ENSMUSG00 Gm14162 | 1058 | 0.03757607 | 0 | 0.12533604 | 0.03410401 | 0.24114544 | 0.02577943 |
| ENSMUSG00 Gm8822 | 2941 | 0.39201256 | 0.65537494 | 0.24047245 | 0.34352168 | 0.46845024 | 4.48858395 |
| ENSMUSG00 Gm11877 | 851 | 0 | 0 | 0.05194106 | 0 | 0 | 0 |
| ENSMUSG00 Gm14253 | 638 | 1.86937996 | 1.58607411 | 1.73204714 | 1.64009293 | 1.99946614 | 1.36800696 |
| ENSMUSG00 Gm9164 | 1006 | 0.07903674 | 0.19159619 | 0.21969107 | 0.17933422 | 0 | 0.0813359 |
| ENSMUSG00 Gm15314 | 2558 | 0 | 0 | 0 | 0 | 0 | 0 |
| ENSMUSG00 Gm14615 | 1813 | 0 | 0.02657829 | 0.0243805 | 0 | 0 | 0 |
| ENSMUSG00 Gm12642 | 875 | 0 | 0 | 0 | 0 | 0 | 0 |
| ENSMUSG00 Gm14366 | 728 | 0 | 0 | 0 | 0 | 0 | 0 |
| ENSMUSG00 Gm13801 | 159 | 0 | 0 | 0 | 0 | 0 | 0 |
| ENSMUSG00 Gm13921 | 462 | 0 | 0 | 0 | 0 | 0 | 0 |
| ENSMUSG00 Gm14287 | 116 | 0 | 0 | 0 | 0 | 0 | 0 |
| ENSMUSG00 Ldha-ps | 994 | 0.15998181 | 0 | 0 | 0.18149922 | 0.05133438 | 0.02743928 |
| ENSMUSG00 Gm14063 | 308 | 0 | 0 | 0 | 0 | 0 | 0 |
| ENSMUSG00 Gm13384 | 1342 | 0 | 0.3590644 | 0 | 0 | 0 | 0 |
| ENSMUSG00 Gm11984 | 765 | 0 | 0 | 0 | 0 | 0 | 0 |
| ENSMUSG00 Gm13244 | 1444 | 0 | 0 | 0.0306107 | 0 | 0 | 0 |
| ENSMUSG00 Gm8401 | 1738 | 0 | 0 | 0 | 0 | 0 | 0 |
| ENSMUSG00 Gm14723 | 214 | 0 | 0 | 0 | 0 | 0 | 0 |
| ENSMUSG00 Gm12327 | 1165 | 0 | 0 | 0 | 0 | 0 | 0 |
| ENSMUSG00 Gm14028 | 478 | 0 | 0 | 0 | 0 | 0 | 0 |
| ENSMUSG00 Gm11321 | 901 | 0 | 0 | 0 | 0 | 0 | 0 |
| ENSMUSG00 Gm14143 | 356 | 0 | 0 | 0.12416248 | 0 | 0 | 0 |
| ENSMUSG00 Gm5395 | 914 | 0.0869923 | 0.10544079 | 0 | 0 | 0.39079281 | 1.01459271 |
| ENSMUSG00 Gm14404 | 1657 | 0.02399244 | 0.0872416 | 0 | 0 | 0 | 0.01646025 |
| ENSMUSG00 4930415H17 | 122 | 0 | 0 | 0 | 0 | 0 | 0 |
| ENSMUSG00 Gm13622 | 432 | 0.09202658 | 0.22308538 | 0.10231908 | 1.08580227 | 0 | 0 |
| ENSMUSG00 Gm6952 | 158 | 0 | 0 | 0 | 0 | 0 | 0 |
| ENSMUSG00 Gm15036 | 399 | 0.29891339 | 0 | 0 | 0 | 0 | 0 |
| ENSMUSG00 Gm12807 | 961 | 0 | 0.05014198 | 0 | 0 | 0 | 0 |
| ENSMUSG00 Gm15064 | 327 | 0 | 0 | 0 | 0 | 0 | 0 |
| ENSMUSG00 Gm14008 | 273 | 0 | 0 | 0 | 0 | 0 | 0 |
| ENSMUSG00 Gm11260 | 941 | 0 | 0 | 0 | 0 | 0 | 0 |
| ENSMUSG00 Gm13228 | 633 | 0 | 0.30449569 | 0 | 0 | 0 | 0 |
| ENSMUSG00 Vmn1r-ps1 | 572 | 0 | 0 | 0 | 0 | 0 | 0 |
| ENSMUSG00 Gm378 | 818 | 0 | 0 | 0 | 0 | 0 | 0 |
| ENSMUSG00 Gm7711 | 398 | 0 | 0 | 0 | 0 | 0 | 0 |
| ENSMUSG00 Gm15565 | 426 | 0.09332272 | 0 | 0 | 0 | 0 | 0 |
| ENSMUSG00 Gm13542 | 636 | 0 | 0.22729454 | 0.27799901 | 0.39712942 | 0 | 0.34307722 |
| ENSMUSG00 Gm14990 | 444 | 0 | 0 | 0 | 0 | 0 | 0 |
| ENSMUSG00 Gm11908 | 346 | 0 | 0 | 0 | 0 | 0 | 0 |
| ENSMUSG00 Gm8814 | 534 | 0.07444847 | 0.09023678 | 0 | 0 | 0.09555501 | 0 |
| ENSMUSG00 Btc | 2789 | 42.1502172 | 65.4983156 | 29.7637365 | 38.410753 | 32.1636317 | 15.0308786 |
| ENSMUSG00 Gm5187 | 990 | 0.28109936 | 0.04867317 | 0.44648326 | 0.18223255 | 0.05154179 | 0.05510028 |
| ENSMUSG00 Mup-ps18 | 525 | 0 | 0 | 0 | 0 | 0 | 0 |
| ENSMUSG00 Gm11955 | 469 | 0 | 0 | 0 | 0 | 0 | 0 |
| ENSMUSG00 Gm15754 | 477 | 0 | 0 | 0 | 0 | 0 | 0 |
| ENSMUSG00 Gm14357 | 867 | 0 | 0 | 0 | 0 | 0 | 0 |
| ENSMUSG00 Gm12363 | 667 | 0.17881026 | 0 | 0 | 0 | 0 | 0 |
| ENSMUSG00 Gm11225 | 1048 | 0.03793462 | 0 | 0 | 0 | 0 | 0.02602542 |
| ENSMUSG00 Gm14208 | 677 | 0 | 0 | 0 | 0 | 0.75371308 | 0 |
| ENSMUSG00 Gm12313 | 420 | 0 | 0 | 0 | 0 | 0 | 0 |
| ENSMUSG00 Gm3712 | 2070 | 0 | 0 | 0 | 0 | 0 | 0 |
| ENSMUSG00 Gm4772 | 842 | 0 | 0 | 0 | 0 | 0 | 0 |
| ENSMUSG00 Gm13754 | 1007 | 0 | 0 | 0 | 0 | 0 | 0 |
| ENSMUSG00 Gm12741 | 555 | 0 | 1.38915869 | 0 | 0 | 0 | 0 |
| ENSMUSG00 Gm14141 | 782 | 0.05083821 | 0.06161949 | 0 | 0 | 0 | 0 |
| ENSMUSG00 Gm14442 | 1733 | 0 | 0 | 0 | 0 | 0 | 0 |
| ENSMUSG00 Gm14415 | 1575 | 0.05048315 | 0.58129676 | 0.14032331 | 0 | 0.0323977 | 0 |
| ENSMUSG00 Olfr408-ps1 | 922 | 0 | 0 | 0 | 0 | 0 | 0 |
| ENSMUSG00 Gm13884 | 828 | 0 | 0 | 0 | 0 | 0 | 0 |
| ENSMUSG00 Gm12218 | 398 | 0 | 0 | 0 | 0 | 0 | 0 |
| ENSMUSG00 Gm12928 | 572 | 0 | 0 | 0 | 0 | 0 | 0.04768294 |
| ENSMUSG00 Gm14970 | 354 | 0.22460724 | 0 | 0 | 0 | 0 | 0 |
| ENSMUSG00 Gm9670 | 335 | 0 | 0 | 0.3958374 | 0 | 0 | 0 |
| ENSMUSG00 Gm12437 | 182 | 0 | 0 | 0 | 0 | 0 | 0 |
| ENSMUSG00 Gm7601 | 401 | 0 | 0 | 0 | 0 | 0 | 0 |
| ENSMUSG00 Gm15724 | 441 | 0 | 0 | 0 | 0 | 0 | 0 |
| ENSMUSG00 Gm12692 | 545 | 0 | 0 | 0 | 0 | 0 | 0 |
| ENSMUSG00 Mkrn1-ps1 | 1393 | 0 | 0 | 0.0317314 | 0.0259024 | 0.03663056 | 0.05873935 |
| ENSMUSG00 Gm12136 | 1208 | 0 | 0 | 0 | 0 | 0 | 0 |
| ENSMUSG00 Gm14938 | 942 | 0 | 0.05115334 | 0 | 0 | 0 | 0 |
| ENSMUSG00 Gm15028 | 369 | 0.10773843 | 0 | 0 | 0.09778332 | 0 | 0 |
| ENSMUSG00 Gm4596 | 441 | 0 | 0 | 0 | 0.81818695 | 0 | 1.48433411 |
| ENSMUSG00 Gm15480 | 518 | 0 | 0 | 0 | 0 | 0 | 0 |
| ENSMUSG00 Zfp389 | 674 | 0.47187514 | 1.21538504 | 1.11488328 | 0 | 0.90848147 | 5.34162074 |
| ENSMUSG00 Gm14803 | 359 | 0 | 0 | 0 | 0 | 0 | 0 |
| ENSMUSG00 Gm11927 | 757 | 0 | 0 | 0 | 0 | 0 | 0.07205981 |
| ENSMUSG00 Gm14036 | 479 | 0 | 0 | 0 | 0 | 0 | 0.17082238 |
| ENSMUSG00 Gm16017 | 207 | 0 | 0 | 0 | 0 | 0 | 0 |
| ENSMUSG00 Gm14328 | 407 | 0 | 0 | 0 | 0 | 0 | 0 |
| ENSMUSG00 Gm15621 | 363 | 0.10951923 | 0 | 0.48707265 | 0.29819872 | 0 | 0 |
| ENSMUSG00 Gm14916 | 203 | 0 | 0 | 0 | 0 | 0 | 0 |
| ENSMUSG00 Gm14799 | 428 | 0 | 0 | 0 | 0.33721537 | 0 | 0 |
| ENSMUSG00 Gm16021 | 218 | 0 | 0 | 0 | 0 | 0 | 0 |
| ENSMUSG00 Gm5401 | 363 | 0 | 0 | 0 | 0 | 0 | 0 |
| ENSMUSG00 Gm13045 | 710 | 0 | 0 | 0 | 0 | 0 | 0 |
| ENSMUSG00 Gm13545 | 481 | 0 | 0 | 0 | 0 | 0 | 0 |
| ENSMUSG00 Gm12011 | 617 | 0 | 0 | 0 | 0 | 0 | 0 |
| ENSMUSG00 Gm12016 | 680 | 0 | 0 | 0 | 0 | 0 | 0 |
| ENSMUSG00 Fau-ps1 | 250 | 0 | 0 | 0 | 0 | 0 | 0 |
| ENSMUSG00 Gm13401 | 290 | 0 | 0 | 0 | 0 | 0.70381208 | 0 |
| ENSMUSG00 Gm13303 | 1283 | 0 | 0 | 0 | 0 | 0 | 0 |
| ENSMUSG00 Gm12486 | 540 | 0 | 0 | 0 | 0.0668186 | 0 | 0.05050859 |
| ENSMUSG00 Gm11414 | 262 | 0 | 0 | 0 | 0 | 0 | 0 |
| ENSMUSG00 Gm13740 | 309 | 0 | 0 | 0 | 0 | 0 | 0 |
| ENSMUSG00 Gm13126 | 3062 | 0 | 0 | 0 | 0 | 0 | 0 |

| | | | | | | | | |
|---|---|---|---|---|---|---|---|---|
| ENSMUSG00 | Gm11425 | 987 | 0 | 0 | 0 | 0 | 0 | 0 |
| ENSMUSG00 | Gm6517 | 520 | 0.68807562 | 0.37066494 | 0.25501063 | 1.11021676 | 0.39251058 | 0.94412213 |
| ENSMUSG00 | Gm15004 | 629 | 0 | 0 | 0 | 0 | 0 | 0 |
| ENSMUSG00 | Gm6973 | 720 | 0 | 0 | 0 | 0 | 0 | 0 |
| ENSMUSG00 | Gm15148 | 155 | 0 | 0 | 0 | 0 | 0 | 0 |
| ENSMUSG00 | Gm13292 | 1997 | 0 | 0.02412942 | 0.02213412 | 0.03613625 | 0.05110303 | 0.02731561 |
| ENSMUSG00 | Mup-ps13 | 539 | 0 | 0 | 0 | 0 | 0 | 0 |
| ENSMUSG00 | Gm14610 | 965 | 0 | 0 | 0 | 0 | 0 | 0 |
| ENSMUSG00 | Olfr1327-ps1 | 866 | 0 | 0 | 0 | 0 | 0 | 0 |
| ENSMUSG00 | Gm13742 | 456 | 0 | 0 | 0 | 0 | 0 | 0 |
| ENSMUSG00 | Gm13171 | 390 | 0 | 0 | 0 | 0 | 0 | 0 |
| ENSMUSG00 | Olfr1315-ps1 | 944 | 0 | 0 | 0 | 0 | 0 | 0 |
| ENSMUSG00 | Gm7110 | 596 | 0.33351913 | 1.21274603 | 0.6674775 | 1.27134721 | 0.59930307 | 0.73220508 |
| ENSMUSG00 | Gm11231 | 1705 | 0 | 0 | 0 | 0 | 0 | 0 |
| ENSMUSG00 | Gm9025 | 1566 | 0 | 0 | 0 | 0 | 0 | 0 |
| ENSMUSG00 | Gm5123 | 836 | 0 | 0 | 0 | 0 | 0 | 0 |
| ENSMUSG00 | Gm13813 | 786 | 0 | 0 | 0 | 0 | 0 | 0 |
| ENSMUSG00 | Gm11688 | 307 | 0.38849004 | 0 | 0.14397995 | 0 | 0 | 0 |
| ENSMUSG00 | Gm15136 | 532 | 0 | 0 | 0 | 0 | 0 | 0 |
| ENSMUSG00 | Gm15682 | 482 | 0.1649605 | 0 | 0 | 0 | 0 | 0 |
| ENSMUSG00 | Gm4914 | 458 | 0 | 0 | 0 | 0 | 0 | 0 |
| ENSMUSG00 | Gm6664 | 463 | 0 | 0 | 0 | 0 | 0 | 0 |
| ENSMUSG00 | Gm12686 | 396 | 0 | 0 | 0 | 0 | 0 | 0 |
| ENSMUSG00 | Gm12604 | 377 | 0 | 0 | 0 | 0 | 0 | 0 |
| ENSMUSG00 | Gm14709 | 303 | 0 | 0 | 0 | 0 | 0 | 0 |
| ENSMUSG00 | Gm13863 | 600 | 1.25892355 | 0 | 0.07366974 | 0 | 1.02052752 | 0 |
| ENSMUSG00 | Gm15364 | 584 | 0 | 0 | 0 | 0 | 0 | 0 |
| ENSMUSG00 | Gm11676 | 489 | 0 | 0 | 0 | 0 | 0 | 0 |
| ENSMUSG00 | Gm15244 | 129 | 0 | 0 | 0 | 0 | 0 | 0 |
| ENSMUSG00 | Mageb7-ps | 1040 | 0 | 0 | 0 | 0 | 0 | 0 |
| ENSMUSG00 | Gm11894 | 327 | 0 | 0 | 0 | 0 | 0 | 0 |
| ENSMUSG00 | Gm15069 | 190 | 0 | 0 | 0 | 0 | 0 | 0 |
| ENSMUSG00 | Gm11502 | 1210 | 0 | 0 | 0 | 0.02981987 | 0 | 0 |
| ENSMUSG00 | Gm12183 | 1847 | 0.90402284 | 0.70440387 | 0.88547276 | 0.46885169 | 0.93930524 | 0.75311673 |
| ENSMUSG00 | Gm13665 | 1063 | 0.11219797 | 0.36264491 | 0.54056817 | 1.08619513 | 1.00804694 | 1.28290871 |
| ENSMUSG00 | Gm11598 | 356 | 2.79181745 | 1.6242621 | 0.99329984 | 1.62166492 | 0.8599951 | 2.45165297 |
| ENSMUSG00 | Gm13776 | 1260 | 0 | 0.34418887 | 0 | 0 | 0 | 0 |
| ENSMUSG00 | Gm9430 | 1526 | 0 | 0 | 0 | 0 | 0 | 0 |
| ENSMUSG00 | Gm11988 | 1067 | 0 | 0 | 0 | 0 | 0 | 0 |
| ENSMUSG00 | Gm13685 | 140 | 0 | 0 | 0 | 0 | 0 | 0 |
| ENSMUSG00 | Gm8844 | 2613 | 0 | 0 | 0 | 0.041426 | 0 | 0 |
| ENSMUSG00 | Gm15147 | 482 | 0 | 0 | 0 | 0 | 0 | 0 |
| ENSMUSG00 | Gm15765 | 909 | 0 | 0 | 0.19450756 | 0 | 0 | 0 |
| ENSMUSG00 | Rab9b-ps1 | 605 | 0 | 0 | 0 | 0 | 0 | 0 |
| ENSMUSG00 | Rps6-ps3 | 749 | 0 | 0 | 0 | 0 | 0 | 0.07282948 |
| ENSMUSG00 | Gm14060 | 446 | 0 | 0 | 0 | 0 | 0 | 0 |
| ENSMUSG00 | Gm6743 | 1065 | 0.03732909 | 0.09049097 | 0 | 0 | 0 | 0.02560999 |
| ENSMUSG00 | Gm12980 | 335 | 0 | 0 | 0 | 0 | 0 | 0 |
| ENSMUSG00 | Gm13079 | 3575 | 0 | 0 | 0 | 0 | 0 | 0 |
| ENSMUSG00 | Defa-ps17 | 170 | 0 | 0 | 0 | 0 | 0 | 0 |
| ENSMUSG00 | Gm13501 | 978 | 0.04064978 | 0.09854078 | 0.31637311 | 0.14757482 | 0 | 0.02788818 |
| ENSMUSG00 | Kif22-ps | 1983 | 0 | 0 | 0 | 0 | 0 | 0 |
| ENSMUSG00 | Gm8213 | 440 | 0 | 0.10951464 | 0.10045873 | 0 | 0.11596904 | 0.7438538 |
| ENSMUSG00 | Gm7206 | 379 | 0 | 0 | 0 | 0 | 0 | 0 |
| ENSMUSG00 | Gm12242 | 737 | 0 | 0 | 0 | 0 | 0 | 0 |
| ENSMUSG00 | Gm11619 | 236 | 0 | 0 | 0 | 0 | 0 | 0.23114101 |
| ENSMUSG00 | Gm4746 | 2405 | 0 | 0 | 0 | 0 | 0 | 0 |
| ENSMUSG00 | Gm11687 | 885 | 0 | 0 | 0 | 0 | 0 | 0 |
| ENSMUSG00 | Gm12334 | 399 | 0 | 0 | 0 | 0 | 0 | 0 |
| ENSMUSG00 | H2afb2-ps | 335 | 0 | 0 | 0 | 0 | 0 | 0 |
| ENSMUSG00 | Gm13486 | 281 | 0 | 0 | 0 | 0 | 0 | 1.55300437 |
| ENSMUSG00 | Gm16177 | 717 | 0.27723487 | 0.47043946 | 0.06164832 | 0.35226543 | 0.07116649 | 0 |
| ENSMUSG00 | Gm15693 | 636 | 0 | 0 | 0 | 0 | 0 | 0 |
| ENSMUSG00 | Gm11919 | 720 | 0 | 0 | 0 | 0 | 0 | 0 |
| ENSMUSG00 | Gm11628 | 170 | 1.16927884 | 0 | 0 | 0 | 0.6003103 | 0.32087811 |
| ENSMUSG00 | 1700119I11R | 331 | 0 | 0 | 0 | 0 | 0 | 0 |
| ENSMUSG00 | Gm14356 | 868 | 0 | 0 | 0 | 0 | 0 | 0 |
| ENSMUSG00 | Gm11880 | 398 | 0 | 0 | 0 | 0 | 0 | 0 |
| ENSMUSG00 | Gm6909 | 1280 | 0.96282804 | 0.41410224 | 0.89784994 | 1.0148075 | 0.83715148 | 1.04410728 |
| ENSMUSG00 | Gm14871 | 193 | 0 | 0 | 0 | 0 | 0 | 0 |
| ENSMUSG00 | Gm14214 | 775 | 0 | 0 | 0 | 0 | 0 | 0 |
| ENSMUSG00 | Gm13687 | 281 | 0 | 0 | 0 | 0 | 0 | 0.19412555 |
| ENSMUSG00 | Gm12570 | 915 | 0 | 0.10532556 | 0 | 0 | 0 | 0 |
| ENSMUSG00 | Gm6798 | 359 | 0 | 0 | 0 | 0 | 0 | 0 |
| ENSMUSG00 | Gm40 | 658 | 0 | 0.07323168 | 0.20152816 | 0.49352341 | 0 | 0.16580328 |
| ENSMUSG00 | Gm12937 | 446 | 0 | 0 | 0 | 0 | 0 | 0 |
| ENSMUSG00 | Gm13615 | 1610 | 0 | 0 | 0 | 0 | 0 | 0 |
| ENSMUSG00 | Gm15195 | 262 | 0 | 0 | 0 | 0 | 0 | 0 |
| ENSMUSG00 | Gm11835 | 662 | 0 | 0 | 0 | 0 | 0 | 0 |
| ENSMUSG00 | Gm12636 | 623 | 0 | 0 | 0 | 0 | 0 | 0 |
| ENSMUSG00 | Gm12229 | 467 | 1.19181312 | 0.10318296 | 0.28395188 | 2.08611392 | 0 | 0.17521182 |
| ENSMUSG00 | Gm15003 | 445 | 0 | 0 | 0 | 0 | 0 | 0 |
| ENSMUSG00 | Gm14816 | 293 | 0 | 0 | 0 | 0 | 0 | 0 |
| ENSMUSG00 | Gm9378 | 470 | 0.76127516 | 0 | 0.3761859 | 0 | 0.7599673 | 0.23212459 |
| ENSMUSG00 | Gm13591 | 605 | 0.91996153 | 0.23894104 | 0.0730609 | 0 | 0 | 0 |
| ENSMUSG00 | Gm15673 | 937 | 0.04242847 | 0 | 0.18869517 | 0 | 0.16337153 | 0 |
| ENSMUSG00 | Gm11793 | 980 | 0.04056682 | 0 | 0 | 0 | 0.05206773 | 0.08349379 |
| ENSMUSG00 | Gm4915 | 1568 | 0 | 0 | 0 | 0 | 0 | 0 |
| ENSMUSG00 | Gm13937 | 987 | 0 | 0.04882112 | 0.22392018 | 0 | 0 | 0 |
| ENSMUSG00 | Gm15374 | 843 | 0 | 0 | 0 | 0.55642536 | 0 | 0 |
| ENSMUSG00 | Gm11452 | 689 | 0.05770026 | 0 | 0.06415362 | 0 | 0 | 0 |
| ENSMUSG00 | Gm12902 | 321 | 0.12384885 | 1.05079469 | 0 | 0 | 1.27168538 | 0 |
| ENSMUSG00 | Gm13342 | 206 | 0 | 0.46782954 | 0.85828821 | 0.17515556 | 0 | 0.26480232 |
| ENSMUSG00 | Gm5757 | 527 | 0 | 0 | 0 | 0 | 0 | 0 |
| ENSMUSG00 | Gm6221 | 1383 | 0 | 0 | 0 | 0 | 0 | 0 |
| ENSMUSG00 | Vamp7-ps | 654 | 1.64128131 | 1.4735915 | 2.70347664 | 3.08959403 | 1.2483517 | 2.00180839 |
| ENSMUSG00 | Gm13349 | 410 | 0 | 0.11752791 | 0.32342812 | 0 | 0.12445458 | 0.6652351 |
| ENSMUSG00 | Gm8402 | 1769 | 0 | 0 | 0 | 0 | 0 | 0 |
| ENSMUSG00 | Gm12667 | 1331 | 0 | 0 | 0 | 0 | 0 | 0 |
| ENSMUSG00 | Gm6272 | 170 | 0 | 0 | 0 | 0 | 0 | 0 |
| ENSMUSG00 | Gm12627 | 381 | 0 | 0 | 0 | 0 | 0 | 0 |
| ENSMUSG00 | Gm14067 | 313 | 0 | 0 | 0 | 0 | 0 | 0 |
| ENSMUSG00 | Gm8200 | 221 | 0 | 0 | 0 | 0 | 0 | 0 |
| ENSMUSG00 | Gm15182 | 1675 | 0 | 0 | 0 | 0 | 0.03046351 | 0 |
| ENSMUSG00 | Gm11199 | 506 | 0.07856814 | 0 | 0 | 0 | 0 | 0 |
| ENSMUSG00 | Gm12168 | 367 | 0 | 0 | 0 | 0 | 0 | 0.07431782 |
| ENSMUSG00 | Gm11556 | 287 | 0 | 0 | 0 | 0 | 0 | 0 |
| ENSMUSG00 | Rpl31-ps6 | 377 | 0 | 0 | 0.23449254 | 0.66995839 | 0.13534848 | 0 |
| ENSMUSG00 | Gm11253 | 200 | 0 | 0 | 0 | 0 | 0 | 0 |
| ENSMUSG00 | Gm13712 | 2292 | 0 | 0 | 0.01928527 | 0.04722781 | 0.02226282 | 0 |
| ENSMUSG00 | Gm7860 | 878 | 0 | 0 | 0 | 0 | 0 | 0 |
| ENSMUSG00 | Gm13456 | 1364 | 0.23316997 | 0.07065461 | 0.29165439 | 0.05290622 | 0.03740937 | 0.05998821 |
| ENSMUSG00 | Gm14981 | 942 | 0 | 0 | 0 | 0 | 0 | 0 |
| ENSMUSG00 | Gm14704 | 873 | 0 | 0 | 0 | 0 | 0 | 0 |
| ENSMUSG00 | Gm15207 | 441 | 0 | 0 | 0 | 0 | 0 | 0 |
| ENSMUSG00 | Gm11311 | 543 | 0 | 0 | 0 | 0 | 0 | 0 |
| ENSMUSG00 | Gm9095 | 880 | 0 | 0 | 0 | 0 | 0 | 0 |
| ENSMUSG00 | Gm13311 | 379 | 0.62937436 | 0 | 0.81639288 | 0 | 0 | 0.28785899 |
| ENSMUSG00 | Gm14336 | 474 | 0 | 0 | 0 | 0.07612246 | 0 | 0 |
| ENSMUSG00 | Gm11667 | 485 | 0 | 0 | 0 | 0 | 0 | 0 |
| ENSMUSG00 | Gm12196 | 261 | 0 | 0 | 0 | 0 | 0 | 0 |
| ENSMUSG00 | Gm14898 | 644 | 0 | 0 | 0 | 0 | 0 | 0 |
| ENSMUSG00 | Olfr1073-ps1 | 593 | 0 | 0 | 0 | 0 | 0 | 0 |
| ENSMUSG00 | Gm12377 | 251 | 0 | 0 | 0 | 0 | 0 | 0 |
| ENSMUSG00 | Gm14813 | 394 | 0 | 0 | 0 | 0 | 0 | 0.13844995 |
| ENSMUSG00 | Gm12344 | 328 | 0 | 0 | 0 | 0 | 0 | 0.49892633 |
| ENSMUSG00 | Gm15065 | 272 | 0 | 0 | 0 | 0 | 0 | 0 |
| ENSMUSG00 | Gm11850 | 478 | 0.24951138 | 0.30242537 | 0 | 0 | 0 | 0 |
| ENSMUSG00 | Gm11404 | 492 | 0 | 0 | 0 | 0 | 0 | 0 |
| ENSMUSG00 | Gm15806 | 954 | 0 | 0 | 0 | 0 | 0 | 0 |
| ENSMUSG00 | Gm11406 | 483 | 0 | 0 | 0 | 0 | 0 | 0 |
| ENSMUSG00 | Gm5387 | 856 | 0 | 0 | 0 | 0 | 0 | 0 |
| ENSMUSG00 | Gm13317 | 487 | 0 | 0 | 0 | 0.14818088 | 0 | 0 |
| ENSMUSG00 | Snrp1c-ps2 | 364 | 0 | 0 | 0 | 0 | 0 | 0 |
| ENSMUSG00 | Gm12812 | 432 | 0 | 0 | 0 | 0 | 0.11811661 | 0 |
| ENSMUSG00 | Gm15157 | 701 | 0 | 1.44353108 | 0 | 0 | 0 | 0 |
| ENSMUSG00 | Gm15580 | 555 | 0 | 0 | 0 | 0 | 0 | 0 |
| ENSMUSG00 | Gm11726 | 423 | 0 | 0 | 0 | 0 | 0 | 0 |
| ENSMUSG00 | Gm15026 | 457 | 0 | 0 | 0 | 0 | 0 | 0 |
| ENSMUSG00 | Gm15586 | 527 | 0 | 0 | 0 | 0 | 0 | 0 |
| ENSMUSG00 | Gm5255 | 640 | 0 | 0 | 0 | 0 | 0 | 0 |
| ENSMUSG00 | Gm11485 | 1174 | 0 | 0 | 0 | 0 | 0 | 0 |
| ENSMUSG00 | Gm14896 | 1350 | 0 | 0 | 0 | 0 | 0 | 0 |
| ENSMUSG00 | Gm14683 | 4287 | 0 | 0 | 0 | 0 | 0 | 0 |
| ENSMUSG00 | Gm12634 | 991 | 0.16046612 | 0.14587218 | 0 | 0.03640973 | 0.05148978 | 0.08256702 |
| ENSMUSG00 | Gm15711 | 534 | 0 | 0 | 0 | 0 | 0 | 0 |
| ENSMUSG00 | Gm8737 | 641 | 0 | 0 | 0 | 0 | 0 | 0 |
| ENSMUSG00 | Gm11903 | 726 | 0 | 0 | 0 | 0 | 0 | 0 |
| ENSMUSG00 | Olfr1268-ps1 | 923 | 0 | 0 | 0 | 0 | 0 | 0 |
| ENSMUSG00 | Gm13050 | 1344 | 0.02957997 | 0.10755902 | 0 | 0.1342338 | 0.15186421 | 0.10146815 |
| ENSMUSG00 | Gm12058 | 436 | 0.82064065 | 0 | 0.20276075 | 0.99308379 | 0.46813189 | 0 |
| ENSMUSG00 | Olfr1003-ps1 | 724 | 0 | 0 | 0 | 0 | 0 | 0 |
| ENSMUSG00 | Gm13527 | 690 | 0 | 0.13967085 | 0 | 0.20917127 | 0.07395127 | 0.03952846 |
| ENSMUSG00 | Gm11979 | 742 | 0 | 0 | 0 | 0 | 0 | 0 |
| ENSMUSG00 | Gm13182 | 768 | 0 | 0 | 0 | 0 | 0 | 0 |
| ENSMUSG00 | Gm13761 | 799 | 0.69659165 | 0 | 0 | 0.045159 | 0.0638628 | 0 |
| ENSMUSG00 | Olfr336-ps1 | 938 | 0 | 0 | 0 | 0 | 0 | 0 |
| ENSMUSG00 | Gm8569 | 646 | 0 | 0 | 0 | 0 | 0 | 0 |
| ENSMUSG00 | Gm15387 | 647 | 0 | 0 | 0 | 0 | 0 | 0.04215555 |
| ENSMUSG00 | Gm15436 | 342 | 0 | 0 | 0 | 0 | 0 | 0 |
| ENSMUSG00 | Gm12129 | 1146 | 0 | 0 | 0 | 0.06297041 | 0 | 0 |
| ENSMUSG00 | Gm15443 | 819 | 1.26207875 | 2.23575678 | 1.40323311 | 1.85036126 | 1.99370455 | 2.33116575 |
| ENSMUSG00 | Gm12433 | 254 | 0 | 0 | 0 | 0 | 0 | 0 |
| ENSMUSG00 | Gm14943 | 542 | 0 | 0 | 0 | 0 | 0 | 0 |
| ENSMUSG00 | Gm11558 | 458 | 0 | 0 | 0 | 0 | 0 | 0 |
| ENSMUSG00 | Gm12950 | 416 | 0 | 0 | 0 | 0 | 0 | 0 |
| ENSMUSG00 | Gm11331 | 287 | 0 | 0 | 0 | 0 | 0 | 0 |
| ENSMUSG00 | Gm15074 | 935 | 0 | 0 | 0 | 0 | 0 | 0 |
| ENSMUSG00 | Gm14793 | 310 | 0 | 0 | 0 | 0.11639369 | 0 | 0 |
| ENSMUSG00 | Gm14227 | 628 | 0.50643924 | 0.23019001 | 0.0703851 | 0 | 0.32500876 | 0.08686191 |
| ENSMUSG00 | Gm12645 | 274 | 0 | 0 | 0 | 0 | 0 | 0 |
| ENSMUSG00 | Gm15212 | 465 | 0 | 0 | 0 | 0 | 0 | 0 |
| ENSMUSG00 | Gm13141 | 455 | 0 | 0 | 0 | 0 | 0 | 0 |
| ENSMUSG00 | Gm9673 | 780 | 0 | 0 | 0.05666903 | 0.23129516 | 0 | 0.31470738 |
| ENSMUSG00 | Gm12700 | 456 | 0 | 0.21134404 | 0 | 0 | 0 | 0.05981281 |
| ENSMUSG00 | Gm15021 | 741 | 0 | 0 | 0 | 0 | 0 | 0 |
| ENSMUSG00 | Gm8482 | 470 | 0 | 0.41009738 | 0.09404647 | 0.30708123 | 0 | 0.1160623 |
| ENSMUSG00 | Gm16148 | 892 | 0.22284462 | 0 | 0 | 0 | 0 | 0 |
| ENSMUSG00 | Gm14638 | 213 | 0 | 0 | 0 | 0 | 0.4791209 | 0 |
| ENSMUSG00 | Gm12045 | 572 | 0 | 0.16848406 | 0.23182785 | 0.0630805 | 0.1784139 | 0.04768294 |
| ENSMUSG00 | Gm7670 | 640 | 0 | 0 | 0.20719614 | 0 | 0 | 0 |
| ENSMUSG00 | Gm15464 | 430 | 0 | 0 | 0 | 0 | 0 | 0 |
| ENSMUSG00 | Gm13829 | 137 | 0 | 1.40690342 | 0 | 0 | 0 | 0 |
| ENSMUSG00 | Gm7746 | 502 | 0 | 0 | 0 | 0 | 0 | 0 |
| ENSMUSG00 | 4930503H13l | 723 | 0 | 0 | 0 | 0 | 0 | 0 |
| ENSMUSG00 | Gm12518 | 778 | 0 | 0 | 0 | 0 | 0 | 0 |
| ENSMUSG00 | Gm12149 | 651 | 0 | 0 | 0 | 0 | 0 | 0 |
| ENSMUSG00 | Gm13239 | 636 | 0 | 0 | 0 | 0 | 0 | 0 |
| ENSMUSG00 | Gm6025 | 200 | 0 | 0 | 0 | 0 | 0 | 0 |
| ENSMUSG00 | Smt3h2-ps | 284 | 0 | 0 | 0 | 0 | 0 | 0 |
| ENSMUSG00 | Gm14852 | 641 | 0 | 0 | 0 | 0 | 0 | 0 |
| ENSMUSG00 | Gm16219 | 273 | 0 | 0 | 0 | 0 | 0 | 0 |
| ENSMUSG00 | Gm8244 | 263 | 0 | 0 | 0 | 0 | 0 | 0 |
| ENSMUSG00 | Rpl7a-ps13 | 562 | 0 | 0 | 0 | 0 | 0 | 0 |
| ENSMUSG00 | Gm12632 | 494 | 0 | 0 | 0 | 0 | 0 | 0 |
| ENSMUSG00 | Gm15493 | 527 | 0 | 0 | 0 | 0 | 0 | 0 |
| ENSMUSG00 | Gm12535 | 483 | 0 | 0 | 0 | 0 | 0 | 0 |
| ENSMUSG00 | Gm15204 | 2280 | 0 | 0.12680643 | 0 | 0 | 0.06713997 | 0.02392512 |
| ENSMUSG00 | Gm12270 | 455 | 0 | 0 | 0 | 0 | 0 | 0 |
| ENSMUSG00 | Gm12566 | 358 | 0 | 0 | 0 | 0 | 0 | 0 |
| ENSMUSG00 | Gm5639 | 2402 | 0 | 0 | 0 | 0 | 0 | 0 |
| ENSMUSG00 | Gm14834 | 793 | 0 | 0 | 0 | 0 | 0 | 0 |
| ENSMUSG00 | Gm13365 | 966 | 0 | 0 | 0.0457576 | 0.03735201 | 0 | 0 |
| ENSMUSG00 | Gm15779 | 411 | 0 | 0 | 0.43018825 | 0 | 0 | 0.26544661 |
| ENSMUSG00 | Gm12900 | 572 | 0 | 0 | 0 | 0 | 0 | 0 |
| ENSMUSG00 | Gm14750 | 512 | 0 | 0 | 0 | 0 | 0 | 0 |
| ENSMUSG00 | Gm14923 | 153 | 0 | 0 | 0 | 0 | 0 | 0 |
| ENSMUSG00 | Gm11248 | 280 | 0 | 0 | 0 | 0 | 0 | 0 |
| ENSMUSG00 | Gm12815 | 134 | 0 | 0 | 0 | 0 | 0 | 0 |
| ENSMUSG00 | Gm2012 | 1311 | 0 | 0 | 0 | 0 | 0 | 0 |
| ENSMUSG00 | Gm11402 | 308 | 0 | 0 | 0 | 0 | 0 | 0 |
| ENSMUSG00 | Rpl30-ps5 | 360 | 0 | 0 | 0 | 0 | 0 | 0 |
| ENSMUSG00 | Gm14362 | 272 | 0 | 0 | 0 | 0 | 0 | 0 |
| ENSMUSG00 | Mup-ps19 | 513 | 0 | 0 | 0 | 0 | 0 | 0 |
| ENSMUSG00 | Gm15481 | 538 | 0 | 0 | 0 | 0 | 0 | 0 |
| ENSMUSG00 | Gm12732 | 179 | 0 | 0 | 0 | 0 | 0 | 0 |
| ENSMUSG00 | Gm14056 | 641 | 0 | 0 | 0 | 0 | 0 | 0 |
| ENSMUSG00 | Gm11366 | 428 | 0.18577327 | 0 | 0 | 0 | 0.23844101 | 0 |
| ENSMUSG00 | Gm12754 | 453 | 0 | 0 | 0 | 0 | 0 | 0 |
| ENSMUSG00 | Gm15358 | 519 | 0 | 0 | 0 | 0 | 0 | 0 |
| ENSMUSG00 | Gm14001 | 246 | 0.64643058 | 0 | 0 | 0 | 0.20742429 | 0 |
| ENSMUSG00 | Gm14725 | 827 | 0 | 0 | 0 | 0 | 0 | 0 |
| ENSMUSG00 | Gm15897 | 1485 | 0 | 0 | 0 | 0 | 0 | 0 |
| ENSMUSG00 | Gm7857 | 357 | 0 | 0 | 0 | 0 | 0 | 0 |
| ENSMUSG00 | Gm11323 | 289 | 0 | 0 | 0 | 0 | 0 | 0 |
| ENSMUSG00 | Gm14736 | 335 | 0 | 0 | 0 | 0 | 0 | 0 |
| ENSMUSG00 | Fau-ps2 | 370 | 0.32234173 | 0 | 0 | 0.29255712 | 0.27581825 | 0.73715241 |
| ENSMUSG00 | Gm11394 | 515 | 0 | 0 | 0 | 0 | 0 | 0 |
| ENSMUSG00 | Gm14612 | 880 | 0 | 0 | 0 | 0 | 0 | 0.06198782 |
| ENSMUSG00 | Gm11590 | 1452 | 0 | 0 | 0 | 0 | 0 | 0 |
| ENSMUSG00 | Gm15516 | 192 | 0 | 0 | 0 | 0 | 0 | 0 |
| ENSMUSG00 | Trav13-3 | 495 | 0 | 0 | 0 | 0 | 0 | 0 |
| ENSMUSG00 | Gm8761 | 954 | 0 | 0 | 0 | 0 | 0 | 0 |
| ENSMUSG00 | Gm11470 | 1473 | 0 | 0 | 0 | 0.02449562 | 0.03464112 | 0.03703278 |
| ENSMUSG00 | Gm13677 | 594 | 0 | 0 | 0 | 0.06074418 | 0.08590299 | 0 |
| ENSMUSG00 | Gm11348 | 317 | 0 | 0 | 0 | 0 | 0 | 0 |
| ENSMUSG00 | Gm13258 | 289 | 0 | 0 | 0 | 0 | 0 | 0 |
| ENSMUSG00 | Gm5393 | 271 | 0 | 0 | 0 | 0 | 0 | 0.10064443 |
| ENSMUSG00 | Gm14050 | 345 | 0 | 0 | 0 | 0 | 0 | 0 |
| ENSMUSG00 | Gm14006 | 952 | 0.04175996 | 0.05061601 | 0.04643051 | 0.22740784 | 0 | 0.11459933 |
| ENSMUSG00 | Gm11876 | 268 | 0 | 0 | 0 | 0 | 0 | 0 |
| ENSMUSG00 | Gm11857 | 323 | 0 | 0 | 0 | 0 | 0 | 0 |
| ENSMUSG00 | Gm11914 | 506 | 0.15713629 | 0.09523012 | 0.08735542 | 0.28523355 | 0.60505584 | 0.75463429 |
| ENSMUSG00 | Gm6382 | 941 | 1.43643607 | 2.04830785 | 1.08038511 | 2.64576097 | 1.46409367 | 0.81157269 |
| ENSMUSG00 | Gm11843 | 424 | 0 | 0 | 0 | 0 | 0 | 0 |
| ENSMUSG00 | Gm15371 | 783 | 0 | 0 | 0 | 0.59906332 | 0 | 0 |
| ENSMUSG00 | Gm12818 | 1348 | 0 | 0.07149324 | 0.19674411 | 0.16060257 | 0.0378534 | 0.02023341 |
| ENSMUSG00 | Gm13469 | 349 | 0 | 0 | 0 | 0 | 0 | 0 |
| ENSMUSG00 | Gm14885 | 495 | 0 | 0 | 0 | 0 | 0 | 0 |
| ENSMUSG00 | Gm12675 | 172 | 0 | 0 | 0 | 0 | 0 | 0 |
| ENSMUSG00 | Gm15349 | 203 | 0 | 0 | 0.21774307 | 0 | 0.25136146 | 0 |
| ENSMUSG00 | Gm12697 | 666 | 0 | 0 | 0 | 0 | 0.15323236 | 0 |
| ENSMUSG00 | Gm12774 | 415 | 0 | 0.34833573 | 0 | 0 | 0.12295512 | 0.5914982 |
| ENSMUSG00 | Gm14689 | 228 | 0 | 0 | 0 | 0 | 0 | 0 |
| ENSMUSG00 | Gm12961 | 1049 | 0.07579691 | 0 | 0 | 0 | 0 | 0 |
| ENSMUSG00 | Gm11474 | 276 | 0.28808319 | 0 | 0 | 0 | 0 | 0 |
| ENSMUSG00 | Gm12124 | 678 | 0 | 0 | 0 | 0 | 0 | 0 |
| ENSMUSG00 | Gm14156 | 482 | 0 | 0 | 0 | 0 | 0 | 0 |
| ENSMUSG00 | Gm14972 | 363 | 0 | 0 | 0 | 0 | 0 | 0 |
| ENSMUSG00 | Dynll1-ps1 | 341 | 0.34975496 | 0.56523686 | 1.03699338 | 0.42324979 | 0.4489124 | 0.63987424 |
| ENSMUSG00 | Gm15652 | 304 | 0 | 0 | 0 | 0 | 0 | 0 |
| ENSMUSG00 | Gm15190 | 275 | 0.28913077 | 0.52567028 | 3.37541347 | 0 | 0.55665137 | 0.39672203 |
| ENSMUSG00 | Gm16187 | 196 | 0 | 0 | 0 | 0 | 0 | 0 |
| ENSMUSG00 | Gm15040 | 1238 | 0 | 0 | 0 | 0 | 0 | 0 |
| ENSMUSG00 | Gm12913 | 544 | 0.07307993 | 0 | 0 | 0 | 0.09379849 | 0 |
| ENSMUSG00 | Gm15316 | 420 | 0 | 0 | 0 | 0 | 0 | 0 |
| ENSMUSG00 | Gm12736 | 235 | 0 | 0 | 0 | 0 | 0 | 0 |
| ENSMUSG00 | Gm13792 | 422 | 0 | 0 | 0 | 0 | 0 | 0 |
| ENSMUSG00 | Gm14460 | 638 | 0 | 0 | 0 | 0 | 0 | 0 |
| ENSMUSG00 | Gm11907 | 1006 | 0.11855511 | 0.19159619 | 0.08787643 | 0.07173369 | 0 | 0.16267181 |
| ENSMUSG00 | Gm12515 | 360 | 0 | 0 | 0 | 0 | 0 | 0 |
| ENSMUSG00 | Gm15616 | 374 | 0.21259615 | 0 | 0 | 0 | 0 | 0.14585369 |
| ENSMUSG00 | Gm11663 | 289 | 0.13756222 | 0 | 0 | 0.12485137 | 0 | 0 |
| ENSMUSG00 | Gm6881 | 832 | 0 | 0 | 0 | 0 | 0 | 0 |
| ENSMUSG00 | Gm15172 | 1045 | 0 | 0.13833428 | 0 | 0 | 0 | 0 |
| ENSMUSG00 | Gm12362 | 341 | 0 | 0 | 0 | 0 | 0 | 0 |
| ENSMUSG00 | Defa-ps14 | 416 | 0 | 0 | 0 | 0 | 0 | 0 |
| ENSMUSG00 | Gm12987 | 488 | 0.65172919 | 1.28365522 | 0.18115509 | 0.59150893 | 1.56843368 | 0.33534393 |
| ENSMUSG00 | Olfr990-ps1 | 895 | 0 | 0 | 0 | 0 | 0 | 0 |
| ENSMUSG00 | Gm14721 | 524 | 0 | 0 | 0 | 0 | 0 | 0 |
| ENSMUSG00 | Gm15120 | 300 | 0 | 0 | 0 | 0 | 0 | 0 |
| ENSMUSG00 | Gm11633 | 443 | 0 | 0.21754601 | 0.19955685 | 0.08144931 | 0 | 0.55411231 |
| ENSMUSG00 | Rpl31-ps5 | 374 | 0 | 0 | 0 | 0 | 0 | 0.21878053 |
| ENSMUSG00 | Gm13087 | 255 | 0 | 0 | 0 | 0 | 0 | 0 |
| ENSMUSG00 | Gm14928 | 476 | 0 | 0 | 0 | 0 | 0 | 0 |
| ENSMUSG00 | Gm12489 | 147 | 0 | 0 | 0 | 0 | 0 | 0 |
| ENSMUSG00 | Gm12087 | 257 | 0 | 0 | 0 | 0 | 0 | 0 |
| ENSMUSG00 | Gm11393 | 286 | 0 | 0 | 0 | 0 | 0 | 0 |
| ENSMUSG00 | Gm14564 | 293 | 0 | 0 | 0 | 0 | 0 | 0 |
| ENSMUSG00 | Gm14416 | 2435 | 0 | 0 | 0 | 0 | 0.02095539 | 0 |
| ENSMUSG00 | Gm12451 | 629 | 0 | 0 | 0.1405464 | 0 | 0 | 0 |
| ENSMUSG00 | Gm11424 | 828 | 0 | 0 | 0 | 0 | 0 | 0 |
| ENSMUSG00 | Gm13260 | 868 | 0.09160249 | 0 | 0 | 0.04156918 | 0 | 0.06284479 |
| ENSMUSG00 | Fam48b1 | 1558 | 0 | 0 | 0 | 0 | 0 | 0 |
| ENSMUSG00 | Gm14845 | 305 | 0 | 0 | 0 | 0 | 0 | 0 |
| ENSMUSG00 | Gm11363 | 508 | 0 | 0 | 0 | 0 | 0 | 0 |
| ENSMUSG00 | Gm14467 | 272 | 0 | 0 | 0 | 0 | 0 | 0 |
| ENSMUSG00 | Rpl23a-ps6 | 467 | 0 | 0 | 0 | 0 | 0 | 0 |
| ENSMUSG00 | Gm15153 | 273 | 0 | 0 | 0 | 0 | 0 | 0.0999071 |
| ENSMUSG00 | Gm15474 | 133 | 0 | 0 | 0 | 0 | 0 | 0 |
| ENSMUSG00 | Gm16394 | 673 | 0 | 0 | 0 | 0 | 0 | 0 |
| ENSMUSG00 | Gm5381 | 1247 | 0 | 0 | 0 | 0 | 0 | 0 |
| ENSMUSG00 | Gm12026 | 1023 | 0 | 0.04710307 | 0 | 0 | 0 | 0 |
| ENSMUSG00 | Gm15422 | 540 | 0 | 0 | 0 | 0 | 0 | 0 |
| ENSMUSG00 | Olfr602-ps1 | 959 | 0 | 0 | 0 | 0 | 0 | 0 |
| ENSMUSG00 | Gm14127 | 676 | 0 | 0.07128172 | 0 | 0 | 0 | 0.0806942 |
| ENSMUSG00 | Gm9703 | 342 | 0 | 0.14089603 | 0 | 0 | 0 | 0 |
| ENSMUSG00 | Gm13354 | 168 | 0 | 0 | 0 | 0 | 0 | 0 |
| ENSMUSG00 | Gm12988 | 477 | 0.25003447 | 0.1010198 | 0.46333169 | 0.2269311 | 0.21394707 | 1.71538612 |
| ENSMUSG00 | Gm15625 | 1092 | 0.03640612 | 0 | 0 | 0 | 0.14018235 | 0 |
| ENSMUSG00 | Gm15181 | 176 | 0 | 0 | 0 | 0 | 0 | 0 |
| ENSMUSG00 | Gm9153 | 400 | 0.1987774 | 1.08419495 | 0.44201843 | 0 | 0.6378297 | 0.75005258 |
| ENSMUSG00 | Gm14265 | 304 | 0 | 0 | 0 | 0.11869094 | 0 | 0 |
| ENSMUSG00 | Gm12616 | 381 | 0 | 0 | 0 | 0 | 0 | 0 |
| ENSMUSG00 | Rpl9-ps2 | 576 | 0 | 0 | 0 | 0.06264244 | 0 | 0 |
| ENSMUSG00 | Gm12156 | 552 | 0 | 0 | 0 | 0 | 0 | 0 |
| ENSMUSG00 | Gm6620 | 713 | 0 | 0 | 0 | 0 | 0 | 0 |
| ENSMUSG00 | Gm14899 | 547 | 0 | 0 | 0 | 0 | 0.09328405 | 0 |
| ENSMUSG00 | Gm12614 | 381 | 0 | 0 | 0 | 0 | 0 | 0 |
| ENSMUSG00 | Olfr1021-ps1 | 948 | 0 | 0 | 0 | 0 | 0 | 0 |
| ENSMUSG00 | Gm8692 | 437 | 0 | 0 | 0 | 0 | 0 | 0 |
| ENSMUSG00 | Gm14888 | 173 | 0 | 0 | 0 | 0 | 0 | 0 |
| ENSMUSG00 | Gm6923 | 405 | 0 | 0 | 0 | 0 | 0 | 0 |
| ENSMUSG00 | Gm13024 | 717 | 0 | 0 | 0 | 0 | 0 | 0 |
| ENSMUSG00 | Gm14826 | 900 | 0 | 0 | 0 | 0.32072929 | 0 | 0 |
| ENSMUSG00 | Gm15546 | 825 | 0 | 0 | 0.42862393 | 0.52482974 | 0 | 0.03306017 |
| ENSMUSG00 | Gm12366 | 872 | 0.0911823 | 0.38681777 | 0.10138037 | 0.04137849 | 0 | 0.06255651 |
| ENSMUSG00 | Gm12457 | 167 | 0 | 0 | 0 | 0 | 0 | 0 |
| ENSMUSG00 | Gm7415 | 631 | 0 | 0 | 0 | 0 | 0 | 0 |
| ENSMUSG00 | Gm14411 | 911 | 1.52737851 | 0.21157604 | 1.11596311 | 4.55481352 | 1.73635307 | 0.17963539 |
| ENSMUSG00 | 1700064H15l | 392 | 0 | 0 | 0 | 0 | 0.13016933 | 0.34789081 |
| ENSMUSG00 | Gm11622 | 690 | 0 | 0 | 0 | 0 | 0 | 0 |
| ENSMUSG00 | Gm12727 | 358 | 0.99943945 | 0 | 0 | 0.20157567 | 0 | 0 |
| ENSMUSG00 | Gm15793 | 655 | 0 | 0 | 0 | 0 | 0 | 0 |
| ENSMUSG00 | Gm14722 | 688 | 0 | 0 | 0 | 0 | 0 | 0 |
| ENSMUSG00 | Gm12803 | 819 | 0 | 0 | 0 | 0 | 0 | 0.19981421 |
| ENSMUSG00 | Gm381 | 2036 | 0 | 0 | 0 | 0 | 0 | 0 |
| ENSMUSG00 | Gm15058 | 457 | 0 | 0 | 0 | 0 | 0 | 0 |
| ENSMUSG00 | Gm8805 | 312 | 0 | 0.30888745 | 0 | 0 | 0 | 0 |
| ENSMUSG00 | Itpa-ps3 | 394 | 0 | 0 | 0 | 0 | 0 | 0 |
| ENSMUSG00 | Gm7061 | 1415 | 0 | 0.03405402 | 0.15619026 | 0.53549324 | 0 | 0 |
| ENSMUSG00 | Gm15861 | 338 | 0.11761977 | 0 | 0 | 0 | 0 | 0 |
| ENSMUSG00 | Gm15191 | 999 | 0 | 0.19293871 | 0.04424609 | 0.10835449 | 0.25538727 | 0.10920777 |
| ENSMUSG00 | Gm13162 | 452 | 0 | 0 | 0 | 0 | 0 | 0 |
| ENSMUSG00 | Gm8329 | 453 | 0 | 0 | 0 | 0 | 0 | 0 |
| ENSMUSG00 | Gm15052 | 268 | 0 | 0 | 0 | 0 | 0 | 0 |
| ENSMUSG00 | Gm5319 | 872 | 0.04559115 | 0.05525968 | 0 | 0.04137849 | 0.05851649 | 0.06255651 |
| ENSMUSG00 | Gm9126 | 412 | 0 | 0 | 0 | 0 | 0 | 0 |
| ENSMUSG00 | Gm15957 | 973 | 0 | 0 | 0 | 0.14833317 | 0.05244232 | 0.02803149 |
| ENSMUSG00 | Gm11296 | 455 | 0 | 0 | 0 | 0 | 0 | 0 |
| ENSMUSG00 | Gm14489 | 900 | 0 | 0 | 0 | 0 | 0 | 0 |
| ENSMUSG00 | Gm13396 | 270 | 0 | 0 | 0 | 0 | 0 | 0 |
| ENSMUSG00 | Gm13284 | 545 | 0 | 0 | 0 | 0 | 0 | 0 |
| ENSMUSG00 | Gm12155 | 159 | 0.50006894 | 3.33365323 | 0.83399704 | 1.81544878 | 0 | 0 |
| ENSMUSG00 | Gm4875 | 785 | 0 | 0.61384003 | 0.95723737 | 1.93050429 | 0.19500526 | 1.21606672 |
| ENSMUSG00 | Gm12176 | 874 | 0 | 0 | 0.20229676 | 0 | 0 | 0 |
| ENSMUSG00 | Gm9308 | 213 | 0 | 0 | 0 | 0 | 0 | 0 |
| ENSMUSG00 | Gm6806 | 1005 | 0 | 0 | 0 | 0 | 0 | 0 |
| ENSMUSG00 | Gm13193 | 224 | 0.53243947 | 1.07559023 | 0 | 2.25512778 | 0.22779632 | 0 |
| ENSMUSG00 | Gm13385 | 219 | 0 | 0 | 0 | 0 | 0 | 0 |
| ENSMUSG00 | Gm12015 | 1261 | 0 | 0 | 0 | 0 | 0 | 0 |
| ENSMUSG00 | Gm11561 | 416 | 0 | 0 | 0 | 0 | 0 | 0 |
| ENSMUSG00 | Gm14554 | 652 | 0 | 0 | 0 | 0 | 0 | 0 |
| ENSMUSG00 | Gm13560 | 328 | 0 | 0 | 0 | 0 | 0 | 0 |
| ENSMUSG00 | Gm15465 | 124 | 0 | 0 | 0 | 0 | 0 | 0 |
| ENSMUSG00 | Gm14423 | 120 | 0 | 0 | 0 | 0 | 0 | 0 |
| ENSMUSG00 | Rpl17-ps1 | 1129 | 0 | 0 | 0 | 0 | 0 | 0 |
| ENSMUSG00 | Gm11395 | 497 | 0 | 0 | 0 | 0 | 0 | 0 |
| ENSMUSG00 | Rpl13-ps1 | 667 | 0 | 0.14448708 | 0 | 0.27048009 | 0 | 0 |
| ENSMUSG00 | Gm14782 | 194 | 0 | 0 | 0 | 0 | 0 | 0 |
| ENSMUSG00 | Gm11391 | 232 | 0 | 0 | 0 | 0 | 0 | 0 |
| ENSMUSG00 | Gm14150 | 1649 | 0.07232653 | 0.02922161 | 0.05361048 | 0.10940584 | 0 | 0.09924065 |
| ENSMUSG00 | Gm8682 | 497 | 0 | 0 | 0 | 0 | 0 | 0 |
| ENSMUSG00 | Gm14860 | 428 | 0 | 0 | 0 | 0 | 0 | 0 |
| ENSMUSG00 | Mup-ps6 | 539 | 0 | 0 | 0 | 0 | 0 | 0 |
| ENSMUSG00 | Gm12956 | 413 | 0 | 0 | 0 | 0 | 0 | 0 |
| ENSMUSG00 | Gm5678 | 831 | 0 | 0 | 0 | 0 | 0 | 0 |
| ENSMUSG00 | Gm11953 | 1002 | 0.07935226 | 0.14427079 | 0.08822723 | 0.18005012 | 0.15277358 | 0.0272202 |
| ENSMUSG00 | Gm16428 | 714 | 0 | 0 | 0 | 0 | 0 | 0 |
| ENSMUSG00 | Gm12175 | 344 | 0 | 0 | 0 | 0 | 0 | 0 |
| ENSMUSG00 | Trav4d-2 | 356 | 0 | 0 | 0 | 0 | 0 | 0 |
| ENSMUSG00 | Gm13803 | 805 | 0.19754276 | 0.65844828 | 0.05490912 | 0.13446725 | 0.31693401 | 0.67763079 |
| ENSMUSG00 | Gm12880 | 491 | 0 | 0 | 0 | 0 | 0 | 0 |
| ENSMUSG00 | Gm14731 | 375 | 0 | 0 | 0 | 0 | 0 | 0 |
| ENSMUSG00 | Gm15828 | 521 | 0 | 0 | 0 | 0 | 0 | 0 |
| ENSMUSG00 | Gm14716 | 717 | 0 | 0 | 0 | 0 | 0 | 0 |
| ENSMUSG00 | Gm15741 | 753 | 0 | 0 | 0 | 0 | 0 | 0 |
| ENSMUSG00 | Gm6097 | 344 | 0.11556826 | 0.14007687 | 0 | 0.20977933 | 0 | 0 |
| ENSMUSG00 | Gm5397 | 1451 | 0 | 0 | 0 | 0 | 0 | 0 |
| ENSMUSG00 | Gm16106 | 293 | 0 | 0 | 0 | 0 | 0 | 0 |
| ENSMUSG00 | Gm15780 | 934 | 0 | 0 | 0 | 0 | 0 | 0 |
| ENSMUSG00 | Gm12591 | 1174 | 2.50588208 | 2.50372485 | 1.39307342 | 3.01195942 | 0.99966494 | 3.22928012 |
| ENSMUSG00 | Gm14786 | 942 | 0 | 0 | 0 | 0 | 0 | 0 |
| ENSMUSG00 | Gm12982 | 189 | 0 | 0 | 0 | 0 | 0 | 0 |
| ENSMUSG00 | Defa-ps1 | 280 | 0 | 0 | 0 | 0 | 0 | 0 |
| ENSMUSG00 | Gm13118 | 548 | 0 | 0 | 0 | 0 | 0 | 0 |
| ENSMUSG00 | Gm13612 | 389 | 0.10219918 | 0 | 0 | 0.0927559 | 0 | 0.21034426 |
| ENSMUSG00 | Gm15373 | 213 | 0 | 0 | 0 | 0 | 0 | 0 |
| ENSMUSG00 | Gm14519 | 311 | 0 | 0 | 0 | 0 | 0 | 0 |
| ENSMUSG00 | Gm14588 | 171 | 0 | 0 | 0 | 0 | 0 | 0 |
| ENSMUSG00 | Gm16421 | 1556 | 0 | 0 | 0 | 0 | 0 | 0 |
| ENSMUSG00 | Gm12994 | 446 | 0 | 0 | 0.09910727 | 0 | 0 | 0 |
| ENSMUSG00 | Gm14748 | 408 | 0 | 0 | 0 | 0 | 0 | 0 |
| ENSMUSG00 | 4930448K20l | 997 | 0.03987511 | 0 | 0 | 0 | 0 | 0 |
| ENSMUSG00 | Gm13537 | 266 | 0.14945669 | 0 | 0 | 0 | 0 | 0 |
| ENSMUSG00 | Gm13879 | 241 | 0 | 0 | 0 | 0 | 0 | 0 |
| ENSMUSG00 | Gm13484 | 411 | 0 | 0 | 0 | 0 | 0 | 0 |
| ENSMUSG00 | Gm5759 | 810 | 0 | 0 | 0 | 0 | 0 | 0 |
| ENSMUSG00 | Gm16066 | 1083 | 0 | 0 | 0.04081426 | 0.09995026 | 0.09423153 | 0.17629037 |
| ENSMUSG00 | Gm14053 | 469 | 0 | 0 | 0 | 0 | 0 | 0 |
| ENSMUSG00 | Gm14299 | 396 | 0 | 0 | 0 | 0 | 0 | 0 |
| ENSMUSG00 | Gm11931 | 446 | 0 | 0.32412405 | 0 | 0 | 0 | 0 |
| ENSMUSG00 | Gm12021 | 1903 | 0 | 0 | 0.11613727 | 0 | 0.0536273 | 0.01433244 |
| ENSMUSG00 | Gm7134 | 2405 | 0 | 0 | 0 | 0 | 0 | 0 |
| ENSMUSG00 | Gm14537 | 597 | 0 | 0 | 0 | 0 | 0 | 0 |
| ENSMUSG00 | Gm15106 | 819 | 0 | 0 | 0 | 0 | 0 | 0 |
| ENSMUSG00 | Gm12293 | 589 | 0 | 0 | 0 | 0 | 0 | 0 |
| ENSMUSG00 | Gm11746 | 820 | 0 | 0 | 0 | 0 | 0 | 0 |
| ENSMUSG00 | Defa-ps9 | 274 | 0 | 0 | 0 | 0 | 0 | 0 |
| ENSMUSG00 | Gm13204 | 820 | 1.55143339 | 0.99898722 | 1.45542654 | 1.49608477 | 2.73800065 | 2.79398744 |
| ENSMUSG00 | Gm11362 | 1478 | 0 | 0.03260246 | 0.05981305 | 0 | 0 | 0 |
| ENSMUSG00 | Gm13806 | 324 | 0 | 0 | 0 | 0 | 0 | 0 |
| ENSMUSG00 | Gm14674 | 341 | 0 | 0 | 0 | 0 | 0 | 0 |
| ENSMUSG00 | Gm13358 | 568 | 0 | 0 | 0 | 0 | 0 | 0 |
| ENSMUSG00 | Gm14745 | 1095 | 0 | 0 | 0 | 0 | 0 | 0 |
| ENSMUSG00 | Gm15730 | 414 | 0 | 0 | 0 | 0.0871547 | 0 | 0 |
| ENSMUSG00 | Mup-ps5 | 98 | 0 | 0 | 0 | 0 | 0 | 0 |
| ENSMUSG00 | Gm11293 | 431 | 0 | 0 | 0 | 0 | 0 | 0 |
| ENSMUSG00 | Gm12165 | 341 | 0 | 0 | 0 | 0 | 0 | 0 |
| ENSMUSG00 | Gm13263 | 795 | 0 | 0 | 0 | 0 | 0 | 0 |
| ENSMUSG00 | Gm15773 | 1447 | 0.02747442 | 0 | 0 | 0 | 0.03526356 | 0.05654728 |

| | | | | | | | |
|---|---|---|---|---|---|---|---|
| ENSMUSG00 Gm15367 | 1015 | 0 | 0 | 0 | 0 | 0 | 0 |
| ENSMUSG00 Rpl15-ps1 | 563 | 0 | 0 | 0 | 0 | 0 | 0.09689037 |
| ENSMUSG00 Gm12598 | 829 | 0.14386784 | 0 | 0 | 0 | 0 | 0 |
| ENSMUSG00 Gm11889 | 1078 | 0 | 0 | 0 | 0 | 0 | 0 |
| ENSMUSG00 Gm13009 | 265 | 0 | 0 | 0 | 0 | 0 | 0 |
| ENSMUSG00 Gm12583 | 852 | 0.18664545 | 0 | 0 | 0.04234982 | 0 | 0.09603746 |
| ENSMUSG00 Gm13525 | 477 | 0 | 0 | 0 | 0 | 0 | 0 |
| ENSMUSG00 Gm14789 | 227 | 0 | 0 | 0 | 0 | 0 | 0 |
| ENSMUSG00 Olfr1185-ps1 | 911 | 0 | 0 | 0 | 0 | 0 | 0 |
| ENSMUSG00 Olfr1293-ps | 1825 | 0 | 0 | 0 | 0 | 0 | 0 |
| ENSMUSG00 Gm13339 | 509 | 2.03073181 | 2.65072766 | 1.30260834 | 2.1975312 | 2.00496565 | 0.69660179 |
| ENSMUSG00 Olfr475-ps1 | 931 | 0 | 0 | 0 | 0 | 0 | 0 |
| ENSMUSG00 Gm5386 | 2449 | 0 | 0 | 0 | 0 | 0 | 0 |
| ENSMUSG00 Gm14607 | 490 | 0 | 0.19667936 | 0 | 0 | 0 | 0 |
| ENSMUSG00 Gm12934 | 342 | 0.23248819 | 0.84537618 | 0 | 0 | 0 | 0.63800326 |
| ENSMUSG00 Gm12831 | 169 | 0 | 0 | 0 | 0 | 0 | 0 |
| ENSMUSG00 Gm15014 | 257 | 0 | 0 | 0 | 0 | 0 | 0 |
| ENSMUSG00 Gm12881 | 201 | 0 | 0 | 0 | 0 | 0 | 0 |
| ENSMUSG00 Olfr1027-ps1 | 1341 | 0 | 0 | 0 | 0 | 0 | 0 |
| ENSMUSG00 Gm11383 | 806 | 0 | 0 | 0 | 0 | 0 | 0.06767901 |
| ENSMUSG00 Gm6480 | 1885 | 0 | 0 | 0 | 0 | 0 | 0 |
| ENSMUSG00 Rpsa-ps9 | 884 | 0.04497226 | 0.2180382 | 0 | 0 | 0 | 0.21597565 |
| ENSMUSG00 Gm5844 | 2190 | 0.03630638 | 0.04400588 | 0.06055047 | 0.03295164 | 0.09319886 | 0 |
| ENSMUSG00 Gm11982 | 313 | 0 | 0 | 0 | 0 | 0 | 0 |
| ENSMUSG00 Gm12041 | 601 | 0 | 0 | 0 | 0 | 0 | 0 |
| ENSMUSG00 Gm12379 | 389 | 0 | 0 | 0 | 0 | 0 | 0 |
| ENSMUSG00 Gm11557 | 1007 | 0.19739563 | 0.19140593 | 0 | 0.03583123 | 0.10134335 | 0.2437654 |
| ENSMUSG00 Gm14323 | 626 | 0 | 0 | 0 | 0.05763905 | 0 | 0 |
| ENSMUSG00 Ccl19-ps1 | 323 | 0 | 0 | 0 | 0 | 0 | 0 |
| ENSMUSG00 Gm14083 | 619 | 0 | 0 | 0 | 0 | 0 | 0 |
| ENSMUSG00 Gm12615 | 381 | 0 | 0 | 0 | 0 | 0 | 0 |
| ENSMUSG00 Gm7823 | 1622 | 0 | 0 | 0 | 0 | 0 | 0 |
| ENSMUSG00 Smt3h1-ps | 279 | 0 | 0 | 0 | 0 | 0 | 0 |
| ENSMUSG00 Gm14832 | 321 | 0 | 0 | 0 | 0 | 0 | 0 |
| ENSMUSG00 Gm13736 | 512 | 0 | 0 | 1.64030277 | 0.14094549 | 0 | 0 |
| ENSMUSG00 Gm12398 | 306 | 0 | 0 | 0 | 0 | 0 | 0 |
| ENSMUSG00 Gm14758 | 137 | 0 | 0 | 0 | 0 | 0 | 0 |
| ENSMUSG00 Gm14119 | 234 | 0 | 0 | 0 | 0 | 0 | 0 |
| ENSMUSG00 Gm8770 | 1108 | 0.14352159 | 0 | 0 | 0 | 0 | 0 |
| ENSMUSG00 Gm14739 | 801 | 0 | 0 | 0 | 0 | 0 | 0 |
| ENSMUSG00 Gm14527 | 631 | 0 | 0 | 0 | 0 | 0 | 0 |
| ENSMUSG00 Gm15240 | 719 | 0 | 0 | 0 | 0 | 0 | 0 |
| ENSMUSG00 Gm11440 | 526 | 0.22674228 | 0 | 0.33613569 | 0 | 0 | 0 |
| ENSMUSG00 Defa-ps16 | 140 | 0 | 0 | 0 | 0 | 0 | 0 |
| ENSMUSG00 Gm12641 | 983 | 0.04044301 | 0 | 0.04496627 | 0.03670605 | 0.05190883 | 0.05549265 |
| ENSMUSG00 Gm13864 | 784 | 0.30425113 | 0.1843869 | 0.28189951 | 0.2761381 | 0.13016933 | 0.06957816 |
| ENSMUSG00 Gm6939 | 858 | 1.11204141 | 1.06706574 | 1.28793249 | 0.8410733 | 1.12995471 | 0.92187009 |
| ENSMUSG00 Gm11462 | 561 | 0 | 0 | 0 | 0 | 0 | 0 |
| ENSMUSG00 Gm13135 | 3120 | 0 | 0 | 0 | 0 | 0 | 0 |
| ENSMUSG00 Gm8172 | 1545 | 0.05146341 | 0 | 0.02860961 | 0.02335407 | 0.03302678 | 0.08826744 |
| ENSMUSG00 Gm5863 | 1007 | 0.27635389 | 0.23925741 | 0.26336749 | 0.28664981 | 0.2026867 | 0.10834018 |
| ENSMUSG00 Gm12785 | 171 | 0 | 0 | 0 | 0 | 0 | 0 |
| ENSMUSG00 Gm6067 | 490 | 0 | 0 | 0 | 0 | 0 | 0 |
| ENSMUSG00 Gm13462 | 960 | 0 | 0 | 0 | 0.03758546 | 0 | 0 |
| ENSMUSG00 Gm13586 | 463 | 0 | 0 | 0 | 0.46758589 | 0 | 0.05890851 |
| ENSMUSG00 Cyp2j8 | 1503 | 0.10580301 | 0.76944419 | 0.14704539 | 0.67218712 | 0 | 0 |
| ENSMUSG00 Gm14811 | 116 | 0 | 0 | 0 | 0 | 0 | 0 |
| ENSMUSG00 Gm13887 | 343 | 0 | 0 | 0 | 0 | 0 | 0 |
| ENSMUSG00 Gm7658 | 1123 | 0.24780798 | 0.25745205 | 0.23616301 | 0.22491034 | 0.04543756 | 0.17001111 |
| ENSMUSG00 Gm16166 | 786 | 0.10115898 | 0.24522362 | 0.56236442 | 0 | 0.06491905 | 0.27760447 |
| ENSMUSG00 Gm15095 | 1429 | 0 | 0 | 0 | 0 | 0 | 0 |
| ENSMUSG00 Gm2810 | 989 | 0.12059296 | 0 | 0 | 0 | 0 | 0.08273399 |
| ENSMUSG00 Gm14881 | 370 | 0 | 0 | 0 | 0 | 0 | 0 |
| ENSMUSG00 Gm14103 | 365 | 0 | 0 | 0 | 0 | 0 | 0 |
| ENSMUSG00 Gm12850 | 435 | 0 | 0 | 0 | 0 | 0.11730201 | 0 |
| ENSMUSG00 Gm13355 | 316 | 0 | 0 | 0 | 0 | 0 | 0 |
| ENSMUSG00 Gm13137 | 619 | 0 | 0 | 0 | 0 | 0 | 0 |
| ENSMUSG00 Gm11540 | 777 | 0 | 0 | 0 | 0 | 0 | 0 |
| ENSMUSG00 Gm12177 | 1995 | 0 | 0 | 0.08862525 | 0 | 0 | 0.06835749 |
| ENSMUSG00 Gm12587 | 426 | 0 | 0 | 0 | 0.16939927 | 0 | 0 |
| ENSMUSG00 Gm11920 | 562 | 0 | 0 | 0 | 0 | 0 | 0 |
| ENSMUSG00 Gm7834 | 436 | 0 | 0 | 0 | 0 | 0 | 0 |
| ENSMUSG00 Gm13359 | 694 | 0 | 0 | 0 | 0.05199142 | 0 | 0 |
| ENSMUSG00 Gm12507 | 414 | 0 | 0 | 0 | 0 | 0 | 0 |
| ENSMUSG00 Gm13217 | 1513 | 0 | 0.03184828 | 0 | 0.04769603 | 0.13490119 | 0.07210744 |
| ENSMUSG00 Gm15504 | 264 | 0 | 0 | 0 | 0 | 0.19328173 | 0 |
| ENSMUSG00 Gm8036 | 1716 | 0.37068047 | 0 | 0 | 0 | 0 | 0.01589431 |
| ENSMUSG00 Naip3 | 4854 | 0 | 0 | 0 | 0 | 0 | 0 |
| ENSMUSG00 Gm15303 | 2623 | 0 | 0 | 0 | 0 | 0 | 0 |
| ENSMUSG00 Gm12731 | 267 | 0 | 0 | 0 | 0 | 0 | 0 |
| ENSMUSG00 Gm2566 | 397 | 0 | 0 | 0 | 0 | 0 | 0 |
| ENSMUSG00 Gm6884 | 614 | 0 | 0.54935683 | 0 | 0 | 0 | 0 |
| ENSMUSG00 Gm15131 | 594 | 0 | 0 | 0 | 0 | 0 | 0.1377507 |
| ENSMUSG00 Gm10601 | 2092 | 0 | 0 | 0 | 0 | 0 | 0.02607518 |
| ENSMUSG00 Gm12465 | 299 | 0 | 0 | 0 | 0 | 0 | 0 |
| ENSMUSG00 Rpl13-ps5 | 629 | 0 | 0 | 0 | 0 | 0.40561507 | 0.52034288 |
| ENSMUSG00 Gm6754 | 912 | 0 | 0 | 0 | 0 | 0 | 0 |
| ENSMUSG00 Gm14614 | 383 | 0 | 0 | 0 | 0 | 0 | 0 |
| ENSMUSG00 Gm16082 | 376 | 0 | 0 | 0 | 0 | 0 | 0 |
| ENSMUSG00 Gm14581 | 197 | 0 | 0 | 0 | 0 | 0 | 0 |
| ENSMUSG00 Gm12630 | 783 | 0.35541298 | 0.30770397 | 0 | 0.09216359 | 0 | 0 |
| ENSMUSG00 Gm15214 | 469 | 0 | 0 | 0 | 0 | 0 | 0 |
| ENSMUSG00 Gm13517 | 1494 | 0 | 0 | 0 | 0 | 0 | 0 |
| ENSMUSG00 Gm13810 | 1346 | 0 | 0 | 0 | 0 | 0 | 0 |
| ENSMUSG00 Gm13672 | 458 | 0 | 0 | 0 | 0 | 0 | 0 |
| ENSMUSG00 Gm15129 | 278 | 0 | 0 | 0 | 0 | 0 | 0 |
| ENSMUSG00 Gm14363 | 407 | 0 | 0 | 0 | 0 | 0 | 0 |
| ENSMUSG00 Gm15056 | 371 | 0 | 0 | 0 | 0 | 0 | 0 |
| ENSMUSG00 Gm11911 | 1022 | 0 | 0 | 0 | 0 | 0 | 0 |
| ENSMUSG00 Rpsa-ps11 | 888 | 0 | 0 | 0.0995537 | 0 | 0 | 0.03071468 |
| ENSMUSG00 Gm14996 | 511 | 0 | 0 | 0 | 0 | 0 | 0 |
| ENSMUSG00 Olfr1274-ps | 1963 | 0 | 0 | 0 | 0 | 0 | 0 |
| ENSMUSG00 Gm11526 | 174 | 0.91391909 | 0.83080073 | 0.25403358 | 0.82947229 | 0.58651007 | 2.0377604 |
| ENSMUSG00 Olfr1149-ps1 | 444 | 0 | 0 | 0 | 0 | 0 | 0 |
| ENSMUSG00 Gm12318 | 272 | 0 | 0 | 0 | 0 | 0 | 0 |
| ENSMUSG00 Gm10599 | 2993 | 0 | 0 | 0 | 0 | 0 | 0 |
| ENSMUSG00 Gm14042 | 832 | 0 | 0.0579164 | 0 | 0 | 0 | 0 |
| ENSMUSG00 Gm11389 | 427 | 0 | 0 | 0 | 0 | 0 | 0 |
| ENSMUSG00 Gm11654 | 427 | 0 | 0 | 0 | 0 | 0 | 0 |
| ENSMUSG00 Gm5903 | 2324 | 0.01710649 | 0 | 0.03803945 | 0.01552584 | 0.13173763 | 0.08215253 |
| ENSMUSG00 Gm13557 | 380 | 0 | 0 | 0 | 0 | 0 | 0 |
| ENSMUSG00 Gm14052 | 1030 | 0.03859755 | 0.32748068 | 0.21457205 | 0.07006222 | 1.53574529 | 0 |
| ENSMUSG00 Gm11806 | 499 | 0 | 0 | 0 | 0 | 0 | 0 |
| ENSMUSG00 Gm11566 | 202 | 0 | 0 | 0 | 0 | 0 | 0 |
| ENSMUSG00 Gm11848 | 972 | 0 | 0 | 0 | 0 | 0 | 0 |
| ENSMUSG00 Gm15126 | 831 | 0 | 0 | 0 | 0 | 0 | 0 |
| ENSMUSG00 Gm6238 | 464 | 1.11383889 | 0 | 0.09526259 | 0 | 0 | 0 |
| ENSMUSG00 Gm13975 | 781 | 0 | 0 | 0 | 0 | 0 | 0 |
| ENSMUSG00 Gm11221 | 1280 | 0 | 0.03764566 | 0.17266345 | 0.0563782 | 0.15945742 | 0.12784987 |
| ENSMUSG00 Gm13238 | 779 | 0 | 0 | 0 | 0 | 0 | 0 |
| ENSMUSG00 Gm15678 | 958 | 0.04149841 | 0 | 0 | 0 | 0 | 0 |
| ENSMUSG00 Gm13250 | 752 | 0 | 0 | 0 | 0 | 0 | 0 |
| ENSMUSG00 Gm13413 | 601 | 0 | 0 | 0.14709432 | 0 | 0.33960982 | 0.09076419 |
| ENSMUSG00 Gm12392 | 937 | 0 | 0 | 0 | 0 | 0 | 0 |
| ENSMUSG00 Gm15715 | 282 | 0 | 0 | 0 | 0 | 0 | 0 |
| ENSMUSG00 Gm13623 | 624 | 0.12742141 | 0 | 0.21250886 | 0 | 0.16354608 | 0 |
| ENSMUSG00 Gm12053 | 250 | 0 | 0 | 0 | 0 | 0 | 0 |
| ENSMUSG00 Gm1335 | 384 | 1.03529897 | 1.12936974 | 0.57554483 | 0.37585463 | 0 | 0.85233248 |
| ENSMUSG00 Gm14687 | 361 | 0 | 0 | 0 | 0 | 0 | 0 |
| ENSMUSG00 Gm12816 | 1211 | 0 | 0 | 0.03650028 | 0.02979525 | 0 | 0.06756723 |
| ENSMUSG00 2810453I06R | 3117 | 0.02550881 | 0.21642932 | 0.01418089 | 0.01157589 | 0.04911105 | 0.05250171 |
| ENSMUSG00 Gm12978 | 521 | 0 | 0 | 0 | 0 | 0 | 0 |
| ENSMUSG00 Gm11764 | 622 | 0 | 0 | 0 | 0 | 0 | 0 |
| ENSMUSG00 Gm16518 | 506 | 0 | 0 | 0.08735542 | 0 | 0 | 0 |
| ENSMUSG00 Gm11703 | 1022 | 0.11669906 | 0 | 0 | 0 | 0 | 0 |
| ENSMUSG00 Gm12163 | 388 | 0 | 0 | 0.11392228 | 0 | 0 | 0.28118185 |
| ENSMUSG00 Gm13555 | 318 | 0 | 0 | 0 | 0 | 0 | 0 |
| ENSMUSG00 Gm6222 | 395 | 0.40258714 | 0 | 0 | 0.1826939 | 0 | 0.20714916 |
| ENSMUSG00 Gm15331 | 384 | 0 | 0 | 0 | 0 | 0 | 0 |
| ENSMUSG00 Gm14370 | 867 | 0 | 0 | 0 | 0 | 0 | 0 |
| ENSMUSG00 Rps15a-ps6 | 392 | 0.50708521 | 0.61462299 | 0.45103921 | 0.2761381 | 0.91118528 | 0.34789081 |
| ENSMUSG00 Gm11804 | 467 | 0 | 0 | 0 | 0 | 0 | 0 |
| ENSMUSG00 Gm14982 | 441 | 0 | 0 | 0 | 0 | 0 | 0 |
| ENSMUSG00 Gm11829 | 281 | 0 | 0 | 0 | 0 | 0 | 0 |
| ENSMUSG00 Gm6373 | 359 | 0 | 0 | 0 | 0 | 0 | 0 |
| ENSMUSG00 Gm13140 | 3149 | 0 | 0.04590642 | 0 | 0 | 0.03240799 | 0 |
| ENSMUSG00 Gm14604 | 441 | 0 | 0 | 0 | 0 | 0 | 0 |
| ENSMUSG00 Gm13720 | 195 | 0 | 0 | 0 | 0 | 0 | 0 |
| ENSMUSG00 Gm14561 | 795 | 0 | 0 | 0 | 0 | 0 | 0 |
| ENSMUSG00 Gm14910 | 1804 | 0 | 0 | 0 | 0 | 0 | 0 |
| ENSMUSG00 Gm8087 | 566 | 0 | 0 | 0 | 0 | 0 | 0 |
| ENSMUSG00 Gm9083 | 818 | 0.29160499 | 0 | 0 | 0 | 0 | 0.03334308 |
| ENSMUSG00 Olfr334-ps1 | 905 | 0 | 0 | 0 | 0 | 0 | 0 |
| ENSMUSG00 Gm15959 | 367 | 0 | 0 | 0 | 0 | 0 | 0 |
| ENSMUSG00 Gm14900 | 732 | 0 | 0 | 0 | 0 | 0 | 0 |
| ENSMUSG00 Gm13233 | 1441 | 0 | 0 | 0 | 0 | 0 | 0 |
| ENSMUSG00 Gm11606 | 616 | 0 | 0 | 0 | 0 | 0 | 0 |
| ENSMUSG00 Gm9835 | 425 | 0 | 0 | 0 | 0 | 0 | 0 |
| ENSMUSG00 Gm11388 | 158 | 0 | 0 | 0 | 0 | 0 | 0 |
| ENSMUSG00 Gm13433 | 239 | 0.16634092 | 0 | 0 | 0 | 0 | 0.11411983 |
| ENSMUSG00 Gm12416 | 1007 | 0.07895825 | 0.04785148 | 0.04389458 | 0.25081858 | 0.15201502 | 0.10834018 |
| ENSMUSG00 Gm16100 | 230 | 0 | 0 | 0 | 0 | 0 | 0 |
| ENSMUSG00 Gm14699 | 227 | 0 | 0 | 0 | 0 | 0 | 0 |
| ENSMUSG00 Gm14082 | 800 | 0 | 0 | 0 | 0 | 0 | 0 |
| ENSMUSG00 Gm14595 | 694 | 0 | 0 | 0 | 0 | 0 | 0 |
| ENSMUSG00 Gm14822 | 290 | 0 | 0.16616015 | 0 | 0.12442084 | 0.17595302 | 0.09405048 |
| ENSMUSG00 Gm11242 | 2456 | 0 | 0 | 0 | 0 | 0 | 0 |
| ENSMUSG00 Gm14625 | 750 | 0 | 0 | 0 | 0 | 0 | 0 |
| ENSMUSG00 Gm12285 | 232 | 0.17135983 | 0 | 0 | 0.77763027 | 0 | 0 |
| ENSMUSG00 Gm15001 | 790 | 0 | 0 | 0 | 0 | 0 | 0 |
| ENSMUSG00 Gm12986 | 966 | 0 | 0 | 0 | 0 | 0 | 0 |
| ENSMUSG00 Gm14140 | 994 | 0 | 0 | 0 | 0 | 0 | 0 |
| ENSMUSG00 Gm2092 | 631 | 0 | 0 | 0 | 0 | 0 | 0 |
| ENSMUSG00 Gm13356 | 334 | 0 | 0 | 0 | 0 | 0 | 0 |
| ENSMUSG00 Gm14795 | 470 | 0 | 0 | 0 | 0 | 0 | 0 |
| ENSMUSG00 Olfr268-ps1 | 1611 | 0 | 0 | 0 | 0 | 0 | 0 |
| ENSMUSG00 Gm14533 | 92 | 0 | 0 | 0 | 0 | 0 | 0 |
| ENSMUSG00 Gm12191 | 347 | 0 | 0 | 0 | 0 | 0 | 0 |
| ENSMUSG00 Gm14980 | 735 | 0 | 0 | 0 | 0 | 0.06942364 | 0 |
| ENSMUSG00 Gm14882 | 217 | 0 | 0 | 0 | 0 | 0 | 0 |
| ENSMUSG00 Gm7785 | 891 | 0.26771367 | 0.27040652 | 0.34726476 | 0.08099224 | 0.2863433 | 1.89789858 |
| ENSMUSG00 Gm13195 | 260 | 0 | 0 | 0 | 0 | 0 | 0 |
| ENSMUSG00 Gm12748 | 503 | 0 | 0 | 0 | 0 | 0 | 0 |
| ENSMUSG00 Gm14751 | 221 | 0 | 0 | 0 | 0 | 0 | 0 |
| ENSMUSG00 Gm11800 | 577 | 0 | 0 | 0 | 0 | 0 | 0 |
| ENSMUSG00 Gm16431 | 427 | 0 | 0 | 0 | 0 | 0 | 0 |
| ENSMUSG00 Gm13350 | 478 | 0 | 0 | 0 | 0 | 0 | 0 |
| ENSMUSG00 Gm13668 | 995 | 0.03995526 | 0.04842859 | 0 | 0.25384353 | 0 | 0.13705849 |
| ENSMUSG00 Gm11845 | 326 | 0 | 0 | 0 | 0 | 0 | 0 |
| ENSMUSG00 Gm14670 | 336 | 0 | 0 | 0 | 0 | 0 | 0 |
| ENSMUSG00 Gm11980 | 985 | 0 | 0 | 0 | 0 | 0 | 0 |
| ENSMUSG00 Gm11408 | 518 | 0.92097638 | 1.02326422 | 0.76798569 | 0.62690811 | 0.49253259 | 0.15796123 |
| ENSMUSG00 Gm14126 | 394 | 0 | 0 | 0 | 0 | 0 | 0 |
| ENSMUSG00 Gm13172 | 255 | 0 | 0.37793288 | 0 | 0 | 0 | 0.53479685 |
| ENSMUSG00 Amy2b | 1564 | 0 | 0 | 0 | 0 | 0 | 0 |
| ENSMUSG00 Gm11484 | 337 | 0 | 0 | 0 | 0 | 0 | 0 |
| ENSMUSG00 Gm13223 | 746 | 0 | 0 | 0 | 0 | 0 | 0 |
| ENSMUSG00 Gm15382 | 536 | 0.37085336 | 0.17980016 | 0.24739838 | 0 | 0.66638924 | 0.50885521 |
| ENSMUSG00 Gm14667 | 1349 | 0 | 0 | 0.03276638 | 0.02674725 | 0.113476 | 0.06065524 |
| ENSMUSG00 Gm12765 | 597 | 0 | 0 | 0 | 0 | 0 | 0 |
| ENSMUSG00 Gm15531 | 251 | 0 | 0 | 0 | 0 | 0 | 0 |
| ENSMUSG00 Gm11249 | 858 | 0 | 0 | 0.2060692 | 0 | 0 | 0.03178862 |
| ENSMUSG00 Gm14153 | 1138 | 0 | 0 | 0 | 0 | 0 | 0 |
| ENSMUSG00 Gm4760 | 1029 | 0 | 0 | 0 | 0 | 0 | 0.05301293 |
| ENSMUSG00 Gm11736 | 504 | 0 | 0 | 0 | 0 | 0 | 0 |
| ENSMUSG00 Gm12438 | 335 | 0 | 0 | 0 | 0 | 0 | 0 |
| ENSMUSG00 Gm15385 | 661 | 0 | 0 | 0 | 0 | 0 | 0 |
| ENSMUSG00 Gm4908 | 1211 | 0.13131455 | 0.07958124 | 0.14600113 | 0.08938574 | 0.16854294 | 0.04504482 |
| ENSMUSG00 Gm11863 | 230 | 0 | 0 | 0 | 0 | 0 | 0 |
| ENSMUSG00 Gm11356 | 340 | 0 | 0 | 0 | 0 | 0 | 0 |
| ENSMUSG00 Gm14494 | 1215 | 0 | 0.03965962 | 0 | 0 | 0 | 0.15713784 |
| ENSMUSG00 Gm14578 | 390 | 0 | 0 | 0 | 0 | 0 | 0 |
| ENSMUSG00 Gm4732 | 467 | 0 | 0 | 0 | 0 | 0 | 0 |
| ENSMUSG00 Gm6044 | 498 | 0.07983028 | 0.3870397 | 0 | 0 | 0 | 0.05476835 |
| ENSMUSG00 Gm13430 | 287 | 0 | 0 | 0 | 0 | 0 | 0 |
| ENSMUSG00 Gm16424 | 398 | 0 | 0 | 0 | 0 | 0 | 0 |
| ENSMUSG00 Gm6429 | 474 | 0 | 0 | 0 | 0 | 0 | 0 |
| ENSMUSG00 Gm12917 | 638 | 0 | 0.67974605 | 0.69281886 | 1.1876535 | 0.39989323 | 0.25650131 |
| ENSMUSG00 Gm12393 | 847 | 0 | 0 | 0 | 0 | 0 | 0 |
| ENSMUSG00 Gm15309 | 239 | 0 | 0 | 0 | 0 | 0 | 0 |
| ENSMUSG00 Gm5303 | 1251 | 0 | 0 | 0 | 0 | 0 | 0 |
| ENSMUSG00 Gm12876 | 440 | 0 | 0 | 0 | 0 | 0 | 0 |
| ENSMUSG00 Gm14073 | 380 | 0 | 0 | 0 | 0 | 0 | 0 |
| ENSMUSG00 Gm14421 | 1955 | 0.06100585 | 0 | 0 | 0 | 0.07830134 | 0 |
| ENSMUSG00 Gm14620 | 382 | 0 | 0 | 0 | 0 | 0 | 0 |
| ENSMUSG00 Rpl10-ps4 | 285 | 0 | 0 | 0 | 0 | 0 | 0 |
| ENSMUSG00 Gm14790 | 1291 | 0 | 0 | 0 | 0 | 0 | 0 |
| ENSMUSG00 Gm13386 | 123 | 0 | 0 | 0 | 0 | 0 | 0 |
| ENSMUSG00 Gm13410 | 1413 | 0 | 0.03410222 | 0 | 0 | 0 | 0.09651323 |
| ENSMUSG00 Gm15171 | 333 | 0 | 0 | 0 | 0 | 0 | 0 |
| ENSMUSG00 Gm11609 | 791 | 0 | 0 | 0 | 0 | 0 | 0 |
| ENSMUSG00 Gm14724 | 536 | 0 | 0 | 0 | 0 | 0 | 0 |
| ENSMUSG00 Gm15854 | 802 | 0 | 0 | 0 | 0 | 0 | 0 |
| ENSMUSG00 Gm6341 | 872 | 0 | 0 | 0 | 0 | 0 | 0 |
| ENSMUSG00 Gm15578 | 474 | 0 | 0 | 0 | 0 | 0 | 0 |
| ENSMUSG00 Gm6826 | 527 | 0 | 0 | 0 | 0 | 0 | 0 |
| ENSMUSG00 Gm11327 | 925 | 0 | 0 | 0 | 0 | 0 | 0 |
| ENSMUSG00 Gm16203 | 495 | 0 | 0 | 0 | 0 | 0 | 0 |
| ENSMUSG00 Gm14109 | 308 | 0 | 0 | 0 | 0 | 0 | 0 |
| ENSMUSG00 Gm15189 | 1964 | 0.0202421 | 0 | 0 | 0 | 0 | 0 |
| ENSMUSG00 Gm12723 | 450 | 0 | 0 | 0 | 0 | 0 | 0 |
| ENSMUSG00 Gm12210 | 152 | 0 | 0 | 0 | 0 | 0 | 0 |
| ENSMUSG00 Gm14560 | 374 | 0 | 0 | 0 | 0 | 0 | 0 |
| ENSMUSG00 Gm14422 | 150 | 0 | 0 | 0 | 0 | 0 | 0 |
| ENSMUSG00 Gm14844 | 272 | 0 | 0 | 0 | 0 | 0 | 0 |
| ENSMUSG00 Gm11504 | 172 | 0 | 0 | 0 | 0 | 0 | 0 |
| ENSMUSG00 Gm15170 | 2021 | 0 | 0 | 0 | 0 | 0 | 0 |
| ENSMUSG00 Gm11792 | 167 | 0 | 0 | 0 | 0 | 0 | 0 |
| ENSMUSG00 Cyp4a29-ps | 1518 | 0 | 0 | 0 | 0 | 0 | 0 |
| ENSMUSG00 Gm12418 | 941 | 0.04224812 | 0 | 0.18789306 | 0.11503309 | 0.27112846 | 0.08695422 |
| ENSMUSG00 Gm5073 | 819 | 0 | 0 | 0 | 0 | 0 | 0 |
| ENSMUSG00 Gm15752 | 393 | 0 | 0 | 0 | 0 | 0 | 0 |
| ENSMUSG00 Gm12288 | 1281 | 0 | 0 | 0 | 0 | 0 | 0 |
| ENSMUSG00 Gm13424 | 404 | 0 | 0 | 0 | 0 | 0 | 0 |
| ENSMUSG00 Gm14334 | 692 | 0 | 0 | 0 | 0 | 0 | 0 |
| ENSMUSG00 Gm12579 | 782 | 0 | 0 | 0 | 0 | 0 | 0 |
| ENSMUSG00 Gm15921 | 482 | 0 | 0.29991562 | 0 | 0 | 0.31759155 | 0 |
| ENSMUSG00 Olfr1103-ps1 | 937 | 0 | 0 | 0 | 0.03850805 | 0 | 0 |
| ENSMUSG00 Gm15102 | 1501 | 0 | 0 | 0 | 0 | 0 | 0 |
| ENSMUSG00 Gm11380 | 999 | 0 | 0 | 0 | 0 | 0 | 0 |
| ENSMUSG00 Gm11852 | 221 | 0 | 0 | 0 | 0 | 0 | 0 |
| ENSMUSG00 Gm4913 | 1289 | 0 | 0 | 0 | 0 | 0 | 0 |
| ENSMUSG00 Apc-ps1 | 469 | 0.67813186 | 0.20548589 | 0.753976 | 0.076934 | 2.17596485 | 0.11630976 |
| ENSMUSG00 Gm12459 | 354 | 0 | 0 | 0 | 0.20385336 | 0 | 0 |
| ENSMUSG00 Gm11892 | 1201 | 0 | 0 | 0 | 0 | 0 | 0 |
| ENSMUSG00 Gm11866 | 334 | 0 | 0 | 0 | 0 | 0 | 0 |
| ENSMUSG00 Gm12201 | 1232 | 0 | 0 | 0.1793906 | 0.02928737 | 0.04141751 | 0 |
| ENSMUSG00 Gm13511 | 362 | 0.65893062 | 1.33111719 | 0 | 0 | 0 | 0.37672154 |
| ENSMUSG00 Gm14655 | 577 | 0 | 0 | 0 | 0 | 0 | 0 |
| ENSMUSG00 Gm12180 | 660 | 0 | 0 | 0.06697249 | 0.05466976 | 0 | 0.04132521 |
| ENSMUSG00 Gm15967 | 1115 | 0 | 0 | 0 | 0 | 0.04576357 | 0 |
| ENSMUSG00 Gm11427 | 1610 | 0 | 0 | 0 | 0 | 0 | 0 |
| ENSMUSG00 Gm4917 | 649 | 0 | 0 | 0 | 0 | 0 | 0 |
| ENSMUSG00 Gm11429 | 143 | 0 | 0 | 0 | 0 | 0 | 0 |
| ENSMUSG00 Gm11344 | 334 | 0 | 0 | 0 | 0 | 0 | 0 |
| ENSMUSG00 Gm8648 | 220 | 0.90353365 | 0 | 1.00458734 | 0.16400929 | 0 | 0.3719269 |
| ENSMUSG00 Gm11287 | 602 | 0 | 0 | 0 | 0.05993695 | 0 | 0 |
| ENSMUSG00 Gm14201 | 288 | 0 | 0 | 0 | 0 | 0 | 0 |
| ENSMUSG00 Gm6580 | 1356 | 0 | 0 | 0.03259723 | 0 | 0 | 0 |
| ENSMUSG00 Gm14405 | 2327 | 0 | 0 | 0 | 0 | 0 | 0 |
| ENSMUSG00 Gm12321 | 803 | 0 | 0 | 0 | 0 | 0 | 0 |
| ENSMUSG00 Gm13409 | 542 | 0 | 0 | 0 | 0 | 0 | 0 |
| ENSMUSG00 Gm13064 | 556 | 0.14300533 | 0 | 0 | 0 | 0 | 0 |
| ENSMUSG00 Gm7190 | 1451 | 0 | 0 | 0 | 0 | 0 | 0 |
| ENSMUSG00 Gm12206 | 600 | 0 | 0 | 0 | 0 | 0 | 0 |
| ENSMUSG00 D11Bhm181e | 281 | 0 | 0 | 0 | 0.25681171 | 0 | 0 |
| ENSMUSG00 Gm7820 | 1051 | 0.03782634 | 0 | 0 | 0.03433116 | 0.14565093 | 0.0778534 |
| ENSMUSG00 Gm12187 | 1374 | 0 | 0 | 0 | 0 | 0 | 0 |
| ENSMUSG00 Gm12693 | 482 | 0 | 0 | 0.09170507 | 0.5240131 | 0 | 0.05658639 |
| ENSMUSG00 Gm15007 | 709 | 0 | 0 | 0 | 0 | 0 | 0 |
| ENSMUSG00 Gm13903 | 192 | 0 | 0 | 0 | 0 | 0 | 0 |
| ENSMUSG00 Gm13828 | 362 | 0 | 0 | 0 | 0 | 0 | 0 |
| ENSMUSG00 Gm15575 | 1580 | 0.0251617 | 0.03049775 | 0 | 0 | 0.03229518 | 0 |
| ENSMUSG00 Gm14969 | 305 | 0 | 0 | 0 | 0 | 0 | 0 |
| ENSMUSG00 Olfr1146-ps1 | 921 | 0 | 0 | 0 | 0 | 0 | 0 |
| ENSMUSG00 Gm11805 | 599 | 0 | 0 | 0.07379273 | 0.60237136 | 0 | 0 |
| ENSMUSG00 Olfr404-ps1 | 1009 | 0 | 0 | 0 | 0 | 0 | 0 |
| ENSMUSG00 Gm15027 | 137 | 0 | 0 | 0 | 0 | 0 | 0 |
| ENSMUSG00 Gm12836 | 1012 | 0.03928407 | 0 | 1.2686536 | 0.53481291 | 0 | 1.4284149 |
| ENSMUSG00 Gm15571 | 636 | 0 | 0 | 0 | 0 | 0 | 0 |
| ENSMUSG00 Gm7689 | 374 | 0 | 0 | 0 | 0.19295211 | 0 | 0 |
| ENSMUSG00 4930595D18 | 814 | 0 | 0.17759131 | 0 | 0.17730734 | 0 | 0.10052078 |
| ENSMUSG00 Gm13567 | 246 | 0 | 0.58763954 | 0 | 0 | 0 | 0 |
| ENSMUSG00 Gm15272 | 232 | 0 | 0 | 0 | 0 | 0 | 0 |
| ENSMUSG00 Gm13581 | 276 | 0 | 0 | 0 | 0 | 0 | 0 |
| ENSMUSG00 Gm9432 | 758 | 0 | 0 | 0 | 0 | 0 | 0 |
| ENSMUSG00 Gm15066 | 2607 | 0 | 0 | 0 | 0 | 0 | 0.01046208 |
| ENSMUSG00 Gm12080 | 1880 | 0.04229306 | 0.02563109 | 0.04702324 | 0.03838515 | 0.05428338 | 0.01450779 |
| ENSMUSG00 Gm12388 | 1886 | 0 | 0 | 0 | 0 | 0 | 0 |
| ENSMUSG00 Gm11421 | 293 | 0 | 0 | 0 | 0 | 0 | 0 |
| ENSMUSG00 Gm13788 | 600 | 0 | 0 | 0 | 0 | 0 | 0 |
| ENSMUSG00 Gm7821 | 787 | 0 | 0 | 0 | 0 | 0 | 0 |
| ENSMUSG00 Gm11338 | 1157 | 0 | 0 | 0 | 0 | 0 | 0 |
| ENSMUSG00 Gm14780 | 2736 | 0 | 0 | 0 | 0 | 0.01864999 | 0 |
| ENSMUSG00 Gm11522 | 175 | 1.36304505 | 0.5507022 | 0 | 0 | 2.04105503 | 0 |
| ENSMUSG00 Gm11969 | 355 | 2.01577084 | 0.67868228 | 0.62256117 | 0.40655825 | 0.71868135 | 2.535389 |
| ENSMUSG00 Gm14869 | 883 | 0 | 0 | 0 | 0 | 0 | 0 |
| ENSMUSG00 Gm13737 | 231 | 0 | 0 | 0 | 0 | 0 | 0 |
| ENSMUSG00 Gm12274 | 824 | 0 | 0 | 0 | 0 | 0 | 0 |
| ENSMUSG00 Gm14875 | 318 | 0 | 0 | 0 | 0 | 0 | 0 |
| ENSMUSG00 Gm11912 | 263 | 0 | 0 | 0 | 0 | 0 | 0 |
| ENSMUSG00 Gm14081 | 566 | 0.07023937 | 0 | 0 | 0.0637492 | 0 | 0 |
| ENSMUSG00 Hbb-bh3 | 298 | 0 | 0 | 0 | 0 | 0 | 0 |
| ENSMUSG00 Gm16425 | 2679 | 0.02967934 | 0 | 0 | 0.02693695 | 0.0190468 | 0.02036181 |
| ENSMUSG00 Gm11410 | 1142 | 2.01910497 | 2.10973915 | 1.1998749 | 1.67456074 | 1.47449247 | 1.8628913 |
| ENSMUSG00 Gm12919 | 1593 | 0 | 0 | 0.16648528 | 0 | 0 | 0 |
| ENSMUSG00 Gm14528 | 230 | 0 | 0 | 0 | 0 | 0 | 0 |
| ENSMUSG00 Gm12427 | 300 | 0 | 0 | 0 | 0 | 0 | 0 |
| ENSMUSG00 Gm15016 | 719 | 0 | 0 | 0 | 0 | 0 | 0 |
| ENSMUSG00 Gm11255 | 1179 | 0 | 0 | 0 | 0 | 0 | 0 |
| ENSMUSG00 Gm15215 | 488 | 0 | 0 | 0 | 0 | 0 | 0 |
| ENSMUSG00 Gm7831 | 551 | 0 | 0 | 0 | 0 | 0 | 0 |
| ENSMUSG00 Gm11546 | 351 | 0 | 0 | 0 | 0 | 0 | 0 |
| ENSMUSG00 Gm11779 | 367 | 0 | 0 | 0 | 0 | 0 | 0 |
| ENSMUSG00 Gm11862 | 658 | 0 | 0 | 0 | 0 | 0 | 0 |
| ENSMUSG00 Gm14158 | 307 | 0 | 0 | 1.2958195 | 0 | 0 | 0 |
| ENSMUSG00 Gm15096 | 877 | 0 | 0 | 0 | 0 | 0 | 0 |
| ENSMUSG00 Gm11796 | 971 | 0 | 0.04962558 | 0.09104396 | 0 | 0 | 0.02808923 |
| ENSMUSG00 Gm13406 | 255 | 0 | 0 | 0 | 0 | 0 | 0 |
| ENSMUSG00 Gm12262 | 304 | 0 | 0 | 0 | 0 | 0 | 0 |
| ENSMUSG00 Gm13246 | 314 | 0 | 0 | 0 | 0 | 0 | 0 |
| ENSMUSG00 Gm7222 | 905 | 0 | 0 | 0 | 0 | 0 | 0 |
| ENSMUSG00 Gm11698 | 454 | 0 | 0 | 0 | 0 | 0 | 0 |
| ENSMUSG00 Hspe1-ps4 | 284 | 0 | 0 | 0 | 0 | 0 | 0 |
| ENSMUSG00 Mup-ps1 | 538 | 0 | 0 | 0 | 0 | 0 | 0 |
| ENSMUSG00 Gm13453 | 356 | 7.70541617 | 0.13535518 | 9.56051099 | 0 | 0 | 4.67346347 |
| ENSMUSG00 Gm13259 | 455 | 0 | 0 | 0 | 0.63440957 | 0 | 0 |
| ENSMUSG00 Gm15659 | 340 | 0 | 0 | 0 | 0 | 0 | 0 |
| ENSMUSG00 Gm14668 | 1722 | 0 | 0 | 0 | 0 | 0 | 0 |
| ENSMUSG00 Gm15180 | 491 | 0 | 0 | 0 | 0 | 0 | 0 |
| ENSMUSG00 Gm11185 | 1006 | 0.19759185 | 0.19159619 | 0.13181464 | 0.50213581 | 0.10144409 | 0.13555984 |
| ENSMUSG00 Gm11839 | 1952 | 0.06109961 | 0.14811406 | 0.06793316 | 0.33272377 | 0.15684337 | 0.13972664 |
| ENSMUSG00 Gm6574 | 361 | 0 | 0 | 0 | 0 | 0 | 0 |
| ENSMUSG00 Gm14623 | 597 | 0 | 0 | 0 | 0 | 0 | 0 |
| ENSMUSG00 Gm5403 | 2794 | 0.27034865 | 0.2242032 | 0.25312437 | 0.28411059 | 0.21915408 | 0.09761861 |
| ENSMUSG00 Gm15300 | 277 | 0 | 0 | 0 | 0 | 0 | 0 |
| ENSMUSG00 Gm13210 | 272 | 0 | 0 | 0 | 0 | 0 | 0 |
| ENSMUSG00 Gm7108 | 3405 | 0 | 0 | 0 | 0 | 0 | 0 |
| ENSMUSG00 Gm15116 | 662 | 0 | 0 | 0 | 0 | 0 | 0 |
| ENSMUSG00 Gm12738 | 535 | 0 | 0 | 0 | 0 | 0 | 0 |
| ENSMUSG00 Kars-ps1 | 1766 | 0 | 0 | 0 | 0 | 0 | 0 |
| ENSMUSG00 Gm15568 | 1212 | 0 | 0 | 0 | 0 | 0.08420194 | 0 |
| ENSMUSG00 Gm5939 | 4090 | 0.00972017 | 0 | 0.0108073 | 0 | 0 | 0 |
| ENSMUSG00 Gm12625 | 381 | 0 | 0 | 0 | 0 | 0 | 0 |
| ENSMUSG00 Gm13460 | 352 | 0.22588341 | 0 | 0 | 0.30751743 | 0 | 0.23245431 |
| ENSMUSG00 Gm7816 | 2371 | 0.05030217 | 0.06096977 | 0.0932135 | 0.04565421 | 0.02152104 | 0.11503433 |
| ENSMUSG00 Gm14470 | 465 | 0 | 0 | 0 | 0 | 0 | 0 |
| ENSMUSG00 Gm14874 | 435 | 0 | 0.22154686 | 0 | 0 | 0 | 0 |
| ENSMUSG00 Gm11461 | 385 | 0 | 0 | 0 | 0 | 0 | 0 |
| ENSMUSG00 Gm12572 | 414 | 0 | 0 | 0 | 0 | 0 | 0 |
| ENSMUSG00 Olfr1127-ps1 | 855 | 0 | 0 | 0 | 0 | 0 | 0 |
| ENSMUSG00 Gm11511 | 539 | 0 | 0 | 0.16401426 | 1.00413853 | 0.2840058 | 0 |
| ENSMUSG00 Gm13254 | 653 | 0 | 0 | 0 | 0 | 0 | 0 |
| ENSMUSG00 Gm13498 | 2165 | 0.01836281 | 0.04451403 | 0 | 0 | 0 | 0.01259799 |
| ENSMUSG00 Gm12384 | 918 | 0 | 0 | 0 | 0 | 0 | 0 |
| ENSMUSG00 Gm12550 | 328 | 0 | 0 | 0 | 0 | 0 | 0 |
| ENSMUSG00 Gm14080 | 785 | 0.91159064 | 0 | 0 | 0 | 0 | 1.45928006 |
| ENSMUSG00 Gm4993 | 1173 | 0.03389214 | 0.12323898 | 0.03768273 | 0 | 0 | 0 |
| ENSMUSG00 Gm13551 | 429 | 0 | 0 | 0 | 0 | 0 | 0 |
| ENSMUSG00 Gm15376 | 782 | 0 | 0 | 0 | 0 | 0 | 0 |
| ENSMUSG00 Gm14589 | 726 | 0 | 0 | 0 | 0 | 0 | 0 |
| ENSMUSG00 Gm13638 | 1156 | 0 | 0.04168377 | 0 | 0 | 0 | 0 |
| ENSMUSG00 Gm11838 | 230 | 0 | 0 | 0.19218193 | 0.62751382 | 0 | 0.11858539 |
| ENSMUSG00 Gm13332 | 1397 | 0 | 0 | 0 | 0.05165647 | 0.03652568 | 0 |
| ENSMUSG00 Ctsf | 1961 | 124.902354 | 89.7874858 | 175.838132 | 78.6592251 | 106.788499 | 106.720197 |
| ENSMUSG00 Gm15361 | 443 | 0 | 0 | 0 | 0 | 0 | 0 |
| ENSMUSG00 Gm13688 | 607 | 0 | 0 | 0 | 0 | 0 | 0.49426859 |
| ENSMUSG00 Gm15815 | 254 | 0 | 0 | 0 | 0 | 0 | 0 |
| ENSMUSG00 Flg | 931 | 0 | 0 | 0 | 0 | 0 | 0 |
| ENSMUSG00 Gm13502 | 3158 | 0.12588816 | 0.01525853 | 0.02799357 | 0.0114256 | 0 | 0.01727336 |
| ENSMUSG00 Csnk2a1-ps | 1174 | 0 | 0 | 0 | 0 | 0 | 0 |
| ENSMUSG00 Gm8812 | 411 | 0 | 0 | 0 | 0 | 0 | 0 |
| ENSMUSG00 Gm15030 | 483 | 0 | 0 | 0 | 0 | 0 | 0 |
| ENSMUSG00 Gm13679 | 483 | 0 | 0 | 0 | 0 | 0 | 0 |
| ENSMUSG00 Gm12010 | 2015 | 0 | 0 | 0 | 0 | 0 | 0 |
| ENSMUSG00 Gm14313 | 288 | 0 | 0 | 0 | 0 | 0 | 0 |
| ENSMUSG00 Gm14957 | 1452 | 0 | 0 | 0 | 0 | 0 | 0 |
| ENSMUSG00 Defa-ps8 | 277 | 0 | 0 | 0 | 0 | 0 | 0 |
| ENSMUSG00 Gm12638 | 297 | 0 | 0 | 0 | 0 | 0 | 0 |
| ENSMUSG00 Gm11316 | 1039 | 0 | 0 | 0 | 0 | 0 | 0 |
| ENSMUSG00 Gm12085 | 939 | 0 | 0 | 0 | 0 | 0 | 0 |
| ENSMUSG00 Gm12332 | 921 | 0 | 0.0523197 | 0.19197326 | 0.19588515 | 0.11080646 | 0.08884247 |
| ENSMUSG00 Gm12309 | 181 | 0 | 0 | 0 | 0 | 0 | 0.90413169 |
| ENSMUSG00 Gm7905 | 1383 | 0 | 0 | 0 | 0 | 0 | 0 |
| ENSMUSG00 Gm12110 | 578 | 0 | 0 | 0 | 0 | 0 | 0 |
| ENSMUSG00 Mup-ps15 | 533 | 0 | 0 | 0 | 0 | 0 | 0 |
| ENSMUSG00 Gm13315 | 988 | 0.24143004 | 0.39017362 | 0 | 0 | 0 | 0.22084728 |
| ENSMUSG00 Gm13868 | 293 | 2.17094774 | 3.45363579 | 1.20687626 | 3.57126038 | 4.87624069 | 6.88847544 |
| ENSMUSG00 AA414768 | 1256 | 0.6013966 | 0.23019001 | 1.61885731 | 0.28727743 | 1.05627848 | 1.12920481 |
| ENSMUSG00 Gm15926 | 548 | 0 | 0 | 0 | 0 | 0 | 0 |
| ENSMUSG00 Gm5760 | 989 | 0.12059296 | 0.04872239 | 0 | 0.07296672 | 0.10318782 | 0.027578 |
| ENSMUSG00 Gm5643 | 965 | 0 | 0 | 0 | 0 | 0 | 0 |
| ENSMUSG00 Olfr1374-ps1 | 935 | 0 | 0 | 0 | 0 | 0 | 0 |
| ENSMUSG00 Gm15090 | 1195 | 0 | 0 | 0 | 0 | 0 | 0 |
| ENSMUSG00 Gm11742 | 232 | 0 | 0 | 0 | 0 | 0 | 0 |
| ENSMUSG00 Gm11871 | 1060 | 0 | 0 | 0 | 0 | 0 | 0 |
| ENSMUSG00 Gm15185 | 745 | 0 | 0 | 0 | 0 | 0 | 0 |
| ENSMUSG00 Gm14402 | 504 | 0 | 0 | 0 | 0 | 0 | 0 |
| ENSMUSG00 Gm15192 | 292 | 0 | 0 | 0 | 0 | 0 | 0 |
| ENSMUSG00 Gm11284 | 1885 | 0 | 0 | 0 | 0 | 0 | 1.60609282 |
| ENSMUSG00 Gm13935 | 796 | 0 | 0 | 0 | 0 | 0 | 0 |
| ENSMUSG00 Gm14931 | 252 | 0 | 0 | 0 | 0 | 0 | 0 |
| ENSMUSG00 Gm15005 | 511 | 0 | 0 | 0 | 0 | 0 | 0 |
| ENSMUSG00 Gm14997 | 635 | 0 | 0 | 0 | 0 | 0 | 0 |
| ENSMUSG00 Gm14121 | 1379 | 0.2882921 | 0.1048291 | 0.44874968 | 0.13082685 | 0.37002448 | 0.53402122 |
| ENSMUSG00 Rpl38-ps1 | 429 | 0 | 0 | 0 | 0 | 0 | 0 |
| ENSMUSG00 Vcp-rs | 2405 | 1.02488141 | 0.70125799 | 1.08436954 | 0.49509666 | 0.7213708 | 0.351565 |
| ENSMUSG00 Gm11826 | 770 | 0 | 0 | 0 | 0.0468598 | 0 | 0.17710805 |
| ENSMUSG00 Gm14220 | 448 | 0 | 0 | 0 | 0 | 0 | 0 |
| ENSMUSG00 Gm11859 | 454 | 0 | 0 | 0 | 0 | 0 | 0 |
| ENSMUSG00 Gm12400 | 1000 | 0 | 0 | 0 | 0 | 0 | 0 |

| | | | | | | | | |
|---|---|---|---|---|---|---|---|---|
| ENSMUSG00 | Gm7599 | 795 | 0 | 0.30305909 | 0 | 0.09077244 | 0 | 0 |
| ENSMUSG00 | Magea7-ps | 1201 | 0 | 0 | 0 | 0 | 0 | 0 |
| ENSMUSG00 | Gm13579 | 515 | 0 | 0 | 0 | 0 | 0 | 0 |
| ENSMUSG00 | Gm13397 | 877 | 0 | 0 | 0 | 0 | 0 | 0 |
| ENSMUSG00 | Olfr1332-ps1 | 953 | 0 | 0 | 0 | 0 | 0 | 0 |
| ENSMUSG00 | Gm11539 | 556 | 0.35751331 | 0.95332889 | 0.31799887 | 0.12979153 | 0.91774057 | 0.53960617 |
| ENSMUSG00 | Gm12475 | 1388 | 0.11456911 | 0 | 0.03184571 | 0 | 0 | 0 |
| ENSMUSG00 | Gm11693 | 563 | 0 | 0 | 0 | 0 | 0.09063299 | 0 |
| ENSMUSG00 | Gm13156 | 320 | 0 | 0 | 0 | 0.22551278 | 0.15945742 | 0.51139949 |
| ENSMUSG00 | Gm14238 | 322 | 0 | 0 | 0.1372728 | 0 | 0 | 1.01644618 |
| ENSMUSG00 | Gm14237 | 429 | 0 | 0 | 0 | 0 | 0 | 0 |
| ENSMUSG00 | Gm14686 | 842 | 0 | 0 | 0 | 0 | 0 | 0 |
| ENSMUSG00 | Gm7363 | 400 | 0 | 0 | 0.22100922 | 0 | 0 | 0 |
| ENSMUSG00 | Gm6491 | 988 | 0 | 0 | 0 | 0 | 0 | 0 |
| ENSMUSG00 | Gm8260 | 352 | 0 | 0 | 0 | 0 | 0 | 0 |
| ENSMUSG00 | Gm13913 | 259 | 0 | 0 | 0 | 0 | 0 | 0 |
| ENSMUSG00 | Gm11289 | 765 | 0 | 0 | 0 | 0 | 0 | 0 |
| ENSMUSG00 | Gm7643 | 398 | 0 | 0 | 0 | 0 | 0 | 0 |
| ENSMUSG00 | Gm9060 | 804 | 0 | 0 | 0 | 0 | 0 | 0 |
| ENSMUSG00 | Gm15783 | 631 | 0 | 0 | 0 | 0 | 0 | 0 |
| ENSMUSG00 | Gm14907 | 853 | 0 | 0 | 0 | 0 | 0 | 0 |
| ENSMUSG00 | Gm12540 | 318 | 0 | 0 | 0 | 0 | 0.64184121 | 0.08576931 |
| ENSMUSG00 | Gm11581 | 761 | 0 | 0 | 0 | 0 | 0 | 0 |
| ENSMUSG00 | Gm11708 | 831 | 0.28704318 | 0.17395828 | 0 | 0 | 0 | 0 |
| ENSMUSG00 | Olfr1004-ps1 | 453 | 0 | 0 | 0 | 0 | 0 | 0 |
| ENSMUSG00 | Gm12656 | 576 | 0 | 0 | 0 | 0 | 0 | 0 |
| ENSMUSG00 | Gm13105 | 2033 | 0 | 0 | 0 | 0 | 0 | 0 |
| ENSMUSG00 | Olfr360 | 1423 | 0 | 0 | 0 | 0 | 0 | 0 |
| ENSMUSG00 | Gm8606 | 887 | 1.0756838 | 1.14082896 | 0.54816265 | 0.77289611 | 0.11505384 | 0.61498623 |
| ENSMUSG00 | Gm12237 | 469 | 0 | 0 | 0 | 0 | 0 | 0 |
| ENSMUSG00 | Gm14535 | 1125 | 0.10601462 | 0 | 0.11787158 | 0.06414586 | 0 | 0 |
| ENSMUSG00 | Gm12895 | 225 | 0 | 0 | 0 | 0 | 0 | 0 |
| ENSMUSG00 | Gm14887 | 661 | 0 | 0 | 0 | 0 | 0 | 0 |
| ENSMUSG00 | Gm8806 | 1406 | 0.02827559 | 0.06854401 | 0 | 0.05132581 | 0 | 0.01939875 |
| ENSMUSG00 | Gm13214 | 407 | 0 | 0 | 0 | 0 | 0 | 0 |
| ENSMUSG00 | Gm15332 | 356 | 0 | 0 | 0 | 0 | 0 | 0 |
| ENSMUSG00 | Gm11422 | 1102 | 0 | 0 | 0 | 0 | 0 | 0 |
| ENSMUSG00 | Gm11235 | 332 | 0.35923627 | 0 | 0 | 0.54340429 | 0 | 0 |
| ENSMUSG00 | Gm13392 | 149 | 0 | 0 | 0 | 0 | 0 | 0 |
| ENSMUSG00 | Gm11449 | 475 | 0 | 0 | 0 | 0 | 0 | 0 |
| ENSMUSG00 | Gm15057 | 533 | 0 | 0 | 0 | 0 | 0 | 0 |
| ENSMUSG00 | Gm11882 | 505 | 0 | 0 | 0 | 0 | 0 | 0 |
| ENSMUSG00 | Gm12653 | 560 | 0 | 0 | 0 | 0 | 0 | 0 |
| ENSMUSG00 | Gm13196 | 378 | 0 | 0 | 0 | 0 | 0 | 0 |
| ENSMUSG00 | Gm4919 | 602 | 0 | 0 | 0.44054993 | 0 | 0 | 0 |
| ENSMUSG00 | Gm3244 | 389 | 0 | 0 | 0 | 0 | 0.1311732 | 0 |
| ENSMUSG00 | Gm12767 | 337 | 0 | 0 | 0 | 0 | 0 | 0 |
| ENSMUSG00 | Gm6433 | 874 | 0.04548682 | 0 | 0 | 0.08256761 | 0 | 0 |
| ENSMUSG00 | Olfr595-ps1 | 926 | 0 | 0 | 0 | 0 | 0 | 0 |
| ENSMUSG00 | Gm12813 | 896 | 0 | 0 | 0 | 0 | 0 | 0 |
| ENSMUSG00 | Gm11411 | 718 | 0 | 0 | 0 | 0 | 0 | 0 |
| ENSMUSG00 | Gm15426 | 555 | 0 | 0 | 0 | 0 | 0 | 0 |
| ENSMUSG00 | Gm11860 | 1009 | 0 | 0 | 0 | 0 | 0 | 0 |
| ENSMUSG00 | Gm11550 | 698 | 0 | 0.06903502 | 0 | 0 | 0 | 0 |
| ENSMUSG00 | Gm13681 | 209 | 0 | 0 | 0 | 0 | 0 | 0 |
| ENSMUSG00 | Gm13121 | 631 | 0.18901179 | 0.38182601 | 0 | 0 | 0 | 0.08644894 |
| ENSMUSG00 | Gm14148 | 1039 | 0.11478965 | 0.09275542 | 0.08508536 | 0.03472767 | 0.04911105 | 0.28875942 |
| ENSMUSG00 | Gm6335 | 1193 | 0 | 0 | 0 | 0.0302448 | 0.08554296 | 0.09144892 |
| ENSMUSG00 | Gm15194 | 433 | 0 | 0 | 0 | 0 | 0 | 0 |
| ENSMUSG00 | Gm12618 | 482 | 0.659842 | 0.09997187 | 0 | 0 | 0 | 0 |
| ENSMUSG00 | Gm11203 | 740 | 0 | 0 | 0 | 0 | 0 | 0 |
| ENSMUSG00 | Gm15542 | 1193 | 0.63315518 | 1.21172948 | 0.629867 | 1.36101593 | 0.8554296 | 1.39459597 |
| ENSMUSG00 | Gm5466 | 1889 | 0 | 0.02550897 | 0 | 0 | 0 | 0.01443867 |
| ENSMUSG00 | Gm13458 | 366 | 0 | 0 | 0 | 0 | 0 | 0 |
| ENSMUSG00 | Gm15033 | 672 | 0 | 0 | 0 | 0 | 0 | 0 |
| ENSMUSG00 | Gm11900 | 351 | 0 | 0 | 0.25186235 | 0 | 0 | 0 |
| ENSMUSG00 | Gm13506 | 1081 | 0 | 0 | 0 | 0 | 0 | 0 |
| ENSMUSG00 | Gm14394 | 294 | 0 | 0 | 0 | 0 | 0 | 0 |
| ENSMUSG00 | Gm13041 | 1521 | 0 | 0 | 0 | 0 | 0 | 0 |
| ENSMUSG00 | Gm13019 | 274 | 0 | 0 | 0 | 0 | 0 | 0 |
| ENSMUSG00 | Gm15725 | 1089 | 0 | 0 | 0 | 0 | 0 | 0 |
| ENSMUSG00 | Gm8410 | 604 | 0 | 0 | 0 | 0 | 0 | 0 |
| ENSMUSG00 | Gm13176 | 476 | 0 | 0 | 0 | 0 | 0 | 0 |
| ENSMUSG00 | Gm11340 | 411 | 0 | 0 | 0 | 0 | 0 | 0 |
| ENSMUSG00 | Gm14300 | 945 | 0 | 0 | 0 | 0 | 0 | 0 |
| ENSMUSG00 | Rpl30-ps10 | 779 | 0 | 0 | 0 | 0 | 0 | 0 |
| ENSMUSG00 | Gm12268 | 441 | 0 | 1.09266309 | 1.30300218 | 0.32727478 | 0.69423641 | 0 |
| ENSMUSG00 | Gm11306 | 1093 | 0 | 0 | 0 | 0 | 0 | 0 |
| ENSMUSG00 | Gm14840 | 593 | 0 | 0 | 0 | 0 | 0 | 0.04599433 |
| ENSMUSG00 | Gm14690 | 374 | 0 | 0 | 0 | 0 | 0 | 0 |
| ENSMUSG00 | Gm15378 | 206 | 0 | 0 | 0 | 0 | 0 | 0 |
| ENSMUSG00 | Olfr269-ps1 | 301 | 0 | 0 | 0 | 0 | 0 | 0 |
| ENSMUSG00 | Olfr1035-ps1 | 924 | 0 | 0 | 0 | 0 | 0 | 0 |
| ENSMUSG00 | Gm14485 | 492 | 0 | 0 | 0 | 0 | 0 | 0 |
| ENSMUSG00 | Gm15665 | 846 | 0 | 0 | 0 | 0 | 0 | 0 |
| ENSMUSG00 | Gm14776 | 885 | 0 | 0 | 0 | 0.04077067 | 0 | 0.0308188 |
| ENSMUSG00 | Gm15604 | 1455 | 0 | 0 | 0 | 0.04959731 | 0 | 0 |
| ENSMUSG00 | Gm13221 | 429 | 0 | 0 | 0 | 0 | 0 | 0 |
| ENSMUSG00 | Rpl35a-ps4 | 332 | 0 | 0.14513989 | 0 | 0 | 0 | 0 |
| ENSMUSG00 | Gm13992 | 155 | 0 | 0 | 0 | 0 | 0 | 0 |
| ENSMUSG00 | Defa-ps10 | 170 | 0 | 0 | 0 | 0 | 0 | 0 |
| ENSMUSG00 | Gm12007 | 596 | 0 | 0 | 0 | 0 | 0 | 0 |
| ENSMUSG00 | Gm14591 | 516 | 0 | 0 | 0 | 0 | 0 | 0 |
| ENSMUSG00 | Gm15198 | 185 | 0 | 0 | 0 | 0.19503808 | 1.37909124 | 0 |
| ENSMUSG00 | Gm13006 | 213 | 0 | 0 | 0 | 0 | 0 | 0 |
| ENSMUSG00 | Gm13437 | 1978 | 0 | 0 | 0 | 0.01824168 | 0 | 0 |
| ENSMUSG00 | Gm14878 | 292 | 0 | 0 | 0 | 0 | 0 | 0 |
| ENSMUSG00 | Gm15405 | 341 | 0 | 0 | 0 | 0 | 0 | 0 |
| ENSMUSG00 | Gm12760 | 480 | 0 | 0 | 0 | 0 | 0 | 0 |
| ENSMUSG00 | Gm15824 | 957 | 0 | 0.35246091 | 0 | 0 | 0 | 0 |
| ENSMUSG00 | Gm11471 | 197 | 0 | 0 | 0 | 0 | 0 | 0 |
| ENSMUSG00 | Gm15187 | 263 | 0 | 0 | 0 | 0 | 0 | 0 |
| ENSMUSG00 | Gm6402 | 1617 | 0 | 0 | 0 | 0 | 0.0315562 | 0 |
| ENSMUSG00 | Gm14491 | 198 | 0 | 0.24336587 | 0 | 0 | 0 | 0 |
| ENSMUSG00 | Rpl7a-ps12 | 792 | 0 | 0 | 0 | 0 | 0 | 0 |
| ENSMUSG00 | Gm15329 | 154 | 0 | 0 | 0 | 0 | 0 | 0 |
| ENSMUSG00 | Gm11187 | 445 | 0 | 0 | 0 | 0 | 0 | 0 |
| ENSMUSG00 | Gm15519 | 1565 | 0 | 0.06158012 | 0.31068388 | 0.04611124 | 0.03260471 | 0.08713942 |
| ENSMUSG00 | Gm14768 | 676 | 0 | 0 | 0 | 0 | 0 | 0 |
| ENSMUSG00 | Gm15038 | 295 | 0 | 0 | 0 | 0 | 0 | 0 |
| ENSMUSG00 | Gm12826 | 378 | 0 | 0 | 0 | 0 | 0 | 0 |
| ENSMUSG00 | Gm11904 | 247 | 0 | 0 | 0 | 0 | 0 | 0 |
| ENSMUSG00 | Gm8749 | 330 | 0 | 0 | 0 | 0 | 0 | 0 |
| ENSMUSG00 | Gm13487 | 863 | 0 | 0.05583597 | 0 | 0.20905008 | 0.05912674 | 0.0632089 |
| ENSMUSG00 | Gm14619 | 367 | 0 | 0 | 0 | 0 | 0 | 0 |
| ENSMUSG00 | Gm11445 | 531 | 0 | 0 | 0 | 0 | 0 | 0 |
| ENSMUSG00 | Gm14905 | 637 | 0 | 0 | 0 | 0 | 0 | 0 |
| ENSMUSG00 | Gm13338 | 360 | 0 | 0 | 0 | 0 | 0 | 0 |
| ENSMUSG00 | Gm13226 | 403 | 0 | 0.11956934 | 0 | 0 | 0 | 0 |
| ENSMUSG00 | Gm15037 | 258 | 0 | 0 | 0 | 0 | 0 | 0 |
| ENSMUSG00 | Gm13108 | 1453 | 0 | 0 | 0 | 0 | 0 | 0 |
| ENSMUSG00 | Gm12839 | 330 | 0 | 0.58407809 | 0 | 0 | 0.15462538 | 0.24795127 |
| ENSMUSG00 | Gm6510 | 485 | 0.08197006 | 0 | 0 | 0 | 0 | 0 |
| ENSMUSG00 | Gm12182 | 1007 | 0.39479127 | 0.04785148 | 0.17557832 | 0.1433249 | 0.15201502 | 0.02708504 |
| ENSMUSG00 | Gm12935 | 244 | 0 | 0 | 0 | 0 | 0 | 0 |
| ENSMUSG00 | Gm12501 | 528 | 0.07529447 | 0 | 0 | 0 | 0 | 0 |
| ENSMUSG00 | Gm4909 | 793 | 0 | 0 | 0 | 0 | 0 | 0 |
| ENSMUSG00 | Gm11652 | 1266 | 0.2826219 | 0.1903098 | 0.59354766 | 0.5700165 | 0.44335714 | 0.0430879 |
| ENSMUSG00 | Gm15225 | 1013 | 0 | 0 | 0 | 0 | 0 | 0 |
| ENSMUSG00 | Gm15840 | 942 | 0 | 0 | 0 | 0 | 0 | 0 |
| ENSMUSG00 | Gm9431 | 1050 | 0 | 0 | 0.04209699 | 0 | 0 | 0 |
| ENSMUSG00 | Gm14281 | 1293 | 0 | 0 | 0 | 0 | 0 | 0 |
| ENSMUSG00 | Nsa2-ps2 | 772 | 0 | 0 | 0.11451255 | 0 | 0 | 0 |
| ENSMUSG00 | Gm12118 | 531 | 0 | 0 | 0.16648528 | 0 | 0 | 0 |
| ENSMUSG00 | Gm13491 | 234 | 0 | 0 | 0 | 0 | 0 | 0 |
| ENSMUSG00 | Gm13207 | 772 | 0 | 0 | 0 | 0.1402152 | 0 | 0 |
| ENSMUSG00 | Gm15267 | 463 | 0 | 0 | 0 | 0 | 0 | 0.11781702 |
| ENSMUSG00 | Gm9440 | 532 | 0 | 0 | 0 | 0 | 0 | 0 |
| ENSMUSG00 | Gm6555 | 1013 | 0.35320763 | 0.90379309 | 0.17453837 | 0.42742797 | 0.10074309 | 0.37694467 |
| ENSMUSG00 | Gm15375 | 848 | 0 | 0 | 0 | 0 | 0 | 0 |
| ENSMUSG00 | Gm11883 | 326 | 0 | 0 | 0 | 0 | 0 | 0 |
| ENSMUSG00 | Rps8-ps2 | 1310 | 0 | 0 | 0 | 0 | 0 | 0 |
| ENSMUSG00 | Gm14706 | 254 | 0 | 0 | 0 | 0 | 0 | 0 |
| ENSMUSG00 | Gm14044 | 951 | 0 | 0 | 0 | 0 | 0 | 0 |
| ENSMUSG00 | Gm14574 | 278 | 0 | 0 | 0 | 0 | 0 | 0 |
| ENSMUSG00 | Gm12308 | 388 | 0 | 0.86934303 | 0 | 0 | 0 | 0.21088639 |
| ENSMUSG00 | Gm14521 | 349 | 0 | 0 | 0 | 0 | 0 | 0 |
| ENSMUSG00 | Gm13888 | 421 | 0 | 0 | 0 | 0 | 0 | 0 |
| ENSMUSG00 | Gm8534 | 765 | 0 | 0 | 0 | 0 | 0 | 0 |
| ENSMUSG00 | Gm11784 | 471 | 0 | 2.25074677 | 0.0938468 | 0 | 0 | 0.92652702 |
| ENSMUSG00 | Gm14940 | 492 | 0 | 0 | 0 | 0 | 0 | 0 |
| ENSMUSG00 | Gm12423 | 134 | 0 | 0 | 0 | 0 | 0 | 0 |
| ENSMUSG00 | Gm14515 | 215 | 0 | 0 | 0 | 0 | 0 | 0 |
| ENSMUSG00 | Gm15044 | 597 | 0 | 0 | 0 | 0 | 0 | 0 |
| ENSMUSG00 | Gm12873 | 418 | 0 | 0 | 0 | 0 | 0 | 0 |
| ENSMUSG00 | Gm13877 | 416 | 0 | 0 | 0 | 0 | 0 | 0 |
| ENSMUSG00 | Gm11263 | 748 | 0 | 0 | 0 | 0 | 0 | 0 |
| ENSMUSG00 | Gm14172 | 643 | 0 | 0 | 0 | 0 | 0 | 0 |
| ENSMUSG00 | Gm14609 | 901 | 0 | 0 | 0 | 0 | 0 | 0 |
| ENSMUSG00 | Gm15470 | 1370 | 1.91522753 | 2.9193246 | 2.09716773 | 1.63291004 | 1.86227649 | 0.79633983 |
| ENSMUSG00 | Gm12255 | 209 | 0 | 0 | 0 | 0 | 0 | 0 |
| ENSMUSG00 | Gm14861 | 454 | 1.22593993 | 0 | 0.09736089 | 0 | 1.12392898 | 0.42053409 |
| ENSMUSG00 | Gm13407 | 359 | 0 | 0 | 0.36937473 | 0 | 0 | 0.3038957 |
| ENSMUSG00 | Gm12450 | 467 | 0 | 0 | 0 | 0 | 0 | 0 |
| ENSMUSG00 | Gm7541 | 534 | 0 | 0 | 0 | 0.06756937 | 0 | 0 |
| ENSMUSG00 | Gm13993 | 1235 | 0 | 0 | 0 | 0 | 0 | 0.04416946 |
| ENSMUSG00 | Gm12842 | 475 | 0 | 0 | 0 | 0 | 0 | 0 |
| ENSMUSG00 | Gm6927 | 755 | 0 | 0 | 0 | 0 | 0 | 0 |
| ENSMUSG00 | Defa-ps4 | 182 | 0 | 0 | 0 | 0 | 0 | 0 |
| ENSMUSG00 | Gm13478 | 412 | 0 | 0 | 0 | 0 | 0 | 0 |
| ENSMUSG00 | Gm12749 | 1158 | 0.30896042 | 0.04161178 | 0.41987934 | 0.3739072 | 0.17625691 | 0 |
| ENSMUSG00 | Olfr647-ps1 | 876 | 0 | 0 | 0 | 0 | 0 | 0 |
| ENSMUSG00 | Gm11881 | 571 | 0 | 0 | 0 | 0 | 0 | 0 |
| ENSMUSG00 | Gm14577 | 255 | 0 | 0 | 0 | 0 | 0 | 0 |
| ENSMUSG00 | Gm15196 | 481 | 0 | 0 | 0 | 0 | 0 | 0 |
| ENSMUSG00 | Gm14998 | 494 | 0 | 0 | 0 | 0 | 0 | 0 |
| ENSMUSG00 | Gm9429 | 867 | 0 | 0 | 0 | 0 | 0 | 0.06291728 |
| ENSMUSG00 | Gm13674 | 822 | 0 | 0 | 0 | 0 | 0 | 0 |
| ENSMUSG00 | Gm6418 | 3575 | 0 | 0 | 0 | 0 | 0 | 0 |
| ENSMUSG00 | Gm12157 | 747 | 0 | 0 | 0 | 0 | 0 | 0 |
| ENSMUSG00 | Gm15321 | 287 | 0.13852084 | 0 | 0 | 0 | 0 | 0 |
| ENSMUSG00 | Mup-ps21 | 443 | 0 | 0 | 0 | 0 | 0 | 0 |
| ENSMUSG00 | Gm15045 | 882 | 0 | 0.21853262 | 0.15034641 | 0.16363739 | 0.05785303 | 0.06184726 |
| ENSMUSG00 | Gm12001 | 911 | 0 | 0 | 0 | 0 | 0 | 0 |
| ENSMUSG00 | Gm13432 | 240 | 0 | 0 | 0 | 0 | 0 | 0 |
| ENSMUSG00 | Gm15035 | 1560 | 0 | 0 | 0 | 0 | 0 | 0 |
| ENSMUSG00 | Gm13867 | 393 | 0 | 0 | 0 | 0 | 0 | 0 |
| ENSMUSG00 | Gm13450 | 27 | 0 | 0 | 0 | 0 | 0 | 0 |
| ENSMUSG00 | H2-M6-ps | 1029 | 0 | 0 | 0 | 0 | 0 | 0 |
| ENSMUSG00 | Gm14381 | 413 | 0 | 0 | 0 | 0 | 0 | 0 |
| ENSMUSG00 | Gm15808 | 603 | 0 | 0 | 0 | 0 | 0 | 0 |
| ENSMUSG00 | Gm14942 | 335 | 0 | 0 | 0 | 0 | 0 | 0 |
| ENSMUSG00 | Gm16016 | 549 | 0 | 1.22879816 | 0.80513375 | 0.78867857 | 0.37177687 | 1.3413757 |
| ENSMUSG00 | Gm12466 | 619 | 0 | 0 | 0 | 0 | 0 | 0 |
| ENSMUSG00 | Gm12585 | 289 | 0 | 0 | 0 | 0 | 0 | 0 |
| ENSMUSG00 | Olfr1379-ps1 | 823 | 0 | 0 | 0 | 0 | 0 | 0 |
| ENSMUSG00 | Gm14663 | 1152 | 0 | 0 | 0 | 0 | 0 | 0 |
| ENSMUSG00 | Oat-rs1 | 1310 | 0 | 0.07356709 | 0.1012256 | 0 | 0.07790286 | 0 |
| ENSMUSG00 | Gm13320 | 329 | 0 | 0 | 0 | 0 | 0 | 0 |
| ENSMUSG00 | Tpt1-ps1 | 435 | 0 | 0 | 0 | 0 | 0 | 0 |
| ENSMUSG00 | Gm13321 | 398 | 0 | 0 | 0 | 0 | 0 | 0 |
| ENSMUSG00 | Gm11514 | 431 | 0 | 0 | 0 | 0 | 0 | 0 |
| ENSMUSG00 | Gm11227 | 470 | 0 | 0 | 0 | 0 | 0 | 0 |
| ENSMUSG00 | Gm13729 | 405 | 0 | 0 | 0 | 0 | 0 | 0 |
| ENSMUSG00 | Gm14516 | 251 | 0 | 0 | 0 | 0 | 0.20329233 | 0.65198341 |
| ENSMUSG00 | Gm14814 | 304 | 0 | 0 | 0 | 0 | 0 | 0 |
| ENSMUSG00 | Gm11989 | 671 | 0.05924811 | 0 | 0 | 0 | 0 | 0 |
| ENSMUSG00 | Gm6645 | 2724 | 0 | 0.1768959 | 0.19472178 | 0.05298391 | 0.0749286 | 0.53067397 |
| ENSMUSG00 | Gm12035 | 622 | 0 | 0 | 0 | 0 | 0 | 0 |
| ENSMUSG00 | Gm9043 | 311 | 0 | 0 | 0 | 0 | 0 | 0 |
| ENSMUSG00 | Gm6877 | 1100 | 0.25298942 | 0.26283514 | 0.20091747 | 0.16400929 | 0.09277523 | 0.3719289 |
| ENSMUSG00 | Gm12896 | 225 | 0 | 0 | 0 | 0 | 0 | 0 |
| ENSMUSG00 | Gm14939 | 678 | 0 | 0 | 0 | 0 | 0 | 0 |
| ENSMUSG00 | Gm6942 | 438 | 0 | 0 | 0 | 0 | 0 | 0 |
| ENSMUSG00 | Gm13234 | 746 | 0 | 0 | 0 | 0 | 0 | 0 |
| ENSMUSG00 | Gm13340 | 1544 | 16.2472203 | 15.6980443 | 8.96060678 | 11.2172159 | 7.6010793 | 7.73723575 |
| ENSMUSG00 | Gm11929 | 944 | 0 | 0 | 0 | 0.03822251 | 0 | 0.08667788 |
| ENSMUSG00 | Gm11941 | 609 | 0 | 0 | 0 | 0 | 0 | 0 |
| ENSMUSG00 | Gm14711 | 215 | 0 | 0 | 0 | 0 | 0 | 0 |
| ENSMUSG00 | Gm11451 | 1246 | 0 | 0.03867291 | 0 | 0.0579166 | 0.04095215 | 0.13133855 |
| ENSMUSG00 | Gm12770 | 461 | 0 | 0 | 0 | 0 | 0 | 0 |
| ENSMUSG00 | Gm13686 | 798 | 0 | 0 | 0 | 0 | 0 | 0 |
| ENSMUSG00 | Gm14671 | 261 | 0 | 0 | 0 | 0 | 0 | 0 |
| ENSMUSG00 | Gm11849 | 260 | 0 | 0 | 0 | 0 | 0 | 0 |
| ENSMUSG00 | Gm7312 | 468 | 0 | 0 | 0 | 0 | 0 | 0 |
| ENSMUSG00 | Gm15709 | 212 | 0 | 0 | 0 | 0 | 0 | 0 |
| ENSMUSG00 | Gm12533 | 137 | 0 | 0 | 0 | 0 | 0 | 0 |
| ENSMUSG00 | Gm16411 | 1099 | 0 | 0 | 0 | 0 | 0 | 0 |
| ENSMUSG00 | Gm12432 | 185 | 0 | 0 | 0 | 0 | 0 | 0 |
| ENSMUSG00 | Gm11247 | 307 | 0 | 0 | 0 | 0 | 0 | 0 |
| ENSMUSG00 | Gm15538 | 312 | 0 | 0 | 0 | 0 | 0 | 0 |
| ENSMUSG00 | Rpsa-ps3 | 874 | 0.04548682 | 0 | 0 | 0 | 0 | 0 |
| ENSMUSG00 | Gm14186 | 136 | 0 | 0 | 0 | 0 | 0 | 0 |
| ENSMUSG00 | Gm15393 | 218 | 0 | 0 | 0 | 0 | 0 | 0 |
| ENSMUSG00 | Gm15394 | 1307 | 0 | 0 | 0 | 0 | 0 | 0 |
| ENSMUSG00 | Gm15885 | 387 | 0 | 0 | 0 | 0 | 0 | 0 |
| ENSMUSG00 | Gm8688 | 964 | 0 | 0 | 0 | 0 | 0 | 0 |
| ENSMUSG00 | Gm14741 | 1008 | 0 | 0 | 0 | 0 | 0 | 0 |
| ENSMUSG00 | Gm8051 | 465 | 0 | 0 | 0 | 0 | 0 | 0 |
| ENSMUSG00 | Gm14043 | 227 | 0 | 0 | 0 | 0 | 0 | 0 |
| ENSMUSG00 | Gm13916 | 627 | 0 | 0 | 0 | 0 | 0 | 0 |
| ENSMUSG00 | Gm13614 | 502 | 0 | 0 | 0 | 0 | 0 | 0 |
| ENSMUSG00 | Gm14365 | 586 | 0 | 0 | 0 | 0 | 0 | 0 |
| ENSMUSG00 | Gm13722 | 3065 | 0 | 0 | 0 | 0 | 0 | 0 |
| ENSMUSG00 | Gm9200 | 1353 | 0.14691604 | 0 | 0.16334754 | 0.05333636 | 0.11314052 | 0.1007932 |
| ENSMUSG00 | Rpl21-ps15 | 1022 | 0 | 0 | 0 | 0 | 0 | 0 |
| ENSMUSG00 | Gm14805 | 339 | 0 | 0 | 0 | 0 | 0 | 0 |
| ENSMUSG00 | Gm14579 | 282 | 0 | 0 | 0 | 0 | 0 | 0 |
| ENSMUSG00 | Gm12821 | 599 | 0 | 0 | 0.14758545 | 0 | 0 | 0 |
| ENSMUSG00 | Gm12028 | 434 | 0 | 0 | 0 | 0 | 0 | 0 |
| ENSMUSG00 | Gm14911 | 988 | 0 | 0 | 0 | 0 | 0 | 0 |
| ENSMUSG00 | Gm14810 | 485 | 0 | 0 | 0 | 0 | 0 | 0 |
| ENSMUSG00 | Gm7398 | 631 | 0 | 0 | 0 | 0 | 0 | 0 |
| ENSMUSG00 | Gm14893 | 280 | 0 | 0 | 0 | 0 | 0 | 0 |
| ENSMUSG00 | Gm15916 | 1169 | 0.06801622 | 0 | 0.37811671 | 0.64818044 | 0 | 0 |
| ENSMUSG00 | Gm12640 | 309 | 0 | 0 | 0 | 0 | 0 | 0 |
| ENSMUSG00 | Gm14863 | 306 | 0 | 0 | 0 | 0 | 0 | 0 |
| ENSMUSG00 | Gm16007 | 285 | 0 | 0 | 0 | 0 | 0 | 0 |
| ENSMUSG00 | Gm9329 | 535 | 1.04033033 | 0.72054493 | 0.74358241 | 1.820963 | 2.09828088 | 0.30588381 |
| ENSMUSG00 | Gm8832 | 394 | 0 | 0 | 0.22437484 | 0 | 0 | 0 |
| ENSMUSG00 | Pou2f3-rs1 | 1045 | 0 | 0 | 0 | 0 | 0 | 0 |
| ENSMUSG00 | Gm12795 | 537 | 0 | 0 | 0 | 0 | 0 | 0.05079076 |
| ENSMUSG00 | Gm12878 | 331 | 0 | 0 | 0 | 0 | 0 | 0 |
| ENSMUSG00 | H2afb3-ps | 347 | 0 | 0 | 0 | 0 | 0 | 0 |
| ENSMUSG00 | Gm5390 | 1229 | 0 | 0 | 0 | 0 | 0 | 0 |
| ENSMUSG00 | Gm11771 | 724 | 0 | 0 | 0 | 0 | 0 | 0 |
| ENSMUSG00 | Gm14414 | 677 | 0.35233808 | 0.85411714 | 0.65290758 | 0.37307875 | 0.52759916 | 1.04747507 |
| ENSMUSG00 | Gm15529 | 993 | 0 | 0 | 0 | 0 | 0 | 0 |
| ENSMUSG00 | Gm14586 | 407 | 0 | 0 | 0 | 0 | 0 | 0 |
| ENSMUSG00 | Gm15570 | 624 | 0 | 0 | 0 | 0 | 0 | 0 |
| ENSMUSG00 | Gm7224 | 2064 | 0.57784129 | 0.37353831 | 0.17132497 | 0.17481611 | 0.61805203 | 0.44929154 |
| ENSMUSG00 | Gm12469 | 206 | 0 | 0 | 0 | 0 | 0 | 0 |
| ENSMUSG00 | Gm15188 | 698 | 0 | 0 | 0 | 0 | 0 | 0 |
| ENSMUSG00 | Gm4459 | 467 | 0.17025902 | 0.72228072 | 0.09465063 | 0 | 0 | 0.11680788 |
| ENSMUSG00 | Gm12502 | 443 | 0 | 0 | 0 | 0 | 0 | 0 |
| ENSMUSG00 | Gm2030 | 1120 | 0 | 0 | 0 | 0 | 0 | 0 |
| ENSMUSG00 | Gm11262 | 831 | 0 | 0 | 0 | 0 | 0 | 0 |
| ENSMUSG00 | Gm15602 | 436 | 0 | 0 | 0 | 0 | 0 | 0 |
| ENSMUSG00 | Gm13366 | 975 | 0 | 0 | 0 | 0 | 0 | 0 |
| ENSMUSG00 | Gm11786 | 154 | 0 | 0 | 0 | 0 | 0 | 0 |
| ENSMUSG00 | Gm13312 | 465 | 0 | 0 | 0 | 0 | 0 | 0 |
| ENSMUSG00 | Gm15068 | 190 | 0 | 0 | 0 | 0 | 0 | 0 |
| ENSMUSG00 | Gm11480 | 658 | 0 | 0 | 0 | 0 | 0 | 0 |
| ENSMUSG00 | Gm15175 | 131 | 0 | 0 | 0 | 0 | 0 | 0 |
| ENSMUSG00 | Gm14754 | 725 | 0 | 0 | 0 | 0 | 0 | 0 |
| ENSMUSG00 | Gm11392 | 706 | 0 | 0 | 0 | 0 | 0 | 0 |
| ENSMUSG00 | Defa-ps6 | 280 | 0 | 0 | 0 | 0 | 0 | 0 |
| ENSMUSG00 | Gm8876 | 1121 | 0 | 0.08597046 | 0 | 0.03218737 | 0 | 0 |
| ENSMUSG00 | Gm9225 | 834 | 0 | 0 | 0 | 0 | 0 | 0 |
| ENSMUSG00 | Gm13148 | 744 | 0 | 0 | 0 | 0 | 0 | 0 |
| ENSMUSG00 | Gm13876 | 260 | 0 | 0 | 0 | 0 | 0 | 0 |
| ENSMUSG00 | Gm12787 | 394 | 0 | 0 | 0 | 0 | 0 | 0 |
| ENSMUSG00 | Gm14086 | 171 | 0 | 0 | 0 | 0 | 0 | 0 |
| ENSMUSG00 | Gm6088 | 2538 | 0 | 0 | 0 | 0 | 0 | 0.01074651 |
| ENSMUSG00 | Gm14229 | 849 | 0.04682624 | 0 | 0 | 0 | 0 | 0 |
| ENSMUSG00 | Rasl2-9-ps | 1005 | 1.78009614 | 0.71920063 | 0.7916748 | 0.07180506 | 1.06622278 | 1.2212525 |
| ENSMUSG00 | Gm13357 | 536 | 0 | 0.08990008 | 0 | 0.06731725 | 0 | 0 |
| ENSMUSG00 | Gm11524 | 806 | 0 | 0 | 0 | 0 | 0 | 0 |
| ENSMUSG00 | Gm11384 | 292 | 0 | 0 | 0 | 0 | 0 | 0 |
| ENSMUSG00 | Gm11875 | 798 | 0 | 0 | 0 | 0 | 0 | 0 |
| ENSMUSG00 | Gm12099 | 749 | 0 | 0 | 0 | 0 | 0 | 0 |
| ENSMUSG00 | Gm12245 | 356 | 0 | 0 | 0 | 0 | 0 | 0 |
| ENSMUSG00 | Gm15798 | 407 | 0 | 0 | 0 | 0 | 0 | 0 |
| ENSMUSG00 | Gm11258 | 1217 | 0 | 0 | 0.07264066 | 0 | 0 | 0 |
| ENSMUSG00 | Gm11853 | 1382 | 0 | 0 | 0 | 0 | 0 | 0 |
| ENSMUSG00 | Gm11473 | 358 | 0 | 0 | 0 | 0 | 0 | 0 |
| ENSMUSG00 | Gm12152 | 683 | 0 | 0 | 0 | 0 | 0 | 0 |
| ENSMUSG00 | Gm14903 | 535 | 0 | 0 | 0 | 0 | 0 | 0 |
| ENSMUSG00 | Gm11433 | 226 | 0 | 0 | 0 | 0 | 0 | 0 |
| ENSMUSG00 | Gm14575 | 559 | 0 | 0 | 0 | 0 | 0 | 0 |
| ENSMUSG00 | Gm12358 | 518 | 0 | 0 | 0 | 0 | 0 | 0 |
| ENSMUSG00 | Gm11334 | 155 | 0 | 0 | 0 | 0 | 0 | 0 |
| ENSMUSG00 | Gm5648 | 1586 | 0 | 0 | 0 | 0 | 0 | 0 |
| ENSMUSG00 | Gm10169 | 347 | 0.34370732 | 1.11092662 | 2.42027383 | 0.10398284 | 1.32345067 | 1.88643038 |
| ENSMUSG00 | Gm6829 | 359 | 0 | 0 | 0 | 0 | 0 | 0 |
| ENSMUSG00 | Gm14304 | 444 | 0 | 0 | 0 | 0 | 0 | 0 |
| ENSMUSG00 | Gm9183 | 1568 | 0 | 0 | 0 | 0 | 0 | 0 |
| ENSMUSG00 | Gm11308 | 1918 | 0 | 0 | 0 | 0 | 0 | 0 |
| ENSMUSG00 | Zfp133-ps | 3107 | 1.151591 | 0.94604859 | 0.49792871 | 0.90582539 | 1.92149532 | 1.16753364 |
| ENSMUSG00 | Gm15143 | 712 | 0 | 0 | 0 | 0 | 0 | 0 |
| ENSMUSG00 | Gm11367 | 122 | 0 | 0 | 0 | 0 | 0 | 0.22356262 |
| ENSMUSG00 | Gm11699 | 400 | 0 | 0 | 0 | 0 | 0 | 0 |
| ENSMUSG00 | Gm12989 | 730 | 0.4356765 | 0.66008825 | 0.42385329 | 0.54370204 | 0.20969744 | 0.26153764 |
| ENSMUSG00 | Gm12892 | 1145 | 0.59025604 | 0 | 0.03860423 | 0.06302541 | 0 | 0 |
| ENSMUSG00 | Gm9119 | 491 | 0 | 0 | 0 | 0 | 0 | 0 |
| ENSMUSG00 | Gm6640 | 1586 | 0 | 0.03038237 | 0 | 0.02275034 | 0 | 0.01719712 |
| ENSMUSG00 | Pcna-ps1 | 736 | 0 | 0 | 0 | 0 | 0 | 0 |
| ENSMUSG00 | Gm6539 | 1062 | 0 | 0 | 0 | 0 | 0 | 0 |
| ENSMUSG00 | Gm14034 | 550 | 0 | 0 | 0 | 0 | 0 | 0 |
| ENSMUSG00 | Gm12412 | 327 | 0 | 0.14735915 | 0 | 0 | 0 | 0 |
| ENSMUSG00 | Gm13504 | 545 | 0 | 0 | 0 | 0 | 0 | 0 |
| ENSMUSG00 | Gm15154 | 365 | 0 | 0 | 0 | 0 | 0 | 0 |
| ENSMUSG00 | Gm13743 | 723 | 0 | 0 | 0 | 0 | 0 | 0.03772426 |
| ENSMUSG00 | Gm14094 | 609 | 0.13055987 | 0.23737164 | 0 | 0.17774406 | 0 | 0 |
| ENSMUSG00 | Defa-ps5 | 277 | 0 | 0 | 0 | 0 | 0 | 0 |
| ENSMUSG00 | Gm14465 | 566 | 0 | 0 | 0 | 0 | 0 | 0 |
| ENSMUSG00 | Gm9575 | 1338 | 0 | 0 | 0.03303576 | 0 | 0.07627261 | 0.08153853 |
| ENSMUSG00 | Gm11971 | 287 | 0 | 0 | 0 | 0 | 0 | 0 |
| ENSMUSG00 | 1700045I19R | 1935 | 0 | 0 | 0.02284333 | 0 | 0 | 0 |
| ENSMUSG00 | Gm12563 | 101 | 0 | 0 | 0 | 0 | 0.50521164 | 0 |
| ENSMUSG00 | Gm13914 | 990 | 0 | 0 | 0 | 0 | 0 | 0 |
| ENSMUSG00 | Gm12687 | 705 | 0.11278151 | 0 | 0 | 0 | 0 | 0 |
| ENSMUSG00 | Gm14932 | 451 | 0 | 0 | 0.09800852 | 0 | 0 | 0 |
| ENSMUSG00 | Gm2986 | 381 | 0 | 0 | 0 | 0 | 0 | 0.07158698 |
| ENSMUSG00 | Gm4912 | 1043 | 0 | 0 | 0 | 0 | 0 | 0 |
| ENSMUSG00 | Gm14428 | 819 | 0.04854149 | 0.11767141 | 0.0539705 | 0.17622488 | 0.12460653 | 0 |
| ENSMUSG00 | Gm13153 | 636 | 0 | 0 | 0 | 0 | 0 | 0 |
| ENSMUSG00 | Gm8624 | 1098 | 0.10862153 | 0 | 0 | 0 | 0 | 0 |
| ENSMUSG00 | Olfr1117-ps1 | 920 | 0 | 0 | 0 | 0 | 0 | 0 |
| ENSMUSG00 | Gm13592 | 1009 | 0.03940087 | 0.2865398 | 0.13142273 | 0.07152041 | 0.20228494 | 0.08109407 |
| ENSMUSG00 | Gm13123 | 491 | 0 | 0 | 0 | 0 | 0 | 0 |
| ENSMUSG00 | Gm14688 | 302 | 0 | 0 | 0 | 0 | 0 | 0 |
| ENSMUSG00 | Gm15078 | 781 | 0 | 0 | 0 | 0 | 0 | 0 |
| ENSMUSG00 | Gm3931 | 454 | 0 | 0 | 0 | 0 | 0 | 0 |
| ENSMUSG00 | Gm5883 | 494 | 0.08047668 | 0 | 0 | 0 | 0.10329226 | 0.11042364 |
| ENSMUSG00 | Gm14499 | 515 | 0 | 0 | 0 | 0 | 0 | 0 |
| ENSMUSG00 | Gm14202 | 310 | 0 | 0 | 0 | 0 | 0 | 0 |
| ENSMUSG00 | Gm13436 | 1181 | 0 | 0 | 0.14970988 | 0 | 0 | 0 |
| ENSMUSG00 | Gm13865 | 864 | 0 | 0 | 0 | 0 | 0 | 0 |
| ENSMUSG00 | Ccnb2-ps | 2917 | 1.30837372 | 1.2719767 | 0.24245097 | 0.98956584 | 1.13702929 | 0.79477008 |
| ENSMUSG00 | Gm15161 | 386 | 0 | 0 | 0 | 0 | 0 | 0 |
| ENSMUSG00 | Gm12901 | 530 | 0 | 0 | 0 | 0 | 0 | 0 |
| ENSMUSG00 | Gm7700 | 969 | 0 | 0 | 0 | 0 | 0 | 0 |
| ENSMUSG00 | Gpr144 | 2644 | 0 | 0 | 0 | 0 | 0 | 0 |
| ENSMUSG00 | Gm11671 | 423 | 0.09398459 | 0 | 0 | 0 | 0 | 0 |
| ENSMUSG00 | Gm12651 | 446 | 0 | 0 | 0 | 0 | 0 | 0 |
| ENSMUSG00 | Gm11865 | 857 | 0.13916738 | 0 | 0 | 0.46313009 | 0 | 0.92294579 |
| ENSMUSG00 | Gm13309 | 320 | 0 | 0 | 0 | 0 | 0 | 0 |
| ENSMUSG00 | Gm11887 | 229 | 0 | 0 | 0 | 0 | 0 | 0 |
| ENSMUSG00 | Gm13854 | 1677 | 0 | 0 | 0 | 0 | 0 | 0 |
| ENSMUSG00 | Sh3d2c-ps1 | 903 | 0 | 0 | 0 | 0 | 0 | 0 |
| ENSMUSG00 | Gm2759 | 631 | 0 | 0 | 0 | 0 | 0 | 0 |
| ENSMUSG00 | Gm16202 | 543 | 0 | 0 | 0.16280605 | 0.06644944 | 0 | 0 |
| ENSMUSG00 | Gm14197 | 1048 | 0.45521543 | 0.32185601 | 0 | 0.03442943 | 0.04868929 | 0 |
| ENSMUSG00 | Gm13035 | 341 | 0 | 0 | 0 | 0 | 0 | 0 |
| ENSMUSG00 | Gm16209 | 626 | 0.82559305 | 0.30790059 | 0.98853962 | 0.28819524 | 1.46721208 | 1.26352163 |
| ENSMUSG00 | Gm12495 | 446 | 0 | 0 | 0 | 0 | 0 | 0 |
| ENSMUSG00 | Gm6039 | 1249 | 0 | 0 | 0 | 0 | 0 | 0 |
| ENSMUSG00 | Prdx6-ps2 | 693 | 0 | 0 | 0 | 0 | 0 | 0 |
| ENSMUSG00 | Gm14592 | 559 | 0 | 0 | 0 | 0 | 0 | 0 |
| ENSMUSG00 | Gm13796 | 312 | 0 | 0 | 0 | 0 | 0 | 0 |
| ENSMUSG00 | Gm11456 | 255 | 0 | 0 | 0 | 0 | 0 | 0 |
| ENSMUSG00 | Gm15297 | 288 | 0 | 0 | 0 | 0 | 0 | 0 |
| ENSMUSG00 | Gm11812 | 916 | 0 | 0 | 0 | 0 | 0 | 0 |
| ENSMUSG00 | Gm12421 | 306 | 0 | 0 | 0 | 0 | 0 | 0 |
| ENSMUSG00 | Gm14824 | 736 | 0 | 0 | 0 | 0 | 0 | 0 |
| ENSMUSG00 | Gm13543 | 795 | 2.10028954 | 1.63652068 | 0.66719763 | 1.54313146 | 3.72267899 | 0.27446178 |
| ENSMUSG00 | Gm14727 | 524 | 0 | 0 | 0 | 0 | 0 | 0 |
| ENSMUSG00 | Gm11662 | 1191 | 0 | 0 | 0 | 0 | 0 | 0 |
| ENSMUSG00 | Gm13100 | 1436 | 0 | 0 | 0 | 0 | 0 | 0 |
| ENSMUSG00 | Gm14108 | 493 | 0 | 0 | 0 | 0 | 0 | 0 |
| ENSMUSG00 | Gm14534 | 356 | 0 | 0.13535518 | 0.24832496 | 1.01354058 | 0 | 0 |
| ENSMUSG00 | Gm12014 | 1617 | 0 | 0 | 0 | 0 | 0 | 0 |
| ENSMUSG00 | Gm12814 | 842 | 0 | 0 | 0 | 0 | 0 | 0 |
| ENSMUSG00 | Gm7803 | 1580 | 0.0251617 | 0 | 0.02797585 | 0.04567347 | 0 | 0.05178729 |
| ENSMUSG00 | Gm14559 | 495 | 0 | 0 | 0 | 0 | 0 | 0 |
| ENSMUSG00 | Gm13468 | 444 | 0 | 0 | 0 | 0 | 0 | 0 |
| ENSMUSG00 | Rps15a-ps4 | 391 | 0.91506779 | 0.36971695 | 0.56524096 | 0 | 1.04401792 | 0.20926833 |
| ENSMUSG00 | Gm14524 | 682 | 0 | 0 | 0 | 0 | 0 | 0.11997642 |
| ENSMUSG00 | Pgam1-ps1 | 748 | 0 | 0.12884075 | 0.05909337 | 0.19295211 | 0 | 0.10939026 |
| ENSMUSG00 | Gm2318 | 631 | 0 | 0 | 0 | 0 | 0 | 0 |
| ENSMUSG00 | Gm9050 | 447 | 0 | 0 | 0 | 0 | 0 | 0 |
| ENSMUSG00 | Gm11516 | 1581 | 0 | 0 | 0 | 0 | 0.03227475 | 0.01725151 |
| ENSMUSG00 | Gm15335 | 409 | 0 | 0 | 0 | 0 | 0 | 0 |
| ENSMUSG00 | Gm11405 | 1555 | 0.02556623 | 0 | 0 | 0.04640777 | 0 | 0 |
| ENSMUSG00 | Gm15362 | 387 | 0 | 0 | 0 | 0 | 0 | 0 |
| ENSMUSG00 | Gm11689 | 301 | 0 | 0 | 0 | 0 | 0 | 0 |
| ENSMUSG00 | Gm15988 | 366 | 0 | 0 | 0 | 0 | 0.13941633 | 0.44712523 |
| ENSMUSG00 | Gm15566 | 589 | 0 | 0 | 0 | 0 | 0 | 0 |
| ENSMUSG00 | Gm13394 | 2249 | 0.01767696 | 0.06427716 | 0.039308 | 0.0160436 | 0.09075389 | 0.01212745 |
| ENSMUSG00 | Gm7180 | 512 | 0.54353196 | 0.09411415 | 0.6906538 | 0.07047274 | 0.19932178 | 0.10654156 |
| ENSMUSG00 | Gm14837 | 205 | 0 | 0 | 0 | 0 | 0 | 0 |
| ENSMUSG00 | Gm13473 | 365 | 0 | 0 | 0 | 0 | 0 | 0 |
| ENSMUSG00 | Gm13361 | 1430 | 0 | 0 | 0 | 0 | 0 | 0 |
| ENSMUSG00 | Gm15094 | 1130 | 0 | 0 | 0 | 0 | 0 | 0 |
| ENSMUSG00 | Gm14775 | 309 | 0 | 0 | 0 | 0 | 0 | 0 |
| ENSMUSG00 | Gm12728 | 492 | 0.16160764 | 0 | 0 | 0 | 0.20742429 | 0.05543626 |
| ENSMUSG00 | Gm14842 | 268 | 0 | 0 | 0 | 0 | 0 | 0 |
| ENSMUSG00 | Gm11660 | 411 | 0 | 0 | 0 | 0 | 0 | 0 |
| ENSMUSG00 | Gm12739 | 381 | 0 | 0 | 0 | 0 | 0 | 0 |
| ENSMUSG00 | Gm14729 | 688 | 0 | 0 | 0 | 0 | 0 | 0 |
| ENSMUSG00 | Gm11888 | 808 | 0 | 0 | 0 | 0 | 0 | 0 |
| ENSMUSG00 | Rps18-ps2 | 397 | 0 | 0 | 0 | 0 | 0 | 0 |
| ENSMUSG00 | Gm11644 | 149 | 0 | 0 | 0 | 0 | 0 | 0 |
| ENSMUSG00 | Gm11499 | 321 | 0 | 0 | 0 | 0 | 0 | 0 |
| ENSMUSG00 | Gm6092 | 2028 | 0 | 0 | 0 | 0 | 0 | 0 |
| ENSMUSG00 | Gm12721 | 267 | 2.97793862 | 2.52662993 | 0 | 2.29735864 | 3.24887037 | 4.59684931 |
| ENSMUSG00 | Rpl12-ps2 | 320 | 0 | 0 | 0 | 0 | 0 | 0 |
| ENSMUSG00 | Gm14274 | 614 | 0 | 0 | 0.07198997 | 0 | 0 | 0 |
| ENSMUSG00 | Gm16123 | 1515 | 0 | 0 | 0 | 0 | 0 | 0.07201225 |
| ENSMUSG00 | Gm15670 | 446 | 0 | 0 | 0 | 0.8899159 | 0 | 0 |

| | | | | | | | |
|---|---|---|---|---|---|---|---|
| ENSMUSG00 Gm13369 | 1570 | 0.17725374 | 0.092076 | 0.36600252 | 0.18385755 | 0.13000351 | 0.36482002 |
| ENSMUSG00 Gm13800 | 165 | 0 | 0 | 0 | 0 | 0 | 0 |
| ENSMUSG00 Gm14584 | 636 | 2.06278437 | 0.90917815 | 1.45949482 | 1.07792271 | 0.88253166 | 2.1871173 |
| ENSMUSG00 Gm11322 | 434 | 0 | 0 | 0 | 0 | 0 | 0 |
| ENSMUSG00 Gm12673 | 178 | 1.34007238 | 1.35355175 | 0 | 0.81083246 | 0.8599951 | 0.61291324 |
| ENSMUSG00 Gm12455 | 415 | 0 | 0 | 0 | 0 | 0 | 0 |
| ENSMUSG00 Gm13676 | 590 | 0 | 0 | 0 | 0 | 0 | 0 |
| ENSMUSG00 Gm14848 | 416 | 0 | 0 | 0 | 0 | 0 | 0 |
| ENSMUSG00 Gm12131 | 357 | 0 | 0 | 0 | 0 | 0 | 0 |
| ENSMUSG00 Gm13466 | 998 | 0.11950545 | 0.14484902 | 0.26574254 | 0.32538918 | 0.25564317 | 0.0546586 |
| ENSMUSG00 Gm3828 | 256 | 0.15529485 | 0 | 0 | 0 | 0 | 0 |
| ENSMUSG00 Gm15110 | 879 | 0 | 0 | 0 | 0 | 0 | 0 |
| ENSMUSG00 Gm14713 | 684 | 0 | 0 | 0 | 0 | 0 | 0 |
| ENSMUSG00 Gm13071 | 375 | 0.42405846 | 0 | 0 | 0.6735315 | 0.68035168 | 0 |
| ENSMUSG00 Gm5054 | 772 | 0 | 0 | 0 | 0 | 0 | 0 |
| ENSMUSG00 Gm15502 | 207 | 0.38411092 | 0.46556949 | 0.42707095 | 0 | 0.49300846 | 0 |
| ENSMUSG00 Gm5133 | 1250 | 0 | 0 | 0 | 0 | 0 | 0 |
| ENSMUSG00 Gm13435 | 1022 | 0 | 0 | 0 | 0 | 0.09985592 | 0 |
| ENSMUSG00 Gm13033 | 1051 | 0.75652675 | 0.22924092 | 0.42056939 | 0.34331156 | 0.87390558 | 3.16603805 |
| ENSMUSG00 Gm14400 | 2132 | 0.14917629 | 0.22601521 | 0.29025601 | 0.4231009 | 0.19146858 | 0.23027369 |
| ENSMUSG00 Gm14908 | 1284 | 0 | 0 | 0 | 0 | 0 | 0 |
| ENSMUSG00 Gm11977 | 1460 | 0.24506803 | 0.09901324 | 0 | 0.02471373 | 0.13979829 | 0.29890016 |
| ENSMUSG00 Gm6415 | 347 | 0 | 0 | 0 | 0.10398284 | 0 | 0 |
| ENSMUSG00 Gm14631 | 593 | 0 | 0 | 0 | 0 | 0 | 0 |
| ENSMUSG00 Hmgb1-ps5 | 556 | 0.07150266 | 0.43333131 | 0.39749859 | 0 | 0.45887029 | 0.34338575 |
| ENSMUSG00 Amd-ps1 | 993 | 0 | 0 | 0.04451344 | 0 | 0 | 0 |
| ENSMUSG00 Gm6947 | 712 | 0 | 0 | 0 | 0 | 0 | 0 |
| ENSMUSG00 Olfr1376-ps1 | 897 | 0 | 0 | 0 | 0 | 0 | 0 |
| ENSMUSG00 Gm13039 | 793 | 0 | 0 | 0 | 0 | 0.064346 | 0 |
| ENSMUSG00 Gm15712 | 1155 | 2.27174174 | 2.16943289 | 2.29619964 | 3.09274667 | 2.87161422 | 2.76288553 |
| ENSMUSG00 Gm15006 | 1121 | 0 | 0 | 0 | 0 | 0 | 0 |
| ENSMUSG00 Gm2199 | 873 | 0.63754493 | 0.38637468 | 0.4556891 | 0.12399328 | 0.23379783 | 0.53112127 |
| ENSMUSG00 Gm8839 | 1644 | 0 | 0 | 0 | 0 | 0 | 0 |
| ENSMUSG00 Gm14248 | 543 | 0 | 0.08874115 | 0 | 0 | 0 | 0 |
| ENSMUSG00 Gm13771 | 237 | 0.67097857 | 0.81327329 | 0.37301133 | 0.30448983 | 1.07650582 | 1.72624299 |
| ENSMUSG00 Gm13841 | 1133 | 0.2456208 | 0.46782954 | 0.1950655 | 0.25477172 | 0.22518259 | 0.24072938 |
| ENSMUSG00 Gm12577 | 573 | 0 | 0 | 0 | 0.12594082 | 0 | 0.14279916 |
| ENSMUSG00 Gm11898 | 384 | 0 | 0 | 0 | 0 | 0 | 0 |
| ENSMUSG00 Ldha-ps2 | 1056 | 0.03764724 | 0.0456311 | 0 | 0.0341686 | 0 | 0 |
| ENSMUSG00 Gm14427 | 634 | 0 | 0 | 0 | 0 | 0 | 0 |
| ENSMUSG00 Gm11957 | 778 | 0 | 0 | 0 | 0 | 0 | 0 |
| ENSMUSG00 Gm14765 | 325 | 0 | 0 | 0 | 0 | 0 | 0 |
| ENSMUSG00 Gm15459 | 3549 | 0 | 0 | 0 | 0 | 0 | 0 |
| ENSMUSG00 Gm1947 | 909 | 0 | 0 | 0 | 0 | 0 | 0 |
| ENSMUSG00 Gm12485 | 290 | 0 | 0 | 0 | 0 | 0 | 0 |
| ENSMUSG00 Gm7071 | 461 | 0 | 0 | 0 | 0 | 0 | 0 |
| ENSMUSG00 2210015D19l | 3185 | 11.2338877 | 12.9657083 | 9.17344372 | 9.30089751 | 9.67658744 | 10.2847227 |
| ENSMUSG00 Gm14571 | 569 | 0 | 0 | 0 | 0 | 0 | 0 |
| ENSMUSG00 Gm13797 | 119 | 0 | 0 | 0 | 0 | 0 | 0 |
| ENSMUSG00 Xlr3d-ps | 650 | 0 | 0 | 0 | 0 | 0 | 0 |
| ENSMUSG00 Gm7309 | 580 | 0 | 0 | 0 | 0 | 0 | 0 |
| ENSMUSG00 Gm13477 | 2686 | 0 | 0.10763911 | 0 | 0.01343338 | 0.01899716 | 0.40617482 |
| ENSMUSG00 Gm12365 | 567 | 0 | 0.08498491 | 0 | 0 | 0 | 0 |
| ENSMUSG00 Gm13007 | 364 | 0 | 0 | 0 | 0 | 0 | 0 |
| ENSMUSG00 Gm15696 | 444 | 0 | 1.62792034 | 0.3982148 | 0 | 0.34477281 | 0 |
| ENSMUSG00 Gm15118 | 350 | 0 | 0.13767555 | 0.12629098 | 0.10309156 | 0 | 0.15585508 |
| ENSMUSG00 Olfr1175-ps | 1882 | 0 | 0 | 0 | 0 | 0 | 0 |
| ENSMUSG00 Gm13886 | 1182 | 0.50451117 | 0.36690184 | 0.71052032 | 0.73263035 | 0.25901714 | 1.47679942 |
| ENSMUSG00 Gm12584 | 1104 | 0 | 0 | 0 | 0 | 0 | 0 |
| ENSMUSG00 Gm14979 | 1034 | 0 | 0 | 0 | 0 | 0 | 0 |
| ENSMUSG00 Gm12003 | 1104 | 0.0360104 | 0 | 0.1201137 | 0.09804903 | 0 | 0 |
| ENSMUSG00 Gm13313 | 1071 | 0 | 0 | 0 | 0 | 0 | 0 |
| ENSMUSG00 Gm9599 | 480 | 0 | 0 | 0 | 0 | 0 | 0 |
| ENSMUSG00 Gm13341 | 671 | 12.9753357 | 6.17590764 | 3.22785441 | 5.75376865 | 4.63876144 | 5.24355956 |
| ENSMUSG00 Gm13719 | 2109 | 0 | 0.02284801 | 0.02095867 | 0.03421721 | 0.04838917 | 0.025865 |
| ENSMUSG00 Gm13472 | 313 | 0 | 0 | 0 | 0 | 0 | 0 |
| ENSMUSG00 Gm12893 | 273 | 0 | 0 | 0 | 0 | 0 | 0 |
| ENSMUSG00 Gm12477 | 570 | 0 | 0 | 0 | 0 | 0 | 0 |
| ENSMUSG00 Gm12442 | 337 | 0 | 0 | 0 | 0 | 0 | 0 |
| ENSMUSG00 Gm11761 | 444 | 0.44769685 | 0 | 0.3982148 | 0 | 0.11492427 | 0.86001115 |
| ENSMUSG00 Gm16479 | 931 | 0 | 0 | 0 | 0 | 0 | 0 |
| ENSMUSG00 Gm11928 | 436 | 0 | 0 | 0 | 0 | 0 | 0 |
| ENSMUSG00 Gm11283 | 516 | 0 | 0 | 0 | 0 | 0 | 0 |
| ENSMUSG00 Gm7494 | 440 | 0 | 0 | 0 | 0 | 0 | 0 |
| ENSMUSG00 Gm13831 | 301 | 0.39623403 | 0 | 2.05589968 | 0.83911732 | 0 | 0.36245368 |
| ENSMUSG00 Gm11439 | 408 | 0 | 0 | 0 | 0 | 0 | 0 |
| ENSMUSG00 Gm14740 | 1149 | 0 | 0 | 0 | 0 | 0 | 0 |
| ENSMUSG00 Gm12034 | 398 | 0 | 0 | 0 | 0 | 0 | 0 |
| ENSMUSG00 Gm14038 | 2386 | 0.08330989 | 0.08078197 | 0.2408315 | 0.15122399 | 0.25662888 | 0.24005341 |
| ENSMUSG00 Hspa1-ps6 | 282 | 0 | 1.87961299 | 0 | 0 | 0 | 0 |
| ENSMUSG00 Gm12605 | 711 | 0 | 0 | 0 | 0 | 0 | 0 |
| ENSMUSG00 Gm11369 | 317 | 0 | 0 | 0 | 0 | 0 | 0 |
| ENSMUSG00 Gm14941 | 443 | 0 | 0 | 0 | 0 | 0 | 0 |
| ENSMUSG00 Gm15388 | 725 | 0 | 0 | 0 | 0 | 0 | 0 |
| ENSMUSG00 Gm13213 | 473 | 0.08404964 | 0.20374817 | 0 | 0.15256678 | 0 | 0.11532617 |
| ENSMUSG00 Olfr409-ps1 | 973 | 0 | 0 | 0 | 0 | 0 | 0 |
| ENSMUSG00 Gm11930 | 189 | 0 | 0 | 0 | 0 | 0 | 0 |
| ENSMUSG00 Gm12441 | 587 | 0 | 0 | 0 | 0 | 0 | 0 |
| ENSMUSG00 E530001F21f | 482 | 0 | 0 | 0 | 0 | 0 | 0 |
| ENSMUSG00 Gm15703 | 879 | 0 | 0 | 0 | 0 | 0 | 0 |
| ENSMUSG00 Gm1848 | 894 | 0.88938435 | 0.916297 | 0.64275611 | 0.36324206 | 1.36983559 | 0.57966236 |
| ENSMUSG00 Gm13347 | 1352 | 0 | 0 | 0 | 0 | 0 | 0 |
| ENSMUSG00 Gm11489 | 497 | 0 | 0 | 0 | 0 | 0 | 0 |
| ENSMUSG00 Gm12074 | 218 | 0 | 0 | 0 | 0 | 0 | 0 |
| ENSMUSG00 Scgb2b1-ps | 327 | 0 | 0 | 0 | 0 | 0 | 0 |
| ENSMUSG00 Gm14843 | 211 | 0 | 0 | 0 | 0.17100495 | 0 | 0 |
| ENSMUSG00 Gm7340 | 2694 | 0 | 0 | 0 | 0 | 0 | 0 |
| ENSMUSG00 Gm12346 | 2255 | 0.88149624 | 1.68812813 | 0.50964431 | 1.58408976 | 0.95038039 | 1.13694727 |
| ENSMUSG00 Gm13294 | 476 | 0 | 0.10123202 | 0 | 0 | 0 | 0 |
| ENSMUSG00 Gm12885 | 327 | 0 | 0 | 0 | 0 | 0 | 0 |
| ENSMUSG00 Gm15975 | 1405 | 0.5376186 | 0.4458532 | 0.28314348 | 0.38521756 | 0.18158853 | 0.95121518 |
| ENSMUSG00 Gm5405 | 608 | 0 | 0.4755241 | 0 | 0 | 0.08392496 | 0 |
| ENSMUSG00 Gm12805 | 1047 | 0 | 0 | 0 | 0 | 0 | 0 |
| ENSMUSG00 Gm15355 | 284 | 0 | 0.16967057 | 0 | 0 | 0 | 0 |
| ENSMUSG00 Gm15186 | 562 | 0 | 0 | 0 | 1.41246438 | 0.7263541 | 0 |
| ENSMUSG00 Plk-ps1 | 1787 | 0.33370577 | 0.45840488 | 0.5441749 | 1.9787579 | 0.37120475 | 0.80892886 |
| ENSMUSG00 Gm6447 | 834 | 0 | 0.05777751 | 0 | 0 | 0 | 0 |
| ENSMUSG00 Gm15842 | 617 | 0 | 0 | 0 | 0.29239906 | 0.99240925 | 1.10513125 |
| ENSMUSG00 Gm13604 | 481 | 0.16530345 | 0 | 0 | 0 | 0 | 0.1701121 |
| ENSMUSG00 Gm4342 | 366 | 0 | 0 | 0 | 0.09858482 | 0 | 0 |
| ENSMUSG00 Gm5391 | 612 | 0 | 0.07873602 | 0 | 0.05895759 | 0 | 0.26739842 |
| ENSMUSG00 Gm14219 | 287 | 0 | 0 | 0 | 0 | 0 | 0 |
| ENSMUSG00 Rps18-ps1 | 968 | 0 | 0 | 0.04566306 | 0 | 0.0527132 | 0 |
| ENSMUSG00 Gm13932 | 609 | 0 | 0.07912388 | 0.07258102 | 0.05924802 | 0 | 0.04478594 |
| ENSMUSG00 Gm15476 | 399 | 0 | 0 | 0 | 0 | 0 | 0 |
| ENSMUSG00 Gm12093 | 1403 | 0 | 0 | 0 | 0 | 0 | 0 |
| ENSMUSG00 Gm14352 | 448 | 0 | 0 | 0 | 0 | 0 | 0 |
| ENSMUSG00 Tpi-rs8 | 707 | 0 | 0 | 0 | 0 | 0 | 0 |
| ENSMUSG00 Gm14987 | 675 | 0 | 0 | 0 | 0 | 0 | 0 |
| ENSMUSG00 Gm15750 | 596 | 0 | 0 | 0.74164266 | 0 | 0 | 0 |
| ENSMUSG00 Gm3835 | 402 | 0 | 0 | 0.659729 | 0 | 0.12693128 | 0.06784736 |
| ENSMUSG00 Gm649 | 1053 | 0 | 0 | 0 | 0 | 0 | 0 |
| ENSMUSG00 Gm12304 | 384 | 0 | 0.12548553 | 0.57554483 | 0 | 1.19593068 | 1.84672037 |
| ENSMUSG00 Gm12012 | 1065 | 0 | 0 | 0 | 0 | 0 | 0 |
| ENSMUSG00 Gm14783 | 662 | 0 | 0 | 0 | 0 | 0 | 0 |
| ENSMUSG00 Gm14749 | 331 | 0 | 0 | 0 | 0 | 0.15415824 | 0 |
| ENSMUSG00 Gm4991 | 1551 | 0.02563216 | 0 | 0 | 0 | 0 | 0 |
| ENSMUSG00 Gm13301 | 2214 | 0 | 0 | 0 | 0 | 0 | 0 |
| ENSMUSG00 Gm13042 | 1953 | 0.02035611 | 0 | 0 | 0 | 0 | 0 |
| ENSMUSG00 Gm14518 | 541 | 0 | 0 | 0 | 0 | 0 | 0 |
| ENSMUSG00 Gm11438 | 188 | 0 | 0 | 0 | 0 | 0 | 0 |
| ENSMUSG00 Gm15515 | 821 | 0 | 0 | 0.05383903 | 0 | 0 | 0.06644248 |
| ENSMUSG00 Gm12368 | 433 | 0 | 0 | 0 | 0 | 0 | 0 |
| ENSMUSG00 Gm12294 | 489 | 0 | 0 | 0 | 0 | 0 | 0.39043451 |
| ENSMUSG00 Gm14829 | 1137 | 0 | 0 | 0 | 0.03173443 | 0.17951232 | 0.0479765 |
| ENSMUSG00 Cct3-ps1 | 1788 | 0.02223461 | 0.02694991 | 0.02472139 | 0 | 0 | 0 |
| ENSMUSG00 Gm12448 | 521 | 0 | 0 | 0 | 0 | 0 | 0 |
| ENSMUSG00 Gm14451 | 1539 | 0 | 0 | 0 | 0 | 0 | 0 |
| ENSMUSG00 Gm11298 | 2033 | 0 | 0 | 0 | 0 | 0 | 0 |
| ENSMUSG00 Gm12889 | 240 | 0 | 0 | 0 | 0 | 0 | 0 |
| ENSMUSG00 Gm12759 | 302 | 0 | 0 | 0 | 0 | 0 | 0 |
| ENSMUSG00 Gm11483 | 246 | 0 | 0 | 0 | 0.29334996 | 0 | 0.22174504 |
| ENSMUSG00 Gm14733 | 132 | 0 | 0 | 0 | 0 | 0 | 0 |
| ENSMUSG00 Gm15242 | 282 | 0 | 0 | 0 | 0 | 0 | 0 |
| ENSMUSG00 Gm11359 | 564 | 0 | 0 | 0 | 0 | 0 | 0 |
| ENSMUSG00 Olfr122 | 1256 | 0 | 0 | 0 | 0 | 0 | 0 |
| ENSMUSG00 Gm7158 | 902 | 0 | 0 | 0 | 0 | 0 | 0 |
| ENSMUSG00 Gm12897 | 247 | 0 | 0 | 0 | 0 | 0 | 0 |
| ENSMUSG00 Gm14466 | 450 | 0 | 0 | 0 | 0 | 0 | 0 |
| ENSMUSG00 Gm13786 | 332 | 0.23949085 | 0 | 0 | 0.43472343 | 0 | 0 |
| ENSMUSG00 Gm14675 | 446 | 0 | 0 | 0 | 0 | 0 | 0 |
| ENSMUSG00 Gm14049 | 635 | 0 | 0 | 0 | 0 | 0 | 0.08590438 |
| ENSMUSG00 Gm12179 | 1271 | 0 | 0 | 0 | 0 | 0 | 0 |
| ENSMUSG00 Gm14785 | 537 | 0 | 0 | 0 | 0 | 0 | 0 |
| ENSMUSG00 Gm12361 | 1041 | 0 | 0 | 0 | 0 | 0 | 0 |
| ENSMUSG00 Gm12742 | 408 | 0 | 0 | 0 | 0 | 0 | 0 |
| ENSMUSG00 Gm12601 | 843 | 0 | 0 | 0 | 0 | 0 | 0 |
| ENSMUSG00 Gm7365 | 630 | 0 | 0 | 0 | 0 | 0 | 0 |
| ENSMUSG00 Gm4918 | 1083 | 0 | 0 | 0 | 0 | 0 | 0 |
| ENSMUSG00 Gm13725 | 206 | 0 | 0 | 0 | 0 | 0 | 0 |
| ENSMUSG00 Gm15707 | 995 | 0.03995526 | 0 | 0 | 0 | 0 | 0 |
| ENSMUSG00 Gm15203 | 454 | 0 | 0 | 0 | 0 | 0 | 0 |
| ENSMUSG00 Gm14122 | 971 | 0.12282847 | 0 | 0.40969782 | 0 | 0.05255034 | 1.1516583 |
| ENSMUSG00 Gm13482 | 253 | 0 | 0 | 0 | 0 | 0 | 0 |
| ENSMUSG00 Gm15115 | 263 | 0 | 0 | 0 | 0 | 0 | 0 |
| ENSMUSG00 Olfr1050-ps1 | 946 | 0 | 0 | 0 | 0 | 0 | 0 |
| ENSMUSG00 Gm15081 | 843 | 0 | 0 | 0 | 0 | 0 | 0 |
| ENSMUSG00 Gm5943 | 458 | 0 | 0 | 0 | 0 | 0 | 0 |
| ENSMUSG00 Gm14802 | 1884 | 0 | 0 | 0 | 0 | 0 | 0 |
| ENSMUSG00 Gm13459 | 645 | 0 | 0 | 0 | 0 | 0 | 0 |
| ENSMUSG00 Gm16053 | 187 | 1.70076922 | 0 | 0 | 0.77180844 | 0 | 0.14585369 |
| ENSMUSG00 Gm14215 | 220 | 0 | 0 | 0 | 0 | 0 | 0 |
| ENSMUSG00 Gm13864 | 344 | 0 | 0 | 0 | 0 | 0 | 0 |
| ENSMUSG00 Mageb8-ps | 1032 | 0 | 0 | 0 | 0 | 0 | 0 |
| ENSMUSG00 Gm5060 | 598 | 0 | 0.16115867 | 0.14783225 | 0 | 0.17065678 | 0 |
| ENSMUSG00 Gm12527 | 563 | 0.77675006 | 0.94147578 | 2.19831546 | 0.5768 | 0.54379797 | 0 |
| ENSMUSG00 Gm15363 | 885 | 0 | 0 | 0 | 0 | 0 | 0 |
| ENSMUSG00 Gm14650 | 888 | 0 | 0 | 0 | 0 | 0 | 0 |
| ENSMUSG00 Gm12375 | 919 | 0 | 0 | 0 | 0 | 0 | 0 |
| ENSMUSG00 Gm12468 | 403 | 0 | 0 | 0 | 0 | 0 | 0.20303702 |
| ENSMUSG00 Gm12213 | 352 | 1.12941706 | 0 | 0 | 0 | 0 | 0.30993908 |
| ENSMUSG00 Gm15311 | 280 | 0 | 0 | 0 | 0 | 0 | 0 |
| ENSMUSG00 Gm15633 | 491 | 0.08096839 | 0.29441818 | 0 | 0.51440797 | 0 | 0.55549163 |
| ENSMUSG00 Gm12607 | 487 | 0 | 0 | 0 | 0 | 0 | 0 |
| ENSMUSG00 Gm12101 | 389 | 0 | 0 | 0 | 0 | 0 | 0 |
| ENSMUSG00 Gm11618 | 265 | 0 | 0 | 0 | 0 | 0 | 0 |
| ENSMUSG00 Gm11478 | 598 | 0.46536516 | 1.12811069 | 0.44349675 | 0.30168934 | 0.42664194 | 0.91219529 |
| ENSMUSG00 Gm14398 | 1072 | 0 | 0 | 0.12369919 | 0 | 0 | 0 |
| ENSMUSG00 Gm16465 | 1756 | 0 | 0 | 0 | 0.02054786 | 0 | 0 |
| ENSMUSG00 Gm12808 | 403 | 4.93244175 | 4.42406541 | 3.07109579 | 6.35688626 | 8.60990957 | 6.42950553 |
| ENSMUSG00 Gm11649 | 374 | 0 | 0 | 0 | 0 | 0 | 0 |
| ENSMUSG00 Gm12920 | 1185 | 0 | 0 | 0 | 0 | 0 | 0 |
| ENSMUSG00 Prdx2-rs3 | 592 | 0 | 0 | 0 | 0 | 0 | 0 |
| ENSMUSG00 Gm7790 | 1309 | 0 | 0 | 0 | 0 | 0 | 0 |
| ENSMUSG00 Gm14653 | 1532 | 0 | 0 | 0 | 0 | 0 | 0 |
| ENSMUSG00 Gm15113 | 834 | 0 | 0 | 0 | 0 | 0 | 0 |
| ENSMUSG00 Gm13146 | 779 | 0 | 0 | 0 | 0 | 0 | 0 |
| ENSMUSG00 Gm15269 | 203 | 0 | 0 | 0 | 0 | 0 | 0 |
| ENSMUSG00 Gm12047 | 943 | 0 | 0 | 0.09374728 | 0.15305215 | 0.05411069 | 0 |
| ENSMUSG00 Gm13701 | 300 | 0 | 0 | 0 | 0 | 0 | 0 |
| ENSMUSG00 Gm14245 | 1842 | 0 | 0 | 0 | 0.03917703 | 0 | 0 |
| ENSMUSG00 Gm9003 | 480 | 0.16564764 | 0.40155369 | 1.47339477 | 0.45102556 | 0.10630495 | 0 |
| ENSMUSG00 Gm14992 | 511 | 0 | 0 | 0 | 0 | 0 | 0 |
| ENSMUSG00 Gm7590 | 800 | 0 | 0 | 0 | 0 | 0 | 0 |
| ENSMUSG00 Gm13302 | 1848 | 0 | 0 | 0 | 0 | 0 | 0 |
| ENSMUSG00 Gm12969 | 1003 | 0 | 0 | 0 | 0.07194824 | 0.25436877 | 0.02719306 |
| ENSMUSG00 Gm11956 | 511 | 0 | 0 | 0 | 0 | 0 | 0 |
| ENSMUSG00 Gm14270 | 845 | 0.51752697 | 0.28512688 | 0.36616911 | 0.42700644 | 0.84540741 | 0.77466431 |
| ENSMUSG00 Gm7209 | 332 | 0 | 0 | 0 | 0 | 0 | 0 |
| ENSMUSG00 Gm14175 | 240 | 0 | 0 | 0 | 0 | 0 | 0 |
| ENSMUSG00 Olfr1096-ps1 | 947 | 0 | 0 | 0 | 0 | 0 | 0 |
| ENSMUSG00 Rpl21-ps9 | 481 | 0.08265173 | 0 | 0 | 0 | 0 | 0.39692822 |
| ENSMUSG00 Gm11895 | 466 | 0 | 0 | 0 | 0 | 0 | 0 |
| ENSMUSG00 Gm8754 | 519 | 0 | 0 | 0 | 0 | 0 | 0 |
| ENSMUSG00 Gm12282 | 4151 | 0.0383093 | 0.01160839 | 0 | 0.13038561 | 0.0245851 | 0.02628248 |
| ENSMUSG00 Gm5832 | 1581 | 0 | 0 | 0 | 0 | 0 | 0 |
| ENSMUSG00 Gm16459 | 685 | 0 | 0 | 0 | 0 | 0 | 0.07963398 |
| ENSMUSG00 Gm15937 | 1234 | 0 | 0.07809796 | 0 | 0 | 0 | 0 |
| ENSMUSG00 Gm6276 | 578 | 0 | 0 | 0 | 0 | 0 | 0 |
| ENSMUSG00 Gm11259 | 714 | 0 | 0 | 0 | 0 | 0 | 0 |
| ENSMUSG00 Gm12376 | 606 | 0 | 0 | 0 | 0 | 0 | 0 |
| ENSMUSG00 Gm13983 | 864 | 0 | 0 | 0 | 0 | 0 | 0 |
| ENSMUSG00 Gm8855 | 877 | 0 | 0 | 0 | 0 | 0 | 0 |
| ENSMUSG00 Gm16423 | 946 | 0 | 0 | 0 | 0 | 0 | 0 |
| ENSMUSG00 Gm7416 | 1485 | 0 | 0 | 0.02976555 | 0 | 0 | 0 |
| ENSMUSG00 Gm12991 | 77 | 0 | 0 | 0 | 0 | 0 | 0 |
| ENSMUSG00 Gm14447 | 1446 | 0.60485517 | 0.63315519 | 0.21397849 | 0.49906009 | 0.74104695 | 0.41496685 |
| ENSMUSG00 Gm12617 | 380 | 0 | 0 | 0 | 0 | 0.40283981 | 0 |
| ENSMUSG00 Ccl19-ps3 | 320 | 0 | 0 | 0 | 0 | 0 | 0 |
| ENSMUSG00 Gm11295 | 801 | 0 | 0 | 0 | 0 | 0 | 0 |
| ENSMUSG00 Gm14551 | 255 | 0 | 0 | 0 | 0 | 0 | 0 |
| ENSMUSG00 Gm14763 | 326 | 0 | 0 | 0 | 0 | 0 | 0 |
| ENSMUSG00 Gm15312 | 270 | 0 | 0 | 0 | 0 | 0 | 0 |
| ENSMUSG00 Gm13732 | 534 | 0 | 0 | 0 | 0 | 0 | 0 |
| ENSMUSG00 Gm14647 | 679 | 0 | 0 | 0 | 0 | 0 | 0 |
| ENSMUSG00 Gm11997 | 312 | 0 | 0 | 0 | 0 | 0 | 0 |
| ENSMUSG00 Gm11833 | 852 | 0 | 0 | 0 | 0 | 0 | 0 |
| ENSMUSG00 Gm14984 | 1262 | 0 | 0 | 0 | 0 | 0 | 0.02161223 |
| ENSMUSG00 Gm12042 | 300 | 0 | 0 | 0 | 0 | 0 | 0.18183093 |
| ENSMUSG00 Gm14439 | 886 | 0 | 0.16315951 | 0.84811663 | 0.20362328 | 0 | 0.09235205 |
| ENSMUSG00 Gm15146 | 446 | 0 | 0 | 0 | 0 | 0 | 0 |
| ENSMUSG00 Gm11745 | 447 | 0.08893844 | 1.07799647 | 0.79108444 | 0.16144092 | 2.16890636 | 1.34237598 |
| ENSMUSG00 Gm14904 | 334 | 0 | 0 | 0 | 0 | 0 | 0 |
| ENSMUSG00 Gm14945 | 1117 | 0 | 0 | 0 | 0 | 0 | 0 |
| ENSMUSG00 Gm12004 | 717 | 0 | 0 | 0 | 0 | 0 | 0 |
| ENSMUSG00 Rhox3b | 644 | 0 | 0 | 0 | 0 | 0 | 0 |
| ENSMUSG00 Gm8916 | 948 | 0 | 0 | 0 | 0 | 0 | 0 |
| ENSMUSG00 Gm15263 | 640 | 0 | 0 | 0 | 0 | 0 | 0 |
| ENSMUSG00 Gm13046 | 500 | 0 | 0 | 0 | 0 | 0 | 0 |
| ENSMUSG00 Gm16412 | 478 | 0 | 0 | 0 | 0 | 0 | 0 |
| ENSMUSG00 Gm4996 | 782 | 0 | 0 | 0 | 0.04614072 | 0 | 0.03487806 |
| ENSMUSG00 Gm14611 | 711 | 0 | 0 | 0 | 0 | 0 | 0 |
| ENSMUSG00 Gm5934 | 2580 | 0 | 0 | 0 | 0 | 0 | 0 |
| ENSMUSG00 Gm12079 | 243 | 0 | 0 | 0 | 0 | 0 | 0 |
| ENSMUSG00 Gm15159 | 763 | 0 | 0 | 0 | 0 | 0 | 0 |
| ENSMUSG00 Xlr4d-ps | 639 | 0 | 0 | 0 | 0 | 0 | 0 |
| ENSMUSG00 Gm14269 | 305 | 0.39103751 | 0.15798834 | 0.14492408 | 0.23660357 | 0 | 0.17885009 |
| ENSMUSG00 Gm11573 | 393 | 0 | 0 | 0 | 0 | 0 | 0 |
| ENSMUSG00 Gm12306 | 170 | 0 | 0 | 0 | 0 | 0.30015515 | 0 |
| ENSMUSG00 Gm6591 | 430 | 0 | 0 | 0 | 0 | 0 | 0 |
| ENSMUSG00 Gm13378 | 335 | 0.11867308 | 0 | 0 | 0 | 0 | 0 |
| ENSMUSG00 Gm11285 | 321 | 0 | 0 | 0 | 0 | 0 | 0 |
| ENSMUSG00 Gm11910 | 430 | 0 | 0 | 0 | 0 | 0 | 0 |
| ENSMUSG00 Gm8310 | 1192 | 0 | 0 | 0 | 0 | 0 | 0 |
| ENSMUSG00 Gm13382 | 131 | 0 | 0 | 0 | 0 | 0 | 0 |
| ENSMUSG00 Gm14600 | 629 | 0 | 0 | 0 | 0 | 0 | 0 |
| ENSMUSG00 Gm11952 | 820 | 0 | 0 | 0 | 0.04400249 | 0 | 0 |
| ENSMUSG00 Olfr349-ps1 | 581 | 0 | 0 | 0 | 0 | 0 | 0 |
| ENSMUSG00 Gm11631 | 155 | 0 | 0 | 0 | 0 | 0 | 0 |
| ENSMUSG00 Gm15227 | 398 | 0 | 0 | 0 | 0 | 0 | 0 |
| ENSMUSG00 Gm12057 | 2019 | 1.10267801 | 0.69212819 | 0.70057404 | 2.28751942 | 0.55600806 | 0.63492226 |
| ENSMUSG00 Gm12872 | 317 | 0 | 0 | 0 | 0 | 0 | 0 |
| ENSMUSG00 Gm15782 | 444 | 0 | 0.21705605 | 0 | 0 | 0 | 0 |
| ENSMUSG00 Gm8383 | 1769 | 0 | 0 | 0 | 0 | 0 | 0 |
| ENSMUSG00 Gm16140 | 1280 | 0 | 0 | 0 | 0 | 0 | 0 |
| ENSMUSG00 Gm5761 | 833 | 0.09545133 | 0.05784687 | 0 | 0.04331578 | 0.12251231 | 0 |
| ENSMUSG00 Gm13650 | 432 | 0 | 0 | 0 | 0 | 0 | 0 |
| ENSMUSG00 Gm12941 | 219 | 0 | 1.10014708 | 0.2018349 | 0.16475819 | 0.9319886 | 1.12087559 |
| ENSMUSG00 Gm14099 | 604 | 0 | 0 | 0 | 0 | 0 | 0 |
| ENSMUSG00 Gm11245 | 1036 | 0 | 0 | 0 | 0.03482823 | 0 | 0.13163436 |
| ENSMUSG00 Gm11995 | 227 | 0 | 0 | 0 | 0 | 0 | 0 |
| ENSMUSG00 Gm5843 | 472 | 0 | 0 | 0 | 0 | 0 | 0 |
| ENSMUSG00 Gm16418 | 170 | 0 | 0 | 0.52002168 | 0.42449464 | 0 | 0.80219527 |
| ENSMUSG00 Gm13961 | 4771 | 0.02499821 | 0.04039945 | 0.01852938 | 0 | 0.04278045 | 0.00571676 |
| ENSMUSG00 Gm11834 | 583 | 0 | 0 | 0 | 0 | 0 | 0 |
| ENSMUSG00 Gm11332 | 594 | 0 | 0 | 0 | 0 | 0 | 0 |
| ENSMUSG00 Gm14922 | 162 | 0 | 0 | 0 | 0 | 0 | 0 |
| ENSMUSG00 Gm13422 | 503 | 0.15807348 | 0.19159619 | 0 | 0.21520106 | 0.10144409 | 0.37956754 |
| ENSMUSG00 Gm12929 | 369 | 0.43095372 | 0.52234626 | 0 | 0.09778332 | 0 | 0.07391501 |
| ENSMUSG00 Mup-ps11 | 539 | 0 | 0 | 0 | 0 | 0 | 0 |
| ENSMUSG00 Gm17014 | 167 | 0 | 0 | 0 | 0 | 0 | 0 |
| ENSMUSG00 Gm15437 | 456 | 0 | 0 | 0 | 0 | 0 | 0 |
| ENSMUSG00 Gm14629 | 143 | 0 | 0 | 0 | 0 | 0 | 0 |
| ENSMUSG00 Gm13578 | 589 | 3.03734571 | 2.04526495 | 1.87613935 | 1.47023611 | 1.99254099 | 2.68578791 |
| ENSMUSG00 Gm13252 | 320 | 1.86353815 | 0.30116526 | 1.38130759 | 0.11275639 | 1.11620197 | 1.27849872 |
| ENSMUSG00 Gm6136 | 1817 | 0.04375947 | 0.02651978 | 0.07298048 | 0.07943213 | 0.05616552 | 0.01501081 |
| ENSMUSG00 Gm13700 | 1127 | 0 | 0 | 0 | 0 | 0 | 0 |
| ENSMUSG00 Gm15794 | 517 | 0 | 0.09320395 | 0.17099359 | 0 | 0 | 0 |
| ENSMUSG00 Gm4992 | 1297 | 0 | 0 | 0 | 0 | 0.03934185 | 0 |
| ENSMUSG00 Gm15710 | 732 | 0 | 0 | 0 | 0 | 0 | 0 |
| ENSMUSG00 Gm15112 | 858 | 0 | 0 | 0 | 0 | 0 | 0 |
| ENSMUSG00 Gm14277 | 359 | 0 | 0 | 0 | 0.10050709 | 0 | 0 |
| ENSMUSG00 Gm8421 | 304 | 0 | 0 | 0 | 0 | 0 | 0 |
| ENSMUSG00 Gm13566 | 1141 | 0 | 0 | 0 | 0 | 0 | 0 |
| ENSMUSG00 Gm12460 | 356 | 0 | 0 | 0 | 0 | 0 | 0 |
| ENSMUSG00 Olfr1063-ps1 | 941 | 0 | 0 | 0 | 0 | 0 | 0 |
| ENSMUSG00 Gm14989 | 206 | 0 | 0 | 0 | 0 | 0 | 0 |
| ENSMUSG00 Gm12351 | 291 | 0 | 0 | 0 | 0 | 0 | 0 |
| ENSMUSG00 Gm11624 | 1040 | 0 | 0 | 0 | 0 | 0 | 0 |
| ENSMUSG00 Gm13106 | 1440 | 0 | 0 | 0 | 0 | 0 | 0 |
| ENSMUSG00 Gm13588 | 134 | 0 | 0 | 0 | 0 | 0 | 0 |
| ENSMUSG00 Gm8757 | 371 | 0.21431526 | 0 | 0 | 0.09725619 | 0 | 0 |
| ENSMUSG00 Gm15844 | 575 | 0 | 0 | 0 | 0 | 0 | 0 |
| ENSMUSG00 Gm13169 | 1457 | 0 | 0 | 0 | 0 | 0 | 0 |
| ENSMUSG00 Esrp2 | 5231 | 19.4407416 | 21.0119049 | 18.8350044 | 23.976522 | 25.1668992 | 18.9686747 |
| ENSMUSG00 Hmgb1-ps1 | 633 | 0 | 0 | 0 | 0.05700165 | 0 | 0 |
| ENSMUSG00 Gm11704 | 213 | 0 | 0 | 0 | 0 | 0 | 0 |
| ENSMUSG00 Rpl3-ps2 | 1213 | 0.03277451 | 0.07945003 | 0 | 0.02974612 | 0 | 0.04497055 |
| ENSMUSG00 Gm15982 | 568 | 0.55993635 | 0.42417643 | 0.38910073 | 0.95287089 | 0 | 0.09603746 |
| ENSMUSG00 Gm12856 | 1292 | 0 | 0 | 0 | 0 | 0 | 0 |
| ENSMUSG00 Gm13930 | 366 | 0 | 0 | 0 | 0 | 0 | 0.37260436 |
| ENSMUSG00 Pom121l12 | 1624 | 0 | 0 | 0 | 0 | 0 | 0 |
| ENSMUSG00 Vmn1r207-ps | 2010 | 0 | 0 | 0 | 0 | 0 | 0 |
| ENSMUSG00 Gm9085 | 1071 | 0 | 0.17996804 | 0.04127156 | 0 | 0.33350572 | 0.33106472 |
| ENSMUSG00 Gm14915 | 203 | 0 | 0 | 0 | 0 | 0 | 0 |
| ENSMUSG00 Gm12322 | 388 | 0.20492516 | 0 | 0.68353366 | 0.55796976 | 0.13151128 | 0 |
| ENSMUSG00 Gm14000 | 232 | 0.17135983 | 0 | 0 | 0 | 0.21994128 | 1.2931941 |
| ENSMUSG00 Olfr1372-ps1 | 905 | 0 | 0 | 0 | 0 | 0.62021009 | 0 |
| ENSMUSG00 Gm9103 | 403 | 0 | 0 | 0 | 0.08953361 | 0 | 0 |
| ENSMUSG00 Gm13400 | 379 | 0 | 0 | 0 | 0 | 0 | 0 |
| ENSMUSG00 Gm12516 | 384 | 0 | 0 | 0 | 0 | 0 | 0 |
| ENSMUSG00 Gm12263 | 1481 | 0.16106204 | 0.09760927 | 0.2686135 | 0.07308989 | 0.17227001 | 0.25782914 |
| ENSMUSG00 Gm12104 | 978 | 0 | 0 | 0 | 0.92234265 | 0 | 1.05975081 |
| ENSMUSG00 Gm15075 | 476 | 0 | 0 | 0 | 0 | 0 | 0 |
| ENSMUSG00 Gm12380 | 543 | 0 | 0 | 0 | 0 | 0 | 0 |
| ENSMUSG00 Gm12944 | 462 | 0 | 0 | 0.28702496 | 0 | 0 | 0 |
| ENSMUSG00 Gm14395 | 562 | 0 | 0 | 0 | 0 | 0 | 0 |
| ENSMUSG00 Gm11577 | 735 | 0 | 0 | 0 | 0 | 0 | 0 |
| ENSMUSG00 Gm11617 | 638 | 0 | 0 | 0 | 0 | 0 | 0 |
| ENSMUSG00 Gm12186 | 1026 | 0 | 0 | 0 | 0 | 0 | 0 |
| ENSMUSG00 Gm15644 | 386 | 1.33891515 | 0.37450603 | 0 | 0.6543376 | 4.09797319 | 2.11979062 |
| ENSMUSG00 Gm7113 | 957 | 0 | 0 | 0 | 0 | 0 | 0 |
| ENSMUSG00 Gm14734 | 446 | 0 | 0 | 0 | 0.08090145 | 0 | 0 |
| ENSMUSG00 Gm15576 | 526 | 0 | 0 | 0 | 0 | 0 | 0 |
| ENSMUSG00 Gm14652 | 234 | 0 | 0 | 0 | 0 | 0 | 0 |
| ENSMUSG00 Gm12696 | 904 | 2.9025019 | 5.4902694 | 4.8895844 | 2.95361869 | 2.59647488 | 2.53436913 |
| ENSMUSG00 Gm13345 | 227 | 0 | 0 | 0 | 0 | 0 | 0 |
| ENSMUSG00 Gm15061 | 1045 | 0 | 0 | 0 | 0 | 0 | 0 |
| ENSMUSG00 Gm11251 | 2797 | 0 | 0 | 0 | 0 | 0 | 0 |
| ENSMUSG00 Gm11901 | 436 | 0 | 0 | 0 | 0 | 0 | 0 |
| ENSMUSG00 Gm13902 | 604 | 0 | 0 | 0 | 0 | 0 | 0.27094013 |
| ENSMUSG00 Gm6451 | 716 | 0 | 0 | 0 | 0.05039392 | 0.07126589 | 0 |
| ENSMUSG00 Gm13924 | 485 | 0 | 0 | 0 | 0 | 0 | 0 |
| ENSMUSG00 Atp5l2 | 695 | 0.22880852 | 0.06933301 | 0.2543991 | 0.10383322 | 0 | 0.31395268 |
| ENSMUSG00 Gm12240 | 228 | 0 | 0 | 0 | 0 | 0 | 0 |
| ENSMUSG00 Gm11954 | 348 | 0 | 0.13846679 | 0 | 0 | 0 | 0.1567508 |
| ENSMUSG00 Gm13457 | 265 | 0 | 0 | 0 | 0 | 0 | 0 |
| ENSMUSG00 Gm16420 | 855 | 0 | 0 | 0 | 0 | 0 | 0 |
| ENSMUSG00 Gm5287 | 597 | 0 | 0 | 0 | 0 | 0 | 0 |
| ENSMUSG00 Sycn | 555 | 0 | 0 | 0 | 0 | 0 | 0 |
| ENSMUSG00 Gm11372 | 569 | 0 | 0 | 0 | 0 | 0 | 0 |
| ENSMUSG00 Gm13508 | 774 | 0 | 0 | 0 | 0 | 0 | 0 |
| ENSMUSG00 Gm11759 | 463 | 0 | 0 | 0 | 0 | 0.22041631 | 0 |
| ENSMUSG00 Gm11890 | 416 | 0 | 0 | 0 | 0 | 0 | 0 |
| ENSMUSG00 Gm15237 | 468 | 0.16989522 | 0.6177749 | 0 | 0.15419677 | 0 | 0 |
| ENSMUSG00 Gm11858 | 837 | 0 | 0 | 0 | 0 | 0 | 0 |
| ENSMUSG00 Taar7c-ps | 1054 | 0 | 0 | 0 | 0 | 0 | 0 |
| ENSMUSG00 Gm12009 | 1099 | 0.1085227 | 0 | 0 | 0.13132682 | 0.04642982 | 0 |
| ENSMUSG00 Gm13474 | 808 | 0 | 0 | 0 | 0 | 0 | 0 |
| ENSMUSG00 Rplp1-ps1 | 344 | 0 | 0 | 0 | 0 | 0 | 0 |
| ENSMUSG00 Rpl36-ps10 | 215 | 0 | 0 | 0 | 0 | 0 | 0 |
| ENSMUSG00 Gm7567 | 401 | 0 | 0 | 0 | 0 | 0 | 0 |
| ENSMUSG00 Gm12851 | 420 | 0 | 0 | 0 | 0 | 0 | 0 |
| ENSMUSG00 Gm8089 | 962 | 0 | 0 | 0 | 0 | 0 | 0 |
| ENSMUSG00 Gm11869 | 327 | 0.24315279 | 0.14735915 | 0.13517383 | 1.87582495 | 0.62417585 | 0.08340868 |
| ENSMUSG00 Gm15167 | 185 | 0 | 0 | 0.47785776 | 0 | 0 | 0.14743048 |
| ENSMUSG00 Gm13282 | 544 | 0 | 0 | 0 | 0 | 0 | 0 |
| ENSMUSG00 Gm6823 | 1838 | 0 | 0 | 0 | 0 | 0 | 0 |
| ENSMUSG00 Gm12407 | 320 | 0 | 0 | 0 | 0 | 0 | 0 |
| ENSMUSG00 Gm15372 | 833 | 0 | 0 | 0 | 0 | 0 | 0 |
| ENSMUSG00 Gm12874 | 911 | 0 | 0 | 0.09704027 | 0 | 0 | 0 |
| ENSMUSG00 Gm9089 | 525 | 0 | 0 | 0 | 0 | 0 | 0 |
| ENSMUSG00 Gm13711 | 451 | 0 | 0 | 0 | 0.32001813 | 0 | 0 |
| ENSMUSG00 Gm12422 | 301 | 0 | 0 | 0 | 0 | 0 | 0 |
| ENSMUSG00 Gm11584 | 324 | 0 | 0 | 0 | 0 | 0 | 0 |
| ENSMUSG00 Gm11926 | 429 | 0 | 0 | 0.515173 | 0 | 0 | 0.19073174 |
| ENSMUSG00 Gm5398 | 983 | 0 | 0 | 0 | 0 | 0 | 0 |
| ENSMUSG00 Gm15593 | 1007 | 0.11843738 | 0.04785148 | 0.04389458 | 0 | 0 | 0.05417009 |
| ENSMUSG00 Gm6584 | 397 | 0 | 0 | 0 | 0 | 0 | 0 |
| ENSMUSG00 Gm8061 | 348 | 0 | 0 | 0 | 0 | 0 | 0 |
| ENSMUSG00 Gm12062 | 258 | 0 | 0 | 0 | 0 | 0 | 0 |
| ENSMUSG00 Gm14070 | 427 | 0 | 0 | 0 | 0 | 0 | 0 |
| ENSMUSG00 Gm12792 | 1018 | 0 | 0 | 0.04342028 | 0.03544405 | 0 | 0.05358475 |
| ENSMUSG00 Gm13811 | 255 | 0 | 0 | 0 | 0 | 0 | 0 |
| ENSMUSG00 Gm15168 | 314 | 0 | 0 | 0 | 0 | 0 | 0 |
| ENSMUSG00 Gm11949 | 371 | 0 | 0 | 0 | 0 | 0 | 0 |
| ENSMUSG00 Gm12225 | 297 | 0 | 0 | 0 | 0 | 0 | 0 |
| ENSMUSG00 Hspa1-ps5 | 296 | 0 | 0 | 0 | 0 | 0 | 0 |
| ENSMUSG00 Gm14825 | 700 | 0 | 0 | 0 | 0 | 0 | 0 |
| ENSMUSG00 Gm11814 | 928 | 0 | 0.05192505 | 0.0476313 | 0.11664454 | 0.05498532 | 0.08817233 |
| ENSMUSG00 Gm7973 | 487 | 0 | 0 | 0 | 0 | 0 | 0 |
| ENSMUSG00 Gm15488 | 677 | 0 | 0 | 0 | 0 | 0 | 0 |
| ENSMUSG00 Gm8145 | 744 | 0 | 0 | 0 | 0 | 0 | 0 |
| ENSMUSG00 Gm14876 | 325 | 0 | 0 | 0 | 0 | 0 | 0 |
| ENSMUSG00 Gm12387 | 954 | 0 | 0 | 0 | 0 | 0 | 0 |
| ENSMUSG00 Hmgb1-ps4 | 634 | 0 | 0 | 0 | 0 | 0 | 0 |
| ENSMUSG00 Gm13973 | 893 | 0 | 0 | 0 | 0 | 0 | 0 |
| ENSMUSG00 Gm12244 | 290 | 0 | 0 | 0.15242015 | 0.24884169 | 0 | 0.84645432 |
| ENSMUSG00 Gm12337 | 399 | 0 | 0 | 0.11078156 | 0 | 0 | 0.06835749 |
| ENSMUSG00 Gm13426 | 794 | 0 | 0 | 0 | 0 | 0 | 0 |
| ENSMUSG00 Gm8080 | 1148 | 0 | 0 | 0 | 0 | 0 | 0 |
| ENSMUSG00 Gm7958 | 547 | 0 | 0 | 0 | 0 | 0 | 0 |
| ENSMUSG00 Gm14388 | 476 | 0.08351992 | 0 | 0 | 0.07580262 | 0.10719827 | 0.05729966 |
| ENSMUSG00 Gm11975 | 480 | 0 | 0 | 0 | 0 | 0.10630495 | 0 |
| ENSMUSG00 4933405O20l | 976 | 0 | 0 | 0 | 0 | 0 | 0 |
| ENSMUSG00 Gm15421 | 398 | 0 | 0 | 0.66635944 | 0.36263361 | 0 | 0.27411698 |
| ENSMUSG00 Olfr637-ps1 | 461 | 0 | 0 | 0 | 0 | 0 | 0 |
| ENSMUSG00 Gm11781 | 471 | 0 | 0 | 0 | 0 | 0 | 0 |
| ENSMUSG00 Gm13076 | 203 | 0 | 0 | 0 | 0 | 0 | 0 |
| ENSMUSG00 Gm14463 | 497 | 0 | 0 | 0 | 0 | 0 | 0 |
| ENSMUSG00 Gm15383 | 1442 | 0 | 0 | 0 | 0 | 0 | 0 |
| ENSMUSG00 Gm13416 | 1611 | 0 | 0.23928711 | 0.1371876 | 0.11198648 | 0.12669491 | 0.13544203 |
| ENSMUSG00 Mup-ps3 | 539 | 0 | 0 | 0 | 0 | 0 | 0 |
| ENSMUSG00 Gm13784 | 517 | 0 | 0 | 0 | 0 | 0 | 0 |
| ENSMUSG00 Gm14020 | 661 | 0 | 0 | 0 | 0 | 0 | 0 |
| ENSMUSG00 Gm14994 | 1091 | 0 | 0 | 0 | 0 | 0 | 0 |
| ENSMUSG00 Gm15162 | 282 | 0 | 0 | 0 | 0 | 0 | 0 |
| ENSMUSG00 Gm14566 | 320 | 0 | 0 | 0 | 0 | 0 | 0 |
| ENSMUSG00 Gm11409 | 402 | 0 | 0 | 0 | 0 | 0 | 0 |
| ENSMUSG00 Gm12267 | 760 | 0.3661689 | 0.63403213 | 0.8724048 | 0 | 0 | 0 |
| ENSMUSG00 Gm13550 | 524 | 0 | 0 | 0 | 0 | 0 | 0 |

| | | | | | | | | |
|---|---|---|---|---|---|---|---|---|
| ENSMUSG00 | Gm12310 | 491 | 0 | 0 | 0 | 0 | 0 | 0 |
| ENSMUSG00 | Gm12036 | 529 | 0.22545641 | 0.5465381 | 0.16711472 | 0.13641605 | 0 | 0 |
| ENSMUSG00 | Gm11256 | 828 | 0 | 0 | 0 | 0 | 0 | 0 |
| ENSMUSG00 | Defa-ps11 | 297 | 0 | 0 | 0 | 0 | 0 | 0 |
| ENSMUSG00 | Gm14976 | 1393 | 0 | 0 | 0 | 0 | 0 | 0 |
| ENSMUSG00 | Gm11597 | 957 | 0 | 0 | 0 | 0 | 0 | 0 |
| ENSMUSG00 | Gm12054 | 377 | 0 | 0 | 0 | 0 | 0 | 0 |
| ENSMUSG00 | Gm12040 | 536 | 0 | 0 | 0.24739838 | 0 | 0 | 0.76328282 |
| ENSMUSG00 | Gm12844 | 581 | 0.47898169 | 0 | 0.91294684 | 0.24841339 | 0 | 0.65722022 |
| ENSMUSG00 | Gm13635 | 295 | 0 | 0 | 0 | 0 | 0 | 0 |
| ENSMUSG00 | Olfr588-ps1 | 941 | 0 | 0 | 0 | 0 | 0 | 0 |
| ENSMUSG00 | Gm14142 | 294 | 0 | 0 | 0 | 0 | 0 | 0 |
| ENSMUSG00 | Gm14541 | 156 | 0.50968565 | 0 | 0 | 0.23129516 | 0 | 1.04902459 |
| ENSMUSG00 | Gm7408 | 631 | 0 | 0 | 0 | 0 | 0 | 0 |
| ENSMUSG00 | Gm12417 | 402 | 0 | 0 | 0 | 0 | 0 | 0 |
| ENSMUSG00 | Olfr1291-ps1 | 1834 | 0 | 0 | 0 | 0 | 0 | 0 |
| ENSMUSG00 | Gm14784 | 662 | 0 | 0 | 0 | 0 | 0 | 0 |
| ENSMUSG00 | Gm15177 | 576 | 0 | 0 | 0 | 0 | 0 | 0 |
| ENSMUSG00 | Gm14846 | 215 | 0 | 0 | 0 | 0 | 0 | 0 |
| ENSMUSG00 | Gm13224 | 93 | 0 | 0 | 0 | 0 | 0 | 0 |
| ENSMUSG00 | Gm12628 | 380 | 0 | 0 | 0 | 0 | 0 | 0 |
| ENSMUSG00 | Gm12504 | 2535 | 0.06273054 | 0.41818609 | 0.01743662 | 0.17080258 | 0.08051499 | 0.08607381 |
| ENSMUSG00 | Gm5396 | 531 | 0 | 0 | 0 | 0.06795112 | 0 | 0 |
| ENSMUSG00 | Gm8317 | 470 | 0 | 0 | 0 | 0 | 0 | 0 |
| ENSMUSG00 | Gm11702 | 68 | 0 | 0 | 0 | 0 | 2.25116364 | 2.40658582 |
| ENSMUSG00 | Gm12189 | 911 | 0 | 0 | 0 | 0 | 0 | 0 |
| ENSMUSG00 | Gm13232 | 631 | 0 | 0 | 0 | 0 | 0 | 0 |
| ENSMUSG00 | Gm11972 | 388 | 0 | 0 | 0 | 0 | 0 | 0 |
| ENSMUSG00 | Gm14770 | 311 | 0 | 0 | 0 | 0 | 0 | 0 |
| ENSMUSG00 | Gm15054 | 280 | 0 | 0 | 0 | 0 | 0 | 0 |
| ENSMUSG00 | Gm14914 | 2030 | 0 | 0 | 0 | 0 | 0 | 0 |
| ENSMUSG00 | Tpt1-ps6 | 522 | 0 | 0 | 0 | 0 | 0 | 0 |
| ENSMUSG00 | Gm13451 | 591 | 0 | 0 | 0 | 0 | 0 | 0 |
| ENSMUSG00 | Gm12658 | 682 | 0.17487748 | 1.27178293 | 0 | 3.1214672 | 1.34673719 | 0.35992926 |
| ENSMUSG00 | Gm14576 | 1110 | 0 | 0 | 0 | 0 | 0 | 0 |
| ENSMUSG00 | Gm5228 | 356 | 0 | 0 | 0 | 0.10135406 | 0 | 0.45968493 |
| ENSMUSG00 | Gm6977 | 510 | 1.48108653 | 0.85034898 | 1.64673533 | 0.56599286 | 0.70036202 | 0.96263433 |
| ENSMUSG00 | Gm7375 | 631 | 0 | 0 | 0 | 0 | 0 | 0 |
| ENSMUSG00 | Gm6654 | 758 | 0 | 0 | 0 | 0 | 0 | 0 |
| ENSMUSG00 | Gm13535 | 1881 | 0 | 0 | 0 | 0 | 0 | 0 |
| ENSMUSG00 | Gm12143 | 331 | 0 | 0 | 0 | 0 | 0 | 0 |
| ENSMUSG00 | Gm13533 | 176 | 0 | 0 | 0 | 0 | 0 | 0 |
| ENSMUSG00 | Olfr1235-ps1 | 290 | 0 | 0 | 0 | 0 | 0 | 0 |
| ENSMUSG00 | Gm12777 | 304 | 0 | 0 | 0 | 0 | 0 | 0 |
| ENSMUSG00 | Gm13755 | 303 | 0 | 0 | 0 | 0 | 0 | 0 |
| ENSMUSG00 | Gm14331 | 407 | 0 | 0 | 0 | 0 | 0 | 0 |
| ENSMUSG00 | Gm6630 | 908 | 0 | 0 | 0 | 0 | 0 | 0 |
| ENSMUSG00 | Gm6954 | 224 | 0 | 0 | 0 | 0 | 0 | 0 |
| ENSMUSG00 | Gm7856 | 1492 | 0 | 0 | 0 | 0 | 0 | 0 |
| ENSMUSG00 | Gm12194 | 1011 | 0 | 0 | 0 | 0 | 0 | 0 |
| ENSMUSG00 | Gm8216 | 600 | 0 | 0 | 0 | 0 | 0 | 0.31820413 |
| ENSMUSG00 | Gm6142 | 324 | 0.9816168 | 0.74361793 | 0.13642544 | 0.22272867 | 0 | 1.85198168 |
| ENSMUSG00 | Gm14645 | 416 | 0 | 0 | 0 | 0 | 0 | 0 |
| ENSMUSG00 | Gm11743 | 1733 | 0 | 0 | 0 | 0 | 0 | 0 |
| ENSMUSG00 | Gm12151 | 289 | 0 | 0 | 0 | 0.24970273 | 0 | 0 |
| ENSMUSG00 | Gm14593 | 293 | 0 | 0 | 0 | 0 | 0 | 0 |
| ENSMUSG00 | Mup-ps20 | 385 | 0 | 0 | 0 | 0 | 0 | 0 |
| ENSMUSG00 | Gm11950 | 837 | 0 | 0 | 0 | 0 | 0 | 0 |
| ENSMUSG00 | Gm11257 | 314 | 0 | 0 | 0 | 0 | 0 | 0 |
| ENSMUSG00 | Gm13122 | 452 | 0 | 0 | 0 | 0 | 0 | 0 |
| ENSMUSG00 | Gm13809 | 594 | 0 | 0.08112196 | 0.07441388 | 0.12148837 | 0 | 0 |
| ENSMUSG00 | Rps15a-ps3 | 394 | 0 | 0 | 0 | 0 | 0 | 0 |
| ENSMUSG00 | Vmn1r-ps128 | 1738 | 0 | 0 | 0 | 0 | 0 | 0 |
| ENSMUSG00 | Gm8668 | 1446 | 0 | 0 | 0 | 0 | 0 | 0 |
| ENSMUSG00 | Gm11987 | 378 | 0 | 0 | 0 | 0 | 0 | 0 |
| ENSMUSG00 | Tpt1-ps3 | 1204 | 0.23113652 | 0.12006589 | 0.25698746 | 0.2397478 | 0.16952284 | 0.31714697 |
| ENSMUSG00 | Gm14125 | 1434 | 0 | 0.33602819 | 0 | 0 | 0 | 0 |
| ENSMUSG00 | Gm14778 | 446 | 0 | 0 | 0 | 0 | 0 | 0 |
| ENSMUSG00 | Gm14438 | 1310 | 0 | 0 | 0 | 0 | 0.03895143 | 0.10410168 |
| ENSMUSG00 | Gm6500 | 450 | 0 | 0 | 0 | 0 | 0 | 0 |
| ENSMUSG00 | Gm15550 | 1073 | 0.1482031 | 0.40417333 | 0.90628196 | 0 | 0.76087793 | 0 |
| ENSMUSG00 | Gm12820 | 341 | 0 | 0 | 0 | 0 | 0 | 0 |
| ENSMUSG00 | Gm14531 | 283 | 0 | 0 | 0 | 0 | 0 | 0 |
| ENSMUSG00 | Gm14046 | 508 | 0.15651764 | 0 | 0 | 0.28411059 | 0 | 0.05369024 |
| ENSMUSG00 | Gm6733 | 1919 | 0.33146831 | 0.25110184 | 0.66797991 | 0.28203787 | 0.07977026 | 0.36953654 |
| ENSMUSG00 | Gm15019 | 429 | 0 | 0 | 0 | 0 | 0 | 0 |
| ENSMUSG00 | Gm8163 | 1059 | 0.07508117 | 0 | 0 | 0 | 0 | 0 |
| ENSMUSG00 | Gm8840 | 385 | 0 | 0 | 0 | 0 | 0 | 0 |
| ENSMUSG00 | Gm11599 | 877 | 0 | 0 | 0 | 0 | 0 | 0 |
| ENSMUSG00 | Gm13494 | 1095 | 0 | 0 | 0 | 0 | 0 | 0 |
| ENSMUSG00 | Gm12619 | 383 | 0 | 0 | 0 | 0 | 0 | 0 |
| ENSMUSG00 | Olfr1267-ps1 | 929 | 0 | 0 | 0 | 0 | 0 | 0 |
| ENSMUSG00 | Gm14986 | 1001 | 0 | 0 | 0 | 0 | 0 | 0 |
| ENSMUSG00 | Gm13531 | 542 | 0 | 0 | 0 | 0 | 0 | 0 |
| ENSMUSG00 | Gm6141 | 693 | 0 | 0 | 0 | 0.05206644 | 0 | 0.07871469 |
| ENSMUSG00 | Gm14766 | 366 | 0.43448613 | 0 | 0 | 0 | 0.41824898 | 0 |
| ENSMUSG00 | Gm3928 | 1306 | 0 | 0 | 0 | 0 | 0 | 0 |
| ENSMUSG00 | Gm11224 | 1110 | 0.03581575 | 0 | 0.03982148 | 0 | 0 | 0.02457175 |
| ENSMUSG00 | Gm16198 | 669 | 0 | 0 | 0 | 0 | 0 | 0 |
| ENSMUSG00 | Gm15305 | 286 | 0 | 0 | 0 | 0 | 0 | 0 |
| ENSMUSG00 | Rpl17-ps4 | 1161 | 0 | 0 | 0 | 0 | 0 | 0 |
| ENSMUSG00 | Cyp4a30b-ps | 1514 | 0.02625857 | 0 | 0.29195405 | 0.02383226 | 0 | 0.09007477 |
| ENSMUSG00 | Akt2-ps | 1448 | 0.10982177 | 0.09983379 | 0.24420908 | 0.09967416 | 0.10571763 | 0.05650823 |
| ENSMUSG00 | Gm14608 | 1552 | 0 | 0 | 0 | 0 | 0 | 0 |
| ENSMUSG00 | Rpl3-ps1 | 1209 | 0.92072246 | 1.91310937 | 1.71835122 | 1.28331507 | 0.92851966 | 0.40607403 |
| ENSMUSG00 | 1700029I01R | 2410 | 0.44539335 | 0.13996062 | 0.18341014 | 0 | 0.10586385 | 0.7129885 |
| ENSMUSG00 | Gm13085 | 369 | 0 | 0 | 0 | 0 | 0 | 0 |
| ENSMUSG00 | Gm12370 | 602 | 0 | 0 | 0 | 0 | 0 | 0 |
| ENSMUSG00 | Gm15452 | 239 | 0 | 0 | 0 | 0 | 0 | 0 |
| ENSMUSG00 | Gm14708 | 307 | 0 | 0 | 0 | 0 | 0 | 0 |
| ENSMUSG00 | Gm14792 | 257 | 0 | 0 | 0 | 0 | 0 | 0 |
| ENSMUSG00 | Olfr590-ps1 | 852 | 0 | 0 | 0 | 0 | 0 | 0 |
| ENSMUSG00 | Gm11268 | 266 | 0 | 0 | 0 | 0 | 0 | 0 |
| ENSMUSG00 | Gm11352 | 516 | 0 | 0 | 0 | 0 | 0 | 0 |
| ENSMUSG00 | Gm13692 | 370 | 0 | 0 | 0 | 0 | 0 | 0 |
| ENSMUSG00 | Gm11797 | 753 | 0 | 0 | 0 | 0 | 0 | 0 |
| ENSMUSG00 | Gm8202 | 309 | 0 | 0 | 0 | 0 | 0 | 0 |
| ENSMUSG00 | 2610024D14 | 293 | 0.13568423 | 0 | 0.15085953 | 0 | 0.34830291 | 0 |
| ENSMUSG00 | Gm2288 | 630 | 0 | 0 | 0 | 0 | 0 | 0 |
| ENSMUSG00 | Gm15801 | 521 | 0 | 0 | 0 | 0 | 0 | 0 |
| ENSMUSG00 | Gm5642 | 398 | 0 | 0 | 0 | 0 | 0 | 0 |
| ENSMUSG00 | Gm15313 | 280 | 0 | 0 | 0 | 0 | 0 | 0 |
| ENSMUSG00 | Gm15142 | 890 | 0 | 0 | 0 | 0 | 0 | 0 |
| ENSMUSG00 | Gm11572 | 119 | 0 | 0 | 0 | 0 | 0 | 0 |
| ENSMUSG00 | Gm14626 | 213 | 0 | 0 | 0 | 0 | 0 | 0 |
| ENSMUSG00 | Gm15442 | 1525 | 0 | 0 | 0.02898482 | 0 | 0 | 0 |
| ENSMUSG00 | Gm14113 | 975 | 0.12232456 | 0.09884398 | 0.04533522 | 0.07401445 | 0.10466949 | 0.08392197 |
| ENSMUSG00 | Gm13988 | 287 | 0 | 0 | 0 | 0 | 0 | 0 |
| ENSMUSG00 | Rps2-ps4 | 632 | 0 | 0.15248874 | 0 | 0 | 0 | 0 |
| ENSMUSG00 | Gm13255 | 546 | 0 | 0 | 0 | 0 | 0 | 0 |
| ENSMUSG00 | Gm11637 | 590 | 0 | 0 | 0 | 0 | 0 | 0 |
| ENSMUSG00 | Gm14855 | 191 | 0 | 0 | 0 | 0 | 0 | 0 |
| ENSMUSG00 | Gm15302 | 2629 | 0 | 0 | 0 | 0 | 0.01940904 | 0 |
| ENSMUSG00 | Gm15238 | 1295 | 0 | 0.03720961 | 0 | 0.02786258 | 0 | 0.0210615 |
| ENSMUSG00 | Gm12957 | 376 | 0.10573266 | 1.02524345 | 0 | 0.19192577 | 0 | 0.07253893 |
| ENSMUSG00 | Gm12483 | 257 | 0 | 0 | 0 | 0 | 0 | 0 |
| ENSMUSG00 | AA413626 | 493 | 0 | 0 | 0 | 0.14637746 | 1.13851954 | 0.9405048 |
| ENSMUSG00 | Gm3267 | 472 | 0 | 0 | 0 | 0 | 0 | 0 |
| ENSMUSG00 | Gm13370 | 482 | 0 | 0 | 0 | 0 | 0.10586385 | 0 |
| ENSMUSG00 | Gm12251 | 1585 | 0.25082322 | 0.03040154 | 0.22310079 | 0.09105879 | 0.06438659 | 0.05162392 |
| ENSMUSG00 | Gm13873 | 426 | 0 | 0 | 0 | 0 | 0 | 0 |
| ENSMUSG00 | Gm13852 | 241 | 0 | 0 | 0 | 0 | 0 | 0 |
| ENSMUSG00 | Olfr379-ps1 | 933 | 0 | 0 | 0 | 0 | 0 | 0 |
| ENSMUSG00 | Gm15425 | 2709 | 0 | 0.03557508 | 0.29369995 | 0.35962171 | 0.15068697 | 0.79538446 |
| ENSMUSG00 | Gm13967 | 378 | 0 | 0 | 0 | 0 | 0 | 0 |
| ENSMUSG00 | Gm15757 | 171 | 0 | 0 | 0 | 0 | 0 | 0 |
| ENSMUSG00 | Gm14101 | 471 | 0 | 0 | 0 | 0 | 0 | 0 |
| ENSMUSG00 | Gm8111 | 986 | 0 | 0 | 0 | 0 | 0 | 0.02766191 |
| ENSMUSG00 | Gm15087 | 1171 | 0 | 0 | 0 | 0 | 0 | 0 |
| ENSMUSG00 | Gm14657 | 301 | 0 | 0 | 0 | 0 | 0 | 0 |
| ENSMUSG00 | Gm9133 | 536 | 0 | 0 | 0 | 0 | 0 | 0 |
| ENSMUSG00 | Gm15686 | 770 | 0.30978297 | 0.25031918 | 0.22961996 | 0 | 0.66268021 | 0 |
| ENSMUSG00 | Gm11576 | 416 | 0 | 0 | 0 | 0 | 0 | 0 |
| ENSMUSG00 | Gm12586 | 1312 | 0.5454258 | 0.62436701 | 0.67380858 | 1.07256078 | 0.50559671 | 0.37419475 |
| ENSMUSG00 | Gm11921 | 319 | 0 | 0 | 0 | 0 | 0 | 0 |
| ENSMUSG00 | Rps15a-ps8 | 386 | 0 | 0 | 0 | 0 | 0 | 0 |
| ENSMUSG00 | Gm12513 | 351 | 0 | 0 | 0 | 0 | 0.14537429 | 0.07770553 |
| ENSMUSG00 | Olfr1169-ps1 | 215 | 0 | 0 | 0 | 0 | 0 | 0 |
| ENSMUSG00 | Gm12894 | 275 | 0 | 0 | 0 | 0 | 0 | 0 |
| ENSMUSG00 | Gm14018 | 451 | 0.08814962 | 0.42737421 | 0.29402556 | 0.08000453 | 0.56570261 | 0.24190367 |
| ENSMUSG00 | Gm11870 | 1321 | 0.72227974 | 0.58363594 | 0.20076537 | 0 | 0.38627082 | 0 |
| ENSMUSG00 | Gm14700 | 446 | 0 | 0 | 0 | 0 | 0 | 0 |
| ENSMUSG00 | Gm14557 | 377 | 0 | 0 | 0 | 0 | 0 | 0 |
| ENSMUSG00 | Gm11510 | 351 | 1.24589825 | 1.78468304 | 1.6371053 | 1.64476556 | 2.76211151 | 1.70952155 |
| ENSMUSG00 | Gm12044 | 212 | 0 | 0 | 0 | 0 | 0 | 0 |
| ENSMUSG00 | Gm12017 | 1002 | 0.43643741 | 0.38472209 | 0.30879531 | 0.39611027 | 0.40739621 | 0.27220199 |
| ENSMUSG00 | Gm11270 | 470 | 0 | 0 | 0.09404647 | 0 | 0 | 0 |
| ENSMUSG00 | Gm12837 | 372 | 0 | 0 | 0 | 0 | 0 | 0 |
| ENSMUSG00 | Rpl10a-ps1 | 1360 | 0.38001562 | 0.17715604 | 0.26001084 | 0.13265458 | 0.18759697 | 0.14038417 |
| ENSMUSG00 | U6 | 98 | 0.40566817 | 0 | 0 | 0 | 0 | 0 |
| ENSMUSG00 | SNORA3 | 128 | 0 | 0 | 0 | 0 | 0 | 0 |
| ENSMUSG00 | 7SK | 309 | 0 | 0 | 0 | 0 | 0.33026781 | 0 |
| ENSMUSG00 | SNORA35 | 128 | 0 | 0 | 0 | 0 | 0 | 0 |
| ENSMUSG00 | SNORD88 | 77 | 0.51630494 | 0 | 0 | 0 | 0 | 0 |
| ENSMUSG00 | AC123043.1 | 113 | 0 | 0 | 0 | 1.27724051 | 0 | 0 |
| ENSMUSG00 | AC102205.1 | 109 | 0 | 0 | 0 | 0 | 0 | 0 |
| ENSMUSG00 | U4 | 162 | 0 | 0 | 0 | 0 | 0 | 0 |
| ENSMUSG00 | SNORA61 | 129 | 0 | 0 | 0 | 0 | 0 | 0 |
| ENSMUSG00 | U4 | 111 | 0 | 0 | 0 | 0 | 0 | 0 |
| ENSMUSG00 | SNORD88 | 90 | 0 | 0 | 0 | 0 | 0 | 0 |
| ENSMUSG00 | n-R5s48 | 109 | 0 | 0 | 0 | 0 | 0 | 0 |
| ENSMUSG00 | 7SK | 317 | 0 | 0 | 0 | 0 | 0 | 0 |
| ENSMUSG00 | SNORA42 | 134 | 3.26350959 | 0 | 0 | 4.30830382 | 1.5231754 | 0 |
| ENSMUSG00 | U4 | 148 | 0 | 0 | 0 | 0 | 0 | 0 |
| ENSMUSG00 | U6 | 101 | 0 | 0 | 0 | 0 | 0 | 0 |
| ENSMUSG00 | 7SK | 313 | 0 | 0 | 0 | 0 | 0 | 0 |
| ENSMUSG00 | U4 | 117 | 0 | 1.2355498 | 0 | 0 | 0 | 0.9324663 |
| ENSMUSG00 | U6 | 102 | 0 | 0 | 0 | 0 | 0 | 0 |
| ENSMUSG00 | U6 | 102 | 0 | 0 | 0 | 0 | 0 | 0 |
| ENSMUSG00 | n-R5s214 | 120 | 0 | 0 | 0 | 0 | 0 | 0 |
| ENSMUSG00 | n-R5s32 | 102 | 0 | 0 | 0 | 0 | 0 | 0 |
| ENSMUSG00 | SNORD33 | 75 | 0 | 0 | 0 | 0 | 0 | 0 |
| ENSMUSG00 | n-R5s207 | 117 | 0 | 0 | 0 | 0 | 0 | 0 |
| ENSMUSG00 | SNORA40 | 121 | 0 | 0 | 0 | 0 | 0 | 0 |
| ENSMUSG00 | U2 | 185 | 0 | 0 | 0 | 0 | 0 | 0 |
| ENSMUSG00 | U6 | 106 | 0 | 0 | 0 | 0 | 0 | 0 |
| ENSMUSG00 | SNORA11 | 109 | 0 | 0 | 0 | 0 | 0 | 0 |
| ENSMUSG00 | U4 | 98 | 0 | 0 | 0 | 0 | 0 | 0 |
| ENSMUSG00 | 7SK | 299 | 0 | 0 | 0 | 0 | 0 | 0 |
| ENSMUSG00 | U6 | 106 | 0 | 0 | 0 | 0 | 0 | 0 |
| ENSMUSG00 | U6 | 105 | 0 | 0 | 0 | 0 | 0 | 0 |
| ENSMUSG00 | U6 | 105 | 0 | 0 | 0 | 0 | 0 | 0 |
| ENSMUSG00 | n-R5s73 | 106 | 0 | 0 | 0 | 0 | 0 | 0 |
| ENSMUSG00 | Mir1907 | 89 | 0 | 0 | 0 | 0 | 0 | 0 |
| ENSMUSG00 | SNORD113 | 73 | 0 | 0 | 0 | 0 | 0 | 0 |
| ENSMUSG00 | SNORA42 | 122 | 0 | 0 | 0 | 0 | 0 | 0 |
| ENSMUSG00 | U2 | 183 | 0 | 0 | 0 | 0 | 0 | 0 |
| ENSMUSG00 | n-R5s154 | 107 | 0 | 0 | 0 | 0 | 0 | 0.25490317 |
| ENSMUSG00 | 7SK | 302 | 0 | 0 | 0 | 0 | 0 | 0 |
| ENSMUSG00 | n-R5s47 | 118 | 0 | 0 | 0 | 0 | 0 | 0 |
| ENSMUSG00 | U6 | 103 | 0 | 0 | 0 | 0 | 0 | 0 |
| ENSMUSG00 | n-R5s24 | 116 | 0 | 0 | 0 | 0 | 0 | 0 |
| ENSMUSG00 | AC158132.1 | 56 | 0 | 0 | 0 | 0 | 0 | 0 |
| ENSMUSG00 | U4 | 79 | 0 | 0 | 0 | 0 | 0 | 0 |
| ENSMUSG00 | n-R5s23 | 114 | 0 | 0 | 0 | 0 | 1.34279936 | 0 |
| ENSMUSG00 | U6 | 80 | 0 | 0 | 0 | 0 | 0 | 0 |
| ENSMUSG00 | U2 | 189 | 0 | 0 | 0.46774437 | 0 | 0 | 0 |
| ENSMUSG00 | 7SK | 288 | 0 | 0 | 0 | 0 | 0 | 0 |
| ENSMUSG00 | U6 | 83 | 0 | 0 | 0 | 0 | 0 | 0 |
| ENSMUSG00 | 7SK | 252 | 1.26207875 | 0 | 0.52621242 | 0 | 0 | 0.86586156 |
| ENSMUSG00 | U4 | 81 | 0 | 1.1897887 | 0 | 0 | 0.62995526 | 0 |
| ENSMUSG00 | U6 | 103 | 0 | 0 | 0 | 0 | 0 | 0 |
| ENSMUSG00 | n-R5s155 | 112 | 0 | 0 | 0 | 0 | 0 | 0 |
| ENSMUSG00 | U2 | 105 | 0 | 0 | 0 | 0 | 0 | 0 |
| ENSMUSG00 | 7SK | 300 | 0 | 0 | 0 | 0 | 0 | 0 |
| ENSMUSG00 | SNORD113 | 72 | 0 | 0 | 0 | 0 | 0 | 0 |
| ENSMUSG00 | n-R5s157 | 120 | 0 | 0 | 0 | 0 | 0 | 0 |
| ENSMUSG00 | U4 | 126 | 0 | 0 | 0 | 0 | 0 | 0 |
| ENSMUSG00 | 7SK | 307 | 0 | 0 | 0 | 0 | 0 | 0 |
| ENSMUSG00 | U6 | 99 | 2.00785255 | 1.94692696 | 0.44648326 | 2.91572077 | 1.03083588 | 6.61203376 |
| ENSMUSG00 | n-R5s98 | 118 | 0 | 0 | 0 | 0 | 0 | 0 |
| ENSMUSG00 | 7SK | 323 | 0 | 0 | 0 | 0 | 0 | 0 |
| ENSMUSG00 | 7SK | 324 | 0 | 0 | 0 | 0 | 0 | 0 |
| ENSMUSG00 | 7SK | 292 | 0 | 0 | 0 | 0 | 0 | 0 |
| ENSMUSG00 | 7SK | 313 | 0 | 0 | 0 | 0 | 0 | 0 |
| ENSMUSG00 | U6 | 102 | 0 | 0 | 0 | 0 | 0.50025859 | 0 |
| ENSMUSG00 | U6 | 103 | 0 | 0 | 0 | 0 | 0 | 0 |
| ENSMUSG00 | U6 | 92 | 0 | 0 | 0 | 0 | 0 | 0 |
| ENSMUSG00 | U6 | 101 | 0 | 0 | 0 | 0 | 0 | 0 |
| ENSMUSG00 | AC123057.1 | 85 | 0 | 0 | 0 | 0 | 0 | 0 |
| ENSMUSG00 | U6 | 106 | 0 | 0 | 0 | 0 | 0 | 0 |
| ENSMUSG00 | 7SK | 191 | 0 | 0 | 0 | 0 | 0 | 0 |
| ENSMUSG00 | 7SK | 358 | 0 | 0 | 0 | 0 | 0 | 0 |
| ENSMUSG00 | n-R5s179 | 117 | 0 | 0 | 0 | 0 | 0 | 0 |
| ENSMUSG00 | Mir1893 | 82 | 0 | 0 | 0 | 0 | 0 | 0 |
| ENSMUSG00 | U4 | 80 | 0 | 0 | 0 | 0 | 0 | 0 |
| ENSMUSG00 | Mir1902 | 79 | 0 | 0 | 0 | 0 | 0 | 0 |
| ENSMUSG00 | SNORA66 | 127 | 0 | 0 | 0 | 0 | 0 | 0 |
| ENSMUSG00 | 7SK | 315 | 0 | 0 | 0 | 0 | 0 | 0 |
| ENSMUSG00 | SNORA67 | 153 | 1.03935897 | 2.8344966 | 0 | 2.12247321 | 1.00051717 | 1.24785931 |
| ENSMUSG00 | SNORA40 | 92 | 0 | 0 | 0 | 0 | 0 | 0 |
| ENSMUSG00 | U6 | 83 | 0 | 0 | 0 | 0 | 0 | 0 |
| ENSMUSG00 | Mir1903 | 79 | 0 | 0 | 0 | 0 | 0 | 0 |
| ENSMUSG00 | Y_RNA | 100 | 0 | 0 | 0 | 0 | 0 | 0 |
| ENSMUSG00 | n-R5s192 | 118 | 0 | 0 | 0 | 0 | 0 | 0 |
| ENSMUSG00 | U6 | 104 | 0 | 0 | 0 | 0 | 0 | 0 |
| ENSMUSG00 | Mir1897 | 78 | 0 | 0 | 0 | 0 | 0 | 0 |
| ENSMUSG00 | 7SK | 313 | 0 | 0 | 0 | 0 | 0 | 0 |
| ENSMUSG00 | n-R5s218 | 108 | 0 | 0 | 0 | 0 | 0 | 0 |
| ENSMUSG00 | Mir1906-1 | 79 | 0 | 0 | 0 | 0 | 0 | 0 |
| ENSMUSG00 | 7SK | 308 | 0 | 0 | 0 | 0 | 0 | 0 |
| ENSMUSG00 | 7SK | 309 | 0 | 0 | 0 | 0 | 0 | 0 |
| ENSMUSG00 | 7SK | 321 | 0 | 0 | 0 | 0 | 0 | 0 |
| ENSMUSG00 | n-R5s84 | 108 | 0 | 0 | 0 | 0 | 0 | 0 |
| ENSMUSG00 | n-R5s45 | 109 | 0 | 0 | 0 | 0 | 0 | 0 |
| ENSMUSG00 | SNORA33 | 130 | 0 | 0 | 0 | 0 | 0 | 1.2588295 |
| ENSMUSG00 | 7SK | 322 | 0 | 0 | 0 | 0 | 0 | 0 |
| ENSMUSG00 | n-R5s80 | 110 | 0 | 0 | 0 | 0 | 0 | 0 |
| ENSMUSG00 | SNORA42 | 133 | 0 | 0 | 0 | 0 | 0 | 0 |
| ENSMUSG00 | 7SK | 246 | 0 | 0 | 0 | 0 | 0 | 0 |
| ENSMUSG00 | n-R5s175 | 116 | 0 | 0 | 0 | 0 | 0 | 0 |
| ENSMUSG00 | SNORD113 | 71 | 0 | 0 | 0 | 0 | 0 | 0 |
| ENSMUSG00 | U6 | 103 | 0 | 0 | 0 | 0 | 0 | 0 |
| ENSMUSG00 | U6 | 106 | 0 | 0 | 0 | 0 | 0 | 0 |
| ENSMUSG00 | U2 | 146 | 0 | 0 | 0.90825705 | 1.48282375 | 1.3979829 | 0 |
| ENSMUSG00 | U4 | 137 | 0 | 0 | 0 | 0 | 0 | 0 |
| ENSMUSG00 | U4 | 132 | 0 | 0 | 0 | 0 | 0 | 0 |
| ENSMUSG00 | U6 | 107 | 0 | 0 | 0 | 0 | 2.38441009 | 0 |
| ENSMUSG00 | Mir1906-2 | 79 | 0 | 0 | 0 | 0 | 0 | 0 |
| ENSMUSG00 | U4 | 132 | 0 | 0 | 0 | 0 | 0 | 0 |
| ENSMUSG00 | n-R5s39 | 107 | 0 | 0 | 2.06550668 | 0 | 0.47688202 | 0.50980634 |
| ENSMUSG00 | Mir1895 | 78 | 0 | 0 | 0 | 0 | 0 | 0 |
| ENSMUSG00 | Mir1901 | 77 | 0 | 0 | 0 | 0 | 0 | 0 |
| ENSMUSG00 | 7SK | 322 | 0 | 0 | 0 | 0 | 0 | 0 |
| ENSMUSG00 | 7SK | 312 | 0 | 0 | 0 | 0.11564758 | 0 | 0 |
| ENSMUSG00 | 7SK | 296 | 0 | 0 | 0 | 0 | 0 | 0 |
| ENSMUSG00 | U6 | 105 | 1.5144945 | 0 | 0 | 0 | 0 | 0 |
| ENSMUSG00 | U6 | 103 | 0 | 0 | 0 | 0 | 0 | 0 |
| ENSMUSG00 | n-R5s201 | 115 | 0 | 0 | 0 | 0 | 0 | 0 |
| ENSMUSG00 | 7SK | 293 | 0 | 0.32891769 | 0.9051572 | 0 | 0 | 0.09308751 |
| ENSMUSG00 | n-R5s30 | 121 | 0 | 0 | 0 | 0 | 0 | 0 |
| ENSMUSG00 | n-R5s198 | 99 | 0 | 0.48673174 | 0 | 0 | 0 | 0 |
| ENSMUSG00 | U6 | 111 | 0 | 0.43411209 | 0.3982148 | 1.62531732 | 0 | 0.49143494 |
| ENSMUSG00 | U6 | 104 | 0 | 0 | 0 | 0 | 0 | 0 |
| ENSMUSG00 | U2 | 86 | 0 | 0 | 0 | 0 | 0 | 0 |
| ENSMUSG00 | U4 | 127 | 0 | 0.3794208 | 1.04413802 | 0 | 0 | 0 |
| ENSMUSG00 | 7SK | 324 | 0 | 0 | 0 | 0 | 0 | 0 |
| ENSMUSG00 | U4 | 116 | 3.08447694 | 0 | 1.14315111 | 0.93315632 | 0 | 1.6458834 |
| ENSMUSG00 | n-R5s15 | 110 | 1.08424038 | 0 | 0 | 0 | 0 | 0 |
| ENSMUSG00 | n-R5s174 | 120 | 0 | 0 | 0 | 0 | 0 | 0 |
| ENSMUSG00 | n-R5s68 | 81 | 0 | 0 | 0 | 0 | 0 | 0 |
| ENSMUSG00 | U4 | 124 | 0 | 0 | 0 | 0 | 0 | 0 |
| ENSMUSG00 | Mir1896 | 80 | 0 | 0 | 0 | 0 | 0 | 0 |
| ENSMUSG00 | 7SK | 277 | 0 | 0 | 0 | 0 | 0 | 0 |
| ENSMUSG00 | Mir1898 | 82 | 0.48482293 | 7.05167446 | 5.92951553 | 2.20012467 | 3.73363726 | 1.99570531 |
| ENSMUSG00 | U6 | 106 | 0 | 0.45458908 | 0 | 0 | 0 | 3.34500293 |
| ENSMUSG00 | AC126443.1 | 109 | 0 | 0 | 0 | 0 | 0 | 0 |
| ENSMUSG00 | n-R5s188 | 104 | 0 | 0 | 0 | 0 | 0.98127646 | 0 |
| ENSMUSG00 | 7SK | 297 | 0 | 0 | 0 | 0 | 0 | 0 |
| ENSMUSG00 | 7SK | 285 | 0 | 0 | 0.46528256 | 0 | 0.89519958 | 0 |
| ENSMUSG00 | 7SK | 273 | 0 | 0 | 0 | 0 | 0 | 0 |
| ENSMUSG00 | n-R5s70 | 115 | 0 | 0 | 0 | 0 | 0 | 0 |
| ENSMUSG00 | CT485616.1 | 58 | 0 | 0 | 0 | 0 | 0 | 0 |
| ENSMUSG00 | U6 | 104 | 0 | 0 | 3.82515949 | 0 | 0.98127646 | 0 |
| ENSMUSG00 | n-R5s34 | 115 | 0 | 0 | 0 | 0 | 0 | 0 |
| ENSMUSG00 | 7SK | 326 | 1.34144259 | 0.88686703 | 3.11853494 | 3.09907131 | 1.8782715 | 3.59757512 |
| ENSMUSG00 | SNORD113 | 70 | 0 | 0 | 0 | 0 | 0 | 0 |
| ENSMUSG00 | 7SK | 285 | 0 | 0 | 0 | 0 | 0 | 0 |
| ENSMUSG00 | U2 | 186 | 0 | 0 | 0 | 0 | 0 | 0 |
| ENSMUSG00 | U6 | 109 | 0 | 0 | 4.46073645 | 0 | 0 | 0.5004521 |
| ENSMUSG00 | 7SK | 306 | 0 | 0 | 0 | 0 | 0 | 0 |
| ENSMUSG00 | Y_RNA | 123 | 0 | 0 | 0 | 0 | 0 | 0 |
| ENSMUSG00 | U2 | 157 | 0 | 0 | 0 | 0 | 0 | 0 |
| ENSMUSG00 | 7SK | 266 | 0 | 0 | 0 | 0 | 0 | 0 |
| ENSMUSG00 | n-R5s6 | 87 | 0 | 0 | 0 | 0 | 0 | 0 |
| ENSMUSG00 | n-R5s164 | 111 | 7.87946461 | 0 | 1.99107401 | 0 | 2.2984854 | 0 |
| ENSMUSG00 | AC151601.1 | 109 | 0 | 0 | 0 | 0 | 0 | 0 |
| ENSMUSG00 | U6 | 103 | 5.78963308 | 5.14612489 | 21.4572054 | 11.5602667 | 8.91723073 | 9.2680813 |
| ENSMUSG00 | U2 | 188 | 2.32611854 | 0.25631086 | 0 | 0 | 0 | 0.43523361 |
| ENSMUSG00 | U6 | 98 | 0 | 0 | 0 | 0 | 0 | 0 |
| ENSMUSG00 | 7SK | 317 | 0 | 0 | 0 | 0 | 0 | 0 |
| ENSMUSG00 | U2 | 133 | 0 | 0 | 0 | 0 | 0 | 0 |
| ENSMUSG00 | 7SK | 322 | 0.74078535 | 0 | 0 | 0.67233623 | 0 | 0 |
| ENSMUSG00 | n-R5s189 | 113 | 0 | 1.27928608 | 0 | 0 | 0 | 0 |
| ENSMUSG00 | SNORA42 | 119 | 0.66815934 | 2.02464043 | 3.34299653 | 0.60642092 | 1.28637922 | 2.52118514 |
| ENSMUSG00 | U6 | 106 | 0 | 0 | 0 | 0 | 0 | 0 |
| ENSMUSG00 | 7SK | 311 | 0 | 0 | 0 | 0 | 0 | 0 |
| ENSMUSG00 | 7SK | 291 | 0 | 0 | 0 | 0 | 0 | 0 |
| ENSMUSG00 | U6 | 103 | 0 | 0 | 0 | 0 | 0 | 0 |
| ENSMUSG00 | n-R5s159 | 119 | 0 | 0 | 0 | 0 | 0 | 0 |
| ENSMUSG00 | 7SK | 293 | 0 | 0 | 0 | 0 | 0 | 0 |
| ENSMUSG00 | U2 | 171 | 19.529008 | 5.35404913 | 8.27168392 | 9.70628098 | 9.25039562 | 4.30652199 |
| ENSMUSG00 | 7SK | 312 | 0 | 0 | 0 | 0 | 0 | 0 |
| ENSMUSG00 | 7SK | 289 | 0 | 0 | 0 | 0 | 0 | 0 |
| ENSMUSG00 | 7SK | 293 | 0 | 0 | 0 | 0 | 0 | 0 |
| ENSMUSG00 | n-R5s14 | 106 | 0 | 0 | 0.83399704 | 0 | 0 | 0 |
| ENSMUSG00 | U6 | 103 | 0 | 0 | 0 | 0 | 1.48620512 | 0 |
| ENSMUSG00 | 7SK | 252 | 0 | 0 | 0 | 0 | 0 | 0 |
| ENSMUSG00 | 7SK | 332 | 0 | 0 | 0 | 0 | 0 | 0 |
| ENSMUSG00 | n-R5s21 | 116 | 0 | 0 | 0 | 0 | 0 | 0 |
| ENSMUSG00 | U6 | 74 | 0 | 1.30233627 | 0 | 0 | 0.68954562 | 0 |
| ENSMUSG00 | 7SK | 349 | 0 | 0 | 0 | 0 | 0 | 0 |
| ENSMUSG00 | Y_RNA | 85 | 0 | 0.56689932 | 0 | 0 | 0 | 0 |
| ENSMUSG00 | 7SK | 323 | 0 | 0 | 0 | 0 | 0 | 0 |
| ENSMUSG00 | U6 | 103 | 0 | 0 | 0 | 0 | 0 | 0 |
| ENSMUSG00 | U6 | 106 | 0 | 0 | 0 | 0 | 0 | 0 |
| ENSMUSG00 | U6 | 105 | 0 | 0 | 0 | 0 | 0 | 0 |
| ENSMUSG00 | n-R5s76 | 112 | 0 | 0 | 0 | 0 | 0 | 0 |
| ENSMUSG00 | 7SK | 314 | 0 | 0 | 0 | 0 | 0 | 0 |
| ENSMUSG00 | Mir1904 | 79 | 0 | 0 | 0 | 0 | 0 | 0 |
| ENSMUSG00 | U4 | 170 | 0 | 0 | 0 | 0 | 0 | 0 |
| ENSMUSG00 | U6 | 81 | 0 | 0 | 0 | 0 | 0 | 0 |
| ENSMUSG00 | n-R5s150 | 107 | 0 | 0 | 0 | 0 | 0 | 0 |
| ENSMUSG00 | U3 | 202 | 0 | 0 | 0 | 0 | 0 | 0 |
| ENSMUSG00 | AC152062.1 | 82 | 0 | 0 | 0 | 0 | 0 | 0 |
| ENSMUSG00 | U4 | 86 | 0 | 0 | 1.54192476 | 3.35646926 | 1.77998985 | 0 |
| ENSMUSG00 | n-R5s61 | 117 | 0 | 0 | 0 | 0 | 0 | 0 |
| ENSMUSG00 | U4 | 127 | 0 | 0 | 0 | 0 | 0 | 0 |
| ENSMUSG00 | Mir1900 | 77 | 0 | 0 | 0 | 0 | 0 | 0 |
| ENSMUSG00 | 7SK | 312 | 0 | 0 | 0 | 0 | 0 | 0 |
| ENSMUSG00 | U4 | 156 | 0 | 0 | 0 | 0 | 0 | 0 |
| ENSMUSG00 | 7SK | 307 | 0 | 0.1569591 | 0 | 0 | 0 | 0 |
| ENSMUSG00 | SNORD44 | 59 | 0 | 0 | 0 | 0 | 0 | 0 |
| ENSMUSG00 | SNORD29 | 64 | 0 | 0 | 0 | 0 | 0 | 0 |
| ENSMUSG00 | U2 | 183 | 0.43448613 | 0 | 0.9661605 | 2.36603571 | 0 | 2.08658442 |
| ENSMUSG00 | 7SK | 312 | 0 | 0 | 0 | 0 | 0 | 0 |
| ENSMUSG00 | n-R5s191 | 115 | 0 | 0 | 0 | 0 | 0 | 0 |
| ENSMUSG00 | 7SK | 267 | 0.44669079 | 0 | 0 | 0 | 0 | 0 |
| ENSMUSG00 | U4 | 122 | 0 | 0 | 0 | 0 | 0 | 0 |
| ENSMUSG00 | U4 | 81 | 0 | 0 | 0 | 0 | 0 | 0 |
| ENSMUSG00 | U2 | 182 | 1.09218353 | 0.52952134 | 0 | 0 | 0.84109411 | 2.09804917 |
| ENSMUSG00 | 7SK | 286 | 0 | 0 | 0 | 0 | 0 | 0 |
| ENSMUSG00 | 7SK | 301 | 0 | 0 | 0 | 0 | 0 | 0 |
| ENSMUSG00 | 7SK | 280 | 0 | 0 | 0 | 0 | 0 | 0 |
| ENSMUSG00 | SNORA19 | 126 | 1.89311812 | 0.76486416 | 2.45565795 | 3.43638519 | 0 | 0.64939617 |
| ENSMUSG00 | n-R5s43 | 109 | 0 | 0 | 0 | 0 | 0 | 0.25022605 |
| ENSMUSG00 | n-R5s18 | 120 | 0 | 0 | 0 | 0 | 0 | 0 |
| ENSMUSG00 | SNORA16 | 134 | 0 | 0 | 0 | 0 | 0 | 0 |
| ENSMUSG00 | SNORD113 | 73 | 0 | 0 | 0 | 0 | 0 | 0 |
| ENSMUSG00 | SNORA33 | 117 | 0 | 0 | 0 | 0 | 0 | 0 |
| ENSMUSG00 | 7SK | 368 | 0 | 0 | 0.1201137 | 0 | 0 | 0 |
| ENSMUSG00 | U6 | 103 | 0 | 0 | 0 | 0 | 0 | 1.32401161 |
| ENSMUSG00 | U2 | 121 | 0 | 0 | 0 | 0 | 0 | 0 |
| ENSMUSG00 | n-R5s87 | 115 | 0 | 0 | 0 | 0 | 0 | 0 |
| ENSMUSG00 | SNORA61 | 129 | 0 | 0 | 0 | 0 | 0 | 0 |
| ENSMUSG00 | U4 | 115 | 0 | 0 | 0 | 0 | 0 | 0 |
| ENSMUSG00 | n-R5s216 | 111 | 0 | 0 | 0 | 0 | 0 | 0 |
| ENSMUSG00 | n-R5s63 | 102 | 0 | 0 | 0 | 0 | 0 | 0 |
| ENSMUSG00 | 7SK | 245 | 0 | 0 | 0.36083137 | 0 | 0.62481277 | 0 |
| ENSMUSG00 | SNORA48 | 143 | 0 | 0 | 0 | 0 | 0 | 0 |
| ENSMUSG00 | 7SK | 288 | 0 | 0 | 0 | 0 | 0 | 0 |
| ENSMUSG00 | Y_RNA | 88 | 1.35530047 | 0 | 0 | 0 | 0 | 0.61987817 |
| ENSMUSG00 | AL808014.1 | 79 | 0 | 0 | 0 | 0 | 0 | 0 |
| ENSMUSG00 | U6 | 80 | 0 | 0 | 0 | 0 | 0 | 0 |
| ENSMUSG00 | U6 | 105 | 0 | 0 | 0 | 0 | 0 | 0 |
| ENSMUSG00 | n-R5s187 | 110 | 0 | 0 | 0 | 0 | 0 | 0 |
| ENSMUSG00 | 7SK | 322 | 0 | 0 | 0 | 0 | 0 | 0 |
| ENSMUSG00 | 7SK | 325 | 0 | 0 | 0 | 0 | 0 | 0 |
| ENSMUSG00 | U4 | 109 | 0 | 0 | 0 | 0 | 0 | 0 |
| ENSMUSG00 | Mir1905 | 81 | 0 | 0 | 0 | 0 | 0 | 0 |
| ENSMUSG00 | 7SK | 287 | 0 | 0 | 0.92408034 | 0 | 0 | 0 |
| ENSMUSG00 | U4 | 121 | 0 | 0 | 0 | 0 | 0 | 0 |
| ENSMUSG00 | 7SK | 304 | 0 | 0 | 0 | 0 | 0 | 0 |
| ENSMUSG00 | 7SK | 283 | 0 | 0 | 0 | 0 | 0 | 0 |
| ENSMUSG00 | SNORA73 | 187 | 0 | 0 | 0 | 0 | 0 | 0 |
| ENSMUSG00 | U6 | 107 | 0 | 0 | 0 | 0 | 0 | 0 |
| ENSMUSG00 | SNORD88 | 52 | 1.52905694 | 0 | 10.2004253 | 4.16331283 | 0 | 3.14707376 |
| ENSMUSG00 | U6 | 106 | 0 | 0 | 0 | 0 | 0 | 0 |
| ENSMUSG00 | U6 | 103 | 0 | 0 | 0 | 0 | 0 | 0 |
| ENSMUSG00 | 7SK | 284 | 0 | 0 | 0 | 0 | 0 | 0 |
| ENSMUSG00 | n-R5s19 | 119 | 0 | 0 | 0 | 0 | 0 | 0 |
| ENSMUSG00 | 7SK | 305 | 0 | 0 | 0 | 0 | 0 | 0 |
| ENSMUSG00 | SNORA48 | 143 | 0 | 0 | 0 | 0 | 0 | 0 |
| ENSMUSG00 | SNORA57 | 143 | 0 | 0 | 0 | 0 | 0 | 0 |
| ENSMUSG00 | 7SK | 321 | 0 | 0 | 0 | 0 | 0 | 0 |
| ENSMUSG00 | Mir1899 | 95 | 0 | 0 | 0 | 0 | 0 | 0 |
| ENSMUSG00 | U4 | 118 | 0 | 0 | 0 | 0 | 0 | 0 |
| ENSMUSG00 | 7SK | 264 | 0 | 0 | 0.33486245 | 0 | 0 | 0 |
| ENSMUSG00 | 7SK | 317 | 0 | 0 | 0 | 0 | 0 | 0 |
| ENSMUSG00 | Mir1894 | 80 | 0 | 0 | 0 | 0 | 0 | 0 |
| ENSMUSG00 | U2 | 150 | 0 | 0 | 0 | 0 | 0 | 0 |
| ENSMUSG00 | U6 | 103 | 0 | 0 | 0 | 0 | 0 | 0 |
| ENSMUSG00 | Gm12532 | 2679 | 0 | 0 | 0 | 0 | 0 | 0 |
| ENSMUSG00 | Gm16276 | 261 | 1.52319849 | 0.18462238 | 0.16935572 | 0.27649076 | 0.78201342 | 0.20900107 |
| ENSMUSG00 | Gm13881 | 3714 | 0 | 0 | 0 | 0 | 0 | 0.03671869 |
| ENSMUSG00 | 1700057H15I | 1242 | 0 | 0 | 0 | 0 | 0 | 0 |
| ENSMUSG00 | Gm12798 | 663 | 0 | 0 | 0 | 0 | 0 | 0 |
| ENSMUSG00 | Gm4726 | 735 | 0 | 0 | 0 | 0 | 0 | 0 |
| ENSMUSG00 | Gm15472 | 845 | 0 | 0 | 0 | 0 | 0 | 0 |
| ENSMUSG00 | Gm12406 | 695 | 0 | 0 | 0 | 0 | 0 | 0 |
| ENSMUSG00 | Gm14074 | 408 | 0 | 0 | 0 | 0 | 0.12506465 | 0 |

| | | | | | | | | |
|---|---|---|---|---|---|---|---|---|
| ENSMUSG00 | Gm5106 | 1331 | 0 | 0 | 0 | 0 | 0 | 0 |
| ENSMUSG00 | 4930564G21 | 719 | 0 | 0 | 0 | 0 | 0 | 0 |
| ENSMUSG00 | Gm16998 | 1227 | 0.09720166 | 0.11781526 | 0 | 0 | 0.16634515 | 0.04445744 |
| ENSMUSG00 | Gm12714 | 665 | 0.35869607 | 0.21738245 | 0 | 0 | 0 | 0 |
| ENSMUSG00 | Gm13605 | 656 | 0.12120573 | 0 | 0 | 0 | 0 | 0 |
| ENSMUSG00 | Gm12061 | 476 | 0 | 0 | 0 | 0 | 0 | 0 |
| ENSMUSG00 | Gm15139 | 6636 | 0.26359873 | 0.61721633 | 0.66609167 | 0.39148692 | 0.66128215 | 0.84257098 |
| ENSMUSG00 | Gm15473 | 1177 | 0 | 0 | 0 | 0 | 0.04335291 | 0 |
| ENSMUSG00 | Gm12159 | 1444 | 0 | 0 | 0 | 0 | 0 | 0 |
| ENSMUSG00 | Gm14110 | 756 | 0 | 0 | 0 | 0 | 0 | 0 |
| ENSMUSG00 | Gm16741 | 1815 | 0.15332692 | 1.03541115 | 0 | 0 | 1.09643452 | 1.09699651 |
| ENSMUSG00 | Gm14951 | 568 | 0 | 0 | 0 | 0 | 0.08983517 | 0 |
| ENSMUSG00 | Gm16742 | 2093 | 0 | 0 | 0 | 0 | 0 | 0 |
| ENSMUSG00 | Gm16743 | 1153 | 0.65512067 | 0 | 0 | 0 | 0 | 0 |
| ENSMUSG00 | 4921504A21 | 2152 | 0.36947473 | 1.45544551 | 0.22593879 | 0.85510422 | 0.28453369 | 0.70974898 |
| ENSMUSG00 | Gm15350 | 615 | 0 | 0 | 0 | 0.11733998 | 0 | 0.04434901 |
| ENSMUSG00 | D930015M05 | 1186 | 0.13408257 | 0 | 0.07453936 | 0.03042331 | 0.04302393 | 0 |
| ENSMUSG00 | Gm11213 | 479 | 0 | 0 | 0 | 0 | 0 | 0 |
| ENSMUSG00 | Gm15419 | 740 | 0 | 0 | 0.05973222 | 0.04875952 | 0 | 0.03685762 |
| ENSMUSG00 | Gm15086 | 571 | 0 | 0 | 0 | 0 | 0 | 0 |
| ENSMUSG00 | Gm13972 | 3162 | 0.07543734 | 0 | 0.08387447 | 0 | 0.01613737 | 0.02587727 |
| ENSMUSG00 | Ubl5 | 4321 | 5.01428764 | 6.3007035 | 6.31162628 | 4.42570785 | 3.88513715 | 10.0488835 |
| ENSMUSG00 | Gm12436 | 2955 | 0 | 0 | 0 | 0 | 0 | 0 |
| ENSMUSG00 | Gm11342 | 813 | 0 | 0 | 0 | 0 | 0 | 0 |
| ENSMUSG00 | Gm12974 | 331 | 0 | 0.29115675 | 0 | 0.87207358 | 0 | 1.48321301 |
| ENSMUSG00 | Gm15879 | 1020 | 0.38975961 | 0 | 0.2166757 | 0.42449464 | 0.20010344 | 0.53479685 |
| ENSMUSG00 | Gm13667 | 614 | 0 | 0 | 0 | 0 | 0 | 0 |
| ENSMUSG00 | 1700056N10 | 1041 | 0 | 0.83319497 | 0.764297 | 0.13864378 | 2.3037845 | 0.36680591 |
| ENSMUSG00 | Gm2568 | 1164 | 0 | 0 | 0 | 0 | 0 | 0 |
| ENSMUSG00 | Gm13662 | 710 | 0 | 0 | 0 | 0 | 0 | 0 |
| ENSMUSG00 | Gm14617 | 924 | 0 | 0 | 0.09567499 | 0 | 0.05522335 | 0.47228813 |
| ENSMUSG00 | A430104N18 | 267 | 0 | 0 | 0 | 0 | 0 | 0 |
| ENSMUSG00 | Gm14321 | 976 | 0 | 0 | 0 | 0.11090792 | 0.47053011 | 0.08383598 |
| ENSMUSG00 | 4930533B01 | 1023 | 0 | 0.09420614 | 0.12962417 | 0.42324979 | 0.14963747 | 0.55988996 |
| ENSMUSG00 | Gm11602 | 1572 | 2.55426433 | 2.42158329 | 1.79956613 | 2.2723425 | 3.18103361 | 2.74134415 |
| ENSMUSG00 | Gm12205 | 573 | 0.55505034 | 1.76599526 | 1.54282175 | 1.3223786 | 0.97956393 | 2.42758569 |
| ENSMUSG00 | Gm16929 | 2288 | 0.05212694 | 0.2737866 | 0 | 0.0630805 | 0 | 0.14304881 |
| ENSMUSG00 | Gm14089 | 316 | 0 | 0 | 0 | 0 | 0 | 0 |
| ENSMUSG00 | 5830444B04 | 5401 | 0 | 0 | 0.04910407 | 0.06012561 | 0.07558063 | 0.97463532 |
| ENSMUSG00 | 1700095J12 | 735 | 0 | 0 | 0 | 0.09818243 | 0.06942364 | 0 |
| ENSMUSG00 | Gm12273 | 1081 | 0 | 0 | 0 | 0 | 0 | 0 |
| ENSMUSG00 | Gm15232 | 2314 | 0 | 0 | 0 | 0 | 0 | 0 |
| ENSMUSG00 | Gm13073 | 484 | 0 | 0 | 0 | 0 | 0 | 0 |
| ENSMUSG00 | 9430091E24 | 1917 | 2.40565245 | 3.97154818 | 2.53636032 | 1.73163698 | 2.76825409 | 1.69310489 |
| ENSMUSG00 | Gm12631 | 635 | 0 | 0 | 0 | 0 | 0 | 0 |
| ENSMUSG00 | Gm16761 | 1785 | 0.2004478 | 0.08098562 | 0.09905175 | 0.02021403 | 0.02858621 | 0.12223928 |
| ENSMUSG00 | 1700092E19 | 1000 | 0 | 0 | 0 | 0 | 0 | 0 |
| ENSMUSG00 | Gm17445 | 3900 | 0.01019371 | 0.0123555 | 0.66869455 | 0.01850361 | 0.35325953 | 0.69235623 |
| ENSMUSG00 | 9430076G02 | 1771 | 0 | 0 | 0 | 0 | 0 | 0 |
| ENSMUSG00 | Gm15525 | 502 | 0 | 0 | 0 | 0 | 0 | 0 |
| ENSMUSG00 | Gm11463 | 824 | 0 | 0 | 0 | 0 | 0 | 0.43030378 |
| ENSMUSG00 | Gm6526 | 877 | 0 | 0.05494463 | 0.20160476 | 0.04114258 | 0 | 0.03109993 |
| ENSMUSG00 | Gm14640 | 435 | 0 | 0 | 0 | 0 | 0 | 0 |
| ENSMUSG00 | Gm11967 | 478 | 0 | 0 | 0 | 0 | 0 | 0 |
| ENSMUSG00 | Gm15880 | 2879 | 0 | 0 | 0 | 0.0626642 | 0 | 0.0568419 |
| ENSMUSG00 | Gm11768 | 1040 | 0 | 0 | 0 | 0 | 0 | 0 |
| ENSMUSG00 | Gm17216 | 726 | 0.32855769 | 0.06637251 | 0.06088408 | 0 | 0.56227411 | 0 |
| ENSMUSG00 | Gm16344 | 1565 | 0 | 0 | 0 | 0 | 0 | 0 |
| ENSMUSG00 | 2410152P15 | 869 | 0 | 0 | 0 | 0 | 0 | 0 |
| ENSMUSG00 | AI847159 | 1228 | 0 | 0 | 0 | 0 | 0 | 0 |
| ENSMUSG00 | B230344G16 | 2021 | 0.2753967 | 0 | 0.21871273 | 0.05356068 | 0.07574425 | 0.18893862 |
| ENSMUSG00 | Gm12367 | 2694 | 0 | 0 | 0.06563006 | 0.0803609 | 0 | 0.01012422 |
| ENSMUSG00 | Gm14266 | 1276 | 0 | 0 | 0 | 0 | 0 | 0 |
| ENSMUSG00 | Gm14539 | 349 | 6.15127779 | 6.48929164 | 4.81280812 | 4.03209094 | 4.24001404 | 13.1293392 |
| ENSMUSG00 | Gm14710 | 1062 | 0 | 0 | 0 | 0.03397556 | 0 | 0 |
| ENSMUSG00 | 4930563E18 | 1550 | 0 | 0 | 0 | 0 | 0 | 0.05278962 |
| ENSMUSG00 | Gm12649 | 578 | 0 | 0 | 0 | 0 | 0 | 0 |
| ENSMUSG00 | 4930565N06 | 2413 | 0 | 0.01996952 | 0.09159106 | 0 | 0 | 0 |
| ENSMUSG00 | Gm12352 | 758 | 0.73427009 | 0.88998706 | 0.46651022 | 1.1424394 | 1.75024508 | 1.3673302 |
| ENSMUSG00 | Gm16274 | 591 | 0 | 0 | 0 | 0 | 0 | 0 |
| ENSMUSG00 | 1700108N11 | 915 | 0 | 0.05266278 | 0 | 0 | 0 | 0.08942505 |
| ENSMUSG00 | Gm10241 | 572 | 0 | 0 | 0 | 0 | 0 | 0 |
| ENSMUSG00 | Gm14097 | 468 | 0 | 0 | 0 | 0 | 0 | 0 |
| ENSMUSG00 | Gm11468 | 380 | 0 | 0 | 0 | 0 | 0 | 0 |
| ENSMUSG00 | Gm16712 | 1957 | 0.040629 | 0.09849043 | 0.11293266 | 0.07374971 | 0.15644264 | 0.11149572 |
| ENSMUSG00 | Pabpc1l2b-ps | 1291 | 0 | 0 | 0 | 0 | 0 | 0 |
| ENSMUSG00 | B230312C02 | 1696 | 0.21096658 | 0.48300089 | 0.31274889 | 0.04254958 | 0 | 0 |
| ENSMUSG00 | Hoxb3os | 2997 | 2.08261943 | 1.39880563 | 1.38637746 | 1.58919916 | 1.02154907 | 3.81317112 |
| ENSMUSG00 | Gm6151 | 5566 | 0.18570652 | 0.27703309 | 0.13500383 | 0.22040775 | 0.41253807 | 0.65172961 |
| ENSMUSG00 | A730011C13 | 3297 | 1.91723427 | 1.87075056 | 1.90374938 | 1.78385601 | 0.74287717 | 2.16741143 |
| ENSMUSG00 | Gm15507 | 1629 | 0 | 0 | 0 | 0 | 0 | 0 |
| ENSMUSG00 | 4933408N05 | 1581 | 0 | 0 | 0 | 0 | 0 | 0 |
| ENSMUSG00 | Gm16105 | 695 | 0 | 0 | 0 | 0 | 0 | 0 |
| ENSMUSG00 | Gm12092 | 597 | 0 | 6.21500175 | 0 | 6.40652717 | 0.68377053 | 0 |
| ENSMUSG00 | Gm11267 | 1067 | 0 | 0 | 0 | 0 | 0 | 0 |
| ENSMUSG00 | Gm16359 | 288 | 0 | 0 | 0 | 0 | 0 | 0 |
| ENSMUSG00 | Gm11791 | 334 | 0 | 0 | 0 | 0 | 0 | 0 |
| ENSMUSG00 | Gm12678 | 690 | 0 | 0 | 0 | 0 | 0 | 0 |
| ENSMUSG00 | Gm9850 | 226 | 0 | 0 | 0 | 0 | 0 | 0 |
| ENSMUSG00 | Gm11528 | 911 | 0 | 0 | 0 | 0 | 0 | 0 |
| ENSMUSG00 | Gm2449 | 582 | 0.40985032 | 0 | 0 | 0.43397648 | 0.08767419 | 0.18745457 |
| ENSMUSG00 | Gm1980 | 2304 | 0 | 0 | 0 | 0 | 0 | 0 |
| ENSMUSG00 | 1700080N15 | 1585 | 0 | 0 | 0 | 0 | 0 | 0 |
| ENSMUSG00 | 2610204G07 | 990 | 0.0803141 | 0 | 0.13394498 | 0 | 0.46387614 | 0 |
| ENSMUSG00 | 1700095J07 | 1974 | 0 | 0 | 0 | 0 | 0 | 0 |
| ENSMUSG00 | Gm16278 | 2069 | 0.1345038 | 0.27947671 | 0.10681934 | 0.47086283 | 1.38109089 | 0.38229316 |
| ENSMUSG00 | Gm12523 | 597 | 0.33296047 | 0.16142862 | 0.14807988 | 0.06043894 | 0.5128279 | 0.13705849 |
| ENSMUSG00 | 1700027A07 | 456 | 0 | 0.21134404 | 0 | 0.31650916 | 0.11189995 | 0.23925122 |
| ENSMUSG00 | Gm15723 | 598 | 0 | 0 | 0 | 0 | 0 | 0 |
| ENSMUSG00 | A930006K02 | 774 | 0.30818202 | 0.06225639 | 1.8845747 | 1.02558783 | 1.5822132 | 0.66953249 |
| ENSMUSG00 | Enox | 3390 | 0.12900008 | 0.12792861 | 0.29989451 | 0.14901139 | 0.33114462 | 0.08045616 |
| ENSMUSG00 | Gm16831 | 1335 | 0 | 0 | 0 | 0 | 0 | 0 |
| ENSMUSG00 | Gm13063 | 667 | 0 | 0 | 0 | 0 | 0 | 0.04089151 |
| ENSMUSG00 | Gm13954 | 541 | 0 | 0 | 0 | 0 | 0 | 0 |
| ENSMUSG00 | Gm2800 | 639 | 0 | 0 | 0 | 0 | 0 | 0 |
| ENSMUSG00 | Gm15252 | 882 | 0 | 0 | 0 | 0 | 0 | 0 |
| ENSMUSG00 | C030047K22 | 648 | 0 | 0 | 0 | 0 | 0 | 0 |
| ENSMUSG00 | Gm12906 | 1475 | 0.05390574 | 0 | 0.17980411 | 0.0244624 | 0.03459415 | 0 |
| ENSMUSG00 | 4930570D08 | 194 | 0 | 0 | 0 | 0 | 0 | 0 |
| ENSMUSG00 | Gm14965 | 1229 | 0 | 0 | 0 | 0 | 0 | 0 |
| ENSMUSG00 | 5430416N02 | 3686 | 1.33740629 | 0.69285986 | 1.15121458 | 0.95931643 | 0.83059755 | 1.24311975 |
| ENSMUSG00 | Gm15817 | 584 | 0.13614891 | 0.24753309 | 0.22706426 | 0 | 0.87373931 | 0 |
| ENSMUSG00 | Gm14872 | 2941 | 0.27035349 | 0.04915312 | 0.43585632 | 0.26990989 | 0.3990502 | 0.28749195 |
| ENSMUSG00 | Gm15585 | 666 | 2.50710237 | 1.01292821 | 1.79196661 | 2.87139393 | 1.07262652 | 1.43335191 |
| ENSMUSG00 | Gm16047 | 997 | 0 | 0 | 0 | 0.07238123 | 0.05117992 | 0 |
| ENSMUSG00 | Ccdc85c | 1541 | 0.54176839 | 0.71920063 | 0.659729 | 0.79609962 | 0.66225017 | 0.24779036 |
| ENSMUSG00 | Gm12289 | 2233 | 0.87237732 | 0.62579795 | 0.41569131 | 1.56737945 | 0.75408437 | 0.24428696 |
| ENSMUSG00 | 3010001F23 | 1807 | 0.46201721 | 0.23999688 | 0.17123016 | 0.27955098 | 0.1976672 | 0.37734697 |
| ENSMUSG00 | 4732487G21 | 2660 | 0.97146851 | 0.83329938 | 0.81424448 | 1.18012702 | 2.32112461 | 1.7226088 |
| ENSMUSG00 | Gm11749 | 182 | 0 | 0 | 0 | 0 | 0 | 0 |
| ENSMUSG00 | Gm16908 | 1775 | 0 | 0.02714729 | 0 | 0 | 0.28747254 | 0 |
| ENSMUSG00 | Gm12148 | 290 | 0 | 0 | 0 | 0 | 0 | 0 |
| ENSMUSG00 | A830036E02 | 5244 | 0 | 0 | 0 | 0 | 0 | 0 |
| ENSMUSG00 | Gm12958 | 672 | 0 | 0 | 0 | 0 | 0 | 0 |
| ENSMUSG00 | Gm14471 | 2356 | 0 | 0 | 0 | 0 | 0 | 0 |
| ENSMUSG00 | Hba-ps4 | 444 | 0 | 0 | 0 | 1.05645626 | 0 | 0 |
| ENSMUSG00 | Gm13834 | 135 | 0 | 0 | 0 | 0 | 0.37797315 | 0 |
| ENSMUSG00 | AA672651 | 487 | 0 | 0 | 0 | 0 | 0 | 0 |
| ENSMUSG00 | Gm11632 | 672 | 0 | 0 | 0 | 0 | 0 | 0 |
| ENSMUSG00 | Gm14226 | 3241 | 0 | 0 | 0 | 0 | 0 | 0 |
| ENSMUSG00 | Gm12371 | 1089 | 0 | 0 | 0 | 0 | 0 | 0 |
| ENSMUSG00 | Gm15344 | 419 | 3.70039079 | 1.15003442 | 3.16480976 | 10.7643331 | 6.94153561 | 6.63964965 |
| ENSMUSG00 | Gm16625 | 1652 | 0 | 0 | 0 | 0 | 0.03088764 | 0 |
| ENSMUSG00 | Gm13266 | 604 | 0 | 0 | 0 | 0 | 0 | 0 |
| ENSMUSG00 | Gm281 | 2493 | 0 | 0 | 0 | 0 | 0 | 0 |
| ENSMUSG00 | Gm16624 | 5828 | 0.22510825 | 0.27284705 | 0.21236301 | 0 | 0.01751077 | 0.00935986 |
| ENSMUSG00 | Gm14827 | 3085 | 0.10309363 | 0.34363103 | 0.28655976 | 0.39766273 | 0.11578108 | 0.06188735 |
| ENSMUSG00 | Gm12081 | 498 | 0 | 0 | 0 | 0 | 0 | 0 |
| ENSMUSG00 | Gm13794 | 465 | 0 | 0 | 0 | 0 | 0 | 0 |
| ENSMUSG00 | Gm14505 | 829 | 0 | 0 | 0 | 0 | 0 | 0 |
| ENSMUSG00 | C79798 | 1791 | 0 | 0 | 0 | 0 | 0 | 0 |
| ENSMUSG00 | 4930528G23 | 1719 | 0 | 0 | 0 | 0 | 0 | 0 |
| ENSMUSG00 | C630043F03 | 2578 | 2.26689513 | 2.28035141 | 5.58952708 | 3.17712339 | 2.25640297 | 3.22682893 |
| ENSMUSG00 | Gm16185 | 3205 | 0.29770095 | 0.87201674 | 1.06194755 | 0.99070824 | 0.76420157 | 1.25097409 |
| ENSMUSG00 | Gm16078 | 409 | 0 | 0 | 0 | 0.44110079 | 0.12475887 | 0.06668616 |
| ENSMUSG00 | Gm7616 | 781 | 0 | 0 | 0 | 0 | 0 | 0 |
| ENSMUSG00 | Gm16786 | 2181 | 0 | 0 | 0 | 0 | 0 | 0 |
| ENSMUSG00 | C230037L18 | 2206 | 0.63075332 | 0.74267409 | 0.1803339 | 1.21036777 | 0.55513736 | 0.90256059 |
| ENSMUSG00 | Gm11656 | 751 | 0 | 0 | 0 | 0 | 0 | 0 |
| ENSMUSG00 | Gm17477 | 2579 | 0.58577288 | 0.7473683 | 0.18853054 | 1.56695967 | 0.55398935 | 0.41245092 |
| ENSMUSG00 | Gm12708 | 1330 | 0 | 0 | 0 | 0 | 0 | 0 |
| ENSMUSG00 | C330008G21 | 627 | 0 | 0 | 0 | 0 | 0 | 0 |
| ENSMUSG00 | Gm15230 | 504 | 0 | 0 | 0.78931863 | 3.15001976 | 0 | 0 |
| ENSMUSG00 | Gm13838 | 529 | 0 | 0 | 0 | 0 | 0 | 0 |
| ENSMUSG00 | 9330179D12 | 1482 | 0 | 0 | 0 | 0 | 0 | 0 |
| ENSMUSG00 | Gm15611 | 404 | 0 | 0 | 0 | 0 | 0 | 0 |
| ENSMUSG00 | Vmn1r-ps103 | 1863 | 0 | 0 | 0 | 0 | 0 | 0 |
| ENSMUSG00 | 1810062O18 | 1612 | 0.36993313 | 0.44838501 | 0.21936399 | 0.47005145 | 0.56977343 | 0.33839503 |
| ENSMUSG00 | Gm5724 | 2435 | 0 | 0 | 0 | 0 | 0 | 0 |
| ENSMUSG00 | Asfx3 | 1146 | 0 | 0 | 0 | 0 | 0 | 0 |
| ENSMUSG00 | 4732418A04 | 2289 | 0 | 0 | 0 | 0 | 0 | 0 |
| ENSMUSG00 | Gm11706 | 313 | 0.25402863 | 0.30790059 | 0 | 0 | 0 | 0.69711538 |
| ENSMUSG00 | 1110019D14 | 1776 | 7.25268901 | 6.75586942 | 7.86474234 | 6.23715522 | 4.9417436 | 9.81334149 |
| ENSMUSG00 | Gm15156 | 415 | 0.19159268 | 0 | 0.10651047 | 0 | 0 | 0.13144405 |
| ENSMUSG00 | Gm13962 | 606 | 0 | 0 | 0 | 0 | 0 | 0 |
| ENSMUSG00 | Gm16035 | 399 | 5.57971656 | 7.48761758 | 3.21266528 | 7.32492633 | 5.75485442 | 15.0386482 |
| ENSMUSG00 | Gm14161 | 337 | 0 | 0 | 0 | 0 | 0 | 0 |
| ENSMUSG00 | Gm11583 | 464 | 0 | 0 | 0 | 0 | 0 | 0 |
| ENSMUSG00 | Gm11228 | 2010 | 0 | 0 | 0 | 0.01795127 | 0 | 0 |
| ENSMUSG00 | BB557941 | 1094 | 0 | 0.5725994 | 0 | 0 | 0 | 0 |
| ENSMUSG00 | Gm830 | 3499 | 0 | 0 | 0 | 0 | 0.6562409 | 0 |
| ENSMUSG00 | Gm15935 | 1068 | 0 | 0 | 0 | 0 | 0 | 0 |
| ENSMUSG00 | Gm11944 | 867 | 2.29270361 | 0.88925383 | 2.03930072 | 2.12247321 | 2.11874225 | 2.13918739 |
| ENSMUSG00 | 7330404K18 | 485 | 0 | 0 | 0 | 0 | 0 | 0.28118185 |
| ENSMUSG00 | Gm13014 | 343 | 0 | 0 | 0.25773669 | 0 | 0 | 0 |
| ENSMUSG00 | Gm14684 | 121 | 0 | 0 | 0 | 0 | 0 | 0 |
| ENSMUSG00 | C030037F17 | 403 | 0 | 0 | 0 | 0 | 0 | 0 |
| ENSMUSG00 | Dlx1as | 1624 | 0.29375971 | 0.94948655 | 0.81653651 | 0.26661609 | 0.94260547 | 0.57102077 |
| ENSMUSG00 | Gm15594 | 2213 | 0.21557423 | 0 | 0 | 0 | 0.20751802 | 0 |
| ENSMUSG00 | 1700061H18 | 1078 | 0 | 0 | 0.04100357 | 0 | 0.1893372 | 0 |
| ENSMUSG00 | Gm16249 | 1255 | 0 | 0 | 0 | 0 | 0 | 0 |
| ENSMUSG00 | Gm5577 | 2096 | 0 | 0 | 0 | 0.3270796 | 0.73033935 | 0.01301271 |
| ENSMUSG00 | Gm15919 | 608 | 0 | 0 | 0 | 0 | 0 | 0 |
| ENSMUSG00 | 7530420F21 | 4284 | 0 | 0 | 0 | 0 | 0 | 0 |
| ENSMUSG00 | Gm16351 | 246 | 0.16160764 | 0 | 0 | 0 | 0 | 0 |
| ENSMUSG00 | Gm14349 | 900 | 0 | 0 | 0 | 0 | 0 | 0 |
| ENSMUSG00 | Gm16194 | 793 | 0 | 0 | 0 | 0 | 0 | 0 |
| ENSMUSG00 | Bbip1 | 4473 | 12.0875148 | 12.3563267 | 15.7419039 | 14.4957375 | 11.3620099 | 15.5184344 |
| ENSMUSG00 | 4930558J18 | 2008 | 0 | 0 | 0 | 0 | 0 | 0.05433195 |
| ENSMUSG00 | 4933407I08 | 1292 | 0 | 0 | 0 | 0 | 0 | 0 |
| ENSMUSG00 | B430010I23 | 3349 | 0.78347618 | 0.21582461 | 0.81830823 | 1.21745925 | 0.33519865 | 0.1140176 |
| ENSMUSG00 | 4933416M07 | 1481 | 0 | 0.09760927 | 0.05969189 | 0 | 0.17227001 | 0.1657473 |
| ENSMUSG00 | Gm11947 | 271 | 0 | 0 | 0 | 0 | 0 | 0 |
| ENSMUSG00 | Gm14936 | 2617 | 0 | 0 | 0 | 0 | 0 | 0 |
| ENSMUSG00 | Gm15503 | 447 | 0 | 0 | 0 | 0 | 0 | 0 |
| ENSMUSG00 | Gm13857 | 3371 | 0.01179338 | 0.02858881 | 0.05244953 | 0.03211099 | 0.03027373 | 0.02427289 |
| ENSMUSG00 | 2810471M01 | 1594 | 0 | 0 | 0 | 0 | 0 | 0 |
| ENSMUSG00 | Gm12296 | 3217 | 0 | 0 | 0 | 0 | 0 | 0 |
| ENSMUSG00 | Gm12743 | 3696 | 0.24739612 | 0.24771169 | 0.22722809 | 0.12691195 | 0.08283503 | 0.04427701 |
| ENSMUSG00 | Gm11205 | 428 | 0 | 0 | 0 | 0 | 0 | 0 |
| ENSMUSG00 | 1700060J05 | 1058 | 0 | 1.22971072 | 0 | 0.03410401 | 1.15749813 | 0.46402978 |
| ENSMUSG00 | 4930579D09 | 467 | 0 | 0 | 0 | 0 | 0 | 0 |
| ENSMUSG00 | 9030407P20 | 3885 | 0.98237481 | 1.46357791 | 1.2970425 | 1.60674227 | 1.93072773 | 2.12019075 |
| ENSMUSG00 | Gm13848 | 2044 | 0 | 0 | 0 | 0 | 0 | 0 |
| ENSMUSG00 | Gm15567 | 2128 | 0.26154921 | 0 | 0.56083166 | 0.30520526 | 0.98312096 | 0 |
| ENSMUSG00 | Gm16890 | 1452 | 0.16427885 | 0 | 0 | 0 | 0 | 0 |
| ENSMUSG00 | Gm16889 | 4781 | 0 | 0 | 0 | 0 | 0 | 0 |
| ENSMUSG00 | 4933417D19 | 1226 | 0.25941586 | 1.25772117 | 0.21632223 | 0.05886141 | 0.91564459 | 0.64515868 |
| ENSMUSG00 | Gm11655 | 760 | 1.77853465 | 0.88764499 | 1.1632064 | 1.47176761 | 0.60425971 | 2.15326099 |
| ENSMUSG00 | Gm16267 | 729 | 0 | 0 | 0 | 0 | 0 | 0 |
| ENSMUSG00 | 4921508A21 | 1979 | 0 | 0 | 0 | 0 | 0 | 0 |
| ENSMUSG00 | Gm15962 | 874 | 0 | 0 | 0 | 0 | 0 | 0 |
| ENSMUSG00 | Gm15275 | 346 | 0 | 0 | 0 | 0 | 0 | 0 |
| ENSMUSG00 | Gm11789 | 1885 | 8.37290491 | 7.92456078 | 16.6255208 | 8.46061734 | 5.38686938 | 11.3005269 |
| ENSMUSG00 | 1700006P19 | 1095 | 0.290451 | 0.13201765 | 0.48440376 | 1.44987211 | 0.27959658 | 0.22417512 |
| ENSMUSG00 | Gm16135 | 566 | 0 | 0 | 0.39047565 | 0 | 0 | 0.48188409 |
| ENSMUSG00 | Gm16224 | 410 | 0 | 0 | 0 | 0 | 0 | 0 |
| ENSMUSG00 | Gm13134 | 1121 | 0.0709286 | 0 | 0 | 0 | 0.09103725 | 0 |
| ENSMUSG00 | E030010N08 | 3127 | 0 | 0 | 0 | 0 | 0 | 0 |
| ENSMUSG00 | Gm14549 | 1033 | 0 | 0 | 0 | 0 | 0 | 0 |
| ENSMUSG00 | Gm16260 | 303 | 0 | 0 | 0 | 0 | 0 | 0 |
| ENSMUSG00 | Gm13842 | 1383 | 0 | 0 | 0 | 0 | 0 | 0 |
| ENSMUSG00 | Gm12305 | 405 | 0 | 0.11897887 | 0.21828071 | 0 | 0 | 0 |
| ENSMUSG00 | Gm16352 | 620 | 0 | 0 | 0 | 0 | 0 | 0 |
| ENSMUSG00 | 4930596M17 | 1021 | 0 | 0 | 0 | 0 | 0 | 0 |
| ENSMUSG00 | Gm11419 | 680 | 0 | 0 | 0 | 0 | 0 | 0 |
| ENSMUSG00 | Gm15471 | 1296 | 0 | 0 | 0 | 0 | 0 | 0 |
| ENSMUSG00 | Gm16279 | 2940 | 1.77141767 | 1.11451635 | 0.88704379 | 2.51592487 | 1.61409964 | 0.89987755 |
| ENSMUSG00 | Gm15839 | 610 | 0 | 0 | 0 | 0 | 0 | 0 |
| ENSMUSG00 | Gm14553 | 1113 | 0 | 0 | 0 | 0 | 0 | 0 |
| ENSMUSG00 | BB114351 | 2114 | 0 | 0.0683819 | 0 | 0 | 0.09654943 | 0.02580382 |
| ENSMUSG00 | Gm12984 | 270 | 0 | 0 | 0 | 0 | 0 | 0 |
| ENSMUSG00 | 4930418C01 | 680 | 0 | 0.28344966 | 0 | 0.15918549 | 0 | 0 |
| ENSMUSG00 | 5430427M07 | 217 | 0 | 0 | 0 | 0 | 0 | 0 |
| ENSMUSG00 | BC021767 | 1924 | 0.0619888 | 0.02504493 | 0 | 0 | 0.10608394 | 0.1842881 |
| ENSMUSG00 | Gm11549 | 2828 | 0 | 0 | 0 | 0 | 0 | 0 |
| ENSMUSG00 | Gm13399 | 4013 | 0 | 0 | 0 | 0 | 0 | 0 |
| ENSMUSG00 | Gm12977 | 1165 | 1.91099306 | 1.36493785 | 0.91059591 | 1.70344416 | 1.05118714 | 1.80270148 |
| ENSMUSG00 | 0610038P03 | 714 | 0 | 0 | 0 | 0 | 0 | 0 |
| ENSMUSG00 | Gm16880 | 1194 | 1.73139446 | 0.64571447 | 0 | 0.36263361 | 2.00857593 | 0.29696006 |
| ENSMUSG00 | Gm15492 | 603 | 0.59336538 | 0.31964472 | 0.659729 | 0 | 0.16924171 | 0 |
| ENSMUSG00 | 4930556M19 | 4027 | 1.92508535 | 1.75897864 | 0.91103873 | 2.13248736 | 3.29447671 | 1.55100382 |
| ENSMUSG00 | Gm13490 | 2402 | 0 | 0 | 0 | 0 | 0 | 0 |
| ENSMUSG00 | Gm14424 | 893 | 0 | 0 | 0.09899629 | 0 | 0 | 0.21379896 |
| ENSMUSG00 | Gm11335 | 1078 | 1.99146192 | 0.17879941 | 1.06609269 | 0.76983954 | 0.2366715 | 0 |
| ENSMUSG00 | Gm13412 | 675 | 0 | 0 | 0 | 0 | 0.37797315 | 0 |
| ENSMUSG00 | Gm15257 | 882 | 0 | 0 | 0 | 0 | 0 | 0 |
| ENSMUSG00 | Gm12747 | 769 | 0 | 0 | 0 | 0 | 0 | 0 |
| ENSMUSG00 | 2310081O03 | 757 | 0.05251715 | 0 | 0 | 0 | 0 | 0 |
| ENSMUSG00 | Gm12051 | 1338 | 0 | 0 | 0 | 0 | 0 | 0.02038463 |
| ENSMUSG00 | Gm5860 | 2817 | 0 | 0.13684471 | 0.54918868 | 0.5123471 | 0.41661578 | 0.16459669 |
| ENSMUSG00 | Gm12744 | 517 | 0 | 0 | 0 | 0 | 0 | 0 |
| ENSMUSG00 | C230035I16 | 910 | 2.92705186 | 4.13026647 | 4.51733121 | 3.13239727 | 4.4297623 | 8.15241964 |
| ENSMUSG00 | Gm13715 | 408 | 0 | 0 | 0 | 0 | 0 | 0 |
| ENSMUSG00 | 4930521C21 | 1269 | 0.06265639 | 0.03797198 | 0 | 0.05686689 | 0.04020991 | 0 |
| ENSMUSG00 | Gm11840 | 694 | 0 | 0 | 0 | 0 | 0 | 0 |
| ENSMUSG00 | Slc2a4rg-ps | 2231 | 1.15827263 | 2.80781599 | 1.22837932 | 4.10795128 | 2.60735403 | 2.02939942 |
| ENSMUSG00 | Gm13569 | 572 | 0 | 0 | 0.23182785 | 0 | 0.3568278 | 0 |
| ENSMUSG00 | 2810455O05 | 1689 | 0.37660609 | 0.08558871 | 0.57574929 | 0 | 0 | 0.74282617 |
| ENSMUSG00 | Gm16982 | 2616 | 0.50150262 | 0.34997798 | 0.25345094 | 0.27585661 | 0.37060441 | 0.13553911 |
| ENSMUSG00 | Gm11217 | 1725 | 0 | 0 | 0 | 0 | 0 | 0 |
| ENSMUSG00 | Gm11646 | 395 | 0.30194036 | 0.24398199 | 1.7904544 | 0.1826939 | 0 | 0 |
| ENSMUSG00 | Gm11240 | 748 | 0 | 0 | 0.70912048 | 0 | 0 | 0 |
| ENSMUSG00 | Gm12031 | 625 | 0 | 0 | 0 | 0 | 0 | 0 |
| ENSMUSG00 | Gm14011 | 300 | 0.79510961 | 0.16062147 | 0.29467895 | 0 | 0 | 1.45464743 |
| ENSMUSG00 | 4933421O10 | 3845 | 2.23333779 | 1.52893263 | 0.68975568 | 0.65689028 | 2.44183437 | 1.7591965 |
| ENSMUSG00 | Gm15927 | 1726 | 0.8522322 | 1.22839134 | 0.99876702 | 1.10796545 | 0.91646446 | 0.20542892 |
| ENSMUSG00 | A230028O05 | 2790 | 0.05699711 | 0 | 0 | 0.02586527 | 0 | 0.17596542 |
| ENSMUSG00 | BB031773 | 3651 | 0 | 0 | 0 | 0 | 0 | 0 |
| ENSMUSG00 | 2010001M06 | 908 | 0.04378357 | 0 | 0 | 0.03973793 | 0 | 0 |
| ENSMUSG00 | Gm13184 | 624 | 0 | 0.46333117 | 0.07083629 | 0.23129516 | 0.57241127 | 0 |
| ENSMUSG00 | 1700109G15 | 381 | 0 | 0.3794208 | 0 | 0 | 0 | 0 |
| ENSMUSG00 | 4921522H14 | 2389 | 0.24961583 | 0.2622117 | 0.05550671 | 0.07551705 | 1.53783971 | 0.76492291 |
| ENSMUSG00 | Gm11467 | 672 | 0 | 0 | 0 | 0 | 0 | 0 |
| ENSMUSG00 | Gm13026 | 606 | 0 | 0 | 0 | 0.05954133 | 0 | 0 |
| ENSMUSG00 | Gm15537 | 482 | 0 | 0 | 0 | 0 | 0 | 0 |
| ENSMUSG00 | A230071N21 | 1280 | 0 | 0 | 0 | 0 | 0 | 0.02130831 |
| ENSMUSG00 | Gm11542 | 1509 | 0 | 0.15966349 | 0.02929214 | 0 | 0 | 0.12652252 |
| ENSMUSG00 | Gm16675 | 1314 | 0.03025531 | 0 | 0.57186555 | 0 | 0 | 0.6019517 |
| ENSMUSG00 | Gm14809 | 691 | 0 | 0 | 0 | 0 | 0 | 0 |
| ENSMUSG00 | Gm15834 | 3974 | 0.98038175 | 0.3758882 | 1.45708139 | 1.22573629 | 0.08988038 | 1.03635393 |
| ENSMUSG00 | Gm15958 | 2390 | 0.41585231 | 0 | 0.46236238 | 0 | 0.27754933 | 0.31953552 |
| ENSMUSG00 | 4930455D15 | 1611 | 0 | 0 | 0 | 0 | 0 | 0 |
| ENSMUSG00 | Gm13415 | 3416 | 0.546987 | 0.87457828 | 0.29761194 | 0.21125319 | 0.17924956 | 0.29542203 |
| ENSMUSG00 | 8030453O22 | 1870 | 0 | 0 | 0 | 0 | 0 | 0 |
| ENSMUSG00 | Gm11750 | 579 | 0 | 0 | 0 | 0 | 0 | 0 |
| ENSMUSG00 | Gm3677 | 1965 | 0 | 0 | 0 | 0 | 0 | 0 |
| ENSMUSG00 | 4933404O12 | 4096 | 0.07764742 | 1.15289827 | 1.44605639 | 0.14975458 | 0.59796534 | 1.0654156 |
| ENSMUSG00 | Gm16024 | 369 | 0 | 0 | 0 | 0 | 0 | 0.1613884 |
| ENSMUSG00 | Asfx4b | 1145 | 0 | 0 | 0 | 0 | 0 | 0 |
| ENSMUSG00 | Gm4989 | 1527 | 0 | 0 | 1.41839575 | 0 | 0 | 1.23244932 |
| ENSMUSG00 | Gm15631 | 742 | 0 | 0 | 0 | 0 | 0 | 0 |
| ENSMUSG00 | Gm15895 | 729 | 1.69056227 | 1.91688179 | 0.06063353 | 1.9798104 | 1.32990554 | 0.41155148 |
| ENSMUSG00 | Gm13111 | 3532 | 3.06157721 | 4.37934539 | 1.87720171 | 3.07494207 | 2.88937575 | 1.90737143 |
| ENSMUSG00 | Gm13211 | 693 | 0 | 0 | 0 | 0 | 0 | 0 |
| ENSMUSG00 | Gm14066 | 2051 | 0 | 0 | 0 | 0.03518483 | 0.14927267 | 0 |
| ENSMUSG00 | C920011F04 | 1050 | 0.9086967 | 0.0917837 | 0.50516392 | 0.27491082 | 0.68035168 | 0.77927541 |
| ENSMUSG00 | Gm14562 | 542 | 0 | 0 | 0 | 0 | 0 | 0 |
| ENSMUSG00 | Gm11494 | 1734 | 0 | 0 | 0 | 0 | 0 | 0 |
| ENSMUSG00 | Gm15669 | 524 | 0 | 0 | 0 | 0 | 0 | 0 |
| ENSMUSG00 | Gm16762 | 1756 | 0 | 0 | 0 | 0 | 0 | 0 |
| ENSMUSG00 | 1700049E15 | 1042 | 0 | 0 | 0 | 0 | 0 | 0.02617528 |
| ENSMUSG00 | C030013C21 | 792 | 0 | 0.60841467 | 0.16743122 | 0.22779069 | 0.19328173 | 0.44768979 |
| ENSMUSG00 | Khdc1b | 2244 | 0 | 0 | 0 | 0 | 0 | 0 |
| ENSMUSG00 | 2310044K18 | 616 | 0 | 0 | 0 | 0 | 0.16567005 | 0 |
| ENSMUSG00 | A430071A18 | 1171 | 0.33950026 | 0.74069681 | 0.52845927 | 0.58544735 | 1.17652617 | 0.279501 |
| ENSMUSG00 | Gm16330 | 296 | 0 | 0 | 0 | 0 | 0 | 0 |
| ENSMUSG00 | Gm11615 | 816 | 0.19487981 | 0 | 0 | 0.39796373 | 0.06253232 | 0.26739842 |
| ENSMUSG00 | 4930570G19 | 4497 | 0.02652134 | 0 | 0.01965837 | 0.00802358 | 0.01134676 | 0 |
| ENSMUSG00 | 1700086P04 | 1007 | 0.71062428 | 2.91894039 | 0.83399704 | 0.10749368 | 0 | 1.73344281 |
| ENSMUSG00 | Gm13528 | 408 | 0.87695913 | 1.18104025 | 0.10833785 | 1.23810937 | 0.12506465 | 0.73534567 |
| ENSMUSG00 | 4931413K12 | 1280 | 0.34164866 | 0 | 0.24172883 | 0.0563782 | 0.11959307 | 0.68186598 |
| ENSMUSG00 | A430092G05 | 1758 | 0 | 0 | 0 | 0 | 0 | 0 |
| ENSMUSG00 | Gm12711 | 338 | 0 | 0 | 0 | 0 | 0 | 0 |
| ENSMUSG00 | E230015J15 | 4061 | 0 | 0 | 0.3156497 | 0.03554006 | 0.03769493 | 0 |
| ENSMUSG00 | Gm16867 | 6098 | 0.53459321 | 0.2054522 | 0.49290346 | 0.34910473 | 0.35144437 | 0.86770745 |
| ENSMUSG00 | Gm12737 | 723 | 0.05498683 | 0 | 0 | 0 | 0 | 0.03772426 |
| ENSMUSG00 | Gm15692 | 351 | 0 | 0 | 0 | 0 | 0 | 0 |
| ENSMUSG00 | Gm15635 | 1051 | 0 | 0 | 0 | 0 | 0 | 0 |
| ENSMUSG00 | B230217O12 | 1639 | 0.89746966 | 0.55859817 | 0.75512606 | 0.57238143 | 1.52549873 | 1.04838455 |
| ENSMUSG00 | Gm13049 | 754 | 0 | 0 | 0 | 0 | 0 | 0 |
| ENSMUSG00 | Gm4221 | 3355 | 0.04739849 | 0 | 0 | 0.19358474 | 0 | 0 |
| ENSMUSG00 | Gm16288 | 366 | 0.43448613 | 0 | 0.24154013 | 0 | 0 | 0.14904175 |
| ENSMUSG00 | Gm13324 | 567 | 0 | 0 | 0 | 0 | 0 | 0 |
| ENSMUSG00 | Gm13227 | 821 | 0 | 0 | 0 | 0 | 0 | 0 |
| ENSMUSG00 | 1700010K24 | 224 | 0.7099193 | 0 | 0 | 0.16108056 | 0.22779632 | 0 |
| ENSMUSG00 | Gm16357 | 586 | 0 | 0 | 0 | 0 | 0 | 0 |
| ENSMUSG00 | Gm11731 | 667 | 0 | 0 | 0 | 0 | 0 | 0 |
| ENSMUSG00 | Gm12758 | 4173 | 0.38107338 | 0.43879339 | 0.13770045 | 0.12969822 | 1.18609117 | 0.7058857 |
| ENSMUSG00 | Gm16000 | 445 | 0 | 0 | 0 | 0 | 0 | 0 |
| ENSMUSG00 | Gm4764 | 1258 | 0 | 0.11491202 | 0 | 0 | 0 | 0 |
| ENSMUSG00 | 4930457A20 | 626 | 0 | 0 | 0 | 0 | 0 | 0 |
| ENSMUSG00 | Gm15356 | 2739 | 0 | 0 | 0 | 0 | 0 | 0 |
| ENSMUSG00 | Gm13920 | 1661 | 0 | 0 | 0 | 0 | 0 | 0 |
| ENSMUSG00 | Ascl4 | 1636 | 1.04491789 | 1.23706025 | 1.05371142 | 0.50726591 | 1.1540195 | 2.93419102 |
| ENSMUSG00 | A530072M11 | 1845 | 0.28011992 | 0.83575401 | 0.83851735 | 0.35201995 | 0.11062629 | 1.52264924 |
| ENSMUSG00 | B130011K05 | 3268 | 0.08515556 | 0 | 0 | 0.12145119 | 0.03122789 | 0 |
| ENSMUSG00 | 1600025M17 | 1148 | 0 | 0 | 0 | 0 | 0 | 0 |
| ENSMUSG00 | Gm11197 | 837 | 0 | 0 | 0 | 0 | 0 | 0 |
| ENSMUSG00 | Gm2735 | 276 | 0 | 0.34917712 | 0 | 0 | 0 | 0 |
| ENSMUSG00 | Gm15774 | 1990 | 0 | 0 | 0.15548387 | 0 | 0 | 0 |
| ENSMUSG00 | Gm13644 | 2143 | 0 | 0.02248551 | 0 | 0.23572031 | 0 | 0.17818243 |
| ENSMUSG00 | Gm16086 | 368 | 0 | 0 | 0 | 0 | 0 | 0 |
| ENSMUSG00 | 5730437C11 | 449 | 0 | 0 | 0 | 0 | 0 | 0 |
| ENSMUSG00 | 4933404K08 | 1430 | 0 | 0 | 0 | 0 | 0 | 0.01907317 |
| ENSMUSG00 | Gm15219 | 385 | 0 | 0 | 0 | 0 | 0 | 0 |
| ENSMUSG00 | Gm12766 | 1820 | 0.02184367 | 0 | 0 | 0 | 0 | 0 |
| ENSMUSG00 | Gm16070 | 8337 | 0 | 0 | 0 | 0.00432794 | 0.16525275 | 0 |
| ENSMUSG00 | Gm12589 | 635 | 0.06260706 | 0.07588416 | 0.0696092 | 0 | 0 | 0.04295219 |
| ENSMUSG00 | 4930444E06 | 1238 | 0 | 0 | 0 | 0 | 0 | 0 |
| ENSMUSG00 | Gm2115 | 1837 | 0.60596269 | 0 | 0 | 0.09820916 | 0 | 0.68297954 |
| ENSMUSG00 | 5031425F14 | 2095 | 0 | 0 | 0 | 0 | 0 | 0 |
| ENSMUSG00 | Gm11417 | 917 | 0 | 0 | 0 | 0 | 0 | 0 |
| ENSMUSG00 | Gm16739 | 408 | 1.46159855 | 2.71639257 | 1.30005421 | 1.14967299 | 0.87545253 | 2.87453306 |
| ENSMUSG00 | Gm12265 | 466 | 0.08531219 | 0 | 0 | 0 | 0 | 0 |
| ENSMUSG00 | B930095G15 | 2791 | 1.55261461 | 0.67333259 | 2.04300887 | 3.1415037 | 2.10248413 | 1.81765779 |
| ENSMUSG00 | Gm11659 | 461 | 0 | 0 | 0 | 0 | 0 | 0 |
| ENSMUSG00 | Gm13713 | 897 | 0 | 0 | 0 | 0 | 0 | 0.12162604 |
| ENSMUSG00 | Gm16740 | 952 | 0.41759959 | 0.55677612 | 0.88217964 | 0.5306183 | 0.75038788 | 1.20329291 |
| ENSMUSG00 | Gm16738 | 3038 | 0 | 0.23791858 | 0 | 0 | 0.01679604 | 0 |
| ENSMUSG00 | A730046J19 | 4704 | 0 | 0 | 0 | 0 | 0 | 0 |
| ENSMUSG00 | Gm2155 | 1972 | 0 | 0 | 0.02241473 | 0.09148591 | 0.02587544 | 0.01383095 |
| ENSMUSG00 | Gm13429 | 1402 | 0 | 0 | 0 | 0 | 0 | 0 |
| ENSMUSG00 | Gm12702 | 610 | 0 | 0 | 0 | 0 | 0 | 0 |
| ENSMUSG00 | Gm11520 | 627 | 0 | 0.69167142 | 0 | 0 | 0.4069089 | 1.08750555 |
| ENSMUSG00 | A930029G22 | 1707 | 1.97962264 | 0.73394698 | 0.49199474 | 0.19023925 | 0.92666529 | 4.33006868 |
| ENSMUSG00 | Gm11614 | 734 | 0.05416278 | 0 | 0 | 0 | 0 | 0.03715891 |
| ENSMUSG00 | Eif2c5 | 1411 | 0 | 0 | 0 | 0 | 0 | 0 |
| ENSMUSG00 | Gm12609 | 606 | 0 | 0 | 0 | 0 | 0 | 0 |
| ENSMUSG00 | 2210403K04 | 2032 | 20.2103402 | 25.3500525 | 22.1226744 | 17.2597182 | 23.0773816 | 19.6640485 |
| ENSMUSG00 | Gm16825 | 2286 | 9.46062178 | 1.62307789 | 6.3421717 | 4.00911606 | 2.09819743 | 0.64428282 |
| ENSMUSG00 | Gm16346 | 934 | 0 | 0 | 0 | 0 | 0 | 0 |
| ENSMUSG00 | 1110018N20 | 1574 | 0 | 0 | 0 | 0 | 0 | 0 |
| ENSMUSG00 | Gm11496 | 579 | 0 | 0 | 0 | 0 | 0 | 0 |
| ENSMUSG00 | B230314M03 | 1203 | 0 | 0 | 0 | 0 | 0 | 0 |
| ENSMUSG00 | C130046K22 | 777 | 1.58612599 | 3.34886471 | 2.90127927 | 1.53244205 | 1.64177528 | 1.19348486 |
| ENSMUSG00 | Gm11780 | 515 | 0 | 0 | 0 | 0 | 0 | 0 |
| ENSMUSG00 | Gm11974 | 1580 | 2.81811001 | 2.71429959 | 2.93746425 | 3.28849013 | 1.93771047 | 3.3489114 |
| ENSMUSG00 | 4930515L19 | 766 | 0 | 0 | 0 | 0 | 0 | 0 |
| ENSMUSG00 | Gm14264 | 1861 | 0 | 0 | 0 | 0 | 0 | 0 |
| ENSMUSG00 | Gm16862 | 4578 | 0 | 0 | 0 | 0 | 0 | 0 |
| ENSMUSG00 | Gm13617 | 673 | 0 | 0 | 0 | 0 | 0 | 0 |
| ENSMUSG00 | Gm7580 | 493 | 0.08063992 | 0.97741262 | 0 | 0.07318873 | 0.51750888 | 0.11064762 |
| ENSMUSG00 | Gm12295 | 3578 | 0 | 0 | 0 | 0 | 0 | 0 |
| ENSMUSG00 | Gm16705 | 2092 | 0 | 0 | 0 | 0 | 0 | 0 |
| ENSMUSG00 | Gm16704 | 3832 | 0 | 0 | 0 | 0 | 0 | 0 |
| ENSMUSG00 | Gm12089 | 426 | 0 | 0 | 1.03760195 | 0.08469964 | 0 | 0 |
| ENSMUSG00 | Gm15522 | 783 | 0 | 0 | 0.28225953 | 0 | 0 | 0.52250267 |
| ENSMUSG00 | Gm12063 | 713 | 0 | 0 | 0 | 0 | 0 | 0 |
| ENSMUSG00 | Gm16252 | 853 | 0 | 0 | 0 | 0 | 0 | 0 |
| ENSMUSG00 | Gm10785 | 1577 | 0.6806582 | 0.55000378 | 0.47649419 | 0.20592162 | 0.32356611 | 1.0031256 |
| ENSMUSG00 | 4930557C09 | 775 | 0 | 0 | 0 | 0 | 0 | 0 |
| ENSMUSG00 | D830026I12 | 2106 | 0.52856289 | 0 | 0 | 0 | 0 | 0.84180985 |
| ENSMUSG00 | Gm6542 | 771 | 0.92814351 | 0.4374904 | 0.28665268 | 1.12317648 | 0.6618207 | 0.38913234 |
| ENSMUSG00 | Gm2869 | 2620 | 0 | 0 | 0 | 0 | 0 | 0 |
| ENSMUSG00 | Gm16206 | 1964 | 0.58702084 | 0 | 0.04501206 | 0.2755757 | 0.10392337 | 0.52771705 |
| ENSMUSG00 | Gm11423 | 987 | 5.71963347 | 5.80971288 | 9.58378359 | 5.19113508 | 5.48003631 | 16.082918 |
| ENSMUSG00 | Gm15397 | 802 | 0 | 0 | 0 | 0 | 0 | 0 |
| ENSMUSG00 | Gm11209 | 429 | 0.46335059 | 0 | 0 | 0 | 0 | 0 |
| ENSMUSG00 | 1700064E03 | 838 | 0.28464545 | 0.69002065 | 0.42197464 | 0.99031864 | 0.12178133 | 0.65094604 |
| ENSMUSG00 | Gm13645 | 697 | 0 | 0 | 0 | 0 | 0 | 0 |
| ENSMUSG00 | AI838599 | 4237 | 6.21150061 | 5.22010313 | 1.70047213 | 7.02565181 | 5.35915441 | 7.04235434 |
| ENSMUSG00 | Gm12709 | 1908 | 0.06250862 | 0 | 0 | 0 | 0 | 0 |
| ENSMUSG00 | Gm13596 | 290 | 0 | 0 | 0 | 0 | 0 | 0 |
| ENSMUSG00 | Gm12603 | 757 | 0 | 0 | 0 | 0 | 0 | 0 |
| ENSMUSG00 | 4933439K11 | 2941 | 0.01351767 | 0.2457656 | 0.3156201 | 0.44167073 | 0.01735001 | 0.62135356 |
| ENSMUSG00 | BC028777 | 1109 | 0.60941674 | 0 | 0.07971478 | 0 | 0.41410044 | 0.44269027 |
| ENSMUSG00 | Gm14377 | 2091 | 0 | 0 | 0 | 0 | 0 | 0 |
| ENSMUSG00 | Gm13860 | 449 | 0.35416909 | 0 | 0 | 0 | 0 | 0 |
| ENSMUSG00 | Gm11368 | 605 | 0 | 0 | 0 | 0 | 0 | 0 |
| ENSMUSG00 | Gm11963 | 497 | 0.31996363 | 0.38781845 | 0 | 0.36299844 | 0.20533753 | 0 |
| ENSMUSG00 | 3010027C24 | 1914 | 0 | 0 | 0 | 0.11310986 | 0 | 0 |
| ENSMUSG00 | Gm16847 | 2865 | 0.52729782 | 0.6391221 | 0.27770791 | 1.96467677 | 0.58773836 | 1.05671377 |
| ENSMUSG00 | Gm12195 | 467 | 0.93642459 | 1.13501256 | 0 | 1.23621566 | 0.76484932 | 1.63531028 |
| ENSMUSG00 | Gm13322 | 244 | 0.65172919 | 0 | 0 | 0.29575446 | 0.41824898 | 0 |
| ENSMUSG00 | Gm16012 | 392 | 0 | 0 | 0 | 0 | 0 | 0 |
| ENSMUSG00 | 6430562O15 | 5689 | 0.27952526 | 0.05929075 | 0.03107881 | 0.12684846 | 0.16144749 | 0.03835421 |
| ENSMUSG00 | Gm14963 | 2556 | 0.04666136 | 0.01885229 | 0 | 0.09881624 | 0.17967034 | 0.01067083 |
| ENSMUSG00 | 1700125H03 | 736 | 0 | 0 | 0 | 0 | 0 | 0 |
| ENSMUSG00 | 4933416E03 | 1119 | 0 | 0 | 0 | 0 | 0 | 0 |
| ENSMUSG00 | Gm15346 | 473 | 0 | 0 | 0 | 0 | 0.10787817 | 0 |
| ENSMUSG00 | 4930552P12 | 2492 | 0.06381297 | 0 | 0.23058746 | 0 | 0 | 0.06566928 |
| ENSMUSG00 | Gm13507 | 1026 | 0.7362126 | 0.14089603 | 0.81855265 | 0.42201222 | 0.9946662 | 1.40892386 |
| ENSMUSG00 | Gm15620 | 268 | 0 | 0 | 0 | 0 | 0 | 0 |
| ENSMUSG00 | Gm12927 | 371 | 0 | 0.12988259 | 0 | 0.09725619 | 0 | 0.51461584 |
| ENSMUSG00 | Gm15327 | 749 | 3.98085586 | 2.57337475 | 1.82944878 | 2.45685483 | 2.9294181 | 6.77314139 |
| ENSMUSG00 | Gm13185 | 942 | 0 | 0 | 0.1876936 | 0 | 0 | 0.08686191 |
| ENSMUSG00 | Gm13380 | 716 | 0 | 0 | 0.61734418 | 0.50393917 | 0.14253178 | 0 |
| ENSMUSG00 | Gm11767 | 490 | 8.03222974 | 6.88377745 | 6.13413332 | 7.29004573 | 6.76880496 | 9.07299224 |
| ENSMUSG00 | 4632419I22 | 6660 | 7.94512681 | 7.69101922 | 6.63027645 | 9.43225819 | 6.59665309 | 6.19617555 |
| ENSMUSG00 | Gm12637 | 365 | 0 | 0 | 0 | 0 | 0 | 0 |

| | | | | | | | | |
|---|---|---|---|---|---|---|---|---|
| ENSMUSG00 | Gm16615 | 3243 | 0 | 0 | 0 | 0 | 0 | 0 |
| ENSMUSG00 | B430219N15 | 833 | 0.143177 | 0.05784687 | 0 | 0 | 0 | 0.19645599 |
| ENSMUSG00 | 4931440J10R | 884 | 0 | 0 | 0 | 0 | 0 | 0 |
| ENSMUSG00 | Gm13091 | 466 | 0.59718533 | 0.93063944 | 1.13824488 | 0.92915136 | 0.43799464 | 0.23411708 |
| ENSMUSG00 | 0610005C13 | 3317 | 2.92443089 | 2.67298925 | 1.31925911 | 1.794856 | 2.56900958 | 3.45352683 |
| ENSMUSG00 | 6030471H07 | 490 | 0 | 0 | 0 | 0 | 0 | 0 |
| ENSMUSG00 | BB218582 | 1472 | 4.67234927 | 6.25245275 | 4.02380908 | 5.41720913 | 5.65033917 | 7.96745576 |
| ENSMUSG00 | Gm16617 | 710 | 0 | 0.20360469 | 0 | 0.50819781 | 0.21560441 | 0.07682997 |
| ENSMUSG00 | Gm12521 | 692 | 0 | 0 | 0 | 0 | 0 | 0 |
| ENSMUSG00 | Gm11309 | 537 | 0 | 0 | 0 | 0 | 0 | 0 |
| ENSMUSG00 | Gm13974 | 1445 | 0 | 0 | 0 | 0 | 0.07062474 | 0 |
| ENSMUSG00 | Gm11210 | 418 | 0 | 0 | 0 | 0 | 0 | 0 |
| ENSMUSG00 | Gm13425 | 671 | 0 | 0 | 0 | 0 | 0 | 0 |
| ENSMUSG00 | Gm14637 | 442 | 0 | 0 | 0 | 0 | 0 | 0 |
| ENSMUSG00 | Gm7855 | 686 | 0 | 0 | 0 | 0 | 0 | 0 |
| ENSMUSG00 | 6330418K02 | 1263 | 2.36077675 | 2.63251347 | 1.43489752 | 0.68564455 | 1.85844283 | 2.4402488 |
| ENSMUSG00 | Gm14376 | 295 | 0 | 0 | 0 | 0 | 0 | 0 |
| ENSMUSG00 | Gm11672 | 643 | 0.9892499 | 0.82434038 | 0.34371573 | 2.02014557 | 1.11099419 | 0.04241779 |
| ENSMUSG00 | Gm16013 | 351 | 0 | 0 | 0 | 0 | 0 | 0 |
| ENSMUSG00 | Gm11962 | 254 | 0.15651764 | 0 | 0 | 0.14205529 | 0.20089124 | 0.32214141 |
| ENSMUSG00 | B230208H11 | 1541 | 0.15479097 | 1.62601881 | 0.659729 | 0.21073225 | 0.69536268 | 1.15045526 |
| ENSMUSG00 | Gm15614 | 1533 | 0 | 0 | 0 | 0 | 0 | 0 |
| ENSMUSG00 | Gm12576 | 3034 | 0.32758306 | 0.31764299 | 0.36422086 | 0.42813237 | 0.33636372 | 0.93492501 |
| ENSMUSG00 | 2610206C17 | 1123 | 3.00909692 | 2.78906388 | 3.0307586 | 2.3133635 | 2.86256605 | 2.0644206 |
| ENSMUSG00 | Gm15406 | 356 | 0.1116727 | 0.5414207 | 0.12416248 | 0.40541623 | 0.14333252 | 0.30645662 |
| ENSMUSG00 | 4930479D17 | 737 | 0.48548077 | 0.45767313 | 0.23990146 | 0 | 0 | 0.29606121 |
| ENSMUSG00 | Gm12150 | 465 | 0 | 0 | 0 | 0 | 0 | 0.17596542 |
| ENSMUSG00 | Gm12119 | 697 | 0 | 0 | 0 | 0 | 0 | 0 |
| ENSMUSG00 | Snhg3 | 981 | 1.37786579 | 1.52271122 | 2.83865047 | 2.61144257 | 1.56043963 | 4.39285729 |
| ENSMUSG00 | Gm11537 | 789 | 0 | 0 | 0.11204523 | 0 | 0 | 0 |
| ENSMUSG00 | 4930480K23 | 4203 | 2.99845047 | 2.06365919 | 2.30316527 | 1.26197015 | 2.78016656 | 1.73265017 |
| ENSMUSG00 | B930086A06 | 4951 | 0 | 0 | 0 | 0 | 0 | 0 |
| ENSMUSG00 | Gm11713 | 521 | 0 | 0 | 0 | 0 | 0 | 0.52350555 |
| ENSMUSG00 | Gm15893 | 1556 | 0.53654569 | 0.1858089 | 0 | 0.11594487 | 0.0655866 | 0.08764344 |
| ENSMUSG00 | 4930545L23 | 2496 | 0 | 0 | 0.01770907 | 0 | 0 | 0 |
| ENSMUSG00 | Gm11587 | 499 | 0 | 0 | 0 | 0 | 0 | 0 |
| ENSMUSG00 | Gm12847 | 409 | 0.09720166 | 0.58907631 | 0 | 0.6175411 | 0.12475887 | 0.6668616 |
| ENSMUSG00 | Gm15234 | 1056 | 1.01647535 | 0.41067991 | 0.75344051 | 0.27334882 | 0.9180882 | 0.72319119 |
| ENSMUSG00 | Gm12326 | 736 | 0.59417158 | 0 | 0.1201137 | 0 | 0.06932932 | 0.66704281 |
| ENSMUSG00 | 8430437L04 | 1656 | 0 | 0 | 0 | 0 | 0 | 0 |
| ENSMUSG00 | Gm16833 | 200 | 0 | 0 | 0 | 0 | 0 | 0.81823918 |
| ENSMUSG00 | Gm15881 | 941 | 0 | 0 | 0 | 0 | 0 | 0 |
| ENSMUSG00 | 5330430P22 | 642 | 0.86694194 | 0.15011353 | 0.13770045 | 0.16860769 | 0.47688202 | 0.63725793 |
| ENSMUSG00 | 3010003L10 | 868 | 0 | 0 | 0 | 0 | 0 | 0 |
| ENSMUSG00 | Gm13264 | 299 | 0 | 0 | 0 | 0 | 0 | 0 |
| ENSMUSG00 | Gm4890 | 2505 | 1.06332024 | 0.4039582 | 0.47642705 | 0.11523208 | 0 | 0.96903908 |
| ENSMUSG00 | 1700013G23 | 524 | 1.28977704 | 0.18391772 | 0 | 0.82630636 | 0.68165006 | 2.02998269 |
| ENSMUSG00 | Gm13814 | 3075 | 0.46543002 | 0 | 0 | 1.14993183 | 0 | 0.42575047 |
| ENSMUSG00 | Gm11574 | 943 | 0.37942664 | 0.15329727 | 0.37498912 | 0 | 0.7034389 | 0.14461633 |
| ENSMUSG00 | 1700125G02 | 516 | 0 | 0 | 0 | 0 | 0 | 0 |
| ENSMUSG00 | Gm15581 | 5635 | 0 | 0 | 0 | 0 | 0 | 0 |
| ENSMUSG00 | 1700081H22 | 1158 | 0 | 0 | 0 | 0 | 0 | 0 |
| ENSMUSG00 | 9330111N05 | 5464 | 0 | 0 | 0 | 0 | 0 | 0.01996679 |
| ENSMUSG00 | Gm14235 | 364 | 0.43687341 | 0.26476067 | 0.48573454 | 0.39650598 | 0.14018235 | 0 |
| ENSMUSG00 | 5033428I22R | 2035 | 0 | 0 | 0 | 0 | 0 | 0 |
| ENSMUSG00 | Gm15777 | 872 | 0.36472918 | 0.22103873 | 0.81104299 | 0.28964944 | 1.22884621 | 0.81323466 |
| ENSMUSG00 | 9530052E02 | 1844 | 0.43118742 | 0.18292033 | 0.43147135 | 0.41091266 | 0.41507356 | 0.29582038 |
| ENSMUSG00 | E130006D01 | 1249 | 0 | 0 | 0 | 0 | 0 | 0 |
| ENSMUSG00 | Gm1078 | 3878 | 0 | 0 | 0.02279621 | 0 | 0.10526328 | 0 |
| ENSMUSG00 | 4930467D21 | 3023 | 0 | 0 | 0 | 0 | 0 | 0 |
| ENSMUSG00 | BC046401 | 845 | 0.70571859 | 0.17107613 | 0.94157772 | 1.70802578 | 1.20772487 | 1.48477326 |
| ENSMUSG00 | Gm14487 | 1256 | 0 | 0 | 0 | 0 | 0 | 0 |
| ENSMUSG00 | Gm15812 | 888 | 0.76108465 | 2.55040854 | 1.04531386 | 4.3070909 | 0.97685629 | 0.43000557 |
| ENSMUSG00 | Gm13943 | 341 | 0 | 0 | 0 | 0 | 0 | 0 |
| ENSMUSG00 | Gm12841 | 336 | 0 | 0 | 0 | 0 | 0 | 0 |
| ENSMUSG00 | Gm15965 | 1426 | 0 | 0 | 0 | 0.17712084 | 0 | 0 |
| ENSMUSG00 | Gm16151 | 636 | 0 | 0 | 0 | 0 | 0 | 0 |
| ENSMUSG00 | Gm13316 | 1121 | 0 | 0 | 0 | 0 | 0 | 0 |
| ENSMUSG00 | Gm15663 | 3361 | 0 | 0 | 0 | 0.13956161 | 0 | 0.1541857 |
| ENSMUSG00 | 9530027J09R | 1680 | 0.02366398 | 0 | 0.23679559 | 0.02147741 | 0.12149137 | 0.08117452 |
| ENSMUSG00 | Gm16189 | 993 | 0 | 0 | 0 | 0.2180184 | 0 | 0 |
| ENSMUSG00 | Gm14968 | 766 | 0 | 0 | 0 | 0 | 0 | 0 |
| ENSMUSG00 | 1700121F15 | 754 | 0.73816542 | 0.31953874 | 0.1758694 | 0.09570834 | 0.27069695 | 0.10851979 |
| ENSMUSG00 | 4833418N02 | 3206 | 0.27280742 | 0.3306618 | 0.2757445 | 0.03376361 | 0.1273272 | 0.33178756 |
| ENSMUSG00 | Gm16356 | 168 | 0 | 0 | 0 | 0 | 0 | 0 |
| ENSMUSG00 | Gm15337 | 3884 | 0 | 0 | 0 | 0 | 0 | 0 |
| ENSMUSG00 | Gm16129 | 484 | 0 | 0 | 0 | 0 | 0 | 0 |
| ENSMUSG00 | Gm16628 | 4352 | 1.49813851 | 0.9522137 | 0.06094004 | 0.5306183 | 0.28139546 | 0.52644065 |
| ENSMUSG00 | Gm12853 | 439 | 0 | 0 | 0 | 0 | 0 | 0 |
| ENSMUSG00 | Gm13817 | 598 | 0 | 0 | 0 | 0 | 0 | 0 |
| ENSMUSG00 | Gm16629 | 1600 | 0.12423588 | 0.12046611 | 0.0552523 | 0.42847428 | 0.12756594 | 0.23865309 |
| ENSMUSG00 | 4930430E12 | 1130 | 0 | 0 | 0 | 0.03193101 | 0 | 0 |
| ENSMUSG00 | Asbx4d | 1145 | 0 | 0 | 0 | 0 | 0 | 0 |
| ENSMUSG00 | Gm11651 | 1522 | 0 | 0 | 0 | 0 | 0 | 0 |
| ENSMUSG00 | F730035M05 | 1291 | 0 | 0 | 0 | 0 | 0 | 0 |
| ENSMUSG00 | Gm16627 | 1793 | 0.4656247 | 0 | 0.17256715 | 0 | 0.7683838 | 0.82143364 |
| ENSMUSG00 | Gm16345 | 157 | 0 | 0 | 0 | 0 | 0 | 0 |
| ENSMUSG00 | Gm12132 | 1383 | 0 | 0 | 0 | 0 | 0 | 0 |
| ENSMUSG00 | Gm14031 | 2419 | 0 | 0 | 0 | 0 | 0 | 0 |
| ENSMUSG00 | Gm16751 | 1684 | 0.37772428 | 1.17318535 | 0.60370688 | 0.47138063 | 0.42420978 | 0.25914147 |
| ENSMUSG00 | Gm12802 | 1809 | 0 | 0.02663706 | 0 | 0 | 0 | 0 |
| ENSMUSG00 | Gm12410 | 403 | 0 | 0 | 0 | 0 | 0 | 0 |
| ENSMUSG00 | Gm16750 | 1749 | 0.86375544 | 0 | 1.44054034 | 0 | 0 | 1.45028099 |
| ENSMUSG00 | Gm11525 | 2802 | 0.07094126 | 0.36114036 | 0.14197594 | 0.07726348 | 0.56453164 | 0.13627586 |
| ENSMUSG00 | Gm16748 | 2969 | 0 | 0 | 0 | 0 | 0 | 0 |
| ENSMUSG00 | Gm16749 | 1330 | 0 | 0 | 0.03323447 | 0 | 0 | 0.14355073 |
| ENSMUSG00 | Gm11491 | 1546 | 0 | 0 | 0 | 0.09335587 | 0.03300542 | 0.10585242 |
| ENSMUSG00 | Gm15445 | 601 | 1.25682884 | 0.32070843 | 2.27996195 | 1.92117375 | 2.54707367 | 0.31767467 |
| ENSMUSG00 | Gm12763 | 1935 | 0 | 0 | 0 | 0 | 0 | 0 |
| ENSMUSG00 | Gm15871 | 266 | 0 | 0 | 0 | 0 | 0 | 0 |
| ENSMUSG00 | Gm14866 | 2115 | 0 | 0 | 0 | 0 | 0 | 0 |
| ENSMUSG00 | A430018G15 | 1710 | 1.37168032 | 0.9299138 | 1.00811221 | 1.22383543 | 0.80567962 | 1.13245578 |
| ENSMUSG00 | D330050G23 | 2477 | 0 | 0 | 0 | 0 | 0 | 0 |
| ENSMUSG00 | Gm14256 | 866 | 0 | 0 | 0 | 0 | 0 | 0 |
| ENSMUSG00 | Gm12524 | 1581 | 0 | 0 | 0 | 0 | 0 | 0 |
| ENSMUSG00 | Gm13330 | 385 | 0 | 0 | 0 | 0 | 0 | 0 |
| ENSMUSG00 | Gm11548 | 2195 | 0 | 0 | 0 | 0 | 0 | 0 |
| ENSMUSG00 | 4930412F09 | 1014 | 0 | 0 | 0 | 0 | 0 | 0 |
| ENSMUSG00 | Gm14261 | 1427 | 0 | 0 | 0 | 0 | 0 | 0 |
| ENSMUSG00 | 9130410C08 | 1086 | 0 | 0 | 0 | 0 | 0 | 0 |
| ENSMUSG00 | Gm11680 | 436 | 0 | 0 | 0 | 0 | 0 | 0 |
| ENSMUSG00 | Gm14974 | 1204 | 0 | 0 | 0 | 0 | 0 | 0 |
| ENSMUSG00 | Gm15642 | 309 | 0 | 0 | 0 | 0 | 0 | 0 |
| ENSMUSG00 | Gm16104 | 399 | 0 | 0 | 0 | 0 | 0.38365696 | 0.13671498 |
| ENSMUSG00 | Gm17131 | 3845 | 0 | 0 | 0 | 0 | 0 | 0 |
| ENSMUSG00 | 2810404F17 | 1702 | 0 | 0 | 0 | 0 | 0 | 0 |
| ENSMUSG00 | Gm13417 | 529 | 0 | 0 | 0 | 0 | 0 | 0 |
| ENSMUSG00 | Gm11274 | 479 | 0 | 0 | 0 | 2.56114721 | 0 | 0 |
| ENSMUSG00 | 1700030A11 | 885 | 0 | 0 | 0 | 0 | 0 | 0 |
| ENSMUSG00 | Gm12940 | 2676 | 2.95640531 | 2.70103375 | 2.44464603 | 2.69671484 | 1.9640197 | 5.32038927 |
| ENSMUSG00 | Gm13684 | 2262 | 1.93329039 | 1.02252397 | 2.52079476 | 2.13748628 | 1.42115901 | 2.41155077 |
| ENSMUSG00 | Gm11732 | 1737 | 0 | 0 | 0 | 0 | 0.0587523 | 0 |
| ENSMUSG00 | Gm15964 | 2743 | 0.07246715 | 0.5094447 | 0.20948741 | 0.05261691 | 0.20462637 | 0.06960353 |
| ENSMUSG00 | 2410004I01R | 1184 | 0.50365896 | 0 | 0 | 0.76186749 | 0.21548301 | 0.27643216 |
| ENSMUSG00 | Gm16714 | 1291 | 0 | 0 | 0 | 0 | 0 | 0 |
| ENSMUSG00 | Gm16335 | 308 | 0.12907624 | 0 | 0 | 0 | 0 | 0 |
| ENSMUSG00 | Gm15863 | 4136 | 0 | 0 | 0 | 0 | 0.06168566 | 0 |
| ENSMUSG00 | Gm12254 | 470 | 0 | 0 | 0 | 0 | 0 | 0 |
| ENSMUSG00 | Gm15463 | 231 | 2.40942306 | 1.46019522 | 0.38269994 | 0 | 2.42982742 | 3.18794485 |
| ENSMUSG00 | Gm16064 | 462 | 0 | 0 | 0 | 0 | 0 | 0 |
| ENSMUSG00 | Gm15099 | 571 | 0 | 0 | 0 | 0 | 0 | 0 |
| ENSMUSG00 | 2210417K05 | 1551 | 1.02528641 | 0.71456362 | 0.85496795 | 0.34895594 | 0.52638428 | 1.03752657 |
| ENSMUSG00 | Gm12300 | 2020 | 0 | 0 | 0 | 0 | 0 | 0 |
| ENSMUSG00 | Gm16956 | 838 | 0 | 0 | 0.26373415 | 0 | 0.24356265 | 0.0325473 |
| ENSMUSG00 | 4930529I22R | 587 | 0 | 0 | 0 | 0 | 0 | 0 |
| ENSMUSG00 | Gm12472 | 1137 | 0 | 0 | 0 | 0 | 0 | 0 |
| ENSMUSG00 | Gm4128 | 457 | 0 | 0 | 0 | 0 | 0 | 0 |
| ENSMUSG00 | 1700092C17 | 390 | 0 | 0 | 0 | 0 | 0 | 0 |
| ENSMUSG00 | Gm2044 | 909 | 0 | 0 | 0 | 0 | 0 | 0 |
| ENSMUSG00 | 3010003L21 | 2101 | 0.3216769 | 0.29815504 | 0.31557718 | 0.3949962 | 0.43716076 | 0.41541573 |
| ENSMUSG00 | Gm13480 | 997 | 0 | 0.24165718 | 0 | 0.07238123 | 0 | 0 |
| ENSMUSG00 | Gm14437 | 357 | 0 | 0 | 0 | 0 | 0 | 0 |
| ENSMUSG00 | Gm12336 | 320 | 0 | 0 | 0 | 0 | 0 | 0 |
| ENSMUSG00 | Gm11647 | 1388 | 0.08592683 | 0.03471646 | 0.31845708 | 0.25995709 | 0.11028756 | 0.15720253 |
| ENSMUSG00 | Gm16305 | 471 | 0 | 0.30692001 | 0.1876936 | 0 | 0 | 0 |
| ENSMUSG00 | C030034L19 | 5953 | 0 | 0 | 0 | 0 | 0 | 0 |
| ENSMUSG00 | Gm15478 | 2170 | 0 | 0 | 0 | 0 | 0 | 0.05027583 |
| ENSMUSG00 | Gm16641 | 1235 | 1.44858026 | 4.48699664 | 2.72011342 | 3.33065026 | 0.86765497 | 3.59981069 |
| ENSMUSG00 | Gm16358 | 209 | 0 | 0 | 0 | 0 | 0 | 0 |
| ENSMUSG00 | Gm15251 | 882 | 0 | 0 | 0 | 0 | 0 | 0 |
| ENSMUSG00 | Gm16188 | 418 | 0 | 0 | 0.10574604 | 0 | 0 | 0.26100133 |
| ENSMUSG00 | 9430093N23 | 513 | 0 | 0 | 0 | 0 | 0 | 0 |
| ENSMUSG00 | 2310002F09 | 2451 | 0.04866032 | 0 | 0 | 0 | 0 | 0 |
| ENSMUSG00 | Gm3160 | 606 | 0 | 0 | 0 | 0.23816531 | 0 | 0.04500766 |
| ENSMUSG00 | Gm16145 | 580 | 0 | 0 | 0 | 0 | 0 | 0 |
| ENSMUSG00 | B130046B21 | 2213 | 0.34132586 | 0.19596836 | 0.35952697 | 0.26087335 | 0.29974825 | 0.23416997 |
| ENSMUSG00 | Gm12506 | 801 | 0 | 0 | 0 | 0 | 0 | 0 |
| ENSMUSG00 | Gm14508 | 899 | 5.88151158 | 7.39680647 | 1.4258659 | 4.41493315 | 8.0597835 | 3.91371353 |
| ENSMUSG00 | 4930551L18 | 697 | 0 | 0.06913406 | 0.50733823 | 0 | 0 | 0.97828692 |
| ENSMUSG00 | Gm11415 | 477 | 0 | 0 | 0 | 0 | 0 | 0 |
| ENSMUSG00 | 2310058D17 | 3199 | 0.13670218 | 0.31632238 | 0.2348957 | 0.22558327 | 0.175458 | 0.36661753 |
| ENSMUSG00 | E130120K24 | 554 | 0 | 0 | 0 | 0 | 0 | 0 |
| ENSMUSG00 | Gm13861 | 493 | 0 | 0 | 0 | 0 | 0 | 0 |
| ENSMUSG00 | Gm14218 | 593 | 0 | 0 | 0 | 0 | 0 | 0 |
| ENSMUSG00 | 9430008C03 | 3170 | 1.63035094 | 2.32571787 | 3.30468038 | 1.70735227 | 2.17304755 | 3.79435833 |
| ENSMUSG00 | Gm13630 | 3642 | 0 | 0 | 0 | 0 | 0 | 0 |
| ENSMUSG00 | A630050E04 | 579 | 0 | 0 | 0 | 0 | 0 | 0 |
| ENSMUSG00 | Gm13336 | 912 | 0 | 0 | 0 | 0 | 0 | 0 |
| ENSMUSG00 | 1700003M07 | 869 | 0 | 0 | 0 | 0 | 0 | 0 |
| ENSMUSG00 | Gm13115 | 618 | 0 | 0 | 0 | 0 | 0 | 0 |
| ENSMUSG00 | Gm16150 | 906 | 0 | 0 | 0 | 0 | 0 | 0 |
| ENSMUSG00 | Gm16606 | 2779 | 0.27180789 | 0.88431398 | 0.39764163 | 0.10387059 | 0.01836142 | 0.17686193 |
| ENSMUSG00 | Gm11280 | 266 | 0 | 0 | 0 | 0 | 0 | 0 |
| ENSMUSG00 | 2210414B05 | 868 | 0 | 0 | 0 | 0 | 0 | 0 |
| ENSMUSG00 | Gm13056 | 562 | 1.90996081 | 0.428705 | 0.86516063 | 0 | 0.7263541 | 0.97062773 |
| ENSMUSG00 | 6720401G13 | 1430 | 18.1540761 | 15.7701083 | 18.3298552 | 19.9839016 | 30.1162666 | 17.2421496 |
| ENSMUSG00 | Gm13524 | 660 | 0 | 0 | 0 | 0 | 0.15462538 | 0 |
| ENSMUSG00 | 4931406G06 | 1250 | 0.31804384 | 0.501139 | 0.10608442 | 0.05773127 | 1.10216972 | 0.50185336 |
| ENSMUSG00 | 9130206I24R | 3219 | 0 | 0 | 0 | 0 | 0 | 0 |
| ENSMUSG00 | Gm11610 | 860 | 0 | 0 | 0.25698746 | 0.50347039 | 0.29666498 | 0 |
| ENSMUSG00 | Slc39a1-ps | 1939 | 0 | 0 | 0 | 0 | 0.02631582 | 0 |
| ENSMUSG00 | Gm12111 | 346 | 0 | 0 | 0 | 0 | 0 | 0 |
| ENSMUSG00 | Gm13068 | 827 | 0.28843154 | 0.05826656 | 0 | 0.39267038 | 0 | 0 |
| ENSMUSG00 | Gm12909 | 617 | 0 | 0 | 0 | 0 | 0 | 0 |
| ENSMUSG00 | 4930441J16R | 511 | 0 | 0 | 0 | 0 | 0 | 0 |
| ENSMUSG00 | 1700095J03R | 1806 | 0 | 0 | 0 | 0 | 0.22603046 | 0 |
| ENSMUSG00 | C530005A16 | 2433 | 1.74839146 | 1.18832163 | 1.27173408 | 2.13555874 | 2.26504258 | 1.99543189 |
| ENSMUSG00 | Gm16155 | 336 | 0 | 0 | 0.26310621 | 0 | 0 | 0 |
| ENSMUSG00 | 2010110K18 | 3073 | 0.12937026 | 0 | 0 | 0 | 0 | 0 |
| ENSMUSG00 | Gm14319 | 701 | 0 | 0 | 0 | 0 | 0.07279084 | 0 |
| ENSMUSG00 | Gm15055 | 672 | 0 | 0 | 0 | 0 | 0.07593211 | 0 |
| ENSMUSG00 | 1700010K10 | 425 | 0 | 0 | 0 | 0 | 0 | 0 |
| ENSMUSG00 | Gm15705 | 822 | 0 | 0 | 0 | 0 | 0 | 0 |
| ENSMUSG00 | Gm12060 | 784 | 0.91275338 | 0 | 0.67655882 | 0.18409206 | 0.32542332 | 0.66099253 |
| ENSMUSG00 | 1700019G24 | 500 | 0 | 0 | 0 | 0 | 0 | 0 |
| ENSMUSG00 | Gm13919 | 2017 | 0 | 0 | 0 | 0 | 0 | 0 |
| ENSMUSG00 | 4930483C13 | 2150 | 0 | 0 | 0.04111799 | 0 | 0.0237332 | 0 |
| ENSMUSG00 | Gm11734 | 723 | 0.71482883 | 4.33211444 | 2.01751151 | 1.89642834 | 1.7643975 | 0.71676092 |
| ENSMUSG00 | 4930453O03 | 1231 | 0 | 0 | 0 | 0 | 0 | 0 |
| ENSMUSG00 | 4732490B19 | 2442 | 0.13023908 | 0 | 0 | 0 | 0.73133626 | 0 |
| ENSMUSG00 | Gm6846 | 1813 | 0 | 0 | 0 | 0 | 0.11257888 | 0.63184493 |
| ENSMUSG00 | Gm15735 | 512 | 0.46588454 | 0 | 0 | 0.35236372 | 0.09966089 | 0.37289546 |
| ENSMUSG00 | Gm13037 | 993 | 0 | 0 | 0 | 0 | 0 | 0 |
| ENSMUSG00 | Gm15648 | 776 | 0 | 0 | 0 | 0 | 0 | 0 |
| ENSMUSG00 | Gm14272 | 482 | 0 | 0 | 0 | 0 | 0 | 0 |
| ENSMUSG00 | 6430710C18 | 2591 | 0 | 0 | 0 | 0 | 0 | 0 |
| ENSMUSG00 | Gm14344 | 576 | 0 | 0 | 0 | 0 | 0 | 0 |
| ENSMUSG00 | Gm13485 | 654 | 0 | 0 | 0.27034766 | 0 | 0 | 0 |
| ENSMUSG00 | Gm15619 | 756 | 0 | 0 | 0 | 0 | 0 | 0 |
| ENSMUSG00 | 4930440I19R | 1508 | 0 | 0 | 0 | 0 | 0 | 0 |
| ENSMUSG00 | A330032B11 | 1916 | 0.14524445 | 0.603588 | 0.76130523 | 0.15065572 | 0.47937096 | 0 |
| ENSMUSG00 | Gm16001 | 1692 | 0.86935743 | 1.05372244 | 1.61968928 | 1.59938141 | 1.26661217 | 2.51468305 |
| ENSMUSG00 | Gm11725 | 1846 | 0.02153601 | 0.05220633 | 0.02394466 | 0.01954607 | 0.08292477 | 0.07387497 |
| ENSMUSG00 | 4930486F22 | 1498 | 0 | 0 | 0 | 0 | 0 | 0.01820737 |
| ENSMUSG00 | Gm11458 | 1270 | 3.72511983 | 2.92154019 | 3.30643708 | 4.43212515 | 2.81247741 | 5.24016691 |
| ENSMUSG00 | 1700047A11 | 949 | 0.41891971 | 0.40620815 | 1.63020496 | 0.60833795 | 0.64522288 | 0.80473119 |
| ENSMUSG00 | 1700020I14R | 8434 | 24.4076213 | 23.8075533 | 23.778013 | 29.3439345 | 29.2460874 | 32.3744858 |
| ENSMUSG00 | Gm13634 | 634 | 0.18811742 | 0 | 0.069719 | 0.28455871 | 0 | 0.51623923 |
| ENSMUSG00 | D330022K07 | 2468 | 0.90206925 | 0.76145512 | 0.37610969 | 2.86551083 | 0.93038368 | 0.07735919 |
| ENSMUSG00 | Gm8289 | 1177 | 0 | 0 | 0 | 0 | 0 | 0 |
| ENSMUSG00 | Gm3362 | 465 | 1.1969392 | 0.62176054 | 1.045635 | 1.93989487 | 0.10973414 | 1.75965415 |
| ENSMUSG00 | Gm15416 | 440 | 0 | 0 | 0 | 0 | 0 | 0 |
| ENSMUSG00 | Gm13936 | 713 | 0.55758037 | 0.27033067 | 0 | 0 | 0 | 0.0765067 |
| ENSMUSG00 | Gm16348 | 2554 | 1.77452027 | 0.20753754 | 0.41536579 | 0.87591494 | 0.99895019 | 0.97180586 |
| ENSMUSG00 | A430046D13 | 1995 | 2.31159686 | 3.42981192 | 1.4401603 | 2.45972835 | 2.40425029 | 3.10343013 |
| ENSMUSG00 | 9530026F06 | 887 | 0 | 0 | 0 | 0 | 0 | 0 |
| ENSMUSG00 | Gm13963 | 888 | 0.67154528 | 0.43411209 | 0.14933055 | 0.81265866 | 0.11492427 | 1.1364433 |
| ENSMUSG00 | Gm15520 | 1197 | 0 | 0 | 0 | 0 | 0 | 0 |
| ENSMUSG00 | Gm13686 | 348 | 0 | 0 | 0 | 0 | 0 | 0 |
| ENSMUSG00 | Gm15137 | 576 | 0.13803986 | 0 | 0.38369655 | 0 | 0 | 0.14205541 |
| ENSMUSG00 | 4930554G24 | 1599 | 0 | 0 | 0 | 0 | 0 | 0 |
| ENSMUSG00 | Gm12018 | 3372 | 0 | 0 | 0 | 0 | 0 | 0 |
| ENSMUSG00 | Gm16320 | 1287 | 0 | 0 | 0 | 0.16821466 | 0 | 0 |
| ENSMUSG00 | 1810059H22 | 2889 | 1.65131799 | 1.78468304 | 0.76500248 | 1.56118227 | 1.7309051 | 1.59550503 |
| ENSMUSG00 | Gm15398 | 689 | 0 | 0 | 0 | 0 | 0 | 0 |
| ENSMUSG00 | 1110046J04R | 395 | 0 | 0 | 0 | 0 | 0 | 0 |
| ENSMUSG00 | G630030J09 | 2132 | 0.22376443 | 0.20341369 | 0.06219772 | 0 | 0 | 0.15351579 |
| ENSMUSG00 | Gm12272 | 330 | 0 | 0 | 0.13394498 | 0 | 0 | 0 |
| ENSMUSG00 | 4833422M21 | 3697 | 0 | 0 | 0.21521049 | 0 | 0 | 0 |
| ENSMUSG00 | Gm16954 | 1726 | 0 | 0.05583597 | 0 | 0 | 0.05912674 | 0.14222002 |
| ENSMUSG00 | 5031434O11 | 3619 | 1.76861906 | 1.55783745 | 1.47787317 | 1.39582377 | 2.32643051 | 1.74093442 |
| ENSMUSG00 | 9430024E24 | 711 | 0 | 0 | 0 | 0 | 0 | 0 |
| ENSMUSG00 | Gm16208 | 468 | 0.16989522 | 0 | 0 | 0 | 0 | 0 |
| ENSMUSG00 | Gm15347 | 382 | 0.72850357 | 0.37842756 | 0 | 0.6611893 | 0.6678845 | 0.07139958 |
| ENSMUSG00 | Hmgn2l6 | 538 | 0 | 0 | 0 | 0 | 0 | 0 |
| ENSMUSG00 | Gm11842 | 553 | 0 | 0 | 0.719379 | 0 | 0 | 0.29592737 |
| ENSMUSG00 | Gm15343 | 636 | 0 | 0.15152969 | 0 | 0 | 0 | 0 |
| ENSMUSG00 | Gm14242 | 373 | 0 | 0 | 0 | 0 | 0 | 0 |
| ENSMUSG00 | 1700012C14 | 619 | 0 | 0 | 0 | 0 | 0 | 0.35249938 |
| ENSMUSG00 | 4933423P22 | 5555 | 0.41508873 | 0.0954187 | 0.12731404 | 0.53911966 | 0.13778499 | 0.51554224 |
| ENSMUSG00 | Gm15691 | 846 | 0 | 0 | 0 | 0 | 0 | 0 |
| ENSMUSG00 | Gm16686 | 395 | 0 | 0 | 0 | 0 | 0 | 0 |
| ENSMUSG00 | Gm12162 | 277 | 0 | 0 | 0 | 0 | 0 | 0 |
| ENSMUSG00 | Gm13335 | 554 | 0 | 0 | 0 | 0 | 0 | 0 |
| ENSMUSG00 | Gm11851 | 690 | 0 | 0 | 0 | 0 | 0 | 0 |
| ENSMUSG00 | 9430073C21 | 2171 | 0 | 0 | 0 | 0 | 0 | 0 |
| ENSMUSG00 | 4930432B10 | 3627 | 0.01096098 | 0 | 0.03656066 | 0 | 0.04220544 | 0.02255967 |
| ENSMUSG00 | Gm16304 | 774 | 0 | 0 | 0 | 0 | 0 | 0 |
| ENSMUSG00 | A230083N12 | 570 | 0.90670394 | 0 | 0.07754709 | 0 | 0.6266397 | 1.43550733 |
| ENSMUSG00 | Gm14198 | 946 | 0.16809928 | 0 | 0 | 0 | 0.43151269 | 0.51896777 |
| ENSMUSG00 | Gm13539 | 593 | 0 | 0 | 0 | 0 | 0 | 0 |
| ENSMUSG00 | Gm14024 | 557 | 0 | 0 | 0 | 0 | 0 | 0 |
| ENSMUSG00 | Gm11634 | 752 | 0 | 0 | 0 | 0.04798144 | 0 | 0 |
| ENSMUSG00 | Gm15654 | 299 | 0 | 0 | 0 | 0 | 0 | 0 |
| ENSMUSG00 | 4930557F10 | 628 | 0 | 0 | 0 | 0 | 0 | 0.04343095 |
| ENSMUSG00 | Gm3669 | 3000 | 0 | 0 | 0 | 0 | 0 | 0 |
| ENSMUSG00 | 4930526L06 | 1019 | 0 | 0 | 0 | 0 | 0 | 0 |
| ENSMUSG00 | 4930527E20 | 349 | 0 | 0 | 0 | 0 | 0.29241476 | 0 |
| ENSMUSG00 | Trmt61b | 4171 | 1.45830461 | 0.98198216 | 1.54722347 | 2.2924339 | 1.00315579 | 1.06587538 |
| ENSMUSG00 | Gm15469 | 958 | 0 | 0 | 0 | 0 | 0 | 0 |
| ENSMUSG00 | 3110045C21 | 1131 | 9.03373872 | 4.04749072 | 8.59805965 | 4.24306978 | 5.86510067 | 11.1172491 |
| ENSMUSG00 | Gm16796 | 3169 | 0 | 0 | 0 | 0 | 0 | 0 |
| ENSMUSG00 | 4930517E11 | 1454 | 0 | 0 | 0 | 0 | 0 | 0 |
| ENSMUSG00 | Gm15985 | 339 | 0 | 0 | 0 | 0 | 0 | 0 |
| ENSMUSG00 | Gm14023 | 1776 | 0.38054233 | 0.10852802 | 0.27377268 | 0.50791166 | 1.00558736 | 1.15180064 |
| ENSMUSG00 | Gm15713 | 1541 | 0.33538043 | 0.06253919 | 0.63104513 | 0.25756164 | 0.96026275 | 0.3008883 |
| ENSMUSG00 | Gm16976 | 3305 | 0 | 0 | 0 | 0 | 0 | 0 |
| ENSMUSG00 | Gm11772 | 530 | 0.52507238 | 0.90917815 | 0.41699852 | 1.42966592 | 1.92552362 | 2.36723284 |
| ENSMUSG00 | Gm12320 | 1178 | 0 | 0 | 0 | 0 | 0 | 0 |
| ENSMUSG00 | Gm14055 | 663 | 0 | 0 | 0 | 0 | 0 | 0 |
| ENSMUSG00 | Gm12868 | 415 | 0.09579634 | 0.23222382 | 0.42604186 | 0.08694469 | 0 | 0.46005416 |
| ENSMUSG00 | Gm16977 | 3906 | 0.14249276 | 0.01233652 | 0.38475747 | 0.02771278 | 0 | 0.10474132 |
| ENSMUSG00 | Gm2709 | 347 | 0 | 0 | 0 | 0 | 0 | 0 |
| ENSMUSG00 | Gm13965 | 573 | 0 | 0.16819003 | 0 | 0 | 0 | 0 |
| ENSMUSG00 | Gm11521 | 325 | 0.48929822 | 0 | 0 | 0.11102168 | 0.78502117 | 0 |
| ENSMUSG00 | Gm12082 | 596 | 0 | 0 | 0 | 0 | 0 | 0 |
| ENSMUSG00 | Gm11738 | 2156 | 0 | 0 | 0 | 0 | 0 | 0 |
| ENSMUSG00 | Gm11958 | 251 | 0 | 0 | 0 | 0 | 0 | 0 |
| ENSMUSG00 | Gm16363 | 876 | 0 | 0 | 0 | 0 | 0.05824929 | 0 |
| ENSMUSG00 | 9530048J24R | 658 | 0.36251198 | 0.8787801 | 0.26870421 | 0.65803121 | 1.16321525 | 0.2072541 |
| ENSMUSG00 | C630028M04 | 3384 | 0 | 0 | 0.06531005 | 0 | 0 | 0 |
| ENSMUSG00 | 4930538E20 | 923 | 0 | 0 | 0 | 0 | 0 | 0 |
| ENSMUSG00 | Gm12963 | 1137 | 0 | 0 | 0 | 0 | 0 | 0 |
| ENSMUSG00 | Gm16360 | 213 | 0 | 0 | 0 | 0 | 0 | 0 |
| ENSMUSG00 | Gm13703 | 447 | 0 | 0 | 0 | 0 | 0 | 0 |
| ENSMUSG00 | Gm16871 | 2497 | 0 | 0 | 0 | 0 | 0 | 0 |
| ENSMUSG00 | 4930526A20 | 2567 | 0 | 0.18771501 | 0.08609631 | 0.05622446 | 0.11926695 | 0.05312552 |
| ENSMUSG00 | 4930556G01 | 884 | 0 | 0 | 0 | 0 | 0 | 0 |
| ENSMUSG00 | Gm15945 | 808 | 0.5412256 | 0 | 0 | 0 | 0 | 0 |
| ENSMUSG00 | Gm14224 | 862 | 0.36896038 | 0.33540447 | 0.05127824 | 0.2092926 | 0 | 0.22148779 |
| ENSMUSG00 | Gm13166 | 1406 | 0.33930709 | 0.27417606 | 0.40869414 | 0.12831453 | 0.10887562 | 0.71775366 |
| ENSMUSG00 | Gm16083 | 2542 | 0 | 0.22747337 | 0.12172026 | 0.01419435 | 0.06021996 | 0.12875518 |
| ENSMUSG00 | Gm15535 | 1855 | 0.15002068 | 0.51953037 | 0.02382849 | 0.75859824 | 0.71519449 | 1.67617729 |
| ENSMUSG00 | 4930401O10 | 782 | 0 | 0 | 0 | 0 | 0 | 0 |
| ENSMUSG00 | Gm15865 | 781 | 0 | 0 | 0 | 0 | 0 | 0 |
| ENSMUSG00 | Gm13901 | 467 | 0 | 0 | 0 | 0 | 0 | 0 |
| ENSMUSG00 | Gm12233 | 518 | 0 | 0 | 0 | 0 | 0 | 0 |
| ENSMUSG00 | B430319H21 | 732 | 0 | 0.59245626 | 0.66423535 | 0 | 0 | 0 |
| ENSMUSG00 | Gm12146 | 514 | 0 | 0 | 0 | 0 | 0 | 0 |
| ENSMUSG00 | Gm2117 | 2386 | 0 | 0 | 0 | 0 | 0 | 0 |
| ENSMUSG00 | Gm16719 | 2882 | 0 | 0 | 0 | 0 | 0 | 0 |
| ENSMUSG00 | Gm14950 | 1151 | 0 | 0 | 0 | 0 | 0 | 0 |
| ENSMUSG00 | Asbx4c | 1145 | 0 | 0 | 0 | 0 | 0 | 0 |
| ENSMUSG00 | Gm11455 | 731 | 0 | 0 | 0 | 0 | 0 | 0 |
| ENSMUSG00 | Gm15561 | 538 | 0 | 0.26869763 | 0 | 0 | 0 | 0 |
| ENSMUSG00 | Gm16036 | 1108 | 0 | 0.04348957 | 0 | 0.26052018 | 0.27631612 | 0 |
| ENSMUSG00 | Gm16010 | 4717 | 1.00294725 | 1.011333 | 0.08433678 | 0.26007834 | 0.03245265 | 0.15033721 |
| ENSMUSG00 | 1700037C06 | 873 | 0 | 0 | 0 | 0 | 0 | 0 |
| ENSMUSG00 | Gm13568 | 444 | 0.08953937 | 0 | 0 | 0.08126587 | 0 | 0 |
| ENSMUSG00 | A930024N18 | 3952 | 0.29172797 | 0.02438585 | 0.08947742 | 0.01826014 | 0.10329226 | 0.60042855 |
| ENSMUSG00 | Gm13553 | 1968 | 0 | 0 | 0 | 0 | 0 | 0 |
| ENSMUSG00 | Gm14252 | 3212 | 0.08664021 | 0.04500602 | 0.233945 | 0.29207134 | 0.07943085 | 0.39060816 |
| ENSMUSG00 | Gm13767 | 852 | 0 | 0 | 0 | 0 | 0 | 0 |
| ENSMUSG00 | Gm11753 | 394 | 0 | 0 | 0 | 0 | 0 | 0 |
| ENSMUSG00 | 1700047F07 | 947 | 0 | 0 | 0 | 0 | 0 | 0 |
| ENSMUSG00 | Gm15658 | 571 | 0 | 0 | 0 | 0 | 0 | 0 |
| ENSMUSG00 | 1810063I02R | 272 | 0 | 0 | 0.6500271 | 0 | 0.18759697 | 0.60164645 |
| ENSMUSG00 | Gm15278 | 382 | 0 | 0 | 0 | 0 | 0 | 0 |
| ENSMUSG00 | Gm14808 | 358 | 0 | 0 | 0 | 0 | 0 | 0 |
| ENSMUSG00 | AI854517 | 3843 | 0 | 0 | 0 | 0 | 0 | 0 |
| ENSMUSG00 | 2610035F20 | 2857 | 0 | 0 | 0 | 0 | 0 | 0 |
| ENSMUSG00 | Gm15020 | 630 | 0 | 0 | 0 | 0 | 0 | 0 |
| ENSMUSG00 | Gm16600 | 1632 | 0.41411959 | 0.02952601 | 0.75836495 | 0.22109096 | 0.4064601 | 0.43452244 |
| ENSMUSG00 | 4930412C18 | 6096 | 1.21953328 | 0.3240886 | 0.57282572 | 0.49127456 | 1.18860652 | 0.40715095 |
| ENSMUSG00 | Gm11959 | 720 | 0 | 0 | 0 | 0 | 0 | 0 |
| ENSMUSG00 | C230038L03 | 4054 | 0 | 0 | 0 | 0 | 0.06293337 | 0 |
| ENSMUSG00 | 1700046C09 | 539 | 0 | 0 | 0 | 0 | 0 | 0 |
| ENSMUSG00 | 2610028E06 | 6967 | 0 | 0 | 0 | 0 | 0 | 0 |
| ENSMUSG00 | 2210411M09 | 640 | 0.55906145 | 0 | 0.13813076 | 0 | 0 | 0.04261662 |
| ENSMUSG00 | Gm12198 | 1807 | 0 | 0 | 0 | 0 | 0 | 0 |
| ENSMUSG00 | Gm15721 | 3304 | 0 | 0 | 0 | 0 | 0 | 0 |
| ENSMUSG00 | A730017L22 | 6007 | 1.72072997 | 2.01345047 | 1.11847514 | 2.2104532 | 1.95373172 | 1.37122375 |
| ENSMUSG00 | Gm14858 | 2270 | 0.03502686 | 0 | 0 | 0.41327452 | 0.1123929 | 0.14418312 |
| ENSMUSG00 | Gm12602 | 1113 | 0 | 0 | 0 | 0 | 0 | 0 |
| ENSMUSG00 | Wscd2 | 188 | 0 | 0 | 0 | 0 | 0 | 0 |
| ENSMUSG00 | Gm16234 | 809 | 0 | 0.23825188 | 0 | 0 | 0 | 0.03371402 |
| ENSMUSG00 | Gm15418 | 674 | 0.23593757 | 0.14298648 | 0 | 0 | 0.07570679 | 0.20233412 |
| ENSMUSG00 | Gm13053 | 456 | 0.95901378 | 0.52836011 | 1.26014026 | 0.71214562 | 2.12609899 | 1.01681769 |
| ENSMUSG00 | Gm15485 | 256 | 0 | 0 | 0 | 0 | 0 | 0 |
| ENSMUSG00 | Dpy19l2 | 3419 | 0 | 0 | 0 | 0 | 0 | 0 |
| ENSMUSG00 | Mageb6-ps | 1031 | 0 | 0 | 0 | 0 | 0 | 0 |
| ENSMUSG00 | Gm11527 | 594 | 0 | 0 | 0 | 0 | 0 | 0 |
| ENSMUSG00 | Gm12066 | 439 | 0 | 0 | 0 | 0 | 0 | 0 |
| ENSMUSG00 | 1700061F12 | 872 | 0 | 0 | 0 | 0 | 0 | 0 |
| ENSMUSG00 | Gm12786 | 640 | 0.37270763 | 2.71048737 | 0.6906538 | 2.31150598 | 1.67430296 | 1.87513145 |
| ENSMUSG00 | 3110099E03 | 3875 | 0.42063863 | 0.65906618 | 0.36502167 | 0 | 0.60573246 | 0.34489221 |
| ENSMUSG00 | Gm15676 | 795 | 0.60008273 | 1.03040191 | 0.44479842 | 0.36308976 | 0.89857769 | 0.20584633 |
| ENSMUSG00 | Rgag1 | 7655 | 0 | 0.16366395 | 0 | 0 | 0.05999182 | 0 |
| ENSMUSG00 | Gm12223 | 525 | 0 | 0 | 0 | 0 | 0 | 0 |
| ENSMUSG00 | Gm11613 | 1967 | 1.1924623 | 2.35175315 | 4.3595107 | 3.30186478 | 0.98576628 | 0.58237664 |
| ENSMUSG00 | Gm14493 | 541 | 0 | 0 | 0 | 0 | 0 | 0 |
| ENSMUSG00 | 3110004A20 | 383 | 0 | 0 | 0 | 0 | 0 | 0 |
| ENSMUSG00 | A430078I02R | 2228 | 0.17843573 | 0.62720234 | 0.23807097 | 0.19433776 | 0.61836272 | 0.69778027 |
| ENSMUSG00 | Gm13789 | 670 | 0 | 0 | 0 | 0 | 0 | 0 |
| ENSMUSG00 | Gm13479 | 4720 | 0 | 0 | 0 | 0 | 0 | 0 |
| ENSMUSG00 | Gm16321 | 572 | 0 | 0 | 0 | 0 | 0 | 0 |
| ENSMUSG00 | 4930589P08 | 1082 | 0 | 0 | 0 | 0 | 0 | 0 |
| ENSMUSG00 | Gm11551 | 406 | 0 | 0 | 0 | 0 | 0 | 0 |
| ENSMUSG00 | Gm16090 | 1701 | 0.21034646 | 0.05665661 | 0.12992899 | 0 | 0.20998509 | 0.17637921 |
| ENSMUSG00 | Gm11476 | 2015 | 0 | 0 | 0 | 0 | 0 | 0 |
| ENSMUSG00 | 4932441J04R | 723 | 0 | 0 | 0.79477726 | 0 | 0 | 0 |
| ENSMUSG00 | Gm14715 | 641 | 0 | 0 | 0 | 0 | 0 | 0 |
| ENSMUSG00 | Gm13449 | 1228 | 0 | 0 | 0 | 0 | 0 | 0 |
| ENSMUSG00 | E330012B07 | 2898 | 0 | 0 | 0 | 0 | 0 | 0 |
| ENSMUSG00 | Gm4969 | 4035 | 0 | 0 | 0 | 0.01788453 | 0 | 0 |
| ENSMUSG00 | 1500002F19 | 1064 | 0.67255512 | 0.27172806 | 0.62314628 | 0.67823392 | 0.81527104 | 0.53831525 |
| ENSMUSG00 | Gm11346 | 3244 | 0.17157112 | 0.17824825 | 0.14988295 | 0.16684053 | 0.23594194 | 0.39516278 |
| ENSMUSG00 | Gm11547 | 1588 | 0 | 0 | 0 | 0 | 0.03213248 | 0 |
| ENSMUSG00 | 5033403F01 | 1369 | 0.17423878 | 0.3167845 | 0.58117836 | 0.8697644 | 0 | 1.73330432 |
| ENSMUSG00 | Gm15792 | 766 | 0 | 0 | 0 | 0 | 0 | 0 |
| ENSMUSG00 | Gm16854 | 2978 | 5.24644185 | 3.05817245 | 4.08176858 | 5.44017394 | 5.72290447 | 4.70757709 |
| ENSMUSG00 | Gm16354 | 149 | 0 | 0 | 0 | 0 | 0 | 0 |
| ENSMUSG00 | 1700016P03 | 860 | 0 | 0 | 0 | 0 | 0.17799899 | 0 |
| ENSMUSG00 | Gm14747 | 983 | 0.04044301 | 0 | 0.13489881 | 0 | 0.05190883 | 0 |
| ENSMUSG00 | Ap3s1-ps1 | 1159 | 0 | 0 | 0 | 0 | 0 | 0 |
| ENSMUSG00 | Gm15868 | 518 | 0.07674803 | 0 | 0 | 0.06965646 | 0.09850652 | 0 |
| ENSMUSG00 | 5730420D15 | 1177 | 0 | 0 | 0 | 0 | 0 | 0 |
| ENSMUSG00 | 1700123M08 | 1363 | 0.58335261 | 0.03535322 | 0.09728946 | 0.3970878 | 0.18718406 | 0.76040814 |
| ENSMUSG00 | A330035P11 | 4245 | 0.58064542 | 0.1929728 | 0.20825368 | 1.08798627 | 0.45677321 | 0.90594208 |
| ENSMUSG00 | Gm13850 | 621 | 0 | 0 | 0 | 0 | 0 | 0 |
| ENSMUSG00 | G730013B05 | 2781 | 0 | 0 | 0 | 0 | 0 | 0 |
| ENSMUSG00 | Gm11466 | 289 | 0 | 0 | 0 | 0 | 0 | 0 |
| ENSMUSG00 | 4930555B12 | 281 | 0 | 0 | 0 | 0 | 0 | 0 |
| ENSMUSG00 | G630018N14 | 767 | 0 | 0 | 0 | 0 | 0 | 0 |
| ENSMUSG00 | 4930467K11 | 721 | 0 | 0 | 1.16481972 | 0 | 0 | 0 |
| ENSMUSG00 | 3110056K07 | 1500 | 19.8777403 | 16.8652548 | 17.9164804 | 21.4086798 | 13.1307874 | 41.3483531 |
| ENSMUSG00 | Gm16041 | 3722 | 0 | 0 | 0 | 0 | 0 | 0 |
| ENSMUSG00 | Gm14573 | 449 | 0.53125364 | 0.21463894 | 0.39378034 | 0.1607218 | 1.59102285 | 0 |
| ENSMUSG00 | Gm8813 | 594 | 0 | 0 | 0 | 0 | 0 | 0 |
| ENSMUSG00 | Gm12898 | 640 | 0 | 0 | 0 | 0 | 0 | 0 |
| ENSMUSG00 | Gm11222 | 737 | 0.05394231 | 0.06538188 | 0.05997536 | 0 | 0 | 0 |
| ENSMUSG00 | D630032N06 | 631 | 1.57509828 | 1.83276484 | 2.45176625 | 1.71546963 | 3.638965 | 2.55024361 |
| ENSMUSG00 | Gm11697 | 523 | 0 | 0 | 0 | 0 | 0 | 0 |
| ENSMUSG00 | Gm11795 | 394 | 0 | 0 | 0 | 0 | 0 | 0 |
| ENSMUSG00 | 9630028H03 | 3636 | 0 | 0 | 0 | 0.09923555 | 0 | 0 |
| ENSMUSG00 | Gm12381 | 2180 | 0 | 0 | 0 | 0 | 0 | 0 |
| ENSMUSG00 | 4930580A09 | 694 | 0.05728455 | 0 | 1.65597683 | 0.31194851 | 0.14705007 | 0.51090823 |
| ENSMUSG00 | Gm15290 | 575 | 0 | 0 | 0 | 0 | 0 | 0 |
| ENSMUSG00 | Gm14565 | 2492 | 0 | 0 | 0 | 0 | 0 | 0 |
| ENSMUSG00 | Gm11769 | 1655 | 0.98487897 | 0.08734703 | 0.37391287 | 1.2645067 | 0.58580129 | 0.57680506 |
| ENSMUSG00 | Gm15980 | 481 | 1.40507935 | 2.70485226 | 0.82706151 | 2.02539543 | 3.81902189 | 2.38156933 |
| ENSMUSG00 | Gm15521 | 832 | 0 | 0 | 0 | 0 | 0 | 0 |
| ENSMUSG00 | Gm15410 | 541 | 1.32273318 | 3.20649153 | 0 | 1.53398711 | 0.94318624 | 0.70581322 |
| ENSMUSG00 | Gm14064 | 722 | 0 | 0 | 0 | 0 | 0 | 0 |
| ENSMUSG00 | 4930465M20 | 1592 | 0 | 0 | 0 | 0 | 0 | 0 |
| ENSMUSG00 | 3110053B16 | 2339 | 0.9178264 | 1.35968584 | 1.30394492 | 0.91014991 | 1.30892798 | 0.43145004 |
| ENSMUSG00 | Gm12519 | 686 | 0 | 0 | 0 | 0 | 0 | 0 |
| ENSMUSG00 | 2310015A10 | 4840 | 1.13352403 | 1.0154994 | 0.96805689 | 0.84241137 | 0.6641863 | 1.94416334 |
| ENSMUSG00 | 0610040B09 | 692 | 14.4199792 | 17.5476639 | 7.53730849 | 17.4674638 | 11.8717435 | 21.8748913 |
| ENSMUSG00 | Gm11493 | 564 | 0 | 0 | 0 | 0 | 0 | 0 |
| ENSMUSG00 | Gm13173 | 439 | 0 | 0 | 0 | 0 | 0 | 0 |
| ENSMUSG00 | A730032A03 | 2148 | 0 | 0 | 0 | 0 | 0 | 0 |
| ENSMUSG00 | 2310010J17R | 447 | 3.37966054 | 3.01839011 | 2.96656665 | 1.21080686 | 2.28305932 | 6.772897 |
| ENSMUSG00 | Gm11695 | 404 | 9.93887013 | 7.75276916 | 8.5340192 | 12.5929908 | 6.31514552 | 15.2575952 |
| ENSMUSG00 | Gm11999 | 809 | 0 | 0 | 0 | 0 | 0 | 0 |
| ENSMUSG00 | Gm15179 | 1021 | 0 | 0 | 0 | 0 | 0 | 0 |
| ENSMUSG00 | Gm12536 | 356 | 0 | 0.40606552 | 0 | 0 | 0.28666503 | 0 |
| ENSMUSG00 | Gm15952 | 946 | 1.76504248 | 2.08841874 | 1.77554972 | 1.10610919 | 1.88786803 | 4.09407904 |
| ENSMUSG00 | Gm13180 | 438 | 0.09076594 | 0.77010296 | 0.2018349 | 0 | 0 | 0 |
| ENSMUSG00 | Gm15984 | 1549 | 0.28231781 | 0.55994575 | 0 | 0.0698813 | 0.09882449 | 0.38737383 |
| ENSMUSG00 | Gm15704 | 2815 | 0 | 0 | 0.03140451 | 0.73061334 | 0 | 0 |

| | | | | | | | | |
|---|---|---|---|---|---|---|---|---|
| ENSMUSG00 | Gm12325 | 318 | 0 | 0 | 0 | 0 | 0 | 0 |
| ENSMUSG00 | Gm12434 | 651 | 0 | 0 | 0 | 0 | 0 | 0 |
| ENSMUSG00 | 1700034P13 | 2844 | 0 | 0 | 0 | 0.08880953 | 0.01794176 | 0.05754143 |
| ENSMUSG00 | Gm16220 | 403 | 0 | 0 | 0 | 0 | 0 | 0 |
| ENSMUSG00 | Gm15718 | 309 | 0 | 2.33914768 | 0.14304804 | 1.40124445 | 0.99080341 | 1.32401161 |
| ENSMUSG00 | 4930556N08 | 2015 | 0.07891907 | 0.55001894 | 0.0438728 | 0.23278738 | 0.05064653 | 0 |
| ENSMUSG00 | Gm12059 | 3103 | 1.37087864 | 0.40375362 | 1.25354888 | 1.24420843 | 0.11510945 | 0.57133469 |
| ENSMUSG00 | Gm9855 | 2434 | 0.11433376 | 0.01979722 | 1.07144977 | 0 | 0 | 0.71716389 |
| ENSMUSG00 | Gm12992 | 5867 | 0.33202975 | 0.11498384 | 0.4897085 | 0.22754997 | 0.46095073 | 0.03719058 |
| ENSMUSG00 | Gm13447 | 710 | 0 | 0 | 0 | 0 | 0 | 0 |
| ENSMUSG00 | Gm14817 | 641 | 0.55818927 | 0 | 0 | 0.28145121 | 0 | 0 |
| ENSMUSG00 | A830052D11 | 1127 | 0 | 0 | 0.50987042 | 0.03201601 | 0 | 0.3146143 |
| ENSMUSG00 | C330046G13 | 1330 | 0.02989134 | 0.61591693 | 0 | 0 | 1.38116506 | 0.32811596 |
| ENSMUSG00 | Gm16193 | 775 | 0 | 0 | 0.22813855 | 0 | 0.06584049 | 0 |
| ENSMUSG00 | Gm16122 | 1156 | 0 | 0 | 0 | 0 | 0 | 0 |
| ENSMUSG00 | Gm12845 | 240 | 0 | 0 | 0 | 0 | 0 | 0 |
| ENSMUSG00 | Gm12022 | 1338 | 0 | 0 | 0 | 0 | 0 | 0 |
| ENSMUSG00 | 4930555O08 | 666 | 0 | 0 | 0 | 0 | 0 | 0 |
| ENSMUSG00 | 4930594O21 | 1152 | 0 | 0 | 0 | 0 | 0 | 0 |
| ENSMUSG00 | Gm16349 | 392 | 0 | 0 | 0 | 0 | 0 | 0 |
| ENSMUSG00 | Gm16328 | 195 | 0 | 0 | 0 | 0 | 0 | 0 |
| ENSMUSG00 | Gm13327 | 2739 | 0 | 0 | 0 | 0 | 0 | 0 |
| ENSMUSG00 | Gm13982 | 752 | 0 | 0 | 0 | 0 | 0 | 0 |
| ENSMUSG00 | Gm14267 | 601 | 0 | 0 | 0 | 0 | 0 | 0 |
| ENSMUSG00 | 9130409I20 | 1499 | 0.37129868 | 0.70720596 | 1.5923279 | 0 | 0 | 0.65502803 |
| ENSMUSG00 | 4930469K13 | 693 | 58.6866617 | 28.8562388 | 123.101814 | 42.382084 | 25.7708969 | 126.573218 |
| ENSMUSG00 | Gm15253 | 272 | 0 | 0 | 0 | 0 | 0 | 0 |
| ENSMUSG00 | Gm9498 | 290 | 0 | 0 | 0 | 0 | 0 | 0 |
| ENSMUSG00 | Gm16153 | 4116 | 1.3039334 | 1.38143833 | 1.18129318 | 2.15650703 | 1.06614877 | 1.58373148 |
| ENSMUSG00 | Gm15395 | 699 | 0 | 0 | 0 | 0 | 0 | 0 |
| ENSMUSG00 | Gm16838 | 813 | 0 | 0 | 0 | 0 | 0 | 0 |
| ENSMUSG00 | Gm12629 | 708 | 0 | 0 | 0 | 0 | 0 | 0 |
| ENSMUSG00 | Gm14216 | 896 | 0 | 0 | 0 | 0 | 0 | 0 |
| ENSMUSG00 | Gm2061 | 2758 | 0 | 0 | 0 | 0 | 0 | 0 |
| ENSMUSG00 | Gm13371 | 467 | 0 | 0 | 0 | 0 | 0 | 0 |
| ENSMUSG00 | Gm14967 | 333 | 0 | 0 | 0 | 0 | 0 | 0 |
| ENSMUSG00 | Gm15243 | 380 | 0 | 0 | 0 | 0 | 0 | 0 |
| ENSMUSG00 | 2700086A05 | 2944 | 0.1890546 | 0.03273536 | 0.09008528 | 0.13481742 | 0.31198192 | 0.24087657 |
| ENSMUSG00 | Gm16022 | 528 | 0 | 0 | 0 | 0 | 0 | 0 |
| ENSMUSG00 | Gm14298 | 236 | 0 | 0 | 0 | 0 | 0 | 0 |
| ENSMUSG00 | Hbb-bh0 | 128 | 0 | 0 | 0 | 0 | 0 | 0 |
| ENSMUSG00 | Gm12968 | 1452 | 0 | 0 | 0 | 0 | 0 | 0 |
| ENSMUSG00 | Gm16722 | 4624 | 0.15475749 | 0.15631415 | 0.14338833 | 0.43697978 | 0.16552674 | 0.27722925 |
| ENSMUSG00 | Gm16723 | 1112 | 0 | 0 | 0 | 0 | 0 | 0 |
| ENSMUSG00 | 4921531C22 | 1964 | 8.1778076 | 5.10324846 | 6.50424268 | 7.10985297 | 5.09224525 | 2.49971236 |
| ENSMUSG00 | Gm16046 | 1152 | 0 | 0.20914254 | 0.11510897 | 0 | 0 | 0 |
| ENSMUSG00 | Gm9428 | 767 | 0 | 0 | 0 | 0 | 0 | 0 |
| ENSMUSG00 | Gm12212 | 376 | 0 | 0 | 0 | 0 | 0 | 0 |
| ENSMUSG00 | Gm16063 | 1697 | 0 | 0 | 0 | 0 | 0 | 0 |
| ENSMUSG00 | Gm14211 | 406 | 0 | 0 | 0 | 0 | 0 | 0 |
| ENSMUSG00 | Gm12676 | 806 | 0 | 0.41849267 | 0 | 0.62673527 | 0.69638974 | 0 |
| ENSMUSG00 | Gm15163 | 2521 | 0.4730918 | 0.22936823 | 0.22793493 | 0.51525331 | 0.30360795 | 0.19474157 |
| ENSMUSG00 | Gm15124 | 951 | 0 | 0 | 0 | 0 | 0 | 0 |
| ENSMUSG00 | Gm15736 | 418 | 0 | 0 | 0 | 0 | 0 | 0 |
| ENSMUSG00 | Gm13008 | 505 | 0 | 0 | 0 | 0 | 0 | 0 |
| ENSMUSG00 | Tsix | 4302 | 0.20330557 | 0.01120094 | 0.03082416 | 0.02516182 | 0.28466597 | 0.06339991 |
| ENSMUSG00 | Gm13388 | 422 | 0 | 0 | 0 | 0 | 0 | 0 |
| ENSMUSG00 | 4930546G22 | 515 | 0 | 0 | 0 | 0 | 0 | 0 |
| ENSMUSG00 | Rpl9-ps3 | 1153 | 0 | 0 | 0 | 0 | 0 | 0 |
| ENSMUSG00 | 4933424L21 | 1182 | 0 | 0 | 0 | 0 | 0 | 0 |
| ENSMUSG00 | Gm7854 | 350 | 1.13587087 | 0.2753511 | 2.39952862 | 1.13400711 | 1.16631716 | 0.23378262 |
| ENSMUSG00 | Gm12796 | 341 | 0 | 0 | 0 | 0 | 0 | 0 |
| ENSMUSG00 | Gm4916 | 1705 | 0 | 0 | 0 | 0 | 0 | 0 |
| ENSMUSG00 | Gm15915 | 902 | 0.52889775 | 0.26710888 | 0.04900426 | 0 | 0.22628105 | 0.03023796 |
| ENSMUSG00 | Gm13096 | 249 | 0 | 0 | 0 | 0 | 0 | 0 |
| ENSMUSG00 | Gm15873 | 759 | 0 | 0 | 0 | 0 | 0 | 0 |
| ENSMUSG00 | 9630001P10 | 5003 | 0 | 0.00963151 | 0 | 0 | 0.01019916 | 0.14174308 |
| ENSMUSG00 | 4930471M09 | 1109 | 0.07169609 | 0 | 0.15942955 | 0 | 0 | 0 |
| ENSMUSG00 | Gm14339 | 421 | 0 | 0 | 0 | 0 | 0 | 0 |
| ENSMUSG00 | Gm13307 | 825 | 0 | 0 | 0 | 0 | 0 | 0 |
| ENSMUSG00 | Gm16229 | 1230 | 0 | 0 | 0 | 0 | 0 | 0.0221745 |
| ENSMUSG00 | Gm4279 | 2327 | 0 | 0.16566031 | 0 | 0 | 0 | 0.05860473 |
| ENSMUSG00 | Gm11641 | 921 | 0 | 0.0523197 | 0 | 0.35259327 | 0 | 0.14807079 |
| ENSMUSG00 | Gm11684 | 755 | 0 | 0 | 0 | 0 | 0 | 0 |
| ENSMUSG00 | 4930550G17 | 665 | 1.43478426 | 1.15937304 | 0.33234468 | 1.08517427 | 0.61385114 | 1.39449283 |
| ENSMUSG00 | Gm16682 | 1273 | 0.40598684 | 0.11355799 | 0.34722579 | 0.19840873 | 0 | 0.38565869 |
| ENSMUSG00 | Gm13442 | 661 | 0.72173338 | 0.43739584 | 0.26748468 | 1.91054699 | 0.92634873 | 0.66020307 |
| ENSMUSG00 | Gm12335 | 613 | 0.06485397 | 0.15721515 | 0.14421482 | 0 | 0.08324042 | 0.0444937 |
| ENSMUSG00 | Gm15862 | 359 | 0 | 0 | 0 | 0 | 0 | 0 |
| ENSMUSG00 | 4930505K14 | 512 | 0 | 0 | 0 | 0 | 0 | 0 |
| ENSMUSG00 | 5430405H02 | 1479 | 3.01055701 | 1.75934272 | 3.3173797 | 2.3420394 | 3.34655744 | 3.81734302 |
| ENSMUSG00 | Gm13536 | 764 | 0 | 0 | 0 | 0 | 0 | 0 |
| ENSMUSG00 | 8430419K02 | 2581 | 0 | 0 | 0 | 0 | 0 | 0 |
| ENSMUSG00 | Gm12259 | 326 | 0.12194933 | 0 | 0.13558848 | 0.66408671 | 0.15652263 | 0.83664538 |
| ENSMUSG00 | Gm14372 | 2190 | 0 | 0 | 0 | 0.46132294 | 0.04659943 | 0.31135433 |
| ENSMUSG00 | 4930455B14 | 982 | 0 | 0 | 0 | 0 | 0 | 0 |
| ENSMUSG00 | Gm11194 | 3622 | 0 | 0 | 0.04881485 | 0 | 0.0704395 | 0.06024216 |
| ENSMUSG00 | Gm12280 | 1622 | 0 | 0 | 0 | 0 | 0 | 0 |
| ENSMUSG00 | Gm12843 | 1012 | 0.39284072 | 0.09523012 | 0.43677711 | 0.42785033 | 0.10084264 | 0 |
| ENSMUSG00 | Gm14009 | 652 | 0 | 0 | 0 | 0 | 0 | 0 |
| ENSMUSG00 | Gm15412 | 1093 | 0 | 0 | 0 | 0 | 0 | 0 |
| ENSMUSG00 | Gm16131 | 657 | 0 | 0 | 0 | 0 | 0 | 0 |
| ENSMUSG00 | Gm15886 | 1452 | 0 | 0.06637251 | 0 | 0 | 0 | 0 |
| ENSMUSG00 | Gm13496 | 1163 | 0 | 0 | 0 | 0 | 0 | 0 |
| ENSMUSG00 | Gm15279 | 287 | 0 | 0 | 0 | 0 | 0 | 0 |
| ENSMUSG00 | 1700019B21 | 774 | 0 | 0 | 0 | 0 | 0 | 0 |
| ENSMUSG00 | 1700061E18 | 586 | 0 | 0 | 0 | 0 | 0 | 0 |
| ENSMUSG00 | Gm14302 | 379 | 0.20979145 | 0 | 0 | 0 | 0 | 0.35982374 |
| ENSMUSG00 | 4930455G09 | 628 | 0.88626867 | 0.38365002 | 1.68924241 | 0.86183227 | 0.32500876 | 3.25732157 |
| ENSMUSG00 | 4930443B20 | 584 | 1.22534015 | 0 | 0.52981661 | 0 | 0 | 0 |
| ENSMUSG00 | 2700081L22 | 624 | 0 | 0.30888745 | 0 | 0.40476653 | 0 | 0.04370936 |
| ENSMUSG00 | Gm16621 | 1159 | 0 | 0 | 0 | 0 | 0.08805242 | 0 |
| ENSMUSG00 | Gm12333 | 2667 | 0.29812884 | 0.43362378 | 0 | 0 | 0.45917999 | 0 |
| ENSMUSG00 | 2810430I11R | 3414 | 0 | 0 | 0 | 0 | 0 | 0 |
| ENSMUSG00 | Gm13563 | 1670 | 1.09506114 | 0.89447887 | 0.6087679 | 1.18833081 | 0.7638679 | 0.86560232 |
| ENSMUSG00 | Gm13707 | 595 | 0 | 0 | 0 | 0.42449464 | 0 | 0.32087811 |
| ENSMUSG00 | D230017M19 | 5327 | 0.31344663 | 0.49751348 | 0.48126655 | 0.18965595 | 0.32567989 | 0.3481651 |
| ENSMUSG00 | Gm11381 | 417 | 0 | 0 | 0 | 0 | 0 | 0 |
| ENSMUSG00 | C230066G23 | 648 | 1.3497231 | 0 | 0 | 0 | 0 | 0 |
| ENSMUSG00 | D630024D03 | 1558 | 5.40960325 | 7.36095843 | 3.20591031 | 6.53089638 | 7.72928414 | 4.77918903 |
| ENSMUSG00 | Gm16233 | 1619 | 0 | 0 | 0 | 0 | 0 | 0 |
| ENSMUSG00 | Gm13055 | 1959 | 0 | 0 | 0 | 0 | 0 | 0 |
| ENSMUSG00 | Gm14364 | 266 | 0 | 0 | 0 | 0 | 0 | 0 |
| ENSMUSG00 | 5430402O13 | 1287 | 1.45183185 | 0.74881806 | 2.16372658 | 1.73821815 | 0.99118834 | 4.11132868 |
| ENSMUSG00 | Gm16892 | 1722 | 1.06199309 | 0.39175969 | 0.77006695 | 1.27816766 | 0.62227288 | 0.22174504 |
| ENSMUSG00 | 2310050B05 | 2695 | 0 | 0 | 0 | 0.01338851 | 0 | 0 |
| ENSMUSG00 | 4930509G22 | 2477 | 0 | 0.33071034 | 0.05353473 | 0.18936883 | 0.24720085 | 0.64965834 |
| ENSMUSG00 | Gm15640 | 525 | 0 | 2.20280878 | 0 | 0.89346015 | 0.29157929 | 0.41561355 |
| ENSMUSG00 | Gm15624 | 2067 | 0 | 0 | 0 | 0.10473743 | 0 | 0 |
| ENSMUSG00 | Gm9816 | 2876 | 0.87086067 | 0.90475239 | 0.737722 | 0.82803023 | 0.81613814 | 0.53107782 |
| ENSMUSG00 | Gm12158 | 629 | 0 | 0 | 0.0702732 | 0.05736414 | 0 | 0 |
| ENSMUSG00 | 4933425H06 | 1236 | 0 | 0.23391477 | 0 | 0 | 0 | 0 |
| ENSMUSG00 | Gm15987 | 2229 | 0 | 0 | 0 | 0 | 0 | 0 |
| ENSMUSG00 | Gm13092 | 597 | 0.59932885 | 0.40357154 | 1.48079876 | 0.18131681 | 0.34188527 | 2.87822826 |
| ENSMUSG00 | 4930470G03 | 2444 | 0.01626656 | 0.01971622 | 0 | 0.14763521 | 0.10439111 | 0.10043852 |
| ENSMUSG00 | 4732419C18 | 3035 | 0.74664329 | 0.03175383 | 0.17476841 | 0.10699783 | 0 | 0.44933508 |
| ENSMUSG00 | Gm12729 | 619 | 0 | 0 | 0 | 0 | 0.08243356 | 0 |
| ENSMUSG00 | Rpl30-ps9 | 777 | 0 | 0 | 0 | 0.04643764 | 0 | 0 |
| ENSMUSG00 | Gm15414 | 666 | 0 | 0 | 0 | 0 | 0 | 0 |
| ENSMUSG00 | Lin52 | 2138 | 12.4212633 | 11.7423463 | 18.5449267 | 14.446319 | 9.02150143 | 10.9073043 |
| ENSMUSG00 | Vax2os1 | 4856 | 0 | 0 | 0 | 0 | 0 | 0 |
| ENSMUSG00 | Zfp703 | 3301 | 0.52991249 | 0.21896293 | 0.45527497 | 0.56839331 | 0.44827776 | 0.47922723 |
| ENSMUSG00 | 4933406D12 | 733 | 0 | 0 | 0 | 0 | 0 | 0 |
| ENSMUSG00 | A530010F05 | 1018 | 0 | 0 | 0 | 0 | 0 | 0 |
| ENSMUSG00 | Gm2018 | 476 | 0 | 0 | 0 | 0 | 0 | 0 |
| ENSMUSG00 | Gm6963 | 378 | 0 | 0 | 0 | 0 | 0 | 0 |
| ENSMUSG00 | 1700066O22 | 2127 | 0 | 0 | 0 | 0 | 0 | 0 |
| ENSMUSG00 | 1700013N06 | 658 | 0 | 0 | 0 | 0 | 0 | 0 |
| ENSMUSG00 | Gm16059 | 3776 | 0.07369925 | 0.02552248 | 0 | 0 | 0 | 0.08667788 |
| ENSMUSG00 | Gm12056 | 1254 | 0 | 0 | 0 | 0 | 0 | 0 |
| ENSMUSG00 | Gm12973 | 603 | 0 | 0 | 0 | 0 | 0 | 0 |
| ENSMUSG00 | Gm14145 | 505 | 0 | 0 | 0 | 0 | 0 | 0 |
| ENSMUSG00 | Gm12023 | 513 | 0 | 0 | 0 | 0 | 0 | 0 |
| ENSMUSG00 | Gm13069 | 315 | 0.12620788 | 0 | 0 | 0.11454617 | 0 | 0 |
| ENSMUSG00 | Gm14718 | 750 | 0 | 0 | 0 | 0 | 0 | 0 |
| ENSMUSG00 | Gm16325 | 3225 | 0 | 0 | 0 | 0 | 0 | 0 |
| ENSMUSG00 | Gm11669 | 4811 | 0 | 0 | 0 | 0.00749991 | 0 | 0 |
| ENSMUSG00 | Gm15843 | 528 | 0 | 0 | 0 | 0 | 0 | 0.72319119 |
| ENSMUSG00 | Gm15870 | 641 | 0 | 0 | 0 | 0 | 0 | 0 |
| ENSMUSG00 | 1810014B01 | 1656 | 3.62504683 | 2.1241608 | 4.59101268 | 6.03546276 | 6.00854063 | 2.91522412 |
| ENSMUSG00 | Gm13548 | 384 | 0 | 0.37645658 | 0.57554483 | 0 | 0.26576237 | 0.07102771 |
| ENSMUSG00 | Gm13052 | 765 | 0.1039359 | 0 | 0 | 0 | 0 | 0 |
| ENSMUSG00 | Gm12668 | 572 | 0 | 0 | 0 | 0 | 0 | 0 |
| ENSMUSG00 | Gm13267 | 868 | 0 | 0 | 0.10184757 | 0.49883011 | 0 | 0 |
| ENSMUSG00 | A930028C08 | 3050 | 0.16944959 | 0.06319533 | 0.08695445 | 0 | 0.20075951 | 0.07154004 |
| ENSMUSG00 | 4933406K04 | 2660 | 0 | 0 | 0 | 0 | 0 | 0.01025362 |
| ENSMUSG00 | Asxl1b | 1146 | 0 | 0 | 0 | 0 | 0 | 0 |
| ENSMUSG00 | Gm15584 | 754 | 0 | 0 | 0 | 0 | 0 | 0 |
| ENSMUSG00 | Gm13012 | 860 | 0 | 0 | 0 | 0 | 0 | 0 |
| ENSMUSG00 | 2410114N07 | 2372 | 0 | 0 | 0 | 0 | 0 | 0 |
| ENSMUSG00 | A930038B10 | 6180 | 0 | 0.07797159 | 0.20741965 | 0.19850963 | 0.10733704 | 0.09709419 |
| ENSMUSG00 | Gm15638 | 4885 | 0.50457314 | 0.60171402 | 0.39813328 | 0.53181314 | 0.57450372 | 0.1116669 |
| ENSMUSG00 | Gm11290 | 393 | 0 | 0 | 0 | 0 | 0 | 0 |
| ENSMUSG00 | Gm15612 | 3115 | 0.03828778 | 0.43313656 | 0 | 0.79924914 | 0.09828515 | 0.08755904 |
| ENSMUSG00 | Gm4285 | 899 | 1.94576323 | 1.82240159 | 2.36005391 | 2.16733082 | 1.92980732 | 5.97675632 |
| ENSMUSG00 | E130003G02 | 2887 | 0 | 0 | 0 | 0 | 0 | 0 |
| ENSMUSG00 | 1700003G13 | 914 | 0 | 0 | 0 | 0 | 0 | 0.05968192 |
| ENSMUSG00 | D430036J16 | 2577 | 0 | 0.29917853 | 0 | 0 | 0 | 0 |
| ENSMUSG00 | Gm13003 | 669 | 0.41597663 | 0.72027567 | 0.39642909 | 1.07868594 | 0.68645349 | 1.06000093 |
| ENSMUSG00 | Gm15622 | 1462 | 0 | 0 | 0 | 0 | 0 | 0 |
| ENSMUSG00 | Gm12707 | 486 | 0 | 0 | 0 | 0 | 0 | 0.22448263 |
| ENSMUSG00 | Gm13074 | 2733 | 0 | 0 | 0 | 0 | 0 | 0 |
| ENSMUSG00 | 4933400F03 | 2382 | 0 | 0 | 0 | 0 | 0 | 0.01145631 |
| ENSMUSG00 | Gm13706 | 634 | 0 | 0 | 0.83662794 | 0.45529394 | 0 | 0.08603987 |
| ENSMUSG00 | Gm15949 | 124 | 0 | 0 | 0 | 0.29098423 | 0 | 0 |
| ENSMUSG00 | Gm11261 | 1133 | 0 | 0 | 0 | 0 | 0 | 0 |
| ENSMUSG00 | Gm4954 | 1599 | 0 | 0 | 0 | 0 | 0 | 0 |
| ENSMUSG00 | 4930448K12 | 1979 | 0.06026601 | 0.02434888 | 0.13401266 | 0.29171941 | 0.15470351 | 0.12403828 |
| ENSMUSG00 | A730085A09 | 2248 | 0 | 0 | 0 | 0 | 0 | 0 |
| ENSMUSG00 | Gm11690 | 1854 | 0 | 0 | 0 | 0 | 0 | 0 |
| ENSMUSG00 | Gm13944 | 960 | 0 | 0 | 0 | 0 | 0 | 0 |
| ENSMUSG00 | Gm15597 | 1047 | 0.1518834 | 0.27614007 | 0.08443523 | 0.20677389 | 0.04873579 | 0.67730718 |
| ENSMUSG00 | Gm14540 | 3259 | 0.24397349 | 0.14785653 | 0.82734349 | 1.55001112 | 0.8141674 | 0.97917545 |
| ENSMUSG00 | Gm16790 | 1079 | 0 | 0 | 0 | 0.03344026 | 0 | 0 |
| ENSMUSG00 | A730020E08 | 2919 | 3.00992161 | 1.7993567 | 4.39141298 | 1.85416467 | 2.62211592 | 3.01805703 |
| ENSMUSG00 | 4833412C05 | 893 | 0 | 0.21584073 | 0 | 0 | 0 | 0 |
| ENSMUSG00 | 4921518K17 | 3385 | 0 | 0 | 0 | 0 | 0 | 0 |
| ENSMUSG00 | Gm13807 | 1289 | 1.01778965 | 0.48597653 | 0.10287473 | 0.22393821 | 0.55420424 | 0.23275487 |
| ENSMUSG00 | Gm12167 | 576 | 0 | 0 | 0 | 0 | 0 | 0 |
| ENSMUSG00 | Gm16520 | 464 | 0 | 0 | 0 | 0 | 0 | 0 |
| ENSMUSG00 | Gm12050 | 613 | 0 | 0 | 0 | 0 | 0 | 0 |
| ENSMUSG00 | Gm15392 | 776 | 0 | 0 | 0 | 0 | 0 | 0 |
| ENSMUSG00 | Gm15831 | 513 | 0 | 0 | 0 | 0 | 0 | 0 |
| ENSMUSG00 | D030068K23 | 5300 | 0 | 0 | 0 | 0 | 0 | 0.00514616 |
| ENSMUSG00 | 2410003L11 | 1140 | 7.63723705 | 3.71965518 | 5.5058436 | 2.18391322 | 5.01311762 | 4.88072492 |
| ENSMUSG00 | 1700016G14 | 861 | 0 | 0 | 0 | 0 | 0 | 0 |
| ENSMUSG00 | Gm13483 | 1326 | 3.38791048 | 3.9246876 | 2.90012092 | 3.12928742 | 2.11647864 | 5.80048889 |
| ENSMUSG00 | Gm12801 | 261 | 0 | 0 | 0 | 0 | 0 | 0 |
| ENSMUSG00 | C330019F10 | 454 | 0 | 0 | 0 | 0 | 0 | 0 |
| ENSMUSG00 | Gm15966 | 680 | 0 | 0 | 0 | 0 | 0 | 0 |
| ENSMUSG00 | Gm15218 | 614 | 0 | 0 | 0 | 0 | 0 | 0 |
| ENSMUSG00 | Gm5834 | 1020 | 0 | 0 | 0 | 0.38912009 | 0.40020687 | 0 |
| ENSMUSG00 | Gm13632 | 382 | 0 | 0 | 0.11571163 | 0 | 0 | 0.07139958 |
| ENSMUSG00 | Gm16079 | 575 | 0 | 0 | 0 | 0 | 0 | 0 |
| ENSMUSG00 | Gm12829 | 2432 | 0 | 0 | 0 | 0 | 0 | 0 |
| ENSMUSG00 | Gm12698 | 606 | 0 | 0.07951558 | 0 | 0 | 0 | 0 |
| ENSMUSG00 | Gm11505 | 735 | 0 | 0 | 0 | 0 | 0 | 0 |
| ENSMUSG00 | Gm12750 | 1038 | 3.06400621 | 2.50680913 | 0.29808565 | 2.71136751 | 2.26128448 | 0.55179906 |
| ENSMUSG00 | Asfx2 | 1146 | 0 | 0 | 0 | 0 | 0 | 0 |
| ENSMUSG00 | Gm12905 | 934 | 2.9795328 | 0.72228072 | 1.65638598 | 1.19758392 | 3.55108611 | 1.98573391 |
| ENSMUSG00 | Gm12409 | 643 | 0.37096871 | 0 | 0 | 0 | 0 | 0.29692453 |
| ENSMUSG00 | Gm16388 | 1531 | 0 | 0 | 0 | 0 | 0 | 0 |
| ENSMUSG00 | Gm12823 | 314 | 0 | 0 | 0 | 0 | 0 | 0 |
| ENSMUSG00 | C430014B12 | 997 | 0.59812659 | 0.14499431 | 0.39901363 | 0.18095308 | 0.9724184 | 0.3009238 |
| ENSMUSG00 | Gm15229 | 1437 | 0 | 0 | 0 | 0 | 0 | 0 |
| ENSMUSG00 | Gm15912 | 575 | 0.48397976 | 0.83802508 | 3.45927467 | 0.50201105 | 0.62119066 | 0.9486831 |
| ENSMUSG00 | 2610507I01R | 1358 | 11.739284 | 14.5481895 | 12.2710566 | 16.2076931 | 20.4406101 | 12.1510727 |
| ENSMUSG00 | Gm3848 | 449 | 0.79688046 | 0.10731947 | 0.29533525 | 0 | 0.56822245 | 0 |
| ENSMUSG00 | Gm15342 | 2098 | 0 | 0.13780679 | 0 | 0.05159492 | 0.5837145 | 0.49401158 |
| ENSMUSG00 | Gm11906 | 675 | 0 | 0 | 0.13096842 | 0.10690976 | 0.07559463 | 0.04040687 |
| ENSMUSG00 | D030047H15 | 2533 | 0.03139004 | 0.0570704 | 0.2094047 | 0.02848957 | 0.18130177 | 0.08614177 |
| ENSMUSG00 | Gm11648 | 2725 | 0 | 0 | 0 | 0 | 0 | 0 |
| ENSMUSG00 | Gm12224 | 1087 | 0 | 0 | 0 | 0 | 0 | 0 |
| ENSMUSG00 | 1700054M17 | 442 | 0 | 0 | 0 | 0 | 0 | 0 |
| ENSMUSG00 | 2010012P19 | 1103 | 0.14417219 | 0 | 0.0801484 | 0.49068964 | 0.13878434 | 0.32145994 |
| ENSMUSG00 | Gm14634 | 3059 | 0.68880041 | 0.26778997 | 0.60689029 | 0.48361027 | 0.71727171 | 0.37448017 |
| ENSMUSG00 | Gm15339 | 502 | 0 | 0 | 0 | 0 | 0 | 0 |
| ENSMUSG00 | Gm12091 | 829 | 0 | 0 | 0 | 0 | 0 | 0 |
| ENSMUSG00 | Gm15832 | 1829 | 0.36951513 | 0.23711207 | 0.24167219 | 0.45373812 | 0.33478213 | 0.29824647 |
| ENSMUSG00 | Gm12861 | 677 | 0 | 0 | 0 | 0 | 0 | 0 |
| ENSMUSG00 | 5330429C05 | 480 | 0 | 0 | 0 | 0 | 0 | 0 |
| ENSMUSG00 | Gm11604 | 741 | 0 | 0 | 0 | 0 | 0 | 0 |
| ENSMUSG00 | Gm6689 | 480 | 0 | 0 | 0 | 0 | 0 | 0 |
| ENSMUSG00 | Gm15338 | 549 | 0 | 0 | 0 | 0 | 0 | 0 |
| ENSMUSG00 | A930041C12 | 534 | 0 | 0.18047357 | 0.33109995 | 0 | 0 | 0 |
| ENSMUSG00 | AI115009 | 1849 | 0 | 0 | 0 | 0 | 0 | 0 |
| ENSMUSG00 | Gm12200 | 1852 | 0 | 0 | 0 | 0 | 0 | 0 |
| ENSMUSG00 | Gm15340 | 702 | 0 | 0 | 0 | 0 | 0 | 0 |
| ENSMUSG00 | Gm16881 | 597 | 0.26636838 | 0.08071431 | 0 | 0.36263361 | 0 | 0.8680371 |
| ENSMUSG00 | Gm14273 | 1781 | 0 | 0 | 0 | 0 | 0 | 0 |
| ENSMUSG00 | Gm16882 | 4955 | 0.88256364 | 0.44734134 | 0.15165113 | 0.53886404 | 0.39132236 | 0.41833957 |
| ENSMUSG00 | Gm14209 | 460 | 0 | 0 | 0 | 0 | 0 | 0 |
| ENSMUSG00 | Gm11846 | 512 | 0 | 0 | 0 | 0 | 0 | 0 |
| ENSMUSG00 | Gm12395 | 534 | 0 | 0 | 0 | 0 | 0 | 0 |
| ENSMUSG00 | Gm14061 | 1002 | 0.23805677 | 0 | 0 | 0 | 0 | 0.40830298 |
| ENSMUSG00 | Gm16197 | 2954 | 0.08074911 | 0 | 0.29926773 | 0.02442928 | 0.39729406 | 0.28622675 |
| ENSMUSG00 | Gm15601 | 4030 | 0.84837998 | 0.05978467 | 0.40582337 | 0 | 0 | 0 |
| ENSMUSG00 | Gm14320 | 1019 | 0.81929842 | 0 | 0.17351067 | 0 | 0.30044971 | 0.05353217 |
| ENSMUSG00 | Gm8091 | 881 | 0 | 0 | 0 | 0 | 0 | 0 |
| ENSMUSG00 | Gm16724 | 1409 | 0 | 0.13679615 | 0 | 0.33290744 | 0.28971683 | 0 |
| ENSMUSG00 | Gm15899 | 237 | 0 | 0 | 0 | 0 | 0 | 0 |
| ENSMUSG00 | Gm13032 | 1998 | 0 | 0.09646935 | 0 | 0 | 0 | 0 |
| ENSMUSG00 | Gm16019 | 259 | 0 | 0 | 0 | 0 | 0 | 0 |
| ENSMUSG00 | Gm12951 | 432 | 0.27607973 | 0 | 0 | 0 | 0 | 0.12627148 |
| ENSMUSG00 | Gm13752 | 487 | 0 | 0 | 0 | 0 | 0 | 0 |
| ENSMUSG00 | Gm13643 | 943 | 0 | 0 | 0 | 0 | 0 | 0 |
| ENSMUSG00 | Gm12781 | 392 | 0 | 0 | 0 | 0 | 0 | 0 |
| ENSMUSG00 | Gm8980 | 2191 | 0 | 0 | 0 | 0 | 0 | 0 |
| ENSMUSG00 | Rtl1 | 6355 | 0 | 0 | 0 | 0 | 0 | 0 |
| ENSMUSG00 | Gm12299 | 2325 | 0 | 0.0414507 | 0 | 0 | 0 | 0 |
| ENSMUSG00 | Gm15888 | 536 | 0 | 0 | 0 | 0 | 0 | 0 |
| ENSMUSG00 | 4933427I22R | 1764 | 0 | 0 | 0 | 0 | 0 | 0 |
| ENSMUSG00 | Gm13421 | 892 | 1.78275697 | 1.35051688 | 0.84241181 | 2.38659263 | 2.11656492 | 1.34538579 |
| ENSMUSG00 | 2310001H17 | 5688 | 2.97047806 | 4.45605988 | 1.78734597 | 3.17811257 | 4.26116887 | 3.52920789 |
| ENSMUSG00 | Gm12648 | 650 | 0 | 0 | 0 | 0 | 0 | 0 |
| ENSMUSG00 | Gm15556 | 660 | 0 | 0 | 0 | 0 | 0 | 0 |
| ENSMUSG00 | Tmem61 | 4003 | 22.3159043 | 17.17763 | 25.154084 | 11.8440686 | 27.1511817 | 22.2734938 |
| ENSMUSG00 | Gm8988 | 1237 | 0 | 0 | 0 | 0 | 0 | 0 |
| ENSMUSG00 | A830009L08 | 2806 | 0.15584828 | 0.01717265 | 0.11026832 | 0.09001223 | 0 | 0.04860057 |
| ENSMUSG00 | 2610307P16 | 3562 | 0.41295699 | 0.17586293 | 0.18613915 | 0.37479945 | 0.35813009 | 0.78868272 |
| ENSMUSG00 | 4930535C22 | 1333 | 0 | 0 | 0 | 0 | 0 | 0 |
| ENSMUSG00 | Gm14371 | 1513 | 0 | 0 | 0 | 0 | 0 | 0 |
| ENSMUSG00 | Cd63-ps | 1631 | 0.56062296 | 0.76814684 | 0.62332458 | 0.79641545 | 0.50056531 | 0.71907387 |
| ENSMUSG00 | 4930405D11 | 929 | 0 | 0 | 0 | 0 | 0 | 0 |
| ENSMUSG00 | Gm11201 | 696 | 0 | 0 | 0 | 0 | 0 | 0 |
| ENSMUSG00 | Gm13364 | 541 | 27.6304264 | 14.7854887 | 17.7297596 | 20.6754784 | 35.9353959 | 10.536783 |
| ENSMUSG00 | Gm14348 | 867 | 0 | 0 | 0 | 0 | 0 | 0 |
| ENSMUSG00 | 1700003D09 | 1386 | 0.02868361 | 0 | 0 | 0 | 0 | 0 |
| ENSMUSG00 | 2310014F06 | 655 | 0.54625851 | 0.29426835 | 0.40490238 | 0.77121927 | 0.54532005 | 0.66625073 |
| ENSMUSG00 | AA387200 | 1175 | 0.0676689 | 0 | 0 | 0.92124369 | 0 | 0 |
| ENSMUSG00 | 2310069G16 | 838 | 1.75531358 | 0 | 0.42197464 | 1.16254797 | 0.73068796 | 1.92029083 |
| ENSMUSG00 | Gm16684 | 1017 | 0.54727309 | 0.47380966 | 0.56501864 | 0.24835232 | 0.5519077 | 0.61683058 |
| ENSMUSG00 | Gm14275 | 428 | 0 | 0 | 0 | 0 | 0 | 0 |
| ENSMUSG00 | Gm13589 | 1673 | 0 | 0 | 0 | 0 | 0 | 0 |
| ENSMUSG00 | Gm12145 | 213 | 0 | 0 | 0.20752039 | 0 | 0.23956045 | 0 |
| ENSMUSG00 | Gm12315 | 908 | 0 | 0 | 0 | 0 | 0 | 0 |
| ENSMUSG00 | Gm15496 | 2452 | 0 | 0 | 0 | 0 | 0 | 0 |
| ENSMUSG00 | 4930478K11 | 712 | 0.2233454 | 0 | 0 | 0 | 0 | 0 |
| ENSMUSG00 | Gm12356 | 351 | 0.22652695 | 0.82369987 | 0.50372471 | 0.82238278 | 0.43612287 | 0 |
| ENSMUSG00 | 4930481B07 | 1515 | 0.55106607 | 0.03180623 | 0.78775562 | 0.19053225 | 0.77465785 | 0.8101378 |
| ENSMUSG00 | Syna | 2025 | 0 | 0 | 0 | 0 | 0 | 0 |
| ENSMUSG00 | A930019D19 | 954 | 3.04208604 | 2.32345528 | 3.47498766 | 7.75347917 | 4.27894137 | 2.54448941 |
| ENSMUSG00 | Gm15904 | 484 | 0 | 0 | 0 | 0 | 0 | 1.07069865 |
| ENSMUSG00 | Gm16984 | 1090 | 0.69298544 | 0.4862852 | 0.52717794 | 0.4634391 | 0.32769232 | 0.82574596 |
| ENSMUSG00 | Gm15249 | 422 | 0 | 0 | 0 | 0 | 0 | 0 |
| ENSMUSG00 | Gm16983 | 1248 | 0 | 0 | 0 | 0 | 0 | 0 |
| ENSMUSG00 | 2310016D23 | 384 | 0 | 0 | 0 | 0 | 0 | 0 |
| ENSMUSG00 | Gm14368 | 413 | 0 | 0 | 0 | 0 | 0 | 0 |
| ENSMUSG00 | Gm12530 | 1754 | 0 | 0 | 0 | 0 | 0 | 0 |
| ENSMUSG00 | Gm13027 | 239 | 0 | 0 | 0 | 0 | 0 | 0 |
| ENSMUSG00 | Gm15600 | 854 | 0 | 0 | 0 | 0 | 0 | 0 |
| ENSMUSG00 | Gm15643 | 381 | 0 | 0 | 0 | 0 | 0 | 0 |
| ENSMUSG00 | Gm15411 | 3484 | 0.15975222 | 0.13830781 | 0.11418387 | 0.26926899 | 0.16110509 | 0.36794146 |
| ENSMUSG00 | 1110028F11 | 973 | 0.32686932 | 0 | 0 | 0 | 0 | 0.30834638 |
| ENSMUSG00 | Gm14742 | 378 | 0 | 0 | 0 | 0 | 0 | 0 |
| ENSMUSG00 | 4930444M15 | 1654 | 0 | 0 | 0 | 0 | 0 | 0 |
| ENSMUSG00 | Gm13572 | 557 | 1.35611154 | 1.73021336 | 2.14263871 | 1.16602657 | 1.92379514 | 1.6159122 |
| ENSMUSG00 | Gm13816 | 713 | 0 | 0 | 0 | 0 | 0 | 0 |
| ENSMUSG00 | Gm5970 | 1215 | 0 | 0 | 0 | 0 | 0 | 0 |
| ENSMUSG00 | Gm15258 | 740 | 0 | 0 | 0 | 0 | 0 | 0 |
| ENSMUSG00 | Gm12055 | 791 | 0 | 0 | 0 | 0 | 0 | 0 |
| ENSMUSG00 | Gm12408 | 993 | 0 | 0 | 0 | 0 | 0 | 0 |
| ENSMUSG00 | Gm11375 | 905 | 0 | 0 | 0 | 0 | 0 | 0 |
| ENSMUSG00 | 9530051G07 | 4410 | 0.38763847 | 0 | 0.26060044 | 1.12909799 | 0.76366005 | 0 |
| ENSMUSG00 | Gm11538 | 1943 | 0 | 0 | 0 | 0 | 0 | 0 |
| ENSMUSG00 | 1700001G11 | 1328 | 0.05987271 | 0.10885491 | 0.23299164 | 0 | 0.03842348 | 0.10269066 |
| ENSMUSG00 | Gm11682 | 513 | 0 | 0 | 0 | 0 | 0 | 0 |
| ENSMUSG00 | Gm13403 | 4162 | 0 | 0 | 0 | 0 | 0 | 0 |
| ENSMUSG00 | Gm16896 | 5482 | 0 | 0 | 0 | 0.11847443 | 0.22339165 | 0.04975308 |
| ENSMUSG00 | Gm7437 | 658 | 0 | 0 | 0 | 0 | 0 | 0 |
| ENSMUSG00 | Gm16731 | 918 | 0 | 0 | 0 | 0 | 0 | 0 |
| ENSMUSG00 | Gm11479 | 385 | 0 | 0 | 0 | 0 | 0 | 0 |
| ENSMUSG00 | Gm11515 | 699 | 0 | 0 | 0 | 0.15485856 | 0 | 0 |
| ENSMUSG00 | Gm15905 | 288 | 0 | 0 | 0 | 0 | 0 | 0 |
| ENSMUSG00 | Gm16733 | 1253 | 0 | 0 | 0.52915215 | 0 | 0 | 0.19590723 |
| ENSMUSG00 | Gm2788 | 1658 | 0 | 0 | 0 | 0 | 0 | 0 |
| ENSMUSG00 | A830012C17 | 1663 | 0 | 0 | 0 | 0 | 0 | 0 |
| ENSMUSG00 | AW822252 | 947 | 0 | 0.10176651 | 0 | 0.07620284 | 0.10776426 | 0.0288011 |
| ENSMUSG00 | Gm13411 | 412 | 0 | 0 | 0 | 0 | 0 | 0 |
| ENSMUSG00 | Gm12493 | 439 | 0 | 0 | 0 | 0 | 0 | 0 |
| ENSMUSG00 | Gm13448 | 1845 | 0 | 0 | 0 | 0 | 0 | 0 |
| ENSMUSG00 | Gm16744 | 2187 | 0.01817809 | 0 | 0 | 0.19798104 | 0.16332173 | 0 |
| ENSMUSG00 | 8230206L02 | 1088 | 0.29231971 | 0.66433514 | 2.23446817 | 2.08930956 | 1.07868258 | 4.28673099 |
| ENSMUSG00 | Gm7805 | 447 | 2.04558401 | 0.32339894 | 0.69219888 | 2.17945235 | 1.02737669 | 0.6101709 |
| ENSMUSG00 | Gm16745 | 2268 | 0 | 0 | 0 | 0 | 0 | 0 |
| ENSMUSG00 | Gm13293 | 666 | 0.77600788 | 0 | 0.46458394 | 0 | 0.07661618 | 0.04095291 |
| ENSMUSG00 | 1700057H21 | 884 | 0 | 0 | 0 | 0 | 0 | 0 |
| ENSMUSG00 | Gm8817 | 1069 | 0 | 0 | 0 | 0 | 0 | 0 |
| ENSMUSG00 | 4930587A21 | 849 | 0 | 0 | 0 | 0 | 0 | 0 |
| ENSMUSG00 | Gm15318 | 1691 | 0 | 0 | 0 | 0 | 0.03017527 | 0 |
| ENSMUSG00 | Olfr396-ps1 | 1570 | 0 | 0 | 0 | 0 | 0 | 0 |
| ENSMUSG00 | Gm15902 | 465 | 0 | 0.10362676 | 0 | 0 | 0 | 0 |
| ENSMUSG00 | Gm15706 | 897 | 4.38772862 | 6.07030988 | 3.10447727 | 5.63153426 | 5.34724563 | 8.17935112 |
| ENSMUSG00 | Gm16095 | 251 | 0 | 0 | 0 | 0 | 0 | 0 |
| ENSMUSG00 | 4833417C18 | 1267 | 0.25102119 | 0.11409576 | 2.5118648 | 0.76891492 | 0.40273383 | 1.07634725 |
| ENSMUSG00 | 1700084C06 | 748 | 1.22242788 | 1.28840754 | 0.47274698 | 0.86828449 | 0.88682204 | 1.45853686 |
| ENSMUSG00 | Gm11872 | 2216 | 0 | 0 | 0 | 0 | 0 | 0 |
| ENSMUSG00 | Gm13179 | 1181 | 0 | 0 | 0 | 0 | 0 | 0 |
| ENSMUSG00 | Gm15762 | 331 | 0 | 0 | 0 | 0 | 0 | 0 |
| ENSMUSG00 | Gm12239 | 495 | 0 | 0 | 0 | 0 | 0 | 0 |
| ENSMUSG00 | Gm15767 | 682 | 0 | 0 | 0 | 0 | 0 | 0 |
| ENSMUSG00 | Rad51ap2 | 3172 | 0.27573158 | 0.22786779 | 0 | 0.27300412 | 0 | 0 |
| ENSMUSG00 | 2810011L19 | 3392 | 0 | 0 | 0 | 0 | 0 | 0 |
| ENSMUSG00 | Gm16639 | 1320 | 0.03011779 | 0.03650488 | 0 | 0 | 0 | 0 |
| ENSMUSG00 | 4933406G16 | 1410 | 0 | 0 | 0 | 0 | 0 | 0.03868743 |
| ENSMUSG00 | Gm12374 | 3291 | 0 | 0 | 0 | 0 | 0 | 0 |
| ENSMUSG00 | Gm11250 | 304 | 0 | 0 | 0 | 0 | 0 | 0 |
| ENSMUSG00 | Gm12243 | 819 | 0 | 0 | 0 | 0 | 0 | 0.29972131 |
| ENSMUSG00 | Pax6os1 | 1914 | 0 | 0 | 0 | 0 | 0 | 0 |
| ENSMUSG00 | Gm14255 | 246 | 0 | 0 | 0 | 0 | 0 | 0 |
| ENSMUSG00 | Gm12348 | 311 | 0 | 0 | 0 | 0 | 0 | 0 |
| ENSMUSG00 | Gm15929 | 1158 | 0.58362968 | 0 | 0 | 0.49854293 | 0.17625691 | 0.23553229 |
| ENSMUSG00 | 2900076A07 | 697 | 16.4839797 | 17.5600521 | 31.5818046 | 20.5517528 | 11.2741203 | 28.9181613 |
| ENSMUSG00 | Gm15201 | 1208 | 0.46074232 | 2.43325577 | 0.29272744 | 0.83633878 | 0.54912491 | 0 |
| ENSMUSG00 | Gm12610 | 577 | 0 | 0 | 0 | 0 | 0 | 0 |
| ENSMUSG00 | Gm14254 | 294 | 0 | 0 | 0 | 0 | 0 | 0 |
| ENSMUSG00 | Gm16085 | 450 | 0 | 0 | 0 | 0 | 0 | 0 |
| ENSMUSG00 | Gm12227 | 559 | 0 | 0 | 0 | 0 | 0 | 0 |
| ENSMUSG00 | Wipf3 | 7104 | 0 | 0.10852802 | 0 | 0.00507912 | 0.00718277 | 0 |
| ENSMUSG00 | Gm12869 | 342 | 0 | 0 | 0 | 0 | 0 | 0 |
| ENSMUSG00 | Gm12670 | 574 | 0 | 0 | 0 | 0 | 0 | 0 |
| ENSMUSG00 | Gm12473 | 577 | 0.06890031 | 0.08351203 | 0 | 0 | 0.08843393 | 0 |
| ENSMUSG00 | A330043C09 | 1007 | 0 | 0 | 0 | 0 | 0 | 0.16251026 |
| ENSMUSG00 | 1700095A21 | 1066 | 0 | 0 | 0 | 0.06769614 | 0 | 0 |
| ENSMUSG00 | 9530046B11 | 1154 | 0.72345329 | 0.75160828 | 0.2681221 | 0.40647017 | 0.79690534 | 1.70171059 |
| ENSMUSG00 | Drr1 | 2901 | 0 | 0 | 0 | 0 | 0 | 0 |
| ENSMUSG00 | Gm12735 | 649 | 0.61256519 | 0.44548329 | 0.06810762 | 0.05559637 | 0.3144923 | 1.21874351 |
| ENSMUSG00 | Gm16045 | 2967 | 0.16079062 | 1.99761793 | 0.44693471 | 0 | 0 | 0.6434866 |
| ENSMUSG00 | Gm11802 | 649 | 0 | 0 | 0 | 0 | 0 | 0 |
| ENSMUSG00 | Gm15178 | 470 | 0 | 0 | 0 | 0 | 0 | 0 |
| ENSMUSG00 | Gm13824 | 1366 | 4.3946395 | 2.29291269 | 4.59492072 | 2.72068125 | 4.18371456 | 5.35110053 |
| ENSMUSG00 | Gm13837 | 335 | 0 | 0 | 0 | 0 | 0 | 0 |
| ENSMUSG00 | Gm4846 | 1736 | 0 | 0 | 0 | 0 | 0 | 0 |
| ENSMUSG00 | Gm15324 | 353 | 0 | 0 | 0 | 0 | 0 | 0 |
| ENSMUSG00 | Gm12575 | 1304 | 0.091462 | 0 | 0 | 0 | 0 | 0 |
| ENSMUSG00 | Gm15797 | 934 | 0 | 1.34137848 | 0.52057845 | 0.23179044 | 1.47506654 | 1.66451225 |
| ENSMUSG00 | Gm16084 | 617 | 0 | 0 | 0 | 0 | 0 | 0 |
| ENSMUSG00 | Gm13075 | 1703 | 9.52450737 | 6.96057825 | 6.48882017 | 11.7166005 | 11.565579 | 10.5703241 |
| ENSMUSG00 | Gm16853 | 1981 | 0 | 0.31621593 | 0 | 0 | 0.05151578 | 0.05507247 |
| ENSMUSG00 | Gm12257 | 582 | 0 | 0 | 0 | 0.06199664 | 0.43837093 | 0 |
| ENSMUSG00 | 4930439A04 | 858 | 0.55602071 | 0 | 0 | 0 | 0 | 0.1271545 |
| ENSMUSG00 | Gm16180 | 310 | 0 | 0 | 0 | 0 | 0 | 0 |
| ENSMUSG00 | 4930423M02 | 1237 | 0 | 0 | 0 | 0 | 0 | 0 |
| ENSMUSG00 | Gm16183 | 1188 | 0 | 0 | 0 | 0 | 0 | 0 |
| ENSMUSG00 | 1700120G11 | 489 | 0 | 0 | 0 | 0 | 0 | 0 |
| ENSMUSG00 | Gm13708 | 388 | 0 | 0 | 0 | 0 | 0 | 0 |
| ENSMUSG00 | 8430436N08 | 1074 | 0 | 0 | 0 | 0 | 0 | 0 |
| ENSMUSG00 | Gm12354 | 608 | 0.13077461 | 0 | 0 | 0 | 0 | 0 |
| ENSMUSG00 | Gm11861 | 379 | 0 | 0 | 0 | 0 | 0 | 0 |
| ENSMUSG00 | Gm13274 | 488 | 0 | 0 | 0 | 0 | 0 | 0 |
| ENSMUSG00 | Gm15728 | 3629 | 0.51488222 | 0.86308039 | 0.31668446 | 0.5369056 | 1.0123723 | 0.45846046 |
| ENSMUSG00 | Ncr3-ps | 522 | 0 | 0 | 0 | 0 | 0 | 0 |
| ENSMUSG00 | Gm14396 | 682 | 0.64121743 | 0.70654607 | 0.12962417 | 2.38078006 | 1.04746226 | 1.23975633 |
| ENSMUSG00 | Gm11657 | 335 | 0 | 0 | 0 | 0 | 0 | 0 |
| ENSMUSG00 | 1700123O12 | 1524 | 0 | 0 | 0 | 0 | 0 | 0 |
| ENSMUSG00 | Gm14759 | 1531 | 0 | 0 | 0 | 0 | 0 | 0 |
| ENSMUSG00 | 1700113A16 | 980 | 2.10947448 | 1.32758565 | 3.20237842 | 2.09864953 | 1.35376099 | 4.53649612 |
| ENSMUSG00 | Gm15829 | 867 | 0.32097851 | 0.61136201 | 0.10196504 | 0 | 0.41197766 | 0.6606314 |
| ENSMUSG00 | 2610528B01 | 2006 | 0.49545714 | 0.55248683 | 0.66104451 | 0 | 0.89029071 | 0.4078959 |
| ENSMUSG00 | 5730422E09 | 1736 | 0.16030436 | 0.02775717 | 0.15277135 | 0.31176882 | 0.08817922 | 0.15711198 |
| ENSMUSG00 | Gm16760 | 1533 | 0 | 0 | 0 | 0 | 0 | 0 |
| ENSMUSG00 | Gm12916 | 944 | 0 | 0 | 0 | 0 | 0 | 0 |
| ENSMUSG00 | Gm11640 | 270 | 0 | 0 | 0 | 0 | 0 | 0 |
| ENSMUSG00 | Gm12531 | 759 | 0 | 0 | 0 | 0 | 0 | 0 |
| ENSMUSG00 | 4930528D03 | 1671 | 0 | 0 | 0 | 0 | 0 | 0 |
| ENSMUSG00 | Gm15323 | 426 | 0 | 0 | 0 | 0.50819781 | 0 | 0 |
| ENSMUSG00 | Gm15341 | 589 | 0 | 0 | 0 | 0 | 0 | 0 |
| ENSMUSG00 | Gm15328 | 2974 | 0 | 0 | 0 | 0 | 0 | 0 |
| ENSMUSG00 | Gm12688 | 699 | 0 | 0 | 0 | 0 | 0 | 0 |
| ENSMUSG00 | Gm16250 | 255 | 0 | 0 | 0 | 0 | 0 | 0 |
| ENSMUSG00 | Gm14291 | 802 | 0 | 0 | 0 | 0 | 0 | 0 |
| ENSMUSG00 | 2610316D01 | 913 | 0 | 0 | 0 | 0 | 0 | 0 |
| ENSMUSG00 | Gm16885 | 1105 | 0.53966716 | 2.04955908 | 1.76007339 | 0.91429615 | 0.8773766 | 2.22146383 |
| ENSMUSG00 | Gm11832 | 429 | 0 | 0 | 0.515173 | 0 | 2.02202422 | 0.38146349 |
| ENSMUSG00 | Gm14707 | 396 | 0 | 0 | 0 | 0 | 0 | 0 |
| ENSMUSG00 | Gm11636 | 772 | 0 | 0 | 0 | 0.0467384 | 0 | 0 |
| ENSMUSG00 | 1110036E04 | 774 | 0 | 0.18676916 | 0 | 0 | 0 | 0 |
| ENSMUSG00 | Gm16312 | 360 | 0 | 0 | 0 | 0 | 0 | 0 |
| ENSMUSG00 | Gm5602 | 1747 | 0.50064143 | 1.29637251 | 1.21447537 | 1.34249164 | 0.554952 | 0.46836816 |
| ENSMUSG00 | Gm13391 | 1113 | 0 | 0 | 0 | 0 | 0 | 0 |
| ENSMUSG00 | Gm15326 | 939 | 0.71974778 | 0 | 0.09414663 | 1.57546733 | 0.54341188 | 0 |

| | | | | | | | | |
|---|---|---|---|---|---|---|---|---|
| [illegible] | | | | | | | | |
| ENSMUSG00 | Gm11934 | 1534 | 0.49649195 | 0.50259653 | 0.37469189 | 0.47043083 | 0.03326361 | 0.81788358 |
| ENSMUSG00 | 1700091E21I | 1705 | 0 | 0 | 0 | 0 | 0 | 0 |
| ENSMUSG00 | Gm13558 | 551 | 0.14430302 | 0 | 0 | 0 | 0 | 0.64350329 |
| ENSMUSG00 | Gm16211 | 1618 | 0 | 0 | 0 | 0 | 0 | 0 |
| ENSMUSG00 | BC048594 | 393 | 0.20231797 | 0.12261181 | 0.22494577 | 0.45905909 | 0 | 0 |
| ENSMUSG00 | Gm13631 | 651 | 0 | 0 | 0 | 0 | 0 | 0 |
| ENSMUSG00 | Gm16331 | 497 | 0.31996363 | 0 | 0 | 0 | 0 | 0 |
| ENSMUSG00 | Gm16159 | 1151 | 0.03453995 | 0 | 0 | 0 | 0 | 0.02369647 |
| ENSMUSG00 | 1700001J04R | 898 | 0 | 0 | 0 | 0 | 0 | 0 |
| ENSMUSG00 | Gm12718 | 779 | 0.20413597 | 0 | 0 | 0.09263683 | 0 | 0 |
| ENSMUSG00 | Gm16248 | 242 | 0 | 0 | 0 | 0 | 0 | 0 |
| ENSMUSG00 | 4930471C06I | 679 | 0 | 0 | 0 | 0 | 0 | 0 |
| ENSMUSG00 | Gm11211 | 356 | 0 | 0 | 0 | 0 | 0 | 0 |
| ENSMUSG00 | Gm13825 | 440 | 1.8070673 | 2.62835139 | 3.51605569 | 0.65603717 | 2.55131879 | 3.78125681 |
| ENSMUSG00 | 9030622O22I | 1670 | 0 | 0 | 0.05293634 | 0 | 0 | 0.0816606 |
| ENSMUSG00 | Smek3-ps | 2444 | 0 | 0 | 0 | 0 | 0 | 0 |
| ENSMUSG00 | Gm16685 | 1589 | 1.37605502 | 0.24260009 | 0.50071314 | 1.56680999 | 0.44957159 | 1.09853802 |
| ENSMUSG00 | Gm11379 | 368 | 0 | 0 | 0 | 0 | 0 | 0 |
| ENSMUSG00 | Gm15479 | 1131 | 0 | 0 | 0 | 0 | 0 | 0 |
| ENSMUSG00 | Gm15729 | 358 | 0 | 0 | 0 | 0 | 0 | 0 |
| ENSMUSG00 | Gm13150 | 2505 | 0.50785444 | 0.09618052 | 0.01764545 | 0.1440401 | 0 | 0 |
| ENSMUSG00 | Gm15271 | 2107 | 0 | 0 | 0 | 0 | 0 | 0 |
| ENSMUSG00 | Gm15598 | 761 | 0.0522411 | 0 | 0 | 0.04741399 | 0 | 0 |
| ENSMUSG00 | BC024582 | 2178 | 0 | 0 | 0 | 0 | 0.18742471 | 0 |
| ENSMUSG00 | Gm5698 | 1139 | 0 | 0 | 0 | 0 | 0 | 0 |
| ENSMUSG00 | Gm11454 | 711 | 0.16774464 | 0 | 0 | 0 | 0.21530116 | 0.61377529 |
| ENSMUSG00 | Gm12923 | 663 | 0 | 0 | 0 | 0.16326717 | 0 | 0 |
| ENSMUSG00 | Gm16196 | 462 | 0.08605082 | 0.31289898 | 0.38269994 | 0 | 0.66268021 | 0.70843219 |
| ENSMUSG00 | 9430041J12R | 2831 | 0 | 0 | 0 | 0 | 0 | 0 |
| ENSMUSG00 | Gm15354 | 1770 | 0 | 0 | 0 | 0 | 0 | 0 |
| ENSMUSG00 | Gm14643 | 1840 | 0 | 0 | 0 | 0 | 0 | 0 |
| ENSMUSG00 | Gm5918 | 707 | 9.72800301 | 13.9038673 | 10.3783677 | 11.4319349 | 9.74336738 | 21.0250048 |
| ENSMUSG00 | Gm13025 | 1159 | 0 | 0 | 0 | 0 | 0 | 0 |
| ENSMUSG00 | 4933438A12I | 1010 | 0 | 0 | 0 | 0 | 0 | 0 |
| ENSMUSG00 | Gm15389 | 663 | 0 | 0 | 0 | 0 | 0 | 0 |
| ENSMUSG00 | Gm16342 | 1014 | 0 | 0 | 0 | 0 | 0 | 0 |
| ENSMUSG00 | Gm14206 | 743 | 0 | 0 | 0 | 0 | 0 | 0 |
| ENSMUSG00 | Gm13029 | 591 | 0 | 0 | 0 | 0 | 0 | 0 |
| ENSMUSG00 | Gm15690 | 430 | 0 | 0 | 0 | 0 | 0.11866599 | 0.25371757 |
| ENSMUSG00 | Gm14342 | 2115 | 0 | 0 | 0 | 0 | 0 | 0 |
| ENSMUSG00 | Gm13827 | 796 | 0.09988814 | 0.24214293 | 0.11105991 | 0 | 0.12820697 | 0.20558773 |
| ENSMUSG00 | Gm15221 | 411 | 0 | 0 | 0 | 0 | 0 | 0 |
| ENSMUSG00 | Gm12144 | 1290 | 0 | 0.03735383 | 0 | 0 | 0 | 0 |
| ENSMUSG00 | C630020P19I | 4601 | 0 | 0 | 0 | 0 | 0 | 0 |
| ENSMUSG00 | 2700068H02I | 1870 | 0 | 0 | 0.21273614 | 0 | 0 | 0 |
| ENSMUSG00 | Kis2 | 2210 | 0 | 0 | 0 | 0 | 0 | 0 |
| ENSMUSG00 | Gm6425 | 789 | 0 | 0.12214561 | 0.28011307 | 0.04573136 | 0.06467221 | 0.13827447 |
| ENSMUSG00 | Gm15802 | 207 | 0 | 0 | 0 | 0 | 0 | 0 |
| ENSMUSG00 | Gm15534 | 843 | 0 | 0 | 0 | 0 | 0 | 0 |
| ENSMUSG00 | Gm14616 | 311 | 0 | 0 | 0 | 0 | 0 | 0 |
| ENSMUSG00 | Gm16950 | 4925 | 0 | 0 | 0 | 0 | 0 | 0 |
| ENSMUSG00 | Gm14317 | 1461 | 0 | 0.06596364 | 0 | 0 | 0 | 0 |
| ENSMUSG00 | Rmst | 5137 | 0 | 0 | 0 | 0 | 0 | 0 |
| ENSMUSG00 | C230034O21 | 2832 | 0.04211386 | 0.01701499 | 0.06243198 | 0 | 0 | 0 |
| ENSMUSG00 | Gm12764 | 615 | 0.38785835 | 0 | 0.64685624 | 0 | 0 | 0.22174504 |
| ENSMUSG00 | 1700036O09I | 347 | 0 | 0 | 0 | 0 | 0 | 0 |
| ENSMUSG00 | 1700024P03I | 1330 | 0 | 0 | 0 | 0 | 0 | 0 |
| ENSMUSG00 | Gm12860 | 953 | 0 | 0 | 0 | 0 | 0 | 0.42929653 |
| ENSMUSG00 | Gm15169 | 4549 | 0 | 0 | 0 | 0 | 0 | 0 |
| ENSMUSG00 | Gm15462 | 4576 | 0.11294171 | 0 | 0.1545519 | 0.02365519 | 0.25646998 | 0.10728661 |
| ENSMUSG00 | 4930579M01 | 929 | 0 | 0 | 0 | 0 | 0 | 0 |
| ENSMUSG00 | 1110035M17 | 535 | 0 | 0 | 0 | 0 | 0 | 0 |
| ENSMUSG00 | Gm13609 | 587 | 0.06772654 | 0 | 0 | 0 | 0.26078216 | 0 |
| ENSMUSG00 | Gm11508 | 438 | 0 | 0 | 0.2018349 | 0.24713729 | 0 | 0 |
| ENSMUSG00 | Gm16329 | 368 | 0 | 0 | 0 | 0 | 0 | 0 |
| ENSMUSG00 | Gm6152 | 339 | 0 | 0 | 0 | 0 | 0 | 0 |
| ENSMUSG00 | Gm13571 | 3093 | 0 | 0 | 0 | 0 | 0 | 0 |
| ENSMUSG00 | Gm13976 | 477 | 0 | 0 | 0 | 0 | 0 | 0 |
| ENSMUSG00 | Gm16598 | 1357 | 1.20116043 | 0.2840763 | 0.16286604 | 0 | 0 | 0 |
| ENSMUSG00 | 8430403D17I | 3612 | 0.38522752 | 0.346857 | 0.23251246 | 0.30967425 | 0.36729949 | 0.95899202 |
| ENSMUSG00 | Gm13659 | 652 | 0 | 0 | 0 | 0 | 0.15652263 | 0 |
| ENSMUSG00 | Gm6270 | 556 | 0 | 0 | 0 | 0 | 0 | 0 |
| ENSMUSG00 | 1700019E08I | 827 | 0 | 0 | 0 | 0 | 0 | 0 |
| ENSMUSG00 | Gm13985 | 618 | 0 | 0 | 0 | 0 | 0 | 0.66200581 |
| ENSMUSG00 | Gm15668 | 673 | 0 | 0 | 0 | 0 | 0 | 0 |
| ENSMUSG00 | Gm12679 | 634 | 0 | 0 | 0 | 0 | 0 | 0 |
| ENSMUSG00 | Gm13783 | 730 | 0 | 0 | 0 | 0 | 0.06989915 | 0 |
| ENSMUSG00 | Gm14717 | 645 | 0 | 0 | 0 | 0 | 0 | 0 |
| ENSMUSG00 | Gm2716 | 1565 | 0 | 0 | 0 | 0 | 0 | 0 |
| ENSMUSG00 | 4933405E24I | 1075 | 0 | 0 | 0 | 0 | 0 | 0 |
| ENSMUSG00 | A230087F16I | 3382 | 0 | 0 | 0 | 0 | 0.03017527 | 0 |
| ENSMUSG00 | Gm12462 | 3054 | 0 | 0 | 0 | 0 | 0 | 0 |
| ENSMUSG00 | Gm13845 | 2939 | 2.51599843 | 2.29537322 | 6.58741315 | 3.15518389 | 3.10776498 | 5.70735051 |
| ENSMUSG00 | A330040F15I | 1104 | 1.40440556 | 0.82929565 | 0.0800758 | 0 | 0.78573224 | 1.16114859 |
| ENSMUSG00 | Gm14062 | 4131 | 0 | 0 | 0 | 0 | 0 | 0 |
| ENSMUSG00 | Gm13610 | 949 | 0 | 0 | 0.04657729 | 0 | 0 | 0 |
| ENSMUSG00 | Gm9530 | 3198 | 0.18647036 | 0 | 0.05528686 | 0.05641345 | 0 | 0 |
| ENSMUSG00 | Gm16755 | 929 | 0.59911381 | 0.25934576 | 0.66612035 | 0.42723627 | 1.20837489 | 1.49731604 |
| ENSMUSG00 | 2410137F16I | 3651 | 0 | 0 | 0 | 0 | 0 | 0 |
| ENSMUSG00 | Gm16599 | 1177 | 0 | 0 | 0 | 0.06131189 | 0 | 0 |
| ENSMUSG00 | 4930412B13I | 1270 | 0.06260706 | 0.1897104 | 0 | 0 | 0 | 0 |
| ENSMUSG00 | Gm11665 | 283 | 0 | 0 | 0.93714155 | 0 | 0 | 0.48188409 |
| ENSMUSG00 | Gm362 | 264 | 0 | 0 | 0 | 0 | 0 | 0 |
| ENSMUSG00 | 2700069I18R | 1134 | 0 | 0 | 0 | 0 | 0 | 0 |
| ENSMUSG00 | Gm8661 | 991 | 0.04011653 | 0 | 0 | 0.18204866 | 0.05148978 | 0 |
| ENSMUSG00 | Gm12660 | 662 | 0 | 0 | 0 | 0 | 0 | 0 |
| ENSMUSG00 | Gm11820 | 660 | 0 | 0 | 0 | 0 | 0 | 0 |
| ENSMUSG00 | Gm514 | 1374 | 0 | 0 | 0.03217019 | 0 | 0 | 0 |
| ENSMUSG00 | Gm4887 | 2101 | 0.03784434 | 0.022935 | 0.02103848 | 0.01717375 | 0.04857342 | 0 |
| ENSMUSG00 | Gm15255 | 272 | 0 | 0 | 0 | 0 | 0 | 0 |
| ENSMUSG00 | 4732447D17I | 3483 | 0.10272734 | 0.09684327 | 0.02538148 | 0 | 0.02930025 | 0.11746184 |
| ENSMUSG00 | Gm7546 | 962 | 0 | 0 | 0 | 0 | 0 | 0 |
| ENSMUSG00 | Gm11816 | 980 | 0 | 0 | 0 | 0 | 0 | 0 |
| ENSMUSG00 | 1700112M01 | 822 | 0 | 0.11724195 | 0.32264119 | 0 | 0 | 0 |
| ENSMUSG00 | Gm16284 | 399 | 0 | 0 | 0 | 0 | 0 | 0 |
| ENSMUSG00 | 5830418P13I | 2381 | 0 | 0 | 0 | 0 | 0 | 0 |
| ENSMUSG00 | Gm15591 | 625 | 0 | 0.15419662 | 0.35361474 | 0.28865636 | 0.40821101 | 0.5673125 |
| ENSMUSG00 | Gm14258 | 2029 | 0 | 0 | 0 | 0 | 0 | 0 |
| ENSMUSG00 | Gm14329 | 589 | 0 | 0 | 0 | 0 | 0 | 0 |
| ENSMUSG00 | Gm7846 | 545 | 0.07294584 | 0.26524647 | 0 | 0 | 0 | 0.05004521 |
| ENSMUSG00 | 4930483K19I | 1233 | 0.03224289 | 0 | 0 | 0.08779086 | 0 | 0 |
| ENSMUSG00 | Gm16718 | 701 | 0 | 0 | 0.50444329 | 0 | 0 | 0 |
| ENSMUSG00 | Gm12121 | 498 | 1053.60006 | 500.926127 | 344.738872 | 694.615584 | 199.289761 | 841.351422 |
| ENSMUSG00 | 4930558G05I | 491 | 0 | 0 | 0 | 0 | 0 | 0 |
| ENSMUSG00 | Gm16170 | 3751 | 0.10598635 | 0.61662203 | 0.09427213 | 0.17314764 | 0.08162044 | 0.15269726 |
| ENSMUSG00 | Gm15277 | 290 | 0 | 0 | 0 | 0 | 0 | 0 |
| ENSMUSG00 | Gm15787 | 3091 | 1.27330721 | 1.32508819 | 0.34320422 | 0.630356 | 1.00699092 | 0.37060332 |
| ENSMUSG00 | Gm16127 | 232 | 0 | 0 | 0 | 0 | 0 | 0 |
| ENSMUSG00 | Gm12724 | 660 | 0 | 0 | 0 | 0 | 0 | 0 |
| ENSMUSG00 | Gm14095 | 513 | 0 | 0 | 0 | 0 | 0 | 0 |
| ENSMUSG00 | Gm12694 | 743 | 0 | 0 | 0 | 0 | 0 | 0 |
| ENSMUSG00 | C030044B11 | 1777 | 31.2092827 | 34.167089 | 48.0325036 | 31.8179875 | 31.4428146 | 58.9236579 |
| ENSMUSG00 | Gm13773 | 579 | 0 | 0 | 0 | 0 | 0 | 0 |
| ENSMUSG00 | 3100003L05I | 660 | 0 | 0 | 0 | 0 | 0 | 0 |
| ENSMUSG00 | Gm11534 | 670 | 0 | 0 | 0 | 0 | 0 | 0 |
| ENSMUSG00 | Gm12052 | 931 | 0 | 0 | 0 | 0 | 0 | 0 |
| ENSMUSG00 | 4930548K13I | 558 | 0 | 0 | 0 | 0 | 0 | 0 |
| ENSMUSG00 | Gm15239 | 601 | 0 | 0.16035422 | 0 | 1.08066024 | 0 | 0.09076419 |
| ENSMUSG00 | 5830403M04 | 1551 | 0.02563216 | 0.06213597 | 0 | 0 | 0.03289902 | 0 |
| ENSMUSG00 | Gm16259 | 552 | 0 | 0 | 0 | 0 | 0 | 0 |
| ENSMUSG00 | Gm3765 | 1446 | 0 | 0 | 0 | 0 | 0 | 0 |
| ENSMUSG00 | A930031H19I | 1043 | 0 | 0 | 0 | 0 | 0 | 0 |
| ENSMUSG00 | 4930519D14 | 1133 | 0 | 0 | 0 | 0 | 0 | 0 |
| ENSMUSG00 | Gm15850 | 2112 | 0 | 0 | 0 | 0 | 0 | 0 |
| ENSMUSG00 | Marcksl1-ps4 | 290 | 20.9744432 | 46.857161 | 9.1452089 | 33.0959443 | 26.041047 | 3.76201921 |
| ENSMUSG00 | Igf2as | 5671 | 0 | 0 | 0 | 0 | 0 | 0 |
| ENSMUSG00 | Gm12496 | 666 | 0 | 0 | 0 | 0 | 0 | 0 |
| ENSMUSG00 | Gm11670 | 508 | 0 | 0 | 0 | 0 | 0 | 0 |
| ENSMUSG00 | Gm16154 | 2910 | 0 | 0 | 0 | 0.11159395 | 0 | 0 |
| ENSMUSG00 | BC039966 | 1368 | 0 | 0 | 0 | 0 | 0 | 0 |
| ENSMUSG00 | 4930564K09I | 2806 | 0 | 0 | 0 | 0 | 0 | 0 |
| ENSMUSG00 | Gm16677 | 2171 | 0 | 0 | 0 | 0 | 0.23503628 | 0.03768951 |
| ENSMUSG00 | 1700121C08I | 673 | 0 | 0.57279575 | 0.06567882 | 0 | 0 | 0 |
| ENSMUSG00 | 2500002B13I | 2615 | 1.03379452 | 0.88449301 | 0.40567657 | 0.95206924 | 1.09272545 | 0.53193369 |
| ENSMUSG00 | 4930558K02I | 1603 | 0.32240876 | 0 | 0 | 0 | 0 | 0.22119171 |
| ENSMUSG00 | Olfr75-ps1 | 1731 | 0 | 0 | 0 | 0 | 0 | 0 |
| ENSMUSG00 | Gm15634 | 489 | 0.1625991 | 0.88686703 | 1.89823866 | 0.66408671 | 1.04348417 | 2.28683069 |
| ENSMUSG00 | 4930540M05 | 2065 | 0.05775615 | 0 | 0.12843151 | 0 | 0.29652131 | 0 |
| ENSMUSG00 | 9530082P21I | 4763 | 0.30882905 | 0.2731543 | 0.30624834 | 0.40150081 | 1.13558594 | 0.41802407 |
| ENSMUSG00 | Gm15686 | 362 | 0 | 0 | 0 | 0 | 0 | 0.52741015 |
| ENSMUSG00 | 2810433D01I | 1201 | 0 | 0 | 0 | 0.09013 | 0 | 0 |
| ENSMUSG00 | 2900093L17I | 412 | 0 | 0 | 0 | 0 | 0 | 0 |
| ENSMUSG00 | D630044L22I | 2957 | 1.80156726 | 3.87838121 | 0.64277283 | 4.20977523 | 4.41756923 | 1.70639441 |
| ENSMUSG00 | Gm15138 | 377 | 0 | 0 | 0 | 0 | 0.27069695 | 0 |
| ENSMUSG00 | Gm15972 | 415 | 0.76637071 | 1.04500718 | 1.17161512 | 0.43472343 | 3.31978831 | 4.27193145 |
| ENSMUSG00 | Gm15265 | 1137 | 0.45454815 | 0.50856403 | 0.42763437 | 0.34907871 | 0.49365887 | 0.79161222 |
| ENSMUSG00 | Gm16933 | 2031 | 0 | 0 | 0 | 0 | 0 | 0 |
| ENSMUSG00 | Snhg12 | 1658 | 1.29481058 | 1.13345672 | 0.5865142 | 1.34926825 | 0.70784478 | 1.11862212 |
| ENSMUSG00 | Gm15513 | 1567 | 0 | 0 | 0 | 0 | 0 | 0 |
| ENSMUSG00 | Gm16052 | 503 | 0.94844089 | 0.67058667 | 1.40602284 | 0.28693475 | 0.91299678 | 0.70491115 |
| ENSMUSG00 | Gm11269 | 592 | 0 | 0 | 0 | 0 | 0 | 0 |
| ENSMUSG00 | Gm16765 | 1094 | 0.03633956 | 1.76184432 | 2.6262521 | 3.2322124 | 0.55970431 | 2.81721594 |
| ENSMUSG00 | 4933433G15I | 1810 | 0.08785742 | 0.26622344 | 0 | 0 | 0.0845741 | 0.18082634 |
| ENSMUSG00 | D030055H07 | 5114 | 0.27208483 | 0.18844913 | 0.04321651 | 0 | 0.05986669 | 0.20799979 |
| ENSMUSG00 | Gm15632 | 244 | 0 | 0 | 0 | 0 | 0 | 0 |
| ENSMUSG00 | Gm11716 | 468 | 0 | 0 | 0.09444838 | 0.15419677 | 0.21806144 | 0 |
| ENSMUSG00 | 4732463B04I | 2929 | 2.59245366 | 1.84256795 | 4.49714893 | 3.07972384 | 2.56090039 | 4.050689 |
| ENSMUSG00 | Gm13790 | 471 | 0 | 0 | 0 | 0 | 0 | 0 |
| ENSMUSG00 | Gm16619 | 1207 | 0 | 0 | 0 | 0 | 0 | 0 |
| ENSMUSG00 | Gm4262 | 1983 | 0.36086669 | 0 | 0 | 0 | 0 | 0 |
| ENSMUSG00 | Gm11754 | 658 | 0 | 0 | 0 | 0 | 0 | 0 |
| ENSMUSG00 | G630016G05 | 3176 | 0.01251747 | 0 | 0 | 0 | 0 | 0 |
| ENSMUSG00 | A930007I19R | 5070 | 0.18035031 | 0.06652961 | 0.27026768 | 0.24197032 | 0.32205997 | 0.09145343 |
| ENSMUSG00 | Gm12278 | 405 | 0 | 0 | 0 | 1.33637202 | 0 | 0 |
| ENSMUSG00 | Gm12122 | 452 | 0 | 0 | 0 | 0 | 0 | 0 |
| ENSMUSG00 | Gm15336 | 1568 | 0 | 0 | 0 | 0 | 0 | 0 |
| ENSMUSG00 | Gm15940 | 815 | 0 | 0 | 0 | 0 | 0.56348145 | 0 |
| ENSMUSG00 | 4930555J06R | 746 | 0 | 0 | 0.1777554 | 0.48367352 | 0.41039981 | 0 |
| ENSMUSG00 | 2210013O21I | 1529 | 27.7170322 | 26.5356339 | 24.485913 | 19.9170998 | 16.3524684 | 38.4235271 |
| ENSMUSG00 | Gm16766 | 2613 | 0 | 0 | 0 | 0 | 0 | 0 |
| ENSMUSG00 | Gm16767 | 2322 | 0.15409101 | 0 | 0 | 0 | 0 | 0 |
| ENSMUSG00 | Gm16087 | 961 | 0 | 0 | 0 | 0 | 0 | 0 |
| ENSMUSG00 | Gm12840 | 625 | 0 | 0 | 0 | 0 | 0 | 0 |
| ENSMUSG00 | Gm11413 | 1458 | 0 | 0 | 0 | 0 | 0 | 0 |
| ENSMUSG00 | E130218I03R | 2090 | 0 | 0 | 0.04229841 | 0 | 0 | 0.0783004 |
| ENSMUSG00 | Gm15564 | 933 | 41.7581682 | 45.9656308 | 42.2120494 | 32.5627883 | 22.0950223 | 36.6877516 |
| ENSMUSG00 | Gm16908 | 1314 | 2.42042499 | 1.28350493 | 1.58104005 | 0.0274597 | 0.54366002 | 1.26617427 |
| ENSMUSG00 | Gm13200 | 256 | 0 | 0 | 0 | 0 | 0 | 0.31962468 |
| ENSMUSG00 | 1700003F17I | 571 | 0 | 0.5063374 | 0 | 0 | 0 | 0 |
| ENSMUSG00 | 2700033N17I | 719 | 0 | 0.13403739 | 0 | 0 | 0 | 0 |
| ENSMUSG00 | Gm14233 | 437 | 0 | 0 | 0 | 0 | 0 | 0 |
| ENSMUSG00 | 1700007J10R | 1291 | 0.06158866 | 0 | 0.13695381 | 0.78256952 | 0.27667284 | 0.29577455 |
| ENSMUSG00 | Gm16310 | 958 | 2.86339056 | 1.307774 | 1.47647075 | 1.28057361 | 3.14254298 | 2.81856919 |
| ENSMUSG00 | 4930480G23I | 1010 | 0 | 0 | 0 | 0 | 0 | 0 |
| ENSMUSG00 | 1700120E14I | 1483 | 0 | 0 | 0.11922277 | 0 | 0.03440754 | 0 |
| ENSMUSG00 | Gm12107 | 886 | 0 | 0 | 0 | 0 | 0 | 0 |
| ENSMUSG00 | Gm12979 | 402 | 0 | 0 | 0 | 0 | 0 | 0 |
| ENSMUSG00 | Gm11535 | 776 | 0 | 0 | 0 | 0 | 0 | 0 |
| ENSMUSG00 | Gm13723 | 561 | 0 | 0 | 0 | 0 | 0 | 0 |
| ENSMUSG00 | Gm11916 | 648 | 0 | 0 | 0 | 0 | 0 | 0 |
| ENSMUSG00 | 1810059C17I | 484 | 0.16427885 | 0.09955877 | 0.18265224 | 0 | 0.21085279 | 0.16905768 |
| ENSMUSG00 | Gm15932 | 509 | 0.85915577 | 0.94668845 | 0.69472445 | 1.13420965 | 1.20297939 | 0.53584753 |
| ENSMUSG00 | Gm12932 | 618 | 0.25731703 | 1.32551702 | 0.7867642 | 0.4086963 | 0.41283476 | 1.19161045 |
| ENSMUSG00 | A530083I20R | 3182 | 0.32484051 | 0.51487713 | 0.41673642 | 0.22678846 | 0.64143779 | 0.28286081 |
| ENSMUSG00 | Gm11345 | 615 | 0 | 0 | 0 | 0 | 0 | 0 |
| ENSMUSG00 | Gm1989 | 628 | 0 | 0 | 0 | 0 | 0 | 0 |
| ENSMUSG00 | Gm12510 | 360 | 0 | 0 | 0 | 0 | 0 | 0 |
| ENSMUSG00 | Gm11626 | 331 | 0.24021438 | 0.5823135 | 0.13354031 | 0.2180184 | 0 | 0.65920578 |
| ENSMUSG00 | 1700017J07R | 471 | 0 | 0 | 0 | 0 | 0 | 0 |
| ENSMUSG00 | Gm13497 | 2035 | 0 | 0 | 0 | 0 | 0 | 0 |
| ENSMUSG00 | B230369F24I | 2558 | 2.54882674 | 2.24166795 | 3.28317051 | 3.27249192 | 1.47613441 | 5.4165429 |
| ENSMUSG00 | 6230400D17I | 1551 | 3.79355971 | 2.08155488 | 1.82393163 | 0.83749426 | 2.33583023 | 1.26613412 |
| ENSMUSG00 | 4930401G09I | 635 | 0 | 0 | 0 | 0 | 0 | 0 |
| ENSMUSG00 | Gm13481 | 1726 | 0.32246624 | 0.86545754 | 0.46096939 | 0.22995509 | 0.20694359 | 2.16490474 |
| ENSMUSG00 | Gm13938 | 488 | 0 | 0 | 0 | 0 | 0 | 0 |
| ENSMUSG00 | 4933428C19I | 811 | 0 | 0 | 0 | 0 | 0 | 0 |
| ENSMUSG00 | Gm13441 | 1164 | 0 | 0 | 0 | 0 | 0 | 0 |
| ENSMUSG00 | 1700112J16R | 2212 | 0.30553489 | 0 | 0.97915475 | 0.89715753 | 0.02306798 | 1.25769132 |
| ENSMUSG00 | Gm13270 | 645 | 1.35600089 | 0.22412299 | 0.13705998 | 0.55941154 | 0.87021726 | 0.63429394 |
| ENSMUSG00 | 9630013K17I | 1068 | 0.1116727 | 0.27071035 | 0.12416248 | 0 | 0.57333007 | 0.0510761 |
| ENSMUSG00 | Gm16214 | 1384 | 0 | 0.87041984 | 0.47906622 | 0.4432043 | 0.22121261 | 0.57150617 |
| ENSMUSG00 | Gm6569 | 807 | 0 | 0 | 0 | 0 | 0 | 0 |
| ENSMUSG00 | A330102I10R | 1825 | 0.19605442 | 0.18482471 | 0.33908263 | 0.03954197 | 0 | 0.1942851 |
| ENSMUSG00 | Gm11751 | 455 | 0 | 0 | 0 | 0 | 0 | 0 |
| ENSMUSG00 | 4930593C16I | 580 | 0 | 0 | 0 | 0 | 0 | 0 |
| ENSMUSG00 | 1700061J23R | 534 | 0 | 0 | 0 | 0 | 0 | 0 |
| ENSMUSG00 | Gm6182 | 1206 | 0 | 0.03995559 | 0 | 0 | 0 | 0 |
| ENSMUSG00 | Gm13830 | 2648 | 1.11099153 | 1.89251888 | 1.38548073 | 2.03029631 | 1.44523345 | 1.56561373 |
| ENSMUSG00 | E330017L17I | 1604 | 0 | 0 | 0 | 0 | 0 | 0 |
| ENSMUSG00 | B230206F22I | 8653 | 1.32778619 | 0.96896347 | 0.98589572 | 2.71876725 | 1.41526987 | 0.92039693 |
| ENSMUSG00 | Gm12478 | 351 | 0 | 0 | 0 | 0 | 0 | 0 |
| ENSMUSG00 | Gm16246 | 1343 | 0 | 0.64583467 | 0 | 0 | 0.26596026 | 0.06092622 |
| ENSMUSG00 | Gm11978 | 1868 | 0 | 0.02579574 | 0 | 0 | 0 | 0 |
| ENSMUSG00 | Gm12127 | 704 | 0 | 0 | 0 | 0 | 0 | 0 |
| ENSMUSG00 | 1700071G01I | 361 | 0 | 0.13348045 | 0.73465667 | 0 | 0 | 0.07555302 |
| ENSMUSG00 | 4930529C04I | 3533 | 0 | 0 | 0 | 0 | 0 | 0 |
| ENSMUSG00 | Gm13753 | 453 | 0 | 0 | 0 | 0 | 0 | 0 |
| ENSMUSG00 | 1700026D11I | 3169 | 0 | 0 | 0 | 0 | 0 | 0 |
| ENSMUSG00 | Gm11195 | 439 | 0 | 0 | 0 | 0 | 0 | 0 |
| ENSMUSG00 | AW064505 | 518 | 0 | 0 | 0 | 0 | 0 | 0 |
| ENSMUSG00 | Gm13705 | 846 | 2.11465322 | 1.02524345 | 0.94046475 | 0.08530034 | 0.48251892 | 2.51468305 |
| ENSMUSG00 | Gm16630 | 4422 | 0 | 0 | 0 | 0 | 0 | 0 |
| ENSMUSG00 | Gm12357 | 1051 | 0 | 0 | 0 | 0 | 0 | 0 |
| ENSMUSG00 | Gm12701 | 657 | 0 | 0 | 0 | 0.71395218 | 0 | 0.62270866 |
| ENSMUSG00 | Gm12108 | 2587 | 0 | 0 | 0 | 0 | 0 | 0 |
| ENSMUSG00 | Gm15563 | 408 | 0.68207932 | 0 | 0 | 0 | 0 | 0 |
| ENSMUSG00 | Gm16631 | 3670 | 0 | 0.01312982 | 0 | 0 | 0 | 0.00743178 |
| ENSMUSG00 | Gm15998 | 1841 | 0 | 0 | 0 | 0 | 0.24944996 | 0 |
| ENSMUSG00 | 1810019D21I | 1732 | 27.7048874 | 30.7146837 | 23.1983114 | 24.5407901 | 25.483727 | 34.1720365 |
| ENSMUSG00 | 1700042O10I | 1071 | 0 | 0 | 0 | 0 | 0 | 0 |
| ENSMUSG00 | A330050B17I | 8235 | 0.22207069 | 0.54418204 | 0.33815618 | 0.30670833 | 0.24785125 | 0.61935125 |
| ENSMUSG00 | Gm16794 | 3057 | 0.63723211 | 0.01576266 | 1.46038147 | 0.82621626 | 1.26856544 | 0.99926712 |
| ENSMUSG00 | Gm11612 | 159 | 0.25003447 | 0.90917815 | 0 | 0.2269311 | 0 | 0.34307722 |
| ENSMUSG00 | A630014C17I | 2982 | 0 | 0 | 0 | 0 | 0 | 0 |
| ENSMUSG00 | 4930555B11I | 1529 | 0 | 0 | 0 | 0 | 0 | 0 |
| ENSMUSG00 | 4933405D12I | 1376 | 0.14446032 | 0.17509608 | 0.25698746 | 0.02622242 | 0.22249873 | 0.37861203 |
| ENSMUSG00 | Gm15639 | 668 | 0.7736845 | 2.81328031 | 0.52936339 | 0.64818044 | 0.45832074 | 0 |
| ENSMUSG00 | Gm16144 | 825 | 0 | 0 | 0 | 0.04373581 | 0 | 0 |
| ENSMUSG00 | Gm16789 | 908 | 0 | 0 | 0 | 0 | 0 | 0.33041964 |
| ENSMUSG00 | Gm15559 | 2258 | 0.95075108 | 0.25608384 | 0.11745397 | 0 | 0.04519608 | 0.39861076 |
| ENSMUSG00 | Gm15322 | 737 | 0 | 0 | 0 | 0 | 0 | 0 |
| ENSMUSG00 | 1500012K07I | 2338 | 0 | 0 | 0 | 0 | 0 | 0 |
| ENSMUSG00 | Gm6787 | 1450 | 0 | 0.29908826 | 0 | 0 | 0 | 0 |
| ENSMUSG00 | 9330198N18I | 1243 | 0 | 0 | 0 | 0 | 0 | 0.02194259 |
| ENSMUSG00 | Gm14123 | 2655 | 0.20963342 | 0 | 0.08324264 | 0.21744358 | 0 | 0.01027293 |
| ENSMUSG00 | Gm11588 | 655 | 0 | 0 | 0 | 0 | 0 | 0 |
| ENSMUSG00 | Gm13898 | 781 | 0 | 0 | 0.11319294 | 0 | 0 | 0 |
| ENSMUSG00 | Gm16158 | 802 | 0 | 0 | 0 | 0 | 0 | 0 |
| ENSMUSG00 | Gm14236 | 870 | 0 | 0 | 0 | 0 | 0 | 0 |
| ENSMUSG00 | Gm16626 | 1565 | 0 | 0.6773813 | 0.19770792 | 0.20750058 | 0 | 0.12199519 |
| ENSMUSG00 | Gm12415 | 510 | 0 | 0 | 0 | 0 | 0 | 0 |
| ENSMUSG00 | Gm15645 | 2203 | 0 | 0 | 0 | 0 | 0 | 0 |
| ENSMUSG00 | Gm4487 | 441 | 0 | 0 | 0 | 0 | 0 | 0 |
| ENSMUSG00 | Gm14002 | 2836 | 0 | 0 | 0.26496368 | 0 | 0 | 0 |
| ENSMUSG00 | Gm12996 | 634 | 1.31682191 | 1.59608089 | 0.27887598 | 1.36588181 | 1.20724864 | 0.68831897 |
| ENSMUSG00 | Gm13362 | 837 | 0.18999035 | 0.17271126 | 0 | 0.6897404 | 0.06096341 | 0 |
| ENSMUSG00 | 4933408J17R | 1855 | 0.12858916 | 0 | 0 | 0 | 0 | 0 |
| ENSMUSG00 | Gm8865 | 429 | 0.09267012 | 0.67393625 | 0 | 0 | 0.1189426 | 0.06357725 |
| ENSMUSG00 | Gm14091 | 872 | 0.18236459 | 0 | 0.25345094 | 0 | 0 | 0.31278256 |
| ENSMUSG00 | 4930421P07I | 398 | 0 | 0 | 0 | 0 | 0 | 0 |
| ENSMUSG00 | Gm11364 | 295 | 0 | 0 | 0 | 0 | 0 | 1.01702045 |
| ENSMUSG00 | Gm15569 | 591 | 0 | 0 | 0 | 0 | 0.51803427 | 0 |
| ENSMUSG00 | F730016J06R | 3794 | 0 | 0 | 0 | 0 | 0 | 0 |
| ENSMUSG00 | Gm15295 | 486 | 0 | 0 | 0 | 0 | 0 | 0 |
| ENSMUSG00 | Hoxa11as | 3375 | 0.10601462 | 0.24271689 | 0.11787158 | 0.17105562 | 0.21166497 | 0.15354612 |
| ENSMUSG00 | Gm15199 | 539 | 0 | 0 | 0 | 0 | 0 | 0 |
| ENSMUSG00 | Gt(ROSA)26S | 1851 | 45.5115414 | 42.9538787 | 51.7717967 | 44.9904694 | 24.5345621 | 98.1061848 |
| ENSMUSG00 | 4930551O13I | 1063 | 0.14959729 | 0 | 0 | 0.0339436 | 0 | 0.20526539 |
| ENSMUSG00 | Gm14929 | 1848 | 0 | 0 | 0 | 0 | 0 | 0 |
| ENSMUSG00 | B430119L08I | 1992 | 0 | 0 | 0 | 0 | 0 | 0 |
| ENSMUSG00 | 1700006D18I | 321 | 0.2476977 | 0 | 0.27540089 | 0 | 0.79480336 | 0 |
| ENSMUSG00 | Gm15200 | 408 | 0 | 0 | 0 | 0 | 0 | 0 |
| ENSMUSG00 | D930007P13I | 1317 | 0.33205033 | 0.65858463 | 0 | 0.38356008 | 0.77488802 | 0.60058052 |
| ENSMUSG00 | Gm13690 | 571 | 0 | 0 | 0.07741128 | 0.25276389 | 0 | 0 |
| ENSMUSG00 | Gm13021 | 277 | 0 | 0 | 0 | 0 | 0 | 0 |
| ENSMUSG00 | 2510003D18I | 494 | 0 | 0 | 0 | 0 | 0 | 0 |
| ENSMUSG00 | Gm12069 | 803 | 0 | 0 | 0 | 0 | 0 | 0 |
| ENSMUSG00 | Gm15046 | 406 | 0 | 0 | 0 | 0 | 0 | 0 |
| ENSMUSG00 | Gm11722 | 532 | 2.76494883 | 0.36230408 | 0.24925851 | 1.42429123 | 0.67139968 | 1.79438416 |
| ENSMUSG00 | 4933421A08I | 1636 | 0 | 0 | 0 | 0 | 0 | 0 |
| ENSMUSG00 | Gm15961 | 264 | 0 | 0 | 0 | 0.13667441 | 0 | 0 |
| ENSMUSG00 | Gm13191 | 414 | 0 | 0 | 0 | 0 | 0 | 0 |
| ENSMUSG00 | 4933426D04I | 984 | 2.14130129 | 0.73454942 | 1.6171406 | 2.49347462 | 1.55568219 | 1.96798718 |
| ENSMUSG00 | Gm13522 | 451 | 0 | 0 | 0 | 0 | 0 | 0 |
| ENSMUSG00 | 9330162O12I | 3834 | 0.89175048 | 0.87977333 | 0.36892514 | 1.47753808 | 1.27765573 | 2.33335464 |
| ENSMUSG00 | 9030204H09I | 650 | 0 | 0 | 0 | 0 | 0 | 0 |
| ENSMUSG00 | Gm14077 | 3122 | 0 | 0 | 0.01415818 | 0 | 0 | 0.0611539 |
| ENSMUSG00 | 4933431K23I | 4433 | 0 | 0 | 0 | 0 | 0 | 0 |
| ENSMUSG00 | Gm14947 | 302 | 0 | 0 | 0 | 0 | 0 | 0 |
| ENSMUSG00 | Gm11457 | 781 | 0.30541983 | 0.43188873 | 0.73575411 | 0.1385994 | 1.11068936 | 0.69845427 |
| ENSMUSG00 | 4930500J02R | 1626 | 0.36674798 | 0 | 0.4893193 | 0 | 0 | 0 |
| ENSMUSG00 | Gm11815 | 706 | 0 | 0 | 0 | 0 | 0 | 0 |
| ENSMUSG00 | Gm16861 | 1644 | 3.02277072 | 0.9672461 | 2.33914863 | 3.66526851 | 2.48303532 | 0.77974942 |
| ENSMUSG00 | D930032P07I | 1460 | 0.10891913 | 0.09901324 | 0 | 0.07414119 | 0.31454615 | 0.69120661 |
| ENSMUSG00 | Gm2639 | 1296 | 0 | 0 | 0 | 0 | 0 | 0 |
| ENSMUSG00 | 1700030C12I | 1319 | 0 | 0 | 0 | 0 | 0 | 0 |
| ENSMUSG00 | Gm12236 | 248 | 0 | 0 | 0 | 0 | 0 | 0 |
| ENSMUSG00 | Gm16899 | 1190 | 0.13363187 | 0 | 0.18572203 | 0.03032105 | 0.12863792 | 0.04583973 |
| ENSMUSG00 | Gm13990 | 2697 | 0 | 0 | 0 | 0 | 0 | 0 |
| ENSMUSG00 | Gm16898 | 5179 | 0 | 0 | 0 | 0 | 0 | 0 |
| ENSMUSG00 | Gm15835 | 354 | 1.45994702 | 0 | 0 | 0.81541344 | 0.43242691 | 0.15409401 |
| ENSMUSG00 | Gm16049 | 614 | 0.12949668 | 0.31391819 | 0.50392981 | 1.05777981 | 1.32967755 | 0.62189731 |
| ENSMUSG00 | 4930571N24I | 1054 | 0 | 0 | 0 | 0 | 0 | 0 |
| ENSMUSG00 | 2900053O20I | 933 | 1.87485653 | 1.70434361 | 2.55830603 | 1.50825267 | 2.62515117 | 2.54329434 |
| ENSMUSG00 | Gm16897 | 2486 | 0.03198349 | 0.44581181 | 0.03556061 | 0.71119074 | 0.41050986 | 0.04388518 |
| ENSMUSG00 | Gm12580 | 1749 | 0 | 0 | 0 | 0.0412602 | 0 | 0 |
| ENSMUSG00 | A530020G20I | 4167 | 2.45192188 | 1.78082844 | 1.83511371 | 1.46337066 | 2.37559801 | 3.69159984 |
| ENSMUSG00 | Gm16172 | 680 | 0.99388701 | 0.56689932 | 1.04004337 | 0.58368013 | 0.45023273 | 0.96263433 |
| ENSMUSG00 | Gm16265 | 107 | 0 | 0 | 0 | 0 | 0 | 0 |
| ENSMUSG00 | 9130204K15I | 1964 | 0 | 0 | 0 | 0 | 0 | 0 |
| ENSMUSG00 | Gm12139 | 372 | 0 | 0 | 0 | 0 | 0 | 0 |
| ENSMUSG00 | 9530059O14I | 6795 | 0 | 0 | 0 | 0 | 0 | 0 |
| ENSMUSG00 | Gm15506 | 1129 | 0.14085201 | 0.68289023 | 0.19575661 | 0.6711452 | 0.22598041 | 0.24158228 |
| ENSMUSG00 | Gm14102 | 462 | 0 | 0 | 0 | 0 | 0 | 0 |
| ENSMUSG00 | Gm16014 | 536 | 0 | 0 | 0 | 0 | 0 | 0 |
| ENSMUSG00 | Gm15287 | 749 | 0 | 0 | 0 | 0 | 0 | 0 |
| ENSMUSG00 | Gm11707 | 488 | 0 | 0.09874271 | 0.18115509 | 0 | 0 | 0.6147972 |
| ENSMUSG00 | 1700012C08I | 361 | 0 | 0 | 0 | 0 | 0 | 0 |
| ENSMUSG00 | 8030443G20I | 897 | 0 | 0 | 0 | 0 | 0 | 0 |
| ENSMUSG00 | A630071L07I | 3786 | 0.07350459 | 0.10182027 | 0.11675077 | 0 | 0.09434354 | 0.15848972 |
| ENSMUSG00 | Gm15458 | 708 | 0 | 0 | 0 | 0 | 0 | 0 |
| ENSMUSG00 | 2310043D23I | 2014 | 1.57916507 | 0 | 1.86551969 | 1.25409291 | 1.01343348 | 0.47398827 |
| ENSMUSG00 | Gm11638 | 1611 | 0 | 0 | 0 | 0 | 0 | 0 |
| ENSMUSG00 | 1600020E01I | 2301 | 13.4418791 | 14.0098782 | 16.4628333 | 18.723147 | 11.730966 | 20.4233826 |
| ENSMUSG00 | Gm16362 | 213 | 0 | 0 | 0 | 0 | 0 | 0 |
| ENSMUSG00 | Gm12292 | 2431 | 0 | 0 | 0 | 0 | 0 | 0 |
| ENSMUSG00 | Gm13291 | 653 | 0 | 0 | 0 | 0 | 0 | 0 |
| ENSMUSG00 | Gm16652 | 1879 | 0.21157786 | 0.05128945 | 0.14114479 | 0.07681116 | 0.02715613 | 0 |
| ENSMUSG00 | Gm12204 | 331 | 0 | 0.72789188 | 0 | 0 | 0 | 0.08240072 |
| ENSMUSG00 | 2210417A02I | 548 | 0 | 0 | 0 | 0 | 0 | 0 |
| ENSMUSG00 | Gm13778 | 740 | 0.16117087 | 0.65116814 | 1.19464441 | 0.19503808 | 1.10327299 | 1.58487769 |
| ENSMUSG00 | Gm14204 | 7395 | 0 | 0 | 0 | 0 | 0 | 0 |
| ENSMUSG00 | Gm15259 | 290 | 0 | 0 | 0 | 0 | 0 | 0 |
| ENSMUSG00 | Gm12681 | 697 | 0 | 0 | 0 | 0 | 0 | 0 |
| ENSMUSG00 | Gm16217 | 614 | 0 | 0 | 0 | 0 | 0 | 0 |
| ENSMUSG00 | Gm15875 | 271 | 0 | 0 | 0 | 0 | 0 | 0 |
| ENSMUSG00 | 4930597A21I | 694 | 0 | 0 | 0 | 0 | 0 | 0 |
| ENSMUSG00 | B130055M24 | 2916 | 3.20388818 | 2.97447174 | 3.25905221 | 3.01921086 | 3.49975143 | 2.534783 |
| ENSMUSG00 | Xist | 16001 | 0 | 0.14455029 | 0.10221037 | 0 | 0 | 0.00170456 |
| ENSMUSG00 | 4930511F01I | 493 | 0.08063992 | 0.19548252 | 0 | 0.36594366 | 0 | 0.11064762 |
| ENSMUSG00 | Gm16058 | 425 | 0 | 0 | 0 | 0 | 0 | 0 |
| ENSMUSG00 | Gm15911 | 806 | 0.14797325 | 0.05978467 | 0 | 0.08953361 | 0.25323263 | 0.10151851 |
| ENSMUSG00 | Gm11206 | 981 | 0 | 0 | 0 | 0 | 0 | 0 |
| ENSMUSG00 | Gm11291 | 191 | 0 | 0 | 0 | 0 | 0 | 0 |
| ENSMUSG00 | Nkx2-2as | 4339 | 0 | 0 | 0 | 0 | 0 | 0 |
| ENSMUSG00 | Gm16881 | 1252 | 0.19052147 | 0.11546272 | 0.31774488 | 0.97986383 | 0.69285015 | 0 |
| ENSMUSG00 | Gm12600 | 706 | 0 | 0 | 0 | 0 | 0 | 0 |
| ENSMUSG00 | 9130208D14 | 7292 | 0 | 0 | 0 | 0 | 0 | 0 |
| ENSMUSG00 | Gm11747 | 1645 | 0 | 0 | 0 | 0 | 0 | 0 |
| ENSMUSG00 | Gm16292 | 606 | 1.24645896 | 1.03370255 | 1.96908905 | 0 | 0.50521164 | 0.90015311 |
| ENSMUSG00 | 4930461G14 | 777 | 5.47469294 | 5.27136111 | 1.02398092 | 11.0521578 | 6.50143012 | 5.65150183 |
| ENSMUSG00 | Gm12534 | 467 | 0 | 0 | 0 | 0 | 0 | 0 |
| ENSMUSG00 | Gm14503 | 1163 | 0 | 0 | 0 | 0 | 0 | 0 |
| ENSMUSG00 | Gm13393 | 394 | 0 | 0 | 0 | 0 | 0 | 0 |
| ENSMUSG00 | 9330185C12I | 1006 | 0 | 0 | 0 | 0 | 0 | 0 |
| ENSMUSG00 | Olfr1343-ps1 | 967 | 0 | 0 | 0 | 0 | 0 | 0 |
| ENSMUSG00 | Gm4473 | 655 | 0 | 0 | 0 | 0 | 0 | 0 |
| ENSMUSG00 | Gm13499 | 600 | 0 | 0 | 0 | 0 | 0 | 0 |
| ENSMUSG00 | Pabpc1l2a-ps | 1114 | 0 | 0 | 0 | 0 | 0 | 0 |
| ENSMUSG00 | Gm15281 | 350 | 0 | 0 | 0 | 0 | 0 | 0 |
| ENSMUSG00 | Gm12762 | 454 | 0.17513428 | 0.21227508 | 0.48680444 | 0.39737935 | 0 | 0.0600763 |
| ENSMUSG00 | Gm15856 | 1160 | 0 | 0 | 0 | 0 | 0 | 0 |
| ENSMUSG00 | Gm15731 | 2189 | 0.32690665 | 0.94655871 | 0 | 0.72526723 | 0.02331036 | 0 |
| ENSMUSG00 | 4930471I20R | 625 | 0 | 1.0793763 | 0 | 0.05773127 | 0 | 0.08727885 |
| ENSMUSG00 | Gm12910 | 722 | 0 | 0 | 0 | 0 | 0 | 0 |
| ENSMUSG00 | Gm11351 | 521 | 0 | 0 | 0 | 0 | 0 | 0 |
| ENSMUSG00 | Gm16266 | 556 | 0.35751331 | 0.08666626 | 0.31799887 | 0.06489576 | 0.09177406 | 0.78488171 |
| ENSMUSG00 | Gm16758 | 3842 | 0 | 0 | 0 | 0 | 0 | 0 |
| ENSMUSG00 | Gm12264 | 498 | 0.95796339 | 2.12871833 | 1.86393314 | 0.86944686 | 2.45910245 | 2.02642902 |
| ENSMUSG00 | Nespas | 2676 | 0 | 0 | 0 | 0 | 0 | 0 |
| ENSMUSG00 | Gm6634 | 1800 | 0 | 0 | 0 | 0 | 0 | 0 |
| ENSMUSG00 | Gm16759 | 1705 | 0 | 0.02826184 | 0.28517318 | 1.18509941 | 0.26934744 | 0.55988996 |
| ENSMUSG00 | Scml1 | 574 | 0 | 0 | 0 | 0 | 0 | 0 |
| ENSMUSG00 | Has2as | 1620 | 0 | 0 | 0 | 0 | 0 | 0 |
| ENSMUSG00 | Gm13782 | 349 | 0 | 0 | 0 | 0 | 0 | 1.0941116 |
| ENSMUSG00 | 4933406C10I | 2559 | 0.01553555 | 0.01883019 | 0 | 0.05640023 | 0 | 0.01065832 |
| ENSMUSG00 | Gm13704 | 3239 | 0.51550793 | 0 | 0.23199485 | 0 | 0.45685857 | 0.15999325 |
| ENSMUSG00 | Gm12 | 900 | 0 | 0 | 0 | 0 | 0 | 0 |
| ENSMUSG00 | Gm13709 | 612 | 0.19487981 | 0 | 0 | 0 | 0 | 0.0445664 |
| ENSMUSG00 | Gm11755 | 885 | 0.26952868 | 0 | 0 | 0 | 0 | 0 |
| ENSMUSG00 | Gm12972 | 1002 | 0 | 0 | 0 | 0 | 0 | 0 |
| ENSMUSG00 | Gm13648 | 1534 | 1.26989475 | 0.75389479 | 2.9391056 | 1.27016324 | 1.1309627 | 2.36475034 |
| ENSMUSG00 | 1810007I06R | 172 | 0 | 0 | 0 | 0 | 0 | 0 |
| ENSMUSG00 | A730090H04 | 1660 | 0.81426888 | 0.02902798 | 0.71894564 | 0.52166811 | 0.09221634 | 0.42719315 |
| ENSMUSG00 | 9230116N13 | 2094 | 0.41767936 | 0.46023345 | 0.90767873 | 1.05110063 | 0.12183948 | 1.54999144 |
| ENSMUSG00 | 9530034E10I | 499 | 0.47802181 | 0.86909415 | 0.62006593 | 1.51848284 | 0.20451453 | 2.84224698 |
| ENSMUSG00 | Gm13446 | 1068 | 0 | 0 | 0 | 0 | 0 | 0 |
| ENSMUSG00 | Gm15444 | 386 | 0 | 0 | 0 | 0 | 0 | 0 |
| ENSMUSG00 | Gm15999 | 1509 | 0 | 0 | 0.14646071 | 0 | 0 | 0 |
| ENSMUSG00 | Gm14223 | 513 | 0 | 0 | 0 | 0 | 0 | 0 |
| ENSMUSG00 | Gm14426 | 347 | 0 | 0 | 0 | 0 | 0 | 0 |
| [illegible] | | | | | | | | |

| | | | | | | | | |
|---|---|---|---|---|---|---|---|---|
| ENSMUSG00 | Gm13133 | 732 | 0 | 0.78994167 | 0 | 0.14787723 | 0 | 0 |
| ENSMUSG00 | 2810429I04R | 7212 | 0.41343054 | 0.10022139 | 0.04903144 | 0.00500306 | 0.13442889 | 0.34414756 |
| ENSMUSG00 | Gm16247 | 384 | 0.82823918 | 0.12548553 | 0.92087173 | 0.28189097 | 0.26576237 | 1.1364433 |
| ENSMUSG00 | 9130020K20R | 1784 | 2.16159283 | 3.4573232 | 3.02277178 | 0.97081734 | 2.94602955 | 6.87981371 |
| ENSMUSG00 | 4930542H20 | 863 | 0.73706569 | 0.44668776 | 0 | 0 | 0.35476043 | 0 |
| ENSMUSG00 | Gm16961 | 1892 | 0 | 0 | 0.09344999 | 0.13349594 | 0.08090863 | 0 |
| ENSMUSG00 | Gm16962 | 1435 | 0.44326668 | 0.36937342 | 0.09240803 | 0.42745279 | 0.74672745 | 0.36112763 |
| ENSMUSG00 | Gm15384 | 484 | 0.08213942 | 0 | 0.45663061 | 0 | 0 | 0 |
| ENSMUSG00 | Gm16960 | 2650 | 0 | 0 | 0 | 0 | 0 | 0 |
| ENSMUSG00 | Gm14104 | 353 | 0.11262176 | 0.68252751 | 0 | 0.30664627 | 0 | 0 |
| ENSMUSG00 | Gm13660 | 720 | 0 | 0.4684793 | 0 | 0.2004558 | 0.70869966 | 0 |
| ENSMUSG00 | Gm7076 | 455 | 0 | 0 | 0 | 0 | 0 | 0 |
| ENSMUSG00 | Gm13583 | 1926 | 0.22705622 | 0 | 0 | 0 | 0 | 0 |
| ENSMUSG00 | A930012L18R | 2359 | 0.50558051 | 0.61279918 | 0 | 2.34020891 | 0.15141358 | 0.0924956 |
| ENSMUSG00 | Gm15280 | 639 | 0 | 0 | 0 | 0 | 0 | 0 |
| ENSMUSG00 | Gm16272 | 386 | 0.20598695 | 0 | 0.34353764 | 0 | 0.66096342 | 0.07065969 |
| ENSMUSG00 | Gm15500 | 1911 | 0.10401748 | 0.10086121 | 0.1387813 | 0.03776248 | 0.0801042 | 0.22835909 |
| ENSMUSG00 | Gm12002 | 2883 | 0 | 0.56827577 | 0.07665946 | 0 | 0.49557354 | 0.44464379 |
| ENSMUSG00 | Gm16126 | 585 | 0 | 0 | 0 | 0 | 0 | 0 |
| ENSMUSG00 | Gm11192 | 149 | 0 | 0.97019682 | 0 | 0 | 2.05475339 | 0 |
| ENSMUSG00 | Gm11837 | 1005 | 0 | 0 | 0 | 0 | 0 | 0.13569472 |
| ENSMUSG00 | Gm11729 | 710 | 0 | 0 | 0 | 0 | 0 | 0 |
| ENSMUSG00 | Gm12915 | 1495 | 0 | 0 | 0 | 0.14481088 | 0 | 0.03648781 |
| ENSMUSG00 | 6030455H03 | 857 | 0 | 0 | 0 | 0 | 0 | 0 |
| ENSMUSG00 | Gm12682 | 1076 | 0 | 0 | 0 | 0 | 0 | 0 |
| ENSMUSG00 | Gm16548 | 1223 | 0 | 0 | 0 | 0 | 0 | 0 |
| ENSMUSG00 | 4930505H01 | 1784 | 0 | 0 | 0.14866091 | 0 | 0.31462452 | 0.27519255 |
| ENSMUSG00 | Gm12214 | 415 | 0 | 0 | 0 | 0 | 0 | 0 |
| ENSMUSG00 | Susd5 | 3869 | 0 | 0 | 0 | 0 | 0 | 0 |
| ENSMUSG00 | F420014N23 | 2393 | 0.48178393 | 0.12081849 | 0.29554095 | 0.54281388 | 0.3624941 | 0.01139768 |
| ENSMUSG00 | Gm13718 | 1087 | 0 | 0 | 0 | 0 | 0 | 0 |
| ENSMUSG00 | C030005K06 | 2686 | 0.87325888 | 0.16145867 | 0.78990635 | 1.46423785 | 0.64590349 | 0.1726243 |
| ENSMUSG00 | Gm16257 | 481 | 0.57856209 | 0.30053914 | 0 | 0.52510252 | 0.74258759 | 0.45363225 |
| ENSMUSG00 | Gm15609 | 2958 | 0 | 0.14661189 | 0 | 0 | 0 | 0.34116351 |
| ENSMUSG00 | Gm13620 | 509 | 0.07810507 | 0.47334423 | 0 | 0.14177621 | 0.20049657 | 0.26792377 |
| ENSMUSG00 | Gm15510 | 648 | 2.14728676 | 0.81797973 | 1.09140353 | 2.67274404 | 3.3072651 | 1.7257102 |
| ENSMUSG00 | Gm14290 | 2066 | 0 | 0 | 0.08557956 | 0 | 0 | 0 |
| ENSMUSG00 | Gm13205 | 560 | 0.49694351 | 0.25814165 | 1.10504608 | 0.45102556 | 2.00460762 | 1.46114139 |
| ENSMUSG00 | 4930511M06 | 4153 | 0.04786357 | 0.04641121 | 0.11707688 | 0 | 0.19658609 | 0.216726 |
| ENSMUSG00 | 4930562D21 | 1748 | 0 | 0 | 0 | 0 | 0 | 0 |
| ENSMUSG00 | 5830432E09 | 4491 | 0 | 0 | 0 | 0 | 0 | 0 |
| ENSMUSG00 | Gm15408 | 1375 | 1.64804537 | 0.80602776 | 0.70722949 | 0.41986379 | 1.70706421 | 1.5075437 |
| ENSMUSG00 | Gm12064 | 529 | 0 | 0 | 0 | 0 | 0 | 0 |
| ENSMUSG00 | 1700007M16 | 433 | 0 | 0 | 0 | 0 | 0 | 0 |
| ENSMUSG00 | Gm14330 | 382 | 0 | 0 | 0 | 0 | 0 | 0 |
| ENSMUSG00 | Gm16862 | 5206 | 0 | 0 | 0 | 0 | 0 | 0 |
| ENSMUSG00 | Gm16863 | 2426 | 0 | 0 | 0 | 0 | 0 | 0 |
| ENSMUSG00 | Gm11635 | 566 | 0 | 0 | 0 | 0 | 0 | 0 |
| ENSMUSG00 | Gm13256 | 518 | 0.07674803 | 0.65116814 | 0 | 0 | 0 | 0 |
| ENSMUSG00 | Gm14333 | 263 | 0 | 0 | 0 | 0 | 0 | 0 |
| ENSMUSG00 | Gm14157 | 1279 | 0.18649952 | 0 | 0 | 0 | 0.71811944 | 0.34119955 |
| ENSMUSG00 | Gm13348 | 684 | 0 | 0 | 0 | 0 | 0 | 0 |
| ENSMUSG00 | Gm16701 | 2122 | 0 | 0.15895622 | 0 | 0 | 0.19237088 | 0.11567943 |
| ENSMUSG00 | Gm16130 | 2794 | 0 | 0 | 0 | 0 | 0 | 0 |
| ENSMUSG00 | Gm11831 | 627 | 0.19021761 | 2.30557139 | 2.39691015 | 3.91320419 | 0.89519958 | 0 |
| ENSMUSG00 | A630072L19R | 3401 | 0 | 0 | 0 | 0 | 0 | 0 |
| ENSMUSG00 | Gm16702 | 2381 | 0 | 0 | 0 | 0 | 0.1285839 | 0 |
| ENSMUSG00 | Gm16157 | 1503 | 0 | 0.06412035 | 0 | 0 | 0 | 0 |
| ENSMUSG00 | 2810403D21 | 7266 | 1.55388886 | 1.33298581 | 1.40526091 | 1.46493299 | 1.22193633 | 2.72521168 |
| ENSMUSG00 | Gm14397 | 571 | 0 | 0 | 0 | 0 | 0 | 0 |
| ENSMUSG00 | Gm12784 | 655 | 0 | 0 | 0 | 0 | 0 | 0 |
| ENSMUSG00 | 1700112J05R | 789 | 0.2015487 | 0 | 0 | 0 | 0 | 0 |
| ENSMUSG00 | Gm16081 | 682 | 0.05829249 | 0.21196382 | 0.45368461 | 0 | 0 | 0.11997642 |
| ENSMUSG00 | 4930448N21 | 2302 | 0.03453995 | 0.14652697 | 0.05760449 | 0.40752961 | 0.3768238 | 0.09478589 |
| ENSMUSG00 | 4932415G12 | 4132 | 18.5499919 | 11.9999635 | 13.8959811 | 12.1903519 | 19.7091229 | 15.511956 |
| ENSMUSG00 | Gm12949 | 360 | 0 | 0 | 0 | 0 | 0 | 0 |
| ENSMUSG00 | Gm14040 | 694 | 0 | 0 | 0 | 0 | 0 | 0 |
| ENSMUSG00 | 4930405A21R | 1133 | 0.28070948 | 1.53107848 | 0.66322271 | 0.57323637 | 0.18014608 | 0.36109408 |
| ENSMUSG00 | Gm17481 | 1509 | 0.15807348 | 0.0319327 | 0.205045 | 0.50213581 | 0 | 0.16267181 |
| ENSMUSG00 | Gm13663 | 1042 | 0 | 0 | 0 | 0 | 0 | 0 |
| ENSMUSG00 | Gm15254 | 272 | 0 | 0 | 0 | 0 | 0 | 0 |
| ENSMUSG00 | Gm15687 | 267 | 0 | 0 | 0 | 0 | 0 | 0 |
| ENSMUSG00 | Gm13470 | 418 | 0.09510881 | 0.69167142 | 0 | 0 | 0 | 0 |
| ENSMUSG00 | Gm15743 | 240 | 0.99388701 | 0 | 0.18417435 | 0 | 1.0630495 | 0.45457732 |
| ENSMUSG00 | 5133400J02R | 1133 | 0 | 0 | 0 | 0 | 0 | 0 |
| ENSMUSG00 | Gm15511 | 554 | 0 | 0 | 0 | 0 | 0 | 0 |
| ENSMUSG00 | Gm15286 | 1093 | 0 | 0 | 0 | 0 | 0 | 0 |
| ENSMUSG00 | A430048G15 | 622 | 0 | 0 | 0 | 0 | 0 | 0 |
| ENSMUSG00 | 1300002E11R | 4033 | 21.8443206 | 18.3163441 | 14.7960546 | 21.9552039 | 17.5106631 | 22.8652505 |
| ENSMUSG00 | Gm14029 | 1346 | 0 | 0 | 0 | 0 | 0 | 0 |
| ENSMUSG00 | 9130015L21R | 602 | 0 | 0 | 0 | 0 | 0 | 0 |
| ENSMUSG00 | Gm13165 | 631 | 0 | 0 | 0 | 0 | 0 | 0 |
| ENSMUSG00 | C330018A13 | 1597 | 0 | 0 | 0 | 0 | 0 | 0 |
| ENSMUSG00 | Gm15701 | 570 | 0 | 0 | 0 | 0 | 0 | 0 |
| ENSMUSG00 | 4921530L18R | 799 | 4.22930644 | 3.13603879 | 7.02582486 | 3.70303836 | 1.91588395 | 5.25693923 |
| ENSMUSG00 | Gm14260 | 258 | 0 | 0 | 0 | 0 | 0 | 0 |
| ENSMUSG00 | Gm14146 | 505 | 0 | 0 | 0 | 0 | 0 | 0 |
| ENSMUSG00 | Gm12594 | 497 | 0 | 0 | 0 | 0 | 0 | 0 |
| ENSMUSG00 | Gm4725 | 586 | 0 | 0 | 0 | 0 | 0 | 0 |
| ENSMUSG00 | BC046251 | 5199 | 0 | 0 | 0 | 0 | 0 | 0 |
| ENSMUSG00 | 9530036O11 | 3652 | 0 | 0.01319454 | 0 | 0.09880078 | 0.09780521 | 0.01493682 |
| ENSMUSG00 | B230209K01 | 2750 | 0 | 0 | 0 | 0 | 0 | 0 |
| ENSMUSG00 | Gm13067 | 402 | 0 | 0 | 0 | 0 | 0 | 0 |
| ENSMUSG00 | Gm12249 | 717 | 0.33268185 | 1.6129353 | 0.49318653 | 1.61035624 | 0.56933195 | 0.72275892 |
| ENSMUSG00 | Gm13434 | 455 | 0 | 0 | 0 | 0 | 0 | 0 |
| ENSMUSG00 | Gm15273 | 867 | 0 | 0 | 0 | 0 | 0 | 0 |
| ENSMUSG00 | AA645442 | 283 | 0 | 0 | 0 | 0 | 0 | 0 |
| ENSMUSG00 | Gm13194 | 310 | 0.12824349 | 0 | 0 | 0.23278738 | 0.82300606 | 0.61587895 |
| ENSMUSG00 | Gm13618 | 380 | 0 | 0 | 0.34896192 | 0 | 0.13427994 | 0.50242757 |
| ENSMUSG00 | Gm14297 | 290 | 0 | 0 | 0 | 0 | 0 | 0.09405048 |
| ENSMUSG00 | Gm16736 | 1739 | 0.02286112 | 0 | 0.10167186 | 0.02074873 | 0 | 0 |
| ENSMUSG00 | Gm12283 | 710 | 0 | 0 | 0.56030505 | 0 | 0.07186814 | 0 |
| ENSMUSG00 | 9430029A11R | 2041 | 0.38956865 | 0 | 1.16947551 | 2.08607607 | 0.02500067 | 0.46771797 |
| ENSMUSG00 | Gm14705 | 1821 | 0 | 0 | 0 | 0 | 0 | 0 |
| ENSMUSG00 | Gm12284 | 1008 | 0.31551969 | 1.57753233 | 1.31553104 | 1.21705309 | 0.10124281 | 4.68106408 |
| ENSMUSG00 | Gm16737 | 1179 | 1.44994543 | 0.40870604 | 0.52487345 | 0.70389061 | 2.12068907 | 1.04101677 |
| ENSMUSG00 | Gm15551 | 521 | 0.15261221 | 0.18497675 | 0 | 0 | 0.1958786 | 0 |
| ENSMUSG00 | Gm11592 | 634 | 1.31682191 | 0.76003852 | 0.76690895 | 0.22764697 | 0.16096649 | 1.07549839 |
| ENSMUSG00 | Gm16178 | 674 | 0.7667971 | 1.85882418 | 0 | 0.5888761 | 0 | 1.13307107 |
| ENSMUSG00 | Gm16023 | 2766 | 1.65288513 | 1.67241448 | 1.18255112 | 1.22621554 | 1.03307196 | 2.76099024 |
| ENSMUSG00 | Gm12867 | 455 | 1.04849619 | 0.52952134 | 1.2629098 | 1.18951795 | 0 | 0.41960984 |
| ENSMUSG00 | Gm12590 | 401 | 0 | 0 | 0 | 0 | 0 | 0 |
| ENSMUSG00 | Gm15015 | 630 | 0 | 0 | 0 | 0 | 0 | 0 |
| ENSMUSG00 | F630206G17 | 681 | 0.11675618 | 0 | 0 | 0.26491956 | 0 | 0 |
| ENSMUSG00 | 4930547E08R | 537 | 0.44419531 | 0 | 0 | 0 | 0.09502118 | 0.10158152 |
| ENSMUSG00 | Gm11560 | 947 | 0.20990222 | 0.10176651 | 0.14002696 | 0.11430426 | 0.32329277 | 0.43201646 |
| ENSMUSG00 | Gm16876 | 756 | 0.47327953 | 0 | 0.05846805 | 0.81136873 | 0 | 0.25254296 |
| ENSMUSG00 | 4933406J10R | 959 | 0 | 0 | 0 | 0.22574793 | 0 | 0.22752567 |
| ENSMUSG00 | Gm15432 | 194 | 0 | 0 | 0 | 0 | 0 | 0 |
| ENSMUSG00 | A730081D07 | 1799 | 0 | 0 | 0 | 0 | 0 | 0 |
| ENSMUSG00 | Gm16237 | 1317 | 0.21130476 | 0 | 0 | 0 | 0 | 0.12425804 |
| ENSMUSG00 | Gm15247 | 401 | 0.5948451 | 0 | 0.11022904 | 0 | 0.12724782 | 0.20404967 |
| ENSMUSG00 | Gm11234 | 843 | 0 | 0 | 0 | 0 | 0 | 0 |
| ENSMUSG00 | 4933427G23 | 2236 | 0 | 0.02155029 | 0.07907306 | 0.24205307 | 0.20538345 | 0.18296941 |
| ENSMUSG00 | 4930511O05 | 388 | 0 | 0 | 0 | 0 | 0 | 0 |
| ENSMUSG00 | Gm16510 | 248 | 0 | 0 | 0.35646648 | 0 | 0 | 0 |
| ENSMUSG00 | Gm15747 | 977 | 0.56967935 | 0.19728328 | 0.09048484 | 0.11079441 | 0.05222761 | 0.36291741 |
| ENSMUSG00 | Gm13595 | 492 | 0 | 0 | 0 | 0 | 0 | 0 |
| ENSMUSG00 | Gm12347 | 466 | 0 | 0 | 0 | 0 | 0 | 0 |
| ENSMUSG00 | 4930507D10 | 1149 | 0 | 0 | 0 | 0 | 0 | 0 |
| ENSMUSG00 | Gm14582 | 1185 | 0 | 0.32530931 | 0 | 0 | 0 | 0 |
| ENSMUSG00 | Gm16894 | 840 | 0 | 0.05736481 | 0 | 0 | 0 | 1.20138292 |
| ENSMUSG00 | Gm15848 | 1144 | 2.18933153 | 1.30575149 | 1.42960506 | 2.4601394 | 0.89206951 | 2.0026833 |
| ENSMUSG00 | Gm12088 | 518 | 0 | 0 | 0 | 0 | 0 | 0 |
| ENSMUSG00 | Gm15577 | 2291 | 0 | 0 | 0 | 0 | 0 | 0 |
| ENSMUSG00 | Gm16263 | 497 | 0 | 0 | 0 | 0 | 0 | 0 |
| ENSMUSG00 | Gm13465 | 976 | 0 | 0 | 0 | 0 | 0 | 0 |
| ENSMUSG00 | Gm15482 | 4527 | 0 | 0 | 0 | 0 | 0 | 0 |
| ENSMUSG00 | AI427809 | 1221 | 0 | 0 | 0.14480538 | 0 | 0 | 0 |
| ENSMUSG00 | 4930407G08 | 695 | 0.5720213 | 0 | 0.19079932 | 0 | 0 | 0 |
| ENSMUSG00 | 0610009E02R | 2076 | 0.44045089 | 0.23211196 | 0.21291832 | 0.34761122 | 0.07373754 | 0.44669448 |
| ENSMUSG00 | B230112J18R | 599 | 0 | 0 | 0 | 0 | 0 | 0 |
| ENSMUSG00 | Gm11629 | 382 | 0.52035969 | 1.76599526 | 0.57855816 | 0.85010053 | 0.5343076 | 2.57038485 |
| ENSMUSG00 | Gm15655 | 438 | 0 | 0 | 0 | 0 | 0 | 0 |
| ENSMUSG00 | Gm16069 | 543 | 0 | 0.08874115 | 0 | 0 | 0 | 0 |
| ENSMUSG00 | Gm16275 | 710 | 0.22397454 | 0.9501552 | 0 | 1.01639562 | 0 | 0.26890489 |
| ENSMUSG00 | Gm13851 | 437 | 0 | 0 | 0.10114838 | 0 | 0 | 0.56172026 |
| ENSMUSG00 | Gm13750 | 464 | 0 | 0.31155027 | 0.09526259 | 0 | 0 | 0 |
| ENSMUSG00 | Gm15499 | 419 | 0 | 0 | 0 | 0 | 0 | 0 |
| ENSMUSG00 | Gm16601 | 1716 | 0 | 0.33696813 | 0 | 0 | 0 | 0.09536587 |
| ENSMUSG00 | Gm12833 | 476 | 0 | 0 | 0.18572203 | 0 | 0 | 0 |
| ENSMUSG00 | A630052C17 | 2201 | 6.21348719 | 3.19637463 | 4.73949793 | 7.57378672 | 7.72002869 | 10.3844378 |
| ENSMUSG00 | Gm9458 | 728 | 0 | 0 | 0 | 0 | 0 | 0 |
| ENSMUSG00 | 4931428L18R | 1542 | 0 | 0 | 0 | 0 | 0 | 0 |
| ENSMUSG00 | 2410133F24R | 2151 | 0.77625764 | 0.98568268 | 0.65758204 | 1.12389446 | 1.02005307 | 1.03975845 |
| ENSMUSG00 | Gm15589 | 269 | 0 | 0 | 0 | 0 | 0 | 0 |
| ENSMUSG00 | Gm11607 | 460 | 0 | 0 | 0 | 0 | 0 | 0 |
| ENSMUSG00 | Gm7518 | 204 | 0 | 0 | 0 | 0 | 0 | 0 |
| ENSMUSG00 | Gm6117 | 1331 | 0 | 0 | 0 | 0 | 0 | 0 |
| ENSMUSG00 | Gm11685 | 1330 | 0 | 0 | 0 | 0 | 0 | 0 |
| ENSMUSG00 | Astx1c | 1146 | 0 | 0 | 0 | 0 | 0 | 0 |
| ENSMUSG00 | Gm13977 | 4046 | 0 | 0 | 0 | 0 | 0 | 0 |
| ENSMUSG00 | Gm14902 | 1004 | 0 | 0 | 0 | 0 | 0 | 0 |
| ENSMUSG00 | Gm13763 | 588 | 0 | 0 | 0 | 0 | 0 | 0 |
| ENSMUSG00 | Gm12154 | 441 | 0 | 0 | 0 | 0 | 0 | 0 |
| ENSMUSG00 | Gm4409 | 1915 | 0 | 0 | 0 | 0 | 0 | 0 |
| ENSMUSG00 | Gm17029 | 5966 | 1.87915614 | 0.83999162 | 1.0742989 | 1.68132893 | 2.8737617 | 1.92467702 |
| ENSMUSG00 | Gm15816 | 1682 | 0.35453758 | 0.20053811 | 0.21023469 | 0 | 0.24269382 | 0.2594496 |
| ENSMUSG00 | Gm16201 | 2946 | 0.06747366 | 0.11449596 | 0.33008844 | 0.01224781 | 0.05196169 | 0.36106956 |
| ENSMUSG00 | Hnrnpa1l2-ps | 964 | 0 | 0 | 0 | 0.03742951 | 0.05293193 | 0 |
| ENSMUSG00 | Gm15997 | 1486 | 0 | 0 | 0 | 0 | 0 | 0 |
| ENSMUSG00 | 2410057H14 | 586 | 0 | 0 | 0 | 0 | 0 | 0 |
| ENSMUSG00 | Gm15222 | 342 | 0 | 0 | 0 | 0 | 0.14919993 | 0 |
| ENSMUSG00 | Gm13261 | 1016 | 0 | 0 | 0 | 0 | 0 | 0 |
| ENSMUSG00 | Gm12037 | 306 | 0.77951923 | 0.31494407 | 0.57780187 | 0 | 0 | 0.17826562 |
| ENSMUSG00 | Gm11770 | 546 | 0 | 0.08825356 | 0 | 0 | 0 | 0 |
| ENSMUSG00 | Astx5 | 1146 | 0 | 0 | 0 | 0 | 0 | 0 |
| ENSMUSG00 | Gm11674 | 3003 | 0 | 0 | 0 | 0 | 0 | 0 |
| ENSMUSG00 | Gm15751 | 1860 | 0.72671309 | 2.69429569 | 0.49905307 | 1.04754323 | 0.8504396 | 0.35193083 |
| ENSMUSG00 | Gm16098 | 794 | 0 | 0 | 0 | 0 | 0 | 0 |
| ENSMUSG00 | Gm11216 | 351 | 0 | 0 | 0 | 0 | 0 | 0 |
| ENSMUSG00 | Gm14051 | 374 | 0.85038461 | 0 | 0.94549397 | 0.67533238 | 1.3643416 | 0.65634159 |
| ENSMUSG00 | Gm2309 | 1937 | 0 | 0 | 0 | 0 | 0 | 0 |
| ENSMUSG00 | Gm13912 | 1857 | 0 | 0 | 0 | 0 | 0 | 0 |
| ENSMUSG00 | 1700025D23 | 844 | 0 | 0 | 0 | 0 | 0 | 0 |
| ENSMUSG00 | Gm11686 | 3003 | 0 | 0 | 0 | 0 | 0 | 0 |
| ENSMUSG00 | Gm14546 | 1645 | 0 | 0 | 0 | 0 | 0 | 0 |
| ENSMUSG00 | B230378P21 | 1975 | 0.08051743 | 0 | 1.3428408 | 0.31057962 | 0 | 0.3452486 |
| ENSMUSG00 | 4930527A07R | 557 | 0 | 0 | 0 | 0 | 0 | 0 |
| ENSMUSG00 | Gm11827 | 1364 | 0.75780241 | 1.66038327 | 0.58330878 | 0.92585891 | 0.48632176 | 0.67986637 |
| ENSMUSG00 | Gm15969 | 234 | 0.67958086 | 0.20592497 | 0 | 0.61678709 | 1.09030718 | 3.26363205 |
| ENSMUSG00 | Gm13070 | 765 | 0 | 0 | 0 | 0 | 0 | 0 |
| ENSMUSG00 | Gm16827 | 1787 | 0.24471756 | 0.3775099 | 0.17314656 | 0.74708206 | 0.42831317 | 0.41209584 |
| ENSMUSG00 | Gm15587 | 3852 | 0 | 0.0625473 | 0 | 0 | 0.03974017 | 0 |
| ENSMUSG00 | Gm15351 | 844 | 0.42393285 | 0 | 0 | 0 | 0.06045779 | 0 |
| ENSMUSG00 | 1700080G11 | 434 | 0 | 0 | 0 | 0 | 0 | 0 |
| ENSMUSG00 | Gm11705 | 108 | 0 | 0 | 0 | 0 | 0 | 0 |
| ENSMUSG00 | Gm16192 | 935 | 1.36061538 | 0.25768151 | 0.52002168 | 1.00335097 | 0.6003103 | 0.32087811 |
| ENSMUSG00 | Gm11915 | 356 | 0 | 0 | 0 | 0 | 0 | 0 |
| ENSMUSG00 | 5730588L14R | 4081 | 0 | 0 | 0 | 0 | 0 | 0.06683323 |
| ENSMUSG00 | Gm15100 | 571 | 0 | 0 | 0 | 0 | 0 | 0 |
| ENSMUSG00 | B230104C08 | 976 | 0 | 0 | 0 | 0 | 0 | 0 |
| ENSMUSG00 | Gm2165 | 2553 | 0 | 0 | 0 | 0 | 0 | 0 |
| ENSMUSG00 | Gm13562 | 1339 | 0.77195108 | 2.19520013 | 0.75925496 | 3.66479317 | 1.33377383 | 0.93699284 |
| ENSMUSG00 | Gm16707 | 1191 | 0.86787783 | 1.61835238 | 1.00205689 | 0.87857203 | 1.11392592 | 1.28243476 |
| ENSMUSG00 | 1700092C02 | 356 | 0 | 0 | 0 | 0 | 0 | 0 |
| ENSMUSG00 | E130102H24 | 739 | 4.46509456 | 3.97750064 | 3.28971768 | 2.6854025 | 4.21192006 | 9.33759639 |
| ENSMUSG00 | Gm14262 | 324 | 0 | 0 | 0 | 0 | 0 | 0 |
| ENSMUSG00 | Isoc2a | 1082 | 0.66136659 | 1.20243432 | 1.22555942 | 0.50021319 | 1.65057593 | 1.53766451 |
| ENSMUSG00 | Gm6081 | 488 | 0 | 0.09874271 | 0 | 0 | 0 | 0 |
| ENSMUSG00 | Gm15908 | 1087 | 0 | 0 | 0 | 0 | 0 | 0 |
| ENSMUSG00 | Gm16706 | 3574 | 0.22247051 | 0.49885237 | 0.66785101 | 0.42401955 | 0.67102397 | 0.21367932 |
| ENSMUSG00 | 1700029M20 | 1579 | 0 | 0 | 0 | 0 | 0 | 0 |
| ENSMUSG00 | Gm12228 | 925 | 0 | 0 | 0 | 0 | 0 | 0 |
| ENSMUSG00 | Gm12911 | 699 | 0 | 0 | 0 | 0 | 0 | 0 |
| ENSMUSG00 | Gm12147 | 180 | 0 | 0 | 0 | 0 | 0 | 0 |
| ENSMUSG00 | Gm12364 | 545 | 0 | 0 | 0 | 0 | 0 | 0 |
| ENSMUSG00 | Gm13764 | 414 | 0 | 0 | 0 | 0 | 0 | 0 |
| ENSMUSG00 | Gm11642 | 438 | 5.3551903 | 6.16082365 | 9.28440538 | 3.37754298 | 5.5919316 | 4.79485667 |
| ENSMUSG00 | Gm11579 | 843 | 0 | 0 | 0 | 0 | 0 | 0 |
| ENSMUSG00 | 4932702P03R | 4270 | 0 | 0.07899417 | 0 | 0 | 0 | 0.02555001 |
| ENSMUSG00 | Gm12965 | 345 | 0 | 0 | 0 | 0 | 0 | 0 |
| ENSMUSG00 | Gm16222 | 1295 | 0 | 0 | 0.17066349 | 0 | 0 | 0.0210615 |
| ENSMUSG00 | 4930502E09R | 752 | 0.05286633 | 0 | 0 | 0 | 0 | 0.25388627 |
| ENSMUSG00 | 4930556L07R | 1488 | 0 | 0 | 0 | 0 | 0 | 0 |
| ENSMUSG00 | Gm15943 | 3819 | 0.04163968 | 0 | 0.02314839 | 0 | 0 | 0 |
| ENSMUSG00 | 2700038G22 | 1257 | 0.72742725 | 3.06675845 | 1.68789854 | 3.4158817 | 1.94850123 | 1.28019389 |
| ENSMUSG00 | Gm13847 | 1187 | 0 | 0 | 0 | 0 | 0 | 0 |
| ENSMUSG00 | 4932443L11R | 2817 | 0.25402863 | 0.42763971 | 0.39227763 | 0.01280868 | 0.86945901 | 0.30982906 |
| ENSMUSG00 | Gm13054 | 817 | 0 | 0 | 0 | 0 | 0 | 0 |
| ENSMUSG00 | C330013F16 | 4018 | 0.94985718 | 0.91144092 | 0.29702582 | 0.71840805 | 1.28264409 | 0.55662529 |
| ENSMUSG00 | Gm11981 | 659 | 0 | 0 | 0 | 0 | 0 | 0 |
| ENSMUSG00 | 2610037D02 | 4983 | 0 | 0 | 0.01774106 | 0 | 0.04096037 | 0.04926184 |
| ENSMUSG00 | Gm13110 | 3412 | 0.09321332 | 0.25420749 | 0.31091566 | 0.21150085 | 0.41873931 | 0.39968698 |
| ENSMUSG00 | Gm8978 | 2217 | 0 | 0 | 0 | 0 | 0 | 0 |
| ENSMUSG00 | Gm13657 | 1935 | 0.18490921 | 0.07470766 | 0.2284333 | 0.09323526 | 0.21096176 | 0.11276337 |
| ENSMUSG00 | Gm15677 | 764 | 0 | 0 | 0 | 0 | 0 | 0 |
| ENSMUSG00 | 3110082J24R | 498 | 0 | 0 | 0 | 0 | 0 | 0 |
| ENSMUSG00 | Gm16726 | 889 | 0.22359663 | 0 | 0 | 0.32469781 | 0 | 0.03068013 |
| ENSMUSG00 | Gm11482 | 715 | 0 | 0 | 0 | 0 | 0 | 0 |
| ENSMUSG00 | 1810035I16R | 319 | 0 | 0 | 0 | 0 | 0 | 0 |
| ENSMUSG00 | Gm13613 | 1439 | 0 | 0 | 0 | 0 | 0 | 0 |
| ENSMUSG00 | Gm11465 | 433 | 1.19358256 | 0.22257017 | 0.61249667 | 0.24999107 | 0.11784382 | 0.25195972 |
| ENSMUSG00 | Gm13377 | 674 | 0 | 0 | 0 | 0 | 0 | 0 |
| ENSMUSG00 | 5330413P13 | 5850 | 0 | 0 | 0 | 0 | 0 | 0 |
| ENSMUSG00 | 1700028P15 | 3524 | 0 | 0 | 0 | 0 | 0 | 0 |
| ENSMUSG00 | 4930448D08 | 1245 | 0 | 0 | 0 | 0 | 0 | 1.29253311 |
| ENSMUSG00 | Gm15675 | 3678 | 0.07566296 | 0 | 0.12017902 | 0.54937316 | 0.08324042 | 0.72673046 |
| ENSMUSG00 | Gm11739 | 720 | 1.15953485 | 2.40932211 | 1.41200332 | 1.00227902 | 2.55131879 | 2.08347939 |
| ENSMUSG00 | 4930461C15 | 1300 | 0 | 0.11119948 | 0 | 0 | 0 | 0.31470738 |
| ENSMUSG00 | Gm15579 | 755 | 0 | 0 | 0 | 0 | 0 | 0 |
| ENSMUSG00 | Gm13467 | 279 | 0 | 0 | 0 | 0 | 0 | 0 |
| ENSMUSG00 | Gm14280 | 1441 | 0 | 0 | 0 | 0 | 0 | 0 |
| ENSMUSG00 | 2810425M01 | 1847 | 0.02152435 | 0 | 0 | 0.01953549 | 0.08287987 | 0 |
| ENSMUSG00 | Gm12440 | 1065 | 0 | 0 | 0 | 0 | 0 | 0 |
| ENSMUSG00 | Gm14547 | 367 | 0 | 0 | 0 | 0 | 0 | 0 |
| ENSMUSG00 | Gm11775 | 770 | 0 | 0 | 0 | 0 | 0 | 0 |
| ENSMUSG00 | Gm13748 | 374 | 0 | 0 | 0 | 0 | 0 | 0 |
| ENSMUSG00 | Gm16618 | 952 | 1.75391826 | 1.46786431 | 0.55716609 | 0.26530915 | 1.98316797 | 0.14324916 |
| ENSMUSG00 | Gm16212 | 664 | 0 | 0 | 0 | 0 | 0 | 0 |
| ENSMUSG00 | Gm11973 | 1213 | 0.03277451 | 0 | 0 | 0 | 0 | 0 |
| ENSMUSG00 | Gm14250 | 563 | 0 | 0 | 0 | 0 | 0 | 0 |
| ENSMUSG00 | 2410006H16 | 457 | 6.4374301 | 4.42851328 | 6.86724476 | 4.18456972 | 3.79627304 | 6.02787432 |
| ENSMUSG00 | Gm15681 | 2703 | 0 | 0 | 0 | 0 | 0 | 0 |
| ENSMUSG00 | E030013I19R | 3315 | 0 | 0 | 0 | 0 | 0 | 0 |
| ENSMUSG00 | B230206H07 | 4996 | 0.15119178 | 0.04822502 | 0.08847447 | 0.18055467 | 0.1327748 | 0.13102309 |
| ENSMUSG00 | Gm13010 | 824 | 0 | 0 | 0 | 0 | 0.24770085 | 0 |
| ENSMUSG00 | Gm13642 | 766 | 0 | 0.06290658 | 0 | 0 | 0 | 0 |
| ENSMUSG00 | Gm16065 | 1342 | 0 | 0.21543864 | 0 | 0 | 0 | 0 |
| ENSMUSG00 | Lce6a | 1228 | 0 | 0 | 0 | 0 | 0 | 0 |
| ENSMUSG00 | Gm13840 | 254 | 0.15651764 | 0 | 0.174023 | 0 | 0 | 0 |
| ENSMUSG00 | Gm13257 | 663 | 0 | 0 | 0 | 0.05442239 | 0.15392572 | 0.04113822 |
| ENSMUSG00 | A430108G06 | 2444 | 0 | 0 | 0 | 0.45766914 | 0 | 0 |
| ENSMUSG00 | Gm11374 | 671 | 0 | 0 | 0 | 0 | 0 | 0 |
| ENSMUSG00 | Gm14720 | 409 | 0 | 0 | 0 | 0 | 0 | 0 |
| ENSMUSG00 | Gm13781 | 490 | 0 | 0 | 0 | 0 | 0 | 0 |
| ENSMUSG00 | Gm13844 | 578 | 0.55024887 | 0.08336755 | 0.53531644 | 0.06242568 | 0 | 0 |
| ENSMUSG00 | Gm16080 | 687 | 0 | 0 | 0 | 0 | 0 | 0 |
| ENSMUSG00 | Gm16913 | 914 | 0 | 0 | 0 | 0 | 0 | 0 |
| ENSMUSG00 | Trav20 | 319 | 0 | 0 | 0 | 0 | 0 | 0 |
| ENSMUSG00 | 2810008D09 | 954 | 21.8363436 | 24.6488299 | 37.9468652 | 26.8913351 | 21.6086539 | 63.0404398 |
| ENSMUSG00 | Gm1720 | 1494 | 0 | 0 | 0 | 0 | 0 | 0.14604894 |
| ENSMUSG00 | Gm15325 | 695 | 0 | 0 | 0 | 0 | 0 | 0 |
| ENSMUSG00 | Gm13546 | 1344 | 0.17747982 | 0.10755902 | 0 | 0.34900787 | 0.03796605 | 0.24352357 |
| ENSMUSG00 | Gm15543 | 359 | 0 | 0 | 0 | 0 | 0 | 0 |
| ENSMUSG00 | A630077J23R | 925 | 0 | 0 | 0 | 0 | 0 | 0 |
| ENSMUSG00 | Gm16978 | 1929 | 0 | 0.29976014 | 0.06874315 | 0.07482021 | 0.02645224 | 0 |
| ENSMUSG00 | 4930512H18 | 1490 | 0 | 0 | 0 | 0 | 0 | 0 |
| ENSMUSG00 | Gm4577 | 714 | 0.5011195 | 0 | 0 | 0 | 0.07146551 | 0.26739842 |
| ENSMUSG00 | Gm15883 | 1686 | 0 | 0 | 0 | 0 | 0.18158853 | 0 |
| ENSMUSG00 | Gm7809 | 2738 | 0.50819643 | 0.10559483 | 0.17758228 | 0.28992147 | 0.20500005 | 0.0498076 |
| ENSMUSG00 | Gm15288 | 647 | 0 | 0 | 0 | 0 | 0 | 0 |
| ENSMUSG00 | Gm16834 | 2246 | 0 | 0 | 0 | 0 | 0 | 0 |
| ENSMUSG00 | Gm14225 | 537 | 3.62759506 | 0.62812867 | 0.98775068 | 3.02363502 | 0.66514829 | 3.96167944 |
| ENSMUSG00 | Gm15672 | 1352 | 1.76429647 | 0.42769032 | 0.91542278 | 0.9607645 | 0.49063823 | 1.10954524 |
| ENSMUSG00 | Gm13175 | 464 | 0 | 0 | 0 | 0 | 0 | 0 |
| ENSMUSG00 | Gm8975 | 1996 | 0 | 0 | 0 | 0 | 0 | 0 |
| ENSMUSG00 | Gm12828 | 286 | 0 | 0 | 0 | 0 | 0 | 0 |
| ENSMUSG00 | A230072C01 | 1575 | 7.39578145 | 9.54550473 | 12.9939386 | 10.6757033 | 8.65018562 | 17.8021138 |
| ENSMUSG00 | Miat | 8753 | 0 | 0 | 0 | 0 | 0 | 0 |
| ENSMUSG00 | Gm14012 | 673 | 0 | 0 | 0 | 0 | 0 | 0 |
| ENSMUSG00 | C430042M11 | 1615 | 0.46771154 | 0 | 0.35580431 | 0 | 0 | 0 |
| ENSMUSG00 | Gm13594 | 474 | 5.36782859 | 8.33605117 | 1.77180383 | 5.70918426 | 3.66011978 | 7.99825919 |
| ENSMUSG00 | Gm13352 | 261 | 0 | 0 | 0 | 0 | 0 | 0 |
| ENSMUSG00 | Gm15992 | 229 | 0 | 0 | 0 | 0 | 0 | 0 |
| ENSMUSG00 | Gm16225 | 274 | 0 | 0 | 0 | 0 | 0 | 0 |
| ENSMUSG00 | Gm13846 | 459 | 0 | 0 | 0 | 0 | 0 | 0 |
| ENSMUSG00 | Gm15135 | 607 | 0 | 0 | 0 | 0 | 0 | 0 |
| ENSMUSG00 | 1700060C16 | 640 | 0 | 0 | 0 | 0 | 0 | 0 |
| ENSMUSG00 | A330033J07R | 4452 | 0.87512064 | 0.63858942 | 0.60564071 | 0.65647925 | 1.04299196 | 1.12112736 |
| ENSMUSG00 | Gm8619 | 495 | 1.60628204 | 3.01773678 | 2.50030627 | 3.20729285 | 4.02025991 | 1.92850985 |
| ENSMUSG00 | 1700021J08R | 573 | 1.38762585 | 0.75685511 | 0.53998761 | 0.31485205 | 1.24671773 | 0.61879635 |
| ENSMUSG00 | Gm15963 | 874 | 1.22814414 | 2.31559562 | 0.45516772 | 4.49993462 | 4.72898906 | 2.37170776 |
| ENSMUSG00 | 4933430M04 | 740 | 0 | 0 | 0 | 0 | 0 | 0 |
| ENSMUSG00 | Gm11365 | 772 | 0 | 0.18725301 | 0 | 0 | 0 | 0 |
| ENSMUSG00 | Gm15708 | 1762 | 0.85738267 | 1.91433085 | 0.3261203 | 2.39591329 | 2.92489441 | 0.72753011 |
| ENSMUSG00 | 9330136K24R | 2475 | 0.61038718 | 1.0902791 | 0.6429359 | 0.67061578 | 1.27823649 | 2.69991378 |
| ENSMUSG00 | Gm12835 | 1999 | 0 | 0 | 0 | 0 | 0 | 0 |
| ENSMUSG00 | 4632428C04 | 2427 | 0.98283017 | 0.93315319 | 0.61922648 | 2.28950756 | 0.50458715 | 1.12380055 |
| ENSMUSG00 | 1700062I23R | 792 | 0 | 0 | 0 | 0 | 0 | 0 |
| ENSMUSG00 | 1600002D24 | 2208 | 0 | 0 | 0 | 0 | 0 | 0.02470529 |
| ENSMUSG00 | Gm16040 | 564 | 0.3524422 | 0.17087391 | 0.39186031 | 0.70372782 | 1.35708446 | 0.33851503 |
| ENSMUSG00 | Gm11728 | 997 | 0 | 0 | 0 | 0 | 0 | 0 |
| ENSMUSG00 | Gm16258 | 2694 | 0 | 0 | 0 | 0 | 0 | 0 |
| ENSMUSG00 | Gm13404 | 1958 | 0 | 0 | 0 | 0 | 0 | 0 |
| ENSMUSG00 | Gm13716 | 806 | 0 | 0 | 0 | 0 | 0 | 0 |
| ENSMUSG00 | Gm15276 | 497 | 0 | 0 | 0 | 0 | 0 | 0 |
| ENSMUSG00 | Gm15298 | 1210 | 0 | 0 | 0 | 0 | 0 | 0 |
| ENSMUSG00 | Gm13598 | 622 | 0 | 0 | 0 | 0 | 0 | 0 |
| ENSMUSG00 | Gm4926 | 1605 | 0 | 0 | 0 | 0 | 0 | 0 |
| ENSMUSG00 | Gm15755 | 656 | 0 | 0 | 0 | 0 | 0 | 0 |
| ENSMUSG00 | Gm12027 | 947 | 0 | 0 | 0 | 0 | 0 | 0 |
| ENSMUSG00 | 4930505O20 | 3414 | 0 | 0 | 0 | 0 | 0 | 0 |
| ENSMUSG00 | Gm14019 | 401 | 0 | 0 | 0 | 0 | 0 | 0.27206623 |
| ENSMUSG00 | Gm16124 | 514 | 6.96107635 | 8.24981889 | 9.45953839 | 6.59866184 | 4.96365524 | 18.1477172 |
| ENSMUSG00 | Gm16365 | 1401 | 0.36889454 | 0.41273184 | 0.82030544 | 0.84989827 | 0.9105349 | 0.7592512 |
| ENSMUSG00 | Gm15903 | 308 | 1.03260988 | 5.94508052 | 2.58322459 | 2.69443839 | 6.1297919 | 4.25059313 |
| ENSMUSG00 | Gm11630 | 380 | 0.31385906 | 0 | 0 | 0 | 0.13427994 | 0 |
| ENSMUSG00 | 4930429F24R | 1517 | 0.15723987 | 0 | 0.37878969 | 0.09514053 | 0.30272735 | 0.07191731 |
| ENSMUSG00 | Gm11497 | 2152 | 0 | 0 | 0 | 0 | 0 | 0 |
| ENSMUSG00 | Gm12207 | 492 | 1.61607644 | 1.17527908 | 2.06634632 | 2.34679964 | 2.59280365 | 0.33261755 |
| ENSMUSG00 | Gm13189 | 250 | 0 | 0 | 0 | 0 | 0 | 0 |
| ENSMUSG00 | Gm13835 | 1346 | 1.53587295 | 0.89499335 | 0.85382461 | 1.8496739 | 0.98565065 | 2.37082674 |
| ENSMUSG00 | 4930406D18 | 1036 | 0 | 0 | 0 | 0 | 0 | 0 |
| ENSMUSG00 | Gm11766 | 3016 | 1.22588186 | 0.97459316 | 0.92331436 | 0.35890628 | 0.60906815 | 0.96763475 |
| ENSMUSG00 | Gm6286 | 761 | 0 | 0 | 0 | 0 | 0 | 0 |
| ENSMUSG00 | Gm6226 | 2278 | 0 | 0.02115296 | 0 | 0 | 0 | 0 |
| ENSMUSG00 | Gm11823 | 2275 | 0 | 0 | 0 | 0 | 0 | 0 |
| ENSMUSG00 | Gm14210 | 501 | 0 | 0 | 0 | 0 | 0.30554716 | 0 |
| ENSMUSG00 | Gm11788 | 559 | 0 | 0 | 0.15814613 | 0.83911732 | 0.73025225 | 0.19516737 |
| ENSMUSG00 | Gm14873 | 1377 | 0.14435541 | 0.06998757 | 0.09630031 | 0.2882371 | 0 | 0.17826562 |
| ENSMUSG00 | Gm14532 | 277 | 0 | 0 | 0.15957344 | 0 | 0 | 0.39385761 |
| ENSMUSG00 | Gm11665 | 725 | 0.16450544 | 0 | 0.42677642 | 0 | 0.14076242 | 0.03762019 |
| ENSMUSG00 | 4933426K07R | 1546 | 0 | 0 | 0 | 0 | 0 | 0 |
| ENSMUSG00 | Gm16801 | 791 | 0 | 0 | 0 | 0 | 0 | 0 |
| ENSMUSG00 | Gm15956 | 500 | 0 | 0 | 0.08840369 | 0 | 0 | 2.40016825 |
| ENSMUSG00 | Gm9697 | 996 | 0 | 0 | 0 | 0 | 0 | 0 |
| ENSMUSG00 | Gm15063 | 893 | 0 | 0 | 0 | 0 | 0 | 0 |
| ENSMUSG00 | 4930481A15R | 2465 | 0.64511936 | 1.95482524 | 1.64972396 | 0.54159661 | 2.31843979 | 6.25159074 |
| ENSMUSG00 | Gm13530 | 388 | 0 | 0 | 0 | 0 | 0 | 0 |
| ENSMUSG00 | Gm12746 | 613 | 0 | 0 | 0 | 0 | 0 | 0 |
| ENSMUSG00 | Gm2020 | 576 | 0 | 0 | 0 | 0 | 0 | 0 |
| ENSMUSG00 | Gm15489 | 969 | 0.08205466 | 1.24320026 | 0.63862312 | 0.96814567 | 0.94785837 | 0.73182727 |
| ENSMUSG00 | 4732414G09 | 2411 | 0 | 0 | 0 | 0 | 0 | 0 |
| ENSMUSG00 | Gm15859 | 708 | 2.69528681 | 0.61253952 | 1.3735036 | 3.82225048 | 1.87384996 | 2.1573161 |
| ENSMUSG00 | 4930562A09R | 732 | 0 | 0 | 0 | 0 | 0 | 0 |
| ENSMUSG00 | Gm15527 | 3563 | 0 | 0 | 0.06202897 | 0 | 0 | 0 |
| ENSMUSG00 | 4930522O17 | 444 | 0 | 0 | 0 | 0 | 0 | 0 |
| ENSMUSG00 | Gm14697 | 644 | 0 | 0 | 0 | 0 | 0 | 0 |
| ENSMUSG00 | Gm13066 | 882 | 0 | 0 | 0 | 0 | 0 | 0 |
| ENSMUSG00 | Gm16907 | 2314 | 8.195058 | 5.93480381 | 7.39244306 | 8.18628927 | 7.80610935 | 5.56336635 |
| ENSMUSG00 | 1700051A21R | 2479 | 0 | 0 | 0 | 0 | 0 | 0 |
| ENSMUSG00 | Gm12596 | 670 | 0 | 0 | 0 | 0 | 0 | 0 |
| ENSMUSG00 | 4933437N03 | 1101 | 0 | 0 | 0.08029399 | 0 | 0 | 0 |
| ENSMUSG00 | 4930583P06R | 703 | 0 | 0 | 0 | 0 | 0 | 0 |
| ENSMUSG00 | 1700021L23R | 724 | 0 | 0 | 0 | 0 | 0 | 0 |
| ENSMUSG00 | 1700125G22 | 858 | 0 | 0.05616135 | 0 | 0.126161 | 0 | 0 |
| ENSMUSG00 | Gm13869 | 1052 | 0 | 0 | 0 | 0 | 0 | 0 |
| ENSMUSG00 | E330010L02R | 962 | 0 | 0 | 0 | 0 | 0 | 0 |
| ENSMUSG00 | Gm13017 | 398 | 0 | 0 | 0 | 0 | 0 | 0 |
| ENSMUSG00 | Gm12946 | 517 | 0 | 0 | 0.0854968 | 0 | 0 | 0 |
| ENSMUSG00 | Gm12248 | 942 | 0.29542289 | 0 | 0 | 0 | 0 | 0 |
| ENSMUSG00 | Gm14135 | 587 | 0 | 0 | 0 | 0 | 0 | 0 |
| ENSMUSG00 | Gm15948 | 400 | 1.19266442 | 1.08419495 | 1.87857833 | 2.07471756 | 0.6378297 | 0.61367938 |
| ENSMUSG00 | Gm16314 | 862 | 1.4758415 | 0.33540447 | 0.46150416 | 0 | 0.47356265 | 0.15820556 |
| ENSMUSG00 | Gm16866 | 2570 | 0 | 0 | 0 | 0 | 0 | 0 |
| ENSMUSG00 | 4930566F21R | 1723 | 0 | 0 | 0 | 0 | 0 | 0 |
| ENSMUSG00 | 4933431E20R | 8844 | 0.62483173 | 0.88265532 | 0.90962635 | 0.77108847 | 0.89428859 | 0.956031 |
| ENSMUSG00 | 4930443O20 | 1440 | 0 | 0 | 0 | 0 | 0.07086997 | 0 |
| ENSMUSG00 | Gm14259 | 1236 | 0 | 0 | 0 | 0 | 0 | 0 |
| ENSMUSG00 | 4930527B05 | 599 | 0 | 0 | 0 | 0 | 0 | 0 |
| ENSMUSG00 | Gm15092 | 572 | 0 | 0 | 0 | 0 | 0 | 0 |
| ENSMUSG00 | Gm15208 | 491 | 0 | 0 | 0 | 0 | 0 | 0 |
| ENSMUSG00 | Gm12316 | 404 | 0 | 0 | 0 | 0 | 0 | 0 |
| ENSMUSG00 | 9430065F17R | 1449 | 2.19491956 | 2.39435737 | 1.46424325 | 0.2988161 | 1.0564467 | 4.36695397 |
| ENSMUSG00 | 4933435F18R | 2600 | 0 | 0 | 0 | 0 | 0 | 0 |
| ENSMUSG00 | Gm13791 | 606 | 0 | 0 | 0 | 0 | 0 | 0 |
| ENSMUSG00 | Gm12171 | 1384 | 0 | 0 | 0 | 0 | 0 | 0 |
| ENSMUSG00 | Gm8327 | 700 | 0 | 0 | 0 | 0 | 0 | 0 |
| ENSMUSG00 | Gm16972 | 588 | 0.33805681 | 0.32779893 | 1.12759804 | 0.12272804 | 0.3471182 | 0.37108353 |
| ENSMUSG00 | Gm6762 | 1315 | 0.27209074 | 0.36643682 | 0.30252212 | 0.21951054 | 0.38803328 | 0.43556458 |
| ENSMUSG00 | Gm16973 | 3109 | 2.74925324 | 4.8976892 | 7.63473454 | 5.25737091 | 6.51575143 | 9.26407818 |
| ENSMUSG00 | Gm12103 | 789 | 0 | 1.77111131 | 0.50420353 | 0.18292545 | 0.45270549 | 0 |
| ENSMUSG00 | Gm12484 | 579 | 0 | 0 | 0 | 0 | 0 | 0 |
| ENSMUSG00 | 1700086O06 | 1927 | 2.92956836 | 2.90068879 | 3.57835366 | 2.88356765 | 2.96572605 | 4.11879607 |
| ENSMUSG00 | Gm16441 | 923 | 0 | 0 | 0 | 0 | 0.22113272 | 0 |
| ENSMUSG00 | 4833428L15R | 1254 | 0.22192055 | 0.03842619 | 0.35248679 | 0.20141492 | 0.36621801 | 0.28275144 |
| ENSMUSG00 | Gm15334 | 323 | 0 | 0 | 0 | 0 | 0 | 0.33776643 |
| ENSMUSG00 | Gm15941 | 3358 | 0.04735614 | 0.20089642 | 0 | 0.41965889 | 0 | 0.0649783 |
| ENSMUSG00 | Gm15623 | 3849 | 0.29953467 | 0.65099896 | 0.03445194 | 0.44997094 | 0.79542285 | 0.84325333 |
| ENSMUSG00 | Gm13402 | 422 | 0 | 0.22837176 | 0 | 0.08550248 | 0 | 0 |
| ENSMUSG00 | Gm13544 | 2936 | 0 | 0 | 0 | 0 | 0 | 0 |
| ENSMUSG00 | 4933416E14R | 2427 | 0 | 0 | 0 | 0 | 0 | 0 |
| ENSMUSG00 | Gm12981 | 620 | 1.09006963 | 0.54404048 | 0.0712933 | 0.17459054 | 0.90530667 | 0.61587895 |
| ENSMUSG00 | Gm16033 | 594 | 0 | 0 | 0 | 0 | 0 | 0 |
| ENSMUSG00 | 1700040N02 | 1166 | 0 | 0 | 0 | 0 | 0 | 0 |
| ENSMUSG00 | Gm13395 | 1837 | 0 | 0 | 0 | 0 | 0 | 0 |
| ENSMUSG00 | Gm15475 | 618 | 0 | 0 | 0 | 0 | 0 | 0 |
| ENSMUSG00 | Gm16277 | 660 | 0 | 0 | 0 | 0 | 0 | 0 |
| ENSMUSG00 | 1700067G03 | 381 | 0 | 0 | 0 | 0 | 0 | 0 |
| ENSMUSG00 | Gm14154 | 575 | 0.48397976 | 0 | 0.15374554 | 0.06275138 | 0 | 0 |
| ENSMUSG00 | Gm13889 | 1380 | 0 | 0 | 0.03203032 | 0 | 0.07395127 | 0.09882116 |
| ENSMUSG00 | Gm2231 | 611 | 0 | 0 | 0 | 0 | 0 | 0 |
| ENSMUSG00 | Gm5530 | 984 | 0 | 0.04896996 | 0 | 0.03666874 | 0 | 0.08315439 |
| ENSMUSG00 | Nphs1as | 1336 | 0 | 0 | 0 | 0 | 0 | 0 |
| ENSMUSG00 | 1700041M05 | 304 | 0 | 0 | 0 | 0 | 0 | 0 |
| ENSMUSG00 | Gm12930 | 674 | 0 | 0 | 0 | 0 | 0 | 0 |
| ENSMUSG00 | Gm13113 | 637 | 0 | 0 | 0 | 0 | 0 | 0 |
| ENSMUSG00 | 2610027K06R | 2568 | 5.51127378 | 6.73634452 | 2.37533269 | 3.40025498 | 4.25219799 | 7.11604685 |
| ENSMUSG00 | Gm16364 | 1311 | 0.36389456 | 0.55133229 | 0 | 0 | 0.89519958 | 0 |
| ENSMUSG00 | Gm14041 | 274 | 0 | 0 | 0 | 0 | 0 | 0 |

[illegible]

| | | |
|---|---|---|
| 0 | 0 | 0 |
| 1.39341229 | 3.00765223 | 3.76966559 |
| 0 | 0 | 0 |
| 0.28145121 | 0.07960433 | 0.17020056 |
| 0.43603679 | 0.57809338 | 0.4326038 |
| 0 | 0 | 0 |
| 0 | 0 | 0 |
| 0 | 0 | 0 |
| 0.03643424 | 0.3434963 | 0.07344231 |
| 0.2466546 | 0 | 0 |
| 0 | 0 | 0 |
| 0.01486693 | 0 | 0.17980809 |
| 0 | 0 | 0 |
| 0 | 0 | 0 |
| 0 | 0 | 0 |
| 0 | 0 | 0 |
| 1.07507023 | 1.04523302 | 0.10158152 |
| 0 | 0.08707573 | 0 |
| 0 | 0 | 0.16530084 |
| 0 | 0 | 0 |
| 0 | 0 | 0 |
| 0.42681217 | 0 | 0.51289889 |
| 0 | 0 | 0 |
| 0 | 0 | 0 |
| 0 | 0 | 0.79811285 |
| 0.12283249 | 0.06514005 | 0 |
| 0 | 0 | 0 |
| 0.53137453 | 0.14544335 | 0.11661366 |
| 0 | 0 | 0.04140363 |
| 0 | 0 | 0 |
| 0 | 0 | 0 |
| 0 | 0 | 0 |
| 0 | 0 | 0 |
| 1.92315486 | 2.20164566 | 0 |
| 0 | 0 | 0 |
| 0 | 0 | 0 |
| 0 | 0 | 0 |
| 0 | 0 | 0 |
| 0 | 0.13716768 | 1.17310276 |
| 0 | 0 | 0 |
| 0 | 0 | 0 |
| 11.6766966 | 10.1656045 | 15.1614127 |
| 1.06198029 | 0 | 0 |
| 0 | 0.20361682 | 0 |
| 0 | 0 | 0 |
| 0 | 0 | 0 |
| 0.30635698 | 0.28882854 | 0 |
| 0.87248361 | 1.48061488 | 0.45433306 |
| 0 | 0 | 0 |
| 0.01050423 | 0 | 0.00794022 |
| 0 | 0.04338978 | 0 |
| 0 | 0 | 0 |
| 0 | 0 | 0 |
| 0 | 0 | 0 |
| 4.50755966 | 4.33098946 | 4.18439375 |
| 0 | 0 | 0 |
| 0 | 0 | 0 |
| 0 | 0 | 0 |
| 0 | 0 | 0 |
| 0 | 0 | 0 |
| 0 | 0 | 0.01122875 |
| 0 | 0 | 0 |
| 0 | 0 | 0 |
| 0 | 0 | 0 |
| 0.02912191 | 0.41183516 | 0.18711415 |
| 0.98719684 | 0.2792141 | 0.41042549 |
| 0 | 0 | 0 |
| 0 | 0.13083686 | 0 |
| 0 | 0 | 0 |
| 0 | 0 | 0 |
| 0 | 0 | 0 |
| 0.05184202 | 0 | 0.09796925 |
| 0 | 0 | 0.18724009 |

[illegible]

| | | | | | | | | |
|---|---|---|---|---|---|---|---|---|
| ENSMUSG00 | Gm2200 | 759 | 0 | 0 | 0 | 0 | 0 | 0 |
| ENSMUSG00 | Gm12882 | 470 | 0 | 0 | 0 | 0 | 0 | 0 |
| ENSMUSG00 | Gm13905 | 576 | 0 | 0 | 0 | 0 | 0 | 0 |
| ENSMUSG00 | A330041J22R | 321 | 0.74309309 | 0 | 0.13770045 | 0.22481025 | 0 | 0 |
| ENSMUSG00 | Gm13601 | 400 | 0 | 0 | 0 | 0 | 0 | 0 |
| ENSMUSG00 | Gm7023 | 867 | 0 | 0 | 0 | 0 | 0 | 0 |
| ENSMUSG00 | Gm16142 | 2847 | 0 | 0 | 0 | 0 | 0 | 0 |
| ENSMUSG00 | A630031M04 | 3562 | 0 | 0 | 0 | 0 | 0 | 0 |
| ENSMUSG00 | Gm16836 | 2826 | 2.78541583 | 2.0120312 | 2.01770621 | 3.40902544 | 1.73338007 | 3.69645677 |
| ENSMUSG00 | A630075F10R | 1910 | 0 | 0 | 0 | 0 | 0 | 0 |
| ENSMUSG00 | Gm13883 | 1814 | 0 | 0 | 0 | 0 | 0 | 0 |
| ENSMUSG00 | Gm12100 | 1931 | 0 | 0 | 0 | 0 | 0 | 0 |
| ENSMUSG00 | 4933406I18R | 1642 | 1.11373453 | 0.17607713 | 0.16151709 | 0.52738677 | 0.90119665 | 1.64445145 |
| ENSMUSG00 | 1810058N05I | 469 | 0.08476648 | 0.20548589 | 0.471235 | 0 | 0 | 0 |
| ENSMUSG00 | Gm13822 | 344 | 0 | 0 | 0.12849373 | 0 | 0 | 0 |
| ENSMUSG00 | 4930506C21I | 1915 | 1.07952219 | 0.85552952 | 0.71553897 | 0.41451957 | 0.77272319 | 0.81182999 |
| ENSMUSG00 | Gm16835 | 2802 | 0.14188251 | 0 | 0 | 0.05150899 | 0.10926419 | 0 |
| ENSMUSG00 | Gm15910 | 2968 | 0 | 0.08117662 | 0.29785609 | 0 | 0 | 0.48704713 |
| ENSMUSG00 | Gm15608 | 3923 | 0.03040185 | 0.4544732 | 0.157743 | 0.11037077 | 0.14307676 | 1.73117134 |
| ENSMUSG00 | Gm12720 | 469 | 0 | 0 | 0 | 0 | 0 | 0 |
| ENSMUSG00 | D830044I16R | 2332 | 0.75010341 | 0.66122048 | 0.72027017 | 1.19138826 | 0.72207136 | 1.00584004 |
| ENSMUSG00 | 2900089D17I | 2893 | 0.23361326 | 0.11659353 | 0.24446232 | 0.06236095 | 0 | 0.73536877 |
| ENSMUSG00 | Gm13383 | 1169 | 0 | 0 | 0 | 0 | 0 | 0 |
| ENSMUSG00 | Gm11723 | 463 | 0 | 0 | 0 | 0 | 0 | 0 |
| ENSMUSG00 | 4933401P06R | 1488 | 0 | 0 | 0 | 0.07274606 | 0 | 0 |
| ENSMUSG00 | Gm16990 | 4324 | 0 | 0 | 0 | 0.0166892 | 0 | 0 |
| ENSMUSG00 | Gm15656 | 613 | 0 | 0 | 0 | 0 | 0 | 0 |
| ENSMUSG00 | A330076H08I | 4519 | 0 | 0.09596769 | 0.01956267 | 0 | 0 | 0 |
| ENSMUSG00 | Gm14552 | 2400 | 0 | 0 | 0 | 0 | 0 | 0 |
| ENSMUSG00 | Gm15700 | 488 | 0 | 0.19748542 | 0.63404283 | 0.07393862 | 1.77755817 | 0.39123458 |
| ENSMUSG00 | 4930550C17I | 1406 | 0 | 0 | 0 | 0 | 0 | 0 |
| ENSMUSG00 | Gm16728 | 1375 | 0 | 0 | 0 | 0 | 0 | 0 |
| ENSMUSG00 | Gm14152 | 1235 | 0 | 0 | 0 | 0 | 0 | 0 |
| ENSMUSG00 | Gm13523 | 486 | 0.0818014 | 0 | 0 | 0 | 0 | 0 |
| ENSMUSG00 | 2810001G20I | 1249 | 8.3712501 | 8.95056411 | 6.54711045 | 7.48218537 | 7.39453485 | 8.42915192 |
| ENSMUSG00 | 2210416O15I | 1501 | 0.23837397 | 0.4494405 | 0.58896526 | 0.31250272 | 0.1699746 | 0.41793251 |
| ENSMUSG00 | Gm16727 | 2492 | 1.53151129 | 4.1186646 | 0.6385499 | 1.28864445 | 3.82902579 | 1.97007828 |
| ENSMUSG00 | Gm12426 | 368 | 0 | 0 | 0 | 0 | 0 | 0 |
| ENSMUSG00 | Gm12116 | 2766 | 0 | 0 | 0 | 0 | 0 | 0 |
| ENSMUSG00 | Gm16091 | 961 | 0.1241066 | 0.85241365 | 0.22997837 | 1.50185409 | 1.00884614 | 0.62439341 |
| ENSMUSG00 | Gm11459 | 419 | 0 | 0 | 0 | 0 | 0 | 0 |
| ENSMUSG00 | Gm13658 | 926 | 0 | 0 | 0 | 0 | 0 | 0 |
| ENSMUSG00 | Gm15241 | 469 | 0.16953297 | 0.82194358 | 0.188494 | 0.076934 | 0.65278946 | 1.51202691 |
| ENSMUSG00 | Gm16141 | 788 | 0 | 0 | 0 | 0 | 0 | 0 |
| ENSMUSG00 | Gm14174 | 159 | 0 | 0 | 0 | 0 | 0 | 0 |
| ENSMUSG00 | Gm2863 | 1907 | 0 | 0 | 0 | 0 | 0 | 0 |
| ENSMUSG00 | 1700112K13R | 699 | 0 | 0 | 0 | 0 | 0 | 0 |
| ENSMUSG00 | A330009N23I | 1389 | 0.22897325 | 0 | 0 | 0 | 0 | 0.17672552 |
| ENSMUSG00 | Gm11985 | 717 | 0 | 0 | 0 | 0 | 0 | 0.22823966 |
| ENSMUSG00 | Gm13236 | 937 | 0 | 0 | 0 | 0 | 0 | 0 |
| ENSMUSG00 | Gm16076 | 478 | 0.08317046 | 0 | 0.09247248 | 0.07548545 | 0 | 0 |
| ENSMUSG00 | Gm13554 | 475 | 0 | 0 | 0 | 0 | 0 | 0 |
| ENSMUSG00 | Gm2301 | 1489 | 0 | 0 | 0 | 0 | 0 | 0 |
| ENSMUSG00 | 4930577H14I | 788 | 0 | 0 | 0 | 0 | 0 | 0 |
| ENSMUSG00 | Gm13561 | 1821 | 0 | 0.02646153 | 0 | 0.07925765 | 0.22416859 | 0 |
| ENSMUSG00 | AV039307 | 1327 | 0.14979458 | 0.4720601 | 0.06661921 | 0.13595345 | 0.30761945 | 0.6782691 |
| ENSMUSG00 | Gm14389 | 225 | 0 | 0 | 0 | 0 | 0 | 0 |
| ENSMUSG00 | AI646519 | 349 | 0 | 0 | 0 | 0 | 0 | 0 |
| ENSMUSG00 | Gm371 | 2425 | 0 | 0 | 0 | 0 | 0 | 0.16870911 |
| ENSMUSG00 | Gm12319 | 355 | 0.55993635 | 1.49310102 | 0.99609787 | 0.8131165 | 1.4373627 | 4.45613824 |
| ENSMUSG00 | Gm14285 | 378 | 0 | 0 | 0 | 0 | 0.13499041 | 0 |
| ENSMUSG00 | Gm13062 | 559 | 1.2801407 | 0 | 0.07907306 | 0.12909497 | 1.00409684 | 0.53671025 |
| ENSMUSG00 | Gm15738 | 578 | 1.03171662 | 0.16673509 | 0 | 0.12485137 | 0.08828093 | 0 |
| ENSMUSG00 | Gm13656 | 343 | 0 | 0 | 0 | 0 | 0 | 0 |
| ENSMUSG00 | 9830144P21I | 4878 | 0.20374888 | 0.19756639 | 0.28996699 | 0.35505087 | 0.88914349 | 0.21806292 |
| ENSMUSG00 | C130060C02 | 1139 | 0.06980769 | 0.04230592 | 0.23284553 | 0.03167871 | 0.49279204 | 0.90995285 |
| ENSMUSG00 | Gm15533 | 463 | 1.63143441 | 1.14481828 | 0.47734172 | 1.55861963 | 1.21228971 | 1.11926165 |
| ENSMUSG00 | Gm15606 | 591 | 0 | 0 | 0 | 0 | 0 | 0 |
| ENSMUSG00 | Gm15938 | 633 | 0.06280487 | 0 | 0.27931654 | 0.05700165 | 0 | 0.0430879 |
| ENSMUSG00 | Gm11418 | 622 | 0 | 0 | 0 | 0 | 0 | 0 |
| ENSMUSG00 | D930049A15I | 3466 | 0.25234292 | 0.12512348 | 0.47186041 | 0.2602571 | 0.2649956 | 0.45641347 |
| ENSMUSG00 | Gm14246 | 678 | 0 | 0 | 0 | 0 | 0 | 0 |
| ENSMUSG00 | Gm13799 | 834 | 0 | 0 | 0 | 0 | 0 | 0 |
| ENSMUSG00 | Gm16341 | 373 | 0 | 0.12918617 | 0 | 0.19346941 | 0 | 0 |
| ENSMUSG00 | Gm13584 | 600 | 0 | 0 | 0 | 0 | 0 | 0 |
| ENSMUSG00 | Gm11412 | 2798 | 0 | 0 | 0 | 0 | 0 | 0 |
| ENSMUSG00 | Gm15830 | 1659 | 0 | 0 | 0 | 0 | 0 | 0 |
| ENSMUSG00 | Gm15878 | 305 | 0 | 0 | 0.86954445 | 0 | 0 | 0 |
| ENSMUSG00 | Gm16576 | 5400 | 1.39144182 | 0.26770246 | 0.62210001 | 2.10478593 | 0.47246644 | 1.5203086 |
| ENSMUSG00 | Gm13923 | 286 | 0 | 0 | 0 | 0 | 0 | 0 |
| ENSMUSG00 | Gm13966 | 830 | 0.6226762 | 0.87083932 | 0.3195314 | 0.9129192 | 1.04511854 | 1.67591157 |
| ENSMUSG00 | Gm2101 | 2553 | 0 | 0 | 0 | 0 | 0 | 0 |
| ENSMUSG00 | 2410019O14I | 627 | 0 | 0 | 0 | 0 | 0 | 0 |
| ENSMUSG00 | 1700028E10R | 2808 | 0.22652695 | 0.20592497 | 0.12593118 | 1.15647579 | 0.18171786 | 0.88390035 |
| ENSMUSG00 | Gm13427 | 790 | 0.15097018 | 0.60995496 | 0.1678551 | 0.09134695 | 0.83967454 | 0.44882318 |
| ENSMUSG00 | Gm14343 | 728 | 0 | 0 | 0 | 0 | 0 | 0 |
| ENSMUSG00 | Gm16912 | 1683 | 0.0236218 | 0 | 0.15758233 | 0.04287825 | 0.27286832 | 0.0486179 |
| ENSMUSG00 | Gm15819 | 478 | 0 | 0 | 0 | 0 | 0 | 0 |
| ENSMUSG00 | 5730412P04R | 1143 | 0 | 0 | 0 | 0 | 0 | 0 |
| ENSMUSG00 | Gm15646 | 764 | 0 | 0 | 0 | 0 | 0 | 0 |
| ENSMUSG00 | Gm13997 | 334 | 0 | 0 | 0 | 0 | 0 | 0 |
| ENSMUSG00 | Gm15764 | 779 | 1.02067986 | 0.30928397 | 1.53202794 | 2.17696546 | 1.76856502 | 3.50123739 |
| ENSMUSG00 | Gm15869 | 732 | 4.61641509 | 2.30399655 | 3.62310189 | 6.45730578 | 5.78577759 | 4.06138754 |
| ENSMUSG00 | 6030443J06R | 7536 | 0.07913113 | 0.12148917 | 0.19942445 | 0.24418581 | 0.31146673 | 0.24610874 |
| ENSMUSG00 | Gm13802 | 661 | 0 | 0 | 0 | 0 | 0 | 0 |
| ENSMUSG00 | Gm15396 | 1086 | 0.2562508 | 0.08874115 | 0.08140303 | 0 | 0 | 0.37672154 |
| ENSMUSG00 | Gm11608 | 877 | 0 | 0 | 0 | 0 | 0 | 0 |
| ENSMUSG00 | Gm16754 | 2074 | 0.28752758 | 0.16263505 | 0.25574837 | 0.17397321 | 0.81189508 | 0.02630148 |
| ENSMUSG00 | Gm14205 | 737 | 0 | 0 | 0 | 0 | 0 | 0 |
| ENSMUSG00 | Gm11817 | 681 | 0 | 0 | 0.06490726 | 0 | 0 | 0 |
| ENSMUSG00 | Gm14662 | 731 | 0 | 0 | 0 | 0 | 0 | 0 |
| ENSMUSG00 | C920006O11 | 1440 | 5.82528221 | 5.75560281 | 6.16984059 | 4.56036952 | 6.55547189 | 10.7015078 |
| ENSMUSG00 | Gm4755 | 2644 | 0 | 0 | 0 | 0.10917411 | 0 | 0.07220971 |
| ENSMUSG00 | Gm15649 | 324 | 0 | 0 | 0 | 0 | 0 | 0 |
| ENSMUSG00 | Gm8464 | 3340 | 0 | 0.01442708 | 0 | 0 | 0 | 0 |
| ENSMUSG00 | Gm13647 | 619 | 0 | 0 | 0 | 0 | 0 | 0 |
| ENSMUSG00 | Gm15202 | 391 | 0 | 0 | 0 | 0 | 0 | 0 |
| ENSMUSG00 | 9530026P05I | 5677 | 0.9173803 | 0.56869678 | 0.68517918 | 1.11227018 | 1.19543914 | 2.26287742 |
| ENSMUSG00 | C030037D09 | 2516 | 2.11734276 | 2.50891253 | 7.08002493 | 7.14183552 | 3.20435905 | 10.4827409 |
| ENSMUSG00 | Gm12976 | 604 | 0 | 0.47867327 | 0.07318186 | 0.53764636 | 0.08448076 | 0.18062675 |
| ENSMUSG00 | Gm16596 | 1490 | 0 | 0.35573884 | 0 | 0.3874582 | 0.23972123 | 0.60406919 |
| ENSMUSG00 | Gm13299 | 421 | 0 | 0 | 0 | 0 | 0 | 0 |
| ENSMUSG00 | Gm15605 | 1542 | 1.88206879 | 1.53121639 | 1.605255 | 1.21677452 | 0.59563863 | 2.51167236 |
| ENSMUSG00 | 1500017E21I | 3963 | 0 | 0 | 0 | 0 | 0 | 0 |
| ENSMUSG00 | Gm16869 | 2683 | 0 | 0 | 0 | 0 | 0 | 0 |
| ENSMUSG00 | Gm11571 | 2393 | 0 | 0 | 0 | 0 | 0 | 0 |
| ENSMUSG00 | Igkv13-2 | 355 | 0 | 0 | 0 | 0 | 0 | 0 |
| ENSMUSG00 | Gm16934 | 1869 | 0 | 0 | 0 | 0 | 0 | 0 |
| ENSMUSG00 | Gm14457 | 1160 | 0 | 0 | 0 | 0 | 0 | 0 |
| ENSMUSG00 | Gm14100 | 647 | 0.12289175 | 0 | 0 | 0 | 0 | 0.12646664 |
| ENSMUSG00 | Gm6938 | 768 | 0 | 0 | 0 | 0 | 0.26576237 | 0 |
| ENSMUSG00 | Gm15245 | 1180 | 0 | 0 | 0 | 0 | 0 | 0 |
| ENSMUSG00 | Gm16355 | 230 | 0 | 0 | 0 | 0 | 0 | 0.83009772 |
| ENSMUSG00 | D430040D24 | 907 | 0 | 0.10625456 | 0.19493646 | 0 | 0 | 0.12028507 |
| ENSMUSG00 | 2410004N09I | 398 | 3.29630869 | 6.78000191 | 5.55299535 | 4.44226176 | 3.07696739 | 8.90880177 |
| ENSMUSG00 | Gm14635 | 1833 | 0 | 0 | 0 | 0 | 0 | 0 |
| ENSMUSG00 | Gm11184 | 265 | 0 | 0 | 0 | 0 | 0 | 0 |
| ENSMUSG00 | Gm16174 | 400 | 0.99388701 | 0 | 0 | 0 | 0 | 0.0681866 |
| ENSMUSG00 | BC039771 | 4708 | 1.78173458 | 2.65086842 | 1.80262401 | 4.41445575 | 0.98627872 | 1.31506863 |
| ENSMUSG00 | 1810012K08I | 572 | 0.83403106 | 0.2527261 | 0.07727595 | 0.37848299 | 0.80286256 | 0.04768294 |
| ENSMUSG00 | Gm11715 | 692 | 0 | 0 | 0 | 0 | 0 | 0.07882844 |
| ENSMUSG00 | Gm15345 | 658 | 0.48334931 | 0.21969503 | 0 | 0 | 1.16321525 | 0 |
| ENSMUSG00 | Zfp111 | 12668 | 1.69152226 | 1.81821277 | 1.4724643 | 3.07044869 | 2.86791756 | 1.48128764 |
| ENSMUSG00 | Gm14454 | 677 | 0.05872301 | 0.14235286 | 0.78348909 | 1.11923624 | 0.97982701 | 0.52373753 |
| ENSMUSG00 | Gm16832 | 2180 | 0 | 0 | 0 | 0 | 0 | 0 |
| ENSMUSG00 | Gm12753 | 651 | 0 | 0 | 0 | 0 | 0 | 0 |
| ENSMUSG00 | Gm11339 | 487 | 0 | 0 | 0 | 0 | 0 | 0 |
| ENSMUSG00 | 4930452L02F | 2862 | 0.43061492 | 0.06734653 | 0.2934434 | 0 | 0.80230151 | 0 |
| ENSMUSG00 | Gm15742 | 657 | 0 | 0.44005883 | 0.2691132 | 0 | 0 | 0 |
| ENSMUSG00 | Gm16121 | 2250 | 0 | 0 | 0 | 0 | 0 | 0.01212206 |
| ENSMUSG00 | Gm16034 | 478 | 0 | 0 | 0 | 0 | 0 | 0 |
| ENSMUSG00 | Gm12914 | 302 | 0 | 0 | 0 | 0 | 0 | 0 |
| ENSMUSG00 | 4930442J19R | 1835 | 0 | 0 | 0 | 0 | 0 | 0 |
| ENSMUSG00 | Gm16253 | 548 | 0 | 0 | 0.24198089 | 0.06584315 | 0.18622765 | 0 |
| ENSMUSG00 | 4930458D05I | 3053 | 0.0520871 | 0.15783309 | 0.05791267 | 0.22455252 | 0.2507028 | 0.45561959 |
| ENSMUSG00 | A230005M16 | 2623 | 0.10609545 | 0.14696589 | 0.3201811 | 0.17882828 | 0.11672065 | 0.04159305 |
| ENSMUSG00 | Gm13855 | 742 | 0.75010341 | 3.63671262 | 1.37013799 | 1.70198323 | 3.30089763 | 1.65412233 |
| ENSMUSG00 | Gm12514 | 1262 | 0 | 0 | 0 | 0 | 0 | 0 |
| ENSMUSG00 | Gm16190 | 3944 | 0 | 0 | 0 | 0 | 0 | 0 |
| ENSMUSG00 | Gm14257 | 763 | 0 | 0.12630784 | 0 | 0 | 0 | 0 |
| ENSMUSG00 | Gm14268 | 1348 | 0 | 0 | 0 | 0 | 0 | 0 |
| ENSMUSG00 | Gm14114 | 285 | 0 | 0 | 0 | 0 | 0 | 0 |
| ENSMUSG00 | 5330434G04 | 7326 | 0 | 0 | 0 | 0 | 0 | 0 |
| ENSMUSG00 | 1700003G18I | 800 | 0 | 0.66256358 | 0 | 0.18041022 | 0.31891485 | 0 |
| ENSMUSG00 | Gm12290 | 2109 | 0 | 0 | 0 | 0 | 0 | 0 |
| ENSMUSG00 | Gm12404 | 1657 | 0 | 0 | 0.02667583 | 0 | 0.03079443 | 0.01646025 |
| ENSMUSG00 | 9230111E07R | 809 | 0 | 0.11912594 | 0 | 0.49060876 | 0.31536697 | 0 |
| ENSMUSG00 | 4930419G24 | 1050 | 0 | 0 | 0 | 0 | 0 | 0 |
| ENSMUSG00 | Gm15050 | 550 | 0.14456538 | 0.08761171 | 0.16073398 | 0.32801859 | 0.18555046 | 0.09918051 |
| ENSMUSG00 | A230059L01F | 2811 | 0 | 0 | 0 | 0 | 0 | 0 |
| ENSMUSG00 | Gm13028 | 275 | 0 | 0 | 0 | 0 | 0 | 0 |
| ENSMUSG00 | Gm12246 | 501 | 0.71417031 | 0 | 1.49986294 | 0.07202005 | 0.40739621 | 0.54440398 |
| ENSMUSG00 | Gm11513 | 748 | 1.75391826 | 0 | 1.24096083 | 0.77180844 | 0.81860496 | 1.0209758 |
| ENSMUSG00 | 1700015O11I | 679 | 0 | 0 | 0 | 0 | 0 | 0 |
| ENSMUSG00 | Gm6058 | 398 | 0 | 0.72642878 | 0 | 0 | 0 | 0 |
| ENSMUSG00 | Gm14455 | 2144 | 0 | 0 | 0.02061653 | 0.06731725 | 0.19039693 | 0.02544276 |
| ENSMUSG00 | Gm13414 | 1454 | 0 | 0 | 0 | 0 | 0 | 0 |
| ENSMUSG00 | Gm4660 | 402 | 0 | 0 | 0 | 0.08975633 | 0 | 0 |
| ENSMUSG00 | Gm11986 | 1437 | 0 | 0 | 0 | 0 | 0 | 0 |
| ENSMUSG00 | Gm11773 | 495 | 0 | 0 | 0 | 0 | 0 | 0 |
| ENSMUSG00 | Gm15512 | 2255 | 0 | 0.02136871 | 0 | 0 | 0.11314052 | 0 |
| ENSMUSG00 | Gm14798 | 1187 | 0.26793921 | 0 | 0 | 0 | 0 | 0 |
| ENSMUSG00 | Gm12811 | 664 | 0 | 0 | 0 | 0 | 0 | 0 |
| ENSMUSG00 | Ighe | 1355 | 0 | 0 | 0 | 0 | 0 | 0 |
| ENSMUSG00 | Gm11314 | 539 | 0 | 0 | 0 | 0 | 0 | 0 |
| ENSMUSG00 | Gm14703 | 1817 | 0.26255683 | 0 | 0.58384383 | 0.07943213 | 0.67398625 | 0.72051881 |
| ENSMUSG00 | F630040L22F | 3152 | 0 | 0.22931365 | 0 | 0 | 0.29139428 | 0.29420613 |
| ENSMUSG00 | Gm1667 | 555 | 0 | 0 | 0 | 0 | 0 | 0 |
| ENSMUSG00 | Gm15407 | 436 | 0 | 0 | 0 | 0 | 0 | 0 |
| ENSMUSG00 | E130018N17I | 1307 | 0.517095 | 0.58988759 | 0.87930216 | 0.38649474 | 0.89793928 | 0.89732937 |
| ENSMUSG00 | Gm15274 | 290 | 0 | 0 | 0 | 0 | 0 | 0 |
| ENSMUSG00 | 1500009L16F | 1187 | 0 | 0 | 0 | 0 | 0 | 0 |
| ENSMUSG00 | Gm15918 | 687 | 0 | 0 | 0 | 0.21008468 | 0.2970968 | 0.63521722 |
| ENSMUSG00 | 1700087M22 | 206 | 0 | 0 | 1.07286027 | 0 | 0 | 0.39720348 |
| ENSMUSG00 | Gm12256 | 443 | 0 | 0 | 0 | 0 | 0 | 0 |
| ENSMUSG00 | Gm5091 | 1174 | 0 | 0 | 0.2259038 | 0.55321704 | 0.08692739 | 0 |
| ENSMUSG00 | Gm13390 | 142 | 0 | 0 | 0 | 0 | 0 | 0 |
| ENSMUSG00 | Gm16764 | 594 | 0 | 0 | 0 | 0 | 0 | 0 |
| ENSMUSG00 | Gm15051 | 1020 | 0.7015673 | 1.8896644 | 0.2166757 | 0.28299643 | 1.20062061 | 0.6952359 |
| ENSMUSG00 | Gm12606 | 1982 | 0 | 0 | 0 | 0 | 0 | 0 |
| ENSMUSG00 | B230398E01I | 1955 | 0 | 0 | 0.06782892 | 0.18456289 | 0 | 0 |
| ENSMUSG00 | Gm6019 | 735 | 0 | 0 | 0 | 0 | 0 | 0 |
| ENSMUSG00 | Gm11373 | 678 | 0 | 0 | 0 | 0 | 0 | 0 |
| ENSMUSG00 | Defa-ps3 | 280 | 0 | 0 | 0 | 0 | 0 | 0 |
| ENSMUSG00 | Gm16361 | 368 | 0 | 0 | 0 | 0 | 0 | 0 |
| ENSMUSG00 | B230359F08I | 1330 | 0.23913071 | 0 | 0.26587575 | 0.16277614 | 0.19182848 | 0.90231889 |
| ENSMUSG00 | Gm13381 | 1015 | 0.23500777 | 0 | 0.08709723 | 0 | 0.15081687 | 0.26871566 |
| ENSMUSG00 | Gm11186 | 1175 | 0 | 0 | 0 | 0 | 0 | 0 |
| ENSMUSG00 | Gm11724 | 470 | 0 | 0 | 0.09404647 | 0 | 0 | 0 |
| ENSMUSG00 | 9530036M11 | 1993 | 0 | 0 | 0 | 0 | 0 | 0 |
| ENSMUSG00 | A330048O09I | 666 | 1.67140158 | 0.14470403 | 2.25655054 | 0.16253173 | 0.68954562 | 1.67906938 |
| ENSMUSG00 | Gm15122 | 460 | 0.43212479 | 0.52376568 | 0.38436385 | 1.09814918 | 0.44370762 | 2.37170776 |
| ENSMUSG00 | Gm16884 | 1091 | 0.10931846 | 0.1766689 | 0.36463482 | 0.59530413 | 0 | 0.17499769 |
| ENSMUSG00 | 4930447M23 | 2053 | 0.07745832 | 0 | 0 | 0 | 0 | 0 |
| ENSMUSG00 | Gm11762 | 1701 | 0 | 0 | 0 | 0 | 0 | 0 |
| ENSMUSG00 | 9230114K14R | 2114 | 3.40385145 | 5.78966713 | 5.12273015 | 6.69071024 | 5.67227924 | 5.57362543 |
| ENSMUSG00 | Gm15226 | 2969 | 0 | 0 | 0 | 0 | 0 | 0 |
| ENSMUSG00 | Gm14120 | 2867 | 0 | 0 | 0 | 0 | 0 | 0 |
| ENSMUSG00 | C330006A16I | 2900 | 6.27862416 | 7.7762948 | 8.12399391 | 7.65188186 | 6.40468993 | 8.69966942 |
| ENSMUSG00 | Gm13565 | 698 | 0 | 0 | 0 | 0 | 0 | 0 |
| ENSMUSG00 | Gm14003 | 311 | 0 | 0 | 0 | 0 | 0 | 0 |
| ENSMUSG00 | Gm11712 | 365 | 0.65351475 | 0 | 0 | 0 | 1.25818461 | 1.19560063 |
| ENSMUSG00 | 1200007C13I | 2453 | 0 | 0.09821941 | 0 | 0 | 0 | 0 |
| ENSMUSG00 | 1700122E12R | 706 | 0.95728494 | 0 | 0.18782653 | 0.61329254 | 0 | 0.19316317 |
| ENSMUSG00 | 1500026H17I | 3024 | 5.04831499 | 4.87600993 | 1.7686584 | 4.20002635 | 2.81792486 | 9.94838859 |
| ENSMUSG00 | 2900053A13R | 1157 | 4.6730729 | 6.37210514 | 7.94639874 | 5.58226964 | 6.52714228 | 14.4270348 |
| ENSMUSG00 | Gm11300 | 563 | 0 | 0 | 0 | 0 | 0 | 0 |
| ENSMUSG00 | Gm15845 | 393 | 0 | 0 | 0 | 0 | 0.30831647 | 0 |
| ENSMUSG00 | Gm16031 | 445 | 0 | 0 | 0 | 0 | 0 | 0 |
| ENSMUSG00 | Gm15674 | 2434 | 0.31033448 | 0 | 0.16344149 | 0 | 0 | 0.11205686 |
| ENSMUSG00 | Gm11940 | 415 | 1.43694508 | 0.69667145 | 0.10651047 | 1.39111497 | 0.73773074 | 0.65722022 |
| ENSMUSG00 | Gm16191 | 741 | 0 | 0 | 0 | 0 | 0 | 0 |
| ENSMUSG00 | A530058N18I | 2518 | 0 | 0 | 0 | 0 | 0 | 0 |
| ENSMUSG00 | Gm16291 | 2591 | 0 | 0 | 0 | 0 | 0 | 0 |
| ENSMUSG00 | Gm16957 | 2364 | 1.31172905 | 2.85368101 | 4.09484079 | 1.90789153 | 2.93552754 | 4.5573107 |
| ENSMUSG00 | Gm2093 | 3523 | 0.07899187 | 0 | 0.53950589 | 0.0204837 | 0.05793514 | 0.43354522 |
| ENSMUSG00 | Gm13031 | 920 | 0 | 0 | 0 | 0 | 0.05546345 | 0 |
| ENSMUSG00 | Gm15283 | 1444 | 0.66075591 | 0.90099303 | 0.24488556 | 1.07446532 | 1.23678889 | 1.69994289 |
| ENSMUSG00 | Gm13493 | 1213 | 0 | 0 | 0.1093203 | 0.05949224 | 0.04206626 | 0.13491165 |
| ENSMUSG00 | Gm13575 | 1518 | 0 | 0 | 0 | 0 | 0 | 0 |
| ENSMUSG00 | Gm15650 | 766 | 0.31140063 | 0 | 0.11540951 | 0.0471045 | 0.13322814 | 0 |
| ENSMUSG00 | Gm14978 | 300 | 0.66259134 | 0 | 0 | 0 | 0 | 0 |
| ENSMUSG00 | SNORA7 | 113 | 0 | 0 | 0 | 0 | 0 | 0 |
| ENSMUSG00 | SNORA17 | 87 | 0 | 0 | 0 | 0 | 0 | 0 |
| ENSMUSG00 | U1 | 138 | 0 | 0 | 0 | 0 | 0 | 0 |
| ENSMUSG00 | U3 | 113 | 0 | 0 | 0 | 0 | 0 | 0 |
| ENSMUSG00 | U6 | 109 | 0 | 0 | 0 | 0 | 0 | 0 |
| ENSMUSG00 | 5S_rRNA | 100 | 0 | 0 | 0 | 0 | 0 | 0 |
| ENSMUSG00 | U6 | 94 | 0 | 0 | 0 | 0 | 0 | 0 |
| ENSMUSG00 | U7 | 61 | 0 | 0 | 0 | 0 | 0 | 0 |
| ENSMUSG00 | U6 | 106 | 0 | 0 | 0 | 3.74436311 | 0.96276181 | 0 |
| ENSMUSG00 | SNORA17 | 136 | 0 | 0 | 0 | 0 | 0 | 0 |
| ENSMUSG00 | SCARNA4 | 118 | 0 | 0 | 0 | 0 | 0 | 0 |
| ENSMUSG00 | SNORA17 | 120 | 0 | 0 | 0 | 0 | 0 | 0 |
| ENSMUSG00 | U6 | 105 | 0 | 0 | 0 | 0 | 0 | 0 |
| ENSMUSG00 | RNaseP_nuc | 304 | 0 | 0 | 0 | 0 | 0 | 0.98691129 |
| ENSMUSG00 | U6 | 101 | 0 | 0 | 0 | 0 | 0 | 1.08018373 |
| ENSMUSG00 | 5S_rRNA | 100 | 0 | 0 | 1.32605529 | 0 | 2.04105503 | 1.36373196 |
| ENSMUSG00 | SNORA70 | 151 | 0 | 0 | 0 | 0 | 0 | 0 |
| ENSMUSG00 | SNORA17 | 131 | 0 | 0 | 0 | 0 | 0 | 0 |
| ENSMUSG00 | SNORA17 | 108 | 0 | 0 | 0 | 0 | 0 | 0 |
| ENSMUSG00 | 7SK | 267 | 0 | 0 | 0 | 0 | 0 | 0 |
| ENSMUSG00 | 7SK | 320 | 0 | 0 | 0 | 0 | 0 | 0 |
| ENSMUSG00 | U6 | 102 | 0 | 0 | 0 | 0 | 0 | 0 |
| ENSMUSG00 | 5S_rRNA | 133 | 0 | 0 | 0 | 0 | 0 | 0 |
| ENSMUSG00 | U6 | 106 | 0 | 0 | 0 | 0 | 0 | 0 |
| ENSMUSG00 | U3 | 89 | 0 | 0 | 0 | 0 | 0 | 0 |
| ENSMUSG00 | U6 | 98 | 0 | 0 | 0 | 0 | 0.5206773 | 1.11325058 |
| ENSMUSG00 | U7 | 59 | 0 | 0 | 0 | 0 | 0 | 0 |
| ENSMUSG00 | SNORA40 | 131 | 0 | 0 | 0 | 0 | 0 | 0 |
| ENSMUSG00 | 5S_rRNA | 112 | 0 | 0 | 0 | 0 | 0 | 0 |
| ENSMUSG00 | SCARNA21 | 138 | 0.86424958 | 0 | 1.60151605 | 0 | 0 | 0.39528463 |
| ENSMUSG00 | U1 | 144 | 0 | 0 | 0 | 0 | 0 | 3.21992269 |
| ENSMUSG00 | SNORA17 | 113 | 0 | 0 | 0 | 0 | 0 | 0 |
| ENSMUSG00 | U1 | 155 | 0 | 0 | 0 | 0 | 0 | 0 |
| ENSMUSG00 | CT025701.1 | 81 | 0 | 0 | 0 | 0 | 0 | 0 |
| ENSMUSG00 | U7 | 59 | 0 | 0 | 0 | 0 | 0 | 0 |
| ENSMUSG00 | SNORA40 | 90 | 0 | 0 | 0 | 0 | 0 | 0 |
| ENSMUSG00 | 5S_rRNA | 118 | 0 | 0 | 0 | 0 | 0 | 0 |
| ENSMUSG00 | U6 | 107 | 0 | 0 | 0 | 0 | 0 | 0 |
| ENSMUSG00 | U1 | 158 | 0 | 0 | 0 | 0 | 0 | 0 |
| ENSMUSG00 | U1 | 119 | 0 | 0 | 0 | 0 | 0 | 0 |
| ENSMUSG00 | U6 | 103 | 0 | 0 | 0 | 0 | 0 | 0 |
| ENSMUSG00 | Y_RNA | 105 | 0 | 0 | 0 | 0 | 0 | 0 |
| ENSMUSG00 | 7SK | 261 | 0 | 0 | 0 | 0 | 0 | 0 |
| ENSMUSG00 | U1 | 157 | 0 | 0 | 0 | 0 | 0 | 0 |
| ENSMUSG00 | U6 | 105 | 0 | 0 | 0 | 0 | 0 | 0 |
| ENSMUSG00 | RNaseP_nuc | 202 | 0 | 0 | 0 | 0 | 0 | 0 |
| ENSMUSG00 | SNORA17 | 123 | 0 | 0 | 0 | 0 | 0 | 0 |
| ENSMUSG00 | U6 | 105 | 0 | 0 | 0 | 0 | 0 | 0 |
| ENSMUSG00 | 5S_rRNA | 113 | 0 | 0 | 0 | 0 | 0 | 0 |
| ENSMUSG00 | U6 | 99 | 0 | 0 | 0 | 0 | 0 | 0 |
| ENSMUSG00 | U7 | 62 | 0 | 0 | 0 | 0 | 0 | 0 |
| ENSMUSG00 | U7 | 59 | 0 | 0 | 0 | 0 | 0 | 0 |
| ENSMUSG00 | SNORA17 | 121 | 0 | 0 | 0 | 0 | 0 | 0 |
| ENSMUSG00 | 7SK | 311 | 0 | 0 | 0 | 0 | 0 | 0 |
| ENSMUSG00 | RNaseP_nuc | 289 | 0 | 0 | 0 | 0 | 0 | 0 |
| ENSMUSG00 | SNORA17 | 110 | 0 | 0 | 0 | 0 | 0 | 0 |
| ENSMUSG00 | U6 | 103 | 0 | 0 | 0 | 0 | 0 | 0 |
| ENSMUSG00 | U1 | 146 | 0 | 0 | 0 | 0 | 0 | 0 |
| ENSMUSG00 | U6 | 102 | 0.38975961 | 0.4724161 | 0 | 3.53745535 | 0.50025859 | 1.0695937 |
| ENSMUSG00 | SNORA48 | 142 | 0 | 0 | 0 | 0 | 0 | 0 |
| ENSMUSG00 | U6 | 111 | 0 | 0 | 0 | 0 | 0 | 0 |
| ENSMUSG00 | U6 | 105 | 0 | 0 | 0 | 0 | 0 | 0 |
| ENSMUSG00 | 7SK | 263 | 0 | 0 | 0 | 0 | 0 | 0 |
| ENSMUSG00 | SNORA25 | 123 | 1.61607644 | 0 | 0 | 1.76009973 | 0 | 0.44349007 |
| ENSMUSG00 | U1 | 159 | 0 | 0 | 0 | 0 | 0 | 0 |
| ENSMUSG00 | U7 | 63 | 0 | 0 | 0 | 0 | 0 | 0 |
| ENSMUSG00 | U6 | 100 | 0 | 0 | 0 | 0 | 0 | 0 |
| ENSMUSG00 | SNORD127 | 85 | 1.87084614 | 5.10209387 | 2.08008673 | 0 | 0 | 0 |
| ENSMUSG00 | SNORA31 | 131 | 0 | 0 | 0 | 0 | 0 | 0 |
| ENSMUSG00 | U6 | 99 | 0 | 1.94692696 | 0 | 0.3644651 | 1.54625381 | 0 |
| ENSMUSG00 | SNORA38 | 146 | 0 | 0 | 0 | 0 | 0 | 0 |
| ENSMUSG00 | 5S_rRNA | 116 | 0 | 0.83080073 | 0 | 0 | 0.43988255 | 0 |
| ENSMUSG00 | SNORA17 | 92 | 0.43212479 | 0 | 0 | 0 | 0.55463452 | 0 |
| ENSMUSG00 | U6 | 96 | 0 | 0 | 0 | 0 | 0 | 0 |
| ENSMUSG00 | snoU13 | 98 | 0 | 0 | 0 | 0 | 0 | 0 |
| ENSMUSG00 | SNORA73 | 167 | 0 | 0 | 0 | 0 | 0 | 0 |
| ENSMUSG00 | 5S_rRNA | 134 | 0 | 0 | 0 | 0 | 0 | 0 |
| ENSMUSG00 | SNORA48 | 130 | 0 | 0 | 0 | 0 | 0 | 0 |
| ENSMUSG00 | 7SK | 217 | 0 | 0 | 0 | 0 | 0 | 0 |
| ENSMUSG00 | SNORA70 | 137 | 0 | 0 | 0 | 0 | 0 | 0 |
| ENSMUSG00 | U6 | 99 | 0 | 0 | 0 | 0 | 0 | 0 |
| ENSMUSG00 | U7 | 65 | 0 | 0 | 0 | 0 | 0 | 0 |
| ENSMUSG00 | SNORA17 | 130 | 0 | 0 | 0 | 0 | 0 | 0 |
| ENSMUSG00 | AL671520.1 | 73 | 0 | 0 | 0 | 0 | 0 | 0 |
| ENSMUSG00 | U1 | 155 | 0 | 0.62176054 | 0 | 0 | 0 | 0 |
| ENSMUSG00 | U3 | 89 | 0 | 0 | 0 | 0 | 0 | 0 |
| ENSMUSG00 | SNORA48 | 142 | 0 | 0 | 0 | 0 | 0 | 0 |
| ENSMUSG00 | SNORA73 | 186 | 0 | 0 | 0 | 0 | 0 | 0 |
| ENSMUSG00 | U1 | 139 | 0 | 0 | 0 | 0 | 0 | 0 |
| ENSMUSG00 | SNORA40 | 118 | 0 | 0 | 0 | 0 | 0 | 0 |
| ENSMUSG00 | 5S_rRNA | 100 | 0 | 0 | 0 | 0 | 0 | 0 |
| ENSMUSG00 | U6 | 101 | 0 | 0 | 0 | 0 | 0 | 0 |
| ENSMUSG00 | SNORA70 | 121 | 0 | 0 | 0 | 0 | 0 | 0 |
| ENSMUSG00 | 7SK | 296 | 0 | 0 | 0 | 0 | 0 | 0 |
| ENSMUSG00 | SNORA48 | 143 | 0 | 0 | 0 | 0 | 0 | 0 |
| ENSMUSG00 | Mir1928 | 66 | 0 | 0 | 0 | 0 | 0 | 0 |
| ENSMUSG00 | U6 | 102 | 0 | 0 | 0 | 0 | 0 | 0 |
| ENSMUSG00 | U6 | 92 | 0 | 0 | 0 | 0 | 0 | 0 |
| ENSMUSG00 | snoU13 | 87 | 0 | 0 | 0 | 0 | 0 | 0 |
| ENSMUSG00 | U7 | 59 | 0 | 2.45015808 | 2.24755134 | 3.66936046 | 3.45941531 | 0 |
| ENSMUSG00 | 7SK | 276 | 0 | 0 | 0 | 0 | 0 | 0 |
| ENSMUSG00 | 7SK | 328 | 0 | 0 | 0 | 0 | 0 | 0 |
| ENSMUSG00 | SNORA72 | 130 | 0 | 0 | 0 | 0 | 0 | 0 |
| ENSMUSG00 | U6 | 104 | 0 | 0 | 0 | 0 | 0 | 0 |
| ENSMUSG00 | 7SK | 323 | 0 | 0 | 0 | 0 | 0 | 0 |
| ENSMUSG00 | 5S_rRNA | 89 | 0 | 0 | 0 | 0 | 0 | 0 |
| ENSMUSG00 | U6 | 101 | 0 | 0 | 0 | 0 | 0 | 0 |
| ENSMUSG00 | 5S_rRNA | 118 | 0 | 0 | 0 | 0 | 0 | 0 |
| ENSMUSG00 | 7SK | 352 | 0 | 0 | 0 | 0 | 0 | 0 |
| ENSMUSG00 | U6 | 99 | 0 | 0 | 0 | 0 | 0 | 0 |
| ENSMUSG00 | U6 | 107 | 0 | 0 | 0 | 0 | 0 | 0 |
| ENSMUSG00 | U6 | 103 | 0 | 0 | 0 | 0 | 0 | 0 |
| ENSMUSG00 | U6 | 106 | 0 | 0 | 0 | 0 | 0 | 0 |
| ENSMUSG00 | Mir1958 | 96 | 0 | 0 | 0 | 0 | 0 | 0 |
| ENSMUSG00 | 7SK | 293 | 0 | 0 | 0 | 0 | 0 | 0 |
| ENSMUSG00 | 5S_rRNA | 95 | 0 | 0 | 0 | 0 | 0 | 0 |
| ENSMUSG00 | SCARNA13 | 274 | 0 | 0 | 0 | 0 | 0 | 0 |
| ENSMUSG00 | 5S_rRNA | 113 | 0 | 0 | 0 | 0 | 0 | 0 |
| ENSMUSG00 | U7 | 61 | 0 | 0 | 0 | 0 | 0 | 0 |
| ENSMUSG00 | SNORA17 | 132 | 0 | 0 | 0 | 0 | 0 | 0 |
| ENSMUSG00 | 7SK | 329 | 0 | 0 | 0 | 0 | 0 | 0 |
| ENSMUSG00 | 7SK | 305 | 0 | 0 | 0 | 0 | 0 | 0 |
| ENSMUSG00 | 5S_rRNA | 128 | 0 | 1.12936974 | 0 | 0 | 0 | 0 |
| ENSMUSG00 | Mir432 | 74 | 0 | 0 | 0 | 0 | 0 | 0 |
| ENSMUSG00 | SNORA17 | 144 | 0 | 0 | 0.30695724 | 0 | 0.35434983 | 0 |
| ENSMUSG00 | SNORA17 | 117 | 0 | 0 | 0 | 0 | 0 | 0 |
| ENSMUSG00 | U6 | 105 | 0 | 1.83567399 | 0 | 0 | 0 | 0 |
| ENSMUSG00 | 7SK | 280 | 0 | 0 | 0 | 0 | 0 | 0 |
| ENSMUSG00 | U7 | 62 | 0 | 0 | 0 | 0 | 0 | 0 |
| ENSMUSG00 | SNORA17 | 128 | 0 | 0 | 0 | 0 | 0 | 0 |
| ENSMUSG00 | 7SK | 326 | 0 | 0 | 0 | 0 | 0 | 0 |
| ENSMUSG00 | U6 | 103 | 0 | 0 | 0 | 0 | 0 | 0 |
| ENSMUSG00 | SNORA17 | 131 | 0 | 0 | 0 | 0 | 0 | 0 |
| ENSMUSG00 | U3 | 89 | 0 | 0 | 0 | 0 | 0 | 0 |
| ENSMUSG00 | 7SK | 307 | 0 | 0 | 0 | 0 | 0 | 0 |
| ENSMUSG00 | SNORA17 | 126 | 0 | 0 | 0 | 0 | 0 | 0 |
| ENSMUSG00 | U6 | 104 | 0 | 0 | 0 | 0 | 0 | 0 |
| ENSMUSG00 | 7SK | 287 | 0 | 0 | 0 | 0 | 0 | 0 |
| ENSMUSG00 | U6 | 108 | 0 | 0 | 0 | 0 | 0 | 0 |
| ENSMUSG00 | U7 | 62 | 0 | 0 | 0 | 0 | 0 | 0 |
| ENSMUSG00 | U6 | 107 | 0 | 0 | 0 | 0 | 0 | 0 |
| ENSMUSG00 | 7SK | 283 | 1.12382984 | 0.34054023 | 0 | 1.01998712 | 1.98335736 | 0 |
| ENSMUSG00 | U6 | 94 | 0 | 0 | 0 | 0 | 0 | 0 |
| ENSMUSG00 | U2 | 189 | 0 | 0 | 0 | 0 | 0 | 0 |
| ENSMUSG00 | RNaseP_nuc | 227 | 0 | 0 | 0 | 0 | 0 | 0 |
| ENSMUSG00 | SNORA17 | 126 | 0 | 0 | 0 | 0 | 0 | 0 |
| ENSMUSG00 | SCARNA21 | 143 | 0 | 0 | 0.92731139 | 0 | 0 | 0 |
| ENSMUSG00 | U7 | 63 | 0 | 0 | 0 | 0 | 0 | 0 |
| ENSMUSG00 | SNORA17 | 131 | 0 | 0 | 0 | 0 | 0 | 0 |
| ENSMUSG00 | SNORA17 | 130 | 0 | 0 | 0 | 0 | 0 | 0 |
| ENSMUSG00 | SNORD113 | 70 | 0 | 0 | 0 | 0 | 0 | 0 |
| ENSMUSG00 | 7SK | 250 | 0 | 0 | 0 | 0 | 0 | 0 |
| ENSMUSG00 | SNORA48 | 143 | 0.55602071 | 0 | 0.3091038 | 0 | 0 | 0 |
| ENSMUSG00 | U7 | 45 | 0 | 0 | 0 | 0 | 0 | 0 |
| ENSMUSG00 | SNORA17 | 135 | 0 | 0 | 0 | 0 | 0 | 0 |
| ENSMUSG00 | 7SK | 323 | 0 | 0 | 0 | 0 | 0 | 0 |
| ENSMUSG00 | SNORA17 | 119 | 0 | 0 | 0 | 0 | 0 | 0 |
| ENSMUSG00 | U6atac | 126 | 0 | 0 | 0 | 0 | 0 | 0 |
| ENSMUSG00 | U1 | 100 | 0 | 0 | 0 | 0 | 0 | 0 |
| ENSMUSG00 | AC100386.1 | 103 | 0 | 0 | 0 | 0 | 0 | 0 |
| ENSMUSG00 | U6 | 83 | 0 | 0 | 0 | 0 | 0 | 0 |
| ENSMUSG00 | Y_RNA | 76 | 0 | 0 | 0 | 0 | 0 | 0 |
| ENSMUSG00 | 7SK | 289 | 1.23805995 | 1.16714566 | 1.07063288 | 0.12485137 | 1.23593298 | 0 |
| ENSMUSG00 | Mir1942 | 62 | 0.64121743 | 0 | 0 | 0 | 0 | 0 |
| ENSMUSG00 | U3 | 127 | 0 | 0 | 0 | 0 | 0 | 0 |
| ENSMUSG00 | 7SK | 241 | 0 | 0 | 0 | 0 | 0 | 0 |
| ENSMUSG00 | SNORA17 | 131 | 0.6069539 | 0 | 1.01225595 | 0.27543545 | 0.38951432 | 0 |
| ENSMUSG00 | 7SK | 214 | 0 | 0 | 0 | 0 | 0 | 0 |
| ENSMUSG00 | 7SK | 295 | 0 | 0 | 0 | 0 | 0 | 0 |
| ENSMUSG00 | SNORA17 | 133 | 0 | 0 | 0 | 0 | 0 | 0 |
| ENSMUSG00 | 7SK | 227 | 0 | 0 | 0 | 0 | 0 | 0 |
| ENSMUSG00 | 7SK | 312 | 0 | 0 | 0 | 0 | 0 | 0 |
| ENSMUSG00 | U6 | 106 | 0 | 0 | 0 | 0 | 0 | 0 |
| ENSMUSG00 | SNORD198 | 78 | 2.54842824 | 2.4710996 | 0 | 1.38777094 | 0.65418431 | 0 |
| ENSMUSG00 | 5S_rRNA | 77 | 0 | 0 | 0 | 0 | 0 | 0 |
| ENSMUSG00 | 7SK | 271 | 0 | 0 | 0 | 0 | 0 | 0 |
| ENSMUSG00 | 7SK | 279 | 0 | 0 | 0 | 0 | 0 | 0 |
| ENSMUSG00 | SNORA70 | 116 | 0 | 0 | 0 | 0 | 0 | 0 |
| ENSMUSG00 | SNORA17 | 135 | 0 | 0 | 0 | 0 | 0 | 0 |
| ENSMUSG00 | SNORA17 | 126 | 0 | 0 | 0 | 0 | 0 | 0 |
| ENSMUSG00 | U7 | 56 | 0 | 0 | 0 | 0 | 0 | 0 |
| ENSMUSG00 | 7SK | 269 | 0 | 0 | 0 | 0 | 0 | 0 |
| ENSMUSG00 | U6 | 104 | 0 | 0 | 0 | 0 | 0 | 0 |
| ENSMUSG00 | U1 | 132 | 0 | 0 | 0 | 0 | 0 | 0 |
| ENSMUSG00 | U6 | 103 | 3.85975539 | 7.01744303 | 0 | 7.70684446 | 8.91723073 | 1.58881394 |
| ENSMUSG00 | U6 | 104 | 0 | 0 | 0 | 0 | 0 | 0 |
| ENSMUSG00 | SNORA70 | 136 | 0 | 0 | 0 | 0 | 0 | 0 |
| ENSMUSG00 | SNORA17 | 133 | 0 | 0 | 0 | 0 | 0 | 0 |
| ENSMUSG00 | SNORA65 | 141 | 0 | 0 | 0 | 0 | 0 | 0 |
| ENSMUSG00 | Mir1971 | 105 | 0 | 0 | 0 | 0 | 0 | 0 |
| ENSMUSG00 | SNORA76 | 112 | 0 | 0 | 0 | 0 | 0 | 0 |
| ENSMUSG00 | Snora81 | 177 | 0 | 0 | 0 | 0 | 0 | 0 |
| ENSMUSG00 | SNORA17 | 124 | 0 | 0 | 0 | 0 | 0 | 0 |
| ENSMUSG00 | U1 | 165 | 0 | 0 | 0 | 0 | 0 | 0 |
| ENSMUSG00 | U6 | 107 | 6.3162913 | 7.20544929 | 9.50133074 | 11.8025379 | 15.2602245 | 21.4118663 |
| ENSMUSG00 | U6 | 104 | 0 | 0 | 0 | 0 | 0 | 0 |
| ENSMUSG00 | SNORA17 | 131 | 0 | 0 | 0 | 0 | 0 | 0 |
| ENSMUSG00 | U7 | 59 | 0 | 0 | 0 | 0 | 0 | 0 |
| ENSMUSG00 | SNORA17 | 129 | 0 | 0 | 0 | 0 | 0 | 0 |

| | | | | | | | | |
|---|---|---|---|---|---|---|---|---|
| ENSMUSG00 | mmu-mir-21 | 56 | 0 | 0 | 0 | 0 | 0 | 0.97409426 |
| ENSMUSG00 | 5S_rRNA | 134 | 0 | 0 | 0 | 0 | 0 | 0 |
| ENSMUSG00 | 7SK | 321 | 0 | 0 | 0 | 0 | 0 | 0 |
| ENSMUSG00 | U7 | 63 | 0 | 0 | 0 | 0 | 0 | 0 |
| ENSMUSG00 | AC124515.1 | 109 | 0 | 0 | 0 | 0 | 0 | 0 |
| ENSMUSG00 | U1 | 113 | 0 | 0 | 0 | 0 | 0 | 0 |
| ENSMUSG00 | Mir1948 | 84 | 0 | 0 | 0 | 0 | 0 | 0 |
| ENSMUSG00 | U6 | 94 | 0 | 0 | 0 | 0 | 0 | 0 |
| ENSMUSG00 | SNORA17 | 133 | 0 | 0 | 0 | 0 | 0 | 0 |
| ENSMUSG00 | 7SK | 83 | 0 | 0 | 0 | 0 | 0 | 0 |
| ENSMUSG00 | 5S_rRNA | 119 | 0 | 0 | 0 | 0 | 0 | 0 |
| ENSMUSG00 | 7SK | 268 | 0 | 0 | 0 | 0 | 0 | 0 |
| ENSMUSG00 | 5S_rRNA | 110 | 0 | 0 | 0 | 0 | 0 | 0 |
| ENSMUSG00 | 7SK | 296 | 0 | 0 | 0 | 0 | 0 | 0 |
| ENSMUSG00 | 7SK | 284 | 0 | 0 | 0 | 0 | 0 | 0 |
| ENSMUSG00 | SNORA64 | 132 | 0 | 0.3650488 | 0 | 0.82004647 | 0 | 0 |
| ENSMUSG00 | SNORA38 | 150 | 0 | 0 | 0 | 0 | 0 | 0 |
| ENSMUSG00 | 7SK | 285 | 0 | 0 | 0 | 0 | 0 | 0 |
| ENSMUSG00 | U6 | 102 | 0 | 0 | 0 | 0 | 0.50025859 | 0 |
| ENSMUSG00 | SNORA70 | 133 | 0 | 0 | 0 | 0 | 0 | 0 |
| ENSMUSG00 | SCARNA6 | 271 | 1.61369109 | 0.3556195 | 1.46795789 | 0.5325763 | 0.37657842 | 3.52255489 |
| ENSMUSG00 | U7 | 51 | 0 | 0 | 0 | 0 | 0 | 0.53479685 |
| ENSMUSG00 | U7 | 65 | 0 | 0 | 0 | 0 | 0 | 0 |
| ENSMUSG00 | SNORA48 | 143 | 0 | 0 | 0 | 0 | 0 | 0 |
| ENSMUSG00 | 7SK | 306 | 0 | 0 | 0 | 0 | 0 | 0 |
| ENSMUSG00 | SNORA32 | 69 | 0.57616638 | 0 | 0 | 0 | 1.47902539 | 0 |
| ENSMUSG00 | U6 | 105 | 3.02898899 | 0 | 0 | 0 | 0 | 0 |
| ENSMUSG00 | U6 | 108 | 0 | 0 | 0 | 0 | 0 | 0 |
| ENSMUSG00 | 7SK | 258 | 0 | 0 | 0 | 0 | 0 | 0 |
| ENSMUSG00 | 5S_rRNA | 92 | 0 | 0 | 0 | 0 | 0 | 0 |
| ENSMUSG00 | 7SK | 277 | 0 | 0 | 0 | 0 | 0 | 0 |
| ENSMUSG00 | snoU13 | 101 | 0 | 0 | 0 | 0 | 0 | 0 |
| ENSMUSG00 | SNORA17 | 124 | 0 | 0 | 0 | 0 | 0 | 0 |
| ENSMUSG00 | U3 | 88 | 0 | 0 | 0 | 0 | 0 | 0 |
| ENSMUSG00 | U6 | 102 | 0 | 0 | 0 | 0 | 0 | 0 |
| ENSMUSG00 | U6 | 99 | 0 | 0 | 0 | 0 | 0 | 0 |
| ENSMUSG00 | U6 | 100 | 0 | 0 | 0 | 0 | 0 | 0 |
| ENSMUSG00 | SCARNA4 | 126 | 0 | 0 | 0 | 0 | 0 | 0 |
| ENSMUSG00 | 7SK | 87 | 0.45695955 | 1.1077343 | 3.04840297 | 0.82947229 | 0 | 1.8810096 |
| ENSMUSG00 | SNORA17 | 116 | 0 | 0 | 0 | 0 | 0 | 0 |
| ENSMUSG00 | Mir1982 | 73 | 4.90136061 | 3.30044124 | 7.87156108 | 2.47137291 | 2.7959658 | 2.61537637 |
| ENSMUSG00 | SNORD113 | 75 | 0 | 0 | 0 | 0 | 0 | 0 |
| ENSMUSG00 | U6 | 96 | 0 | 0 | 0 | 0 | 0 | 0 |
| ENSMUSG00 | U7 | 61 | 0 | 0 | 0 | 0 | 0 | 0 |
| ENSMUSG00 | U1 | 129 | 0 | 0 | 0 | 0 | 0 | 0 |
| ENSMUSG00 | U6 | 102 | 0 | 0 | 0.4333514 | 0 | 0 | 0 |
| ENSMUSG00 | U6 | 105 | 0 | 0 | 0 | 0 | 0 | 0 |
| ENSMUSG00 | U6 | 98 | 3.24534535 | 0 | 0 | 0 | 0 | 0 |
| ENSMUSG00 | U6 | 73 | 0 | 0 | 0 | 0 | 0 | 0 |
| ENSMUSG00 | 5S_rRNA | 108 | 0 | 1.33851228 | 0 | 0 | 0 | 0.50508591 |
| ENSMUSG00 | SNORA70 | 122 | 0 | 0.39497084 | 0 | 0 | 0 | 0 |
| ENSMUSG00 | snoR38 | 71 | 0 | 0 | 3.11280585 | 0 | 4.3120881 | 0.7682997 |
| ENSMUSG00 | SCARNA20 | 128 | 3.7270763 | 1.50582632 | 0 | 0.84567292 | 3.18914849 | 0.42616624 |
| ENSMUSG00 | SNORA17 | 133 | 0 | 0 | 0 | 0 | 0 | 0 |
| ENSMUSG00 | 5S_rRNA | 99 | 0 | 0 | 0 | 0 | 0 | 0 |
| ENSMUSG00 | SNORA17 | 113 | 0 | 0 | 0 | 0 | 0 | 0 |
| ENSMUSG00 | U3 | 130 | 0 | 0 | 0 | 0 | 0 | 0 |
| ENSMUSG00 | U7 | 61 | 0 | 0 | 0 | 0 | 0 | 0 |
| ENSMUSG00 | 7SK | 273 | 0 | 0 | 0.32382303 | 0 | 0.74763921 | 0 |
| ENSMUSG00 | Mir1932 | 89 | 0.89338158 | 0 | 0.99329984 | 0 | 0 | 0 |
| ENSMUSG00 | U6 | 101 | 0 | 0 | 0 | 0 | 0 | 0 |
| ENSMUSG00 | 7SK | 269 | 0 | 0 | 0 | 0 | 0 | 0 |
| ENSMUSG00 | U6 | 105 | 0.37862362 | 0 | 0 | 0 | 0 | 0.51951694 |
| ENSMUSG00 | SNORA17 | 118 | 0 | 0 | 0 | 0 | 0 | 0 |
| ENSMUSG00 | SNORA17 | 136 | 0 | 0 | 0 | 0 | 0 | 0 |
| ENSMUSG00 | U1 | 147 | 0 | 0 | 0 | 0 | 0 | 0 |
| ENSMUSG00 | 7SK | 287 | 0 | 0 | 0 | 0 | 0 | 0 |
| ENSMUSG00 | U7 | 61 | 0 | 0 | 0 | 0 | 0 | 0 |
| ENSMUSG00 | SNORA5 | 91 | 0 | 0 | 0 | 0 | 0 | 0 |
| ENSMUSG00 | RNaseP_nuc | 289 | 0 | 0 | 0 | 0 | 0 | 0 |
| ENSMUSG00 | 7SK | 202 | 0 | 0 | 0 | 0 | 0 | 0 |
| ENSMUSG00 | SNORA17 | 101 | 0 | 0 | 0 | 0 | 0 | 0 |
| ENSMUSG00 | 7SK | 276 | 0 | 0 | 0 | 0 | 0 | 0 |
| ENSMUSG00 | U7 | 58 | 0 | 0 | 0 | 0 | 0 | 0 |
| ENSMUSG00 | 5S_rRNA | 107 | 0 | 0 | 0 | 0 | 0 | 0 |
| ENSMUSG00 | 7SK | 322 | 0 | 0 | 0 | 0 | 0 | 0 |
| ENSMUSG00 | U6 | 106 | 0 | 0 | 0 | 0 | 0 | 0 |
| ENSMUSG00 | AL671963.1 | 90 | 0 | 1.07080983 | 0 | 0 | 0 | 0 |
| ENSMUSG00 | SNORA70 | 129 | 0 | 0.74707682 | 0 | 1.9579404 | 0 | 0.21143131 |
| ENSMUSG00 | snoU13 | 109 | 0 | 0 | 0 | 0 | 0 | 0 |
| ENSMUSG00 | SNORA17 | 120 | 0 | 0 | 0 | 0 | 0 | 0 |
| ENSMUSG00 | SNORA17 | 107 | 0 | 0 | 0 | 0 | 0 | 0 |
| ENSMUSG00 | SNORA17 | 133 | 0 | 0 | 0 | 0 | 0 | 0 |
| ENSMUSG00 | U3 | 89 | 0 | 0 | 0 | 0 | 0 | 0 |
| ENSMUSG00 | SNORA17 | 131 | 0 | 0 | 0 | 0 | 0 | 0 |
| ENSMUSG00 | U6 | 104 | 0 | 0 | 0 | 0 | 0 | 0 |
| ENSMUSG00 | SNORA17 | 77 | 0 | 0 | 0 | 0 | 0 | 0 |
| ENSMUSG00 | SNORA17 | 130 | 0 | 0 | 0 | 0 | 0 | 0 |
| ENSMUSG00 | SNORA17 | 128 | 0 | 0 | 0 | 0 | 0 | 0 |
| ENSMUSG00 | U6 | 107 | 0 | 0 | 0 | 0 | 0 | 0 |
| ENSMUSG00 | 7SK | 274 | 0 | 0 | 0 | 0 | 0 | 0 |
| ENSMUSG00 | U7 | 59 | 0 | 0 | 0 | 0 | 0 | 0 |
| ENSMUSG00 | U1 | 150 | 0 | 0 | 0 | 0 | 0 | 0 |
| ENSMUSG00 | 5S_rRNA | 100 | 0 | 0 | 0 | 0 | 0 | 0 |
| ENSMUSG00 | Mir1968 | 67 | 0 | 0.71920083 | 0 | 5.92391776 | 0 | 0 |
| ENSMUSG00 | SNORA17 | 67 | 0 | 2.15760189 | 0 | 0 | 0 | 0 |
| ENSMUSG00 | SNORA48 | 143 | 0 | 0 | 0 | 0 | 0 | 0 |
| ENSMUSG00 | U6 | 107 | 0 | 0 | 0 | 0 | 0 | 0 |
| ENSMUSG00 | SNORA70 | 126 | 0 | 0 | 0 | 0 | 0 | 0 |
| ENSMUSG00 | SNORA74 | 140 | 9.93887013 | 7.22798632 | 1.2629098 | 5.92778446 | 6.92500815 | 2.53264507 |
| ENSMUSG00 | SNORA48 | 142 | 0 | 0 | 0 | 0 | 0 | 0 |
| ENSMUSG00 | 5S_rRNA | 130 | 0 | 0 | 0 | 0 | 0 | 0 |
| ENSMUSG00 | U6 | 102 | 0 | 0 | 0 | 0 | 0 | 0 |
| ENSMUSG00 | U1 | 157 | 0 | 0 | 0 | 0 | 0 | 0 |
| ENSMUSG00 | U7 | 63 | 0 | 0 | 0 | 0 | 0 | 0 |
| ENSMUSG00 | U6 | 103 | 1.54390216 | 0 | 0 | 0.70062222 | 0 | 0.52960465 |
| ENSMUSG00 | U6 | 102 | 0 | 0 | 0 | 0 | 0 | 0 |
| ENSMUSG00 | U6 | 100 | 0 | 0 | 0 | 0 | 0 | 0 |
| ENSMUSG00 | 7SK | 266 | 0 | 0.90576019 | 0 | 0 | 0.19182848 | 0 |
| ENSMUSG00 | 7SK | 302 | 0 | 0 | 0 | 0 | 0 | 0 |
| ENSMUSG00 | U6 | 103 | 0 | 0.46782954 | 0 | 0 | 3.46781195 | 0 |
| ENSMUSG00 | RNaseP_nuc | 355 | 0 | 0 | 0 | 0 | 0 | 0 |
| ENSMUSG00 | 5S_rRNA | 116 | 0 | 0 | 0 | 0 | 0 | 0 |
| ENSMUSG00 | U7 | 61 | 0 | 0 | 0 | 0 | 0 | 0 |
| ENSMUSG00 | U6 | 103 | 0 | 0 | 0 | 0 | 0 | 0 |
| ENSMUSG00 | SCARNA4 | 123 | 6.46430577 | 0 | 0 | 4.98694924 | 0.41484858 | 1.99570531 |
| ENSMUSG00 | U3 | 197 | 0 | 0 | 0 | 0 | 0 | 0 |
| ENSMUSG00 | 7SK | 231 | 0 | 0 | 0 | 0 | 0 | 0 |
| ENSMUSG00 | SNORA17 | 121 | 0 | 0 | 0 | 0 | 0 | 0 |
| ENSMUSG00 | U6 | 106 | 0 | 0 | 0 | 0 | 0 | 0 |
| ENSMUSG00 | 7SK | 267 | 0 | 0 | 0.6621999 | 0 | 0 | 0 |
| ENSMUSG00 | RNase_MRP | 270 | 0 | 0 | 0 | 0 | 0 | 0 |
| ENSMUSG00 | SNORA17 | 113 | 0 | 0 | 0 | 0 | 0 | 0 |
| ENSMUSG00 | 5S_rRNA | 110 | 0 | 0 | 0 | 0 | 0 | 0 |
| ENSMUSG00 | 5S_rRNA | 119 | 0 | 0 | 0 | 0 | 0 | 0 |
| ENSMUSG00 | SNORA17 | 131 | 0 | 0 | 0 | 0 | 0 | 0 |
| ENSMUSG00 | RNaseP_nuc | 153 | 0 | 0 | 0 | 0 | 0 | 0 |
| ENSMUSG00 | SNORA17 | 118 | 0 | 0 | 0 | 0 | 0 | 0 |
| ENSMUSG00 | SNORA17 | 127 | 0 | 0 | 0 | 0 | 0 | 0 |
| ENSMUSG00 | U4 | 106 | 0 | 0 | 0 | 0 | 0 | 0 |
| ENSMUSG00 | 7SK | 102 | 0 | 0 | 0 | 0 | 0 | 0 |
| ENSMUSG00 | SNORA17 | 130 | 0 | 0 | 0 | 0 | 0 | 0 |
| ENSMUSG00 | 7SK | 297 | 0 | 0 | 0 | 0 | 0 | 0 |
| ENSMUSG00 | U3 | 121 | 0 | 0 | 0 | 0 | 0 | 0 |
| ENSMUSG00 | U7 | 64 | 0 | 0 | 0 | 0 | 0 | 0 |
| ENSMUSG00 | 7SK | 275 | 0 | 0 | 0 | 0 | 0 | 0 |
| ENSMUSG00 | SNORA48 | 143 | 0 | 0 | 0 | 0 | 0 | 0 |
| ENSMUSG00 | SNORA17 | 94 | 0 | 0 | 0 | 0 | 0 | 0 |
| ENSMUSG00 | SNORA17 | 130 | 0 | 0 | 0 | 0 | 0 | 0 |
| ENSMUSG00 | U6 | 110 | 0 | 0 | 0.40183494 | 0 | 0 | 0 |
| ENSMUSG00 | U7 | 61 | 0 | 0 | 0 | 0 | 0 | 0 |
| ENSMUSG00 | Snora47 | 137 | 0.58037198 | 0.35172586 | 0 | 0 | 0.3724553 | 0.59725487 |
| ENSMUSG00 | U6 | 73 | 0 | 0 | 0 | 0 | 0 | 0 |
| ENSMUSG00 | SNORA17 | 105 | 0 | 0 | 0 | 0 | 0 | 0 |
| ENSMUSG00 | 7SK | 263 | 0 | 0 | 0 | 0 | 0 | 0 |
| ENSMUSG00 | Mir1960 | 77 | 0 | 0 | 0 | 0 | 0 | 0 |
| ENSMUSG00 | SNORA72 | 119 | 0 | 0 | 0 | 0 | 0 | 0 |
| ENSMUSG00 | 7SK | 265 | 0 | 0 | 0 | 0 | 0 | 0 |
| ENSMUSG00 | AC141566.1 | 104 | 0 | 0 | 0 | 0 | 0 | 0 |
| ENSMUSG00 | 7SK | 311 | 0 | 0 | 0.99489679 | 0.69611662 | 0 | 0 |
| ENSMUSG00 | U6 | 94 | 2.96051451 | 2.56310863 | 9.40464745 | 4.22236691 | 4.88550407 | 9.28498357 |
| ENSMUSG00 | U1 | 170 | 0 | 0 | 0 | 0 | 0 | 0 |
| ENSMUSG00 | 7SK | 277 | 0 | 0 | 0 | 0 | 0 | 0 |
| ENSMUSG00 | 7SK | 278 | 0 | 0 | 0 | 0 | 0 | 0 |
| ENSMUSG00 | U6 | 108 | 0 | 0 | 0 | 0 | 0 | 0 |
| ENSMUSG00 | SNORA17 | 136 | 0 | 0 | 0 | 0 | 0 | 0 |
| ENSMUSG00 | SNORA17 | 111 | 0 | 0 | 0 | 0 | 0 | 0 |
| ENSMUSG00 | U7 | 64 | 0 | 0 | 0.6906538 | 0 | 0 | 0 |
| ENSMUSG00 | U6 | 104 | 0 | 0 | 0 | 0.69388547 | 0 | 0 |
| ENSMUSG00 | U7 | 62 | 0 | 0 | 0 | 0 | 0 | 0 |
| ENSMUSG00 | U7 | 59 | 0 | 0 | 0 | 0 | 0 | 0 |
| ENSMUSG00 | SCARNA15 | 127 | 0 | 0 | 0 | 0 | 0 | 0 |
| ENSMUSG00 | 5S_rRNA | 121 | 0 | 0 | 0 | 0 | 0 | 0 |
| ENSMUSG00 | 7SK | 307 | 0 | 0 | 0 | 0 | 0 | 0 |
| ENSMUSG00 | SNORA17 | 115 | 0 | 0 | 0 | 0 | 0 | 0 |
| ENSMUSG00 | SNORA81 | 139 | 0 | 0 | 0 | 0 | 0 | 0 |
| ENSMUSG00 | 7SK | 297 | 0 | 0 | 0 | 0 | 0 | 0 |
| ENSMUSG00 | snoU13 | 103 | 0 | 0 | 0 | 0 | 0 | 0 |
| ENSMUSG00 | U6 | 102 | 0 | 0 | 0 | 0 | 0 | 0 |
| ENSMUSG00 | U7 | 62 | 0 | 0 | 0 | 0 | 0 | 0 |
| ENSMUSG00 | Mir1962 | 120 | 0 | 0 | 0 | 0 | 0 | 0 |
| ENSMUSG00 | U1 | 165 | 0 | 0 | 0 | 0 | 0 | 0 |
| ENSMUSG00 | SNORA22 | 120 | 0 | 0.40155369 | 0 | 0 | 0 | 0 |
| ENSMUSG00 | SNORA17 | 109 | 0 | 0 | 0 | 0 | 0 | 0 |
| ENSMUSG00 | SCARNA3 | 135 | 0 | 0 | 0 | 0 | 0 | 0 |
| ENSMUSG00 | SNORA17 | 123 | 0 | 0 | 0 | 0 | 0 | 0 |
| ENSMUSG00 | Mir1983 | 131 | 0 | 0 | 0 | 0 | 0 | 0 |
| ENSMUSG00 | U6 | 103 | 0 | 0 | 0 | 0 | 0 | 0 |
| ENSMUSG00 | 7SK | 328 | 0 | 0 | 0 | 0 | 0 | 0 |
| ENSMUSG00 | SCARNA11 | 130 | 0.30581139 | 2.59465458 | 3.74015595 | 2.4979877 | 1.17753175 | 0.83921967 |
| ENSMUSG00 | U2 | 171 | 0 | 0 | 0 | 0 | 0 | 0 |
| ENSMUSG00 | 7SK | 239 | 1.16438646 | 0.80646765 | 0 | 1.20776718 | 0.21349948 | 0.45647932 |
| ENSMUSG00 | U6 | 106 | 0 | 0 | 0 | 0 | 0 | 0 |
| ENSMUSG00 | SNORA17 | 129 | 0 | 0 | 0 | 0 | 0 | 0 |
| ENSMUSG00 | 7SK | 287 | 0 | 0 | 0 | 0 | 0 | 0 |
| ENSMUSG00 | RNaseP_nuc | 231 | 0 | 0 | 0 | 0 | 0 | 0 |
| ENSMUSG00 | Scarna3b | 138 | 0 | 0 | 0 | 0 | 0 | 0 |
| ENSMUSG00 | SNORA17 | 119 | 0 | 0 | 0 | 0 | 0 | 0 |
| ENSMUSG00 | U6 | 100 | 0 | 0 | 0 | 0 | 0 | 0 |
| ENSMUSG00 | 7SK | 329 | 0 | 0 | 0 | 0 | 0 | 0 |
| ENSMUSG00 | 7SK | 223 | 0 | 0 | 0 | 0 | 0 | 0 |
| ENSMUSG00 | SNORA17 | 96 | 0 | 0 | 0 | 0 | 0 | 0 |
| ENSMUSG00 | 7SK | 244 | 0 | 0 | 0 | 0 | 0 | 0 |
| ENSMUSG00 | 5S_rRNA | 133 | 0 | 0 | 0 | 0 | 0 | 0 |
| ENSMUSG00 | U6 | 105 | 0 | 0 | 0 | 0 | 0 | 0 |
| ENSMUSG00 | U1 | 163 | 0 | 0 | 0 | 0 | 0 | 0 |
| ENSMUSG00 | SNORD124 | 103 | 0 | 0 | 0 | 0 | 0 | 0 |
| ENSMUSG00 | 7SK | 329 | 0 | 0 | 0 | 3.72884351 | 0 | 0 |
| ENSMUSG00 | U6 | 94 | 0 | 0 | 0.47023237 | 1.15155461 | 2.17133514 | 0.58031147 |
| ENSMUSG00 | U6 | 100 | 0 | 0 | 0 | 0 | 0 | 0 |
| ENSMUSG00 | U7 | 56 | 0 | 0 | 0 | 0 | 0 | 0 |
| ENSMUSG00 | U1 | 136 | 0 | 0 | 0 | 0 | 0 | 0 |
| ENSMUSG00 | 7SK | 321 | 1.11463964 | 0 | 0 | 0 | 1.27168538 | 0.50980634 |
| ENSMUSG00 | 7SK | 298 | 0 | 0 | 0 | 0 | 0 | 0 |
| ENSMUSG00 | U6 | 100 | 0 | 0 | 0 | 0 | 0 | 0 |
| ENSMUSG00 | U7 | 58 | 0 | 0 | 0 | 0 | 0 | 0 |
| ENSMUSG00 | 7SK | 312 | 0 | 0 | 0 | 0 | 0 | 0 |
| ENSMUSG00 | 5S_rRNA | 113 | 0 | 0 | 0 | 0 | 0 | 0 |
| ENSMUSG00 | U6 | 95 | 0 | 0 | 1.86113023 | 0 | 0 | 0 |
| ENSMUSG00 | U6 | 106 | 0 | 0 | 0 | 0 | 0 | 0 |
| ENSMUSG00 | Scarna2 | 413 | 0.57756146 | 0 | 0.53513127 | 0.3494629 | 0.37065164 | 0.72644318 |
| ENSMUSG00 | 7SK | 318 | 0 | 0 | 0 | 0 | 0 | 0 |
| ENSMUSG00 | U12 | 171 | 0 | 0 | 0 | 0 | 0 | 0 |
| ENSMUSG00 | SNORA61 | 118 | 0 | 5.71703551 | 0 | 0 | 0 | 0 |
| ENSMUSG00 | U3 | 88 | 0 | 0 | 0 | 0 | 0 | 0 |
| ENSMUSG00 | SNORA17 | 130 | 0 | 0 | 0 | 0 | 0 | 0 |
| ENSMUSG00 | SNORA17 | 123 | 0 | 0 | 0 | 0 | 0 | 0 |
| ENSMUSG00 | 5S_rRNA | 108 | 0 | 0 | 0 | 0 | 0 | 0 |
| ENSMUSG00 | 5S_rRNA | 107 | 0 | 0 | 0 | 0 | 0 | 0 |
| ENSMUSG00 | U6 | 106 | 0 | 0 | 0 | 0 | 0 | 0 |
| ENSMUSG00 | SNORA70 | 133 | 0 | 0 | 0 | 0 | 0 | 0 |
| ENSMUSG00 | SNORA17 | 113 | 0 | 0 | 0 | 0 | 0 | 0 |
| ENSMUSG00 | SNORA17 | 127 | 0 | 0 | 0 | 0 | 0 | 0 |
| ENSMUSG00 | U1 | 148 | 0 | 0 | 0 | 0 | 0 | 0 |
| ENSMUSG00 | U6 | 62 | 0 | 0 | 0 | 0 | 0 | 0 |
| ENSMUSG00 | 7SK | 342 | 0 | 0 | 0 | 0 | 0 | 0 |
| ENSMUSG00 | AC117688.1 | 103 | 0 | 0 | 0 | 0 | 0 | 0 |
| ENSMUSG00 | 5S_rRNA | 112 | 0 | 0 | 0 | 0 | 0 | 0 |
| ENSMUSG00 | 5S_rRNA | 113 | 1.75909206 | 0 | 0 | 0 | 0 | 0 |
| ENSMUSG00 | SCARNA11 | 135 | 0 | 0 | 0 | 1.87092083 | 0 | 0 |
| ENSMUSG00 | 5S_rRNA | 114 | 0 | 0 | 0 | 0 | 0 | 0 |
| ENSMUSG00 | U6 | 104 | 0 | 0 | 0 | 0 | 0 | 0 |
| ENSMUSG00 | U6 | 101 | 0 | 0 | 0 | 0 | 0 | 0 |
| ENSMUSG00 | U1 | 160 | 0 | 0 | 0 | 1.80410223 | 0 | 0.1704665 |
| ENSMUSG00 | 7SK | 324 | 0 | 0 | 0 | 0 | 0 | 0 |
| ENSMUSG00 | U7 | 62 | 0 | 0 | 0 | 0 | 0 | 0 |
| ENSMUSG00 | U6 | 105 | 0 | 0 | 0 | 0 | 0 | 0 |
| ENSMUSG00 | U6 | 105 | 0.37862362 | 0.4589185 | 0 | 1.37455408 | 0.97193097 | 0 |
| ENSMUSG00 | U6 | 102 | 0 | 0 | 0 | 0 | 0 | 0 |
| ENSMUSG00 | snoU83B | 92 | 2.16062394 | 2.61882838 | 0.48045482 | 1.17658841 | 1.10926904 | 0.88939041 |
| ENSMUSG00 | 7SK | 249 | 0 | 0 | 0 | 0 | 0 | 0 |
| ENSMUSG00 | U6 | 94 | 0 | 0 | 0 | 0 | 0 | 0 |
| ENSMUSG00 | SNORA17 | 131 | 0 | 0 | 0 | 0 | 0 | 0 |
| ENSMUSG00 | 7SK | 268 | 0 | 0 | 0 | 0 | 0 | 0 |
| ENSMUSG00 | SNORA17 | 104 | 0 | 0 | 0 | 0 | 0 | 0 |
| ENSMUSG00 | SNORA40 | 126 | 0.63103937 | 0 | 0 | 0 | 1.21491371 | 0 |
| ENSMUSG00 | 7SK | 315 | 0.63103937 | 1.98864682 | 0.56129325 | 0 | 2.42982742 | 1.81830928 |
| ENSMUSG00 | U6 | 79 | 0 | 0 | 0 | 0 | 0 | 0 |
| ENSMUSG00 | SNORA70 | 95 | 0 | 0 | 0 | 0 | 0 | 0 |
| ENSMUSG00 | 7SK | 310 | 0 | 0 | 0 | 0 | 0 | 0 |
| ENSMUSG00 | Y_RNA | 102 | 0 | 0 | 0 | 0 | 0 | 0 |
| ENSMUSG00 | U1 | 169 | 0 | 0 | 0 | 0 | 0 | 0 |
| ENSMUSG00 | SNORA17 | 131 | 0 | 0 | 0 | 0 | 0 | 0.20820335 |
| ENSMUSG00 | SNORA63 | 113 | 0 | 1.27928608 | 0 | 0 | 0 | 0 |
| ENSMUSG00 | U6 | 106 | 0 | 0 | 0 | 0 | 0 | 0 |
| ENSMUSG00 | U6 | 95 | 0 | 0 | 0 | 0 | 0 | 0 |
| ENSMUSG00 | U7 | 57 | 0 | 0 | 0 | 0 | 0 | 0 |
| ENSMUSG00 | U6 | 73 | 0 | 0 | 0 | 0 | 0 | 0 |
| ENSMUSG00 | 7SK | 288 | 0 | 0 | 0 | 0 | 0 | 0 |
| ENSMUSG00 | SNORA70 | 135 | 0 | 0 | 0 | 0 | 0 | 0 |
| ENSMUSG00 | SNORA17 | 118 | 0 | 0 | 0 | 0 | 0 | 0 |
| ENSMUSG00 | SCARNA17 | 134 | 0 | 0 | 0 | 0.26926899 | 0 | 0 |
| ENSMUSG00 | U1 | 152 | 0 | 0 | 0 | 0 | 0 | 0 |
| ENSMUSG00 | SNORA40 | 120 | 0 | 0 | 0 | 0 | 0 | 0 |
| ENSMUSG00 | SNORA17 | 139 | 0 | 0 | 0 | 0 | 0 | 0 |
| ENSMUSG00 | mmu-mir-21 | 55 | 2.89130767 | 0 | 0 | 0 | 0 | 0 |
| ENSMUSG00 | U7 | 56 | 0 | 0 | 0 | 0 | 0 | 0 |
| ENSMUSG00 | Mir1969 | 93 | 0 | 0 | 0 | 0 | 0 | 0 |
| ENSMUSG00 | 5S_rRNA | 97 | 0 | 0 | 1.82275641 | 0 | 0 | 0 |
| ENSMUSG00 | U4 | 128 | 0 | 1.12936974 | 0 | 0 | 0 | 0 |
| ENSMUSG00 | U3 | 89 | 0 | 0 | 0 | 0 | 0 | 0 |
| ENSMUSG00 | Snord13 | 103 | 0.38597554 | 0.46782954 | 0.42914411 | 1.05093334 | 0.99080341 | 3.17762787 |
| ENSMUSG00 | U6 | 99 | 0 | 0 | 0 | 0 | 0 | 0 |
| ENSMUSG00 | SCARNA14 | 134 | 0 | 0 | 0 | 0 | 0 | 0 |
| ENSMUSG00 | SNORA48 | 143 | 0 | 0 | 0 | 0 | 0 | 0 |
| ENSMUSG00 | snoMe28S-C | 78 | 0 | 0 | 0 | 0 | 0 | 0 |
| ENSMUSG00 | SNORA17 | 127 | 0 | 0 | 0 | 0 | 0 | 0 |
| ENSMUSG00 | U1 | 161 | 0 | 0 | 0 | 0 | 0 | 0 |
| ENSMUSG00 | 7SK | 95 | 0 | 0 | 0 | 0 | 0 | 0 |
| ENSMUSG00 | SNORA48 | 142 | 0 | 0 | 0 | 0 | 0 | 0 |
| ENSMUSG00 | SNORA17 | 116 | 0 | 0 | 0 | 0 | 0 | 0 |
| ENSMUSG00 | 7SK | 207 | 0 | 0 | 0 | 0 | 0 | 0.26352309 |
| ENSMUSG00 | snoU13 | 103 | 2.31585323 | 2.80697721 | 3.86229696 | 3.50311112 | 3.96321366 | 4.23683717 |
| ENSMUSG00 | U6 | 99 | 0 | 0 | 0 | 0 | 0 | 0 |
| ENSMUSG00 | U7 | 61 | 0.65172919 | 0 | 0 | 0 | 0 | 2.6827514 |
| ENSMUSG00 | SNORA48 | 131 | 0 | 0 | 0 | 0 | 0 | 0 |
| ENSMUSG00 | U6 | 85 | 0 | 0 | 0 | 0 | 0 | 0.96263433 |
| ENSMUSG00 | 5S_rRNA | 106 | 0 | 0 | 0 | 0 | 0 | 0 |
| ENSMUSG00 | U6 | 102 | 0 | 0 | 0 | 0 | 0 | 0 |
| ENSMUSG00 | SNORA5 | 91 | 0 | 0 | 0 | 0 | 0 | 0 |
| ENSMUSG00 | SNORA48 | 143 | 0 | 0 | 0 | 1.26160995 | 0 | 0 |
| ENSMUSG00 | U6 | 106 | 0 | 0 | 0 | 0 | 0 | 0 |
| ENSMUSG00 | U6 | 105 | 0 | 0 | 0 | 0 | 0 | 0 |
| ENSMUSG00 | U1 | 172 | 0 | 0 | 0.25698746 | 0 | 0 | 0 |
| ENSMUSG00 | 7SK | 286 | 0 | 0 | 0 | 0 | 0 | 0 |
| ENSMUSG00 | SNORA25 | 128 | 0 | 0 | 0 | 0 | 0 | 0 |
| ENSMUSG00 | U6 | 101 | 0 | 0 | 0 | 0 | 0 | 0 |
| ENSMUSG00 | SNORA17 | 122 | 0 | 0 | 0 | 0 | 0 | 0 |
| ENSMUSG00 | U6 | 83 | 0 | 0 | 0 | 0 | 0 | 0 |
| ENSMUSG00 | U6 | 98 | 0 | 0 | 0 | 0 | 0 | 0 |
| ENSMUSG00 | U6 | 104 | 0 | 0 | 0 | 0 | 0 | 0 |
| ENSMUSG00 | U6 | 103 | 0 | 0 | 0 | 0 | 0 | 0 |
| ENSMUSG00 | 7SK | 285 | 0 | 0 | 0 | 0 | 0 | 0 |
| ENSMUSG00 | SCARNA4 | 126 | 1.89311812 | 0 | 0.35080828 | 0 | 0 | 0 |
| ENSMUSG00 | U6 | 54 | 0 | 0 | 0 | 0 | 0 | 0 |
| ENSMUSG00 | U6 | 106 | 0 | 0 | 0 | 0 | 0 | 0 |
| ENSMUSG00 | AF357425 | 72 | 0 | 0 | 0 | 0 | 0 | 0 |
| ENSMUSG00 | 7SK | 260 | 0 | 0 | 0 | 0 | 0 | 0 |
| ENSMUSG00 | U7 | 62 | 0 | 0 | 0 | 0 | 0 | 0 |
| ENSMUSG00 | SNORA17 | 126 | 0 | 0 | 0 | 0 | 0 | 0 |
| ENSMUSG00 | 7SK | 291 | 0 | 0 | 0 | 0 | 0 | 0 |
| ENSMUSG00 | 7SK | 268 | 0 | 0.17980016 | 0 | 0 | 0 | 0 |
| ENSMUSG00 | 5S_rRNA | 93 | 0 | 0 | 0 | 0 | 0 | 0 |
| ENSMUSG00 | 7SK | 307 | 0.38849004 | 1.41263185 | 2.30367912 | 0 | 2.32693571 | 2.75411667 |
| ENSMUSG00 | SNORA70 | 103 | 0 | 0 | 0 | 0 | 0 | 0 |
| ENSMUSG00 | U6 | 99 | 0 | 0 | 0 | 0 | 0 | 0 |
| ENSMUSG00 | SNORD26 | 79 | 0 | 0 | 0 | 0 | 0 | 0 |
| ENSMUSG00 | SNORA17 | 125 | 0 | 0 | 0 | 0 | 0 | 0 |
| ENSMUSG00 | snoU109 | 133 | 0 | 0.72460815 | 0.66468937 | 0.54258714 | 1.53462784 | 0.41014495 |
| ENSMUSG00 | U3 | 221 | 0 | 0 | 0.80003336 | 0.81633585 | 0 | 0 |
| ENSMUSG00 | SCARNA17 | 138 | 0 | 0 | 0 | 0 | 0 | 0 |
| ENSMUSG00 | SNORA17 | 130 | 0 | 0 | 0 | 0 | 0 | 0 |
| ENSMUSG00 | U6 | 105 | 0 | 0 | 0 | 0 | 0 | 0 |
| ENSMUSG00 | SNORA17 | 115 | 0 | 0 | 0 | 0 | 0 | 0 |
| ENSMUSG00 | SNORA18 | 124 | 0 | 0 | 0 | 0 | 0 | 0 |
| ENSMUSG00 | U6 | 103 | 0 | 0 | 0 | 0 | 0 | 0 |
| ENSMUSG00 | 7SK | 265 | 0 | 0 | 0 | 0 | 0 | 0 |
| ENSMUSG00 | U6 | 100 | 0 | 0 | 0 | 0 | 0 | 0 |
| ENSMUSG00 | SNORA17 | 109 | 0 | 0 | 0 | 0 | 0 | 0 |
| ENSMUSG00 | U6 | 106 | 0 | 0 | 0 | 0 | 0 | 0 |
| ENSMUSG00 | snoU109 | 134 | 2.37346152 | 0.71920083 | 4.2882385 | 0.80780697 | 0.38079385 | 3.25667334 |
| ENSMUSG00 | 5S_rRNA | 111 | 0 | 0 | 0 | 0 | 0 | 0 |
| ENSMUSG00 | SNORA17 | 128 | 0 | 0 | 0 | 0 | 0 | 0 |
| ENSMUSG00 | 7SK | 274 | 0 | 0 | 0 | 0 | 0 | 0 |
| ENSMUSG00 | U6 | 106 | 0 | 0 | 0 | 0 | 0 | 0 |
| ENSMUSG00 | U7 | 59 | 0 | 0 | 0 | 0 | 0 | 0 |
| ENSMUSG00 | RNaseP_nuc | 295 | 0 | 0 | 0 | 0 | 0 | 0.0924564 |
| ENSMUSG00 | U1 | 164 | 0 | 0 | 0 | 0 | 0 | 0 |
| ENSMUSG00 | 7SK | 269 | 0 | 0 | 0 | 0 | 0 | 0 |
| ENSMUSG00 | U1 | 104 | 0 | 0 | 0 | 0 | 0 | 0 |
| ENSMUSG00 | U7 | 62 | 0 | 0 | 0 | 0 | 0 | 0 |
| ENSMUSG00 | HAR1F | 131 | 0 | 0 | 0 | 0 | 0 | 0 |
| ENSMUSG00 | SNORA25 | 133 | 0 | 0 | 0 | 0 | 0 | 1.02536238 |
| ENSMUSG00 | snoZ178 | 107 | 0 | 0 | 0 | 0 | 0 | 0 |
| ENSMUSG00 | U6 | 103 | 0 | 0 | 0 | 0 | 0 | 0 |
| ENSMUSG00 | 7SK | 306 | 0 | 0.4724161 | 0 | 0 | 0.33350572 | 0.89132808 |
| ENSMUSG00 | U2 | 81 | 0 | 0 | 0 | 0 | 0 | 0 |
| ENSMUSG00 | SNORA75 | 128 | 0 | 0 | 0 | 0 | 0 | 0 |
| ENSMUSG00 | U6 | 104 | 0 | 0 | 0 | 0 | 0 | 0 |
| ENSMUSG00 | 5S_rRNA | 113 | 0 | 0 | 0 | 0 | 0 | 0 |
| ENSMUSG00 | U7 | 61 | 0 | 0 | 0 | 0 | 0 | 0 |
| ENSMUSG00 | U6 | 106 | 0 | 0 | 0 | 0 | 0 | 0 |
| ENSMUSG00 | SNORA17 | 131 | 0 | 0 | 0 | 0 | 0 | 0 |
| ENSMUSG00 | U6 | 96 | 0 | 0 | 0 | 0 | 0 | 0 |
| ENSMUSG00 | U6 | 98 | 0 | 0 | 0 | 0 | 0 | 0 |
| ENSMUSG00 | 7SK | 231 | 0 | 0 | 0 | 0 | 0 | 0 |
| ENSMUSG00 | snoU54 | 67 | 0 | 0 | 0 | 0 | 0 | 0 |
| ENSMUSG00 | SNORA17 | 98 | 0 | 0 | 0 | 0 | 0 | 0 |
| ENSMUSG00 | U1 | 156 | 0 | 0 | 0 | 0 | 0 | 0 |
| ENSMUSG00 | AC164075.1 | 59 | 0 | 0 | 0 | 0 | 0 | 0 |
| ENSMUSG00 | 7SK | 281 | 0 | 0 | 0 | 0 | 0 | 0 |
| ENSMUSG00 | 7SK | 267 | 0 | 0 | 0.16554997 | 0 | 0 | 0 |
| ENSMUSG00 | SNORA17 | 137 | 0 | 0 | 0 | 0 | 0 | 0 |
| ENSMUSG00 | 5S_rRNA | 92 | 0 | 2.61882838 | 1.44136445 | 2.74537295 | 2.21853808 | 0.59292694 |
| ENSMUSG00 | U6 | 97 | 0 | 0 | 0 | 0 | 0 | 0 |
| ENSMUSG00 | 7SK | 267 | 0 | 0 | 0 | 0 | 0 | 0 |
| ENSMUSG00 | SNORA17 | 86 | 0 | 0 | 0 | 0 | 0 | 0 |
| ENSMUSG00 | U1 | 129 | 0 | 0 | 0 | 0 | 0 | 0 |
| ENSMUSG00 | U7 | 61 | 0 | 0 | 0 | 0 | 0 | 0 |
| ENSMUSG00 | 7SK | 241 | 0 | 0 | 0 | 0 | 0 | 0 |
| ENSMUSG00 | 7SK | 247 | 0 | 0 | 0 | 0 | 0 | 0 |
| ENSMUSG00 | 7SK | 305 | 0 | 0 | 0 | 0 | 0 | 0 |
| ENSMUSG00 | 7SK | 174 | 0 | 0 | 0 | 0 | 0 | 0 |
| ENSMUSG00 | 7SK | 255 | 0 | 0 | 0 | 0 | 0 | 0 |
| ENSMUSG00 | U6 | 102 | 0 | 0 | 0 | 0 | 0 | 0 |
| ENSMUSG00 | 5S_rRNA | 114 | 1.74366143 | 0 | 0 | 0 | 0 | 0 |
| ENSMUSG00 | SNORA17 | 133 | 0 | 0 | 0 | 0 | 0 | 0 |
| ENSMUSG00 | Mir1956 | 64 | 0 | 0 | 0 | 0 | 0 | 2.13083119 |
| ENSMUSG00 | SNORA17 | 123 | 0 | 0 | 0 | 0 | 0 | 0 |
| ENSMUSG00 | RNase_MRP | 239 | 0 | 0 | 0 | 0 | 0 | 0 |
| ENSMUSG00 | snoU13 | 102 | 0 | 0 | 0 | 0 | 0 | 0 |
| ENSMUSG00 | SNORA40 | 105 | 0 | 0 | 0 | 0 | 0 | 0 |
| ENSMUSG00 | U1 | 162 | 0 | 0 | 0 | 0 | 0 | 0 |
| ENSMUSG00 | SNORA17 | 95 | 0 | 0 | 0 | 0 | 0 | 0 |
| ENSMUSG00 | U6 | 92 | 0 | 0 | 0 | 0 | 0 | 0 |
| ENSMUSG00 | snoU13 | 97 | 0 | 0 | 0 | 0 | 0 | 0 |
| ENSMUSG00 | RNaseP_nuc | 240 | 0 | 0 | 0 | 0 | 0 | 0 |
| ENSMUSG00 | 5S_rRNA | 96 | 0 | 0 | 0 | 0 | 0 | 0 |
| ENSMUSG00 | U6 | 103 | 0 | 0 | 0.42914411 | 5.25466668 | 0 | 0 |
| ENSMUSG00 | U1 | 145 | 0 | 0 | 0 | 0 | 0 | 0 |
| ENSMUSG00 | U1 | 139 | 0 | 0 | 0 | 0 | 0 | 0 |
| ENSMUSG00 | U7 | 60 | 0 | 0 | 0 | 0 | 0 | 0 |
| ENSMUSG00 | 7SK | 284 | 0 | 0 | 0 | 0 | 0 | 0 |
| ENSMUSG00 | U7 | 63 | 0 | 0 | 0 | 0 | 0 | 0 |
| ENSMUSG00 | SNORA17 | 116 | 0 | 0 | 0 | 0 | 0 | 0 |
| ENSMUSG00 | SNORA17 | 125 | 0 | 0 | 0 | 0 | 0 | 0 |
| ENSMUSG00 | AC090489.1 | 68 | 0 | 0 | 0 | 0 | 0 | 0 |
| ENSMUSG00 | U1 | 164 | 0 | 0 | 0 | 0 | 0 | 0 |
| ENSMUSG00 | 7SK | 307 | 0 | 0 | 0 | 0 | 0 | 0 |
| ENSMUSG00 | SNORA13 | 76 | 0 | 0 | 0 | 0 | 0 | 0 |
| ENSMUSG00 | SNORA40 | 118 | 0 | 0 | 0 | 0 | 0 | 0 |
| ENSMUSG00 | U6 | 106 | 0 | 0 | 0 | 0 | 0 | 0 |
| ENSMUSG00 | U2 | 83 | 0 | 0 | 0 | 0 | 0 | 0 |
| ENSMUSG00 | 7SK | 301 | 0 | 0 | 0 | 0 | 0 | 0 |
| ENSMUSG00 | 5S_rRNA | 108 | 5.52159452 | 0 | 0 | 0 | 0 | 0 |
| ENSMUSG00 | U6 | 113 | 0 | 0 | 0 | 0 | 0 | 0 |
| ENSMUSG00 | 7SK | 270 | 0 | 0 | 0 | 0 | 0 | 0 |
| ENSMUSG00 | U7 | 61 | 0 | 0 | 0 | 0 | 0 | 0 |
| ENSMUSG00 | U6 | 102 | 0 | 0 | 0 | 0 | 0 | 0 |
| ENSMUSG00 | U4 | 84 | 0 | 0 | 0 | 0 | 0 | 0 |
| ENSMUSG00 | SNORA31 | 134 | 0 | 0 | 1.319458 | 0 | 0 | 0.61062625 |
| ENSMUSG00 | SNORA79 | 144 | 0 | 0 | 0 | 0 | 0 | 0 |
| ENSMUSG00 | Mir1967 | 81 | 0 | 0 | 0 | 0 | 0 | 0 |
| ENSMUSG00 | SNORA61 | 131 | 0 | 0 | 0 | 0 | 0 | 0 |
| ENSMUSG00 | 7SK | 254 | 0 | 0 | 0 | 0 | 0 | 0 |
| ENSMUSG00 | 5S_rRNA | 97 | 0 | 0 | 0 | 0 | 0 | 0 |
| ENSMUSG00 | SNORA17 | 131 | 0 | 0 | 0 | 0 | 0 | 0 |
| ENSMUSG00 | SNORA70 | 133 | 0 | 0 | 0 | 0 | 0 | 0 |
| ENSMUSG00 | SNORA70 | 102 | 0 | 0 | 0 | 0 | 0 | 0 |
| ENSMUSG00 | 7SK | 336 | 0 | 0 | 0 | 0 | 0 | 0 |
| ENSMUSG00 | 7SK | 294 | 0 | 0 | 0 | 0 | 0 | 0 |
| ENSMUSG00 | SNORA48 | 146 | 0 | 0 | 0 | 0 | 0 | 0 |
| ENSMUSG00 | SNORA17 | 118 | 0 | 0 | 0 | 0 | 0 | 0 |
| ENSMUSG00 | U6 | 61 | 0 | 0 | 0 | 0 | 0 | 0 |
| ENSMUSG00 | 5S_rRNA | 99 | 0 | 0 | 0 | 0 | 0 | 0 |
| ENSMUSG00 | U6 | 101 | 0.39361862 | 0 | 0 | 0 | 0 | 0 |
| ENSMUSG00 | U7 | 59 | 0 | 0 | 0 | 0 | 0 | 0 |
| ENSMUSG00 | U6 | 104 | 0 | 0 | 0 | 0 | 0 | 0 |
| ENSMUSG00 | SNORA17 | 135 | 0 | 0 | 0 | 0 | 0 | 0 |
| ENSMUSG00 | U3 | 89 | 0 | 0 | 0 | 0 | 0 | 0 |
| ENSMUSG00 | SNORA48 | 143 | 0 | 0 | 0 | 0 | 0 | 0 |

| | | | | | | | | |
|---|---|---|---|---|---|---|---|---|
| ENSMUSG00 | U6 | 96 | 0 | 0 | 0 | 0 | 0 | 0 |
| ENSMUSG00 | U6 | 107 | 0 | 0 | 0 | 0 | 0 | 0 |
| ENSMUSG00 | SNORA17 | 131 | 0 | 0 | 0 | 0 | 0 | 0 |
| ENSMUSG00 | 7SK | 287 | 0.83112503 | 2.68635218 | 0 | 0.50288564 | 4.08922176 | 2.56590683 |
| ENSMUSG00 | SNORA17 | 134 | 0 | 0 | 0 | 0 | 0 | 0 |
| ENSMUSG00 | U6 | 102 | 0 | 0 | 0 | 0 | 0 | 0 |
| ENSMUSG00 | 7SK | 294 | 0 | 0 | 0 | 0 | 0 | 0 |
| ENSMUSG00 | SNORA17 | 109 | 0 | 0 | 0 | 0 | 0 | 0 |
| ENSMUSG00 | SNORA17 | 128 | 0 | 0 | 0 | 0 | 0 | 0 |
| ENSMUSG00 | SCARNA24 | 137 | 0 | 0 | 0 | 0 | 0 | 0 |
| ENSMUSG00 | AL732470.1 | 91 | 0 | 0 | 0 | 0 | 0 | 0 |
| ENSMUSG00 | SNORA40 | 121 | 0 | 0 | 0 | 0 | 0 | 0 |
| ENSMUSG00 | 5S_rRNA | 79 | 0 | 0 | 0 | 0 | 0 | 0 |
| ENSMUSG00 | U6 | 102 | 0 | 0 | 0 | 0 | 0 | 0 |
| ENSMUSG00 | U6 | 99 | 0 | 0 | 0 | 0 | 0 | 0 |
| ENSMUSG00 | SNORA70 | 117 | 0 | 0 | 0 | 0 | 0 | 0 |
| ENSMUSG00 | U7 | 55 | 0 | 0 | 0 | 0 | 0 | 0 |
| ENSMUSG00 | 7SK | 246 | 0 | 0 | 0 | 0 | 0 | 0 |
| ENSMUSG00 | U6 | 103 | 0 | 0 | 0 | 0 | 0 | 0 |
| ENSMUSG00 | 7SK | 235 | 0 | 0 | 0 | 0 | 0 | 0 |
| ENSMUSG00 | U6 | 102 | 0 | 0 | 0 | 0 | 0 | 0 |
| ENSMUSG00 | Mir1941 | 82 | 0.96964587 | 0 | 0.53904687 | 2.20012467 | 0.62227288 | 0.33261755 |
| ENSMUSG00 | 5S_rRNA | 86 | 0 | 0 | 0 | 0 | 0 | 0 |
| ENSMUSG00 | 7SK | 107 | 0 | 0 | 0 | 0 | 0 | 0 |
| ENSMUSG00 | 7SK | 342 | 0 | 0 | 0 | 0 | 0 | 0 |
| ENSMUSG00 | Mir1970 | 103 | 0 | 0 | 1.71657643 | 0 | 0 | 0 |
| ENSMUSG00 | SNORA17 | 84 | 0 | 0 | 0 | 0 | 0 | 0 |
| ENSMUSG00 | snoU13 | 99 | 0 | 0 | 0 | 0 | 0 | 0 |
| ENSMUSG00 | U6 | 103 | 0 | 0 | 0 | 0 | 0 | 0 |
| ENSMUSG00 | SNORA17 | 101 | 0 | 0 | 0 | 0 | 0 | 0 |
| ENSMUSG00 | 7SK | 296 | 0 | 0 | 0 | 0 | 0 | 0 |
| ENSMUSG00 | U6 | 102 | 0 | 0 | 0 | 0 | 0 | 0 |
| ENSMUSG00 | SNORA17 | 116 | 0 | 0 | 0 | 0 | 0 | 1.6458834 |
| ENSMUSG00 | U1 | 147 | 0 | 0 | 0.30069281 | 0 | 0 | 0 |
| ENSMUSG00 | 7SK | 292 | 0 | 0 | 0 | 0 | 0 | 0 |
| ENSMUSG00 | 5S_rRNA | 113 | 0 | 0 | 0 | 0.95793039 | 2.25780424 | 0.72410547 |
| ENSMUSG00 | SNORA17 | 115 | 0 | 0 | 0 | 0 | 0 | 0 |
| ENSMUSG00 | SNORA70 | 132 | 0 | 0 | 0 | 0 | 0 | 0 |
| ENSMUSG00 | Mir1946b | 132 | 0 | 0 | 0 | 0 | 0 | 0 |
| ENSMUSG00 | SNORA17 | 115 | 2.07419898 | 0.41901254 | 0.76872771 | 0.31375691 | 1.33112285 | 2.13453699 |
| ENSMUSG00 | U6 | 63 | 0 | 0 | 0 | 0 | 0 | 0 |
| ENSMUSG00 | SNORA17 | 76 | 0 | 0 | 0 | 0 | 0 | 0 |
| ENSMUSG00 | U6 | 100 | 0 | 0 | 0 | 0 | 0 | 0 |
| ENSMUSG00 | 7SK | 257 | 0 | 0 | 0 | 0 | 0 | 0 |
| ENSMUSG00 | SNORA17 | 116 | 0 | 0 | 0 | 0 | 0 | 0 |
| ENSMUSG00 | U6 | 103 | 0 | 0 | 1.71657643 | 1.40124445 | 0.49540171 | 1.05920929 |
| ENSMUSG00 | U7 | 61 | 0 | 0 | 0 | 0 | 0 | 0 |
| ENSMUSG00 | 7SK | 96 | 0 | 0 | 0 | 0 | 0 | 0 |
| ENSMUSG00 | U6 | 106 | 0 | 0 | 0 | 0 | 0 | 0 |
| ENSMUSG00 | U6 | 106 | 0 | 0 | 0 | 0 | 0 | 0 |
| ENSMUSG00 | Mir1953 | 88 | 0 | 0 | 0 | 1.2300697 | 0 | 0 |
| ENSMUSG00 | SNORA40 | 90 | 2.20863781 | 4.2832393 | 4.91131589 | 4.81093927 | 2.26783893 | 11.2129073 |
| ENSMUSG00 | U6 | 73 | 0 | 0 | 1.8165141 | 0 | 0.69899145 | 0 |
| ENSMUSG00 | SNORA17 | 108 | 0 | 0 | 0 | 0 | 0 | 0 |
| ENSMUSG00 | 5S_rRNA | 93 | 0 | 0 | 0 | 0 | 0 | 0 |
| ENSMUSG00 | U2 | 90 | 0 | 0 | 0 | 0 | 0 | 0 |
| ENSMUSG00 | U6 | 104 | 0 | 0 | 0 | 0 | 0 | 0 |
| ENSMUSG00 | 7SK | 331 | 0 | 0 | 0 | 0 | 0 | 0 |
| ENSMUSG00 | U7 | 60 | 0 | 0 | 0 | 0 | 0 | 0 |
| ENSMUSG00 | U6 | 72 | 0 | 0 | 0 | 0 | 0 | 0 |
| ENSMUSG00 | SNORA17 | 131 | 0 | 0 | 0 | 0 | 0 | 0 |
| ENSMUSG00 | SNORA17 | 139 | 0 | 0 | 0.31799887 | 0 | 0.36709623 | 3.33574725 |
| ENSMUSG00 | SCARNA13 | 263 | 0 | 0 | 0 | 0 | 0 | 0 |
| ENSMUSG00 | U1 | 151 | 0 | 0 | 0 | 0 | 0 | 0 |
| ENSMUSG00 | SNORA17 | 129 | 0 | 0 | 0 | 0 | 0 | 0 |
| ENSMUSG00 | SNORA17 | 98 | 0 | 0 | 0 | 0 | 0 | 0 |
| ENSMUSG00 | U6 | 109 | 0 | 0 | 0 | 0 | 0 | 0 |
| ENSMUSG00 | SNORA17 | 125 | 0 | 0 | 0 | 0 | 0 | 0 |
| ENSMUSG00 | 7SK | 301 | 0 | 0 | 0 | 0 | 0 | 0 |
| ENSMUSG00 | SNORA17 | 108 | 0 | 0 | 0 | 0 | 0 | 0 |
| ENSMUSG00 | U6 | 99 | 0 | 0 | 0 | 0 | 0 | 0 |
| ENSMUSG00 | U6 | 103 | 0 | 0 | 0 | 0 | 0 | 0 |
| ENSMUSG00 | SNORA9 | 114 | 0 | 0 | 0 | 0 | 0 | 0 |
| ENSMUSG00 | U6 | 116 | 0 | 0 | 0 | 0 | 0 | 0 |
| ENSMUSG00 | U6 | 73 | 0 | 0 | 0 | 0 | 0 | 0 |
| ENSMUSG00 | SNORA17 | 130 | 0 | 0 | 0 | 0 | 0 | 0 |
| ENSMUSG00 | SNORA61 | 127 | 0 | 0 | 0 | 0 | 0 | 0 |
| ENSMUSG00 | 5S_rRNA | 106 | 0 | 0 | 0 | 0 | 0 | 0 |
| ENSMUSG00 | Snord2 | 68 | 0 | 0 | 0 | 0 | 0 | 0 |
| ENSMUSG00 | SNORA17 | 115 | 0 | 0 | 0 | 0 | 0 | 0 |
| ENSMUSG00 | U7 | 61 | 0 | 0 | 0 | 0 | 0 | 0 |
| ENSMUSG00 | 7SK | 262 | 0 | 0 | 0.50612797 | 0.68858864 | 0 | 0 |
| ENSMUSG00 | U6 | 103 | 0 | 0 | 0 | 0 | 0 | 0 |
| ENSMUSG00 | SCARNA16 | 192 | 0 | 0 | 0 | 0 | 0 | 0 |
| ENSMUSG00 | SNORA17 | 138 | 0 | 0 | 0 | 0 | 0 | 0 |
| ENSMUSG00 | 7SK | 100 | 0 | 0 | 0 | 0 | 0 | 0 |
| ENSMUSG00 | U1 | 111 | 0 | 0 | 0 | 0 | 0 | 0 |
| ENSMUSG00 | SNORA17 | 88 | 0 | 0 | 0 | 0 | 0 | 0 |
| ENSMUSG00 | U6 | 104 | 0 | 0 | 0 | 0 | 0 | 0 |
| ENSMUSG00 | SNORA48 | 142 | 0 | 0 | 0 | 0 | 0 | 0 |
| ENSMUSG00 | 5S_rRNA | 96 | 0 | 0 | 0 | 0 | 0 | 0 |
| ENSMUSG00 | SNORA17 | 137 | 0 | 0 | 0 | 0 | 0 | 0 |
| ENSMUSG00 | SNORA17 | 135 | 0 | 0 | 0 | 0 | 0 | 0 |
| ENSMUSG00 | U7 | 57 | 0 | 0 | 0 | 0 | 0 | 0 |
| ENSMUSG00 | SNORA48 | 143 | 0 | 0 | 0 | 0 | 0 | 0 |
| ENSMUSG00 | U6 | 108 | 0 | 0 | 0 | 0 | 0 | 0 |
| ENSMUSG00 | U1 | 183 | 0 | 0 | 0 | 0 | 0 | 0 |
| ENSMUSG00 | Mir1945 | 77 | 0 | 0 | 0 | 0 | 0 | 0 |
| ENSMUSG00 | U6 | 100 | 0.79510961 | 0 | 0 | 0 | 0 | 0.27274639 |
| ENSMUSG00 | 7SK | 294 | 0 | 0 | 0 | 0 | 0 | 0 |
| ENSMUSG00 | U6 | 105 | 0 | 0 | 0 | 0 | 0 | 0 |
| ENSMUSG00 | U6 | 89 | 0 | 0 | 0 | 0 | 0 | 0 |
| ENSMUSG00 | U1 | 145 | 0 | 0.66464058 | 0 | 0.49768337 | 0.35190604 | 0.37620192 |
| ENSMUSG00 | SNORA17 | 122 | 0 | 0 | 0 | 0 | 0 | 0 |
| ENSMUSG00 | U7 | 62 | 0 | 0 | 0 | 0 | 0 | 0 |
| ENSMUSG00 | Mir1957 | 65 | 1.22324555 | 0.74132988 | 0 | 2.77554189 | 0 | 0 |
| ENSMUSG00 | U6 | 102 | 0 | 0 | 0 | 0 | 0 | 0 |
| ENSMUSG00 | 7SK | 277 | 0 | 0 | 0 | 0 | 0 | 0 |
| ENSMUSG00 | U4 | 84 | 1.41983859 | 6.31012933 | 1.05242483 | 4.72502964 | 1.82237057 | 2.2728866 |
| ENSMUSG00 | SNORA17 | 97 | 0 | 0 | 0 | 0 | 0 | 0 |
| ENSMUSG00 | Mir1981 | 81 | 0 | 0 | 1.09140353 | 0 | 0 | 0.67344788 |
| ENSMUSG00 | snoU13 | 104 | 0 | 0 | 0 | 1.04082821 | 0 | 0 |
| ENSMUSG00 | 7SK | 332 | 0.47898169 | 0.87083932 | 0.13313808 | 0.21736171 | 0.61477561 | 0.08215253 |
| ENSMUSG00 | U7 | 63 | 0 | 0 | 0 | 0 | 0 | 0 |
| ENSMUSG00 | 5S_rRNA | 119 | 0 | 0 | 0 | 0 | 0 | 0 |
| ENSMUSG00 | 7SK | 307 | 0 | 0 | 0 | 0 | 0 | 0 |
| ENSMUSG00 | U6 | 101 | 0 | 0 | 0 | 0 | 0 | 0 |
| ENSMUSG00 | U6 | 99 | 0 | 0 | 0 | 0 | 0 | 0 |
| ENSMUSG00 | 7SK | 324 | 0 | 0 | 0 | 0 | 0 | 0 |
| ENSMUSG00 | 7SK | 314 | 0 | 0 | 0 | 0 | 0 | 0 |
| ENSMUSG00 | 7SK | 287 | 0 | 0.16789701 | 0 | 0.25144282 | 0 | 0 |
| ENSMUSG00 | 7SK | 290 | 0 | 0 | 0 | 0 | 0 | 0 |
| ENSMUSG00 | U6 | 101 | 0 | 0 | 1.75056804 | 0 | 0 | 0 |
| ENSMUSG00 | 7SK | 285 | 0.83695748 | 0 | 0 | 0.63301833 | 0 | 0 |
| ENSMUSG00 | SNORA70 | 108 | 0 | 0 | 0 | 0 | 0 | 0 |
| ENSMUSG00 | SNORA17 | 113 | 0 | 0 | 0 | 0 | 0 | 0 |
| ENSMUSG00 | SNORA17 | 126 | 0 | 0 | 0 | 0 | 0 | 0 |
| ENSMUSG00 | 5S_rRNA | 117 | 0 | 0 | 0 | 0 | 0 | 0 |
| ENSMUSG00 | 7SK | 238 | 0 | 0 | 0 | 0 | 0 | 0 |
| ENSMUSG00 | SNORA17 | 128 | 0 | 0 | 0 | 0 | 0 | 0 |
| ENSMUSG00 | U6 | 100 | 0 | 0 | 0 | 0 | 0 | 0 |
| ENSMUSG00 | U7 | 59 | 0 | 0 | 0 | 0 | 0 | 0 |
| ENSMUSG00 | SNORA66 | 90 | 0 | 0 | 0 | 0 | 0 | 0 |
| ENSMUSG00 | SNORD113 | 72 | 0 | 0 | 0 | 0 | 0 | 0 |
| ENSMUSG00 | SNORA5 | 91 | 0 | 0 | 0 | 0 | 0 | 0 |
| ENSMUSG00 | U6 | 105 | 0 | 0 | 0 | 0 | 0 | 0 |
| ENSMUSG00 | 5S_rRNA | 106 | 0 | 0 | 0 | 0 | 0 | 0 |
| ENSMUSG00 | U7 | 65 | 0 | 0 | 0 | 0 | 0 | 0 |
| ENSMUSG00 | SNORA17 | 141 | 0 | 0 | 0 | 0 | 0 | 0 |
| ENSMUSG00 | 5_8S_rRNA | 142 | 3.35961807 | 3.73275256 | 3.73536702 | 4.57378029 | 1.4373627 | 3.26527371 |
| ENSMUSG00 | U6 | 62 | 0 | 0 | 0 | 0 | 0 | 0 |
| ENSMUSG00 | U6 | 106 | 0 | 0 | 0 | 0 | 0 | 0 |
| ENSMUSG00 | SNORA17 | 49 | 0 | 0 | 0 | 0 | 0 | 0 |
| ENSMUSG00 | 5S_rRNA | 93 | 0 | 0 | 4.27759771 | 0 | 0.54867071 | 0 |
| ENSMUSG00 | 7SK | 215 | 0 | 0 | 0 | 0 | 0 | 0 |
| ENSMUSG00 | U6atac | 122 | 0 | 0 | 0 | 0 | 0 | 0 |
| ENSMUSG00 | U1 | 144 | 0 | 0 | 0 | 0 | 0 | 0 |
| ENSMUSG00 | SCARNA15 | 126 | 1.26207875 | 0.76486416 | 1.05242483 | 2.29092346 | 0 | 0.86586156 |
| ENSMUSG00 | 5S_rRNA | 107 | 0 | 0 | 0 | 0 | 0 | 0 |
| ENSMUSG00 | Mir1951 | 88 | 0 | 0 | 0 | 0 | 0 | 0 |
| ENSMUSG00 | 5S_rRNA | 126 | 0 | 0 | 0 | 0 | 0 | 0 |
| ENSMUSG00 | mmu-mir-21 | 56 | 0 | 0.86047218 | 0.78931863 | 0 | 0 | 0 |
| ENSMUSG00 | 7SK | 273 | 0 | 0 | 0 | 0 | 0 | 0 |
| ENSMUSG00 | U7 | 64 | 0 | 0 | 0 | 0 | 0 | 0 |
| ENSMUSG00 | U7 | 61 | 0 | 0 | 0 | 0 | 0 | 0 |
| ENSMUSG00 | U6 | 102 | 0 | 0 | 0 | 0 | 0 | 0 |
| ENSMUSG00 | U6 | 108 | 0.7362126 | 0 | 0 | 0 | 0 | 0.75762887 |
| ENSMUSG00 | U6 | 106 | 0 | 0 | 0 | 0 | 0 | 0 |
| ENSMUSG00 | 7SK | 305 | 0 | 0 | 0 | 0 | 0 | 0 |
| ENSMUSG00 | 5S_rRNA | 117 | 0 | 0 | 0 | 0 | 0 | 0 |
| ENSMUSG00 | Mir1965 | 125 | 0 | 0 | 0 | 0 | 0 | 0 |
| ENSMUSG00 | U6 | 106 | 0 | 0 | 0 | 0 | 0 | 0 |
| ENSMUSG00 | SNORA17 | 133 | 0 | 0 | 0 | 0 | 0 | 0 |
| ENSMUSG00 | U6 | 113 | 1.75909206 | 0 | 0 | 0 | 0 | 0 |
| ENSMUSG00 | U6atac | 118 | 0 | 1.22507904 | 0 | 0 | 0 | 0.92456404 |
| ENSMUSG00 | SNORA42 | 128 | 0 | 0 | 0 | 0 | 0 | 0 |
| ENSMUSG00 | U6 | 107 | 0 | 1.80136232 | 0 | 0 | 0 | 0 |
| ENSMUSG00 | SNORA28 | 145 | 1.37087864 | 0 | 0 | 0 | 0 | 0 |
| ENSMUSG00 | U6 | 105 | 0 | 0 | 0 | 0 | 0 | 0 |
| ENSMUSG00 | SNORA40 | 108 | 0 | 0 | 0 | 0 | 0 | 0 |
| ENSMUSG00 | 7SK | 272 | 0 | 0 | 0 | 0 | 0 | 0 |
| ENSMUSG00 | SNORA17 | 129 | 0 | 0 | 0 | 0 | 0 | 0 |
| ENSMUSG00 | 5S_rRNA | 109 | 0 | 0 | 0 | 0 | 0 | 0 |
| ENSMUSG00 | 7SK | 280 | 0.14198386 | 0 | 0 | 1.15978 | 0.36447411 | 0.19481885 |
| ENSMUSG00 | SNORA17 | 128 | 0 | 0 | 0 | 0 | 0 | 0 |
| ENSMUSG00 | SNORD125 | 96 | 4.14119589 | 0 | 0.46043587 | 0 | 0 | 1.1364433 |
| ENSMUSG00 | SNORA48 | 142 | 0 | 0 | 0 | 0 | 0 | 0 |
| ENSMUSG00 | 7SK | 215 | 0 | 0 | 0 | 0 | 0 | 0 |
| ENSMUSG00 | SNORA17 | 134 | 0 | 0 | 0 | 0 | 0 | 0 |
| ENSMUSG00 | Mir1955 | 97 | 0 | 1.98706978 | 0 | 0 | 0 | 0.28118185 |
| ENSMUSG00 | U7 | 61 | 0 | 0 | 0 | 0 | 0 | 0 |
| ENSMUSG00 | AC127677.1 | 103 | 0 | 0 | 0 | 0 | 0 | 0 |
| ENSMUSG00 | U6 | 105 | 0 | 0 | 0 | 0 | 0 | 2.59758469 |
| ENSMUSG00 | U6 | 63 | 0 | 0 | 0 | 0 | 0 | 0 |
| ENSMUSG00 | 5S_rRNA | 109 | 0 | 0 | 0 | 0 | 0 | 0 |
| ENSMUSG00 | U6 | 93 | 0 | 0 | 0 | 0 | 0 | 0 |
| ENSMUSG00 | U7 | 60 | 0 | 0 | 0 | 0 | 0 | 0 |
| ENSMUSG00 | SNORA17 | 126 | 0 | 0 | 0 | 0 | 0 | 0 |
| ENSMUSG00 | U6 | 97 | 0 | 0 | 0 | 0 | 0 | 0 |
| ENSMUSG00 | SNORA40 | 120 | 0 | 0 | 0 | 0 | 0 | 0 |
| ENSMUSG00 | U7 | 64 | 0 | 0 | 0 | 0 | 0 | 0 |
| ENSMUSG00 | U6 | 102 | 0 | 0 | 0 | 0 | 0 | 0 |
| ENSMUSG00 | SNORA17 | 118 | 0 | 0 | 0 | 0 | 0 | 0 |
| ENSMUSG00 | U7 | 59 | 0 | 0 | 0 | 0 | 0 | 0 |
| ENSMUSG00 | U6 | 106 | 0 | 0 | 0 | 0 | 0 | 0 |
| ENSMUSG00 | U6 | 107 | 0 | 0 | 0 | 0 | 0 | 0 |
| ENSMUSG00 | U6 | 107 | 0 | 0 | 0 | 0 | 0 | 0 |
| ENSMUSG00 | 5S_rRNA | 108 | 0 | 0 | 0 | 0 | 0 | 0 |
| ENSMUSG00 | 7SK | 245 | 0 | 0 | 0 | 0 | 0 | 0 |
| ENSMUSG00 | U6 | 109 | 0.72945836 | 0.44207745 | 2.02760748 | 0 | 3.27692322 | 0.5004521 |
| ENSMUSG00 | U3 | 88 | 0 | 0 | 0 | 0 | 0 | 0 |
| ENSMUSG00 | U6 | 104 | 0 | 0 | 0 | 0 | 0 | 0 |
| ENSMUSG00 | U7 | 60 | 0 | 0 | 0 | 0 | 0 | 0 |
| ENSMUSG00 | 5S_rRNA | 125 | 0 | 0 | 0 | 0 | 0 | 0 |
| ENSMUSG00 | U7 | 56 | 0 | 0 | 0 | 0 | 0 | 0 |
| ENSMUSG00 | U6 | 102 | 0 | 0 | 0 | 0 | 0 | 0 |
| ENSMUSG00 | SNORA17 | 104 | 0 | 0 | 0 | 0 | 0 | 0 |
| ENSMUSG00 | SNORA31 | 79 | 0 | 0 | 0 | 0 | 0 | 0 |
| ENSMUSG00 | 5S_rRNA | 110 | 0 | 0 | 0 | 0 | 0 | 0 |
| ENSMUSG00 | U6atac | 88 | 0 | 0 | 0 | 0 | 0 | 0 |
| ENSMUSG00 | 7SK | 282 | 0.56390753 | 0 | 0.31348825 | 0 | 0.36188919 | 0 |
| ENSMUSG00 | RNaseP_nuc | 326 | 0 | 0 | 0 | 0 | 0 | 0 |
| ENSMUSG00 | SNORA2 | 100 | 0 | 0 | 0 | 0 | 0 | 0 |
| ENSMUSG00 | SNORA17 | 115 | 0 | 0 | 0 | 0 | 0 | 0 |
| ENSMUSG00 | U7 | 61 | 0 | 0 | 0 | 0 | 0 | 0 |
| ENSMUSG00 | 5S_rRNA | 120 | 0 | 0 | 0 | 0 | 0 | 0 |
| ENSMUSG00 | U3 | 88 | 0 | 0 | 0 | 0 | 0 | 0 |
| ENSMUSG00 | 5S_rRNA | 103 | 0 | 0 | 0 | 0 | 0 | 0 |
| ENSMUSG00 | 7SK | 322 | 0 | 0 | 0 | 0 | 0 | 0 |
| ENSMUSG00 | SNORA17 | 87 | 0 | 0 | 0 | 0 | 0 | 0 |
| ENSMUSG00 | SNORA70 | 128 | 0 | 0 | 0 | 0 | 0 | 0 |
| ENSMUSG00 | 7SK | 309 | 0 | 0 | 0 | 0 | 0 | 0 |
| ENSMUSG00 | SNORA70 | 130 | 0 | 0 | 0 | 0 | 0 | 0 |
| ENSMUSG00 | SNORA70 | 88 | 0 | 0 | 0 | 0 | 0 | 0.30993908 |
| ENSMUSG00 | Scarna17 | 78 | 0 | 0 | 0 | 0 | 0 | 0 |
| ENSMUSG00 | SNORA68 | 135 | 0 | 0 | 0 | 0 | 0 | 0 |
| ENSMUSG00 | U1 | 161 | 0 | 0 | 0 | 0 | 0 | 0 |
| ENSMUSG00 | SNORA17 | 114 | 0 | 0 | 0 | 0 | 0 | 0 |
| ENSMUSG00 | U6 | 103 | 0 | 0 | 0 | 0 | 0 | 0 |
| ENSMUSG00 | SNORA70 | 132 | 1.50588941 | 0.3650488 | 1.00458734 | 0.27334882 | 0.38656345 | 1.23975633 |
| ENSMUSG00 | U6 | 100 | 0 | 0 | 0 | 0 | 0 | 0 |
| ENSMUSG00 | SNORA17 | 125 | 0 | 0 | 0 | 0 | 0 | 0 |
| ENSMUSG00 | U7 | 64 | 0 | 0 | 0 | 0 | 0 | 0 |
| ENSMUSG00 | AC152987.1 | 111 | 0 | 0 | 0 | 1.30025386 | 0 | 0 |
| ENSMUSG00 | 5S_rRNA | 112 | 0 | 0 | 0 | 0 | 0 | 0 |
| ENSMUSG00 | 5S_rRNA | 78 | 0 | 0 | 0 | 0 | 0 | 0 |
| ENSMUSG00 | U1 | 143 | 0 | 0 | 0 | 0 | 0 | 0 |
| ENSMUSG00 | 5S_rRNA | 110 | 0 | 0 | 0 | 0 | 0 | 0 |
| ENSMUSG00 | SNORA17 | 127 | 0 | 0 | 0 | 0 | 0 | 0 |
| ENSMUSG00 | U6 | 93 | 0 | 0 | 0 | 0.77595795 | 0.54867071 | 0 |
| ENSMUSG00 | U6 | 90 | 0 | 0 | 0 | 0 | 0 | 0 |
| ENSMUSG00 | 5S_rRNA | 102 | 0 | 0 | 0 | 0 | 0 | 0 |
| ENSMUSG00 | U6 | 105 | 0 | 0 | 0 | 0.34363852 | 0 | 0 |
| ENSMUSG00 | 7SK | 314 | 0 | 0 | 0 | 0 | 0 | 0 |
| ENSMUSG00 | U1 | 170 | 0 | 0 | 0 | 0 | 0 | 0 |
| ENSMUSG00 | 5S_rRNA | 93 | 0 | 0 | 0 | 0 | 0 | 0 |
| ENSMUSG00 | SCARNA23 | 128 | 0 | 0 | 0 | 0.56378195 | 0 | 2.77008055 |
| ENSMUSG00 | SNORA19 | 128 | 0 | 0 | 0 | 0 | 0 | 0 |
| ENSMUSG00 | 7SK | 178 | 0 | 0 | 0 | 0 | 0 | 0 |
| ENSMUSG00 | SNORA17 | 115 | 0 | 0 | 0 | 0 | 0 | 0 |
| ENSMUSG00 | SNORA17 | 126 | 0 | 0 | 0 | 0 | 0 | 0 |
| ENSMUSG00 | SNORA70 | 109 | 0 | 0 | 0 | 0 | 0 | 0 |
| ENSMUSG00 | SNORA17 | 120 | 0 | 0 | 0 | 0 | 0 | 0 |
| ENSMUSG00 | U6 | 90 | 0 | 0 | 0 | 0 | 0 | 0 |
| ENSMUSG00 | U6 | 111 | 0 | 0 | 0 | 0 | 0 | 0 |
| ENSMUSG00 | U1 | 155 | 0 | 0 | 0.57034636 | 0.23278738 | 0 | 0 |
| ENSMUSG00 | U1 | 164 | 0 | 0 | 0 | 0 | 0 | 0 |
| ENSMUSG00 | U6 | 104 | 0 | 0 | 0 | 0 | 0 | 0 |
| ENSMUSG00 | SNORA48 | 143 | 0 | 0 | 0 | 0 | 0 | 0 |
| ENSMUSG00 | SNORA32 | 69 | 0 | 0 | 0 | 0 | 0 | 0 |
| ENSMUSG00 | U7 | 59 | 0 | 0 | 0 | 0 | 0 | 0 |
| ENSMUSG00 | SNORA17 | 129 | 0 | 0 | 0 | 0 | 0 | 0 |
| ENSMUSG00 | SNORA71 | 133 | 0 | 0 | 0 | 0 | 0 | 0 |
| ENSMUSG00 | 7SK | 312 | 0 | 0 | 0 | 0 | 0 | 0 |
| ENSMUSG00 | U1 | 159 | 0 | 0 | 0 | 0 | 0 | 0 |
| ENSMUSG00 | SNORA17 | 129 | 0 | 0 | 0 | 0 | 0 | 0 |
| ENSMUSG00 | snoU109 | 157 | 0 | 0 | 0 | 0 | 0 | 0 |
| ENSMUSG00 | U6 | 102 | 0 | 0 | 0 | 0 | 0 | 0 |
| ENSMUSG00 | 5S_rRNA | 89 | 0 | 0 | 0 | 0 | 0 | 0 |
| ENSMUSG00 | U7 | 62 | 0 | 0 | 0 | 0 | 0 | 0 |
| ENSMUSG00 | SNORA17 | 137 | 0 | 0 | 0 | 0 | 0 | 0 |
| ENSMUSG00 | SNORA17 | 136 | 0 | 0 | 0 | 0 | 0 | 0 |
| ENSMUSG00 | U3 | 88 | 0 | 0 | 0 | 0 | 0 | 0 |
| ENSMUSG00 | AC109196.1 | 75 | 0 | 0 | 0 | 0 | 0 | 0 |
| ENSMUSG00 | 7SK | 92 | 4.32124788 | 1.57129703 | 0.48045482 | 0 | 0 | 0 |
| ENSMUSG00 | SNORA17 | 126 | 0 | 0 | 0 | 0 | 0 | 0 |
| ENSMUSG00 | U3 | 89 | 0 | 0 | 0 | 0 | 0 | 0 |
| ENSMUSG00 | SNORA17 | 113 | 0 | 0 | 0 | 0 | 0 | 0 |
| ENSMUSG00 | 7SK | 270 | 0 | 0 | 0 | 0 | 0 | 0 |
| ENSMUSG00 | U6 | 86 | 0 | 0 | 0 | 0 | 0 | 0 |
| ENSMUSG00 | 7SK | 275 | 0 | 0 | 0 | 0 | 0 | 0 |
| ENSMUSG00 | U6 | 109 | 0 | 0 | 0 | 0 | 0 | 0 |
| ENSMUSG00 | SNORA17 | 122 | 0 | 0 | 0 | 0 | 0 | 0 |
| ENSMUSG00 | Mir1934 | 82 | 0 | 0 | 1.07809373 | 0 | 0 | 1.99570531 |
| ENSMUSG00 | SNORA61 | 133 | 0 | 0 | 0 | 0 | 0 | 0 |
| ENSMUSG00 | U6 | 103 | 0 | 0 | 0 | 0 | 0 | 0 |
| ENSMUSG00 | U7 | 59 | 0 | 0 | 0 | 0 | 0 | 0 |
| ENSMUSG00 | SNORA17 | 125 | 0 | 0 | 0 | 0 | 0 | 0 |
| ENSMUSG00 | snR78 | 68 | 0 | 0 | 0 | 0 | 0 | 0 |
| ENSMUSG00 | U6 | 105 | 0 | 0 | 0 | 0 | 0 | 0 |
| ENSMUSG00 | SNORA17 | 113 | 0 | 0 | 0 | 0 | 0 | 0 |
| ENSMUSG00 | U6 | 96 | 0 | 0 | 0 | 0 | 0 | 0 |
| ENSMUSG00 | U6 | 102 | 0 | 0 | 0 | 0 | 0 | 0 |
| ENSMUSG00 | U3 | 89 | 0 | 0 | 0 | 0 | 0 | 0 |
| ENSMUSG00 | U6 | 101 | 0 | 0.47709349 | 0 | 0 | 0 | 0 |
| ENSMUSG00 | SNORA17 | 124 | 0 | 0 | 0 | 0 | 0 | 0 |
| ENSMUSG00 | SNORA17 | 122 | 0 | 0 | 0 | 0 | 0 | 0 |
| ENSMUSG00 | 7SK | 274 | 0 | 0 | 0 | 0 | 0 | 0 |
| ENSMUSG00 | U6 | 99 | 0 | 0 | 0 | 0 | 0 | 0 |
| ENSMUSG00 | 7SK | 282 | 2.53758386 | 1.70873908 | 2.50790599 | 3.19876281 | 0.72377838 | 0.77374863 |
| ENSMUSG00 | SNORA17 | 122 | 0 | 0 | 0 | 0 | 0 | 0 |
| ENSMUSG00 | SNORA81 | 160 | 0 | 0 | 0 | 0 | 0 | 0 |
| ENSMUSG00 | 5S_rRNA | 107 | 0 | 0 | 0 | 0 | 0 | 0 |
| ENSMUSG00 | SNORA48 | 143 | 0 | 0 | 0 | 0 | 0 | 0 |
| ENSMUSG00 | 7SK | 310 | 3.46257411 | 0 | 0.28517318 | 0 | 0 | 1.75965415 |
| ENSMUSG00 | U6 | 105 | 0 | 0 | 0 | 0 | 0 | 0 |
| ENSMUSG00 | SCARNA4 | 143 | 0 | 1.68484063 | 0 | 0 | 0 | 0.57219523 |
| ENSMUSG00 | snoU13 | 108 | 0 | 0 | 0 | 0 | 0 | 0 |
| ENSMUSG00 | U6 | 115 | 0 | 0 | 0 | 0 | 0 | 0 |
| ENSMUSG00 | SNORA5 | 91 | 0 | 0 | 0 | 0 | 0 | 0 |
| ENSMUSG00 | SNORA17 | 114 | 0 | 0.84537618 | 0 | 0 | 0 | 0 |
| ENSMUSG00 | SNORA17 | 134 | 0 | 0 | 0 | 0 | 0 | 0 |
| ENSMUSG00 | AC134903.1 | 100 | 0 | 0 | 0 | 0 | 0 | 0 |
| ENSMUSG00 | SNORA55 | 93 | 0 | 0 | 0 | 0 | 0 | 0 |
| ENSMUSG00 | 7SK | 254 | 0 | 0 | 0 | 0 | 0 | 0 |
| ENSMUSG00 | U6 | 100 | 0 | 0 | 0 | 0 | 1.53079128 | 0 |
| ENSMUSG00 | Scarna13 | 274 | 2.46658091 | 0.35172586 | 2.25848833 | 0.79011776 | 2.23473179 | 1.99084958 |
| ENSMUSG00 | SNORA18 | 126 | 0 | 0 | 0 | 0 | 0 | 0 |
| ENSMUSG00 | U6 | 94 | 0 | 0 | 0 | 0 | 0 | 0 |
| ENSMUSG00 | SNORA17 | 116 | 0 | 0 | 0 | 0 | 0 | 0 |
| ENSMUSG00 | U6 | 116 | 0.34271966 | 0 | 0.38105037 | 0.31105211 | 0.43988255 | 0 |
| ENSMUSG00 | 7SK | 298 | 0 | 0 | 0 | 0 | 0 | 0 |
| ENSMUSG00 | Mir1935 | 59 | 0 | 0 | 0 | 0 | 0 | 0 |
| ENSMUSG00 | SNORA17 | 133 | 0 | 0 | 0 | 0 | 0 | 0 |
| ENSMUSG00 | 7SK | 198 | 0 | 0 | 0 | 0 | 0 | 0 |
| ENSMUSG00 | U6atac | 103 | 0 | 0 | 0 | 0 | 0 | 0 |
| ENSMUSG00 | 5S_rRNA | 118 | 0 | 0 | 0 | 0 | 0 | 0 |
| ENSMUSG00 | SNORA17 | 71 | 0 | 0 | 0 | 0 | 0 | 0 |
| ENSMUSG00 | U6 | 105 | 0 | 0 | 0 | 0 | 0 | 0 |
| ENSMUSG00 | 7SK | 259 | 0 | 0 | 0 | 0 | 0 | 0 |
| ENSMUSG00 | U6 | 93 | 0 | 0 | 0 | 0 | 0 | 0 |
| ENSMUSG00 | SNORA84 | 132 | 0 | 0 | 0 | 0 | 0 | 0 |
| ENSMUSG00 | SNORA17 | 135 | 0 | 0 | 0 | 0 | 0 | 0 |
| ENSMUSG00 | AL806518.1 | 102 | 0 | 0 | 0 | 0 | 0 | 0 |
| ENSMUSG00 | SNORA40 | 129 | 0 | 0 | 0 | 0 | 0 | 0 |
| ENSMUSG00 | U6 | 106 | 0 | 0 | 0 | 0 | 0 | 0 |
| ENSMUSG00 | U6 | 103 | 0.38597554 | 0 | 0 | 0 | 0 | 0 |
| ENSMUSG00 | 7SK | 326 | 0 | 0 | 0 | 0 | 0 | 0 |
| ENSMUSG00 | 5S_rRNA | 100 | 0 | 0 | 0 | 0 | 0 | 0 |
| ENSMUSG00 | SCARNA3 | 130 | 0 | 0 | 0 | 0 | 0 | 0 |
| ENSMUSG00 | SNORA17 | 128 | 0 | 0 | 0 | 0 | 0 | 0 |
| ENSMUSG00 | 7SK | 300 | 0 | 0 | 0 | 0 | 0 | 0 |
| ENSMUSG00 | 7SK | 247 | 0 | 0 | 0 | 0 | 0 | 0 |
| ENSMUSG00 | SNORD113 | 67 | 0 | 0 | 0 | 0 | 0 | 0 |
| ENSMUSG00 | U6 | 101 | 0 | 0 | 0 | 0 | 0 | 0 |
| ENSMUSG00 | SNORA70 | 122 | 0 | 0 | 0 | 0 | 0 | 0 |
| ENSMUSG00 | SNORA17 | 125 | 0 | 0 | 0 | 0 | 0 | 0 |
| ENSMUSG00 | Y_RNA | 109 | 0 | 0 | 0 | 1.32411173 | 0 | 0 |
| ENSMUSG00 | U3 | 89 | 0 | 0 | 0 | 0 | 0 | 0 |
| ENSMUSG00 | U7 | 62 | 0 | 0 | 0 | 0 | 0 | 0 |
| ENSMUSG00 | SNORA17 | 111 | 0 | 0 | 0 | 0 | 0 | 0 |
| ENSMUSG00 | U6 | 103 | 0 | 0 | 0 | 0 | 0 | 0 |
| ENSMUSG00 | U7 | 63 | 0 | 0 | 0 | 0 | 0 | 0 |
| ENSMUSG00 | U6 | 105 | 0 | 0 | 0 | 0 | 0 | 0 |
| ENSMUSG00 | AL772216.39 | 95 | 0 | 0 | 0 | 0 | 0 | 0 |
| ENSMUSG00 | snoU89 | 270 | 0.14724252 | 0 | 0 | 0.1336372 | 0 | 0.30305155 |
| ENSMUSG00 | SNORA48 | 136 | 0 | 0 | 0 | 0 | 0 | 0 |
| ENSMUSG00 | U6 | 106 | 0 | 0 | 0 | 0 | 0 | 0 |
| ENSMUSG00 | U7 | 57 | 0 | 0 | 0 | 0 | 0 | 0 |
| ENSMUSG00 | SNORA17 | 96 | 0 | 0 | 0 | 0 | 0 | 0 |
| ENSMUSG00 | 7SK | 282 | 0 | 0 | 0 | 0 | 0 | 0 |
| ENSMUSG00 | U6atac | 114 | 0 | 0.42268809 | 0.38773547 | 0 | 0 | 0 |
| ENSMUSG00 | U6 | 104 | 0.38226424 | 0 | 0 | 0 | 2.45319115 | 1.04902459 |
| ENSMUSG00 | SNORA17 | 128 | 0 | 0 | 0 | 0 | 0 | 0 |
| ENSMUSG00 | SNORA17 | 124 | 0 | 0 | 0 | 0 | 0 | 0 |
| ENSMUSG00 | 7SK | 254 | 0.15651764 | 0 | 0.34804601 | 0.14205529 | 1.00445622 | 0 |
| ENSMUSG00 | SNORA26 | 136 | 0 | 0 | 0 | 0 | 0 | 0 |
| ENSMUSG00 | SNORA4 | 124 | 0 | 0 | 0 | 0 | 0 | 0 |
| ENSMUSG00 | SNORA17 | 130 | 0 | 0 | 0 | 0 | 0 | 0 |
| ENSMUSG00 | SNORA17 | 116 | 0 | 0 | 0 | 0 | 0 | 0 |
| ENSMUSG00 | U1 | 124 | 0 | 0 | 0 | 0 | 0 | 0 |
| ENSMUSG00 | U6 | 106 | 0 | 0 | 0 | 0 | 0 | 0 |
| ENSMUSG00 | SNORA48 | 143 | 0 | 0 | 0 | 0 | 0 | 0 |
| ENSMUSG00 | SNORA40 | 120 | 0 | 0 | 0 | 0 | 0 | 0 |
| ENSMUSG00 | SNORA17 | 131 | 0 | 0 | 0 | 0 | 0 | 0 |
| ENSMUSG00 | Mir1947 | 84 | 0 | 0 | 0 | 0 | 0 | 0 |
| ENSMUSG00 | 5S_rRNA | 129 | 0 | 0 | 0 | 0 | 0 | 0 |
| ENSMUSG00 | U6 | 70 | 0 | 0 | 0 | 0 | 0 | 0 |
| ENSMUSG00 | U7 | 61 | 0 | 0 | 0 | 0 | 0 | 0 |
| ENSMUSG00 | 7SK | 292 | 0 | 0 | 0 | 0 | 0 | 0 |
| ENSMUSG00 | 7SK | 310 | 0 | 0 | 0 | 0 | 0 | 0 |
| ENSMUSG00 | SNORD19 | 59 | 0 | 0 | 0 | 0 | 0 | 0 |
| ENSMUSG00 | 5S_rRNA | 114 | 0 | 0 | 0 | 0 | 0 | 0 |
| ENSMUSG00 | SNORA17 | 122 | 0 | 0 | 0 | 0 | 0 | 0 |
| ENSMUSG00 | snoU13 | 99 | 0 | 0 | 0 | 0 | 0 | 0 |
| ENSMUSG00 | SNORA17 | 111 | 0 | 0 | 0 | 0 | 0 | 0 |
| ENSMUSG00 | AC154485.1 | 103 | 0 | 0 | 0 | 0 | 0 | 0 |
| ENSMUSG00 | U6 | 91 | 0 | 0 | 0 | 0 | 0 | 0 |
| ENSMUSG00 | RNaseP_nuc | 328 | 0 | 0 | 0 | 0 | 0 | 0 |
| ENSMUSG00 | SNORA70 | 128 | 0 | 0.75291316 | 0.3453269 | 2.81890973 | 1.19593068 | 1.0654156 |
| ENSMUSG00 | SNORA70 | 108 | 0 | 0 | 0 | 0 | 0 | 0 |
| ENSMUSG00 | SNORA17 | 125 | 0 | 0 | 0 | 0 | 0 | 0 |
| ENSMUSG00 | SNORA17 | 113 | 0 | 0 | 0 | 0 | 0 | 0 |
| ENSMUSG00 | U6 | 102 | 0 | 0 | 0 | 0 | 0 | 0 |
| ENSMUSG00 | 7SK | 257 | 0 | 0 | 0 | 0 | 0 | 0 |
| ENSMUSG00 | 7SK | 202 | 0 | 0 | 0 | 0 | 0 | 0 |
| ENSMUSG00 | SNORA17 | 131 | 0 | 0 | 0 | 0 | 0 | 0 |
| ENSMUSG00 | SNORA17 | 116 | 0 | 0 | 0 | 0 | 0 | 0 |
| ENSMUSG00 | 5S_rRNA | 132 | 0 | 0 | 0 | 0 | 0 | 0 |
| ENSMUSG00 | U7 | 57 | 0 | 0 | 0 | 0 | 0 | 0 |
| ENSMUSG00 | SNORA17 | 132 | 0 | 0 | 0 | 0 | 0 | 0 |
| ENSMUSG00 | U6 | 106 | 0 | 0 | 0 | 0 | 0 | 0 |
| ENSMUSG00 | SNORA17 | 135 | 0 | 0 | 0 | 0 | 0 | 0 |
| ENSMUSG00 | SNORA76 | 123 | 0 | 0 | 1.07809373 | 0 | 0.41484858 | 0 |
| ENSMUSG00 | AL772376.1 | 67 | 0 | 0 | 0 | 0 | 0 | 0 |
| ENSMUSG00 | SNORA44 | 128 | 0 | 4.89393553 | 0.6906538 | 1.40945486 | 0 | 0 |
| ENSMUSG00 | SNORA81 | 177 | 0 | 0 | 0 | 0 | 0 | 0 |
| ENSMUSG00 | SNORA17 | 150 | 0 | 0 | 0 | 0 | 0 | 0 |
| ENSMUSG00 | SNORA17 | 133 | 0 | 0 | 0 | 0 | 0 | 0 |
| ENSMUSG00 | 7SK | 304 | 0 | 0 | 0 | 0 | 0 | 0 |
| ENSMUSG00 | SNORA17 | 122 | 0 | 0 | 0 | 0 | 0 | 0 |
| ENSMUSG00 | SNORA17 | 144 | 0 | 0 | 0 | 0 | 0 | 0 |
| ENSMUSG00 | Mir1943 | 72 | 0 | 2.67702456 | 0 | 0 | 0 | 0 |
| ENSMUSG00 | 7SK | 266 | 0 | 0 | 0.16617234 | 1.22082106 | 0 | 0 |
| ENSMUSG00 | SCARNA16 | 166 | 0 | 0 | 0 | 0 | 0 | 0 |
| ENSMUSG00 | U1 | 153 | 0 | 0 | 0 | 0 | 0 | 0 |
| ENSMUSG00 | SNORA17 | 130 | 0 | 0 | 0 | 0 | 0 | 0 |
| ENSMUSG00 | SNORA17 | 109 | 0 | 0 | 0 | 0 | 0 | 0 |
| ENSMUSG00 | SNORA17 | 131 | 0 | 0 | 0 | 0 | 0 | 0 |
| ENSMUSG00 | 7SK | 285 | 0 | 0 | 0 | 0 | 0 | 0 |
| ENSMUSG00 | SNORD86 | 65 | 0 | 0 | 0 | 0 | 0 | 0 |
| ENSMUSG00 | U6 | 102 | 0 | 0 | 0 | 0 | 0 | 0 |
| ENSMUSG00 | SNORA17 | 120 | 0 | 0 | 0 | 0 | 0 | 0 |
| ENSMUSG00 | AC107678.1 | 59 | 0 | 0 | 0 | 0 | 0 | 0 |
| ENSMUSG00 | U1 | 149 | 0 | 0 | 0 | 0 | 0 | 0 |
| ENSMUSG00 | U6 | 107 | 0 | 0 | 0 | 0 | 0 | 0 |
| ENSMUSG00 | U6 | 102 | 0 | 0 | 0 | 0 | 0 | 0 |
| ENSMUSG00 | U6 | 104 | 0 | 0 | 0 | 0.69388547 | 0 | 0 |
| ENSMUSG00 | SNORA17 | 131 | 0 | 0 | 0 | 0 | 0 | 0 |
| ENSMUSG00 | U6 | 100 | 0 | 0 | 0 | 0 | 0 | 0 |
| ENSMUSG00 | 7SK | 276 | 0 | 0 | 0 | 0 | 0 | 0 |

| | | | | | | | | |
|---|---|---|---|---|---|---|---|---|
| ENSMUSG00 | snoU13 | 87 | 0 | 0 | 0 | 0 | 0 | 0 |
| ENSMUSG00 | U6 | 105 | 0 | 0 | 0 | 0 | 0 | 0 |
| ENSMUSG00 | SNORA17 | 130 | 0 | 0 | 0 | 0 | 0 | 0 |
| ENSMUSG00 | SNORA17 | 59 | 0 | 0 | 0 | 0 | 0 | 0 |
| ENSMUSG00 | SNORD70 | 67 | 0 | 0 | 0 | 0 | 0 | 0 |
| ENSMUSG00 | 5S_rRNA | 104 | 0 | 0 | 0 | 0 | 0 | 0 |
| ENSMUSG00 | U6 | 105 | 0 | 0 | 0 | 0 | 0 | 0 |
| ENSMUSG00 | SNORD5 | 70 | 5.67935436 | 5.50702196 | 1.8943647 | 0 | 5.83158581 | 5.06529015 |
| ENSMUSG00 | SNORA70 | 136 | 0 | 0 | 0 | 0 | 0 | 0 |
| ENSMUSG00 | U6 | 98 | 0 | 0 | 0 | 0 | 0 | 0 |
| ENSMUSG00 | U6 | 103 | 0 | 0 | 0 | 0 | 0 | 0 |
| ENSMUSG00 | U3 | 89 | 0 | 0 | 0 | 0 | 0 | 0 |
| ENSMUSG00 | U6 | 102 | 0 | 0 | 0 | 0 | 0 | 0 |
| ENSMUSG00 | 5S_rRNA | 137 | 0 | 0 | 0 | 0 | 0 | 0 |
| ENSMUSG00 | U6 | 85 | 0 | 0 | 0 | 0 | 0 | 0 |
| ENSMUSG00 | 7SK | 305 | 0 | 0 | 0 | 0 | 0 | 0 |
| ENSMUSG00 | 7SK | 82 | 0 | 0 | 0 | 0 | 0 | 0 |
| ENSMUSG00 | SNORA17 | 135 | 0 | 0 | 0 | 0 | 0 | 0 |
| ENSMUSG00 | U2 | 171 | 0.92995276 | 5.35404913 | 3.61886434 | 1.26603665 | 1.19359943 | 2.2330114 |
| ENSMUSG00 | 7SK | 302 | 0 | 0 | 0 | 0 | 0 | 0 |
| ENSMUSG00 | U6 | 105 | 0 | 0 | 0 | 0 | 0 | 0 |
| ENSMUSG00 | U3 | 88 | 0 | 0 | 0 | 0 | 0 | 0 |
| ENSMUSG00 | U6 | 106 | 0 | 0 | 0 | 0 | 0 | 0 |
| ENSMUSG00 | 5S_rRNA | 131 | 0 | 0 | 0 | 0 | 0 | 0 |
| ENSMUSG00 | snoU97 | 167 | 3.094738 | 0.28854157 | 0.2646817 | 0.43212029 | 0.30554716 | 0.48996358 |
| ENSMUSG00 | 5S_rRNA | 106 | 0 | 0 | 0 | 0 | 0 | 0 |
| ENSMUSG00 | 5S_rRNA | 136 | 0 | 0 | 0 | 0 | 0 | 0 |
| ENSMUSG00 | U6atac | 107 | 0 | 0 | 0 | 0 | 0 | 0 |
| ENSMUSG00 | snoU13 | 101 | 0 | 1.90837395 | 2.18821005 | 1.42899186 | 1.01042328 | 1.35022967 |
| ENSMUSG00 | SNORA17 | 113 | 0 | 0 | 0 | 0 | 0 | 0 |
| ENSMUSG00 | RNaseP_nuc | 432 | 0 | 0 | 0 | 0 | 0 | 0 |
| ENSMUSG00 | SNORA61 | 127 | 0 | 0 | 0 | 0 | 0 | 0 |
| ENSMUSG00 | SNORA25 | 125 | 0 | 0 | 0 | 0 | 0 | 0 |
| ENSMUSG00 | U6 | 105 | 0 | 0 | 0 | 0 | 0 | 0 |
| ENSMUSG00 | Scarna8 | 130 | 0 | 0 | 0 | 0.27755419 | 0 | 0 |
| ENSMUSG00 | U6 | 109 | 0 | 0 | 0 | 0 | 0 | 0 |
| ENSMUSG00 | U6 | 110 | 0 | 0 | 0 | 0 | 0 | 0 |
| ENSMUSG00 | U6 | 101 | 0 | 0 | 0 | 0 | 0 | 0 |
| ENSMUSG00 | U6 | 104 | 0 | 0 | 0 | 0 | 0 | 0 |
| ENSMUSG00 | U6 | 87 | 0 | 0 | 0 | 0 | 0 | 0 |
| ENSMUSG00 | snoU13 | 90 | 0 | 0 | 0 | 0 | 0 | 0 |
| ENSMUSG00 | SNORA25 | 116 | 0 | 0 | 0 | 0 | 0 | 0 |
| ENSMUSG00 | U6 | 103 | 0 | 0 | 0 | 0 | 0 | 0 |
| ENSMUSG00 | U6 | 109 | 0 | 0 | 0 | 0 | 0 | 0 |
| ENSMUSG00 | U7 | 59 | 0 | 0 | 0 | 0 | 0 | 0 |
| ENSMUSG00 | 5S_rRNA | 47 | 0 | 0 | 0 | 0 | 0 | 0 |
| ENSMUSG00 | U6 | 107 | 0 | 0 | 0 | 0 | 0 | 0 |
| ENSMUSG00 | AL683804.1 | 105 | 0 | 0 | 0 | 0 | 0 | 0 |
| ENSMUSG00 | U6 | 103 | 0 | 0 | 0 | 0 | 0 | 0 |
| ENSMUSG00 | SNORA17 | 115 | 0 | 0 | 0 | 0 | 0 | 0 |
| ENSMUSG00 | 7SK | 267 | 0 | 0 | 0 | 0 | 0 | 0 |
| ENSMUSG00 | 5S_rRNA | 99 | 0 | 0 | 0 | 0 | 0 | 0 |
| ENSMUSG00 | SNORA72 | 89 | 0 | 0 | 0 | 0 | 0 | 0 |
| ENSMUSG00 | Mir669m-2 | 97 | 0 | 0 | 0 | 0 | 0 | 0 |
| ENSMUSG00 | U6 | 104 | 0 | 0 | 0 | 0 | 0 | 0 |
| ENSMUSG00 | U6 | 105 | 0.37862362 | 0.4589185 | 0 | 1.7181926 | 0.97193097 | 0 |
| ENSMUSG00 | Mir1952 | 78 | 0 | 0 | 0 | 0 | 0 | 0 |
| ENSMUSG00 | SNORA17 | 132 | 0 | 0 | 0 | 0 | 1.93281727 | 0 |
| ENSMUSG00 | Mir1946a | 133 | 0 | 0 | 0 | 0 | 0 | 0 |
| ENSMUSG00 | AC113291.1 | 104 | 0 | 0 | 0 | 0 | 0 | 0 |
| ENSMUSG00 | U7 | 61 | 0 | 0 | 0 | 0 | 0 | 0 |
| ENSMUSG00 | 5S_rRNA | 94 | 0 | 0 | 0 | 0 | 0 | 0 |
| ENSMUSG00 | U1 | 123 | 0 | 0 | 0 | 0 | 0 | 0 |
| ENSMUSG00 | SCARNA1 | 164 | 0.48482293 | 0 | 0 | 0 | 0 | 0 |
| ENSMUSG00 | U2 | 152 | 0 | 0 | 0 | 0 | 0 | 0 |
| ENSMUSG00 | U6 | 97 | 0 | 0 | 0 | 0 | 0 | 0 |
| ENSMUSG00 | U6 | 66 | 1.20471153 | 0.73009761 | 0 | 0 | 0 | 0 |
| ENSMUSG00 | 7SK | 283 | 0 | 0 | 0 | 0 | 0 | 0 |
| ENSMUSG00 | U6 | 91 | 0 | 0 | 0 | 0 | 0 | 0 |
| ENSMUSG00 | SNORA25 | 125 | 0 | 0 | 0 | 0 | 0 | 0 |
| ENSMUSG00 | U6 | 98 | 0 | 0 | 0 | 0.36818413 | 0 | 0 |
| ENSMUSG00 | SNORA17 | 106 | 0 | 0 | 0 | 0 | 0 | 0 |
| ENSMUSG00 | U6 | 103 | 0 | 0 | 0 | 0 | 0 | 0 |
| ENSMUSG00 | SNORD19B | 76 | 0 | 0 | 0 | 0 | 0 | 0 |
| ENSMUSG00 | SNORA70 | 134 | 0 | 0 | 0 | 0 | 0 | 0 |
| ENSMUSG00 | U6 | 107 | 0 | 0 | 0 | 0 | 0 | 0 |
| ENSMUSG00 | 5S_rRNA | 110 | 0 | 0 | 0 | 0 | 0 | 0 |
| ENSMUSG00 | U6 | 98 | 0 | 0 | 0 | 0 | 0 | 0 |
| ENSMUSG00 | U3 | 89 | 0 | 0 | 0 | 0 | 0 | 0 |
| ENSMUSG00 | SNORA68 | 128 | 0 | 0 | 0 | 0 | 0 | 0 |
| ENSMUSG00 | U7 | 63 | 0 | 0 | 0 | 0 | 0 | 0 |
| ENSMUSG00 | U6 | 105 | 0 | 0 | 0 | 0 | 0 | 0 |
| ENSMUSG00 | SNORA48 | 142 | 0 | 0 | 0 | 0.50819781 | 0.71868135 | 0.38414985 |
| ENSMUSG00 | AL591146.1 | 77 | 0 | 0 | 0 | 0 | 0 | 0 |
| ENSMUSG00 | SNORA36 | 117 | 0 | 0 | 0 | 0 | 0 | 0 |
| ENSMUSG00 | U6 | 98 | 7.7076952 | 4.4252855 | 8.11870586 | 7.73186668 | 8.85151417 | 9.46262994 |
| ENSMUSG00 | 7SK | 280 | 0 | 0 | 0 | 0 | 0 | 0 |
| ENSMUSG00 | SNORA17 | 126 | 0 | 0 | 0 | 0 | 0 | 0 |
| ENSMUSG00 | U6 | 107 | 0 | 0 | 0 | 0 | 0 | 0 |
| ENSMUSG00 | U6 | 104 | 0 | 0 | 0 | 0 | 0 | 0 |
| ENSMUSG00 | U1 | 163 | 0 | 0 | 0 | 0 | 0 | 0 |
| ENSMUSG00 | SNORA31 | 125 | 0 | 0 | 0 | 0 | 0 | 0 |
| ENSMUSG00 | SNORA17 | 130 | 0 | 0 | 0 | 0 | 0 | 0 |
| ENSMUSG00 | SNORA17 | 127 | 0 | 0 | 0 | 0 | 0 | 0 |
| ENSMUSG00 | U6 | 99 | 0 | 0 | 3.12538284 | 0.72893019 | 0 | 0 |
| ENSMUSG00 | SNORA17 | 127 | 0 | 0 | 0 | 0 | 0 | 0 |
| ENSMUSG00 | U7 | 64 | 0 | 0 | 0 | 0 | 1.59457424 | 0 |
| ENSMUSG00 | SNORA17 | 101 | 0 | 0 | 0 | 0 | 0 | 0 |
| ENSMUSG00 | 7SK | 327 | 0 | 0 | 0 | 0 | 0 | 0 |
| ENSMUSG00 | U7 | 58 | 0 | 0 | 0 | 0 | 0 | 0 |
| ENSMUSG00 | U7 | 47 | 0 | 0 | 0 | 0 | 0 | 0 |
| ENSMUSG00 | SNORA70 | 134 | 0 | 0 | 0 | 0 | 0 | 0 |
| ENSMUSG00 | ACA64 | 140 | 0 | 0 | 0 | 0 | 0 | 0 |
| ENSMUSG00 | U6 | 103 | 0 | 0 | 0 | 0 | 0 | 0 |
| ENSMUSG00 | SNORA17 | 125 | 0 | 0 | 0 | 0 | 0 | 0 |
| ENSMUSG00 | SNORA17 | 129 | 0 | 0 | 0 | 0 | 0 | 0 |
| ENSMUSG00 | 5S_rRNA | 105 | 0 | 0 | 0 | 0 | 0 | 0 |
| ENSMUSG00 | U6 | 102 | 0 | 0 | 0 | 0 | 0 | 0 |
| ENSMUSG00 | RNaseP_nuc | 218 | 0 | 0 | 0 | 0 | 0 | 0 |
| ENSMUSG00 | SNORA61 | 135 | 0 | 0 | 0 | 0 | 0 | 0 |
| ENSMUSG00 | 7SK | 220 | 0 | 0 | 0 | 0 | 0 | 0 |
| ENSMUSG00 | U1 | 147 | 0 | 0 | 0 | 0 | 0 | 0 |
| ENSMUSG00 | SNORA70 | 134 | 0 | 0 | 0 | 0 | 0 | 0 |
| ENSMUSG00 | SNORA17 | 133 | 0 | 0 | 0 | 0 | 0 | 0 |
| ENSMUSG00 | 5S_rRNA | 130 | 0 | 0 | 0 | 0 | 0 | 0 |
| ENSMUSG00 | 7SK | 252 | 0.63103937 | 0.19121604 | 0.35080828 | 0 | 0.20248562 | 0 |
| ENSMUSG00 | 7SK | 287 | 0 | 0 | 0 | 0 | 0 | 0 |
| ENSMUSG00 | SNORA25 | 71 | 0 | 0 | 0 | 0 | 0 | 0 |
| ENSMUSG00 | U6 | 103 | 0 | 0 | 0 | 0 | 0 | 0 |
| ENSMUSG00 | SNORA40 | 124 | 0 | 0 | 0 | 0 | 0 | 0 |
| ENSMUSG00 | U7 | 63 | 0 | 0 | 0 | 0 | 0 | 0 |
| ENSMUSG00 | U6 | 102 | 0 | 0 | 0 | 0 | 0 | 0 |
| ENSMUSG00 | U6 | 103 | 0 | 0 | 0 | 0 | 0 | 0 |
| ENSMUSG00 | U7 | 61 | 0 | 0 | 0 | 0 | 0 | 0 |
| ENSMUSG00 | U6 | 97 | 0 | 0 | 0 | 0 | 0 | 0 |
| ENSMUSG00 | U3 | 200 | 0 | 0 | 0 | 0 | 0 | 0 |
| ENSMUSG00 | U1 | 175 | 0 | 0 | 0 | 0 | 0 | 0 |
| ENSMUSG00 | 7SK | 303 | 0 | 0 | 0 | 0 | 0 | 0 |
| ENSMUSG00 | U6 | 98 | 0 | 0 | 0 | 0 | 5.20677304 | 0 |
| ENSMUSG00 | 7SK | 331 | 0 | 0 | 0 | 0 | 0 | 0 |
| ENSMUSG00 | 7SK | 310 | 0 | 0 | 0 | 0 | 0 | 0 |
| ENSMUSG00 | U7 | 60 | 0 | 0 | 0 | 0 | 0 | 0 |
| ENSMUSG00 | AC099599.1 | 117 | 0 | 0 | 0 | 0 | 0 | 0 |
| ENSMUSG00 | U7 | 61 | 0 | 0 | 0 | 0 | 0 | 0 |
| ENSMUSG00 | SNORA44 | 135 | 0 | 2.85549287 | 0.65484212 | 1.33637202 | 0 | 1.21220619 |
| ENSMUSG00 | U6 | 101 | 0 | 0 | 0 | 0.35724797 | 0 | 0 |
| ENSMUSG00 | U6 | 103 | 0 | 0 | 0 | 0 | 0 | 0 |
| ENSMUSG00 | SNORA4 | 123 | 2.26250702 | 0.39175969 | 1.07809373 | 0 | 2.4890915 | 0 |
| ENSMUSG00 | 5S_rRNA | 102 | 0 | 0 | 0 | 0 | 0 | 0 |
| ENSMUSG00 | U6 | 103 | 0 | 0 | 0 | 0 | 0 | 0 |
| ENSMUSG00 | 7SK | 299 | 0 | 0 | 0 | 0 | 0 | 0 |
| ENSMUSG00 | U3 | 89 | 0 | 0 | 0 | 0 | 0 | 0 |
| ENSMUSG00 | U7 | 61 | 0 | 0 | 0 | 0 | 0 | 0 |
| ENSMUSG00 | U6 | 104 | 0 | 0 | 0 | 0 | 0 | 0 |
| ENSMUSG00 | SNORA17 | 134 | 0 | 0 | 0 | 0 | 0 | 0 |
| ENSMUSG00 | U6 | 99 | 0 | 0 | 0 | 0 | 0 | 0 |
| ENSMUSG00 | U6 | 103 | 0 | 0 | 0 | 0 | 0 | 0 |
| ENSMUSG00 | 7SK | 275 | 0 | 0 | 0 | 0 | 0 | 0 |
| ENSMUSG00 | U7 | 61 | 0 | 0 | 0 | 0 | 0 | 0 |
| ENSMUSG00 | 7SK | 295 | 0 | 0 | 0 | 0 | 0 | 0 |
| ENSMUSG00 | U1 | 177 | 0.89842894 | 0.81671936 | 2.49727927 | 1.01926679 | 0.57656922 | 4.62282021 |
| ENSMUSG00 | U7 | 64 | 0 | 0 | 0 | 0 | 0 | 0 |
| ENSMUSG00 | SNORA17 | 131 | 0 | 0 | 0 | 0 | 0 | 0 |
| ENSMUSG00 | AC112941.1 | 84 | 0 | 0 | 0 | 0 | 0 | 0 |
| ENSMUSG00 | 5S_rRNA | 127 | 0 | 0 | 0 | 0 | 0 | 0 |
| ENSMUSG00 | 7SK | 289 | 0 | 0 | 0 | 0 | 0 | 0 |
| ENSMUSG00 | U7 | 65 | 0 | 0 | 0 | 0 | 0 | 0 |
| ENSMUSG00 | U6 | 87 | 0.91391909 | 3.32320291 | 1.01613432 | 0.41473614 | 4.10557047 | 0.6270032 |
| ENSMUSG00 | Snord11 | 85 | 0 | 0 | 0 | 0 | 0 | 0.32087811 |
| ENSMUSG00 | 5S_rRNA | 36 | 0 | 0 | 0 | 0 | 0 | 0 |
| ENSMUSG00 | 5S_rRNA | 103 | 0 | 0 | 0 | 0 | 0 | 0 |
| ENSMUSG00 | U6 | 94 | 0 | 0 | 0 | 0 | 0 | 0 |
| ENSMUSG00 | SNORA17 | 127 | 0 | 0 | 0 | 0 | 0 | 0 |
| ENSMUSG00 | SNORA17 | 137 | 0 | 0 | 0 | 0 | 0 | 0 |
| ENSMUSG00 | SNORA5 | 91 | 0 | 0 | 0 | 0 | 0 | 0 |
| ENSMUSG00 | U6 | 98 | 0 | 0 | 0 | 0 | 0.5206773 | 0 |
| ENSMUSG00 | 5S_rRNA | 116 | 1.37087864 | 5.40020472 | 8.38310816 | 3.42157319 | 3.5190604 | 3.52689301 |
| ENSMUSG00 | 5S_rRNA | 107 | 0 | 0 | 0 | 0 | 0 | 0 |
| ENSMUSG00 | U6 | 66 | 0 | 0 | 0 | 0 | 0 | 0 |
| ENSMUSG00 | U6 | 107 | 0 | 0 | 0 | 0 | 0 | 0 |
| ENSMUSG00 | U7 | 63 | 0 | 0 | 0 | 0 | 0 | 0 |
| ENSMUSG00 | RNaseP_nuc | 398 | 0 | 0 | 0 | 0 | 0 | 0 |
| ENSMUSG00 | SNORA74 | 124 | 0 | 0 | 0 | 0 | 0 | 0 |
| ENSMUSG00 | RNaseP_nuc | 301 | 0 | 0 | 0 | 0 | 0 | 0 |
| ENSMUSG00 | SNORA17 | 125 | 0 | 0 | 0 | 0 | 0 | 0 |
| ENSMUSG00 | U6 | 100 | 0 | 0 | 0 | 0 | 0 | 0 |
| ENSMUSG00 | SNORA17 | 142 | 0 | 0 | 0 | 0 | 0 | 0 |
| ENSMUSG00 | U6 | 102 | 0 | 0 | 0 | 0 | 0 | 0 |
| ENSMUSG00 | 7SK | 317 | 0.50164644 | 0 | 0 | 0.11382348 | 0 | 0 |
| ENSMUSG00 | SNORA17 | 129 | 0 | 0 | 0 | 0 | 0 | 0 |
| ENSMUSG00 | 5S_rRNA | 99 | 0 | 0 | 0 | 0 | 0 | 0 |
| ENSMUSG00 | SNORA17 | 108 | 0 | 0 | 0 | 0 | 0 | 0 |
| ENSMUSG00 | U6 | 112 | 0 | 0 | 0 | 0 | 0 | 0 |
| ENSMUSG00 | U6 | 104 | 0 | 0 | 0 | 0 | 0.98127646 | 1.57353688 |
| ENSMUSG00 | SNORA40 | 75 | 0 | 0 | 0 | 0 | 0 | 0 |
| ENSMUSG00 | SNORA17 | 123 | 0 | 0 | 0 | 0 | 0 | 0 |
| ENSMUSG00 | 7SK | 306 | 0 | 0 | 0 | 0 | 0 | 0.62392966 |
| ENSMUSG00 | SNORA17 | 91 | 0 | 0 | 0 | 0 | 0 | 0 |
| ENSMUSG00 | SNORA18 | 128 | 0 | 0 | 0 | 0 | 0 | 0 |
| ENSMUSG00 | 5S_rRNA | 109 | 0 | 0 | 0 | 0 | 0 | 0 |
| ENSMUSG00 | SNORA17 | 136 | 0 | 0 | 0 | 0 | 0 | 0 |
| ENSMUSG00 | AC147047.1 | 94 | 0 | 0 | 0 | 0 | 0 | 0 |
| ENSMUSG00 | U6atac | 101 | 0 | 0 | 0 | 0 | 0 | 0 |
| ENSMUSG00 | U6 | 93 | 0 | 0 | 0 | 0 | 0 | 0 |
| ENSMUSG00 | U6 | 98 | 0 | 0 | 0 | 0 | 0 | 0 |
| ENSMUSG00 | Mir1964 | 82 | 0 | 0.58763954 | 0 | 0.88004987 | 0.62227288 | 0 |
| ENSMUSG00 | 7SK | 236 | 0 | 0 | 0 | 0 | 0 | 0 |
| ENSMUSG00 | U7 | 65 | 2.44649111 | 0 | 0 | 0.55510838 | 0 | 0 |
| ENSMUSG00 | U6 | 100 | 0 | 0 | 0 | 0 | 0 | 0 |
| ENSMUSG00 | SNORA17 | 116 | 0 | 0 | 0 | 0 | 0 | 0 |
| ENSMUSG00 | 7SK | 346 | 0 | 0 | 0 | 0 | 0 | 0 |
| ENSMUSG00 | 7SK | 303 | 0 | 0 | 0 | 0 | 0 | 0 |
| ENSMUSG00 | U7 | 62 | 0 | 0 | 0 | 0 | 0 | 0 |
| ENSMUSG00 | 7SK | 264 | 0 | 0 | 0 | 0 | 0 | 0 |
| ENSMUSG00 | 7SK | 311 | 0 | 0 | 0 | 0 | 0 | 0 |
| ENSMUSG00 | U6atac | 94 | 0 | 0 | 0 | 0 | 0 | 0 |
| ENSMUSG00 | Mir1950 | 73 | 0 | 0 | 0 | 0 | 2.09697435 | 0 |
| ENSMUSG00 | SNORA17 | 86 | 0 | 0 | 0 | 0 | 0 | 0 |
| ENSMUSG00 | U6 | 102 | 0 | 0 | 0 | 0 | 0 | 0 |
| ENSMUSG00 | SNORA17 | 138 | 0 | 0 | 0 | 0 | 0 | 0 |
| ENSMUSG00 | U7 | 62 | 0 | 0 | 0 | 0 | 0 | 0 |
| ENSMUSG00 | SNORA17 | 137 | 0 | 0 | 0 | 0 | 0 | 0 |
| ENSMUSG00 | 7SK | 288 | 0 | 0 | 0 | 0 | 0 | 0 |
| ENSMUSG00 | SNORA48 | 150 | 0 | 0 | 0 | 0 | 0 | 0 |
| ENSMUSG00 | U6 | 96 | 0 | 0 | 0 | 0 | 0 | 0 |
| ENSMUSG00 | U6 | 106 | 0 | 0 | 0 | 0 | 0 | 0 |
| ENSMUSG00 | SNORA17 | 131 | 0 | 0 | 0 | 0 | 0 | 0 |
| ENSMUSG00 | U6 | 98 | 0 | 0 | 0 | 0 | 0 | 0 |
| ENSMUSG00 | 7SK | 311 | 0 | 0 | 0 | 0 | 0.16407195 | 0 |
| ENSMUSG00 | Y_RNA | 106 | 0 | 0 | 0 | 0.68079329 | 0 | 0 |
| ENSMUSG00 | SNORA17 | 106 | 0 | 0 | 0 | 0 | 0 | 0 |
| ENSMUSG00 | snoU13 | 104 | 0 | 0.46333117 | 1.70007089 | 0.34694274 | 0 | 0 |
| ENSMUSG00 | SNORA40 | 121 | 0 | 0 | 0 | 0 | 0 | 0 |
| ENSMUSG00 | SCARNA14 | 141 | 0 | 0 | 0 | 0 | 0 | 0 |
| ENSMUSG00 | snoU13 | 101 | 0 | 0 | 0 | 0 | 0 | 0 |
| ENSMUSG00 | SNORA25 | 127 | 0 | 0.3794208 | 0 | 2.27288469 | 0.80356497 | 0 |
| ENSMUSG00 | SNORA17 | 129 | 0 | 0 | 0 | 0 | 0 | 0 |
| ENSMUSG00 | SNORA17 | 131 | 0 | 0 | 0 | 0 | 0 | 0 |
| ENSMUSG00 | U3 | 89 | 0 | 0 | 0 | 0 | 0 | 0 |
| ENSMUSG00 | U1 | 156 | 0 | 0 | 0 | 0 | 0 | 0 |
| ENSMUSG00 | U2 | 81 | 0 | 0 | 0 | 0 | 0 | 0 |
| ENSMUSG00 | SNORA70 | 90 | 0 | 0 | 0 | 0 | 0 | 0 |
| ENSMUSG00 | U1 | 154 | 0 | 0 | 0 | 0 | 0 | 0 |
| ENSMUSG00 | 7SK | 242 | 0 | 0.19911753 | 0.18265224 | 0 | 0 | 0 |
| ENSMUSG00 | AC154701.1 | 104 | 0 | 0 | 0 | 0 | 0 | 0 |
| ENSMUSG00 | SNORA70 | 123 | 0 | 0 | 0 | 0 | 0 | 0 |
| ENSMUSG00 | U6 | 75 | 0 | 0 | 0 | 0 | 0 | 0 |
| ENSMUSG00 | SNORA48 | 143 | 0 | 0 | 0 | 0 | 0 | 0 |
| ENSMUSG00 | SNORA17 | 114 | 0 | 0 | 0 | 0 | 0 | 0 |
| ENSMUSG00 | SNORA17 | 126 | 0 | 0 | 0 | 0 | 0 | 0 |
| ENSMUSG00 | U6 | 97 | 0 | 0 | 0 | 0 | 0 | 0 |
| ENSMUSG00 | SNORA17 | 123 | 0 | 0 | 0 | 0 | 0 | 0 |
| ENSMUSG00 | U6 | 55 | 0 | 0 | 0 | 0 | 0 | 0 |
| ENSMUSG00 | U7 | 60 | 0 | 0 | 0 | 0 | 0 | 0 |
| ENSMUSG00 | SNORA70 | 128 | 0 | 0 | 0 | 0 | 0 | 0 |
| ENSMUSG00 | Mir1927 | 108 | 0 | 0 | 0.40927632 | 0 | 0 | 0 |
| ENSMUSG00 | U7 | 62 | 0 | 0 | 0 | 0 | 0 | 0 |
| ENSMUSG00 | U6 | 106 | 0 | 0 | 0 | 0 | 0 | 0 |
| ENSMUSG00 | 5S_rRNA | 109 | 0 | 0 | 0 | 0 | 0 | 0 |
| ENSMUSG00 | 5S_rRNA | 72 | 0 | 0 | 0 | 0 | 0 | 0 |
| ENSMUSG00 | SCARNA20 | 138 | 0 | 0 | 0 | 0 | 0 | 0 |
| ENSMUSG00 | SCARNA21 | 143 | 0 | 0 | 0 | 0 | 0 | 0 |
| ENSMUSG00 | U6 | 106 | 0 | 0 | 0 | 0 | 0 | 0 |
| ENSMUSG00 | AC163703.1 | 56 | 0 | 0 | 0 | 0 | 0 | 0 |
| ENSMUSG00 | Mir1931 | 118 | 0 | 0 | 0 | 0.30578004 | 0 | 0 |
| ENSMUSG00 | SCARNA3 | 87 | 0 | 0 | 0 | 0 | 0 | 0 |
| ENSMUSG00 | 7SK | 294 | 0 | 0 | 0 | 0 | 0 | 0 |
| ENSMUSG00 | U6 | 102 | 0 | 0 | 0 | 0 | 0 | 0 |
| ENSMUSG00 | Mir1963 | 58 | 0 | 0 | 0 | 0 | 0 | 0 |
| ENSMUSG00 | SNORA17 | 112 | 0 | 0 | 0 | 0 | 0 | 0 |
| ENSMUSG00 | Y_RNA | 81 | 0 | 0 | 0 | 0 | 0 | 0 |
| ENSMUSG00 | U6 | 72 | 0 | 0 | 0 | 0 | 0 | 0 |
| ENSMUSG00 | U6 | 90 | 0 | 0 | 0 | 0 | 0 | 0 |
| ENSMUSG00 | 7SK | 191 | 0 | 0 | 0.69426979 | 0.94455614 | 0 | 0 |
| ENSMUSG00 | SNORD79 | 83 | 0 | 0 | 0 | 0 | 0 | 0 |
| ENSMUSG00 | 5S_rRNA | 83 | 0 | 0 | 0 | 0 | 0 | 0 |
| ENSMUSG00 | 7SK | 248 | 0 | 0 | 0 | 0 | 0 | 0 |
| ENSMUSG00 | U6 | 103 | 0 | 0 | 0 | 0 | 0 | 0 |
| ENSMUSG00 | RNaseP_nuc | 211 | 0 | 0 | 0 | 0 | 0 | 0 |
| ENSMUSG00 | U7 | 57 | 0 | 0 | 0 | 0 | 0 | 0 |
| ENSMUSG00 | SNORA17 | 124 | 0 | 0 | 0 | 0 | 0 | 0 |
| ENSMUSG00 | snoU13 | 95 | 0 | 0 | 0 | 0 | 0 | 0 |
| ENSMUSG00 | 7SK | 310 | 0 | 0 | 0 | 0 | 0 | 0 |
| ENSMUSG00 | 7SK | 316 | 0 | 0 | 0 | 0 | 0 | 0 |
| ENSMUSG00 | SCARNA17 | 139 | 0.28601065 | 0 | 0 | 0.51916611 | 0 | 0.98110213 |
| ENSMUSG00 | U3 | 89 | 0 | 0 | 0 | 0 | 0 | 0 |
| ENSMUSG00 | SNORA17 | 115 | 0 | 0 | 0 | 0 | 0 | 0 |
| ENSMUSG00 | SNORD63 | 66 | 0 | 3.65048804 | 0.66972489 | 4.37358115 | 3.86563453 | 0 |
| ENSMUSG00 | U7 | 55 | 0 | 0 | 0 | 0 | 0 | 0 |
| ENSMUSG00 | 7SK | 301 | 0 | 0 | 0 | 1.31861292 | 0 | 0 |
| ENSMUSG00 | U1 | 145 | 3.29010873 | 1.32928116 | 0 | 0 | 0 | 0 |
| ENSMUSG00 | U7 | 61 | 0 | 0 | 0 | 0 | 0 | 0 |
| ENSMUSG00 | AC102477.1 | 104 | 0 | 0 | 0 | 0 | 0 | 0 |
| ENSMUSG00 | AC151572.1 | 104 | 1.14679271 | 0 | 0 | 3.12248462 | 0.49063823 | 0.52451229 |
| ENSMUSG00 | U7 | 63 | 0 | 0 | 0 | 0 | 0 | 0 |
| ENSMUSG00 | SNORD75 | 74 | 0 | 0 | 0 | 0 | 0 | 0 |
| ENSMUSG00 | 7SK | 289 | 0 | 0 | 0 | 0 | 0 | 0 |
| ENSMUSG00 | U6 | 108 | 0 | 0 | 0.81855265 | 0 | 0 | 0 |
| ENSMUSG00 | 7SK | 295 | 0 | 0 | 0 | 0 | 0 | 0 |
| ENSMUSG00 | SNORA17 | 124 | 0 | 0 | 0 | 0 | 0 | 1.53969738 |
| ENSMUSG00 | U6 | 103 | 0 | 0 | 0 | 0 | 0.99080341 | 0 |
| ENSMUSG00 | Snora78 | 127 | 0.62607056 | 1.13826241 | 0 | 0.28411059 | 0 | 1.28856563 |
| ENSMUSG00 | 7SK | 352 | 0 | 0 | 0 | 0 | 0 | 0 |
| ENSMUSG00 | U6 | 117 | 0 | 0 | 0 | 0 | 0 | 0 |
| ENSMUSG00 | SNORA17 | 127 | 0 | 0 | 0 | 0 | 0 | 0 |
| ENSMUSG00 | 5S_rRNA | 92 | 0 | 0 | 0 | 0 | 0 | 0 |
| ENSMUSG00 | Mir1929 | 92 | 0 | 0 | 0 | 0.39219614 | 0 | 0 |
| ENSMUSG00 | Mir669n | 53 | 0 | 0 | 1.66799408 | 0 | 0.96276181 | 1.02923167 |
| ENSMUSG00 | SNORA17 | 115 | 0 | 0 | 0 | 0 | 0 | 0 |
| ENSMUSG00 | AC161364.1 | 104 | 0 | 0 | 0 | 0 | 0 | 0 |
| ENSMUSG00 | SNORA55 | 57 | 0 | 0 | 0 | 0 | 0 | 0 |
| ENSMUSG00 | SNORD121A | 76 | 0 | 0 | 0 | 1.89905498 | 2.68559873 | 0 |
| ENSMUSG00 | U1 | 163 | 0 | 0 | 0 | 0 | 0 | 0 |
| ENSMUSG00 | SNORA48 | 143 | 0 | 0 | 0 | 0 | 0 | 0 |
| ENSMUSG00 | Mir1961 | 111 | 0 | 0 | 0 | 0 | 0 | 0 |
| ENSMUSG00 | U6 | 105 | 0 | 0 | 0 | 0 | 0 | 0 |
| ENSMUSG00 | snoR9_plant | 91 | 0 | 0 | 0 | 0 | 0 | 0 |
| ENSMUSG00 | U6 | 102 | 0 | 0 | 0 | 0 | 0 | 0 |
| ENSMUSG00 | SNORA17 | 138 | 0 | 0 | 0 | 0 | 0 | 0 |
| ENSMUSG00 | 7SK | 278 | 0 | 0 | 0 | 0 | 0 | 0 |
| ENSMUSG00 | U1 | 148 | 0 | 0 | 0 | 0 | 0 | 0 |
| ENSMUSG00 | SNORA17 | 107 | 0 | 0 | 0 | 0 | 0 | 0 |
| ENSMUSG00 | 7SK | 84 | 0 | 0 | 0 | 0 | 0 | 0 |
| ENSMUSG00 | U6 | 101 | 0 | 0 | 0 | 0 | 0 | 0 |
| ENSMUSG00 | SNORA17 | 130 | 0 | 0 | 0 | 0 | 0 | 0 |
| ENSMUSG00 | SNORA17 | 72 | 0 | 0 | 0 | 0 | 0 | 0 |
| ENSMUSG00 | 7SK | 270 | 0 | 0 | 0 | 0 | 0 | 0 |
| ENSMUSG00 | Scarna6 | 264 | 0 | 0 | 0 | 0 | 0 | 0 |
| ENSMUSG00 | 7SK | 321 | 0 | 0.45034058 | 0.55080178 | 0 | 0 | 2.20916081 |
| ENSMUSG00 | SNORA17 | 131 | 0 | 0 | 0 | 0 | 0 | 0 |
| ENSMUSG00 | 7SK | 293 | 0 | 0 | 0 | 0 | 0 | 0 |
| ENSMUSG00 | U6 | 103 | 1.54390216 | 0 | 0 | 0 | 0 | 0.52980465 |
| ENSMUSG00 | SNORA38 | 132 | 0 | 0 | 0 | 0 | 0 | 0 |
| ENSMUSG00 | U6 | 107 | 0 | 0 | 0 | 0 | 0 | 0 |
| ENSMUSG00 | SNORA17 | 130 | 0 | 0 | 0 | 0 | 0 | 0 |
| ENSMUSG00 | U6 | 101 | 0 | 0 | 0 | 0 | 0 | 0 |
| ENSMUSG00 | SNORA18 | 126 | 0 | 0 | 0 | 0 | 0 | 0 |
| ENSMUSG00 | U6 | 109 | 3.28256261 | 0 | 2.02760748 | 3.31027931 | 0 | 3.25293863 |
| ENSMUSG00 | U6 | 97 | 0 | 0 | 0 | 0 | 0 | 2.53063663 |
| ENSMUSG00 | Mir2139 | 94 | 0 | 0 | 0 | 0 | 0 | 0 |
| ENSMUSG00 | U3 | 85 | 0 | 1.70069796 | 0 | 0 | 0 | 0 |
| ENSMUSG00 | U6 | 102 | 15.5903845 | 16.0621474 | 3.03345982 | 7.42865623 | 12.5064647 | 15.2417102 |
| ENSMUSG00 | SNORA62 | 106 | 0 | 0 | 0 | 0 | 0 | 0 |
| ENSMUSG00 | 7SK | 287 | 0 | 0 | 0 | 0 | 0 | 0.47516793 |
| ENSMUSG00 | SNORA17 | 84 | 0 | 0 | 0 | 0 | 1.82237057 | 1.29879235 |
| ENSMUSG00 | 7SK | 356 | 0.2233454 | 0 | 0.12416248 | 0 | 0 | 0 |
| ENSMUSG00 | SNORA17 | 123 | 0 | 0 | 0 | 0 | 0 | 0 |
| ENSMUSG00 | SNORA17 | 126 | 0 | 0 | 0 | 0 | 0 | 0 |
| ENSMUSG00 | 7SK | 90 | 0 | 0 | 0 | 0 | 0 | 0 |
| ENSMUSG00 | U6 | 53 | 0 | 0 | 0 | 0 | 0 | 0 |
| ENSMUSG00 | SNORA17 | 75 | 0 | 0 | 0 | 0 | 0 | 0 |
| ENSMUSG00 | SNORD121A | 80 | 0 | 0.60233053 | 0 | 0 | 0 | 0 |
| ENSMUSG00 | SNORA17 | 129 | 0 | 0 | 0 | 0 | 0 | 0 |
| ENSMUSG00 | SNORA17 | 109 | 0 | 0 | 0 | 0 | 0 | 0 |
| ENSMUSG00 | SNORA17 | 124 | 0 | 0 | 0 | 0 | 0 | 0 |
| ENSMUSG00 | 7SK | 300 | 0 | 0 | 0 | 0 | 0 | 0 |
| ENSMUSG00 | U6 | 133 | 0 | 0.36230408 | 0 | 0 | 0 | 0 |
| ENSMUSG00 | U6 | 84 | 0 | 0 | 0 | 0 | 0 | 0 |
| ENSMUSG00 | 7SK | 323 | 0 | 0 | 0 | 0 | 0 | 0 |
| ENSMUSG00 | SNORD123 | 70 | 2.83967718 | 1.37675549 | 0 | 0.51545778 | 0 | 0.3896377 |
| ENSMUSG00 | 7SK | 303 | 0 | 0 | 0 | 0 | 0 | 0 |
| ENSMUSG00 | SNORA17 | 129 | 0 | 0 | 0 | 0 | 0 | 0 |
| ENSMUSG00 | U6 | 99 | 0 | 0 | 0 | 0 | 0 | 0 |
| ENSMUSG00 | SNORA17 | 84 | 0 | 0 | 0 | 0 | 0 | 0 |
| ENSMUSG00 | AL772326.1 | 89 | 0 | 0 | 0 | 0 | 0 | 0 |
| ENSMUSG00 | Mir1936 | 92 | 0 | 0 | 0 | 0 | 0 | 0 |
| ENSMUSG00 | 5S_rRNA | 93 | 0 | 0 | 0 | 0 | 0 | 0 |
| ENSMUSG00 | SNORA17 | 124 | 0 | 0 | 0 | 0 | 0 | 0 |
| ENSMUSG00 | SNORA17 | 123 | 0 | 0 | 0 | 0 | 0 | 0 |
| ENSMUSG00 | U7 | 63 | 0 | 0 | 0 | 0 | 0 | 0 |
| ENSMUSG00 | SNORA17 | 110 | 0 | 0 | 0 | 0 | 0 | 0 |
| ENSMUSG00 | SNORA17 | 100 | 0 | 0 | 0 | 0 | 0 | 0 |
| ENSMUSG00 | SNORA17 | 129 | 0 | 0 | 0 | 0 | 0 | 0 |
| ENSMUSG00 | SNORA17 | 141 | 0 | 0 | 0 | 0 | 0 | 0 |
| ENSMUSG00 | SNORA73 | 193 | 0 | 0 | 1.14512547 | 0 | 0 | 0 |
| ENSMUSG00 | 7SK | 85 | 0 | 0 | 0 | 0 | 0 | 0 |
| ENSMUSG00 | U3 | 89 | 0 | 0 | 0 | 0 | 0 | 0 |
| ENSMUSG00 | 7SK | 259 | 0 | 0 | 0 | 0 | 0 | 0 |
| ENSMUSG00 | U1 | 163 | 0 | 0 | 0 | 0 | 0 | 0 |
| ENSMUSG00 | 7SK | 327 | 0 | 0 | 0 | 0 | 0.46813189 | 0.83408683 |
| ENSMUSG00 | 7SK | 104 | 0 | 0 | 0 | 0 | 0 | 0 |
| ENSMUSG00 | 7SK | 274 | 0 | 0 | 0 | 0 | 0 | 0 |
| ENSMUSG00 | U6 | 101 | 0 | 0 | 0 | 0.35724797 | 0 | 0.54009187 |
| ENSMUSG00 | 7SK | 317 | 0 | 0 | 0 | 0 | 0 | 0 |
| ENSMUSG00 | SNORA17 | 142 | 0 | 0 | 0 | 0 | 0 | 0 |
| ENSMUSG00 | SNORD22 | 124 | 0 | 0 | 0 | 0 | 0 | 0 |
| ENSMUSG00 | U7 | 62 | 0 | 0 | 0 | 0 | 0 | 0 |
| ENSMUSG00 | U1 | 140 | 0 | 0 | 0 | 0 | 0 | 0 |
| ENSMUSG00 | 5S_rRNA | 117 | 0 | 0 | 0 | 0 | 0 | 0 |
| ENSMUSG00 | Mir2137 | 65 | 0 | 0 | 0 | 0 | 0 | 0 |
| ENSMUSG00 | 7SK | 307 | 0 | 0 | 0 | 0 | 0 | 0 |
| ENSMUSG00 | SNORA17 | 129 | 0 | 0 | 0 | 0 | 0 | 0 |
| ENSMUSG00 | U7 | 65 | 0 | 0 | 0 | 0 | 0 | 0 |
| ENSMUSG00 | 5S_rRNA | 105 | 0 | 0 | 0 | 0 | 0 | 0 |
| ENSMUSG00 | SNORA17 | 127 | 0 | 0 | 0 | 0 | 0 | 0 |
| ENSMUSG00 | SNORA17 | 123 | 0 | 0 | 0 | 0 | 0 | 0 |
| ENSMUSG00 | 7SK | 275 | 0 | 0 | 0 | 0 | 0 | 0 |
| ENSMUSG00 | snoMe18S-Um | 81 | 0 | 0 | 0 | 0 | 0 | 0 |
| ENSMUSG00 | U7 | 58 | 0 | 0 | 0 | 0 | 0 | 0 |
| ENSMUSG00 | U6 | 108 | 0 | 0 | 0 | 0 | 0 | 0 |
| ENSMUSG00 | U6 | 119 | 0 | 0 | 0 | 0 | 0 | 0 |
| ENSMUSG00 | Mir1938 | 99 | 0 | 0 | 0 | 0 | 0 | 0 |
| ENSMUSG00 | U6 | 104 | 0 | 0 | 0 | 0 | 0 | 0 |
| ENSMUSG00 | SNORA17 | 127 | 0 | 0 | 0 | 0 | 0 | 0 |
| ENSMUSG00 | Snord107 | 69 | 0 | 0 | 0 | 0 | 0 | 0 |
| ENSMUSG00 | 7SK | 267 | 0 | 0 | 0 | 0 | 0 | 0 |
| ENSMUSG00 | 5S_rRNA | 107 | 0 | 0 | 0 | 0 | 0 | 0 |
| ENSMUSG00 | AC113092.1 | 89 | 0 | 0 | 0 | 0 | 0 | 0 |
| ENSMUSG00 | SNORA17 | 149 | 0 | 0 | 0 | 0 | 0 | 0 |
| ENSMUSG00 | U6 | 103 | 0 | 0 | 0 | 0 | 0 | 0 |
| ENSMUSG00 | 7SK | 258 | 0 | 0 | 0 | 0 | 0 | 0 |
| ENSMUSG00 | U6 | 74 | 0 | 0 | 0 | 0 | 0 | 0 |
| ENSMUSG00 | SNORA17 | 125 | 0 | 0 | 0 | 0 | 0 | 0 |
| ENSMUSG00 | SNORA17 | 106 | 0 | 0 | 0 | 0 | 0 | 0 |
| ENSMUSG00 | SNORA64 | 132 | 0 | 0 | 0 | 0 | 0 | 0 |
| ENSMUSG00 | SNORA17 | 110 | 0 | 0 | 0 | 0 | 0 | 0 |
| ENSMUSG00 | SCARNA20 | 136 | 1.16927884 | 0.35431208 | 0.32501355 | 0 | 2.62635758 | 1.00274409 |
| ENSMUSG00 | U1 | 168 | 0 | 0 | 0 | 0 | 0 | 0 |
| ENSMUSG00 | 7SK | 252 | 0 | 0 | 0 | 0 | 0 | 0 |
| ENSMUSG00 | 5S_rRNA | 108 | 0 | 0 | 0 | 0 | 0 | 0 |
| ENSMUSG00 | U2 | 179 | 0 | 0 | 0 | 0 | 0 | 0 |
| ENSMUSG00 | U6 | 110 | 0 | 0 | 0 | 0 | 0 | 0 |
| ENSMUSG00 | 7SK | 291 | 0 | 0 | 0 | 0 | 0 | 0 |
| ENSMUSG00 | U1 | 157 | 1.01287848 | 0 | 0 | 0 | 0 | 0 |
| ENSMUSG00 | U6 | 105 | 0 | 0 | 0 | 0 | 0 | 0 |
| ENSMUSG00 | 7SK | 264 | 0 | 0 | 0 | 0 | 0 | 0 |
| ENSMUSG00 | U6 | 101 | 0 | 0 | 0 | 0 | 0 | 0 |
| ENSMUSG00 | U1 | 164 | 0 | 0 | 0 | 0 | 0 | 0 |
| ENSMUSG00 | 5S_rRNA | 84 | 0 | 0 | 0 | 0 | 0 | 0 |
| ENSMUSG00 | U7 | 65 | 0 | 0 | 0 | 0 | 0 | 0 |
| ENSMUSG00 | U7 | 62 | 0 | 0 | 0 | 0 | 0 | 0 |
| ENSMUSG00 | Y_RNA | 115 | 0 | 0 | 0 | 0 | 0 | 0 |
| ENSMUSG00 | 5S_rRNA | 91 | 0.43687341 | 0 | 0 | 0 | 0 | 0 |
| ENSMUSG00 | Mir2136 | 74 | 0 | 0 | 0 | 0 | 0 | 0 |
| ENSMUSG00 | U6 | 103 | 1.54390216 | 0 | 1.28743232 | 0 | 0 | 0.79440697 |
| ENSMUSG00 | RNaseP_nuc | 269 | 0 | 0 | 0 | 0 | 0 | 0 |
| ENSMUSG00 | SNORA17 | 118 | 0 | 0 | 0 | 0 | 0 | 0 |
| ENSMUSG00 | 7SK | 289 | 0 | 0 | 0 | 0 | 0 | 0 |
| ENSMUSG00 | U3 | 88 | 0 | 0 | 0 | 0 | 0 | 0 |
| ENSMUSG00 | snoU13 | 104 | 0 | 0 | 0 | 0 | 0 | 0 |
| ENSMUSG00 | U6 | 102 | 0 | 0 | 0 | 0 | 0 | 0 |

| | | | | | | | | |
|---|---|---|---|---|---|---|---|---|
| ENSMUSG00 | snoZ159 | 86 | 0 | 0 | 0 | 0 | 0 | 0 |
| ENSMUSG00 | SNORA17 | 122 | 0 | 0 | 0 | 0 | 0 | 0 |
| ENSMUSG00 | U4 | 140 | 0 | 0 | 0 | 0 | 0 | 0 |
| ENSMUSG00 | snoU90 | 342 | 0 | 0 | 0 | 0.42201222 | 0 | 0 |
| ENSMUSG00 | Y_RNA | 109 | 0 | 0 | 0 | 0 | 0 | 0 |
| ENSMUSG00 | snoU13 | 102 | 0 | 0 | 0 | 0 | 0 | 0 |
| ENSMUSG00 | U7 | 61 | 0 | 0 | 0 | 0 | 0 | 0 |
| ENSMUSG00 | 7SK | 254 | 0 | 0 | 0 | 0 | 0 | 0 |
| ENSMUSG00 | U6 | 98 | 0 | 0 | 0 | 0 | 0 | 0 |
| ENSMUSG00 | U6atac | 119 | 0 | 0 | 0 | 0 | 0 | 0 |
| ENSMUSG00 | Mir1930 | 83 | 0 | 0 | 0 | 0 | 0 | 0 |
| ENSMUSG00 | SNORA17 | 80 | 0 | 0 | 0 | 0 | 0 | 0 |
| ENSMUSG00 | U6 | 104 | 0 | 0 | 0 | 0 | 0 | 0 |
| ENSMUSG00 | SNORA17 | 133 | 0 | 0 | 0 | 0 | 0 | 0 |
| ENSMUSG00 | 5S_rRNA | 115 | 0 | 0 | 0 | 0 | 0.88741523 | 0 |
| ENSMUSG00 | U3 | 88 | 0 | 0 | 0 | 0 | 0 | 0 |
| ENSMUSG00 | U6 | 106 | 0 | 0.90917815 | 0 | 0 | 0 | 0 |
| ENSMUSG00 | Mir1966 | 107 | 0.74309309 | 1.35102174 | 0.82620267 | 2.36050759 | 0 | 1.78432219 |
| ENSMUSG00 | U6 | 111 | 0 | 0 | 0 | 0 | 0 | 0 |
| ENSMUSG00 | 5S_rRNA | 90 | 0 | 0 | 0 | 0 | 0 | 0 |
| ENSMUSG00 | U6 | 106 | 0 | 0 | 0 | 1.02118994 | 0.96276181 | 3.08769501 |
| ENSMUSG00 | U6 | 104 | 0 | 0 | 0 | 0 | 0 | 0 |
| ENSMUSG00 | SNORA17 | 128 | 0 | 0 | 0 | 0 | 0 | 0 |
| ENSMUSG00 | SCARNA3 | 87 | 0 | 0 | 0 | 0 | 0 | 0 |
| ENSMUSG00 | 5S_rRNA | 105 | 0 | 0 | 0 | 0 | 0 | 0 |
| ENSMUSG00 | SNORA17 | 122 | 0 | 0 | 0 | 0 | 0 | 0 |
| ENSMUSG00 | U6 | 103 | 0 | 0 | 0 | 0 | 0 | 0 |
| ENSMUSG00 | SNORA17 | 139 | 0 | 0 | 0 | 0 | 0 | 0 |
| ENSMUSG00 | SNORA17 | 117 | 0 | 0 | 0 | 0 | 0 | 0 |
| ENSMUSG00 | 7SK | 273 | 0 | 0 | 0 | 0 | 0 | 0 |
| ENSMUSG00 | SNORA50 | 139 | 0 | 0 | 0 | 0 | 0 | 0 |
| ENSMUSG00 | U7 | 62 | 0 | 0 | 0 | 0 | 0 | 0 |
| ENSMUSG00 | SNORD112 | 76 | 0 | 0 | 0 | 0 | 0 | 0 |
| ENSMUSG00 | U6 | 97 | 0 | 0 | 0 | 0 | 0 | 0 |
| ENSMUSG00 | U6 | 96 | 0 | 0 | 0 | 0 | 0 | 0 |
| ENSMUSG00 | U6 | 89 | 0 | 0 | 0 | 0 | 0 | 0 |
| ENSMUSG00 | 7SK | 257 | 0.92814351 | 0 | 0 | 0 | 0 | 0 |
| ENSMUSG00 | SNORA17 | 128 | 0 | 0 | 0 | 0 | 0 | 0 |
| ENSMUSG00 | SNORA17 | 117 | 0 | 0 | 0 | 0 | 0 | 0 |
| ENSMUSG00 | snoU13 | 101 | 0 | 0 | 0 | 0 | 0 | 0 |
| ENSMUSG00 | snoU85 | 312 | 0 | 0 | 0 | 0 | 0 | 0 |
| ENSMUSG00 | 5S_rRNA | 92 | 0 | 0 | 0 | 0 | 0 | 0 |
| ENSMUSG00 | SNORA70 | 134 | 0 | 0 | 0 | 0 | 0 | 0 |
| ENSMUSG00 | SNORA17 | 127 | 0 | 0 | 0 | 0 | 0 | 0 |
| ENSMUSG00 | 7SK | 229 | 0 | 0 | 0 | 0 | 0 | 0 |
| ENSMUSG00 | SNORA17 | 142 | 0 | 0 | 0 | 0 | 0 | 0 |
| ENSMUSG00 | 5S_rRNA | 129 | 0 | 0 | 0 | 0 | 0 | 0 |
| ENSMUSG00 | snoU13 | 109 | 0 | 0 | 0 | 0 | 0 | 0 |
| ENSMUSG00 | AC103664.1 | 101 | 0 | 0 | 0 | 0 | 0 | 0 |
| ENSMUSG00 | U7 | 61 | 0 | 0 | 0 | 0 | 0 | 0 |
| ENSMUSG00 | 7SK | 339 | 0 | 0 | 0 | 0 | 0 | 0 |
| ENSMUSG00 | U4 | 123 | 0 | 0 | 0 | 0 | 0 | 0 |
| ENSMUSG00 | U6 | 105 | 0 | 0 | 0 | 0 | 0 | 0 |
| ENSMUSG00 | 5S_rRNA | 130 | 0 | 0 | 0 | 0 | 0 | 0 |
| ENSMUSG00 | U8 | 131 | 0 | 0 | 0 | 0 | 0 | 0 |
| ENSMUSG00 | U6 | 73 | 0 | 0 | 0 | 0 | 0 | 0 |
| ENSMUSG00 | 7SK | 244 | 0 | 0 | 0 | 0 | 0 | 0 |
| ENSMUSG00 | U6 | 106 | 0.75010341 | 0 | 0.41699852 | 0 | 0 | 1.02923167 |
| ENSMUSG00 | 5S_rRNA | 120 | 0 | 0 | 0 | 0 | 0 | 0 |
| ENSMUSG00 | U6 | 104 | 0 | 0 | 0 | 0 | 0 | 0 |
| ENSMUSG00 | 7SK | 291 | 0 | 0 | 0 | 0 | 0 | 0 |
| ENSMUSG00 | 5S_rRNA | 122 | 0 | 0 | 0 | 0 | 0 | 0 |
| ENSMUSG00 | 5S_rRNA | 105 | 0 | 0 | 0 | 0 | 0 | 0 |
| ENSMUSG00 | U7 | 62 | 0 | 0 | 0 | 0 | 0 | 0 |
| ENSMUSG00 | U6 | 117 | 0 | 0 | 0 | 0 | 0 | 0 |
| ENSMUSG00 | 7SK | 261 | 0 | 0 | 0 | 0 | 0 | 0 |
| ENSMUSG00 | U6 | 99 | 0 | 0 | 0 | 0 | 0 | 0 |
| ENSMUSG00 | U1 | 129 | 0 | 0 | 0 | 0 | 0 | 0 |
| ENSMUSG00 | 7SK | 252 | 0 | 0 | 0 | 0 | 0 | 0 |
| ENSMUSG00 | U7 | 62 | 0 | 0 | 0 | 0 | 0 | 0 |
| ENSMUSG00 | 7SK | 314 | 0 | 0 | 0 | 0 | 0 | 0 |
| ENSMUSG00 | SNORA19 | 82 | 0 | 0 | 0 | 0 | 0 | 0.99785266 |
| ENSMUSG00 | U1 | 185 | 0 | 0.52093451 | 0.95571553 | 0.39007616 | 0 | 0 |
| ENSMUSG00 | U6 | 103 | 0 | 0 | 0 | 0 | 0 | 0 |
| ENSMUSG00 | U6 | 102 | 0 | 0 | 0 | 0 | 0 | 0 |
| ENSMUSG00 | U6 | 100 | 0 | 0 | 0 | 0 | 0 | 0 |
| ENSMUSG00 | U6 | 102 | 0 | 0 | 0 | 0 | 0 | 0 |
| ENSMUSG00 | U7 | 63 | 0 | 0 | 0 | 0 | 0 | 0 |
| ENSMUSG00 | SNORA17 | 121 | 0 | 0 | 0 | 0 | 0 | 0 |
| ENSMUSG00 | U7 | 56 | 0 | 0 | 0 | 0 | 0 | 0 |
| ENSMUSG00 | U7 | 61 | 0 | 0 | 0 | 0 | 0 | 0 |
| ENSMUSG00 | 7SK | 273 | 0 | 0 | 0 | 0 | 0 | 0 |
| ENSMUSG00 | SNORA17 | 127 | 0 | 0 | 0 | 0 | 0 | 0 |
| ENSMUSG00 | 5S_rRNA | 115 | 0 | 1.25703762 | 0.76872771 | 0.31375691 | 0 | 1.42302466 |
| ENSMUSG00 | SNORD113 | 74 | 0 | 0 | 0 | 0 | 0 | 0 |
| ENSMUSG00 | U6 | 105 | 0 | 0 | 0 | 0 | 0 | 0 |
| ENSMUSG00 | SNORD11 | 74 | 3.76065356 | 0.65116814 | 4.18125542 | 0 | 0 | 1.10572862 |
| ENSMUSG00 | U6 | 103 | 0 | 0 | 0 | 0 | 0 | 0 |
| ENSMUSG00 | SNORA17 | 136 | 0 | 0 | 0 | 0 | 0 | 0 |
| ENSMUSG00 | U1 | 144 | 0 | 0 | 0 | 0 | 0 | 0 |
| ENSMUSG00 | SNORD113 | 72 | 0 | 0 | 0 | 0 | 0 | 0 |
| ENSMUSG00 | U7 | 56 | 0 | 0 | 0 | 0 | 0 | 0 |
| ENSMUSG00 | SNORA17 | 84 | 0 | 0 | 0 | 0 | 0 | 0 |
| ENSMUSG00 | snosnR60_Z1 | 86 | 0 | 0 | 1.02794984 | 0 | 0 | 0 |
| ENSMUSG00 | 7SK | 350 | 0 | 0 | 0 | 0 | 0 | 0 |
| ENSMUSG00 | SNORA70 | 107 | 0 | 0 | 0 | 0 | 0 | 0 |
| ENSMUSG00 | 7SK | 251 | 0 | 0 | 0 | 0 | 0 | 0 |
| ENSMUSG00 | U6 | 99 | 0 | 0 | 0 | 0 | 0 | 0 |
| ENSMUSG00 | SNORA17 | 113 | 0 | 0 | 0 | 0 | 0 | 0 |
| ENSMUSG00 | 7SK | 307 | 0 | 0 | 0 | 0 | 0 | 0 |
| ENSMUSG00 | U6 | 103 | 0 | 0 | 0 | 0 | 0 | 0.52960465 |
| ENSMUSG00 | SNORA73 | 158 | 0 | 0 | 0 | 0 | 0 | 0 |
| ENSMUSG00 | SNORA17 | 113 | 0 | 0 | 0 | 0 | 0 | 0 |
| ENSMUSG00 | 5S_rRNA | 115 | 0 | 0 | 0 | 0 | 0 | 0 |
| ENSMUSG00 | SNORA17 | 110 | 0 | 0 | 0 | 0 | 0 | 0 |
| ENSMUSG00 | snoMe28S-C | 82 | 0 | 0 | 0 | 0 | 0 | 0 |
| ENSMUSG00 | Scarna3a | 139 | 0 | 0.34666505 | 0 | 2.07666443 | 0 | 0 |
| ENSMUSG00 | SNORA17 | 131 | 0 | 0 | 0 | 0 | 0 | 0 |
| ENSMUSG00 | U6 | 101 | 0 | 0 | 0 | 0 | 0 | 0 |
| ENSMUSG00 | snoU13 | 107 | 0 | 0 | 0 | 0 | 0 | 0 |
| ENSMUSG00 | U6 | 100 | 0 | 0 | 0 | 0 | 0 | 0 |
| ENSMUSG00 | snoU6-77 | 143 | 6.39423812 | 2.69574502 | 1.85462278 | 3.02786388 | 8.20703947 | 4.76829358 |
| ENSMUSG00 | SNORA17 | 122 | 0.32586459 | 0 | 0 | 0 | 0 | 0 |
| ENSMUSG00 | AL603828.1 | 107 | 0 | 0 | 0 | 0 | 0 | 0 |
| ENSMUSG00 | 7SK | 119 | 0 | 0 | 0 | 0 | 0 | 0 |
| ENSMUSG00 | U6 | 102 | 0 | 0 | 0 | 0 | 0 | 0 |
| ENSMUSG00 | SNORA17 | 131 | 0 | 0 | 0 | 0 | 0 | 0 |
| ENSMUSG00 | U1 | 162 | 0 | 0 | 0 | 0 | 0 | 1.34689577 |
| ENSMUSG00 | SNORA17 | 126 | 0 | 0 | 0 | 0 | 0 | 0 |
| ENSMUSG00 | U6 | 95 | 0 | 0 | 0 | 0 | 0 | 0 |
| ENSMUSG00 | SNORA17 | 130 | 0 | 0 | 0 | 0 | 0 | 0 |
| ENSMUSG00 | SNORA17 | 120 | 0 | 0 | 0 | 0 | 0 | 0 |
| ENSMUSG00 | Mir1933 | 87 | 0 | 0 | 0 | 0 | 0 | 0 |
| ENSMUSG00 | SNORA70 | 128 | 0 | 0 | 0 | 0 | 0 | 0 |
| ENSMUSG00 | 5S_rRNA | 110 | 0 | 0 | 0 | 0 | 0 | 0 |
| ENSMUSG00 | U6 | 103 | 0 | 0 | 0 | 0 | 0 | 0 |
| ENSMUSG00 | U6 | 103 | 0 | 0 | 0 | 0 | 0 | 0 |
| ENSMUSG00 | U6 | 96 | 0 | 0 | 0 | 0 | 0 | 0 |
| ENSMUSG00 | U2 | 166 | 0.23949085 | 0 | 0 | 0 | 0 | 0 |
| ENSMUSG00 | U6 | 102 | 0 | 0 | 0 | 0 | 0 | 0 |
| ENSMUSG00 | 5S_rRNA | 99 | 0 | 0 | 0 | 0 | 0 | 0 |
| ENSMUSG00 | U3 | 107 | 0 | 0 | 0 | 0 | 0 | 0 |
| ENSMUSG00 | snoU13 | 103 | 0 | 0 | 0 | 0 | 0 | 0 |
| ENSMUSG00 | U7 | 59 | 0 | 0 | 0 | 0 | 0 | 0 |
| ENSMUSG00 | snoU13 | 95 | 0 | 0 | 0 | 0 | 0 | 0 |
| ENSMUSG00 | Mir669m-1 | 97 | 0 | 0 | 0 | 0 | 0 | 0 |
| ENSMUSG00 | snoU105B | 79 | 7.54850896 | 6.09954964 | 3.357102 | 5.48081689 | 10.9803594 | 1.72624299 |
| ENSMUSG00 | snoCD11 | 69 | 0 | 0 | 0 | 0 | 0 | 0 |
| ENSMUSG00 | snoU109 | 138 | 0 | 0 | 0 | 0 | 0 | 0 |
| ENSMUSG00 | AC161366.1 | 70 | 0 | 0 | 0 | 0 | 0 | 0 |
| ENSMUSG00 | SNORA17 | 135 | 0 | 0 | 0 | 0 | 0 | 0 |
| ENSMUSG00 | 7SK | 176 | 0 | 0 | 0 | 0 | 0 | 0 |
| ENSMUSG00 | 7SK | 285 | 0 | 0 | 0 | 0 | 0 | 0 |
| ENSMUSG00 | SNORA17 | 122 | 0 | 0 | 0 | 0 | 0 | 0 |
| ENSMUSG00 | U6 | 96 | 0 | 0 | 0 | 0 | 0 | 0 |
| ENSMUSG00 | 5S_rRNA | 114 | 0 | 0 | 0 | 0 | 0 | 0 |
| ENSMUSG00 | U6 | 117 | 0 | 0 | 0 | 0 | 0 | 0 |
| ENSMUSG00 | SNORA40 | 94 | 0 | 0 | 0 | 0 | 0 | 0 |
| ENSMUSG00 | 5S_rRNA | 110 | 0 | 0.43805857 | 0 | 0 | 0.92775229 | 0 |
| ENSMUSG00 | SNORA48 | 138 | 0 | 0 | 0 | 0 | 0 | 0 |
| ENSMUSG00 | 7SK | 105 | 0 | 0 | 0 | 0 | 0 | 2.33782622 |
| ENSMUSG00 | SNORA48 | 142 | 0 | 0 | 0 | 0 | 0 | 0 |
| ENSMUSG00 | U7 | 62 | 0 | 0 | 0 | 0 | 0 | 0 |
| ENSMUSG00 | SNORA17 | 124 | 0 | 0 | 0 | 0 | 0 | 0 |
| ENSMUSG00 | U6 | 103 | 0 | 0 | 0 | 0 | 0 | 0 |
| ENSMUSG00 | U7 | 64 | 0 | 0 | 0 | 0 | 0 | 0 |
| ENSMUSG00 | NRON | 369 | 0 | 0 | 0 | 0 | 0 | 0 |
| ENSMUSG00 | 5S_rRNA | 134 | 0 | 0 | 0 | 0 | 0 | 0 |
| ENSMUSG00 | SNORA19 | 118 | 0.33691085 | 0.40835968 | 1.12377567 | 0 | 1.72970766 | 3.00483314 |
| ENSMUSG00 | 5S_rRNA | 125 | 0 | 0 | 1.06084423 | 0 | 0 | 0.21819711 |
| ENSMUSG00 | 5S_rRNA | 116 | 5.14079489 | 9.13880799 | 12.5746622 | 10.8868238 | 7.47800335 | 5.64302881 |
| ENSMUSG00 | U6 | 106 | 0 | 0 | 0 | 0 | 0 | 0 |
| ENSMUSG00 | 5S_rRNA | 110 | 0 | 0 | 0 | 0 | 0 | 0 |
| ENSMUSG00 | SNORA17 | 126 | 0 | 0 | 0 | 0 | 0 | 0 |
| ENSMUSG00 | U6 | 98 | 0 | 0 | 0 | 0 | 0 | 0 |
| ENSMUSG00 | U6 | 114 | 0 | 0 | 0 | 0 | 0 | 0 |
| ENSMUSG00 | AL845355.1 | 149 | 0 | 0 | 0 | 0 | 0 | 0 |
| ENSMUSG00 | 7SK | 295 | 0 | 0 | 0 | 0 | 0 | 0 |
| ENSMUSG00 | U6 | 102 | 0 | 0 | 0 | 0 | 0 | 0 |
| ENSMUSG00 | U6 | 94 | 13.9567112 | 4.1009738 | 3.29162661 | 3.83851538 | 7.59967299 | 7.25389342 |
| ENSMUSG00 | SCARNA14 | 129 | 0 | 0 | 0 | 0 | 0 | 0 |
| ENSMUSG00 | SNORA17 | 130 | 0 | 0 | 0 | 0 | 0 | 0 |
| ENSMUSG00 | 7SK | 248 | 0 | 0 | 0 | 0 | 0 | 0 |
| ENSMUSG00 | Scarna10 | 324 | 0 | 0 | 0 | 0 | 0 | 0 |
| ENSMUSG00 | 7SK | 324 | 0 | 0 | 0 | 0 | 0 | 0 |
| ENSMUSG00 | 7SK | 258 | 0 | 0 | 0 | 0 | 0 | 0 |
| ENSMUSG00 | U7 | 62 | 0 | 0 | 0 | 0 | 0 | 0 |
| ENSMUSG00 | 7SK | 298 | 1.33407653 | 0.32339894 | 0.444985 | 2.05837167 | 0 | 0 |
| ENSMUSG00 | 5S_rRNA | 127 | 0 | 0 | 0 | 0 | 0 | 0 |
| ENSMUSG00 | U6 | 112 | 0 | 0 | 0 | 0 | 0 | 0 |
| ENSMUSG00 | SCARNA18 | 75 | 0 | 0 | 0 | 0 | 0 | 0 |
| ENSMUSG00 | Gm16228 | 808 | 0 | 0 | 0 | 0 | 0 | 0 |
| ENSMUSG00 | Gm9727 | 554 | 0.21528239 | 0.17395828 | 0 | 0 | 0.27631612 | 0 |
| ENSMUSG00 | Gm6903 | 1365 | 0 | 0 | 0 | 0 | 0 | 0 |
| ENSMUSG00 | Gm16326 | 413 | 1.25138316 | 0.46669678 | 0.42810502 | 1.13575443 | 2.1003593 | 1.84912809 |
| ENSMUSG00 | A230009B12I | 3237 | 0 | 0 | 0 | 0 | 0 | 0 |
| ENSMUSG00 | Nat8b | 698 | 0 | 0 | 0.12665285 | 0 | 0.36551845 | 0 |
| ENSMUSG00 | Gm16559 | 1561 | 0.15280774 | 1.0804135 | 0.25484727 | 1.6180289 | 0.62107696 | 0.3669234 |
| ENSMUSG00 | 1700058P15I | 411 | 0.58037198 | 0.46896781 | 0.86037651 | 0 | 0.12415177 | 0.53089322 |
| ENSMUSG00 | Hmga1-rs1 | 1564 | 0 | 0 | 0 | 0 | 0 | 0 |
| ENSMUSG00 | Gm2862 | 327 | 1.09418754 | 0.14735915 | 0.13517383 | 1.98616759 | 0.31208793 | 0.16681737 |
| ENSMUSG00 | Gm6397 | 1068 | 0 | 0.04511839 | 0.08277499 | 0 | 0 | 0 |
| ENSMUSG00 | Gm16566 | 924 | 0 | 0 | 0 | 0 | 0 | 0 |
| ENSMUSG00 | Gm16301 | 855 | 0.18599055 | 0.73265935 | 0.46528256 | 0.12660367 | 0.77583963 | 0.31900163 |
| ENSMUSG00 | Gm5766 | 2774 | 0 | 0.03474149 | 0 | 0 | 0.01839451 | 0.00983224 |
| ENSMUSG00 | Gm15796 | 203 | 0 | 0 | 0 | 0 | 1.00544583 | 0 |
| ENSMUSG00 | Gm2245 | 3369 | 0 | 0 | 0 | 0 | 0 | 0 |
| ENSMUSG00 | Gm15790 | 1250 | 0.15902192 | 0.03854915 | 0.03536147 | 0.05773127 | 0.0408211 | 0.06545913 |
| ENSMUSG00 | Gm16353 | 1222 | 0.09759938 | 0.23659464 | 1.48304056 | 0.17716225 | 0 | 0.20087705 |
| ENSMUSG00 | Gm16025 | 2606 | 0 | 0 | 0 | 0 | 0 | 0 |
| ENSMUSG00 | Gm15841 | 514 | 0 | 0 | 0 | 0 | 0 | 0 |
| ENSMUSG00 | Ear-ps12 | 450 | 0 | 0 | 0 | 0 | 0 | 0 |
| ENSMUSG00 | Gm16004 | 308 | 0 | 0 | 0 | 0 | 0 | 0 |
| ENSMUSG00 | Gm16271 | 217 | 0 | 0 | 0 | 0 | 0 | 0 |
| ENSMUSG00 | Gm16585 | 609 | 0 | 0 | 0 | 0 | 0 | 0 |
| ENSMUSG00 | Gm13695 | 2725 | 0 | 0 | 0 | 0 | 0 | 0 |
| ENSMUSG00 | Gm3470 | 1119 | 0.03552769 | 0 | 0 | 0 | 0 | 0.07312236 |
| ENSMUSG00 | Vmn2r-ps110 | 863 | 0 | 0 | 0 | 0 | 0 | 0 |
| ENSMUSG00 | Mia1 | 2265 | 0.07020835 | 0.04254874 | 0 | 0.06372105 | 0.09011281 | 0.04816713 |
| ENSMUSG00 | Gm14057 | 481 | 0.08265173 | 2.30413341 | 1.10274868 | 1.42527827 | 1.16692336 | 0.85056048 |
| ENSMUSG00 | Gm15651 | 2505 | 1.72987919 | 1.76972163 | 1.25282669 | 2.29823756 | 1.38514713 | 3.17931923 |
| ENSMUSG00 | Gm12494 | 623 | 0 | 0 | 0 | 0 | 0 | 0 |
| ENSMUSG00 | 2400009B08I | 364 | 3.05811389 | 0.52952134 | 1.94293815 | 0.99126496 | 0.70091176 | 5.09526228 |
| ENSMUSG00 | Gm16269 | 408 | 0 | 0 | 0 | 0.08843638 | 0 | 0 |
| ENSMUSG00 | Gm15914 | 308 | 0 | 0 | 0 | 0 | 0 | 0 |
| ENSMUSG00 | Tnfsf13 | 2043 | 0 | 0.44813627 | 0.04327151 | 0.0176613 | 0 | 0.20025433 |
| ENSMUSG00 | Gm16581 | 990 | 0 | 0.04867317 | 0 | 0 | 0.20616718 | 0 |
| ENSMUSG00 | Gm16537 | 699 | 0 | 0 | 0.06323583 | 0 | 0.07299911 | 0 |
| ENSMUSG00 | Gp49a | 2173 | 0 | 0 | 0 | 0 | 0 | 0 |
| ENSMUSG00 | Gm16546 | 354 | 0 | 0 | 0 | 0 | 0 | 0 |
| ENSMUSG00 | Ugt1a8 | 2222 | 0 | 0 | 0 | 0 | 0 | 0 |
| ENSMUSG00 | B230217C12 | 2230 | 0 | 0 | 0 | 0 | 0 | 0 |
| ENSMUSG00 | Agpt2 | 3294 | 0.42241707 | 0.0877713 | 0.05367558 | 0 | 0.34079547 | 0.62100727 |
| ENSMUSG00 | Gm16299 | 2764 | 0.17259977 | 0.66247641 | 2.07895788 | 0.86158283 | 1.77226197 | 1.42096529 |
| ENSMUSG00 | Gm15722 | 659 | 0 | 0 | 0 | 0.32851634 | 0 | 0 |
| ENSMUSG00 | Mup17 | 920 | 0 | 0 | 0 | 0 | 0 | 0 |
| ENSMUSG00 | Bcl2l2 | 4012 | 8.48222615 | 6.11338461 | 7.1062285 | 6.55628376 | 5.83776334 | 5.66975302 |
| ENSMUSG00 | 4930570N18I | 329 | 0 | 0 | 0 | 0 | 0 | 0 |
| ENSMUSG00 | Gm14163 | 216 | 0 | 0 | 0 | 0 | 0 | 0 |
| ENSMUSG00 | Gm9017 | 877 | 0 | 0 | 0 | 0 | 0 | 0 |
| ENSMUSG00 | Gm3833 | 603 | 0 | 0 | 0 | 0.05983755 | 0 | 0 |
| ENSMUSG00 | Rab42-ps | 759 | 1.72849915 | 1.90460246 | 3.61069073 | 1.33108992 | 2.62190864 | 1.9045532 |
| ENSMUSG00 | 1600010F14I | 255 | 0 | 0 | 0 | 0 | 0 | 0 |
| ENSMUSG00 | Gm4982 | 1097 | 0 | 0 | 0 | 0 | 0 | 0 |
| ENSMUSG00 | Gm16569 | 311 | 0 | 0 | 0 | 0 | 0.16407195 | 0 |
| ENSMUSG00 | Gm4477 | 1819 | 0 | 0 | 0 | 0 | 0 | 0 |
| ENSMUSG00 | Gm15946 | 425 | 0.18708461 | 0.90703891 | 0.41601735 | 0 | 0 | 0.12835124 |
| ENSMUSG00 | Gm4778 | 1206 | 0 | 0 | 0 | 0 | 0 | 0 |
| ENSMUSG00 | Gm15947 | 2740 | 0 | 0 | 0 | 0 | 0 | 0 |
| ENSMUSG00 | Gm2541 | 710 | 0 | 0 | 0 | 0 | 0 | 0 |
| ENSMUSG00 | Gm1992 | 248 | 0 | 0 | 0 | 0 | 0 | 0 |
| ENSMUSG00 | Gm2343 | 1041 | 0 | 0 | 0 | 0 | 0 | 0 |
| ENSMUSG00 | Gm15978 | 736 | 0 | 0.13094142 | 0 | 0 | 0.06932932 | 0.07411587 |
| ENSMUSG00 | Gm16568 | 1908 | 0 | 0 | 0 | 0 | 0 | 0 |
| ENSMUSG00 | Gm15833 | 956 | 0 | 0 | 0 | 0 | 0 | 0 |
| ENSMUSG00 | Galnt2 | 4406 | 0.09925336 | 0.16404826 | 0.1404507 | 0.22930033 | 0.20846 | 0.16713919 |
| ENSMUSG00 | Gm15970 | 833 | 0.04772567 | 0 | 0.05306344 | 0 | 0 | 0 |
| ENSMUSG00 | B230216N24 | 4715 | 0.56492411 | 0.68472781 | 0.90934863 | 1.43869022 | 1.50427704 | 1.45773258 |
| ENSMUSG00 | 4921525B02I | 3098 | 0 | 0 | 0 | 0 | 0 | 0 |
| ENSMUSG00 | 5330439A09I | 1115 | 0 | 0 | 0 | 0 | 0 | 0 |
| ENSMUSG00 | Gm16262 | 1749 | 0 | 0 | 0.07581791 | 0.1650408 | 0 | 0 |
| ENSMUSG00 | Gm15810 | 677 | 0.4110611 | 0.42705857 | 0.65290758 | 0.63956357 | 0.82908439 | 0.64460004 |
| ENSMUSG00 | Gm16240 | 612 | 0.12991987 | 0 | 0 | 0.05895759 | 0.08337643 | 0.84676368 |
| ENSMUSG00 | Gm15889 | 386 | 0 | 0 | 0 | 0 | 0 | 0 |
| ENSMUSG00 | Gm16564 | 3008 | 0.21146532 | 0 | 0.5730957 | 0.0599768 | 0 | 1.87694492 |
| ENSMUSG00 | Gm16092 | 2403 | 0 | 0 | 0 | 0 | 0 | 0 |
| ENSMUSG00 | Cbx6 | 10455 | 6.10306516 | 7.1300264 | 4.37156439 | 5.72204972 | 7.2281265 | 5.81493744 |
| ENSMUSG00 | Gm6264 | 1460 | 0 | 0 | 0 | 0 | 0 | 0.03736252 |
| ENSMUSG00 | Olfr275 | 959 | 0 | 0 | 0 | 0 | 0 | 0 |
| ENSMUSG00 | 2310075C17 | 324 | 0 | 0 | 0 | 0 | 0 | 0 |
| ENSMUSG00 | Gm15758 | 1211 | 0 | 0 | 0 | 0 | 0 | 0 |
| ENSMUSG00 | Gm13734 | 811 | 0 | 0 | 0 | 0 | 0 | 0 |
| ENSMUSG00 | Gm11711 | 848 | 0 | 0 | 0 | 0 | 0 | 0 |
| ENSMUSG00 | Gm16117 | 322 | 0 | 0 | 0.54909122 | 0 | 0 | 0.42351924 |
| ENSMUSG00 | Krtap4-8 | 982 | 0 | 0 | 0 | 0 | 0 | 0 |
| ENSMUSG00 | Mir17hg | 1008 | 5.28495475 | 5.35404913 | 2.98187036 | 6.80117903 | 4.25219799 | 5.79044921 |
| ENSMUSG00 | Klra8 | 1170 | 0 | 0 | 0 | 0 | 0 | 0 |
| ENSMUSG00 | Clec2f-ps | 371 | 0 | 0 | 0 | 0 | 0 | 0 |
| ENSMUSG00 | Gm16552 | 683 | 0.6402786 | 0.3527558 | 0.06471719 | 0.31697257 | 1.3447654 | 0 |
| ENSMUSG00 | 1700007P06I | 530 | 0 | 0 | 0 | 0 | 0 | 0 |
| ENSMUSG00 | Gm10013 | 600 | 0 | 0 | 0 | 0 | 0 | 0 |
| ENSMUSG00 | Olfr1118 | 995 | 0 | 0 | 0 | 0 | 0 | 0 |
| ENSMUSG00 | Gm15628 | 545 | 0 | 0 | 0 | 0 | 0 | 0 |
| ENSMUSG00 | Gm16108 | 639 | 0 | 0 | 0 | 0 | 0 | 0 |
| ENSMUSG00 | Gm8428 | 1783 | 0 | 0 | 0 | 0 | 0.08585481 | 0 |
| ENSMUSG00 | Gm14378 | 1864 | 1.57827551 | 2.14564093 | 0.68768962 | 2.67131016 | 2.57321852 | 1.90220124 |
| ENSMUSG00 | Gm15688 | 2416 | 0 | 0 | 0 | 0 | 0 | 0 |
| ENSMUSG00 | Gm20431 | 2359 | 0 | 0 | 0 | 0 | 0 | 0 |
| ENSMUSG00 | Gm15898 | 624 | 2.03874259 | 0.84944049 | 0.5666903 | 0.40476653 | 0.89950342 | 0 |
| ENSMUSG00 | Gm16226 | 1249 | 0 | 0 | 0 | 0 | 0 | 0 |
| ENSMUSG00 | Gm16146 | 437 | 1.09168368 | 0 | 0 | 0 | 0 | 0 |
| ENSMUSG00 | 1810008B01I | 827 | 0 | 0 | 0 | 0 | 0 | 0 |
| ENSMUSG00 | Gm3513 | 635 | 0 | 0 | 0 | 0 | 0 | 0 |
| ENSMUSG00 | Matr3-ps2 | 1024 | 0 | 0 | 0 | 0 | 0 | 0 |
| ENSMUSG00 | Gm11357 | 953 | 0 | 0 | 0 | 0 | 0 | 0 |
| ENSMUSG00 | Olfr1213 | 2219 | 0 | 0 | 0 | 0 | 0 | 0 |
| ENSMUSG00 | Gm11709 | 1212 | 0 | 0 | 0 | 0 | 0 | 0 |
| ENSMUSG00 | Ear-ps7 | 441 | 0 | 0 | 0 | 0 | 0 | 0 |
| ENSMUSG00 | 0610012D04 | 529 | 0 | 0.63762778 | 0 | 0 | 0 | 0 |
| ENSMUSG00 | Gm8898 | 3543 | 0 | 0 | 0 | 0 | 0 | 0 |
| ENSMUSG00 | Gm16334 | 691 | 0 | 0 | 0 | 0 | 0 | 0 |
| ENSMUSG00 | 3632454L22I | 6679 | 0 | 0 | 0.06618033 | 0 | 0.07639823 | 0 |
| ENSMUSG00 | D030046N08 | 1568 | 0 | 0 | 0 | 0 | 0 | 0 |
| ENSMUSG00 | Vmn2r-ps111 | 865 | 0 | 0 | 0 | 0 | 0 | 0 |
| ENSMUSG00 | Ier5l | 1551 | 3.28091651 | 7.17670415 | 1.7954327 | 4.39684488 | 3.68468994 | 1.8464456 |
| ENSMUSG00 | Gm2635 | 913 | 0 | 0 | 0 | 0 | 0 | 0 |
| ENSMUSG00 | Gm16580 | 1043 | 0.49551414 | 0.13859955 | 0.97472904 | 0.62270067 | 1.02737669 | 0.62760436 |
| ENSMUSG00 | Gm16538 | 440 | 0 | 0 | 0 | 0 | 0 | 0 |
| ENSMUSG00 | 4930556I23R | 558 | 0 | 0 | 0 | 0 | 0 | 0 |
| ENSMUSG00 | Tmsb15b1 | 436 | 9.93887013 | 6.41012304 | 3.64969346 | 3.06200837 | 2.45769241 | 6.38076423 |
| ENSMUSG00 | Gm16574 | 744 | 0 | 0 | 0 | 0 | 0 | 0 |
| ENSMUSG00 | Ifna14 | 569 | 0 | 0 | 0 | 0 | 0 | 0 |
| ENSMUSG00 | Gm16311 | 699 | 0 | 0 | 0 | 0 | 0 | 0 |
| ENSMUSG00 | Skint1 | 3067 | 0 | 0 | 0 | 0 | 0 | 0 |
| ENSMUSG00 | Slc5a3 | 10911 | 25.7676435 | 18.6059464 | 49.0145815 | 46.4757633 | 47.8790244 | 42.4105517 |
| ENSMUSG00 | Gm16270 | 3237 | 0 | 0 | 0 | 0 | 0 | 0 |
| ENSMUSG00 | Gm15684 | 1812 | 0 | 0 | 0 | 0 | 0 | 0 |
| ENSMUSG00 | Gm6619 | 597 | 0 | 0 | 0 | 0 | 0 | 0 |
| ENSMUSG00 | Gm15641 | 668 | 0 | 0 | 0 | 0 | 0 | 0 |
| ENSMUSG00 | Gm16306 | 174 | 0 | 0.27693358 | 0.25403358 | 0 | 0 | 0 |
| ENSMUSG00 | Gm15756 | 680 | 0.87695913 | 0.92121139 | 0.84503523 | 0.58368013 | 0.75038788 | 0.92252456 |
| ENSMUSG00 | Gm3531 | 488 | 0.97759378 | 0.49371355 | 2.80790396 | 1.10907924 | 1.04562246 | 1.50904766 |
| ENSMUSG00 | Gm454 | 749 | 0 | 0 | 0 | 0 | 0 | 0 |
| ENSMUSG00 | Gm8429 | 825 | 0.38550769 | 0.05840781 | 2.08954167 | 0.21867906 | 0.80405198 | 0.26448135 |
| ENSMUSG00 | Gm15825 | 915 | 0 | 0 | 0 | 0 | 0 | 0 |
| ENSMUSG00 | Gm16586 | 2638 | 21.8820916 | 21.5359421 | 26.0720499 | 27.4103174 | 20.7741955 | 49.7725979 |
| ENSMUSG00 | Rdh1 | 3926 | 0.07088343 | 0 | 0 | 0 | 0.09097927 | 0 |
| ENSMUSG00 | Gm16588 | 354 | 0 | 0 | 0 | 0 | 0 | 0 |
| ENSMUSG00 | Gm3555 | 1394 | 0.028519 | 0 | 0 | 0 | 0 | 0 |
| ENSMUSG00 | Gm16571 | 287 | 0 | 0 | 0 | 0 | 0 | 0 |
| ENSMUSG00 | Gm16112 | 381 | 0 | 0 | 0 | 0 | 0 | 0 |
| ENSMUSG00 | Gm16337 | 915 | 0.04344861 | 0 | 0 | 0 | 0 | 0 |
| ENSMUSG00 | Gm16543 | 600 | 0 | 0 | 0 | 0 | 0 | 0 |
| ENSMUSG00 | Gm16575 | 1613 | 0.17252843 | 0 | 0.0822105 | 0.35791241 | 0.50615128 | 0.52418712 |
| ENSMUSG00 | Gm16118 | 677 | 0 | 0 | 0 | 0 | 0 | 0 |
| ENSMUSG00 | 1700028K03I | 5315 | 0.05983892 | 0.43517389 | 0.70689683 | 0.06109848 | 0.23041073 | 0.08210616 |
| ENSMUSG00 | Rps19-ps10 | 434 | 0 | 0 | 0 | 0 | 0 | 0 |
| ENSMUSG00 | Gm16107 | 861 | 0 | 0 | 0 | 0.37716423 | 0 | 0 |
| ENSMUSG00 | Gm10171 | 1256 | 0 | 0 | 0 | 0 | 0 | 0.02171548 |
| ENSMUSG00 | Gm16136 | 2033 | 16.6022641 | 15.7619203 | 20.2854498 | 20.019944 | 18.5482006 | 28.6162349 |
| ENSMUSG00 | Gm16570 | 435 | 0.27417573 | 0 | 0 | 0 | 0.23460403 | 0.12540064 |
| ENSMUSG00 | Gm16298 | 419 | 0.09488182 | 0 | 2.32086049 | 1.29171997 | 0 | 0.32547302 |
| ENSMUSG00 | Gm7241 | 718 | 0.05536975 | 0 | 0 | 0 | 0 | 0 |
| ENSMUSG00 | A930011G23 | 421331 | 0.00434042 | 0.00160114 | 0.00514059 | 0 | 0.00326991 | 0.00576137 |
| ENSMUSG00 | Gm16536 | 1588 | 4.08069479 | 5.12815411 | 6.6803793 | 4.77155501 | 3.34177776 | 7.09346726 |
| ENSMUSG00 | Gm16138 | 677 | 0.29361507 | 0.85411714 | 0.58761682 | 0.90604839 | 0.45222785 | 0.16115001 |
| ENSMUSG00 | Gm15867 | 601 | 0 | 0 | 0 | 0 | 0.08490246 | 0 |
| ENSMUSG00 | Gm15310 | 371 | 0 | 0 | 0 | 0 | 0 | 0 |
| ENSMUSG00 | Gm5083 | 1364 | 0 | 0 | 0 | 0 | 0 | 0 |
| ENSMUSG00 | Ear-ps8 | 353 | 0 | 0 | 0 | 0 | 0 | 0 |
| ENSMUSG00 | Gm7162 | 1023 | 0 | 0 | 0 | 0 | 0.99758311 | 0.02666143 |
| ENSMUSG00 | Gm15950 | 1972 | 0.64511936 | 0 | 0 | 0 | 0 | 0 |
| ENSMUSG00 | Gm15679 | 1933 | 0 | 0 | 0 | 0 | 0 | 0 |
| ENSMUSG00 | Gm15775 | 536 | 0.8158774 | 1.70810149 | 0 | 0.13463449 | 0 | 0 |
| ENSMUSG00 | Gm16535 | 330 | 0.12047115 | 0 | 0.13394496 | 0.43735812 | 0 | 0.33060169 |
| ENSMUSG00 | Gm15759 | 3717 | 0 | 0 | 0 | 0 | 0 | 0 |
| ENSMUSG00 | Rbm12 | 8677 | 2.84524068 | 2.36017494 | 3.84607485 | 4.23736353 | 2.117033 | 2.68440036 |
| ENSMUSG00 | Gm16560 | 403 | 0 | 0 | 0 | 0 | 0 | 0 |
| ENSMUSG00 | 1700023H06 | 1042 | 2.63256061 | 3.60704653 | 1.44228662 | 1.86989482 | 3.03611833 | 5.47063302 |
| ENSMUSG00 | Gm16300 | 4113 | 0 | 0 | 0 | 0 | 0 | 0 |
| ENSMUSG00 | Gm16565 | 2178 | 0 | 0 | 0 | 0 | 0 | 0 |
| ENSMUSG00 | Gm15749 | 610 | 0 | 0 | 0 | 0 | 0 | 0 |
| ENSMUSG00 | Gm15508 | 324 | 0 | 0 | 0 | 0 | 0 | 0 |
| ENSMUSG00 | Shkbp1 | 4534 | 5.36620072 | 5.50520005 | 3.85084429 | 4.12229799 | 4.51291943 | 4.00637621 |
| ENSMUSG00 | Gm16542 | 1507 | 0.02638054 | 0 | 0.02933102 | 0 | 0.03385957 | 0 |
| ENSMUSG00 | Gm16303 | 389 | 0 | 0 | 0 | 0 | 0 | 0 |
| ENSMUSG00 | Gm7097 | 489 | 0 | 0 | 0 | 0 | 0 | 0 |
| ENSMUSG00 | Gm16290 | 293 | 0 | 0 | 0 | 0 | 0 | 0 |
| ENSMUSG00 | Npcd | 1444 | 0 | 0 | 0 | 0 | 0 | 0 |
| ENSMUSG00 | Gm2962 | 335 | 0.94938461 | 1.58224138 | 2.3750244 | 1.83102913 | 1.5231754 | 2.76817234 |
| ENSMUSG00 | Rps19-ps11 | 432 | 0 | 0 | 0 | 0 | 0.11811661 | 0 |
| ENSMUSG00 | Gm4491 | 1521 | 0 | 0 | 0 | 0 | 0 | 0 |
| ENSMUSG00 | Gm16338 | 1567 | 0 | 0 | 0 | 0 | 0 | 0 |
| ENSMUSG00 | A530032D15I | 1587 | 0 | 0 | 0.11140981 | 0 | 0.16076363 | 0.08593144 |
| ENSMUSG00 | Fxc1 | 2575 | 8.67673012 | 5.65138079 | 9.33817577 | 7.51067024 | 7.3715774 | 9.79768594 |
| ENSMUSG00 | 9530072K05I | 816 | 0 | 0 | 0 | 0 | 0 | 0 |
| ENSMUSG00 | Gm16579 | 2564 | 0 | 0 | 0 | 0 | 0 | 0 |
| ENSMUSG00 | Gm15853 | 554 | 0.28704318 | 0.17395828 | 0 | 0 | 0 | 0.1969288 |
| ENSMUSG00 | Gm16133 | 942 | 0 | 0.25576668 | 0.1876936 | 0 | 0.37917689 | 0.02895397 |
| ENSMUSG00 | Gm15662 | 158 | 0 | 0 | 0 | 0 | 0 | 0 |
| ENSMUSG00 | Fzd10 | 2428 | 0 | 0 | 0 | 0 | 0 | 0 |
| ENSMUSG00 | Zfp882 | 3646 | 7.98162692 | 7.38788293 | 4.43715703 | 10.3119831 | 5.05225498 | 9.00676513 |
| ENSMUSG00 | Gm13769 | 1164 | 0 | 0 | 0 | 0 | 0 | 0 |
| ENSMUSG00 | Gm16139 | 419 | 0.47440908 | 0.69002065 | 0.10549366 | 0 | 0.12178133 | 0.13018921 |
| ENSMUSG00 | Gm16039 | 10558 | 8.12581236 | 9.30137991 | 12.7983552 | 8.99487982 | 8.51567287 | 9.51434896 |
| ENSMUSG00 | Gm20510 | 4367 | 0 | 0 | 0 | 0 | 0 | 0 |
| ENSMUSG00 | Gm15699 | 278 | 0 | 0 | 0 | 0 | 0 | 0 |
| ENSMUSG00 | Gm16134 | 335 | 0 | 0 | 0 | 0 | 0 | 0 |
| ENSMUSG00 | Gm15562 | 424 | 0.09376293 | 0 | 0 | 0 | 0 | 0 |
| ENSMUSG00 | Speer7-ps1 | 1078 | 0 | 0 | 0 | 0.03347128 | 0 | 0 |
| ENSMUSG00 | Rps6kc1 | 9500 | 1.12570782 | 1.61297775 | 1.33070812 | 2.32064518 | 2.27201652 | 1.83457837 |
| ENSMUSG00 | Mup13 | 1227 | 0 | 0 | 0 | 0 | 0 | 0 |
| ENSMUSG00 | 9230117E06I | 495 | 0 | 0.38938539 | 0 | 0 | 0 | 0 |
| ENSMUSG00 | Etohd2 | 3301 | 3.82982212 | 0.91964431 | 2.75843068 | 3.07151 | 2.05589457 | 2.1730476 |
| ENSMUSG00 | Tmem102 | 1931 | 5.53817932 | 4.79119466 | 4.23477005 | 4.82090497 | 3.27668079 | 2.81079918 |
| ENSMUSG00 | Gm16102 | 419 | 0.09488182 | 0.46001377 | 0.21098732 | 0.51668799 | 0 | 0 |
| ENSMUSG00 | 4930448H16 | 576 | 0 | 0.50194211 | 0 | 0.50113951 | 0 | 0 |
| ENSMUSG00 | Gm16280 | 621 | 0 | 0 | 0 | 0 | 0 | 0 |
| ENSMUSG00 | Gm3355 | 486 | 0 | 0.09914906 | 0 | 0 | 0 | 0 |
| ENSMUSG00 | Gm13747 | 549 | 0 | 0 | 0 | 0 | 0 | 0 |
| ENSMUSG00 | Gm16296 | 324 | 0.1227021 | 0.14872359 | 0.40927632 | 0 | 0.31497763 | 0 |
| ENSMUSG00 | Gm16184 | 572 | 0.20850777 | 0 | 0 | 0 | 0.08920695 | 0 |
| ENSMUSG00 | 4930428N03 | 1122 | 0 | 0 | 0 | 0 | 0 | 0 |
| ENSMUSG00 | Gm16547 | 292 | 0 | 0 | 0 | 0 | 0 | 0 |
| ENSMUSG00 | 0610040B10 | 706 | 1.01359582 | 1.09204402 | 1.87826528 | 1.2776928 | 0.65047788 | 2.89744751 |
| ENSMUSG00 | Gm16055 | 624 | 0 | 0.38610931 | 0 | 0.34694274 | 0.40886519 | 0.08741872 |
| ENSMUSG00 | Gm16561 | 191 | 0 | 0 | 0 | 0 | 0 | 0 |
| ENSMUSG00 | Olfr1199 | 932 | 0 | 0 | 0 | 0 | 0 | 0 |
| ENSMUSG00 | Gm9141 | 1006 | 0 | 0 | 0 | 0 | 0 | 0 |
| ENSMUSG00 | Gm15524 | 461 | 0 | 0 | 0 | 0 | 0 | 0 |
| ENSMUSG00 | Wnk1 | 1301 | 2.26126484 | 2.81488824 | 1.08720905 | 7.3772666 | 3.33377552 | 2.80922495 |
| ENSMUSG00 | Gm14199 | 493 | 0 | 0 | 0 | 0 | 0 | 0 |
| ENSMUSG00 | 0610038L08I | 1812 | 0.83372421 | 1.86150715 | 1.04893998 | 3.46483209 | 2.92866816 | 4.51566875 |
| ENSMUSG00 | Gm8113 | 2719 | 2.3247964 | 0.67344053 | 2.90994112 | 0.65024648 | 2.72115649 | 1.52473158 |
| ENSMUSG00 | Gm13625 | 478 | 0 | 0 | 0.18494495 | 0 | 0 | 0.17117974 |
| ENSMUSG00 | Gm15664 | 183 | 0 | 0 | 0 | 0 | 0 | 0 |
| ENSMUSG00 | Gm16563 | 422 | 0 | 0 | 0 | 0 | 0 | 0 |
| ENSMUSG00 | Gm6428 | 344 | 0 | 0 | 0 | 0 | 0.14833249 | 0.07928674 |
| ENSMUSG00 | Gm16169 | 2926 | 0 | 0 | 0 | 0 | 0 | 0 |
| ENSMUSG00 | Gm16558 | 531 | 0.44921447 | 0 | 0 | 0 | 0 | 0 |
| ENSMUSG00 | Gm16164 | 467 | 0 | 0 | 0 | 0 | 0 | 0 |
| ENSMUSG00 | Gm16113 | 371 | 0 | 0 | 0 | 0 | 0 | 0 |
| ENSMUSG00 | Hiat1 | 4073 | 58.4082483 | 58.7985803 | 66.9484826 | 63.7127582 | 53.8827504 | 59.2903648 |
| ENSMUSG00 | 4930486I03R | 534 | 0.74448465 | 0 | 0.16554997 | 0.33784686 | 0.09555501 | 0.0510761 |
| ENSMUSG00 | Gm16160 | 779 | 0 | 0 | 0 | 0 | 0 | 0 |
| ENSMUSG00 | Uckl1 | 4698 | 18.9807456 | 18.380184 | 18.1398794 | 18.7245689 | 18.1166443 | 19.4080713 |
| ENSMUSG00 | 3110039M20 | 4709 | 0 | 0 | 0 | 0 | 0 | 0 |
| ENSMUSG00 | Gm15386 | 196 | 0 | 0 | 0 | 0 | 0 | 0 |
| ENSMUSG00 | Gm15689 | 1156 | 0 | 0 | 0 | 0 | 0 | 0 |
| ENSMUSG00 | Ang-ps2 | 432 | 0 | 0 | 0 | 0 | 0 | 0 |
| ENSMUSG00 | Gm15734 | 1038 | 0 | 0 | 0 | 0 | 0 | 0.39414219 |
| ENSMUSG00 | Gm16539 | 325 | 0 | 0 | 0 | 0 | 0 | 0 |
| ENSMUSG00 | Bcl2a1b | 815 | 0 | 0 | 0 | 0 | 0 | 0 |
| ENSMUSG00 | Gm8459 | 759 | 0 | 0.06348675 | 0.05823695 | 0.04753893 | 0.06722843 | 0 |
| ENSMUSG00 | Gm16114 | 299 | 0.26592295 | 0 | 0 | 1.20675734 | 0 | 0 |
| ENSMUSG00 | Gm16101 | 429 | 0 | 0.11232271 | 0 | 0 | 0 | 0 |
| ENSMUSG00 | 4930473D10 | 485 | 0 | 0 | 0 | 0 | 0 | 0 |
| ENSMUSG00 | Gm16343 | 408 | 0 | 0 | 0 | 0.44218192 | 0 | 0 |
| ENSMUSG00 | Gm16199 | 287 | 0 | 0 | 0 | 0 | 0 | 0 |
| ENSMUSG00 | Gm7281 | 538 | 0 | 0 | 0 | 0 | 0 | 0 |
| ENSMUSG00 | Gm15744 | 212 | 1.31268096 | 0 | 0 | 0 | 0.24069045 | 0 |

| | | | | | | | | |
|---|---|---|---|---|---|---|---|---|
| ENSMUSG00 | Gm4117 | 53120 | 0 | 0.00725699 | 0.02413128 | 0.04483085 | 0.02977819 | 0.03748209 |
| ENSMUSG00 | Gm16168 | 1776 | 0 | 0 | 0 | 0 | 0 | 0 |
| ENSMUSG00 | Pira2 | 4736 | 0 | 0 | 0 | 0 | 0 | 0 |
| ENSMUSG00 | Ugt1a5 | 2198 | 0 | 0 | 0 | 0 | 0 | 0 |
| ENSMUSG00 | Gm8292 | 622 | 0.06391556 | 0.30988066 | 0.63957651 | 0.40606803 | 0 | 0.30694932 |
| ENSMUSG00 | Gm20459 | 8529 | 0 | 0 | 0 | 0 | 0 | 0 |
| ENSMUSG00 | Gm15509 | 2159 | 0 | 0 | 0 | 0 | 0 | 0 |
| ENSMUSG00 | Gm16583 | 552 | 0 | 0 | 0 | 0 | 0 | 0 |
| ENSMUSG00 | Gm16137 | 662 | 2.40214384 | 1.96530806 | 1.8027942 | 0 | 1.00202853 | 0.57680506 |
| ENSMUSG00 | Gm16215 | 429 | 0 | 0 | 0 | 0 | 0 | 0 |
| ENSMUSG00 | Gm14435 | 1504 | 0 | 0 | 0 | 0 | 0 | 0 |
| ENSMUSG00 | 4933413C19I | 2215 | 0.23332788 | 0.45684663 | 0.01995569 | 0 | 0 | 0.04925443 |
| ENSMUSG00 | Gm757 | 2019 | 0 | 0 | 0 | 0 | 0 | 0 |
| ENSMUSG00 | Gm8783 | 521 | 0 | 0 | 0 | 0 | 0 | 0 |
| ENSMUSG00 | C230030N03 | 2671 | 0.07442059 | 0.2164872 | 0.0661952 | 0.10807052 | 0.09551924 | 0.03063419 |
| ENSMUSG00 | Gm16553 | 713 | 0 | 0 | 0 | 0 | 0 | 0 |
| ENSMUSG00 | A830011K09I | 2637 | 0 | 0 | 0 | 0 | 0 | 0 |
| ENSMUSG00 | Gm16165 | 794 | 0 | 0.06068821 | 0 | 0 | 0 | 0 |
| ENSMUSG00 | Gm16268 | 1154 | 0.03445016 | 0 | 0.11490947 | 0 | 0 | 0.18907896 |
| ENSMUSG00 | Ugt1a1 | 2219 | 0 | 0 | 0 | 0.08130249 | 0.02299521 | 0 |
| ENSMUSG00 | Gm16567 | 573 | 0 | 0 | 0 | 0 | 0 | 0 |
| ENSMUSG00 | Gm16273 | 267 | 0 | 0 | 0 | 0 | 0 | 0 |
| ENSMUSG00 | Gm14555 | 230 | 0 | 0 | 0 | 0 | 0 | 0 |
| ENSMUSG00 | Gm16582 | 1926 | 0 | 0 | 0 | 0 | 0 | 0 |
| ENSMUSG00 | Gm16236 | 378 | 0 | 0.12747736 | 0 | 0 | 0 | 0 |
| ENSMUSG00 | Gm16030 | 602 | 0 | 0 | 0 | 0 | 0 | 0 |
| ENSMUSG00 | Gm16550 | 125 | 0 | 0 | 0 | 0 | 0 | 0 |
| ENSMUSG00 | 4921518J05R | 1704 | 0 | 0 | 0 | 0 | 0 | 0.03201249 |
| ENSMUSG00 | Gm16318 | 631 | 0 | 0 | 0 | 0 | 0 | 0 |
| ENSMUSG00 | Vmn1r190-ps | 1906 | 0 | 0 | 0 | 0 | 0 | 0 |
| ENSMUSG00 | Gm7420 | 845 | 1.50553299 | 0.6843045 | 0 | 0.93941418 | 0.90579365 | 0.03227768 |
| ENSMUSG00 | Gm2389 | 187 | 0 | 0 | 0 | 0 | 0 | 0 |
| ENSMUSG00 | Gm16245 | 894 | 0 | 0 | 0 | 0 | 0 | 0 |
| ENSMUSG00 | Gm15954 | 464 | 0.85679915 | 0.10385009 | 0.57157556 | 0.62210422 | 0.21994128 | 0.3526893 |
| ENSMUSG00 | Gm11565 | 502 | 0 | 0 | 0 | 0 | 0 | 0 |
| ENSMUSG00 | Gm7568 | 1010 | 0.03936186 | 0.1908374 | 0.61269881 | 0.42869756 | 0.05052116 | 0.10801837 |
| ENSMUSG00 | 2010320O07I | 523 | 0.83615733 | 0.8292122 | 2.36644666 | 0.5519242 | 1.36590681 | 5.68438944 |
| ENSMUSG00 | Fbxo24 | 3114 | 0.56173447 | 0.12379304 | 0.25550198 | 0.25491489 | 0.57351418 | 0.35910733 |
| ENSMUSG00 | Gpx2-ps1 | 302 | 0 | 0 | 0 | 0 | 0 | 0 |
| ENSMUSG00 | Gm16238 | 587 | 0.06772654 | 0.16417868 | 0.2259038 | 0 | 0.78234648 | 0 |
| ENSMUSG00 | Flt3l | 3926 | 0.32403856 | 0.28229449 | 0.20265746 | 0.52386056 | 0.63685492 | 0.65998236 |
| ENSMUSG00 | Gm15852 | 913 | 0 | 0 | 0.04841385 | 0 | 0 | 0 |
| ENSMUSG00 | Gm16332 | 3406 | 0 | 0 | 0 | 0 | 0 | 0 |
| ENSMUSG00 | G6pd2 | 1634 | 0.46227303 | 1.09112507 | 0.9467959 | 0.88328138 | 0.68701363 | 0.41729864 |
| ENSMUSG00 | Gm5822 | 1377 | 0.05774217 | 0 | 0 | 0.02620337 | 0 | 0.05942187 |
| ENSMUSG00 | Gm16239 | 2631 | 0 | 0 | 0 | 0 | 0 | 0 |
| ENSMUSG00 | Gm15716 | 398 | 0 | 0 | 0 | 0.36263361 | 0 | 0 |
| ENSMUSG00 | Tmsb15b2 | 447 | 0.35575374 | 1.29359576 | 0.39554222 | 0.64576366 | 0.45661186 | 1.09830762 |
| ENSMUSG00 | 1810020O05I | 1042 | 0 | 0 | 0.33936156 | 0 | 0 | 0 |
| ENSMUSG00 | A130049A11I | 4356 | 0.17340545 | 0.68584927 | 0.27397837 | 0.99399572 | 0.93712352 | 0.54474142 |
| ENSMUSG00 | Gm6485 | 438 | 0.54459562 | 0.66008825 | 0.8073396 | 1.23568646 | 1.16498575 | 0.31135433 |
| ENSMUSG00 | Ier3ip1 | 2061 | 34.4316025 | 33.0127396 | 40.9848239 | 36.4671998 | 26.5654058 | 39.8731141 |
| ENSMUSG00 | Gm9719 | 542 | 0 | 0 | 0.16310643 | 0 | 0 | 0.05032221 |
| ENSMUSG00 | Gm16006 | 564 | 0 | 0.08543695 | 0.6269765 | 0 | 0.1809446 | 0.87046721 |
| ENSMUSG00 | Gm15882 | 853 | 0 | 0 | 0 | 0 | 0 | 0 |
| ENSMUSG00 | Gm16125 | 338 | 1.999536 | 1.99588814 | 3.66169114 | 3.52280316 | 4.07607144 | 2.42082597 |
| ENSMUSG00 | Gm16540 | 424 | 0 | 0 | 0.31274889 | 0 | 1.08310703 | 0 |
| ENSMUSG00 | Gm16227 | 561 | 0 | 0 | 0 | 0 | 0 | 0 |
| ENSMUSG00 | Rpl30-ps2 | 290 | 0 | 0 | 0 | 0 | 0 | 0 |
| ENSMUSG00 | Gm16111 | 284 | 0 | 0 | 0 | 0 | 0 | 0 |
| ENSMUSG00 | Gm16282 | 448 | 0 | 0 | 0 | 0 | 0 | 0 |
| ENSMUSG00 | Vmn2r-ps12 | 597 | 0 | 0 | 0 | 0 | 0 | 0 |
| ENSMUSG00 | Gm16555 | 1197 | 0 | 0 | 0 | 0 | 0 | 0 |
| ENSMUSG00 | Gm6507 | 650 | 0 | 0 | 0 | 0 | 0 | 0 |
| ENSMUSG00 | Gm15446 | 3775 | 29.3506024 | 20.9212129 | 18.8048106 | 33.2623881 | 11.5840011 | 36.7322559 |
| ENSMUSG00 | Gm16578 | 781 | 0 | 0 | 0 | 0.0923996 | 0 | 0 |
| ENSMUSG00 | Gm16213 | 637 | 0 | 0 | 0 | 0 | 0 | 0 |
| ENSMUSG00 | Gm13741 | 537 | 0 | 0 | 0 | 0 | 0 | 0 |
| ENSMUSG00 | Gimap1 | 1738 | 0.36598831 | 0.77630632 | 0.66124736 | 1.49476824 | 2.90656571 | 0.5178729 |
| ENSMUSG00 | Gm6493 | 1784 | 0.15599124 | 0.27010338 | 0.04955364 | 0.2629297 | 0.25742006 | 0.25990407 |
| ENSMUSG00 | Gm16302 | 426 | 0 | 0.22622743 | 0 | 0 | 0.11978023 | 0 |
| ENSMUSG00 | Gm16350 | 444 | 0.62677559 | 1.4108643 | 0.99553701 | 1.13772213 | 1.1492427 | 1.04429925 |
| ENSMUSG00 | Gm16088 | 500 | 0 | 0 | 0 | 0 | 0 | 0 |
| ENSMUSG00 | Gm15996 | 760 | 0 | 0.31701607 | 0 | 0 | 0 | 0 |
| ENSMUSG00 | Gm15740 | 1291 | 0 | 0 | 0.03423845 | 0.16769347 | 0.07904938 | 0 |
| ENSMUSG00 | Gm17018 | 651 | 0 | 0 | 0 | 0 | 0 | 0 |
| ENSMUSG00 | A430072P03I | 1226 | 0 | 0 | 0 | 0 | 0 | 0 |
| ENSMUSG00 | 4732440D04 | 411 | 12.3812689 | 12.6621308 | 12.3679123 | 15.7145644 | 17.0087919 | 15.4622651 |
| ENSMUSG00 | Gm11503 | 301 | 0 | 0 | 0 | 0 | 0 | 0 |
| ENSMUSG00 | Gm16243 | 492 | 0.96964587 | 2.25261823 | 0.08984114 | 3.00683704 | 1.55568219 | 0.72067136 |
| ENSMUSG00 | Gm15917 | 484 | 0.16427885 | 0 | 0.27397837 | 0.29819872 | 1.26511676 | 0.45082048 |
| ENSMUSG00 | Galnt4 | 5088 | 5.00068937 | 7.0650719 | 6.99341266 | 4.64499591 | 6.87973542 | 5.12471602 |
| ENSMUSG00 | Gm15900 | 341 | 0 | 0 | 0 | 0 | 0 | 0.07998428 |
| ENSMUSG00 | Krtap4-9 | 688 | 0 | 0 | 0 | 0 | 0 | 0 |
| ENSMUSG00 | Gm16573 | 1296 | 0.52148393 | 0.59489435 | 0.23874452 | 0 | 0.15748881 | 1.19957904 |
| ENSMUSG00 | Gm15894 | 2064 | 0 | 0 | 0 | 0 | 0 | 0 |
| ENSMUSG00 | Gm16232 | 166 | 0 | 0 | 0 | 0 | 0 | 0 |
| ENSMUSG00 | Gm16027 | 757 | 0 | 0 | 0 | 0 | 0 | 0 |
| ENSMUSG00 | Gm8550 | 652 | 0 | 0 | 0 | 0 | 0 | 0 |
| ENSMUSG00 | Gm16167 | 1064 | 0.07472835 | 0 | 0 | 0 | 0 | 0 |
| ENSMUSG00 | Gm16557 | 190 | 0 | 0 | 0 | 0 | 0 | 0 |
| ENSMUSG00 | Tmem229a | 893 | 0 | 0 | 0 | 0 | 0 | 0 |
| ENSMUSG00 | Gm16297 | 458 | 0.08680236 | 0.10521057 | 0 | 0.07878176 | 0.1114113 | 0 |
| ENSMUSG00 | Gm16317 | 736 | 0 | 0 | 0 | 0 | 0 | 0 |
| ENSMUSG00 | Gm16339 | 927 | 0 | 0 | 0 | 0 | 0 | 0 |
| ENSMUSG00 | Gm8194 | 858 | 0 | 0 | 0 | 0 | 0 | 0 |
| ENSMUSG00 | Palm2 | 6583 | 0 | 0.02927932 | 0 | 0.08769751 | 0 | 0.20301646 |
| ENSMUSG00 | Gm15629 | 473 | 0 | 0 | 0 | 0 | 0 | 0 |
| ENSMUSG00 | Gm16333 | 316 | 0 | 0 | 0 | 0 | 0 | 0 |
| ENSMUSG00 | Olfr1062 | 955 | 0 | 0 | 0 | 0 | 0 | 0 |
| ENSMUSG00 | 3110047P20I | 10362 | 0 | 0 | 0 | 0 | 0 | 0 |
| ENSMUSG00 | 1810009N23I | 1384 | 0 | 0 | 0 | 0 | 0 | 0 |
| ENSMUSG00 | Dlx6as1 | 8889 | 0 | 0 | 0 | 0 | 0 | 0 |
| ENSMUSG00 | Gm11554 | 867 | 0 | 0 | 0 | 0 | 0 | 0 |
| ENSMUSG00 | 1110002E22I | 9645 | 0 | 0.40467619 | 0 | 0 | 0 | 0 |
| ENSMUSG00 | Gm2976 | 512 | 0 | 0 | 0 | 0 | 0 | 0 |
| ENSMUSG00 | D030054H15 | 4878 | 0 | 0 | 0.01812294 | 0.00739689 | 0.01046051 | 0 |
| ENSMUSG00 | Gm16577 | 469 | 0 | 0 | 0 | 2.30801991 | 0 | 1.39571715 |
| ENSMUSG00 | Cdk5r2 | 3353 | 0 | 0 | 0 | 0 | 0 | 0 |
| ENSMUSG00 | Gm13538 | 545 | 0 | 0 | 0 | 0 | 0 | 0 |
| ENSMUSG00 | Gm6921 | 434 | 0 | 0 | 0 | 0 | 0 | 0 |
| ENSMUSG00 | Gm16251 | 336 | 0 | 0 | 0 | 0 | 0.30372843 | 0.08117452 |
| ENSMUSG00 | Lime1 | 1822 | 0.65459079 | 0.63472811 | 0.21834061 | 0.55449904 | 0.78415945 | 0.11975692 |
| ENSMUSG00 | Gm16147 | 278 | 0 | 0 | 0 | 0 | 0 | 0 |
| ENSMUSG00 | Gm15849 | 332 | 0 | 0 | 0 | 0 | 0 | 0.16430506 |
| ENSMUSG00 | Gm15872 | 637 | 0 | 0 | 0 | 0 | 0 | 0 |
| ENSMUSG00 | Gm16587 | 861 | 0 | 0 | 0 | 0 | 0 | 0 |
| ENSMUSG00 | Rnf8 | 8382 | 5.90498368 | 6.19720647 | 6.89763907 | 6.99514702 | 7.1590334 | 6.25735281 |
| ENSMUSG00 | Srpx | 3186 | 0 | 0 | 0.11099019 | 0 | 0.32031623 | 0 |
| ENSMUSG00 | AI480526 | 4686 | 1.04351773 | 0.73009761 | 1.41491175 | 0.73919682 | 1.03446558 | 1.6704698 |
| ENSMUSG00 | Gm15939 | 2142 | 0 | 0 | 0 | 0 | 0 | 0 |
| ENSMUSG00 | Gm15934 | 523 | 0 | 0 | 0 | 0 | 0 | 0 |
| ENSMUSG00 | Mageа9-ps | 1629 | 0 | 0 | 0 | 0 | 0 | 0 |
| ENSMUSG00 | Mup-ps16 | 590 | 0 | 0 | 0 | 0 | 0 | 0 |
| ENSMUSG00 | Ang-ps3 | 432 | 0 | 0 | 0 | 0 | 0 | 0 |
| ENSMUSG00 | Gm14399 | 1788 | 3.95776036 | 6.79137775 | 4.3756858 | 6.09439455 | 3.76704788 | 1.96780115 |
| ENSMUSG00 | Rpl29-ps1 | 506 | 0 | 0 | 0 | 0 | 0 | 0 |
| ENSMUSG00 | Gm13757 | 286 | 0 | 0 | 0 | 0 | 0 | 0 |
| ENSMUSG00 | Alms1-ps2 | 2411 | 0 | 0 | 0.01833341 | 0 | 0 | 0 |
| ENSMUSG00 | Gm8837 | 1884 | 0 | 0 | 0.0938468 | 0.01915183 | 0.08125219 | 0 |
| ENSMUSG00 | Ttbk2 | 11388 | 0.88322239 | 1.64598929 | 1.42060718 | 1.50817116 | 0.8378936 | 1.76274451 |
| ENSMUSG00 | Snhg9 | 181 | 2.63572247 | 0.79867031 | 2.1978817 | 2.59152806 | 7.32975564 | 9.19200549 |
| ENSMUSG00 | Gm4985 | 1232 | 0 | 0 | 0 | 0 | 0 | 0 |
| ENSMUSG00 | Gm16094 | 425 | 0 | 0 | 0 | 0 | 0 | 0 |
| ENSMUSG00 | Gm16241 | 2192 | 0 | 0 | 0 | 0 | 0.02327846 | 0 |
| ENSMUSG00 | Gm15890 | 714 | 0.05567995 | 0.13497603 | 0.06190734 | 0.35374554 | 0 | 0.03819978 |
| ENSMUSG00 | Gm15838 | 179 | 0 | 0 | 0 | 0 | 0 | 0 |
| ENSMUSG00 | Gm11492 | 1970 | 0 | 0 | 0 | 0 | 0 | 0 |
| ENSMUSG00 | Gm15717 | 344 | 0 | 1.54084553 | 0.77096238 | 0.52444832 | 2.52165229 | 0.9514409 |
| ENSMUSG00 | Ear-ps10 | 459 | 0 | 0 | 0 | 0 | 0 | 0 |
| ENSMUSG00 | Mtcp1 | 1455 | 8.36094642 | 8.27945741 | 7.86823185 | 9.42348929 | 7.64518895 | 19.9264203 |
| ENSMUSG00 | Gm16584 | 188 | 0 | 0 | 0 | 0 | 0 | 0 |
| ENSMUSG00 | Shprh | 14133 | 2.79607639 | 4.00275123 | 4.26599546 | 3.77083279 | 4.24587971 | 2.92373017 |
| ENSMUSG00 | Nhlrc4 | 2478 | 9.30515686 | 7.50603982 | 2.42592843 | 6.33401508 | 6.77468832 | 3.45610844 |
| ENSMUSG00 | Gm3076 | 450 | 0.88345512 | 1.07080983 | 0.19645264 | 0.48109393 | 0.22678389 | 0.30305155 |
| ENSMUSG00 | Usp49 | 8574 | 0.24574766 | 0.17422203 | 0.31963078 | 0.65649626 | 0.57132401 | 0.89706651 |
| ENSMUSG00 | Gm15680 | 463 | 0 | 0 | 0 | 0.07793098 | 0 | 0 |
| ENSMUSG00 | Gm16541 | 857 | 0 | 0.11245377 | 0 | 0 | 0 | 0 |
| ENSMUSG00 | Gm16163 | 1117 | 0 | 0 | 0.15828771 | 0.16151318 | 0.31977138 | 0.19534209 |
| ENSMUSG00 | Gm8185 | 668 | 0.17854258 | 0 | 0.19851127 | 0 | 0 | 0 |
| ENSMUSG00 | A730090N16I | 623 | 0 | 0 | 0.70949989 | 0.40541623 | 0 | 0 |
| ENSMUSG00 | Abhd12b | 3390 | 0 | 0 | 0.19558338 | 0 | 0 | 0 |
| ENSMUSG00 | Kcne1l | 2693 | 0 | 0 | 0.03282721 | 0 | 0.05684335 | 0.10127976 |
| ENSMUSG00 | Ugt1a7c | 2792 | 0 | 0.84567896 | 0.03166321 | 0 | 0.29241476 | 0.13676395 |
| ENSMUSG00 | Pou3f1 | 3006 | 0 | 0 | 0 | 0 | 0 | 0 |
| ENSMUSG00 | 4930519F09I | 1393 | 0 | 0.03459185 | 0 | 0.0259024 | 0 | 0.01957978 |
| ENSMUSG00 | Gm11567 | 556 | 0 | 0 | 0 | 0 | 0 | 0 |
| ENSMUSG00 | Gm16038 | 191 | 0 | 0 | 0 | 0 | 0 | 0 |
| ENSMUSG00 | Olfr287 | 3193 | 0 | 0 | 0 | 0 | 0 | 0 |
| ENSMUSG00 | Gm16162 | 294 | 0 | 0 | 0 | 0 | 0 | 0 |
| ENSMUSG00 | Cypt2-ps | 695 | 0 | 0 | 0 | 0 | 0 | 0 |
| ENSMUSG00 | Gm15809 | 659 | 0.84457774 | 0.36560275 | 0.8048894 | 0.65703268 | 0.07743001 | 0.2069396 |
| ENSMUSG00 | Gm10177 | 664 | 0 | 0.14513989 | 0.06656904 | 0.10868086 | 0 | 0.28753385 |
| ENSMUSG00 | Uba52 | 1758 | 1.67343888 | 2.00091597 | 2.38860926 | 2.09349746 | 1.3061359 | 2.38924599 |
| ENSMUSG00 | Gm3687 | 320 | 0 | 0 | 0 | 0 | 0 | 0 |
| ENSMUSG00 | Gm614 | 768 | 0 | 0 | 0 | 0 | 0 | 0 |
| ENSMUSG00 | Gm15795 | 302 | 1.8429693 | 1.11690429 | 0 | 0 | 0 | 0.81282038 |
| ENSMUSG00 | Ugt1a6b | 2376 | 0 | 0 | 0 | 0 | 0 | 0 |
| ENSMUSG00 | 4930405H06 | 789 | 0 | 0 | 0 | 0 | 0 | 0 |
| ENSMUSG00 | Gm3329 | 555 | 0 | 0 | 0 | 0 | 0 | 0 |
| ENSMUSG00 | Acad11 | 4054 | 47.5614407 | 34.5886894 | 51.8450329 | 36.0820445 | 51.7186429 | 32.6030838 |
| ENSMUSG00 | Gm16043 | 614 | 0.12949668 | 0 | 0 | 0 | 0 | 0 |
| ENSMUSG00 | Gm16285 | 346 | 0 | 0 | 0 | 0 | 0 | 0 |
| ENSMUSG00 | Gm16534 | 2045 | 0.01944033 | 0 | 0 | 0.07057613 | 0.02495177 | 0 |
| ENSMUSG00 | Gm8580 | 463 | 0.17172994 | 1.35296706 | 0 | 0.77930982 | 0.44083262 | 0.29454254 |
| ENSMUSG00 | 4930480K15I | 2196 | 0 | 0 | 0 | 0 | 0 | 0 |
| ENSMUSG00 | Gm6808 | 806 | 0 | 0 | 0 | 0 | 0 | 0 |
| ENSMUSG00 | Gm16231 | 317 | 0 | 0 | 0 | 0 | 0 | 0 |
| ENSMUSG00 | BC035044 | 2611 | 0 | 0 | 0 | 0 | 0 | 0 |
| ENSMUSG00 | Ugt1a10 | 1917 | 0 | 0.02513638 | 0 | 0 | 0 | 0 |
| ENSMUSG00 | Ear10 | 1146 | 0 | 0 | 0 | 0 | 0 | 0 |
| ENSMUSG00 | D630014O11 | 3017 | 0 | 0.17568806 | 0.26371666 | 0 | 0 | 0 |
| ENSMUSG00 | Ugt1a2 | 2206 | 0.21625828 | 0 | 0 | 0 | 0 | 0 |
| ENSMUSG00 | Fbxw10 | 4683 | 0 | 0.22637235 | 0 | 0.12327839 | 0 | 0.22714306 |
| ENSMUSG00 | Gm10612 | 1678 | 0 | 0 | 0 | 0 | 0 | 0 |
| ENSMUSG00 | Ugt1a9 | 2185 | 0 | 0 | 0 | 0 | 0 | 0 |
| ENSMUSG00 | Cd200r2 | 1744 | 0 | 0 | 0 | 0 | 0 | 0 |
| ENSMUSG00 | Gm16307 | 209 | 0 | 0 | 0 | 0 | 0 | 0 |
| ENSMUSG00 | Ang-ps1 | 435 | 0 | 0 | 0 | 0 | 0 | 0 |
| ENSMUSG00 | Gm15523 | 435 | 0 | 0 | 0 | 0 | 0 | 0 |
| ENSMUSG00 | Gm7592 | 1761 | 0 | 0 | 0 | 0 | 0 | 0.17036969 |
| ENSMUSG00 | AK157302 | 389 | 0 | 0 | 0 | 0 | 0 | 0 |
| ENSMUSG00 | Gm7418 | 800 | 0 | 0 | 0 | 0.13530767 | 0 | 0 |
| ENSMUSG00 | Gm2666 | 890 | 0 | 0 | 0 | 0 | 0 | 0 |
| ENSMUSG00 | 9230105E05I | 2247 | 0 | 0 | 0.01967149 | 0.256926 | 0 | 0 |
| ENSMUSG00 | Gm16556 | 798 | 0 | 0 | 0 | 0 | 0 | 0 |
| ENSMUSG00 | Gm15995 | 572 | 0 | 0 | 0 | 0 | 0 | 0 |
| ENSMUSG00 | Gm16161 | 288 | 0 | 0 | 1.07435035 | 0.50113951 | 0.17717492 | 1.04173969 |
| ENSMUSG00 | Dnaja1-ps | 641 | 0 | 0 | 0 | 0.05629024 | 0 | 0 |
| ENSMUSG00 | Gm16029 | 2425 | 0 | 0 | 0 | 0 | 0 | 0 |
| ENSMUSG00 | 1700025N21I | 684 | 0 | 0 | 0 | 0 | 0 | 0 |
| ENSMUSG00 | 4930503B20I | 2571 | 0 | 0 | 0 | 0 | 0 | 0 |
| ENSMUSG00 | AU015336 | 846 | 0 | 0.11391594 | 0 | 0 | 0 | 1.74093442 |
| ENSMUSG00 | Tepp | 1568 | 0.35495965 | 0.73754758 | 0.25370956 | 0.43721865 | 0.6183043 | 0.97409426 |
| ENSMUSG00 | 4930524O07I | 1339 | 0.74226065 | 0.14394755 | 0.39613302 | 0.08084103 | 0.60972518 | 0.54997406 |
| ENSMUSG00 | Gm15851 | 659 | 0 | 0 | 0 | 0 | 0 | 0 |
| ENSMUSG00 | Itga10 | 5157 | 0.0462542 | 0.00934389 | 0.0257137 | 0.10495068 | 0.04947293 | 0.11106601 |
| ENSMUSG00 | Gm16050 | 2361 | 0 | 0 | 0.01872166 | 0 | 0 | 0 |
| ENSMUSG00 | Gm15647 | 689 | 0 | 0 | 0 | 0 | 0 | 0 |
| ENSMUSG00 | Tmem189 | 4212 | 16.6969243 | 14.7693962 | 11.4492428 | 14.6915257 | 11.6178287 | 9.96573357 |
| ENSMUSG00 | Gm15657 | 1004 | 0 | 0 | 0 | 0 | 0 | 0 |
| ENSMUSG00 | Trim34b | 2037 | 0.23420018 | 0.3075227 | 0.23869429 | 0.30112654 | 0.1502966 | 0.62931176 |
| ENSMUSG00 | Gm16549 | 501 | 0.31740903 | 0.09618052 | 0 | 0.86424059 | 0.10184905 | 0.48996358 |
| ENSMUSG00 | Gm16116 | 428 | 0 | 0 | 0 | 0 | 0.1192205 | 0.31862896 |
| ENSMUSG00 | 4930516K23I | 1070 | 0 | 0 | 0 | 0 | 0 | 0 |
| ENSMUSG00 | 4930568K20I | 1612 | 0 | 0 | 0 | 0 | 0 | 0.20303702 |
| ENSMUSG00 | Gm4245 | 1753 | 0 | 0 | 0 | 0.02058303 | 0 | 0 |
| ENSMUSG00 | Gm16340 | 2157 | 0.29489462 | 0.29041435 | 0.32787644 | 0 | 0.09462471 | 0 |
| ENSMUSG00 | Pcp4 | 638 | 2.80406994 | 1.73712879 | 4.50332256 | 3.56296051 | 4.95867602 | 5.21552663 |
| ENSMUSG00 | Gm15974 | 530 | 0.07501034 | 0.63642471 | 0.16679941 | 0.40847598 | 0.09627618 | 0.05146158 |
| ENSMUSG00 | Gm11559 | 1095 | 0 | 0 | 0 | 0 | 0 | 0 |
| ENSMUSG00 | Gm16110 | 890 | 0 | 0 | 0 | 0 | 0 | 0 |
| ENSMUSG00 | Gm16554 | 458 | 0 | 0 | 0 | 0 | 0.1114113 | 0 |
| ENSMUSG00 | Gm16289 | 182 | 0 | 0 | 0 | 0 | 0 | 0 |
| ENSMUSG00 | Gm16315 | 781 | 0 | 0 | 0 | 2.81818786 | 0 | 0 |
| ENSMUSG00 | Cfb | 3011 | 0.07922049 | 0.03200694 | 0.0734006 | 0 | 0.03389331 | 0.027175 |
| ENSMUSG00 | Gm16132 | 596 | 0.93385357 | 0.16169947 | 0 | 0.12108069 | 0 | 0.82373072 |
| ENSMUSG00 | Gm16042 | 1246 | 0 | 0 | 0 | 0 | 0 | 0 |
| ENSMUSG00 | Gm16244 | 2474 | 0 | 0.13633997 | 0.1071993 | 0 | 0 | 0 |
| ENSMUSG00 | Car15 | 1536 | 7.42827012 | 10.6976411 | 6.96409245 | 8.33927461 | 7.60744796 | 2.85886518 |
| ENSMUSG00 | Gm16308 | 936 | 0 | 0 | 0 | 0 | 0 | 0 |
| ENSMUSG00 | Gm16207 | 728 | 0 | 0 | 0 | 0 | 0 | 0 |
| ENSMUSG00 | Gm16103 | 1278 | 0.06221515 | 0.03770457 | 0 | 0.16939927 | 0.4791209 | 0.02134166 |
| ENSMUSG00 | Gm16572 | 277 | 3.58803976 | 0 | 0 | 0 | 0.18421074 | 0 |
| ENSMUSG00 | Gm17017 | 2403 | 0.03308821 | 0 | 0.12876109 | 0 | 0.02123445 | 0.1248527 |
| ENSMUSG00 | Bloc1s1 | 1281 | 2.6689862 | 5.04058021 | 4.03717067 | 4.16872958 | 2.27049447 | 7.17529542 |
| ENSMUSG00 | Gm14027 | 617 | 1.22423684 | 1.32766534 | 0.7163994 | 0.40935869 | 0.90970848 | 1.1493365 |
| ENSMUSG00 | Gm3287 | 1323 | 0 | 0 | 0 | 0 | 0 | 0 |
| ENSMUSG00 | Gm16028 | 888 | 0 | 0 | 0 | 0 | 0 | 0 |
| ENSMUSG00 | Krtap1-4 | 520 | 0 | 0 | 0 | 0 | 0 | 0 |
| ENSMUSG00 | Gm1965 | 2434 | 0 | 0 | 0 | 0 | 0 | 0 |
| ENSMUSG00 | 4921534H16I | 3949 | 0.07047059 | 0 | 0 | 0 | 0 | 0 |
| ENSMUSG00 | Alms1-ps1 | 2157 | 0 | 0 | 0 | 0 | 0 | 0 |
| ENSMUSG00 | Gm4524 | 2150 | 0.98001882 | 0.76201816 | 0.16447197 | 0.93981139 | 1.35279229 | 0.32983285 |
| ENSMUSG00 | Churc1 | 5772 | 2.75505755 | 2.63806579 | 3.26229819 | 2.6567687 | 1.77690602 | 6.9084412 |
| ENSMUSG00 | Gm16545 | 462 | 0 | 0 | 0 | 0 | 0 | 0 |
| ENSMUSG00 | 2810442N19I | 1087 | 0 | 0.17731901 | 0 | 0 | 0 | 0.02509166 |
| ENSMUSG00 | Vmn2r-ps10 | 743 | 0 | 0 | 0 | 0 | 0 | 0 |
| ENSMUSG00 | Mpv17 | 4399 | 26.8410496 | 26.5304758 | 31.470828 | 30.5700796 | 26.7601384 | 29.6555125 |
| ENSMUSG00 | D730045A05I | 1266 | 0.25121947 | 0.72317725 | 0 | 0 | 0 | 0 |
| ENSMUSG00 | Eif4ebp3 | 741 | 1.18032466 | 0.7153183 | 0.29825805 | 0.43824346 | 0.27544602 | 0.47850244 |
| ENSMUSG00 | Gm9550 | 861 | 0 | 0 | 0 | 0 | 0 | 0 |
| ENSMUSG00 | 1110005A03I | 2163 | 10.641897 | 11.606828 | 10.9533924 | 14.8465186 | 6.1571355 | 9.88595339 |
| ENSMUSG00 | Gm5286 | 1591 | 0 | 0 | 0 | 0 | 0 | 0 |
| ENSMUSG00 | Gm17226 | 1165 | 0 | 0 | 0 | 0 | 0.04379946 | 0 |
| ENSMUSG00 | Mndal | 1897 | 0.44009757 | 1.19386546 | 0.27961103 | 1.86401706 | 1.80219672 | 0 |
| ENSMUSG00 | Prr22 | 1958 | 0.28425778 | 0.12305016 | 0.04514999 | 0.38698822 | 0.13030229 | 0.2089477 |
| ENSMUSG00 | Gm17671 | 52 | 0.76452847 | 0 | 0 | 0 | 0 | 1.04902459 |
| ENSMUSG00 | Cbln1 | 1815 | 0 | 0 | 0 | 0 | 0 | 0 |
| ENSMUSG00 | Gm8338 | 362 | 0 | 0 | 0 | 0 | 0 | 0 |
| ENSMUSG00 | D130062J21I | 2742 | 0.15948588 | 0 | 0 | 0.51320195 | 0 | 0 |
| ENSMUSG00 | Gm9603 | 2058 | 0 | 0 | 0 | 0 | 0 | 0 |
| ENSMUSG00 | Vmn2r20 | 2556 | 0 | 0 | 0 | 0 | 0 | 0 |
| ENSMUSG00 | Gm17423 | 2108 | 0 | 0 | 0 | 0 | 0 | 0 |
| ENSMUSG00 | Gm17688 | 1595 | 0.04985013 | 0.12084374 | 0.08313826 | 0.1583538 | 0 | 0 |
| ENSMUSG00 | Gm17613 | 409 | 0 | 0 | 0 | 0 | 0 | 0 |
| ENSMUSG00 | Gm17615 | 2274 | 0 | 0 | 0 | 0 | 0 | 0 |
| ENSMUSG00 | Gm17614 | 1212 | 0 | 0 | 0 | 0 | 0 | 0 |
| ENSMUSG00 | Gm17296 | 5030 | 0.54535351 | 0.65142705 | 0.60634735 | 1.11904552 | 1.23761786 | 0.62899765 |
| ENSMUSG00 | Lrrc10b | 1358 | 0 | 0 | 0 | 0 | 0 | 0 |
| ENSMUSG00 | Gm17152 | 534 | 0 | 0 | 0 | 0 | 0 | 0 |
| ENSMUSG00 | Gm17034 | 2227 | 0 | 0 | 0 | 0 | 0 | 0 |
| ENSMUSG00 | Gm17298 | 2769 | 0 | 0 | 0 | 0 | 0 | 0 |
| ENSMUSG00 | Gm6541 | 476 | 0 | 0 | 0 | 0 | 0 | 0 |
| ENSMUSG00 | Gm17297 | 172 | 0 | 0 | 0 | 0 | 0 | 0 |
| ENSMUSG00 | Gm4794 | 876 | 0 | 0 | 0 | 0 | 0 | 0 |
| ENSMUSG00 | Gm17295 | 1499 | 0 | 0 | 0 | 0 | 0 | 0 |
| ENSMUSG00 | Gm17119 | 279 | 0 | 0 | 0 | 0 | 0 | 0 |
| ENSMUSG00 | Gm7714 | 682 | 0 | 0 | 0 | 0 | 0 | 0 |
| ENSMUSG00 | Gm9479 | 1572 | 0.17702822 | 0 | 0 | 0 | 0.19475716 | 0 |
| ENSMUSG00 | Vmn2r99 | 2565 | 0 | 0 | 0 | 0 | 0 | 0 |
| ENSMUSG00 | Gm5459 | 812 | 0.04895995 | 0.23737164 | 0.38105037 | 0.22218008 | 0 | 0.20153674 |
| ENSMUSG00 | Adh6-ps1 | 2785 | 0 | 0 | 0 | 0 | 0 | 0 |
| ENSMUSG00 | 1700071M16 | 3298 | 0.15670747 | 0.4091026 | 0 | 0 | 0 | 0 |
| ENSMUSG00 | Gm5174 | 1442 | 0 | 0.06683279 | 0 | 0 | 0 | 0 |
| ENSMUSG00 | Gm17506 | 443 | 0 | 0 | 0 | 0 | 0 | 0 |
| ENSMUSG00 | Gm5156 | 1333 | 0 | 0 | 0 | 0 | 0 | 0 |
| ENSMUSG00 | Gm17327 | 5282 | 7.22553224 | 4.74383755 | 5.08798193 | 6.43492729 | 6.43384443 | 3.51131289 |
| ENSMUSG00 | Gm17326 | 172 | 0 | 0 | 0 | 0 | 0 | 0 |
| ENSMUSG00 | Gm17325 | 2655 | 0 | 0 | 0 | 0 | 0 | 0 |
| ENSMUSG00 | 2310050C09I | 1065 | 0 | 0 | 0 | 0 | 0 | 0 |
| ENSMUSG00 | Vmn2r104 | 2577 | 0 | 0 | 0 | 0 | 0 | 0 |
| ENSMUSG00 | Gm3662 | 1865 | 0 | 0 | 0 | 0 | 0 | 0 |
| ENSMUSG00 | Gm17324 | 921 | 0.08633112 | 0.20927879 | 0.09598663 | 0.15670812 | 0.16620969 | 0 |
| ENSMUSG00 | Gm17223 | 480 | 0 | 0 | 0 | 0 | 0 | 0 |
| ENSMUSG00 | Gm4462 | 491 | 0 | 0.29441818 | 0 | 0 | 0 | 0 |
| ENSMUSG00 | Gm17385 | 1499 | 0 | 0 | 0 | 0 | 0 | 0 |
| ENSMUSG00 | Gm17090 | 487 | 0 | 0 | 0.09076354 | 0 | 0 | 0 |
| ENSMUSG00 | Gm5263 | 361 | 0 | 0 | 0 | 0 | 0 | 0.07555302 |
| ENSMUSG00 | 4930505N22I | 943 | 0.37942664 | 0.76648635 | 0.18749456 | 0.49741949 | 0.32466411 | 0.34707919 |
| ENSMUSG00 | Gm17384 | 2418 | 0 | 0 | 0 | 0 | 0 | 0 |
| ENSMUSG00 | Gm17111 | 409 | 1.0692183 | 0.70689157 | 0.75651076 | 0.70576126 | 0.99807092 | 0.3334308 |
| ENSMUSG00 | Gm17073 | 555 | 0 | 0 | 0 | 0 | 0 | 0 |
| ENSMUSG00 | Gm17160 | 258 | 0 | 0 | 0 | 0 | 0 | 0.84572525 |
| ENSMUSG00 | 9130221H12I | 1463 | 1.33152327 | 1.28453263 | 1.11788667 | 0.41927188 | 0.76731392 | 1.13722009 |
| ENSMUSG00 | Gm17199 | 415 | 0.09579634 | 0.11611191 | 0 | 0 | 0 | 0 |
| ENSMUSG00 | Gm3216 | 483 | 0 | 0 | 0 | 0 | 0 | 0 |
| ENSMUSG00 | Gm3025 | 486 | 0 | 0 | 0 | 0 | 0 | 0 |
| ENSMUSG00 | Gm17149 | 374 | 0.31889423 | 0 | 0 | 0.09647606 | 0.13643416 | 0.65634159 |
| ENSMUSG00 | Gm17698 | 1656 | 0 | 0 | 0 | 0 | 0 | 0 |
| ENSMUSG00 | 4932443I19R | 777 | 0 | 0 | 0 | 0 | 0 | 0 |
| ENSMUSG00 | Gm8122 | 1399 | 0 | 0 | 0 | 0 | 0 | 0 |
| ENSMUSG00 | Gm17081 | 477 | 0 | 0 | 0 | 0 | 0 | 0 |
| ENSMUSG00 | Gm6669 | 1004 | 0 | 0 | 0 | 0 | 0 | 0 |
| ENSMUSG00 | C030048H21 | 374 | 0 | 0 | 0 | 0 | 0 | 0 |
| ENSMUSG00 | Gm17217 | 468 | 0 | 0 | 0 | 0 | 0 | 0 |
| ENSMUSG00 | Vmn2r46 | 2424 | 0 | 0 | 0 | 0 | 0 | 0 |
| ENSMUSG00 | Olfr723 | 1005 | 0 | 0 | 0 | 0 | 0 | 0 |
| ENSMUSG00 | Gm17237 | 61 | 0 | 0 | 0 | 0 | 0 | 0 |
| ENSMUSG00 | Gm5798 | 1746 | 0 | 0 | 0 | 0 | 0 | 0 |
| ENSMUSG00 | Vmn1r5 | 1055 | 0 | 0 | 0.12569245 | 0 | 0 | 0 |
| ENSMUSG00 | Gm17236 | 3653 | 0 | 0 | 0 | 0 | 0 | 0 |
| ENSMUSG00 | Gm17261 | 43 | 0 | 0 | 0 | 0 | 0 | 0 |
| ENSMUSG00 | Gm17148 | 248 | 0 | 0 | 0 | 0 | 0 | 0 |
| ENSMUSG00 | Gm17204 | 475 | 0 | 0 | 0 | 0 | 0 | 0 |
| ENSMUSG00 | Gm3972 | 2869 | 0 | 0 | 0 | 0 | 0 | 0 |
| ENSMUSG00 | Gm17555 | 1499 | 0 | 0 | 0 | 0 | 0 | 0 |
| ENSMUSG00 | Gm17556 | 3023 | 0 | 0 | 0 | 0 | 0 | 0 |
| ENSMUSG00 | Vmn2r-ps128 | 2567 | 0 | 0 | 0 | 0 | 0 | 0 |
| ENSMUSG00 | 2310042E22I | 1247 | 0 | 0 | 0 | 0 | 0 | 0 |
| ENSMUSG00 | Gm17077 | 277 | 0 | 0 | 0 | 0.26052018 | 0.18421074 | 0 |
| ENSMUSG00 | Gm2822 | 2908 | 0 | 0 | 0 | 0.59557708 | 0.05264069 | 0 |
| ENSMUSG00 | Gm17621 | 1130 | 0 | 0 | 0 | 0 | 0 | 0 |
| ENSMUSG00 | Vmn2r79 | 2550 | 0 | 0 | 0 | 0 | 0 | 0 |
| ENSMUSG00 | Gm3512 | 1855 | 0 | 0 | 0 | 0 | 0 | 0 |
| ENSMUSG00 | Gm3043 | 816 | 0 | 0 | 0 | 0 | 0 | 0 |
| ENSMUSG00 | Gm17070 | 783 | 0 | 0 | 0 | 0 | 0 | 0 |
| ENSMUSG00 | Gm3623 | 4849 | 0 | 0 | 0 | 0 | 0 | 0 |
| ENSMUSG00 | 4933411K16I | 1379 | 0 | 0.38437336 | 0.03205355 | 0 | 0.62904162 | 0.01977856 |
| ENSMUSG00 | Gm20533 | 1606 | 0 | 0 | 0 | 0 | 0 | 0 |
| ENSMUSG00 | Gm17435 | 2992 | 0.11958534 | 1.07904132 | 0.87162725 | 0.02411901 | 0.18759697 | 0.78396356 |
| ENSMUSG00 | Gm17232 | 449 | 0 | 0 | 0 | 0 | 0 | 0 |
| ENSMUSG00 | Gm6882 | 1355 | 0 | 0 | 0 | 0 | 0 | 0 |
| ENSMUSG00 | Gm8281 | 1947 | 0 | 0 | 0 | 0 | 0 | 0 |
| ENSMUSG00 | Gm10925 | 770 | 14.2500164 | 14.0178741 | 9.29960853 | 13.3550425 | 9.87393506 | 7.72191085 |
| ENSMUSG00 | Gm8229 | 1398 | 0 | 0 | 0 | 0 | 0 | 0 |
| ENSMUSG00 | 2610020F03I | 586 | 0 | 0 | 0 | 0 | 0 | 0 |
| ENSMUSG00 | Gm6158 | 563 | 0 | 0.42794354 | 0 | 0.19226867 | 0 | 0.09689037 |
| ENSMUSG00 | Gm17078 | 476 | 0 | 0 | 0 | 0 | 0 | 0 |
| ENSMUSG00 | Vmn2r58 | 2583 | 0 | 0 | 0 | 0 | 0 | 0 |
| ENSMUSG00 | Gm17492 | 6842 | 0 | 0 | 0 | 0 | 0 | 0 |
| ENSMUSG00 | Gm7316 | 442 | 0 | 0 | 0 | 0 | 0 | 0 |
| ENSMUSG00 | 2810055G20I | 2171 | 2.72849682 | 1.30953482 | 1.95457252 | 2.84202193 | 1.5277358 | 3.45487139 |
| ENSMUSG00 | Gm17056 | 2297 | 0.74422536 | 0.46151577 | 0.01924329 | 0.76970839 | 0.24435792 | 0.02374805 |
| ENSMUSG00 | Mir706 | 2231 | 0.05345874 | 0 | 0.41606396 | 0.42049895 | 0.16010069 | 0.75796846 |
| ENSMUSG00 | Gm7236 | 329 | 0 | 0 | 0 | 0 | 0 | 0 |
| ENSMUSG00 | Gm7360 | 506 | 0 | 0 | 0 | 0 | 0 | 0 |
| ENSMUSG00 | Gm14478 | 499 | 0 | 0 | 0 | 0 | 0 | 0 |
| ENSMUSG00 | Gm13278 | 1555 | 0 | 0 | 0 | 0 | 0 | 0 |
| ENSMUSG00 | Vmn1r91 | 923 | 0 | 0 | 0 | 0 | 0 | 0 |
| ENSMUSG00 | 4930523C07I | 5773 | 13.2495314 | 8.4052914 | 10.6963407 | 7.06265552 | 9.62545007 | 11.6175884 |
| ENSMUSG00 | Gm17309 | 277 | 0 | 0 | 0 | 0 | 0 | 0 |
| ENSMUSG00 | Gm6536 | 611 | 0 | 0 | 0 | 0 | 0 | 0 |
| ENSMUSG00 | Gm17096 | 731 | 2.06663237 | 4.21878563 | 0.12093528 | 1.33271574 | 0.76783876 | 4.10425488 |
| ENSMUSG00 | Gm17609 | 1089 | 0 | 0 | 0 | 0 | 0 | 0 |
| ENSMUSG00 | Nav1 | 1643 | 1.23404109 | 1.37843139 | 1.45276903 | 1.38354766 | 2.01869411 | 3.00469246 |
| ENSMUSG00 | A330094K24I | 1025 | 0 | 0 | 0 | 0 | 0 | 0 |
| ENSMUSG00 | 0610007N19I | 1480 | 15.6604359 | 19.6652777 | 5.34603372 | 14.4084381 | 13.618526 | 13.7478925 |
| ENSMUSG00 | Gm17400 | 850 | 0 | 0 | 0.26001084 | 0 | 0 | 0 |
| ENSMUSG00 | Lca5 | 58 | 2.05631796 | 0 | 0.76210074 | 0 | 0 | 0 |
| ENSMUSG00 | Gm8362 | 2139 | 0 | 0 | 0 | 0 | 0 | 0 |
| ENSMUSG00 | Gm16501 | 335 | 0 | 0 | 0 | 0 | 0 | 0 |
| ENSMUSG00 | Gm17058 | 926 | 0.04293248 | 0.10407439 | 0.04773417 | 0 | 0 | 0.14727127 |
| ENSMUSG00 | Gm16042 | 1250 | 0 | 0 | 0 | 0 | 0 | 0 |
| ENSMUSG00 | Gm17402 | 226 | 0 | 0 | 0 | 0 | 0 | 0 |
| ENSMUSG00 | Gm17401 | 2835 | 1.17794016 | 0.40792755 | 1.9333434 | 0.59818557 | 0.48596548 | 1.0871373 |
| ENSMUSG00 | Gm17582 | 341 | 0 | 0 | 0 | 0 | 0 | 0 |
| ENSMUSG00 | Vmn1r174 | 941 | 0 | 0 | 0 | 0 | 0 | 0 |
| ENSMUSG00 | Gm16484 | 326 | 0 | 0 | 0 | 0 | 0 | 0 |
| ENSMUSG00 | Gm17583 | 66 | 1.8070673 | 0 | 0.66972489 | 0 | 0 | 0 |
| ENSMUSG00 | Gm6713 | 1445 | 0 | 0 | 0 | 0 | 0 | 0 |
| ENSMUSG00 | Gm17584 | 172 | 0 | 0 | 0 | 0 | 0 | 0 |
| ENSMUSG00 | Gm4242 | 107 | 0 | 0 | 0 | 0 | 0 | 0.76470951 |
| ENSMUSG00 | Vmn2r94 | 2451 | 0 | 0 | 0 | 0 | 0 | 0 |
| ENSMUSG00 | Gm17585 | 2978 | 0.02669945 | 0 | 0.04452838 | 0 | 0 | 0.02747613 |
| ENSMUSG00 | Gm7980 | 1398 | 0 | 0 | 0 | 0 | 0 | 0 |
| ENSMUSG00 | 1700040D17I | 960 | 0.49694351 | 1.05407842 | 0.59856662 | 0.15034185 | 0.4252198 | 0.51139949 |
| ENSMUSG00 | Gm4303 | 869 | 0 | 0 | 0 | 0 | 0 | 0 |
| ENSMUSG00 | Gm17272 | 2213 | 0 | 0.91451901 | 0 | 0.27717793 | 0.25363314 | 0.30811838 |
| ENSMUSG00 | Gm2871 | 1203 | 0 | 0 | 0 | 0 | 0 | 0 |
| ENSMUSG00 | Efna5 | 2415 | 0 | 0 | 0 | 0 | 0 | 0 |
| ENSMUSG00 | Gm17392 | 1506 | 0 | 0 | 0 | 0 | 0 | 0 |
| ENSMUSG00 | Gm17225 | 495 | 0 | 0 | 0 | 0 | 0 | 0 |
| ENSMUSG00 | Gm17271 | 376 | 0 | 0 | 0 | 0 | 0 | 0.43523361 |
| ENSMUSG00 | Snhg4 | 1147 | 44.3653139 | 30.9199838 | 58.7687974 | 42.8454618 | 31.6301249 | 88.1729401 |
| ENSMUSG00 | Gm17456 | 3375 | 0 | 0 | 0 | 0 | 0 | 0 |
| ENSMUSG00 | 9430037G07I | 4892 | 0.32506525 | 0.29550149 | 0.03614215 | 0.37616195 | 0.40679245 | 0.01115071 |
| ENSMUSG00 | Gm6410 | 1103 | 0 | 0 | 0 | 0.32712642 | 0 | 0 |
| ENSMUSG00 | Vmn2r75 | 2556 | 0 | 0 | 0 | 0 | 0 | 0 |
| ENSMUSG00 | Gm17454 | 617 | 0 | 0 | 0 | 0 | 0 | 0 |
| ENSMUSG00 | Gm17455 | 853 | 0 | 0 | 0 | 0 | 0 | 0 |
| ENSMUSG00 | Gm9732 | 1462 | 1.2236639 | 1.48316682 | 0.27210437 | 0.76507755 | 0.52352643 | 0.18655704 |
| ENSMUSG00 | Gm17651 | 34 | 0 | 0 | 1.30005421 | 0 | 0 | 0 |
| ENSMUSG00 | Psg-ps1 | 1917 | 0 | 0 | 0 | 0 | 0 | 0 |
| ENSMUSG00 | Gm16478 | 395 | 0 | 0 | 0 | 0 | 0 | 0 |
| ENSMUSG00 | D930048G16 | 2092 | 8.28555904 | 9.19043519 | 8.19804737 | 11.9353608 | 10.5857778 | 8.13545645 |
| ENSMUSG00 | Gm17653 | 2139 | 0.65051043 | 0.58571645 | 0.04132945 | 0.33737302 | 0.42939447 | 0.05100447 |
| ENSMUSG00 | A930017M01 | 836 | 0 | 0.46111428 | 1.32182545 | 0 | 0.85450869 | 1.2723815 |
| ENSMUSG00 | Gm17652 | 2359 | 0.58984392 | 0.16341312 | 0.28106301 | 0.26002321 | 0.43261022 | 0.53184968 |
| ENSMUSG00 | Gm17192 | 510 | 0 | 0 | 0 | 0 | 0 | 0 |
| ENSMUSG00 | Gm17076 | 1400 | 0 | 0 | 0 | 0 | 0 | 0 |
| ENSMUSG00 | Trav5d-3 | 269 | 0 | 0 | 0 | 0 | 0 | 0 |
| ENSMUSG00 | Gm6133 | 947 | 0.04198044 | 0 | 0 | 0 | 0 | 0 |
| ENSMUSG00 | 1700022N22I | 707 | 0.33738739 | 0.68156212 | 1.37544632 | 0.56138966 | 1.51563493 | 1.11876172 |
| ENSMUSG00 | Gm17343 | 52 | 0 | 0 | 0 | 0 | 0 | 0 |
| ENSMUSG00 | Gm17341 | 1955 | 0 | 0 | 0 | 0 | 0 | 0 |
| ENSMUSG00 | Gm2431 | 514 | 0 | 0 | 0 | 0 | 0 | 0 |
| ENSMUSG00 | Vmn2r-ps100 | 660 | 0 | 0 | 0 | 0 | 0 | 0 |
| ENSMUSG00 | 4930571K23I | 2152 | 0 | 0 | 0 | 0 | 0 | 0 |
| ENSMUSG00 | Gm17122 | 370 | 0 | 0 | 0 | 0 | 0 | 0 |
| ENSMUSG00 | Gm17210 | 389 | 0.30659754 | 0 | 0.90903533 | 0 | 0 | 0.21034426 |
| ENSMUSG00 | Gm17511 | 496 | 0 | 0.29145026 | 0.53469971 | 0 | 0.30862727 | 0.76984869 |
| ENSMUSG00 | Ifitfb | 1106 | 0 | 0 | 0 | 0 | 0 | 0 |
| ENSMUSG00 | Gm3578 | 433 | 0 | 0 | 0 | 0 | 0 | 0 |
| ENSMUSG00 | Vmn1r-ps73 | 4211 | 0 | 0 | 0 | 0 | 0 | 0 |
| ENSMUSG00 | Gm17140 | 625 | 0 | 0 | 0 | 0 | 0 | 0 |
| ENSMUSG00 | Gm3091 | 483 | 0 | 0 | 0 | 0 | 0 | 0 |
| ENSMUSG00 | Gm17043 | 1279 | 0 | 0 | 0 | 0 | 0 | 0 |
| ENSMUSG00 | Gm4613 | 348 | 0 | 0 | 0 | 0 | 0 | 0 |
| ENSMUSG00 | Gm5331 | 398 | 0 | 0 | 0 | 0 | 0 | 0.06852924 |
| ENSMUSG00 | AC166053.1 | 846 | 0 | 0 | 0 | 0 | 0 | 0 |

| | | | | | | | | |
|---|---|---|---|---|---|---|---|---|
| ENSMUSG00 | 2410012M07 | 1813 | 0 | 0 | 0.19504399 | 0 | 0 | 0 |
| ENSMUSG00 | Gm4553 | 713 | 0 | 0 | 0 | 0 | 0 | 0 |
| ENSMUSG00 | Gm3047 | 817 | 0 | 0 | 0 | 0 | 0 | 0 |
| ENSMUSG00 | Gm17262 | 1648 | 0.62721025 | 0.58478692 | 0.75100219 | 1.29177223 | 1.14561645 | 0.26480232 |
| ENSMUSG00 | Gm9372 | 446 | 0 | 0 | 0 | 0 | 0 | 0 |
| ENSMUSG00 | Gm6245 | 767 | 0.05183244 | 0.06282457 | 0 | 0.09408617 | 0.06652722 | 1.20904529 |
| ENSMUSG00 | Gm17166 | 428 | 0 | 0 | 0 | 0 | 0 | 0 |
| ENSMUSG00 | Gm8068 | 1397 | 0 | 0 | 0 | 0 | 0 | 0 |
| ENSMUSG00 | Gm5656 | 1092 | 0 | 0 | 0 | 0 | 0 | 0 |
| ENSMUSG00 | Gm2663 | 865 | 0 | 0 | 0 | 0 | 0 | 0 |
| ENSMUSG00 | Gm2966 | 478 | 0 | 0 | 0 | 0 | 0 | 0 |
| ENSMUSG00 | Gm3675 | 1648 | 0 | 0 | 0 | 0 | 0 | 0 |
| ENSMUSG00 | Tmem29 | 1253 | 0 | 0 | 0 | 0 | 0 | 0 |
| ENSMUSG00 | Gm17417 | 3282 | 0.82369673 | 0.29364072 | 0.41750674 | 0.85752574 | 0.23321013 | 0.21606966 |
| ENSMUSG00 | Gm17416 | 200 | 0 | 0 | 0 | 0 | 0 | 0 |
| ENSMUSG00 | BC035947 | 2086 | 0.13340765 | 0 | 0 | 0 | 0 | 0 |
| ENSMUSG00 | Gm2888 | 1753 | 0 | 0 | 0 | 0 | 0 | 0 |
| ENSMUSG00 | A430105J06R | 1273 | 0.15614878 | 1.05987461 | 0.31250321 | 0.11337642 | 0.40083563 | 0.81416834 |
| ENSMUSG00 | Gm17415 | 800 | 0 | 0 | 0 | 0 | 0.06378297 | 0 |
| ENSMUSG00 | Gm17553 | 28 | 1.41983859 | 0 | 4.73591175 | 0 | 0 | 4.8704713 |
| ENSMUSG00 | Fryl | 1637 | 0 | 0 | 0 | 0 | 0 | 0 |
| ENSMUSG00 | Gm17176 | 164 | 0 | 0 | 0 | 0 | 0 | 0 |
| ENSMUSG00 | Gm3089 | 1845 | 0 | 0 | 0 | 0 | 0 | 0 |
| ENSMUSG00 | Traj36 | 62 | 0 | 0 | 0 | 0 | 0 | 0 |
| ENSMUSG00 | Vmn1r151 | 917 | 0 | 0 | 0 | 0 | 0 | 0 |
| ENSMUSG00 | Gm13693 | 2725 | 0 | 0 | 0 | 0 | 0 | 0 |
| ENSMUSG00 | Gm17611 | 230 | 0 | 0 | 0 | 0 | 0 | 0 |
| ENSMUSG00 | Kcnb2 | 4064 | 0 | 0 | 0 | 0 | 0 | 0 |
| ENSMUSG00 | Gm17057 | 589 | 0.74246229 | 0.57267419 | 1.95118492 | 0.85763773 | 1.73264434 | 1.20397389 |
| ENSMUSG00 | Gm3233 | 692 | 0 | 0 | 0 | 0 | 0 | 0 |
| ENSMUSG00 | Gm17735 | 1097 | 0 | 0 | 0 | 0 | 0 | 0 |
| ENSMUSG00 | Gm7352 | 1191 | 0.16689958 | 0 | 0 | 0 | 0 | 0 |
| ENSMUSG00 | Gm3486 | 878 | 0 | 0 | 0 | 0 | 0 | 0 |
| ENSMUSG00 | Gm17734 | 948 | 0 | 0 | 0 | 0 | 0.16147587 | 0 |
| ENSMUSG00 | Gm6476 | 795 | 0 | 0 | 0.0555998 | 0 | 0 | 0 |
| ENSMUSG00 | Gm7957 | 395 | 0 | 0 | 0 | 0.09134695 | 0 | 0 |
| ENSMUSG00 | Sfta2 | 1281 | 0 | 0.15046508 | 0.03450573 | 0.11266837 | 0.03983324 | 0.21291678 |
| ENSMUSG00 | Gm17554 | 3524 | 0 | 0 | 0 | 0 | 0.01447968 | 0 |
| ENSMUSG00 | Gm3510 | 1453 | 0 | 0 | 0 | 0 | 0 | 0 |
| ENSMUSG00 | Gm3424 | 1753 | 0 | 0 | 0 | 0 | 0 | 0 |
| ENSMUSG00 | Gm9140 | 305 | 0 | 0 | 0 | 0 | 0 | 0 |
| ENSMUSG00 | Gm7782 | 517 | 0 | 0 | 0 | 0 | 0 | 0 |
| ENSMUSG00 | Krtap27-1 | 608 | 0 | 0 | 0 | 0 | 0 | 0 |
| ENSMUSG00 | Gm6202 | 474 | 1.42582947 | 1.72820573 | 1.678551 | 1.59857159 | 1.29180698 | 2.58936449 |
| ENSMUSG00 | 6330403N20I | 1187 | 0 | 0 | 0.03723828 | 0 | 0 | 0 |
| ENSMUSG00 | Vmn2r35 | 2580 | 0 | 0 | 0 | 0 | 0 | 0 |
| ENSMUSG00 | Gm17061 | 496 | 0 | 0 | 0 | 0 | 0 | 0 |
| ENSMUSG00 | Gm17639 | 1226 | 0 | 0 | 0 | 0 | 0 | 0 |
| ENSMUSG00 | Gm5919 | 3836 | 0 | 0 | 0 | 0 | 0 | 0 |
| ENSMUSG00 | Gypc | 4203 | 0.19863552 | 0 | 0 | 0 | 0 | 0 |
| ENSMUSG00 | Gm13691 | 2725 | 0 | 0 | 0 | 0 | 0 | 0 |
| ENSMUSG00 | Gm7742 | 1526 | 0 | 0 | 0 | 0 | 0 | 0 |
| ENSMUSG00 | Gm17380 | 48 | 0 | 0 | 0 | 0 | 0 | 2.2728866 |
| ENSMUSG00 | Gm5538 | 1298 | 0 | 0 | 0 | 0 | 0 | 0 |
| ENSMUSG00 | Gm17381 | 1352 | 0 | 0 | 0 | 0 | 0 | 0 |
| ENSMUSG00 | Gm6803 | 493 | 0 | 0 | 0 | 0 | 0 | 0 |
| ENSMUSG00 | Gm17697 | 281 | 0 | 0 | 0 | 0.12840585 | 0 | 0 |
| ENSMUSG00 | Gm17696 | 2261 | 0 | 0 | 0 | 0 | 0 | 0 |
| ENSMUSG00 | Gm8214 | 416 | 0 | 0 | 0 | 0 | 0 | 0 |
| ENSMUSG00 | Gm4675 | 3126 | 0 | 0 | 0 | 0 | 0 | 0 |
| ENSMUSG00 | Gm8540 | 727 | 0 | 0 | 0 | 0 | 0 | 0 |
| ENSMUSG00 | Scn8a | 1069 | 0.3347047 | 0.27045711 | 0 | 0.23627157 | 0 | 0.30616995 |
| ENSMUSG00 | Gm17222 | 846 | 0 | 0 | 0 | 0 | 0 | 0 |
| ENSMUSG00 | Gm4775 | 404 | 0 | 0 | 0 | 0 | 0 | 0 |
| ENSMUSG00 | Gm9602 | 2057 | 0 | 0 | 0 | 0 | 0 | 0 |
| ENSMUSG00 | Gm17487 | 3072 | 0.1035299 | 0 | 0.2733838 | 0.1644364 | 0.06644059 | 0.21308312 |
| ENSMUSG00 | Gm10814 | 1928 | 0.8248025 | 0.12496484 | 0.1375576 | 0.44915408 | 0.31759155 | 0.65074347 |
| ENSMUSG00 | Gm17694 | 782 | 0 | 0 | 0 | 0 | 0 | 0 |
| ENSMUSG00 | Gm6576 | 954 | 0 | 0.1010198 | 0.18533268 | 0 | 0 | 0 |
| ENSMUSG00 | Gm17488 | 57 | 0 | 0 | 0 | 0 | 0 | 0 |
| ENSMUSG00 | Cdr1 | 2312 | 0 | 0 | 0 | 0 | 0 | 0 |
| ENSMUSG00 | Gm8265 | 471 | 0 | 0 | 0 | 0 | 0 | 0 |
| ENSMUSG00 | Gm17489 | 2859 | 0 | 0.75844348 | 0 | 0.55530254 | 0.07139052 | 0.14309884 |
| ENSMUSG00 | Rpl28-ps4 | 392 | 0 | 0 | 0 | 0 | 0 | 0 |
| ENSMUSG00 | Prlh | 259 | 0 | 0.74419216 | 0 | 0 | 0 | 0 |
| ENSMUSG00 | A730015C16I | 1220 | 0.71690211 | 0.2369825 | 0.07246204 | 0.41405625 | 0.46007388 | 0.38005645 |
| ENSMUSG00 | Gm17158 | 492 | 0 | 0 | 0 | 0 | 0 | 0 |
| ENSMUSG00 | Snrpe | 1329 | 26.982275 | 19.7604296 | 13.7361634 | 14.1993298 | 14.5899344 | 39.0546565 |
| ENSMUSG00 | Gm17565 | 1316 | 0.36251198 | 0.32954254 | 0.10076408 | 0.13706984 | 0.34896458 | 0.10362705 |
| ENSMUSG00 | Gm8893 | 1238 | 0 | 0 | 0 | 0 | 0 | 0 |
| ENSMUSG00 | Olfr753-ps1 | 1801 | 0 | 0 | 0 | 0 | 0 | 0 |
| ENSMUSG00 | Gm17202 | 948 | 0.25161697 | 0 | 0.2797585 | 0.03806123 | 0 | 0.08631215 |
| ENSMUSG00 | Gm17137 | 5326 | 0.3209699 | 0.75998144 | 0.18258366 | 0.51487709 | 0.01916124 | 0.17411523 |
| ENSMUSG00 | Gm17190 | 848 | 0 | 0 | 0.10424963 | 0 | 0 | 0 |
| ENSMUSG00 | Gm17207 | 681 | 0 | 0.56606887 | 0 | 0.15895174 | 0 | 0.68086471 |
| ENSMUSG00 | Gm7358 | 1526 | 0 | 0 | 0 | 0 | 0 | 0 |
| ENSMUSG00 | Ighd6-1 | 28 | 0 | 0 | 0 | 0 | 0 | 0 |
| ENSMUSG00 | A430057M04 | 5891 | 0.1754613 | 0.3108275 | 0.30763462 | 0.17762337 | 0.22520553 | 0 |
| ENSMUSG00 | 4930405O22I | 1008 | 0.07887992 | 1.33851228 | 1.0085738 | 0.50113951 | 0.60745686 | 1.16350148 |
| ENSMUSG00 | Gm17465 | 2399 | 0 | 0 | 0 | 0 | 0 | 0 |
| ENSMUSG00 | Gm17466 | 1280 | 0 | 0 | 0 | 0 | 0 | 0 |
| ENSMUSG00 | Gm17041 | 477 | 0 | 0 | 0 | 0 | 0 | 0 |
| ENSMUSG00 | Gm17434 | 1685 | 0 | 0 | 0 | 0 | 0 | 0 |
| ENSMUSG00 | Vmn2r109 | 2571 | 0 | 0 | 0 | 0 | 0 | 0 |
| ENSMUSG00 | Gm2636 | 645 | 0 | 0 | 0 | 0 | 0 | 0 |
| ENSMUSG00 | Traj61 | 71 | 0 | 0 | 0 | 0 | 0 | 0 |
| ENSMUSG00 | Gm13289 | 1555 | 0 | 0 | 0 | 0 | 0 | 0 |
| ENSMUSG00 | Gm17055 | 2230 | 0 | 0.04321654 | 0.25767891 | 0 | 0.18305426 | 0 |
| ENSMUSG00 | Gm3027 | 448 | 0 | 0 | 0 | 0 | 0 | 0 |
| ENSMUSG00 | Gm8323 | 1200 | 0 | 0 | 0 | 0 | 0 | 0 |
| ENSMUSG00 | Gm7137 | 690 | 0 | 0 | 0 | 0 | 0 | 0 |
| ENSMUSG00 | Gm4144 | 920 | 0 | 0 | 0 | 0 | 0 | 0 |
| ENSMUSG00 | Vmn2r6 | 2736 | 0 | 0 | 0 | 0 | 0 | 0 |
| ENSMUSG00 | Gm17024 | 632 | 0.69194665 | 1.21990993 | 0.13987925 | 1.94112265 | 1.37254492 | 0.56102897 |
| ENSMUSG00 | Gm17169 | 350 | 0 | 0 | 0 | 0 | 0 | 0 |
| ENSMUSG00 | 4933406F09I | 1291 | 0 | 0 | 0 | 0 | 0 | 0 |
| ENSMUSG00 | Gm17260 | 631 | 0.25201572 | 0.1527304 | 0 | 0 | 0.16173178 | 0.08644894 |
| ENSMUSG00 | Olfr1381 | 937 | 0 | 0 | 0 | 0 | 0 | 0 |
| ENSMUSG00 | Gm9573 | 5518 | 0 | 0 | 0 | 0 | 0 | 0 |
| ENSMUSG00 | Gm17180 | 431 | 0 | 0 | 0 | 0 | 0 | 0 |
| ENSMUSG00 | Gm17570 | 1274 | 0 | 0 | 0 | 0 | 0 | 0 |
| ENSMUSG00 | Gm17569 | 1514 | 0 | 0 | 0 | 0 | 0 | 0 |
| ENSMUSG00 | Gm17571 | 73 | 0 | 0 | 0 | 0 | 0 | 0 |
| ENSMUSG00 | 4930539J05R | 1404 | 1.44410934 | 0.24024579 | 0.34631074 | 0.48828978 | 0.36343573 | 0.48565953 |
| ENSMUSG00 | Gm3327 | 1384 | 0 | 0 | 0 | 0 | 0 | 0 |
| ENSMUSG00 | Gm17145 | 477 | 0 | 0 | 0 | 0 | 0 | 0 |
| ENSMUSG00 | Gm17121 | 846 | 0 | 0 | 0 | 0 | 0 | 0 |
| ENSMUSG00 | Gm8164 | 1191 | 0 | 0 | 0 | 0 | 0 | 0 |
| ENSMUSG00 | AL626783.1 | 2287 | 0.01738325 | 0 | 0 | 0 | 0.4239183 | 0.27429677 |
| ENSMUSG00 | Gm3395 | 2695 | 0 | 0 | 0 | 0 | 0 | 0 |
| ENSMUSG00 | Gm2877 | 1203 | 0 | 0 | 0 | 0 | 0 | 0 |
| ENSMUSG00 | Gm5611 | 854 | 0 | 0.05642441 | 0 | 0 | 0.05974986 | 0 |
| ENSMUSG00 | Gm17186 | 453 | 0 | 0 | 0 | 0 | 0 | 0 |
| ENSMUSG00 | Gm17378 | 925 | 0 | 0 | 0 | 0 | 0 | 0 |
| ENSMUSG00 | E230013L22F | 1854 | 2.61605643 | 2.67702456 | 1.4543217 | 3.38634075 | 4.65127158 | 3.32474028 |
| ENSMUSG00 | Gm17052 | 288 | 0 | 0 | 0 | 0 | 0 | 0 |
| ENSMUSG00 | Gm17200 | 2019 | 0.29536018 | 0.38186383 | 0.19703645 | 0.10722747 | 0.12636547 | 0 |
| ENSMUSG00 | Gm3571 | 1140 | 0.31385906 | 0 | 0.11632064 | 0 | 0 | 0.04785024 |
| ENSMUSG00 | C330013E15I | 876 | 2.04223359 | 4.51060303 | 2.6238537 | 2.7185102 | 2.50471936 | 3.48716849 |
| ENSMUSG00 | D930030I03F | 918 | 1.21258546 | 0.20996271 | 1.15560374 | 1.80803273 | 0.27792144 | 1.01017182 |
| ENSMUSG00 | Gm17630 | 4427 | 0.8531219 | 0.94696645 | 1.37787539 | 2.05391354 | 1.70587387 | 1.95302928 |
| ENSMUSG00 | Gm17629 | 2536 | 0 | 0.22801156 | 0 | 0.55488949 | 0.14084568 | 0 |
| ENSMUSG00 | Gm17628 | 1594 | 0 | 0 | 0 | 0.09054465 | 0 | 0 |
| ENSMUSG00 | Gm17395 | 2970 | 0.04015705 | 0 | 0 | 0 | 0 | 0.02755014 |
| ENSMUSG00 | Gm8379 | 1191 | 0 | 0 | 0 | 0 | 0 | 0 |
| ENSMUSG00 | Vmn2r60 | 2556 | 0 | 0 | 0 | 0 | 0 | 0 |
| ENSMUSG00 | Gm6724 | 1109 | 0.03584804 | 0.13035106 | 0.03985739 | 0.22774961 | 0.09202232 | 0.27053294 |
| ENSMUSG00 | Gm4419 | 1727 | 2.1408568 | 5.18973841 | 0.74224288 | 4.15768782 | 2.12733009 | 2.66902955 |
| ENSMUSG00 | A930033H14I | 2967 | 0.46897264 | 0.12992637 | 0.47673036 | 0 | 0.82550254 | 0.05515599 |
| ENSMUSG00 | Cfhr3 | 965 | 0 | 0 | 0 | 0 | 0 | 0 |
| ENSMUSG00 | Gm17060 | 823 | 0 | 0 | 0 | 0 | 0 | 0 |
| ENSMUSG00 | AL593857.1 | 1471 | 0.10810464 | 0 | 0 | 0 | 0 | 0 |
| ENSMUSG00 | Tex9 | 1467 | 0 | 0 | 0 | 0 | 0 | 0 |
| ENSMUSG00 | Gm8356 | 2368 | 0.01678863 | 0 | 0 | 0 | 0 | 0 |
| ENSMUSG00 | Gm17083 | 1365 | 0 | 0 | 0 | 0.68727704 | 0.74763921 | 0 |
| ENSMUSG00 | Olfr180 | 1140 | 0 | 0 | 0 | 0 | 0 | 0 |
| ENSMUSG00 | Gm17403 | 3268 | 1.83692704 | 1.12061493 | 0.78448803 | 0.65142002 | 0.93683677 | 2.19499086 |
| ENSMUSG00 | Olfr456 | 1051 | 0 | 0 | 0.08411388 | 0 | 0.33985217 | 0 |
| ENSMUSG00 | Gm17404 | 170 | 0 | 0 | 0 | 0 | 0 | 0 |
| ENSMUSG00 | Vmn1r-ps79 | 3054 | 0 | 0 | 0 | 0 | 0 | 0 |
| ENSMUSG00 | Gm8126 | 1759 | 0 | 0.24654803 | 0.15077377 | 0.02051282 | 0 | 0 |
| ENSMUSG00 | Gm2951 | 487 | 0 | 0 | 0 | 0 | 0 | 0 |
| ENSMUSG00 | Gm17069 | 489 | 0 | 0 | 0 | 0 | 0 | 0 |
| ENSMUSG00 | Gm6189 | 2084 | 0 | 0 | 0 | 0 | 0 | 0 |
| ENSMUSG00 | Gm17028 | 702 | 0 | 0 | 0 | 0 | 0 | 0 |
| ENSMUSG00 | Gm20425 | 3005 | 0 | 0 | 0 | 0 | 0 | 0 |
| ENSMUSG00 | Gm17447 | 1946 | 0 | 0 | 0 | 0 | 0 | 0 |
| ENSMUSG00 | Zfp712 | 3966 | 2.35565757 | 6.63383697 | 4.87045018 | 3.37529968 | 8.47866406 | 5.86617934 |
| ENSMUSG00 | Gm17182 | 857 | 0 | 0 | 0 | 0 | 0.11908139 | 0 |
| ENSMUSG00 | Gm3453 | 1751 | 0 | 0 | 0 | 0 | 0 | 0 |
| ENSMUSG00 | B230364G03 | 1372 | 0 | 0.14048525 | 0 | 0 | 0 | 0 |
| ENSMUSG00 | Gm17448 | 43 | 0 | 0 | 0 | 0 | 0 | 0 |
| ENSMUSG00 | 2310010G23I | 1018 | 0 | 0 | 0 | 0 | 0 | 0 |
| ENSMUSG00 | Gm17446 | 3803 | 0 | 0 | 0 | 0 | 0 | 0 |
| ENSMUSG00 | Psmd14 | 336 | 0 | 0 | 0 | 0 | 0 | 0 |
| ENSMUSG00 | Gm11168 | 668 | 0 | 0 | 0 | 0 | 0 | 0 |
| ENSMUSG00 | Vmn1r117 | 923 | 0 | 0 | 0 | 0 | 0 | 0 |
| ENSMUSG00 | Gm4064 | 1721 | 0 | 0 | 0 | 0 | 0 | 0 |
| ENSMUSG00 | Gm17645 | 1162 | 0 | 0.29027977 | 0.34235507 | 0 | 0 | 0 |
| ENSMUSG00 | Vmn2r120 | 2564 | 0 | 0 | 0 | 0 | 0 | 0 |
| ENSMUSG00 | Gm17559 | 1785 | 0.15590385 | 0.21596165 | 0.04952588 | 0.28299643 | 0.02858621 | 0.07639955 |
| ENSMUSG00 | Gm8114 | 481 | 0 | 0 | 0 | 0 | 0 | 0 |
| ENSMUSG00 | Prss47 | 868 | 0 | 0 | 0 | 0 | 0 | 0 |
| ENSMUSG00 | Zfp493 | 3270 | 1.61696603 | 3.86080974 | 1.91946841 | 0.94894674 | 5.07142879 | 3.20289342 |
| ENSMUSG00 | Gm17030 | 390 | 0 | 0 | 0 | 0 | 0 | 0 |
| ENSMUSG00 | Gm17724 | 1831 | 0.17369953 | 0 | 0.26554903 | 0.82766022 | 0.19507626 | 0.20854448 |
| ENSMUSG00 | Vmn2r45 | 3613 | 0 | 0 | 0 | 0 | 0 | 0 |
| ENSMUSG00 | Srcap | 3051 | 4.22182094 | 5.30666816 | 4.60707508 | 3.58336922 | 4.44871058 | 5.24752974 |
| ENSMUSG00 | Gm17568 | 2380 | 0 | 0 | 0.07428881 | 0 | 0.08575862 | 0 |
| ENSMUSG00 | Gad1-ps | 3277 | 0 | 0 | 0.18883912 | 0 | 0 | 0 |
| ENSMUSG00 | Gm17567 | 65 | 0 | 0 | 0 | 7.77151728 | 0 | 0 |
| ENSMUSG00 | Gm765 | 822 | 0 | 0 | 0 | 0 | 0 | 0 |
| ENSMUSG00 | 4930448E22I | 1581 | 0.12572891 | 0 | 0 | 0 | 0.03227475 | 0 |
| ENSMUSG00 | Gm2990 | 1107 | 0 | 0 | 0 | 0 | 0 | 0 |
| ENSMUSG00 | Gm5067 | 1008 | 0 | 0 | 0 | 0 | 0 | 0 |
| ENSMUSG00 | Gm17256 | 2594 | 0.27586687 | 1.02168632 | 0.54528102 | 0.4590237 | 0.62946956 | 3.37515775 |
| ENSMUSG00 | Gm340 | 3727 | 1.32269374 | 1.96521041 | 1.67224574 | 1.02621323 | 2.03995975 | 2.71502312 |
| ENSMUSG00 | Gm17082 | 1090 | 0 | 0.04420775 | 0 | 0.03310279 | 0 | 0.10009042 |
| ENSMUSG00 | Olfr111 | 1040 | 0 | 0 | 0 | 0 | 0 | 0 |
| ENSMUSG00 | Gm6721 | 1430 | 0 | 0 | 0 | 0 | 0 | 0 |
| ENSMUSG00 | D930019O06 | 3661 | 0 | 0 | 0 | 0 | 0 | 0 |
| ENSMUSG00 | 6030442K20I | 1538 | 0.87885978 | 0.81459525 | 0.51731676 | 0.11730184 | 0.76307324 | 2.39406781 |
| ENSMUSG00 | Gm17255 | 622 | 0 | 0 | 0 | 0 | 0 | 0 |
| ENSMUSG00 | Vmn2r101 | 2568 | 0 | 0 | 0 | 0 | 0 | 0 |
| ENSMUSG00 | Gm3221 | 437 | 0 | 0 | 0 | 0 | 0 | 0 |
| ENSMUSG00 | Gm3852 | 263 | 0 | 0 | 0 | 0 | 0 | 0 |
| ENSMUSG00 | Gm4169 | 1151 | 0 | 0 | 0 | 0 | 0 | 0 |
| ENSMUSG00 | Gm17221 | 480 | 0 | 0 | 0 | 0 | 0 | 0 |
| ENSMUSG00 | Gm9047 | 680 | 0 | 0.56689932 | 0.32501355 | 0.79592745 | 0.52527152 | 0.08021953 |
| ENSMUSG00 | Gm3015 | 1398 | 0 | 0 | 0 | 0 | 0 | 0 |
| ENSMUSG00 | 9330151L19F | 2544 | 16.002206 | 12.6716705 | 9.59096594 | 6.70865057 | 6.79950527 | 10.4852976 |
| ENSMUSG00 | Vmn2r12 | 2490 | 0 | 0 | 0 | 0 | 0 | 0 |
| ENSMUSG00 | Gm17695 | 1685 | 0 | 0 | 0 | 0 | 0 | 0 |
| ENSMUSG00 | Gm8020 | 1401 | 0 | 0 | 0 | 0 | 0 | 0 |
| ENSMUSG00 | Gm3667 | 2182 | 0 | 0 | 0.04051498 | 0 | 0 | 0 |
| ENSMUSG00 | Gm17172 | 188 | 0 | 0 | 0 | 0 | 0 | 0 |
| ENSMUSG00 | Trim43c | 1832 | 0.06510177 | 0.1841185 | 0 | 0.33482247 | 0 | 0.10421533 |
| ENSMUSG00 | Gm17379 | 40 | 0 | 0 | 0 | 0 | 0 | 0 |
| ENSMUSG00 | Nsun4 | 3914 | 0.01015725 | 0 | 0 | 0 | 0 | 0.09755875 |
| ENSMUSG00 | Apold1 | 3282 | 0.29071649 | 0.13213832 | 0.12121164 | 0.85752574 | 0.21766279 | 0.96400309 |
| ENSMUSG00 | Gm9071 | 471 | 0 | 0 | 0 | 0 | 0 | 0 |
| ENSMUSG00 | Cyp4f40 | 1846 | 0 | 0 | 0 | 0 | 0 | 0 |
| ENSMUSG00 | Gm6486 | 431 | 0 | 0 | 0 | 0 | 0 | 0 |
| ENSMUSG00 | Gm17552 | 3875 | 0.01025948 | 0.02487042 | 0.05703464 | 0 | 0.4213791 | 0 |
| ENSMUSG00 | Gm17048 | 846 | 0 | 0 | 0 | 0 | 0 | 0 |
| ENSMUSG00 | Trp53-ps | 140 | 0 | 0 | 0 | 0 | 0 | 0 |
| ENSMUSG00 | Gm17233 | 696 | 0 | 0 | 0 | 0 | 0 | 0 |
| ENSMUSG00 | Gm8237 | 2180 | 0 | 0 | 0 | 0 | 0 | 0 |
| ENSMUSG00 | Gm17196 | 342 | 0 | 0 | 0 | 0.10550305 | 0.14919993 | 0 |
| ENSMUSG00 | Gm17173 | 2862 | 0 | 0 | 0 | 0 | 0 | 0 |
| ENSMUSG00 | Gm9513 | 297 | 0 | 0 | 0 | 0 | 0 | 0 |
| ENSMUSG00 | Gm17027 | 1398 | 0 | 0 | 0 | 0 | 0 | 0 |
| ENSMUSG00 | Gm8159 | 1945 | 0 | 0 | 0 | 0 | 0 | 0 |
| ENSMUSG00 | Gm8127 | 1398 | 0.11374959 | 0.3791494 | 0 | 0.36133664 | 0.03649955 | 0.05852927 |
| ENSMUSG00 | Zscan4d | 1988 | 0 | 0 | 0 | 0 | 0 | 0 |
| ENSMUSG00 | Vmn1r167 | 947 | 0 | 0 | 0 | 0 | 0 | 0 |
| ENSMUSG00 | Gm3371 | 1434 | 0 | 0 | 0 | 0 | 0 | 0 |
| ENSMUSG00 | Gm17376 | 82 | 0 | 0 | 0 | 0 | 0 | 0 |
| ENSMUSG00 | Gm14476 | 499 | 0 | 0 | 0 | 0 | 0 | 0 |
| ENSMUSG00 | Gm17632 | 3939 | 0.25231963 | 0.62389148 | 0 | 0.10992245 | 0.03886243 | 0.18003062 |
| ENSMUSG00 | Gm8378 | 2103 | 0 | 0 | 0 | 0 | 0 | 0 |
| ENSMUSG00 | Gm9625 | 1096 | 0.79801147 | 0.87931464 | 1.41155521 | 1.02056878 | 0.3724553 | 0.97053917 |
| ENSMUSG00 | 4930542C12I | 1202 | 0 | 0 | 0 | 0 | 0 | 0 |
| ENSMUSG00 | Gm17633 | 1249 | 0 | 0 | 0 | 0 | 0 | 0 |
| ENSMUSG00 | Gm17421 | 267 | 0 | 0 | 0 | 0 | 0 | 0 |
| ENSMUSG00 | Gm3956 | 2554 | 0 | 0 | 0 | 0 | 0 | 0 |
| ENSMUSG00 | Gm17420 | 150 | 0 | 0 | 0 | 0 | 0 | 0 |
| ENSMUSG00 | Gm2959 | 617 | 0 | 0 | 0 | 0 | 0 | 0 |
| ENSMUSG00 | 4930579O11I | 1293 | 0 | 0 | 0 | 0 | 0 | 0 |
| ENSMUSG00 | Rps27 | 341 | 0.46633995 | 0 | 1.16661756 | 1.05812447 | 1.79564959 | 1.51970131 |
| ENSMUSG00 | Gm9593 | 466 | 0 | 0 | 0 | 0 | 0 | 0 |
| ENSMUSG00 | Gm6788 | 269 | 0 | 0 | 0 | 0.134134 | 0 | 0.10139271 |
| ENSMUSG00 | Gm17727 | 327 | 0 | 0 | 0 | 0 | 0 | 0 |
| ENSMUSG00 | D930016D06 | 4504 | 5.89623912 | 9.29707332 | 4.75974553 | 5.96026668 | 3.41006641 | 11.6874009 |
| ENSMUSG00 | Gm17079 | 1347 | 0 | 0 | 0 | 0 | 0 | 0 |
| ENSMUSG00 | 1700091H14I | 1397 | 0 | 0 | 0 | 0 | 0 | 0 |
| ENSMUSG00 | Gm3008 | 567 | 0 | 0 | 0 | 0 | 0 | 0 |
| ENSMUSG00 | Gm17213 | 1359 | 0 | 0 | 0.71555596 | 0.55755919 | 0 | 0 |
| ENSMUSG00 | Gm6871 | 2273 | 1.48667657 | 1.01757555 | 0.40837602 | 1.15881621 | 1.41428142 | 1.87190661 |
| ENSMUSG00 | Vmn1r126 | 965 | 0 | 0 | 0 | 0 | 0 | 0 |
| ENSMUSG00 | Gm17654 | 1398 | 0 | 0 | 0 | 0.02580976 | 0 | 0 |
| ENSMUSG00 | Gm17495 | 616 | 0 | 0 | 0 | 0 | 0 | 0 |
| ENSMUSG00 | Olfr247 | 962 | 0 | 0 | 0 | 0 | 0 | 0 |
| ENSMUSG00 | Gm3495 | 385 | 0 | 0 | 0 | 0 | 0 | 0 |
| ENSMUSG00 | Vmn2r63 | 2574 | 0 | 0 | 0 | 0 | 0 | 0 |
| ENSMUSG00 | Gm6401 | 1399 | 0 | 0 | 0 | 0 | 0 | 0 |
| ENSMUSG00 | Rpl31-ps4 | 389 | 0 | 0 | 0 | 0 | 0 | 0 |
| ENSMUSG00 | C230091D08 | 52 | 0 | 0 | 0 | 2.77554189 | 2.94382937 | 40.3874466 |
| ENSMUSG00 | Gm17635 | 367 | 143.072119 | 131.574956 | 101.373089 | 175.87296 | 77.914526 | 231.916094 |
| ENSMUSG00 | Gm13277 | 1555 | 0 | 0 | 0 | 0 | 0 | 0 |
| ENSMUSG00 | Gm17178 | 354 | 0 | 0 | 0 | 0 | 0 | 0 |
| ENSMUSG00 | Gm17634 | 1277 | 2.92640185 | 1.58483208 | 1.66146317 | 1.44102136 | 2.23764843 | 4.80563338 |
| ENSMUSG00 | Gm10634 | 3668 | 0 | 0 | 0.01205067 | 0.01967396 | 0 | 0.02974334 |
| ENSMUSG00 | Gm17201 | 388 | 0.20492516 | 0.86934303 | 0.68353366 | 0.55796976 | 0.13151128 | 0.63265916 |
| ENSMUSG00 | Vmn2r56 | 2334 | 0 | 0 | 0 | 0 | 0 | 0 |
| ENSMUSG00 | Gm3127 | 1944 | 0 | 0 | 0 | 0 | 0 | 0 |
| ENSMUSG00 | Gm17686 | 293 | 0 | 0 | 0 | 0 | 0 | 0 |
| ENSMUSG00 | Gm17684 | 250 | 0 | 0 | 0 | 0 | 0 | 0 |
| ENSMUSG00 | Gm17685 | 2730 | 0 | 0 | 0 | 0 | 0 | 0 |
| ENSMUSG00 | Gm2824 | 5459 | 0 | 0 | 0 | 0 | 0 | 0 |
| ENSMUSG00 | Gm7970 | 1398 | 0 | 0 | 0 | 0 | 0 | 0 |
| ENSMUSG00 | Gm17501 | 1318 | 0.15081745 | 0.21936165 | 0 | 0 | 0.27100503 | 0.18624564 |
| ENSMUSG00 | Gm2930 | 471 | 0 | 0 | 0 | 0 | 0 | 0 |
| ENSMUSG00 | Vmn2r-ps69 | 2658 | 0 | 0 | 0 | 0 | 0 | 0 |
| ENSMUSG00 | Vmn2r74 | 2556 | 0 | 0 | 0 | 0 | 0 | 0 |
| ENSMUSG00 | Vmn1r61 | 902 | 0 | 0 | 0 | 0 | 0 | 0 |
| ENSMUSG00 | Gm7873 | 742 | 0 | 0 | 0 | 0 | 0 | 0 |
| ENSMUSG00 | Gm3235 | 693 | 0.91787545 | 0.76486416 | 0.8291832 | 1.04132885 | 2.06167175 | 1.6136511 |
| ENSMUSG00 | Gm17110 | 732 | 0.81466149 | 1.18491251 | 1.02654553 | 1.52806473 | 0.7667898 | 1.3413757 |
| ENSMUSG00 | Gm17532 | 58 | 2.05631796 | 0.83080073 | 1.52420148 | 4.35472951 | 4.3988255 | 0 |
| ENSMUSG00 | Igkv2-95-2 | 122 | 0 | 0 | 0 | 0 | 0 | 0 |
| ENSMUSG00 | Traj60 | 53 | 0 | 0 | 0 | 0 | 0 | 0 |
| ENSMUSG00 | Gm17485 | 40 | 0 | 0 | 0 | 0 | 0 | 1.36373196 |
| ENSMUSG00 | Gm17531 | 1416 | 0 | 0.30626976 | 0 | 0.56059674 | 0 | 0.15409401 |
| ENSMUSG00 | Gm17116 | 1009 | 0 | 0 | 0 | 0 | 0 | 0 |
| ENSMUSG00 | Gm9597 | 598 | 0 | 0 | 0 | 0 | 0 | 0 |
| ENSMUSG00 | Gm17377 | 206 | 1.54390216 | 0 | 0 | 0 | 0.49540171 | 4.104436 |
| ENSMUSG00 | Vmn1r158 | 886 | 0 | 0 | 0 | 0 | 0 | 0 |
| ENSMUSG00 | Gm8221 | 481 | 0 | 0 | 0 | 0 | 0 | 0 |
| ENSMUSG00 | Gm6650 | 767 | 0 | 0 | 0 | 0 | 0 | 0 |
| ENSMUSG00 | Vmn1r60 | 1834 | 0 | 0 | 0 | 0 | 0 | 0 |
| ENSMUSG00 | Gm7346 | 473 | 0 | 0 | 0 | 0 | 0 | 0 |
| ENSMUSG00 | Gm3141 | 496 | 0 | 0 | 0 | 0 | 0 | 0 |
| ENSMUSG00 | Gm17335 | 932 | 0 | 0 | 0 | 0 | 0 | 0 |
| ENSMUSG00 | Gm8046 | 612 | 0.12991987 | 0.31494407 | 0 | 0.58957589 | 0 | 0.17826562 |
| ENSMUSG00 | Klhl33 | 1599 | 0 | 0 | 0 | 0 | 0 | 0 |
| ENSMUSG00 | 4833407H14I | 1483 | 0 | 0 | 0 | 0 | 0 | 0 |
| ENSMUSG00 | Gm4862 | 437 | 0 | 0 | 0 | 0 | 0 | 0 |
| ENSMUSG00 | Gm8074 | 256 | 0 | 0 | 0 | 0 | 0 | 0 |
| ENSMUSG00 | Gm17413 | 70 | 0 | 0 | 0 | 0 | 0 | 0 |
| ENSMUSG00 | Gm17729 | 57 | 0 | 0 | 0 | 0 | 0 | 0 |
| ENSMUSG00 | Vmn2r70 | 2559 | 0 | 0 | 0 | 0 | 0 | 0 |
| ENSMUSG00 | Gm1758 | 468 | 0 | 0 | 0 | 0 | 0 | 0 |
| ENSMUSG00 | Gm17412 | 172 | 0 | 0 | 0 | 0 | 0 | 0 |
| ENSMUSG00 | Gm3261 | 480 | 0 | 0 | 0 | 0 | 0 | 0 |
| ENSMUSG00 | Samd15 | 1193 | 0.2332677 | 0.96938358 | 0.37051 | 0 | 0.38494332 | 0.06858669 |
| ENSMUSG00 | Gm4651 | 1281 | 0.09310417 | 0.82755795 | 0.48308025 | 0.08450128 | 0 | 0.10645839 |
| ENSMUSG00 | Vmn1r88 | 950 | 0 | 0 | 0 | 0 | 0 | 0 |
| ENSMUSG00 | Gm17601 | 80 | 0 | 0 | 0 | 0 | 0 | 0 |
| ENSMUSG00 | Gm6729 | 1451 | 0 | 0 | 0 | 0 | 0 | 0 |
| ENSMUSG00 | Gm4450 | 1684 | 0.02360777 | 0.02861428 | 0 | 0 | 0.0303007 | 0 |
| ENSMUSG00 | Trbj2-6 | 47 | 0 | 0 | 0 | 0 | 0 | 0 |
| ENSMUSG00 | Gm17603 | 1170 | 0.10193713 | 0 | 0 | 0 | 0.3488983 | 0 |
| ENSMUSG00 | Gm8161 | 478 | 0 | 0 | 0 | 0 | 0 | 0 |
| ENSMUSG00 | 2700012I20R | 682 | 0.17487748 | 0 | 2.78691972 | 1.32265559 | 0.14963747 | 1.27974847 |
| ENSMUSG00 | Olfr1344 | 1060 | 0 | 0 | 0 | 0 | 0 | 0 |
| ENSMUSG00 | Gm2221 | 445 | 0 | 0.10828414 | 0 | 0 | 0.11466601 | 0.18387397 |
| ENSMUSG00 | Gm17285 | 37 | 0 | 0 | 0 | 2.92557118 | 0 | 0 |
| ENSMUSG00 | Gm8297 | 1945 | 0 | 0 | 0 | 0 | 0 | 0 |
| ENSMUSG00 | Sox2ot | 8689 | 0 | 0 | 0 | 0.00415261 | 0.01174505 | 0 |
| ENSMUSG00 | Vmn1r135 | 923 | 0 | 0 | 0 | 0 | 0 | 0 |
| ENSMUSG00 | Vmn1r163 | 923 | 0 | 0 | 0 | 0 | 0 | 0 |
| ENSMUSG00 | Gm17474 | 57 | 0 | 0 | 0 | 0 | 0 | 0 |
| ENSMUSG00 | Gm17476 | 3613 | 0 | 0 | 0 | 0 | 0 | 0 |
| ENSMUSG00 | Vmn1r116 | 923 | 0 | 0 | 0 | 0 | 0 | 0 |
| ENSMUSG00 | Gm17475 | 3214 | 0 | 0 | 0 | 0 | 0 | 0 |
| ENSMUSG00 | Gm7233 | 1756 | 0 | 0 | 0.05034379 | 0 | 0 | 0 |
| ENSMUSG00 | Gm7084 | 317 | 0 | 0 | 0 | 0 | 0 | 0 |
| ENSMUSG00 | Gm17094 | 3323 | 0 | 0 | 0 | 0 | 0 | 0 |
| ENSMUSG00 | 1700092M07 | 378 | 0 | 0 | 0.70161656 | 1.05000659 | 0.94493289 | 0 |
| ENSMUSG00 | Myl6 | 659 | 49.4681245 | 50.0875764 | 69.0863404 | 52.0698397 | 34.3789239 | 87.0387957 |
| ENSMUSG00 | Gm17674 | 86 | 0 | 0 | 0 | 0.41955866 | 0 | 0 |
| ENSMUSG00 | Gm17673 | 1698 | 0.49167555 | 0.42567528 | 0.44253906 | 0.14874812 | 0 | 0.94770537 |
| ENSMUSG00 | Gm3302 | 1845 | 0 | 0 | 0 | 0 | 0 | 0 |
| ENSMUSG00 | Lsm3 | 627 | 10.3985627 | 12.7574951 | 23.8986041 | 16.0556466 | 6.51054237 | 36.9316886 |
| ENSMUSG00 | Gm17672 | 1868 | 0 | 0 | 0 | 0.01931587 | 0 | 0 |
| ENSMUSG00 | Vmn1r165 | 923 | 0 | 0 | 0 | 0 | 0 | 0 |
| ENSMUSG00 | Vmn2r-ps36 | 3460 | 0 | 0 | 0 | 0 | 0 | 0 |
| ENSMUSG00 | Orai2-ps | 3033 | 0 | 0 | 0 | 0 | 0 | 0 |
| ENSMUSG00 | Vmn2r98 | 2568 | 0 | 0 | 0 | 0 | 0 | 0 |
| ENSMUSG00 | Tnfrsf22 | 1779 | 0 | 0 | 0 | 0 | 0 | 0 |
| ENSMUSG00 | Gm17364 | 70 | 0 | 0 | 0 | 0 | 0 | 0 |
| ENSMUSG00 | Gm4340 | 797 | 0 | 0 | 0 | 0 | 0 | 0 |
| ENSMUSG00 | Gm17219 | 592 | 0.53723622 | 0 | 0.14933055 | 0 | 0.0861932 | 0.1842881 |
| ENSMUSG00 | Gm17365 | 294 | 0 | 0 | 0 | 0 | 0 | 0 |
| ENSMUSG00 | Gm3238 | 692 | 0 | 0 | 0 | 0 | 0 | 0 |
| ENSMUSG00 | Gm17134 | 435 | 0 | 0 | 0 | 0 | 0 | 0 |
| ENSMUSG00 | 4930570N19I | 564 | 0.14097688 | 0 | 0 | 0.31987628 | 0 | 0.38687432 |
| ENSMUSG00 | Rps13 | 1443 | 3.14076561 | 2.40431312 | 2.02170592 | 2.40046866 | 1.59125912 | 5.74600855 |
| ENSMUSG00 | A530084C06 | 3199 | 0.23612195 | 0.24100753 | 0.34543485 | 0.40604989 | 1.0048958 | 0.75028704 |
| ENSMUSG00 | Vmn2r40 | 3066 | 0 | 0 | 0 | 0 | 0 | 0 |
| ENSMUSG00 | Gm17074 | 555 | 0 | 0 | 0 | 0 | 0 | 0 |
| ENSMUSG00 | Snhg1 | 3501 | 3.497483 | 1.03227168 | 2.71448051 | 6.33831973 | 3.01698366 | 3.40446083 |
| ENSMUSG00 | Trav9d-4 | 305 | 0 | 0 | 0 | 0 | 0 | 0 |
| ENSMUSG00 | Gm17540 | 1526 | 0 | 0 | 0 | 0 | 0 | 0 |
| ENSMUSG00 | Gm17722 | 50 | 0 | 0 | 0.88403686 | 0.72164089 | 0 | 0 |
| ENSMUSG00 | Gm3239 | 1746 | 0 | 0 | 0 | 0 | 0 | 0 |
| ENSMUSG00 | Gm17112 | 428 | 0 | 0 | 0 | 0 | 0 | 0.31862896 |
| ENSMUSG00 | Olfr733 | 993 | 0 | 0 | 0 | 0 | 0 | 0 |
| ENSMUSG00 | Gm14482 | 499 | 0 | 0 | 0 | 0 | 0 | 0 |
| ENSMUSG00 | Hspa1b | 5006 | 46.5375781 | 50.2367242 | 46.9920512 | 80.6692454 | 52.1883828 | 72.9975263 |
| ENSMUSG00 | Gm17640 | 1194 | 0 | 0.12107146 | 0 | 0 | 0.08547132 | 0.2512739 |
| ENSMUSG00 | Gm17721 | 622 | 0 | 0.69723148 | 0.21319217 | 0 | 0 | 0 |
| ENSMUSG00 | Vmn1r129 | 923 | 0 | 0 | 0 | 0 | 0 | 0 |
| ENSMUSG00 | Gm6904 | 892 | 0 | 0.48618608 | 0 | 0 | 0 | 0 |
| ENSMUSG00 | Gm17065 | 238 | 0 | 0 | 0 | 0 | 0 | 0 |
| ENSMUSG00 | Gm17301 | 76 | 0 | 0 | 0 | 0 | 0 | 0 |
| ENSMUSG00 | Vmn2r102 | 2568 | 0 | 0 | 0 | 0 | 0 | 0 |
| ENSMUSG00 | Gm3944 | 335 | 0 | 0 | 0 | 0 | 0 | 0 |
| ENSMUSG00 | Gm17427 | 3194 | 0 | 0 | 0 | 0 | 0 | 0 |
| ENSMUSG00 | Gm17426 | 34 | 0 | 0 | 0 | 0 | 0 | 0.80219527 |
| ENSMUSG00 | Gm17429 | 3218 | 0 | 0 | 0 | 0 | 0 | 0 |
| ENSMUSG00 | Gm17428 | 92 | 0 | 0 | 0 | 0 | 0 | 0 |
| ENSMUSG00 | D6Ertd527e | 1403 | 0 | 0.44648877 | 0 | 0 | 0 | 0 |
| ENSMUSG00 | Vmn2r41 | 3611 | 0 | 0 | 0 | 0 | 0 | 0 |
| ENSMUSG00 | Gm4840 | 1574 | 0 | 0 | 0.02808249 | 0 | 0.22692797 | 0 |
| ENSMUSG00 | Olfr110 | 1101 | 0 | 0 | 0 | 0 | 0 | 0 |
| ENSMUSG00 | Vmn2r33 | 2580 | 0 | 0 | 0 | 0 | 0 | 0 |
| ENSMUSG00 | Gm17616 | 70 | 0 | 0 | 0 | 0 | 0 | 0.3896377 |
| ENSMUSG00 | Gm5494 | 1224 | 0 | 0 | 0 | 0 | 0 | 0 |
| ENSMUSG00 | Gm7732 | 1526 | 0 | 0 | 0 | 0 | 0 | 0 |
| ENSMUSG00 | Gm3326 | 1042 | 0 | 0 | 0 | 0 | 0 | 0 |
| ENSMUSG00 | Abl1 | 483 | 0 | 0 | 0 | 0 | 0 | 0 |
| ENSMUSG00 | Gm17246 | 1455 | 0 | 0.23182481 | 0.39493056 | 0.59516775 | 0 | 0.61860007 |
| ENSMUSG00 | Gm17247 | 43 | 0 | 0 | 0 | 0 | 0 | 0 |
| ENSMUSG00 | Gm17248 | 2597 | 0.19900703 | 0.46386641 | 0.42550869 | 0.375131 | 0.4519086 | 0.88219857 |
| ENSMUSG00 | Gm17249 | 2507 | 1.30033881 | 0.42285669 | 0.6347293 | 1.39607432 | 0.89555666 | 1.6536678 |
| ENSMUSG00 | Mcart1 | 572 | 0 | 0 | 0 | 0.126161 | 0.3568278 | 0 |
| ENSMUSG00 | C030017D09 | 921 | 0 | 0 | 0.04799332 | 0.58765545 | 0 | 0 |
| ENSMUSG00 | Gm17323 | 64 | 0 | 0 | 0 | 0 | 0 | 0 |
| ENSMUSG00 | Gm17706 | 34 | 1.16927884 | 0 | 1.30005421 | 4.24494642 | 12.0062061 | 17.648296 |
| ENSMUSG00 | 4931403G20I | 1697 | 0.11713459 | 0.14197538 | 0.46884689 | 0.44650733 | 0.12027431 | 0 |
| ENSMUSG00 | 9530052C20I | 3093 | 1.11824339 | 2.36803725 | 1.75778425 | 1.79651951 | 0.82486867 | 3.51845492 |
| ENSMUSG00 | Gm17563 | 1352 | 0 | 0 | 0 | 0 | 0 | 0 |
| ENSMUSG00 | Gm17562 | 28 | 0 | 0 | 0 | 0 | 0 | 0 |
| ENSMUSG00 | 1700007L15F | 735 | 3.84032533 | 2.29459248 | 5.47260914 | 4.81093927 | 2.2215565 | 6.15998655 |
| ENSMUSG00 | Pabpc4l | 5216 | 0.16768032 | 0.16628757 | 0.23727983 | 0 | 0.01956533 | 0 |
| ENSMUSG00 | 9130213A22F | 1579 | 0.0755329 | 0.18310238 | 1.2877041 | 0.0914048 | 0.3554719 | 0.24182707 |
| ENSMUSG00 | Gm17391 | 335 | 0 | 0 | 0 | 0 | 0 | 0 |
| ENSMUSG00 | Gm17390 | 2234 | 1.1567172 | 1.61771851 | 2.730463 | 3.10105307 | 3.12919136 | 3.98009507 |
| ENSMUSG00 | 1810064F22F | 1205 | 1.8145655 | 0.91974122 | 0.03668203 | 2.00622156 | 3.13356997 | 3.66679797 |
| ENSMUSG00 | Gm17047 | 429 | 0 | 0 | 0 | 0 | 0 | 0 |
| ENSMUSG00 | C530050E15I | 3835 | 0.87078497 | 0.288993 | 1.65973022 | 1.45833557 | 0.27941431 | 2.93015681 |
| ENSMUSG00 | Gm17704 | 956 | 0.12475569 | 0 | 0 | 0 | 0 | 0.05705992 |
| ENSMUSG00 | 4632411P08F | 3235 | 0.18433762 | 0.05958138 | 0.17762719 | 0.02230729 | 0.12618578 | 0.28665772 |
| ENSMUSG00 | Vmn1r101 | 923 | 0 | 0 | 0 | 0 | 0 | 0 |
| ENSMUSG00 | Gm17703 | 2237 | 0.05331535 | 0.2800285 | 0.01975943 | 0.37098213 | 0.02281018 | 0.04877003 |
| ENSMUSG00 | 4930598A11F | 625 | 0 | 0 | 0 | 0 | 0 | 0 |
| ENSMUSG00 | Synj2bp | 6020 | 23.5693206 | 24.4454143 | 28.231908 | 27.8407138 | 20.2240752 | 32.4169508 |
| ENSMUSG00 | Gm17705 | 1553 | 1.61274647 | 1.24111892 | 3.35854313 | 1.18492226 | 2.33282208 | 5.86589151 |
| ENSMUSG00 | Gm5895 | 279 | 0 | 0 | 0 | 0 | 0 | 0 |
| ENSMUSG00 | Gm6905 | 512 | 0 | 0 | 0 | 0 | 0 | 0 |
| ENSMUSG00 | Gm17147 | 431 | 0 | 0 | 0 | 0 | 0 | 0 |
| ENSMUSG00 | AW121686 | 2198 | 0 | 0 | 0 | 0 | 0 | 0 |
| ENSMUSG00 | Gm17212 | 258 | 0.92454606 | 0.18676916 | 0.85662487 | 2.23764617 | 0 | 1.05715656 |
| ENSMUSG00 | F830016B08I | 3295 | 0 | 0.01462411 | 0.99269693 | 0.60227995 | 5.42010062 | 4.37056435 |
| ENSMUSG00 | Gm3594 | 2136 | 0 | 0 | 0 | 0 | 0 | 0 |
| ENSMUSG00 | Gm17513 | 3521 | 0 | 0.01368544 | 0.01255378 | 0 | 0.01449201 | 0.03098511 |
| ENSMUSG00 | Gm17514 | 1098 | 0 | 0 | 0 | 0 | 0 | 0 |
| ENSMUSG00 | 2010109K11I | 4239 | 6.04913539 | 5.50182543 | 8.71732502 | 10.8356789 | 6.69277305 | 11.1504954 |
| ENSMUSG00 | Gm3102 | 486 | 0 | 0 | 0 | 0 | 0 | 0 |
| ENSMUSG00 | Gm17512 | 1438 | 0.19352459 | 0.03350935 | 0.27664575 | 0 | 0.46129547 | 0.18967065 |
| ENSMUSG00 | Vmn2r77 | 2559 | 0 | 0 | 0 | 0 | 0 | 0 |
| ENSMUSG00 | Olfr181 | 1389 | 0 | 0 | 0.09546834 | 0 | 0 | 0 |
| ENSMUSG00 | Gm17251 | 923 | 6.54694804 | 5.89931524 | 2.63391264 | 2.81463403 | 5.14133581 | 3.3095987 |
| ENSMUSG00 | Gm17250 | 1008 | 0 | 0 | 0 | 0 | 0 | 0 |
| ENSMUSG00 | Gm17345 | 92 | 3.45699831 | 4.1901254 | 0 | 0 | 0 | 7.70805022 |
| ENSMUSG00 | Gm17097 | 712 | 0 | 0 | 0.18624372 | 0 | 0 | 0 |
| ENSMUSG00 | Gm7535 | 908 | 0 | 0.05306877 | 0.04868044 | 0.07947587 | 0.05619645 | 0 |
| ENSMUSG00 | Lrrc32 | 3815 | 0.26052084 | 0.83363176 | 0.23172657 | 0.97416791 | 1.75215078 | 0.31456989 |
| ENSMUSG00 | Ighv1-62-1 | 89 | 0 | 0 | 0 | 0 | 0 | 0 |

| | | | | | | | | |
|---|---|---|---|---|---|---|---|---|
| ENSMUSG00 | Vmn2r21 | 2556 | 0 | 0 | 0 | 0 | 0 | 0 |
| ENSMUSG00 | Vmn2r8 | 2541 | 0 | 0 | 0 | 0 | 0 | 0 |
| ENSMUSG00 | Dio3os | 1277 | 0 | 0 | 0 | 0 | 0 | 0 |
| ENSMUSG00 | Gm17655 | 254 | 2.66079988 | 5.69131206 | 3.82850609 | 4.54576939 | 3.61604238 | 9.01995944 |
| ENSMUSG00 | Gm17206 | 574 | 0 | 0 | 0 | 0 | 0.17779225 | 0 |
| ENSMUSG00 | Gm17203 | 4015 | 0.28715042 | 0.07200963 | 0 | 0 | 0.01270894 | 0.00679319 |
| ENSMUSG00 | Vmn2r116 | 2565 | 0 | 0 | 0 | 0 | 0 | 0 |
| ENSMUSG00 | Vmn2r61 | 2592 | 0 | 0 | 0 | 0 | 0 | 0 |
| ENSMUSG00 | Gm3978 | 2506 | 0 | 0 | 0 | 0 | 0 | 0 |
| ENSMUSG00 | Gm17656 | 1092 | 0 | 0 | 0 | 0 | 0 | 0 |
| ENSMUSG00 | Gm3209 | 254 | 0.46955292 | 0.1897104 | 0 | 0.14205529 | 0 | 0.10738047 |
| ENSMUSG00 | Vmn1r142 | 917 | 0 | 0 | 0 | 0 | 0 | 0 |
| ENSMUSG00 | Slco6b1 | 2477 | 0 | 0 | 0 | 0 | 0 | 0 |
| ENSMUSG00 | Gm17162 | 470 | 0 | 0 | 0 | 0 | 0 | 0 |
| ENSMUSG00 | Gm14479 | 499 | 0 | 0 | 0 | 0 | 0 | 0 |
| ENSMUSG00 | Gm17459 | 448 | 0 | 0 | 0 | 0 | 0 | 0 |
| ENSMUSG00 | Gm8922 | 1119 | 0 | 0 | 0 | 0 | 0 | 0 |
| ENSMUSG00 | Gm17274 | 3682 | 0 | 0 | 0 | 0 | 0 | 0.05926049 |
| ENSMUSG00 | Olfr718-ps1 | 989 | 0 | 0 | 0 | 0 | 0 | 0 |
| ENSMUSG00 | 9430076C15l | 610 | 0.39103751 | 0 | 0 | 0 | 0 | 0 |
| ENSMUSG00 | AL451076.1 | 628 | 0 | 0 | 0 | 0 | 0 | 0.04343095 |
| ENSMUSG00 | Gm17276 | 2780 | 1.2298458 | 0.50266433 | 0.41339853 | 0 | 0.36709623 | 0.45130698 |
| ENSMUSG00 | Vmn1r170 | 914 | 0 | 0 | 0 | 0 | 0 | 0 |
| ENSMUSG00 | Gm17275 | 2548 | 0.39006555 | 0.35931805 | 0.55512519 | 0.2548967 | 0.90117226 | 2.42988348 |
| ENSMUSG00 | Gm4781 | 785 | 0 | 0 | 0 | 0 | 0 | 0 |
| ENSMUSG00 | Gm8693 | 920 | 0 | 0 | 0 | 0 | 0 | 0 |
| ENSMUSG00 | Gm17042 | 1212 | 0 | 0 | 0 | 0 | 0 | 0 |
| ENSMUSG00 | Gm17197 | 449 | 0 | 0 | 0 | 0 | 0 | 0 |
| ENSMUSG00 | Gm6647 | 637 | 0 | 0 | 0 | 0 | 0 | 0 |
| ENSMUSG00 | Gm17588 | 4243 | 0 | 0 | 0 | 0 | 0 | 0 |
| ENSMUSG00 | Gm17587 | 1814 | 0 | 0 | 0 | 0 | 0 | 0 |
| ENSMUSG00 | Gm6153 | 1057 | 0.03761162 | 0.27352758 | 0 | 0 | 0 | 0 |
| ENSMUSG00 | Gm17589 | 2142 | 0.14847985 | 0.02249601 | 0 | 0.01684503 | 0.04764368 | 0 |
| ENSMUSG00 | Gm20458 | 1363 | 0 | 0 | 0 | 0 | 0 | 0 |
| ENSMUSG00 | Gm17586 | 906 | 1.57968797 | 2.76566776 | 3.02485018 | 0.75668747 | 1.18273057 | 3.73295283 |
| ENSMUSG00 | Gm8266 | 1191 | 0 | 0 | 0 | 0 | 0 | 0 |
| ENSMUSG00 | Tcerg1l | 3973 | 0.01000641 | 0.3274689 | 0 | 0 | 0 | 0 |
| ENSMUSG00 | Gm7244 | 377 | 0 | 0 | 0 | 0 | 0 | 0 |
| ENSMUSG00 | Gm6872 | 699 | 0 | 0 | 0 | 0 | 0 | 0 |
| ENSMUSG00 | Traj55 | 62 | 0 | 0 | 0 | 0 | 0 | 0 |
| ENSMUSG00 | Vmn2r69 | 2547 | 0 | 0 | 0 | 0 | 0 | 0 |
| ENSMUSG00 | D630036H23 | 939 | 0.59273347 | 2.87373883 | 1.08268625 | 1.6523194 | 1.95628278 | 3.34034453 |
| ENSMUSG00 | Gm17472 | 350 | 0 | 0 | 0 | 0 | 0 | 0 |
| ENSMUSG00 | Gm17473 | 2195 | 0 | 0.24148103 | 0 | 0 | 0 | 0 |
| ENSMUSG00 | Gm17107 | 552 | 0 | 0 | 0 | 0 | 0 | 0 |
| ENSMUSG00 | Gm17187 | 408 | 0 | 0 | 0 | 0 | 0 | 0 |
| ENSMUSG00 | Vmn1r34 | 929 | 0 | 0 | 0 | 0 | 0 | 0 |
| ENSMUSG00 | Vmn1r1 | 1175 | 0 | 0 | 0 | 0 | 0 | 0 |
| ENSMUSG00 | Gm5244 | 500 | 0 | 0 | 0 | 0 | 0 | 0 |
| ENSMUSG00 | Gm17284 | 848 | 1.82837705 | 0.96600179 | 1.45949482 | 1.95728072 | 1.26362487 | 3.34500293 |
| ENSMUSG00 | Gm2878 | 483 | 0 | 0 | 0 | 0 | 0 | 0 |
| ENSMUSG00 | Fam71a | 2233 | 0 | 0 | 0 | 0 | 0 | 0 |
| ENSMUSG00 | Rplp2-ps1 | 441 | 0.18029696 | 0.43706524 | 0.70161656 | 0.40909348 | 0.11570607 | 0.30923627 |
| ENSMUSG00 | Gm7502 | 732 | 0 | 0 | 0 | 0 | 0 | 0 |
| ENSMUSG00 | Gm5828 | 3144 | 0.49315004 | 0.62838554 | 0.77325107 | 0.44758261 | 0.45443337 | 0.45978241 |
| ENSMUSG00 | Gm17300 | 1525 | 1.40773505 | 0.66355101 | 1.01446853 | 1.25399892 | 0.70265829 | 1.28772067 |
| ENSMUSG00 | Gm3115 | 1792 | 0 | 0 | 0 | 0 | 0 | 0 |
| ENSMUSG00 | Gm17351 | 1764 | 0.04507424 | 0 | 0 | 0 | 0 | 0 |
| ENSMUSG00 | Gm8305 | 1194 | 0 | 0 | 0 | 0 | 0 | 0 |
| ENSMUSG00 | Gm6900 | 692 | 0.05745012 | 0.13926717 | 0 | 0.15642505 | 0.14747508 | 0.23648531 |
| ENSMUSG00 | Gm11107 | 504 | 0 | 0 | 0 | 0 | 0 | 0 |
| ENSMUSG00 | Gm10722 | 647 | 0.06144587 | 0 | 0 | 0 | 0 | 0 |
| ENSMUSG00 | Gm8138 | 1399 | 0 | 0 | 0 | 0 | 0 | 0 |
| ENSMUSG00 | Gm17063 | 453 | 0 | 0 | 0 | 0 | 0 | 0 |
| ENSMUSG00 | Gm17299 | 40 | 1.98777403 | 0 | 0 | 0 | 2.55131879 | 0.68186598 |
| ENSMUSG00 | 4930477E14l | 619 | 0 | 0 | 0 | 0 | 0 | 0 |
| ENSMUSG00 | Gm17660 | 510 | 0 | 0 | 0 | 0 | 0 | 0 |
| ENSMUSG00 | Gm17659 | 1535 | 0.49208738 | 0.09417546 | 0.2591639 | 0.11753109 | 0.4321452 | 0.78181376 |
| ENSMUSG00 | Gm17174 | 476 | 0 | 0 | 0 | 0 | 0 | 0 |
| ENSMUSG00 | Gm17661 | 268 | 0 | 0 | 0 | 0 | 0 | 0 |
| ENSMUSG00 | Gm17662 | 22 | 0 | 0 | 0 | 0 | 0 | 0 |
| ENSMUSG00 | Krtap20-2 | 527 | 0 | 0 | 0 | 0 | 0 | 0 |
| ENSMUSG00 | Gm9359 | 1220 | 0 | 0 | 0 | 0 | 0.0418249 | 0 |
| ENSMUSG00 | Gm17720 | 25 | 3.18043844 | 19.2745769 | 3.53614744 | 11.5462543 | 0 | 12.0008413 |
| ENSMUSG00 | Gm3727 | 990 | 0 | 0 | 0 | 0 | 0 | 0 |
| ENSMUSG00 | Glyatl3 | 868 | 0 | 0 | 0 | 0 | 0 | 0 |
| ENSMUSG00 | Gm3170 | 843 | 0 | 0 | 0 | 0 | 0 | 0 |
| ENSMUSG00 | Vmn2r55 | 2341 | 0 | 0 | 0 | 0 | 0 | 0 |
| ENSMUSG00 | 4930532G15l | 697 | 0.11407598 | 0.89874282 | 0.06341728 | 0 | 0.2928343 | 0.39131477 |
| ENSMUSG00 | Gm8207 | 499 | 0 | 0 | 0 | 0 | 0 | 0 |
| ENSMUSG00 | Gm1979 | 4317 | 0 | 0 | 0 | 0 | 0 | 0 |
| ENSMUSG00 | 9330020H09l | 2145 | 0.68575887 | 0.62900717 | 1.15398751 | 0.9420021 | 1.16563749 | 0.82650422 |
| ENSMUSG00 | Ighd5-8 | 28 | 0 | 0 | 0 | 0 | 0 | 0 |
| ENSMUSG00 | Gm5551 | 617 | 0 | 0 | 0 | 0 | 0 | 0.13261575 |
| ENSMUSG00 | Gm9742 | 587 | 0 | 0 | 0 | 0 | 0 | 0 |
| ENSMUSG00 | Siglec15 | 1150 | 0 | 0 | 0 | 0 | 0 | 0 |
| ENSMUSG00 | Gm17536 | 3633 | 0.67845852 | 0.25200727 | 0.6570051 | 0.98324316 | 0.64608128 | 0.71320967 |
| ENSMUSG00 | Gm17535 | 668 | 0 | 0 | 0 | 0 | 0.38193395 | 0 |
| ENSMUSG00 | Gm17538 | 326 | 0 | 0.44343352 | 0.67794238 | 0 | 0.15652263 | 0.08366454 |
| ENSMUSG00 | Vmn2r14 | 2541 | 0 | 0 | 0 | 0 | 0 | 0 |
| ENSMUSG00 | 7420461P10l | 731 | 0 | 0 | 0 | 0 | 0 | 0 |
| ENSMUSG00 | 4930445B16l | 1118 | 0 | 0 | 0 | 0 | 0 | 0 |
| ENSMUSG00 | Ighd5-6 | 11 | 0 | 0 | 0 | 0 | 0 | 0 |
| ENSMUSG00 | Gm13698 | 2725 | 0 | 0 | 0 | 0 | 0 | 0 |
| ENSMUSG00 | Gm996 | 2824 | 0 | 0 | 0 | 0 | 0 | 0 |
| ENSMUSG00 | Gm17599 | 1272 | 0.09376293 | 0 | 0 | 0 | 0.08023015 | 0.30019257 |
| ENSMUSG00 | Gm17598 | 994 | 0.07999091 | 0.29086384 | 0 | 0 | 0.05133438 | 0.05487855 |
| ENSMUSG00 | Gm7889 | 424 | 0 | 0 | 0 | 0 | 0 | 0 |
| ENSMUSG00 | Vmn1r175 | 914 | 0 | 0 | 0 | 0 | 0 | 0 |
| ENSMUSG00 | Gm3764 | 1086 | 0 | 0 | 0 | 0 | 0 | 0 |
| ENSMUSG00 | 1700030C10l | 1470 | 0 | 0.78671742 | 0 | 0.02454561 | 0 | 0 |
| ENSMUSG00 | Gm17411 | 34 | 0 | 0 | 0 | 0 | 0 | 0 |
| ENSMUSG00 | Gm17410 | 3372 | 0.01178988 | 0 | 0.01310849 | 0.03210146 | 0.01513238 | 0.01617713 |
| ENSMUSG00 | Vmn2r115 | 2568 | 0 | 0 | 0 | 0 | 0 | 0 |
| ENSMUSG00 | Gm17218 | 408 | 0 | 0 | 0 | 0 | 0 | 0 |
| ENSMUSG00 | Gm17105 | 650 | 0 | 0.22239896 | 0.40801701 | 0 | 0 | 0 |
| ENSMUSG00 | Prr23a | 1058 | 0 | 0 | 0.04177868 | 0 | 0 | 0 |
| ENSMUSG00 | Gm17679 | 172 | 0 | 0 | 0 | 0 | 0 | 0 |
| ENSMUSG00 | Gm17130 | 2680 | 0.02966827 | 0 | 0 | 0 | 0.01903969 | 0 |
| ENSMUSG00 | Gm4535 | 557 | 0 | 0 | 0 | 0 | 0 | 0 |
| ENSMUSG00 | BC065403 | 1018 | 0 | 0 | 0.13026083 | 0.24810836 | 0.15037242 | 0.21433901 |
| ENSMUSG00 | Gm5428 | 887 | 0 | 0 | 0 | 0 | 0 | 0 |
| ENSMUSG00 | Ighv8-14 | 119 | 0 | 0 | 0 | 0 | 0 | 0 |
| ENSMUSG00 | 4921514A10l | 2114 | 0.4137278 | 0.25073362 | 0 | 0 | 0 | 0.23223439 |
| ENSMUSG00 | A930018M24 | 1420 | 0 | 0 | 0 | 0 | 0 | 0 |
| ENSMUSG00 | Gm5269 | 748 | 0 | 0 | 0 | 0 | 0 | 0 |
| ENSMUSG00 | Kcnmb3 | 716 | 0 | 0 | 0 | 0 | 0 | 0 |
| ENSMUSG00 | Gm17302 | 785 | 0 | 0 | 0 | 0 | 0 | 0 |
| ENSMUSG00 | Gm9222 | 335 | 2.4921346 | 3.02064264 | 0.659729 | 1.29249115 | 2.13244556 | 0.89558517 |
| ENSMUSG00 | Gm17305 | 501 | 0.07935226 | 0 | 0 | 0 | 0 | 0.27220199 |
| ENSMUSG00 | Gm17304 | 67 | 0 | 0 | 0 | 0 | 0 | 0 |
| ENSMUSG00 | Gm17303 | 74 | 11.2819607 | 4.55817696 | 16.1276995 | 7.31392794 | 13.7909124 | 19.1659627 |
| ENSMUSG00 | Cldn21 | 653 | 0 | 0 | 0 | 0 | 0 | 0 |
| ENSMUSG00 | Gm17164 | 503 | 0 | 0 | 0 | 0 | 0 | 0 |
| ENSMUSG00 | Gm17626 | 3795 | 0.27236956 | 0.0507894 | 0.17471084 | 0.00950779 | 0.0941198 | 0.10061791 |
| ENSMUSG00 | Gm4302 | 884 | 0 | 0 | 0 | 0 | 0 | 0 |
| ENSMUSG00 | 5830462I19R | 3382 | 0.19983535 | 0.29920618 | 0.02613947 | 0.06401309 | 0.10561343 | 0.06451718 |
| ENSMUSG00 | Gm17623 | 58 | 0 | 0 | 0 | 0 | 0 | 0 |
| ENSMUSG00 | Gm5633 | 548 | 0 | 0 | 0 | 0 | 0 | 0 |
| ENSMUSG00 | Gm17625 | 3197 | 0 | 0 | 0 | 0 | 0 | 0 |
| ENSMUSG00 | Gm17624 | 199 | 0 | 0 | 0 | 0 | 0 | 0 |
| ENSMUSG00 | Gm13276 | 1555 | 0 | 0 | 0 | 0 | 0 | 0 |
| ENSMUSG00 | Gm17622 | 43 | 0 | 0 | 3.08384951 | 0 | 0 | 0 |
| ENSMUSG00 | Gm2832 | 1755 | 0 | 0 | 0 | 0 | 0 | 0 |
| ENSMUSG00 | Gm8324 | 1203 | 0 | 0 | 0 | 0 | 0 | 0 |
| ENSMUSG00 | Gm17310 | 40 | 0 | 0 | 0 | 0 | 0 | 0 |
| ENSMUSG00 | Gm17311 | 1743 | 0 | 0 | 0 | 0 | 0 | 0 |
| ENSMUSG00 | Gm3138 | 1755 | 0 | 0 | 0 | 0 | 0 | 0 |
| ENSMUSG00 | BC027582 | 873 | 0 | 0 | 0 | 0 | 0 | 0 |
| ENSMUSG00 | Gm7991 | 587 | 0 | 0 | 0 | 0 | 0 | 0 |
| ENSMUSG00 | Gm9064 | 565 | 0 | 0 | 0 | 0 | 0 | 0 |
| ENSMUSG00 | Gm20603 | 3483 | 0 | 0 | 0 | 0.08287578 | 0 | 0.00783079 |
| ENSMUSG00 | Ccdc152 | 789 | 0.05038717 | 0.48858243 | 0.28011307 | 0.04573136 | 0.45270549 | 0.5876665 |
| ENSMUSG00 | Gm4545 | 1510 | 0 | 0 | 0 | 0 | 0 | 0 |
| ENSMUSG00 | Vmn1r104 | 923 | 0 | 0 | 0 | 0 | 0 | 0 |
| ENSMUSG00 | Gm8247 | 1400 | 0 | 0.06883777 | 0 | 0 | 0 | 0 |
| ENSMUSG00 | Gm17181 | 122 | 0 | 0 | 0 | 0 | 0 | 0 |
| ENSMUSG00 | Gm17239 | 49 | 0 | 0 | 0 | 0 | 0 | 0 |
| ENSMUSG00 | Gm17238 | 696 | 9.42479064 | 13.6389786 | 16.9567415 | 14.8786592 | 18.1084983 | 25.5895682 |
| ENSMUSG00 | Iqcf6 | 472 | 0 | 0 | 0 | 0 | 0 | 0 |
| ENSMUSG00 | Gm14474 | 524 | 0 | 0 | 0 | 0 | 0 | 0 |
| ENSMUSG00 | Gm7945 | 1423 | 0 | 0 | 0 | 0 | 0 | 0 |
| ENSMUSG00 | Gm17109 | 357 | 0 | 0 | 0 | 0 | 0.14293103 | 0 |
| ENSMUSG00 | Gm17108 | 2875 | 0.02765599 | 0.11732351 | 0.26136742 | 0 | 0.21297966 | 0.21819711 |
| ENSMUSG00 | Gm17436 | 59 | 0 | 0 | 0 | 0 | 0 | 0 |
| ENSMUSG00 | Gm17209 | 518 | 0 | 0 | 0 | 0 | 0 | 0 |
| ENSMUSG00 | Gm17437 | 1966 | 0 | 0 | 0 | 0 | 0 | 0 |
| ENSMUSG00 | Gm1153 | 506 | 0 | 0 | 0 | 0 | 0 | 0 |
| ENSMUSG00 | E530011L22f | 2005 | 3.31130436 | 1.75441909 | 2.35890135 | 1.24172622 | 3.10484681 | 2.14932319 |
| ENSMUSG00 | Gm6190 | 772 | 0.05149674 | 0 | 0.17176882 | 0 | 0.13219268 | 0.14131937 |
| ENSMUSG00 | Gm17124 | 1374 | 0 | 0 | 0 | 0.10504234 | 0 | 0 |
| ENSMUSG00 | Gm17367 | 58 | 0 | 0 | 0 | 0 | 0.8797651 | 0 |
| ENSMUSG00 | Gm17175 | 1398 | 0 | 0 | 0 | 0 | 0 | 0 |
| ENSMUSG00 | Gm6907 | 3448 | 0.72639074 | 0.18167742 | 0.93582788 | 0.2092926 | 1.33189496 | 0.74356615 |
| ENSMUSG00 | Gm4308 | 915 | 0 | 0 | 0 | 0 | 0 | 0 |
| ENSMUSG00 | Gm3077 | 624 | 0 | 0 | 0 | 0 | 0 | 0 |
| ENSMUSG00 | 4921509A06l | 390 | 0 | 0 | 0 | 0 | 0 | 0 |
| ENSMUSG00 | Gm3182 | 1939 | 0 | 0 | 0 | 0 | 0 | 0 |
| ENSMUSG00 | Gm17020 | 273 | 0 | 0 | 0 | 0 | 0.1869098 | 0 |
| ENSMUSG00 | Gm17544 | 73 | 0 | 0 | 0 | 0 | 0 | 0 |
| ENSMUSG00 | Vmn1r224 | 896 | 0 | 0 | 0 | 0 | 0 | 0 |
| ENSMUSG00 | Gm13696 | 2725 | 0 | 0 | 0 | 0 | 0 | 0 |
| ENSMUSG00 | Olfr92 | 938 | 0 | 0 | 0 | 0 | 0 | 0 |
| ENSMUSG00 | 1700024N20l | 615 | 0.19392917 | 0 | 0.28749166 | 0.17600997 | 0.16593943 | 0.44349007 |
| ENSMUSG00 | Serpina3 | 1393 | 0.22831575 | 0.31132662 | 0.12692561 | 0.1036096 | 0.14652226 | 0.07831914 |
| ENSMUSG00 | Gm3500 | 2040 | 0 | 0 | 0 | 0 | 0 | 0 |
| ENSMUSG00 | Serpina3l-ps | 1260 | 0 | 0 | 0 | 0 | 0 | 0 |
| ENSMUSG00 | Mirg | 3166 | 0 | 0 | 0 | 0 | 0 | 0 |
| ENSMUSG00 | Gm17545 | 70 | 0 | 0 | 0 | 0 | 0 | 0 |
| ENSMUSG00 | Gm17133 | 287 | 0 | 0 | 0 | 0 | 0 | 0 |
| ENSMUSG00 | Gm6164 | 923 | 0 | 0 | 0 | 0 | 0 | 0 |
| ENSMUSG00 | Gm7677 | 1782 | 0 | 0 | 0 | 0 | 0.14317165 | 0 |
| ENSMUSG00 | Gm3962 | 2611 | 0 | 0 | 0 | 0 | 0 | 0 |
| ENSMUSG00 | Gm17716 | 2541 | 0 | 0 | 0.10437271 | 0 | 0 | 0 |
| ENSMUSG00 | Gm17036 | 252 | 0 | 0.38243208 | 1.57863725 | 0 | 0.20248562 | 0.64939617 |
| ENSMUSG00 | Tstd1 | 547 | 0.21803737 | 1.14519881 | 0.16161551 | 0.32961759 | 2.70523748 | 0.69807121 |
| ENSMUSG00 | Gm14477 | 499 | 0 | 0 | 0 | 0 | 0 | 0 |
| ENSMUSG00 | 1700026J14R | 792 | 0 | 0 | 0 | 0 | 0 | 0 |
| ENSMUSG00 | Gm17493 | 1132 | 0.52679524 | 0.34054023 | 0.11714269 | 0.92436333 | 0.67614456 | 1.46974646 |
| ENSMUSG00 | Gm4307 | 869 | 0 | 0 | 0 | 0 | 0 | 0 |
| ENSMUSG00 | Gm10060 | 4008 | 0 | 0.02404513 | 0 | 0.12603509 | 0.01273113 | 0.10207575 |
| ENSMUSG00 | Gm17120 | 741 | 0.05365112 | 0 | 0 | 0 | 0.06886151 | 0 |
| ENSMUSG00 | Gm7354 | 506 | 0 | 0 | 0 | 0 | 0 | 0 |
| ENSMUSG00 | Gm17677 | 348 | 0 | 0 | 0 | 0 | 0 | 0 |
| ENSMUSG00 | 9130204L05R | 488 | 0 | 0 | 0 | 0 | 0 | 0 |
| ENSMUSG00 | Gm17678 | 242 | 0 | 0 | 0 | 0 | 0.21085279 | 0 |
| ENSMUSG00 | Gm15494 | 449 | 0.08854227 | 0 | 0 | 0.1607218 | 0 | 0.1822359 |
| ENSMUSG00 | Vmn2r44 | 2580 | 0 | 0 | 0 | 0 | 0 | 0 |
| ENSMUSG00 | Gm3597 | 1754 | 0 | 0 | 0 | 0 | 0 | 0 |
| ENSMUSG00 | Trav5n-3 | 269 | 0 | 0 | 0 | 0 | 0 | 0 |
| ENSMUSG00 | Gm17647 | 34 | 0 | 0 | 0 | 0 | 0 | 0 |
| ENSMUSG00 | Gm17649 | 49 | 0 | 0 | 0 | 0 | 0 | 0 |
| ENSMUSG00 | Gm3604 | 1674 | 7.50461878 | 4.86470055 | 2.71970719 | 10.4538779 | 5.79152414 | 20.284902 |
| ENSMUSG00 | Gm17059 | 517 | 0 | 0 | 0 | 0 | 0 | 0 |
| ENSMUSG00 | Gm3278 | 1495 | 0 | 0 | 0 | 0 | 0 | 0 |
| ENSMUSG00 | Gm2078 | 8379 | 0 | 0 | 0 | 0 | 0 | 0 |
| ENSMUSG00 | Gm8082 | 1398 | 0 | 0 | 0 | 0 | 0 | 0 |
| ENSMUSG00 | 2700016F22R | 3603 | 0.3751558 | 0.05349591 | 0.88330078 | 0.04005778 | 1.1612997 | 0.32550916 |
| ENSMUSG00 | Ear-ps3 | 444 | 0 | 0 | 0 | 0 | 0 | 0 |
| ENSMUSG00 | 2310031A07R | 988 | 0.32190673 | 0.09754341 | 0.49212578 | 0.21912173 | 0.51646129 | 0.69014775 |
| ENSMUSG00 | Gm17334 | 906 | 0 | 0 | 0 | 0.03982566 | 0.0563205 | 0.18062675 |
| ENSMUSG00 | Gm17071 | 263 | 0 | 0.73287365 | 0 | 0 | 0.97008319 | 0 |
| ENSMUSG00 | Gm17333 | 376 | 0 | 0 | 0 | 0 | 0 | 0 |
| ENSMUSG00 | Gm19840 | 880 | 0 | 0 | 0.35160557 | 0 | 0 | 0 |
| ENSMUSG00 | Gm17332 | 86 | 0.46227303 | 0 | 0 | 0.41955866 | 0 | 0 |
| ENSMUSG00 | Gm17330 | 335 | 0 | 0 | 0 | 0 | 0 | 0 |
| ENSMUSG00 | Vmn1r95 | 923 | 0 | 0 | 0 | 0 | 0 | 0 |
| ENSMUSG00 | Gm5218 | 464 | 0 | 0 | 0 | 0 | 0 | 0 |
| ENSMUSG00 | Gm2619 | 490 | 0 | 0 | 0 | 0 | 0 | 0 |
| ENSMUSG00 | Vmn1r77 | 920 | 0 | 0 | 0 | 0 | 0 | 0 |
| ENSMUSG00 | Gm3685 | 979 | 0 | 0 | 0 | 0 | 0 | 0 |
| ENSMUSG00 | Gm8011 | 1746 | 0 | 0 | 0 | 0 | 0 | 0 |
| ENSMUSG00 | Gm7271 | 516 | 0 | 0 | 0 | 0 | 0 | 0 |
| ENSMUSG00 | Vmn2r71 | 2562 | 0 | 0 | 0 | 0 | 0 | 0 |
| ENSMUSG00 | Vmn2r91 | 2565 | 0 | 0 | 0 | 0 | 0 | 0 |
| ENSMUSG00 | Purb | 1125 | 4.87667228 | 8.73780818 | 5.73641696 | 5.83727298 | 3.80996939 | 9.64916126 |
| ENSMUSG00 | Gm3494 | 887 | 0 | 0 | 0 | 0 | 0 | 0 |
| ENSMUSG00 | Gm3264 | 1004 | 0 | 0 | 0 | 0 | 0 | 0 |
| ENSMUSG00 | 1110054M08 | 1601 | 1.04292953 | 1.26410404 | 2.48480067 | 2.18610763 | 1.24299104 | 3.56052442 |
| ENSMUSG00 | Gm4211 | 2218 | 0 | 0 | 0.25907302 | 0 | 0 | 0.01229695 |
| ENSMUSG00 | Krtap11-1 | 1007 | 0 | 0 | 0 | 0 | 0 | 0 |
| ENSMUSG00 | Gm17253 | 64 | 11.1812289 | 6.77621843 | 4.14392278 | 7.32916529 | 3.18914849 | 4.26166238 |
| ENSMUSG00 | 4732491K20R | 3140 | 3.82361628 | 3.60631016 | 3.81487244 | 2.02243307 | 3.3638407 | 3.30943871 |
| ENSMUSG00 | Gm17252 | 383 | 0 | 0 | 0 | 0 | 0 | 0 |
| ENSMUSG00 | Gm17208 | 527 | 0 | 0 | 0 | 0 | 0 | 0 |
| ENSMUSG00 | 5430400D12l | 1656 | 2.08860314 | 3.3462807 | 1.65489992 | 2.22244477 | 4.00569375 | 3.60697222 |
| ENSMUSG00 | Gm17254 | 4474 | 0.29323443 | 1.12011399 | 0.28651172 | 0.74196426 | 1.17472434 | 0.88395679 |
| ENSMUSG00 | Olfr724 | 1005 | 0 | 0 | 0 | 0 | 0 | 0 |
| ENSMUSG00 | Gm3084 | 597 | 0 | 0 | 0 | 0 | 0 | 0 |
| ENSMUSG00 | Gm17723 | 66 | 0 | 0 | 0 | 0 | 0 | 0 |
| ENSMUSG00 | Gm8775 | 512 | 0 | 0 | 0 | 0 | 0 | 0 |
| ENSMUSG00 | Vmn1r122 | 917 | 0 | 0 | 0 | 0 | 0 | 0 |
| ENSMUSG00 | Vmn1r176 | 914 | 0 | 0 | 0 | 0 | 0 | 0 |
| ENSMUSG00 | Gm3755 | 1457 | 0 | 0 | 0 | 0 | 0 | 0 |
| ENSMUSG00 | Gm20390 | 983 | 0 | 0 | 0 | 0 | 0 | 0 |
| ENSMUSG00 | Gm7177 | 539 | 0 | 0.17879941 | 0 | 0 | 0 | 0 |
| ENSMUSG00 | Gm6970 | 665 | 0.17934803 | 0 | 0.06646894 | 0 | 0.84404531 | 0 |
| ENSMUSG00 | Gm4033 | 2552 | 0 | 0 | 0 | 0 | 0 | 0 |
| ENSMUSG00 | Gm17394 | 43 | 0 | 0 | 0 | 0 | 0 | 0 |
| ENSMUSG00 | Gm6434 | 407 | 0 | 0 | 0 | 0 | 0 | 0.06701386 |
| ENSMUSG00 | Gm17393 | 2797 | 0.95231219 | 0.99921832 | 1.12203463 | 0.50311038 | 0.8756761 | 1.89176976 |
| ENSMUSG00 | Gm4207 | 1869 | 0 | 0 | 0 | 0 | 0 | 0 |
| ENSMUSG00 | Vmn1r159 | 917 | 0 | 0 | 0 | 0 | 0 | 0 |
| ENSMUSG00 | Gm17114 | 509 | 0 | 0 | 0 | 0 | 0 | 0 |
| ENSMUSG00 | Gm17103 | 547 | 0 | 0 | 0 | 0 | 0 | 0 |
| ENSMUSG00 | Vmn2r76 | 2556 | 0 | 0 | 0 | 0 | 0 | 0 |
| ENSMUSG00 | Gm9182 | 1040 | 0.03822642 | 0 | 0.04250177 | 0.06938855 | 0 | 0.07867684 |
| ENSMUSG00 | Ighv1-62-2 | 301 | 0 | 0 | 0 | 0 | 0 | 0 |
| ENSMUSG00 | Vgll3 | 3318 | 0 | 0 | 0 | 0 | 0 | 0 |
| ENSMUSG00 | Gm17690 | 3465 | 1.50302105 | 0.7926774 | 1.33944979 | 0.85388965 | 1.2664555 | 1.8655381 |
| ENSMUSG00 | Gm3250 | 692 | 0 | 0 | 0 | 0 | 0 | 0 |
| ENSMUSG00 | Gm3834 | 2375 | 0 | 0 | 0 | 0 | 0 | 0.12632465 |
| ENSMUSG00 | Gm16529 | 1685 | 0 | 0 | 0 | 0 | 0 | 0 |
| ENSMUSG00 | Gm17689 | 327 | 0 | 0 | 0 | 0 | 0 | 0 |
| ENSMUSG00 | Kiss1 | 776 | 0.4610816 | 0.86934303 | 0.4556891 | 0.41847732 | 1.38086842 | 1.37076151 |
| ENSMUSG00 | Gm6346 | 1443 | 0 | 0 | 0 | 0 | 0 | 0 |
| ENSMUSG00 | Gm17505 | 82 | 0 | 0 | 0 | 0 | 0 | 0 |
| ENSMUSG00 | Speer4e | 1598 | 0 | 0.09046266 | 0 | 0.18063602 | 0.0638628 | 0 |
| ENSMUSG00 | Gm3348 | 846 | 0 | 0 | 0 | 0 | 0 | 0 |
| ENSMUSG00 | Vmn2r110 | 2571 | 0 | 0 | 0 | 0 | 0 | 0 |
| ENSMUSG00 | Vmn2r19 | 2559 | 0 | 0 | 0 | 0 | 0 | 0 |
| ENSMUSG00 | Gm8453 | 920 | 0 | 0 | 0 | 0 | 0 | 0 |
| ENSMUSG00 | D430020J02R | 37 | 0 | 0 | 0 | 0 | 0 | 0 |
| ENSMUSG00 | BC024659 | 4462 | 5.80918272 | 6.8575506 | 7.46933878 | 14.3131373 | 7.29601698 | 8.55159577 |
| ENSMUSG00 | Gm17320 | 93 | 0 | 0 | 0 | 0 | 0 | 0 |
| ENSMUSG00 | Gm17321 | 2775 | 0.37248378 | 0.41674761 | 0.76457242 | 0.4030787 | 0.31259401 | 0.97304118 |
| ENSMUSG00 | Gm17322 | 2167 | 0 | 0 | 0.50994281 | 0 | 0.42384622 | 0 |
| ENSMUSG00 | Igkv17-134 | 344 | 0 | 0 | 0 | 0 | 0 | 0 |
| ENSMUSG00 | Gm6682 | 296 | 0.13430906 | 0 | 0 | 0 | 0 | 0 |
| ENSMUSG00 | Gm17642 | 2446 | 0.87767619 | 1.00470505 | 1.40954365 | 2.5224978 | 1.83578131 | 2.74307492 |
| ENSMUSG00 | Gm17104 | 321 | 0.12384885 | 0.15011353 | 0 | 0.11240512 | 0 | 0.08496772 |
| ENSMUSG00 | Gm17641 | 2349 | 0 | 0 | 0 | 0 | 0 | 0 |
| ENSMUSG00 | Gm6673 | 695 | 0 | 0 | 0 | 0 | 0 | 0 |
| ENSMUSG00 | Igkv4-53 | 291 | 0 | 0 | 0 | 0 | 0 | 0 |
| ENSMUSG00 | Gm3248 | 2048 | 0 | 0.23528536 | 0 | 0 | 0 | 0 |
| ENSMUSG00 | Vmn1r149 | 923 | 0 | 0 | 0 | 0 | 0 | 0 |
| ENSMUSG00 | Gm1818 | 2397 | 0.01658552 | 0.08041125 | 0.18440485 | 0.16558302 | 0.0212876 | 0.12516522 |
| ENSMUSG00 | Gm17643 | 1592 | 0.19977628 | 0.84750024 | 2.0268433 | 1.13323004 | 0.06410349 | 0.770954 |
| ENSMUSG00 | Gm17032 | 336 | 0 | 0 | 0 | 0 | 0 | 0 |
| ENSMUSG00 | Gm8065 | 601 | 0 | 0 | 0 | 0 | 0 | 0 |
| ENSMUSG00 | Gm17431 | 2024 | 0 | 0 | 0 | 0 | 0 | 0 |
| ENSMUSG00 | Gm17234 | 390 | 0 | 0 | 0.11333806 | 0 | 0 | 0 |
| ENSMUSG00 | Gm17430 | 473 | 0 | 0 | 0 | 0 | 0 | 0 |
| ENSMUSG00 | Gm3111 | 487 | 0 | 0 | 0 | 0 | 0 | 0 |
| ENSMUSG00 | Gm17075 | 669 | 0 | 0 | 0 | 0 | 0 | 0 |
| ENSMUSG00 | Gm17051 | 486 | 0 | 0 | 0 | 0 | 0 | 0 |
| ENSMUSG00 | Gm3275 | 486 | 0 | 0 | 0 | 0 | 0 | 0 |
| ENSMUSG00 | Gm17731 | 34 | 0 | 2.8344966 | 2.60010841 | 1.0612366 | 1.50077576 | 3.20878109 |
| ENSMUSG00 | Gm3104 | 470 | 0 | 0 | 0 | 0 | 0 | 0 |
| ENSMUSG00 | Gm6047 | 573 | 0 | 0 | 0 | 0 | 0 | 0.19039888 |
| ENSMUSG00 | Glis3 | 1271 | 0 | 0 | 0 | 0 | 0 | 0 |
| ENSMUSG00 | Flg | 935 | 0 | 0 | 0 | 0 | 0 | 0 |
| ENSMUSG00 | Gm4181 | 893 | 0 | 0 | 0 | 0 | 0 | 0 |
| ENSMUSG00 | Gm8439 | 417 | 0 | 0 | 0 | 0.08652769 | 0 | 0.06540681 |
| ENSMUSG00 | Gm17730 | 2757 | 0 | 0 | 0 | 0 | 0 | 0 |
| ENSMUSG00 | Snhg7 | 1529 | 2.2100823 | 1.92241529 | 2.57289995 | 1.60469524 | 2.10246022 | 2.83627707 |
| ENSMUSG00 | Ighv1-71 | 301 | 0 | 0 | 0 | 0 | 0 | 0 |
| ENSMUSG00 | 2310043M15 | 1551 | 1.89677986 | 1.33592328 | 1.31095086 | 2.39616414 | 2.96091156 | 1.26613412 |
| ENSMUSG00 | 4831440E17R | 1026 | 2.40237796 | 2.34826716 | 4.00659981 | 1.54737813 | 2.63586542 | 10.6865546 |
| ENSMUSG00 | Speer6-ps1 | 541 | 0 | 0 | 0 | 0 | 0 | 0 |
| ENSMUSG00 | Gm17100 | 271 | 0 | 0 | 0 | 0 | 0 | 0 |
| ENSMUSG00 | Gm17606 | 156 | 0 | 0 | 1.41672574 | 0.23129516 | 0 | 0.52451229 |
| ENSMUSG00 | Gm8895 | 1219 | 0 | 0 | 0 | 0 | 0 | 0 |
| ENSMUSG00 | Olfr1380 | 937 | 0 | 0 | 0 | 0 | 0 | 0 |
| ENSMUSG00 | Gm17605 | 709 | 0 | 0 | 0 | 0 | 0 | 0 |
| ENSMUSG00 | Gm17288 | 1634 | 0 | 0 | 0 | 0 | 0 | 0 |
| ENSMUSG00 | Gm4934 | 4347 | 0 | 0 | 0 | 0 | 0 | 0 |
| ENSMUSG00 | Gm17490 | 61 | 0 | 0 | 0 | 0 | 0 | 0 |
| ENSMUSG00 | Fth-ps2 | 515 | 0.30878043 | 0.93565907 | 0.51497293 | 1.61143112 | 0.39632137 | 0.47664418 |
| ENSMUSG00 | Gm17368 | 158 | 0 | 0 | 0 | 0 | 0 | 0 |
| ENSMUSG00 | Gm17214 | 540 | 0.66259134 | 0.35693661 | 0 | 1.47000922 | 0.28347987 | 0.10101718 |
| ENSMUSG00 | Gm17289 | 1850 | 2.64320222 | 1.61489698 | 2.31761015 | 1.83335794 | 0.99294569 | 2.58003344 |
| ENSMUSG00 | Gm5415 | 3747 | 0 | 0 | 0 | 0 | 0 | 0 |
| ENSMUSG00 | Gm17287 | 1053 | 0 | 0 | 0 | 0 | 0 | 0 |
| ENSMUSG00 | Vmn1r123 | 923 | 0 | 0 | 0 | 0 | 0 | 0 |
| ENSMUSG00 | Gm17480 | 1900 | 0 | 0.12680643 | 0.09305651 | 0 | 0.10742395 | 0.11484059 |
| ENSMUSG00 | 4930480E11l | 1468 | 0 | 0 | 0 | 0 | 0 | 0 |
| ENSMUSG00 | Gm12166 | 811 | 0 | 0 | 0 | 0 | 0 | 0 |
| ENSMUSG00 | Gm4673 | 3658 | 0 | 0 | 0 | 0 | 0 | 0 |
| ENSMUSG00 | Gm3760 | 1750 | 0 | 0 | 0 | 0 | 0 | 0 |
| ENSMUSG00 | Traj54 | 53 | 0 | 0 | 0 | 0 | 0 | 0 |
| ENSMUSG00 | Raxos1 | 2545 | 0 | 0 | 0 | 0 | 0 | 0 |
| ENSMUSG00 | Gm3542 | 2058 | 0 | 0 | 0 | 0 | 0 | 0 |
| ENSMUSG00 | 1700112D23 | 543 | 0.07321451 | 0 | 0 | 0.26579775 | 0.09397123 | 0 |
| ENSMUSG00 | Gm20520 | 588 | 0 | 0 | 0 | 0 | 0 | 0 |
| ENSMUSG00 | 4930500F10l | 716 | 0.16657324 | 0 | 0 | 0 | 0 | 0 |
| ENSMUSG00 | BC051077 | 1934 | 0 | 0 | 0 | 0 | 0 | 0.01410271 |
| ENSMUSG00 | Ighd5-3 | 9 | 0 | 0 | 0 | 0 | 0 | 0 |
| ENSMUSG00 | Gm8766 | 785 | 0 | 0 | 0 | 0 | 0 | 0 |
| ENSMUSG00 | Gm17157 | 347 | 0 | 0 | 0 | 0 | 0 | 0 |
| ENSMUSG00 | Eid1 | 1690 | 40.3671033 | 33.8445603 | 30.6274309 | 35.847191 | 25.2414498 | 59.633013 |
| ENSMUSG00 | Ighv1-78 | 293 | 0 | 0 | 0 | 0 | 0 | 0 |
| ENSMUSG00 | Gm2866 | 480 | 0 | 0 | 0 | 0 | 0 | 0 |
| ENSMUSG00 | Flg | 1487 | 0 | 0 | 0 | 0.12132497 | 0 | 0 |
| ENSMUSG00 | Gm8104 | 1756 | 0 | 0.13720513 | 0 | 0.16438289 | 0 | 0 |
| ENSMUSG00 | Gm17329 | 138 | 0 | 0 | 0 | 0 | 0 | 0 |
| ENSMUSG00 | Gm17328 | 1395 | 0 | 0 | 0.19011545 | 0 | 0.43893657 | 0.07820685 |
| ENSMUSG00 | 1700037F24l | 971 | 0 | 0 | 0 | 0 | 0 | 0 |
| ENSMUSG00 | Col6a5 | 9252 | 0 | 0 | 0 | 0 | 0 | 0 |
| ENSMUSG00 | Gm17353 | 254 | 0 | 0 | 0 | 0 | 0 | 0 |
| ENSMUSG00 | Gm7349 | 506 | 0 | 0 | 0 | 0 | 0 | 0 |
| ENSMUSG00 | Vmn2r92 | 2577 | 0 | 0 | 0 | 0 | 0 | 0 |
| ENSMUSG00 | Gm17135 | 561 | 0.42519231 | 2.57681509 | 0.07879116 | 1.22203003 | 0.45478053 | 1.0695937 |
| ENSMUSG00 | Gm17519 | 1735 | 0 | 0 | 0 | 0 | 0 | 0 |
| ENSMUSG00 | Gm13285 | 1555 | 0 | 0 | 0 | 0 | 0 | 0 |
| ENSMUSG00 | 6030419C18 | 535 | 0 | 0 | 0 | 0 | 0 | 0 |
| ENSMUSG00 | Gm17142 | 642 | 0 | 0 | 0 | 0 | 0 | 0 |
| ENSMUSG00 | Speer5-ps1 | 772 | 0.15449021 | 0 | 0 | 0.6543376 | 0.85925244 | 0 |
| ENSMUSG00 | Gm3129 | 1865 | 0 | 0 | 0 | 0 | 0 | 0.04387341 |
| ENSMUSG00 | Gm6277 | 587 | 0 | 0 | 0 | 0 | 0 | 0.04646446 |
| ENSMUSG00 | Gm7147 | 282 | 0 | 0 | 0 | 0 | 0 | 0 |
| ENSMUSG00 | Gm7701 | 1478 | 0.43037056 | 0.48903696 | 1.04672835 | 0.31736575 | 0.75952657 | 0.44288995 |
| ENSMUSG00 | Gm17045 | 1691 | 0 | 0 | 0 | 0 | 0 | 0 |
| ENSMUSG00 | Gm17040 | 1434 | 0.08317046 | 0.26882255 | 0.24659327 | 0 | 0.92516442 | 0 |
| ENSMUSG00 | Gm17711 | 235 | 0 | 0 | 0 | 0 | 0 | 0 |
| ENSMUSG00 | Gm17085 | 501 | 0 | 0 | 0 | 0 | 0 | 0 |
| ENSMUSG00 | 5730435O14l | 694 | 0 | 0 | 0 | 0 | 0 | 0 |
| ENSMUSG00 | Gm17397 | 1262 | 0 | 0 | 0 | 0 | 0 | 0 |
| ENSMUSG00 | Gm14475 | 499 | 0 | 0 | 0 | 0 | 0 | 0 |
| ENSMUSG00 | Gm8810 | 1031 | 0 | 0 | 0 | 0 | 0 | 0 |
| ENSMUSG00 | Vmn2r15 | 2562 | 0 | 0 | 0 | 0 | 0 | 0 |
| ENSMUSG00 | Aadacl2 | 1585 | 0 | 0 | 0 | 0 | 0 | 0 |
| ENSMUSG00 | Gm17398 | 1475 | 0 | 0 | 0 | 0 | 0 | 0 |
| ENSMUSG00 | Gm4219 | 2141 | 0.35280436 | 0 | 0.57807174 | 0.69096863 | 0 | 1.5796615 |
| ENSMUSG00 | Gm3159 | 1984 | 0 | 0 | 0 | 0 | 0 | 0 |
| ENSMUSG00 | Gm17575 | 3238 | 0 | 0 | 0 | 0 | 0 | 0 |
| ENSMUSG00 | Vmn2r83 | 2604 | 0 | 0 | 0 | 0 | 0 | 0 |
| ENSMUSG00 | Vmn1r18 | 1086 | 0 | 0 | 0 | 0 | 0 | 0 |
| ENSMUSG00 | Hist1h2al | 377 | 0 | 0 | 0 | 0 | 0 | 0.07234652 |
| ENSMUSG00 | Gm17574 | 5749 | 0.06915199 | 0.01676342 | 0.00768861 | 0 | 0.03550278 | 0.12335026 |
| ENSMUSG00 | Gm17573 | 245 | 0 | 0 | 0 | 0 | 0 | 0 |
| ENSMUSG00 | Olfr443-ps1 | 914 | 0 | 0 | 0 | 0 | 0 | 0 |
| ENSMUSG00 | Gcnt4 | 5090 | 6.45928927 | 8.47286163 | 1.87575601 | 7.22349772 | 12.1300422 | 0.69124331 |
| ENSMUSG00 | Gm17576 | 190 | 0 | 0 | 0 | 0 | 0 | 0 |
| ENSMUSG00 | Speer1-ps1 | 1093 | 0 | 0 | 0 | 0 | 0 | 0 |
| ENSMUSG00 | Gm17168 | 341 | 0 | 0 | 0 | 1.16393692 | 1.64601212 | 0.55988996 |
| ENSMUSG00 | Gm5373 | 471 | 0 | 0 | 0 | 0 | 0 | 0 |
| ENSMUSG00 | Gm17263 | 2459 | 0.11317136 | 0 | 0 | 0 | 0.08300346 | 0 |
| ENSMUSG00 | 5330438I03R | 2844 | 0 | 0 | 0 | 0 | 0 | 0 |
| ENSMUSG00 | Ighd5-2 | 9 | 0 | 0 | 0 | 0 | 0 | 0 |
| ENSMUSG00 | Gm6297 | 2144 | 2.67014421 | 2.29245201 | 2.7007656 | 5.82294189 | 5.99750313 | 4.248941 |
| ENSMUSG00 | Spanxn4 | 422 | 0 | 0 | 0 | 0 | 0 | 0 |
| ENSMUSG00 | Gm17265 | 2718 | 1.00924509 | 0.53185919 | 1.17090975 | 0.67703616 | 0.54443153 | 2.09727726 |
| ENSMUSG00 | Gm3164 | 2191 | 0 | 0 | 0 | 0 | 0 | 0 |
| ENSMUSG00 | Gm6356 | 1752 | 0 | 0 | 0 | 0.02059477 | 0 | 0 |
| ENSMUSG00 | Gm9252 | 1601 | 0 | 0 | 0 | 0 | 0 | 0 |
| ENSMUSG00 | BC048943 | 758 | 0 | 0 | 0 | 0 | 0 | 0 |
| ENSMUSG00 | 9530085L02R | 3626 | 0 | 0 | 0 | 0 | 0 | 0 |
| ENSMUSG00 | Gm3140 | 480 | 0 | 0 | 0 | 0 | 0 | 0 |
| ENSMUSG00 | Hist2h4 | 1607 | 0.19791154 | 0.71964817 | 1.15524419 | 0.83076269 | 0.92082446 | 0.15275156 |
| ENSMUSG00 | Gm17650 | 64 | 0 | 0 | 0 | 0 | 0 | 0 |
| ENSMUSG00 | Vmn2r117 | 2568 | 0 | 0 | 0 | 0 | 0 | 0 |
| ENSMUSG00 | Gm6728 | 1096 | 0 | 0.08793146 | 1.20990446 | 0.03292157 | 0.04655691 | 0.69679735 |
| ENSMUSG00 | Gm17150 | 380 | 0 | 0 | 0 | 0 | 0 | 0 |
| ENSMUSG00 | Gm5916 | 1139 | 0 | 0 | 0 | 0 | 0 | 0 |
| ENSMUSG00 | Gm2895 | 3192 | 0.01245472 | 0.03019201 | 0 | 0 | 0.01598571 | 0 |
| ENSMUSG00 | Olfr772 | 1003 | 0 | 0 | 0 | 0 | 0 | 0 |
| ENSMUSG00 | Akd1 | 5637 | 0 | 0 | 0 | 0.01920279 | 0 | 0 |
| ENSMUSG00 | Gm6327 | 1409 | 0 | 0 | 0 | 0 | 0 | 0 |
| ENSMUSG00 | Gpr173 | 398 | 0 | 0 | 0 | 0 | 0 | 0 |
| ENSMUSG00 | Gm17143 | 1948 | 0 | 0 | 0 | 0 | 0 | 0 |
| ENSMUSG00 | Gm17450 | 463 | 0 | 0 | 0 | 0 | 0.77145709 | 0.17672552 |
| ENSMUSG00 | Gm17510 | 338 | 0.70571859 | 0.85538063 | 0.65387342 | 0.74726128 | 0.45289683 | 1.53318978 |
| ENSMUSG00 | 4933402N22l | 1314 | 0 | 0 | 0 | 0 | 0 | 0 |
| ENSMUSG00 | Gm17509 | 2374 | 0.41865502 | 0.34505877 | 0.14895314 | 0.12159071 | 0 | 1.06846733 |
| ENSMUSG00 | Gm17118 | 333 | 0 | 0 | 0 | 0 | 0 | 0 |
| ENSMUSG00 | Gm17228 | 490 | 0 | 0.39335871 | 0.54124706 | 0 | 0.41654184 | 0 |
| ENSMUSG00 | Gm17508 | 679 | 0 | 0 | 0 | 0 | 0.1502986 | 0 |
| ENSMUSG00 | Gm17507 | 34 | 0 | 2.8344966 | 0 | 2.12247321 | 0 | 0 |
| ENSMUSG00 | Gm17093 | 1399 | 0 | 0 | 0 | 0 | 0 | 0 |
| ENSMUSG00 | Gm17337 | 1464 | 0 | 0 | 0 | 0 | 0 | 0 |
| ENSMUSG00 | Gm17336 | 2385 | 0.01666897 | 0 | 0.20386594 | 0 | 0 | 0 |
| ENSMUSG00 | Gm17338 | 4274 | 0.30695621 | 0.41714983 | 0.33094501 | 0 | 1.03867447 | 0.06381525 |
| ENSMUSG00 | Vmn1r119 | 923 | 0 | 0 | 0 | 0 | 0 | 0 |
| ENSMUSG00 | 1010001B22l | 587 | 1.4222574 | 2.62685886 | 2.33433924 | 2.33580527 | 1.9124025 | 4.46058836 |
| ENSMUSG00 | Vmn1r115 | 887 | 0 | 0 | 0 | 0 | 0 | 0 |
| ENSMUSG00 | Vmn1r155 | 923 | 0 | 0 | 0 | 0 | 0 | 0 |
| ENSMUSG00 | C1galt1c1 | 533 | 0.96964587 | 0.36162433 | 0.6634423 | 0.47387301 | 1.05307717 | 1.63750179 |
| ENSMUSG00 | Gm17088 | 579 | 0 | 0 | 0 | 0 | 0 | 0 |
| ENSMUSG00 | Gm17387 | 70 | 0 | 0 | 0 | 0 | 0 | 0 |
| ENSMUSG00 | Gm17031 | 1668 | 0 | 0 | 0 | 0 | 0 | 0 |

| | | | | | | | | |
|---|---|---|---|---|---|---|---|---|
| ENSMUSG00 | Gm17023 | 445 | 0 | 0 | 0.09932998 | 0 | 0 | 0 |
| ENSMUSG00 | 2010300F17 | 2804 | 0 | 0 | 0.07881926 | 0 | 0.12738396 | 0.10699752 |
| ENSMUSG00 | Gm17386 | 739 | 0 | 0 | 0 | 0 | 0 | 0 |
| ENSMUSG00 | Gm17388 | 1917 | 1.67980904 | 0.527864 | 0.46115642 | 1.93868054 | 2.58192929 | 1.25204395 |
| ENSMUSG00 | Gm10269 | 444 | 1.07447245 | 2.27908848 | 1.09509071 | 1.46278559 | 0.80446989 | 3.07146838 |
| ENSMUSG00 | Vmn2r11 | 2580 | 0 | 0 | 0 | 0 | 0 | 0 |
| ENSMUSG00 | C430039J16 | 1031 | 0 | 0 | 0 | 0 | 0 | 0 |
| ENSMUSG00 | Gm6541 | 718 | 0 | 0 | 0 | 0 | 0 | 0 |
| ENSMUSG00 | Dynll1b | 802 | 5.00661288 | 5.94820171 | 7.22000179 | 4.49900805 | 4.00830633 | 5.30529143 |
| ENSMUSG00 | Otogl | 6919 | 0 | 0 | 0 | 0 | 0 | 0 |
| ENSMUSG00 | A930011O12 | 4049 | 0 | 0 | 0 | 0 | 0 | 0 |
| ENSMUSG00 | Gm17171 | 529 | 0 | 0 | 0 | 0 | 0 | 0 |
| ENSMUSG00 | Gm17444 | 841 | 0 | 0 | 0 | 0 | 0 | 0 |
| ENSMUSG00 | Gm17054 | 258 | 0 | 0 | 0 | 0 | 0 | 0 |
| ENSMUSG00 | Gm7808 | 158 | 0 | 0 | 0 | 0 | 0 | 0 |
| ENSMUSG00 | 2210008F06 | 1080 | 0.11043189 | 0 | 0 | 0 | 0.56695973 | 0.0252543 |
| ENSMUSG00 | Gm17084 | 4640 | 0.04283996 | 0 | 0.17147267 | 0.10886824 | 0.03299119 | 0.07053786 |
| ENSMUSG00 | Ms4a12 | 783 | 0 | 0 | 0 | 0 | 0 | 0 |
| ENSMUSG00 | Gm17183 | 490 | 0 | 0 | 0 | 0 | 0 | 0 |
| ENSMUSG00 | Gm17240 | 2027 | 0 | 0 | 0 | 0 | 0 | 0 |
| ENSMUSG00 | Vmn2r82 | 3039 | 0 | 0 | 0 | 0 | 0 | 0 |
| ENSMUSG00 | 4930525G20 | 2866 | 0.97099918 | 0.30263641 | 0.81741022 | 2.54311689 | 0.07121616 | 0.75181315 |
| ENSMUSG00 | Gm17159 | 479 | 0 | 0 | 0 | 0 | 0 | 0 |
| ENSMUSG00 | Gm20538 | 223 | 0 | 0 | 0 | 0 | 0 | 0 |
| ENSMUSG00 | Gm3739 | 3181 | 0 | 0 | 0 | 0 | 0 | 0 |
| ENSMUSG00 | 1700045I11R | 490 | 0 | 0 | 0 | 0 | 0 | 0 |
| ENSMUSG00 | 2610021A01 | 3998 | 8.561648 | 8.01500351 | 8.5462793 | 12.4094075 | 10.4911658 | 11.2427728 |
| ENSMUSG00 | 2810468N07 | 1233 | 0 | 0 | 0 | 0 | 0 | 0 |
| ENSMUSG00 | Gm16432 | 1655 | 0.09608575 | 0.08734703 | 0 | 0.17441472 | 0 | 0 |
| ENSMUSG00 | Gm5799 | 1310 | 0 | 0.03678354 | 0 | 0.13771773 | 0 | 0 |
| ENSMUSG00 | Gm10039 | 943 | 5.22765597 | 5.62089993 | 11.2028001 | 7.57608146 | 2.8678663 | 10.788378 |
| ENSMUSG00 | Gm17035 | 417 | 0 | 0 | 0 | 0 | 0 | 0 |
| ENSMUSG00 | Gm17665 | 58 | 0 | 0 | 0 | 0 | 0 | 0 |
| ENSMUSG00 | Gm17154 | 428 | 0 | 0 | 0 | 0 | 0 | 0 |
| ENSMUSG00 | Gm8223 | 261 | 0 | 0 | 0 | 0 | 0 | 0 |
| ENSMUSG00 | Gm8715 | 527 | 0 | 0 | 0 | 0 | 0 | 0 |
| ENSMUSG00 | Gm7953 | 466 | 0 | 0 | 0 | 0 | 0 | 0 |
| ENSMUSG00 | Gm6586 | 639 | 0 | 0 | 0 | 0 | 0 | 0 |
| ENSMUSG00 | Gm6225 | 1698 | 0.65556741 | 0.25540517 | 0.13015855 | 0.33999571 | 0.24040695 | 0.36944447 |
| ENSMUSG00 | Gm17664 | 54 | 0 | 0 | 0 | 0 | 0 | 0 |
| ENSMUSG00 | Trim12 | 1531 | 0 | 0 | 0 | 0 | 0 | 0 |
| ENSMUSG00 | Vmn2r97 | 2580 | 0 | 0 | 0 | 0 | 0 | 0 |
| ENSMUSG00 | Gm17025 | 578 | 0 | 0.91704302 | 0 | 0.18727705 | 0.35312371 | 1.17969893 |
| ENSMUSG00 | Gm4498 | 920 | 0 | 0 | 0 | 0 | 0 | 0 |
| ENSMUSG00 | Gm3269 | 1753 | 0 | 0 | 0 | 0 | 0 | 0 |
| ENSMUSG00 | Gm8398 | 363 | 0 | 0 | 0 | 0 | 0 | 0 |
| ENSMUSG00 | Gm44 | 992 | 0 | 0 | 0 | 0 | 0 | 0 |
| ENSMUSG00 | Gm3452 | 329 | 56.0685196 | 60.4893635 | 70.2661517 | 61.1969023 | 32.8802179 | 56.2073113 |
| ENSMUSG00 | Gm5327 | 1004 | 0 | 0 | 0 | 0 | 0 | 0 |
| ENSMUSG00 | Gm17292 | 2107 | 0 | 0 | 0.02097857 | 0 | 0 | 0 |
| ENSMUSG00 | Gm17293 | 506 | 0 | 0 | 0 | 0 | 0 | 0 |
| ENSMUSG00 | Gm7138 | 692 | 0 | 0 | 0 | 0 | 0 | 0 |
| ENSMUSG00 | Vmn2r118 | 2556 | 0 | 0 | 0 | 0 | 0 | 0 |
| ENSMUSG00 | Gm8720 | 923 | 0 | 0 | 0 | 0 | 0 | 0 |
| ENSMUSG00 | Gm17294 | 3549 | 0.24644141 | 0.85538063 | 0.84692176 | 2.10453176 | 0.80514991 | 0.23824002 |
| ENSMUSG00 | Gm17194 | 347 | 0 | 0.13886583 | 0.3821485 | 3.01550228 | 0 | 0.07860127 |
| ENSMUSG00 | Gm7951 | 1403 | 0 | 0 | 0 | 0 | 0 | 0 |
| ENSMUSG00 | Gm17066 | 1502 | 1.4822283 | 0.80203799 | 1.41257554 | 1.12906531 | 0.74739033 | 3.15964529 |
| ENSMUSG00 | Mtag2 | 711 | 0 | 0 | 0 | 0 | 0 | 0 |
| ENSMUSG00 | Vmn2r87 | 3030 | 0 | 0 | 0.07294034 | 0 | 0 | 0 |
| ENSMUSG00 | Lamtor3 | 1288 | 48.8918332 | 45.6798493 | 63.9348086 | 45.6908499 | 32.7234367 | 54.8669179 |
| ENSMUSG00 | 4731419I09R | 2818 | 0.04232308 | 0 | 0 | 0.07682479 | 0.14485841 | 0.20325317 |
| ENSMUSG00 | Gm17484 | 2291 | 0.53793972 | 0.44169153 | 0.86821604 | 0.55123158 | 0.33408801 | 2.41674019 |
| ENSMUSG00 | Zfp422-rs1 | 432 | 1.56445178 | 1.33851228 | 0.92087173 | 1.83751153 | 2.71668205 | 4.36695877 |
| ENSMUSG00 | Ighd5-1 | 9 | 0 | 0 | 0 | 0 | 0 | 0 |
| ENSMUSG00 | Gm4066 | 239 | 0 | 0 | 0 | 0 | 0 | 0 |
| ENSMUSG00 | Skor2 | 3132 | 0 | 0 | 0 | 0 | 0 | 0 |
| ENSMUSG00 | Gm17419 | 2647 | 0 | 0 | 0 | 0 | 0 | 0 |
| ENSMUSG00 | AU040972 | 1038 | 0.30640062 | 0 | 0 | 0 | 0 | 0 |
| ENSMUSG00 | 2610020C07 | 2923 | 0.5576376 | 0.51104335 | 0.28731954 | 1.91334824 | 0.17456851 | 0.66250407 |
| ENSMUSG00 | Gm6494 | 284 | 0 | 0 | 0 | 0 | 0 | 0 |
| ENSMUSG00 | Gm8330 | 3272 | 0.71686227 | 0.11781526 | 2.75586063 | 1.42255004 | 0.8577172 | 0.10836501 |
| ENSMUSG00 | Gm9268 | 2587 | 0 | 0 | 0 | 0 | 0 | 0 |
| ENSMUSG00 | Gm3485 | 446 | 0 | 0 | 0 | 0 | 0 | 0 |
| ENSMUSG00 | Cldn20 | 659 | 0 | 0 | 0 | 0 | 0 | 0 |
| ENSMUSG00 | Olfr102 | 1026 | 0 | 0 | 0 | 0 | 0 | 0 |
| ENSMUSG00 | Gm6465 | 1093 | 0 | 0 | 0 | 0 | 0 | 0 |
| ENSMUSG00 | Gm17195 | 3528 | 1.50998707 | 1.93947698 | 1.36564651 | 0.78750494 | 1.35954629 | 2.58985378 |
| ENSMUSG00 | Gm17612 | 3613 | 0 | 0 | 0 | 0 | 0 | 0 |
| ENSMUSG00 | Ccdc72 | 633 | 14.3823144 | 16.8995105 | 24.9988307 | 16.8154868 | 11.0436232 | 25.2495081 |
| ENSMUSG00 | Vmn1r238 | 1010 | 0 | 0 | 0 | 0 | 0 | 0 |
| ENSMUSG00 | Gm17548 | 2667 | 0 | 0.16260892 | 0 | 0 | 0.0573975 | 0 |
| ENSMUSG00 | Vmn1r9 | 1050 | 0 | 0 | 0 | 0 | 0 | 0 |
| ENSMUSG00 | Gm17167 | 5052 | 0 | 0 | 0 | 0 | 0 | 0 |
| ENSMUSG00 | Gm17549 | 1207 | 7.01567303 | 4.63100852 | 5.0171106 | 8.5795748 | 7.05998738 | 7.77338517 |
| ENSMUSG00 | Vmn2r48 | 3115 | 0 | 0 | 0 | 0 | 0 | 0 |
| ENSMUSG00 | Vmn2r47 | 2553 | 0 | 0 | 0 | 0 | 0 | 0 |
| ENSMUSG00 | Gm2366 | 3980 | 0 | 0.09685717 | 0.2443318 | 0 | 0.25641395 | 0 |
| ENSMUSG00 | Gm17547 | 2009 | 0.15830953 | 0.69557333 | 0.74806504 | 0.21552242 | 1.01595572 | 0.35298189 |
| ENSMUSG00 | Gm17572 | 419 | 0 | 0 | 0 | 0 | 0 | 0 |
| ENSMUSG00 | Gm6548 | 1164 | 1.67355545 | 1.36611047 | 3.15184963 | 1.8598992 | 2.54255137 | 2.85868212 |
| ENSMUSG00 | Gm6811 | 1532 | 0 | 0 | 0 | 0 | 0 | 0 |
| ENSMUSG00 | Gm8711 | 877 | 0 | 0 | 0 | 0 | 0 | 0 |
| ENSMUSG00 | Serpina3e-ps | 1224 | 0 | 0 | 0 | 0 | 0 | 0 |
| ENSMUSG00 | Gm8653 | 923 | 0 | 0 | 0 | 0 | 0 | 0 |
| ENSMUSG00 | Gm17725 | 1670 | 0 | 0 | 0.13234085 | 0 | 0 | 0.34297451 |
| ENSMUSG00 | Gm14569 | 5711 | 0 | 0 | 0 | 0 | 0 | 0 |
| ENSMUSG00 | Gm8584 | 738 | 0 | 0 | 0 | 0 | 0 | 0 |
| ENSMUSG00 | Gm17227 | 1122 | 0.67322115 | 0.30062843 | 0.23637349 | 0.80396712 | 0 | 0.5104879 |
| ENSMUSG00 | Gm17726 | 3345 | 0.01188505 | 0.30251578 | 0.39642909 | 0 | 0.30509044 | 0.05707697 |
| ENSMUSG00 | Gm6665 | 252 | 0.15775984 | 0 | 0.17540414 | 0.14318272 | 0.80994247 | 0.21646539 |
| ENSMUSG00 | Gm9581 | 6258 | 0.33034276 | 0.41579864 | 0.62156635 | 0.73801561 | 0.47291943 | 0.62324599 |
| ENSMUSG00 | Gm8108 | 2082 | 0 | 0 | 0 | 0 | 0 | 0 |
| ENSMUSG00 | Gm17683 | 781 | 0 | 0 | 0 | 0 | 0 | 0 |
| ENSMUSG00 | Gm3317 | 685 | 0 | 0 | 0 | 0 | 0 | 0 |
| ENSMUSG00 | Gm13279 | 1555 | 0 | 0 | 0 | 0 | 0 | 0 |
| ENSMUSG00 | Gm8989 | 2132 | 0 | 0 | 0 | 0 | 0 | 0 |
| ENSMUSG00 | Gm8206 | 2332 | 0 | 0 | 0 | 0 | 0 | 0 |
| ENSMUSG00 | Gm8232 | 1397 | 0 | 0 | 0 | 0 | 0 | 0 |
| ENSMUSG00 | Gm7707 | 1155 | 0.03442033 | 0 | 0.03826999 | 0 | 0 | 0 |
| ENSMUSG00 | Gm6026 | 335 | 0 | 0 | 0 | 0 | 0 | 0 |
| ENSMUSG00 | Vmn2r3 | 2742 | 0 | 0 | 0 | 0 | 0 | 0 |
| ENSMUSG00 | Serpina3d-ps | 1268 | 0 | 0 | 0 | 0 | 0 | 0 |
| ENSMUSG00 | Gm14296 | 1264 | 0 | 0 | 0 | 0 | 0 | 0 |
| ENSMUSG00 | 2010016I18R | 1424 | 1.67509047 | 0.43990432 | 1.48994976 | 0.91218652 | 1.39749203 | 2.60488128 |
| ENSMUSG00 | Trim30b | 2769 | 0 | 0.45245486 | 0 | 0.05212285 | 0 | 0 |
| ENSMUSG00 | Gm6211 | 5205 | 0 | 0.03703089 | 0.01698438 | 0.33967727 | 0.37252686 | 0.0681211 |
| ENSMUSG00 | Gstm2-ps1 | 255 | 2.80626921 | 4.53519456 | 1.56006505 | 5.80142677 | 1.00051717 | 0.96263433 |
| ENSMUSG00 | CT030161.2 | 152 | 0 | 0 | 0 | 0 | 0 | 0 |
| ENSMUSG00 | Gm6467 | 607 | 0 | 0 | 0 | 0.17832971 | 0 | 0 |
| ENSMUSG00 | Gm17277 | 982 | 0 | 0 | 0 | 0 | 0 | 0 |
| ENSMUSG00 | Gm3187 | 596 | 0 | 0 | 0 | 0 | 0 | 0 |
| ENSMUSG00 | Gm17033 | 590 | 0 | 0 | 0 | 0 | 0 | 0 |
| ENSMUSG00 | Gm8180 | 1387 | 0 | 0 | 0 | 0 | 0 | 0 |
| ENSMUSG00 | Cyp4f17 | 1680 | 2.34273367 | 1.77830917 | 2.05222843 | 1.22421223 | 1.82237057 | 1.70466495 |
| ENSMUSG00 | Gm17191 | 415 | 1.2453524 | 1.97390245 | 2.76927209 | 3.91251085 | 2.82796782 | 2.30027078 |
| ENSMUSG00 | Gm17279 | 1015 | 0 | 0 | 0 | 0 | 0 | 0.02687157 |
| ENSMUSG00 | Gm17278 | 517 | 0 | 0 | 0 | 0 | 0 | 0 |
| ENSMUSG00 | Gm17461 | 1744 | 0.18236459 | 0.02762984 | 0 | 0 | 0 | 0.14075215 |
| ENSMUSG00 | Gm17460 | 2925 | 0 | 0 | 0 | 0 | 0 | 0 |
| ENSMUSG00 | 4631405K08 | 1739 | 0 | 0 | 0 | 0 | 0 | 0 |
| ENSMUSG00 | Gm17067 | 3669 | 0.02167102 | 0.0131334 | 0.01204738 | 0.0196686 | 0.02781487 | 0 |
| ENSMUSG00 | Gm3344 | 599 | 0 | 0 | 0 | 0 | 0 | 0 |
| ENSMUSG00 | 3110083C13 | 854 | 4.05003139 | 2.1441274 | 5.38289423 | 2.02803061 | 1.43399651 | 1.5330008 |
| ENSMUSG00 | Gm10310 | 962 | 0 | 0 | 0 | 0 | 0 | 0 |
| ENSMUSG00 | Gm10716 | 665 | 0 | 0 | 0 | 0 | 0 | 0 |
| ENSMUSG00 | Gm3991 | 2554 | 0 | 0 | 0 | 0 | 0 | 0 |
| ENSMUSG00 | Olfr93 | 1044 | 0 | 0 | 0 | 0 | 0 | 0 |
| ENSMUSG00 | Gm8879 | 1120 | 0 | 0 | 0 | 0 | 0 | 0 |
| ENSMUSG00 | Gm17346 | 546 | 0 | 0 | 0 | 0 | 0 | 0 |
| ENSMUSG00 | Gm17349 | 293 | 0.4070527 | 0 | 0.60343813 | 0 | 0 | 0.37235002 |
| ENSMUSG00 | Gm8256 | 450 | 0 | 0 | 0 | 0 | 0 | 0 |
| ENSMUSG00 | Ighv2-1 | 351 | 0 | 0 | 0 | 0 | 0 | 0 |
| ENSMUSG00 | Gm17348 | 1523 | 0 | 0 | 0 | 0 | 0 | 0 |
| ENSMUSG00 | Vmn1r111 | 920 | 0 | 0 | 0 | 0 | 0 | 0 |
| ENSMUSG00 | Acer2 | 2939 | 0 | 0 | 0 | 0 | 0 | 0 |
| ENSMUSG00 | Gm9364 | 787 | 0 | 0 | 0 | 0 | 0 | 0 |
| ENSMUSG00 | Gm17087 | 1098 | 0 | 0.0877713 | 0 | 0 | 0 | 0 |
| ENSMUSG00 | Gm17046 | 849 | 0 | 0 | 0 | 0 | 0 | 0 |
| ENSMUSG00 | Gm17657 | 509 | 0 | 0 | 0 | 0 | 0 | 0 |
| ENSMUSG00 | Gm17044 | 739 | 0 | 0 | 0 | 0 | 0 | 0 |
| ENSMUSG00 | Vmn1r169 | 914 | 0 | 0 | 0 | 0 | 0 | 0 |
| ENSMUSG00 | Gm3752 | 1946 | 0 | 0 | 0 | 0 | 0 | 0 |
| ENSMUSG00 | H60c | 1068 | 0 | 0 | 0 | 0 | 0 | 0 |
| ENSMUSG00 | Gm3625 | 101 | 10.6277027 | 3.33965441 | 0.87528402 | 7.14495931 | 12.1250794 | 1.08018373 |
| ENSMUSG00 | Vmn2r23 | 2596 | 0 | 0 | 0 | 0 | 0 | 0 |
| ENSMUSG00 | Gm17463 | 1358 | 0 | 0 | 0 | 0 | 0 | 0 |
| ENSMUSG00 | Gm17464 | 1347 | 0 | 1.21628733 | 0 | 1.39292228 | 0 | 0 |
| ENSMUSG00 | Gm17092 | 696 | 0.22847977 | 0 | 0 | 0.10368404 | 0 | 0.0391877 |
| ENSMUSG00 | Vmn2r9 | 2541 | 0 | 0 | 0 | 0 | 0 | 0 |
| ENSMUSG00 | Lsm5 | 994 | 1.39984086 | 2.32691069 | 2.31237006 | 1.01639562 | 1.33469394 | 4.36284471 |
| ENSMUSG00 | Gm17462 | 1865 | 0 | 0 | 0 | 0 | 0 | 0 |
| ENSMUSG00 | Gm5929 | 484 | 0 | 0 | 0 | 0 | 0 | 0 |
| ENSMUSG00 | Ighd5-4 | 9 | 0 | 0 | 0 | 0 | 0 | 0 |
| ENSMUSG00 | Vmn2r95 | 2532 | 0 | 0 | 0 | 0 | 0 | 0 |
| ENSMUSG00 | Gm17280 | 1489 | 0.3737923 | 0 | 0.05937118 | 0.121162 | 0 | 0.03663484 |
| ENSMUSG00 | Gm13290 | 1555 | 0 | 0 | 0 | 0 | 0 | 0 |
| ENSMUSG00 | Vmn2r13 | 2490 | 0 | 0 | 0 | 0 | 0 | 0 |
| ENSMUSG00 | A330050F15 | 2294 | 0 | 0 | 0 | 0 | 0 | 0 |
| ENSMUSG00 | Gm17281 | 3587 | 0.33249635 | 0.3224072 | 0.83794963 | 0 | 0.29873262 | 0.38779108 |
| ENSMUSG00 | Vmn1r113 | 1561 | 0 | 0 | 0 | 0 | 0 | 0 |
| ENSMUSG00 | Gm3756 | 296 | 0 | 0.16279203 | 0.14933055 | 0 | 0 | 0 |
| ENSMUSG00 | Gm17592 | 199 | 0 | 0 | 0 | 0.18131681 | 0.5128279 | 0 |
| ENSMUSG00 | Gm17593 | 1408 | 0 | 0 | 0 | 0 | 0 | 0 |
| ENSMUSG00 | Mocs3 | 1056 | 0 | 0 | 0 | 0 | 0 | 0 |
| ENSMUSG00 | Zscan4b | 1741 | 0 | 0 | 0 | 0 | 0 | 0 |
| ENSMUSG00 | Gm17590 | 107 | 0 | 0 | 0 | 0 | 0 | 0 |
| ENSMUSG00 | Gm17591 | 1806 | 0 | 0.02668131 | 0.12237498 | 0 | 0.02825381 | 0 |
| ENSMUSG00 | Olfr819 | 962 | 0 | 0 | 0 | 0 | 0 | 0 |
| ENSMUSG00 | Gm4971 | 1049 | 0 | 0 | 0 | 0 | 0 | 0 |
| ENSMUSG00 | C2cd4d | 1314 | 0 | 0.14668628 | 0.10091745 | 0.0274597 | 0.11649858 | 0.08302782 |
| ENSMUSG00 | Gm4902 | 1154 | 0.41340188 | 0 | 0 | 0 | 0 | 0 |
| ENSMUSG00 | Apol11a | 1128 | 0 | 0 | 0 | 0 | 0 | 0 |
| ENSMUSG00 | Vmn2r36 | 2555 | 0 | 0 | 0 | 0 | 0 | 0 |
| ENSMUSG00 | Vmn1r57 | 932 | 0 | 0 | 0 | 0 | 0 | 0 |
| ENSMUSG00 | Gm17406 | 1840 | 0 | 0 | 0 | 0 | 0 | 0 |
| ENSMUSG00 | Gm8741 | 769 | 0.1550929 | 0 | 0 | 0 | 0 | 0 |
| ENSMUSG00 | Vmn2r106 | 2583 | 0 | 0 | 0 | 0 | 0 | 0 |
| ENSMUSG00 | Gm3072 | 1390 | 0 | 0 | 0 | 0 | 0 | 0 |
| ENSMUSG00 | Gm17405 | 103 | 0 | 0 | 0 | 0 | 0 | 0.26480232 |
| ENSMUSG00 | Gm17099 | 410 | 0 | 0 | 0 | 0 | 0 | 0 |
| ENSMUSG00 | Gm17021 | 459 | 0 | 0 | 0 | 0 | 0 | 0 |
| ENSMUSG00 | Vmn1r69 | 2389 | 0 | 0 | 0 | 0 | 0 | 0 |
| ENSMUSG00 | Gm2783 | 1865 | 0 | 0 | 0 | 0 | 0 | 0 |
| ENSMUSG00 | Gm10662 | 1355 | 0 | 0 | 0 | 0 | 0 | 0 |
| ENSMUSG00 | Gm17101 | 1587 | 0 | 0 | 0 | 0 | 0 | 0 |
| ENSMUSG00 | Gm17717 | 1055 | 0 | 0 | 0.50276978 | 1.33383861 | 0.24183117 | 0 |
| ENSMUSG00 | Gm17718 | 2437 | 0 | 0.03954571 | 0.10882686 | 0.94757934 | 0 | 0 |
| ENSMUSG00 | Gm9307 | 728 | 0.10921835 | 0 | 0 | 0 | 0 | 0 |
| ENSMUSG00 | Gm17117 | 298 | 0 | 0 | 0 | 0 | 0 | 0 |
| ENSMUSG00 | Vmn2r105 | 2577 | 0 | 0 | 0 | 0 | 0 | 0 |
| ENSMUSG00 | Trbv10 | 288 | 0 | 0 | 0 | 0 | 0 | 0 |
| ENSMUSG00 | Gm17129 | 504 | 0 | 0 | 0 | 0 | 0 | 0 |
| ENSMUSG00 | Gm17525 | 1046 | 0 | 0 | 0.29580583 | 0 | 0.19512954 | 0.07822554 |
| ENSMUSG00 | C630004L07 | 1268 | 0 | 0 | 0 | 0.08536761 | 0 | 0 |
| ENSMUSG00 | Gm8374 | 668 | 0 | 0 | 0 | 0 | 0 | 0 |
| ENSMUSG00 | Gm7356 | 1463 | 0 | 0 | 0 | 0 | 0 | 0 |
| ENSMUSG00 | Gm17527 | 193 | 0 | 0 | 0 | 0 | 0 | 0 |
| ENSMUSG00 | Vmn2r96 | 2338 | 0 | 0 | 0 | 0 | 0 | 0 |
| ENSMUSG00 | Klhdc7b | 1817 | 0 | 0 | 0 | 0 | 0 | 0 |
| ENSMUSG00 | Gm17037 | 395 | 0 | 0 | 0 | 0 | 0 | 0 |
| ENSMUSG00 | Gm17356 | 516 | 0 | 0 | 0 | 0 | 0 | 0 |
| ENSMUSG00 | Gm17357 | 2524 | 0.69304324 | 0.190913 | 0.47284063 | 0.50034531 | 0.14151531 | 0.74562207 |
| ENSMUSG00 | Gm17358 | 2912 | 0.04095688 | 0.16547542 | 0.0151792 | 0.04956325 | 0.14018235 | 0 |
| ENSMUSG00 | Gm17359 | 773 | 0 | 0 | 0 | 0 | 0 | 0.10585242 |
| ENSMUSG00 | 4930471C04 | 861 | 0 | 0 | 0.35936458 | 0 | 0 | 0 |
| ENSMUSG00 | Gm4983 | 335 | 0 | 0 | 0.1319458 | 0 | 0.45695262 | 0.16283367 |
| ENSMUSG00 | Hivep3 | 1147 | 0 | 0 | 0 | 0 | 0 | 0 |
| ENSMUSG00 | Gm17355 | 53 | 0 | 0 | 0 | 0 | 0 | 0 |
| ENSMUSG00 | Gm17668 | 1603 | 0 | 0 | 0 | 0 | 0 | 0 |
| ENSMUSG00 | Gm17667 | 1814 | 0 | 0 | 0 | 0 | 0 | 0 |
| ENSMUSG00 | 4930433I11R | 2422 | 0 | 0 | 0 | 0 | 0 | 0 |
| ENSMUSG00 | Apol11b | 2032 | 0 | 0 | 0 | 0 | 0 | 0 |
| ENSMUSG00 | Gm17669 | 464 | 0.17135983 | 0 | 0 | 0 | 0.21994128 | 0 |
| ENSMUSG00 | Gm17670 | 3427 | 1.09046255 | 2.16536682 | 1.3027097 | 3.29550042 | 1.80163159 | 1.86234771 |
| ENSMUSG00 | Eif3s6-ps2 | 218 | 0.54709377 | 17.4620593 | 1.41932523 | 19.365134 | 14.5120885 | 0 |
| ENSMUSG00 | Gm6526 | 1718 | 0 | 0.81339163 | 0 | 0 | 0.1782062 | 0 |
| ENSMUSG00 | Gm7876 | 1931 | 0 | 0 | 0 | 0 | 0 | 0 |
| ENSMUSG00 | Vmn1r94 | 917 | 0 | 0 | 0 | 0 | 0 | 0 |
| ENSMUSG00 | Gm14483 | 524 | 0 | 0 | 0 | 0 | 0 | 0 |
| ENSMUSG00 | Gm4222 | 428 | 0 | 0 | 0 | 0 | 0 | 0 |
| ENSMUSG00 | Ighd6-2 | 28 | 0 | 0 | 0 | 0 | 0 | 0 |
| ENSMUSG00 | H2-Q2 | 2173 | 3.23825129 | 9.64615846 | 2.84779476 | 7.22305079 | 7.09156259 | 6.92848636 |
| ENSMUSG00 | AA543186 | 1694 | 2.98048762 | 1.93428457 | 2.40057235 | 4.1321822 | 2.50011168 | 1.56177096 |
| ENSMUSG00 | Gm8224 | 494 | 0 | 0 | 0 | 0 | 0 | 0 |
| ENSMUSG00 | Gm17189 | 1627 | 1.29504638 | 0.62195162 | 0.16300618 | 0.48789489 | 0.12544899 | 0 |
| ENSMUSG00 | 4930426I24R | 635 | 0.37564234 | 0.07588416 | 0.20882761 | 1.02279811 | 0.0803565 | 0.25771313 |
| ENSMUSG00 | Gm2885 | 1440 | 0.6073754 | 0 | 0 | 0.55125346 | 2.23240394 | 0.94703609 |
| ENSMUSG00 | Sec14l5 | 2321 | 0 | 0.06228321 | 0.05713293 | 0 | 0 | 0 |
| ENSMUSG00 | Gm17126 | 522 | 0 | 0 | 0 | 0 | 0 | 0 |
| ENSMUSG00 | Gm17675 | 3326 | 0 | 0 | 0 | 0 | 0 | 0 |
| ENSMUSG00 | Gm4305 | 869 | 0 | 0 | 0 | 0 | 0 | 0 |
| ENSMUSG00 | Gm17676 | 37 | 1.07447245 | 6.51168137 | 3.58393322 | 1.95038079 | 4.13727372 | 1.47430482 |
| ENSMUSG00 | Olfr773 | 1003 | 0 | 0 | 0 | 0 | 0 | 0 |
| ENSMUSG00 | Gm16506 | 1400 | 0 | 0.24093221 | 0 | 0 | 0 | 0 |
| ENSMUSG00 | Vmn2r22 | 2556 | 0 | 0 | 0 | 0 | 0 | 0 |
| ENSMUSG00 | Gm8890 | 1120 | 0 | 0 | 0 | 0 | 0 | 0 |
| ENSMUSG00 | Gm5549 | 652 | 0 | 0 | 0 | 0 | 0 | 0 |
| ENSMUSG00 | Siah3 | 903 | 1.14467607 | 0 | 0.53844992 | 0.63932748 | 0.3955533 | 0.69470288 |
| ENSMUSG00 | Gm17366 | 187 | 0 | 0 | 0 | 0 | 0 | 0 |
| ENSMUSG00 | A430010J10R | 2035 | 0.68375519 | 0.80508061 | 0.49957857 | 0.39007616 | 0.7271572 | 0.34847205 |
| ENSMUSG00 | Gm8220 | 1560 | 0 | 0 | 0 | 0 | 0 | 0 |
| ENSMUSG00 | 1810019N24 | 1472 | 0.24307019 | 0.49103032 | 0.03002843 | 0 | 0.45064055 | 0.1111738 |
| ENSMUSG00 | Gm8705 | 810 | 0 | 0 | 0 | 0 | 0 | 0 |
| ENSMUSG00 | Gm13697 | 2725 | 0 | 0 | 0 | 0 | 0 | 0 |
| ENSMUSG00 | Gm17230 | 3941 | 0.09078897 | 0 | 0 | 0.03662222 | 0.16831842 | 0.01384148 |
| ENSMUSG00 | Gm17542 | 97 | 0 | 0 | 0 | 0 | 0 | 0 |
| ENSMUSG00 | Gm17541 | 524 | 0 | 0.09195886 | 0 | 0.06885886 | 0.09737858 | 0 |
| ENSMUSG00 | Gm8094 | 1403 | 0 | 0.03434529 | 0.0945157 | 0.02571778 | 0 | 0 |
| ENSMUSG00 | Vmn1r29 | 1022 | 0 | 0 | 0 | 0 | 0 | 0 |
| ENSMUSG00 | Gpr62 | 1967 | 0.24253471 | 0 | 0 | 0 | 0 | 0.20799166 |
| ENSMUSG00 | Yy2 | 2866 | 0 | 0 | 0 | 0 | 0 | 0 |
| ENSMUSG00 | Gm17543 | 1475 | 0 | 0 | 0.0599347 | 0.07338721 | 0 | 0.18491281 |
| ENSMUSG00 | Gm8723 | 871 | 0 | 0 | 0 | 0 | 0 | 0 |
| ENSMUSG00 | Gm8701 | 914 | 0 | 0 | 0 | 0 | 0 | 0 |
| ENSMUSG00 | Gm7929 | 620 | 0 | 0 | 0 | 0 | 0 | 0 |
| ENSMUSG00 | Gm17161 | 521 | 0 | 0 | 0 | 0 | 0 | 0 |
| ENSMUSG00 | Gm5093 | 521 | 0.30522442 | 0 | 0.08484039 | 0 | 0 | 0 |
| ENSMUSG00 | Gm5705 | 626 | 0 | 0 | 0 | 0 | 0 | 0 |
| ENSMUSG00 | Gm17732 | 820 | 0 | 0 | 0 | 0 | 0 | 0 |
| ENSMUSG00 | Gm17098 | 626 | 0.38104294 | 0 | 0.07060997 | 0 | 0.24453535 | 0 |
| ENSMUSG00 | Gm4511 | 1865 | 0 | 0 | 0 | 0 | 0 | 0 |
| ENSMUSG00 | D17H6S56E-5 | 5355 | 5.72389831 | 23.0898993 | 5.99263082 | 10.571938 | 16.9802057 | 5.85729881 |
| ENSMUSG00 | Gm17733 | 981 | 0.48630557 | 0 | 2.16278131 | 0.73561763 | 0.15604396 | 1.4457505 |
| ENSMUSG00 | Tspy-ps | 698 | 0 | 0 | 0 | 0 | 0 | 0 |
| ENSMUSG00 | 2810029C07 | 1948 | 8.46946838 | 7.34670088 | 3.83475948 | 4.9455369 | 6.78430767 | 5.30651349 |
| ENSMUSG00 | C030010L15 | 713 | 0.50182234 | 0.202748 | 1.05390089 | 2.4796917 | 0.14313149 | 3.63406834 |
| ENSMUSG00 | Gm3208 | 426 | 0 | 0 | 0 | 0 | 0 | 0 |
| ENSMUSG00 | Gm17432 | 1722 | 1.17742712 | 0.81150222 | 0.43637127 | 1.7810533 | 0.68153696 | 1.86899386 |
| ENSMUSG00 | Gm3636 | 1945 | 0.10219918 | 0.02477452 | 0 | 0.03710236 | 0 | 0.08413771 |
| ENSMUSG00 | Vmn1r152 | 923 | 0 | 0 | 0 | 0 | 0 | 0 |
| ENSMUSG00 | Gm3095 | 1564 | 0 | 0 | 0 | 0 | 0 | 0 |
| ENSMUSG00 | Gm17064 | 449 | 0 | 0 | 0 | 0 | 0 | 0 |
| ENSMUSG00 | Rgs21 | 456 | 0 | 0 | 0 | 0 | 0 | 0 |
| ENSMUSG00 | Gm17433 | 3681 | 0.03240056 | 0.09163409 | 0.1681135 | 0 | 0.23565563 | 0 |
| ENSMUSG00 | Gm17619 | 1447 | 0 | 0 | 0 | 0.04987152 | 0 | 0 |
| ENSMUSG00 | Gm4349 | 2557 | 0.10883393 | 0 | 0 | 0 | 0.09977782 | 0 |
| ENSMUSG00 | Gm3191 | 345 | 0 | 0 | 0 | 0 | 0 | 0.07905693 |
| ENSMUSG00 | Vmn1r107 | 923 | 0 | 0 | 0 | 0 | 0 | 0 |
| ENSMUSG00 | Zfp964 | 1413 | 14.9118221 | 9.13939596 | 10.9800757 | 24.871841 | 9.24469371 | 15.1332747 |
| ENSMUSG00 | Gm5891 | 1355 | 0 | 0 | 0 | 0 | 0 | 0 |
| ENSMUSG00 | Gm4220 | 920 | 0 | 0 | 0 | 0 | 0 | 0 |
| ENSMUSG00 | Gm17618 | 633 | 0 | 0 | 0 | 0 | 0 | 0 |
| ENSMUSG00 | Gm17617 | 37 | 0 | 0 | 0 | 0 | 0 | 0 |
| ENSMUSG00 | 4921507G05 | 1651 | 0.36119455 | 0.26267594 | 0.85672864 | 1.68280886 | 0.52540787 | 0.1817208 |
| ENSMUSG00 | Gm17095 | 578 | 0 | 0 | 0 | 0 | 0 | 0 |
| ENSMUSG00 | Vmn2r103 | 2565 | 0 | 0 | 0 | 0 | 0 | 0 |
| ENSMUSG00 | Gm17307 | 101 | 0.78723724 | 0 | 0 | 0 | 0 | 0 |
| ENSMUSG00 | Gm17308 | 1995 | 0 | 0 | 0 | 0 | 0 | 0 |
| ENSMUSG00 | Gm7954 | 1398 | 0 | 0 | 0 | 0 | 0 | 0 |
| ENSMUSG00 | Gm16494 | 260 | 0 | 0 | 0 | 0 | 0 | 0 |
| ENSMUSG00 | Gm8340 | 1190 | 0 | 0 | 0 | 0 | 0 | 0 |
| ENSMUSG00 | Gm17564 | 82 | 0 | 0 | 0 | 0 | 0 | 0 |
| ENSMUSG00 | Gm17306 | 3971 | 0 | 0.01213459 | 0 | 0 | 0.06424877 | 0 |
| ENSMUSG00 | Gm6763 | 862 | 0 | 0 | 0 | 0 | 0 | 0 |
| ENSMUSG00 | Sco2 | 767 | 0 | 0 | 0 | 0 | 0 | 0 |
| ENSMUSG00 | Gm9590 | 323 | 0.12308198 | 0 | 0 | 0 | 0 | 0 |
| ENSMUSG00 | Gm8212 | 1399 | 0 | 0 | 0 | 0 | 0 | 0 |
| ENSMUSG00 | Gm17022 | 911 | 0 | 0.47604608 | 0.63076176 | 1.26742637 | 0.72814806 | 0.26945308 |
| ENSMUSG00 | Rpl29-ps5 | 476 | 0 | 0 | 0 | 0 | 0 | 0 |
| ENSMUSG00 | Gm8425 | 768 | 0 | 0 | 0 | 0 | 0 | 0 |
| ENSMUSG00 | Gm8032 | 570 | 0 | 0 | 0 | 0 | 0 | 0 |
| ENSMUSG00 | Vmn1r143 | 923 | 0 | 0 | 0 | 0 | 0 | 0 |
| ENSMUSG00 | Gm16500 | 359 | 0 | 0 | 0 | 0.10050709 | 0 | 0 |
| ENSMUSG00 | Gm3573 | 893 | 0 | 0 | 0 | 0 | 0 | 0 |
| ENSMUSG00 | Rian | 6489 | 0 | 0 | 0 | 0 | 0 | 0 |
| ENSMUSG00 | Vmn2r18 | 2380 | 0 | 0 | 0 | 0 | 0 | 0 |
| ENSMUSG00 | Gm17692 | 2465 | 0.11289589 | 0.01954825 | 0.05379535 | 0.14637746 | 0.02070036 | 0.04425905 |
| ENSMUSG00 | Gm17693 | 172 | 0 | 0 | 0 | 0 | 0 | 0 |
| ENSMUSG00 | Vmn2r53 | 2416 | 0 | 0 | 0 | 0 | 0 | 0 |
| ENSMUSG00 | Gm17691 | 1958 | 0 | 0 | 0.022575 | 0 | 0.05212092 | 0 |
| ENSMUSG00 | Igkv1-115 | 341 | 0 | 0 | 0 | 0 | 0 | 0 |
| ENSMUSG00 | Gm17637 | 1968 | 0 | 0 | 0 | 0 | 0 | 0 |
| ENSMUSG00 | Gm17638 | 4341 | 0.43959066 | 0.73262041 | 0.67203908 | 0.28260528 | 0.4466718 | 0.8607753 |
| ENSMUSG00 | Cox16 | 3740 | 1.50943268 | 1.99703169 | 1.99735601 | 2.12247321 | 1.00961278 | 2.53056145 |
| ENSMUSG00 | Gm17636 | 103 | 0 | 0 | 0 | 0 | 0 | 0 |
| ENSMUSG00 | Vmn2r108 | 2562 | 0 | 0 | 0 | 0 | 0 | 0 |
| ENSMUSG00 | Gm5258 | 326 | 0 | 0 | 0 | 0.44272447 | 0 | 0 |
| ENSMUSG00 | Olfr1438-ps1 | 959 | 0 | 0 | 0 | 0 | 0 | 0 |
| ENSMUSG00 | Gm9671 | 544 | 0 | 0 | 0 | 0 | 0 | 0 |
| ENSMUSG00 | Olfr721-ps1 | 950 | 0 | 0 | 0 | 0 | 0 | 0 |
| ENSMUSG00 | Gm17442 | 1487 | 0.08020608 | 2.30076489 | 1.30792273 | 0.97059972 | 1.26965427 | 2.53120391 |
| ENSMUSG00 | Prr24 | 1678 | 5.54397046 | 5.77203508 | 4.5571626 | 5.09621249 | 6.02099071 | 6.09534548 |
| ENSMUSG00 | Ear-ps5 | 462 | 0 | 0 | 0 | 0 | 0 | 0 |
| ENSMUSG00 | Ces2h | 1594 | 0 | 0 | 0 | 0 | 0 | 0 |
| ENSMUSG00 | Gm8050 | 1590 | 0 | 0 | 0 | 0 | 0 | 0 |
| ENSMUSG00 | Gm17153 | 487 | 0 | 0 | 0 | 0 | 0 | 0 |
| ENSMUSG00 | Gm17141 | 3143 | 0 | 0 | 0 | 0 | 0 | 0 |
| ENSMUSG00 | Gm17091 | 354 | 0 | 0 | 0 | 0 | 0 | 0 |
| ENSMUSG00 | Rpl19-ps7 | 681 | 8.11455476 | 8.06645287 | 7.91868553 | 5.82823039 | 4.42078733 | 5.60712114 |
| ENSMUSG00 | Gm17443 | 4609 | 0.03450248 | 0.35546518 | 0.12467432 | 0.32097284 | 0.04428412 | 0.56218067 |
| ENSMUSG00 | Gm1815 | 302 | 0.26328133 | 0 | 0 | 0 | 0 | 0 |
| ENSMUSG00 | Gm17503 | 1082 | 0 | 0 | 0 | 0 | 0 | 0 |
| ENSMUSG00 | Gm17502 | 1130 | 0.03518184 | 0.5543573 | 0.19558338 | 0.15965506 | 0.13546825 | 1.11029505 |
| ENSMUSG00 | Gm8897 | 1452 | 0 | 0 | 0 | 0 | 0 | 0 |
| ENSMUSG00 | Gm6553 | 917 | 0 | 0 | 0 | 0 | 0 | 0 |
| ENSMUSG00 | Gm5778 | 1071 | 0.48255952 | 0.26995206 | 0.20635781 | 0.50535076 | 0.1905747 | 0.56026337 |
| ENSMUSG00 | Gm17504 | 2035 | 0 | 0 | 0 | 0 | 0 | 0 |
| ENSMUSG00 | Gm3495 | 1000 | 0 | 0 | 0.17680737 | 0.03608205 | 0 | 0.21819711 |
| ENSMUSG00 | 5033417F24 | 1783 | 0 | 2.62147218 | 0.86767499 | 1.49751615 | 0.2861827 | 1.6367843 |
| ENSMUSG00 | 9330159M07 | 3902 | 0.66225173 | 0.34577662 | 0.19257594 | 1.06341238 | 0.30077054 | 0.36347546 |
| ENSMUSG00 | Gm4707 | 1598 | 1.31854848 | 1.80925315 | 1.63198294 | 0.51932855 | 2.90575732 | 1.9798862 |
| ENSMUSG00 | Gm17155 | 470 | 0 | 0 | 0 | 0 | 0 | 0 |
| ENSMUSG00 | Gm17317 | 3766 | 0.39058757 | 0.23031226 | 0.07042248 | 0.105391 | 0.17613991 | 0.73871832 |
| ENSMUSG00 | Gm17177 | 478 | 0 | 0 | 0 | 0 | 0 | 0.17117974 |
| ENSMUSG00 | Gm2075 | 995 | 0 | 0 | 0 | 0 | 0 | 0 |
| ENSMUSG00 | Gm17316 | 1499 | 0 | 0 | 0 | 0 | 0 | 0 |
| ENSMUSG00 | Gm17315 | 2502 | 0 | 0 | 0 | 0 | 0 | 0.01090114 |
| ENSMUSG00 | Gm3005 | 1954 | 0 | 0 | 0 | 0 | 0 | 0 |
| ENSMUSG00 | 4930519A11 | 1394 | 0 | 0 | 0 | 0 | 0 | 0 |
| ENSMUSG00 | Gm16490 | 353 | 0 | 0 | 0 | 0 | 0 | 0 |
| ENSMUSG00 | Ighv1-76 | 293 | 0 | 0 | 0 | 0 | 0 | 0 |
| ENSMUSG00 | Gm8251 | 5984 | 0 | 0.1368933 | 0 | 0.00602975 | 0 | 0 |
| ENSMUSG00 | Gm4604 | 389 | 0 | 0 | 0 | 0.0927559 | 0 | 0.07011475 |
| ENSMUSG00 | Gm2855 | 453 | 0 | 0 | 0 | 0 | 0 | 0 |
| ENSMUSG00 | Gm7697 | 471 | 0 | 0 | 0 | 0 | 0 | 0 |
| ENSMUSG00 | Gm17188 | 2874 | 0 | 0 | 0 | 0 | 0 | 0 |
| ENSMUSG00 | Gm3460 | 1804 | 0 | 0 | 0 | 0 | 0 | 0 |
| ENSMUSG00 | Gm8024 | 1403 | 0 | 0 | 0 | 0 | 0 | 0 |
| ENSMUSG00 | 4930534D22 | 2713 | 0 | 0 | 0.04887782 | 0 | 0 | 0 |
| ENSMUSG00 | Gm17702 | 964 | 0.24744075 | 0.74978904 | 0.87119815 | 0.22457704 | 0.37052348 | 2.15028276 |
| ENSMUSG00 | Mier2 | 1397 | 0 | 0 | 0 | 0 | 0 | 0 |
| ENSMUSG00 | Gm17701 | 1348 | 0 | 0 | 0 | 0 | 0 | 0 |
| ENSMUSG00 | Ighd5-5 | 9 | 0 | 0 | 0 | 0 | 0 | 0 |
| ENSMUSG00 | Vmn2r100 | 2547 | 0 | 0 | 0 | 0 | 0 | 0 |
| ENSMUSG00 | Flg | 935 | 0 | 0 | 0 | 0 | 0 | 0 |
| ENSMUSG00 | C230036H16 | 1301 | 0 | 0 | 0 | 0 | 0 | 0 |
| ENSMUSG00 | Gm7157 | 635 | 0 | 0 | 0 | 0 | 0 | 0 |
| ENSMUSG00 | Gm17102 | 904 | 1.09943254 | 0.21321435 | 0.68454182 | 0.1197413 | 1.18534723 | 1.90077685 |
| ENSMUSG00 | Gm17244 | 1287 | 0 | 0 | 0 | 0 | 0.07929507 | 0 |
| ENSMUSG00 | Gm6257 | 909 | 0.0437354 | 0.15903116 | 0.58352268 | 0 | 0.16840388 | 0.15002552 |
| ENSMUSG00 | Cyp2a22 | 2217 | 0 | 0 | 0 | 0 | 0 | 0 |
| ENSMUSG00 | 9330104G04 | 2233 | 0.05341086 | 0.12947544 | 0.0791793 | 0.08079276 | 0.04570208 | 0.20764392 |
| ENSMUSG00 | 1700101I11R | 882 | 0.9916333 | 1.80289409 | 0.80184749 | 5.52276192 | 3.00835776 | 2.01003577 |
| ENSMUSG00 | Gm17560 | 468 | 0.42473804 | 0 | 0 | 0.07709839 | 0.10903072 | 0 |
| ENSMUSG00 | Gm17561 | 2038 | 0 | 0 | 0 | 0 | 0 | 0 |
| ENSMUSG00 | Olfr732 | 1043 | 0 | 0 | 0 | 0 | 0 | 0 |
| ENSMUSG00 | Vmn1r56 | 1830 | 0 | 0 | 0 | 0 | 0 | 0 |
| ENSMUSG00 | Mir155 | 1259 | 0.15788515 | 0.68892451 | 1.68521721 | 0.88843795 | 0.85111509 | 0.56325705 |
| ENSMUSG00 | Gm3851 | 317 | 0 | 0 | 0 | 0 | 0 | 0 |
| ENSMUSG00 | Gm17699 | 3465 | 0 | 0 | 0 | 0 | 0 | 0 |
| ENSMUSG00 | C430002E04 | 984 | 0 | 0 | 0 | 0 | 0 | 0 |
| ENSMUSG00 | Vmn2r17 | 2544 | 0 | 0 | 0 | 0 | 0 | 0 |
| ENSMUSG00 | Zscan4e | 1741 | 0 | 0 | 0 | 0 | 0 | 0 |
| ENSMUSG00 | Gm17146 | 370 | 0 | 0.26046726 | 0.71678664 | 0.19503808 | 0 | 0.07371524 |
| ENSMUSG00 | Gm6337 | 2135 | 0 | 0 | 0 | 0 | 0 | 0 |
| ENSMUSG00 | 1700039M10 | 520 | 0 | 0 | 0 | 0 | 0 | 0 |
| ENSMUSG00 | Vmn1r103 | 917 | 0 | 0 | 0 | 0 | 0 | 0 |
| ENSMUSG00 | Gm8926 | 1093 | 0 | 0 | 0 | 0 | 0 | 0 |
| ENSMUSG00 | AW047730 | 2577 | 0 | 0 | 0 | 0 | 0 | 0 |
| ENSMUSG00 | Gm17053 | 493 | 0 | 0 | 0 | 0 | 0 | 0 |
| ENSMUSG00 | Vmn2r80 | 2684 | 0 | 0 | 0 | 0 | 0 | 0 |
| ENSMUSG00 | Gm4216 | 917 | 0 | 0 | 0 | 0 | 0 | 0 |
| ENSMUSG00 | A830073O21 | 2024 | 0.41248275 | 1.38083678 | 0.37126054 | 0.12478968 | 0.70589848 | 1.6440247 |
| ENSMUSG00 | Gm17422 | 969 | 0.20513664 | 0.04972801 | 0.13684781 | 0.14894549 | 0 | 0.19703042 |
| ENSMUSG00 | Taar8c | 1108 | 0 | 0 | 0 | 0 | 0 | 0 |
| ENSMUSG00 | Gm3030 | 512 | 0 | 0 | 0 | 0 | 0 | 0 |
| ENSMUSG00 | Fgfr1 | 756 | 0 | 0 | 0 | 0 | 0 | 0 |
| ENSMUSG00 | Ube2d2 | 2877 | 12.7820019 | 10.3340406 | 12.2603652 | 14.1970366 | 9.45327018 | 21.2452091 |
| ENSMUSG00 | Gm17019 | 1157 | 0.03436083 | 0 | 0 | 0.09355759 | 0 | 0 |
| ENSMUSG00 | Tnnc1 | 1044 | 0 | 0 | 0 | 0 | 0 | 0 |
| ENSMUSG00 | Gm4353 | 1542 | 0.02578176 | 0.15624657 | 0.05733054 | 0.18719608 | 0.09927311 | 0.0530635 |
| ENSMUSG00 | Vmn2r67 | 2550 | 0 | 0 | 0 | 0 | 0 | 0 |
| ENSMUSG00 | Gm6918 | 1218 | 0 | 0 | 0 | 0 | 0 | 0 |
| ENSMUSG00 | Gm8871 | 1224 | 0 | 0 | 0 | 0 | 0 | 0 |
| ENSMUSG00 | Gm6395 | 721 | 0 | 0 | 0 | 0.10008889 | 0 | 0 |
| ENSMUSG00 | 1700099I09R | 874 | 0 | 0 | 0 | 0 | 0 | 0 |
| ENSMUSG00 | Gm9107 | 1022 | 0 | 0 | 0 | 0 | 0 | 0 |
| ENSMUSG00 | Gm17231 | 1100 | 0.18070673 | 0 | 0.04018349 | 0.03280186 | 0 | 0.07438538 |
| ENSMUSG00 | Gm17282 | 5126 | 3.20308495 | 2.65091235 | 1.58664438 | 3.21683464 | 4.23063007 | 1.45791088 |
| ENSMUSG00 | Gm17469 | 172 | 0 | 0 | 0 | 0 | 0 | 0 |
| ENSMUSG00 | Gm9587 | 451 | 0 | 0 | 0 | 0 | 0 | 0 |
| ENSMUSG00 | Gm17468 | 1995 | 1.03623308 | 0.16907524 | 0.1772505 | 0.56067337 | 0.6650054 | 0.10937199 |
| ENSMUSG00 | Gm17558 | 36 | 1.1043189 | 0 | 1.22782897 | 0 | 0 | 0 |
| ENSMUSG00 | 2010015M23 | 1197 | 1.72705513 | 0.60384013 | 1.84635936 | 1.26603665 | 1.27885654 | 3.23558795 |
| ENSMUSG00 | Gm4312 | 915 | 0 | 0 | 0 | 0 | 0 | 0 |
| ENSMUSG00 | Gm17467 | 172 | 0 | 0 | 0 | 0 | 0 | 0 |
| ENSMUSG00 | Gm4301 | 915 | 0 | 0 | 0 | 0 | 0 | 0 |
| ENSMUSG00 | Gm17470 | 1728 | 0 | 0 | 0 | 0 | 0 | 0 |
| ENSMUSG00 | Gm5861 | 1091 | 0 | 0 | 0 | 0 | 0 | 0 |
| ENSMUSG00 | B930041F14 | 2136 | 0 | 0 | 0 | 0 | 0 | 0 |

| | | | | | | | | |
|---|---|---|---|---|---|---|---|---|
| ENSMUSG00 | Gm17409 | 961 | 0 | 0 | 0 | 0 | 0.63716598 | 0 |
| ENSMUSG00 | Gm8267 | 2534 | 0 | 0 | 0 | 0 | 0 | 0 |
| ENSMUSG00 | Vmn1r185 | 1143 | 0 | 0 | 0 | 0 | 0 | 0 |
| ENSMUSG00 | Vmn2r62 | 2582 | 0 | 0 | 0 | 0 | 0 | 0 |
| ENSMUSG00 | Gm17408 | 2437 | 0.24469931 | 1.06773405 | 1.05199298 | 1.14005639 | 0.02093819 | 1.56686479 |
| ENSMUSG00 | Gm8979 | 2132 | 0 | 0 | 0 | 0 | 0 | 0 |
| ENSMUSG00 | Gm8582 | 784 | 0 | 0 | 0 | 0 | 0 | 0 |
| ENSMUSG00 | Vmn2r52 | 2553 | 0 | 0 | 0 | 0 | 0 | 0 |
| ENSMUSG00 | Moap1 | 7408 | 0.6117879 | 0.53988589 | 0.81148092 | 1.14461129 | 0.48904869 | 0.58908508 |
| ENSMUSG00 | Gm17596 | 2503 | 1.50889758 | 0.48128688 | 0.98893456 | 0.63428284 | 0.97853218 | 1.65631049 |
| ENSMUSG00 | Gm8857 | 1172 | 0 | 0 | 0 | 0 | 0 | 0 |
| ENSMUSG00 | Gm17597 | 1113 | 0.32147289 | 1.77506211 | 0.75456875 | 1.10223676 | 1.55875721 | 4.19044323 |
| ENSMUSG00 | Vmn1r118 | 920 | 0 | 0 | 0 | 0 | 0 | 0 |
| ENSMUSG00 | Gm3053 | 612 | 0 | 0 | 0 | 0 | 0 | 0 |
| ENSMUSG00 | Gm3646 | 161 | 0 | 0 | 0 | 0 | 0 | 0 |
| ENSMUSG00 | Gm2442 | 703 | 0 | 0 | 0 | 0 | 0 | 0 |
| ENSMUSG00 | Gm17595 | 4310 | 0 | 0 | 0 | 0 | 0 | 0 |
| ENSMUSG00 | Vmn1r-ps70 | 3054 | 0 | 0 | 0 | 0 | 0 | 0 |
| ENSMUSG00 | Gm7399 | 857 | 0 | 0 | 0 | 0.08420547 | 0 | 0 |
| ENSMUSG00 | Gm17138 | 458 | 0 | 0 | 0.09651057 | 0 | 0 | 0 |
| ENSMUSG00 | A730099G02I | 2219 | 0 | 0 | 0 | 0 | 0 | 0 |
| ENSMUSG00 | Gm17517 | 6618 | 0 | 0 | 0 | 0 | 0 | 0 |
| ENSMUSG00 | Vmn2r114 | 2565 | 0 | 0 | 0 | 0 | 0 | 0 |
| ENSMUSG00 | Gm17518 | 1344 | 0.29579971 | 0.28682406 | 0.49332414 | 0.1342338 | 0.41762659 | 0 |
| ENSMUSG00 | 6530401F13I | 1088 | 0.14615986 | 0.35431208 | 0.20313347 | 0.03316364 | 0 | 0.10027441 |
| ENSMUSG00 | Olfr421 | 1049 | 0 | 0 | 0 | 0 | 0 | 0 |
| ENSMUSG00 | Gm17709 | 1433 | 0.69357084 | 0.77340417 | 0.06169134 | 0.73020188 | 0.64094541 | 0.36163164 |
| ENSMUSG00 | Gm17710 | 1553 | 0 | 0 | 0 | 0.74348063 | 0.0657133 | 1.22937846 |
| ENSMUSG00 | Gm17179 | 392 | 0 | 0.2458492 | 0 | 0 | 0 | 0 |
| ENSMUSG00 | Gm3788 | 474 | 0 | 0.60995496 | 0.2797585 | 0 | 0.43060233 | 0.05754143 |
| ENSMUSG00 | C2cd4b | 1324 | 0.0600536 | 0 | 0.13354031 | 0 | 0.07707912 | 0 |
| ENSMUSG00 | Gm8841 | 950 | 0 | 0.05072257 | 0 | 0.2658677 | 0.05371198 | 0.11484059 |
| ENSMUSG00 | Gm17708 | 4867 | 0 | 0 | 0 | 0 | 0 | 0.01681198 |
| ENSMUSG00 | Gm17038 | 491 | 0 | 0 | 0 | 0 | 0 | 0 |
| ENSMUSG00 | Gm3123 | 477 | 0 | 0 | 0 | 0 | 0 | 0 |
| ENSMUSG00 | Vmn2r78 | 2605 | 0 | 0 | 0 | 0 | 0 | 0 |
| ENSMUSG00 | Gm17658 | 46 | 0 | 0 | 1.92181926 | 3.13756909 | 0 | 0 |
| ENSMUSG00 | BC037034 | 3577 | 0 | 0 | 0 | 0 | 0 | 0 |
| ENSMUSG00 | Gm2670 | 2802 | 0 | 0 | 0 | 0 | 0 | 0 |
| ENSMUSG00 | Vmn2r-ps5 | 3197 | 0 | 0 | 0 | 0 | 0 | 0 |
| ENSMUSG00 | Gm17115 | 883 | 0 | 0 | 0 | 0 | 0.05778752 | 0 |
| ENSMUSG00 | Pcdha3 | 2437 | 0 | 0 | 0 | 0 | 0 | 0 |
| ENSMUSG00 | Gm4868 | 788 | 0 | 0 | 0 | 0 | 0 | 0 |
| ENSMUSG00 | Hspa1a | 2925 | 29.8879664 | 27.9069515 | 32.1880088 | 39.5113807 | 44.4670879 | 37.5877165 |
| ENSMUSG00 | Tmem207 | 1533 | 0.12966563 | 0 | 0.25950201 | 0.49427458 | 0 | 0 |
| ENSMUSG00 | Gm20474 | 1481 | 0 | 0 | 0 | 0 | 0 | 0 |
| ENSMUSG00 | Gm7822 | 519 | 0 | 0 | 0 | 0 | 0 | 0 |
| ENSMUSG00 | Gm2104 | 1649 | 0 | 0 | 0 | 0 | 0 | 0 |
| ENSMUSG00 | Gm17350 | 22 | 0 | 0 | 0 | 0 | 0 | 0 |
| ENSMUSG00 | Gm725 | 905 | 0 | 0 | 0 | 0 | 0 | 0 |
| ENSMUSG00 | Gm3719 | 312 | 0 | 0 | 0 | 0 | 0 | 0 |
| ENSMUSG00 | Gm8038 | 2063 | 0 | 0 | 0 | 0 | 0 | 0 |
| ENSMUSG00 | Gm17259 | 4606 | 0.45745559 | 0.48123672 | 1.03643054 | 1.69208025 | 0.94165044 | 1.18430913 |
| ENSMUSG00 | Olfr457 | 1038 | 0 | 0 | 0 | 0 | 0 | 0 |
| ENSMUSG00 | Gm17258 | 1264 | 0 | 0 | 0 | 0 | 0 | 0 |
| ENSMUSG00 | Gm17354 | 3037 | 1.91119531 | 2.96702821 | 1.41178096 | 2.34052116 | 1.32732423 | 2.64933111 |
| ENSMUSG00 | Rps2-ps5 | 665 | 0 | 0 | 0 | 0 | 0 | 0.12304349 |
| ENSMUSG00 | Gm3376 | 1496 | 0 | 0 | 0 | 0 | 0 | 0 |
| ENSMUSG00 | Vmn1r120 | 917 | 0 | 0 | 0 | 0 | 0 | 0 |
| ENSMUSG00 | Gm17257 | 467 | 0 | 0 | 0 | 0.30905392 | 0 | 0.29201969 |
| ENSMUSG00 | Gm9672 | 422 | 0 | 0 | 0 | 0 | 0 | 0 |
| ENSMUSG00 | Gm17458 | 1815 | 0.24094231 | 0 | 0 | 1.01387563 | 0.81529747 | 0.88661362 |
| ENSMUSG00 | B930036N10 | 1315 | 0.03023231 | 0.84280469 | 0 | 0 | 0.07760686 | 0.24889405 |
| ENSMUSG00 | E130317F20I | 3263 | 0.4142465 | 0.31011808 | 0.47412335 | 0.1548111 | 0.12510298 | 0.36778551 |
| ENSMUSG00 | Gm3339 | 471 | 0 | 0 | 0 | 0 | 0 | 0 |
| ENSMUSG00 | BC049352 | 2678 | 1.11339098 | 1.61940993 | 2.47583139 | 2.78901539 | 0.93364168 | 4.16554423 |
| ENSMUSG00 | Gm6611 | 1271 | 0 | 0.30329783 | 0.03477722 | 0.31227576 | 0 | 0 |
| ENSMUSG00 | Gm17072 | 1351 | 0 | 0 | 0 | 0 | 0 | 0 |
| ENSMUSG00 | Gm2858 | 245 | 0 | 0 | 0 | 0 | 0 | 0 |
| ENSMUSG00 | Gm13272 | 1555 | 0 | 0 | 0 | 0 | 0 | 0 |
| ENSMUSG00 | Tmem200a | 2883 | 0 | 0 | 0 | 0 | 0 | 0 |
| ENSMUSG00 | Gm17375 | 3219 | 1.0003709 | 1.63166269 | 1.41434912 | 1.60289915 | 1.53760747 | 1.28789847 |
| ENSMUSG00 | Gm3906 | 2136 | 0.27918175 | 0.31582874 | 0.28971245 | 0 | 0 | 0 |
| ENSMUSG00 | Gm17482 | 46 | 0 | 0 | 0 | 0 | 0 | 0 |
| ENSMUSG00 | C1rb | 2155 | 0 | 0 | 0 | 0 | 0 | 0 |
| ENSMUSG00 | Gm17139 | 569 | 0 | 0 | 0 | 0 | 0 | 0 |
| ENSMUSG00 | Cyp2c69 | 1761 | 0 | 0 | 0 | 0 | 0 | 0 |
| ENSMUSG00 | Myh15 | 6299 | 0 | 0 | 0 | 0 | 0 | 0 |
| ENSMUSG00 | Gm9793 | 3132 | 0.17770649 | 0.18462238 | 0.19758167 | 0.20736807 | 0.4561745 | 0.67054509 |
| ENSMUSG00 | Gm4782 | 2072 | 0 | 0 | 0 | 0 | 0 | 0 |
| ENSMUSG00 | Vmn1r130 | 923 | 0 | 0 | 0 | 0 | 0 | 0 |
| ENSMUSG00 | Gm4468 | 293 | 0 | 0 | 0 | 0.24629382 | 0 | 0 |
| ENSMUSG00 | Gm17551 | 878 | 0.22639795 | 0 | 0 | 0 | 0 | 0.18638706 |
| ENSMUSG00 | Gm3252 | 2077 | 0 | 0 | 0 | 0 | 0 | 0 |
| ENSMUSG00 | Gm9396 | 490 | 0 | 0 | 0 | 0 | 0 | 0 |
| ENSMUSG00 | Gm17550 | 4568 | 0 | 0 | 0 | 0 | 0 | 0 |
| ENSMUSG00 | Gm4027 | 996 | 0 | 0 | 0 | 0 | 0 | 0 |
| ENSMUSG00 | Gm5176 | 1224 | 0.19487981 | 2.12587245 | 0.07222523 | 0.29478795 | 0.20844108 | 0.20054882 |
| ENSMUSG00 | Gbp11 | 1322 | 0 | 0 | 0 | 0.10917411 | 0 | 0.61894038 |
| ENSMUSG00 | Gm17497 | 46 | 0 | 0 | 0 | 0 | 0 | 2.37170776 |
| ENSMUSG00 | Gm17496 | 1237 | 0 | 0 | 0 | 0 | 0 | 0 |
| ENSMUSG00 | Gm17499 | 690 | 0 | 0 | 0 | 0.67980664 | 0.29580508 | 0.59292694 |
| ENSMUSG00 | Gm17498 | 34 | 0 | 0 | 0 | 0 | 0 | 0 |
| ENSMUSG00 | Gm17062 | 458 | 0 | 0 | 0 | 0 | 0 | 0 |
| ENSMUSG00 | Gm17500 | 2984 | 0 | 0 | 0 | 0 | 0 | 0 |
| ENSMUSG00 | Gm9027 | 660 | 0 | 0.07300976 | 0.20091747 | 0.16400929 | 0 | 0.04132521 |
| ENSMUSG00 | Gm4904 | 1194 | 0 | 0 | 0 | 0 | 0 | 0 |
| ENSMUSG00 | Gm2108 | 2175 | 0 | 0 | 0 | 0 | 0 | 0 |
| ENSMUSG00 | Vmn2r59 | 2592 | 0 | 0 | 0 | 0 | 0 | 0 |
| ENSMUSG00 | A730056A06I | 2661 | 0 | 0 | 0 | 0.0677979 | 0 | 0 |
| ENSMUSG00 | Gm17370 | 2650 | 0 | 0 | 0 | 0 | 0 | 0 |
| ENSMUSG00 | Peg10 | 6662 | 0 | 0 | 0 | 0 | 0.01531864 | 0 |
| ENSMUSG00 | Gm2244 | 1496 | 0 | 0 | 0 | 0 | 0 | 0 |
| ENSMUSG00 | Gm7850 | 745 | 0 | 0 | 0 | 0.09686455 | 0 | 0 |
| ENSMUSG00 | Gm17369 | 991 | 0.56163141 | 0.09724812 | 0.8028589 | 0.1092292 | 0.61787741 | 0.82567021 |
| ENSMUSG00 | Gm5501 | 860 | 0 | 0 | 0 | 0 | 0 | 0 |
| ENSMUSG00 | Gm17198 | 1150 | 0 | 0 | 0 | 0 | 0 | 0 |
| ENSMUSG00 | Gm17163 | 430 | 3.05100199 | 0.11206149 | 0.51397492 | 0.50347039 | 0.35599797 | 1.77602302 |
| ENSMUSG00 | Gm5494 | 655 | 0 | 0 | 0 | 0 | 0 | 0 |
| ENSMUSG00 | 4930556M19 | 2461 | 0.37154655 | 0.25454033 | 0.50290598 | 0.16127692 | 0.29027601 | 0.19948944 |
| ENSMUSG00 | Gm17086 | 483 | 0 | 0 | 0 | 0 | 0 | 0 |
| ENSMUSG00 | Gm6450 | 692 | 0.3447007 | 0.13926717 | 0.19162649 | 0.15642505 | 0.07373754 | 1.0641839 |
| ENSMUSG00 | Gm17631 | 67 | 0 | 0 | 0 | 0 | 0 | 0 |
| ENSMUSG00 | Vmn2r85 | 3036 | 0 | 0 | 0 | 0.01188473 | 0 | 0 |
| ENSMUSG00 | Vmn2r4 | 2736 | 0 | 0 | 0 | 0 | 0 | 0 |
| ENSMUSG00 | 5330439K02I | 2310 | 0.10326099 | 0.08343973 | 0.26788996 | 0.39049832 | 0.15462538 | 0.29518008 |
| ENSMUSG00 | Gm17229 | 1757 | 0 | 0 | 0.12578783 | 0.123217 | 0.05808352 | 0.01552341 |
| ENSMUSG00 | Gm5101 | 3843 | 5.4621113 | 8.72697469 | 9.88014133 | 5.90570024 | 6.61231724 | 5.01064151 |
| ENSMUSG00 | Kif4-ps | 3668 | 0 | 0 | 0 | 0 | 0 | 0 |
| ENSMUSG00 | Gm7925 | 398 | 0 | 0 | 0 | 0 | 0 | 0 |
| ENSMUSG00 | Gm7289 | 921 | 0.04316556 | 0 | 0 | 0 | 0.05540323 | 0 |
| ENSMUSG00 | Gm10668 | 1333 | 0 | 0 | 0 | 0 | 0 | 0 |
| ENSMUSG00 | Gm3109 | 281 | 0 | 0 | 0 | 0 | 0 | 0.29118832 |
| ENSMUSG00 | Bend4 | 7930 | 0.05013302 | 0.01215295 | 0.005574 | 0.26390398 | 0.36033759 | 0.0412731 |
| ENSMUSG00 | Gm17243 | 2250 | 0 | 0 | 0 | 0 | 0 | 0 |
| ENSMUSG00 | Gm7664 | 749 | 0 | 0.19300311 | 0 | 0.04817362 | 0.068126 | 0 |
| ENSMUSG00 | Gm17242 | 2082 | 0 | 0 | 0 | 0 | 0 | 0 |
| ENSMUSG00 | Gm9312 | 335 | 0 | 0 | 0 | 0.1077076 | 0.91390524 | 0 |
| ENSMUSG00 | Gm17241 | 338 | 0.35285929 | 0.57025375 | 0.7846481 | 0.42700644 | 0.60386244 | 0.403471 |
| ENSMUSG00 | 1700121N20I | 1023 | 0 | 0 | 0 | 0 | 0 | 0 |
| ENSMUSG00 | A230107N01I | 2346 | 4.57543893 | 4.55984235 | 6.66984332 | 0.72287131 | 1.56602688 | 5.60374089 |
| ENSMUSG00 | Gm14484 | 524 | 0 | 0 | 0 | 0 | 0 | 0 |
| ENSMUSG00 | Gm17441 | 1041 | 0 | 0 | 0 | 0 | 0 | 0 |
| ENSMUSG00 | A230065N10I | 1027 | 2.32261814 | 1.45450799 | 2.92670431 | 2.1080065 | 3.03077792 | 3.9305225 |
| ENSMUSG00 | Gm4540 | 293 | 6.78421169 | 9.20969543 | 9.50415055 | 6.77308003 | 4.17963488 | 17.5004511 |
| ENSMUSG00 | Gm7919 | 841 | 0 | 0 | 0 | 0 | 0 | 0 |
| ENSMUSG00 | Dynlt1a | 737 | 0.16182692 | 1.30763751 | 0.35985218 | 0.63645397 | 0 | 0.51810712 |
| ENSMUSG00 | Serpina4-ps1 | 1283 | 0 | 0 | 0 | 0 | 0 | 0 |
| ENSMUSG00 | Gm4482 | 194 | 0.20492516 | 0.24838372 | 0.22784455 | 0.37197984 | 0 | 0 |
| ENSMUSG00 | Olfr101 | 1038 | 0 | 0 | 0 | 0 | 0 | 0 |
| ENSMUSG00 | Gm17439 | 1729 | 0.29891339 | 0 | 0.25564976 | 0 | 0 | 0 |
| ENSMUSG00 | Gm17440 | 1104 | 0 | 0 | 0 | 0 | 0 | 0 |
| ENSMUSG00 | Vmn2r16 | 2547 | 0 | 0 | 0 | 0 | 0 | 0 |
| ENSMUSG00 | Gm17360 | 4229 | 0.00940068 | 0.10254859 | 0.11497287 | 0.00853205 | 0.01206583 | 0.00644943 |
| ENSMUSG00 | Gm17361 | 172 | 0 | 0 | 0 | 0 | 0 | 0 |
| ENSMUSG00 | Kcnb2 | 5713 | 0 | 0 | 0 | 0 | 0 | 0 |
| ENSMUSG00 | Gm2056 | 434 | 0 | 0 | 0 | 0 | 0 | 0 |
| ENSMUSG00 | Gm17224 | 498 | 0 | 0.09675992 | 0.08875872 | 0.07245391 | 0 | 0 |
| ENSMUSG00 | Gm6793 | 1603 | 0.32240876 | 0.21042115 | 0.0551489 | 0.04501815 | 0.1273272 | 0.17014747 |
| ENSMUSG00 | Gm10772 | 86 | 0 | 0 | 0 | 0 | 0 | 0 |
| ENSMUSG00 | Gm8906 | 1094 | 0 | 0 | 0 | 0 | 0 | 0 |
| ENSMUSG00 | Gm3294 | 608 | 0 | 0 | 0 | 0 | 0 | 0 |
| ENSMUSG00 | 1700066J03I | 442 | 0.26983358 | 0.4360764 | 0 | 0 | 0 | 0 |
| ENSMUSG00 | Gm17215 | 274 | 0 | 0 | 0 | 0 | 0 | 0 |
| ENSMUSG00 | Gm17528 | 237 | 0 | 0 | 0 | 0 | 0 | 0 |
| ENSMUSG00 | Zfp804b | 3921 | 0 | 0 | 0 | 0 | 0 | 0 |
| ENSMUSG00 | Gm4695 | 279 | 0 | 0 | 0 | 0 | 0 | 0 |
| ENSMUSG00 | Gm5819 | 464 | 0 | 0 | 0 | 0 | 0 | 0.2351262 |
| ENSMUSG00 | Gm17529 | 3085 | 0.24484737 | 0.56230532 | 0.30088775 | 0 | 0.14886139 | 0.1237747 |
| ENSMUSG00 | Gm17530 | 113 | 4.57363935 | 4.69071561 | 2.34700051 | 0.95793039 | 1.35468254 | 0 |
| ENSMUSG00 | 4930544I03I | 1271 | 0 | 0 | 0 | 0 | 0 | 0 |
| ENSMUSG00 | Gm2956 | 2083 | 0 | 0 | 0 | 0 | 0 | 0 |
| ENSMUSG00 | Gm17712 | 1481 | 1.9327445 | 0.7158013 | 1.07445398 | 1.16943831 | 0.99916604 | 1.32597841 |
| ENSMUSG00 | Gm17039 | 609 | 0.52223948 | 0 | 0 | 0.53323219 | 0 | 0.40307349 |
| ENSMUSG00 | Gm17713 | 744 | 0.16030436 | 0 | 0 | 0 | 0 | 0 |
| ENSMUSG00 | Gm17714 | 172 | 0 | 0 | 0 | 0 | 0 | 0 |
| ENSMUSG00 | Gm17715 | 3087 | 0 | 0.01560947 | 0 | 0.04675354 | 0 | 0 |
| ENSMUSG00 | Ear-ps4 | 449 | 0 | 0 | 0 | 0 | 0 | 0 |
| ENSMUSG00 | Gm2963 | 521 | 0 | 0 | 0 | 0 | 0 | 0 |
| ENSMUSG00 | Gm9376 | 610 | 0 | 0 | 0 | 0 | 0 | 0 |
| ENSMUSG00 | Gm17438 | 70 | 0 | 0 | 0 | 0 | 0 | 0 |
| ENSMUSG00 | Vmn2r113 | 2565 | 0 | 0 | 0 | 0 | 0 | 0 |
| ENSMUSG00 | Gm6996 | 506 | 0 | 0 | 0 | 0 | 0 | 0 |
| ENSMUSG00 | Taar8b | 1125 | 0 | 0 | 0 | 0 | 0 | 0.02424412 |
| ENSMUSG00 | Gm17522 | 172 | 0 | 0 | 0 | 0 | 0 | 0 |
| ENSMUSG00 | Gm17521 | 467 | 0 | 0 | 0 | 0 | 0 | 0 |
| ENSMUSG00 | Gm10320 | 525 | 0 | 0 | 0 | 0 | 0 | 0 |
| ENSMUSG00 | Gm17520 | 952 | 1.29455871 | 3.13819266 | 1.48577624 | 2.31197974 | 1.28637922 | 6.41756218 |
| ENSMUSG00 | Fancf | 1690 | 3.34040132 | 3.45003521 | 2.17085975 | 2.4979877 | 3.20047091 | 5.43878901 |
| ENSMUSG00 | Gm17523 | 2044 | 0.01944984 | 0.07072374 | 0.17300134 | 0 | 0.09985592 | 0 |
| ENSMUSG00 | Vmn2r90 | 2586 | 0 | 0 | 0 | 0 | 0 | 0 |
| ENSMUSG00 | Gm17581 | 56 | 0 | 0 | 0 | 0 | 0 | 0 |
| ENSMUSG00 | B930094E09I | 1239 | 0.35295423 | 0 | 0.60648211 | 0.72804771 | 0.16473406 | 1.05664462 |
| ENSMUSG00 | AA543401 | 1268 | 0 | 0 | 0 | 0 | 0 | 0 |
| ENSMUSG00 | Gm17579 | 72 | 1.65647836 | 2.00776842 | 1.84174346 | 4.00911606 | 1.41739933 | 0 |
| ENSMUSG00 | Gm17580 | 1432 | 0 | 0 | 0 | 0 | 0 | 0 |
| ENSMUSG00 | Gm17577 | 172 | 0 | 0 | 0 | 0 | 0 | 0 |
| ENSMUSG00 | Gm17578 | 1398 | 0.31281136 | 0.06893626 | 0.31617914 | 0.59362448 | 0 | 0.05852927 |
| ENSMUSG00 | D030025P21I | 2363 | 0 | 0 | 0 | 0 | 0 | 0 |
| ENSMUSG00 | BC050972 | 1201 | 0.62893766 | 0.16048773 | 0.0368042 | 0 | 0.25491945 | 0.04541988 |
| ENSMUSG00 | AU022751 | 1828 | 0 | 0 | 0 | 0 | 0 | 0 |
| ENSMUSG00 | Gm3642 | 1757 | 0 | 0 | 0 | 0 | 0 | 0 |
| ENSMUSG00 | Gm17089 | 1390 | 0.31461172 | 0.31199855 | 0.54059808 | 0.20766644 | 0 | 0.66714945 |
| ENSMUSG00 | Gm17399 | 1179 | 2.69757289 | 0.53131785 | 0.78731018 | 0.67328667 | 0.34623495 | 0.87908082 |
| ENSMUSG00 | Vmn1r128 | 923 | 0 | 0 | 0 | 0 | 0 | 0 |
| ENSMUSG00 | Gcom1 | 1712 | 0 | 0 | 0 | 0 | 0 | 0 |
| ENSMUSG00 | Gm8607 | 995 | 0 | 0 | 0 | 0 | 0 | 0 |
| ENSMUSG00 | Gm17451 | 2289 | 0 | 0 | 0 | 0 | 0 | 0 |
| ENSMUSG00 | Gm6482 | 1405 | 0 | 0 | 0 | 0 | 0 | 0 |
| ENSMUSG00 | Gm17452 | 3016 | 0 | 0 | 0 | 0 | 0 | 0 |
| ENSMUSG00 | Gm3370 | 497 | 0 | 0 | 0 | 0 | 0 | 0 |
| ENSMUSG00 | Vmn1r160 | 923 | 0 | 0 | 0 | 0 | 0 | 0 |
| ENSMUSG00 | Gm13694 | 2725 | 0 | 0 | 0 | 0 | 0 | 0 |
| ENSMUSG00 | Gm7052 | 1511 | 0 | 0 | 0 | 0 | 0 | 0 |
| ENSMUSG00 | Gm8005 | 1396 | 0 | 0 | 0 | 0 | 0 | 0 |
| ENSMUSG00 | Gm17453 | 3585 | 0.23287729 | 0.56452736 | 0.06164832 | 0.26168289 | 0.22773278 | 0.89013467 |
| ENSMUSG00 | Gm17267 | 172 | 0 | 0 | 0 | 0 | 0 | 0 |
| ENSMUSG00 | Gm17026 | 1374 | 0 | 0 | 0 | 0 | 0 | 0 |
| ENSMUSG00 | Gm17266 | 2029 | 0 | 0 | 0 | 0 | 0 | 0 |
| ENSMUSG00 | Gm17270 | 3338 | 0.08336979 | 0.43307168 | 0.45022848 | 0.58371193 | 0.21401116 | 0.76806953 |
| ENSMUSG00 | Gm17269 | 209 | 0 | 0 | 0 | 0 | 0 | 0 |
| ENSMUSG00 | Ighv1-77 | 293 | 0 | 0 | 0 | 0 | 0 | 0 |
| ENSMUSG00 | Gm17268 | 64 | 0 | 0 | 0 | 0 | 0 | 0 |
| ENSMUSG00 | Gm2135 | 1649 | 0 | 0 | 0 | 0 | 0 | 0 |
| ENSMUSG00 | Ighd5-7 | 28 | 0 | 0 | 0 | 0 | 0 | 0 |
| ENSMUSG00 | Gm17340 | 82 | 0 | 0 | 0 | 0 | 0 | 0 |
| ENSMUSG00 | Vmn2r86 | 3033 | 0 | 0 | 0 | 0 | 0 | 0 |
| ENSMUSG00 | Flg | 935 | 0 | 0 | 0 | 0 | 0 | 0 |
| ENSMUSG00 | Rergl | 677 | 0 | 0 | 0 | 0 | 0 | 0 |
| ENSMUSG00 | Gm5624 | 1743 | 0 | 0 | 0 | 0 | 0 | 0 |
| ENSMUSG00 | Gm7942 | 1496 | 0 | 0 | 0 | 0 | 0 | 0 |
| ENSMUSG00 | Gm3696 | 1939 | 0.14352159 | 0.02485118 | 0.11398103 | 0 | 0.10526328 | 0.40792395 |
| ENSMUSG00 | Gm9075 | 789 | 0 | 0 | 0 | 0 | 0 | 0 |
| ENSMUSG00 | B930082K07I | 2476 | 0 | 0 | 0.03570424 | 0 | 0.02060839 | 0 |
| ENSMUSG00 | Vmn1r110 | 923 | 0 | 0 | 0 | 0 | 0 | 0 |
| ENSMUSG00 | 4833427F10I | 1472 | 0 | 0 | 0 | 0 | 0 | 0 |
| ENSMUSG00 | Gm20523 | 1123 | 0 | 0 | 0 | 0 | 0 | 0 |
| ENSMUSG00 | Vmn2r-ps38 | 2567 | 0 | 0 | 0 | 0 | 0 | 0 |
| ENSMUSG00 | Gm7029 | 448 | 0 | 0 | 0 | 0 | 0 | 0 |
| ENSMUSG00 | Gm3584 | 2559 | 0 | 0 | 0 | 0 | 0 | 0 |
| ENSMUSG00 | Gm20460 | 1560 | 0 | 0 | 0 | 0 | 0 | 0 |
| ENSMUSG00 | Vax2os2 | 1429 | 0 | 0 | 0 | 0 | 0 | 0 |
| ENSMUSG00 | Gm3912 | 2438 | 0 | 0.09882371 | 0 | 0 | 0 | 0 |
| ENSMUSG00 | Vmn2r-ps75 | 211 | 0 | 0 | 0 | 0 | 0 | 0 |
| ENSMUSG00 | Gm20505 | 1707 | 0 | 0 | 0 | 0 | 0 | 0 |
| ENSMUSG00 | Gm20432 | 249 | 0 | 0 | 0 | 0 | 0.2049252 | 0 |
| ENSMUSG00 | Gm20407 | 596 | 0 | 0 | 0 | 0 | 0.3424589 | 0 |
| ENSMUSG00 | 4930515G01I | 1246 | 0.44669079 | 0.65743942 | 0.21284997 | 0 | 0.69618651 | 1.02881866 |
| ENSMUSG00 | Gm18489 | 550 | 0 | 0 | 0 | 0 | 0 | 0 |
| ENSMUSG00 | Gm20416 | 480 | 0 | 0.20077684 | 0 | 0.60136741 | 0.74413465 | 0 |
| ENSMUSG00 | Gm5967 | 493 | 0 | 0 | 0.08965891 | 0 | 0 | 0 |
| ENSMUSG00 | Gm20457 | 680 | 0.35078365 | 0.35431208 | 0.13000542 | 0 | 0.45023273 | 0.80219527 |
| ENSMUSG00 | Gm20471 | 624 | 0 | 0 | 0 | 0 | 0 | 0 |
| ENSMUSG00 | Gm3644 | 298 | 0 | 0 | 0 | 0 | 0 | 0 |
| ENSMUSG00 | Gm20470 | 329 | 2.65842119 | 2.0504869 | 2.1496337 | 2.63212483 | 1.55095367 | 1.24352459 |
| ENSMUSG00 | Gm7211 | 1116 | 0 | 0 | 0 | 0 | 0 | 0 |
| ENSMUSG00 | Dyx1c1 | 1966 | 1.07173981 | 1.22549446 | 0.89932539 | 1.0828284 | 0.70076915 | 0.04161949 |
| ENSMUSG00 | Cd9-ps | 649 | 0.12251304 | 0.22274164 | 0.20432285 | 0.33357822 | 0 | 0.08405128 |
| ENSMUSG00 | Mir196a-2 | 109 | 0 | 0 | 0 | 0 | 0 | 0 |
| ENSMUSG00 | Vmn2r-ps87 | 875 | 0 | 0 | 0 | 0 | 0 | 0 |
| ENSMUSG00 | Gm20446 | 659 | 0 | 0 | 0 | 0 | 0 | 0 |
| ENSMUSG00 | Gm20480 | 249 | 0 | 0 | 0 | 0 | 0 | 0 |
| ENSMUSG00 | Gm20527 | 509 | 0 | 0 | 0 | 0 | 0 | 0 |
| ENSMUSG00 | Gm18358 | 1273 | 0 | 0 | 0 | 0 | 0 | 0 |
| ENSMUSG00 | Tnxa | 2957 | 0 | 0 | 0 | 0 | 0 | 0 |
| ENSMUSG00 | A530058N18I | 2311 | 0 | 0 | 0 | 0 | 0 | 0 |
| ENSMUSG00 | Stk19-ps1 | 512 | 0 | 0 | 0 | 0 | 0 | 0 |
| ENSMUSG00 | 1110038B12I | 2042 | 1.83007599 | 2.14738797 | 2.98719605 | 2.70350285 | 1.77417859 | 1.8833125 |
| ENSMUSG00 | Vmn2r-ps49 | 3545 | 0 | 0 | 0 | 0 | 0 | 0 |
| ENSMUSG00 | Mir10a | 109 | 0 | 0 | 0 | 0 | 0 | 0 |
| ENSMUSG00 | Runx2 | 3002 | 0 | 0 | 0 | 0 | 0 | 0 |
| ENSMUSG00 | Vmn1r-ps75 | 919 | 0 | 0 | 0 | 0 | 0 | 0 |
| ENSMUSG00 | Vmn1r-ps58 | 839 | 0 | 0 | 0 | 0 | 0 | 0 |
| ENSMUSG00 | A930009A15I | 996 | 0 | 0 | 0 | 0 | 0 | 0 |
| ENSMUSG00 | Vmn2r-ps67 | 384 | 0 | 0 | 0 | 0 | 0 | 0 |
| ENSMUSG00 | Slc22a13b | 2201 | 0.37931172 | 0.19703679 | 0.2209088 | 0.11475434 | 0.32456577 | 0.38414985 |
| ENSMUSG00 | Gm18522 | 649 | 0 | 0 | 0 | 0 | 0 | 0 |
| ENSMUSG00 | Gm18560 | 1122 | 0 | 0 | 0 | 0 | 0 | 0 |
| ENSMUSG00 | Gm20419 | 541 | 0 | 0 | 0 | 0 | 0 | 0 |
| ENSMUSG00 | Gm19345 | 986 | 0 | 0 | 0 | 0 | 0 | 0 |
| ENSMUSG00 | Vmn2r-ps62 | 614 | 0 | 0 | 0 | 0 | 0 | 0 |
| ENSMUSG00 | Gm20529 | 201 | 0 | 0 | 0 | 0 | 0 | 0 |
| ENSMUSG00 | Gm20443 | 731 | 0 | 0 | 0 | 0 | 0 | 0 |
| ENSMUSG00 | Gm20528 | 1029 | 0 | 0 | 0 | 0 | 0 | 0 |
| ENSMUSG00 | Gm7230 | 810 | 0 | 0 | 0 | 0 | 0 | 0 |
| ENSMUSG00 | Gm20506 | 587 | 0 | 0 | 0 | 0 | 0 | 0 |
| ENSMUSG00 | Gm19807 | 666 | 0 | 0 | 0 | 0.37924071 | 0.07661618 | 0 |
| ENSMUSG00 | Gm8815 | 616 | 0 | 0 | 0 | 0 | 0 | 0 |
| ENSMUSG00 | Gm2381 | 4069 | 0.4885166 | 0.17763496 | 0.07604151 | 0.60299313 | 0.18810411 | 0.40888498 |
| ENSMUSG00 | Gm9282 | 988 | 0 | 0 | 0.44738708 | 0 | 0 | 0.16563546 |
| ENSMUSG00 | Gm7651 | 896 | 0 | 0 | 0 | 0 | 0 | 0 |
| ENSMUSG00 | Vmn1r-ps28 | 941 | 0 | 0 | 0 | 0 | 0 | 0 |
| ENSMUSG00 | Gm4433 | 503 | 0 | 0 | 0 | 0 | 0 | 0 |
| ENSMUSG00 | Gm18889 | 928 | 0.04283996 | 0.05192505 | 0 | 0 | 0 | 0 |
| ENSMUSG00 | Gm9612 | 367 | 0 | 0 | 0 | 0 | 0 | 0 |
| ENSMUSG00 | Gm20521 | 1236 | 0 | 0 | 0 | 0 | 0 | 0 |
| ENSMUSG00 | Gm20403 | 371 | 0 | 0 | 0 | 0.19451237 | 0 | 0 |
| ENSMUSG00 | Gm3595 | 871 | 0 | 0 | 0 | 0 | 0 | 0 |
| ENSMUSG00 | Vmn1r-ps78 | 923 | 0 | 0 | 0 | 0 | 0 | 0 |
| ENSMUSG00 | Vmn1r-ps27 | 984 | 0 | 0 | 0 | 0 | 0 | 0 |
| ENSMUSG00 | Gm6004 | 1114 | 0 | 0 | 0 | 0 | 0 | 0 |
| ENSMUSG00 | 1700031A10I | 653 | 0 | 0 | 0 | 0 | 0 | 0 |
| ENSMUSG00 | Gm4246 | 215 | 0 | 0 | 0 | 0 | 0 | 0 |
| ENSMUSG00 | Gm20522 | 782 | 0.76257316 | 0.80105339 | 0.90438554 | 1.01509588 | 1.56602688 | 0.76731722 |
| ENSMUSG00 | Gm20515 | 4822 | 0 | 0 | 0 | 0 | 0 | 0 |
| ENSMUSG00 | Gm7030 | 908 | 0 | 0 | 0 | 0 | 0 | 0 |
| ENSMUSG00 | Gm20410 | 1407 | 0 | 0 | 0 | 0 | 0 | 0 |
| ENSMUSG00 | Gm17973 | 519 | 0 | 0 | 0 | 0 | 0 | 0 |
| ENSMUSG00 | Gm20493 | 212 | 0 | 0 | 2.29349185 | 0.85099162 | 0 | 0 |
| ENSMUSG00 | Gm20426 | 438 | 0 | 0 | 0 | 0.24713729 | 0 | 0 |
| ENSMUSG00 | Vmn1r-ps63 | 988 | 0 | 0 | 0 | 0 | 0 | 0 |
| ENSMUSG00 | Gm5708 | 885 | 0 | 0 | 0 | 0 | 0 | 0 |
| ENSMUSG00 | Gm20467 | 992 | 0 | 0 | 0 | 0 | 0 | 0 |
| ENSMUSG00 | Trav9n-1 | 581 | 0 | 0 | 0 | 0 | 0 | 0 |
| ENSMUSG00 | Gm20499 | 442 | 0 | 0 | 0 | 0 | 0 | 0 |
| ENSMUSG00 | H2-Q3 | 605 | 0 | 0 | 0 | 0 | 0 | 0 |
| ENSMUSG00 | A930015D03I | 1064 | 0 | 0 | 0 | 0 | 0 | 0 |
| ENSMUSG00 | Gm20455 | 1136 | 0 | 0 | 0 | 0 | 0 | 0 |
| ENSMUSG00 | Gm18359 | 1455 | 0 | 0 | 0 | 0 | 0 | 0 |
| ENSMUSG00 | Gm2308 | 1003 | 0.07927314 | 0.38433852 | 0.1322089 | 0.10792237 | 0.05087376 | 0.19035142 |
| ENSMUSG00 | Gm20444 | 727 | 0 | 0 | 0 | 0 | 0 | 0 |
| ENSMUSG00 | Vmn2r-ps68 | 860 | 0 | 0 | 0 | 0 | 0 | 0 |
| ENSMUSG00 | Zfp963 | 2607 | 4.68160051 | 3.10522527 | 6.52769834 | 6.14515833 | 6.22416092 | 3.33740312 |
| ENSMUSG00 | Gm20519 | 358 | 0 | 0 | 0 | 0 | 0 | 0.30474457 |
| ENSMUSG00 | Gm20434 | 1534 | 0 | 0 | 0 | 0 | 0 | 0 |
| ENSMUSG00 | Vmn2r-ps79 | 376 | 0 | 0 | 0 | 0 | 0 | 0 |
| ENSMUSG00 | Gm3585 | 870 | 0 | 0 | 0 | 0 | 0 | 0 |
| ENSMUSG00 | Gm5682 | 573 | 0 | 0 | 0 | 0 | 0 | 0 |
| ENSMUSG00 | Gm20500 | 319 | 0 | 0 | 0 | 0 | 0 | 0 |
| ENSMUSG00 | Gm20417 | 423 | 0 | 0 | 0 | 0 | 0 | 0 |
| ENSMUSG00 | Gm20540 | 797 | 2.24466327 | 0 | 1.77472895 | 1.62960377 | 2.88103753 | 2.01907618 |
| ENSMUSG00 | Gm9577 | 2619 | 0 | 0 | 0 | 0 | 0 | 0 |
| ENSMUSG00 | Gm7031 | 918 | 0 | 0 | 0 | 0 | 0 | 0 |
| ENSMUSG00 | Vmn1r-ps46 | 948 | 0 | 0 | 0 | 0 | 0 | 0 |
| ENSMUSG00 | Gm19301 | 2208 | 0 | 0.02182357 | 0 | 0 | 0 | 0 |
| ENSMUSG00 | Rnf169 | 713 | 0 | 0 | 0.18598251 | 0.40484762 | 0.42939447 | 0 |
| ENSMUSG00 | Neat1 | 23411 | 69.9129953 | 62.6479322 | 30.7436081 | 80.0922441 | 62.9378338 | 44.4658988 |
| ENSMUSG00 | Gm20465 | 1925 | 0 | 0 | 0 | 0 | 0 | 0 |
| ENSMUSG00 | 1700116B05I | 465 | 0 | 0 | 0 | 0 | 0 | 0 |
| ENSMUSG00 | Gm19684 | 1608 | 0 | 0 | 0 | 0 | 0.06346564 | 0.28835129 |
| ENSMUSG00 | Gm8752 | 1692 | 0 | 0 | 0 | 0.02132509 | 0 | 0 |
| ENSMUSG00 | 1500011B03I | 577 | 9.71494411 | 12.2762686 | 11.4909471 | 7.62913246 | 7.78218557 | 25.147501 |
| ENSMUSG00 | Gm20411 | 1675 | 0 | 0 | 0 | 0 | 0 | 0 |
| ENSMUSG00 | Hmg1l1 | 647 | 0 | 0 | 0 | 0 | 0 | 0 |
| ENSMUSG00 | Gm4441 | 857 | 0 | 0 | 0 | 0 | 0 | 0 |
| ENSMUSG00 | Gm20412 | 631 | 0 | 0 | 0 | 0 | 0 | 0 |
| ENSMUSG00 | Gm8801 | 1058 | 0 | 0 | 0 | 0 | 0 | 0 |
| ENSMUSG00 | Vmn1r-ps56 | 964 | 0 | 0 | 0 | 0 | 0 | 0 |
| ENSMUSG00 | Gm20448 | 1659 | 1.34195715 | 0.75518234 | 1.41211433 | 1.58769696 | 0.92271927 | 2.13725323 |
| ENSMUSG00 | Vmn2r-ps77 | 319 | 0 | 0 | 0 | 0 | 0 | 0 |
| ENSMUSG00 | Gm9289 | 707 | 0 | 0 | 0 | 0 | 0 | 0 |
| ENSMUSG00 | Gm6659 | 546 | 0 | 0.08825356 | 0 | 0.06608433 | 0 | 0 |
| ENSMUSG00 | Gm20452 | 510 | 0 | 0 | 0 | 0 | 0 | 0 |
| ENSMUSG00 | Olfr113 | 994 | 0 | 0 | 0 | 0 | 0 | 0 |
| ENSMUSG00 | Gm17919 | 310 | 0 | 0 | 0 | 0 | 0 | 0 |
| ENSMUSG00 | Gm18200 | 829 | 0 | 0 | 0 | 0 | 0 | 0 |
| ENSMUSG00 | Gm20543 | 723 | 0 | 3.26574781 | 0 | 0 | 0 | 0 |
| ENSMUSG00 | Gm20454 | 220 | 0 | 0 | 0 | 0 | 0 | 0 |
| ENSMUSG00 | Gm4214 | 959 | 0 | 0 | 0 | 0 | 0 | 0 |
| ENSMUSG00 | Gm8877 | 652 | 0 | 0 | 0 | 0 | 0 | 0 |
| ENSMUSG00 | Psmb9 | 3976 | 0 | 0 | 0 | 0 | 0 | 0 |
| ENSMUSG00 | Cdk3-ps | 1542 | 0.12890882 | 0.12499726 | 0.02866527 | 0 | 0.43018345 | 0 |
| ENSMUSG00 | Gm6887 | 481 | 0 | 0 | 0 | 0 | 0 | 0 |
| ENSMUSG00 | Gm20524 | 86 | 0 | 0 | 0 | 0 | 0 | 0 |
| ENSMUSG00 | Vmn1r-ps29 | 783 | 0 | 0 | 0 | 0 | 0 | 0 |
| ENSMUSG00 | Gm20530 | 257 | 0 | 0 | 0 | 0 | 0 | 0 |
| ENSMUSG00 | Prps1l1 | 1675 | 0 | 0 | 0 | 0 | 0 | 0 |
| ENSMUSG00 | Gm20447 | 342 | 0 | 0 | 0 | 0 | 0 | 0 |
| ENSMUSG00 | Vmn2r-ps76 | 738 | 0 | 0 | 0 | 0 | 0 | 0 |
| ENSMUSG00 | Gm3623 | 2741 | 0 | 0 | 0 | 0 | 0 | 0 |
| ENSMUSG00 | 4930518C09I | 475 | 0.08369575 | 1.31878684 | 0.55833907 | 0.60769759 | 0.85939159 | 2.64133349 |
| ENSMUSG00 | Gm20509 | 383 | 0 | 0 | 0 | 0 | 0 | 0 |
| ENSMUSG00 | Vmn1r-ps26 | 839 | 0 | 0 | 0 | 0 | 0 | 0 |
| ENSMUSG00 | Zfp419 | 912 | 0 | 0 | 0 | 0 | 0 | 0 |
| ENSMUSG00 | Gm20415 | 420 | 0 | 0 | 0 | 0 | 0 | 0 |
| ENSMUSG00 | Gm3563 | 1623 | 0 | 0 | 0 | 0 | 0 | 0 |
| ENSMUSG00 | Gm20440 | 620 | 0 | 0 | 0 | 0 | 0 | 0 |
| ENSMUSG00 | Gm6726 | 680 | 0 | 0 | 0 | 0 | 0 | 0 |
| ENSMUSG00 | Gm6633 | 269 | 0 | 0 | 0 | 0 | 0 | 0 |
| ENSMUSG00 | Vmn1r-ps80 | 850 | 0 | 0 | 0 | 0 | 0 | 0 |
| ENSMUSG00 | Znhit2-ps | 383 | 0 | 0 | 0 | 0 | 0 | 0 |
| ENSMUSG00 | Gm20436 | 485 | 0 | 0 | 0 | 0 | 0 | 0 |
| ENSMUSG00 | Gm20408 | 1113 | 0 | 0 | 0 | 0 | 0 | 0 |
| ENSMUSG00 | BB365896 | 2996 | 1.97715841 | 1.80136232 | 1.99173859 | 3.7816295 | 2.60581959 | 5.96291345 |
| ENSMUSG00 | Gm3616 | 2508 | 0 | 0 | 0 | 0 | 0 | 0 |
| ENSMUSG00 | Gm18284 | 623 | 1.27625941 | 0.92814977 | 1.63184974 | 0.69499925 | 0.49142577 | 1.88251924 |
| ENSMUSG00 | Vmn1r-ps55 | 391 | 0 | 0 | 0 | 0 | 0 | 0 |
| ENSMUSG00 | Gm20488 | 2987 | 0 | 0 | 0 | 0 | 0 | 0 |
| ENSMUSG00 | Vmn2r-ps57 | 2592 | 0 | 0 | 0 | 0 | 0 | 0 |
| ENSMUSG00 | Gm20388 | 3937 | 0 | 0 | 0 | 0 | 0 | 0 |
| ENSMUSG00 | Vmn2r-ps82 | 1635 | 0 | 0 | 0 | 0 | 0 | 0 |
| ENSMUSG00 | Vmn1r-ps66 | 611 | 0 | 0 | 0 | 0 | 0 | 0 |
| ENSMUSG00 | Gm8868 | 574 | 0 | 0 | 0 | 0 | 0 | 0 |
| ENSMUSG00 | Gm7704 | 814 | 0 | 0 | 0 | 0 | 0 | 0 |
| ENSMUSG00 | Gm7221 | 1421 | 10.8551207 | 8.91839147 | 5.97238132 | 0.48244817 | 8.76174222 | 12.9175455 |
| ENSMUSG00 | Vmn1r-ps40 | 342 | 0 | 0 | 0 | 0 | 0 | 0 |
| ENSMUSG00 | Vmn1r-ps76 | 923 | 0 | 0 | 0 | 0 | 0 | 0 |
| ENSMUSG00 | Mir219-1 | 104 | 0 | 0 | 0 | 0 | 0 | 0 |
| ENSMUSG00 | Gm4831 | 593 | 0 | 0 | 0 | 0 | 0 | 0 |
| ENSMUSG00 | Olfr754-ps1 | 783 | 0 | 0 | 0 | 0 | 0 | 0 |
| ENSMUSG00 | Malat1 | 8810 | 707.864155 | 688.376964 | 575.843147 | 762.658107 | 541.215512 | 1096.34762 |
| ENSMUSG00 | Gm20516 | 1305 | 0 | 0 | 0 | 0 | 0 | 0 |
| ENSMUSG00 | Gm20525 | 47 | 0 | 0 | 0 | 0 | 0 | 0 |
| ENSMUSG00 | Hspe1-ps3 | 276 | 0.28808319 | 0 | 0 | 0 | 0 | 0 |
| ENSMUSG00 | Gm20503 | 1291 | 0 | 0 | 0 | 0 | 0 | 0 |
| ENSMUSG00 | Gm20424 | 618 | 0 | 0 | 0 | 0 | 0 | 0 |
| ENSMUSG00 | Gm4133 | 923 | 0 | 0 | 0 | 0 | 0 | 0 |
| ENSMUSG00 | Alx1 | 4126 | 0 | 0 | 0 | 0 | 0 | 0 |
| ENSMUSG00 | BC033916 | 804 | 0 | 0 | 0 | 0 | 0 | 0 |
| ENSMUSG00 | Vmn1r-ps62 | 908 | 0 | 0 | 0 | 0 | 0 | 0 |
| ENSMUSG00 | Vmn1r-ps64 | 654 | 0 | 0 | 0 | 0 | 0 | 0 |
| ENSMUSG00 | Vmn2r-ps74 | 916 | 0 | 0 | 0 | 0 | 0 | 0 |
| ENSMUSG00 | Gm20539 | 1379 | 0.20180447 | 0.1048291 | 0.16026774 | 0.91578793 | 0 | 0.17800707 |
| ENSMUSG00 | Gm20548 | 1651 | 0.04815927 | 0.26267594 | 0.18740939 | 1.31127963 | 1.32897284 | 0.94164412 |
| ENSMUSG00 | Gm20514 | 249 | 0 | 0 | 0 | 0 | 0 | 0 |
| ENSMUSG00 | Gm20532 | 773 | 0 | 0 | 0 | 0 | 0 | 0 |
| ENSMUSG00 | Vmn2r-ps83 | 2363 | 0 | 0 | 0 | 0 | 0 | 0 |
| ENSMUSG00 | Gm20430 | 206 | 0 | 0 | 0 | 0.52546667 | 0 | 0 |
| ENSMUSG00 | Vmn1r-ps68 | 398 | 0 | 0 | 0 | 0 | 0 | 0 |
| ENSMUSG00 | Gm20441 | 750 | 0 | 0 | 0 | 0 | 0 | 0 |
| ENSMUSG00 | Gm20433 | 420 | 0 | 0 | 0 | 0 | 0 | 0 |
| ENSMUSG00 | Gm20392 | 342 | 0 | 0 | 0 | 0 | 0 | 0 |
| ENSMUSG00 | Gm20391 | 105 | 0 | 0 | 0 | 0 | 0 | 0 |
| ENSMUSG00 | Gm20476 | 466 | 0 | 0 | 0.09485374 | 0 | 0 | 0 |
| ENSMUSG00 | BC023719 | 978 | 0 | 0 | 0 | 0 | 0 | 0 |
| ENSMUSG00 | Gm17782 | 276 | 0 | 0 | 0 | 0 | 0 | 0 |
| ENSMUSG00 | Gm20479 | 549 | 0 | 0 | 0 | 0 | 0 | 0 |
| ENSMUSG00 | A930015D03I | 1109 | 0 | 0 | 0 | 0 | 0 | 0 |
| ENSMUSG00 | 1700039E22I | 767 | 0.15549732 | 0 | 0 | 0 | 0 | 0.35560156 |
| ENSMUSG00 | Vmn2r-ps58 | 241 | 0 | 0 | 0 | 0 | 0 | 0 |
| ENSMUSG00 | Gm20511 | 454 | 0.17513428 | 0 | 0 | 0 | 0 | 0 |
| ENSMUSG00 | Olfr296-ps1 | 925 | 0 | 0 | 0 | 0 | 0 | 0 |
| ENSMUSG00 | Gm20535 | 1266 | 0.97347543 | 0.22837176 | 0.06982914 | 1.140033 | 0.04030519 | 0 |
| ENSMUSG00 | Gm6623 | 543 | 0 | 0 | 0 | 0 | 0 | 0 |
| ENSMUSG00 | A730060N03I | 2122 | 0 | 0 | 0 | 0 | 0 | 0 |
| ENSMUSG00 | Gm19203 | 938 | 0 | 0 | 0 | 0 | 0 | 0 |
| ENSMUSG00 | Gm20549 | 428 | 0 | 0 | 0 | 0 | 0 | 0 |
| ENSMUSG00 | Gm9285 | 1118 | 0 | 0 | 0 | 0 | 0 | 0 |
| ENSMUSG00 | Alx3 | 1968 | 0 | 0 | 0 | 0 | 0 | 0 |
| ENSMUSG00 | Gm6838 | 832 | 0 | 0 | 0 | 0 | 0 | 0 |
| ENSMUSG00 | Gm20512 | 616 | 0.32269059 | 0 | 0.21526872 | 0 | 0 | 0 |

| | | | | | | | | |
|---|---|---|---|---|---|---|---|---|
| ENSMUSG00 | Vmn2r-ps59 | 134 | 0 | 0 | 0 | 0 | 0 | 0 |
| ENSMUSG00 | Gm9566 | 528 | 0 | 0 | 0 | 0 | 0 | 0 |
| ENSMUSG00 | Gm4189 | 428 | 0.27865991 | 0 | 1.34257934 | 0.42151921 | 1.07298454 | 0.50980634 |
| ENSMUSG00 | Gm20401 | 209 | 0 | 0 | 0 | 0 | 0 | 0 |
| ENSMUSG00 | Gm20536 | 404 | 0 | 0 | 0 | 0 | 0 | 0 |
| ENSMUSG00 | Gm7648 | 1654 | 0 | 0 | 0 | 0 | 0 | 0 |
| ENSMUSG00 | Olfr306-ps1 | 627 | 0 | 0 | 0 | 0 | 0 | 0 |
| ENSMUSG00 | Gm20483 | 287 | 0 | 0 | 0 | 0 | 0 | 0 |
| ENSMUSG00 | Gm20541 | 1403 | 1.19011417 | 0.65256051 | 0.75612561 | 1.28588897 | 3.05503604 | 1.71074002 |
| ENSMUSG00 | Gm4175 | 944 | 0 | 0 | 0 | 0 | 0 | 0 |
| ENSMUSG00 | Gm20546 | 1885 | 0 | 0 | 0 | 0 | 0 | 0 |
| ENSMUSG00 | Gm20399 | 137 | 0 | 0 | 0 | 0 | 0 | 0 |
| ENSMUSG00 | Gm20445 | 564 | 0 | 0 | 0 | 0 | 0 | 0 |
| ENSMUSG00 | Gm20463 | 364 | 0 | 0 | 0 | 0 | 0 | 0 |
| ENSMUSG00 | Olfr758-ps1 | 838 | 0 | 0 | 0 | 0 | 0 | 0 |
| ENSMUSG00 | C130080G10 | 2860 | 0 | 0 | 0 | 0 | 0 | 0 |
| ENSMUSG00 | Vmn1r-ps59 | 198 | 0 | 0 | 0 | 0 | 0 | 0 |
| ENSMUSG00 | 4930556N13I | 1029 | 0 | 0 | 0 | 0.52597733 | 0 | 0 |
| ENSMUSG00 | Gm20469 | 3838 | 0 | 0 | 0 | 0 | 0 | 0 |
| ENSMUSG00 | Olfr300-ps1 | 905 | 0 | 0 | 0 | 0 | 0 | 0 |
| ENSMUSG00 | Gm20485 | 392 | 2.02834084 | 1.59801977 | 1.35311764 | 0.92046032 | 0.78101596 | 1.66987587 |
| ENSMUSG00 | Gm20497 | 668 | 0 | 0 | 0 | 0 | 0 | 0 |
| ENSMUSG00 | Gm9921 | 4397 | 0 | 0 | 0 | 0 | 0 | 0 |
| ENSMUSG00 | Gm20402 | 685 | 0 | 0 | 0 | 0 | 0 | 0 |
| ENSMUSG00 | Gm20462 | 407 | 0 | 0 | 0 | 0 | 0.50148772 | 0 |
| ENSMUSG00 | Gm4040 | 2355 | 0 | 0 | 0 | 0 | 0 | 0 |
| ENSMUSG00 | Vhl-ps1 | 596 | 0 | 0 | 0 | 0 | 0 | 0 |
| ENSMUSG00 | Gm9613 | 195 | 0 | 0 | 0 | 0 | 0 | 0 |
| ENSMUSG00 | Gm18354 | 1066 | 0 | 0 | 0 | 0 | 0 | 0 |
| ENSMUSG00 | Gm5594 | 1425 | 0 | 0 | 0 | 0 | 0 | 0 |
| ENSMUSG00 | Gm20507 | 2850 | 0.25108725 | 0.11835267 | 0.23264128 | 0.1519244 | 0.6266397 | 0.55506283 |
| ENSMUSG00 | Olfr115 | 1424 | 0 | 0 | 0 | 0 | 0 | 0 |
| ENSMUSG00 | Gm20453 | 225 | 0 | 0 | 0 | 0 | 0 | 0 |
| ENSMUSG00 | Gm20513 | 534 | 0 | 0 | 0 | 0 | 0 | 0 |
| ENSMUSG00 | Zfp141 | 4932 | 5.11855842 | 5.73508547 | 6.65895567 | 9.2546201 | 9.43553422 | 8.83162994 |
| ENSMUSG00 | Gpank1 | 1943 | 2.59853115 | 3.24880284 | 3.70813197 | 4.15974162 | 3.78167685 | 2.24598162 |
| ENSMUSG00 | Gm20406 | 421 | 0 | 0 | 0 | 0 | 0 | 0 |
| ENSMUSG00 | Vmn2r-ps86 | 870 | 0 | 0 | 0 | 0 | 0 | 0 |
| ENSMUSG00 | Gm4187 | 923 | 0 | 0 | 0 | 0 | 0 | 0 |
| ENSMUSG00 | Gm20464 | 654 | 0 | 0 | 0 | 0 | 0 | 0 |
| ENSMUSG00 | Vmn2r-ps65 | 219 | 0 | 0 | 0 | 0 | 0 | 0 |
| ENSMUSG00 | Vmn1r63 | 902 | 0 | 0 | 0 | 0 | 0 | 0 |
| ENSMUSG00 | Mir615 | 91 | 0 | 0 | 0 | 0 | 0 | 0 |
| ENSMUSG00 | Gm8018 | 786 | 0 | 0 | 0 | 0 | 0 | 0 |
| ENSMUSG00 | Gm18779 | 576 | 0 | 0 | 0 | 0 | 0 | 0 |
| ENSMUSG00 | Vmn1r-ps69 | 200 | 0 | 0 | 0 | 0 | 0 | 0 |
| ENSMUSG00 | Gm20545 | 209 | 0 | 0 | 0 | 0.17264136 | 0.24414534 | 0.13050067 |
| ENSMUSG00 | Gm18958 | 759 | 0 | 0 | 0 | 0 | 0 | 0.3234147 |
| ENSMUSG00 | Gm20437 | 1968 | 0 | 0 | 0 | 0 | 0 | 0 |
| ENSMUSG00 | Vmn2r-ps70 | 301 | 0 | 0 | 0 | 0 | 0 | 0 |
| ENSMUSG00 | Vmn1r-ps72 | 923 | 0 | 0 | 0 | 0 | 0 | 0 |
| ENSMUSG00 | Vmn2r-ps40 | 2552 | 0 | 0 | 0 | 0 | 0 | 0 |
| ENSMUSG00 | Gm20526 | 156 | 0 | 0 | 0 | 0 | 0 | 0 |
| ENSMUSG00 | Gm3603 | 2506 | 0 | 0 | 0 | 0 | 0 | 0 |
| ENSMUSG00 | Gm5227 | 572 | 0 | 0 | 0 | 0 | 0 | 0 |
| ENSMUSG00 | Gm18734 | 802 | 0 | 0 | 0 | 0 | 0 | 0 |
| ENSMUSG00 | Vmn2r-ps37 | 1329 | 0 | 0 | 0 | 0 | 0 | 0 |
| ENSMUSG00 | Vmn2r-ps78 | 307 | 0 | 0 | 0 | 0 | 0 | 0 |
| ENSMUSG00 | Vmn2r-ps46 | 2550 | 0 | 0 | 0 | 0 | 0 | 0 |
| ENSMUSG00 | Gm3589 | 324 | 0 | 0 | 0 | 0 | 0 | 0 |
| ENSMUSG00 | CT009512.1 | 452 | 0 | 0 | 0 | 0 | 0 | 0 |
| ENSMUSG00 | Gm18733 | 416 | 0 | 0 | 0 | 0 | 0 | 0 |
| ENSMUSG00 | Gm4437 | 921 | 0 | 0 | 0 | 0 | 0 | 0 |
| ENSMUSG00 | Gsx2 | 2184 | 0 | 0 | 0 | 0 | 0 | 0 |
| ENSMUSG00 | Gm7696 | 902 | 0 | 0 | 0 | 0 | 0 | 0 |
| ENSMUSG00 | Gm20387 | 1780 | 0 | 0 | 0 | 0 | 0 | 0 |
| ENSMUSG00 | Mir196a-1 | 101 | 0 | 0 | 0 | 0 | 0 | 0 |
| ENSMUSG00 | Vmn1r-ps31 | 237 | 0 | 0 | 0 | 0 | 0 | 0 |
| ENSMUSG00 | Vmn2r-ps56 | 2589 | 0 | 0 | 0 | 0 | 0 | 0 |
| ENSMUSG00 | Vmn2r-ps45 | 2429 | 0 | 0 | 0 | 0 | 0 | 0 |
| ENSMUSG00 | Gm5726 | 923 | 0 | 0 | 0 | 0 | 0 | 0 |
| ENSMUSG00 | Gm2991 | 479 | 0 | 0 | 0 | 0 | 0 | 0 |
| ENSMUSG00 | Gm20456 | 383 | 0.20760042 | 0 | 0 | 0 | 1.19905322 | 0.9969842 |
| ENSMUSG00 | V1rd19 | 1156 | 0 | 0 | 0 | 0 | 0 | 0 |
| ENSMUSG00 | Gm8835 | 908 | 0 | 0 | 0 | 0 | 0 | 0 |
| ENSMUSG00 | Gm18737 | 936 | 0 | 0 | 0 | 0.11564758 | 0.10903072 | 0 |
| ENSMUSG00 | Gm2302 | 2239 | 0 | 0 | 0 | 0 | 0 | 0 |
| ENSMUSG00 | Gm8878 | 1723 | 0.02307341 | 0 | 0 | 0 | 0 | 0 |
| ENSMUSG00 | Gm18833 | 861 | 0 | 0 | 0 | 0 | 0 | 0 |
| ENSMUSG00 | Vmn1r-ps83 | 704 | 0 | 0 | 0 | 0 | 0 | 0 |
| ENSMUSG00 | Gm20489 | 1338 | 0 | 0 | 0 | 0 | 0 | 0 |
| ENSMUSG00 | Gm8902 | 1094 | 0 | 0 | 0 | 0.1649088 | 0.04664203 | 0.07479334 |
| ENSMUSG00 | Gm4425 | 1234 | 0 | 0 | 0 | 0 | 0 | 0 |
| ENSMUSG00 | Vmn1r-ps57 | 287 | 0 | 0 | 0 | 0 | 0 | 0 |
| ENSMUSG00 | Gm18360 | 1522 | 0 | 0 | 0 | 0 | 0 | 0 |
| ENSMUSG00 | Gm18520 | 657 | 0 | 0 | 0 | 0 | 0 | 0 |
| ENSMUSG00 | Gm20395 | 687 | 1.21523303 | 1.75350954 | 1.73719034 | 0.52521171 | 0.1485484 | 1.31013551 |
| ENSMUSG00 | Gm20518 | 3795 | 12.2147273 | 22.9441109 | 5.26462004 | 17.4087546 | 14.144861 | 5.63460268 |
| ENSMUSG00 | Cyp21a2-ps | 1374 | 0 | 0 | 0 | 0 | 0 | 0 |
| ENSMUSG00 | Gm20473 | 111 | 0 | 0 | 0 | 0 | 0 | 0 |
| ENSMUSG00 | Vmn1r180 | 969 | 0 | 0 | 0 | 0 | 0 | 0 |
| ENSMUSG00 | Gm20478 | 1028 | 0 | 0 | 0 | 0 | 0 | 0 |
| ENSMUSG00 | 9930038K12I | 2285 | 0.60894609 | 1.09658424 | 0.6770523 | 0.28423492 | 0.96023377 | 1.27719317 |
| ENSMUSG00 | Vmn1r-ps65 | 950 | 0 | 0 | 0 | 0 | 0 | 0 |
| ENSMUSG00 | Gm18808 | 459 | 0 | 0 | 0 | 0 | 0 | 0 |
| ENSMUSG00 | Gm8909 | 1530 | 0 | 0 | 0 | 0 | 0 | 0 |
| ENSMUSG00 | Gm8677 | 923 | 0 | 0 | 0 | 0 | 0 | 0 |
| ENSMUSG00 | Gm20495 | 234 | 0 | 0 | 0 | 0 | 0 | 0 |
| ENSMUSG00 | Gm20435 | 1707 | 0 | 0 | 0 | 0 | 0 | 0 |
| ENSMUSG00 | Gm20531 | 1371 | 0 | 0.10544079 | 0 | 0.02631805 | 0.11165509 | 0 |
| ENSMUSG00 | Gm20421 | 670 | 0 | 0 | 0 | 0 | 0 | 0 |
| ENSMUSG00 | Gm20487 | 414 | 0 | 0 | 0.10676774 | 0 | 0 | 0 |
| ENSMUSG00 | Rpl9-ps5 | 570 | 0 | 0 | 0 | 0 | 0 | 0 |
| ENSMUSG00 | 2610524H06I | 945 | 4.4593449 | 8.66846049 | 7.10971443 | 7.06368067 | 8.63938638 | 13.7383368 |
| ENSMUSG00 | Vmn1r-ps71 | 923 | 0 | 0 | 0 | 0 | 0 | 0 |
| ENSMUSG00 | Gm18159 | 1069 | 0 | 0 | 0 | 0 | 0 | 0 |
| ENSMUSG00 | Vmn1r-ps53 | 921 | 0 | 0 | 0 | 0 | 0 | 0 |
| ENSMUSG00 | Gm20482 | 1715 | 0 | 0 | 0 | 0 | 0.02975299 | 0 |
| ENSMUSG00 | Nrap | 5658 | 0 | 0 | 0 | 0 | 0 | 0 |
| ENSMUSG00 | B230208B08I | 402 | 0 | 0 | 0 | 0 | 0 | 0 |
| ENSMUSG00 | Gm20451 | 271 | 0 | 0 | 0 | 0 | 0 | 0 |
| ENSMUSG00 | Gm20420 | 637 | 0.12482098 | 0 | 0 | 0 | 0 | 0 |
| ENSMUSG00 | Gm20405 | 227 | 0 | 0 | 0 | 0.47685521 | 0.67435739 | 0.1201526 |
| ENSMUSG00 | Vmn1r-ps84 | 922 | 0 | 0 | 0 | 0 | 0 | 0 |
| ENSMUSG00 | Gm9286 | 832 | 0 | 0 | 0 | 0 | 0 | 0 |
| ENSMUSG00 | Vmn1r109 | 893 | 0 | 0 | 0 | 0 | 0 | 0 |
| ENSMUSG00 | 1700092C10I | 582 | 0 | 0 | 0 | 0 | 0 | 0 |
| ENSMUSG00 | Gm20400 | 662 | 0 | 0 | 0 | 0 | 0.15415824 | 0.04120036 |
| ENSMUSG00 | Gm4172 | 920 | 0 | 0 | 0 | 0 | 0 | 0 |
| ENSMUSG00 | Gm4252 | 312 | 0 | 0 | 0 | 0 | 0 | 0 |
| ENSMUSG00 | Gm20484 | 229 | 0 | 0 | 0 | 0 | 0 | 0 |
| ENSMUSG00 | Gm20396 | 743 | 0 | 0 | 0 | 0 | 0 | 0 |
| ENSMUSG00 | Gm19246 | 254 | 0 | 0.1897104 | 0.52206901 | 0.14205529 | 0.20089124 | 0 |
| ENSMUSG00 | Vmn1r-ps77 | 920 | 0 | 0 | 0 | 0 | 0 | 0 |
| ENSMUSG00 | Gm4558 | 547 | 0 | 0 | 0 | 0 | 0 | 0 |
| ENSMUSG00 | Gm4256 | 934 | 0 | 0 | 0 | 0 | 0 | 0 |
| ENSMUSG00 | Gm20394 | 940 | 0 | 0 | 0.28213942 | 0 | 0.05428338 | 0.17409344 |
| ENSMUSG00 | Gm18343 | 603 | 0.06592949 | 0 | 0 | 0 | 0.08462086 | 0 |
| ENSMUSG00 | Gm20547 | 4027 | 0 | 0 | 0 | 0 | 0 | 0 |
| ENSMUSG00 | Gm4152 | 258 | 0 | 0.37353831 | 0 | 0 | 0 | 0 |
| ENSMUSG00 | Gm20542 | 371 | 0 | 0 | 0 | 0 | 0 | 0 |
| ENSMUSG00 | Vmn1r-ps86 | 546 | 0 | 0 | 0 | 0 | 0 | 0 |
| ENSMUSG00 | C87198 | 1040 | 0 | 0 | 0 | 0 | 0 | 0 |
| ENSMUSG00 | Gm20397 | 361 | 0 | 0 | 0 | 0 | 0 | 0 |
| ENSMUSG00 | Art2a-ps | 1253 | 0 | 0 | 0 | 0 | 0 | 0 |
| ENSMUSG00 | Fam71e2 | 2733 | 0 | 0 | 0 | 0 | 0 | 0 |
| ENSMUSG00 | Actl9 | 1371 | 0 | 0 | 0 | 0 | 0 | 0 |
| ENSMUSG00 | Gm3611 | 872 | 0 | 0 | 0 | 0 | 0 | 0 |
| ENSMUSG00 | Gm4271 | 673 | 0 | 0 | 0 | 0 | 0 | 0 |
| ENSMUSG00 | Gm20389 | 1004 | 0 | 0 | 0 | 0 | 0 | 0 |
| ENSMUSG00 | Gm18191 | 820 | 0 | 0 | 0 | 0 | 0 | 0 |
| ENSMUSG00 | Olfr756-ps1 | 927 | 0 | 0 | 0 | 0 | 0 | 1.14747673 |
| ENSMUSG00 | Gm20461 | 303 | 0 | 0 | 0 | 0 | 0.33680776 | 0 |
| ENSMUSG00 | Gm17907 | 709 | 2.07468657 | 0.54371162 | 0 | 1.01782918 | 2.44696302 | 0 |
| ENSMUSG00 | Nlrp1c-ps | 3444 | 0 | 0 | 0 | 0 | 0 | 0 |
| ENSMUSG00 | Vmn2r-ps80 | 912 | 0 | 0 | 0 | 0 | 0 | 0 |
| ENSMUSG00 | Vmn2r-ps64 | 522 | 0 | 0 | 0 | 0 | 0 | 0 |
| ENSMUSG00 | Vmn2r-ps84 | 896 | 0 | 0 | 0 | 0 | 0 | 0 |
| ENSMUSG00 | Vmn2r-ps72 | 906 | 0 | 0 | 0 | 0 | 0 | 0 |
| ENSMUSG00 | Gm20418 | 688 | 0 | 0 | 0 | 0 | 0 | 0 |
| ENSMUSG00 | Vmn1r-ps61 | 821 | 0 | 0 | 0 | 0 | 0 | 0 |
| ENSMUSG00 | Gm20501 | 559 | 0 | 0 | 0 | 0 | 0 | 0 |
| ENSMUSG00 | Gm20504 | 1707 | 0 | 0 | 0 | 0 | 0 | 0 |
| ENSMUSG00 | Vmn2r-ps55 | 239 | 0 | 0 | 0 | 0 | 0 | 0 |
| ENSMUSG00 | Gm20468 | 449 | 0.17708455 | 0 | 0 | 0 | 0 | 0.54670769 |
| ENSMUSG00 | Gm8738 | 328 | 0.7272344 | 1.02836919 | 0.8085703 | 1.6500935 | 0.93340931 | 0.49892633 |
| ENSMUSG00 | Gm20537 | 1000 | 0 | 0 | 0 | 0 | 0 | 0 |
| ENSMUSG00 | Olfr757-ps1 | 368 | 0 | 0 | 0 | 0 | 0 | 0 |
| ENSMUSG00 | Skiv2l-ps1 | 653 | 0 | 0 | 0 | 0 | 0 | 0 |
| ENSMUSG00 | Gm20422 | 1760 | 0 | 0 | 0 | 0 | 0 | 0 |
| ENSMUSG00 | Gm20319 | 1430 | 0 | 0 | 0 | 0 | 0 | 0 |
| ENSMUSG00 | Gm20439 | 1707 | 0 | 0 | 0 | 0 | 0 | 0 |
| ENSMUSG00 | Vmn1r-ps80 | 965 | 0 | 0 | 0 | 0 | 0 | 0 |
| ENSMUSG00 | Vmn2r-ps41 | 978 | 0 | 0 | 0 | 0 | 0 | 0 |
| ENSMUSG00 | Gm20491 | 1240 | 0 | 0 | 0 | 0 | 0 | 0 |
| ENSMUSG00 | Gm20496 | 660 | 0 | 0 | 0 | 0 | 0 | 0 |
| ENSMUSG00 | Gm20287 | 2361 | 0 | 0 | 0 | 0 | 0 | 0 |
| ENSMUSG00 | Gm4287 | 584 | 0 | 0 | 0 | 0 | 0.08737393 | 0 |
| ENSMUSG00 | Gm20428 | 282 | 0.98683817 | 0 | 0.15674412 | 0 | 0 | 0 |
| ENSMUSG00 | Gm18128 | 510 | 0 | 0 | 0 | 0 | 0 | 0 |
| ENSMUSG00 | Gm318 | 1073 | 0 | 0 | 0 | 0 | 0 | 0 |
| ENSMUSG00 | Olfr755-ps1 | 906 | 0 | 0 | 0 | 0 | 0 | 0 |
| ENSMUSG00 | Gm8696 | 384 | 0.62117938 | 0 | 1.95685243 | 2.53701876 | 2.65762374 | 1.56260954 |
| ENSMUSG00 | Med20 | 6497 | 10.3167216 | 8.34381213 | 10.3139902 | 9.76331142 | 10.9168328 | 7.64882141 |
| ENSMUSG00 | Gm20475 | 261 | 0 | 0 | 0 | 0 | 0.97751678 | 0.73150374 |
| ENSMUSG00 | Gm8750 | 1533 | 0 | 0 | 0 | 0 | 0 | 0 |
| ENSMUSG00 | Gm4011 | 2549 | 0 | 0 | 0 | 0 | 0 | 0 |
| ENSMUSG00 | Gm20414 | 645 | 0 | 0 | 0 | 0 | 0 | 0 |
| ENSMUSG00 | Gm3617 | 554 | 0.50232557 | 0 | 0 | 0.06513004 | 0.18421074 | 0.34462541 |
| ENSMUSG00 | BC051226 | 1378 | 0.20195092 | 0.87420976 | 0.32076809 | 0.23565922 | 0.3332637 | 0.85109542 |
| ENSMUSG00 | Gm20477 | 279 | 0 | 0 | 0 | 0 | 0 | 0 |
| ENSMUSG00 | Gm20409 | 129 | 0 | 0 | 0 | 0 | 0 | 0 |
| ENSMUSG00 | Vmn2r-ps73 | 1191 | 0 | 0 | 0 | 0 | 0 | 0 |
| ENSMUSG00 | Olfr759-ps1 | 720 | 0 | 0 | 0 | 0 | 0 | 0 |
| ENSMUSG00 | Gm20544 | 688 | 0.34670477 | 0.14007687 | 0.83520924 | 1.20623114 | 0.51916371 | 0.79286742 |
| ENSMUSG00 | 2210019I11R | 1948 | 0 | 0 | 0 | 0 | 0 | 0 |
| ENSMUSG00 | Gm4134 | 560 | 0 | 0 | 0 | 0 | 0 | 0.09740943 |
| ENSMUSG00 | Serpinb10-ps | 3546 | 0 | 0 | 0 | 0 | 0 | 0 |
| ENSMUSG00 | Vmn2r-ps43 | 2468 | 0 | 0 | 0 | 0 | 0 | 0 |
| ENSMUSG00 | Gm18202 | 826 | 0 | 0 | 0 | 0 | 0 | 0 |
| ENSMUSG00 | Gm5583 | 1525 | 0 | 0 | 0 | 0 | 0 | 0 |
| ENSMUSG00 | Gm7235 | 856 | 0 | 0 | 0 | 0 | 0 | 0 |
| ENSMUSG00 | Gm7393 | 738 | 0 | 0 | 0 | 0.04889166 | 0.62227288 | 0 |
| ENSMUSG00 | 4930488B22I | 1719 | 0 | 0 | 0 | 0 | 0 | 0 |
| ENSMUSG00 | Vmn1r62 | 1756 | 0 | 0 | 0 | 0 | 0 | 0 |
| ENSMUSG00 | Vmn1r-ps45 | 954 | 0 | 0 | 0 | 0 | 0 | 0 |
| ENSMUSG00 | Gm3211 | 392 | 0 | 0 | 0 | 0 | 0 | 0 |
| ENSMUSG00 | Vmn1r158 | 923 | 0 | 0 | 0 | 0 | 0 | 0 |
| ENSMUSG00 | Vmn2r-ps53 | 2348 | 0 | 0 | 0 | 0 | 0 | 0 |
| ENSMUSG00 | Gm18199 | 831 | 0 | 0 | 0 | 0 | 0 | 0 |
| ENSMUSG00 | Gm20398 | 613 | 0 | 0 | 0 | 0 | 0 | 0 |
| ENSMUSG00 | Ly6g6c | 1356 | 0 | 0 | 0 | 0 | 0.2634105 | 0 |
| ENSMUSG00 | Vmn2r-ps85 | 939 | 0 | 0 | 0 | 0 | 0 | 0 |
| ENSMUSG00 | Gm20534 | 171 | 0 | 0 | 0 | 0 | 0 | 0 |
| ENSMUSG00 | Gm16451 | 923 | 0 | 0 | 0 | 0 | 0 | 0 |
| ENSMUSG00 | Gm20429 | 882 | 0 | 0 | 0 | 0 | 0 | 0 |
| ENSMUSG00 | Gm20449 | 3839 | 0.04142275 | 0.01255182 | 0.44904191 | 0.16917864 | 0.05316632 | 0.05683697 |
| ENSMUSG00 | Gm20492 | 284 | 0.13998409 | 1.52703514 | 0.77820146 | 0 | 0.89835169 | 2.20886163 |
| ENSMUSG00 | Gm4141 | 920 | 0 | 0 | 0 | 0 | 0 | 0 |
| ENSMUSG00 | Gm20427 | 3692 | 0.20459213 | 0.23492848 | 0.1197233 | 0.45933264 | 0.20731193 | 0.11081246 |
| ENSMUSG00 | Gm20490 | 196 | 0 | 0 | 0 | 0 | 0 | 0 |
| ENSMUSG00 | Gm20486 | 417 | 0 | 0 | 0 | 0 | 0 | 0 |
| ENSMUSG00 | Gm4201 | 920 | 0 | 0 | 0 | 0 | 0 | 0 |
| ENSMUSG00 | 1700010K23I | 373 | 0 | 0 | 0 | 0 | 0 | 0 |
| ENSMUSG00 | Gm20442 | 490 | 0 | 0 | 0 | 0 | 0 | 0 |
| ENSMUSG00 | Olfr760-ps1 | 556 | 0 | 0 | 0 | 0 | 0 | 0 |
| ENSMUSG00 | 4931413I07R | 762 | 0 | 0 | 0 | 0 | 0 | 0 |
| ENSMUSG00 | Gm20386 | 131 | 0.6069539 | 0 | 0 | 0 | 0 | 0 |
| ENSMUSG00 | Gm20508 | 485 | 0 | 0 | 0 | 0 | 0 | 0 |
| ENSMUSG00 | Gm20450 | 146 | 0 | 0 | 0 | 0 | 0 | 0 |
| ENSMUSG00 | Scnm1 | 1533 | 16.0266712 | 11.535828 | 19.9528215 | 20.85368 | 15.5442384 | 22.7911369 |
| ENSMUSG00 | Vmn2r-ps81 | 837 | 0 | 0 | 0 | 0 | 0 | 0 |
| ENSMUSG00 | Gm20481 | 562 | 0.07073929 | 0 | 0 | 0 | 0 | 0 |
| ENSMUSG00 | Vmn1r-ps67 | 690 | 0 | 0 | 0 | 0 | 0 | 0 |
| ENSMUSG00 | Vmn2r-ps42 | 2077 | 0 | 0 | 0 | 0 | 0 | 0 |
| ENSMUSG00 | Gm18604 | 307 | 0 | 0 | 0 | 0 | 0 | 0 |
| ENSMUSG00 | Gm16456 | 919 | 0 | 0 | 0 | 0 | 0 | 0 |
| ENSMUSG00 | Vmn2r-ps47 | 2550 | 0 | 0 | 0 | 0 | 0 | 0 |
| ENSMUSG00 | Olfr762-ps1 | 887 | 0 | 0 | 0 | 0 | 0 | 0 |
| ENSMUSG00 | Vmn1r-ps74 | 569 | 0 | 0 | 0 | 0 | 0 | 0 |
| ENSMUSG00 | Gm8412 | 439 | 0 | 0.10976411 | 0 | 0.08219145 | 0 | 0 |
| ENSMUSG00 | Btnl6 | 1742 | 0 | 0 | 0 | 0 | 0 | 0 |
| ENSMUSG00 | Gm20438 | 327 | 0 | 0 | 0 | 0 | 0 | 0 |
| ENSMUSG00 | Gm18201 | 870 | 0 | 0 | 0 | 0 | 0 | 0 |
| ENSMUSG00 | Vmn2r-ps66 | 1253 | 0 | 0 | 0 | 0 | 0 | 0 |
| ENSMUSG00 | 2410004A20I | 1547 | 0 | 0 | 0.1428631 | 0 | 0 | 0.38787464 |
| ENSMUSG00 | Gm20413 | 1785 | 0 | 0 | 0 | 0 | 0 | 0 |
| ENSMUSG00 | Gm3654 | 904 | 0 | 0 | 0 | 0 | 0.22578042 | 0 |
| ENSMUSG00 | Gm20404 | 616 | 1.22622424 | 1.17337116 | 0.35878119 | 2.5772889 | 0.66268021 | 2.83372875 |
| ENSMUSG00 | 9130230N09I | 661 | 2.52606684 | 1.82248268 | 1.40429456 | 1.69219876 | 0.77195727 | 3.42480342 |
| ENSMUSG00 | D130058E05I | 1370 | 0 | 0 | 0 | 0 | 0 | 0 |
| ENSMUSG00 | Vmn2r-ps71 | 874 | 0 | 0 | 0 | 0 | 0 | 0 |
| ENSMUSG00 | Vmn1r-ps85 | 889 | 0 | 0 | 0 | 0 | 0 | 0 |
| ENSMUSG00 | Gm9574 | 246 | 0 | 0 | 0 | 0 | 0 | 0 |
| ENSMUSG00 | Vmn1r-ps54 | 177 | 0 | 0 | 0 | 0 | 0 | 0 |
| ENSMUSG00 | AC154909.1 | 70 | 0 | 0 | 0 | 0 | 0 | 0 |
| ENSMUSG00 | Mir3962 | 63 | 0 | 0 | 0 | 0 | 0 | 0 |
| ENSMUSG00 | AC164093.1 | 125 | 0 | 0.38549154 | 0.70722949 | 0.57731271 | 1.22463302 | 1.09098557 |
| ENSMUSG00 | AL929466.1 | 268 | 0.74170673 | 0.71920063 | 0.16493225 | 0.13463449 | 0.19039693 | 0.61062625 |
| ENSMUSG00 | AC124193.1 | 91 | 0 | 0 | 0 | 0.39650598 | 0.56072941 | 0.29972131 |
| ENSMUSG00 | BX000454.1 | 101 | 0 | 0 | 0 | 0 | 0 | 0 |
| ENSMUSG00 | Mir5112 | 59 | 0 | 0 | 0 | 0 | 0 | 0 |
| ENSMUSG00 | Mir3964 | 73 | 0 | 0 | 0 | 0 | 0 | 0 |
| ENSMUSG00 | AC161342.1 | 77 | 0 | 0 | 0 | 0 | 0 | 0 |
| ENSMUSG00 | AL596258.1 | 248 | 0 | 0 | 0 | 0 | 0 | 0 |
| ENSMUSG00 | AC130216.1 | 125 | 0 | 0 | 0 | 0 | 0 | 0 |
| ENSMUSG00 | AC124746.1 | 61 | 0 | 0 | 0 | 0 | 0 | 0 |
| ENSMUSG00 | AL645623.1 | 66 | 1.8070673 | 0.73009761 | 2.00917468 | 4.37358115 | 3.09250763 | 0.82650422 |
| ENSMUSG00 | AC134325.1 | 70 | 1.70380631 | 2.06513324 | 1.8943647 | 1.03091556 | 1.45789645 | 0.77927541 |
| ENSMUSG00 | AC117644.1 | 87 | 0 | 0 | 0 | 0 | 0 | 0 |
| ENSMUSG00 | AC161253.1 | 64 | 0 | 0 | 0 | 0 | 0 | 0 |
| ENSMUSG00 | AC126547.1 | 107 | 0 | 0 | 0 | 0 | 0 | 0 |
| ENSMUSG00 | AC123616.1 | 256 | 0 | 0 | 0.51799035 | 0 | 0 | 0 |
| ENSMUSG00 | AC154824.1 | 83 | 0 | 0 | 0 | 0 | 0 | 0 |
| ENSMUSG00 | AL935155.1 | 288 | 0 | 0 | 0 | 0 | 0 | 0 |
| ENSMUSG00 | Mir3104 | 62 | 0 | 0 | 0 | 0 | 0 | 0 |
| ENSMUSG00 | AC159204.1 | 220 | 0 | 0 | 0 | 0 | 0 | 0 |
| ENSMUSG00 | AL831731.1 | 71 | 0 | 0 | 0 | 0.50819781 | 0 | 0 |
| ENSMUSG00 | AC108409.1 | 88 | 0 | 0 | 0 | 0 | 0 | 0 |
| ENSMUSG00 | AC102525.1 | 85 | 0 | 0 | 0 | 0 | 0 | 0 |
| ENSMUSG00 | AC122925.1 | 124 | 0 | 0 | 0 | 0 | 0 | 0 |
| ENSMUSG00 | AC166781.1 | 70 | 0 | 0 | 0 | 0 | 0 | 0 |
| ENSMUSG00 | Mir3100 | 64 | 0 | 0 | 0 | 0 | 0 | 0 |
| ENSMUSG00 | AL845291.1 | 83 | 2.87389016 | 0 | 0 | 0.86944686 | 0.61477561 | 0 |
| ENSMUSG00 | AL591611.1 | 109 | 0 | 0 | 0 | 0 | 0 | 0 |
| ENSMUSG00 | AC087556.1 | 256 | 0 | 0 | 0 | 0 | 0 | 0 |
| ENSMUSG00 | AC140340.1 | 92 | 0 | 0 | 0 | 0 | 0 | 0 |
| ENSMUSG00 | AC099934.1 | 296 | 0 | 0 | 0 | 0 | 0 | 0 |
| ENSMUSG00 | AC165162.1 | 64 | 0 | 0 | 0 | 0 | 0 | 0 |
| ENSMUSG00 | AC159897.1 | 285 | 0 | 0 | 0 | 0 | 0.17903992 | 0 |
| ENSMUSG00 | AC126033.1 | 116 | 0 | 0 | 0 | 0 | 0 | 0 |
| ENSMUSG00 | AL596111.1 | 267 | 0 | 0 | 0 | 0 | 0 | 0 |
| ENSMUSG00 | AC111135.1 | 125 | 0 | 0 | 0 | 0 | 0 | 0 |
| ENSMUSG00 | AC093451.1 | 76 | 0 | 0 | 0 | 0 | 0 | 0 |
| ENSMUSG00 | AL606976.2 | 258 | 0 | 0 | 0 | 0 | 0 | 0 |
| ENSMUSG00 | Mir3067 | 83 | 0 | 0 | 0 | 0 | 0 | 0 |
| ENSMUSG00 | AC122546.1 | 290 | 0 | 0 | 0 | 0 | 0 | 0 |
| ENSMUSG00 | AL732626.1 | 132 | 0 | 0 | 0 | 0 | 0 | 0 |
| ENSMUSG00 | AC141876.1 | 77 | 0 | 0 | 0 | 0 | 0 | 0 |
| ENSMUSG00 | Mir5106 | 72 | 1.65647836 | 0 | 3.06957243 | 0.50113951 | 0 | 3.40932991 |
| ENSMUSG00 | AC087117.1 | 69 | 0 | 0 | 0 | 0 | 0 | 0 |
| ENSMUSG00 | Mir5129 | 77 | 0 | 0 | 0 | 0 | 0 | 0 |
| ENSMUSG00 | Snord55 | 78 | 3.56779953 | 1.2355498 | 0 | 0.92518063 | 1.30836861 | 3.84642348 |
| ENSMUSG00 | AL645484.1 | 69 | 0 | 0 | 0 | 0 | 0 | 0 |
| ENSMUSG00 | AL671866.1 | 124 | 0 | 0 | 0 | 0 | 0 | 0 |
| ENSMUSG00 | AC073553.1 | 77 | 0 | 0 | 0 | 0 | 0 | 0 |
| ENSMUSG00 | AC158995.1 | 318 | 0 | 0 | 0 | 0 | 0 | 0 |
| ENSMUSG00 | AC160633.1 | 100 | 0 | 0 | 0 | 0 | 0 | 0 |
| ENSMUSG00 | AC152453.1 | 132 | 0 | 0 | 0 | 0 | 0 | 0 |
| ENSMUSG00 | AC087117.2 | 62 | 0 | 0 | 0 | 0 | 0 | 0 |
| ENSMUSG00 | AC102320.1 | 101 | 0 | 0 | 0 | 0 | 0 | 0 |
| ENSMUSG00 | AL844842.1 | 125 | 0 | 0 | 0 | 0 | 0 | 0 |
| ENSMUSG00 | Mir5111 | 66 | 0 | 0 | 0 | 0.54669764 | 0 | 0 |
| ENSMUSG00 | CT030723.1 | 91 | 0 | 0 | 0 | 0 | 0 | 0 |
| ENSMUSG00 | AC129776.1 | 109 | 0 | 0 | 0 | 0 | 0 | 0 |
| ENSMUSG00 | AL450399.1 | 69 | 0 | 0 | 0 | 0 | 0 | 0 |
| ENSMUSG00 | Mir3092 | 95 | 0 | 0 | 0 | 0 | 0 | 0 |
| ENSMUSG00 | Mir5133 | 76 | 1.04619686 | 1.26806427 | 1.1632064 | 0 | 4.02839809 | 4.66539882 |
| ENSMUSG00 | AL772216.40 | 101 | 0 | 0 | 0 | 0 | 0 | 0 |
| ENSMUSG00 | AC022236.1 | 67 | 0 | 0 | 0 | 0 | 0 | 0 |
| ENSMUSG00 | AC161805.1 | 70 | 0 | 0 | 1.2629098 | 0 | 0 | 0 |
| ENSMUSG00 | AC098732.1 | 108 | 0 | 6.24639065 | 4.09276324 | 0 | 0 | 0 |
| ENSMUSG00 | AC149599.1 | 287 | 0 | 0 | 0 | 0 | 0 | 0.19006717 |
| ENSMUSG00 | AC112688.1 | 64 | 0 | 0 | 0 | 0 | 0 | 0 |
| ENSMUSG00 | AL669947.1 | 137 | 0 | 0 | 0 | 0 | 0 | 0 |
| ENSMUSG00 | CT025708.1 | 93 | 0 | 0 | 0 | 0 | 0 | 0 |
| ENSMUSG00 | AL683847.1 | 234 | 0 | 0 | 1.32227736 | 0 | 0 | 0.23311658 |
| ENSMUSG00 | AC103368.1 | 275 | 0 | 0 | 0.16073398 | 0 | 0 | 0 |
| ENSMUSG00 | AC102862.1 | 133 | 0 | 2.53612853 | 0 | 0 | 0 | 0 |
| ENSMUSG00 | AC160138.1 | 70 | 0 | 0 | 0 | 0 | 0 | 0 |
| ENSMUSG00 | AL824704.1 | 59 | 2.02146511 | 0 | 0 | 0 | 0 | 0 |
| ENSMUSG00 | CT010429.1 | 75 | 2.12029229 | 0 | 0 | 0 | 0 | 1.81830928 |
| ENSMUSG00 | AC162941.1 | 83 | 0 | 0 | 0 | 0 | 0 | 0 |
| ENSMUSG00 | AC119801.1 | 130 | 0 | 0 | 0 | 0 | 0 | 0 |
| ENSMUSG00 | AL772272.1 | 92 | 0 | 0 | 0 | 0 | 0 | 0 |
| ENSMUSG00 | AC158354.1 | 109 | 0 | 0 | 0 | 0 | 0 | 0 |
| ENSMUSG00 | AL683823.1 | 73 | 0 | 0 | 0 | 0 | 0 | 0 |
| ENSMUSG00 | AL833780.1 | 121 | 0 | 0 | 0 | 0 | 0 | 0 |
| ENSMUSG00 | AC134596.1 | 117 | 0 | 0 | 0 | 0 | 0 | 0 |
| ENSMUSG00 | AL807764.1 | 332 | 0 | 0 | 0 | 0 | 0 | 0 |
| ENSMUSG00 | AC121953.1 | 66 | 0 | 0 | 0 | 0 | 0 | 0 |
| ENSMUSG00 | AL626784.1 | 84 | 0 | 0 | 0 | 0 | 0 | 0 |
| ENSMUSG00 | AL773548.1 | 71 | 0 | 0 | 0 | 0 | 0 | 0 |
| ENSMUSG00 | AL808023.1 | 72 | 0 | 0 | 0 | 0 | 0 | 0 |
| ENSMUSG00 | AC141881.1 | 118 | 0 | 0 | 0 | 0 | 0 | 0 |
| ENSMUSG00 | Mir3062 | 104 | 0 | 0 | 0 | 0 | 0 | 0 |
| ENSMUSG00 | AC118632.1 | 197 | 0 | 0 | 0 | 0 | 0 | 0 |
| ENSMUSG00 | AC166252.1 | 100 | 0 | 0 | 0 | 0 | 0 | 0 |
| ENSMUSG00 | Snora24 | 129 | 0.61636404 | 0 | 0.34264995 | 0.83911732 | 0.3955533 | 0 |
| ENSMUSG00 | AC165951.1 | 97 | 0 | 0 | 0 | 0 | 0 | 0 |
| ENSMUSG00 | AC154226.1 | 75 | 0 | 0 | 0 | 0 | 0 | 0 |
| ENSMUSG00 | CT572991.1 | 107 | 0 | 0 | 0 | 0 | 0 | 0 |
| ENSMUSG00 | Mir5100 | 63 | 0 | 0 | 1.40323311 | 0 | 0 | 0 |
| ENSMUSG00 | AC166064.1 | 101 | 0 | 0 | 0 | 0 | 0 | 0 |
| ENSMUSG00 | AC127313.1 | 248 | 0 | 0.19430017 | 0 | 0 | 0 | 0 |
| ENSMUSG00 | AC073553.2 | 69 | 0 | 0 | 0 | 0 | 0 | 0 |
| ENSMUSG00 | Mir5135 | 78 | 0 | 1.8533247 | 2.26676118 | 0 | 0 | 0.69934972 |
| ENSMUSG00 | AC122362.1 | 101 | 0 | 0 | 0 | 0 | 0 | 0 |
| ENSMUSG00 | AC134382.1 | 110 | 0 | 0 | 0 | 0 | 0 | 0 |
| ENSMUSG00 | Mir3074-1 | 116 | 11.3097488 | 13.708212 | 14.4799141 | 13.0641885 | 16.2756544 | 12.6968148 |
| ENSMUSG00 | AC167016.1 | 100 | 0 | 0 | 0 | 0 | 0 | 0 |
| ENSMUSG00 | AC101659.1 | 157 | 0 | 0 | 0 | 0 | 0 | 0 |
| ENSMUSG00 | Mir5122 | 88 | 0 | 0 | 0 | 0 | 0 | 0 |
| ENSMUSG00 | AC156953.1 | 281 | 0 | 0 | 0 | 0 | 0 | 0 |
| ENSMUSG00 | AL671173.1 | 58 | 0 | 0 | 0 | 0 | 0 | 0 |
| ENSMUSG00 | AC073553.3 | 77 | 0 | 0 | 0 | 0 | 0 | 0 |
| ENSMUSG00 | AC152826.1 | 249 | 0 | 0 | 0.17751744 | 0 | 0 | 0 |
| ENSMUSG00 | AC158898.1 | 280 | 0 | 0 | 0.15786373 | 0 | 0 | 0 |
| ENSMUSG00 | Mir3080 | 78 | 0 | 0 | 0 | 0 | 0 | 0 |
| ENSMUSG00 | Mir3473 | 77 | 0 | 0 | 0 | 0 | 0 | 0 |
| ENSMUSG00 | AL805972.1 | 84 | 0 | 0 | 0 | 0 | 0 | 0 |
| ENSMUSG00 | AC121599.1 | 106 | 1.87525852 | 0.45458908 | 0.83399704 | 0 | 1.44414271 | 3.60231084 |
| ENSMUSG00 | AC133195.1 | 272 | 0 | 0 | 0 | 0 | 0 | 0 |
| ENSMUSG00 | AC190403.1 | 285 | 0 | 1.85982759 | 0 | 0 | 0 | 0 |
| ENSMUSG00 | AL807778.1 | 78 | 0 | 0 | 0 | 0 | 0 | 0 |
| ENSMUSG00 | AL672100.1 | 187 | 0 | 0.51536302 | 1.65461444 | 1.9295211 | 1.63720992 | 0.72926843 |
| ENSMUSG00 | AC155162.1 | 110 | 0 | 0 | 0 | 0 | 0 | 0 |
| ENSMUSG00 | AC153910.1 | 124 | 0 | 0 | 0.71293295 | 1.45492115 | 0 | 0 |
| ENSMUSG00 | Mir5120 | 77 | 0 | 0 | 0 | 0 | 0 | 0 |
| ENSMUSG00 | AC132957.1 | 65 | 0 | 0 | 0 | 0 | 0 | 0 |
| ENSMUSG00 | BX005053.1 | 88 | 0 | 0 | 0 | 0 | 0 | 0 |
| ENSMUSG00 | AC122546.2 | 97 | 0 | 0 | 0 | 0 | 0 | 0 |
| ENSMUSG00 | AL808027.1 | 81 | 0 | 0 | 0 | 0 | 0 | 0 |
| ENSMUSG00 | AC122448.1 | 60 | 0 | 0 | 0 | 0 | 0 | 0 |
| ENSMUSG00 | AC162038.1 | 72 | 0 | 0 | 0 | 0 | 0 | 0 |
| ENSMUSG00 | AL603706.1 | 76 | 0 | 0 | 0 | 0 | 0 | 0 |
| ENSMUSG00 | AC162387.1 | 95 | 0 | 0 | 0 | 0 | 0 | 0 |
| ENSMUSG00 | Mirlet7a-2 | 95 | 0 | 0 | 0 | 0 | 0 | 0 |
| ENSMUSG00 | AC149593.1 | 262 | 0 | 0.18391772 | 0 | 0 | 0 | 0 |
| ENSMUSG00 | AL596130.1 | 70 | 0 | 0 | 0 | 0 | 0 | 0 |
| ENSMUSG00 | AC157822.1 | 296 | 0 | 0.4883761 | 0 | 0 | 0 | 0 |
| ENSMUSG00 | AC126690.2 | 114 | 0 | 0 | 0 | 0 | 0 | 0 |
| ENSMUSG00 | AC147101.1 | 280 | 0 | 0 | 0 | 0 | 0 | 0 |
| ENSMUSG00 | CT033779.1 | 100 | 0 | 0 | 0 | 0 | 0 | 0 |
| ENSMUSG00 | AL671889.1 | 70 | 0 | 0 | 0 | 0 | 0 | 0 |
| ENSMUSG00 | AC104325.1 | 81 | 0 | 0 | 0 | 0 | 0 | 0 |
| ENSMUSG00 | AC122218.1 | 82 | 0.48482293 | 0 | 0 | 0 | 0 | 0 |
| ENSMUSG00 | AL662835.1 | 70 | 0 | 0 | 0 | 0 | 0 | 0 |
| ENSMUSG00 | Mir3057 | 90 | 4.85900317 | 5.88945404 | 4.4201843 | 4.00911606 | 7.93743624 | 8.18239178 |
| ENSMUSG00 | AC141881.2 | 108 | 0 | 0 | 0 | 0 | 0 | 0 |
| ENSMUSG00 | Mir3077 | 63 | 0 | 0 | 0 | 0 | 0 | 0 |
| ENSMUSG00 | AL672076.1 | 294 | 0 | 0 | 0 | 0 | 0 | 0 |
| ENSMUSG00 | AC134576.1 | 66 | 1.20471153 | 0 | 3.34862447 | 0 | 0 | 0 |
| ENSMUSG00 | Mir5116 | 62 | 0 | 0 | 0 | 0 | 0 | 0 |
| ENSMUSG00 | AL591905.1 | 124 | 0 | 0 | 0 | 0 | 0 | 0 |
| ENSMUSG00 | AL732624.1 | 71 | 0 | 0 | 0 | 0 | 0 | 0 |
| ENSMUSG00 | AC122246.1 | 441 | 0 | 0 | 0 | 0 | 0 | 0 |
| ENSMUSG00 | AL591584.1 | 124 | 0 | 0 | 0 | 0 | 0 | 0 |
| ENSMUSG00 | AC151406.1 | 69 | 0 | 0 | 0 | 0 | 0 | 0 |
| ENSMUSG00 | Mir466m | 83 | 0 | 0 | 0 | 0 | 0 | 0 |
| ENSMUSG00 | Mir3110 | 79 | 0 | 0 | 0 | 0 | 0 | 0 |
| ENSMUSG00 | AL928551.1 | 120 | 0 | 0.80310737 | 0 | 0 | 0 | 0 |
| ENSMUSG00 | AL589902.1 | 124 | 0 | 0 | 0 | 0 | 0 | 0 |
| ENSMUSG00 | AC165333.1 | 259 | 0.46048819 | 0 | 0 | 0 | 0 | 0.73715241 |
| ENSMUSG00 | AL627204.1 | 276 | 0 | 0 | 0 | 0 | 0 | 0 |
| ENSMUSG00 | AC165361.1 | 116 | 0 | 0 | 0 | 0 | 0 | 0 |
| ENSMUSG00 | AL671914.1 | 244 | 0 | 0 | 0 | 0 | 0 | 0 |
| ENSMUSG00 | AL844882.1 | 125 | 0 | 0 | 0 | 0 | 0 | 0 |
| ENSMUSG00 | AC124394.1 | 125 | 0 | 0 | 0 | 0 | 0 | 0 |
| ENSMUSG00 | AC084162.1 | 77 | 0 | 0 | 0 | 0 | 0 | 0 |
| ENSMUSG00 | AC118293.1 | 70 | 0 | 0 | 0 | 0 | 0 | 0 |
| ENSMUSG00 | CT025673.1 | 298 | 0.53363061 | 0.48509841 | 0.59331333 | 0 | 0.3424589 | 0 |
| ENSMUSG00 | AC164649.1 | 100 | 0 | 0 | 0 | 0 | 0 | 0 |
| ENSMUSG00 | Mir199b | 109 | 0 | 0 | 0 | 0 | 0 | 0 |
| ENSMUSG00 | AC083946.1 | 100 | 0 | 0 | 0 | 0 | 0 | 0 |
| ENSMUSG00 | Mir5123 | 82 | 0 | 0 | 0 | 0 | 0 | 0 |
| ENSMUSG00 | AC164408.1 | 83 | 0 | 0 | 0 | 0 | 0 | 0 |
| ENSMUSG00 | AL928693.1 | 94 | 0 | 0 | 0 | 0 | 0 | 0 |
| ENSMUSG00 | AL669982.1 | 90 | 0 | 0 | 0 | 0 | 0 | 0 |
| ENSMUSG00 | Mir3086 | 86 | 0 | 0 | 0 | 0 | 0 | 0 |
| ENSMUSG00 | AL928557.1 | 125 | 1.27217538 | 0.38549154 | 1.06084423 | 0.86596907 | 0 | 0 |
| ENSMUSG00 | Mir3099 | 83 | 0 | 0 | 0 | 0 | 0 | 0 |
| ENSMUSG00 | AC102618.1 | 93 | 0 | 0 | 0 | 0 | 0 | 0 |
| ENSMUSG00 | AL670236.2 | 125 | 0 | 0 | 0 | 0 | 0 | 0 |
| ENSMUSG00 | CU062626.1 | 47 | 0 | 0 | 4.70232373 | 0 | 0 | 0 |
| ENSMUSG00 | AC150277.1 | 131 | 1.2139078 | 0 | 0.6748373 | 0.82630636 | 1.16854296 | 0.20820335 |
| ENSMUSG00 | AC124472.1 | 244 | 0 | 0 | 0 | 0.4436317 | 0 | 0 |
| ENSMUSG00 | AC123838.1 | 100 | 0 | 0 | 0 | 0 | 0 | 0 |
| ENSMUSG00 | AL807379.1 | 109 | 0 | 3.09454216 | 0 | 0 | 0 | 0.75067814 |
| ENSMUSG00 | AL672228.1 | 71 | 0 | 0 | 0 | 0 | 0 | 0 |
| ENSMUSG00 | CT010479.2 | 89 | 0 | 0.5414207 | 0 | 0 | 0 | 0 |

| | | | | | | | |
|---|---|---|---|---|---|---|---|
| ENSMUSG00 AL928905.1 | 110 | 0 | 0 | 0 | 0 | 0 | 0 |
| ENSMUSG00 AC161765.1 | 108 | 0 | 0 | 0 | 0 | 0 | 0 |
| ENSMUSG00 Mir3091 | 75 | 0 | 0 | 2.35743163 | 0 | 0.68035168 | 1.09098557 |
| ENSMUSG00 AC154187.1 | 82 | 0 | 0 | 0 | 0 | 0 | 0 |
| ENSMUSG00 AL954675.1 | 76 | 0 | 0 | 0 | 0 | 0 | 0 |
| ENSMUSG00 Mir3963 | 70 | 0 | 0 | 0 | 0 | 0 | 0 |
| ENSMUSG00 AC114668.1 | 288 | 0 | 0 | 0 | 0 | 0 | 0 |
| ENSMUSG00 Mir3088 | 73 | 0 | 0 | 0 | 0 | 0 | 0 |
| ENSMUSG00 AL808119.1 | 90 | 0 | 0 | 0 | 0 | 0 | 0 |
| ENSMUSG00 AC114406.1 | 160 | 0 | 0 | 0 | 0 | 0 | 0 |
| ENSMUSG00 AC154848.1 | 165 | 0 | 0 | 0 | 0 | 0 | 0 |
| ENSMUSG00 AC103359.1 | 225 | 0 | 0 | 0 | 0 | 0 | 0 |
| ENSMUSG00 AC027184.1 | 318 | 0 | 0 | 0.13899951 | 0 | 0 | 0.17153861 |
| ENSMUSG00 AL671885.1 | 77 | 0 | 0 | 0 | 0 | 0 | 0 |
| ENSMUSG00 AL606908.1 | 294 | 0 | 0.16389946 | 0 | 0 | 0 | 0 |
| ENSMUSG00 CT033785.1 | 66 | 0 | 0 | 0 | 0 | 0 | 0 |
| ENSMUSG00 AC124992.1 | 75 | 0 | 0 | 0 | 0 | 0 | 0 |
| ENSMUSG00 AC102287.1 | 83 | 0 | 0 | 0 | 0 | 0 | 0 |
| ENSMUSG00 AL935159.1 | 101 | 0 | 0 | 0 | 0 | 0 | 0 |
| ENSMUSG00 AL592465.1 | 78 | 0 | 0 | 0 | 0 | 0 | 0 |
| ENSMUSG00 AL732585.1 | 66 | 0 | 0 | 0 | 0 | 0 | 0.82650422 |
| ENSMUSG00 AC107711.1 | 70 | 0 | 0 | 0 | 0 | 0 | 0 |
| ENSMUSG00 Mir344d-1 | 68 | 0 | 0 | 0 | 0 | 0 | 0 |
| ENSMUSG00 AL671011.1 | 75 | 0 | 0 | 0 | 0 | 0 | 0 |
| ENSMUSG00 AL831766.1 | 77 | 0 | 0 | 0 | 0 | 0 | 0 |
| ENSMUSG00 Mir344g | 70 | 0 | 0 | 0 | 0 | 0 | 0 |
| ENSMUSG00 AL844840.1 | 124 | 0 | 0 | 0 | 0 | 0 | 0 |
| ENSMUSG00 Mir5103 | 78 | 0 | 0 | 0 | 0 | 0 | 1.04902459 |
| ENSMUSG00 AL772185.1 | 101 | 0 | 0 | 0 | 0 | 0 | 0 |
| ENSMUSG00 AC124745.1 | 74 | 0 | 0 | 0 | 0 | 0.68954562 | 0 |
| ENSMUSG00 AC152394.1 | 70 | 0 | 0 | 0 | 0 | 0 | 0 |
| ENSMUSG00 AC099728.1 | 204 | 0 | 0 | 0 | 0 | 0 | 0 |
| ENSMUSG00 AC158898.2 | 79 | 0 | 0 | 0 | 0 | 0 | 0 |
| ENSMUSG00 AL844147.1 | 66 | 0 | 0 | 0 | 0 | 0 | 0 |
| ENSMUSG00 AC139317.1 | 86 | 0 | 0 | 0 | 0 | 0 | 0 |
| ENSMUSG00 CR956641.1 | 294 | 0 | 0 | 0 | 0 | 0 | 0 |
| ENSMUSG00 Mir3094 | 63 | 0 | 0 | 2.10484967 | 2.29092346 | 0 | 0.43293078 |
| ENSMUSG00 BX000694.1 | 271 | 0 | 0 | 0 | 0 | 0 | 0 |
| ENSMUSG00 AC133502.1 | 70 | 0 | 0 | 0 | 0 | 0 | 0 |
| ENSMUSG00 AL833775.1 | 82 | 0 | 0 | 0 | 0 | 0 | 0 |
| ENSMUSG00 AC144794.1 | 184 | 0 | 0 | 0 | 0 | 0 | 0 |
| ENSMUSG00 Mir3058 | 90 | 0 | 0.53540491 | 0.49113159 | 0.80182321 | 0 | 0.90915464 |
| ENSMUSG00 Mir5113 | 81 | 0 | 0 | 0 | 0 | 0 | 0 |
| ENSMUSG00 AL808027.2 | 84 | 0 | 0 | 0 | 0.8590963 | 0 | 0.32469809 |
| ENSMUSG00 Mir3061 | 90 | 0 | 0 | 0 | 0 | 0 | 0 |
| ENSMUSG00 AC138790.1 | 114 | 0 | 0 | 0 | 0 | 0 | 0 |
| ENSMUSG00 AL672094.1 | 69 | 0 | 0 | 0 | 0 | 0 | 0 |
| ENSMUSG00 AC110520.1 | 105 | 0 | 0 | 0 | 0 | 0 | 0 |
| ENSMUSG00 AL645950.1 | 125 | 0 | 0 | 0 | 0 | 0 | 0 |
| ENSMUSG00 AC149277.1 | 298 | 0 | 0 | 0 | 0 | 0 | 0 |
| ENSMUSG00 AC044864.1 | 107 | 0 | 0 | 0 | 0 | 0 | 0 |
| ENSMUSG00 AC166251.1 | 95 | 0 | 0 | 0 | 0 | 0 | 0 |
| ENSMUSG00 AL590633.1 | 65 | 0 | 0 | 0 | 0 | 0 | 0 |
| ENSMUSG00 BX005219.1 | 104 | 0 | 0 | 0 | 0 | 0 | 0 |
| ENSMUSG00 Mir3471-1 | 130 | 0 | 0 | 0 | 0 | 0 | 0 |
| ENSMUSG00 AC148768.1 | 100 | 0 | 0 | 0 | 0 | 0 | 0 |
| ENSMUSG00 AL772216.41 | 126 | 0 | 0 | 0 | 0 | 0 | 0 |
| ENSMUSG00 AC121802.1 | 70 | 0 | 0 | 0 | 0 | 0 | 0 |
| ENSMUSG00 AC125484.1 | 70 | 0 | 0 | 0 | 0 | 0 | 0 |
| ENSMUSG00 AL646020.1 | 52 | 55.0460499 | 135.292703 | 68.8528709 | 74.2457455 | 75.5582873 | 56.1228154 |
| ENSMUSG00 BX470220.1 | 69 | 0 | 0 | 0 | 0 | 0 | 0 |
| ENSMUSG00 Snord53 | 77 | 3.09782965 | 0.62579795 | 0 | 1.40579394 | 1.98804062 | 1.41686438 |
| ENSMUSG00 AC162464.1 | 199 | 0 | 0 | 0 | 0 | 0 | 0 |
| ENSMUSG00 AL669944.1 | 53 | 0 | 11.819316 | 20.0159289 | 4.76555305 | 12.5159035 | 10.2923167 |
| ENSMUSG00 AC120002.1 | 70 | 0 | 0 | 0 | 0 | 0 | 0 |
| ENSMUSG00 AC163023.1 | 73 | 0 | 0 | 0 | 0 | 0 | 0 |
| ENSMUSG00 Mir344d-3 | 79 | 0 | 0 | 0 | 0 | 0 | 0 |
| ENSMUSG00 AC124489.1 | 259 | 1.53496064 | 0.37209608 | 0 | 1.11450331 | 1.57610427 | 0.10530749 |
| ENSMUSG00 AL606479.2 | 76 | 0 | 0 | 0 | 0 | 0 | 0 |
| ENSMUSG00 AC127086.1 | 100 | 0 | 0 | 0 | 0 | 0 | 0 |
| ENSMUSG00 CT030685.1 | 80 | 0 | 0 | 0 | 0 | 0 | 0 |
| ENSMUSG00 AC156551.1 | 83 | 0 | 0 | 0 | 0 | 0 | 0 |
| ENSMUSG00 AL672284.1 | 108 | 0 | 0 | 0 | 0 | 0 | 0 |
| ENSMUSG00 AC113276.1 | 90 | 0 | 0 | 0 | 0 | 0 | 0 |
| ENSMUSG00 AC134897.1 | 101 | 0 | 0 | 0 | 0 | 0 | 0 |
| ENSMUSG00 Mir5130 | 83 | 0.47898169 | 1.74167863 | 1.59765698 | 5.21668114 | 0.61477561 | 4.60054156 |
| ENSMUSG00 AC132606.1 | 125 | 0 | 0 | 0 | 0 | 0 | 0 |
| ENSMUSG00 AC142212.1 | 302 | 0 | 0 | 0 | 0 | 0 | 0 |
| ENSMUSG00 AC124426.2 | 109 | 0 | 0 | 0 | 0 | 0 | 0 |
| ENSMUSG00 AC122537.1 | 212 | 0 | 0 | 0 | 0 | 0 | 0 |
| ENSMUSG00 AC098743.1 | 100 | 0 | 0 | 0 | 0 | 0 | 0 |
| ENSMUSG00 Mir5132 | 70 | 0 | 0 | 0 | 0 | 0 | 0 |
| ENSMUSG00 CT030729.1 | 124 | 0 | 0 | 0 | 0 | 0 | 0 |
| ENSMUSG00 AL807811.1 | 59 | 0 | 0 | 0 | 0 | 0 | 0 |
| ENSMUSG00 AC146595.1 | 101 | 0 | 0 | 0 | 0 | 0 | 0 |
| ENSMUSG00 AC137107.1 | 72 | 0 | 0 | 0 | 0.50113951 | 0.70869966 | 0.37881443 |
| ENSMUSG00 AC125186.1 | 97 | 0.40985032 | 0 | 0 | 0 | 0 | 0 |
| ENSMUSG00 AL589878.1 | 115 | 0 | 0 | 0 | 0 | 0 | 0 |
| ENSMUSG00 AC153950.1 | 136 | 0 | 0 | 0 | 0 | 0 | 0 |
| ENSMUSG00 CT010508.1 | 119 | 0 | 0.80985617 | 0 | 0.90963138 | 0 | 0 |
| ENSMUSG00 AC125115.1 | 82 | 0 | 0 | 0 | 0 | 0 | 0 |
| ENSMUSG00 Mir3472 | 70 | 0 | 0 | 0 | 0 | 0 | 0 |
| ENSMUSG00 AL928590.1 | 124 | 0 | 0 | 0 | 0 | 0 | 0 |
| ENSMUSG00 AC124226.1 | 225 | 0 | 0 | 0 | 0 | 0 | 0 |
| ENSMUSG00 AC110211.1 | 284 | 0 | 0 | 0 | 0 | 0 | 0 |
| ENSMUSG00 Mir344b | 62 | 0 | 0 | 0 | 0 | 0 | 0 |
| ENSMUSG00 Mir5101 | 82 | 0 | 2.35055815 | 0 | 0 | 0.62227288 | 5.32188083 |
| ENSMUSG00 BX005039.1 | 94 | 0 | 1.02524345 | 0 | 0 | 0 | 0 |
| ENSMUSG00 AL806523.1 | 107 | 1.11463964 | 0 | 0 | 0 | 0 | 0 |
| ENSMUSG00 Mir3960 | 72 | 0 | 0 | 0 | 0 | 0 | 0 |
| ENSMUSG00 AC154229.1 | 125 | 0 | 0 | 0 | 0 | 0 | 0 |
| ENSMUSG00 AC166290.1 | 92 | 0 | 0 | 0 | 0 | 0 | 0 |
| ENSMUSG00 AC141881.3 | 118 | 0 | 0 | 0 | 0 | 0 | 0 |
| ENSMUSG00 Mir3471-2 | 101 | 0 | 0 | 0 | 0 | 0 | 0 |
| ENSMUSG00 AC132354.1 | 157 | 0 | 0 | 0 | 0 | 0 | 0 |
| ENSMUSG00 AC161342.2 | 247 | 0 | 0 | 0 | 0 | 0 | 0 |
| ENSMUSG00 AC114657.1 | 74 | 0 | 0 | 0 | 0 | 0 | 0 |
| ENSMUSG00 AC134918.1 | 101 | 0 | 0 | 0 | 0 | 0 | 0 |
| ENSMUSG00 AC125110.1 | 62 | 0 | 0 | 0 | 0 | 0 | 0 |
| ENSMUSG00 AC153877.1 | 76 | 0 | 0 | 0 | 0 | 0 | 0 |
| ENSMUSG00 AC158384.1 | 77 | 1.03260988 | 0 | 0 | 0 | 0 | 0 |
| ENSMUSG00 CAAA01010SI | 100 | 0 | 0 | 0 | 0 | 0 | 0 |
| ENSMUSG00 AC015535.1 | 125 | 0 | 0 | 0 | 0 | 0 | 0 |
| ENSMUSG00 AC117662.1 | 69 | 0 | 0 | 0 | 0 | 0 | 0 |
| ENSMUSG00 AC093346.1 | 295 | 0 | 0 | 0 | 0 | 0.34594153 | 0 |
| ENSMUSG00 AL591712.1 | 299 | 0 | 0 | 0 | 0 | 0 | 0 |
| ENSMUSG00 AC159649.1 | 276 | 0 | 0 | 0 | 0 | 0 | 0 |
| ENSMUSG00 AC119249.1 | 79 | 0 | 0 | 0 | 0 | 0 | 0 |
| ENSMUSG00 AC140327.1 | 249 | 0 | 0 | 0 | 0 | 0 | 0 |
| ENSMUSG00 Mir3070a | 88 | 0 | 0 | 0 | 0 | 0 | 0 |
| ENSMUSG00 AC159140.1 | 259 | 0 | 0 | 0 | 0 | 0 | 0 |
| ENSMUSG00 AC122189.1 | 71 | 0 | 0 | 0 | 0 | 0 | 0 |
| ENSMUSG00 AC159624.1 | 90 | 0 | 0 | 0 | 0 | 0 | 0 |
| ENSMUSG00 AL662901.1 | 80 | 0 | 0 | 0 | 0 | 0 | 0 |
| ENSMUSG00 AL928889.1 | 287 | 0 | 0 | 0 | 0 | 0 | 0 |
| ENSMUSG00 Mir3059 | 80 | 0 | 0 | 0 | 0 | 0 | 0 |
| ENSMUSG00 AC162893.1 | 83 | 0 | 0 | 0 | 0 | 0 | 0 |
| ENSMUSG00 AL808021.1 | 70 | 0 | 0 | 0.6314549 | 2.5772889 | 3.64474113 | 1.55855081 |
| ENSMUSG00 AL732525.1 | 255 | 0 | 0 | 0 | 0 | 0 | 0 |
| ENSMUSG00 AC044807.1 | 88 | 0 | 0 | 0 | 0 | 0 | 0 |
| ENSMUSG00 AC084822.1 | 70 | 0 | 0 | 0 | 0 | 0 | 0 |
| ENSMUSG00 Mir5128 | 83 | 0 | 0 | 0 | 0 | 0 | 0 |
| ENSMUSG00 CT030685.2 | 66 | 0 | 0 | 0 | 0 | 0 | 0 |
| ENSMUSG00 AC113329.1 | 70 | 0 | 0 | 0 | 0.51545778 | 0.72894823 | 0 |
| ENSMUSG00 AL772216.42 | 102 | 0 | 0 | 0 | 0 | 0 | 0 |
| ENSMUSG00 AC141881.4 | 118 | 0 | 0 | 0 | 0 | 0 | 0 |
| ENSMUSG00 AL672070.1 | 87 | 0 | 0 | 0 | 0 | 0 | 0 |
| ENSMUSG00 Mir3470a | 76 | 0 | 0 | 0 | 0 | 0 | 0 |
| ENSMUSG00 AC091260.1 | 117 | 0 | 0 | 0 | 0 | 0 | 0 |
| ENSMUSG00 AC126679.1 | 81 | 0 | 0 | 0 | 0 | 0 | 0 |
| ENSMUSG00 AC167020.1 | 87 | 0 | 0 | 0 | 1.24420843 | 0 | 0 |
| ENSMUSG00 AC118255.1 | 70 | 0 | 0 | 0 | 0 | 0 | 0 |
| ENSMUSG00 AC160534.1 | 109 | 0 | 0 | 0 | 0 | 0 | 0 |
| ENSMUSG00 AC091763.1 | 125 | 0 | 0.38549154 | 0 | 0 | 0.40821101 | 0 |
| ENSMUSG00 AL833775.2 | 82 | 0 | 0 | 0 | 0 | 0 | 0 |
| ENSMUSG00 Mir3078 | 86 | 0 | 0 | 0 | 0 | 0 | 0 |
| ENSMUSG00 AC117949.1 | 69 | 0 | 0 | 0 | 0 | 0 | 0 |
| ENSMUSG00 AL589670.1 | 232 | 0 | 0 | 0 | 0 | 0 | 0 |
| ENSMUSG00 AC122252.1 | 125 | 0 | 0 | 0 | 0 | 0 | 0 |
| ENSMUSG00 AC155948.1 | 100 | 0 | 0 | 0 | 0 | 0 | 0 |
| ENSMUSG00 AC108434.1 | 251 | 0 | 0 | 0 | 0 | 0 | 0 |
| ENSMUSG00 AC113178.1 | 101 | 0 | 0 | 0 | 0 | 0 | 0 |
| ENSMUSG00 AC103598.1 | 85 | 0 | 0 | 0 | 0 | 0 | 0 |
| ENSMUSG00 AC161246.1 | 295 | 0 | 0 | 0 | 0 | 0 | 0 |
| ENSMUSG00 Mir5125 | 78 | 4.07748518 | 9.88439839 | 3.40014177 | 8.78921597 | 7.85021166 | 9.09154642 |
| ENSMUSG00 AC123853.1 | 55 | 0 | 0 | 0 | 0 | 0 | 0 |
| ENSMUSG00 Mir5115 | 73 | 18.5162512 | 20.4627357 | 8.47706578 | 8.4026679 | 11.1838632 | 12.7032566 |
| ENSMUSG00 Mir378b | 91 | 0 | 0.52952134 | 0.97146908 | 0.39650598 | 3.92510583 | 0 |
| ENSMUSG00 AC162799.1 | 246 | 0 | 0 | 0 | 0 | 0 | 0 |
| ENSMUSG00 BX470216.1 | 206 | 0 | 0 | 0 | 0 | 0 | 0 |
| ENSMUSG00 Mir3093 | 86 | 0 | 0 | 0 | 0 | 0 | 0 |
| ENSMUSG00 AL596108.1 | 296 | 0 | 0 | 0 | 0 | 0 | 0 |
| ENSMUSG00 Mir466q | 75 | 0 | 0 | 0 | 0 | 0 | 0 |
| ENSMUSG00 AC158379.1 | 285 | 0 | 0 | 0 | 0 | 0 | 0 |
| ENSMUSG00 AL731688.1 | 69 | 0 | 0 | 0 | 0 | 0 | 0 |
| ENSMUSG00 AL772336.1 | 100 | 0 | 0 | 0 | 0 | 0 | 0 |
| ENSMUSG00 AC113124.1 | 105 | 0 | 0 | 0 | 0.34363852 | 0.48596548 | 3.3768601 |
| ENSMUSG00 Mir16-1 | 92 | 0 | 0 | 0 | 0 | 0 | 0 |
| ENSMUSG00 AC119249.2 | 79 | 0 | 0 | 0 | 0 | 0 | 0 |
| ENSMUSG00 AC091333.1 | 97 | 0 | 0 | 0 | 0 | 0 | 0 |
| ENSMUSG00 Mir5099 | 73 | 0 | 0 | 0 | 0 | 0 | 0 |
| ENSMUSG00 AC160471.1 | 106 | 0 | 0 | 0 | 0 | 0 | 0 |
| ENSMUSG00 AC122183.1 | 109 | 0 | 0 | 0 | 0 | 0 | 0 |
| ENSMUSG00 AL954388.2 | 312 | 0 | 0 | 0 | 0 | 0 | 0 |
| ENSMUSG00 AC127694.1 | 129 | 0 | 0 | 0 | 0 | 0 | 0 |
| ENSMUSG00 Mir3473c | 80 | 0.99388701 | 0 | 0 | 0 | 0 | 0 |
| ENSMUSG00 AL596123.1 | 70 | 0 | 0 | 0 | 0 | 0 | 0 |
| ENSMUSG00 AC138118.1 | 101 | 0 | 0 | 0 | 0 | 0 | 0 |
| ENSMUSG00 AL935177.1 | 95 | 0 | 0 | 0 | 0 | 0 | 0 |
| ENSMUSG00 Mir15a | 83 | 0 | 0 | 0 | 0 | 0 | 0 |
| ENSMUSG00 Snord52 | 65 | 0 | 0 | 0 | 0 | 0 | 0 |
| ENSMUSG00 AC139323.1 | 123 | 0 | 0 | 0 | 0 | 0 | 0 |
| ENSMUSG00 Mir100 | 79 | 0 | 0 | 0 | 0 | 0 | 0 |
| ENSMUSG00 AC122247.1 | 292 | 0.13614891 | 0 | 0 | 0 | 0 | 0 |
| ENSMUSG00 AL591209.1 | 69 | 0 | 0 | 0 | 0 | 0 | 0 |
| ENSMUSG00 AL732400.1 | 63 | 0 | 0 | 0 | 0 | 0 | 0 |
| ENSMUSG00 AC103663.1 | 100 | 0 | 0 | 0 | 0 | 0 | 0 |
| ENSMUSG00 AC122266.1 | 283 | 0 | 0 | 0 | 0.50999356 | 0 | 0.86739135 |
| ENSMUSG00 Mir5108 | 84 | 0 | 0 | 0 | 0 | 0 | 0 |
| ENSMUSG00 AL672099.1 | 269 | 0 | 0 | 0 | 0 | 0 | 0 |
| ENSMUSG00 AC119249.3 | 68 | 0 | 0 | 0 | 0 | 0 | 0 |
| ENSMUSG00 AC149222.1 | 280 | 0 | 0 | 0 | 0 | 0 | 0 |
| ENSMUSG00 AC138604.1 | 73 | 0 | 0 | 0 | 0 | 0 | 0 |
| ENSMUSG00 AC127294.1 | 100 | 0 | 0 | 0 | 0 | 0 | 0 |
| ENSMUSG00 AC142099.1 | 263 | 0 | 0 | 0 | 0.41158226 | 0.38803328 | 0 |
| ENSMUSG00 AL773580.1 | 101 | 0 | 0 | 0 | 0 | 0 | 0 |
| ENSMUSG00 AC079245.1 | 83 | 0 | 0 | 0 | 0 | 0 | 0 |
| ENSMUSG00 AC116595.1 | 112 | 0.35495965 | 0 | 0 | 0 | 0 | 0 |
| ENSMUSG00 AC101777.1 | 120 | 0 | 0 | 0 | 0 | 0 | 0 |
| ENSMUSG00 AL596246.1 | 229 | 0 | 0 | 0 | 0 | 0 | 0 |
| ENSMUSG00 AC122043.1 | 116 | 0 | 0 | 0 | 0 | 0 | 0 |
| ENSMUSG00 AC140381.1 | 124 | 0 | 0 | 0 | 0 | 0 | 0 |
| ENSMUSG00 BX537327.1 | 253 | 0 | 0 | 0 | 0 | 0.20168528 | 0 |
| ENSMUSG00 AC125372.1 | 122 | 0 | 0 | 0 | 0 | 0 | 0 |
| ENSMUSG00 AL683878.1 | 103 | 0 | 0 | 0 | 0 | 0 | 0 |
| ENSMUSG00 AC154327.1 | 118 | 0 | 0 | 1.12377567 | 0 | 0 | 0 |
| ENSMUSG00 AC133597.1 | 70 | 0 | 0 | 0 | 0 | 0 | 0 |
| ENSMUSG00 Mir5127 | 70 | 0 | 0 | 0 | 0 | 0 | 0 |
| ENSMUSG00 Mir3066 | 82 | 0 | 0 | 0 | 0 | 0 | 0 |
| ENSMUSG00 AC131596.1 | 100 | 0 | 0 | 0 | 0 | 0 | 0 |
| ENSMUSG00 Mir1912 | 86 | 0 | 0 | 0 | 0 | 0 | 0 |
| ENSMUSG00 AL663077.1 | 125 | 0 | 0 | 0 | 0 | 0 | 0 |
| ENSMUSG00 AC132338.1 | 91 | 0 | 0 | 0 | 0 | 0 | 0 |
| ENSMUSG00 Mir3109 | 87 | 0 | 0 | 0 | 0 | 0 | 0 |
| ENSMUSG00 AC159379.1 | 311 | 0 | 0 | 0 | 0 | 0 | 0 |
| ENSMUSG00 Snord8 | 109 | 1.8236459 | 2.65246471 | 2.02760748 | 5.95850277 | 9.36263777 | 1.50135629 |
| ENSMUSG00 AC162893.2 | 70 | 0 | 0 | 0 | 0 | 0 | 0 |
| ENSMUSG00 Mir3081 | 83 | 0 | 0 | 0 | 0 | 0 | 0 |
| ENSMUSG00 AL645683.1 | 100 | 0 | 0 | 0 | 0 | 0 | 0 |
| ENSMUSG00 AC135118.1 | 103 | 0 | 0 | 0 | 0 | 0 | 0 |
| ENSMUSG00 AC122782.1 | 74 | 0 | 0 | 0 | 0 | 0 | 0 |
| ENSMUSG00 AC156026.1 | 69 | 0 | 0 | 0 | 0 | 0 | 0 |
| ENSMUSG00 CT030727.1 | 83 | 0 | 2.90279772 | 0.53255233 | 0 | 1.22955123 | 0 |
| ENSMUSG00 AC121954.1 | 148 | 0 | 0.32558407 | 0 | 0 | 1.72386405 | 0 |
| ENSMUSG00 AC124134.1 | 69 | 0 | 0 | 0 | 0 | 0 | 0 |
| ENSMUSG00 AC115933.1 | 99 | 0 | 0 | 0 | 0 | 0 | 0 |
| ENSMUSG00 AL844869.1 | 277 | 0 | 0 | 0 | 0 | 0 | 0 |
| ENSMUSG00 AC156988.1 | 99 | 0 | 0 | 0 | 0 | 0 | 0 |
| ENSMUSG00 AC134453.1 | 69 | 0 | 0 | 0 | 0 | 0 | 0 |
| ENSMUSG00 AL589767.1 | 267 | 0 | 0 | 0 | 0 | 0 | 0 |
| ENSMUSG00 Mir5104 | 93 | 0 | 0 | 0 | 0 | 0 | 0 |
| ENSMUSG00 Mir5098 | 81 | 0 | 0 | 0 | 0.44545734 | 0 | 0 |
| ENSMUSG00 AL732357.1 | 115 | 0 | 0 | 0 | 0 | 0 | 0 |
| ENSMUSG00 AC124613.1 | 285 | 0 | 0 | 0 | 0 | 0 | 0 |
| ENSMUSG00 AL806517.1 | 64 | 0 | 0 | 0 | 0 | 0 | 0 |
| ENSMUSG00 AL592465.2 | 78 | 0 | 0 | 0 | 0 | 0 | 0 |
| ENSMUSG00 AC168054.1 | 85 | 0 | 0 | 0 | 0 | 0 | 0 |
| ENSMUSG00 AC132118.1 | 78 | 0 | 0 | 0 | 0 | 0 | 0 |
| ENSMUSG00 AC123530.1 | 125 | 0 | 0 | 0 | 0 | 0 | 0 |
| ENSMUSG00 AC115293.1 | 73 | 0 | 0.66008825 | 0 | 0 | 1.3979829 | 0 |
| ENSMUSG00 AL954710.1 | 63 | 0 | 0 | 0 | 0 | 0 | 0 |
| ENSMUSG00 AC110817.1 | 99 | 0 | 0 | 0 | 0 | 0 | 0 |
| ENSMUSG00 Mir124a-2 | 108 | 0 | 0 | 0 | 0 | 0 | 0 |
| ENSMUSG00 CAAA010375 | 113 | 0 | 0 | 0 | 0 | 0 | 0 |
| ENSMUSG00 AL807824.1 | 111 | 0 | 1.30233627 | 0 | 0.65012693 | 0 | 0 |
| ENSMUSG00 Mir3082 | 63 | 0 | 0 | 0 | 0 | 0 | 0 |
| ENSMUSG00 Mir5105 | 81 | 24.5404201 | 28.5549287 | 25.1022812 | 31.6274711 | 22.048434 | 31.3153266 |
| ENSMUSG00 AC150681.1 | 70 | 0 | 0 | 0 | 0 | 0 | 0 |
| ENSMUSG00 AC140192.1 | 82 | 0 | 0 | 0 | 0 | 0 | 0 |
| ENSMUSG00 Mir3060 | 82 | 1.4544688 | 6.46403492 | 1.07809373 | 2.20012467 | 3.11136438 | 0 |
| ENSMUSG00 Mir3075 | 84 | 0 | 0 | 0 | 0 | 0 | 0 |
| ENSMUSG00 AC105995.1 | 69 | 0 | 0 | 0 | 0 | 0 | 0 |
| ENSMUSG00 AL627205.1 | 76 | 0 | 0 | 0 | 0 | 0 | 0 |
| ENSMUSG00 Mir1186b | 63 | 0 | 0 | 0 | 0 | 0 | 0 |
| ENSMUSG00 AC153815.1 | 73 | 0.54459562 | 0 | 0 | 0 | 2.7959658 | 2.24175117 |
| ENSMUSG00 AC127419.1 | 80 | 0.99388701 | 4.21631369 | 0 | 1.80410223 | 6.37829698 | 0.34093299 |
| ENSMUSG00 Mir3473d | 80 | 0 | 1.20466105 | 0 | 0 | 0.6378297 | 1.70466495 |
| ENSMUSG00 AC160635.1 | 125 | 0 | 0 | 0 | 0 | 0 | 0 |
| ENSMUSG00 AC121784.1 | 79 | 0 | 0 | 0 | 0 | 0 | 0 |
| ENSMUSG00 Mir3073 | 84 | 0 | 0 | 0 | 0 | 0 | 0 |
| ENSMUSG00 Mir3089 | 83 | 0 | 0 | 0 | 0 | 0 | 0 |
| ENSMUSG00 AC169127.1 | 100 | 0 | 0 | 0 | 0 | 0 | 0 |
| ENSMUSG00 AC132327.1 | 247 | 0 | 0 | 0 | 0 | 0 | 0 |
| ENSMUSG00 AC140313.1 | 100 | 0 | 0 | 0 | 0 | 0 | 0 |
| ENSMUSG00 AC170189.1 | 80 | 0 | 0 | 0 | 0 | 0 | 0 |
| ENSMUSG00 Mir3968 | 99 | 0 | 0 | 0 | 0 | 0 | 0 |
| ENSMUSG00 Mir3097 | 66 | 0 | 0 | 0 | 0 | 0 | 0 |
| ENSMUSG00 Mir5102 | 86 | 90.6055137 | 136.715022 | 99.7111342 | 91.4637873 | 53.3996956 | 72.6266557 |
| ENSMUSG00 Rn5s | 118 | 0 | 0 | 0 | 0 | 0 | 0 |
| ENSMUSG00 Mir3971 | 82 | 0 | 4.11347677 | 2.69523433 | 1.3200748 | 0.62227288 | 1.33047021 |
| ENSMUSG00 AC119249.4 | 78 | 0 | 0 | 0 | 0 | 0 | 0 |
| ENSMUSG00 Mir3475 | 64 | 0 | 0 | 0 | 0 | 0 | 0 |
| ENSMUSG00 AC141881.5 | 118 | 0 | 0 | 0 | 0 | 0 | 0 |
| ENSMUSG00 AL845455.1 | 152 | 1.8308445 | 0.63403213 | 0.5816032 | 0 | 1.67849921 | 0.71775366 |
| ENSMUSG00 AC153918.1 | 124 | 4.16791328 | 7.7720068 | 0.35646648 | 3.78279499 | 2.46901819 | 0.43991354 |
| ENSMUSG00 AC136976.1 | 124 | 0 | 0 | 0 | 0 | 0 | 0 |
| ENSMUSG00 AC167122.1 | 126 | 7.57247248 | 10.3256662 | 17.8912222 | 13.7455408 | 9.31433844 | 19.4818852 |
| ENSMUSG00 Mir344f | 67 | 0 | 0 | 0 | 0 | 0 | 0 |
| ENSMUSG00 AC165159.1 | 284 | 0 | 0 | 0 | 0.38114836 | 0 | 0 |
| ENSMUSG00 AC115767.1 | 85 | 0 | 0 | 0 | 0 | 1.20062061 | 0 |
| ENSMUSG00 AC120136.1 | 190 | 0.20923937 | 0 | 0 | 0 | 0 | 0.28710147 |
| ENSMUSG00 AC069308.1 | 275 | 0 | 0 | 0 | 0 | 0 | 0 |
| ENSMUSG00 AL662785.1 | 68 | 0 | 0 | 0 | 0 | 0 | 0 |
| ENSMUSG00 AC133174.1 | 67 | 0 | 0 | 0 | 0 | 0 | 0 |
| ENSMUSG00 AL772159.1 | 100 | 0 | 0 | 0 | 0 | 0 | 0 |
| ENSMUSG00 AC130536.1 | 107 | 0 | 0 | 0 | 0 | 0 | 0 |
| ENSMUSG00 AC166574.1 | 104 | 0 | 0 | 0 | 0 | 0 | 0 |
| ENSMUSG00 Mir3083 | 63 | 0 | 0 | 0 | 0 | 0 | 0 |
| ENSMUSG00 AC122463.1 | 124 | 0 | 0 | 0 | 0 | 0 | 0 |
| ENSMUSG00 Mir3474 | 57 | 2.09239371 | 0 | 0 | 0 | 0 | 0 |
| ENSMUSG00 AC167242.1 | 237 | 0 | 0 | 0 | 0 | 0 | 0.46033146 |
| ENSMUSG00 Mir1264 | 85 | 0 | 0 | 0 | 0 | 0 | 0 |
| ENSMUSG00 AL671229.1 | 125 | 0 | 0 | 0 | 0 | 0 | 0 |
| ENSMUSG00 AC114917.1 | 85 | 2.80626921 | 5.10209387 | 0 | 2.12247321 | 4.80248243 | 2.24614676 |
| ENSMUSG00 AC130474.1 | 124 | 0 | 0 | 0 | 0 | 0 | 0 |
| ENSMUSG00 AC161763.1 | 70 | 0 | 0 | 0 | 0 | 0 | 0 |
| ENSMUSG00 AC167724.1 | 125 | 0 | 0 | 0 | 0 | 0 | 0 |
| ENSMUSG00 AC134912.1 | 95 | 0 | 0 | 0 | 0 | 0 | 0 |
| ENSMUSG00 AC107770.1 | 105 | 0 | 0 | 0 | 0 | 0 | 0 |
| ENSMUSG00 AC140983.1 | 106 | 0 | 0 | 0 | 0 | 0 | 0 |
| ENSMUSG00 AC144905.1 | 79 | 0 | 0 | 0 | 0 | 0 | 0.3452486 |
| ENSMUSG00 AL603664.1 | 100 | 0 | 0 | 0 | 0 | 0 | 0 |
| ENSMUSG00 CT025520.1 | 116 | 0 | 0 | 0 | 0 | 0 | 0 |
| ENSMUSG00 AL671140.1 | 109 | 0 | 0 | 0 | 0 | 0 | 0 |
| ENSMUSG00 AC122057.1 | 141 | 0 | 0 | 0 | 0 | 0 | 0 |
| ENSMUSG00 AL845277.1 | 52 | 0 | 0 | 0.85003544 | 0 | 4.90638229 | 0 |
| ENSMUSG00 AC161198.1 | 132 | 0 | 0 | 0 | 0 | 0 | 0 |
| ENSMUSG00 Mir28b | 79 | 0 | 0 | 0 | 0 | 0 | 0 |
| ENSMUSG00 AC163282.1 | 89 | 0 | 0 | 0 | 0 | 0 | 0 |
| ENSMUSG00 Mir3106 | 81 | 0 | 0 | 0 | 0 | 0 | 0 |
| ENSMUSG00 AC134910.1 | 274 | 0 | 0 | 0.1613206 | 0 | 0 | 0 |
| ENSMUSG00 AC125186.2 | 99 | 0 | 0 | 0 | 0 | 0 | 0 |
| ENSMUSG00 AC140457.1 | 73 | 0.54459562 | 0 | 3.63302819 | 0 | 0 | 0 |
| ENSMUSG00 AC131793.1 | 129 | 0 | 0 | 0 | 0 | 0 | 0 |
| ENSMUSG00 AL772271.1 | 104 | 0 | 0 | 0 | 0 | 0 | 0 |
| ENSMUSG00 AC122359.1 | 82 | 0 | 0 | 0 | 0 | 0 | 0 |
| ENSMUSG00 AC163694.1 | 83 | 0 | 0.58055954 | 0 | 0 | 0 | 0 |
| ENSMUSG00 AC160932.1 | 70 | 0 | 0 | 0 | 0 | 0 | 0 |
| ENSMUSG00 AC123662.1 | 94 | 0 | 0.51262173 | 0 | 1.15155461 | 4.34267028 | 0 |
| ENSMUSG00 AC073947.1 | 83 | 0 | 0 | 0 | 0 | 0 | 0 |
| ENSMUSG00 AC149058.1 | 69 | 0 | 0 | 0 | 0 | 0 | 0 |
| ENSMUSG00 AC126942.1 | 57 | 0 | 0 | 7.7547093 | 4.43112828 | 4.47599788 | 20.0971026 |
| ENSMUSG00 AC117610.1 | 76 | 0 | 0 | 0 | 0.94952749 | 0 | 1.0766305 |
| ENSMUSG00 Mir3076 | 59 | 0 | 0 | 0 | 0.61156008 | 0 | 0 |
| ENSMUSG00 Mir3098 | 86 | 2.77363818 | 2.80153733 | 4.11179935 | 6.71293852 | 5.33996956 | 3.80576362 |
| ENSMUSG00 AC124466.1 | 49 | 0 | 0 | 0 | 0 | 0 | 0 |
| ENSMUSG00 AC026231.1 | 109 | 0 | 0 | 0 | 0 | 0 | 0 |
| ENSMUSG00 AL845460.1 | 77 | 0 | 0 | 0 | 0 | 0 | 0 |
| ENSMUSG00 AC153489.1 | 72 | 0 | 0 | 0 | 0 | 0 | 0 |
| ENSMUSG00 AC158662.1 | 61 | 0 | 0 | 0 | 0 | 0 | 0 |
| ENSMUSG00 AL607123.1 | 81 | 0 | 0 | 0 | 0 | 0 | 0 |
| ENSMUSG00 AL645971.1 | 251 | 0 | 0 | 0 | 0 | 0 | 0 |
| ENSMUSG00 AC127366.1 | 125 | 0 | 0 | 0 | 0 | 0 | 0 |
| ENSMUSG00 AC160757.1 | 320 | 0 | 0 | 0 | 0 | 0 | 0 |
| ENSMUSG00 Mir5131 | 92 | 0 | 0 | 0 | 0 | 0 | 0 |
| ENSMUSG00 AL663049.1 | 88 | 6.32473554 | 0 | 13.0596354 | 11.8906738 | 15.6558199 | 15.8068932 |
| ENSMUSG00 Mir3079 | 89 | 0 | 0 | 0 | 0 | 0 | 0 |
| ENSMUSG00 AC091333.2 | 69 | 0 | 0 | 0 | 0 | 0 | 0 |
| ENSMUSG00 AC132452.1 | 72 | 0 | 0 | 0 | 0 | 0 | 0 |
| ENSMUSG00 Mir3112 | 81 | 0 | 0 | 0 | 0 | 0 | 0 |
| ENSMUSG00 AC139568.1 | 76 | 0 | 0 | 0 | 0 | 0 | 0 |
| ENSMUSG00 BX323066.1 | 277 | 0 | 0 | 0 | 0 | 0 | 0 |
| ENSMUSG00 AC133646.1 | 324 | 0 | 0 | 0 | 0 | 0 | 0 |
| ENSMUSG00 Mir5119 | 56 | 0.7099193 | 4.30236091 | 0 | 4.51025557 | 2.73355585 | 5.35751842 |
| ENSMUSG00 AC110195.1 | 70 | 0 | 0 | 0 | 0 | 0 | 0 |
| ENSMUSG00 Snord87 | 86 | 3.2359112 | 0.56030747 | 2.56987459 | 0.83911732 | 1.1866599 | 15.5402014 |
| ENSMUSG00 AL672246.1 | 66 | 0 | 0 | 0 | 0 | 0 | 0 |
| ENSMUSG00 AL670285.1 | 76 | 0 | 0 | 0 | 0 | 0 | 0 |
| ENSMUSG00 AC157798.1 | 195 | 0 | 0 | 0 | 0 | 0.78502117 | 0 |
| ENSMUSG00 AC136021.1 | 316 | 0 | 0 | 0 | 0 | 0 | 0 |
| ENSMUSG00 AL772349.1 | 278 | 0 | 0 | 0 | 0 | 0 | 0 |
| ENSMUSG00 AL808020.1 | 84 | 0 | 0 | 0 | 0 | 0 | 0 |
| ENSMUSG00 AL772376.2 | 124 | 0 | 0 | 0 | 0 | 0 | 0 |
| ENSMUSG00 AL713978.1 | 279 | 0 | 0 | 0 | 0 | 0 | 0 |
| ENSMUSG00 Mir344-2 | 79 | 0 | 0 | 0 | 0 | 0 | 0 |
| ENSMUSG00 AL732564.1 | 251 | 0 | 0 | 0 | 0 | 0 | 0 |
| ENSMUSG00 AC161829.1 | 281 | 0 | 0 | 0 | 0.25681171 | 0 | 0.97062773 |
| ENSMUSG00 AL732525.2 | 125 | 0 | 0 | 0 | 0 | 0 | 0 |
| ENSMUSG00 AC113598.1 | 300 | 0 | 0 | 0 | 0 | 0 | 0 |
| ENSMUSG00 AC125201.1 | 104 | 0 | 0 | 0 | 0 | 0 | 0 |
| ENSMUSG00 AC105256.2 | 70 | 0 | 0 | 0 | 0 | 0 | 0 |
| ENSMUSG00 AC126881.1 | 85 | 0 | 0 | 0 | 0 | 0 | 0 |
| ENSMUSG00 AC115706.1 | 122 | 0 | 0 | 0 | 0 | 0 | 0 |
| ENSMUSG00 AC156032.1 | 228 | 0 | 0.21134404 | 0 | 0 | 0.67139968 | 0.95700489 |
| ENSMUSG00 AL929170.1 | 68 | 0 | 0 | 0 | 0 | 0 | 0 |
| ENSMUSG00 Mir344d-2 | 79 | 0 | 0 | 0 | 0 | 0 | 0 |
| ENSMUSG00 AC141881.6 | 118 | 0 | 0 | 0 | 0 | 0 | 0 |
| ENSMUSG00 AC153589.1 | 327 | 0 | 0 | 0 | 0 | 0 | 0 |
| ENSMUSG00 AC122454.1 | 109 | 0 | 0 | 0 | 0 | 0 | 0 |
| ENSMUSG00 AL844481.1 | 125 | 0 | 0 | 0 | 0 | 0 | 0 |
| ENSMUSG00 AC166574.2 | 106 | 0 | 0 | 0 | 0 | 0 | 0 |
| ENSMUSG00 AC122417.1 | 268 | 0.14834135 | 1.43840126 | 0 | 0.67317247 | 0.19039693 | 0.20354208 |
| ENSMUSG00 AL672290.1 | 90 | 0 | 0 | 0 | 0 | 0 | 0 |
| ENSMUSG00 AC134475.1 | 100 | 0 | 0 | 0 | 0 | 0 | 0 |
| ENSMUSG00 AC122860.1 | 61 | 0 | 0 | 0 | 0 | 0 | 0 |
| ENSMUSG00 Mir3070b | 88 | 0 | 0 | 0 | 0 | 0 | 0 |
| ENSMUSG00 AL662894.1 | 315 | 0 | 0 | 0 | 0 | 0 | 0 |
| ENSMUSG00 AC157995.1 | 75 | 0 | 0 | 0 | 0 | 0 | 0 |
| ENSMUSG00 AC123823.1 | 77 | 0 | 0 | 0 | 0 | 0 | 0 |
| ENSMUSG00 AC101833.1 | 101 | 0 | 0 | 0 | 0 | 0 | 0 |
| ENSMUSG00 AC154353.1 | 97 | 0 | 0 | 0 | 0 | 0 | 0 |
| ENSMUSG00 Mir3103 | 66 | 0 | 0 | 0 | 0 | 0 | 0 |
| ENSMUSG00 AC122376.1 | 89 | 0 | 0 | 0 | 0 | 0 | 0 |
| ENSMUSG00 AL671117.1 | 68 | 0 | 0 | 0 | 0 | 0 | 0 |
| ENSMUSG00 AC121784.2 | 119 | 0 | 0 | 0 | 0 | 0 | 0 |
| ENSMUSG00 Mir3113 | 75 | 0 | 0 | 0 | 0 | 0 | 0 |
| ENSMUSG00 AL627182.1 | 252 | 0 | 0 | 0 | 0 | 0 | 0 |
| ENSMUSG00 Mir3105 | 84 | 0 | 0 | 0 | 0 | 0 | 0 |
| ENSMUSG00 AC154683.1 | 106 | 0 | 0 | 0 | 0 | 0 | 0 |
| ENSMUSG00 AC117806.1 | 125 | 0 | 0 | 0 | 0 | 0 | 0 |
| ENSMUSG00 AC121784.3 | 111 | 0 | 0 | 0 | 0 | 0 | 0 |
| ENSMUSG00 AL645531.1 | 73 | 0 | 0 | 0 | 0 | 0 | 0 |
| ENSMUSG00 AC163349.1 | 110 | 0 | 0 | 0 | 0 | 0 | 0 |
| ENSMUSG00 Mir1251 | 83 | 0 | 0 | 0 | 0 | 0 | 0 |
| ENSMUSG00 AC167297.1 | 69 | 0 | 0 | 0 | 0 | 0 | 0 |
| ENSMUSG00 AC139241.1 | 102 | 0 | 0 | 0 | 0 | 0 | 0 |
| ENSMUSG00 AC161253.2 | 66 | 0 | 0 | 0 | 0 | 0 | 0 |
| ENSMUSG00 AC147162.1 | 96 | 0 | 2.00776842 | 1.84174346 | 0 | 0 | 0 |
| ENSMUSG00 AL591070.1 | 280 | 0 | 0 | 0 | 0.90205111 | 0 | 0 |
| ENSMUSG00 Mir3085 | 88 | 0 | 0 | 0 | 0 | 0 | 0 |
| ENSMUSG00 AC132271.1 | 77 | 0 | 0.62579795 | 0 | 0 | 0 | 0 |
| ENSMUSG00 AC157986.1 | 76 | 0 | 0 | 0 | 0 | 0 | 0 |
| ENSMUSG00 AC132575.1 | 275 | 0 | 0 | 0 | 0 | 0 | 0 |
| ENSMUSG00 Mir9-3 | 89 | 0 | 0 | 0 | 0 | 0 | 0 |
| ENSMUSG00 AC160118.1 | 126 | 0.31551969 | 0 | 0 | 0 | 0 | 0 |
| ENSMUSG00 AL807771.1 | 111 | 0 | 0 | 0 | 0 | 0 | 0 |
| ENSMUSG00 Mir3108 | 78 | 0 | 0 | 0 | 0 | 0 | 0 |
| ENSMUSG00 AL772404.1 | 73 | 0 | 0 | 0 | 0 | 0 | 0 |
| ENSMUSG00 AL583889.1 | 76 | [illegible] | [illegible] | [illegible] | [illegible] | [illegible] | [illegible] |

| | | | | | | | | |
|---|---|---|---|---|---|---|---|---|
| ENSMUSG00 | AL671848.1 | 82 | 0 | 0 | 0 | 0 | 0 | 0 |
| ENSMUSG00 | AC122525.1 | 275 | 0.28913077 | 0 | 0.32146795 | 0 | 0 | 0.49590253 |
| ENSMUSG00 | AC154368.1 | 125 | 0 | 0 | 0 | 0 | 0 | 0 |
| ENSMUSG00 | Mir5124 | 71 | 0 | 3.39341142 | 0 | 0 | 1.4373627 | 0 |
| ENSMUSG00 | AC117577.2 | 77 | 0 | 0 | 0 | 0 | 0 | 0 |
| ENSMUSG00 | AC027654.1 | 69 | 0 | 0 | 0 | 0 | 0 | 0 |
| ENSMUSG00 | Mir3064 | 66 | 0 | 0 | 0 | 0 | 0 | 0 |
| ENSMUSG00 | AC122304.1 | 100 | 0 | 0 | 0 | 0 | 0 | 0 |
| ENSMUSG00 | AC158667.1 | 243 | 0 | 0 | 0 | 0 | 0 | 0 |
| ENSMUSG00 | AC171917.1 | 124 | 0 | 0 | 0 | 0 | 0 | 0 |
| ENSMUSG00 | AL772271.2 | 136 | 0 | 0 | 0 | 0 | 0.37519394 | 2.60713463 |
| ENSMUSG00 | Mir5110 | 86 | 0.46227303 | 0 | 0 | 0 | 0 | 0 |
| ENSMUSG00 | AC090443.1 | 116 | 0 | 0 | 0.38105037 | 0.31105211 | 2.19941275 | 0 |
| ENSMUSG00 | Mir2861 | 81 | 0 | 0 | 0 | 0 | 0 | 0 |
| ENSMUSG00 | AC134552.1 | 68 | 0 | 0 | 0 | 0 | 0 | 0 |
| ENSMUSG00 | AC153974.1 | 84 | 0.47327953 | 0 | 0 | 0 | 0 | 0 |
| ENSMUSG00 | AC102240.1 | 341 | 0.11658499 | 0 | 0.38887252 | 0 | 0 | 0 |
| ENSMUSG00 | AC131316.1 | 290 | 0 | 0 | 0 | 0 | 0 | 0 |
| ENSMUSG00 | AL807811.2 | 70 | 0 | 0 | 0 | 0 | 0 | 0 |
| ENSMUSG00 | AC116712.1 | 123 | 0 | 0 | 0 | 0 | 0.41484858 | 0 |
| ENSMUSG00 | AC153624.1 | 303 | 0 | 0 | 0 | 0 | 0 | 0 |
| ENSMUSG00 | AC153909.1 | 125 | 0.63608769 | 0 | 0 | 0 | 0 | 0 |
| ENSMUSG00 | Snord92 | 84 | 0 | 0 | 0 | 1.28864445 | 0 | 0 |
| ENSMUSG00 | Mir3572 | 85 | 0 | 0 | 0 | 0 | 0 | 0 |
| ENSMUSG00 | AC121784.4 | 116 | 0 | 0 | 0 | 0 | 0 | 0 |
| ENSMUSG00 | Mir5118 | 66 | 0 | 0 | 0 | 0 | 0 | 0 |
| ENSMUSG00 | Mir3095 | 86 | 0 | 0 | 0 | 0 | 0 | 0 |
| ENSMUSG00 | AL671011.2 | 75 | 0 | 0 | 0 | 0 | 0 | 0 |
| ENSMUSG00 | AC166252.2 | 100 | 0 | 0 | 0 | 0 | 0 | 0 |
| ENSMUSG00 | Mir3102 | 103 | 0 | 0 | 0 | 0 | 0 | 0 |
| ENSMUSG00 | AL645683.2 | 100 | 0 | 0 | 0 | 0 | 0 | 0 |
| ENSMUSG00 | Mir3473b | 54 | 0 | 0 | 0 | 0 | 0 | 0 |
| ENSMUSG00 | AL844901.1 | 74 | 0 | 0 | 0 | 0 | 0 | 0 |
| ENSMUSG00 | AC159245.1 | 119 | 0 | 0 | 0 | 0 | 0 | 0 |
| ENSMUSG00 | CAAA010391 | 101 | 0 | 0 | 0 | 0 | 0 | 0 |
| ENSMUSG00 | AC171500.1 | 66 | 0 | 0 | 0 | 0 | 0 | 0 |
| ENSMUSG00 | AC100729.1 | 63 | 0 | 0 | 0 | 0 | 0 | 0 |
| ENSMUSG00 | Mir3470b | 125 | 0 | 0 | 0 | 0 | 0 | 0 |
| ENSMUSG00 | AL731857.1 | 101 | 0 | 0 | 0 | 0 | 0 | 0 |
| ENSMUSG00 | AC137708.1 | 87 | 0 | 0 | 0 | 0 | 0 | 0 |
| ENSMUSG00 | AL772216.43 | 126 | 0 | 0 | 0 | 0 | 0 | 0 |
| ENSMUSG00 | AC165289.1 | 250 | 0 | 0.38549154 | 0.17680737 | 0 | 0 | 0 |
| ENSMUSG00 | AC159324.1 | 79 | 0 | 2.43981986 | 0 | 0 | 0 | 0 |
| ENSMUSG00 | Mir3965 | 68 | 0 | 0 | 0 | 0 | 0 | 0 |
| ENSMUSG00 | AC155932.1 | 251 | 0 | 0 | 0 | 0 | 0 | 0 |
| ENSMUSG00 | Mir5136 | 75 | 0 | 0 | 0.58935791 | 0 | 0 | 2.545633 |
| ENSMUSG00 | Mir5114 | 60 | 0 | 0 | 0 | 0 | 0 | 0 |
| ENSMUSG00 | AC110235.1 | 233 | 0 | 0 | 0 | 0 | 0 | 0 |
| ENSMUSG00 | AC163724.1 | 110 | 0 | 0 | 0 | 0 | 0 | 0 |
| ENSMUSG00 | AC122199.1 | 100 | 0 | 0 | 0 | 0 | 0 | 0 |
| ENSMUSG00 | AC155810.1 | 226 | 1.75909206 | 0.85285738 | 0.39116675 | 0.47896519 | 0 | 0.36205273 |
| ENSMUSG00 | AC163629.1 | 70 | 0 | 0 | 0 | 0 | 0 | 0 |
| ENSMUSG00 | Snord93 | 70 | 0.56793544 | 0 | 0 | 0 | 0 | 0 |
| ENSMUSG00 | AC155256.1 | 109 | 0 | 0 | 0 | 0 | 0 | 0 |
| ENSMUSG00 | AC166172.1 | 81 | 0 | 0 | 0 | 0 | 0 | 0 |
| ENSMUSG00 | AC123830.1 | 81 | 0 | 0 | 0 | 0 | 0 | 0 |
| ENSMUSG00 | AL669920.1 | 107 | 0 | 0 | 0 | 0 | 0 | 0 |
| ENSMUSG00 | AL772393.1 | 100 | 0 | 0 | 0 | 0 | 0 | 0 |
| ENSMUSG00 | Mir5107 | 87 | 0 | 0 | 0 | 0 | 0 | 0 |
| ENSMUSG00 | AL646096.2 | 265 | 0 | 0 | 0 | 0 | 0 | 0 |
| ENSMUSG00 | AL627403.1 | 100 | 0 | 0 | 0 | 0 | 0 | 0 |
| ENSMUSG00 | AC166903.1 | 76 | 0 | 0 | 0 | 0.94952749 | 0 | 0 |
| ENSMUSG00 | AC144543.1 | 69 | 0 | 0 | 0 | 0 | 0 | 0 |
| ENSMUSG00 | AL805896.1 | 124 | 0.96182614 | 0 | 0 | 0.58196846 | 0 | 0 |
| ENSMUSG00 | AC120136.3 | 190 | 0 | 0 | 0.23264128 | 0 | 1.34279936 | 0.71775366 |
| ENSMUSG00 | AC165250.1 | 165 | 0.48188461 | 0 | 0.26788996 | 0.21867906 | 0 | 0 |
| ENSMUSG00 | AC154106.1 | 98 | 0 | 0.49169839 | 0 | 0 | 0 | 0 |
| ENSMUSG00 | Mir1954 | 85 | 0 | 0 | 0 | 0 | 0 | 0 |
| ENSMUSG00 | Mir5109 | 86 | 7.39636847 | 2.80153733 | 2.05589968 | 5.03470389 | 2.96664976 | 2.53717574 |
| ENSMUSG00 | Mir3090 | 86 | 0 | 0 | 0 | 0 | 0 | 0 |
| ENSMUSG00 | AC124681.1 | 125 | 0 | 0 | 0 | 0 | 0 | 0 |
| ENSMUSG00 | AC117657.1 | 94 | 0 | 0 | 0 | 0 | 0 | 0 |
| ENSMUSG00 | AL607091.1 | 76 | 0 | 0 | 0 | 0 | 0 | 0 |
| ENSMUSG00 | AC122826.1 | 103 | 0 | 0 | 0 | 0 | 0 | 0 |
| ENSMUSG00 | AC151577.1 | 72 | 0 | 0 | 0 | 0 | 0.70869966 | 0 |
| ENSMUSG00 | AC161570.1 | 78 | 0 | 0 | 0 | 0 | 0 | 0 |
| ENSMUSG00 | AC124584.1 | 231 | 0.86050824 | 0 | 0.19134997 | 0 | 0 | 0 |
| ENSMUSG00 | AC109138.1 | 86 | 0 | 0 | 0 | 0 | 0 | 0 |
| ENSMUSG00 | AC102240.2 | 70 | 0 | 0 | 0 | 0 | 0 | 0 |
| ENSMUSG00 | Mir3101 | 87 | 0 | 0 | 0 | 0 | 0 | 0 |
| ENSMUSG00 | AC154460.1 | 104 | 0 | 0 | 0 | 0 | 0 | 0 |
| ENSMUSG00 | Mir3072 | 82 | 0 | 0 | 0 | 0 | 0 | 0 |
| ENSMUSG00 | AC159243.1 | 77 | 0 | 0 | 0 | 0 | 0 | 0 |
| ENSMUSG00 | Mir125b-1 | 76 | 0 | 0 | 0 | 0 | 0 | 0 |
| ENSMUSG00 | Snora26 | 120 | 0 | 0 | 0 | 0 | 0 | 0 |
| ENSMUSG00 | AC142112.1 | 107 | 0 | 0 | 0 | 0 | 0 | 0 |
| ENSMUSG00 | AC121874.1 | 169 | 2.82287436 | 1.14050751 | 1.5692962 | 2.98904511 | 0 | 0 |
| ENSMUSG00 | Mir3063 | 91 | 0 | 0 | 0 | 0 | 0 | 0 |
| ENSMUSG00 | AC160540.1 | 100 | 0 | 0 | 0 | 0 | 0 | 0 |
| ENSMUSG00 | AC091522.1 | 70 | 0.56793544 | 0.68837775 | 0 | 0.51545778 | 2.18684468 | 0 |
| ENSMUSG00 | Mir5097 | 82 | 0 | 0 | 0 | 0 | 0 | 0 |
| ENSMUSG00 | AC123985.1 | 88 | 0 | 0 | 0 | 0 | 0 | 0 |
| ENSMUSG00 | BX649381.1 | 124 | 0 | 0 | 0 | 0 | 0 | 0 |
| ENSMUSG00 | AC131766.1 | 70 | 0 | 0 | 0 | 0 | 0 | 0 |
| ENSMUSG00 | AL663080.1 | 83 | 0 | 0 | 0 | 0 | 0 | 0 |
| ENSMUSG00 | AC150660.1 | 100 | 0 | 0 | 0 | 0 | 0 | 0 |
| ENSMUSG00 | AC141881.7 | 118 | 0 | 0 | 0 | 0 | 0 | 0 |
| ENSMUSG00 | AL845495.1 | 93 | 0 | 0 | 0 | 0 | 0 | 0 |
| ENSMUSG00 | AC127354.1 | 273 | 0 | 0 | 0 | 0 | 0 | 0 |
| ENSMUSG00 | AC126428.1 | 77 | 0 | 0 | 0 | 0 | 0 | 0 |
| ENSMUSG00 | AC152058.1 | 101 | 0 | 0 | 0 | 0 | 0 | 0 |
| ENSMUSG00 | AC141881.8 | 118 | 0 | 0 | 0 | 0 | 0 | 0 |
| ENSMUSG00 | AC102038.1 | 69 | 0 | 0 | 0 | 0 | 0 | 0 |
| ENSMUSG00 | AL669940.1 | 80 | 0 | 0 | 0 | 0 | 0 | 0 |
| ENSMUSG00 | Vmn2r-ps8 | 896 | 0 | 0 | 0 | 0 | 0 | 0 |
| ENSMUSG00 | Vmn1r23 | 996 | 0 | 0 | 0 | 0 | 0 | 0 |
| ENSMUSG00 | RP24-84K14. | 423 | 0 | 0 | 0 | 0 | 0 | 0 |
| ENSMUSG00 | RP23-466I14. | 791 | 0 | 0 | 0 | 0 | 0 | 0 |
| ENSMUSG00 | Vmn1r31 | 1543 | 0 | 0 | 0 | 0 | 0 | 0 |
| ENSMUSG00 | RP23-120A1 | 486 | 0 | 0 | 0 | 0 | 0 | 0 |
| ENSMUSG00 | RP24-106J5. | 700 | 0 | 0 | 0 | 0 | 0 | 0 |
| ENSMUSG00 | Vmn1r-ps14 | 912 | 0 | 0 | 0 | 0 | 0 | 0 |
| ENSMUSG00 | RP23-263O1 | 1508 | 0 | 0 | 0 | 0 | 0 | 0 |
| ENSMUSG00 | RP24-399B3. | 278 | 0 | 0 | 0 | 0.12979153 | 0.55064434 | 0.09811021 |
| ENSMUSG00 | A330044P14 | 2698 | 0.53046601 | 0.92872313 | 0.54064523 | 0.04012088 | 0 | 0.26283937 |
| ENSMUSG00 | 1700120O09 | 975 | 0 | 0 | 0 | 0 | 0 | 0 |
| ENSMUSG00 | Mir30d | 683 | 0.34924287 | 0.21165348 | 0 | 0.21131505 | 0 | 0 |
| ENSMUSG00 | AC167240.1 | 95 | 0 | 0 | 0 | 0 | 0 | 0 |
| ENSMUSG00 | RP23-349I13. | 723 | 0 | 0 | 0 | 0 | 0 | 0 |
| ENSMUSG00 | RP23-398K14 | 380 | 0 | 0 | 0.11632064 | 0 | 0 | 0 |
| ENSMUSG00 | AC109305.1 | 70 | 3.40761262 | 0 | 0.6314549 | 0 | 1.45789645 | 5.84456555 |
| ENSMUSG00 | RP23-71G16. | 491 | 0 | 0 | 0 | 0 | 0 | 0 |
| ENSMUSG00 | AL591496.1 | 81 | 0 | 0 | 0 | 0 | 0 | 0 |
| ENSMUSG00 | RP23-455J20 | 1834 | 0.23844618 | 0 | 0.26511465 | 0.13771773 | 0.13911226 | 0.02974334 |
| ENSMUSG00 | Vmn2r-ps6 | 1099 | 0 | 0 | 0 | 0 | 0 | 0 |
| ENSMUSG00 | RP23-285E1 | 878 | 0 | 0 | 0 | 0 | 0 | 0 |
| ENSMUSG00 | Gm4014 | 1131 | 0 | 0 | 0 | 0 | 0 | 0 |
| ENSMUSG00 | Vmn1r-ps11 | 852 | 0 | 0 | 0 | 0 | 0 | 0 |
| ENSMUSG00 | AC091333.3 | 59 | 0 | 0 | 0 | 0 | 0 | 0 |
| ENSMUSG00 | RP23-193A2. | 853 | 0.09321332 | 0 | 0 | 0 | 0 | 0 |
| ENSMUSG00 | Vmn2r-ps126 | 1145 | 0 | 0 | 0 | 0 | 0 | 0 |
| ENSMUSG00 | Gm18588 | 491 | 2.59098855 | 1.66836969 | 1.98053064 | 1.17578964 | 0.51961686 | 3.22185148 |
| ENSMUSG00 | AL645645.1 | 59 | 0 | 0 | 0 | 0 | 0 | 0 |
| ENSMUSG00 | AC139376.1 | 94 | 2.53758386 | 0.51262173 | 2.35116186 | 1.91925769 | 0 | 0.58031147 |
| ENSMUSG00 | RP23-181K4. | 378 | 0 | 0 | 0 | 0 | 0 | 0 |
| ENSMUSG00 | Rpl48-ps1 | 555 | 0.71631496 | 0 | 0.3982148 | 1.49529194 | 0 | 1.6708788 |
| ENSMUSG00 | AC114613.1 | 70 | 0 | 0 | 0 | 0 | 0 | 0 |
| ENSMUSG00 | AL671885.2 | 56 | 0 | 0 | 0 | 0 | 0 | 0 |
| ENSMUSG00 | AC084273.1 | 96 | 4.96943506 | 2.00776842 | 4.60435865 | 2.25512778 | 2.65762374 | 2.2728866 |
| ENSMUSG00 | RP23-393E8. | 548 | 0 | 0 | 0 | 0 | 0 | 0 |
| ENSMUSG00 | Vmn1r17 | 979 | 0 | 0 | 0 | 0 | 0 | 0 |
| ENSMUSG00 | RP23-103L13 | 748 | 0 | 0 | 0 | 0 | 0 | 0 |
| ENSMUSG00 | Snora15 | 130 | 0 | 0 | 0 | 0 | 0.78502117 | 0 |
| ENSMUSG00 | RP24-549B2 | 232 | 0 | 0 | 0 | 0 | 0 | 0 |
| ENSMUSG00 | Gm3989 | 1134 | 0 | 0 | 0 | 0 | 0 | 0 |
| ENSMUSG00 | Gm18294 | 1744 | 0 | 0 | 0 | 0 | 0 | 0 |
| ENSMUSG00 | RP23-460G2 | 201 | 0 | 0 | 0 | 0 | 0 | 0 |
| ENSMUSG00 | Vmn1r-ps50 | 373 | 0 | 0 | 0 | 0 | 0 | 0 |
| ENSMUSG00 | Gm19898 | 250 | 0.15902192 | 0 | 0 | 0 | 0 | 0 |
| ENSMUSG00 | Gm4023 | 1136 | 0 | 0 | 0 | 0 | 0 | 0 |
| ENSMUSG00 | RP23-163B8. | 1002 | 0 | 0 | 0 | 0 | 0 | 0 |
| ENSMUSG00 | AC102352.1 | 78 | 0 | 0 | 0 | 0 | 0 | 0 |
| ENSMUSG00 | RP23-113F14 | 883 | 0 | 0 | 0 | 0 | 0 | 0 |
| ENSMUSG00 | RP24-372P1 | 555 | 0.42978898 | 0.26046726 | 0.31857184 | 0.52010154 | 0.64357591 | 0.63886542 |
| ENSMUSG00 | Shox2 | 5584 | 0 | 0 | 0 | 0 | 0 | 0 |
| ENSMUSG00 | Vmn2r-ps123 | 2404 | 0 | 0 | 0 | 0 | 0 | 0 |
| ENSMUSG00 | Gm18513 | 559 | 0 | 0 | 0 | 0 | 0 | 0 |
| ENSMUSG00 | AL954850.1 | 95 | 0 | 0 | 0 | 0 | 0 | 1.43550733 |
| ENSMUSG00 | Vmn1r-ps145 | 423 | 0.28195376 | 0.68349563 | 0 | 2.47370991 | 0 | 0 |
| ENSMUSG00 | Isl2 | 2124 | 0 | 0 | 0 | 0 | 0 | 0 |
| ENSMUSG00 | AL627405.1 | 70 | 0 | 0 | 0 | 0 | 2.18684468 | 0 |
| ENSMUSG00 | AC102096.1 | 69 | 0 | 0 | 0 | 0 | 0 | 0 |
| ENSMUSG00 | Gm18415 | 646 | 0 | 0 | 0 | 0 | 0 | 0 |
| ENSMUSG00 | Vmn2r-ps11 | 3462 | 0 | 0 | 0 | 0 | 0 | 0 |
| ENSMUSG00 | Gm10678 | 1359 | 0 | 0 | 0 | 0 | 0 | 0 |
| ENSMUSG00 | RP23-114H2. | 658 | 1.3292106 | 0.95201178 | 0.87328869 | 1.42573428 | 1.39585831 | 0.78756557 |
| ENSMUSG00 | RP24-262G2 | 459 | 0 | 0 | 0 | 0.15722024 | 0.22233715 | 0 |
| ENSMUSG00 | Gm3685 | 2456 | 0 | 0 | 0 | 0 | 0 | 0 |
| ENSMUSG00 | AC102542.1 | 66 | 0 | 0 | 0 | 0 | 0 | 0 |
| ENSMUSG00 | RP24-312B1 | 437 | 0 | 0 | 0 | 0 | 0 | 0 |
| ENSMUSG00 | AC159892.1 | 68 | 0 | 0 | 0 | 0 | 0 | 0 |
| ENSMUSG00 | Vmn1r25 | 1052 | 0 | 0 | 0 | 0 | 0 | 0 |
| ENSMUSG00 | AC158612.1 | 53 | 0 | 0 | 0 | 0 | 0 | 0 |
| ENSMUSG00 | Vmn1r-ps152 | 698 | 0 | 0 | 0 | 0 | 0 | 0 |
| ENSMUSG00 | Lrch4 | 4190 | 0.28464545 | 0.33350998 | 0.29538224 | 0.68030585 | 0.51148157 | 0.54679468 |
| ENSMUSG00 | RP23-230G1 | 403 | 0 | 0 | 0 | 0 | 0 | 0 |
| ENSMUSG00 | Gm5139 | 248 | 0 | 0.19430017 | 0 | 0 | 0 | 0 |
| ENSMUSG00 | AC119886.1 | 70 | 0 | 0 | 0 | 0 | 0 | 0 |
| ENSMUSG00 | Vmn1r-ps149 | 182 | 0 | 0 | 0 | 0 | 0 | 0 |
| ENSMUSG00 | Six3 | 4505 | 0 | 0 | 0 | 0 | 0 | 0 |
| ENSMUSG00 | Vmn1r24 | 945 | 0 | 0 | 0 | 0 | 0 | 0 |
| ENSMUSG00 | RP23-171A1 | 787 | 0 | 0.12245602 | 0 | 0 | 0.06483656 | 0 |
| ENSMUSG00 | Vmn1r-ps146 | 524 | 0 | 0 | 0 | 0 | 0 | 0 |
| ENSMUSG00 | Vmn1r-ps143 | 603 | 0 | 0 | 0 | 0 | 0 | 0 |
| ENSMUSG00 | AC127411.1 | 277 | 0 | 0 | 0 | 0 | 0 | 0 |
| ENSMUSG00 | RP24-484G1 | 1842 | 0 | 0 | 0 | 0 | 0 | 0 |
| ENSMUSG00 | AC117644.2 | 84 | 0 | 0 | 0 | 0 | 0 | 0 |
| ENSMUSG00 | RP23-89G22. | 510 | 0 | 0 | 0 | 0 | 0 | 0 |
| ENSMUSG00 | 5730480H06 | 1731 | 0.5512025 | 2.0321261 | 1.73640978 | 4.8151082 | 1.56233271 | 0.97690793 |
| ENSMUSG00 | Six3os1 | 10321 | 0 | 0 | 0 | 0 | 0 | 0 |
| ENSMUSG00 | Vmn1r-ps21 | 924 | 0 | 0 | 0 | 0 | 0 | 0 |
| ENSMUSG00 | AC117644.3 | 87 | 0 | 0 | 0 | 0 | 0 | 0 |
| ENSMUSG00 | AC087184.1 | 70 | 0 | 0 | 0 | 0 | 0 | 0 |
| ENSMUSG00 | Gm4897 | 1134 | 0 | 0 | 0 | 0 | 0 | 0 |
| ENSMUSG00 | RP23-127O4. | 3515 | 0.38454803 | 0.86365458 | 1.63477656 | 0.97519039 | 0.116134 | 1.34239334 |
| ENSMUSG00 | AL591496.2 | 81 | 0 | 0 | 0 | 0 | 0 | 0 |
| ENSMUSG00 | RP23-398F7. | 602 | 0 | 0 | 0 | 0 | 0 | 0 |
| ENSMUSG00 | AC105169.1 | 81 | 0 | 0 | 0 | 0 | 0 | 0 |
| ENSMUSG00 | Vmn2r-ps127 | 1027 | 0 | 0 | 0 | 0 | 0 | 0 |
| ENSMUSG00 | RP23-164N1 | 3505 | 0.28356263 | 0.64615201 | 0.41616571 | 0.89561708 | 0.34939601 | 0.00778164 |
| ENSMUSG00 | AC188461.1 | 70 | 0 | 0 | 0 | 0 | 0 | 0 |
| ENSMUSG00 | 4930438E09 | 770 | 0 | 0 | 0 | 0 | 0 | 0 |
| ENSMUSG00 | RP23-117L24 | 167 | 0 | 0 | 0 | 0 | 0 | 0 |
| ENSMUSG00 | Vmn2r-ps12 | 621 | 0 | 0 | 0 | 0 | 0 | 0 |
| ENSMUSG00 | AC121819.1 | 59 | 0 | 0 | 0 | 0 | 0 | 0 |
| ENSMUSG00 | AL589870.2 | 95 | 0 | 1.01445141 | 0 | 0 | 2.68559873 | 0 |
| ENSMUSG00 | Alkbh1 | 2497 | 0 | 0 | 0 | 0 | 0 | 0 |
| ENSMUSG00 | Vmn2r-ps51 | 2456 | 0 | 0 | 0 | 0 | 0 | 0 |
| ENSMUSG00 | RP24-465G1 | 509 | 0 | 0 | 0 | 0 | 0 | 0 |
| ENSMUSG00 | AC131739.1 | 60 | 0 | 0 | 0 | 0 | 0 | 0 |
| ENSMUSG00 | RP23-5H10.4 | 850 | 0 | 0 | 0 | 0 | 0 | 0 |
| ENSMUSG00 | RP23-360N5. | 2883 | 0 | 0 | 0 | 0 | 0 | 0 |
| ENSMUSG00 | RP23-381B1 | 628 | 0.31652453 | 0.84403004 | 12.1766224 | 9.76743244 | 0.8937741 | 29.0987393 |
| ENSMUSG00 | RP23-445D1 | 511 | 0 | 0 | 0 | 0 | 0 | 0 |
| ENSMUSG00 | RP23-268N2 | 1777 | 0 | 0 | 0 | 0 | 0.0574298 | 0 |
| ENSMUSG00 | Lhx8 | 3142 | 0 | 0 | 0 | 0 | 0 | 0 |
| ENSMUSG00 | Vmn1r-ps48 | 895 | 0 | 0 | 0 | 0 | 0 | 0 |
| ENSMUSG00 | AC124379.1 | 70 | 0 | 0 | 0 | 0 | 0 | 0 |
| ENSMUSG00 | RP23-425D1 | 695 | 0 | 0 | 0 | 0 | 0 | 0 |
| ENSMUSG00 | Gm19932 | 314 | 0 | 0 | 0 | 0 | 0 | 0 |
| ENSMUSG00 | Gm3683 | 2507 | 0 | 0 | 0 | 0 | 0 | 0 |
| ENSMUSG00 | RP23-356F8. | 328 | 0 | 0 | 0 | 0 | 0 | 0 |
| ENSMUSG00 | Vmn2r-ps133 | 2563 | 0 | 0 | 0 | 0 | 0 | 0 |
| ENSMUSG00 | Gm3984 | 1137 | 0 | 0 | 0 | 0 | 0 | 0 |
| ENSMUSG00 | AC164703.1 | 95 | 0 | 0 | 0 | 0 | 0 | 0 |
| ENSMUSG00 | Zhx2 | 4415 | 0 | 0 | 0 | 0 | 0 | 0 |
| ENSMUSG00 | RP24-308C1. | 431 | 0.18448019 | 0 | 0 | 0 | 0.11839066 | 0 |
| ENSMUSG00 | AC155932.2 | 59 | 1.34764341 | 0 | 0 | 0 | 0 | 0 |
| ENSMUSG00 | AC158547.1 | 91 | 0 | 0 | 0 | 0 | 0 | 0 |
| ENSMUSG00 | RP23-354F1 | 1703 | 0 | 0 | 0 | 0 | 0 | 0 |
| ENSMUSG00 | Vmn1r22 | 1095 | 0 | 0 | 0 | 0 | 0 | 0 |
| ENSMUSG00 | RP23-14C20. | 2203 | 0 | 0 | 0 | 0 | 0 | 0 |
| ENSMUSG00 | Vmn2r-ps4 | 1012 | 0 | 0 | 0 | 0 | 0 | 0 |
| ENSMUSG00 | Vmn1r-ps24 | 899 | 0 | 0 | 0 | 0 | 0 | 0 |
| ENSMUSG00 | RP24-399B3. | 332 | 0 | 0 | 0 | 0 | 0.1536939 | 0 |
| ENSMUSG00 | RP23-352J18 | 476 | 0 | 0 | 0 | 0 | 0 | 0 |
| ENSMUSG00 | RP24-201C1 | 1159 | 0.10290461 | 0.12472763 | 0 | 0 | 0.57234071 | 0.28239488 |
| ENSMUSG00 | AC163731.1 | 88 | 0 | 0 | 0 | 0 | 0 | 0 |
| ENSMUSG00 | Gm8064 | 1001 | 0 | 0 | 0 | 0 | 0 | 0 |
| ENSMUSG00 | Vmn2r-ps21 | 2942 | 0 | 0 | 0 | 0 | 0 | 0 |
| ENSMUSG00 | RP23-432N1. | 787 | 0 | 0 | 0 | 0 | 0 | 0 |
| ENSMUSG00 | AC159810.1 | 109 | 0 | 0 | 0 | 0 | 0 | 0 |
| ENSMUSG00 | Gm4018 | 1134 | 0 | 0 | 0 | 0 | 0 | 0 |
| ENSMUSG00 | RP23-85E24. | 439 | 0 | 0.43905642 | 0.20137514 | 0.24657434 | 1.04609882 | 0.12425804 |
| ENSMUSG00 | AC016814.1 | 80 | 0 | 0 | 0 | 0 | 0 | 0 |
| ENSMUSG00 | Gm19553 | 492 | 0 | 0 | 0 | 0 | 0 | 0 |
| ENSMUSG00 | Vmn2r-ps131 | 1044 | 0 | 0 | 0 | 0 | 0 | 0 |
| ENSMUSG00 | Vmn1r-ps22 | 899 | 0 | 0 | 0 | 0 | 0 | 0 |
| ENSMUSG00 | Gm20345 | 2408 | 0 | 0 | 0 | 0 | 0 | 0 |
| ENSMUSG00 | Vmn1r-ps9 | 357 | 0 | 0 | 0 | 0 | 0 | 0 |
| ENSMUSG00 | AL772249.1 | 80 | 0 | 0 | 0 | 0 | 0 | 0 |
| ENSMUSG00 | Vmn1r-ps19 | 925 | 0 | 0 | 0 | 0 | 0 | 0 |
| ENSMUSG00 | Vmn1r16 | 1012 | 0 | 0 | 0 | 0 | 0 | 0 |
| ENSMUSG00 | AC110575.2 | 2252 | 0 | 0 | 0 | 0 | 0 | 0 |
| ENSMUSG00 | Gm8974 | 478 | 0.33268185 | 0 | 0.3698899 | 0.07548545 | 0.10674974 | 0 |
| ENSMUSG00 | A930033C23 | 1216 | 0 | 0 | 0 | 0 | 0 | 0 |
| ENSMUSG00 | Gm3998 | 1137 | 0 | 0 | 0 | 0 | 0 | 0 |
| ENSMUSG00 | 4930529F24 | 922 | 0 | 0 | 0 | 0 | 0 | 0 |
| ENSMUSG00 | AC133578.1 | 91 | 0 | 0 | 0 | 0 | 0 | 0 |
| ENSMUSG00 | Vmn1r-ps12 | 501 | 0 | 0 | 0 | 0 | 0 | 0 |
| ENSMUSG00 | Vmn2r-ps117 | 2183 | 0 | 0 | 0 | 0 | 0 | 0 |
| ENSMUSG00 | Gm3994 | 1134 | 0 | 0 | 0 | 0 | 0 | 0 |
| ENSMUSG00 | Vmn2r-ps119 | 2400 | 0 | 0 | 0 | 0 | 0 | 0 |
| ENSMUSG00 | Vmn1r-ps51 | 319 | 0 | 0 | 0 | 0 | 0 | 0 |
| ENSMUSG00 | 4930533I22R | 2557 | 0 | 0 | 0 | 0 | 0 | 0 |
| ENSMUSG00 | AC188461.4 | 3690 | 0.02154769 | 0.01305866 | 0.03593646 | 0.01955666 | 0.02765657 | 0.02956601 |
| ENSMUSG00 | Gm7584 | 939 | 0 | 0 | 0.09414663 | 0 | 0 | 0 |
| ENSMUSG00 | RP23-45N11. | 372 | 0 | 0 | 0 | 0 | 0 | 0 |
| ENSMUSG00 | AL645852.1 | 62 | 0 | 0 | 0 | 0 | 0 | 0 |
| ENSMUSG00 | 4930505M18 | 1167 | 0 | 0 | 0 | 0.03091863 | 0 | 0 |
| ENSMUSG00 | RP24-171M1 | 288 | 0 | 0 | 0 | 0 | 0 | 0 |
| ENSMUSG00 | RP24-292K9. | 335 | 0 | 0 | 0 | 0 | 0 | 0 |
| ENSMUSG00 | Gm8473 | 1884 | 0 | 0 | 0 | 0 | 0 | 0 |
| ENSMUSG00 | RP23-67O10. | 344 | 0 | 0 | 0 | 0 | 0 | 0 |
| ENSMUSG00 | Gm5871 | 717 | 0 | 0 | 0 | 0 | 0 | 0 |
| ENSMUSG00 | Vmn2r-ps7 | 1096 | 0 | 0 | 0 | 0 | 0 | 0 |
| ENSMUSG00 | RP24-247F6. | 368 | 0 | 0 | 0 | 0 | 0 | 0 |
| ENSMUSG00 | Gm6407 | 548 | 0 | 1.05517757 | 0.64528238 | 0.46090203 | 0.27934147 | 0 |
| ENSMUSG00 | RP24-194K24 | 720 | 0 | 0 | 0 | 0 | 0 | 0 |
| ENSMUSG00 | Higd1c | 691 | 0 | 0 | 0 | 0 | 0 | 0 |
| ENSMUSG00 | 4930505G20 | 557 | 0 | 0 | 0 | 0 | 0 | 0 |
| ENSMUSG00 | Gm5777 | 447 | 0 | 0 | 0 | 0 | 0 | 0 |
| ENSMUSG00 | RP24-201C1 | 2830 | 1.41883517 | 0 | 1.21828401 | 1.26223407 | 1.6768385 | 2.59253638 |
| ENSMUSG00 | RP24-370G2 | 723 | 0 | 0 | 0 | 0 | 0 | 0 |
| ENSMUSG00 | AC091333.4 | 59 | 0 | 0 | 0 | 0 | 0 | 0 |
| ENSMUSG00 | RP23-352G2. | 613 | 0 | 0 | 0 | 0 | 0 | 0 |
| ENSMUSG00 | 4930555M17 | 400 | 0 | 0 | 0 | 0 | 0 | 0 |
| ENSMUSG00 | AC102862.2 | 295 | 0 | 0 | 0 | 0 | 0 | 0 |
| ENSMUSG00 | RP23-333F8. | 1919 | 0 | 0.02511018 | 0.11516895 | 0.22563029 | 0.10636035 | 0 |
| ENSMUSG00 | Gm6695 | 1593 | 0 | 0 | 0 | 0 | 0 | 0 |
| ENSMUSG00 | RP23-152I2.3 | 2163 | 0.42273511 | 0.11138799 | 0.04087087 | 0.9008 | 0.07077167 | 0.16392525 |
| ENSMUSG00 | AL807823.1 | 77 | 0 | 0 | 0 | 0 | 0 | 0 |
| ENSMUSG00 | RP24-189B1 | 550 | 0 | 0 | 0 | 0 | 0 | 0 |
| ENSMUSG00 | Gm18396 | 360 | 0 | 0 | 0 | 0 | 0 | 0 |
| ENSMUSG00 | 2610019E17 | 665 | 0.5380441 | 0.14492163 | 0.26587575 | 0.70536328 | 0.46038835 | 1.6405798 |
| ENSMUSG00 | AC101752.1 | 77 | 0 | 0 | 0 | 0 | 0 | 0 |
| ENSMUSG00 | Gm3959 | 713 | 0 | 0 | 0 | 0 | 0 | 0 |
| ENSMUSG00 | RP23-425D1 | 1128 | 0 | 0 | 0 | 0 | 0 | 0 |
| ENSMUSG00 | Vmn2r-ps129 | 2562 | 0 | 0 | 0 | 0 | 0 | 0 |
| ENSMUSG00 | RP23-353M1 | 1028 | 0 | 0 | 0 | 0 | 0 | 0 |
| ENSMUSG00 | AC149067.1 | 370 | 0 | 0 | 0 | 0 | 0 | 0 |
| ENSMUSG00 | AL713994.1 | 95 | 0 | 0 | 0 | 0 | 0 | 0 |
| ENSMUSG00 | RP23-425D1 | 584 | 0 | 0 | 0 | 0 | 0 | 0 |
| ENSMUSG00 | RP23-118L1. | 1719 | 0 | 0 | 0 | 0 | 0 | 0 |
| ENSMUSG00 | RP23-312H1 | 1078 | 0 | 0 | 0 | 0 | 0 | 0 |
| ENSMUSG00 | AC165293.1 | 82 | 0 | 0 | 0 | 0 | 0 | 0 |
| ENSMUSG00 | RP24-201C1 | 2426 | 2.26144118 | 0.57601271 | 2.42326674 | 2.84075454 | 2.33467754 | 1.91124842 |
| ENSMUSG00 | A430110C17 | 4347 | 0 | 0.24386973 | 0 | 0 | 0 | 0 |
| ENSMUSG00 | Gm20036 | 842 | 3.21065638 | 0 | 3.46475254 | 4.67095351 | 0.90902095 | 4.34061955 |
| ENSMUSG00 | RP23-313A2. | 2022 | 0 | 0 | 0 | 0 | 0 | 0 |
| ENSMUSG00 | Gm18006 | 660 | 0 | 0 | 0 | 0 | 0 | 0 |
| ENSMUSG00 | Vmn2r-ps115 | 1805 | 0 | 0 | 0 | 0 | 0 | 0 |
| ENSMUSG00 | AL714025.1 | 86 | 0 | 0 | 0 | 0 | 0 | 0 |
| ENSMUSG00 | Gm8040 | 1001 | 0 | 0 | 0 | 0 | 0 | 0 |
| ENSMUSG00 | Zeb1 | 8983 | 0 | 0 | 0 | 0 | 0 | 0 |
| ENSMUSG00 | AC159323.1 | 70 | 0 | 0 | 0 | 0 | 0 | 0.3896377 |
| ENSMUSG00 | Gm20554 | 10403 | 0.04585848 | 0.01389593 | 0.02549371 | 0 | 0 | 0 |
| ENSMUSG00 | Vmn1r-ps23 | 294 | 0 | 0 | 0 | 0 | 0 | 0 |
| ENSMUSG00 | Gm18367 | 599 | 0.1327395 | 0 | 0 | 0 | 0 | 0 |
| ENSMUSG00 | RP24-553E2 | 320 | 0 | 0 | 0 | 0 | 0 | 0 |
| ENSMUSG00 | Gm3996 | 1137 | 0 | 0 | 0 | 0 | 0 | 0 |
| ENSMUSG00 | Vmn2r-ps122 | 905 | 0 | 0 | 0 | 0 | 0 | 0 |
| ENSMUSG00 | RP23-356J2.1 | 2773 | 0.01433663 | 0 | 0.11158056 | 0.01301192 | 0.01840115 | 0.05901473 |
| ENSMUSG00 | RP23-406N5. | 2512 | 2.0415832 | 2.2443526 | 2.92098166 | 2.12585294 | 3.33133982 | 3.20303287 |
| ENSMUSG00 | Vmn1r-ps49 | 876 | 0 | 0 | 0 | 0 | 0 | 0 |
| ENSMUSG00 | Atxn3 | 5288 | 0 | 0 | 0 | 0 | 0 | 0 |
| ENSMUSG00 | RP23-393F2 | 978 | 0.69104619 | 0.09854078 | 0 | 0.36893706 | 0.31304525 | 0 |
| ENSMUSG00 | A730085K08 | 2010 | 0.11867308 | 0 | 0 | 0 | 0.15231754 | 0.18997261 |
| ENSMUSG00 | AC149056.1 | 55 | 5.78261535 | 1.75223426 | 0.80366987 | 1.96811152 | 5.56651373 | 10.4139532 |
| ENSMUSG00 | AC152922.1 | 70 | 0 | 1.37675549 | 0 | 0 | 0 | 0.77927541 |
| ENSMUSG00 | Gm19459 | 453 | 0 | 0 | 0 | 0 | 0 | 0 |
| ENSMUSG00 | Vmn2r-ps116 | 125 | 0 | 0 | 0 | 0 | 0 | 0 |
| ENSMUSG00 | Snrl2 | 2551 | 0 | 0 | 0 | 0 | 0 | 0 |
| ENSMUSG00 | AC131761.1 | 88 | 0 | 0 | 0.50229367 | 0 | 0 | 0 |
| ENSMUSG00 | RP23-90I17.3 | 538 | 0 | 0 | 0 | 0 | 0 | 0 |
| ENSMUSG00 | B130034C11 | 2573 | 1.96227984 | 1.62931227 | 1.83816448 | 1.52854367 | 2.35994509 | 2.74548448 |
| ENSMUSG00 | Vmn2r-ps118 | 906 | 0 | 0 | 0 | 0 | 0 | 0 |
| ENSMUSG00 | AL954710.2 | 95 | 0 | 0 | 0 | 0 | 0 | 0 |
| ENSMUSG00 | AC122438.1 | 70 | 0 | 0 | 0 | 0 | 2.18684468 | 0 |
| ENSMUSG00 | Cyp2c53-ps | 1951 | 0 | 0 | 0 | 0 | 0 | 0 |
| ENSMUSG00 | Tdh | 1543 | 0 | 0 | 0 | 0 | 0 | 0 |
| ENSMUSG00 | Gm4001 | 1137 | 0 | 0 | 0 | 0 | 0 | 0 |
| ENSMUSG00 | Gm3980 | 1134 | 0 | 0 | 0 | 0 | 0 | 0 |
| ENSMUSG00 | RP23-325I3.4 | 210 | 0 | 0 | 0 | 0 | 0 | 0 |
| ENSMUSG00 | CT030639.1 | 50 | 0 | 0 | 0 | 0 | 0 | 0 |
| ENSMUSG00 | 1700055C04 | 352 | 0 | 0 | 0 | 0 | 0 | 0 |
| ENSMUSG00 | 4933438K21 | 1690 | 0 | 0 | 0 | 0 | 0 | 0 |
| ENSMUSG00 | CT010444.1 | 60 | 0 | 0 | 0 | 0 | 0.8504396 | 0 |
| ENSMUSG00 | 4930535L15 | 532 | 0 | 0 | 0 | 0 | 0 | 0 |
| ENSMUSG00 | CT025759.1 | 2124 | 0 | 0 | 0 | 0 | 0 | 0 |
| ENSMUSG00 | Vmn1r-ps10 | 888 | 0 | 0 | 0 | 0 | 0 | 0 |
| ENSMUSG00 | RP23-349B1 | 383 | 0.20760042 | 0 | 0.11540951 | 0 | 0.13322814 | 1.28183683 |
| ENSMUSG00 | Gm3981 | 1124 | 0 | 0 | 0 | 0 | 0 | 0 |
| ENSMUSG00 | RP23-260N2 | 290 | 0 | 0 | 0 | 0 | 0 | 0 |
| ENSMUSG00 | RP24-113P1 | 297 | 0 | 0 | 0 | 0 | 0 | 0 |
| ENSMUSG00 | RP23-138I21. | 427 | 0 | 0 | 0 | 0 | 0 | 0 |
| ENSMUSG00 | Vmn2r-ps121 | 800 | 0 | 0 | 0 | 0 | 0 | 0 |
| ENSMUSG00 | RP23-397G2 | 1193 | 0.06664791 | 0 | 0 | 0 | 0 | 0 |
| ENSMUSG00 | Gm19357 | 1131 | 0.45695955 | 0 | 0.19541045 | 0.19141668 | 0.13534848 | 1.03696683 |
| ENSMUSG00 | Gm3968 | 723 | 0 | 0 | 0 | 0 | 0 | 0 |
| ENSMUSG00 | Eif2c2 | 840 | 16.091504 | 7.17060151 | 4.10445685 | 6.6150415 | 2.85504722 | 12.208648 |
| ENSMUSG00 | Vmn1r38 | 1035 | 0 | 0 | 0 | 0 | 0 | 0 |
| ENSMUSG00 | CT009530.1 | 81 | 0 | 0 | 0 | 0 | 0 | 0 |
| ENSMUSG00 | Gm10860 | 1883 | 0 | 0 | 0 | 0 | 0 | 0 |
| ENSMUSG00 | Vmn1r11 | 1026 | 0 | 0 | 0 | 0 | 0 | 0 |
| ENSMUSG00 | Gm4003 | 1134 | 0 | 0 | 0 | 0 | 0 | 0 |
| ENSMUSG00 | RP23-459I1.1 | 704 | 0.39529597 | 0.34223325 | 0.37672025 | 0.92255228 | 0.07248065 | 0.07748477 |
| ENSMUSG00 | AC149605.1 | 97 | 0 | 0 | 0 | 2.23187904 | 1.57813534 | 0 |
| ENSMUSG00 | RP23-49P22. | 2208 | 0 | 0 | 0 | 0 | 0 | 0 |
| ENSMUSG00 | AC135509.1 | 109 | 0 | 0 | 0 | 0 | 0 | 0 |
| ENSMUSG00 | RP23-344J14 | 2575 | 0.04631707 | 0 | 0 | 0 | 0 | 0.06355256 |
| ENSMUSG00 | Vmn2r-ps124 | 902 | 0 | 0 | 0 | 0 | 0 | 0 |
| ENSMUSG00 | Gm10679 | 1359 | 0 | 0 | 0 | 0 | 0 | 0 |
| ENSMUSG00 | Gm19560 | 513 | 0 | 0 | 0 | 0 | 0 | 0 |
| ENSMUSG00 | AL833804.1 | 85 | 0 | 0 | 0 | 0 | 0 | 0 |
| ENSMUSG00 | Vmn2r-ps10 | 896 | 0 | 0 | 0 | 0 | 0 | 0 |
| ENSMUSG00 | AC124460.1 | 68 | 0 | 0 | 0 | 0 | 0 | 0 |
| ENSMUSG00 | Vmn1r-ps148 | 454 | 0 | 0 | 0 | 0 | 0 | 0 |
| ENSMUSG00 | RP24-106J5.3 | 627 | 0 | 0 | 0 | 0 | 0 | 0 |
| ENSMUSG00 | RP23-454G7. | 7321 | 0 | 0 | 0 | 0 | 0 | 0 |
| ENSMUSG00 | Gm5873 | 1065 | 0.18664545 | 0.09049097 | 0.04150408 | 0.06775971 | 0 | 0.07682997 |
| ENSMUSG00 | Vmn2r-ps31 | 759 | 0 | 0 | 0 | 0 | 0 | 0 |
| ENSMUSG00 | Vmn2r-ps134 | 1044 | 0 | 0 | 0 | 0 | 0 | 0 |
| ENSMUSG00 | AL845457.2 | 70 | 2.83967718 | 0.68837775 | 1.8943647 | 1.03091556 | 2.18684468 | 0.77927541 |
| ENSMUSG00 | AC117644.4 | 87 | 0 | 0.55386715 | 0 | 0 | 0 | 0 |
| ENSMUSG00 | RP24-201C1 | 3214 | 0.04947789 | 0.10494869 | 0.23379942 | 0.04490609 | 0.06350514 | 0 |
| ENSMUSG00 | AC137678.1 | 124 | 0 | 0 | 0 | 0 | 0 | 0 |
| ENSMUSG00 | RP23-120A1 | 1370 | 0 | 0 | 0.38716943 | 0 | 0 | 0 |
| ENSMUSG00 | Irx2 | 3137 | 0 | 0 | 0 | 0 | 0 | 0 |
| ENSMUSG00 | AL929450.1 | 165 | 0 | 0 | 0 | 0 | 0 | 0 |
| ENSMUSG00 | Eif4e3 | 2508 | 9.74865252 | 5.43730587 | 10.1516195 | 7.55305956 | 9.88788622 | 11.6145593 |
| ENSMUSG00 | RP23-342N1 | 2736 | 0 | 0 | 0 | 0 | 0 | 0 |
| ENSMUSG00 | Gm7171 | 1346 | 0 | 0 | 0 | 0 | 0 | 0 |
| ENSMUSG00 | RP24-370G2 | 216 | 0 | 0 | 0 | 0 | 0 | 0 |
| ENSMUSG00 | Gm8057 | 991 | 0 | 0 | 0 | 0 | 0 | 0 |
| ENSMUSG00 | Vmn1r-ps15 | 909 | 0 | 0 | 0 | 0 | 0 | 0 |
| ENSMUSG00 | AC130680.1 | 70 | 0 | 3.44188873 | 0 | 3.60820445 | 0.72894823 | 3.11710163 |
| ENSMUSG00 | Pou5f2 | 3383 | 1.53945254 | 0.74067248 | 0.19598807 | 0.90658403 | 0.54299425 | 1.53183017 |
| ENSMUSG00 | Gm9399 | 638 | 0 | 0 | 0 | 0 | 0 | 0 |
| ENSMUSG00 | AL805897.1 | 95 | 0 | 0 | 0 | 0 | 0 | 0 |
| ENSMUSG00 | RP23-445D1 | 514 | 0 | 0 | 0 | 0 | 0 | 0 |
| ENSMUSG00 | RP24-309I5.3 | 470 | 0 | 0 | 0 | 0.38385154 | 0 | 0 |
| ENSMUSG00 | RP23-333M1 | 2062 | 0.01928006 | 0 | 0 | 0 | 0 | 0.03968182 |
| ENSMUSG00 | Rpl41 | 1456 | 5.16056718 | 6.48663644 | 6.314549 | 5.55108377 | 3.71483231 | 9.34755837 |
| ENSMUSG00 | RP24-129K7. | 955 | 0 | 0 | 0 | 0 | 0 | 0 |
| ENSMUSG00 | AC133950.1 | 87 | 0 | 0 | 0 | 0 | 0 | 0 |
| ENSMUSG00 | RP23-102L5.1 | 1490 | 1.38743959 | 0.87317714 | 0.65264466 | 0.67805184 | 1.09586847 | 2.05017423 |
| ENSMUSG00 | RP24-370G2 | 402 | 0 | 0 | 0 | 0 | 0 | 0 |
| ENSMUSG00 | Vmn1r20 | 1051 | 0 | 0 | 0 | 0 | 0 | 0 |
| ENSMUSG00 | AL591496.3 | 81 | 0 | 0 | 0 | 0 | 0 | 0 |
| ENSMUSG00 | Vmn2r-ps52 | 611 | 0 | 0 | 0 | 0 | 0 | 0 |
| ENSMUSG00 | RP23-354I24. | 500 | 0 | 0.19274577 | 0.35361474 | 0.36082045 | 0.30615826 | 1.03643629 |
| ENSMUSG00 | RP24-370G2 | 366 | 0 | 0 | 0 | 0 | 0 | 0 |
| ENSMUSG00 | AC170999.1 | 73 | 0 | 0 | 0 | 0 | 0 | 0 |
| ENSMUSG00 | Gm18416 | 1001 | 0 | 0 | 0 | 0 | 0 | 0 |
| ENSMUSG00 | Gm4705 | 353 | 0 | 0 | 0 | 0 | 0 | 0 |
| ENSMUSG00 | RP23-260N2 | 461 | 0 | 0 | 0 | 0 | 0 | 0 |
| ENSMUSG00 | Gm8488 | 1278 | 0 | 0 | 0 | 0 | 0 | 0 |
| ENSMUSG00 | Vmn1r-ps52 | 861 | 0 | 0 | 0 | 0 | 0 | 0 |
| ENSMUSG00 | AL714024.1 | 60 | 0 | 0 | 0 | 0 | 0 | 0.45457732 |
| ENSMUSG00 | Ppp1r3e | 1145 | 0 | 0 | 0 | 0 | 0 | 0 |
| ENSMUSG00 | Vmn1r14 | 1528 | 0 | 0 | 0 | 0 | 0 | 0 |
| ENSMUSG00 | RP24-91B1.5 | 812 | 0 | 0 | 0 | 0 | 0 | 0 |
| ENSMUSG00 | Vmn2r-ps34 | 903 | 0 | 0 | 0 | 0 | 0 | 0 |
| ENSMUSG00 | RP23-133P6. | 528 | 0 | 0 | 0 | 0 | 0 | 0 |
| ENSMUSG00 | Vmn1r7 | 1218 | 0 | 0 | 0 | 0 | 0 | 0 |
| ENSMUSG00 | RP24-302G1 | 533 | 0 | 0 | 0 | 0 | 0 | 0 |
| ENSMUSG00 | AC125145.1 | 53 | 0 | 0 | 0 | 0 | 0 | 0 |
| ENSMUSG00 | Vmn2r-ps9 | 1097 | 0 | 0 | 0 | 0 | 0 | 0 |
| ENSMUSG00 | AL663090.1 | 109 | 0 | 0 | 0 | 0 | 0 | 0 |
| ENSMUSG00 | RP23-190G1 | 878 | 0 | 0 | 0 | 0 | 0 | 0 |

| | | | | | | | |
|---|---|---|---|---|---|---|---|
| ENSMUSG00 Vmn1r-ps20 | 914 | 0 | 0 | 0 | 0 | 0 | 0 |
| ENSMUSG00 Gm18397 | 586 | 0 | 0 | 0 | 0 | 0 | 0 |
| ENSMUSG00 AL691456.1 | 70 | 0 | 0 | 0 | 0 | 0 | 0 |
| ENSMUSG00 Vmn1r-ps142 | 851 | 0 | 0 | 0 | 0 | 0 | 0 |
| ENSMUSG00 AL591496.4 | 81 | 0 | 0 | 0 | 0 | 0 | 0 |
| ENSMUSG00 Gm3965 | 1133 | 0 | 0 | 0 | 0 | 0 | 0 |
| ENSMUSG00 RP24-91B1.4 | 258 | 0 | 0 | 0 | 0 | 0 | 0 |
| ENSMUSG00 RP23-278M1 | 2430 | 0 | 0 | 0 | 0 | 0 | 0 |
| ENSMUSG00 AC105298.1 | 79 | 0 | 0 | 0 | 0 | 0 | 0 |
| ENSMUSG00 Vmn2r-ps125 | 930 | 0 | 0 | 0 | 0 | 0 | 0 |
| ENSMUSG00 AC162914.1 | 70 | 0 | 0 | 0 | 0 | 0 | 0 |
| ENSMUSG00 Vmn1r235 | 986 | 0 | 0 | 0 | 0 | 0 | 0 |
| ENSMUSG00 Vmn2r-ps132 | 650 | 0 | 0 | 0 | 0 | 0 | 0 |
| ENSMUSG00 AC150683.1 | 68 | 0 | 0 | 0 | 0 | 0 | 0 |
| ENSMUSG00 Gm19815 | 334 | 0 | 0 | 0 | 0.43212029 | 0 | 0.89826656 |
| ENSMUSG00 RP23-345J21 | 1607 | 0 | 0 | 0 | 0 | 0 | 0 |
| ENSMUSG00 RP23-322O2 | 1529 | 0 | 0 | 0 | 0 | 0 | 0 |
| ENSMUSG00 Gm4016 | 947 | 0 | 0 | 0 | 0 | 0 | 0 |
| ENSMUSG00 RP23-372N1 | 1285 | 0.21656682 | 0 | 0.20638993 | 0.02807941 | 0 | 0.55186041 |
| ENSMUSG00 Gm3896 | 4092 | 0 | 0 | 0 | 0 | 0 | 0 |
| ENSMUSG00 RP24-166B2. | 235 | 0 | 0 | 0 | 0 | 0 | 0 |
| ENSMUSG00 Gm3987 | 1134 | 0 | 0 | 0 | 0 | 0 | 0 |
| ENSMUSG00 Vmn2r-ps32 | 756 | 0 | 0 | 0 | 0 | 0 | 0 |
| ENSMUSG00 Gm18595 | 1066 | 0 | 0 | 0 | 0 | 0 | 0 |
| ENSMUSG00 RP23-358B2 | 1929 | 0 | 0 | 0 | 0 | 0.10580897 | 0 |
| ENSMUSG00 Gm8053 | 1000 | 0 | 0 | 0 | 0 | 0 | 0 |
| ENSMUSG00 Vmn1r-ps8 | 837 | 0 | 0 | 0 | 0 | 0 | 0 |
| ENSMUSG00 Vmn1r-ps141 | 752 | 0 | 0 | 0 | 0 | 0 | 0 |
| ENSMUSG00 RP24-399B3. | 314 | 0 | 0 | 0 | 0 | 0 | 0 |
| ENSMUSG00 Vmn1r-ps18 | 874 | 0 | 0 | 0 | 0 | 0 | 0 |
| ENSMUSG00 RP23-60H16. | 1523 | 0 | 0 | 0 | 0 | 0 | 0 |
| ENSMUSG00 AL663090.2 | 109 | 0 | 0 | 0 | 0 | 0 | 0 |
| ENSMUSG00 AL645907.1 | 254 | 1.25214112 | 2.08681442 | 0.87011502 | 2.84110587 | 7.83475849 | 2.36237033 |
| ENSMUSG00 Vmn2r-ps33 | 903 | 0 | 0 | 0 | 0 | 0 | 0 |
| ENSMUSG00 RP23-349I13. | 723 | 0 | 0 | 0 | 0 | 0 | 0 |
| ENSMUSG00 RP24-356D2 | 663 | 0 | 0 | 0 | 0 | 0 | 0 |
| ENSMUSG00 AI606473 | 1728 | 0 | 0 | 0 | 0 | 0 | 0 |
| ENSMUSG00 RP23-329D1 | 679 | 0.58550045 | 0 | 0.19529533 | 0 | 2.17932975 | 0 |
| ENSMUSG00 Gm18417 | 647 | 0 | 0 | 0 | 0 | 0 | 0 |
| ENSMUSG00 RP24-97C18. | 260 | 0 | 0 | 0 | 0 | 0 | 0 |
| ENSMUSG00 Gm3977 | 1359 | 0 | 0 | 0 | 0 | 0 | 0 |
| ENSMUSG00 Vmn1r-ps17 | 893 | 0 | 0 | 0 | 0 | 0 | 0 |
| ENSMUSG00 RP23-177N1 | 336 | 0 | 0 | 0 | 0 | 0 | 0 |
| ENSMUSG00 AC136903.1 | 58 | 0 | 0 | 0 | 0 | 0 | 0 |
| ENSMUSG00 AL833801.1 | 59 | 0 | 0 | 0 | 0 | 0 | 0 |
| ENSMUSG00 RP24-97C18. | 429 | 0 | 0 | 0 | 0 | 0 | 0 |
| ENSMUSG00 Gm19031 | 1364 | 0 | 0 | 0 | 0 | 0 | 0 |
| ENSMUSG00 4921523L03F | 1184 | 0 | 0 | 0 | 0 | 0 | 0 |
| ENSMUSG00 AC139849.1 | 1617 | 0 | 0 | 0 | 0 | 0 | 0 |
| ENSMUSG00 Gm3307 | 631 | 0 | 0.0763652 | 0 | 0 | 0 | 0 |
| ENSMUSG00 RP23-391M1 | 5276 | 0 | 0 | 0 | 0 | 0 | 0 |
| ENSMUSG00 RP23-309A11 | 738 | 0 | 0 | 0 | 0 | 0 | 0 |
| ENSMUSG00 Vmn2r-ps130 | 2562 | 0 | 0 | 0 | 0 | 0 | 0 |
| ENSMUSG00 Vmn1r39 | 1231 | 0 | 0 | 0 | 0 | 0 | 0 |
| ENSMUSG00 Gm18594 | 1402 | 0 | 0 | 0 | 0 | 0 | 0 |
| ENSMUSG00 AC140469.1 | 69 | 0 | 0 | 0 | 0 | 0 | 0 |
| ENSMUSG00 RP24-262G2 | 346 | 0 | 0 | 0 | 0 | 0 | 0 |
| ENSMUSG00 2600006L11F | 302 | 0 | 0 | 0 | 0 | 0 | 0 |
| ENSMUSG00 RP23-152I2.1 | 1049 | 0 | 0 | 0 | 0 | 0.14592863 | 0 |
| ENSMUSG00 Gm5496 | 1041 | 0 | 0 | 0 | 0 | 0 | 0 |
| ENSMUSG00 AC147634.1 | 93 | 0 | 0 | 0 | 0 | 0 | 0 |
| ENSMUSG00 AC157086.1 | 59 | 0 | 0 | 0 | 0 | 0 | 0 |
| ENSMUSG00 Vmn1r36 | 1028 | 0 | 0 | 0 | 0 | 0 | 0 |
| ENSMUSG00 RP24-386J17 | 3151 | 0 | 0 | 0 | 0 | 0 | 0 |
| ENSMUSG00 RP24-347O5. | 1398 | 0 | 0 | 0 | 0 | 0 | 0 |
| ENSMUSG00 Vmn2r-ps13 | 605 | 0 | 0 | 0 | 0 | 0 | 0 |
| ENSMUSG00 RP23-142A1 | 196 | 0 | 0 | 0 | 0 | 0 | 0 |
| ENSMUSG00 Hist2h3c1 | 2364 | 0 | 0 | 0 | 0 | 0 | 0 |
| ENSMUSG00 Gm8072 | 1002 | 0 | 0 | 0 | 0 | 0 | 0 |
| ENSMUSG00 AU023070 | 385 | 0 | 0 | 0 | 0 | 0 | 0 |
| ENSMUSG00 4931403E22F | 664 | 0.17961814 | 0 | 0.26827616 | 0.05434043 | 0.07684695 | 0 |
| ENSMUSG00 Gm18355 | 408 | 0 | 0 | 0 | 0 | 0 | 0 |
| ENSMUSG00 Vmn1r-ps151 | 981 | 0 | 0 | 0 | 0 | 0 | 0 |
| ENSMUSG00 Vmn2r-ps114 | 1279 | 0 | 0 | 0 | 0 | 0 | 0 |
| ENSMUSG00 Vmn2r-ps120 | 2403 | 0 | 0 | 0 | 0 | 0 | 0 |
| ENSMUSG00 AC130205.1 | 68 | 0 | 0 | 0 | 0 | 0 | 0 |
| ENSMUSG00 RP23-71K9.1 | 1628 | 1.09889228 | 2.10149717 | 1.60190954 | 1.90605395 | 1.66117808 | 1.77586718 |
| ENSMUSG00 RP24-63G9.3 | 327 | 0 | 0 | 0 | 0 | 0 | 0 |
| ENSMUSG00 AC132271.2 | 95 | 0 | 0 | 0 | 0 | 0 | 0 |
| ENSMUSG00 Vmn1r-ps47 | 906 | 0 | 0 | 0 | 0 | 0 | 0 |
| ENSMUSG00 AC138214.1 | 52 | 0 | 0 | 0 | 0 | 0 | 0 |
| ENSMUSG00 AC132307.1 | 70 | 0 | 0 | 0 | 0 | 0 | 0 |
| ENSMUSG00 AC158612.2 | 75 | 1.06014615 | 0 | 0 | 0.48109393 | 0 | 0 |
| ENSMUSG00 Vmn1r10 | 971 | 0 | 0 | 0 | 0 | 0 | 0 |
| ENSMUSG00 AL591496.5 | 81 | 0 | 0 | 0 | 0 | 0 | 0 |
| ENSMUSG00 RP23-286K8. | 147 | 2.43400901 | 4.26138604 | 2.10484967 | 0.24545609 | 0.69423641 | 0 |
| ENSMUSG00 Vmn1r-ps147 | 512 | 0 | 0 | 0 | 0 | 0 | 0 |
| ENSMUSG00 RP24-73K9.3 | 1109 | 0 | 0 | 0 | 0 | 0 | 0 |

Suppl. Table 4. Upregulated genes in ICs after Treg depletion (FC>2, p<005)

| ID | Name | FC: [B1Foxp3-DTR / B1WT] | P-value: B1Foxp3-DTR / B1W |
|---|---|---|---|
| **ENSMUSG00000090754** | C230091D08Rik | 70.7652638 | 0.008142127 |
| **ENSMUSG00000065200** | SNORA65 | 20.53744846 | 3.59500036069428E-04 |
| **ENSMUSG00000064967** | U1 | 18.82317073 | 0.001495821 |
| **ENSMUSG00000077557** | SNORD86 | 17.57244833 | 2.71843108460866E-05 |
| **ENSMUSG00000090911** | Gm17706 | 17.38397491 | 0.041030272 |
| **ENSMUSG00000079361** | Gm11102 | 16.38127189 | 0.014160369 |
| **ENSMUSG00000089437** | U6 | 15.60261161 | 0.001542784 |
| **ENSMUSG00000065817** | SNORA3 | 14.57138924 | 0.001517763 |
| **ENSMUSG00000065701** | Y_RNA | 14.05580077 | 2.0351602833862E-04 |
| **ENSMUSG00000092954** | AL808021.1 | 13.14879906 | 0.021601162 |
| **ENSMUSG00000088475** | 5S_rRNA | 12.71498572 | 0.001311684 |
| **ENSMUSG00000070167** | SNORA57 | 12.67941127 | 0.004551907 |
| **ENSMUSG00000091390** | Gm17168 | 11.33482251 | 0.00105742 |
| **ENSMUSG00000064952** | SNORD62 | 11.06455507 | 1.29100455839877E-05 |
| **ENSMUSG00000065431** | Mir186 | 10.92228985 | 0.034084232 |
| **ENSMUSG00000065374** | U1 | 10.70413064 | 0.003756012 |
| **ENSMUSG00000038642** | Ctss | 9.643240737 | 0.005309522 |
| **ENSMUSG00000064602** | Snora41 | 9.488576248 | 0.023977911 |
| **ENSMUSG00000034459** | Ifit1 | 7.167671856 | 0.045123216 |
| **ENSMUSG00000088171** | U6 | 6.973684851 | 0.045458602 |
| **ENSMUSG00000091992** | Gm17458 | 6.657150742 | 0.010092041 |
| **ENSMUSG00000041827** | Oasl1 | 5.73423128 | 0.0181689 |
| **ENSMUSG00000024810** | Il33 | 5.360673425 | 0.011603432 |
| **ENSMUSG00000092354** | Gm20548 | 5.142835401 | 0.004688602 |
| **ENSMUSG00000072974** | Gm4787 | 5.089014313 | 0.017499702 |
| **ENSMUSG00000028268** | Gbp3 | 4.932727084 | 0.011566234 |
| **ENSMUSG00000081137** | BC022960 | 4.227283917 | 0.004059146 |
| **ENSMUSG00000030604** | Zfp626 | 4.194691663 | 3.31740190235084E-04 |
| **ENSMUSG00000040253** | Gbp7 | 4.181107687 | 0.04103462 |
| **ENSMUSG00000078920** | Ifi47 | 4.097940382 | 0.022155636 |
| **ENSMUSG00000056155** | Nanos3 | 3.798319297 | 0.039449599 |
| **ENSMUSG00000026989** | Dapl1 | 3.773666834 | 0.026357574 |
| **ENSMUSG00000064514** | SNORD18 | 3.519946933 | 0.029813147 |
| **ENSMUSG00000069874** | Irgm2 | 3.329939451 | 0.046484544 |
| **ENSMUSG00000084899** | Gm15344 | 3.215386779 | 0.037162026 |
| **ENSMUSG00000019850** | Tnfaip3 | 3.110228086 | 0.017189198 |
| **ENSMUSG00000082455** | Gm13665 | 3.06891233 | 0.028450877 |
| **ENSMUSG00000088252** | Snord13 | 3.043102985 | 0.036815164 |
| **ENSMUSG00000077649** | SNORD86 | 2.873879283 | 0.003271857 |
| **ENSMUSG00000089900** | 0610038L08Rik | 2.871704536 | 0.014010572 |
| **ENSMUSG00000085501** | Gm11772 | 2.861885112 | 0.012303995 |
| **ENSMUSG00000064936** | SNORA33 | 2.783059405 | 0.047002834 |
| **ENSMUSG00000064927** | U1 | 2.761610127 | 0.041285893 |
| **ENSMUSG00000086465** | Gm16049 | 2.751401924 | 0.043208112 |
| **ENSMUSG00000031323** | Dmrtc1a | 2.724858559 | 0.048672124 |
| **ENSMUSG00000074608** | Zfp850 | 2.629095846 | 0.010807136 |
| **ENSMUSG00000026121** | Sema4c | 2.530089367 | 0.016525802 |
| **ENSMUSG00000073366** | Gm10492 | 2.399110318 | 0.005381287 |
| **ENSMUSG00000039167** | Eltd1 | 2.373498908 | 0.040819151 |
| **ENSMUSG00000040428** | Plekha4 | 2.335494156 | 0.03491477 |
| **ENSMUSG00000054181** | A930012O16Rik | 2.298122306 | 0.016333991 |
| **ENSMUSG00000064647** | SNORD58 | 2.287877937 | 0.049414138 |
| **ENSMUSG00000073177** | Gm773 | 2.284145789 | 0.018017241 |
| **ENSMUSG00000087252** | Gm14379 | 2.263633087 | 0.0402788 |
| **ENSMUSG00000057596** | Trim30d | 2.250843768 | 0.042235678 |
| **ENSMUSG00000025395** | Prim1 | 2.250019872 | 0.013360671 |
| **ENSMUSG00000037259** | 6330439K17Rik | 2.215631359 | 0.018981505 |
| **ENSMUSG00000053070** | 9230110C19Rik | 2.189517563 | 0.003330091 |
| **ENSMUSG00000021068** | Nin | 2.175598594 | 0.030074055 |
| **ENSMUSG00000070883** | 4930578N16Rik | 2.13840733 | 0.033334607 |
| **ENSMUSG00000063047** | Zfp780b | 2.134701552 | 0.03099 |
| **ENSMUSG00000091515** | Zfp422-rs1 | 2.119897517 | 0.049752983 |
| **ENSMUSG00000026980** | Ly75 | 2.113637449 | 0.02799894 |
| **ENSMUSG00000026611** | Spata17 | 2.108916259 | 0.009192125 |
| **ENSMUSG00000091270** | Gm17642 | 2.068668459 | 0.014056583 |
| **ENSMUSG00000087938** | U6 | 2.059764641 | 0.02569841 |
| **ENSMUSG00000026814** | Eng | 2.048768766 | 0.012673181 |
| **ENSMUSG00000091395** | Gm6297 | 2.037746775 | 0.003982654 |
| **ENSMUSG00000087385** | AA415398 | 2.019073829 | 0.048951696 |
| **ENSMUSG00000027811** | 4930579G24Rik | 2.0149098 | 0.047910456 |
| **ENSMUSG00000005980** | Dnase1 | 2.013023436 | 0.019762035 |

Suppl. Table 5 downregulated genes in ICs after Treg depletion (FC<0.5, p<0.05)

| ID | Name | FC: [B1Foxp3-DTR / B1WT] | P-value: B1Foxp3-DTR / B1WT |
|---|---|---|---|
| ENSMUSG00000087790 | SNORD127 | 0.03550028 | 4.11688425961459E-04 |
| ENSMUSG00000065623 | U6 | 0.039946403 | 0.007317081 |
| ENSMUSG00000088743 | 7SK | 0.061547586 | 0.008936245 |
| ENSMUSG00000037852 | Cpe | 0.070910089 | 0.025018526 |
| ENSMUSG00000052616 | Ccdc79 | 0.087533737 | 9.81073226328484E-04 |
| ENSMUSG00000077440 | SNORD101 | 0.094327358 | 0.022541107 |
| ENSMUSG00000055691 | Gja6 | 0.096147382 | 1.02227208456009E-04 |
| ENSMUSG00000053337 | Gm9903 | 0.103104033 | 0.005891144 |
| ENSMUSG00000093787 | RP23-286K8.2 | 0.104034406 | 0.023740328 |
| ENSMUSG00000080410 | AC084287.1 | 0.109391773 | 0.039214591 |
| ENSMUSG00000025359 | Pmel | 0.110055642 | 0.004630337 |
| ENSMUSG00000064525 | SNORA65 | 0.110453679 | 0.006951702 |
| ENSMUSG00000051037 | Zfp455 | 0.131630842 | 0.026735015 |
| ENSMUSG00000025666 | Tmem47 | 0.151912073 | 0.039588657 |
| ENSMUSG00000015134 | Aldh1a3 | 0.165349235 | 0.049189796 |
| ENSMUSG00000005220 | Corin | 0.177978227 | 0.035156974 |
| ENSMUSG00000065226 | SNORA8 | 0.217922402 | 0.049273728 |
| ENSMUSG00000059187 | Fam19a1 | 0.250308648 | 0.037837324 |
| ENSMUSG00000074171 | Gm6658 | 0.26823019 | 0.011318246 |
| ENSMUSG00000027386 | Fbln7 | 0.295265396 | 0.017312021 |
| ENSMUSG00000024030 | Abcg1 | 0.30404678 | 0.006286639 |
| ENSMUSG00000007908 | Hmgcll1 | 0.327230269 | 0.038622181 |
| ENSMUSG00000022193 | Psmb5 | 0.327792657 | 0.019269724 |
| ENSMUSG00000055313 | Pgbd1 | 0.337662695 | 0.016344145 |
| ENSMUSG00000045822 | Zswim3 | 0.345399731 | 0.032668972 |
| ENSMUSG00000080465 | SNORD94 | 0.356790125 | 0.012038169 |
| ENSMUSG00000080665 | Mir1199 | 0.360946513 | 0.007135555 |
| ENSMUSG00000073016 | Uprt | 0.366571973 | 0.020293715 |
| ENSMUSG00000020930 | Ccdc103 | 0.368512458 | 0.002628392 |
| ENSMUSG00000056069 | Fam105a | 0.377765392 | 0.025917782 |
| ENSMUSG00000027487 | Cdk5rap1 | 0.379814535 | 0.034402535 |
| ENSMUSG00000039633 | Lonrf1 | 0.391469461 | 0.016856825 |
| ENSMUSG00000041939 | Mvk | 0.393718991 | 0.018185132 |
| ENSMUSG00000002781 | Tmem143 | 0.397630546 | 0.035852562 |
| ENSMUSG00000029314 | Agpat9 | 0.403940349 | 0.003646606 |
| ENSMUSG00000054000 | Tusc1 | 0.406794893 | 0.024515996 |
| ENSMUSG00000022235 | Cmbl | 0.416874413 | 0.023334583 |
| ENSMUSG00000007670 | Khsrp | 0.429713798 | 0.041426671 |
| ENSMUSG00000026315 | Serpinb8 | 0.450750363 | 0.034736727 |
| ENSMUSG00000029727 | Cyp3a13 | 0.471686336 | 0.028838591 |
| ENSMUSG00000039206 | Daglb | 0.482305335 | 0.043785608 |
